# Lysine-specific demethylase 1a is obligatory for gene regulation during kidney development

**DOI:** 10.1101/2025.02.25.640014

**Authors:** Savithri Balasubramanian Kota, Satya K. Kota

**Author notes:** Address for Correspondence: Satya K. Kota Harvard School of Dental Medicine 188 Longwood Ave, REB Room 314 Boston, MA-02215, USA.

## Abstract

Histone methyltransferases and demethylases play crucial roles in gene regulation and are vital for proper functioning of multiple tissues. Lysine-specific histone demethylase 1A (Kdm1a), is responsible for the demethylation of specific lysines, namely K4 and K9, on histone H3. In this study, we investigated the functions of Kdm1a during mouse kidney development upon targeted deletion in renal progenitor cells. Loss of *Kdm1*a in Six2-positive nephron progenitors resulted in significant reduction in renal mass, tissue structural changes and impaired function. To further understand the molecular function of Kdm1a during kidney development, we conducted multi-omics analyses that included transcriptome profiling, Chromatin immunoprecipitation (ChIP) sequencing, and methylome assessments. These omic analyses identified Kdm1a as a critical gene regulator required for sustained expression of several nephron segment marker genes, as well as vast number of solute carrier (Slc) genes and a few imprinted genes. Absence of Kdm1a in kidneys led to an increase in global H3K9 methylation peaks, which correlated with the transcriptional downregulation of numerous genes. Among these were markers of nephron progenitors and presumptive tubular precursors. We also observed that specific gene bodies exhibited altered DNA methylation patterns at intragenic differentially methylated regions (DMRs) upon Kdm1a deletion, while the overall global levels of DNA methylation remained unchanged. Our data point to a key regulatory role for Kdm1a in the renal progenitor epigenome, influencing kidney specific gene expression in the developing nephrons. Together the study highlights an indispensable role for Kdm1a for proper development of mouse kidneys, and its absence leading to significant developmental and functional impairment.

## Introduction

Histone tails undergo post-translational modifications that establish tissue and cell type-specific histone codes, forming one of the core gene regulatory mechanisms in eukaryotes during both development and disease^1,2^. Histone codes, covalent modifications on DNA, non-coding RNAs along with transcription factors confer spatio-temporal gene regulation crucial for tissue-specific gene expression programs during development^3^. Various enzymatic complexes coordinate the establishment and maintenance of histone post-translational modifications. The discovery of lysine-specific demethylases has expanded our understanding of the gene regulatory messages embedded within histone codes, particularly those required for gene expression during early embryonic development. Kdm1a (Lysine-specific histone demethylase 1a, also known as Lysine-specific demethylase 1 (Lsd1)), a pioneering member of histone demethylases, targets histone H3 at lysine residues K4 and K9^4–6^. Kdm1a plays crucial roles in both gene repression and activation in normal and malignant cells^7–12^. Kdm1a demethylation function is required for gene regulation, retroviral silencing, erasure of transcriptional memory and/or enhancer decommissioning during stem cell differentiation in multiple stem cell lineages including embryonic stem cells^13–18^. Furthermore, Kdm1a is essential for demethylation of several non-histone proteins^19^ including DNA methyl transferase, Dnmt1 and its stability through lysine demethylation dependent and independent functions, thereby regulating global DNA methylation levels^20,21^. In mammalian embryonic development, Kdm1a is indispensable. Loss of Kdm1a activity during early mouse embryonic development led to lethality, affecting gastrulation with growth arrest and resorption of homozygous knockout embryos by 7.5 days post coitum (dpc)^20,22^. Conditional deletion of Kdm1a in various somatic cell lineages resulted in developmental phenotypes in multiple tissues, including heart and brain^23,24^. The role of *Kdm1a* in kidney development remains largely unexplored. In this study, we investigated the expression dynamics and molecular gene regulatory functions of Kdm1a in embryonic kidneys. We specifically targeted Kdm1a in the nephron progenitor cells of the developing mouse kidneys during early renal morphogenesis. We analyzed kidney phenotypes as well as molecular changes including genome-wide gene expression, histone H3K9 trimethylation, and DNA methylation in the Kdm1a conditional knockout and littermate control kidneys to elucidate essential gene regulatory mechanisms mediated by Lysine-specific histone demethylase 1a during kidney development.

### Kdm1a expression dynamics and cell type abundance in developing kidney

*Kdm1a* is known to be expressed in a variety of tissues throughout development, and to delve deeper into its role in kidney development, we examined the expression dynamics of Kdm1a using poly A+ RNA sequencing datasets^25,26^ derived from embryonic mouse kidneys at different developmental stages: E14.5, E15.5, E16.5, and postnatal day zero (P0). Our analysis revealed *Kdm1a* expression across these embryonic and neonatal stages (Fig 1A). Notably, the highest levels of Kdm1a transcripts per million (TPM) were detected at E14.5, (Fig 1B) indicating its expression during early phases of nephrogenesis. In addition, we also evaluated the expression of *Kdm1b* (Lysine-specific histone demethylase 1b, also known as Lsd2), a homolog of *Kdm1a* that demethylates histone H3 K4/K9^17,27,28^ and is highly expressed in oocytes. In the embryonic kidneys, *Kdm1b* expression was detected in the same developmental time points (Suppl Fig 1A). However, Kdm1a kidney expression was 50% or more across all embryonic and neonatal stages (Suppl Fig 1B).

**Figure 1:**
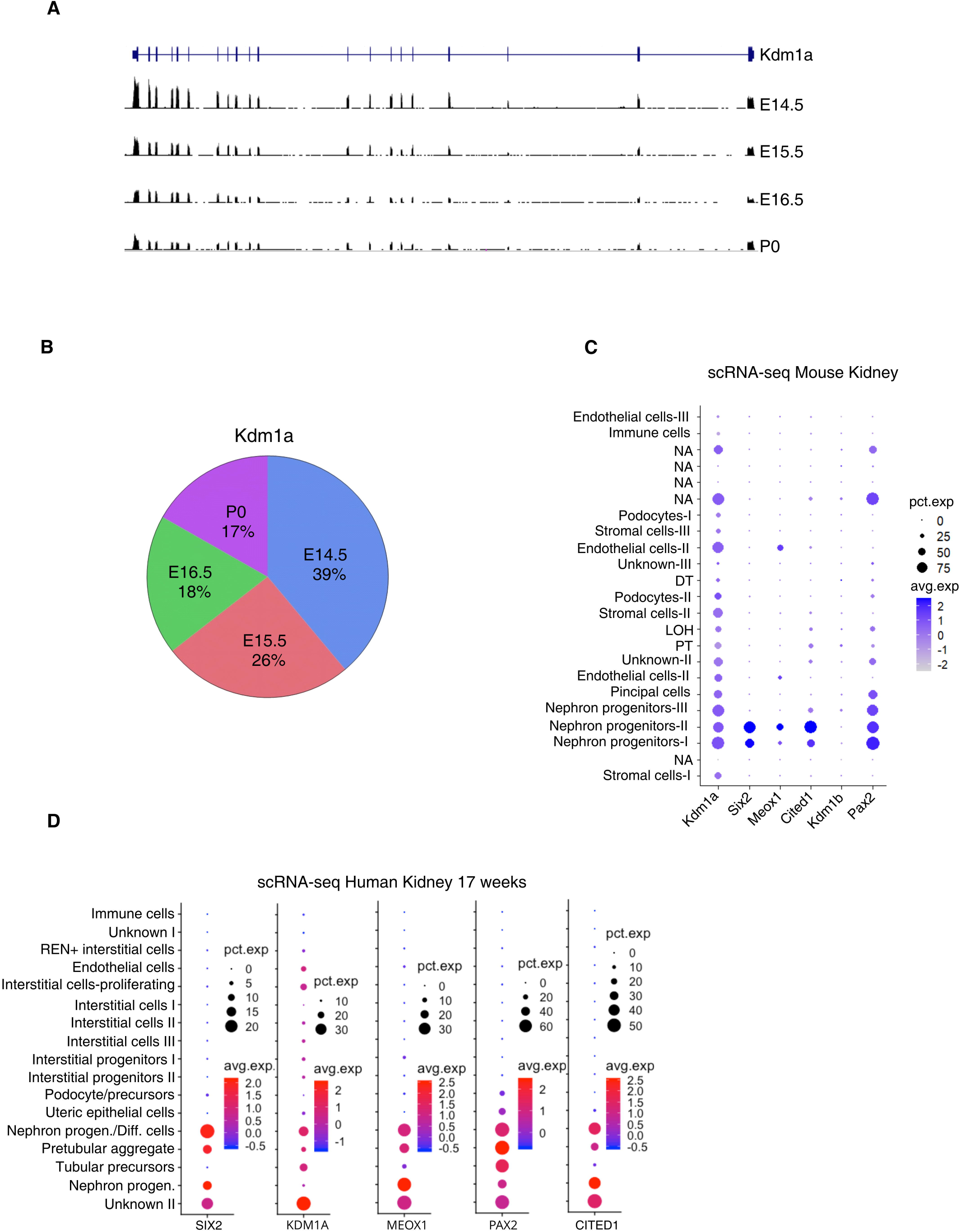
Analysis of *Kdm1a* gene expression during embryonic kidney development. **A.** RNA-sequencing tracks ENCSR504GEG (E14.5), ENCSR062VTB (E15.5), ENCSR537GNQ (E16.5) and ENCSR173PJN (P0) prepared from developing mouse kidneys (Mus musculus strain B6NCrl) shows expression of *Kdm1a* in embryonic and neonatal kidney, of embryonic stages, E14.5, E15.5, E16.5 and P0. Annotated *Kdm1a* protein coding gene and its exon-intron structure is shown on the top, **B.** In the analyzed time-points, highest expression of Kdm1a was seen in E14.5 kidney (39%) and lowest in P0 kidney (17%) with a gradual decline in *Kdm1a* expression during the developmental period. Dot plots showing expression patterns of genes upon analysis of single cell RNA (scRNA) sequencing data (GSE149134) from embryonic and neonatal mouse kidneys, **C**, and Human 17-week-old kidney (GSE112570), **D**, for *Kdm1a, Kdm1b, Pax2, Cited1, Meox1* and *Six2*. Cell cluster overlap is seen for *Kdm1a*, and nephron progenitor cell marker genes in developing kidneys in both mice and humans.

Analysis of single-cell RNA sequencing (scRNA-seq) datasets from developing mouse kidneys^29^ indicated that *Kdm1a* was expressed across a diverse array of cell types, including those enriched for renal progenitor marker genes^30^ such as *Pax2*^31^, *Six2*^32^, *Cited1*^33^, and *Meox1*^34^ (Fig 1C), as well as in highly proliferative cells (Suppl Fig 2). In contrast, *Kdm1b* had a more limited expression pattern, noted in fewer cell clusters and with significantly lower expression levels in clusters expressing nephron progenitor markers (Suppl Fig 2). Analysis of scRNA-seq data from 17-week-old human kidneys^35,36^ revealed co-expression of *KDM1A* in nephron progenitor cell clusters (Fig 1D). *KDM1A* was also prominently expressed in tubular precursor cluster overlapping with PAX2, a gene critical for mesenchymal to epithelial differentiation that is eventually downregulated as epithelial differentiation proceeds ^37^. Unlike *KDM1A*, the expression pattern of *KDM1B* was broader in human kidneys (Suppl Fig 3). Together, these data highlighted significant expression of *Kdm1a* in the nephrogenic zone and pointed to its potential involvement in kidney development.

### Six2-cre mediated deletion of Kdm1a in mouse kidneys: phenotype and functional analyses

To investigate the role of *Kdm1a* in renal progenitor cells within the murine kidney, we utilized a *Six2-Cre* BAC transgenic mouse line (*Six2-TGC^tg/+^*)^32^, in which Cre recombinase expression is driven by *Six2*, which is specifically expressed in multipotent nephron progenitor cells during early kidney organogenesis. By crossing *Six2-cre* line with mice carrying Kdm1a floxed alleles (*Kdm1a^fl/fl^*), we generated conditional knockout mice that exhibit a targeted deletion of Kdm1a expression in the developing renal tissues (Suppl Fig 4).

Upon comparing *Six2-cre;Kdm1a^fl/fl^* mice, referred hereafter as KKO, with those of control mice (Kdm1a^fl/fl^, termed KWT), we observed that the KKO mice (both male and female) were significantly smaller in size (Fig 2A) and lower in total body weight (Fig 2B) when analyzed postnatally at 4- and 7 weeks of age, while KWT and KKO mice appeared similar at birth (Fig 2C). Examination of gross morphology of the postnatal kidneys from KKO and KWT mice revealed distinct differences in size and coloration (Fig 2D). Kidneys from KKO mice (both male and female) were significantly smaller, suggesting that Kdm1a plays a crucial role in early nephrogenesis. Mice and kidneys from strains carrying either a single Kdm1a floxed allele with and without the background of Six-Cre allele were indistinguishable from *Kdm1a^fl/fl^* (KWT) (data not shown). Histomorphometric analyses of Periodic Acid Schiff (PAS) stained KKO mouse kidney sections revealed reduction in number of glomeruli and cortex region, loss of brush border, vacuolation and enlargement of tubules, luminal casts, glomerulosclerosis and atrophy. (Fig 2E). Nephron numbers were significantly reduced in KKO kidneys compared to wildtype, as deduced by glomerular density (Fig 2F).

**Figure 2.**
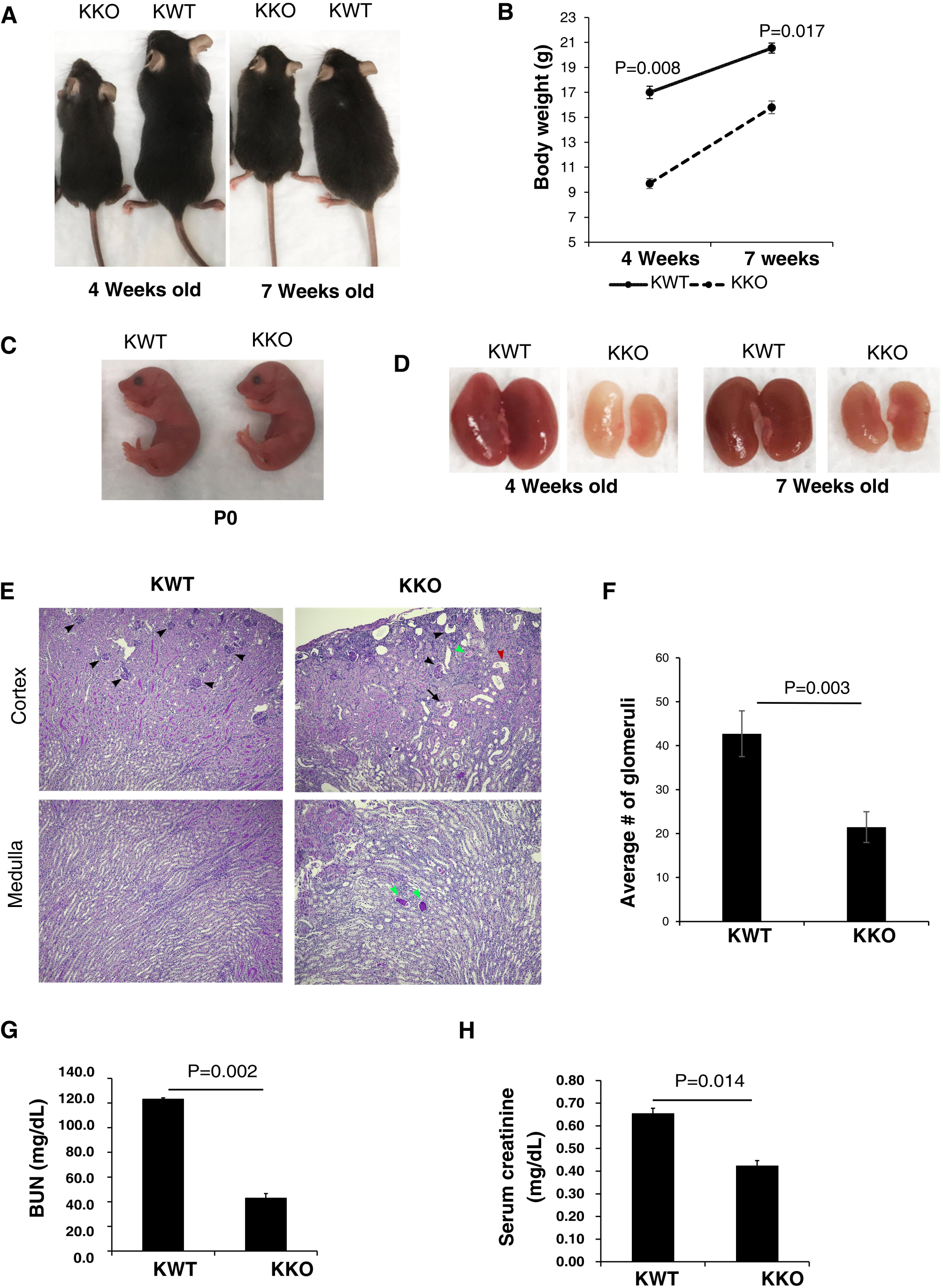
Morphological and functional phenotypes of control and *Kdm1a* conditional knockout mice and kidneys. Representative images of Control (KWT) and *Kdm1a* conditional knockout (KKO) male mice at 4 and 7 weeks of age, **A** and at P0, **C**. **B.** KKO mice weighed significantly lower than control littermates. **D.** KKO kidneys were visibly smaller at 4 weeks and 7 weeks of age. **E.** Histological analyses of Periodic acid Schiff (PAS) stained kidney sections from 4 weeks old mice revealed significant structural alterations in the glomerular and tubulointerstitial compartments in the *Kdm1a* knockout genotype. Black arrowhead: Glomerulus; green arrowhead: tubular cast; red arrowhead>: dilated tubule; black arrow: vacuolated tubule. **F.** Significant reduction in glomerular density was seen in KKO kidneys compared to littermate controls (N=3 each). Renal function impairment was observed in KKO mice, as deduced by BUN, **G** and serum creatinine, **H** levels (N=3).

To assess functional implications of the anatomical and histopathological changes observed in postnatal kidneys upon deletion of Kdm1a in nephron progenitor cells, we measured blood urea nitrogen (BUN) and creatinine levels in KKO and KWT mice at 7 weeks of age. Significant elevation of BUN levels was seen in the KKO mice (123.3+0.9 mg/dL vs. 43.0+3.58 mg/dL; KKO vs. KWT) (Fig 2G). Similarly, a 1.5-fold increase in the levels of serum creatinine was seen in KKO mice compared to littermate controls (0.65+0.02 mg/dL vs. 0.42+0.02 mg/dL; KKO vs. KWT) (Fig 2H). Interestingly, the phenotypes observed in KKO mice viz., renal hypoplasia, pathological tissue structural changes and impaired kidney function recapitulate the observations made in heterozygous Brachyrhine mice (Br/+), a mutant strain carrying a functionally inactive copy of *Six2* gene^38^, a gene critical for the maintenance of nephron progenitor cells and kidney development^32,39^.

### Genome wide gene expression changes underlying Kdm1a knockout kidney phenotypes

To explore transcriptomic alterations linked to the notably reduced size, structural and functional impairment observed in KKO kidneys, we conducted bulk RNA sequencing of kidney samples from postnatal day zero to one (P0/P1) KWT (N=4) and KKO (N=4) mice (Suppl Table-1). Our analysis uncovered a total of 1,153 genes exhibiting significant expression changes (p-value < 0.05 and log2 fold change > 1) when comparing KWT and KKO kidneys. Interestingly, we found that a larger number of genes were downregulated in KKO kidneys (667 genes) compared to those that were upregulated (486 genes) (Fig 3A and Fig 3B). Among the most significantly differentially expressed genes, several of them are unique to proximal tubular segments including Kap (log2 fold change of -7.6), Acmsd (log2 fold change of -5.6). On the other hand, we observed that Cyp11b2 and Phox2b were markedly upregulated, with log2 fold changes of 10 and 4.7, respectively (Suppl Table-2).

**Figure 3.**
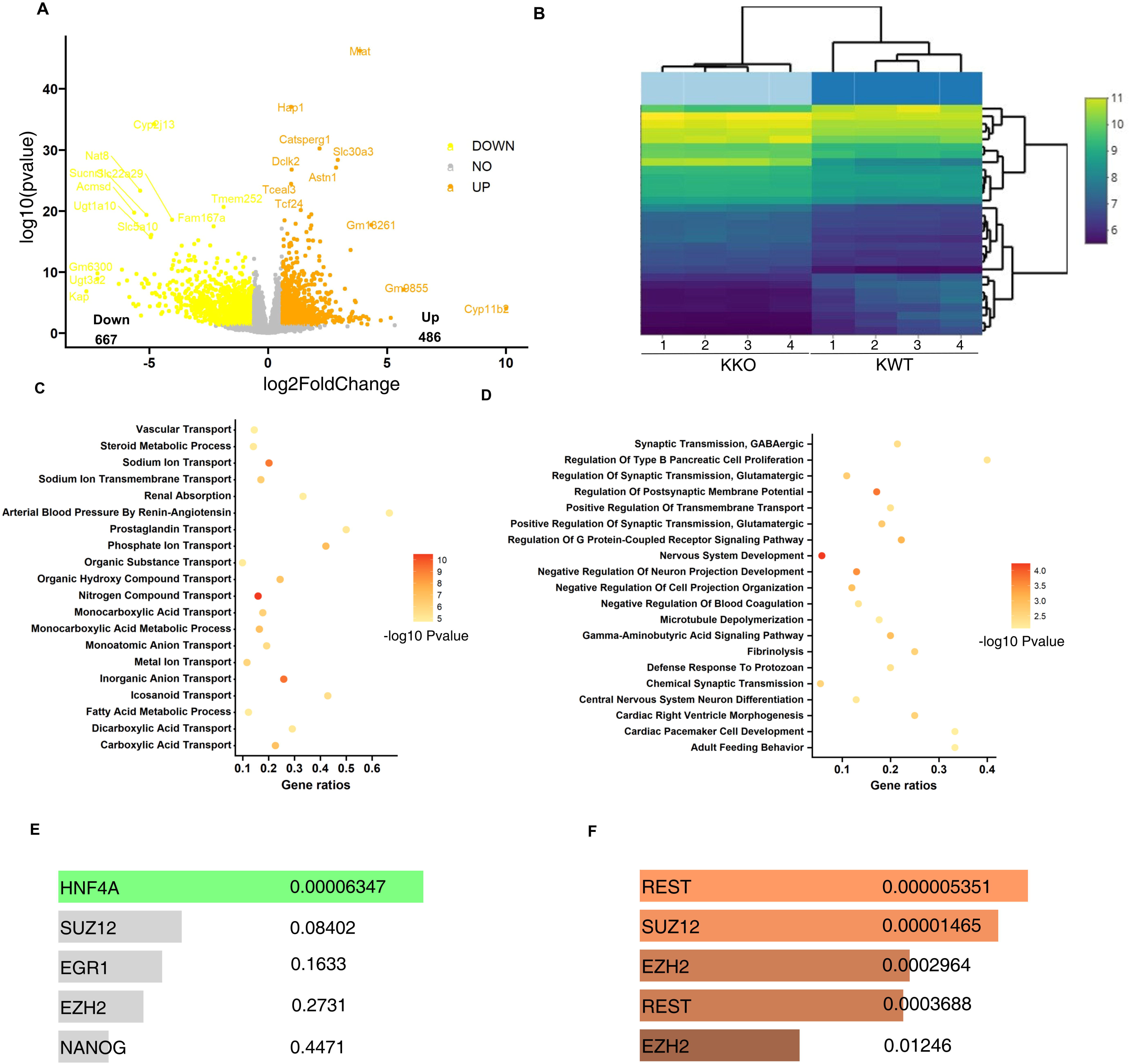
Genome wide analysis of gene expression differences between control (KWT) and *Kdm1a* conditional knockout (KKO) mice kidneys via bulk RNA sequencing. Control and knockout kidneys collected from P0/P1 pups were analyzed bulk RNA sequencing. **A.** Volcano plot showing global transcriptional changes between KWT (n=4) and KKO (n=4) kidneys, the log2 fold change of each gene is on the x-axis and the log10 of its adjusted p-value is on the y-axis. Genes upregulated in KKO kidneys (with an adjusted p-value less than 0.05 and a log2 fold change greater than 1 are indicated by orange dots. Genes downregulated in KKO kidneys (with an adjusted p-value less than 0.05 and a log2 fold change less than -1 are indicated by yellow dots. **B.** Bi-clustering heatmap showing top 30 differentially expressed genes in individual samples, by plotting their log2 transformed expression sorted by their adjusted p-value. **C**. Gene ontology (GO) analysis showing top 20 enrichment terms for genes involved in multiple solute and other transport pathways downregulated in *Kdm1a* knockout kidneys. **D.** GO analysis showing top 20 enrichment terms for genes involved in pathways involved in nervous system development, synaptic transmission that were upregulated in *Kdm1a* knockout kidneys. **E.** Bar graph showing top 5 Encode and ChIP enrichment analyses (ChEA) consensus transcription factors for genes with log2 fold change <1.5 **F.** Bar graph showing top 5 Encode and ChEA consensus transcription factors for genes with log2 fold >1.5.

Gene ontology (GO) analysis of the top downregulated genes (Fig 3C) revealed significant enrichment in pathways critical to various kidney functions. These include transport mechanisms, renal absorption processes, blood pressure regulation, and monocarboxylic acid metabolism, implicating defective tubular development corroborating renal hypoplastic phenotype observed in KKO mice. Analysis of upregulated genes (Fig 3D) revealed enrichment of neuronal pathways, membrane potential regulation, and microtubule polymerization. Comprehensive analysis of cell lineages in human kidney organoids and human fetal kidney samples previously revealed the consistent presence of neural clusters among committed renal cell lineages^40^. Similarly, neural precursors were reported in developing mouse kidneys^41^. Together, our data suggest potential de-repression and expansion of neural lineage concurrent to impaired renal tubular development due to the absence of functional Kdm1a in the cap mesenchyme during early nephrogenesis.

Encode-ChIA transcription factor and regulatory factor enrichment analysis highlighted Hepatocyte nuclear factor 4 alpha (*Hnf4a*) as a potential trans-regulatory factor associated with downregulated genes (Fig 3E), while REST and polycomb family proteins were linked to upregulated genes (Fig 3F). This suggests that these transcription factors may play a crucial role in mediating the downstream effects of Kdm1a loss on kidney gene expression. Indeed, Kdm1 and Hnf4a were previously found in protein interaction complexes^42^. Consistent with this, around 600 of the downregulated genes in Kdm1a knockout kidneys are also potential transcriptional targets of Hnf4a (Suppl. Table 3). *Hnf4a* expression was found in the metanephric mesenchyme very early on, subsequently widely distributed in the developing nephron and finally restricted to proximal tubules in the fully developed kidneys^43^. Overall, global transcriptomic analysis revealed significant alterations in gene expression in KKO kidneys, implicating dysregulation of cell lineage commitment and modulation of a variety of pathways critical for renal function, thus highlighting the importance of *Kdm1a* in regulating kidney development at the molecular level.

### Reduced expression of progenitor cell markers and solute carrier gene family in KKO knockout kidneys

We found a significant relationship between the observed reduction in renal mass in KKO kidneys (as shown in Figure 2) and changed expression of various renal cell type specific marker genes. Specifically, decreased expression levels of markers of nephron progenitors (*Cited1*, *Meox1*) and presumptive nephron precursors (*Lhx1*) (Fig 4A), the Loop of Henle (*Umod*), podocyte (*Nphs1*, *Nphs2*), principal cell (*Aqp2* and *Fxyd4*), the proximal tubule (*Kap*) (Suppl Fig 5A-D) was observed in KKO kidneys. In contrast, there was an increase in the expression of stromal cell markers, including *Foxd1* and *Col3a1* (Suppl Fig 5E). These data further corroborate the consequence of Kdm1a deletion within the cap mesenchyme and importantly the developmental impairment appears to be wide spread along the entire nephron segment. Further investigation into the downregulated genes revealed significant alterations in the expression of solute carrier (Slc) superfamily genes following the loss of Kdm1a. Out of 346 differentially expressed Slc family genes (Suppl Table-4) identified in the RNA sequencing data from P0 kidneys, 86 demonstrated significant changes in expression (p-< 0.05 and a log2 FC > 1). Intriguingly, a striking ∼88% (76 out of 86) of these altered genes were downregulated, while only 12% (10 out of 86) exhibited increased expression as a result of Kdm1a loss during kidney development (Fig 4B and Fig 4C, Suppl Table-4). The Slc genes differentially expressed in KKO kidneys mapped predominantly to the proximal and distal tubular segments of the kidney^44^. Interestingly, *Hnf4a* and closely related *Hnf1a* were shown to regulate majority of the solute carrier genes in the developing proximal tubules^45^. Together, the data suggest that *Hnf4a* may intersect with *Kdm1a* mediated gene regulation in the proximal tubules resulting in marked changes in the proximal tubular transport/reabsorption processes.

**Figure 4.**
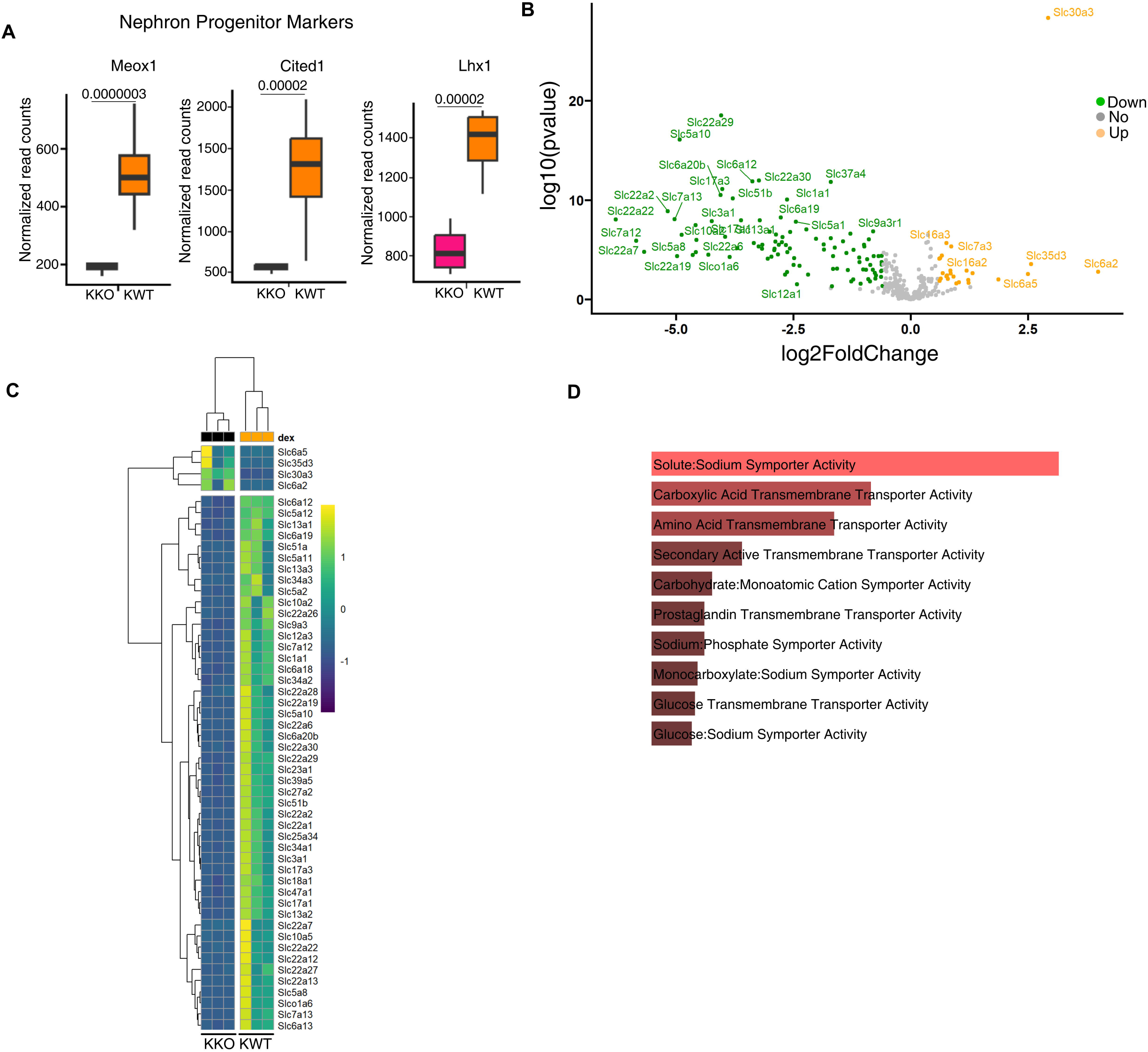
Altered expression of renal progenitor markers and the solute-carrier gene (SLC) superfamily genes in KKO kidneys. **A.** Box plots showing normalized RNA expression counts for nephron progenitor cell marker genes, *Meox1* and *Cited1* and tubular precursor marker, *Lhx1* in KWT (n=4 kidneys) and KKO (n=4 kidneys). **B.** Volcano plot showing transcriptional changes in Slc super family genes between KWT(n=4) and KKO(n=4) kidneys; the log2 fold change of each Slc gene is plotted on the x-axis and the log10 of its adjusted p-value is on the y-axis. Genes upregulated in KKO kidneys (with an adjusted p-value less than 0.05 and a log2 fold change greater than 1) are indicated by orange dots. Slc genes downregulated in KKO kidneys (with an adjusted p-value less than 0.05 and a log2 fold change less than -1 are indicated by yellow dots. **C.** Bi-clustering heatmap showing differentially expressed (log 2-fold lesser or greater than 2) Slc family genes between individual samples, (KWT (n=3) and KKO (n=3) kidneys), by plotting their log2 transformed expression sorted by their adjusted p-value. **D.** Gene Ontology analysis showing top 10 enrichment terms for Slc genes downregulated in *Kdm1a* knockout kidneys.

Pathway analysis of the Slc genes downregulated in KKO kidneys highlighted their involvement in various transport processes crucial for kidney function. These include sodium symporter activity, carboxylic acid transmembrane transporter activity, amino acid transmembrane transporter activity, and cation symporter activity (Fig 4D). Among the downregulated Slc family genes, several are particularly vital for the transport of essential cellular metabolites including *Slc22a12* (Urate transporter 1), *Slc5a2* (Sodium-glucose cotransporter 2), *Slc22a6* (Organic anion transporter 1), and *Slc22a8* (Organic anion transporter 3). Also downregulated were Slc3a1 (log2 FC, -4.2) and *Slc7a9* (log2 FC, - 2.2), that constitutes a transporter system expressed in the proximal tubule segment essential for the reabsorption of neutral and dibasic cations. Two major electron neutral chloride transporters, *Slc12a1* (sodium-, potassium-, chloride symporter) expressed in thick ascending limb and *Slc12a3* (thiazide sensitive sodium chloride co-transporter) expressed in distal convoluted tubules and play a key role in salt reabsorption, were significantly downregulated in the KKO kidneys.

These data show that the loss of *Kdm1a* in the nephron progenitors significantly disrupted the expression of genes in differentiated nephrons that are essential for renal transport / reabsorption functions, particularly those involved in the handling of metabolites, electrolytes and amino acids. The profound dysregulation of genes observed in the differentiated nephrons likely contributes to the renal function impairment observed in KKO mice.

### Perturbation of H3K9 trimethylation levels upon loss of Kdm1a in the nephron progenitors

To explore the mechanisms underlying Kdm1a-mediated downregulation of gene expression in the kidney, we conducted chromatin immunoprecipitation followed by sequencing (ChIP-seq) to evaluate the presence of gene-repressive histone modification namely trimethylation of histone H3 at lysine 9 (H3K9me3), in both KWT and KKO kidneys. Our analysis revealed a significant increase in both the total and differential peaks of H3K9me3 in the P0/P1 KKO kidneys compared to their wildtype counterparts (KWT) (Fig 5A and Suppl Fig-6, Supplementary Table-5). Genome-wide distribution of differential H3K9me3 peaks in KWT vs KKO was characterized by approximately 56% residing in intergenic regions, 26% in intronic regions, and 13% within promoter-transcription start site (TSS) regions (Fig 5B). Elevated levels of H3K9me3 observed in the genic regions corresponded to the downregulation of several genes that are markers of nephron progenitors and tubular precursors, including Meox1, Lhx1 and Esrrg (Fig 5C). About 4% of the significantly downregulated genes in the KKO kidneys were found to be associated with increased H3K9me3 in their genic regions.

**Figure 5.**
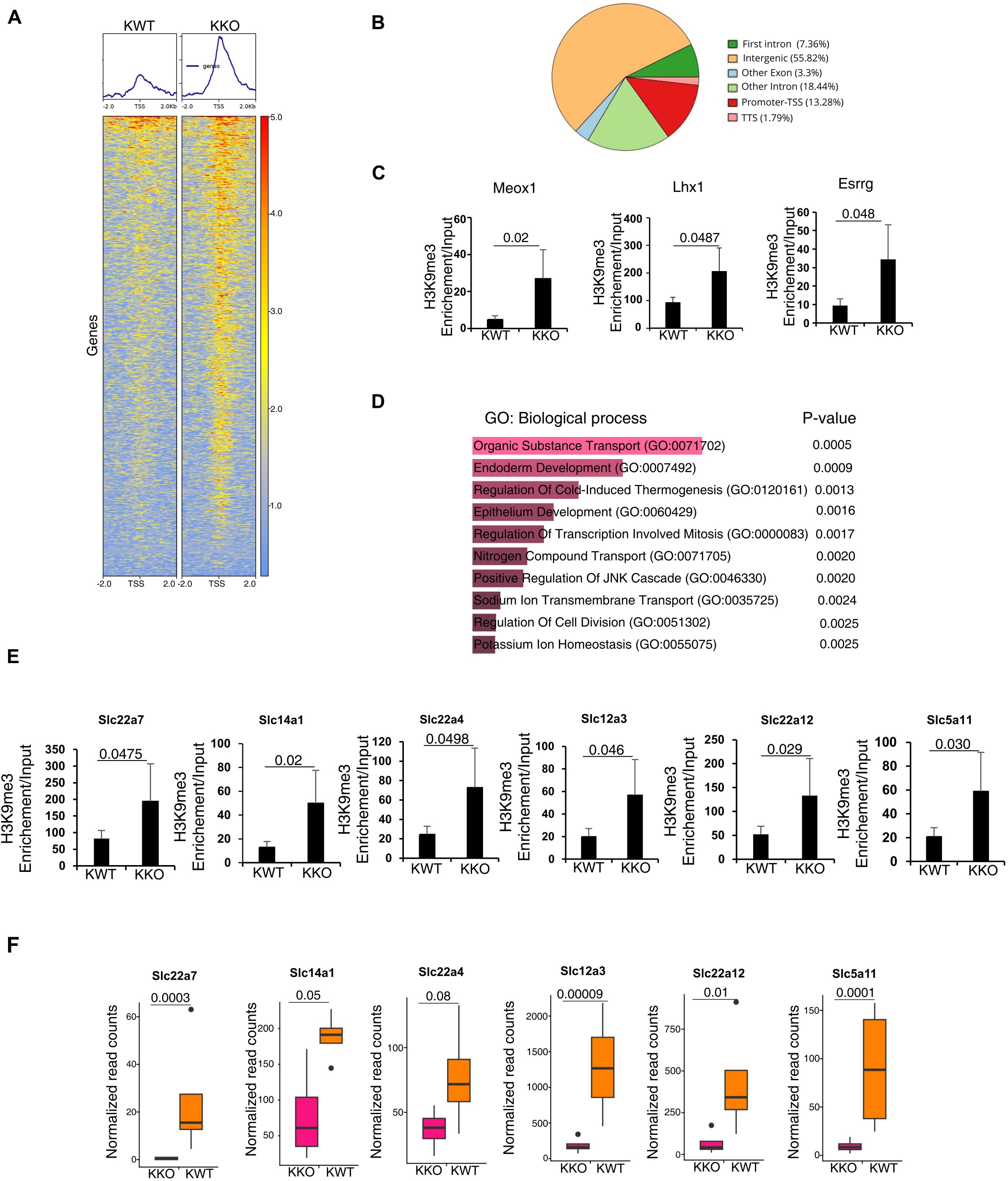
Differences in repressive H3K9 trimethylation enrichment between control (KWT) and *Kdm1a* conditional knockout (KKO) mice kidneys as assessed by ChIP-sequencing. Heatmap depicting H3K9 trimethylation enrichment (peaks) at regions spanning 2kb upstream and 2kb downstream of TSS. **A.** Differential peaks, **B.** Distribution of differential H3K9me3 peaks between control and KKO kidneys across genome; majority of the H3K9 differential peaks were found in intergenic regions with ∼13% located near promoter-TSS regions. **C.** H3K9me3 enrichment at *Meox1, Lhx1 and Esrrg* genes (H3K9me3/input reads) KWT(n=2) vs KKO (n=2) kidneys. **D.** Gene Ontology analysis showing top 10 enrichment terms for Slc genes that are downregulated in *Kdm1a* knockout kidneys coinciding with H3K9me3 enrichment. **E.** H3K9me3 enrichment at Slc gene family genes *Slc22a4, Slc5a11, Slc12a3, Slc14a1, Slc22a12, Slc22a7* (H3K9me3/input reads) KWT(n=2) vs KKO (n=2) kidneys. **F.** Box plots showing normalized RNA expression counts of *Slc22a4, Slc5a11, Slc12a3, Slc14a1, Slc22a12, Slc22a7* genes in KWT (n=4) and KKO (n=4) kidneys.

Gene Ontology (GO) analysis of the biological pathways highlighted that numerous genes with increased H3K9me3 peaks in KKO kidneys were linked to organic substrate transport processes (Fig 5D). Furthermore, approximately 8% of the Slc superfamily genes, which were downregulated in KKO kidneys, *Slc22a4*, *Slc5a11*, *Slc12a3*, *Slc14a1*, *Slc22a12*, *Slc22a7* (p-< 0.05 and a log2 FC > 1), *Slc22a23* and *Slc25a3* (p-< 0.05 and a log2 FC > 0.5) demonstrated H3K9me3 peak enrichment in their genic regions compared to KWT kidneys (Fig 5E, 5F, and Suppl Fig-7,8A, 8B). This data suggests that the loss of Kdm1a and subsequent establishment of a repressive chromatin landscape characterized by increased H3K9 trimethylation play a critical role in the altered expression of essential genes including those that belong to Slc family involved in substrate transport during kidney development.

### Specific gene body and promoter associated DNA methylation changes in KKO kidneys

Kdm1a has been established as a regulator of DNA methylation in mouse embryonic stem cells and early embryos^20,21^; however, its role in the regulation of DNA methylation within solid organs, including kidneys, remains less understood. To investigate whether the increased H3K9 trimethylation observed in KKO kidneys might influence DNA methylation patterns and, subsequently, gene expression, we employed Reduced Representation Bisulfite Sequencing (RRBS-seq) to assess global methylation levels in both KWT and KKO kidneys. The overall distribution and levels of DNA methylation did not significantly differ between the two genotypes (Suppl Fig 9A, Suppl Fig 9B). However, detailed analysis of genomic regions known for varying methylation levels across multiple tissues and during development, namely differentially methylation regions (DMRs) within the RRBS data revealed that 467 DMRs were hypomethylated and 553 DMRs were hypermethylated in KKO kidneys compared to KWT kidneys (Fig 6A, Fig 6B).

**Figure 6.**
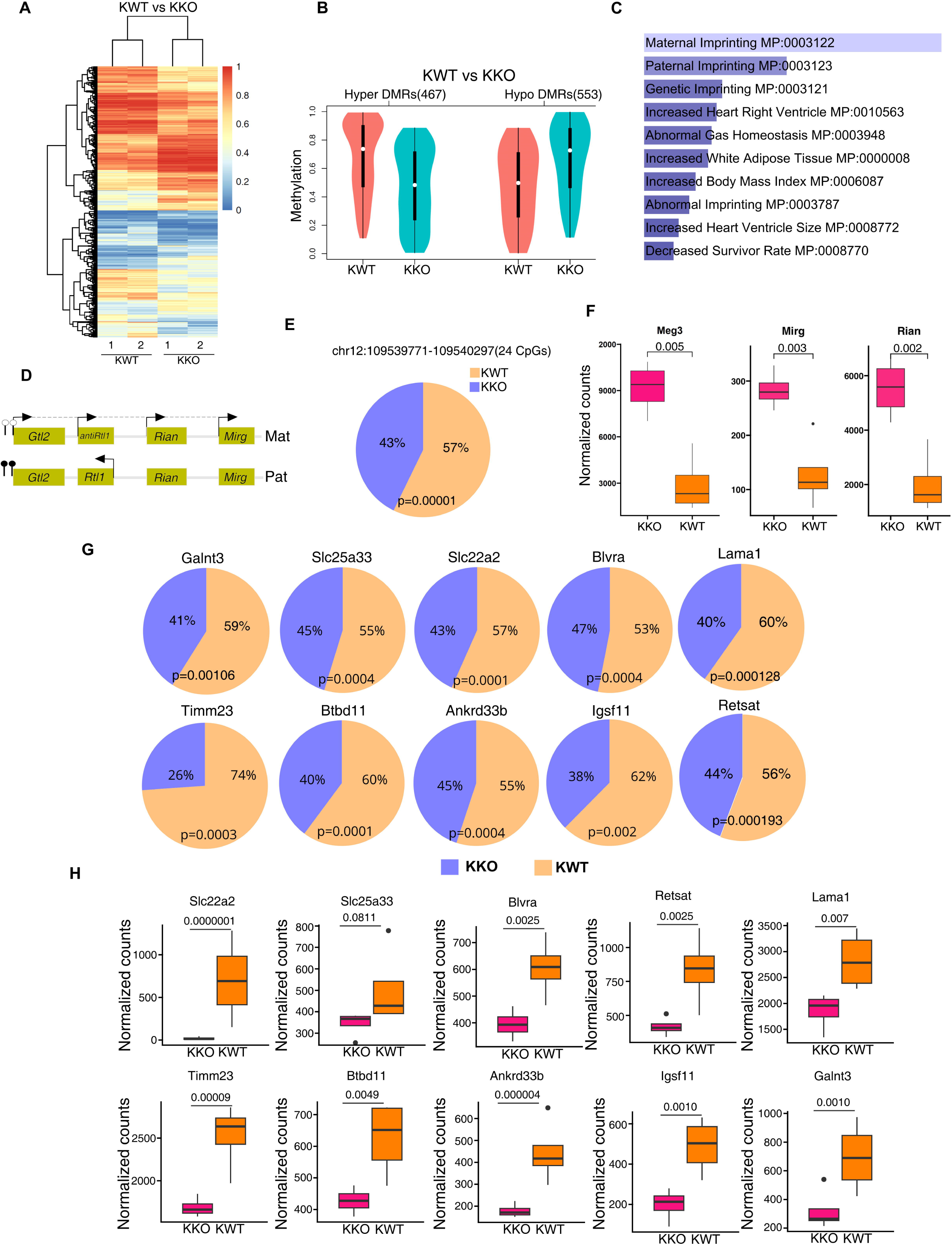
Global DNA methylation analysis by reduced representation bisulfite sequencing (RRBS) **A.** Heatmap showing methylation levels at Differentially methylated regions (DMRs) between control (n=2) and KKO (n=2) kidney samples. **B.** Violin plots showing average methylation levels at hyper and hypo methylated DMRs between control and KKO kidneys. **C.** Top 10 MGI mammalian phenotyping gene ontology terms for genes associated with differentially methylated promoters detected in KKO (N=2) kidneys versus KWT (N=2) kidneys identified by RRBS analysis. **D.** Cartoon showing imprinted Meg3 locus wherein the promoter DMR is methylated on paternal allele and unmethylated on maternal allele. **E.** CpG methylation levels calculated at 24 CpGs spanning *Meg3* promoter regions (chr12:109539771-109540297) expressed as percentage (KWT vs KKO). *Meg3* promoter region is significantly hypomethylated in KKO kidneys. **F.** Box plots showing normalized RNA expression counts of non-coding RNAs, *Meg3, Mirg* and *Rian* from paternally methylated Dlk-Dio3 imprinted domain in KWT (n=4) and KKO (n=4) kidneys. **G.** Pie-charts showing gene body CpG methylation levels at individual gene loci, expressed as percentages between KWT (N=2) and KKO (N=2) kidneys. **H.** Box plots showing normalized RNA expression counts of genes shown in G, KWT (n=4) and KKO (n=4) kidneys.

Gene Ontology analysis of the MGI mammalian phenotype for genes associated with hypo- and hypermethylated promoter DMRs between KWT vs KKO kidneys, revealed significant enrichment of imprinted genes, whose expression is dependent on the parent of origin (Fig 6C, Suppl Table-6). The 1-megabase Dlk1-Dio3 imprinted domain, on the maternal chromosome encompasses several non-coding RNA (ncRNA) genes, such as *Meg3*, *Mirg* locus (>50 miRNAs), and *Rian* locus (>40 C/D box containing snoRNAs). On the paternal chromosome, three key protein-coding genes are transcribed from the imprinted region namely, *Dlk1*, *Rtl1* (*Peg11*), and *Dio3*. The *Meg3* promoter-associated DMR controls non-coding RNA transcription in conjunction with imprinting control region, IG-DMR^46,47^. So, we analyzed a few of these DMRs in the KKO and KWT kidneys. In the KKO kidneys, Meg3 promoter associated DMR was hypomethylated, with increased maternally encoded gene expression from the Dlk1-Dio3 imprinted cluster (Fig 6D, 6E). Specifically, non-coding RNA genes including *Meg3*, *Rian* and *Mirg* that are under the control of Meg3 DMR showed increased expression in KKO kidneys (Fig 6F). At the *Plagl1 (Zac1)* loci maternal ICR regulates paternal expression of Plagl1. In majority of the tissues, Plagl1 expression is paternal and in few adult tissues such as liver and kidney, it is bi-allelic^48^. In KKO kidneys, the promoter-Exon1 spanning DMR was hypermethylated and correlated with increased expression of *Plagl1* transcript levels (Supp Fig 10). At *Nespas-Gnas* and *Igf2R-Airn* imprinted loci, DMR hypermethylation did not lead to significant changes in transcript levels of the genes at these loci (Supp Fig 11).

Interestingly, most of the DMRs (around 811) with significant methylation changes were concentrated within intragenic or gene body regions (Supp Fig 12). Gene body CpG methylation including on active X chromosome has been previously reported to be closely associated with gene expression^49^. Consistent with this, among the hyper and hypomethylated DMR-associated genes, several exhibited reduced expressions in KKO kidneys, including Lama1^50^ essential for proper glomerular development in mice (Fig 6G, 6H). Together, these findings suggest that Kdm1a plays a key role in regulating DNA methylation at DMRs in the kidneys, significantly impacting the expression of associated genes, including at specific imprinted loci.

## Discussion

Lysine-specific demethylases, including Kdm1a, are pivotal in the regulation of gene expression, influencing both gene activation and repression. *Kdm1a* is expressed in nephron progenitor cells in both murine and human embryonic kidneys, overlapping with key transcription factors such as *Six2*, *Pax2*, *Cited 1* and *Meox1*^34^. In our study, we investigated the role of Kdm1a during kidney development by employing *Six2-Cre* to delete *Kdm1a* in nephron progenitor cells. This targeted deletion of *Kdm1a* led to pronounced renal phenotypic changes in adult mice, including reduced kidney size, pathological tissue alterations and impaired kidney function. Interestingly, while the kidney phenotypes resulting from Kdm1a deletion in multipotent nephron progenitor cells mirrors that of *Six-2* mutant, unlike *Six2*, *Kdm1a* is expressed more broadly in developing kidneys.

Subsequent dissection of molecular mechanisms underlying the *Kdm1a* conditional knockout kidney phenotypes revealed significant alterations in the renal tissue transcriptome. Significant downregulation of marker genes associated with various renal cell types, and especially the solute carrier gene family was observed. Out of the 196 solute carrier gene family members that showed significant changes in gene expression (p<0.05), 145 of them were downregulated. The Slc family genes carryout important metabolic and transport functions within kidney and are markers of preserved renal tubule cell state and function^51^. Indeed, many Slc genes that are hallmarks of differentiated tubular epithelium (*Slc5a12, Slc7a13, Slc34a1, Slc22a30*) were highly downregulated upon Kdm1a deletion. Our study also revealed an interesting overlap between Kdm1a and Hnf4a bound Slc genes. Further, similar to our observation in Kdm1a conditional knockout kidneys, downregulation of renal solute carriers was reported upon deletion of hepatocyte nuclear factors, *Hnf1a* and *Hnf4a*^52,53^. In the developing mouse kidneys, Hnf4a is first detected the comma-shaped body, subsequently widely distributed in the developing nephron and finally restricted to proximal tubules in the fully developed kidneys^43^. In contrast to *Hnf4a*, *Kdm1a* is expressed more broadly both in embryonic and adult kidneys. Disruption of solute carrier expression that we observed upon Kdm1a deletion was not just limited to proximal tubules but spanned all the tubular segments derived from metanephric mesenchyme. Taken together, it is tempting to postulate that sustenance of Kdm1a function not just in multipotent nephron progenitors but also in the committed tubular precursors might be crucial for the generation of fully differentiated nephrons including the maintenance of Slc gene expression.

Kdm1a exerts its regulatory effects primarily through the demethylation of histones H3K9 and H3K4, allowing for the activation or repression of target genes. In our analysis, we found that the number of significantly downregulated genes exceeded that of upregulated genes, prompting us to focus on the H3K9 methylation mediated repressive mechanisms underlying Kdm1a loss-of-function. In KKO kidneys, we identified genome-wide increase in H3K9me3 levels primarily localized to intergenic and genic regions. This elevated trimethylation correlated with reduced expression of several Slc family genes and markers pertaining to multiple renal cell types including progenitor cells. Our findings suggest that Kdm1a plays a crucial role in H3K9 demethylation to promote gene expression; about one tenth of down-regulated Slc genes in KKO kidneys exhibited increased H3K9me3 levels. Previous studies have highlighted the involvement of Kdm1a in regulating DNA methylation in mice and embryonic stem cells by controlling the stability of the DNA methyltransferase, Dnmt1 both by enzyme dependent and independent functions^20,21^. Of note, deletion of Dnmt1 was previously shown to impair proper kidney development in mice^54^. However, whether Kdm1a influences DNA methylation levels during organ development remains unclear. In our investigation, we found no global alterations in DNA methylation levels in *Kdm1a* conditional knockout kidneys. Nonetheless, we identified >1,000 differentially methylated regions (DMRs), through Gene Ontology (GO) analysis of promoter DMR associated genes revealing an enrichment for genes involved in nuclear pathways and imprinted gene regions.

The changes in DNA methylation at imprinted genes in KKO kidneys were restricted to specific imprinted loci and those DMRs with altered methylation patterns were not at known germline parental DMRs or imprint control regions (ICRs). Instead, changes were seen within the promoter DMR regions of genes characterized by imprinted expression. For instance, several maternally expressed non-coding RNAs from the *Dlk1-Dio3* imprinted domain exhibited increased expression and decreased DNA methylation at the *Meg3* upstream region DMR in KKO kidneys, while at the imprint control region (ICR) IG-DMR, no changes in methylation were noted. Similarly, alterations were seen at the *Igf2R-Airn* locus, with changes observed at the DMR1 region associated with the *Igf2R* upstream CpG island but not at the DMR2 imprinted control region of the Airn locus. Moreover, only a small number of downregulated genes (5% of hypomethylated, 9% of hypermethylated) correlated with methylation changes at the associated DMRs. Most of the changes in gene body DMR methylation in KKO kidneys did not lead to notable changes in gene expression. These results point to the existence of localized, chromatin-restricted changes in DNA methylation that are contingent upon Kdm1a function.

Genetic ablation of *Kdm1a* in mice led to severe phenotypic outcomes, including embryonic lethality, contingent upon the specific cell types in which the deletion occurs. In contrast, de novo mutations in Kdm1a in humans have been linked to a rare autosomal dominant syndrome characterized by cleft palate, psychomotor retardation, and distinctive facial features (CPRF, OMIM#616728). However, due to limited number of documented patients with this disorder and individuals carrying other rare Kdm1a variants, the impact of Kdm1a mutations on kidney function in affected individuals remains unknown. Importantly, Kdm1a is considered to be an evolutionarily constrained gene^55^. Interestingly, postnatal deletion of *Kdm1a* using tamoxifen inducible cre did not reveal significant structural or functional impairment in kidney^56^ and in light of our current findings, it appears that Kdm1a plays an indispensable role specifically during kidney development. Kidneys are exposed to significant metabolic and physiological pressures, and it would be interesting to see whether Kdm1a is required for mediating stress responses in adult kidneys. Future research would be required to dissect significant genome-wide association (GWAS) observed between *Kdm1a* SNP (rs10917347-G) and elevated serum gamma glutamyl transferase levels^57^ and any causal relationship to kidney calculus, a condition arising from solute carrier dysfunction^58^.

In summary, our study highlights the complex role of Kdm1a in mouse kidney development and points to a likely role in human kidneys. It influences both developing and differentiated nephrons by maintaining expression levels of critical kidney developmental genes, the Slc gene family among others, through histone modification and localized DNA methylation changes (Figure-7). These insights add to our growing understanding of the complex molecular mechanisms involved in kidney development.

**Figure 7.**
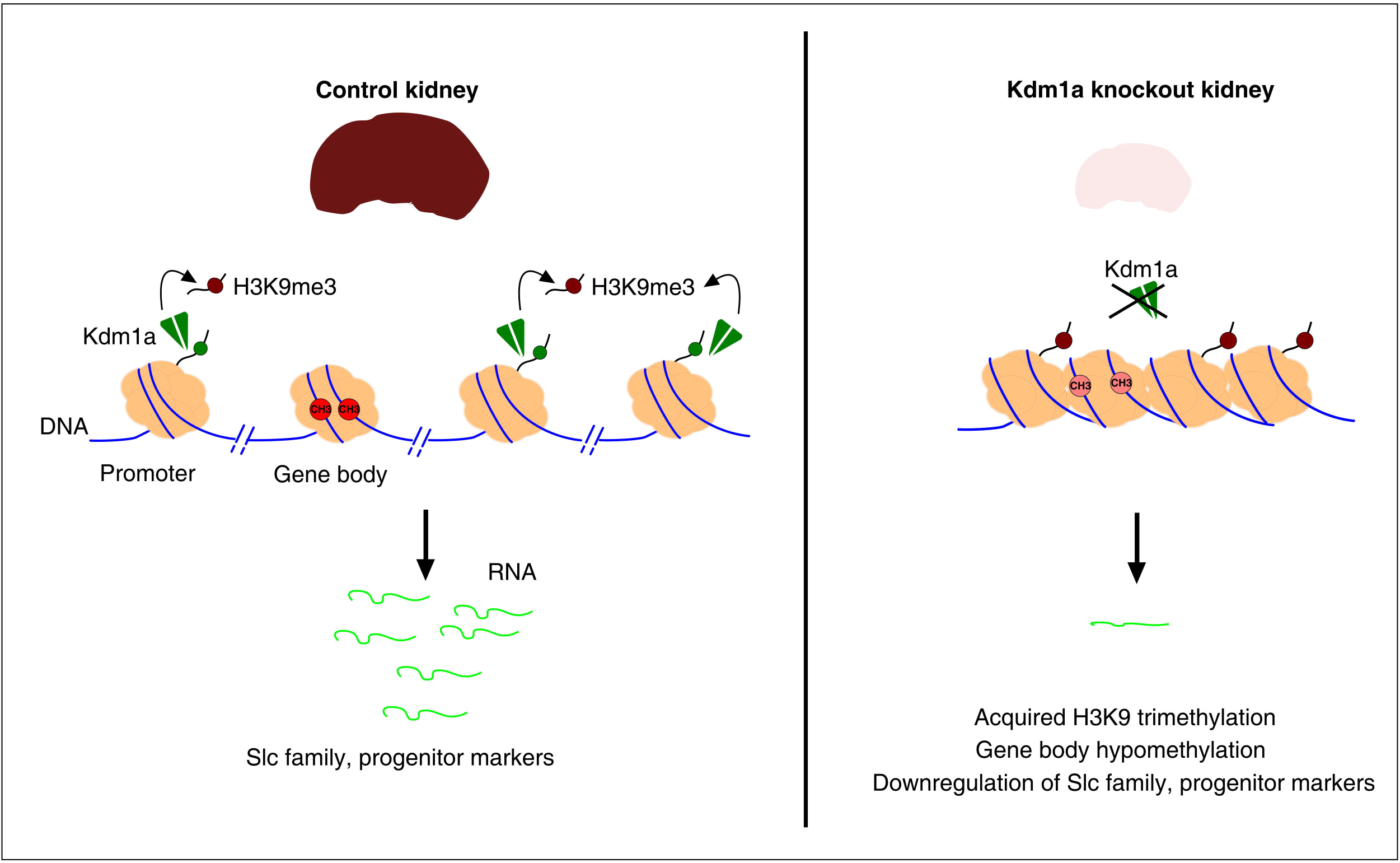
Kdm1a is obligatory for kidney development and gene regulation via H3K9me3 and DNA methylation. The loss of *Kdm1a* in *Six2*-expressing renal progenitor cells within the developing mouse kidney results in hypoplasia and histopathological changes, leading to a significant decline in renal function. At the molecular level, deletion of *Kdm1a* led to significant downregulation of several genes. This encompasses various renal cell type marker genes, with over a quarter belonging to the Slc family. ChIP-sequencing in these knockout kidneys revealed a global increase in the localized enrichment of H3K9me3 peaks at specific gene loci, such as renal progenitor marker genes and Slc family genes. RRBS analysis of CpG methylation levels in Kdm1a knockout kidneys indicated altered DNA methylation predominantly in differentially methylated regions (DMRs) within gene bodies. Both hypermethylated and hypomethylated gene bodies contribute to downregulated expression. Specific promoter DMR-associated methylation changes are primarily enriched at certain imprinted loci.

### Study limitations

First, in our study, we explored gene expression, chromatin structure, and DNA methylation changes in kidneys just after birth (P0/P1) to understand the functions of the *Kdm1a* gene. We used control and *Kdm1a* conditional knockout kidneys, targeting nephron progenitors with a cre line, to reveal targets of *Kdm1a* regulation. Future analysis at additional timepoints during nephrogenesis could help identify direct and indirect gene targets. Second, our study was mainly focused on dissecting the molecular role of Kdm1a during kidney development. While we captured salient phenotypes upon *Kdm1a* deletion, we suspect that detailed phenotypic characterization may further reveal additional pathological alterations in the kidneys. Finally, our findings showed more genes were downregulated than upregulated in *Kdm1a* conditional knockout kidneys. This prompted us to further examine the connections between repressive chromatin and epigenetic modifications, focusing on H3K9me3 and DNA methylation patterns. Kdm1a is known to function in gene repression through demethylation of H3K4 methylation, polycomb proteins, and various histone post-translational modifications, which may also coordinate gene expression changes observed in the absence of *Kdm1a*. However, these were not analyzed in our study.

## Methods

### Mouse strains

*Six2 Cre* (*Six2-TGCtg/+)* strain (Kobayashi et al.2009) was kindly provided by Dr. M. Todd Valerius (Brigham and Women’s Hospital, Boston). *Kdm1a* floxed mice^59^ (*Kdm1afl/fl*; Strain #023969) were purchased from Jackson laboratory. Six2-Cre; Kdm1afl/fl (KKO) mice were maintained by breeding *Kdm1af/fl; Six2-TGCtg/+* and *Kdm1af/fl* mice such that only a single copy of *Six2-TGCtg/+* allele is retained. Animals were housed in an AALAC-approved facility in a pathogen free environment. All the animal studies were approved by the Institutional Animal Care and Use Committee at BIDMC.

### Renal histopathology

Transverse sections of the kidneys were fixed in 10% formalin and processed for paraffin embedded sections. Renal morphology was assessed by PAS (Periodic acid-Schiff reagent) staining of 5 μm sections. Glomerular density was deduced by counting total number of glomeruli per section from KWT and KKO (N=3).

### Analysis of kidney function

Blood plasma was collected and stored at -80 deg C until further processing. Plasma Creatinine levels were quantified using QuantiChrom™ Creatinine Assay Kit (Bioassays systems). Blood Urea Nitrogen levels were measured using Infinity Urea liquid stable reagent (Thermo fisher).

### RNA sequencing and data analysis

Total RNA was extracted from fresh frozen tissue samples using Qiagen RNeasy plus Universal mini kit following manufacturer’s instructions (Qiagen, Hilden, Germany). RNA samples were quantified using Qubit 2.0 Fluorometer (Life Technologies, Carlsbad, CA, USA) and RNA integrity was checked using Agilent TapeStation 4200 (Agilent Technologies, Palo Alto, CA, USA). RNA sequencing libraries were prepared using the NEBNext Ultra RNA Library Prep Kit for Illumina using manufacturer’s instructions (NEB, Ipswich, MA, USA). Briefly, mRNAs were initially enriched with Oligo d(T) beads. Enriched mRNAs were fragmented for 15 minutes at 94 °C. First strand and second strand cDNA were subsequently synthesized. cDNA fragments were end repaired and adenylated at 3’ends, and universal adapters were ligated to cDNA fragments, followed by index addition and library enrichment by PCR with limited cycles. The sequencing library was validated on the Agilent TapeStation (Agilent Technologies, Palo Alto, CA, USA), and quantified by using Qubit 2.0 Fluorometer (Invitrogen, Carlsbad, CA) as well as by quantitative PCR (KAPA Biosystems, Wilmington, MA, USA). The sequencing libraries were clustered on 3 lanes of a flow cell. After clustering, the flow cell was loaded on the Illumina instrument (4000 or equivalent) according to manufacturer’s instructions. The samples were sequenced using a 2x150bp Paired End (PE) configuration. Image analysis and base calling were conducted by the Control software. Raw sequence data (.bcl files) generated by the sequencer were converted into fastq files and de-multiplexed using Illumina’s bcl2fastq 2.17 software. One mismatch was allowed for index sequence identification.

After investigating the quality of the raw data, sequence reads were trimmed to remove possible adapter sequences and nucleotides with poor quality. The trimmed reads were mapped to the reference genome available on ENSEMBL using the STAR aligner v.2.5.2b. Unique gene hit counts were calculated using feature Counts of Subread package v.1.5.2. Unique reads that fell within the exon regions were counted. Gene hit counts table was used for downstream differential expression analysis, using DESeq2. Comparison of gene expression between the groups of samples was performed, and wald test was used to generate p-values and log2 fold changes. Genes with adjusted p-values < 0.05 and absolute log2 fold changes > 1 were called as differentially expressed genes for each comparison. A gene ontology analysis was performed on the statistically significant set of genes by implementing the software GeneSCF. The mgi GO list was used to cluster the set of genes based on their biological process and determine their statistical significance. A PCA analysis was performed using the “plotPCA” function within the DESeq2 R package. The plot shows the samples in a 2D plane spanned by their first two principal components. The top 500 genes, selected by highest row variance, were used to generate the plot.

### ChIP-sequencing and data analysis

For ChIP on cross-linked chromatin, cells obtained from minced frozen tissue in presence of ice-cold PBS containing protease inhibitor cocktail were fixed with 1% formaldehyde for 10 min and quenched with 125 mM of glycine for 5 minutes at room temperature. Subsequently, cells were washed in PBS, centrifuged to remove the supernatant. Cross liked cells were re-suspended in SDS lysis buffer (100 mM NaCl, 1% SDS, 5 mM ethylenediaminetetraacetic acid (EDTA), 50 mM Tris–Cl and ‘Complete protease inhibitor cocktail’ from Roche). After 10 min on ice with occasional vortexing, cells were diluted in ChIP dilution buffer (0.01% SDS, 1.1% Triton X-100, 1.2 mM EDTA, 16.7 mM Tris–Cl, 167 mM NaCl and ‘Complete protease inhibitor cocktail’ from Roche), and sonicated. Sheared chromatin was incubated with anti-H3K9 tri methyl antibody (Abcam, ab8898) overnight at 4*c with gentle rotation. Genomic DNA was quantified with Qubit 2.0 DNA HS Assay (ThermoFisher, Massachusetts, USA) and quality assessed by TapeStation genomic DNA Assay (Agilent Technologies, California, USA). Library preparation was performed using KAPA Hyper Prep kit (Roche, Basel, Switzerland) per manufacturer’s recommendations. Library quality and quantity were assessed with Qubit 2.0 DNA HS Assay (ThermoFisher, Massachusetts, USA), Tapestation High Sensitivity D1000 Assay (Agilent Technologies, California, USA), and QuantStudio ® 5 System (Applied Biosystems, California, USA). Illumina® 8-nt dual-indices were used. Equimolar pooling of libraries was performed based on QC values and sequenced on an Illumina® NovaSeq S4 (Illumina, California, USA) with a read length configuration of 150 PE for 20M PE reads (10M in each direction) per sample.

Raw sequencing data were processed with Trimmomatic (v0.36)^60^ to filter out low-quality reads and trim adaptor sequences. Reads were aligned to the reference genome using STAR software (v2.5.3a)^61^ with default parameters. Distribution of reads was analyzed using RSeQC (v2.6)^62^. H3K9 trimethylation peak calling was performed with MACS2 (v2.1.1)^63^, followed by annotation and distribution analysis of peaks using bedtools (v2.25.0)^64^. Differential peak analysis was performed using Python script employing the Fisher test. For GO analysis, list of genes with H3K9me3 peaks that were downregulated in expression were input into Enrichr^65^ and top 10 GO biological process terms based on p-value ranking were used.

### Reduced-representation bisulfite sequencing (*RRBS*-Seq) and data analysis

Genomic DNA from control and Kdm1a conditional knockout frozen kidney tissues was isolated and quantified using the Qubit High Sensitivity assay (Life Technologies). For RRBS library preparation, 1 µg of genomic DNA was digested with MspI (NEB), followed by end preparation and adaptor ligation using the Premium RRBS kit (Diagenode). Size selection was performed using AMPure XP beads (Beckman Coulter, Inc.) to obtain DNA fractions of MspI-digested products enriched for the most CpG-rich regions within the 150-350 bp range. Bisulfite treatment was performed using the ZYMO EZ DNA Methylation-Gold Kit. The converted DNAs were amplified through twelve cycles of PCR using 25 μl KAPA HiFi HotStart Uracil+ ReadyMix (2X) and 8-bp index primers, each with a final concentration of 1 μM, followed by clean-up using AMPure XP beads. The constructed RRBS libraries were quantified with a Qubit fluorometer using the Quant-iT dsDNA HS Assay Kit (Invitrogen) and sequenced on the Illumina Novaseq6000 platform with a paired-end 150 bp strategy at CD Genomics (Shirley, NY).

FastQC tool (https://www.bioinformatics.babraham.ac.uk/projects/fastqc/) is used to perform basic statistics on the quality of the raw reads. Sequencing adapters and low-quality data of the sequencing data are removed by Trimmomatic (v0.36)^60^. The BSMAP software is used to map the bisulfite sequence to the reference genome with parameters ‘-n 0 -g 0 -v 0.08 -m 50 -x 1000^66^.

The alignment’s statistical information is compiled, retaining only uniquely mapped read s for subsequent analysis. Methylated cytosines with a sequencing depth of at least 5 ar e considered. If the base in the alignment is C, methylation has occurred. Conversely, if the base is T, no methylation is present. Methylation levels of individual cytosines were calculated as the ratio of the sequenced depth of the ascertained methylated CpG cytosines to the total sequenced depth of individual CpG cytosines, i.e.,ML = mC / (mC + umC). Where ML is the methylation level, mC and umC represent the number of reads supporting methylation C and the number of reads supporting unmethylated C, respectively. The software metilene (v0.2-7) was used to identify DMR (differentially methylated regions) by a binary segmentation algorithm combined with a two-dimensional statistical test (parameters: -M 300 -m 5 -d 0.1 -t 1 -f 1 -v 0.7). Gene Ontology(GO) (http://www.geneontology.org/)^67,68^ enrichment analysis of DMR-related genes are applied to uncover biological processes of interest, we choose to deem pathways with a Q value ≤ 0.05 as significantly enriched with DMR-related genes. Based on the results of the DMR annotation and the database of Kyoto Encyclopedia of Genes and Genomes (KEGG)^69^ functional enrichment analysis is performed on genes whose gene body and its upstream and downstream regions (upstream 2k, gene body, and downstream 2k) that overlap with DMR.

### Data Availability

RNA sequencing, ChIP sequencing and RRBS data generated and described in this study is deposited in NCBI GEO, Accession codes, GSE289719, GSE289720 and GSE290356). Other NGS data used for analysis are publicly available from GEO accession codes, (GSE112570, GSE149134).

## Acknowledgements

This work was supported by grants K01AR069197 (NIH/NIAMS), and R01GM135377 (NIH/NIGMS) to S.K.K, and R21AG047412 (NIH/NIA) to S.B.K.

## Notes

### Competing Interest Statement

The authors have declared no competing interest.

