## Supplementary figures and images for "Lysine-specific demethylase 1a is obligatory for gene regulation during kidney development"

### Suppl Fig 1-3

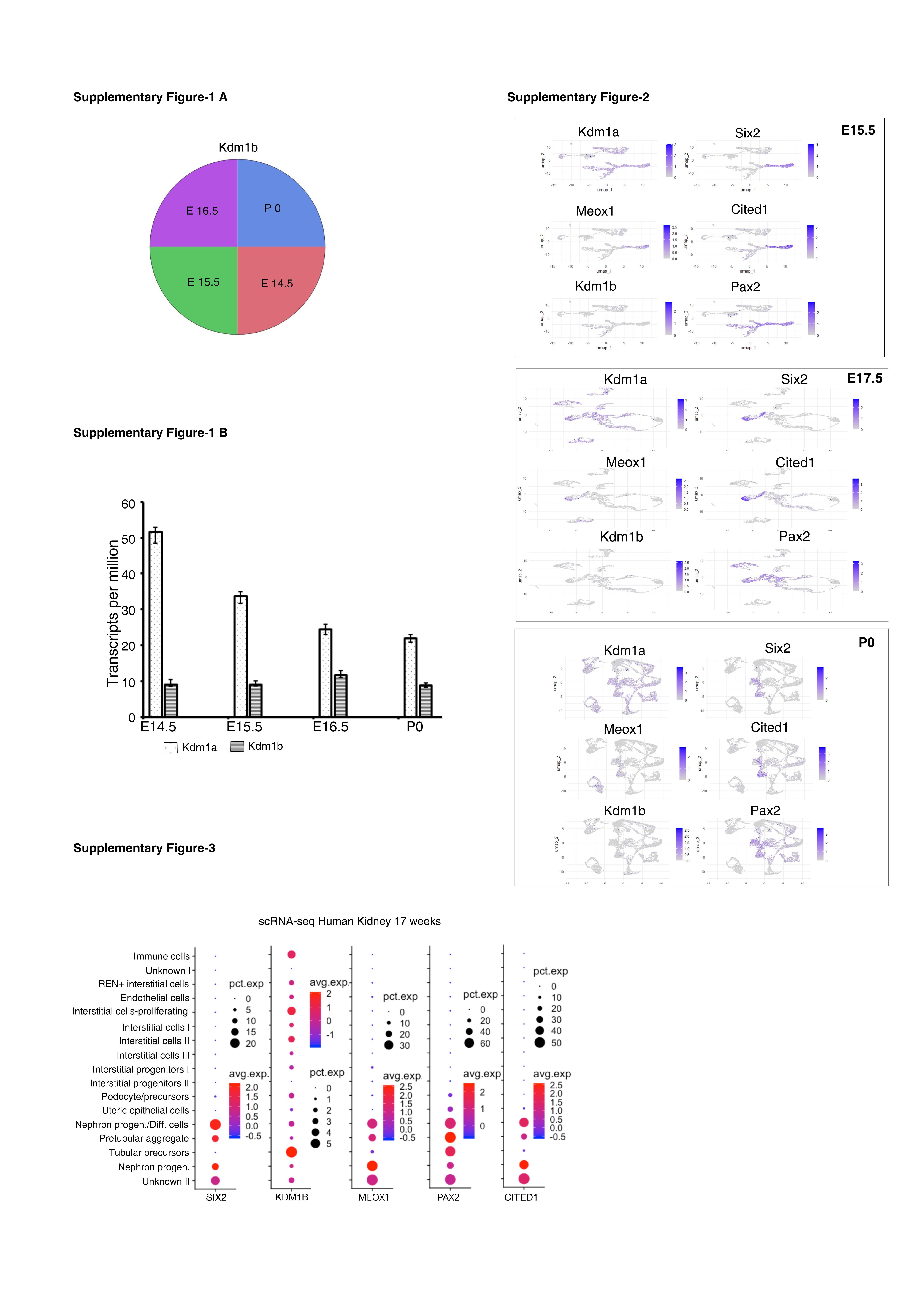

### Suppl Fig 6-8

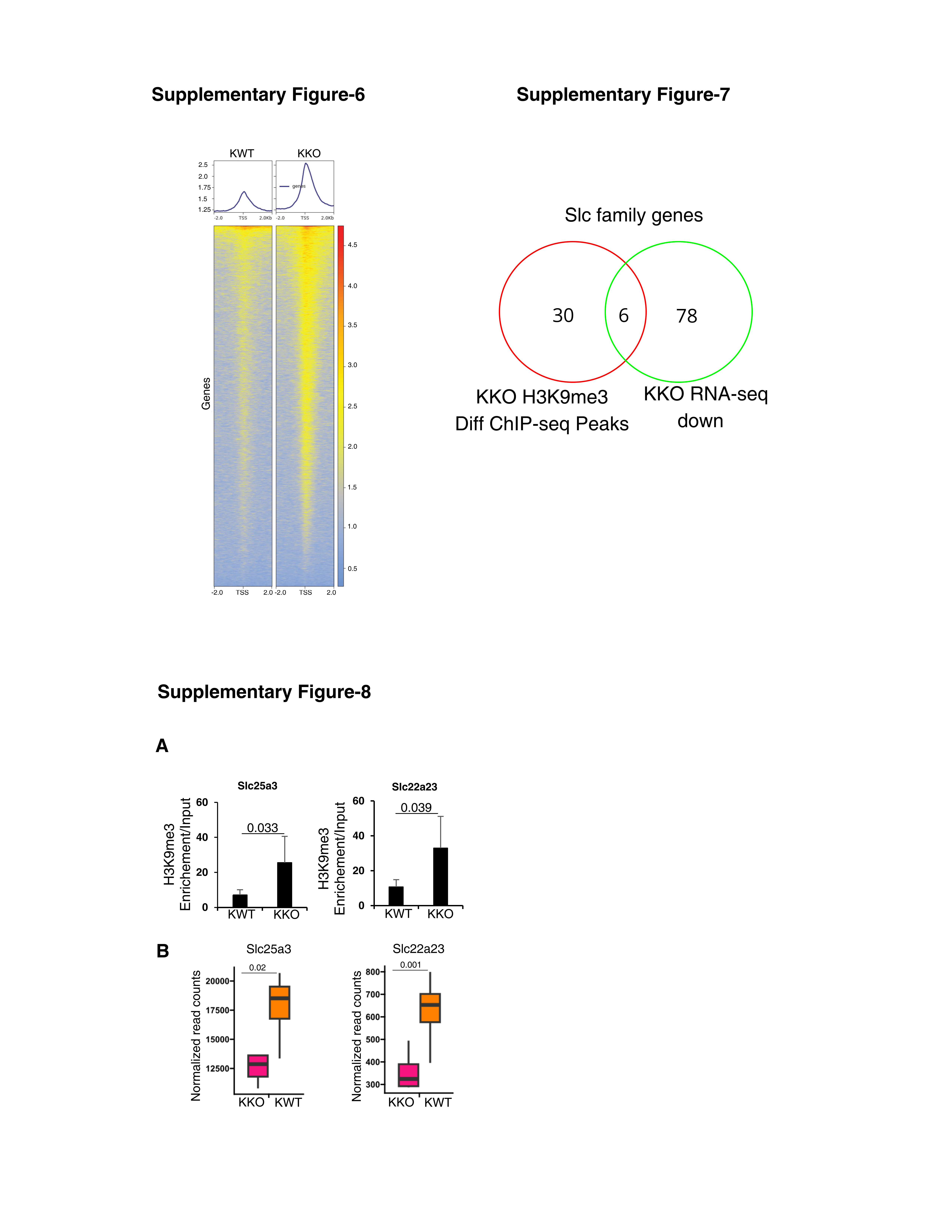

### Suppl Fig 9-12

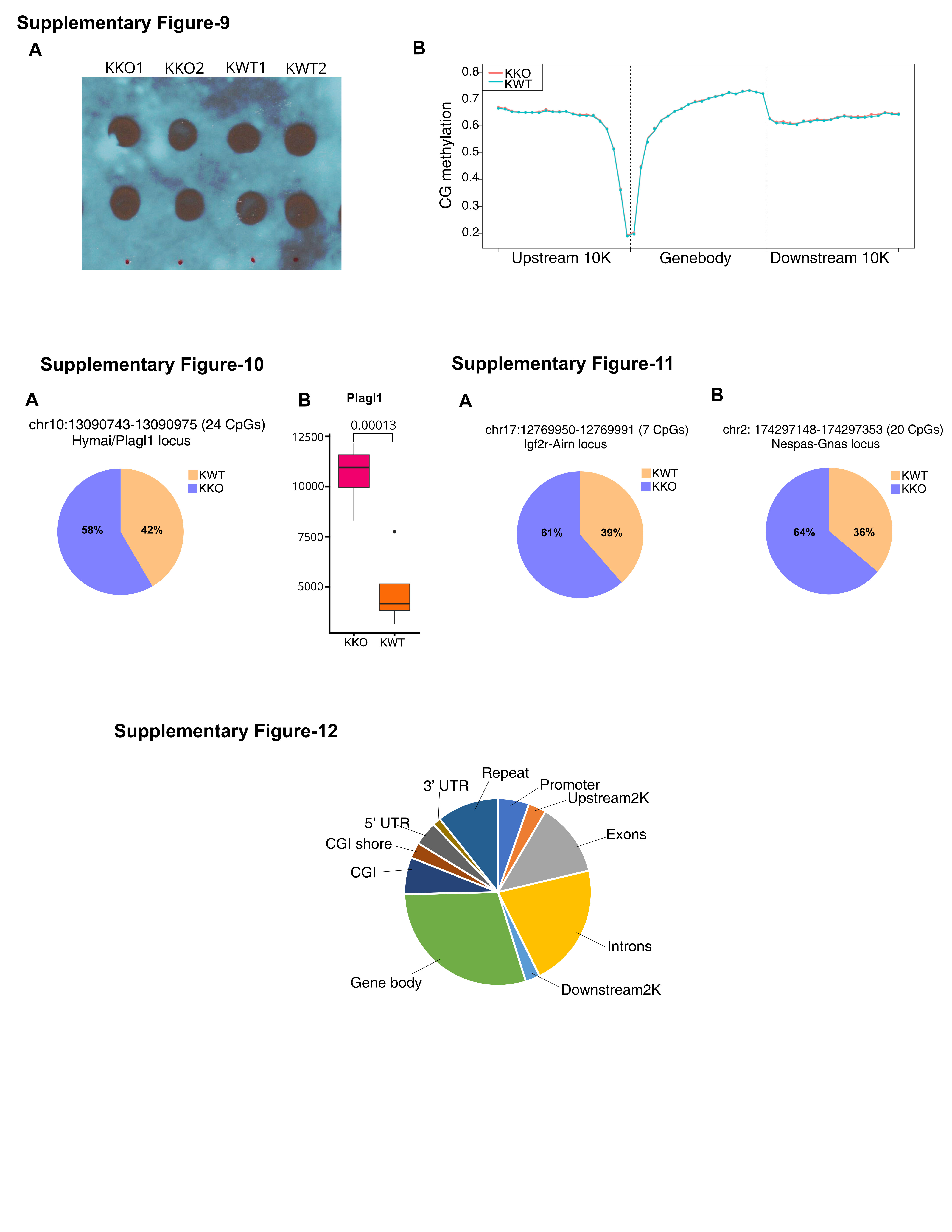

### Suppl Fig-4-5

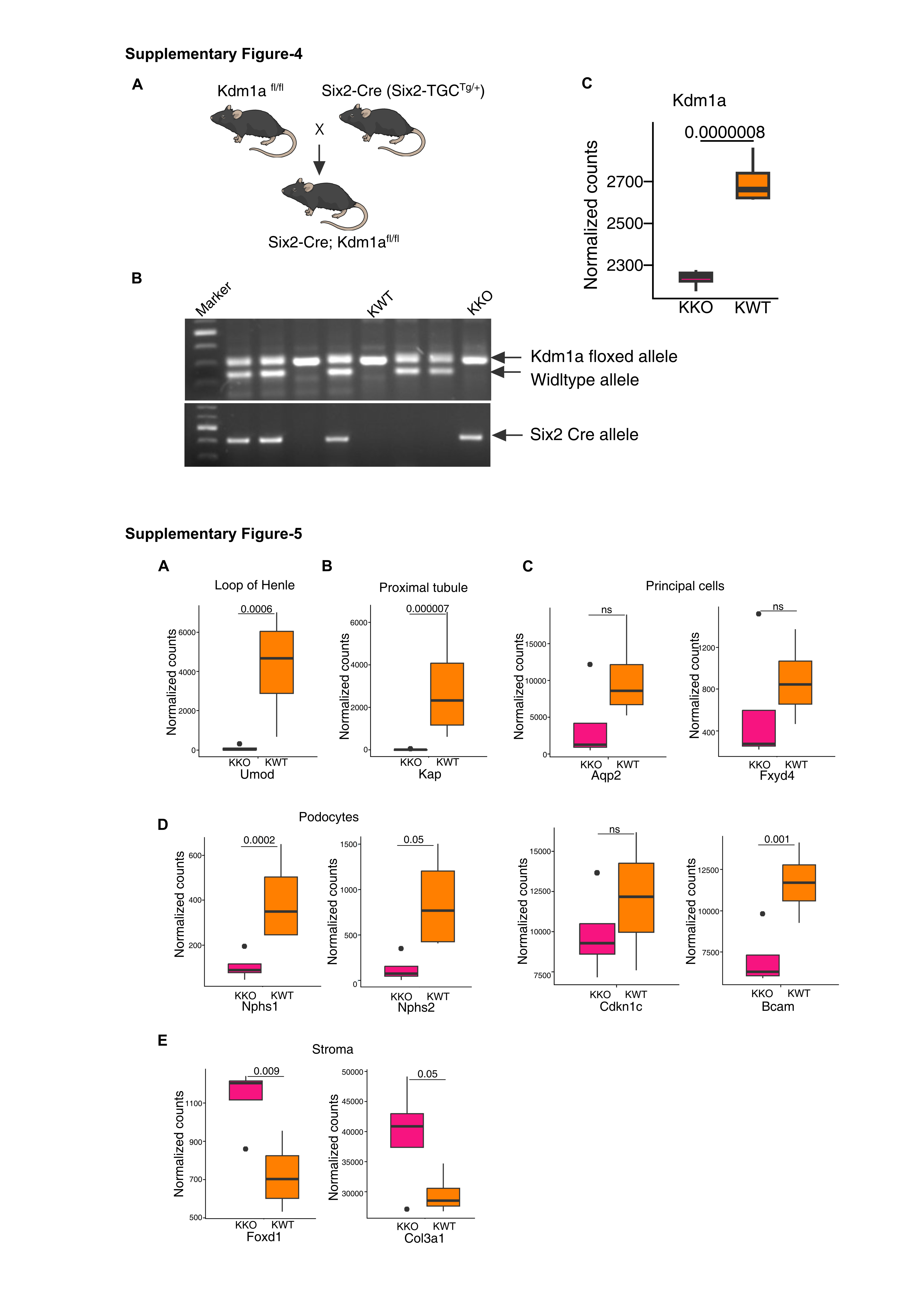
