## Supplementary material for "Lysine-specific demethylase 1a is obligatory for gene regulation during kidney development": Suppl Table-1

**Supplementary Table-1**

| ID | log2FoldChange | pvalue | padj | KKO-1 | KKO-2 | KKO-3 | KKO-4 | KWT-1 | KWT-2 | KWT-3 | KWT-4 |
|---|---|---|---|---|---|---|---|---|---|---|---|
| ENSMUSG00000000001 | -0.120649107 | 0.015826 | 0.059901 | 5200.295 | 5227.622 | 5167.385 | 5110.182 | 5925.751 | 5327.64 | 5632.413 | 5620.671 |
| ENSMUSG00000000028 | -0.17307832 | 0.231663 | 0.392183 | 534.2149 | 670.7002 | 482.7862 | 470.1298 | 701.105 | 553.1002 | 598.8236 | 574.2481 |
| ENSMUSG00000000031 | -0.058614427 | 0.627044 | 0.75698 | 49109.54 | 57909.39 | 52936.72 | 50490.22 | 54264.94 | 67166.47 | 46661.95 | 51078.72 |
| ENSMUSG00000000037 | 0.486799679 | 0.085277 | 0.197374 | 205.2351 | 221.967 | 152.3716 | 237.2215 | 118.9636 | 226.8269 | 143.6791 | 93.53283 |
| ENSMUSG00000000049 | 2.874240897 | 0.008411 | 0.03765 | 117.7084 | 1.199822 | 3.312427 | 115.5915 | 4.875557 | 4.469496 | 15.42863 | 7.613138 |
| ENSMUSG00000000056 | -0.008433724 | 0.917109 | 0.950961 | 1616.729 | 1410.99 | 1467.405 | 1611.381 | 1582.606 | 1400.07 | 1563.113 | 1597.671 |
| ENSMUSG00000000058 | -0.258552366 | 0.301556 | 0.469941 | 581.4994 | 427.1365 | 872.8244 | 614.1879 | 602.6188 | 540.8091 | 911.2533 | 933.1532 |
| ENSMUSG00000000078 | 0.362229802 | 0.0879 | 0.201777 | 2430.627 | 1986.904 | 1857.443 | 2445.538 | 1253.018 | 2344.251 | 1922.793 | 1264.868 |
| ENSMUSG00000000085 | 0.197737876 | 0.055656 | 0.146914 | 991.9696 | 1010.25 | 957.2913 | 1018.758 | 825.9193 | 1029.102 | 837.003 | 777.6276 |
| ENSMUSG00000000088 | -0.830541157 | 2.50E-05 | 0.000467 | 1620.754 | 1643.756 | 2309.589 | 1732.148 | 3114.506 | 2243.687 | 3348.012 | 4289.459 |
| ENSMUSG00000000093 | -0.356058951 | 0.056311 | 0.147981 | 2207.283 | 2148.88 | 3478.876 | 2266.112 | 2980.915 | 2569.96 | 3707.692 | 3672.795 |
| ENSMUSG00000000120 | 0.411055201 | 0.048073 | 0.132462 | 447.6942 | 417.5379 | 553.1752 | 514.9862 | 314.961 | 252.5265 | 399.2157 | 488.3284 |
| ENSMUSG00000000126 | 0.309697915 | 0.245951 | 0.40887 | 112.6781 | 131.9804 | 108.482 | 150.9591 | 91.66046 | 156.4324 | 81.00029 | 78.30656 |
| ENSMUSG00000000127 | -0.058418568 | 0.56705 | 0.711356 | 429.5852 | 466.7306 | 471.1927 | 425.2734 | 443.6756 | 462.5929 | 433.9301 | 527.4817 |
| ENSMUSG00000000131 | 0.042443946 | 0.569053 | 0.712996 | 2204.265 | 2073.292 | 2249.138 | 2178.124 | 2074.062 | 2354.307 | 2011.507 | 2017.481 |
| ENSMUSG00000000134 | 0.000622956 | 0.995601 | 0.997547 | 899.4126 | 785.8831 | 998.6966 | 794.4762 | 907.8286 | 764.2839 | 883.2889 | 923.3648 |
| ENSMUSG00000000142 | 0.607823597 | 0.000163 | 0.001932 | 1344.089 | 1406.191 | 1231.395 | 1227.513 | 934.1566 | 1095.027 | 714.5383 | 674.3065 |
| ENSMUSG00000000148 | 0.045542212 | 0.606294 | 0.742598 | 581.4994 | 649.1034 | 578.8465 | 620.2263 | 638.6979 | 555.3349 | 563.1449 | 593.8247 |
| ENSMUSG00000000149 | -0.114871479 | 0.140774 | 0.27731 | 2821.982 | 3209.523 | 3356.316 | 3027.808 | 3519.177 | 3237.033 | 3185.047 | 3504.218 |
| ENSMUSG00000000154 | -1.335082452 | 0.001083 | 0.008295 | 78.47224 | 76.78858 | 136.6376 | 64.69676 | 239.8774 | 90.5073 | 190.9293 | 379.5693 |
| ENSMUSG00000000157 | -0.169089026 | 0.510103 | 0.66585 | 41.24823 | 40.79393 | 54.65504 | 48.30692 | 48.75557 | 45.81234 | 46.28588 | 68.51824 |
| ENSMUSG00000000159 | -1.090125269 | 2.70E-10 | 4.14E-08 | 404.4339 | 392.3416 | 484.4424 | 359.714 | 942.9326 | 621.26 | 913.1818 | 1016.898 |
| ENSMUSG00000000167 | -0.337731281 | 0.253706 | 0.417294 | 29.17558 | 62.39072 | 47.20208 | 60.38364 | 77.03379 | 56.98608 | 63.64308 | 53.29196 |
| ENSMUSG00000000168 | -0.587486043 | 0.000337 | 0.003383 | 1392.379 | 1264.612 | 1565.95 | 1465.597 | 2087.713 | 1525.216 | 2277.651 | 2659.16 |
| ENSMUSG00000000171 | -0.771417016 | 6.59E-06 | 0.000167 | 2766.649 | 2636.008 | 3445.752 | 2855.284 | 4990.62 | 3505.203 | 5220.662 | 6263.437 |
| ENSMUSG00000000184 | 0.572329327 | 3.10E-06 | 9.31E-05 | 6136.932 | 7438.894 | 6620.713 | 5896.032 | 4528.417 | 5068.409 | 3600.656 | 4350.364 |
| ENSMUSG00000000194 | -0.028954401 | 0.735565 | 0.834534 | 1558.378 | 1546.57 | 1499.701 | 1478.537 | 1541.651 | 1365.431 | 1566.006 | 1731.445 |
| ENSMUSG00000000197 | 0.492791706 | 0.188144 | 0.34043 | 43.26034 | 28.79572 | 72.87338 | 73.323 | 30.22845 | 36.87335 | 53.0359 | 35.89051 |
| ENSMUSG00000000202 | 1.245229166 | 9.48E-05 | 0.001281 | 54.32694 | 103.1847 | 62.9361 | 56.07053 | 31.20356 | 35.75597 | 23.14294 | 26.10219 |
| ENSMUSG00000000204 | 0.606980581 | 0.343304 | 0.51199 | 12.07265 | 19.19714 | 34.78048 | 8.626235 | 5.850668 | 8.938993 | 10.60718 | 23.927 |
| ENSMUSG00000000214 | 3.826258112 | 0.004683 | 0.024588 | 610.675 | 8.398751 | 4.140533 | 444.2511 | 8.776002 | 22.34748 | 11.57147 | 32.62773 |
| ENSMUSG00000000216 | -0.429161691 | 0.123886 | 0.253569 | 681.0988 | 603.5102 | 1286.05 | 690.9614 | 993.6384 | 702.8283 | 1275.755 | 1421.482 |
| ENSMUSG00000000223 | 1.067523913 | 0.001226 | 0.009106 | 91.55095 | 109.1838 | 96.88848 | 87.12497 | 35.10401 | 81.56831 | 33.75012 | 33.71532 |
| ENSMUSG00000000244 | 0.899755876 | 0.031444 | 0.097617 | 19.11503 | 22.79661 | 16.56213 | 25.01608 | 10.72622 | 14.52586 | 7.714313 | 11.9635 |
| ENSMUSG00000000247 | -0.039287055 | 0.939425 | 0.964398 | 14.08476 | 9.598572 | 11.59349 | 7.763611 | 12.67645 | 5.58687 | 10.60718 | 15.22628 |
| ENSMUSG00000000253 | -0.43740379 | 0.028706 | 0.091587 | 723.3531 | 631.1061 | 691.469 | 731.5047 | 1043.369 | 565.3913 | 1038.539 | 1113.693 |
| ENSMUSG00000000266 | 0.004726537 | 0.967325 | 0.981738 | 720.3349 | 569.9152 | 635.1578 | 697.8624 | 584.0917 | 649.1943 | 645.1095 | 739.5619 |
| ENSMUSG00000000275 | 0.288085013 | 0.052718 | 0.141215 | 1704.256 | 1606.561 | 1763.867 | 1169.717 | 1290.072 | 1081.618 | 1323.005 | 1418.219 |
| ENSMUSG00000000276 | -0.025259162 | 0.889009 | 0.933305 | 708.2623 | 609.5093 | 693.9534 | 728.0542 | 764.4873 | 915.1294 | 597.8593 | 512.2554 |
| ENSMUSG00000000278 | -0.124355635 | 0.336421 | 0.504685 | 2962.83 | 2502.828 | 2632.551 | 2823.367 | 3205.191 | 2290.617 | 3030.761 | 3378.058 |
| ENSMUSG00000000282 | -0.059277545 | 0.617211 | 0.750007 | 565.4025 | 571.1151 | 720.4528 | 537.4144 | 673.8019 | 594.443 | 618.1094 | 610.1386 |
| ENSMUSG00000000290 | 0.54004058 | 0.1455 | 0.283834 | 75.45408 | 37.19447 | 40.57722 | 43.13117 | 26.32801 | 22.34748 | 39.53586 | 46.76642 |
| ENSMUSG00000000295 | -0.065747095 | 0.379045 | 0.547176 | 880.2976 | 920.2631 | 963.9161 | 919.5566 | 1018.991 | 1002.285 | 935.3605 | 900.5254 |
| ENSMUSG00000000296 | 0.417276517 | 0.000179 | 0.002079 | 990.9635 | 1057.043 | 1137.819 | 1163.679 | 813.2428 | 944.1811 | 801.3243 | 700.4087 |
| ENSMUSG00000000301 | -0.194334358 | 0.212069 | 0.369694 | 152.9203 | 130.7805 | 134.9814 | 119.042 | 170.6445 | 134.0849 | 148.5005 | 160.9635 |
| ENSMUSG00000000303 | -0.336276794 | 0.077802 | 0.185446 | 3855.2 | 3196.325 | 5396.771 | 3916.311 | 5043.276 | 3755.494 | 5811.771 | 6052.444 |
| ENSMUSG00000000305 | 0.410885109 | 0.028352 | 0.090844 | 973.8606 | 1144.63 | 946.5259 | 998.918 | 680.6277 | 1098.379 | 626.788 | 651.4671 |
| ENSMUSG00000000308 | -1.404995927 | 0.023916 | 0.080457 | 155.9384 | 131.9804 | 548.2066 | 172.5247 | 553.8632 | 274.874 | 734.7883 | 1109.343 |
| ENSMUSG00000000317 | -0.021967118 | 0.849498 | 0.909614 | 616.7113 | 573.5147 | 741.1554 | 582.2709 | 665.0259 | 584.3867 | 617.1451 | 688.4452 |
| ENSMUSG00000000318 | 0.57748542 | 0.190406 | 0.343009 | 42.25428 | 44.3934 | 47.20208 | 31.91707 | 26.32801 | 48.04709 | 27.0001 | 9.78832 |
| ENSMUSG00000000320 | -0.312566969 | 0.180664 | 0.330853 | 123.7447 | 113.983 | 168.1056 | 113.0037 | 158.9431 | 112.8548 | 164.8934 | 208.8175 |
| ENSMUSG00000000325 | -0.290737159 | 0.001906 | 0.012687 | 798.8072 | 747.4888 | 907.6049 | 753.9329 | 951.7086 | 922.951 | 1014.432 | 1038.649 |
| ENSMUSG00000000326 | -0.282556584 | 0.047246 | 0.130749 | 1520.148 | 1624.558 | 1912.926 | 1820.998 | 2214.478 | 1721.873 | 1871.685 | 2561.277 |
| ENSMUSG00000000339 | -0.364013649 | 8.73E-05 | 0.001213 | 1017.121 | 1043.845 | 1117.116 | 1141.251 | 1582.606 | 1280.511 | 1323.969 | 1373.628 |
| ENSMUSG00000000340 | -0.417944352 | 0.000161 | 0.001913 | 1145.896 | 904.6654 | 987.9312 | 1083.455 | 1274.47 | 1244.755 | 1495.612 | 1494.35 |
| ENSMUSG00000000346 | -0.189652969 | 0.025591 | 0.084402 | 5830.085 | 5141.235 | 6356.547 | 6045.265 | 6593.703 | 6162.318 | 6948.668 | 6958.408 |
| ENSMUSG00000000355 | 0.308562295 | 0.000703 | 0.005967 | 780.6982 | 868.6708 | 756.0614 | 880.7386 | 676.7272 | 705.0631 | 614.2522 | 656.905 |
| ENSMUSG00000000359 | 0.156887486 | 0.185425 | 0.337031 | 952.7335 | 1067.841 | 784.217 | 877.2881 | 889.3015 | 735.2322 | 832.1815 | 840.7079 |
| ENSMUSG00000000365 | 0.697048561 | 0.175932 | 0.324376 | 44.26639 | 58.79126 | 60.45178 | 68.14726 | 22.42756 | 80.45093 | 13.50005 | 27.18978 |
| ENSMUSG00000000374 | -0.265878144 | 0.016281 | 0.060951 | 1228.392 | 1069.041 | 1295.159 | 1218.887 | 1428.538 | 1221.29 | 1517.791 | 1620.511 |
| ENSMUSG00000000378 | -0.059436172 | 0.482178 | 0.640887 | 1009.073 | 1055.843 | 958.1194 | 967.8636 | 1082.374 | 935.2421 | 1012.504 | 1124.569 |
| ENSMUSG00000000384 | -0.530290068 | 0.000256 | 0.002753 | 902.4308 | 863.8715 | 996.2123 | 924.7324 | 1449.015 | 981.0545 | 1345.183 | 1550.905 |

| | | | | | | | | | | | |
|---|---|---|---|---|---|---|---|---|---|---|---|
| ENSMUSG00000000385 | -1.535046527 | 0.001534 | 0.010701 | 908.4671 | 554.3175 | 2042.111 | 952.3363 | 3530.878 | 1622.427 | 3598.727 | 4167.649 |
| ENSMUSG00000000386 | 1.210412926 | 0.059355 | 0.153577 | 11.0666 | 21.59679 | 41.40533 | 8.626235 | 13.65156 | 8.938993 | 7.714313 | 5.437955 |
| ENSMUSG00000000392 | 0.819540097 | 0.001601 | 0.011069 | 494.9787 | 856.6726 | 466.224 | 701.3129 | 388.0943 | 491.6446 | 297.0011 | 250.146 |
| ENSMUSG00000000399 | -0.678506725 | 9.12E-06 | 0.000213 | 2535.257 | 2406.842 | 2766.704 | 2542.151 | 4282.689 | 2957.689 | 4082.8 | 5084.488 |
| ENSMUSG00000000402 | -0.081300411 | 0.616551 | 0.749474 | 574.457 | 688.6976 | 772.6235 | 708.2139 | 713.7815 | 538.5743 | 798.4314 | 852.6714 |
| ENSMUSG00000000409 | -0.007049328 | 0.952963 | 0.973191 | 182.0958 | 219.5673 | 203.7142 | 204.4418 | 202.8232 | 208.949 | 206.3579 | 194.6788 |
| ENSMUSG00000000416 | 0.022077975 | 0.890797 | 0.934016 | 653.9353 | 565.1159 | 447.1776 | 596.0728 | 471.9539 | 684.9503 | 538.0734 | 532.9196 |
| ENSMUSG00000000420 | -0.203702059 | 0.002605 | 0.015894 | 2566.445 | 2661.204 | 2602.739 | 2641.353 | 2941.911 | 2801.257 | 3007.618 | 3308.452 |
| ENSMUSG00000000439 | -0.015697397 | 0.800257 | 0.877926 | 882.3097 | 875.8697 | 846.325 | 896.2658 | 892.2268 | 895.0167 | 873.646 | 877.686 |
| ENSMUSG00000000440 | -0.376118805 | 0.165723 | 0.311164 | 92.557 | 62.39072 | 129.1846 | 87.9876 | 150.1671 | 88.27255 | 134.0362 | 112.0219 |
| ENSMUSG00000000441 | -0.167588631 | 0.03222 | 0.099525 | 1483.93 | 1648.555 | 1605.699 | 1483.712 | 1804.931 | 1573.263 | 1816.721 | 1789.087 |
| ENSMUSG00000000486 | 0.398585918 | 0.003088 | 0.018171 | 273.6468 | 236.3648 | 235.1823 | 276.0395 | 192.0969 | 223.4748 | 185.1435 | 175.1022 |
| ENSMUSG00000000489 | -0.473768942 | 0.011065 | 0.046079 | 382.3007 | 327.5513 | 517.5666 | 407.1583 | 500.2321 | 460.3581 | 611.3593 | 701.4963 |
| ENSMUSG00000000530 | -0.158174215 | 0.213722 | 0.371838 | 705.2441 | 704.2952 | 838.872 | 770.3228 | 815.1931 | 724.0584 | 811.9315 | 1020.16 |
| ENSMUSG00000000531 | -0.321066775 | 0.023974 | 0.080535 | 414.4944 | 425.9366 | 515.9104 | 455.4652 | 640.6481 | 459.2408 | 635.4666 | 528.5693 |
| ENSMUSG00000000532 | -0.3052487 | 0.031597 | 0.097993 | 963.8001 | 682.6985 | 795.8105 | 839.3327 | 929.2811 | 863.7302 | 1104.111 | 1159.372 |
| ENSMUSG00000000538 | -0.394824433 | 0.007046 | 0.033083 | 1284.731 | 1073.84 | 1300.956 | 1125.724 | 1370.031 | 1273.806 | 1802.256 | 1846.73 |
| ENSMUSG00000000544 | -0.207234519 | 0.745199 | 0.840853 | 10.06054 | 8.398751 | 34.78048 | 8.626235 | 31.20356 | 11.17374 | 15.42863 | 14.13868 |
| ENSMUSG00000000552 | 0.166552086 | 0.095688 | 0.214115 | 760.5771 | 886.6681 | 846.325 | 876.4255 | 831.7699 | 783.2792 | 675.0024 | 712.3722 |
| ENSMUSG00000000555 | 0.145875332 | 0.117995 | 0.246712 | 1192.174 | 1177.025 | 1410.266 | 1286.172 | 1224.74 | 1034.688 | 1141.718 | 1180.036 |
| ENSMUSG00000000560 | 1.130104087 | 0.001341 | 0.009746 | 216.3017 | 263.9607 | 175.5586 | 197.5408 | 98.48624 | 168.7235 | 78.10742 | 44.59123 |
| ENSMUSG00000000561 | 0.113452722 | 0.261537 | 0.425729 | 1535.239 | 1542.97 | 1456.64 | 1673.49 | 1359.305 | 1681.648 | 1394.362 | 1305.109 |
| ENSMUSG00000000563 | -0.476256165 | 0.000127 | 0.001609 | 5810.97 | 5667.957 | 6740.788 | 6391.177 | 8477.618 | 6837.212 | 8889.782 | 10035.2 |
| ENSMUSG00000000567 | 0.903174906 | 0.008697 | 0.038501 | 1574.475 | 1679.75 | 1026.024 | 1284.446 | 561.6641 | 1429.121 | 581.4664 | 403.4963 |
| ENSMUSG00000000568 | 0.039782959 | 0.644931 | 0.770145 | 8632.952 | 9140.24 | 8078.18 | 8935.917 | 8006.639 | 9680.929 | 8250.458 | 7902.437 |
| ENSMUSG00000000579 | 0.004610127 | 0.973662 | 0.985571 | 284.7134 | 225.5664 | 223.5888 | 242.3972 | 278.8818 | 222.3574 | 227.5722 | 243.6204 |
| ENSMUSG00000000581 | 0.042109396 | 0.597972 | 0.736677 | 2051.345 | 2297.658 | 2133.203 | 2245.409 | 2087.713 | 2346.486 | 1951.721 | 2091.438 |
| ENSMUSG00000000594 | 0.215816242 | 0.000208 | 0.002346 | 2693.208 | 2571.218 | 2538.147 | 2708.638 | 2401.699 | 2231.396 | 2189.901 | 2227.387 |
| ENSMUSG00000000600 | 0.151555909 | 0.120616 | 0.249718 | 1348.113 | 1277.81 | 1149.412 | 1471.636 | 1216.939 | 1255.928 | 1140.754 | 1110.431 |
| ENSMUSG00000000605 | -0.015176596 | 0.768969 | 0.856611 | 2075.49 | 1968.907 | 1949.363 | 2056.494 | 2018.48 | 2085.02 | 2019.222 | 2014.219 |
| ENSMUSG00000000617 | 0.077022543 | 0.803933 | 0.880139 | 35.2119 | 31.19536 | 24.01509 | 31.05445 | 24.37778 | 24.58223 | 28.92867 | 36.9781 |
| ENSMUSG00000000627 | 0.883962958 | 0.000314 | 0.003219 | 124.7507 | 111.5834 | 124.216 | 133.7066 | 65.33246 | 99.44629 | 54.00019 | 50.02919 |
| ENSMUSG00000000628 | 0.134833343 | 0.511496 | 0.667009 | 1056.357 | 847.074 | 763.5143 | 873.8376 | 743.0348 | 912.8946 | 1040.468 | 527.4817 |
| ENSMUSG00000000631 | -0.339558876 | 0.023943 | 0.080488 | 1995.006 | 1830.928 | 2384.947 | 1880.519 | 2350.993 | 2019.095 | 2766.546 | 3103.985 |
| ENSMUSG00000000632 | 0.569248594 | 0.182448 | 0.33303 | 16.09687 | 25.19625 | 18.21835 | 22.42821 | 7.80089 | 13.40849 | 14.46434 | 19.57664 |
| ENSMUSG00000000673 | -3.119130414 | 1.80E-06 | 5.98E-05 | 10.06054 | 5.999108 | 6.624853 | 12.07673 | 77.03379 | 20.11273 | 36.64299 | 169.6642 |
| ENSMUSG00000000682 | -0.490234591 | 0.261347 | 0.425537 | 28.16952 | 20.39697 | 47.20208 | 21.56559 | 37.05423 | 20.11273 | 54.00019 | 54.37955 |
| ENSMUSG00000000686 | -1.332481725 | 0.021999 | 0.075423 | 6.036326 | 3.599465 | 14.90592 | 11.21411 | 37.05423 | 12.29112 | 17.3572 | 25.0146 |
| ENSMUSG00000000690 | 0.178637541 | 0.004981 | 0.025691 | 2086.557 | 1968.907 | 1931.973 | 1901.222 | 1672.316 | 1693.939 | 1758.863 | 1843.467 |
| ENSMUSG00000000693 | 0.157746468 | 0.085892 | 0.198451 | 870.237 | 878.2694 | 874.4806 | 899.7163 | 763.5122 | 781.0445 | 901.6104 | 710.197 |
| ENSMUSG00000000708 | 0.245453753 | 0.03501 | 0.106117 | 781.7042 | 605.9099 | 655.8605 | 678.0221 | 501.2072 | 583.2693 | 583.3949 | 629.7152 |
| ENSMUSG00000000711 | -0.105122369 | 0.314889 | 0.483212 | 1997.018 | 1860.923 | 2076.063 | 1777.004 | 2182.299 | 1724.108 | 2217.865 | 2168.657 |
| ENSMUSG00000000724 | 2.141414653 | NA | NA | 2.012109 | 0 | 54.65504 | 0.862623 | 7.80089 | 2.234748 | 1.928578 | 1.087591 |
| ENSMUSG00000000730 | 1.421697775 | 0.030466 | 0.095511 | 30.18163 | 7.198929 | 17.39024 | 13.80198 | 11.70134 | 5.58687 | 2.892867 | 5.437955 |
| ENSMUSG00000000731 | -0.312420158 | 0.259759 | 0.42377 | 46.2785 | 58.79126 | 45.54586 | 44.85642 | 68.25779 | 72.62932 | 63.64308 | 36.9781 |
| ENSMUSG00000000732 | -0.341487082 | 0.079758 | 0.188683 | 244.4712 | 208.7689 | 352.7734 | 239.8093 | 329.5876 | 267.0524 | 354.8584 | 376.3065 |
| ENSMUSG00000000738 | -0.352851306 | 0.001315 | 0.009604 | 1528.197 | 1516.574 | 1620.605 | 1500.102 | 2009.704 | 1582.202 | 2131.079 | 2150.168 |
| ENSMUSG00000000739 | 1.1428241 | 8.01E-08 | 4.77E-06 | 339.0403 | 439.1347 | 336.2113 | 474.4429 | 142.3663 | 239.1181 | 194.7864 | 143.562 |
| ENSMUSG00000000740 | -0.153226088 | 0.065571 | 0.164783 | 36696.84 | 45154.08 | 43408.52 | 41530.15 | 49690.7 | 43620.05 | 46957.03 | 45207.9 |
| ENSMUSG00000000743 | -0.133171992 | 0.134739 | 0.268691 | 2422.579 | 2513.626 | 2729.439 | 2630.139 | 3036.497 | 2436.993 | 2865.867 | 2952.81 |
| ENSMUSG00000000751 | -0.048533351 | 0.466532 | 0.627229 | 2093.599 | 2451.235 | 2362.588 | 2289.403 | 2324.665 | 2421.35 | 2386.616 | 2377.474 |
| ENSMUSG00000000753 | 0.520456188 | 0.000755 | 0.006303 | 1419.543 | 1976.106 | 1676.916 | 1774.417 | 1206.213 | 1505.103 | 1027.932 | 1034.299 |
| ENSMUSG00000000759 | -0.056361073 | 0.448393 | 0.611056 | 1631.82 | 1784.135 | 1581.684 | 1498.377 | 1751.3 | 1657.066 | 1703.899 | 1634.649 |
| ENSMUSG00000000776 | -0.199019254 | 0.028441 | 0.091004 | 859.1704 | 825.4772 | 809.8883 | 790.1631 | 1057.021 | 859.2607 | 887.146 | 964.6933 |
| ENSMUSG00000000782 | 0.307961907 | 0.011138 | 0.046317 | 906.455 | 929.8617 | 1059.976 | 907.4799 | 870.7744 | 850.3217 | 673.0738 | 679.7444 |
| ENSMUSG00000000787 | 0.045671635 | 0.672589 | 0.790252 | 12757.78 | 10652.02 | 10959.16 | 13454.34 | 11205 | 13255.41 | 10721.93 | 11154.33 |
| ENSMUSG00000000794 | -0.220558173 | 0.3674 | 0.535942 | 72.43591 | 83.98751 | 101.029 | 106.1027 | 78.98402 | 106.1505 | 143.6791 | 95.70802 |
| ENSMUSG00000000804 | 0.052060618 | 0.628348 | 0.75801 | 1261.592 | 1017.449 | 1114.632 | 1201.635 | 1086.274 | 1273.806 | 1060.718 | 1014.722 |
| ENSMUSG00000000805 | -2.427067873 | 5.31E-06 | 0.000141 | 146.8839 | 56.39161 | 278.2438 | 196.6782 | 865.8988 | 497.2315 | 841.8244 | 1447.584 |
| ENSMUSG00000000811 | -0.045208932 | 0.695487 | 0.807789 | 1061.387 | 1265.812 | 1410.266 | 1341.38 | 1397.335 | 1379.957 | 1323.005 | 1141.971 |
| ENSMUSG00000000823 | 0.267768347 | 0.000185 | 0.002131 | 2694.214 | 2757.19 | 2781.61 | 3004.518 | 2256.408 | 2542.026 | 2207.258 | 2332.883 |
| ENSMUSG00000000826 | -0.154481873 | 0.029968 | 0.094508 | 3725.419 | 3787.837 | 4105.753 | 3762.764 | 4357.772 | 3892.931 | 4306.515 | 4564.62 |
| ENSMUSG00000000827 | -0.083204095 | 0.09992 | 0.2207 | 2567.451 | 2625.21 | 2650.769 | 2679.309 | 2903.881 | 2737.567 | 2687.474 | 2820.124 |
| ENSMUSG00000000838 | 0.14003816 | 0.197055 | 0.351414 | 2441.694 | 2496.829 | 2150.593 | 2538.701 | 2069.186 | 2599.012 | 1983.543 | 2086 |

| | | | | | | | | | | | |
|---|---|---|---|---|---|---|---|---|---|---|---|
| ENSMUSG00000000861 | 1.515011403 | 1.82E-10 | 3.00E-08 | 297.7921 | 329.9509 | 227.7293 | 265.688 | 79.95913 | 145.2586 | 75.21455 | 92.44524 |
| ENSMUSG00000000869 | 0.629692636 | 0.073429 | 0.178433 | 133.8052 | 160.7761 | 130.8408 | 177.7004 | 90.68535 | 174.3104 | 75.21455 | 50.02919 |
| ENSMUSG00000000876 | -0.493172809 | 0.003992 | 0.021812 | 819.9343 | 790.6824 | 799.1229 | 809.1408 | 1073.598 | 796.6877 | 1182.219 | 1478.036 |
| ENSMUSG00000000881 | 0.303648237 | 0.000487 | 0.004486 | 1256.562 | 1244.215 | 1118.772 | 1205.948 | 935.1317 | 1096.144 | 967.182 | 910.3137 |
| ENSMUSG00000000884 | -0.251869645 | 0.0453 | 0.12706 | 197.1867 | 207.5691 | 248.432 | 229.4578 | 266.2054 | 251.4092 | 285.4296 | 250.146 |
| ENSMUSG00000000889 | 3.693565762 | 0.002777 | 0.016703 | 444.676 | 11.99822 | 8.281066 | 556.3922 | 18.52711 | 10.05637 | 31.82154 | 18.48905 |
| ENSMUSG00000000901 | 0.105564826 | 0.358325 | 0.526574 | 1053.339 | 1214.219 | 1373.001 | 1337.929 | 1185.735 | 1295.037 | 1048.182 | 1101.73 |
| ENSMUSG00000000902 | -0.113295561 | 0.119983 | 0.248921 | 2060.399 | 2166.878 | 2305.449 | 2015.951 | 2173.523 | 2307.378 | 2302.723 | 2465.569 |
| ENSMUSG00000000903 | 0.928624157 | 0.001882 | 0.012567 | 52.31483 | 55.19179 | 37.2648 | 49.16954 | 29.25334 | 30.1691 | 18.32149 | 23.927 |
| ENSMUSG00000000915 | -0.661326628 | 0.008845 | 0.039024 | 1133.823 | 830.2765 | 1757.242 | 1116.235 | 1805.906 | 1239.168 | 2258.365 | 2350.284 |
| ENSMUSG00000000916 | -0.124147618 | 0.161667 | 0.305616 | 458.7608 | 469.1302 | 451.3181 | 418.3724 | 509.9832 | 453.6539 | 504.3232 | 487.2408 |
| ENSMUSG00000000934 | -0.021733443 | 0.884882 | 0.931253 | 353.1251 | 305.9545 | 313.0243 | 285.5284 | 270.1058 | 300.5736 | 309.5368 | 396.9707 |
| ENSMUSG00000000938 | 0.029763386 | 0.731297 | 0.831495 | 3713.347 | 3804.634 | 3622.138 | 4140.593 | 3736.627 | 3300.723 | 3825.335 | 4105.656 |
| ENSMUSG00000000942 | 0.0780952 | 0.712161 | 0.819637 | 105.6357 | 112.7832 | 94.40416 | 128.5309 | 98.48624 | 135.2023 | 104.1432 | 80.48174 |
| ENSMUSG00000000948 | 0.553567462 | 0.003977 | 0.021776 | 140.8476 | 145.1784 | 121.7317 | 125.943 | 108.2374 | 67.04245 | 98.35749 | 88.09488 |
| ENSMUSG00000000957 | 0.535167007 | 0.00014 | 0.001735 | 6004.132 | 6780.191 | 5800.059 | 6229.867 | 3938.475 | 5595.809 | 3941.05 | 3648.868 |
| ENSMUSG00000000958 | -0.737487383 | 0.003997 | 0.021824 | 320.9313 | 463.1311 | 445.5214 | 452.8773 | 910.754 | 434.6585 | 536.1448 | 923.3648 |
| ENSMUSG00000000959 | -0.069280861 | 0.423104 | 0.588279 | 3173.095 | 2725.995 | 3142.665 | 3227.074 | 3346.582 | 2875.004 | 3377.905 | 3275.824 |
| ENSMUSG00000000976 | 0.101252745 | 0.319642 | 0.487789 | 740.456 | 829.0767 | 693.1253 | 683.1978 | 700.1299 | 749.758 | 669.2167 | 623.1897 |
| ENSMUSG00000001016 | 0.051047473 | 0.441022 | 0.604313 | 4842.14 | 5068.046 | 4651.475 | 5220.597 | 4876.532 | 5069.526 | 4628.588 | 4518.941 |
| ENSMUSG00000001017 | 0.015387074 | 0.734038 | 0.833414 | 5694.268 | 6066.298 | 5725.529 | 5930.537 | 5890.647 | 5772.355 | 5629.52 | 5871.904 |
| ENSMUSG00000001018 | 0.24428188 | 0.009568 | 0.04124 | 1442.682 | 1527.373 | 1389.563 | 1680.391 | 1344.678 | 1369.901 | 1218.862 | 1165.898 |
| ENSMUSG00000001020 | -0.583771345 | 0.042402 | 0.12154 | 33.19979 | 34.79482 | 37.2648 | 58.6584 | 71.18313 | 58.10345 | 48.21446 | 69.60583 |
| ENSMUSG00000001025 | 0.237864165 | 0.133953 | 0.267557 | 2257.586 | 2333.653 | 3449.892 | 2368.764 | 2596.721 | 2138.654 | 2133.972 | 1959.839 |
| ENSMUSG00000001027 | 0.479801951 | 0.01187 | 0.048577 | 98.59333 | 123.5816 | 96.06037 | 119.9047 | 81.90935 | 89.38993 | 66.53595 | 76.13138 |
| ENSMUSG00000001029 | -0.664308665 | 8.86E-05 | 0.001224 | 355.1372 | 297.5557 | 397.4912 | 366.615 | 471.9539 | 464.8276 | 610.395 | 701.4963 |
| ENSMUSG00000001034 | -0.102683206 | 0.255288 | 0.419218 | 948.7093 | 1013.849 | 1034.305 | 984.2534 | 1223.765 | 1014.576 | 1008.646 | 1026.686 |
| ENSMUSG00000001036 | 0.127356751 | 0.092752 | 0.209295 | 1341.07 | 1279.01 | 1260.378 | 1296.523 | 1187.686 | 1255.928 | 1232.362 | 1062.576 |
| ENSMUSG00000001039 | -0.237345858 | 0.149666 | 0.289574 | 387.3309 | 511.124 | 591.2681 | 489.9701 | 659.1752 | 464.8276 | 635.4666 | 574.2481 |
| ENSMUSG00000001052 | 0.177111518 | 0.051346 | 0.138565 | 1941.685 | 1970.107 | 1841.709 | 1893.459 | 1614.784 | 1965.461 | 1549.613 | 1634.649 |
| ENSMUSG00000001053 | -0.200406584 | 0.127227 | 0.258402 | 494.9787 | 490.727 | 643.4389 | 473.5803 | 651.3744 | 532.9874 | 620.0379 | 613.4014 |
| ENSMUSG00000001054 | -0.206278849 | 0.018268 | 0.066232 | 811.8859 | 767.8858 | 905.9487 | 869.5245 | 1018.016 | 908.4251 | 929.5748 | 1020.16 |
| ENSMUSG00000001056 | -0.440818522 | 0.000268 | 0.002845 | 1730.413 | 2007.301 | 2029.689 | 1739.049 | 2898.031 | 2096.194 | 2444.473 | 2748.343 |
| ENSMUSG00000001062 | 0.032720816 | 0.676195 | 0.792674 | 790.7587 | 704.2952 | 739.4992 | 821.2176 | 756.6864 | 744.1711 | 755.0384 | 734.124 |
| ENSMUSG00000001065 | 0.069063051 | 0.4151 | 0.581002 | 680.0927 | 711.4942 | 631.8454 | 627.1273 | 642.5984 | 615.6731 | 595.9307 | 668.8685 |
| ENSMUSG00000001076 | 1.268371504 | 4.48E-07 | 1.92E-05 | 179.0777 | 272.3595 | 198.7456 | 220.8316 | 115.0631 | 82.68568 | 107.0361 | 55.46715 |
| ENSMUSG00000001082 | 0.056706183 | 0.40273 | 0.569693 | 1313.907 | 1315.004 | 1294.331 | 1337.929 | 1298.848 | 1150.895 | 1316.255 | 1289.883 |
| ENSMUSG00000001089 | -0.091595272 | 0.165145 | 0.310557 | 1858.182 | 2052.895 | 2055.361 | 1876.206 | 2055.535 | 2112.954 | 2007.65 | 2180.62 |
| ENSMUSG00000001095 | -2.568567286 | 1.66E-07 | 8.51E-06 | 9.054489 | 10.79839 | 24.01509 | 13.80198 | 102.3867 | 35.75597 | 69.42882 | 138.1241 |
| ENSMUSG00000001098 | -0.328149025 | 0.029133 | 0.092653 | 1480.912 | 1498.577 | 1473.202 | 1223.2 | 1921.944 | 1289.45 | 1926.65 | 1984.854 |
| ENSMUSG00000001100 | -0.39274569 | 2.79E-06 | 8.54E-05 | 2516.142 | 2552.02 | 2590.318 | 2498.158 | 3493.824 | 2852.656 | 3467.584 | 3518.357 |
| ENSMUSG00000001105 | -0.066051577 | 0.411846 | 0.577629 | 1586.548 | 1791.334 | 1565.95 | 1544.959 | 1603.083 | 1843.667 | 1666.292 | 1675.978 |
| ENSMUSG00000001119 | 0.190786139 | 0.148215 | 0.287688 | 4475.936 | 5975.111 | 6034.413 | 5455.231 | 3925.798 | 5364.513 | 4840.732 | 5093.189 |
| ENSMUSG00000001120 | -0.249477007 | 0.025031 | 0.083147 | 589.5479 | 555.5174 | 685.6723 | 682.3352 | 730.3584 | 677.1287 | 831.2173 | 752.613 |
| ENSMUSG00000001123 | 0.143696973 | 0.509783 | 0.665571 | 919.5337 | 1378.595 | 1667.807 | 833.2943 | 1095.05 | 916.2468 | 1167.754 | 1164.81 |
| ENSMUSG00000001127 | 0.148506667 | 0.013457 | 0.053275 | 3218.368 | 3121.936 | 3063.166 | 3109.758 | 2641.577 | 2945.398 | 2746.296 | 2958.248 |
| ENSMUSG00000001128 | 0.640265319 | 0.004901 | 0.025356 | 338.0343 | 341.9491 | 289.0092 | 297.6051 | 209.6489 | 286.0478 | 128.2505 | 189.2408 |
| ENSMUSG00000001131 | 0.440401028 | 0.001265 | 0.009324 | 259.562 | 244.7636 | 215.3077 | 277.7648 | 183.3209 | 201.1273 | 185.1435 | 165.3138 |
| ENSMUSG00000001134 | -0.538119084 | 7.61E-05 | 0.001103 | 275.6589 | 255.562 | 355.2577 | 301.0556 | 455.377 | 392.1983 | 410.7872 | 470.9269 |
| ENSMUSG00000001138 | 0.196064142 | 0.053968 | 0.143808 | 960.7819 | 1011.45 | 939.0729 | 975.6272 | 810.3175 | 1000.05 | 821.5744 | 761.3138 |
| ENSMUSG00000001143 | -0.096788373 | 0.11671 | 0.244967 | 838.0433 | 866.2711 | 829.7628 | 836.7448 | 895.1522 | 896.134 | 888.1103 | 923.3648 |
| ENSMUSG00000001150 | 0.043503425 | 0.508535 | 0.664274 | 1392.379 | 1372.596 | 1518.748 | 1385.373 | 1369.056 | 1439.178 | 1374.112 | 1322.511 |
| ENSMUSG00000001151 | 0.168388897 | 0.051671 | 0.139116 | 1248.513 | 1052.243 | 1141.959 | 1061.89 | 1002.414 | 1019.045 | 1055.897 | 930.978 |
| ENSMUSG00000001155 | 0.434729498 | 0.44775 | 0.610316 | 20.12109 | 10.79839 | 14.07781 | 22.42821 | 2.925334 | 14.52586 | 15.42863 | 17.40146 |
| ENSMUSG00000001156 | 0.185887005 | 0.249222 | 0.412686 | 592.566 | 424.7368 | 385.0696 | 469.2672 | 427.0988 | 427.9543 | 444.5373 | 344.7664 |
| ENSMUSG00000001157 | -0.055843017 | 0.526289 | 0.679434 | 1126.781 | 1072.64 | 971.3691 | 1041.187 | 1081.398 | 1210.116 | 1080.968 | 1003.847 |
| ENSMUSG00000001158 | -0.139379983 | 0.066909 | 0.166964 | 1486.948 | 1546.57 | 1440.077 | 1383.648 | 1597.232 | 1771.038 | 1544.791 | 1535.679 |
| ENSMUSG00000001166 | -0.163553785 | 0.590905 | 0.73091 | 45.27245 | 71.98929 | 79.49824 | 50.03216 | 73.13335 | 71.51194 | 87.75031 | 43.50364 |
| ENSMUSG00000001173 | 0.313332059 | 0.004789 | 0.024962 | 1787.759 | 1526.173 | 1427.656 | 1657.962 | 1194.511 | 1479.403 | 1306.612 | 1170.248 |
| ENSMUSG00000001175 | -0.152108772 | 0.044913 | 0.126301 | 8658.104 | 9834.937 | 9834.594 | 9012.69 | 9880.803 | 9882.057 | 10402.75 | 11326.17 |
| ENSMUSG00000001211 | -0.572596495 | 2.40E-05 | 0.000452 | 2623.79 | 2499.228 | 2929.841 | 2579.244 | 3796.108 | 3054.901 | 4246.729 | 4715.795 |
| ENSMUSG00000001227 | -0.267899413 | 0.261585 | 0.425771 | 237.4288 | 139.1793 | 158.9965 | 171.6621 | 145.2916 | 219.0053 | 268.0724 | 219.6934 |
| ENSMUSG00000001228 | -0.071334148 | 0.512199 | 0.667488 | 2179.114 | 2678.002 | 2129.89 | 1968.507 | 2398.774 | 2372.185 | 2159.043 | 2474.27 |
| ENSMUSG00000001229 | -0.118896188 | 0.100178 | 0.221186 | 1145.896 | 1065.442 | 1161.834 | 1126.586 | 1286.172 | 1139.722 | 1185.111 | 1277.92 |

| | | | | | | | | | | | |
|---|---|---|---|---|---|---|---|---|---|---|---|
| ENSMUSG00000001240 | -0.132511989 | 0.052621 | 0.141057 | 1980.921 | 1817.73 | 1909.614 | 1981.446 | 2024.331 | 2213.518 | 2211.115 | 1983.766 |
| ENSMUSG00000001247 | -0.337175267 | 0.041574 | 0.11992 | 583.5115 | 572.3149 | 787.5294 | 598.6607 | 726.4579 | 640.2554 | 901.6104 | 945.1167 |
| ENSMUSG00000001248 | 0.331713597 | 0.00161 | 0.011105 | 2651.959 | 2352.85 | 2216.013 | 2734.516 | 2174.498 | 2012.391 | 1998.971 | 1723.832 |
| ENSMUSG00000001249 | -1.718618889 | 1.24E-07 | 6.69E-06 | 418.5186 | 315.5531 | 731.2182 | 458.0531 | 1624.535 | 751.9928 | 1705.828 | 2252.401 |
| ENSMUSG00000001260 | 1.036964455 | 0.009726 | 0.041725 | 173.0413 | 290.3568 | 132.4971 | 205.3044 | 96.53602 | 177.6625 | 69.42882 | 46.76642 |
| ENSMUSG00000001270 | -0.168495612 | 0.349936 | 0.518737 | 1208.271 | 1587.364 | 2014.783 | 1514.767 | 1735.698 | 1406.774 | 1767.542 | 2200.197 |
| ENSMUSG00000001280 | 0.296119178 | 0.008802 | 0.038874 | 3036.272 | 3119.536 | 2738.549 | 2896.69 | 2247.632 | 2926.403 | 2295.008 | 2133.854 |
| ENSMUSG00000001281 | 0.988180273 | 0.000832 | 0.006764 | 42.25428 | 47.99286 | 47.20208 | 48.30692 | 29.25334 | 26.81698 | 22.17865 | 15.22628 |
| ENSMUSG00000001285 | -0.087148757 | 0.282563 | 0.449168 | 1275.677 | 1267.012 | 1217.317 | 1261.156 | 1472.418 | 1192.238 | 1325.898 | 1339.912 |
| ENSMUSG00000001288 | 0.416165103 | 2.20E-05 | 0.000424 | 1151.932 | 1197.422 | 1198.27 | 1091.219 | 914.6544 | 744.1711 | 852.4316 | 963.6057 |
| ENSMUSG00000001289 | -0.049295455 | 0.528043 | 0.680556 | 4121.805 | 4566.521 | 5052.279 | 4667.656 | 5048.151 | 4602.464 | 4711.517 | 4688.605 |
| ENSMUSG00000001300 | 0.054323005 | 0.602482 | 0.740217 | 2549.342 | 2404.442 | 2861.937 | 2503.333 | 2125.743 | 2716.336 | 2401.08 | 2699.401 |
| ENSMUSG00000001305 | -0.090108118 | 0.312934 | 0.481091 | 1183.12 | 1349.799 | 1153.553 | 1233.552 | 1454.866 | 1278.276 | 1215.969 | 1283.357 |
| ENSMUSG00000001313 | 0.302811936 | 0.001228 | 0.009116 | 491.9606 | 521.9224 | 467.8802 | 483.0692 | 385.169 | 440.2454 | 385.7157 | 380.6569 |
| ENSMUSG00000001323 | -0.220164881 | 0.088075 | 0.201952 | 624.7598 | 687.4977 | 693.9534 | 596.0728 | 774.2384 | 596.6778 | 836.0387 | 822.2189 |
| ENSMUSG00000001333 | 0.657275409 | 0.004796 | 0.024968 | 60.36326 | 64.79036 | 62.9361 | 70.73513 | 48.75557 | 42.46022 | 31.82154 | 41.32846 |
| ENSMUSG00000001334 | 0.176362584 | 0.35498 | 0.523535 | 325.9616 | 484.7279 | 398.3193 | 375.2412 | 302.2845 | 464.8276 | 291.2153 | 343.6788 |
| ENSMUSG00000001348 | 0.357771551 | 0.044448 | 0.125491 | 116.7023 | 118.7823 | 114.2787 | 109.5532 | 92.63557 | 73.74669 | 82.92887 | 108.7591 |
| ENSMUSG00000001349 | 0.253809808 | 0.42605 | 0.591039 | 246.4833 | 547.1186 | 513.4261 | 533.1013 | 423.1983 | 560.9218 | 215.0365 | 344.7664 |
| ENSMUSG00000001366 | -0.355457754 | 0.020812 | 0.072472 | 756.5529 | 745.0892 | 640.9545 | 611.6001 | 843.4713 | 683.8329 | 909.3247 | 1084.328 |
| ENSMUSG00000001376 | 0.145030672 | 0.190016 | 0.342541 | 971.8485 | 935.8608 | 857.9185 | 986.8413 | 757.6615 | 1003.402 | 795.5386 | 838.5327 |
| ENSMUSG00000001380 | -0.131262097 | 0.069136 | 0.170721 | 1772.668 | 1904.117 | 1938.598 | 1973.683 | 2194.976 | 1919.649 | 2032.722 | 2165.394 |
| ENSMUSG00000001383 | -0.137574344 | 0.050604 | 0.137208 | 2805.886 | 2932.364 | 3050.745 | 2980.364 | 3523.077 | 2980.037 | 3191.797 | 3251.897 |
| ENSMUSG00000001403 | 0.000334469 | 0.997014 | 0.998149 | 2502.057 | 2259.264 | 2348.51 | 2706.913 | 2704.959 | 2420.232 | 2441.58 | 2249.138 |
| ENSMUSG00000001415 | -0.200408485 | 0.069848 | 0.172067 | 1741.48 | 1616.16 | 1870.693 | 1544.096 | 2154.996 | 1706.23 | 1789.721 | 2131.679 |
| ENSMUSG00000001416 | -0.283973135 | 3.90E-06 | 0.000111 | 5945.781 | 6371.052 | 6499.809 | 6264.372 | 7947.157 | 7083.034 | 7581.241 | 7925.276 |
| ENSMUSG00000001418 | -0.213592153 | 0.024091 | 0.08084 | 1725.383 | 1655.754 | 1923.692 | 1774.417 | 2100.39 | 1767.686 | 2112.758 | 2230.649 |
| ENSMUSG00000001419 | 0.109622261 | 0.188859 | 0.341288 | 1345.095 | 1107.435 | 1158.521 | 1179.206 | 1097.975 | 1125.196 | 1153.29 | 1063.664 |
| ENSMUSG00000001420 | -0.674062552 | 0.02344 | 0.079105 | 69.41775 | 93.58608 | 124.216 | 79.36136 | 161.8685 | 82.68568 | 150.4291 | 190.3284 |
| ENSMUSG00000001424 | -0.178177686 | 0.031271 | 0.097259 | 3432.657 | 3681.052 | 3820.884 | 3433.242 | 4142.273 | 3574.48 | 4186.944 | 4351.452 |
| ENSMUSG00000001435 | -0.421755813 | 0.052245 | 0.140292 | 4813.97 | 4254.567 | 8297.628 | 4811.714 | 6247.538 | 6448.366 | 7835.814 | 9180.356 |
| ENSMUSG00000001436 | -0.993473969 | 3.27E-06 | 9.68E-05 | 486.9303 | 401.9402 | 569.7374 | 493.4206 | 961.4597 | 619.0252 | 1015.396 | 1293.146 |
| ENSMUSG00000001440 | -0.08271695 | 0.109481 | 0.234988 | 7574.583 | 7944.018 | 8009.447 | 7434.089 | 8113.901 | 8370.249 | 7853.171 | 8452.758 |
| ENSMUSG00000001441 | 0.466678162 | 0.008095 | 0.0366 | 3186.174 | 2956.36 | 2431.321 | 3190.844 | 1811.757 | 2955.454 | 1912.185 | 1834.766 |
| ENSMUSG00000001445 | -0.175779303 | 0.020635 | 0.072021 | 1255.556 | 1222.618 | 1401.156 | 1318.951 | 1449.015 | 1376.605 | 1559.256 | 1491.087 |
| ENSMUSG00000001467 | -0.964107289 | 2.88E-09 | 3.13E-07 | 1061.387 | 999.4513 | 1175.083 | 1002.369 | 2052.609 | 1461.525 | 2217.865 | 2537.35 |
| ENSMUSG00000001472 | -0.029011725 | 0.535502 | 0.686367 | 6825.073 | 6699.803 | 6884.05 | 6810.412 | 7093.935 | 6546.695 | 7095.24 | 7035.627 |
| ENSMUSG00000001473 | -0.056679429 | 0.582238 | 0.722932 | 1716.329 | 1604.161 | 2045.423 | 1772.691 | 2031.157 | 1748.69 | 1707.756 | 1941.35 |
| ENSMUSG00000001482 | -0.292562885 | 0.026651 | 0.087056 | 1102.636 | 1178.225 | 1275.284 | 1121.411 | 1535.8 | 1124.078 | 1385.684 | 1683.591 |
| ENSMUSG00000001493 | -1.45138218 | 2.97E-09 | 3.18E-07 | 159.9626 | 206.3693 | 192.9488 | 200.9913 | 756.6864 | 319.569 | 485.0374 | 516.6058 |
| ENSMUSG00000001494 | -0.641911844 | 0.003674 | 0.020516 | 121.7326 | 151.1775 | 107.6539 | 139.745 | 259.3796 | 138.5544 | 211.1793 | 200.1168 |
| ENSMUSG00000001504 | -1.848453805 | 0.006141 | 0.029934 | 113.6841 | 27.5959 | 219.4483 | 104.3774 | 241.8276 | 198.8926 | 595.9307 | 640.5912 |
| ENSMUSG00000001506 | 0.683635857 | 0.002966 | 0.017615 | 10767.8 | 21939.94 | 18489.14 | 17594.07 | 9573.643 | 14995.16 | 8727.781 | 9532.736 |
| ENSMUSG00000001507 | -0.077065541 | 0.551188 | 0.698868 | 1674.074 | 1311.405 | 1849.99 | 1539.783 | 1515.323 | 1543.094 | 1892.9 | 1777.124 |
| ENSMUSG00000001510 | 0.804185905 | 0.05582 | 0.147195 | 21.12714 | 17.99732 | 18.21835 | 16.38985 | 9.751113 | 11.17374 | 13.50005 | 7.613138 |
| ENSMUSG00000001517 | -0.040687078 | 0.58206 | 0.722865 | 1058.369 | 1133.831 | 1038.446 | 1023.071 | 1146.731 | 1119.609 | 1094.468 | 1009.285 |
| ENSMUSG00000001518 | -0.027115509 | 0.753736 | 0.846886 | 831.0009 | 853.0731 | 936.5886 | 897.9911 | 973.1611 | 917.3641 | 883.2889 | 813.5181 |
| ENSMUSG00000001520 | -0.44654686 | 0.025491 | 0.084148 | 263.5862 | 195.5709 | 335.3832 | 301.0556 | 383.2187 | 282.6956 | 412.7158 | 417.635 |
| ENSMUSG00000001521 | -0.021275894 | 0.763693 | 0.853191 | 1192.174 | 1210.62 | 1094.757 | 1167.13 | 1114.552 | 1157.6 | 1242.004 | 1215.927 |
| ENSMUSG00000001524 | -0.219869302 | 0.006505 | 0.031321 | 610.675 | 609.5093 | 618.5957 | 627.1273 | 698.1797 | 693.8893 | 791.6814 | 687.3576 |
| ENSMUSG00000001525 | 0.022332008 | 0.724728 | 0.827662 | 29321.45 | 32777.92 | 32175.26 | 32639.95 | 33288.35 | 30827.23 | 31205.36 | 29642.3 |
| ENSMUSG00000001542 | -0.311370365 | 0.043221 | 0.123138 | 298.7981 | 201.57 | 237.6666 | 224.2821 | 310.0854 | 263.7003 | 327.8583 | 292.562 |
| ENSMUSG00000001552 | -0.583773453 | 0.00191 | 0.012711 | 1896.412 | 1724.144 | 2619.301 | 1950.392 | 3025.77 | 2164.354 | 3299.798 | 3788.08 |
| ENSMUSG00000001555 | 0.233072852 | 0.011199 | 0.046491 | 3440.706 | 4345.754 | 4191.048 | 3660.974 | 3321.229 | 3392.348 | 3154.19 | 3436.788 |
| ENSMUSG00000001558 | 0.412188592 | 0.263024 | 0.427321 | 32.19374 | 19.19714 | 19.87456 | 30.19182 | 14.62667 | 20.11273 | 22.17865 | 19.57664 |
| ENSMUSG00000001569 | 0.012988339 | 0.847331 | 0.908255 | 1290.768 | 1189.023 | 1213.176 | 1220.612 | 1239.366 | 1287.215 | 1197.647 | 1146.321 |
| ENSMUSG00000001576 | -0.105830123 | 0.110809 | 0.236866 | 3311.931 | 3352.301 | 3735.589 | 3457.395 | 3818.536 | 3514.142 | 3901.514 | 3680.408 |
| ENSMUSG00000001583 | -0.492749535 | 0.029977 | 0.094508 | 124.7507 | 151.1775 | 197.9175 | 108.6906 | 214.5245 | 157.5497 | 233.358 | 214.2554 |
| ENSMUSG00000001588 | -0.52130848 | 0.377778 | 0.545818 | 19.11503 | 10.79839 | 17.39024 | 6.038364 | 19.50223 | 35.75597 | 9.642892 | 11.9635 |
| ENSMUSG00000001604 | -0.03612399 | 0.884644 | 0.931133 | 71.42986 | 93.58608 | 101.029 | 71.59775 | 71.18313 | 98.32892 | 66.53595 | 110.9343 |
| ENSMUSG00000001627 | -0.060832863 | 0.52319 | 0.676427 | 1094.587 | 1024.648 | 892.6989 | 998.0554 | 985.8375 | 1157.6 | 1033.718 | 1003.847 |
| ENSMUSG00000001630 | -0.002223669 | 0.983344 | 0.990839 | 478.8819 | 515.9233 | 417.3657 | 470.9924 | 490.481 | 452.5365 | 432.0015 | 507.905 |
| ENSMUSG00000001632 | -0.063891162 | 0.354605 | 0.523232 | 1416.525 | 1388.194 | 1418.547 | 1319.814 | 1525.074 | 1433.591 | 1493.684 | 1338.825 |
| ENSMUSG00000001642 | -1.185603713 | 3.00E-05 | 0.000535 | 1831.019 | 1677.351 | 4328.513 | 1770.103 | 6270.941 | 4631.516 | 5394.234 | 5559.766 |

| | | | | | | | | | | | |
|---|---|---|---|---|---|---|---|---|---|---|---|
| ENSMUSG00000001656 | 0.330110763 | 0.067108 | 0.167299 | 433.6094 | 473.9295 | 415.7095 | 488.2449 | 412.4721 | 441.3628 | 335.5726 | 251.2335 |
| ENSMUSG00000001657 | 0.90605905 | 0.085243 | 0.197322 | 1927.6 | 2544.821 | 1439.249 | 1960.743 | 822.0188 | 2244.805 | 586.2878 | 548.1459 |
| ENSMUSG00000001661 | 1.010336101 | 4.37E-05 | 0.000721 | 1097.605 | 1129.032 | 787.5294 | 1071.378 | 418.3228 | 813.4483 | 401.1443 | 395.8832 |
| ENSMUSG00000001663 | -0.134862239 | 0.502714 | 0.658721 | 255.5378 | 206.3693 | 305.5713 | 233.771 | 201.848 | 261.4655 | 289.2867 | 349.1167 |
| ENSMUSG00000001665 | -0.135410174 | 0.303948 | 0.472512 | 755.5468 | 632.3059 | 913.4016 | 751.3451 | 838.5957 | 740.819 | 846.6459 | 930.978 |
| ENSMUSG00000001666 | -0.649098418 | 0.001154 | 0.008721 | 358.1554 | 334.7502 | 464.5678 | 436.4875 | 547.0374 | 459.2408 | 674.0381 | 822.2189 |
| ENSMUSG00000001670 | 2.988764174 | 0.004445 | 0.023594 | 82.49646 | 8.398751 | 0.828107 | 68.14726 | 4.875557 | 3.352122 | 9.642892 | 2.175182 |
| ENSMUSG00000001672 | -0.035514208 | 0.839295 | 0.902882 | 212.2775 | 214.7681 | 221.1045 | 186.3267 | 233.0516 | 153.0803 | 217.9294 | 250.146 |
| ENSMUSG00000001674 | -0.146046819 | 0.044595 | 0.12583 | 1775.686 | 1936.512 | 1738.196 | 1700.231 | 2122.817 | 1964.344 | 1944.007 | 1873.919 |
| ENSMUSG00000001687 | 0.004793462 | 0.952475 | 0.972978 | 1982.933 | 1959.309 | 2117.469 | 2153.971 | 1819.558 | 2119.659 | 2185.079 | 2066.423 |
| ENSMUSG00000001700 | -0.095594855 | 0.471754 | 0.631194 | 623.7537 | 632.3059 | 857.0904 | 710.8018 | 765.4624 | 654.7812 | 787.8242 | 812.4305 |
| ENSMUSG00000001707 | -0.230102724 | 0.050348 | 0.136691 | 805.8495 | 745.0892 | 651.7199 | 720.2906 | 986.8126 | 740.819 | 819.6458 | 877.686 |
| ENSMUSG00000001729 | -0.145277351 | 0.161171 | 0.304864 | 5456.839 | 5875.526 | 6922.971 | 5915.872 | 6780.924 | 5837.162 | 6741.346 | 7374.955 |
| ENSMUSG00000001739 | 0.055212421 | 0.812299 | 0.88583 | 209.2593 | 343.149 | 202.058 | 237.2215 | 171.6196 | 250.2918 | 299.8939 | 230.5693 |
| ENSMUSG00000001741 | 0.130471961 | 0.53163 | 0.683254 | 104.6297 | 145.1784 | 140.7781 | 101.7896 | 91.66046 | 115.0895 | 107.0361 | 135.9489 |
| ENSMUSG00000001750 | 0.415915399 | 1.99E-08 | 1.51E-06 | 1789.771 | 1865.722 | 1873.177 | 1746.813 | 1449.991 | 1443.647 | 1293.112 | 1265.956 |
| ENSMUSG00000001751 | -0.387188229 | 0.004789 | 0.024962 | 421.5368 | 415.1383 | 420.6782 | 321.7586 | 500.2321 | 440.2454 | 542.8948 | 579.686 |
| ENSMUSG00000001755 | -0.288950805 | 0.036907 | 0.110452 | 1008.066 | 891.4674 | 949.8383 | 894.5406 | 1184.76 | 872.6692 | 1150.397 | 1366.014 |
| ENSMUSG00000001761 | -0.073171664 | 0.352755 | 0.521562 | 4690.225 | 5457.988 | 5236.946 | 4917.817 | 5749.256 | 4885.16 | 5409.662 | 5310.707 |
| ENSMUSG00000001763 | -0.483179793 | 0.017176 | 0.06335 | 393.3673 | 287.9572 | 487.7548 | 367.4776 | 476.8294 | 419.0153 | 578.5735 | 676.4817 |
| ENSMUSG00000001767 | -0.015775195 | 0.847834 | 0.908531 | 947.7032 | 839.8751 | 831.4191 | 894.5406 | 962.4349 | 851.4391 | 864.0031 | 873.3356 |
| ENSMUSG00000001768 | 0.458488814 | 0.00017 | 0.001998 | 1977.903 | 1877.721 | 1763.039 | 1876.206 | 1192.561 | 1697.291 | 1245.862 | 1320.336 |
| ENSMUSG00000001773 | -0.069597337 | 0.853684 | 0.912204 | 46.2785 | 35.99465 | 22.35888 | 43.13117 | 53.63112 | 42.46022 | 36.64299 | 21.75182 |
| ENSMUSG00000001774 | 0.286917886 | 0.041032 | 0.118883 | 1958.788 | 1940.111 | 1320.002 | 1832.212 | 1302.749 | 1525.216 | 1383.755 | 1566.131 |
| ENSMUSG00000001783 | -0.130352968 | 0.079814 | 0.188683 | 4237.501 | 4764.491 | 4829.518 | 4537.4 | 5450.872 | 4687.384 | 5062.518 | 4902.861 |
| ENSMUSG00000001785 | -0.238427938 | 0.021493 | 0.074131 | 1217.326 | 1415.789 | 1285.221 | 1128.312 | 1576.755 | 1297.271 | 1498.505 | 1575.919 |
| ENSMUSG00000001786 | 0.032918739 | 0.719703 | 0.824664 | 1154.95 | 1203.421 | 1166.802 | 1172.305 | 1135.03 | 1257.046 | 1214.04 | 984.2699 |
| ENSMUSG00000001794 | 0.090922386 | 0.103587 | 0.226379 | 5917.612 | 6301.463 | 6532.105 | 6139.291 | 5916 | 5658.382 | 5686.413 | 6111.174 |
| ENSMUSG00000001802 | -0.102327081 | 0.391805 | 0.559211 | 516.1059 | 695.8965 | 654.2042 | 633.1656 | 702.0801 | 669.3071 | 596.895 | 714.5473 |
| ENSMUSG00000001819 | 1.125643112 | 0.086735 | 0.199942 | 10.06054 | 14.39786 | 12.4216 | 18.11509 | 5.850668 | 6.704245 | 0.964289 | 11.9635 |
| ENSMUSG00000001823 | 0.972288803 | 0.001513 | 0.010619 | 272.6407 | 371.9447 | 323.7897 | 322.6212 | 161.8685 | 280.4609 | 121.5004 | 94.62042 |
| ENSMUSG00000001827 | -1.67406505 | 0.00042 | 0.004005 | 333.004 | 313.1534 | 823.138 | 456.3278 | 1646.963 | 746.4059 | 1378.934 | 2375.299 |
| ENSMUSG00000001829 | -0.197076944 | 0.032212 | 0.099517 | 739.45 | 740.2899 | 828.1066 | 766.8723 | 843.4713 | 813.4483 | 881.3603 | 989.7079 |
| ENSMUSG00000001833 | 0.068135615 | 0.158823 | 0.301808 | 5844.17 | 6102.292 | 5864.651 | 6191.049 | 5718.053 | 5741.068 | 5505.127 | 5931.722 |
| ENSMUSG00000001844 | 0.024635896 | 0.73807 | 0.836054 | 1130.805 | 1057.043 | 1157.693 | 1063.615 | 1092.125 | 998.9324 | 1128.218 | 1115.868 |
| ENSMUSG00000001847 | -0.007228192 | 0.904807 | 0.942937 | 7017.229 | 7572.074 | 7036.422 | 7549.681 | 6872.584 | 7723.29 | 7348.848 | 7377.13 |
| ENSMUSG00000001855 | -0.016500816 | 0.887938 | 0.932608 | 1173.059 | 1021.048 | 1090.616 | 988.5665 | 1017.041 | 1305.093 | 1014.432 | 987.5327 |
| ENSMUSG00000001864 | -1.233598652 | 1.05E-05 | 0.000236 | 1395.397 | 1166.227 | 2216.841 | 1418.153 | 3621.563 | 1926.353 | 3914.05 | 5112.766 |
| ENSMUSG00000001865 | -0.248854115 | 0.700266 | 0.810746 | 56.33904 | 133.1802 | 72.87338 | 69.00988 | 83.85957 | 251.4092 | 36.64299 | 21.75182 |
| ENSMUSG00000001870 | 0.477742009 | 0.000314 | 0.003219 | 5894.472 | 5918.72 | 6501.465 | 6240.218 | 3965.778 | 5679.613 | 4207.194 | 3782.642 |
| ENSMUSG00000001891 | -0.081224088 | 0.295986 | 0.463846 | 1657.978 | 1973.706 | 1822.663 | 1902.085 | 1942.422 | 2063.79 | 1848.542 | 1927.211 |
| ENSMUSG00000001901 | 0.154712764 | 0.755674 | 0.848334 | 14.08476 | 17.99732 | 9.109173 | 7.763611 | 9.751113 | 8.938993 | 12.53576 | 11.9635 |
| ENSMUSG00000001909 | -0.017244161 | 0.819673 | 0.890713 | 2512.118 | 2469.233 | 2545.6 | 2653.43 | 2818.072 | 2622.477 | 2540.902 | 2322.007 |
| ENSMUSG00000001910 | -0.33095315 | 0.00049 | 0.004504 | 2134.847 | 2282.061 | 2604.395 | 2213.492 | 2988.716 | 2629.181 | 2812.831 | 3187.729 |
| ENSMUSG00000001911 | 0.745280934 | 0.000184 | 0.002126 | 1473.87 | 1943.711 | 1733.227 | 1622.595 | 1012.166 | 1441.413 | 749.2527 | 838.5327 |
| ENSMUSG00000001918 | 0.089454044 | 0.317598 | 0.485914 | 2416.543 | 2300.058 | 2379.978 | 2339.435 | 2274.935 | 1873.836 | 2377.937 | 2340.496 |
| ENSMUSG00000001924 | -0.197756966 | 0.057524 | 0.150314 | 7173.168 | 6979.362 | 7596.222 | 6803.511 | 7946.182 | 6861.794 | 8508.888 | 9430.502 |
| ENSMUSG00000001930 | 0.242560239 | 0.63537 | 0.76294 | 67.40564 | 74.38894 | 158.9965 | 415.7845 | 160.8934 | 113.9722 | 179.3578 | 152.2628 |
| ENSMUSG00000001942 | 0.039097975 | 0.692031 | 0.805049 | 415.5004 | 387.5424 | 423.9906 | 366.615 | 416.3725 | 389.9636 | 376.0728 | 368.6934 |
| ENSMUSG00000001943 | -0.114203734 | 0.531788 | 0.683379 | 110.666 | 88.78679 | 135.8095 | 122.4925 | 139.4409 | 113.9722 | 120.5361 | 123.9854 |
| ENSMUSG00000001946 | -0.346985858 | 0.06007 | 0.155078 | 711.2804 | 649.1034 | 1045.899 | 727.1916 | 845.4215 | 884.9603 | 1014.432 | 1244.204 |
| ENSMUSG00000001948 | -0.179956133 | 0.304093 | 0.472636 | 105.6357 | 131.9804 | 93.57605 | 95.75121 | 118.9636 | 110.62 | 121.5004 | 129.4233 |
| ENSMUSG00000001962 | -0.125523833 | 0.098495 | 0.218725 | 1071.448 | 1007.85 | 1073.226 | 1100.708 | 1073.598 | 1174.36 | 1138.826 | 1258.343 |
| ENSMUSG00000001964 | -0.123332798 | 0.198424 | 0.353044 | 1524.172 | 1612.56 | 1801.96 | 1838.251 | 1889.766 | 1650.362 | 1996.079 | 1847.817 |
| ENSMUSG00000001983 | -0.646181092 | 0.001288 | 0.009446 | 286.7255 | 254.3622 | 279.9 | 231.1831 | 443.6756 | 269.2872 | 416.5729 | 516.6058 |
| ENSMUSG00000001985 | 0.577632968 | 0.079785 | 0.188683 | 27.16347 | 32.39518 | 34.78048 | 24.15346 | 18.52711 | 21.23011 | 23.14294 | 16.31387 |
| ENSMUSG00000001986 | 0.116774939 | 0.259013 | 0.422872 | 345.0766 | 349.1481 | 363.5388 | 337.2858 | 292.5334 | 349.7381 | 338.4655 | 306.7007 |
| ENSMUSG00000001988 | -0.745772163 | 0.103196 | 0.225862 | 11.0666 | 16.7975 | 17.39024 | 10.35148 | 22.42756 | 13.40849 | 20.25007 | 36.9781 |
| ENSMUSG00000001995 | -0.193496501 | 0.034232 | 0.104289 | 1258.574 | 1048.644 | 1214.004 | 1187.833 | 1229.615 | 1373.253 | 1317.219 | 1470.423 |
| ENSMUSG00000001998 | 0.261571287 | 0.060712 | 0.156363 | 698.2017 | 632.3059 | 506.8013 | 609.8748 | 537.2863 | 581.0345 | 490.8232 | 430.6861 |
| ENSMUSG00000001999 | -0.614148642 | 0.000234 | 0.00257 | 378.2764 | 331.1507 | 462.0835 | 408.8835 | 621.1459 | 465.945 | 596.895 | 738.4743 |
| ENSMUSG00000002006 | 0.382443868 | 0.000141 | 0.001737 | 641.8627 | 746.289 | 727.9057 | 687.5109 | 508.033 | 611.2036 | 527.4662 | 504.6423 |
| ENSMUSG00000002007 | 0.187155241 | 0.572618 | 0.715181 | 78.47224 | 52.79215 | 60.45178 | 66.42201 | 67.28268 | 79.33356 | 52.07161 | 28.27737 |
| ENSMUSG00000002010 | -0.396848918 | 0.001855 | 0.012439 | 3656.002 | 3376.298 | 3877.195 | 3758.451 | 5049.126 | 3668.339 | 5201.376 | 5394.452 |

| | | | | | | | | | | | |
|---|---|---|---|---|---|---|---|---|---|---|---|
| ENSMUSG00000002012 | 0.161049166 | 0.112175 | 0.238651 | 423.5489 | 447.5334 | 427.303 | 509.8105 | 387.1192 | 401.1373 | 426.2158 | 403.4963 |
| ENSMUSG00000002014 | -0.19965193 | 0.0072 | 0.033572 | 2107.684 | 2224.469 | 2307.105 | 2322.182 | 2638.651 | 2317.434 | 2723.153 | 2612.394 |
| ENSMUSG00000002015 | -0.078641662 | 0.220482 | 0.379329 | 2782.746 | 3030.749 | 3066.479 | 2940.683 | 3010.169 | 2964.393 | 3222.654 | 3285.613 |
| ENSMUSG00000002017 | -0.216385399 | 0.069278 | 0.170936 | 984.9272 | 1054.643 | 940.7291 | 900.5789 | 1077.498 | 1072.679 | 991.2893 | 1366.014 |
| ENSMUSG00000002020 | 0.862533052 | 0.138236 | 0.273684 | 81.4904 | 106.7841 | 66.24853 | 132.844 | 41.92979 | 128.498 | 29.89296 | 13.05109 |
| ENSMUSG00000002028 | 0.612273196 | 0.000197 | 0.002236 | 2399.44 | 2338.452 | 1984.972 | 2247.134 | 1350.529 | 2007.921 | 1302.755 | 1207.226 |
| ENSMUSG00000002031 | 0.173992274 | 0.009361 | 0.040574 | 1882.328 | 1964.108 | 1787.054 | 1871.893 | 1632.336 | 1796.738 | 1593.97 | 1628.124 |
| ENSMUSG00000002032 | -1.225891688 | 0.000174 | 0.002036 | 186.1201 | 166.7752 | 247.6039 | 255.3366 | 497.3068 | 224.5922 | 495.6446 | 786.3284 |
| ENSMUSG00000002043 | -0.436961532 | 0.001381 | 0.009969 | 445.6821 | 491.9268 | 470.3646 | 473.5803 | 629.9219 | 511.7573 | 623.8951 | 780.8904 |
| ENSMUSG00000002052 | -0.115886716 | 0.080654 | 0.190036 | 2719.365 | 2584.416 | 2773.329 | 2515.41 | 2957.513 | 2655.998 | 2877.439 | 2987.613 |
| ENSMUSG00000002055 | 0.089195191 | 0.415338 | 0.581203 | 1526.184 | 1383.394 | 1235.535 | 1400.901 | 1510.447 | 1310.68 | 1201.504 | 1188.737 |
| ENSMUSG00000002058 | 0.127753998 | 0.146307 | 0.285106 | 1078.49 | 1183.024 | 1295.159 | 1255.98 | 1141.855 | 1172.125 | 1040.468 | 1053.876 |
| ENSMUSG00000002059 | -0.088343745 | 0.127717 | 0.259113 | 2359.197 | 2373.247 | 2532.35 | 2604.26 | 2595.746 | 2583.369 | 2614.188 | 2705.927 |
| ENSMUSG00000002064 | -0.068920485 | 0.234674 | 0.395501 | 2031.224 | 2055.294 | 2187.858 | 2220.393 | 2279.81 | 2152.063 | 2255.472 | 2227.387 |
| ENSMUSG00000002068 | -0.250727142 | 0.065799 | 0.1652 | 314.895 | 464.3309 | 401.6317 | 391.6311 | 496.3317 | 444.7149 | 432.0015 | 495.9415 |
| ENSMUSG00000002076 | -1.692100809 | 3.88E-06 | 0.000111 | 20.12109 | 5.999108 | 16.56213 | 17.25247 | 39.00445 | 49.16446 | 51.10733 | 57.64233 |
| ENSMUSG00000002083 | 0.279815123 | 0.009211 | 0.04015 | 401.4157 | 370.7449 | 449.6619 | 414.0593 | 366.6419 | 312.8647 | 343.2869 | 326.2773 |
| ENSMUSG00000002102 | -0.25726861 | 0.00549 | 0.027513 | 4820.006 | 5060.847 | 5526.784 | 5083.44 | 6397.705 | 5209.198 | 6291.987 | 6592.977 |
| ENSMUSG00000002103 | 0.126915133 | 0.062446 | 0.15938 | 1765.625 | 1749.34 | 1620.605 | 1658.825 | 1572.855 | 1664.887 | 1487.898 | 1494.35 |
| ENSMUSG00000002104 | -0.806037877 | 0.038172 | 0.112828 | 31.18769 | 16.7975 | 59.62368 | 31.05445 | 56.55646 | 49.16446 | 59.78593 | 79.39415 |
| ENSMUSG00000002105 | -0.05343208 | 0.649482 | 0.773046 | 836.0312 | 867.471 | 934.1043 | 931.6334 | 915.6295 | 749.758 | 1005.754 | 1033.212 |
| ENSMUSG00000002107 | 0.970946262 | 0.004012 | 0.021877 | 1382.319 | 1982.105 | 1204.895 | 1722.659 | 640.6481 | 1533.037 | 567.9663 | 468.7518 |
| ENSMUSG00000002108 | -0.600661056 | 0.012638 | 0.050872 | 68.4117 | 63.59054 | 58.79557 | 58.6584 | 83.85957 | 65.92507 | 108.0004 | 119.635 |
| ENSMUSG00000002109 | 0.214321732 | 0.130892 | 0.263472 | 452.7245 | 417.5379 | 332.0708 | 435.6249 | 363.7165 | 321.8037 | 404.0372 | 319.7518 |
| ENSMUSG00000002111 | -0.163703139 | 0.557011 | 0.703689 | 76.46013 | 105.5843 | 118.4192 | 63.83414 | 102.3867 | 68.15982 | 121.5004 | 115.2847 |
| ENSMUSG00000002129 | -0.011673987 | 0.853152 | 0.911952 | 1971.867 | 2171.677 | 2154.733 | 1991.798 | 2061.385 | 2156.532 | 2006.686 | 2132.766 |
| ENSMUSG00000002147 | -0.00321221 | 0.983377 | 0.990839 | 1232.417 | 992.2524 | 1343.189 | 1167.13 | 1149.656 | 920.7163 | 1228.504 | 1448.671 |
| ENSMUSG00000002204 | -2.948546453 | 3.03E-08 | 2.15E-06 | 396.3854 | 397.1409 | 775.1078 | 420.9603 | 3322.204 | 1191.121 | 4097.265 | 6751.765 |
| ENSMUSG00000002205 | -0.135817746 | 0.049033 | 0.134366 | 956.7577 | 940.6601 | 1016.087 | 977.3524 | 1057.021 | 1011.224 | 1134.004 | 1074.54 |
| ENSMUSG00000002210 | 0.130772611 | 0.072647 | 0.177093 | 742.4681 | 753.4879 | 735.3587 | 728.9169 | 699.1548 | 706.1804 | 632.5737 | 665.6057 |
| ENSMUSG00000002221 | -0.086362941 | 0.259602 | 0.423619 | 1147.908 | 1147.029 | 1214.004 | 1191.283 | 1226.69 | 1127.43 | 1322.04 | 1314.898 |
| ENSMUSG00000002222 | 0.119328799 | 0.110332 | 0.23618 | 3928.642 | 3918.617 | 3848.212 | 4089.698 | 3552.33 | 4012.49 | 3681.656 | 3286.7 |
| ENSMUSG00000002227 | 0.130919715 | 0.280113 | 0.446478 | 789.7527 | 845.8742 | 822.3099 | 653.8686 | 728.4081 | 808.9788 | 642.2166 | 661.2554 |
| ENSMUSG00000002228 | -0.113212752 | 0.503917 | 0.659736 | 98.59333 | 101.9848 | 107.6539 | 100.0643 | 109.2125 | 92.74205 | 129.2147 | 109.8467 |
| ENSMUSG00000002233 | 0.081066789 | 0.21116 | 0.368562 | 2506.081 | 2690 | 2731.096 | 2744.005 | 2548.941 | 2508.505 | 2360.58 | 2673.299 |
| ENSMUSG00000002249 | 0.106230878 | 0.237864 | 0.399097 | 1234.429 | 1168.626 | 1410.266 | 1419.878 | 1191.586 | 1261.515 | 1252.612 | 1161.547 |
| ENSMUSG00000002250 | -0.499890779 | 0.013662 | 0.053809 | 324.9556 | 268.76 | 377.6166 | 297.6051 | 431.9743 | 301.691 | 532.2876 | 529.6569 |
| ENSMUSG00000002257 | -0.040362893 | 0.793143 | 0.873246 | 223.3441 | 269.9598 | 318.8211 | 284.6658 | 299.3592 | 241.3528 | 270.001 | 318.6642 |
| ENSMUSG00000002265 | 0.353685459 | 0.038224 | 0.112947 | 20480.25 | 16190.39 | 15308.38 | 18998.42 | 12414.14 | 17712.61 | 15044.84 | 10374.53 |
| ENSMUSG00000002266 | 0.265154104 | 0.120913 | 0.250136 | 2538.275 | 1648.555 | 1728.259 | 2444.675 | 1606.008 | 1916.297 | 1945.936 | 1488.912 |
| ENSMUSG00000002274 | -0.590153396 | 0.000855 | 0.006921 | 117.7084 | 170.3747 | 187.9802 | 163.0358 | 270.1058 | 210.0663 | 251.6795 | 230.5693 |
| ENSMUSG00000002279 | -0.254109011 | 0.006976 | 0.032896 | 405.4399 | 453.5325 | 462.0835 | 439.0754 | 520.7094 | 486.0577 | 520.7161 | 572.0729 |
| ENSMUSG00000002280 | -0.334641775 | 0.010452 | 0.044072 | 671.0383 | 704.2952 | 667.4539 | 690.0988 | 814.2179 | 686.0677 | 916.0747 | 1028.861 |
| ENSMUSG00000002289 | -1.764245601 | 0.013848 | 0.054358 | 159.9626 | 171.5745 | 155.684 | 114.7289 | 515.8339 | 88.27255 | 260.3581 | 1180.036 |
| ENSMUSG00000002297 | -0.083172935 | 0.398135 | 0.565316 | 1165.011 | 1231.017 | 1165.146 | 1337.066 | 1303.724 | 1420.182 | 1347.112 | 1118.044 |
| ENSMUSG00000002307 | -0.165413043 | 0.007229 | 0.033674 | 922.5518 | 917.8635 | 904.2924 | 885.0517 | 998.514 | 1049.214 | 1032.754 | 988.6203 |
| ENSMUSG00000002308 | -0.346967938 | 9.75E-06 | 0.000222 | 866.2128 | 890.2676 | 825.6223 | 937.6717 | 1134.054 | 1059.271 | 1200.54 | 1079.978 |
| ENSMUSG00000002319 | -0.175732542 | 0.014512 | 0.056278 | 2054.363 | 2208.871 | 2223.466 | 2034.066 | 2547.966 | 2214.635 | 2365.401 | 2493.846 |
| ENSMUSG00000002320 | -0.313757684 | 0.011777 | 0.048271 | 1754.559 | 1835.727 | 2186.202 | 1927.964 | 2463.131 | 1929.705 | 2452.187 | 2733.116 |
| ENSMUSG00000002324 | 0.98480065 | 6.06E-09 | 5.56E-07 | 822.9525 | 1211.82 | 899.3238 | 940.2596 | 443.6756 | 622.3774 | 417.5372 | 474.1897 |
| ENSMUSG00000002325 | 0.346789718 | 0.185547 | 0.337199 | 911.4852 | 1506.976 | 1658.698 | 682.3352 | 909.7789 | 763.1665 | 1044.325 | 1024.511 |
| ENSMUSG00000002326 | -0.052626491 | 0.435705 | 0.599856 | 1142.878 | 1149.429 | 1108.007 | 1136.938 | 1181.835 | 1083.853 | 1240.076 | 1196.35 |
| ENSMUSG00000002329 | 0.033325233 | 0.621462 | 0.753063 | 1146.902 | 1127.832 | 1119.6 | 1089.493 | 1125.278 | 1087.205 | 1018.289 | 1150.671 |
| ENSMUSG00000002332 | -0.213779375 | 0.081521 | 0.19142 | 1458.779 | 1397.792 | 1597.418 | 1590.678 | 1699.619 | 1410.126 | 1871.685 | 2030.533 |
| ENSMUSG00000002342 | 0.011844488 | 0.877354 | 0.92652 | 1081.508 | 1120.633 | 1150.24 | 1176.618 | 1176.959 | 1024.632 | 1099.29 | 1192 |
| ENSMUSG00000002343 | -0.216795 | 0.092144 | 0.20843 | 274.6528 | 266.3604 | 264.9941 | 263.1002 | 307.1601 | 258.1134 | 313.394 | 363.2554 |
| ENSMUSG00000002345 | -0.132285687 | 0.233695 | 0.394285 | 662.9898 | 646.7038 | 813.2007 | 693.5493 | 814.2179 | 689.4198 | 750.217 | 836.3575 |
| ENSMUSG00000002346 | -0.459993402 | 0.091529 | 0.207539 | 147.89 | 183.5727 | 129.1846 | 137.1571 | 210.624 | 129.6154 | 170.6792 | 309.9635 |
| ENSMUSG00000002365 | -0.268707004 | 0.041169 | 0.119094 | 1171.047 | 1282.609 | 1406.125 | 1435.405 | 1665.49 | 1241.403 | 1696.185 | 1777.124 |
| ENSMUSG00000002372 | -0.051275697 | 0.494592 | 0.651408 | 2468.857 | 2324.054 | 2440.43 | 2385.154 | 2637.676 | 2241.452 | 2424.223 | 2663.511 |
| ENSMUSG00000002379 | -0.467834492 | 0.00297 | 0.017621 | 1110.684 | 1247.814 | 1386.251 | 1080.005 | 1472.418 | 1303.976 | 1936.293 | 1959.839 |
| ENSMUSG00000002384 | 0.418877892 | 0.259025 | 0.422872 | 27.16347 | 25.19625 | 37.2648 | 31.05445 | 30.22845 | 12.29112 | 27.0001 | 20.66423 |
| ENSMUSG00000002393 | -0.49430546 | 5.13E-05 | 0.000807 | 1962.812 | 2162.078 | 2447.883 | 2205.728 | 3127.182 | 2524.148 | 3127.19 | 3589.051 |
| ENSMUSG00000002395 | -0.147932299 | 0.056676 | 0.148729 | 2843.11 | 3198.724 | 3189.039 | 2894.102 | 3624.489 | 3231.446 | 3444.441 | 3130.087 |

| | | | | | | | | | | | |
|---|---|---|---|---|---|---|---|---|---|---|---|
| ENSMUSG00000002396 | -0.159777461 | 0.125583 | 0.256106 | 354.1311 | 301.1552 | 347.8048 | 307.094 | 360.7912 | 350.8555 | 364.5013 | 389.3576 |
| ENSMUSG00000002409 | -0.259505252 | 0.028141 | 0.090419 | 395.3794 | 326.3515 | 315.5086 | 307.9566 | 412.4721 | 372.0856 | 426.2158 | 396.9707 |
| ENSMUSG00000002413 | 0.310146303 | 0.028708 | 0.091587 | 1543.287 | 1451.784 | 1358.923 | 1598.441 | 1049.22 | 1547.563 | 1165.826 | 1039.737 |
| ENSMUSG00000002416 | -0.54267292 | 0.000254 | 0.002747 | 949.7153 | 783.4835 | 939.0729 | 907.4799 | 1251.068 | 1004.519 | 1356.755 | 1604.197 |
| ENSMUSG00000002428 | 0.488167881 | 0.005425 | 0.027214 | 1055.351 | 1030.647 | 787.5294 | 1019.621 | 597.7432 | 940.829 | 632.5737 | 604.7006 |
| ENSMUSG00000002455 | 0.045279353 | 0.502506 | 0.658496 | 2688.177 | 2749.991 | 2601.911 | 2639.628 | 2641.577 | 2334.194 | 2719.295 | 2649.372 |
| ENSMUSG00000002458 | -0.130103829 | 0.209665 | 0.366924 | 433.6094 | 478.7288 | 426.4749 | 450.2895 | 460.2525 | 438.0106 | 529.3948 | 527.4817 |
| ENSMUSG00000002459 | 0.099929179 | 0.828558 | 0.896148 | 17.10292 | 13.19804 | 11.59349 | 21.56559 | 10.72622 | 23.46486 | 13.50005 | 11.9635 |
| ENSMUSG00000002475 | -0.599136127 | 0.026901 | 0.08756 | 103.6236 | 73.18911 | 137.4657 | 67.28463 | 145.2916 | 115.0895 | 139.8219 | 179.4525 |
| ENSMUSG00000002477 | -0.087557937 | 0.321291 | 0.489294 | 2376.3 | 2701.998 | 2510.819 | 2613.749 | 2723.486 | 3070.544 | 2545.723 | 2501.46 |
| ENSMUSG00000002486 | -0.020208745 | 0.839586 | 0.903048 | 625.7658 | 657.5022 | 615.2832 | 537.4144 | 653.3246 | 559.8044 | 610.395 | 642.7663 |
| ENSMUSG00000002489 | 0.083742316 | 0.4536 | 0.61591 | 656.9535 | 483.5281 | 563.1125 | 570.1941 | 541.1868 | 538.5743 | 507.2161 | 561.197 |
| ENSMUSG00000002496 | -0.099122337 | 0.272069 | 0.437357 | 1653.953 | 1710.946 | 2032.174 | 1883.107 | 1884.89 | 1862.663 | 1945.936 | 2109.927 |
| ENSMUSG00000002504 | -0.785032401 | 4.45E-05 | 0.000729 | 1229.398 | 1131.432 | 1688.509 | 1335.341 | 2120.867 | 1643.657 | 2629.617 | 2887.554 |
| ENSMUSG00000002524 | -0.206294564 | 0.000419 | 0.003994 | 3878.34 | 4156.182 | 4145.502 | 4182.861 | 4809.249 | 4394.632 | 4858.089 | 4814.766 |
| ENSMUSG00000002546 | -0.126755196 | 0.258981 | 0.422872 | 1753.553 | 1815.33 | 1777.945 | 1424.191 | 1916.094 | 1592.258 | 1865.9 | 2015.306 |
| ENSMUSG00000002550 | -0.22674243 | 0.077452 | 0.184946 | 1121.751 | 1015.049 | 1141.131 | 1102.433 | 1395.384 | 1017.928 | 1215.969 | 1497.613 |
| ENSMUSG00000002565 | -1.360899612 | 0.039155 | 0.114759 | 336.0222 | 311.9536 | 1372.173 | 428.7239 | 1388.559 | 555.3349 | 1816.721 | 2530.824 |
| ENSMUSG00000002578 | 0.579823917 | 0.023949 | 0.080488 | 360.1675 | 313.1534 | 252.5725 | 257.0618 | 195.9974 | 301.691 | 151.3934 | 142.4744 |
| ENSMUSG00000002580 | -0.380272029 | 0.004272 | 0.022922 | 499.003 | 470.33 | 545.7223 | 496.0085 | 655.2748 | 517.3442 | 682.7167 | 763.4889 |
| ENSMUSG00000002602 | 0.25163595 | 0.003554 | 0.020039 | 4621.814 | 5751.944 | 5304.851 | 5160.214 | 4670.783 | 4378.989 | 4407.766 | 4042.576 |
| ENSMUSG00000002603 | -0.194160312 | 0.307587 | 0.476044 | 336.0222 | 393.5415 | 568.9093 | 431.3117 | 462.2028 | 398.9026 | 555.4306 | 564.4598 |
| ENSMUSG00000002608 | 0.005572327 | 0.945423 | 0.968566 | 686.1291 | 724.6922 | 777.5921 | 797.9267 | 729.3833 | 778.8097 | 724.1812 | 747.1751 |
| ENSMUSG00000002617 | 0.52299649 | 0.000514 | 0.004676 | 321.9374 | 382.7431 | 348.6329 | 393.3563 | 210.624 | 307.2779 | 262.2867 | 227.3065 |
| ENSMUSG00000002625 | 0.144173677 | 0.061241 | 0.157157 | 1689.165 | 1786.534 | 1637.995 | 1806.334 | 1647.938 | 1630.249 | 1571.791 | 1409.518 |
| ENSMUSG00000002633 | -0.079427711 | 0.89256 | 0.934962 | 145.8779 | 23.99643 | 225.245 | 86.26235 | 61.43201 | 78.21619 | 221.7865 | 147.9124 |
| ENSMUSG00000002635 | -0.184450922 | 0.076137 | 0.182871 | 528.1785 | 495.5263 | 544.0661 | 503.7721 | 653.3246 | 506.1705 | 584.3592 | 610.1386 |
| ENSMUSG00000002658 | -0.229791759 | 0.000858 | 0.006939 | 1823.977 | 1899.317 | 1911.27 | 1783.905 | 2189.125 | 1995.63 | 2184.115 | 2328.533 |
| ENSMUSG00000002660 | -0.174908005 | 0.01175 | 0.04818 | 1338.052 | 1421.789 | 1459.124 | 1367.258 | 1652.814 | 1461.525 | 1556.363 | 1634.649 |
| ENSMUSG00000002661 | -0.4011776 | 0.000735 | 0.006171 | 519.124 | 405.5397 | 467.8802 | 464.0914 | 666.001 | 529.6353 | 599.7879 | 657.9926 |
| ENSMUSG00000002664 | -0.166765216 | 0.675943 | 0.792479 | 19.11503 | 11.99822 | 14.90592 | 15.52722 | 19.50223 | 16.76061 | 12.53576 | 20.66423 |
| ENSMUSG00000002668 | 0.581138273 | 1.23E-06 | 4.42E-05 | 379.2825 | 403.14 | 388.382 | 400.2573 | 260.3547 | 222.3574 | 260.3581 | 306.7007 |
| ENSMUSG00000002679 | -0.016344282 | 0.837248 | 0.901479 | 800.8193 | 799.0811 | 780.0764 | 896.2658 | 822.0188 | 877.1387 | 800.36 | 816.7809 |
| ENSMUSG00000002688 | 0.169597923 | 0.255017 | 0.418922 | 443.67 | 543.5192 | 436.4122 | 541.7276 | 398.8205 | 530.7527 | 446.4659 | 370.8686 |
| ENSMUSG00000002699 | -0.029075926 | 0.933062 | 0.960274 | 42.25428 | 32.39518 | 50.5145 | 46.58167 | 24.37778 | 61.45558 | 37.60728 | 53.29196 |
| ENSMUSG00000002718 | -0.185292742 | 0.001672 | 0.011474 | 2439.682 | 2494.429 | 2245.825 | 2374.802 | 2820.022 | 2708.515 | 2613.224 | 2715.715 |
| ENSMUSG00000002728 | -0.092136775 | 0.244842 | 0.407651 | 1699.226 | 1685.749 | 1812.725 | 1981.446 | 1933.646 | 1798.972 | 2021.15 | 1902.197 |
| ENSMUSG00000002731 | -0.123394599 | 0.066062 | 0.165658 | 1226.38 | 1197.422 | 1233.879 | 1329.303 | 1400.26 | 1296.154 | 1330.719 | 1408.43 |
| ENSMUSG00000002732 | 0.400253484 | 6.89E-05 | 0.001018 | 918.5276 | 1117.034 | 876.9649 | 1010.995 | 725.4828 | 779.9271 | 761.7884 | 702.5838 |
| ENSMUSG00000002733 | 0.10848723 | 0.300798 | 0.469207 | 876.2733 | 800.281 | 751.0927 | 794.4762 | 672.8268 | 827.9742 | 804.2172 | 683.0072 |
| ENSMUSG00000002741 | -0.190068501 | 0.017723 | 0.064912 | 3031.242 | 3245.517 | 3331.473 | 3375.446 | 3851.69 | 3289.549 | 3685.513 | 3986.021 |
| ENSMUSG00000002748 | 0.149851144 | 0.035362 | 0.106904 | 4241.525 | 4360.151 | 3802.666 | 3935.288 | 3537.704 | 3910.809 | 3738.549 | 3535.759 |
| ENSMUSG00000002763 | -0.035836711 | 0.658116 | 0.779672 | 1451.736 | 1565.767 | 1408.609 | 1507.003 | 1398.31 | 1452.586 | 1563.113 | 1666.19 |
| ENSMUSG00000002767 | -0.322276882 | 0.006107 | 0.029849 | 1369.24 | 1309.005 | 1516.263 | 1364.67 | 1859.537 | 1396.718 | 1743.435 | 1952.226 |
| ENSMUSG00000002768 | -0.265518988 | 0.000112 | 0.001463 | 1135.835 | 1149.429 | 1183.364 | 1165.404 | 1365.156 | 1296.154 | 1411.719 | 1498.701 |
| ENSMUSG00000002769 | -1.259737685 | 0.000167 | 0.001967 | 53.32088 | 33.595 | 39.74912 | 60.38364 | 134.5654 | 58.10345 | 113.7861 | 142.4744 |
| ENSMUSG00000002771 | 0.369793334 | 0.036331 | 0.109187 | 100.6054 | 118.7823 | 102.6852 | 120.7673 | 86.78491 | 75.98144 | 98.35749 | 80.48174 |
| ENSMUSG00000002778 | -0.049871375 | 0.471548 | 0.631172 | 3200.259 | 3387.096 | 3558.374 | 3349.567 | 3313.428 | 3444.864 | 3400.084 | 3815.27 |
| ENSMUSG00000002781 | -0.28313904 | 0.09466 | 0.212459 | 247.4894 | 220.7672 | 255.0568 | 182.0136 | 265.2303 | 221.2401 | 315.3226 | 299.0875 |
| ENSMUSG00000002797 | -0.582850527 | 0.017304 | 0.063687 | 102.6175 | 141.5789 | 135.8095 | 108.6906 | 171.6196 | 119.559 | 190.9293 | 249.0584 |
| ENSMUSG00000002799 | -0.488192694 | 0.017474 | 0.064192 | 246.4833 | 184.7725 | 365.195 | 228.5952 | 332.513 | 357.5597 | 363.537 | 388.27 |
| ENSMUSG00000002803 | -0.499433973 | 4.13E-06 | 0.000116 | 365.1977 | 445.1338 | 469.5365 | 424.4108 | 617.2455 | 550.8654 | 605.5736 | 636.2408 |
| ENSMUSG00000002804 | -0.371022035 | 0.013904 | 0.054484 | 469.8274 | 427.1365 | 540.7536 | 420.0976 | 610.4197 | 462.5929 | 660.5381 | 669.9561 |
| ENSMUSG00000002808 | -0.11147823 | 0.27331 | 0.439009 | 550.3117 | 550.7181 | 460.4273 | 593.485 | 578.241 | 600.0299 | 569.8949 | 578.5985 |
| ENSMUSG00000002812 | -0.081444461 | 0.290806 | 0.457906 | 2157.987 | 1943.711 | 2109.188 | 1985.759 | 2129.643 | 1990.043 | 2187.008 | 2367.686 |
| ENSMUSG00000002814 | -0.027421542 | 0.780521 | 0.864611 | 464.7971 | 519.5227 | 467.0521 | 433.8996 | 512.9085 | 444.7149 | 464.7874 | 494.8539 |
| ENSMUSG00000002820 | -0.538280066 | 0.000116 | 0.001505 | 650.9172 | 603.5102 | 645.9232 | 621.0889 | 830.7948 | 721.8237 | 1070.361 | 1038.649 |
| ENSMUSG00000002825 | -0.139011328 | 0.070322 | 0.17287 | 698.2017 | 769.0856 | 722.109 | 743.5815 | 852.2473 | 748.6406 | 813.8601 | 811.343 |
| ENSMUSG00000002831 | 1.01935656 | 0.005085 | 0.026035 | 28.16952 | 28.79572 | 29.81184 | 41.40593 | 18.52711 | 12.29112 | 20.25007 | 11.9635 |
| ENSMUSG00000002833 | -0.069425639 | 0.226532 | 0.386474 | 2148.932 | 2285.66 | 2219.326 | 2147.07 | 2192.05 | 2312.964 | 2306.58 | 2422.065 |
| ENSMUSG00000002835 | -0.224860379 | 0.037845 | 0.112203 | 729.3894 | 971.8554 | 857.9185 | 790.1631 | 987.7878 | 912.8946 | 960.432 | 1049.525 |
| ENSMUSG00000002844 | 0.06134092 | 0.339692 | 0.508231 | 2018.145 | 2198.073 | 2135.687 | 2050.456 | 2082.838 | 1957.639 | 1901.578 | 2108.839 |
| ENSMUSG00000002845 | 0.133124148 | 0.225799 | 0.385543 | 1345.095 | 1322.203 | 1114.632 | 1360.357 | 1208.163 | 1315.149 | 1148.468 | 1015.81 |
| ENSMUSG00000002846 | 0.007414378 | 0.944479 | 0.967814 | 709.2683 | 718.6931 | 660.8291 | 695.2745 | 635.7726 | 649.1943 | 663.4309 | 821.1313 |

| | | | | | | | | | | | |
|---|---|---|---|---|---|---|---|---|---|---|---|
| ENSMUSG00000002847 | -2.438215362 | 0.001814 | 0.012234 | 199.1988 | 31.19536 | 355.2577 | 93.16334 | 828.8446 | 289.3999 | 1191.861 | 1371.452 |
| ENSMUSG00000002870 | -0.270292127 | 0.021064 | 0.073131 | 2063.417 | 2529.224 | 2330.292 | 1964.194 | 2787.843 | 2326.373 | 2571.759 | 3028.941 |
| ENSMUSG00000002871 | 0.128488964 | 0.209946 | 0.367312 | 944.685 | 899.8662 | 812.3726 | 852.272 | 788.865 | 729.6453 | 765.6456 | 924.4524 |
| ENSMUSG00000002881 | -0.113554055 | 0.147554 | 0.28672 | 1114.708 | 971.8554 | 1002.837 | 1083.455 | 1064.822 | 1118.491 | 1197.647 | 1135.445 |
| ENSMUSG00000002885 | -0.251558477 | 0.015513 | 0.059117 | 932.6124 | 975.4549 | 1096.413 | 898.8537 | 1203.287 | 1005.637 | 1192.826 | 1245.292 |
| ENSMUSG00000002897 | -0.057484501 | 0.647802 | 0.771952 | 236.4228 | 218.3675 | 197.0894 | 203.5791 | 213.5494 | 213.4185 | 214.0722 | 247.9708 |
| ENSMUSG00000002900 | -0.081558299 | 0.526481 | 0.679534 | 12198.41 | 11280.72 | 14825.59 | 12772 | 12302 | 11519.01 | 14280.16 | 15949.52 |
| ENSMUSG00000002908 | -1.483810538 | 1.83E-05 | 0.000365 | 85.51462 | 61.1909 | 100.2009 | 70.73513 | 221.3503 | 98.32892 | 235.2866 | 334.9781 |
| ENSMUSG00000002910 | -0.233973372 | 0.148282 | 0.287728 | 303.8284 | 220.7672 | 266.6503 | 259.6497 | 286.6827 | 346.386 | 353.8941 | 250.146 |
| ENSMUSG00000002944 | -0.121467133 | 0.482193 | 0.640887 | 472.8455 | 334.7502 | 313.8524 | 300.193 | 361.7663 | 421.25 | 343.2869 | 419.8102 |
| ENSMUSG00000002948 | 0.066428961 | 0.406623 | 0.573149 | 1405.458 | 1213.02 | 1194.13 | 1274.958 | 1151.606 | 1269.337 | 1195.719 | 1243.117 |
| ENSMUSG00000002949 | -0.100006831 | 0.130937 | 0.263472 | 2306.883 | 2045.696 | 2167.155 | 2299.754 | 2361.72 | 2239.218 | 2426.152 | 2428.591 |
| ENSMUSG00000002957 | -0.08029529 | 0.309916 | 0.478464 | 3287.786 | 3202.324 | 3484.673 | 3233.975 | 3513.326 | 3165.521 | 3401.048 | 3887.051 |
| ENSMUSG00000002963 | -0.151711694 | 0.197841 | 0.352244 | 705.2441 | 706.6949 | 693.1253 | 609.8748 | 804.4668 | 617.9079 | 833.1458 | 756.9634 |
| ENSMUSG00000002968 | -0.198405153 | 0.047579 | 0.131415 | 1693.189 | 1785.334 | 1859.099 | 1915.024 | 2203.752 | 1758.747 | 2057.793 | 2302.43 |
| ENSMUSG00000002980 | -0.720950359 | 0.000117 | 0.001515 | 6135.926 | 5929.518 | 9817.204 | 6502.456 | 11062.64 | 9264.149 | 12347.72 | 14115.84 |
| ENSMUSG00000002981 | -0.44805409 | 0.001583 | 0.01097 | 2135.853 | 2028.898 | 2433.805 | 2041.83 | 2888.28 | 2253.744 | 3085.725 | 3560.773 |
| ENSMUSG00000002983 | 0.406856394 | 0.000909 | 0.00726 | 364.1917 | 382.7431 | 340.3518 | 360.5766 | 252.5538 | 299.4563 | 303.7511 | 234.9197 |
| ENSMUSG00000002984 | -0.571482567 | 4.34E-05 | 0.000718 | 1772.668 | 1546.57 | 1984.143 | 1726.972 | 2630.85 | 2035.856 | 2663.367 | 3120.299 |
| ENSMUSG00000002985 | -0.235534292 | 0.223788 | 0.383494 | 5006.126 | 4475.334 | 6932.081 | 5596.701 | 5707.326 | 4657.215 | 7920.671 | 7630.539 |
| ENSMUSG00000002996 | 0.17435963 | 0.032109 | 0.099278 | 4922.624 | 5063.247 | 4597.648 | 4862.609 | 4260.261 | 4709.732 | 4459.837 | 3800.043 |
| ENSMUSG00000002997 | 0.36008995 | 0.043394 | 0.123365 | 1258.574 | 763.0865 | 737.0149 | 804.8277 | 672.8268 | 719.5889 | 722.2526 | 661.2554 |
| ENSMUSG00000003031 | 0.297010859 | 0.004457 | 0.023645 | 3053.375 | 2823.18 | 2431.321 | 3166.691 | 2266.159 | 2466.045 | 2480.152 | 2126.241 |
| ENSMUSG00000003032 | 0.43969494 | 0.004325 | 0.023159 | 636.8324 | 701.8956 | 703.0625 | 722.0159 | 498.2819 | 635.7859 | 524.5733 | 379.5693 |
| ENSMUSG00000003033 | -0.059969556 | 0.325312 | 0.493178 | 2186.156 | 2213.671 | 2220.154 | 2278.189 | 2370.496 | 2182.232 | 2469.545 | 2251.314 |
| ENSMUSG00000003037 | -0.326415961 | 0.004462 | 0.023665 | 1871.261 | 2240.067 | 2172.124 | 2012.501 | 2593.796 | 2181.114 | 2595.866 | 3030.029 |
| ENSMUSG00000003038 | -1.055901554 | 3.01E-09 | 3.21E-07 | 4549.378 | 4956.463 | 4748.363 | 4908.328 | 6324.572 | 12207.31 | 10783.65 | 10524.62 |
| ENSMUSG00000003039 | -0.03192878 | 0.652028 | 0.775356 | 3270.683 | 3117.136 | 3510.344 | 3452.219 | 3648.867 | 3441.512 | 3172.511 | 3392.197 |
| ENSMUSG00000003051 | -0.55692773 | 0.041729 | 0.120224 | 429.5852 | 277.1588 | 657.5167 | 347.6373 | 599.6935 | 438.0106 | 716.4668 | 766.7517 |
| ENSMUSG00000003062 | 0.071504188 | 0.399436 | 0.566906 | 839.0493 | 838.6753 | 773.4516 | 854.8599 | 719.6321 | 822.3873 | 758.8956 | 846.1459 |
| ENSMUSG00000003068 | -0.026762083 | 0.635506 | 0.763054 | 2913.533 | 3143.532 | 3016.792 | 3051.099 | 3176.913 | 2896.234 | 3130.083 | 3144.226 |
| ENSMUSG00000003070 | 0.341951422 | 0.021647 | 0.074563 | 382.3007 | 397.1409 | 346.1486 | 375.2412 | 283.7574 | 378.7898 | 255.5366 | 266.4598 |
| ENSMUSG00000003072 | -0.667963932 | 4.46E-05 | 0.000729 | 4325.028 | 4364.951 | 5389.318 | 4403.693 | 7436.199 | 5240.485 | 7510.848 | 9180.356 |
| ENSMUSG00000003099 | -0.215218981 | 0.053008 | 0.141745 | 1925.588 | 1918.515 | 2076.063 | 1967.644 | 2268.109 | 1849.254 | 2459.902 | 2579.766 |
| ENSMUSG00000003119 | 0.147763271 | 0.120751 | 0.249926 | 897.4005 | 839.8751 | 828.9347 | 766.0097 | 700.1299 | 848.0869 | 729.0026 | 730.8612 |
| ENSMUSG00000003123 | -0.051069259 | 0.704871 | 0.814341 | 492.9666 | 398.3407 | 366.8512 | 375.2412 | 431.9743 | 458.1234 | 433.9301 | 366.5182 |
| ENSMUSG00000003131 | 0.005919596 | 0.922421 | 0.953784 | 3395.433 | 3262.315 | 3044.12 | 3425.478 | 3262.722 | 3397.935 | 3197.583 | 3214.919 |
| ENSMUSG00000003134 | 0.362960004 | 0.049793 | 0.135722 | 858.1644 | 885.4683 | 636.814 | 820.3549 | 626.9966 | 802.2746 | 609.4308 | 449.1751 |
| ENSMUSG00000003135 | -0.028773688 | 0.620467 | 0.752486 | 1176.078 | 1252.614 | 1243.816 | 1254.255 | 1274.47 | 1282.745 | 1236.219 | 1233.328 |
| ENSMUSG00000003153 | 0.523154285 | 0.000436 | 0.004116 | 393.3673 | 323.9518 | 370.1637 | 453.74 | 286.6827 | 305.0431 | 248.7866 | 233.8321 |
| ENSMUSG00000003154 | 0.109846619 | 0.289205 | 0.456248 | 647.899 | 673.0999 | 722.109 | 649.5555 | 592.8677 | 725.1758 | 610.395 | 568.8101 |
| ENSMUSG00000003161 | -0.144288061 | 0.048008 | 0.132362 | 3297.846 | 3364.3 | 3556.718 | 3540.207 | 3670.319 | 3495.146 | 3896.693 | 4146.985 |
| ENSMUSG00000003166 | 0.022862053 | 0.813082 | 0.886475 | 2248.531 | 2427.239 | 2365.901 | 2406.72 | 2115.992 | 2721.923 | 2331.651 | 2131.679 |
| ENSMUSG00000003184 | 0.030050441 | 0.717498 | 0.823362 | 1563.408 | 1581.365 | 1364.72 | 1523.393 | 1464.617 | 1374.37 | 1454.148 | 1611.81 |
| ENSMUSG00000003190 | 0.061478017 | 0.624084 | 0.754985 | 559.3662 | 549.5183 | 431.4436 | 427.8613 | 456.3521 | 487.1751 | 485.0374 | 453.5255 |
| ENSMUSG00000003198 | 0.140895033 | 0.224668 | 0.384466 | 374.2522 | 339.5495 | 322.1335 | 395.0816 | 314.961 | 362.0292 | 314.3583 | 307.7883 |
| ENSMUSG00000003199 | -0.269425361 | 0.018777 | 0.067462 | 1654.959 | 1828.528 | 1929.488 | 1958.155 | 2240.806 | 1807.911 | 2381.794 | 2454.693 |
| ENSMUSG00000003200 | -0.18651329 | 0.05567 | 0.146914 | 1853.152 | 2002.502 | 2106.703 | 1963.331 | 2434.853 | 1906.24 | 2261.258 | 2416.627 |
| ENSMUSG00000003206 | -0.154558577 | 0.704959 | 0.814392 | 15.09082 | 23.99643 | 30.63995 | 18.11509 | 16.57689 | 18.99536 | 30.85725 | 31.54014 |
| ENSMUSG00000003208 | -0.334805844 | 0.006388 | 0.030873 | 452.7245 | 495.5263 | 512.598 | 417.5098 | 604.569 | 511.7573 | 570.8592 | 680.832 |
| ENSMUSG00000003226 | 0.068167188 | 0.252509 | 0.416216 | 2875.303 | 2752.391 | 2594.458 | 2793.175 | 2498.235 | 2620.242 | 2668.188 | 2718.978 |
| ENSMUSG00000003227 | 0.330318836 | 0.220499 | 0.379329 | 59.35721 | 113.983 | 81.98256 | 93.16334 | 69.2329 | 84.92043 | 72.32169 | 50.02919 |
| ENSMUSG00000003228 | 0.161199843 | 0.20369 | 0.359742 | 443.67 | 511.124 | 503.4888 | 432.1744 | 438.8001 | 348.6207 | 475.3946 | 425.2481 |
| ENSMUSG00000003233 | -0.140846525 | 0.287874 | 0.454754 | 863.1946 | 740.2899 | 900.98 | 862.6235 | 999.4891 | 730.7627 | 926.6819 | 1057.139 |
| ENSMUSG00000003234 | -0.069265699 | 0.431339 | 0.596045 | 1383.325 | 1339.001 | 1235.535 | 1293.073 | 1360.28 | 1210.116 | 1456.077 | 1479.124 |
| ENSMUSG00000003235 | -0.232579157 | 0.000961 | 0.007604 | 1730.413 | 1845.326 | 1845.85 | 1707.132 | 2175.473 | 1936.409 | 2047.186 | 2214.335 |
| ENSMUSG00000003269 | -0.005815232 | 0.941922 | 0.966348 | 2704.274 | 2591.615 | 2568.787 | 2905.316 | 2828.798 | 2433.641 | 2873.582 | 2676.562 |
| ENSMUSG00000003271 | -0.327888713 | 0.323979 | 0.491966 | 37.22401 | 26.39607 | 31.46805 | 37.09281 | 51.6809 | 24.58223 | 36.64299 | 53.29196 |
| ENSMUSG00000003273 | 0.817901329 | 1.13E-07 | 6.24E-06 | 418.5186 | 489.5272 | 358.5702 | 460.6409 | 228.176 | 300.5736 | 224.6794 | 226.2189 |
| ENSMUSG00000003279 | 0.764533784 | 0.038479 | 0.113374 | 523.1483 | 531.5209 | 356.0859 | 414.0593 | 230.1263 | 518.4616 | 199.6079 | 126.1606 |
| ENSMUSG00000003282 | 0.282046789 | 0.027024 | 0.08784 | 279.6831 | 298.7556 | 289.0092 | 299.3304 | 205.7485 | 275.9914 | 255.5366 | 222.9562 |
| ENSMUSG00000003283 | 0.255064289 | 0.33658 | 0.504829 | 91.55095 | 80.38804 | 141.6062 | 100.9269 | 92.63557 | 59.22083 | 87.75031 | 108.7591 |
| ENSMUSG00000003299 | -0.266704659 | 0.007175 | 0.033491 | 1219.338 | 1222.618 | 1381.282 | 1244.766 | 1485.095 | 1320.736 | 1591.077 | 1702.08 |
| ENSMUSG00000003308 | -0.11899469 | 0.133046 | 0.26641 | 1528.197 | 1600.562 | 1474.03 | 1454.383 | 1705.47 | 1503.986 | 1578.541 | 1785.825 |

| | | | | | | | | | | | |
|---|---|---|---|---|---|---|---|---|---|---|---|
| ENSMUSG00000003309 | -0.65274449 | 0.001425 | 0.010206 | 559.3662 | 575.9143 | 850.4655 | 591.7597 | 941.9575 | 716.2368 | 1137.861 | 1258.343 |
| ENSMUSG00000003316 | 0.23397146 | 0.004431 | 0.023569 | 6302.931 | 6047.101 | 6120.536 | 5823.571 | 4678.584 | 5859.51 | 5148.34 | 4972.466 |
| ENSMUSG00000003341 | 0.266341678 | 0.618516 | 0.751043 | 12.07265 | 11.99822 | 9.109173 | 20.70296 | 5.850668 | 12.29112 | 9.642892 | 17.40146 |
| ENSMUSG00000003344 | -0.096087687 | 0.328879 | 0.49675 | 1438.658 | 1749.34 | 1755.586 | 1592.403 | 1703.519 | 1575.497 | 1796.471 | 1909.81 |
| ENSMUSG00000003345 | -0.260642576 | 0.001448 | 0.010317 | 4161.041 | 4904.87 | 4855.189 | 4830.692 | 6108.097 | 5242.719 | 5590.949 | 5521.7 |
| ENSMUSG00000003346 | -0.348666516 | 0.002328 | 0.014615 | 1594.596 | 1767.337 | 1925.348 | 1856.366 | 2393.898 | 1881.658 | 2253.544 | 2568.89 |
| ENSMUSG00000003348 | -0.473932408 | 0.005575 | 0.027854 | 127.7689 | 118.7823 | 145.7468 | 146.646 | 190.1467 | 148.6108 | 221.7865 | 189.2408 |
| ENSMUSG00000003352 | 0.378161167 | 0.000368 | 0.003615 | 1597.614 | 2120.085 | 1810.241 | 1869.305 | 1522.149 | 1460.408 | 1427.148 | 1279.007 |
| ENSMUSG00000003354 | -0.252277807 | 0.593393 | 0.732847 | 23.13925 | 15.59768 | 19.04645 | 37.95543 | 21.45245 | 49.16446 | 27.96439 | 16.31387 |
| ENSMUSG00000003355 | 0.609371575 | 1.04E-06 | 3.86E-05 | 328.9798 | 299.9554 | 284.0406 | 357.9887 | 218.4249 | 217.8879 | 202.5007 | 194.6788 |
| ENSMUSG00000003360 | -0.148172067 | 0.070156 | 0.172649 | 3563.445 | 3313.907 | 3298.349 | 3293.496 | 3921.898 | 3245.972 | 3801.228 | 3954.481 |
| ENSMUSG00000003363 | -0.234146414 | 0.13468 | 0.268622 | 1477.894 | 1370.196 | 1711.696 | 1545.821 | 1704.495 | 1340.849 | 1899.65 | 2238.262 |
| ENSMUSG00000003378 | 0.389946426 | 1.23E-08 | 1.00E-06 | 1642.887 | 1689.349 | 1593.277 | 1670.039 | 1206.213 | 1374.37 | 1244.897 | 1208.314 |
| ENSMUSG00000003379 | 0.083838827 | 0.789488 | 0.8709 | 60.36326 | 83.98751 | 55.48314 | 76.77349 | 88.73513 | 65.92507 | 70.39311 | 34.80291 |
| ENSMUSG00000003380 | -0.277570103 | 0.036793 | 0.1102 | 3044.32 | 2823.18 | 3302.489 | 3426.341 | 3798.059 | 2939.811 | 4266.98 | 4265.532 |
| ENSMUSG00000003382 | -0.171756039 | 0.021051 | 0.073099 | 787.7406 | 759.487 | 766.8267 | 688.3735 | 829.8197 | 857.0259 | 862.0745 | 830.9196 |
| ENSMUSG00000003402 | -0.109268744 | 0.241294 | 0.403491 | 2646.929 | 2994.755 | 3163.367 | 2825.955 | 3144.734 | 2834.778 | 3049.082 | 3518.357 |
| ENSMUSG00000003410 | 1.882440649 | 0.007633 | 0.034999 | 82.49646 | 25.19625 | 20.70267 | 251.8861 | 32.17867 | 31.28647 | 20.25007 | 19.57664 |
| ENSMUSG00000003411 | 0.037488658 | 0.917505 | 0.951211 | 185.114 | 89.98662 | 115.1068 | 253.6113 | 115.0631 | 109.5027 | 186.1078 | 217.5182 |
| ENSMUSG00000003418 | -0.873710443 | 0.001916 | 0.012726 | 92.557 | 101.9848 | 190.4645 | 105.2401 | 178.4454 | 183.2494 | 247.8223 | 291.4744 |
| ENSMUSG00000003420 | 0.220587235 | 0.02495 | 0.08294 | 3020.175 | 3783.037 | 3419.252 | 3602.316 | 3082.327 | 3171.108 | 2996.046 | 2613.481 |
| ENSMUSG00000003421 | -0.139703587 | 0.087638 | 0.201326 | 1476.888 | 1484.179 | 1610.667 | 1629.496 | 1798.105 | 1531.92 | 1691.363 | 1813.014 |
| ENSMUSG00000003423 | 0.123512664 | 0.035324 | 0.106824 | 1819.952 | 1938.912 | 1812.725 | 1936.59 | 1778.603 | 1657.066 | 1699.078 | 1754.284 |
| ENSMUSG00000003429 | -0.071783689 | 0.394343 | 0.561649 | 26024.61 | 29983.54 | 28255 | 29007.44 | 33086.5 | 27913.12 | 30673.07 | 27372.49 |
| ENSMUSG00000003435 | -0.112509037 | 0.12307 | 0.252685 | 3385.373 | 3251.516 | 3368.738 | 3050.237 | 3745.403 | 3234.798 | 3442.512 | 3690.197 |
| ENSMUSG00000003437 | 0.008039691 | 0.903984 | 0.942489 | 2336.058 | 2444.036 | 2297.996 | 2345.473 | 2217.403 | 2214.635 | 2426.152 | 2510.16 |
| ENSMUSG00000003438 | -0.313183852 | 0.000335 | 0.003372 | 1384.331 | 1445.785 | 1568.434 | 1392.274 | 1887.815 | 1582.202 | 1820.578 | 1904.372 |
| ENSMUSG00000003444 | -0.371481173 | 0.003683 | 0.020561 | 365.1977 | 368.3452 | 448.0057 | 362.3019 | 488.5308 | 442.4801 | 487.9303 | 580.7736 |
| ENSMUSG00000003452 | 0.626505445 | 0.003808 | 0.021065 | 533.2088 | 633.5058 | 459.5992 | 593.485 | 255.4792 | 516.2268 | 350.037 | 316.489 |
| ENSMUSG00000003458 | -0.096004555 | 0.359211 | 0.527582 | 3025.205 | 2550.821 | 3060.682 | 2840.619 | 2861.952 | 2697.341 | 3404.905 | 3305.189 |
| ENSMUSG00000003464 | -0.323162162 | 0.004005 | 0.021854 | 2318.955 | 2069.692 | 2236.716 | 2143.619 | 2671.805 | 2244.805 | 2902.51 | 3151.839 |
| ENSMUSG00000003469 | 0.143321566 | 0.665194 | 0.785063 | 62.37537 | 58.79126 | 53.82693 | 31.05445 | 41.92979 | 30.1691 | 55.92877 | 57.64233 |
| ENSMUSG00000003477 | 0.15818501 | 0.432117 | 0.596508 | 205.2351 | 251.9625 | 230.2136 | 221.6942 | 149.192 | 168.7235 | 253.6081 | 242.5328 |
| ENSMUSG00000003484 | -0.385036927 | 0.493802 | 0.650647 | 12.07265 | 10.79839 | 7.45296 | 14.6646 | 18.52711 | 4.469496 | 21.21436 | 14.13868 |
| ENSMUSG00000003500 | -0.177841376 | 0.111731 | 0.238257 | 771.6437 | 628.7065 | 702.2344 | 683.1978 | 858.098 | 716.2368 | 711.6454 | 866.8101 |
| ENSMUSG00000003518 | -0.504462092 | 0.000226 | 0.002492 | 942.6729 | 820.6779 | 988.7593 | 879.0133 | 1392.459 | 1033.571 | 1195.719 | 1531.328 |
| ENSMUSG00000003526 | -0.690795294 | 5.60E-05 | 0.000865 | 655.9474 | 615.5084 | 616.9394 | 627.1273 | 1110.652 | 687.1851 | 1242.969 | 1017.985 |
| ENSMUSG00000003527 | -0.124018842 | 0.281376 | 0.447795 | 872.2491 | 755.8876 | 838.0439 | 957.5121 | 1032.643 | 794.453 | 971.0392 | 934.2407 |
| ENSMUSG00000003528 | -0.583101661 | 0.000298 | 0.003083 | 1120.745 | 1035.446 | 1291.018 | 1067.065 | 1704.495 | 1231.346 | 1726.078 | 2102.314 |
| ENSMUSG00000003531 | -0.332793816 | 0.003279 | 0.018947 | 1085.533 | 1143.43 | 1209.864 | 1182.657 | 1554.327 | 1163.186 | 1540.934 | 1561.781 |
| ENSMUSG00000003534 | -0.268617169 | 0.119048 | 0.247986 | 1948.727 | 1984.505 | 2463.617 | 1798.57 | 2343.192 | 1773.273 | 2747.26 | 3009.365 |
| ENSMUSG00000003541 | -0.676728804 | 0.007515 | 0.034606 | 321.9374 | 245.9634 | 400.8036 | 377.8291 | 431.9743 | 348.6207 | 667.2881 | 706.9342 |
| ENSMUSG00000003545 | 0.137334164 | 0.742413 | 0.839119 | 15.09082 | 10.79839 | 14.90592 | 18.97772 | 13.65156 | 13.40849 | 11.57147 | 16.31387 |
| ENSMUSG00000003546 | -0.504354811 | 0.000321 | 0.003267 | 1058.369 | 883.0686 | 964.7442 | 986.8413 | 1439.264 | 1051.449 | 1395.326 | 1636.825 |
| ENSMUSG00000003549 | -0.013282996 | 0.856698 | 0.91426 | 831.0009 | 751.0883 | 840.5282 | 823.8054 | 862.9735 | 802.2746 | 801.3243 | 813.5181 |
| ENSMUSG00000003559 | -0.360633482 | 0.007498 | 0.034561 | 757.5589 | 687.4977 | 791.6699 | 754.7956 | 914.6544 | 772.1055 | 982.6107 | 1173.511 |
| ENSMUSG00000003573 | -0.127385667 | 0.058347 | 0.151649 | 1394.391 | 1558.568 | 1540.278 | 1522.53 | 1636.237 | 1572.145 | 1625.792 | 1736.883 |
| ENSMUSG00000003575 | 0.197287017 | 0.028854 | 0.091974 | 971.8485 | 947.859 | 950.6664 | 1013.583 | 823.9691 | 972.1155 | 792.6457 | 801.5546 |
| ENSMUSG00000003581 | 0.078462262 | 0.280452 | 0.446902 | 817.9222 | 855.4728 | 861.2309 | 828.1186 | 852.2473 | 770.9881 | 756.0027 | 804.8174 |
| ENSMUSG00000003585 | -0.195671813 | 0.202356 | 0.3579 | 102.6175 | 117.5825 | 122.5598 | 103.5148 | 124.8142 | 124.0285 | 138.8576 | 122.8978 |
| ENSMUSG00000003604 | -0.165361579 | 0.252523 | 0.416216 | 381.2946 | 597.5111 | 474.5051 | 463.2288 | 505.1077 | 582.1519 | 524.5733 | 535.0948 |
| ENSMUSG00000003617 | -0.036157794 | 0.872418 | 0.92356 | 358.1554 | 184.7725 | 250.9163 | 273.4516 | 239.8774 | 220.1227 | 323.0369 | 312.1386 |
| ENSMUSG00000003623 | -0.015245318 | 0.869008 | 0.921481 | 1625.784 | 1652.154 | 1507.982 | 1766.653 | 1633.311 | 1458.173 | 1717.399 | 1811.927 |
| ENSMUSG00000003644 | -0.15260325 | 0.33698 | 0.505306 | 1188.15 | 1144.63 | 1496.389 | 1138.663 | 1546.527 | 1022.397 | 1384.719 | 1569.394 |
| ENSMUSG00000003657 | -0.158387849 | 0.530997 | 0.682647 | 35.2119 | 56.39161 | 47.20208 | 43.13117 | 45.83023 | 44.69496 | 53.0359 | 58.72992 |
| ENSMUSG00000003660 | -0.206712021 | 0.002136 | 0.013757 | 4709.34 | 4586.918 | 4763.269 | 4433.022 | 5542.533 | 4809.178 | 5489.698 | 5497.773 |
| ENSMUSG00000003662 | -0.123844028 | 0.234442 | 0.395292 | 1137.847 | 1139.83 | 1070.742 | 1086.043 | 1164.283 | 1017.928 | 1282.505 | 1364.927 |
| ENSMUSG00000003665 | 0.776783021 | 0.068488 | 0.169531 | 36.21796 | 32.39518 | 24.01509 | 32.77969 | 21.45245 | 25.6996 | 19.28578 | 6.525547 |
| ENSMUSG00000003680 | -0.045935439 | 0.552355 | 0.69976 | 906.455 | 997.0517 | 870.3401 | 959.2373 | 947.8082 | 920.7163 | 1022.147 | 958.1677 |
| ENSMUSG00000003721 | 0.053789063 | 0.608966 | 0.744309 | 1058.369 | 920.2631 | 847.1531 | 1014.445 | 905.8784 | 826.8568 | 983.5749 | 982.0948 |
| ENSMUSG00000003731 | -0.127552694 | 0.124819 | 0.255099 | 1312.901 | 1118.234 | 1185.849 | 1119.685 | 1312.5 | 1196.708 | 1316.255 | 1348.613 |
| ENSMUSG00000003746 | 0.068602063 | 0.41656 | 0.582046 | 1337.046 | 1286.209 | 1428.484 | 1293.073 | 1141.855 | 1386.661 | 1313.362 | 1258.343 |
| ENSMUSG00000003752 | -0.274920354 | 0.165498 | 0.310915 | 178.0716 | 152.3773 | 193.777 | 162.1732 | 204.7734 | 146.376 | 216.9651 | 263.197 |
| ENSMUSG00000003762 | 0.174821953 | 0.048214 | 0.132712 | 816.9161 | 761.8867 | 748.6084 | 847.9589 | 677.7024 | 762.0491 | 725.1455 | 649.2919 |

| | | | | | | | | | | | |
|---|---|---|---|---|---|---|---|---|---|---|---|
| ENSMUSG00000003778 | 0.115558341 | 0.219146 | 0.377925 | 3572.499 | 3101.539 | 2996.09 | 3123.56 | 3202.266 | 3161.051 | 2754.01 | 2690.7 |
| ENSMUSG00000003779 | -0.122304746 | 0.123214 | 0.25284 | 1565.421 | 1492.578 | 1493.076 | 1707.132 | 1846.861 | 1603.432 | 1659.542 | 1703.168 |
| ENSMUSG00000003808 | -0.23652901 | 0.005008 | 0.025762 | 1338.052 | 1422.988 | 1423.515 | 1403.488 | 1671.341 | 1440.295 | 1672.077 | 1798.876 |
| ENSMUSG00000003809 | -0.654465774 | 0.000264 | 0.00282 | 982.9151 | 927.462 | 969.7129 | 1021.346 | 1660.615 | 998.9324 | 1572.756 | 1908.722 |
| ENSMUSG00000003810 | -0.001911293 | 0.975588 | 0.986739 | 1619.748 | 1611.36 | 1598.246 | 1535.47 | 1568.954 | 1707.348 | 1520.684 | 1577.007 |
| ENSMUSG00000003812 | 0.03787535 | 0.705242 | 0.814617 | 326.9677 | 328.7511 | 322.1335 | 312.2697 | 339.3387 | 322.9211 | 290.251 | 303.4379 |
| ENSMUSG00000003813 | -0.141267667 | 0.106547 | 0.230776 | 984.9272 | 979.0544 | 1131.194 | 1047.225 | 1141.855 | 1030.219 | 1196.683 | 1202.876 |
| ENSMUSG00000003814 | -0.234148341 | 0.041885 | 0.120481 | 10841.24 | 12791.3 | 12106.92 | 10159.98 | 12519.45 | 11800.59 | 14114.3 | 15550.38 |
| ENSMUSG00000003824 | -0.026618354 | 0.847484 | 0.908344 | 471.8395 | 590.3122 | 530.8164 | 446.839 | 530.4606 | 612.321 | 473.466 | 460.051 |
| ENSMUSG00000003847 | 0.238479295 | 0.092938 | 0.209613 | 5918.618 | 5663.158 | 8500.515 | 6153.956 | 5369.938 | 5803.641 | 4960.303 | 6108.999 |
| ENSMUSG00000003848 | -0.109050889 | 0.275592 | 0.441808 | 1712.305 | 1794.933 | 1623.089 | 1635.534 | 2144.27 | 1654.831 | 1786.828 | 1706.43 |
| ENSMUSG00000003849 | 0.106997935 | 0.411206 | 0.576995 | 281.6952 | 254.3622 | 307.2276 | 236.3588 | 240.8525 | 232.4138 | 260.3581 | 269.7226 |
| ENSMUSG00000003863 | -0.196065341 | 0.310068 | 0.47852 | 83.50251 | 100.785 | 90.26362 | 115.5915 | 131.64 | 92.74205 | 108.0004 | 114.1971 |
| ENSMUSG00000003865 | -0.687171995 | 6.23E-05 | 0.000944 | 672.0443 | 566.3158 | 795.8105 | 640.0666 | 979.9869 | 821.27 | 1177.397 | 1330.124 |
| ENSMUSG00000003868 | -0.198738072 | 0.011021 | 0.045926 | 1740.474 | 2012.101 | 1845.85 | 1921.063 | 2257.383 | 1977.752 | 2120.472 | 2270.89 |
| ENSMUSG00000003873 | -0.166109893 | 0.057407 | 0.150093 | 1997.018 | 2042.096 | 2328.636 | 2209.179 | 2523.588 | 2143.124 | 2410.723 | 2549.314 |
| ENSMUSG00000003923 | -0.084487084 | 0.337347 | 0.505774 | 1725.383 | 1760.138 | 1628.058 | 1789.944 | 1809.807 | 2088.372 | 1749.221 | 1672.715 |
| ENSMUSG00000003929 | 0.148569988 | 0.267418 | 0.431699 | 720.3349 | 784.6833 | 669.9383 | 626.2647 | 602.6188 | 766.5186 | 619.0736 | 537.27 |
| ENSMUSG00000003934 | 0.267888744 | 0.053316 | 0.142362 | 605.6447 | 697.0963 | 700.5782 | 753.9329 | 509.9832 | 717.3542 | 544.8234 | 519.8685 |
| ENSMUSG00000003948 | 0.161055406 | 0.102873 | 0.225315 | 846.0917 | 897.4665 | 881.1055 | 1017.033 | 789.8402 | 878.256 | 858.2174 | 731.9488 |
| ENSMUSG00000003949 | 0.236808491 | 0.091119 | 0.206901 | 337.0282 | 353.9474 | 286.5249 | 386.4553 | 264.2552 | 337.447 | 278.6796 | 277.3357 |
| ENSMUSG00000003955 | 0.064315588 | 0.198831 | 0.353494 | 2248.531 | 2402.043 | 2335.261 | 2319.595 | 2189.125 | 2286.147 | 2240.044 | 2182.795 |
| ENSMUSG00000003970 | -0.200708717 | 0.018217 | 0.066143 | 31662.54 | 35919.06 | 36801.06 | 34981.11 | 43215.96 | 35389.47 | 41677.54 | 39880.88 |
| ENSMUSG00000003974 | 1.682097861 | 0.00031 | 0.003189 | 27.16347 | 21.59679 | 24.01509 | 16.38985 | 5.850668 | 5.58687 | 9.642892 | 6.525547 |
| ENSMUSG00000003992 | 0.274991149 | 0.003507 | 0.01981 | 1385.337 | 1437.386 | 1424.343 | 1661.413 | 1187.686 | 1339.732 | 1242.969 | 1114.781 |
| ENSMUSG00000004018 | -0.078013483 | 0.492386 | 0.649523 | 481.9 | 543.5192 | 510.9418 | 530.5134 | 603.5939 | 602.2646 | 494.6803 | 480.7153 |
| ENSMUSG00000004031 | 1.596443018 | 0.002171 | 0.013939 | 33.19979 | 16.7975 | 29.81184 | 64.69676 | 9.751113 | 18.99536 | 8.678602 | 10.87591 |
| ENSMUSG00000004032 | -0.149174559 | 0.267129 | 0.431449 | 1504.051 | 1442.185 | 1647.932 | 1802.02 | 1914.143 | 1382.192 | 1790.685 | 2007.693 |
| ENSMUSG00000004035 | -0.38930515 | 0.016778 | 0.062231 | 364.1917 | 227.9661 | 269.9628 | 261.3749 | 393.945 | 375.4377 | 325.9297 | 377.3941 |
| ENSMUSG00000004040 | 0.056916643 | 0.623646 | 0.754953 | 1733.432 | 1428.987 | 1800.304 | 1532.882 | 1346.629 | 1495.047 | 1690.399 | 1715.131 |
| ENSMUSG00000004043 | -0.262140742 | 0.163264 | 0.307971 | 153.9263 | 123.5816 | 201.2299 | 172.5247 | 178.4454 | 177.6625 | 195.7507 | 232.7445 |
| ENSMUSG00000004044 | 0.400609364 | 0.001021 | 0.00796 | 4338.106 | 5286.414 | 5024.123 | 5146.412 | 3484.073 | 4576.764 | 3267.012 | 3669.532 |
| ENSMUSG00000004054 | -0.585536368 | 3.81E-05 | 0.000647 | 1068.43 | 1003.051 | 1247.129 | 905.7547 | 1641.112 | 1269.337 | 1603.613 | 1826.065 |
| ENSMUSG00000004056 | -0.006328129 | 0.8927 | 0.935055 | 3465.857 | 3528.675 | 3607.232 | 3561.772 | 3616.688 | 3419.165 | 3665.263 | 3523.795 |
| ENSMUSG00000004069 | -0.47378426 | 0.0092 | 0.040114 | 1088.551 | 937.0606 | 1040.93 | 969.5888 | 1317.375 | 965.4112 | 1461.862 | 1860.868 |
| ENSMUSG00000004070 | -0.159521474 | 0.122567 | 0.252114 | 1399.422 | 1498.577 | 1659.526 | 1483.712 | 1705.47 | 1433.591 | 1737.649 | 1871.744 |
| ENSMUSG00000004071 | 0.06415014 | 0.310438 | 0.479011 | 2750.553 | 2543.622 | 2434.633 | 2569.755 | 2432.903 | 2349.838 | 2505.223 | 2560.189 |
| ENSMUSG00000004085 | 0.165156436 | 0.055337 | 0.146424 | 698.2017 | 661.1017 | 615.2832 | 636.6161 | 603.5939 | 610.0863 | 577.6092 | 535.0948 |
| ENSMUSG00000004096 | -0.088672905 | 0.071585 | 0.175199 | 2508.094 | 2607.212 | 2642.488 | 2574.931 | 2710.809 | 2841.482 | 2772.331 | 2664.598 |
| ENSMUSG00000004098 | 1.379957252 | 0.000627 | 0.005456 | 445.6821 | 470.33 | 311.3681 | 367.4776 | 163.8187 | 289.3999 | 103.1789 | 56.55474 |
| ENSMUSG00000004099 | -0.179588484 | 0.046799 | 0.129943 | 2440.688 | 2875.972 | 2630.895 | 2333.397 | 3007.243 | 2765.501 | 2736.653 | 3130.087 |
| ENSMUSG00000004100 | -0.138539757 | 0.076599 | 0.183719 | 1354.149 | 1371.396 | 1229.738 | 1362.945 | 1579.68 | 1460.408 | 1344.219 | 1467.16 |
| ENSMUSG00000004105 | -0.047094263 | 0.774793 | 0.860668 | 1078.49 | 1530.972 | 1768.836 | 1327.578 | 1533.85 | 1298.389 | 1343.255 | 1720.569 |
| ENSMUSG00000004110 | 0.552760852 | 0.39623 | 0.563522 | 62.37537 | 21.59679 | 34.78048 | 72.46037 | 49.73068 | 60.3382 | 8.678602 | 11.9635 |
| ENSMUSG00000004113 | 0.862289855 | 0.010714 | 0.044909 | 70.4238 | 53.99197 | 48.85829 | 80.22398 | 30.22845 | 55.8687 | 28.92867 | 25.0146 |
| ENSMUSG00000004127 | 0.105608282 | 0.349982 | 0.518737 | 389.343 | 415.1383 | 456.2868 | 472.7177 | 378.3432 | 448.067 | 373.1799 | 415.4598 |
| ENSMUSG00000004151 | 0.570287898 | 0.001238 | 0.009169 | 429.5852 | 461.9313 | 326.274 | 473.5803 | 323.737 | 291.6346 | 306.644 | 215.343 |
| ENSMUSG00000004187 | 0.111464718 | 0.490675 | 0.647961 | 244.4712 | 176.3738 | 171.4181 | 221.6942 | 191.1218 | 201.1273 | 171.6435 | 190.3284 |
| ENSMUSG00000004207 | -0.050677125 | 0.643423 | 0.769406 | 12446.9 | 11817.04 | 14679.02 | 13035.97 | 12958.25 | 11589.4 | 13891.55 | 15401.38 |
| ENSMUSG00000004221 | -0.003289557 | 0.967259 | 0.981728 | 1164.005 | 1065.442 | 1043.414 | 1035.148 | 1096.025 | 997.8151 | 1161.004 | 1059.314 |
| ENSMUSG00000004231 | 0.106916244 | 0.242852 | 0.405434 | 2911.521 | 2740.392 | 2925.701 | 2657.743 | 3023.82 | 2510.74 | 2485.937 | 2411.189 |
| ENSMUSG00000004233 | -0.097215815 | 0.31339 | 0.481484 | 364.1917 | 371.9447 | 348.6329 | 342.4615 | 403.6961 | 394.4331 | 378.9656 | 346.9416 |
| ENSMUSG00000004263 | 0.469605259 | 0.001481 | 0.010472 | 528.1785 | 448.7333 | 464.5678 | 495.1459 | 320.8116 | 450.3018 | 309.5368 | 319.7518 |
| ENSMUSG00000004264 | -0.455354227 | 1.77E-06 | 5.93E-05 | 5841.152 | 5753.144 | 6393.811 | 6147.055 | 8588.78 | 6971.297 | 8393.173 | 9141.203 |
| ENSMUSG00000004266 | -0.414445851 | 0.042199 | 0.12109 | 264.5923 | 253.1623 | 304.7432 | 207.0296 | 344.2143 | 233.5312 | 381.8585 | 412.197 |
| ENSMUSG00000004267 | 0.361660572 | 0.009974 | 0.042491 | 601.6205 | 423.537 | 633.5016 | 565.0184 | 402.721 | 429.0717 | 447.4302 | 454.6131 |
| ENSMUSG00000004268 | -0.058238909 | 0.343073 | 0.511905 | 2796.831 | 2914.367 | 2779.954 | 2908.766 | 3202.266 | 2963.276 | 2858.153 | 2842.963 |
| ENSMUSG00000004270 | -0.553902464 | 0.00101 | 0.007894 | 1534.233 | 1312.605 | 1657.041 | 1371.571 | 2314.914 | 1501.751 | 2189.901 | 2620.007 |
| ENSMUSG00000004285 | -0.371707777 | 0.008913 | 0.039209 | 2355.173 | 2172.877 | 2664.019 | 2416.208 | 3062.825 | 2335.312 | 3402.976 | 3632.554 |
| ENSMUSG00000004317 | -0.627419231 | 0.008741 | 0.038657 | 967.8243 | 815.8786 | 1621.433 | 830.7064 | 1469.493 | 1322.971 | 1625.792 | 2128.416 |
| ENSMUSG00000004319 | -0.357893753 | 6.76E-05 | 0.001006 | 1191.168 | 1118.234 | 1172.599 | 1140.388 | 1379.783 | 1344.201 | 1528.398 | 1672.715 |
| ENSMUSG00000004328 | -0.061993972 | 0.840324 | 0.903737 | 1410.488 | 629.9063 | 818.1694 | 1293.073 | 709.881 | 1201.177 | 1613.256 | 810.2554 |
| ENSMUSG00000004341 | -3.257660957 | 4.44E-07 | 1.91E-05 | 48.29061 | 33.595 | 110.9663 | 37.95543 | 442.7005 | 144.1413 | 857.2531 | 767.8393 |
| ENSMUSG00000004347 | 0.233445951 | 0.234819 | 0.39571 | 141.8537 | 123.5816 | 175.5586 | 115.5915 | 104.3369 | 140.7891 | 115.7147 | 114.1971 |

| | | | | | | | | | | | |
|---|---|---|---|---|---|---|---|---|---|---|---|
| ENSMUSG00000004356 | -0.036536025 | 0.657793 | 0.779519 | 1181.108 | 1161.427 | 1146.928 | 1044.637 | 1095.05 | 1206.764 | 1092.54 | 1256.168 |
| ENSMUSG00000004360 | 0.983914958 | 0.067534 | 0.167798 | 36.21796 | 15.59768 | 16.56213 | 62.10889 | 15.60178 | 22.34748 | 16.39292 | 11.9635 |
| ENSMUSG00000004364 | 0.013554329 | 0.865833 | 0.919615 | 3190.198 | 3121.936 | 3013.48 | 3349.567 | 2982.865 | 3519.728 | 3136.833 | 2919.094 |
| ENSMUSG00000004383 | 0.042429098 | 0.695034 | 0.807475 | 665.0019 | 733.091 | 761.03 | 772.9107 | 787.8899 | 663.7202 | 626.788 | 770.0145 |
| ENSMUSG00000004393 | -0.295610943 | 0.006972 | 0.032892 | 1168.029 | 1216.619 | 1274.456 | 1211.986 | 1667.44 | 1210.116 | 1553.47 | 1546.555 |
| ENSMUSG00000004394 | -0.132051561 | 0.100394 | 0.221451 | 2485.96 | 2411.641 | 2581.208 | 2683.622 | 2770.291 | 2491.744 | 2832.117 | 3045.255 |
| ENSMUSG00000004415 | 0.577747318 | 4.91E-05 | 0.000785 | 2568.457 | 3186.726 | 3110.369 | 3026.946 | 1768.852 | 2574.43 | 1800.328 | 1826.065 |
| ENSMUSG00000004446 | -0.732364761 | 6.26E-05 | 0.000947 | 320.9313 | 269.9598 | 369.3356 | 315.7202 | 587.017 | 374.3203 | 525.5376 | 635.1532 |
| ENSMUSG00000004451 | -0.131985438 | 0.070969 | 0.174105 | 1934.643 | 1914.915 | 2037.97 | 1996.111 | 2351.968 | 1996.748 | 2066.472 | 2225.211 |
| ENSMUSG00000004455 | -0.081510041 | 0.301729 | 0.470132 | 4901.497 | 5136.436 | 5114.387 | 5287.019 | 5854.568 | 4745.488 | 5612.163 | 5412.941 |
| ENSMUSG00000004460 | -0.234460876 | 0.014518 | 0.056289 | 1824.983 | 1926.913 | 1932.801 | 1792.532 | 2054.56 | 1965.461 | 2232.329 | 2543.876 |
| ENSMUSG00000004462 | -0.069835217 | 0.524403 | 0.677583 | 568.4207 | 475.1293 | 563.1125 | 543.4528 | 575.3157 | 505.0531 | 541.9305 | 637.3284 |
| ENSMUSG00000004473 | 0.23172273 | 0.357809 | 0.5262 | 117.7084 | 196.7707 | 209.511 | 174.2499 | 146.2667 | 205.5968 | 129.2147 | 114.1971 |
| ENSMUSG00000004500 | -0.066198774 | 0.700987 | 0.811326 | 375.2583 | 356.347 | 348.6329 | 277.7648 | 345.1894 | 282.6956 | 340.3941 | 452.4379 |
| ENSMUSG00000004508 | 0.121351975 | 0.119446 | 0.24843 | 1555.36 | 1540.571 | 1716.665 | 1770.966 | 1492.895 | 1530.803 | 1603.613 | 1429.095 |
| ENSMUSG00000004530 | -0.358341631 | 0.000842 | 0.006838 | 1651.941 | 1560.968 | 1995.737 | 1764.065 | 2190.1 | 2041.442 | 2174.472 | 2538.438 |
| ENSMUSG00000004535 | -0.149716325 | 0.014727 | 0.056909 | 4671.11 | 4172.979 | 4568.664 | 4696.985 | 4943.814 | 4824.821 | 5211.019 | 5116.028 |
| ENSMUSG00000004552 | -0.852210834 | 0.22824 | 0.38824 | 131.7931 | 88.78679 | 581.3309 | 101.7896 | 629.9219 | 234.6486 | 279.6439 | 488.3284 |
| ENSMUSG00000004558 | -0.144177468 | 0.434823 | 0.59894 | 2610.711 | 1654.554 | 2329.464 | 2545.602 | 2611.348 | 1830.259 | 2986.404 | 2674.386 |
| ENSMUSG00000004561 | 0.188934347 | 0.046954 | 0.130236 | 925.57 | 965.8563 | 871.1682 | 901.4416 | 791.7904 | 898.3688 | 819.6458 | 702.5838 |
| ENSMUSG00000004562 | 0.231770694 | 0.000871 | 0.007006 | 2194.205 | 2444.036 | 2443.743 | 2366.176 | 2132.568 | 2069.377 | 1938.221 | 1905.46 |
| ENSMUSG00000004565 | 0.095396715 | 0.275156 | 0.441292 | 1257.568 | 1139.83 | 1256.238 | 1319.814 | 1065.797 | 1112.905 | 1211.147 | 1269.219 |
| ENSMUSG00000004567 | -0.123670272 | 0.02921 | 0.092788 | 1089.557 | 1042.645 | 1079.851 | 1068.791 | 1162.333 | 1191.121 | 1144.611 | 1169.16 |
| ENSMUSG00000004568 | -0.4135422 | 0.002572 | 0.015735 | 891.3642 | 782.2836 | 863.7152 | 772.9107 | 1090.174 | 848.0869 | 1193.79 | 1276.832 |
| ENSMUSG00000004591 | 0.049455142 | 0.560783 | 0.70652 | 2307.889 | 2038.497 | 1926.176 | 2139.306 | 1933.646 | 2229.161 | 2010.543 | 1955.489 |
| ENSMUSG00000004609 | 0.319017457 | 0.433719 | 0.597888 | 80.48435 | 37.19447 | 63.76421 | 103.5148 | 44.85512 | 88.27255 | 64.60737 | 31.54014 |
| ENSMUSG00000004610 | -0.577995014 | 0.001676 | 0.011495 | 769.6316 | 856.6726 | 885.246 | 678.8847 | 1187.686 | 843.6174 | 1161.004 | 1569.394 |
| ENSMUSG00000004626 | -0.380468856 | 0.008304 | 0.037316 | 1734.438 | 1540.571 | 2001.534 | 1609.655 | 2196.926 | 1765.451 | 2295.973 | 2708.102 |
| ENSMUSG00000004630 | -0.362017724 | 0.277808 | 0.444068 | 17.10292 | 20.39697 | 21.53077 | 27.60395 | 24.37778 | 27.93435 | 25.07152 | 34.80291 |
| ENSMUSG00000004631 | 0.493929096 | 1.96E-08 | 1.49E-06 | 1323.968 | 1345 | 1381.282 | 1553.585 | 974.1362 | 1071.562 | 1022.147 | 913.5765 |
| ENSMUSG00000004633 | -0.625003506 | 9.44E-07 | 3.55E-05 | 272.6407 | 326.3515 | 342.008 | 258.787 | 470.9788 | 419.0153 | 490.8232 | 467.6642 |
| ENSMUSG00000004637 | -0.124469103 | 0.205539 | 0.361948 | 434.6155 | 410.339 | 416.5376 | 431.3117 | 491.4561 | 401.1373 | 475.3946 | 476.3649 |
| ENSMUSG00000004642 | 0.000246384 | 0.997449 | 0.998477 | 1835.043 | 1841.726 | 1555.184 | 1661.413 | 1720.096 | 1674.944 | 1730.899 | 1759.722 |
| ENSMUSG00000004655 | -1.993536476 | 0.000652 | 0.005628 | 884.3218 | 932.2613 | 2998.574 | 1168.855 | 6283.617 | 2153.18 | 7025.811 | 8369.013 |
| ENSMUSG00000004661 | 0.042197926 | 0.815275 | 0.887529 | 177.0656 | 182.3729 | 168.1056 | 163.0358 | 216.4747 | 144.1413 | 135.9648 | 172.927 |
| ENSMUSG00000004665 | -0.076077111 | 0.500804 | 0.657105 | 5124.841 | 5558.773 | 6606.635 | 5166.252 | 5796.062 | 5337.696 | 5768.378 | 6772.43 |
| ENSMUSG00000004667 | -0.241914722 | 0.000168 | 0.001979 | 2469.863 | 2622.81 | 2601.083 | 2534.388 | 3219.818 | 2803.492 | 2963.261 | 3106.16 |
| ENSMUSG00000004668 | -3.058995119 | 0.003404 | 0.019397 | 16.09687 | 9.598572 | 224.4169 | 31.05445 | 527.5352 | 160.9019 | 885.2175 | 773.2773 |
| ENSMUSG00000004677 | 0.150147257 | 0.017776 | 0.065041 | 1691.177 | 1613.76 | 1762.211 | 1599.304 | 1564.079 | 1520.746 | 1469.577 | 1455.197 |
| ENSMUSG00000004698 | 0.274052415 | 0.188777 | 0.341288 | 271.6347 | 356.347 | 290.6654 | 337.2858 | 251.5787 | 356.4423 | 247.8223 | 182.7153 |
| ENSMUSG00000004730 | 0.02609961 | 0.901698 | 0.940849 | 70.4238 | 106.7841 | 112.6225 | 87.12497 | 92.63557 | 89.38993 | 105.1075 | 82.65692 |
| ENSMUSG00000004748 | -0.231950026 | 0.447643 | 0.61026 | 184.1079 | 217.1677 | 174.7305 | 163.8985 | 205.7485 | 98.32892 | 270.001 | 293.6496 |
| ENSMUSG00000004768 | 0.239565074 | 0.009388 | 0.040662 | 885.3278 | 938.2604 | 820.6537 | 870.3871 | 765.4624 | 810.0962 | 740.5741 | 657.9926 |
| ENSMUSG00000004771 | -0.413257312 | 0.001 | 0.007842 | 3433.664 | 2676.802 | 3694.184 | 3483.274 | 4097.418 | 3976.734 | 4544.695 | 5082.313 |
| ENSMUSG00000004788 | -0.354540855 | 0.000647 | 0.005588 | 1315.919 | 1373.796 | 1503.842 | 1410.389 | 2004.829 | 1500.633 | 1828.292 | 1831.503 |
| ENSMUSG00000004789 | -0.603481693 | 0.000318 | 0.003241 | 3356.197 | 2861.574 | 3497.922 | 3089.917 | 4642.505 | 3523.081 | 4906.303 | 6386.335 |
| ENSMUSG00000004791 | 0.022951826 | 0.9086 | 0.945375 | 81.4904 | 124.7814 | 130.0127 | 106.9653 | 114.088 | 119.559 | 96.42892 | 106.5839 |
| ENSMUSG00000004798 | 0.150450129 | 0.074831 | 0.180746 | 1365.216 | 1328.202 | 1300.127 | 1526.844 | 1165.258 | 1346.436 | 1217.897 | 1247.467 |
| ENSMUSG00000004814 | 1.788560466 | 0.000902 | 0.007205 | 20.12109 | 19.19714 | 27.32752 | 23.29083 | 1.950223 | 12.29112 | 6.750024 | 5.437955 |
| ENSMUSG00000004815 | -0.242107483 | 0.110789 | 0.236865 | 368.2159 | 311.9536 | 370.9918 | 370.0655 | 468.0534 | 317.3342 | 422.3587 | 474.1897 |
| ENSMUSG00000004837 | -0.436432527 | 0.094887 | 0.212854 | 117.7084 | 129.5807 | 206.1986 | 95.75121 | 191.1218 | 143.0239 | 214.0722 | 195.7664 |
| ENSMUSG00000004843 | 0.035889609 | 0.631437 | 0.760118 | 2048.327 | 1931.713 | 2224.294 | 2315.281 | 2034.082 | 2097.311 | 2101.186 | 2084.912 |
| ENSMUSG00000004846 | -0.142982615 | 0.064223 | 0.16275 | 1484.936 | 1515.375 | 1647.932 | 1650.199 | 1683.042 | 1614.606 | 1860.114 | 1799.963 |
| ENSMUSG00000004849 | -0.253505712 | 0.027827 | 0.089707 | 1918.546 | 2033.698 | 2238.372 | 1978.858 | 2503.111 | 2001.217 | 2410.723 | 2824.474 |
| ENSMUSG00000004864 | 0.028578712 | 0.859267 | 0.91552 | 1067.424 | 682.6985 | 755.2332 | 725.4664 | 766.4375 | 699.4762 | 901.6104 | 799.3794 |
| ENSMUSG00000004865 | -0.101075821 | 0.156174 | 0.298288 | 2681.135 | 2469.233 | 2395.712 | 2484.356 | 2836.599 | 2716.336 | 2459.902 | 2745.08 |
| ENSMUSG00000004872 | 2.817048786 | 0.002123 | 0.013707 | 36.21796 | 53.99197 | 53.82693 | 39.68068 | 2.925334 | 22.34748 | 0 | 1.087591 |
| ENSMUSG00000004880 | -0.043763426 | 0.541844 | 0.691151 | 1613.711 | 1526.173 | 1432.624 | 1468.185 | 1590.407 | 1647.009 | 1522.613 | 1463.898 |
| ENSMUSG00000004885 | 1.082293785 | 1.10E-10 | 2.00E-08 | 483.9121 | 729.4915 | 639.2983 | 590.0345 | 281.8072 | 354.2076 | 243.9652 | 274.073 |
| ENSMUSG00000004891 | -0.459376619 | 0.001503 | 0.010581 | 1617.735 | 2042.096 | 2302.136 | 1728.697 | 2365.62 | 2454.871 | 2555.366 | 3199.693 |
| ENSMUSG00000004892 | 1.983511599 | 0.037371 | 0.111263 | 59.35721 | 3.599465 | 2.48432 | 23.29083 | 7.80089 | 3.352122 | 5.785735 | 5.437955 |
| ENSMUSG00000004895 | 0.0073129 | 0.895963 | 0.93709 | 2118.75 | 2313.256 | 2214.357 | 2192.789 | 2102.34 | 2257.096 | 2216.901 | 2216.511 |
| ENSMUSG00000004896 | 0.298159656 | 0.03302 | 0.101332 | 1123.763 | 1019.848 | 800.7791 | 936.8091 | 744.985 | 936.3595 | 781.0742 | 692.7955 |
| ENSMUSG00000004897 | -0.057751012 | 0.245547 | 0.408521 | 7913.624 | 8066.4 | 7390.024 | 8142.303 | 8087.573 | 8146.775 | 8350.744 | 8209.138 |

| | | | | | | | | | | | |
|---|---|---|---|---|---|---|---|---|---|---|---|
| ENSMUSG00000004902 | 0.047795798 | 0.917704 | 0.951364 | 19.11503 | 25.19625 | 14.07781 | 18.97772 | 26.32801 | 25.6996 | 12.53576 | 9.78832 |
| ENSMUSG00000004929 | -0.318674712 | 2.51E-05 | 0.000469 | 861.1825 | 914.264 | 911.7454 | 920.4193 | 1131.129 | 1020.163 | 1184.147 | 1162.635 |
| ENSMUSG00000004931 | -0.372438192 | 0.006909 | 0.032677 | 701.2199 | 681.4986 | 645.0951 | 710.8018 | 861.0233 | 683.8329 | 940.1819 | 1059.314 |
| ENSMUSG00000004933 | 0.503761806 | 0.000705 | 0.005981 | 190.1443 | 157.1766 | 171.4181 | 166.4863 | 137.4907 | 120.6764 | 108.9647 | 116.3722 |
| ENSMUSG00000004934 | -0.067950305 | 0.575044 | 0.717289 | 926.5761 | 718.6931 | 809.0602 | 701.3129 | 751.8108 | 785.514 | 918.9676 | 851.5838 |
| ENSMUSG00000004936 | -0.238442428 | 0.118081 | 0.246817 | 1216.32 | 1333.002 | 1528.685 | 1247.354 | 1452.916 | 1222.407 | 1656.649 | 1951.138 |
| ENSMUSG00000004937 | -0.150903143 | 0.102396 | 0.224557 | 2477.912 | 2699.598 | 2911.623 | 2550.778 | 2996.517 | 2581.134 | 3024.975 | 3210.569 |
| ENSMUSG00000004945 | -0.150116474 | 0.063771 | 0.161915 | 2236.459 | 2206.472 | 2243.341 | 2186.751 | 2653.278 | 2209.049 | 2323.937 | 2659.16 |
| ENSMUSG00000004947 | -0.141087492 | 0.16841 | 0.314592 | 566.4086 | 527.9215 | 516.7385 | 582.2709 | 615.2952 | 553.1002 | 566.0377 | 685.1824 |
| ENSMUSG00000004951 | 0.195690857 | 0.091043 | 0.206817 | 1236.441 | 1046.244 | 1346.501 | 1304.287 | 1182.81 | 1023.515 | 1157.147 | 946.2042 |
| ENSMUSG00000004952 | 1.084677924 | 4.00E-16 | 2.64E-13 | 1173.059 | 1202.221 | 1086.476 | 1221.475 | 567.5148 | 679.3635 | 517.8233 | 443.7372 |
| ENSMUSG00000004961 | 0.618827315 | 0.000788 | 0.006508 | 214.2896 | 188.372 | 179.6991 | 200.9913 | 126.7645 | 131.8501 | 158.1434 | 92.44524 |
| ENSMUSG00000004980 | 0.095747124 | 0.128914 | 0.260595 | 42276.42 | 43193.58 | 37100.83 | 41161.81 | 38740.2 | 39295.81 | 37402.85 | 37774.21 |
| ENSMUSG00000004988 | -0.618999717 | 0.400526 | 0.56788 | 265.5983 | 220.7672 | 1521.232 | 291.5667 | 722.5575 | 465.945 | 971.0392 | 1372.54 |
| ENSMUSG00000004994 | -0.027897503 | 0.820969 | 0.891475 | 278.6771 | 271.1597 | 280.7281 | 248.4356 | 314.961 | 241.3528 | 267.1081 | 275.1605 |
| ENSMUSG00000004996 | -0.097235524 | 0.211426 | 0.368886 | 968.8303 | 961.057 | 929.9637 | 942.8475 | 1039.469 | 931.89 | 984.5392 | 1110.431 |
| ENSMUSG00000005034 | 0.193929149 | 0.010782 | 0.045121 | 2730.432 | 2890.37 | 2844.546 | 3093.368 | 2458.256 | 2777.792 | 2510.045 | 2361.16 |
| ENSMUSG00000005043 | -0.110807504 | 0.581306 | 0.722532 | 217.3077 | 227.9661 | 249.2601 | 169.9368 | 223.3005 | 188.8362 | 210.215 | 311.0511 |
| ENSMUSG00000005045 | 0.63375119 | 0.070726 | 0.173673 | 116.7023 | 58.79126 | 52.17072 | 131.1188 | 55.58134 | 62.57295 | 66.53595 | 46.76642 |
| ENSMUSG00000005054 | -0.176358673 | 0.090204 | 0.205506 | 1538.257 | 1323.403 | 1628.058 | 1453.521 | 1688.893 | 1455.938 | 1714.506 | 1859.781 |
| ENSMUSG00000005057 | -0.169167557 | 0.290661 | 0.457794 | 120.7265 | 111.5834 | 128.3565 | 143.1955 | 155.0427 | 120.6764 | 149.4648 | 142.4744 |
| ENSMUSG00000005069 | 0.084154471 | 0.304207 | 0.472636 | 1590.572 | 1470.981 | 1498.873 | 1511.316 | 1542.626 | 1259.281 | 1449.327 | 1474.774 |
| ENSMUSG00000005078 | 0.034851365 | 0.714087 | 0.820631 | 1391.373 | 1419.389 | 1197.442 | 1428.505 | 1182.81 | 1432.474 | 1353.862 | 1336.649 |
| ENSMUSG00000005087 | 0.900373993 | 0.001331 | 0.009698 | 445.6821 | 429.5361 | 391.6944 | 415.7845 | 205.7485 | 383.2593 | 170.6792 | 142.4744 |
| ENSMUSG00000005089 | 0.073189052 | 0.775181 | 0.860918 | 60.36326 | 65.99018 | 81.98256 | 79.36136 | 56.55646 | 97.21155 | 64.60737 | 56.55474 |
| ENSMUSG00000005102 | 0.123559438 | 0.038033 | 0.112563 | 1222.356 | 1213.02 | 1194.13 | 1244.766 | 1114.552 | 1149.778 | 1143.647 | 1065.839 |
| ENSMUSG00000005103 | -0.309480151 | 0.022603 | 0.076883 | 3437.688 | 3269.514 | 4051.926 | 3149.438 | 4391.901 | 3377.822 | 4480.087 | 4987.693 |
| ENSMUSG00000005107 | -0.478126972 | 0.063684 | 0.161739 | 203.223 | 165.5754 | 209.511 | 201.8539 | 213.5494 | 168.7235 | 381.8585 | 323.0146 |
| ENSMUSG00000005124 | 0.126486719 | 0.223585 | 0.383289 | 649.9111 | 777.4844 | 722.109 | 618.501 | 599.6935 | 673.7766 | 601.7164 | 659.0802 |
| ENSMUSG00000005125 | -1.136274736 | 6.49E-08 | 4.02E-06 | 3672.098 | 2484.83 | 3699.152 | 3913.723 | 8975.9 | 4749.957 | 8140.529 | 8404.904 |
| ENSMUSG00000005131 | 0.601372266 | 7.52E-05 | 0.001092 | 169.0171 | 215.9679 | 178.871 | 191.5024 | 121.8889 | 139.6718 | 109.929 | 126.1606 |
| ENSMUSG00000005142 | -0.150820089 | 0.094681 | 0.21248 | 1415.518 | 1406.191 | 1566.778 | 1449.207 | 1655.739 | 1444.765 | 1568.898 | 1814.102 |
| ENSMUSG00000005148 | 0.766322856 | 0.000163 | 0.001932 | 394.3733 | 560.3167 | 395.835 | 401.9825 | 258.4045 | 339.6817 | 241.0723 | 190.3284 |
| ENSMUSG00000005150 | -0.097959623 | 0.209855 | 0.367187 | 1025.169 | 1090.638 | 1076.539 | 1027.385 | 1212.063 | 1017.928 | 1159.076 | 1123.482 |
| ENSMUSG00000005161 | -0.177176996 | 0.003434 | 0.019521 | 10797.98 | 10897.98 | 10696.65 | 11233.08 | 13109.4 | 11305.59 | 12395.94 | 12512.74 |
| ENSMUSG00000005198 | 0.235386181 | 0.000213 | 0.002383 | 2392.397 | 2428.439 | 2265.7 | 2416.208 | 2003.854 | 2161.002 | 1896.757 | 2010.956 |
| ENSMUSG00000005204 | -0.013421761 | 0.832329 | 0.898647 | 2372.276 | 2362.449 | 2559.678 | 2488.669 | 2566.493 | 2583.369 | 2347.08 | 2382.912 |
| ENSMUSG00000005220 | -1.541595778 | 0.18331 | 0.334206 | 85.51462 | 14.39786 | 1006.15 | 129.3935 | 641.6232 | 243.5876 | 1249.719 | 1462.81 |
| ENSMUSG00000005225 | -0.444919011 | 5.45E-06 | 0.000143 | 576.4691 | 512.3238 | 541.5817 | 544.3154 | 644.5486 | 793.3356 | 741.5384 | 784.1532 |
| ENSMUSG00000005232 | 0.355969236 | 0.064101 | 0.16253 | 131.7931 | 106.7841 | 108.482 | 120.7673 | 78.0089 | 111.7374 | 78.10742 | 98.97079 |
| ENSMUSG00000005233 | -0.192440029 | 0.014244 | 0.05553 | 1431.615 | 1518.974 | 1406.125 | 1421.604 | 1550.427 | 1640.305 | 1570.827 | 1840.204 |
| ENSMUSG00000005237 | 0.487774921 | 0.046315 | 0.129162 | 70.4238 | 76.78858 | 64.59232 | 72.46037 | 40.95467 | 70.39457 | 49.17875 | 42.41605 |
| ENSMUSG00000005251 | -0.46905026 | 0.037657 | 0.111861 | 293.7679 | 242.3639 | 397.4912 | 255.3366 | 411.497 | 284.9304 | 527.4662 | 423.0729 |
| ENSMUSG00000005262 | -0.022159226 | 0.72671 | 0.828973 | 1670.05 | 1588.564 | 1639.651 | 1717.483 | 1663.54 | 1578.85 | 1770.435 | 1706.43 |
| ENSMUSG00000005267 | 0.042775032 | 0.794145 | 0.873694 | 225.3562 | 194.3711 | 190.4645 | 182.8762 | 212.5743 | 187.7188 | 218.8936 | 149 |
| ENSMUSG00000005268 | 0.537739585 | 0.373539 | 0.542368 | 437.6336 | 41.99375 | 104.3414 | 169.0742 | 78.0089 | 102.7984 | 93.53605 | 244.708 |
| ENSMUSG00000005299 | -0.682312774 | 0.000292 | 0.003043 | 986.9393 | 1009.05 | 1355.611 | 1065.34 | 1673.291 | 1248.107 | 1871.685 | 2296.992 |
| ENSMUSG00000005312 | -0.2674888 | 0.004938 | 0.025514 | 2645.923 | 2728.394 | 2702.94 | 2519.723 | 2789.793 | 2988.976 | 3364.405 | 3611.89 |
| ENSMUSG00000005320 | -0.403501704 | 0.038938 | 0.114251 | 618.7234 | 401.9402 | 557.3158 | 501.1842 | 595.793 | 525.1658 | 752.1455 | 878.7736 |
| ENSMUSG00000005338 | 0.431513446 | 0.010259 | 0.043457 | 237.4288 | 237.5647 | 224.4169 | 220.8316 | 166.744 | 223.4748 | 142.7148 | 150.0876 |
| ENSMUSG00000005354 | -0.415509661 | 0.002097 | 0.013581 | 2460.809 | 1899.317 | 2389.088 | 2118.603 | 2974.089 | 2384.476 | 2984.475 | 3486.817 |
| ENSMUSG00000005357 | 0.174863798 | 0.613047 | 0.747022 | 31.18769 | 33.595 | 24.01509 | 17.25247 | 21.45245 | 24.58223 | 23.14294 | 23.927 |
| ENSMUSG00000005360 | 0.849322123 | 0.002528 | 0.015553 | 1241.471 | 1456.583 | 938.2448 | 1343.967 | 580.1912 | 1179.947 | 547.7162 | 456.7883 |
| ENSMUSG00000005362 | -0.032602828 | 0.71045 | 0.818711 | 1252.538 | 1096.637 | 1029.337 | 1156.778 | 1211.088 | 1074.914 | 1198.611 | 1151.759 |
| ENSMUSG00000005364 | -2.494827935 | 0.006599 | 0.031619 | 4.024217 | 5.999108 | 9.109173 | 0 | 27.30312 | 3.352122 | 45.32159 | 31.54014 |
| ENSMUSG00000005370 | -0.078814334 | 0.308774 | 0.477081 | 1578.499 | 1878.921 | 1684.369 | 1629.496 | 1813.707 | 1874.954 | 1745.363 | 1712.956 |
| ENSMUSG00000005371 | 0.408118874 | 0.0086 | 0.03826 | 2509.1 | 2566.418 | 2257.419 | 2762.12 | 1795.18 | 2546.496 | 1641.22 | 1625.949 |
| ENSMUSG00000005373 | 0.002719378 | 0.992042 | 0.995436 | 216.3017 | 129.5807 | 195.4332 | 150.9591 | 127.7396 | 124.0285 | 199.6079 | 240.3576 |
| ENSMUSG00000005374 | -0.094676468 | 0.335016 | 0.503219 | 949.7153 | 946.6592 | 883.5898 | 859.173 | 1033.618 | 850.3217 | 926.6819 | 1072.365 |
| ENSMUSG00000005378 | -0.089566576 | 0.199857 | 0.354806 | 1058.369 | 1055.843 | 1042.586 | 1079.142 | 1057.996 | 1091.674 | 1142.683 | 1215.927 |
| ENSMUSG00000005397 | 0.350257196 | 0.000299 | 0.003086 | 7157.071 | 8104.794 | 9481.821 | 8807.386 | 6220.235 | 6696.423 | 6881.167 | 6524.459 |
| ENSMUSG00000005410 | -0.327148782 | 0.008085 | 0.036563 | 1967.842 | 2693.599 | 2367.557 | 2055.632 | 3072.576 | 2495.096 | 2773.296 | 3051.781 |
| ENSMUSG00000005413 | -0.432801031 | 0.149269 | 0.289229 | 1430.609 | 958.6574 | 1479.827 | 1075.691 | 2406.575 | 845.8522 | 1248.754 | 2174.095 |
| ENSMUSG00000005417 | -0.107188133 | 0.21539 | 0.373588 | 7357.276 | 8168.385 | 9333.59 | 8026.712 | 8432.763 | 8573.611 | 9153.033 | 9266.276 |

| | | | | | | | | | | | |
|---|---|---|---|---|---|---|---|---|---|---|---|
| ENSMUSG00000005442 | 0.194432687 | 0.015381 | 0.058734 | 1664.014 | 1690.549 | 1867.38 | 1725.247 | 1356.38 | 1581.084 | 1554.434 | 1583.533 |
| ENSMUSG00000005447 | -0.1921788 | 0.007537 | 0.034673 | 1443.688 | 1684.549 | 1592.449 | 1586.365 | 1878.064 | 1838.08 | 1711.613 | 1776.036 |
| ENSMUSG00000005465 | -0.005691174 | 0.9873 | 0.992833 | 22.1332 | 38.39429 | 46.37397 | 36.23019 | 29.25334 | 27.93435 | 33.75012 | 53.29196 |
| ENSMUSG00000005469 | -0.065676077 | 0.375972 | 0.544226 | 1816.934 | 1847.725 | 1772.976 | 1745.95 | 1795.18 | 1759.864 | 1876.507 | 2084.912 |
| ENSMUSG00000005470 | -0.126891853 | 0.2785 | 0.444752 | 323.9495 | 417.5379 | 380.1009 | 357.9887 | 436.8499 | 408.9589 | 363.537 | 404.5839 |
| ENSMUSG00000005481 | -0.270482031 | 0.006538 | 0.031449 | 1485.942 | 1440.986 | 1617.292 | 1482.85 | 1964.849 | 1541.976 | 1784.899 | 1979.416 |
| ENSMUSG00000005483 | -0.325533454 | 0.007786 | 0.035526 | 1190.162 | 1271.811 | 1389.563 | 1407.802 | 1667.44 | 1340.849 | 1663.399 | 1920.686 |
| ENSMUSG00000005501 | 0.036485552 | 0.544412 | 0.69299 | 1311.895 | 1376.195 | 1256.238 | 1258.568 | 1278.371 | 1273.806 | 1257.433 | 1257.255 |
| ENSMUSG00000005505 | -0.117040263 | 0.482491 | 0.641014 | 271.6347 | 301.1552 | 325.4459 | 350.2251 | 354.9405 | 253.6439 | 339.4298 | 406.7591 |
| ENSMUSG00000005506 | 0.127058645 | 0.094464 | 0.212121 | 4704.31 | 4733.296 | 4403.871 | 4389.891 | 3898.495 | 4667.272 | 4061.586 | 4066.503 |
| ENSMUSG00000005510 | -0.402252969 | 1.83E-05 | 0.000365 | 2240.483 | 2076.891 | 2396.541 | 2173.811 | 2891.205 | 2563.256 | 3016.297 | 3278 |
| ENSMUSG00000005514 | -0.069097887 | 0.396665 | 0.563836 | 2059.393 | 1866.922 | 1995.737 | 1942.628 | 2058.46 | 1857.076 | 2048.15 | 2288.292 |
| ENSMUSG00000005533 | 0.53392832 | 0.012002 | 0.048987 | 6468.93 | 6205.477 | 4947.109 | 6561.114 | 3390.462 | 6291.934 | 3903.443 | 3117.036 |
| ENSMUSG00000005534 | 0.490956889 | 0.001559 | 0.010831 | 2410.506 | 2114.086 | 1696.79 | 2174.674 | 1332.002 | 1887.245 | 1479.22 | 1275.744 |
| ENSMUSG00000005553 | -0.525835295 | 0.067356 | 0.167583 | 75.45408 | 59.99108 | 94.40416 | 71.59775 | 91.66046 | 70.39457 | 116.679 | 156.6131 |
| ENSMUSG00000005566 | -0.070835475 | 0.149579 | 0.289495 | 7992.096 | 8811.489 | 8552.685 | 8262.208 | 8784.778 | 8887.594 | 8648.71 | 8986.765 |
| ENSMUSG00000005575 | -0.328732958 | 0.006986 | 0.032918 | 2357.185 | 2010.901 | 2418.071 | 2159.147 | 2654.253 | 2346.486 | 2910.225 | 3325.854 |
| ENSMUSG00000005580 | -0.42750327 | 0.120059 | 0.248995 | 81.4904 | 41.99375 | 71.21717 | 46.58167 | 75.08357 | 71.51194 | 80.036 | 98.97079 |
| ENSMUSG00000005583 | 0.279966448 | 0.025161 | 0.083399 | 952.7335 | 922.6628 | 893.5271 | 891.9527 | 655.2748 | 934.1247 | 675.9667 | 751.5254 |
| ENSMUSG00000005609 | -0.029746222 | 0.60352 | 0.740729 | 2319.961 | 2231.668 | 2175.436 | 2130.68 | 2193.025 | 2195.64 | 2336.473 | 2313.306 |
| ENSMUSG00000005610 | -0.027910596 | 0.529533 | 0.681668 | 24584.95 | 26123.71 | 26399.21 | 26015 | 25799.49 | 26957.77 | 25723.38 | 26661.21 |
| ENSMUSG00000005611 | 0.432864988 | 4.41E-05 | 0.000726 | 964.8061 | 1190.223 | 1005.321 | 1099.845 | 775.2135 | 877.1387 | 703.9311 | 799.3794 |
| ENSMUSG00000005615 | -0.369119498 | 0.002712 | 0.016409 | 1273.665 | 1111.035 | 1265.347 | 1170.58 | 1614.784 | 1224.642 | 1637.363 | 1749.934 |
| ENSMUSG00000005621 | 0.113035634 | 0.116946 | 0.245141 | 1182.114 | 1130.232 | 1236.363 | 1243.903 | 1053.12 | 1071.562 | 1181.254 | 1128.92 |
| ENSMUSG00000005625 | -0.146413424 | 0.029733 | 0.093932 | 3101.666 | 3106.338 | 3377.019 | 3171.867 | 3531.853 | 3276.141 | 3514.834 | 3800.043 |
| ENSMUSG00000005628 | 0.89440866 | 0.026872 | 0.087531 | 25.15136 | 37.19447 | 55.48314 | 43.9938 | 23.40267 | 33.52122 | 14.46434 | 16.31387 |
| ENSMUSG00000005640 | -1.007781074 | 0.016695 | 0.061995 | 44.26639 | 23.99643 | 60.45178 | 44.85642 | 57.53157 | 46.92971 | 106.0718 | 140.2993 |
| ENSMUSG00000005649 | -0.102743304 | 0.794448 | 0.873694 | 13.07871 | 14.39786 | 26.49941 | 18.11509 | 18.52711 | 20.11273 | 22.17865 | 17.40146 |
| ENSMUSG00000005656 | -0.078167896 | 0.215106 | 0.373295 | 2642.905 | 2787.185 | 2858.624 | 2870.811 | 2945.811 | 2813.548 | 2863.939 | 3160.54 |
| ENSMUSG00000005667 | -0.213695769 | 0.23129 | 0.391871 | 574.457 | 434.3354 | 504.3169 | 542.5902 | 468.0534 | 494.9967 | 746.3598 | 675.3941 |
| ENSMUSG00000005672 | -0.049510737 | 0.745759 | 0.841098 | 1047.303 | 967.0562 | 1335.736 | 1018.758 | 906.8535 | 1307.328 | 1248.754 | 1061.489 |
| ENSMUSG00000005674 | -0.419119921 | 0.005957 | 0.029304 | 284.7134 | 307.1543 | 282.3844 | 283.8031 | 337.3885 | 312.8647 | 437.7873 | 458.9634 |
| ENSMUSG00000005677 | -0.385749249 | 0.301717 | 0.470132 | 38.23007 | 19.19714 | 35.60859 | 23.29083 | 38.02934 | 24.58223 | 35.6787 | 54.37955 |
| ENSMUSG00000005681 | 0.225555155 | 0.764725 | 0.853982 | 95.57516 | 9.598572 | 13.24971 | 103.5148 | 39.00445 | 17.87799 | 47.25017 | 85.9197 |
| ENSMUSG00000005682 | 0.164782125 | 0.060927 | 0.156655 | 2749.547 | 2434.438 | 2364.244 | 2746.593 | 2329.541 | 2469.397 | 2312.365 | 2072.949 |
| ENSMUSG00000005683 | -0.558112915 | 0.002176 | 0.013962 | 3769.686 | 3487.881 | 4906.532 | 3894.745 | 5765.833 | 4121.993 | 6237.987 | 7520.692 |
| ENSMUSG00000005686 | -0.478001185 | 0.072073 | 0.176159 | 288.7376 | 213.5682 | 422.3344 | 385.5927 | 354.9405 | 349.7381 | 477.3231 | 646.0291 |
| ENSMUSG00000005687 | 0.119440003 | 0.106091 | 0.230107 | 2904.479 | 3065.544 | 2820.531 | 3130.461 | 2863.902 | 2936.459 | 2628.652 | 2543.876 |
| ENSMUSG00000005698 | -0.015686748 | 0.767044 | 0.855485 | 3448.754 | 3526.275 | 3333.129 | 3366.819 | 3408.014 | 3660.518 | 3390.441 | 3362.832 |
| ENSMUSG00000005699 | -0.242376799 | 0.118537 | 0.247389 | 141.8537 | 165.5754 | 150.7154 | 193.2277 | 191.1218 | 217.8879 | 173.572 | 189.2408 |
| ENSMUSG00000005705 | -0.09591809 | 0.80188 | 0.879143 | 28.16952 | 16.7975 | 14.90592 | 25.8787 | 29.25334 | 23.46486 | 18.32149 | 20.66423 |
| ENSMUSG00000005718 | 0.1525132 | 0.273006 | 0.438635 | 768.6255 | 893.867 | 688.1566 | 762.5592 | 773.2633 | 784.3966 | 684.6453 | 555.759 |
| ENSMUSG00000005732 | -0.272800476 | 0.001201 | 0.00898 | 3701.274 | 4630.111 | 4116.518 | 4022.413 | 5285.103 | 4889.629 | 4719.231 | 4999.656 |
| ENSMUSG00000005763 | 0.7057799 | 0.15229 | 0.293022 | 13.07871 | 22.79661 | 30.63995 | 15.52722 | 15.60178 | 6.704245 | 10.60718 | 17.40146 |
| ENSMUSG00000005774 | 0.219883086 | 0.04149 | 0.119779 | 783.7163 | 849.4736 | 664.1415 | 729.7795 | 656.2499 | 708.4152 | 591.1093 | 640.5912 |
| ENSMUSG00000005779 | -0.212733671 | 0.002406 | 0.01498 | 4901.497 | 5202.426 | 5329.694 | 5118.808 | 6105.172 | 5401.386 | 5905.307 | 6405.912 |
| ENSMUSG00000005800 | -0.008277645 | 0.983789 | 0.990959 | 49.29666 | 16.7975 | 28.15563 | 49.16954 | 45.83023 | 29.05173 | 41.46443 | 28.27737 |
| ENSMUSG00000005802 | -0.082132436 | 0.368248 | 0.536797 | 941.6669 | 916.6637 | 956.4632 | 1044.637 | 1003.39 | 1119.609 | 1054.932 | 910.3137 |
| ENSMUSG00000005803 | -0.793516981 | 0.00025 | 0.002712 | 397.3915 | 275.959 | 481.13 | 311.4071 | 650.3992 | 459.2408 | 650.8952 | 783.0656 |
| ENSMUSG00000005804 | -0.040686331 | 0.502275 | 0.658427 | 953.7395 | 951.4585 | 996.2123 | 961.8252 | 995.5886 | 1019.045 | 965.2535 | 997.321 |
| ENSMUSG00000005813 | -0.119544846 | 0.13093 | 0.263472 | 1492.985 | 1450.584 | 1415.234 | 1414.703 | 1502.647 | 1473.816 | 1530.327 | 1765.16 |
| ENSMUSG00000005823 | -0.044547489 | 0.623238 | 0.754607 | 910.4792 | 814.6788 | 880.2773 | 830.7064 | 920.5051 | 778.8097 | 891.9675 | 952.7298 |
| ENSMUSG00000005836 | 0.281993313 | 0.318485 | 0.486689 | 488.9424 | 407.9393 | 350.2891 | 472.7177 | 231.1014 | 595.5604 | 276.751 | 312.1386 |
| ENSMUSG00000005846 | -0.140151385 | 0.021817 | 0.074942 | 3287.786 | 3389.496 | 3266.053 | 3390.973 | 3757.104 | 3896.283 | 3648.87 | 3391.109 |
| ENSMUSG00000005864 | 0.952905902 | 0.098051 | 0.218027 | 20.12109 | 11.99822 | 19.87456 | 16.38985 | 5.850668 | 17.87799 | 8.678602 | 3.262773 |
| ENSMUSG00000005871 | 0.126172273 | 0.077599 | 0.185201 | 2818.964 | 2468.033 | 2629.239 | 2673.27 | 2351.968 | 2635.885 | 2363.473 | 2355.722 |
| ENSMUSG00000005873 | 0.02085104 | 0.811485 | 0.885204 | 2647.935 | 2303.657 | 2610.192 | 2335.122 | 2373.421 | 2295.086 | 2362.508 | 2726.591 |
| ENSMUSG00000005881 | -0.104417373 | 0.250034 | 0.413794 | 3539.299 | 3641.458 | 4043.645 | 3901.646 | 4141.298 | 3519.728 | 4170.551 | 4431.934 |
| ENSMUSG00000005882 | -0.433055022 | 0.001431 | 0.010239 | 947.7032 | 889.0678 | 1074.054 | 929.9081 | 1363.206 | 1002.285 | 1317.219 | 1504.138 |
| ENSMUSG00000005883 | 1.22593528 | 0.101918 | 0.223849 | 12.07265 | 16.7975 | 18.21835 | 16.38985 | 5.850668 | 15.64324 | 5.785735 | 0 |
| ENSMUSG00000005886 | 0.169240473 | 0.17299 | 0.32066 | 1837.055 | 1599.362 | 1601.558 | 1823.586 | 1439.264 | 1873.836 | 1471.505 | 1319.248 |
| ENSMUSG00000005893 | 0.089208865 | 0.372887 | 0.541843 | 888.346 | 808.6797 | 824.7942 | 695.2745 | 787.8899 | 698.3588 | 777.2171 | 758.051 |
| ENSMUSG00000005897 | 0.324551063 | 0.000868 | 0.006989 | 673.0504 | 737.8902 | 728.7338 | 714.2523 | 552.8881 | 648.077 | 510.109 | 569.8977 |
| ENSMUSG00000005899 | -0.056624776 | 0.498354 | 0.654775 | 1443.688 | 1692.948 | 1580.856 | 1476.811 | 1613.809 | 1756.512 | 1533.22 | 1536.766 |

| | | | | | | | | | | | |
|---|---|---|---|---|---|---|---|---|---|---|---|
| ENSMUSG00000005907 | -0.21873612 | 0.100391 | 0.221451 | 563.3904 | 478.7288 | 423.9906 | 476.1682 | 545.0872 | 616.7905 | 477.3231 | 621.0145 |
| ENSMUSG00000005936 | -0.027836719 | 0.634604 | 0.762368 | 2811.922 | 2892.77 | 2748.486 | 2916.53 | 3089.153 | 2914.112 | 2803.189 | 2782.058 |
| ENSMUSG00000005947 | 0.867738562 | 0.058131 | 0.151252 | 21.12714 | 34.79482 | 19.87456 | 13.80198 | 13.65156 | 15.64324 | 10.60718 | 8.700729 |
| ENSMUSG00000005949 | -0.142018434 | 0.224337 | 0.384078 | 594.5781 | 518.3229 | 601.2054 | 536.5518 | 622.121 | 526.2832 | 709.7168 | 625.3649 |
| ENSMUSG00000005950 | -1.569345178 | 0.006202 | 0.030184 | 9.054489 | 14.39786 | 6.624853 | 8.626235 | 35.10401 | 7.821619 | 27.96439 | 42.41605 |
| ENSMUSG00000005951 | -0.356364849 | 0.091975 | 0.208232 | 124.7507 | 113.983 | 95.23226 | 90.57547 | 133.5902 | 98.32892 | 147.5362 | 162.0511 |
| ENSMUSG00000005958 | 0.261708047 | 0.044738 | 0.126005 | 923.5579 | 1199.822 | 979.6501 | 884.1891 | 765.4624 | 743.0538 | 875.5746 | 938.5911 |
| ENSMUSG00000005968 | 0.12760477 | 0.334791 | 0.503039 | 496.9909 | 379.1436 | 391.6944 | 353.6756 | 374.4427 | 369.8508 | 348.1084 | 391.5328 |
| ENSMUSG00000005973 | 0.153226781 | 0.057171 | 0.149688 | 7588.668 | 8024.406 | 6711.804 | 7467.732 | 6426.959 | 7056.217 | 7074.025 | 6228.634 |
| ENSMUSG00000005980 | -0.435139765 | 0.013306 | 0.052892 | 137.8294 | 147.578 | 130.8408 | 179.4257 | 218.4249 | 200.01 | 226.608 | 159.8759 |
| ENSMUSG00000005981 | -0.446566627 | 0.000123 | 0.001567 | 2101.648 | 2091.289 | 2125.75 | 1954.705 | 2931.185 | 2249.274 | 2829.224 | 3263.861 |
| ENSMUSG00000005982 | -0.134991093 | 0.047871 | 0.132023 | 798.8072 | 881.8688 | 871.1682 | 870.3871 | 961.4597 | 909.5425 | 931.5033 | 954.905 |
| ENSMUSG00000005983 | -0.416050412 | 0.040095 | 0.116921 | 76.46013 | 106.7841 | 101.029 | 75.04824 | 116.0382 | 98.32892 | 134.0362 | 129.4233 |
| ENSMUSG00000005986 | 0.563913707 | 3.07E-05 | 0.000546 | 303.8284 | 262.7609 | 262.5098 | 259.6497 | 203.7983 | 200.01 | 174.5363 | 157.7007 |
| ENSMUSG00000006005 | -0.011594617 | 0.851069 | 0.910569 | 4427.645 | 4261.766 | 4212.578 | 3951.678 | 4005.757 | 4259.43 | 4274.694 | 4448.248 |
| ENSMUSG00000006007 | 0.078818628 | 0.632748 | 0.761079 | 199.1988 | 291.5566 | 215.3077 | 259.6497 | 241.8276 | 248.057 | 193.8221 | 229.4817 |
| ENSMUSG00000006010 | 0.100637389 | 0.186684 | 0.338586 | 1117.726 | 1149.429 | 1127.053 | 1052.401 | 1000.464 | 1140.839 | 1012.504 | 992.9707 |
| ENSMUSG00000006014 | 1.586974992 | 5.39E-06 | 0.000143 | 97.58727 | 87.58697 | 57.96746 | 100.9269 | 19.50223 | 45.81234 | 29.89296 | 19.57664 |
| ENSMUSG00000006019 | -0.136846633 | 0.290692 | 0.457805 | 418.5186 | 514.7234 | 578.8465 | 432.1744 | 542.1619 | 522.9311 | 568.9306 | 503.5547 |
| ENSMUSG00000006021 | -0.082037065 | 0.406723 | 0.573149 | 472.8455 | 519.5227 | 448.0057 | 490.8328 | 483.6552 | 540.8091 | 469.6088 | 549.2335 |
| ENSMUSG00000006024 | -0.258324451 | 0.002503 | 0.015454 | 2541.293 | 2577.217 | 2808.938 | 2536.113 | 3150.585 | 2727.51 | 3299.798 | 3337.817 |
| ENSMUSG00000006050 | -0.002611789 | 0.960888 | 0.977573 | 2086.557 | 2129.683 | 1984.972 | 2095.312 | 2124.768 | 2003.452 | 2071.293 | 2106.664 |
| ENSMUSG00000006057 | -0.94268455 | 1.45E-05 | 0.000303 | 2312.919 | 2298.858 | 3171.648 | 2354.962 | 4729.29 | 3002.384 | 5168.59 | 6588.627 |
| ENSMUSG00000006058 | -0.077194188 | 0.307575 | 0.476044 | 1112.696 | 1186.623 | 1208.208 | 1084.318 | 1125.278 | 1250.342 | 1192.826 | 1276.832 |
| ENSMUSG00000006095 | -0.264601338 | 0.011307 | 0.046782 | 1275.677 | 1458.983 | 1541.106 | 1418.153 | 1679.142 | 1483.873 | 1747.292 | 1930.474 |
| ENSMUSG00000006127 | -0.068958381 | 0.473117 | 0.632492 | 1358.173 | 1183.024 | 1427.656 | 1483.712 | 1567.004 | 1392.248 | 1421.362 | 1343.175 |
| ENSMUSG00000006134 | 0.0340529 | 0.736214 | 0.834773 | 1169.035 | 1190.223 | 1083.163 | 1151.602 | 1215.964 | 939.7116 | 1117.611 | 1210.489 |
| ENSMUSG00000006143 | -0.733286415 | 0.074721 | 0.180551 | 22.1332 | 43.19358 | 61.27989 | 33.64232 | 109.2125 | 43.57759 | 54.00019 | 59.81751 |
| ENSMUSG00000006154 | -0.088039814 | 0.803028 | 0.879572 | 106.6418 | 314.3532 | 255.0568 | 300.193 | 191.1218 | 173.193 | 341.3584 | 331.7153 |
| ENSMUSG00000006169 | -0.190381041 | 0.016131 | 0.060548 | 4144.944 | 4001.405 | 4354.185 | 4230.306 | 4759.518 | 4206.913 | 4959.339 | 5167.145 |
| ENSMUSG00000006191 | -0.157654963 | 0.18417 | 0.335278 | 428.5792 | 514.7234 | 427.303 | 446.839 | 526.5601 | 576.565 | 462.8588 | 458.9634 |
| ENSMUSG00000006204 | 0.238034542 | 0.483945 | 0.642244 | 28.16952 | 22.79661 | 33.12427 | 24.15346 | 15.60178 | 24.58223 | 25.07152 | 27.18978 |
| ENSMUSG00000006205 | 0.372749712 | 0.062503 | 0.159484 | 229.3804 | 388.7422 | 279.0719 | 371.7907 | 222.3254 | 288.2825 | 216.0008 | 253.4087 |
| ENSMUSG00000006215 | -0.25972525 | 8.02E-05 | 0.001139 | 1039.254 | 1093.037 | 1132.022 | 1095.532 | 1300.798 | 1251.459 | 1285.397 | 1384.503 |
| ENSMUSG00000006216 | -2.861607153 | 0.00012 | 0.001538 | 176.0595 | 83.98751 | 642.6107 | 156.9975 | 1367.106 | 725.1758 | 2425.187 | 3188.817 |
| ENSMUSG00000006218 | -0.283356165 | 0.296481 | 0.464307 | 70.4238 | 75.58876 | 75.3577 | 43.9938 | 68.25779 | 74.86406 | 108.9647 | 69.60583 |
| ENSMUSG00000006219 | 0.287232647 | 0.002655 | 0.016119 | 2144.908 | 2268.863 | 2654.91 | 2301.479 | 1985.327 | 1756.512 | 1846.614 | 2092.525 |
| ENSMUSG00000006221 | -0.214450346 | 0.581632 | 0.722577 | 19.11503 | 41.99375 | 48.85829 | 46.58167 | 38.02934 | 53.63396 | 28.92867 | 61.99269 |
| ENSMUSG00000006235 | -0.334598594 | 0.038745 | 0.113787 | 266.6044 | 221.967 | 268.3065 | 251.8861 | 396.8703 | 296.1041 | 324.9654 | 254.4963 |
| ENSMUSG00000006241 | -0.600766532 | 0.005194 | 0.026408 | 61.36932 | 45.59322 | 57.96746 | 51.75741 | 91.66046 | 92.74205 | 69.42882 | 76.13138 |
| ENSMUSG00000006262 | 0.154077841 | 0.174235 | 0.322358 | 1209.277 | 1228.617 | 1189.161 | 1224.063 | 947.8082 | 1328.558 | 1062.647 | 1022.336 |
| ENSMUSG00000006269 | -1.745236847 | 0.027611 | 0.089135 | 137.8294 | 49.19268 | 524.1915 | 116.4542 | 523.6348 | 278.2261 | 732.8598 | 1242.029 |
| ENSMUSG00000006273 | -0.498778004 | 0.000113 | 0.001483 | 2424.591 | 2127.284 | 2487.632 | 2482.63 | 3295.876 | 2653.763 | 3511.941 | 3995.81 |
| ENSMUSG00000006276 | -0.107682864 | 0.133905 | 0.267491 | 1687.153 | 1684.549 | 1537.794 | 1498.377 | 1666.465 | 1730.812 | 1676.899 | 1826.065 |
| ENSMUSG00000006281 | 0.177975837 | 0.175899 | 0.324376 | 638.8445 | 575.9143 | 683.188 | 564.1558 | 534.361 | 445.8323 | 633.538 | 562.2846 |
| ENSMUSG00000006288 | 0.245328268 | 0.0011 | 0.008398 | 2034.242 | 2135.682 | 1815.21 | 2052.181 | 1662.565 | 1794.503 | 1696.185 | 1623.773 |
| ENSMUSG00000006289 | -0.073866895 | 0.472883 | 0.632292 | 1280.707 | 1538.171 | 1658.698 | 1413.84 | 1686.943 | 1546.446 | 1426.184 | 1542.204 |
| ENSMUSG00000006299 | -0.25437459 | 0.004885 | 0.025341 | 4224.422 | 4086.592 | 4477.573 | 4415.77 | 5084.23 | 4419.215 | 5308.412 | 5712.028 |
| ENSMUSG00000006301 | 0.295056478 | 0.000213 | 0.002383 | 981.9091 | 1023.448 | 997.0404 | 916.9688 | 814.2179 | 848.0869 | 801.3243 | 728.686 |
| ENSMUSG00000006304 | -0.147361923 | 0.001803 | 0.012174 | 5771.734 | 6097.493 | 6064.225 | 5785.616 | 6500.092 | 6535.521 | 6405.773 | 6827.897 |
| ENSMUSG00000006307 | 0.220053561 | 0.006553 | 0.031487 | 1517.13 | 1458.983 | 1534.482 | 1481.125 | 1292.998 | 1430.239 | 1156.183 | 1268.131 |
| ENSMUSG00000006310 | -1.12500474 | 7.67E-05 | 0.001108 | 18.10898 | 19.19714 | 19.87456 | 22.42821 | 45.83023 | 39.10809 | 42.42872 | 46.76642 |
| ENSMUSG00000006313 | -1.051265912 | 0.122262 | 0.251777 | 255.5378 | 141.5789 | 617.7675 | 316.5828 | 1214.989 | 155.315 | 672.1095 | 717.8101 |
| ENSMUSG00000006315 | -0.425352883 | 0.000464 | 0.00433 | 1359.179 | 1307.805 | 1531.997 | 1433.68 | 1911.218 | 1539.742 | 1886.15 | 2229.562 |
| ENSMUSG00000006333 | -0.036570181 | 0.658885 | 0.780312 | 32330.56 | 39058.99 | 36422.61 | 37776.87 | 40340.35 | 35479.98 | 38289.99 | 35212.94 |
| ENSMUSG00000006335 | -0.047111593 | 0.534986 | 0.685864 | 655.9474 | 649.1034 | 708.0312 | 667.6706 | 665.0259 | 682.7156 | 689.4668 | 736.2992 |
| ENSMUSG00000006342 | -1.557815914 | 1.33E-06 | 4.72E-05 | 195.1745 | 154.777 | 323.7897 | 244.9851 | 571.4152 | 405.6068 | 606.5379 | 1125.657 |
| ENSMUSG00000006344 | -0.088176447 | 0.496365 | 0.652999 | 168.0111 | 223.1668 | 192.9488 | 194.0903 | 209.6489 | 203.3621 | 202.5007 | 209.9051 |
| ENSMUSG00000006345 | -2.890728516 | 2.61E-05 | 0.000482 | 518.118 | 161.9759 | 1337.392 | 565.0184 | 4682.485 | 1748.69 | 4379.801 | 8347.262 |
| ENSMUSG00000006356 | -0.236266674 | 0.039977 | 0.116688 | 1737.456 | 1732.542 | 2122.437 | 1861.541 | 2282.736 | 1842.55 | 2187.972 | 2469.919 |
| ENSMUSG00000006360 | 0.257662806 | 0.079678 | 0.188618 | 803.8374 | 1244.215 | 1030.165 | 851.4094 | 827.8695 | 878.256 | 778.1814 | 800.467 |
| ENSMUSG00000006362 | 0.208748611 | 0.278589 | 0.444855 | 181.0898 | 231.5656 | 278.2438 | 207.8923 | 152.1174 | 227.9443 | 186.1078 | 213.1679 |
| ENSMUSG00000006369 | 0.324403526 | 0.004898 | 0.025356 | 2789.789 | 3831.03 | 3683.418 | 3471.197 | 2552.841 | 3040.375 | 2642.152 | 2766.832 |
| ENSMUSG00000006373 | -0.032276773 | 0.634565 | 0.762368 | 5742.558 | 5150.834 | 5562.392 | 5659.673 | 5721.953 | 5222.607 | 5644.949 | 6029.605 |

| | | | | | | | | | | | |
|---|---|---|---|---|---|---|---|---|---|---|---|
| ENSMUSG00000006378 | -1.044095857 | 1.50E-11 | 3.37E-09 | 396.3854 | 371.9447 | 503.4888 | 450.2895 | 937.082 | 695.0067 | 884.2532 | 1039.737 |
| ENSMUSG00000006386 | -0.433659403 | 0.016059 | 0.060359 | 1214.308 | 895.0669 | 1473.202 | 1159.366 | 1486.07 | 1257.046 | 1718.363 | 1946.788 |
| ENSMUSG00000006390 | -0.724099778 | 0.001521 | 0.010658 | 844.0796 | 640.7047 | 952.3226 | 723.7411 | 1276.421 | 779.9271 | 1467.648 | 1698.817 |
| ENSMUSG00000006392 | -0.133784191 | 0.184772 | 0.336042 | 957.7637 | 849.4736 | 934.9324 | 1009.269 | 1061.896 | 886.0777 | 1052.039 | 1118.044 |
| ENSMUSG00000006395 | 0.061683724 | 0.452494 | 0.614875 | 719.3289 | 801.4808 | 698.922 | 703.9008 | 724.5077 | 681.5982 | 666.3238 | 724.3357 |
| ENSMUSG00000006398 | -0.153982921 | 0.080769 | 0.190212 | 2381.331 | 2451.235 | 2186.202 | 2574.069 | 2900.956 | 2377.772 | 2749.188 | 2641.759 |
| ENSMUSG00000006403 | -0.394380393 | 0.175878 | 0.324376 | 181.0898 | 173.9741 | 241.8071 | 118.1794 | 239.8774 | 132.9675 | 297.0011 | 269.7226 |
| ENSMUSG00000006411 | -0.395178286 | 0.309266 | 0.477681 | 151.9142 | 85.18733 | 235.1823 | 98.33908 | 234.0267 | 87.15518 | 225.6437 | 204.4671 |
| ENSMUSG00000006412 | -0.175730512 | 0.005241 | 0.026581 | 2044.302 | 1866.922 | 2002.362 | 2119.466 | 2373.421 | 2231.396 | 2231.365 | 2242.613 |
| ENSMUSG00000006418 | -0.054873661 | 0.393308 | 0.560748 | 2310.907 | 2103.287 | 2299.652 | 2284.227 | 2439.728 | 2239.218 | 2424.223 | 2245.876 |
| ENSMUSG00000006423 | 0.15563776 | 0.056799 | 0.148882 | 1112.696 | 1226.218 | 1087.304 | 1204.222 | 1008.265 | 1134.135 | 977.7892 | 1036.474 |
| ENSMUSG00000006435 | 0.229812008 | 0.103323 | 0.225992 | 223.3441 | 209.9688 | 255.0568 | 207.8923 | 182.3458 | 169.8409 | 218.8936 | 193.5912 |
| ENSMUSG00000006442 | -0.469995476 | 0.000157 | 0.001887 | 1027.181 | 1136.231 | 945.6978 | 947.1606 | 1549.452 | 1148.661 | 1348.076 | 1567.219 |
| ENSMUSG00000006445 | -0.143830309 | 0.1527 | 0.29368 | 726.3712 | 652.7029 | 725.4214 | 597.7981 | 762.537 | 673.7766 | 775.2885 | 773.2773 |
| ENSMUSG00000006456 | -0.174876396 | 0.055571 | 0.146791 | 838.0433 | 955.0579 | 977.1658 | 877.2881 | 940.0073 | 1019.045 | 1032.754 | 1126.744 |
| ENSMUSG00000006457 | 1.16603983 | 2.72E-12 | 6.98E-10 | 517.1119 | 611.909 | 431.4436 | 533.9639 | 287.6578 | 205.5968 | 242.0366 | 195.7664 |
| ENSMUSG00000006463 | -0.110051127 | 0.249158 | 0.412654 | 362.1796 | 323.9518 | 336.2113 | 335.5605 | 375.4179 | 340.7991 | 365.4656 | 383.9197 |
| ENSMUSG00000006464 | -0.208542833 | 0.04189 | 0.120481 | 333.004 | 382.7431 | 384.2415 | 385.5927 | 463.1779 | 396.6678 | 441.6444 | 414.3722 |
| ENSMUSG00000006469 | -3.852462647 | 5.41E-05 | 0.000842 | 1.006054 | 1.199822 | 4.96864 | 0 | 49.73068 | 12.29112 | 8.678602 | 35.89051 |
| ENSMUSG00000006471 | 0.065400899 | 0.513135 | 0.668037 | 1147.908 | 1059.442 | 1047.555 | 1054.989 | 1179.885 | 1064.858 | 922.8247 | 950.5546 |
| ENSMUSG00000006476 | 0.051749895 | 0.353613 | 0.522353 | 1859.188 | 1948.51 | 1926.176 | 1935.727 | 1950.223 | 1782.212 | 1822.507 | 1842.379 |
| ENSMUSG00000006494 | 0.085209541 | 0.333473 | 0.501623 | 2116.738 | 2019.3 | 1862.412 | 2274.738 | 1973.625 | 2095.076 | 1940.15 | 1790.175 |
| ENSMUSG00000006498 | -0.089932574 | 0.122003 | 0.251472 | 12170.24 | 12588.53 | 13648.03 | 12740.09 | 13918.74 | 12863.21 | 13716.05 | 13941.83 |
| ENSMUSG00000006517 | -0.919679386 | 1.84E-06 | 6.08E-05 | 455.7426 | 385.1427 | 546.5504 | 436.4875 | 927.3309 | 574.3303 | 928.6105 | 1022.336 |
| ENSMUSG00000006519 | -1.199217546 | 8.76E-07 | 3.35E-05 | 901.4247 | 973.0553 | 1369.688 | 1029.972 | 2678.631 | 1465.995 | 2177.365 | 3494.43 |
| ENSMUSG00000006522 | 1.788279331 | 0.057002 | 0.149308 | 147.89 | 2.399643 | 6.624853 | 121.6299 | 15.60178 | 18.99536 | 22.17865 | 23.927 |
| ENSMUSG00000006526 | -0.055030371 | 0.540041 | 0.689654 | 568.4207 | 637.1052 | 671.5945 | 594.3476 | 626.9966 | 680.4808 | 608.4665 | 653.6422 |
| ENSMUSG00000006527 | 0.298212979 | 0.019052 | 0.068193 | 962.794 | 1064.242 | 928.3075 | 1086.043 | 802.5166 | 1007.871 | 767.5742 | 709.1094 |
| ENSMUSG00000006538 | -3.672258131 | 9.94E-06 | 0.000226 | 15.09082 | 9.598572 | 87.7793 | 16.38985 | 377.3681 | 122.9112 | 614.2522 | 535.0948 |
| ENSMUSG00000006542 | -0.245388586 | 0.622937 | 0.754426 | 7.04238 | 11.99822 | 10.76539 | 7.763611 | 11.70134 | 7.821619 | 10.60718 | 14.13868 |
| ENSMUSG00000006567 | -0.04501129 | 0.921482 | 0.953139 | 27.16347 | 14.39786 | 36.43669 | 40.5433 | 33.15378 | 13.40849 | 40.50014 | 35.89051 |
| ENSMUSG00000006574 | -0.189456979 | 0.755316 | 0.848145 | 349.1009 | 23.99643 | 102.6852 | 111.2784 | 148.2169 | 98.32892 | 152.3577 | 270.8102 |
| ENSMUSG00000006575 | -0.222785532 | 0.421311 | 0.586625 | 219.3198 | 97.18554 | 128.3565 | 212.2054 | 194.0472 | 151.9629 | 205.3936 | 216.4306 |
| ENSMUSG00000006576 | 0.42567303 | 4.63E-05 | 0.000749 | 1741.48 | 1564.567 | 1310.893 | 1681.253 | 1198.412 | 1201.177 | 1104.111 | 1184.387 |
| ENSMUSG00000006585 | -0.175337764 | 0.118625 | 0.24744 | 676.0685 | 843.4745 | 840.5282 | 780.6743 | 989.738 | 925.1858 | 826.3958 | 804.8174 |
| ENSMUSG00000006586 | 1.284872196 | 0.007912 | 0.035933 | 124.7507 | 195.5709 | 95.23226 | 134.5693 | 37.05423 | 127.3806 | 27.96439 | 33.71532 |
| ENSMUSG00000006589 | -0.196329835 | 0.020547 | 0.071883 | 2805.886 | 2914.367 | 3088.838 | 3020.045 | 3726.875 | 3001.267 | 3279.547 | 3546.635 |
| ENSMUSG00000006599 | -0.003018659 | 0.979076 | 0.988557 | 1511.094 | 1684.549 | 1517.091 | 1657.962 | 1559.203 | 1927.47 | 1378.934 | 1519.365 |
| ENSMUSG00000006611 | -0.262739315 | 0.091406 | 0.207374 | 301.8163 | 238.7645 | 315.5086 | 285.5284 | 330.5627 | 273.7567 | 369.3228 | 398.0583 |
| ENSMUSG00000006638 | -0.519865805 | 0.218277 | 0.37692 | 15.09082 | 9.598572 | 22.35888 | 32.77969 | 31.20356 | 29.05173 | 31.82154 | 23.927 |
| ENSMUSG00000006641 | -0.763781821 | 0.002098 | 0.013581 | 298.7981 | 239.9643 | 275.7595 | 251.0234 | 506.0828 | 271.5219 | 383.7871 | 648.2043 |
| ENSMUSG00000006642 | -0.368596154 | 0.293545 | 0.461117 | 89.53884 | 45.59322 | 35.60859 | 79.36136 | 73.13335 | 64.8077 | 111.8575 | 72.8686 |
| ENSMUSG00000006649 | -1.92680269 | 1.16E-05 | 0.000258 | 90.54489 | 45.59322 | 194.6051 | 87.9876 | 246.7032 | 244.7049 | 455.1445 | 650.3795 |
| ENSMUSG00000006651 | -0.391253961 | 0.077956 | 0.185669 | 436.6276 | 266.3604 | 411.569 | 505.4974 | 422.2232 | 459.2408 | 569.8949 | 676.4817 |
| ENSMUSG00000006673 | -0.100219559 | 0.15154 | 0.291884 | 2712.323 | 2922.765 | 2885.124 | 2878.575 | 3158.386 | 3296.254 | 2931.439 | 2833.175 |
| ENSMUSG00000006675 | 0.274337575 | 0.067151 | 0.167344 | 236.4228 | 241.1641 | 273.2752 | 296.7425 | 239.8774 | 245.8223 | 189.965 | 192.5036 |
| ENSMUSG00000006676 | -0.002235752 | 0.970184 | 0.983398 | 3312.937 | 3144.732 | 3216.366 | 3133.911 | 3191.539 | 2982.271 | 3383.691 | 3268.211 |
| ENSMUSG00000006678 | -0.101545431 | 0.1716 | 0.318778 | 675.0625 | 733.091 | 669.1102 | 683.1978 | 769.3628 | 750.8754 | 736.7169 | 700.4087 |
| ENSMUSG00000006699 | 0.032900669 | 0.482399 | 0.640976 | 11048.49 | 11350.31 | 11542.98 | 11736.86 | 10924.17 | 11694.44 | 10879.11 | 11157.6 |
| ENSMUSG00000006705 | -0.024430391 | 0.795931 | 0.874595 | 849.1099 | 1012.649 | 963.9161 | 888.5022 | 935.1317 | 1031.336 | 858.2174 | 952.7298 |
| ENSMUSG00000006711 | -0.224385058 | 0.565965 | 0.710478 | 33.19979 | 13.19804 | 25.67131 | 25.01608 | 25.35289 | 24.58223 | 23.14294 | 41.32846 |
| ENSMUSG00000006715 | -0.352672321 | 0.000136 | 0.001702 | 720.3349 | 862.6717 | 746.9522 | 681.4726 | 992.6633 | 927.4205 | 964.2892 | 953.8174 |
| ENSMUSG00000006717 | -0.91615035 | 5.91E-05 | 0.000903 | 434.6155 | 332.3506 | 514.2542 | 463.2288 | 842.4962 | 507.2878 | 864.9674 | 1079.978 |
| ENSMUSG00000006720 | 0.216390894 | 0.190217 | 0.342803 | 378.2764 | 473.9295 | 348.6329 | 386.4553 | 350.065 | 417.8979 | 327.8583 | 268.635 |
| ENSMUSG00000006724 | -3.970560773 | 1.24E-12 | 3.62E-10 | 55.33299 | 26.39607 | 130.0127 | 105.2401 | 1611.859 | 604.4994 | 1074.218 | 1693.379 |
| ENSMUSG00000006728 | -0.200533032 | 0.009636 | 0.041456 | 9020.283 | 10788.8 | 10467.27 | 10001.26 | 12471.67 | 11050.83 | 11323.65 | 11434.93 |
| ENSMUSG00000006731 | -0.486211596 | 0.007422 | 0.034305 | 420.5307 | 319.1525 | 524.1915 | 336.4232 | 571.4152 | 462.5929 | 615.2165 | 594.9123 |
| ENSMUSG00000006732 | 0.040666411 | 0.710705 | 0.818775 | 956.7577 | 1137.431 | 924.167 | 897.9911 | 902.9531 | 1043.627 | 863.0388 | 995.1458 |
| ENSMUSG00000006736 | -0.173152605 | 0.032114 | 0.099278 | 1819.952 | 1848.925 | 1799.476 | 1833.075 | 2185.224 | 1792.268 | 2067.436 | 2184.97 |
| ENSMUSG00000006740 | 0.120212057 | 0.252237 | 0.415948 | 4848.176 | 4423.742 | 4229.969 | 4400.242 | 3576.708 | 4798.004 | 3837.871 | 4260.094 |
| ENSMUSG00000006763 | -0.101012569 | 0.246473 | 0.40948 | 801.8253 | 893.867 | 757.7176 | 884.1891 | 956.5842 | 882.7255 | 860.1459 | 876.5984 |
| ENSMUSG00000006777 | 0.401051928 | 0.020613 | 0.072009 | 1720.353 | 2291.659 | 2013.127 | 2019.402 | 1767.877 | 1824.672 | 1424.255 | 1074.54 |
| ENSMUSG00000006782 | 0.215182975 | 0.00682 | 0.032358 | 698.2017 | 771.4852 | 719.6247 | 782.3995 | 667.9512 | 642.4901 | 643.1809 | 604.7006 |
| ENSMUSG00000006784 | -0.001158402 | 0.996851 | 0.998149 | 37.22401 | 39.59411 | 27.32752 | 38.81806 | 40.95467 | 25.6996 | 40.50014 | 34.80291 |

| | | | | | | | | | | | |
|---|---|---|---|---|---|---|---|---|---|---|---|
| ENSMUSG00000006800 | -0.175992086 | 0.074444 | 0.180141 | 2652.965 | 3520.276 | 3001.058 | 2787.137 | 3509.426 | 3247.089 | 3374.048 | 3376.97 |
| ENSMUSG00000006818 | -0.956041662 | 1.67E-05 | 0.000339 | 1314.913 | 969.4558 | 1522.06 | 1211.123 | 2408.525 | 1534.155 | 2487.866 | 3306.277 |
| ENSMUSG00000006850 | -0.15958873 | 0.127465 | 0.258772 | 435.6215 | 437.9349 | 380.9291 | 421.8229 | 509.9832 | 432.4238 | 492.7518 | 432.8613 |
| ENSMUSG00000006906 | 0.018140593 | 0.840379 | 0.903743 | 556.3481 | 526.7217 | 527.5039 | 582.2709 | 524.6099 | 498.3488 | 581.4664 | 561.197 |
| ENSMUSG00000006920 | 0.160600984 | 0.095339 | 0.213618 | 1799.831 | 1577.765 | 1522.06 | 1745.95 | 1511.423 | 1604.549 | 1526.47 | 1302.934 |
| ENSMUSG00000006930 | 0.968892546 | 9.51E-38 | 8.77E-34 | 1064.406 | 1054.643 | 1115.46 | 1166.267 | 579.2161 | 555.3349 | 533.2519 | 582.9488 |
| ENSMUSG00000006931 | 0.08738949 | 0.294271 | 0.461826 | 1780.716 | 2183.675 | 1941.91 | 1957.293 | 1788.354 | 1988.926 | 1827.328 | 1794.525 |
| ENSMUSG00000006932 | -0.135860935 | 0.00583 | 0.02881 | 23175.47 | 23577.69 | 24526.86 | 23825.66 | 26638.09 | 24550.94 | 26469.74 | 26839.57 |
| ENSMUSG00000006941 | -0.016682638 | 0.824536 | 0.893455 | 1429.603 | 1554.969 | 1503.014 | 1638.122 | 1447.065 | 1632.484 | 1513.934 | 1606.372 |
| ENSMUSG00000006958 | 1.502001172 | 2.08E-08 | 1.56E-06 | 583.5115 | 826.677 | 609.4865 | 684.0604 | 229.1512 | 378.7898 | 204.4293 | 142.4744 |
| ENSMUSG00000006998 | -0.229433718 | 0.009622 | 0.041433 | 5594.668 | 5631.962 | 6021.163 | 5501.813 | 6788.725 | 5705.312 | 6854.167 | 7322.751 |
| ENSMUSG00000007021 | 1.302463351 | 0.001983 | 0.013056 | 74.44802 | 44.3934 | 57.96746 | 107.8279 | 26.32801 | 14.52586 | 27.0001 | 47.85401 |
| ENSMUSG00000007029 | -0.172966543 | 0.004076 | 0.022128 | 2590.59 | 2783.586 | 2699.628 | 2451.576 | 2925.334 | 3049.314 | 2970.975 | 2915.832 |
| ENSMUSG00000007030 | 0.469173527 | 0.276531 | 0.44271 | 13.07871 | 40.79393 | 29.81184 | 19.84034 | 19.50223 | 17.87799 | 17.3572 | 19.57664 |
| ENSMUSG00000007033 | -0.0035202 | 0.986323 | 0.992283 | 75.45408 | 62.39072 | 65.42042 | 79.36136 | 80.93424 | 71.51194 | 72.32169 | 58.72992 |
| ENSMUSG00000007034 | -1.370332491 | 2.21E-05 | 0.000425 | 159.9626 | 94.7859 | 196.2613 | 142.3329 | 405.6463 | 208.949 | 330.7512 | 591.6496 |
| ENSMUSG00000007035 | -0.028440175 | 0.873924 | 0.924677 | 94.56911 | 103.1847 | 120.9036 | 87.9876 | 108.2374 | 107.2679 | 108.0004 | 91.35765 |
| ENSMUSG00000007036 | -0.407201713 | 0.005603 | 0.027941 | 1276.683 | 1383.394 | 1581.684 | 1369.846 | 2044.808 | 1382.192 | 1856.257 | 2158.868 |
| ENSMUSG00000007038 | -1.023860878 | 1.54E-05 | 0.000318 | 813.898 | 595.1115 | 998.6966 | 870.3871 | 1665.49 | 994.4629 | 1836.971 | 2171.919 |
| ENSMUSG00000007039 | 0.039993667 | 0.436873 | 0.600681 | 4348.167 | 4609.714 | 4668.865 | 4659.892 | 4638.604 | 4459.44 | 4319.051 | 4371.029 |
| ENSMUSG00000007041 | -0.198609113 | 0.022022 | 0.075449 | 3993.03 | 4258.167 | 4768.238 | 4114.714 | 5064.728 | 4405.806 | 4950.661 | 5243.277 |
| ENSMUSG00000007050 | -0.19009863 | 0.014032 | 0.054854 | 788.7466 | 843.4745 | 799.1229 | 772.048 | 929.2811 | 986.6413 | 868.8245 | 867.8977 |
| ENSMUSG00000007080 | -0.012585723 | 0.919401 | 0.952207 | 869.231 | 1075.04 | 843.0126 | 792.751 | 960.4846 | 969.8807 | 890.0389 | 786.3284 |
| ENSMUSG00000007097 | 0.94993706 | 0.003041 | 0.017946 | 748.5044 | 1030.647 | 624.3924 | 898.8537 | 361.7663 | 770.9881 | 326.894 | 250.146 |
| ENSMUSG00000007122 | 0.116884757 | 0.653955 | 0.776716 | 80.48435 | 157.1766 | 117.5911 | 115.5915 | 111.1627 | 143.0239 | 83.89316 | 95.70802 |
| ENSMUSG00000007207 | 0.385838477 | 0.015721 | 0.059687 | 570.4328 | 761.8867 | 616.9394 | 684.9231 | 525.585 | 616.7905 | 480.216 | 392.6204 |
| ENSMUSG00000007216 | -0.174078921 | 0.167349 | 0.313301 | 295.78 | 290.3568 | 258.3693 | 288.1162 | 335.4383 | 262.5829 | 343.2869 | 333.8905 |
| ENSMUSG00000007279 | -1.095619436 | 0.019699 | 0.069716 | 38.23007 | 44.3934 | 120.9036 | 45.71905 | 170.6445 | 52.51658 | 135.9648 | 175.1022 |
| ENSMUSG00000007338 | -0.069675216 | 0.56817 | 0.712423 | 1044.284 | 1347.4 | 1103.866 | 1086.906 | 1250.093 | 1011.224 | 1209.219 | 1334.474 |
| ENSMUSG00000007379 | 0.460647654 | 0.000521 | 0.004732 | 297.7921 | 291.5566 | 277.4157 | 279.49 | 192.0969 | 234.6486 | 230.4651 | 175.1022 |
| ENSMUSG00000007411 | -0.148684842 | 0.154156 | 0.295564 | 3381.349 | 2684.001 | 2779.954 | 3045.061 | 3046.248 | 3037.023 | 3643.084 | 3455.277 |
| ENSMUSG00000007415 | -0.378521875 | 4.44E-05 | 0.000728 | 2200.241 | 2040.896 | 2039.627 | 2161.734 | 2877.553 | 2312.964 | 2860.082 | 2923.445 |
| ENSMUSG00000007458 | -0.265514778 | 0.032815 | 0.100851 | 3270.683 | 3124.335 | 3685.903 | 3280.557 | 3781.482 | 3282.845 | 4190.801 | 4808.24 |
| ENSMUSG00000007476 | 0.068979523 | 0.625295 | 0.75573 | 412.4823 | 367.1454 | 344.4924 | 375.2412 | 277.9067 | 391.0809 | 367.3942 | 393.708 |
| ENSMUSG00000007564 | -0.187353933 | 0.044704 | 0.126002 | 6586.638 | 7050.151 | 7833.061 | 6700.859 | 8217.263 | 6956.771 | 8385.459 | 8518.013 |
| ENSMUSG00000007570 | -0.276817233 | 0.001438 | 0.010272 | 842.0675 | 919.0633 | 956.4632 | 907.4799 | 1069.697 | 988.8761 | 1169.683 | 1163.722 |
| ENSMUSG00000007589 | -0.182056036 | 0.038599 | 0.113583 | 602.6266 | 586.7127 | 627.7048 | 615.0506 | 706.9557 | 617.9079 | 751.1813 | 683.0072 |
| ENSMUSG00000007591 | 0.396273652 | 0.188839 | 0.341288 | 55.33299 | 79.18822 | 66.24853 | 56.07053 | 38.02934 | 74.86406 | 45.32159 | 36.9781 |
| ENSMUSG00000007603 | 0.087845833 | 0.212579 | 0.370462 | 1906.473 | 1840.526 | 1795.335 | 1903.81 | 1876.114 | 1597.845 | 1736.685 | 1793.438 |
| ENSMUSG00000007610 | -0.073542102 | 0.48905 | 0.646694 | 692.1654 | 674.2997 | 674.9069 | 693.5493 | 791.7904 | 592.2083 | 738.6455 | 753.7006 |
| ENSMUSG00000007613 | -0.035190029 | 0.673247 | 0.790599 | 2327.004 | 2196.873 | 2505.851 | 2241.096 | 2201.801 | 2606.834 | 2457.973 | 2237.175 |
| ENSMUSG00000007617 | -0.109173544 | 0.53961 | 0.689303 | 320.9313 | 262.7609 | 251.7444 | 243.2598 | 223.3005 | 358.6771 | 270.001 | 312.1386 |
| ENSMUSG00000007646 | 0.194510223 | 0.180247 | 0.330351 | 184.1079 | 220.7672 | 164.7932 | 211.3428 | 170.6445 | 182.132 | 162.9649 | 165.3138 |
| ENSMUSG00000007653 | 0.793973578 | 0.029669 | 0.093812 | 40.24217 | 55.19179 | 44.71776 | 31.91707 | 24.37778 | 39.10809 | 18.32149 | 17.40146 |
| ENSMUSG00000007655 | 0.198859416 | 0.178368 | 0.327787 | 2069.454 | 2743.992 | 2370.869 | 2514.547 | 1845.886 | 2686.167 | 2096.365 | 1821.715 |
| ENSMUSG00000007656 | -0.148955728 | 0.014927 | 0.057503 | 2961.824 | 3181.927 | 3076.416 | 3156.339 | 3228.594 | 3371.118 | 3465.655 | 3657.569 |
| ENSMUSG00000007659 | -0.289972561 | 0.040196 | 0.117074 | 1686.147 | 1544.17 | 1912.926 | 1705.407 | 1867.338 | 1701.761 | 2564.045 | 2241.525 |
| ENSMUSG00000007670 | 0.112253848 | 0.072036 | 0.176092 | 3072.49 | 3215.522 | 2899.201 | 3056.275 | 2833.673 | 3005.736 | 2686.51 | 2798.372 |
| ENSMUSG00000007682 | -0.25806423 | 0.423815 | 0.588965 | 25.15136 | 29.99554 | 29.81184 | 26.74133 | 24.37778 | 45.81234 | 27.0001 | 36.9781 |
| ENSMUSG00000007721 | -0.46875973 | 0.001333 | 0.009701 | 1156.963 | 1226.218 | 1435.937 | 1292.21 | 1844.911 | 1330.793 | 1791.649 | 2107.752 |
| ENSMUSG00000007739 | -0.089810642 | 0.081737 | 0.191829 | 6804.952 | 7190.53 | 7028.141 | 7206.357 | 7898.402 | 7461.824 | 7530.134 | 7149.824 |
| ENSMUSG00000007777 | -0.272956091 | 0.003664 | 0.02048 | 504.0332 | 542.3193 | 494.3797 | 458.0531 | 572.3903 | 583.2693 | 596.895 | 659.0802 |
| ENSMUSG00000007783 | 0.637682766 | 1.24E-06 | 4.43E-05 | 946.6971 | 1011.45 | 856.2623 | 1017.033 | 624.0712 | 740.819 | 594.9664 | 502.4671 |
| ENSMUSG00000007805 | 0.468846472 | 0.179904 | 0.329821 | 55.33299 | 119.9822 | 79.49824 | 64.69676 | 64.35735 | 82.68568 | 36.64299 | 46.76642 |
| ENSMUSG00000007812 | -0.196661601 | 0.006029 | 0.02956 | 755.5468 | 805.0802 | 744.4679 | 758.2461 | 915.6295 | 898.3688 | 865.9317 | 826.5692 |
| ENSMUSG00000007815 | -0.143749615 | 0.061033 | 0.156753 | 8421.681 | 9528.983 | 8580.013 | 7692.014 | 9275.259 | 9199.341 | 9524.284 | 9802.458 |
| ENSMUSG00000007817 | 0.140738458 | 0.301243 | 0.469613 | 3037.278 | 2727.194 | 3027.558 | 3061.451 | 2446.554 | 3494.029 | 2446.402 | 2367.686 |
| ENSMUSG00000007827 | 0.161883141 | 0.195121 | 0.34895 | 633.8142 | 584.3131 | 473.677 | 540.0023 | 448.5512 | 549.7481 | 473.466 | 522.0437 |
| ENSMUSG00000007833 | -0.26906167 | 0.00155 | 0.010786 | 1307.871 | 1075.04 | 1229.738 | 1227.513 | 1435.364 | 1358.727 | 1521.648 | 1520.452 |
| ENSMUSG00000007836 | -0.080746591 | 0.139077 | 0.274966 | 13628.01 | 14755.41 | 14356.88 | 14191.88 | 14068.91 | 15271.15 | 15343.77 | 15525.36 |
| ENSMUSG00000007837 | -0.228207236 | 0.103687 | 0.226466 | 452.7245 | 527.9215 | 623.5643 | 482.2065 | 672.8268 | 517.3442 | 591.1093 | 663.4306 |
| ENSMUSG00000007850 | 0.25416333 | 0.023817 | 0.080199 | 24221.76 | 24039.62 | 19432.35 | 24159.5 | 19111.21 | 22511.74 | 17986.89 | 17405.81 |
| ENSMUSG00000007867 | 0.029872497 | 0.753626 | 0.846815 | 694.1775 | 695.8965 | 708.0312 | 802.2398 | 740.1095 | 653.6638 | 676.931 | 772.1897 |
| ENSMUSG00000007872 | -0.06275007 | 0.727497 | 0.829154 | 4991.036 | 5497.582 | 7409.07 | 5773.539 | 5939.403 | 4515.309 | 6395.166 | 7875.247 |

| | | | | | | | | | | | |
|---|---|---|---|---|---|---|---|---|---|---|---|
| ENSMUSG00000007877 | -1.861138297 | 2.78E-05 | 0.000507 | 22.1332 | 8.398751 | 7.45296 | 12.07673 | 61.43201 | 30.1691 | 48.21446 | 41.32846 |
| ENSMUSG00000007880 | 0.505925424 | 0.000318 | 0.003241 | 4232.471 | 4223.372 | 3545.124 | 3719.633 | 2690.332 | 3547.663 | 2377.937 | 2454.693 |
| ENSMUSG00000007888 | 0.263661833 | 0.249789 | 0.413494 | 898.4065 | 1358.198 | 888.5584 | 966.1383 | 936.1069 | 1131.9 | 816.7529 | 538.3576 |
| ENSMUSG00000007891 | -0.429944748 | 0.011729 | 0.048149 | 5681.189 | 5390.798 | 7987.917 | 5365.518 | 8111.951 | 6294.168 | 8865.675 | 9636.057 |
| ENSMUSG00000007892 | -0.202534987 | 0.014004 | 0.054769 | 33535.82 | 39421.34 | 38583.14 | 36311.27 | 46699.06 | 38933.78 | 42229.12 | 42270.32 |
| ENSMUSG00000007908 | 1.078411305 | 4.82E-08 | 3.16E-06 | 167.005 | 171.5745 | 138.2938 | 196.6782 | 73.13335 | 96.09417 | 64.60737 | 85.9197 |
| ENSMUSG00000007946 | 3.483160398 | 0.002265 | 0.014359 | 82.49646 | 8.398751 | 5.796746 | 118.1794 | 0.975111 | 14.52586 | 3.857157 | 0 |
| ENSMUSG00000007950 | -0.077403331 | 0.384168 | 0.552197 | 754.5408 | 865.0713 | 850.4655 | 826.3933 | 942.9326 | 800.0399 | 831.2173 | 902.7006 |
| ENSMUSG00000007987 | -0.207188745 | 0.021784 | 0.074842 | 814.904 | 799.0811 | 882.7617 | 792.751 | 973.1611 | 844.7348 | 1038.539 | 940.7663 |
| ENSMUSG00000007989 | -0.196074342 | 0.046289 | 0.129107 | 284.7134 | 281.9581 | 308.0557 | 318.3081 | 356.8907 | 335.2122 | 344.2512 | 332.8029 |
| ENSMUSG00000008028 | 0.573558049 | 0.123593 | 0.253279 | 21.12714 | 22.79661 | 33.95237 | 21.56559 | 18.52711 | 20.11273 | 14.46434 | 14.13868 |
| ENSMUSG00000008035 | -0.055155101 | 0.574162 | 0.716582 | 851.122 | 772.6851 | 931.62 | 894.5406 | 834.6953 | 834.6784 | 936.3248 | 983.1823 |
| ENSMUSG00000008036 | -0.126894697 | 0.010248 | 0.04344 | 2230.423 | 2309.656 | 2395.712 | 2290.265 | 2509.937 | 2464.927 | 2546.688 | 2554.751 |
| ENSMUSG00000008090 | 0.122993078 | 0.197452 | 0.351754 | 953.7395 | 1018.648 | 838.0439 | 952.3363 | 903.9282 | 922.951 | 783.9671 | 841.7955 |
| ENSMUSG00000008129 | -0.104756262 | 0.669975 | 0.788387 | 53.32088 | 68.38983 | 66.24853 | 55.2079 | 44.85512 | 75.98144 | 73.28598 | 67.43065 |
| ENSMUSG00000008136 | 0.249809869 | 0.036088 | 0.108634 | 2572.481 | 3168.729 | 3057.37 | 3346.979 | 2925.334 | 2680.58 | 2377.937 | 2230.649 |
| ENSMUSG00000008140 | -0.207978414 | 0.061032 | 0.156753 | 2764.637 | 3166.329 | 3411.799 | 2959.661 | 3510.401 | 2976.685 | 3846.549 | 3876.175 |
| ENSMUSG00000008153 | 0.865962112 | 0.000107 | 0.001417 | 362.1796 | 199.1704 | 318.8211 | 270.0012 | 137.4907 | 127.3806 | 185.1435 | 181.6277 |
| ENSMUSG00000008167 | -0.172685579 | 0.040137 | 0.116969 | 681.0988 | 766.686 | 787.5294 | 762.5592 | 877.6002 | 890.5472 | 790.7171 | 822.2189 |
| ENSMUSG00000008200 | 0.276283359 | 0.005486 | 0.027498 | 3334.064 | 3214.322 | 2943.091 | 3147.713 | 2815.146 | 2896.234 | 2490.759 | 2232.825 |
| ENSMUSG00000008206 | 0.681200899 | 1.01E-05 | 0.000229 | 1478.9 | 1688.149 | 1377.141 | 1471.636 | 887.3513 | 1241.403 | 838.9316 | 784.1532 |
| ENSMUSG00000008226 | -0.15965332 | 0.165325 | 0.310719 | 434.6155 | 381.5432 | 395.835 | 417.5098 | 429.049 | 424.6022 | 431.0373 | 537.27 |
| ENSMUSG00000008301 | 0.003952263 | 0.960836 | 0.977573 | 1780.716 | 2110.486 | 1829.288 | 2034.066 | 2022.381 | 1914.062 | 1964.257 | 1828.241 |
| ENSMUSG00000008305 | 0.355305597 | 0.005086 | 0.026035 | 3535.275 | 3269.514 | 2918.248 | 3866.278 | 2480.683 | 2918.581 | 2979.654 | 2243.7 |
| ENSMUSG00000008307 | 0.260488955 | 0.294167 | 0.461764 | 141.8537 | 127.1811 | 139.1219 | 183.7388 | 131.64 | 174.3104 | 101.2504 | 88.09488 |
| ENSMUSG00000008318 | -0.720703071 | 0.009737 | 0.041733 | 31.18769 | 29.99554 | 54.65504 | 50.89479 | 66.30757 | 67.04245 | 81.00029 | 63.08028 |
| ENSMUSG00000008333 | 0.125746605 | 0.218588 | 0.377351 | 2632.844 | 2710.397 | 2582.036 | 2938.958 | 2571.369 | 2870.534 | 2349.008 | 2167.569 |
| ENSMUSG00000008348 | -0.188581695 | 0.130209 | 0.262638 | 2317.949 | 2458.434 | 3127.759 | 2475.729 | 2784.918 | 2712.984 | 2876.475 | 3458.54 |
| ENSMUSG00000008373 | -0.219628486 | 0.00054 | 0.004867 | 1893.394 | 1920.914 | 1964.269 | 1785.631 | 2241.781 | 2055.968 | 2232.329 | 2275.241 |
| ENSMUSG00000008384 | -0.414817023 | 0.007256 | 0.03375 | 280.6892 | 323.9518 | 305.5713 | 250.1608 | 388.0943 | 306.1605 | 391.5014 | 458.9634 |
| ENSMUSG00000008393 | 0.33530067 | 4.91E-09 | 4.72E-07 | 2752.565 | 2923.965 | 2924.873 | 2894.964 | 2409.5 | 2201.227 | 2298.865 | 2200.197 |
| ENSMUSG00000008398 | 0.026229124 | 0.674398 | 0.791543 | 2704.274 | 2717.596 | 2978.7 | 2640.491 | 2692.282 | 2812.431 | 2613.224 | 2728.766 |
| ENSMUSG00000008429 | 0.157256358 | 0.026211 | 0.08598 | 1380.307 | 1421.789 | 1295.159 | 1425.054 | 1144.781 | 1263.75 | 1260.326 | 1282.27 |
| ENSMUSG00000008435 | -0.152542276 | 0.188886 | 0.341305 | 762.5892 | 795.4817 | 762.6862 | 795.3389 | 940.9824 | 690.5372 | 876.5389 | 953.8174 |
| ENSMUSG00000008450 | -0.182950092 | 0.016476 | 0.061457 | 1208.271 | 1205.821 | 1393.703 | 1267.194 | 1526.049 | 1407.891 | 1395.326 | 1436.708 |
| ENSMUSG00000008461 | 0.591962032 | 0.278104 | 0.444297 | 11.0666 | 32.39518 | 22.35888 | 18.11509 | 14.62667 | 18.99536 | 17.3572 | 4.350364 |
| ENSMUSG00000008475 | -0.110586978 | 0.124972 | 0.255312 | 4025.223 | 3967.81 | 4251.499 | 4073.308 | 4309.992 | 3976.734 | 4610.266 | 4722.321 |
| ENSMUSG00000008489 | 0.590676077 | 0.389168 | 0.556608 | 94.56911 | 50.3925 | 25.67131 | 148.3712 | 41.92979 | 130.7328 | 22.17865 | 17.40146 |
| ENSMUSG00000008496 | 0.985749769 | 0.004248 | 0.022832 | 47.28455 | 61.1909 | 45.54586 | 102.6522 | 29.25334 | 29.05173 | 36.64299 | 34.80291 |
| ENSMUSG00000008540 | 0.368785442 | 0.001049 | 0.008103 | 3225.41 | 2543.622 | 2337.745 | 2520.586 | 2001.904 | 2201.227 | 2085.757 | 1940.262 |
| ENSMUSG00000008575 | 0.889930889 | 0.069603 | 0.171539 | 4715.377 | 6069.897 | 4254.812 | 5099.83 | 1792.255 | 5680.73 | 1869.757 | 1525.89 |
| ENSMUSG00000008601 | -0.669791986 | 0.004264 | 0.022899 | 264.5923 | 245.9634 | 434.756 | 273.4516 | 574.3406 | 331.8601 | 496.6089 | 538.3576 |
| ENSMUSG00000008604 | 0.028576335 | 0.640942 | 0.767484 | 2354.167 | 2201.673 | 2348.51 | 2232.47 | 2228.129 | 2391.181 | 2200.508 | 2141.467 |
| ENSMUSG00000008658 | 1.302031086 | 0.00148 | 0.010472 | 623.7537 | 773.8849 | 467.0521 | 691.824 | 193.072 | 559.8044 | 148.5005 | 135.9489 |
| ENSMUSG00000008668 | -0.236637519 | 0.005278 | 0.026705 | 29684.64 | 33700.59 | 31758.72 | 32338.03 | 42376.39 | 33827.38 | 37506.03 | 36490.86 |
| ENSMUSG00000008682 | -0.095292751 | 0.203761 | 0.35981 | 28052.82 | 33182.26 | 32592.62 | 32414.8 | 35705.65 | 32911.14 | 34286.27 | 31957.78 |
| ENSMUSG00000008683 | -0.145963327 | 0.089373 | 0.204103 | 20591.92 | 24056.42 | 25046.91 | 23498.73 | 27898.91 | 23929.68 | 26645.24 | 24641.55 |
| ENSMUSG00000008690 | -0.123159004 | 0.113238 | 0.240221 | 3355.191 | 3646.258 | 3678.45 | 3392.698 | 4157.875 | 3454.921 | 3942.978 | 3767.416 |
| ENSMUSG00000008730 | 0.124515937 | 0.140711 | 0.277216 | 2644.917 | 2220.87 | 2476.867 | 2640.491 | 2136.469 | 2369.95 | 2439.652 | 2215.423 |
| ENSMUSG00000008734 | 0.425190363 | 4.37E-05 | 0.000721 | 1482.924 | 1488.979 | 1335.736 | 1399.175 | 1115.527 | 1147.543 | 1107.968 | 876.5984 |
| ENSMUSG00000008763 | 0.127737662 | 0.093623 | 0.210617 | 3224.404 | 3589.866 | 3219.679 | 3116.659 | 2821.972 | 3267.202 | 2926.618 | 3018.065 |
| ENSMUSG00000008822 | -0.058252965 | 0.576341 | 0.71847 | 379.2825 | 329.9509 | 344.4924 | 401.9825 | 375.4179 | 365.3813 | 400.18 | 376.3065 |
| ENSMUSG00000008845 | 1.19771114 | 1.23E-06 | 4.42E-05 | 427.5731 | 303.5548 | 240.979 | 327.7969 | 116.0382 | 177.6625 | 171.6435 | 101.146 |
| ENSMUSG00000008855 | 0.014783149 | 0.80898 | 0.883359 | 1976.897 | 2051.695 | 1922.035 | 2066.846 | 2105.265 | 1953.17 | 1978.721 | 1894.584 |
| ENSMUSG00000008859 | -0.010456884 | 0.843727 | 0.905813 | 2207.283 | 2262.863 | 2257.419 | 2204.866 | 2265.184 | 2343.133 | 2133.972 | 2256.752 |
| ENSMUSG00000008892 | -0.151786704 | 0.004432 | 0.023569 | 3178.126 | 3465.085 | 3156.742 | 3340.941 | 3585.484 | 3685.1 | 3599.691 | 3723.912 |
| ENSMUSG00000008932 | 0.159832738 | 0.731917 | 0.831995 | 19.11503 | 26.39607 | 9.93728 | 37.95543 | 21.45245 | 23.46486 | 22.17865 | 16.31387 |
| ENSMUSG00000008958 | -0.133246301 | 0.012619 | 0.050816 | 1644.899 | 1679.75 | 1729.087 | 1733.011 | 1852.711 | 1907.358 | 1797.435 | 1891.321 |
| ENSMUSG00000008976 | 0.204886473 | 0.012547 | 0.050639 | 2350.143 | 2601.213 | 2122.437 | 2372.215 | 1929.745 | 2167.706 | 2063.579 | 2030.533 |
| ENSMUSG00000008999 | 0.257446631 | 4.45E-06 | 0.000124 | 3380.343 | 3510.678 | 3358.801 | 3474.647 | 2786.868 | 2886.177 | 2751.117 | 3058.306 |
| ENSMUSG00000009013 | -0.229387071 | 0.000243 | 0.002646 | 4327.04 | 4343.354 | 4339.279 | 4295.002 | 4847.278 | 4755.544 | 5203.304 | 5480.371 |
| ENSMUSG00000009030 | 0.012988359 | 0.825441 | 0.893982 | 1594.596 | 1559.768 | 1491.42 | 1645.886 | 1578.705 | 1557.619 | 1580.47 | 1517.19 |
| ENSMUSG00000009035 | 0.032566318 | 0.723068 | 0.826722 | 1116.72 | 1100.236 | 1252.925 | 1129.174 | 1163.308 | 1039.158 | 1047.218 | 1249.642 |
| ENSMUSG00000009073 | -0.103210897 | 0.241906 | 0.40433 | 2555.378 | 2897.569 | 2909.967 | 2830.268 | 2816.121 | 2737.567 | 3159.011 | 3310.627 |

| | | | | | | | | | | | |
|---|---|---|---|---|---|---|---|---|---|---|---|
| ENSMUSG00000009076 | 0.045886017 | 0.755994 | 0.848334 | 481.9 | 424.7368 | 436.4122 | 432.1744 | 435.8748 | 331.8601 | 425.2515 | 526.3941 |
| ENSMUSG00000009079 | 0.104890111 | 0.106485 | 0.230667 | 13277.91 | 13790.75 | 13223.21 | 12917.79 | 12164.51 | 13611.85 | 11685.26 | 12016.79 |
| ENSMUSG00000009090 | -0.335580426 | 0.01259 | 0.050767 | 1565.421 | 1625.758 | 1958.472 | 1597.579 | 2113.066 | 1738.634 | 2094.436 | 2569.978 |
| ENSMUSG00000009092 | -0.308755989 | 0.339714 | 0.508231 | 37.22401 | 46.79304 | 54.65504 | 35.36756 | 39.97956 | 80.45093 | 40.50014 | 55.46715 |
| ENSMUSG00000009097 | 0.121849918 | 0.806526 | 0.881775 | 19.11503 | 7.198929 | 19.04645 | 17.25247 | 17.552 | 13.40849 | 19.28578 | 7.613138 |
| ENSMUSG00000009108 | -0.126021316 | 0.616247 | 0.749338 | 68.4117 | 58.79126 | 50.5145 | 43.9938 | 68.25779 | 45.81234 | 67.50024 | 58.72992 |
| ENSMUSG00000009112 | -0.223781945 | 0.041395 | 0.119578 | 1422.561 | 1225.018 | 1423.515 | 1304.287 | 1694.743 | 1296.154 | 1683.649 | 1603.109 |
| ENSMUSG00000009145 | -1.021159934 | 0.00288 | 0.017225 | 108.6539 | 41.99375 | 74.5296 | 105.2401 | 157.968 | 110.62 | 157.1791 | 246.8832 |
| ENSMUSG00000009207 | 0.244022455 | 0.166703 | 0.312415 | 577.4752 | 559.1168 | 473.677 | 648.6929 | 405.6463 | 641.3727 | 423.3229 | 438.2992 |
| ENSMUSG00000009246 | 0.886996445 | 0.000184 | 0.002124 | 570.4328 | 896.2667 | 674.9069 | 684.0604 | 444.6508 | 525.1658 | 280.6081 | 277.3357 |
| ENSMUSG00000009248 | -0.058285042 | 0.9021 | 0.941162 | 11.0666 | 29.99554 | 16.56213 | 11.21411 | 20.47734 | 17.87799 | 20.25007 | 11.9635 |
| ENSMUSG00000009281 | 0.425267853 | 0.04653 | 0.129488 | 836.0312 | 1629.358 | 1222.285 | 1123.998 | 844.4464 | 1163.186 | 841.8244 | 733.0364 |
| ENSMUSG00000009291 | -0.403582387 | 0.002258 | 0.01434 | 3601.675 | 3268.314 | 4332.654 | 3939.601 | 5116.409 | 4001.317 | 5223.554 | 5692.452 |
| ENSMUSG00000009292 | 0.195018212 | 0.720506 | 0.825136 | 18.10898 | 15.59768 | 5.796746 | 17.25247 | 8.776002 | 7.821619 | 17.3572 | 15.22628 |
| ENSMUSG00000009293 | -0.201561891 | 0.004982 | 0.025691 | 1349.119 | 1267.012 | 1265.347 | 1364.67 | 1585.531 | 1382.192 | 1515.863 | 1548.73 |
| ENSMUSG00000009356 | -5.537024401 | 2.24E-10 | 3.57E-08 | 0 | 0 | 3.312427 | 1.725247 | 35.10401 | 25.6996 | 126.3219 | 67.43065 |
| ENSMUSG00000009376 | -0.213130884 | 0.009072 | 0.039721 | 1548.318 | 1653.354 | 1697.619 | 1550.134 | 1702.544 | 1811.263 | 1904.471 | 2058.81 |
| ENSMUSG00000009378 | -2.493031591 | 0.000333 | 0.003357 | 133.8052 | 32.39518 | 345.3205 | 127.6683 | 902.9531 | 404.4894 | 1020.218 | 1274.657 |
| ENSMUSG00000009394 | 0.588930891 | 0.153558 | 0.294877 | 141.8537 | 43.19358 | 137.4657 | 212.2054 | 75.08357 | 69.27719 | 111.8575 | 100.0584 |
| ENSMUSG00000009406 | -0.037613801 | 0.697407 | 0.80906 | 587.5357 | 698.2961 | 611.9708 | 643.5171 | 673.8019 | 613.4384 | 708.7525 | 607.9634 |
| ENSMUSG00000009418 | 0.56651538 | 5.55E-05 | 0.000859 | 2484.954 | 2385.245 | 2578.724 | 2349.786 | 1595.282 | 2150.945 | 1509.113 | 1362.752 |
| ENSMUSG00000009470 | 0.059749567 | 0.513429 | 0.668226 | 5571.529 | 4715.299 | 4641.538 | 5183.505 | 4528.417 | 5263.949 | 5026.839 | 4477.613 |
| ENSMUSG00000009535 | 0.075289109 | 0.504585 | 0.660282 | 1473.87 | 1345 | 1214.832 | 1396.587 | 1319.326 | 1434.708 | 1329.755 | 1069.102 |
| ENSMUSG00000009545 | -0.437958743 | 0.088676 | 0.202915 | 232.3986 | 242.3639 | 472.8489 | 247.5729 | 361.7663 | 326.2732 | 475.3946 | 457.8758 |
| ENSMUSG00000009549 | -0.199567066 | 0.002309 | 0.014546 | 1851.14 | 1761.338 | 1849.99 | 1818.41 | 2120.867 | 1907.358 | 2167.722 | 2165.394 |
| ENSMUSG00000009555 | -0.042554912 | 0.492301 | 0.649457 | 2533.245 | 2693.599 | 2704.596 | 2696.561 | 2766.391 | 2621.36 | 2928.546 | 2627.62 |
| ENSMUSG00000009563 | -0.400612328 | 0.000594 | 0.005237 | 958.7698 | 1075.04 | 1108.007 | 940.2596 | 1374.907 | 1124.078 | 1345.183 | 1543.292 |
| ENSMUSG00000009566 | -0.394161326 | 0.000446 | 0.004197 | 819.9343 | 969.4558 | 922.5108 | 938.5344 | 1213.038 | 1017.928 | 1184.147 | 1381.241 |
| ENSMUSG00000009569 | 0.260122751 | 0.009252 | 0.040236 | 1444.694 | 1207.02 | 1274.456 | 1423.329 | 1143.806 | 1246.989 | 1043.361 | 1035.387 |
| ENSMUSG00000009575 | 0.103877497 | 0.16419 | 0.309151 | 5275.749 | 5865.927 | 5260.133 | 5258.553 | 4916.511 | 5558.936 | 4775.16 | 4902.861 |
| ENSMUSG00000009585 | 0.48996388 | 0.000712 | 0.006032 | 1427.591 | 1114.634 | 1187.505 | 1163.679 | 795.6908 | 1014.576 | 974.8963 | 699.3211 |
| ENSMUSG00000009596 | -4.030283203 | NA | NA | 5.030272 | 3.599465 | 0.828107 | 6.900988 | 3.900445 | 261.4655 | 1.928578 | 0 |
| ENSMUSG00000009614 | -0.831950298 | 2.10E-05 | 0.000408 | 1038.248 | 1136.231 | 1391.219 | 1080.867 | 2222.279 | 1439.178 | 1838.899 | 2772.27 |
| ENSMUSG00000009621 | 0.080897067 | 0.458236 | 0.619893 | 878.2855 | 815.8786 | 833.9034 | 791.8884 | 814.2179 | 915.1294 | 716.4668 | 693.8831 |
| ENSMUSG00000009628 | -0.879510707 | NA | NA | 53.32088 | 35.99465 | 33.12427 | 61.24627 | 7.80089 | 286.0478 | 17.3572 | 27.18978 |
| ENSMUSG00000009630 | -0.011424012 | 0.868524 | 0.92134 | 2621.778 | 2652.805 | 2519.928 | 2839.757 | 2473.857 | 2737.567 | 2800.296 | 2708.102 |
| ENSMUSG00000009633 | -0.647325099 | 0.001431 | 0.010239 | 224.3501 | 189.5718 | 303.9151 | 289.8415 | 355.9156 | 309.5126 | 470.5731 | 445.9123 |
| ENSMUSG00000009640 | -0.034404936 | 0.66362 | 0.783707 | 1217.326 | 1105.036 | 1247.129 | 1275.82 | 1288.122 | 1252.576 | 1285.397 | 1139.795 |
| ENSMUSG00000009647 | 0.020315783 | 0.823527 | 0.892957 | 841.0614 | 886.6681 | 1026.024 | 859.173 | 936.1069 | 892.7819 | 847.6102 | 888.5619 |
| ENSMUSG00000009654 | 0.514603341 | 0.26305 | 0.427326 | 55.33299 | 15.59768 | 46.37397 | 28.46658 | 28.27823 | 14.52586 | 27.96439 | 31.54014 |
| ENSMUSG00000009681 | -0.086135286 | 0.375143 | 0.54362 | 734.4197 | 631.1061 | 750.2646 | 619.3637 | 684.5281 | 732.9974 | 743.4669 | 744.9999 |
| ENSMUSG00000009687 | -0.147769783 | 0.280517 | 0.446928 | 293.7679 | 327.5513 | 336.2113 | 272.589 | 341.289 | 279.3435 | 376.0728 | 364.343 |
| ENSMUSG00000009731 | 0.85301092 | 2.82E-06 | 8.61E-05 | 157.9505 | 194.3711 | 163.9651 | 154.4096 | 109.2125 | 103.9158 | 79.07171 | 78.30656 |
| ENSMUSG00000009733 | 0.178894085 | 0.052664 | 0.141129 | 1020.139 | 1108.635 | 949.0102 | 1052.401 | 913.6793 | 808.9788 | 931.5033 | 990.7955 |
| ENSMUSG00000009739 | 0.101067395 | 0.291006 | 0.458127 | 755.5468 | 740.2899 | 922.5108 | 841.9205 | 785.9397 | 735.2322 | 754.0741 | 768.9269 |
| ENSMUSG00000009741 | 0.175361905 | 0.137411 | 0.272519 | 4166.071 | 3659.456 | 3456.517 | 4285.514 | 3572.808 | 3964.443 | 3334.512 | 2914.744 |
| ENSMUSG00000009772 | -1.136624507 | 1.52E-10 | 2.64E-08 | 110.666 | 93.58608 | 134.1533 | 109.5532 | 231.1014 | 203.3621 | 285.4296 | 268.635 |
| ENSMUSG00000009828 | 0.162131888 | 0.149777 | 0.289697 | 1310.889 | 1408.59 | 1325.799 | 1469.91 | 1328.102 | 1413.478 | 1144.611 | 1043 |
| ENSMUSG00000009863 | -0.841322774 | 2.63E-06 | 8.11E-05 | 2769.668 | 2439.237 | 3184.07 | 2655.155 | 5027.674 | 3297.371 | 5360.483 | 6111.174 |
| ENSMUSG00000009876 | -0.521850405 | 0.162969 | 0.307479 | 33.19979 | 50.3925 | 95.23226 | 63.83414 | 86.78491 | 50.28183 | 107.0361 | 105.4963 |
| ENSMUSG00000009894 | -0.113397746 | 0.174726 | 0.322927 | 1343.083 | 1420.589 | 1321.658 | 1414.703 | 1487.045 | 1322.971 | 1638.327 | 1497.613 |
| ENSMUSG00000009905 | 0.248237604 | 0.00732 | 0.033955 | 946.6971 | 964.6565 | 843.0126 | 892.8153 | 740.1095 | 853.6738 | 774.3242 | 700.4087 |
| ENSMUSG00000009907 | -0.007629622 | 0.901663 | 0.940849 | 2873.291 | 2642.007 | 2746.002 | 2817.328 | 2902.906 | 2624.712 | 2895.76 | 2713.54 |
| ENSMUSG00000009927 | -0.175051725 | 0.044729 | 0.126005 | 19498.34 | 22422.26 | 22443.35 | 20906.54 | 26791.18 | 22712.86 | 24788.98 | 21974.78 |
| ENSMUSG00000009941 | -1.079374665 | 0.207847 | 0.364745 | 9.054489 | 1.199822 | 7.45296 | 12.93935 | 7.80089 | 43.57759 | 6.750024 | 7.613138 |
| ENSMUSG00000009995 | 0.059284634 | 0.485455 | 0.643832 | 1659.99 | 1578.965 | 1373.829 | 1476.811 | 1352.479 | 1479.403 | 1525.505 | 1483.474 |
| ENSMUSG00000010021 | -0.104872523 | 0.830992 | 0.897886 | 11.0666 | 10.79839 | 10.76539 | 14.6646 | 11.70134 | 21.23011 | 9.642892 | 8.700729 |
| ENSMUSG00000010025 | -0.048805568 | 0.469671 | 0.629685 | 1490.973 | 1479.38 | 1539.45 | 1593.266 | 1625.511 | 1459.291 | 1571.791 | 1658.576 |
| ENSMUSG00000010044 | -1.98536623 | 1.13E-08 | 9.52E-07 | 13.07871 | 25.19625 | 27.32752 | 25.8787 | 89.71024 | 55.8687 | 92.57176 | 125.073 |
| ENSMUSG00000010045 | -0.202893466 | 0.053561 | 0.142932 | 941.6669 | 955.0579 | 1006.978 | 984.2534 | 1105.776 | 931.89 | 1179.326 | 1258.343 |
| ENSMUSG00000010047 | -0.264501818 | 0.063511 | 0.161389 | 1168.029 | 1184.224 | 1160.177 | 1239.59 | 1556.278 | 1052.566 | 1410.755 | 1687.941 |
| ENSMUSG00000010048 | -0.188377271 | 0.049437 | 0.135042 | 796.795 | 755.8876 | 699.7501 | 742.7188 | 909.7789 | 782.1619 | 785.8957 | 933.1532 |
| ENSMUSG00000010051 | 0.240222998 | 0.135708 | 0.270154 | 171.0292 | 171.5745 | 123.3879 | 154.4096 | 129.6898 | 135.2023 | 131.1433 | 127.2482 |
| ENSMUSG00000010054 | -0.212351721 | 0.148904 | 0.288674 | 494.9787 | 466.7306 | 480.3018 | 441.6632 | 561.6641 | 432.4238 | 507.2161 | 680.832 |

| | | | | | | | | | | | |
|---|---|---|---|---|---|---|---|---|---|---|---|
| ENSMUSG00000010057 | -0.237606865 | 0.006124 | 0.029884 | 560.3723 | 607.1097 | 564.7687 | 602.1112 | 664.0508 | 630.199 | 726.1097 | 729.7736 |
| ENSMUSG00000010064 | -1.086796083 | 0.000177 | 0.002063 | 293.7679 | 256.7618 | 342.8361 | 324.3464 | 617.2455 | 332.9775 | 604.6093 | 1033.212 |
| ENSMUSG00000010066 | -0.29614433 | 0.144805 | 0.282897 | 134.8113 | 98.38537 | 129.1846 | 142.3329 | 135.5405 | 124.0285 | 171.6435 | 190.3284 |
| ENSMUSG00000010067 | -0.010847328 | 0.932355 | 0.960082 | 871.2431 | 755.8876 | 758.5457 | 889.3648 | 1002.414 | 725.1758 | 809.0386 | 762.4014 |
| ENSMUSG00000010080 | -0.52209388 | 0.022934 | 0.077806 | 841.0614 | 821.8778 | 1232.223 | 821.2176 | 1250.093 | 831.3263 | 1543.827 | 1712.956 |
| ENSMUSG00000010086 | 0.519096286 | 0.45247 | 0.614875 | 44.26639 | 50.3925 | 23.18699 | 47.44429 | 8.776002 | 75.98144 | 23.14294 | 7.613138 |
| ENSMUSG00000010095 | -0.429953573 | 0.007365 | 0.03411 | 1890.376 | 1706.146 | 2308.761 | 1810.647 | 2429.002 | 1991.161 | 2758.831 | 3218.182 |
| ENSMUSG00000010097 | 0.044349036 | 0.271341 | 0.436529 | 5620.826 | 5881.525 | 5651.828 | 5693.315 | 5564.96 | 5655.03 | 5435.698 | 5496.685 |
| ENSMUSG00000010110 | 0.05971213 | 0.222972 | 0.382486 | 2849.146 | 2851.976 | 2696.315 | 2841.482 | 2771.266 | 2704.045 | 2676.867 | 2626.532 |
| ENSMUSG00000010122 | -2.90255208 | 1.10E-05 | 0.000246 | 98.59333 | 35.99465 | 278.2438 | 122.4925 | 1127.229 | 414.5458 | 858.2174 | 1608.547 |
| ENSMUSG00000010142 | -0.363994823 | 0.28543 | 0.452218 | 26.15741 | 20.39697 | 19.04645 | 20.70296 | 18.52711 | 31.28647 | 30.85725 | 30.45255 |
| ENSMUSG00000010154 | -0.025224992 | 0.887711 | 0.932533 | 162.9808 | 151.1775 | 198.7456 | 141.4703 | 169.6694 | 131.8501 | 183.2149 | 181.6277 |
| ENSMUSG00000010175 | 0.626759732 | 0.008279 | 0.037224 | 368.2159 | 205.1695 | 258.3693 | 313.995 | 215.4996 | 174.3104 | 130.179 | 222.9562 |
| ENSMUSG00000010277 | -0.228092104 | 0.016127 | 0.060546 | 5690.243 | 5481.985 | 5980.586 | 5499.225 | 6845.281 | 5619.274 | 6624.667 | 7443.473 |
| ENSMUSG00000010290 | -0.01285786 | 0.876545 | 0.92607 | 861.1825 | 982.6538 | 835.5596 | 899.7163 | 945.858 | 869.317 | 929.5748 | 860.2845 |
| ENSMUSG00000010307 | -0.135563174 | 0.154448 | 0.295939 | 799.8132 | 819.4781 | 1010.29 | 860.0356 | 953.6589 | 991.1108 | 950.7891 | 942.9415 |
| ENSMUSG00000010311 | 0.42272508 | 0.436591 | 0.600382 | 27.16347 | 17.99732 | 18.21835 | 18.97772 | 11.70134 | 29.05173 | 4.821446 | 16.31387 |
| ENSMUSG00000010342 | -0.250550878 | 0.561133 | 0.706672 | 21.12714 | 17.99732 | 28.15563 | 22.42821 | 19.50223 | 46.92971 | 16.39292 | 25.0146 |
| ENSMUSG00000010358 | -0.063763711 | 0.797139 | 0.875505 | 276.6649 | 357.5468 | 505.9732 | 227.7326 | 350.065 | 290.5173 | 419.4658 | 369.781 |
| ENSMUSG00000010362 | 0.123117742 | 0.122878 | 0.252404 | 553.3299 | 543.5192 | 554.0033 | 607.2869 | 510.9583 | 530.7527 | 518.7876 | 515.5182 |
| ENSMUSG00000010376 | -0.191663948 | 0.004076 | 0.022128 | 3046.333 | 3125.535 | 3326.504 | 2935.508 | 3509.426 | 3348.77 | 3585.227 | 3757.627 |
| ENSMUSG00000010392 | 0.010693668 | 0.877193 | 0.926493 | 1297.81 | 1205.821 | 1246.3 | 1274.095 | 1231.566 | 1183.299 | 1348.076 | 1223.54 |
| ENSMUSG00000010406 | -0.141567973 | 0.067592 | 0.167851 | 1453.749 | 1604.161 | 1594.105 | 1431.955 | 1732.773 | 1578.85 | 1607.47 | 1790.175 |
| ENSMUSG00000010453 | -0.031262906 | 0.669561 | 0.7881 | 2498.033 | 2208.871 | 2127.406 | 2124.642 | 2327.591 | 2257.096 | 2237.151 | 2330.708 |
| ENSMUSG00000010461 | -0.202106112 | 0.326098 | 0.493883 | 748.5044 | 727.0918 | 510.9418 | 624.5394 | 959.5095 | 815.6831 | 720.324 | 505.7299 |
| ENSMUSG00000010476 | 1.022481864 | 0.000263 | 0.002808 | 312.8829 | 513.5236 | 327.1021 | 446.839 | 198.9227 | 303.9258 | 152.3577 | 132.6861 |
| ENSMUSG00000010492 | -0.294045792 | 0.259526 | 0.423578 | 32.19374 | 47.99286 | 35.60859 | 34.50494 | 48.75557 | 51.39921 | 39.53586 | 43.50364 |
| ENSMUSG00000010505 | 1.853997916 | 0.00137 | 0.009903 | 42.25428 | 16.7975 | 15.73403 | 67.28463 | 9.751113 | 11.17374 | 7.714313 | 10.87591 |
| ENSMUSG00000010517 | -0.309116382 | 0.001347 | 0.009765 | 987.9454 | 1097.837 | 1177.568 | 1046.362 | 1363.206 | 1164.304 | 1418.469 | 1393.204 |
| ENSMUSG00000010529 | -0.236449705 | 0.02821 | 0.090562 | 1151.932 | 1204.621 | 1130.366 | 1118.823 | 1321.276 | 1121.844 | 1453.184 | 1526.978 |
| ENSMUSG00000010538 | 0.277621076 | 0.246674 | 0.40974 | 77.46619 | 104.3845 | 82.81066 | 62.10889 | 54.60623 | 72.62932 | 79.07171 | 61.99269 |
| ENSMUSG00000010554 | -0.028295825 | 0.792303 | 0.872948 | 740.456 | 823.0776 | 612.7989 | 709.9391 | 701.105 | 720.7063 | 743.4669 | 773.2773 |
| ENSMUSG00000010592 | -3.463379984 | NA | NA | 4.024217 | 1.199822 | 2.48432 | 9.488858 | 1.950223 | 183.2494 | 0 | 5.437955 |
| ENSMUSG00000010601 | 1.284317836 | 0.080932 | 0.190425 | 16.09687 | 32.39518 | 9.109173 | 3.450494 | 6.825779 | 4.469496 | 5.785735 | 7.613138 |
| ENSMUSG00000010607 | -0.401389651 | 0.001315 | 0.009604 | 477.8758 | 496.7261 | 623.5643 | 564.1558 | 733.2837 | 651.4291 | 664.3952 | 811.343 |
| ENSMUSG00000010608 | 0.387352929 | 0.047175 | 0.130651 | 10246.66 | 10626.82 | 8457.453 | 8916.076 | 6126.624 | 10820.65 | 6294.88 | 5999.152 |
| ENSMUSG00000010609 | -0.010868053 | 0.923002 | 0.954016 | 713.2925 | 579.5138 | 631.8454 | 749.6198 | 622.121 | 643.6075 | 722.2526 | 709.1094 |
| ENSMUSG00000010651 | -2.429228994 | 4.02E-08 | 2.69E-06 | 91.55095 | 55.19179 | 200.4018 | 100.0643 | 620.1708 | 244.7049 | 516.859 | 1033.212 |
| ENSMUSG00000010660 | -0.050553239 | 0.703151 | 0.812914 | 628.784 | 608.3095 | 780.9046 | 714.2523 | 828.8446 | 610.0863 | 664.3952 | 728.686 |
| ENSMUSG00000010663 | 0.163099392 | 0.043613 | 0.123854 | 3361.228 | 3017.551 | 2800.657 | 2874.261 | 2742.013 | 2819.135 | 2694.224 | 2506.897 |
| ENSMUSG00000010721 | -0.011029275 | 0.898783 | 0.938869 | 616.7113 | 591.512 | 539.0974 | 583.1335 | 549.9628 | 623.4947 | 572.7878 | 601.4379 |
| ENSMUSG00000010751 | -0.071212477 | 0.604778 | 0.741291 | 290.7497 | 269.9598 | 313.0243 | 318.3081 | 371.5174 | 275.9914 | 326.894 | 278.4233 |
| ENSMUSG00000010755 | -0.044910552 | 0.591601 | 0.731496 | 1276.683 | 1225.018 | 1200.755 | 1066.203 | 1190.611 | 1324.088 | 1213.076 | 1189.825 |
| ENSMUSG00000010760 | -0.698178841 | 0.158762 | 0.301763 | 34.20585 | 38.39429 | 91.91984 | 50.03216 | 83.85957 | 25.6996 | 117.6433 | 121.8102 |
| ENSMUSG00000010797 | 0.317153937 | 0.435142 | 0.599282 | 30.18163 | 68.38983 | 43.06154 | 29.3292 | 37.05423 | 50.28183 | 22.17865 | 27.18978 |
| ENSMUSG00000010803 | 0.176839255 | 0.654037 | 0.77673 | 52.31483 | 85.18733 | 53.82693 | 64.69676 | 35.10401 | 105.0332 | 47.25017 | 39.15328 |
| ENSMUSG00000010825 | -0.28117257 | 0.562906 | 0.708373 | 12.07265 | 8.398751 | 10.76539 | 6.900988 | 14.62667 | 11.17374 | 9.642892 | 10.87591 |
| ENSMUSG00000010830 | 0.305269316 | 0.006905 | 0.032666 | 548.2996 | 599.9108 | 673.2507 | 574.5072 | 490.481 | 508.4052 | 419.4658 | 523.1313 |
| ENSMUSG00000010911 | -0.153606985 | 0.13431 | 0.268006 | 462.785 | 448.7333 | 476.1613 | 428.7239 | 565.5646 | 488.2925 | 451.2873 | 515.5182 |
| ENSMUSG00000010914 | -0.359683631 | 0.002895 | 0.017287 | 722.347 | 665.901 | 667.4539 | 625.402 | 773.2633 | 781.0445 | 841.8244 | 1044.087 |
| ENSMUSG00000010936 | -0.295745336 | 0.000608 | 0.005325 | 707.2562 | 656.3024 | 721.2809 | 616.7758 | 844.4464 | 768.7534 | 853.3959 | 849.4086 |
| ENSMUSG00000011008 | 0.017195127 | 0.957526 | 0.975954 | 27.16347 | 22.79661 | 28.98373 | 19.84034 | 23.40267 | 21.23011 | 27.96439 | 25.0146 |
| ENSMUSG00000011034 | -2.442223861 | 1.55E-08 | 1.22E-06 | 59.35721 | 35.99465 | 110.1382 | 52.62003 | 325.6872 | 151.9629 | 318.2154 | 612.3138 |
| ENSMUSG00000011096 | -0.192020238 | 0.004719 | 0.024695 | 2218.35 | 2357.649 | 2562.99 | 2435.186 | 2819.047 | 2582.252 | 2764.617 | 2774.445 |
| ENSMUSG00000011114 | 0.014387223 | 0.849422 | 0.909599 | 2665.038 | 2977.957 | 2778.298 | 2901.865 | 2922.409 | 2511.857 | 2930.475 | 2841.876 |
| ENSMUSG00000011148 | -0.763751358 | 0.000341 | 0.00341 | 244.4712 | 160.7761 | 202.8861 | 194.9529 | 382.2436 | 223.4748 | 380.8942 | 377.3941 |
| ENSMUSG00000011158 | -0.047736064 | 0.458827 | 0.62042 | 1706.268 | 1587.364 | 1583.34 | 1613.106 | 1778.603 | 1638.07 | 1699.078 | 1590.058 |
| ENSMUSG00000011171 | 0.021381296 | 0.94524 | 0.968433 | 39.23612 | 77.9884 | 105.9976 | 63.83414 | 66.30757 | 68.15982 | 67.50024 | 81.56933 |
| ENSMUSG00000011179 | -0.299610041 | 0.000115 | 0.0015 | 6072.544 | 7201.329 | 7281.542 | 6776.77 | 8853.036 | 8005.985 | 8203.208 | 8577.831 |
| ENSMUSG00000011254 | -0.176351134 | 0.11212 | 0.238643 | 786.7345 | 843.4745 | 794.1543 | 732.3673 | 1053.12 | 858.1433 | 865.9317 | 786.3284 |
| ENSMUSG00000011256 | 0.445862418 | 2.90E-05 | 0.000524 | 1202.235 | 1240.615 | 1334.08 | 1348.281 | 960.4846 | 987.7587 | 780.1099 | 1037.562 |
| ENSMUSG00000011257 | -0.057916013 | 0.444425 | 0.607449 | 4951.8 | 5397.997 | 5258.477 | 5080.852 | 5435.27 | 5983.538 | 5138.697 | 4978.992 |
| ENSMUSG00000011263 | -0.842429582 | 0.008117 | 0.036662 | 237.4288 | 121.182 | 274.1033 | 205.3044 | 353.9654 | 206.7142 | 398.2514 | 545.9707 |
| ENSMUSG00000011267 | 0.290592927 | 0.5441 | 0.692833 | 15.09082 | 11.99822 | 19.87456 | 12.07673 | 8.776002 | 16.76061 | 15.42863 | 7.613138 |

| | | | | | | | | | | | |
|---|---|---|---|---|---|---|---|---|---|---|---|
| ENSMUSG00000011305 | -0.222557477 | 0.713759 | 0.820362 | 24.1453 | 5.999108 | 8.281066 | 13.80198 | 6.825779 | 12.29112 | 20.25007 | 21.75182 |
| ENSMUSG00000011306 | 0.081122726 | 0.161098 | 0.304774 | 1327.992 | 1403.791 | 1389.563 | 1407.802 | 1337.853 | 1279.393 | 1269.969 | 1339.912 |
| ENSMUSG00000011382 | 0.458395246 | 0.03114 | 0.096968 | 324.9556 | 214.7681 | 177.2148 | 204.4418 | 156.0178 | 167.6061 | 156.2148 | 190.3284 |
| ENSMUSG00000011427 | -0.10178287 | 0.513696 | 0.668338 | 278.6771 | 275.959 | 218.6202 | 249.2982 | 295.4587 | 214.5358 | 276.751 | 307.7883 |
| ENSMUSG00000011463 | 0.547798888 | 0.057834 | 0.150759 | 80.48435 | 100.785 | 70.38906 | 69.8725 | 74.10846 | 63.69032 | 34.71441 | 46.76642 |
| ENSMUSG00000011589 | 0.103975471 | 0.513208 | 0.668037 | 212.2775 | 231.5656 | 156.5122 | 213.068 | 191.1218 | 203.3621 | 186.1078 | 174.0146 |
| ENSMUSG00000011658 | -0.062627999 | 0.486223 | 0.64438 | 369.2219 | 388.7422 | 399.9755 | 408.8835 | 390.0445 | 400.0199 | 418.5015 | 429.5985 |
| ENSMUSG00000011751 | 1.609701475 | 3.06E-17 | 2.35E-14 | 152.9203 | 193.1713 | 159.8246 | 178.5631 | 57.53157 | 67.04245 | 55.92877 | 43.50364 |
| ENSMUSG00000011752 | -0.852169425 | 4.65E-05 | 0.000752 | 1650.935 | 1444.585 | 1708.384 | 1299.974 | 3061.85 | 1664.887 | 2762.688 | 3529.233 |
| ENSMUSG00000011831 | 0.028102217 | 0.74305 | 0.839394 | 2077.502 | 1961.708 | 1894.708 | 2103.939 | 1821.508 | 2217.988 | 1859.15 | 1987.029 |
| ENSMUSG00000011832 | -0.035715777 | 0.727666 | 0.829154 | 657.9595 | 635.9054 | 640.1264 | 658.1817 | 637.7228 | 584.3867 | 664.3952 | 771.1021 |
| ENSMUSG00000011837 | -0.204523367 | 0.029638 | 0.093744 | 615.7053 | 691.0972 | 650.0637 | 604.6991 | 783.9895 | 650.3117 | 755.0384 | 758.051 |
| ENSMUSG00000011877 | -0.091141399 | 0.33544 | 0.503669 | 1609.687 | 1820.129 | 2081.032 | 1802.883 | 2034.082 | 1829.141 | 1901.578 | 2028.357 |
| ENSMUSG00000011884 | -0.350066788 | 0.01827 | 0.066232 | 695.1836 | 731.8911 | 884.4179 | 772.9107 | 999.4891 | 751.9928 | 1013.468 | 1168.073 |
| ENSMUSG00000011958 | 0.088974608 | 0.246968 | 0.410044 | 2620.772 | 2459.634 | 2272.325 | 2452.439 | 2421.201 | 2435.876 | 2133.972 | 2226.299 |
| ENSMUSG00000011960 | 0.210991813 | 0.12122 | 0.250475 | 2287.768 | 1990.504 | 1606.527 | 1850.327 | 1601.133 | 1994.513 | 1525.505 | 1560.693 |
| ENSMUSG00000012017 | 0.401975929 | 2.05E-06 | 6.56E-05 | 3078.526 | 3701.449 | 3476.392 | 3522.092 | 2733.237 | 2783.379 | 2447.366 | 2463.394 |
| ENSMUSG00000012076 | -0.046126947 | 0.612973 | 0.746981 | 795.789 | 725.892 | 684.8442 | 728.0542 | 790.8153 | 674.894 | 767.5742 | 793.9415 |
| ENSMUSG00000012114 | 0.091439819 | 0.179197 | 0.328916 | 1209.277 | 1077.44 | 1160.177 | 1125.724 | 1108.702 | 1064.858 | 1092.54 | 1026.686 |
| ENSMUSG00000012117 | -0.174138889 | 0.008277 | 0.037224 | 1172.053 | 1178.225 | 1139.475 | 1150.74 | 1345.654 | 1204.529 | 1358.683 | 1322.511 |
| ENSMUSG00000012123 | -0.083124354 | 0.761196 | 0.851536 | 78.47224 | 86.38715 | 88.60741 | 58.6584 | 106.2871 | 52.51658 | 75.21455 | 95.70802 |
| ENSMUSG00000012126 | 0.091855129 | 0.610363 | 0.745326 | 279.6831 | 309.554 | 205.3704 | 239.8093 | 291.5583 | 234.6486 | 242.0366 | 199.0292 |
| ENSMUSG00000012187 | -0.06454105 | 0.9105 | 0.946981 | 19.11503 | 9.598572 | 9.93728 | 12.93935 | 14.62667 | 6.704245 | 7.714313 | 25.0146 |
| ENSMUSG00000012296 | -0.005495729 | 0.95883 | 0.976771 | 949.7153 | 919.0633 | 801.6072 | 835.8822 | 918.5549 | 750.8754 | 953.682 | 891.8247 |
| ENSMUSG00000012350 | 0.015707151 | 0.908409 | 0.945287 | 1545.299 | 1273.011 | 1463.264 | 1375.022 | 1177.934 | 1403.422 | 1746.328 | 1268.131 |
| ENSMUSG00000012405 | -0.12817674 | 0.129472 | 0.261379 | 15078.74 | 18351.27 | 18113.18 | 18429.95 | 20368.12 | 18209.85 | 19224.07 | 18671.76 |
| ENSMUSG00000012422 | 0.143533964 | 0.151524 | 0.291883 | 2650.953 | 3061.945 | 2611.848 | 2799.213 | 2429.977 | 2928.638 | 2298.865 | 2412.277 |
| ENSMUSG00000012428 | 0.133141091 | 0.593663 | 0.733018 | 288.7376 | 136.7797 | 180.5272 | 257.9244 | 191.1218 | 159.7845 | 226.608 | 210.9927 |
| ENSMUSG00000012429 | 0.044174293 | 0.654915 | 0.777385 | 707.2562 | 776.2845 | 643.4389 | 739.2683 | 695.2544 | 760.9318 | 693.3239 | 627.5401 |
| ENSMUSG00000012443 | 0.002962476 | 0.979425 | 0.988724 | 2407.488 | 2246.066 | 1848.334 | 2240.233 | 2376.346 | 2366.598 | 2080.936 | 1897.846 |
| ENSMUSG00000012483 | -0.135878017 | 0.13289 | 0.266242 | 799.8132 | 951.4585 | 862.8871 | 984.2534 | 942.9326 | 1030.219 | 1002.861 | 977.7444 |
| ENSMUSG00000012519 | 0.130107767 | 0.756218 | 0.848356 | 86.52067 | 32.39518 | 58.79557 | 37.95543 | 51.6809 | 24.58223 | 68.46453 | 52.20437 |
| ENSMUSG00000012520 | 4.735000042 | 0.008486 | 0.037894 | 112.6781 | 3.599465 | 5.796746 | 194.0903 | 10.72622 | 0 | 0 | 1.087591 |
| ENSMUSG00000012535 | -0.045156368 | 0.524452 | 0.677583 | 3080.538 | 3263.515 | 2824.672 | 2972.601 | 3320.254 | 3058.253 | 3181.19 | 2959.335 |
| ENSMUSG00000012609 | 0.160807803 | 0.199794 | 0.354729 | 425.561 | 597.5111 | 504.3169 | 471.8551 | 412.4721 | 454.7713 | 450.323 | 468.7518 |
| ENSMUSG00000012640 | 0.089621098 | 0.31481 | 0.483132 | 734.4197 | 675.4995 | 610.3146 | 732.3673 | 649.4241 | 635.7859 | 648.9666 | 651.4671 |
| ENSMUSG00000012819 | -0.012887414 | 0.979656 | 0.988775 | 14.08476 | 32.39518 | 25.67131 | 8.626235 | 19.50223 | 29.05173 | 18.32149 | 14.13868 |
| ENSMUSG00000012848 | -0.170178795 | 0.05769 | 0.15054 | 25514.54 | 31281.75 | 30449.48 | 30728.37 | 36277.07 | 31656.33 | 33625.73 | 31183.41 |
| ENSMUSG00000013033 | 0.400124449 | 0.00743 | 0.034333 | 3244.525 | 3237.118 | 3165.852 | 3188.256 | 2300.288 | 3253.793 | 2214.008 | 1959.839 |
| ENSMUSG00000013076 | 0.374039786 | 0.006241 | 0.030325 | 3831.055 | 3796.235 | 3415.112 | 3616.98 | 2547.966 | 3683.982 | 2610.331 | 2469.919 |
| ENSMUSG00000013089 | 0.366722177 | 0.038169 | 0.112828 | 3666.062 | 4590.517 | 3127.759 | 3718.77 | 3005.293 | 3811.363 | 2672.045 | 2223.036 |
| ENSMUSG00000013150 | -0.187158923 | 0.021076 | 0.073158 | 470.8334 | 436.735 | 453.8024 | 462.3662 | 509.9832 | 505.0531 | 513.0018 | 550.3211 |
| ENSMUSG00000013155 | -0.353330032 | 0.00026 | 0.002783 | 282.7013 | 310.7538 | 316.3367 | 276.9021 | 384.1939 | 374.3203 | 378.0014 | 378.4817 |
| ENSMUSG00000013160 | -0.191912014 | 0.034746 | 0.105507 | 2012.109 | 2001.302 | 2166.327 | 2212.629 | 2424.127 | 2069.377 | 2469.545 | 2625.445 |
| ENSMUSG00000013236 | 0.442455191 | 0.000167 | 0.001976 | 7718.449 | 8313.563 | 7475.319 | 8037.063 | 5902.349 | 7098.678 | 5198.483 | 5013.795 |
| ENSMUSG00000013275 | 0.177205847 | 0.00672 | 0.032025 | 993.9817 | 938.2604 | 958.9475 | 915.2435 | 839.5708 | 824.6221 | 833.1458 | 868.9853 |
| ENSMUSG00000013338 | -0.669499375 | 0.263974 | 0.428225 | 43.26034 | 10.79839 | 53.82693 | 30.19182 | 80.93424 | 12.29112 | 60.75022 | 66.34306 |
| ENSMUSG00000013367 | 0.389169197 | 0.02583 | 0.084918 | 198.1927 | 177.5736 | 149.0592 | 156.9975 | 151.1423 | 105.0332 | 134.0362 | 128.3357 |
| ENSMUSG00000013415 | 0.098686225 | 0.442766 | 0.606004 | 1147.908 | 964.6565 | 908.433 | 893.6779 | 835.6704 | 1095.027 | 856.2888 | 868.9853 |
| ENSMUSG00000013418 | 0.687784623 | 0.000257 | 0.002766 | 176.0595 | 161.9759 | 125.8722 | 134.5693 | 96.53602 | 105.0332 | 91.60747 | 77.21897 |
| ENSMUSG00000013419 | 0.304616538 | 0.000127 | 0.001605 | 1265.616 | 1542.97 | 1459.124 | 1415.565 | 1177.934 | 1127.43 | 1145.576 | 1148.496 |
| ENSMUSG00000013465 | -0.128154661 | 0.049856 | 0.135799 | 2028.206 | 2080.491 | 2028.861 | 1908.123 | 2257.383 | 2038.09 | 2306.58 | 2184.97 |
| ENSMUSG00000013483 | -1.215782467 | 0.007839 | 0.035703 | 10.06054 | 22.79661 | 9.93728 | 13.80198 | 34.1289 | 23.46486 | 22.17865 | 50.02919 |
| ENSMUSG00000013495 | 0.152009295 | 0.113598 | 0.240708 | 569.4268 | 685.0981 | 674.9069 | 691.824 | 603.5939 | 549.7481 | 610.395 | 594.9123 |
| ENSMUSG00000013523 | 0.381554965 | 0.302675 | 0.470931 | 26.15741 | 45.59322 | 25.67131 | 21.56559 | 22.42756 | 21.23011 | 27.96439 | 18.48905 |
| ENSMUSG00000013539 | -0.148853526 | 0.401167 | 0.56844 | 344.0706 | 297.5557 | 295.6341 | 288.9789 | 359.8161 | 241.3528 | 335.5726 | 421.9853 |
| ENSMUSG00000013584 | 0.777284184 | 2.94E-05 | 0.000529 | 12635.04 | 14944.98 | 11094.97 | 14163.42 | 7518.108 | 10682.1 | 6957.346 | 5671.788 |
| ENSMUSG00000013593 | -0.465060307 | 0.002584 | 0.015783 | 5629.88 | 4511.329 | 5501.112 | 5287.882 | 7038.353 | 5244.954 | 8029.636 | 8580.006 |
| ENSMUSG00000013611 | -0.912691871 | 0.153684 | 0.295027 | 161.9748 | 109.1838 | 472.8489 | 222.5569 | 811.2926 | 154.1976 | 337.5012 | 517.6934 |
| ENSMUSG00000013622 | -0.077366853 | 0.347774 | 0.516479 | 2047.321 | 2067.293 | 2177.92 | 2408.445 | 2302.238 | 2105.133 | 2433.866 | 2341.584 |
| ENSMUSG00000013629 | -0.221772255 | 0.051818 | 0.139412 | 930.6003 | 1102.636 | 1166.802 | 939.397 | 1130.154 | 1099.496 | 1248.754 | 1348.613 |
| ENSMUSG00000013646 | -0.041470846 | 0.739919 | 0.837336 | 1057.363 | 1107.435 | 947.354 | 995.4675 | 1024.842 | 855.9086 | 1105.075 | 1238.766 |
| ENSMUSG00000013662 | -0.0555602 | 0.271168 | 0.436327 | 2552.36 | 2562.819 | 2538.147 | 2636.177 | 2560.642 | 2714.102 | 2649.867 | 2772.27 |
| ENSMUSG00000013663 | 0.161938208 | 0.079991 | 0.188964 | 2206.277 | 2315.656 | 2254.106 | 2524.036 | 1879.039 | 2372.185 | 2041.4 | 2022.919 |

| | | | | | | | | | | | |
|---|---|---|---|---|---|---|---|---|---|---|---|
| ENSMUSG00000013698 | 0.050608444 | 0.494719 | 0.651529 | 2950.757 | 3135.134 | 3511.172 | 3181.355 | 3237.37 | 3089.539 | 2920.832 | 3094.197 |
| ENSMUSG00000013701 | -0.582716715 | 3.28E-06 | 9.69E-05 | 1632.826 | 1580.165 | 1845.85 | 1689.879 | 2695.208 | 1971.048 | 2581.402 | 2861.452 |
| ENSMUSG00000013707 | 0.146336844 | 0.626915 | 0.756874 | 39.23612 | 34.79482 | 61.27989 | 32.77969 | 35.10401 | 39.10809 | 40.50014 | 38.06569 |
| ENSMUSG00000013736 | 0.098850625 | 0.342214 | 0.511012 | 1461.797 | 1482.979 | 1315.033 | 1718.346 | 1396.359 | 1538.624 | 1326.862 | 1321.423 |
| ENSMUSG00000013766 | -0.13676043 | 0.791811 | 0.872614 | 13.07871 | 4.799286 | 11.59349 | 12.93935 | 16.57689 | 8.938993 | 9.642892 | 11.9635 |
| ENSMUSG00000013787 | 0.026505658 | 0.623939 | 0.754981 | 5060.453 | 5027.252 | 5483.722 | 5165.389 | 5133.961 | 4871.751 | 5267.912 | 5088.839 |
| ENSMUSG00000013822 | -0.255550275 | 0.000555 | 0.004958 | 1327.992 | 1446.985 | 1389.563 | 1503.553 | 1652.814 | 1577.732 | 1744.399 | 1791.263 |
| ENSMUSG00000013833 | -0.242865015 | 0.003432 | 0.019512 | 1532.221 | 1575.366 | 1723.29 | 1447.482 | 1974.6 | 1706.23 | 1888.078 | 1859.781 |
| ENSMUSG00000013846 | 0.201117194 | 0.042542 | 0.121823 | 710.2744 | 790.6824 | 741.1554 | 713.3896 | 720.6073 | 626.8469 | 566.0377 | 655.8174 |
| ENSMUSG00000013858 | -0.227028614 | 0.008223 | 0.037017 | 3082.551 | 3030.749 | 3246.178 | 3097.681 | 3668.369 | 3180.047 | 3655.62 | 4077.379 |
| ENSMUSG00000013878 | -0.065026949 | 0.52138 | 0.674748 | 651.9232 | 704.2952 | 723.7652 | 744.4441 | 702.0801 | 854.7912 | 716.4668 | 684.0948 |
| ENSMUSG00000013921 | 0.466630606 | 1.35E-06 | 4.77E-05 | 1617.735 | 1828.528 | 1612.324 | 1870.168 | 1260.819 | 1411.243 | 1187.04 | 1155.022 |
| ENSMUSG00000013973 | -0.216858344 | 0.017914 | 0.065432 | 597.5963 | 599.9108 | 575.5341 | 601.2486 | 746.9353 | 602.2646 | 695.2525 | 712.3722 |
| ENSMUSG00000013997 | -0.356483652 | 0.002834 | 0.016992 | 911.4852 | 856.6726 | 871.1682 | 930.7707 | 1154.532 | 909.5425 | 1201.504 | 1305.109 |
| ENSMUSG00000014039 | 0.092314129 | 0.558473 | 0.704826 | 559.3662 | 577.1142 | 470.3646 | 568.4689 | 620.1708 | 545.2786 | 486.0017 | 387.1824 |
| ENSMUSG00000014074 | 0.071922587 | 0.595722 | 0.734838 | 1191.168 | 1443.385 | 1023.54 | 1218.887 | 1152.582 | 1368.783 | 1094.468 | 1021.248 |
| ENSMUSG00000014075 | 0.017095672 | 0.844868 | 0.906563 | 382.3007 | 406.7395 | 391.6944 | 413.1967 | 406.6214 | 392.1983 | 375.1085 | 401.3211 |
| ENSMUSG00000014077 | -0.186319547 | 0.058016 | 0.151063 | 2283.743 | 1941.311 | 2284.746 | 2413.621 | 2446.554 | 2278.326 | 2736.653 | 2696.138 |
| ENSMUSG00000014158 | -0.079611226 | 0.563032 | 0.708435 | 1192.174 | 1012.649 | 1324.143 | 1123.998 | 1121.378 | 1016.81 | 1361.576 | 1419.306 |
| ENSMUSG00000014164 | -0.986386813 | 0.000347 | 0.003462 | 99.59938 | 76.78858 | 87.7793 | 83.67448 | 170.6445 | 92.74205 | 209.2507 | 216.4306 |
| ENSMUSG00000014177 | 0.016567056 | 0.877076 | 0.926493 | 1168.029 | 1078.64 | 1035.961 | 1137.8 | 969.2606 | 983.2892 | 1187.04 | 1230.066 |
| ENSMUSG00000014195 | -0.041889963 | 0.401612 | 0.568792 | 6001.114 | 5931.918 | 5984.727 | 6248.845 | 6558.599 | 6098.628 | 6215.808 | 6005.678 |
| ENSMUSG00000014198 | -0.822045101 | 0.030378 | 0.095285 | 13.07871 | 21.59679 | 38.92101 | 21.56559 | 47.78045 | 33.52122 | 41.46443 | 46.76642 |
| ENSMUSG00000014226 | -0.157435666 | 0.01434 | 0.055766 | 2124.787 | 2393.644 | 2171.296 | 2285.09 | 2381.222 | 2523.031 | 2599.724 | 2501.46 |
| ENSMUSG00000014232 | 0.057627121 | 0.62736 | 0.757172 | 617.7174 | 701.8956 | 537.4412 | 576.2325 | 550.9379 | 658.1333 | 588.2164 | 537.27 |
| ENSMUSG00000014243 | -0.26811166 | 0.045596 | 0.127697 | 233.4046 | 242.3639 | 284.0406 | 251.8861 | 252.5538 | 319.569 | 325.9297 | 323.0146 |
| ENSMUSG00000014245 | -0.138983658 | 0.248512 | 0.411905 | 287.7315 | 271.1597 | 290.6654 | 332.9727 | 325.6872 | 328.508 | 290.251 | 361.0802 |
| ENSMUSG00000014294 | -0.330030987 | 0.002274 | 0.0144 | 1686.147 | 1665.352 | 1938.598 | 1637.259 | 2126.718 | 1904.005 | 2145.543 | 2534.087 |
| ENSMUSG00000014301 | -0.155278667 | 0.066789 | 0.166804 | 558.3602 | 596.3113 | 608.6584 | 552.9417 | 614.3201 | 619.0252 | 648.0023 | 698.2335 |
| ENSMUSG00000014303 | -0.173799266 | 0.092803 | 0.209385 | 1741.48 | 1809.331 | 1996.565 | 1728.697 | 1957.048 | 1914.062 | 1918.935 | 2419.89 |
| ENSMUSG00000014313 | -0.563085408 | 1.20E-06 | 4.38E-05 | 4485.996 | 4471.735 | 5352.881 | 4473.565 | 6740.944 | 5785.763 | 7266.883 | 7961.167 |
| ENSMUSG00000014329 | -0.120785741 | 0.249268 | 0.41271 | 4751.595 | 4606.115 | 5782.669 | 4623.662 | 4746.842 | 5690.786 | 5308.412 | 5749.006 |
| ENSMUSG00000014349 | -0.05654038 | 0.320464 | 0.488405 | 3120.781 | 3150.731 | 3274.334 | 3365.094 | 3241.27 | 3264.967 | 3384.655 | 3541.197 |
| ENSMUSG00000014353 | 0.191232776 | 0.074273 | 0.179869 | 1479.906 | 1268.211 | 1139.475 | 1354.319 | 1056.046 | 1265.985 | 1171.611 | 1097.379 |
| ENSMUSG00000014355 | -0.048972204 | 0.382703 | 0.550691 | 3626.826 | 3729.045 | 3644.497 | 3374.583 | 3699.572 | 3843.767 | 3753.978 | 3569.474 |
| ENSMUSG00000014361 | -0.002514492 | 0.986709 | 0.992387 | 920.5397 | 742.6895 | 894.3552 | 767.7349 | 640.6481 | 797.8051 | 912.2175 | 982.0948 |
| ENSMUSG00000014402 | -0.077114822 | 0.219469 | 0.378199 | 1963.818 | 1854.924 | 1906.301 | 2012.501 | 2010.68 | 2146.476 | 2087.686 | 1920.686 |
| ENSMUSG00000014418 | -0.137316051 | 0.079068 | 0.187617 | 1088.551 | 1043.845 | 1048.383 | 1048.95 | 1092.125 | 1092.792 | 1183.183 | 1284.445 |
| ENSMUSG00000014426 | -0.064035095 | 0.375387 | 0.543895 | 1316.925 | 1469.781 | 1371.345 | 1311.188 | 1433.414 | 1366.549 | 1384.719 | 1529.153 |
| ENSMUSG00000014444 | -0.222029535 | 0.148571 | 0.28824 | 1176.078 | 1107.435 | 1543.591 | 1223.2 | 1530.925 | 1138.604 | 1515.863 | 1707.518 |
| ENSMUSG00000014470 | 0.058398755 | 0.476076 | 0.635594 | 996.9999 | 989.8528 | 953.1507 | 966.1383 | 937.082 | 1046.98 | 863.0388 | 904.8758 |
| ENSMUSG00000014496 | -0.216606736 | 0.0002 | 0.002266 | 1269.641 | 1249.014 | 1276.94 | 1299.974 | 1463.642 | 1462.643 | 1435.827 | 1562.868 |
| ENSMUSG00000014498 | -0.01098999 | 0.919347 | 0.952207 | 954.7456 | 831.4763 | 887.7303 | 826.3933 | 1006.315 | 757.5796 | 866.896 | 895.0875 |
| ENSMUSG00000014503 | 0.387321056 | 0.010028 | 0.042676 | 451.7184 | 418.7377 | 409.9128 | 484.7944 | 312.0356 | 434.6585 | 307.6082 | 296.9124 |
| ENSMUSG00000014504 | 0.123716047 | 0.214253 | 0.372375 | 1824.983 | 1670.152 | 1456.64 | 1754.576 | 1500.696 | 1716.287 | 1505.255 | 1432.357 |
| ENSMUSG00000014547 | -0.494520017 | 0.027701 | 0.089348 | 118.7144 | 81.58786 | 87.7793 | 97.47645 | 167.7191 | 100.5637 | 121.5004 | 153.3503 |
| ENSMUSG00000014550 | -0.02982092 | 0.636992 | 0.76434 | 1200.223 | 1261.012 | 1296.815 | 1244.766 | 1285.197 | 1321.854 | 1296.005 | 1205.051 |
| ENSMUSG00000014551 | -0.352729141 | 0.02722 | 0.088212 | 595.5842 | 739.0901 | 751.9208 | 645.2424 | 808.3673 | 688.3024 | 883.2889 | 1108.255 |
| ENSMUSG00000014554 | 0.116115366 | 0.259895 | 0.423918 | 779.6921 | 1007.85 | 814.8569 | 941.9849 | 843.4713 | 808.9788 | 820.6101 | 793.9415 |
| ENSMUSG00000014592 | 0.209229116 | 0.181928 | 0.332463 | 1300.828 | 1343.8 | 1035.961 | 1244.766 | 989.738 | 1392.248 | 980.6821 | 897.2626 |
| ENSMUSG00000014599 | 0.520039869 | 0.000287 | 0.003003 | 386.3249 | 515.9233 | 524.1915 | 474.4429 | 290.5832 | 378.7898 | 349.0727 | 307.7883 |
| ENSMUSG00000014601 | -0.178771498 | 0.031897 | 0.098719 | 1125.775 | 1145.83 | 1300.127 | 1109.334 | 1349.554 | 1251.459 | 1285.397 | 1414.956 |
| ENSMUSG00000014602 | 0.790549068 | 1.46E-09 | 1.79E-07 | 527.1725 | 423.537 | 494.3797 | 602.9738 | 298.3841 | 312.8647 | 282.5367 | 292.562 |
| ENSMUSG00000014606 | -0.537461199 | 0.000254 | 0.002745 | 1918.546 | 2072.092 | 2382.463 | 1952.117 | 2943.861 | 2293.969 | 3199.511 | 3646.693 |
| ENSMUSG00000014633 | -0.424228952 | 0.002885 | 0.017243 | 355.1372 | 383.9429 | 388.382 | 361.4392 | 478.7797 | 400.0199 | 504.3232 | 614.489 |
| ENSMUSG00000014668 | -0.005038947 | 0.957599 | 0.975954 | 1410.488 | 1167.426 | 1138.647 | 1371.571 | 1230.59 | 1286.098 | 1266.112 | 1324.686 |
| ENSMUSG00000014704 | 0.127229817 | 0.565383 | 0.710019 | 220.3259 | 310.7538 | 269.1347 | 385.5927 | 294.4836 | 345.2686 | 232.3937 | 214.2554 |
| ENSMUSG00000014747 | 1.031186079 | 0.002256 | 0.014333 | 33.19979 | 44.3934 | 51.34261 | 55.2079 | 26.32801 | 25.6996 | 25.07152 | 13.05109 |
| ENSMUSG00000014748 | -0.212061798 | 0.031242 | 0.097202 | 2031.224 | 1900.517 | 2088.485 | 1854.641 | 2388.048 | 1916.297 | 2346.116 | 2471.007 |
| ENSMUSG00000014763 | 0.046231412 | 0.593336 | 0.732826 | 1138.854 | 1124.233 | 991.2436 | 1164.542 | 1142.83 | 1017.928 | 1108.933 | 1006.022 |
| ENSMUSG00000014767 | 0.052435816 | 0.623032 | 0.754457 | 1300.828 | 1354.599 | 1124.569 | 1237.865 | 1201.337 | 1404.539 | 1119.54 | 1111.518 |
| ENSMUSG00000014769 | -0.119684203 | 0.047688 | 0.131618 | 4465.875 | 4990.058 | 5114.387 | 4906.602 | 5320.207 | 5137.686 | 5292.019 | 5412.941 |
| ENSMUSG00000014773 | -0.938396894 | 8.83E-05 | 0.001222 | 433.6094 | 208.7689 | 327.1021 | 291.5667 | 529.4854 | 454.7713 | 743.4669 | 690.6203 |
| ENSMUSG00000014776 | 0.073957998 | 0.889043 | 0.933305 | 14.08476 | 13.19804 | 14.90592 | 13.80198 | 6.825779 | 6.704245 | 22.17865 | 17.40146 |

| | | | | | | | | | | | |
|---|---|---|---|---|---|---|---|---|---|---|---|
| ENSMUSG00000014778 | 0.285500274 | 0.000184 | 0.002125 | 736.4318 | 736.6904 | 735.3587 | 722.8785 | 620.1708 | 563.1565 | 579.5378 | 641.6787 |
| ENSMUSG00000014782 | -0.039242294 | 0.905626 | 0.943335 | 27.16347 | 32.39518 | 48.85829 | 29.3292 | 44.85512 | 29.05173 | 39.53586 | 28.27737 |
| ENSMUSG00000014786 | 0.611667747 | 7.91E-05 | 0.001131 | 301.8163 | 437.9349 | 377.6166 | 360.5766 | 241.8276 | 283.813 | 214.0722 | 227.3065 |
| ENSMUSG00000014791 | -0.280182932 | 0.01381 | 0.054229 | 435.6215 | 447.5334 | 510.1137 | 445.9763 | 524.6099 | 490.5272 | 608.4665 | 611.2262 |
| ENSMUSG00000014813 | 0.135061834 | 0.416993 | 0.582503 | 515.0998 | 446.3336 | 609.4865 | 458.0531 | 377.3681 | 553.1002 | 406.93 | 513.343 |
| ENSMUSG00000014837 | -0.286372826 | 0.074689 | 0.180498 | 197.1867 | 199.1704 | 180.5272 | 182.8762 | 292.5334 | 215.6532 | 193.8221 | 222.9562 |
| ENSMUSG00000014846 | 0.805771242 | 0.000614 | 0.005368 | 405.4399 | 590.3122 | 395.0069 | 575.3699 | 313.9858 | 379.9072 | 202.5007 | 228.3941 |
| ENSMUSG00000014850 | 0.155151213 | 0.103654 | 0.226423 | 1372.258 | 1424.188 | 1281.081 | 1302.561 | 1266.67 | 1358.727 | 1114.718 | 1089.766 |
| ENSMUSG00000014856 | -0.143727479 | 0.064861 | 0.163984 | 773.6558 | 843.4745 | 826.4504 | 784.9874 | 931.2313 | 821.27 | 876.5389 | 935.3283 |
| ENSMUSG00000014859 | -0.152168726 | 0.10161 | 0.223539 | 1270.647 | 1262.212 | 1204.895 | 1242.178 | 1463.642 | 1176.595 | 1383.755 | 1507.401 |
| ENSMUSG00000014867 | -0.146267432 | 0.036239 | 0.108983 | 5411.566 | 5663.158 | 5839.808 | 5098.967 | 6268.991 | 5624.861 | 6141.558 | 6324.342 |
| ENSMUSG00000014873 | -0.134740853 | 0.073907 | 0.179265 | 686.1291 | 667.1008 | 656.6886 | 626.2647 | 744.0099 | 681.5982 | 715.5026 | 750.4379 |
| ENSMUSG00000014905 | 0.018718427 | 0.850936 | 0.910548 | 708.2623 | 609.5093 | 646.7513 | 725.4664 | 591.8926 | 688.3024 | 705.8597 | 672.1313 |
| ENSMUSG00000014907 | -0.012088615 | 0.912202 | 0.948044 | 394.3733 | 423.537 | 388.382 | 428.7239 | 361.7663 | 464.8276 | 400.18 | 423.0729 |
| ENSMUSG00000014932 | 0.156720425 | 0.134458 | 0.268248 | 1136.841 | 1101.436 | 1011.946 | 1218.024 | 933.1815 | 1158.717 | 928.6105 | 989.7079 |
| ENSMUSG00000014956 | 0.184736412 | 0.023806 | 0.08019 | 6668.128 | 7214.527 | 6264.627 | 6987.25 | 5627.367 | 6690.836 | 5795.378 | 5758.795 |
| ENSMUSG00000014959 | -0.199004258 | 0.00355 | 0.020023 | 2746.528 | 2760.789 | 3008.511 | 2815.603 | 3225.668 | 3010.206 | 3290.155 | 3484.642 |
| ENSMUSG00000014980 | -0.114796319 | 0.334749 | 0.503039 | 703.232 | 854.2729 | 739.4992 | 792.751 | 704.0304 | 887.195 | 808.0743 | 946.2042 |
| ENSMUSG00000015002 | 0.041529385 | 0.635141 | 0.762769 | 1810.898 | 1697.747 | 1672.775 | 1825.311 | 1533.85 | 1816.85 | 1600.72 | 1859.781 |
| ENSMUSG00000015013 | -0.066780555 | 0.406379 | 0.572907 | 725.3652 | 617.9081 | 708.0312 | 714.2523 | 737.1841 | 711.7673 | 731.8955 | 719.9853 |
| ENSMUSG00000015016 | -0.313550445 | 0.005923 | 0.029156 | 449.7063 | 473.9295 | 546.5504 | 499.459 | 586.0419 | 555.3349 | 602.6807 | 706.9342 |
| ENSMUSG00000015023 | -0.166467244 | 0.0202 | 0.070958 | 1289.762 | 1274.21 | 1390.391 | 1298.248 | 1518.248 | 1360.962 | 1560.22 | 1456.284 |
| ENSMUSG00000015027 | -0.38192282 | 0.004443 | 0.023594 | 452.7245 | 529.1213 | 451.3181 | 449.4268 | 639.673 | 518.4616 | 559.2877 | 733.0364 |
| ENSMUSG00000015053 | -0.3215083 | 0.305441 | 0.473876 | 708.2623 | 406.7395 | 1407.781 | 744.4441 | 897.1024 | 883.8429 | 993.2178 | 1311.635 |
| ENSMUSG00000015083 | -0.513492139 | 0.000694 | 0.005912 | 88.53278 | 106.7841 | 109.3101 | 100.9269 | 156.0178 | 141.9065 | 141.7505 | 139.2117 |
| ENSMUSG00000015085 | 0.15620388 | 0.348865 | 0.517452 | 119.7205 | 157.1766 | 163.137 | 167.349 | 154.0676 | 132.9675 | 131.1433 | 127.2482 |
| ENSMUSG00000015087 | -0.143660608 | 0.019851 | 0.070105 | 2223.38 | 2318.055 | 2270.668 | 2315.281 | 2701.058 | 2407.941 | 2406.866 | 2565.627 |
| ENSMUSG00000015090 | 0.335448812 | 0.124235 | 0.254143 | 117.7084 | 119.9822 | 171.4181 | 120.7673 | 107.2622 | 134.0849 | 87.75031 | 92.44524 |
| ENSMUSG00000015092 | -0.388067851 | 0.004501 | 0.023803 | 1948.727 | 2199.273 | 2325.323 | 2073.747 | 2871.703 | 2186.701 | 2719.295 | 3407.423 |
| ENSMUSG00000015093 | 0.048890733 | 0.827859 | 0.895602 | 49.29666 | 45.59322 | 52.17072 | 61.24627 | 48.75557 | 48.04709 | 50.14304 | 55.46715 |
| ENSMUSG00000015094 | -0.00980131 | 0.919198 | 0.952207 | 756.5529 | 917.8635 | 891.0427 | 954.0616 | 841.5211 | 889.4298 | 944.0391 | 868.9853 |
| ENSMUSG00000015095 | -0.215623942 | 0.037158 | 0.110754 | 1576.487 | 1619.759 | 1700.103 | 1636.397 | 2050.659 | 1555.385 | 1915.078 | 2064.248 |
| ENSMUSG00000015112 | -0.987453102 | 4.82E-06 | 0.000131 | 775.6679 | 703.0954 | 858.7466 | 640.9293 | 1486.07 | 888.3124 | 1600.72 | 1930.474 |
| ENSMUSG00000015120 | -0.097344281 | 0.189867 | 0.342466 | 6570.541 | 6808.987 | 6486.559 | 6975.174 | 7936.431 | 7138.903 | 7123.204 | 6513.583 |
| ENSMUSG00000015126 | -0.099039642 | 0.125349 | 0.2559 | 866.2128 | 884.2685 | 937.4167 | 935.9465 | 968.2855 | 982.1718 | 953.682 | 982.0948 |
| ENSMUSG00000015127 | 0.031382098 | 0.854473 | 0.912783 | 302.8224 | 251.9625 | 298.9465 | 267.4133 | 336.4134 | 202.2447 | 270.9653 | 287.124 |
| ENSMUSG00000015133 | -0.016166451 | 0.868997 | 0.921481 | 761.5831 | 688.6976 | 853.7779 | 693.5493 | 726.4579 | 743.0538 | 801.3243 | 763.4889 |
| ENSMUSG00000015134 | 0.926709503 | 1.37E-08 | 1.10E-06 | 5152.004 | 6026.704 | 6617.4 | 5498.362 | 3168.137 | 3864.997 | 2218.829 | 3003.927 |
| ENSMUSG00000015143 | 0.096826704 | 0.104125 | 0.227075 | 3182.15 | 3312.707 | 3427.533 | 3256.404 | 2999.442 | 3195.69 | 2911.189 | 3221.445 |
| ENSMUSG00000015149 | -0.015371486 | 0.888602 | 0.932987 | 1786.753 | 1539.371 | 1704.243 | 1980.584 | 1831.259 | 1515.159 | 1860.114 | 1881.533 |
| ENSMUSG00000015165 | 0.030170043 | 0.650482 | 0.773836 | 10078.65 | 10724 | 10236.23 | 9881.352 | 10500 | 10676.51 | 9538.748 | 9355.459 |
| ENSMUSG00000015176 | -0.372799628 | 4.42E-07 | 1.91E-05 | 1992.994 | 1932.912 | 1938.598 | 1851.19 | 2645.477 | 2327.49 | 2345.151 | 2671.124 |
| ENSMUSG00000015189 | 0.178817079 | 0.037458 | 0.111414 | 1879.31 | 1775.736 | 1603.214 | 1757.164 | 1382.708 | 1647.009 | 1570.827 | 1596.584 |
| ENSMUSG00000015202 | 0.030552899 | 0.681154 | 0.796463 | 1029.194 | 1089.438 | 1070.742 | 1075.691 | 981.9371 | 985.524 | 1088.682 | 1119.131 |
| ENSMUSG00000015214 | 0.176661391 | 0.03852 | 0.11341 | 697.1957 | 663.5013 | 631.0173 | 717.7027 | 570.4401 | 636.9032 | 617.1451 | 573.1605 |
| ENSMUSG00000015217 | -0.10098212 | 0.136641 | 0.2714 | 3682.159 | 4134.585 | 3796.041 | 4007.749 | 4375.324 | 4384.576 | 4023.979 | 3966.445 |
| ENSMUSG00000015222 | 0.294571401 | 0.082387 | 0.192898 | 502.0211 | 362.3461 | 427.303 | 479.6187 | 275.9565 | 346.386 | 407.8943 | 415.4598 |
| ENSMUSG00000015224 | 0.124521417 | 0.268304 | 0.432776 | 207.2472 | 225.5664 | 230.2136 | 215.6559 | 197.9476 | 200.01 | 199.6079 | 208.8175 |
| ENSMUSG00000015243 | 0.501021345 | 0.009344 | 0.040521 | 2165.029 | 1380.995 | 1826.803 | 2221.255 | 1336.878 | 1573.263 | 1495.612 | 961.4305 |
| ENSMUSG00000015247 | -0.060511689 | 0.394396 | 0.561649 | 748.5044 | 760.6869 | 801.6072 | 816.9044 | 832.7451 | 811.2136 | 781.0742 | 840.7079 |
| ENSMUSG00000015289 | -0.094580846 | 0.245839 | 0.408795 | 1021.145 | 1191.423 | 1111.319 | 1093.807 | 1210.113 | 1077.149 | 1183.183 | 1243.117 |
| ENSMUSG00000015290 | -0.346077399 | 0.002036 | 0.013289 | 1080.502 | 1221.418 | 1364.72 | 1251.667 | 1679.142 | 1331.91 | 1539.006 | 1703.168 |
| ENSMUSG00000015291 | 0.135359978 | 0.165869 | 0.311327 | 3398.452 | 2807.582 | 2930.669 | 3332.315 | 2700.083 | 2574.43 | 3024.011 | 3055.043 |
| ENSMUSG00000015305 | 0.194694216 | 0.034379 | 0.104663 | 1421.555 | 1458.983 | 1590.793 | 1450.933 | 1254.968 | 1487.225 | 1239.112 | 1196.35 |
| ENSMUSG00000015312 | -0.146472381 | 0.606795 | 0.742837 | 319.9253 | 206.3693 | 346.9767 | 260.5123 | 265.2303 | 195.5405 | 488.8946 | 305.6131 |
| ENSMUSG00000015335 | -0.204990498 | 0.16924 | 0.315546 | 316.9071 | 212.3684 | 284.0406 | 264.8254 | 315.9361 | 272.6393 | 329.7869 | 326.2773 |
| ENSMUSG00000015337 | -1.320942895 | 2.18E-06 | 6.88E-05 | 143.8658 | 145.1784 | 219.4483 | 144.0581 | 340.3138 | 251.4092 | 434.8944 | 605.7882 |
| ENSMUSG00000015340 | 0.042635549 | 0.881989 | 0.929373 | 46.2785 | 80.38804 | 78.67013 | 44.85642 | 53.63112 | 52.51658 | 66.53595 | 69.60583 |
| ENSMUSG00000015341 | -0.106681259 | 0.046582 | 0.129599 | 2843.11 | 2658.805 | 2890.092 | 2885.476 | 3113.53 | 2950.985 | 2994.118 | 3090.934 |
| ENSMUSG00000015342 | 0.653500967 | 0.000869 | 0.006991 | 138.8355 | 121.182 | 141.6062 | 165.6237 | 86.78491 | 116.2069 | 81.96458 | 77.21897 |
| ENSMUSG00000015354 | -0.363156736 | 0.386538 | 0.554049 | 14.08476 | 22.79661 | 29.81184 | 10.35148 | 25.35289 | 26.81698 | 24.10723 | 22.83941 |
| ENSMUSG00000015355 | 0.146646466 | 0.774392 | 0.8603 | 11.0666 | 21.59679 | 12.4216 | 14.6646 | 21.45245 | 15.64324 | 7.714313 | 8.700729 |
| ENSMUSG00000015357 | -0.158962822 | 0.049536 | 0.135201 | 1445.7 | 1268.211 | 1354.782 | 1384.511 | 1527.999 | 1368.783 | 1613.256 | 1579.182 |
| ENSMUSG00000015363 | -0.423725942 | 0.000124 | 0.001573 | 1204.247 | 1090.638 | 1230.566 | 1165.404 | 1559.203 | 1296.154 | 1663.399 | 1774.949 |

| | | | | | | | | | | | |
|---|---|---|---|---|---|---|---|---|---|---|---|
| ENSMUSG00000015365 | -5.304014643 | NA | NA | 3.018163 | 0 | 0 | 1.725247 | 0 | 187.7188 | 0 | 0 |
| ENSMUSG00000015377 | 0.211289865 | 0.02143 | 0.073978 | 688.1412 | 705.4951 | 638.4702 | 658.1817 | 634.7975 | 606.7341 | 536.1448 | 543.7955 |
| ENSMUSG00000015396 | 0.389111885 | 0.043908 | 0.124422 | 1034.224 | 1617.359 | 1202.411 | 1275.82 | 1001.439 | 1303.976 | 786.86 | 824.394 |
| ENSMUSG00000015401 | -3.221421816 | 2.92E-07 | 1.35E-05 | 160.9687 | 81.58786 | 523.3634 | 176.8378 | 2451.43 | 1103.966 | 1915.078 | 3330.204 |
| ENSMUSG00000015405 | -1.301904198 | 0.004586 | 0.02415 | 844.0796 | 319.1525 | 880.2773 | 615.0506 | 1350.529 | 833.5611 | 1698.113 | 2674.386 |
| ENSMUSG00000015451 | 1.646745325 | 0.013137 | 0.052422 | 240.447 | 29.99554 | 27.32752 | 52.62003 | 32.17867 | 32.40385 | 21.21436 | 26.10219 |
| ENSMUSG00000015452 | 0.143331052 | 0.458084 | 0.619779 | 340.0464 | 375.5441 | 295.6341 | 275.1769 | 293.5085 | 388.8462 | 249.7509 | 231.6569 |
| ENSMUSG00000015461 | -0.150093149 | 0.081371 | 0.191131 | 1459.785 | 1311.405 | 1509.638 | 1306.875 | 1470.468 | 1455.938 | 1611.327 | 1665.102 |
| ENSMUSG00000015467 | 0.120888625 | 0.598077 | 0.736699 | 126.7628 | 68.38983 | 105.9976 | 102.6522 | 83.85957 | 82.68568 | 97.39321 | 108.7591 |
| ENSMUSG00000015468 | -0.391134215 | 0.010287 | 0.043496 | 567.4147 | 415.1383 | 466.224 | 462.3662 | 605.5441 | 500.5836 | 737.6812 | 662.343 |
| ENSMUSG00000015474 | -0.788337243 | 3.08E-06 | 9.26E-05 | 637.8385 | 514.7234 | 590.44 | 576.2325 | 935.1317 | 735.2322 | 1061.682 | 1274.657 |
| ENSMUSG00000015476 | 0.828238038 | 0.001931 | 0.012809 | 48.29061 | 63.59054 | 53.82693 | 48.30692 | 32.17867 | 35.75597 | 23.14294 | 29.36496 |
| ENSMUSG00000015478 | -0.439126998 | 0.001142 | 0.008647 | 1398.416 | 1380.995 | 1445.046 | 1545.821 | 1922.919 | 1507.338 | 2032.722 | 2361.16 |
| ENSMUSG00000015484 | -0.165385126 | 0.811329 | 0.885135 | 47.28455 | 11.99822 | 7.45296 | 43.9938 | 30.22845 | 11.17374 | 23.14294 | 59.81751 |
| ENSMUSG00000015488 | 0.005357368 | 0.942023 | 0.966348 | 2010.097 | 2106.887 | 1950.191 | 2034.066 | 2186.2 | 1919.649 | 1868.792 | 2093.613 |
| ENSMUSG00000015501 | 0.768442837 | 2.23E-06 | 7.03E-05 | 926.5761 | 959.8572 | 783.3889 | 1040.324 | 499.257 | 707.2978 | 530.359 | 441.562 |
| ENSMUSG00000015522 | 0.178035744 | 0.006248 | 0.030345 | 2202.253 | 2142.881 | 2015.612 | 2273.876 | 1937.546 | 1986.691 | 1889.042 | 1818.452 |
| ENSMUSG00000015533 | 0.374733305 | 0.316126 | 0.484426 | 128.775 | 45.59322 | 69.56096 | 52.62003 | 64.35735 | 39.10809 | 58.82164 | 66.34306 |
| ENSMUSG00000015536 | -0.097568944 | 0.24687 | 0.409947 | 1362.198 | 1358.198 | 1304.268 | 1416.428 | 1426.588 | 1319.619 | 1438.719 | 1636.825 |
| ENSMUSG00000015542 | -0.165202978 | 0.107393 | 0.231872 | 435.6215 | 405.5397 | 422.3344 | 417.5098 | 455.377 | 455.8886 | 540.0019 | 432.8613 |
| ENSMUSG00000015568 | -0.049990146 | 0.810679 | 0.884586 | 1370.246 | 2740.392 | 1612.324 | 1674.352 | 1576.755 | 1996.748 | 2153.258 | 1928.299 |
| ENSMUSG00000015575 | -0.544098278 | 0.004006 | 0.021854 | 1710.292 | 1412.19 | 2126.578 | 1705.407 | 2459.231 | 1743.104 | 2857.189 | 3083.321 |
| ENSMUSG00000015597 | 0.169381009 | 0.118604 | 0.247423 | 1304.852 | 1259.813 | 1146.928 | 1343.967 | 1060.921 | 1335.262 | 1060.718 | 1039.737 |
| ENSMUSG00000015599 | 0.226885061 | 0.514193 | 0.668674 | 26.15741 | 23.99643 | 19.87456 | 26.74133 | 18.52711 | 15.64324 | 21.21436 | 27.18978 |
| ENSMUSG00000015605 | -0.213439924 | 0.128715 | 0.260363 | 1431.615 | 1341.4 | 1501.357 | 1165.404 | 1662.565 | 1221.29 | 1599.756 | 1822.803 |
| ENSMUSG00000015619 | 0.552508979 | 0.000447 | 0.004199 | 2847.134 | 2398.443 | 2686.378 | 2185.888 | 1583.581 | 2281.678 | 1553.47 | 1481.299 |
| ENSMUSG00000015627 | -0.510676345 | 0.256713 | 0.420325 | 22.1332 | 41.99375 | 93.57605 | 37.09281 | 88.73513 | 43.57759 | 64.60737 | 81.56933 |
| ENSMUSG00000015647 | -0.033721599 | 0.814291 | 0.887216 | 3605.699 | 3256.316 | 5050.622 | 3726.533 | 3784.407 | 3625.879 | 4238.051 | 4364.503 |
| ENSMUSG00000015652 | -0.302559459 | 0.372334 | 0.541342 | 119.7205 | 44.3934 | 85.29498 | 56.93315 | 83.85957 | 106.1505 | 70.39311 | 118.5474 |
| ENSMUSG00000015653 | -0.629061331 | 3.30E-06 | 9.74E-05 | 669.0261 | 613.1088 | 804.9196 | 689.2362 | 1000.464 | 957.5896 | 1028.897 | 1311.635 |
| ENSMUSG00000015656 | -0.204757434 | 0.008921 | 0.039228 | 34188.75 | 36500.97 | 35161.41 | 35078.58 | 40405.69 | 35788.37 | 41407.54 | 44816.37 |
| ENSMUSG00000015659 | 0.278956394 | 0.046941 | 0.130236 | 316.9071 | 279.5584 | 238.4947 | 243.2598 | 228.176 | 197.7752 | 227.5722 | 232.7445 |
| ENSMUSG00000015668 | -0.075604878 | 0.257313 | 0.421045 | 811.8859 | 836.2756 | 834.7315 | 807.4156 | 834.6953 | 843.6174 | 879.4317 | 909.2261 |
| ENSMUSG00000015671 | -0.227233343 | 0.002534 | 0.015568 | 3121.787 | 3105.138 | 3401.034 | 3426.341 | 3833.163 | 3456.038 | 3947.8 | 4048.014 |
| ENSMUSG00000015672 | 0.028549416 | 0.794779 | 0.873746 | 746.4923 | 788.2827 | 656.6886 | 656.4565 | 711.8313 | 601.1473 | 736.7169 | 737.3868 |
| ENSMUSG00000015697 | 0.025466115 | 0.694842 | 0.807303 | 2063.417 | 1982.105 | 1934.457 | 2091.862 | 1939.496 | 2140.889 | 1938.221 | 1914.16 |
| ENSMUSG00000015702 | -0.271695674 | 0.287004 | 0.454071 | 154.9324 | 178.7734 | 317.1648 | 149.2339 | 234.0267 | 223.4748 | 273.8581 | 236.0073 |
| ENSMUSG00000015709 | 0.250231582 | 0.027059 | 0.087907 | 2313.925 | 2680.401 | 2102.563 | 2313.556 | 1993.128 | 2290.617 | 1873.614 | 1752.109 |
| ENSMUSG00000015711 | -0.136610088 | 0.342136 | 0.510962 | 835.0251 | 679.099 | 814.0288 | 790.1631 | 826.8944 | 661.4855 | 948.8605 | 991.8831 |
| ENSMUSG00000015714 | -0.150375899 | 0.089378 | 0.204103 | 2965.848 | 2699.598 | 3039.151 | 2844.932 | 3221.768 | 2816.9 | 3208.19 | 3573.824 |
| ENSMUSG00000015733 | 0.042571342 | 0.398537 | 0.565843 | 3831.055 | 4148.983 | 3922.741 | 4028.452 | 3833.163 | 3994.612 | 3886.085 | 3751.102 |
| ENSMUSG00000015745 | -0.045267139 | 0.600708 | 0.738618 | 1030.2 | 1000.651 | 1174.255 | 1174.893 | 1166.233 | 1092.792 | 1080.968 | 1185.474 |
| ENSMUSG00000015747 | -0.256569607 | 0.026959 | 0.087692 | 404.4339 | 379.1436 | 486.9267 | 404.5704 | 460.2525 | 526.2832 | 480.216 | 539.4452 |
| ENSMUSG00000015748 | 0.002707153 | 0.980312 | 0.988885 | 1100.623 | 1210.62 | 953.1507 | 1048.95 | 1183.785 | 1145.308 | 1002.861 | 969.0437 |
| ENSMUSG00000015749 | -0.046366992 | 0.38685 | 0.55432 | 5952.824 | 6273.867 | 5891.151 | 6372.2 | 6627.832 | 6171.257 | 6167.593 | 6319.992 |
| ENSMUSG00000015750 | -0.274752605 | 0.002421 | 0.015053 | 1760.595 | 1704.946 | 1920.379 | 1710.582 | 2175.473 | 1860.428 | 2317.187 | 2232.825 |
| ENSMUSG00000015755 | 0.114955236 | 0.141482 | 0.278289 | 3634.874 | 3541.873 | 3812.603 | 3812.796 | 3145.709 | 3834.828 | 3384.655 | 3308.452 |
| ENSMUSG00000015757 | 9.92E-05 | 0.999084 | 0.999528 | 1317.931 | 1421.789 | 1159.349 | 1290.485 | 1293.973 | 1385.544 | 1283.469 | 1221.365 |
| ENSMUSG00000015759 | -0.045461606 | 0.572354 | 0.715045 | 1700.232 | 1701.347 | 1756.414 | 1921.063 | 1714.246 | 2016.86 | 1801.292 | 1778.211 |
| ENSMUSG00000015766 | -0.92966305 | 0.000485 | 0.004472 | 647.899 | 499.1258 | 864.5433 | 456.3278 | 1049.22 | 753.1101 | 1216.933 | 1683.591 |
| ENSMUSG00000015776 | -0.187317879 | 0.027611 | 0.089135 | 1477.894 | 1574.166 | 1811.897 | 1634.672 | 1969.725 | 1746.456 | 1829.257 | 1857.606 |
| ENSMUSG00000015790 | 0.09127919 | 0.439643 | 0.602962 | 945.6911 | 991.0526 | 1031.821 | 1230.101 | 1119.428 | 887.195 | 999.0036 | 936.4159 |
| ENSMUSG00000015804 | -0.323673612 | 0.000471 | 0.004373 | 1401.434 | 1534.572 | 1633.854 | 1553.585 | 2023.356 | 1669.357 | 1915.078 | 2056.635 |
| ENSMUSG00000015806 | -0.591478733 | 0.00015 | 0.001828 | 2100.641 | 2244.866 | 2400.681 | 2441.224 | 3658.618 | 2622.477 | 3125.261 | 4438.459 |
| ENSMUSG00000015812 | 0.703316908 | 0.077216 | 0.184574 | 24.1453 | 35.99465 | 28.98373 | 43.9938 | 12.67645 | 15.64324 | 23.14294 | 30.45255 |
| ENSMUSG00000015829 | 0.424410853 | 0.205522 | 0.361948 | 22.1332 | 22.79661 | 26.49941 | 26.74133 | 15.60178 | 21.23011 | 18.32149 | 18.48905 |
| ENSMUSG00000015837 | -0.103700682 | 0.3339 | 0.502081 | 7108.78 | 6450.241 | 7392.508 | 6561.977 | 7223.625 | 6129.914 | 7938.993 | 8272.218 |
| ENSMUSG00000015839 | 0.000560856 | 0.996113 | 0.997718 | 1961.806 | 1779.335 | 2136.515 | 1966.782 | 1924.87 | 1617.958 | 2056.829 | 2243.7 |
| ENSMUSG00000015843 | 0.559160284 | 0.10126 | 0.222987 | 23.13925 | 34.79482 | 28.98373 | 27.60395 | 22.42756 | 23.46486 | 16.39292 | 15.22628 |
| ENSMUSG00000015846 | -0.011153019 | 0.920982 | 0.952855 | 1900.437 | 1922.114 | 2573.755 | 2033.204 | 2120.867 | 2001.217 | 2184.115 | 2192.584 |
| ENSMUSG00000015850 | 0.834068606 | 5.01E-05 | 0.000794 | 356.1432 | 351.5477 | 426.4749 | 420.0976 | 217.4498 | 292.752 | 217.9294 | 144.6496 |
| ENSMUSG00000015852 | 0.290793386 | 0.338528 | 0.507092 | 94.56911 | 151.1775 | 231.0418 | 134.5693 | 167.7191 | 111.7374 | 92.57176 | 128.3357 |
| ENSMUSG00000015869 | -7.60E-05 | 0.999075 | 0.999528 | 2365.234 | 2514.826 | 2606.052 | 2689.66 | 2480.683 | 2588.956 | 2660.474 | 2449.255 |
| ENSMUSG00000015879 | 0.383992368 | 0.004087 | 0.022161 | 272.6407 | 295.1561 | 288.1811 | 251.0234 | 202.8232 | 252.5265 | 200.5721 | 192.5036 |

| | | | | | | | | | | | |
|---|---|---|---|---|---|---|---|---|---|---|---|
| ENSMUSG00000015880 | 0.076386411 | 0.591418 | 0.731386 | 1055.351 | 987.4531 | 769.3111 | 971.3141 | 951.7086 | 1051.449 | 823.5029 | 760.2262 |
| ENSMUSG00000015882 | 0.226692824 | 0.111045 | 0.237177 | 936.6366 | 1005.45 | 981.3064 | 1044.637 | 734.2588 | 1103.966 | 751.1813 | 803.7298 |
| ENSMUSG00000015889 | -0.104365139 | 0.119288 | 0.248293 | 2000.036 | 1985.705 | 2043.767 | 2180.712 | 2285.661 | 2049.264 | 2301.758 | 2190.408 |
| ENSMUSG00000015890 | -1.194369962 | 0.018188 | 0.066063 | 34.20585 | 7.198929 | 19.04645 | 22.42821 | 31.20356 | 30.1691 | 51.10733 | 78.30656 |
| ENSMUSG00000015932 | -0.006232069 | 0.932609 | 0.960099 | 8230.531 | 8654.313 | 9670.629 | 9539.753 | 8773.076 | 8730.044 | 9379.641 | 9373.948 |
| ENSMUSG00000015937 | -0.02519508 | 0.621824 | 0.75339 | 5594.668 | 6005.107 | 5838.98 | 5721.782 | 5993.034 | 6120.975 | 5821.414 | 5630.459 |
| ENSMUSG00000015942 | 0.372720811 | 0.004193 | 0.022606 | 947.7032 | 944.2595 | 889.3865 | 962.6878 | 648.449 | 911.7773 | 704.8954 | 627.5401 |
| ENSMUSG00000015943 | -0.490294815 | 0.000904 | 0.007221 | 283.7073 | 351.5477 | 351.9453 | 313.1323 | 522.6597 | 359.7945 | 452.2516 | 490.5036 |
| ENSMUSG00000015944 | -0.148038795 | 0.512931 | 0.668016 | 1944.703 | 1081.039 | 1229.738 | 1491.476 | 1163.308 | 1509.572 | 2187.972 | 1507.401 |
| ENSMUSG00000015947 | 0.519283588 | 0.135183 | 0.269373 | 75.45408 | 112.7832 | 175.5586 | 64.69676 | 89.71024 | 64.8077 | 58.82164 | 85.9197 |
| ENSMUSG00000015950 | 0.403619517 | 0.091598 | 0.207631 | 172.0353 | 171.5745 | 178.871 | 187.1893 | 157.968 | 150.8455 | 153.322 | 73.95619 |
| ENSMUSG00000015957 | -0.021533165 | 0.869351 | 0.921582 | 2458.797 | 3630.66 | 3412.627 | 3123.56 | 3604.987 | 3171.108 | 3013.404 | 3024.591 |
| ENSMUSG00000015961 | -0.033801481 | 0.578062 | 0.719775 | 3149.956 | 3439.888 | 3244.522 | 3577.3 | 3342.682 | 3506.32 | 3454.084 | 3427 |
| ENSMUSG00000015962 | -3.782088492 | 6.25E-06 | 0.00016 | 3.018163 | 1.199822 | 12.4216 | 0 | 37.05423 | 27.93435 | 101.2504 | 68.51824 |
| ENSMUSG00000015966 | 0.394820771 | 0.036431 | 0.109396 | 165.999 | 159.5763 | 115.1068 | 152.6844 | 101.4116 | 135.2023 | 101.2504 | 113.1095 |
| ENSMUSG00000015968 | 0.603624449 | 9.17E-05 | 0.001252 | 848.1038 | 692.297 | 729.5619 | 755.6582 | 402.721 | 639.138 | 499.5018 | 451.3503 |
| ENSMUSG00000015970 | -0.52924795 | 0.000799 | 0.006572 | 609.6689 | 488.3274 | 640.1264 | 603.8364 | 887.3513 | 643.6075 | 829.2887 | 1022.336 |
| ENSMUSG00000015971 | -0.013236429 | 0.878479 | 0.927104 | 1132.817 | 1178.225 | 1119.6 | 1067.065 | 1165.258 | 987.7587 | 1189.933 | 1192 |
| ENSMUSG00000015980 | -0.061326594 | 0.764855 | 0.853994 | 67.40564 | 88.78679 | 68.73285 | 75.04824 | 79.95913 | 82.68568 | 62.6788 | 87.00729 |
| ENSMUSG00000015981 | -0.537130707 | 0.020827 | 0.072513 | 69.41775 | 65.99018 | 86.9512 | 68.14726 | 91.66046 | 80.45093 | 136.9291 | 113.1095 |
| ENSMUSG00000015994 | -0.070848067 | 0.238035 | 0.399162 | 2257.586 | 2326.454 | 2350.167 | 2411.895 | 2626.95 | 2432.523 | 2414.58 | 2342.671 |
| ENSMUSG00000016018 | -0.151654363 | 0.032551 | 0.100245 | 2036.254 | 1870.522 | 1964.269 | 2040.967 | 2154.996 | 2020.212 | 2287.294 | 2328.533 |
| ENSMUSG00000016024 | 0.662782767 | 5.39E-08 | 3.49E-06 | 1707.274 | 1300.607 | 1329.111 | 1533.745 | 794.7157 | 958.707 | 991.2893 | 964.6933 |
| ENSMUSG00000016028 | 0.034071289 | 0.773341 | 0.859443 | 940.6608 | 863.8715 | 1120.428 | 916.9688 | 844.4464 | 907.3078 | 926.6819 | 1077.803 |
| ENSMUSG00000016087 | 0.1120531 | 0.312765 | 0.481048 | 679.0867 | 509.9242 | 656.6886 | 591.7597 | 548.9877 | 593.3256 | 554.4663 | 562.2846 |
| ENSMUSG00000016128 | 0.576156816 | 3.24E-08 | 2.28E-06 | 849.1099 | 943.0597 | 756.8895 | 959.2373 | 589.9423 | 627.9642 | 562.1806 | 572.0729 |
| ENSMUSG00000016150 | 1.010314642 | 0.006548 | 0.031473 | 38.23007 | 31.19536 | 20.70267 | 34.50494 | 14.62667 | 14.52586 | 19.28578 | 13.05109 |
| ENSMUSG00000016179 | 0.438817807 | 0.00868 | 0.038465 | 235.4167 | 235.165 | 226.9012 | 251.0234 | 166.744 | 155.315 | 228.5365 | 147.9124 |
| ENSMUSG00000016181 | 0.14744217 | 0.248938 | 0.412369 | 780.6982 | 698.2961 | 622.7362 | 632.303 | 729.3833 | 550.8654 | 569.8949 | 615.5766 |
| ENSMUSG00000016194 | -1.559891389 | 5.07E-05 | 0.0008 | 42.25428 | 27.5959 | 27.32752 | 25.8787 | 118.9636 | 39.10809 | 89.67889 | 114.1971 |
| ENSMUSG00000016200 | -0.115641854 | 0.534198 | 0.685327 | 251.5136 | 217.1677 | 178.0429 | 169.0742 | 240.8525 | 220.1227 | 174.5363 | 246.8832 |
| ENSMUSG00000016206 | 0.061145911 | 0.814819 | 0.887383 | 40.24217 | 47.99286 | 52.99882 | 40.5433 | 36.07912 | 45.81234 | 38.57157 | 54.37955 |
| ENSMUSG00000016239 | 0.129377019 | 0.746403 | 0.841465 | 46.2785 | 53.99197 | 60.45178 | 43.9938 | 33.15378 | 86.03781 | 43.39301 | 25.0146 |
| ENSMUSG00000016252 | -0.308761532 | 0.004445 | 0.023594 | 2426.603 | 2825.58 | 2987.809 | 2484.356 | 3304.652 | 2867.182 | 3406.834 | 3704.335 |
| ENSMUSG00000016253 | -0.180189605 | 0.005998 | 0.02947 | 1637.856 | 1737.342 | 1641.307 | 1544.096 | 1838.085 | 1758.747 | 1896.757 | 1933.737 |
| ENSMUSG00000016256 | 0.104078517 | 0.293124 | 0.460495 | 2320.967 | 2508.827 | 2689.69 | 2526.624 | 2468.982 | 1982.222 | 2324.901 | 2571.065 |
| ENSMUSG00000016257 | -0.289566612 | 0.062144 | 0.15892 | 1291.774 | 1088.238 | 1351.47 | 1282.721 | 1501.671 | 1138.604 | 1597.827 | 1892.408 |
| ENSMUSG00000016262 | 0.934285763 | 1.30E-05 | 0.000281 | 4089.611 | 4484.933 | 3071.447 | 4872.96 | 1847.836 | 2845.952 | 2473.402 | 1476.949 |
| ENSMUSG00000016308 | -0.073264583 | 0.427073 | 0.592191 | 676.0685 | 655.1026 | 746.9522 | 798.7894 | 727.433 | 795.5704 | 766.6099 | 742.8247 |
| ENSMUSG00000016319 | -1.285689171 | 2.48E-07 | 1.20E-05 | 4903.509 | 5234.821 | 7362.696 | 4583.119 | 13273.22 | 7408.19 | 15104.63 | 18056.19 |
| ENSMUSG00000016344 | -0.025511052 | 0.784296 | 0.867232 | 1608.681 | 1971.307 | 1904.645 | 1876.206 | 2072.112 | 1857.076 | 1806.114 | 1755.372 |
| ENSMUSG00000016346 | 1.083498592 | 0.039053 | 0.114497 | 79.47829 | 27.5959 | 25.67131 | 113.8663 | 33.15378 | 36.87335 | 27.0001 | 19.57664 |
| ENSMUSG00000016349 | 0.85457236 | 0.020018 | 0.07042 | 158.9566 | 83.98751 | 87.7793 | 260.5123 | 85.8098 | 90.5073 | 76.17884 | 75.04379 |
| ENSMUSG00000016356 | 0.054487805 | 0.855616 | 0.913423 | 62.37537 | 34.79482 | 49.6864 | 51.75741 | 36.07912 | 60.3382 | 38.57157 | 57.64233 |
| ENSMUSG00000016382 | 0.181677025 | 0.04424 | 0.125039 | 5273.737 | 6281.066 | 6037.725 | 5798.555 | 4553.77 | 5636.035 | 5205.233 | 5229.138 |
| ENSMUSG00000016386 | 0.46327306 | 0.018023 | 0.065724 | 789.7527 | 783.4835 | 693.9534 | 730.6421 | 404.6712 | 786.6314 | 486.0017 | 498.1167 |
| ENSMUSG00000016409 | 0.373777489 | 0.01451 | 0.056278 | 859.1704 | 869.8706 | 746.9522 | 796.2015 | 554.8383 | 810.0962 | 648.9666 | 511.1678 |
| ENSMUSG00000016427 | -0.480969927 | 0.003831 | 0.021163 | 755.5468 | 668.3006 | 856.2623 | 728.0542 | 953.6589 | 793.3356 | 1124.361 | 1329.036 |
| ENSMUSG00000016458 | 0.042672701 | 0.763812 | 0.853273 | 4824.031 | 4978.06 | 4018.801 | 4957.497 | 4616.177 | 5599.162 | 4455.016 | 3559.686 |
| ENSMUSG00000016477 | 0.159211309 | 0.1539 | 0.29538 | 1258.574 | 1207.02 | 1024.368 | 1048.088 | 993.6384 | 1145.308 | 1025.039 | 897.2626 |
| ENSMUSG00000016481 | 0.030351945 | 0.687858 | 0.801763 | 1442.682 | 1505.776 | 1483.967 | 1659.688 | 1510.447 | 1382.192 | 1552.506 | 1520.452 |
| ENSMUSG00000016487 | 0.262532375 | 0.001754 | 0.011909 | 4868.297 | 5254.018 | 4669.693 | 4977.338 | 4126.671 | 4616.99 | 4032.657 | 3702.16 |
| ENSMUSG00000016493 | 0.628769898 | 0.000583 | 0.005152 | 179.0777 | 215.9679 | 183.0116 | 202.7165 | 119.9387 | 165.3714 | 118.6076 | 101.146 |
| ENSMUSG00000016494 | -0.136617226 | 0.243854 | 0.406473 | 1933.636 | 2618.011 | 2617.645 | 2631.864 | 2732.262 | 2727.51 | 2858.153 | 2456.868 |
| ENSMUSG00000016495 | 0.013590496 | 0.883151 | 0.930174 | 671.0383 | 635.9054 | 660.001 | 726.329 | 626.9966 | 680.4808 | 736.7169 | 625.3649 |
| ENSMUSG00000016496 | 0.641993057 | 0.132055 | 0.265176 | 18.10898 | 28.79572 | 31.46805 | 18.97772 | 13.65156 | 12.29112 | 24.10723 | 11.9635 |
| ENSMUSG00000016503 | -0.219723507 | 0.057615 | 0.150422 | 733.4136 | 941.8599 | 853.7779 | 836.7448 | 1064.822 | 889.4298 | 1076.147 | 885.2991 |
| ENSMUSG00000016510 | -0.242196858 | 0.037146 | 0.110737 | 474.8577 | 469.1302 | 430.6154 | 397.6694 | 572.3903 | 444.7149 | 532.2876 | 542.708 |
| ENSMUSG00000016520 | -0.326108273 | 0.035861 | 0.108075 | 493.9727 | 436.735 | 549.0347 | 450.2895 | 609.4446 | 457.006 | 636.4308 | 717.8101 |
| ENSMUSG00000016526 | 0.114137084 | 0.44149 | 0.604865 | 206.2411 | 262.7609 | 224.4169 | 251.0234 | 235.9769 | 207.8316 | 243.9652 | 182.7153 |
| ENSMUSG00000016528 | -0.177974448 | 0.00689 | 0.03263 | 3693.226 | 3798.635 | 4153.783 | 3761.038 | 4391.901 | 4076.181 | 4566.873 | 4396.043 |
| ENSMUSG00000016534 | -0.279007159 | 0.014204 | 0.055412 | 5529.275 | 4998.457 | 6342.469 | 5649.321 | 6736.069 | 5908.674 | 6699.881 | 7984.006 |
| ENSMUSG00000016541 | -0.090560158 | 0.238176 | 0.399326 | 3951.782 | 4108.189 | 4614.21 | 4402.83 | 4674.684 | 4177.862 | 4560.123 | 4774.525 |
| ENSMUSG00000016552 | 0.252649399 | 0.133131 | 0.266493 | 353.1251 | 418.7377 | 283.2125 | 312.2697 | 308.1352 | 330.7427 | 268.0724 | 238.1824 |

| | | | | | | | | | | | |
|---|---|---|---|---|---|---|---|---|---|---|---|
| ENSMUSG00000016554 | -0.116445832 | 0.098285 | 0.218364 | 3570.487 | 3697.85 | 3917.772 | 3876.63 | 4305.116 | 3739.851 | 3999.871 | 4286.196 |
| ENSMUSG00000016559 | 0.174233359 | 0.065157 | 0.164392 | 39158.65 | 46101.94 | 47055.5 | 50267.66 | 40541.23 | 44255.84 | 39291.89 | 37726.36 |
| ENSMUSG00000016619 | -0.211831058 | 0.028228 | 0.090603 | 1403.446 | 1590.963 | 1300.956 | 1310.325 | 1674.266 | 1470.464 | 1592.041 | 1748.846 |
| ENSMUSG00000016624 | -0.503485817 | 0.035641 | 0.107552 | 63.38142 | 64.79036 | 66.24853 | 50.03216 | 79.95913 | 62.57295 | 94.50034 | 108.7591 |
| ENSMUSG00000016637 | 0.071549 | 0.60871 | 0.744176 | 481.9 | 659.9018 | 573.8779 | 601.2486 | 557.7637 | 634.6685 | 553.502 | 457.8758 |
| ENSMUSG00000016664 | -0.446076718 | 0.001029 | 0.007993 | 2349.137 | 2206.472 | 2793.204 | 2323.045 | 3118.406 | 2662.702 | 3435.762 | 3962.094 |
| ENSMUSG00000016756 | -1.560195727 | 0.000411 | 0.003947 | 19.11503 | 37.19447 | 24.01509 | 24.15346 | 48.75557 | 42.46022 | 75.21455 | 140.2993 |
| ENSMUSG00000016757 | -0.285710245 | 0.000822 | 0.006712 | 770.6376 | 815.8786 | 769.3111 | 740.9936 | 907.8286 | 858.1433 | 976.8249 | 1028.861 |
| ENSMUSG00000016758 | 0.282492029 | 0.065467 | 0.164684 | 176.0595 | 178.7734 | 163.137 | 173.3873 | 151.1423 | 149.7281 | 155.2506 | 110.9343 |
| ENSMUSG00000016763 | -0.124461455 | 0.662734 | 0.782975 | 190.1443 | 184.7725 | 396.6631 | 188.0519 | 273.0312 | 210.0663 | 235.2866 | 329.5401 |
| ENSMUSG00000016831 | 0.011382864 | 0.845568 | 0.906998 | 1788.765 | 1806.931 | 1652.901 | 1767.516 | 1748.375 | 1733.047 | 1683.649 | 1792.35 |
| ENSMUSG00000016833 | -0.100208029 | 0.246205 | 0.409107 | 903.4368 | 887.8679 | 936.5886 | 891.9527 | 1019.966 | 842.5001 | 1018.289 | 998.4086 |
| ENSMUSG00000016918 | 0.537197454 | 0.092969 | 0.209658 | 1339.058 | 1769.737 | 1142.787 | 1623.457 | 885.4011 | 1843.667 | 761.7884 | 557.9342 |
| ENSMUSG00000016921 | 0.142658409 | 0.089112 | 0.203774 | 12191.37 | 12168.59 | 11085.86 | 12502.86 | 10907.6 | 12259.83 | 10268.72 | 9998.225 |
| ENSMUSG00000016933 | 0.116917192 | 0.169262 | 0.315546 | 4086.593 | 3622.261 | 3622.967 | 4255.322 | 3447.018 | 3794.602 | 3780.978 | 3353.043 |
| ENSMUSG00000016940 | 0.046950002 | 0.632935 | 0.761254 | 1366.222 | 1131.432 | 1079.851 | 1139.526 | 1122.353 | 1203.412 | 1177.397 | 1061.489 |
| ENSMUSG00000016946 | -0.174653879 | 0.066777 | 0.166804 | 1303.846 | 1523.773 | 1615.636 | 1371.571 | 1719.121 | 1497.281 | 1717.399 | 1628.124 |
| ENSMUSG00000016984 | 0.035265961 | 0.804867 | 0.880483 | 641.8627 | 897.4665 | 775.9359 | 779.8116 | 811.2926 | 869.317 | 708.7525 | 629.7152 |
| ENSMUSG00000016995 | 1.929493761 | 1.30E-05 | 0.000281 | 33.19979 | 73.18911 | 53.82693 | 73.323 | 16.57689 | 25.6996 | 11.57147 | 7.613138 |
| ENSMUSG00000017002 | 2.368217345 | 3.67E-06 | 0.000107 | 178.0716 | 387.5424 | 219.4483 | 396.8068 | 78.98402 | 71.51194 | 69.42882 | 8.700729 |
| ENSMUSG00000017009 | -0.350520265 | 0.00031 | 0.003189 | 3432.657 | 3382.297 | 4078.425 | 3697.204 | 4301.216 | 4347.703 | 4712.481 | 5247.627 |
| ENSMUSG00000017057 | -0.136531813 | 0.197248 | 0.351608 | 632.8082 | 724.6922 | 794.1543 | 763.4218 | 755.7113 | 749.758 | 810.9672 | 890.7371 |
| ENSMUSG00000017119 | 0.009193388 | 0.899078 | 0.939124 | 4191.222 | 4019.402 | 3629.591 | 4115.577 | 3804.884 | 3814.715 | 4250.587 | 3980.583 |
| ENSMUSG00000017132 | -0.05934752 | 0.496154 | 0.652768 | 1281.713 | 1078.64 | 1088.96 | 1171.443 | 1281.296 | 1112.905 | 1208.254 | 1211.576 |
| ENSMUSG00000017144 | 0.200499962 | 0.030228 | 0.094975 | 1037.242 | 1075.04 | 949.0102 | 942.8475 | 875.65 | 955.3549 | 871.7174 | 778.7152 |
| ENSMUSG00000017146 | -0.106610358 | 0.413975 | 0.579867 | 580.4934 | 754.6877 | 548.2066 | 626.2647 | 742.0597 | 718.4715 | 612.3236 | 625.3649 |
| ENSMUSG00000017167 | 0.196971478 | 0.283598 | 0.450166 | 112.6781 | 113.983 | 122.5598 | 103.5148 | 106.2871 | 119.559 | 85.82174 | 83.74451 |
| ENSMUSG00000017176 | -0.152339951 | 0.1374 | 0.272519 | 1471.858 | 1770.937 | 1549.388 | 1555.31 | 1995.078 | 1545.328 | 1775.256 | 1733.62 |
| ENSMUSG00000017188 | -0.287213315 | 0.00519 | 0.026403 | 979.8969 | 1065.442 | 1163.49 | 1063.615 | 1362.23 | 1149.778 | 1237.183 | 1466.073 |
| ENSMUSG00000017195 | -0.416309912 | 0.413123 | 0.57898 | 11.0666 | 9.598572 | 9.109173 | 12.93935 | 18.52711 | 5.58687 | 16.39292 | 16.31387 |
| ENSMUSG00000017210 | -0.109937413 | 0.095896 | 0.214476 | 1797.819 | 1748.14 | 1869.037 | 1695.055 | 2037.008 | 1830.259 | 1857.221 | 1948.963 |
| ENSMUSG00000017221 | -0.535382336 | 1.69E-05 | 0.000342 | 3842.122 | 3408.693 | 4240.734 | 3652.348 | 5417.718 | 4501.9 | 5511.877 | 6520.109 |
| ENSMUSG00000017264 | -0.185884996 | 0.004034 | 0.02196 | 1989.976 | 1965.308 | 2066.954 | 1896.909 | 2268.109 | 2111.837 | 2228.472 | 2400.314 |
| ENSMUSG00000017286 | -0.185725516 | 0.0136 | 0.053642 | 1662.002 | 1667.752 | 1730.743 | 1779.592 | 2126.718 | 1796.738 | 1994.15 | 1861.956 |
| ENSMUSG00000017288 | -0.389155084 | 0.000648 | 0.005596 | 891.3642 | 883.0686 | 984.6188 | 805.6903 | 1175.984 | 979.9371 | 1197.647 | 1314.898 |
| ENSMUSG00000017291 | 0.193756119 | 0.103214 | 0.225862 | 2420.567 | 2362.449 | 2260.731 | 2291.128 | 1840.035 | 2544.261 | 1813.828 | 1965.277 |
| ENSMUSG00000017299 | -0.084883726 | 0.202998 | 0.358726 | 1482.924 | 1473.381 | 1560.153 | 1532.882 | 1716.196 | 1496.164 | 1624.827 | 1579.182 |
| ENSMUSG00000017300 | 0.499312416 | 0.380669 | 0.548577 | 46.2785 | 11.99822 | 24.8432 | 21.56559 | 10.72622 | 26.81698 | 27.96439 | 8.700729 |
| ENSMUSG00000017307 | -0.458307659 | 0.000853 | 0.006913 | 300.8103 | 319.1525 | 334.5551 | 307.9566 | 405.6463 | 377.6724 | 412.7158 | 539.4452 |
| ENSMUSG00000017309 | -0.435967393 | 0.119589 | 0.248522 | 34.20585 | 49.19268 | 32.29616 | 57.79577 | 54.60623 | 67.04245 | 62.6788 | 50.02919 |
| ENSMUSG00000017310 | 0.240943631 | 0.538099 | 0.688091 | 23.13925 | 20.39697 | 19.04645 | 31.91707 | 13.65156 | 29.05173 | 18.32149 | 19.57664 |
| ENSMUSG00000017314 | -0.34600768 | 0.083922 | 0.195292 | 263.5862 | 218.3675 | 295.6341 | 272.589 | 301.3094 | 240.2354 | 356.787 | 438.2992 |
| ENSMUSG00000017344 | 0.299597031 | 0.113102 | 0.240016 | 196.1806 | 169.1748 | 152.3716 | 200.9913 | 147.2418 | 186.6015 | 132.1076 | 118.5474 |
| ENSMUSG00000017376 | 0.032763932 | 0.771887 | 0.858471 | 946.6971 | 825.4772 | 820.6537 | 793.6136 | 703.0553 | 930.7726 | 811.9315 | 865.7225 |
| ENSMUSG00000017386 | -0.275297485 | 0.03783 | 0.112177 | 1978.909 | 2004.902 | 2359.276 | 2210.041 | 2732.262 | 2015.743 | 2579.474 | 3025.678 |
| ENSMUSG00000017390 | -0.796079796 | 6.63E-06 | 0.000168 | 197.1867 | 199.1704 | 245.1196 | 209.6175 | 329.5876 | 283.813 | 422.3587 | 443.7372 |
| ENSMUSG00000017400 | 0.954190322 | 6.11E-09 | 5.58E-07 | 234.4107 | 183.5727 | 208.6829 | 207.0296 | 119.9387 | 109.5027 | 115.7147 | 84.8321 |
| ENSMUSG00000017404 | -0.186159455 | 0.029298 | 0.092968 | 20543.63 | 24136.81 | 24934.29 | 23463.36 | 28442.05 | 24150.92 | 27158.24 | 26145.69 |
| ENSMUSG00000017405 | 0.110933567 | 0.144757 | 0.282892 | 756.5529 | 716.2935 | 693.1253 | 740.131 | 669.9015 | 630.199 | 703.9311 | 685.1824 |
| ENSMUSG00000017412 | 0.748228138 | 0.035233 | 0.106584 | 716.3107 | 763.0865 | 549.0347 | 735.8178 | 312.0356 | 817.9178 | 292.1796 | 224.0438 |
| ENSMUSG00000017417 | 1.335890433 | 3.26E-09 | 3.46E-07 | 149.9021 | 166.7752 | 147.403 | 106.9653 | 42.9049 | 56.98608 | 59.78593 | 66.34306 |
| ENSMUSG00000017418 | 0.248478384 | 0.132985 | 0.266363 | 1293.786 | 1313.805 | 1093.929 | 1328.44 | 1019.966 | 1417.948 | 963.3249 | 833.0948 |
| ENSMUSG00000017421 | 0.09770708 | 0.155662 | 0.297649 | 6377.379 | 6378.251 | 5738.779 | 6661.179 | 5913.075 | 6221.539 | 5788.628 | 5583.693 |
| ENSMUSG00000017428 | -0.032244323 | 0.628675 | 0.758208 | 1719.347 | 1642.556 | 1641.307 | 1594.128 | 1646.963 | 1584.436 | 1702.935 | 1810.839 |
| ENSMUSG00000017446 | 0.409056312 | 0.001495 | 0.010537 | 516.1059 | 513.5236 | 518.3948 | 447.7016 | 436.8499 | 313.9821 | 397.2871 | 352.3795 |
| ENSMUSG00000017453 | -1.659787862 | 1.50E-06 | 5.18E-05 | 113.6841 | 65.99018 | 173.0743 | 129.3935 | 360.7912 | 201.1273 | 405.9657 | 560.1094 |
| ENSMUSG00000017466 | 0.295987391 | 0.002368 | 0.014796 | 5289.834 | 6421.445 | 6304.376 | 5945.201 | 4726.365 | 5546.645 | 4792.517 | 4451.51 |
| ENSMUSG00000017478 | -0.274973295 | 0.001414 | 0.010154 | 1689.165 | 1542.97 | 1509.638 | 1572.563 | 1900.492 | 1678.296 | 2049.114 | 2009.868 |
| ENSMUSG00000017485 | 0.124043824 | 0.133619 | 0.267122 | 5284.804 | 5819.134 | 5010.045 | 5400.886 | 4699.061 | 5542.176 | 4708.624 | 4790.839 |
| ENSMUSG00000017491 | 0.141389059 | 0.135919 | 0.270344 | 1228.392 | 1590.963 | 1519.576 | 1447.482 | 1323.226 | 1269.337 | 1293.112 | 1359.489 |
| ENSMUSG00000017493 | -0.08631993 | 0.304199 | 0.472636 | 10694.36 | 11005.96 | 12981.4 | 10942.38 | 11320.07 | 12069.87 | 12123.04 | 12929.28 |
| ENSMUSG00000017499 | -0.285729385 | 0.163805 | 0.30871 | 312.8829 | 512.3238 | 397.4912 | 277.7648 | 524.6099 | 415.6632 | 395.3586 | 490.5036 |
| ENSMUSG00000017548 | 0.135011838 | 0.202563 | 0.358173 | 2506.081 | 2627.609 | 2371.697 | 2507.646 | 2070.161 | 2704.045 | 2321.044 | 2022.919 |
| ENSMUSG00000017550 | 0.05382153 | 0.685169 | 0.79974 | 659.9717 | 676.6993 | 561.4563 | 611.6001 | 641.6232 | 715.1194 | 551.5734 | 507.905 |

| | | | | | | | | | | | |
|---|---|---|---|---|---|---|---|---|---|---|---|
| ENSMUSG00000017561 | 0.068858616 | 0.454846 | 0.61698 | 683.1109 | 835.0758 | 707.2031 | 746.1693 | 715.7317 | 738.5843 | 712.6097 | 662.343 |
| ENSMUSG00000017607 | 0.531309041 | 0.122574 | 0.252114 | 23.13925 | 31.19536 | 30.63995 | 25.01608 | 17.552 | 17.87799 | 25.07152 | 15.22628 |
| ENSMUSG00000017615 | -0.211125558 | 6.16E-05 | 0.000936 | 2621.778 | 2747.591 | 2550.568 | 2503.333 | 3087.202 | 2983.389 | 2966.153 | 3020.24 |
| ENSMUSG00000017631 | 0.212663078 | 0.003212 | 0.018674 | 3392.415 | 3217.921 | 3224.647 | 3685.99 | 2930.209 | 3088.422 | 2911.189 | 2740.73 |
| ENSMUSG00000017639 | -0.541575119 | 0.000324 | 0.003286 | 741.4621 | 639.5049 | 844.6688 | 684.0604 | 1003.39 | 863.7302 | 1053.968 | 1317.073 |
| ENSMUSG00000017652 | -0.30904825 | 0.04181 | 0.120362 | 169.0171 | 178.7734 | 219.4483 | 174.2499 | 247.6783 | 193.3057 | 227.5722 | 251.2335 |
| ENSMUSG00000017664 | -0.296237196 | 0.001519 | 0.010649 | 780.6982 | 724.6922 | 732.0463 | 730.6421 | 881.5006 | 839.1479 | 879.4317 | 1045.175 |
| ENSMUSG00000017667 | 0.153064383 | 0.234659 | 0.395501 | 995.9938 | 977.8546 | 794.1543 | 897.9911 | 849.3219 | 947.5332 | 822.5387 | 675.3941 |
| ENSMUSG00000017670 | -0.215937695 | 0.021238 | 0.073568 | 1415.518 | 1478.18 | 1586.652 | 1491.476 | 1672.316 | 1537.507 | 1755.006 | 1972.89 |
| ENSMUSG00000017677 | 0.281168728 | 0.000142 | 0.001749 | 8637.983 | 7656.061 | 8173.412 | 7767.925 | 6390.88 | 7186.95 | 6164.701 | 6788.744 |
| ENSMUSG00000017679 | -0.002810023 | 0.973828 | 0.985583 | 883.3157 | 943.0597 | 832.2472 | 826.3933 | 840.5459 | 941.9464 | 811.9315 | 895.0875 |
| ENSMUSG00000017686 | -0.283015924 | 0.000225 | 0.002484 | 1335.034 | 1348.599 | 1319.174 | 1349.143 | 1560.178 | 1496.164 | 1651.827 | 1803.226 |
| ENSMUSG00000017688 | -1.670519181 | 0.028767 | 0.09173 | 10.06054 | 0 | 14.90592 | 3.450494 | 13.65156 | 23.46486 | 20.25007 | 34.80291 |
| ENSMUSG00000017692 | 0.695950065 | 8.45E-05 | 0.001184 | 419.5247 | 493.1267 | 420.6782 | 514.9862 | 330.5627 | 349.7381 | 232.3937 | 228.3941 |
| ENSMUSG00000017697 | -0.220537458 | 0.225019 | 0.384774 | 118.7144 | 101.9848 | 138.2938 | 136.2945 | 139.4409 | 125.1459 | 137.8934 | 177.2773 |
| ENSMUSG00000017707 | -0.233327888 | 0.125479 | 0.256017 | 10248.68 | 7232.524 | 10235.4 | 9336.174 | 10013.42 | 9241.801 | 13543.44 | 10760.63 |
| ENSMUSG00000017713 | -0.310457689 | 0.080014 | 0.188964 | 147.89 | 122.3818 | 158.1684 | 107.8279 | 148.2169 | 163.1366 | 189.0007 | 165.3138 |
| ENSMUSG00000017715 | -0.184787886 | 0.017731 | 0.064918 | 782.7103 | 767.8858 | 790.8418 | 724.6037 | 845.4215 | 830.209 | 861.1102 | 948.3794 |
| ENSMUSG00000017716 | -0.051002561 | 0.637943 | 0.765283 | 1533.227 | 1698.947 | 1499.701 | 1701.094 | 1759.101 | 1891.714 | 1599.756 | 1412.781 |
| ENSMUSG00000017718 | -0.14786778 | 0.46501 | 0.625928 | 109.6599 | 92.38626 | 96.88848 | 106.1027 | 140.416 | 82.68568 | 109.929 | 115.2847 |
| ENSMUSG00000017721 | -0.210444378 | 0.065509 | 0.164695 | 2153.962 | 2171.677 | 2809.766 | 2299.754 | 2694.233 | 2400.12 | 2919.868 | 2906.043 |
| ENSMUSG00000017723 | -0.385757725 | 0.008684 | 0.03847 | 3544.329 | 3013.952 | 3989.818 | 3143.4 | 4159.825 | 3529.785 | 4929.446 | 5271.554 |
| ENSMUSG00000017724 | 0.228039668 | 0.172842 | 0.320504 | 3513.142 | 4489.732 | 3545.124 | 3723.083 | 3599.136 | 4066.124 | 2993.154 | 2378.562 |
| ENSMUSG00000017734 | -0.368928395 | 0.00295 | 0.01755 | 210.2654 | 202.7698 | 214.4796 | 219.969 | 269.1307 | 234.6486 | 303.7511 | 287.124 |
| ENSMUSG00000017737 | -0.366415662 | 0.264181 | 0.428371 | 31.18769 | 26.39607 | 43.88965 | 43.9938 | 55.58134 | 59.22083 | 29.89296 | 44.59123 |
| ENSMUSG00000017740 | 0.100368664 | 0.369372 | 0.538266 | 262.5802 | 283.1579 | 287.353 | 253.6113 | 253.5289 | 263.7003 | 230.4651 | 266.4598 |
| ENSMUSG00000017747 | -0.328133018 | 0.006668 | 0.031865 | 256.5439 | 231.5656 | 244.2915 | 251.0234 | 284.7325 | 275.9914 | 327.8583 | 346.9416 |
| ENSMUSG00000017754 | 0.272911539 | 0.118964 | 0.247866 | 655.9474 | 809.8795 | 820.6537 | 720.2906 | 688.4286 | 422.3674 | 656.6809 | 719.9853 |
| ENSMUSG00000017756 | -0.283410581 | 0.002935 | 0.017499 | 896.3944 | 820.6779 | 899.3238 | 781.5369 | 1071.647 | 949.768 | 972.9678 | 1141.971 |
| ENSMUSG00000017760 | -0.390653441 | 0.000662 | 0.005695 | 3592.62 | 3271.913 | 3858.977 | 3514.328 | 4947.715 | 3815.833 | 4553.373 | 5350.948 |
| ENSMUSG00000017764 | -0.440884969 | 0.000233 | 0.002557 | 207.2472 | 245.9634 | 231.0418 | 215.6559 | 318.8614 | 269.2872 | 303.7511 | 327.3649 |
| ENSMUSG00000017765 | 0.089169141 | 0.376402 | 0.544682 | 1011.085 | 1125.433 | 1197.442 | 1133.487 | 1129.179 | 992.2282 | 931.5033 | 1148.496 |
| ENSMUSG00000017774 | -0.588919053 | 0.001962 | 0.012949 | 2310.907 | 2019.3 | 2585.349 | 2130.68 | 3388.512 | 2177.762 | 3796.406 | 4244.868 |
| ENSMUSG00000017776 | -0.122340308 | 0.013508 | 0.053408 | 4129.853 | 4102.19 | 4205.954 | 4119.027 | 4524.516 | 4246.022 | 4588.088 | 4663.591 |
| ENSMUSG00000017778 | -0.408067976 | 9.20E-05 | 0.001252 | 3709.322 | 3619.862 | 3964.146 | 3792.956 | 4698.086 | 4267.252 | 5236.09 | 5818.612 |
| ENSMUSG00000017781 | -0.124173967 | 0.099125 | 0.219744 | 3389.397 | 3228.72 | 3483.845 | 3558.322 | 3688.846 | 3371.118 | 3769.406 | 4062.153 |
| ENSMUSG00000017801 | -0.135990893 | 0.253119 | 0.41677 | 474.8577 | 454.7324 | 447.1776 | 452.0147 | 548.0126 | 407.8415 | 497.5732 | 554.6715 |
| ENSMUSG00000017802 | -0.118602924 | 0.042326 | 0.121356 | 2509.1 | 2434.438 | 2572.927 | 2418.796 | 2629.875 | 2563.256 | 2836.939 | 2757.043 |
| ENSMUSG00000017817 | -0.13898131 | 0.577586 | 0.719439 | 183.1019 | 265.1606 | 393.3507 | 241.5346 | 316.9112 | 223.4748 | 291.2153 | 362.1678 |
| ENSMUSG00000017830 | 0.508735828 | 0.145721 | 0.284175 | 93.56306 | 128.3809 | 157.3403 | 66.42201 | 79.95913 | 42.46022 | 102.2147 | 88.09488 |
| ENSMUSG00000017831 | -0.102974859 | 0.176985 | 0.325959 | 1876.291 | 1684.549 | 1787.054 | 1908.123 | 1926.82 | 1811.263 | 2112.758 | 1943.525 |
| ENSMUSG00000017832 | -0.040637435 | 0.940837 | 0.965583 | 9.054489 | 19.19714 | 19.04645 | 6.900988 | 10.72622 | 22.34748 | 9.642892 | 13.05109 |
| ENSMUSG00000017837 | -0.287595193 | 0.064586 | 0.163444 | 1084.527 | 1151.829 | 1222.285 | 1094.669 | 1603.083 | 969.8807 | 1503.327 | 1480.211 |
| ENSMUSG00000017843 | 0.104610827 | 0.058158 | 0.151285 | 3252.574 | 3491.481 | 3295.036 | 3407.363 | 3192.514 | 3262.732 | 3002.796 | 3046.343 |
| ENSMUSG00000017858 | -0.033971165 | 0.642438 | 0.768726 | 1210.283 | 1199.822 | 1154.381 | 1237.002 | 1339.803 | 1150.895 | 1231.397 | 1190.912 |
| ENSMUSG00000017861 | -0.175700648 | 0.158835 | 0.301808 | 368.2159 | 464.3309 | 390.0382 | 380.417 | 521.6845 | 414.5458 | 422.3587 | 448.0875 |
| ENSMUSG00000017868 | -2.559350758 | 0.000658 | 0.005671 | 79.47829 | 15.59768 | 159.8246 | 50.89479 | 480.7299 | 141.9065 | 435.8587 | 747.1751 |
| ENSMUSG00000017897 | -0.45923017 | 0.002261 | 0.014356 | 311.8769 | 317.9527 | 298.1184 | 314.8576 | 533.3859 | 366.4987 | 447.4302 | 358.9051 |
| ENSMUSG00000017929 | 0.126907792 | 0.143579 | 0.281335 | 1050.321 | 943.0597 | 933.2762 | 954.9242 | 904.9033 | 803.392 | 877.5031 | 967.9561 |
| ENSMUSG00000017943 | 0.621596441 | 0.168677 | 0.314901 | 57.3451 | 37.19447 | 16.56213 | 58.6584 | 37.05423 | 30.1691 | 21.21436 | 21.75182 |
| ENSMUSG00000017950 | -2.26204967 | 0.000792 | 0.006528 | 705.2441 | 79.18822 | 843.0126 | 656.4565 | 2614.273 | 1533.037 | 2914.082 | 3896.839 |
| ENSMUSG00000017969 | 0.915819445 | 1.76E-06 | 5.90E-05 | 1901.443 | 2565.218 | 1712.525 | 2359.275 | 1238.391 | 1387.779 | 1107.968 | 790.6787 |
| ENSMUSG00000017978 | 0.316624646 | 0.000267 | 0.002837 | 1262.598 | 1258.613 | 1081.507 | 1257.705 | 1027.767 | 982.1718 | 995.1464 | 893.9999 |
| ENSMUSG00000017999 | -0.017373191 | 0.820157 | 0.89103 | 1704.256 | 1882.52 | 1721.634 | 1754.576 | 1840.035 | 1610.136 | 1790.685 | 1903.284 |
| ENSMUSG00000018001 | 0.127202269 | 0.331613 | 0.499409 | 6073.55 | 5201.226 | 5249.368 | 6399.804 | 5303.63 | 4892.981 | 6389.38 | 4402.569 |
| ENSMUSG00000018008 | -0.052248522 | 0.824329 | 0.893455 | 135.8173 | 107.9839 | 164.7932 | 81.94923 | 131.64 | 116.2069 | 124.3933 | 137.0365 |
| ENSMUSG00000018012 | -0.256512143 | 0.163605 | 0.308459 | 203.223 | 185.9723 | 237.6666 | 285.5284 | 274.0063 | 214.5358 | 297.0011 | 306.7007 |
| ENSMUSG00000018040 | -0.270567914 | 0.000625 | 0.005449 | 1657.978 | 1595.763 | 1760.555 | 1593.266 | 2102.34 | 1782.212 | 1999.936 | 2087.087 |
| ENSMUSG00000018042 | -0.035873759 | 0.516945 | 0.670985 | 4005.102 | 4237.77 | 4239.906 | 3837.812 | 4133.497 | 4148.81 | 4136.801 | 4310.123 |
| ENSMUSG00000018068 | 0.091769435 | 0.45455 | 0.616715 | 1126.781 | 1160.227 | 955.6351 | 1098.982 | 1030.693 | 1220.173 | 905.4675 | 916.8393 |
| ENSMUSG00000018076 | 0.281036568 | 0.149641 | 0.289554 | 1523.166 | 1271.811 | 1332.424 | 1601.892 | 982.9122 | 1692.822 | 1134.004 | 907.051 |
| ENSMUSG00000018102 | -1.209706855 | 2.87E-08 | 2.05E-06 | 369.2219 | 359.9465 | 470.3646 | 347.6373 | 999.4891 | 574.3303 | 819.6458 | 1186.562 |
| ENSMUSG00000018126 | -1.879179527 | 2.50E-06 | 7.74E-05 | 164.9929 | 89.98662 | 346.9767 | 179.4257 | 576.2908 | 377.6724 | 878.4674 | 1047.35 |
| ENSMUSG00000018143 | -0.352817259 | 0.074656 | 0.180461 | 482.9061 | 272.3595 | 356.0859 | 362.3019 | 446.601 | 368.7335 | 540.0019 | 527.4817 |

| | | | | | | | | | | | |
|---|---|---|---|---|---|---|---|---|---|---|---|
| ENSMUSG00000018160 | 0.02186738 | 0.794066 | 0.873694 | 1748.522 | 1776.936 | 1519.576 | 1726.11 | 1736.673 | 1781.094 | 1593.006 | 1555.255 |
| ENSMUSG00000018166 | -0.22385775 | 0.045076 | 0.126643 | 873.2552 | 802.6806 | 945.6978 | 784.1248 | 909.7789 | 890.5472 | 1069.397 | 1109.343 |
| ENSMUSG00000018167 | -0.153334742 | 0.077056 | 0.184335 | 986.9393 | 1073.84 | 915.8859 | 973.0393 | 1128.204 | 988.8761 | 1164.861 | 1103.905 |
| ENSMUSG00000018168 | 0.508647829 | 0.126691 | 0.257823 | 34.20585 | 37.19447 | 43.88965 | 35.36756 | 26.32801 | 40.22547 | 19.28578 | 20.66423 |
| ENSMUSG00000018169 | -0.288858508 | 0.0785 | 0.186652 | 247.4894 | 256.7618 | 380.1009 | 274.3143 | 393.945 | 334.0949 | 342.3227 | 348.0291 |
| ENSMUSG00000018171 | -0.546679014 | 3.04E-07 | 1.39E-05 | 2023.175 | 1920.914 | 2017.268 | 1956.43 | 2832.698 | 2380.007 | 3041.368 | 3311.715 |
| ENSMUSG00000018189 | -0.137998053 | 0.140331 | 0.276733 | 769.6316 | 790.6824 | 720.4528 | 795.3389 | 841.5211 | 739.7017 | 923.789 | 876.5984 |
| ENSMUSG00000018196 | -0.102184675 | 0.103363 | 0.226011 | 855.1462 | 881.8688 | 835.5596 | 882.4638 | 906.8535 | 919.5989 | 945.9677 | 934.2407 |
| ENSMUSG00000018199 | 0.463074135 | 0.033321 | 0.101953 | 1584.536 | 1796.133 | 1450.015 | 1684.704 | 956.5842 | 1830.259 | 1028.897 | 911.4013 |
| ENSMUSG00000018209 | -0.114704717 | 0.216238 | 0.374591 | 1107.666 | 1093.037 | 955.6351 | 976.4898 | 1072.622 | 1023.515 | 1214.04 | 1159.372 |
| ENSMUSG00000018211 | -0.618807452 | 0.001106 | 0.008427 | 537.233 | 527.9215 | 809.0602 | 559.8426 | 849.3219 | 724.0584 | 1092.54 | 1074.54 |
| ENSMUSG00000018217 | 0.275394141 | 0.010763 | 0.045083 | 2033.236 | 1962.908 | 1928.66 | 2212.629 | 1508.497 | 1946.466 | 1772.363 | 1497.613 |
| ENSMUSG00000018238 | 0.422759919 | 0.412849 | 0.57864 | 24.1453 | 15.59768 | 7.45296 | 15.52722 | 11.70134 | 15.64324 | 11.57147 | 7.613138 |
| ENSMUSG00000018239 | 0.059020572 | 0.528557 | 0.68098 | 384.3128 | 419.9375 | 419.022 | 373.516 | 386.1441 | 402.2547 | 377.0371 | 366.5182 |
| ENSMUSG00000018286 | -0.231789818 | 0.007083 | 0.033196 | 3768.68 | 3934.215 | 4197.673 | 4001.71 | 4802.423 | 4086.237 | 4623.767 | 5162.795 |
| ENSMUSG00000018287 | -0.262356491 | 0.000641 | 0.005555 | 1795.807 | 1906.516 | 2112.5 | 1977.133 | 2446.554 | 2168.823 | 2407.83 | 2325.27 |
| ENSMUSG00000018293 | -0.073049322 | 0.210263 | 0.36776 | 11921.74 | 12293.37 | 13017.84 | 12055.16 | 13097.7 | 12459.84 | 12512.62 | 13780.87 |
| ENSMUSG00000018322 | 0.223395735 | 0.057887 | 0.150833 | 1483.93 | 1470.981 | 1431.796 | 1473.361 | 1254.968 | 1506.22 | 1236.219 | 1022.336 |
| ENSMUSG00000018326 | 0.003862609 | 0.951744 | 0.972667 | 6914.612 | 7458.091 | 7152.357 | 6918.24 | 6682.438 | 7692.003 | 6942.882 | 7049.765 |
| ENSMUSG00000018334 | -0.193811205 | 0.221152 | 0.380196 | 262.5802 | 231.5656 | 289.8373 | 270.0012 | 259.3796 | 282.6956 | 282.5367 | 383.9197 |
| ENSMUSG00000018339 | -2.534035817 | 1.87E-07 | 9.48E-06 | 3185.168 | 1402.591 | 4210.922 | 3283.145 | 23916.56 | 6982.471 | 16035.16 | 23047.14 |
| ENSMUSG00000018340 | 0.283582569 | 0.001745 | 0.011852 | 6165.101 | 7030.954 | 7760.187 | 7108.88 | 5635.168 | 5307.527 | 5892.771 | 6223.196 |
| ENSMUSG00000018347 | -0.027047166 | 0.833011 | 0.898985 | 422.5428 | 433.1356 | 390.8663 | 357.9887 | 472.929 | 386.6114 | 421.3944 | 350.2043 |
| ENSMUSG00000018362 | -0.185819498 | 0.015273 | 0.058419 | 2667.05 | 2559.219 | 2519.928 | 2767.296 | 3223.718 | 2682.815 | 3068.368 | 2982.175 |
| ENSMUSG00000018363 | 0.082251514 | 0.316636 | 0.484845 | 1766.631 | 1814.13 | 1627.23 | 1773.554 | 1519.223 | 1830.259 | 1667.256 | 1577.007 |
| ENSMUSG00000018372 | 0.497433576 | 4.65E-06 | 0.000126 | 837.0372 | 860.272 | 686.5004 | 848.8215 | 602.6188 | 603.382 | 566.0377 | 515.5182 |
| ENSMUSG00000018377 | 0.256867402 | 0.010772 | 0.045094 | 3133.859 | 3223.92 | 3023.417 | 3434.967 | 2578.194 | 3161.051 | 2592.009 | 2395.963 |
| ENSMUSG00000018378 | 0.227000467 | 0.001607 | 0.011092 | 1506.063 | 1680.95 | 1599.074 | 1564.799 | 1319.326 | 1469.347 | 1349.041 | 1287.708 |
| ENSMUSG00000018379 | 0.11976386 | 0.172086 | 0.319423 | 12285.94 | 12253.78 | 10618.81 | 12269.96 | 10854.94 | 12285.53 | 10170.36 | 10338.64 |
| ENSMUSG00000018381 | -1.070204867 | 0.001384 | 0.009976 | 46.2785 | 26.39607 | 52.99882 | 27.60395 | 82.88446 | 51.39921 | 87.75031 | 101.146 |
| ENSMUSG00000018387 | 0.237303502 | 0.458642 | 0.62026 | 108.6539 | 74.38894 | 55.48314 | 69.8725 | 91.66046 | 69.27719 | 41.46443 | 58.72992 |
| ENSMUSG00000018395 | 0.341998995 | 0.001182 | 0.008877 | 1601.639 | 1802.132 | 1447.53 | 1672.627 | 1216.939 | 1483.873 | 1242.004 | 1202.876 |
| ENSMUSG00000018398 | 0.286910486 | 0.000215 | 0.0024 | 2055.369 | 2398.443 | 2423.868 | 2251.447 | 1772.752 | 1973.283 | 1863.007 | 1875.007 |
| ENSMUSG00000018401 | -0.197331018 | 0.001513 | 0.010619 | 1345.095 | 1426.588 | 1357.267 | 1385.373 | 1509.472 | 1600.08 | 1672.077 | 1537.854 |
| ENSMUSG00000018405 | 0.10953533 | 0.432705 | 0.596995 | 334.01 | 334.7502 | 307.2276 | 281.2153 | 270.1058 | 242.4702 | 331.7155 | 318.6642 |
| ENSMUSG00000018411 | 0.252343564 | 0.465912 | 0.626786 | 402.4217 | 155.9768 | 229.3855 | 508.0852 | 195.9974 | 250.2918 | 297.9654 | 344.7664 |
| ENSMUSG00000018412 | 0.198531513 | 0.000479 | 0.004437 | 2380.325 | 2285.66 | 2314.558 | 2352.374 | 1924.87 | 2026.917 | 2030.793 | 2153.43 |
| ENSMUSG00000018415 | 0.026702099 | 0.692116 | 0.805049 | 1372.258 | 1406.191 | 1268.659 | 1376.747 | 1355.405 | 1320.736 | 1386.648 | 1256.168 |
| ENSMUSG00000018417 | 0.078564166 | 0.313382 | 0.481484 | 2788.783 | 2651.606 | 3074.76 | 2916.53 | 2511.887 | 2939.811 | 2700.01 | 2680.912 |
| ENSMUSG00000018425 | 0.273889979 | 0.000327 | 0.003318 | 2549.342 | 2514.826 | 2406.478 | 2651.705 | 2194.976 | 2222.457 | 2079.007 | 1875.007 |
| ENSMUSG00000018427 | 0.388481101 | 0.013533 | 0.053434 | 1198.211 | 818.2783 | 1048.383 | 1065.34 | 671.8517 | 944.1811 | 849.5388 | 691.7079 |
| ENSMUSG00000018428 | -0.153049903 | 0.212951 | 0.370828 | 1252.538 | 1120.633 | 1063.289 | 1162.816 | 1221.814 | 1053.684 | 1325.898 | 1511.752 |
| ENSMUSG00000018433 | -0.28654482 | 2.91E-05 | 0.000526 | 1137.847 | 1142.23 | 1052.524 | 1082.592 | 1407.086 | 1264.867 | 1323.005 | 1384.503 |
| ENSMUSG00000018442 | -0.018914569 | 0.779302 | 0.863832 | 1005.048 | 1052.243 | 1035.961 | 1058.439 | 1016.066 | 1101.731 | 1092.54 | 996.2334 |
| ENSMUSG00000018446 | -0.400794231 | 5.85E-05 | 0.000895 | 2290.786 | 2360.049 | 2626.754 | 2467.103 | 3318.304 | 2764.384 | 3141.654 | 3643.43 |
| ENSMUSG00000018449 | 0.319630089 | 0.003659 | 0.020463 | 596.5902 | 610.7092 | 491.8953 | 634.8909 | 447.5761 | 500.5836 | 459.9659 | 461.1386 |
| ENSMUSG00000018451 | 0.533695933 | 3.97E-05 | 0.000669 | 1216.32 | 1528.573 | 1414.406 | 1584.639 | 1100.901 | 1110.67 | 911.2533 | 845.0583 |
| ENSMUSG00000018459 | -2.898627159 | 8.10E-06 | 0.000195 | 17.10292 | 9.598572 | 57.96746 | 30.19182 | 317.8863 | 52.51658 | 101.2504 | 390.4452 |
| ENSMUSG00000018470 | 0.719332164 | 4.03E-07 | 1.76E-05 | 201.2109 | 212.3684 | 217.792 | 184.6014 | 125.7894 | 126.2633 | 108.0004 | 135.9489 |
| ENSMUSG00000018474 | 0.202472205 | 0.001347 | 0.009765 | 8062.52 | 7975.214 | 8327.44 | 7867.126 | 6506.918 | 7572.444 | 6825.239 | 7111.758 |
| ENSMUSG00000018476 | 0.258202129 | 0.030765 | 0.096237 | 1230.404 | 1057.043 | 1146.928 | 1174.031 | 854.1975 | 1178.83 | 916.0747 | 907.051 |
| ENSMUSG00000018481 | 0.179432876 | 0.088919 | 0.203358 | 2646.929 | 2141.681 | 2243.341 | 2399.819 | 1926.82 | 2410.176 | 1966.186 | 2028.357 |
| ENSMUSG00000018486 | 0.271394859 | 0.139141 | 0.275033 | 1068.43 | 987.4531 | 1503.014 | 846.2336 | 794.7157 | 861.4954 | 931.5033 | 1063.664 |
| ENSMUSG00000018500 | -1.073103297 | 0.038666 | 0.113652 | 14.08476 | 20.39697 | 36.43669 | 18.11509 | 23.40267 | 24.58223 | 79.07171 | 60.9051 |
| ENSMUSG00000018501 | 0.124612658 | 0.28996 | 0.457144 | 5730.486 | 5318.809 | 5034.06 | 5176.604 | 4280.739 | 6036.055 | 4581.338 | 4603.773 |
| ENSMUSG00000018507 | 0.111966737 | 0.56324 | 0.708577 | 69.41775 | 61.1909 | 67.07664 | 63.83414 | 60.4569 | 60.3382 | 62.6788 | 58.72992 |
| ENSMUSG00000018509 | 0.163741123 | 0.136046 | 0.270539 | 516.1059 | 609.5093 | 493.5516 | 584.8587 | 512.9085 | 526.2832 | 481.1803 | 444.8248 |
| ENSMUSG00000018537 | 0.253026992 | 0.01912 | 0.068323 | 1521.154 | 1654.554 | 1388.735 | 1550.997 | 1386.608 | 1439.178 | 1178.361 | 1125.657 |
| ENSMUSG00000018541 | -0.056298439 | 0.413931 | 0.57985 | 943.679 | 920.2631 | 929.1356 | 941.1222 | 1038.494 | 959.8243 | 972.9678 | 910.3137 |
| ENSMUSG00000018547 | 0.133222697 | 0.097175 | 0.216524 | 1157.969 | 1166.227 | 1161.005 | 1047.225 | 1136.005 | 1000.05 | 995.1464 | 998.4086 |
| ENSMUSG00000018548 | -0.167119469 | 0.014604 | 0.056542 | 1161.993 | 1244.215 | 1161.005 | 1141.251 | 1404.16 | 1261.515 | 1274.79 | 1341 |
| ENSMUSG00000018554 | -0.593510951 | 0.007219 | 0.033635 | 252.5196 | 237.5647 | 300.6027 | 220.8316 | 274.0063 | 513.9921 | 306.644 | 433.9488 |
| ENSMUSG00000018559 | -0.031184062 | 0.671964 | 0.789942 | 1678.099 | 1718.144 | 1649.588 | 1735.598 | 1729.847 | 1920.766 | 1642.184 | 1637.912 |
| ENSMUSG00000018565 | -0.285053271 | 1.45E-05 | 0.000303 | 1234.429 | 1250.214 | 1281.081 | 1232.689 | 1618.685 | 1441.413 | 1462.827 | 1567.219 |

| | | | | | | | | | | | |
|---|---|---|---|---|---|---|---|---|---|---|---|
| ENSMUSG00000018566 | -0.236037337 | 0.054802 | 0.145465 | 497.9969 | 479.9286 | 479.4737 | 525.3377 | 649.4241 | 496.1141 | 660.5381 | 527.4817 |
| ENSMUSG00000018567 | -0.173865605 | 0.04181 | 0.120362 | 4342.131 | 4561.721 | 5227.009 | 4948.008 | 5290.954 | 5009.188 | 5348.912 | 5878.43 |
| ENSMUSG00000018569 | -0.20503188 | 0.16634 | 0.311956 | 1376.282 | 1558.568 | 2018.924 | 1568.25 | 1764.951 | 1573.263 | 2005.721 | 2176.27 |
| ENSMUSG00000018570 | 0.01764708 | 0.904176 | 0.942582 | 458.7608 | 532.7208 | 418.1938 | 487.3823 | 500.2321 | 512.8747 | 500.4661 | 357.8175 |
| ENSMUSG00000018572 | -0.137197891 | 0.250249 | 0.413857 | 1313.907 | 1455.384 | 1206.551 | 1262.881 | 1459.742 | 1171.008 | 1490.791 | 1635.737 |
| ENSMUSG00000018574 | -0.81949796 | 0.00074 | 0.006199 | 1206.259 | 1229.817 | 1430.14 | 1134.35 | 2113.066 | 1231.346 | 2224.615 | 3256.248 |
| ENSMUSG00000018581 | -0.128650967 | 0.687373 | 0.801308 | 44.26639 | 63.59054 | 44.71776 | 43.9938 | 40.95467 | 80.45093 | 55.92877 | 36.9781 |
| ENSMUSG00000018583 | -0.008232963 | 0.87082 | 0.922508 | 6588.65 | 6925.37 | 6530.449 | 6441.21 | 6755.571 | 6814.865 | 6321.88 | 6740.89 |
| ENSMUSG00000018585 | -0.278339114 | 0.018334 | 0.066372 | 1309.883 | 1307.805 | 1526.201 | 1242.178 | 1754.225 | 1343.084 | 1634.47 | 1801.051 |
| ENSMUSG00000018593 | 0.395009494 | 0.001973 | 0.013013 | 30772.18 | 36037.84 | 31186.5 | 32556.27 | 23207.65 | 31468.61 | 21969.4 | 22638.21 |
| ENSMUSG00000018599 | 0.045875292 | 0.661623 | 0.782137 | 353.1251 | 337.1499 | 380.9291 | 334.6979 | 358.841 | 313.9821 | 325.9297 | 364.343 |
| ENSMUSG00000018604 | 0.323047115 | 0.065899 | 0.165361 | 2441.694 | 1847.725 | 3135.212 | 1972.82 | 1554.327 | 1985.574 | 1949.793 | 2025.095 |
| ENSMUSG00000018648 | -1.29564148 | 2.98E-05 | 0.000533 | 112.6781 | 63.59054 | 178.0429 | 105.2401 | 242.8027 | 221.2401 | 273.8581 | 394.7956 |
| ENSMUSG00000018651 | 0.129908939 | 0.176889 | 0.325815 | 947.7032 | 998.2515 | 876.9649 | 1018.758 | 825.9193 | 960.9417 | 926.6819 | 796.1167 |
| ENSMUSG00000018654 | 0.075295337 | 0.75254 | 0.846016 | 62.37537 | 50.3925 | 67.90474 | 49.16954 | 51.6809 | 63.69032 | 58.82164 | 44.59123 |
| ENSMUSG00000018659 | -0.523734017 | 0.000249 | 0.0027 | 437.6336 | 358.7466 | 399.1474 | 376.1038 | 549.9628 | 443.5975 | 620.0379 | 646.0291 |
| ENSMUSG00000018661 | -0.260941561 | 0.009909 | 0.042314 | 838.0433 | 765.4861 | 896.0114 | 824.6681 | 1035.568 | 848.0869 | 1044.325 | 1057.139 |
| ENSMUSG00000018666 | 0.165338458 | 0.096236 | 0.214897 | 6912.599 | 7875.629 | 6558.605 | 7276.229 | 6449.386 | 7327.739 | 5963.164 | 5781.634 |
| ENSMUSG00000018669 | -0.196777389 | 0.001028 | 0.007993 | 1754.559 | 1907.716 | 1854.959 | 1745.95 | 2172.548 | 2015.743 | 2062.615 | 2068.598 |
| ENSMUSG00000018672 | 0.159269681 | 0.075063 | 0.181045 | 1130.805 | 1434.987 | 1223.942 | 1295.66 | 1148.681 | 1131.9 | 1168.718 | 1100.642 |
| ENSMUSG00000018677 | -0.620762052 | 0.000178 | 0.002074 | 2956.794 | 2346.851 | 3024.245 | 2574.069 | 4324.619 | 2974.45 | 4329.658 | 5136.693 |
| ENSMUSG00000018678 | -0.134808994 | 0.282678 | 0.449248 | 365.1977 | 374.3443 | 445.5214 | 350.2251 | 396.8703 | 481.5882 | 423.3229 | 386.0948 |
| ENSMUSG00000018697 | -0.224964838 | 0.001722 | 0.01173 | 744.4802 | 761.8867 | 697.2658 | 747.8946 | 858.098 | 854.7912 | 826.3958 | 908.1386 |
| ENSMUSG00000018698 | -0.726790705 | 7.32E-07 | 2.88E-05 | 989.9575 | 869.8706 | 703.8906 | 753.0703 | 1340.778 | 1115.139 | 1535.148 | 1493.263 |
| ENSMUSG00000018707 | 0.002097096 | 0.98281 | 0.990592 | 6358.264 | 5990.709 | 7438.882 | 6339.42 | 5915.025 | 6175.727 | 6923.596 | 7079.13 |
| ENSMUSG00000018727 | -0.466861383 | 0.13888 | 0.274693 | 29.17558 | 39.59411 | 46.37397 | 37.95543 | 38.02934 | 46.92971 | 49.17875 | 78.30656 |
| ENSMUSG00000018733 | -0.141806477 | 0.149434 | 0.289398 | 336.0222 | 351.5477 | 372.648 | 397.6694 | 422.2232 | 393.3157 | 409.8229 | 385.0072 |
| ENSMUSG00000018736 | -0.217998156 | 0.010044 | 0.042724 | 1291.774 | 1174.625 | 1332.424 | 1324.99 | 1456.816 | 1403.422 | 1439.684 | 1665.102 |
| ENSMUSG00000018740 | -0.631538379 | 0.001463 | 0.0104 | 154.9324 | 154.777 | 163.137 | 144.0581 | 213.5494 | 169.8409 | 301.8225 | 269.7226 |
| ENSMUSG00000018750 | 0.176409711 | 0.102154 | 0.224277 | 630.7961 | 536.3202 | 602.8616 | 646.105 | 494.3814 | 604.4994 | 498.5375 | 544.8831 |
| ENSMUSG00000018752 | 0.081243441 | 0.570197 | 0.713761 | 124.7507 | 149.9777 | 137.4657 | 129.3935 | 121.8889 | 126.2633 | 131.1433 | 131.5985 |
| ENSMUSG00000018761 | -0.011086716 | 0.842994 | 0.905412 | 2216.338 | 2261.664 | 2382.463 | 2327.358 | 2380.247 | 2202.344 | 2306.58 | 2372.036 |
| ENSMUSG00000018765 | 0.008680747 | 0.868798 | 0.921473 | 2362.216 | 2357.649 | 2187.03 | 2321.32 | 2253.482 | 2270.504 | 2300.794 | 2343.759 |
| ENSMUSG00000018770 | -0.671401734 | 4.20E-05 | 0.000699 | 7259.688 | 6730.999 | 8814.367 | 7126.133 | 11305.44 | 8730.044 | 12629.3 | 15006.58 |
| ENSMUSG00000018774 | 0.296299115 | 0.138898 | 0.274699 | 157.9505 | 194.3711 | 214.4796 | 138.8824 | 166.744 | 158.6671 | 128.2505 | 120.7226 |
| ENSMUSG00000018796 | -1.666557315 | 3.57E-08 | 2.42E-06 | 639.8506 | 482.3283 | 949.8383 | 714.2523 | 1993.128 | 1070.444 | 2774.26 | 3011.54 |
| ENSMUSG00000018800 | 0.272322186 | 0.03704 | 0.110598 | 415.5004 | 343.149 | 384.2415 | 448.5642 | 295.4587 | 375.4377 | 338.4655 | 311.0511 |
| ENSMUSG00000018809 | 0.077906583 | 0.479472 | 0.638512 | 381.2946 | 374.3443 | 343.6643 | 331.2474 | 315.9361 | 374.3203 | 314.3583 | 350.2043 |
| ENSMUSG00000018819 | -0.144860798 | 0.261447 | 0.425621 | 7699.334 | 10113.3 | 8864.881 | 8112.111 | 9226.503 | 11742.48 | 8768.281 | 8725.743 |
| ENSMUSG00000018820 | 0.04381588 | 0.625116 | 0.755691 | 1264.61 | 1089.438 | 1085.648 | 1098.12 | 1019.966 | 1204.529 | 1107.004 | 1071.277 |
| ENSMUSG00000018821 | -0.667417332 | 0.002169 | 0.013927 | 254.5318 | 172.7743 | 266.6503 | 192.365 | 361.7663 | 240.2354 | 399.2157 | 407.8467 |
| ENSMUSG00000018822 | 0.570378345 | 0.241546 | 0.403838 | 27.16347 | 21.59679 | 14.90592 | 51.75741 | 25.35289 | 21.23011 | 19.28578 | 11.9635 |
| ENSMUSG00000018830 | 0.666386833 | 0.145715 | 0.284175 | 1650.935 | 3709.848 | 2991.949 | 3217.586 | 2058.46 | 3192.338 | 957.5391 | 1082.153 |
| ENSMUSG00000018841 | 0.112557308 | 0.445966 | 0.608649 | 369.2219 | 419.9375 | 362.7107 | 394.2189 | 367.617 | 271.5219 | 369.3228 | 419.8102 |
| ENSMUSG00000018844 | 0.275186608 | 0.621204 | 0.752895 | 20.12109 | 10.79839 | 9.109173 | 9.488858 | 9.751113 | 16.76061 | 5.785735 | 8.700729 |
| ENSMUSG00000018845 | 0.045563061 | 0.83577 | 0.900729 | 163.9869 | 152.3773 | 149.8873 | 136.2945 | 116.0382 | 208.949 | 140.7862 | 118.5474 |
| ENSMUSG00000018846 | -0.111670275 | 0.19338 | 0.346575 | 2147.926 | 1857.324 | 1893.052 | 1783.905 | 1914.143 | 2164.354 | 2190.865 | 2030.533 |
| ENSMUSG00000018848 | -0.166233296 | 0.025897 | 0.085107 | 1988.969 | 2072.092 | 2026.377 | 1955.567 | 2362.695 | 2000.1 | 2292.115 | 2366.598 |
| ENSMUSG00000018849 | -0.847412182 | 6.92E-05 | 0.001021 | 1377.288 | 1087.038 | 2003.19 | 1312.913 | 2391.948 | 1907.358 | 2913.118 | 3192.08 |
| ENSMUSG00000018858 | -0.220543342 | 0.006083 | 0.029755 | 1558.378 | 1672.551 | 1581.684 | 1649.336 | 1982.401 | 1678.296 | 1838.899 | 2027.27 |
| ENSMUSG00000018861 | 1.118573933 | 0.12673 | 0.257848 | 1712.305 | 193.1713 | 219.4483 | 483.9318 | 271.0809 | 226.8269 | 324.0012 | 379.5693 |
| ENSMUSG00000018865 | 0.797447766 | 0.068463 | 0.169512 | 63.38142 | 44.3934 | 31.46805 | 100.0643 | 18.52711 | 45.81234 | 34.71441 | 39.15328 |
| ENSMUSG00000018882 | -0.081260052 | 0.199772 | 0.354724 | 1444.694 | 1557.368 | 1500.529 | 1614.831 | 1641.112 | 1658.183 | 1623.863 | 1548.73 |
| ENSMUSG00000018899 | 0.153312396 | 0.199348 | 0.354242 | 443.67 | 441.5343 | 491.0672 | 433.8996 | 420.273 | 476.0014 | 373.1799 | 359.9926 |
| ENSMUSG00000018900 | -0.361100222 | 0.02318 | 0.078469 | 929.5942 | 796.6815 | 872.8244 | 971.3141 | 1180.86 | 883.8429 | 1050.111 | 1472.598 |
| ENSMUSG00000018906 | 0.085248621 | 0.409665 | 0.575883 | 418.5186 | 388.7422 | 359.3983 | 436.4875 | 365.6667 | 374.3203 | 369.3228 | 402.4087 |
| ENSMUSG00000018909 | -0.084465507 | 0.532372 | 0.683958 | 1535.239 | 1129.032 | 1642.135 | 1263.743 | 1467.543 | 1336.379 | 1469.577 | 1635.737 |
| ENSMUSG00000018919 | 0.505803025 | 0.375957 | 0.544226 | 8.048435 | 21.59679 | 21.53077 | 18.97772 | 7.80089 | 24.58223 | 8.678602 | 8.700729 |
| ENSMUSG00000018920 | -0.312807285 | 0.285862 | 0.452787 | 199.1988 | 163.1757 | 339.5237 | 164.7611 | 287.6578 | 168.7235 | 329.7869 | 291.4744 |
| ENSMUSG00000018921 | -0.105424194 | 0.219389 | 0.378197 | 1065.412 | 1141.03 | 926.6513 | 1017.033 | 1071.647 | 1126.313 | 1093.504 | 1168.073 |
| ENSMUSG00000018923 | -0.405980168 | 0.002217 | 0.014159 | 433.6094 | 383.9429 | 441.3808 | 506.36 | 582.1415 | 506.1705 | 567.9663 | 686.27 |
| ENSMUSG00000018924 | -1.980865676 | 8.71E-11 | 1.62E-08 | 24.1453 | 11.99822 | 17.39024 | 21.56559 | 66.30757 | 97.21155 | 68.46453 | 67.43065 |
| ENSMUSG00000018927 | 1.497839986 | 2.97E-10 | 4.42E-08 | 334.01 | 296.3559 | 186.324 | 262.2375 | 113.1129 | 106.1505 | 94.50034 | 67.43065 |
| ENSMUSG00000018931 | -0.18210774 | 0.187469 | 0.339609 | 798.8072 | 898.6663 | 1059.976 | 792.751 | 1143.806 | 833.5611 | 1000.932 | 1049.525 |

| | | | | | | | | | | | |
|---|---|---|---|---|---|---|---|---|---|---|---|
| ENSMUSG00000018932 | -0.174081397 | 0.01977 | 0.069885 | 1443.688 | 1529.772 | 1366.376 | 1334.479 | 1680.117 | 1492.812 | 1590.113 | 1631.387 |
| ENSMUSG00000018965 | -0.022081791 | 0.785527 | 0.868177 | 3235.471 | 3501.079 | 3402.69 | 3702.38 | 3641.066 | 3133.117 | 3510.977 | 3769.591 |
| ENSMUSG00000018973 | 0.623116965 | 0.345888 | 0.514649 | 19.11503 | 10.79839 | 19.04645 | 9.488858 | 5.850668 | 22.34748 | 5.785735 | 4.350364 |
| ENSMUSG00000018974 | -0.126782755 | 0.071193 | 0.174541 | 1735.444 | 1749.34 | 1907.13 | 1712.308 | 2039.933 | 1849.254 | 1853.364 | 2016.394 |
| ENSMUSG00000018983 | 0.054025457 | 0.741706 | 0.838783 | 347.0888 | 338.3497 | 306.3995 | 226.0074 | 330.5627 | 291.6346 | 278.6796 | 269.7226 |
| ENSMUSG00000018986 | -1.006414176 | 0.016032 | 0.060347 | 44.26639 | 50.3925 | 48.03018 | 19.84034 | 106.2871 | 34.6386 | 90.64318 | 93.53283 |
| ENSMUSG00000018995 | -0.677255631 | 3.57E-08 | 2.42E-06 | 228.3743 | 233.9652 | 279.0719 | 270.0012 | 375.4179 | 389.9636 | 399.2157 | 457.8758 |
| ENSMUSG00000018999 | -0.193012028 | 0.220515 | 0.379329 | 1454.755 | 1437.386 | 1743.164 | 1468.185 | 1597.232 | 1390.013 | 1699.078 | 2292.642 |
| ENSMUSG00000019027 | 0.435293956 | 0.000251 | 0.002716 | 565.4025 | 509.9242 | 522.5353 | 573.6446 | 431.9743 | 443.5975 | 401.1443 | 329.5401 |
| ENSMUSG00000019039 | -0.092540694 | 0.215508 | 0.373712 | 867.2189 | 898.6663 | 915.0578 | 948.8858 | 979.9869 | 887.195 | 1012.504 | 990.7955 |
| ENSMUSG00000019054 | -0.162272176 | 0.041089 | 0.118954 | 2588.578 | 2688.8 | 3083.041 | 2818.191 | 3089.153 | 2940.929 | 3130.083 | 3354.131 |
| ENSMUSG00000019055 | 0.224528155 | 0.009319 | 0.040442 | 1801.843 | 2174.077 | 1908.786 | 1883.107 | 1768.852 | 1522.981 | 1690.399 | 1660.752 |
| ENSMUSG00000019066 | -0.29418795 | 0.014512 | 0.056278 | 412.4823 | 333.5504 | 422.3344 | 413.1967 | 517.7841 | 420.1327 | 496.6089 | 507.905 |
| ENSMUSG00000019080 | -0.179273382 | 0.277647 | 0.444018 | 188.1322 | 165.5754 | 149.0592 | 125.943 | 163.8187 | 194.4231 | 171.6435 | 180.5401 |
| ENSMUSG00000019082 | 0.248490833 | 0.015327 | 0.05858 | 525.1604 | 567.5156 | 501.8326 | 522.7498 | 502.1823 | 398.9026 | 433.9301 | 443.7372 |
| ENSMUSG00000019087 | -0.350860122 | 0.01332 | 0.052912 | 2406.482 | 2195.673 | 2855.312 | 2687.935 | 3041.372 | 2576.665 | 3478.191 | 3845.722 |
| ENSMUSG00000019088 | 0.022591006 | 0.888342 | 0.93282 | 214.2896 | 250.7627 | 255.8849 | 276.9021 | 221.3503 | 201.1273 | 270.001 | 290.3868 |
| ENSMUSG00000019122 | 1.705873676 | 9.12E-08 | 5.25E-06 | 428.5792 | 243.5638 | 212.8234 | 351.9504 | 89.71024 | 128.498 | 110.8933 | 50.02919 |
| ENSMUSG00000019124 | 0.417803371 | 0.040282 | 0.117282 | 462.785 | 291.5566 | 295.6341 | 461.5036 | 244.7529 | 334.0949 | 278.6796 | 275.1605 |
| ENSMUSG00000019132 | 0.221130648 | 0.120539 | 0.249599 | 5257.64 | 5839.531 | 4841.111 | 5449.193 | 4072.065 | 6011.473 | 4216.837 | 4048.014 |
| ENSMUSG00000019139 | -0.179870568 | 0.030311 | 0.095156 | 3194.223 | 3337.904 | 3255.287 | 3564.36 | 4110.094 | 3334.244 | 3911.157 | 3767.416 |
| ENSMUSG00000019143 | 0.07600313 | 0.402443 | 0.569517 | 1539.263 | 1374.995 | 1247.957 | 1547.547 | 1373.932 | 1395.6 | 1347.112 | 1299.671 |
| ENSMUSG00000019146 | 1.109629322 | 0.120053 | 0.248995 | 13.07871 | 4.799286 | 7.45296 | 31.91707 | 7.80089 | 7.821619 | 6.750024 | 4.350364 |
| ENSMUSG00000019158 | -0.482900349 | 2.65E-05 | 0.000487 | 424.5549 | 418.7377 | 430.6154 | 390.7684 | 546.0623 | 499.4662 | 621.0022 | 659.0802 |
| ENSMUSG00000019173 | -0.315704791 | 0.019024 | 0.068124 | 3807.916 | 3996.606 | 4627.46 | 4122.478 | 5371.888 | 3990.143 | 5104.947 | 6138.364 |
| ENSMUSG00000019178 | 0.053023611 | 0.928061 | 0.957316 | 11.0666 | 8.398751 | 14.07781 | 15.52722 | 5.850668 | 21.23011 | 5.785735 | 15.22628 |
| ENSMUSG00000019179 | -0.765923458 | 4.46E-06 | 0.000124 | 4777.752 | 4337.355 | 5771.903 | 4622.799 | 8312.824 | 5859.51 | 8897.496 | 10108.07 |
| ENSMUSG00000019188 | -0.004721437 | 0.928036 | 0.957316 | 4227.44 | 4380.548 | 4403.871 | 4092.286 | 4479.661 | 4278.425 | 4149.336 | 4250.306 |
| ENSMUSG00000019189 | 0.139756357 | 0.120433 | 0.249434 | 3781.758 | 3427.89 | 3686.731 | 4049.155 | 3555.256 | 3663.87 | 3369.226 | 2980 |
| ENSMUSG00000019194 | -0.09476387 | 0.691153 | 0.804404 | 58.35115 | 52.79215 | 87.7793 | 66.42201 | 66.30757 | 61.45558 | 80.036 | 77.21897 |
| ENSMUSG00000019210 | -0.702910529 | 0.000965 | 0.007629 | 2236.459 | 1511.775 | 2551.397 | 1949.529 | 3157.41 | 2255.978 | 3788.692 | 4228.554 |
| ENSMUSG00000019214 | -0.088829456 | 0.477354 | 0.636748 | 415.5004 | 482.3283 | 353.6015 | 407.1583 | 489.5059 | 407.8415 | 432.9658 | 428.5109 |
| ENSMUSG00000019230 | 0.102921778 | 0.905665 | 0.943335 | 8.048435 | 19.19714 | 14.90592 | 11.21411 | 4.875557 | 37.99072 | 4.821446 | 2.175182 |
| ENSMUSG00000019232 | -3.604779915 | 6.84E-11 | 1.33E-08 | 20.12109 | 3.599465 | 13.24971 | 16.38985 | 161.8685 | 44.69496 | 200.5721 | 247.9708 |
| ENSMUSG00000019235 | 0.143256808 | 0.692271 | 0.805177 | 50.30272 | 20.39697 | 36.43669 | 49.16954 | 25.35289 | 37.99072 | 34.71441 | 44.59123 |
| ENSMUSG00000019254 | -0.043703514 | 0.430735 | 0.595479 | 1777.698 | 1840.526 | 1761.383 | 1921.925 | 1884.89 | 1898.419 | 1896.757 | 1845.642 |
| ENSMUSG00000019256 | 0.040078834 | 0.676401 | 0.792815 | 561.3783 | 521.9224 | 616.1113 | 535.6892 | 514.8588 | 556.4523 | 587.2521 | 517.6934 |
| ENSMUSG00000019261 | -0.148391947 | 0.066393 | 0.166099 | 709.2683 | 836.2756 | 809.0602 | 743.5815 | 850.2971 | 865.9649 | 834.1101 | 882.0364 |
| ENSMUSG00000019278 | -0.388590319 | 0.142393 | 0.279604 | 569.4268 | 376.744 | 531.6445 | 666.808 | 881.5006 | 378.7898 | 726.1097 | 822.2189 |
| ENSMUSG00000019295 | -0.221874588 | 0.201026 | 0.356523 | 487.9364 | 334.7502 | 357.7421 | 401.9825 | 482.6801 | 357.5597 | 455.1445 | 550.3211 |
| ENSMUSG00000019297 | -0.228454395 | 0.008142 | 0.036751 | 900.4186 | 931.0615 | 920.0265 | 823.8054 | 1125.278 | 938.5942 | 1053.968 | 1066.927 |
| ENSMUSG00000019302 | -0.190947256 | 0.127071 | 0.258228 | 1037.242 | 913.0642 | 1238.019 | 1092.944 | 1310.55 | 1023.515 | 1271.897 | 1284.445 |
| ENSMUSG00000019303 | -0.223817846 | 0.06119 | 0.157046 | 290.7497 | 349.1481 | 362.7107 | 288.1162 | 396.8703 | 385.4941 | 367.3942 | 356.7299 |
| ENSMUSG00000019312 | -0.276640744 | 0.012927 | 0.051821 | 1704.256 | 1641.356 | 1854.959 | 1524.256 | 1961.924 | 1731.93 | 2141.686 | 2311.131 |
| ENSMUSG00000019320 | 0.23175524 | 0.363556 | 0.532101 | 58.35115 | 41.99375 | 44.71776 | 52.62003 | 39.97956 | 52.51658 | 40.50014 | 35.89051 |
| ENSMUSG00000019326 | 0.394175846 | 0.015624 | 0.059457 | 346.0827 | 343.149 | 308.8838 | 462.3662 | 265.2303 | 278.2261 | 321.1083 | 246.8832 |
| ENSMUSG00000019338 | -0.09222168 | 0.201337 | 0.356815 | 1343.083 | 1232.217 | 1373.829 | 1343.967 | 1324.201 | 1371.018 | 1467.648 | 1484.562 |
| ENSMUSG00000019359 | 0.965435374 | 0.000151 | 0.001837 | 106.6418 | 105.5843 | 86.12309 | 117.3168 | 56.55646 | 74.86406 | 40.50014 | 41.32846 |
| ENSMUSG00000019362 | -0.324504701 | 0.002866 | 0.017155 | 2035.248 | 2188.474 | 2411.447 | 2092.725 | 2887.305 | 2319.669 | 2625.759 | 3097.459 |
| ENSMUSG00000019370 | -0.260741338 | 0.022022 | 0.075449 | 8334.154 | 9548.18 | 9722.8 | 8375.212 | 11058.74 | 8863.011 | 10857.9 | 12326.76 |
| ENSMUSG00000019373 | -0.201146175 | 0.000429 | 0.004066 | 1893.394 | 2063.693 | 1946.879 | 1922.788 | 2219.353 | 2161.002 | 2294.044 | 2317.657 |
| ENSMUSG00000019428 | -0.169800958 | 0.01606 | 0.060359 | 4821.012 | 4792.087 | 5323.898 | 5135.198 | 5714.152 | 5202.494 | 5616.02 | 6051.357 |
| ENSMUSG00000019432 | -0.046666371 | 0.453986 | 0.616176 | 10793.96 | 11634.67 | 11578.59 | 10739.66 | 12183.04 | 11960.37 | 10988.08 | 11084.73 |
| ENSMUSG00000019433 | -0.528205299 | 0.000993 | 0.007793 | 2064.424 | 2279.661 | 2907.482 | 2253.173 | 3408.014 | 2561.021 | 3730.835 | 4008.861 |
| ENSMUSG00000019437 | -0.586721751 | 0.001952 | 0.012907 | 546.2875 | 428.3363 | 640.9545 | 551.2164 | 760.5868 | 582.1519 | 953.682 | 960.3429 |
| ENSMUSG00000019461 | 0.154217042 | 0.026602 | 0.086957 | 1884.34 | 1968.907 | 2108.359 | 2022.852 | 1870.263 | 1834.728 | 1656.649 | 1817.365 |
| ENSMUSG00000019464 | -0.176602575 | 0.675781 | 0.792479 | 225.3562 | 235.165 | 820.6537 | 224.2821 | 453.4268 | 270.4045 | 466.716 | 512.2554 |
| ENSMUSG00000019467 | 0.589682223 | 1.53E-06 | 5.29E-05 | 2933.655 | 3349.902 | 2722.815 | 3366.819 | 2040.908 | 2405.706 | 2053.936 | 1720.569 |
| ENSMUSG00000019470 | -0.064960304 | 0.296716 | 0.464438 | 1892.388 | 1713.345 | 1684.369 | 1775.279 | 1891.716 | 1858.193 | 1850.471 | 1789.087 |
| ENSMUSG00000019471 | -0.245657183 | 0.003702 | 0.02065 | 6086.629 | 6992.56 | 6821.942 | 6308.366 | 8232.865 | 6865.146 | 7782.778 | 8190.648 |
| ENSMUSG00000019478 | -0.451960161 | 0.031313 | 0.09733 | 337.0282 | 220.7672 | 388.382 | 263.9628 | 380.2934 | 329.6254 | 445.5016 | 502.4671 |
| ENSMUSG00000019487 | -0.061045328 | 0.523989 | 0.677271 | 1198.211 | 1075.04 | 1127.053 | 1175.756 | 1267.645 | 1016.81 | 1208.254 | 1281.182 |
| ENSMUSG00000019494 | -0.257060031 | 0.028011 | 0.090126 | 2716.347 | 3019.951 | 3340.582 | 3145.988 | 3696.647 | 2967.746 | 3863.907 | 4079.554 |
| ENSMUSG00000019505 | -0.092561426 | 0.27394 | 0.439753 | 21891.74 | 24265.19 | 26738.74 | 24751.26 | 27406.48 | 23576.59 | 25894.06 | 27241.98 |

| | | | | | | | | | | | |
|---|---|---|---|---|---|---|---|---|---|---|---|
| ENSMUSG00000019518 | 0.176401783 | 0.052843 | 0.141445 | 1451.736 | 1600.562 | 1332.424 | 1419.878 | 1335.902 | 1346.436 | 1309.505 | 1139.795 |
| ENSMUSG00000019528 | -0.200243924 | 0.015715 | 0.059687 | 839.0493 | 801.4808 | 944.8697 | 825.5307 | 1009.24 | 917.3641 | 996.1107 | 999.4962 |
| ENSMUSG00000019539 | 0.311176352 | 0.001509 | 0.010607 | 5267.701 | 6991.36 | 6011.226 | 6163.445 | 5056.927 | 5248.306 | 4634.374 | 4751.685 |
| ENSMUSG00000019558 | -0.327204521 | 0.018048 | 0.065766 | 1440.67 | 1148.229 | 1478.998 | 1266.331 | 1628.436 | 1338.614 | 1777.185 | 1950.051 |
| ENSMUSG00000019564 | -0.186134841 | 0.118556 | 0.247389 | 540.2512 | 487.1275 | 589.6119 | 439.938 | 585.0668 | 534.1048 | 590.145 | 631.8904 |
| ENSMUSG00000019577 | -0.390896687 | 0.351359 | 0.520189 | 502.0211 | 260.3613 | 285.6968 | 526.2003 | 425.1485 | 236.8833 | 357.7513 | 1045.175 |
| ENSMUSG00000019578 | -0.043920975 | 0.506938 | 0.662721 | 2184.144 | 2339.652 | 2264.044 | 2453.301 | 2518.713 | 2311.847 | 2418.437 | 2276.328 |
| ENSMUSG00000019579 | -0.305051593 | 0.010523 | 0.044254 | 1381.313 | 1445.785 | 1623.917 | 1455.246 | 1901.467 | 1454.821 | 1897.721 | 2043.584 |
| ENSMUSG00000019590 | -0.164720316 | 0.537622 | 0.687763 | 598.6023 | 377.9438 | 588.7838 | 843.6458 | 636.7477 | 431.3064 | 770.467 | 863.5473 |
| ENSMUSG00000019647 | 0.534903825 | 0.005708 | 0.028373 | 1179.096 | 1330.602 | 1077.367 | 1204.222 | 721.5824 | 1191.121 | 622.9308 | 772.1897 |
| ENSMUSG00000019659 | -0.219697011 | 0.01877 | 0.067448 | 1218.332 | 1289.808 | 1472.374 | 1211.123 | 1488.995 | 1469.347 | 1433.898 | 1656.401 |
| ENSMUSG00000019689 | -0.66012041 | 0.003509 | 0.019815 | 392.3612 | 341.9491 | 483.6143 | 376.9665 | 584.0917 | 396.6678 | 689.4668 | 851.5838 |
| ENSMUSG00000019699 | 0.098242566 | 0.39953 | 0.566948 | 1082.514 | 1115.834 | 883.5898 | 985.116 | 1050.195 | 974.3502 | 962.3606 | 808.0802 |
| ENSMUSG00000019710 | 0.021907821 | 0.710934 | 0.818885 | 2392.397 | 2432.038 | 2328.636 | 2452.439 | 2499.21 | 2338.664 | 2399.151 | 2219.773 |
| ENSMUSG00000019715 | 0.10000212 | 0.338323 | 0.506825 | 1553.348 | 1662.953 | 1668.635 | 1681.253 | 1514.348 | 1825.789 | 1414.612 | 1373.628 |
| ENSMUSG00000019718 | -0.149195588 | 0.276175 | 0.442378 | 513.0877 | 496.7261 | 515.0823 | 593.485 | 586.0419 | 476.0014 | 581.4664 | 706.9342 |
| ENSMUSG00000019726 | -0.552849468 | 0.000135 | 0.001684 | 343.0645 | 238.7645 | 364.3669 | 304.5061 | 497.3068 | 444.7149 | 440.6801 | 456.7883 |
| ENSMUSG00000019731 | -0.357375802 | 0.000155 | 0.001873 | 811.8859 | 770.2854 | 814.0288 | 756.5208 | 1053.12 | 921.8336 | 1132.075 | 929.8904 |
| ENSMUSG00000019732 | 0.017380239 | 0.949407 | 0.971355 | 59.35721 | 65.99018 | 75.3577 | 56.07053 | 85.8098 | 46.92971 | 72.32169 | 47.85401 |
| ENSMUSG00000019734 | -0.002192492 | 0.984467 | 0.991234 | 679.0867 | 664.7011 | 872.8244 | 696.9998 | 718.657 | 690.5372 | 771.4313 | 740.6495 |
| ENSMUSG00000019737 | -0.64659471 | 7.78E-06 | 0.00019 | 405.4399 | 359.9465 | 464.5678 | 402.8452 | 601.6437 | 537.4569 | 636.4308 | 784.1532 |
| ENSMUSG00000019738 | -0.193548338 | 0.006004 | 0.029494 | 715.3046 | 754.6877 | 790.0137 | 747.0319 | 877.6002 | 842.5001 | 832.1815 | 888.5619 |
| ENSMUSG00000019761 | 0.010354434 | 0.917204 | 0.951006 | 298.7981 | 304.7547 | 323.7897 | 335.5605 | 332.513 | 315.0995 | 305.6797 | 302.3503 |
| ENSMUSG00000019762 | -2.400344786 | 1.82E-09 | 2.17E-07 | 21.12714 | 10.79839 | 31.46805 | 25.8787 | 119.9387 | 60.3382 | 129.2147 | 167.489 |
| ENSMUSG00000019763 | -0.156122593 | 0.089255 | 0.203974 | 768.6255 | 805.0802 | 656.6886 | 690.0988 | 781.0642 | 796.6877 | 835.0744 | 835.27 |
| ENSMUSG00000019767 | -1.610796836 | 0.010858 | 0.045379 | 13.07871 | 3.599465 | 10.76539 | 9.488858 | 17.552 | 20.11273 | 14.46434 | 61.99269 |
| ENSMUSG00000019768 | 0.452071743 | 0.326573 | 0.494239 | 222.338 | 387.5424 | 305.5713 | 243.2598 | 107.2622 | 488.2925 | 122.4647 | 129.4233 |
| ENSMUSG00000019773 | 0.048741731 | 0.72269 | 0.826557 | 945.6911 | 1053.443 | 904.2924 | 970.4514 | 893.202 | 1190.003 | 868.8245 | 792.8539 |
| ENSMUSG00000019774 | 0.06980732 | 0.6905 | 0.80399 | 405.4399 | 343.149 | 260.8536 | 383.0048 | 315.9361 | 335.2122 | 393.43 | 280.5985 |
| ENSMUSG00000019775 | 1.163688735 | 0.003314 | 0.019073 | 98.59333 | 131.9804 | 86.9512 | 131.1188 | 38.02934 | 90.5073 | 50.14304 | 21.75182 |
| ENSMUSG00000019777 | 0.060899411 | 0.467576 | 0.62788 | 4720.407 | 5100.441 | 4391.449 | 5038.584 | 4415.304 | 5160.034 | 4415.48 | 4463.474 |
| ENSMUSG00000019779 | 0.318512598 | 0.000554 | 0.004956 | 863.1946 | 1094.237 | 968.0567 | 931.6334 | 758.6366 | 778.8097 | 756.0027 | 797.2043 |
| ENSMUSG00000019782 | -0.041060464 | 0.539021 | 0.688884 | 1520.148 | 1625.758 | 1641.307 | 1676.94 | 1648.913 | 1782.212 | 1593.006 | 1630.299 |
| ENSMUSG00000019787 | 2.0410957 | 0.025271 | 0.083584 | 22.1332 | 25.19625 | 14.07781 | 8.626235 | 0.975111 | 12.29112 | 3.857157 | 0 |
| ENSMUSG00000019789 | -0.400469114 | 0.109334 | 0.234746 | 97.58727 | 63.59054 | 89.43552 | 138.0198 | 132.6151 | 109.5027 | 126.3219 | 146.8248 |
| ENSMUSG00000019790 | 0.358586159 | 0.002724 | 0.016472 | 879.2915 | 925.0624 | 823.9661 | 936.8091 | 658.2001 | 841.3827 | 679.8239 | 601.4379 |
| ENSMUSG00000019791 | -0.466099913 | 6.65E-05 | 0.000992 | 220.3259 | 227.9661 | 211.9953 | 181.1509 | 292.5334 | 281.5783 | 294.1082 | 290.3868 |
| ENSMUSG00000019792 | 0.086879824 | 0.444509 | 0.607468 | 379.2825 | 457.132 | 376.7885 | 429.5865 | 382.2436 | 431.3064 | 366.4299 | 365.4306 |
| ENSMUSG00000019794 | 0.12535691 | 0.131983 | 0.265115 | 1201.229 | 1127.832 | 1113.803 | 1160.229 | 1148.681 | 1071.562 | 1061.682 | 936.4159 |
| ENSMUSG00000019795 | -0.056470898 | 0.321473 | 0.489489 | 1930.618 | 1854.924 | 1853.303 | 1909.848 | 1937.546 | 1998.982 | 2052.007 | 1860.868 |
| ENSMUSG00000019796 | 0.21491472 | 0.436083 | 0.600086 | 102.6175 | 68.38983 | 54.65504 | 113.0037 | 65.33246 | 71.51194 | 79.07171 | 76.13138 |
| ENSMUSG00000019797 | -0.194470357 | 0.058367 | 0.151661 | 540.2512 | 574.7145 | 645.0951 | 664.2201 | 668.9264 | 646.9596 | 711.6454 | 750.4379 |
| ENSMUSG00000019802 | 0.111709655 | 0.027196 | 0.088176 | 3217.362 | 3235.919 | 3195.663 | 3150.301 | 2915.583 | 3071.661 | 3038.475 | 2817.949 |
| ENSMUSG00000019804 | -0.161957254 | 0.075137 | 0.181153 | 2370.264 | 2355.25 | 2732.752 | 2420.522 | 2756.64 | 2440.345 | 2837.903 | 3020.24 |
| ENSMUSG00000019806 | -0.39162111 | 0.008767 | 0.038736 | 478.8819 | 440.3345 | 526.6758 | 439.938 | 548.0126 | 564.2739 | 567.9663 | 796.1167 |
| ENSMUSG00000019808 | 0.065457826 | 0.670509 | 0.788764 | 249.5015 | 232.7654 | 270.7909 | 309.6818 | 302.2845 | 216.7706 | 240.108 | 257.7591 |
| ENSMUSG00000019809 | 0.115906726 | 0.141123 | 0.27785 | 808.8677 | 849.4736 | 765.1705 | 824.6681 | 761.5619 | 758.697 | 785.8957 | 687.3576 |
| ENSMUSG00000019810 | -0.434588556 | 0.010077 | 0.042832 | 2041.284 | 1964.108 | 2544.772 | 2045.28 | 3086.227 | 2033.621 | 2967.118 | 3531.408 |
| ENSMUSG00000019813 | 0.285254531 | 0.054651 | 0.145212 | 410.4702 | 381.5432 | 272.4471 | 359.714 | 282.7823 | 312.8647 | 287.3582 | 283.8613 |
| ENSMUSG00000019814 | -0.067380272 | 0.371866 | 0.540904 | 1150.926 | 1280.21 | 1116.288 | 1241.315 | 1180.86 | 1255.928 | 1272.862 | 1305.109 |
| ENSMUSG00000019817 | 1.1387554 | 4.95E-06 | 0.000134 | 10512.26 | 11387.51 | 8303.425 | 12150.91 | 4282.689 | 7751.224 | 4050.014 | 3151.839 |
| ENSMUSG00000019818 | -0.076305356 | 0.167849 | 0.313863 | 5456.839 | 5106.44 | 5538.377 | 5701.079 | 5517.18 | 5767.885 | 5831.057 | 5879.517 |
| ENSMUSG00000019820 | 0.097254773 | 0.16435 | 0.309357 | 3210.319 | 2815.981 | 2923.216 | 3037.297 | 2623.049 | 2980.037 | 2802.224 | 2803.81 |
| ENSMUSG00000019822 | -0.257852519 | 0.009725 | 0.041725 | 966.8182 | 958.6574 | 1042.586 | 1136.075 | 1261.794 | 1097.261 | 1188.004 | 1363.839 |
| ENSMUSG00000019823 | -0.045671804 | 0.657584 | 0.779321 | 527.1725 | 628.7065 | 678.2193 | 655.5939 | 661.1255 | 613.4384 | 631.6094 | 665.6057 |
| ENSMUSG00000019826 | 0.164695545 | 0.0874 | 0.201003 | 733.4136 | 709.0945 | 647.5794 | 740.9936 | 683.553 | 664.8376 | 610.395 | 565.5474 |
| ENSMUSG00000019831 | 0.064082094 | 0.656839 | 0.778842 | 337.0282 | 283.1579 | 306.3995 | 322.6212 | 266.2054 | 373.2029 | 271.9295 | 286.0365 |
| ENSMUSG00000019832 | 0.268327548 | 0.075492 | 0.181771 | 336.0222 | 428.3363 | 365.195 | 434.7622 | 321.7867 | 385.4941 | 267.1081 | 325.1897 |
| ENSMUSG00000019837 | -0.12700749 | 0.135749 | 0.27018 | 1507.069 | 1757.739 | 1565.95 | 1461.284 | 1795.18 | 1594.493 | 1678.827 | 1796.7 |
| ENSMUSG00000019838 | -0.115940742 | 0.579092 | 0.720342 | 391.3551 | 284.3577 | 355.2577 | 354.5383 | 345.1894 | 280.4609 | 344.2512 | 532.9196 |
| ENSMUSG00000019841 | 0.128598667 | 0.309716 | 0.478216 | 1958.788 | 2224.469 | 1861.584 | 1995.248 | 1730.823 | 2297.321 | 1684.613 | 1641.175 |
| ENSMUSG00000019842 | 0.349058289 | 7.82E-05 | 0.001122 | 499.003 | 550.7181 | 494.3797 | 486.5197 | 402.721 | 413.4284 | 378.9656 | 396.9707 |
| ENSMUSG00000019843 | -0.063459926 | 0.446265 | 0.608934 | 1697.214 | 1500.977 | 1845.85 | 1666.589 | 1639.162 | 1822.437 | 1785.864 | 1771.686 |
| ENSMUSG00000019845 | 0.117610308 | 0.518756 | 0.672487 | 306.8466 | 266.3604 | 203.7142 | 272.589 | 246.7032 | 293.8694 | 193.8221 | 232.7445 |

| | | | | | | | | | | | |
|---|---|---|---|---|---|---|---|---|---|---|---|
| ENSMUSG00000019846 | 0.42359499 | 0.001543 | 0.01075 | 3790.813 | 5320.009 | 4215.891 | 4367.463 | 3428.491 | 3633.701 | 3436.727 | 2690.7 |
| ENSMUSG00000019848 | -0.317762719 | 0.428016 | 0.592975 | 13.07871 | 25.19625 | 16.56213 | 12.93935 | 22.42756 | 24.58223 | 21.21436 | 15.22628 |
| ENSMUSG00000019849 | -0.190473297 | 0.014204 | 0.055412 | 1679.105 | 1514.175 | 1662.01 | 1579.464 | 1823.458 | 1660.418 | 1897.721 | 1963.102 |
| ENSMUSG00000019850 | -0.196547311 | 0.313529 | 0.481526 | 163.9869 | 152.3773 | 183.0116 | 174.2499 | 153.0925 | 173.193 | 261.3224 | 184.8905 |
| ENSMUSG00000019851 | -0.410252243 | 0.142706 | 0.280131 | 355.1372 | 277.1588 | 654.2042 | 401.1199 | 654.2997 | 376.5551 | 522.6447 | 691.7079 |
| ENSMUSG00000019852 | 0.464638845 | 0.029681 | 0.093817 | 339.0403 | 208.7689 | 290.6654 | 264.8254 | 146.2667 | 255.8787 | 195.7507 | 203.3795 |
| ENSMUSG00000019853 | 0.436480953 | 0.007645 | 0.03501 | 292.7618 | 322.752 | 358.5702 | 350.2251 | 274.9814 | 177.6625 | 261.3224 | 264.2846 |
| ENSMUSG00000019854 | 0.251625283 | 0.050364 | 0.136713 | 1753.553 | 1808.131 | 1833.428 | 1917.612 | 1500.696 | 1941.996 | 1386.648 | 1314.898 |
| ENSMUSG00000019856 | 0.470464065 | 0.000471 | 0.004373 | 582.5055 | 442.7341 | 449.6619 | 573.6446 | 363.7165 | 389.9636 | 390.5371 | 334.9781 |
| ENSMUSG00000019857 | -0.144444088 | 0.036436 | 0.109396 | 1190.162 | 1294.607 | 1178.396 | 1243.04 | 1417.812 | 1388.896 | 1269.969 | 1343.175 |
| ENSMUSG00000019861 | -0.064236741 | 0.463154 | 0.624014 | 963.8001 | 934.661 | 975.5096 | 1061.027 | 982.9122 | 938.5942 | 1079.04 | 1115.868 |
| ENSMUSG00000019863 | -0.416987647 | 0.004272 | 0.022922 | 387.3309 | 346.7484 | 438.0684 | 378.6917 | 517.7841 | 413.4284 | 521.6804 | 619.9269 |
| ENSMUSG00000019864 | -0.682421061 | 0.001557 | 0.010826 | 211.2714 | 175.1739 | 246.7758 | 193.2277 | 305.2098 | 227.9443 | 358.7156 | 436.124 |
| ENSMUSG00000019865 | 1.058982831 | 0.132274 | 0.265382 | 25.15136 | 29.99554 | 15.73403 | 12.07673 | 3.900445 | 16.76061 | 1.928578 | 17.40146 |
| ENSMUSG00000019866 | 0.153639626 | 0.307903 | 0.476373 | 785.7285 | 693.4968 | 1068.258 | 796.2015 | 655.2748 | 737.4669 | 805.1815 | 811.343 |
| ENSMUSG00000019868 | 0.002930869 | 0.970922 | 0.983692 | 1036.236 | 1154.228 | 1150.24 | 1092.081 | 1030.693 | 1052.566 | 1160.04 | 1180.036 |
| ENSMUSG00000019872 | -0.619690405 | 0.000372 | 0.003649 | 746.4923 | 849.4736 | 1023.54 | 919.5566 | 1379.783 | 956.4722 | 1469.577 | 1633.562 |
| ENSMUSG00000019873 | 0.1150352 | 0.014773 | 0.057015 | 2476.906 | 2332.453 | 2427.181 | 2392.918 | 2228.129 | 2212.401 | 2252.579 | 2200.197 |
| ENSMUSG00000019874 | 0.734002842 | 0.168451 | 0.314619 | 75.45408 | 21.59679 | 43.88965 | 100.9269 | 39.97956 | 60.3382 | 16.39292 | 29.36496 |
| ENSMUSG00000019876 | 0.008029105 | 0.968635 | 0.982421 | 72.43591 | 83.98751 | 94.40416 | 69.8725 | 74.10846 | 70.39457 | 80.036 | 94.62042 |
| ENSMUSG00000019877 | 0.058449232 | 0.304182 | 0.472636 | 5175.144 | 5156.833 | 4838.627 | 5296.508 | 4697.111 | 5184.616 | 4881.232 | 4891.985 |
| ENSMUSG00000019878 | 0.385915903 | 0.001277 | 0.009386 | 788.7466 | 817.0785 | 783.3889 | 789.3005 | 540.2117 | 725.1758 | 643.1809 | 524.2189 |
| ENSMUSG00000019880 | 1.208492374 | 0.018685 | 0.06728 | 1700.232 | 2920.366 | 1395.36 | 2538.701 | 840.5459 | 1805.677 | 675.0024 | 380.6569 |
| ENSMUSG00000019883 | -0.333951102 | 0.091976 | 0.208232 | 181.0898 | 223.1668 | 189.6364 | 190.6398 | 251.5787 | 166.4887 | 261.3224 | 307.7883 |
| ENSMUSG00000019888 | 1.985029152 | 0.00167 | 0.011463 | 111.672 | 14.39786 | 24.8432 | 37.09281 | 10.72622 | 12.29112 | 12.53576 | 11.9635 |
| ENSMUSG00000019889 | -0.103061858 | 0.361244 | 0.529809 | 1740.474 | 1606.561 | 2018.096 | 1867.58 | 1825.408 | 1687.235 | 2134.936 | 2124.065 |
| ENSMUSG00000019890 | -1.293369875 | 0.312438 | 0.480769 | 11.0666 | 13.19804 | 605.3459 | 60.38364 | 319.8365 | 87.15518 | 591.1093 | 693.8831 |
| ENSMUSG00000019891 | -0.170335952 | 0.035719 | 0.107717 | 755.5468 | 712.694 | 714.656 | 748.7572 | 781.0642 | 811.2136 | 796.5028 | 912.4889 |
| ENSMUSG00000019892 | -0.537517082 | 0.281254 | 0.447678 | 114.6902 | 137.9795 | 478.6456 | 146.646 | 306.185 | 118.4417 | 387.6442 | 463.3138 |
| ENSMUSG00000019893 | 0.663129712 | 0.000376 | 0.003675 | 1147.908 | 1172.226 | 972.1972 | 1162.816 | 729.3833 | 938.5942 | 662.4667 | 482.8904 |
| ENSMUSG00000019894 | -0.33041047 | 0.240966 | 0.403112 | 137.8294 | 140.3791 | 93.57605 | 147.5086 | 133.5902 | 110.62 | 159.1077 | 249.0584 |
| ENSMUSG00000019897 | 0.068377154 | 0.501733 | 0.657987 | 1464.815 | 1449.384 | 1403.641 | 1509.591 | 1297.873 | 1649.244 | 1363.505 | 1248.555 |
| ENSMUSG00000019899 | 0.306774212 | 0.002105 | 0.013614 | 1281.713 | 1177.025 | 1445.874 | 1449.207 | 1032.643 | 1005.637 | 1129.183 | 1164.81 |
| ENSMUSG00000019906 | 0.402503164 | 0.102142 | 0.224277 | 125.7568 | 129.5807 | 116.763 | 130.2561 | 98.48624 | 138.5544 | 78.10742 | 65.25547 |
| ENSMUSG00000019907 | 0.207634358 | 0.115861 | 0.243805 | 2141.89 | 2013.301 | 1714.181 | 1737.324 | 1657.689 | 1997.865 | 1423.291 | 1507.401 |
| ENSMUSG00000019913 | -0.022806854 | 0.851514 | 0.910624 | 695.1836 | 577.1142 | 547.3785 | 618.501 | 546.0623 | 575.4477 | 672.1095 | 683.0072 |
| ENSMUSG00000019916 | 0.186939902 | 0.034485 | 0.104905 | 1208.271 | 1485.379 | 1310.893 | 1381.06 | 1129.179 | 1185.534 | 1142.683 | 1272.482 |
| ENSMUSG00000019917 | 0.092102217 | 0.278899 | 0.445274 | 1259.58 | 1342.6 | 1269.487 | 1282.721 | 1295.923 | 1190.003 | 1285.397 | 1060.401 |
| ENSMUSG00000019920 | 0.010362134 | 0.889555 | 0.933475 | 2700.25 | 2767.988 | 2817.219 | 3011.419 | 2674.73 | 3104.065 | 2698.081 | 2743.992 |
| ENSMUSG00000019923 | -0.151484037 | 0.089889 | 0.205017 | 3144.926 | 2700.798 | 2800.657 | 2826.817 | 3253.946 | 2776.675 | 3321.976 | 3390.021 |
| ENSMUSG00000019927 | 0.099434354 | 0.373402 | 0.542211 | 895.3884 | 944.2595 | 833.9034 | 919.5566 | 821.0437 | 1000.05 | 790.7171 | 741.7371 |
| ENSMUSG00000019929 | 0.55353806 | 0.20662 | 0.363213 | 3742.522 | 7867.23 | 5195.541 | 5679.513 | 3406.064 | 6962.358 | 2374.08 | 2577.591 |
| ENSMUSG00000019933 | 0.570576323 | 0.118773 | 0.24758 | 52.31483 | 89.98662 | 35.60859 | 56.93315 | 35.10401 | 53.63396 | 42.42872 | 26.10219 |
| ENSMUSG00000019935 | -0.256202037 | 0.850384 | 0.910207 | 7.04238 | 4.799286 | 427.303 | 18.11509 | 122.864 | 45.81234 | 175.5006 | 202.2919 |
| ENSMUSG00000019942 | -0.037131169 | 0.639772 | 0.766629 | 2423.585 | 2367.248 | 2218.498 | 2504.196 | 2631.825 | 2558.787 | 2330.687 | 2239.35 |
| ENSMUSG00000019943 | -0.05508409 | 0.626402 | 0.756502 | 4060.435 | 3701.449 | 4427.886 | 3607.491 | 3594.26 | 4753.309 | 3812.799 | 4255.744 |
| ENSMUSG00000019944 | -0.181846436 | 0.046053 | 0.128548 | 598.6023 | 639.5049 | 603.6897 | 621.0889 | 746.9353 | 690.5372 | 616.1808 | 739.5619 |
| ENSMUSG00000019945 | -0.104952724 | 0.506493 | 0.66231 | 157.9505 | 116.3827 | 144.9187 | 148.3712 | 146.2667 | 139.6718 | 173.572 | 152.2628 |
| ENSMUSG00000019947 | 0.263298664 | 0.008986 | 0.039441 | 524.1543 | 437.9349 | 514.2542 | 477.0308 | 379.3183 | 405.6068 | 436.823 | 407.8467 |
| ENSMUSG00000019948 | 0.114591353 | 0.354465 | 0.523068 | 426.567 | 428.3363 | 335.3832 | 404.5704 | 377.3681 | 396.6678 | 372.2156 | 324.1021 |
| ENSMUSG00000019951 | 0.040863063 | 0.745717 | 0.841098 | 2523.184 | 1722.944 | 1889.739 | 2122.916 | 1841.01 | 2014.625 | 2057.793 | 2116.452 |
| ENSMUSG00000019952 | 0.069590904 | 0.37984 | 0.547896 | 811.8859 | 841.0749 | 893.5271 | 838.47 | 822.0188 | 870.4344 | 785.8957 | 749.3503 |
| ENSMUSG00000019960 | 0.351868002 | 0.001485 | 0.010486 | 3614.753 | 3294.71 | 2810.594 | 3579.025 | 2652.303 | 2825.839 | 2684.581 | 2256.752 |
| ENSMUSG00000019961 | 0.108086886 | 0.228974 | 0.389058 | 9192.319 | 10348.46 | 8893.865 | 10290.24 | 9133.868 | 9943.512 | 8756.71 | 8093.853 |
| ENSMUSG00000019966 | 0.484916722 | 0.019975 | 0.070342 | 1878.303 | 1838.127 | 1383.766 | 1988.347 | 1156.482 | 1849.254 | 1089.647 | 970.1312 |
| ENSMUSG00000019969 | -0.261651466 | 0.017744 | 0.06495 | 1405.458 | 1439.786 | 1523.716 | 1439.719 | 1715.221 | 1429.121 | 1811.899 | 2007.693 |
| ENSMUSG00000019970 | -1.066864105 | 0.037251 | 0.11096 | 2865.243 | 787.0829 | 2352.651 | 2774.197 | 2578.194 | 3688.452 | 8358.458 | 3767.416 |
| ENSMUSG00000019971 | 0.301682596 | 0.049997 | 0.136097 | 410.4702 | 503.925 | 358.5702 | 389.0432 | 298.3841 | 400.0199 | 302.7868 | 345.854 |
| ENSMUSG00000019975 | 0.457015268 | 1.07E-07 | 5.97E-06 | 1494.997 | 1552.569 | 1399.5 | 1344.83 | 1090.174 | 1139.722 | 1024.075 | 962.5181 |
| ENSMUSG00000019977 | -0.027135744 | 0.683452 | 0.798241 | 1955.77 | 1965.308 | 1882.286 | 2118.603 | 2146.22 | 2004.569 | 1970.043 | 1951.138 |
| ENSMUSG00000019978 | -0.206606279 | 0.297614 | 0.465488 | 2953.776 | 3021.151 | 5043.997 | 3127.873 | 4102.293 | 3178.929 | 4128.122 | 4918.087 |
| ENSMUSG00000019979 | 0.41337429 | 5.10E-05 | 0.000804 | 1132.817 | 1105.036 | 1031.821 | 1167.13 | 799.5913 | 966.5286 | 739.6098 | 827.6568 |
| ENSMUSG00000019982 | -0.030416249 | 0.862084 | 0.917537 | 245.4773 | 239.9643 | 158.1684 | 205.3044 | 213.5494 | 236.8833 | 224.6794 | 189.2408 |
| ENSMUSG00000019984 | 0.20386235 | 0.013259 | 0.052784 | 1523.166 | 1539.371 | 1487.28 | 1467.323 | 1309.574 | 1463.76 | 1274.79 | 1175.686 |

| | | | | | | | | | | | |
|---|---|---|---|---|---|---|---|---|---|---|---|
| ENSMUSG00000019986 | 0.516582732 | 0.000855 | 0.006925 | 796.795 | 757.0874 | 529.1601 | 742.7188 | 451.4765 | 566.5087 | 509.1447 | 446.9999 |
| ENSMUSG00000019987 | 1.533319166 | 0.003129 | 0.018333 | 148.896 | 33.595 | 34.78048 | 90.57547 | 29.25334 | 26.81698 | 32.78583 | 17.40146 |
| ENSMUSG00000019988 | 0.129135657 | 0.260417 | 0.424448 | 1295.798 | 1160.227 | 955.6351 | 1170.58 | 1063.846 | 1157.6 | 972.0035 | 995.1458 |
| ENSMUSG00000019989 | -0.85220791 | 9.78E-05 | 0.001314 | 89.53884 | 121.182 | 109.3101 | 130.2561 | 174.5449 | 248.057 | 152.3577 | 239.27 |
| ENSMUSG00000019990 | -0.182438796 | 0.339123 | 0.507612 | 251.5136 | 190.7716 | 293.1497 | 175.9752 | 229.1512 | 288.2825 | 263.2509 | 255.5839 |
| ENSMUSG00000019992 | 0.098268195 | 0.638247 | 0.765498 | 152.9203 | 170.3747 | 130.0127 | 163.0358 | 162.8436 | 166.4887 | 149.4648 | 95.70802 |
| ENSMUSG00000019996 | 0.097172133 | 0.327956 | 0.495681 | 1658.984 | 1616.16 | 1720.806 | 1648.473 | 1409.036 | 1413.478 | 1566.97 | 1823.89 |
| ENSMUSG00000019997 | 0.346026191 | 0.188123 | 0.340426 | 3096.635 | 1433.787 | 1454.155 | 2197.965 | 1423.663 | 1387.779 | 2230.401 | 1395.379 |
| ENSMUSG00000019998 | -0.279102004 | 0.016058 | 0.060359 | 2481.936 | 2318.055 | 2470.242 | 2499.02 | 2692.282 | 2471.632 | 3253.512 | 3437.875 |
| ENSMUSG00000020000 | 0.303844377 | 0.25639 | 0.420094 | 1225.374 | 1564.567 | 1103.038 | 1423.329 | 1055.07 | 1745.338 | 850.503 | 655.8174 |
| ENSMUSG00000020003 | -0.426509463 | 0.003314 | 0.019073 | 608.6629 | 488.3274 | 544.8942 | 542.5902 | 686.4784 | 588.8561 | 767.5742 | 893.9999 |
| ENSMUSG00000020009 | -0.125904183 | 0.185609 | 0.337199 | 1332.016 | 1227.417 | 1444.218 | 1299.974 | 1407.086 | 1338.614 | 1649.899 | 1393.204 |
| ENSMUSG00000020014 | 0.14097282 | 0.647844 | 0.771952 | 47.28455 | 55.19179 | 60.45178 | 82.81186 | 49.73068 | 80.45093 | 38.57157 | 55.46715 |
| ENSMUSG00000020015 | 0.044576106 | 0.691365 | 0.804479 | 946.6971 | 853.0731 | 789.1856 | 933.3586 | 863.9486 | 983.2892 | 816.7529 | 751.5254 |
| ENSMUSG00000020017 | 3.682129046 | 9.04E-06 | 0.000212 | 12.07265 | 22.79661 | 15.73403 | 54.34528 | 2.925334 | 0 | 1.928578 | 3.262773 |
| ENSMUSG00000020018 | -0.161484126 | 0.023267 | 0.078692 | 2168.047 | 2376.846 | 2375.838 | 2347.199 | 2778.092 | 2460.458 | 2680.724 | 2443.817 |
| ENSMUSG00000020019 | -0.91472632 | 0.001183 | 0.008877 | 254.5318 | 202.7698 | 389.2101 | 296.7425 | 421.2481 | 346.386 | 615.2165 | 775.4524 |
| ENSMUSG00000020020 | 0.002040249 | 0.99663 | 0.998036 | 20.12109 | 9.598572 | 15.73403 | 17.25247 | 23.40267 | 7.821619 | 18.32149 | 13.05109 |
| ENSMUSG00000020021 | 0.551758893 | 0.000459 | 0.004286 | 743.4742 | 638.3051 | 504.3169 | 616.7758 | 446.601 | 496.1141 | 415.6086 | 348.0291 |
| ENSMUSG00000020022 | -0.575032397 | 7.92E-05 | 0.001131 | 2064.424 | 2021.699 | 2289.715 | 2031.478 | 3158.386 | 2287.265 | 3332.583 | 3746.751 |
| ENSMUSG00000020023 | -0.011267789 | 0.930097 | 0.958773 | 425.561 | 451.1329 | 588.7838 | 484.7944 | 468.0534 | 464.8276 | 535.1805 | 500.2919 |
| ENSMUSG00000020024 | 0.027425991 | 0.742014 | 0.838976 | 857.1583 | 758.2872 | 741.1554 | 861.7609 | 779.1139 | 774.3402 | 800.36 | 804.8174 |
| ENSMUSG00000020027 | 0.239897277 | 0.060238 | 0.155468 | 2417.549 | 2630.009 | 2269.012 | 2746.593 | 2081.863 | 2586.721 | 2095.4 | 1757.547 |
| ENSMUSG00000020029 | -0.253477546 | 0.072314 | 0.176516 | 8751.667 | 8434.745 | 10809.28 | 8872.945 | 9455.654 | 9321.135 | 11792.29 | 13383.9 |
| ENSMUSG00000020032 | 0.269882054 | 0.045707 | 0.127852 | 1272.659 | 1554.969 | 1271.972 | 1363.808 | 1340.778 | 1202.295 | 913.1818 | 1073.452 |
| ENSMUSG00000020034 | 0.542619405 | 7.97E-05 | 0.001136 | 1075.472 | 961.057 | 889.3865 | 1052.401 | 620.1708 | 679.3635 | 848.5745 | 581.8612 |
| ENSMUSG00000020038 | -0.239687387 | 0.015503 | 0.059098 | 716.3107 | 649.1034 | 695.6096 | 718.5654 | 761.5619 | 735.2322 | 894.8603 | 891.8247 |
| ENSMUSG00000020042 | -0.550275073 | 0.00055 | 0.004924 | 440.6518 | 476.3291 | 414.8814 | 377.8291 | 722.5575 | 719.5889 | 583.3949 | 475.2773 |
| ENSMUSG00000020044 | -0.672860863 | 0.001712 | 0.011683 | 2629.826 | 2026.499 | 3061.51 | 2331.671 | 3649.842 | 2540.909 | 4921.732 | 4910.474 |
| ENSMUSG00000020048 | -0.252248319 | 0.046414 | 0.129262 | 14970.09 | 14932.98 | 16175.41 | 13117.05 | 15681.74 | 14859.96 | 18975.28 | 20988.33 |
| ENSMUSG00000020051 | -4.371501582 | 3.06E-07 | 1.39E-05 | 33.19979 | 4.799286 | 33.95237 | 42.26855 | 1150.631 | 135.2023 | 234.3223 | 851.5838 |
| ENSMUSG00000020052 | -0.627545183 | 0.361403 | 0.529959 | 8.048435 | 7.198929 | 15.73403 | 4.313117 | 23.40267 | 16.76061 | 3.857157 | 10.87591 |
| ENSMUSG00000020053 | 0.364316254 | 0.201916 | 0.357508 | 974.8667 | 1668.952 | 1597.418 | 1372.434 | 825.9193 | 1847.019 | 757.9313 | 930.978 |
| ENSMUSG00000020056 | -0.082354259 | 0.228185 | 0.388181 | 730.3955 | 707.8947 | 683.188 | 725.4664 | 756.6864 | 732.9974 | 774.3242 | 748.2627 |
| ENSMUSG00000020057 | -0.591911139 | 0.024229 | 0.081141 | 85.51462 | 77.9884 | 165.6213 | 103.5148 | 168.6943 | 132.9675 | 180.3221 | 172.927 |
| ENSMUSG00000020059 | -1.692811754 | NA | NA | 9.054489 | 28.79572 | 36.43669 | 25.8787 | 18.52711 | 262.5829 | 27.0001 | 16.31387 |
| ENSMUSG00000020061 | 0.458124508 | 0.313275 | 0.481484 | 241.453 | 254.3622 | 245.1196 | 200.9913 | 120.9138 | 393.3157 | 92.57176 | 79.39415 |
| ENSMUSG00000020062 | -4.876815617 | 3.08E-07 | 1.40E-05 | 6.036326 | 7.198929 | 73.70149 | 13.80198 | 628.9468 | 205.5968 | 648.9666 | 1490 |
| ENSMUSG00000020063 | 0.03692059 | 0.765089 | 0.854012 | 1022.151 | 933.4612 | 835.5596 | 1023.934 | 846.3966 | 1103.966 | 941.1462 | 827.6568 |
| ENSMUSG00000020064 | 0.235051485 | 0.049533 | 0.135201 | 1411.494 | 1285.009 | 1017.743 | 1253.392 | 1031.668 | 1178.83 | 1030.825 | 977.7444 |
| ENSMUSG00000020069 | 0.232488507 | 0.022198 | 0.075895 | 3658.014 | 3789.036 | 3407.659 | 3772.253 | 2992.617 | 3708.565 | 2923.725 | 2825.562 |
| ENSMUSG00000020070 | 0.311502526 | 0.005121 | 0.026168 | 1182.114 | 1066.641 | 957.2913 | 1130.899 | 939.0322 | 955.3549 | 845.6816 | 753.7006 |
| ENSMUSG00000020072 | -0.336930399 | 0.224805 | 0.384523 | 364.1917 | 202.7698 | 446.3495 | 345.912 | 442.7005 | 282.6956 | 394.3943 | 599.2627 |
| ENSMUSG00000020074 | 0.286721683 | 0.045734 | 0.12787 | 5267.701 | 5287.613 | 4109.065 | 4847.081 | 3463.595 | 5101.93 | 3780.014 | 3648.868 |
| ENSMUSG00000020075 | -0.216613347 | 0.00517 | 0.026332 | 2574.493 | 2574.817 | 2486.804 | 2385.154 | 3016.994 | 2611.303 | 2852.367 | 3160.54 |
| ENSMUSG00000020076 | 0.170932297 | 0.038292 | 0.112999 | 3468.875 | 3423.091 | 3233.756 | 3513.465 | 3042.347 | 3413.578 | 2917.939 | 2741.817 |
| ENSMUSG00000020077 | -0.185260511 | 0.307563 | 0.476044 | 276.6649 | 196.7707 | 321.3054 | 295.8799 | 269.1307 | 296.1041 | 371.2513 | 306.7007 |
| ENSMUSG00000020078 | -0.150935485 | 0.004743 | 0.024772 | 1748.522 | 1733.742 | 1775.461 | 1680.391 | 1969.725 | 1928.588 | 1956.543 | 1846.73 |
| ENSMUSG00000020079 | -0.27863299 | 0.001466 | 0.0104 | 632.8082 | 728.2917 | 720.4528 | 740.131 | 867.8491 | 788.8661 | 858.2174 | 908.1386 |
| ENSMUSG00000020080 | 0.407999149 | 0.003458 | 0.019605 | 207.2472 | 178.7734 | 180.5272 | 185.4641 | 154.0676 | 136.3196 | 125.3576 | 151.1752 |
| ENSMUSG00000020083 | -0.111877046 | 0.318852 | 0.486986 | 208.2533 | 243.5638 | 240.979 | 226.87 | 250.6036 | 258.1134 | 236.2508 | 249.0584 |
| ENSMUSG00000020085 | 0.24556086 | 0.213305 | 0.371183 | 1018.127 | 641.9045 | 568.9093 | 590.8971 | 568.4899 | 530.7527 | 609.4308 | 668.8685 |
| ENSMUSG00000020086 | -0.00688906 | 0.931104 | 0.959275 | 3281.749 | 3268.314 | 3337.27 | 3420.302 | 3456.77 | 3124.178 | 3727.942 | 3060.481 |
| ENSMUSG00000020087 | -0.58605573 | 4.61E-05 | 0.000747 | 728.3834 | 788.2827 | 760.2019 | 738.4057 | 1065.797 | 919.5989 | 1080.968 | 1459.547 |
| ENSMUSG00000020088 | 0.042427826 | 0.443868 | 0.606906 | 5420.621 | 6061.498 | 5484.55 | 5812.357 | 5462.574 | 5644.974 | 5535.02 | 5471.671 |
| ENSMUSG00000020089 | -0.447747529 | 2.25E-05 | 0.000431 | 926.5761 | 1105.036 | 1031.821 | 992.017 | 1293.973 | 1222.407 | 1446.434 | 1566.131 |
| ENSMUSG00000020090 | 0.465782364 | 0.292789 | 0.460125 | 15.09082 | 15.59768 | 21.53077 | 18.11509 | 13.65156 | 13.40849 | 17.3572 | 6.525547 |
| ENSMUSG00000020091 | -0.433044838 | 0.009324 | 0.040449 | 759.571 | 661.1017 | 876.1368 | 703.0381 | 1178.91 | 724.0584 | 1045.289 | 1102.817 |
| ENSMUSG00000020092 | -0.040855292 | 0.620698 | 0.752618 | 1454.755 | 1647.355 | 1465.749 | 1669.176 | 1625.511 | 1502.868 | 1721.256 | 1562.868 |
| ENSMUSG00000020097 | 0.03384712 | 0.696692 | 0.808587 | 2691.195 | 2766.788 | 3286.755 | 2719.852 | 2900.956 | 2619.125 | 2827.296 | 2853.839 |
| ENSMUSG00000020098 | -0.91692294 | 2.69E-05 | 0.000493 | 926.5761 | 909.4647 | 1324.971 | 1082.592 | 2149.145 | 1244.755 | 1999.936 | 2621.095 |
| ENSMUSG00000020099 | -0.198921462 | 0.3562 | 0.524872 | 1140.866 | 1243.015 | 2163.843 | 1287.034 | 1580.655 | 1331.91 | 1802.256 | 1984.854 |
| ENSMUSG00000020100 | -0.21877757 | 0.115043 | 0.242689 | 512.0817 | 555.5174 | 585.4714 | 552.079 | 625.0463 | 536.3396 | 598.8236 | 806.9926 |
| ENSMUSG00000020101 | -0.206287752 | 0.121512 | 0.250741 | 222.338 | 274.7591 | 314.6805 | 283.8031 | 322.7618 | 296.1041 | 315.3226 | 331.7153 |

| | | | | | | | | | | | |
|---|---|---|---|---|---|---|---|---|---|---|---|
| ENSMUSG00000020102 | -0.601962203 | 0.043311 | 0.123203 | 462.785 | 442.7341 | 808.2321 | 477.8934 | 627.9717 | 492.762 | 1015.396 | 1192 |
| ENSMUSG00000020105 | 0.414234364 | 0.001653 | 0.011368 | 2358.191 | 2562.819 | 2120.781 | 2574.069 | 1940.472 | 2121.893 | 1702.935 | 1449.759 |
| ENSMUSG00000020107 | -0.034340619 | 0.530551 | 0.682263 | 1635.844 | 1636.557 | 1652.073 | 1739.049 | 1675.241 | 1731.93 | 1751.149 | 1668.365 |
| ENSMUSG00000020108 | -0.459295414 | 0.264094 | 0.428305 | 1947.721 | 1217.819 | 1377.969 | 2006.462 | 1254.968 | 2233.631 | 4000.836 | 1516.102 |
| ENSMUSG00000020109 | -0.348221399 | 7.26E-06 | 0.00018 | 887.3399 | 860.272 | 978.822 | 885.9143 | 1196.462 | 1063.74 | 1166.79 | 1174.598 |
| ENSMUSG00000020111 | -0.040555316 | 0.682171 | 0.797147 | 1724.377 | 2060.094 | 2259.075 | 1954.705 | 2180.349 | 1900.653 | 2094.436 | 2051.197 |
| ENSMUSG00000020114 | -0.139140939 | 0.062274 | 0.159098 | 3189.192 | 3573.069 | 3113.681 | 2941.546 | 3508.45 | 3426.986 | 3528.334 | 3643.43 |
| ENSMUSG00000020115 | 0.089107743 | 0.293619 | 0.461155 | 1331.01 | 1138.631 | 1190.817 | 1227.513 | 1217.914 | 1162.069 | 1176.433 | 1038.649 |
| ENSMUSG00000020116 | -0.355763915 | 0.000474 | 0.004395 | 723.3531 | 889.0678 | 887.7303 | 816.0418 | 1051.17 | 978.8197 | 1036.611 | 1176.774 |
| ENSMUSG00000020120 | 0.547305285 | 0.002294 | 0.014477 | 147.89 | 116.3827 | 173.0743 | 134.5693 | 105.312 | 92.74205 | 93.53605 | 101.146 |
| ENSMUSG00000020121 | 0.329883762 | 0.024234 | 0.081141 | 834.0191 | 706.6949 | 816.5131 | 829.8438 | 480.7299 | 759.8144 | 634.5023 | 663.4306 |
| ENSMUSG00000020122 | 0.801546275 | 0.000158 | 0.001891 | 762.5892 | 1051.044 | 861.2309 | 1004.956 | 485.6054 | 776.575 | 431.0373 | 418.7226 |
| ENSMUSG00000020123 | 0.055896973 | 0.845202 | 0.906816 | 153.9263 | 235.165 | 231.8699 | 145.7834 | 131.64 | 138.5544 | 190.9293 | 276.2481 |
| ENSMUSG00000020124 | -0.135502421 | 0.107837 | 0.23245 | 1476.888 | 1352.199 | 1430.14 | 1271.507 | 1497.771 | 1403.422 | 1511.041 | 1662.927 |
| ENSMUSG00000020128 | -0.057600378 | 0.521722 | 0.675049 | 1727.395 | 1592.163 | 1545.247 | 1789.944 | 1629.411 | 1936.409 | 1621.934 | 1741.233 |
| ENSMUSG00000020130 | 0.096256189 | 0.221203 | 0.380229 | 1721.359 | 1784.135 | 1535.31 | 1751.988 | 1653.789 | 1620.192 | 1615.184 | 1460.635 |
| ENSMUSG00000020131 | -0.230671182 | 0.173191 | 0.320975 | 192.1564 | 146.3782 | 168.1056 | 213.9306 | 234.0267 | 179.8972 | 200.5721 | 232.7445 |
| ENSMUSG00000020132 | -0.183660587 | 0.127922 | 0.259325 | 1041.266 | 1006.65 | 971.3691 | 1021.346 | 1031.668 | 973.2328 | 1209.219 | 1374.715 |
| ENSMUSG00000020133 | -0.261013047 | 0.007553 | 0.03472 | 821.9464 | 947.859 | 949.0102 | 918.694 | 1183.785 | 957.5896 | 1062.647 | 1153.934 |
| ENSMUSG00000020134 | 0.15860056 | 0.283453 | 0.449975 | 1843.092 | 1551.369 | 1708.384 | 1847.74 | 1383.683 | 2012.391 | 1548.648 | 1284.445 |
| ENSMUSG00000020135 | 0.554306604 | 0.006868 | 0.032534 | 116.7023 | 109.1838 | 135.8095 | 166.4863 | 77.03379 | 94.9768 | 104.1432 | 84.8321 |
| ENSMUSG00000020137 | -0.102414509 | 0.55829 | 0.704774 | 323.9495 | 279.5584 | 261.6817 | 237.2215 | 303.2596 | 219.0053 | 311.4654 | 348.0291 |
| ENSMUSG00000020140 | 0.450427503 | 0.052221 | 0.14027 | 580.4934 | 607.1097 | 461.2554 | 510.6731 | 313.0107 | 604.4994 | 369.3228 | 293.6496 |
| ENSMUSG00000020142 | -0.140719609 | 0.231315 | 0.391871 | 355.1372 | 284.3577 | 335.3832 | 319.1707 | 393.945 | 364.264 | 348.1084 | 321.927 |
| ENSMUSG00000020143 | 0.017042088 | 0.953306 | 0.973347 | 46.2785 | 65.99018 | 51.34261 | 46.58167 | 39.00445 | 40.22547 | 54.96448 | 72.8686 |
| ENSMUSG00000020149 | -0.231446177 | 0.003417 | 0.01944 | 4905.521 | 5132.837 | 5202.166 | 5140.373 | 5951.104 | 5303.057 | 6033.557 | 6639.744 |
| ENSMUSG00000020150 | -0.07772781 | 0.603115 | 0.740579 | 231.3925 | 195.5709 | 222.7607 | 204.4418 | 213.5494 | 191.071 | 223.7151 | 274.073 |
| ENSMUSG00000020151 | -0.119270267 | 0.726238 | 0.828695 | 67.40564 | 34.79482 | 88.60741 | 87.12497 | 97.51113 | 87.15518 | 65.57166 | 53.29196 |
| ENSMUSG00000020152 | -0.188076921 | 0.016824 | 0.062382 | 4159.029 | 3898.22 | 4482.541 | 4256.184 | 4766.344 | 4381.224 | 4741.41 | 5250.89 |
| ENSMUSG00000020153 | -0.592493774 | 1.91E-06 | 6.24E-05 | 1545.299 | 1590.963 | 1702.587 | 1612.243 | 2338.317 | 1960.992 | 2485.937 | 2943.021 |
| ENSMUSG00000020154 | -0.458143643 | 0.024741 | 0.082421 | 683.1109 | 413.9384 | 801.6072 | 672.8463 | 800.5664 | 757.5796 | 898.7175 | 1079.978 |
| ENSMUSG00000020155 | -0.255965989 | 0.594803 | 0.734023 | 13.07871 | 14.39786 | 36.43669 | 11.21411 | 20.47734 | 21.23011 | 27.0001 | 21.75182 |
| ENSMUSG00000020156 | 0.032613137 | 0.711602 | 0.819194 | 2271.671 | 2542.422 | 2220.154 | 2400.681 | 2429.977 | 2490.627 | 2263.187 | 2037.058 |
| ENSMUSG00000020159 | -1.808603259 | 3.58E-05 | 0.000614 | 39.23612 | 39.59411 | 23.18699 | 23.29083 | 72.15824 | 210.0663 | 63.64308 | 91.35765 |
| ENSMUSG00000020160 | 0.832834083 | 3.20E-05 | 0.000562 | 4052.387 | 5214.424 | 3768.713 | 4722.864 | 2576.244 | 3464.977 | 2166.758 | 1761.898 |
| ENSMUSG00000020163 | -0.810022334 | 3.66E-06 | 0.000107 | 1515.118 | 1329.402 | 1647.932 | 1337.929 | 2484.584 | 1794.503 | 2648.902 | 3295.401 |
| ENSMUSG00000020166 | 0.025005126 | 0.613377 | 0.747226 | 2672.08 | 2725.995 | 2637.52 | 2668.957 | 2757.615 | 2616.89 | 2567.902 | 2574.328 |
| ENSMUSG00000020167 | 0.028351194 | 0.591705 | 0.731496 | 4528.251 | 4574.92 | 4634.913 | 4499.444 | 4568.396 | 4713.084 | 4360.516 | 4241.605 |
| ENSMUSG00000020170 | 0.269976891 | 0.004363 | 0.023299 | 1659.99 | 1415.789 | 1465.749 | 1588.952 | 1265.694 | 1436.943 | 1208.254 | 1175.686 |
| ENSMUSG00000020171 | -0.007015016 | 0.896039 | 0.937117 | 1750.535 | 1720.544 | 1795.335 | 1841.701 | 1799.08 | 1838.08 | 1747.292 | 1764.073 |
| ENSMUSG00000020173 | -0.909743413 | 3.61E-05 | 0.000618 | 264.5923 | 271.1597 | 434.756 | 286.391 | 564.5894 | 419.0153 | 703.9311 | 676.4817 |
| ENSMUSG00000020175 | 0.355529431 | 0.033346 | 0.101995 | 411.4762 | 502.7252 | 481.13 | 526.2003 | 438.8001 | 443.5975 | 324.9654 | 294.7372 |
| ENSMUSG00000020176 | 0.323885459 | 0.00644 | 0.031081 | 15196.45 | 14501.04 | 12203.81 | 14604.22 | 10970.98 | 13665.49 | 10315.97 | 10190.73 |
| ENSMUSG00000020178 | 0.007697771 | 0.96597 | 0.98085 | 305.8405 | 243.5638 | 366.0231 | 272.589 | 337.3885 | 225.7096 | 307.6082 | 312.1386 |
| ENSMUSG00000020180 | -0.20946827 | 0.001942 | 0.012855 | 3198.247 | 3427.89 | 3578.249 | 3366.819 | 4072.065 | 3613.588 | 3915.014 | 4091.518 |
| ENSMUSG00000020181 | -0.72050962 | 0.046981 | 0.130272 | 170.0232 | 146.3782 | 404.9441 | 171.6621 | 360.7912 | 204.4795 | 432.9658 | 475.2773 |
| ENSMUSG00000020182 | 0.037896904 | 0.929321 | 0.958295 | 326.9677 | 87.58697 | 139.95 | 397.6694 | 211.5992 | 166.4887 | 230.4651 | 319.7518 |
| ENSMUSG00000020183 | 0.514653413 | 0.000266 | 0.002832 | 2156.981 | 2002.502 | 1617.292 | 2082.373 | 1409.036 | 1413.478 | 1591.077 | 1085.416 |
| ENSMUSG00000020184 | 0.089873898 | 0.482293 | 0.640923 | 1989.976 | 1823.729 | 1623.917 | 1889.145 | 1648.913 | 2080.551 | 1730.899 | 1423.657 |
| ENSMUSG00000020185 | 0.0625839 | 0.594808 | 0.734023 | 342.0585 | 362.3461 | 343.6643 | 316.5828 | 283.7574 | 368.7335 | 339.4298 | 314.3138 |
| ENSMUSG00000020186 | 0.009972371 | 0.900765 | 0.940407 | 2392.397 | 2234.068 | 2534.006 | 2630.139 | 2255.432 | 2383.359 | 2595.866 | 2493.846 |
| ENSMUSG00000020189 | 0.061696595 | 0.592926 | 0.732467 | 764.6013 | 622.7074 | 717.9685 | 778.0864 | 664.0508 | 709.5326 | 611.3593 | 781.978 |
| ENSMUSG00000020190 | -0.22743356 | 0.011735 | 0.048149 | 2852.164 | 2939.563 | 3290.896 | 3095.093 | 3656.667 | 3087.305 | 3824.371 | 3690.197 |
| ENSMUSG00000020191 | 0.327833574 | 0.430166 | 0.594915 | 33.19979 | 19.19714 | 27.32752 | 25.01608 | 14.62667 | 35.75597 | 16.39292 | 17.40146 |
| ENSMUSG00000020193 | 0.144987454 | 0.560172 | 0.706138 | 51.30877 | 81.58786 | 53.82693 | 64.69676 | 53.63112 | 65.92507 | 46.28588 | 60.9051 |
| ENSMUSG00000020196 | -0.032488829 | 0.68588 | 0.800266 | 1340.064 | 1287.409 | 1512.951 | 1382.785 | 1447.065 | 1507.338 | 1344.219 | 1356.226 |
| ENSMUSG00000020198 | -0.06713955 | 0.219432 | 0.378197 | 3446.742 | 3346.302 | 3512.828 | 3384.935 | 3545.505 | 3391.23 | 3625.727 | 3782.642 |
| ENSMUSG00000020205 | 0.388201103 | 0.052276 | 0.140316 | 584.5176 | 549.5183 | 520.8791 | 491.6954 | 343.2392 | 601.1473 | 334.6083 | 362.1678 |
| ENSMUSG00000020211 | -0.104864336 | 0.123336 | 0.252965 | 978.8909 | 982.6538 | 978.822 | 947.1606 | 1087.249 | 975.4676 | 1025.039 | 1090.854 |
| ENSMUSG00000020212 | 0.012351002 | 0.926502 | 0.956348 | 501.0151 | 483.5281 | 461.2554 | 563.2931 | 510.9583 | 595.5604 | 465.7517 | 420.8978 |
| ENSMUSG00000020216 | -0.704505862 | 0.00163 | 0.01122 | 80.48435 | 58.79126 | 73.70149 | 67.28463 | 136.5156 | 109.5027 | 128.2505 | 82.65692 |
| ENSMUSG00000020218 | 0.139271374 | 0.619448 | 0.751645 | 79.47829 | 101.9848 | 86.9512 | 103.5148 | 82.88446 | 122.9112 | 81.96458 | 50.02919 |
| ENSMUSG00000020219 | -0.480981853 | 0.000712 | 0.006035 | 1538.257 | 1636.557 | 1632.198 | 1534.607 | 2247.632 | 1654.831 | 2227.508 | 2720.065 |
| ENSMUSG00000020220 | -0.008018528 | 0.897429 | 0.937817 | 1575.481 | 1466.182 | 1565.95 | 1431.955 | 1483.144 | 1538.624 | 1496.577 | 1557.43 |

| | | | | | | | | | | | |
|---|---|---|---|---|---|---|---|---|---|---|---|
| ENSMUSG00000020224 | -0.097890401 | 0.204077 | 0.360071 | 1009.073 | 1017.449 | 1056.664 | 1144.701 | 1194.511 | 1154.247 | 1120.504 | 1058.226 |
| ENSMUSG00000020225 | -0.725443047 | 8.24E-05 | 0.001163 | 1461.797 | 1180.624 | 1546.075 | 1350.868 | 2299.312 | 1575.497 | 2276.687 | 3009.365 |
| ENSMUSG00000020227 | 0.892666743 | 7.21E-06 | 0.000179 | 195.1745 | 154.777 | 149.8873 | 144.9207 | 91.66046 | 108.3853 | 71.3574 | 76.13138 |
| ENSMUSG00000020228 | 0.147956972 | 0.257766 | 0.421524 | 341.0524 | 376.744 | 370.1637 | 410.6088 | 379.3183 | 343.0338 | 276.751 | 354.5547 |
| ENSMUSG00000020230 | 0.309290197 | 0.000219 | 0.00243 | 1096.599 | 1127.832 | 980.4783 | 1127.449 | 891.2517 | 798.9225 | 907.3961 | 895.0875 |
| ENSMUSG00000020231 | 0.027630697 | 0.824572 | 0.893455 | 1190.162 | 1059.442 | 1216.489 | 1172.305 | 1128.204 | 920.7163 | 1145.576 | 1357.314 |
| ENSMUSG00000020232 | -0.23518911 | 0.013568 | 0.053553 | 1447.712 | 1610.16 | 1602.386 | 1547.547 | 1834.184 | 1559.854 | 1900.614 | 2010.956 |
| ENSMUSG00000020235 | -0.160599423 | 0.027718 | 0.089387 | 1840.073 | 1697.747 | 1815.21 | 1821.861 | 2082.838 | 1804.559 | 2100.222 | 2032.708 |
| ENSMUSG00000020238 | -0.358236361 | 0.001779 | 0.012064 | 1201.229 | 1238.216 | 1431.796 | 1293.935 | 1714.246 | 1350.905 | 1735.72 | 1821.715 |
| ENSMUSG00000020241 | 0.052703075 | 0.759238 | 0.84986 | 2831.037 | 4264.166 | 4533.884 | 3542.795 | 2820.022 | 3862.762 | 3786.764 | 4158.948 |
| ENSMUSG00000020246 | 0.030287938 | 0.740775 | 0.837968 | 591.56 | 644.3042 | 544.8942 | 546.9033 | 547.0374 | 576.565 | 552.5377 | 599.2627 |
| ENSMUSG00000020248 | 0.04042496 | 0.659737 | 0.780834 | 1277.689 | 1347.4 | 1200.755 | 1412.977 | 1175.984 | 1419.065 | 1284.433 | 1214.839 |
| ENSMUSG00000020250 | -0.194196247 | 0.096036 | 0.214632 | 2778.722 | 2686.4 | 3040.808 | 2663.781 | 3443.118 | 2586.721 | 3102.118 | 3647.781 |
| ENSMUSG00000020251 | 0.344299767 | 0.343244 | 0.511966 | 73.44197 | 187.1722 | 128.3565 | 166.4863 | 101.4116 | 178.7799 | 78.10742 | 79.39415 |
| ENSMUSG00000020253 | 0.223620752 | 0.010206 | 0.043294 | 1311.895 | 1201.021 | 1074.054 | 1239.59 | 1018.991 | 970.9981 | 1084.825 | 1056.051 |
| ENSMUSG00000020255 | -0.116393697 | 0.144802 | 0.282897 | 1500.027 | 1454.184 | 1458.296 | 1601.892 | 1793.23 | 1497.281 | 1645.077 | 1583.533 |
| ENSMUSG00000020256 | 0.384079602 | 0.204725 | 0.36081 | 30.18163 | 34.79482 | 29.81184 | 29.3292 | 27.30312 | 25.6996 | 23.14294 | 18.48905 |
| ENSMUSG00000020257 | 0.020141783 | 0.778852 | 0.863592 | 3338.088 | 3863.425 | 3383.644 | 3417.714 | 3538.679 | 3558.836 | 3237.119 | 3469.416 |
| ENSMUSG00000020258 | -1.265166811 | 1.42E-05 | 0.000298 | 84.50857 | 75.58876 | 110.9663 | 99.2017 | 199.8978 | 139.6718 | 201.5364 | 351.2919 |
| ENSMUSG00000020260 | -0.279448528 | 0.000794 | 0.006544 | 1598.62 | 1517.774 | 1680.228 | 1651.061 | 2013.605 | 1714.052 | 2031.757 | 2068.598 |
| ENSMUSG00000020261 | 0.151505312 | 0.190857 | 0.343486 | 561.3783 | 491.9268 | 539.9255 | 622.8142 | 497.3068 | 560.9218 | 442.6087 | 498.1167 |
| ENSMUSG00000020262 | -0.040788521 | 0.790684 | 0.871818 | 512.0817 | 352.7475 | 453.8024 | 497.7338 | 430.9992 | 557.5697 | 459.0016 | 424.1605 |
| ENSMUSG00000020263 | 0.404306045 | 0.004506 | 0.023821 | 2156.981 | 2012.101 | 1615.636 | 2090.999 | 1590.407 | 1703.995 | 1485.005 | 1170.248 |
| ENSMUSG00000020264 | 0.787567172 | 0.148953 | 0.288738 | 30.18163 | 32.39518 | 11.59349 | 26.74133 | 12.67645 | 25.6996 | 14.46434 | 5.437955 |
| ENSMUSG00000020265 | 0.014879823 | 0.806723 | 0.88179 | 5522.232 | 5960.713 | 6185.957 | 6129.803 | 6218.285 | 5706.43 | 5839.735 | 5791.423 |
| ENSMUSG00000020267 | -0.238954194 | 0.001109 | 0.008441 | 5520.22 | 5833.532 | 6019.507 | 6156.544 | 7307.484 | 6227.126 | 7146.347 | 7087.831 |
| ENSMUSG00000020268 | -0.700777044 | 0.003147 | 0.018406 | 52.31483 | 65.99018 | 63.76421 | 52.62003 | 95.56091 | 71.51194 | 88.7146 | 125.073 |
| ENSMUSG00000020271 | -0.124786278 | 0.135676 | 0.270123 | 2371.27 | 2260.464 | 2552.225 | 2622.375 | 2648.402 | 2420.232 | 2739.546 | 2888.642 |
| ENSMUSG00000020272 | -0.022065824 | 0.857504 | 0.914643 | 239.4409 | 269.9598 | 293.9779 | 244.9851 | 243.7778 | 261.4655 | 270.9653 | 289.2992 |
| ENSMUSG00000020273 | 0.099851012 | 0.418365 | 0.583933 | 456.7487 | 481.1284 | 392.5225 | 445.9763 | 462.2028 | 439.128 | 402.1086 | 351.2919 |
| ENSMUSG00000020275 | 0.260552806 | 0.203849 | 0.35982 | 105.6357 | 89.98662 | 108.482 | 89.71284 | 78.0089 | 106.1505 | 78.10742 | 67.43065 |
| ENSMUSG00000020277 | -0.394211803 | 0.018162 | 0.066009 | 2478.918 | 2625.21 | 2765.876 | 2458.477 | 3432.392 | 2290.617 | 3691.299 | 4158.948 |
| ENSMUSG00000020280 | 0.114813149 | 0.188852 | 0.341288 | 923.5579 | 925.0624 | 838.0439 | 1033.423 | 882.4757 | 881.6082 | 844.7173 | 826.5692 |
| ENSMUSG00000020282 | -0.101671061 | 0.235044 | 0.395908 | 1747.516 | 1529.772 | 1818.522 | 1740.774 | 2002.879 | 1697.291 | 1801.292 | 1838.029 |
| ENSMUSG00000020283 | 0.01537519 | 0.839334 | 0.902882 | 911.4852 | 991.0526 | 872.8244 | 1009.269 | 920.5051 | 945.2985 | 929.5748 | 947.2918 |
| ENSMUSG00000020284 | 0.050126977 | 0.596775 | 0.735694 | 670.0322 | 649.1034 | 676.5631 | 766.0097 | 697.2046 | 695.0067 | 677.8953 | 599.2627 |
| ENSMUSG00000020287 | 0.236886322 | 0.00132 | 0.009628 | 860.1765 | 799.0811 | 840.5282 | 857.4478 | 739.1344 | 701.7109 | 741.5384 | 666.6933 |
| ENSMUSG00000020288 | 0.18246171 | 0.082083 | 0.192406 | 837.0372 | 868.6708 | 705.5469 | 829.8438 | 705.9806 | 802.2746 | 684.6453 | 661.2554 |
| ENSMUSG00000020289 | -0.022276471 | 0.733516 | 0.833185 | 787.7406 | 829.0767 | 799.1229 | 792.751 | 817.1433 | 819.0352 | 783.9671 | 837.4451 |
| ENSMUSG00000020290 | 0.216885897 | 0.119601 | 0.248522 | 4960.854 | 5397.997 | 4588.539 | 5118.808 | 3904.346 | 5642.739 | 3877.407 | 3841.372 |
| ENSMUSG00000020297 | 0.009920091 | 0.972476 | 0.984801 | 392.3612 | 208.7689 | 356.914 | 562.4305 | 332.513 | 275.9914 | 456.1088 | 446.9999 |
| ENSMUSG00000020300 | -0.040506584 | 0.833091 | 0.898985 | 919.5337 | 489.5272 | 719.6247 | 724.6037 | 648.449 | 692.7719 | 704.8954 | 890.7371 |
| ENSMUSG00000020303 | -0.351344487 | 0.033295 | 0.101905 | 249.5015 | 185.9723 | 255.0568 | 274.3143 | 249.6285 | 319.569 | 340.3941 | 325.1897 |
| ENSMUSG00000020305 | 0.289195392 | 0.041537 | 0.119857 | 572.4449 | 536.3202 | 462.0835 | 489.1075 | 420.273 | 519.579 | 369.3228 | 376.3065 |
| ENSMUSG00000020307 | -0.564536624 | 3.72E-06 | 0.000107 | 689.1472 | 692.297 | 737.843 | 718.5654 | 1080.423 | 827.9742 | 1092.54 | 1196.35 |
| ENSMUSG00000020308 | -0.276600024 | 0.0049 | 0.025356 | 815.9101 | 850.6735 | 977.9939 | 985.116 | 1125.278 | 1003.402 | 1092.54 | 1180.036 |
| ENSMUSG00000020309 | -0.694462276 | 0.005229 | 0.026536 | 117.7084 | 118.7823 | 91.09173 | 96.61383 | 145.2916 | 122.9112 | 167.7863 | 249.0584 |
| ENSMUSG00000020310 | 0.360460193 | 0.31996 | 0.488031 | 23.13925 | 20.39697 | 24.8432 | 27.60395 | 26.32801 | 17.87799 | 14.46434 | 16.31387 |
| ENSMUSG00000020311 | 0.010763502 | 0.887008 | 0.932224 | 1159.981 | 1221.418 | 1228.082 | 1300.836 | 1142.83 | 1314.032 | 1241.04 | 1178.949 |
| ENSMUSG00000020312 | 0.080360949 | 0.448667 | 0.611295 | 467.8153 | 557.917 | 540.7536 | 593.485 | 509.9832 | 555.3349 | 481.1803 | 498.1167 |
| ENSMUSG00000020315 | 0.110780213 | 0.043223 | 0.123138 | 11447.89 | 10948.37 | 11331.81 | 12099.16 | 10055.35 | 10784.89 | 10680.47 | 10925.94 |
| ENSMUSG00000020319 | -0.103550941 | 0.312775 | 0.481048 | 292.7618 | 287.9572 | 327.1021 | 311.4071 | 331.5378 | 301.691 | 339.4298 | 339.3284 |
| ENSMUSG00000020321 | -1.1230685 | 4.63E-07 | 1.98E-05 | 5155.023 | 4426.142 | 6289.47 | 4979.063 | 10855.91 | 6752.292 | 12210.79 | 15596.06 |
| ENSMUSG00000020325 | -0.924741856 | 0.028166 | 0.090484 | 109.6599 | 79.18822 | 167.2775 | 108.6906 | 213.5494 | 68.15982 | 277.7153 | 324.1021 |
| ENSMUSG00000020326 | -0.115527914 | 0.320577 | 0.488526 | 1197.205 | 1247.814 | 1286.05 | 1230.101 | 1340.778 | 1076.031 | 1409.791 | 1547.642 |
| ENSMUSG00000020328 | -0.211159987 | 0.001688 | 0.011553 | 1588.56 | 1706.146 | 1523.716 | 1608.793 | 1969.725 | 1778.86 | 1799.364 | 1885.883 |
| ENSMUSG00000020329 | -0.273552516 | 0.079442 | 0.188213 | 605.6447 | 515.9233 | 595.4087 | 597.7981 | 671.8517 | 525.1658 | 738.6455 | 863.5473 |
| ENSMUSG00000020330 | 0.000475628 | 0.996908 | 0.998149 | 863.1946 | 865.0713 | 631.0173 | 771.1854 | 819.0935 | 722.941 | 759.8599 | 823.3065 |
| ENSMUSG00000020331 | -0.367788484 | 0.257487 | 0.421256 | 28.16952 | 27.5959 | 21.53077 | 34.50494 | 37.05423 | 24.58223 | 35.6787 | 46.76642 |
| ENSMUSG00000020333 | 0.334153842 | 0.14227 | 0.279433 | 80.48435 | 71.98929 | 68.73285 | 98.33908 | 56.55646 | 55.8687 | 79.07171 | 61.99269 |
| ENSMUSG00000020334 | -1.065579167 | 0.026728 | 0.087216 | 34.20585 | 41.99375 | 55.48314 | 16.38985 | 77.03379 | 33.52122 | 66.53595 | 132.6861 |
| ENSMUSG00000020335 | -0.017884318 | 0.936332 | 0.962298 | 57.3451 | 67.19001 | 47.20208 | 56.93315 | 54.60623 | 51.39921 | 57.85735 | 66.34306 |
| ENSMUSG00000020340 | -0.736382413 | 0.006686 | 0.03192 | 1060.381 | 707.8947 | 1277.769 | 1074.829 | 1378.807 | 992.2282 | 2285.365 | 2211.073 |
| ENSMUSG00000020346 | -0.233095 | 0.059484 | 0.153824 | 1858.182 | 1768.537 | 2095.11 | 1932.277 | 2235.93 | 1834.728 | 2236.187 | 2691.788 |

| | | | | | | | | | | | |
|---|---|---|---|---|---|---|---|---|---|---|---|
| ENSMUSG00000020349 | -0.251077093 | 0.000189 | 0.002171 | 6318.021 | 6206.677 | 6116.396 | 6045.265 | 7056.881 | 6762.348 | 7605.349 | 7952.466 |
| ENSMUSG00000020354 | -0.04086631 | 0.886983 | 0.932224 | 30.18163 | 44.3934 | 35.60859 | 34.50494 | 43.88001 | 27.93435 | 33.75012 | 42.41605 |
| ENSMUSG00000020357 | -0.104365766 | 0.438352 | 0.601907 | 423.5489 | 346.7484 | 505.9732 | 408.0209 | 466.1032 | 427.9543 | 429.1087 | 491.5912 |
| ENSMUSG00000020358 | -0.275170675 | 2.24E-05 | 0.00043 | 15521.41 | 17540.19 | 17136.01 | 17114.45 | 20994.15 | 19014.35 | 20213.43 | 21233.04 |
| ENSMUSG00000020359 | -0.026664924 | 0.742245 | 0.839031 | 674.0564 | 629.9063 | 663.3134 | 634.0283 | 698.1797 | 693.8893 | 612.3236 | 647.1167 |
| ENSMUSG00000020361 | -0.184750065 | 0.001014 | 0.007919 | 5548.39 | 5450.789 | 5454.738 | 5242.163 | 6045.69 | 5931.022 | 6052.843 | 6629.955 |
| ENSMUSG00000020362 | 0.125443305 | 0.206538 | 0.363138 | 5494.063 | 5747.145 | 5025.779 | 5776.127 | 4804.373 | 5948.9 | 4763.588 | 4690.78 |
| ENSMUSG00000020363 | 0.443540504 | 0.089226 | 0.203932 | 437.6336 | 478.7288 | 322.1335 | 442.5259 | 339.3387 | 425.7195 | 301.8225 | 168.5766 |
| ENSMUSG00000020364 | 0.051868511 | 0.741275 | 0.838346 | 288.7376 | 363.5459 | 236.0104 | 332.11 | 279.8569 | 297.2215 | 321.1083 | 276.2481 |
| ENSMUSG00000020366 | 0.088371931 | 0.361057 | 0.529619 | 1766.631 | 1562.168 | 1459.952 | 1753.714 | 1577.73 | 1687.235 | 1429.077 | 1460.635 |
| ENSMUSG00000020368 | -0.040437201 | 0.426735 | 0.591766 | 9954.908 | 10436.05 | 10116.15 | 9868.413 | 9849.599 | 10302.19 | 10467.36 | 10902.01 |
| ENSMUSG00000020372 | -0.182589034 | 0.041168 | 0.119094 | 35735.05 | 43134.78 | 41927.04 | 41368.83 | 50278.69 | 41952.93 | 47764.14 | 44046.35 |
| ENSMUSG00000020375 | -0.282742249 | 0.010941 | 0.04565 | 901.4247 | 892.6672 | 1040.102 | 955.7868 | 1180.86 | 975.4676 | 1140.754 | 1315.985 |
| ENSMUSG00000020376 | -0.018505615 | 0.736348 | 0.834871 | 1760.595 | 1749.34 | 1738.196 | 1859.816 | 1763.976 | 1860.428 | 1811.899 | 1766.248 |
| ENSMUSG00000020377 | -0.317509552 | 0.351491 | 0.520274 | 18.10898 | 26.39607 | 33.95237 | 36.23019 | 41.92979 | 39.10809 | 37.60728 | 25.0146 |
| ENSMUSG00000020380 | 0.153067503 | 0.181142 | 0.331497 | 1215.314 | 1304.206 | 993.728 | 1067.065 | 1002.414 | 1141.956 | 1035.647 | 935.3283 |
| ENSMUSG00000020381 | -0.435990206 | 0.011732 | 0.048149 | 223.3441 | 159.5763 | 225.245 | 192.365 | 270.1058 | 216.7706 | 279.6439 | 318.6642 |
| ENSMUSG00000020385 | 0.247231027 | 0.093915 | 0.211016 | 2863.231 | 2474.032 | 1999.049 | 2389.467 | 1919.019 | 2523.031 | 2014.4 | 1736.883 |
| ENSMUSG00000020386 | -0.270250728 | 0.003263 | 0.018892 | 1313.907 | 1149.429 | 1243.816 | 1216.299 | 1559.203 | 1281.628 | 1570.827 | 1525.89 |
| ENSMUSG00000020387 | -0.226887376 | 0.076114 | 0.182864 | 192.1564 | 158.3764 | 173.9024 | 175.9752 | 196.9725 | 214.5358 | 218.8936 | 190.3284 |
| ENSMUSG00000020388 | 0.405974259 | 0.000991 | 0.007787 | 988.9514 | 1268.211 | 1032.649 | 1113.647 | 842.4962 | 950.8854 | 823.5029 | 704.759 |
| ENSMUSG00000020389 | 0.201912262 | 0.087679 | 0.20137 | 264.5923 | 299.9554 | 241.8071 | 282.9405 | 235.9769 | 222.3574 | 248.7866 | 237.0949 |
| ENSMUSG00000020390 | 0.326946474 | 0.016286 | 0.060954 | 3944.739 | 3577.868 | 3107.056 | 4067.27 | 2783.943 | 3622.527 | 2779.081 | 2531.912 |
| ENSMUSG00000020392 | -0.041033332 | 0.469333 | 0.629429 | 1354.149 | 1396.592 | 1312.549 | 1420.741 | 1422.687 | 1417.948 | 1391.469 | 1408.43 |
| ENSMUSG00000020393 | 0.009336453 | 0.939059 | 0.964188 | 1447.712 | 1514.175 | 1596.59 | 1272.37 | 1527.024 | 1168.773 | 1573.72 | 1521.54 |
| ENSMUSG00000020396 | 0.846740838 | 0.00511 | 0.026125 | 47.28455 | 43.19358 | 37.2648 | 63.83414 | 24.37778 | 27.93435 | 23.14294 | 31.54014 |
| ENSMUSG00000020397 | -0.090926719 | 0.38526 | 0.552791 | 697.1957 | 607.1097 | 644.267 | 754.7956 | 677.7024 | 766.5186 | 667.2881 | 772.1897 |
| ENSMUSG00000020399 | -0.243596142 | 0.706673 | 0.815556 | 17.10292 | 3.599465 | 7.45296 | 13.80198 | 6.825779 | 21.23011 | 14.46434 | 7.613138 |
| ENSMUSG00000020400 | -0.120540242 | 0.314138 | 0.48222 | 636.8324 | 643.1043 | 688.1566 | 502.0469 | 721.5824 | 620.1426 | 665.3595 | 676.4817 |
| ENSMUSG00000020402 | -0.370561749 | 7.98E-06 | 0.000193 | 8874.405 | 8259.571 | 9116.626 | 9011.828 | 11133.82 | 10160.28 | 11558.93 | 12740.04 |
| ENSMUSG00000020407 | -0.955355533 | 0.010253 | 0.04344 | 138.8355 | 86.38715 | 183.8397 | 100.9269 | 207.6987 | 109.5027 | 333.6441 | 339.3284 |
| ENSMUSG00000020409 | 0.010069851 | 0.882636 | 0.929757 | 2077.502 | 2080.491 | 2152.249 | 2248.859 | 2174.498 | 1967.696 | 2266.08 | 2091.438 |
| ENSMUSG00000020411 | -0.432695309 | 0.384228 | 0.552197 | 27.16347 | 17.99732 | 70.38906 | 26.74133 | 68.25779 | 25.6996 | 40.50014 | 58.72992 |
| ENSMUSG00000020412 | 0.145503268 | 0.076503 | 0.183536 | 1372.258 | 1366.597 | 1250.441 | 1177.481 | 1105.776 | 1176.595 | 1141.718 | 1244.204 |
| ENSMUSG00000020413 | -0.197241273 | 0.006287 | 0.030453 | 592.566 | 615.5084 | 590.44 | 611.6001 | 696.2295 | 672.6592 | 669.2167 | 724.3357 |
| ENSMUSG00000020415 | 0.064620384 | 0.573317 | 0.715844 | 832.007 | 874.6699 | 971.3691 | 818.6297 | 766.4375 | 730.7627 | 910.289 | 936.4159 |
| ENSMUSG00000020420 | -0.071549475 | 0.735393 | 0.834456 | 130.7871 | 109.1838 | 89.43552 | 77.63611 | 118.9636 | 97.21155 | 105.1075 | 104.4087 |
| ENSMUSG00000020422 | 0.127134553 | 0.0422 | 0.12109 | 4471.912 | 4139.384 | 4335.138 | 4167.334 | 3619.613 | 3977.852 | 4113.658 | 3961.007 |
| ENSMUSG00000020423 | 0.271359612 | 0.151854 | 0.292335 | 1277.689 | 835.0758 | 862.059 | 1092.944 | 721.5824 | 883.8429 | 1074.218 | 690.6203 |
| ENSMUSG00000020424 | -0.031312235 | 0.855994 | 0.913721 | 124.7507 | 112.7832 | 132.4971 | 153.547 | 146.2667 | 122.9112 | 150.4291 | 116.3722 |
| ENSMUSG00000020427 | 0.143063708 | 0.201959 | 0.35754 | 5708.352 | 4894.072 | 5369.443 | 6045.265 | 4138.372 | 4953.319 | 5382.662 | 5467.32 |
| ENSMUSG00000020429 | 0.875839606 | 0.331132 | 0.499012 | 653.9353 | 22.79661 | 118.4192 | 535.6892 | 111.1627 | 170.9582 | 259.3938 | 183.8029 |
| ENSMUSG00000020430 | -0.265752913 | 0.002737 | 0.016518 | 2783.752 | 2922.765 | 3127.759 | 2978.639 | 3800.984 | 3049.314 | 3622.834 | 3729.35 |
| ENSMUSG00000020431 | -0.896424057 | 0.00275 | 0.016573 | 366.2038 | 211.1686 | 472.0208 | 340.7363 | 525.585 | 394.4331 | 731.8955 | 938.5911 |
| ENSMUSG00000020432 | -1.470661687 | 6.38E-07 | 2.57E-05 | 1632.826 | 1135.031 | 1994.909 | 1546.684 | 5249.999 | 2045.912 | 3941.05 | 6252.561 |
| ENSMUSG00000020434 | -0.044195738 | 0.900821 | 0.940413 | 17.10292 | 35.99465 | 27.32752 | 35.36756 | 24.37778 | 27.93435 | 27.96439 | 39.15328 |
| ENSMUSG00000020435 | 0.619488468 | 0.000415 | 0.003965 | 219.3198 | 272.3595 | 238.4947 | 270.0012 | 165.7689 | 208.949 | 140.7862 | 135.9489 |
| ENSMUSG00000020436 | -0.173586469 | 0.628009 | 0.7577 | 19.11503 | 16.7975 | 15.73403 | 18.97772 | 20.47734 | 18.99536 | 16.39292 | 23.927 |
| ENSMUSG00000020437 | -0.1294716 | 0.754523 | 0.847512 | 34.20585 | 29.99554 | 42.23344 | 18.11509 | 32.17867 | 18.99536 | 52.07161 | 32.62773 |
| ENSMUSG00000020439 | -0.277895417 | 0.036968 | 0.110508 | 1682.123 | 1508.176 | 1864.068 | 1494.064 | 1918.044 | 1607.901 | 2071.293 | 2343.759 |
| ENSMUSG00000020440 | 0.023202104 | 0.743563 | 0.839647 | 3544.329 | 3627.06 | 4191.048 | 3770.527 | 3687.871 | 3813.598 | 3606.441 | 3790.255 |
| ENSMUSG00000020441 | -0.035591004 | 0.70371 | 0.813356 | 643.8748 | 706.6949 | 610.3146 | 626.2647 | 626.0215 | 734.1148 | 663.4309 | 625.3649 |
| ENSMUSG00000020444 | -0.273000248 | 0.009017 | 0.039559 | 985.9333 | 1052.243 | 1113.803 | 1029.972 | 1314.45 | 1068.21 | 1244.897 | 1425.832 |
| ENSMUSG00000020448 | -0.222125911 | 0.004742 | 0.024772 | 1108.672 | 1109.835 | 1197.442 | 1149.877 | 1253.018 | 1263.75 | 1342.291 | 1470.423 |
| ENSMUSG00000020451 | -0.034098222 | 0.592675 | 0.732341 | 1503.045 | 1469.781 | 1536.138 | 1409.527 | 1561.153 | 1577.732 | 1476.327 | 1445.409 |
| ENSMUSG00000020453 | -0.065459069 | 0.422161 | 0.587233 | 1716.329 | 1718.144 | 1848.334 | 1862.404 | 1675.241 | 1878.306 | 1880.364 | 2047.934 |
| ENSMUSG00000020454 | -0.075300925 | 0.25833 | 0.422222 | 1861.201 | 2087.689 | 2051.22 | 2053.907 | 2107.216 | 2257.096 | 2062.615 | 2059.898 |
| ENSMUSG00000020455 | 0.144543543 | 0.145351 | 0.283633 | 690.1533 | 617.9081 | 545.7223 | 594.3476 | 564.5894 | 562.0392 | 563.1449 | 522.0437 |
| ENSMUSG00000020456 | -0.586187719 | 0.00393 | 0.021566 | 2906.491 | 2628.809 | 3517.797 | 2492.982 | 4000.882 | 2870.534 | 4671.017 | 5792.51 |
| ENSMUSG00000020457 | -0.085626786 | 0.295121 | 0.462844 | 1984.945 | 2244.866 | 1828.459 | 2130.68 | 2180.349 | 2080.551 | 2159.043 | 2264.365 |
| ENSMUSG00000020458 | 0.043116048 | 0.517279 | 0.67123 | 5944.775 | 5499.982 | 5948.29 | 5845.999 | 5239.273 | 5566.758 | 5635.306 | 6118.787 |
| ENSMUSG00000020459 | 0.180521045 | 0.024251 | 0.08117 | 1375.276 | 1562.168 | 1288.534 | 1405.214 | 1192.561 | 1284.98 | 1225.612 | 1261.606 |
| ENSMUSG00000020460 | -0.04982265 | 0.547521 | 0.695844 | 22950.11 | 27651.09 | 25923.88 | 25960.65 | 28084.18 | 27033.75 | 27071.45 | 23894.38 |
| ENSMUSG00000020462 | -0.023411467 | 0.737582 | 0.835859 | 1176.078 | 1340.201 | 1274.456 | 1275.82 | 1214.014 | 1320.736 | 1296.005 | 1318.16 |

| | | | | | | | | | | | |
|---|---|---|---|---|---|---|---|---|---|---|---|
| ENSMUSG00000020463 | 0.068341178 | 0.320602 | 0.488526 | 2259.598 | 2349.251 | 2018.924 | 2286.815 | 2090.639 | 2123.011 | 2079.972 | 2204.547 |
| ENSMUSG00000020464 | 0.007683249 | 0.920564 | 0.952564 | 1113.702 | 1042.645 | 941.5572 | 975.6272 | 1010.215 | 1012.341 | 1022.147 | 1002.759 |
| ENSMUSG00000020467 | 0.12791229 | 0.600665 | 0.738618 | 239.4409 | 481.1284 | 539.9255 | 345.0494 | 344.2143 | 334.0949 | 352.9298 | 438.2992 |
| ENSMUSG00000020469 | 2.392814541 | 1.10E-08 | 9.47E-07 | 33.19979 | 65.99018 | 49.6864 | 77.63611 | 8.776002 | 10.05637 | 15.42863 | 8.700729 |
| ENSMUSG00000020471 | -0.304393061 | 0.000315 | 0.003224 | 1101.63 | 1257.413 | 1218.973 | 1160.229 | 1550.427 | 1315.149 | 1440.648 | 1542.204 |
| ENSMUSG00000020472 | -0.101731858 | 0.216313 | 0.374654 | 1419.543 | 1557.368 | 1567.606 | 1420.741 | 1699.619 | 1679.413 | 1580.47 | 1439.971 |
| ENSMUSG00000020473 | 0.431112811 | 0.000296 | 0.003067 | 2172.071 | 2216.07 | 2061.986 | 1759.752 | 1356.38 | 1797.855 | 1415.576 | 1519.365 |
| ENSMUSG00000020474 | 0.150054988 | 0.114124 | 0.241462 | 502.0211 | 460.7315 | 549.0347 | 477.8934 | 461.2276 | 465.945 | 444.5373 | 424.1605 |
| ENSMUSG00000020475 | -1.82864481 | 2.05E-07 | 1.02E-05 | 126.7628 | 85.18733 | 150.7154 | 118.1794 | 318.8614 | 198.8926 | 511.0733 | 681.9196 |
| ENSMUSG00000020476 | -0.322887072 | 0.018806 | 0.067514 | 1447.712 | 1625.758 | 1816.038 | 1426.779 | 1976.551 | 1574.38 | 2021.15 | 2328.533 |
| ENSMUSG00000020477 | -0.291903878 | 0.002192 | 0.014021 | 971.8485 | 1133.831 | 1213.176 | 1084.318 | 1363.206 | 1262.633 | 1296.005 | 1470.423 |
| ENSMUSG00000020482 | -0.183053589 | 0.01202 | 0.049025 | 953.7395 | 898.6663 | 868.6839 | 960.9626 | 1044.344 | 1098.379 | 986.4678 | 1052.788 |
| ENSMUSG00000020483 | -0.277290736 | 0.004325 | 0.023159 | 4437.706 | 4951.663 | 5580.611 | 4976.475 | 6084.695 | 5358.926 | 6117.45 | 6614.729 |
| ENSMUSG00000020484 | -0.091070354 | 0.279784 | 0.446185 | 2595.62 | 2342.052 | 2583.693 | 2679.309 | 2495.31 | 2549.848 | 2891.903 | 2932.146 |
| ENSMUSG00000020485 | -0.053562245 | 0.600881 | 0.738753 | 1971.867 | 2176.476 | 2116.641 | 2264.387 | 2540.165 | 2116.307 | 2279.58 | 1914.16 |
| ENSMUSG00000020486 | -1.107751129 | 8.54E-06 | 0.000203 | 924.564 | 675.4995 | 1392.047 | 935.9465 | 2076.987 | 1345.318 | 2383.723 | 2663.511 |
| ENSMUSG00000020491 | 0.112706697 | 0.425916 | 0.590898 | 396.3854 | 519.5227 | 408.2566 | 439.938 | 450.5014 | 455.8886 | 377.0371 | 345.854 |
| ENSMUSG00000020492 | 0.192218001 | 0.122173 | 0.251727 | 1485.942 | 1596.962 | 1276.112 | 1372.434 | 1437.314 | 1365.431 | 1124.361 | 1086.503 |
| ENSMUSG00000020493 | 0.247656449 | 0.072674 | 0.177135 | 661.9838 | 698.2961 | 556.4877 | 645.2424 | 512.9085 | 669.3071 | 497.5732 | 477.4525 |
| ENSMUSG00000020495 | -0.273794734 | 0.016182 | 0.06069 | 527.1725 | 490.727 | 462.9116 | 492.558 | 679.6526 | 515.1095 | 627.7522 | 560.1094 |
| ENSMUSG00000020496 | -0.20969006 | 0.005853 | 0.028904 | 4233.477 | 4955.263 | 5026.607 | 4740.116 | 5404.067 | 5263.949 | 5398.091 | 5855.59 |
| ENSMUSG00000020513 | -0.341481156 | 0.1098 | 0.235472 | 154.9324 | 119.9822 | 150.7154 | 127.6683 | 143.3414 | 132.9675 | 212.1436 | 213.1679 |
| ENSMUSG00000020514 | -0.291059355 | 0.015836 | 0.059928 | 597.5963 | 556.7172 | 719.6247 | 607.2869 | 779.1139 | 655.8986 | 756.0027 | 848.321 |
| ENSMUSG00000020515 | 0.134167618 | 0.029271 | 0.092936 | 1875.285 | 1918.515 | 2009.815 | 1921.063 | 1822.483 | 1636.953 | 1787.792 | 1791.263 |
| ENSMUSG00000020516 | 0.111216354 | 0.250248 | 0.413857 | 1990.982 | 1960.508 | 1711.696 | 1802.02 | 1726.922 | 1972.165 | 1616.149 | 1594.409 |
| ENSMUSG00000020519 | -0.246889719 | 0.000101 | 0.001352 | 754.5408 | 799.0811 | 794.1543 | 764.2844 | 926.3557 | 903.9556 | 936.3248 | 924.4524 |
| ENSMUSG00000020520 | 0.07106902 | 0.47837 | 0.637735 | 1464.815 | 1457.783 | 1368.86 | 1343.105 | 1463.642 | 1119.609 | 1358.683 | 1418.219 |
| ENSMUSG00000020521 | 0.126043476 | 0.083956 | 0.195322 | 1096.599 | 1040.245 | 1020.227 | 1154.19 | 1028.742 | 989.9934 | 990.325 | 941.8539 |
| ENSMUSG00000020522 | 0.152871409 | 0.006995 | 0.032932 | 1848.122 | 1932.912 | 1796.163 | 1828.762 | 1615.759 | 1718.521 | 1623.863 | 1700.992 |
| ENSMUSG00000020523 | -0.002042718 | 0.978266 | 0.988064 | 2225.392 | 2348.051 | 1997.393 | 2082.373 | 2106.24 | 2292.852 | 2173.508 | 2087.087 |
| ENSMUSG00000020524 | 1.194789825 | 0.034463 | 0.104856 | 328.9798 | 1057.043 | 522.5353 | 598.6607 | 270.1058 | 535.2222 | 164.8934 | 125.073 |
| ENSMUSG00000020525 | -0.09595923 | 0.242701 | 0.405292 | 877.2794 | 937.0606 | 928.3075 | 961.8252 | 1069.697 | 889.4298 | 997.075 | 1001.671 |
| ENSMUSG00000020526 | -0.048502429 | 0.646753 | 0.77119 | 341.0524 | 356.347 | 313.8524 | 350.2251 | 352.9903 | 374.3203 | 366.4299 | 312.1386 |
| ENSMUSG00000020527 | -0.838072548 | 0.000677 | 0.005796 | 410.4702 | 325.1516 | 446.3495 | 335.5605 | 617.2455 | 402.2547 | 731.8955 | 962.5181 |
| ENSMUSG00000020528 | -0.103743469 | 0.304276 | 0.472686 | 731.4015 | 944.2595 | 912.5735 | 791.0257 | 904.9033 | 895.0167 | 928.6105 | 900.5254 |
| ENSMUSG00000020530 | 0.051812107 | 0.460463 | 0.621841 | 3192.21 | 3049.946 | 2974.559 | 2863.91 | 2939.961 | 3138.704 | 2874.546 | 2699.401 |
| ENSMUSG00000020532 | -0.029125978 | 0.742832 | 0.839394 | 1067.424 | 1184.224 | 1077.367 | 1054.989 | 1228.64 | 1134.135 | 1109.897 | 996.2334 |
| ENSMUSG00000020534 | -0.667981739 | 2.00E-05 | 0.000392 | 844.0796 | 747.4888 | 782.5608 | 744.4441 | 1198.412 | 897.2514 | 1356.755 | 1501.963 |
| ENSMUSG00000020536 | -0.06253879 | 0.461937 | 0.623028 | 1609.687 | 1896.918 | 1863.24 | 1790.806 | 2027.256 | 1784.446 | 1917.971 | 1745.584 |
| ENSMUSG00000020537 | -0.276924541 | 0.000875 | 0.007031 | 1262.598 | 1331.802 | 1448.359 | 1354.319 | 1637.212 | 1489.46 | 1623.863 | 1791.263 |
| ENSMUSG00000020538 | -0.085536177 | 0.551057 | 0.69875 | 2079.514 | 1653.354 | 2023.065 | 2124.642 | 2180.349 | 2247.039 | 2354.794 | 1581.357 |
| ENSMUSG00000020541 | 0.05769524 | 0.486169 | 0.644362 | 1318.937 | 1131.432 | 1157.693 | 1298.248 | 1236.441 | 1107.318 | 1172.576 | 1198.525 |
| ENSMUSG00000020542 | 0.413110589 | 0.342292 | 0.511039 | 63.38142 | 76.78858 | 69.56096 | 62.10889 | 22.42756 | 102.7984 | 42.42872 | 36.9781 |
| ENSMUSG00000020544 | -0.387427697 | 0.015724 | 0.059687 | 377.2704 | 337.1499 | 387.5539 | 325.2091 | 429.049 | 360.9118 | 506.2518 | 570.9853 |
| ENSMUSG00000020546 | 0.127557307 | 0.144421 | 0.28259 | 579.4873 | 626.3068 | 621.9081 | 635.7535 | 567.5148 | 611.2036 | 513.9661 | 564.4598 |
| ENSMUSG00000020547 | -0.111249154 | 0.030088 | 0.094697 | 2964.842 | 3250.317 | 3113.681 | 3016.594 | 3377.786 | 3373.352 | 3298.833 | 3280.175 |
| ENSMUSG00000020549 | -0.301475352 | 0.001172 | 0.008818 | 784.7224 | 781.0838 | 745.296 | 829.8438 | 1000.464 | 835.7958 | 1028.897 | 1003.847 |
| ENSMUSG00000020553 | -0.381801923 | 0.001785 | 0.012097 | 362.1796 | 319.1525 | 361.0545 | 339.011 | 468.0534 | 379.9072 | 437.7873 | 515.5182 |
| ENSMUSG00000020561 | -0.069481487 | 0.556152 | 0.702989 | 776.674 | 806.2801 | 711.3436 | 698.725 | 740.1095 | 943.0637 | 757.9313 | 698.2335 |
| ENSMUSG00000020564 | 0.191080104 | 0.068409 | 0.169426 | 476.8698 | 616.7083 | 587.9557 | 554.6669 | 484.6303 | 512.8747 | 481.1803 | 479.6277 |
| ENSMUSG00000020566 | -1.561837548 | 0.007788 | 0.035527 | 41.24823 | 33.595 | 217.792 | 43.9938 | 168.6943 | 134.0849 | 314.3583 | 379.5693 |
| ENSMUSG00000020570 | -0.272073468 | 2.28E-07 | 1.13E-05 | 2618.759 | 2591.615 | 2437.946 | 2552.503 | 2993.592 | 3011.323 | 3186.011 | 3121.386 |
| ENSMUSG00000020571 | -0.301504429 | 0.000233 | 0.002561 | 7607.783 | 8099.995 | 8385.408 | 7534.154 | 9412.749 | 8738.983 | 10141.43 | 10684.49 |
| ENSMUSG00000020572 | -0.132315196 | 0.147302 | 0.286518 | 1458.779 | 1310.205 | 1383.766 | 1388.824 | 1406.111 | 1405.657 | 1731.863 | 1530.241 |
| ENSMUSG00000020573 | 0.11160109 | 0.50208 | 0.658262 | 102.6175 | 134.38 | 113.4506 | 117.3168 | 121.8889 | 107.2679 | 101.2504 | 101.146 |
| ENSMUSG00000020576 | -0.144119642 | 0.168209 | 0.314313 | 597.5963 | 674.2997 | 679.8755 | 610.7374 | 666.9761 | 627.9642 | 756.967 | 778.7152 |
| ENSMUSG00000020577 | -0.219800537 | 0.018797 | 0.067505 | 1244.489 | 1280.21 | 1398.672 | 1375.022 | 1452.916 | 1441.413 | 1507.184 | 1772.773 |
| ENSMUSG00000020580 | 0.192336039 | 0.057342 | 0.149964 | 2566.445 | 2145.281 | 2138.171 | 2205.728 | 1798.105 | 2241.452 | 2009.579 | 1877.182 |
| ENSMUSG00000020581 | 0.522273499 | 0.315799 | 0.484206 | 22.1332 | 8.398751 | 46.37397 | 36.23019 | 22.42756 | 24.58223 | 17.3572 | 15.22628 |
| ENSMUSG00000020585 | 0.046821837 | 0.507399 | 0.663071 | 13825.2 | 15609.68 | 15066.57 | 15644.54 | 14734.91 | 15560.55 | 14463.37 | 13466.55 |
| ENSMUSG00000020589 | 0.411033901 | 0.001329 | 0.009688 | 364.1917 | 407.9393 | 318.8211 | 376.9665 | 279.8569 | 313.9821 | 256.5009 | 252.3211 |
| ENSMUSG00000020590 | -0.024168718 | 0.725517 | 0.828291 | 1039.254 | 935.8608 | 1046.727 | 1029.972 | 1012.166 | 1021.28 | 1074.218 | 1016.898 |
| ENSMUSG00000020592 | 0.104833933 | 0.354149 | 0.522769 | 1313.907 | 1536.971 | 1839.225 | 1517.355 | 1373.932 | 1491.694 | 1379.898 | 1530.241 |
| ENSMUSG00000020593 | -0.234447884 | 0.234601 | 0.395487 | 521.1362 | 405.5397 | 374.3042 | 397.6694 | 428.0739 | 563.1565 | 637.3951 | 368.6934 |

| | | | | | | | | | | | |
|---|---|---|---|---|---|---|---|---|---|---|---|
| ENSMUSG00000020594 | 0.287695771 | 0.011622 | 0.047793 | 5412.572 | 5429.192 | 4381.512 | 5579.449 | 3902.395 | 4942.146 | 3901.514 | 4295.985 |
| ENSMUSG00000020598 | -0.319468719 | 0.104046 | 0.226955 | 128.775 | 130.7805 | 194.6051 | 144.0581 | 165.7689 | 160.9019 | 202.5007 | 219.6934 |
| ENSMUSG00000020599 | 0.133234594 | 0.501713 | 0.657987 | 149.9021 | 157.1766 | 147.403 | 212.2054 | 153.0925 | 176.5451 | 116.679 | 163.1387 |
| ENSMUSG00000020601 | 0.178115224 | 0.136141 | 0.270698 | 3080.538 | 2986.356 | 2621.786 | 3527.267 | 2473.857 | 3126.413 | 2727.974 | 2469.919 |
| ENSMUSG00000020604 | -0.081513949 | 0.458493 | 0.62015 | 287.7315 | 316.7529 | 308.0557 | 322.6212 | 365.6667 | 299.4563 | 310.5011 | 330.6277 |
| ENSMUSG00000020605 | 0.232526031 | 0.048043 | 0.132413 | 386.3249 | 377.9438 | 472.0208 | 387.3179 | 321.7867 | 331.8601 | 357.7513 | 373.0437 |
| ENSMUSG00000020607 | 1.04107406 | 0.000116 | 0.001503 | 817.9222 | 1243.015 | 804.0915 | 935.0839 | 440.7503 | 717.3542 | 416.5729 | 271.8978 |
| ENSMUSG00000020608 | 0.147530015 | 0.146951 | 0.285998 | 2935.667 | 3131.534 | 2625.098 | 2769.021 | 2379.272 | 3019.145 | 2476.295 | 2471.007 |
| ENSMUSG00000020609 | 1.734187592 | 6.24E-05 | 0.000944 | 59.35721 | 31.19536 | 25.67131 | 48.30692 | 15.60178 | 8.938993 | 9.642892 | 15.22628 |
| ENSMUSG00000020610 | 0.123357545 | 0.049124 | 0.134456 | 2179.114 | 2280.861 | 2031.346 | 2185.025 | 2084.788 | 1952.053 | 1998.971 | 1922.861 |
| ENSMUSG00000020611 | -0.096450541 | 0.147357 | 0.286577 | 2066.436 | 1836.927 | 1896.364 | 1985.759 | 2138.419 | 1955.405 | 2148.436 | 2081.649 |
| ENSMUSG00000020612 | -0.118401016 | 0.070043 | 0.172453 | 8525.305 | 7783.242 | 8516.249 | 8542.56 | 8485.419 | 8764.682 | 9411.462 | 9565.364 |
| ENSMUSG00000020614 | 0.105095676 | 0.517088 | 0.671078 | 128.775 | 134.38 | 161.4808 | 141.4703 | 143.3414 | 108.3853 | 135.9648 | 139.2117 |
| ENSMUSG00000020620 | 1.090761413 | 5.07E-05 | 0.0008 | 276.6649 | 243.5638 | 154.0278 | 247.5729 | 91.66046 | 158.6671 | 103.1789 | 79.39415 |
| ENSMUSG00000020621 | -0.094745137 | 0.256033 | 0.419812 | 766.6134 | 758.2872 | 712.1717 | 793.6136 | 759.6117 | 766.5186 | 873.646 | 835.27 |
| ENSMUSG00000020622 | 0.045785898 | 0.935915 | 0.962083 | 13.07871 | 11.99822 | 7.45296 | 11.21411 | 6.825779 | 16.76061 | 4.821446 | 14.13868 |
| ENSMUSG00000020623 | -0.173714392 | 0.158473 | 0.301338 | 301.8163 | 341.9491 | 274.9314 | 301.9182 | 303.2596 | 382.1419 | 330.7512 | 358.9051 |
| ENSMUSG00000020627 | 0.63526467 | 0.004262 | 0.022891 | 344.0706 | 310.7538 | 287.353 | 321.7586 | 204.7734 | 297.2215 | 168.7506 | 143.562 |
| ENSMUSG00000020628 | -0.100911455 | 0.325751 | 0.49356 | 970.8425 | 1097.837 | 996.2123 | 947.1606 | 985.8375 | 970.9981 | 1128.218 | 1214.839 |
| ENSMUSG00000020629 | -0.160894835 | 0.093699 | 0.210713 | 900.4186 | 992.2524 | 783.3889 | 935.0839 | 1007.29 | 957.5896 | 1080.004 | 986.4451 |
| ENSMUSG00000020630 | 0.034769362 | 0.726535 | 0.828888 | 563.3904 | 519.5227 | 541.5817 | 634.8909 | 590.9175 | 563.1565 | 537.1091 | 516.6058 |
| ENSMUSG00000020634 | -0.218249967 | 0.02704 | 0.087861 | 592.566 | 561.5165 | 605.3459 | 551.2164 | 693.3041 | 577.6824 | 692.3596 | 724.3357 |
| ENSMUSG00000020635 | 0.299569652 | 0.12789 | 0.259292 | 144.8718 | 136.7797 | 111.7944 | 159.5853 | 112.1378 | 90.5073 | 137.8934 | 107.6715 |
| ENSMUSG00000020638 | 0.898167436 | 2.78E-15 | 1.31E-12 | 1153.944 | 1151.829 | 929.1356 | 1183.519 | 586.0419 | 574.3303 | 665.3595 | 542.708 |
| ENSMUSG00000020639 | 0.12418749 | 0.629928 | 0.759023 | 38.23007 | 65.99018 | 57.13936 | 45.71905 | 44.85512 | 51.39921 | 44.3573 | 48.9416 |
| ENSMUSG00000020640 | -0.083603871 | 0.511134 | 0.66663 | 1044.284 | 886.6681 | 871.9963 | 758.2461 | 864.9237 | 884.9603 | 928.6105 | 1094.117 |
| ENSMUSG00000020641 | 0.185614263 | 0.107085 | 0.231398 | 2078.508 | 1874.121 | 2421.384 | 1813.235 | 1695.719 | 1756.512 | 2022.114 | 1726.007 |
| ENSMUSG00000020642 | 0.383256683 | 0.000443 | 0.004176 | 2156.981 | 2283.26 | 2078.548 | 2555.091 | 1892.691 | 1905.123 | 1651.827 | 1507.401 |
| ENSMUSG00000020644 | 0.178377966 | 0.079958 | 0.188951 | 4387.403 | 4528.126 | 5453.91 | 5090.341 | 4026.235 | 3964.443 | 4405.837 | 4803.89 |
| ENSMUSG00000020646 | -0.435545396 | 0.003257 | 0.018873 | 579.4873 | 537.52 | 749.4365 | 586.584 | 782.0393 | 692.7719 | 962.3606 | 883.124 |
| ENSMUSG00000020647 | 0.31150136 | 0.009293 | 0.040366 | 1503.045 | 1631.757 | 1581.684 | 1540.646 | 1154.532 | 1573.263 | 1182.219 | 1133.27 |
| ENSMUSG00000020648 | 0.172378683 | 0.218904 | 0.377684 | 280.6892 | 323.9518 | 274.9314 | 339.8737 | 287.6578 | 309.5126 | 249.7509 | 234.9197 |
| ENSMUSG00000020649 | -0.171528385 | 0.025315 | 0.083681 | 1527.191 | 1785.334 | 1676.088 | 1555.31 | 1844.911 | 1817.968 | 1753.078 | 1951.138 |
| ENSMUSG00000020650 | 0.157487746 | 0.087622 | 0.201314 | 557.3541 | 526.7217 | 532.4726 | 607.2869 | 484.6303 | 540.8091 | 495.6446 | 475.2773 |
| ENSMUSG00000020651 | -1.649642411 | 0.000861 | 0.006955 | 31.18769 | 27.5959 | 43.88965 | 39.68068 | 102.3867 | 29.05173 | 110.8933 | 205.5547 |
| ENSMUSG00000020652 | 0.24166223 | 0.045954 | 0.12833 | 398.3975 | 459.5316 | 351.9453 | 385.5927 | 331.5378 | 373.2029 | 330.7512 | 311.0511 |
| ENSMUSG00000020653 | -0.188152581 | 0.562449 | 0.707942 | 501.0151 | 614.3086 | 722.109 | 735.8178 | 855.1726 | 422.3674 | 428.1444 | 1226.803 |
| ENSMUSG00000020654 | 0.042328946 | 0.632523 | 0.760958 | 399.4036 | 440.3345 | 426.4749 | 433.8996 | 419.2979 | 421.25 | 385.7157 | 425.2481 |
| ENSMUSG00000020656 | -0.537153705 | 0.054917 | 0.145666 | 108.6539 | 106.7841 | 172.2462 | 114.7289 | 224.2756 | 102.7984 | 200.5721 | 202.2919 |
| ENSMUSG00000020657 | 0.110108657 | 0.227789 | 0.387722 | 528.1785 | 476.3291 | 491.0672 | 475.3055 | 497.3068 | 450.3018 | 431.0373 | 446.9999 |
| ENSMUSG00000020658 | 0.573358276 | 2.30E-05 | 0.000438 | 363.1856 | 424.7368 | 433.0998 | 401.9825 | 310.0854 | 301.691 | 231.4294 | 247.9708 |
| ENSMUSG00000020659 | 0.201394922 | 0.046917 | 0.130211 | 550.3117 | 503.925 | 467.8802 | 500.3216 | 424.1734 | 473.7666 | 399.2157 | 462.2262 |
| ENSMUSG00000020660 | 0.297277174 | 0.185109 | 0.336588 | 57.3451 | 56.39161 | 74.5296 | 61.24627 | 49.73068 | 58.10345 | 44.3573 | 52.20437 |
| ENSMUSG00000020661 | 0.240936285 | 0.005756 | 0.028549 | 5188.222 | 5033.251 | 4622.491 | 4681.458 | 3723.95 | 4616.99 | 3931.407 | 4250.306 |
| ENSMUSG00000020664 | -0.426846258 | 0.000154 | 0.001867 | 3380.343 | 3088.341 | 3379.503 | 3409.951 | 4311.942 | 3622.527 | 4795.41 | 5094.277 |
| ENSMUSG00000020668 | 0.066236478 | 0.526974 | 0.679938 | 586.5297 | 491.9268 | 503.4888 | 510.6731 | 468.0534 | 461.4755 | 538.0734 | 530.7445 |
| ENSMUSG00000020669 | -0.170346622 | 0.13136 | 0.264232 | 518.118 | 530.3211 | 544.8942 | 597.7981 | 596.7681 | 535.2222 | 622.9308 | 712.3722 |
| ENSMUSG00000020671 | 0.087023116 | 0.172565 | 0.320022 | 3469.881 | 3479.482 | 3573.28 | 3721.358 | 3098.904 | 3548.78 | 3324.869 | 3443.313 |
| ENSMUSG00000020672 | 0.756252863 | 0.005747 | 0.028515 | 122.7386 | 217.1677 | 160.6527 | 200.1287 | 74.10846 | 149.7281 | 103.1789 | 88.09488 |
| ENSMUSG00000020674 | 0.241899777 | 0.002445 | 0.015178 | 6183.21 | 6848.581 | 6896.472 | 6862.17 | 5477.2 | 6376.854 | 5405.805 | 5397.715 |
| ENSMUSG00000020676 | 0.199368597 | 0.613922 | 0.747693 | 43.26034 | 82.78769 | 69.56096 | 59.52102 | 41.92979 | 30.1691 | 52.07161 | 97.8832 |
| ENSMUSG00000020677 | 0.044197377 | 0.600538 | 0.738607 | 1762.607 | 1682.15 | 1542.763 | 1650.199 | 1685.967 | 1752.043 | 1491.755 | 1506.314 |
| ENSMUSG00000020679 | -0.758167394 | 0.000823 | 0.006715 | 1270.647 | 807.4799 | 1424.343 | 1060.164 | 1714.246 | 1365.431 | 1993.186 | 2647.197 |
| ENSMUSG00000020680 | 0.069739701 | 0.099537 | 0.220245 | 4254.604 | 4202.975 | 4210.094 | 4056.918 | 3942.375 | 4014.725 | 3916.943 | 4061.065 |
| ENSMUSG00000020681 | -2.449873437 | 0.000145 | 0.001776 | 179.0777 | 111.5834 | 509.2856 | 242.3972 | 1413.911 | 489.4099 | 1252.612 | 2542.788 |
| ENSMUSG00000020682 | -0.508305462 | 0.090247 | 0.205554 | 49.29666 | 47.99286 | 98.54469 | 68.14726 | 104.3369 | 65.92507 | 107.0361 | 100.0584 |
| ENSMUSG00000020684 | 0.230992848 | 0.540823 | 0.690376 | 73.44197 | 21.59679 | 47.20208 | 59.52102 | 34.1289 | 41.34284 | 47.25017 | 50.02919 |
| ENSMUSG00000020687 | 0.204948714 | 0.03958 | 0.115854 | 2036.254 | 1857.324 | 1595.761 | 1897.772 | 1569.929 | 1779.977 | 1523.577 | 1534.591 |
| ENSMUSG00000020689 | 0.788276372 | 0.000633 | 0.005498 | 243.4652 | 259.1615 | 175.5586 | 167.349 | 142.3663 | 137.437 | 123.429 | 84.8321 |
| ENSMUSG00000020691 | 0.12050498 | 0.163689 | 0.308553 | 1020.139 | 1025.847 | 920.8546 | 984.2534 | 961.4597 | 977.7023 | 832.1815 | 861.3721 |
| ENSMUSG00000020692 | -0.069217058 | 0.542932 | 0.692109 | 569.4268 | 598.7109 | 614.4551 | 558.1174 | 719.6321 | 558.687 | 545.7877 | 630.8028 |
| ENSMUSG00000020694 | 0.01883186 | 0.807467 | 0.882437 | 2059.393 | 1853.724 | 1841.709 | 1827.899 | 1741.549 | 1968.813 | 1977.757 | 1795.613 |
| ENSMUSG00000020695 | 0.424102595 | 8.70E-05 | 0.00121 | 2091.587 | 2655.205 | 2525.725 | 2437.774 | 1821.508 | 2068.259 | 1665.327 | 1682.503 |
| ENSMUSG00000020696 | -0.083744674 | 0.528282 | 0.680769 | 855.1462 | 856.6726 | 812.3726 | 816.9044 | 947.8082 | 667.0723 | 918.9676 | 1004.934 |

| | | | | | | | | | | | |
|---|---|---|---|---|---|---|---|---|---|---|---|
| ENSMUSG00000020697 | -0.278161317 | 0.046453 | 0.129332 | 1002.03 | 899.8662 | 959.7756 | 800.5146 | 1189.636 | 845.8522 | 1180.29 | 1223.54 |
| ENSMUSG00000020698 | -1.158056308 | 0.031901 | 0.098719 | 14.08476 | 7.198929 | 4.96864 | 6.038364 | 17.552 | 25.6996 | 17.3572 | 10.87591 |
| ENSMUSG00000020700 | -0.174811364 | 0.070487 | 0.173155 | 1254.55 | 1124.233 | 1228.082 | 1303.424 | 1368.081 | 1201.177 | 1509.113 | 1466.073 |
| ENSMUSG00000020701 | -0.244037408 | 0.5227 | 0.675983 | 17.10292 | 14.39786 | 14.90592 | 18.11509 | 22.42756 | 22.34748 | 13.50005 | 18.48905 |
| ENSMUSG00000020704 | -1.023430711 | 6.21E-09 | 5.65E-07 | 280.6892 | 227.9661 | 222.7607 | 266.5507 | 539.2366 | 446.9496 | 642.2166 | 399.1459 |
| ENSMUSG00000020705 | 0.034875482 | 0.643887 | 0.769677 | 4141.926 | 3797.435 | 3609.717 | 3967.205 | 3821.461 | 4107.467 | 3696.12 | 3520.532 |
| ENSMUSG00000020706 | -0.162410897 | 0.099133 | 0.219744 | 1954.764 | 2066.093 | 1907.958 | 2039.242 | 2409.5 | 1888.362 | 2157.115 | 2460.131 |
| ENSMUSG00000020707 | 0.004171121 | 0.979739 | 0.988793 | 220.3259 | 251.9625 | 277.4157 | 275.1769 | 267.1805 | 191.071 | 286.3939 | 277.3357 |
| ENSMUSG00000020708 | -0.341182622 | 0.003008 | 0.017797 | 2879.328 | 2796.784 | 3144.321 | 2681.896 | 3695.672 | 2984.506 | 3657.549 | 4233.992 |
| ENSMUSG00000020709 | -0.604109983 | 0.010297 | 0.043528 | 66.39959 | 47.99286 | 79.49824 | 49.16954 | 101.4116 | 78.21619 | 94.50034 | 96.79561 |
| ENSMUSG00000020712 | 0.367690322 | 0.288272 | 0.455197 | 51.30877 | 55.19179 | 61.27989 | 61.24627 | 69.2329 | 35.75597 | 49.17875 | 22.83941 |
| ENSMUSG00000020715 | 0.259472193 | 0.124416 | 0.254458 | 812.8919 | 545.9188 | 508.4575 | 531.3761 | 473.9041 | 464.8276 | 513.9661 | 550.3211 |
| ENSMUSG00000020716 | 0.407255785 | 0.001125 | 0.008541 | 2100.641 | 1934.112 | 1706.728 | 1871.03 | 1379.783 | 1759.864 | 1350.969 | 1249.642 |
| ENSMUSG00000020717 | -0.447251078 | 0.003642 | 0.020416 | 1884.34 | 1552.569 | 2516.616 | 2143.619 | 2742.988 | 2359.894 | 2918.903 | 3023.503 |
| ENSMUSG00000020718 | 0.124845294 | 0.322763 | 0.490873 | 316.9071 | 278.3586 | 317.9929 | 328.6596 | 263.2801 | 337.447 | 273.8581 | 267.5474 |
| ENSMUSG00000020719 | -0.017460768 | 0.742197 | 0.839029 | 25130.23 | 25149.46 | 23917.38 | 26334.17 | 25170.55 | 26725.35 | 25369.48 | 24490.38 |
| ENSMUSG00000020720 | -0.210913023 | 0.000354 | 0.003512 | 2447.73 | 2368.448 | 2281.434 | 2406.72 | 2763.465 | 2593.425 | 2737.617 | 2903.868 |
| ENSMUSG00000020721 | 0.394971087 | 0.049649 | 0.13543 | 1782.728 | 1569.367 | 1277.769 | 1557.898 | 898.0775 | 1699.526 | 1010.575 | 1098.467 |
| ENSMUSG00000020723 | 1.133639439 | 6.71E-08 | 4.07E-06 | 353.1251 | 303.5548 | 325.4459 | 385.5927 | 118.9636 | 224.5922 | 148.5005 | 132.6861 |
| ENSMUSG00000020728 | 0.142058377 | 0.183097 | 0.33395 | 470.8334 | 519.5227 | 462.0835 | 444.2511 | 469.0285 | 415.6632 | 450.323 | 379.5693 |
| ENSMUSG00000020732 | -0.665132525 | 0.053975 | 0.143808 | 40.24217 | 43.19358 | 48.03018 | 19.84034 | 42.9049 | 61.45558 | 55.92877 | 79.39415 |
| ENSMUSG00000020733 | -0.80028617 | 1.46E-07 | 7.71E-06 | 1469.845 | 1534.572 | 1902.989 | 1505.278 | 2607.448 | 2214.635 | 2862.01 | 3485.729 |
| ENSMUSG00000020736 | -0.270284137 | 0.01705 | 0.062999 | 879.2915 | 843.4745 | 871.9963 | 779.8116 | 1097 | 845.8522 | 981.6464 | 1144.146 |
| ENSMUSG00000020737 | 0.216378938 | 0.002266 | 0.014359 | 4003.09 | 4444.139 | 4093.331 | 4449.412 | 3857.54 | 3818.067 | 3476.262 | 3470.503 |
| ENSMUSG00000020738 | -0.251211645 | 0.001035 | 0.008024 | 4171.101 | 4451.338 | 4637.397 | 4494.268 | 5490.852 | 4675.093 | 5444.377 | 5520.612 |
| ENSMUSG00000020739 | -0.048866623 | 0.483128 | 0.64153 | 1642.887 | 1782.935 | 1692.65 | 1555.31 | 1725.947 | 1804.559 | 1739.578 | 1629.211 |
| ENSMUSG00000020740 | 0.165249732 | 0.091384 | 0.207351 | 793.7769 | 675.4995 | 687.3285 | 688.3735 | 645.5237 | 696.1241 | 617.1451 | 578.5985 |
| ENSMUSG00000020741 | -0.841380106 | 4.47E-06 | 0.000124 | 1651.941 | 1325.803 | 1847.506 | 1522.53 | 2698.133 | 1995.63 | 3024.011 | 3658.656 |
| ENSMUSG00000020743 | -0.253432854 | 0.065176 | 0.164396 | 783.7163 | 681.4986 | 753.577 | 684.0604 | 842.4962 | 687.1851 | 880.396 | 1050.613 |
| ENSMUSG00000020744 | -0.343649152 | 0.020263 | 0.071112 | 641.8627 | 697.0963 | 647.5794 | 631.4404 | 885.4011 | 664.8376 | 729.9669 | 1040.825 |
| ENSMUSG00000020745 | 0.058148987 | 0.26455 | 0.428819 | 5133.895 | 5165.232 | 5132.605 | 5192.131 | 4672.733 | 5162.268 | 4844.589 | 5133.43 |
| ENSMUSG00000020747 | -0.103465735 | 0.28518 | 0.4519 | 1242.477 | 1112.235 | 1215.661 | 1099.845 | 1203.287 | 1115.139 | 1290.219 | 1409.518 |
| ENSMUSG00000020752 | 0.019217513 | 0.84049 | 0.90381 | 558.3602 | 581.9134 | 528.332 | 526.2003 | 508.033 | 510.64 | 537.1091 | 607.9634 |
| ENSMUSG00000020755 | -0.203039671 | 0.016342 | 0.061103 | 1528.197 | 1576.565 | 1542.763 | 1408.664 | 1880.99 | 1556.502 | 1832.149 | 1696.642 |
| ENSMUSG00000020758 | 0.415850413 | 0.001599 | 0.01106 | 375.2583 | 382.7431 | 410.7409 | 378.6917 | 311.0605 | 311.7474 | 310.5011 | 226.2189 |
| ENSMUSG00000020766 | -0.388858882 | 0.005263 | 0.02664 | 1609.687 | 1594.563 | 1922.864 | 1551.86 | 2221.304 | 1700.643 | 2220.758 | 2603.693 |
| ENSMUSG00000020770 | 0.118357013 | 0.12494 | 0.255284 | 1379.301 | 1450.584 | 1383.766 | 1262.881 | 1318.35 | 1319.619 | 1232.362 | 1171.336 |
| ENSMUSG00000020773 | -0.71663566 | 0.009315 | 0.040432 | 348.0948 | 211.1686 | 354.4296 | 345.912 | 460.2525 | 297.2215 | 590.145 | 724.3357 |
| ENSMUSG00000020774 | -2.481020269 | 0.000196 | 0.00223 | 96.58122 | 17.99732 | 174.7305 | 103.5148 | 481.705 | 261.4655 | 571.8235 | 882.0364 |
| ENSMUSG00000020775 | -0.354099641 | 0.003042 | 0.017946 | 1224.368 | 1145.83 | 1031.821 | 1056.714 | 1443.165 | 1204.529 | 1366.398 | 1682.503 |
| ENSMUSG00000020776 | -0.083313191 | 0.183945 | 0.335001 | 1331.01 | 1317.404 | 1209.864 | 1257.705 | 1354.43 | 1308.445 | 1390.505 | 1360.576 |
| ENSMUSG00000020777 | -0.839526884 | 3.45E-05 | 0.000598 | 2266.64 | 1606.561 | 2039.627 | 1823.586 | 3828.287 | 2195.64 | 3408.762 | 4412.357 |
| ENSMUSG00000020778 | -0.110101287 | 0.257578 | 0.421292 | 573.451 | 677.8992 | 685.6723 | 626.2647 | 672.8268 | 642.4901 | 695.2525 | 755.8758 |
| ENSMUSG00000020780 | -0.202423095 | 0.012963 | 0.051885 | 2545.318 | 2495.629 | 2671.472 | 2507.646 | 3065.75 | 2554.317 | 3011.475 | 3127.912 |
| ENSMUSG00000020781 | -0.540584312 | 6.53E-05 | 0.000976 | 545.2815 | 545.9188 | 551.519 | 587.4466 | 874.6748 | 613.4384 | 841.8244 | 913.5765 |
| ENSMUSG00000020782 | -1.068767259 | 1.46E-06 | 5.06E-05 | 751.5226 | 641.9045 | 1048.383 | 660.7696 | 1525.074 | 1084.97 | 1873.614 | 2027.27 |
| ENSMUSG00000020783 | -0.05150487 | 0.460396 | 0.621813 | 1429.603 | 1570.566 | 1568.434 | 1473.361 | 1575.78 | 1450.352 | 1599.756 | 1633.562 |
| ENSMUSG00000020785 | -0.526676216 | 0.004891 | 0.025347 | 167.005 | 130.7805 | 197.9175 | 180.2883 | 285.7076 | 203.3621 | 273.8581 | 213.1679 |
| ENSMUSG00000020787 | -0.859125607 | 0.025417 | 0.08399 | 22.1332 | 14.39786 | 33.95237 | 14.6646 | 40.95467 | 27.93435 | 44.3573 | 42.41605 |
| ENSMUSG00000020788 | 0.023732255 | 0.914593 | 0.949313 | 439.6458 | 719.8929 | 789.1856 | 660.7696 | 847.3717 | 471.5319 | 620.0379 | 627.5401 |
| ENSMUSG00000020790 | 0.003067285 | 0.968305 | 0.982153 | 1732.426 | 1790.134 | 1992.425 | 1656.237 | 1781.528 | 1797.855 | 1728.006 | 1851.08 |
| ENSMUSG00000020792 | -0.186468495 | 0.050483 | 0.136958 | 1360.185 | 1340.201 | 1440.906 | 1367.258 | 1541.651 | 1334.145 | 1676.899 | 1716.219 |
| ENSMUSG00000020793 | 0.559208774 | 0.099877 | 0.2207 | 29.17558 | 28.79572 | 21.53077 | 30.19182 | 20.47734 | 21.23011 | 18.32149 | 14.13868 |
| ENSMUSG00000020794 | -0.035094304 | 0.659659 | 0.780834 | 1963.818 | 1728.943 | 1909.614 | 2111.702 | 1924.87 | 2091.724 | 1964.257 | 1929.387 |
| ENSMUSG00000020798 | -1.098455515 | 0.030101 | 0.094705 | 18.10898 | 9.598572 | 14.07781 | 12.93935 | 55.58134 | 15.64324 | 23.14294 | 22.83941 |
| ENSMUSG00000020799 | 0.277430001 | 0.134026 | 0.267618 | 156.9445 | 241.1641 | 170.59 | 186.3267 | 168.6943 | 168.7235 | 156.2148 | 127.2482 |
| ENSMUSG00000020801 | 0.00677776 | 0.951876 | 0.972676 | 529.1846 | 433.1356 | 482.7862 | 478.756 | 495.3565 | 512.8747 | 497.5732 | 410.0218 |
| ENSMUSG00000020802 | 0.133811447 | 0.182896 | 0.333682 | 945.6911 | 868.6708 | 963.9161 | 1078.279 | 813.2428 | 981.0545 | 872.6817 | 852.6714 |
| ENSMUSG00000020803 | -0.349457212 | 0.010495 | 0.044181 | 1642.887 | 1583.764 | 2008.159 | 1682.116 | 2244.706 | 1785.564 | 2128.186 | 2656.985 |
| ENSMUSG00000020805 | 0.944498709 | 0.086744 | 0.199942 | 12.07265 | 25.19625 | 15.73403 | 11.21411 | 5.850668 | 12.29112 | 10.60718 | 4.350364 |
| ENSMUSG00000020806 | -0.049390532 | 0.693865 | 0.806523 | 391.3551 | 340.7493 | 452.9743 | 341.5989 | 394.9201 | 417.8979 | 372.2156 | 398.0583 |
| ENSMUSG00000020807 | 0.248073356 | 0.045159 | 0.126859 | 676.0685 | 679.099 | 592.9243 | 629.7152 | 519.7343 | 661.4855 | 507.2161 | 481.8029 |
| ENSMUSG00000020808 | -0.044615013 | 0.631219 | 0.760032 | 1354.149 | 1315.004 | 1337.392 | 1458.696 | 1606.008 | 1402.304 | 1369.291 | 1259.43 |
| ENSMUSG00000020810 | 0.111045144 | 0.406826 | 0.573203 | 985.9333 | 1372.596 | 1328.283 | 1209.398 | 1164.283 | 1259.281 | 1153.29 | 955.9926 |

| | | | | | | | | | | | |
|---|---|---|---|---|---|---|---|---|---|---|---|
| ENSMUSG00000020811 | -1.182433202 | 1.93E-06 | 6.28E-05 | 247.4894 | 262.7609 | 438.0684 | 281.2153 | 636.7477 | 487.1751 | 701.0382 | 969.0437 |
| ENSMUSG00000020812 | -0.035332853 | 0.732042 | 0.832065 | 698.2017 | 650.3033 | 696.4377 | 803.9651 | 798.6162 | 678.2461 | 758.8956 | 685.1824 |
| ENSMUSG00000020814 | -0.152167193 | 0.269787 | 0.434674 | 562.3844 | 545.9188 | 637.6421 | 582.2709 | 674.777 | 536.3396 | 587.2521 | 790.6787 |
| ENSMUSG00000020817 | 0.029946332 | 0.807156 | 0.882202 | 1577.493 | 1371.396 | 1106.35 | 1218.024 | 1176.959 | 1338.614 | 1313.362 | 1333.387 |
| ENSMUSG00000020818 | -0.103700578 | 0.231463 | 0.392023 | 751.5226 | 782.2836 | 809.0602 | 771.1854 | 857.1228 | 748.6406 | 821.5744 | 919.0145 |
| ENSMUSG00000020821 | -0.317351189 | 0.026394 | 0.086387 | 2964.842 | 2837.578 | 3632.904 | 3007.968 | 3889.719 | 2975.567 | 4051.943 | 4589.634 |
| ENSMUSG00000020823 | -0.818162495 | 0.000116 | 0.001505 | 2359.197 | 2092.489 | 3271.021 | 2382.566 | 3981.379 | 2957.689 | 5602.52 | 5278.08 |
| ENSMUSG00000020826 | 0.087581667 | 0.78803 | 0.869849 | 38.23007 | 43.19358 | 24.01509 | 31.05445 | 36.07912 | 39.10809 | 24.10723 | 28.27737 |
| ENSMUSG00000020827 | 0.005643968 | 0.925597 | 0.955735 | 1714.317 | 1641.356 | 1735.712 | 1799.433 | 1736.673 | 1624.662 | 1757.899 | 1747.759 |
| ENSMUSG00000020828 | 0.281615391 | 0.00907 | 0.039721 | 550.3117 | 525.5218 | 637.6421 | 656.4565 | 510.9583 | 507.2878 | 478.2874 | 456.7883 |
| ENSMUSG00000020829 | -0.62033744 | 4.11E-05 | 0.000689 | 296.786 | 259.1615 | 254.2287 | 244.1224 | 368.5921 | 331.8601 | 432.9658 | 486.1532 |
| ENSMUSG00000020831 | -0.062023665 | 0.632522 | 0.760958 | 201.2109 | 199.1704 | 197.0894 | 184.6014 | 234.0267 | 182.132 | 200.5721 | 197.9416 |
| ENSMUSG00000020832 | -0.079823063 | 0.470724 | 0.630616 | 722.347 | 653.9027 | 634.3297 | 746.1693 | 728.4081 | 840.2653 | 649.9309 | 697.1459 |
| ENSMUSG00000020834 | -0.245127738 | 0.068505 | 0.169548 | 228.3743 | 242.3639 | 246.7758 | 233.771 | 323.737 | 233.5312 | 268.0724 | 301.2627 |
| ENSMUSG00000020836 | 0.275477184 | 0.467308 | 0.62788 | 31.18769 | 32.39518 | 26.49941 | 44.85642 | 24.37778 | 40.22547 | 31.82154 | 15.22628 |
| ENSMUSG00000020838 | -0.281912994 | 0.445912 | 0.608619 | 61.36932 | 44.3934 | 53.82693 | 49.16954 | 78.98402 | 34.6386 | 41.46443 | 98.97079 |
| ENSMUSG00000020839 | -1.730609818 | 0.007033 | 0.033064 | 21.12714 | 16.7975 | 134.1533 | 28.46658 | 149.192 | 77.09881 | 147.5362 | 294.7372 |
| ENSMUSG00000020840 | -0.205356497 | 0.0023 | 0.014505 | 3678.135 | 3841.829 | 3830.821 | 3817.972 | 4576.197 | 3948.8 | 4348.944 | 4613.561 |
| ENSMUSG00000020841 | 0.214610579 | 0.016338 | 0.061103 | 5207.337 | 5220.423 | 5177.323 | 5771.814 | 4143.248 | 5228.193 | 4673.91 | 4379.729 |
| ENSMUSG00000020843 | -0.27744777 | 0.023915 | 0.080457 | 808.8677 | 871.0704 | 836.3877 | 744.4441 | 1070.672 | 834.6784 | 899.6818 | 1145.233 |
| ENSMUSG00000020844 | -0.051813878 | 0.54916 | 0.697351 | 1489.967 | 1547.77 | 1684.369 | 1499.24 | 1488.02 | 1569.911 | 1586.256 | 1807.576 |
| ENSMUSG00000020846 | -0.263368467 | 0.093247 | 0.21005 | 1288.756 | 1632.957 | 2158.874 | 1666.589 | 2071.136 | 1854.841 | 1878.435 | 2296.992 |
| ENSMUSG00000020847 | -0.254803923 | 0.158314 | 0.301192 | 275.6589 | 209.9688 | 259.1974 | 270.8638 | 280.8321 | 250.2918 | 281.5724 | 401.3211 |
| ENSMUSG00000020848 | 1.191005608 | 5.89E-14 | 2.31E-11 | 325.9616 | 296.3559 | 278.2438 | 394.2189 | 145.2916 | 147.4934 | 150.4291 | 123.9854 |
| ENSMUSG00000020849 | -0.062900686 | 0.065249 | 0.16449 | 15160.23 | 15339.72 | 14741.13 | 15170.96 | 15728.55 | 15725.92 | 15595.45 | 16049.58 |
| ENSMUSG00000020850 | -0.104959312 | 0.098977 | 0.219531 | 6291.864 | 6080.696 | 6584.276 | 6163.445 | 6811.152 | 6179.079 | 6979.525 | 7047.59 |
| ENSMUSG00000020857 | -0.549179062 | 0.000137 | 0.001703 | 2504.069 | 3136.333 | 3303.317 | 2388.604 | 4599.6 | 3411.343 | 3995.05 | 4574.408 |
| ENSMUSG00000020859 | 0.048299036 | 0.428747 | 0.593664 | 3805.904 | 3653.457 | 3411.799 | 3847.301 | 3447.994 | 3663.87 | 3497.477 | 3624.941 |
| ENSMUSG00000020863 | 0.53781398 | 0.000507 | 0.00463 | 11315.09 | 11575.88 | 8584.153 | 10950.14 | 6932.066 | 9488.741 | 6574.524 | 6227.547 |
| ENSMUSG00000020864 | -0.05674825 | 0.469208 | 0.629339 | 1755.565 | 1632.957 | 1535.31 | 1636.397 | 1669.391 | 1588.906 | 1698.113 | 1865.219 |
| ENSMUSG00000020865 | -0.595780904 | 0.033191 | 0.101653 | 254.5318 | 231.5656 | 423.1625 | 265.688 | 368.5921 | 273.7567 | 572.7878 | 562.2846 |
| ENSMUSG00000020866 | 0.181855391 | 0.040983 | 0.118778 | 869.231 | 1000.651 | 1065.773 | 919.5566 | 817.1433 | 878.256 | 837.9673 | 866.8101 |
| ENSMUSG00000020868 | -0.146423677 | 0.060153 | 0.155271 | 1000.018 | 1076.24 | 1185.849 | 1028.247 | 1162.333 | 1167.656 | 1197.647 | 1223.54 |
| ENSMUSG00000020869 | -0.333757788 | 0.001506 | 0.010593 | 3231.447 | 3258.715 | 3673.481 | 3182.218 | 4299.266 | 3608.001 | 4055.8 | 4858.269 |
| ENSMUSG00000020873 | -0.325066964 | 3.73E-05 | 0.000636 | 1014.103 | 982.6538 | 1094.757 | 1038.599 | 1213.038 | 1238.05 | 1318.183 | 1409.518 |
| ENSMUSG00000020875 | 0.375126105 | 2.54E-05 | 0.000473 | 3731.456 | 3904.219 | 3593.983 | 3568.673 | 2806.37 | 3262.732 | 2799.331 | 2540.613 |
| ENSMUSG00000020876 | -0.013118789 | 0.915828 | 0.950114 | 772.6497 | 651.5031 | 618.5957 | 693.5493 | 783.9895 | 626.8469 | 743.4669 | 604.7006 |
| ENSMUSG00000020877 | -0.750792502 | 4.12E-08 | 2.76E-06 | 436.6276 | 418.7377 | 448.8338 | 415.7845 | 705.9806 | 594.443 | 695.2525 | 899.4378 |
| ENSMUSG00000020878 | -0.06059355 | 0.880983 | 0.928526 | 21.12714 | 15.59768 | 23.18699 | 21.56559 | 23.40267 | 13.40849 | 31.82154 | 16.31387 |
| ENSMUSG00000020882 | 0.205402662 | 0.051616 | 0.139029 | 481.9 | 482.3283 | 427.303 | 454.6026 | 446.601 | 356.4423 | 404.0372 | 390.4452 |
| ENSMUSG00000020883 | 0.365440038 | 0.000677 | 0.005792 | 2411.512 | 2063.693 | 2081.86 | 2380.841 | 1821.508 | 1766.568 | 1892.9 | 1456.284 |
| ENSMUSG00000020884 | 0.7945009 | 0.020601 | 0.071986 | 98.59333 | 154.777 | 135.8095 | 131.9814 | 67.28268 | 129.6154 | 63.64308 | 40.24087 |
| ENSMUSG00000020886 | 0.334239355 | 0.000153 | 0.00185 | 898.4065 | 1009.05 | 876.9649 | 996.3301 | 788.865 | 783.2792 | 732.8598 | 691.7079 |
| ENSMUSG00000020887 | -1.723158226 | 0.004949 | 0.025556 | 5.030272 | 5.999108 | 11.59349 | 7.763611 | 12.67645 | 11.17374 | 40.50014 | 36.9781 |
| ENSMUSG00000020888 | 0.093051251 | 0.179935 | 0.329845 | 1390.367 | 1534.572 | 1348.986 | 1377.61 | 1349.554 | 1280.511 | 1371.219 | 1289.883 |
| ENSMUSG00000020889 | 0.152453077 | 0.410746 | 0.576758 | 223.3441 | 161.9759 | 250.9163 | 195.8155 | 228.176 | 174.3104 | 163.9292 | 183.8029 |
| ENSMUSG00000020890 | 0.302695548 | 0.023348 | 0.078879 | 162.9808 | 173.9741 | 173.9024 | 173.3873 | 140.416 | 154.1976 | 129.2147 | 131.5985 |
| ENSMUSG00000020891 | -0.476545872 | 0.407436 | 0.573711 | 6.036326 | 9.598572 | 27.32752 | 15.52722 | 19.50223 | 16.76061 | 12.53576 | 33.71532 |
| ENSMUSG00000020892 | 0.654446631 | 0.042153 | 0.121028 | 47.28455 | 45.59322 | 32.29616 | 37.09281 | 26.32801 | 26.81698 | 32.78583 | 16.31387 |
| ENSMUSG00000020893 | 0.063269046 | 0.901195 | 0.940696 | 1601.639 | 847.074 | 818.9975 | 1846.877 | 644.5486 | 1711.817 | 1909.293 | 629.7152 |
| ENSMUSG00000020894 | 0.213191771 | 0.011611 | 0.047758 | 2254.568 | 2045.696 | 2057.017 | 2365.314 | 1717.171 | 2034.738 | 1871.685 | 1904.372 |
| ENSMUSG00000020895 | -0.190915029 | 0.168768 | 0.315007 | 320.9313 | 421.1374 | 318.8211 | 364.0271 | 461.2276 | 350.8555 | 398.2514 | 412.197 |
| ENSMUSG00000020897 | -0.062922626 | 0.355526 | 0.524172 | 1164.005 | 1097.837 | 1108.835 | 1192.146 | 1212.063 | 1217.938 | 1225.612 | 1111.518 |
| ENSMUSG00000020898 | 0.171543032 | 0.047942 | 0.132201 | 1558.378 | 1602.962 | 1452.499 | 1476.811 | 1311.525 | 1516.277 | 1356.755 | 1221.365 |
| ENSMUSG00000020899 | 0.060863924 | 0.488249 | 0.646025 | 1347.107 | 1216.619 | 1106.35 | 1220.612 | 1194.511 | 1259.281 | 1107.968 | 1125.657 |
| ENSMUSG00000020900 | 0.12042989 | 0.075815 | 0.182312 | 12209.48 | 13827.94 | 13678.67 | 12387.27 | 12443.4 | 12314.58 | 11839.54 | 11330.52 |
| ENSMUSG00000020901 | 0.207679831 | 0.520047 | 0.673542 | 45.27245 | 37.19447 | 68.73285 | 37.09281 | 40.95467 | 54.75133 | 34.71441 | 33.71532 |
| ENSMUSG00000020902 | 1.301761483 | 8.30E-09 | 7.38E-07 | 5648.995 | 8017.207 | 6619.056 | 7033.832 | 2581.12 | 4269.486 | 2247.758 | 1983.766 |
| ENSMUSG00000020903 | -0.314111039 | 0.002779 | 0.016707 | 549.3057 | 547.1186 | 622.7362 | 615.0506 | 763.5122 | 657.016 | 675.9667 | 809.1678 |
| ENSMUSG00000020904 | -2.226759968 | 2.24E-05 | 0.00043 | 30.18163 | 9.598572 | 52.17072 | 24.15346 | 129.6898 | 56.98608 | 243.0009 | 117.4598 |
| ENSMUSG00000020905 | -0.48462955 | 0.024973 | 0.083001 | 161.9748 | 115.1829 | 219.4483 | 172.5247 | 224.2756 | 236.8833 | 282.5367 | 195.7664 |
| ENSMUSG00000020907 | -0.052880085 | 0.720833 | 0.82531 | 113.6841 | 131.9804 | 119.2474 | 132.844 | 123.8391 | 134.0849 | 120.5361 | 138.1241 |
| ENSMUSG00000020908 | 1.644335357 | 0.012735 | 0.051195 | 15.09082 | 8.398751 | 38.09291 | 24.15346 | 2.925334 | 5.58687 | 12.53576 | 6.525547 |
| ENSMUSG00000020910 | 0.125248684 | 0.291539 | 0.458729 | 472.8455 | 524.322 | 390.0382 | 506.36 | 459.2774 | 426.8369 | 419.4658 | 427.4233 |

| | | | | | | | | | | | |
|---|---|---|---|---|---|---|---|---|---|---|---|
| ENSMUSG00000020911 | -0.38346767 | 0.08951 | 0.204279 | 1214.308 | 1300.607 | 2461.133 | 1335.341 | 2193.025 | 1743.104 | 2364.437 | 1934.825 |
| ENSMUSG00000020914 | 0.056022884 | 0.481038 | 0.640274 | 6561.487 | 6585.82 | 5963.196 | 6092.71 | 6249.488 | 6650.611 | 5835.878 | 5504.298 |
| ENSMUSG00000020916 | 0.48691959 | 0.107054 | 0.231384 | 39.23612 | 65.99018 | 38.09291 | 52.62003 | 28.27823 | 32.40385 | 43.39301 | 34.80291 |
| ENSMUSG00000020917 | -0.243489203 | 0.15372 | 0.295066 | 5523.238 | 3550.272 | 3649.466 | 3674.776 | 5014.997 | 3842.65 | 5086.625 | 5467.32 |
| ENSMUSG00000020918 | -0.11450137 | 0.114032 | 0.241406 | 1555.36 | 1643.756 | 1483.139 | 1620.007 | 1824.433 | 1747.573 | 1596.863 | 1650.963 |
| ENSMUSG00000020919 | 0.104820538 | 0.149619 | 0.289543 | 995.9938 | 945.4594 | 891.8708 | 935.9465 | 880.5255 | 841.3827 | 926.6819 | 852.6714 |
| ENSMUSG00000020921 | 0.095292348 | 0.284816 | 0.451421 | 762.5892 | 819.4781 | 692.2971 | 761.6965 | 771.313 | 676.0113 | 709.7168 | 679.7444 |
| ENSMUSG00000020922 | -0.149518581 | 0.004044 | 0.022004 | 1397.41 | 1431.387 | 1398.672 | 1369.846 | 1549.452 | 1536.389 | 1532.255 | 1587.883 |
| ENSMUSG00000020923 | -0.187636861 | 0.000484 | 0.004463 | 5952.824 | 6325.459 | 6410.373 | 5927.949 | 7231.425 | 6988.058 | 6702.774 | 7112.846 |
| ENSMUSG00000020925 | -0.127909273 | 0.051925 | 0.139617 | 786.7345 | 765.4861 | 795.8105 | 799.652 | 892.2268 | 843.6174 | 874.6103 | 829.832 |
| ENSMUSG00000020926 | 0.454220968 | 0.000672 | 0.005761 | 617.7174 | 631.1061 | 611.9708 | 658.1817 | 546.0623 | 507.2878 | 390.5371 | 394.7956 |
| ENSMUSG00000020928 | -0.917695647 | 0.212947 | 0.370828 | 5.030272 | 2.399643 | 23.18699 | 4.313117 | 19.50223 | 10.05637 | 21.21436 | 16.31387 |
| ENSMUSG00000020929 | -0.260194118 | 0.005423 | 0.027212 | 2983.957 | 3448.287 | 3467.282 | 3005.38 | 3957.977 | 3383.409 | 4040.372 | 4070.853 |
| ENSMUSG00000020930 | -0.088991714 | 0.794435 | 0.873694 | 34.20585 | 59.99108 | 54.65504 | 34.50494 | 55.58134 | 30.1691 | 64.60737 | 43.50364 |
| ENSMUSG00000020932 | 0.228138673 | 0.366246 | 0.53485 | 45.27245 | 51.59233 | 66.24853 | 55.2079 | 41.92979 | 44.69496 | 58.82164 | 41.32846 |
| ENSMUSG00000020935 | -0.095078745 | 0.314024 | 0.482085 | 1782.728 | 1960.508 | 1948.535 | 1628.633 | 2040.908 | 1814.616 | 1834.078 | 2127.328 |
| ENSMUSG00000020936 | -0.177228421 | 0.006609 | 0.03165 | 2345.113 | 2394.844 | 2507.507 | 2403.269 | 2777.117 | 2534.204 | 2675.902 | 2926.708 |
| ENSMUSG00000020937 | 0.26594446 | 0.019934 | 0.070247 | 463.7911 | 436.735 | 426.4749 | 399.3947 | 394.9201 | 326.2732 | 388.6085 | 323.0146 |
| ENSMUSG00000020941 | -0.019501607 | 0.90547 | 0.943292 | 220.3259 | 238.7645 | 215.3077 | 182.0136 | 201.848 | 269.2872 | 212.1436 | 183.8029 |
| ENSMUSG00000020946 | -0.054067129 | 0.472213 | 0.631533 | 2807.898 | 2959.96 | 3259.428 | 2785.411 | 3221.768 | 2895.116 | 3040.404 | 3107.248 |
| ENSMUSG00000020948 | -0.185023497 | 0.266698 | 0.431164 | 133.8052 | 116.3827 | 108.482 | 131.1188 | 125.7894 | 131.8501 | 165.8577 | 132.6861 |
| ENSMUSG00000020949 | -0.033572476 | 0.608593 | 0.744128 | 2817.958 | 3107.538 | 2784.923 | 2963.974 | 3057.949 | 3140.939 | 2830.189 | 2916.919 |
| ENSMUSG00000020952 | -0.094790973 | 0.20273 | 0.358389 | 1437.652 | 1451.784 | 1416.062 | 1442.306 | 1617.71 | 1371.018 | 1526.47 | 1620.511 |
| ENSMUSG00000020953 | 0.410175343 | 0.121071 | 0.25028 | 64.38748 | 55.19179 | 76.18581 | 48.30692 | 39.00445 | 41.34284 | 58.82164 | 44.59123 |
| ENSMUSG00000020954 | 0.199045447 | 0.031032 | 0.096765 | 2640.893 | 2526.824 | 2379.978 | 2616.337 | 1991.177 | 2540.909 | 2102.15 | 2221.949 |
| ENSMUSG00000020955 | -0.135974374 | 0.130575 | 0.263031 | 481.9 | 513.5236 | 472.8489 | 474.4429 | 563.6143 | 547.5133 | 480.216 | 541.6204 |
| ENSMUSG00000020956 | -0.173742369 | 0.133503 | 0.267007 | 632.8082 | 505.1249 | 620.2519 | 583.9961 | 693.3041 | 565.3913 | 673.0738 | 712.3722 |
| ENSMUSG00000020961 | -0.429250483 | 0.155516 | 0.297462 | 107.6478 | 88.78679 | 131.669 | 62.97152 | 143.3414 | 83.80306 | 122.4647 | 177.2773 |
| ENSMUSG00000020962 | 0.238751538 | 0.113087 | 0.24001 | 1418.537 | 1412.19 | 1344.845 | 1466.46 | 1097 | 1605.667 | 1048.182 | 1032.124 |
| ENSMUSG00000020963 | -1.350650666 | 0.00338 | 0.019307 | 57.3451 | 11.99822 | 27.32752 | 32.77969 | 66.30757 | 52.51658 | 127.2862 | 84.8321 |
| ENSMUSG00000020964 | 0.020012483 | 0.782762 | 0.866262 | 3082.551 | 2602.413 | 2833.781 | 2778.51 | 2653.278 | 2763.266 | 2796.439 | 2932.146 |
| ENSMUSG00000020973 | -0.145134033 | 0.287834 | 0.454754 | 440.6518 | 574.7145 | 467.0521 | 440.8006 | 628.9468 | 513.9921 | 494.6803 | 485.0656 |
| ENSMUSG00000020974 | -0.056364473 | 0.701388 | 0.811639 | 305.8405 | 422.3372 | 289.0092 | 351.9504 | 371.5174 | 376.5551 | 319.1797 | 353.4671 |
| ENSMUSG00000020978 | 0.016760026 | 0.80427 | 0.880231 | 3765.661 | 4100.99 | 3670.169 | 4050.017 | 4062.314 | 3699.626 | 3965.157 | 3673.883 |
| ENSMUSG00000020982 | 0.153789048 | 0.224722 | 0.384523 | 2087.563 | 2164.478 | 1746.477 | 1968.507 | 1568.954 | 2193.405 | 1747.292 | 1650.963 |
| ENSMUSG00000020986 | 0.182219059 | 0.057798 | 0.150724 | 2721.377 | 2566.418 | 2105.875 | 2713.814 | 2270.059 | 2264.917 | 2167.722 | 2203.46 |
| ENSMUSG00000020988 | -0.525686908 | 9.25E-07 | 3.49E-05 | 558.3602 | 555.5174 | 595.4087 | 643.5171 | 833.7202 | 729.6453 | 891.0032 | 935.3283 |
| ENSMUSG00000020990 | -1.088149786 | 0.030604 | 0.095865 | 403.4278 | 295.1561 | 1122.084 | 553.8043 | 1175.009 | 654.7812 | 1445.469 | 1774.949 |
| ENSMUSG00000020992 | -0.969962454 | 0.048621 | 0.133594 | 63.38142 | 23.99643 | 143.2624 | 66.42201 | 133.5902 | 70.39457 | 174.5363 | 205.5547 |
| ENSMUSG00000020993 | 0.033729012 | 0.675637 | 0.792479 | 1339.058 | 1287.409 | 1382.938 | 1418.153 | 1317.375 | 1184.417 | 1391.469 | 1410.606 |
| ENSMUSG00000020994 | 0.142029969 | 0.055223 | 0.146226 | 6915.618 | 6522.23 | 6219.081 | 6496.418 | 6546.897 | 5644.974 | 5544.663 | 5962.174 |
| ENSMUSG00000021000 | -0.348417067 | 0.000108 | 0.00142 | 1745.504 | 1638.956 | 1735.712 | 1607.068 | 2058.46 | 1878.306 | 2287.294 | 2340.496 |
| ENSMUSG00000021003 | -0.825019685 | 3.11E-06 | 9.31E-05 | 564.3965 | 500.3256 | 614.4551 | 606.4243 | 983.8873 | 719.5889 | 1038.539 | 1309.46 |
| ENSMUSG00000021007 | 0.128580682 | 0.345558 | 0.514314 | 418.5186 | 467.9304 | 415.7095 | 485.657 | 405.6463 | 500.5836 | 375.1085 | 354.5547 |
| ENSMUSG00000021009 | 0.138776393 | 0.173364 | 0.321149 | 531.1967 | 500.3256 | 463.7397 | 476.1682 | 412.4721 | 503.9357 | 436.823 | 437.2116 |
| ENSMUSG00000021010 | 0.041425062 | 0.875109 | 0.925241 | 93.56306 | 85.18733 | 60.45178 | 91.43809 | 68.25779 | 99.44629 | 96.42892 | 56.55474 |
| ENSMUSG00000021012 | -0.245710129 | 3.57E-05 | 0.000613 | 2699.244 | 2849.576 | 2846.202 | 2808.702 | 3438.242 | 3077.248 | 3328.726 | 3437.875 |
| ENSMUSG00000021013 | 0.127585351 | 0.176045 | 0.324519 | 626.7719 | 716.2935 | 687.3285 | 744.4441 | 573.3654 | 662.6028 | 663.4309 | 641.6787 |
| ENSMUSG00000021018 | -0.000752571 | 0.993347 | 0.996207 | 954.7456 | 1141.03 | 1112.147 | 991.1544 | 1122.353 | 1061.505 | 983.5749 | 1032.124 |
| ENSMUSG00000021022 | 0.275252437 | 0.004795 | 0.024968 | 875.2673 | 1096.637 | 958.9475 | 1042.912 | 840.5459 | 878.256 | 785.8957 | 777.6276 |
| ENSMUSG00000021023 | -0.079304826 | 0.294133 | 0.461764 | 504.0332 | 524.322 | 526.6758 | 546.9033 | 546.0623 | 555.3349 | 567.9663 | 552.4963 |
| ENSMUSG00000021024 | -0.163623633 | 0.008534 | 0.038027 | 3911.539 | 4132.185 | 4101.612 | 4117.302 | 4813.149 | 4177.862 | 4634.374 | 4587.459 |
| ENSMUSG00000021025 | 0.012188518 | 0.951999 | 0.972676 | 1837.055 | 1288.608 | 1470.717 | 1965.919 | 1271.545 | 1793.385 | 2144.579 | 1298.584 |
| ENSMUSG00000021027 | 0.203812699 | 0.027239 | 0.08826 | 1797.819 | 1840.526 | 1575.059 | 1706.269 | 1376.857 | 1689.47 | 1503.327 | 1436.708 |
| ENSMUSG00000021028 | 0.345144515 | 0.026792 | 0.087374 | 747.4984 | 928.6619 | 761.8581 | 818.6297 | 645.5237 | 824.6221 | 559.2877 | 534.0072 |
| ENSMUSG00000021032 | 0.131050717 | 0.791444 | 0.87239 | 14.08476 | 15.59768 | 11.59349 | 21.56559 | 5.850668 | 20.11273 | 15.42863 | 16.31387 |
| ENSMUSG00000021033 | -0.640640928 | 3.92E-06 | 0.000111 | 1012.091 | 965.8563 | 1038.446 | 1051.538 | 1781.528 | 1240.285 | 1439.684 | 1881.533 |
| ENSMUSG00000021036 | -0.00365486 | 0.950287 | 0.97177 | 2968.866 | 2765.589 | 3006.855 | 2727.615 | 2817.097 | 2832.543 | 2872.617 | 2978.912 |
| ENSMUSG00000021037 | -0.279772982 | 0.000681 | 0.005816 | 2462.821 | 2624.01 | 2428.837 | 2555.091 | 3028.696 | 2687.285 | 3167.69 | 3338.905 |
| ENSMUSG00000021038 | -0.148892895 | 0.084857 | 0.196764 | 1496.003 | 1457.783 | 1453.327 | 1539.783 | 1686.943 | 1417.948 | 1729.935 | 1757.547 |
| ENSMUSG00000021039 | -0.179831919 | 0.018879 | 0.067695 | 1965.83 | 2138.082 | 1914.583 | 1771.829 | 2298.337 | 2098.429 | 2223.651 | 2195.846 |
| ENSMUSG00000021040 | -0.168752932 | 0.088342 | 0.202413 | 709.2683 | 769.0856 | 798.2948 | 660.7696 | 843.4713 | 876.0213 | 736.7169 | 846.1459 |
| ENSMUSG00000021044 | -0.57008625 | 8.22E-07 | 3.16E-05 | 386.3249 | 403.14 | 428.1311 | 425.2734 | 570.4401 | 539.6917 | 614.2522 | 716.7225 |
| ENSMUSG00000021047 | 0.941292 | 0.002328 | 0.014615 | 391.3551 | 484.7279 | 366.0231 | 423.5481 | 174.5449 | 392.1983 | 162.0006 | 139.2117 |

| | | | | | | | | | | | |
|---|---|---|---|---|---|---|---|---|---|---|---|
| ENSMUSG00000021048 | -1.008424864 | 5.00E-07 | 2.11E-05 | 1453.749 | 1409.79 | 1734.055 | 1456.108 | 2961.413 | 1899.536 | 3332.583 | 3986.021 |
| ENSMUSG00000021051 | -0.007558167 | 0.931097 | 0.959275 | 1570.451 | 1654.554 | 1434.281 | 1571.7 | 1430.488 | 1740.869 | 1585.291 | 1505.226 |
| ENSMUSG00000021054 | -0.382246962 | 0.003818 | 0.021109 | 815.9101 | 566.3158 | 674.0788 | 699.5877 | 783.0144 | 897.2514 | 921.8604 | 992.9707 |
| ENSMUSG00000021055 | -0.774914139 | 0.068583 | 0.169697 | 11.0666 | 14.39786 | 21.53077 | 18.97772 | 38.02934 | 16.76061 | 35.6787 | 22.83941 |
| ENSMUSG00000021057 | 0.625487556 | 0.013239 | 0.052726 | 132.7992 | 109.1838 | 89.43552 | 106.1027 | 83.85957 | 87.15518 | 46.28588 | 66.34306 |
| ENSMUSG00000021061 | 0.691109958 | 0.056758 | 0.148796 | 644.8808 | 202.7698 | 230.2136 | 235.4962 | 239.8774 | 163.1366 | 220.8222 | 189.2408 |
| ENSMUSG00000021062 | -0.069579178 | 0.73476 | 0.833942 | 515.0998 | 527.9215 | 932.4481 | 596.9355 | 668.9264 | 558.687 | 735.7526 | 738.4743 |
| ENSMUSG00000021065 | 0.07698931 | 0.2844 | 0.451019 | 971.8485 | 1063.042 | 1030.165 | 1130.037 | 1002.414 | 1003.402 | 991.2893 | 981.0072 |
| ENSMUSG00000021066 | 0.451444358 | 0.011971 | 0.048893 | 257.5499 | 191.9714 | 163.137 | 224.2821 | 135.5405 | 163.1366 | 161.0363 | 152.2628 |
| ENSMUSG00000021067 | -0.096159234 | 0.170945 | 0.317902 | 1415.518 | 1430.187 | 1529.513 | 1601.892 | 1667.44 | 1518.511 | 1593.006 | 1613.985 |
| ENSMUSG00000021068 | -0.090498636 | 0.233936 | 0.394655 | 1348.113 | 1138.631 | 1334.08 | 1276.683 | 1350.529 | 1331.91 | 1341.326 | 1409.518 |
| ENSMUSG00000021069 | -0.063823493 | 0.604582 | 0.741111 | 1577.493 | 1339.001 | 1658.698 | 1563.936 | 1504.597 | 1337.497 | 1832.149 | 1744.496 |
| ENSMUSG00000021070 | -0.801048835 | 0.097768 | 0.21756 | 24.1453 | 13.19804 | 9.109173 | 15.52722 | 35.10401 | 12.29112 | 33.75012 | 26.10219 |
| ENSMUSG00000021071 | -0.387411492 | 0.19897 | 0.353673 | 107.6478 | 51.59233 | 52.99882 | 104.3774 | 90.68535 | 102.7984 | 123.429 | 97.8832 |
| ENSMUSG00000021072 | -0.052114988 | 0.391557 | 0.559029 | 2299.84 | 2256.864 | 2112.5 | 2317.007 | 2267.134 | 2251.509 | 2347.08 | 2449.255 |
| ENSMUSG00000021076 | -0.06170884 | 0.308285 | 0.476744 | 1821.964 | 1799.732 | 1893.052 | 1927.101 | 1997.028 | 1817.968 | 1994.15 | 1959.839 |
| ENSMUSG00000021079 | -0.178452135 | 0.227351 | 0.387335 | 526.1664 | 554.3175 | 565.5968 | 522.7498 | 613.345 | 493.8794 | 563.1449 | 784.1532 |
| ENSMUSG00000021087 | 0.672924263 | 0.081033 | 0.190588 | 386.3249 | 152.3773 | 283.2125 | 661.6322 | 246.7032 | 149.7281 | 275.7867 | 258.8467 |
| ENSMUSG00000021090 | 1.154653472 | 0.02292 | 0.077773 | 28.16952 | 19.19714 | 10.76539 | 22.42821 | 11.70134 | 11.17374 | 7.714313 | 5.437955 |
| ENSMUSG00000021091 | 2.438610958 | 9.67E-06 | 0.000221 | 268.6165 | 41.99375 | 80.32634 | 191.5024 | 26.32801 | 15.64324 | 42.42872 | 22.83941 |
| ENSMUSG00000021094 | 0.736877356 | 9.71E-05 | 0.001308 | 481.9 | 434.3354 | 352.7734 | 413.1967 | 256.4543 | 336.3296 | 220.8222 | 195.7664 |
| ENSMUSG00000021096 | -0.093575529 | 0.128935 | 0.260609 | 2620.772 | 2436.838 | 2445.399 | 2682.759 | 2749.814 | 2738.684 | 2801.26 | 2579.766 |
| ENSMUSG00000021097 | -0.361187638 | 0.02311 | 0.078304 | 734.4197 | 637.1052 | 933.2762 | 636.6161 | 813.2428 | 902.8383 | 942.1105 | 1123.482 |
| ENSMUSG00000021098 | 1.713430075 | 8.29E-05 | 0.001166 | 269.6226 | 289.157 | 208.6829 | 301.9182 | 64.35735 | 178.7799 | 44.3573 | 39.15328 |
| ENSMUSG00000021102 | -0.614217798 | 2.59E-05 | 0.000479 | 1770.656 | 1582.565 | 1830.116 | 1603.617 | 2718.61 | 1912.944 | 2650.831 | 3107.248 |
| ENSMUSG00000021103 | -0.077386396 | 0.290179 | 0.457309 | 1226.38 | 1267.012 | 1088.96 | 1214.574 | 1276.421 | 1296.154 | 1273.826 | 1209.401 |
| ENSMUSG00000021108 | -0.515851039 | 0.001593 | 0.011029 | 400.4096 | 293.9563 | 457.943 | 338.1484 | 509.0081 | 463.7102 | 613.2879 | 548.1459 |
| ENSMUSG00000021109 | -0.150904073 | 0.012399 | 0.050216 | 3098.647 | 3205.923 | 3313.255 | 3200.333 | 3335.856 | 3554.367 | 3798.335 | 3544.459 |
| ENSMUSG00000021111 | 0.052957128 | 0.388336 | 0.555851 | 5461.869 | 5472.386 | 4827.862 | 5169.703 | 5115.434 | 5136.569 | 4857.125 | 5062.737 |
| ENSMUSG00000021112 | -0.254939666 | 0.002021 | 0.013241 | 1039.254 | 979.0544 | 985.4469 | 957.5121 | 1120.403 | 1114.022 | 1321.076 | 1169.16 |
| ENSMUSG00000021113 | 0.063816148 | 0.677272 | 0.793634 | 451.7184 | 381.5432 | 332.8989 | 370.9281 | 399.7956 | 289.3999 | 368.3585 | 411.1094 |
| ENSMUSG00000021114 | -0.522753973 | 0.000194 | 0.002211 | 1563.408 | 1380.995 | 1767.18 | 1501.828 | 2156.946 | 1748.69 | 2380.83 | 2642.846 |
| ENSMUSG00000021115 | 0.110923281 | 0.193771 | 0.347106 | 1431.615 | 1342.6 | 1286.05 | 1430.23 | 1332.002 | 1372.135 | 1240.076 | 1139.795 |
| ENSMUSG00000021116 | -0.275525398 | 2.44E-05 | 0.000457 | 1850.134 | 1871.722 | 1771.32 | 1877.931 | 2336.367 | 2048.147 | 2232.329 | 2301.343 |
| ENSMUSG00000021118 | -1.046031369 | 0.000759 | 0.006322 | 40.24217 | 37.19447 | 67.07664 | 33.64232 | 100.4365 | 62.57295 | 114.7504 | 91.35765 |
| ENSMUSG00000021120 | -0.272005835 | 0.01788 | 0.065359 | 339.0403 | 345.5486 | 336.2113 | 301.9182 | 419.2979 | 337.447 | 416.5729 | 420.8978 |
| ENSMUSG00000021123 | -0.333828308 | 0.054459 | 0.144809 | 164.9929 | 122.3818 | 162.3089 | 194.0903 | 202.8232 | 175.4277 | 223.7151 | 212.0803 |
| ENSMUSG00000021124 | -0.197183214 | 0.02643 | 0.086487 | 1472.864 | 1491.378 | 1578.371 | 1393.137 | 1798.105 | 1482.755 | 1706.792 | 1816.277 |
| ENSMUSG00000021125 | -0.806187114 | 0.001681 | 0.011514 | 301.8163 | 429.5361 | 510.9418 | 389.9058 | 588.9672 | 499.4662 | 1094.468 | 671.0437 |
| ENSMUSG00000021127 | 0.207340627 | 0.019332 | 0.068816 | 2995.024 | 2872.373 | 2548.912 | 2630.139 | 2457.28 | 2567.726 | 2132.043 | 2409.014 |
| ENSMUSG00000021130 | 0.335744876 | 0.06687 | 0.166901 | 361.1735 | 400.7404 | 475.3332 | 503.7721 | 334.4632 | 452.5365 | 317.2511 | 277.3357 |
| ENSMUSG00000021131 | -0.118933464 | 0.114556 | 0.242071 | 4217.38 | 5047.649 | 4877.548 | 4664.205 | 5313.382 | 5232.663 | 5057.697 | 4816.941 |
| ENSMUSG00000021133 | -0.030441283 | 0.824137 | 0.893455 | 1074.466 | 892.6672 | 1058.32 | 1161.091 | 945.858 | 909.5425 | 1176.433 | 1246.379 |
| ENSMUSG00000021134 | 0.08674061 | 0.28965 | 0.456865 | 24032.63 | 20944.08 | 20699.35 | 24087.9 | 22214.99 | 21497.16 | 21458.33 | 19355.86 |
| ENSMUSG00000021135 | -1.700370793 | 0.073673 | 0.178862 | 17.10292 | 1.199822 | 1.656213 | 7.763611 | 33.15378 | 6.704245 | 5.785735 | 44.59123 |
| ENSMUSG00000021136 | 0.824228556 | 0.146832 | 0.285828 | 325.9616 | 53.99197 | 43.06154 | 149.2339 | 75.08357 | 108.3853 | 54.00019 | 85.9197 |
| ENSMUSG00000021139 | -0.151970664 | 0.408726 | 0.574958 | 138.8355 | 111.5834 | 91.91984 | 140.6076 | 139.4409 | 132.9675 | 130.179 | 133.7737 |
| ENSMUSG00000021140 | 0.137236163 | 0.103374 | 0.226011 | 2351.149 | 2192.074 | 2052.048 | 2260.936 | 1876.114 | 2226.927 | 2062.615 | 1886.971 |
| ENSMUSG00000021143 | -0.242035087 | 0.005141 | 0.026247 | 1370.246 | 1385.794 | 1657.869 | 1490.613 | 1693.768 | 1673.826 | 1754.042 | 1867.394 |
| ENSMUSG00000021144 | -0.013071557 | 0.850481 | 0.910257 | 2694.214 | 2613.211 | 2623.442 | 2739.692 | 2784.918 | 2872.769 | 2677.831 | 2432.941 |
| ENSMUSG00000021147 | 0.085025046 | 0.408418 | 0.574721 | 937.6427 | 871.0704 | 890.2146 | 891.0901 | 790.8153 | 994.4629 | 842.7887 | 758.051 |
| ENSMUSG00000021149 | -0.067819151 | 0.386528 | 0.554049 | 1890.376 | 1960.508 | 1710.868 | 1860.679 | 2051.634 | 2060.438 | 1840.828 | 1822.803 |
| ENSMUSG00000021156 | 0.183087331 | 0.013514 | 0.053419 | 4005.102 | 4042.199 | 3854.836 | 4238.069 | 3362.184 | 3958.856 | 3397.191 | 3500.956 |
| ENSMUSG00000021171 | 0.047446971 | 0.581836 | 0.722782 | 1657.978 | 1718.144 | 1727.43 | 1824.449 | 1514.348 | 1881.658 | 1708.72 | 1602.022 |
| ENSMUSG00000021175 | -0.028104091 | 0.8101 | 0.884215 | 1056.357 | 1371.396 | 1066.601 | 1168.855 | 1326.151 | 1197.825 | 1173.54 | 1052.788 |
| ENSMUSG00000021176 | -0.111928459 | 0.468506 | 0.628746 | 144.8718 | 140.3791 | 112.6225 | 124.2178 | 143.3414 | 153.0803 | 131.1433 | 134.8613 |
| ENSMUSG00000021177 | -0.028594092 | 0.726971 | 0.829129 | 743.4742 | 872.2703 | 804.0915 | 769.4602 | 834.6953 | 801.1572 | 843.753 | 768.9269 |
| ENSMUSG00000021178 | -0.121723776 | 0.019055 | 0.068193 | 5255.628 | 5321.209 | 5200.51 | 5521.653 | 5710.252 | 5507.537 | 6029.7 | 5925.196 |
| ENSMUSG00000021179 | 0.073673953 | 0.422105 | 0.587229 | 1317.931 | 1214.219 | 1311.721 | 1129.174 | 1242.292 | 1271.572 | 1142.683 | 1069.102 |
| ENSMUSG00000021180 | -0.095479035 | 0.56544 | 0.710019 | 241.453 | 309.554 | 211.1672 | 286.391 | 273.0312 | 327.3906 | 278.6796 | 239.27 |
| ENSMUSG00000021182 | 0.030386033 | 0.778504 | 0.86331 | 804.8435 | 713.8938 | 835.5596 | 688.3735 | 705.0055 | 749.758 | 686.5739 | 840.7079 |
| ENSMUSG00000021185 | -0.381795066 | 0.168895 | 0.315147 | 80.48435 | 137.9795 | 144.0906 | 106.1027 | 161.8685 | 98.32892 | 144.6434 | 205.5547 |
| ENSMUSG00000021186 | 0.248752462 | 0.083704 | 0.194883 | 2509.1 | 3689.451 | 3833.306 | 3725.671 | 2660.104 | 2991.21 | 2734.724 | 3194.255 |
| ENSMUSG00000021187 | 0.573247304 | 0.111595 | 0.238021 | 32.19374 | 28.79572 | 38.92101 | 62.97152 | 25.35289 | 32.40385 | 21.21436 | 31.54014 |

| | | | | | | | | | | | |
|---|---|---|---|---|---|---|---|---|---|---|---|
| ENSMUSG00000021188 | 0.163462188 | 0.051346 | 0.138565 | 1445.7 | 1269.411 | 1311.721 | 1274.958 | 1182.81 | 1299.506 | 1156.183 | 1096.292 |
| ENSMUSG00000021189 | 0.123037025 | 0.382066 | 0.549989 | 718.3228 | 617.9081 | 596.2368 | 755.6582 | 611.3948 | 734.1148 | 618.1094 | 505.7299 |
| ENSMUSG00000021190 | -0.973376849 | 6.61E-05 | 0.000988 | 2394.409 | 1840.526 | 2813.906 | 2169.498 | 5403.092 | 2466.045 | 4361.48 | 5870.817 |
| ENSMUSG00000021192 | -0.048132088 | 0.462575 | 0.623614 | 1570.451 | 1602.962 | 1707.556 | 1532.019 | 1691.818 | 1600.08 | 1606.506 | 1733.62 |
| ENSMUSG00000021193 | -0.049821804 | 0.541405 | 0.690735 | 3030.236 | 2748.791 | 2872.702 | 2711.226 | 3039.422 | 2586.721 | 3023.047 | 3111.598 |
| ENSMUSG00000021194 | 3.036095014 | 0.0057 | 0.028348 | 1591.578 | 64.79036 | 44.71776 | 1147.289 | 74.10846 | 45.81234 | 76.17884 | 151.1752 |
| ENSMUSG00000021196 | -0.683356375 | 0.0099 | 0.042286 | 908.4671 | 614.3086 | 1038.446 | 890.2274 | 1168.183 | 783.2792 | 1789.721 | 1803.226 |
| ENSMUSG00000021198 | 0.162759379 | 0.703033 | 0.812828 | 33.19979 | 15.59768 | 20.70267 | 48.30692 | 23.40267 | 34.6386 | 24.10723 | 23.927 |
| ENSMUSG00000021200 | 0.553396515 | 0.186782 | 0.338697 | 30.18163 | 83.98751 | 42.23344 | 39.68068 | 39.00445 | 46.92971 | 23.14294 | 23.927 |
| ENSMUSG00000021203 | -0.060453797 | 0.685327 | 0.799822 | 190.1443 | 184.7725 | 177.2148 | 144.9207 | 192.0969 | 198.8926 | 163.9292 | 170.7518 |
| ENSMUSG00000021207 | -2.83811215 | 4.81E-07 | 2.04E-05 | 6.036326 | 3.599465 | 9.109173 | 8.626235 | 33.15378 | 21.23011 | 57.85735 | 87.00729 |
| ENSMUSG00000021209 | -0.504175514 | 0.010402 | 0.043891 | 125.7568 | 149.9777 | 132.4971 | 111.2784 | 149.192 | 157.5497 | 234.3223 | 193.5912 |
| ENSMUSG00000021211 | -0.600995806 | 0.248586 | 0.411988 | 27.16347 | 14.39786 | 87.7793 | 46.58167 | 61.43201 | 37.99072 | 81.00029 | 88.09488 |
| ENSMUSG00000021213 | -0.166786203 | 0.615042 | 0.748514 | 89.53884 | 79.18822 | 173.0743 | 57.79577 | 106.2871 | 101.681 | 116.679 | 125.073 |
| ENSMUSG00000021214 | 0.099910725 | 0.781186 | 0.865089 | 28.16952 | 38.39429 | 35.60859 | 34.50494 | 30.22845 | 51.39921 | 20.25007 | 26.10219 |
| ENSMUSG00000021215 | -0.028305069 | 0.778691 | 0.863465 | 1995.006 | 1720.544 | 1752.274 | 1821.861 | 1579.68 | 1949.818 | 2040.436 | 1865.219 |
| ENSMUSG00000021217 | -0.070205454 | 0.738305 | 0.836217 | 178.0716 | 169.1748 | 273.2752 | 219.1064 | 179.4205 | 213.4185 | 212.1436 | 279.5109 |
| ENSMUSG00000021218 | -0.04852823 | 0.374949 | 0.543518 | 8270.773 | 9064.652 | 8807.742 | 9205.055 | 8820.857 | 9442.928 | 9116.39 | 9179.269 |
| ENSMUSG00000021219 | -0.124082613 | 0.603817 | 0.740916 | 82.49646 | 82.78769 | 124.216 | 85.39973 | 88.73513 | 87.15518 | 98.35749 | 135.9489 |
| ENSMUSG00000021221 | -0.395808525 | 0.00353 | 0.019915 | 135.8173 | 163.1757 | 149.8873 | 138.8824 | 201.848 | 197.7752 | 199.6079 | 171.8394 |
| ENSMUSG00000021222 | 0.155319404 | 0.120428 | 0.249434 | 773.6558 | 752.2881 | 683.188 | 690.0988 | 693.3041 | 702.8283 | 571.8235 | 634.0656 |
| ENSMUSG00000021223 | -0.520078127 | 0.001817 | 0.01224 | 187.1261 | 181.1731 | 207.0267 | 158.7227 | 255.4792 | 204.4795 | 292.1796 | 300.1751 |
| ENSMUSG00000021224 | -0.051506919 | 0.665522 | 0.7853 | 1126.781 | 949.0588 | 1069.086 | 1043.774 | 951.7086 | 1293.919 | 985.5035 | 1113.693 |
| ENSMUSG00000021226 | -0.279552776 | 0.111136 | 0.237288 | 626.7719 | 680.2988 | 798.2948 | 688.3735 | 873.6997 | 611.2036 | 811.9315 | 1095.204 |
| ENSMUSG00000021228 | -3.045649416 | 1.40E-05 | 0.000296 | 13.07871 | 7.198929 | 36.43669 | 6.900988 | 148.2169 | 34.6386 | 61.71451 | 284.9489 |
| ENSMUSG00000021234 | -0.005870843 | 0.976731 | 0.987311 | 68.4117 | 76.78858 | 59.62368 | 81.08661 | 64.35735 | 74.86406 | 70.39311 | 77.21897 |
| ENSMUSG00000021235 | -0.200564941 | 0.054902 | 0.145647 | 578.4813 | 577.1142 | 676.5631 | 677.1594 | 722.5575 | 641.3727 | 752.1455 | 771.1021 |
| ENSMUSG00000021236 | -1.439738138 | 8.12E-07 | 3.13E-05 | 868.2249 | 705.4951 | 1110.491 | 775.4985 | 2090.639 | 1187.769 | 2372.151 | 3736.963 |
| ENSMUSG00000021238 | -1.06674792 | 4.96E-09 | 4.74E-07 | 1885.346 | 1382.194 | 1538.622 | 1907.261 | 3684.946 | 2414.645 | 3687.442 | 4277.496 |
| ENSMUSG00000021239 | 0.003379961 | 0.991051 | 0.994824 | 40.24217 | 61.1909 | 57.96746 | 51.75741 | 47.78045 | 79.33356 | 40.50014 | 43.50364 |
| ENSMUSG00000021240 | -0.506660611 | 0.000298 | 0.003078 | 387.3309 | 387.5424 | 385.8977 | 445.1137 | 509.0081 | 472.6492 | 657.6452 | 642.7663 |
| ENSMUSG00000021241 | 0.014191023 | 0.886871 | 0.932224 | 934.6245 | 772.6851 | 819.8256 | 916.1062 | 845.4215 | 812.331 | 806.1457 | 948.3794 |
| ENSMUSG00000021242 | -0.147970644 | 0.08472 | 0.196531 | 4958.842 | 5453.189 | 5446.457 | 5476.797 | 6297.269 | 5094.109 | 6069.236 | 6177.517 |
| ENSMUSG00000021243 | -0.05186444 | 0.360002 | 0.528534 | 1424.573 | 1390.593 | 1454.155 | 1405.214 | 1538.726 | 1431.356 | 1439.684 | 1473.686 |
| ENSMUSG00000021244 | 0.303633753 | 0.060699 | 0.156363 | 1877.297 | 2068.492 | 1794.507 | 1912.436 | 1326.151 | 2135.302 | 1369.291 | 1370.365 |
| ENSMUSG00000021245 | -0.201801907 | 0.089482 | 0.204238 | 460.7729 | 458.3318 | 409.9128 | 397.6694 | 481.705 | 459.2408 | 459.0016 | 584.0364 |
| ENSMUSG00000021248 | -0.145030482 | 0.002904 | 0.017327 | 4899.485 | 5102.841 | 5312.304 | 5128.297 | 5681.974 | 5450.551 | 5672.913 | 5802.298 |
| ENSMUSG00000021250 | 0.744676104 | 0.05095 | 0.13788 | 232.3986 | 140.3791 | 105.1695 | 201.8539 | 102.3867 | 42.46022 | 150.4291 | 109.8467 |
| ENSMUSG00000021252 | -0.158752472 | 0.013399 | 0.053134 | 949.7153 | 1033.046 | 931.62 | 988.5665 | 1092.125 | 1074.914 | 1100.254 | 1084.328 |
| ENSMUSG00000021253 | 0.804158688 | 0.000808 | 0.006628 | 1786.753 | 2775.187 | 1886.427 | 2191.926 | 1270.57 | 1833.611 | 1017.325 | 826.5692 |
| ENSMUSG00000021254 | -0.22694099 | 0.102411 | 0.224557 | 503.0272 | 544.719 | 525.0196 | 537.4144 | 643.5735 | 465.945 | 642.2166 | 716.7225 |
| ENSMUSG00000021255 | -2.603649274 | 7.50E-06 | 0.000185 | 55.33299 | 71.98929 | 207.0267 | 69.00988 | 527.5352 | 259.2308 | 1003.825 | 664.5182 |
| ENSMUSG00000021256 | -0.106234394 | 0.634378 | 0.762319 | 153.9263 | 140.3791 | 186.324 | 173.3873 | 172.5947 | 120.6764 | 171.6435 | 240.3576 |
| ENSMUSG00000021257 | -0.257796731 | 0.054832 | 0.145525 | 155.9384 | 182.3729 | 196.2613 | 183.7388 | 216.4747 | 213.4185 | 194.7864 | 236.0073 |
| ENSMUSG00000021258 | 0.182135279 | 0.058482 | 0.151849 | 719.3289 | 680.2988 | 665.7977 | 656.4565 | 669.9015 | 573.2129 | 537.1091 | 617.7517 |
| ENSMUSG00000021259 | 0.321849896 | 0.019881 | 0.070169 | 159.9626 | 196.7707 | 180.5272 | 190.6398 | 153.0925 | 154.1976 | 139.8219 | 134.8613 |
| ENSMUSG00000021260 | -0.258218267 | 0.392595 | 0.560025 | 33.19979 | 28.79572 | 29.81184 | 27.60395 | 29.25334 | 29.05173 | 48.21446 | 35.89051 |
| ENSMUSG00000021262 | 0.002973898 | 0.972556 | 0.984801 | 1913.515 | 2296.458 | 2172.124 | 2191.926 | 2167.672 | 2338.664 | 2052.972 | 1996.817 |
| ENSMUSG00000021263 | 0.054840387 | 0.701883 | 0.812145 | 254.5318 | 220.7672 | 264.9941 | 238.0841 | 250.6036 | 245.8223 | 189.0007 | 258.8467 |
| ENSMUSG00000021264 | -0.148982596 | 0.037002 | 0.110558 | 2126.799 | 2025.299 | 2011.471 | 2274.738 | 2167.672 | 2383.359 | 2430.973 | 2377.474 |
| ENSMUSG00000021265 | -0.44431449 | 0.039707 | 0.116046 | 89.53884 | 121.182 | 122.5598 | 139.745 | 156.0178 | 186.6015 | 184.1792 | 117.4598 |
| ENSMUSG00000021266 | -0.235783269 | 0.02015 | 0.070849 | 1449.724 | 1630.557 | 1825.975 | 1533.745 | 1951.198 | 1682.765 | 1902.543 | 2047.934 |
| ENSMUSG00000021268 | 1.658559114 | 0.000624 | 0.005443 | 8713.437 | 10857.19 | 7024 | 10075.44 | 2831.723 | 5572.345 | 1811.899 | 1399.73 |
| ENSMUSG00000021270 | -0.113071531 | 0.144101 | 0.282148 | 11396.58 | 12193.79 | 10967.44 | 10822.47 | 11611.63 | 13454.3 | 11395.01 | 12617.14 |
| ENSMUSG00000021271 | 0.069158879 | 0.633892 | 0.761959 | 591.56 | 533.9206 | 416.5376 | 487.3823 | 532.4108 | 479.3535 | 515.8947 | 403.4963 |
| ENSMUSG00000021273 | -0.69245379 | 2.04E-08 | 1.54E-06 | 913.4974 | 766.686 | 966.4004 | 885.9143 | 1371.982 | 1217.938 | 1680.756 | 1441.058 |
| ENSMUSG00000021275 | -0.018280988 | 0.898048 | 0.93831 | 347.0888 | 285.5575 | 285.6968 | 265.688 | 315.9361 | 245.8223 | 309.5368 | 326.2773 |
| ENSMUSG00000021276 | -0.086716269 | 0.339007 | 0.50752 | 629.79 | 707.8947 | 681.5318 | 600.386 | 725.4828 | 651.4291 | 671.1453 | 730.8612 |
| ENSMUSG00000021277 | 0.241906358 | 0.006832 | 0.032405 | 933.6184 | 928.6619 | 990.4155 | 914.3809 | 887.3513 | 735.2322 | 738.6455 | 824.394 |
| ENSMUSG00000021278 | -2.47375294 | 0.000385 | 0.003746 | 141.8537 | 35.99465 | 336.2113 | 147.5086 | 948.7833 | 385.4941 | 777.2171 | 1567.219 |
| ENSMUSG00000021279 | -0.298862596 | 0.041046 | 0.118888 | 1936.655 | 1844.126 | 2153.905 | 1748.538 | 2537.24 | 1726.343 | 2508.116 | 2679.824 |
| ENSMUSG00000021280 | -0.862212669 | 0.002516 | 0.015496 | 104.6297 | 152.3773 | 208.6829 | 119.9047 | 299.3592 | 160.9019 | 270.9653 | 333.8905 |
| ENSMUSG00000021281 | -1.201068579 | 0.000112 | 0.001469 | 391.3551 | 250.7627 | 565.5968 | 324.3464 | 987.7878 | 440.2454 | 968.1463 | 1128.92 |
| ENSMUSG00000021282 | -0.107199919 | 0.158028 | 0.300708 | 8300.954 | 8127.591 | 7360.212 | 8026.712 | 8146.08 | 9513.323 | 8148.243 | 8461.459 |

| | | | | | | | | | | | |
|---|---|---|---|---|---|---|---|---|---|---|---|
| ENSMUSG00000021285 | -0.133480327 | 0.083981 | 0.195355 | 826.9767 | 889.0678 | 849.6374 | 774.6359 | 940.9824 | 862.6128 | 903.5389 | 952.7298 |
| ENSMUSG00000021286 | -0.295971858 | 0.013916 | 0.054508 | 1070.442 | 1130.232 | 988.7593 | 1129.174 | 1443.165 | 1046.98 | 1417.505 | 1391.029 |
| ENSMUSG00000021287 | -0.06548485 | 0.464149 | 0.62496 | 493.9727 | 547.1186 | 475.3332 | 545.178 | 535.3361 | 566.5087 | 516.859 | 537.27 |
| ENSMUSG00000021288 | 0.040620454 | 0.604501 | 0.74111 | 2460.809 | 2278.461 | 2360.932 | 2634.452 | 2350.018 | 2171.058 | 2403.009 | 2542.788 |
| ENSMUSG00000021290 | -0.336889305 | 0.000355 | 0.003518 | 1800.837 | 1802.132 | 2095.11 | 1864.129 | 2351.968 | 2095.076 | 2528.366 | 2578.678 |
| ENSMUSG00000021294 | -0.338792577 | 0.077989 | 0.1857 | 299.8042 | 217.1677 | 369.3356 | 301.9182 | 387.1192 | 291.6346 | 387.6442 | 439.3868 |
| ENSMUSG00000021301 | 0.861676114 | 0.06335 | 0.161111 | 34.20585 | 22.79661 | 17.39024 | 26.74133 | 7.80089 | 23.46486 | 10.60718 | 14.13868 |
| ENSMUSG00000021302 | 0.05226631 | 0.634233 | 0.762269 | 901.4247 | 829.0767 | 732.0463 | 876.4255 | 879.5504 | 819.0352 | 825.4315 | 693.8831 |
| ENSMUSG00000021303 | 0.491493857 | 0.37828 | 0.546457 | 24.1453 | 14.39786 | 20.70267 | 56.93315 | 9.751113 | 25.6996 | 16.39292 | 31.54014 |
| ENSMUSG00000021306 | -0.645979473 | 0.007382 | 0.034147 | 309.8647 | 254.3622 | 303.087 | 311.4071 | 524.6099 | 292.752 | 371.2513 | 656.905 |
| ENSMUSG00000021311 | -0.270798756 | 0.037022 | 0.110565 | 516.1059 | 409.1391 | 413.2252 | 379.5543 | 513.8837 | 454.7713 | 547.7162 | 554.6715 |
| ENSMUSG00000021313 | 0.763459885 | 0.008365 | 0.037499 | 58.35115 | 57.59143 | 38.09291 | 74.18562 | 30.22845 | 34.6386 | 34.71441 | 34.80291 |
| ENSMUSG00000021314 | 0.417967031 | 0.013277 | 0.052818 | 1033.218 | 1235.816 | 963.9161 | 1214.574 | 1004.365 | 924.0684 | 796.5028 | 602.5255 |
| ENSMUSG00000021318 | 0.345414233 | 0.147311 | 0.286518 | 1064.406 | 1131.432 | 1088.96 | 1120.548 | 791.7904 | 1391.131 | 699.1096 | 586.2116 |
| ENSMUSG00000021319 | 1.584667682 | 0.015002 | 0.057719 | 109.6599 | 184.7725 | 125.8722 | 147.5086 | 19.50223 | 135.2023 | 24.10723 | 10.87591 |
| ENSMUSG00000021322 | -1.435435224 | 0.000269 | 0.002849 | 27.16347 | 29.99554 | 32.29616 | 27.60395 | 100.4365 | 35.75597 | 60.75022 | 119.635 |
| ENSMUSG00000021326 | -0.115143198 | 0.153423 | 0.294729 | 1650.935 | 1934.112 | 1817.694 | 1754.576 | 2068.211 | 1785.564 | 1890.007 | 2004.43 |
| ENSMUSG00000021327 | 0.067253165 | 0.50664 | 0.662455 | 2101.648 | 2034.897 | 1646.276 | 1790.806 | 1760.076 | 1709.582 | 1782.006 | 1972.89 |
| ENSMUSG00000021335 | -3.610677438 | 1.12E-08 | 9.52E-07 | 52.31483 | 17.99732 | 119.2474 | 56.07053 | 894.1771 | 313.9821 | 600.7521 | 1200.701 |
| ENSMUSG00000021336 | -1.009108673 | 0.000973 | 0.007672 | 65.39353 | 38.39429 | 76.18581 | 42.26855 | 144.3165 | 82.68568 | 92.57176 | 129.4233 |
| ENSMUSG00000021337 | -0.863678722 | 0.076109 | 0.182864 | 10.06054 | 9.598572 | 10.76539 | 7.763611 | 18.52711 | 12.29112 | 11.57147 | 27.18978 |
| ENSMUSG00000021338 | -0.122636533 | 0.308178 | 0.476719 | 1122.757 | 952.6583 | 1238.848 | 1105.883 | 1032.643 | 1158.717 | 1259.362 | 1366.014 |
| ENSMUSG00000021339 | -0.606103494 | 0.004483 | 0.023742 | 226.3622 | 230.3657 | 232.698 | 214.7932 | 348.1147 | 214.5358 | 360.6441 | 452.4379 |
| ENSMUSG00000021340 | -0.113984313 | 0.243525 | 0.40608 | 425.561 | 397.1409 | 457.1149 | 439.0754 | 436.8499 | 449.1844 | 463.8231 | 514.4306 |
| ENSMUSG00000021357 | 0.080171737 | 0.276826 | 0.442894 | 1064.406 | 1066.641 | 1038.446 | 963.5504 | 1038.494 | 939.7116 | 937.2891 | 990.7955 |
| ENSMUSG00000021359 | 0.34259221 | 0.029726 | 0.093926 | 481.9 | 421.1374 | 433.9279 | 392.4937 | 325.6872 | 435.7759 | 328.8226 | 274.073 |
| ENSMUSG00000021360 | -0.306125157 | 0.027926 | 0.089945 | 473.8516 | 430.7359 | 543.238 | 484.7944 | 527.5352 | 516.2268 | 712.6097 | 635.1532 |
| ENSMUSG00000021361 | -0.370919763 | 0.000322 | 0.003278 | 1650.935 | 1568.167 | 1835.084 | 1695.918 | 2207.652 | 1844.785 | 2240.044 | 2439.467 |
| ENSMUSG00000021363 | 0.282885001 | 0.26546 | 0.429616 | 70.4238 | 58.79126 | 83.63877 | 56.07053 | 62.40712 | 39.10809 | 59.78593 | 59.81751 |
| ENSMUSG00000021364 | -1.236996503 | 1.65E-05 | 0.000336 | 207.2472 | 137.9795 | 251.7444 | 203.5791 | 493.4063 | 263.7003 | 445.5016 | 687.3576 |
| ENSMUSG00000021365 | -0.022365396 | 0.807705 | 0.882476 | 1424.573 | 1293.408 | 1340.705 | 1277.545 | 1249.118 | 1233.581 | 1490.791 | 1445.409 |
| ENSMUSG00000021366 | 0.39862528 | 6.24E-08 | 3.88E-06 | 1431.615 | 1353.399 | 1516.263 | 1465.597 | 1102.851 | 1179.947 | 1038.539 | 1058.226 |
| ENSMUSG00000021367 | -0.050376643 | 0.85753 | 0.914643 | 147.89 | 125.9813 | 125.0441 | 219.969 | 120.9138 | 207.8316 | 196.715 | 116.3722 |
| ENSMUSG00000021368 | 0.03560403 | 0.720914 | 0.825333 | 458.7608 | 495.5263 | 539.0974 | 524.4751 | 546.0623 | 465.945 | 479.2517 | 478.5401 |
| ENSMUSG00000021371 | -0.745184402 | 3.17E-05 | 0.000559 | 808.8677 | 764.2863 | 953.9788 | 734.9552 | 1253.993 | 993.3456 | 1442.577 | 1779.299 |
| ENSMUSG00000021373 | 0.147627906 | 0.627327 | 0.757172 | 207.2472 | 316.7529 | 223.5888 | 228.5952 | 185.2711 | 376.5551 | 173.572 | 145.7372 |
| ENSMUSG00000021374 | 0.02435599 | 0.69051 | 0.80399 | 2078.508 | 2078.091 | 1941.91 | 2196.239 | 2080.888 | 2044.795 | 1971.007 | 2058.81 |
| ENSMUSG00000021375 | 0.070784232 | 0.290742 | 0.457845 | 1392.379 | 1425.388 | 1522.06 | 1339.654 | 1369.056 | 1372.135 | 1370.255 | 1296.409 |
| ENSMUSG00000021376 | -1.192754244 | 1.52E-05 | 0.000315 | 122.7386 | 145.1784 | 142.4343 | 132.844 | 322.7618 | 162.0192 | 310.5011 | 445.9123 |
| ENSMUSG00000021377 | 0.162620268 | 0.23684 | 0.398021 | 6662.092 | 8403.55 | 6243.096 | 6865.62 | 6166.604 | 7745.637 | 5790.556 | 5466.233 |
| ENSMUSG00000021379 | -0.066975668 | 0.711295 | 0.818994 | 1057.363 | 1295.807 | 1876.49 | 1216.299 | 1202.312 | 1409.009 | 1536.113 | 1559.606 |
| ENSMUSG00000021384 | -2.278960502 | 1.88E-09 | 2.20E-07 | 102.6175 | 47.99286 | 153.1997 | 112.1411 | 415.3974 | 268.1698 | 537.1091 | 804.8174 |
| ENSMUSG00000021385 | -0.063939096 | 0.614253 | 0.748023 | 321.9374 | 298.7556 | 298.1184 | 357.9887 | 359.8161 | 280.4609 | 354.8584 | 339.3284 |
| ENSMUSG00000021388 | 1.118140677 | 7.18E-07 | 2.84E-05 | 1882.328 | 2223.269 | 1411.922 | 2299.754 | 947.8082 | 1239.168 | 805.1815 | 609.051 |
| ENSMUSG00000021390 | 1.274779245 | 1.75E-08 | 1.35E-06 | 3975.927 | 4632.511 | 2990.293 | 4987.689 | 1525.074 | 2507.387 | 1568.898 | 1253.993 |
| ENSMUSG00000021391 | 0.163962969 | 0.236863 | 0.398021 | 301.8163 | 316.7529 | 261.6817 | 282.9405 | 288.6329 | 287.1651 | 243.9652 | 216.4306 |
| ENSMUSG00000021392 | 0.159795325 | 0.219883 | 0.378701 | 733.4136 | 763.0865 | 545.7223 | 597.7981 | 583.1166 | 632.4337 | 580.5021 | 563.3722 |
| ENSMUSG00000021395 | 0.090866974 | 0.247408 | 0.410589 | 4822.019 | 5377.6 | 5132.605 | 5268.904 | 4861.905 | 5377.922 | 4662.338 | 4441.722 |
| ENSMUSG00000021396 | -0.292612714 | 0.336948 | 0.505299 | 25.15136 | 53.99197 | 53.82693 | 40.5433 | 54.60623 | 62.57295 | 44.3573 | 51.11678 |
| ENSMUSG00000021400 | 0.059342857 | 0.453507 | 0.615843 | 1457.773 | 1349.799 | 1351.47 | 1447.482 | 1312.5 | 1274.924 | 1496.577 | 1295.321 |
| ENSMUSG00000021408 | -0.370066958 | 0.032266 | 0.099617 | 405.4399 | 380.3434 | 430.6154 | 347.6373 | 510.9583 | 357.5597 | 538.0734 | 614.489 |
| ENSMUSG00000021411 | 0.120588622 | 0.449979 | 0.612494 | 270.6286 | 263.9607 | 279.9 | 278.6274 | 222.3254 | 196.6578 | 286.3939 | 300.1751 |
| ENSMUSG00000021413 | 0.002682823 | 0.974192 | 0.985749 | 3034.26 | 2773.987 | 2439.602 | 2708.638 | 2863.902 | 2597.895 | 2692.295 | 2777.708 |
| ENSMUSG00000021414 | -0.483395279 | 0.013413 | 0.053173 | 106.6418 | 83.98751 | 135.8095 | 99.2017 | 150.1671 | 134.0849 | 139.8219 | 174.0146 |
| ENSMUSG00000021416 | -2.701804457 | 1.05E-07 | 5.87E-06 | 13.07871 | 20.39697 | 17.39024 | 19.84034 | 122.864 | 32.40385 | 94.50034 | 209.9051 |
| ENSMUSG00000021417 | -0.111183809 | 0.142749 | 0.280155 | 2925.606 | 3100.339 | 2951.372 | 3127.01 | 3346.582 | 2927.52 | 3242.904 | 3555.335 |
| ENSMUSG00000021418 | -0.040644578 | 0.734079 | 0.833414 | 194.1685 | 207.5691 | 195.4332 | 208.7549 | 227.2009 | 207.8316 | 201.5364 | 191.416 |
| ENSMUSG00000021420 | -0.413760274 | 0.000241 | 0.002634 | 692.1654 | 675.4995 | 768.483 | 756.5208 | 1013.141 | 791.1009 | 1003.825 | 1047.35 |
| ENSMUSG00000021423 | -0.014170976 | 0.964355 | 0.979965 | 32.19374 | 61.1909 | 86.9512 | 55.2079 | 57.53157 | 54.75133 | 57.85735 | 68.51824 |
| ENSMUSG00000021427 | -0.03028487 | 0.672692 | 0.790252 | 5382.391 | 5997.908 | 5880.385 | 5631.206 | 5593.238 | 6439.427 | 5611.199 | 5734.868 |
| ENSMUSG00000021428 | 0.118205372 | 0.084164 | 0.195609 | 1494.997 | 1490.178 | 1390.391 | 1379.335 | 1329.077 | 1396.718 | 1331.683 | 1240.941 |
| ENSMUSG00000021431 | 0.264657979 | 0.061601 | 0.157881 | 861.1825 | 904.6654 | 755.2332 | 818.6297 | 709.881 | 855.9086 | 659.5738 | 553.5839 |
| ENSMUSG00000021432 | 0.067703171 | 0.507623 | 0.663318 | 982.9151 | 1021.048 | 1046.727 | 1037.736 | 952.6837 | 1155.365 | 885.2175 | 910.3137 |
| ENSMUSG00000021451 | -0.089114322 | 0.370484 | 0.53963 | 709.2683 | 733.091 | 766.8267 | 832.4317 | 726.4579 | 803.392 | 802.2886 | 907.051 |

| | | | | | | | | | | | |
|---|---|---|---|---|---|---|---|---|---|---|---|
| ENSMUSG00000021453 | -0.97940059 | 0.003977 | 0.021776 | 448.7002 | 254.3622 | 462.0835 | 409.7462 | 507.0579 | 505.0531 | 1432.934 | 661.2554 |
| ENSMUSG00000021456 | -2.757720064 | 2.95E-07 | 1.35E-05 | 16.09687 | 13.19804 | 64.59232 | 18.11509 | 186.2463 | 93.85942 | 162.0006 | 321.927 |
| ENSMUSG00000021457 | 0.291886174 | 0.001819 | 0.012246 | 476.8698 | 547.1186 | 533.3007 | 489.1075 | 387.1192 | 423.4848 | 433.9301 | 426.3357 |
| ENSMUSG00000021458 | -0.017968082 | 0.835386 | 0.900426 | 1503.045 | 1676.151 | 1606.527 | 1689.879 | 1737.648 | 1489.46 | 1548.648 | 1780.387 |
| ENSMUSG00000021460 | -0.122622772 | 0.168469 | 0.314619 | 1143.884 | 1048.644 | 1014.431 | 1231.826 | 1173.059 | 1145.308 | 1280.576 | 1234.416 |
| ENSMUSG00000021461 | 0.030122935 | 0.765419 | 0.854292 | 531.1967 | 566.3158 | 496.864 | 609.8748 | 550.9379 | 509.5226 | 579.5378 | 516.6058 |
| ENSMUSG00000021464 | 0.309419113 | 0.014094 | 0.055042 | 767.6195 | 747.4888 | 835.5596 | 749.6198 | 536.3112 | 703.9457 | 715.5026 | 547.0583 |
| ENSMUSG00000021466 | 0.095640969 | 0.32368 | 0.491634 | 1992.994 | 1665.352 | 2044.595 | 1675.215 | 1624.535 | 1734.165 | 1711.613 | 1838.029 |
| ENSMUSG00000021468 | -0.035143915 | 0.611499 | 0.746072 | 1375.276 | 1415.789 | 1454.155 | 1457.834 | 1439.264 | 1345.318 | 1510.077 | 1549.817 |
| ENSMUSG00000021469 | -1.294015603 | 0.000876 | 0.007032 | 19.11503 | 11.99822 | 27.32752 | 16.38985 | 57.53157 | 43.57759 | 27.96439 | 56.55474 |
| ENSMUSG00000021470 | 0.158727562 | 0.010302 | 0.043538 | 1310.889 | 1342.6 | 1238.848 | 1316.363 | 1131.129 | 1217.938 | 1159.076 | 1156.109 |
| ENSMUSG00000021474 | -0.363791897 | 1.91E-05 | 0.00038 | 1986.957 | 1968.907 | 2034.658 | 1993.523 | 2562.593 | 2247.039 | 2589.116 | 2875.591 |
| ENSMUSG00000021476 | -0.332019146 | 0.007781 | 0.035513 | 1063.399 | 1083.439 | 1200.755 | 1091.219 | 1394.409 | 1117.374 | 1422.327 | 1654.226 |
| ENSMUSG00000021477 | -0.263928702 | 0.021761 | 0.074822 | 8599.753 | 7682.457 | 9258.232 | 9186.94 | 11419.53 | 8414.944 | 10831.86 | 11034.7 |
| ENSMUSG00000021478 | 0.307499212 | 0.47897 | 0.638266 | 25.15136 | 16.7975 | 11.59349 | 25.01608 | 18.52711 | 12.29112 | 19.28578 | 13.05109 |
| ENSMUSG00000021481 | 0.255723773 | 0.006484 | 0.031237 | 910.4792 | 764.2863 | 800.7791 | 728.9169 | 675.7521 | 659.2507 | 695.2525 | 652.5547 |
| ENSMUSG00000021482 | -0.093924236 | 0.463107 | 0.624014 | 502.0211 | 494.3265 | 479.4737 | 583.9961 | 613.345 | 451.4191 | 590.145 | 542.708 |
| ENSMUSG00000021483 | 0.044005048 | 0.76063 | 0.850954 | 193.1624 | 253.1623 | 220.2764 | 212.2054 | 247.6783 | 204.4795 | 205.3936 | 192.5036 |
| ENSMUSG00000021484 | -0.270614528 | 0.017286 | 0.063666 | 3451.772 | 3540.673 | 3739.73 | 3296.084 | 4206.63 | 3472.799 | 4257.337 | 4985.518 |
| ENSMUSG00000021485 | 0.023601243 | 0.835392 | 0.900426 | 439.6458 | 531.5209 | 472.0208 | 408.8835 | 495.3565 | 453.6539 | 430.073 | 439.3868 |
| ENSMUSG00000021486 | -0.407369759 | 0.000162 | 0.001922 | 4342.131 | 4512.529 | 5169.87 | 5001.491 | 6771.173 | 5241.602 | 6375.88 | 6847.473 |
| ENSMUSG00000021488 | 0.196736383 | 0.054929 | 0.145678 | 5338.124 | 5176.03 | 4880.032 | 4920.404 | 4170.551 | 5315.349 | 4001.8 | 4238.342 |
| ENSMUSG00000021490 | -3.04464571 | 8.27E-05 | 0.001166 | 535.2209 | 105.5843 | 1284.393 | 628.8525 | 6439.635 | 1949.818 | 3478.191 | 9211.897 |
| ENSMUSG00000021493 | -0.023545859 | 0.799485 | 0.877351 | 1434.634 | 1685.749 | 1816.866 | 1729.56 | 1739.599 | 1765.451 | 1573.72 | 1700.992 |
| ENSMUSG00000021494 | -0.226179842 | 0.009045 | 0.039653 | 1633.832 | 1874.121 | 1691.822 | 1594.128 | 2185.224 | 1828.024 | 1945.936 | 1981.591 |
| ENSMUSG00000021495 | 0.292602973 | 0.000516 | 0.00469 | 1961.806 | 1998.903 | 1685.197 | 1852.053 | 1405.135 | 1625.779 | 1535.148 | 1553.08 |
| ENSMUSG00000021496 | 0.133565397 | 0.264746 | 0.429059 | 355.1372 | 365.9456 | 378.4447 | 400.2573 | 340.3138 | 296.1041 | 332.6798 | 399.1459 |
| ENSMUSG00000021497 | 0.036708312 | 0.584122 | 0.724628 | 1286.744 | 1318.604 | 1322.486 | 1433.68 | 1277.396 | 1255.928 | 1360.612 | 1334.474 |
| ENSMUSG00000021500 | 0.023307384 | 0.718872 | 0.824245 | 4200.277 | 3903.019 | 3764.573 | 4095.736 | 3909.221 | 4203.561 | 3826.299 | 3769.591 |
| ENSMUSG00000021501 | 0.045539098 | 0.586618 | 0.726928 | 1031.206 | 907.0651 | 870.3401 | 967.8636 | 896.1273 | 877.1387 | 912.2175 | 973.394 |
| ENSMUSG00000021504 | 0.162263927 | 0.051984 | 0.139735 | 535.2209 | 536.3202 | 554.8314 | 582.2709 | 479.7548 | 505.0531 | 522.6447 | 467.6642 |
| ENSMUSG00000021508 | -0.594900317 | 0.187089 | 0.339035 | 782.7103 | 959.8572 | 2085.173 | 919.5566 | 1804.931 | 886.0777 | 2073.222 | 2406.839 |
| ENSMUSG00000021509 | 0.104702334 | 0.597634 | 0.736409 | 223.3441 | 172.7743 | 227.7293 | 195.8155 | 158.9431 | 197.7752 | 251.6795 | 154.4379 |
| ENSMUSG00000021510 | -0.044963426 | 0.734369 | 0.833601 | 257.5499 | 289.157 | 227.7293 | 266.5507 | 293.5085 | 264.8177 | 281.5724 | 230.5693 |
| ENSMUSG00000021514 | 0.31382446 | 0.009529 | 0.04111 | 272.6407 | 238.7645 | 221.9326 | 237.2215 | 190.1467 | 197.7752 | 201.5364 | 190.3284 |
| ENSMUSG00000021518 | -0.093937217 | 0.276728 | 0.442803 | 1591.578 | 1821.329 | 1957.644 | 1811.509 | 1958.999 | 1791.151 | 1874.578 | 2042.496 |
| ENSMUSG00000021519 | 0.0404503 | 0.681268 | 0.796487 | 783.7163 | 825.4772 | 843.8407 | 766.8723 | 800.5664 | 889.4298 | 697.1811 | 744.9999 |
| ENSMUSG00000021520 | -0.61906208 | 8.23E-05 | 0.001162 | 2413.524 | 2178.876 | 2991.949 | 2411.895 | 3795.133 | 2889.529 | 4031.693 | 4639.664 |
| ENSMUSG00000021532 | -0.262568153 | 0.030487 | 0.095562 | 374.2522 | 413.9384 | 353.6015 | 315.7202 | 470.0037 | 459.2408 | 424.2872 | 390.4452 |
| ENSMUSG00000021534 | 1.911850251 | 0.000579 | 0.005135 | 12.07265 | 16.7975 | 16.56213 | 16.38985 | 4.875557 | 3.352122 | 4.821446 | 3.262773 |
| ENSMUSG00000021536 | 0.844714749 | 1.37E-07 | 7.36E-06 | 895.3884 | 763.0865 | 705.5469 | 808.2782 | 450.5014 | 557.5697 | 421.3944 | 337.1532 |
| ENSMUSG00000021537 | 0.029288027 | 0.647591 | 0.77184 | 2535.257 | 2775.187 | 2615.161 | 2809.565 | 2617.199 | 2794.553 | 2572.723 | 2535.175 |
| ENSMUSG00000021540 | 0.326861197 | 0.016021 | 0.060331 | 5774.752 | 6844.982 | 5560.736 | 5877.054 | 4740.991 | 6050.581 | 4187.908 | 4200.277 |
| ENSMUSG00000021546 | -0.017322972 | 0.758436 | 0.849288 | 21812.26 | 22740.22 | 21424.77 | 23859.3 | 22831.26 | 23682.74 | 22614.51 | 21793.15 |
| ENSMUSG00000021548 | 0.148074194 | 0.095485 | 0.213737 | 977.8848 | 988.6529 | 872.8244 | 1042.049 | 835.6704 | 941.9464 | 827.3601 | 898.3502 |
| ENSMUSG00000021549 | 0.081126124 | 0.355956 | 0.524597 | 1988.969 | 2141.681 | 1827.631 | 2046.143 | 1771.777 | 2123.011 | 1864.935 | 1805.401 |
| ENSMUSG00000021550 | -0.155481265 | 0.176075 | 0.324542 | 813.898 | 875.8697 | 949.0102 | 839.3327 | 1002.414 | 797.8051 | 980.6821 | 1093.029 |
| ENSMUSG00000021552 | -0.017631048 | 0.84304 | 0.905412 | 1604.657 | 1662.953 | 1443.39 | 1710.582 | 1518.248 | 1806.794 | 1586.256 | 1588.971 |
| ENSMUSG00000021555 | -0.10935232 | 0.059707 | 0.154271 | 2034.242 | 2078.091 | 1966.753 | 1904.673 | 2180.349 | 2222.457 | 2149.401 | 2054.46 |
| ENSMUSG00000021556 | 0.279108482 | 0.031721 | 0.098292 | 1554.354 | 2106.887 | 1782.085 | 1739.049 | 1545.551 | 1702.878 | 1421.362 | 1247.467 |
| ENSMUSG00000021557 | 0.193823646 | 0.026096 | 0.085668 | 777.68 | 893.867 | 772.6235 | 874.7002 | 753.761 | 749.758 | 712.6097 | 683.0072 |
| ENSMUSG00000021559 | -0.151286045 | 0.123688 | 0.253358 | 1937.661 | 1677.351 | 1857.443 | 1902.085 | 1780.553 | 1964.344 | 2230.401 | 2217.598 |
| ENSMUSG00000021565 | -2.762476486 | 5.62E-09 | 5.29E-07 | 31.18769 | 14.39786 | 51.34261 | 21.56559 | 284.7325 | 72.62932 | 190.9293 | 261.0219 |
| ENSMUSG00000021567 | -0.652894523 | 0.070665 | 0.173569 | 186.1201 | 241.1641 | 549.8628 | 182.0136 | 459.2774 | 298.3389 | 479.2517 | 587.2992 |
| ENSMUSG00000021569 | 0.040456323 | 0.730708 | 0.831219 | 547.2936 | 691.0972 | 584.6433 | 579.683 | 600.6686 | 640.2554 | 592.0735 | 500.2919 |
| ENSMUSG00000021572 | -0.181744134 | 0.308222 | 0.476744 | 412.4823 | 351.5477 | 284.0406 | 376.9665 | 430.9992 | 443.5975 | 445.5016 | 294.7372 |
| ENSMUSG00000021573 | -0.437949458 | 0.105599 | 0.229366 | 38.23007 | 51.59233 | 50.5145 | 60.38364 | 90.68535 | 70.39457 | 58.82164 | 52.20437 |
| ENSMUSG00000021575 | -0.331148203 | 0.231064 | 0.391599 | 51.30877 | 34.79482 | 74.5296 | 48.30692 | 75.08357 | 69.27719 | 60.75022 | 59.81751 |
| ENSMUSG00000021576 | -0.163557652 | 0.043663 | 0.123943 | 2264.628 | 2002.502 | 2295.512 | 2310.106 | 2535.289 | 2240.335 | 2558.259 | 2608.043 |
| ENSMUSG00000021577 | -0.566088172 | 0.000393 | 0.003808 | 6892.478 | 6240.272 | 7323.775 | 6793.16 | 10236.72 | 6971.297 | 10862.72 | 12273.47 |
| ENSMUSG00000021578 | 0.116038664 | 0.200713 | 0.356086 | 1422.561 | 1503.376 | 1330.767 | 1511.316 | 1276.421 | 1515.159 | 1268.04 | 1262.693 |
| ENSMUSG00000021583 | -0.366354082 | 0.023441 | 0.079105 | 541.2572 | 617.9081 | 655.8605 | 414.0593 | 722.5575 | 611.2036 | 801.3243 | 736.2992 |
| ENSMUSG00000021585 | 0.131062982 | 0.115899 | 0.243849 | 1252.538 | 1121.833 | 1094.757 | 1135.213 | 996.5638 | 982.1718 | 1111.825 | 1112.606 |
| ENSMUSG00000021589 | 0.360370746 | 0.001497 | 0.010545 | 2610.711 | 3029.549 | 2388.26 | 2988.99 | 2144.27 | 2457.106 | 2056.829 | 1922.861 |

| | | | | | | | | | | | |
|---|---|---|---|---|---|---|---|---|---|---|---|
| ENSMUSG00000021590 | 0.287133161 | 0.461511 | 0.622645 | 26.15741 | 26.39607 | 20.70267 | 14.6646 | 19.50223 | 17.87799 | 12.53576 | 21.75182 |
| ENSMUSG00000021591 | -0.358099394 | 0.01653 | 0.061559 | 333.004 | 387.5424 | 466.224 | 369.2029 | 558.7388 | 405.6068 | 522.6447 | 507.905 |
| ENSMUSG00000021592 | 0.378484174 | 0.001653 | 0.011368 | 534.2149 | 557.917 | 503.4888 | 614.1879 | 361.7663 | 476.0014 | 446.4659 | 416.5474 |
| ENSMUSG00000021594 | 0.021883059 | 0.882127 | 0.929407 | 401.4157 | 425.9366 | 426.4749 | 477.0308 | 450.5014 | 458.1234 | 477.3231 | 318.6642 |
| ENSMUSG00000021595 | -0.213325027 | 0.004823 | 0.025082 | 2626.808 | 2848.376 | 2596.942 | 2536.976 | 3306.602 | 2800.139 | 3027.868 | 3158.365 |
| ENSMUSG00000021596 | 0.48559027 | 0.376641 | 0.544772 | 21.12714 | 23.99643 | 16.56213 | 22.42821 | 14.62667 | 31.28647 | 6.750024 | 7.613138 |
| ENSMUSG00000021597 | 0.232415964 | 0.189016 | 0.341505 | 831.0009 | 862.6717 | 658.3448 | 816.9044 | 611.3948 | 924.0684 | 583.3949 | 578.5985 |
| ENSMUSG00000021598 | -0.174617366 | 0.283046 | 0.449522 | 211.2714 | 209.9688 | 189.6364 | 177.7004 | 256.4543 | 178.7799 | 200.5721 | 252.3211 |
| ENSMUSG00000021606 | -0.536942648 | 0.00055 | 0.004927 | 1170.041 | 1180.624 | 1414.406 | 1166.267 | 1771.777 | 1317.384 | 1886.15 | 2180.62 |
| ENSMUSG00000021607 | -0.564506108 | 0.003221 | 0.018713 | 372.2401 | 415.1383 | 538.2693 | 408.8835 | 606.5192 | 476.0014 | 648.0023 | 836.3575 |
| ENSMUSG00000021608 | 0.180641256 | 0.175726 | 0.324255 | 1382.319 | 1667.752 | 1370.516 | 1514.767 | 1412.936 | 1526.333 | 1250.683 | 1045.175 |
| ENSMUSG00000021610 | -0.260635925 | 0.014044 | 0.054885 | 2323.986 | 2471.632 | 2648.285 | 2170.361 | 2709.834 | 2516.326 | 3092.475 | 3198.605 |
| ENSMUSG00000021611 | 0.168107505 | 0.27173 | 0.436965 | 244.4712 | 259.1615 | 237.6666 | 238.0841 | 214.5245 | 165.3714 | 241.0723 | 249.0584 |
| ENSMUSG00000021612 | -3.691801868 | 7.18E-06 | 0.000179 | 106.6418 | 17.99732 | 364.3669 | 153.547 | 1953.148 | 835.7958 | 2301.758 | 3220.357 |
| ENSMUSG00000021613 | 0.330827469 | 0.0753 | 0.181451 | 193.1624 | 244.7636 | 230.2136 | 225.1447 | 125.7894 | 213.4185 | 179.3578 | 192.5036 |
| ENSMUSG00000021614 | 0.6524453 | 0.004239 | 0.02279 | 12075.67 | 14305.47 | 10738.89 | 13098.08 | 7696.554 | 12393.91 | 6322.844 | 5535.839 |
| ENSMUSG00000021615 | 0.213418065 | 0.072831 | 0.177423 | 394.3733 | 358.7466 | 380.1009 | 407.1583 | 347.1396 | 273.7567 | 351.9655 | 355.6423 |
| ENSMUSG00000021619 | -0.054338386 | 0.715615 | 0.821867 | 244.4712 | 188.372 | 212.8234 | 195.8155 | 213.5494 | 186.6015 | 222.7508 | 251.2335 |
| ENSMUSG00000021620 | -0.995872697 | 0.11492 | 0.24259 | 8.048435 | 3.599465 | 6.624853 | 9.488858 | 17.552 | 7.821619 | 6.750024 | 23.927 |
| ENSMUSG00000021621 | 0.051812238 | 0.46314 | 0.624014 | 908.4671 | 940.6601 | 870.3401 | 955.7868 | 913.6793 | 841.3827 | 874.6103 | 913.5765 |
| ENSMUSG00000021624 | 0.291580678 | 0.677179 | 0.793626 | 12.07265 | 1.199822 | 24.8432 | 12.93935 | 9.751113 | 12.29112 | 11.57147 | 8.700729 |
| ENSMUSG00000021629 | -0.142326968 | 0.013912 | 0.054502 | 1902.449 | 1952.11 | 1824.319 | 2017.676 | 2097.464 | 2144.241 | 2184.115 | 2066.423 |
| ENSMUSG00000021635 | 0.055221493 | 0.599708 | 0.737734 | 729.3894 | 628.7065 | 599.5492 | 600.386 | 636.7477 | 559.8044 | 596.895 | 666.6933 |
| ENSMUSG00000021636 | -0.596115882 | 0.000354 | 0.003517 | 355.1372 | 352.7475 | 478.6456 | 433.037 | 553.8632 | 493.8794 | 673.0738 | 730.8612 |
| ENSMUSG00000021638 | -0.309398757 | 0.031331 | 0.09733 | 717.3168 | 760.6869 | 882.7617 | 720.2906 | 861.0233 | 772.1055 | 1093.504 | 1091.941 |
| ENSMUSG00000021639 | -0.162359655 | 0.055665 | 0.146914 | 879.2915 | 908.2649 | 820.6537 | 857.4478 | 972.186 | 941.9464 | 890.0389 | 1072.365 |
| ENSMUSG00000021643 | -0.076542625 | 0.299975 | 0.468126 | 1002.03 | 1091.838 | 1045.899 | 1127.449 | 1107.726 | 1202.295 | 1121.468 | 1069.102 |
| ENSMUSG00000021645 | -0.159529347 | 0.090641 | 0.206247 | 717.3168 | 713.8938 | 806.5759 | 687.5109 | 838.5957 | 882.7255 | 735.7526 | 813.5181 |
| ENSMUSG00000021646 | -0.512777885 | 4.69E-05 | 0.000756 | 878.2855 | 884.2685 | 873.6525 | 904.892 | 1270.57 | 974.3502 | 1383.755 | 1422.569 |
| ENSMUSG00000021647 | 1.923403309 | 0.003663 | 0.02048 | 62.37537 | 19.19714 | 10.76539 | 61.24627 | 12.67645 | 4.469496 | 14.46434 | 8.700729 |
| ENSMUSG00000021650 | -0.170546982 | 0.033417 | 0.102193 | 687.1351 | 760.6869 | 700.5782 | 659.907 | 754.7362 | 789.9835 | 775.2885 | 837.4451 |
| ENSMUSG00000021660 | -0.054301149 | 0.405952 | 0.572627 | 14208.51 | 14868.19 | 14319.62 | 15813.61 | 16575.92 | 15130.36 | 15202.02 | 14571.55 |
| ENSMUSG00000021661 | 0.260814153 | 0.055959 | 0.147372 | 818.9282 | 749.8885 | 724.5933 | 851.4094 | 560.689 | 823.5047 | 625.8237 | 616.6641 |
| ENSMUSG00000021662 | -0.088675721 | 0.375881 | 0.544226 | 1777.698 | 1523.773 | 1777.117 | 1602.754 | 1592.357 | 1673.826 | 1996.079 | 1844.554 |
| ENSMUSG00000021665 | 0.240742044 | 0.00988 | 0.042258 | 961.788 | 1065.442 | 1138.647 | 1194.734 | 984.8624 | 886.0777 | 884.2532 | 937.5035 |
| ENSMUSG00000021666 | -0.130667305 | 0.068056 | 0.168801 | 929.5942 | 907.0651 | 862.8871 | 844.5084 | 924.4055 | 977.7023 | 954.6463 | 1021.248 |
| ENSMUSG00000021668 | 0.551634114 | 0.002251 | 0.014305 | 515.0998 | 386.3425 | 317.9929 | 440.8006 | 242.8027 | 337.447 | 288.3225 | 264.2846 |
| ENSMUSG00000021669 | 0.120829541 | 0.255375 | 0.419287 | 1978.909 | 1478.18 | 1609.011 | 1692.467 | 1552.377 | 1690.587 | 1496.577 | 1478.036 |
| ENSMUSG00000021670 | -0.757413939 | 1.11E-07 | 6.15E-06 | 1474.876 | 1306.606 | 1555.184 | 1414.703 | 2350.018 | 1890.597 | 3026.904 | 2455.781 |
| ENSMUSG00000021671 | -0.059834304 | 0.563776 | 0.70908 | 616.7113 | 518.3229 | 536.6131 | 555.5295 | 617.2455 | 511.7573 | 618.1094 | 573.1605 |
| ENSMUSG00000021676 | 0.310014198 | 0.024452 | 0.081621 | 1663.008 | 1215.419 | 1218.145 | 1272.37 | 993.6384 | 956.4722 | 1169.683 | 1210.489 |
| ENSMUSG00000021678 | -0.289805454 | 0.193309 | 0.346547 | 742.4681 | 523.1222 | 919.1984 | 727.1916 | 816.1682 | 619.0252 | 1181.254 | 945.1167 |
| ENSMUSG00000021681 | -0.075799439 | 0.343324 | 0.51199 | 1715.323 | 1715.745 | 1753.102 | 1755.439 | 1931.696 | 1595.61 | 1846.614 | 1939.175 |
| ENSMUSG00000021684 | 0.116867235 | 0.374127 | 0.54288 | 737.4378 | 620.3077 | 742.8116 | 593.485 | 514.8588 | 686.0677 | 619.0736 | 666.6933 |
| ENSMUSG00000021686 | 0.037704513 | 0.543435 | 0.692451 | 2365.234 | 2380.446 | 2427.181 | 2378.253 | 2243.731 | 2175.527 | 2453.152 | 2431.854 |
| ENSMUSG00000021687 | -0.133982119 | 0.077211 | 0.184574 | 1342.077 | 1267.012 | 1421.031 | 1450.07 | 1398.31 | 1493.929 | 1567.934 | 1559.606 |
| ENSMUSG00000021690 | 0.286174559 | 0.026503 | 0.086711 | 1563.408 | 1498.577 | 1289.362 | 1416.428 | 1044.344 | 1469.347 | 1132.075 | 1084.328 |
| ENSMUSG00000021692 | 0.182108927 | 0.049121 | 0.134456 | 539.2451 | 488.3274 | 456.2868 | 532.2387 | 450.5014 | 435.7759 | 448.3945 | 441.562 |
| ENSMUSG00000021693 | 0.389020301 | 0.008883 | 0.039117 | 1531.215 | 1702.547 | 1334.908 | 1581.189 | 1065.797 | 1528.568 | 1039.504 | 1062.576 |
| ENSMUSG00000021694 | -0.133981647 | 0.170156 | 0.316796 | 303.8284 | 304.7547 | 336.2113 | 313.995 | 346.1645 | 326.2732 | 343.2869 | 367.6058 |
| ENSMUSG00000021696 | -0.989208878 | 0.000122 | 0.001563 | 365.1977 | 267.5602 | 501.0045 | 344.1868 | 612.3699 | 478.2361 | 870.7531 | 975.5692 |
| ENSMUSG00000021697 | -0.205213504 | 0.109934 | 0.235678 | 228.3743 | 193.1713 | 196.2613 | 205.3044 | 227.2009 | 214.5358 | 266.1438 | 240.3576 |
| ENSMUSG00000021699 | -0.171942923 | 0.310247 | 0.478756 | 240.447 | 160.7761 | 217.792 | 248.4356 | 249.6285 | 213.4185 | 274.8224 | 241.4452 |
| ENSMUSG00000021700 | 1.795172055 | 0.000511 | 0.004657 | 121.7326 | 39.59411 | 44.71776 | 197.5408 | 35.10401 | 25.6996 | 24.10723 | 31.54014 |
| ENSMUSG00000021701 | -0.204801269 | 0.012405 | 0.05023 | 1190.162 | 1033.046 | 1136.162 | 1050.675 | 1314.45 | 1352.023 | 1188.969 | 1230.066 |
| ENSMUSG00000021702 | 0.83917722 | 2.20E-09 | 2.49E-07 | 203.223 | 185.9723 | 231.8699 | 207.0296 | 113.1129 | 120.6764 | 120.5361 | 109.8467 |
| ENSMUSG00000021703 | 0.388503627 | 0.000341 | 0.003409 | 2552.36 | 2315.656 | 2133.203 | 2718.989 | 1988.252 | 1933.057 | 1901.578 | 1602.022 |
| ENSMUSG00000021704 | 0.309612044 | 0.00408 | 0.022136 | 1678.099 | 1584.964 | 1511.295 | 1632.084 | 1248.142 | 1553.15 | 1215.004 | 1153.934 |
| ENSMUSG00000021706 | 0.015374715 | 0.906292 | 0.943776 | 754.5408 | 863.8715 | 593.7525 | 739.2683 | 683.553 | 812.331 | 728.0383 | 692.7955 |
| ENSMUSG00000021707 | -0.265406278 | 0.019689 | 0.069694 | 665.0019 | 886.6681 | 727.0776 | 718.5654 | 943.9077 | 822.3873 | 865.9317 | 965.7809 |
| ENSMUSG00000021708 | 0.584130105 | 0.00086 | 0.006949 | 220.3259 | 261.5611 | 249.2601 | 203.5791 | 153.0925 | 197.7752 | 127.2862 | 145.7372 |
| ENSMUSG00000021709 | 0.170179606 | 0.037734 | 0.112019 | 5721.431 | 6073.497 | 5864.651 | 6375.65 | 4913.586 | 6036.055 | 5184.983 | 5229.138 |
| ENSMUSG00000021710 | -0.088562939 | 0.281055 | 0.447438 | 617.7174 | 664.7011 | 601.2054 | 617.6384 | 692.329 | 612.321 | 654.7523 | 696.0583 |
| ENSMUSG00000021711 | -0.025420522 | 0.789131 | 0.8707 | 1133.823 | 1160.227 | 966.4004 | 1100.708 | 1055.07 | 1015.693 | 1188.969 | 1174.598 |

| | | | | | | | | | | | |
|---|---|---|---|---|---|---|---|---|---|---|---|
| ENSMUSG00000021712 | 0.344229787 | 0.006851 | 0.032461 | 1236.441 | 1118.234 | 983.7907 | 1201.635 | 923.4304 | 1034.688 | 880.396 | 737.3868 |
| ENSMUSG00000021713 | 0.078233906 | 0.401374 | 0.568602 | 705.2441 | 727.0918 | 606.1741 | 620.2263 | 656.2499 | 616.7905 | 630.6451 | 609.051 |
| ENSMUSG00000021714 | -0.003580697 | 0.981328 | 0.989687 | 320.9313 | 404.3399 | 304.7432 | 403.7078 | 365.6667 | 423.4848 | 330.7512 | 316.489 |
| ENSMUSG00000021715 | 0.00739309 | 0.946712 | 0.969349 | 1037.242 | 1067.841 | 809.0602 | 1049.813 | 913.6793 | 1024.632 | 981.6464 | 1021.248 |
| ENSMUSG00000021716 | 0.062850835 | 0.606479 | 0.742598 | 1087.545 | 1143.43 | 952.3226 | 1010.995 | 860.0482 | 1200.06 | 949.8248 | 1004.934 |
| ENSMUSG00000021719 | 0.057848795 | 0.789496 | 0.8709 | 91.55095 | 81.58786 | 138.2938 | 123.3552 | 95.56091 | 109.5027 | 101.2504 | 114.1971 |
| ENSMUSG00000021720 | 0.224873093 | 0.150978 | 0.291195 | 183.1019 | 250.7627 | 192.1207 | 184.6014 | 169.6694 | 163.1366 | 163.9292 | 194.6788 |
| ENSMUSG00000021725 | 0.211533947 | 0.022011 | 0.075439 | 953.7395 | 853.0731 | 953.1507 | 891.0901 | 765.4624 | 888.3124 | 721.2883 | 781.978 |
| ENSMUSG00000021728 | -0.039294109 | 0.809802 | 0.884031 | 1326.986 | 1066.641 | 1319.174 | 1199.909 | 1133.079 | 941.9464 | 1492.72 | 1481.299 |
| ENSMUSG00000021730 | -0.113292081 | 0.608229 | 0.743951 | 51.30877 | 73.18911 | 77.01392 | 75.91087 | 68.25779 | 88.27255 | 67.50024 | 77.21897 |
| ENSMUSG00000021731 | -0.06726737 | 0.540824 | 0.690376 | 691.1593 | 722.2926 | 678.2193 | 805.6903 | 674.777 | 874.9039 | 746.3598 | 741.7371 |
| ENSMUSG00000021732 | 0.665433293 | 0.028918 | 0.0921 | 352.119 | 464.3309 | 279.9 | 383.0048 | 181.3707 | 402.2547 | 194.7864 | 154.4379 |
| ENSMUSG00000021733 | 0.277600489 | 0.214017 | 0.37205 | 1487.954 | 1330.602 | 1216.489 | 1454.383 | 978.0366 | 1743.104 | 1024.075 | 784.1532 |
| ENSMUSG00000021737 | -0.18698117 | 0.002249 | 0.014301 | 2498.033 | 2717.596 | 2659.879 | 2767.296 | 3011.144 | 2873.886 | 3055.832 | 3174.678 |
| ENSMUSG00000021738 | 0.174678692 | 0.045617 | 0.127735 | 921.5458 | 860.272 | 786.7013 | 925.595 | 727.433 | 778.8097 | 768.5385 | 821.1313 |
| ENSMUSG00000021745 | 0.37533537 | 8.06E-05 | 0.001142 | 3257.604 | 3346.302 | 3197.32 | 3239.151 | 2141.344 | 2878.356 | 2539.938 | 2494.934 |
| ENSMUSG00000021747 | -0.575687121 | 8.02E-06 | 0.000194 | 212.2775 | 220.7672 | 264.166 | 200.9913 | 353.9654 | 311.7474 | 352.9298 | 320.8394 |
| ENSMUSG00000021748 | -0.540014532 | 0.000111 | 0.001452 | 3607.711 | 3180.727 | 3831.649 | 3539.344 | 5055.952 | 3870.584 | 5651.699 | 6011.116 |
| ENSMUSG00000021749 | 0.544625842 | 0.029137 | 0.092653 | 71.42986 | 73.18911 | 105.9976 | 74.18562 | 65.33246 | 51.39921 | 63.64308 | 42.41605 |
| ENSMUSG00000021750 | 0.804965036 | 6.03E-06 | 0.000155 | 1499.021 | 1726.543 | 1246.3 | 1451.795 | 694.2793 | 1158.717 | 792.6457 | 744.9999 |
| ENSMUSG00000021751 | -6.148751376 | 4.19E-11 | 8.40E-09 | 0 | 2.399643 | 3.312427 | 0 | 119.9387 | 15.64324 | 82.92887 | 185.9781 |
| ENSMUSG00000021752 | 0.095975346 | 0.416473 | 0.582046 | 644.8808 | 641.9045 | 741.9835 | 741.8562 | 667.9512 | 725.1758 | 540.9662 | 662.343 |
| ENSMUSG00000021754 | 0.069178667 | 0.529407 | 0.6816 | 1819.952 | 1598.162 | 2129.062 | 1681.253 | 1566.029 | 1858.193 | 1727.042 | 1743.409 |
| ENSMUSG00000021756 | 0.149956302 | 0.012493 | 0.050507 | 3618.778 | 3246.717 | 3603.92 | 3546.245 | 3101.829 | 3313.014 | 3105.011 | 3119.211 |
| ENSMUSG00000021758 | -0.854658196 | NA | NA | 36.21796 | 16.7975 | 30.63995 | 35.36756 | 12.67645 | 169.8409 | 19.28578 | 14.13868 |
| ENSMUSG00000021759 | -0.245434016 | 0.178069 | 0.327335 | 1533.227 | 1045.045 | 1358.095 | 1221.475 | 1386.608 | 1122.961 | 1714.506 | 1891.321 |
| ENSMUSG00000021760 | 0.322730936 | 0.000118 | 0.001515 | 1887.358 | 1973.706 | 1973.378 | 2068.571 | 1482.169 | 1798.972 | 1504.291 | 1536.766 |
| ENSMUSG00000021763 | 0.156502341 | 0.071427 | 0.174927 | 737.4378 | 873.4701 | 829.7628 | 740.131 | 711.8313 | 734.1148 | 691.3953 | 714.5473 |
| ENSMUSG00000021764 | -0.306858575 | 0.001055 | 0.008137 | 1485.942 | 1518.974 | 1815.21 | 1632.084 | 2007.754 | 1807.911 | 2020.186 | 2150.168 |
| ENSMUSG00000021765 | 1.284222812 | 0.011319 | 0.046796 | 126.7628 | 266.3604 | 166.4494 | 172.5247 | 75.08357 | 147.4934 | 61.71451 | 16.31387 |
| ENSMUSG00000021767 | 0.274420552 | 0.065491 | 0.164684 | 1633.832 | 1437.386 | 1396.188 | 1448.345 | 1144.781 | 1614.606 | 1088.682 | 1044.087 |
| ENSMUSG00000021770 | 0.148269398 | 0.142969 | 0.280438 | 683.1109 | 607.1097 | 654.2042 | 658.1817 | 511.9334 | 631.3164 | 638.3594 | 568.8101 |
| ENSMUSG00000021771 | -0.242162327 | 0.001359 | 0.009839 | 6088.641 | 6216.275 | 6527.965 | 6811.275 | 7757.01 | 6763.465 | 7893.671 | 7918.751 |
| ENSMUSG00000021772 | -0.05417294 | 0.650059 | 0.773564 | 320.9313 | 279.5584 | 271.619 | 325.2091 | 297.4089 | 288.2825 | 313.394 | 344.7664 |
| ENSMUSG00000021773 | -0.391418699 | 0.055378 | 0.146491 | 121.7326 | 117.5825 | 101.8571 | 97.47645 | 118.9636 | 126.2633 | 137.8934 | 191.416 |
| ENSMUSG00000021774 | -0.124947627 | 0.157317 | 0.299756 | 1417.531 | 1631.757 | 1539.45 | 1738.186 | 1812.732 | 1750.925 | 1755.971 | 1579.182 |
| ENSMUSG00000021775 | 0.142631215 | 0.282243 | 0.448864 | 1170.041 | 1174.625 | 1103.038 | 1137.8 | 1026.792 | 1268.22 | 1047.218 | 811.343 |
| ENSMUSG00000021779 | -0.170427254 | 0.346188 | 0.514997 | 119.7205 | 103.1847 | 107.6539 | 117.3168 | 102.3867 | 112.8548 | 144.6434 | 144.6496 |
| ENSMUSG00000021782 | 0.097941813 | 0.18652 | 0.338393 | 1830.013 | 1728.943 | 1874.005 | 1880.519 | 1624.535 | 1890.597 | 1646.042 | 1678.153 |
| ENSMUSG00000021785 | 0.123046115 | 0.265805 | 0.430023 | 1101.63 | 1123.033 | 864.5433 | 1149.877 | 969.2606 | 970.9981 | 916.0747 | 1034.299 |
| ENSMUSG00000021786 | -0.302457918 | 0.040284 | 0.117282 | 224.3501 | 231.5656 | 213.6515 | 244.9851 | 254.5041 | 240.2354 | 288.3225 | 344.7664 |
| ENSMUSG00000021792 | 0.165683957 | 0.089693 | 0.204645 | 2985.969 | 3552.672 | 2813.078 | 3316.787 | 3086.227 | 2725.275 | 2854.296 | 2623.27 |
| ENSMUSG00000021794 | -0.696283077 | 3.96E-06 | 0.000112 | 5166.089 | 4926.467 | 6238.955 | 5511.302 | 8753.574 | 6626.028 | 9102.89 | 10912.89 |
| ENSMUSG00000021795 | -0.430882651 | 0.31636 | 0.484663 | 18.10898 | 10.79839 | 28.15563 | 17.25247 | 22.42756 | 34.6386 | 16.39292 | 28.27737 |
| ENSMUSG00000021796 | 0.223049649 | 0.003763 | 0.02093 | 4192.229 | 4469.335 | 4263.921 | 4468.39 | 3817.561 | 4124.228 | 3393.334 | 3568.386 |
| ENSMUSG00000021798 | 0.606325095 | 0.113075 | 0.24001 | 38.23007 | 29.99554 | 29.81184 | 38.81806 | 18.52711 | 37.99072 | 14.46434 | 19.57664 |
| ENSMUSG00000021803 | -0.22001425 | 0.061679 | 0.158037 | 242.4591 | 298.7556 | 293.9779 | 276.9021 | 349.0898 | 301.691 | 305.6797 | 338.2408 |
| ENSMUSG00000021806 | -0.261367995 | 0.061251 | 0.157159 | 1839.067 | 2309.656 | 2890.92 | 2242.821 | 2497.26 | 2765.501 | 2815.724 | 3050.693 |
| ENSMUSG00000021807 | 0.006973498 | 0.919729 | 0.952288 | 4057.417 | 4064.995 | 4421.261 | 4472.703 | 4370.449 | 4499.665 | 4075.086 | 3994.722 |
| ENSMUSG00000021809 | 0.179237657 | 0.126219 | 0.257062 | 460.7729 | 460.7315 | 356.0859 | 445.1137 | 398.8205 | 382.1419 | 362.5727 | 375.2189 |
| ENSMUSG00000021810 | -0.175137832 | 0.025435 | 0.08399 | 1229.398 | 1445.785 | 1219.801 | 1218.024 | 1450.966 | 1436.943 | 1450.291 | 1426.92 |
| ENSMUSG00000021811 | -0.156065935 | 0.062453 | 0.15938 | 1574.475 | 1884.92 | 1647.104 | 1617.419 | 2004.829 | 1900.653 | 1808.042 | 1772.773 |
| ENSMUSG00000021814 | -0.047097564 | 0.558556 | 0.70487 | 1732.426 | 1496.177 | 1702.587 | 1606.205 | 1652.814 | 1594.493 | 1843.721 | 1665.102 |
| ENSMUSG00000021815 | 0.919654551 | 9.94E-05 | 0.001331 | 155.9384 | 115.1829 | 95.23226 | 125.0804 | 80.93424 | 60.3382 | 66.53595 | 51.11678 |
| ENSMUSG00000021816 | 0.18472583 | 0.041207 | 0.119148 | 2323.986 | 2351.65 | 1983.315 | 2273.013 | 1822.483 | 2188.936 | 1940.15 | 1905.46 |
| ENSMUSG00000021819 | 0.059747705 | 0.392012 | 0.55942 | 2645.923 | 2568.818 | 2763.392 | 2733.654 | 2633.776 | 2347.603 | 2537.045 | 2761.394 |
| ENSMUSG00000021820 | 0.13016882 | 0.151511 | 0.291883 | 1387.349 | 1333.002 | 1275.284 | 1341.38 | 1275.446 | 1367.666 | 1106.04 | 1127.832 |
| ENSMUSG00000021822 | -2.429339789 | 0.009252 | 0.040236 | 419.5247 | 206.3693 | 3204.773 | 536.5518 | 4974.043 | 1752.043 | 6910.096 | 9890.553 |
| ENSMUSG00000021823 | -0.521445348 | 0.004358 | 0.023284 | 1208.271 | 1043.845 | 1303.44 | 984.2534 | 1734.723 | 1082.735 | 1776.221 | 1922.861 |
| ENSMUSG00000021824 | -0.040039982 | 0.577936 | 0.71973 | 1509.082 | 1419.389 | 1387.079 | 1330.165 | 1440.239 | 1379.957 | 1430.041 | 1551.992 |
| ENSMUSG00000021830 | 0.084439351 | 0.406711 | 0.573149 | 1308.877 | 1088.238 | 1193.302 | 1338.792 | 1278.371 | 1067.092 | 1209.219 | 1095.204 |
| ENSMUSG00000021831 | -0.393115387 | 0.003347 | 0.019176 | 660.9777 | 577.1142 | 571.3936 | 592.6223 | 750.8357 | 635.7859 | 825.4315 | 941.8539 |
| ENSMUSG00000021832 | -0.010199603 | 0.851447 | 0.910624 | 2923.594 | 2986.356 | 2755.111 | 2992.441 | 2892.18 | 3044.844 | 2855.26 | 2946.284 |
| ENSMUSG00000021835 | -0.125653924 | 0.370781 | 0.539807 | 841.0614 | 925.0624 | 843.8407 | 765.147 | 1018.991 | 692.7719 | 944.0391 | 1023.423 |

| | | | | | | | | | | | |
|---|---|---|---|---|---|---|---|---|---|---|---|
| ENSMUSG00000021838 | -0.010468573 | 0.936711 | 0.962366 | 499.003 | 424.7368 | 503.4888 | 442.5259 | 412.4721 | 570.9782 | 459.0016 | 443.7372 |
| ENSMUSG00000021840 | 0.11502367 | 0.109836 | 0.235494 | 2706.286 | 2807.582 | 2678.925 | 2806.114 | 2421.201 | 2832.543 | 2478.223 | 2425.328 |
| ENSMUSG00000021843 | 0.114140088 | 0.243942 | 0.406525 | 4052.387 | 4207.774 | 3597.295 | 3870.592 | 3594.26 | 4205.796 | 3455.048 | 3274.737 |
| ENSMUSG00000021846 | 0.083907063 | 0.461232 | 0.622349 | 770.6376 | 748.6886 | 764.3424 | 795.3389 | 642.5984 | 697.2414 | 878.4674 | 686.27 |
| ENSMUSG00000021848 | -0.087299912 | 0.778461 | 0.86331 | 26.15741 | 35.99465 | 47.20208 | 31.91707 | 29.25334 | 46.92971 | 35.6787 | 39.15328 |
| ENSMUSG00000021850 | -0.835869269 | 7.81E-05 | 0.001122 | 711.2804 | 689.8974 | 1147.756 | 790.1631 | 1308.599 | 1118.491 | 1643.149 | 1893.496 |
| ENSMUSG00000021866 | -0.405364854 | 0.072404 | 0.176687 | 633.8142 | 562.7163 | 955.6351 | 690.0988 | 953.6589 | 610.0863 | 1026.004 | 1176.774 |
| ENSMUSG00000021868 | -0.513341424 | 0.002357 | 0.014753 | 854.1401 | 679.099 | 922.5108 | 704.7634 | 1204.262 | 845.8522 | 1093.504 | 1369.277 |
| ENSMUSG00000021870 | -0.087076525 | 0.292241 | 0.459577 | 1860.195 | 1704.946 | 1923.692 | 2003.874 | 1862.463 | 2185.584 | 1944.007 | 1973.978 |
| ENSMUSG00000021871 | 0.20703787 | 0.065444 | 0.164684 | 1934.643 | 2442.837 | 2095.938 | 2119.466 | 1809.807 | 1583.319 | 2018.257 | 2029.445 |
| ENSMUSG00000021874 | 0.022328026 | 0.967173 | 0.981694 | 21.12714 | 16.7975 | 24.01509 | 27.60395 | 15.60178 | 48.04709 | 15.42863 | 9.78832 |
| ENSMUSG00000021876 | 1.039620679 | 4.18E-12 | 1.01E-09 | 2857.194 | 3467.484 | 2913.279 | 3397.011 | 1633.311 | 1866.015 | 1492.72 | 1153.934 |
| ENSMUSG00000021877 | -0.005322292 | 0.920348 | 0.952447 | 5546.378 | 6067.497 | 5864.651 | 5694.178 | 5564.96 | 6029.351 | 5840.699 | 5821.875 |
| ENSMUSG00000021879 | -0.861857048 | 0.046962 | 0.130236 | 7.04238 | 15.59768 | 28.15563 | 16.38985 | 29.25334 | 33.52122 | 27.0001 | 33.71532 |
| ENSMUSG00000021884 | -0.920401674 | 1.35E-05 | 0.00029 | 553.3299 | 356.347 | 484.4424 | 511.5357 | 922.4553 | 611.2036 | 890.0389 | 1185.474 |
| ENSMUSG00000021886 | -0.010192417 | 0.985051 | 0.991442 | 17.10292 | 27.5959 | 26.49941 | 53.48266 | 9.751113 | 51.39921 | 35.6787 | 29.36496 |
| ENSMUSG00000021890 | -0.150854364 | 0.075057 | 0.181045 | 1135.835 | 1132.632 | 1039.274 | 932.496 | 1176.959 | 1190.003 | 1152.326 | 1183.299 |
| ENSMUSG00000021891 | 0.076904691 | 0.483024 | 0.641437 | 764.6013 | 800.281 | 640.9545 | 770.3228 | 732.3086 | 688.3024 | 626.788 | 772.1897 |
| ENSMUSG00000021892 | -0.340690573 | 0.009036 | 0.039624 | 1474.876 | 1256.213 | 1500.529 | 1451.795 | 1963.874 | 1424.652 | 1736.685 | 2074.036 |
| ENSMUSG00000021893 | -0.04355348 | 0.491201 | 0.648331 | 1990.982 | 2036.097 | 2036.314 | 2216.942 | 2240.806 | 2147.593 | 2066.472 | 2081.649 |
| ENSMUSG00000021895 | -0.153011652 | 0.117986 | 0.246712 | 809.8738 | 783.4835 | 861.2309 | 815.1792 | 936.1069 | 772.1055 | 936.3248 | 991.8831 |
| ENSMUSG00000021898 | 0.945294808 | 0.010649 | 0.044686 | 49.29666 | 49.19268 | 40.57722 | 47.44429 | 22.42756 | 41.34284 | 20.25007 | 13.05109 |
| ENSMUSG00000021900 | -0.397363117 | 0.004842 | 0.025154 | 853.1341 | 785.8831 | 992.0717 | 944.5727 | 1123.328 | 944.1811 | 1232.362 | 1412.781 |
| ENSMUSG00000021901 | -0.168086737 | 0.017512 | 0.064269 | 2686.165 | 2838.778 | 2851.171 | 2722.44 | 3212.992 | 2790.083 | 3264.119 | 3199.693 |
| ENSMUSG00000021902 | -0.236751422 | 0.013505 | 0.053408 | 431.5973 | 429.5361 | 430.6154 | 445.1137 | 536.3112 | 450.3018 | 551.5734 | 506.8174 |
| ENSMUSG00000021903 | 1.713988677 | 4.71E-07 | 2.00E-05 | 613.6932 | 341.9491 | 237.6666 | 585.7214 | 127.7396 | 179.8972 | 155.2506 | 79.39415 |
| ENSMUSG00000021904 | -1.33109632 | 8.19E-06 | 0.000196 | 371.2341 | 226.7663 | 555.6596 | 327.7969 | 646.4988 | 689.4198 | 1116.647 | 1279.007 |
| ENSMUSG00000021905 | -0.084062599 | 0.302285 | 0.4706 | 1339.058 | 1457.783 | 1490.592 | 1588.952 | 1627.461 | 1552.033 | 1617.113 | 1433.445 |
| ENSMUSG00000021906 | -0.178778249 | 0.066154 | 0.165775 | 806.8556 | 791.8822 | 683.188 | 892.8153 | 886.3762 | 902.8383 | 917.039 | 886.3867 |
| ENSMUSG00000021908 | -0.156221798 | 0.647869 | 0.771952 | 25.15136 | 23.99643 | 22.35888 | 27.60395 | 33.15378 | 35.75597 | 23.14294 | 18.48905 |
| ENSMUSG00000021910 | 0.03163721 | 0.52655 | 0.679534 | 10761.76 | 10446.85 | 10985.66 | 10423.08 | 10697.95 | 10101.06 | 10038.25 | 10858.51 |
| ENSMUSG00000021911 | 0.018517752 | 0.808854 | 0.883359 | 1611.699 | 1485.379 | 1464.921 | 1585.502 | 1434.389 | 1667.122 | 1509.113 | 1460.635 |
| ENSMUSG00000021913 | -0.379213711 | 0.002323 | 0.014609 | 260.5681 | 229.1659 | 276.5876 | 244.9851 | 356.8907 | 291.6346 | 357.7513 | 309.9635 |
| ENSMUSG00000021916 | 0.166403917 | 0.029007 | 0.092317 | 1693.189 | 1768.537 | 1676.916 | 1852.053 | 1563.103 | 1699.526 | 1529.363 | 1437.795 |
| ENSMUSG00000021917 | -0.114791947 | 0.074746 | 0.180589 | 2078.508 | 2159.679 | 2287.231 | 2222.981 | 2328.566 | 2223.574 | 2480.152 | 2442.73 |
| ENSMUSG00000021918 | -0.189698274 | 0.21259 | 0.370462 | 405.4399 | 554.3175 | 414.0533 | 366.615 | 562.6392 | 450.3018 | 473.466 | 493.7664 |
| ENSMUSG00000021922 | 3.645006374 | 5.45E-06 | 0.000143 | 60.36326 | 19.19714 | 13.24971 | 38.81806 | 0 | 3.352122 | 1.928578 | 5.437955 |
| ENSMUSG00000021928 | 0.266991796 | 0.021201 | 0.073454 | 617.7174 | 772.6851 | 669.1102 | 661.6322 | 646.4988 | 574.3303 | 524.5733 | 513.343 |
| ENSMUSG00000021929 | 0.085891472 | 0.230417 | 0.390753 | 1474.876 | 1565.767 | 1443.39 | 1621.732 | 1467.543 | 1520.746 | 1370.255 | 1394.292 |
| ENSMUSG00000021930 | -0.166744492 | 0.026892 | 0.08756 | 587.5357 | 592.7118 | 572.2217 | 614.1879 | 637.7228 | 648.077 | 699.1096 | 671.0437 |
| ENSMUSG00000021932 | -0.093927502 | 0.210835 | 0.368293 | 963.8001 | 1058.243 | 938.2448 | 973.9019 | 1109.677 | 1072.679 | 1017.325 | 994.0583 |
| ENSMUSG00000021936 | 0.072160948 | 0.451625 | 0.614001 | 1479.906 | 1414.59 | 1391.219 | 1450.07 | 1347.604 | 1586.671 | 1313.362 | 1209.401 |
| ENSMUSG00000021938 | -0.104014065 | 0.13469 | 0.268622 | 1832.025 | 2163.278 | 2061.986 | 2084.961 | 2195.951 | 2129.715 | 2219.794 | 2203.46 |
| ENSMUSG00000021939 | -0.15182515 | 0.160724 | 0.304207 | 7912.618 | 7422.096 | 8469.047 | 7283.993 | 8582.93 | 7251.758 | 8686.317 | 10017.8 |
| ENSMUSG00000021943 | 0.279131349 | 0.259621 | 0.423619 | 152.9203 | 243.5638 | 216.9639 | 264.8254 | 136.5156 | 236.8833 | 142.7148 | 208.8175 |
| ENSMUSG00000021944 | -0.296321615 | NA | NA | 14.08476 | 25.19625 | 13.24971 | 27.60395 | 2.925334 | 92.74205 | 2.892867 | 0 |
| ENSMUSG00000021945 | 0.194553529 | 0.022384 | 0.076377 | 2423.585 | 2297.658 | 2041.283 | 2443.812 | 2119.892 | 2105.133 | 1921.828 | 1896.759 |
| ENSMUSG00000021947 | -1.374064737 | 3.75E-07 | 1.66E-05 | 466.8092 | 435.5352 | 626.8767 | 494.2833 | 1515.323 | 695.0067 | 1137.861 | 1898.934 |
| ENSMUSG00000021948 | -0.065250678 | 0.589251 | 0.729391 | 1724.377 | 1347.4 | 1613.152 | 1455.246 | 1606.983 | 1335.262 | 1714.506 | 1768.423 |
| ENSMUSG00000021950 | 1.301099415 | 1.97E-05 | 0.000389 | 265.5983 | 334.7502 | 235.1823 | 345.912 | 187.2214 | 73.74669 | 137.8934 | 79.39415 |
| ENSMUSG00000021951 | -0.057315291 | 0.568817 | 0.712804 | 407.452 | 493.1267 | 422.3344 | 429.5865 | 464.153 | 422.3674 | 450.323 | 482.8904 |
| ENSMUSG00000021952 | -0.120733356 | 0.358186 | 0.526533 | 503.0272 | 419.9375 | 385.8977 | 390.7684 | 468.0534 | 401.1373 | 458.0374 | 518.7809 |
| ENSMUSG00000021957 | -0.309819976 | 0.003582 | 0.02017 | 6290.858 | 6431.043 | 7435.569 | 5996.959 | 8465.916 | 6868.499 | 8207.065 | 8879.094 |
| ENSMUSG00000021958 | -0.086309035 | 0.535527 | 0.686367 | 330.9919 | 327.5513 | 308.0557 | 351.0878 | 290.5832 | 402.2547 | 315.3226 | 392.6204 |
| ENSMUSG00000021959 | 0.085700432 | 0.30083 | 0.469207 | 1264.61 | 1225.018 | 1376.313 | 1372.434 | 1191.586 | 1133.017 | 1307.576 | 1307.284 |
| ENSMUSG00000021962 | 0.183574633 | 0.175782 | 0.324294 | 1283.725 | 1253.814 | 1112.975 | 1067.065 | 1066.772 | 1272.689 | 940.1819 | 873.3356 |
| ENSMUSG00000021963 | -0.206677903 | 0.000189 | 0.002171 | 2895.424 | 3049.946 | 2939.779 | 3057.138 | 3508.45 | 3239.268 | 3453.119 | 3578.175 |
| ENSMUSG00000021965 | 0.115605516 | 0.357687 | 0.526139 | 527.1725 | 527.9215 | 438.0684 | 527.063 | 530.4606 | 498.3488 | 416.5729 | 417.635 |
| ENSMUSG00000021967 | -0.372179621 | 0.001838 | 0.012341 | 785.7285 | 723.4924 | 782.5608 | 722.8785 | 1053.12 | 777.6924 | 987.4321 | 1083.241 |
| ENSMUSG00000021969 | 0.381409151 | 0.008297 | 0.037296 | 3341.107 | 3067.944 | 2801.485 | 2922.568 | 2124.768 | 3051.549 | 2157.115 | 1981.591 |
| ENSMUSG00000021972 | 0.246422909 | 0.044643 | 0.125859 | 1428.597 | 1407.391 | 1428.484 | 1420.741 | 1097.975 | 1498.399 | 1167.754 | 1029.949 |
| ENSMUSG00000021973 | -0.155711921 | 0.03876 | 0.113789 | 2259.598 | 2418.84 | 2461.133 | 2530.075 | 2762.49 | 2464.927 | 2635.402 | 2910.394 |
| ENSMUSG00000021974 | 0.010001456 | 0.95561 | 0.974791 | 346.0827 | 399.5406 | 307.2276 | 268.2759 | 271.0809 | 317.3342 | 308.5725 | 413.2846 |
| ENSMUSG00000021975 | 0.122149304 | 0.169647 | 0.316093 | 842.0675 | 826.677 | 832.2472 | 755.6582 | 813.2428 | 676.0113 | 775.2885 | 724.3357 |

| | | | | | | | | | | | |
|---|---|---|---|---|---|---|---|---|---|---|---|
| ENSMUSG00000021978 | -0.004028843 | 0.958379 | 0.976487 | 1724.377 | 1838.127 | 1792.023 | 1605.342 | 1681.092 | 1625.779 | 1843.721 | 1824.978 |
| ENSMUSG00000021981 | -0.623936199 | 0.000421 | 0.004009 | 678.0806 | 752.2881 | 1098.069 | 673.7089 | 1241.317 | 1032.454 | 1354.826 | 1308.372 |
| ENSMUSG00000021982 | -0.140716651 | 0.055682 | 0.146915 | 940.6608 | 927.462 | 941.5572 | 1016.17 | 1075.548 | 992.2282 | 1028.897 | 1123.482 |
| ENSMUSG00000021983 | -0.229916695 | 0.308402 | 0.476744 | 65.39353 | 101.9848 | 59.62368 | 75.91087 | 97.51113 | 86.03781 | 90.64318 | 78.30656 |
| ENSMUSG00000021986 | 0.178061492 | 0.416525 | 0.582046 | 77.46619 | 85.18733 | 94.40416 | 112.1411 | 75.08357 | 92.74205 | 64.60737 | 95.70802 |
| ENSMUSG00000021987 | 0.018916071 | 0.828435 | 0.896119 | 1373.264 | 1289.808 | 1242.988 | 1467.323 | 1366.131 | 1197.825 | 1403.041 | 1335.562 |
| ENSMUSG00000021990 | 0.132277249 | 0.250207 | 0.413857 | 2134.847 | 1799.732 | 2121.609 | 2028.028 | 1551.402 | 1750.925 | 1963.293 | 2113.189 |
| ENSMUSG00000021991 | 1.055022453 | 8.90E-05 | 0.001228 | 283.7073 | 434.3354 | 382.5853 | 368.3402 | 175.52 | 268.1698 | 159.1077 | 104.4087 |
| ENSMUSG00000021993 | -0.37951518 | 0.002365 | 0.014782 | 543.2694 | 610.7092 | 554.8314 | 574.5072 | 732.3086 | 612.321 | 737.6812 | 886.3867 |
| ENSMUSG00000021994 | 0.243467911 | 0.066231 | 0.1659 | 1284.731 | 1430.187 | 1743.993 | 1330.165 | 1043.369 | 1280.511 | 1182.219 | 1386.679 |
| ENSMUSG00000021996 | -0.296209025 | 0.000667 | 0.005722 | 4006.108 | 4069.795 | 4523.118 | 4368.325 | 5139.812 | 4690.736 | 5142.554 | 5865.379 |
| ENSMUSG00000021998 | -0.030561849 | 0.873378 | 0.924258 | 375.2583 | 367.1454 | 597.893 | 356.2635 | 415.3974 | 423.4848 | 396.3228 | 500.2919 |
| ENSMUSG00000021999 | -1.973204701 | 0.000151 | 0.001836 | 47.28455 | 11.99822 | 76.18581 | 69.00988 | 147.2418 | 92.74205 | 226.608 | 340.416 |
| ENSMUSG00000022000 | 0.227746397 | 0.147411 | 0.286613 | 3118.769 | 3145.932 | 2641.66 | 2819.916 | 2168.648 | 3383.409 | 2299.83 | 2162.131 |
| ENSMUSG00000022003 | -0.753413966 | 0.007975 | 0.036144 | 297.7921 | 265.1606 | 419.8501 | 276.0395 | 537.2863 | 267.0524 | 606.5379 | 712.3722 |
| ENSMUSG00000022008 | 0.116062248 | 0.132773 | 0.266153 | 1385.337 | 1456.583 | 1373.001 | 1237.002 | 1268.62 | 1190.003 | 1322.04 | 1244.204 |
| ENSMUSG00000022009 | 0.005920773 | 0.955724 | 0.974844 | 756.5529 | 727.0918 | 746.1241 | 787.5752 | 744.985 | 855.9086 | 773.3599 | 631.8904 |
| ENSMUSG00000022010 | 0.197224934 | 0.181662 | 0.3322 | 27539.73 | 26111.72 | 21450.45 | 26644.71 | 20114.6 | 28850.6 | 20938.57 | 18842.52 |
| ENSMUSG00000022012 | 0.031508418 | 0.860972 | 0.916754 | 330.9919 | 371.9447 | 394.1788 | 354.5383 | 296.4338 | 489.4099 | 306.644 | 329.5401 |
| ENSMUSG00000022013 | -0.154617897 | 0.294365 | 0.461894 | 866.2128 | 689.8974 | 745.296 | 741.8562 | 758.6366 | 680.4808 | 938.2534 | 1010.372 |
| ENSMUSG00000022014 | 0.292803466 | 0.261349 | 0.425537 | 62.37537 | 119.9822 | 123.3879 | 85.39973 | 76.05868 | 81.56831 | 90.64318 | 70.69342 |
| ENSMUSG00000022015 | 0.095394115 | 0.886404 | 0.932078 | 16.09687 | 16.7975 | 4.140533 | 6.900988 | 17.552 | 6.704245 | 10.60718 | 5.437955 |
| ENSMUSG00000022016 | 0.162751189 | 0.034374 | 0.104663 | 2001.042 | 2043.296 | 1824.319 | 1963.331 | 1658.664 | 1933.057 | 1714.506 | 1689.029 |
| ENSMUSG00000022018 | -0.62732517 | 0.003163 | 0.018476 | 252.5196 | 179.9732 | 239.3228 | 255.3366 | 285.7076 | 312.8647 | 503.3589 | 331.7153 |
| ENSMUSG00000022019 | 0.049203781 | 0.645627 | 0.770506 | 1075.472 | 1216.619 | 1045.071 | 1116.235 | 957.5593 | 1252.576 | 1085.79 | 1007.109 |
| ENSMUSG00000022020 | 0.40130274 | 0.005526 | 0.027645 | 578.4813 | 670.7002 | 473.677 | 615.9132 | 421.2481 | 525.1658 | 398.2514 | 425.2481 |
| ENSMUSG00000022021 | -0.005827566 | 0.952078 | 0.972685 | 316.9071 | 309.554 | 347.8048 | 327.7969 | 325.6872 | 343.0338 | 312.4297 | 329.5401 |
| ENSMUSG00000022022 | 0.044790766 | 0.772716 | 0.858991 | 272.6407 | 278.3586 | 235.1823 | 289.8415 | 281.8072 | 197.7752 | 286.3939 | 275.1605 |
| ENSMUSG00000022023 | 0.167928045 | 0.115046 | 0.242689 | 1855.164 | 2054.094 | 1628.886 | 1746.813 | 1611.859 | 1850.372 | 1485.005 | 1534.591 |
| ENSMUSG00000022024 | -0.069548504 | 0.346205 | 0.514997 | 1966.836 | 1944.911 | 2012.299 | 2096.175 | 2261.283 | 1895.066 | 2144.579 | 2115.365 |
| ENSMUSG00000022025 | 1.16589502 | 5.00E-05 | 0.000794 | 91.55095 | 100.785 | 97.71658 | 79.36136 | 29.25334 | 46.92971 | 59.78593 | 28.27737 |
| ENSMUSG00000022026 | 0.111517996 | 0.611855 | 0.746201 | 105.6357 | 76.78858 | 65.42042 | 87.9876 | 87.76002 | 73.74669 | 68.46453 | 80.48174 |
| ENSMUSG00000022031 | -0.007816956 | 0.903523 | 0.942315 | 2725.401 | 2771.588 | 2634.207 | 2831.13 | 2925.334 | 2539.791 | 2754.974 | 2798.372 |
| ENSMUSG00000022032 | 1.41588078 | 0.023082 | 0.078235 | 400.4096 | 520.7225 | 307.2276 | 509.8105 | 215.4996 | 302.8084 | 98.35749 | 34.80291 |
| ENSMUSG00000022033 | -0.04371783 | 0.61919 | 0.751531 | 1114.708 | 1293.408 | 1034.305 | 1207.673 | 1236.441 | 1198.942 | 1190.897 | 1161.547 |
| ENSMUSG00000022034 | 0.249478768 | 0.158319 | 0.301192 | 485.9243 | 568.7154 | 398.3193 | 467.5419 | 352.9903 | 534.1048 | 365.4656 | 362.1678 |
| ENSMUSG00000022035 | -0.16307089 | 0.123461 | 0.25312 | 827.9827 | 801.4808 | 789.1856 | 760.8339 | 815.1931 | 783.2792 | 939.2176 | 1021.248 |
| ENSMUSG00000022037 | -0.079932033 | 0.69609 | 0.808244 | 3476.924 | 2632.408 | 4412.98 | 3050.237 | 3200.315 | 2719.689 | 3578.477 | 4849.569 |
| ENSMUSG00000022039 | 0.184155058 | 0.613254 | 0.747176 | 24.1453 | 47.99286 | 38.92101 | 41.40593 | 47.78045 | 37.99072 | 23.14294 | 25.0146 |
| ENSMUSG00000022040 | -1.005135463 | 0.00033 | 0.003338 | 138.8355 | 104.3845 | 214.4796 | 133.7066 | 280.8321 | 202.2447 | 287.3582 | 419.8102 |
| ENSMUSG00000022043 | 0.088877205 | 0.037404 | 0.111326 | 4503.099 | 4424.942 | 4414.636 | 4323.469 | 4269.037 | 4062.772 | 4185.979 | 4088.255 |
| ENSMUSG00000022044 | 2.68129849 | 0.000217 | 0.002418 | 84.50857 | 17.99732 | 10.76539 | 105.2401 | 10.72622 | 7.821619 | 5.785735 | 9.78832 |
| ENSMUSG00000022048 | 0.480986077 | 0.015845 | 0.059937 | 1390.367 | 1412.19 | 1165.974 | 1525.981 | 901.978 | 1433.591 | 851.4673 | 750.4379 |
| ENSMUSG00000022051 | 0.07913347 | 0.466534 | 0.627229 | 5000.09 | 5543.175 | 4835.315 | 5613.954 | 4912.611 | 5870.684 | 4757.803 | 4330.788 |
| ENSMUSG00000022052 | 0.238287433 | 0.001279 | 0.009394 | 2379.319 | 2214.871 | 2018.924 | 2304.067 | 1849.786 | 1921.883 | 1815.756 | 1971.803 |
| ENSMUSG00000022053 | 1.536601284 | 0.005677 | 0.028246 | 55.33299 | 142.7788 | 43.88965 | 87.9876 | 28.27823 | 53.63396 | 24.10723 | 7.613138 |
| ENSMUSG00000022054 | 0.135078375 | 0.885009 | 0.93128 | 291.7558 | 134.38 | 125.8722 | 607.2869 | 99.46135 | 781.0445 | 89.67889 | 85.9197 |
| ENSMUSG00000022055 | 0.477234986 | 0.318466 | 0.486689 | 211.2714 | 93.58608 | 82.81066 | 307.9566 | 88.73513 | 227.9443 | 76.17884 | 107.6715 |
| ENSMUSG00000022057 | 0.137162635 | 0.890459 | 0.933768 | 5.030272 | 21.59679 | 14.90592 | 2.58787 | 4.875557 | 30.1691 | 2.892867 | 2.175182 |
| ENSMUSG00000022061 | -0.980939652 | 0.010118 | 0.042989 | 29.17558 | 45.59322 | 50.5145 | 67.28463 | 133.5902 | 46.92971 | 83.89316 | 116.3722 |
| ENSMUSG00000022064 | 0.127216442 | 0.342924 | 0.511724 | 388.337 | 405.5397 | 334.5551 | 382.1422 | 398.8205 | 307.2779 | 372.2156 | 301.2627 |
| ENSMUSG00000022066 | -0.088027929 | 0.846693 | 0.907835 | 12.07265 | 9.598572 | 9.93728 | 11.21411 | 9.751113 | 13.40849 | 10.60718 | 11.9635 |
| ENSMUSG00000022070 | 0.33032385 | 0.035446 | 0.107087 | 1296.804 | 1336.601 | 1109.663 | 1267.194 | 1013.141 | 1299.506 | 871.7174 | 800.467 |
| ENSMUSG00000022074 | 0.424134854 | 0.008818 | 0.038935 | 211.2714 | 275.959 | 227.7293 | 184.6014 | 177.4703 | 172.0756 | 149.4648 | 169.6642 |
| ENSMUSG00000022075 | 0.079326281 | 0.38726 | 0.554697 | 695.1836 | 710.2943 | 596.2368 | 613.3253 | 607.4943 | 615.6731 | 644.1452 | 602.5255 |
| ENSMUSG00000022089 | -0.15428427 | 0.1237 | 0.253358 | 898.4065 | 767.8858 | 969.7129 | 897.9911 | 1039.469 | 883.8429 | 991.2893 | 1022.336 |
| ENSMUSG00000022090 | 0.163735241 | 0.121414 | 0.250705 | 1244.489 | 1424.188 | 1460.78 | 1300.836 | 1329.077 | 1022.397 | 1246.826 | 1247.467 |
| ENSMUSG00000022091 | 0.127470407 | 0.096351 | 0.215129 | 1302.84 | 1456.583 | 1364.72 | 1256.842 | 1308.599 | 1194.473 | 1242.969 | 1174.598 |
| ENSMUSG00000022092 | -0.342999991 | 0.003781 | 0.02097 | 231.3925 | 215.9679 | 228.5574 | 263.9628 | 320.8116 | 282.6956 | 310.5011 | 279.5109 |
| ENSMUSG00000022094 | 0.178193666 | 0.109815 | 0.235476 | 530.1906 | 455.9322 | 453.8024 | 549.4912 | 485.6054 | 426.8369 | 434.8944 | 411.1094 |
| ENSMUSG00000022095 | 0.057688846 | 0.442585 | 0.605825 | 886.3339 | 767.8858 | 818.1694 | 798.7894 | 797.641 | 776.575 | 766.6099 | 803.7298 |
| ENSMUSG00000022096 | 0.808353572 | 0.001563 | 0.010856 | 157.9505 | 233.9652 | 174.7305 | 210.4801 | 117.9885 | 159.7845 | 87.75031 | 78.30656 |
| ENSMUSG00000022098 | 0.425014643 | 6.97E-06 | 0.000175 | 4172.107 | 4888.073 | 4125.627 | 4626.25 | 3266.623 | 3759.964 | 3156.118 | 3083.321 |
| ENSMUSG00000022099 | -1.287046801 | 0.000813 | 0.00665 | 395.3794 | 195.5709 | 590.44 | 316.5828 | 792.7655 | 405.6068 | 1022.147 | 1437.795 |

| | | | | | | | | | | | |
|---|---|---|---|---|---|---|---|---|---|---|---|
| ENSMUSG00000022100 | -0.074285613 | 0.286381 | 0.453452 | 2140.884 | 2186.075 | 1863.24 | 2012.501 | 2179.374 | 2156.532 | 2112.758 | 2180.62 |
| ENSMUSG00000022102 | -0.111687478 | 0.55843 | 0.704826 | 93.56306 | 146.3782 | 126.7003 | 134.5693 | 112.1378 | 146.376 | 143.6791 | 139.2117 |
| ENSMUSG00000022103 | 1.306774131 | 2.76E-10 | 4.18E-08 | 282.7013 | 488.3274 | 378.4447 | 383.8675 | 183.3209 | 173.193 | 140.7862 | 121.8102 |
| ENSMUSG00000022105 | -0.238930364 | 0.068674 | 0.169854 | 438.6397 | 428.3363 | 512.598 | 505.4974 | 493.4063 | 493.8794 | 590.145 | 650.3795 |
| ENSMUSG00000022106 | 0.069453023 | 0.327596 | 0.495339 | 2620.772 | 2827.979 | 2435.462 | 2612.024 | 2605.497 | 2595.66 | 2378.901 | 2417.715 |
| ENSMUSG00000022108 | -0.217652947 | 0.090831 | 0.206527 | 17300.11 | 17236.64 | 22248.74 | 18158.22 | 20982.45 | 17892.51 | 23660.76 | 24614.36 |
| ENSMUSG00000022109 | 0.040631171 | 0.665761 | 0.785381 | 1202.235 | 1367.797 | 1226.426 | 1413.84 | 1412.936 | 1255.928 | 1197.647 | 1197.438 |
| ENSMUSG00000022110 | -0.39334929 | 0.001166 | 0.008786 | 2485.96 | 2072.092 | 2456.164 | 2421.384 | 3041.372 | 2518.561 | 3265.083 | 3570.562 |
| ENSMUSG00000022111 | -0.281106702 | 0.002668 | 0.016181 | 674.0564 | 760.6869 | 782.5608 | 811.7287 | 901.978 | 843.6174 | 945.0034 | 991.8831 |
| ENSMUSG00000022112 | 0.20057036 | 0.186912 | 0.338866 | 329.9858 | 356.347 | 334.5551 | 348.4999 | 227.2009 | 332.9775 | 282.5367 | 350.2043 |
| ENSMUSG00000022114 | 0.327026378 | 0.000137 | 0.001702 | 1729.407 | 1934.112 | 1706.728 | 1996.973 | 1532.875 | 1538.624 | 1450.291 | 1349.701 |
| ENSMUSG00000022119 | 0.326504501 | 0.021767 | 0.074822 | 2285.755 | 2549.621 | 2057.845 | 2428.285 | 1733.748 | 2384.476 | 1566.97 | 1748.846 |
| ENSMUSG00000022120 | -0.095543731 | 0.414281 | 0.580208 | 672.0443 | 657.5022 | 561.4563 | 578.8204 | 768.3877 | 614.5558 | 648.9666 | 602.5255 |
| ENSMUSG00000022122 | -0.7222507 | 2.03E-05 | 0.000397 | 1124.769 | 833.876 | 1268.659 | 1127.449 | 1736.673 | 1364.314 | 2057.793 | 2029.445 |
| ENSMUSG00000022123 | -1.504916161 | 1.78E-06 | 5.95E-05 | 68.4117 | 40.79393 | 115.9349 | 73.323 | 181.3707 | 157.5497 | 247.8223 | 265.3722 |
| ENSMUSG00000022124 | 0.146277622 | 0.062034 | 0.15866 | 2671.074 | 2403.243 | 2332.776 | 2601.672 | 2123.792 | 2462.693 | 2254.508 | 2204.547 |
| ENSMUSG00000022125 | -0.299179841 | 0.005227 | 0.02653 | 706.2502 | 707.8947 | 703.0625 | 621.9515 | 777.1637 | 746.4059 | 908.3604 | 936.4159 |
| ENSMUSG00000022130 | 0.17559622 | 0.120301 | 0.249311 | 811.8859 | 700.6958 | 609.4865 | 623.6768 | 620.1708 | 594.443 | 595.9307 | 617.7517 |
| ENSMUSG00000022131 | 0.02887606 | 0.774885 | 0.860693 | 1017.121 | 1286.209 | 1146.928 | 1201.635 | 1204.262 | 1097.261 | 1037.575 | 1219.19 |
| ENSMUSG00000022132 | -2.356029569 | 0.002477 | 0.015326 | 123.7447 | 99.58519 | 845.4969 | 199.266 | 1317.375 | 724.0584 | 2124.329 | 2329.62 |
| ENSMUSG00000022136 | -0.221153893 | 0.0053 | 0.026792 | 2012.109 | 2100.888 | 2114.984 | 2022.852 | 2151.096 | 2318.551 | 2564.045 | 2584.116 |
| ENSMUSG00000022139 | 0.137967486 | 0.120624 | 0.249718 | 3109.714 | 2947.962 | 2704.596 | 3010.556 | 2330.516 | 2862.712 | 2820.546 | 2685.262 |
| ENSMUSG00000022141 | 0.248621263 | 0.068549 | 0.169635 | 3657.008 | 3379.897 | 3051.573 | 2987.265 | 2473.857 | 3494.029 | 2618.045 | 2419.89 |
| ENSMUSG00000022142 | 0.104575377 | 0.294675 | 0.462262 | 1493.991 | 1427.788 | 1332.424 | 1440.581 | 1319.326 | 1545.328 | 1238.147 | 1194.175 |
| ENSMUSG00000022144 | 0.619399571 | 0.000457 | 0.004277 | 1884.34 | 2765.589 | 2072.751 | 2318.732 | 1727.897 | 1755.395 | 1243.933 | 1157.197 |
| ENSMUSG00000022146 | 0.803380621 | 1.50E-05 | 0.000312 | 633.8142 | 469.1302 | 440.5527 | 460.6409 | 283.7574 | 332.9775 | 321.1083 | 209.9051 |
| ENSMUSG00000022148 | 0.486167005 | 0.062644 | 0.159691 | 59.35721 | 67.19001 | 73.70149 | 63.83414 | 34.1289 | 60.3382 | 38.57157 | 56.55474 |
| ENSMUSG00000022150 | -0.082221694 | 0.495058 | 0.651883 | 2708.298 | 2244.866 | 2636.692 | 2401.544 | 2532.364 | 2272.739 | 2583.331 | 3190.992 |
| ENSMUSG00000022151 | -0.057575863 | 0.602846 | 0.74031 | 721.341 | 734.2908 | 649.2356 | 750.4824 | 798.6162 | 631.3164 | 822.5387 | 715.6349 |
| ENSMUSG00000022159 | 0.288602608 | 0.230597 | 0.391023 | 240.447 | 206.3693 | 202.8861 | 191.5024 | 165.7689 | 99.44629 | 216.9651 | 205.5547 |
| ENSMUSG00000022160 | 0.086863925 | 0.343253 | 0.511966 | 866.2128 | 943.0597 | 763.5143 | 914.3809 | 801.5415 | 797.8051 | 812.8958 | 867.8977 |
| ENSMUSG00000022174 | -0.221189459 | 1.57E-05 | 0.000324 | 2608.699 | 2616.811 | 2823.844 | 2723.302 | 3178.863 | 3081.718 | 3110.797 | 3193.167 |
| ENSMUSG00000022175 | -0.352412991 | 0.007643 | 0.03501 | 1747.516 | 1739.741 | 2236.716 | 1912.436 | 2481.658 | 1972.165 | 2493.652 | 2804.897 |
| ENSMUSG00000022176 | -0.367674334 | 0.050179 | 0.136381 | 89.53884 | 91.18644 | 85.29498 | 111.2784 | 142.3663 | 127.3806 | 100.2861 | 117.4598 |
| ENSMUSG00000022177 | 0.328439487 | 5.55E-07 | 2.28E-05 | 1123.763 | 1223.818 | 1199.098 | 1209.398 | 940.0073 | 993.3456 | 940.1819 | 914.6641 |
| ENSMUSG00000022178 | -0.110955609 | 0.268836 | 0.433406 | 2122.775 | 2264.063 | 2155.562 | 2348.061 | 2584.045 | 1992.278 | 2435.794 | 2587.379 |
| ENSMUSG00000022179 | 0.181210441 | 0.119409 | 0.248401 | 373.2462 | 349.1481 | 337.0394 | 341.5989 | 317.8863 | 258.1134 | 324.0012 | 333.8905 |
| ENSMUSG00000022180 | -0.865765938 | 0.015853 | 0.059945 | 512.0817 | 263.9607 | 835.5596 | 339.8737 | 841.5211 | 547.5133 | 816.7529 | 1352.963 |
| ENSMUSG00000022184 | -0.114821252 | 0.316706 | 0.484882 | 312.8829 | 292.7565 | 305.5713 | 362.3019 | 318.8614 | 334.0949 | 368.3585 | 359.9926 |
| ENSMUSG00000022185 | 0.198989158 | 0.02023 | 0.071048 | 8310.009 | 7792.841 | 6951.127 | 7546.23 | 6738.994 | 7373.552 | 6185.915 | 6358.057 |
| ENSMUSG00000022186 | -0.735849532 | 2.51E-07 | 1.21E-05 | 4478.954 | 4454.937 | 5125.98 | 4765.995 | 7768.712 | 5824.871 | 8276.494 | 9483.794 |
| ENSMUSG00000022191 | 0.070113486 | 0.381165 | 0.549078 | 2113.72 | 2370.847 | 2145.624 | 2223.843 | 2021.406 | 2353.19 | 1989.329 | 2069.686 |
| ENSMUSG00000022193 | -0.28272667 | 0.009303 | 0.040391 | 2588.578 | 2577.217 | 2769.189 | 2737.967 | 3298.802 | 2625.829 | 3384.655 | 3674.97 |
| ENSMUSG00000022194 | 0.273169089 | 0.027093 | 0.08797 | 6202.325 | 6305.062 | 5203.822 | 5996.959 | 4654.206 | 6062.872 | 4565.909 | 4335.138 |
| ENSMUSG00000022195 | 0.096239458 | 0.38673 | 0.554239 | 947.7032 | 863.8715 | 699.7501 | 818.6297 | 817.1433 | 785.514 | 722.2526 | 787.4159 |
| ENSMUSG00000022197 | 0.451918925 | 0.028307 | 0.090811 | 1162.999 | 1246.615 | 1120.428 | 1327.578 | 729.3833 | 1333.027 | 809.0386 | 680.832 |
| ENSMUSG00000022199 | 0.228524485 | 0.004496 | 0.023786 | 2583.548 | 2905.968 | 2658.222 | 2897.552 | 2507.986 | 2500.683 | 2231.365 | 2186.058 |
| ENSMUSG00000022200 | -0.062085211 | 0.435932 | 0.600013 | 2346.119 | 2217.27 | 2255.762 | 2574.069 | 2291.512 | 2358.777 | 2527.402 | 2631.97 |
| ENSMUSG00000022201 | 0.202849533 | 0.036917 | 0.110463 | 3526.221 | 3372.698 | 3003.543 | 3255.541 | 2657.178 | 3317.484 | 2784.867 | 2672.211 |
| ENSMUSG00000022203 | 0.359293174 | 0.00135 | 0.009783 | 893.3763 | 1124.233 | 1028.508 | 1003.231 | 890.2766 | 824.6221 | 729.9669 | 710.197 |
| ENSMUSG00000022204 | -0.114650511 | 0.122389 | 0.251874 | 1186.138 | 1159.028 | 1325.799 | 1167.992 | 1314.45 | 1262.633 | 1287.326 | 1379.065 |
| ENSMUSG00000022205 | 0.068321407 | 0.497617 | 0.654039 | 6464.905 | 6691.405 | 5867.135 | 6775.045 | 6238.762 | 7155.664 | 5701.842 | 5508.649 |
| ENSMUSG00000022206 | -0.765249412 | 0.009359 | 0.040574 | 320.9313 | 178.7734 | 489.411 | 347.6373 | 477.8045 | 404.4894 | 689.4668 | 703.6714 |
| ENSMUSG00000022208 | 1.782806275 | 8.38E-18 | 6.72E-15 | 149.9021 | 172.7743 | 144.0906 | 169.9368 | 42.9049 | 55.8687 | 33.75012 | 53.29196 |
| ENSMUSG00000022210 | -0.302290225 | 0.015263 | 0.058395 | 1699.226 | 1842.926 | 1942.738 | 1770.966 | 2162.797 | 1896.184 | 2114.686 | 2774.445 |
| ENSMUSG00000022211 | 0.424531711 | 0.004138 | 0.022371 | 259.562 | 315.5531 | 266.6503 | 309.6818 | 204.7734 | 258.1134 | 212.1436 | 182.7153 |
| ENSMUSG00000022212 | -0.188540209 | 0.687249 | 0.801255 | 12.07265 | 10.79839 | 9.93728 | 12.93935 | 8.776002 | 18.99536 | 10.60718 | 14.13868 |
| ENSMUSG00000022214 | -0.352350163 | 0.000768 | 0.006382 | 2275.695 | 2230.468 | 2412.275 | 2381.703 | 3078.426 | 2410.176 | 3246.762 | 3137.7 |
| ENSMUSG00000022215 | -2.420550806 | 0.001798 | 0.012158 | 4.024217 | 11.99822 | 24.01509 | 2.58787 | 78.0089 | 14.52586 | 29.89296 | 106.5839 |
| ENSMUSG00000022216 | -0.050216447 | 0.624447 | 0.755179 | 2317.949 | 2606.012 | 2868.561 | 2294.578 | 2654.253 | 2308.495 | 2706.76 | 2774.445 |
| ENSMUSG00000022217 | -0.298256943 | 0.20319 | 0.358963 | 112.6781 | 95.98572 | 112.6225 | 93.16334 | 117.0134 | 89.38993 | 126.3219 | 177.2773 |
| ENSMUSG00000022218 | -1.550549858 | 1.87E-06 | 6.11E-05 | 41.24823 | 43.19358 | 57.13936 | 56.93315 | 134.5654 | 90.5073 | 121.5004 | 237.0949 |
| ENSMUSG00000022219 | -1.897865831 | 7.94E-05 | 0.001132 | 64.38748 | 25.19625 | 95.23226 | 52.62003 | 202.8232 | 82.68568 | 203.465 | 399.1459 |
| ENSMUSG00000022220 | 0.00453262 | 0.977221 | 0.987494 | 426.567 | 356.347 | 443.037 | 357.1261 | 355.9156 | 316.2169 | 455.1445 | 451.3503 |

| | | | | | | | | | | | |
|---|---|---|---|---|---|---|---|---|---|---|---|
| ENSMUSG00000022221 | -0.032181 | 0.736197 | 0.834773 | 445.6821 | 463.1311 | 431.4436 | 458.0531 | 515.8339 | 433.5411 | 446.4659 | 440.4744 |
| ENSMUSG00000022223 | -0.115170708 | 0.118749 | 0.247558 | 1030.2 | 1155.428 | 1079.023 | 1152.465 | 1150.631 | 1268.22 | 1213.076 | 1151.759 |
| ENSMUSG00000022225 | -0.066585172 | 0.94508 | 0.968362 | 65.39353 | 133.1802 | 77.84202 | 100.0643 | 96.53602 | 261.4655 | 30.85725 | 5.437955 |
| ENSMUSG00000022228 | 0.202609153 | 0.017362 | 0.063858 | 1421.555 | 1394.193 | 1170.943 | 1259.43 | 1133.079 | 1127.43 | 1111.825 | 1182.212 |
| ENSMUSG00000022229 | -1.526759539 | 0.080831 | 0.190282 | 74.44802 | 57.59143 | 373.4761 | 70.73513 | 321.7867 | 139.6718 | 198.6436 | 1001.671 |
| ENSMUSG00000022231 | -0.052604989 | 0.758508 | 0.849288 | 1881.322 | 2340.852 | 3366.253 | 2398.093 | 2258.358 | 2610.186 | 2606.474 | 2885.379 |
| ENSMUSG00000022234 | -0.113837647 | 0.140618 | 0.277122 | 8910.623 | 10313.67 | 9955.498 | 9852.886 | 11524.84 | 9742.385 | 10622.61 | 10344.08 |
| ENSMUSG00000022235 | -0.967001195 | 0.005017 | 0.025791 | 304.8345 | 127.1811 | 218.6202 | 221.6942 | 476.8294 | 239.1181 | 315.3226 | 675.3941 |
| ENSMUSG00000022236 | -0.463163345 | 0.109977 | 0.235741 | 37.22401 | 28.79572 | 29.81184 | 24.15346 | 35.10401 | 44.69496 | 35.6787 | 50.02919 |
| ENSMUSG00000022237 | -1.314140885 | 8.43E-08 | 4.95E-06 | 150.9082 | 163.1757 | 223.5888 | 177.7004 | 420.273 | 297.2215 | 414.6443 | 649.2919 |
| ENSMUSG00000022240 | 0.803701352 | 0.003629 | 0.020376 | 636.8324 | 754.6877 | 520.8791 | 670.2585 | 356.8907 | 601.1473 | 300.8582 | 220.781 |
| ENSMUSG00000022241 | -0.134976388 | 0.09777 | 0.21756 | 1842.086 | 1878.921 | 1947.707 | 1970.232 | 2059.435 | 1867.132 | 2148.436 | 2314.394 |
| ENSMUSG00000022244 | -1.147290803 | 7.25E-06 | 0.00018 | 288.7376 | 319.1525 | 335.3832 | 288.1162 | 648.449 | 394.4331 | 638.3594 | 1046.263 |
| ENSMUSG00000022246 | -0.237455714 | 0.004904 | 0.025366 | 3100.66 | 2531.623 | 2729.439 | 2732.791 | 3208.116 | 3149.878 | 3558.227 | 3163.802 |
| ENSMUSG00000022247 | 0.037091888 | 0.55663 | 0.703449 | 1786.753 | 1720.544 | 1613.152 | 1758.027 | 1754.225 | 1681.648 | 1648.934 | 1616.16 |
| ENSMUSG00000022248 | -0.058527803 | 0.557406 | 0.703981 | 402.4217 | 478.7288 | 420.6782 | 410.6088 | 428.0739 | 478.2361 | 453.2159 | 420.8978 |
| ENSMUSG00000022253 | -0.37445307 | 8.08E-06 | 0.000195 | 1314.913 | 1309.005 | 1262.035 | 1339.654 | 1686.943 | 1481.638 | 1829.257 | 1773.861 |
| ENSMUSG00000022255 | 0.14864588 | 0.174209 | 0.322358 | 5206.331 | 5141.235 | 4747.535 | 5022.194 | 4066.214 | 5541.058 | 4342.194 | 4199.189 |
| ENSMUSG00000022257 | 0.046663226 | 0.581416 | 0.722532 | 3256.598 | 2907.168 | 3537.672 | 2946.722 | 3062.825 | 2910.76 | 3033.654 | 3242.109 |
| ENSMUSG00000022261 | -0.10794132 | 0.543691 | 0.692599 | 2275.695 | 2309.656 | 3624.623 | 2593.046 | 2580.145 | 2426.937 | 3271.833 | 3366.094 |
| ENSMUSG00000022262 | 0.109992513 | 0.807741 | 0.882476 | 40.24217 | 27.5959 | 12.4216 | 32.77969 | 17.552 | 39.10809 | 24.10723 | 23.927 |
| ENSMUSG00000022263 | 0.21818354 | 0.070169 | 0.172649 | 2423.585 | 2277.261 | 2099.25 | 2499.883 | 1814.682 | 2439.228 | 1968.114 | 1773.861 |
| ENSMUSG00000022265 | -0.115634179 | 0.424519 | 0.589668 | 894.3823 | 892.6672 | 755.2332 | 883.3265 | 1138.93 | 735.2322 | 974.8963 | 859.197 |
| ENSMUSG00000022269 | 0.987516917 | 0.049196 | 0.134578 | 23.13925 | 25.19625 | 20.70267 | 40.5433 | 15.60178 | 24.58223 | 6.750024 | 8.700729 |
| ENSMUSG00000022270 | -1.266684644 | 5.99E-08 | 3.75E-06 | 577.4752 | 350.3479 | 658.3448 | 587.4466 | 1240.342 | 887.195 | 1686.542 | 1420.394 |
| ENSMUSG00000022272 | 0.045590445 | 0.674592 | 0.7917 | 2860.213 | 2500.428 | 2399.025 | 2191.926 | 2264.208 | 2740.919 | 2385.651 | 2250.226 |
| ENSMUSG00000022280 | -0.235689482 | 0.003134 | 0.018355 | 1442.682 | 1340.201 | 1426.828 | 1324.127 | 1690.843 | 1459.291 | 1741.506 | 1622.686 |
| ENSMUSG00000022283 | 0.121405254 | 0.247032 | 0.410092 | 24082.93 | 26918 | 24333.91 | 26567.08 | 24134 | 27205.82 | 22466.97 | 19870.29 |
| ENSMUSG00000022285 | -0.011251058 | 0.825312 | 0.893947 | 10883.5 | 10629.22 | 10573.27 | 10681 | 10520.48 | 11472.08 | 10825.11 | 10285.35 |
| ENSMUSG00000022286 | -0.112684508 | 0.322324 | 0.490583 | 534.2149 | 557.917 | 596.2368 | 459.7783 | 548.0126 | 539.6917 | 635.4666 | 598.1751 |
| ENSMUSG00000022292 | -0.004230231 | 0.976879 | 0.987311 | 412.4823 | 440.3345 | 385.8977 | 333.8353 | 372.4925 | 326.2732 | 414.6443 | 461.1386 |
| ENSMUSG00000022295 | -0.372405421 | 0.000544 | 0.004894 | 1063.399 | 957.4576 | 1004.493 | 894.5406 | 1229.615 | 1092.792 | 1314.326 | 1436.708 |
| ENSMUSG00000022296 | 0.34340566 | 0.47911 | 0.638368 | 15.09082 | 9.598572 | 12.4216 | 13.80198 | 6.825779 | 13.40849 | 12.53576 | 7.613138 |
| ENSMUSG00000022297 | -0.209391518 | 0.065264 | 0.164507 | 1044.284 | 851.8733 | 1106.35 | 921.2819 | 1117.478 | 1004.519 | 1190.897 | 1226.803 |
| ENSMUSG00000022299 | -0.171760321 | 0.104984 | 0.228462 | 278.6771 | 290.3568 | 321.3054 | 295.8799 | 346.1645 | 344.1512 | 302.7868 | 345.854 |
| ENSMUSG00000022300 | -0.042074823 | 0.51216 | 0.667488 | 1093.581 | 1138.631 | 1112.975 | 1109.334 | 1144.781 | 1076.031 | 1154.254 | 1209.401 |
| ENSMUSG00000022305 | 0.149690888 | 0.122343 | 0.251807 | 695.1836 | 765.4861 | 619.4238 | 750.4824 | 658.2001 | 651.4291 | 597.8593 | 641.6787 |
| ENSMUSG00000022306 | 0.905092253 | 0.000514 | 0.004679 | 309.8647 | 488.3274 | 358.5702 | 429.5865 | 174.5449 | 334.0949 | 163.9292 | 175.1022 |
| ENSMUSG00000022307 | 0.070861795 | 0.298276 | 0.466248 | 2134.847 | 2034.897 | 1941.91 | 2123.779 | 1896.591 | 2120.776 | 1898.685 | 1926.124 |
| ENSMUSG00000022309 | -0.827153814 | 5.48E-06 | 0.000144 | 237.4288 | 215.9679 | 349.461 | 256.1992 | 443.6756 | 391.0809 | 526.5019 | 522.0437 |
| ENSMUSG00000022312 | -0.157575536 | 0.049928 | 0.13595 | 9033.362 | 9897.328 | 10068.95 | 10058.19 | 11872.96 | 9682.047 | 11136.58 | 10872.65 |
| ENSMUSG00000022313 | 0.083960289 | 0.435331 | 0.599453 | 344.0706 | 346.7484 | 304.7432 | 320.8959 | 320.8116 | 284.9304 | 335.5726 | 296.9124 |
| ENSMUSG00000022314 | -0.077463266 | 0.27911 | 0.445456 | 7679.213 | 6999.759 | 7385.883 | 8077.606 | 7597.092 | 7947.882 | 8601.459 | 7662.079 |
| ENSMUSG00000022321 | 0.909854393 | 0.016336 | 0.061103 | 64.38748 | 77.9884 | 54.65504 | 48.30692 | 31.20356 | 55.8687 | 26.03581 | 17.40146 |
| ENSMUSG00000022322 | 0.020610627 | 0.853521 | 0.912083 | 781.7042 | 889.0678 | 735.3587 | 784.1248 | 811.2926 | 901.7209 | 727.074 | 702.5838 |
| ENSMUSG00000022323 | -1.417347786 | 1.88E-08 | 1.44E-06 | 863.1946 | 686.2979 | 1153.553 | 983.3908 | 2526.513 | 1499.516 | 2206.294 | 3618.416 |
| ENSMUSG00000022324 | 0.751918649 | 0.00182 | 0.012248 | 1763.613 | 2486.03 | 1998.221 | 2391.192 | 1159.407 | 2035.856 | 990.325 | 945.1167 |
| ENSMUSG00000022325 | -0.224741382 | 0.02573 | 0.084708 | 349.1009 | 361.1463 | 359.3983 | 349.3625 | 389.0694 | 383.2593 | 421.3944 | 464.4014 |
| ENSMUSG00000022329 | 0.153163334 | 0.106794 | 0.231039 | 950.7214 | 974.2551 | 978.822 | 979.0777 | 752.7859 | 981.0545 | 861.1102 | 899.4378 |
| ENSMUSG00000022330 | -1.42592984 | 1.38E-05 | 0.000294 | 428.5792 | 116.3827 | 313.0243 | 306.2313 | 542.1619 | 812.331 | 857.2531 | 920.1021 |
| ENSMUSG00000022332 | -0.086760002 | 0.281503 | 0.447843 | 1035.23 | 1237.016 | 1168.458 | 1176.618 | 1152.582 | 1268.22 | 1218.862 | 1263.781 |
| ENSMUSG00000022335 | 0.067398791 | 0.614475 | 0.748118 | 221.332 | 206.3693 | 190.4645 | 190.6398 | 196.9725 | 216.7706 | 170.6792 | 187.0657 |
| ENSMUSG00000022336 | 0.032954184 | 0.647306 | 0.771749 | 8111.816 | 8797.091 | 8365.533 | 9148.122 | 8707.744 | 8857.424 | 8433.673 | 7645.765 |
| ENSMUSG00000022337 | -0.222234884 | 0.012853 | 0.051581 | 969.8364 | 999.4513 | 1121.256 | 1084.318 | 1174.034 | 1127.43 | 1227.54 | 1345.35 |
| ENSMUSG00000022338 | -0.061426123 | 0.232333 | 0.392957 | 2710.31 | 2937.163 | 2829.64 | 2935.508 | 2909.732 | 3021.38 | 2962.296 | 3015.89 |
| ENSMUSG00000022339 | -0.114262018 | 0.201051 | 0.356523 | 695.1836 | 709.0945 | 729.5619 | 746.1693 | 765.4624 | 708.4152 | 768.5385 | 876.5984 |
| ENSMUSG00000022340 | -0.824847903 | 0.02288 | 0.077667 | 18.10898 | 17.99732 | 18.21835 | 25.01608 | 23.40267 | 48.04709 | 40.50014 | 29.36496 |
| ENSMUSG00000022346 | -0.411774766 | 0.031046 | 0.096788 | 850.1159 | 704.2952 | 536.6131 | 503.7721 | 988.7629 | 895.0167 | 879.4317 | 685.1824 |
| ENSMUSG00000022350 | -0.222404505 | 0.004892 | 0.025347 | 1742.486 | 1582.565 | 1746.477 | 1632.084 | 2097.464 | 1758.747 | 2009.579 | 1955.489 |
| ENSMUSG00000022351 | -1.061273975 | 1.60E-12 | 4.33E-10 | 889.3521 | 892.6672 | 924.167 | 817.7671 | 1766.902 | 1347.553 | 2165.793 | 2071.861 |
| ENSMUSG00000022353 | 0.048519325 | 0.584255 | 0.724713 | 1470.851 | 1436.186 | 1628.058 | 1580.326 | 1346.629 | 1638.07 | 1409.791 | 1524.803 |
| ENSMUSG00000022354 | -0.619112427 | 0.000429 | 0.004065 | 2156.981 | 1876.521 | 2603.567 | 2109.114 | 3381.686 | 2374.42 | 3432.869 | 4247.043 |
| ENSMUSG00000022358 | 0.44385527 | 0.022842 | 0.07758 | 763.5953 | 1007.85 | 763.5143 | 916.9688 | 601.6437 | 864.8476 | 615.2165 | 455.7007 |
| ENSMUSG00000022359 | 0.176815052 | 0.118596 | 0.247423 | 290.7497 | 296.3559 | 286.5249 | 276.9021 | 249.6285 | 290.5173 | 244.9294 | 232.7445 |

| | | | | | | | | | | | |
|---|---|---|---|---|---|---|---|---|---|---|---|
| ENSMUSG00000022360 | 0.170007292 | 0.192677 | 0.345886 | 1273.665 | 1428.987 | 992.8999 | 1180.932 | 1079.448 | 1231.346 | 996.1107 | 1024.511 |
| ENSMUSG00000022361 | 0.021990376 | 0.777442 | 0.862807 | 1578.499 | 1520.174 | 1532.825 | 1776.142 | 1576.755 | 1674.944 | 1518.755 | 1544.379 |
| ENSMUSG00000022362 | -0.540301785 | 0.23158 | 0.392114 | 13.07871 | 32.39518 | 12.4216 | 17.25247 | 25.35289 | 31.28647 | 34.71441 | 16.31387 |
| ENSMUSG00000022364 | -0.019178056 | 0.847068 | 0.908105 | 429.5852 | 435.5352 | 377.6166 | 448.5642 | 439.7752 | 453.6539 | 416.5729 | 402.4087 |
| ENSMUSG00000022365 | -0.092499852 | 0.092119 | 0.208429 | 2329.016 | 2331.253 | 2283.09 | 2277.326 | 2550.891 | 2321.903 | 2415.544 | 2539.525 |
| ENSMUSG00000022366 | -6.283077507 | 8.78E-09 | 7.60E-07 | 0 | 3.599465 | 0 | 0 | 48.75557 | 20.11273 | 38.57157 | 130.5109 |
| ENSMUSG00000022367 | -0.107355527 | 0.56789 | 0.712168 | 421.5368 | 567.5156 | 382.5853 | 466.6793 | 586.0419 | 581.0345 | 428.1444 | 382.8321 |
| ENSMUSG00000022369 | 0.089309563 | 0.32863 | 0.496455 | 684.117 | 742.6895 | 649.2356 | 671.9837 | 691.3539 | 657.016 | 653.7881 | 576.4233 |
| ENSMUSG00000022370 | -0.060397114 | 0.506997 | 0.662734 | 1881.322 | 2210.071 | 2047.908 | 1952.117 | 2385.122 | 1997.865 | 1978.721 | 2071.861 |
| ENSMUSG00000022371 | 0.24874122 | 0.189966 | 0.342514 | 1438.658 | 2718.796 | 2153.905 | 1923.65 | 1455.841 | 2019.095 | 1624.827 | 1830.416 |
| ENSMUSG00000022372 | -0.604237351 | 0.116975 | 0.245141 | 27.16347 | 37.19447 | 35.60859 | 62.10889 | 33.15378 | 72.62932 | 84.85745 | 56.55474 |
| ENSMUSG00000022376 | 1.378351721 | 0.001178 | 0.008849 | 196.1806 | 239.9643 | 194.6051 | 213.9306 | 64.35735 | 173.193 | 53.0359 | 34.80291 |
| ENSMUSG00000022377 | -0.075760292 | 0.237049 | 0.398088 | 1941.685 | 1880.12 | 2058.673 | 2109.977 | 2139.394 | 2139.771 | 2145.543 | 2002.255 |
| ENSMUSG00000022378 | 0.048043868 | 0.510799 | 0.666363 | 1468.839 | 1401.392 | 1545.247 | 1558.761 | 1369.056 | 1558.737 | 1457.041 | 1399.73 |
| ENSMUSG00000022382 | -1.4662388 | 0.042403 | 0.12154 | 139.8416 | 117.5825 | 720.4528 | 160.448 | 673.8019 | 320.6864 | 1011.539 | 1140.883 |
| ENSMUSG00000022383 | -1.500917255 | 0.000748 | 0.006253 | 52.31483 | 16.7975 | 54.65504 | 18.97772 | 96.53602 | 59.22083 | 130.179 | 119.635 |
| ENSMUSG00000022385 | -0.149144731 | 0.206782 | 0.363428 | 391.3551 | 344.3488 | 327.1021 | 393.3563 | 443.6756 | 360.9118 | 386.68 | 423.0729 |
| ENSMUSG00000022386 | 0.051057317 | 0.58221 | 0.722932 | 711.2804 | 693.4968 | 639.2983 | 650.4181 | 699.1548 | 688.3024 | 586.2878 | 625.3649 |
| ENSMUSG00000022387 | 0.00098334 | 0.986498 | 0.992305 | 2217.344 | 2048.095 | 2101.735 | 2147.932 | 2065.286 | 2251.509 | 2096.365 | 2100.138 |
| ENSMUSG00000022389 | 0.175021038 | 0.175381 | 0.323752 | 2393.403 | 2243.666 | 2083.516 | 2555.091 | 2296.387 | 2297.321 | 1996.079 | 1625.949 |
| ENSMUSG00000022390 | -0.024903058 | 0.681109 | 0.796461 | 4123.817 | 4021.802 | 4239.078 | 4128.516 | 4461.134 | 3928.687 | 4340.266 | 4069.766 |
| ENSMUSG00000022391 | -0.261987056 | 0.006395 | 0.030891 | 2113.72 | 1946.111 | 2229.263 | 2183.3 | 2569.418 | 2233.631 | 2470.509 | 2889.73 |
| ENSMUSG00000022394 | -0.340641473 | 0.001988 | 0.01308 | 432.6034 | 395.9411 | 419.8501 | 360.5766 | 550.9379 | 491.6446 | 454.1802 | 540.5328 |
| ENSMUSG00000022400 | -0.090578774 | 0.03402 | 0.103765 | 3288.792 | 3154.331 | 3307.458 | 3308.161 | 3472.371 | 3433.691 | 3499.405 | 3505.306 |
| ENSMUSG00000022401 | -0.020364551 | 0.847309 | 0.908255 | 441.6579 | 529.1213 | 409.9128 | 435.6249 | 464.153 | 468.1797 | 451.2873 | 453.5255 |
| ENSMUSG00000022403 | -0.105454351 | 0.006564 | 0.031504 | 5417.603 | 5376.4 | 5310.648 | 5477.659 | 5723.903 | 5698.608 | 5895.664 | 5900.182 |
| ENSMUSG00000022404 | -0.224337736 | 0.010375 | 0.043785 | 2164.023 | 1949.71 | 1927.832 | 2160.872 | 2452.405 | 2101.781 | 2538.973 | 2488.408 |
| ENSMUSG00000022407 | -0.29287684 | 0.005087 | 0.026035 | 1479.906 | 1785.334 | 1768.836 | 1687.292 | 2239.831 | 1755.395 | 2083.829 | 2153.43 |
| ENSMUSG00000022408 | -0.356788391 | 0.304149 | 0.472636 | 84.50857 | 70.78947 | 164.7932 | 56.93315 | 129.6898 | 91.62468 | 113.7861 | 149 |
| ENSMUSG00000022412 | 0.03145528 | 0.766883 | 0.855443 | 961.788 | 895.0669 | 859.5747 | 880.7386 | 926.3557 | 995.5803 | 847.6102 | 749.3503 |
| ENSMUSG00000022414 | -0.07582247 | 0.451652 | 0.614001 | 1079.496 | 1247.814 | 1205.723 | 978.215 | 1284.222 | 1174.36 | 1166.79 | 1125.657 |
| ENSMUSG00000022415 | -0.089480813 | 0.570597 | 0.713919 | 319.9253 | 284.3577 | 332.0708 | 313.1323 | 327.6374 | 246.9397 | 370.287 | 385.0072 |
| ENSMUSG00000022416 | 0.283179253 | 0.578546 | 0.719984 | 12.07265 | 16.7975 | 21.53077 | 20.70296 | 15.60178 | 24.58223 | 5.785735 | 13.05109 |
| ENSMUSG00000022419 | 0.027043208 | 0.888313 | 0.93282 | 937.6427 | 639.5049 | 698.922 | 753.9329 | 522.6597 | 879.3734 | 876.5389 | 696.0583 |
| ENSMUSG00000022420 | -0.166608533 | 0.081491 | 0.191373 | 785.7285 | 946.6592 | 910.9173 | 930.7707 | 1019.966 | 919.5989 | 1094.468 | 975.5692 |
| ENSMUSG00000022421 | 0.315724833 | 0.059455 | 0.153771 | 268.6165 | 244.7636 | 321.3054 | 313.995 | 220.3752 | 292.752 | 199.6079 | 213.1679 |
| ENSMUSG00000022422 | -0.194717266 | 0.121495 | 0.250741 | 249.5015 | 254.3622 | 230.2136 | 225.1447 | 311.0605 | 281.5783 | 241.0723 | 262.1095 |
| ENSMUSG00000022425 | -0.819572217 | 3.67E-07 | 1.63E-05 | 1766.631 | 1422.988 | 2196.967 | 1713.17 | 2920.458 | 2512.974 | 3507.12 | 3594.489 |
| ENSMUSG00000022426 | -0.096696112 | 0.175912 | 0.324376 | 1198.211 | 1228.617 | 1197.442 | 1230.964 | 1330.052 | 1177.712 | 1298.898 | 1383.416 |
| ENSMUSG00000022427 | -0.110612938 | 0.029306 | 0.092968 | 2382.337 | 2492.029 | 2594.458 | 2511.097 | 2697.158 | 2655.998 | 2751.117 | 2673.299 |
| ENSMUSG00000022428 | -0.151229783 | 0.088561 | 0.202802 | 653.9353 | 729.4915 | 680.7037 | 743.5815 | 796.6659 | 707.2978 | 774.3242 | 838.5327 |
| ENSMUSG00000022429 | 1.601642035 | 2.51E-09 | 2.79E-07 | 227.3683 | 266.3604 | 222.7607 | 216.5185 | 68.25779 | 119.559 | 75.21455 | 44.59123 |
| ENSMUSG00000022432 | -1.802038054 | NA | NA | 26.15741 | 33.595 | 19.04645 | 33.64232 | 15.60178 | 346.386 | 19.28578 | 10.87591 |
| ENSMUSG00000022433 | 0.105953865 | 0.1969 | 0.351214 | 3066.454 | 2874.772 | 2808.938 | 3079.566 | 2682.531 | 3079.483 | 2728.938 | 2502.547 |
| ENSMUSG00000022434 | -0.006506187 | 0.956043 | 0.97505 | 423.5489 | 441.5343 | 549.0347 | 545.178 | 495.3565 | 509.5226 | 486.966 | 480.7153 |
| ENSMUSG00000022435 | -0.806162358 | 0.205626 | 0.362018 | 362.1796 | 283.1579 | 1082.335 | 390.7684 | 1464.617 | 274.874 | 873.646 | 1091.941 |
| ENSMUSG00000022436 | 0.010621308 | 0.935904 | 0.962083 | 806.8556 | 769.0856 | 929.1356 | 784.1248 | 909.7789 | 681.5982 | 738.6455 | 936.4159 |
| ENSMUSG00000022437 | -0.50573354 | 0.000641 | 0.005555 | 3215.35 | 3198.724 | 3868.914 | 3396.149 | 5095.932 | 3535.372 | 5084.697 | 5707.678 |
| ENSMUSG00000022438 | -0.684891027 | 4.66E-05 | 0.000752 | 213.2835 | 229.1659 | 298.1184 | 251.8861 | 437.825 | 305.0431 | 417.5372 | 437.2116 |
| ENSMUSG00000022439 | -0.243284946 | 0.356347 | 0.524955 | 39.23612 | 33.595 | 40.57722 | 33.64232 | 39.97956 | 35.75597 | 44.3573 | 54.37955 |
| ENSMUSG00000022440 | 0.092914239 | 0.239746 | 0.401703 | 1087.545 | 1268.211 | 1225.598 | 1179.206 | 1161.358 | 1125.196 | 1035.647 | 1140.883 |
| ENSMUSG00000022441 | -1.005673397 | 0.095074 | 0.213129 | 3.018163 | 5.999108 | 15.73403 | 6.038364 | 13.65156 | 11.17374 | 17.3572 | 20.66423 |
| ENSMUSG00000022442 | 0.009538751 | 0.908336 | 0.945278 | 513.0877 | 503.925 | 520.051 | 542.5902 | 522.6597 | 543.0438 | 479.2517 | 524.2189 |
| ENSMUSG00000022443 | -0.228142889 | 0.126816 | 0.257937 | 7267.737 | 6390.249 | 9359.261 | 7102.842 | 8219.213 | 7258.462 | 9493.427 | 10312.54 |
| ENSMUSG00000022445 | -3.863399547 | 4.62E-09 | 4.56E-07 | 67.40564 | 21.59679 | 163.137 | 78.49874 | 1295.923 | 476.0014 | 1086.754 | 1964.19 |
| ENSMUSG00000022449 | 0.604681074 | 0.002863 | 0.017148 | 331.9979 | 315.5531 | 322.1335 | 363.1645 | 183.3209 | 312.8647 | 215.0365 | 166.4014 |
| ENSMUSG00000022450 | -0.34250976 | 0.000581 | 0.005147 | 2497.027 | 2288.06 | 2531.522 | 2549.052 | 3133.033 | 2651.529 | 3178.297 | 3548.81 |
| ENSMUSG00000022451 | -0.030527851 | 0.738513 | 0.836401 | 2003.054 | 1974.906 | 2336.917 | 2124.642 | 2210.577 | 1911.827 | 2174.472 | 2326.357 |
| ENSMUSG00000022452 | -0.409138964 | 0.008485 | 0.037894 | 1773.674 | 1854.924 | 2385.775 | 1935.727 | 2563.568 | 2092.842 | 2580.438 | 3322.591 |
| ENSMUSG00000022453 | -0.28304115 | 0.045051 | 0.126612 | 1320.949 | 1277.81 | 1682.713 | 1333.616 | 1588.456 | 1420.182 | 1784.899 | 2041.408 |
| ENSMUSG00000022454 | 0.114762261 | 0.408642 | 0.574884 | 507.0514 | 539.9197 | 436.4122 | 508.9479 | 495.3565 | 357.5597 | 505.2875 | 478.5401 |
| ENSMUSG00000022456 | 0.299971178 | 0.075006 | 0.18103 | 328.9798 | 327.5513 | 324.6178 | 325.2091 | 245.728 | 359.7945 | 229.5008 | 227.3065 |
| ENSMUSG00000022462 | 0.365686723 | 1.62E-06 | 5.53E-05 | 8515.244 | 8414.348 | 7226.058 | 8278.598 | 6328.472 | 6675.193 | 6270.772 | 5894.744 |
| ENSMUSG00000022463 | -0.564912845 | 5.21E-06 | 0.000139 | 2154.968 | 1973.706 | 2386.603 | 1980.584 | 3017.969 | 2553.2 | 3481.084 | 3517.27 |

| | | | | | | | | | | | |
|---|---|---|---|---|---|---|---|---|---|---|---|
| ENSMUSG00000022464 | 0.265948754 | 0.025683 | 0.084625 | 1891.382 | 1989.304 | 2529.038 | 2035.791 | 1527.024 | 1810.146 | 1769.471 | 1920.686 |
| ENSMUSG00000022466 | -0.195150128 | 0.008069 | 0.036501 | 995.9938 | 959.8572 | 1056.664 | 1043.774 | 1118.453 | 1096.144 | 1222.719 | 1210.489 |
| ENSMUSG00000022468 | -0.623101943 | 0.030859 | 0.096451 | 37.22401 | 49.19268 | 66.24853 | 40.5433 | 61.43201 | 63.69032 | 72.32169 | 101.146 |
| ENSMUSG00000022469 | -0.299487521 | 0.071424 | 0.174927 | 881.3036 | 679.099 | 929.1356 | 728.9169 | 868.8242 | 791.1009 | 1146.54 | 1156.109 |
| ENSMUSG00000022471 | -0.106868488 | 0.149921 | 0.289823 | 1141.872 | 1079.839 | 1050.867 | 1004.956 | 1181.835 | 1083.853 | 1129.183 | 1208.314 |
| ENSMUSG00000022472 | 0.003108022 | 0.977781 | 0.987843 | 1021.145 | 841.0749 | 1002.009 | 804.8277 | 838.5957 | 893.8993 | 992.2536 | 937.5035 |
| ENSMUSG00000022474 | 0.26856034 | 0.004447 | 0.0236 | 893.3763 | 904.6654 | 913.4016 | 1029.972 | 842.4962 | 820.1526 | 736.7169 | 708.0218 |
| ENSMUSG00000022475 | 0.087791766 | 0.217243 | 0.375767 | 2012.109 | 1809.331 | 2058.673 | 2018.539 | 1844.911 | 1927.47 | 1926.65 | 1737.971 |
| ENSMUSG00000022476 | -0.329074245 | 0.001333 | 0.009701 | 593.5721 | 662.3015 | 587.1276 | 687.5109 | 885.4011 | 787.7487 | 788.7885 | 714.5473 |
| ENSMUSG00000022477 | -0.951421931 | 1.61E-05 | 0.000331 | 5039.326 | 4676.904 | 6461.716 | 5075.677 | 10118.73 | 6040.524 | 11120.18 | 13822.2 |
| ENSMUSG00000022479 | -0.669035098 | 0.00184 | 0.012349 | 398.3975 | 328.7511 | 399.1474 | 484.7944 | 449.5263 | 553.1002 | 689.4668 | 872.2481 |
| ENSMUSG00000022483 | -0.204780062 | 0.240815 | 0.403048 | 909.4731 | 1081.039 | 914.2297 | 1014.445 | 1170.134 | 1517.394 | 953.682 | 875.5108 |
| ENSMUSG00000022484 | 0.367639763 | 0.016091 | 0.060435 | 1539.263 | 1955.709 | 1522.888 | 1848.602 | 1364.181 | 1666.005 | 1202.469 | 1088.679 |
| ENSMUSG00000022485 | 0.949888554 | 0.001888 | 0.012591 | 378.2764 | 509.9242 | 373.4761 | 500.3216 | 175.52 | 406.7242 | 170.6792 | 159.8759 |
| ENSMUSG00000022488 | -0.06586016 | 0.758203 | 0.849269 | 83.50251 | 68.38983 | 119.2474 | 91.43809 | 102.3867 | 92.74205 | 97.39321 | 89.18247 |
| ENSMUSG00000022489 | 0.585155179 | 5.08E-06 | 0.000136 | 378.2764 | 455.9322 | 404.9441 | 400.2573 | 273.0312 | 321.8037 | 260.3581 | 237.0949 |
| ENSMUSG00000022490 | -0.481636589 | 0.012768 | 0.051295 | 555.342 | 716.2935 | 660.001 | 665.9453 | 817.1433 | 612.321 | 1083.861 | 1112.606 |
| ENSMUSG00000022494 | 0.48355842 | 0.152785 | 0.293791 | 59.35721 | 73.18911 | 56.31125 | 67.28463 | 33.15378 | 73.74669 | 49.17875 | 27.18978 |
| ENSMUSG00000022498 | -0.299860561 | 0.010759 | 0.045077 | 767.6195 | 664.7011 | 797.4667 | 739.2683 | 916.6046 | 759.8144 | 941.1462 | 1039.737 |
| ENSMUSG00000022500 | -0.043531726 | 0.582904 | 0.723622 | 2161.005 | 2320.455 | 2439.602 | 2355.825 | 2526.513 | 2289.5 | 2558.259 | 2186.058 |
| ENSMUSG00000022503 | -0.302620492 | 0.018074 | 0.065832 | 741.4621 | 847.074 | 783.3889 | 791.8884 | 1022.892 | 753.1101 | 1040.468 | 1083.241 |
| ENSMUSG00000022504 | -0.802143989 | 0.068138 | 0.168913 | 15.09082 | 15.59768 | 14.90592 | 12.07673 | 30.22845 | 15.64324 | 16.39292 | 38.06569 |
| ENSMUSG00000022505 | 0.126647288 | 0.146552 | 0.285493 | 1261.592 | 1316.204 | 1480.655 | 1306.875 | 1351.504 | 1202.295 | 1146.54 | 1215.927 |
| ENSMUSG00000022507 | -0.148924852 | 0.20606 | 0.362636 | 1030.2 | 869.8706 | 812.3726 | 786.7126 | 947.8082 | 1076.031 | 990.325 | 863.5473 |
| ENSMUSG00000022508 | -0.430445104 | 0.154054 | 0.295521 | 98.59333 | 56.39161 | 77.01392 | 43.9938 | 121.8889 | 78.21619 | 78.10742 | 93.53283 |
| ENSMUSG00000022510 | 0.37049257 | 0.098804 | 0.219228 | 157.9505 | 134.38 | 114.2787 | 165.6237 | 114.088 | 147.4934 | 94.50034 | 87.00729 |
| ENSMUSG00000022512 | 1.017838586 | 2.33E-05 | 0.000443 | 6673.159 | 6888.175 | 4421.261 | 7740.321 | 3095.978 | 4508.604 | 3111.761 | 1987.029 |
| ENSMUSG00000022514 | 0.018068756 | 0.901334 | 0.940735 | 411.4762 | 458.3318 | 511.7699 | 397.6694 | 380.2934 | 406.7242 | 434.8944 | 536.1824 |
| ENSMUSG00000022515 | -0.033538518 | 0.650368 | 0.7738 | 1352.137 | 1237.016 | 1177.568 | 1213.711 | 1338.828 | 1212.351 | 1247.79 | 1295.321 |
| ENSMUSG00000022516 | -0.290245143 | 0.006116 | 0.029877 | 1321.955 | 1391.793 | 1549.388 | 1399.175 | 1728.872 | 1470.464 | 1758.863 | 1967.452 |
| ENSMUSG00000022517 | -0.260563356 | 0.063225 | 0.160933 | 2071.466 | 1829.728 | 2435.462 | 2019.402 | 2278.835 | 2043.677 | 2816.689 | 2873.416 |
| ENSMUSG00000022518 | -0.085807885 | 0.876258 | 0.925873 | 39.23612 | 17.99732 | 26.49941 | 32.77969 | 16.57689 | 69.27719 | 22.17865 | 16.31387 |
| ENSMUSG00000022519 | -0.031635966 | 0.784426 | 0.867324 | 319.9253 | 299.9554 | 361.8826 | 326.0717 | 332.513 | 303.9258 | 326.894 | 376.3065 |
| ENSMUSG00000022521 | 0.639612255 | 0.008275 | 0.037224 | 2874.297 | 2626.409 | 2240.857 | 2471.416 | 1255.943 | 2697.341 | 1288.29 | 1314.898 |
| ENSMUSG00000022523 | 1.01071381 | 5.53E-06 | 0.000144 | 335.0161 | 338.3497 | 279.0719 | 331.2474 | 135.5405 | 240.2354 | 136.9291 | 125.073 |
| ENSMUSG00000022526 | -0.032945752 | 0.785704 | 0.868321 | 847.0978 | 781.0838 | 628.5329 | 737.5431 | 746.9353 | 679.3635 | 787.8242 | 846.1459 |
| ENSMUSG00000022528 | 0.197465827 | 0.238208 | 0.399343 | 2089.575 | 1941.311 | 1623.917 | 1438.856 | 1331.027 | 1995.63 | 1379.898 | 1479.124 |
| ENSMUSG00000022529 | 0.138134386 | 0.197665 | 0.351998 | 1160.987 | 1031.847 | 932.4481 | 1075.691 | 985.8375 | 1064.858 | 920.8962 | 845.0583 |
| ENSMUSG00000022533 | 0.225620749 | 0.055712 | 0.146951 | 2439.682 | 2282.061 | 2030.517 | 2284.227 | 1669.391 | 2333.077 | 1859.15 | 1867.394 |
| ENSMUSG00000022536 | -0.168850401 | 0.013522 | 0.053429 | 4151.986 | 3923.416 | 4173.657 | 4219.954 | 4653.231 | 4180.096 | 4853.267 | 4828.904 |
| ENSMUSG00000022537 | 0.656730695 | 5.51E-15 | 2.37E-12 | 1089.557 | 1130.232 | 1174.255 | 1239.59 | 794.7157 | 687.1851 | 699.1096 | 759.1386 |
| ENSMUSG00000022538 | -0.173233117 | 0.003302 | 0.019044 | 1332.016 | 1466.182 | 1369.688 | 1356.907 | 1587.481 | 1521.864 | 1572.756 | 1541.117 |
| ENSMUSG00000022540 | -0.334592552 | 0.103349 | 0.226011 | 628.784 | 515.9233 | 846.325 | 575.3699 | 812.2677 | 589.9735 | 795.5386 | 1040.825 |
| ENSMUSG00000022543 | 0.086645878 | 0.759156 | 0.84986 | 35.2119 | 35.99465 | 51.34261 | 39.68068 | 30.22845 | 48.04709 | 36.64299 | 39.15328 |
| ENSMUSG00000022544 | -0.127087714 | 0.132614 | 0.265977 | 634.8203 | 742.6895 | 672.4226 | 623.6768 | 711.8313 | 735.2322 | 750.217 | 717.8101 |
| ENSMUSG00000022545 | 0.073746254 | 0.445607 | 0.608384 | 623.7537 | 550.7181 | 561.4563 | 571.0568 | 553.8632 | 506.1705 | 610.395 | 519.8685 |
| ENSMUSG00000022546 | 0.5292746 | 0.006902 | 0.032666 | 142.8597 | 152.3773 | 115.1068 | 169.9368 | 113.1129 | 91.62468 | 83.89316 | 113.1095 |
| ENSMUSG00000022548 | 1.032783165 | 0.05291 | 0.141564 | 20.12109 | 92.38626 | 38.09291 | 58.6584 | 12.67645 | 39.10809 | 31.82154 | 18.48905 |
| ENSMUSG00000022550 | 0.104296573 | 0.324541 | 0.492495 | 891.3642 | 733.091 | 684.8442 | 750.4824 | 750.8357 | 662.6028 | 707.7882 | 723.2481 |
| ENSMUSG00000022551 | -0.90225481 | 1.93E-05 | 0.000381 | 2667.05 | 2594.014 | 3479.704 | 2687.072 | 5148.588 | 3298.488 | 5791.521 | 7121.546 |
| ENSMUSG00000022552 | 0.03533335 | 0.632017 | 0.760703 | 1290.768 | 1313.805 | 1250.441 | 1336.204 | 1288.122 | 1369.901 | 1247.79 | 1159.372 |
| ENSMUSG00000022553 | 0.070458862 | 0.354123 | 0.522769 | 3934.679 | 3801.035 | 3873.055 | 4428.709 | 3944.325 | 3768.903 | 4012.407 | 3548.81 |
| ENSMUSG00000022554 | -0.401761167 | 0.007883 | 0.035845 | 359.1614 | 416.3381 | 395.0069 | 351.0878 | 572.3903 | 382.1419 | 487.9303 | 565.5474 |
| ENSMUSG00000022555 | -0.144736811 | 0.221823 | 0.380951 | 815.9101 | 710.2943 | 682.3599 | 707.3513 | 836.6455 | 699.4762 | 750.217 | 936.4159 |
| ENSMUSG00000022556 | -0.095651576 | 0.210776 | 0.368274 | 903.4368 | 873.4701 | 828.1066 | 929.0455 | 954.634 | 876.0213 | 953.682 | 990.7955 |
| ENSMUSG00000022557 | -0.277223188 | 0.008469 | 0.037839 | 1073.46 | 1165.027 | 1208.208 | 1023.934 | 1434.389 | 1173.243 | 1303.719 | 1505.226 |
| ENSMUSG00000022558 | -0.551354971 | 0.006596 | 0.031614 | 852.128 | 628.7065 | 1011.118 | 746.1693 | 1147.706 | 825.7395 | 1285.397 | 1488.912 |
| ENSMUSG00000022559 | 0.039532981 | 0.578302 | 0.719943 | 1306.865 | 1316.204 | 1229.738 | 1374.159 | 1333.952 | 1204.529 | 1227.54 | 1318.16 |
| ENSMUSG00000022560 | -0.526053809 | 0.007551 | 0.03472 | 303.8284 | 341.9491 | 336.2113 | 331.2474 | 441.7254 | 335.2122 | 456.1088 | 657.9926 |
| ENSMUSG00000022561 | -0.225721386 | 0.016954 | 0.062705 | 936.6366 | 883.0686 | 1044.242 | 879.0133 | 1096.025 | 996.6977 | 1090.611 | 1196.35 |
| ENSMUSG00000022562 | -0.525998383 | 0.001728 | 0.011767 | 650.9172 | 497.9259 | 795.8105 | 663.3575 | 886.3762 | 767.636 | 1100.254 | 1004.934 |
| ENSMUSG00000022564 | -0.514344762 | 0.000362 | 0.003573 | 1885.346 | 1756.539 | 2227.607 | 2050.456 | 2966.289 | 2127.48 | 2923.725 | 3297.576 |
| ENSMUSG00000022565 | 0.021791625 | 0.833886 | 0.899486 | 1405.458 | 1334.202 | 1745.649 | 1351.731 | 1392.459 | 1414.596 | 1470.541 | 1475.861 |
| ENSMUSG00000022568 | -0.155748256 | 0.040449 | 0.117619 | 1667.032 | 1634.157 | 1759.727 | 1515.629 | 1914.143 | 1718.521 | 1777.185 | 1916.335 |

| | | | | | | | | | | | |
|---|---|---|---|---|---|---|---|---|---|---|---|
| ENSMUSG00000022570 | -0.06464027 | 0.476466 | 0.635931 | 1089.557 | 1377.395 | 1252.097 | 1192.146 | 1342.728 | 1211.234 | 1269.005 | 1309.46 |
| ENSMUSG00000022571 | -0.474027402 | 0.000536 | 0.004837 | 814.904 | 965.8563 | 1040.102 | 868.6619 | 1302.749 | 1022.397 | 1301.79 | 1497.613 |
| ENSMUSG00000022574 | 0.234079673 | 0.066505 | 0.166249 | 238.4349 | 296.3559 | 276.5876 | 312.2697 | 241.8276 | 256.996 | 226.608 | 230.5693 |
| ENSMUSG00000022575 | 1.002502513 | 3.45E-09 | 3.58E-07 | 530.1906 | 758.2872 | 613.627 | 634.0283 | 306.185 | 406.7242 | 266.1438 | 287.124 |
| ENSMUSG00000022577 | 0.774845365 | 0.002432 | 0.015116 | 566.4086 | 632.3059 | 617.7675 | 659.907 | 294.4836 | 603.382 | 287.3582 | 263.197 |
| ENSMUSG00000022579 | -0.116353502 | 0.141364 | 0.278147 | 675.0625 | 700.6958 | 650.0637 | 719.428 | 690.3788 | 755.3449 | 781.0742 | 748.2627 |
| ENSMUSG00000022580 | -1.391387438 | 7.30E-06 | 0.000181 | 173.0413 | 182.3729 | 271.619 | 203.5791 | 366.6419 | 328.508 | 609.4308 | 876.5984 |
| ENSMUSG00000022587 | 0.097171937 | 0.418748 | 0.584292 | 6072.544 | 5525.178 | 6968.517 | 5146.412 | 5356.286 | 4843.817 | 6263.058 | 5705.503 |
| ENSMUSG00000022589 | 10.00473679 | 9.26E-05 | 0.001256 | 1670.05 | 0 | 1.656213 | 439.938 | 0 | 0 | 0.964289 | 1.087591 |
| ENSMUSG00000022594 | 0.240511225 | 0.211596 | 0.369104 | 309.8647 | 202.7698 | 306.3995 | 217.3811 | 228.176 | 181.0146 | 218.8936 | 250.146 |
| ENSMUSG00000022595 | -0.75237187 | 0.244078 | 0.406635 | 3.018163 | 7.198929 | 17.39024 | 6.900988 | 23.40267 | 8.938993 | 15.42863 | 10.87591 |
| ENSMUSG00000022598 | -0.982257001 | 0.178794 | 0.328373 | 303.8284 | 106.7841 | 698.0939 | 180.2883 | 950.7335 | 164.254 | 816.7529 | 615.5766 |
| ENSMUSG00000022601 | -0.023810805 | 0.823056 | 0.892656 | 579.4873 | 484.7279 | 464.5678 | 548.6285 | 528.5103 | 566.5087 | 523.609 | 493.7664 |
| ENSMUSG00000022602 | -0.631008644 | 0.101221 | 0.222928 | 24.1453 | 14.39786 | 18.21835 | 16.38985 | 25.35289 | 18.99536 | 40.50014 | 28.27737 |
| ENSMUSG00000022604 | 0.431346437 | 0.002969 | 0.017618 | 715.3046 | 781.0838 | 543.238 | 640.9293 | 516.809 | 547.5133 | 506.2518 | 414.3722 |
| ENSMUSG00000022607 | -0.186033287 | 0.001041 | 0.008061 | 2138.872 | 2086.49 | 2270.668 | 2153.108 | 2440.704 | 2358.777 | 2485.937 | 2559.102 |
| ENSMUSG00000022610 | -0.049556009 | 0.655865 | 0.778183 | 505.0393 | 547.1186 | 501.8326 | 558.98 | 640.6481 | 535.2222 | 510.109 | 499.2043 |
| ENSMUSG00000022613 | -5.937722768 | 2.44E-09 | 2.73E-07 | 4.024217 | 3.599465 | 26.49941 | 18.97772 | 1627.461 | 177.6625 | 183.2149 | 1289.883 |
| ENSMUSG00000022614 | -0.131832716 | 0.071961 | 0.175931 | 1116.72 | 1257.413 | 1264.519 | 1167.992 | 1373.932 | 1264.867 | 1274.79 | 1351.876 |
| ENSMUSG00000022615 | -0.562032074 | 0.009892 | 0.042279 | 111.672 | 86.38715 | 120.0755 | 82.81186 | 168.6943 | 109.5027 | 143.6791 | 170.7518 |
| ENSMUSG00000022617 | 0.090506498 | 0.250274 | 0.413857 | 1429.603 | 1299.407 | 1300.127 | 1496.652 | 1229.615 | 1352.023 | 1340.362 | 1270.306 |
| ENSMUSG00000022619 | 0.664052302 | 0.007378 | 0.034135 | 148.896 | 118.7823 | 95.23226 | 184.6014 | 90.68535 | 72.62932 | 99.32178 | 82.65692 |
| ENSMUSG00000022620 | -0.111351376 | 0.492927 | 0.64992 | 358.1554 | 385.1427 | 516.7385 | 374.3786 | 426.1236 | 368.7335 | 513.9661 | 457.8758 |
| ENSMUSG00000022621 | -0.008127043 | 0.9327 | 0.960099 | 747.4984 | 778.6842 | 815.685 | 781.5369 | 904.9033 | 735.2322 | 772.3956 | 727.5984 |
| ENSMUSG00000022622 | 0.388862889 | 0.374064 | 0.542874 | 30.18163 | 15.59768 | 19.04645 | 20.70296 | 12.67645 | 25.6996 | 16.39292 | 10.87591 |
| ENSMUSG00000022623 | 0.09915927 | 0.453913 | 0.616123 | 513.0877 | 445.1338 | 538.2693 | 508.9479 | 377.3681 | 460.3581 | 523.609 | 513.343 |
| ENSMUSG00000022629 | -0.863192189 | 1.82E-05 | 0.000365 | 896.3944 | 586.7127 | 815.685 | 742.7188 | 1272.52 | 950.8854 | 1611.327 | 1699.905 |
| ENSMUSG00000022634 | -0.079031063 | 0.281165 | 0.447574 | 1101.63 | 1142.23 | 1187.505 | 1286.172 | 1251.068 | 1287.215 | 1242.004 | 1207.226 |
| ENSMUSG00000022635 | 0.136402697 | 0.226884 | 0.386817 | 1722.365 | 1863.323 | 1500.529 | 1707.132 | 1469.493 | 1833.611 | 1451.255 | 1424.744 |
| ENSMUSG00000022636 | 0.738530779 | 0.001188 | 0.008902 | 12117.92 | 16150.8 | 10939.29 | 13665.68 | 7235.326 | 12171.56 | 6627.559 | 5655.474 |
| ENSMUSG00000022637 | 0.040161163 | 0.852355 | 0.911279 | 707.2562 | 609.5093 | 672.4226 | 775.4985 | 540.2117 | 1004.519 | 642.2166 | 503.5547 |
| ENSMUSG00000022639 | 0.706323555 | 0.001164 | 0.008778 | 515.0998 | 541.1195 | 317.1648 | 505.4974 | 290.5832 | 376.5551 | 256.5009 | 227.3065 |
| ENSMUSG00000022641 | 0.35931676 | 0.037018 | 0.110565 | 2235.453 | 1977.306 | 1801.132 | 1958.155 | 1488.02 | 2144.241 | 1336.505 | 1246.379 |
| ENSMUSG00000022651 | -0.376924177 | 0.413195 | 0.579038 | 25.15136 | 9.598572 | 33.95237 | 35.36756 | 26.32801 | 23.46486 | 35.6787 | 51.11678 |
| ENSMUSG00000022656 | 0.077927765 | 0.352699 | 0.521546 | 1797.819 | 1761.338 | 1940.254 | 1952.117 | 1593.332 | 1845.902 | 1706.792 | 1919.598 |
| ENSMUSG00000022658 | 1.545818987 | 0.000126 | 0.0016 | 184.1079 | 93.58608 | 93.57605 | 207.0296 | 52.65601 | 48.04709 | 74.25027 | 22.83941 |
| ENSMUSG00000022661 | -0.138778343 | 0.434643 | 0.598818 | 495.9848 | 434.3354 | 466.224 | 600.386 | 419.2979 | 489.4099 | 660.5381 | 630.8028 |
| ENSMUSG00000022663 | -0.322228563 | 8.41E-05 | 0.00118 | 832.007 | 796.6815 | 838.0439 | 841.0579 | 1070.672 | 940.829 | 997.075 | 1128.92 |
| ENSMUSG00000022664 | 0.021871614 | 0.846194 | 0.907479 | 556.3481 | 562.7163 | 467.0521 | 485.657 | 506.0828 | 451.4191 | 515.8947 | 563.3722 |
| ENSMUSG00000022665 | 0.211319927 | 0.037131 | 0.110724 | 2750.553 | 3629.46 | 3415.94 | 3229.662 | 2787.843 | 3068.309 | 2671.081 | 2723.328 |
| ENSMUSG00000022667 | 0.240730548 | 0.683115 | 0.797997 | 13.07871 | 13.19804 | 15.73403 | 6.038364 | 12.67645 | 3.352122 | 14.46434 | 9.78832 |
| ENSMUSG00000022668 | -0.084470495 | 0.372517 | 0.541523 | 587.5357 | 683.8983 | 684.8442 | 685.7857 | 730.3584 | 749.758 | 644.1452 | 678.6568 |
| ENSMUSG00000022671 | -0.104498192 | 0.243522 | 0.40608 | 642.8687 | 692.297 | 616.9394 | 697.8624 | 731.3335 | 657.016 | 769.5028 | 687.3576 |
| ENSMUSG00000022672 | -0.189088078 | 0.14946 | 0.289418 | 464.7971 | 423.537 | 419.8501 | 388.1806 | 389.0694 | 547.5133 | 488.8946 | 508.9926 |
| ENSMUSG00000022673 | -0.048911596 | 0.607419 | 0.743305 | 1884.34 | 2420.04 | 1901.333 | 2041.83 | 2155.971 | 2156.532 | 2160.008 | 2053.372 |
| ENSMUSG00000022674 | -0.05467202 | 0.486395 | 0.644476 | 581.4994 | 596.3113 | 595.4087 | 540.0023 | 625.0463 | 596.6778 | 600.7521 | 577.5109 |
| ENSMUSG00000022676 | 0.311971981 | 0.000961 | 0.007604 | 1062.393 | 1033.046 | 976.3377 | 1143.839 | 751.8108 | 842.5001 | 918.9676 | 883.124 |
| ENSMUSG00000022677 | 0.032111086 | 0.692632 | 0.805496 | 1227.386 | 1444.585 | 1238.019 | 1353.456 | 1292.022 | 1358.727 | 1222.719 | 1271.394 |
| ENSMUSG00000022678 | -0.006780747 | 0.930161 | 0.958785 | 2615.741 | 2695.999 | 2531.522 | 2771.609 | 2969.214 | 2467.162 | 2650.831 | 2574.328 |
| ENSMUSG00000022679 | -1.557217989 | 7.81E-10 | 1.04E-07 | 201.2109 | 158.3764 | 174.7305 | 168.2116 | 477.8045 | 287.1651 | 676.931 | 625.3649 |
| ENSMUSG00000022680 | -0.136614774 | 0.013615 | 0.05367 | 2426.603 | 2571.218 | 2475.211 | 2435.186 | 2776.142 | 2557.669 | 2767.51 | 2785.321 |
| ENSMUSG00000022681 | 0.106192003 | 0.167812 | 0.313825 | 1325.98 | 1479.38 | 1440.077 | 1469.048 | 1365.156 | 1426.887 | 1230.433 | 1287.708 |
| ENSMUSG00000022682 | -0.094198652 | 0.165393 | 0.31075 | 1598.62 | 1460.183 | 1551.872 | 1623.457 | 1683.042 | 1712.934 | 1549.613 | 1715.131 |
| ENSMUSG00000022684 | 0.299594647 | 0.00365 | 0.020442 | 1650.935 | 1716.945 | 1406.953 | 1607.068 | 1202.312 | 1491.694 | 1245.862 | 1244.204 |
| ENSMUSG00000022685 | -0.106911977 | 0.188085 | 0.34039 | 1403.446 | 1468.582 | 1355.611 | 1318.089 | 1572.855 | 1345.318 | 1456.077 | 1593.321 |
| ENSMUSG00000022686 | 0.136165406 | 0.514223 | 0.668674 | 259.562 | 277.1588 | 332.0708 | 275.1769 | 253.5289 | 367.6161 | 227.5722 | 193.5912 |
| ENSMUSG00000022687 | 0.68506338 | 1.54E-07 | 8.06E-06 | 2096.617 | 2494.429 | 2098.422 | 2354.1 | 1518.248 | 1634.718 | 1352.898 | 1118.044 |
| ENSMUSG00000022696 | -0.05012262 | 0.890339 | 0.933695 | 76.46013 | 35.99465 | 85.29498 | 66.42201 | 99.46135 | 39.10809 | 61.71451 | 73.95619 |
| ENSMUSG00000022698 | -0.082222786 | 0.229073 | 0.389155 | 3034.26 | 3361.9 | 3145.149 | 3112.346 | 3231.519 | 3552.132 | 3129.118 | 3482.467 |
| ENSMUSG00000022701 | 0.305189881 | 0.096769 | 0.215853 | 695.1836 | 563.9161 | 525.8477 | 577.9577 | 416.3725 | 653.6638 | 459.0016 | 383.9197 |
| ENSMUSG00000022702 | -0.066534918 | 0.439574 | 0.602911 | 897.4005 | 1093.037 | 1025.196 | 919.5566 | 1028.742 | 1023.515 | 1009.611 | 1056.051 |
| ENSMUSG00000022704 | 0.198510192 | 0.048547 | 0.133411 | 1050.321 | 980.2542 | 823.138 | 1004.956 | 882.4757 | 872.6692 | 816.7529 | 788.5035 |
| ENSMUSG00000022706 | -0.19417472 | 0.040931 | 0.118646 | 687.1351 | 777.4844 | 838.872 | 758.2461 | 862.9735 | 808.9788 | 876.5389 | 955.9926 |
| ENSMUSG00000022707 | -0.137792854 | 0.162306 | 0.30651 | 696.1896 | 699.496 | 708.8593 | 711.6644 | 814.2179 | 652.5465 | 786.86 | 843.9707 |

| | | | | | | | | | | | |
|---|---|---|---|---|---|---|---|---|---|---|---|
| ENSMUSG00000022708 | 0.954910609 | 0.018562 | 0.067001 | 5791.855 | 6030.303 | 3695.012 | 4969.574 | 2111.116 | 4914.211 | 2006.686 | 1536.766 |
| ENSMUSG00000022710 | 0.039535637 | 0.552232 | 0.699721 | 5095.665 | 5009.255 | 4726.005 | 4948.008 | 4499.164 | 5258.362 | 4736.588 | 4750.598 |
| ENSMUSG00000022711 | -0.136030658 | 0.193497 | 0.346682 | 1099.617 | 1005.45 | 1075.711 | 959.2373 | 1135.03 | 975.4676 | 1146.54 | 1292.058 |
| ENSMUSG00000022718 | 0.082062713 | 0.14303 | 0.280497 | 1610.693 | 1725.343 | 1644.62 | 1677.803 | 1623.56 | 1516.277 | 1584.327 | 1561.781 |
| ENSMUSG00000022721 | 0.006385809 | 0.924767 | 0.955253 | 1263.604 | 1298.207 | 1181.708 | 1239.59 | 1308.599 | 1238.05 | 1170.647 | 1239.854 |
| ENSMUSG00000022722 | -0.129989619 | 0.23922 | 0.40093 | 477.8758 | 611.909 | 583.8152 | 516.7115 | 548.0126 | 621.26 | 634.5023 | 591.6496 |
| ENSMUSG00000022723 | 0.557536231 | 0.000577 | 0.00512 | 1421.555 | 1508.176 | 1136.99 | 1692.467 | 923.4304 | 1207.881 | 984.5392 | 797.2043 |
| ENSMUSG00000022724 | -0.334493218 | 0.011512 | 0.047468 | 445.6821 | 477.529 | 442.2089 | 395.0816 | 611.3948 | 449.1844 | 534.2162 | 622.1021 |
| ENSMUSG00000022742 | -1.020684549 | 2.03E-05 | 0.000397 | 518.118 | 423.537 | 591.2681 | 479.6187 | 1100.901 | 597.7951 | 951.7534 | 1434.533 |
| ENSMUSG00000022744 | -0.063327257 | 0.470829 | 0.630643 | 1491.979 | 1376.195 | 1432.624 | 1672.627 | 1544.576 | 1440.295 | 1645.077 | 1613.985 |
| ENSMUSG00000022747 | -0.114678059 | 0.205547 | 0.361948 | 1620.754 | 1502.177 | 1549.388 | 1651.924 | 1709.37 | 1496.164 | 1913.15 | 1728.182 |
| ENSMUSG00000022748 | -0.142231927 | 0.083134 | 0.194017 | 709.2683 | 653.9027 | 645.9232 | 701.3129 | 790.8153 | 714.002 | 780.1099 | 704.759 |
| ENSMUSG00000022749 | 0.490642944 | 3.84E-05 | 0.00065 | 1451.736 | 1618.559 | 1361.407 | 1608.793 | 1024.842 | 1299.506 | 1013.468 | 961.4305 |
| ENSMUSG00000022750 | -0.042303855 | 0.545863 | 0.694406 | 1690.171 | 1906.516 | 1693.478 | 1719.209 | 1914.143 | 1750.925 | 1744.399 | 1802.138 |
| ENSMUSG00000022751 | -0.302687762 | 0.012515 | 0.050565 | 1093.581 | 1162.627 | 1315.033 | 1376.747 | 1706.445 | 1286.098 | 1478.255 | 1634.649 |
| ENSMUSG00000022752 | -0.260069358 | 2.00E-06 | 6.44E-05 | 2633.85 | 2682.801 | 2529.866 | 2531.8 | 3155.46 | 3068.309 | 2963.261 | 3236.671 |
| ENSMUSG00000022754 | 0.46000669 | 0.000265 | 0.002824 | 386.3249 | 447.5334 | 435.5841 | 496.0085 | 273.0312 | 338.5644 | 340.3941 | 332.8029 |
| ENSMUSG00000022756 | -0.235082494 | 0.331574 | 0.499409 | 150.9082 | 133.1802 | 186.324 | 123.3552 | 145.2916 | 126.2633 | 189.965 | 238.1824 |
| ENSMUSG00000022757 | -0.203538586 | 0.015193 | 0.058236 | 1734.438 | 1700.147 | 1826.803 | 1847.74 | 2082.838 | 1788.916 | 2111.793 | 2204.547 |
| ENSMUSG00000022758 | 0.238178169 | 0.419623 | 0.585157 | 35.2119 | 55.19179 | 59.62368 | 55.2079 | 39.97956 | 60.3382 | 39.53586 | 34.80291 |
| ENSMUSG00000022759 | -0.106372881 | 0.848686 | 0.909191 | 14.08476 | 9.598572 | 13.24971 | 10.35148 | 15.60178 | 22.34748 | 6.750024 | 6.525547 |
| ENSMUSG00000022760 | -0.389128168 | 0.008318 | 0.037341 | 338.0343 | 491.9268 | 458.7711 | 407.1583 | 627.9717 | 479.3535 | 535.1805 | 576.4233 |
| ENSMUSG00000022761 | -0.028447339 | 0.766018 | 0.854851 | 2151.95 | 2319.255 | 2266.528 | 2461.065 | 2539.19 | 1995.63 | 2356.723 | 2489.496 |
| ENSMUSG00000022762 | 0.18644832 | 0.736908 | 0.835233 | 29.17558 | 13.19804 | 11.59349 | 35.36756 | 17.552 | 35.75597 | 11.57147 | 14.13868 |
| ENSMUSG00000022763 | -1.141427363 | 1.92E-05 | 0.00038 | 32.19374 | 25.19625 | 33.12427 | 28.46658 | 78.98402 | 71.51194 | 49.17875 | 64.16787 |
| ENSMUSG00000022765 | 0.027729669 | 0.829442 | 0.896882 | 677.0746 | 658.702 | 682.3599 | 636.6161 | 691.3539 | 502.8183 | 667.2881 | 741.7371 |
| ENSMUSG00000022768 | 0.213287211 | 0.565797 | 0.710316 | 36.21796 | 23.99643 | 38.92101 | 31.91707 | 40.95467 | 15.64324 | 25.07152 | 31.54014 |
| ENSMUSG00000022769 | -0.438266903 | 0.000287 | 0.003001 | 330.9919 | 358.7466 | 279.0719 | 289.8415 | 435.8748 | 382.1419 | 430.073 | 451.3503 |
| ENSMUSG00000022770 | -0.008925145 | 0.905156 | 0.943178 | 3035.266 | 2950.361 | 3173.305 | 2874.261 | 2814.171 | 3308.545 | 3107.904 | 2879.941 |
| ENSMUSG00000022771 | 0.029811285 | 0.554073 | 0.701081 | 2213.32 | 2300.058 | 2289.715 | 2285.09 | 2271.034 | 2167.706 | 2167.722 | 2295.905 |
| ENSMUSG00000022772 | -0.281214688 | 0.046598 | 0.129599 | 425.561 | 343.149 | 391.6944 | 382.1422 | 536.3112 | 368.7335 | 471.5374 | 498.1167 |
| ENSMUSG00000022773 | 0.412771694 | 0.002881 | 0.017226 | 1182.114 | 1361.797 | 1069.914 | 1320.677 | 1020.942 | 1074.914 | 850.503 | 759.1386 |
| ENSMUSG00000022774 | 0.152251265 | 0.019579 | 0.069438 | 2760.613 | 2994.755 | 2654.082 | 2936.37 | 2535.289 | 2690.637 | 2511.973 | 2468.832 |
| ENSMUSG00000022779 | -0.139703484 | 0.117299 | 0.245573 | 1703.25 | 1714.545 | 1640.479 | 1628.633 | 1802.981 | 1600.08 | 2049.114 | 1910.898 |
| ENSMUSG00000022780 | 1.141599418 | 0.000409 | 0.003927 | 111.672 | 169.1748 | 147.403 | 90.57547 | 51.6809 | 87.15518 | 34.71441 | 61.99269 |
| ENSMUSG00000022781 | 0.009653173 | 0.885884 | 0.93183 | 2526.202 | 2506.427 | 2312.902 | 2392.918 | 2240.806 | 2391.181 | 2484.973 | 2553.664 |
| ENSMUSG00000022783 | 0.187025668 | 0.7428 | 0.839394 | 21.12714 | 14.39786 | 25.67131 | 6.900988 | 23.40267 | 15.64324 | 6.750024 | 14.13868 |
| ENSMUSG00000022787 | 0.00569253 | 0.965545 | 0.980631 | 323.9495 | 433.1356 | 322.1335 | 333.8353 | 331.5378 | 376.5551 | 359.6799 | 336.0656 |
| ENSMUSG00000022788 | 0.260083164 | 0.064747 | 0.163784 | 648.9051 | 782.2836 | 664.9696 | 659.0443 | 470.0037 | 698.3588 | 533.2519 | 599.2627 |
| ENSMUSG00000022789 | 0.078480348 | 0.17714 | 0.326146 | 2995.024 | 2879.572 | 2844.546 | 3019.182 | 2691.307 | 2896.234 | 2648.902 | 2884.292 |
| ENSMUSG00000022790 | -1.298935786 | 6.99E-05 | 0.00103 | 279.6831 | 89.98662 | 229.3855 | 196.6782 | 571.4152 | 320.6864 | 436.823 | 631.8904 |
| ENSMUSG00000022791 | -0.006156199 | 0.950347 | 0.971777 | 1094.587 | 1111.035 | 1384.594 | 1137.8 | 1234.491 | 1173.243 | 1124.361 | 1220.277 |
| ENSMUSG00000022792 | -0.301196583 | 0.019228 | 0.068535 | 437.6336 | 383.9429 | 471.1927 | 376.1038 | 485.6054 | 472.6492 | 494.6803 | 605.7882 |
| ENSMUSG00000022793 | -0.106665593 | 0.214832 | 0.372961 | 645.8869 | 675.4995 | 647.5794 | 613.3253 | 637.7228 | 758.697 | 676.931 | 706.9342 |
| ENSMUSG00000022797 | -0.295022913 | 0.027594 | 0.089113 | 1699.226 | 1439.786 | 1479.827 | 1610.518 | 1954.123 | 1688.352 | 1628.684 | 2373.124 |
| ENSMUSG00000022799 | 0.022421432 | 0.803763 | 0.880109 | 828.9888 | 761.8867 | 835.5596 | 784.1248 | 709.881 | 860.3781 | 761.7884 | 833.0948 |
| ENSMUSG00000022800 | -0.019562626 | 0.761734 | 0.851959 | 2478.918 | 2584.416 | 2337.745 | 2631.864 | 2648.402 | 2600.13 | 2462.795 | 2455.781 |
| ENSMUSG00000022801 | 0.328647968 | 0.008671 | 0.038451 | 2018.145 | 1790.134 | 1803.616 | 2118.603 | 1419.762 | 1854.841 | 1577.577 | 1305.109 |
| ENSMUSG00000022802 | -0.163309359 | 0.30133 | 0.469688 | 244.4712 | 172.7743 | 183.8397 | 182.8762 | 206.7236 | 250.2918 | 206.3579 | 215.343 |
| ENSMUSG00000022803 | 0.551654874 | 0.27319 | 0.438854 | 21.12714 | 47.99286 | 25.67131 | 33.64232 | 30.22845 | 30.1691 | 20.25007 | 6.525547 |
| ENSMUSG00000022805 | 0.063037191 | 0.866212 | 0.919627 | 16.09687 | 23.99643 | 21.53077 | 24.15346 | 16.57689 | 16.76061 | 28.92867 | 19.57664 |
| ENSMUSG00000022807 | -0.229492669 | 0.014695 | 0.056798 | 878.2855 | 914.264 | 881.9336 | 915.2435 | 1052.145 | 898.3688 | 1108.933 | 1147.409 |
| ENSMUSG00000022808 | -0.039252706 | 0.511646 | 0.667063 | 2686.165 | 2546.021 | 2696.315 | 2794.9 | 2824.897 | 2584.486 | 2837.903 | 2774.445 |
| ENSMUSG00000022809 | -0.823892977 | 0.141598 | 0.278458 | 33.19979 | 4.799286 | 17.39024 | 8.626235 | 25.35289 | 24.58223 | 27.96439 | 35.89051 |
| ENSMUSG00000022811 | 0.190227523 | 0.080806 | 0.190273 | 2232.435 | 2502.828 | 2245.825 | 2457.614 | 1961.924 | 2490.627 | 1867.828 | 1953.314 |
| ENSMUSG00000022812 | 0.25072701 | 0.049853 | 0.135799 | 2953.776 | 2944.362 | 2897.545 | 3190.844 | 2254.457 | 3218.037 | 2267.044 | 2337.233 |
| ENSMUSG00000022814 | -0.291638021 | 0.007178 | 0.033494 | 1278.695 | 1347.4 | 1356.439 | 1166.267 | 1635.262 | 1366.549 | 1485.97 | 1813.014 |
| ENSMUSG00000022816 | 0.20655076 | 0.004376 | 0.023353 | 35472.47 | 37318.05 | 39838.55 | 37161.82 | 29681.41 | 34796.15 | 33882.23 | 31453.13 |
| ENSMUSG00000022817 | -0.083013332 | 0.4675 | 0.62788 | 3056.393 | 2650.406 | 3052.401 | 2825.955 | 2872.678 | 2586.721 | 3234.226 | 3579.262 |
| ENSMUSG00000022820 | -0.464290648 | 8.20E-05 | 0.001159 | 701.2199 | 613.1088 | 808.2321 | 685.7857 | 927.3309 | 857.0259 | 1021.182 | 1073.452 |
| ENSMUSG00000022821 | -3.49123003 | 2.86E-07 | 1.33E-05 | 75.45408 | 10.79839 | 63.76421 | 41.40593 | 620.1708 | 183.2494 | 435.8587 | 917.9269 |
| ENSMUSG00000022822 | 0.193169667 | 0.001418 | 0.010173 | 2284.749 | 2061.293 | 2206.076 | 2207.454 | 1872.214 | 1946.466 | 1997.043 | 1848.905 |
| ENSMUSG00000022827 | 0.001680038 | 0.988829 | 0.993729 | 441.6579 | 410.339 | 458.7711 | 497.7338 | 415.3974 | 521.8137 | 466.716 | 405.6715 |
| ENSMUSG00000022828 | -0.088805044 | 0.397804 | 0.56502 | 609.6689 | 647.9036 | 579.6746 | 596.9355 | 692.329 | 577.6824 | 593.0378 | 723.2481 |

| | | | | | | | | | | | |
|---|---|---|---|---|---|---|---|---|---|---|---|
| ENSMUSG00000022829 | -1.511696289 | 0.002873 | 0.017193 | 9.054489 | 3.599465 | 11.59349 | 6.900988 | 18.52711 | 18.99536 | 18.32149 | 34.80291 |
| ENSMUSG00000022831 | 0.034136294 | 0.86446 | 0.918772 | 167.005 | 149.9777 | 180.5272 | 143.1955 | 136.5156 | 116.2069 | 187.0721 | 185.9781 |
| ENSMUSG00000022833 | 0.224239943 | 0.126288 | 0.257145 | 455.7426 | 473.9295 | 408.2566 | 452.0147 | 378.3432 | 458.1234 | 401.1443 | 293.6496 |
| ENSMUSG00000022836 | 0.219558748 | 0.049648 | 0.13543 | 6353.233 | 7659.661 | 8698.432 | 7139.072 | 6471.814 | 7089.739 | 5838.771 | 6238.422 |
| ENSMUSG00000022837 | 0.275505346 | 0.119835 | 0.248762 | 968.8303 | 958.6574 | 683.188 | 925.595 | 681.6028 | 945.2985 | 717.4311 | 576.4233 |
| ENSMUSG00000022838 | -1.149652862 | 0.000518 | 0.004709 | 89.53884 | 38.39429 | 102.6852 | 77.63611 | 151.1423 | 115.0895 | 179.3578 | 241.4452 |
| ENSMUSG00000022840 | 0.323592787 | 0.090726 | 0.206365 | 220.3259 | 268.76 | 330.4145 | 385.5927 | 211.5992 | 241.3528 | 247.8223 | 264.2846 |
| ENSMUSG00000022841 | -0.421817636 | 0.002613 | 0.015928 | 5485.008 | 5492.783 | 5894.463 | 5399.16 | 7804.791 | 5412.56 | 7867.635 | 8749.67 |
| ENSMUSG00000022842 | -0.323998776 | 0.003907 | 0.021472 | 325.9616 | 374.3443 | 323.7897 | 326.9343 | 457.3272 | 365.3813 | 431.0373 | 432.8613 |
| ENSMUSG00000022843 | 0.496868898 | 0.000544 | 0.004893 | 1229.398 | 1581.365 | 1144.443 | 1304.287 | 1077.498 | 1019.045 | 806.1457 | 822.2189 |
| ENSMUSG00000022844 | -0.4274662 | 0.013256 | 0.052784 | 443.67 | 436.735 | 605.3459 | 508.0852 | 575.3157 | 555.3349 | 716.4668 | 837.4451 |
| ENSMUSG00000022847 | 1.044031186 | 0.024013 | 0.080612 | 31.18769 | 43.19358 | 28.98373 | 38.81806 | 11.70134 | 32.40385 | 17.3572 | 7.613138 |
| ENSMUSG00000022848 | -0.120068287 | 0.295518 | 0.463271 | 791.7648 | 857.8724 | 1006.978 | 1017.033 | 957.5593 | 948.6506 | 1123.397 | 965.7809 |
| ENSMUSG00000022849 | -0.15876678 | 0.197255 | 0.351608 | 214.2896 | 197.9706 | 195.4332 | 177.7004 | 203.7983 | 231.2964 | 227.5722 | 213.1679 |
| ENSMUSG00000022853 | -1.949806543 | 1.70E-07 | 8.72E-06 | 170.0232 | 111.5834 | 207.0267 | 158.7227 | 767.4126 | 253.6439 | 464.7874 | 1017.985 |
| ENSMUSG00000022855 | 0.105046929 | 0.33882 | 0.507282 | 1737.456 | 1505.776 | 1371.345 | 1636.397 | 1470.468 | 1645.892 | 1399.184 | 1296.409 |
| ENSMUSG00000022856 | 0.184262841 | 0.117716 | 0.246326 | 314.895 | 305.9545 | 356.0859 | 293.292 | 262.3049 | 282.6956 | 265.1795 | 309.9635 |
| ENSMUSG00000022858 | -0.061918529 | 0.390103 | 0.557515 | 6200.313 | 6980.562 | 5970.649 | 6784.534 | 6717.542 | 7115.438 | 6461.702 | 6775.692 |
| ENSMUSG00000022860 | 1.871222642 | 7.90E-05 | 0.001131 | 86.52067 | 209.9688 | 122.5598 | 204.4418 | 35.10401 | 82.68568 | 37.60728 | 15.22628 |
| ENSMUSG00000022861 | 0.087664383 | 0.824628 | 0.893455 | 37.22401 | 19.19714 | 18.21835 | 25.8787 | 18.52711 | 22.34748 | 20.25007 | 33.71532 |
| ENSMUSG00000022863 | 0.336976128 | 0.045427 | 0.12732 | 629.79 | 443.934 | 455.4586 | 554.6669 | 332.513 | 493.8794 | 432.0015 | 392.6204 |
| ENSMUSG00000022864 | 0.02815952 | 0.836031 | 0.900957 | 230.3864 | 218.3675 | 193.777 | 188.9145 | 226.2258 | 204.4795 | 201.5364 | 180.5401 |
| ENSMUSG00000022865 | 0.140424802 | 0.117994 | 0.246712 | 1735.444 | 1700.147 | 1728.259 | 1767.516 | 1347.604 | 1737.517 | 1570.827 | 1635.737 |
| ENSMUSG00000022867 | 0.047716009 | 0.550423 | 0.698381 | 1653.953 | 1426.588 | 1583.34 | 1645.023 | 1420.737 | 1488.342 | 1607.47 | 1591.146 |
| ENSMUSG00000022868 | 3.439510106 | 0.00744 | 0.034372 | 990.9635 | 16.7975 | 15.73403 | 916.1062 | 12.67645 | 41.34284 | 70.39311 | 54.37955 |
| ENSMUSG00000022875 | 1.47312742 | 0.096515 | 0.215443 | 197.1867 | 3.599465 | 14.90592 | 113.8663 | 24.37778 | 14.52586 | 48.21446 | 31.54014 |
| ENSMUSG00000022876 | 0.111999997 | 0.737296 | 0.835587 | 65.39353 | 52.79215 | 33.95237 | 36.23019 | 49.73068 | 50.28183 | 27.96439 | 45.67883 |
| ENSMUSG00000022881 | -0.086383075 | 0.442197 | 0.605393 | 587.5357 | 755.8876 | 696.4377 | 669.3958 | 778.1388 | 736.3495 | 731.8955 | 627.5401 |
| ENSMUSG00000022883 | 0.409700088 | 0.003237 | 0.018784 | 2930.636 | 2935.963 | 2256.591 | 2639.628 | 1921.944 | 2520.796 | 1898.685 | 1759.722 |
| ENSMUSG00000022884 | 0.220994292 | 0.006483 | 0.031237 | 11776.87 | 11549.48 | 10566.64 | 12336.38 | 9660.428 | 11060.89 | 9313.105 | 9630.619 |
| ENSMUSG00000022885 | 0.368415222 | 8.11E-08 | 4.80E-06 | 1968.848 | 1876.521 | 1758.07 | 1907.261 | 1375.882 | 1529.685 | 1494.648 | 1417.131 |
| ENSMUSG00000022887 | -0.434262007 | 0.141201 | 0.277974 | 147.89 | 99.58519 | 92.74794 | 69.00988 | 158.9431 | 84.92043 | 154.2863 | 153.3503 |
| ENSMUSG00000022889 | -0.18194345 | 0.035399 | 0.106981 | 1184.126 | 1088.238 | 1202.411 | 1210.261 | 1442.19 | 1174.36 | 1340.362 | 1359.489 |
| ENSMUSG00000022890 | -0.363939137 | 0.0004 | 0.003866 | 4420.603 | 4771.69 | 5173.182 | 4757.369 | 6203.658 | 5182.381 | 6253.415 | 6971.459 |
| ENSMUSG00000022892 | -0.069423158 | 0.486253 | 0.64438 | 17201.52 | 18567.24 | 19985.53 | 18980.3 | 18864.5 | 16852.24 | 20387 | 22316.28 |
| ENSMUSG00000022893 | -0.287397964 | 0.161852 | 0.30586 | 2668.056 | 1440.986 | 2549.74 | 2272.15 | 2041.883 | 2669.407 | 3165.761 | 3025.678 |
| ENSMUSG00000022894 | 0.95534059 | 4.69E-09 | 4.58E-07 | 1477.894 | 1678.55 | 1335.736 | 1681.253 | 787.8899 | 1052.566 | 675.0024 | 668.8685 |
| ENSMUSG00000022895 | 0.035696827 | 0.736613 | 0.834974 | 3308.913 | 2875.972 | 3184.07 | 3319.375 | 2864.877 | 3279.493 | 3544.727 | 2690.7 |
| ENSMUSG00000022897 | 0.220137277 | 0.003749 | 0.020871 | 2316.943 | 2144.081 | 2244.997 | 2415.346 | 1807.856 | 2127.48 | 1911.221 | 1989.204 |
| ENSMUSG00000022898 | -0.162483612 | 0.099721 | 0.220467 | 947.7032 | 946.6592 | 924.9951 | 948.8858 | 1094.075 | 878.256 | 1148.468 | 1094.117 |
| ENSMUSG00000022899 | -0.094595712 | 0.738062 | 0.836054 | 772.6497 | 914.264 | 635.9859 | 522.7498 | 418.3228 | 625.7295 | 947.8962 | 1045.175 |
| ENSMUSG00000022900 | -0.938475933 | 0.00023 | 0.002529 | 256.5439 | 260.3613 | 363.5388 | 217.3811 | 462.2028 | 330.7427 | 724.1812 | 587.2992 |
| ENSMUSG00000022901 | -0.070587842 | 0.874346 | 0.924965 | 15.09082 | 26.39607 | 28.15563 | 25.01608 | 11.70134 | 34.6386 | 33.75012 | 19.57664 |
| ENSMUSG00000022902 | 0.312260722 | 0.625757 | 0.755971 | 60.36326 | 10.79839 | 26.49941 | 34.50494 | 37.05423 | 5.58687 | 37.60728 | 26.10219 |
| ENSMUSG00000022905 | -0.067111779 | 0.439513 | 0.602873 | 1096.599 | 1034.246 | 1001.181 | 961.8252 | 964.3851 | 1073.797 | 1082.897 | 1166.985 |
| ENSMUSG00000022906 | 0.454232082 | 0.007731 | 0.035307 | 500.009 | 680.2988 | 778.4202 | 499.459 | 430.9992 | 405.6068 | 473.466 | 483.978 |
| ENSMUSG00000022911 | 0.256837573 | 0.019648 | 0.069614 | 747.4984 | 683.8983 | 583.8152 | 642.6545 | 584.0917 | 528.5179 | 596.895 | 511.1678 |
| ENSMUSG00000022912 | 0.196349164 | 0.065617 | 0.164856 | 1698.22 | 1348.599 | 1618.12 | 1659.688 | 1240.342 | 1339.732 | 1542.863 | 1399.73 |
| ENSMUSG00000022913 | -0.302024509 | 3.59E-05 | 0.000615 | 776.674 | 802.6806 | 751.0927 | 774.6359 | 981.9371 | 887.195 | 945.0034 | 1010.372 |
| ENSMUSG00000022914 | 0.071461582 | 0.501911 | 0.65809 | 2378.312 | 2166.878 | 2058.673 | 2215.217 | 1894.641 | 2491.744 | 2078.043 | 1929.387 |
| ENSMUSG00000022935 | 1.265022336 | 0.000217 | 0.002413 | 152.9203 | 124.7814 | 157.3403 | 210.4801 | 53.63112 | 120.6764 | 45.32159 | 50.02919 |
| ENSMUSG00000022940 | 0.036019032 | 0.663249 | 0.783419 | 1135.835 | 1281.409 | 1295.987 | 1336.204 | 1146.731 | 1312.915 | 1257.433 | 1210.489 |
| ENSMUSG00000022941 | -0.783983207 | 0.002034 | 0.013285 | 113.6841 | 81.58786 | 148.2311 | 107.8279 | 166.744 | 154.1976 | 185.1435 | 274.073 |
| ENSMUSG00000022945 | -0.160822987 | 0.146722 | 0.285743 | 760.5771 | 977.8546 | 830.591 | 746.1693 | 1012.166 | 882.7255 | 896.7889 | 909.2261 |
| ENSMUSG00000022946 | -0.062626707 | 0.713709 | 0.820362 | 505.0393 | 481.1284 | 737.0149 | 576.2325 | 530.4606 | 541.9264 | 729.0026 | 602.5255 |
| ENSMUSG00000022947 | 0.243836297 | 0.157194 | 0.299616 | 251.5136 | 182.3729 | 296.4622 | 239.8093 | 224.2756 | 181.0146 | 208.2865 | 207.7299 |
| ENSMUSG00000022948 | 0.048751124 | 0.659937 | 0.780969 | 370.228 | 377.9438 | 343.6643 | 389.9058 | 366.6419 | 405.6068 | 329.7869 | 330.6277 |
| ENSMUSG00000022949 | -0.378825331 | 0.205091 | 0.361387 | 141.8537 | 88.78679 | 247.6039 | 149.2339 | 191.1218 | 160.9019 | 213.1079 | 253.4087 |
| ENSMUSG00000022951 | -0.809660994 | 1.02E-05 | 0.000231 | 551.3178 | 489.5272 | 638.4702 | 602.9738 | 983.8873 | 687.1851 | 1080.004 | 1251.817 |
| ENSMUSG00000022952 | 0.234302844 | 0.336429 | 0.504685 | 193.1624 | 153.5772 | 168.9338 | 143.1955 | 117.9885 | 210.0663 | 127.2862 | 105.4963 |
| ENSMUSG00000022956 | -0.771426588 | 2.19E-06 | 6.90E-05 | 4448.772 | 4320.557 | 5546.658 | 4683.183 | 7989.087 | 5813.697 | 8624.602 | 10005.84 |
| ENSMUSG00000022957 | 0.27285243 | 0.000722 | 0.006102 | 2281.731 | 2387.645 | 1990.768 | 2313.556 | 1849.786 | 1983.339 | 1801.292 | 1790.175 |
| ENSMUSG00000022960 | 0.120494009 | 0.380832 | 0.548644 | 377.2704 | 464.3309 | 338.6956 | 408.8835 | 364.6916 | 420.1327 | 344.2512 | 330.6277 |
| ENSMUSG00000022961 | 0.321216506 | 0.011175 | 0.04641 | 15265.87 | 15022.97 | 13576.81 | 15327.09 | 11392.23 | 14996.28 | 10747 | 10242.93 |

| | | | | | | | | | | | |
|---|---|---|---|---|---|---|---|---|---|---|---|
| ENSMUSG00000022962 | -0.199849857 | 0.012954 | 0.051884 | 2028.206 | 2043.296 | 1884.771 | 1915.887 | 2308.088 | 1986.691 | 2384.687 | 2356.81 |
| ENSMUSG00000022964 | -0.363035224 | 0.041729 | 0.120224 | 744.4802 | 535.1204 | 855.4342 | 776.3611 | 878.5753 | 753.1101 | 1154.254 | 961.4305 |
| ENSMUSG00000022965 | -0.107391263 | 0.116045 | 0.244043 | 1291.774 | 1262.212 | 1341.533 | 1368.983 | 1410.986 | 1343.084 | 1522.613 | 1396.467 |
| ENSMUSG00000022967 | -0.091534876 | 0.162723 | 0.307141 | 2237.465 | 2333.653 | 2404.822 | 2182.437 | 2621.099 | 2343.133 | 2389.509 | 2402.489 |
| ENSMUSG00000022969 | -0.152153434 | 0.311468 | 0.480077 | 573.451 | 475.1293 | 606.1741 | 503.7721 | 614.3201 | 465.945 | 609.4308 | 710.197 |
| ENSMUSG00000022971 | 0.11876706 | 0.141077 | 0.27779 | 910.4792 | 831.4763 | 818.1694 | 892.8153 | 854.1975 | 767.636 | 751.1813 | 806.9926 |
| ENSMUSG00000022972 | -0.482234674 | 9.57E-06 | 0.000219 | 608.6629 | 548.3184 | 655.0323 | 591.7597 | 914.6544 | 712.8847 | 850.503 | 882.0364 |
| ENSMUSG00000022973 | 0.06171652 | 0.48134 | 0.640445 | 514.0938 | 465.5308 | 467.0521 | 491.6954 | 430.0241 | 472.6492 | 480.216 | 475.2773 |
| ENSMUSG00000022974 | 0.230626742 | 0.101148 | 0.222793 | 2343.101 | 2028.898 | 1774.633 | 2015.088 | 1734.723 | 2140.889 | 1650.863 | 1429.095 |
| ENSMUSG00000022978 | 0.049136358 | 0.631443 | 0.760118 | 675.0625 | 589.1124 | 560.6282 | 565.881 | 562.6392 | 627.9642 | 590.145 | 528.5693 |
| ENSMUSG00000022982 | -0.140057856 | 0.030266 | 0.095064 | 7050.429 | 6509.032 | 6653.009 | 6779.358 | 7718.981 | 6777.991 | 7474.205 | 7771.926 |
| ENSMUSG00000022983 | 0.066969774 | 0.459375 | 0.620842 | 1139.86 | 1053.443 | 928.3075 | 969.5888 | 978.0366 | 940.829 | 946.932 | 1036.474 |
| ENSMUSG00000022986 | 1.115598787 | 0.01456 | 0.056443 | 27.16347 | 33.595 | 32.29616 | 48.30692 | 8.776002 | 27.93435 | 9.642892 | 19.57664 |
| ENSMUSG00000022987 | -0.347894156 | 0.028639 | 0.091448 | 250.5075 | 207.5691 | 289.8373 | 257.9244 | 299.3592 | 267.0524 | 339.4298 | 377.3941 |
| ENSMUSG00000022992 | -0.116627036 | 0.060281 | 0.155535 | 1952.752 | 2002.502 | 1933.629 | 2002.149 | 2272.984 | 2138.654 | 1987.4 | 2155.606 |
| ENSMUSG00000022993 | 0.378014472 | 0.433349 | 0.59757 | 44.26639 | 16.7975 | 31.46805 | 17.25247 | 13.65156 | 27.93435 | 28.92867 | 14.13868 |
| ENSMUSG00000022994 | -0.020298175 | 0.797731 | 0.875947 | 3575.517 | 3129.135 | 3205.601 | 3442.73 | 3243.22 | 3126.413 | 3557.263 | 3616.24 |
| ENSMUSG00000022995 | 0.111983116 | 0.287707 | 0.454656 | 3359.215 | 3373.898 | 3356.316 | 3171.867 | 2777.117 | 3721.973 | 2879.367 | 2894.08 |
| ENSMUSG00000022996 | 0.55619951 | 0.474548 | 0.634014 | 7.04238 | 14.39786 | 11.59349 | 14.6646 | 19.50223 | 7.821619 | 3.857157 | 1.087591 |
| ENSMUSG00000022999 | 0.225094417 | 0.021459 | 0.074029 | 1046.297 | 963.4567 | 934.1043 | 1032.56 | 889.3015 | 926.3031 | 856.2888 | 729.7736 |
| ENSMUSG00000023000 | -0.72283179 | 0.046401 | 0.129251 | 30.18163 | 25.19625 | 44.71776 | 31.05445 | 59.48179 | 39.10809 | 36.64299 | 82.65692 |
| ENSMUSG00000023004 | -0.196537694 | 0.035824 | 0.108 | 14300.06 | 17337.42 | 17938.45 | 16268.22 | 20175.05 | 16961.74 | 19255.89 | 19060.03 |
| ENSMUSG00000023007 | 0.29416345 | 0.002445 | 0.015178 | 1541.275 | 1544.17 | 1324.971 | 1640.71 | 1208.163 | 1375.488 | 1192.826 | 1158.285 |
| ENSMUSG00000023008 | -0.154177661 | 0.135357 | 0.269633 | 2134.847 | 2285.66 | 2782.438 | 2320.457 | 2621.099 | 2404.589 | 2844.653 | 2729.854 |
| ENSMUSG00000023009 | 0.269225793 | 0.008007 | 0.036248 | 649.9111 | 580.7136 | 627.7048 | 586.584 | 548.0126 | 509.5226 | 439.7159 | 532.9196 |
| ENSMUSG00000023010 | -0.613704504 | 3.31E-05 | 0.000577 | 9480.05 | 8471.94 | 11509.85 | 8957.482 | 14280.51 | 11456.44 | 15474.91 | 17579.82 |
| ENSMUSG00000023011 | 0.316279882 | 0.336228 | 0.504507 | 78.47224 | 56.39161 | 52.99882 | 107.8279 | 56.55646 | 86.03781 | 51.10733 | 44.59123 |
| ENSMUSG00000023013 | -1.433176993 | 0.151709 | 0.292126 | 1077.484 | 533.9206 | 12160.75 | 1550.997 | 7266.529 | 5257.245 | 18921.28 | 9935.145 |
| ENSMUSG00000023015 | -0.041536295 | 0.579576 | 0.72075 | 2942.709 | 2890.37 | 2772.501 | 3005.38 | 3297.826 | 2882.825 | 3019.189 | 2747.255 |
| ENSMUSG00000023017 | -0.11092258 | 0.473136 | 0.632492 | 184.1079 | 154.777 | 170.59 | 184.6014 | 223.3005 | 158.6671 | 175.5006 | 192.5036 |
| ENSMUSG00000023018 | -0.01244684 | 0.871322 | 0.922824 | 1735.444 | 1700.147 | 1881.458 | 1804.608 | 1983.376 | 1729.695 | 1783.935 | 1687.941 |
| ENSMUSG00000023019 | -1.97574196 | 6.57E-08 | 4.03E-06 | 260.5681 | 167.975 | 334.5551 | 244.1224 | 1024.842 | 427.9543 | 772.3956 | 1740.146 |
| ENSMUSG00000023020 | -0.439009519 | 0.002126 | 0.013714 | 1480.912 | 1261.012 | 1445.046 | 1414.703 | 1943.397 | 1441.413 | 1892.9 | 2317.657 |
| ENSMUSG00000023021 | 0.141789646 | 0.017447 | 0.064118 | 2134.847 | 1988.104 | 2065.298 | 2161.734 | 1860.512 | 2002.334 | 1866.864 | 1843.467 |
| ENSMUSG00000023022 | 0.078405281 | 0.527994 | 0.680556 | 2885.364 | 2811.182 | 3392.753 | 2926.882 | 2792.719 | 2290.617 | 2999.904 | 3298.664 |
| ENSMUSG00000023025 | -0.150149725 | 0.040742 | 0.118267 | 1648.923 | 1463.782 | 1415.234 | 1453.521 | 1579.68 | 1710.7 | 1670.149 | 1675.978 |
| ENSMUSG00000023026 | 0.118217292 | 0.264707 | 0.429037 | 1122.757 | 1007.85 | 1108.835 | 1034.286 | 907.8286 | 1159.834 | 982.6107 | 889.6495 |
| ENSMUSG00000023027 | 0.03266309 | 0.698648 | 0.809838 | 1534.233 | 1569.367 | 1396.188 | 1509.591 | 1517.273 | 1617.958 | 1376.041 | 1361.664 |
| ENSMUSG00000023030 | -0.615793788 | 0.000315 | 0.003228 | 1137.847 | 927.462 | 1068.258 | 953.199 | 1632.336 | 1102.848 | 1550.577 | 1977.241 |
| ENSMUSG00000023031 | -0.710782169 | 0.173604 | 0.321497 | 333.004 | 46.79304 | 148.2311 | 136.2945 | 409.5467 | 127.3806 | 183.2149 | 367.6058 |
| ENSMUSG00000023032 | 0.122683986 | 0.291818 | 0.459069 | 657.9595 | 571.1151 | 549.8628 | 508.9479 | 517.7841 | 505.0531 | 486.0017 | 591.6496 |
| ENSMUSG00000023033 | 0.474003788 | 0.019734 | 0.0698 | 1012.091 | 892.6672 | 798.2948 | 800.5146 | 536.3112 | 928.5379 | 509.1447 | 549.2335 |
| ENSMUSG00000023034 | 0.931205025 | 0.157188 | 0.299616 | 1296.804 | 161.9759 | 221.1045 | 458.9157 | 246.7032 | 284.9304 | 310.5011 | 279.5109 |
| ENSMUSG00000023036 | 0.420136923 | 0.050505 | 0.136996 | 90.54489 | 100.785 | 86.12309 | 76.77349 | 60.4569 | 80.45093 | 52.07161 | 71.78101 |
| ENSMUSG00000023039 | -1.058915386 | 0.041686 | 0.120199 | 1653.953 | 1342.6 | 4112.378 | 1489.751 | 4732.215 | 2043.677 | 5124.233 | 6015.466 |
| ENSMUSG00000023043 | -0.196803528 | 0.393358 | 0.560777 | 4039.308 | 2980.357 | 6711.804 | 3854.202 | 5751.206 | 3928.687 | 5020.089 | 5458.62 |
| ENSMUSG00000023044 | -0.525159911 | 0.000661 | 0.005694 | 1449.724 | 1415.789 | 1471.545 | 1381.06 | 1786.404 | 1667.122 | 2029.829 | 2746.167 |
| ENSMUSG00000023045 | 0.604382431 | 0.075869 | 0.182394 | 21.12714 | 34.79482 | 37.2648 | 29.3292 | 16.57689 | 21.23011 | 21.21436 | 21.75182 |
| ENSMUSG00000023046 | 1.38809093 | 0.000964 | 0.007629 | 2826.007 | 3606.664 | 2178.749 | 3216.723 | 1033.618 | 1997.865 | 944.0391 | 543.7955 |
| ENSMUSG00000023047 | -1.443761582 | NA | NA | 18.10898 | 16.7975 | 16.56213 | 16.38985 | 10.72622 | 145.2586 | 11.57147 | 17.40146 |
| ENSMUSG00000023048 | 0.10050767 | 0.298083 | 0.466056 | 948.7093 | 887.8679 | 920.8546 | 860.8982 | 771.313 | 782.1619 | 857.2531 | 964.6933 |
| ENSMUSG00000023050 | 0.238685182 | 0.041378 | 0.119549 | 654.9414 | 621.5076 | 592.0962 | 667.6706 | 581.1663 | 548.6307 | 437.7873 | 582.9488 |
| ENSMUSG00000023051 | -0.135973169 | 0.051812 | 0.139412 | 1017.121 | 1120.633 | 1096.413 | 1110.196 | 1235.466 | 1152.013 | 1139.79 | 1246.379 |
| ENSMUSG00000023052 | 0.812773689 | 0.000727 | 0.006118 | 100.6054 | 80.38804 | 74.5296 | 95.75121 | 42.9049 | 67.04245 | 49.17875 | 41.32846 |
| ENSMUSG00000023055 | 0.396712867 | 2.39E-08 | 1.76E-06 | 3588.596 | 3243.118 | 3224.647 | 3498.801 | 2663.029 | 2500.683 | 2725.081 | 2405.751 |
| ENSMUSG00000023057 | 1.3573964 | 0.541965 | 0.691162 | 15.09082 | 98.38537 | 33.12427 | 0 | 37.05423 | 20.11273 | 0 | 0 |
| ENSMUSG00000023064 | 0.641663317 | 0.199487 | 0.354456 | 92.557 | 27.5959 | 39.74912 | 140.6076 | 42.9049 | 34.6386 | 49.17875 | 66.34306 |
| ENSMUSG00000023066 | 0.336987065 | 0.013371 | 0.053057 | 303.8284 | 380.3434 | 293.9779 | 288.9789 | 248.6534 | 269.2872 | 260.3581 | 221.8686 |
| ENSMUSG00000023067 | -0.463257659 | 0.060834 | 0.156502 | 1088.551 | 959.8572 | 1255.41 | 1224.925 | 1175.984 | 1091.674 | 2508.116 | 1468.248 |
| ENSMUSG00000023068 | -0.393265945 | 0.002115 | 0.013661 | 2081.526 | 1950.91 | 2122.437 | 2134.993 | 2725.436 | 2149.828 | 2701.938 | 3311.715 |
| ENSMUSG00000023070 | 0.360845965 | 0.004184 | 0.022581 | 294.7739 | 291.5566 | 252.5725 | 263.9628 | 210.624 | 240.2354 | 189.965 | 217.5182 |
| ENSMUSG00000023072 | -0.025136312 | 0.720461 | 0.825136 | 1184.126 | 1143.43 | 1047.555 | 1173.168 | 1191.586 | 1175.478 | 1111.825 | 1147.409 |
| ENSMUSG00000023073 | -4.577577047 | 3.35E-08 | 2.33E-06 | 2.012109 | 4.799286 | 2.48432 | 0.862623 | 31.20356 | 10.05637 | 91.60747 | 102.2336 |
| ENSMUSG00000023074 | 0.016616138 | 0.827046 | 0.894992 | 579.4873 | 634.7056 | 578.0184 | 608.1496 | 594.8179 | 586.6214 | 610.395 | 577.5109 |

| | | | | | | | | | | | |
|---|---|---|---|---|---|---|---|---|---|---|---|
| ENSMUSG00000023075 | -0.179861541 | 0.053016 | 0.141745 | 969.8364 | 848.2738 | 823.9661 | 900.5789 | 989.738 | 924.0684 | 986.4678 | 1112.606 |
| ENSMUSG00000023078 | 1.229665392 | 0.00014 | 0.001734 | 50.30272 | 85.18733 | 89.43552 | 56.07053 | 21.45245 | 35.75597 | 26.03581 | 36.9781 |
| ENSMUSG00000023079 | 0.247563578 | 0.000246 | 0.002673 | 1760.595 | 1991.704 | 1892.224 | 1891.733 | 1650.863 | 1630.249 | 1567.934 | 1496.525 |
| ENSMUSG00000023084 | 0.271935745 | 0.559433 | 0.705627 | 16.09687 | 33.595 | 43.88965 | 39.68068 | 33.15378 | 37.99072 | 11.57147 | 28.27737 |
| ENSMUSG00000023087 | -0.71299858 | 0.002017 | 0.013217 | 151.9142 | 139.1793 | 165.6213 | 127.6683 | 183.3209 | 215.6532 | 351.0013 | 207.7299 |
| ENSMUSG00000023088 | -0.359501472 | 0.018339 | 0.066379 | 934.6245 | 881.8688 | 1111.319 | 941.9849 | 1189.636 | 933.0074 | 1432.934 | 1410.606 |
| ENSMUSG00000023089 | -0.395569876 | 0.002479 | 0.015329 | 1445.7 | 1418.189 | 1760.555 | 1429.367 | 1881.965 | 1649.244 | 2087.686 | 2347.022 |
| ENSMUSG00000023092 | -0.121796716 | 0.206192 | 0.362677 | 2136.859 | 1870.522 | 1979.175 | 2385.154 | 2144.27 | 2283.913 | 2215.937 | 2469.919 |
| ENSMUSG00000023094 | 0.006054295 | 0.954821 | 0.9743 | 572.4449 | 521.9224 | 568.0811 | 586.584 | 534.361 | 501.701 | 555.4306 | 650.3795 |
| ENSMUSG00000023104 | -0.154953568 | 0.121846 | 0.251178 | 1657.978 | 2058.894 | 1854.959 | 1584.639 | 2002.879 | 1849.254 | 2035.614 | 2075.124 |
| ENSMUSG00000023106 | -0.052469595 | 0.530658 | 0.682354 | 2911.521 | 3245.517 | 2990.293 | 2963.112 | 3242.245 | 3485.09 | 2860.082 | 2970.211 |
| ENSMUSG00000023110 | -0.207476999 | 0.002854 | 0.017097 | 2709.304 | 2704.398 | 2600.255 | 2418.796 | 3209.091 | 2820.252 | 3043.297 | 2966.948 |
| ENSMUSG00000023118 | -0.089082045 | 0.101641 | 0.223559 | 2480.93 | 2633.608 | 2443.743 | 2434.323 | 2687.407 | 2557.669 | 2736.653 | 2638.496 |
| ENSMUSG00000023120 | -5.4525277 | 2.11E-05 | 0.000411 | 0 | 0 | 10.76539 | 1.725247 | 114.088 | 30.1691 | 212.1436 | 204.4671 |
| ENSMUSG00000023122 | -3.438552607 | 1.92E-06 | 6.28E-05 | 13.07871 | 2.399643 | 34.78048 | 3.450494 | 148.2169 | 46.92971 | 143.6791 | 250.146 |
| ENSMUSG00000023143 | -0.198491048 | 0.263813 | 0.428038 | 319.9253 | 251.9625 | 294.806 | 268.2759 | 396.8703 | 235.7659 | 318.2154 | 351.2919 |
| ENSMUSG00000023147 | -0.134021742 | 0.080933 | 0.190425 | 768.6255 | 778.6842 | 716.3122 | 719.428 | 877.6002 | 793.3356 | 812.8958 | 784.1532 |
| ENSMUSG00000023150 | -0.32681925 | 0.036621 | 0.109808 | 12827.19 | 9653.764 | 14076.16 | 12938.49 | 13054.79 | 13388.38 | 18826.78 | 16813.07 |
| ENSMUSG00000023153 | -2.469702097 | 1.22E-05 | 0.000268 | 11.0666 | 1.199822 | 14.90592 | 6.038364 | 34.1289 | 34.6386 | 45.32159 | 73.95619 |
| ENSMUSG00000023156 | -0.095072471 | 0.302031 | 0.470419 | 1011.085 | 974.2551 | 1004.493 | 910.0678 | 1159.407 | 920.7163 | 1045.289 | 1037.562 |
| ENSMUSG00000023169 | 0.177365437 | 0.00101 | 0.007894 | 3140.902 | 2915.566 | 2903.342 | 2991.578 | 2531.389 | 2685.05 | 2666.26 | 2687.438 |
| ENSMUSG00000023170 | 0.130467449 | 0.046528 | 0.129488 | 1431.615 | 1607.761 | 1580.027 | 1526.844 | 1415.862 | 1440.295 | 1418.469 | 1338.825 |
| ENSMUSG00000023175 | -0.364896895 | 0.000865 | 0.00697 | 10073.62 | 10986.77 | 12445.61 | 11027.78 | 14390.69 | 12028.53 | 14685.16 | 16247.52 |
| ENSMUSG00000023186 | 0.202728187 | 0.019863 | 0.070134 | 1099.617 | 1027.047 | 958.9475 | 1099.845 | 901.0028 | 836.9132 | 907.3961 | 990.7955 |
| ENSMUSG00000023191 | 0.129926742 | 0.194837 | 0.348638 | 1736.45 | 2279.661 | 2114.984 | 2208.316 | 1940.472 | 2009.039 | 1857.221 | 1814.102 |
| ENSMUSG00000023206 | -0.440298327 | 0.050023 | 0.136148 | 78.47224 | 58.79126 | 52.17072 | 59.52102 | 70.20801 | 79.33356 | 87.75031 | 100.0584 |
| ENSMUSG00000023216 | -1.446166419 | 0.016768 | 0.06223 | 31.18769 | 5.999108 | 12.4216 | 13.80198 | 65.33246 | 22.34748 | 20.25007 | 65.25547 |
| ENSMUSG00000023224 | 0.717163089 | 3.41E-08 | 2.35E-06 | 3624.814 | 4547.324 | 4236.594 | 4240.657 | 2449.48 | 3114.122 | 2441.58 | 2122.978 |
| ENSMUSG00000023232 | 0.011279961 | 0.951954 | 0.972676 | 660.9777 | 769.0856 | 1064.117 | 767.7349 | 635.7726 | 725.1758 | 917.039 | 960.3429 |
| ENSMUSG00000023235 | 0.154707875 | 0.3121 | 0.48057 | 169.0171 | 177.5736 | 132.4971 | 149.2339 | 142.3663 | 149.7281 | 132.1076 | 138.1241 |
| ENSMUSG00000023236 | -0.409908919 | 0.167552 | 0.313498 | 386.3249 | 223.1668 | 487.7548 | 366.615 | 367.617 | 311.7474 | 687.5382 | 579.686 |
| ENSMUSG00000023243 | -0.821450747 | 0.000505 | 0.00462 | 267.6105 | 184.7725 | 263.3379 | 200.1287 | 379.3183 | 260.3482 | 441.6444 | 538.3576 |
| ENSMUSG00000023247 | -0.518479384 | 0.440548 | 0.603843 | 9.054489 | 19.19714 | 99.3728 | 26.74133 | 34.1289 | 36.87335 | 58.82164 | 92.44524 |
| ENSMUSG00000023249 | 0.670640772 | 4.23E-05 | 0.000701 | 286.7255 | 260.3613 | 361.8826 | 265.688 | 204.7734 | 162.0192 | 167.7863 | 204.4671 |
| ENSMUSG00000023259 | -0.309842487 | 0.050063 | 0.136196 | 208.2533 | 181.1731 | 177.2148 | 163.0358 | 230.1263 | 182.132 | 226.608 | 264.2846 |
| ENSMUSG00000023262 | -0.672907457 | 0.000368 | 0.003618 | 550.3117 | 508.7243 | 571.3936 | 511.5357 | 893.202 | 565.3913 | 841.8244 | 1114.781 |
| ENSMUSG00000023266 | -0.247498437 | 0.065165 | 0.164392 | 174.0474 | 152.3773 | 167.2775 | 161.3106 | 190.1467 | 177.6625 | 188.0364 | 222.9562 |
| ENSMUSG00000023267 | 0.472893344 | 0.259692 | 0.423699 | 20.12109 | 27.5959 | 24.01509 | 12.07673 | 16.57689 | 17.87799 | 12.53576 | 13.05109 |
| ENSMUSG00000023272 | -0.435214767 | 0.000959 | 0.007599 | 829.9948 | 926.2622 | 927.4794 | 883.3265 | 1055.07 | 1006.754 | 1330.719 | 1430.182 |
| ENSMUSG00000023274 | 0.618836426 | 0.323587 | 0.491571 | 13.07871 | 16.7975 | 18.21835 | 26.74133 | 5.850668 | 29.05173 | 6.750024 | 7.613138 |
| ENSMUSG00000023277 | 0.025435815 | 0.793554 | 0.873442 | 844.0796 | 801.4808 | 751.0927 | 759.9713 | 848.3468 | 688.3024 | 737.6812 | 824.394 |
| ENSMUSG00000023279 | 0.15041453 | 0.6815 | 0.796515 | 20.12109 | 25.19625 | 19.04645 | 18.97772 | 12.67645 | 22.34748 | 20.25007 | 19.57664 |
| ENSMUSG00000023284 | 0.110295042 | 0.475087 | 0.634588 | 178.0716 | 197.9706 | 163.9651 | 141.4703 | 167.7191 | 158.6671 | 142.7148 | 159.8759 |
| ENSMUSG00000023286 | -0.095115091 | 0.14192 | 0.278943 | 1853.152 | 1944.911 | 2076.063 | 2108.252 | 2082.838 | 2166.588 | 2105.043 | 2178.445 |
| ENSMUSG00000023307 | -0.03700507 | 0.471284 | 0.630997 | 2109.696 | 2202.872 | 2115.812 | 2256.623 | 2250.557 | 2261.565 | 2214.008 | 2183.883 |
| ENSMUSG00000023328 | -0.807126307 | 0.094913 | 0.212854 | 111.672 | 52.79215 | 310.54 | 144.9207 | 191.1218 | 151.9629 | 401.1443 | 342.5912 |
| ENSMUSG00000023330 | 0.026873737 | 0.81058 | 0.884531 | 295.78 | 359.9465 | 326.274 | 364.0271 | 321.7867 | 338.5644 | 350.037 | 309.9635 |
| ENSMUSG00000023333 | -1.461732817 | 0.013089 | 0.052309 | 10.06054 | 7.198929 | 7.45296 | 1.725247 | 15.60178 | 11.17374 | 20.25007 | 25.0146 |
| ENSMUSG00000023336 | 0.395216323 | 0.035191 | 0.106518 | 147.89 | 152.3773 | 130.8408 | 191.5024 | 114.088 | 136.3196 | 125.3576 | 97.8832 |
| ENSMUSG00000023341 | 0.43857502 | 0.485824 | 0.644184 | 33.19979 | 57.59143 | 82.81066 | 6.900988 | 31.20356 | 27.93435 | 52.07161 | 21.75182 |
| ENSMUSG00000023345 | -0.106250741 | 0.382377 | 0.550307 | 328.9798 | 301.1552 | 302.2589 | 295.8799 | 313.0107 | 284.9304 | 346.1798 | 377.3941 |
| ENSMUSG00000023348 | -0.129177785 | 0.042455 | 0.12165 | 2063.417 | 2362.449 | 2291.371 | 2191.064 | 2458.256 | 2348.72 | 2484.009 | 2448.168 |
| ENSMUSG00000023349 | 0.305508555 | 0.265204 | 0.429351 | 68.4117 | 43.19358 | 68.73285 | 60.38364 | 50.70579 | 33.52122 | 59.78593 | 51.11678 |
| ENSMUSG00000023353 | -0.040551434 | 0.490766 | 0.648034 | 1439.664 | 1472.181 | 1439.249 | 1542.371 | 1470.468 | 1564.324 | 1483.077 | 1546.555 |
| ENSMUSG00000023367 | -0.454678896 | 0.000726 | 0.006116 | 2144.908 | 1974.906 | 2467.758 | 2153.971 | 2976.04 | 2441.462 | 2884.189 | 3681.496 |
| ENSMUSG00000023393 | 0.589514073 | 0.161502 | 0.305367 | 43.26034 | 27.5959 | 28.98373 | 32.77969 | 20.47734 | 27.93435 | 30.85725 | 8.700729 |
| ENSMUSG00000023403 | -2.860642093 | NA | NA | 0 | 2.399643 | 5.796746 | 2.58787 | 0.975111 | 77.09881 | 0.964289 | 0 |
| ENSMUSG00000023411 | 0.25050526 | 0.000301 | 0.003111 | 1676.087 | 1733.742 | 1735.712 | 1707.132 | 1524.099 | 1495.047 | 1433.898 | 1306.197 |
| ENSMUSG00000023439 | -0.154092429 | 0.315976 | 0.484317 | 127.7689 | 112.7832 | 139.1219 | 144.9207 | 158.9431 | 136.3196 | 155.2506 | 134.8613 |
| ENSMUSG00000023452 | -0.171657645 | 0.01187 | 0.048577 | 788.7466 | 787.0829 | 761.8581 | 817.7671 | 911.7291 | 840.2653 | 906.4318 | 893.9999 |
| ENSMUSG00000023456 | -0.700451851 | 0.000367 | 0.003611 | 6127.877 | 5918.72 | 7751.906 | 6282.487 | 10695.02 | 6766.818 | 10975.54 | 13946.18 |
| ENSMUSG00000023460 | 0.11346068 | 0.127042 | 0.25821 | 2169.053 | 2242.466 | 2215.185 | 2329.083 | 2013.605 | 2306.26 | 2025.007 | 1935.912 |
| ENSMUSG00000023467 | 0.48324764 | 0.107689 | 0.232293 | 42.25428 | 64.79036 | 43.88965 | 43.13117 | 31.20356 | 48.04709 | 30.85725 | 28.27737 |
| ENSMUSG00000023473 | 0.524947127 | 0.090675 | 0.206299 | 50.30272 | 37.19447 | 34.78048 | 64.69676 | 28.27823 | 37.99072 | 33.75012 | 30.45255 |

| | | | | | | | | | | | |
|---|---|---|---|---|---|---|---|---|---|---|---|
| ENSMUSG00000023484 | 3.549787245 | 0.000353 | 0.003511 | 637.8385 | 75.58876 | 58.79557 | 1006.682 | 29.25334 | 69.27719 | 41.46443 | 11.9635 |
| ENSMUSG00000023495 | -0.158393675 | 0.057253 | 0.149817 | 2905.485 | 3037.948 | 3096.291 | 3101.994 | 3882.893 | 3244.854 | 3160.94 | 3261.686 |
| ENSMUSG00000023505 | -0.150091443 | 0.01366 | 0.053809 | 1936.655 | 2003.702 | 1980.831 | 1961.606 | 2355.869 | 2149.828 | 2160.972 | 2077.299 |
| ENSMUSG00000023571 | -0.604924021 | 1.75E-10 | 2.94E-08 | 1061.387 | 1163.827 | 1223.942 | 1216.299 | 2030.182 | 1662.653 | 1693.292 | 1710.781 |
| ENSMUSG00000023572 | 0.168341212 | 0.054244 | 0.144381 | 1402.44 | 1287.409 | 1458.296 | 1612.243 | 1340.778 | 1299.506 | 1262.255 | 1227.89 |
| ENSMUSG00000023707 | -0.25971763 | 0.026514 | 0.086731 | 548.2996 | 425.9366 | 460.4273 | 474.4429 | 570.4401 | 502.8183 | 583.3949 | 629.7152 |
| ENSMUSG00000023723 | -0.338362124 | 0.000776 | 0.006435 | 1241.471 | 1235.816 | 1286.05 | 1349.143 | 1677.191 | 1359.844 | 1618.077 | 1809.752 |
| ENSMUSG00000023755 | 0.316697281 | 0.081316 | 0.19111 | 429.5852 | 401.9402 | 337.0394 | 418.3724 | 388.0943 | 372.0856 | 271.9295 | 241.4452 |
| ENSMUSG00000023764 | 0.264008785 | 0.042292 | 0.121297 | 827.9827 | 814.6788 | 799.951 | 687.5109 | 633.8224 | 800.0399 | 581.4664 | 591.6496 |
| ENSMUSG00000023781 | 0.723233846 | 0.015243 | 0.058358 | 43.26034 | 43.19358 | 44.71776 | 43.9938 | 21.45245 | 32.40385 | 33.75012 | 18.48905 |
| ENSMUSG00000023791 | -0.063691504 | 0.33228 | 0.500209 | 1161.993 | 1185.424 | 1220.629 | 1309.462 | 1248.142 | 1283.863 | 1303.719 | 1265.956 |
| ENSMUSG00000023795 | -0.016030162 | 0.958392 | 0.976487 | 41.24823 | 58.79126 | 48.03018 | 64.69676 | 42.9049 | 35.75597 | 65.57166 | 70.69342 |
| ENSMUSG00000023800 | 0.362263832 | 0.031486 | 0.097721 | 339.0403 | 262.7609 | 245.9477 | 241.5346 | 209.6489 | 240.2354 | 223.7151 | 172.927 |
| ENSMUSG00000023805 | 0.396890657 | 6.40E-05 | 0.000959 | 533.2088 | 562.7163 | 485.2705 | 467.5419 | 395.8952 | 403.372 | 385.7157 | 367.6058 |
| ENSMUSG00000023806 | -0.349467617 | 0.005167 | 0.026328 | 292.7618 | 311.9536 | 332.0708 | 316.5828 | 431.9743 | 324.0385 | 417.5372 | 423.0729 |
| ENSMUSG00000023809 | -0.26923589 | 0.011247 | 0.046617 | 604.6387 | 533.9206 | 568.9093 | 583.1335 | 807.3922 | 644.7249 | 662.4667 | 646.0291 |
| ENSMUSG00000023826 | -0.275716005 | 0.17823 | 0.327597 | 64.38748 | 49.19268 | 75.3577 | 63.83414 | 76.05868 | 75.98144 | 81.00029 | 75.04379 |
| ENSMUSG00000023827 | 0.331434123 | 0.001598 | 0.011054 | 1110.684 | 913.0642 | 854.606 | 1064.477 | 799.5913 | 767.636 | 791.6814 | 774.3649 |
| ENSMUSG00000023828 | 0.185533683 | 0.550091 | 0.698052 | 49.29666 | 87.58697 | 91.91984 | 61.24627 | 85.8098 | 70.39457 | 44.3573 | 54.37955 |
| ENSMUSG00000023829 | -3.163770175 | 8.10E-06 | 0.000195 | 60.36326 | 10.79839 | 107.6539 | 47.44429 | 612.3699 | 203.3621 | 372.2156 | 845.0583 |
| ENSMUSG00000023830 | -0.051082969 | 0.583562 | 0.724151 | 2948.745 | 2752.391 | 3460.658 | 3094.23 | 2893.155 | 3152.112 | 3262.19 | 3396.547 |
| ENSMUSG00000023832 | -0.638980433 | 9.25E-08 | 5.30E-06 | 1016.115 | 1156.628 | 1073.226 | 1138.663 | 1935.596 | 1390.013 | 1630.613 | 1869.569 |
| ENSMUSG00000023845 | 0.067059473 | 0.697678 | 0.809274 | 298.7981 | 236.3648 | 245.1196 | 183.7388 | 219.4 | 205.5968 | 243.9652 | 250.146 |
| ENSMUSG00000023852 | -0.048774748 | 0.498946 | 0.655273 | 2847.134 | 2556.82 | 2375.01 | 2625.826 | 2793.694 | 2659.35 | 2642.152 | 2664.598 |
| ENSMUSG00000023861 | -0.587089808 | 0.000194 | 0.002211 | 773.6558 | 734.2908 | 853.7779 | 875.5628 | 1310.55 | 867.0823 | 1262.255 | 1424.744 |
| ENSMUSG00000023868 | 0.081680311 | 0.729697 | 0.83034 | 362.1796 | 172.7743 | 245.9477 | 234.6336 | 220.3752 | 241.3528 | 296.0368 | 202.2919 |
| ENSMUSG00000023873 | 0.204947465 | 0.395236 | 0.56256 | 104.6297 | 99.58519 | 60.45178 | 85.39973 | 90.68535 | 74.86406 | 72.32169 | 64.16787 |
| ENSMUSG00000023882 | 0.091183539 | 0.69908 | 0.810259 | 103.6236 | 103.1847 | 75.3577 | 69.00988 | 88.73513 | 61.45558 | 82.92887 | 94.62042 |
| ENSMUSG00000023883 | -0.07914669 | 0.2289 | 0.388968 | 1998.024 | 2034.897 | 2003.19 | 2000.424 | 1926.82 | 2225.809 | 2197.615 | 2140.379 |
| ENSMUSG00000023885 | -0.509450182 | 0.067221 | 0.167403 | 555.342 | 867.471 | 1250.441 | 844.5084 | 1155.507 | 817.9178 | 1189.933 | 1845.642 |
| ENSMUSG00000023886 | 0.824790335 | 5.42E-06 | 0.000143 | 2181.126 | 4138.184 | 3564.999 | 3208.959 | 1659.639 | 2077.198 | 1762.721 | 1892.408 |
| ENSMUSG00000023892 | 0.053976893 | 0.664791 | 0.784688 | 287.7315 | 285.5575 | 240.1509 | 248.4356 | 273.0312 | 251.4092 | 228.5365 | 267.5474 |
| ENSMUSG00000023903 | -0.384102276 | 0.524427 | 0.677583 | 22.1332 | 10.79839 | 36.43669 | 6.038364 | 18.52711 | 23.46486 | 40.50014 | 16.31387 |
| ENSMUSG00000023904 | -0.3414463 | 0.018304 | 0.066318 | 737.4378 | 665.901 | 923.3389 | 718.5654 | 937.082 | 832.4437 | 927.6462 | 1164.81 |
| ENSMUSG00000023905 | -0.358636589 | 0.117828 | 0.246482 | 576.4691 | 291.5566 | 499.3483 | 401.1199 | 547.0374 | 435.7759 | 561.2163 | 725.4233 |
| ENSMUSG00000023906 | -0.668086503 | 1.66E-06 | 5.65E-05 | 1210.283 | 1067.841 | 1330.767 | 1299.974 | 1762.026 | 1615.723 | 2033.686 | 2392.7 |
| ENSMUSG00000023908 | -0.159336986 | 0.342441 | 0.511165 | 319.9253 | 427.1365 | 430.6154 | 364.0271 | 419.2979 | 481.5882 | 492.7518 | 327.3649 |
| ENSMUSG00000023909 | -0.200178178 | 0.101536 | 0.223462 | 280.6892 | 320.3523 | 379.2728 | 303.6435 | 352.9903 | 378.7898 | 378.9656 | 366.5182 |
| ENSMUSG00000023911 | 0.419412716 | 0.001872 | 0.012518 | 351.113 | 415.1383 | 469.5365 | 461.5036 | 298.3841 | 318.4516 | 359.6799 | 293.6496 |
| ENSMUSG00000023912 | 0.683734941 | 0.002296 | 0.014489 | 746.4923 | 697.0963 | 505.145 | 880.7386 | 437.825 | 593.3256 | 419.4658 | 311.0511 |
| ENSMUSG00000023913 | 0.657672315 | 3.05E-10 | 4.47E-08 | 3139.896 | 3535.874 | 3049.089 | 3610.079 | 2255.432 | 2301.791 | 2093.472 | 1801.051 |
| ENSMUSG00000023914 | -3.657771973 | 3.28E-05 | 0.000573 | 169.0171 | 64.79036 | 857.0904 | 257.9244 | 4263.187 | 1259.281 | 3851.371 | 7656.641 |
| ENSMUSG00000023915 | -0.778853112 | 7.54E-05 | 0.001094 | 961.788 | 803.8804 | 1344.017 | 1027.385 | 1494.846 | 1387.779 | 2102.15 | 2117.54 |
| ENSMUSG00000023918 | 0.932173212 | 0.108548 | 0.233384 | 15.09082 | 8.398751 | 33.12427 | 12.93935 | 9.751113 | 5.58687 | 11.57147 | 9.78832 |
| ENSMUSG00000023919 | -0.036151297 | 0.724098 | 0.827323 | 520.1301 | 525.5218 | 455.4586 | 470.9924 | 524.6099 | 539.6917 | 507.2161 | 446.9999 |
| ENSMUSG00000023921 | -0.590991273 | 6.43E-06 | 0.000163 | 1097.605 | 969.4558 | 964.7442 | 1026.522 | 1488.995 | 1191.121 | 1678.827 | 1753.197 |
| ENSMUSG00000023923 | -0.041716884 | 0.705627 | 0.814807 | 519.124 | 634.7056 | 645.0951 | 547.7659 | 603.5939 | 543.0438 | 644.1452 | 623.1897 |
| ENSMUSG00000023927 | 0.725975577 | 8.93E-06 | 0.000211 | 956.7577 | 916.6637 | 842.1844 | 1052.401 | 545.0872 | 750.8754 | 537.1091 | 445.9123 |
| ENSMUSG00000023931 | 1.072475961 | 0.01052 | 0.044254 | 26.15741 | 47.99286 | 24.8432 | 50.03216 | 18.52711 | 25.6996 | 12.53576 | 14.13868 |
| ENSMUSG00000023932 | 0.026568292 | 0.711247 | 0.818994 | 2100.641 | 2405.642 | 2177.092 | 2177.262 | 2326.616 | 2071.612 | 2181.222 | 2113.189 |
| ENSMUSG00000023935 | 0.682525978 | 0.104678 | 0.227957 | 34.20585 | 25.19625 | 31.46805 | 19.84034 | 12.67645 | 22.34748 | 24.10723 | 9.78832 |
| ENSMUSG00000023938 | -0.258407792 | 0.036024 | 0.108478 | 435.6215 | 417.5379 | 478.6456 | 456.3278 | 520.7094 | 472.6492 | 503.3589 | 644.9415 |
| ENSMUSG00000023939 | -0.387447939 | 0.001719 | 0.011727 | 761.5831 | 718.6931 | 705.5469 | 664.2201 | 1076.523 | 750.8754 | 917.039 | 981.0072 |
| ENSMUSG00000023940 | -0.033640574 | 0.723961 | 0.827323 | 554.3359 | 539.9197 | 523.3634 | 549.4912 | 582.1415 | 575.4477 | 580.5021 | 478.5401 |
| ENSMUSG00000023942 | -0.161089038 | 0.255729 | 0.419644 | 1262.598 | 1396.592 | 1657.869 | 1312.913 | 1564.079 | 1231.346 | 1733.792 | 1766.248 |
| ENSMUSG00000023944 | -0.036076361 | 0.506764 | 0.662569 | 80699.64 | 82235.77 | 80865.44 | 81895.75 | 88563.51 | 79041.93 | 86262.42 | 80070.63 |
| ENSMUSG00000023945 | 0.540108428 | 0.366498 | 0.535049 | 15.09082 | 19.19714 | 10.76539 | 10.35148 | 4.875557 | 7.821619 | 5.785735 | 19.57664 |
| ENSMUSG00000023947 | -0.178736732 | 0.51282 | 0.66798 | 56.33904 | 45.59322 | 62.108 | 84.5371 | 82.88446 | 50.28183 | 74.25027 | 75.04379 |
| ENSMUSG00000023951 | -0.325914041 | 0.038048 | 0.112572 | 2801.861 | 1943.711 | 2579.552 | 2543.014 | 2949.712 | 2447.049 | 3203.369 | 3771.766 |
| ENSMUSG00000023952 | 0.089471683 | 0.229555 | 0.38976 | 2276.701 | 2111.686 | 2098.422 | 2301.479 | 2145.245 | 1901.771 | 2203.401 | 2007.693 |
| ENSMUSG00000023953 | 0.022470528 | 0.889329 | 0.933378 | 511.0756 | 500.3256 | 390.0382 | 408.0209 | 485.6054 | 359.7945 | 405.0014 | 528.5693 |
| ENSMUSG00000023959 | -1.272839763 | 4.46E-06 | 0.000124 | 275.6589 | 175.1739 | 341.1799 | 254.4739 | 534.361 | 393.3157 | 694.2882 | 910.3137 |
| ENSMUSG00000023960 | -0.359915899 | 0.0174 | 0.06397 | 893.3763 | 728.2917 | 1094.757 | 918.694 | 1153.557 | 954.2375 | 1224.647 | 1336.649 |
| ENSMUSG00000023961 | 0.180644248 | 0.143635 | 0.281384 | 327.9737 | 349.1481 | 310.54 | 347.6373 | 244.7529 | 296.1041 | 317.2511 | 319.7518 |

| | | | | | | | | | | | |
|---|---|---|---|---|---|---|---|---|---|---|---|
| ENSMUSG00000023963 | 0.476302695 | 0.000294 | 0.003051 | 763.5953 | 647.9036 | 684.0161 | 892.8153 | 517.7841 | 560.9218 | 592.0735 | 478.5401 |
| ENSMUSG00000023965 | 0.12794936 | 0.251854 | 0.415496 | 927.5821 | 968.256 | 914.2297 | 976.4898 | 758.6366 | 1042.51 | 847.6102 | 817.8685 |
| ENSMUSG00000023966 | -0.033918053 | 0.829623 | 0.896882 | 194.1685 | 160.7761 | 209.511 | 244.1224 | 227.2009 | 196.6578 | 204.4293 | 202.2919 |
| ENSMUSG00000023967 | -0.305163561 | 0.000944 | 0.007498 | 977.8848 | 965.8563 | 1016.087 | 1020.484 | 1286.172 | 1048.097 | 1259.362 | 1324.686 |
| ENSMUSG00000023971 | -0.194885896 | 0.031173 | 0.097037 | 703.232 | 754.6877 | 809.8883 | 795.3389 | 937.082 | 784.3966 | 893.8961 | 891.8247 |
| ENSMUSG00000023972 | -0.023300428 | 0.766832 | 0.855443 | 6403.536 | 6844.982 | 7677.377 | 6703.447 | 7473.253 | 6938.893 | 7113.561 | 6554.911 |
| ENSMUSG00000023973 | -0.225366098 | 0.009222 | 0.040161 | 1215.314 | 1199.822 | 1455.811 | 1211.123 | 1470.468 | 1443.647 | 1545.756 | 1485.649 |
| ENSMUSG00000023977 | 0.140318239 | 0.059382 | 0.153623 | 3011.121 | 2615.611 | 2615.989 | 2849.245 | 2532.364 | 2495.096 | 2660.474 | 2375.299 |
| ENSMUSG00000023978 | 0.069563885 | 0.558484 | 0.704826 | 738.4439 | 895.0669 | 851.2936 | 815.1792 | 735.2339 | 916.2468 | 679.8239 | 814.6057 |
| ENSMUSG00000023979 | -0.011983664 | 0.932576 | 0.960099 | 319.9253 | 313.1534 | 370.1637 | 308.8192 | 319.8365 | 350.8555 | 270.001 | 385.0072 |
| ENSMUSG00000023980 | 0.073292983 | 0.468711 | 0.62887 | 1059.375 | 1001.851 | 957.2913 | 1055.851 | 987.7878 | 1076.031 | 989.3607 | 818.9561 |
| ENSMUSG00000023982 | 0.285572621 | 0.081338 | 0.19111 | 151.9142 | 201.57 | 212.8234 | 162.1732 | 159.9183 | 146.376 | 137.8934 | 153.3503 |
| ENSMUSG00000023984 | -0.220827352 | 0.611664 | 0.746175 | 7.04238 | 15.59768 | 14.90592 | 15.52722 | 15.60178 | 14.52586 | 14.46434 | 17.40146 |
| ENSMUSG00000023988 | -0.618242748 | 0.000163 | 0.001932 | 409.4641 | 417.5379 | 438.0684 | 370.0655 | 594.8179 | 463.7102 | 673.0738 | 777.6276 |
| ENSMUSG00000023990 | -0.543891593 | 0.000827 | 0.006733 | 329.9858 | 298.7556 | 395.835 | 333.8353 | 445.6259 | 419.0153 | 486.0017 | 632.978 |
| ENSMUSG00000023991 | 0.566640485 | 1.65E-07 | 8.49E-06 | 1671.056 | 1658.153 | 1777.117 | 1717.483 | 1124.303 | 1391.131 | 1075.182 | 1019.073 |
| ENSMUSG00000023992 | 0.70866292 | 0.013728 | 0.054032 | 65.39353 | 85.18733 | 71.21717 | 42.26855 | 39.00445 | 33.52122 | 41.46443 | 46.76642 |
| ENSMUSG00000023994 | 0.054504843 | 0.653261 | 0.776084 | 1405.458 | 1466.182 | 1089.788 | 1330.165 | 1286.172 | 1109.552 | 1278.647 | 1417.131 |
| ENSMUSG00000023995 | 1.047624625 | 0.02543 | 0.08399 | 48.29061 | 20.39697 | 23.18699 | 31.91707 | 12.67645 | 22.34748 | 17.3572 | 7.613138 |
| ENSMUSG00000023999 | 0.067097875 | 0.841823 | 0.90461 | 55.33299 | 52.79215 | 42.23344 | 26.74133 | 44.85512 | 44.69496 | 51.10733 | 27.18978 |
| ENSMUSG00000024002 | 0.475386865 | 0.005965 | 0.029335 | 2873.291 | 2622.81 | 2302.965 | 2515.41 | 1703.519 | 2585.604 | 1621.934 | 1508.489 |
| ENSMUSG00000024006 | 0.293425552 | 0.000468 | 0.004349 | 2793.813 | 2745.192 | 2411.447 | 2710.363 | 2153.046 | 2405.706 | 2014.4 | 2125.153 |
| ENSMUSG00000024007 | -0.290718711 | 0.007535 | 0.034672 | 1315.919 | 1682.15 | 1596.59 | 1426.779 | 1882.94 | 1603.432 | 1894.828 | 1981.591 |
| ENSMUSG00000024008 | -0.067688989 | 0.767636 | 0.855731 | 201.2109 | 165.5754 | 164.7932 | 119.9047 | 183.3209 | 119.559 | 167.7863 | 210.9927 |
| ENSMUSG00000024011 | 1.336768093 | 0.00018 | 0.002091 | 348.0948 | 601.1106 | 332.0708 | 475.3055 | 202.8232 | 290.5173 | 111.8575 | 90.27006 |
| ENSMUSG00000024012 | -0.174849745 | 0.092204 | 0.208519 | 7599.735 | 7873.229 | 7876.95 | 7314.185 | 8666.789 | 7166.837 | 8781.781 | 9998.225 |
| ENSMUSG00000024013 | -0.374151209 | 0.08128 | 0.191071 | 125.7568 | 177.5736 | 176.3867 | 123.3552 | 186.2463 | 144.1413 | 223.7151 | 226.2189 |
| ENSMUSG00000024014 | 0.388986793 | 0.001933 | 0.012811 | 1548.318 | 1309.005 | 1153.553 | 1453.521 | 1138.93 | 1005.637 | 1129.183 | 897.2626 |
| ENSMUSG00000024018 | 0.04391162 | 0.723678 | 0.827317 | 278.6771 | 297.5557 | 318.8211 | 351.0878 | 315.9361 | 267.0524 | 298.9296 | 328.4525 |
| ENSMUSG00000024019 | -0.192597978 | 0.200582 | 0.355956 | 410.4702 | 435.5352 | 412.3971 | 359.714 | 531.4357 | 347.5033 | 498.5375 | 468.7518 |
| ENSMUSG00000024026 | -0.185641163 | 0.00605 | 0.02964 | 1956.776 | 2057.694 | 2112.5 | 2128.955 | 2368.545 | 2183.349 | 2322.008 | 2517.773 |
| ENSMUSG00000024028 | -0.662594402 | NA | NA | 10.06054 | 1.199822 | 83.63877 | 6.038364 | 38.02934 | 55.8687 | 29.89296 | 36.9781 |
| ENSMUSG00000024029 | -0.100030861 | 0.911317 | 0.947671 | 11.0666 | 0 | 9.93728 | 17.25247 | 6.825779 | 22.34748 | 4.821446 | 7.613138 |
| ENSMUSG00000024030 | -0.080474066 | 0.722976 | 0.826667 | 382.3007 | 365.9456 | 403.2879 | 431.3117 | 384.1939 | 486.0577 | 554.4663 | 249.0584 |
| ENSMUSG00000024033 | -0.371349356 | 0.116094 | 0.244118 | 64.38748 | 59.99108 | 43.88965 | 81.94923 | 79.95913 | 78.21619 | 84.85745 | 80.48174 |
| ENSMUSG00000024036 | -0.375344471 | 0.150816 | 0.290973 | 95.57516 | 88.78679 | 101.029 | 87.12497 | 117.9885 | 77.09881 | 176.4649 | 110.9343 |
| ENSMUSG00000024037 | 0.011472818 | 0.884667 | 0.931133 | 1125.775 | 1262.212 | 1164.318 | 1174.893 | 1286.172 | 1169.891 | 1118.575 | 1111.518 |
| ENSMUSG00000024038 | -0.298079898 | 0.035218 | 0.106555 | 1529.203 | 1730.143 | 1801.132 | 1611.381 | 2064.311 | 1591.141 | 2004.757 | 2542.788 |
| ENSMUSG00000024039 | -0.422135281 | 0.204716 | 0.36081 | 122.7386 | 40.79393 | 101.8571 | 106.9653 | 132.6151 | 88.27255 | 126.3219 | 153.3503 |
| ENSMUSG00000024041 | 1.31851946 | NA | NA | 9.054489 | 3.599465 | 57.13936 | 3.450494 | 5.850668 | 4.469496 | 11.57147 | 7.613138 |
| ENSMUSG00000024042 | -0.58856919 | 0.004705 | 0.024657 | 608.6629 | 356.347 | 522.5353 | 561.5679 | 630.897 | 640.2554 | 1008.646 | 803.7298 |
| ENSMUSG00000024043 | 0.322074826 | 0.012351 | 0.050054 | 1125.775 | 1157.828 | 1026.024 | 1117.96 | 786.9148 | 1111.787 | 858.2174 | 785.2408 |
| ENSMUSG00000024044 | 0.536047894 | 0.00369 | 0.020592 | 1016.115 | 980.2542 | 799.951 | 1061.89 | 665.0259 | 901.7209 | 593.0378 | 501.3795 |
| ENSMUSG00000024045 | -0.017766893 | 0.78193 | 0.865498 | 3909.527 | 3985.807 | 3675.137 | 3897.333 | 4086.691 | 4124.228 | 3661.406 | 3784.817 |
| ENSMUSG00000024048 | -0.411411432 | 0.002237 | 0.014252 | 7888.472 | 8641.115 | 10258.58 | 8412.304 | 12005.57 | 9118.89 | 12362.19 | 13330.6 |
| ENSMUSG00000024049 | 0.698787725 | 0.011992 | 0.048962 | 310.8708 | 398.3407 | 270.7909 | 438.2127 | 193.072 | 350.8555 | 172.6078 | 157.7007 |
| ENSMUSG00000024050 | 0.026790974 | 0.657949 | 0.779604 | 1685.141 | 1630.557 | 1763.867 | 1685.566 | 1692.793 | 1700.643 | 1689.435 | 1560.693 |
| ENSMUSG00000024052 | 0.106836439 | 0.243144 | 0.405701 | 3320.985 | 3591.066 | 2895.889 | 3289.183 | 3208.116 | 3163.286 | 3064.511 | 2721.153 |
| ENSMUSG00000024053 | 0.875496452 | 1.66E-08 | 1.29E-06 | 321.9374 | 301.1552 | 328.7583 | 342.4615 | 159.9183 | 229.0617 | 164.8934 | 153.3503 |
| ENSMUSG00000024054 | -0.178901097 | 0.007371 | 0.034119 | 1839.067 | 1899.317 | 1752.274 | 1789.081 | 1904.392 | 2074.964 | 2183.151 | 2074.036 |
| ENSMUSG00000024055 | 0.144615085 | 0.224255 | 0.383974 | 538.2391 | 447.5334 | 462.0835 | 497.7338 | 488.5308 | 472.6492 | 418.5015 | 380.6569 |
| ENSMUSG00000024056 | 0.066273091 | 0.525206 | 0.678439 | 880.2976 | 923.8626 | 789.1856 | 852.272 | 865.8988 | 926.3031 | 746.3598 | 750.4379 |
| ENSMUSG00000024059 | 0.334605393 | 0.103497 | 0.226212 | 558.3602 | 608.3095 | 399.9755 | 517.5741 | 397.8454 | 572.0955 | 346.1798 | 336.0656 |
| ENSMUSG00000024063 | 0.07501316 | 0.645832 | 0.770604 | 3637.893 | 3624.661 | 4945.453 | 3628.194 | 3744.427 | 3107.417 | 3368.262 | 4815.853 |
| ENSMUSG00000024064 | -1.483637194 | 2.30E-05 | 0.000438 | 168.0111 | 87.58697 | 206.1986 | 217.3811 | 449.5263 | 235.7659 | 512.0375 | 705.8466 |
| ENSMUSG00000024065 | -0.580691811 | 0.010452 | 0.044072 | 632.8082 | 391.1418 | 494.3797 | 498.5964 | 565.5646 | 563.1565 | 921.8604 | 966.8685 |
| ENSMUSG00000024066 | 1.49932047 | 2.50E-07 | 1.21E-05 | 870.237 | 634.7056 | 343.6643 | 721.1532 | 263.2801 | 274.874 | 224.6794 | 145.7372 |
| ENSMUSG00000024067 | -0.242447389 | 0.00033 | 0.003338 | 985.9333 | 911.8644 | 1021.055 | 975.6272 | 1115.527 | 1129.665 | 1173.54 | 1194.175 |
| ENSMUSG00000024068 | 0.220743419 | 0.019655 | 0.069615 | 1806.874 | 1742.141 | 1608.183 | 1836.525 | 1484.119 | 1724.108 | 1383.755 | 1410.606 |
| ENSMUSG00000024069 | -0.127159494 | 0.335452 | 0.503669 | 577.4752 | 517.1231 | 479.4737 | 530.5134 | 629.9219 | 458.1234 | 566.0377 | 642.7663 |
| ENSMUSG00000024070 | 0.101962341 | 0.211908 | 0.369482 | 2806.892 | 3115.937 | 2698.8 | 2966.562 | 2646.452 | 2996.797 | 2617.081 | 2535.175 |
| ENSMUSG00000024072 | 0.071964004 | 0.338174 | 0.506643 | 1432.621 | 1426.588 | 1312.549 | 1489.751 | 1249.118 | 1341.966 | 1431.005 | 1362.752 |
| ENSMUSG00000024073 | -0.022114358 | 0.700885 | 0.811311 | 3047.339 | 2872.373 | 2963.794 | 2893.239 | 3006.268 | 3169.99 | 2956.511 | 2827.737 |
| ENSMUSG00000024074 | -0.177300225 | 0.138293 | 0.273768 | 1185.132 | 893.867 | 1065.773 | 1014.445 | 1030.693 | 1277.159 | 1286.362 | 1111.518 |

| | | | | | | | | | | | |
|---|---|---|---|---|---|---|---|---|---|---|---|
| ENSMUSG00000024076 | 1.222536486 | 9.85E-08 | 5.59E-06 | 150.9082 | 135.5798 | 102.6852 | 162.1732 | 60.4569 | 69.27719 | 61.71451 | 44.59123 |
| ENSMUSG00000024077 | 0.038449884 | 0.63856 | 0.765743 | 1162.999 | 1004.251 | 1096.413 | 1180.069 | 1018.991 | 1148.661 | 1070.361 | 1094.117 |
| ENSMUSG00000024078 | -0.087210445 | 0.384598 | 0.552354 | 773.6558 | 921.4629 | 753.577 | 778.949 | 785.9397 | 938.5942 | 862.0745 | 838.5327 |
| ENSMUSG00000024079 | 0.371081378 | 0.04249 | 0.121695 | 953.7395 | 1195.022 | 1236.363 | 674.5716 | 771.313 | 775.4576 | 835.0744 | 755.8758 |
| ENSMUSG00000024081 | 0.168347295 | 0.097952 | 0.217858 | 2344.107 | 2492.029 | 2167.983 | 2527.487 | 2078.937 | 2471.632 | 2020.186 | 1910.898 |
| ENSMUSG00000024082 | -0.306916474 | 0.000342 | 0.003422 | 737.4378 | 713.8938 | 763.5143 | 664.2201 | 852.2473 | 832.4437 | 950.7891 | 925.54 |
| ENSMUSG00000024083 | 0.045608537 | 0.468815 | 0.628949 | 3161.023 | 2959.96 | 2837.921 | 3176.18 | 2823.922 | 3031.436 | 2896.725 | 3007.189 |
| ENSMUSG00000024084 | 0.543043818 | 0.043285 | 0.123188 | 101.6115 | 137.9795 | 91.91984 | 178.5631 | 81.90935 | 112.8548 | 75.21455 | 80.48174 |
| ENSMUSG00000024085 | 0.218306191 | 0.035245 | 0.106602 | 3395.433 | 2757.19 | 2871.046 | 3112.346 | 2270.059 | 2852.656 | 2761.724 | 2549.314 |
| ENSMUSG00000024087 | 0.194463659 | 0.423769 | 0.588965 | 1416.525 | 925.0624 | 898.4957 | 1488.888 | 839.5708 | 890.5472 | 1523.577 | 878.7736 |
| ENSMUSG00000024091 | -0.137588968 | 0.022439 | 0.076468 | 2724.395 | 2592.814 | 2702.94 | 2651.705 | 2715.685 | 3070.544 | 2938.189 | 3020.24 |
| ENSMUSG00000024095 | -0.1064225 | 0.036537 | 0.109646 | 3079.532 | 2939.563 | 2832.125 | 3001.93 | 3153.51 | 3114.122 | 3253.512 | 3236.671 |
| ENSMUSG00000024096 | 0.193726863 | 0.06299 | 0.160461 | 3071.484 | 3153.131 | 2572.927 | 2918.255 | 2256.408 | 2889.529 | 2632.509 | 2463.394 |
| ENSMUSG00000024097 | 0.032192446 | 0.770045 | 0.857226 | 5460.863 | 6062.698 | 4582.742 | 5548.394 | 5892.598 | 5547.762 | 4833.982 | 4898.51 |
| ENSMUSG00000024098 | 0.252758443 | 0.003275 | 0.018935 | 4929.666 | 5858.729 | 5502.769 | 5474.209 | 4317.793 | 5116.456 | 4499.373 | 4334.05 |
| ENSMUSG00000024099 | -0.382248248 | 7.11E-05 | 0.001044 | 3070.478 | 3093.14 | 3449.064 | 3347.842 | 4311.942 | 3583.419 | 4348.944 | 4650.539 |
| ENSMUSG00000024101 | -0.012925212 | 0.86329 | 0.917949 | 1512.1 | 1333.002 | 1544.419 | 1405.214 | 1527.999 | 1410.126 | 1498.505 | 1412.781 |
| ENSMUSG00000024104 | -0.116672345 | 0.165634 | 0.311107 | 2521.172 | 2295.259 | 2275.637 | 2556.816 | 2741.038 | 2318.551 | 2764.617 | 2636.321 |
| ENSMUSG00000024109 | 0.974808244 | 0.000535 | 0.004832 | 162.9808 | 88.78679 | 104.3414 | 164.7611 | 61.43201 | 88.27255 | 61.71451 | 54.37955 |
| ENSMUSG00000024112 | 0.055893804 | 0.630348 | 0.759331 | 494.9787 | 440.3345 | 587.9557 | 542.5902 | 491.4561 | 508.4052 | 530.359 | 461.1386 |
| ENSMUSG00000024114 | 1.039774439 | 0.000104 | 0.001383 | 111.672 | 152.3773 | 101.8571 | 112.1411 | 55.58134 | 34.6386 | 69.42882 | 71.78101 |
| ENSMUSG00000024118 | 0.103341972 | 0.313176 | 0.481393 | 390.3491 | 436.735 | 354.4296 | 380.417 | 372.4925 | 344.1512 | 368.3585 | 364.343 |
| ENSMUSG00000024120 | -0.336508754 | 0.000576 | 0.00512 | 1781.722 | 1593.363 | 1733.227 | 1755.439 | 2308.088 | 1814.616 | 2246.794 | 2298.08 |
| ENSMUSG00000024121 | -0.666727072 | 0.030807 | 0.096321 | 859.1704 | 685.0981 | 1888.911 | 1305.149 | 1517.273 | 1352.023 | 2587.188 | 2068.598 |
| ENSMUSG00000024122 | 0.062244815 | 0.408433 | 0.574721 | 1317.931 | 1244.215 | 1258.722 | 1263.743 | 1160.382 | 1122.961 | 1296.005 | 1289.883 |
| ENSMUSG00000024127 | -0.340996 | 0.017337 | 0.063778 | 969.8364 | 922.6628 | 848.8093 | 900.5789 | 1198.412 | 881.6082 | 1117.611 | 1413.868 |
| ENSMUSG00000024130 | -0.732587216 | 0.003311 | 0.019067 | 1279.701 | 1129.032 | 1165.974 | 986.8413 | 2153.046 | 1031.336 | 1678.827 | 2715.715 |
| ENSMUSG00000024131 | -4.233535956 | 1.35E-08 | 1.09E-06 | 66.39959 | 15.59768 | 103.5133 | 56.07053 | 1446.09 | 279.3435 | 729.0026 | 2099.051 |
| ENSMUSG00000024132 | -0.679274313 | 0.000346 | 0.003454 | 861.1825 | 788.2827 | 1040.93 | 906.6173 | 1641.112 | 984.4066 | 1310.469 | 1826.065 |
| ENSMUSG00000024134 | 0.465404119 | 0.021361 | 0.073787 | 4448.772 | 5237.221 | 3230.444 | 4472.703 | 3855.59 | 3818.067 | 2601.652 | 2317.657 |
| ENSMUSG00000024135 | -0.062611373 | 0.500501 | 0.656847 | 781.7042 | 794.2819 | 719.6247 | 704.7634 | 865.8988 | 736.3495 | 800.36 | 725.4233 |
| ENSMUSG00000024136 | -0.771428666 | 0.055644 | 0.146914 | 95.57516 | 94.7859 | 266.6503 | 91.43809 | 235.0018 | 149.7281 | 353.8941 | 199.0292 |
| ENSMUSG00000024137 | -0.062859368 | 0.462879 | 0.623888 | 796.795 | 896.2667 | 888.5584 | 905.7547 | 992.6633 | 896.134 | 912.2175 | 840.7079 |
| ENSMUSG00000024140 | -0.322484467 | 0.067447 | 0.16764 | 1151.932 | 880.669 | 1499.701 | 1225.788 | 1415.862 | 1201.177 | 1607.47 | 1729.27 |
| ENSMUSG00000024142 | -0.224255828 | 0.013382 | 0.053091 | 777.68 | 793.082 | 836.3877 | 786.7126 | 1024.842 | 819.0352 | 921.8604 | 964.6933 |
| ENSMUSG00000024143 | 0.168508606 | 0.028673 | 0.091535 | 2010.097 | 1950.91 | 1818.522 | 2162.597 | 1766.902 | 1860.428 | 1772.363 | 1667.277 |
| ENSMUSG00000024145 | -0.224392406 | 0.108746 | 0.233672 | 270.6286 | 229.1659 | 300.6027 | 302.7808 | 306.185 | 290.5173 | 323.0369 | 373.0437 |
| ENSMUSG00000024146 | -0.082808309 | 0.222812 | 0.382247 | 1036.236 | 959.8572 | 995.3842 | 1048.95 | 1019.966 | 1121.844 | 1058.79 | 1083.241 |
| ENSMUSG00000024150 | -0.322226836 | 0.043128 | 0.123006 | 2448.736 | 1757.739 | 1962.613 | 2079.785 | 2728.361 | 1955.405 | 2494.616 | 3135.525 |
| ENSMUSG00000024151 | 0.04955324 | 0.574246 | 0.716633 | 2460.809 | 2987.556 | 2444.571 | 2543.014 | 2673.755 | 2451.519 | 2571.759 | 2379.649 |
| ENSMUSG00000024155 | -0.172481197 | 0.80642 | 0.881775 | 23.13925 | 27.5959 | 7.45296 | 3.450494 | 8.776002 | 29.05173 | 12.53576 | 18.48905 |
| ENSMUSG00000024158 | -0.724028944 | 9.06E-05 | 0.001242 | 528.1785 | 463.1311 | 548.2066 | 568.4689 | 819.0935 | 589.9735 | 1019.254 | 1054.963 |
| ENSMUSG00000024160 | -0.367527852 | 0.019439 | 0.069097 | 265.5983 | 267.5602 | 316.3367 | 317.4454 | 332.513 | 309.5126 | 446.4659 | 418.7226 |
| ENSMUSG00000024163 | 0.168062277 | 0.086126 | 0.198843 | 2219.356 | 1883.72 | 1729.915 | 1933.139 | 1844.911 | 1794.503 | 1597.827 | 1673.803 |
| ENSMUSG00000024164 | 1.834888712 | 6.03E-06 | 0.000155 | 528.1785 | 1391.793 | 507.6294 | 846.2336 | 213.5494 | 353.0902 | 259.3938 | 91.35765 |
| ENSMUSG00000024165 | -0.300261597 | 0.003498 | 0.019783 | 2278.713 | 2734.393 | 2932.326 | 2608.573 | 3226.643 | 2879.473 | 3504.227 | 3385.671 |
| ENSMUSG00000024168 | -0.209637466 | 0.148825 | 0.288612 | 823.9585 | 885.4683 | 741.9835 | 844.5084 | 1008.265 | 700.5936 | 1032.754 | 1066.927 |
| ENSMUSG00000024169 | 0.109796295 | 0.189131 | 0.341613 | 1156.963 | 1043.845 | 1088.96 | 1117.097 | 1075.548 | 1082.735 | 1018.289 | 908.1386 |
| ENSMUSG00000024170 | 0.016055167 | 0.870606 | 0.922435 | 844.0796 | 779.884 | 822.3099 | 839.3327 | 766.4375 | 889.4298 | 883.2889 | 711.2846 |
| ENSMUSG00000024172 | -0.333039665 | 0.301879 | 0.470286 | 30.18163 | 33.595 | 18.21835 | 26.74133 | 29.25334 | 42.46022 | 33.75012 | 30.45255 |
| ENSMUSG00000024173 | 0.193717797 | 0.820767 | 0.891378 | 43.26034 | 58.79126 | 14.07781 | 5.175741 | 46.80534 | 15.64324 | 2.892867 | 40.24087 |
| ENSMUSG00000024174 | 0.40574698 | 8.74E-05 | 0.001214 | 687.1351 | 681.4986 | 587.9557 | 712.527 | 471.9539 | 548.6307 | 523.609 | 469.8393 |
| ENSMUSG00000024176 | -0.561644939 | 0.006532 | 0.031425 | 75.45408 | 106.7841 | 69.56096 | 87.9876 | 142.3663 | 136.3196 | 115.7147 | 104.4087 |
| ENSMUSG00000024177 | -0.093761563 | 0.330288 | 0.498145 | 641.8627 | 777.4844 | 800.7791 | 773.7733 | 841.5211 | 763.1665 | 813.8601 | 776.54 |
| ENSMUSG00000024180 | -0.852247674 | 0.000195 | 0.00222 | 371.2341 | 226.7663 | 459.5992 | 335.5605 | 644.5486 | 469.2971 | 620.0379 | 785.2408 |
| ENSMUSG00000024181 | -0.195208364 | 0.01803 | 0.065724 | 1387.349 | 1279.01 | 1291.018 | 1249.079 | 1553.352 | 1352.023 | 1434.862 | 1619.423 |
| ENSMUSG00000024182 | -0.11134383 | 0.20825 | 0.365349 | 1513.106 | 1697.747 | 1941.91 | 1707.995 | 1809.807 | 1811.263 | 1895.792 | 1897.846 |
| ENSMUSG00000024186 | -0.198341214 | 0.511928 | 0.667336 | 171.0292 | 140.3791 | 330.4145 | 232.0457 | 209.6489 | 196.6578 | 233.358 | 365.4306 |
| ENSMUSG00000024187 | -0.433847052 | 0.019097 | 0.068265 | 1302.84 | 1156.628 | 1825.975 | 1310.325 | 1841.985 | 1412.361 | 1976.793 | 2330.708 |
| ENSMUSG00000024188 | 0.229641649 | 0.046943 | 0.130236 | 4885.4 | 4699.701 | 3999.755 | 4692.672 | 4069.139 | 4555.534 | 3591.977 | 3370.445 |
| ENSMUSG00000024190 | -0.420309284 | 0.121051 | 0.25028 | 843.0736 | 395.9411 | 501.8326 | 837.6074 | 693.3041 | 674.894 | 1134.004 | 949.467 |
| ENSMUSG00000024191 | 0.084649075 | 0.365561 | 0.534244 | 930.6003 | 884.2685 | 934.9324 | 981.6655 | 874.6748 | 993.3456 | 873.646 | 779.8028 |
| ENSMUSG00000024193 | 0.218192873 | 0.094373 | 0.211942 | 604.6387 | 579.5138 | 593.7525 | 742.7188 | 492.4312 | 633.5511 | 550.6091 | 492.6788 |
| ENSMUSG00000024194 | -0.188581841 | 0.021765 | 0.074822 | 1017.121 | 974.2551 | 964.7442 | 897.1284 | 1074.573 | 1007.871 | 1116.647 | 1189.825 |

| | | | | | | | | | | | |
|---|---|---|---|---|---|---|---|---|---|---|---|
| ENSMUSG00000024197 | 0.055390795 | 0.436432 | 0.600273 | 2728.419 | 2454.835 | 2640.832 | 2498.158 | 2388.048 | 2479.453 | 2369.258 | 2700.489 |
| ENSMUSG00000024201 | -0.134306156 | 0.015871 | 0.059989 | 1242.477 | 1243.015 | 1318.346 | 1278.408 | 1411.961 | 1375.488 | 1413.648 | 1380.153 |
| ENSMUSG00000024205 | -0.146601828 | 0.678191 | 0.794458 | 17.10292 | 15.59768 | 21.53077 | 18.11509 | 19.50223 | 23.46486 | 19.28578 | 18.48905 |
| ENSMUSG00000024206 | -0.184475468 | 0.384601 | 0.552354 | 103.6236 | 124.7814 | 119.2474 | 165.6237 | 136.5156 | 174.3104 | 158.1434 | 115.2847 |
| ENSMUSG00000024208 | -0.293632769 | 0.050196 | 0.136381 | 770.6376 | 862.6717 | 946.5259 | 709.0765 | 964.3851 | 852.5564 | 950.7891 | 1263.781 |
| ENSMUSG00000024209 | 0.56020413 | 0.0611 | 0.15688 | 34.20585 | 49.19268 | 56.31125 | 55.2079 | 34.1289 | 43.57759 | 27.96439 | 27.18978 |
| ENSMUSG00000024211 | 0.346734833 | 0.44069 | 0.603986 | 24.1453 | 13.19804 | 22.35888 | 31.05445 | 17.552 | 29.05173 | 12.53576 | 13.05109 |
| ENSMUSG00000024212 | -0.261984701 | 0.002519 | 0.015502 | 1090.563 | 1233.417 | 1327.455 | 1125.724 | 1434.389 | 1347.553 | 1486.934 | 1459.547 |
| ENSMUSG00000024213 | -0.114868733 | 0.095971 | 0.214618 | 2950.757 | 3057.145 | 3218.022 | 2995.029 | 3297.826 | 3068.309 | 3275.69 | 3593.401 |
| ENSMUSG00000024215 | -0.159011379 | 0.744083 | 0.839946 | 18.10898 | 7.198929 | 18.21835 | 23.29083 | 12.67645 | 26.81698 | 13.50005 | 22.83941 |
| ENSMUSG00000024217 | -0.124531865 | 0.117718 | 0.246326 | 1723.371 | 1775.736 | 1937.77 | 1668.314 | 2071.136 | 1781.094 | 1933.4 | 1959.839 |
| ENSMUSG00000024218 | -0.166217628 | 0.02977 | 0.094017 | 1077.484 | 1245.415 | 1134.506 | 1179.206 | 1393.434 | 1262.633 | 1297.933 | 1244.204 |
| ENSMUSG00000024219 | -0.26680115 | 0.015402 | 0.058767 | 861.1825 | 783.4835 | 985.4469 | 815.1792 | 1008.265 | 924.0684 | 1056.861 | 1159.372 |
| ENSMUSG00000024220 | -0.125742868 | 0.161301 | 0.305048 | 1434.634 | 1571.766 | 1548.559 | 1463.009 | 1737.648 | 1430.239 | 1611.327 | 1784.737 |
| ENSMUSG00000024222 | -0.427425291 | 0.115045 | 0.242689 | 2258.592 | 1277.81 | 1426 | 2161.734 | 1931.696 | 1794.503 | 3817.621 | 2037.058 |
| ENSMUSG00000024227 | 0.22840324 | 0.481103 | 0.640315 | 28.16952 | 40.79393 | 32.29616 | 42.26855 | 36.07912 | 35.75597 | 19.28578 | 31.54014 |
| ENSMUSG00000024228 | -0.028423453 | 0.874691 | 0.925069 | 278.6771 | 225.5664 | 183.0116 | 206.167 | 216.4747 | 189.9536 | 232.3937 | 270.8102 |
| ENSMUSG00000024231 | -0.053987043 | 0.405094 | 0.572028 | 1113.702 | 1117.034 | 1033.477 | 1115.372 | 1142.83 | 1168.773 | 1082.897 | 1150.671 |
| ENSMUSG00000024232 | -0.222811741 | 0.032484 | 0.100055 | 660.9777 | 641.9045 | 749.4365 | 672.8463 | 688.4286 | 858.1433 | 785.8957 | 852.6714 |
| ENSMUSG00000024234 | 0.272828011 | 0.041971 | 0.1206 | 1022.151 | 1118.234 | 987.9312 | 1058.439 | 795.6908 | 1070.444 | 891.9675 | 706.9342 |
| ENSMUSG00000024235 | 0.474775824 | 0.041471 | 0.11974 | 147.89 | 79.18822 | 122.5598 | 115.5915 | 92.63557 | 80.45093 | 92.57176 | 69.60583 |
| ENSMUSG00000024236 | 0.451887904 | 4.30E-07 | 1.86E-05 | 2157.987 | 2404.442 | 2189.514 | 2243.684 | 1454.866 | 1776.625 | 1568.898 | 1777.124 |
| ENSMUSG00000024238 | 0.490115066 | 0.000331 | 0.003338 | 2183.138 | 2247.266 | 1805.272 | 2272.15 | 1505.572 | 1862.663 | 1427.148 | 1261.606 |
| ENSMUSG00000024240 | 0.079871128 | 0.275846 | 0.442082 | 1892.388 | 1816.53 | 1796.163 | 1827.899 | 1842.96 | 1621.31 | 1849.507 | 1620.511 |
| ENSMUSG00000024241 | 0.271882298 | 0.038762 | 0.113789 | 1050.321 | 993.4522 | 857.0904 | 972.1767 | 687.4535 | 978.8197 | 730.9312 | 811.343 |
| ENSMUSG00000024242 | -0.258393409 | 0.009464 | 0.040932 | 1828.001 | 1622.159 | 1939.426 | 1931.414 | 1919.994 | 2235.866 | 2430.009 | 2176.27 |
| ENSMUSG00000024245 | -1.126471319 | 0.000225 | 0.002482 | 269.6226 | 197.9706 | 490.2391 | 285.5284 | 556.7886 | 439.128 | 810.9672 | 911.4013 |
| ENSMUSG00000024246 | 0.114745813 | 0.400351 | 0.567711 | 358.1554 | 434.3354 | 416.5376 | 389.9058 | 336.4134 | 417.8979 | 411.7515 | 309.9635 |
| ENSMUSG00000024247 | 0.413320051 | 0.009303 | 0.040391 | 6499.111 | 7316.512 | 6388.843 | 6836.291 | 4287.564 | 6970.18 | 4376.909 | 4671.204 |
| ENSMUSG00000024248 | -0.108739939 | 0.1837 | 0.334719 | 6430.699 | 7016.556 | 7253.386 | 7424.6 | 8028.091 | 7611.552 | 7944.778 | 6743.065 |
| ENSMUSG00000024251 | -0.333866887 | 0.002159 | 0.013884 | 456.7487 | 399.5406 | 381.7572 | 349.3625 | 508.033 | 472.6492 | 512.0375 | 504.6423 |
| ENSMUSG00000024253 | -0.167690341 | 0.117171 | 0.245433 | 787.7406 | 933.4612 | 958.9475 | 935.0839 | 1090.174 | 871.5518 | 1040.468 | 1058.226 |
| ENSMUSG00000024254 | 0.396629049 | 0.244809 | 0.407641 | 35.2119 | 70.78947 | 46.37397 | 37.09281 | 41.92979 | 30.1691 | 44.3573 | 26.10219 |
| ENSMUSG00000024258 | -0.19390801 | 0.033849 | 0.103328 | 1531.215 | 1696.548 | 1791.195 | 1739.049 | 2088.688 | 1696.174 | 1948.828 | 1996.817 |
| ENSMUSG00000024259 | 0.032850958 | 0.704615 | 0.814096 | 1496.003 | 1664.152 | 1455.811 | 1584.639 | 1462.667 | 1705.113 | 1434.862 | 1457.372 |
| ENSMUSG00000024260 | -0.002424925 | 0.972855 | 0.98497 | 1175.071 | 1078.64 | 1230.566 | 1156.778 | 1191.586 | 1095.027 | 1189.933 | 1175.686 |
| ENSMUSG00000024261 | 0.52011285 | 0.214964 | 0.373119 | 200.2048 | 70.78947 | 80.32634 | 233.771 | 81.90935 | 136.3196 | 119.5719 | 70.69342 |
| ENSMUSG00000024268 | 1.117413524 | 0.003045 | 0.01796 | 1025.169 | 827.8769 | 661.6572 | 964.4131 | 289.6081 | 824.6221 | 284.4653 | 205.5547 |
| ENSMUSG00000024269 | -0.131443151 | 0.375167 | 0.54362 | 379.2825 | 488.3274 | 499.3483 | 454.6026 | 442.7005 | 543.0438 | 426.2158 | 585.124 |
| ENSMUSG00000024270 | 0.222866348 | 0.006233 | 0.030311 | 1607.675 | 1712.145 | 1544.419 | 1709.72 | 1416.837 | 1545.328 | 1389.541 | 1280.095 |
| ENSMUSG00000024271 | 0.025591313 | 0.707186 | 0.815963 | 2920.576 | 2946.762 | 2703.768 | 2919.981 | 2938.01 | 2950.985 | 2586.224 | 2812.511 |
| ENSMUSG00000024273 | -0.000669149 | 0.995884 | 0.99765 | 452.7245 | 531.5209 | 456.2868 | 401.9825 | 532.4108 | 460.3581 | 436.823 | 410.0218 |
| ENSMUSG00000024274 | -0.532901323 | 0.131386 | 0.264232 | 21.12714 | 34.79482 | 19.87456 | 25.01608 | 46.80534 | 32.40385 | 24.10723 | 41.32846 |
| ENSMUSG00000024276 | 0.029098469 | 0.755635 | 0.848334 | 1356.161 | 1508.176 | 1162.662 | 1323.264 | 1340.778 | 1318.501 | 1333.612 | 1244.204 |
| ENSMUSG00000024277 | 0.175561761 | 0.005539 | 0.027694 | 1602.645 | 1713.345 | 1575.059 | 1736.461 | 1522.149 | 1458.173 | 1458.005 | 1426.92 |
| ENSMUSG00000024283 | 0.051282519 | 0.417275 | 0.582809 | 3086.575 | 3309.108 | 2943.091 | 3268.48 | 3069.65 | 3176.695 | 2988.332 | 2928.883 |
| ENSMUSG00000024286 | -0.090552012 | 0.107415 | 0.231883 | 2879.328 | 2819.581 | 3067.307 | 3075.253 | 3050.148 | 3155.464 | 3258.333 | 3151.839 |
| ENSMUSG00000024287 | 0.185926563 | 0.080074 | 0.189056 | 1808.886 | 2030.098 | 1739.024 | 1983.171 | 1596.257 | 1959.874 | 1513.934 | 1577.007 |
| ENSMUSG00000024290 | 0.470583616 | 0.021841 | 0.074979 | 2504.069 | 2420.04 | 1842.537 | 2012.501 | 1255.943 | 2333.077 | 1372.183 | 1374.715 |
| ENSMUSG00000024293 | -0.007985721 | 0.936195 | 0.962264 | 572.4449 | 688.6976 | 587.9557 | 610.7374 | 599.6935 | 581.0345 | 603.645 | 686.27 |
| ENSMUSG00000024294 | 0.20179064 | 0.144589 | 0.28269 | 3768.68 | 4043.399 | 3312.427 | 4012.062 | 2867.802 | 4214.735 | 3095.368 | 2983.262 |
| ENSMUSG00000024295 | -0.321518403 | 0.376207 | 0.544443 | 14.08476 | 20.39697 | 17.39024 | 18.97772 | 28.27823 | 21.23011 | 21.21436 | 17.40146 |
| ENSMUSG00000024298 | -0.303926248 | 0.085712 | 0.198135 | 316.9071 | 277.1588 | 275.7595 | 232.0457 | 378.3432 | 243.5876 | 365.4656 | 370.8686 |
| ENSMUSG00000024299 | 0.564771614 | 3.09E-13 | 1.06E-10 | 3346.137 | 3297.11 | 3085.525 | 3682.54 | 2288.586 | 2406.824 | 2134.936 | 2238.262 |
| ENSMUSG00000024300 | 0.036228626 | 0.908525 | 0.945354 | 29.17558 | 47.99286 | 52.17072 | 34.50494 | 33.15378 | 31.28647 | 46.28588 | 48.9416 |
| ENSMUSG00000024301 | 0.113312374 | 0.196641 | 0.351004 | 459.7668 | 494.3265 | 513.4261 | 482.2065 | 468.0534 | 470.4145 | 442.6087 | 421.9853 |
| ENSMUSG00000024302 | 0.940707057 | 1.74E-05 | 0.000352 | 901.4247 | 925.0624 | 650.0637 | 929.0455 | 455.377 | 626.8469 | 379.9299 | 312.1386 |
| ENSMUSG00000024304 | 0.195531343 | 0.215675 | 0.37387 | 1344.089 | 1452.984 | 1189.989 | 1520.805 | 1285.197 | 1489.46 | 1124.361 | 910.3137 |
| ENSMUSG00000024306 | -0.97963573 | 0.048111 | 0.132529 | 9.054489 | 7.198929 | 4.96864 | 7.763611 | 10.72622 | 13.40849 | 16.39292 | 16.31387 |
| ENSMUSG00000024308 | 0.086261801 | 0.602038 | 0.739909 | 1012.091 | 1407.391 | 1582.512 | 1000.643 | 1045.319 | 1223.525 | 1217.897 | 1225.715 |
| ENSMUSG00000024309 | -0.165562904 | 0.008219 | 0.037007 | 1832.025 | 1822.529 | 1777.945 | 1849.465 | 2184.249 | 1910.71 | 2006.686 | 2063.16 |
| ENSMUSG00000024312 | -0.229474864 | 0.019094 | 0.068265 | 886.3339 | 975.4549 | 992.0717 | 906.6173 | 1143.806 | 966.5286 | 1065.54 | 1232.241 |
| ENSMUSG00000024313 | -5.228780073 | 3.48E-08 | 2.39E-06 | 7.04238 | 7.198929 | 63.76421 | 12.07673 | 776.1886 | 175.4277 | 687.5382 | 1755.372 |
| ENSMUSG00000024317 | 0.167822284 | 0.263753 | 0.428017 | 889.3521 | 1051.044 | 836.3877 | 1062.752 | 822.9939 | 1086.088 | 753.1098 | 755.8758 |

| | | | | | | | | | | | |
|---|---|---|---|---|---|---|---|---|---|---|---|
| ENSMUSG00000024319 | -0.077847676 | 0.327325 | 0.495051 | 1100.623 | 1096.637 | 1268.659 | 1124.861 | 1239.366 | 1130.783 | 1245.862 | 1232.241 |
| ENSMUSG00000024325 | -0.150546472 | 0.035838 | 0.108022 | 886.3339 | 923.8626 | 924.9951 | 985.116 | 1028.742 | 970.9981 | 1082.897 | 1047.35 |
| ENSMUSG00000024327 | -0.517139706 | 0.001532 | 0.010697 | 702.2259 | 667.1008 | 757.7176 | 668.5332 | 1026.792 | 700.5936 | 1058.79 | 1214.839 |
| ENSMUSG00000024330 | -1.049741062 | 4.80E-05 | 0.000771 | 145.8779 | 213.5682 | 149.8873 | 153.547 | 511.9334 | 238.0007 | 341.3584 | 277.3357 |
| ENSMUSG00000024331 | 0.170252324 | 0.304882 | 0.473405 | 326.9677 | 315.5531 | 433.9279 | 286.391 | 313.0107 | 293.8694 | 261.3224 | 344.7664 |
| ENSMUSG00000024334 | 1.024371697 | 0.04923 | 0.134647 | 23.13925 | 10.79839 | 16.56213 | 12.07673 | 7.80089 | 4.469496 | 8.678602 | 9.78832 |
| ENSMUSG00000024335 | 0.020302933 | 0.626588 | 0.756629 | 6264.7 | 6225.874 | 6329.219 | 5977.118 | 6019.362 | 6152.262 | 6231.237 | 6045.919 |
| ENSMUSG00000024338 | -0.32709363 | 0.227851 | 0.387732 | 104.6297 | 110.3836 | 184.6678 | 80.22398 | 140.416 | 130.7328 | 178.3935 | 153.3503 |
| ENSMUSG00000024339 | -0.104666999 | 0.651906 | 0.775328 | 210.2654 | 196.7707 | 310.54 | 164.7611 | 274.9814 | 182.132 | 222.7508 | 269.7226 |
| ENSMUSG00000024346 | -0.40189938 | 0.000286 | 0.002998 | 1629.808 | 1695.348 | 1792.023 | 1620.007 | 2388.048 | 1824.672 | 2170.615 | 2517.773 |
| ENSMUSG00000024347 | 1.286324004 | 0.00049 | 0.004507 | 106.6418 | 80.38804 | 86.9512 | 138.8824 | 23.40267 | 72.62932 | 42.42872 | 31.54014 |
| ENSMUSG00000024349 | 0.453308302 | 0.00072 | 0.006096 | 557.3541 | 721.0927 | 650.8918 | 627.9899 | 558.7388 | 426.8369 | 468.6445 | 411.1094 |
| ENSMUSG00000024350 | 0.132261342 | 0.114422 | 0.241901 | 938.6487 | 865.0713 | 835.5596 | 807.4156 | 835.6704 | 770.9881 | 737.6812 | 798.2919 |
| ENSMUSG00000024352 | -0.066845111 | 0.497569 | 0.654039 | 545.2815 | 568.7154 | 520.051 | 523.6125 | 599.6935 | 612.321 | 550.6091 | 494.8539 |
| ENSMUSG00000024354 | -2.629234026 | 1.80E-05 | 0.000363 | 151.9142 | 53.99197 | 332.0708 | 144.0581 | 1004.365 | 453.6539 | 1006.718 | 1759.722 |
| ENSMUSG00000024357 | -0.272544999 | 0.003296 | 0.019031 | 730.3955 | 853.0731 | 778.4202 | 768.5975 | 1001.439 | 832.4437 | 974.8963 | 967.9561 |
| ENSMUSG00000024359 | -0.551496366 | 1.17E-05 | 0.000259 | 8148.034 | 7107.743 | 8840.866 | 7624.729 | 11143.57 | 9401.586 | 12420.04 | 13528.55 |
| ENSMUSG00000024360 | -0.260402634 | 1.00E-05 | 0.000228 | 3100.66 | 3065.544 | 3100.431 | 2890.651 | 3788.307 | 3413.578 | 3612.227 | 3744.576 |
| ENSMUSG00000024365 | 4.342487068 | NA | NA | 16528.47 | 14.39786 | 45.54586 | 1408.664 | 1.950223 | 2.234748 | 9.642892 | 873.3356 |
| ENSMUSG00000024366 | 1.863140234 | 0.000364 | 0.003587 | 46.2785 | 23.99643 | 28.15563 | 95.75121 | 12.67645 | 15.64324 | 15.42863 | 9.78832 |
| ENSMUSG00000024369 | -0.188767951 | 0.022255 | 0.076034 | 1187.144 | 1427.788 | 1351.47 | 1279.271 | 1527.024 | 1384.427 | 1549.613 | 1513.927 |
| ENSMUSG00000024370 | -0.012900315 | 0.849727 | 0.909714 | 2147.926 | 2250.865 | 2059.501 | 2177.262 | 2363.67 | 2174.41 | 2067.436 | 2103.401 |
| ENSMUSG00000024371 | 0.586981482 | 0.030971 | 0.096667 | 116.7023 | 269.9598 | 134.9814 | 158.7227 | 117.0134 | 113.9722 | 104.1432 | 116.3722 |
| ENSMUSG00000024376 | -0.485811557 | 0.000155 | 0.001874 | 577.4752 | 581.9134 | 592.0962 | 615.0506 | 955.6091 | 648.077 | 825.4315 | 884.2116 |
| ENSMUSG00000024378 | -0.607266354 | 0.000217 | 0.002419 | 1218.332 | 1094.237 | 1565.122 | 1051.538 | 1783.479 | 1491.694 | 2109.865 | 2126.241 |
| ENSMUSG00000024379 | -0.027931761 | 0.919497 | 0.952207 | 43.26034 | 47.99286 | 43.88965 | 43.13117 | 28.27823 | 50.28183 | 51.10733 | 52.20437 |
| ENSMUSG00000024381 | 0.166129797 | 0.00625 | 0.030347 | 2023.175 | 2061.293 | 2119.953 | 2038.379 | 1761.051 | 1963.226 | 1859.15 | 1765.16 |
| ENSMUSG00000024382 | -0.116835581 | 0.16392 | 0.308862 | 1115.714 | 1121.833 | 1173.427 | 1112.784 | 1237.416 | 1072.679 | 1316.255 | 1277.92 |
| ENSMUSG00000024383 | 0.218157087 | 0.098432 | 0.218611 | 951.7274 | 851.8733 | 826.4504 | 893.6779 | 667.9512 | 953.1201 | 685.6096 | 724.3357 |
| ENSMUSG00000024384 | 0.001597319 | 0.981778 | 0.989807 | 1497.009 | 1402.591 | 1409.437 | 1463.872 | 1368.081 | 1566.558 | 1386.648 | 1448.671 |
| ENSMUSG00000024386 | -2.232133332 | 0.000177 | 0.002063 | 21.12714 | 4.799286 | 38.92101 | 22.42821 | 131.64 | 33.52122 | 80.036 | 168.5766 |
| ENSMUSG00000024387 | -0.256466631 | 0.001069 | 0.008221 | 1800.837 | 1900.517 | 2066.126 | 2010.775 | 2407.55 | 2102.898 | 2370.223 | 2413.365 |
| ENSMUSG00000024388 | -1.875016164 | 6.60E-08 | 4.03E-06 | 51.30877 | 26.39607 | 64.59232 | 41.40593 | 147.2418 | 106.1505 | 166.822 | 257.7591 |
| ENSMUSG00000024391 | -2.463233726 | 5.34E-10 | 7.35E-08 | 63.38142 | 26.39607 | 28.98373 | 37.95543 | 235.9769 | 102.7984 | 192.8578 | 332.8029 |
| ENSMUSG00000024392 | -0.072313156 | 0.343189 | 0.511957 | 3812.946 | 3905.419 | 4298.702 | 3954.266 | 4309.017 | 3763.316 | 4436.694 | 4284.021 |
| ENSMUSG00000024393 | -0.007950135 | 0.91669 | 0.950634 | 4960.854 | 4972.06 | 5642.719 | 4685.771 | 5320.207 | 4853.873 | 4989.232 | 5211.736 |
| ENSMUSG00000024395 | -0.193964353 | 0.395988 | 0.563264 | 164.9929 | 115.1829 | 134.1533 | 198.4034 | 154.0676 | 148.6108 | 229.5008 | 169.6642 |
| ENSMUSG00000024397 | 0.376798173 | 0.157146 | 0.299602 | 50.30272 | 64.79036 | 86.12309 | 55.2079 | 55.58134 | 42.46022 | 55.92877 | 43.50364 |
| ENSMUSG00000024399 | 0.444727609 | 0.269363 | 0.434105 | 14.08476 | 23.99643 | 22.35888 | 26.74133 | 19.50223 | 16.76061 | 10.60718 | 17.40146 |
| ENSMUSG00000024400 | -0.032564532 | 0.589587 | 0.729659 | 1965.83 | 1910.116 | 1811.897 | 1820.136 | 1885.865 | 1877.188 | 1888.078 | 2025.095 |
| ENSMUSG00000024403 | 0.037972816 | 0.785034 | 0.867736 | 236.4228 | 223.1668 | 207.0267 | 228.5952 | 188.1965 | 232.4138 | 251.6795 | 199.0292 |
| ENSMUSG00000024404 | -0.096509141 | 0.130256 | 0.262661 | 2275.695 | 2224.469 | 2272.325 | 2337.71 | 2315.889 | 2310.73 | 2603.581 | 2511.248 |
| ENSMUSG00000024410 | -0.008116537 | 0.946268 | 0.969217 | 558.3602 | 448.7333 | 470.3646 | 458.0531 | 497.3068 | 430.189 | 548.6805 | 468.7518 |
| ENSMUSG00000024411 | -1.145454943 | 0.148901 | 0.288674 | 127.7689 | 97.18554 | 874.4806 | 179.4257 | 666.9761 | 303.9258 | 1040.468 | 818.9561 |
| ENSMUSG00000024413 | 0.172571483 | 0.088046 | 0.201935 | 1178.09 | 983.8537 | 965.5723 | 936.8091 | 870.7744 | 882.7255 | 982.6107 | 867.8977 |
| ENSMUSG00000024414 | -0.274518495 | 0.006451 | 0.031119 | 1214.308 | 1210.62 | 1294.331 | 1231.826 | 1610.884 | 1235.816 | 1560.22 | 1581.357 |
| ENSMUSG00000024420 | 0.332392219 | 0.050726 | 0.137433 | 1077.484 | 1464.982 | 1241.332 | 1252.529 | 1072.622 | 1284.98 | 843.753 | 798.2919 |
| ENSMUSG00000024421 | -0.233909491 | 0.112877 | 0.239757 | 152.9203 | 151.1775 | 175.5586 | 135.4319 | 198.9227 | 160.9019 | 187.0721 | 176.1898 |
| ENSMUSG00000024422 | -0.11000816 | 0.203804 | 0.35981 | 1580.511 | 1559.768 | 1727.43 | 1678.665 | 1947.297 | 1564.324 | 1792.614 | 1761.898 |
| ENSMUSG00000024423 | 0.303745318 | 0.024887 | 0.082775 | 1316.925 | 1348.599 | 1152.724 | 1362.082 | 985.8375 | 1330.793 | 968.1463 | 912.4889 |
| ENSMUSG00000024424 | 0.152989169 | 0.322587 | 0.490822 | 357.1493 | 287.9572 | 263.3379 | 356.2635 | 255.4792 | 268.1698 | 280.6081 | 333.8905 |
| ENSMUSG00000024425 | -0.014784143 | 0.858007 | 0.914917 | 4055.405 | 3531.075 | 3873.055 | 4168.197 | 3830.237 | 3621.409 | 4101.122 | 4239.43 |
| ENSMUSG00000024426 | 0.401229806 | 0.001004 | 0.007863 | 960.7819 | 929.8617 | 822.3099 | 885.0517 | 672.8268 | 817.9178 | 655.7166 | 577.5109 |
| ENSMUSG00000024427 | 0.278573695 | 0.065129 | 0.164345 | 946.6971 | 775.0847 | 610.3146 | 936.8091 | 709.881 | 701.7109 | 642.2166 | 640.5912 |
| ENSMUSG00000024429 | -0.179050005 | 0.037055 | 0.110607 | 1365.216 | 1545.37 | 1704.243 | 1459.559 | 1688.893 | 1660.418 | 1739.578 | 1790.175 |
| ENSMUSG00000024430 | 0.401623143 | 0.232452 | 0.393014 | 57.3451 | 45.59322 | 81.15445 | 58.6584 | 28.27823 | 53.63396 | 65.57166 | 36.9781 |
| ENSMUSG00000024431 | 0.043614397 | 0.674413 | 0.791543 | 1521.154 | 1672.551 | 1815.21 | 1677.803 | 1400.26 | 1619.075 | 1627.72 | 1843.467 |
| ENSMUSG00000024436 | -0.338436548 | 0.013189 | 0.052586 | 481.9 | 447.5334 | 501.8326 | 496.8711 | 676.7272 | 463.7102 | 653.7881 | 643.8539 |
| ENSMUSG00000024440 | -0.428012148 | 0.034617 | 0.105219 | 315.9011 | 319.1525 | 438.8965 | 306.2313 | 424.1734 | 357.5597 | 449.3588 | 627.5401 |
| ENSMUSG00000024442 | -0.239238712 | 0.015677 | 0.059619 | 1216.32 | 1166.227 | 1216.489 | 1117.097 | 1483.144 | 1160.952 | 1467.648 | 1453.022 |
| ENSMUSG00000024446 | -0.234420954 | 0.091059 | 0.206817 | 371.2341 | 428.3363 | 407.4285 | 389.9058 | 444.6508 | 374.3203 | 510.109 | 548.1459 |
| ENSMUSG00000024451 | -0.042046419 | 0.791751 | 0.8726 | 884.3218 | 703.0954 | 1005.321 | 816.9044 | 896.1273 | 672.6592 | 918.0033 | 1025.598 |
| ENSMUSG00000024454 | 0.06883781 | 0.307865 | 0.476354 | 2732.444 | 2938.363 | 2955.513 | 3150.301 | 2957.513 | 2783.379 | 2815.724 | 2672.211 |
| ENSMUSG00000024456 | -0.069716981 | 0.405394 | 0.572277 | 1118.732 | 1089.438 | 1268.659 | 1224.063 | 1173.059 | 1249.224 | 1187.04 | 1331.211 |

| | | | | | | | | | | | |
|---|---|---|---|---|---|---|---|---|---|---|---|
| ENSMUSG00000024457 | -0.019449164 | 0.755916 | 0.848334 | 1318.937 | 1390.593 | 1469.889 | 1362.945 | 1400.26 | 1397.835 | 1450.291 | 1370.365 |
| ENSMUSG00000024459 | 1.122664977 | 0.000623 | 0.005433 | 111.672 | 113.983 | 80.32634 | 125.943 | 41.92979 | 83.80306 | 41.46443 | 31.54014 |
| ENSMUSG00000024462 | 0.496110287 | 0.00033 | 0.003338 | 2112.714 | 1841.726 | 1652.073 | 2142.757 | 1418.787 | 1605.667 | 1369.291 | 1100.642 |
| ENSMUSG00000024472 | 0.185155794 | 0.025562 | 0.084334 | 1591.578 | 1857.324 | 1580.856 | 1512.179 | 1469.493 | 1457.056 | 1388.576 | 1433.445 |
| ENSMUSG00000024474 | -0.045122058 | 0.301044 | 0.469406 | 5019.205 | 4898.871 | 4886.657 | 4900.564 | 4872.631 | 5215.902 | 5143.518 | 5100.802 |
| ENSMUSG00000024477 | -0.254389816 | 0.048081 | 0.132465 | 487.9364 | 451.1329 | 483.6143 | 448.5642 | 603.5939 | 434.6585 | 588.2164 | 604.7006 |
| ENSMUSG00000024479 | 0.020708695 | 0.856429 | 0.914026 | 2559.402 | 2725.995 | 3298.349 | 2666.369 | 2716.66 | 2981.154 | 3007.618 | 2386.175 |
| ENSMUSG00000024480 | 0.092844847 | 0.454101 | 0.616212 | 1338.052 | 1414.59 | 1204.067 | 1500.102 | 1250.093 | 1549.798 | 1177.397 | 1139.795 |
| ENSMUSG00000024481 | 1.951193469 | 0.003952 | 0.021671 | 60.36326 | 107.9839 | 37.2648 | 96.61383 | 19.50223 | 40.22547 | 17.3572 | 1.087591 |
| ENSMUSG00000024483 | 0.060039836 | 0.548764 | 0.697048 | 2595.62 | 2362.449 | 2153.905 | 2058.22 | 2077.962 | 2453.754 | 2111.793 | 2151.255 |
| ENSMUSG00000024485 | -1.675977914 | 0.048155 | 0.132611 | 148.896 | 69.58965 | 772.6235 | 121.6299 | 779.1139 | 339.6817 | 1079.04 | 1359.489 |
| ENSMUSG00000024486 | -0.767357754 | 0.003969 | 0.021743 | 222.338 | 153.5772 | 345.3205 | 207.8923 | 345.1894 | 296.1041 | 429.1087 | 514.4306 |
| ENSMUSG00000024487 | 0.031976045 | 0.700142 | 0.810722 | 1484.936 | 1695.348 | 1641.307 | 1659.688 | 1590.407 | 1735.282 | 1443.541 | 1571.569 |
| ENSMUSG00000024491 | 0.281154369 | 0.063015 | 0.160504 | 1191.168 | 1136.231 | 938.2448 | 1060.164 | 776.1886 | 1154.247 | 786.86 | 842.8831 |
| ENSMUSG00000024493 | -0.04170635 | 0.556685 | 0.70347 | 1944.703 | 2028.898 | 1900.505 | 1883.107 | 1924.87 | 1841.433 | 2114.686 | 2099.051 |
| ENSMUSG00000024498 | 0.215009337 | 0.080901 | 0.190399 | 3535.275 | 3586.267 | 3209.741 | 3539.344 | 2905.832 | 3732.029 | 2740.51 | 2572.153 |
| ENSMUSG00000024500 | 0.111201924 | 0.564869 | 0.70973 | 171.0292 | 207.5691 | 225.245 | 214.7932 | 145.2916 | 244.7049 | 169.7149 | 200.1168 |
| ENSMUSG00000024501 | 0.772071275 | 7.80E-06 | 0.00019 | 3085.569 | 2695.999 | 2654.082 | 3473.785 | 1649.888 | 2376.655 | 1431.969 | 1517.19 |
| ENSMUSG00000024502 | 0.994054381 | 0.005305 | 0.026803 | 50.30272 | 50.3925 | 37.2648 | 47.44429 | 24.37778 | 33.52122 | 24.10723 | 10.87591 |
| ENSMUSG00000024503 | -2.621590653 | 0.000306 | 0.003155 | 102.6175 | 27.5959 | 313.8524 | 99.2017 | 784.9646 | 368.7335 | 959.4677 | 1234.416 |
| ENSMUSG00000024505 | 0.438732977 | 0.009955 | 0.042431 | 245.4773 | 219.5673 | 216.9639 | 211.3428 | 179.4205 | 204.4795 | 142.7148 | 132.6861 |
| ENSMUSG00000024507 | -0.32084478 | 0.007318 | 0.033955 | 2313.925 | 2288.06 | 2484.32 | 2366.176 | 2991.641 | 2326.373 | 3048.118 | 3441.138 |
| ENSMUSG00000024511 | -0.279579198 | 0.251372 | 0.415004 | 181.0898 | 215.9679 | 361.8826 | 231.1831 | 392.9699 | 251.4092 | 271.9295 | 287.124 |
| ENSMUSG00000024513 | 0.013156015 | 0.850869 | 0.910548 | 2012.109 | 1812.93 | 1980.003 | 2040.105 | 1862.463 | 1974.4 | 1869.757 | 2074.036 |
| ENSMUSG00000024515 | 0.01300157 | 0.80673 | 0.88179 | 2594.614 | 2566.418 | 2512.476 | 2600.81 | 2487.509 | 2702.928 | 2508.116 | 2485.146 |
| ENSMUSG00000024516 | 0.178291338 | 0.07901 | 0.187542 | 677.0746 | 663.5013 | 567.253 | 602.1112 | 506.0828 | 603.382 | 566.0377 | 540.5328 |
| ENSMUSG00000024517 | 0.70999683 | 0.027562 | 0.089052 | 53.32088 | 94.7859 | 66.24853 | 55.2079 | 49.73068 | 33.52122 | 53.0359 | 27.18978 |
| ENSMUSG00000024518 | 0.681536469 | 0.151992 | 0.29251 | 11.0666 | 11.99822 | 14.07781 | 15.52722 | 6.825779 | 8.938993 | 9.642892 | 7.613138 |
| ENSMUSG00000024519 | 0.333228688 | 0.118697 | 0.247522 | 81.4904 | 134.38 | 125.8722 | 114.7289 | 97.51113 | 98.32892 | 75.21455 | 91.35765 |
| ENSMUSG00000024521 | 0.458538186 | 0.095446 | 0.213737 | 186.1201 | 106.7841 | 271.619 | 176.8378 | 146.2667 | 131.8501 | 110.8933 | 152.2628 |
| ENSMUSG00000024524 | 0.151340415 | 0.196699 | 0.351025 | 263.5862 | 242.3639 | 271.619 | 276.9021 | 225.2507 | 269.2872 | 229.5008 | 228.3941 |
| ENSMUSG00000024525 | -0.784421919 | 0.000336 | 0.003377 | 297.7921 | 290.3568 | 390.0382 | 337.2858 | 568.4899 | 370.9682 | 549.6448 | 778.7152 |
| ENSMUSG00000024527 | -0.519128502 | 0.000219 | 0.002428 | 1789.771 | 1576.565 | 1934.457 | 1645.886 | 2439.728 | 1924.118 | 2605.509 | 2987.613 |
| ENSMUSG00000024528 | -0.181692936 | 0.016467 | 0.061447 | 523.1483 | 526.7217 | 497.6921 | 524.4751 | 596.7681 | 570.9782 | 572.7878 | 607.9634 |
| ENSMUSG00000024529 | 0.881759288 | 4.50E-09 | 4.48E-07 | 5439.736 | 4424.942 | 4403.871 | 6300.602 | 2303.213 | 3137.586 | 3065.475 | 2658.073 |
| ENSMUSG00000024530 | 0.42603666 | 0.021197 | 0.073453 | 140.8476 | 137.9795 | 146.5749 | 127.6683 | 103.3618 | 74.86406 | 120.5361 | 112.0219 |
| ENSMUSG00000024533 | -0.455519084 | 6.36E-06 | 0.000162 | 1089.557 | 923.8626 | 1064.117 | 962.6878 | 1282.271 | 1271.572 | 1552.506 | 1435.62 |
| ENSMUSG00000024534 | -0.569293352 | 3.48E-09 | 3.59E-07 | 1672.062 | 1572.966 | 1553.528 | 1667.451 | 2748.839 | 2273.856 | 2461.83 | 2107.752 |
| ENSMUSG00000024535 | 0.104750085 | 0.198743 | 0.353372 | 655.9474 | 699.496 | 665.7977 | 659.907 | 612.3699 | 654.7812 | 570.8592 | 655.8174 |
| ENSMUSG00000024537 | -0.067154393 | 0.402147 | 0.569261 | 745.4863 | 695.8965 | 659.1729 | 738.4057 | 725.4828 | 767.636 | 773.3599 | 706.9342 |
| ENSMUSG00000024538 | 0.099662453 | 0.139208 | 0.275107 | 4965.884 | 5777.141 | 5224.525 | 5616.542 | 5147.613 | 5089.639 | 5086.625 | 4815.853 |
| ENSMUSG00000024539 | 0.087540052 | 0.220179 | 0.378964 | 1227.386 | 1337.801 | 1238.019 | 1230.964 | 1119.428 | 1272.689 | 1190.897 | 1152.847 |
| ENSMUSG00000024542 | 0.22189432 | 0.056191 | 0.147729 | 1165.011 | 1037.846 | 855.4342 | 937.6717 | 899.0526 | 902.8383 | 834.1101 | 787.4159 |
| ENSMUSG00000024544 | -0.295046915 | 0.093613 | 0.210617 | 216.3017 | 185.9723 | 174.7305 | 175.1126 | 299.3592 | 212.3011 | 198.6436 | 210.9927 |
| ENSMUSG00000024548 | 0.492031201 | 0.001803 | 0.012174 | 1449.724 | 1323.403 | 1480.655 | 1383.648 | 903.9282 | 1356.492 | 928.6105 | 821.1313 |
| ENSMUSG00000024552 | -1.56598976 | 0.000823 | 0.006715 | 88.53278 | 99.58519 | 384.2415 | 108.6906 | 378.3432 | 311.7474 | 743.4669 | 586.2116 |
| ENSMUSG00000024556 | 0.037315948 | 0.762363 | 0.852273 | 1145.896 | 1388.194 | 1201.583 | 1273.232 | 1264.719 | 1451.469 | 1045.289 | 1119.131 |
| ENSMUSG00000024558 | -0.194787312 | 0.30586 | 0.474366 | 222.338 | 223.1668 | 228.5574 | 176.8378 | 256.4543 | 168.7235 | 261.3224 | 286.0365 |
| ENSMUSG00000024560 | -0.018063604 | 0.776995 | 0.862569 | 2168.047 | 2270.062 | 2191.17 | 2354.962 | 2352.944 | 2106.25 | 2323.937 | 2312.219 |
| ENSMUSG00000024561 | 0.008192609 | 0.891816 | 0.934606 | 1229.398 | 1249.014 | 1168.458 | 1197.321 | 1218.889 | 1173.243 | 1174.504 | 1246.379 |
| ENSMUSG00000024563 | -0.182735383 | 0.00765 | 0.035025 | 1225.374 | 1189.023 | 1186.677 | 1164.542 | 1450.966 | 1278.276 | 1380.862 | 1295.321 |
| ENSMUSG00000024565 | -1.224547991 | 0.000522 | 0.004735 | 72.43591 | 25.19625 | 57.96746 | 47.44429 | 92.63557 | 83.80306 | 136.9291 | 163.1387 |
| ENSMUSG00000024566 | 0.008983422 | 0.873101 | 0.924072 | 1918.546 | 1938.912 | 1840.053 | 1783.905 | 1840.035 | 1904.005 | 1809.971 | 1877.182 |
| ENSMUSG00000024570 | -0.240002403 | 0.008429 | 0.037706 | 1011.085 | 934.661 | 1042.586 | 1008.407 | 1141.855 | 1074.914 | 1165.826 | 1341 |
| ENSMUSG00000024571 | -0.03724658 | 0.581886 | 0.722795 | 2846.128 | 2839.978 | 2932.326 | 3118.384 | 3218.842 | 2799.022 | 2985.439 | 3041.992 |
| ENSMUSG00000024575 | 0.126479622 | 0.31658 | 0.4848 | 386.3249 | 508.7243 | 501.0045 | 442.5259 | 427.0988 | 441.3628 | 370.287 | 445.9123 |
| ENSMUSG00000024576 | 0.040580685 | 0.482022 | 0.640751 | 11195.37 | 11310.72 | 10853.99 | 11132.16 | 10501.95 | 11720.14 | 10753.75 | 10282.09 |
| ENSMUSG00000024579 | 0.133298247 | 0.292781 | 0.460125 | 404.4339 | 465.5308 | 343.6643 | 360.5766 | 343.2392 | 365.3813 | 340.3941 | 382.8321 |
| ENSMUSG00000024580 | -0.19957825 | 0.038028 | 0.112563 | 860.1765 | 739.0901 | 775.9359 | 740.9936 | 854.1975 | 830.209 | 891.9675 | 1002.759 |
| ENSMUSG00000024581 | -0.011510465 | 0.91918 | 0.952207 | 1022.151 | 847.074 | 780.0764 | 920.4193 | 793.7406 | 931.89 | 964.2892 | 908.1386 |
| ENSMUSG00000024583 | -0.210502364 | 4.76E-05 | 0.000766 | 2271.671 | 2415.241 | 2391.572 | 2461.927 | 2691.307 | 2837.013 | 2770.403 | 2742.905 |
| ENSMUSG00000024587 | -0.159176468 | 0.044602 | 0.12583 | 4048.363 | 3748.242 | 3969.943 | 3946.502 | 4564.496 | 3857.175 | 4378.837 | 4746.248 |
| ENSMUSG00000024588 | -0.083670323 | 0.466197 | 0.627123 | 1422.561 | 1350.999 | 1554.356 | 1552.722 | 1683.042 | 1253.694 | 1644.113 | 1652.051 |
| ENSMUSG00000024589 | -0.534201083 | 0.00487 | 0.025275 | 1208.271 | 963.4567 | 1794.507 | 1267.194 | 1734.723 | 1603.432 | 2151.329 | 2093.613 |

| | | | | | | | | | | | |
|---|---|---|---|---|---|---|---|---|---|---|---|
| ENSMUSG00000024590 | -0.007317721 | 0.917285 | 0.951037 | 5834.109 | 6702.203 | 6119.708 | 6194.499 | 6230.961 | 6751.174 | 5946.771 | 6045.919 |
| ENSMUSG00000024592 | 0.042198805 | 0.772673 | 0.858991 | 273.6468 | 245.9634 | 309.7119 | 253.6113 | 242.8027 | 284.9304 | 224.6794 | 302.3503 |
| ENSMUSG00000024593 | 1.426343265 | 3.20E-08 | 2.26E-06 | 526.1664 | 441.5343 | 382.5853 | 433.8996 | 134.5654 | 269.2872 | 119.5719 | 141.3868 |
| ENSMUSG00000024594 | 0.108207611 | 0.175318 | 0.323697 | 1930.618 | 1913.715 | 1848.334 | 1945.216 | 1760.076 | 2000.1 | 1683.649 | 1643.35 |
| ENSMUSG00000024597 | 0.642559254 | 0.008216 | 0.037001 | 7048.417 | 8055.602 | 5841.464 | 7535.016 | 3912.147 | 7451.768 | 3517.727 | 3362.832 |
| ENSMUSG00000024598 | 0.575995777 | 0.010162 | 0.043136 | 7367.336 | 9046.654 | 6688.617 | 7811.918 | 4864.83 | 7902.07 | 4505.159 | 3466.153 |
| ENSMUSG00000024600 | 0.387652748 | 0.014634 | 0.056634 | 637.8385 | 782.2836 | 626.0486 | 803.9651 | 610.4197 | 604.4994 | 553.502 | 408.9342 |
| ENSMUSG00000024601 | 0.227286153 | 0.005278 | 0.026705 | 2699.244 | 3244.317 | 2948.06 | 3142.537 | 2562.593 | 2752.092 | 2547.652 | 2416.627 |
| ENSMUSG00000024603 | 0.000430825 | 0.993525 | 0.996332 | 2157.987 | 2176.476 | 2052.048 | 2203.14 | 2139.394 | 2176.645 | 2193.758 | 2074.036 |
| ENSMUSG00000024604 | -0.045964678 | 0.497793 | 0.65413 | 1610.693 | 1748.14 | 1840.053 | 1773.554 | 1890.741 | 1792.268 | 1789.721 | 1727.095 |
| ENSMUSG00000024608 | -0.163705063 | 0.043784 | 0.124184 | 47109.5 | 56543.99 | 52168.23 | 54059.75 | 63888.32 | 55044.08 | 59676.96 | 56487.31 |
| ENSMUSG00000024610 | -1.066020768 | 0.000695 | 0.005913 | 43.26034 | 47.99286 | 95.23226 | 68.14726 | 181.3707 | 93.85942 | 132.1076 | 128.3357 |
| ENSMUSG00000024613 | 0.042913002 | 0.718907 | 0.824245 | 1282.719 | 1255.013 | 1153.553 | 949.7485 | 1056.046 | 1254.811 | 1055.897 | 1136.533 |
| ENSMUSG00000024614 | 0.093076373 | 0.302342 | 0.470649 | 1849.128 | 1917.315 | 1828.459 | 1768.378 | 1561.153 | 1966.578 | 1765.613 | 1609.635 |
| ENSMUSG00000024617 | 0.275098807 | 0.260699 | 0.424778 | 84.50857 | 124.7814 | 151.5435 | 119.042 | 125.7894 | 103.9158 | 77.14313 | 90.27006 |
| ENSMUSG00000024620 | -0.057679641 | 0.646745 | 0.77119 | 2442.7 | 2075.691 | 2655.738 | 2377.39 | 2418.276 | 2019.095 | 2796.439 | 2709.189 |
| ENSMUSG00000024621 | -0.395558 | 0.114502 | 0.241985 | 428.5792 | 644.3042 | 724.5933 | 501.1842 | 622.121 | 499.4662 | 850.503 | 1051.701 |
| ENSMUSG00000024622 | -0.191236911 | 0.164278 | 0.309253 | 662.9898 | 662.3015 | 681.5318 | 603.8364 | 822.0188 | 558.687 | 805.1815 | 792.8539 |
| ENSMUSG00000024639 | 0.324933548 | 0.045627 | 0.127741 | 3784.776 | 3873.024 | 3514.485 | 3397.874 | 2389.023 | 4006.904 | 2697.117 | 2539.525 |
| ENSMUSG00000024640 | -0.023345256 | 0.826452 | 0.894552 | 3229.434 | 2853.176 | 3000.23 | 2886.338 | 3136.933 | 2916.346 | 3527.37 | 2581.941 |
| ENSMUSG00000024642 | 0.292693824 | 0.004239 | 0.02279 | 794.7829 | 914.264 | 828.9347 | 911.793 | 670.8766 | 812.331 | 668.2524 | 665.6057 |
| ENSMUSG00000024644 | -0.998696424 | 7.00E-06 | 0.000175 | 1937.661 | 2076.891 | 2611.02 | 1850.327 | 4527.442 | 2559.904 | 4068.336 | 5781.634 |
| ENSMUSG00000024645 | -0.041964536 | 0.619384 | 0.751617 | 1055.351 | 1033.046 | 893.5271 | 1049.813 | 1095.05 | 994.4629 | 1052.039 | 1004.934 |
| ENSMUSG00000024646 | -0.632759192 | 5.94E-06 | 0.000153 | 2042.29 | 1859.723 | 2317.87 | 2178.124 | 3376.81 | 2495.096 | 3287.262 | 3865.299 |
| ENSMUSG00000024647 | 0.280520223 | 0.48263 | 0.64109 | 52.31483 | 21.59679 | 43.88965 | 47.44429 | 17.552 | 40.22547 | 34.71441 | 44.59123 |
| ENSMUSG00000024650 | -4.559778162 | 1.03E-06 | 3.82E-05 | 60.36326 | 9.598572 | 227.7293 | 56.07053 | 2274.935 | 611.2036 | 1394.362 | 4074.116 |
| ENSMUSG00000024654 | -0.112967444 | 0.150677 | 0.290798 | 764.6013 | 757.0874 | 768.483 | 711.6644 | 871.7495 | 751.9928 | 823.5029 | 796.1167 |
| ENSMUSG00000024659 | 0.123483468 | 0.318417 | 0.486684 | 1033.218 | 1324.603 | 1114.632 | 1205.948 | 1092.125 | 1243.637 | 1048.182 | 909.2261 |
| ENSMUSG00000024660 | 0.016972468 | 0.788244 | 0.86993 | 1990.982 | 1917.315 | 1845.022 | 1942.628 | 1919.994 | 1971.048 | 1950.757 | 1761.898 |
| ENSMUSG00000024661 | -0.374566068 | 0.002969 | 0.017618 | 11101.81 | 10629.22 | 11999.27 | 11941.3 | 16360.42 | 11586.05 | 14178.91 | 17087.14 |
| ENSMUSG00000024663 | -0.141181493 | 0.236308 | 0.397348 | 431.5973 | 489.5272 | 546.5504 | 414.0593 | 562.6392 | 503.9357 | 518.7876 | 489.416 |
| ENSMUSG00000024664 | -1.365913954 | 1.01E-05 | 0.000229 | 400.4096 | 281.9581 | 464.5678 | 401.9825 | 1027.767 | 496.1141 | 851.4673 | 1619.423 |
| ENSMUSG00000024665 | 0.158860311 | 0.531654 | 0.683254 | 1802.849 | 759.487 | 844.6688 | 985.9787 | 1012.166 | 836.9132 | 979.7178 | 1106.08 |
| ENSMUSG00000024666 | 0.030837102 | 0.746175 | 0.841259 | 425.561 | 478.7288 | 439.7246 | 426.136 | 473.9041 | 413.4284 | 413.6801 | 428.5109 |
| ENSMUSG00000024667 | -0.074133347 | 0.468343 | 0.628589 | 397.3915 | 391.1418 | 419.022 | 411.4714 | 483.6552 | 410.0763 | 421.3944 | 389.3576 |
| ENSMUSG00000024668 | -0.057492156 | 0.410191 | 0.576403 | 979.8969 | 1120.633 | 1014.431 | 1004.956 | 1071.647 | 1062.623 | 1050.111 | 1098.467 |
| ENSMUSG00000024672 | 0.148463308 | 0.762059 | 0.852086 | 12.07265 | 50.3925 | 62.9361 | 25.01608 | 37.05423 | 35.75597 | 35.6787 | 27.18978 |
| ENSMUSG00000024675 | -0.236388616 | 0.718428 | 0.824127 | 7.04238 | 10.79839 | 20.70267 | 2.58787 | 11.70134 | 10.05637 | 10.60718 | 16.31387 |
| ENSMUSG00000024677 | 0.504039583 | 0.01788 | 0.065359 | 136.8234 | 141.5789 | 161.4808 | 145.7834 | 78.98402 | 129.6154 | 123.429 | 81.56933 |
| ENSMUSG00000024678 | 2.103231877 | 0.002266 | 0.014359 | 72.43591 | 61.1909 | 33.95237 | 83.67448 | 13.65156 | 37.99072 | 3.857157 | 3.262773 |
| ENSMUSG00000024679 | 1.113360627 | 0.000133 | 0.001673 | 271.6347 | 145.1784 | 223.5888 | 181.1509 | 91.66046 | 84.92043 | 140.7862 | 61.99269 |
| ENSMUSG00000024680 | -2.330851848 | 8.58E-07 | 3.29E-05 | 22.1332 | 28.79572 | 4.96864 | 12.93935 | 62.40712 | 63.69032 | 104.1432 | 109.8467 |
| ENSMUSG00000024683 | -0.587194412 | 0.000108 | 0.001425 | 546.2875 | 574.7145 | 644.267 | 513.261 | 934.1566 | 644.7249 | 846.6459 | 997.321 |
| ENSMUSG00000024687 | -0.11608487 | 0.133821 | 0.267352 | 2262.616 | 2042.096 | 2269.84 | 2228.156 | 2299.312 | 2197.875 | 2446.402 | 2600.43 |
| ENSMUSG00000024691 | 0.10897047 | 0.499307 | 0.655685 | 493.9727 | 761.8867 | 549.8628 | 508.0852 | 492.4312 | 559.8044 | 493.7161 | 597.0875 |
| ENSMUSG00000024694 | -2.737531005 | 5.27E-05 | 0.000824 | 316.9071 | 34.79482 | 410.7409 | 339.011 | 1937.546 | 1055.919 | 2195.686 | 2161.043 |
| ENSMUSG00000024695 | 0.02502273 | 0.777549 | 0.862822 | 4439.718 | 4460.936 | 4271.374 | 4572.767 | 3987.23 | 5038.24 | 4364.373 | 4051.277 |
| ENSMUSG00000024696 | 0.276529012 | 0.408256 | 0.574587 | 32.19374 | 44.3934 | 42.23344 | 39.68068 | 39.00445 | 42.46022 | 31.82154 | 17.40146 |
| ENSMUSG00000024697 | -0.392629809 | 0.241157 | 0.403297 | 169.0171 | 99.58519 | 246.7758 | 125.943 | 209.6489 | 125.1459 | 206.3579 | 302.3503 |
| ENSMUSG00000024712 | 0.075525941 | 0.416958 | 0.582497 | 2344.107 | 2330.053 | 2054.533 | 2157.421 | 1945.347 | 2416.88 | 2033.686 | 2035.971 |
| ENSMUSG00000024713 | 0.366889402 | 0.009219 | 0.040161 | 1571.457 | 1186.623 | 1344.017 | 1110.196 | 830.7948 | 1054.801 | 1095.432 | 1061.489 |
| ENSMUSG00000024725 | -0.540948862 | 0.00099 | 0.007785 | 1559.384 | 1286.209 | 1927.004 | 1602.754 | 2233.98 | 1773.273 | 2556.331 | 2715.715 |
| ENSMUSG00000024726 | -0.135586343 | 0.138828 | 0.274647 | 684.117 | 695.8965 | 597.0649 | 728.0542 | 761.5619 | 701.7109 | 775.2885 | 729.7736 |
| ENSMUSG00000024727 | 0.224830507 | 0.104863 | 0.228252 | 243.4652 | 205.1695 | 207.0267 | 221.6942 | 159.9183 | 204.4795 | 183.2149 | 204.4671 |
| ENSMUSG00000024732 | -0.113686644 | 0.340409 | 0.50896 | 470.8334 | 447.5334 | 576.3622 | 458.0531 | 587.017 | 498.3488 | 530.359 | 499.2043 |
| ENSMUSG00000024735 | -0.263838445 | 0.00173 | 0.011775 | 5101.702 | 5906.721 | 5861.339 | 5518.202 | 7051.03 | 6014.825 | 6613.095 | 7199.853 |
| ENSMUSG00000024736 | -0.252555987 | 0.07292 | 0.177571 | 1077.484 | 1294.607 | 1441.734 | 1133.487 | 1457.791 | 1190.003 | 1534.184 | 1711.868 |
| ENSMUSG00000024737 | -0.205923799 | 0.516686 | 0.670791 | 38.23007 | 41.99375 | 52.17072 | 43.9938 | 53.63112 | 26.81698 | 57.85735 | 65.25547 |
| ENSMUSG00000024740 | -0.190879998 | 0.026032 | 0.085505 | 14141.1 | 13925.13 | 14008.25 | 13279.23 | 16268.76 | 13411.84 | 16786.35 | 16714.1 |
| ENSMUSG00000024742 | -0.522223643 | 0.001054 | 0.008137 | 150.9082 | 196.7707 | 167.2775 | 195.8155 | 258.4045 | 219.0053 | 239.1437 | 303.4379 |
| ENSMUSG00000024743 | 0.240820428 | 0.142451 | 0.279688 | 292.7618 | 250.7627 | 233.5261 | 310.5445 | 208.6738 | 274.874 | 193.8221 | 244.708 |
| ENSMUSG00000024747 | -0.022845224 | 0.88942 | 0.933421 | 153.9263 | 208.7689 | 188.8083 | 164.7611 | 208.6738 | 150.8455 | 182.2507 | 183.8029 |
| ENSMUSG00000024750 | 0.155180283 | 0.096212 | 0.21487 | 3728.437 | 2899.969 | 2946.403 | 3160.652 | 2791.744 | 2840.365 | 2914.082 | 2890.817 |
| ENSMUSG00000024754 | -0.101888028 | 0.240228 | 0.402328 | 1236.441 | 1136.231 | 1085.648 | 999.7806 | 1171.109 | 1149.778 | 1199.576 | 1260.518 |

| | | | | | | | | | | | |
|---|---|---|---|---|---|---|---|---|---|---|---|
| ENSMUSG00000024757 | -4.972972177 | 4.41E-05 | 0.000725 | 14.08476 | 0 | 78.67013 | 22.42821 | 980.962 | 203.3621 | 743.4669 | 1699.905 |
| ENSMUSG00000024758 | -0.16899725 | 0.057655 | 0.150505 | 5750.607 | 5340.406 | 6243.924 | 6100.473 | 6116.873 | 6051.698 | 6975.668 | 7208.554 |
| ENSMUSG00000024759 | 0.202002529 | 0.00183 | 0.012301 | 2204.265 | 2157.279 | 2002.362 | 2140.169 | 1843.935 | 1947.583 | 1734.756 | 1866.306 |
| ENSMUSG00000024764 | -0.165486132 | 0.039611 | 0.115854 | 1195.193 | 1347.4 | 1384.594 | 1217.162 | 1409.036 | 1401.187 | 1409.791 | 1549.817 |
| ENSMUSG00000024766 | 0.215064463 | 0.150924 | 0.291135 | 376.2643 | 476.3291 | 414.8814 | 433.037 | 366.6419 | 426.8369 | 387.6442 | 282.7737 |
| ENSMUSG00000024767 | -0.230076102 | 0.009521 | 0.041098 | 2892.406 | 3003.153 | 3369.566 | 3071.802 | 3793.183 | 3154.347 | 3699.013 | 3825.058 |
| ENSMUSG00000024769 | -0.396893235 | 0.017315 | 0.063708 | 647.899 | 608.3095 | 758.5457 | 601.2486 | 859.0731 | 616.7905 | 946.932 | 1022.336 |
| ENSMUSG00000024772 | -0.344301864 | 0.003796 | 0.021024 | 1712.305 | 1851.325 | 2004.846 | 1842.564 | 2542.115 | 1885.01 | 2325.865 | 2655.897 |
| ENSMUSG00000024773 | 0.049837611 | 0.487305 | 0.645251 | 727.3773 | 733.091 | 719.6247 | 691.824 | 653.3246 | 700.5936 | 702.0025 | 717.8101 |
| ENSMUSG00000024776 | -0.037051541 | 0.814644 | 0.887294 | 136.8234 | 182.3729 | 181.3554 | 142.3329 | 157.968 | 169.8409 | 176.4649 | 154.4379 |
| ENSMUSG00000024777 | -0.416450147 | 0.010064 | 0.042797 | 453.7305 | 321.5522 | 462.0835 | 443.3885 | 565.5646 | 443.5975 | 618.1094 | 618.8393 |
| ENSMUSG00000024778 | 0.026472331 | 0.904876 | 0.942939 | 150.9082 | 116.3827 | 115.1068 | 149.2339 | 113.1129 | 111.7374 | 180.3221 | 116.3722 |
| ENSMUSG00000024780 | -0.43503686 | 0.001259 | 0.009302 | 768.6255 | 634.7056 | 631.0173 | 699.5877 | 841.5211 | 797.8051 | 1129.183 | 926.6276 |
| ENSMUSG00000024781 | -0.406428345 | 0.032373 | 0.099811 | 1011.085 | 741.4897 | 837.2158 | 885.9143 | 1327.126 | 808.9788 | 1019.254 | 1451.934 |
| ENSMUSG00000024782 | -0.166209555 | 0.10162 | 0.223539 | 2653.971 | 2240.067 | 2513.304 | 2723.302 | 2872.678 | 2436.993 | 3035.582 | 3024.591 |
| ENSMUSG00000024784 | 0.800554914 | 0.305846 | 0.474366 | 38.23007 | 9.598572 | 4.140533 | 8.626235 | 3.900445 | 13.40849 | 7.714313 | 9.78832 |
| ENSMUSG00000024785 | -0.35140008 | 0.002043 | 0.013322 | 930.6003 | 1031.847 | 872.8244 | 800.5146 | 1306.649 | 1022.397 | 1134.004 | 1169.16 |
| ENSMUSG00000024787 | -0.185886844 | 0.087573 | 0.201227 | 495.9848 | 407.9393 | 450.49 | 439.938 | 507.0579 | 478.2361 | 479.2517 | 578.5985 |
| ENSMUSG00000024789 | 0.044068147 | 0.654126 | 0.776731 | 1402.44 | 1442.185 | 1643.792 | 1573.425 | 1385.633 | 1631.366 | 1546.72 | 1319.248 |
| ENSMUSG00000024790 | -0.466455729 | 0.0007 | 0.005944 | 306.8466 | 341.9491 | 340.3518 | 276.9021 | 451.4765 | 363.1466 | 427.1801 | 505.7299 |
| ENSMUSG00000024791 | -0.036937198 | 0.756614 | 0.84857 | 483.9121 | 609.5093 | 503.4888 | 521.8872 | 616.2703 | 484.9404 | 552.5377 | 515.5182 |
| ENSMUSG00000024792 | 0.036586389 | 0.662566 | 0.782896 | 761.5831 | 829.0767 | 843.0126 | 916.1062 | 789.8402 | 858.1433 | 800.36 | 821.1313 |
| ENSMUSG00000024793 | 0.469318521 | 0.138217 | 0.273676 | 48.29061 | 39.59411 | 35.60859 | 42.26855 | 17.552 | 37.99072 | 30.85725 | 33.71532 |
| ENSMUSG00000024795 | 0.288336407 | 0.027138 | 0.088078 | 1040.26 | 951.4585 | 783.3889 | 814.3166 | 718.657 | 846.9696 | 647.038 | 725.4233 |
| ENSMUSG00000024797 | -0.132259242 | 0.167519 | 0.313498 | 1278.695 | 1234.616 | 1264.519 | 1190.42 | 1449.015 | 1140.839 | 1416.541 | 1436.708 |
| ENSMUSG00000024799 | -0.196751265 | 0.077679 | 0.185298 | 394.3733 | 400.7404 | 334.5551 | 344.1868 | 446.601 | 389.9636 | 399.2157 | 449.1751 |
| ENSMUSG00000024800 | 0.001037281 | 0.994234 | 0.996664 | 530.1906 | 488.3274 | 433.9279 | 533.9639 | 488.5308 | 610.0863 | 474.4303 | 412.197 |
| ENSMUSG00000024803 | -0.797827907 | 0.317788 | 0.486138 | 7.04238 | 2.399643 | 32.29616 | 3.450494 | 12.67645 | 15.64324 | 24.10723 | 27.18978 |
| ENSMUSG00000024805 | -0.757377126 | 5.56E-05 | 0.00086 | 129.781 | 71.98929 | 105.9976 | 101.7896 | 169.6694 | 177.6625 | 170.6792 | 177.2773 |
| ENSMUSG00000024807 | -0.014287561 | 0.863194 | 0.917928 | 753.5347 | 754.6877 | 715.4841 | 696.1372 | 705.0055 | 682.7156 | 767.5742 | 790.6787 |
| ENSMUSG00000024810 | 1.184438856 | 8.45E-08 | 4.95E-06 | 6305.949 | 6181.481 | 4281.311 | 6681.019 | 2542.115 | 3657.165 | 2399.151 | 1719.482 |
| ENSMUSG00000024811 | 0.227738521 | 0.002451 | 0.015197 | 7595.71 | 6968.563 | 6587.588 | 7174.44 | 5753.157 | 6631.615 | 6023.914 | 5781.634 |
| ENSMUSG00000024812 | 0.396596194 | 2.41E-08 | 1.76E-06 | 2716.347 | 2982.756 | 3019.277 | 2921.706 | 2071.136 | 2387.828 | 2243.901 | 2141.467 |
| ENSMUSG00000024816 | -0.29032014 | 0.026885 | 0.087556 | 946.6971 | 850.6735 | 929.1356 | 873.8376 | 1103.826 | 846.9696 | 1258.397 | 1193.087 |
| ENSMUSG00000024817 | 0.233393577 | 0.019901 | 0.070188 | 2851.158 | 2962.359 | 2404.822 | 2913.08 | 2230.08 | 2687.285 | 2254.508 | 2295.905 |
| ENSMUSG00000024818 | -0.857583124 | 0.000346 | 0.003454 | 202.2169 | 171.5745 | 270.7909 | 189.7772 | 380.2934 | 244.7049 | 378.9656 | 510.0802 |
| ENSMUSG00000024824 | -0.054476821 | 0.641899 | 0.768381 | 395.3794 | 531.5209 | 502.6607 | 502.0469 | 528.5103 | 465.945 | 501.4304 | 508.9926 |
| ENSMUSG00000024826 | -0.117115444 | 0.079103 | 0.187628 | 2139.878 | 2314.456 | 2299.652 | 2189.338 | 2603.547 | 2288.382 | 2458.937 | 2344.846 |
| ENSMUSG00000024827 | -0.389729274 | 0.033151 | 0.101583 | 529.1846 | 519.5227 | 520.8791 | 464.9541 | 893.202 | 479.3535 | 689.4668 | 601.4379 |
| ENSMUSG00000024829 | -0.080069382 | 0.223393 | 0.383102 | 1021.145 | 1024.648 | 987.9312 | 938.5344 | 1061.896 | 1040.275 | 1011.539 | 1082.153 |
| ENSMUSG00000024830 | 0.014465484 | 0.852367 | 0.911279 | 699.2078 | 677.8992 | 649.2356 | 694.4119 | 713.7815 | 643.6075 | 642.2166 | 692.7955 |
| ENSMUSG00000024831 | -0.0408104 | 0.639585 | 0.766541 | 429.5852 | 419.9375 | 413.2252 | 406.2957 | 441.7254 | 454.7713 | 407.8943 | 412.197 |
| ENSMUSG00000024833 | -0.095449535 | 0.426194 | 0.59115 | 786.7345 | 1031.847 | 959.7756 | 771.1854 | 997.5389 | 873.7865 | 932.4676 | 985.3575 |
| ENSMUSG00000024835 | -0.126044792 | 0.106935 | 0.231236 | 3342.113 | 3301.909 | 3750.495 | 3531.581 | 3914.097 | 3433.691 | 3793.514 | 4059.978 |
| ENSMUSG00000024841 | -0.279170596 | 0.036606 | 0.109783 | 1141.872 | 1064.242 | 1297.643 | 1179.206 | 1549.452 | 1091.674 | 1462.827 | 1580.27 |
| ENSMUSG00000024842 | 0.360412362 | 0.253975 | 0.417583 | 25.15136 | 34.79482 | 34.78048 | 36.23019 | 30.22845 | 30.1691 | 22.17865 | 19.57664 |
| ENSMUSG00000024843 | 0.064177386 | 0.610737 | 0.745684 | 2442.7 | 1950.91 | 2539.803 | 2189.338 | 1874.164 | 2576.665 | 2164.829 | 2114.277 |
| ENSMUSG00000024844 | -0.198964287 | 0.000536 | 0.004837 | 3040.296 | 3091.94 | 3286.755 | 3217.586 | 3761.979 | 3496.264 | 3498.441 | 3753.277 |
| ENSMUSG00000024845 | -0.153805287 | 0.064249 | 0.162793 | 1597.614 | 1565.767 | 1794.507 | 1718.346 | 1973.625 | 1671.592 | 1921.828 | 1863.044 |
| ENSMUSG00000024847 | -0.159763983 | 0.108307 | 0.233029 | 1412.5 | 1511.775 | 1551.872 | 1494.927 | 1733.748 | 1387.779 | 1725.113 | 1822.803 |
| ENSMUSG00000024851 | -0.10987716 | 0.355113 | 0.523689 | 609.6689 | 501.5254 | 588.7838 | 634.8909 | 648.449 | 535.2222 | 688.5025 | 649.2919 |
| ENSMUSG00000024853 | 0.015334759 | 0.713154 | 0.820112 | 8172.18 | 8619.518 | 8338.206 | 8052.59 | 8098.299 | 8239.517 | 8331.458 | 8155.846 |
| ENSMUSG00000024854 | -0.285911893 | 0.051327 | 0.138554 | 419.5247 | 511.124 | 476.9894 | 401.1199 | 606.5192 | 429.0717 | 609.4308 | 556.8466 |
| ENSMUSG00000024855 | -0.030853804 | 0.701328 | 0.811619 | 1192.174 | 1082.239 | 1189.989 | 1039.461 | 1150.631 | 1232.464 | 1110.861 | 1109.343 |
| ENSMUSG00000024856 | -0.293732365 | 0.005792 | 0.028665 | 573.451 | 569.9152 | 597.893 | 552.079 | 692.329 | 591.0909 | 766.6099 | 760.2262 |
| ENSMUSG00000024858 | -0.228907481 | 0.015738 | 0.059717 | 1302.84 | 1313.805 | 1407.781 | 1321.539 | 1549.452 | 1338.614 | 1728.006 | 1648.788 |
| ENSMUSG00000024862 | -0.588846732 | 0.0001 | 0.001338 | 549.3057 | 496.7261 | 602.0335 | 554.6669 | 794.7157 | 658.1333 | 810.9672 | 1051.701 |
| ENSMUSG00000024866 | -1.058892735 | 3.90E-05 | 0.00066 | 442.6639 | 429.5361 | 553.1752 | 478.756 | 1182.81 | 551.9828 | 828.3244 | 1405.168 |
| ENSMUSG00000024867 | 0.113042653 | 0.600268 | 0.738323 | 352.119 | 233.9652 | 284.0406 | 267.4133 | 239.8774 | 358.6771 | 192.8578 | 262.1095 |
| ENSMUSG00000024868 | -0.620155936 | 0.005353 | 0.026934 | 588.5418 | 373.1445 | 324.6178 | 331.2474 | 735.2339 | 484.9404 | 689.4668 | 574.2481 |
| ENSMUSG00000024869 | -0.055309409 | 0.79891 | 0.876929 | 111.672 | 94.7859 | 89.43552 | 67.28463 | 104.3369 | 102.7984 | 79.07171 | 90.27006 |
| ENSMUSG00000024870 | -0.413102065 | 0.000526 | 0.004767 | 3391.409 | 3648.657 | 4034.536 | 3434.104 | 5187.592 | 3898.518 | 4763.588 | 5469.496 |
| ENSMUSG00000024871 | 0.123290245 | 0.615026 | 0.748514 | 152.9203 | 190.7716 | 97.71658 | 125.943 | 161.8685 | 127.3806 | 122.4647 | 106.5839 |
| ENSMUSG00000024873 | -0.010591792 | 0.965496 | 0.980631 | 79.47829 | 74.38894 | 115.9349 | 69.8725 | 80.93424 | 73.74669 | 80.036 | 108.7591 |

| | | | | | | | | | | | |
|---|---|---|---|---|---|---|---|---|---|---|---|
| ENSMUSG00000024875 | -0.409947789 | 0.001959 | 0.012939 | 803.8374 | 770.2854 | 815.685 | 752.2077 | 1057.021 | 827.9742 | 1018.289 | 1271.394 |
| ENSMUSG00000024878 | -0.178159462 | 0.114083 | 0.241457 | 426.567 | 397.1409 | 351.9453 | 351.0878 | 398.8205 | 408.9589 | 462.8588 | 453.5255 |
| ENSMUSG00000024883 | -0.45070525 | 0.023274 | 0.078699 | 101.6115 | 122.3818 | 140.7781 | 85.39973 | 152.1174 | 137.437 | 150.4291 | 175.1022 |
| ENSMUSG00000024885 | 0.66064886 | 0.123047 | 0.252685 | 343.0645 | 83.98751 | 111.7944 | 112.1411 | 111.1627 | 90.5073 | 82.92887 | 127.2482 |
| ENSMUSG00000024887 | -0.153550152 | 0.224081 | 0.383871 | 417.5126 | 359.9465 | 325.4459 | 313.995 | 358.841 | 397.7852 | 393.43 | 424.1605 |
| ENSMUSG00000024889 | -0.287495986 | 0.000348 | 0.003464 | 879.2915 | 929.8617 | 914.2297 | 982.5282 | 1133.079 | 1045.862 | 1110.861 | 1234.416 |
| ENSMUSG00000024891 | -0.056003821 | 0.721169 | 0.825522 | 260.5681 | 196.7707 | 260.0255 | 213.9306 | 258.4045 | 197.7752 | 249.7509 | 263.197 |
| ENSMUSG00000024892 | -2.037307432 | 2.85E-06 | 8.66E-05 | 753.5347 | 427.1365 | 565.5968 | 508.9479 | 2866.827 | 991.1108 | 1683.649 | 3716.299 |
| ENSMUSG00000024896 | -0.421547744 | 1.03E-07 | 5.78E-06 | 1270.647 | 1280.21 | 1383.766 | 1318.951 | 1723.997 | 1626.897 | 1734.756 | 1954.401 |
| ENSMUSG00000024897 | 0.481903535 | 7.89E-05 | 0.00113 | 478.8819 | 358.7466 | 406.6004 | 405.433 | 294.4836 | 281.5783 | 284.4653 | 321.927 |
| ENSMUSG00000024899 | -1.091619428 | 8.81E-08 | 5.11E-06 | 554.3359 | 554.3175 | 438.8965 | 565.881 | 1094.075 | 763.1665 | 1087.718 | 1557.43 |
| ENSMUSG00000024900 | -0.570420933 | 0.042708 | 0.122167 | 2024.181 | 2271.262 | 2579.552 | 1798.57 | 3166.186 | 1716.287 | 2718.331 | 5279.167 |
| ENSMUSG00000024901 | -0.329129466 | 0.095406 | 0.213692 | 69.41775 | 69.58965 | 75.3577 | 68.14726 | 83.85957 | 74.86406 | 86.78602 | 109.8467 |
| ENSMUSG00000024902 | -0.229456611 | 0.000109 | 0.001434 | 1407.47 | 1463.782 | 1551.872 | 1441.444 | 1737.648 | 1661.535 | 1743.435 | 1734.708 |
| ENSMUSG00000024905 | 0.436744583 | 0.211716 | 0.369218 | 41.24823 | 74.38894 | 37.2648 | 43.13117 | 28.27823 | 36.87335 | 27.96439 | 51.11678 |
| ENSMUSG00000024906 | 0.063616784 | 0.574071 | 0.716559 | 541.2572 | 622.7074 | 573.0498 | 571.9194 | 509.0081 | 486.0577 | 576.6449 | 636.2408 |
| ENSMUSG00000024908 | -0.0050556 | 0.922984 | 0.954016 | 3547.348 | 3479.482 | 3233.756 | 3305.573 | 3398.263 | 3393.465 | 3357.655 | 3458.54 |
| ENSMUSG00000024909 | -0.079626614 | 0.432681 | 0.596995 | 1690.171 | 2088.889 | 2204.42 | 1966.782 | 2126.718 | 1909.592 | 2136.865 | 2228.474 |
| ENSMUSG00000024911 | -0.044016798 | 0.557677 | 0.704145 | 1659.99 | 1817.73 | 1789.538 | 1592.403 | 1893.666 | 1743.104 | 1675.935 | 1756.46 |
| ENSMUSG00000024912 | -0.032319127 | 0.841872 | 0.90461 | 95.57516 | 98.38537 | 100.2009 | 88.85022 | 106.2871 | 93.85942 | 98.35749 | 92.44524 |
| ENSMUSG00000024913 | -0.040537163 | 0.765081 | 0.854012 | 1505.057 | 1365.397 | 1560.153 | 1353.456 | 1480.219 | 1137.487 | 1599.756 | 1731.445 |
| ENSMUSG00000024914 | -0.174272147 | 0.024873 | 0.082752 | 3359.215 | 3454.286 | 3574.936 | 3510.015 | 4136.422 | 3457.155 | 3888.014 | 4201.364 |
| ENSMUSG00000024921 | 0.437575833 | 0.000987 | 0.007773 | 4739.522 | 5005.655 | 4233.281 | 4938.52 | 3333.906 | 4415.862 | 3288.226 | 2929.97 |
| ENSMUSG00000024922 | -0.599264608 | 0.0084 | 0.037618 | 198.1927 | 173.9741 | 310.54 | 182.8762 | 305.2098 | 259.2308 | 347.1441 | 402.4087 |
| ENSMUSG00000024924 | -0.06845067 | 0.220845 | 0.379783 | 3181.144 | 3379.897 | 3122.79 | 3223.624 | 3441.168 | 3187.868 | 3483.977 | 3413.948 |
| ENSMUSG00000024925 | -0.31065769 | 0.000764 | 0.006351 | 1190.162 | 1222.618 | 1218.145 | 1237.865 | 1590.407 | 1299.506 | 1475.362 | 1672.715 |
| ENSMUSG00000024926 | -0.182133837 | 0.001576 | 0.010933 | 1341.07 | 1313.805 | 1397.016 | 1381.923 | 1600.158 | 1512.925 | 1558.291 | 1496.525 |
| ENSMUSG00000024927 | -0.086201577 | 0.399721 | 0.567044 | 1167.023 | 1055.843 | 1207.379 | 1283.584 | 1394.409 | 1121.844 | 1180.29 | 1310.547 |
| ENSMUSG00000024935 | -2.629325979 | 8.99E-11 | 1.66E-08 | 48.29061 | 25.19625 | 103.5133 | 49.16954 | 271.0809 | 221.2401 | 390.5371 | 525.3065 |
| ENSMUSG00000024937 | -0.05419462 | 0.709949 | 0.818517 | 585.5236 | 547.1186 | 717.9685 | 547.7659 | 628.9468 | 510.64 | 617.1451 | 735.2116 |
| ENSMUSG00000024939 | 0.147145954 | 0.49168 | 0.648814 | 89.53884 | 95.98572 | 98.54469 | 128.5309 | 115.0631 | 92.74205 | 74.25027 | 91.35765 |
| ENSMUSG00000024940 | 0.366259556 | 6.24E-05 | 0.000944 | 2506.081 | 2885.571 | 2880.983 | 2738.83 | 1956.073 | 2428.054 | 2099.258 | 2060.985 |
| ENSMUSG00000024941 | -0.082115071 | 0.350332 | 0.519044 | 1582.524 | 1733.742 | 1951.847 | 1665.726 | 1877.089 | 1703.995 | 1830.221 | 1930.474 |
| ENSMUSG00000024942 | -0.429641017 | 0.041897 | 0.120482 | 721.341 | 701.8956 | 1083.992 | 672.8463 | 1156.482 | 736.3495 | 1085.79 | 1306.197 |
| ENSMUSG00000024943 | 0.41908118 | 0.003771 | 0.020945 | 2021.163 | 1984.505 | 1620.605 | 2116.015 | 1371.982 | 1830.259 | 1360.612 | 1227.89 |
| ENSMUSG00000024944 | -0.284658155 | 0.005037 | 0.025859 | 882.3097 | 921.4629 | 1040.93 | 1018.758 | 1299.823 | 1034.688 | 1158.111 | 1215.927 |
| ENSMUSG00000024947 | -0.174703893 | 0.069537 | 0.171437 | 1243.483 | 1171.026 | 1238.019 | 1223.2 | 1345.654 | 1176.595 | 1441.612 | 1540.029 |
| ENSMUSG00000024948 | 0.035034179 | 0.727191 | 0.829154 | 962.794 | 837.4754 | 1099.726 | 977.3524 | 888.3264 | 960.9417 | 946.932 | 994.0583 |
| ENSMUSG00000024949 | 0.054495587 | 0.556776 | 0.703537 | 5116.792 | 5022.453 | 4889.97 | 5203.345 | 5137.861 | 5474.016 | 4691.267 | 4179.613 |
| ENSMUSG00000024952 | -0.193345487 | 0.190372 | 0.342982 | 478.8819 | 505.1249 | 665.7977 | 546.0407 | 656.2499 | 524.0485 | 606.5379 | 726.5108 |
| ENSMUSG00000024953 | -0.89032967 | 0.000142 | 0.00175 | 2872.285 | 2922.765 | 4591.851 | 3024.358 | 5844.817 | 3848.236 | 6916.846 | 8251.554 |
| ENSMUSG00000024955 | -1.075674739 | 0.000135 | 0.001689 | 537.233 | 386.3425 | 788.3575 | 513.261 | 1119.428 | 651.4291 | 1327.826 | 1594.409 |
| ENSMUSG00000024957 | 0.245637578 | 0.732651 | 0.832561 | 12.07265 | 5.999108 | 25.67131 | 9.488858 | 20.47734 | 2.234748 | 12.53576 | 9.78832 |
| ENSMUSG00000024958 | -0.19956016 | 0.175825 | 0.324341 | 452.7245 | 477.529 | 525.8477 | 533.1013 | 564.5894 | 438.0106 | 600.7521 | 681.9196 |
| ENSMUSG00000024959 | -0.318128507 | 0.008512 | 0.037966 | 738.4439 | 757.0874 | 903.4643 | 817.7671 | 981.9371 | 841.3827 | 1043.361 | 1146.321 |
| ENSMUSG00000024960 | 0.188755994 | 0.064671 | 0.163614 | 1979.915 | 1866.922 | 2222.638 | 2005.6 | 1743.499 | 1520.746 | 1848.542 | 1973.978 |
| ENSMUSG00000024962 | -0.245835542 | 0.038117 | 0.112702 | 739.45 | 622.7074 | 758.5457 | 729.7795 | 828.8446 | 710.6499 | 950.7891 | 891.8247 |
| ENSMUSG00000024963 | -0.097919307 | 0.505641 | 0.661429 | 401.4157 | 404.3399 | 463.7397 | 438.2127 | 524.6099 | 345.2686 | 459.9659 | 498.1167 |
| ENSMUSG00000024965 | 0.054389915 | 0.772905 | 0.859063 | 150.9082 | 109.1838 | 139.95 | 104.3774 | 113.1129 | 136.3196 | 128.2505 | 108.7591 |
| ENSMUSG00000024966 | -0.311504657 | 0.001171 | 0.008818 | 3669.08 | 3591.066 | 3612.201 | 3504.839 | 4453.333 | 3767.785 | 4532.159 | 5087.751 |
| ENSMUSG00000024968 | 0.512104957 | 9.24E-06 | 0.000214 | 717.3168 | 752.2881 | 764.3424 | 714.2523 | 601.6437 | 547.5133 | 440.6801 | 477.4525 |
| ENSMUSG00000024969 | 0.046601617 | 0.415 | 0.580938 | 1380.307 | 1350.999 | 1391.219 | 1347.418 | 1298.848 | 1378.84 | 1278.647 | 1343.175 |
| ENSMUSG00000024970 | 0.094362822 | 0.181013 | 0.331328 | 954.7456 | 922.6628 | 907.6049 | 904.0294 | 870.7744 | 889.4298 | 894.8603 | 798.2919 |
| ENSMUSG00000024972 | -1.350193183 | 0.000631 | 0.005483 | 30.18163 | 27.5959 | 50.5145 | 29.3292 | 55.58134 | 71.51194 | 71.3574 | 154.4379 |
| ENSMUSG00000024974 | 0.174255856 | 0.071604 | 0.175223 | 4688.213 | 4998.457 | 4220.86 | 4361.424 | 3922.873 | 4657.215 | 3902.478 | 3705.423 |
| ENSMUSG00000024975 | 0.229892185 | 0.041696 | 0.120204 | 4125.829 | 4394.946 | 3770.369 | 4121.615 | 3450.919 | 4176.744 | 3404.905 | 2961.511 |
| ENSMUSG00000024976 | -0.016972404 | 0.764993 | 0.854012 | 1440.67 | 1360.598 | 1437.593 | 1484.575 | 1419.762 | 1445.882 | 1464.755 | 1466.073 |
| ENSMUSG00000024978 | -0.249491147 | 0.058596 | 0.152065 | 838.0433 | 700.6958 | 820.6537 | 665.0827 | 937.082 | 725.1758 | 968.1463 | 964.6933 |
| ENSMUSG00000024981 | -0.485286148 | 0.001585 | 0.010978 | 1416.525 | 1448.185 | 1699.275 | 1350.868 | 2007.754 | 1546.446 | 2205.329 | 2521.036 |
| ENSMUSG00000024982 | -0.274364097 | 0.000585 | 0.005158 | 1228.392 | 1141.03 | 1123.741 | 1190.42 | 1442.19 | 1273.806 | 1417.505 | 1530.241 |
| ENSMUSG00000024983 | -0.191507502 | 0.20777 | 0.364644 | 694.1775 | 655.1026 | 753.577 | 624.5394 | 741.0846 | 594.443 | 824.4672 | 954.905 |
| ENSMUSG00000024985 | 0.441253124 | 0.06676 | 0.166804 | 1007.06 | 939.4603 | 746.1241 | 923.8698 | 545.0872 | 1065.975 | 550.6091 | 502.4671 |
| ENSMUSG00000024986 | 0.119719797 | 0.570561 | 0.713919 | 231.3925 | 146.3782 | 209.511 | 163.0358 | 156.9929 | 135.2023 | 196.715 | 202.2919 |
| ENSMUSG00000024987 | 0.415421354 | 0.350758 | 0.519466 | 66.39959 | 21.59679 | 58.79557 | 59.52102 | 28.27823 | 21.23011 | 49.17875 | 56.55474 |

| | | | | | | | | | | | |
|---|---|---|---|---|---|---|---|---|---|---|---|
| ENSMUSG00000024989 | 0.063975967 | 0.618068 | 0.750712 | 599.6084 | 603.5102 | 501.0045 | 541.7276 | 600.6686 | 598.9125 | 471.5374 | 475.2773 |
| ENSMUSG00000024990 | 0.404091988 | 0.252746 | 0.41636 | 147.89 | 81.58786 | 55.48314 | 167.349 | 79.95913 | 73.74669 | 102.2147 | 85.9197 |
| ENSMUSG00000024991 | -0.063518518 | 0.303425 | 0.471858 | 11141.05 | 11159.54 | 10564.98 | 9882.215 | 10918.32 | 10999.43 | 10989.04 | 11760.12 |
| ENSMUSG00000024992 | -0.239670469 | 0.489484 | 0.647036 | 17.10292 | 25.19625 | 25.67131 | 17.25247 | 23.40267 | 23.46486 | 22.17865 | 31.54014 |
| ENSMUSG00000024993 | 0.074546644 | 0.519728 | 0.673278 | 507.0514 | 502.7252 | 479.4737 | 552.9417 | 481.705 | 568.7434 | 464.7874 | 425.2481 |
| ENSMUSG00000024997 | -0.17358064 | 0.328998 | 0.496849 | 2591.596 | 1594.563 | 1941.91 | 1806.334 | 2280.785 | 1730.812 | 2253.544 | 2684.175 |
| ENSMUSG00000024998 | 0.077493433 | 0.533484 | 0.684747 | 446.6881 | 417.5379 | 494.3797 | 460.6409 | 379.3183 | 513.9921 | 404.0372 | 430.6861 |
| ENSMUSG00000024999 | 0.210253357 | 0.055981 | 0.147408 | 589.5479 | 661.1017 | 501.0045 | 591.7597 | 489.5059 | 528.5179 | 476.3588 | 528.5693 |
| ENSMUSG00000025001 | 0.047323307 | 0.749023 | 0.843439 | 1020.139 | 1373.796 | 1012.774 | 989.4291 | 1056.046 | 1278.276 | 935.3605 | 982.0948 |
| ENSMUSG00000025006 | -0.359993545 | 0.030579 | 0.095812 | 258.556 | 219.5673 | 256.7131 | 213.068 | 275.9565 | 241.3528 | 367.3942 | 331.7153 |
| ENSMUSG00000025007 | 0.069797267 | 0.492934 | 0.64992 | 1333.022 | 1263.412 | 1422.687 | 1211.123 | 1265.694 | 1060.388 | 1355.791 | 1300.759 |
| ENSMUSG00000025008 | -0.17875804 | 0.139949 | 0.276098 | 327.9737 | 424.7368 | 362.7107 | 312.2697 | 405.6463 | 396.6678 | 415.6086 | 393.708 |
| ENSMUSG00000025010 | 0.136628237 | 0.22477 | 0.384523 | 502.0211 | 560.3167 | 481.9581 | 533.9639 | 515.8339 | 519.579 | 426.2158 | 427.4233 |
| ENSMUSG00000025016 | -0.002368453 | 0.958213 | 0.976412 | 8380.433 | 8709.505 | 8196.599 | 8630.548 | 8227.014 | 8770.269 | 8520.459 | 8452.758 |
| ENSMUSG00000025017 | -0.027481545 | 0.918179 | 0.951643 | 50.30272 | 49.19268 | 59.62368 | 34.50494 | 57.53157 | 40.22547 | 49.17875 | 50.02919 |
| ENSMUSG00000025019 | -0.281527788 | 0.130612 | 0.263048 | 205.2351 | 149.9777 | 136.6376 | 132.844 | 188.1965 | 170.9582 | 180.3221 | 218.6058 |
| ENSMUSG00000025020 | 0.462808643 | 0.002395 | 0.01494 | 574.457 | 457.132 | 406.6004 | 554.6669 | 402.721 | 376.5551 | 361.6084 | 304.5255 |
| ENSMUSG00000025024 | 0.056815393 | 0.556323 | 0.703109 | 1478.9 | 1436.186 | 1547.731 | 1533.745 | 1434.389 | 1683.883 | 1319.148 | 1331.211 |
| ENSMUSG00000025025 | 0.12124205 | 0.246283 | 0.4092 | 2689.183 | 2489.63 | 2143.968 | 2745.731 | 2357.819 | 2282.795 | 2531.259 | 2082.737 |
| ENSMUSG00000025026 | 0.135367459 | 0.08588 | 0.198449 | 3086.575 | 3247.917 | 3199.804 | 3269.343 | 2759.565 | 3285.08 | 2694.224 | 2921.27 |
| ENSMUSG00000025027 | -0.368793946 | 0.005171 | 0.026332 | 2273.683 | 2174.077 | 2498.398 | 2164.322 | 3131.082 | 2285.03 | 2870.689 | 3478.116 |
| ENSMUSG00000025034 | -0.042057109 | 0.671982 | 0.789942 | 1770.656 | 1611.36 | 2059.501 | 1711.445 | 1755.2 | 1824.672 | 1761.756 | 2028.357 |
| ENSMUSG00000025035 | -0.243225068 | 0.001284 | 0.009421 | 1632.826 | 1665.352 | 1856.615 | 1710.582 | 2115.992 | 1855.958 | 2095.4 | 2060.985 |
| ENSMUSG00000025036 | -0.523630512 | 1.06E-10 | 1.93E-08 | 1104.648 | 1001.851 | 1088.132 | 1066.203 | 1505.572 | 1398.952 | 1537.077 | 1686.854 |
| ENSMUSG00000025037 | -0.073415174 | 0.530884 | 0.682549 | 1073.46 | 1004.251 | 1189.161 | 1153.328 | 1057.996 | 1029.102 | 1377.969 | 1187.649 |
| ENSMUSG00000025038 | 0.28839036 | 0.486344 | 0.644455 | 19.11503 | 19.19714 | 12.4216 | 15.52722 | 12.67645 | 12.29112 | 11.57147 | 17.40146 |
| ENSMUSG00000025040 | -0.074300325 | 0.328809 | 0.496686 | 1038.248 | 1101.436 | 1006.978 | 1011.857 | 1092.125 | 1193.356 | 1035.647 | 1054.963 |
| ENSMUSG00000025041 | -0.083513658 | 0.154823 | 0.296474 | 1494.997 | 1484.179 | 1508.81 | 1627.771 | 1638.187 | 1603.432 | 1646.042 | 1595.496 |
| ENSMUSG00000025044 | 0.705123239 | 0.000274 | 0.002884 | 132.7992 | 130.7805 | 149.0592 | 120.7673 | 85.8098 | 67.04245 | 71.3574 | 103.3212 |
| ENSMUSG00000025047 | 0.003934737 | 0.964786 | 0.980188 | 1612.705 | 1487.779 | 1488.108 | 1392.274 | 1352.479 | 1419.065 | 1555.398 | 1636.825 |
| ENSMUSG00000025049 | 0.01143135 | 0.935129 | 0.96165 | 456.7487 | 387.5424 | 354.4296 | 444.2511 | 426.1236 | 459.2408 | 335.5726 | 410.0218 |
| ENSMUSG00000025050 | -0.024401786 | 0.792867 | 0.873246 | 759.571 | 796.6815 | 669.1102 | 708.2139 | 736.209 | 777.6924 | 784.9314 | 679.7444 |
| ENSMUSG00000025056 | 1.493748171 | 0.009868 | 0.042218 | 141.8537 | 34.79482 | 20.70267 | 72.46037 | 18.52711 | 35.75597 | 23.14294 | 18.48905 |
| ENSMUSG00000025059 | -1.588480945 | 2.81E-09 | 3.08E-07 | 588.5418 | 353.9474 | 734.5306 | 612.4627 | 1533.85 | 1125.196 | 1716.435 | 2515.598 |
| ENSMUSG00000025060 | -0.025685361 | 0.625449 | 0.755733 | 3584.572 | 3335.504 | 3541.812 | 3685.99 | 3494.799 | 3621.409 | 3642.12 | 3651.043 |
| ENSMUSG00000025064 | 0.791235122 | 0.03869 | 0.113668 | 26.15741 | 53.99197 | 49.6864 | 42.26855 | 20.47734 | 37.99072 | 16.39292 | 25.0146 |
| ENSMUSG00000025066 | 0.289354459 | 0.004212 | 0.022677 | 4636.905 | 5047.649 | 4316.092 | 4896.251 | 3760.029 | 4524.248 | 3710.585 | 3467.24 |
| ENSMUSG00000025068 | -0.397958735 | 0.002692 | 0.016303 | 1193.18 | 1029.447 | 1334.908 | 1196.459 | 1711.32 | 1231.346 | 1631.577 | 1692.292 |
| ENSMUSG00000025069 | 0.014692985 | 0.971503 | 0.984087 | 19.11503 | 21.59679 | 23.18699 | 30.19182 | 13.65156 | 17.87799 | 35.6787 | 26.10219 |
| ENSMUSG00000025075 | -0.434830495 | 0.080021 | 0.188964 | 668.0201 | 566.3158 | 1094.757 | 703.0381 | 997.5389 | 699.4762 | 998.0393 | 1406.255 |
| ENSMUSG00000025076 | 0.020680804 | 0.857365 | 0.914643 | 513.0877 | 680.2988 | 620.2519 | 525.3377 | 588.9672 | 591.0909 | 555.4306 | 567.7225 |
| ENSMUSG00000025077 | -0.148561884 | 0.232207 | 0.392814 | 418.5186 | 533.9206 | 467.0521 | 420.9603 | 559.7139 | 449.1844 | 486.0017 | 541.6204 |
| ENSMUSG00000025078 | -0.330187476 | 0.002828 | 0.016964 | 561.3783 | 428.3363 | 481.9581 | 505.4974 | 667.9512 | 591.0909 | 647.038 | 580.7736 |
| ENSMUSG00000025081 | -0.810623872 | NA | NA | 54.32694 | 47.99286 | 54.65504 | 71.59775 | 50.70579 | 265.935 | 49.17875 | 35.89051 |
| ENSMUSG00000025082 | 0.230444742 | 0.042465 | 0.12166 | 1490.973 | 1225.018 | 1526.201 | 1272.37 | 1053.12 | 1139.722 | 1334.576 | 1174.598 |
| ENSMUSG00000025083 | -0.110347018 | 0.571296 | 0.714304 | 569.4268 | 547.1186 | 718.7966 | 597.7981 | 595.793 | 483.823 | 639.3237 | 909.2261 |
| ENSMUSG00000025085 | 0.307166913 | 1.51E-09 | 1.85E-07 | 3117.762 | 3009.152 | 2907.482 | 2982.089 | 2334.416 | 2451.519 | 2455.08 | 2469.919 |
| ENSMUSG00000025086 | 0.147918167 | 0.056143 | 0.147645 | 1457.773 | 1426.588 | 1339.048 | 1436.268 | 1288.122 | 1405.657 | 1227.54 | 1186.562 |
| ENSMUSG00000025089 | 0.568485606 | 0.002632 | 0.016007 | 8132.943 | 9647.765 | 6927.94 | 8583.966 | 5880.896 | 7347.852 | 5436.662 | 3783.729 |
| ENSMUSG00000025092 | -0.198551478 | 0.049369 | 0.134927 | 878.2855 | 727.0918 | 834.7315 | 785.85 | 831.7699 | 895.0167 | 952.7177 | 1025.598 |
| ENSMUSG00000025094 | 1.027606431 | 0.01887 | 0.067676 | 293.7679 | 128.3809 | 90.26362 | 393.3563 | 88.73513 | 159.7845 | 107.0361 | 89.18247 |
| ENSMUSG00000025102 | -0.190208303 | 0.049099 | 0.134427 | 423.5489 | 350.3479 | 398.3193 | 395.9442 | 468.0534 | 422.3674 | 447.4302 | 453.5255 |
| ENSMUSG00000025103 | -0.165672489 | 0.006579 | 0.031552 | 2500.045 | 2560.419 | 2547.256 | 2649.979 | 2675.705 | 2843.717 | 2964.225 | 3024.591 |
| ENSMUSG00000025104 | 0.273221836 | 0.057274 | 0.14985 | 1741.48 | 1955.709 | 1600.73 | 1854.641 | 1472.418 | 1887.245 | 1330.719 | 1227.89 |
| ENSMUSG00000025105 | 0.570351224 | NA | NA | 27.16347 | 134.38 | 19.87456 | 27.60395 | 13.65156 | 113.9722 | 8.678602 | 4.350364 |
| ENSMUSG00000025127 | -0.375086335 | 0.252717 | 0.41636 | 85.51462 | 68.38983 | 115.1068 | 82.81186 | 86.78491 | 65.92507 | 172.6078 | 131.5985 |
| ENSMUSG00000025128 | 0.560585994 | 0.000747 | 0.006244 | 446.6881 | 609.5093 | 467.8802 | 489.1075 | 343.2392 | 421.25 | 330.7512 | 268.635 |
| ENSMUSG00000025130 | -0.239462025 | 0.007483 | 0.034516 | 13189.37 | 15327.72 | 15878.12 | 14448.08 | 17135.63 | 15523.68 | 17906.85 | 18901.25 |
| ENSMUSG00000025132 | -0.343890363 | 0.005337 | 0.026896 | 6297.9 | 6979.362 | 8005.307 | 6741.403 | 9331.815 | 7176.894 | 8951.496 | 10108.07 |
| ENSMUSG00000025133 | -0.065275428 | 0.421117 | 0.586533 | 1401.434 | 1416.989 | 1246.3 | 1378.472 | 1419.762 | 1548.68 | 1312.398 | 1412.781 |
| ENSMUSG00000025134 | -0.26269696 | 5.60E-05 | 0.000864 | 2079.514 | 2294.059 | 2076.891 | 2050.456 | 2507.011 | 2630.299 | 2421.33 | 2635.233 |
| ENSMUSG00000025135 | 0.003356411 | 0.971264 | 0.983952 | 1280.707 | 1641.356 | 1576.715 | 1552.722 | 1559.203 | 1471.582 | 1463.791 | 1542.204 |
| ENSMUSG00000025137 | -0.551537979 | 0.001203 | 0.008984 | 1221.35 | 941.8599 | 1275.284 | 1108.471 | 1569.929 | 1210.116 | 1938.221 | 1947.876 |
| ENSMUSG00000025138 | -0.165154106 | 0.154938 | 0.296601 | 672.0443 | 633.5058 | 750.2646 | 644.3797 | 768.3877 | 640.2554 | 755.0384 | 865.7225 |

| | | | | | | | | | | | |
|---|---|---|---|---|---|---|---|---|---|---|---|
| ENSMUSG00000025139 | -0.123852912 | 0.041949 | 0.120575 | 1588.56 | 1560.968 | 1625.573 | 1552.722 | 1739.599 | 1630.249 | 1697.149 | 1829.328 |
| ENSMUSG00000025140 | -0.294637092 | 0.024002 | 0.080601 | 223.3441 | 274.7591 | 235.1823 | 228.5952 | 325.6872 | 311.7474 | 281.5724 | 257.7591 |
| ENSMUSG00000025142 | -0.081115003 | 0.265289 | 0.429407 | 1732.426 | 1682.15 | 1796.991 | 1667.451 | 1857.587 | 1653.714 | 1811.899 | 1954.401 |
| ENSMUSG00000025144 | -0.228804989 | 0.031857 | 0.098632 | 616.7113 | 760.6869 | 762.6862 | 708.2139 | 856.1477 | 749.758 | 815.7886 | 915.7517 |
| ENSMUSG00000025145 | 0.316216723 | 0.000188 | 0.00216 | 1421.555 | 1570.566 | 1329.939 | 1462.147 | 1261.794 | 1145.308 | 1158.111 | 1075.628 |
| ENSMUSG00000025147 | -0.115657309 | 0.144562 | 0.28269 | 1144.89 | 988.6529 | 1136.162 | 1189.558 | 1169.158 | 1241.403 | 1234.29 | 1193.087 |
| ENSMUSG00000025150 | 0.829945377 | 0.002065 | 0.013422 | 90.54489 | 145.1784 | 110.9663 | 126.8057 | 75.08357 | 90.5073 | 54.96448 | 45.67883 |
| ENSMUSG00000025151 | 0.157610075 | 0.010099 | 0.042917 | 14283.96 | 16039.21 | 14778.39 | 14811.25 | 13168.88 | 14343.73 | 13137.48 | 13059.79 |
| ENSMUSG00000025153 | -0.127768563 | 0.09223 | 0.208526 | 2920.576 | 2965.959 | 2909.139 | 2687.072 | 3171.062 | 3050.431 | 3440.584 | 2878.854 |
| ENSMUSG00000025154 | -0.006858871 | 0.929754 | 0.958527 | 1093.581 | 1000.651 | 910.0892 | 988.5665 | 1017.041 | 996.6977 | 1002.861 | 991.8831 |
| ENSMUSG00000025155 | -0.428782557 | 2.82E-07 | 1.31E-05 | 1117.726 | 1141.03 | 1127.053 | 1062.752 | 1579.68 | 1345.318 | 1434.862 | 1625.949 |
| ENSMUSG00000025156 | -0.369504502 | 0.000254 | 0.002742 | 2200.241 | 2445.236 | 2478.523 | 2346.336 | 3277.349 | 2568.843 | 3018.225 | 3369.357 |
| ENSMUSG00000025157 | -0.263820813 | 0.03133 | 0.09733 | 840.0554 | 872.2703 | 883.5898 | 897.9911 | 1159.407 | 813.4483 | 1096.397 | 1124.569 |
| ENSMUSG00000025158 | -0.215765719 | 0.041868 | 0.120472 | 505.0393 | 556.7172 | 443.8652 | 444.2511 | 571.4152 | 573.2129 | 529.3948 | 585.124 |
| ENSMUSG00000025159 | 0.032165876 | 0.660713 | 0.781535 | 2088.569 | 1964.108 | 1989.94 | 1928.826 | 1999.953 | 2119.659 | 1876.507 | 1799.963 |
| ENSMUSG00000025161 | 0.758619556 | 2.16E-06 | 6.84E-05 | 608.6629 | 403.14 | 427.303 | 540.8649 | 313.0107 | 275.9914 | 311.4654 | 269.7226 |
| ENSMUSG00000025162 | -0.055276379 | 0.488404 | 0.646164 | 5850.206 | 5285.214 | 5776.872 | 5432.803 | 5877.971 | 5136.569 | 5934.236 | 6269.963 |
| ENSMUSG00000025169 | -0.383263369 | 0.005065 | 0.025964 | 402.4217 | 493.1267 | 461.2554 | 452.8773 | 668.9264 | 468.1797 | 579.5378 | 641.6787 |
| ENSMUSG00000025170 | 0.092007917 | 0.606436 | 0.742598 | 89.53884 | 92.38626 | 94.40416 | 83.67448 | 79.95913 | 74.86406 | 83.89316 | 98.97079 |
| ENSMUSG00000025171 | -0.475844679 | 4.11E-05 | 0.000689 | 520.1301 | 587.9126 | 692.2971 | 636.6161 | 829.8197 | 763.1665 | 877.5031 | 922.2772 |
| ENSMUSG00000025173 | -0.065546191 | 0.296524 | 0.464315 | 1853.152 | 1853.724 | 2052.048 | 1927.964 | 2103.315 | 1939.761 | 2003.793 | 2001.168 |
| ENSMUSG00000025175 | -0.297770073 | 0.210649 | 0.368171 | 70.4238 | 82.78769 | 86.12309 | 92.30071 | 72.15824 | 88.27255 | 119.5719 | 128.3357 |
| ENSMUSG00000025176 | -1.804755162 | 5.59E-07 | 2.29E-05 | 334.01 | 166.7752 | 438.0684 | 332.9727 | 1129.179 | 513.9921 | 1004.789 | 1799.963 |
| ENSMUSG00000025178 | -0.201425092 | 0.005618 | 0.028002 | 1238.453 | 1108.635 | 1146.1 | 1059.302 | 1314.45 | 1262.633 | 1346.148 | 1309.46 |
| ENSMUSG00000025184 | 0.076188008 | 0.449765 | 0.612329 | 589.5479 | 613.1088 | 569.7374 | 627.9899 | 582.1415 | 486.0577 | 589.1807 | 617.7517 |
| ENSMUSG00000025185 | 0.355985918 | 0.161686 | 0.30562 | 149.9021 | 153.5772 | 147.403 | 88.85022 | 135.5405 | 86.03781 | 83.89316 | 115.2847 |
| ENSMUSG00000025188 | 0.133233359 | 0.357963 | 0.526294 | 453.7305 | 409.1391 | 539.0974 | 357.1261 | 424.1734 | 372.0856 | 415.6086 | 392.6204 |
| ENSMUSG00000025189 | -1.300722857 | 0.02002 | 0.07042 | 15.09082 | 20.39697 | 93.57605 | 18.11509 | 88.73513 | 75.98144 | 102.2147 | 97.8832 |
| ENSMUSG00000025190 | -0.747304215 | 0.012164 | 0.049482 | 1975.891 | 812.2792 | 1385.422 | 1230.101 | 2237.88 | 1217.938 | 2783.903 | 2832.087 |
| ENSMUSG00000025192 | -0.107153264 | 0.354066 | 0.522769 | 265.5983 | 242.3639 | 237.6666 | 225.1447 | 249.6285 | 287.1651 | 248.7866 | 259.9343 |
| ENSMUSG00000025193 | -0.356019279 | 0.00713 | 0.033333 | 317.9132 | 321.5522 | 364.3669 | 369.2029 | 465.1281 | 351.9728 | 470.5731 | 470.9269 |
| ENSMUSG00000025194 | -3.111950633 | 3.41E-13 | 1.14E-10 | 48.29061 | 21.59679 | 50.5145 | 30.19182 | 447.5761 | 130.7328 | 238.1794 | 490.5036 |
| ENSMUSG00000025195 | -0.08904298 | 0.330669 | 0.498551 | 1042.272 | 962.2569 | 1089.788 | 898.8537 | 1057.996 | 976.585 | 1087.718 | 1125.657 |
| ENSMUSG00000025196 | -1.29065985 | 4.94E-05 | 0.000787 | 577.4752 | 250.7627 | 755.2332 | 556.3922 | 1195.486 | 845.8522 | 1313.362 | 1884.795 |
| ENSMUSG00000025197 | -0.664054379 | 0.250654 | 0.414226 | 10.06054 | 7.198929 | 8.281066 | 12.07673 | 19.50223 | 6.704245 | 8.678602 | 25.0146 |
| ENSMUSG00000025198 | -0.54932203 | 6.45E-05 | 0.000965 | 1099.617 | 1202.221 | 1123.741 | 1112.784 | 1949.248 | 1258.163 | 1606.506 | 1824.978 |
| ENSMUSG00000025199 | -0.161513859 | 0.08783 | 0.201642 | 1518.136 | 1355.798 | 1393.703 | 1410.389 | 1666.465 | 1362.079 | 1581.434 | 1740.146 |
| ENSMUSG00000025200 | -0.02295019 | 0.819386 | 0.890665 | 609.6689 | 549.5183 | 523.3634 | 518.4367 | 597.7432 | 583.2693 | 551.5734 | 501.3795 |
| ENSMUSG00000025202 | 0.28138049 | 0.576903 | 0.718897 | 24.1453 | 19.19714 | 9.109173 | 22.42821 | 15.60178 | 6.704245 | 19.28578 | 19.57664 |
| ENSMUSG00000025203 | 0.059256594 | 0.743065 | 0.839394 | 18215.62 | 14694.21 | 15167.6 | 17373.24 | 15141.53 | 19404.32 | 17883.71 | 10387.58 |
| ENSMUSG00000025204 | -0.761916575 | 8.69E-06 | 0.000205 | 2245.513 | 2207.672 | 2779.126 | 2441.224 | 4182.252 | 2825.839 | 4295.908 | 5101.89 |
| ENSMUSG00000025207 | 0.276164893 | 0.021677 | 0.074641 | 1487.954 | 1438.586 | 1334.908 | 1440.581 | 1451.941 | 1026.867 | 1140.754 | 1086.503 |
| ENSMUSG00000025208 | -0.398006681 | 2.98E-05 | 0.000533 | 999.012 | 955.0579 | 1141.959 | 1067.928 | 1416.837 | 1206.764 | 1416.541 | 1450.847 |
| ENSMUSG00000025209 | -0.088911422 | 0.32919 | 0.497016 | 941.6669 | 929.8617 | 852.9498 | 923.8698 | 1097.975 | 908.4251 | 966.2177 | 903.7882 |
| ENSMUSG00000025212 | 0.237588008 | 0.016435 | 0.061389 | 1217.326 | 1289.808 | 1152.724 | 1265.469 | 1098.95 | 1182.182 | 951.7534 | 944.0291 |
| ENSMUSG00000025213 | -0.001647514 | 0.994964 | 0.997071 | 638.8445 | 705.4951 | 463.7397 | 589.1718 | 882.4757 | 372.0856 | 680.7882 | 462.2262 |
| ENSMUSG00000025217 | 0.124769252 | 0.084051 | 0.1954 | 1345.095 | 1491.378 | 1513.779 | 1451.795 | 1275.446 | 1429.121 | 1298.898 | 1320.336 |
| ENSMUSG00000025218 | -0.053710715 | 0.532571 | 0.684146 | 454.7366 | 455.9322 | 462.9116 | 440.8006 | 478.7797 | 430.189 | 478.2874 | 494.8539 |
| ENSMUSG00000025219 | -0.160484908 | 0.444479 | 0.607468 | 220.3259 | 265.1606 | 254.2287 | 258.787 | 374.4427 | 184.3667 | 270.001 | 286.0365 |
| ENSMUSG00000025220 | 0.294557555 | 0.028951 | 0.092173 | 5220.416 | 4925.267 | 5200.51 | 5417.276 | 4149.099 | 5440.494 | 3834.978 | 3506.394 |
| ENSMUSG00000025221 | 0.332279354 | 0.297561 | 0.465485 | 104.6297 | 71.98929 | 79.49824 | 123.3552 | 47.78045 | 116.2069 | 63.64308 | 75.04379 |
| ENSMUSG00000025223 | 0.163771927 | 0.002513 | 0.015492 | 6287.84 | 6318.26 | 6069.194 | 6667.217 | 5634.193 | 5917.613 | 5436.662 | 5636.985 |
| ENSMUSG00000025224 | -0.060403041 | 0.416057 | 0.581855 | 1306.865 | 1267.012 | 1403.641 | 1238.727 | 1260.819 | 1390.013 | 1375.076 | 1417.131 |
| ENSMUSG00000025225 | -0.272918129 | 0.03094 | 0.09662 | 426.567 | 310.7538 | 368.5075 | 334.6979 | 436.8499 | 396.6678 | 461.8945 | 445.9123 |
| ENSMUSG00000025226 | -1.097455621 | 9.65E-06 | 0.000221 | 175.0535 | 133.1802 | 164.7932 | 164.7611 | 391.9947 | 210.0663 | 294.1082 | 469.8393 |
| ENSMUSG00000025227 | -0.390292786 | 0.030036 | 0.094633 | 222.338 | 203.9697 | 255.8849 | 200.1287 | 291.5583 | 212.3011 | 299.8939 | 353.4671 |
| ENSMUSG00000025228 | -0.264998543 | 0.038685 | 0.113668 | 2883.352 | 2796.784 | 3410.143 | 3023.495 | 3672.269 | 2883.943 | 3744.335 | 4257.919 |
| ENSMUSG00000025231 | -0.207877367 | 0.047566 | 0.13141 | 953.7395 | 992.2524 | 1089.788 | 997.1928 | 1080.423 | 1055.919 | 1159.076 | 1364.927 |
| ENSMUSG00000025232 | -0.012499529 | 0.894638 | 0.936266 | 1298.816 | 1571.766 | 1604.043 | 1518.217 | 1556.278 | 1453.704 | 1406.898 | 1629.211 |
| ENSMUSG00000025234 | 0.109686523 | 0.116977 | 0.245141 | 3411.53 | 3287.511 | 3018.449 | 3348.704 | 2831.723 | 3216.92 | 2968.082 | 3093.109 |
| ENSMUSG00000025235 | 0.149760496 | 0.185668 | 0.337272 | 536.227 | 503.925 | 490.2391 | 631.4404 | 508.033 | 513.9921 | 472.5017 | 455.7007 |
| ENSMUSG00000025236 | -0.153824394 | 0.065453 | 0.164684 | 896.3944 | 873.4701 | 849.6374 | 879.876 | 948.7833 | 1036.923 | 870.7531 | 1038.649 |
| ENSMUSG00000025237 | 0.245410591 | 0.040355 | 0.117437 | 723.3531 | 676.6993 | 614.4551 | 809.1408 | 574.3406 | 674.894 | 543.8591 | 590.562 |
| ENSMUSG00000025239 | -0.026850812 | 0.681706 | 0.796654 | 2207.283 | 2174.077 | 2142.312 | 2199.69 | 2013.605 | 2281.678 | 2311.401 | 2281.766 |

| | | | | | | | | | | | |
|---|---|---|---|---|---|---|---|---|---|---|---|
| ENSMUSG00000025240 | -0.683530881 | 0.000874 | 0.007029 | 592.566 | 458.3318 | 544.0661 | 529.6508 | 932.2064 | 519.579 | 927.6462 | 1033.212 |
| ENSMUSG00000025241 | 0.027284696 | 0.767334 | 0.855545 | 783.7163 | 760.6869 | 814.0288 | 700.4503 | 724.5077 | 716.2368 | 723.2169 | 838.5327 |
| ENSMUSG00000025243 | -4.050333588 | 3.12E-11 | 6.39E-09 | 10.06054 | 7.198929 | 35.60859 | 6.900988 | 285.7076 | 81.56831 | 176.4649 | 457.8758 |
| ENSMUSG00000025245 | 0.204276629 | 0.065997 | 0.165539 | 1033.218 | 1161.427 | 956.4632 | 1042.912 | 805.4419 | 1061.505 | 878.4674 | 893.9999 |
| ENSMUSG00000025246 | 0.113498116 | 0.176842 | 0.325762 | 3951.782 | 4204.175 | 3923.569 | 4095.736 | 3690.796 | 4273.956 | 3570.763 | 3416.124 |
| ENSMUSG00000025255 | 1.245736482 | 0.002753 | 0.016576 | 114.6902 | 113.983 | 51.34261 | 89.71284 | 36.07912 | 69.27719 | 29.89296 | 20.66423 |
| ENSMUSG00000025257 | -0.31196934 | 0.230092 | 0.390383 | 81.4904 | 62.39072 | 79.49824 | 56.93315 | 87.76002 | 120.6764 | 66.53595 | 73.95619 |
| ENSMUSG00000025260 | -0.174155092 | 0.018659 | 0.067222 | 2413.524 | 2627.609 | 2587.005 | 2680.171 | 2866.827 | 2671.641 | 2925.653 | 3167.065 |
| ENSMUSG00000025261 | 0.164023536 | 0.002346 | 0.014699 | 6676.177 | 6510.232 | 6723.398 | 6426.545 | 5542.533 | 6227.126 | 5835.878 | 5904.532 |
| ENSMUSG00000025262 | 0.226874952 | 0.162872 | 0.307328 | 253.5257 | 196.7707 | 212.8234 | 256.1992 | 195.0223 | 166.4887 | 188.0364 | 237.0949 |
| ENSMUSG00000025264 | 0.015419571 | 0.854859 | 0.913001 | 904.4429 | 1011.45 | 935.7605 | 930.7707 | 1025.817 | 926.3031 | 854.3602 | 933.1532 |
| ENSMUSG00000025265 | 0.036298304 | 0.661067 | 0.781793 | 736.4318 | 856.6726 | 818.1694 | 855.7225 | 839.5708 | 805.6267 | 763.717 | 776.54 |
| ENSMUSG00000025266 | 0.179766144 | 0.025226 | 0.083539 | 1571.457 | 1398.992 | 1478.17 | 1405.214 | 1421.712 | 1220.173 | 1241.04 | 1284.445 |
| ENSMUSG00000025268 | 0.233385295 | 0.003625 | 0.020376 | 7940.787 | 9944.121 | 9374.167 | 8810.836 | 7737.508 | 7842.849 | 7431.777 | 7668.605 |
| ENSMUSG00000025269 | 0.248265484 | 0.021106 | 0.073206 | 390.3491 | 361.1463 | 324.6178 | 342.4615 | 300.3343 | 319.569 | 291.2153 | 281.6861 |
| ENSMUSG00000025270 | 0.243975485 | 0.419788 | 0.58528 | 390.3491 | 209.9688 | 252.5725 | 565.881 | 289.6081 | 384.3767 | 244.9294 | 280.5985 |
| ENSMUSG00000025271 | 0.604144968 | 9.19E-05 | 0.001252 | 128.775 | 118.7823 | 131.669 | 126.8057 | 79.95913 | 84.92043 | 82.92887 | 85.9197 |
| ENSMUSG00000025272 | 0.478438841 | 9.82E-08 | 5.59E-06 | 1285.737 | 1563.367 | 1386.251 | 1492.339 | 1055.07 | 1099.496 | 968.1463 | 987.5327 |
| ENSMUSG00000025277 | -0.088121971 | 0.650059 | 0.773564 | 671.0383 | 560.3167 | 603.6897 | 476.1682 | 570.4401 | 455.8886 | 594.9664 | 835.27 |
| ENSMUSG00000025278 | -0.080518227 | 0.490499 | 0.647775 | 3099.653 | 2626.409 | 3590.67 | 2982.089 | 2850.25 | 3165.521 | 3501.334 | 3492.255 |
| ENSMUSG00000025280 | -0.114764526 | 0.202851 | 0.358535 | 705.2441 | 585.5129 | 624.3924 | 627.1273 | 715.7317 | 638.0206 | 702.0025 | 697.1459 |
| ENSMUSG00000025283 | -0.551249934 | 0.003445 | 0.019571 | 2786.771 | 1720.544 | 2674.784 | 2670.682 | 3525.027 | 2659.35 | 4240.944 | 4015.386 |
| ENSMUSG00000025287 | -0.475651472 | 0.002787 | 0.016749 | 689.1472 | 577.1142 | 780.9046 | 691.824 | 879.5504 | 736.3495 | 1128.218 | 1066.927 |
| ENSMUSG00000025289 | 0.13670288 | 0.147469 | 0.286645 | 2439.682 | 3070.343 | 2701.284 | 2735.379 | 2646.452 | 2637.003 | 2393.366 | 2277.416 |
| ENSMUSG00000025290 | -0.059384783 | 0.45439 | 0.616543 | 33534.81 | 38735.04 | 38113.61 | 36097.34 | 40424.21 | 39496.94 | 38549.39 | 34163.41 |
| ENSMUSG00000025314 | 0.173960723 | 0.115063 | 0.242698 | 1296.804 | 1433.787 | 1602.386 | 1547.547 | 1129.179 | 1477.169 | 1312.398 | 1297.496 |
| ENSMUSG00000025316 | 0.073062969 | 0.604556 | 0.741111 | 548.2996 | 603.5102 | 472.0208 | 572.782 | 450.5014 | 641.3727 | 486.966 | 508.9926 |
| ENSMUSG00000025318 | 0.679492082 | 0.13268 | 0.26608 | 29.17558 | 15.59768 | 14.07781 | 30.19182 | 9.751113 | 13.40849 | 17.3572 | 15.22628 |
| ENSMUSG00000025321 | 0.270712189 | 0.160685 | 0.304203 | 571.4389 | 465.5308 | 370.9918 | 615.9132 | 344.2143 | 412.311 | 404.0372 | 517.6934 |
| ENSMUSG00000025323 | 0.201327412 | 0.272541 | 0.43804 | 731.4015 | 718.6931 | 584.6433 | 669.3958 | 570.4401 | 796.6877 | 551.5734 | 432.8613 |
| ENSMUSG00000025324 | 0.44359571 | 0.078088 | 0.185889 | 497.9969 | 315.5531 | 278.2438 | 322.6212 | 288.6329 | 256.996 | 326.894 | 166.4014 |
| ENSMUSG00000025326 | 0.349126046 | 0.099602 | 0.220361 | 2178.108 | 2024.099 | 1365.548 | 1714.896 | 1156.482 | 2051.499 | 1284.433 | 1224.628 |
| ENSMUSG00000025329 | 0.355443981 | 0.298243 | 0.466235 | 28.16952 | 22.79661 | 23.18699 | 19.84034 | 16.57689 | 20.11273 | 19.28578 | 17.40146 |
| ENSMUSG00000025330 | 0.096632165 | 0.827111 | 0.895002 | 14.08476 | 22.79661 | 19.87456 | 15.52722 | 16.57689 | 11.17374 | 12.53576 | 27.18978 |
| ENSMUSG00000025332 | -0.133478841 | 0.353056 | 0.521776 | 1973.879 | 1265.812 | 1420.203 | 1565.662 | 1524.099 | 1741.986 | 1703.899 | 1860.868 |
| ENSMUSG00000025337 | -0.208673291 | 0.13413 | 0.267766 | 1519.142 | 1102.636 | 1288.534 | 1289.622 | 1505.572 | 1208.999 | 1563.113 | 1732.533 |
| ENSMUSG00000025340 | -0.061724786 | 0.507951 | 0.663698 | 845.0857 | 884.2685 | 851.2936 | 770.3228 | 980.962 | 804.5093 | 836.0387 | 872.2481 |
| ENSMUSG00000025347 | -4.33948442 | 1.42E-06 | 4.95E-05 | 5.030272 | 2.399643 | 31.46805 | 12.07673 | 258.4045 | 71.51194 | 162.9649 | 549.2335 |
| ENSMUSG00000025348 | 0.116962871 | 0.628927 | 0.75846 | 105.6357 | 151.1775 | 160.6527 | 126.8057 | 113.1129 | 174.3104 | 92.57176 | 122.8978 |
| ENSMUSG00000025350 | -0.045330471 | 0.750872 | 0.844855 | 118.7144 | 122.3818 | 122.5598 | 113.0037 | 123.8391 | 124.0285 | 116.679 | 127.2482 |
| ENSMUSG00000025351 | -0.573887534 | 0.000967 | 0.007642 | 9262.742 | 8763.496 | 11085.86 | 9713.141 | 14912.38 | 9975.916 | 14238.69 | 18667.41 |
| ENSMUSG00000025352 | 0.153744253 | 0.247107 | 0.410164 | 643.8748 | 650.3033 | 603.6897 | 538.2771 | 519.7343 | 483.823 | 512.0375 | 673.2189 |
| ENSMUSG00000025353 | -0.420697369 | 0.020602 | 0.071986 | 374.2522 | 403.14 | 478.6456 | 429.5865 | 598.7183 | 386.6114 | 583.3949 | 688.4452 |
| ENSMUSG00000025354 | -0.140442235 | 0.212848 | 0.370806 | 1628.802 | 1751.739 | 1819.35 | 1764.065 | 2049.684 | 1530.803 | 1990.293 | 2104.489 |
| ENSMUSG00000025355 | 0.619844224 | 0.001959 | 0.012938 | 129.781 | 115.1829 | 96.88848 | 103.5148 | 87.76002 | 71.51194 | 68.46453 | 60.9051 |
| ENSMUSG00000025357 | 0.22799192 | 0.038562 | 0.113508 | 356.1432 | 341.9491 | 325.4459 | 395.0816 | 290.5832 | 324.0385 | 311.4654 | 286.0365 |
| ENSMUSG00000025358 | 0.078324697 | 0.256694 | 0.420325 | 1722.365 | 1802.132 | 1604.871 | 1814.96 | 1668.415 | 1717.404 | 1566.006 | 1623.773 |
| ENSMUSG00000025359 | 0.167905631 | 0.510436 | 0.66605 | 49.29666 | 65.99018 | 63.76421 | 43.13117 | 53.63112 | 56.98608 | 45.32159 | 41.32846 |
| ENSMUSG00000025362 | -0.156402293 | 0.083409 | 0.194467 | 23619.14 | 28488.56 | 27452.56 | 26734.43 | 33123.56 | 28222.63 | 29848.61 | 27268.08 |
| ENSMUSG00000025364 | -0.223026379 | 0.00015 | 0.001828 | 4174.12 | 4344.554 | 4167.861 | 4133.692 | 5235.373 | 4630.398 | 4753.946 | 5007.269 |
| ENSMUSG00000025366 | 0.018098904 | 0.859662 | 0.915887 | 1035.23 | 977.8546 | 1143.615 | 1069.653 | 1059.946 | 884.9603 | 1126.29 | 1103.905 |
| ENSMUSG00000025369 | 0.176818063 | 0.000322 | 0.003279 | 3835.079 | 3900.62 | 3637.872 | 3661.837 | 3367.059 | 3376.705 | 3239.047 | 3312.802 |
| ENSMUSG00000025371 | -0.718297413 | 0.000118 | 0.001515 | 827.9827 | 787.0829 | 1134.506 | 876.4255 | 1588.456 | 1060.388 | 1452.219 | 1867.394 |
| ENSMUSG00000025372 | -0.653590457 | 2.65E-05 | 0.000487 | 575.4631 | 400.7404 | 628.5329 | 577.9577 | 828.8446 | 744.1711 | 957.5391 | 908.1386 |
| ENSMUSG00000025373 | -0.201652381 | 0.082183 | 0.192582 | 1222.356 | 995.8519 | 1023.54 | 1137.8 | 1362.23 | 1044.745 | 1324.933 | 1304.022 |
| ENSMUSG00000025374 | -0.277089915 | 0.010186 | 0.043225 | 1078.49 | 1083.439 | 1127.881 | 1018.758 | 1442.19 | 1071.562 | 1395.326 | 1309.46 |
| ENSMUSG00000025375 | 0.711574132 | 0.000576 | 0.00512 | 227.3683 | 163.1757 | 216.9639 | 310.5445 | 148.2169 | 137.437 | 135.0005 | 141.3868 |
| ENSMUSG00000025377 | -0.164433889 | 0.150358 | 0.290364 | 374.2522 | 358.7466 | 309.7119 | 299.3304 | 377.3681 | 365.3813 | 358.7156 | 399.1459 |
| ENSMUSG00000025380 | -0.338946112 | 0.296458 | 0.464307 | 22.1332 | 45.59322 | 47.20208 | 27.60395 | 41.92979 | 52.51658 | 44.3573 | 41.32846 |
| ENSMUSG00000025381 | -0.073799935 | 0.270937 | 0.436077 | 3016.151 | 3436.289 | 3150.118 | 3198.608 | 3478.222 | 3162.169 | 3319.083 | 3508.569 |
| ENSMUSG00000025384 | -0.177791901 | 0.086678 | 0.199841 | 519.124 | 561.5165 | 591.2681 | 569.3315 | 627.9717 | 550.8654 | 648.0023 | 709.1094 |
| ENSMUSG00000025386 | 0.190356308 | 0.18832 | 0.340615 | 174.0474 | 236.3648 | 230.2136 | 204.4418 | 178.4454 | 202.2447 | 178.3935 | 181.6277 |
| ENSMUSG00000025393 | -0.893256864 | 8.18E-06 | 0.000196 | 22494.37 | 20947.68 | 27961.02 | 23100.19 | 43840.03 | 27155.54 | 48102.6 | 56430.75 |
| ENSMUSG00000025395 | 0.072181996 | 0.522678 | 0.675983 | 1210.283 | 1503.376 | 1149.412 | 1142.114 | 1245.217 | 1229.112 | 1205.361 | 1075.628 |

| | | | | | | | | | | | |
|---|---|---|---|---|---|---|---|---|---|---|---|
| ENSMUSG00000025400 | -1.716211352 | 0.001921 | 0.012745 | 5.030272 | 4.799286 | 8.281066 | 6.038364 | 34.1289 | 13.40849 | 18.32149 | 14.13868 |
| ENSMUSG00000025401 | 0.07453149 | 0.867395 | 0.920566 | 17.10292 | 14.39786 | 9.109173 | 12.07673 | 12.67645 | 14.52586 | 11.57147 | 10.87591 |
| ENSMUSG00000025402 | 0.021155783 | 0.879314 | 0.927685 | 1459.785 | 1473.381 | 1218.145 | 1205.085 | 1607.959 | 1092.792 | 1343.255 | 1230.066 |
| ENSMUSG00000025403 | -0.551751296 | 6.06E-05 | 0.000923 | 1275.677 | 1214.219 | 1225.598 | 1176.618 | 1833.209 | 1339.732 | 1869.757 | 2127.328 |
| ENSMUSG00000025404 | 0.050835601 | 0.459581 | 0.621026 | 2024.181 | 1836.927 | 2008.987 | 1897.772 | 1829.309 | 2028.034 | 1833.114 | 1813.014 |
| ENSMUSG00000025407 | 0.51569492 | 0.000346 | 0.003455 | 1669.044 | 1168.626 | 1696.79 | 1861.541 | 1187.686 | 1105.083 | 1142.683 | 1040.825 |
| ENSMUSG00000025408 | 0.195806295 | 0.104397 | 0.22748 | 344.0706 | 329.9509 | 334.5551 | 293.292 | 306.185 | 288.2825 | 243.0009 | 299.0875 |
| ENSMUSG00000025409 | 0.383097584 | 0.000785 | 0.006494 | 987.9454 | 1016.249 | 1026.024 | 1040.324 | 788.865 | 940.829 | 677.8953 | 715.6349 |
| ENSMUSG00000025410 | -0.18092316 | 0.030657 | 0.096012 | 3527.227 | 3544.273 | 4028.739 | 3742.061 | 4212.481 | 3777.842 | 4228.408 | 4610.299 |
| ENSMUSG00000025413 | -0.046845974 | 0.533264 | 0.684646 | 1629.808 | 1616.16 | 1414.406 | 1499.24 | 1674.266 | 1616.84 | 1537.077 | 1529.153 |
| ENSMUSG00000025417 | -0.3374632 | 0.016145 | 0.060588 | 1622.766 | 1608.961 | 2129.062 | 1638.122 | 2181.324 | 1778.86 | 2407.83 | 2477.532 |
| ENSMUSG00000025418 | -2.450628883 | 0.001517 | 0.01064 | 87.52673 | 44.3934 | 316.3367 | 65.55939 | 638.6979 | 230.1791 | 608.4665 | 1334.474 |
| ENSMUSG00000025420 | 0.037702958 | 0.83917 | 0.902811 | 114.6902 | 97.18554 | 91.09173 | 80.22398 | 85.8098 | 102.7984 | 91.60747 | 92.44524 |
| ENSMUSG00000025421 | -0.163917148 | 0.193264 | 0.346501 | 937.6427 | 926.2622 | 1012.774 | 997.1928 | 1037.518 | 863.7302 | 1182.219 | 1257.255 |
| ENSMUSG00000025422 | -0.889513898 | 0.00318 | 0.018551 | 62.37537 | 67.19001 | 128.3565 | 65.55939 | 156.0178 | 105.0332 | 190.9293 | 149 |
| ENSMUSG00000025423 | -0.014367644 | 0.784613 | 0.867441 | 2496.021 | 2516.026 | 2486.804 | 2600.81 | 2487.509 | 2585.604 | 2665.295 | 2462.306 |
| ENSMUSG00000025425 | 0.678884659 | 0.270941 | 0.436077 | 27.16347 | 7.198929 | 15.73403 | 10.35148 | 5.850668 | 5.58687 | 15.42863 | 10.87591 |
| ENSMUSG00000025427 | 1.263683315 | 8.40E-10 | 1.11E-07 | 190.1443 | 202.7698 | 146.5749 | 168.2116 | 70.20801 | 96.09417 | 69.42882 | 58.72992 |
| ENSMUSG00000025428 | -0.706531922 | 2.64E-05 | 0.000486 | 23140.26 | 22926.19 | 27570.15 | 23974.89 | 39731.89 | 27082.91 | 43356.37 | 49119.96 |
| ENSMUSG00000025429 | -0.020784886 | 0.945118 | 0.968362 | 29.17558 | 29.99554 | 32.29616 | 30.19182 | 32.17867 | 21.23011 | 30.85725 | 39.15328 |
| ENSMUSG00000025431 | 0.018207728 | NA | NA | 6.036326 | 4.799286 | 295.6341 | 9.488858 | 81.90935 | 60.3382 | 73.28598 | 96.79561 |
| ENSMUSG00000025432 | 0.774591877 | 0.132994 | 0.266363 | 28.16952 | 31.19536 | 33.12427 | 12.07673 | 6.825779 | 24.58223 | 12.53576 | 17.40146 |
| ENSMUSG00000025436 | -0.301239331 | 0.043505 | 0.123608 | 239.4409 | 203.9697 | 216.9639 | 246.7103 | 345.1894 | 254.7613 | 260.3581 | 257.7591 |
| ENSMUSG00000025437 | -0.117221054 | 0.195971 | 0.35013 | 904.4429 | 836.2756 | 881.9336 | 856.5851 | 948.7833 | 820.1526 | 975.8606 | 1028.861 |
| ENSMUSG00000025439 | -0.206771844 | 0.012273 | 0.04983 | 1990.982 | 2316.855 | 2336.089 | 2242.821 | 2774.192 | 2375.537 | 2595.866 | 2509.073 |
| ENSMUSG00000025450 | 0.756034734 | 0.000433 | 0.004096 | 535.2209 | 495.5263 | 447.1776 | 611.6001 | 296.4338 | 442.4801 | 281.5724 | 217.5182 |
| ENSMUSG00000025451 | 0.084326432 | 0.290614 | 0.457766 | 2786.771 | 2800.383 | 2444.571 | 2722.44 | 2374.396 | 2739.801 | 2619.974 | 2406.839 |
| ENSMUSG00000025453 | -0.465920144 | 0.005256 | 0.026624 | 1276.683 | 1325.803 | 1277.769 | 1396.587 | 1265.694 | 1843.667 | 2325.865 | 1853.255 |
| ENSMUSG00000025464 | -0.060870085 | 0.485714 | 0.644129 | 731.4015 | 752.2881 | 679.8755 | 808.2782 | 792.7655 | 786.6314 | 803.2529 | 715.6349 |
| ENSMUSG00000025465 | -0.351664107 | 0.000321 | 0.003269 | 3094.623 | 2764.389 | 3060.682 | 3140.812 | 3858.515 | 3249.324 | 4085.693 | 4198.102 |
| ENSMUSG00000025466 | -0.937780042 | 3.39E-05 | 0.000589 | 314.895 | 307.1543 | 390.0382 | 332.9727 | 663.0757 | 373.2029 | 729.9669 | 811.343 |
| ENSMUSG00000025467 | -1.342591806 | 0.000563 | 0.005013 | 11.0666 | 14.39786 | 13.24971 | 7.763611 | 32.17867 | 35.75597 | 24.10723 | 25.0146 |
| ENSMUSG00000025468 | 0.865175683 | 0.019959 | 0.070312 | 46.2785 | 35.99465 | 41.40533 | 75.91087 | 31.20356 | 37.99072 | 19.28578 | 21.75182 |
| ENSMUSG00000025470 | -0.116182002 | 0.213021 | 0.370828 | 552.3238 | 518.3229 | 537.4412 | 613.3253 | 617.2455 | 611.2036 | 629.6808 | 551.4087 |
| ENSMUSG00000025473 | 0.746936189 | 0.013162 | 0.05249 | 134.8113 | 92.38626 | 83.63877 | 106.1027 | 47.78045 | 94.9768 | 43.39301 | 63.08028 |
| ENSMUSG00000025474 | -0.073942871 | 0.406175 | 0.572897 | 1520.148 | 1734.942 | 1748.133 | 1601.029 | 1894.641 | 1709.582 | 1789.721 | 1555.255 |
| ENSMUSG00000025475 | 1.092632264 | 0.001504 | 0.010583 | 79.47829 | 67.19001 | 57.13936 | 67.28463 | 29.25334 | 53.63396 | 26.03581 | 18.48905 |
| ENSMUSG00000025477 | -0.0409371 | 0.565493 | 0.71003 | 1467.833 | 1252.614 | 1395.36 | 1351.731 | 1399.285 | 1379.957 | 1462.827 | 1385.591 |
| ENSMUSG00000025478 | 0.220356669 | 0.127339 | 0.258573 | 336.0222 | 418.7377 | 352.7734 | 345.912 | 363.7165 | 251.4092 | 307.6082 | 321.927 |
| ENSMUSG00000025479 | -3.424689237 | 1.23E-07 | 6.68E-06 | 14.08476 | 2.399643 | 7.45296 | 25.01608 | 156.0178 | 40.22547 | 93.53605 | 241.4452 |
| ENSMUSG00000025484 | 0.131542546 | 0.099446 | 0.220148 | 881.3036 | 896.2667 | 908.433 | 914.3809 | 820.0686 | 912.8946 | 765.6456 | 790.6787 |
| ENSMUSG00000025485 | 0.041831376 | 0.586883 | 0.727191 | 1461.797 | 1463.782 | 1687.681 | 1447.482 | 1519.223 | 1415.713 | 1507.184 | 1447.584 |
| ENSMUSG00000025486 | -0.420275196 | 0.000646 | 0.005588 | 457.7547 | 466.7306 | 487.7548 | 494.2833 | 666.9761 | 522.9311 | 617.1451 | 744.9999 |
| ENSMUSG00000025487 | -0.199427937 | 0.040135 | 0.116969 | 2685.159 | 2856.775 | 3078.072 | 2937.233 | 3285.15 | 2889.529 | 3299.798 | 3797.868 |
| ENSMUSG00000025489 | 0.857766568 | 0.009573 | 0.041244 | 36.21796 | 32.39518 | 27.32752 | 33.64232 | 16.57689 | 23.46486 | 17.3572 | 14.13868 |
| ENSMUSG00000025491 | 0.451489672 | 3.06E-05 | 0.000545 | 1527.191 | 1569.367 | 1703.415 | 1756.301 | 1217.914 | 1386.661 | 1168.718 | 1023.423 |
| ENSMUSG00000025492 | 0.242801602 | 0.33191 | 0.499775 | 2318.955 | 4334.955 | 4731.801 | 2093.587 | 2780.042 | 2654.881 | 3206.261 | 2749.43 |
| ENSMUSG00000025494 | -1.877582383 | 3.27E-08 | 2.29E-06 | 155.9384 | 101.9848 | 251.7444 | 138.0198 | 609.4446 | 324.0385 | 532.2876 | 919.0145 |
| ENSMUSG00000025495 | -0.46272182 | 0.001837 | 0.012338 | 1254.55 | 1303.006 | 1449.187 | 1275.82 | 1966.8 | 1319.619 | 1853.364 | 2140.379 |
| ENSMUSG00000025496 | 0.869892787 | 1.56E-07 | 8.15E-06 | 257.5499 | 239.9643 | 269.9628 | 266.5507 | 154.0676 | 175.4277 | 126.3219 | 110.9343 |
| ENSMUSG00000025497 | -2.314805506 | 4.11E-05 | 0.000689 | 117.7084 | 89.98662 | 269.1347 | 100.0643 | 674.777 | 272.6393 | 648.9666 | 1276.832 |
| ENSMUSG00000025498 | 0.30118489 | 0.528023 | 0.680556 | 161.9748 | 334.7502 | 462.0835 | 74.18562 | 184.296 | 128.498 | 297.9654 | 227.3065 |
| ENSMUSG00000025499 | -0.477366493 | 0.001704 | 0.011646 | 1186.138 | 1033.046 | 1301.784 | 1132.625 | 1596.257 | 1208.999 | 1715.47 | 1959.839 |
| ENSMUSG00000025500 | 0.197474775 | 0.31892 | 0.487005 | 224.3501 | 193.1713 | 133.3252 | 201.8539 | 167.7191 | 191.071 | 154.2863 | 142.4744 |
| ENSMUSG00000025503 | -0.395068987 | 0.007473 | 0.034499 | 3075.508 | 2917.966 | 3372.05 | 2649.979 | 3806.835 | 3053.783 | 4021.086 | 4919.174 |
| ENSMUSG00000025504 | -1.419302796 | 1.33E-05 | 0.000286 | 458.7608 | 367.1454 | 767.6548 | 373.516 | 1260.819 | 629.0816 | 1572.756 | 1801.051 |
| ENSMUSG00000025505 | 0.017295243 | 0.826865 | 0.894894 | 842.0675 | 969.4558 | 867.8558 | 860.8982 | 905.8784 | 824.6221 | 902.5747 | 859.197 |
| ENSMUSG00000025507 | 0.134190384 | 0.263681 | 0.427975 | 293.7679 | 377.9438 | 321.3054 | 300.193 | 312.0356 | 289.3999 | 295.0725 | 278.4233 |
| ENSMUSG00000025508 | -0.168717874 | 0.055281 | 0.146317 | 16885.62 | 18874.39 | 20668.71 | 18229.82 | 22302.75 | 18924.97 | 22111.15 | 20581.57 |
| ENSMUSG00000025509 | -0.32158507 | 0.049758 | 0.135689 | 1107.666 | 874.6699 | 1067.429 | 1161.091 | 1237.416 | 1024.632 | 1696.185 | 1305.109 |
| ENSMUSG00000025510 | -0.239440971 | 0.012141 | 0.049419 | 2636.868 | 2409.242 | 2587.005 | 2567.168 | 3078.426 | 2543.143 | 3062.582 | 3358.481 |
| ENSMUSG00000025511 | -0.088928057 | 0.462157 | 0.623214 | 1290.768 | 1333.002 | 1674.432 | 1454.383 | 1610.884 | 1283.863 | 1563.113 | 1662.927 |
| ENSMUSG00000025512 | -0.357731019 | 0.006814 | 0.032357 | 632.8082 | 760.6869 | 760.2019 | 634.0283 | 946.8331 | 734.1148 | 863.0388 | 1026.686 |
| ENSMUSG00000025515 | 0.079268672 | 0.848143 | 0.908809 | 20.12109 | 15.59768 | 23.18699 | 24.15346 | 10.72622 | 29.05173 | 21.21436 | 18.48905 |

| | | | | | | | | | | | |
|---|---|---|---|---|---|---|---|---|---|---|---|
| ENSMUSG00000025521 | -0.185146337 | 0.20912 | 0.366213 | 446.6881 | 357.5468 | 385.0696 | 439.0754 | 560.689 | 377.6724 | 455.1445 | 457.8758 |
| ENSMUSG00000025525 | -0.179222343 | 0.041688 | 0.120199 | 668.0201 | 605.9099 | 624.3924 | 680.6099 | 767.4126 | 659.2507 | 763.717 | 729.7736 |
| ENSMUSG00000025529 | 0.552558897 | 0.016559 | 0.061653 | 453.7305 | 476.3291 | 490.2391 | 491.6954 | 252.5538 | 511.7573 | 289.2867 | 251.2335 |
| ENSMUSG00000025531 | 0.133618408 | 0.103611 | 0.226381 | 880.2976 | 790.6824 | 798.2948 | 900.5789 | 736.209 | 816.8005 | 765.6456 | 755.8758 |
| ENSMUSG00000025532 | -0.29435844 | 0.042448 | 0.121648 | 614.6992 | 523.1222 | 634.3297 | 572.782 | 842.4962 | 549.7481 | 718.3954 | 765.6641 |
| ENSMUSG00000025533 | -0.628027165 | 0.003308 | 0.019066 | 1022.151 | 735.4906 | 939.0729 | 1028.247 | 1721.071 | 872.6692 | 1459.934 | 1704.255 |
| ENSMUSG00000025534 | 0.124918606 | 0.179184 | 0.328916 | 1536.245 | 1568.167 | 1691.822 | 1582.051 | 1648.913 | 1289.45 | 1400.148 | 1510.664 |
| ENSMUSG00000025537 | 1.123614683 | 0.012727 | 0.051175 | 23.13925 | 28.79572 | 13.24971 | 21.56559 | 8.776002 | 12.29112 | 7.714313 | 10.87591 |
| ENSMUSG00000025538 | -0.061444113 | 0.536906 | 0.687419 | 809.8738 | 830.2765 | 684.8442 | 822.9428 | 832.7451 | 756.4623 | 796.5028 | 896.1751 |
| ENSMUSG00000025544 | -0.202323428 | 0.003186 | 0.018573 | 3808.922 | 3720.647 | 3988.162 | 4014.65 | 4103.268 | 4391.28 | 4581.338 | 4800.627 |
| ENSMUSG00000025545 | -0.463542413 | 0.002168 | 0.013927 | 363.1856 | 296.3559 | 414.0533 | 339.011 | 487.5557 | 394.4331 | 513.0018 | 555.759 |
| ENSMUSG00000025551 | 1.252970614 | 0.060993 | 0.156745 | 26.15741 | 13.19804 | 8.281066 | 17.25247 | 8.776002 | 12.29112 | 1.928578 | 4.350364 |
| ENSMUSG00000025555 | 0.08440396 | 0.418806 | 0.584292 | 3408.512 | 3739.844 | 3884.648 | 3426.341 | 3108.655 | 4051.598 | 3191.797 | 3287.788 |
| ENSMUSG00000025557 | -1.079661193 | 0.090925 | 0.206612 | 12.07265 | 2.399643 | 19.04645 | 6.038364 | 14.62667 | 13.40849 | 23.14294 | 33.71532 |
| ENSMUSG00000025558 | -0.164701058 | 0.075321 | 0.181454 | 1610.693 | 1347.4 | 1403.641 | 1507.866 | 1818.583 | 1496.164 | 1626.756 | 1637.912 |
| ENSMUSG00000025571 | 0.499947542 | 0.004835 | 0.025125 | 1820.958 | 1616.16 | 1323.314 | 1658.825 | 968.2855 | 1563.206 | 988.3964 | 1020.16 |
| ENSMUSG00000025572 | -0.12942789 | 0.093583 | 0.210604 | 1235.435 | 1262.212 | 1176.74 | 1229.238 | 1428.538 | 1206.764 | 1403.041 | 1320.336 |
| ENSMUSG00000025574 | -0.074270179 | 0.556892 | 0.703587 | 944.685 | 1395.392 | 1180.052 | 1059.302 | 1220.839 | 1274.924 | 1160.04 | 1162.635 |
| ENSMUSG00000025575 | -0.06522038 | 0.326645 | 0.494306 | 1381.313 | 1373.796 | 1409.437 | 1366.396 | 1421.712 | 1335.262 | 1512.97 | 1516.102 |
| ENSMUSG00000025576 | 0.46814843 | 0.154419 | 0.295923 | 57.3451 | 64.79036 | 68.73285 | 56.07053 | 32.17867 | 74.86406 | 40.50014 | 31.54014 |
| ENSMUSG00000025577 | -0.032988362 | 0.784627 | 0.867441 | 791.7648 | 837.4754 | 731.2182 | 832.4317 | 972.186 | 722.941 | 842.7887 | 725.4233 |
| ENSMUSG00000025578 | -8.50E-05 | 0.999492 | 0.999733 | 261.5741 | 263.9607 | 221.1045 | 232.9083 | 269.1307 | 244.7049 | 208.2865 | 255.5839 |
| ENSMUSG00000025579 | -0.127331325 | 0.288622 | 0.455594 | 1757.577 | 1883.72 | 2254.934 | 1927.964 | 2210.577 | 1774.39 | 2164.829 | 2398.138 |
| ENSMUSG00000025580 | -0.291793604 | 0.000412 | 0.003954 | 2940.697 | 3125.535 | 3283.443 | 3108.895 | 3954.076 | 3321.953 | 3940.086 | 4034.963 |
| ENSMUSG00000025582 | 0.224014923 | 0.549126 | 0.697351 | 30.18163 | 50.3925 | 47.20208 | 68.14726 | 28.27823 | 52.51658 | 57.85735 | 29.36496 |
| ENSMUSG00000025583 | -0.103304499 | 0.160222 | 0.303599 | 1041.266 | 1025.847 | 983.7907 | 981.6655 | 1021.917 | 1080.501 | 1050.111 | 1178.949 |
| ENSMUSG00000025584 | 0.064551558 | 0.690361 | 0.803922 | 700.2138 | 482.3283 | 688.1566 | 674.5716 | 495.3565 | 587.7388 | 638.3594 | 715.6349 |
| ENSMUSG00000025586 | 0.207816774 | NA | NA | 52.31483 | 53.99197 | 50.5145 | 58.6584 | 21.45245 | 137.437 | 9.642892 | 18.48905 |
| ENSMUSG00000025591 | -0.117906752 | 0.228436 | 0.388537 | 730.3955 | 712.694 | 737.843 | 768.5975 | 865.8988 | 882.7255 | 722.2526 | 731.9488 |
| ENSMUSG00000025592 | 0.536141484 | 0.012617 | 0.050816 | 186.1201 | 226.7663 | 231.0418 | 194.0903 | 91.66046 | 157.5497 | 174.5363 | 154.4379 |
| ENSMUSG00000025597 | 0.334616024 | 0.017902 | 0.065414 | 396.3854 | 548.3184 | 602.8616 | 534.8266 | 409.5467 | 393.3157 | 417.5372 | 431.7737 |
| ENSMUSG00000025602 | 0.026585966 | 0.903559 | 0.942315 | 326.9677 | 287.9572 | 293.1497 | 322.6212 | 342.2641 | 402.2547 | 274.8224 | 189.2408 |
| ENSMUSG00000025607 | -0.101331712 | 0.165196 | 0.310588 | 2568.457 | 2736.793 | 2489.289 | 2593.909 | 2913.633 | 2963.276 | 2733.76 | 2529.737 |
| ENSMUSG00000025608 | -1.136143995 | 2.00E-05 | 0.000392 | 1864.219 | 1397.792 | 3039.151 | 2085.824 | 3096.954 | 3794.602 | 5057.697 | 6489.656 |
| ENSMUSG00000025609 | 0.04549953 | 0.681312 | 0.796487 | 2448.736 | 2318.055 | 2283.918 | 2489.531 | 2086.738 | 2824.722 | 2178.329 | 2156.693 |
| ENSMUSG00000025610 | -0.983204584 | 0.000149 | 0.001818 | 110.666 | 115.1829 | 103.5133 | 163.8985 | 198.9227 | 194.4231 | 219.8579 | 363.2554 |
| ENSMUSG00000025612 | 0.148123705 | 0.016028 | 0.060346 | 1398.416 | 1298.207 | 1426.828 | 1419.016 | 1214.989 | 1261.515 | 1277.683 | 1252.905 |
| ENSMUSG00000025613 | -0.105937387 | 0.058501 | 0.151878 | 7216.428 | 7782.042 | 7438.882 | 7839.522 | 8539.05 | 7761.28 | 8264.922 | 8014.459 |
| ENSMUSG00000025616 | 0.018869073 | 0.782849 | 0.866307 | 1582.524 | 1503.376 | 1447.53 | 1557.898 | 1386.608 | 1546.446 | 1531.291 | 1548.73 |
| ENSMUSG00000025626 | 0.182326629 | 0.123686 | 0.253358 | 1293.786 | 1487.779 | 1113.803 | 1368.983 | 1145.756 | 1331.91 | 1091.575 | 1068.014 |
| ENSMUSG00000025630 | -0.215945964 | 0.001333 | 0.009701 | 1531.215 | 1449.384 | 1570.918 | 1451.795 | 1700.594 | 1676.061 | 1722.22 | 1877.182 |
| ENSMUSG00000025644 | 0.019079079 | 0.930909 | 0.959196 | 76.46013 | 79.18822 | 110.9663 | 75.91087 | 95.56091 | 90.5073 | 68.46453 | 84.8321 |
| ENSMUSG00000025645 | -0.147020984 | 0.445884 | 0.608619 | 206.2411 | 235.165 | 185.4959 | 202.7165 | 221.3503 | 178.7799 | 207.3222 | 309.9635 |
| ENSMUSG00000025646 | -0.015304344 | 0.957709 | 0.975959 | 38.23007 | 34.79482 | 62.108 | 38.81806 | 37.05423 | 48.04709 | 44.3573 | 47.85401 |
| ENSMUSG00000025647 | -0.107020078 | 0.470159 | 0.630173 | 1545.299 | 1650.954 | 1981.659 | 1453.521 | 1685.967 | 1421.3 | 1945.936 | 2089.262 |
| ENSMUSG00000025648 | -0.419695637 | 0.00505 | 0.025911 | 183.1019 | 171.5745 | 217.792 | 182.8762 | 266.2054 | 206.7142 | 256.5009 | 282.7737 |
| ENSMUSG00000025650 | -0.671462055 | 0.012008 | 0.048998 | 161.9748 | 134.38 | 283.2125 | 156.1349 | 287.6578 | 213.4185 | 367.3942 | 305.6131 |
| ENSMUSG00000025651 | -0.742675562 | 6.24E-05 | 0.000944 | 3666.062 | 3454.286 | 4882.517 | 3930.113 | 6541.047 | 4630.398 | 6888.882 | 8602.845 |
| ENSMUSG00000025656 | 0.307055806 | 0.017113 | 0.063141 | 467.8153 | 545.9188 | 523.3634 | 580.5456 | 372.4925 | 486.0577 | 387.6442 | 467.6642 |
| ENSMUSG00000025658 | 0.466163879 | 0.034536 | 0.105006 | 136.8234 | 139.1793 | 121.7317 | 128.5309 | 74.10846 | 135.2023 | 81.00029 | 91.35765 |
| ENSMUSG00000025665 | -0.043707706 | 0.64517 | 0.770248 | 597.5963 | 733.091 | 673.2507 | 730.6421 | 680.6277 | 721.8237 | 736.7169 | 678.6568 |
| ENSMUSG00000025666 | 0.578013302 | 6.08E-07 | 2.47E-05 | 885.3278 | 1082.239 | 1097.241 | 1071.378 | 593.8428 | 769.8708 | 722.2526 | 686.27 |
| ENSMUSG00000025701 | 1.65192938 | 0.000223 | 0.002465 | 29.17558 | 35.99465 | 25.67131 | 24.15346 | 10.72622 | 13.40849 | 3.857157 | 8.700729 |
| ENSMUSG00000025702 | -0.204779265 | 0.001848 | 0.012395 | 2373.282 | 2211.271 | 2404.822 | 2345.473 | 2788.818 | 2470.514 | 2790.653 | 2710.277 |
| ENSMUSG00000025722 | -0.165121417 | 0.101589 | 0.223524 | 930.6003 | 1077.44 | 944.8697 | 884.1891 | 1217.914 | 1005.637 | 1044.325 | 1028.861 |
| ENSMUSG00000025723 | -0.096743472 | 0.64509 | 0.770202 | 117.7084 | 85.18733 | 81.98256 | 112.1411 | 117.0134 | 102.7984 | 119.5719 | 84.8321 |
| ENSMUSG00000025724 | -0.081587382 | 0.179662 | 0.329508 | 2297.828 | 2332.453 | 2403.165 | 2479.18 | 2607.448 | 2349.838 | 2517.759 | 2592.817 |
| ENSMUSG00000025726 | -0.811674875 | 0.165568 | 0.311015 | 19.11503 | 5.999108 | 4.96864 | 10.35148 | 19.50223 | 8.938993 | 17.3572 | 25.0146 |
| ENSMUSG00000025727 | -0.034337377 | 0.904036 | 0.942489 | 101.6115 | 82.78769 | 173.9024 | 93.16334 | 93.61069 | 99.44629 | 126.3219 | 144.6496 |
| ENSMUSG00000025728 | -0.358441805 | 0.002278 | 0.01441 | 1898.425 | 1762.538 | 2004.846 | 1853.778 | 2407.55 | 1928.588 | 2541.866 | 2763.569 |
| ENSMUSG00000025730 | -0.093232036 | 0.17627 | 0.324836 | 688.1412 | 677.8992 | 702.2344 | 648.6929 | 738.1593 | 720.7063 | 721.2883 | 717.8101 |
| ENSMUSG00000025731 | -0.338468369 | 0.024426 | 0.081566 | 838.0433 | 752.2881 | 925.8232 | 811.7287 | 999.4891 | 803.392 | 1125.325 | 1281.182 |
| ENSMUSG00000025732 | -1.469895001 | 8.06E-06 | 0.000194 | 347.0888 | 154.777 | 278.2438 | 166.4863 | 595.793 | 378.7898 | 653.7881 | 995.1458 |
| ENSMUSG00000025733 | -0.238042054 | 0.008467 | 0.037836 | 901.4247 | 893.867 | 933.2762 | 853.9973 | 1056.046 | 966.5286 | 1004.789 | 1198.525 |

| | | | | | | | | | | | |
|---|---|---|---|---|---|---|---|---|---|---|---|
| ENSMUSG00000025735 | 0.175160259 | 0.644889 | 0.770145 | 42.25428 | 47.99286 | 37.2648 | 31.91707 | 44.85512 | 23.46486 | 20.25007 | 52.20437 |
| ENSMUSG00000025736 | -0.127294965 | 0.125927 | 0.256641 | 1558.378 | 1677.351 | 1523.716 | 1644.16 | 1880.99 | 1554.267 | 1718.363 | 1836.941 |
| ENSMUSG00000025737 | -0.098275486 | 0.258181 | 0.422053 | 896.3944 | 929.8617 | 847.9812 | 883.3265 | 901.0028 | 877.1387 | 965.2535 | 1062.576 |
| ENSMUSG00000025738 | 0.409769078 | 0.115706 | 0.243691 | 99.59938 | 87.58697 | 56.31125 | 111.2784 | 68.25779 | 71.51194 | 71.3574 | 55.46715 |
| ENSMUSG00000025739 | -1.074908309 | 0.07922 | 0.187783 | 4.024217 | 14.39786 | 13.24971 | 8.626235 | 8.776002 | 36.87335 | 25.07152 | 14.13868 |
| ENSMUSG00000025742 | -0.084166657 | 0.321975 | 0.490173 | 1096.599 | 1180.624 | 1224.77 | 1132.625 | 1101.876 | 1210.116 | 1267.076 | 1335.562 |
| ENSMUSG00000025743 | 0.148904944 | 0.177948 | 0.327177 | 2085.551 | 1719.344 | 2088.485 | 1719.209 | 1612.834 | 1554.267 | 1906.4 | 1793.438 |
| ENSMUSG00000025745 | -0.392100833 | 0.003802 | 0.021044 | 4265.67 | 4306.159 | 4873.408 | 4439.06 | 5678.073 | 4595.76 | 5916.878 | 7280.335 |
| ENSMUSG00000025747 | -0.042955229 | 0.665154 | 0.785063 | 965.8122 | 1239.416 | 1051.695 | 976.4898 | 1133.079 | 1073.797 | 1051.075 | 1098.467 |
| ENSMUSG00000025757 | 0.774274525 | 0.000148 | 0.001805 | 964.8061 | 1009.05 | 891.0427 | 978.215 | 466.1032 | 848.0869 | 461.8945 | 472.0145 |
| ENSMUSG00000025758 | -0.086985226 | 0.33602 | 0.504276 | 934.6245 | 1033.046 | 870.3401 | 898.8537 | 1068.722 | 1027.984 | 927.6462 | 939.6787 |
| ENSMUSG00000025759 | 0.011591233 | 0.926409 | 0.956313 | 270.6286 | 271.1597 | 231.8699 | 261.3749 | 267.1805 | 286.0478 | 246.858 | 225.1314 |
| ENSMUSG00000025762 | -1.003961148 | 2.31E-08 | 1.71E-06 | 409.4641 | 335.95 | 402.4598 | 381.2796 | 623.0961 | 632.4337 | 777.2171 | 1036.474 |
| ENSMUSG00000025764 | -0.035688153 | 0.701968 | 0.812145 | 1515.118 | 1318.604 | 1281.081 | 1413.84 | 1253.993 | 1524.098 | 1431.005 | 1459.547 |
| ENSMUSG00000025766 | -0.011890473 | 0.947246 | 0.969627 | 226.3622 | 307.1543 | 223.5888 | 209.6175 | 211.5992 | 292.752 | 212.1436 | 256.6715 |
| ENSMUSG00000025776 | 0.990753461 | 0.008413 | 0.03765 | 617.7174 | 663.5013 | 516.7385 | 763.4218 | 214.5245 | 650.3117 | 268.0724 | 156.6131 |
| ENSMUSG00000025777 | 0.982628659 | 0.025924 | 0.085178 | 45.27245 | 31.19536 | 18.21835 | 60.38364 | 18.52711 | 24.58223 | 20.25007 | 15.22628 |
| ENSMUSG00000025779 | 0.307543551 | 0.110251 | 0.236138 | 129.781 | 100.785 | 123.3879 | 106.1027 | 99.46135 | 99.44629 | 102.2147 | 70.69342 |
| ENSMUSG00000025780 | 0.851680216 | 0.003864 | 0.021297 | 1198.211 | 1410.99 | 1051.695 | 1297.386 | 524.6099 | 1241.403 | 535.1805 | 446.9999 |
| ENSMUSG00000025781 | -0.533665064 | 7.81E-05 | 0.001122 | 5532.293 | 5569.572 | 6729.194 | 5972.805 | 8671.665 | 6667.371 | 8898.46 | 10223.36 |
| ENSMUSG00000025782 | 0.468149148 | 0.004911 | 0.025395 | 1123.763 | 998.2515 | 905.1205 | 876.4255 | 581.1663 | 939.7116 | 630.6451 | 671.0437 |
| ENSMUSG00000025783 | 0.000110764 | 0.999679 | 0.999733 | 140.8476 | 73.18911 | 156.5122 | 101.7896 | 117.9885 | 87.15518 | 138.8576 | 129.4233 |
| ENSMUSG00000025784 | 1.93106553 | 0.012499 | 0.050512 | 404.4339 | 1009.05 | 708.0312 | 866.074 | 203.7983 | 444.7149 | 108.9647 | 26.10219 |
| ENSMUSG00000025785 | -0.105374388 | 0.159081 | 0.302057 | 1013.097 | 1072.64 | 1035.961 | 1021.346 | 1144.781 | 1061.505 | 1042.397 | 1207.226 |
| ENSMUSG00000025786 | -0.159878379 | 0.047077 | 0.130438 | 1830.013 | 2038.497 | 2180.405 | 1973.683 | 2357.819 | 2058.203 | 2272.83 | 2274.153 |
| ENSMUSG00000025787 | -0.275725897 | 0.466582 | 0.627229 | 26.15741 | 25.19625 | 24.01509 | 35.36756 | 22.42756 | 45.81234 | 22.17865 | 44.59123 |
| ENSMUSG00000025789 | 0.645310419 | 0.000153 | 0.00185 | 298.7981 | 424.7368 | 341.1799 | 355.4009 | 199.8978 | 277.1088 | 190.9293 | 240.3576 |
| ENSMUSG00000025790 | 0.329164854 | 2.06E-06 | 6.57E-05 | 1182.114 | 1123.033 | 1053.352 | 1117.96 | 876.6251 | 858.1433 | 921.8604 | 903.7882 |
| ENSMUSG00000025791 | -0.223411785 | 0.067214 | 0.167403 | 1139.86 | 1237.016 | 1235.535 | 1195.596 | 1471.443 | 1100.613 | 1426.184 | 1613.985 |
| ENSMUSG00000025792 | -1.217626256 | 1.06E-05 | 0.000238 | 893.3763 | 782.2836 | 1283.565 | 873.8376 | 2289.561 | 1202.295 | 2049.114 | 3375.883 |
| ENSMUSG00000025793 | -0.1757161 | 0.079167 | 0.18773 | 1027.181 | 1022.248 | 1132.022 | 974.7645 | 1180.86 | 1015.693 | 1188.004 | 1310.547 |
| ENSMUSG00000025794 | -0.111419517 | 0.18109 | 0.331435 | 23691.57 | 28830.51 | 26821.55 | 26074.52 | 30537.56 | 28379.07 | 28978.82 | 25982.55 |
| ENSMUSG00000025795 | 0.066439279 | 0.506952 | 0.662721 | 801.8253 | 899.8662 | 972.1972 | 888.5022 | 878.5753 | 777.6924 | 803.2529 | 944.0291 |
| ENSMUSG00000025804 | 0.28814019 | 0.359609 | 0.528083 | 94.56911 | 61.1909 | 87.7793 | 56.07053 | 77.03379 | 37.99072 | 52.07161 | 78.30656 |
| ENSMUSG00000025809 | -0.015152446 | 0.741986 | 0.838976 | 16152.2 | 16663.12 | 17037.47 | 17345.63 | 16365.29 | 16928.22 | 17506.67 | 17112.16 |
| ENSMUSG00000025810 | 0.092978455 | 0.245874 | 0.408808 | 3060.417 | 2889.17 | 2898.373 | 2842.344 | 2500.185 | 3033.671 | 2583.331 | 2846.226 |
| ENSMUSG00000025812 | -0.107445141 | 0.171563 | 0.31877 | 1774.68 | 1833.327 | 1970.066 | 1918.475 | 1847.836 | 2126.363 | 1940.15 | 2167.569 |
| ENSMUSG00000025813 | -0.071434382 | 0.628677 | 0.758208 | 288.7376 | 260.3613 | 345.3205 | 244.9851 | 303.2596 | 286.0478 | 272.8938 | 337.1532 |
| ENSMUSG00000025815 | 0.462780204 | 0.018275 | 0.06624 | 378.2764 | 275.959 | 261.6817 | 368.3402 | 251.5787 | 263.7003 | 246.858 | 169.6642 |
| ENSMUSG00000025816 | 0.340950746 | 0.005321 | 0.026853 | 1655.965 | 1552.569 | 1385.422 | 1603.617 | 1154.532 | 1505.103 | 1119.54 | 1114.781 |
| ENSMUSG00000025817 | 0.041620041 | 0.615545 | 0.74883 | 926.5761 | 950.2587 | 871.9963 | 916.9688 | 851.2722 | 884.9603 | 834.1101 | 990.7955 |
| ENSMUSG00000025821 | -0.152595963 | 0.04424 | 0.125039 | 1201.229 | 1142.23 | 1197.442 | 1102.433 | 1356.38 | 1168.773 | 1305.648 | 1329.036 |
| ENSMUSG00000025823 | -0.167237987 | 0.037431 | 0.11137 | 4479.96 | 4866.476 | 4693.708 | 4225.992 | 4883.357 | 4718.671 | 5366.269 | 5539.101 |
| ENSMUSG00000025825 | -0.231815064 | 0.137247 | 0.272281 | 1727.395 | 1178.225 | 1328.283 | 1419.878 | 1725.947 | 1296.154 | 1689.435 | 1928.299 |
| ENSMUSG00000025854 | -0.186401736 | 0.030305 | 0.095156 | 1080.502 | 1037.846 | 1163.49 | 996.3301 | 1176.959 | 1136.369 | 1330.719 | 1224.628 |
| ENSMUSG00000025855 | -0.051848753 | 0.757148 | 0.848808 | 540.2512 | 458.3318 | 596.2368 | 600.386 | 544.1121 | 422.3674 | 636.4308 | 674.3065 |
| ENSMUSG00000025856 | -0.180399166 | 0.089369 | 0.204103 | 1787.759 | 1563.367 | 1895.536 | 1674.352 | 1694.743 | 1987.809 | 1942.078 | 2223.036 |
| ENSMUSG00000025857 | -0.431241819 | 0.000725 | 0.006116 | 469.8274 | 429.5361 | 433.9279 | 450.2895 | 636.7477 | 476.0014 | 607.5022 | 684.0948 |
| ENSMUSG00000025858 | -0.288158563 | 0.001489 | 0.010503 | 1638.863 | 1544.17 | 1890.567 | 1801.158 | 2169.623 | 1935.292 | 2057.793 | 2238.262 |
| ENSMUSG00000025860 | 0.26625032 | 1.45E-05 | 0.000303 | 3412.536 | 3435.089 | 3246.178 | 3510.015 | 2697.158 | 3039.258 | 2761.724 | 2814.686 |
| ENSMUSG00000025862 | 0.1882031 | 0.057426 | 0.150121 | 3967.878 | 4054.197 | 3709.918 | 3912.86 | 3130.107 | 4054.951 | 3425.155 | 3121.386 |
| ENSMUSG00000025867 | -0.05758661 | 0.43003 | 0.594816 | 986.9393 | 884.2685 | 953.1507 | 969.5888 | 938.0571 | 1000.05 | 969.1106 | 1046.263 |
| ENSMUSG00000025868 | -0.324076213 | 0.00813 | 0.036706 | 1162.999 | 1011.45 | 1236.363 | 1158.503 | 1366.131 | 1187.769 | 1506.22 | 1662.927 |
| ENSMUSG00000025869 | -0.121449389 | 0.047565 | 0.13141 | 1215.314 | 1263.412 | 1201.583 | 1274.958 | 1409.036 | 1296.154 | 1321.076 | 1361.664 |
| ENSMUSG00000025870 | -0.416736322 | 0.058789 | 0.152432 | 309.8647 | 261.5611 | 276.5876 | 234.6336 | 428.0739 | 215.6532 | 392.4657 | 407.8467 |
| ENSMUSG00000025871 | -0.739717112 | 4.81E-05 | 0.000772 | 2130.823 | 1692.948 | 2175.436 | 1869.305 | 3358.283 | 2251.509 | 3261.226 | 4269.883 |
| ENSMUSG00000025872 | -0.107868096 | 0.082732 | 0.193427 | 1487.954 | 1659.353 | 1657.869 | 1628.633 | 1779.578 | 1726.343 | 1735.72 | 1691.204 |
| ENSMUSG00000025873 | 0.033332996 | 0.598552 | 0.736956 | 1975.891 | 1947.31 | 1803.616 | 1962.468 | 1878.064 | 1773.273 | 1895.792 | 1963.102 |
| ENSMUSG00000025875 | -0.02826612 | 0.894322 | 0.936192 | 198.1927 | 147.578 | 230.2136 | 256.1992 | 222.3254 | 170.9582 | 195.7507 | 262.1095 |
| ENSMUSG00000025876 | -0.38045553 | 0.11367 | 0.240805 | 62.37537 | 43.19358 | 48.85829 | 42.26855 | 69.2329 | 56.98608 | 55.92877 | 73.95619 |
| ENSMUSG00000025877 | 0.522344958 | 0.119552 | 0.248501 | 24.1453 | 29.99554 | 43.06154 | 32.77969 | 17.552 | 25.6996 | 25.07152 | 22.83941 |
| ENSMUSG00000025878 | 0.254468798 | 0.00917 | 0.040005 | 1267.628 | 1208.22 | 1097.241 | 1255.117 | 954.634 | 1138.604 | 1041.432 | 912.4889 |
| ENSMUSG00000025880 | -0.232244927 | 0.044811 | 0.126129 | 827.9827 | 725.892 | 894.3552 | 701.3129 | 805.4419 | 931.89 | 989.3607 | 975.5692 |
| ENSMUSG00000025885 | -0.390216156 | 0.024829 | 0.082652 | 1729.407 | 1605.361 | 2232.575 | 1662.275 | 2149.145 | 1781.094 | 2592.974 | 2953.897 |

| | | | | | | | | | | | |
|---|---|---|---|---|---|---|---|---|---|---|---|
| ENSMUSG00000025887 | 0.735722186 | 0.000221 | 0.002447 | 348.0948 | 490.727 | 351.9453 | 483.9318 | 255.4792 | 326.2732 | 223.7151 | 200.1168 |
| ENSMUSG00000025888 | 0.484494486 | 0.221391 | 0.380447 | 24.1453 | 28.79572 | 27.32752 | 22.42821 | 10.72622 | 13.40849 | 24.10723 | 25.0146 |
| ENSMUSG00000025889 | 0.515426179 | 0.164476 | 0.3095 | 193.1624 | 74.38894 | 139.1219 | 284.6658 | 135.5405 | 147.4934 | 89.67889 | 112.0219 |
| ENSMUSG00000025892 | 0.85738589 | 7.33E-08 | 4.39E-06 | 1083.521 | 1480.58 | 1223.113 | 1426.779 | 723.5326 | 906.1904 | 656.6809 | 591.6496 |
| ENSMUSG00000025893 | -0.045661498 | 0.787537 | 0.869461 | 127.7689 | 159.5763 | 160.6527 | 114.7289 | 156.9929 | 146.376 | 134.0362 | 142.4744 |
| ENSMUSG00000025894 | 0.091984339 | 0.372149 | 0.541244 | 859.1704 | 853.0731 | 698.922 | 830.7064 | 709.881 | 803.392 | 713.574 | 813.5181 |
| ENSMUSG00000025898 | 0.37997939 | 0.041956 | 0.120575 | 1349.119 | 1466.182 | 1156.865 | 1276.683 | 851.2722 | 1445.882 | 895.8246 | 840.7079 |
| ENSMUSG00000025899 | 0.22893609 | 0.012773 | 0.051303 | 1115.714 | 1139.83 | 986.275 | 1105.021 | 839.5708 | 1024.632 | 942.1105 | 901.613 |
| ENSMUSG00000025900 | 0.137883471 | 0.374491 | 0.543238 | 213.2835 | 266.3604 | 296.4622 | 303.6435 | 259.3796 | 259.2308 | 209.2507 | 255.5839 |
| ENSMUSG00000025902 | -0.343008045 | 0.034931 | 0.105913 | 588.5418 | 511.124 | 782.5608 | 544.3154 | 693.3041 | 720.7063 | 745.3955 | 922.2772 |
| ENSMUSG00000025903 | -0.225151203 | 0.002078 | 0.013498 | 1422.561 | 1541.771 | 1521.232 | 1516.492 | 1670.366 | 1722.991 | 1691.363 | 1932.649 |
| ENSMUSG00000025907 | 0.225300083 | 0.053374 | 0.142496 | 1742.486 | 1683.35 | 1373.829 | 1594.128 | 1233.516 | 1605.667 | 1303.719 | 1325.774 |
| ENSMUSG00000025909 | 0.485268158 | 0.549395 | 0.69758 | 13.07871 | 20.39697 | 18.21835 | 23.29083 | 3.900445 | 39.10809 | 8.678602 | 2.175182 |
| ENSMUSG00000025911 | -1.679507065 | 2.36E-08 | 1.74E-06 | 243.4652 | 179.9732 | 294.806 | 235.4962 | 881.5006 | 358.6771 | 729.0026 | 1088.679 |
| ENSMUSG00000025912 | 0.383442837 | 0.085133 | 0.197167 | 210.2654 | 307.1543 | 203.7142 | 217.3811 | 186.2463 | 239.1181 | 158.1434 | 134.8613 |
| ENSMUSG00000025915 | -0.328687185 | 0.025129 | 0.083355 | 529.1846 | 410.339 | 523.3634 | 480.4813 | 526.5601 | 596.6778 | 564.1092 | 756.9634 |
| ENSMUSG00000025917 | -0.146869428 | 0.018723 | 0.067349 | 2778.722 | 2724.795 | 2754.283 | 2707.775 | 3231.519 | 2790.083 | 3102.118 | 3013.715 |
| ENSMUSG00000025920 | 0.157222319 | 0.043928 | 0.124457 | 1178.09 | 1335.401 | 1205.723 | 1274.958 | 1087.249 | 1205.647 | 1052.039 | 1133.27 |
| ENSMUSG00000025921 | 0.286538925 | 0.22192 | 0.381072 | 2307.889 | 3001.953 | 2212.701 | 2411.895 | 1829.309 | 3224.742 | 1547.684 | 1543.292 |
| ENSMUSG00000025922 | -0.206386724 | 0.674719 | 0.791798 | 11.0666 | 8.398751 | 9.93728 | 12.07673 | 18.52711 | 8.938993 | 10.60718 | 9.78832 |
| ENSMUSG00000025925 | 0.161856248 | 0.4056 | 0.572441 | 781.7042 | 789.4826 | 643.4389 | 724.6037 | 700.1299 | 895.0167 | 564.1092 | 467.6642 |
| ENSMUSG00000025927 | -0.275620867 | 0.183819 | 0.334837 | 815.9101 | 512.3238 | 779.2483 | 780.6743 | 678.6775 | 749.758 | 894.8603 | 1175.686 |
| ENSMUSG00000025930 | -0.039917593 | 0.933789 | 0.960915 | 69.41775 | 113.983 | 52.99882 | 65.55939 | 122.864 | 22.34748 | 105.1075 | 58.72992 |
| ENSMUSG00000025931 | 0.850957943 | 2.87E-07 | 1.33E-05 | 509.0635 | 409.1391 | 351.9453 | 370.9281 | 231.1014 | 254.7613 | 239.1437 | 183.8029 |
| ENSMUSG00000025932 | 0.193801727 | 0.392649 | 0.560025 | 2453.767 | 2620.41 | 1730.743 | 2297.166 | 2670.83 | 2302.908 | 1731.863 | 1250.73 |
| ENSMUSG00000025933 | 0.036275354 | 0.803357 | 0.879822 | 179.0777 | 195.5709 | 158.1684 | 208.7549 | 168.6943 | 201.1273 | 171.6435 | 181.6277 |
| ENSMUSG00000025934 | -0.554868319 | 0.05469 | 0.145273 | 209.2593 | 152.3773 | 393.3507 | 202.7165 | 323.737 | 283.813 | 351.0013 | 451.3503 |
| ENSMUSG00000025935 | 0.109790896 | 0.213939 | 0.37205 | 2555.378 | 2604.813 | 2474.383 | 2650.842 | 2213.503 | 2755.445 | 2216.901 | 2348.109 |
| ENSMUSG00000025937 | -1.639609521 | 1.82E-09 | 2.17E-07 | 503.0272 | 393.5415 | 703.0625 | 523.6125 | 1618.685 | 939.7116 | 1585.291 | 2476.445 |
| ENSMUSG00000025938 | 0.846388894 | 0.001291 | 0.009465 | 131.7931 | 112.7832 | 91.91984 | 123.3552 | 52.65601 | 96.09417 | 51.10733 | 56.55474 |
| ENSMUSG00000025939 | 0.095699595 | 0.457465 | 0.619168 | 1086.539 | 1087.038 | 887.7303 | 1057.576 | 873.6997 | 1182.182 | 896.7889 | 901.613 |
| ENSMUSG00000025940 | -0.406387555 | 8.98E-05 | 0.001235 | 896.3944 | 879.4692 | 822.3099 | 946.298 | 1193.536 | 1006.754 | 1168.718 | 1327.949 |
| ENSMUSG00000025949 | 0.088694825 | 0.245652 | 0.408625 | 820.9404 | 895.0669 | 787.5294 | 878.1507 | 817.1433 | 812.331 | 762.7527 | 785.2408 |
| ENSMUSG00000025950 | -0.73885964 | 2.56E-05 | 0.000474 | 2788.783 | 1954.509 | 2498.398 | 2293.716 | 4008.683 | 3009.088 | 3682.62 | 5214.999 |
| ENSMUSG00000025955 | 3.340408278 | 0.082772 | 0.193494 | 936.6366 | 0 | 8.281066 | 169.0742 | 0.975111 | 34.6386 | 4.821446 | 69.60583 |
| ENSMUSG00000025956 | 0.180833738 | 0.159403 | 0.302421 | 480.894 | 391.1418 | 351.9453 | 421.8229 | 375.4179 | 392.1983 | 351.9655 | 331.7153 |
| ENSMUSG00000025958 | 0.276117998 | 0.005865 | 0.028934 | 2400.446 | 2366.048 | 2050.392 | 2296.304 | 1784.454 | 2190.053 | 1732.828 | 1818.452 |
| ENSMUSG00000025959 | 0.454094835 | 0.027571 | 0.089052 | 685.123 | 487.1275 | 563.1125 | 603.8364 | 331.5378 | 612.321 | 393.43 | 371.9562 |
| ENSMUSG00000025962 | -0.199226155 | 0.025616 | 0.084467 | 515.0998 | 557.917 | 464.5678 | 493.4206 | 556.7886 | 585.504 | 597.8593 | 586.2116 |
| ENSMUSG00000025964 | 0.651663048 | 0.008545 | 0.038064 | 367.2098 | 203.9697 | 195.4332 | 328.6596 | 159.9183 | 202.2447 | 192.8578 | 142.4744 |
| ENSMUSG00000025967 | -0.111892994 | 0.135792 | 0.270228 | 17830.3 | 20333.38 | 18727.63 | 19630.72 | 22552.37 | 19728.36 | 21084.18 | 19323.23 |
| ENSMUSG00000025968 | -0.563469311 | 0.000331 | 0.003341 | 2471.876 | 2284.46 | 2808.938 | 2368.764 | 3557.206 | 2669.407 | 3942.014 | 4513.503 |
| ENSMUSG00000025969 | 0.201460836 | 0.042837 | 0.122459 | 783.7163 | 704.2952 | 737.0149 | 844.5084 | 715.7317 | 680.4808 | 593.0378 | 683.0072 |
| ENSMUSG00000025971 | -0.007627899 | 0.921367 | 0.953074 | 737.4378 | 669.5004 | 746.1241 | 697.8624 | 721.5824 | 758.697 | 691.3953 | 698.2335 |
| ENSMUSG00000025977 | 3.223396968 | 5.56E-08 | 3.55E-06 | 28.16952 | 17.99732 | 33.95237 | 25.8787 | 1.950223 | 3.352122 | 1.928578 | 4.350364 |
| ENSMUSG00000025978 | 0.764275815 | 1.03E-12 | 3.10E-10 | 524.1543 | 562.7163 | 499.3483 | 559.8426 | 313.0107 | 313.9821 | 354.8584 | 279.5109 |
| ENSMUSG00000025979 | 0.347359179 | 0.056831 | 0.148904 | 3058.405 | 1846.525 | 1680.228 | 2180.712 | 1687.918 | 1943.114 | 1583.363 | 1675.978 |
| ENSMUSG00000025980 | -0.311428574 | 0.055857 | 0.147209 | 9553.492 | 6169.482 | 6484.075 | 6680.156 | 8963.223 | 7381.373 | 8950.532 | 10551.81 |
| ENSMUSG00000025981 | -0.182704033 | 0.339665 | 0.508231 | 303.8284 | 189.5718 | 223.5888 | 200.1287 | 248.6534 | 217.8879 | 307.6082 | 266.4598 |
| ENSMUSG00000025982 | 0.156250166 | 0.073763 | 0.179032 | 10545.46 | 10440.85 | 10186.54 | 11148.55 | 9371.795 | 10924.57 | 9128.926 | 8553.904 |
| ENSMUSG00000025983 | -0.664080463 | 0.104938 | 0.228388 | 16.09687 | 10.79839 | 24.01509 | 18.97772 | 20.47734 | 40.22547 | 29.89296 | 21.75182 |
| ENSMUSG00000025986 | -0.064343401 | 0.459041 | 0.620528 | 1675.081 | 1977.306 | 1604.871 | 1653.649 | 1707.42 | 1877.188 | 1839.864 | 1795.613 |
| ENSMUSG00000025991 | 2.616224666 | 0.000498 | 0.004571 | 126.7628 | 17.99732 | 10.76539 | 148.3712 | 12.67645 | 13.40849 | 11.57147 | 11.9635 |
| ENSMUSG00000025993 | 0.451487548 | 0.000826 | 0.006728 | 3917.576 | 5684.754 | 5250.196 | 4720.276 | 3946.275 | 3881.758 | 3091.511 | 3393.284 |
| ENSMUSG00000025995 | -0.03107148 | 0.734974 | 0.834031 | 1230.404 | 1288.608 | 1174.255 | 1177.481 | 1321.276 | 1368.783 | 1180.29 | 1103.905 |
| ENSMUSG00000025997 | 0.84521697 | 3.94E-05 | 0.000666 | 403.4278 | 386.3425 | 383.4134 | 364.8897 | 191.1218 | 316.2169 | 182.2507 | 167.489 |
| ENSMUSG00000026000 | 0.099174283 | 0.241741 | 0.404128 | 1644.899 | 1556.169 | 1522.06 | 1778.73 | 1544.576 | 1640.305 | 1494.648 | 1392.117 |
| ENSMUSG00000026003 | -0.572191284 | 0.00363 | 0.020376 | 2209.295 | 1838.127 | 2259.075 | 1982.309 | 3064.775 | 1986.691 | 3109.833 | 4163.299 |
| ENSMUSG00000026004 | -0.044833018 | 0.687058 | 0.801184 | 626.7719 | 679.099 | 703.8906 | 630.5778 | 562.6392 | 721.8237 | 729.0026 | 711.2846 |
| ENSMUSG00000026005 | -0.049615801 | 0.55 | 0.697988 | 715.3046 | 695.8965 | 675.735 | 637.4788 | 658.2001 | 747.5233 | 681.7524 | 731.9488 |
| ENSMUSG00000026014 | 0.502112526 | 0.026352 | 0.086328 | 1280.707 | 1244.215 | 1083.992 | 1202.497 | 656.2499 | 1344.201 | 722.2526 | 675.3941 |
| ENSMUSG00000026017 | 0.281512156 | 0.175215 | 0.323572 | 514.0938 | 499.1258 | 395.835 | 474.4429 | 332.513 | 567.626 | 318.2154 | 331.7153 |
| ENSMUSG00000026018 | 0.104982389 | 0.679253 | 0.794996 | 54.32694 | 43.19358 | 62.108 | 58.6584 | 48.75557 | 67.04245 | 46.28588 | 42.41605 |
| ENSMUSG00000026019 | -0.101572977 | 0.135593 | 0.270044 | 1096.599 | 1108.635 | 1083.163 | 1113.647 | 1273.495 | 1108.435 | 1162.933 | 1175.686 |

| | | | | | | | | | | | |
|---|---|---|---|---|---|---|---|---|---|---|---|
| ENSMUSG00000026020 | 0.028695732 | 0.770311 | 0.857419 | 3832.061 | 4525.727 | 3653.606 | 3886.119 | 3935.549 | 4371.167 | 3602.584 | 3671.708 |
| ENSMUSG00000026021 | -0.003555207 | 0.960088 | 0.977364 | 3252.574 | 3462.685 | 3232.928 | 3334.902 | 3357.308 | 3666.104 | 3156.118 | 3135.525 |
| ENSMUSG00000026023 | 0.349995314 | 0.396739 | 0.563898 | 16.09687 | 19.19714 | 33.95237 | 28.46658 | 16.57689 | 13.40849 | 21.21436 | 26.10219 |
| ENSMUSG00000026024 | -0.160493375 | 0.149387 | 0.289368 | 793.7769 | 800.281 | 859.5747 | 715.9775 | 910.754 | 757.5796 | 868.8245 | 1004.934 |
| ENSMUSG00000026027 | -0.07664105 | 0.493487 | 0.650418 | 513.0877 | 449.9331 | 409.9128 | 442.5259 | 443.6756 | 515.1095 | 445.5016 | 510.0802 |
| ENSMUSG00000026028 | -0.499038311 | 0.001878 | 0.012551 | 1218.332 | 1124.233 | 1325.799 | 1191.283 | 1587.481 | 1271.572 | 1817.685 | 2192.584 |
| ENSMUSG00000026029 | 0.072074331 | 0.51308 | 0.668035 | 954.7456 | 975.4549 | 1012.774 | 996.3301 | 931.2313 | 797.8051 | 906.4318 | 1112.606 |
| ENSMUSG00000026031 | -0.088780341 | 0.243685 | 0.406273 | 1461.797 | 1256.213 | 1276.94 | 1240.453 | 1395.384 | 1395.6 | 1353.862 | 1422.569 |
| ENSMUSG00000026032 | -0.272648528 | 0.00363 | 0.020376 | 1521.154 | 1518.974 | 1678.572 | 1598.441 | 1863.438 | 1695.057 | 1892.9 | 2182.795 |
| ENSMUSG00000026034 | 0.391703838 | 0.043992 | 0.124553 | 7420.657 | 5802.337 | 4214.235 | 5414.688 | 4253.436 | 5552.232 | 4288.194 | 3323.678 |
| ENSMUSG00000026035 | -0.014224761 | 0.87102 | 0.922556 | 535.2209 | 507.5245 | 486.9267 | 511.5357 | 485.6054 | 540.8091 | 545.7877 | 488.3284 |
| ENSMUSG00000026036 | -0.263495559 | 0.003506 | 0.01981 | 855.1462 | 854.2729 | 734.5306 | 732.3673 | 981.9371 | 959.8243 | 889.0746 | 976.6568 |
| ENSMUSG00000026037 | -0.123361865 | 0.21925 | 0.378035 | 888.346 | 910.6645 | 793.3262 | 942.8475 | 1055.07 | 1021.28 | 902.5747 | 870.0729 |
| ENSMUSG00000026039 | 0.188080579 | 0.282823 | 0.449379 | 355.1372 | 380.3434 | 286.5249 | 298.4677 | 282.7823 | 367.6161 | 277.7153 | 229.4817 |
| ENSMUSG00000026042 | 0.519628366 | 0.003504 | 0.019809 | 3722.401 | 5087.243 | 4124.799 | 5167.977 | 2769.316 | 4305.242 | 2863.939 | 2689.613 |
| ENSMUSG00000026043 | 0.416024257 | 0.015646 | 0.059526 | 27069.9 | 49190.28 | 40866.23 | 40954.78 | 26703.42 | 34681.06 | 27881.46 | 29213.78 |
| ENSMUSG00000026047 | 0.12307894 | 0.159707 | 0.302809 | 1227.386 | 1269.411 | 1057.492 | 1285.309 | 1078.473 | 1146.426 | 1152.326 | 1063.664 |
| ENSMUSG00000026048 | 0.115467281 | 0.25653 | 0.420248 | 931.6063 | 903.4656 | 817.3412 | 946.298 | 730.3584 | 915.1294 | 794.5743 | 883.124 |
| ENSMUSG00000026049 | 0.074950981 | 0.555829 | 0.702687 | 364.1917 | 394.7413 | 313.0243 | 389.9058 | 313.9858 | 389.9636 | 360.6441 | 321.927 |
| ENSMUSG00000026051 | 0.715985544 | 0.000667 | 0.005722 | 437.6336 | 551.9179 | 382.5853 | 442.5259 | 260.3547 | 391.0809 | 242.0366 | 210.9927 |
| ENSMUSG00000026058 | 1.309505671 | 0.00123 | 0.009119 | 148.896 | 179.9732 | 157.3403 | 191.5024 | 47.78045 | 143.0239 | 46.28588 | 36.9781 |
| ENSMUSG00000026062 | -0.551014086 | 0.090707 | 0.206346 | 248.4954 | 172.7743 | 289.8373 | 200.9913 | 264.2552 | 174.3104 | 352.9298 | 545.9707 |
| ENSMUSG00000026064 | 0.085441692 | 0.408868 | 0.57507 | 1239.459 | 1160.227 | 1257.066 | 1309.462 | 1022.892 | 1363.196 | 1170.647 | 1127.832 |
| ENSMUSG00000026065 | 0.530065781 | 0.057305 | 0.149911 | 97.58727 | 50.3925 | 106.8258 | 84.5371 | 57.53157 | 56.98608 | 50.14304 | 71.78101 |
| ENSMUSG00000026068 | -0.367220765 | 0.533285 | 0.684646 | 9.054489 | 5.999108 | 9.109173 | 10.35148 | 7.80089 | 11.17374 | 20.25007 | 5.437955 |
| ENSMUSG00000026069 | 0.1784399 | 0.802585 | 0.87942 | 9.054489 | 34.79482 | 9.109173 | 20.70296 | 14.62667 | 32.40385 | 14.46434 | 3.262773 |
| ENSMUSG00000026070 | 0.879409447 | 0.021838 | 0.074979 | 43.26034 | 64.79036 | 57.96746 | 68.14726 | 36.07912 | 52.51658 | 17.3572 | 21.75182 |
| ENSMUSG00000026072 | 0.100537025 | 0.515883 | 0.66989 | 406.446 | 364.7457 | 515.0823 | 430.4491 | 322.7618 | 436.8933 | 398.2514 | 446.9999 |
| ENSMUSG00000026073 | 0.318278256 | 0.537459 | 0.687697 | 57.3451 | 10.79839 | 42.23344 | 40.5433 | 32.17867 | 17.87799 | 47.25017 | 23.927 |
| ENSMUSG00000026074 | 0.222805996 | 0.000837 | 0.006802 | 6994.09 | 6589.42 | 6468.341 | 7215.846 | 5597.139 | 6294.168 | 5642.056 | 5836.014 |
| ENSMUSG00000026077 | 0.788250336 | 0.001432 | 0.01024 | 192.1564 | 110.3836 | 123.3879 | 125.0804 | 78.0089 | 98.32892 | 81.96458 | 60.9051 |
| ENSMUSG00000026078 | -0.155336808 | 0.037727 | 0.112016 | 1962.812 | 1890.919 | 2111.672 | 2105.664 | 2270.059 | 2045.912 | 2340.33 | 2336.146 |
| ENSMUSG00000026080 | 0.201253411 | 0.250139 | 0.413857 | 250.5075 | 231.5656 | 206.1986 | 216.5185 | 210.624 | 243.5876 | 183.2149 | 149 |
| ENSMUSG00000026082 | -0.198054272 | 0.052988 | 0.141745 | 658.9656 | 605.9099 | 571.3936 | 624.5394 | 757.6615 | 762.0491 | 687.5382 | 614.489 |
| ENSMUSG00000026083 | 0.256365584 | 0.073506 | 0.178552 | 6015.199 | 6013.506 | 5079.606 | 5222.323 | 4017.459 | 6084.102 | 4308.444 | 4285.109 |
| ENSMUSG00000026087 | -0.078644202 | 0.356455 | 0.524955 | 1602.645 | 1913.715 | 1788.71 | 1932.277 | 2044.808 | 1911.827 | 1888.078 | 1796.7 |
| ENSMUSG00000026088 | 0.35958185 | 0.010219 | 0.043338 | 466.8092 | 629.9063 | 446.3495 | 444.2511 | 382.2436 | 364.264 | 406.93 | 391.5328 |
| ENSMUSG00000026090 | 0.613737548 | 0.001607 | 0.011092 | 235.4167 | 249.5629 | 256.7131 | 277.7648 | 182.3458 | 204.4795 | 169.7149 | 109.8467 |
| ENSMUSG00000026094 | 0.12760587 | 0.238248 | 0.399374 | 614.6992 | 518.3229 | 599.5492 | 614.1879 | 503.1574 | 534.1048 | 612.3236 | 500.2919 |
| ENSMUSG00000026095 | 0.115595634 | 0.450996 | 0.613562 | 458.7608 | 448.7333 | 428.9592 | 530.5134 | 410.5219 | 550.8654 | 401.1443 | 362.1678 |
| ENSMUSG00000026096 | -0.103604566 | 0.312585 | 0.480916 | 355.1372 | 301.1552 | 346.9767 | 349.3625 | 371.5174 | 335.2122 | 378.9656 | 369.781 |
| ENSMUSG00000026097 | -0.036454495 | 0.64883 | 0.772469 | 471.8395 | 465.5308 | 448.0057 | 445.9763 | 460.2525 | 462.5929 | 466.716 | 487.2408 |
| ENSMUSG00000026098 | -0.046048915 | 0.667758 | 0.786832 | 273.6468 | 253.1623 | 284.8687 | 268.2759 | 271.0809 | 292.752 | 295.0725 | 257.7591 |
| ENSMUSG00000026102 | 0.409046425 | 0.00107 | 0.008228 | 816.9161 | 801.4808 | 742.8116 | 669.3958 | 533.3859 | 688.3024 | 515.8947 | 544.8831 |
| ENSMUSG00000026103 | -0.138089605 | 0.168983 | 0.315281 | 2590.59 | 2421.24 | 2614.333 | 2711.226 | 2523.588 | 2835.895 | 2680.724 | 3339.992 |
| ENSMUSG00000026104 | 0.285071876 | 0.372669 | 0.541659 | 401.4157 | 571.1151 | 730.39 | 214.7932 | 386.1441 | 307.2779 | 432.0015 | 448.0875 |
| ENSMUSG00000026107 | -1.429503907 | 1.87E-05 | 0.000372 | 303.8284 | 232.7654 | 450.49 | 235.4962 | 822.0188 | 357.5597 | 1071.325 | 1044.087 |
| ENSMUSG00000026109 | 0.984908743 | 9.07E-05 | 0.001242 | 119.7205 | 105.5843 | 171.4181 | 135.4319 | 47.78045 | 87.15518 | 62.6788 | 72.8686 |
| ENSMUSG00000026110 | -0.588668863 | 7.51E-13 | 2.31E-10 | 685.123 | 659.9018 | 656.6886 | 708.2139 | 1097.975 | 948.6506 | 1061.682 | 965.7809 |
| ENSMUSG00000026111 | -0.099736476 | 0.154107 | 0.295563 | 1419.543 | 1380.995 | 1416.89 | 1445.757 | 1496.796 | 1387.779 | 1621.934 | 1561.781 |
| ENSMUSG00000026112 | -0.23874506 | 0.003025 | 0.017883 | 1748.522 | 1574.166 | 1695.134 | 1535.47 | 1916.094 | 1786.681 | 1921.828 | 2108.839 |
| ENSMUSG00000026113 | 0.249040318 | 0.087988 | 0.201878 | 700.2138 | 647.9036 | 540.7536 | 678.8847 | 495.3565 | 670.4245 | 529.3948 | 465.489 |
| ENSMUSG00000026116 | 0.014824213 | 0.843368 | 0.90548 | 1491.979 | 1474.581 | 1515.435 | 1476.811 | 1333.952 | 1598.962 | 1528.398 | 1438.883 |
| ENSMUSG00000026117 | -0.524786594 | 0.308648 | 0.477006 | 11.0666 | 8.398751 | 13.24971 | 4.313117 | 15.60178 | 12.29112 | 14.46434 | 10.87591 |
| ENSMUSG00000026121 | -0.020004911 | 0.745635 | 0.841093 | 1450.73 | 1402.591 | 1509.638 | 1421.604 | 1500.696 | 1402.304 | 1528.398 | 1434.533 |
| ENSMUSG00000026123 | -0.409810865 | 0.064044 | 0.162451 | 1006.054 | 803.8804 | 1117.116 | 972.1767 | 1371.006 | 789.9835 | 1247.79 | 1772.773 |
| ENSMUSG00000026126 | -0.756397089 | 0.00035 | 0.003484 | 173.0413 | 157.1766 | 253.4006 | 208.7549 | 277.9067 | 278.2261 | 357.7513 | 428.5109 |
| ENSMUSG00000026127 | -0.188448361 | 0.018092 | 0.065859 | 1330.004 | 1436.186 | 1475.686 | 1337.929 | 1668.415 | 1426.887 | 1611.327 | 1649.876 |
| ENSMUSG00000026131 | 0.199726576 | 0.021572 | 0.074347 | 4474.93 | 3816.632 | 3603.092 | 4215.641 | 3366.084 | 3612.47 | 3674.906 | 3373.708 |
| ENSMUSG00000026134 | 0.096990863 | 0.437328 | 0.601127 | 672.0443 | 779.884 | 692.2971 | 642.6545 | 694.2793 | 754.2275 | 600.7521 | 554.6715 |
| ENSMUSG00000026135 | -0.069007481 | 0.711047 | 0.818913 | 367.2098 | 368.3452 | 316.3367 | 247.5729 | 418.3228 | 269.2872 | 318.2154 | 354.5547 |
| ENSMUSG00000026142 | -0.10756231 | 0.370609 | 0.539684 | 296.786 | 261.5611 | 272.4471 | 238.9467 | 274.9814 | 264.8177 | 300.8582 | 311.0511 |
| ENSMUSG00000026147 | -0.006345385 | 0.976148 | 0.986896 | 385.3188 | 409.1391 | 336.2113 | 389.0432 | 355.9156 | 547.5133 | 359.6799 | 263.197 |
| ENSMUSG00000026150 | -0.014925247 | 0.814924 | 0.887383 | 3183.156 | 3403.894 | 3574.936 | 3513.465 | 3367.059 | 3677.278 | 3370.191 | 3408.51 |

| | | | | | | | | | | | |
|---|---|---|---|---|---|---|---|---|---|---|---|
| ENSMUSG00000026153 | 0.182411521 | 0.032226 | 0.099525 | 1424.573 | 1454.184 | 1257.894 | 1341.38 | 1196.462 | 1320.736 | 1196.683 | 1110.431 |
| ENSMUSG00000026154 | -0.188950779 | 0.11211 | 0.238643 | 397.3915 | 407.9393 | 395.0069 | 381.2796 | 379.3183 | 431.3064 | 474.4303 | 517.6934 |
| ENSMUSG00000026155 | -0.031206178 | 0.720732 | 0.82531 | 910.4792 | 833.876 | 787.5294 | 766.0097 | 783.9895 | 862.6128 | 841.8244 | 879.8612 |
| ENSMUSG00000026156 | -0.957716852 | 0.003806 | 0.021062 | 127.7689 | 82.78769 | 193.777 | 150.0965 | 180.3956 | 211.1837 | 254.5723 | 433.9488 |
| ENSMUSG00000026158 | 0.27840421 | 0.153575 | 0.294878 | 1851.14 | 1964.108 | 1624.745 | 1813.235 | 1332.002 | 2185.584 | 1347.112 | 1115.868 |
| ENSMUSG00000026159 | 0.072940347 | 0.309074 | 0.477465 | 2099.635 | 2044.496 | 1999.878 | 2223.843 | 1932.671 | 2175.527 | 1930.507 | 1919.598 |
| ENSMUSG00000026162 | -0.553625353 | 0.003385 | 0.019325 | 104.6297 | 74.38894 | 81.98256 | 106.1027 | 153.0925 | 120.6764 | 140.7862 | 125.073 |
| ENSMUSG00000026163 | -0.126669033 | 0.518289 | 0.672184 | 55.33299 | 74.38894 | 66.24853 | 70.73513 | 78.0089 | 71.51194 | 69.42882 | 71.78101 |
| ENSMUSG00000026167 | -2.445566701 | 0.000133 | 0.001669 | 3.018163 | 2.399643 | 11.59349 | 1.725247 | 30.22845 | 18.99536 | 23.14294 | 32.62773 |
| ENSMUSG00000026170 | 0.481481131 | 0.005542 | 0.027702 | 434.6155 | 487.1275 | 453.8024 | 547.7659 | 367.617 | 401.1373 | 369.3228 | 239.27 |
| ENSMUSG00000026171 | 0.04435432 | 0.471917 | 0.631304 | 1029.194 | 1049.844 | 1049.211 | 1006.682 | 999.4891 | 987.7587 | 973.9321 | 1048.438 |
| ENSMUSG00000026172 | -0.170180241 | 0.111476 | 0.237821 | 337.0282 | 301.1552 | 310.54 | 323.4838 | 398.8205 | 325.1559 | 357.7513 | 349.1167 |
| ENSMUSG00000026173 | 0.056514434 | 0.874663 | 0.925069 | 68.4117 | 39.59411 | 38.92101 | 33.64232 | 36.07912 | 65.92507 | 33.75012 | 38.06569 |
| ENSMUSG00000026174 | -0.081967507 | 0.19288 | 0.346082 | 1553.348 | 1612.56 | 1464.921 | 1459.559 | 1596.257 | 1564.324 | 1605.541 | 1673.803 |
| ENSMUSG00000026175 | -2.319144863 | 9.71E-05 | 0.001308 | 112.6781 | 27.5959 | 212.8234 | 109.5532 | 619.1957 | 319.569 | 540.0019 | 834.1824 |
| ENSMUSG00000026176 | -0.069215085 | 0.292427 | 0.459782 | 4167.077 | 4626.512 | 4811.3 | 4568.454 | 4881.407 | 4499.665 | 4823.374 | 4862.62 |
| ENSMUSG00000026177 | 0.390389881 | 0.230724 | 0.391167 | 53.32088 | 46.79304 | 81.98256 | 50.89479 | 44.85512 | 36.87335 | 64.60737 | 31.54014 |
| ENSMUSG00000026179 | -0.372889125 | 0.006902 | 0.032666 | 1080.502 | 974.2551 | 1139.475 | 1116.235 | 1381.733 | 1086.088 | 1420.398 | 1695.555 |
| ENSMUSG00000026181 | -0.142222477 | 0.218612 | 0.377357 | 1095.593 | 1180.624 | 1318.346 | 1087.768 | 1370.031 | 1077.149 | 1291.183 | 1429.095 |
| ENSMUSG00000026185 | 0.214636998 | 0.344304 | 0.513037 | 14981.16 | 25162.66 | 20180.13 | 17505.22 | 16331.16 | 24691.73 | 12992.83 | 13054.36 |
| ENSMUSG00000026187 | -0.105314555 | 0.107639 | 0.232214 | 1700.232 | 1808.131 | 1657.041 | 1800.295 | 1991.177 | 1881.658 | 1816.721 | 1799.963 |
| ENSMUSG00000026188 | 0.011855151 | 0.980268 | 0.988885 | 101.6115 | 25.19625 | 110.9663 | 115.5915 | 95.56091 | 39.10809 | 91.60747 | 125.073 |
| ENSMUSG00000026189 | -0.48339445 | 0.007105 | 0.033261 | 563.3904 | 519.5227 | 557.3158 | 611.6001 | 849.3219 | 537.4569 | 764.6813 | 997.321 |
| ENSMUSG00000026192 | -0.165187491 | 0.047861 | 0.132018 | 2491.997 | 2678.002 | 2698.8 | 2574.069 | 3196.415 | 2556.552 | 2962.296 | 2991.963 |
| ENSMUSG00000026193 | 0.101205547 | 0.569377 | 0.71316 | 7052.441 | 8530.731 | 13118.04 | 9167.962 | 7863.298 | 8369.132 | 9631.32 | 9442.466 |
| ENSMUSG00000026196 | 0.410579877 | 0.052712 | 0.141215 | 297.7921 | 363.5459 | 247.6039 | 274.3143 | 186.2463 | 301.691 | 229.5008 | 171.8394 |
| ENSMUSG00000026197 | -0.384157456 | 0.006498 | 0.031296 | 387.3309 | 345.5486 | 438.0684 | 439.938 | 568.4899 | 420.1327 | 539.0376 | 577.5109 |
| ENSMUSG00000026198 | -0.383710866 | 0.00451 | 0.023837 | 179.0777 | 209.9688 | 195.4332 | 166.4863 | 259.3796 | 220.1227 | 231.4294 | 266.4598 |
| ENSMUSG00000026199 | 0.132539889 | 0.096018 | 0.214632 | 1143.884 | 1043.845 | 958.9475 | 1042.049 | 986.8126 | 952.0027 | 912.2175 | 967.9561 |
| ENSMUSG00000026200 | -0.425504358 | 0.000598 | 0.005253 | 789.7527 | 706.6949 | 895.1833 | 817.7671 | 1039.469 | 935.2421 | 1058.79 | 1281.182 |
| ENSMUSG00000026201 | -0.193889446 | 0.014841 | 0.057243 | 1502.039 | 1490.178 | 1498.873 | 1560.486 | 1761.051 | 1549.798 | 1701.97 | 1909.81 |
| ENSMUSG00000026202 | -1.537182333 | 6.24E-06 | 0.00016 | 527.1725 | 419.9375 | 908.433 | 569.3315 | 1485.095 | 804.5093 | 2008.614 | 2742.905 |
| ENSMUSG00000026203 | -0.237662966 | 0.019883 | 0.070169 | 632.8082 | 617.9081 | 640.9545 | 660.7696 | 719.6321 | 648.077 | 816.7529 | 825.4816 |
| ENSMUSG00000026204 | 1.616973253 | 0.002396 | 0.014942 | 177.0656 | 40.79393 | 50.5145 | 200.9913 | 50.70579 | 40.22547 | 23.14294 | 39.15328 |
| ENSMUSG00000026205 | -1.702891512 | 7.21E-07 | 2.85E-05 | 48.29061 | 35.99465 | 66.24853 | 46.58167 | 155.0427 | 79.33356 | 179.3578 | 230.5693 |
| ENSMUSG00000026207 | 0.013210327 | 0.889974 | 0.933475 | 933.6184 | 919.0633 | 1022.712 | 969.5888 | 921.4802 | 1002.285 | 837.003 | 1053.876 |
| ENSMUSG00000026208 | 0.167876971 | 0.441917 | 0.605226 | 1034.224 | 1334.202 | 1168.458 | 1192.146 | 885.4011 | 1619.075 | 803.2529 | 902.7006 |
| ENSMUSG00000026209 | -0.209350804 | 0.039784 | 0.116162 | 1906.473 | 1925.714 | 2057.017 | 1890.871 | 2341.242 | 1854.841 | 2312.365 | 2486.233 |
| ENSMUSG00000026211 | 0.147771682 | 0.02171 | 0.074712 | 3001.06 | 3132.734 | 2841.234 | 2959.661 | 2654.253 | 2898.468 | 2654.688 | 2562.365 |
| ENSMUSG00000026213 | 0.087412858 | 0.188642 | 0.341096 | 1253.544 | 1231.017 | 1308.408 | 1353.456 | 1196.462 | 1172.125 | 1281.54 | 1196.35 |
| ENSMUSG00000026219 | 0.144663044 | 0.035337 | 0.106844 | 6114.798 | 6152.685 | 6191.753 | 6056.48 | 5323.133 | 6200.309 | 5282.376 | 5373.788 |
| ENSMUSG00000026220 | -0.576721812 | 0.223032 | 0.382554 | 14.08476 | 5.999108 | 23.18699 | 12.93935 | 18.52711 | 21.23011 | 17.3572 | 28.27737 |
| ENSMUSG00000026222 | 0.464199305 | 0.110796 | 0.236865 | 312.8829 | 715.0936 | 638.4702 | 282.9405 | 343.2392 | 351.9728 | 354.8584 | 362.1678 |
| ENSMUSG00000026223 | 0.116887454 | 0.103638 | 0.226413 | 8190.289 | 7330.91 | 8306.738 | 7679.937 | 7642.922 | 6645.024 | 7348.848 | 7420.634 |
| ENSMUSG00000026224 | -0.42767314 | 0.125575 | 0.256106 | 38.23007 | 40.79393 | 36.43669 | 43.13117 | 54.60623 | 65.92507 | 58.82164 | 33.71532 |
| ENSMUSG00000026227 | -0.679609675 | 0.058844 | 0.152512 | 17.10292 | 26.39607 | 24.8432 | 22.42821 | 30.22845 | 54.75133 | 26.03581 | 34.80291 |
| ENSMUSG00000026228 | 0.846179062 | 0.003297 | 0.019035 | 113.6841 | 208.7689 | 130.8408 | 146.646 | 99.46135 | 110.62 | 56.89306 | 66.34306 |
| ENSMUSG00000026229 | -0.170415809 | 0.004547 | 0.023986 | 3773.71 | 4094.991 | 4189.391 | 3999.123 | 4249.535 | 4544.36 | 4621.838 | 4657.065 |
| ENSMUSG00000026234 | -0.116398084 | 0.011311 | 0.046782 | 21169.4 | 22812.21 | 21104.3 | 21762.27 | 23780.04 | 24086.12 | 23335.8 | 22936.21 |
| ENSMUSG00000026235 | 0.286080257 | 0.012958 | 0.051885 | 1401.434 | 1267.012 | 1403.641 | 1238.727 | 1009.24 | 1112.905 | 945.0034 | 1290.971 |
| ENSMUSG00000026238 | -0.126724832 | 0.083236 | 0.194185 | 55684.1 | 67245.2 | 61791.66 | 62795.54 | 70894.49 | 65293.76 | 65222.59 | 68827.11 |
| ENSMUSG00000026239 | 0.009304218 | 0.912255 | 0.948044 | 943.679 | 965.8563 | 891.8708 | 1015.308 | 968.2855 | 850.3217 | 1000.932 | 970.1312 |
| ENSMUSG00000026240 | 0.037473337 | 0.626357 | 0.756498 | 883.3157 | 808.6797 | 825.6223 | 803.9651 | 872.7246 | 789.9835 | 772.3956 | 800.467 |
| ENSMUSG00000026241 | 0.505040496 | 0.122867 | 0.252404 | 79.47829 | 76.78858 | 78.67013 | 54.34528 | 82.88446 | 34.6386 | 46.28588 | 39.15328 |
| ENSMUSG00000026245 | -0.150216202 | 0.050174 | 0.136381 | 1472.864 | 1306.606 | 1270.316 | 1387.961 | 1488.995 | 1415.713 | 1549.613 | 1579.182 |
| ENSMUSG00000026247 | 3.47232237 | 2.62E-14 | 1.05E-11 | 191.1503 | 95.98572 | 62.108 | 159.5853 | 8.776002 | 18.99536 | 9.642892 | 8.700729 |
| ENSMUSG00000026248 | -0.140276493 | 0.055844 | 0.147209 | 804.8435 | 801.4808 | 882.7617 | 864.3487 | 908.8037 | 898.3688 | 979.7178 | 912.4889 |
| ENSMUSG00000026249 | 0.170033724 | 0.045813 | 0.128013 | 4603.705 | 5714.75 | 5318.101 | 5267.179 | 4847.278 | 4338.764 | 4875.446 | 4514.591 |
| ENSMUSG00000026251 | 0.742360842 | 0.119329 | 0.248319 | 27.16347 | 46.79304 | 24.8432 | 36.23019 | 29.25334 | 29.05173 | 14.46434 | 7.613138 |
| ENSMUSG00000026254 | -0.23288945 | 0.001521 | 0.010658 | 2407.488 | 2501.628 | 2572.099 | 2559.404 | 3055.024 | 2640.355 | 2951.689 | 3152.927 |
| ENSMUSG00000026255 | -0.853343324 | 0.130765 | 0.263271 | 818.9282 | 181.1731 | 537.4412 | 414.9219 | 693.3041 | 368.7335 | 1195.719 | 1270.306 |
| ENSMUSG00000026259 | -0.413989097 | 0.001889 | 0.012598 | 167.005 | 152.3773 | 178.0429 | 167.349 | 197.9476 | 210.0663 | 234.3223 | 245.7956 |
| ENSMUSG00000026260 | -0.50120646 | 1.81E-06 | 5.99E-05 | 2693.208 | 2701.998 | 2992.777 | 2738.83 | 3839.988 | 3287.315 | 4155.122 | 4467.824 |
| ENSMUSG00000026269 | -0.320588412 | 5.47E-06 | 0.000143 | 1141.872 | 1113.434 | 1170.943 | 1128.312 | 1391.484 | 1325.206 | 1431.005 | 1542.204 |

| | | | | | | | | | | | |
|---|---|---|---|---|---|---|---|---|---|---|---|
| ENSMUSG00000026270 | -0.232956004 | 0.063499 | 0.161378 | 717.3168 | 796.6815 | 726.2495 | 643.5171 | 861.0233 | 686.0677 | 880.396 | 958.1677 |
| ENSMUSG00000026271 | 0.611657821 | 0.111865 | 0.238407 | 30.18163 | 34.79482 | 25.67131 | 37.95543 | 9.751113 | 23.46486 | 26.03581 | 25.0146 |
| ENSMUSG00000026273 | 0.026688559 | 0.74789 | 0.842597 | 502.0211 | 539.9197 | 493.5516 | 480.4813 | 506.0828 | 483.823 | 508.1804 | 476.3649 |
| ENSMUSG00000026274 | 0.047106754 | 0.660963 | 0.781719 | 511.0756 | 629.9063 | 546.5504 | 529.6508 | 545.0872 | 587.7388 | 527.4662 | 482.8904 |
| ENSMUSG00000026275 | -0.048741195 | 0.536763 | 0.687331 | 1155.956 | 1299.407 | 1253.753 | 1205.085 | 1213.038 | 1205.647 | 1269.005 | 1394.292 |
| ENSMUSG00000026276 | 0.128664202 | 0.020602 | 0.071986 | 7002.138 | 7384.902 | 6889.847 | 7354.728 | 6479.615 | 6951.184 | 6282.344 | 6475.517 |
| ENSMUSG00000026277 | -0.185380449 | 0.003035 | 0.01793 | 3905.503 | 3838.229 | 4000.583 | 4128.516 | 4650.306 | 4132.049 | 4644.017 | 4624.437 |
| ENSMUSG00000026278 | -0.641014591 | 0.004658 | 0.02448 | 745.4863 | 854.2729 | 1359.751 | 912.6557 | 1599.183 | 1011.224 | 1573.72 | 1856.518 |
| ENSMUSG00000026279 | -0.47417036 | 0.000293 | 0.003043 | 710.2744 | 722.2926 | 896.8395 | 788.4379 | 1027.767 | 972.1155 | 1018.289 | 1317.073 |
| ENSMUSG00000026280 | -0.099229256 | 0.176785 | 0.325688 | 1651.941 | 1664.152 | 1602.386 | 1683.841 | 1922.919 | 1622.427 | 1701.97 | 1822.803 |
| ENSMUSG00000026281 | -0.129436307 | 0.108647 | 0.233542 | 2129.817 | 2508.827 | 2170.467 | 2310.106 | 2704.959 | 2454.871 | 2447.366 | 2362.248 |
| ENSMUSG00000026283 | 0.079928683 | 0.480012 | 0.639046 | 799.8132 | 739.0901 | 741.9835 | 696.1372 | 729.3833 | 814.5657 | 672.1095 | 600.3503 |
| ENSMUSG00000026288 | -0.179001925 | 0.293068 | 0.460446 | 190.1443 | 173.9741 | 226.0731 | 210.4801 | 196.9725 | 222.3574 | 286.3939 | 202.2919 |
| ENSMUSG00000026289 | -0.037452937 | 0.517831 | 0.671804 | 1051.327 | 1070.241 | 1019.399 | 1041.187 | 1071.647 | 1089.44 | 1050.111 | 1078.89 |
| ENSMUSG00000026295 | -3.781394294 | 1.99E-06 | 6.43E-05 | 115.6963 | 28.79572 | 411.569 | 138.0198 | 3226.643 | 744.1711 | 1914.114 | 3667.357 |
| ENSMUSG00000026303 | -0.045010689 | 0.89424 | 0.93619 | 43.26034 | 73.18911 | 116.763 | 59.52102 | 57.53157 | 80.45093 | 69.42882 | 95.70802 |
| ENSMUSG00000026304 | -0.801752044 | 0.014878 | 0.057337 | 146.8839 | 83.98751 | 225.245 | 131.9814 | 241.8276 | 155.315 | 273.8581 | 356.7299 |
| ENSMUSG00000026305 | -0.46698287 | 0.02665 | 0.087056 | 1341.07 | 959.8572 | 1744.821 | 1218.024 | 1793.23 | 1295.037 | 1923.757 | 2266.54 |
| ENSMUSG00000026307 | -0.852374962 | 0.001755 | 0.011914 | 327.9737 | 197.9706 | 390.8663 | 287.2536 | 468.0534 | 360.9118 | 550.6091 | 797.2043 |
| ENSMUSG00000026308 | -0.239723236 | 0.700357 | 0.810801 | 15.09082 | 7.198929 | 5.796746 | 12.93935 | 8.776002 | 14.52586 | 4.821446 | 20.66423 |
| ENSMUSG00000026309 | 0.046524694 | 0.599149 | 0.737465 | 2200.241 | 2100.888 | 2007.33 | 2032.341 | 1904.392 | 2317.434 | 1990.293 | 1864.131 |
| ENSMUSG00000026311 | -0.123787048 | 0.330071 | 0.497965 | 502.0211 | 471.5299 | 476.9894 | 484.7944 | 600.6686 | 414.5458 | 553.502 | 538.3576 |
| ENSMUSG00000026313 | 0.162649735 | 0.407819 | 0.57412 | 930.6003 | 740.2899 | 843.0126 | 720.2906 | 645.5237 | 1032.454 | 655.7166 | 556.8466 |
| ENSMUSG00000026315 | 0.023305201 | 0.934515 | 0.961287 | 45.27245 | 64.79036 | 81.98256 | 47.44429 | 48.75557 | 53.63396 | 59.78593 | 73.95619 |
| ENSMUSG00000026317 | -0.061888021 | 0.661655 | 0.782137 | 714.2986 | 655.1026 | 697.2658 | 634.0283 | 693.3041 | 541.9264 | 731.8955 | 851.5838 |
| ENSMUSG00000026319 | 0.193331257 | 0.088612 | 0.202855 | 1138.854 | 1161.427 | 1247.129 | 1205.085 | 964.3851 | 1271.572 | 955.6106 | 967.9561 |
| ENSMUSG00000026321 | -0.332707436 | 0.054005 | 0.143868 | 109.6599 | 79.18822 | 100.2009 | 100.0643 | 115.0631 | 135.2023 | 111.8575 | 130.5109 |
| ENSMUSG00000026333 | 0.064436033 | 0.58151 | 0.722532 | 320.9313 | 291.5566 | 331.2427 | 332.11 | 298.3841 | 327.3906 | 331.7155 | 264.2846 |
| ENSMUSG00000026335 | 0.084127415 | 0.431578 | 0.596141 | 3237.483 | 2597.614 | 2803.969 | 3185.669 | 2450.455 | 2700.693 | 3006.654 | 2999.576 |
| ENSMUSG00000026339 | 0.005164794 | 0.953888 | 0.973726 | 752.5287 | 704.2952 | 631.8454 | 659.907 | 713.7815 | 638.0206 | 696.2168 | 686.27 |
| ENSMUSG00000026341 | -0.190239373 | 0.0281 | 0.09032 | 4942.745 | 5216.824 | 5829.043 | 5452.643 | 5945.254 | 5539.941 | 6224.487 | 6757.203 |
| ENSMUSG00000026342 | -0.122652429 | 0.236433 | 0.397487 | 1482.924 | 1298.207 | 1651.245 | 1480.262 | 1612.834 | 1410.126 | 1703.899 | 1714.044 |
| ENSMUSG00000026343 | -0.305481781 | 0.364301 | 0.532986 | 51.30877 | 41.99375 | 77.01392 | 48.30692 | 79.95913 | 35.75597 | 70.39311 | 84.8321 |
| ENSMUSG00000026344 | 0.630379005 | 0.099843 | 0.220685 | 25.15136 | 68.38983 | 51.34261 | 53.48266 | 30.22845 | 21.23011 | 31.82154 | 44.59123 |
| ENSMUSG00000026347 | -1.267475405 | 0.041523 | 0.119837 | 10.06054 | 8.398751 | 25.67131 | 5.175741 | 22.42756 | 13.40849 | 32.78583 | 51.11678 |
| ENSMUSG00000026348 | -5.6311638 | 2.01E-20 | 3.09E-17 | 3.018163 | 3.599465 | 5.796746 | 5.175741 | 307.1601 | 77.09881 | 105.1075 | 394.7956 |
| ENSMUSG00000026349 | 0.234075909 | 0.068118 | 0.168908 | 2871.279 | 2811.182 | 2399.853 | 2638.765 | 2199.851 | 2848.187 | 2052.972 | 2014.219 |
| ENSMUSG00000026353 | 0.09359795 | 0.196343 | 0.350591 | 3595.638 | 3683.452 | 3400.206 | 3742.923 | 3271.498 | 3725.325 | 3348.976 | 3170.328 |
| ENSMUSG00000026354 | 0.326334958 | 0.556262 | 0.70308 | 36.21796 | 45.59322 | 29.81184 | 17.25247 | 27.30312 | 45.81234 | 6.750024 | 22.83941 |
| ENSMUSG00000026355 | -0.103754402 | 0.318944 | 0.487005 | 3635.88 | 4764.491 | 3996.443 | 3619.568 | 4441.632 | 4068.359 | 4176.336 | 4518.941 |
| ENSMUSG00000026356 | -0.065753242 | 0.326886 | 0.494591 | 3022.187 | 3161.53 | 2849.515 | 3049.374 | 3285.15 | 2906.29 | 3160.94 | 3287.788 |
| ENSMUSG00000026358 | 1.418812179 | 0.143387 | 0.281048 | 31.18769 | 9.598572 | 3.312427 | 33.64232 | 0.975111 | 4.469496 | 20.25007 | 3.262773 |
| ENSMUSG00000026360 | 0.502748549 | 0.119119 | 0.248104 | 2144.908 | 991.0526 | 747.7803 | 1158.503 | 674.777 | 1110.67 | 1156.183 | 616.6641 |
| ENSMUSG00000026361 | 0.038255678 | 0.617428 | 0.750131 | 1274.671 | 1264.612 | 1145.271 | 1156.778 | 1118.453 | 1269.337 | 1164.861 | 1159.372 |
| ENSMUSG00000026365 | 0.26587678 | 0.10323 | 0.225871 | 6814.006 | 7849.232 | 9355.121 | 8373.486 | 6487.416 | 4887.394 | 7962.136 | 7603.349 |
| ENSMUSG00000026368 | -3.498695397 | 2.65E-05 | 0.000487 | 8.048435 | 0 | 2.48432 | 7.763611 | 65.33246 | 11.17374 | 29.89296 | 103.3212 |
| ENSMUSG00000026374 | 0.104527777 | 0.284446 | 0.451047 | 3678.135 | 3981.008 | 3696.668 | 3820.559 | 3501.625 | 4162.219 | 3252.547 | 3199.693 |
| ENSMUSG00000026377 | -0.204708914 | 0.006946 | 0.032801 | 1270.647 | 1374.995 | 1413.578 | 1395.725 | 1619.66 | 1508.455 | 1689.435 | 1468.248 |
| ENSMUSG00000026380 | -0.436064211 | 0.090093 | 0.205304 | 3387.385 | 2044.496 | 4547.962 | 2827.68 | 3117.431 | 3506.32 | 4914.018 | 5792.51 |
| ENSMUSG00000026383 | -0.527811547 | 1.25E-09 | 1.58E-07 | 1387.349 | 1196.222 | 1327.455 | 1264.606 | 1808.831 | 1683.883 | 1996.079 | 1975.065 |
| ENSMUSG00000026384 | 0.188014687 | 0.151505 | 0.291883 | 761.5831 | 879.4692 | 809.8883 | 866.9366 | 648.449 | 908.4251 | 648.0023 | 709.1094 |
| ENSMUSG00000026385 | -0.788191034 | 0.000135 | 0.001687 | 1857.176 | 1769.737 | 2373.354 | 1853.778 | 3528.928 | 2114.072 | 3446.369 | 4475.437 |
| ENSMUSG00000026387 | 0.661465827 | 0.106754 | 0.231007 | 50.30272 | 22.79661 | 28.98373 | 43.9938 | 31.20356 | 13.40849 | 27.96439 | 19.57664 |
| ENSMUSG00000026388 | 0.337176452 | 0.112444 | 0.239087 | 92.557 | 110.3836 | 93.57605 | 105.2401 | 74.10846 | 107.2679 | 68.46453 | 68.51824 |
| ENSMUSG00000026389 | 0.737298026 | 0.003258 | 0.018873 | 196.1806 | 307.1543 | 275.7595 | 237.2215 | 139.4409 | 224.5922 | 107.0361 | 139.2117 |
| ENSMUSG00000026393 | 0.037652606 | 0.603371 | 0.740659 | 2000.036 | 1936.512 | 1904.645 | 2222.118 | 1891.716 | 1995.63 | 2033.686 | 1937 |
| ENSMUSG00000026394 | -2.060048061 | 0.005861 | 0.028931 | 98.59333 | 38.39429 | 279.9 | 56.07053 | 450.5014 | 156.4324 | 583.3949 | 784.1532 |
| ENSMUSG00000026395 | 0.926462944 | 1.34E-07 | 7.23E-06 | 884.3218 | 968.256 | 838.0439 | 999.7806 | 394.9201 | 633.5511 | 536.1448 | 377.3941 |
| ENSMUSG00000026398 | -0.014839328 | 0.961241 | 0.977878 | 24.1453 | 34.79482 | 44.71776 | 37.09281 | 32.17867 | 44.69496 | 34.71441 | 31.54014 |
| ENSMUSG00000026399 | 0.688991946 | 0.000786 | 0.006496 | 464.7971 | 829.0767 | 558.972 | 601.2486 | 380.2934 | 483.823 | 327.8583 | 329.5401 |
| ENSMUSG00000026404 | -0.115462257 | 0.277831 | 0.444068 | 422.5428 | 355.1472 | 411.569 | 355.4009 | 438.8001 | 431.3064 | 388.6085 | 416.5474 |
| ENSMUSG00000026405 | -1.061998961 | 0.201796 | 0.357355 | 9.054489 | 2.399643 | 102.6852 | 18.97772 | 74.10846 | 42.46022 | 61.71451 | 101.146 |
| ENSMUSG00000026409 | 0.361501521 | 0.000556 | 0.004966 | 551.3178 | 494.3265 | 529.9882 | 593.485 | 465.1281 | 403.372 | 387.6442 | 433.9488 |
| ENSMUSG00000026411 | -0.406912207 | 0.099341 | 0.219967 | 3051.363 | 3258.715 | 5193.057 | 3358.193 | 6250.463 | 3096.244 | 4122.336 | 6236.247 |

| | | | | | | | | | | | |
|---|---|---|---|---|---|---|---|---|---|---|---|
| ENSMUSG00000026413 | -0.867406023 | 0.025193 | 0.083462 | 16.09687 | 28.79572 | 33.95237 | 37.09281 | 45.83023 | 31.28647 | 76.17884 | 58.72992 |
| ENSMUSG00000026414 | 0.829847935 | 0.251514 | 0.415163 | 237.4288 | 443.934 | 285.6968 | 389.9058 | 243.7778 | 413.4284 | 61.71451 | 44.59123 |
| ENSMUSG00000026418 | -0.173717965 | 0.418291 | 0.583873 | 100.6054 | 75.58876 | 75.3577 | 82.81186 | 95.56091 | 88.27255 | 75.21455 | 118.5474 |
| ENSMUSG00000026421 | -0.298689948 | 0.094342 | 0.211925 | 5251.604 | 5396.797 | 8442.547 | 5980.569 | 7672.176 | 6032.703 | 7998.779 | 9137.94 |
| ENSMUSG00000026425 | 0.165336086 | 0.069607 | 0.171539 | 1697.214 | 1613.76 | 1521.232 | 1518.217 | 1344.678 | 1598.962 | 1434.862 | 1283.357 |
| ENSMUSG00000026426 | 0.240163738 | 0.013266 | 0.052802 | 1171.047 | 1023.448 | 1205.723 | 1292.21 | 933.1815 | 1074.914 | 954.6463 | 1015.81 |
| ENSMUSG00000026427 | 0.058886894 | 0.374781 | 0.543403 | 1087.545 | 1208.22 | 1152.724 | 1165.404 | 1161.358 | 1098.379 | 1072.29 | 1095.204 |
| ENSMUSG00000026429 | -0.198725402 | 0.183405 | 0.334314 | 225.3562 | 266.3604 | 202.8861 | 237.2215 | 256.4543 | 236.8833 | 320.144 | 252.3211 |
| ENSMUSG00000026430 | -0.336742735 | 0.099714 | 0.220467 | 165.999 | 127.1811 | 149.0592 | 144.9207 | 186.2463 | 127.3806 | 197.6793 | 230.5693 |
| ENSMUSG00000026433 | 0.318904093 | 0.001031 | 0.008003 | 336.0222 | 338.3497 | 335.3832 | 336.4232 | 255.4792 | 282.6956 | 259.3938 | 282.7737 |
| ENSMUSG00000026434 | 0.251926521 | 0.126978 | 0.258124 | 6845.194 | 6901.373 | 6095.693 | 6434.309 | 4968.192 | 7710.999 | 4696.088 | 4691.868 |
| ENSMUSG00000026435 | -0.24678013 | 0.277674 | 0.444018 | 456.7487 | 425.9366 | 403.2879 | 425.2734 | 405.6463 | 402.2547 | 791.6814 | 429.5985 |
| ENSMUSG00000026436 | 0.365246667 | 0.024238 | 0.081141 | 2001.042 | 1859.723 | 1830.116 | 1934.002 | 1288.122 | 2048.147 | 1330.719 | 1253.993 |
| ENSMUSG00000026437 | -1.235844007 | 0.000176 | 0.002056 | 192.1564 | 158.3764 | 244.2915 | 152.6844 | 533.3859 | 183.2494 | 445.5016 | 599.2627 |
| ENSMUSG00000026439 | -0.003124344 | 0.975226 | 0.986504 | 1188.15 | 1005.45 | 938.2448 | 992.8796 | 1103.826 | 945.2985 | 1027.932 | 1053.876 |
| ENSMUSG00000026442 | 0.527518717 | 0.018904 | 0.067758 | 101.6115 | 76.78858 | 85.29498 | 119.042 | 53.63112 | 74.86406 | 66.53595 | 71.78101 |
| ENSMUSG00000026443 | 0.040725276 | 0.861815 | 0.917409 | 78.47224 | 75.58876 | 83.63877 | 94.88858 | 94.5858 | 53.63396 | 95.46463 | 79.39415 |
| ENSMUSG00000026447 | 0.332286009 | 0.021317 | 0.073719 | 477.8758 | 447.5334 | 438.8965 | 355.4009 | 379.3183 | 375.4377 | 286.3939 | 324.1021 |
| ENSMUSG00000026452 | 0.88578278 | 2.43E-05 | 0.000457 | 112.6781 | 110.3836 | 89.43552 | 97.47645 | 41.92979 | 56.98608 | 61.71451 | 60.9051 |
| ENSMUSG00000026455 | -0.184052995 | 0.039616 | 0.115854 | 873.2552 | 889.0678 | 804.0915 | 980.8029 | 1075.548 | 954.2375 | 1026.004 | 972.3064 |
| ENSMUSG00000026456 | 0.032068864 | 0.767262 | 0.855545 | 876.2733 | 841.0749 | 714.656 | 787.5752 | 867.8491 | 808.9788 | 686.5739 | 783.0656 |
| ENSMUSG00000026457 | -0.178384328 | 0.047697 | 0.131622 | 3061.423 | 2613.211 | 3020.105 | 3026.083 | 3338.781 | 2918.581 | 3489.762 | 3519.445 |
| ENSMUSG00000026458 | 0.62190585 | 0.000115 | 0.001497 | 284.7134 | 250.7627 | 264.9941 | 259.6497 | 162.8436 | 217.8879 | 136.9291 | 172.927 |
| ENSMUSG00000026463 | 0.448284997 | 0.000108 | 0.001424 | 1220.344 | 1479.38 | 1236.363 | 1409.527 | 868.8242 | 1139.722 | 918.9676 | 990.7955 |
| ENSMUSG00000026466 | -0.026986308 | 0.606083 | 0.742507 | 1386.343 | 1409.79 | 1409.437 | 1368.983 | 1409.036 | 1422.417 | 1451.255 | 1395.379 |
| ENSMUSG00000026468 | 0.415719246 | 0.402642 | 0.569693 | 10.06054 | 14.39786 | 14.90592 | 21.56559 | 6.825779 | 17.87799 | 10.60718 | 10.87591 |
| ENSMUSG00000026469 | 0.1503485 | 0.151305 | 0.291643 | 1675.081 | 1350.999 | 1382.938 | 1543.233 | 1185.735 | 1384.427 | 1461.862 | 1332.299 |
| ENSMUSG00000026470 | -0.068066453 | 0.51021 | 0.665943 | 1458.779 | 1592.163 | 1891.396 | 1642.435 | 1830.284 | 1541.976 | 1740.542 | 1792.35 |
| ENSMUSG00000026471 | 0.329194207 | 0.002737 | 0.016518 | 558.3602 | 640.7047 | 746.9522 | 642.6545 | 528.5103 | 491.6446 | 508.1804 | 534.0072 |
| ENSMUSG00000026473 | -0.378582656 | 0.004209 | 0.022673 | 7373.372 | 6049.5 | 6002.117 | 7673.036 | 8613.158 | 7642.839 | 10657.32 | 8315.721 |
| ENSMUSG00000026475 | -0.125195429 | 0.54106 | 0.690581 | 107.6478 | 148.7779 | 118.4192 | 98.33908 | 114.088 | 108.3853 | 134.0362 | 157.7007 |
| ENSMUSG00000026478 | 0.212593075 | 0.016151 | 0.060598 | 14355.39 | 12500.94 | 14252.54 | 14223.8 | 10371.28 | 12578.28 | 12679.44 | 12126.64 |
| ENSMUSG00000026479 | -0.470140274 | 0.000287 | 0.003003 | 3229.434 | 2904.768 | 3781.135 | 3057.138 | 4136.422 | 3801.307 | 5276.59 | 4758.211 |
| ENSMUSG00000026480 | -0.281074282 | 0.121844 | 0.251178 | 159.9626 | 205.1695 | 188.8083 | 204.4418 | 214.5245 | 175.4277 | 282.5367 | 247.9708 |
| ENSMUSG00000026482 | 0.302115301 | 0.002357 | 0.014753 | 2211.307 | 2809.982 | 2394.884 | 2468.828 | 2092.589 | 2166.588 | 1812.864 | 1943.525 |
| ENSMUSG00000026483 | -0.229872483 | 0.43977 | 0.603068 | 845.0857 | 434.3354 | 1157.693 | 652.1434 | 711.8313 | 671.5418 | 1269.969 | 971.2188 |
| ENSMUSG00000026484 | 0.156282964 | 0.023463 | 0.079164 | 2381.331 | 2338.452 | 2249.138 | 2415.346 | 1996.053 | 2253.744 | 2192.794 | 1978.328 |
| ENSMUSG00000026489 | -0.42447474 | 0.000945 | 0.007498 | 470.8334 | 449.9331 | 432.2717 | 522.7498 | 702.0801 | 508.4052 | 663.4309 | 642.7663 |
| ENSMUSG00000026490 | 0.013770418 | 0.886184 | 0.932038 | 2065.43 | 1767.337 | 2025.549 | 2090.137 | 1742.524 | 2174.41 | 2042.364 | 1918.511 |
| ENSMUSG00000026491 | 0.019358577 | 0.815353 | 0.887534 | 2173.077 | 1960.508 | 1878.974 | 2156.559 | 2042.858 | 2190.053 | 1892.9 | 1935.912 |
| ENSMUSG00000026492 | -0.022889509 | 0.802756 | 0.879425 | 674.0564 | 611.909 | 595.4087 | 635.7535 | 663.0757 | 566.5087 | 662.4667 | 663.4306 |
| ENSMUSG00000026494 | 0.211349344 | 0.024725 | 0.082398 | 2927.618 | 3383.497 | 3728.964 | 3127.873 | 3081.352 | 2832.543 | 2734.724 | 2725.503 |
| ENSMUSG00000026495 | 0.071593466 | 0.577009 | 0.718915 | 261.5741 | 295.1561 | 291.4935 | 232.9083 | 244.7529 | 265.935 | 238.1794 | 279.5109 |
| ENSMUSG00000026496 | -0.235079518 | 0.003479 | 0.019693 | 3534.269 | 3943.813 | 3425.049 | 3446.181 | 4381.175 | 3822.537 | 4185.015 | 4492.839 |
| ENSMUSG00000026499 | -0.079854801 | 0.302446 | 0.470753 | 1678.099 | 1548.97 | 1618.12 | 1482.85 | 1683.042 | 1520.746 | 1703.899 | 1779.299 |
| ENSMUSG00000026500 | 0.007014183 | 0.946986 | 0.969522 | 342.0585 | 380.3434 | 341.1799 | 332.11 | 341.289 | 385.4941 | 339.4298 | 320.8394 |
| ENSMUSG00000026502 | 0.147157365 | 0.067254 | 0.16744 | 1354.149 | 1215.419 | 1162.662 | 1360.357 | 1099.926 | 1186.651 | 1165.826 | 1147.409 |
| ENSMUSG00000026504 | 0.158330618 | 0.063731 | 0.161834 | 565.4025 | 563.9161 | 597.893 | 571.9194 | 491.4561 | 487.1751 | 518.7876 | 564.4598 |
| ENSMUSG00000026509 | -0.28056398 | 0.004408 | 0.023464 | 3352.173 | 3427.89 | 3669.34 | 3435.829 | 4072.065 | 3571.128 | 4495.516 | 4727.758 |
| ENSMUSG00000026510 | -0.322156241 | 1.17E-06 | 4.30E-05 | 1426.585 | 1383.394 | 1407.781 | 1349.143 | 1701.569 | 1616.84 | 1814.792 | 1824.978 |
| ENSMUSG00000026511 | -0.423513036 | 0.002246 | 0.014293 | 2073.478 | 1930.513 | 1917.895 | 2021.127 | 2190.1 | 2239.218 | 3153.226 | 3070.27 |
| ENSMUSG00000026514 | 0.339426305 | 0.500842 | 0.657109 | 17.10292 | 10.79839 | 9.109173 | 11.21411 | 7.80089 | 8.938993 | 13.50005 | 7.613138 |
| ENSMUSG00000026516 | 0.026295268 | 0.715452 | 0.82173 | 1774.68 | 1619.759 | 1508.81 | 1567.387 | 1542.626 | 1577.732 | 1595.899 | 1634.649 |
| ENSMUSG00000026519 | 0.112784635 | 0.384445 | 0.552315 | 870.237 | 593.9117 | 758.5457 | 740.9936 | 752.7859 | 634.6685 | 672.1095 | 683.0072 |
| ENSMUSG00000026520 | -0.227207572 | 0.033602 | 0.102657 | 892.3702 | 1106.235 | 934.1043 | 882.4638 | 1106.751 | 1012.341 | 1102.183 | 1239.854 |
| ENSMUSG00000026525 | -0.52257277 | 0.030206 | 0.09495 | 105.6357 | 73.18911 | 104.3414 | 81.94923 | 113.1129 | 94.9768 | 164.8934 | 152.2628 |
| ENSMUSG00000026526 | -0.700747477 | 1.61E-07 | 8.34E-06 | 2286.762 | 2138.082 | 2523.241 | 2245.409 | 3841.939 | 2851.539 | 3858.121 | 4392.78 |
| ENSMUSG00000026527 | 0.3571374 | 0.400077 | 0.567462 | 37.22401 | 19.19714 | 18.21835 | 50.03216 | 27.30312 | 20.11273 | 24.10723 | 26.10219 |
| ENSMUSG00000026532 | 0.649567615 | 0.338753 | 0.507264 | 54.32694 | 9.598572 | 23.18699 | 18.97772 | 30.22845 | 22.34748 | 9.642892 | 5.437955 |
| ENSMUSG00000026535 | 2.138205585 | 0.008938 | 0.039268 | 45.27245 | 93.58608 | 76.18581 | 31.05445 | 28.27823 | 24.58223 | 1.928578 | 1.087591 |
| ENSMUSG00000026536 | 0.8408145 | 0.095334 | 0.213618 | 26.15741 | 56.39161 | 50.5145 | 22.42821 | 19.50223 | 7.821619 | 31.82154 | 27.18978 |
| ENSMUSG00000026542 | -2.117172233 | 0.011081 | 0.046135 | 8.048435 | 0 | 3.312427 | 17.25247 | 18.52711 | 15.64324 | 30.85725 | 60.9051 |
| ENSMUSG00000026544 | -0.293066657 | 0.053666 | 0.143131 | 245.4773 | 260.3613 | 240.1509 | 277.7648 | 282.7823 | 262.5829 | 383.7871 | 324.1021 |
| ENSMUSG00000026546 | -0.038442046 | 0.891761 | 0.934602 | 44.26639 | 59.99108 | 51.34261 | 60.38364 | 34.1289 | 63.69032 | 55.92877 | 68.51824 |

| | | | | | | | | | | | |
|---|---|---|---|---|---|---|---|---|---|---|---|
| ENSMUSG00000026547 | -0.295356054 | 0.021497 | 0.074131 | 2457.791 | 2463.234 | 2835.437 | 2331.671 | 2851.225 | 2556.552 | 3268.94 | 3704.335 |
| ENSMUSG00000026548 | -0.208705326 | 0.592477 | 0.732205 | 13.07871 | 19.19714 | 29.81184 | 18.11509 | 26.32801 | 17.87799 | 26.03581 | 22.83941 |
| ENSMUSG00000026553 | -0.119666488 | 0.232563 | 0.393087 | 7612.813 | 7880.428 | 7939.886 | 7601.438 | 7853.546 | 7171.307 | 9192.569 | 9500.108 |
| ENSMUSG00000026554 | 0.033108169 | 0.591307 | 0.731321 | 6062.484 | 5657.159 | 5255.993 | 5679.513 | 5622.492 | 5425.969 | 5709.556 | 5378.138 |
| ENSMUSG00000026556 | 0.239230714 | 0.014574 | 0.056473 | 1154.95 | 1089.438 | 1035.961 | 990.2918 | 853.2224 | 973.2328 | 810.0029 | 982.0948 |
| ENSMUSG00000026558 | -0.261681349 | 8.99E-06 | 0.000211 | 1982.933 | 2150.08 | 2138.171 | 2098.763 | 2439.728 | 2407.941 | 2592.009 | 2593.905 |
| ENSMUSG00000026563 | 0.089966905 | 0.382144 | 0.550015 | 822.9525 | 794.2819 | 715.4841 | 759.9713 | 817.1433 | 632.4337 | 737.6812 | 714.5473 |
| ENSMUSG00000026565 | 0.316122721 | 0.076945 | 0.184236 | 1121.751 | 965.8563 | 906.7768 | 1036.873 | 685.5032 | 1128.548 | 763.717 | 661.2554 |
| ENSMUSG00000026566 | 0.100516507 | 0.212663 | 0.370555 | 5787.831 | 7129.34 | 6350.75 | 6508.494 | 6183.181 | 6301.99 | 5904.343 | 5648.948 |
| ENSMUSG00000026568 | -0.796401197 | 8.94E-05 | 0.001232 | 1398.416 | 1174.625 | 1963.441 | 1648.473 | 2468.007 | 1944.231 | 2893.832 | 3440.051 |
| ENSMUSG00000026571 | 0.107120462 | 0.27062 | 0.435788 | 1862.207 | 1599.362 | 1795.335 | 2072.884 | 1734.723 | 1781.094 | 1735.72 | 1557.43 |
| ENSMUSG00000026574 | 0.876227745 | 0.331133 | 0.499012 | 503.0272 | 1076.24 | 616.9394 | 862.6235 | 391.0196 | 1109.552 | 122.4647 | 43.50364 |
| ENSMUSG00000026575 | 0.088884959 | 0.349984 | 0.518737 | 938.6487 | 874.6699 | 774.2797 | 839.3327 | 797.641 | 863.7302 | 828.3244 | 729.7736 |
| ENSMUSG00000026576 | -1.726853917 | 0.001014 | 0.007919 | 5303.919 | 2560.419 | 11402.2 | 5633.794 | 15996.7 | 10829.59 | 26881.49 | 28716.76 |
| ENSMUSG00000026577 | -0.064192821 | 0.443143 | 0.60632 | 775.6679 | 805.0802 | 716.3122 | 698.725 | 727.433 | 787.7487 | 810.9672 | 801.5546 |
| ENSMUSG00000026578 | -0.019507561 | 0.849535 | 0.909614 | 411.4762 | 502.7252 | 479.4737 | 476.1682 | 442.7005 | 512.8747 | 484.0732 | 455.7007 |
| ENSMUSG00000026579 | 0.253302313 | 0.551849 | 0.699466 | 33.19979 | 15.59768 | 55.48314 | 37.95543 | 19.50223 | 31.28647 | 32.78583 | 36.9781 |
| ENSMUSG00000026581 | -0.911885451 | 0.162757 | 0.307141 | 15.09082 | 4.799286 | 7.45296 | 4.313117 | 14.62667 | 7.821619 | 27.0001 | 9.78832 |
| ENSMUSG00000026582 | -1.722261117 | 0.034212 | 0.104245 | 8.048435 | 1.199822 | 8.281066 | 2.58787 | 30.22845 | 3.352122 | 19.28578 | 14.13868 |
| ENSMUSG00000026584 | -0.009989247 | 0.915015 | 0.949629 | 999.012 | 886.6681 | 882.7617 | 1042.912 | 1048.245 | 910.6599 | 963.3249 | 915.7517 |
| ENSMUSG00000026585 | 0.151084134 | 0.121141 | 0.250368 | 1684.135 | 1750.54 | 1598.246 | 1718.346 | 1442.19 | 1779.977 | 1490.791 | 1367.102 |
| ENSMUSG00000026586 | 0.367080638 | 0.136893 | 0.271724 | 424.5549 | 568.7154 | 529.1601 | 619.3637 | 297.4089 | 636.9032 | 410.7872 | 316.489 |
| ENSMUSG00000026587 | 2.85634451 | 8.62E-28 | 2.65E-24 | 405.4399 | 311.9536 | 284.8687 | 452.8773 | 36.07912 | 75.98144 | 47.25017 | 42.41605 |
| ENSMUSG00000026589 | 0.660479921 | 5.86E-06 | 0.000152 | 229.3804 | 197.9706 | 190.4645 | 177.7004 | 128.7147 | 120.6764 | 134.0362 | 118.5474 |
| ENSMUSG00000026594 | 0.247287488 | 0.013522 | 0.053429 | 1618.741 | 1454.184 | 1503.014 | 1479.399 | 1206.213 | 1483.873 | 1133.04 | 1281.182 |
| ENSMUSG00000026596 | 0.218656658 | 0.044313 | 0.125167 | 1197.205 | 1057.043 | 1136.162 | 1188.695 | 935.1317 | 1127.43 | 1034.682 | 839.6203 |
| ENSMUSG00000026600 | 0.573635258 | 2.53E-05 | 0.000472 | 3075.508 | 2358.849 | 2949.716 | 2527.487 | 1700.594 | 2216.87 | 1854.328 | 1561.781 |
| ENSMUSG00000026601 | -0.367277603 | 0.394205 | 0.561508 | 13.07871 | 17.99732 | 12.4216 | 11.21411 | 12.67645 | 16.76061 | 25.07152 | 15.22628 |
| ENSMUSG00000026602 | -2.768061758 | 0.013476 | 0.053323 | 61.36932 | 2.399643 | 349.461 | 93.16334 | 430.9992 | 407.8415 | 1107.004 | 1506.314 |
| ENSMUSG00000026603 | -0.236958618 | 0.009152 | 0.039984 | 1025.169 | 1165.027 | 1066.601 | 1224.925 | 1389.534 | 1364.314 | 1192.826 | 1335.562 |
| ENSMUSG00000026604 | 0.208970158 | 0.024121 | 0.080911 | 3449.76 | 2998.354 | 3362.941 | 3172.729 | 2513.837 | 3187.868 | 2715.438 | 2820.124 |
| ENSMUSG00000026605 | 0.283296525 | 0.015394 | 0.058751 | 2680.129 | 2642.007 | 2312.074 | 2625.826 | 2038.958 | 2572.195 | 1954.614 | 1865.219 |
| ENSMUSG00000026608 | -0.112580181 | 0.044237 | 0.125039 | 1631.82 | 1679.75 | 1668.635 | 1638.985 | 1726.922 | 1825.789 | 1733.792 | 1871.744 |
| ENSMUSG00000026609 | 0.146790822 | 0.645587 | 0.770506 | 25.15136 | 38.39429 | 53.82693 | 37.95543 | 37.05423 | 27.93435 | 36.64299 | 39.15328 |
| ENSMUSG00000026610 | -0.948790173 | 0.000784 | 0.006485 | 398.3975 | 274.7591 | 616.1113 | 381.2796 | 651.3744 | 544.1612 | 903.5389 | 1128.92 |
| ENSMUSG00000026611 | -0.631989846 | 0.196719 | 0.351025 | 9.054489 | 8.398751 | 6.624853 | 12.93935 | 14.62667 | 8.938993 | 16.39292 | 17.40146 |
| ENSMUSG00000026614 | 1.316142293 | 0.002327 | 0.014615 | 62.37537 | 76.78858 | 55.48314 | 44.85642 | 12.67645 | 46.92971 | 17.3572 | 19.57664 |
| ENSMUSG00000026615 | -0.094378213 | 0.174523 | 0.322656 | 5046.369 | 5125.638 | 4844.424 | 4834.142 | 5060.828 | 5887.444 | 5140.626 | 5102.977 |
| ENSMUSG00000026616 | -0.167556228 | 0.667448 | 0.786689 | 66.39959 | 39.59411 | 97.71658 | 52.62003 | 46.80534 | 58.10345 | 119.5719 | 64.16787 |
| ENSMUSG00000026617 | -1.535396696 | 1.13E-07 | 6.24E-06 | 603.6326 | 505.1249 | 808.2321 | 556.3922 | 1687.918 | 849.2043 | 2122.4 | 2512.335 |
| ENSMUSG00000026618 | -0.283994638 | 0.005761 | 0.028566 | 2133.841 | 1826.128 | 1878.146 | 1958.155 | 2350.993 | 2007.921 | 2519.688 | 2613.481 |
| ENSMUSG00000026620 | -0.155053191 | 0.385076 | 0.552598 | 410.4702 | 386.3425 | 587.1276 | 433.8996 | 461.2276 | 427.9543 | 509.1447 | 628.6276 |
| ENSMUSG00000026621 | 1.334457232 | 0.049318 | 0.134807 | 22.1332 | 13.19804 | 4.140533 | 27.60395 | 5.850668 | 4.469496 | 9.642892 | 6.525547 |
| ENSMUSG00000026622 | -0.23684394 | 0.006028 | 0.02956 | 1029.194 | 1065.442 | 993.728 | 1043.774 | 1373.932 | 1112.905 | 1209.219 | 1169.16 |
| ENSMUSG00000026623 | 0.012187025 | 0.865536 | 0.91951 | 2020.157 | 2109.286 | 1856.615 | 1884.832 | 1927.795 | 1926.353 | 1862.042 | 2083.825 |
| ENSMUSG00000026626 | -0.215832391 | 0.011162 | 0.046389 | 2257.586 | 1978.506 | 2410.618 | 2336.847 | 2417.301 | 2583.369 | 2687.474 | 2752.693 |
| ENSMUSG00000026627 | -0.263762746 | 0.077365 | 0.18481 | 383.3067 | 429.5361 | 352.7734 | 374.3786 | 472.929 | 362.0292 | 451.2873 | 560.1094 |
| ENSMUSG00000026628 | -0.223151047 | 0.365189 | 0.53394 | 67.40564 | 73.18911 | 97.71658 | 69.00988 | 71.18313 | 77.09881 | 119.5719 | 91.35765 |
| ENSMUSG00000026630 | -0.16642981 | 0.43936 | 0.602708 | 170.0232 | 97.18554 | 190.4645 | 149.2339 | 158.9431 | 191.071 | 167.7863 | 166.4014 |
| ENSMUSG00000026632 | 0.111898139 | 0.395819 | 0.56311 | 299.8042 | 269.9598 | 231.8699 | 302.7808 | 256.4543 | 277.1088 | 237.2151 | 251.2335 |
| ENSMUSG00000026634 | 0.1262041 | 0.253823 | 0.417444 | 2240.483 | 2627.609 | 2081.032 | 2380.841 | 2216.428 | 2399.002 | 2087.686 | 1842.379 |
| ENSMUSG00000026637 | 0.131597866 | 0.219794 | 0.378619 | 387.3309 | 350.3479 | 339.5237 | 331.2474 | 299.3592 | 316.2169 | 351.9655 | 316.489 |
| ENSMUSG00000026638 | -0.337814474 | 0.173807 | 0.321743 | 436.6276 | 357.5468 | 751.9208 | 430.4491 | 642.5984 | 453.6539 | 636.4308 | 767.8393 |
| ENSMUSG00000026639 | 0.188764651 | 0.4247 | 0.589675 | 105.6357 | 158.3764 | 167.2775 | 119.9047 | 159.9183 | 110.62 | 117.6433 | 94.62042 |
| ENSMUSG00000026640 | -0.256216212 | 0.069304 | 0.170977 | 436.6276 | 332.3506 | 525.0196 | 433.8996 | 509.9832 | 511.7573 | 520.7161 | 526.3941 |
| ENSMUSG00000026641 | 0.062324118 | 0.397222 | 0.564489 | 1680.111 | 1727.743 | 1777.945 | 1703.681 | 1745.449 | 1735.282 | 1485.97 | 1633.562 |
| ENSMUSG00000026643 | 0.179541735 | 0.05321 | 0.14212 | 1599.626 | 1474.581 | 1534.482 | 1430.23 | 1291.047 | 1529.685 | 1306.612 | 1206.139 |
| ENSMUSG00000026646 | -0.457727704 | 0.000211 | 0.002369 | 225.3562 | 263.9607 | 232.698 | 273.4516 | 370.5423 | 364.264 | 316.2868 | 315.4014 |
| ENSMUSG00000026648 | 0.186137761 | 0.157155 | 0.299602 | 311.8769 | 283.1579 | 275.7595 | 259.6497 | 285.7076 | 213.4185 | 249.7509 | 242.5328 |
| ENSMUSG00000026649 | -1.337510853 | 9.22E-05 | 0.001253 | 22.1332 | 19.19714 | 19.87456 | 18.11509 | 45.83023 | 30.1691 | 58.82164 | 65.25547 |
| ENSMUSG00000026655 | -0.370301044 | 0.000174 | 0.002041 | 1630.814 | 1544.17 | 1931.145 | 1768.378 | 2067.236 | 2108.485 | 2260.294 | 2456.868 |
| ENSMUSG00000026656 | -0.09610446 | 0.714597 | 0.821054 | 120.7265 | 111.5834 | 180.5272 | 93.16334 | 116.0382 | 103.9158 | 163.9292 | 157.7007 |
| ENSMUSG00000026657 | -0.124956227 | 0.25111 | 0.414793 | 1601.639 | 1644.955 | 1607.355 | 1589.815 | 1693.768 | 1450.352 | 1847.578 | 2033.795 |
| ENSMUSG00000026659 | 0.218928353 | 0.086829 | 0.200089 | 521.1362 | 567.5156 | 544.8942 | 569.3315 | 451.4765 | 586.6214 | 414.6443 | 441.562 |

| | | | | | | | | | | | |
|---|---|---|---|---|---|---|---|---|---|---|---|
| ENSMUSG00000026662 | 0.074841731 | 0.262227 | 0.426418 | 3977.939 | 4159.781 | 3668.512 | 4193.213 | 3758.079 | 4010.256 | 3667.192 | 3753.277 |
| ENSMUSG00000026663 | -0.232711252 | 0.019769 | 0.069885 | 864.2007 | 831.4763 | 896.8395 | 776.3611 | 935.1317 | 877.1387 | 1071.325 | 1074.54 |
| ENSMUSG00000026664 | -0.973913165 | 1.62E-05 | 0.000332 | 776.674 | 566.3158 | 879.4492 | 752.2077 | 1485.095 | 878.256 | 1611.327 | 1870.657 |
| ENSMUSG00000026667 | -0.091129833 | 0.326318 | 0.494054 | 1823.977 | 1491.378 | 1553.528 | 1496.652 | 1588.456 | 1764.334 | 1632.542 | 1796.7 |
| ENSMUSG00000026669 | -0.122875792 | 0.308014 | 0.476506 | 520.1301 | 620.3077 | 605.3459 | 457.1905 | 611.3948 | 593.3256 | 627.7522 | 563.3722 |
| ENSMUSG00000026670 | -0.107871027 | 0.185595 | 0.337199 | 1095.593 | 1033.046 | 1020.227 | 1042.912 | 1100.901 | 1014.576 | 1182.219 | 1218.102 |
| ENSMUSG00000026672 | -0.761079581 | 0.001163 | 0.008774 | 443.67 | 266.3604 | 515.9104 | 380.417 | 688.4286 | 460.3581 | 795.5386 | 780.8904 |
| ENSMUSG00000026674 | 0.410167986 | 0.002286 | 0.014443 | 1344.089 | 1612.56 | 1497.217 | 1507.866 | 977.0615 | 1421.3 | 1073.254 | 1015.81 |
| ENSMUSG00000026675 | -0.548937137 | 0.000104 | 0.001387 | 375.2583 | 404.3399 | 463.7397 | 360.5766 | 610.4197 | 498.3488 | 694.2882 | 542.708 |
| ENSMUSG00000026676 | -0.942664854 | 0.004435 | 0.023569 | 92.557 | 119.9822 | 264.9941 | 152.6844 | 307.1601 | 197.7752 | 323.0369 | 386.0948 |
| ENSMUSG00000026678 | -0.647544715 | 0.001202 | 0.008983 | 2182.132 | 2454.835 | 2962.137 | 2791.45 | 3634.24 | 2671.641 | 4967.053 | 5005.094 |
| ENSMUSG00000026679 | 0.968693459 | 0.012416 | 0.050265 | 66.39959 | 62.39072 | 53.82693 | 55.2079 | 22.42756 | 56.98608 | 25.07152 | 17.40146 |
| ENSMUSG00000026683 | 0.015665927 | 0.871567 | 0.92303 | 1193.18 | 1162.627 | 1026.852 | 1178.344 | 1226.69 | 1208.999 | 1053.004 | 1021.248 |
| ENSMUSG00000026687 | -0.491962574 | 0.000152 | 0.001844 | 1131.811 | 1118.234 | 1205.723 | 1075.691 | 1633.311 | 1305.093 | 1476.327 | 1958.752 |
| ENSMUSG00000026688 | -0.839714704 | 0.005734 | 0.028477 | 542.2633 | 309.554 | 744.4679 | 421.8229 | 967.3104 | 505.0531 | 923.789 | 1218.102 |
| ENSMUSG00000026691 | 1.753408163 | 0.016461 | 0.061438 | 18.10898 | 21.59679 | 7.45296 | 28.46658 | 1.950223 | 5.58687 | 12.53576 | 2.175182 |
| ENSMUSG00000026692 | -0.13844858 | 0.729047 | 0.829805 | 31.18769 | 19.19714 | 25.67131 | 22.42821 | 15.60178 | 27.93435 | 23.14294 | 42.41605 |
| ENSMUSG00000026694 | -0.140828368 | 0.103808 | 0.226596 | 699.2078 | 704.2952 | 713.8279 | 634.0283 | 769.3628 | 806.7441 | 769.5028 | 686.27 |
| ENSMUSG00000026696 | 0.206705528 | 0.065092 | 0.164345 | 971.8485 | 1073.84 | 876.9649 | 1128.312 | 853.2224 | 983.2892 | 874.6103 | 798.2919 |
| ENSMUSG00000026697 | 1.714318542 | 0.019525 | 0.0693 | 24.1453 | 45.59322 | 33.95237 | 50.03216 | 14.62667 | 24.58223 | 7.714313 | 0 |
| ENSMUSG00000026698 | 0.011570126 | 0.911853 | 0.948028 | 918.5276 | 819.4781 | 794.1543 | 986.8413 | 786.9148 | 910.6599 | 932.4676 | 862.4597 |
| ENSMUSG00000026701 | -0.045879529 | 0.529002 | 0.681459 | 4158.023 | 3898.22 | 3845.727 | 4109.538 | 4431.881 | 3888.462 | 4356.658 | 3848.985 |
| ENSMUSG00000026705 | 0.128430457 | 0.129271 | 0.261061 | 696.1896 | 677.8992 | 637.6421 | 690.0988 | 594.8179 | 671.5418 | 630.6451 | 574.2481 |
| ENSMUSG00000026707 | 0.251882564 | 0.030995 | 0.096726 | 267.6105 | 250.7627 | 248.432 | 248.4356 | 205.7485 | 221.2401 | 233.358 | 191.416 |
| ENSMUSG00000026708 | 0.186967528 | 0.163426 | 0.308183 | 456.7487 | 496.7261 | 464.5678 | 522.7498 | 484.6303 | 459.2408 | 425.2515 | 334.9781 |
| ENSMUSG00000026709 | -0.266472866 | 0.008314 | 0.037339 | 476.8698 | 464.3309 | 462.9116 | 455.4652 | 635.7726 | 512.8747 | 515.8947 | 570.9853 |
| ENSMUSG00000026712 | 0.809938042 | 5.78E-17 | 4.27E-14 | 689.1472 | 692.297 | 627.7048 | 616.7758 | 365.6667 | 394.4331 | 397.2871 | 338.2408 |
| ENSMUSG00000026715 | 1.410246822 | 0.066838 | 0.1669 | 42.25428 | 5.999108 | 10.76539 | 47.44429 | 2.925334 | 13.40849 | 7.714313 | 16.31387 |
| ENSMUSG00000026718 | 0.111494477 | 0.24041 | 0.402559 | 1223.362 | 1087.038 | 982.1345 | 1183.519 | 1045.319 | 1063.74 | 957.5391 | 1076.715 |
| ENSMUSG00000026721 | 0.210707048 | 0.034766 | 0.105537 | 752.5287 | 770.2854 | 870.3401 | 784.1248 | 639.673 | 785.514 | 670.181 | 653.6422 |
| ENSMUSG00000026723 | -0.139608615 | 0.317178 | 0.485393 | 241.453 | 271.1597 | 346.1486 | 275.1769 | 310.0854 | 332.9775 | 317.2511 | 291.4744 |
| ENSMUSG00000026725 | -0.234055692 | 0.489456 | 0.647036 | 136.8234 | 145.1784 | 91.09173 | 124.2178 | 224.2756 | 119.559 | 168.7506 | 70.69342 |
| ENSMUSG00000026726 | -3.124019044 | 3.18E-06 | 9.51E-05 | 72.43591 | 51.59233 | 303.9151 | 144.9207 | 1235.466 | 524.0485 | 1069.397 | 2171.919 |
| ENSMUSG00000026727 | 0.131597999 | 0.079286 | 0.187911 | 2019.151 | 2081.69 | 2298.824 | 2386.879 | 1998.003 | 2062.673 | 2007.65 | 1957.664 |
| ENSMUSG00000026728 | 0.318978796 | 4.90E-05 | 0.000784 | 15651.19 | 17620.58 | 18313.58 | 19284.81 | 13937.27 | 14982.87 | 14189.52 | 13705.82 |
| ENSMUSG00000026730 | -0.920270833 | 0.000686 | 0.005846 | 386.3249 | 362.3461 | 501.0045 | 361.4392 | 786.9148 | 451.4191 | 620.0379 | 1192 |
| ENSMUSG00000026736 | -0.229361394 | 0.51872 | 0.672487 | 20.12109 | 25.19625 | 28.15563 | 14.6646 | 22.42756 | 22.34748 | 28.92867 | 29.36496 |
| ENSMUSG00000026737 | -0.079296014 | 0.54459 | 0.69317 | 668.0201 | 530.3211 | 675.735 | 653.006 | 564.5894 | 631.3164 | 747.3241 | 729.7736 |
| ENSMUSG00000026739 | 0.068845038 | 0.535193 | 0.686082 | 2003.054 | 2262.863 | 1953.504 | 2178.124 | 1996.053 | 2385.594 | 1808.042 | 1816.277 |
| ENSMUSG00000026740 | 0.217926598 | 0.067436 | 0.16764 | 1347.107 | 1343.8 | 1094.757 | 1179.206 | 941.9575 | 1248.107 | 1008.646 | 1069.102 |
| ENSMUSG00000026743 | 0.195892358 | 0.151881 | 0.292357 | 2214.326 | 2033.698 | 1757.242 | 2014.226 | 1602.108 | 2220.222 | 1622.899 | 1556.343 |
| ENSMUSG00000026748 | 0.536872635 | 0.000166 | 0.001963 | 2199.235 | 2661.204 | 2030.517 | 2385.154 | 1601.133 | 1909.592 | 1629.649 | 1251.817 |
| ENSMUSG00000026749 | 0.049491198 | 0.550508 | 0.698381 | 911.4852 | 915.4638 | 968.8848 | 897.1284 | 942.9326 | 948.6506 | 801.3243 | 877.686 |
| ENSMUSG00000026750 | -0.315978195 | 0.001081 | 0.008283 | 3624.814 | 3377.498 | 3848.212 | 3673.913 | 4824.851 | 3792.368 | 4625.695 | 4839.78 |
| ENSMUSG00000026751 | 3.07893511 | 0.014752 | 0.056968 | 1183.12 | 11.99822 | 20.70267 | 238.0841 | 36.07912 | 36.87335 | 24.10723 | 75.04379 |
| ENSMUSG00000026753 | 0.022904783 | 0.769203 | 0.856805 | 1693.189 | 1520.174 | 1471.545 | 1551.86 | 1506.547 | 1642.54 | 1565.041 | 1423.657 |
| ENSMUSG00000026754 | 0.057462031 | 0.516284 | 0.670315 | 907.461 | 776.2845 | 741.1554 | 797.9267 | 791.7904 | 740.819 | 785.8957 | 776.54 |
| ENSMUSG00000026755 | -0.21750807 | 0.000747 | 0.006245 | 1470.851 | 1454.184 | 1627.23 | 1503.553 | 1778.603 | 1706.23 | 1732.828 | 1829.328 |
| ENSMUSG00000026761 | 0.122433604 | 0.337446 | 0.50584 | 1009.073 | 957.4576 | 779.2483 | 984.2534 | 758.6366 | 1003.402 | 794.5743 | 870.0729 |
| ENSMUSG00000026764 | 1.027960553 | 6.44E-05 | 0.000965 | 1412.5 | 1129.032 | 946.5259 | 1364.67 | 529.4854 | 947.5332 | 513.0018 | 390.4452 |
| ENSMUSG00000026765 | -0.10769443 | 0.514919 | 0.669147 | 275.6589 | 263.9607 | 395.835 | 308.8192 | 289.6081 | 328.508 | 380.8942 | 344.7664 |
| ENSMUSG00000026766 | -0.133809495 | 0.017073 | 0.063045 | 1963.818 | 2147.681 | 2048.736 | 2040.105 | 2247.632 | 2168.823 | 2241.008 | 2336.146 |
| ENSMUSG00000026767 | -0.016651901 | 0.818995 | 0.890502 | 855.1462 | 835.0758 | 820.6537 | 879.0133 | 833.7202 | 841.3827 | 829.2887 | 926.6276 |
| ENSMUSG00000026768 | 0.266049589 | 0.012183 | 0.049536 | 3254.586 | 4261.766 | 3535.187 | 3611.805 | 3431.417 | 2831.426 | 2981.582 | 2945.197 |
| ENSMUSG00000026771 | 0.329360476 | 0.033636 | 0.102746 | 921.5458 | 802.6806 | 755.2332 | 839.3327 | 632.8472 | 868.1997 | 574.7163 | 566.635 |
| ENSMUSG00000026773 | -0.140394187 | 0.355808 | 0.524431 | 589.5479 | 443.934 | 586.2995 | 516.7115 | 634.7975 | 481.5882 | 695.2525 | 543.7955 |
| ENSMUSG00000026775 | -0.074955593 | 0.205438 | 0.361894 | 3083.557 | 3119.536 | 2847.859 | 2901.865 | 3104.754 | 3049.314 | 3119.475 | 3310.627 |
| ENSMUSG00000026778 | -0.917633227 | 0.000116 | 0.001503 | 145.8779 | 97.18554 | 140.7781 | 96.61383 | 182.3458 | 178.7799 | 258.4295 | 289.2992 |
| ENSMUSG00000026779 | 0.081717245 | 0.502313 | 0.65843 | 388.337 | 442.7341 | 386.7258 | 393.3563 | 385.169 | 430.189 | 388.6085 | 316.489 |
| ENSMUSG00000026781 | -0.039671858 | 0.721562 | 0.825853 | 1167.023 | 1025.847 | 920.0265 | 1031.698 | 969.2606 | 969.8807 | 1162.933 | 1156.109 |
| ENSMUSG00000026782 | 0.364591959 | 0.011305 | 0.046782 | 1749.529 | 1961.708 | 1562.637 | 1729.56 | 1242.292 | 1754.277 | 1269.005 | 1173.511 |
| ENSMUSG00000026784 | -0.392539789 | 0.008099 | 0.036608 | 273.6468 | 334.7502 | 272.4471 | 305.3687 | 356.8907 | 362.0292 | 351.9655 | 485.0656 |
| ENSMUSG00000026785 | 0.03469976 | 0.792984 | 0.873246 | 312.8829 | 262.7609 | 236.0104 | 262.2375 | 277.9067 | 255.8787 | 236.2508 | 277.3357 |
| ENSMUSG00000026786 | 0.053466147 | 0.804634 | 0.880385 | 107.6478 | 148.7779 | 124.216 | 95.75121 | 146.2667 | 110.62 | 100.2861 | 100.0584 |

| | | | | | | | | | | | |
|---|---|---|---|---|---|---|---|---|---|---|---|
| ENSMUSG00000026787 | -0.212709502 | 0.644054 | 0.769762 | 11.0666 | 9.598572 | 8.281066 | 14.6646 | 10.72622 | 12.29112 | 12.53576 | 15.22628 |
| ENSMUSG00000026788 | -0.124656011 | 0.115922 | 0.243868 | 685.123 | 649.1034 | 658.3448 | 644.3797 | 778.1388 | 681.5982 | 686.5739 | 727.5984 |
| ENSMUSG00000026790 | 0.099742363 | 0.299619 | 0.467712 | 2493.003 | 2334.853 | 2300.48 | 2612.024 | 2518.713 | 2439.228 | 2103.115 | 2029.445 |
| ENSMUSG00000026791 | -0.07161116 | 0.630502 | 0.759418 | 608.6629 | 555.5174 | 811.5445 | 664.2201 | 612.3699 | 630.199 | 787.8242 | 747.1751 |
| ENSMUSG00000026792 | 0.240725196 | 0.027504 | 0.088898 | 417.5126 | 398.3407 | 353.6015 | 387.3179 | 313.9858 | 299.4563 | 343.2869 | 358.9051 |
| ENSMUSG00000026796 | -0.405451761 | 0.021522 | 0.074204 | 1732.426 | 1770.937 | 2533.178 | 1890.008 | 2842.449 | 1878.306 | 2932.403 | 2847.313 |
| ENSMUSG00000026797 | 0.051366419 | 0.584147 | 0.724628 | 737.4378 | 760.6869 | 833.0753 | 796.2015 | 742.0597 | 673.7766 | 766.6099 | 837.4451 |
| ENSMUSG00000026798 | -0.410041481 | 0.001172 | 0.008818 | 311.8769 | 363.5459 | 378.4447 | 336.4232 | 475.8543 | 383.2593 | 475.3946 | 512.2554 |
| ENSMUSG00000026799 | -0.261834796 | 0.03798 | 0.112482 | 320.9313 | 317.9527 | 354.4296 | 309.6818 | 397.8454 | 359.7945 | 345.2155 | 461.1386 |
| ENSMUSG00000026803 | 0.16479389 | 0.085978 | 0.19855 | 726.3712 | 705.4951 | 622.7362 | 684.0604 | 548.9877 | 640.2554 | 653.7881 | 598.1751 |
| ENSMUSG00000026806 | -0.058068776 | 0.69561 | 0.807828 | 338.0343 | 339.5495 | 242.6352 | 290.7041 | 301.3094 | 292.752 | 303.7511 | 359.9926 |
| ENSMUSG00000026807 | 0.070158578 | 0.811639 | 0.88532 | 56.33904 | 65.99018 | 47.20208 | 45.71905 | 65.33246 | 31.28647 | 58.82164 | 47.85401 |
| ENSMUSG00000026809 | -0.034190909 | 0.919482 | 0.952207 | 24.1453 | 22.79661 | 31.46805 | 19.84034 | 25.35289 | 29.05173 | 27.96439 | 18.48905 |
| ENSMUSG00000026810 | -0.019086895 | 0.819718 | 0.890713 | 840.0554 | 965.8563 | 831.4191 | 828.9812 | 895.1522 | 912.8946 | 823.5029 | 876.5984 |
| ENSMUSG00000026811 | 0.293645614 | 0.00415 | 0.022424 | 698.2017 | 788.2827 | 844.6688 | 784.1248 | 702.0801 | 659.2507 | 605.5736 | 575.3357 |
| ENSMUSG00000026812 | 0.071940555 | 0.41594 | 0.581824 | 1508.075 | 1420.589 | 1364.72 | 1450.07 | 1281.296 | 1446.999 | 1502.363 | 1232.241 |
| ENSMUSG00000026814 | -0.646367697 | 0.00364 | 0.020411 | 1085.533 | 928.6619 | 1618.12 | 955.7868 | 1801.031 | 1253.694 | 1839.864 | 2289.379 |
| ENSMUSG00000026816 | -0.080564577 | 0.348842 | 0.517452 | 928.5882 | 1071.441 | 1076.539 | 902.3042 | 1058.971 | 1051.449 | 1033.718 | 1061.489 |
| ENSMUSG00000026817 | -0.06422535 | 0.566084 | 0.710579 | 547.2936 | 621.5076 | 605.3459 | 609.8748 | 603.5939 | 548.6307 | 608.4665 | 731.9488 |
| ENSMUSG00000026819 | -0.306365924 | 0.056154 | 0.147653 | 486.9303 | 421.1374 | 388.382 | 365.7524 | 548.0126 | 393.3157 | 506.2518 | 605.7882 |
| ENSMUSG00000026820 | -0.783117775 | 0.00018 | 0.002091 | 431.5973 | 373.1445 | 524.1915 | 465.8167 | 767.4126 | 503.9357 | 799.3957 | 1020.16 |
| ENSMUSG00000026821 | 0.116488572 | 0.146492 | 0.285408 | 1153.944 | 1022.248 | 1164.318 | 1230.101 | 1081.398 | 1020.163 | 1090.611 | 1027.774 |
| ENSMUSG00000026822 | 0.246060837 | 0.234489 | 0.395334 | 823.9585 | 1130.232 | 804.9196 | 916.1062 | 877.6002 | 983.2892 | 730.9312 | 505.7299 |
| ENSMUSG00000026825 | 0.634315525 | 2.43E-05 | 0.000457 | 775.6679 | 1004.251 | 856.2623 | 891.9527 | 663.0757 | 651.4291 | 479.2517 | 478.5401 |
| ENSMUSG00000026826 | 0.583547431 | 0.056706 | 0.148787 | 324.9556 | 167.975 | 127.5284 | 185.4641 | 141.3911 | 165.3714 | 135.0005 | 95.70802 |
| ENSMUSG00000026827 | -0.110313306 | 0.251873 | 0.415496 | 541.2572 | 602.3104 | 583.8152 | 520.162 | 624.0712 | 555.3349 | 584.3592 | 660.1678 |
| ENSMUSG00000026828 | 0.217624403 | 0.318166 | 0.486502 | 117.7084 | 135.5798 | 146.5749 | 83.67448 | 108.2374 | 89.38993 | 112.8218 | 104.4087 |
| ENSMUSG00000026829 | 0.72298555 | 2.95E-07 | 1.35E-05 | 186.1201 | 178.7734 | 162.3089 | 169.0742 | 111.1627 | 106.1505 | 98.35749 | 105.4963 |
| ENSMUSG00000026832 | -0.203082925 | 0.591565 | 0.731496 | 160.9687 | 71.98929 | 125.0441 | 170.7995 | 88.73513 | 99.44629 | 181.2864 | 240.3576 |
| ENSMUSG00000026833 | 0.106696951 | 0.280432 | 0.446902 | 723.3531 | 677.8992 | 847.1531 | 805.6903 | 712.8064 | 665.955 | 750.217 | 711.2846 |
| ENSMUSG00000026834 | 2.393758305 | 0.00579 | 0.028665 | 112.6781 | 14.39786 | 8.281066 | 39.68068 | 12.67645 | 7.821619 | 11.57147 | 1.087591 |
| ENSMUSG00000026836 | 0.403187906 | 0.00535 | 0.026934 | 817.9222 | 765.4861 | 728.7338 | 937.6717 | 555.8134 | 764.2839 | 611.3593 | 527.4817 |
| ENSMUSG00000026837 | 0.375088635 | 0.018156 | 0.066009 | 3059.411 | 3829.83 | 4251.499 | 3814.521 | 2348.068 | 3720.856 | 2639.259 | 2825.562 |
| ENSMUSG00000026839 | -0.071435511 | 0.73074 | 0.831219 | 105.6357 | 98.38537 | 78.67013 | 118.1794 | 89.71024 | 101.681 | 129.2147 | 100.0584 |
| ENSMUSG00000026840 | 0.731191607 | 0.000584 | 0.005158 | 272.6407 | 232.7654 | 337.8675 | 248.4356 | 142.3663 | 119.559 | 192.8578 | 203.3795 |
| ENSMUSG00000026841 | 0.001758575 | 0.993968 | 0.99656 | 63.38142 | 80.38804 | 86.9512 | 76.77349 | 58.50668 | 69.27719 | 81.00029 | 98.97079 |
| ENSMUSG00000026842 | 0.067277492 | 0.138432 | 0.273954 | 3713.347 | 3859.826 | 3944.272 | 3842.988 | 3675.195 | 3557.719 | 3698.049 | 3730.437 |
| ENSMUSG00000026843 | 0.026126309 | 0.604141 | 0.741013 | 2268.653 | 2301.258 | 2235.06 | 2311.831 | 2336.367 | 2168.823 | 2222.687 | 2221.949 |
| ENSMUSG00000026848 | -0.09299469 | 0.136297 | 0.27095 | 1462.803 | 1562.168 | 1530.341 | 1599.304 | 1714.246 | 1579.967 | 1593.006 | 1677.065 |
| ENSMUSG00000026849 | -0.172515311 | 0.022404 | 0.076393 | 814.904 | 790.6824 | 881.9336 | 883.3265 | 940.9824 | 901.7209 | 966.2177 | 995.1458 |
| ENSMUSG00000026851 | 0.006066553 | 0.927815 | 0.957316 | 1538.257 | 1419.389 | 1420.203 | 1548.409 | 1519.223 | 1426.887 | 1422.327 | 1534.591 |
| ENSMUSG00000026853 | -0.787704561 | 0.000391 | 0.003796 | 697.1957 | 581.9134 | 932.4481 | 747.8946 | 1271.545 | 826.8568 | 1294.076 | 1719.482 |
| ENSMUSG00000026854 | -0.326204508 | 0.057576 | 0.150407 | 550.3117 | 452.3327 | 556.4877 | 545.178 | 589.9423 | 506.1705 | 683.681 | 860.2845 |
| ENSMUSG00000026856 | -0.315626486 | 0.006664 | 0.031853 | 342.0585 | 338.3497 | 372.648 | 312.2697 | 459.2774 | 366.4987 | 418.5015 | 454.6131 |
| ENSMUSG00000026857 | -0.255226172 | 0.000341 | 0.00341 | 884.3218 | 841.0749 | 850.4655 | 816.9044 | 971.2109 | 970.9981 | 1029.861 | 1076.715 |
| ENSMUSG00000026858 | -0.576763791 | 9.96E-05 | 0.001333 | 470.8334 | 502.7252 | 482.7862 | 438.2127 | 674.777 | 546.3959 | 732.8598 | 870.0729 |
| ENSMUSG00000026860 | -0.154158091 | 0.206197 | 0.362677 | 1083.521 | 1131.432 | 1348.158 | 1148.152 | 1381.733 | 1141.956 | 1198.611 | 1522.628 |
| ENSMUSG00000026864 | -0.283277201 | 0.004681 | 0.024587 | 10700.39 | 11367.11 | 11157.08 | 10061.64 | 11710.11 | 11894.45 | 14077.66 | 14993.53 |
| ENSMUSG00000026866 | -1.972272197 | 4.65E-05 | 0.000751 | 23.13925 | 11.99822 | 46.37397 | 24.15346 | 109.2125 | 48.04709 | 84.85745 | 176.1898 |
| ENSMUSG00000026867 | -2.30E-05 | 0.999648 | 0.999733 | 2511.112 | 2386.445 | 2346.026 | 2342.023 | 2418.276 | 2460.458 | 2332.615 | 2373.124 |
| ENSMUSG00000026869 | -0.213848532 | 0.006693 | 0.031944 | 1608.681 | 1895.718 | 1696.79 | 1707.132 | 2074.062 | 1848.137 | 1985.471 | 2099.051 |
| ENSMUSG00000026870 | 0.092202799 | 0.461903 | 0.623027 | 485.9243 | 598.7109 | 444.6933 | 535.6892 | 532.4108 | 498.3488 | 436.823 | 466.5766 |
| ENSMUSG00000026872 | -0.093611335 | 0.699815 | 0.810529 | 1334.028 | 945.4594 | 2086.829 | 1170.58 | 1229.615 | 1328.558 | 1397.255 | 1955.489 |
| ENSMUSG00000026873 | 0.221402778 | 0.259973 | 0.424007 | 567.4147 | 821.8778 | 520.051 | 671.9837 | 729.3833 | 446.9496 | 531.3233 | 503.5547 |
| ENSMUSG00000026875 | -0.669595321 | 0.020236 | 0.071055 | 347.0888 | 482.3283 | 306.3995 | 393.3563 | 899.0526 | 306.1605 | 638.3594 | 586.2116 |
| ENSMUSG00000026878 | 0.089538572 | 0.353887 | 0.522675 | 4566.481 | 4712.899 | 4456.87 | 4770.308 | 3886.794 | 5134.334 | 4261.194 | 4112.182 |
| ENSMUSG00000026879 | 0.427759373 | 0.000996 | 0.007811 | 4633.886 | 6435.843 | 6132.13 | 5724.37 | 4189.078 | 5039.357 | 3943.943 | 3871.824 |
| ENSMUSG00000026880 | -0.192794161 | 0.067191 | 0.167398 | 717.3168 | 728.2917 | 823.9661 | 866.074 | 1016.066 | 863.7302 | 862.0745 | 845.0583 |
| ENSMUSG00000026883 | 0.120762144 | 0.162072 | 0.306162 | 1726.389 | 1656.954 | 2066.954 | 1767.516 | 1662.565 | 1674.944 | 1700.042 | 1605.284 |
| ENSMUSG00000026885 | -0.669226715 | 0.014738 | 0.056929 | 58.35115 | 80.38804 | 76.18581 | 58.6584 | 105.312 | 65.92507 | 141.7505 | 120.7226 |
| ENSMUSG00000026887 | -0.171103226 | 0.075354 | 0.181486 | 523.1483 | 608.3095 | 573.8779 | 646.9676 | 710.8561 | 644.7249 | 640.288 | 652.5547 |
| ENSMUSG00000026888 | 0.354871663 | 0.006387 | 0.030873 | 669.0261 | 717.4933 | 806.5759 | 835.8822 | 478.7797 | 649.1943 | 627.7522 | 615.5766 |
| ENSMUSG00000026889 | 0.153268173 | 0.062572 | 0.15955 | 1668.038 | 1682.15 | 1517.091 | 1665.726 | 1429.513 | 1623.545 | 1480.184 | 1339.912 |
| ENSMUSG00000026890 | -0.14230224 | 0.644959 | 0.770145 | 79.47829 | 67.19001 | 112.6225 | 69.8725 | 90.68535 | 55.8687 | 83.89316 | 133.7737 |

| | | | | | | | | | | | |
|---|---|---|---|---|---|---|---|---|---|---|---|
| ENSMUSG00000026893 | 0.000930539 | 0.995561 | 0.997547 | 543.2694 | 472.7297 | 586.2995 | 577.0951 | 414.4223 | 483.823 | 616.1808 | 665.6057 |
| ENSMUSG00000026895 | -0.557363375 | 8.30E-05 | 0.001166 | 2230.423 | 2136.882 | 2335.261 | 2247.997 | 3292.951 | 2447.049 | 3405.869 | 4026.262 |
| ENSMUSG00000026896 | 0.497734857 | 0.072823 | 0.177423 | 302.8224 | 554.3175 | 521.7072 | 222.5569 | 256.4543 | 271.5219 | 300.8582 | 304.5255 |
| ENSMUSG00000026904 | 0.060443166 | 0.8314 | 0.897939 | 45.27245 | 55.19179 | 33.12427 | 39.68068 | 34.1289 | 52.51658 | 40.50014 | 38.06569 |
| ENSMUSG00000026914 | -0.20505356 | 0.007046 | 0.033083 | 1612.705 | 1872.921 | 1685.197 | 1563.074 | 1912.193 | 1899.536 | 1923.757 | 2020.744 |
| ENSMUSG00000026915 | -0.004126619 | 0.962202 | 0.978478 | 2096.617 | 1952.11 | 2081.032 | 2110.84 | 1825.408 | 2307.378 | 2005.721 | 2130.591 |
| ENSMUSG00000026917 | -0.114607568 | 0.103466 | 0.226171 | 1964.824 | 1970.107 | 2051.22 | 1828.762 | 2118.917 | 2262.683 | 2109.865 | 1969.627 |
| ENSMUSG00000026918 | 0.413159784 | 0.015093 | 0.057923 | 4894.454 | 5143.635 | 4146.33 | 4610.723 | 3365.109 | 4875.103 | 3015.332 | 2859.277 |
| ENSMUSG00000026921 | -0.365658505 | 0.016192 | 0.060702 | 1795.807 | 1636.557 | 2572.099 | 1992.66 | 2523.588 | 2305.143 | 2500.402 | 2980 |
| ENSMUSG00000026922 | -0.352404101 | 0.002508 | 0.015472 | 706.2502 | 583.1133 | 737.0149 | 579.683 | 861.9984 | 770.9881 | 789.7528 | 907.051 |
| ENSMUSG00000026923 | 0.025492624 | 0.768054 | 0.856029 | 1123.763 | 955.0579 | 1098.898 | 1035.148 | 957.5593 | 1022.397 | 1114.718 | 1047.35 |
| ENSMUSG00000026924 | 0.063783359 | 0.262838 | 0.427169 | 2084.545 | 2026.499 | 2057.845 | 1950.392 | 1863.438 | 2038.09 | 1917.007 | 1951.138 |
| ENSMUSG00000026925 | -0.082993895 | 0.372913 | 0.541843 | 757.5589 | 663.5013 | 772.6235 | 795.3389 | 821.0437 | 716.2368 | 830.253 | 801.5546 |
| ENSMUSG00000026926 | -0.163659563 | 0.042906 | 0.122582 | 2037.26 | 1944.911 | 2069.438 | 2053.044 | 2396.824 | 1997.865 | 2242.937 | 2441.642 |
| ENSMUSG00000026927 | -0.008544521 | 0.897252 | 0.937817 | 1809.892 | 1619.759 | 1628.058 | 1746.813 | 1727.897 | 1626.897 | 1742.471 | 1747.759 |
| ENSMUSG00000026928 | 0.529770618 | 0.042993 | 0.122716 | 54.32694 | 53.99197 | 50.5145 | 37.09281 | 32.17867 | 31.28647 | 34.71441 | 36.9781 |
| ENSMUSG00000026930 | 0.277329538 | 0.000286 | 0.002998 | 1223.362 | 1317.404 | 1228.91 | 1243.903 | 1102.851 | 1084.97 | 997.075 | 949.467 |
| ENSMUSG00000026932 | -0.063336784 | 0.655939 | 0.778221 | 698.2017 | 755.8876 | 604.5178 | 664.2201 | 723.5326 | 865.9649 | 573.7521 | 680.832 |
| ENSMUSG00000026933 | 0.155819586 | 0.105497 | 0.2292 | 1563.408 | 1610.16 | 1533.653 | 1489.751 | 1382.708 | 1612.371 | 1341.326 | 1225.715 |
| ENSMUSG00000026938 | 1.479339163 | 0.00099 | 0.007785 | 118.7144 | 341.9491 | 156.5122 | 171.6621 | 73.13335 | 129.6154 | 45.32159 | 34.80291 |
| ENSMUSG00000026939 | -0.117903191 | 0.494308 | 0.651128 | 352.119 | 278.3586 | 374.3042 | 262.2375 | 342.2641 | 283.813 | 335.5726 | 414.3722 |
| ENSMUSG00000026940 | 0.317391589 | 0.624554 | 0.755259 | 20.12109 | 19.19714 | 12.4216 | 12.07673 | 1.950223 | 22.34748 | 9.642892 | 17.40146 |
| ENSMUSG00000026941 | 0.271924532 | 0.218106 | 0.37673 | 249.5015 | 289.157 | 156.5122 | 226.0074 | 192.0969 | 236.8833 | 173.572 | 158.7883 |
| ENSMUSG00000026942 | -0.160328976 | 0.114746 | 0.242317 | 544.2754 | 526.7217 | 537.4412 | 459.7783 | 597.7432 | 517.3442 | 620.0379 | 573.1605 |
| ENSMUSG00000026944 | -0.063258113 | 0.599497 | 0.737591 | 1762.607 | 1617.359 | 1966.753 | 1518.217 | 1644.038 | 1600.08 | 2069.365 | 1859.781 |
| ENSMUSG00000026946 | 0.016802991 | 0.919882 | 0.952339 | 396.3854 | 544.719 | 616.1113 | 401.1199 | 472.929 | 444.7149 | 521.6804 | 495.9415 |
| ENSMUSG00000026950 | 0.346497108 | 0.377373 | 0.545463 | 143.8658 | 86.38715 | 89.43552 | 108.6906 | 42.9049 | 157.5497 | 66.53595 | 70.69342 |
| ENSMUSG00000026955 | -0.145161228 | 0.111951 | 0.238489 | 508.0574 | 476.3291 | 458.7711 | 499.459 | 588.9672 | 529.6353 | 501.4304 | 527.4817 |
| ENSMUSG00000026956 | -1.192911422 | 5.06E-05 | 0.000799 | 380.2885 | 279.5584 | 664.9696 | 360.5766 | 1109.677 | 568.7434 | 971.0392 | 1207.226 |
| ENSMUSG00000026958 | -0.165685584 | 0.215396 | 0.373588 | 471.8395 | 469.1302 | 547.3785 | 511.5357 | 650.3992 | 445.8323 | 577.6092 | 569.8977 |
| ENSMUSG00000026959 | -0.011851137 | 0.972582 | 0.984801 | 28.16952 | 22.79661 | 19.87456 | 34.50494 | 19.50223 | 25.6996 | 30.85725 | 30.45255 |
| ENSMUSG00000026960 | 0.019236359 | 0.814347 | 0.887225 | 789.7527 | 869.8706 | 737.0149 | 844.5084 | 793.7406 | 768.7534 | 825.4315 | 805.905 |
| ENSMUSG00000026965 | -0.017598015 | 0.759176 | 0.84986 | 3026.212 | 3263.515 | 3334.785 | 3359.919 | 3286.125 | 3182.281 | 3281.476 | 3396.547 |
| ENSMUSG00000026966 | -0.248898055 | 0.005014 | 0.025782 | 1124.769 | 1275.41 | 1206.551 | 1136.075 | 1448.04 | 1255.928 | 1388.576 | 1540.029 |
| ENSMUSG00000026970 | 0.342203989 | 0.015023 | 0.057753 | 4226.434 | 4093.791 | 3994.786 | 4269.124 | 2845.375 | 4310.829 | 3111.761 | 2815.773 |
| ENSMUSG00000026971 | -1.661881406 | 0.001615 | 0.011128 | 65.39353 | 43.19358 | 253.4006 | 73.323 | 274.0063 | 186.6015 | 320.144 | 600.3503 |
| ENSMUSG00000026972 | -0.180301283 | 0.054527 | 0.144964 | 721.341 | 616.7083 | 729.5619 | 669.3958 | 764.4873 | 738.5843 | 854.3602 | 746.0875 |
| ENSMUSG00000026974 | -0.339423859 | 9.28E-06 | 0.000214 | 1065.412 | 1131.432 | 1076.539 | 987.7039 | 1407.086 | 1279.393 | 1413.648 | 1284.445 |
| ENSMUSG00000026975 | -0.062705382 | 0.532738 | 0.684218 | 618.7234 | 659.9018 | 616.1113 | 597.7981 | 723.5326 | 679.3635 | 567.9663 | 630.8028 |
| ENSMUSG00000026976 | 0.032416751 | 0.692685 | 0.805506 | 3210.319 | 3059.545 | 3294.208 | 2748.318 | 2779.067 | 3048.197 | 3186.976 | 3024.591 |
| ENSMUSG00000026977 | 0.161934884 | 0.078862 | 0.187333 | 2821.982 | 2986.356 | 2625.926 | 3076.115 | 2371.471 | 2921.933 | 2455.08 | 2540.613 |
| ENSMUSG00000026979 | -0.015552556 | 0.929721 | 0.958527 | 173.0413 | 177.5736 | 187.1521 | 154.4096 | 202.8232 | 128.498 | 192.8578 | 174.0146 |
| ENSMUSG00000026980 | -0.356427529 | 0.055222 | 0.146226 | 89.53884 | 130.7805 | 104.3414 | 127.6683 | 121.8889 | 147.4934 | 161.0363 | 147.9124 |
| ENSMUSG00000026981 | 1.159544463 | 0.009734 | 0.041731 | 55.33299 | 69.58965 | 72.87338 | 67.28463 | 55.58134 | 16.76061 | 33.75012 | 11.9635 |
| ENSMUSG00000026986 | 0.33236157 | 0.125558 | 0.256106 | 77.46619 | 93.58608 | 78.67013 | 102.6522 | 69.2329 | 74.86406 | 81.96458 | 53.29196 |
| ENSMUSG00000026987 | 0.424828546 | 0.014826 | 0.057195 | 2930.636 | 2598.813 | 2250.794 | 2660.331 | 1788.354 | 2664.937 | 1821.542 | 1503.051 |
| ENSMUSG00000026988 | 0.183998059 | 0.037136 | 0.110724 | 880.2976 | 957.4576 | 871.1682 | 1038.599 | 859.0731 | 813.4483 | 779.1456 | 847.2335 |
| ENSMUSG00000026989 | -0.277397695 | 0.276141 | 0.442363 | 34.20585 | 35.99465 | 40.57722 | 30.19182 | 43.88001 | 42.46022 | 46.28588 | 38.06569 |
| ENSMUSG00000026991 | -0.400651537 | 0.00016 | 0.001911 | 2401.452 | 2153.68 | 2727.783 | 2437.774 | 3055.999 | 2822.487 | 3522.548 | 3435.7 |
| ENSMUSG00000026994 | -1.110557889 | 0.000722 | 0.006102 | 264.5923 | 214.7681 | 539.9255 | 265.688 | 574.3406 | 423.4848 | 974.8963 | 804.8174 |
| ENSMUSG00000026999 | -0.113666259 | 0.428328 | 0.593351 | 417.5126 | 358.7466 | 287.353 | 313.1323 | 361.7663 | 349.7381 | 365.4656 | 410.0218 |
| ENSMUSG00000027001 | -0.03781136 | 0.768982 | 0.856611 | 229.3804 | 249.5629 | 245.1196 | 223.4195 | 250.6036 | 202.2447 | 250.7152 | 267.5474 |
| ENSMUSG00000027002 | -0.035104468 | 0.489344 | 0.646944 | 4745.558 | 5057.248 | 4721.864 | 4702.161 | 5043.276 | 4745.488 | 4861.946 | 5042.072 |
| ENSMUSG00000027004 | 0.157379548 | 0.445524 | 0.60836 | 1616.729 | 854.2729 | 885.246 | 1189.558 | 1064.822 | 1005.637 | 1140.754 | 864.6349 |
| ENSMUSG00000027006 | 0.05314436 | 0.306089 | 0.474682 | 3929.648 | 4178.978 | 3811.775 | 3928.387 | 3866.316 | 3913.044 | 3791.585 | 3697.81 |
| ENSMUSG00000027007 | 0.182756881 | 0.05896 | 0.152725 | 3408.512 | 3497.48 | 2787.407 | 3434.104 | 2899.006 | 2775.557 | 3161.904 | 2724.416 |
| ENSMUSG00000027009 | 0.404024961 | 0.029488 | 0.09342 | 215.2956 | 250.7627 | 351.1172 | 248.4356 | 183.3209 | 214.5358 | 182.2507 | 227.3065 |
| ENSMUSG00000027010 | -0.28201544 | 0.056072 | 0.147526 | 1548.318 | 1183.024 | 1427.656 | 1457.834 | 1688.893 | 1295.037 | 1931.471 | 1915.248 |
| ENSMUSG00000027011 | 0.052114998 | 0.581516 | 0.722532 | 2773.692 | 3318.706 | 3118.65 | 3376.308 | 2991.641 | 3406.874 | 2911.189 | 2832.087 |
| ENSMUSG00000027012 | 0.129427767 | 0.151364 | 0.291726 | 4730.468 | 4624.112 | 4379.028 | 4799.637 | 3852.665 | 4811.413 | 4399.087 | 3880.525 |
| ENSMUSG00000027014 | -0.085291899 | 0.201255 | 0.356773 | 1174.065 | 1148.229 | 1222.285 | 1130.037 | 1323.226 | 1208.999 | 1196.683 | 1231.153 |
| ENSMUSG00000027015 | 0.796971248 | 2.71E-05 | 0.000495 | 388.337 | 375.5441 | 346.9767 | 339.011 | 165.7689 | 293.8694 | 191.8935 | 183.8029 |
| ENSMUSG00000027016 | -0.63996932 | 0.002199 | 0.014063 | 70.4238 | 76.78858 | 79.49824 | 79.36136 | 98.48624 | 112.8548 | 108.0004 | 158.7883 |
| ENSMUSG00000027018 | -0.103522513 | 0.11104 | 0.237177 | 1506.063 | 1604.161 | 1466.577 | 1441.444 | 1601.133 | 1667.122 | 1536.113 | 1657.489 |

| | | | | | | | | | | | |
|---|---|---|---|---|---|---|---|---|---|---|---|
| ENSMUSG00000027030 | -0.431529531 | 0.008852 | 0.039036 | 1132.817 | 788.2827 | 1092.273 | 895.4032 | 1109.677 | 1127.43 | 1530.327 | 1506.314 |
| ENSMUSG00000027035 | -0.013196299 | 0.902427 | 0.94145 | 921.5458 | 854.2729 | 952.3226 | 935.9465 | 832.7451 | 1099.496 | 902.5747 | 866.8101 |
| ENSMUSG00000027067 | -0.13998615 | 0.073808 | 0.179118 | 6266.713 | 7681.257 | 7009.923 | 6637.888 | 7922.779 | 7285.279 | 7757.706 | 7435.86 |
| ENSMUSG00000027070 | -2.241107802 | 7.91E-06 | 0.000192 | 1172.053 | 447.5334 | 2086.001 | 1198.184 | 5810.688 | 2947.633 | 5414.484 | 9015.043 |
| ENSMUSG00000027071 | -0.894160036 | 0.093121 | 0.209847 | 18.10898 | 4.799286 | 9.93728 | 9.488858 | 11.70134 | 17.87799 | 29.89296 | 19.57664 |
| ENSMUSG00000027074 | 0.100432241 | 0.439248 | 0.602643 | 2110.702 | 1960.508 | 1758.07 | 2306.655 | 1821.508 | 1945.348 | 2234.258 | 1586.795 |
| ENSMUSG00000027075 | -0.056832771 | 0.684231 | 0.798746 | 516.1059 | 488.3274 | 403.2879 | 498.5964 | 510.9583 | 449.1844 | 590.145 | 429.5985 |
| ENSMUSG00000027076 | -0.500490351 | 0.000763 | 0.006351 | 425.561 | 412.7386 | 501.8326 | 425.2734 | 660.1504 | 474.884 | 638.3594 | 725.4233 |
| ENSMUSG00000027078 | 0.646826733 | 4.67E-06 | 0.000127 | 475.8637 | 585.5129 | 643.4389 | 513.261 | 349.0898 | 417.8979 | 319.1797 | 331.7153 |
| ENSMUSG00000027079 | -0.158965055 | 0.122202 | 0.251736 | 443.67 | 428.3363 | 433.0998 | 433.037 | 521.6845 | 412.311 | 511.0733 | 493.7664 |
| ENSMUSG00000027080 | -0.091795584 | 0.316571 | 0.4848 | 789.7527 | 974.2551 | 862.8871 | 852.272 | 870.7744 | 988.8761 | 946.932 | 898.3502 |
| ENSMUSG00000027082 | 0.404312358 | 1.97E-05 | 0.000388 | 1053.339 | 1109.835 | 977.9939 | 1023.071 | 705.9806 | 883.8429 | 804.2172 | 751.5254 |
| ENSMUSG00000027086 | -0.195488228 | 0.135658 | 0.27012 | 339.0403 | 271.1597 | 271.619 | 340.7363 | 342.2641 | 315.0995 | 370.287 | 373.0437 |
| ENSMUSG00000027087 | -0.010955081 | 0.862642 | 0.917774 | 2619.766 | 2379.246 | 2619.301 | 2715.539 | 2499.21 | 2642.59 | 2597.795 | 2680.912 |
| ENSMUSG00000027088 | -0.132486966 | 0.216152 | 0.374544 | 453.7305 | 394.7413 | 374.3042 | 416.6471 | 456.3521 | 421.25 | 424.2872 | 494.8539 |
| ENSMUSG00000027091 | -8.16E-05 | 0.999037 | 0.999528 | 3737.492 | 4147.783 | 3489.641 | 3794.681 | 3831.212 | 3872.819 | 3746.263 | 3713.036 |
| ENSMUSG00000027099 | -0.18236282 | 0.003767 | 0.020938 | 1895.406 | 1874.121 | 2051.22 | 1957.293 | 2294.437 | 2068.259 | 2229.437 | 2237.175 |
| ENSMUSG00000027102 | -0.301912607 | 0.031552 | 0.09787 | 2722.383 | 2342.052 | 2760.908 | 2461.927 | 2424.127 | 3010.206 | 3703.835 | 3545.547 |
| ENSMUSG00000027104 | 0.288538641 | 0.008548 | 0.038069 | 2166.035 | 2219.67 | 1942.738 | 2254.898 | 1627.461 | 2093.959 | 1735.72 | 1570.482 |
| ENSMUSG00000027107 | 0.894556369 | 0.114232 | 0.241664 | 26.15741 | 25.19625 | 14.90592 | 6.900988 | 9.751113 | 5.58687 | 11.57147 | 11.9635 |
| ENSMUSG00000027108 | -0.072190697 | 0.469045 | 0.629175 | 1534.233 | 1942.511 | 1516.263 | 1712.308 | 1695.719 | 1842.55 | 1675.935 | 1831.503 |
| ENSMUSG00000027109 | 0.173675301 | 0.06433 | 0.162952 | 2920.576 | 3393.095 | 2953.856 | 3206.372 | 2660.104 | 3168.873 | 2660.474 | 2568.89 |
| ENSMUSG00000027111 | -0.388413766 | 0.01719 | 0.063383 | 3186.174 | 2206.472 | 2986.153 | 2688.797 | 3113.53 | 3014.675 | 3827.264 | 4534.167 |
| ENSMUSG00000027115 | 0.409654345 | 0.007561 | 0.034739 | 433.6094 | 430.7359 | 296.4622 | 417.5098 | 309.1103 | 281.5783 | 273.8581 | 321.927 |
| ENSMUSG00000027122 | 0.434124766 | 0.000289 | 0.003017 | 1841.079 | 1990.504 | 1606.527 | 1814.097 | 1303.724 | 1619.075 | 1194.754 | 1249.642 |
| ENSMUSG00000027130 | 0.382076441 | 0.008534 | 0.038027 | 1387.349 | 1262.212 | 1079.023 | 1446.62 | 945.858 | 1235.816 | 936.3248 | 853.759 |
| ENSMUSG00000027131 | -0.259668429 | 0.001467 | 0.010407 | 1048.309 | 1107.435 | 1126.225 | 1167.13 | 1366.131 | 1193.356 | 1335.54 | 1432.357 |
| ENSMUSG00000027132 | 0.085315253 | 0.442173 | 0.605393 | 748.5044 | 814.6788 | 701.4063 | 801.3772 | 749.8606 | 831.3263 | 667.2881 | 640.5912 |
| ENSMUSG00000027133 | -0.375716294 | 0.001433 | 0.010245 | 1218.332 | 1274.21 | 1103.038 | 1123.998 | 1647.938 | 1230.229 | 1548.648 | 1692.292 |
| ENSMUSG00000027134 | -0.038870961 | 0.658692 | 0.780183 | 487.9364 | 512.3238 | 525.0196 | 475.3055 | 495.3565 | 502.8183 | 497.5732 | 560.1094 |
| ENSMUSG00000027160 | 0.27382381 | 0.086879 | 0.200155 | 2200.241 | 2363.648 | 1888.911 | 2167.773 | 1622.585 | 2399.002 | 1474.398 | 1634.649 |
| ENSMUSG00000027162 | 0.010719761 | 0.845521 | 0.906998 | 2514.13 | 2488.43 | 2317.042 | 2425.697 | 2372.446 | 2521.913 | 2398.187 | 2377.474 |
| ENSMUSG00000027163 | -0.172441493 | 0.040078 | 0.116897 | 849.1099 | 842.2747 | 842.1844 | 805.6903 | 949.7584 | 827.9742 | 993.2178 | 989.7079 |
| ENSMUSG00000027164 | -0.100408546 | 0.253669 | 0.417294 | 1043.278 | 884.2685 | 1102.21 | 1075.691 | 1090.174 | 1076.031 | 1132.075 | 1109.343 |
| ENSMUSG00000027165 | 0.213660985 | 0.037966 | 0.112471 | 737.4378 | 878.2694 | 782.5608 | 885.9143 | 715.7317 | 784.3966 | 646.0737 | 686.27 |
| ENSMUSG00000027166 | -0.026026369 | 0.869145 | 0.921481 | 309.8647 | 298.7556 | 269.1347 | 332.11 | 304.2347 | 232.4138 | 346.1798 | 348.0291 |
| ENSMUSG00000027167 | 0.045832818 | 0.579001 | 0.720277 | 470.8334 | 523.1222 | 522.5353 | 524.4751 | 501.2072 | 494.9967 | 477.3231 | 504.6423 |
| ENSMUSG00000027168 | 0.177305118 | 0.492517 | 0.649627 | 80.48435 | 95.98572 | 77.84202 | 98.33908 | 58.50668 | 115.0895 | 71.3574 | 67.43065 |
| ENSMUSG00000027170 | -0.214172536 | 7.73E-05 | 0.001114 | 5489.033 | 5868.327 | 5788.465 | 5933.987 | 6990.573 | 6441.662 | 6843.56 | 6495.094 |
| ENSMUSG00000027171 | -0.145030011 | 0.487486 | 0.645366 | 107.6478 | 79.18822 | 95.23226 | 78.49874 | 107.2622 | 80.45093 | 91.60747 | 119.635 |
| ENSMUSG00000027173 | -0.485635002 | 0.00255 | 0.015655 | 255.5378 | 285.5575 | 366.8512 | 305.3687 | 397.8454 | 359.7945 | 433.9301 | 510.0802 |
| ENSMUSG00000027175 | 0.006572266 | 0.953244 | 0.973337 | 265.5983 | 334.7502 | 321.3054 | 298.4677 | 302.2845 | 290.5173 | 309.5368 | 311.0511 |
| ENSMUSG00000027176 | 0.100615448 | 0.304956 | 0.473481 | 1187.144 | 1177.025 | 1059.148 | 1132.625 | 998.514 | 1238.05 | 1009.611 | 1002.759 |
| ENSMUSG00000027177 | 0.063010174 | 0.487895 | 0.645722 | 2236.459 | 2074.491 | 2346.854 | 2435.186 | 1951.198 | 2391.181 | 2267.044 | 2100.138 |
| ENSMUSG00000027180 | 0.081582891 | 0.256333 | 0.42008 | 2515.136 | 2481.231 | 2174.608 | 2493.845 | 2298.337 | 2382.242 | 2268.972 | 2180.62 |
| ENSMUSG00000027184 | -0.043014195 | 0.364418 | 0.533108 | 9308.015 | 9869.732 | 9198.608 | 9431.063 | 9832.047 | 9455.22 | 9561.891 | 10096.11 |
| ENSMUSG00000027185 | 0.101661213 | 0.197317 | 0.35165 | 1272.659 | 1141.03 | 1081.507 | 1179.206 | 1065.797 | 1162.069 | 1062.647 | 1065.839 |
| ENSMUSG00000027186 | -0.557199872 | 0.177837 | 0.327072 | 308.8587 | 176.3738 | 790.0137 | 232.0457 | 474.8792 | 398.9026 | 614.2522 | 731.9488 |
| ENSMUSG00000027187 | -0.065272622 | 0.529865 | 0.681904 | 2544.311 | 2514.826 | 2452.852 | 2644.804 | 2867.802 | 2153.18 | 2756.903 | 2847.313 |
| ENSMUSG00000027188 | 0.12877096 | 0.316737 | 0.484882 | 998.0059 | 722.2926 | 910.0892 | 1036.011 | 791.7904 | 834.6784 | 922.8247 | 806.9926 |
| ENSMUSG00000027189 | -0.022374785 | 0.67303 | 0.790437 | 4847.17 | 4840.08 | 5029.092 | 5114.495 | 4790.722 | 5242.719 | 4946.803 | 5168.233 |
| ENSMUSG00000027193 | -0.147170467 | 0.020671 | 0.072118 | 3509.118 | 3678.653 | 3564.999 | 3561.772 | 4167.626 | 4132.049 | 3626.692 | 3925.116 |
| ENSMUSG00000027194 | 0.010102653 | 0.886926 | 0.932224 | 778.6861 | 837.4754 | 738.6711 | 784.9874 | 771.313 | 775.4576 | 779.1456 | 787.4159 |
| ENSMUSG00000027195 | -0.214947054 | 0.000106 | 0.001401 | 1970.86 | 2117.685 | 2005.674 | 1965.056 | 2386.097 | 2238.1 | 2390.473 | 2331.795 |
| ENSMUSG00000027196 | 0.15795642 | 0.728608 | 0.829664 | 26.15741 | 23.99643 | 24.8432 | 29.3292 | 25.35289 | 41.34284 | 17.3572 | 9.78832 |
| ENSMUSG00000027198 | -0.287717855 | 0.009246 | 0.040236 | 2757.595 | 2220.87 | 3066.479 | 2617.2 | 3256.872 | 2959.924 | 3491.691 | 3311.715 |
| ENSMUSG00000027199 | -3.002615965 | 1.86E-09 | 2.20E-07 | 205.2351 | 153.5772 | 469.5365 | 313.995 | 2908.757 | 1078.266 | 1644.113 | 3531.408 |
| ENSMUSG00000027200 | 0.04881122 | 0.728715 | 0.829692 | 802.8314 | 782.2836 | 1091.445 | 890.2274 | 898.0775 | 762.0491 | 799.3957 | 991.8831 |
| ENSMUSG00000027201 | 0.159512411 | 0.022614 | 0.076904 | 3246.537 | 3085.941 | 2809.766 | 3240.014 | 2867.802 | 2794.553 | 2638.295 | 2784.233 |
| ENSMUSG00000027202 | -2.418862845 | 0.030377 | 0.095285 | 567.4147 | 67.19001 | 5578.126 | 899.7163 | 7565.889 | 3592.358 | 12942.69 | 13935.3 |
| ENSMUSG00000027203 | -0.041875118 | 0.668462 | 0.787337 | 2026.193 | 2399.643 | 2138.171 | 2102.213 | 2354.894 | 2482.805 | 2016.329 | 2065.335 |
| ENSMUSG00000027204 | 0.523964074 | 7.97E-06 | 0.000193 | 4061.441 | 5171.231 | 4678.802 | 4848.807 | 3200.315 | 3858.293 | 3145.511 | 2842.963 |
| ENSMUSG00000027206 | 0.047439401 | 0.510599 | 0.666216 | 3589.602 | 3675.053 | 3403.518 | 3838.675 | 3421.666 | 3824.772 | 3501.334 | 3289.963 |
| ENSMUSG00000027207 | -0.217007601 | 0.002265 | 0.014359 | 1432.621 | 1605.361 | 1371.345 | 1468.185 | 1730.823 | 1701.761 | 1754.042 | 1636.825 |

| | | | | | | | | | | | |
|---|---|---|---|---|---|---|---|---|---|---|---|
| ENSMUSG00000027208 | 1.080370626 | 0.020012 | 0.070419 | 48.29061 | 71.98929 | 44.71776 | 72.46037 | 28.27823 | 50.28183 | 25.07152 | 8.700729 |
| ENSMUSG00000027210 | 0.407964636 | 0.009373 | 0.040618 | 6201.319 | 7268.519 | 5602.141 | 6796.611 | 4933.088 | 6337.746 | 4302.658 | 3922.941 |
| ENSMUSG00000027215 | -0.526082389 | 0.018781 | 0.067463 | 1144.89 | 1145.83 | 1803.616 | 1111.922 | 1613.809 | 1311.797 | 2186.044 | 2387.262 |
| ENSMUSG00000027217 | -0.215636189 | 0.009113 | 0.039867 | 1638.863 | 1467.382 | 1844.193 | 1588.952 | 1907.318 | 1874.954 | 1910.257 | 1907.635 |
| ENSMUSG00000027219 | -0.380377493 | 0.377674 | 0.545752 | 12.07265 | 26.39607 | 13.24971 | 15.52722 | 21.45245 | 25.6996 | 13.50005 | 26.10219 |
| ENSMUSG00000027220 | -0.083280661 | 0.539465 | 0.689165 | 207.2472 | 253.1623 | 243.4633 | 246.7103 | 221.3503 | 279.3435 | 273.8581 | 232.7445 |
| ENSMUSG00000027221 | -0.495346367 | 0.003652 | 0.020442 | 191.1503 | 195.5709 | 219.4483 | 167.349 | 297.4089 | 203.3621 | 304.7154 | 283.8613 |
| ENSMUSG00000027222 | -0.270358913 | 0.151233 | 0.291565 | 603.6326 | 433.1356 | 462.0835 | 467.5419 | 675.7521 | 415.6632 | 571.8235 | 708.0218 |
| ENSMUSG00000027223 | -0.348464255 | 0.00021 | 0.002365 | 1302.84 | 1602.962 | 1396.188 | 1433.68 | 1957.048 | 1928.588 | 1743.435 | 1669.452 |
| ENSMUSG00000027227 | -1.142881087 | 2.26E-05 | 0.000433 | 1100.623 | 861.4719 | 1184.192 | 1178.344 | 2807.345 | 1234.698 | 2033.686 | 3475.941 |
| ENSMUSG00000027230 | -0.21039835 | 0.317362 | 0.485594 | 268.6165 | 320.3523 | 491.8953 | 319.1707 | 334.4632 | 407.8415 | 388.6085 | 491.5912 |
| ENSMUSG00000027236 | 0.031401813 | 0.667466 | 0.786689 | 2054.363 | 2070.892 | 1830.944 | 1882.244 | 1821.508 | 2015.743 | 1856.257 | 1972.89 |
| ENSMUSG00000027238 | 1.130964371 | 0.001471 | 0.010426 | 386.3249 | 130.7805 | 147.403 | 179.4257 | 101.4116 | 118.4417 | 73.28598 | 92.44524 |
| ENSMUSG00000027239 | 0.283345555 | 0.003483 | 0.019708 | 12260.78 | 14504.64 | 13351.56 | 13553.54 | 11619.43 | 12433.02 | 9961.107 | 10086.32 |
| ENSMUSG00000027242 | 0.211924716 | 0.134784 | 0.268723 | 549.3057 | 555.5174 | 460.4273 | 462.3662 | 419.2979 | 538.5743 | 380.8942 | 411.1094 |
| ENSMUSG00000027243 | -0.145373525 | 0.277704 | 0.444018 | 361.1735 | 311.9536 | 292.3216 | 340.7363 | 409.5467 | 311.7474 | 388.6085 | 332.8029 |
| ENSMUSG00000027244 | 0.094527838 | 0.249166 | 0.412654 | 1320.949 | 1270.611 | 1287.706 | 1318.089 | 1313.475 | 1192.238 | 1283.469 | 1076.715 |
| ENSMUSG00000027245 | 0.168475944 | 0.076854 | 0.184138 | 734.4197 | 743.8894 | 681.5318 | 822.9428 | 688.4286 | 705.0631 | 655.7166 | 604.7006 |
| ENSMUSG00000027246 | -0.439630614 | 0.130313 | 0.262675 | 41.24823 | 23.99643 | 38.92101 | 35.36756 | 53.63112 | 56.98608 | 36.64299 | 43.50364 |
| ENSMUSG00000027247 | -0.080620516 | 0.305316 | 0.47378 | 2122.775 | 2498.028 | 2419.728 | 2290.265 | 2603.547 | 2273.856 | 2476.295 | 2510.16 |
| ENSMUSG00000027248 | -0.06929634 | 0.306977 | 0.475531 | 11870.44 | 13557.98 | 13265.44 | 12419.19 | 12587.71 | 13469.94 | 13400.73 | 14169.14 |
| ENSMUSG00000027249 | 2.723884424 | 0.009283 | 0.040329 | 63.38142 | 4.799286 | 4.140533 | 59.52102 | 1.950223 | 1.117374 | 13.50005 | 3.262773 |
| ENSMUSG00000027253 | 0.541155084 | 1.11E-05 | 0.000247 | 2338.07 | 2088.889 | 1942.738 | 2154.833 | 1486.07 | 1706.23 | 1489.827 | 1175.686 |
| ENSMUSG00000027254 | 0.607643762 | 0.006136 | 0.029933 | 317.9132 | 374.3443 | 383.4134 | 394.2189 | 201.848 | 364.264 | 189.0007 | 210.9927 |
| ENSMUSG00000027255 | -0.213356475 | 0.093656 | 0.210665 | 1435.64 | 1396.592 | 1497.217 | 1382.785 | 1780.553 | 1252.576 | 1799.364 | 1789.087 |
| ENSMUSG00000027257 | -0.000172336 | 0.998269 | 0.999081 | 1005.048 | 982.6538 | 1006.978 | 997.1928 | 1010.215 | 882.7255 | 1042.397 | 1056.051 |
| ENSMUSG00000027259 | -0.209476062 | 0.083934 | 0.195295 | 364.1917 | 391.1418 | 374.3042 | 350.2251 | 446.601 | 369.8508 | 396.3228 | 497.0291 |
| ENSMUSG00000027263 | 0.053404047 | 0.56198 | 0.707497 | 601.6205 | 577.1142 | 502.6607 | 527.063 | 547.0374 | 510.64 | 533.2519 | 532.9196 |
| ENSMUSG00000027270 | -1.42194908 | 1.13E-06 | 4.15E-05 | 27.16347 | 40.79393 | 43.88965 | 40.5433 | 135.5405 | 65.92507 | 108.9647 | 97.8832 |
| ENSMUSG00000027272 | 0.183179428 | 0.044273 | 0.125111 | 1529.203 | 1440.986 | 1312.549 | 1388.824 | 1264.719 | 1363.196 | 1261.29 | 1103.905 |
| ENSMUSG00000027273 | 0.724684708 | 0.008514 | 0.037966 | 315.9011 | 245.9634 | 197.0894 | 499.459 | 216.4747 | 186.6015 | 179.3578 | 179.4525 |
| ENSMUSG00000027274 | -0.318163729 | 0.002093 | 0.013564 | 643.8748 | 641.9045 | 593.7525 | 638.3414 | 769.3628 | 667.0723 | 859.1816 | 840.7079 |
| ENSMUSG00000027276 | -0.254173542 | 0.040289 | 0.117282 | 2293.804 | 1835.727 | 2447.883 | 1926.238 | 2384.147 | 2241.452 | 2733.76 | 2785.321 |
| ENSMUSG00000027281 | 0.029199712 | 0.865758 | 0.919615 | 338.0343 | 365.9456 | 317.1648 | 275.1769 | 313.0107 | 402.2547 | 246.858 | 307.7883 |
| ENSMUSG00000027282 | -0.285276313 | 0.008942 | 0.039277 | 2385.355 | 2177.676 | 2562.162 | 2396.368 | 3000.417 | 2402.354 | 2913.118 | 3289.963 |
| ENSMUSG00000027284 | 0.255142806 | 0.052671 | 0.141129 | 825.9706 | 803.8804 | 752.7489 | 842.7832 | 694.2793 | 834.6784 | 587.2521 | 587.2992 |
| ENSMUSG00000027285 | 0.082815054 | 0.467062 | 0.627647 | 771.6437 | 872.2703 | 888.5584 | 972.1767 | 945.858 | 850.3217 | 756.0027 | 758.051 |
| ENSMUSG00000027286 | 0.104044702 | 0.418813 | 0.584292 | 333.004 | 298.7556 | 277.4157 | 330.3848 | 245.728 | 289.3999 | 297.9654 | 320.8394 |
| ENSMUSG00000027287 | -0.227676211 | 0.009141 | 0.03996 | 1391.373 | 1298.207 | 1375.485 | 1440.581 | 1709.37 | 1395.6 | 1673.042 | 1669.452 |
| ENSMUSG00000027288 | -0.049336954 | 0.466706 | 0.627305 | 3512.136 | 3555.071 | 3987.333 | 3693.754 | 3557.206 | 3917.514 | 3900.55 | 3892.489 |
| ENSMUSG00000027291 | 0.112053795 | 0.142151 | 0.279278 | 1308.877 | 1118.234 | 1141.959 | 1214.574 | 1077.498 | 1088.322 | 1108.933 | 1152.847 |
| ENSMUSG00000027293 | -0.602870151 | 0.011515 | 0.047468 | 996.9999 | 865.0713 | 1763.039 | 1013.583 | 1763.001 | 1250.342 | 2010.543 | 2024.007 |
| ENSMUSG00000027296 | 0.089577027 | 0.801132 | 0.878636 | 26.15741 | 26.39607 | 30.63995 | 14.6646 | 22.42756 | 22.34748 | 24.10723 | 22.83941 |
| ENSMUSG00000027297 | 0.183465171 | 0.61511 | 0.748547 | 33.19979 | 22.79661 | 19.87456 | 39.68068 | 26.32801 | 18.99536 | 25.07152 | 31.54014 |
| ENSMUSG00000027298 | -0.063740359 | 0.533873 | 0.685151 | 701.2199 | 745.0892 | 788.3575 | 810.0035 | 880.5255 | 688.3024 | 791.6814 | 822.2189 |
| ENSMUSG00000027300 | -0.275496173 | 0.026796 | 0.087374 | 316.9071 | 392.3416 | 360.2264 | 349.3625 | 469.0285 | 374.3203 | 471.5374 | 399.1459 |
| ENSMUSG00000027303 | -0.143264748 | 0.081091 | 0.190701 | 2957.8 | 3298.309 | 2766.704 | 2809.565 | 3502.6 | 3060.488 | 3311.369 | 3184.467 |
| ENSMUSG00000027304 | 0.262944037 | 0.082058 | 0.192385 | 3456.803 | 3731.445 | 3195.663 | 3456.532 | 2744.938 | 3850.471 | 2538.973 | 2400.314 |
| ENSMUSG00000027305 | -0.329698811 | 0.012781 | 0.051324 | 462.785 | 440.3345 | 469.5365 | 444.2511 | 564.5894 | 470.4145 | 548.6805 | 700.4087 |
| ENSMUSG00000027306 | 0.13387871 | 0.1873 | 0.339336 | 2402.458 | 2285.66 | 2029.689 | 2308.38 | 2084.788 | 2349.838 | 1986.436 | 1804.314 |
| ENSMUSG00000027309 | 0.036457445 | 0.757886 | 0.849068 | 1725.383 | 1396.592 | 1647.932 | 1650.199 | 1289.097 | 1712.934 | 1553.47 | 1708.606 |
| ENSMUSG00000027312 | -0.100494482 | 0.253344 | 0.417064 | 1394.391 | 1258.613 | 1369.688 | 1308.6 | 1365.156 | 1489.46 | 1285.397 | 1580.27 |
| ENSMUSG00000027313 | -0.843354112 | 0.118684 | 0.247522 | 17.10292 | 8.398751 | 11.59349 | 6.900988 | 23.40267 | 7.821619 | 19.28578 | 28.27737 |
| ENSMUSG00000027314 | -0.736528561 | 0.001047 | 0.008098 | 224.3501 | 140.3791 | 269.9628 | 240.672 | 344.2143 | 306.1605 | 339.4298 | 473.1021 |
| ENSMUSG00000027315 | -0.8601025 | 0.001218 | 0.009052 | 731.4015 | 647.9036 | 1273.628 | 737.5431 | 1561.153 | 915.1294 | 1758.863 | 1921.773 |
| ENSMUSG00000027316 | -0.304062793 | 0.066209 | 0.165867 | 186.1201 | 247.1632 | 276.5876 | 238.0841 | 288.6329 | 347.5033 | 284.4653 | 251.2335 |
| ENSMUSG00000027318 | 1.215480718 | 0.004127 | 0.022339 | 271.6347 | 473.9295 | 221.1045 | 339.011 | 122.864 | 270.4045 | 115.7147 | 53.29196 |
| ENSMUSG00000027322 | 0.528649188 | 0.178523 | 0.327959 | 25.15136 | 37.19447 | 59.62368 | 24.15346 | 21.45245 | 27.93435 | 24.10723 | 28.27737 |
| ENSMUSG00000027323 | -0.010438142 | 0.919853 | 0.952339 | 648.9051 | 847.074 | 699.7501 | 779.8116 | 730.3584 | 779.9271 | 721.2883 | 763.4889 |
| ENSMUSG00000027324 | -0.330514367 | 0.075914 | 0.182475 | 114.6902 | 98.38537 | 120.0755 | 113.8663 | 152.1174 | 108.3853 | 133.0719 | 169.6642 |
| ENSMUSG00000027326 | 0.315942583 | 0.049165 | 0.134527 | 546.2875 | 622.7074 | 428.9592 | 529.6508 | 411.497 | 527.4006 | 363.537 | 405.6715 |
| ENSMUSG00000027327 | -0.315275721 | 8.62E-05 | 0.001202 | 988.9514 | 1033.046 | 1052.524 | 983.3908 | 1344.678 | 1122.961 | 1290.219 | 1288.795 |
| ENSMUSG00000027329 | 0.290668362 | 0.135889 | 0.270342 | 580.4934 | 724.6922 | 591.2681 | 621.9515 | 455.377 | 738.5843 | 456.1088 | 408.9342 |
| ENSMUSG00000027330 | 0.262160551 | 0.005323 | 0.026853 | 2055.369 | 2286.86 | 2138.999 | 2215.217 | 1873.189 | 2007.921 | 1821.542 | 1547.642 |

| | | | | | | | | | | | |
|---|---|---|---|---|---|---|---|---|---|---|---|
| ENSMUSG00000027331 | 0.105869521 | 0.377296 | 0.545462 | 2548.336 | 2648.006 | 2198.623 | 2543.877 | 2579.169 | 2538.674 | 2238.115 | 1877.182 |
| ENSMUSG00000027332 | -0.398000423 | 0.000312 | 0.003201 | 2483.948 | 2068.492 | 2245.825 | 2546.465 | 3329.03 | 2616.89 | 3354.762 | 3013.715 |
| ENSMUSG00000027333 | -0.028893094 | 0.823174 | 0.892731 | 970.8425 | 1336.601 | 1194.958 | 1181.794 | 1411.961 | 1087.205 | 1140.754 | 1136.533 |
| ENSMUSG00000027335 | -1.023232549 | 0.032339 | 0.099758 | 34.20585 | 9.598572 | 13.24971 | 16.38985 | 47.78045 | 25.6996 | 27.0001 | 48.9416 |
| ENSMUSG00000027338 | -0.35521402 | 0.27162 | 0.436865 | 22.1332 | 34.79482 | 43.06154 | 43.9938 | 58.50668 | 35.75597 | 39.53586 | 51.11678 |
| ENSMUSG00000027339 | 1.071096548 | 6.30E-05 | 0.00095 | 1196.199 | 1364.197 | 876.9649 | 1242.178 | 493.4063 | 901.7209 | 482.1446 | 350.2043 |
| ENSMUSG00000027340 | 0.445402942 | 0.008345 | 0.037423 | 1668.038 | 1100.236 | 991.2436 | 1135.213 | 896.1273 | 1010.106 | 849.5388 | 838.5327 |
| ENSMUSG00000027341 | 0.001736868 | 0.984848 | 0.991442 | 769.6316 | 770.2854 | 765.1705 | 917.8314 | 831.7699 | 797.8051 | 826.3958 | 764.5765 |
| ENSMUSG00000027342 | -0.2897432 | 0.008601 | 0.03826 | 3495.033 | 4686.503 | 4291.249 | 3899.921 | 5267.551 | 4535.421 | 4778.053 | 5430.342 |
| ENSMUSG00000027344 | 0.289403908 | 0.491078 | 0.648215 | 20.12109 | 26.39607 | 31.46805 | 13.80198 | 14.62667 | 18.99536 | 26.03581 | 15.22628 |
| ENSMUSG00000027346 | 0.165374808 | 0.058056 | 0.151147 | 1370.246 | 1385.794 | 1332.424 | 1460.422 | 1300.798 | 1363.196 | 1111.825 | 1173.511 |
| ENSMUSG00000027347 | -1.176272411 | 0.013859 | 0.054385 | 20.12109 | 20.39697 | 59.62368 | 32.77969 | 98.48624 | 30.1691 | 80.036 | 93.53283 |
| ENSMUSG00000027349 | -0.032861604 | 0.536649 | 0.68728 | 1667.032 | 1606.561 | 1600.73 | 1658.825 | 1646.963 | 1652.596 | 1648.934 | 1736.883 |
| ENSMUSG00000027350 | 2.306723415 | 0.006567 | 0.031504 | 1306.865 | 122.3818 | 123.3879 | 1159.366 | 117.9885 | 120.6764 | 105.1075 | 204.4671 |
| ENSMUSG00000027351 | 0.552580796 | 0.00097 | 0.007653 | 3502.075 | 3808.234 | 3019.277 | 3579.887 | 2103.315 | 3261.615 | 2167.722 | 1951.138 |
| ENSMUSG00000027353 | -0.18217348 | 0.191784 | 0.344927 | 263.5862 | 321.5522 | 295.6341 | 271.7264 | 301.3094 | 384.3767 | 341.3584 | 279.5109 |
| ENSMUSG00000027356 | -0.018417371 | 0.876153 | 0.925815 | 485.9243 | 607.1097 | 637.6421 | 540.0023 | 522.6597 | 627.9642 | 562.1806 | 588.3868 |
| ENSMUSG00000027357 | 0.008444908 | 0.932763 | 0.960099 | 851.122 | 920.2631 | 768.483 | 892.8153 | 802.5166 | 967.646 | 814.8243 | 826.5692 |
| ENSMUSG00000027358 | -0.245576775 | 0.090041 | 0.205211 | 363.1856 | 241.1641 | 296.4622 | 275.1769 | 340.3138 | 378.7898 | 345.2155 | 331.7153 |
| ENSMUSG00000027359 | -3.949421069 | 5.19E-07 | 2.17E-05 | 125.7568 | 28.79572 | 423.9906 | 170.7995 | 2991.641 | 1130.783 | 2665.295 | 4798.452 |
| ENSMUSG00000027360 | -2.02097751 | 0.000867 | 0.006982 | 517.1119 | 165.5754 | 1163.49 | 693.5493 | 1994.103 | 1582.202 | 2419.402 | 4315.561 |
| ENSMUSG00000027361 | 0.106878188 | 0.258623 | 0.422514 | 1283.725 | 1223.818 | 1157.693 | 1314.638 | 1164.283 | 1317.384 | 1069.397 | 1074.54 |
| ENSMUSG00000027363 | -0.161015298 | 0.100894 | 0.222341 | 1939.673 | 1809.331 | 1689.338 | 1718.346 | 2118.917 | 1698.409 | 2023.079 | 2157.781 |
| ENSMUSG00000027365 | 0.069860703 | 0.428916 | 0.593809 | 3475.918 | 3191.525 | 2851.999 | 3240.014 | 2795.644 | 3261.615 | 3185.047 | 2912.569 |
| ENSMUSG00000027366 | 0.147937968 | 0.049944 | 0.135973 | 1987.963 | 1923.314 | 1748.961 | 1951.254 | 1610.884 | 1851.489 | 1732.828 | 1673.803 |
| ENSMUSG00000027367 | -0.576285394 | 5.55E-07 | 2.28E-05 | 2306.883 | 2403.243 | 2688.034 | 2460.202 | 3687.871 | 2975.567 | 3838.835 | 4198.102 |
| ENSMUSG00000027368 | -0.584716424 | 0.029186 | 0.092758 | 99.59938 | 92.38626 | 135.8095 | 82.81186 | 147.2418 | 103.9158 | 216.9651 | 147.9124 |
| ENSMUSG00000027371 | -0.174727008 | 0.174244 | 0.322358 | 525.1604 | 530.3211 | 491.0672 | 488.2449 | 570.4401 | 452.5365 | 621.0022 | 650.3795 |
| ENSMUSG00000027374 | -0.217109418 | 7.61E-05 | 0.001103 | 1504.051 | 1485.379 | 1516.263 | 1487.163 | 1702.544 | 1674.944 | 1785.864 | 1803.226 |
| ENSMUSG00000027375 | -1.086662009 | 0.010353 | 0.043725 | 1749.529 | 1481.78 | 3863.946 | 1962.468 | 4602.525 | 2667.172 | 5406.769 | 6562.525 |
| ENSMUSG00000027376 | -0.379897567 | 0.150197 | 0.290155 | 137.8294 | 93.58608 | 183.8397 | 104.3774 | 195.9974 | 118.4417 | 189.0007 | 174.0146 |
| ENSMUSG00000027377 | -0.240869287 | 0.482497 | 0.641014 | 22.1332 | 52.79215 | 67.07664 | 46.58167 | 51.6809 | 56.98608 | 64.60737 | 50.02919 |
| ENSMUSG00000027378 | -0.244259013 | 0.003983 | 0.0218 | 1205.253 | 1340.201 | 1342.361 | 1333.616 | 1659.639 | 1372.135 | 1548.648 | 1603.109 |
| ENSMUSG00000027379 | -0.057827958 | 0.543491 | 0.692451 | 978.8909 | 909.4647 | 827.2785 | 987.7039 | 1055.07 | 962.0591 | 952.7177 | 883.124 |
| ENSMUSG00000027380 | -0.831973739 | 0.011133 | 0.04631 | 26.15741 | 37.19447 | 19.04645 | 22.42821 | 42.9049 | 52.51658 | 54.96448 | 33.71532 |
| ENSMUSG00000027381 | -0.078299485 | 0.704284 | 0.813867 | 1169.035 | 1057.043 | 1010.29 | 1082.592 | 1153.557 | 1658.183 | 953.682 | 795.0291 |
| ENSMUSG00000027384 | -0.249456571 | 0.019287 | 0.068706 | 282.7013 | 310.7538 | 273.2752 | 281.2153 | 352.0152 | 324.0385 | 316.2868 | 369.781 |
| ENSMUSG00000027386 | -0.050709392 | 0.793149 | 0.873246 | 169.0171 | 161.9759 | 223.5888 | 205.3044 | 184.296 | 170.9582 | 175.5006 | 258.8467 |
| ENSMUSG00000027387 | -0.06875502 | 0.549922 | 0.697988 | 260.5681 | 274.7591 | 245.9477 | 257.9244 | 254.5041 | 301.691 | 249.7509 | 283.8613 |
| ENSMUSG00000027394 | 0.17355964 | 0.011565 | 0.047632 | 1151.932 | 1219.019 | 1182.536 | 1231.826 | 1128.204 | 1002.285 | 1081.932 | 1027.774 |
| ENSMUSG00000027395 | -0.050101241 | 0.442956 | 0.606154 | 1098.611 | 1095.437 | 1029.337 | 1090.356 | 1097.975 | 1101.731 | 1080.968 | 1184.387 |
| ENSMUSG00000027397 | -0.226714176 | 0.004025 | 0.021935 | 1096.599 | 958.6574 | 1172.599 | 1067.928 | 1277.396 | 1252.576 | 1250.683 | 1252.905 |
| ENSMUSG00000027400 | -2.97412148 | 5.47E-06 | 0.000143 | 2.012109 | 4.799286 | 2.48432 | 6.038364 | 49.73068 | 11.17374 | 32.78583 | 26.10219 |
| ENSMUSG00000027404 | -0.318958445 | 0.000584 | 0.005158 | 5025.242 | 5389.598 | 5783.497 | 5417.276 | 7228.5 | 5752.242 | 6792.453 | 7191.152 |
| ENSMUSG00000027405 | -0.115150617 | 0.228861 | 0.388938 | 3673.104 | 4108.189 | 3345.551 | 3703.243 | 4549.869 | 3978.969 | 3736.621 | 3791.343 |
| ENSMUSG00000027406 | -0.480321087 | 3.79E-05 | 0.000644 | 4062.447 | 3971.409 | 4234.937 | 4200.114 | 6130.525 | 4497.431 | 5826.235 | 6521.196 |
| ENSMUSG00000027408 | 0.25081966 | 0.083397 | 0.194462 | 2601.657 | 3211.922 | 2485.148 | 2888.926 | 2833.673 | 2490.627 | 2226.544 | 1848.905 |
| ENSMUSG00000027411 | 0.102849706 | 0.253417 | 0.417148 | 2080.52 | 2004.902 | 1930.317 | 2187.613 | 2154.996 | 1906.24 | 1866.864 | 1709.693 |
| ENSMUSG00000027412 | -0.382230822 | 0.132167 | 0.265309 | 197.1867 | 82.78769 | 145.7468 | 136.2945 | 176.4951 | 154.1976 | 206.3579 | 196.854 |
| ENSMUSG00000027419 | -1.144682399 | 0.000813 | 0.00665 | 37.22401 | 19.19714 | 38.92101 | 34.50494 | 55.58134 | 49.16446 | 88.7146 | 95.70802 |
| ENSMUSG00000027420 | 0.579849638 | 0.002326 | 0.014615 | 152.9203 | 147.578 | 129.1846 | 156.9975 | 80.93424 | 87.15518 | 97.39321 | 127.2482 |
| ENSMUSG00000027422 | 0.164333811 | 0.127966 | 0.259325 | 6265.707 | 7328.51 | 6971.002 | 6184.148 | 5185.642 | 6789.165 | 5559.127 | 6336.306 |
| ENSMUSG00000027423 | -0.069175788 | 0.298826 | 0.466909 | 3712.341 | 4103.39 | 3508.688 | 3938.739 | 4074.99 | 3868.349 | 4020.122 | 4042.576 |
| ENSMUSG00000027424 | -0.039638665 | 0.719982 | 0.824828 | 497.9969 | 519.5227 | 400.8036 | 461.5036 | 454.4019 | 474.884 | 488.8946 | 510.0802 |
| ENSMUSG00000027425 | -0.295822805 | 0.002083 | 0.013514 | 970.8425 | 1082.239 | 1202.411 | 1028.247 | 1330.052 | 1203.412 | 1298.898 | 1428.007 |
| ENSMUSG00000027427 | 0.041938988 | 0.672742 | 0.790252 | 1129.799 | 1155.428 | 1128.709 | 1067.928 | 1053.12 | 1276.041 | 975.8606 | 1049.525 |
| ENSMUSG00000027428 | -0.056232806 | 0.420628 | 0.585984 | 1695.202 | 1864.523 | 1667.807 | 1876.206 | 1794.205 | 1783.329 | 1905.435 | 1900.022 |
| ENSMUSG00000027429 | -0.573384247 | 0.00014 | 0.001734 | 1642.887 | 1468.582 | 1830.116 | 1541.508 | 2213.503 | 1890.597 | 2513.902 | 3031.116 |
| ENSMUSG00000027430 | 0.05266494 | 0.568487 | 0.712674 | 657.9595 | 801.4808 | 722.9371 | 787.5752 | 759.6117 | 710.6499 | 693.3239 | 698.2335 |
| ENSMUSG00000027433 | 0.041845411 | 0.478513 | 0.637833 | 2982.951 | 3013.952 | 2993.605 | 3171.004 | 3055.024 | 2975.567 | 3037.511 | 2745.08 |
| ENSMUSG00000027435 | -0.258517782 | 0.038723 | 0.113747 | 1619.748 | 1196.222 | 1734.883 | 1512.179 | 1661.59 | 1765.451 | 1872.65 | 1958.752 |
| ENSMUSG00000027438 | 0.401680559 | 0.019225 | 0.068535 | 278.6771 | 235.165 | 201.2299 | 281.2153 | 174.5449 | 230.1791 | 180.3221 | 169.6642 |
| ENSMUSG00000027439 | -0.234920226 | 0.007181 | 0.033504 | 1044.284 | 982.6538 | 967.2285 | 955.7868 | 1222.79 | 1015.693 | 1169.683 | 1237.679 |
| ENSMUSG00000027447 | 0.009851996 | 0.886474 | 0.932078 | 5533.299 | 5299.612 | 5573.986 | 5969.355 | 5729.754 | 5085.17 | 5582.27 | 5829.488 |

| | | | | | | | | | | | |
|---|---|---|---|---|---|---|---|---|---|---|---|
| ENSMUSG00000027452 | -1.769342405 | 3.73E-06 | 0.000107 | 306.8466 | 166.7752 | 511.7699 | 276.0395 | 1050.195 | 441.3628 | 1324.933 | 1487.825 |
| ENSMUSG00000027454 | 0.055935553 | 0.691934 | 0.80504 | 561.3783 | 687.4977 | 525.0196 | 492.558 | 607.4943 | 578.7998 | 525.5376 | 464.4014 |
| ENSMUSG00000027455 | -0.188937828 | 0.001195 | 0.008947 | 3673.104 | 3611.463 | 3691.699 | 3712.732 | 4238.809 | 3939.861 | 4077.015 | 4490.664 |
| ENSMUSG00000027456 | -0.019154551 | 0.954648 | 0.974178 | 52.31483 | 46.79304 | 71.21717 | 35.36756 | 71.18313 | 42.46022 | 59.78593 | 34.80291 |
| ENSMUSG00000027457 | 0.659431754 | 0.040773 | 0.118301 | 46.2785 | 50.3925 | 36.43669 | 31.91707 | 18.52711 | 34.6386 | 26.03581 | 25.0146 |
| ENSMUSG00000027459 | -0.105385134 | 0.353081 | 0.521776 | 372.2401 | 335.95 | 428.9592 | 343.3242 | 420.273 | 391.0809 | 381.8585 | 402.4087 |
| ENSMUSG00000027460 | -0.772270078 | 0.222276 | 0.381505 | 12.07265 | 17.99732 | 55.48314 | 12.93935 | 36.07912 | 13.40849 | 60.75022 | 58.72992 |
| ENSMUSG00000027463 | -0.899372924 | 0.009247 | 0.040236 | 171.0292 | 127.1811 | 262.5098 | 168.2116 | 283.7574 | 203.3621 | 277.7153 | 597.0875 |
| ENSMUSG00000027465 | -0.002007978 | 0.974941 | 0.986285 | 2019.151 | 2021.699 | 2163.843 | 2241.096 | 2093.564 | 2051.499 | 2096.365 | 2221.949 |
| ENSMUSG00000027466 | -0.074813173 | 0.388466 | 0.555993 | 1791.783 | 2019.3 | 2059.501 | 1865.855 | 2079.912 | 1806.794 | 2075.15 | 2184.97 |
| ENSMUSG00000027469 | 0.004389737 | 0.963185 | 0.978992 | 2991 | 2935.963 | 2396.541 | 2977.776 | 3005.293 | 2896.234 | 2728.938 | 2633.058 |
| ENSMUSG00000027470 | -0.832579492 | 0.237493 | 0.398614 | 9.054489 | 1.199822 | 4.96864 | 16.38985 | 20.47734 | 7.821619 | 13.50005 | 15.22628 |
| ENSMUSG00000027472 | 0.096972504 | 0.21659 | 0.375024 | 1442.682 | 1298.207 | 1229.738 | 1331.891 | 1306.649 | 1222.407 | 1164.861 | 1262.693 |
| ENSMUSG00000027474 | -0.471493798 | 0.001811 | 0.012223 | 231.3925 | 227.9661 | 291.4935 | 265.688 | 320.8116 | 307.2779 | 360.6441 | 424.1605 |
| ENSMUSG00000027475 | -0.239592071 | 0.113492 | 0.240595 | 847.0978 | 823.0776 | 1017.743 | 854.8599 | 1118.453 | 798.9225 | 1000.932 | 1265.956 |
| ENSMUSG00000027478 | -0.275446177 | 0.017131 | 0.063195 | 440.6518 | 356.347 | 342.8361 | 388.1806 | 467.0783 | 426.8369 | 499.5018 | 454.6131 |
| ENSMUSG00000027479 | -0.006499235 | 0.906824 | 0.944222 | 4738.516 | 5083.644 | 4643.194 | 4820.34 | 4828.751 | 5084.052 | 4644.981 | 4811.503 |
| ENSMUSG00000027487 | -0.161073725 | 0.374811 | 0.543403 | 249.5015 | 227.9661 | 241.8071 | 306.2313 | 282.7823 | 307.2779 | 212.1436 | 346.9416 |
| ENSMUSG00000027488 | -0.177193283 | 0.062539 | 0.159511 | 615.7053 | 591.512 | 663.3134 | 672.8463 | 720.6073 | 665.955 | 688.5025 | 804.8174 |
| ENSMUSG00000027489 | 1.130753899 | 0.002727 | 0.016482 | 24.1453 | 39.59411 | 34.78048 | 36.23019 | 12.67645 | 22.34748 | 10.60718 | 16.31387 |
| ENSMUSG00000027490 | -0.12545128 | 0.27937 | 0.445756 | 560.3723 | 727.0918 | 624.3924 | 572.782 | 723.5326 | 715.1194 | 608.4665 | 660.1678 |
| ENSMUSG00000027495 | -0.106273228 | 0.355815 | 0.524431 | 482.9061 | 451.1329 | 468.7084 | 401.1199 | 554.8383 | 433.5411 | 469.6088 | 481.8029 |
| ENSMUSG00000027496 | -0.010181713 | 0.916913 | 0.950812 | 940.6608 | 1035.446 | 848.8093 | 1005.819 | 1057.996 | 910.6599 | 904.5032 | 981.0072 |
| ENSMUSG00000027498 | -0.176897341 | 0.036302 | 0.109153 | 1212.295 | 1337.801 | 1257.066 | 1179.206 | 1511.423 | 1254.811 | 1423.291 | 1442.146 |
| ENSMUSG00000027499 | 0.28891662 | 0.030099 | 0.094705 | 829.9948 | 1105.036 | 867.0276 | 985.9787 | 699.1548 | 917.3641 | 713.574 | 770.0145 |
| ENSMUSG00000027500 | 1.933319499 | 0.002689 | 0.0163 | 1583.53 | 1313.805 | 892.6989 | 2166.91 | 228.176 | 948.6506 | 226.608 | 156.6131 |
| ENSMUSG00000027502 | -0.121596093 | 0.184729 | 0.336008 | 1929.612 | 1980.905 | 2105.875 | 1907.261 | 2215.453 | 1833.611 | 2268.008 | 2302.43 |
| ENSMUSG00000027505 | -0.013413169 | 0.980284 | 0.988885 | 8.048435 | 10.79839 | 9.93728 | 11.21411 | 4.875557 | 7.821619 | 12.53576 | 15.22628 |
| ENSMUSG00000027506 | 0.092119215 | 0.265005 | 0.429179 | 3117.762 | 3273.113 | 3353.832 | 3407.363 | 2887.305 | 2796.787 | 3316.19 | 3338.905 |
| ENSMUSG00000027508 | 0.282572518 | 0.039599 | 0.115854 | 497.9969 | 493.1267 | 453.8024 | 558.98 | 386.1441 | 503.9357 | 405.9657 | 352.3795 |
| ENSMUSG00000027509 | -0.232168266 | 1.05E-05 | 0.000236 | 1524.172 | 1528.573 | 1591.621 | 1585.502 | 1880.015 | 1783.329 | 1810.935 | 1846.73 |
| ENSMUSG00000027510 | -0.134534952 | 0.403706 | 0.570586 | 351.113 | 221.967 | 308.0557 | 309.6818 | 354.9405 | 287.1651 | 318.2154 | 349.1167 |
| ENSMUSG00000027513 | -3.860321624 | 1.10E-13 | 4.07E-11 | 263.5862 | 101.9848 | 246.7758 | 288.9789 | 4061.339 | 1080.501 | 3388.512 | 4568.97 |
| ENSMUSG00000027514 | 0.88957551 | 0.116176 | 0.244263 | 18.10898 | 38.39429 | 74.5296 | 13.80198 | 16.57689 | 15.64324 | 22.17865 | 23.927 |
| ENSMUSG00000027519 | 0.106928326 | 0.057339 | 0.149964 | 1956.776 | 1828.528 | 1808.585 | 1898.634 | 1766.902 | 1747.573 | 1764.649 | 1677.065 |
| ENSMUSG00000027520 | 1.061992503 | 2.06E-06 | 6.57E-05 | 402.4217 | 365.9456 | 221.1045 | 443.3885 | 176.4951 | 189.9536 | 172.6078 | 146.8248 |
| ENSMUSG00000027522 | 0.325086253 | 0.001203 | 0.008983 | 2358.191 | 2445.236 | 2140.656 | 2371.352 | 1775.678 | 2192.288 | 1754.042 | 1714.044 |
| ENSMUSG00000027523 | -0.052831182 | 0.451064 | 0.613563 | 13745.72 | 14365.46 | 14469.51 | 15122.65 | 15056.69 | 13517.99 | 15214.55 | 16066.98 |
| ENSMUSG00000027524 | -0.176149624 | 0.552039 | 0.699659 | 238.4349 | 385.1427 | 562.2844 | 371.7907 | 394.9201 | 264.8177 | 577.6092 | 523.1313 |
| ENSMUSG00000027525 | 0.568422189 | 0.301342 | 0.469688 | 78.47224 | 26.39607 | 43.88965 | 70.73513 | 25.35289 | 78.21619 | 30.85725 | 14.13868 |
| ENSMUSG00000027530 | -0.14506627 | 0.765786 | 0.85465 | 7.04238 | 15.59768 | 21.53077 | 13.80198 | 9.751113 | 20.11273 | 17.3572 | 17.40146 |
| ENSMUSG00000027531 | -0.100454788 | 0.383844 | 0.55186 | 1610.693 | 1436.186 | 1480.655 | 1384.511 | 1634.287 | 1299.506 | 1577.577 | 1826.065 |
| ENSMUSG00000027533 | -0.192250437 | 0.091727 | 0.207872 | 1618.741 | 1694.148 | 1830.944 | 1946.079 | 2345.143 | 1722.991 | 1941.114 | 2092.525 |
| ENSMUSG00000027534 | 0.355307219 | 0.057498 | 0.150268 | 147.89 | 244.7636 | 192.1207 | 223.4195 | 139.4409 | 173.193 | 161.0363 | 157.7007 |
| ENSMUSG00000027536 | -0.405422356 | 0.077741 | 0.185374 | 207.2472 | 183.5727 | 303.087 | 203.5791 | 299.3592 | 210.0663 | 294.1082 | 387.1824 |
| ENSMUSG00000027540 | -0.260042236 | 0.024849 | 0.082691 | 1272.659 | 1035.446 | 1292.674 | 1163.679 | 1485.095 | 1188.886 | 1529.363 | 1504.138 |
| ENSMUSG00000027544 | 0.494581653 | 0.005324 | 0.026853 | 203.223 | 185.9723 | 208.6829 | 168.2116 | 112.1378 | 168.7235 | 118.6076 | 145.7372 |
| ENSMUSG00000027546 | 0.688969172 | 3.74E-19 | 4.06E-16 | 2131.829 | 2423.639 | 2422.212 | 2537.838 | 1474.368 | 1415.713 | 1456.077 | 1557.43 |
| ENSMUSG00000027550 | 0.188362629 | 0.123261 | 0.25288 | 1297.81 | 1360.598 | 1199.098 | 1157.641 | 968.2855 | 1272.689 | 956.5749 | 1203.963 |
| ENSMUSG00000027551 | -0.277853623 | 0.001559 | 0.010831 | 496.9909 | 476.3291 | 516.7385 | 455.4652 | 599.6935 | 545.2786 | 593.0378 | 621.0145 |
| ENSMUSG00000027552 | -0.065221054 | 0.569253 | 0.713058 | 821.9464 | 812.2792 | 711.3436 | 938.5344 | 764.4873 | 937.4769 | 837.003 | 898.3502 |
| ENSMUSG00000027555 | 0.279798039 | 0.144964 | 0.283118 | 100.6054 | 86.38715 | 105.1695 | 89.71284 | 63.38224 | 87.15518 | 87.75031 | 77.21897 |
| ENSMUSG00000027559 | 1.215920924 | 0.038356 | 0.113119 | 75.45408 | 320.3523 | 87.7793 | 72.46037 | 39.97956 | 125.1459 | 32.78583 | 41.32846 |
| ENSMUSG00000027560 | 0.420292639 | 0.417831 | 0.583496 | 13.07871 | 33.595 | 31.46805 | 14.6646 | 19.50223 | 8.938993 | 13.50005 | 27.18978 |
| ENSMUSG00000027562 | -1.194662199 | 7.09E-06 | 0.000177 | 978.8909 | 867.471 | 1546.903 | 1026.522 | 2453.38 | 1488.342 | 2453.152 | 3724.999 |
| ENSMUSG00000027566 | -0.230429924 | 0.00639 | 0.030874 | 3984.981 | 4466.936 | 4667.209 | 4524.46 | 5357.262 | 4593.525 | 5253.447 | 5495.598 |
| ENSMUSG00000027568 | 0.682416865 | 0.170109 | 0.316741 | 734.4197 | 1167.426 | 819.8256 | 1022.209 | 427.0988 | 1246.989 | 365.4656 | 293.6496 |
| ENSMUSG00000027569 | 0.079029595 | 0.430422 | 0.595225 | 1120.745 | 1237.016 | 1167.63 | 1280.133 | 1074.573 | 1324.088 | 1107.968 | 1044.087 |
| ENSMUSG00000027570 | 0.717936442 | 0.003527 | 0.019909 | 328.9798 | 380.3434 | 324.6178 | 421.8229 | 220.3752 | 335.2122 | 183.2149 | 146.8248 |
| ENSMUSG00000027573 | -0.231268584 | 0.00213 | 0.013732 | 1015.109 | 1069.041 | 1109.663 | 1021.346 | 1308.599 | 1176.595 | 1166.79 | 1296.409 |
| ENSMUSG00000027574 | 0.592418739 | 0.000578 | 0.005125 | 310.8708 | 533.9206 | 420.6782 | 475.3055 | 278.8818 | 292.752 | 276.751 | 305.6131 |
| ENSMUSG00000027575 | 0.150367879 | 0.021155 | 0.073349 | 2278.713 | 2236.467 | 2013.127 | 2103.076 | 1991.177 | 1990.043 | 1880.364 | 1910.898 |
| ENSMUSG00000027577 | -0.33066456 | 0.248178 | 0.411541 | 76.46013 | 57.59143 | 115.9349 | 73.323 | 99.46135 | 115.0895 | 69.42882 | 125.073 |
| ENSMUSG00000027580 | 0.019671346 | 0.926165 | 0.956215 | 376.2643 | 297.5557 | 445.5214 | 248.4356 | 326.6623 | 263.7003 | 399.2157 | 359.9926 |

| | | | | | | | | | | | |
|---|---|---|---|---|---|---|---|---|---|---|---|
| ENSMUSG00000027581 | 2.458883764 | 0.00092 | 0.007332 | 206.2411 | 34.79482 | 26.49941 | 314.8576 | 27.30312 | 32.40385 | 38.57157 | 7.613138 |
| ENSMUSG00000027582 | -0.156867355 | 0.2753 | 0.441437 | 971.8485 | 932.2613 | 926.6513 | 879.0133 | 1152.582 | 775.4576 | 987.4321 | 1219.19 |
| ENSMUSG00000027583 | -0.374805146 | 3.79E-05 | 0.000644 | 647.899 | 609.5093 | 560.6282 | 653.006 | 788.865 | 836.9132 | 842.7887 | 734.124 |
| ENSMUSG00000027584 | 0.506398295 | 0.225745 | 0.385543 | 22.1332 | 16.7975 | 14.90592 | 27.60395 | 17.552 | 14.52586 | 15.42863 | 9.78832 |
| ENSMUSG00000027589 | 0.381816771 | 0.003382 | 0.019315 | 2727.413 | 2658.805 | 2246.653 | 2969.15 | 1915.119 | 2373.303 | 2150.365 | 1697.73 |
| ENSMUSG00000027593 | -0.094065755 | 0.17057 | 0.317375 | 7169.143 | 8216.378 | 8270.301 | 7820.545 | 8150.955 | 8068.558 | 8522.388 | 8855.167 |
| ENSMUSG00000027596 | 0.425854189 | 0.400345 | 0.567711 | 17.10292 | 14.39786 | 14.07781 | 17.25247 | 15.60178 | 17.87799 | 4.821446 | 8.700729 |
| ENSMUSG00000027597 | -0.17909878 | 0.093097 | 0.209847 | 2996.03 | 3155.531 | 2645.801 | 2585.283 | 3647.891 | 2821.37 | 3232.297 | 3179.029 |
| ENSMUSG00000027598 | 0.100166728 | 0.264459 | 0.428723 | 2085.551 | 1857.324 | 1672.775 | 1798.57 | 1659.639 | 1636.953 | 1839.864 | 1777.124 |
| ENSMUSG00000027599 | -0.158499707 | 0.037855 | 0.112206 | 1174.065 | 1238.216 | 1176.74 | 1245.628 | 1226.69 | 1375.488 | 1335.54 | 1459.547 |
| ENSMUSG00000027601 | -0.163782444 | 0.102174 | 0.224277 | 951.7274 | 806.2801 | 779.2483 | 866.074 | 1006.315 | 851.4391 | 942.1105 | 1011.46 |
| ENSMUSG00000027602 | 0.199668349 | 0.050605 | 0.137208 | 2000.036 | 1553.769 | 1782.085 | 1636.397 | 1437.314 | 1442.53 | 1532.255 | 1660.752 |
| ENSMUSG00000027603 | 0.993235584 | 2.18E-05 | 0.000422 | 181.0898 | 125.9813 | 135.8095 | 175.1126 | 86.78491 | 100.5637 | 60.75022 | 63.08028 |
| ENSMUSG00000027605 | -0.570566921 | 0.023949 | 0.080488 | 449.7063 | 279.5584 | 397.4912 | 329.5222 | 543.137 | 312.8647 | 585.3235 | 722.1605 |
| ENSMUSG00000027610 | -1.458421354 | 2.67E-07 | 1.26E-05 | 673.0504 | 469.1302 | 751.0927 | 579.683 | 1552.377 | 902.8383 | 1679.792 | 2663.511 |
| ENSMUSG00000027611 | 0.308941433 | 0.37997 | 0.548037 | 309.8647 | 79.18822 | 165.6213 | 182.0136 | 148.2169 | 125.1459 | 144.6434 | 177.2773 |
| ENSMUSG00000027612 | 1.602478155 | 0.012115 | 0.049352 | 22.1332 | 23.99643 | 5.796746 | 24.15346 | 2.925334 | 8.938993 | 7.714313 | 5.437955 |
| ENSMUSG00000027613 | -0.405760334 | 0.000355 | 0.003521 | 2289.78 | 2283.26 | 2514.96 | 2323.908 | 3176.913 | 2501.801 | 3224.583 | 3566.211 |
| ENSMUSG00000027615 | -0.256550281 | 0.002693 | 0.016308 | 477.8758 | 407.9393 | 448.8338 | 433.8996 | 527.5352 | 531.8701 | 518.7876 | 537.27 |
| ENSMUSG00000027618 | -0.527389441 | 0.002084 | 0.013522 | 746.4923 | 655.1026 | 886.9022 | 812.5913 | 1103.826 | 805.6267 | 1195.719 | 1367.102 |
| ENSMUSG00000027620 | 0.245762599 | 0.071957 | 0.175931 | 15965.08 | 15584.48 | 12651.81 | 14831.09 | 11420.5 | 15794.08 | 11742.15 | 10829.14 |
| ENSMUSG00000027624 | -0.076081466 | 0.601169 | 0.739038 | 867.2189 | 904.6654 | 1222.285 | 957.5121 | 945.858 | 917.3641 | 1082.897 | 1222.452 |
| ENSMUSG00000027628 | -0.041016385 | 0.679456 | 0.795133 | 800.8193 | 869.8706 | 817.3412 | 703.0381 | 795.6908 | 746.4059 | 875.5746 | 861.3721 |
| ENSMUSG00000027630 | 0.271516924 | 0.114804 | 0.242399 | 1520.148 | 1502.177 | 1334.908 | 1521.668 | 1174.034 | 1690.587 | 1012.504 | 994.0583 |
| ENSMUSG00000027634 | -0.094969064 | 0.37075 | 0.539805 | 1350.125 | 1345 | 1435.937 | 1519.943 | 1538.726 | 1246.989 | 1645.077 | 1605.284 |
| ENSMUSG00000027635 | -0.007134982 | 0.923884 | 0.954715 | 627.7779 | 632.3059 | 587.1276 | 642.6545 | 643.5735 | 627.9642 | 624.8594 | 603.6131 |
| ENSMUSG00000027637 | -0.572487369 | 0.001142 | 0.008647 | 1139.86 | 923.8626 | 1011.946 | 988.5665 | 1648.913 | 1017.928 | 1522.613 | 1854.343 |
| ENSMUSG00000027639 | 0.041622946 | 0.537265 | 0.687647 | 1151.932 | 1154.228 | 1191.645 | 1057.576 | 1075.548 | 1093.909 | 1134.004 | 1121.306 |
| ENSMUSG00000027641 | 0.041660341 | 0.719777 | 0.82468 | 913.4974 | 1036.646 | 847.1531 | 854.8599 | 940.0073 | 1003.402 | 782.0385 | 820.0437 |
| ENSMUSG00000027642 | -0.311910298 | 0.001008 | 0.007893 | 4214.362 | 4537.725 | 4937.172 | 4131.967 | 5424.544 | 4901.92 | 5680.627 | 6115.525 |
| ENSMUSG00000027646 | 0.009991474 | 0.879481 | 0.927685 | 1754.559 | 1638.956 | 1849.162 | 1687.292 | 1748.375 | 1631.366 | 1740.542 | 1765.16 |
| ENSMUSG00000027649 | -0.289125429 | 0.012353 | 0.050054 | 1395.397 | 1653.354 | 1524.544 | 1574.288 | 1963.874 | 1508.455 | 2037.543 | 1998.992 |
| ENSMUSG00000027650 | -0.095903974 | 0.479223 | 0.638386 | 671.0383 | 677.8992 | 520.8791 | 536.5518 | 704.0304 | 545.2786 | 641.2523 | 676.4817 |
| ENSMUSG00000027651 | -0.029148154 | 0.710163 | 0.818662 | 1081.508 | 1077.44 | 1059.976 | 933.3586 | 1106.751 | 1033.571 | 1073.254 | 1019.073 |
| ENSMUSG00000027652 | 0.021537433 | 0.788413 | 0.870064 | 1719.347 | 1486.579 | 1516.263 | 1576.013 | 1465.592 | 1505.103 | 1546.72 | 1689.029 |
| ENSMUSG00000027654 | -0.037871994 | 0.771449 | 0.858295 | 602.6266 | 620.3077 | 534.1288 | 673.7089 | 744.985 | 551.9828 | 630.6451 | 565.5474 |
| ENSMUSG00000027655 | -0.145483025 | 0.113729 | 0.240848 | 718.3228 | 664.7011 | 772.6235 | 666.808 | 848.3468 | 735.2322 | 796.5028 | 742.8247 |
| ENSMUSG00000027660 | 0.223128938 | 0.018758 | 0.067439 | 1592.584 | 1642.556 | 1417.719 | 1661.413 | 1291.047 | 1536.389 | 1256.469 | 1325.774 |
| ENSMUSG00000027661 | -0.262985269 | 0.144408 | 0.28259 | 183.1019 | 153.5772 | 172.2462 | 135.4319 | 211.5992 | 150.8455 | 225.6437 | 183.8029 |
| ENSMUSG00000027663 | 0.186660888 | 0.075922 | 0.182475 | 431.5973 | 459.5316 | 502.6607 | 486.5197 | 407.5965 | 412.311 | 374.1442 | 461.1386 |
| ENSMUSG00000027665 | 0.333792256 | 0.026309 | 0.086252 | 2839.085 | 2688.8 | 1994.909 | 2427.423 | 1872.214 | 2451.519 | 1874.578 | 1695.555 |
| ENSMUSG00000027667 | -0.021736638 | 0.833133 | 0.898985 | 1124.769 | 1039.045 | 1079.023 | 1215.437 | 1108.702 | 1318.501 | 1054.932 | 1047.35 |
| ENSMUSG00000027668 | -0.28249617 | 0.003627 | 0.020376 | 1437.652 | 1395.392 | 1448.359 | 1568.25 | 1768.852 | 1530.803 | 1815.756 | 2001.168 |
| ENSMUSG00000027669 | 0.646995597 | 5.37E-12 | 1.29E-09 | 1571.457 | 1544.17 | 1430.14 | 1623.457 | 979.0118 | 1072.679 | 1035.647 | 851.5838 |
| ENSMUSG00000027671 | -0.281891683 | 1.85E-05 | 0.000369 | 1936.655 | 2207.672 | 2148.109 | 2073.747 | 2617.199 | 2406.824 | 2552.473 | 2591.73 |
| ENSMUSG00000027673 | -0.37664037 | 5.63E-05 | 0.000866 | 2126.799 | 2102.087 | 2288.059 | 2177.262 | 2931.185 | 2371.068 | 2976.761 | 3009.365 |
| ENSMUSG00000027674 | 0.323243603 | 0.14004 | 0.276218 | 81.4904 | 82.78769 | 86.9512 | 123.3552 | 68.25779 | 87.15518 | 70.39311 | 75.04379 |
| ENSMUSG00000027676 | 0.698573041 | 0.000303 | 0.003123 | 419.5247 | 449.9331 | 351.9453 | 450.2895 | 257.4294 | 346.386 | 239.1437 | 187.0657 |
| ENSMUSG00000027677 | 0.364396475 | 0.002321 | 0.014607 | 2593.608 | 2488.43 | 1998.221 | 2329.083 | 1709.37 | 2162.119 | 1753.078 | 1683.591 |
| ENSMUSG00000027678 | 0.353777293 | 0.000241 | 0.002628 | 2029.212 | 1982.105 | 1715.837 | 1962.468 | 1330.052 | 1673.826 | 1499.47 | 1513.927 |
| ENSMUSG00000027679 | -0.559942555 | 6.90E-07 | 2.75E-05 | 868.2249 | 941.8599 | 1050.039 | 935.0839 | 1471.443 | 1235.816 | 1282.505 | 1607.46 |
| ENSMUSG00000027680 | 0.021762128 | 0.813152 | 0.886498 | 4315.973 | 4255.767 | 3545.953 | 4261.36 | 4074.015 | 4244.904 | 4191.765 | 3619.503 |
| ENSMUSG00000027684 | 0.189796554 | 0.00725 | 0.03375 | 4576.541 | 4398.546 | 4189.391 | 4249.283 | 3534.778 | 4152.162 | 3884.157 | 3695.635 |
| ENSMUSG00000027690 | -2.365064103 | 0.000406 | 0.003904 | 101.6115 | 16.7975 | 180.5272 | 135.4319 | 566.5397 | 357.5597 | 413.6801 | 903.7882 |
| ENSMUSG00000027692 | 0.993318007 | 1.80E-11 | 3.90E-09 | 256.5439 | 203.9697 | 230.2136 | 251.8861 | 129.6898 | 126.2633 | 113.7861 | 104.4087 |
| ENSMUSG00000027694 | 0.00704345 | 0.973565 | 0.985571 | 90.54489 | 123.5816 | 102.6852 | 94.88858 | 120.9138 | 73.74669 | 101.2504 | 112.0219 |
| ENSMUSG00000027695 | -0.068197567 | 0.593583 | 0.732983 | 264.5923 | 314.3532 | 329.5864 | 332.9727 | 286.6827 | 320.6864 | 361.6084 | 333.8905 |
| ENSMUSG00000027698 | -0.608075633 | 0.021081 | 0.073162 | 961.788 | 428.3363 | 716.3122 | 587.4466 | 825.9193 | 757.5796 | 1167.754 | 1356.226 |
| ENSMUSG00000027699 | 0.115646733 | 0.253709 | 0.417294 | 1310.889 | 1166.227 | 977.9939 | 1152.465 | 1057.996 | 1096.144 | 1025.039 | 1071.277 |
| ENSMUSG00000027702 | 0.656670159 | 0.092481 | 0.208888 | 62.37537 | 145.1784 | 72.87338 | 106.9653 | 34.1289 | 98.32892 | 49.17875 | 64.16787 |
| ENSMUSG00000027703 | 0.700343344 | 0.322714 | 0.490873 | 5.030272 | 34.79482 | 12.4216 | 16.38985 | 3.900445 | 8.938993 | 9.642892 | 19.57664 |
| ENSMUSG00000027706 | 0.084726653 | 0.37966 | 0.547789 | 6192.265 | 6699.803 | 5558.252 | 6159.132 | 5113.484 | 6543.343 | 5737.521 | 5809.912 |
| ENSMUSG00000027708 | 0.01663468 | 0.847795 | 0.908531 | 1342.077 | 1377.395 | 1180.88 | 1258.568 | 1215.964 | 1388.896 | 1304.683 | 1186.562 |
| ENSMUSG00000027709 | -1.194343334 | 2.84E-06 | 8.64E-05 | 949.7153 | 848.2738 | 958.1194 | 853.9973 | 2163.772 | 1051.449 | 2041.4 | 3005.014 |

| | | | | | | | | | | | |
|---|---|---|---|---|---|---|---|---|---|---|---|
| ENSMUSG00000027710 | -0.377321967 | 0.00012 | 0.001543 | 1002.03 | 941.8599 | 1078.195 | 1051.538 | 1404.16 | 1147.543 | 1293.112 | 1449.759 |
| ENSMUSG00000027712 | 0.04943084 | 0.527978 | 0.680556 | 5618.814 | 5616.365 | 5672.53 | 5487.148 | 5648.82 | 4687.384 | 5537.913 | 5764.233 |
| ENSMUSG00000027713 | -0.022050242 | 0.942065 | 0.966348 | 28.16952 | 45.59322 | 65.42042 | 49.16954 | 51.6809 | 45.81234 | 52.07161 | 42.41605 |
| ENSMUSG00000027714 | -0.070317108 | 0.330391 | 0.498218 | 908.4671 | 886.6681 | 939.901 | 910.0678 | 1023.867 | 975.4676 | 894.8603 | 935.3283 |
| ENSMUSG00000027715 | 0.004244798 | 0.960503 | 0.977472 | 2845.122 | 2959.96 | 2657.394 | 3091.643 | 2924.359 | 3202.394 | 2695.188 | 2698.313 |
| ENSMUSG00000027716 | 0.345977099 | 0.032428 | 0.099944 | 301.8163 | 201.57 | 247.6039 | 240.672 | 194.0472 | 181.0146 | 181.2864 | 225.1314 |
| ENSMUSG00000027722 | -0.238896561 | 0.002983 | 0.017679 | 669.0261 | 589.1124 | 634.3297 | 642.6545 | 787.8899 | 762.0491 | 701.0382 | 743.9123 |
| ENSMUSG00000027737 | -0.224560633 | 0.472786 | 0.632209 | 50.30272 | 31.19536 | 35.60859 | 31.91707 | 38.02934 | 54.75133 | 30.85725 | 51.11678 |
| ENSMUSG00000027739 | 0.283795474 | 0.015811 | 0.059881 | 719.3289 | 757.0874 | 640.1264 | 764.2844 | 596.7681 | 689.4198 | 515.8947 | 564.4598 |
| ENSMUSG00000027742 | -0.064890912 | 0.422779 | 0.587871 | 1214.308 | 1082.239 | 1122.084 | 1114.51 | 1170.134 | 1076.031 | 1220.79 | 1274.657 |
| ENSMUSG00000027746 | -0.100860758 | 0.112606 | 0.239294 | 1629.808 | 1715.745 | 1753.102 | 1748.538 | 1760.076 | 1781.094 | 1843.721 | 1960.927 |
| ENSMUSG00000027748 | 0.284012973 | 0.399683 | 0.567034 | 56.33904 | 41.99375 | 25.67131 | 44.85642 | 25.35289 | 43.57759 | 33.75012 | 35.89051 |
| ENSMUSG00000027750 | 0.626623787 | 0.021771 | 0.074822 | 6210.374 | 10319.67 | 7024 | 7783.452 | 5145.662 | 8219.404 | 3387.548 | 3544.459 |
| ENSMUSG00000027751 | 0.169791339 | 0.074344 | 0.179922 | 2628.82 | 2815.981 | 2448.711 | 2614.612 | 2450.455 | 2629.181 | 2090.579 | 2169.744 |
| ENSMUSG00000027752 | -0.052583712 | 0.454975 | 0.61711 | 1222.356 | 1319.804 | 1180.88 | 1208.536 | 1320.301 | 1336.379 | 1216.933 | 1236.591 |
| ENSMUSG00000027761 | -2.224328388 | 0.000158 | 0.001895 | 20.12109 | 10.79839 | 100.2009 | 44.85642 | 180.3956 | 102.7984 | 200.5721 | 343.6788 |
| ENSMUSG00000027762 | -5.116213779 | 4.84E-20 | 6.38E-17 | 3.018163 | 3.599465 | 6.624853 | 5.175741 | 209.6489 | 65.92507 | 104.1432 | 270.8102 |
| ENSMUSG00000027763 | 0.410619065 | 0.021751 | 0.074822 | 3061.423 | 2663.604 | 2091.797 | 2706.05 | 1697.669 | 2705.163 | 1752.113 | 1761.898 |
| ENSMUSG00000027765 | -0.374395842 | 0.182494 | 0.333081 | 36.21796 | 49.19268 | 36.43669 | 56.07053 | 48.75557 | 77.09881 | 54.96448 | 50.02919 |
| ENSMUSG00000027770 | 0.213586755 | 0.061369 | 0.157418 | 2392.397 | 2324.054 | 1944.394 | 2180.712 | 1760.076 | 2263.8 | 1737.649 | 1863.044 |
| ENSMUSG00000027774 | -0.507650561 | 0.000646 | 0.005588 | 1129.799 | 997.0517 | 1129.537 | 1085.18 | 1481.194 | 1160.952 | 1615.184 | 1916.335 |
| ENSMUSG00000027775 | -0.482119246 | 0.00016 | 0.001913 | 2065.43 | 1954.509 | 2350.167 | 2220.393 | 3218.842 | 2352.072 | 3003.761 | 3427 |
| ENSMUSG00000027777 | -0.023966412 | 0.913175 | 0.948699 | 135.8173 | 154.777 | 198.7456 | 144.9207 | 187.2214 | 113.9722 | 149.4648 | 194.6788 |
| ENSMUSG00000027778 | 0.354002703 | 0.00334 | 0.019158 | 1202.235 | 1286.209 | 971.3691 | 1198.184 | 900.0277 | 1044.745 | 884.2532 | 813.5181 |
| ENSMUSG00000027782 | -0.002779329 | 0.959492 | 0.977106 | 1810.898 | 1750.54 | 1666.979 | 1794.257 | 1736.673 | 1768.803 | 1749.221 | 1780.387 |
| ENSMUSG00000027784 | 0.030476418 | 0.76832 | 0.856029 | 688.1412 | 667.1008 | 728.7338 | 663.3575 | 566.5397 | 703.9457 | 693.3239 | 728.686 |
| ENSMUSG00000027787 | -0.249768649 | 0.002508 | 0.015472 | 949.7153 | 951.4585 | 1006.978 | 1021.346 | 1209.138 | 1043.627 | 1161.968 | 1259.43 |
| ENSMUSG00000027792 | 0.877827731 | 1.02E-05 | 0.000231 | 395.3794 | 526.7217 | 381.7572 | 480.4813 | 251.5787 | 317.3342 | 226.608 | 175.1022 |
| ENSMUSG00000027793 | 0.804597467 | 0.004195 | 0.022612 | 40.24217 | 46.79304 | 39.74912 | 45.71905 | 28.27823 | 21.23011 | 27.96439 | 20.66423 |
| ENSMUSG00000027796 | -0.090399286 | 0.626013 | 0.756182 | 220.3259 | 143.9786 | 246.7758 | 213.068 | 223.3005 | 240.2354 | 226.608 | 190.3284 |
| ENSMUSG00000027797 | -0.093533084 | 0.525407 | 0.678628 | 687.1351 | 778.6842 | 1042.586 | 822.9428 | 783.9895 | 850.3217 | 1003.825 | 919.0145 |
| ENSMUSG00000027799 | 0.17943111 | 0.083293 | 0.19427 | 1430.609 | 1187.823 | 1372.173 | 1337.929 | 1024.842 | 1322.971 | 1169.683 | 1192 |
| ENSMUSG00000027800 | -0.412747801 | 0.012294 | 0.049892 | 2083.539 | 1656.954 | 2009.815 | 2006.462 | 2423.152 | 1850.372 | 2993.154 | 3060.481 |
| ENSMUSG00000027801 | 1.171654156 | 0.128664 | 0.260297 | 39.23612 | 4.799286 | 16.56213 | 33.64232 | 6.825779 | 4.469496 | 5.785735 | 25.0146 |
| ENSMUSG00000027803 | 0.15647781 | 0.007871 | 0.035814 | 2582.542 | 2766.788 | 2761.736 | 2817.328 | 2311.989 | 2467.162 | 2507.152 | 2521.036 |
| ENSMUSG00000027804 | 0.293128442 | 0.016501 | 0.061505 | 2988.987 | 2983.956 | 2521.585 | 3099.406 | 2480.683 | 2802.374 | 2114.686 | 2064.248 |
| ENSMUSG00000027805 | -0.414250174 | 0.1577 | 0.300269 | 1870.255 | 1956.909 | 4530.571 | 2310.106 | 3314.403 | 2375.537 | 3864.871 | 4663.591 |
| ENSMUSG00000027806 | -0.052426509 | 0.720519 | 0.825136 | 898.4065 | 827.8769 | 1239.676 | 937.6717 | 902.9531 | 997.8151 | 992.2536 | 1159.372 |
| ENSMUSG00000027808 | -0.229163802 | 0.011494 | 0.047437 | 3478.936 | 3444.688 | 3538.5 | 3380.621 | 3693.722 | 3648.226 | 4307.48 | 4576.583 |
| ENSMUSG00000027809 | -0.516896228 | 0.001383 | 0.009973 | 1512.1 | 1323.403 | 1557.669 | 1497.514 | 2146.22 | 1477.169 | 2215.937 | 2590.642 |
| ENSMUSG00000027810 | 0.136461607 | 0.22725 | 0.387197 | 3244.525 | 3343.903 | 2861.108 | 3645.447 | 3100.854 | 3376.705 | 2880.332 | 2554.751 |
| ENSMUSG00000027811 | 0.111737853 | 0.306414 | 0.475001 | 302.8224 | 317.9527 | 273.2752 | 309.6818 | 270.1058 | 302.8084 | 278.6796 | 261.0219 |
| ENSMUSG00000027820 | -0.497330569 | 0.001873 | 0.012519 | 2760.613 | 2879.572 | 3304.145 | 2838.031 | 3559.156 | 3221.39 | 4515.766 | 5336.809 |
| ENSMUSG00000027822 | -0.327832947 | 0.000982 | 0.007744 | 601.6205 | 663.5013 | 606.1741 | 598.6607 | 744.0099 | 700.5936 | 766.6099 | 886.3867 |
| ENSMUSG00000027823 | -0.079154785 | 0.286935 | 0.454019 | 2518.154 | 3012.752 | 2731.096 | 2561.129 | 2870.728 | 2936.459 | 2819.582 | 2801.635 |
| ENSMUSG00000027827 | -0.072364033 | 0.747765 | 0.842597 | 101.6115 | 80.38804 | 116.763 | 130.2561 | 103.3618 | 147.4934 | 96.42892 | 106.5839 |
| ENSMUSG00000027828 | 0.090317646 | 0.09821 | 0.218244 | 9706.412 | 9635.767 | 9122.423 | 9891.704 | 8876.438 | 9581.483 | 8757.674 | 8813.838 |
| ENSMUSG00000027829 | 0.230551309 | 0.044627 | 0.125859 | 2592.602 | 2422.44 | 2145.624 | 2643.078 | 2090.639 | 2444.815 | 2008.614 | 1811.927 |
| ENSMUSG00000027831 | -0.283688802 | 0.035017 | 0.106121 | 776.674 | 724.6922 | 1019.399 | 753.9329 | 945.858 | 896.134 | 1082.897 | 1064.752 |
| ENSMUSG00000027832 | -0.084524992 | 0.799126 | 0.877076 | 103.6236 | 97.18554 | 213.6515 | 99.2017 | 137.4907 | 97.21155 | 116.679 | 194.6788 |
| ENSMUSG00000027833 | 3.654136851 | 0.002727 | 0.016482 | 18.10898 | 5.999108 | 2.48432 | 50.89479 | 0 | 0 | 2.892867 | 3.262773 |
| ENSMUSG00000027834 | -0.172484862 | 0.54848 | 0.696823 | 144.8718 | 76.78858 | 76.18581 | 121.6299 | 94.5858 | 99.44629 | 159.1077 | 119.635 |
| ENSMUSG00000027835 | -0.141580623 | 0.095023 | 0.213065 | 1337.046 | 1167.426 | 1191.645 | 1317.226 | 1340.778 | 1321.854 | 1522.613 | 1345.35 |
| ENSMUSG00000027840 | 1.06620754 | 0.001677 | 0.011496 | 210.2654 | 266.3604 | 154.8559 | 215.6559 | 77.03379 | 183.2494 | 77.14313 | 67.43065 |
| ENSMUSG00000027843 | -0.80546883 | 0.242197 | 0.404634 | 19.11503 | 10.79839 | 13.24971 | 24.15346 | 7.80089 | 74.86406 | 18.32149 | 17.40146 |
| ENSMUSG00000027845 | -0.307971979 | 0.014759 | 0.056975 | 421.5368 | 397.1409 | 337.0394 | 384.7301 | 442.7005 | 425.7195 | 553.502 | 481.8029 |
| ENSMUSG00000027848 | 0.26701623 | 0.042925 | 0.122598 | 2352.155 | 2556.82 | 3199.804 | 2661.193 | 1965.824 | 2023.564 | 2343.223 | 2620.007 |
| ENSMUSG00000027852 | 0.167896659 | 0.120012 | 0.248953 | 3243.519 | 3220.321 | 2947.232 | 3230.525 | 2722.511 | 3401.287 | 2510.045 | 2620.007 |
| ENSMUSG00000027854 | 0.075298771 | 0.26217 | 0.426384 | 1579.505 | 1684.549 | 1743.164 | 1621.732 | 1556.278 | 1673.826 | 1489.827 | 1574.832 |
| ENSMUSG00000027855 | -3.702191385 | NA | NA | 3.018163 | 0 | 3.312427 | 18.11509 | 1.950223 | 316.2169 | 0 | 1.087591 |
| ENSMUSG00000027858 | -0.012612935 | 0.935264 | 0.961682 | 317.9132 | 295.1561 | 349.461 | 408.8835 | 318.8614 | 367.6161 | 295.0725 | 405.6715 |
| ENSMUSG00000027859 | -0.473625329 | 0.127835 | 0.259223 | 22.1332 | 37.19447 | 50.5145 | 38.81806 | 43.88001 | 45.81234 | 56.89306 | 60.9051 |
| ENSMUSG00000027860 | 0.04053298 | 0.606419 | 0.742598 | 688.1412 | 749.8885 | 711.3436 | 725.4664 | 736.209 | 679.3635 | 650.8952 | 727.5984 |
| ENSMUSG00000027861 | -0.65376641 | 0.127734 | 0.259118 | 15.09082 | 9.598572 | 12.4216 | 8.626235 | 23.40267 | 16.76061 | 15.42863 | 16.31387 |

| | | | | | | | | | | | |
|---|---|---|---|---|---|---|---|---|---|---|---|
| ENSMUSG00000027864 | 0.233438321 | 0.004727 | 0.024729 | 1975.891 | 2304.857 | 2240.028 | 2033.204 | 1669.391 | 1885.01 | 1915.078 | 1805.401 |
| ENSMUSG00000027865 | -0.259695526 | 0.029286 | 0.092965 | 383.3067 | 377.9438 | 334.5551 | 389.0432 | 416.3725 | 400.0199 | 441.6444 | 518.7809 |
| ENSMUSG00000027867 | 0.600242712 | 0.247042 | 0.410092 | 54.32694 | 16.7975 | 23.18699 | 25.01608 | 15.60178 | 31.28647 | 22.17865 | 9.78832 |
| ENSMUSG00000027869 | -1.375607579 | 0.316289 | 0.484635 | 12.07265 | 1.199822 | 224.4169 | 7.763611 | 117.0134 | 41.34284 | 172.6078 | 306.7007 |
| ENSMUSG00000027870 | -2.333951908 | 0.000626 | 0.005449 | 472.8455 | 56.39161 | 439.7246 | 269.1385 | 1461.692 | 688.3024 | 1488.862 | 2605.868 |
| ENSMUSG00000027871 | 4.176439867 | 0.038415 | 0.113239 | 11014.28 | 9.598572 | 31.46805 | 1269.782 | 1.950223 | 32.40385 | 6.750024 | 640.5912 |
| ENSMUSG00000027875 | 0.286740591 | 0.305775 | 0.474313 | 624.7598 | 807.4799 | 728.7338 | 774.6359 | 541.1868 | 476.0014 | 366.4299 | 1023.423 |
| ENSMUSG00000027878 | 0.229563744 | 0.092005 | 0.208271 | 2792.807 | 3059.545 | 3089.666 | 2827.68 | 2273.96 | 3263.85 | 2206.294 | 2295.905 |
| ENSMUSG00000027879 | 0.087105107 | 0.132862 | 0.266242 | 2242.495 | 2237.667 | 2038.799 | 2195.377 | 2048.709 | 2105.133 | 1992.221 | 2054.46 |
| ENSMUSG00000027881 | 0.283598001 | 0.006337 | 0.030664 | 4796.867 | 4368.55 | 3854.008 | 4408.869 | 3429.466 | 4156.632 | 3299.798 | 3432.437 |
| ENSMUSG00000027882 | 0.054819509 | 0.636791 | 0.76415 | 789.7527 | 722.2926 | 674.0788 | 803.9651 | 668.9264 | 630.199 | 822.5387 | 755.8758 |
| ENSMUSG00000027883 | 0.02050077 | 0.79502 | 0.873958 | 1109.678 | 1150.629 | 1081.507 | 1188.695 | 1217.914 | 1139.722 | 1053.968 | 1053.876 |
| ENSMUSG00000027884 | -0.058980324 | 0.376626 | 0.544772 | 1046.297 | 1082.239 | 980.4783 | 1084.318 | 1110.652 | 1082.735 | 1115.683 | 1054.963 |
| ENSMUSG00000027887 | -0.634543777 | 0.02736 | 0.088541 | 99.59938 | 116.3827 | 91.91984 | 73.323 | 135.5405 | 119.559 | 105.1075 | 230.5693 |
| ENSMUSG00000027889 | 0.066971981 | 0.389269 | 0.556667 | 1590.572 | 1367.797 | 1438.421 | 1337.066 | 1374.907 | 1341.966 | 1334.576 | 1422.569 |
| ENSMUSG00000027890 | -0.342509511 | 0.041767 | 0.120306 | 279.6831 | 322.752 | 307.2276 | 317.4454 | 405.6463 | 278.2261 | 399.2157 | 472.0145 |
| ENSMUSG00000027893 | -0.014723491 | 0.790027 | 0.871324 | 5699.298 | 5676.356 | 5186.432 | 5716.606 | 5701.476 | 5647.209 | 5743.306 | 5409.678 |
| ENSMUSG00000027894 | -0.889202686 | 9.36E-07 | 3.53E-05 | 162.9808 | 170.3747 | 236.0104 | 191.5024 | 351.0401 | 275.9914 | 363.537 | 421.9853 |
| ENSMUSG00000027895 | 1.074702826 | 0.000287 | 0.003003 | 144.8718 | 103.1847 | 78.67013 | 121.6299 | 44.85512 | 77.09881 | 53.0359 | 38.06569 |
| ENSMUSG00000027896 | -0.440994896 | 0.022521 | 0.076682 | 295.78 | 320.3523 | 281.5563 | 364.8897 | 429.049 | 353.0902 | 346.1798 | 586.2116 |
| ENSMUSG00000027900 | -0.211576898 | 0.002402 | 0.014968 | 1244.489 | 1190.223 | 1295.159 | 1334.479 | 1413.911 | 1422.417 | 1472.47 | 1561.781 |
| ENSMUSG00000027901 | -0.255450201 | 0.18027 | 0.330361 | 216.3017 | 179.9732 | 279.0719 | 198.4034 | 268.1556 | 200.01 | 278.6796 | 298 |
| ENSMUSG00000027905 | -0.066905924 | 0.404302 | 0.571173 | 840.0554 | 754.6877 | 790.0137 | 834.1569 | 827.8695 | 895.0167 | 782.0385 | 871.1605 |
| ENSMUSG00000027907 | 0.323124154 | 2.92E-05 | 0.000527 | 8112.822 | 8595.521 | 9847.016 | 8611.57 | 7053.955 | 7363.495 | 7050.882 | 6645.182 |
| ENSMUSG00000027931 | 0.193763129 | 0.290271 | 0.457376 | 1870.255 | 1037.846 | 1345.673 | 1377.61 | 1131.129 | 1044.745 | 1306.612 | 1442.146 |
| ENSMUSG00000027932 | -0.125195497 | 0.477761 | 0.63706 | 557.3541 | 428.3363 | 362.7107 | 398.5321 | 488.5308 | 392.1983 | 444.5373 | 578.5985 |
| ENSMUSG00000027933 | -0.056082314 | 0.55099 | 0.698714 | 1534.233 | 1584.964 | 1454.983 | 1449.207 | 1463.642 | 1384.427 | 1692.327 | 1718.394 |
| ENSMUSG00000027935 | 0.132445905 | 0.055868 | 0.147215 | 1273.665 | 1250.214 | 1242.16 | 1186.107 | 1118.453 | 1125.196 | 1060.718 | 1213.752 |
| ENSMUSG00000027936 | 0.189848672 | 0.023951 | 0.080488 | 1112.696 | 1094.237 | 1050.867 | 1091.219 | 1054.095 | 850.3217 | 964.2892 | 940.7663 |
| ENSMUSG00000027937 | -0.018738607 | 0.804515 | 0.880306 | 1375.276 | 1228.617 | 1193.302 | 1343.967 | 1347.604 | 1243.637 | 1332.648 | 1283.357 |
| ENSMUSG00000027938 | 0.495267262 | 0.000828 | 0.006734 | 149.9021 | 135.5798 | 147.403 | 164.7611 | 109.2125 | 111.7374 | 105.1075 | 98.97079 |
| ENSMUSG00000027939 | 0.547627202 | 0.134249 | 0.267976 | 81.4904 | 76.78858 | 52.99882 | 80.22398 | 69.2329 | 69.27719 | 35.6787 | 25.0146 |
| ENSMUSG00000027940 | -0.108123335 | 0.227193 | 0.387145 | 6146.992 | 6270.267 | 7131.654 | 6067.694 | 6935.967 | 6211.483 | 6804.989 | 7659.904 |
| ENSMUSG00000027942 | -0.058246111 | 0.434413 | 0.598591 | 1651.941 | 1430.187 | 1438.421 | 1601.029 | 1640.137 | 1614.606 | 1582.399 | 1537.854 |
| ENSMUSG00000027944 | -0.169183358 | 0.092901 | 0.209563 | 1523.166 | 1446.985 | 1270.316 | 1497.514 | 1636.237 | 1406.774 | 1623.863 | 1782.562 |
| ENSMUSG00000027947 | 0.605309501 | 0.00629 | 0.030455 | 163.9869 | 159.5763 | 116.763 | 188.0519 | 83.85957 | 106.1505 | 131.1433 | 91.35765 |
| ENSMUSG00000027950 | -0.039792122 | 0.865653 | 0.919563 | 64.38748 | 56.39161 | 64.59232 | 77.63611 | 53.63112 | 58.10345 | 79.07171 | 80.48174 |
| ENSMUSG00000027951 | 0.00903547 | 0.955405 | 0.974735 | 694.1775 | 914.264 | 1005.321 | 622.8142 | 787.8899 | 750.8754 | 862.0745 | 814.6057 |
| ENSMUSG00000027952 | -0.560786514 | 0.000209 | 0.002355 | 636.8324 | 550.7181 | 669.9383 | 606.4243 | 878.5753 | 681.5982 | 1016.361 | 1059.314 |
| ENSMUSG00000027953 | 0.210043588 | 0.044495 | 0.125603 | 1147.908 | 1240.615 | 932.4481 | 1164.542 | 972.186 | 1021.28 | 948.8605 | 932.0656 |
| ENSMUSG00000027954 | -0.209891164 | 0.055564 | 0.146791 | 1095.593 | 1007.85 | 1284.393 | 1080.005 | 1373.932 | 1120.726 | 1297.933 | 1377.978 |
| ENSMUSG00000027955 | 0.054841548 | 0.643251 | 0.76925 | 479.8879 | 415.1383 | 477.8175 | 442.5259 | 391.9947 | 444.7149 | 404.0372 | 510.0802 |
| ENSMUSG00000027956 | -0.333916962 | 0.26702 | 0.431421 | 241.453 | 119.9822 | 96.06037 | 128.5309 | 184.296 | 141.9065 | 182.2507 | 229.4817 |
| ENSMUSG00000027957 | -0.187760517 | 0.087507 | 0.201124 | 1018.127 | 824.2774 | 897.6676 | 873.8376 | 877.6002 | 1062.623 | 1055.897 | 1122.394 |
| ENSMUSG00000027959 | -0.227654707 | 0.045356 | 0.127178 | 257.5499 | 225.5664 | 221.1045 | 227.7326 | 292.5334 | 274.874 | 254.5723 | 268.635 |
| ENSMUSG00000027961 | 0.535759544 | 0.055009 | 0.145849 | 100.6054 | 94.7859 | 78.67013 | 111.2784 | 69.2329 | 98.32892 | 54.00019 | 44.59123 |
| ENSMUSG00000027962 | -0.06697888 | 0.821689 | 0.891697 | 546.2875 | 201.57 | 409.9128 | 464.0914 | 274.0063 | 459.2408 | 497.5732 | 469.8393 |
| ENSMUSG00000027963 | -0.030900576 | 0.757725 | 0.848991 | 690.1533 | 754.6877 | 820.6537 | 705.626 | 681.6028 | 745.2885 | 777.2171 | 833.0948 |
| ENSMUSG00000027966 | 0.569358977 | 0.000125 | 0.00159 | 502.0211 | 565.1159 | 549.8628 | 558.98 | 300.3343 | 471.5319 | 348.1084 | 348.0291 |
| ENSMUSG00000027968 | 0.141320502 | 0.306766 | 0.475352 | 2925.606 | 3360.7 | 2921.56 | 3083.016 | 2660.104 | 3597.945 | 2521.616 | 2364.423 |
| ENSMUSG00000027977 | -0.437125981 | 0.324757 | 0.492702 | 39.23612 | 14.39786 | 34.78048 | 31.91707 | 28.27823 | 69.27719 | 40.50014 | 26.10219 |
| ENSMUSG00000027978 | -0.499153062 | 0.086576 | 0.19963 | 91.55095 | 76.78858 | 149.8873 | 98.33908 | 120.9138 | 102.7984 | 165.8577 | 201.2044 |
| ENSMUSG00000027981 | 0.126259512 | 0.108862 | 0.233894 | 806.8556 | 772.6851 | 758.5457 | 773.7733 | 725.4828 | 772.1055 | 693.3239 | 660.1678 |
| ENSMUSG00000027983 | -0.770468943 | 0.005901 | 0.02906 | 36.21796 | 39.59411 | 52.99882 | 50.03216 | 71.18313 | 54.75133 | 75.21455 | 105.4963 |
| ENSMUSG00000027984 | -0.250712959 | 0.04757 | 0.13141 | 3478.936 | 3012.752 | 3552.577 | 3342.666 | 4131.547 | 3188.986 | 3855.228 | 4753.861 |
| ENSMUSG00000027985 | 0.499255885 | 0.00291 | 0.017357 | 391.3551 | 362.3461 | 442.2089 | 367.4776 | 211.5992 | 319.569 | 248.7866 | 328.4525 |
| ENSMUSG00000027993 | 0.532440382 | 0.000667 | 0.005722 | 3319.979 | 2844.777 | 2563.818 | 2697.424 | 1615.759 | 2575.547 | 1762.721 | 1946.788 |
| ENSMUSG00000027994 | 0.994941146 | 0.000448 | 0.004211 | 47.28455 | 55.19179 | 39.74912 | 59.52102 | 26.32801 | 21.23011 | 25.07152 | 28.27737 |
| ENSMUSG00000027995 | 0.235088136 | 0.1732 | 0.320975 | 137.8294 | 143.9786 | 198.7456 | 156.9975 | 127.7396 | 130.7328 | 133.0719 | 152.2628 |
| ENSMUSG00000027996 | -0.308681943 | 0.003777 | 0.020951 | 1403.446 | 1370.196 | 1801.132 | 1590.678 | 1872.214 | 2069.377 | 1883.257 | 1818.452 |
| ENSMUSG00000027997 | 0.067225112 | 0.455359 | 0.617359 | 1245.495 | 1141.03 | 1122.084 | 1254.255 | 1274.47 | 1125.196 | 1040.468 | 1106.08 |
| ENSMUSG00000027998 | -0.136235467 | 0.05009 | 0.13625 | 1447.712 | 1451.784 | 1541.935 | 1550.134 | 1787.379 | 1577.732 | 1632.542 | 1588.971 |
| ENSMUSG00000027999 | -0.364850864 | 7.46E-06 | 0.000184 | 836.0312 | 729.4915 | 811.5445 | 832.4317 | 1088.224 | 954.2375 | 1053.968 | 1039.737 |
| ENSMUSG00000028001 | 1.072700605 | 0.405129 | 0.572034 | 761.5831 | 1.199822 | 49.6864 | 406.2957 | 149.192 | 77.09881 | 172.6078 | 180.5401 |

| | | | | | | | | | | | |
|---|---|---|---|---|---|---|---|---|---|---|---|
| ENSMUSG00000028004 | 0.151548669 | 0.745364 | 0.840858 | 25.15136 | 17.99732 | 24.01509 | 25.8787 | 13.65156 | 20.11273 | 11.57147 | 39.15328 |
| ENSMUSG00000028005 | 0.120016924 | 0.296299 | 0.464124 | 818.9282 | 855.4728 | 850.4655 | 872.975 | 832.7451 | 643.6075 | 748.2884 | 901.613 |
| ENSMUSG00000028007 | -0.681607015 | 0.000215 | 0.002402 | 705.2441 | 557.917 | 691.469 | 732.3673 | 1004.365 | 797.8051 | 1054.932 | 1455.197 |
| ENSMUSG00000028010 | -0.146950646 | 0.043466 | 0.123545 | 1596.608 | 1614.96 | 1594.105 | 1495.789 | 1902.442 | 1670.474 | 1762.721 | 1636.825 |
| ENSMUSG00000028011 | 1.272993394 | 0.116411 | 0.244577 | 24.1453 | 32.39518 | 11.59349 | 22.42821 | 5.850668 | 26.81698 | 3.857157 | 1.087591 |
| ENSMUSG00000028012 | -0.245760009 | 0.560364 | 0.706282 | 10.06054 | 21.59679 | 10.76539 | 18.11509 | 19.50223 | 17.87799 | 17.3572 | 16.31387 |
| ENSMUSG00000028013 | -0.45847679 | 7.67E-05 | 0.001108 | 720.3349 | 805.0802 | 852.9498 | 751.3451 | 1118.453 | 892.7819 | 1074.218 | 1214.839 |
| ENSMUSG00000028015 | 0.511115758 | 0.000126 | 0.001592 | 980.903 | 919.0633 | 891.0427 | 993.7423 | 549.9628 | 825.7395 | 613.2879 | 668.8685 |
| ENSMUSG00000028016 | -0.035992791 | 0.718951 | 0.824245 | 791.7648 | 657.5022 | 671.5945 | 686.6483 | 724.5077 | 645.8422 | 729.9669 | 777.6276 |
| ENSMUSG00000028017 | -0.566778318 | 0.003246 | 0.018828 | 80.48435 | 113.983 | 92.74794 | 86.26235 | 128.7147 | 116.2069 | 139.8219 | 166.4014 |
| ENSMUSG00000028018 | -0.057202954 | 0.46937 | 0.629429 | 867.2189 | 867.471 | 869.512 | 934.2212 | 851.2722 | 1003.402 | 919.9319 | 910.3137 |
| ENSMUSG00000028019 | 0.1035373 | 0.508814 | 0.664543 | 2638.881 | 3117.136 | 2285.574 | 3149.438 | 3025.77 | 2860.478 | 2533.188 | 1994.642 |
| ENSMUSG00000028020 | 0.544644586 | 0.006181 | 0.030108 | 317.9132 | 419.9375 | 250.0882 | 312.2697 | 192.0969 | 270.4045 | 239.1437 | 188.1533 |
| ENSMUSG00000028023 | -3.374290411 | 1.23E-06 | 4.42E-05 | 5.030272 | 8.398751 | 39.74912 | 6.038364 | 114.088 | 46.92971 | 176.4649 | 282.7737 |
| ENSMUSG00000028024 | -1.193570204 | 0.004381 | 0.023367 | 1554.354 | 795.4817 | 2146.452 | 1437.993 | 2656.203 | 1909.592 | 3600.656 | 5408.59 |
| ENSMUSG00000028028 | 0.212967807 | 0.296611 | 0.464315 | 189.1382 | 110.3836 | 166.4494 | 125.0804 | 127.7396 | 120.6764 | 134.0362 | 128.3357 |
| ENSMUSG00000028029 | 0.015411358 | 0.842079 | 0.904759 | 2773.692 | 2622.81 | 2889.264 | 2939.821 | 3024.795 | 2818.017 | 2736.653 | 2529.737 |
| ENSMUSG00000028030 | 0.078350832 | 0.56872 | 0.712804 | 590.5539 | 729.4915 | 589.6119 | 580.5456 | 533.3859 | 720.7063 | 568.9306 | 534.0072 |
| ENSMUSG00000028031 | 0.13478147 | 0.311819 | 0.480258 | 530.1906 | 665.901 | 480.3018 | 518.4367 | 445.6259 | 474.884 | 542.8948 | 531.832 |
| ENSMUSG00000028032 | -0.315462764 | 0.014299 | 0.055669 | 2897.437 | 2925.165 | 3414.284 | 3136.499 | 3756.129 | 3010.206 | 4249.622 | 4382.992 |
| ENSMUSG00000028033 | 0.713582772 | 0.011521 | 0.047483 | 118.7144 | 140.3791 | 109.3101 | 110.4158 | 95.56091 | 87.15518 | 70.39311 | 38.06569 |
| ENSMUSG00000028034 | 0.130671164 | 0.071903 | 0.175884 | 6522.25 | 6295.464 | 5539.205 | 6218.653 | 5469.399 | 6004.768 | 5454.984 | 5516.262 |
| ENSMUSG00000028035 | 0.032831357 | 0.743864 | 0.839833 | 1004.042 | 843.4745 | 862.8871 | 891.9527 | 846.3966 | 798.9225 | 987.4321 | 887.4743 |
| ENSMUSG00000028036 | 0.335050763 | 0.193196 | 0.346447 | 195.1745 | 154.777 | 149.8873 | 192.365 | 81.90935 | 124.0285 | 184.1792 | 158.7883 |
| ENSMUSG00000028037 | -0.227337684 | 0.508074 | 0.663765 | 232.3986 | 283.1579 | 475.3332 | 151.8217 | 337.3885 | 204.4795 | 443.573 | 352.3795 |
| ENSMUSG00000028039 | -0.243423243 | 0.221285 | 0.3803 | 94.56911 | 101.9848 | 135.8095 | 101.7896 | 108.2374 | 116.2069 | 138.8576 | 152.2628 |
| ENSMUSG00000028040 | 0.162522557 | 0.137136 | 0.272118 | 1096.599 | 1263.412 | 1229.738 | 1187.833 | 1107.726 | 1203.412 | 1069.397 | 887.4743 |
| ENSMUSG00000028041 | -0.510542029 | 0.002875 | 0.017201 | 1010.079 | 867.471 | 1225.598 | 960.9626 | 1395.384 | 1081.618 | 1503.327 | 1811.927 |
| ENSMUSG00000028042 | -0.63770121 | 0.008973 | 0.039391 | 276.6649 | 245.9634 | 437.2403 | 302.7808 | 450.5014 | 326.2732 | 575.6806 | 614.489 |
| ENSMUSG00000028044 | -0.122611542 | 0.121816 | 0.251178 | 1898.425 | 2096.088 | 1994.081 | 1997.836 | 2304.188 | 2321.903 | 2115.65 | 1951.138 |
| ENSMUSG00000028047 | 0.800255611 | 2.79E-05 | 0.000508 | 865.2067 | 1103.836 | 978.822 | 1001.506 | 537.2863 | 806.7441 | 484.0732 | 440.4744 |
| ENSMUSG00000028048 | -0.10450629 | 0.411647 | 0.577482 | 707.2562 | 753.4879 | 856.2623 | 769.4602 | 974.1362 | 678.2461 | 844.7173 | 821.1313 |
| ENSMUSG00000028049 | -0.084960626 | 0.397629 | 0.564936 | 1438.658 | 1379.795 | 1580.027 | 1390.549 | 1530.925 | 1315.149 | 1581.434 | 1714.044 |
| ENSMUSG00000028051 | -0.201549832 | 0.479454 | 0.638512 | 65.39353 | 37.19447 | 37.2648 | 66.42201 | 57.53157 | 59.22083 | 66.53595 | 54.37955 |
| ENSMUSG00000028053 | 0.281105774 | 0.004938 | 0.025514 | 2395.415 | 2164.478 | 1948.535 | 2101.351 | 1570.904 | 1977.752 | 1743.435 | 1793.438 |
| ENSMUSG00000028057 | -0.092477007 | 0.45106 | 0.613563 | 711.2804 | 739.0901 | 821.4818 | 706.4886 | 810.3175 | 642.4901 | 815.7886 | 907.051 |
| ENSMUSG00000028059 | 0.080733914 | 0.192289 | 0.34549 | 2779.728 | 2797.984 | 2762.564 | 2594.771 | 2645.477 | 2457.106 | 2735.688 | 2496.022 |
| ENSMUSG00000028060 | 0.139434563 | 0.180892 | 0.331139 | 3778.74 | 3300.709 | 3454.861 | 3630.782 | 3164.236 | 3810.246 | 3028.832 | 2859.277 |
| ENSMUSG00000028062 | -0.280499865 | 0.029678 | 0.093817 | 1352.137 | 1334.202 | 1411.922 | 1505.278 | 1812.732 | 1321.854 | 1687.506 | 1984.854 |
| ENSMUSG00000028063 | -0.249204004 | 0.078679 | 0.187031 | 1642.887 | 1473.381 | 2101.735 | 1791.669 | 2071.136 | 1710.7 | 2155.186 | 2398.138 |
| ENSMUSG00000028064 | -0.272254098 | 0.158406 | 0.301271 | 598.6023 | 585.5129 | 724.5933 | 540.8649 | 845.4215 | 526.2832 | 643.1809 | 944.0291 |
| ENSMUSG00000028066 | -0.245740378 | 0.001312 | 0.009598 | 1126.781 | 1233.417 | 1217.317 | 1242.178 | 1519.223 | 1465.995 | 1429.077 | 1299.671 |
| ENSMUSG00000028068 | -0.039208145 | 0.59463 | 0.733932 | 813.898 | 811.0794 | 747.7803 | 761.6965 | 772.2882 | 845.8522 | 817.7172 | 781.978 |
| ENSMUSG00000028069 | -0.202846529 | 0.007245 | 0.033739 | 984.9272 | 1027.047 | 956.4632 | 909.2052 | 1198.412 | 1061.505 | 1107.004 | 1089.766 |
| ENSMUSG00000028070 | -0.147803339 | 0.027071 | 0.087924 | 1673.068 | 1882.52 | 1797.819 | 1741.637 | 2010.68 | 1848.137 | 1948.828 | 2049.022 |
| ENSMUSG00000028072 | 0.667738534 | 0.249808 | 0.413494 | 100.6054 | 11.99822 | 77.84202 | 162.1732 | 50.70579 | 39.10809 | 59.78593 | 72.8686 |
| ENSMUSG00000028073 | 0.196752095 | 0.021312 | 0.073719 | 1971.867 | 1696.548 | 1962.613 | 1897.772 | 1627.461 | 1634.718 | 1801.292 | 1507.401 |
| ENSMUSG00000028076 | 0.670560497 | 5.80E-06 | 0.000151 | 772.6497 | 615.5084 | 605.3459 | 693.5493 | 439.7752 | 510.64 | 392.4657 | 345.854 |
| ENSMUSG00000028078 | 0.985693334 | 1.68E-27 | 4.42E-24 | 665.0019 | 655.1026 | 664.9696 | 658.1817 | 309.1103 | 367.6161 | 312.4297 | 348.0291 |
| ENSMUSG00000028080 | -0.292660694 | 0.049776 | 0.135713 | 1571.457 | 1377.395 | 1927.004 | 1504.415 | 1781.528 | 1600.08 | 2151.329 | 2285.029 |
| ENSMUSG00000028081 | -0.026833678 | 0.692934 | 0.805613 | 45349.91 | 49971.37 | 47292.34 | 48909.03 | 52230.86 | 49420.34 | 49007.1 | 44456.37 |
| ENSMUSG00000028082 | 0.35801962 | 0.000121 | 0.001551 | 1828.001 | 2055.294 | 1672.775 | 1908.986 | 1408.061 | 1595.61 | 1340.362 | 1479.124 |
| ENSMUSG00000028085 | -0.471819396 | 3.17E-05 | 0.000559 | 355.1372 | 332.3506 | 356.0859 | 325.2091 | 502.1823 | 406.7242 | 461.8945 | 527.4817 |
| ENSMUSG00000028086 | 0.240836016 | 0.222303 | 0.381516 | 747.4984 | 808.6797 | 676.5631 | 768.5975 | 514.8588 | 931.89 | 528.4305 | 565.5474 |
| ENSMUSG00000028088 | -0.095178248 | 0.715912 | 0.822105 | 277.671 | 266.3604 | 363.5388 | 361.4392 | 531.4357 | 234.6486 | 268.0724 | 321.927 |
| ENSMUSG00000028089 | 0.015390821 | 0.851252 | 0.910607 | 820.9404 | 891.4674 | 854.606 | 794.4762 | 890.2766 | 789.9835 | 861.1102 | 779.8028 |
| ENSMUSG00000028093 | 0.166789136 | 0.107318 | 0.231764 | 1255.556 | 1297.007 | 1006.978 | 1212.849 | 1158.432 | 1002.285 | 1046.254 | 1039.737 |
| ENSMUSG00000028096 | -0.443438299 | 0.000143 | 0.001761 | 463.7911 | 484.7279 | 530.8164 | 517.5741 | 712.8064 | 556.4523 | 708.7525 | 738.4743 |
| ENSMUSG00000028098 | -0.126453865 | 0.090869 | 0.206558 | 1083.521 | 998.2515 | 1135.334 | 1170.58 | 1173.059 | 1213.468 | 1167.754 | 1243.117 |
| ENSMUSG00000028099 | -0.016197092 | 0.802705 | 0.87942 | 1436.646 | 1295.807 | 1351.47 | 1412.977 | 1441.215 | 1371.018 | 1414.612 | 1333.387 |
| ENSMUSG00000028100 | 0.128567924 | 0.59913 | 0.737465 | 67.40564 | 61.1909 | 62.9361 | 98.33908 | 59.48179 | 80.45093 | 65.57166 | 60.9051 |
| ENSMUSG00000028101 | 0.037421171 | 0.727857 | 0.829218 | 1017.121 | 1136.231 | 1032.649 | 1157.641 | 1134.054 | 1193.356 | 974.8963 | 929.8904 |
| ENSMUSG00000028102 | 0.004970114 | 0.95055 | 0.971873 | 784.7224 | 785.8831 | 741.9835 | 762.5592 | 811.2926 | 792.2182 | 767.5742 | 690.6203 |
| ENSMUSG00000028104 | 0.078268626 | 0.437277 | 0.6011 | 1062.393 | 963.4567 | 919.1984 | 1038.599 | 930.2562 | 939.7116 | 1067.468 | 834.1824 |

| | | | | | | | | | | | |
|---|---|---|---|---|---|---|---|---|---|---|---|
| ENSMUSG00000028106 | 0.177298925 | 0.098855 | 0.219314 | 1744.498 | 1512.975 | 1416.89 | 1601.892 | 1478.269 | 1519.629 | 1358.683 | 1193.087 |
| ENSMUSG00000028107 | 0.063123726 | 0.591326 | 0.731321 | 1120.745 | 958.6574 | 889.3865 | 884.1891 | 970.2358 | 1021.28 | 807.11 | 888.5619 |
| ENSMUSG00000028108 | 0.745939306 | 0.003565 | 0.020088 | 4221.404 | 5024.853 | 4231.625 | 4572.767 | 1989.227 | 4600.229 | 2182.186 | 1992.467 |
| ENSMUSG00000028111 | 0.777041073 | 0.000246 | 0.002673 | 482.9061 | 791.8822 | 560.6282 | 696.9998 | 354.9405 | 502.8183 | 313.394 | 306.7007 |
| ENSMUSG00000028114 | 0.016584669 | 0.857136 | 0.914569 | 1160.987 | 1145.83 | 1050.039 | 1220.612 | 1178.91 | 1248.107 | 1042.397 | 1056.051 |
| ENSMUSG00000028115 | 0.096647221 | 0.630967 | 0.759778 | 143.8658 | 175.1739 | 123.3879 | 109.5532 | 151.1423 | 113.9722 | 115.7147 | 132.6861 |
| ENSMUSG00000028121 | -0.34465889 | 0.036208 | 0.108926 | 612.6871 | 465.5308 | 595.4087 | 600.386 | 739.1344 | 549.7481 | 893.8961 | 705.8466 |
| ENSMUSG00000028124 | -0.700069221 | 3.32E-05 | 0.000578 | 1107.666 | 844.6744 | 1180.052 | 1108.471 | 1462.667 | 1386.661 | 1910.257 | 2133.854 |
| ENSMUSG00000028125 | 0.067482535 | 0.570835 | 0.714034 | 277.671 | 362.3461 | 337.0394 | 344.1868 | 304.2347 | 325.1559 | 298.9296 | 332.8029 |
| ENSMUSG00000028126 | 0.025611805 | 0.703531 | 0.813207 | 1464.815 | 1341.4 | 1408.609 | 1504.415 | 1369.056 | 1401.187 | 1484.041 | 1367.102 |
| ENSMUSG00000028127 | -0.253723782 | 0.020771 | 0.072357 | 2134.847 | 2176.476 | 2442.086 | 2194.514 | 2613.298 | 2337.547 | 2532.223 | 3187.729 |
| ENSMUSG00000028128 | -0.376143034 | 0.082785 | 0.1935 | 295.78 | 267.5602 | 398.3193 | 227.7326 | 501.2072 | 320.6864 | 327.8583 | 394.7956 |
| ENSMUSG00000028132 | -0.026047167 | 0.900296 | 0.940077 | 239.4409 | 248.3631 | 144.0906 | 195.8155 | 254.5041 | 207.8316 | 184.1792 | 193.5912 |
| ENSMUSG00000028133 | 0.283359997 | 0.087956 | 0.201829 | 303.8284 | 217.1677 | 226.9012 | 270.0012 | 205.7485 | 224.5922 | 234.3223 | 171.8394 |
| ENSMUSG00000028134 | 0.341714419 | 0.132806 | 0.26616 | 3577.529 | 3562.27 | 3189.039 | 3855.927 | 2535.289 | 4412.51 | 2319.115 | 1927.211 |
| ENSMUSG00000028136 | 0.02700147 | 0.754477 | 0.847512 | 1378.294 | 1372.596 | 1427.656 | 1371.571 | 1291.047 | 1213.468 | 1438.719 | 1504.138 |
| ENSMUSG00000028137 | 1.243281001 | 0.001275 | 0.009377 | 158.9566 | 64.79036 | 81.15445 | 197.5408 | 45.83023 | 64.8077 | 61.71451 | 40.24087 |
| ENSMUSG00000028138 | -0.241214585 | 0.001973 | 0.013013 | 4575.535 | 4109.389 | 4342.591 | 4352.798 | 5165.165 | 4548.83 | 5344.091 | 5485.809 |
| ENSMUSG00000028139 | 0.541885929 | 0.155258 | 0.297153 | 36.21796 | 29.99554 | 39.74912 | 37.95543 | 20.47734 | 36.87335 | 29.89296 | 11.9635 |
| ENSMUSG00000028140 | -0.168434866 | 0.119258 | 0.248282 | 2240.483 | 2159.679 | 2379.15 | 2495.57 | 2815.146 | 2126.363 | 2770.403 | 2712.452 |
| ENSMUSG00000028141 | 0.565417008 | 0.280694 | 0.447018 | 22.1332 | 11.99822 | 15.73403 | 25.8787 | 4.875557 | 20.11273 | 11.57147 | 15.22628 |
| ENSMUSG00000028145 | -0.486124181 | 0.015749 | 0.059733 | 173.0413 | 121.182 | 159.8246 | 170.7995 | 249.6285 | 154.1976 | 233.358 | 239.27 |
| ENSMUSG00000028149 | 0.103136807 | 0.130602 | 0.263048 | 1364.21 | 1456.583 | 1330.767 | 1388.824 | 1221.814 | 1279.393 | 1273.826 | 1381.241 |
| ENSMUSG00000028150 | -0.300916705 | 0.079001 | 0.187542 | 771.6437 | 555.5174 | 780.0764 | 834.1569 | 770.3379 | 763.1665 | 1076.147 | 1016.898 |
| ENSMUSG00000028152 | 0.368279996 | 0.000596 | 0.005242 | 730.3955 | 784.6833 | 713.8279 | 816.0418 | 604.569 | 673.7766 | 564.1092 | 516.6058 |
| ENSMUSG00000028156 | -0.207697083 | 2.91E-05 | 0.000526 | 2442.7 | 2463.234 | 2483.492 | 2620.65 | 2881.454 | 2888.412 | 2835.01 | 2960.423 |
| ENSMUSG00000028158 | 0.43028784 | 0.000231 | 0.002539 | 280.6892 | 283.1579 | 284.0406 | 253.6113 | 206.7236 | 219.0053 | 209.2507 | 181.6277 |
| ENSMUSG00000028159 | 0.497569791 | 7.11E-06 | 0.000177 | 392.3612 | 466.7306 | 460.4273 | 434.7622 | 304.2347 | 278.2261 | 320.144 | 339.3284 |
| ENSMUSG00000028161 | 0.437435545 | 0.030341 | 0.095218 | 4888.418 | 5153.233 | 4606.757 | 5206.795 | 2953.612 | 5546.645 | 3339.333 | 2823.386 |
| ENSMUSG00000028163 | 0.063096278 | 0.459646 | 0.621026 | 1833.031 | 1581.365 | 1483.967 | 1640.71 | 1487.045 | 1606.784 | 1629.649 | 1534.591 |
| ENSMUSG00000028164 | -0.399667923 | 0.032213 | 0.099517 | 1136.841 | 854.2729 | 1364.72 | 1102.433 | 1377.832 | 1086.088 | 1625.792 | 1794.525 |
| ENSMUSG00000028165 | -0.153713992 | 0.011307 | 0.046782 | 1565.421 | 1640.156 | 1671.119 | 1688.154 | 1858.562 | 1816.85 | 1728.97 | 1902.197 |
| ENSMUSG00000028167 | -0.181422614 | 0.339091 | 0.507605 | 519.124 | 502.7252 | 510.9418 | 645.2424 | 625.0463 | 426.8369 | 617.1451 | 801.5546 |
| ENSMUSG00000028172 | 1.942226275 | 1.17E-08 | 9.73E-07 | 232.3986 | 467.9304 | 279.0719 | 372.6533 | 92.63557 | 144.1413 | 50.14304 | 65.25547 |
| ENSMUSG00000028173 | -0.166056345 | 0.238025 | 0.399162 | 3326.016 | 3447.087 | 4809.643 | 3792.093 | 3839.013 | 3944.331 | 4523.48 | 4947.452 |
| ENSMUSG00000028175 | 0.2896188 | 0.030954 | 0.09664 | 562.3844 | 569.9152 | 490.2391 | 675.4342 | 458.3023 | 539.6917 | 415.6086 | 467.6642 |
| ENSMUSG00000028176 | 1.513727494 | 0.002573 | 0.015735 | 65.39353 | 57.59143 | 33.12427 | 66.42201 | 10.72622 | 41.34284 | 16.39292 | 9.78832 |
| ENSMUSG00000028177 | -0.268070428 | 0.630544 | 0.759418 | 14.08476 | 15.59768 | 6.624853 | 16.38985 | 9.751113 | 30.1691 | 11.57147 | 11.9635 |
| ENSMUSG00000028179 | -1.188040851 | 6.01E-07 | 2.44E-05 | 470.8334 | 353.9474 | 578.8465 | 602.9738 | 1079.448 | 727.4105 | 1272.862 | 1496.525 |
| ENSMUSG00000028180 | 0.217313097 | 0.025431 | 0.08399 | 3942.727 | 4402.145 | 3631.248 | 4179.411 | 3332.93 | 3987.908 | 3274.726 | 3299.751 |
| ENSMUSG00000028182 | 0.079593737 | 0.814651 | 0.887294 | 34.20585 | 31.19536 | 19.87456 | 25.01608 | 27.30312 | 18.99536 | 30.85725 | 26.10219 |
| ENSMUSG00000028184 | 0.420428802 | 0.004138 | 0.022371 | 4279.755 | 4438.14 | 3553.406 | 4490.818 | 3004.318 | 4027.016 | 2883.225 | 2610.219 |
| ENSMUSG00000028187 | -0.143103907 | 0.071262 | 0.174639 | 1165.011 | 1107.435 | 1158.521 | 1164.542 | 1398.31 | 1148.661 | 1269.969 | 1257.255 |
| ENSMUSG00000028188 | 0.529621409 | 0.004135 | 0.022371 | 190.1443 | 214.7681 | 175.5586 | 176.8378 | 122.864 | 173.193 | 108.9647 | 119.635 |
| ENSMUSG00000028189 | -0.216393304 | 0.061503 | 0.157675 | 403.4278 | 358.7466 | 370.9918 | 402.8452 | 425.1485 | 387.7288 | 465.7517 | 506.8174 |
| ENSMUSG00000028191 | 0.069071353 | 0.375007 | 0.543559 | 1194.187 | 1107.435 | 1016.087 | 1174.031 | 1033.618 | 1090.557 | 1099.29 | 1057.139 |
| ENSMUSG00000028194 | -0.826026347 | 8.55E-07 | 3.28E-05 | 1142.878 | 1009.05 | 1310.893 | 1233.552 | 1905.367 | 1537.507 | 2274.758 | 2611.306 |
| ENSMUSG00000028195 | 0.399267933 | 0.139406 | 0.27535 | 2157.987 | 913.0642 | 1028.508 | 1642.435 | 1049.22 | 808.9788 | 1355.791 | 1139.795 |
| ENSMUSG00000028197 | 1.154947292 | 1.39E-05 | 0.000294 | 614.6992 | 658.702 | 469.5365 | 821.2176 | 309.1103 | 417.8979 | 250.7152 | 174.0146 |
| ENSMUSG00000028199 | -1.003841333 | 2.47E-07 | 1.20E-05 | 921.5458 | 970.6556 | 1212.348 | 1016.17 | 2002.879 | 1378.84 | 2140.722 | 2742.905 |
| ENSMUSG00000028207 | -0.124219962 | 0.085751 | 0.198199 | 1593.59 | 1691.748 | 1655.385 | 1645.023 | 1713.271 | 1680.531 | 1812.864 | 1970.715 |
| ENSMUSG00000028211 | 0.645382145 | 0.009648 | 0.041489 | 2255.574 | 1733.742 | 1430.968 | 2275.601 | 961.4597 | 1640.305 | 1533.22 | 785.2408 |
| ENSMUSG00000028212 | 0.134757596 | 0.534479 | 0.6855 | 325.9616 | 481.1284 | 317.1648 | 389.0432 | 353.9654 | 470.4145 | 275.7867 | 277.3357 |
| ENSMUSG00000028214 | 0.80422157 | 0.018393 | 0.066535 | 860.1765 | 393.5415 | 334.5551 | 653.006 | 221.3503 | 384.3767 | 458.0374 | 219.6934 |
| ENSMUSG00000028217 | 0.316421646 | 0.577049 | 0.718916 | 13.07871 | 8.398751 | 13.24971 | 27.60395 | 7.80089 | 16.76061 | 18.32149 | 7.613138 |
| ENSMUSG00000028218 | 0.166601287 | 0.216977 | 0.375514 | 1502.039 | 1982.105 | 1699.275 | 1890.008 | 1437.314 | 1927.47 | 1533.22 | 1404.08 |
| ENSMUSG00000028221 | 0.103936801 | 0.266184 | 0.430523 | 1568.439 | 1775.736 | 1623.917 | 1725.247 | 1546.527 | 1785.564 | 1455.112 | 1441.058 |
| ENSMUSG00000028222 | 0.249273639 | 0.057543 | 0.150343 | 13594.81 | 15991.22 | 11061.02 | 14247.95 | 9641.901 | 12219.6 | 12498.15 | 11822.12 |
| ENSMUSG00000028223 | -0.372634793 | 0.00144 | 0.010278 | 1337.046 | 1250.214 | 1356.439 | 1245.628 | 1696.694 | 1412.361 | 1592.041 | 2018.569 |
| ENSMUSG00000028224 | -0.116204143 | 0.118821 | 0.247643 | 776.674 | 890.2676 | 799.1229 | 822.9428 | 877.6002 | 888.3124 | 865.9317 | 929.8904 |
| ENSMUSG00000028226 | 0.572465902 | 0.001979 | 0.013039 | 447.6942 | 545.9188 | 482.7862 | 463.2288 | 332.513 | 444.7149 | 264.2152 | 263.197 |
| ENSMUSG00000028228 | 0.204399653 | 0.087222 | 0.200669 | 1700.232 | 1841.726 | 1503.014 | 1626.908 | 1448.04 | 1722.991 | 1378.934 | 1238.766 |
| ENSMUSG00000028229 | 0.202014183 | 0.206566 | 0.363153 | 487.9364 | 583.1133 | 431.4436 | 571.9194 | 358.841 | 543.0438 | 444.5373 | 456.7883 |
| ENSMUSG00000028232 | 0.241601059 | 0.044552 | 0.125726 | 624.7598 | 645.504 | 517.5666 | 570.1941 | 458.3023 | 579.9172 | 486.966 | 467.6642 |

| | | | | | | | | | | | |
|---|---|---|---|---|---|---|---|---|---|---|---|
| ENSMUSG00000028233 | -0.026028765 | 0.781329 | 0.865154 | 1085.533 | 1052.243 | 916.714 | 973.0393 | 978.0366 | 1136.369 | 1028.897 | 954.905 |
| ENSMUSG00000028234 | -0.080015988 | 0.301546 | 0.469941 | 19220.67 | 22704.22 | 22592.41 | 21175.68 | 23886.33 | 22532.97 | 23176.69 | 20981.81 |
| ENSMUSG00000028238 | -1.015297224 | 0.009938 | 0.04238 | 185.114 | 131.9804 | 443.037 | 170.7995 | 366.6419 | 275.9914 | 597.8593 | 643.8539 |
| ENSMUSG00000028243 | 0.299938062 | 0.019671 | 0.069657 | 694.1775 | 730.6913 | 608.6584 | 679.7473 | 476.8294 | 663.7202 | 501.4304 | 562.2846 |
| ENSMUSG00000028245 | 0.225079728 | 0.020777 | 0.072365 | 2511.112 | 2608.412 | 2262.387 | 2672.408 | 2217.403 | 2368.833 | 2153.258 | 1860.868 |
| ENSMUSG00000028246 | -0.0030038 | 0.981264 | 0.989683 | 278.6771 | 295.1561 | 333.727 | 253.6113 | 317.8863 | 279.3435 | 295.0725 | 270.8102 |
| ENSMUSG00000028247 | -0.037107219 | 0.769987 | 0.857214 | 389.343 | 341.9491 | 328.7583 | 288.1162 | 344.2143 | 312.8647 | 351.0013 | 373.0437 |
| ENSMUSG00000028248 | 0.354502828 | 0.002573 | 0.015735 | 8473.996 | 8095.196 | 7003.298 | 8181.984 | 5978.407 | 7559.036 | 5799.235 | 5499.948 |
| ENSMUSG00000028249 | 0.033435127 | 0.679332 | 0.795038 | 5228.464 | 4963.662 | 6084.099 | 5492.324 | 5250.974 | 5076.231 | 5657.485 | 5290.043 |
| ENSMUSG00000028251 | -0.390014236 | 0.034193 | 0.104221 | 175.0535 | 115.1829 | 189.6364 | 144.0581 | 228.176 | 185.4841 | 196.715 | 209.9051 |
| ENSMUSG00000028252 | 0.140714242 | 0.338724 | 0.507262 | 625.7658 | 638.3051 | 458.7711 | 645.2424 | 498.2819 | 612.321 | 473.466 | 563.3722 |
| ENSMUSG00000028256 | 0.161928205 | 0.229282 | 0.389418 | 774.6619 | 623.9072 | 544.0661 | 693.5493 | 567.5148 | 672.6592 | 558.3234 | 557.9342 |
| ENSMUSG00000028261 | -0.469335354 | 0.000292 | 0.003041 | 586.5297 | 566.3158 | 652.548 | 640.9293 | 765.4624 | 716.2368 | 899.6818 | 1008.197 |
| ENSMUSG00000028262 | -0.205709969 | 0.677878 | 0.794142 | 21.12714 | 14.39786 | 31.46805 | 16.38985 | 36.07912 | 8.938993 | 23.14294 | 28.27737 |
| ENSMUSG00000028264 | 0.515710579 | 0.048768 | 0.133838 | 57.3451 | 63.59054 | 69.56096 | 96.61383 | 58.50668 | 53.63396 | 39.53586 | 50.02919 |
| ENSMUSG00000028266 | 0.193637594 | 0.021866 | 0.075029 | 2371.27 | 2486.03 | 2630.067 | 2538.701 | 2040.908 | 2490.627 | 2165.793 | 2072.949 |
| ENSMUSG00000028268 | 0.853904332 | 0.190273 | 0.34287 | 153.9263 | 553.1177 | 692.2971 | 105.2401 | 177.4703 | 140.7891 | 300.8582 | 213.1679 |
| ENSMUSG00000028270 | 1.120092563 | 0.00199 | 0.013086 | 108.6539 | 280.7582 | 188.8083 | 119.042 | 53.63112 | 118.4417 | 80.036 | 68.51824 |
| ENSMUSG00000028271 | -0.05968494 | 0.353456 | 0.522236 | 1287.75 | 1303.006 | 1310.893 | 1308.6 | 1392.459 | 1246.989 | 1389.541 | 1399.73 |
| ENSMUSG00000028273 | 0.210970535 | 0.010464 | 0.044093 | 3208.307 | 3022.35 | 2972.903 | 3224.487 | 2340.267 | 2860.478 | 2748.224 | 2790.759 |
| ENSMUSG00000028274 | -0.013175258 | 0.922666 | 0.953883 | 518.118 | 609.5093 | 499.3483 | 508.0852 | 577.2659 | 439.128 | 512.0375 | 623.1897 |
| ENSMUSG00000028277 | 0.103276095 | 0.088035 | 0.201935 | 1472.864 | 1474.581 | 1396.188 | 1490.613 | 1338.828 | 1434.708 | 1331.683 | 1325.774 |
| ENSMUSG00000028278 | 0.346291415 | 7.31E-05 | 0.001067 | 1809.892 | 1624.558 | 1797.819 | 1774.417 | 1362.23 | 1254.811 | 1554.434 | 1339.912 |
| ENSMUSG00000028280 | 0.04160083 | 0.935081 | 0.96165 | 8.048435 | 22.79661 | 20.70267 | 19.84034 | 13.65156 | 23.46486 | 8.678602 | 23.927 |
| ENSMUSG00000028282 | -0.072188611 | 0.468593 | 0.628788 | 736.4318 | 809.8795 | 665.7977 | 636.6161 | 726.4579 | 788.8661 | 733.8241 | 740.6495 |
| ENSMUSG00000028284 | -0.042506481 | 0.436194 | 0.600149 | 2572.481 | 2674.402 | 2519.928 | 2618.925 | 2731.287 | 2783.379 | 2546.688 | 2633.058 |
| ENSMUSG00000028289 | 0.158727365 | 0.313472 | 0.481504 | 807.8616 | 736.6904 | 767.6548 | 695.2745 | 534.361 | 871.5518 | 594.0021 | 696.0583 |
| ENSMUSG00000028291 | 0.081926733 | 0.481769 | 0.6406 | 1548.318 | 1527.373 | 1271.972 | 1506.141 | 1149.656 | 1543.094 | 1377.005 | 1460.635 |
| ENSMUSG00000028292 | -0.282017083 | 0.015995 | 0.060312 | 704.238 | 903.4656 | 792.498 | 779.8116 | 935.1317 | 858.1433 | 952.7177 | 1118.044 |
| ENSMUSG00000028293 | -0.044585711 | 0.488945 | 0.646694 | 1038.248 | 1061.842 | 1070.742 | 1060.164 | 1064.822 | 1127.43 | 1033.718 | 1140.883 |
| ENSMUSG00000028294 | -0.40977848 | 0.327805 | 0.495558 | 13.07871 | 13.19804 | 11.59349 | 12.93935 | 10.72622 | 20.11273 | 19.28578 | 17.40146 |
| ENSMUSG00000028295 | -0.083880864 | 0.464076 | 0.624924 | 318.9192 | 410.339 | 385.8977 | 418.3724 | 407.5965 | 387.7288 | 417.5372 | 412.197 |
| ENSMUSG00000028300 | 0.182616489 | 0.237047 | 0.398088 | 437.6336 | 347.9482 | 332.0708 | 387.3179 | 338.3636 | 388.8462 | 335.5726 | 263.197 |
| ENSMUSG00000028307 | -3.063767691 | 3.50E-05 | 0.000604 | 1417.531 | 223.1668 | 2804.797 | 1874.481 | 15074.25 | 5379.039 | 11604.26 | 20792.57 |
| ENSMUSG00000028309 | 0.040754095 | 0.590892 | 0.73091 | 2508.094 | 2195.673 | 2212.701 | 2408.445 | 2174.498 | 2453.754 | 2208.222 | 2231.737 |
| ENSMUSG00000028312 | 0.215716905 | 0.150452 | 0.290485 | 2841.098 | 3233.519 | 2414.759 | 2593.909 | 2313.939 | 3048.197 | 2145.543 | 2034.883 |
| ENSMUSG00000028318 | -0.299091519 | 0.030042 | 0.094633 | 466.8092 | 411.5388 | 469.5365 | 444.2511 | 626.9966 | 425.7195 | 555.4306 | 597.0875 |
| ENSMUSG00000028322 | -0.128335355 | 0.156104 | 0.298241 | 1051.327 | 939.4603 | 873.6525 | 904.892 | 1095.05 | 948.6506 | 1013.468 | 1059.314 |
| ENSMUSG00000028327 | 1.201172437 | 1.12E-06 | 4.15E-05 | 250.5075 | 134.38 | 177.2148 | 162.1732 | 66.30757 | 96.09417 | 66.53595 | 87.00729 |
| ENSMUSG00000028328 | -0.77227148 | 0.000155 | 0.001874 | 118.7144 | 94.7859 | 70.38906 | 100.0643 | 191.1218 | 139.6718 | 163.9292 | 158.7883 |
| ENSMUSG00000028329 | 0.014663754 | 0.881493 | 0.928904 | 794.7829 | 681.4986 | 749.4365 | 751.3451 | 791.7904 | 738.5843 | 775.2885 | 641.6787 |
| ENSMUSG00000028330 | -0.295090466 | 6.46E-07 | 2.59E-05 | 1921.564 | 1811.731 | 1888.083 | 1797.707 | 2281.76 | 2205.696 | 2207.258 | 2410.102 |
| ENSMUSG00000028331 | -0.210489423 | 0.13478 | 0.268723 | 162.9808 | 170.3747 | 152.3716 | 149.2339 | 202.8232 | 163.1366 | 168.7506 | 197.9416 |
| ENSMUSG00000028333 | 0.020492195 | 0.801395 | 0.878873 | 10083.68 | 10610.02 | 9636.677 | 9776.975 | 9892.504 | 11183.8 | 9289.962 | 9173.831 |
| ENSMUSG00000028334 | -0.025989753 | 0.784036 | 0.8671 | 1320.949 | 1623.359 | 1409.437 | 1382.785 | 1412.936 | 1330.793 | 1526.47 | 1566.131 |
| ENSMUSG00000028337 | -0.006146968 | 0.977284 | 0.987503 | 459.7668 | 382.7431 | 532.4726 | 480.4813 | 380.2934 | 431.3064 | 683.681 | 368.6934 |
| ENSMUSG00000028339 | -0.934291121 | 0.169501 | 0.315895 | 662.9898 | 788.2827 | 3781.963 | 919.5566 | 2704.959 | 1258.163 | 3581.37 | 4214.415 |
| ENSMUSG00000028341 | 2.83029428 | 0.002088 | 0.013543 | 184.1079 | 7.198929 | 7.45296 | 53.48266 | 9.751113 | 5.58687 | 13.50005 | 6.525547 |
| ENSMUSG00000028343 | -0.016780672 | 0.827053 | 0.894992 | 1681.117 | 1762.538 | 1637.167 | 1839.113 | 1641.112 | 1714.052 | 1728.97 | 1917.423 |
| ENSMUSG00000028344 | 0.092834124 | 0.407203 | 0.573603 | 551.3178 | 517.1231 | 499.3483 | 523.6125 | 477.8045 | 578.7998 | 434.8944 | 470.9269 |
| ENSMUSG00000028345 | 0.081956247 | 0.244814 | 0.407641 | 1172.053 | 1192.623 | 1174.255 | 1199.909 | 1198.412 | 1152.013 | 1093.504 | 1032.124 |
| ENSMUSG00000028347 | 0.426033535 | 0.000805 | 0.006603 | 2124.787 | 2055.294 | 1632.198 | 2145.345 | 1473.393 | 1725.226 | 1444.505 | 1279.007 |
| ENSMUSG00000028348 | 0.225894136 | 0.535978 | 0.68669 | 43.26034 | 61.1909 | 63.76421 | 74.18562 | 46.80534 | 88.27255 | 42.42872 | 30.45255 |
| ENSMUSG00000028351 | 0.442099328 | 0.036957 | 0.1105 | 168.0111 | 118.7823 | 120.0755 | 114.7289 | 84.83468 | 121.7938 | 90.64318 | 87.00729 |
| ENSMUSG00000028354 | 0.138418291 | 0.619918 | 0.751969 | 71.42986 | 55.19179 | 51.34261 | 69.00988 | 52.65601 | 81.56831 | 47.25017 | 43.50364 |
| ENSMUSG00000028356 | 0.173639246 | 0.699385 | 0.810259 | 125.7568 | 23.99643 | 97.71658 | 115.5915 | 63.38224 | 62.57295 | 84.85745 | 112.0219 |
| ENSMUSG00000028357 | -0.710008544 | 0.006258 | 0.030366 | 1106.66 | 752.2881 | 1430.14 | 886.777 | 1543.601 | 1130.783 | 1640.256 | 2518.861 |
| ENSMUSG00000028358 | 0.109607966 | 0.364617 | 0.533315 | 2199.235 | 1901.717 | 1989.94 | 1806.334 | 1641.112 | 2234.748 | 1702.935 | 1742.321 |
| ENSMUSG00000028360 | -0.553857707 | 0.109485 | 0.234988 | 23.13925 | 14.39786 | 24.8432 | 31.91707 | 33.15378 | 45.81234 | 30.85725 | 30.45255 |
| ENSMUSG00000028364 | 0.225820457 | 0.022817 | 0.077509 | 3260.622 | 3465.085 | 3319.879 | 3245.19 | 2580.145 | 3386.761 | 2646.974 | 2752.693 |
| ENSMUSG00000028367 | -0.19559404 | 0.017075 | 0.063045 | 5240.537 | 5429.192 | 5511.878 | 5679.513 | 6527.395 | 5449.433 | 6261.13 | 6797.444 |
| ENSMUSG00000028369 | 0.508322431 | 0.002752 | 0.016576 | 535.2209 | 680.2988 | 618.5957 | 702.1755 | 394.9201 | 595.5604 | 391.5014 | 402.4087 |
| ENSMUSG00000028370 | 0.929026597 | 0.002041 | 0.013319 | 1225.374 | 731.8911 | 728.7338 | 1080.005 | 348.1147 | 701.7109 | 641.2523 | 287.124 |
| ENSMUSG00000028373 | 0.427217949 | 0.023211 | 0.078544 | 130.7871 | 105.5843 | 104.3414 | 123.3552 | 72.15824 | 100.5637 | 93.53605 | 79.39415 |

| | | | | | | | | | | | |
|---|---|---|---|---|---|---|---|---|---|---|---|
| ENSMUSG00000028378 | 0.688354467 | 4.06E-07 | 1.77E-05 | 1614.717 | 1793.733 | 1764.695 | 1869.305 | 1359.305 | 1136.369 | 905.4675 | 969.0437 |
| ENSMUSG00000028381 | 0.261208099 | 0.151714 | 0.292126 | 1197.205 | 1096.637 | 992.8999 | 1263.743 | 793.7406 | 1334.145 | 878.4674 | 791.7663 |
| ENSMUSG00000028382 | 0.054516114 | 0.5024 | 0.658496 | 5640.947 | 4982.859 | 5458.051 | 5331.876 | 4769.269 | 5779.059 | 4933.303 | 5143.218 |
| ENSMUSG00000028383 | -0.291861851 | 0.023527 | 0.079323 | 722.347 | 608.3095 | 703.0625 | 717.7027 | 825.9193 | 707.2978 | 819.6458 | 1017.985 |
| ENSMUSG00000028385 | 0.022267451 | 0.794298 | 0.873694 | 538.2391 | 602.3104 | 612.7989 | 609.8748 | 599.6935 | 576.565 | 547.7162 | 604.7006 |
| ENSMUSG00000028389 | 0.021350402 | 0.846195 | 0.907479 | 414.4944 | 416.3381 | 381.7572 | 419.235 | 457.3272 | 411.1937 | 380.8942 | 356.7299 |
| ENSMUSG00000028391 | -0.099074266 | 0.348801 | 0.517452 | 311.8769 | 315.5531 | 317.9929 | 280.3526 | 331.5378 | 358.6771 | 313.394 | 308.8759 |
| ENSMUSG00000028392 | -0.181194564 | 0.307304 | 0.475766 | 952.7335 | 1031.847 | 1589.965 | 977.3524 | 1182.81 | 1181.064 | 1367.362 | 1432.357 |
| ENSMUSG00000028393 | -0.336323746 | 0.000226 | 0.002494 | 908.4671 | 887.8679 | 1003.665 | 849.6841 | 1146.731 | 1057.036 | 1136.897 | 1269.219 |
| ENSMUSG00000028394 | -0.178292546 | 0.013531 | 0.053434 | 1991.988 | 2249.665 | 2167.155 | 2210.904 | 2604.522 | 2260.448 | 2447.366 | 2438.379 |
| ENSMUSG00000028397 | 0.220208696 | 0.007351 | 0.034073 | 812.8919 | 805.0802 | 731.2182 | 794.4762 | 654.2997 | 649.1943 | 734.7883 | 656.905 |
| ENSMUSG00000028398 | -0.166454227 | 0.048922 | 0.13412 | 800.8193 | 904.6654 | 869.512 | 910.9304 | 1033.618 | 891.6645 | 962.3606 | 1023.423 |
| ENSMUSG00000028399 | 0.464820013 | 0.005369 | 0.026978 | 3678.135 | 4025.401 | 3323.192 | 3840.4 | 2639.626 | 3678.396 | 2298.865 | 2155.606 |
| ENSMUSG00000028402 | 0.208700671 | 0.0643 | 0.162898 | 2347.125 | 2357.649 | 2288.059 | 2399.819 | 1958.999 | 2498.448 | 1889.042 | 1782.562 |
| ENSMUSG00000028403 | 0.128103519 | 0.170956 | 0.317902 | 1105.654 | 1156.628 | 1035.133 | 1044.637 | 925.3806 | 1138.604 | 966.2177 | 941.8539 |
| ENSMUSG00000028405 | -0.535675194 | 0.000213 | 0.002383 | 2962.83 | 2711.597 | 2978.7 | 2749.181 | 4192.004 | 3031.436 | 4266.98 | 5038.81 |
| ENSMUSG00000028407 | -0.111685091 | 0.560986 | 0.706583 | 110.666 | 103.1847 | 110.1382 | 94.02596 | 114.088 | 125.1459 | 83.89316 | 129.4233 |
| ENSMUSG00000028409 | -0.092400104 | 0.157825 | 0.300382 | 2305.877 | 2405.642 | 2569.615 | 2644.804 | 2682.531 | 2550.965 | 2656.617 | 2697.226 |
| ENSMUSG00000028410 | -0.112631385 | 0.052655 | 0.141126 | 7783.843 | 8044.803 | 7635.143 | 8464.924 | 8801.355 | 8297.62 | 8395.101 | 9027.006 |
| ENSMUSG00000028411 | -0.051508209 | 0.514786 | 0.669147 | 689.1472 | 706.6949 | 662.4853 | 663.3575 | 765.4624 | 668.1897 | 683.681 | 699.3211 |
| ENSMUSG00000028412 | 0.090708499 | 0.173302 | 0.321099 | 2817.958 | 2945.562 | 3150.946 | 2895.827 | 2672.78 | 2918.581 | 2869.725 | 2631.97 |
| ENSMUSG00000028413 | 0.146018939 | 0.029436 | 0.093297 | 2132.835 | 2178.876 | 2152.249 | 2100.488 | 1918.044 | 2054.851 | 2003.793 | 1761.898 |
| ENSMUSG00000028414 | 0.15467301 | 0.137607 | 0.272753 | 1013.097 | 951.4585 | 872.8244 | 982.5282 | 849.3219 | 986.6413 | 756.967 | 839.6203 |
| ENSMUSG00000028416 | -0.28469523 | 0.000377 | 0.003682 | 3584.572 | 3631.86 | 3898.726 | 3813.658 | 4408.478 | 4056.068 | 4805.053 | 4918.087 |
| ENSMUSG00000028417 | 0.27211032 | 0.512374 | 0.667588 | 15.09082 | 23.99643 | 17.39024 | 21.56559 | 15.60178 | 10.05637 | 23.14294 | 15.22628 |
| ENSMUSG00000028419 | -0.068801786 | 0.513783 | 0.668338 | 2061.405 | 2165.678 | 2060.329 | 2137.581 | 2204.727 | 1834.728 | 2232.329 | 2563.452 |
| ENSMUSG00000028420 | -0.121211858 | 0.320372 | 0.488405 | 383.3067 | 351.5477 | 449.6619 | 383.0048 | 403.6961 | 394.4331 | 486.0017 | 423.0729 |
| ENSMUSG00000028423 | 0.004701566 | 0.949565 | 0.971453 | 3007.096 | 2817.181 | 2625.098 | 2618.925 | 2719.585 | 2968.863 | 2686.51 | 2654.81 |
| ENSMUSG00000028426 | -0.217413176 | 0.013535 | 0.053434 | 4282.773 | 4490.932 | 5080.434 | 4512.383 | 5005.246 | 4916.446 | 5795.378 | 5639.16 |
| ENSMUSG00000028427 | -1.204203717 | 0.009659 | 0.041508 | 30.18163 | 38.39429 | 46.37397 | 21.56559 | 78.98402 | 31.28647 | 61.71451 | 142.4744 |
| ENSMUSG00000028430 | -0.007774379 | 0.893403 | 0.935548 | 1400.428 | 1386.994 | 1483.967 | 1383.648 | 1395.384 | 1391.131 | 1444.505 | 1457.372 |
| ENSMUSG00000028431 | 0.015316808 | 0.818954 | 0.890502 | 1786.753 | 1994.103 | 1836.741 | 1741.637 | 1837.11 | 1890.597 | 1785.864 | 1762.985 |
| ENSMUSG00000028433 | -0.066959908 | 0.373355 | 0.542186 | 1971.867 | 1958.109 | 1896.364 | 1978.858 | 1973.625 | 1972.165 | 1935.328 | 2295.905 |
| ENSMUSG00000028434 | -0.569249823 | 0.012198 | 0.049577 | 494.9787 | 457.132 | 834.7315 | 545.178 | 819.0935 | 619.0252 | 988.3964 | 1036.474 |
| ENSMUSG00000028435 | -0.796529023 | 0.133815 | 0.267352 | 915.5095 | 601.1106 | 2386.603 | 847.0963 | 1640.137 | 1096.144 | 2800.296 | 2715.715 |
| ENSMUSG00000028436 | -0.068179151 | 0.266966 | 0.431387 | 2022.169 | 2049.295 | 1895.536 | 1952.98 | 2155.971 | 1941.996 | 2107.936 | 2090.35 |
| ENSMUSG00000028437 | 0.002988815 | 0.959818 | 0.977262 | 1265.616 | 1240.615 | 1284.393 | 1290.485 | 1326.151 | 1211.234 | 1258.397 | 1275.744 |
| ENSMUSG00000028438 | -0.196075767 | 0.232166 | 0.392783 | 106.6418 | 154.777 | 120.9036 | 124.2178 | 153.0925 | 150.8455 | 135.0005 | 139.2117 |
| ENSMUSG00000028439 | 0.165821532 | 0.206889 | 0.363479 | 321.9374 | 273.5593 | 297.2903 | 290.7041 | 222.3254 | 252.5265 | 297.0011 | 283.8613 |
| ENSMUSG00000028443 | -0.366540897 | 0.003343 | 0.019166 | 326.9677 | 381.5432 | 368.5075 | 358.8514 | 519.7343 | 379.9072 | 466.716 | 482.8904 |
| ENSMUSG00000028444 | 0.542970835 | 1.20E-05 | 0.000264 | 1289.762 | 1599.362 | 1361.407 | 1426.779 | 941.9575 | 1177.712 | 849.5388 | 927.7152 |
| ENSMUSG00000028445 | -0.349284299 | 0.009679 | 0.041562 | 292.7618 | 363.5459 | 331.2427 | 331.2474 | 499.257 | 382.1419 | 427.1801 | 368.6934 |
| ENSMUSG00000028447 | -0.209832989 | 0.022069 | 0.075574 | 1224.368 | 1070.241 | 1271.144 | 1211.123 | 1410.986 | 1221.29 | 1450.291 | 1445.409 |
| ENSMUSG00000028452 | -0.117447655 | 0.028402 | 0.090909 | 12529.4 | 12908.88 | 13023.63 | 12478.71 | 13382.43 | 13239.77 | 13877.09 | 14761.87 |
| ENSMUSG00000028453 | 0.245474449 | 0.020911 | 0.072735 | 276.6649 | 277.1588 | 249.2601 | 263.1002 | 222.3254 | 220.1227 | 224.6794 | 230.5693 |
| ENSMUSG00000028454 | -0.226701482 | 0.009672 | 0.041544 | 718.3228 | 706.6949 | 787.5294 | 703.0381 | 822.0188 | 781.0445 | 896.7889 | 913.5765 |
| ENSMUSG00000028455 | -0.273026861 | 0.004813 | 0.025038 | 2317.949 | 2594.014 | 2352.651 | 2342.023 | 3135.958 | 2436.993 | 3024.975 | 3005.014 |
| ENSMUSG00000028456 | 0.301284977 | 0.001384 | 0.009977 | 1602.645 | 1448.185 | 1424.343 | 1733.011 | 1179.885 | 1312.915 | 1347.112 | 1199.613 |
| ENSMUSG00000028457 | -0.031558664 | 0.928663 | 0.957777 | 34.20585 | 23.99643 | 19.04645 | 21.56559 | 18.52711 | 26.81698 | 26.03581 | 29.36496 |
| ENSMUSG00000028458 | -0.323270158 | 0.008406 | 0.037638 | 462.785 | 458.3318 | 505.9732 | 445.1137 | 583.1166 | 488.2925 | 584.3592 | 687.3576 |
| ENSMUSG00000028459 | 0.744426946 | 0.000588 | 0.005181 | 89.53884 | 80.38804 | 100.2009 | 102.6522 | 49.73068 | 58.10345 | 69.42882 | 45.67883 |
| ENSMUSG00000028461 | -0.402927579 | 0.000504 | 0.004619 | 578.4813 | 591.512 | 592.9243 | 622.8142 | 767.4126 | 672.6592 | 774.3242 | 940.7663 |
| ENSMUSG00000028463 | 0.327768955 | 0.252535 | 0.416216 | 75.45408 | 93.58608 | 70.38906 | 87.9876 | 61.43201 | 100.5637 | 50.14304 | 48.9416 |
| ENSMUSG00000028464 | 0.044451033 | 0.727159 | 0.829154 | 2368.252 | 3243.118 | 3097.947 | 3009.693 | 3271.498 | 2756.562 | 2464.723 | 2870.153 |
| ENSMUSG00000028465 | -0.131940162 | 0.236782 | 0.397964 | 4826.043 | 4244.969 | 5256.821 | 4497.719 | 4883.357 | 4458.323 | 5364.341 | 5925.196 |
| ENSMUSG00000028466 | -0.129201706 | 0.400195 | 0.567586 | 1383.325 | 977.8546 | 1049.211 | 1016.17 | 1289.097 | 978.8197 | 1166.79 | 1406.255 |
| ENSMUSG00000028467 | -0.038184993 | 0.694367 | 0.806903 | 923.5579 | 951.4585 | 785.0451 | 945.4353 | 1009.24 | 854.7912 | 917.039 | 916.8393 |
| ENSMUSG00000028468 | 0.030964178 | 0.756326 | 0.848356 | 1503.045 | 1293.408 | 1377.969 | 1256.842 | 1429.513 | 1176.595 | 1275.755 | 1433.445 |
| ENSMUSG00000028469 | 0.408152353 | 0.001827 | 0.012287 | 1263.604 | 1422.988 | 1469.889 | 1529.431 | 928.306 | 1331.91 | 1049.147 | 977.7444 |
| ENSMUSG00000028470 | -0.547289616 | 7.51E-05 | 0.001091 | 861.1825 | 769.0856 | 896.8395 | 772.9107 | 1251.068 | 946.4159 | 1177.397 | 1448.671 |
| ENSMUSG00000028475 | -0.723591336 | 0.062517 | 0.159499 | 33.19979 | 11.99822 | 27.32752 | 23.29083 | 55.58134 | 27.93435 | 38.57157 | 36.9781 |
| ENSMUSG00000028476 | 0.223635995 | 0.006737 | 0.032087 | 1497.009 | 1763.738 | 1748.961 | 1812.372 | 1470.468 | 1528.568 | 1393.398 | 1451.934 |
| ENSMUSG00000028478 | -0.193983853 | 0.034887 | 0.105812 | 6001.114 | 6066.298 | 7035.594 | 6561.977 | 7609.769 | 6409.258 | 7389.348 | 7953.554 |
| ENSMUSG00000028479 | -0.420269648 | 0.002089 | 0.013545 | 518.118 | 545.9188 | 600.3773 | 498.5964 | 729.3833 | 566.5087 | 769.5028 | 828.7444 |

| | | | | | | | | | | | |
|---|---|---|---|---|---|---|---|---|---|---|---|
| ENSMUSG00000028480 | -0.227785132 | 0.202097 | 0.357578 | 578.4813 | 508.7243 | 863.7152 | 609.8748 | 700.1299 | 676.0113 | 735.7526 | 890.7371 |
| ENSMUSG00000028483 | 0.327425164 | 0.000667 | 0.005722 | 799.8132 | 725.892 | 725.4214 | 789.3005 | 585.0668 | 679.3635 | 545.7877 | 615.5766 |
| ENSMUSG00000028484 | 0.343912406 | 0.013414 | 0.053173 | 6258.664 | 6583.421 | 5238.603 | 5756.287 | 4457.234 | 5990.243 | 4295.908 | 4037.138 |
| ENSMUSG00000028487 | 0.747564872 | 0.008872 | 0.03909 | 204.229 | 243.5638 | 176.3867 | 178.5631 | 95.56091 | 198.8926 | 92.57176 | 91.35765 |
| ENSMUSG00000028488 | -0.710156599 | 0.000548 | 0.004912 | 368.2159 | 333.5504 | 500.1764 | 392.4937 | 765.4624 | 431.3064 | 672.1095 | 741.7371 |
| ENSMUSG00000028494 | 0.014270277 | 0.905212 | 0.943183 | 1360.185 | 1273.011 | 1406.125 | 1311.188 | 1379.783 | 1109.552 | 1221.754 | 1587.883 |
| ENSMUSG00000028495 | -0.192537124 | 0.02676 | 0.087305 | 24714.73 | 29455.62 | 29148.53 | 27713.5 | 34539.42 | 28622.65 | 32404.94 | 31315.01 |
| ENSMUSG00000028496 | 0.187489385 | 0.074664 | 0.180461 | 1428.597 | 1243.015 | 1401.156 | 1294.798 | 1033.618 | 1368.783 | 1147.504 | 1166.985 |
| ENSMUSG00000028497 | 0.068927473 | 0.429599 | 0.594488 | 943.679 | 1078.64 | 1021.884 | 910.0678 | 903.9282 | 1022.397 | 913.1818 | 928.8028 |
| ENSMUSG00000028514 | -0.265913827 | 0.002802 | 0.01682 | 1802.849 | 1649.755 | 1944.394 | 1597.579 | 1953.148 | 2052.616 | 2179.294 | 2228.474 |
| ENSMUSG00000028517 | -0.078767727 | 0.45134 | 0.613849 | 2081.526 | 1839.326 | 2315.386 | 2062.533 | 1947.297 | 2125.246 | 2470.509 | 2225.211 |
| ENSMUSG00000028518 | -0.874574445 | 0.004212 | 0.022677 | 380.2885 | 196.7707 | 532.4726 | 330.3848 | 562.6392 | 446.9496 | 696.2168 | 937.5035 |
| ENSMUSG00000028519 | 0.064796424 | 0.553651 | 0.700643 | 493.9727 | 393.5415 | 510.9418 | 444.2511 | 443.6756 | 423.4848 | 446.4659 | 451.3503 |
| ENSMUSG00000028521 | -0.028710432 | 0.765122 | 0.854012 | 545.2815 | 501.5254 | 461.2554 | 546.9033 | 488.5308 | 547.5133 | 525.5376 | 535.0948 |
| ENSMUSG00000028522 | 0.226644047 | 0.021334 | 0.073719 | 2433.645 | 2294.059 | 2042.111 | 2335.984 | 1880.99 | 2241.452 | 1873.614 | 1785.825 |
| ENSMUSG00000028523 | 1.944708517 | 0.001235 | 0.009155 | 17.10292 | 25.19625 | 12.4216 | 18.11509 | 0.975111 | 6.704245 | 4.821446 | 6.525547 |
| ENSMUSG00000028524 | 0.807515052 | 0.004945 | 0.025544 | 357.1493 | 403.14 | 334.5551 | 374.3786 | 137.4907 | 364.264 | 167.7863 | 170.7518 |
| ENSMUSG00000028525 | -0.135980213 | 0.313488 | 0.481504 | 266.6044 | 268.76 | 271.619 | 338.1484 | 317.8863 | 272.6393 | 321.1083 | 348.0291 |
| ENSMUSG00000028527 | -1.776205685 | 4.86E-09 | 4.69E-07 | 275.6589 | 285.5575 | 349.461 | 320.8959 | 1029.718 | 562.0392 | 817.7172 | 1810.839 |
| ENSMUSG00000028528 | -0.639372618 | 0.014413 | 0.056002 | 81.4904 | 77.9884 | 101.8571 | 109.5532 | 170.6445 | 106.1505 | 108.0004 | 194.6788 |
| ENSMUSG00000028530 | -0.039109933 | 0.452046 | 0.614447 | 4288.81 | 4334.955 | 4504.9 | 4303.629 | 4713.688 | 4268.369 | 4487.802 | 4440.634 |
| ENSMUSG00000028532 | 0.089910699 | 0.210991 | 0.368412 | 1199.217 | 1310.205 | 1396.188 | 1325.852 | 1184.76 | 1265.985 | 1218.862 | 1249.642 |
| ENSMUSG00000028536 | -2.09854898 | 8.32E-09 | 7.38E-07 | 72.43591 | 74.38894 | 110.1382 | 69.8725 | 337.3885 | 143.0239 | 357.7513 | 563.3722 |
| ENSMUSG00000028538 | -0.570873862 | 0.000483 | 0.004461 | 411.4762 | 349.1481 | 541.5817 | 383.0048 | 609.4446 | 526.2832 | 683.681 | 687.3576 |
| ENSMUSG00000028539 | -0.447366257 | 0.124945 | 0.255284 | 27.16347 | 31.19536 | 48.03018 | 33.64232 | 51.6809 | 36.87335 | 50.14304 | 53.29196 |
| ENSMUSG00000028540 | -0.094562891 | 0.532255 | 0.683893 | 306.8466 | 308.3541 | 281.5563 | 248.4356 | 362.7414 | 249.1744 | 287.3582 | 320.8394 |
| ENSMUSG00000028541 | -0.007306049 | 0.940292 | 0.965131 | 1295.798 | 1629.358 | 1570.918 | 1423.329 | 1626.486 | 1442.53 | 1406.898 | 1471.511 |
| ENSMUSG00000028542 | -0.0079559 | 0.946459 | 0.969279 | 325.9616 | 310.7538 | 346.9767 | 330.3848 | 379.3183 | 298.3389 | 343.2869 | 300.1751 |
| ENSMUSG00000028544 | -1.652801749 | 4.55E-05 | 0.00074 | 44.26639 | 20.39697 | 63.76421 | 36.23019 | 124.8142 | 78.21619 | 106.0718 | 212.0803 |
| ENSMUSG00000028545 | -0.194816063 | 0.10617 | 0.230174 | 458.7608 | 620.3077 | 641.7826 | 609.8748 | 663.0757 | 665.955 | 654.7523 | 685.1824 |
| ENSMUSG00000028546 | 0.358468365 | 0.169418 | 0.315772 | 266.6044 | 344.3488 | 170.59 | 307.9566 | 297.4089 | 196.6578 | 182.2507 | 171.8394 |
| ENSMUSG00000028549 | 0.096583539 | 0.397868 | 0.565068 | 247.4894 | 291.5566 | 263.3379 | 293.292 | 257.4294 | 270.4045 | 238.1794 | 258.8467 |
| ENSMUSG00000028550 | 0.370880722 | 0.005841 | 0.028856 | 457.7547 | 431.9358 | 346.1486 | 379.5543 | 274.0063 | 345.2686 | 324.9654 | 303.4379 |
| ENSMUSG00000028551 | 0.223935099 | 0.151461 | 0.291852 | 434.6155 | 659.9018 | 642.6107 | 597.7981 | 533.3859 | 557.5697 | 457.0731 | 451.3503 |
| ENSMUSG00000028552 | 0.183806526 | 0.040632 | 0.11802 | 1799.831 | 1492.578 | 1651.245 | 1866.717 | 1507.522 | 1585.554 | 1465.72 | 1441.058 |
| ENSMUSG00000028553 | -0.322201628 | 0.342508 | 0.511186 | 37.22401 | 38.39429 | 37.2648 | 29.3292 | 32.17867 | 71.51194 | 39.53586 | 34.80291 |
| ENSMUSG00000028555 | -0.168758728 | 0.570598 | 0.713919 | 55.33299 | 44.3934 | 81.15445 | 53.48266 | 67.28268 | 44.69496 | 64.60737 | 88.09488 |
| ENSMUSG00000028556 | 0.522650931 | 0.000159 | 0.001899 | 4301.888 | 4915.669 | 4008.036 | 4307.079 | 2864.877 | 3887.344 | 2912.153 | 2539.525 |
| ENSMUSG00000028557 | 0.090642165 | 0.167187 | 0.313068 | 1978.909 | 1934.112 | 1943.566 | 1972.82 | 1763.976 | 1860.428 | 1734.756 | 1996.817 |
| ENSMUSG00000028559 | -0.131801207 | 0.040761 | 0.118283 | 2449.742 | 2277.261 | 2353.479 | 2473.142 | 2618.174 | 2433.641 | 2648.902 | 2769.007 |
| ENSMUSG00000028560 | -0.064805194 | 0.463161 | 0.624014 | 2432.639 | 2892.77 | 2715.362 | 2956.211 | 2912.657 | 3133.117 | 2741.474 | 2715.715 |
| ENSMUSG00000028563 | -0.089902819 | 0.360939 | 0.52953 | 987.9454 | 886.6681 | 968.0567 | 1078.279 | 1119.428 | 933.0074 | 1009.611 | 1113.693 |
| ENSMUSG00000028565 | 0.970883337 | 0.026324 | 0.086279 | 2820.976 | 3598.265 | 2312.074 | 3022.633 | 1066.772 | 2978.919 | 1078.075 | 873.3356 |
| ENSMUSG00000028567 | -0.035082216 | 0.61638 | 0.749401 | 1361.192 | 1552.569 | 1464.093 | 1428.505 | 1447.065 | 1574.38 | 1496.577 | 1429.095 |
| ENSMUSG00000028568 | 0.097241881 | 0.225035 | 0.384774 | 2053.357 | 2176.476 | 2147.28 | 2340.298 | 2050.659 | 2249.274 | 1951.721 | 1900.022 |
| ENSMUSG00000028571 | -4.791852713 | 6.04E-35 | 3.71E-31 | 9.054489 | 5.999108 | 15.73403 | 6.900988 | 290.5832 | 151.9629 | 256.5009 | 365.4306 |
| ENSMUSG00000028572 | -0.236629683 | 0.004737 | 0.024762 | 1137.847 | 1069.041 | 1074.882 | 1101.57 | 1216.939 | 1193.356 | 1308.54 | 1446.496 |
| ENSMUSG00000028573 | -0.9514527 | 2.84E-05 | 0.000517 | 218.3138 | 218.3675 | 276.5876 | 213.9306 | 427.0988 | 296.1041 | 434.8944 | 636.2408 |
| ENSMUSG00000028575 | 0.630513147 | 0.347632 | 0.516412 | 10.06054 | 7.198929 | 14.07781 | 18.97772 | 4.875557 | 11.17374 | 14.46434 | 2.175182 |
| ENSMUSG00000028576 | 0.120797037 | 0.092079 | 0.208389 | 1337.046 | 1496.177 | 1384.594 | 1440.581 | 1238.391 | 1394.483 | 1308.54 | 1261.606 |
| ENSMUSG00000028577 | -0.129810641 | 0.036016 | 0.108471 | 2127.805 | 2085.29 | 1955.16 | 2121.191 | 2130.618 | 2296.204 | 2283.437 | 2358.985 |
| ENSMUSG00000028578 | 0.034919557 | 0.790957 | 0.871985 | 444.676 | 508.7243 | 491.0672 | 505.4974 | 434.8996 | 588.8561 | 422.3587 | 458.9634 |
| ENSMUSG00000028580 | 0.17599213 | 0.016266 | 0.060915 | 3195.229 | 3217.921 | 2914.935 | 3449.631 | 2767.366 | 3007.971 | 2813.796 | 2721.153 |
| ENSMUSG00000028581 | -0.018298152 | 0.922966 | 0.954016 | 157.9505 | 188.372 | 236.8385 | 169.9368 | 151.1423 | 186.6015 | 214.0722 | 212.0803 |
| ENSMUSG00000028582 | -0.164862944 | 0.048202 | 0.132708 | 1283.725 | 1461.383 | 1337.392 | 1346.555 | 1580.655 | 1358.727 | 1516.827 | 1625.949 |
| ENSMUSG00000028583 | 0.64549996 | 2.01E-12 | 5.39E-10 | 1779.71 | 1984.505 | 1864.896 | 1973.683 | 1368.081 | 1250.342 | 1147.504 | 1093.029 |
| ENSMUSG00000028587 | -0.041302064 | 0.784146 | 0.867163 | 159.9626 | 195.5709 | 163.137 | 154.4096 | 166.744 | 194.4231 | 180.3221 | 149 |
| ENSMUSG00000028591 | -1.542443402 | 0.058902 | 0.152617 | 2.012109 | 4.799286 | 41.40533 | 12.93935 | 30.22845 | 91.62468 | 20.25007 | 38.06569 |
| ENSMUSG00000028597 | 0.097521751 | 0.349225 | 0.517902 | 748.5044 | 980.2542 | 908.433 | 938.5344 | 891.2517 | 814.5657 | 833.1458 | 801.5546 |
| ENSMUSG00000028599 | 0.369912499 | 0.027978 | 0.090054 | 142.8597 | 188.372 | 158.1684 | 154.4096 | 140.416 | 101.681 | 132.1076 | 121.8102 |
| ENSMUSG00000028600 | 0.170372077 | 0.323355 | 0.491365 | 148.896 | 158.3764 | 144.9187 | 165.6237 | 116.0382 | 126.2633 | 132.1076 | 175.1022 |
| ENSMUSG00000028601 | 0.14287584 | 0.229683 | 0.389904 | 912.4913 | 933.4612 | 899.3238 | 1031.698 | 894.1771 | 682.7156 | 904.5032 | 938.5911 |
| ENSMUSG00000028603 | -0.43871412 | 0.002131 | 0.01374 | 2589.584 | 2560.419 | 2851.999 | 2796.625 | 3665.443 | 2791.2 | 3586.191 | 4595.072 |
| ENSMUSG00000028607 | -0.68808178 | 0.002657 | 0.016124 | 808.8677 | 725.892 | 828.9347 | 709.9391 | 1248.142 | 782.1619 | 1072.29 | 1849.992 |

| | | | | | | | | | | | |
|---|---|---|---|---|---|---|---|---|---|---|---|
| ENSMUSG00000028608 | 0.039905535 | 0.547724 | 0.696007 | 1454.755 | 1582.565 | 1388.735 | 1411.252 | 1446.09 | 1382.192 | 1427.148 | 1414.956 |
| ENSMUSG00000028609 | -0.222776356 | 6.73E-05 | 0.001002 | 2089.575 | 2038.497 | 2271.496 | 2143.619 | 2527.489 | 2501.801 | 2468.58 | 2479.708 |
| ENSMUSG00000028613 | 0.30819277 | 0.054198 | 0.144298 | 157.9505 | 155.9768 | 164.7932 | 159.5853 | 159.9183 | 119.559 | 123.429 | 112.0219 |
| ENSMUSG00000028614 | -0.080412029 | 0.325028 | 0.492991 | 1401.434 | 1428.987 | 1426 | 1327.578 | 1401.235 | 1664.887 | 1446.434 | 1392.117 |
| ENSMUSG00000028617 | 0.028280299 | 0.753858 | 0.84692 | 1816.934 | 2087.689 | 1860.756 | 2090.999 | 1994.103 | 2115.189 | 1789.721 | 1804.314 |
| ENSMUSG00000028618 | -0.07782477 | 0.288403 | 0.455361 | 5785.819 | 5225.223 | 6042.694 | 6112.55 | 5936.478 | 5821.519 | 6484.845 | 6213.408 |
| ENSMUSG00000028619 | 0.096207057 | 0.318816 | 0.486971 | 794.7829 | 770.2854 | 712.9998 | 778.0864 | 765.4624 | 613.4384 | 738.6455 | 738.4743 |
| ENSMUSG00000028621 | -0.393685552 | 0.024397 | 0.081541 | 258.556 | 323.9518 | 313.0243 | 278.6274 | 385.169 | 281.5783 | 397.2871 | 477.4525 |
| ENSMUSG00000028622 | -0.272279417 | 3.68E-06 | 0.000107 | 1519.142 | 1504.576 | 1420.203 | 1421.604 | 1761.051 | 1700.643 | 1785.864 | 1830.416 |
| ENSMUSG00000028626 | -0.551718742 | 0.165369 | 0.310736 | 112.6781 | 140.3791 | 447.1776 | 154.4096 | 271.0809 | 389.9636 | 287.3582 | 306.7007 |
| ENSMUSG00000028629 | -0.026344244 | 0.777513 | 0.862822 | 540.2512 | 614.3086 | 578.8465 | 514.1236 | 613.345 | 566.5087 | 546.752 | 559.0218 |
| ENSMUSG00000028630 | -0.415316744 | 0.004695 | 0.024631 | 407.452 | 400.7404 | 552.3471 | 471.8551 | 681.6028 | 518.4616 | 580.5021 | 666.6933 |
| ENSMUSG00000028631 | 0.527738054 | 0.060271 | 0.155531 | 74.44802 | 89.98662 | 83.63877 | 119.042 | 46.80534 | 92.74205 | 62.6788 | 53.29196 |
| ENSMUSG00000028633 | -0.265838425 | 0.00133 | 0.00969 | 1298.816 | 1274.21 | 1165.974 | 1308.6 | 1673.291 | 1410.126 | 1453.184 | 1529.153 |
| ENSMUSG00000028634 | 0.273245158 | 0.215043 | 0.373221 | 129.781 | 160.7761 | 190.4645 | 144.0581 | 132.6151 | 170.9582 | 97.39321 | 117.4598 |
| ENSMUSG00000028635 | 0.671195841 | 0.210497 | 0.367996 | 15.09082 | 5.999108 | 22.35888 | 19.84034 | 6.825779 | 11.17374 | 12.53576 | 9.78832 |
| ENSMUSG00000028636 | -0.385554635 | 0.06637 | 0.166069 | 444.676 | 343.149 | 360.2264 | 333.8353 | 539.2366 | 315.0995 | 459.9659 | 621.0145 |
| ENSMUSG00000028637 | 0.452661332 | 0.001465 | 0.0104 | 264.5923 | 242.3639 | 243.4633 | 275.1769 | 198.9227 | 166.4887 | 163.9292 | 220.781 |
| ENSMUSG00000028639 | -0.412818581 | 5.52E-05 | 0.000855 | 17053.63 | 18320.08 | 18665.52 | 18301.42 | 24674.22 | 19726.12 | 24820.8 | 27084.28 |
| ENSMUSG00000028640 | 0.095945747 | 0.861259 | 0.916954 | 28.16952 | 11.99822 | 16.56213 | 16.38985 | 30.22845 | 7.821619 | 20.25007 | 9.78832 |
| ENSMUSG00000028641 | 0.241977187 | 0.00626 | 0.030366 | 1612.705 | 1858.524 | 1870.693 | 1649.336 | 1491.92 | 1328.558 | 1536.113 | 1553.08 |
| ENSMUSG00000028642 | 0.548852292 | 0.286047 | 0.453002 | 14.08476 | 9.598572 | 13.24971 | 11.21411 | 5.850668 | 11.17374 | 10.60718 | 5.437955 |
| ENSMUSG00000028643 | 0.058609616 | 0.546704 | 0.695045 | 1226.38 | 1431.387 | 1482.311 | 1475.086 | 1389.534 | 1482.755 | 1281.54 | 1239.854 |
| ENSMUSG00000028644 | 1.458942459 | 0.001383 | 0.009973 | 80.48435 | 25.19625 | 32.29616 | 35.36756 | 16.57689 | 15.64324 | 13.50005 | 17.40146 |
| ENSMUSG00000028645 | -0.927202588 | 9.17E-05 | 0.001252 | 1324.974 | 892.6672 | 1589.137 | 1157.641 | 2124.768 | 1588.906 | 3354.762 | 2374.211 |
| ENSMUSG00000028646 | -0.171139734 | 0.033447 | 0.10227 | 1660.996 | 1542.97 | 1856.615 | 1608.793 | 1865.388 | 1776.625 | 1975.828 | 1895.671 |
| ENSMUSG00000028647 | -0.070101266 | 0.495962 | 0.652653 | 497.9969 | 485.9277 | 412.3971 | 431.3117 | 445.6259 | 503.9357 | 472.5017 | 493.7664 |
| ENSMUSG00000028648 | -0.486456987 | 0.001039 | 0.008051 | 940.6608 | 927.462 | 1189.161 | 935.9465 | 1358.33 | 1100.613 | 1462.827 | 1674.89 |
| ENSMUSG00000028649 | 0.082005633 | 0.33228 | 0.500209 | 5667.104 | 5218.024 | 5927.587 | 5692.452 | 4717.589 | 5760.063 | 5633.377 | 5155.182 |
| ENSMUSG00000028651 | -0.053214789 | 0.409338 | 0.575555 | 1322.961 | 1382.194 | 1257.894 | 1376.747 | 1452.916 | 1348.671 | 1357.719 | 1376.89 |
| ENSMUSG00000028653 | 0.065162133 | 0.559016 | 0.70529 | 660.9777 | 583.1133 | 551.519 | 604.6991 | 582.1415 | 659.2507 | 522.6447 | 530.7445 |
| ENSMUSG00000028654 | -0.327572038 | 0.119206 | 0.248203 | 468.8213 | 308.3541 | 569.7374 | 510.6731 | 626.0215 | 469.2971 | 718.3954 | 519.8685 |
| ENSMUSG00000028655 | 0.32119337 | 0.096623 | 0.215606 | 133.8052 | 148.7779 | 117.5911 | 98.33908 | 109.2125 | 101.681 | 102.2147 | 83.74451 |
| ENSMUSG00000028656 | -0.139977447 | 0.093821 | 0.210858 | 4419.597 | 4141.784 | 4603.445 | 4434.747 | 4841.428 | 4260.547 | 4956.446 | 5336.809 |
| ENSMUSG00000028657 | -0.109476064 | 0.080121 | 0.189093 | 2079.514 | 1959.309 | 2057.017 | 2212.629 | 2350.018 | 2212.401 | 2226.544 | 2178.445 |
| ENSMUSG00000028661 | -1.572085602 | 0.006508 | 0.03133 | 10.06054 | 10.79839 | 32.29616 | 16.38985 | 40.95467 | 16.76061 | 61.71451 | 89.18247 |
| ENSMUSG00000028664 | -0.083245777 | 0.378969 | 0.54711 | 930.6003 | 1097.837 | 1015.259 | 999.7806 | 999.4891 | 1213.468 | 1022.147 | 1048.438 |
| ENSMUSG00000028668 | -0.050966177 | 0.603887 | 0.740916 | 1365.216 | 1360.598 | 1305.924 | 1400.901 | 1407.086 | 1182.182 | 1474.398 | 1562.868 |
| ENSMUSG00000028669 | -0.034508046 | 0.746138 | 0.841259 | 863.1946 | 914.264 | 811.5445 | 935.9465 | 1039.469 | 794.453 | 859.1816 | 914.6641 |
| ENSMUSG00000028670 | -0.268420139 | 0.014566 | 0.056454 | 1342.077 | 1317.404 | 1482.311 | 1293.935 | 1659.639 | 1359.844 | 1663.399 | 1865.219 |
| ENSMUSG00000028671 | -0.878069075 | 0.000671 | 0.005755 | 132.7992 | 163.1757 | 202.058 | 149.2339 | 290.5832 | 191.071 | 273.8581 | 435.0364 |
| ENSMUSG00000028672 | -0.536848207 | 0.000169 | 0.00199 | 876.2733 | 813.479 | 934.9324 | 806.553 | 1418.787 | 957.5896 | 1168.718 | 1433.445 |
| ENSMUSG00000028673 | -0.231771421 | 0.064441 | 0.163145 | 1799.831 | 1992.904 | 2196.967 | 1982.309 | 2568.443 | 1828.024 | 2494.616 | 2469.919 |
| ENSMUSG00000028675 | -0.074852662 | 0.307024 | 0.475531 | 3132.853 | 3317.507 | 2895.889 | 2870.811 | 3023.82 | 3243.737 | 3212.047 | 3382.408 |
| ENSMUSG00000028676 | 0.153256078 | 0.230801 | 0.391225 | 5107.738 | 5573.171 | 4793.909 | 5516.477 | 4507.94 | 5955.604 | 4260.23 | 4152.423 |
| ENSMUSG00000028677 | -0.102731566 | 0.077822 | 0.185448 | 2331.028 | 2343.251 | 2456.164 | 2435.186 | 2744.938 | 2483.923 | 2534.152 | 2510.16 |
| ENSMUSG00000028678 | 0.07363004 | 0.509807 | 0.665571 | 1233.423 | 1103.836 | 992.0717 | 1168.855 | 1203.287 | 920.7163 | 1063.611 | 1084.328 |
| ENSMUSG00000028680 | -0.780404514 | 0.010494 | 0.044181 | 133.8052 | 61.1909 | 118.4192 | 134.5693 | 173.5698 | 166.4887 | 285.4296 | 145.7372 |
| ENSMUSG00000028681 | 0.226260895 | 0.070797 | 0.173776 | 395.3794 | 380.3434 | 481.13 | 376.9665 | 385.169 | 353.0902 | 343.2869 | 316.489 |
| ENSMUSG00000028683 | -0.228782181 | 0.013688 | 0.053888 | 709.2683 | 589.1124 | 603.6897 | 639.204 | 772.2882 | 745.2885 | 688.5025 | 773.2773 |
| ENSMUSG00000028684 | -0.09448143 | 0.252038 | 0.415689 | 1775.686 | 1671.351 | 1661.182 | 1837.388 | 1914.143 | 1634.718 | 1904.471 | 1962.014 |
| ENSMUSG00000028687 | 0.045344678 | 0.749312 | 0.843611 | 253.5257 | 248.3631 | 223.5888 | 240.672 | 265.2303 | 211.1837 | 260.3581 | 196.854 |
| ENSMUSG00000028688 | -0.09127237 | 0.326199 | 0.493995 | 603.6326 | 673.0999 | 683.188 | 592.6223 | 711.8313 | 616.7905 | 702.9668 | 685.1824 |
| ENSMUSG00000028689 | 0.239889285 | 0.015996 | 0.060312 | 900.4186 | 931.0615 | 811.5445 | 921.2819 | 800.5664 | 779.9271 | 788.7885 | 646.0291 |
| ENSMUSG00000028690 | -0.163499341 | 0.107885 | 0.2325 | 349.1009 | 376.744 | 352.7734 | 361.4392 | 385.169 | 401.1373 | 373.1799 | 453.5255 |
| ENSMUSG00000028691 | -0.279350138 | 0.000988 | 0.00778 | 8043.405 | 8342.359 | 9233.389 | 7871.439 | 9930.534 | 9198.224 | 10416.25 | 11102.13 |
| ENSMUSG00000028692 | -0.402154323 | 0.000437 | 0.004124 | 11382.5 | 11386.31 | 12946.62 | 12003.41 | 17403.79 | 12697.84 | 15421.88 | 17537.41 |
| ENSMUSG00000028693 | -0.146167023 | 0.093453 | 0.210389 | 3778.74 | 4570.12 | 4162.892 | 3949.953 | 4556.695 | 5013.658 | 4251.551 | 4392.78 |
| ENSMUSG00000028696 | -0.45356964 | 0.013317 | 0.052912 | 221.332 | 188.372 | 187.9802 | 277.7648 | 311.0605 | 233.5312 | 327.8583 | 327.3649 |
| ENSMUSG00000028698 | -0.044447455 | 0.733803 | 0.833293 | 689.1472 | 781.0838 | 780.0764 | 614.1879 | 656.2499 | 728.5279 | 690.431 | 878.7736 |
| ENSMUSG00000028699 | -1.455859072 | 0.000272 | 0.002873 | 47.28455 | 28.79572 | 88.60741 | 31.05445 | 132.6151 | 82.68568 | 141.7505 | 182.7153 |
| ENSMUSG00000028700 | -0.235883966 | 0.020469 | 0.071685 | 1297.81 | 1345 | 1353.126 | 1268.919 | 1495.821 | 1328.558 | 1581.434 | 1793.438 |
| ENSMUSG00000028701 | -0.203362814 | 0.19665 | 0.351004 | 233.4046 | 217.1677 | 274.1033 | 255.3366 | 291.5583 | 216.7706 | 301.8225 | 319.7518 |
| ENSMUSG00000028702 | -0.10862678 | 0.323033 | 0.491133 | 445.6821 | 466.7306 | 392.5225 | 453.74 | 442.7005 | 539.6917 | 467.6802 | 444.8248 |

| | | | | | | | | | | | |
|---|---|---|---|---|---|---|---|---|---|---|---|
| ENSMUSG00000028703 | -0.42330986 | 0.000795 | 0.006545 | 924.564 | 837.4754 | 885.246 | 843.6458 | 1213.038 | 909.5425 | 1223.683 | 1334.474 |
| ENSMUSG00000028706 | 0.003133505 | 0.967963 | 0.981932 | 651.9232 | 645.504 | 614.4551 | 650.4181 | 607.4943 | 629.0816 | 630.6451 | 689.5327 |
| ENSMUSG00000028708 | 0.0493789 | 0.55478 | 0.70188 | 665.0019 | 668.3006 | 662.4853 | 699.5877 | 649.4241 | 584.3867 | 681.7524 | 688.4452 |
| ENSMUSG00000028709 | 0.066395418 | 0.684011 | 0.798636 | 439.6458 | 454.7324 | 379.2728 | 322.6212 | 376.393 | 458.1234 | 312.4297 | 376.3065 |
| ENSMUSG00000028710 | -0.596159281 | 2.90E-05 | 0.000524 | 526.1664 | 466.7306 | 514.2542 | 538.2771 | 777.1637 | 591.0909 | 807.11 | 917.9269 |
| ENSMUSG00000028712 | -4.550915923 | 1.20E-08 | 9.91E-07 | 18.10898 | 5.999108 | 62.9361 | 31.05445 | 676.7272 | 198.8926 | 547.7162 | 1358.401 |
| ENSMUSG00000028713 | -0.156717645 | 0.717259 | 0.823139 | 112.6781 | 26.39607 | 80.32634 | 45.71905 | 51.6809 | 70.39457 | 74.25027 | 100.0584 |
| ENSMUSG00000028715 | -2.440454421 | 0.002272 | 0.014388 | 21.12714 | 0 | 5.796746 | 12.93935 | 74.10846 | 20.11273 | 29.89296 | 93.53283 |
| ENSMUSG00000028716 | -1.70893299 | 0.005006 | 0.025762 | 899.4126 | 513.5236 | 2466.102 | 949.7485 | 4156.9 | 1633.601 | 4023.979 | 5974.138 |
| ENSMUSG00000028717 | -0.19535617 | 0.459155 | 0.62059 | 142.8597 | 76.78858 | 121.7317 | 88.85022 | 102.3867 | 98.32892 | 134.0362 | 158.7883 |
| ENSMUSG00000028718 | 0.032236785 | 0.770554 | 0.857548 | 706.2502 | 668.3006 | 568.0811 | 678.8847 | 658.2001 | 716.2368 | 601.7164 | 586.2116 |
| ENSMUSG00000028719 | -0.178883753 | 0.009215 | 0.040157 | 2068.448 | 2014.5 | 2005.674 | 2163.46 | 2206.677 | 2220.222 | 2412.651 | 2503.635 |
| ENSMUSG00000028729 | -0.110461653 | 0.099856 | 0.220685 | 2275.695 | 2366.048 | 2179.577 | 2276.463 | 2672.78 | 2365.481 | 2435.794 | 2341.584 |
| ENSMUSG00000028730 | 0.352196177 | 0.64758 | 0.77184 | 3.018163 | 17.99732 | 17.39024 | 14.6646 | 6.825779 | 17.87799 | 1.928578 | 15.22628 |
| ENSMUSG00000028737 | -1.304137114 | 9.89E-08 | 5.60E-06 | 717.3168 | 569.9152 | 780.0764 | 718.5654 | 1771.777 | 964.2938 | 1669.185 | 2476.445 |
| ENSMUSG00000028741 | -0.148292203 | 0.06677 | 0.166804 | 1053.339 | 1257.413 | 1185.021 | 1145.564 | 1316.4 | 1193.356 | 1303.719 | 1326.861 |
| ENSMUSG00000028743 | -0.700727164 | 4.50E-07 | 1.93E-05 | 1143.884 | 1137.431 | 1301.784 | 1155.053 | 1912.193 | 1497.281 | 1927.614 | 2365.511 |
| ENSMUSG00000028744 | -0.595945297 | 0.001536 | 0.010706 | 204.229 | 172.7743 | 178.0429 | 184.6014 | 331.5378 | 204.4795 | 247.8223 | 333.8905 |
| ENSMUSG00000028745 | -0.085897741 | 0.177062 | 0.326069 | 6146.992 | 6565.423 | 6589.244 | 6299.739 | 6969.121 | 6227.126 | 6801.131 | 7172.663 |
| ENSMUSG00000028750 | -0.44746459 | 0.445367 | 0.608236 | 13.07871 | 7.198929 | 18.21835 | 12.93935 | 25.35289 | 5.58687 | 12.53576 | 27.18978 |
| ENSMUSG00000028755 | -1.115561313 | 1.84E-06 | 6.06E-05 | 109.6599 | 121.182 | 149.8873 | 135.4319 | 318.8614 | 191.071 | 237.2151 | 373.0437 |
| ENSMUSG00000028756 | -0.084600119 | 0.351837 | 0.520605 | 4855.218 | 4165.78 | 5216.244 | 5039.446 | 5158.339 | 4820.352 | 5546.591 | 4920.262 |
| ENSMUSG00000028757 | -0.173379962 | 0.02604 | 0.085514 | 5321.021 | 6093.894 | 6271.252 | 5691.59 | 6583.952 | 6057.285 | 6901.418 | 6819.196 |
| ENSMUSG00000028758 | 0.002588662 | 0.992458 | 0.995642 | 54.32694 | 43.19358 | 32.29616 | 51.75741 | 43.88001 | 43.57759 | 54.00019 | 39.15328 |
| ENSMUSG00000028759 | 0.033463793 | 0.552169 | 0.699721 | 8024.29 | 7668.059 | 7253.386 | 7755.848 | 7834.044 | 7455.12 | 7564.848 | 7138.948 |
| ENSMUSG00000028760 | 0.32838899 | 0.007055 | 0.033098 | 5644.971 | 5474.786 | 5381.037 | 5494.912 | 3831.212 | 5538.823 | 4182.122 | 3967.532 |
| ENSMUSG00000028763 | 0.194326836 | 0.128634 | 0.260297 | 5121.823 | 4993.657 | 6329.219 | 5604.465 | 4037.936 | 5792.467 | 4713.445 | 4731.021 |
| ENSMUSG00000028766 | -0.877454974 | 0.002977 | 0.017657 | 391.3551 | 491.9268 | 554.8314 | 426.9986 | 946.8331 | 416.7805 | 741.5384 | 1321.423 |
| ENSMUSG00000028771 | 0.047343666 | 0.465464 | 0.62632 | 1676.087 | 1554.969 | 1484.795 | 1652.787 | 1503.622 | 1538.624 | 1564.077 | 1556.343 |
| ENSMUSG00000028772 | -0.221345527 | 0.00346 | 0.019606 | 941.6669 | 911.8644 | 857.9185 | 890.2274 | 1010.215 | 1036.923 | 1007.682 | 1143.058 |
| ENSMUSG00000028773 | -2.655731342 | 0.000366 | 0.003599 | 246.4833 | 113.983 | 739.4992 | 213.068 | 1864.413 | 568.7434 | 2356.723 | 3487.905 |
| ENSMUSG00000028776 | -1.208813419 | 0.000159 | 0.001898 | 1255.556 | 686.2979 | 1970.894 | 1146.427 | 2638.651 | 1631.366 | 3873.55 | 3554.248 |
| ENSMUSG00000028778 | 0.68038527 | 0.154142 | 0.295564 | 69.41775 | 29.99554 | 28.15563 | 36.23019 | 31.20356 | 18.99536 | 39.53586 | 11.9635 |
| ENSMUSG00000028779 | -0.192988737 | 0.02832 | 0.090822 | 1230.404 | 1249.014 | 1251.269 | 1222.337 | 1533.85 | 1220.173 | 1497.541 | 1407.343 |
| ENSMUSG00000028780 | 0.050108301 | 0.712366 | 0.819818 | 2633.85 | 2909.567 | 3643.669 | 2698.286 | 2543.09 | 2685.05 | 2864.903 | 3388.934 |
| ENSMUSG00000028782 | 0.74945905 | 3.54E-11 | 7.17E-09 | 485.9243 | 449.9331 | 544.0661 | 512.3984 | 319.8365 | 312.8647 | 261.3224 | 293.6496 |
| ENSMUSG00000028785 | 0.673766246 | 0.01347 | 0.053316 | 67.40564 | 58.79126 | 61.27989 | 92.30071 | 37.05423 | 45.81234 | 56.89306 | 35.89051 |
| ENSMUSG00000028786 | -0.73743174 | 0.085427 | 0.197648 | 49.29666 | 19.19714 | 54.65504 | 32.77969 | 75.08357 | 30.1691 | 65.57166 | 90.27006 |
| ENSMUSG00000028788 | 0.207945223 | 0.114713 | 0.242317 | 8552.468 | 9236.226 | 8296.8 | 8803.073 | 6476.689 | 9735.68 | 7119.347 | 6874.663 |
| ENSMUSG00000028789 | 0.359074865 | 2.32E-05 | 0.000442 | 651.9232 | 628.7065 | 624.3924 | 624.5394 | 444.6508 | 501.701 | 500.4661 | 526.3941 |
| ENSMUSG00000028790 | 0.078396922 | 0.406243 | 0.572907 | 5587.626 | 6216.275 | 5618.703 | 5855.488 | 5268.526 | 6461.774 | 5069.268 | 5247.627 |
| ENSMUSG00000028792 | -0.341301371 | 9.97E-05 | 0.001334 | 2408.494 | 2639.607 | 2614.333 | 2444.675 | 3259.797 | 2774.44 | 3266.047 | 3502.043 |
| ENSMUSG00000028793 | -0.353910784 | 0.000286 | 0.002998 | 513.0877 | 464.3309 | 473.677 | 552.079 | 680.6277 | 584.3867 | 656.6809 | 639.5036 |
| ENSMUSG00000028794 | 0.147607045 | 0.614632 | 0.748195 | 34.20585 | 31.19536 | 38.92101 | 31.91707 | 24.37778 | 30.1691 | 28.92867 | 40.24087 |
| ENSMUSG00000028795 | -0.361958563 | 6.39E-05 | 0.000959 | 551.3178 | 502.7252 | 522.5353 | 545.178 | 688.4286 | 629.0816 | 753.1098 | 655.8174 |
| ENSMUSG00000028796 | 0.002146647 | 0.974956 | 0.986285 | 2879.328 | 3111.137 | 3421.737 | 3201.196 | 3233.469 | 3101.83 | 3132.011 | 3131.175 |
| ENSMUSG00000028797 | -0.056529336 | 0.312859 | 0.481091 | 3816.97 | 4218.573 | 4199.329 | 4169.059 | 4177.377 | 4429.271 | 4238.051 | 4216.591 |
| ENSMUSG00000028798 | -0.18694574 | 0.007368 | 0.034115 | 4431.669 | 4912.069 | 4833.658 | 4665.068 | 5606.89 | 4863.929 | 5431.841 | 5543.452 |
| ENSMUSG00000028799 | 0.111774749 | 0.118507 | 0.247362 | 1945.709 | 1971.307 | 2049.564 | 1841.701 | 1819.558 | 1920.766 | 1658.577 | 1829.328 |
| ENSMUSG00000028800 | -0.230608219 | 0.005342 | 0.026914 | 3394.427 | 3802.234 | 3868.914 | 3598.865 | 4238.809 | 3851.589 | 4440.552 | 4674.466 |
| ENSMUSG00000028801 | -0.573947169 | 0.426348 | 0.591319 | 7.04238 | 2.399643 | 22.35888 | 11.21411 | 7.80089 | 8.938993 | 22.17865 | 26.10219 |
| ENSMUSG00000028803 | -0.284966963 | 0.045724 | 0.127861 | 682.1049 | 597.5111 | 657.5167 | 725.4664 | 716.7068 | 667.0723 | 974.8963 | 886.3867 |
| ENSMUSG00000028804 | 0.27069559 | 0.621472 | 0.753063 | 18.10898 | 13.19804 | 8.281066 | 32.77969 | 9.751113 | 21.23011 | 17.3572 | 11.9635 |
| ENSMUSG00000028807 | -0.206958758 | 0.09143 | 0.207403 | 308.8587 | 344.3488 | 340.3518 | 293.292 | 354.9405 | 429.0717 | 371.2513 | 329.5401 |
| ENSMUSG00000028809 | 0.211007301 | 0.033196 | 0.101653 | 5564.487 | 5230.022 | 4821.237 | 5316.349 | 4381.175 | 5281.827 | 4406.801 | 4014.299 |
| ENSMUSG00000028811 | -0.317384187 | 0.000155 | 0.001872 | 1420.549 | 1498.577 | 1560.153 | 1350.006 | 1864.413 | 1688.352 | 1729.935 | 1980.503 |
| ENSMUSG00000028820 | -0.010664711 | 0.9131 | 0.948699 | 9797.963 | 10938.77 | 9705.41 | 9827.869 | 9059.759 | 11877.69 | 9740.285 | 9890.553 |
| ENSMUSG00000028821 | -0.02707398 | 0.652589 | 0.775593 | 2363.222 | 2177.676 | 2142.312 | 2346.336 | 2226.179 | 2301.791 | 2352.866 | 2320.919 |
| ENSMUSG00000028822 | 0.057374645 | 0.402451 | 0.569517 | 3657.008 | 3185.526 | 3343.895 | 3724.808 | 3370.96 | 3382.291 | 3320.048 | 3299.751 |
| ENSMUSG00000028825 | 1.537624473 | 0.019073 | 0.068209 | 72.43591 | 15.59768 | 14.90592 | 21.56559 | 9.751113 | 10.05637 | 5.785735 | 17.40146 |
| ENSMUSG00000028826 | 0.001385637 | 0.983693 | 0.990959 | 2222.374 | 2120.085 | 2088.485 | 2222.118 | 2220.328 | 2318.551 | 2015.364 | 2092.525 |
| ENSMUSG00000028830 | -0.163418213 | 0.216812 | 0.375339 | 909.4731 | 829.0767 | 891.8708 | 976.4898 | 979.0118 | 806.7441 | 1042.397 | 1212.664 |
| ENSMUSG00000028832 | -0.211165767 | 0.003991 | 0.02181 | 6896.503 | 8031.605 | 7578.832 | 7914.571 | 8960.298 | 9142.355 | 8991.032 | 8121.043 |
| ENSMUSG00000028833 | -0.109387124 | 0.30214 | 0.470457 | 746.4923 | 752.2881 | 838.872 | 715.9775 | 938.0571 | 725.1758 | 796.5028 | 834.1824 |

| | | | | | | | | | | | |
|---|---|---|---|---|---|---|---|---|---|---|---|
| ENSMUSG00000028834 | -0.091431889 | 0.64235 | 0.768671 | 181.0898 | 212.3684 | 132.4971 | 175.9752 | 231.1014 | 166.4887 | 186.1078 | 160.9635 |
| ENSMUSG00000028836 | 0.261027976 | 0.507987 | 0.663699 | 270.6286 | 91.18644 | 115.1068 | 104.3774 | 136.5156 | 78.21619 | 97.39321 | 172.927 |
| ENSMUSG00000028837 | -0.245995306 | 0.000365 | 0.003599 | 1935.649 | 2007.301 | 2226.779 | 2034.929 | 2389.998 | 2331.96 | 2449.294 | 2563.452 |
| ENSMUSG00000028838 | -0.124197577 | 0.672465 | 0.790252 | 68.4117 | 38.39429 | 59.62368 | 42.26855 | 76.05868 | 48.04709 | 48.21446 | 55.46715 |
| ENSMUSG00000028840 | 0.341404043 | 0.18194 | 0.332463 | 96.58122 | 136.7797 | 79.49824 | 106.9653 | 88.73513 | 107.2679 | 61.71451 | 72.8686 |
| ENSMUSG00000028841 | -0.016039337 | 0.904227 | 0.942582 | 311.8769 | 269.9598 | 378.4447 | 306.2313 | 339.3387 | 306.1605 | 336.5369 | 301.2627 |
| ENSMUSG00000028842 | 0.418150938 | 0.000114 | 0.001489 | 577.4752 | 519.5227 | 452.1462 | 498.5964 | 358.841 | 403.372 | 380.8942 | 388.27 |
| ENSMUSG00000028843 | -0.458655194 | 0.000791 | 0.006525 | 1044.284 | 963.4567 | 1218.145 | 1057.576 | 1151.606 | 1455.938 | 1664.363 | 1618.336 |
| ENSMUSG00000028845 | 1.257637144 | 4.22E-06 | 0.000118 | 109.6599 | 146.3782 | 139.1219 | 164.7611 | 44.85512 | 84.92043 | 64.60737 | 40.24087 |
| ENSMUSG00000028847 | -0.13669397 | 0.087221 | 0.200669 | 1874.279 | 1943.711 | 1988.284 | 2021.127 | 2244.706 | 1881.658 | 2240.044 | 2238.262 |
| ENSMUSG00000028848 | -0.149802149 | 0.209098 | 0.366209 | 355.1372 | 371.9447 | 404.9441 | 386.4553 | 467.0783 | 357.5597 | 457.0731 | 402.4087 |
| ENSMUSG00000028849 | -0.064976324 | 0.313656 | 0.48164 | 1846.11 | 1800.932 | 2051.22 | 1861.541 | 2012.63 | 1924.118 | 1961.364 | 2015.306 |
| ENSMUSG00000028850 | -0.361249466 | 0.005869 | 0.028941 | 237.4288 | 202.7698 | 190.4645 | 178.5631 | 274.0063 | 252.5265 | 262.2867 | 247.9708 |
| ENSMUSG00000028851 | -0.308444301 | 0.000132 | 0.001655 | 3810.934 | 3928.216 | 4483.369 | 4139.73 | 5357.262 | 4605.816 | 5056.732 | 5246.539 |
| ENSMUSG00000028854 | 0.131020322 | 0.174486 | 0.322656 | 809.8738 | 712.694 | 854.606 | 714.2523 | 720.6073 | 652.5465 | 707.7882 | 743.9123 |
| ENSMUSG00000028857 | -0.092430453 | 0.194742 | 0.348542 | 1223.362 | 1258.613 | 1408.609 | 1316.363 | 1445.115 | 1339.732 | 1373.148 | 1397.555 |
| ENSMUSG00000028859 | -0.151052449 | 0.572784 | 0.715292 | 41.24823 | 31.19536 | 43.06154 | 37.09281 | 44.85512 | 34.6386 | 38.57157 | 52.20437 |
| ENSMUSG00000028860 | 0.036460523 | 0.890018 | 0.933475 | 56.33904 | 75.58876 | 42.23344 | 47.44429 | 49.73068 | 51.39921 | 50.14304 | 63.08028 |
| ENSMUSG00000028861 | -0.163555797 | 0.00433 | 0.02316 | 1112.696 | 1093.037 | 1081.507 | 1076.554 | 1224.74 | 1243.637 | 1177.397 | 1242.029 |
| ENSMUSG00000028862 | 0.22188291 | 0.511327 | 0.666835 | 347.0888 | 187.1722 | 161.4808 | 332.9727 | 150.1671 | 263.7003 | 318.2154 | 150.0876 |
| ENSMUSG00000028863 | 0.126103675 | 0.091961 | 0.208232 | 1303.846 | 1319.804 | 1230.566 | 1322.402 | 1262.769 | 1245.872 | 1129.183 | 1103.905 |
| ENSMUSG00000028864 | 0.77669241 | 0.010629 | 0.044634 | 105.6357 | 151.1775 | 105.9976 | 83.67448 | 83.85957 | 82.68568 | 53.0359 | 40.24087 |
| ENSMUSG00000028865 | -0.526044488 | 0.004232 | 0.022765 | 82.49646 | 101.9848 | 120.0755 | 119.042 | 128.7147 | 155.315 | 162.0006 | 166.4014 |
| ENSMUSG00000028868 | 0.261045627 | 0.000734 | 0.006166 | 3279.737 | 3346.302 | 3426.705 | 3278.832 | 3018.945 | 2964.393 | 2567.902 | 2574.328 |
| ENSMUSG00000028869 | 0.081342032 | 0.226942 | 0.386817 | 2317.949 | 2424.839 | 2307.105 | 2389.467 | 2274.935 | 2358.777 | 2264.151 | 2021.832 |
| ENSMUSG00000028871 | 0.023156795 | 0.897422 | 0.937817 | 1896.412 | 2237.667 | 1633.026 | 2138.444 | 2361.72 | 2266.035 | 1755.971 | 1394.292 |
| ENSMUSG00000028873 | -0.039986614 | 0.627776 | 0.757468 | 1453.749 | 1536.971 | 1406.953 | 1502.69 | 1648.913 | 1574.38 | 1487.898 | 1351.876 |
| ENSMUSG00000028874 | -0.257909128 | 0.608616 | 0.744128 | 25.15136 | 8.398751 | 10.76539 | 12.93935 | 22.42756 | 10.05637 | 17.3572 | 18.48905 |
| ENSMUSG00000028878 | 0.030873498 | 0.668828 | 0.78764 | 907.461 | 1033.046 | 936.5886 | 982.5282 | 929.2811 | 946.4159 | 981.6464 | 916.8393 |
| ENSMUSG00000028879 | 0.018892245 | 0.742754 | 0.839394 | 2100.641 | 2021.699 | 2174.608 | 2150.52 | 2057.485 | 1985.574 | 2140.722 | 2157.781 |
| ENSMUSG00000028882 | -0.109382869 | 0.203183 | 0.358963 | 1506.063 | 1451.784 | 1380.454 | 1448.345 | 1676.216 | 1353.14 | 1607.47 | 1602.022 |
| ENSMUSG00000028883 | 0.506965346 | 0.00108 | 0.008282 | 1066.418 | 1595.763 | 1734.883 | 1324.99 | 1023.867 | 1101.731 | 967.182 | 934.2407 |
| ENSMUSG00000028884 | -0.116854835 | 0.29235 | 0.459709 | 1042.272 | 1395.392 | 1164.318 | 1074.829 | 1367.106 | 1193.356 | 1251.647 | 1253.993 |
| ENSMUSG00000028885 | -0.161634021 | 0.408573 | 0.574874 | 238.4349 | 273.5593 | 304.7432 | 245.8477 | 351.0401 | 210.0663 | 358.7156 | 267.5474 |
| ENSMUSG00000028886 | -0.016772843 | 0.868511 | 0.92134 | 1054.345 | 890.2676 | 789.1856 | 941.1222 | 921.4802 | 937.4769 | 928.6105 | 928.8028 |
| ENSMUSG00000028889 | -0.126825947 | 0.156136 | 0.298246 | 524.1543 | 542.3193 | 561.4563 | 558.1174 | 614.3201 | 530.7527 | 602.6807 | 639.5036 |
| ENSMUSG00000028890 | 0.187256954 | 0.02101 | 0.072998 | 485.9243 | 499.1258 | 488.5829 | 489.1075 | 446.601 | 441.3628 | 429.1087 | 405.6715 |
| ENSMUSG00000028893 | -0.581233792 | 0.00199 | 0.013086 | 271.6347 | 305.9545 | 330.4145 | 244.1224 | 386.1441 | 318.4516 | 489.8589 | 528.5693 |
| ENSMUSG00000028894 | -0.263068128 | 0.040751 | 0.118273 | 799.8132 | 760.6869 | 746.1241 | 739.2683 | 1010.215 | 712.8847 | 892.9318 | 1037.562 |
| ENSMUSG00000028896 | -0.129994931 | 0.224953 | 0.384741 | 1168.029 | 1156.628 | 1204.067 | 1110.196 | 1383.683 | 1043.627 | 1253.576 | 1394.292 |
| ENSMUSG00000028898 | -0.04303223 | 0.526541 | 0.679534 | 1008.066 | 1045.045 | 1010.29 | 948.8858 | 1071.647 | 993.3456 | 1060.718 | 1002.759 |
| ENSMUSG00000028899 | -0.072484936 | 0.537746 | 0.687843 | 442.6639 | 524.322 | 549.0347 | 570.1941 | 533.3859 | 487.1751 | 590.145 | 584.0364 |
| ENSMUSG00000028901 | 0.065201347 | 0.325681 | 0.49356 | 1104.648 | 1136.231 | 1077.367 | 1109.334 | 1125.278 | 1070.444 | 1024.075 | 1009.285 |
| ENSMUSG00000028902 | -0.148569326 | 0.079152 | 0.187719 | 2195.211 | 2454.835 | 2422.212 | 2301.479 | 2782.968 | 2285.03 | 2733.76 | 2585.204 |
| ENSMUSG00000028906 | -0.455079564 | 8.29E-05 | 0.001166 | 2631.838 | 2412.841 | 2888.436 | 2750.044 | 3687.871 | 2958.807 | 4035.55 | 3965.357 |
| ENSMUSG00000028907 | 0.071468803 | 0.457386 | 0.619143 | 1664.014 | 1517.774 | 1288.534 | 1569.975 | 1471.443 | 1496.164 | 1350.005 | 1429.095 |
| ENSMUSG00000028909 | 0.128948599 | 0.200664 | 0.356032 | 1582.524 | 1319.804 | 1527.029 | 1576.876 | 1226.69 | 1303.976 | 1485.005 | 1480.211 |
| ENSMUSG00000028910 | -0.445984175 | 0.000334 | 0.003362 | 544.2754 | 459.5316 | 454.6305 | 449.4268 | 637.7228 | 556.4523 | 648.9666 | 754.7882 |
| ENSMUSG00000028911 | -0.054855329 | 0.534687 | 0.685626 | 1025.169 | 1153.028 | 1194.958 | 1112.784 | 1228.64 | 1178.83 | 1214.04 | 1037.562 |
| ENSMUSG00000028914 | -0.033617686 | 0.728308 | 0.829425 | 774.6619 | 785.8831 | 977.9939 | 833.2943 | 876.6251 | 872.6692 | 838.9316 | 867.8977 |
| ENSMUSG00000028917 | -0.048073418 | 0.606609 | 0.742658 | 980.903 | 881.8688 | 984.6188 | 980.8029 | 1093.1 | 869.317 | 1007.682 | 988.6203 |
| ENSMUSG00000028919 | 0.313764859 | 0.000502 | 0.004607 | 621.7416 | 610.7092 | 692.2971 | 667.6706 | 503.1574 | 527.4006 | 491.7875 | 567.7225 |
| ENSMUSG00000028920 | 0.028742943 | 0.788114 | 0.86989 | 771.6437 | 736.6904 | 645.9232 | 625.402 | 666.9761 | 641.3727 | 756.0027 | 655.8174 |
| ENSMUSG00000028923 | -0.157398016 | 0.090405 | 0.205863 | 1532.221 | 1490.178 | 1752.274 | 1512.179 | 1818.583 | 1544.211 | 1778.149 | 1872.832 |
| ENSMUSG00000028926 | 0.489796129 | 0.001026 | 0.007991 | 1332.016 | 1436.186 | 1184.192 | 1401.763 | 1063.846 | 1144.191 | 868.8245 | 735.2116 |
| ENSMUSG00000028927 | 0.014509358 | 0.877778 | 0.926842 | 722.347 | 782.2836 | 761.03 | 643.5171 | 773.2633 | 677.1287 | 723.2169 | 702.5838 |
| ENSMUSG00000028931 | -0.272462689 | 0.060955 | 0.156706 | 164.9929 | 200.3702 | 216.1358 | 163.8985 | 246.7032 | 212.3011 | 218.8936 | 221.8686 |
| ENSMUSG00000028932 | -0.244617815 | 0.000169 | 0.00199 | 2797.837 | 2977.957 | 3006.855 | 2902.728 | 3472.371 | 3190.103 | 3486.87 | 3694.547 |
| ENSMUSG00000028933 | -0.206144738 | 0.072152 | 0.176275 | 238.4349 | 286.7573 | 258.3693 | 263.1002 | 287.6578 | 336.3296 | 290.251 | 292.562 |
| ENSMUSG00000028936 | -0.070230442 | 0.342818 | 0.511608 | 15944.96 | 17506.6 | 17170.79 | 17340.46 | 19775.26 | 17353.94 | 17774.74 | 16446.55 |
| ENSMUSG00000028937 | -0.519446158 | 0.009155 | 0.039984 | 630.7961 | 890.2676 | 780.0764 | 703.0381 | 1097 | 706.1804 | 1134.004 | 1367.102 |
| ENSMUSG00000028943 | -0.331197396 | 7.09E-05 | 0.001042 | 532.2028 | 517.1231 | 521.7072 | 496.8711 | 692.329 | 636.9032 | 600.7521 | 671.0437 |
| ENSMUSG00000028944 | -0.019576145 | 0.851307 | 0.910613 | 1067.424 | 947.859 | 1032.649 | 1054.126 | 937.082 | 1082.735 | 945.0034 | 1197.438 |
| ENSMUSG00000028945 | -0.096183736 | 0.073101 | 0.177872 | 2904.479 | 2907.168 | 2720.33 | 2932.92 | 2951.662 | 3121.943 | 3154.19 | 3024.591 |

| | | | | | | | | | | | |
|---|---|---|---|---|---|---|---|---|---|---|---|
| ENSMUSG00000028948 | -0.204087051 | 0.018915 | 0.067785 | 963.8001 | 969.4558 | 990.4155 | 1068.791 | 1247.167 | 1024.632 | 1134.968 | 1193.087 |
| ENSMUSG00000028949 | -0.178992712 | 0.064928 | 0.164084 | 528.1785 | 647.9036 | 623.5643 | 553.8043 | 689.4037 | 625.7295 | 682.7167 | 663.4306 |
| ENSMUSG00000028950 | 0.161858109 | 0.663708 | 0.783755 | 20.12109 | 25.19625 | 24.8432 | 21.56559 | 25.35289 | 11.17374 | 22.17865 | 22.83941 |
| ENSMUSG00000028952 | 0.043172143 | 0.675834 | 0.792479 | 435.6215 | 431.9358 | 395.835 | 372.6533 | 401.7459 | 426.8369 | 394.3943 | 362.1678 |
| ENSMUSG00000028953 | -0.317165118 | 0.000115 | 0.001503 | 1641.881 | 1619.759 | 1707.556 | 1748.538 | 2134.519 | 1860.428 | 2072.257 | 2304.606 |
| ENSMUSG00000028954 | -0.030752977 | 0.576906 | 0.718897 | 1926.594 | 1960.508 | 1993.253 | 1887.42 | 1955.098 | 1978.87 | 2082.865 | 1916.335 |
| ENSMUSG00000028955 | -0.113981463 | 0.164171 | 0.309145 | 2368.252 | 2446.436 | 2396.541 | 2593.909 | 2708.859 | 2844.834 | 2735.688 | 2322.007 |
| ENSMUSG00000028957 | 0.273769698 | 0.102316 | 0.224482 | 565.4025 | 608.3095 | 524.1915 | 540.0023 | 491.4561 | 587.7388 | 432.9658 | 338.2408 |
| ENSMUSG00000028958 | -0.320115976 | 0.008687 | 0.038475 | 230.3864 | 247.1632 | 245.1196 | 229.4578 | 269.1307 | 269.2872 | 332.6798 | 316.489 |
| ENSMUSG00000028959 | -0.264242319 | 0.000752 | 0.006277 | 1603.651 | 1528.573 | 1666.979 | 1664.001 | 1829.309 | 1849.254 | 1925.685 | 2163.219 |
| ENSMUSG00000028960 | 0.311746038 | 0.002005 | 0.013164 | 3341.107 | 3286.311 | 2853.655 | 3298.672 | 2528.464 | 2967.746 | 2519.688 | 2279.591 |
| ENSMUSG00000028961 | -0.202067117 | 0.079774 | 0.188683 | 2922.588 | 2441.637 | 2931.497 | 2497.295 | 3128.157 | 2574.43 | 3296.905 | 3417.211 |
| ENSMUSG00000028962 | -0.210627883 | 0.113729 | 0.240848 | 1421.555 | 1478.18 | 1465.749 | 1357.769 | 1508.497 | 1326.323 | 1767.542 | 2019.657 |
| ENSMUSG00000028964 | -0.156950622 | 0.017261 | 0.063586 | 3991.018 | 4490.932 | 4436.995 | 4256.184 | 4858.98 | 4491.844 | 4750.088 | 5046.423 |
| ENSMUSG00000028967 | -0.215805708 | 0.457203 | 0.618995 | 1906.473 | 1179.425 | 1151.068 | 1928.826 | 1209.138 | 2119.659 | 2666.26 | 1165.898 |
| ENSMUSG00000028969 | 0.124878464 | 0.090569 | 0.206135 | 1206.259 | 1198.622 | 1066.601 | 1184.382 | 1105.776 | 1023.515 | 1101.218 | 1034.299 |
| ENSMUSG00000028970 | 2.409277944 | 0.002602 | 0.015881 | 848.1038 | 141.5789 | 66.24853 | 184.6014 | 47.78045 | 93.85942 | 46.28588 | 45.67883 |
| ENSMUSG00000028973 | -0.484659327 | 5.83E-06 | 0.000151 | 973.8606 | 980.2542 | 996.2123 | 851.4094 | 1302.749 | 1140.839 | 1374.112 | 1499.788 |
| ENSMUSG00000028974 | -0.617008323 | 0.003647 | 0.020436 | 647.899 | 477.529 | 589.6119 | 551.2164 | 869.7993 | 560.9218 | 857.2531 | 1188.737 |
| ENSMUSG00000028975 | -0.371302032 | 0.003848 | 0.021239 | 740.456 | 794.2819 | 877.793 | 744.4441 | 1009.24 | 844.7348 | 1005.754 | 1224.628 |
| ENSMUSG00000028976 | -0.482626019 | 0.17005 | 0.316662 | 100.6054 | 103.1847 | 100.2009 | 106.1027 | 109.2125 | 74.86406 | 129.2147 | 259.9343 |
| ENSMUSG00000028977 | -0.318393426 | 0.021987 | 0.075411 | 404.4339 | 285.5575 | 349.461 | 309.6818 | 399.7956 | 388.8462 | 423.3229 | 472.0145 |
| ENSMUSG00000028978 | -0.542672999 | 0.019656 | 0.069615 | 254.5318 | 223.1668 | 390.8663 | 312.2697 | 403.6961 | 298.3389 | 479.2517 | 541.6204 |
| ENSMUSG00000028980 | -0.337933992 | 0.031684 | 0.098197 | 1829.007 | 1818.929 | 2152.249 | 1809.784 | 2468.007 | 1693.939 | 2658.545 | 2798.372 |
| ENSMUSG00000028982 | -0.545764576 | 0.02422 | 0.081141 | 377.2704 | 269.9598 | 381.7572 | 357.1261 | 391.9947 | 463.7102 | 778.1814 | 390.4452 |
| ENSMUSG00000028986 | 0.239099537 | 0.032647 | 0.100491 | 1770.656 | 1745.74 | 1523.716 | 1822.723 | 1467.543 | 1682.765 | 1426.184 | 1237.679 |
| ENSMUSG00000028988 | -0.19393864 | 0.002214 | 0.014142 | 1027.181 | 1088.238 | 1079.023 | 1059.302 | 1253.993 | 1262.633 | 1186.076 | 1162.635 |
| ENSMUSG00000028989 | 0.903944257 | 0.06919 | 0.17081 | 94.56911 | 140.3791 | 116.763 | 125.943 | 78.98402 | 120.6764 | 40.50014 | 15.22628 |
| ENSMUSG00000028990 | -0.231249473 | 0.008887 | 0.039124 | 636.8324 | 675.4995 | 741.1554 | 711.6644 | 856.1477 | 784.3966 | 754.0741 | 854.8466 |
| ENSMUSG00000028991 | -0.439131289 | 0.000955 | 0.007573 | 1192.174 | 1028.247 | 1142.787 | 1070.516 | 1529.95 | 1166.539 | 1538.041 | 1777.124 |
| ENSMUSG00000028992 | -0.334349954 | 0.050938 | 0.137866 | 252.5196 | 181.1731 | 212.8234 | 157.8601 | 245.728 | 232.4138 | 257.4652 | 278.4233 |
| ENSMUSG00000028995 | 0.349675759 | 0.031969 | 0.098898 | 1758.583 | 1420.589 | 1226.426 | 1400.901 | 1134.054 | 1455.938 | 956.5749 | 1010.372 |
| ENSMUSG00000028998 | -0.159449353 | 0.015935 | 0.060156 | 1269.641 | 1307.805 | 1422.687 | 1255.117 | 1481.194 | 1439.178 | 1487.898 | 1462.81 |
| ENSMUSG00000028999 | 0.076167147 | 0.441951 | 0.605227 | 722.347 | 794.2819 | 621.9081 | 681.4726 | 687.4535 | 641.3727 | 646.0737 | 694.9707 |
| ENSMUSG00000029001 | -0.188592929 | 0.299985 | 0.468126 | 483.9121 | 406.7395 | 508.4575 | 423.5481 | 490.481 | 363.1466 | 609.4308 | 614.489 |
| ENSMUSG00000029003 | -0.025584064 | 0.822797 | 0.89248 | 872.2491 | 937.0606 | 891.0427 | 984.2534 | 1118.453 | 912.8946 | 906.4318 | 811.343 |
| ENSMUSG00000029004 | 0.478917608 | 0.004329 | 0.02316 | 2835.061 | 2701.998 | 2112.5 | 2434.323 | 1726.922 | 2417.998 | 1621.934 | 1468.248 |
| ENSMUSG00000029005 | 0.095998019 | 0.742987 | 0.839394 | 79.47829 | 124.7814 | 95.23226 | 131.1188 | 149.192 | 67.04245 | 85.82174 | 100.0584 |
| ENSMUSG00000029007 | 0.014018291 | 0.860276 | 0.91625 | 1128.793 | 1193.822 | 1172.599 | 1061.027 | 1205.238 | 1051.449 | 1079.04 | 1173.511 |
| ENSMUSG00000029009 | 0.099581548 | 0.257556 | 0.421292 | 1130.805 | 1001.851 | 1069.914 | 1019.621 | 998.514 | 869.317 | 1045.289 | 1026.686 |
| ENSMUSG00000029012 | 0.078176804 | 0.389187 | 0.556608 | 798.8072 | 803.8804 | 726.2495 | 819.4923 | 744.985 | 823.5047 | 738.6455 | 674.3065 |
| ENSMUSG00000029014 | -0.041282827 | 0.656151 | 0.778423 | 2146.92 | 2276.061 | 1909.614 | 2035.791 | 2031.157 | 2441.462 | 2058.757 | 2077.299 |
| ENSMUSG00000029016 | -0.0757407 | 0.49691 | 0.653568 | 873.2552 | 679.099 | 781.7327 | 843.6458 | 859.0731 | 874.9039 | 884.2532 | 733.0364 |
| ENSMUSG00000029017 | -0.2277045 | 0.001424 | 0.010201 | 2049.333 | 1845.326 | 1952.675 | 1935.727 | 2240.806 | 2093.959 | 2362.508 | 2417.715 |
| ENSMUSG00000029020 | -0.468262054 | 0.005103 | 0.026094 | 1199.217 | 943.0597 | 1293.503 | 1096.394 | 1496.796 | 1182.182 | 1615.184 | 1978.328 |
| ENSMUSG00000029022 | -0.02746767 | 0.799765 | 0.877502 | 1198.211 | 1040.245 | 1004.493 | 1079.142 | 1250.093 | 1110.67 | 1096.397 | 946.2042 |
| ENSMUSG00000029026 | 0.778120318 | 0.07901 | 0.187542 | 45.27245 | 16.7975 | 24.8432 | 26.74133 | 14.62667 | 24.58223 | 15.42863 | 11.9635 |
| ENSMUSG00000029027 | -0.022708602 | 0.860693 | 0.916562 | 389.343 | 452.3327 | 471.1927 | 485.657 | 477.8045 | 530.7527 | 404.0372 | 416.5474 |
| ENSMUSG00000029028 | -0.316465757 | 0.00054 | 0.004867 | 1338.052 | 1304.206 | 1341.533 | 1327.578 | 1733.748 | 1393.365 | 1734.756 | 1751.022 |
| ENSMUSG00000029029 | -0.225708831 | 0.009112 | 0.039867 | 426.567 | 459.5316 | 494.3797 | 450.2895 | 557.7637 | 519.579 | 538.0734 | 526.3941 |
| ENSMUSG00000029030 | -0.190381071 | 0.040428 | 0.117586 | 2852.164 | 2662.404 | 2821.359 | 2957.073 | 3295.876 | 2723.041 | 3483.977 | 3383.496 |
| ENSMUSG00000029032 | -1.001127846 | 7.61E-07 | 2.98E-05 | 487.9364 | 460.7315 | 689.8128 | 512.3984 | 988.7629 | 754.2275 | 1227.54 | 1337.737 |
| ENSMUSG00000029033 | -0.166988728 | 0.264467 | 0.428723 | 405.4399 | 398.3407 | 428.9592 | 376.9665 | 517.7841 | 344.1512 | 428.1444 | 516.6058 |
| ENSMUSG00000029034 | -0.189983026 | 0.019355 | 0.068879 | 1077.484 | 979.0544 | 982.9626 | 960.0999 | 1191.586 | 1040.275 | 1119.54 | 1209.401 |
| ENSMUSG00000029036 | -0.563404388 | 3.57E-06 | 0.000104 | 1773.674 | 1760.138 | 1985.8 | 1758.889 | 2848.3 | 2128.598 | 2693.26 | 3086.584 |
| ENSMUSG00000029038 | -0.35549056 | 0.001079 | 0.008281 | 1457.773 | 1482.979 | 1632.198 | 1422.466 | 2017.505 | 1569.911 | 1979.686 | 2103.401 |
| ENSMUSG00000029047 | -0.479457932 | 0.002778 | 0.016707 | 294.7739 | 254.3622 | 288.1811 | 251.0234 | 391.0196 | 286.0478 | 384.7514 | 455.7007 |
| ENSMUSG00000029048 | -0.256040692 | 0.000886 | 0.007095 | 2350.143 | 2422.44 | 2589.489 | 2534.388 | 3087.202 | 2695.106 | 2845.617 | 3193.167 |
| ENSMUSG00000029049 | -0.068481289 | 0.656449 | 0.778543 | 132.7992 | 143.9786 | 166.4494 | 131.9814 | 148.2169 | 139.6718 | 170.6792 | 144.6496 |
| ENSMUSG00000029050 | 0.162909488 | 0.149343 | 0.289343 | 3129.835 | 3229.92 | 3796.041 | 3574.712 | 2863.902 | 3669.457 | 2862.01 | 2873.416 |
| ENSMUSG00000029053 | -1.123616355 | 4.65E-06 | 0.000126 | 218.3138 | 196.7707 | 313.8524 | 269.1385 | 545.0872 | 329.6254 | 593.0378 | 710.197 |
| ENSMUSG00000029055 | -0.085056097 | 0.660522 | 0.781448 | 117.7084 | 172.7743 | 197.0894 | 137.1571 | 177.4703 | 166.4887 | 149.4648 | 169.6642 |
| ENSMUSG00000029056 | -0.14864148 | 0.080095 | 0.189063 | 1175.071 | 1060.642 | 1103.866 | 1224.925 | 1297.873 | 1137.487 | 1334.576 | 1290.971 |
| ENSMUSG00000029059 | -0.386656736 | 0.001883 | 0.012567 | 384.3128 | 328.7511 | 319.6492 | 320.0333 | 469.0285 | 381.0246 | 424.2872 | 492.6788 |

| | | | | | | | | | | | |
|---|---|---|---|---|---|---|---|---|---|---|---|
| ENSMUSG00000029060 | -0.05470352 | 0.576152 | 0.718319 | 979.8969 | 955.0579 | 1019.399 | 1098.12 | 940.9824 | 989.9934 | 1119.54 | 1161.547 |
| ENSMUSG00000029061 | -0.255103462 | 0.054795 | 0.145465 | 634.8203 | 853.0731 | 978.822 | 793.6136 | 973.1611 | 1008.989 | 954.6463 | 955.9926 |
| ENSMUSG00000029062 | 0.12087385 | 0.132064 | 0.265176 | 3031.242 | 3207.123 | 3078.072 | 3230.525 | 2877.553 | 3261.615 | 2724.117 | 2676.562 |
| ENSMUSG00000029063 | -0.3449776 | 0.002262 | 0.014356 | 2064.424 | 1889.719 | 2053.704 | 2031.478 | 2570.393 | 2052.616 | 2673.974 | 2914.744 |
| ENSMUSG00000029064 | -0.07903457 | 0.130273 | 0.262661 | 10430.77 | 10951.97 | 11552.92 | 10907.01 | 11519.96 | 11372.63 | 11345.83 | 12077.7 |
| ENSMUSG00000029066 | -0.431999975 | 0.00056 | 0.004992 | 1023.157 | 958.6574 | 1264.519 | 1085.18 | 1540.676 | 1219.055 | 1470.541 | 1617.248 |
| ENSMUSG00000029068 | 0.297278203 | 0.002556 | 0.015675 | 6971.957 | 6493.434 | 5960.712 | 6819.901 | 5323.133 | 6180.196 | 5069.268 | 4786.488 |
| ENSMUSG00000029070 | 0.248732057 | 0.017238 | 0.063542 | 1872.267 | 2282.061 | 2095.938 | 2456.752 | 1894.641 | 1914.062 | 1873.614 | 1645.525 |
| ENSMUSG00000029071 | -0.391651499 | 0.002737 | 0.016518 | 1258.574 | 1076.24 | 1204.067 | 1105.021 | 1581.631 | 1198.942 | 1518.755 | 1793.438 |
| ENSMUSG00000029072 | -0.748961001 | 0.069261 | 0.170916 | 12.07265 | 8.398751 | 15.73403 | 12.07673 | 21.45245 | 14.52586 | 23.14294 | 22.83941 |
| ENSMUSG00000029073 | -0.309025736 | 0.119824 | 0.248762 | 258.556 | 200.3702 | 279.0719 | 230.3205 | 274.9814 | 214.5358 | 357.7513 | 353.4671 |
| ENSMUSG00000029074 | -0.507069446 | 0.012692 | 0.051056 | 195.1745 | 134.38 | 172.2462 | 185.4641 | 212.5743 | 185.4841 | 285.4296 | 294.7372 |
| ENSMUSG00000029075 | 0.149944558 | 0.552856 | 0.700117 | 78.47224 | 61.1909 | 81.15445 | 53.48266 | 55.58134 | 80.45093 | 60.75022 | 51.11678 |
| ENSMUSG00000029076 | -0.20452513 | 0.008025 | 0.036321 | 3763.649 | 3976.209 | 4300.358 | 3836.949 | 4744.892 | 4127.58 | 4605.445 | 4818.029 |
| ENSMUSG00000029082 | -2.109104783 | 6.70E-07 | 2.67E-05 | 38.23007 | 26.39607 | 71.21717 | 30.19182 | 186.2463 | 73.74669 | 232.3937 | 227.3065 |
| ENSMUSG00000029084 | 0.184088841 | 0.120187 | 0.249148 | 630.7961 | 484.7279 | 640.9545 | 608.1496 | 487.5557 | 558.687 | 492.7518 | 547.0583 |
| ENSMUSG00000029086 | -0.164533616 | 0.180848 | 0.331124 | 2548.336 | 2372.047 | 3123.618 | 2427.423 | 2878.529 | 2489.509 | 3210.119 | 3160.54 |
| ENSMUSG00000029088 | 0.199934645 | 0.529662 | 0.681737 | 69.41775 | 27.5959 | 42.23344 | 48.30692 | 42.9049 | 41.34284 | 41.46443 | 38.06569 |
| ENSMUSG00000029089 | 0.17690856 | 0.066312 | 0.166014 | 569.4268 | 550.7181 | 546.5504 | 640.0666 | 531.4357 | 537.4569 | 505.2875 | 467.6642 |
| ENSMUSG00000029090 | -0.375290316 | 0.000825 | 0.006724 | 3184.162 | 3069.143 | 3338.926 | 3146.851 | 3766.855 | 3476.151 | 4487.802 | 4794.102 |
| ENSMUSG00000029093 | 0.551632965 | 0.000295 | 0.003061 | 1015.109 | 1556.169 | 1194.13 | 1213.711 | 864.9237 | 989.9934 | 811.9315 | 728.686 |
| ENSMUSG00000029094 | 0.408686739 | 0.012535 | 0.0506 | 1223.362 | 1245.415 | 1058.32 | 1241.315 | 760.5868 | 1205.647 | 739.6098 | 887.4743 |
| ENSMUSG00000029095 | 0.097194983 | 0.577655 | 0.719477 | 191.1503 | 238.7645 | 183.0116 | 182.0136 | 182.3458 | 231.2964 | 173.572 | 154.4379 |
| ENSMUSG00000029096 | 0.166442637 | 0.395607 | 0.562932 | 312.8829 | 579.5138 | 554.0033 | 505.4974 | 419.2979 | 401.1373 | 418.5015 | 500.2919 |
| ENSMUSG00000029097 | 0.044044556 | 0.791733 | 0.8726 | 273.6468 | 231.5656 | 206.1986 | 173.3873 | 215.4996 | 206.7142 | 232.3937 | 201.2044 |
| ENSMUSG00000029098 | -0.181870981 | 0.149565 | 0.289495 | 722.347 | 647.9036 | 637.6421 | 610.7374 | 761.5619 | 588.8561 | 803.2529 | 814.6057 |
| ENSMUSG00000029101 | 0.093933142 | 0.226155 | 0.385973 | 1435.64 | 1534.572 | 1516.263 | 1626.908 | 1418.787 | 1535.272 | 1465.72 | 1309.46 |
| ENSMUSG00000029102 | -0.88646197 | 0.006848 | 0.032457 | 25.15136 | 37.19447 | 22.35888 | 25.8787 | 48.75557 | 32.40385 | 54.96448 | 66.34306 |
| ENSMUSG00000029103 | -0.631170026 | 0.00026 | 0.002785 | 2488.978 | 2525.624 | 3216.366 | 2744.868 | 4194.929 | 3176.695 | 3964.193 | 5666.35 |
| ENSMUSG00000029104 | -0.058347514 | 0.504023 | 0.65977 | 1135.835 | 1048.644 | 1021.055 | 988.5665 | 1074.573 | 1217.938 | 1034.682 | 1039.737 |
| ENSMUSG00000029106 | -0.005905431 | 0.925068 | 0.95551 | 5135.907 | 5376.4 | 5659.281 | 5114.495 | 5282.178 | 5075.113 | 5334.448 | 5682.663 |
| ENSMUSG00000029108 | 0.922198837 | 9.67E-05 | 0.001303 | 929.5942 | 1029.447 | 1200.755 | 1167.992 | 449.5263 | 895.0167 | 405.9657 | 535.0948 |
| ENSMUSG00000029110 | -0.284767295 | 6.78E-05 | 0.001007 | 3138.89 | 3321.106 | 3456.517 | 3259.854 | 4266.112 | 3622.527 | 4068.336 | 4093.693 |
| ENSMUSG00000029111 | -0.010129765 | 0.885724 | 0.931739 | 992.9756 | 967.0562 | 889.3865 | 925.595 | 988.7629 | 914.012 | 939.2176 | 954.905 |
| ENSMUSG00000029119 | -0.255336935 | 0.232985 | 0.39359 | 555.342 | 500.3256 | 690.6409 | 458.0531 | 747.9104 | 432.4238 | 624.8594 | 826.5692 |
| ENSMUSG00000029120 | 0.766619258 | 0.090473 | 0.205991 | 63.38142 | 27.5959 | 22.35888 | 67.28463 | 25.35289 | 34.6386 | 18.32149 | 28.27737 |
| ENSMUSG00000029121 | 0.861233214 | 0.028339 | 0.090843 | 276.6649 | 110.3836 | 115.1068 | 362.3019 | 114.088 | 88.27255 | 147.5362 | 126.1606 |
| ENSMUSG00000029122 | 0.245523956 | 0.000444 | 0.004187 | 1534.233 | 1327.003 | 1473.202 | 1435.405 | 1191.586 | 1230.229 | 1188.004 | 1261.606 |
| ENSMUSG00000029123 | -0.96496679 | 0.003932 | 0.02157 | 143.8658 | 82.78769 | 171.4181 | 155.2722 | 210.624 | 145.2586 | 349.0727 | 377.3941 |
| ENSMUSG00000029125 | -0.412995659 | 0.00257 | 0.015735 | 703.232 | 579.5138 | 689.8128 | 559.8426 | 880.5255 | 673.7766 | 882.3246 | 935.3283 |
| ENSMUSG00000029126 | 0.460618629 | 3.85E-06 | 0.00011 | 1043.278 | 1058.243 | 952.3226 | 1138.663 | 864.9237 | 714.002 | 752.1455 | 713.4598 |
| ENSMUSG00000029127 | 0.107053674 | 0.58227 | 0.722932 | 321.9374 | 263.9607 | 252.5725 | 254.4739 | 314.961 | 274.874 | 248.7866 | 175.1022 |
| ENSMUSG00000029128 | -0.04899538 | 0.49654 | 0.653183 | 1091.569 | 1083.439 | 1127.053 | 1177.481 | 1129.179 | 1260.398 | 1128.218 | 1121.306 |
| ENSMUSG00000029130 | 0.33804222 | 0.329034 | 0.496849 | 50.30272 | 43.19358 | 42.23344 | 62.10889 | 24.37778 | 58.10345 | 46.28588 | 28.27737 |
| ENSMUSG00000029131 | -0.11209304 | 0.131514 | 0.264403 | 2533.245 | 2678.002 | 2667.331 | 2948.447 | 3095.003 | 2743.153 | 3014.368 | 2849.489 |
| ENSMUSG00000029135 | 0.151228641 | 0.428024 | 0.592975 | 2006.072 | 1177.025 | 1816.038 | 1615.694 | 1193.536 | 1472.699 | 1899.65 | 1392.117 |
| ENSMUSG00000029136 | -0.472549523 | 0.066865 | 0.166901 | 55.33299 | 53.99197 | 81.15445 | 66.42201 | 89.71024 | 61.45558 | 112.8218 | 93.53283 |
| ENSMUSG00000029141 | -0.041997982 | 0.675475 | 0.792434 | 891.3642 | 935.8608 | 779.2483 | 742.7188 | 842.4962 | 815.6831 | 879.4317 | 904.8758 |
| ENSMUSG00000029145 | -0.279525854 | 0.019403 | 0.069024 | 1076.478 | 1103.836 | 1101.382 | 1106.746 | 1358.33 | 1050.332 | 1376.041 | 1541.117 |
| ENSMUSG00000029146 | -0.182167914 | 0.010904 | 0.04552 | 3230.441 | 3180.727 | 3237.897 | 3216.723 | 3775.631 | 3238.15 | 3692.263 | 3890.313 |
| ENSMUSG00000029147 | -0.333457169 | 0.001669 | 0.011462 | 2725.401 | 2947.962 | 3347.207 | 2813.015 | 3698.597 | 3229.211 | 3773.264 | 4211.153 |
| ENSMUSG00000029148 | -0.054406566 | 0.322715 | 0.490873 | 2760.613 | 2859.175 | 2864.421 | 2826.817 | 3111.58 | 2889.529 | 2792.581 | 2951.722 |
| ENSMUSG00000029149 | -0.234234827 | 0.051319 | 0.138554 | 223.3441 | 226.7663 | 248.432 | 248.4356 | 301.3094 | 294.9868 | 243.9652 | 276.2481 |
| ENSMUSG00000029151 | 2.927377171 | 4.72E-29 | 1.74E-25 | 286.7255 | 202.7698 | 213.6515 | 248.4356 | 27.30312 | 45.81234 | 33.75012 | 18.48905 |
| ENSMUSG00000029152 | -0.056878261 | 0.359775 | 0.528285 | 3968.884 | 4415.343 | 4302.842 | 4402.83 | 4319.743 | 4262.782 | 4526.373 | 4667.941 |
| ENSMUSG00000029153 | 0.274126476 | 0.020468 | 0.071685 | 503.0272 | 412.7386 | 538.2693 | 558.98 | 418.3228 | 395.5504 | 419.4658 | 435.0364 |
| ENSMUSG00000029154 | -1.985956642 | 6.28E-05 | 0.000949 | 35.2119 | 9.598572 | 47.20208 | 25.8787 | 76.05868 | 60.3382 | 152.3577 | 181.6277 |
| ENSMUSG00000029156 | 0.241041971 | 0.01431 | 0.055672 | 858.1644 | 839.8751 | 744.4679 | 934.2212 | 699.1548 | 774.3402 | 711.6454 | 672.1313 |
| ENSMUSG00000029161 | -2.198293983 | 8.78E-08 | 5.11E-06 | 122.7386 | 70.78947 | 181.3554 | 132.844 | 659.1752 | 220.1227 | 465.7517 | 989.7079 |
| ENSMUSG00000029162 | -0.560064904 | 0.00037 | 0.003629 | 668.0201 | 873.4701 | 752.7489 | 652.1434 | 1218.889 | 836.9132 | 1032.754 | 1251.817 |
| ENSMUSG00000029163 | 0.176102107 | 0.133595 | 0.267103 | 5193.253 | 7095.745 | 7111.78 | 5740.759 | 5672.222 | 5233.78 | 5454.02 | 5891.481 |
| ENSMUSG00000029165 | 0.090286325 | 0.251644 | 0.415267 | 944.685 | 1006.65 | 997.8685 | 1031.698 | 997.5389 | 943.0637 | 952.7177 | 845.0583 |
| ENSMUSG00000029166 | 0.323644014 | 0.007624 | 0.034974 | 707.2562 | 532.7208 | 543.238 | 646.9676 | 498.2819 | 451.4191 | 508.1804 | 483.978 |
| ENSMUSG00000029167 | -0.814914747 | 0.000323 | 0.003286 | 599.6084 | 419.9375 | 631.0173 | 581.4082 | 749.8606 | 703.9457 | 1313.362 | 1161.547 |

| | | | | | | | | | | | |
|---|---|---|---|---|---|---|---|---|---|---|---|
| ENSMUSG00000029168 | 1.172004869 | 3.62E-09 | 3.71E-07 | 379.2825 | 321.5522 | 260.8536 | 467.5419 | 165.7689 | 139.6718 | 168.7506 | 159.8759 |
| ENSMUSG00000029169 | -0.113909389 | 0.015968 | 0.06023 | 5850.206 | 5773.541 | 5501.112 | 5952.965 | 6425.984 | 6105.332 | 6225.451 | 6213.408 |
| ENSMUSG00000029171 | -0.274167281 | 0.005126 | 0.026178 | 879.2915 | 713.8938 | 847.9812 | 803.1025 | 987.7878 | 886.0777 | 1058.79 | 992.9707 |
| ENSMUSG00000029173 | -0.082714092 | 0.302041 | 0.470419 | 638.8445 | 623.9072 | 616.1113 | 558.98 | 664.0508 | 619.0252 | 646.0737 | 649.2919 |
| ENSMUSG00000029174 | -0.397679277 | 0.009506 | 0.04105 | 1187.144 | 1075.04 | 1498.873 | 1184.382 | 1502.647 | 1300.623 | 1842.757 | 1871.744 |
| ENSMUSG00000029175 | -0.240042333 | 0.014047 | 0.054885 | 503.0272 | 473.9295 | 566.4249 | 522.7498 | 618.2206 | 555.3349 | 610.395 | 660.1678 |
| ENSMUSG00000029176 | 0.081470299 | 0.219017 | 0.377809 | 2424.591 | 2679.201 | 2337.745 | 2605.123 | 2446.554 | 2365.481 | 2297.901 | 2380.737 |
| ENSMUSG00000029177 | 0.083165682 | 0.40373 | 0.570586 | 3127.823 | 2899.969 | 2654.082 | 3320.238 | 3005.293 | 2928.638 | 2913.118 | 2481.883 |
| ENSMUSG00000029178 | 0.114836323 | 0.311945 | 0.480371 | 1812.91 | 1811.731 | 1753.102 | 1748.538 | 1596.257 | 2023.564 | 1490.791 | 1471.511 |
| ENSMUSG00000029179 | 0.265944203 | 0.050561 | 0.137126 | 791.7648 | 686.2979 | 678.2193 | 754.7956 | 556.7886 | 753.1101 | 597.8593 | 514.4306 |
| ENSMUSG00000029185 | 0.268891629 | 0.013067 | 0.052236 | 1066.418 | 1392.993 | 1208.208 | 1164.542 | 953.6589 | 1115.139 | 932.4676 | 1008.197 |
| ENSMUSG00000029186 | 0.457277119 | 0.000935 | 0.007436 | 1300.828 | 1463.782 | 1157.693 | 1406.076 | 952.6837 | 1205.647 | 896.7889 | 825.4816 |
| ENSMUSG00000029188 | -2.867176583 | 3.06E-07 | 1.39E-05 | 2.012109 | 8.398751 | 24.01509 | 10.35148 | 69.2329 | 51.39921 | 91.60747 | 119.635 |
| ENSMUSG00000029189 | 0.951558197 | 1.75E-15 | 8.96E-13 | 3313.943 | 3509.478 | 2886.78 | 3286.596 | 1679.142 | 1991.161 | 1652.792 | 1396.467 |
| ENSMUSG00000029190 | -0.121618053 | 0.056561 | 0.148469 | 1538.257 | 1440.986 | 1546.075 | 1519.08 | 1595.282 | 1689.47 | 1564.077 | 1733.62 |
| ENSMUSG00000029191 | -0.00073196 | 0.991879 | 0.995385 | 2289.78 | 2580.816 | 2452.852 | 2277.326 | 2422.176 | 2553.2 | 2257.401 | 2370.949 |
| ENSMUSG00000029192 | -0.28491406 | 0.003327 | 0.019111 | 2118.75 | 2232.868 | 2401.509 | 1940.04 | 2536.265 | 2401.237 | 2739.546 | 2913.657 |
| ENSMUSG00000029193 | -2.869818902 | 0.000204 | 0.002312 | 5.030272 | 1.199822 | 38.92101 | 12.07673 | 72.15824 | 48.04709 | 91.60747 | 212.0803 |
| ENSMUSG00000029196 | -0.530910358 | 0.002418 | 0.015044 | 368.2159 | 346.7484 | 471.1927 | 378.6917 | 565.5646 | 416.7805 | 590.145 | 690.6203 |
| ENSMUSG00000029198 | -0.685821403 | 9.80E-07 | 3.67E-05 | 1334.028 | 1267.012 | 1429.312 | 1225.788 | 2193.025 | 1582.202 | 2192.794 | 2487.321 |
| ENSMUSG00000029199 | -0.092622194 | 0.181535 | 0.332054 | 1198.211 | 1171.026 | 1244.644 | 1094.669 | 1242.292 | 1308.445 | 1210.183 | 1261.606 |
| ENSMUSG00000029201 | 0.077998111 | 0.162197 | 0.306335 | 2367.246 | 2433.238 | 2548.084 | 2626.689 | 2347.093 | 2383.359 | 2386.616 | 2338.321 |
| ENSMUSG00000029202 | 0.091586599 | 0.388113 | 0.555703 | 3487.99 | 3654.656 | 3164.195 | 3302.123 | 2994.567 | 3825.889 | 2946.868 | 3003.927 |
| ENSMUSG00000029203 | -0.079738663 | 0.130993 | 0.263557 | 3011.121 | 3250.317 | 2982.84 | 3111.483 | 3237.37 | 3375.587 | 3188.904 | 3251.897 |
| ENSMUSG00000029204 | 0.806301777 | 0.046326 | 0.129173 | 19.11503 | 26.39607 | 19.87456 | 27.60395 | 8.776002 | 17.87799 | 11.57147 | 15.22628 |
| ENSMUSG00000029206 | 1.236128888 | 3.42E-05 | 0.000593 | 113.6841 | 99.58519 | 96.88848 | 95.75121 | 23.40267 | 65.92507 | 43.39301 | 40.24087 |
| ENSMUSG00000029207 | 0.027830821 | 0.780394 | 0.864523 | 1424.573 | 1186.623 | 1440.077 | 1400.038 | 1182.81 | 1435.826 | 1409.791 | 1323.598 |
| ENSMUSG00000029208 | 0.300379589 | 0.037793 | 0.11212 | 2585.56 | 2600.013 | 2200.279 | 2692.248 | 1940.472 | 2638.12 | 1912.185 | 1693.379 |
| ENSMUSG00000029209 | 0.030193706 | 0.76176 | 0.851959 | 579.4873 | 560.3167 | 611.9708 | 595.2102 | 552.8881 | 614.5558 | 628.7165 | 503.5547 |
| ENSMUSG00000029211 | -0.372009395 | 0.433741 | 0.597888 | 49.29666 | 25.19625 | 28.98373 | 40.5433 | 31.20356 | 92.74205 | 24.10723 | 39.15328 |
| ENSMUSG00000029212 | -0.372031052 | 0.571589 | 0.714525 | 12.07265 | 13.19804 | 17.39024 | 26.74133 | 6.825779 | 49.16446 | 9.642892 | 25.0146 |
| ENSMUSG00000029213 | 0.012713702 | 0.875501 | 0.925381 | 1183.12 | 1398.992 | 1382.938 | 1406.076 | 1287.147 | 1398.952 | 1327.826 | 1311.635 |
| ENSMUSG00000029217 | -0.018317138 | 0.88961 | 0.933475 | 288.7376 | 243.5638 | 275.7595 | 286.391 | 275.9565 | 264.8177 | 243.9652 | 326.2773 |
| ENSMUSG00000029219 | 1.270118792 | 0.059255 | 0.153404 | 49.29666 | 8.398751 | 18.21835 | 92.30071 | 20.47734 | 21.23011 | 17.3572 | 10.87591 |
| ENSMUSG00000029221 | -0.153840595 | 0.037324 | 0.11116 | 2231.429 | 2016.9 | 1972.55 | 2149.658 | 2172.548 | 2267.152 | 2399.151 | 2474.27 |
| ENSMUSG00000029223 | 0.687179152 | 1.27E-05 | 0.000277 | 1969.854 | 1451.784 | 1540.278 | 2132.405 | 1227.665 | 1238.05 | 958.5034 | 983.1823 |
| ENSMUSG00000029227 | 0.186395262 | 0.042571 | 0.121868 | 2848.14 | 3160.33 | 2877.671 | 3000.205 | 2368.545 | 2986.741 | 2660.474 | 2429.678 |
| ENSMUSG00000029228 | -0.400392244 | 0.141334 | 0.278117 | 65.39353 | 81.58786 | 90.26362 | 93.16334 | 119.9387 | 61.45558 | 125.3576 | 129.4233 |
| ENSMUSG00000029229 | 0.262621883 | 0.015693 | 0.059655 | 1761.601 | 1833.327 | 1553.528 | 1839.113 | 1353.454 | 1658.183 | 1536.113 | 1275.744 |
| ENSMUSG00000029231 | 0.897516923 | 6.84E-15 | 2.87E-12 | 7911.611 | 8686.708 | 7814.842 | 8450.26 | 4113.995 | 5183.498 | 4698.017 | 3645.605 |
| ENSMUSG00000029233 | -0.029859846 | 0.734358 | 0.833601 | 680.0927 | 788.2827 | 702.2344 | 797.0641 | 798.6162 | 724.0584 | 762.7527 | 741.7371 |
| ENSMUSG00000029234 | 0.077789556 | 0.267457 | 0.431699 | 2852.164 | 3121.936 | 2986.981 | 3132.186 | 2620.124 | 3038.14 | 2860.082 | 2941.934 |
| ENSMUSG00000029238 | 0.169453055 | 0.28786 | 0.454754 | 956.7577 | 859.0722 | 920.0265 | 910.9304 | 785.9397 | 1089.44 | 708.7525 | 660.1678 |
| ENSMUSG00000029245 | 0.990279638 | 0.097147 | 0.216513 | 48.29061 | 19.19714 | 12.4216 | 51.75741 | 11.70134 | 31.28647 | 10.60718 | 13.05109 |
| ENSMUSG00000029246 | -0.536246453 | 1.17E-05 | 0.000259 | 961.788 | 953.8581 | 929.1356 | 911.793 | 1493.871 | 1055.919 | 1400.148 | 1495.438 |
| ENSMUSG00000029247 | -0.091508827 | 0.219924 | 0.378736 | 7636.959 | 8207.979 | 7686.486 | 8219.939 | 9328.89 | 7869.666 | 8638.102 | 7989.444 |
| ENSMUSG00000029249 | 0.261169529 | 0.14971 | 0.289628 | 2369.258 | 2440.437 | 1955.988 | 2064.258 | 1770.802 | 2572.195 | 1591.077 | 1433.445 |
| ENSMUSG00000029250 | -0.288794234 | 0.000511 | 0.004657 | 2287.768 | 2120.085 | 2226.779 | 2027.165 | 2693.257 | 2331.96 | 2808.01 | 2746.167 |
| ENSMUSG00000029253 | 0.326462677 | 0.009221 | 0.040161 | 1354.149 | 1315.004 | 979.6501 | 1225.788 | 1048.245 | 1040.275 | 894.8603 | 901.613 |
| ENSMUSG00000029260 | -0.561087438 | 0.248815 | 0.41222 | 102.6175 | 27.5959 | 199.5737 | 119.9047 | 126.7645 | 118.4417 | 158.1434 | 262.1095 |
| ENSMUSG00000029263 | -0.212019128 | 0.091051 | 0.206817 | 376.2643 | 361.1463 | 365.195 | 329.5222 | 441.7254 | 334.0949 | 432.9658 | 448.0875 |
| ENSMUSG00000029265 | -0.084017677 | 0.156247 | 0.298334 | 1266.622 | 1389.393 | 1288.534 | 1315.501 | 1358.33 | 1392.248 | 1404.969 | 1416.044 |
| ENSMUSG00000029267 | 0.333829712 | 0.037858 | 0.112206 | 2077.502 | 2078.091 | 1520.404 | 1883.107 | 1451.941 | 1937.527 | 1331.683 | 1275.744 |
| ENSMUSG00000029269 | -1.669342357 | 7.65E-07 | 2.98E-05 | 18.10898 | 14.39786 | 26.49941 | 22.42821 | 57.53157 | 45.81234 | 68.46453 | 90.27006 |
| ENSMUSG00000029270 | 0.009944276 | 0.931552 | 0.95963 | 315.9011 | 326.3515 | 339.5237 | 356.2635 | 310.0854 | 297.2215 | 376.0728 | 345.854 |
| ENSMUSG00000029272 | 2.505406315 | 4.24E-07 | 1.84E-05 | 66.39959 | 140.3791 | 60.45178 | 123.3552 | 16.57689 | 32.40385 | 14.46434 | 5.437955 |
| ENSMUSG00000029273 | -2.864320882 | 3.21E-06 | 9.56E-05 | 438.6397 | 81.58786 | 647.5794 | 523.6125 | 3008.218 | 1648.127 | 3447.334 | 4218.766 |
| ENSMUSG00000029276 | 0.064589987 | 0.528169 | 0.680671 | 468.8213 | 541.1195 | 520.051 | 541.7276 | 527.5352 | 536.3396 | 463.8231 | 453.5255 |
| ENSMUSG00000029279 | 0.638378185 | 0.007903 | 0.035924 | 297.7921 | 337.1499 | 236.8385 | 287.2536 | 168.6943 | 279.3435 | 164.8934 | 131.5985 |
| ENSMUSG00000029283 | 0.234692096 | 0.103729 | 0.226532 | 763.5953 | 974.2551 | 639.2983 | 686.6483 | 671.8517 | 680.4808 | 593.0378 | 654.7298 |
| ENSMUSG00000029287 | 0.419165433 | 0.019407 | 0.069024 | 1231.411 | 928.6619 | 970.541 | 1146.427 | 580.1912 | 1020.163 | 851.4673 | 748.2627 |
| ENSMUSG00000029290 | 0.082237752 | 0.538772 | 0.688661 | 2144.908 | 2547.221 | 1936.941 | 2130.68 | 1879.039 | 2562.139 | 1930.507 | 1901.109 |
| ENSMUSG00000029291 | -0.287033899 | 0.003775 | 0.020951 | 1000.018 | 853.0731 | 864.5433 | 949.7485 | 1135.03 | 1067.092 | 1018.289 | 1256.168 |
| ENSMUSG00000029298 | 0.441479205 | 0.108137 | 0.232788 | 188.1322 | 433.1356 | 441.3808 | 246.7103 | 231.1014 | 274.874 | 237.2151 | 220.781 |

| | | | | | | | | | | | |
|---|---|---|---|---|---|---|---|---|---|---|---|
| ENSMUSG00000029299 | 0.234103743 | 0.485163 | 0.64349 | 24.1453 | 25.19625 | 25.67131 | 19.84034 | 19.50223 | 23.46486 | 21.21436 | 16.31387 |
| ENSMUSG00000029304 | -2.506112048 | 0.001029 | 0.007993 | 799.8132 | 382.7431 | 3035.011 | 802.2398 | 8444.464 | 2699.576 | 7180.097 | 10196.17 |
| ENSMUSG00000029309 | 0.07321417 | 0.578239 | 0.719913 | 3558.414 | 4206.574 | 4164.548 | 4754.781 | 4042.811 | 4786.831 | 3573.656 | 3457.452 |
| ENSMUSG00000029310 | -0.162039952 | 0.060924 | 0.156655 | 1284.731 | 1268.211 | 1334.908 | 1252.529 | 1514.348 | 1250.342 | 1552.506 | 1432.357 |
| ENSMUSG00000029311 | 0.041331318 | 0.685223 | 0.799751 | 2936.673 | 2711.597 | 3105.4 | 3045.061 | 3025.77 | 2373.303 | 2985.439 | 3082.233 |
| ENSMUSG00000029312 | 0.107306035 | 0.384292 | 0.552246 | 398.3975 | 475.1293 | 379.2728 | 376.9665 | 384.1939 | 408.9589 | 387.6442 | 328.4525 |
| ENSMUSG00000029313 | -0.058534616 | 0.612775 | 0.746981 | 1465.821 | 1222.618 | 1418.547 | 1412.977 | 1198.412 | 1396.718 | 1527.434 | 1629.211 |
| ENSMUSG00000029314 | 0.399747768 | 0.061073 | 0.156833 | 68.4117 | 63.59054 | 72.87338 | 81.08661 | 48.75557 | 58.10345 | 48.21446 | 63.08028 |
| ENSMUSG00000029319 | -0.16649012 | 0.003902 | 0.021455 | 1435.64 | 1539.371 | 1493.076 | 1519.943 | 1671.341 | 1760.982 | 1645.077 | 1643.35 |
| ENSMUSG00000029321 | 1.234948999 | 0.021852 | 0.075006 | 77.46619 | 62.39072 | 36.43669 | 85.39973 | 12.67645 | 62.57295 | 22.17865 | 14.13868 |
| ENSMUSG00000029322 | 0.008284051 | 0.969965 | 0.983284 | 1269.641 | 1252.614 | 2313.73 | 1323.264 | 1259.844 | 1367.666 | 1847.578 | 1650.963 |
| ENSMUSG00000029326 | -0.076108934 | 0.32561 | 0.493509 | 606.6508 | 653.9027 | 631.0173 | 637.4788 | 695.2544 | 639.138 | 633.538 | 697.1459 |
| ENSMUSG00000029328 | 0.188003451 | 0.062325 | 0.159186 | 10984.1 | 11772.65 | 11460.17 | 12271.68 | 10940.75 | 11660.92 | 9120.247 | 9087.911 |
| ENSMUSG00000029330 | -0.690171663 | 0.011222 | 0.046557 | 742.4681 | 569.9152 | 1291.018 | 719.428 | 1347.604 | 844.7348 | 1545.756 | 1625.949 |
| ENSMUSG00000029334 | -0.190759577 | 0.621056 | 0.75282 | 140.8476 | 127.1811 | 88.60741 | 81.94923 | 72.15824 | 231.2964 | 82.92887 | 114.1971 |
| ENSMUSG00000029335 | -0.356176084 | 0.015301 | 0.058517 | 587.5357 | 589.1124 | 864.5433 | 602.9738 | 834.6953 | 786.6314 | 906.4318 | 860.2845 |
| ENSMUSG00000029338 | 0.132821176 | 0.069969 | 0.172293 | 969.8364 | 938.2604 | 928.3075 | 959.2373 | 842.4962 | 946.4159 | 823.5029 | 851.5838 |
| ENSMUSG00000029343 | 0.027343468 | 0.912383 | 0.948085 | 147.89 | 106.7841 | 98.54469 | 117.3168 | 89.71024 | 151.9629 | 133.0719 | 87.00729 |
| ENSMUSG00000029344 | 0.094165366 | 0.311294 | 0.47993 | 1498.015 | 1309.005 | 1194.958 | 1363.808 | 1174.034 | 1217.938 | 1324.933 | 1308.372 |
| ENSMUSG00000029345 | -0.173065312 | 0.027472 | 0.088826 | 1084.527 | 1045.045 | 1112.147 | 1029.972 | 1202.312 | 1082.735 | 1248.754 | 1282.27 |
| ENSMUSG00000029346 | -0.051442524 | 0.653296 | 0.776084 | 310.8708 | 346.7484 | 394.1788 | 389.0432 | 389.0694 | 374.3203 | 349.0727 | 383.9197 |
| ENSMUSG00000029348 | 0.307815315 | 0.091486 | 0.20748 | 140.8476 | 134.38 | 159.8246 | 156.9975 | 139.4409 | 106.1505 | 137.8934 | 94.62042 |
| ENSMUSG00000029352 | 1.196450267 | 0.001056 | 0.008143 | 42.25428 | 55.19179 | 52.17072 | 66.42201 | 16.57689 | 39.10809 | 15.42863 | 23.927 |
| ENSMUSG00000029359 | 0.970991287 | 2.01E-09 | 2.33E-07 | 646.893 | 479.9286 | 597.893 | 502.9095 | 256.4543 | 239.1181 | 290.251 | 351.2919 |
| ENSMUSG00000029360 | -0.079559673 | 0.84082 | 0.904006 | 19.11503 | 17.99732 | 13.24971 | 25.01608 | 24.37778 | 21.23011 | 13.50005 | 20.66423 |
| ENSMUSG00000029361 | 0.273941807 | 0.399491 | 0.566937 | 137.8294 | 172.7743 | 134.1533 | 111.2784 | 71.18313 | 196.6578 | 86.78602 | 105.4963 |
| ENSMUSG00000029363 | -0.187983295 | 0.031913 | 0.098741 | 1035.23 | 1217.819 | 1173.427 | 1213.711 | 1443.165 | 1297.271 | 1232.362 | 1312.722 |
| ENSMUSG00000029364 | -0.598629387 | 0.000427 | 0.004053 | 1381.313 | 1029.447 | 1340.705 | 1175.756 | 1822.483 | 1379.957 | 1891.935 | 2368.773 |
| ENSMUSG00000029366 | 0.068879407 | 0.605797 | 0.742255 | 551.3178 | 685.0981 | 490.2391 | 522.7498 | 583.1166 | 565.3913 | 512.0375 | 479.6277 |
| ENSMUSG00000029368 | 2.989708023 | 0.01069 | 0.044829 | 2127.805 | 40.79393 | 56.31125 | 1977.133 | 83.85957 | 131.8501 | 143.6791 | 169.6642 |
| ENSMUSG00000029369 | -1.126771517 | 0.000132 | 0.001655 | 135.8173 | 146.3782 | 239.3228 | 163.0358 | 355.9156 | 211.1837 | 381.8585 | 548.1459 |
| ENSMUSG00000029370 | -1.411976013 | 0.000264 | 0.002818 | 143.8658 | 75.58876 | 250.9163 | 160.448 | 325.6872 | 217.8879 | 536.1448 | 602.5255 |
| ENSMUSG00000029372 | 1.421349775 | 0.051014 | 0.137971 | 49.29666 | 5.999108 | 8.281066 | 37.95543 | 6.825779 | 7.821619 | 13.50005 | 9.78832 |
| ENSMUSG00000029373 | 1.154920441 | 1.29E-18 | 1.33E-15 | 541.2572 | 527.9215 | 505.145 | 505.4974 | 210.624 | 292.752 | 221.7865 | 209.9051 |
| ENSMUSG00000029376 | 0.215637336 | 0.020667 | 0.072117 | 870.237 | 957.4576 | 824.7942 | 960.9626 | 751.8108 | 858.1433 | 774.3242 | 726.5108 |
| ENSMUSG00000029381 | 0.277339707 | 0.000207 | 0.002341 | 1647.917 | 1542.97 | 1702.587 | 1464.735 | 1318.35 | 1337.497 | 1242.969 | 1349.701 |
| ENSMUSG00000029385 | 0.311672284 | 0.013586 | 0.053608 | 2805.886 | 2498.028 | 2041.283 | 2552.503 | 2075.037 | 2287.265 | 1849.507 | 1761.898 |
| ENSMUSG00000029386 | -0.146516582 | 0.15402 | 0.295488 | 574.457 | 607.1097 | 608.6584 | 501.1842 | 642.5984 | 589.9735 | 692.3596 | 607.9634 |
| ENSMUSG00000029387 | -0.159164426 | 0.039726 | 0.116065 | 1147.908 | 1215.419 | 1072.398 | 1133.487 | 1342.728 | 1194.473 | 1218.862 | 1341 |
| ENSMUSG00000029388 | -0.212329626 | 0.016438 | 0.061389 | 1221.35 | 1199.822 | 1253.753 | 1136.075 | 1376.857 | 1225.759 | 1435.827 | 1534.591 |
| ENSMUSG00000029389 | 0.054432846 | 0.66362 | 0.783707 | 735.4257 | 718.6931 | 581.3309 | 629.7152 | 628.9468 | 751.9928 | 601.7164 | 581.8612 |
| ENSMUSG00000029390 | 0.000594764 | 0.993759 | 0.996513 | 3068.466 | 3196.325 | 3342.238 | 3434.104 | 3051.123 | 3606.884 | 3172.511 | 3210.569 |
| ENSMUSG00000029392 | -0.084159544 | 0.373875 | 0.542642 | 750.5165 | 819.4781 | 762.6862 | 747.0319 | 779.1139 | 721.8237 | 887.146 | 873.3356 |
| ENSMUSG00000029394 | -0.250641373 | 0.000188 | 0.002166 | 1312.901 | 1257.413 | 1236.363 | 1162.816 | 1550.427 | 1448.117 | 1432.934 | 1476.949 |
| ENSMUSG00000029397 | 0.07481003 | 0.340867 | 0.509511 | 1828.001 | 1937.712 | 1712.525 | 1985.759 | 1698.644 | 1926.353 | 1738.613 | 1722.744 |
| ENSMUSG00000029401 | -0.151978575 | 0.208922 | 0.366165 | 629.79 | 635.9054 | 644.267 | 655.5939 | 746.9353 | 560.9218 | 798.4314 | 742.8247 |
| ENSMUSG00000029402 | -0.343940092 | 0.005561 | 0.027793 | 351.113 | 406.7395 | 421.5063 | 331.2474 | 517.7841 | 417.8979 | 486.966 | 492.6788 |
| ENSMUSG00000029403 | 0.042637198 | 0.856143 | 0.913826 | 124.7507 | 62.39072 | 84.46688 | 98.33908 | 89.71024 | 86.03781 | 88.7146 | 95.70802 |
| ENSMUSG00000029404 | -0.107698723 | 0.067042 | 0.167252 | 1663.008 | 1838.127 | 1848.334 | 1768.378 | 1942.422 | 1901.771 | 1935.328 | 1889.146 |
| ENSMUSG00000029405 | 0.025504747 | 0.611752 | 0.746183 | 2036.254 | 2140.482 | 2078.548 | 2113.428 | 2086.738 | 2107.368 | 1998.971 | 2028.357 |
| ENSMUSG00000029406 | -0.308943334 | 0.003897 | 0.021439 | 573.451 | 460.7315 | 566.4249 | 495.1459 | 668.9264 | 586.6214 | 674.0381 | 668.8685 |
| ENSMUSG00000029407 | -0.059328172 | 0.438453 | 0.601998 | 2195.211 | 1972.507 | 1989.94 | 2045.28 | 2040.908 | 1975.517 | 2267.044 | 2263.277 |
| ENSMUSG00000029408 | -0.424987479 | 0.139895 | 0.276031 | 61.36932 | 68.38983 | 67.90474 | 69.8725 | 104.3369 | 45.81234 | 105.1075 | 103.3212 |
| ENSMUSG00000029410 | 0.242616183 | 0.367242 | 0.535809 | 35.2119 | 50.3925 | 38.92101 | 42.26855 | 40.95467 | 34.6386 | 28.92867 | 35.89051 |
| ENSMUSG00000029413 | -0.125645576 | 0.114013 | 0.241393 | 552.3238 | 629.9063 | 619.4238 | 603.8364 | 644.5486 | 686.0677 | 647.038 | 647.1167 |
| ENSMUSG00000029414 | 0.034017524 | 0.772747 | 0.858991 | 598.6023 | 713.8938 | 623.5643 | 584.8587 | 686.4784 | 662.6028 | 542.8948 | 567.7225 |
| ENSMUSG00000029415 | -0.126956122 | 0.093257 | 0.21005 | 1844.098 | 1703.747 | 1652.073 | 1695.918 | 1996.053 | 1974.4 | 1731.863 | 1827.153 |
| ENSMUSG00000029416 | -0.060166294 | 0.649641 | 0.773185 | 284.7134 | 334.7502 | 330.4145 | 257.9244 | 303.2596 | 282.6956 | 327.8583 | 343.6788 |
| ENSMUSG00000029420 | 1.668182668 | 0.015928 | 0.060154 | 47.28455 | 11.99822 | 9.93728 | 73.323 | 14.62667 | 11.17374 | 12.53576 | 6.525547 |
| ENSMUSG00000029422 | 0.068595347 | 0.405035 | 0.572028 | 3850.17 | 3741.044 | 3181.586 | 3383.209 | 3287.1 | 3577.832 | 3400.084 | 3229.058 |
| ENSMUSG00000029426 | 0.162295513 | 0.015505 | 0.059098 | 1992.994 | 2058.894 | 2181.233 | 2101.351 | 1980.451 | 1919.649 | 1777.185 | 1772.773 |
| ENSMUSG00000029427 | 0.056160273 | 0.432126 | 0.596508 | 1919.552 | 1896.918 | 1726.602 | 1833.938 | 1754.225 | 1862.663 | 1638.327 | 1839.117 |
| ENSMUSG00000029428 | -0.176000573 | 0.075029 | 0.181045 | 1026.175 | 938.2604 | 1024.368 | 986.8413 | 1120.403 | 979.9371 | 1105.075 | 1287.708 |
| ENSMUSG00000029430 | -0.291168453 | 7.28E-05 | 0.001063 | 8476.008 | 9093.447 | 9293.013 | 9117.068 | 11618.45 | 9965.86 | 10689.15 | 11752.51 |

| | | | | | | | | | | | |
|---|---|---|---|---|---|---|---|---|---|---|---|
| ENSMUSG00000029432 | -0.392395505 | 0.003306 | 0.01906 | 3051.363 | 2791.985 | 3471.423 | 3222.761 | 3912.147 | 3240.385 | 4637.267 | 4669.029 |
| ENSMUSG00000029433 | -0.088604762 | 0.202056 | 0.357575 | 1352.137 | 1345 | 1324.143 | 1449.207 | 1409.036 | 1573.263 | 1426.184 | 1411.693 |
| ENSMUSG00000029434 | -0.23721973 | 0.03968 | 0.116004 | 1020.139 | 1035.446 | 1172.599 | 980.8029 | 1270.57 | 1022.397 | 1278.647 | 1389.941 |
| ENSMUSG00000029436 | 0.044837517 | 0.757244 | 0.848812 | 157.9505 | 171.5745 | 162.3089 | 175.9752 | 169.6694 | 178.7799 | 165.8577 | 132.6861 |
| ENSMUSG00000029438 | -0.026948884 | 0.643131 | 0.769207 | 2267.647 | 2082.89 | 2148.937 | 2108.252 | 2120.867 | 2174.41 | 2173.508 | 2303.518 |
| ENSMUSG00000029439 | 0.194048237 | 0.062411 | 0.159339 | 3024.199 | 3214.322 | 2950.544 | 2924.294 | 2589.896 | 3165.521 | 2475.33 | 2357.897 |
| ENSMUSG00000029440 | -0.141844655 | 0.082325 | 0.192817 | 1127.787 | 1121.833 | 1216.489 | 1216.299 | 1365.156 | 1154.247 | 1355.791 | 1292.058 |
| ENSMUSG00000029442 | 1.119962521 | 0.000308 | 0.003171 | 236.4228 | 303.5548 | 212.8234 | 266.5507 | 100.4365 | 204.4795 | 91.60747 | 72.8686 |
| ENSMUSG00000029445 | 1.66302343 | 0.11712 | 0.245381 | 79.47829 | 4.799286 | 0.828107 | 53.48266 | 9.751113 | 2.234748 | 15.42863 | 16.31387 |
| ENSMUSG00000029446 | -0.18885346 | 0.033013 | 0.101329 | 595.5842 | 661.1017 | 606.1741 | 579.683 | 704.0304 | 696.1241 | 634.5023 | 747.1751 |
| ENSMUSG00000029447 | -0.172952317 | 6.76E-05 | 0.001006 | 6963.908 | 7040.553 | 6695.242 | 6951.883 | 7972.51 | 7560.153 | 7676.706 | 7957.904 |
| ENSMUSG00000029449 | 0.354038033 | 0.152925 | 0.293979 | 117.7084 | 110.3836 | 149.0592 | 145.7834 | 107.2622 | 61.45558 | 125.3576 | 115.2847 |
| ENSMUSG00000029452 | -1.659184745 | 6.38E-05 | 0.000959 | 75.45408 | 38.39429 | 149.8873 | 71.59775 | 239.8774 | 140.7891 | 284.4653 | 398.0583 |
| ENSMUSG00000029454 | -0.166500992 | 0.101873 | 0.223803 | 429.5852 | 406.7395 | 348.6329 | 395.0816 | 449.5263 | 430.189 | 424.2872 | 466.5766 |
| ENSMUSG00000029455 | 0.006508939 | 0.945723 | 0.968767 | 5092.647 | 5256.418 | 5477.925 | 5077.402 | 5307.531 | 4333.177 | 5469.448 | 5698.977 |
| ENSMUSG00000029456 | -0.81510773 | 6.89E-05 | 0.001018 | 136.8234 | 173.9741 | 139.1219 | 185.4641 | 274.9814 | 196.6578 | 304.7154 | 340.416 |
| ENSMUSG00000029458 | -0.183436255 | 0.080379 | 0.189533 | 1364.21 | 1232.217 | 1491.42 | 1264.606 | 1548.477 | 1303.976 | 1566.97 | 1660.752 |
| ENSMUSG00000029461 | 0.340105783 | 0.000879 | 0.007046 | 2340.082 | 2525.624 | 2275.637 | 2623.238 | 1790.304 | 2271.622 | 1855.292 | 1797.788 |
| ENSMUSG00000029462 | -0.363145744 | 0.000929 | 0.007403 | 1423.567 | 1352.199 | 1487.28 | 1495.789 | 1833.209 | 1549.798 | 1862.042 | 2164.306 |
| ENSMUSG00000029463 | -0.033928643 | 0.658066 | 0.779672 | 758.565 | 777.4844 | 808.2321 | 840.1953 | 875.65 | 778.8097 | 801.3243 | 805.905 |
| ENSMUSG00000029464 | -0.023051213 | 0.822734 | 0.89248 | 837.0372 | 713.8938 | 667.4539 | 725.4664 | 799.5913 | 674.894 | 744.4312 | 770.0145 |
| ENSMUSG00000029465 | -0.217749388 | 0.065737 | 0.165067 | 2887.376 | 2701.998 | 3450.72 | 2989.853 | 3615.713 | 2869.417 | 3693.228 | 3814.182 |
| ENSMUSG00000029466 | -0.175030578 | 0.049407 | 0.13499 | 2065.43 | 2022.899 | 2224.294 | 2098.763 | 2587.945 | 2053.734 | 2351.901 | 2503.635 |
| ENSMUSG00000029467 | -0.352608958 | 0.001244 | 0.009208 | 5019.205 | 4733.296 | 5434.036 | 4754.781 | 5734.63 | 5527.65 | 7102.954 | 7098.707 |
| ENSMUSG00000029468 | 1.059594237 | 4.03E-10 | 5.67E-08 | 168.0111 | 157.1766 | 165.6213 | 184.6014 | 64.35735 | 89.38993 | 90.64318 | 80.48174 |
| ENSMUSG00000029469 | 0.13254555 | 0.229292 | 0.389418 | 1223.362 | 1216.619 | 1068.258 | 1389.686 | 1152.582 | 1221.29 | 1111.825 | 982.0948 |
| ENSMUSG00000029470 | -0.16190112 | 0.329218 | 0.497018 | 538.2391 | 467.9304 | 655.0323 | 466.6793 | 556.7886 | 482.7056 | 622.9308 | 719.9853 |
| ENSMUSG00000029471 | 0.249165831 | 0.001319 | 0.009624 | 1623.772 | 1599.362 | 1792.023 | 1513.042 | 1404.16 | 1357.61 | 1300.826 | 1432.357 |
| ENSMUSG00000029472 | 0.016105341 | 0.762104 | 0.852086 | 10328.15 | 11628.67 | 10910.3 | 10888.9 | 11136.75 | 10807.24 | 10650.57 | 10670.36 |
| ENSMUSG00000029474 | -0.264315586 | 0.001308 | 0.009572 | 1134.829 | 1060.642 | 971.3691 | 1114.51 | 1318.35 | 1184.417 | 1346.148 | 1289.883 |
| ENSMUSG00000029475 | 0.00138662 | 0.985016 | 0.991442 | 2531.233 | 2811.182 | 2438.774 | 2521.448 | 2773.217 | 2561.021 | 2501.366 | 2450.343 |
| ENSMUSG00000029477 | 1.021000659 | 0.058379 | 0.151667 | 25.15136 | 16.7975 | 18.21835 | 24.15346 | 8.776002 | 20.11273 | 9.642892 | 3.262773 |
| ENSMUSG00000029478 | 0.242664376 | 0.114891 | 0.242557 | 1693.189 | 1655.754 | 1955.988 | 1710.582 | 1355.405 | 1986.691 | 1298.898 | 1290.971 |
| ENSMUSG00000029480 | -0.21414519 | 0.034741 | 0.105507 | 455.7426 | 471.5299 | 537.4412 | 486.5197 | 583.1166 | 516.2268 | 545.7877 | 621.0145 |
| ENSMUSG00000029482 | -0.440381078 | 0.001786 | 0.012103 | 536.227 | 533.9206 | 603.6897 | 542.5902 | 787.8899 | 564.2739 | 850.503 | 804.8174 |
| ENSMUSG00000029484 | 0.19128552 | 0.015877 | 0.059997 | 2029.212 | 1812.93 | 1983.315 | 2038.379 | 1820.533 | 1554.267 | 1706.792 | 1807.576 |
| ENSMUSG00000029486 | 0.043398231 | 0.671086 | 0.789191 | 696.1896 | 700.6958 | 563.1125 | 667.6706 | 672.8268 | 615.6731 | 599.7879 | 657.9926 |
| ENSMUSG00000029490 | 0.53025416 | 0.024707 | 0.082353 | 58.35115 | 79.18822 | 55.48314 | 64.69676 | 46.80534 | 50.28183 | 42.42872 | 38.06569 |
| ENSMUSG00000029491 | 0.057662258 | 0.645965 | 0.770635 | 298.7981 | 383.9429 | 337.0394 | 366.615 | 334.4632 | 347.5033 | 291.2153 | 358.9051 |
| ENSMUSG00000029499 | -1.715113853 | 2.60E-07 | 1.24E-05 | 118.7144 | 49.19268 | 116.763 | 125.943 | 317.8863 | 207.8316 | 327.8583 | 499.2043 |
| ENSMUSG00000029500 | -0.325268716 | 0.002098 | 0.013581 | 589.5479 | 557.917 | 589.6119 | 586.584 | 812.2677 | 621.26 | 698.1454 | 779.8028 |
| ENSMUSG00000029501 | 0.117778668 | 0.233347 | 0.393914 | 1573.469 | 1565.767 | 1327.455 | 1552.722 | 1340.778 | 1559.854 | 1393.398 | 1251.817 |
| ENSMUSG00000029502 | 0.199246861 | 0.030047 | 0.094633 | 1500.027 | 1337.801 | 1226.426 | 1243.903 | 1104.801 | 1120.726 | 1243.933 | 1150.671 |
| ENSMUSG00000029503 | -0.403229514 | 0.397653 | 0.564936 | 15.09082 | 19.19714 | 8.281066 | 8.626235 | 15.60178 | 23.46486 | 13.50005 | 14.13868 |
| ENSMUSG00000029504 | -0.073980672 | 0.432323 | 0.596728 | 560.3723 | 585.5129 | 533.3007 | 552.9417 | 663.0757 | 548.6307 | 572.7878 | 561.197 |
| ENSMUSG00000029505 | 0.171831214 | 0.100251 | 0.221292 | 2657.996 | 2477.631 | 2627.582 | 2492.119 | 2021.406 | 2730.862 | 2144.579 | 2209.985 |
| ENSMUSG00000029507 | -0.415137707 | 0.00429 | 0.023003 | 760.5771 | 671.9001 | 708.0312 | 690.0988 | 970.2358 | 699.4762 | 988.3964 | 1115.868 |
| ENSMUSG00000029510 | 0.448788439 | 0.013123 | 0.05238 | 286.7255 | 361.1463 | 247.6039 | 291.5667 | 260.3547 | 187.7188 | 179.3578 | 240.3576 |
| ENSMUSG00000029512 | 0.152935479 | 0.14098 | 0.277628 | 1609.687 | 1469.781 | 1368.032 | 1594.128 | 1234.491 | 1565.441 | 1371.219 | 1263.781 |
| ENSMUSG00000029513 | -0.246559597 | 0.093089 | 0.209847 | 1382.319 | 1425.388 | 1816.866 | 1375.022 | 1866.363 | 1440.295 | 1685.577 | 2127.328 |
| ENSMUSG00000029516 | 0.271962907 | 0.053174 | 0.142096 | 538.2391 | 428.3363 | 447.1776 | 423.5481 | 441.7254 | 406.7242 | 343.2869 | 329.5401 |
| ENSMUSG00000029518 | -0.148697106 | 0.137217 | 0.272251 | 1386.343 | 1270.611 | 1425.172 | 1362.945 | 1527.999 | 1291.684 | 1503.327 | 1715.131 |
| ENSMUSG00000029521 | 0.19345862 | 0.151271 | 0.291608 | 554.3359 | 494.3265 | 414.8814 | 477.8934 | 416.3725 | 491.6446 | 366.4299 | 423.0729 |
| ENSMUSG00000029524 | 0.333270802 | 0.009276 | 0.04031 | 504.0332 | 448.7333 | 407.4285 | 522.7498 | 364.6916 | 429.0717 | 366.4299 | 334.9781 |
| ENSMUSG00000029528 | -0.264700314 | 0.005248 | 0.02661 | 2302.858 | 2265.263 | 2659.879 | 2543.877 | 2834.649 | 2604.599 | 3175.404 | 3129 |
| ENSMUSG00000029534 | -0.141912784 | 0.141727 | 0.278593 | 469.8274 | 479.9286 | 484.4424 | 483.9318 | 509.9832 | 569.8608 | 568.9306 | 467.6642 |
| ENSMUSG00000029535 | -0.154011502 | 0.169954 | 0.316547 | 617.7174 | 640.7047 | 622.7362 | 673.7089 | 787.8899 | 591.0909 | 688.5025 | 774.3649 |
| ENSMUSG00000029536 | -0.056773634 | 0.507374 | 0.663071 | 930.6003 | 1040.245 | 1062.461 | 1031.698 | 1157.457 | 997.8151 | 994.1821 | 1078.89 |
| ENSMUSG00000029538 | -0.155441265 | 0.031318 | 0.09733 | 2565.439 | 2809.982 | 2403.994 | 2650.842 | 3072.576 | 2950.985 | 2779.081 | 2808.16 |
| ENSMUSG00000029544 | -0.020115181 | 0.914436 | 0.949313 | 82.49646 | 106.7841 | 92.74794 | 99.2017 | 111.1627 | 105.0332 | 83.89316 | 85.9197 |
| ENSMUSG00000029545 | -0.457729612 | 0.001654 | 0.011369 | 1463.809 | 1370.196 | 1474.03 | 1410.389 | 2088.688 | 1432.474 | 1969.078 | 2363.335 |
| ENSMUSG00000029546 | 0.046930483 | 0.595424 | 0.734618 | 2835.061 | 3034.349 | 2561.334 | 2955.348 | 3028.696 | 2602.364 | 2836.939 | 2548.226 |
| ENSMUSG00000029547 | -0.148011929 | 0.168822 | 0.315075 | 1146.902 | 1329.402 | 1244.644 | 1100.708 | 1320.301 | 1182.182 | 1300.826 | 1536.766 |
| ENSMUSG00000029550 | 0.157098608 | 0.060523 | 0.156029 | 2265.634 | 2397.243 | 2558.021 | 2436.911 | 2102.34 | 2429.171 | 2152.293 | 1980.503 |

| | | | | | | | | | | | |
|---|---|---|---|---|---|---|---|---|---|---|---|
| ENSMUSG00000029551 | -0.152847604 | 0.243005 | 0.405543 | 398.3975 | 385.1427 | 417.3657 | 370.0655 | 449.5263 | 344.1512 | 462.8588 | 489.416 |
| ENSMUSG00000029552 | 0.164220321 | 0.019728 | 0.069796 | 2083.539 | 2400.843 | 2328.636 | 2389.467 | 2007.754 | 1984.456 | 2148.436 | 2070.773 |
| ENSMUSG00000029553 | -1.924467955 | 3.82E-07 | 1.69E-05 | 20.12109 | 20.39697 | 42.23344 | 28.46658 | 89.71024 | 64.8077 | 106.0718 | 165.3138 |
| ENSMUSG00000029554 | -0.202230312 | 0.044138 | 0.124863 | 764.6013 | 787.0829 | 934.9324 | 759.1087 | 934.1566 | 852.5564 | 965.2535 | 984.2699 |
| ENSMUSG00000029556 | -1.601848662 | 1.71E-06 | 5.78E-05 | 126.7628 | 57.59143 | 93.57605 | 67.28463 | 222.3254 | 167.6061 | 244.9294 | 414.3722 |
| ENSMUSG00000029557 | -0.183147597 | 0.107784 | 0.23239 | 235.4167 | 278.3586 | 254.2287 | 266.5507 | 313.0107 | 282.6956 | 269.0367 | 308.8759 |
| ENSMUSG00000029559 | -0.114336893 | 0.148806 | 0.288612 | 1207.265 | 1246.615 | 1269.487 | 1223.2 | 1254.968 | 1283.863 | 1314.326 | 1503.051 |
| ENSMUSG00000029560 | -0.525966747 | 1.66E-05 | 0.000338 | 769.6316 | 752.2881 | 841.3563 | 770.3228 | 1220.839 | 937.4769 | 1041.432 | 1313.81 |
| ENSMUSG00000029561 | 0.906872791 | 0.083471 | 0.194528 | 436.6276 | 1428.987 | 1247.957 | 321.7586 | 374.4427 | 439.128 | 593.0378 | 425.2481 |
| ENSMUSG00000029563 | 1.218192309 | 0.003718 | 0.020731 | 371.2341 | 607.1097 | 373.4761 | 589.1718 | 151.1423 | 433.5411 | 162.9649 | 87.00729 |
| ENSMUSG00000029569 | -0.082825041 | 0.311712 | 0.480258 | 1025.169 | 1093.037 | 1010.29 | 1004.094 | 1061.896 | 1217.938 | 1017.325 | 1078.89 |
| ENSMUSG00000029570 | -0.151941854 | 0.240651 | 0.402912 | 744.4802 | 676.6993 | 815.685 | 723.7411 | 809.3424 | 669.3071 | 837.003 | 975.5692 |
| ENSMUSG00000029571 | -0.289782986 | 0.010816 | 0.045254 | 2302.858 | 2210.071 | 2691.347 | 2484.356 | 2731.287 | 2575.547 | 3129.118 | 3411.773 |
| ENSMUSG00000029575 | -0.432685773 | 0.002013 | 0.013203 | 690.1533 | 653.9027 | 744.4679 | 775.4985 | 1009.24 | 728.5279 | 1048.182 | 1081.066 |
| ENSMUSG00000029576 | 0.077653444 | 0.776235 | 0.861881 | 35.2119 | 53.99197 | 65.42042 | 43.9938 | 52.65601 | 43.57759 | 44.3573 | 47.85401 |
| ENSMUSG00000029577 | -0.224298443 | 0.060326 | 0.155584 | 1658.984 | 1683.35 | 1896.364 | 1513.042 | 2055.535 | 1596.728 | 2123.365 | 2111.014 |
| ENSMUSG00000029578 | 0.041899997 | 0.438305 | 0.601907 | 2898.443 | 2995.954 | 2771.673 | 2857.009 | 2913.633 | 2812.431 | 2775.224 | 2685.262 |
| ENSMUSG00000029580 | 0.037526009 | 0.606968 | 0.742971 | 43035.99 | 44987.31 | 49301.33 | 41400.75 | 42460.25 | 44198.85 | 41584.01 | 45896.34 |
| ENSMUSG00000029581 | 0.215009675 | 0.003631 | 0.020377 | 4536.299 | 5027.252 | 4994.311 | 5026.507 | 4624.953 | 4196.857 | 3916.943 | 4133.934 |
| ENSMUSG00000029587 | -0.045395052 | 0.624664 | 0.755344 | 923.5579 | 1001.851 | 841.3563 | 954.9242 | 1028.742 | 860.3781 | 991.2893 | 954.905 |
| ENSMUSG00000029591 | -0.445159743 | 0.000258 | 0.002772 | 594.5781 | 815.8786 | 703.8906 | 605.5617 | 973.1611 | 953.1201 | 851.4673 | 921.1897 |
| ENSMUSG00000029592 | -0.150561272 | 0.049047 | 0.134366 | 850.1159 | 891.4674 | 937.4167 | 973.9019 | 1055.07 | 1039.158 | 1007.682 | 955.9926 |
| ENSMUSG00000029594 | -0.271437143 | 0.037412 | 0.111332 | 554.3359 | 617.9081 | 679.0474 | 580.5456 | 774.2384 | 615.6731 | 684.6453 | 861.3721 |
| ENSMUSG00000029596 | 0.286075849 | 0.078986 | 0.187542 | 301.8163 | 281.9581 | 220.2764 | 205.3044 | 206.7236 | 211.1837 | 216.9651 | 190.3284 |
| ENSMUSG00000029598 | -0.257747426 | 0.03245 | 0.099966 | 1109.678 | 1131.432 | 1305.096 | 1073.104 | 1485.095 | 1134.135 | 1357.719 | 1546.555 |
| ENSMUSG00000029599 | -0.290055823 | 0.025824 | 0.084911 | 1646.911 | 1635.357 | 1994.909 | 1754.576 | 2356.844 | 1692.822 | 2140.722 | 2409.014 |
| ENSMUSG00000029600 | -0.262662902 | 0.119486 | 0.24843 | 241.453 | 268.76 | 269.9628 | 287.2536 | 320.8116 | 230.1791 | 371.2513 | 357.8175 |
| ENSMUSG00000029601 | 0.322963648 | 0.209068 | 0.366191 | 62.37537 | 82.78769 | 60.45178 | 64.69676 | 52.65601 | 64.8077 | 61.71451 | 35.89051 |
| ENSMUSG00000029602 | 0.609041021 | 0.127601 | 0.25899 | 43.26034 | 15.59768 | 30.63995 | 25.01608 | 19.50223 | 21.23011 | 17.3572 | 17.40146 |
| ENSMUSG00000029603 | 0.360689766 | 0.009772 | 0.041855 | 349.1009 | 443.934 | 484.4424 | 470.1298 | 360.7912 | 315.0995 | 374.1442 | 311.0511 |
| ENSMUSG00000029605 | 0.683646288 | 0.10756 | 0.232097 | 81.4904 | 148.7779 | 155.684 | 48.30692 | 65.33246 | 55.8687 | 106.0718 | 42.41605 |
| ENSMUSG00000029607 | 0.367596982 | 0.315266 | 0.48367 | 31.18769 | 22.79661 | 28.98373 | 30.19182 | 17.552 | 32.40385 | 24.10723 | 14.13868 |
| ENSMUSG00000029608 | -0.222560522 | 0.352139 | 0.520922 | 39.23612 | 44.3934 | 52.99882 | 55.2079 | 55.58134 | 64.8077 | 57.85735 | 46.76642 |
| ENSMUSG00000029610 | -0.456822324 | 2.55E-05 | 0.000473 | 504.0332 | 535.1204 | 501.0045 | 508.9479 | 738.1593 | 584.3867 | 710.6811 | 776.54 |
| ENSMUSG00000029613 | -0.24938486 | 0.017512 | 0.064269 | 1398.416 | 1289.808 | 1358.923 | 1373.297 | 1690.843 | 1310.68 | 1717.399 | 1723.832 |
| ENSMUSG00000029614 | -0.092733532 | 0.222449 | 0.381724 | 34986.55 | 41335.05 | 38427.46 | 39705.7 | 43887.81 | 41505.98 | 41369.93 | 37941.7 |
| ENSMUSG00000029616 | -0.251443568 | 0.016912 | 0.062633 | 2609.705 | 2829.179 | 3366.253 | 2878.575 | 3352.433 | 3186.751 | 3444.441 | 3928.379 |
| ENSMUSG00000029617 | 0.038164591 | 0.637484 | 0.764831 | 1617.735 | 1811.731 | 1792.023 | 1701.956 | 1722.047 | 1825.789 | 1675.935 | 1518.277 |
| ENSMUSG00000029618 | -1.62801185 | 5.87E-06 | 0.000152 | 36.21796 | 59.99108 | 34.78048 | 46.58167 | 210.624 | 146.376 | 119.5719 | 69.60583 |
| ENSMUSG00000029621 | -0.127999851 | 0.06191 | 0.158433 | 2819.97 | 2829.179 | 3117.821 | 2941.546 | 3414.84 | 2967.746 | 3207.226 | 3207.306 |
| ENSMUSG00000029622 | 0.058028825 | 0.521797 | 0.675051 | 4139.914 | 4272.564 | 4553.758 | 4040.528 | 4397.752 | 3488.442 | 4191.765 | 4256.832 |
| ENSMUSG00000029623 | -0.128835399 | 0.075182 | 0.181239 | 4163.053 | 4932.466 | 4759.129 | 4409.731 | 5147.613 | 5005.836 | 4752.017 | 5062.737 |
| ENSMUSG00000029624 | -0.283988692 | 0.030047 | 0.094633 | 541.2572 | 526.7217 | 481.9581 | 511.5357 | 644.5486 | 498.3488 | 626.788 | 738.4743 |
| ENSMUSG00000029625 | 0.012744036 | 0.867988 | 0.921001 | 1119.738 | 1015.049 | 1019.399 | 1087.768 | 1101.876 | 985.524 | 1011.539 | 1106.08 |
| ENSMUSG00000029627 | -0.00878997 | 0.930097 | 0.958773 | 319.9253 | 290.3568 | 317.1648 | 325.2091 | 296.4338 | 313.9821 | 336.5369 | 315.4014 |
| ENSMUSG00000029629 | 0.280416114 | 0.025686 | 0.084625 | 3021.181 | 3468.684 | 3039.151 | 3274.519 | 2536.265 | 3304.075 | 2371.187 | 2330.708 |
| ENSMUSG00000029632 | -0.400012249 | 0.000265 | 0.002824 | 4398.47 | 4428.541 | 5152.479 | 4558.103 | 6439.635 | 5080.7 | 6028.736 | 6913.817 |
| ENSMUSG00000029634 | -0.368463971 | 2.83E-06 | 8.64E-05 | 1213.302 | 1019.848 | 1100.554 | 1038.599 | 1464.617 | 1358.727 | 1417.505 | 1404.08 |
| ENSMUSG00000029635 | -0.053155103 | 0.630744 | 0.75956 | 1868.243 | 1593.363 | 1565.122 | 1859.816 | 1696.694 | 2091.724 | 1698.113 | 1660.752 |
| ENSMUSG00000029636 | 0.309061255 | 0.028944 | 0.092167 | 640.8566 | 543.5192 | 474.5051 | 460.6409 | 429.049 | 478.2361 | 374.1442 | 428.5109 |
| ENSMUSG00000029638 | 0.461068063 | 0.000266 | 0.00283 | 266.6044 | 230.3657 | 227.7293 | 225.1447 | 176.4951 | 164.254 | 162.9649 | 185.9781 |
| ENSMUSG00000029640 | -0.183807087 | 0.086297 | 0.199186 | 1611.699 | 1390.593 | 1766.351 | 1727.835 | 1862.463 | 1632.484 | 1845.649 | 2043.584 |
| ENSMUSG00000029641 | -0.134267231 | 0.446465 | 0.609149 | 564.3965 | 475.1293 | 602.8616 | 545.178 | 623.0961 | 429.0717 | 584.3592 | 765.6641 |
| ENSMUSG00000029642 | -0.173739492 | 0.024609 | 0.082043 | 2172.071 | 2036.097 | 2335.261 | 2309.243 | 2501.161 | 2283.913 | 2638.295 | 2566.715 |
| ENSMUSG00000029647 | 0.345681245 | 0.020976 | 0.072905 | 2380.325 | 2250.865 | 2017.268 | 2371.352 | 1609.909 | 2350.955 | 1638.327 | 1499.788 |
| ENSMUSG00000029648 | -0.302590702 | 0.140472 | 0.276915 | 712.2865 | 392.3416 | 743.6398 | 543.4528 | 621.1459 | 696.1241 | 809.0386 | 826.5692 |
| ENSMUSG00000029649 | -0.162943432 | 0.008831 | 0.038974 | 2652.965 | 2667.203 | 2730.268 | 2561.992 | 3005.293 | 2867.182 | 2817.653 | 3192.08 |
| ENSMUSG00000029650 | -0.233951681 | 0.374543 | 0.543269 | 54.32694 | 79.18822 | 62.108 | 56.93315 | 51.6809 | 65.92507 | 94.50034 | 83.74451 |
| ENSMUSG00000029651 | 0.596438933 | 0.003238 | 0.018789 | 180.0837 | 166.7752 | 150.7154 | 185.4641 | 109.2125 | 149.7281 | 82.92887 | 110.9343 |
| ENSMUSG00000029655 | 0.178766004 | 0.07211 | 0.176227 | 2124.787 | 2072.092 | 1753.102 | 2114.29 | 1693.768 | 2022.447 | 1698.113 | 1709.693 |
| ENSMUSG00000029657 | -0.348788551 | 0.000213 | 0.002383 | 906.455 | 1117.034 | 988.7593 | 961.8252 | 1227.665 | 1197.825 | 1235.254 | 1396.467 |
| ENSMUSG00000029659 | 1.414425017 | 0.016729 | 0.062112 | 98.59333 | 131.9804 | 100.2009 | 100.0643 | 40.95467 | 94.9768 | 18.32149 | 7.613138 |
| ENSMUSG00000029661 | 0.544583333 | 0.002249 | 0.014301 | 20535.58 | 34260.9 | 31050.69 | 30162.49 | 17351.13 | 25506.3 | 17994.6 | 18683.73 |
| ENSMUSG00000029663 | 0.088833959 | 0.611745 | 0.746183 | 104.6297 | 141.5789 | 151.5435 | 127.6683 | 112.1378 | 138.5544 | 123.429 | 120.7226 |

| | | | | | | | | | | | |
|---|---|---|---|---|---|---|---|---|---|---|---|
| ENSMUSG00000029664 | -0.422882072 | 0.166462 | 0.312079 | 25.15136 | 33.595 | 28.98373 | 31.91707 | 41.92979 | 30.1691 | 33.75012 | 54.37955 |
| ENSMUSG00000029669 | -0.222308122 | 0.125257 | 0.255808 | 1118.732 | 853.0731 | 1078.195 | 862.6235 | 1120.403 | 962.0591 | 1109.897 | 1373.628 |
| ENSMUSG00000029670 | 0.116673328 | 0.173851 | 0.321792 | 1178.09 | 1168.626 | 1014.431 | 1161.091 | 1059.946 | 1059.271 | 1098.325 | 949.467 |
| ENSMUSG00000029671 | -0.070845846 | 0.889097 | 0.933305 | 36.21796 | 7.198929 | 40.57722 | 26.74133 | 20.47734 | 37.99072 | 23.14294 | 35.89051 |
| ENSMUSG00000029672 | 0.143278122 | 0.003991 | 0.02181 | 2427.609 | 2470.433 | 2479.351 | 2417.071 | 2154.996 | 2264.917 | 2290.187 | 2157.781 |
| ENSMUSG00000029673 | 0.419764457 | 0.051459 | 0.138728 | 1086.539 | 1078.64 | 958.1194 | 1051.538 | 641.6232 | 1210.116 | 630.6451 | 639.5036 |
| ENSMUSG00000029674 | -0.17614392 | 0.064874 | 0.163993 | 500.009 | 458.3318 | 559.8001 | 481.3439 | 538.2614 | 594.443 | 568.9306 | 562.2846 |
| ENSMUSG00000029675 | 0.928533926 | 0.038749 | 0.113787 | 3756.607 | 6169.482 | 5774.388 | 15775.66 | 3655.692 | 4372.285 | 4274.694 | 4235.08 |
| ENSMUSG00000029676 | 0.190024728 | 0.157771 | 0.300343 | 790.7587 | 775.0847 | 581.3309 | 721.1532 | 632.8472 | 667.0723 | 681.7524 | 529.6569 |
| ENSMUSG00000029681 | -0.12678307 | 0.160995 | 0.304657 | 809.8738 | 873.4701 | 881.1055 | 804.8277 | 950.7335 | 829.0916 | 883.2889 | 1014.722 |
| ENSMUSG00000029684 | 0.086012116 | 0.406312 | 0.572907 | 2057.381 | 2022.899 | 2158.046 | 2130.68 | 1680.117 | 2315.199 | 1935.328 | 1957.664 |
| ENSMUSG00000029685 | -0.900238619 | 0.323321 | 0.491365 | 3.018163 | 3.599465 | 51.34261 | 5.175741 | 23.40267 | 23.46486 | 14.46434 | 57.64233 |
| ENSMUSG00000029686 | -0.025727421 | 0.752549 | 0.846016 | 4225.428 | 4763.291 | 4406.355 | 4223.405 | 4371.424 | 4977.902 | 4487.802 | 4095.868 |
| ENSMUSG00000029687 | 0.099836958 | 0.21519 | 0.373372 | 2651.959 | 2681.601 | 2256.591 | 2460.202 | 2433.878 | 2454.871 | 2230.401 | 2254.576 |
| ENSMUSG00000029695 | -0.452947364 | 0.074082 | 0.179534 | 65.39353 | 38.39429 | 40.57722 | 61.24627 | 75.08357 | 72.62932 | 72.32169 | 61.99269 |
| ENSMUSG00000029699 | 0.098396383 | 0.315158 | 0.483544 | 369.2219 | 428.3363 | 423.9906 | 382.1422 | 379.3183 | 366.4987 | 391.5014 | 358.9051 |
| ENSMUSG00000029700 | -3.211759482 | 1.12E-08 | 9.52E-07 | 5.030272 | 5.999108 | 16.56213 | 18.97772 | 172.5947 | 36.87335 | 91.60747 | 137.0365 |
| ENSMUSG00000029701 | 0.111047348 | 0.16096 | 0.304622 | 1873.273 | 2206.472 | 1938.598 | 1842.564 | 1802.006 | 1839.198 | 1739.578 | 1893.496 |
| ENSMUSG00000029703 | -0.067086203 | 0.264768 | 0.429059 | 1069.436 | 1010.25 | 1035.133 | 1036.873 | 1100.901 | 1043.627 | 1104.111 | 1100.642 |
| ENSMUSG00000029705 | -0.100681141 | 0.122602 | 0.252118 | 3186.174 | 3309.108 | 3455.689 | 3001.067 | 3455.794 | 3362.179 | 3432.869 | 3636.905 |
| ENSMUSG00000029708 | -0.099781663 | 0.407289 | 0.573635 | 576.4691 | 428.3363 | 494.3797 | 475.3055 | 530.4606 | 499.4662 | 588.2164 | 498.1167 |
| ENSMUSG00000029710 | -0.074396336 | 0.51304 | 0.668031 | 2736.468 | 2903.568 | 3356.316 | 2627.551 | 2961.413 | 2654.881 | 3162.868 | 3460.715 |
| ENSMUSG00000029712 | 0.982312009 | 0.207111 | 0.36373 | 24.1453 | 5.999108 | 3.312427 | 28.46658 | 7.80089 | 12.29112 | 3.857157 | 7.613138 |
| ENSMUSG00000029713 | -0.137810423 | 0.077507 | 0.185055 | 7092.683 | 7207.328 | 8406.939 | 7246.9 | 8154.856 | 7746.755 | 8326.637 | 8732.269 |
| ENSMUSG00000029714 | 0.438230882 | 4.24E-05 | 0.000703 | 1813.916 | 1736.142 | 1573.403 | 1692.467 | 1352.479 | 1429.121 | 1147.504 | 1100.642 |
| ENSMUSG00000029715 | -0.340384966 | 0.014049 | 0.054885 | 339.0403 | 357.5468 | 400.8036 | 375.2412 | 451.4765 | 370.9682 | 511.0733 | 531.832 |
| ENSMUSG00000029718 | -0.023006735 | 0.877555 | 0.92666 | 2236.459 | 1961.708 | 2474.383 | 1928.826 | 1993.128 | 1771.038 | 2244.865 | 2732.029 |
| ENSMUSG00000029720 | 0.600936771 | 8.03E-05 | 0.001139 | 156.9445 | 154.777 | 166.4494 | 153.547 | 98.48624 | 124.0285 | 99.32178 | 95.70802 |
| ENSMUSG00000029722 | -0.619052887 | 2.93E-05 | 0.000527 | 501.0151 | 476.3291 | 651.7199 | 447.7016 | 850.2971 | 755.3449 | 891.9675 | 693.8831 |
| ENSMUSG00000029723 | -0.100015561 | 0.429711 | 0.594554 | 1231.411 | 1095.437 | 1481.483 | 1176.618 | 1185.735 | 1246.989 | 1384.719 | 1529.153 |
| ENSMUSG00000029725 | 0.034821836 | 0.735598 | 0.834534 | 584.5176 | 472.7297 | 474.5051 | 510.6731 | 474.8792 | 486.0577 | 522.6447 | 510.0802 |
| ENSMUSG00000029726 | -0.119007014 | 0.119493 | 0.24843 | 1460.791 | 1479.38 | 1395.36 | 1302.561 | 1519.223 | 1421.3 | 1637.363 | 1538.941 |
| ENSMUSG00000029729 | -0.04682927 | 0.674415 | 0.791543 | 1655.965 | 1619.759 | 1413.578 | 1488.026 | 1519.223 | 1898.419 | 1542.863 | 1419.306 |
| ENSMUSG00000029730 | -0.068304363 | 0.455131 | 0.61714 | 4631.874 | 5821.534 | 4858.502 | 5149.862 | 5859.444 | 5146.625 | 5298.769 | 5143.218 |
| ENSMUSG00000029735 | -0.739093903 | 2.77E-06 | 8.51E-05 | 382.3007 | 332.3506 | 390.8663 | 426.9986 | 633.8224 | 499.4662 | 632.5737 | 795.0291 |
| ENSMUSG00000029736 | -3.331034427 | NA | NA | 2.012109 | 2.399643 | 2.48432 | 2.58787 | 1.950223 | 90.5073 | 0 | 3.262773 |
| ENSMUSG00000029752 | -0.963553382 | 4.35E-13 | 1.43E-10 | 214.2896 | 172.7743 | 183.8397 | 170.7995 | 364.6916 | 311.7474 | 391.5014 | 377.3941 |
| ENSMUSG00000029757 | -0.080832155 | 0.761998 | 0.852086 | 55.33299 | 76.78858 | 42.23344 | 65.55939 | 48.75557 | 72.62932 | 60.75022 | 70.69342 |
| ENSMUSG00000029759 | -0.00582171 | 0.983804 | 0.990959 | 175.0535 | 85.18733 | 158.9965 | 122.4925 | 124.8142 | 90.5073 | 153.322 | 176.1898 |
| ENSMUSG00000029761 | 0.368260687 | 5.61E-05 | 0.000865 | 7539.371 | 7568.474 | 6882.394 | 7739.458 | 5231.472 | 6666.254 | 5563.948 | 5571.729 |
| ENSMUSG00000029762 | 0.522072536 | 0.03236 | 0.099797 | 533.2088 | 277.1588 | 454.6305 | 350.2251 | 331.5378 | 204.4795 | 247.8223 | 341.5036 |
| ENSMUSG00000029763 | 0.006087236 | 0.939234 | 0.964314 | 1469.845 | 1635.357 | 1529.513 | 1394 | 1494.846 | 1616.84 | 1482.112 | 1406.255 |
| ENSMUSG00000029765 | 0.664022718 | 0.0253 | 0.083665 | 302.8224 | 224.3666 | 240.1509 | 315.7202 | 132.6151 | 290.5173 | 113.7861 | 147.9124 |
| ENSMUSG00000029767 | 0.032308602 | 0.558037 | 0.704551 | 11294.97 | 12734.91 | 12540.02 | 11932.67 | 11942.19 | 11884.39 | 12005.4 | 11593.72 |
| ENSMUSG00000029769 | 0.596514022 | 3.50E-12 | 8.73E-10 | 1194.187 | 1233.417 | 1252.925 | 1260.293 | 795.6908 | 930.7726 | 791.6814 | 751.5254 |
| ENSMUSG00000029771 | 0.267488245 | 0.169241 | 0.315546 | 134.8113 | 152.3773 | 173.0743 | 111.2784 | 109.2125 | 121.7938 | 137.8934 | 105.4963 |
| ENSMUSG00000029772 | -0.634956564 | 0.001154 | 0.008721 | 1178.09 | 742.6895 | 961.4318 | 871.2497 | 1548.477 | 1001.167 | 1533.22 | 1746.671 |
| ENSMUSG00000029775 | 0.226852764 | 0.013237 | 0.052726 | 5063.472 | 5637.961 | 4904.876 | 5305.134 | 4435.781 | 5114.221 | 4255.408 | 4062.153 |
| ENSMUSG00000029776 | -0.399905756 | 0.000411 | 0.003942 | 2141.89 | 1923.314 | 2076.891 | 2204.003 | 2680.581 | 2305.143 | 2773.296 | 3255.16 |
| ENSMUSG00000029777 | -0.257434804 | 0.001493 | 0.010524 | 2379.319 | 2416.441 | 2606.052 | 2348.924 | 2799.545 | 2639.238 | 3035.582 | 3182.292 |
| ENSMUSG00000029778 | 0.406590136 | 0.119629 | 0.248522 | 210.2654 | 119.9822 | 139.1219 | 219.1064 | 105.312 | 172.0756 | 111.8575 | 131.5985 |
| ENSMUSG00000029780 | -0.17303033 | 0.02321 | 0.078544 | 823.9585 | 807.4799 | 801.6072 | 794.4762 | 910.754 | 987.7587 | 837.9673 | 903.7882 |
| ENSMUSG00000029781 | 0.012912752 | 0.862748 | 0.917798 | 5454.827 | 6681.806 | 6279.533 | 5976.256 | 6022.287 | 5866.214 | 6115.522 | 6167.729 |
| ENSMUSG00000029782 | 0.018679468 | 0.757393 | 0.848813 | 1356.161 | 1352.199 | 1407.781 | 1270.644 | 1329.077 | 1356.492 | 1316.255 | 1315.985 |
| ENSMUSG00000029787 | 0.105818028 | 0.335431 | 0.503669 | 1310.889 | 1130.232 | 1083.992 | 1228.376 | 1034.593 | 1290.567 | 1082.897 | 1010.372 |
| ENSMUSG00000029788 | -1.198163163 | 0.055476 | 0.146683 | 7.04238 | 7.198929 | 2.48432 | 8.626235 | 23.40267 | 7.821619 | 16.39292 | 9.78832 |
| ENSMUSG00000029790 | 0.153232792 | 0.276833 | 0.442894 | 348.0948 | 433.1356 | 339.5237 | 454.6026 | 319.8365 | 334.0949 | 379.9299 | 381.7445 |
| ENSMUSG00000029797 | 0.194088762 | 0.626781 | 0.756762 | 30.18163 | 14.39786 | 24.8432 | 31.05445 | 14.62667 | 23.46486 | 20.25007 | 30.45255 |
| ENSMUSG00000029798 | 0.406685864 | 0.144482 | 0.282595 | 114.6902 | 194.3711 | 171.4181 | 85.39973 | 115.0631 | 101.681 | 118.6076 | 90.27006 |
| ENSMUSG00000029802 | -0.683442716 | 0.000197 | 0.002239 | 1405.458 | 1049.844 | 1300.127 | 1411.252 | 2216.428 | 1423.535 | 2047.186 | 2612.394 |
| ENSMUSG00000029804 | -0.147855988 | 0.288459 | 0.455376 | 626.7719 | 539.9197 | 582.9871 | 646.105 | 571.4152 | 567.626 | 726.1097 | 790.6787 |
| ENSMUSG00000029810 | -0.355166137 | 0.016052 | 0.060359 | 4694.25 | 4230.571 | 5592.204 | 5164.527 | 6124.674 | 4933.207 | 6348.88 | 7771.926 |
| ENSMUSG00000029811 | -1.621771111 | 0.036959 | 0.1105 | 36.21796 | 39.59411 | 235.1823 | 45.71905 | 289.6081 | 93.85942 | 277.7153 | 438.2992 |
| ENSMUSG00000029814 | 0.183584422 | 0.083011 | 0.193881 | 594.5781 | 505.1249 | 640.1264 | 591.7597 | 471.9539 | 522.9311 | 508.1804 | 554.6715 |

| | | | | | | | | | | | |
|---|---|---|---|---|---|---|---|---|---|---|---|
| ENSMUSG00000029815 | -0.055414523 | 0.549947 | 0.697988 | 538.2391 | 537.52 | 556.4877 | 564.1558 | 535.3361 | 644.7249 | 569.8949 | 535.0948 |
| ENSMUSG00000029816 | 0.279702857 | 0.362928 | 0.531351 | 40.24217 | 33.595 | 35.60859 | 33.64232 | 39.97956 | 20.11273 | 27.96439 | 29.36496 |
| ENSMUSG00000029817 | 0.312204797 | 0.007302 | 0.033914 | 4456.821 | 4402.145 | 3322.364 | 4558.965 | 3310.503 | 3716.386 | 3199.511 | 3255.16 |
| ENSMUSG00000029819 | 2.446354074 | 2.68E-07 | 1.26E-05 | 822.9525 | 148.7779 | 206.1986 | 669.3958 | 103.3618 | 75.98144 | 87.75031 | 71.78101 |
| ENSMUSG00000029821 | 0.632040069 | 0.023957 | 0.080492 | 1397.41 | 1518.974 | 1170.943 | 1626.045 | 682.5779 | 1619.075 | 721.2883 | 664.5182 |
| ENSMUSG00000029822 | 0.107745167 | 0.208839 | 0.366165 | 1254.55 | 1023.448 | 1089.788 | 1152.465 | 1012.166 | 1052.566 | 1026.968 | 1106.08 |
| ENSMUSG00000029823 | 0.294248878 | 0.005885 | 0.028998 | 7801.952 | 7666.86 | 6952.783 | 7657.509 | 5955.98 | 7402.603 | 5632.413 | 5539.101 |
| ENSMUSG00000029826 | 0.48474748 | 0.025702 | 0.084646 | 441.6579 | 496.7261 | 683.188 | 313.1323 | 353.9654 | 329.6254 | 350.037 | 349.1167 |
| ENSMUSG00000029828 | 0.6386478 | 0.006027 | 0.02956 | 230.3864 | 215.9679 | 205.3704 | 240.672 | 163.8187 | 200.01 | 101.2504 | 108.7591 |
| ENSMUSG00000029829 | -2.220610276 | 0.000118 | 0.001515 | 161.9748 | 76.78858 | 392.5225 | 136.2945 | 700.1299 | 460.3581 | 1089.647 | 1331.211 |
| ENSMUSG00000029830 | -0.031669136 | 0.851038 | 0.910569 | 489.9485 | 307.1543 | 362.7107 | 469.2672 | 419.2979 | 385.4941 | 465.7517 | 395.8832 |
| ENSMUSG00000029832 | 0.665650466 | 3.96E-07 | 1.74E-05 | 399.4036 | 406.7395 | 360.2264 | 464.0914 | 257.4294 | 293.8694 | 250.7152 | 226.2189 |
| ENSMUSG00000029833 | 0.324271789 | 0.002602 | 0.015881 | 2982.951 | 2973.158 | 2577.068 | 3028.671 | 2227.154 | 2671.641 | 2349.008 | 1985.941 |
| ENSMUSG00000029836 | 0.085928468 | 0.302141 | 0.470457 | 4737.51 | 5396.797 | 4536.368 | 5054.111 | 4717.589 | 5070.644 | 4389.444 | 4403.656 |
| ENSMUSG00000029838 | 0.25620477 | 0.142856 | 0.280305 | 12317.12 | 18949.98 | 13961.88 | 15087.28 | 13699.34 | 15778.44 | 11016.04 | 10006.93 |
| ENSMUSG00000029840 | -0.04558836 | 0.335909 | 0.504269 | 4351.185 | 4637.31 | 4272.202 | 4408.869 | 4452.358 | 4587.938 | 4604.481 | 4585.284 |
| ENSMUSG00000029843 | 0.191211658 | 0.766983 | 0.855469 | 18.10898 | 13.19804 | 7.45296 | 6.038364 | 7.80089 | 10.05637 | 3.857157 | 17.40146 |
| ENSMUSG00000029844 | 0.695464993 | 0.055509 | 0.146732 | 23.13925 | 43.19358 | 24.8432 | 43.13117 | 20.47734 | 23.46486 | 20.25007 | 18.48905 |
| ENSMUSG00000029847 | 0.472360022 | 0.154773 | 0.296408 | 27.16347 | 34.79482 | 28.98373 | 26.74133 | 28.27823 | 20.11273 | 16.39292 | 19.57664 |
| ENSMUSG00000029851 | 0.112858192 | 0.447368 | 0.609996 | 269.6226 | 268.76 | 327.1021 | 317.4454 | 307.1601 | 219.0053 | 270.9653 | 298 |
| ENSMUSG00000029859 | -0.328556913 | 0.015073 | 0.057883 | 313.889 | 417.5379 | 445.5214 | 362.3019 | 516.809 | 440.2454 | 504.3232 | 470.9269 |
| ENSMUSG00000029860 | -0.114510404 | 0.180351 | 0.330477 | 1769.65 | 1875.321 | 2177.92 | 2038.379 | 2236.905 | 2177.762 | 2060.686 | 2040.321 |
| ENSMUSG00000029861 | 0.805176696 | 0.037091 | 0.110672 | 56.33904 | 75.58876 | 56.31125 | 98.33908 | 48.75557 | 64.8077 | 28.92867 | 21.75182 |
| ENSMUSG00000029862 | 0.580011432 | 0.082658 | 0.193291 | 33.19979 | 33.595 | 32.29616 | 23.29083 | 19.50223 | 25.6996 | 21.21436 | 15.22628 |
| ENSMUSG00000029863 | 0.108328363 | 0.213972 | 0.37205 | 1737.456 | 2027.698 | 1950.191 | 2003.012 | 1967.775 | 1774.39 | 1786.828 | 1629.211 |
| ENSMUSG00000029864 | -0.346733017 | 0.005189 | 0.026403 | 307.8526 | 333.5504 | 313.0243 | 300.193 | 412.4721 | 325.1559 | 443.573 | 411.1094 |
| ENSMUSG00000029866 | -0.18295063 | 0.648642 | 0.772344 | 102.6175 | 45.59322 | 35.60859 | 32.77969 | 51.6809 | 53.63396 | 71.3574 | 68.51824 |
| ENSMUSG00000029868 | -0.164413092 | 0.339759 | 0.508231 | 219.3198 | 203.9697 | 222.7607 | 169.0742 | 203.7983 | 186.6015 | 250.7152 | 271.8978 |
| ENSMUSG00000029869 | 0.343856421 | 0.004138 | 0.022371 | 619.7295 | 593.9117 | 651.7199 | 749.6198 | 520.7094 | 576.565 | 521.6804 | 443.7372 |
| ENSMUSG00000029875 | 0.050583776 | 0.865875 | 0.919615 | 40.24217 | 63.59054 | 52.99882 | 58.6584 | 75.08357 | 36.87335 | 45.32159 | 50.02919 |
| ENSMUSG00000029910 | -0.132661812 | 0.049622 | 0.135397 | 1813.916 | 1895.718 | 1653.729 | 1845.152 | 2050.659 | 2007.921 | 1925.685 | 1913.073 |
| ENSMUSG00000029911 | 0.083964508 | 0.331245 | 0.499089 | 2179.114 | 2228.069 | 1996.565 | 2227.294 | 2091.614 | 2272.739 | 1899.65 | 1878.27 |
| ENSMUSG00000029913 | 0.004299535 | 0.970506 | 0.983563 | 653.9353 | 699.496 | 573.0498 | 711.6644 | 673.8019 | 702.8283 | 693.3239 | 557.9342 |
| ENSMUSG00000029915 | 0.004305948 | 0.994475 | 0.996743 | 6.036326 | 14.39786 | 19.87456 | 5.175741 | 7.80089 | 14.52586 | 15.42863 | 7.613138 |
| ENSMUSG00000029916 | 0.077430232 | 0.378813 | 0.546971 | 786.7345 | 724.6922 | 681.5318 | 720.2906 | 691.3539 | 738.5843 | 629.6808 | 701.4963 |
| ENSMUSG00000029918 | -0.108517058 | 0.33397 | 0.502097 | 789.7527 | 916.6637 | 923.3389 | 841.0579 | 1087.249 | 817.9178 | 945.9677 | 888.5619 |
| ENSMUSG00000029919 | 0.058907689 | 0.85786 | 0.914865 | 67.40564 | 69.58965 | 149.8873 | 63.83414 | 99.46135 | 84.92043 | 66.53595 | 87.00729 |
| ENSMUSG00000029920 | 0.176299511 | 0.046384 | 0.129251 | 1392.379 | 1503.376 | 1223.942 | 1327.578 | 1222.79 | 1288.332 | 1169.683 | 1135.445 |
| ENSMUSG00000029922 | 0.046983253 | 0.462872 | 0.623888 | 2308.895 | 2177.676 | 2254.106 | 2496.432 | 2244.706 | 2286.147 | 2253.544 | 2161.043 |
| ENSMUSG00000029923 | 0.290810564 | 0.251587 | 0.41521 | 67.40564 | 94.7859 | 101.8571 | 81.94923 | 84.83468 | 58.10345 | 53.0359 | 87.00729 |
| ENSMUSG00000029924 | 0.040393693 | 0.569257 | 0.713058 | 1996.012 | 2216.07 | 1976.691 | 2049.593 | 2068.211 | 1898.419 | 1918.935 | 2120.803 |
| ENSMUSG00000029925 | -0.275721564 | 0.428531 | 0.59347 | 67.40564 | 93.58608 | 183.0116 | 84.5371 | 112.1378 | 113.9722 | 106.0718 | 188.1533 |
| ENSMUSG00000029992 | 0.047201007 | 0.567134 | 0.711413 | 1963.818 | 1894.518 | 1643.792 | 1801.158 | 1761.051 | 1710.7 | 1685.577 | 1907.635 |
| ENSMUSG00000029993 | -0.079181496 | 0.246034 | 0.408918 | 1269.641 | 1347.4 | 1209.036 | 1276.683 | 1344.678 | 1310.68 | 1296.969 | 1434.533 |
| ENSMUSG00000029994 | -0.228241916 | 0.124785 | 0.25507 | 1180.102 | 1195.022 | 1623.089 | 1326.715 | 1412.936 | 1303.976 | 1710.649 | 1813.014 |
| ENSMUSG00000029998 | -0.142923373 | 0.146971 | 0.286007 | 3341.107 | 3156.73 | 3592.327 | 3309.024 | 3558.181 | 3162.169 | 3985.407 | 4090.43 |
| ENSMUSG00000029999 | 0.316973282 | 0.009248 | 0.040236 | 688.1412 | 585.5129 | 595.4087 | 602.1112 | 458.3023 | 518.4616 | 576.6449 | 429.5985 |
| ENSMUSG00000030000 | 0.5679396 | 0.04721 | 0.130728 | 226.3622 | 93.58608 | 124.216 | 190.6398 | 94.5858 | 108.3853 | 103.1789 | 122.8978 |
| ENSMUSG00000030002 | 0.197560766 | 0.012924 | 0.051818 | 3263.64 | 3013.952 | 2647.457 | 3020.907 | 2538.215 | 2685.05 | 2671.081 | 2519.949 |
| ENSMUSG00000030004 | -5.380759626 | 5.21E-24 | 1.07E-20 | 8.048435 | 1.199822 | 8.281066 | 5.175741 | 290.5832 | 87.15518 | 218.8936 | 373.0437 |
| ENSMUSG00000030007 | -0.237578978 | 7.66E-05 | 0.001108 | 6740.564 | 6828.184 | 7102.671 | 6831.115 | 8246.516 | 7409.308 | 8342.066 | 8427.743 |
| ENSMUSG00000030008 | -0.642076281 | 5.99E-08 | 3.75E-06 | 201.2109 | 235.165 | 254.2287 | 235.4962 | 347.1396 | 347.5033 | 353.8941 | 399.1459 |
| ENSMUSG00000030016 | 0.261506168 | 0.007046 | 0.033083 | 2181.126 | 2324.054 | 1778.773 | 2107.389 | 1660.615 | 1843.667 | 1715.47 | 1777.124 |
| ENSMUSG00000030019 | 0.268556832 | 0.044647 | 0.125859 | 1458.779 | 1536.971 | 1183.364 | 1596.716 | 1118.453 | 1436.943 | 1070.361 | 1169.16 |
| ENSMUSG00000030020 | 0.367957598 | 0.011574 | 0.04765 | 598.6023 | 777.4844 | 737.843 | 678.0221 | 433.9245 | 646.9596 | 525.5376 | 557.9342 |
| ENSMUSG00000030022 | -0.002523588 | 0.981505 | 0.989764 | 2168.047 | 1811.731 | 1862.412 | 1971.957 | 1640.137 | 2071.612 | 2162.901 | 1954.401 |
| ENSMUSG00000030029 | 0.207864312 | 0.163369 | 0.308107 | 1111.69 | 1286.209 | 1122.913 | 1242.178 | 1032.643 | 1334.145 | 902.5747 | 854.8466 |
| ENSMUSG00000030030 | 0.215599646 | 0.470123 | 0.630173 | 60.36326 | 74.38894 | 57.96746 | 49.16954 | 60.4569 | 30.1691 | 49.17875 | 67.43065 |
| ENSMUSG00000030031 | -0.551078451 | 0.106102 | 0.230107 | 57.3451 | 22.79661 | 27.32752 | 35.36756 | 54.60623 | 61.45558 | 51.10733 | 42.41605 |
| ENSMUSG00000030032 | 0.214461897 | 0.214922 | 0.373081 | 178.0716 | 175.1739 | 134.1533 | 157.8601 | 165.7689 | 139.6718 | 119.5719 | 129.4233 |
| ENSMUSG00000030034 | -0.058233848 | 0.518968 | 0.672618 | 1179.096 | 1222.618 | 1238.019 | 1359.495 | 1424.638 | 1194.473 | 1366.398 | 1219.19 |
| ENSMUSG00000030035 | -0.27096245 | 0.003005 | 0.017786 | 1714.317 | 2098.488 | 2066.126 | 1931.414 | 2498.235 | 2216.87 | 2221.722 | 2486.233 |
| ENSMUSG00000030036 | -0.238408199 | 0.001896 | 0.012638 | 1340.064 | 1355.798 | 1242.988 | 1218.024 | 1529.95 | 1386.661 | 1564.077 | 1596.584 |
| ENSMUSG00000030037 | 0.104694663 | 0.666554 | 0.785864 | 51.30877 | 71.98929 | 62.108 | 63.83414 | 59.48179 | 73.74669 | 44.3573 | 54.37955 |

| | | | | | | | | | | | |
|---|---|---|---|---|---|---|---|---|---|---|---|
| ENSMUSG00000030042 | 0.028475686 | 0.733124 | 0.832956 | 1123.763 | 1330.602 | 1217.317 | 1270.644 | 1162.333 | 1301.741 | 1236.219 | 1144.146 |
| ENSMUSG00000030043 | 1.912931415 | 8.31E-10 | 1.10E-07 | 61.36932 | 67.19001 | 57.13936 | 56.93315 | 14.62667 | 24.58223 | 13.50005 | 11.9635 |
| ENSMUSG00000030045 | -0.197267937 | 0.045552 | 0.127594 | 813.898 | 771.4852 | 667.4539 | 703.9008 | 847.3717 | 767.636 | 860.1459 | 910.3137 |
| ENSMUSG00000030047 | -0.172379394 | 0.371446 | 0.540434 | 135.8173 | 105.5843 | 162.3089 | 107.8279 | 152.1174 | 124.0285 | 149.4648 | 152.2628 |
| ENSMUSG00000030051 | 0.004548562 | 0.959894 | 0.977262 | 482.9061 | 489.5272 | 453.8024 | 507.2226 | 491.4561 | 510.64 | 440.6801 | 485.0656 |
| ENSMUSG00000030055 | -0.255112906 | 0.233657 | 0.394256 | 117.7084 | 101.9848 | 137.4657 | 87.9876 | 119.9387 | 106.1505 | 153.322 | 152.2628 |
| ENSMUSG00000030056 | -0.046006117 | 0.526089 | 0.679403 | 1141.872 | 1262.212 | 1124.569 | 1198.184 | 1267.645 | 1182.182 | 1268.04 | 1156.109 |
| ENSMUSG00000030057 | -0.11338572 | 0.043365 | 0.1233 | 12061.59 | 13188.44 | 12255.15 | 13006.64 | 14192.75 | 13968.29 | 13425.8 | 13051.09 |
| ENSMUSG00000030058 | -0.04693594 | 0.452259 | 0.614691 | 4897.473 | 5189.228 | 5101.965 | 4812.576 | 5182.717 | 4786.831 | 5198.483 | 5491.247 |
| ENSMUSG00000030059 | -0.031271458 | 0.676307 | 0.792754 | 1319.943 | 1130.232 | 1132.022 | 1202.497 | 1216.939 | 1238.05 | 1196.683 | 1238.766 |
| ENSMUSG00000030060 | -0.122630582 | 0.080098 | 0.189063 | 761.5831 | 695.8965 | 781.7327 | 747.0319 | 804.4668 | 810.0962 | 823.5029 | 817.8685 |
| ENSMUSG00000030061 | -0.062722652 | 0.395249 | 0.56256 | 2012.109 | 1982.105 | 1748.961 | 1907.261 | 2050.659 | 1850.372 | 2027.9 | 2055.547 |
| ENSMUSG00000030062 | -0.316597916 | 0.003839 | 0.021201 | 4404.506 | 5000.856 | 5442.317 | 4703.886 | 5989.134 | 5167.855 | 6299.701 | 6893.152 |
| ENSMUSG00000030064 | -0.260680095 | 0.000812 | 0.00665 | 806.8556 | 827.8769 | 776.764 | 783.2621 | 913.6793 | 978.8197 | 896.7889 | 1037.562 |
| ENSMUSG00000030067 | 0.424434629 | 0.052069 | 0.139923 | 2039.272 | 2097.288 | 1649.588 | 1990.935 | 1213.038 | 2241.452 | 1238.147 | 1102.817 |
| ENSMUSG00000030068 | -0.183010258 | 0.696324 | 0.808286 | 26.15741 | 8.398751 | 13.24971 | 11.21411 | 15.60178 | 17.87799 | 17.3572 | 16.31387 |
| ENSMUSG00000030074 | 0.44662159 | 0.130062 | 0.262456 | 1148.914 | 1394.193 | 881.9336 | 1086.043 | 630.897 | 1469.347 | 642.2166 | 567.7225 |
| ENSMUSG00000030075 | 0.643393527 | 0.007907 | 0.035924 | 398.3975 | 514.7234 | 373.4761 | 456.3278 | 305.2098 | 398.9026 | 235.2866 | 176.1898 |
| ENSMUSG00000030077 | 1.195972264 | 0.009451 | 0.040887 | 38.23007 | 20.39697 | 20.70267 | 55.2079 | 11.70134 | 17.87799 | 16.39292 | 13.05109 |
| ENSMUSG00000030079 | -0.244058907 | 0.000105 | 0.001395 | 1802.849 | 1895.718 | 1813.554 | 1795.119 | 2246.656 | 1994.513 | 2229.437 | 2177.357 |
| ENSMUSG00000030082 | -0.140311838 | 0.02938 | 0.09315 | 4605.717 | 4835.281 | 5102.793 | 4746.154 | 5431.37 | 4956.671 | 5239.947 | 5633.722 |
| ENSMUSG00000030083 | 0.118272375 | 0.204457 | 0.360476 | 1580.511 | 1534.572 | 1363.892 | 1472.498 | 1449.991 | 1334.145 | 1485.97 | 1208.314 |
| ENSMUSG00000030084 | 0.062170147 | 0.39323 | 0.560681 | 1726.389 | 1678.55 | 1777.945 | 1605.342 | 1571.879 | 1641.423 | 1528.398 | 1762.985 |
| ENSMUSG00000030086 | -0.522646808 | 9.28E-08 | 5.30E-06 | 495.9848 | 587.9126 | 602.8616 | 541.7276 | 844.4464 | 726.2932 | 813.8601 | 815.6933 |
| ENSMUSG00000030087 | -0.974181119 | 0.005515 | 0.027606 | 403.4278 | 194.3711 | 260.8536 | 492.558 | 521.6845 | 612.321 | 1112.79 | 408.9342 |
| ENSMUSG00000030088 | -2.06857743 | 0.000797 | 0.006558 | 458.7608 | 91.18644 | 407.4285 | 269.1385 | 1173.059 | 442.4801 | 1617.113 | 1914.16 |
| ENSMUSG00000030089 | 0.184831102 | 0.110042 | 0.235853 | 233.4046 | 223.1668 | 248.432 | 246.7103 | 190.1467 | 217.8879 | 213.1079 | 218.6058 |
| ENSMUSG00000030091 | 0.065873142 | 0.325184 | 0.493053 | 1523.166 | 1420.589 | 1541.106 | 1458.696 | 1365.156 | 1353.14 | 1493.684 | 1468.248 |
| ENSMUSG00000030092 | 0.073766306 | 0.895119 | 0.936579 | 16.09687 | 16.7975 | 57.96746 | 19.84034 | 14.62667 | 26.81698 | 21.21436 | 43.50364 |
| ENSMUSG00000030094 | -0.249642036 | 0.019493 | 0.069213 | 1310.889 | 1235.816 | 1405.297 | 1288.76 | 1670.366 | 1277.159 | 1674.97 | 1608.547 |
| ENSMUSG00000030095 | 0.093151535 | 0.11874 | 0.247558 | 1323.968 | 1400.192 | 1359.751 | 1312.913 | 1315.425 | 1217.938 | 1271.897 | 1249.642 |
| ENSMUSG00000030096 | 0.130375827 | 0.211268 | 0.368646 | 5621.832 | 4356.552 | 4731.801 | 4843.631 | 4126.671 | 4551.065 | 5007.554 | 4179.613 |
| ENSMUSG00000030098 | 0.398039779 | 0.233012 | 0.3936 | 46.2785 | 27.5959 | 42.23344 | 64.69676 | 27.30312 | 42.46022 | 32.78583 | 35.89051 |
| ENSMUSG00000030101 | -0.068324112 | 0.405782 | 0.572519 | 1095.593 | 1143.43 | 1141.959 | 1180.069 | 1177.934 | 1069.327 | 1225.612 | 1309.46 |
| ENSMUSG00000030102 | -0.175377493 | 0.219056 | 0.377841 | 2307.889 | 1944.911 | 2472.726 | 2128.092 | 2249.582 | 2079.433 | 2537.045 | 3134.438 |
| ENSMUSG00000030103 | -0.071559637 | 0.586211 | 0.726603 | 955.7516 | 793.082 | 955.6351 | 847.0963 | 897.1024 | 764.2839 | 1103.147 | 967.9561 |
| ENSMUSG00000030104 | 0.020681425 | 0.795979 | 0.874596 | 1093.581 | 1065.442 | 914.2297 | 1007.544 | 979.0118 | 1015.693 | 1004.789 | 1019.073 |
| ENSMUSG00000030105 | 0.030666737 | 0.622201 | 0.753747 | 2274.689 | 2321.655 | 2147.28 | 2126.367 | 2163.772 | 2297.321 | 2132.043 | 2086 |
| ENSMUSG00000030107 | 0.471993754 | 0.427209 | 0.59233 | 68.4117 | 194.3711 | 203.7142 | 18.97772 | 74.10846 | 56.98608 | 128.2505 | 90.27006 |
| ENSMUSG00000030108 | -2.570414224 | 7.44E-06 | 0.000184 | 131.7931 | 53.99197 | 219.4483 | 110.4158 | 692.329 | 291.6346 | 738.6455 | 1344.263 |
| ENSMUSG00000030109 | -3.373332539 | 1.37E-12 | 3.88E-10 | 25.15136 | 4.799286 | 33.95237 | 8.626235 | 210.624 | 126.2633 | 198.6436 | 225.1314 |
| ENSMUSG00000030110 | 0.818457736 | 0.000401 | 0.003866 | 4017.175 | 4967.261 | 3801.009 | 4266.536 | 2340.267 | 3711.917 | 2058.757 | 1558.518 |
| ENSMUSG00000030111 | 0.653640876 | 0.284404 | 0.451019 | 112.6781 | 15.59768 | 26.49941 | 70.73513 | 17.552 | 56.98608 | 37.60728 | 31.54014 |
| ENSMUSG00000030116 | 1.167913664 | 1.31E-06 | 4.64E-05 | 1191.168 | 1762.538 | 1416.89 | 1649.336 | 723.5326 | 992.2282 | 508.1804 | 455.7007 |
| ENSMUSG00000030120 | -0.293498792 | 0.007881 | 0.035842 | 4646.965 | 3917.417 | 4634.085 | 4435.61 | 5411.868 | 4515.309 | 5497.413 | 6190.568 |
| ENSMUSG00000030122 | 0.042485179 | 0.436447 | 0.600273 | 12789.97 | 13194.44 | 12519.32 | 12489.06 | 11785.2 | 12107.87 | 12400.76 | 13216.41 |
| ENSMUSG00000030123 | -0.207014825 | 0.134083 | 0.267701 | 3510.124 | 2943.162 | 4293.733 | 3746.374 | 4002.832 | 3601.297 | 4228.408 | 4901.773 |
| ENSMUSG00000030124 | 0.459736382 | 0.061625 | 0.157923 | 150.9082 | 112.7832 | 131.669 | 148.3712 | 116.0382 | 132.9675 | 73.28598 | 73.95619 |
| ENSMUSG00000030125 | 0.416095881 | 0.250359 | 0.413922 | 54.32694 | 26.39607 | 28.15563 | 35.36756 | 35.10401 | 23.46486 | 21.21436 | 28.27737 |
| ENSMUSG00000030126 | 0.289389447 | 0.03861 | 0.113596 | 1151.932 | 1341.4 | 1090.616 | 1253.392 | 890.2766 | 1260.398 | 883.2889 | 924.4524 |
| ENSMUSG00000030127 | -0.256957729 | 0.02504 | 0.083164 | 1979.915 | 1974.906 | 2162.186 | 2056.494 | 2500.185 | 2013.508 | 2360.58 | 2894.08 |
| ENSMUSG00000030134 | -0.362579896 | 0.24895 | 0.412369 | 64.38748 | 45.59322 | 104.3414 | 46.58167 | 88.73513 | 69.27719 | 99.32178 | 79.39415 |
| ENSMUSG00000030137 | -0.332858957 | 0.544123 | 0.692833 | 14.08476 | 7.198929 | 9.109173 | 6.900988 | 11.70134 | 5.58687 | 16.39292 | 13.05109 |
| ENSMUSG00000030138 | -0.094645234 | 0.075627 | 0.181937 | 2084.545 | 2190.874 | 2176.264 | 2081.51 | 2371.471 | 2282.795 | 2207.258 | 2248.051 |
| ENSMUSG00000030144 | 1.420885469 | 0.029392 | 0.093173 | 61.36932 | 5.999108 | 27.32752 | 31.91707 | 13.65156 | 6.704245 | 9.642892 | 17.40146 |
| ENSMUSG00000030145 | 0.175117227 | 0.224228 | 0.383974 | 316.9071 | 302.355 | 241.8071 | 281.2153 | 268.1556 | 288.2825 | 226.608 | 227.3065 |
| ENSMUSG00000030148 | 0.408328883 | 0.183062 | 0.33392 | 36.21796 | 52.79215 | 66.24853 | 39.68068 | 31.20356 | 33.52122 | 46.28588 | 35.89051 |
| ENSMUSG00000030157 | 0.230912168 | 0.156983 | 0.299425 | 615.7053 | 571.1151 | 756.8895 | 565.881 | 456.3521 | 531.8701 | 678.8596 | 472.0145 |
| ENSMUSG00000030159 | 0.105754623 | 0.824281 | 0.893455 | 71.42986 | 19.19714 | 32.29616 | 44.85642 | 18.52711 | 50.28183 | 36.64299 | 51.11678 |
| ENSMUSG00000030160 | -2.1323115 | 6.02E-13 | 1.91E-10 | 516.1059 | 277.1588 | 593.7525 | 428.7239 | 1553.352 | 1168.773 | 2448.33 | 2796.197 |
| ENSMUSG00000030161 | -0.210689243 | 0.039983 | 0.116688 | 3934.679 | 3712.248 | 4153.783 | 4188.9 | 4505.989 | 3978.969 | 5384.591 | 4635.313 |
| ENSMUSG00000030166 | 0.037163033 | 0.84175 | 0.904584 | 525.1604 | 566.3158 | 357.7421 | 424.4108 | 431.9743 | 572.0955 | 436.823 | 382.8321 |
| ENSMUSG00000030168 | -0.277521957 | 0.011514 | 0.047468 | 1975.891 | 1875.321 | 1866.552 | 1899.497 | 2359.769 | 1853.724 | 2415.544 | 2602.605 |
| ENSMUSG00000030170 | 0.540133327 | 0.002435 | 0.015131 | 598.6023 | 890.2676 | 719.6247 | 659.907 | 576.2908 | 575.4477 | 401.1443 | 418.7226 |

| | | | | | | | | | | | |
|---|---|---|---|---|---|---|---|---|---|---|---|
| ENSMUSG00000030172 | 0.080086125 | 0.447383 | 0.609996 | 794.7829 | 729.4915 | 933.2762 | 788.4379 | 730.3584 | 830.209 | 797.4671 | 716.7225 |
| ENSMUSG00000030177 | 0.177257305 | 0.121242 | 0.250492 | 644.8808 | 682.6985 | 532.4726 | 604.6991 | 539.2366 | 556.4523 | 595.9307 | 483.978 |
| ENSMUSG00000030180 | 0.239269507 | 0.12864 | 0.260297 | 3455.797 | 2982.756 | 2557.193 | 2615.474 | 2261.283 | 3154.347 | 2394.33 | 2026.182 |
| ENSMUSG00000030188 | 0.115263595 | 0.271861 | 0.437137 | 456.7487 | 536.3202 | 468.7084 | 496.0085 | 502.1823 | 434.6585 | 457.0731 | 410.0218 |
| ENSMUSG00000030189 | 0.157313574 | 0.102203 | 0.224314 | 5489.033 | 6968.563 | 5718.076 | 6271.273 | 5679.048 | 5889.679 | 5256.34 | 5093.189 |
| ENSMUSG00000030199 | 0.0247166 | 0.860912 | 0.916743 | 1067.424 | 1019.848 | 1218.145 | 1145.564 | 1020.942 | 1402.304 | 1022.147 | 933.1532 |
| ENSMUSG00000030200 | -0.373765752 | 0.157215 | 0.299624 | 115.6963 | 100.785 | 204.5423 | 126.8057 | 197.9476 | 129.6154 | 174.5363 | 209.9051 |
| ENSMUSG00000030201 | 0.163278509 | 0.045543 | 0.127587 | 4191.222 | 4030.201 | 3629.591 | 3987.908 | 3292.951 | 3936.509 | 3422.262 | 3492.255 |
| ENSMUSG00000030203 | -0.030420567 | 0.710477 | 0.818711 | 878.2855 | 793.082 | 825.6223 | 810.0035 | 811.2926 | 783.2792 | 869.7888 | 913.5765 |
| ENSMUSG00000030204 | 0.074162893 | 0.282916 | 0.449379 | 2940.697 | 2794.384 | 2821.359 | 3139.087 | 2981.89 | 2639.238 | 2764.617 | 2724.416 |
| ENSMUSG00000030206 | -0.121391795 | 0.777119 | 0.862655 | 11.0666 | 14.39786 | 16.56213 | 13.80198 | 14.62667 | 20.11273 | 16.39292 | 9.78832 |
| ENSMUSG00000030207 | 0.199070141 | 0.017907 | 0.065417 | 1381.313 | 1397.792 | 1553.528 | 1311.188 | 1131.129 | 1240.285 | 1306.612 | 1239.854 |
| ENSMUSG00000030208 | 0.363727138 | 3.72E-06 | 0.000107 | 6855.254 | 7663.26 | 6633.134 | 8035.338 | 5470.374 | 5667.321 | 5712.449 | 5831.663 |
| ENSMUSG00000030209 | 1.75399373 | 5.63E-05 | 0.000866 | 45.27245 | 46.79304 | 48.03018 | 40.5433 | 10.72622 | 26.81698 | 6.750024 | 9.78832 |
| ENSMUSG00000030213 | 0.263432277 | 8.98E-06 | 0.000211 | 2997.036 | 2914.367 | 2948.06 | 2726.753 | 2444.604 | 2509.622 | 2403.009 | 2293.73 |
| ENSMUSG00000030214 | -0.339011702 | 0.264022 | 0.428237 | 112.6781 | 56.39161 | 80.32634 | 93.16334 | 84.83468 | 87.15518 | 98.35749 | 164.2263 |
| ENSMUSG00000030216 | -0.031159474 | 0.6429 | 0.76908 | 1925.588 | 1916.115 | 2017.268 | 2002.149 | 1957.048 | 1967.696 | 2195.686 | 1913.073 |
| ENSMUSG00000030217 | 0.231384645 | 0.194435 | 0.348161 | 486.9303 | 429.5361 | 650.8918 | 515.8488 | 526.5601 | 410.0763 | 490.8232 | 348.0291 |
| ENSMUSG00000030218 | 0.132600265 | 0.411143 | 0.576974 | 4936.709 | 4907.27 | 5723.045 | 5525.103 | 6178.305 | 4121.993 | 5264.055 | 3674.97 |
| ENSMUSG00000030219 | 1.136880713 | 0.034767 | 0.105537 | 24.1453 | 7.198929 | 21.53077 | 14.6646 | 7.80089 | 8.938993 | 7.714313 | 6.525547 |
| ENSMUSG00000030220 | 0.025990799 | 0.830615 | 0.897636 | 1796.813 | 1917.315 | 2316.214 | 1754.576 | 1859.537 | 1649.244 | 2052.007 | 2086 |
| ENSMUSG00000030222 | 0.049077168 | 0.854777 | 0.913001 | 1000.018 | 440.3345 | 650.8918 | 911.793 | 654.2997 | 567.626 | 1013.468 | 667.7809 |
| ENSMUSG00000030223 | -1.445003849 | 0.000819 | 0.006695 | 120.7265 | 62.39072 | 266.6503 | 147.5086 | 266.2054 | 233.5312 | 509.1447 | 621.0145 |
| ENSMUSG00000030224 | -0.125524258 | 0.016772 | 0.062231 | 4343.137 | 4572.52 | 4499.103 | 4506.345 | 5010.122 | 5120.925 | 4714.41 | 4704.919 |
| ENSMUSG00000030225 | -0.39785714 | 0.015815 | 0.059885 | 1045.29 | 1277.81 | 807.404 | 1140.388 | 1277.396 | 1175.478 | 1547.684 | 1622.686 |
| ENSMUSG00000030226 | 1.508075603 | 0.00045 | 0.004225 | 44.26639 | 46.79304 | 23.18699 | 52.62003 | 9.751113 | 23.46486 | 11.57147 | 14.13868 |
| ENSMUSG00000030228 | -1.264002827 | 0.060973 | 0.156731 | 2.012109 | 10.79839 | 11.59349 | 9.488858 | 15.60178 | 11.17374 | 12.53576 | 42.41605 |
| ENSMUSG00000030231 | 0.083054303 | 0.346735 | 0.515618 | 2546.324 | 2462.034 | 2979.528 | 2492.982 | 2457.28 | 2674.994 | 2318.151 | 2449.255 |
| ENSMUSG00000030232 | 0.149582712 | 0.036329 | 0.109187 | 2755.583 | 2632.408 | 2424.696 | 2728.478 | 2296.387 | 2556.552 | 2288.258 | 2362.248 |
| ENSMUSG00000030235 | 0.178979098 | 0.783383 | 0.866794 | 4.024217 | 17.99732 | 15.73403 | 20.70296 | 9.751113 | 25.6996 | 6.750024 | 9.78832 |
| ENSMUSG00000030237 | 1.192736343 | 0.001313 | 0.009599 | 528.1785 | 551.9179 | 401.6317 | 527.9256 | 163.8187 | 446.9496 | 150.4291 | 118.5474 |
| ENSMUSG00000030243 | -0.128387914 | 0.245754 | 0.408689 | 519.124 | 506.3247 | 576.3622 | 512.3984 | 565.5646 | 528.5179 | 542.8948 | 676.4817 |
| ENSMUSG00000030245 | 0.010015223 | 0.901151 | 0.940696 | 1104.648 | 1208.22 | 1085.648 | 1045.5 | 1139.905 | 1173.243 | 1032.754 | 1063.664 |
| ENSMUSG00000030246 | -0.769365297 | 4.92E-05 | 0.000787 | 7741.588 | 7364.505 | 10386.11 | 8606.395 | 13524.79 | 10006.09 | 15805.66 | 18788.14 |
| ENSMUSG00000030247 | -0.190327672 | 0.197257 | 0.351608 | 1152.938 | 941.8599 | 1064.945 | 1212.849 | 1205.238 | 1002.285 | 1555.398 | 1226.803 |
| ENSMUSG00000030249 | 0.006698478 | 0.954946 | 0.97432 | 1098.611 | 915.4638 | 1021.055 | 1255.117 | 1019.966 | 1027.984 | 1202.469 | 1022.336 |
| ENSMUSG00000030254 | 0.031890345 | 0.799139 | 0.877076 | 739.45 | 676.6993 | 588.7838 | 699.5877 | 712.8064 | 716.2368 | 674.0381 | 540.5328 |
| ENSMUSG00000030255 | 0.094920028 | 0.287611 | 0.454542 | 792.7708 | 938.2604 | 859.5747 | 869.5245 | 854.1975 | 840.2653 | 732.8598 | 811.343 |
| ENSMUSG00000030256 | -0.233035687 | 0.45676 | 0.61851 | 73.44197 | 73.18911 | 75.3577 | 75.04824 | 60.4569 | 148.6108 | 71.3574 | 69.60583 |
| ENSMUSG00000030257 | 0.598980001 | 3.14E-05 | 0.000556 | 806.8556 | 1004.251 | 1064.945 | 921.2819 | 508.033 | 750.8754 | 610.395 | 639.5036 |
| ENSMUSG00000030259 | 0.284291093 | 0.000105 | 0.001389 | 2113.72 | 2279.661 | 2282.262 | 2189.338 | 1807.856 | 1962.109 | 1857.221 | 1652.051 |
| ENSMUSG00000030263 | 0.064011429 | 0.816915 | 0.888921 | 33.19979 | 47.99286 | 38.92101 | 33.64232 | 38.02934 | 29.05173 | 36.64299 | 42.41605 |
| ENSMUSG00000030264 | -0.206220596 | 0.000753 | 0.006287 | 851.122 | 843.4745 | 816.5131 | 822.9428 | 972.186 | 943.0637 | 974.8963 | 952.7298 |
| ENSMUSG00000030265 | -0.032239422 | 0.64672 | 0.77119 | 1956.776 | 1991.704 | 1912.926 | 1916.749 | 1861.487 | 2140.889 | 1874.578 | 2078.387 |
| ENSMUSG00000030268 | 0.520858964 | 0.023732 | 0.079984 | 569.4268 | 607.1097 | 484.4424 | 556.3922 | 376.393 | 511.7573 | 435.8587 | 220.781 |
| ENSMUSG00000030269 | -0.144842186 | 0.194595 | 0.34838 | 1085.533 | 1023.448 | 962.2599 | 1154.19 | 1249.118 | 1125.196 | 1302.755 | 992.9707 |
| ENSMUSG00000030271 | 0.011472447 | 0.937566 | 0.962923 | 341.0524 | 501.5254 | 436.4122 | 487.3823 | 430.0241 | 492.762 | 400.18 | 429.5985 |
| ENSMUSG00000030272 | 0.350147899 | 4.47E-06 | 0.000124 | 2138.872 | 2193.274 | 2445.399 | 2146.207 | 1798.105 | 1752.043 | 1619.042 | 1834.766 |
| ENSMUSG00000030275 | -0.061690035 | 0.59781 | 0.736533 | 5438.73 | 4603.715 | 4829.518 | 4833.279 | 4173.476 | 5013.658 | 5875.414 | 5504.298 |
| ENSMUSG00000030276 | 0.46408579 | 0.043113 | 0.122982 | 125.7568 | 160.7761 | 105.9976 | 112.1411 | 117.9885 | 70.39457 | 92.57176 | 82.65692 |
| ENSMUSG00000030279 | 0.282198412 | 0.00249 | 0.015382 | 2178.108 | 2062.493 | 1772.148 | 1964.194 | 1615.759 | 1787.799 | 1661.47 | 1492.175 |
| ENSMUSG00000030281 | -0.466177593 | 0.000278 | 0.002921 | 306.8466 | 373.1445 | 391.6944 | 295.8799 | 492.4312 | 444.7149 | 504.3232 | 445.9123 |
| ENSMUSG00000030282 | -0.033216752 | 0.550539 | 0.698381 | 1641.881 | 1637.756 | 1539.45 | 1676.94 | 1638.187 | 1690.587 | 1658.577 | 1658.576 |
| ENSMUSG00000030283 | 0.581197872 | 0.000595 | 0.005238 | 586.5297 | 831.4763 | 643.4389 | 721.1532 | 527.5352 | 553.1002 | 383.7871 | 394.7956 |
| ENSMUSG00000030284 | 0.196889219 | 0.048388 | 0.133111 | 657.9595 | 692.297 | 658.3448 | 597.7981 | 577.2659 | 492.762 | 595.9307 | 604.7006 |
| ENSMUSG00000030286 | -0.080158112 | 0.328456 | 0.496233 | 2168.047 | 1941.311 | 2261.559 | 2076.335 | 2190.1 | 2058.203 | 2242.937 | 2442.73 |
| ENSMUSG00000030287 | -0.497554445 | 0.005055 | 0.025924 | 436.6276 | 371.9447 | 616.1113 | 453.74 | 543.137 | 645.8422 | 756.0027 | 711.2846 |
| ENSMUSG00000030291 | 0.007213814 | 0.936528 | 0.962308 | 834.0191 | 883.0686 | 898.4957 | 826.3933 | 851.2722 | 962.0591 | 840.8602 | 771.1021 |
| ENSMUSG00000030298 | -0.314900775 | 0.003654 | 0.020445 | 2162.011 | 2387.645 | 2550.568 | 2342.885 | 3012.119 | 2463.81 | 2907.332 | 3363.919 |
| ENSMUSG00000030301 | -0.323963062 | 0.009055 | 0.039688 | 582.5055 | 607.1097 | 639.2983 | 590.0345 | 727.433 | 615.6731 | 798.4314 | 886.3867 |
| ENSMUSG00000030302 | 0.140019915 | 0.750556 | 0.844651 | 57.3451 | 37.19447 | 43.88965 | 62.10889 | 29.25334 | 88.27255 | 24.10723 | 41.32846 |
| ENSMUSG00000030303 | 0.916260853 | 1.52E-05 | 0.000315 | 450.7124 | 575.9143 | 462.9116 | 588.3092 | 235.0018 | 401.1373 | 245.8937 | 219.6934 |
| ENSMUSG00000030304 | 0.084613154 | 0.137395 | 0.272519 | 2113.72 | 2036.097 | 2035.486 | 2131.543 | 1955.098 | 1966.578 | 2058.757 | 1861.956 |
| ENSMUSG00000030306 | 1.138671599 | 1.61E-05 | 0.000331 | 2258.592 | 1940.111 | 1348.158 | 2245.409 | 723.5326 | 1339.732 | 925.7176 | 550.3211 |

| | | | | | | | | | | | |
|---|---|---|---|---|---|---|---|---|---|---|---|
| ENSMUSG00000030307 | -0.105024344 | 0.690014 | 0.803616 | 35.2119 | 39.59411 | 61.27989 | 47.44429 | 48.75557 | 45.81234 | 53.0359 | 51.11678 |
| ENSMUSG00000030309 | 0.624847308 | 0.005162 | 0.026322 | 1109.678 | 1069.041 | 905.1205 | 1167.13 | 603.5939 | 1052.566 | 616.1808 | 485.0656 |
| ENSMUSG00000030310 | 0.437625082 | 0.016689 | 0.061987 | 116.7023 | 103.1847 | 99.3728 | 93.16334 | 73.13335 | 89.38993 | 72.32169 | 69.60583 |
| ENSMUSG00000030313 | 0.22594446 | 0.116449 | 0.244614 | 963.8001 | 955.0579 | 783.3889 | 840.1953 | 722.5575 | 954.2375 | 710.6811 | 640.5912 |
| ENSMUSG00000030314 | 0.221984001 | 0.015412 | 0.058792 | 1368.234 | 1282.609 | 1164.318 | 1186.97 | 958.5344 | 1130.783 | 1125.325 | 1072.365 |
| ENSMUSG00000030315 | -0.068595128 | 0.560478 | 0.706328 | 2016.133 | 1864.523 | 1849.99 | 1699.368 | 1942.422 | 1566.558 | 2110.829 | 2169.744 |
| ENSMUSG00000030316 | -0.24645591 | 0.007007 | 0.032971 | 539.2451 | 635.9054 | 631.0173 | 601.2486 | 746.9353 | 654.7812 | 735.7526 | 716.7225 |
| ENSMUSG00000030317 | 0.899957728 | 0.134005 | 0.267618 | 44.26639 | 22.79661 | 21.53077 | 83.67448 | 15.60178 | 41.34284 | 27.0001 | 8.700729 |
| ENSMUSG00000030319 | 0.226963115 | 0.025316 | 0.083681 | 1066.418 | 1249.014 | 1030.993 | 1092.944 | 892.2268 | 1055.919 | 865.9317 | 977.7444 |
| ENSMUSG00000030321 | 0.542044059 | 0.004984 | 0.025697 | 130.7871 | 137.9795 | 171.4181 | 121.6299 | 94.5858 | 118.4417 | 85.82174 | 88.09488 |
| ENSMUSG00000030322 | 0.43590439 | 0.004218 | 0.022693 | 652.9293 | 613.1088 | 448.0057 | 523.6125 | 449.5263 | 442.4801 | 412.7158 | 346.9416 |
| ENSMUSG00000030323 | -0.063374367 | 0.444914 | 0.607797 | 970.8425 | 955.0579 | 981.3064 | 976.4898 | 1133.079 | 941.9464 | 1030.825 | 950.5546 |
| ENSMUSG00000030324 | 0.156428773 | 0.344045 | 0.512751 | 1981.927 | 3329.505 | 2475.211 | 2750.044 | 2398.774 | 2688.402 | 1954.614 | 2411.189 |
| ENSMUSG00000030327 | 0.008014308 | 0.928859 | 0.957925 | 959.7759 | 843.4745 | 829.7628 | 761.6965 | 855.1726 | 842.5001 | 834.1101 | 841.7955 |
| ENSMUSG00000030329 | 2.045833463 | 1.28E-10 | 2.26E-08 | 215.2956 | 249.5629 | 240.1509 | 210.4801 | 71.18313 | 84.92043 | 38.57157 | 27.18978 |
| ENSMUSG00000030330 | 0.298671276 | 0.016602 | 0.061763 | 317.9132 | 293.9563 | 281.5563 | 317.4454 | 247.6783 | 283.813 | 239.1437 | 214.2554 |
| ENSMUSG00000030335 | -0.484932218 | 1.45E-05 | 0.000304 | 1351.131 | 1196.222 | 1292.674 | 1183.519 | 1811.757 | 1450.352 | 1770.435 | 1997.905 |
| ENSMUSG00000030337 | 0.179955897 | 0.049782 | 0.135713 | 533.2088 | 530.3211 | 469.5365 | 538.2771 | 438.8001 | 478.2361 | 436.823 | 474.1897 |
| ENSMUSG00000030339 | -0.337331497 | 0.003364 | 0.019253 | 1388.355 | 1485.379 | 1817.694 | 1482.85 | 1845.886 | 1735.282 | 2125.293 | 2096.876 |
| ENSMUSG00000030340 | -2.223041826 | 0.001066 | 0.008203 | 305.8405 | 75.58876 | 494.3797 | 169.9368 | 879.5504 | 449.1844 | 1942.078 | 1613.985 |
| ENSMUSG00000030341 | -0.106064583 | 0.226267 | 0.386128 | 1934.643 | 1643.756 | 1905.473 | 1778.73 | 2060.41 | 1769.921 | 1907.364 | 2081.649 |
| ENSMUSG00000030342 | -0.093036735 | 0.391642 | 0.559104 | 2536.263 | 2207.672 | 3070.619 | 2702.599 | 2828.798 | 2603.482 | 2895.76 | 2895.167 |
| ENSMUSG00000030346 | -0.056727498 | 0.631358 | 0.760118 | 529.1846 | 706.6949 | 581.3309 | 568.4689 | 675.7521 | 633.5511 | 604.6093 | 563.3722 |
| ENSMUSG00000030347 | -0.187817483 | 0.175129 | 0.323477 | 393.3673 | 357.5468 | 401.6317 | 355.4009 | 450.5014 | 331.8601 | 453.2159 | 481.8029 |
| ENSMUSG00000030350 | 0.054580652 | 0.936558 | 0.962308 | 5.030272 | 22.79661 | 10.76539 | 8.626235 | 12.67645 | 21.23011 | 5.785735 | 5.437955 |
| ENSMUSG00000030351 | 0.962539407 | 3.54E-06 | 0.000104 | 423.5489 | 662.3015 | 547.3785 | 605.5617 | 300.3343 | 384.3767 | 238.1794 | 226.2189 |
| ENSMUSG00000030352 | -0.035012445 | 0.768214 | 0.856029 | 2728.419 | 3195.125 | 3855.664 | 2883.75 | 3342.682 | 3054.901 | 3117.547 | 3460.715 |
| ENSMUSG00000030353 | -0.149108148 | 0.535664 | 0.686453 | 96.58122 | 82.78769 | 101.8571 | 73.323 | 104.3369 | 65.92507 | 124.3933 | 97.8832 |
| ENSMUSG00000030357 | -0.192782437 | 0.01876 | 0.067439 | 7011.193 | 8006.409 | 7830.576 | 7330.574 | 9062.684 | 7699.825 | 8494.423 | 9233.648 |
| ENSMUSG00000030359 | 1.28484339 | 0.092142 | 0.20843 | 118.7144 | 22.79661 | 11.59349 | 119.042 | 7.80089 | 39.10809 | 18.32149 | 46.76642 |
| ENSMUSG00000030364 | -2.585756858 | 0.00072 | 0.006093 | 8.048435 | 34.79482 | 8.281066 | 0.862623 | 115.0631 | 32.40385 | 55.92877 | 103.3212 |
| ENSMUSG00000030374 | -0.171428694 | 0.107702 | 0.232295 | 2301.852 | 2416.441 | 2897.545 | 2349.786 | 2942.886 | 2431.406 | 2890.939 | 2959.335 |
| ENSMUSG00000030376 | 0.189255755 | 0.217297 | 0.375825 | 122.7386 | 167.975 | 147.403 | 156.1349 | 135.5405 | 127.3806 | 131.1433 | 126.1606 |
| ENSMUSG00000030380 | 0.175773127 | 0.364306 | 0.532986 | 136.8234 | 159.5763 | 120.9036 | 165.6237 | 119.9387 | 162.0192 | 106.0718 | 128.3357 |
| ENSMUSG00000030386 | 0.061920606 | 0.533298 | 0.684646 | 871.2431 | 1065.442 | 871.9963 | 936.8091 | 819.0935 | 968.7633 | 884.2532 | 913.5765 |
| ENSMUSG00000030393 | 0.378188869 | 0.003217 | 0.018695 | 340.0464 | 385.1427 | 318.8211 | 326.9343 | 252.5538 | 310.63 | 245.8937 | 244.708 |
| ENSMUSG00000030397 | -0.263888402 | 0.014404 | 0.055978 | 680.0927 | 652.7029 | 706.375 | 727.1916 | 875.65 | 690.5372 | 840.8602 | 915.7517 |
| ENSMUSG00000030399 | 1.029538709 | 0.103197 | 0.225862 | 142.8597 | 100.785 | 108.482 | 74.18562 | 33.15378 | 130.7328 | 37.60728 | 7.613138 |
| ENSMUSG00000030400 | 0.093499939 | 0.340231 | 0.508746 | 724.3591 | 808.6797 | 682.3599 | 727.1916 | 755.7113 | 616.7905 | 705.8597 | 674.3065 |
| ENSMUSG00000030401 | -0.196258144 | 0.427375 | 0.592477 | 150.9082 | 92.38626 | 214.4796 | 154.4096 | 174.5449 | 150.8455 | 194.7864 | 183.8029 |
| ENSMUSG00000030402 | 0.713425587 | 0.217596 | 0.37606 | 8.048435 | 22.79661 | 8.281066 | 20.70296 | 10.72622 | 7.821619 | 5.785735 | 11.9635 |
| ENSMUSG00000030403 | -0.297645441 | 0.073582 | 0.178711 | 1902.449 | 1632.957 | 2428.837 | 1883.107 | 2486.534 | 1786.681 | 2595.866 | 2778.795 |
| ENSMUSG00000030406 | 0.523892485 | 0.358939 | 0.527337 | 12.07265 | 15.59768 | 11.59349 | 30.19182 | 8.776002 | 20.11273 | 5.785735 | 14.13868 |
| ENSMUSG00000030407 | -0.130642643 | 0.28483 | 0.451421 | 425.561 | 407.9393 | 423.1625 | 370.0655 | 480.7299 | 364.264 | 480.216 | 453.5255 |
| ENSMUSG00000030409 | 0.566373197 | 3.47E-05 | 0.0006 | 493.9727 | 621.5076 | 659.1729 | 615.9132 | 453.4268 | 432.4238 | 342.3227 | 387.1824 |
| ENSMUSG00000030410 | -0.135241922 | 0.014669 | 0.056722 | 1644.899 | 1592.163 | 1605.699 | 1648.473 | 1820.533 | 1707.348 | 1751.149 | 1851.08 |
| ENSMUSG00000030411 | -0.099464566 | 0.729946 | 0.830572 | 62.37537 | 34.79482 | 34.78048 | 50.89479 | 55.58134 | 41.34284 | 52.07161 | 46.76642 |
| ENSMUSG00000030413 | 0.075942535 | 0.80761 | 0.882437 | 42.25428 | 46.79304 | 33.12427 | 39.68068 | 46.80534 | 25.6996 | 49.17875 | 30.45255 |
| ENSMUSG00000030417 | -0.021567199 | 0.680101 | 0.795588 | 2082.533 | 2103.287 | 2195.311 | 2094.45 | 2199.851 | 2171.058 | 2060.686 | 2175.182 |
| ENSMUSG00000030421 | 0.102257722 | 0.306788 | 0.475352 | 2281.731 | 2459.634 | 2284.746 | 2538.701 | 2268.109 | 2575.547 | 2112.758 | 1954.401 |
| ENSMUSG00000030423 | -0.037598176 | 0.642236 | 0.768658 | 1532.221 | 1521.374 | 1426 | 1563.936 | 1602.108 | 1705.113 | 1466.684 | 1429.095 |
| ENSMUSG00000030424 | -0.05619743 | 0.766167 | 0.854869 | 68.4117 | 79.18822 | 69.56096 | 87.12497 | 78.98402 | 80.45093 | 84.85745 | 71.78101 |
| ENSMUSG00000030428 | 0.467012905 | 0.047243 | 0.130749 | 224.3501 | 167.975 | 208.6829 | 243.2598 | 120.9138 | 225.7096 | 131.1433 | 134.8613 |
| ENSMUSG00000030431 | -0.82007744 | 5.02E-05 | 0.000796 | 217.3077 | 261.5611 | 317.9929 | 262.2375 | 477.8045 | 317.3342 | 500.4661 | 575.3357 |
| ENSMUSG00000030432 | -0.148308066 | 0.107551 | 0.232097 | 29271.15 | 33882.96 | 32508.15 | 31804.07 | 40094.63 | 31489.84 | 36114.56 | 33564.15 |
| ENSMUSG00000030435 | -0.150950756 | 0.05583 | 0.147199 | 5922.642 | 6219.875 | 6715.117 | 6235.043 | 7055.905 | 6213.717 | 7050.882 | 7541.357 |
| ENSMUSG00000030443 | 0.233343485 | 0.302727 | 0.470971 | 81.4904 | 103.1847 | 67.90474 | 109.5532 | 78.98402 | 86.03781 | 78.10742 | 64.16787 |
| ENSMUSG00000030446 | 0.444633187 | 0.0075 | 0.034561 | 219.3198 | 226.7663 | 179.6991 | 166.4863 | 127.7396 | 164.254 | 137.8934 | 151.1752 |
| ENSMUSG00000030447 | -0.069454229 | 0.14325 | 0.28087 | 2756.589 | 2760.789 | 2723.643 | 2822.504 | 2938.985 | 2776.675 | 2950.725 | 2940.846 |
| ENSMUSG00000030451 | 0.039424674 | 0.613605 | 0.747357 | 1970.86 | 1750.54 | 1984.143 | 1927.101 | 1681.092 | 1902.888 | 1891.935 | 1956.576 |
| ENSMUSG00000030452 | -0.140908197 | 0.170614 | 0.317421 | 788.7466 | 707.8947 | 693.1253 | 664.2201 | 877.6002 | 701.7109 | 807.11 | 756.9634 |
| ENSMUSG00000030465 | 0.534225463 | 0.002824 | 0.016943 | 961.788 | 793.082 | 732.0463 | 839.3327 | 508.033 | 778.8097 | 559.2877 | 451.3503 |
| ENSMUSG00000030468 | -4.124328764 | 5.32E-09 | 5.06E-07 | 5.030272 | 5.999108 | 7.45296 | 12.93935 | 250.6036 | 12.29112 | 138.8576 | 150.0876 |
| ENSMUSG00000030469 | -0.066003942 | 0.767376 | 0.855545 | 261.5741 | 302.355 | 207.8548 | 172.5247 | 294.4836 | 256.996 | 254.5723 | 179.4525 |

| | | | | | | | | | | | |
|---|---|---|---|---|---|---|---|---|---|---|---|
| ENSMUSG00000030471 | -0.273921197 | 0.003727 | 0.020766 | 870.237 | 909.4647 | 824.7942 | 753.0703 | 1028.742 | 910.6599 | 1082.897 | 1031.036 |
| ENSMUSG00000030474 | 0.066267581 | 0.8875 | 0.932519 | 18.10898 | 15.59768 | 16.56213 | 10.35148 | 15.60178 | 7.821619 | 12.53576 | 21.75182 |
| ENSMUSG00000030486 | -0.127672102 | 0.599601 | 0.737651 | 94.56911 | 94.7859 | 92.74794 | 77.63611 | 111.1627 | 59.22083 | 114.7504 | 106.5839 |
| ENSMUSG00000030491 | -0.966010494 | 0.111867 | 0.238407 | 4.024217 | 7.198929 | 14.07781 | 4.313117 | 14.62667 | 22.34748 | 9.642892 | 11.9635 |
| ENSMUSG00000030492 | -2.223035781 | 8.86E-08 | 5.13E-06 | 64.38748 | 25.19625 | 101.029 | 70.73513 | 334.4632 | 159.7845 | 254.5723 | 477.4525 |
| ENSMUSG00000030493 | -0.202648004 | 0.197441 | 0.351754 | 165.999 | 170.3747 | 165.6213 | 190.6398 | 187.2214 | 177.6625 | 184.1792 | 249.0584 |
| ENSMUSG00000030494 | -0.195903298 | 0.14215 | 0.279278 | 512.0817 | 619.1079 | 697.2658 | 515.8488 | 760.5868 | 644.7249 | 649.9309 | 629.7152 |
| ENSMUSG00000030495 | 0.866537673 | 0.106667 | 0.230871 | 28.16952 | 70.78947 | 48.03018 | 43.13117 | 20.47734 | 53.63396 | 8.678602 | 21.75182 |
| ENSMUSG00000030498 | -0.7328296 | 7.32E-07 | 2.88E-05 | 1206.259 | 1252.614 | 1322.486 | 1297.386 | 2206.677 | 1586.671 | 2003.793 | 2643.934 |
| ENSMUSG00000030499 | 0.008118607 | 0.929556 | 0.958482 | 1641.881 | 1934.112 | 1967.581 | 1798.57 | 1883.915 | 1702.878 | 1716.435 | 1997.905 |
| ENSMUSG00000030500 | -0.219323555 | 0.613332 | 0.747221 | 8.048435 | 15.59768 | 14.07781 | 11.21411 | 14.62667 | 14.52586 | 13.50005 | 14.13868 |
| ENSMUSG00000030505 | -0.118727072 | 0.204091 | 0.360071 | 1157.969 | 1245.415 | 1100.554 | 1085.18 | 1376.857 | 1204.529 | 1111.825 | 1285.533 |
| ENSMUSG00000030509 | 0.150629303 | 0.119728 | 0.248615 | 686.1291 | 745.0892 | 670.7664 | 671.1211 | 569.465 | 706.1804 | 628.7165 | 592.7371 |
| ENSMUSG00000030510 | -0.620179024 | 0.141274 | 0.278088 | 68.4117 | 32.39518 | 128.3565 | 47.44429 | 147.2418 | 71.51194 | 108.9647 | 98.97079 |
| ENSMUSG00000030512 | 0.064963139 | 0.458085 | 0.619779 | 1506.063 | 1623.359 | 1377.141 | 1434.543 | 1473.393 | 1521.864 | 1284.433 | 1396.467 |
| ENSMUSG00000030513 | -0.718119387 | 0.000862 | 0.006958 | 253.5257 | 224.3666 | 371.8199 | 260.5123 | 348.1147 | 392.1983 | 548.6805 | 540.5328 |
| ENSMUSG00000030515 | -0.433404631 | 0.000646 | 0.005588 | 464.7971 | 425.9366 | 409.9128 | 370.0655 | 647.4739 | 477.1187 | 567.9663 | 560.1094 |
| ENSMUSG00000030516 | 0.310819111 | 3.04E-10 | 4.47E-08 | 4850.188 | 5092.043 | 4875.892 | 5165.389 | 4100.343 | 3946.565 | 3917.907 | 4144.81 |
| ENSMUSG00000030518 | 0.473802342 | 0.134294 | 0.268006 | 39.23612 | 52.79215 | 53.82693 | 36.23019 | 22.42756 | 27.93435 | 42.42872 | 38.06569 |
| ENSMUSG00000030519 | 0.905880671 | 7.98E-05 | 0.001136 | 133.8052 | 100.785 | 84.46688 | 129.3935 | 65.33246 | 53.63396 | 66.53595 | 53.29196 |
| ENSMUSG00000030521 | -0.218656572 | 0.015478 | 0.059023 | 880.2976 | 859.0722 | 790.8418 | 889.3648 | 1105.776 | 972.1155 | 903.5389 | 996.2334 |
| ENSMUSG00000030522 | -0.059458489 | 0.514759 | 0.669147 | 628.784 | 575.9143 | 629.361 | 583.9961 | 612.3699 | 693.8893 | 641.2523 | 574.2481 |
| ENSMUSG00000030523 | 0.209422313 | 0.414425 | 0.580277 | 68.4117 | 70.78947 | 108.482 | 89.71284 | 73.13335 | 96.09417 | 55.92877 | 68.51824 |
| ENSMUSG00000030525 | -0.791976798 | 0.039847 | 0.116325 | 22.1332 | 17.99732 | 22.35888 | 15.52722 | 32.17867 | 17.87799 | 42.42872 | 42.41605 |
| ENSMUSG00000030527 | 0.255518192 | 0.02416 | 0.081029 | 882.3097 | 1017.449 | 1082.335 | 1045.5 | 827.8695 | 988.8761 | 773.3599 | 786.3284 |
| ENSMUSG00000030528 | -0.125377313 | 0.242907 | 0.405453 | 583.5115 | 649.1034 | 568.0811 | 575.3699 | 702.0801 | 710.6499 | 568.9306 | 607.9634 |
| ENSMUSG00000030530 | -0.338946447 | 0.009635 | 0.041456 | 1865.225 | 1658.153 | 2343.542 | 1889.145 | 2443.629 | 2071.612 | 2601.652 | 2697.226 |
| ENSMUSG00000030532 | -0.525957094 | 0.024426 | 0.081566 | 110.666 | 122.3818 | 110.1382 | 93.16334 | 178.4454 | 101.681 | 146.572 | 200.1168 |
| ENSMUSG00000030533 | -0.198959075 | 0.071917 | 0.175895 | 914.5034 | 967.0562 | 1219.801 | 1064.477 | 1268.62 | 1107.318 | 1165.826 | 1244.204 |
| ENSMUSG00000030534 | -0.012010788 | 0.890968 | 0.934089 | 496.9909 | 500.3256 | 463.7397 | 518.4367 | 499.257 | 462.5929 | 528.4305 | 503.5547 |
| ENSMUSG00000030536 | -0.097419114 | 0.22223 | 0.381505 | 5795.879 | 5338.006 | 6096.521 | 5498.362 | 5616.641 | 5787.998 | 6262.094 | 6653.882 |
| ENSMUSG00000030538 | -0.882616171 | 6.05E-05 | 0.000923 | 562.3844 | 520.7225 | 771.7954 | 582.2709 | 1165.258 | 729.6453 | 1075.182 | 1525.89 |
| ENSMUSG00000030539 | 0.010218182 | 0.938135 | 0.963453 | 1043.278 | 844.6744 | 1099.726 | 939.397 | 857.1228 | 920.7163 | 1171.611 | 951.6422 |
| ENSMUSG00000030541 | -0.591406725 | 0.000483 | 0.004461 | 7439.772 | 7552.877 | 9288.044 | 8051.728 | 11774.47 | 8378.071 | 14051.62 | 14512.82 |
| ENSMUSG00000030543 | 1.103111331 | 0.00992 | 0.042337 | 25.15136 | 16.7975 | 29.81184 | 31.91707 | 11.70134 | 17.87799 | 11.57147 | 7.613138 |
| ENSMUSG00000030545 | -0.641572539 | 0.000338 | 0.00339 | 374.2522 | 250.7627 | 250.9163 | 286.391 | 388.0943 | 412.311 | 554.4663 | 457.8758 |
| ENSMUSG00000030546 | 2.269785103 | 0.013361 | 0.053047 | 152.9203 | 20.39697 | 7.45296 | 17.25247 | 9.751113 | 7.821619 | 2.892867 | 20.66423 |
| ENSMUSG00000030549 | -1.208312366 | 0.025776 | 0.084815 | 899.4126 | 320.3523 | 1444.218 | 721.1532 | 1340.778 | 1019.045 | 2885.153 | 2578.678 |
| ENSMUSG00000030551 | 0.628534032 | 0.001061 | 0.008167 | 12686.35 | 13418.8 | 10557.53 | 13383.6 | 6736.069 | 11846.4 | 7236.026 | 6553.824 |
| ENSMUSG00000030553 | 0.191403286 | 0.63888 | 0.76591 | 23.13925 | 29.99554 | 20.70267 | 37.09281 | 22.42756 | 12.29112 | 33.75012 | 28.27737 |
| ENSMUSG00000030554 | 0.291963504 | 0.119723 | 0.248615 | 156.9445 | 176.3738 | 131.669 | 166.4863 | 117.9885 | 166.4887 | 114.7504 | 116.3722 |
| ENSMUSG00000030555 | 0.156658156 | 0.055089 | 0.146018 | 1117.726 | 1280.21 | 1126.225 | 1301.699 | 1071.647 | 1128.548 | 1076.147 | 1051.701 |
| ENSMUSG00000030556 | -0.181375648 | 0.074035 | 0.179481 | 291.7558 | 326.3515 | 342.008 | 323.4838 | 349.0898 | 382.1419 | 347.1441 | 379.5693 |
| ENSMUSG00000030557 | 0.195426837 | 0.088455 | 0.202604 | 3090.599 | 3034.349 | 2895.061 | 3117.521 | 2316.864 | 3176.695 | 2782.939 | 2324.182 |
| ENSMUSG00000030559 | -0.666575469 | 0.001123 | 0.008534 | 51.30877 | 58.79126 | 55.48314 | 61.24627 | 89.71024 | 71.51194 | 98.35749 | 100.0584 |
| ENSMUSG00000030560 | 0.025064697 | 0.86898 | 0.921481 | 747.4984 | 1035.446 | 1146.1 | 913.5183 | 816.1682 | 891.6645 | 1059.754 | 1009.285 |
| ENSMUSG00000030562 | -0.330717595 | 0.007967 | 0.036131 | 241.453 | 302.355 | 232.698 | 244.1224 | 327.6374 | 318.4516 | 295.0725 | 338.2408 |
| ENSMUSG00000030579 | -0.087353559 | 0.673269 | 0.790599 | 112.6781 | 181.1731 | 203.7142 | 161.3106 | 152.1174 | 182.132 | 167.7863 | 199.0292 |
| ENSMUSG00000030583 | -0.211679775 | 0.068452 | 0.169509 | 523.1483 | 488.3274 | 654.2042 | 518.4367 | 590.9175 | 665.955 | 633.538 | 643.8539 |
| ENSMUSG00000030584 | 1.321155163 | 0.001207 | 0.009002 | 64.38748 | 52.79215 | 31.46805 | 86.26235 | 28.27823 | 23.46486 | 28.92867 | 13.05109 |
| ENSMUSG00000030587 | -0.306223663 | 0.35316 | 0.521852 | 117.7084 | 86.38715 | 197.0894 | 121.6299 | 195.9974 | 87.15518 | 159.1077 | 205.5547 |
| ENSMUSG00000030588 | -0.40583339 | 0.003136 | 0.018364 | 788.7466 | 920.2631 | 969.7129 | 818.6297 | 1241.317 | 889.4298 | 1242.004 | 1259.43 |
| ENSMUSG00000030589 | 0.384250373 | 0.125726 | 0.25637 | 81.4904 | 62.39072 | 91.09173 | 65.55939 | 45.83023 | 48.04709 | 68.46453 | 68.51824 |
| ENSMUSG00000030590 | -0.157833402 | 0.299544 | 0.467675 | 276.6649 | 208.7689 | 259.1974 | 306.2313 | 332.513 | 261.4655 | 280.6081 | 300.1751 |
| ENSMUSG00000030591 | -0.204427975 | 0.035639 | 0.107552 | 3765.661 | 3480.682 | 4169.517 | 3788.642 | 4458.209 | 3756.612 | 4518.659 | 4788.664 |
| ENSMUSG00000030592 | 0.981915016 | 2.25E-05 | 0.000431 | 72.43591 | 80.38804 | 81.15445 | 82.81186 | 31.20356 | 37.99072 | 39.53586 | 52.20437 |
| ENSMUSG00000030595 | -0.321379553 | 0.026206 | 0.08598 | 399.4036 | 304.7547 | 412.3971 | 346.7746 | 473.9041 | 374.3203 | 492.7518 | 489.416 |
| ENSMUSG00000030598 | 0.017225821 | 0.910069 | 0.946586 | 232.3986 | 214.7681 | 298.1184 | 255.3366 | 281.8072 | 213.4185 | 250.7152 | 244.708 |
| ENSMUSG00000030600 | -0.05455084 | 0.718981 | 0.824245 | 242.4591 | 271.1597 | 289.0092 | 312.2697 | 264.2552 | 243.5876 | 307.6082 | 343.6788 |
| ENSMUSG00000030602 | -0.349828684 | 0.002446 | 0.015178 | 1206.259 | 1137.431 | 1312.549 | 1056.714 | 1476.319 | 1260.398 | 1612.291 | 1657.489 |
| ENSMUSG00000030603 | -0.286935864 | 0.00391 | 0.021475 | 2299.84 | 2218.47 | 2371.697 | 2321.32 | 2785.893 | 2390.063 | 2812.831 | 3250.81 |
| ENSMUSG00000030604 | 0.291054021 | 0.084942 | 0.196921 | 506.0453 | 490.727 | 390.0382 | 467.5419 | 401.7459 | 463.7102 | 367.3942 | 281.6861 |
| ENSMUSG00000030605 | -0.071294027 | 0.439105 | 0.602537 | 4948.781 | 5010.455 | 5356.194 | 5127.434 | 5852.618 | 4551.065 | 5751.985 | 5321.583 |
| ENSMUSG00000030606 | 0.15099453 | 0.481718 | 0.640579 | 148.896 | 130.7805 | 158.1684 | 115.5915 | 129.6898 | 165.3714 | 104.1432 | 100.0584 |

| | | | | | | | | | | | |
|---|---|---|---|---|---|---|---|---|---|---|---|
| ENSMUSG00000030607 | 1.063384916 | 0.197275 | 0.351608 | 16.09687 | 11.99822 | 12.4216 | 62.97152 | 0.975111 | 12.29112 | 21.21436 | 15.22628 |
| ENSMUSG00000030609 | -0.333855927 | 7.98E-05 | 0.001136 | 737.4378 | 715.0936 | 774.2797 | 740.131 | 1001.439 | 878.256 | 864.9674 | 997.321 |
| ENSMUSG00000030610 | -0.126826867 | 0.260187 | 0.424206 | 368.2159 | 353.9474 | 360.2264 | 410.6088 | 441.7254 | 355.325 | 434.8944 | 398.0583 |
| ENSMUSG00000030611 | -0.184411626 | 0.060632 | 0.156243 | 777.68 | 736.6904 | 723.7652 | 784.1248 | 890.2766 | 746.4059 | 836.0387 | 961.4305 |
| ENSMUSG00000030612 | -0.442472102 | 8.46E-06 | 0.000202 | 549.3057 | 574.7145 | 624.3924 | 631.4404 | 807.3922 | 709.5326 | 850.503 | 868.9853 |
| ENSMUSG00000030613 | 0.082896806 | 0.280989 | 0.447372 | 1076.478 | 1055.843 | 1042.586 | 1074.829 | 938.0571 | 1115.139 | 982.6107 | 978.832 |
| ENSMUSG00000030614 | -0.14177097 | 0.103212 | 0.225862 | 636.8324 | 679.099 | 602.0335 | 589.1718 | 714.7566 | 633.5511 | 695.2525 | 716.7225 |
| ENSMUSG00000030615 | -0.114817587 | 0.201392 | 0.356878 | 563.3904 | 611.909 | 573.0498 | 590.0345 | 566.5397 | 625.7295 | 682.7167 | 655.8174 |
| ENSMUSG00000030616 | -0.721145121 | 0.020297 | 0.071217 | 256.5439 | 185.9723 | 482.7862 | 290.7041 | 460.2525 | 318.4516 | 537.1091 | 691.7079 |
| ENSMUSG00000030619 | -0.070019803 | 0.224091 | 0.383871 | 1186.138 | 1205.821 | 1143.615 | 1203.36 | 1207.188 | 1272.689 | 1255.504 | 1237.679 |
| ENSMUSG00000030621 | -1.160659653 | 1.13E-05 | 0.000252 | 316.9071 | 166.7752 | 339.5237 | 263.9628 | 494.3814 | 427.9543 | 714.5383 | 797.2043 |
| ENSMUSG00000030629 | 0.203113724 | 0.02949 | 0.09342 | 3054.381 | 3171.128 | 2892.576 | 3271.068 | 2489.459 | 3115.239 | 2654.688 | 2503.635 |
| ENSMUSG00000030630 | -1.652129095 | 1.83E-07 | 9.33E-06 | 294.7739 | 183.5727 | 381.7572 | 318.3081 | 1010.215 | 435.7759 | 946.932 | 1314.898 |
| ENSMUSG00000030638 | 0.615712388 | 5.24E-05 | 0.000821 | 513.0877 | 577.1142 | 459.5992 | 617.6384 | 390.0445 | 360.9118 | 385.7157 | 276.2481 |
| ENSMUSG00000030641 | 0.128506107 | 0.405093 | 0.572028 | 183.1019 | 225.5664 | 206.1986 | 163.0358 | 200.8729 | 165.3714 | 175.5006 | 167.489 |
| ENSMUSG00000030643 | 0.553025868 | 0.030168 | 0.094851 | 194.1685 | 239.9643 | 198.7456 | 251.8861 | 120.9138 | 235.7659 | 135.0005 | 112.0219 |
| ENSMUSG00000030647 | -0.322100845 | 0.000134 | 0.00168 | 2431.633 | 2373.247 | 2457.82 | 2536.976 | 3119.381 | 2640.355 | 3189.869 | 3301.927 |
| ENSMUSG00000030649 | -0.316590695 | 0.018724 | 0.067349 | 333.004 | 368.3452 | 479.4737 | 408.8835 | 507.0579 | 449.1844 | 508.1804 | 518.7809 |
| ENSMUSG00000030650 | -2.493651911 | 0.00014 | 0.001735 | 4.024217 | 2.399643 | 19.87456 | 5.175741 | 36.07912 | 32.40385 | 33.75012 | 79.39415 |
| ENSMUSG00000030652 | -0.373180925 | 0.000402 | 0.003873 | 1030.2 | 1154.228 | 1300.956 | 1224.925 | 1521.174 | 1357.61 | 1527.434 | 1697.73 |
| ENSMUSG00000030653 | 0.986367988 | 0.120177 | 0.249148 | 21.12714 | 23.99643 | 25.67131 | 31.05445 | 4.875557 | 30.1691 | 3.857157 | 13.05109 |
| ENSMUSG00000030654 | -0.16190093 | 0.005486 | 0.027498 | 6575.571 | 6324.259 | 6405.405 | 6737.089 | 7660.474 | 6751.174 | 7335.348 | 7388.006 |
| ENSMUSG00000030655 | 0.220929367 | 0.024773 | 0.082484 | 3230.441 | 3046.347 | 2832.953 | 3049.374 | 2373.421 | 3038.14 | 2397.223 | 2625.445 |
| ENSMUSG00000030657 | 0.055584378 | 0.784035 | 0.8671 | 559.3662 | 565.1159 | 472.0208 | 481.3439 | 572.3903 | 646.9596 | 448.3945 | 330.6277 |
| ENSMUSG00000030659 | 0.000369682 | 0.996984 | 0.998149 | 631.8021 | 650.3033 | 745.296 | 711.6644 | 676.7272 | 648.077 | 761.7884 | 653.6422 |
| ENSMUSG00000030660 | 0.038693772 | 0.569905 | 0.713532 | 1363.204 | 1305.406 | 1230.566 | 1237.002 | 1211.088 | 1215.703 | 1255.504 | 1314.898 |
| ENSMUSG00000030662 | -0.150473288 | 0.005055 | 0.025924 | 5841.152 | 6399.848 | 6449.294 | 6282.487 | 6998.374 | 6660.667 | 7005.561 | 7053.028 |
| ENSMUSG00000030663 | -0.11785934 | 0.015109 | 0.057966 | 4189.21 | 4258.167 | 4151.299 | 4391.616 | 4792.672 | 4498.548 | 4658.481 | 4484.138 |
| ENSMUSG00000030669 | 0.863511175 | 0.161823 | 0.305849 | 23.13925 | 16.7975 | 19.87456 | 18.11509 | 1.950223 | 20.11273 | 14.46434 | 6.525547 |
| ENSMUSG00000030670 | 0.281380934 | 0.548364 | 0.696723 | 16.09687 | 17.99732 | 29.81184 | 31.05445 | 23.40267 | 22.34748 | 25.07152 | 7.613138 |
| ENSMUSG00000030671 | 0.141035275 | 0.221163 | 0.380196 | 778.6861 | 705.4951 | 651.7199 | 803.1025 | 660.1504 | 745.2885 | 684.6453 | 575.3357 |
| ENSMUSG00000030672 | 0.25786165 | 0.30106 | 0.469406 | 124.7507 | 64.79036 | 116.763 | 107.8279 | 72.15824 | 94.9768 | 101.2504 | 79.39415 |
| ENSMUSG00000030674 | -1.973134879 | 1.10E-07 | 6.14E-06 | 129.781 | 86.38715 | 217.792 | 139.745 | 448.5512 | 306.1605 | 504.3232 | 998.4086 |
| ENSMUSG00000030677 | 0.032987359 | 0.616561 | 0.749474 | 1642.887 | 1662.953 | 1515.435 | 1605.342 | 1572.855 | 1668.24 | 1493.684 | 1544.379 |
| ENSMUSG00000030678 | -0.042768144 | 0.453247 | 0.615723 | 5039.326 | 5378.8 | 5428.239 | 5199.894 | 5505.478 | 5729.894 | 5175.34 | 5270.466 |
| ENSMUSG00000030681 | -0.328379992 | 0.03295 | 0.101169 | 834.0191 | 659.9018 | 1004.493 | 682.3352 | 915.6295 | 924.0684 | 1031.789 | 1125.657 |
| ENSMUSG00000030682 | -0.323865892 | 0.031099 | 0.096889 | 1755.565 | 1766.137 | 2408.962 | 1795.982 | 2385.122 | 1926.353 | 2574.652 | 2787.496 |
| ENSMUSG00000030683 | 0.248491096 | 0.366019 | 0.534678 | 149.9021 | 79.18822 | 76.18581 | 124.2178 | 78.0089 | 88.27255 | 84.85745 | 110.9343 |
| ENSMUSG00000030685 | -0.147318283 | 0.281305 | 0.44772 | 329.9858 | 251.9625 | 288.1811 | 279.49 | 341.289 | 264.8177 | 336.5369 | 330.6277 |
| ENSMUSG00000030688 | -0.844704953 | 3.40E-05 | 0.00059 | 1250.526 | 1012.649 | 1687.681 | 1289.622 | 2302.238 | 1641.423 | 2458.937 | 3012.627 |
| ENSMUSG00000030689 | 0.159421919 | 0.101847 | 0.223792 | 927.5821 | 991.0526 | 1022.712 | 924.7324 | 917.5797 | 743.0538 | 860.1459 | 939.6787 |
| ENSMUSG00000030691 | 0.100070418 | 0.466571 | 0.627229 | 989.9575 | 1045.045 | 1081.507 | 1081.73 | 884.426 | 1269.337 | 882.3246 | 883.124 |
| ENSMUSG00000030693 | 0.695606095 | 0.141526 | 0.278346 | 19.11503 | 35.99465 | 32.29616 | 26.74133 | 15.60178 | 29.05173 | 19.28578 | 6.525547 |
| ENSMUSG00000030695 | -0.77889383 | 0.000446 | 0.004196 | 8434.76 | 6653.01 | 10593.14 | 7682.525 | 13132.8 | 9118.89 | 15532.77 | 19463.53 |
| ENSMUSG00000030697 | -0.085174994 | 0.128171 | 0.259634 | 2741.498 | 2916.766 | 2902.514 | 3063.176 | 3175.938 | 3038.14 | 3019.189 | 3099.635 |
| ENSMUSG00000030699 | 1.342301035 | 0.000175 | 0.002044 | 40.24217 | 31.19536 | 43.06154 | 55.2079 | 11.70134 | 22.34748 | 19.28578 | 14.13868 |
| ENSMUSG00000030701 | 0.466350974 | 0.007456 | 0.034437 | 277.671 | 280.7582 | 294.806 | 397.6694 | 208.6738 | 270.4045 | 194.7864 | 233.8321 |
| ENSMUSG00000030703 | -0.161945879 | 0.45388 | 0.616122 | 70.4238 | 76.78858 | 72.87338 | 52.62003 | 78.0089 | 69.27719 | 88.7146 | 67.43065 |
| ENSMUSG00000030704 | 0.093483906 | 0.455011 | 0.617113 | 3090.599 | 3202.324 | 2803.141 | 3119.247 | 2742.988 | 3599.062 | 2569.831 | 2537.35 |
| ENSMUSG00000030706 | -0.174857817 | 0.035592 | 0.107475 | 1441.676 | 1680.95 | 1463.264 | 1526.844 | 1845.886 | 1744.221 | 1731.863 | 1572.657 |
| ENSMUSG00000030707 | -0.156189241 | 0.440027 | 0.603263 | 155.9384 | 182.3729 | 229.3855 | 169.0742 | 173.5698 | 167.6061 | 258.4295 | 221.8686 |
| ENSMUSG00000030708 | 0.402580327 | 0.216171 | 0.374544 | 25.15136 | 38.39429 | 32.29616 | 46.58167 | 29.25334 | 26.81698 | 30.85725 | 20.66423 |
| ENSMUSG00000030711 | 0.466630118 | 0.090779 | 0.206447 | 1008.066 | 704.2952 | 542.4098 | 1054.126 | 446.601 | 668.1897 | 855.3245 | 424.1605 |
| ENSMUSG00000030714 | -0.254731258 | 0.009744 | 0.041752 | 329.9858 | 386.3425 | 335.3832 | 354.5383 | 421.2481 | 446.9496 | 397.2871 | 410.0218 |
| ENSMUSG00000030717 | -0.069313227 | 0.776716 | 0.862312 | 123.7447 | 254.3622 | 209.511 | 185.4641 | 190.1467 | 246.9397 | 155.2506 | 218.6058 |
| ENSMUSG00000030718 | -0.149816425 | 0.087533 | 0.20116 | 1176.078 | 1123.033 | 1158.521 | 1174.893 | 1214.989 | 1147.543 | 1419.434 | 1357.314 |
| ENSMUSG00000030720 | 0.030808276 | 0.727688 | 0.829154 | 455.7426 | 472.7297 | 451.3181 | 511.5357 | 437.825 | 464.8276 | 475.3946 | 474.1897 |
| ENSMUSG00000030722 | 0.195193815 | 0.156353 | 0.298443 | 296.786 | 337.1499 | 245.9477 | 290.7041 | 287.6578 | 249.1744 | 243.0009 | 239.27 |
| ENSMUSG00000030725 | -0.45359738 | 8.71E-05 | 0.001211 | 238.4349 | 259.1615 | 265.8222 | 245.8477 | 389.0694 | 347.5033 | 309.5368 | 336.0656 |
| ENSMUSG00000030726 | 0.03343739 | 0.669783 | 0.788262 | 1185.132 | 1346.2 | 1124.569 | 1224.925 | 1192.561 | 1238.05 | 1188.969 | 1144.146 |
| ENSMUSG00000030727 | -0.234003465 | 0.030582 | 0.095812 | 794.7829 | 705.4951 | 878.6211 | 816.0418 | 969.2606 | 819.0352 | 933.4319 | 1039.737 |
| ENSMUSG00000030729 | 0.883990463 | 0.001707 | 0.011655 | 2202.253 | 2463.234 | 1633.854 | 2060.808 | 973.1611 | 1966.578 | 819.6458 | 771.1021 |
| ENSMUSG00000030730 | 1.272268705 | 7.68E-06 | 0.000188 | 133.8052 | 182.3729 | 133.3252 | 150.9591 | 58.50668 | 99.44629 | 45.32159 | 45.67883 |
| ENSMUSG00000030731 | 0.543253121 | 0.030808 | 0.096321 | 95.57516 | 75.58876 | 77.84202 | 69.8725 | 38.02934 | 50.28183 | 69.42882 | 60.9051 |

| | | | | | | | | | | | |
|---|---|---|---|---|---|---|---|---|---|---|---|
| ENSMUSG00000030733 | -0.02545165 | 0.742983 | 0.839394 | 1617.735 | 1694.148 | 1669.463 | 1689.879 | 1653.789 | 1520.746 | 1776.221 | 1838.029 |
| ENSMUSG00000030735 | -0.317802668 | 0.362459 | 0.531001 | 17.10292 | 34.79482 | 25.67131 | 28.46658 | 37.05423 | 22.34748 | 31.82154 | 40.24087 |
| ENSMUSG00000030737 | 0.326224131 | 0.107423 | 0.231883 | 511.0756 | 394.7413 | 348.6329 | 421.8229 | 312.0356 | 298.3389 | 460.9302 | 264.2846 |
| ENSMUSG00000030738 | -0.277299805 | 0.00339 | 0.019347 | 7546.414 | 7519.282 | 8046.712 | 7424.6 | 9665.303 | 7679.712 | 9788.499 | 9874.239 |
| ENSMUSG00000030739 | -0.462785228 | 0.067569 | 0.167851 | 163.9869 | 143.9786 | 258.3693 | 172.5247 | 284.7325 | 164.254 | 262.2867 | 308.8759 |
| ENSMUSG00000030741 | -0.185411258 | 0.04566 | 0.127761 | 1565.421 | 1496.177 | 1616.464 | 1589.815 | 1776.653 | 1517.394 | 1911.221 | 1922.861 |
| ENSMUSG00000030742 | -0.300709992 | 0.375649 | 0.544104 | 28.16952 | 17.99732 | 34.78048 | 30.19182 | 28.27823 | 48.04709 | 31.82154 | 30.45255 |
| ENSMUSG00000030744 | -0.16314003 | 0.075535 | 0.181836 | 36574.1 | 43791.09 | 41734.92 | 43554.72 | 51485.88 | 42138.41 | 47509.56 | 44350.88 |
| ENSMUSG00000030745 | 0.502537228 | 0.231577 | 0.392114 | 12.07265 | 17.99732 | 26.49941 | 18.97772 | 13.65156 | 13.40849 | 12.53576 | 14.13868 |
| ENSMUSG00000030747 | -1.696218784 | 9.33E-10 | 1.21E-07 | 240.447 | 197.9706 | 332.8989 | 221.6942 | 913.6793 | 426.8369 | 789.7528 | 1090.854 |
| ENSMUSG00000030748 | 0.034291867 | 0.864982 | 0.919168 | 853.1341 | 471.5299 | 707.2031 | 509.8105 | 720.6073 | 531.8701 | 630.6451 | 599.2627 |
| ENSMUSG00000030750 | -0.158609547 | 0.008433 | 0.03771 | 1213.302 | 1225.018 | 1259.55 | 1208.536 | 1433.414 | 1388.896 | 1347.112 | 1307.284 |
| ENSMUSG00000030751 | -0.110111369 | 0.033206 | 0.101667 | 3360.222 | 3509.478 | 3435.814 | 3631.645 | 3821.461 | 3595.71 | 3787.728 | 3837.021 |
| ENSMUSG00000030752 | -0.121267129 | 0.334909 | 0.5031 | 269.6226 | 307.1543 | 294.806 | 289.8415 | 375.4179 | 289.3999 | 300.8582 | 295.8248 |
| ENSMUSG00000030753 | -0.086989751 | 0.219447 | 0.378197 | 3337.082 | 3945.013 | 3830.821 | 3749.824 | 4067.189 | 4046.012 | 3805.085 | 3867.474 |
| ENSMUSG00000030754 | -0.089497541 | 0.154107 | 0.295563 | 3859.225 | 3786.637 | 3805.978 | 4112.126 | 3999.907 | 3924.218 | 4229.372 | 4409.094 |
| ENSMUSG00000030757 | 0.564368541 | 0.008522 | 0.037989 | 370.228 | 485.9277 | 388.382 | 461.5036 | 322.7618 | 392.1983 | 225.6437 | 213.1679 |
| ENSMUSG00000030759 | 0.052249473 | 0.434189 | 0.598416 | 2787.777 | 2606.012 | 2851.999 | 2899.278 | 2523.588 | 2847.069 | 2628.652 | 2757.043 |
| ENSMUSG00000030760 | 0.131048518 | 0.243281 | 0.405862 | 466.8092 | 379.1436 | 423.1625 | 462.3662 | 360.7912 | 433.5411 | 385.7157 | 404.5839 |
| ENSMUSG00000030761 | -0.255359216 | 0.168274 | 0.314398 | 556.3481 | 387.5424 | 491.8953 | 365.7524 | 518.7592 | 429.0717 | 534.2162 | 668.8685 |
| ENSMUSG00000030763 | 0.078942715 | 0.291889 | 0.459113 | 1147.908 | 1291.008 | 1134.506 | 1198.184 | 1143.806 | 1196.708 | 1092.54 | 1081.066 |
| ENSMUSG00000030766 | 0.019401968 | 0.817751 | 0.889464 | 780.6982 | 685.0981 | 709.6874 | 792.751 | 705.0055 | 774.3402 | 715.5026 | 737.3868 |
| ENSMUSG00000030768 | 0.261809269 | 0.026985 | 0.087728 | 957.7637 | 1118.234 | 870.3401 | 927.3203 | 862.9735 | 896.134 | 707.7882 | 761.3138 |
| ENSMUSG00000030769 | -3.234341524 | 4.63E-06 | 0.000126 | 2.012109 | 7.198929 | 19.04645 | 9.488858 | 134.5654 | 24.58223 | 42.42872 | 157.7007 |
| ENSMUSG00000030770 | 0.166597382 | 0.007817 | 0.035624 | 6487.038 | 7196.53 | 6932.909 | 6972.586 | 6011.561 | 6626.028 | 5930.378 | 6013.291 |
| ENSMUSG00000030771 | 0.232319164 | 0.712798 | 0.819906 | 10.06054 | 31.19536 | 9.93728 | 28.46658 | 8.776002 | 34.6386 | 13.50005 | 10.87591 |
| ENSMUSG00000030772 | 0.258491138 | 0.048753 | 0.133817 | 1293.786 | 1280.21 | 1608.183 | 1315.501 | 1052.145 | 1064.858 | 1073.254 | 1408.43 |
| ENSMUSG00000030774 | -0.315662148 | 0.003991 | 0.02181 | 503.0272 | 422.3372 | 553.1752 | 480.4813 | 593.8428 | 568.7434 | 657.6452 | 622.1021 |
| ENSMUSG00000030779 | 0.262954643 | 0.024747 | 0.082429 | 5064.478 | 4865.276 | 4491.65 | 4731.49 | 3619.613 | 4963.376 | 3723.12 | 3656.481 |
| ENSMUSG00000030780 | -0.14773819 | 0.07534 | 0.181476 | 924.564 | 934.661 | 818.1694 | 881.6012 | 981.9371 | 902.8383 | 1046.254 | 1006.022 |
| ENSMUSG00000030781 | -2.882209946 | 1.55E-06 | 5.33E-05 | 21.12714 | 7.198929 | 51.34261 | 28.46658 | 333.4881 | 62.57295 | 114.7504 | 292.562 |
| ENSMUSG00000030782 | 0.157138347 | 0.157293 | 0.299743 | 1003.036 | 1163.827 | 1291.846 | 1155.053 | 969.2606 | 1045.862 | 949.8248 | 1175.686 |
| ENSMUSG00000030786 | 0.89175144 | 0.008642 | 0.038378 | 89.53884 | 46.79304 | 55.48314 | 112.1411 | 43.88001 | 49.16446 | 37.60728 | 33.71532 |
| ENSMUSG00000030787 | 1.152302911 | 1.82E-05 | 0.000365 | 1072.454 | 1201.021 | 799.1229 | 1144.701 | 365.6667 | 708.4152 | 544.8234 | 278.4233 |
| ENSMUSG00000030788 | 0.045585877 | 0.538034 | 0.688091 | 1086.539 | 981.454 | 1025.196 | 1098.12 | 981.9371 | 1034.688 | 1074.218 | 972.3064 |
| ENSMUSG00000030790 | 0.255728214 | 0.239395 | 0.401187 | 246.4833 | 147.578 | 157.3403 | 250.1608 | 160.8934 | 151.9629 | 175.5006 | 183.8029 |
| ENSMUSG00000030792 | 0.484556486 | 0.328277 | 0.496085 | 47.28455 | 51.59233 | 29.81184 | 37.95543 | 32.17867 | 41.34284 | 38.57157 | 6.525547 |
| ENSMUSG00000030793 | -0.189971954 | 0.459477 | 0.620935 | 191.1503 | 220.7672 | 375.1323 | 210.4801 | 238.9023 | 232.4138 | 293.1439 | 375.2189 |
| ENSMUSG00000030795 | -0.010437583 | 0.855301 | 0.913351 | 16952.02 | 17228.24 | 18413.78 | 16884.99 | 18133.17 | 16377.35 | 17766.06 | 17707.07 |
| ENSMUSG00000030796 | 0.061014424 | 0.43235 | 0.596728 | 5001.096 | 5122.038 | 5033.232 | 5262.003 | 5457.698 | 4884.042 | 4848.446 | 4380.817 |
| ENSMUSG00000030798 | 0.322413943 | 0.337712 | 0.506116 | 39.23612 | 33.595 | 27.32752 | 43.13117 | 32.17867 | 20.11273 | 23.14294 | 39.15328 |
| ENSMUSG00000030800 | -1.341099451 | 4.64E-06 | 0.000126 | 798.8072 | 622.7074 | 1195.786 | 806.553 | 1936.571 | 1143.074 | 2355.758 | 3242.109 |
| ENSMUSG00000030801 | 0.021138311 | 0.793117 | 0.873246 | 760.5771 | 838.6753 | 711.3436 | 759.1087 | 768.3877 | 721.8237 | 778.1814 | 750.4379 |
| ENSMUSG00000030802 | -0.254438025 | 0.000318 | 0.003246 | 1708.28 | 1760.138 | 1876.49 | 1721.796 | 2207.652 | 1925.236 | 2146.508 | 2150.168 |
| ENSMUSG00000030805 | 0.145694279 | 0.031073 | 0.096857 | 1873.273 | 2038.497 | 1793.679 | 1850.327 | 1759.101 | 1705.113 | 1614.22 | 1746.671 |
| ENSMUSG00000030806 | 0.417389327 | 0.001205 | 0.008988 | 358.1554 | 421.1374 | 389.2101 | 419.235 | 269.1307 | 357.5597 | 281.5724 | 281.6861 |
| ENSMUSG00000030811 | 0.00968601 | 0.880606 | 0.928341 | 1568.439 | 1614.96 | 1551.044 | 1716.621 | 1550.427 | 1643.657 | 1553.47 | 1662.927 |
| ENSMUSG00000030814 | -0.129062108 | 0.102353 | 0.22451 | 1633.832 | 1929.313 | 1873.177 | 1745.95 | 1966.8 | 1966.578 | 1839.864 | 2080.562 |
| ENSMUSG00000030815 | 0.193476603 | 0.046738 | 0.129849 | 877.2794 | 735.4906 | 719.6247 | 777.2238 | 733.2837 | 688.3024 | 659.5738 | 637.3284 |
| ENSMUSG00000030816 | -0.005777838 | 0.926415 | 0.956313 | 2118.75 | 2097.288 | 2095.938 | 2120.329 | 2199.851 | 2024.682 | 1994.15 | 2248.051 |
| ENSMUSG00000030822 | 0.04423997 | 0.471704 | 0.631172 | 1097.605 | 1155.428 | 1103.038 | 1073.966 | 1061.896 | 1078.266 | 1042.397 | 1111.518 |
| ENSMUSG00000030823 | -0.458835186 | 0.048456 | 0.133246 | 113.6841 | 145.1784 | 104.3414 | 72.46037 | 173.5698 | 126.2633 | 147.5362 | 147.9124 |
| ENSMUSG00000030824 | -0.156134246 | 0.166488 | 0.312079 | 4075.526 | 4351.753 | 5065.528 | 4544.301 | 4936.989 | 4201.327 | 5338.305 | 5623.934 |
| ENSMUSG00000030825 | -0.553009501 | 0.045425 | 0.12732 | 158.9566 | 104.3845 | 210.3391 | 154.4096 | 238.9023 | 138.5544 | 281.5724 | 264.2846 |
| ENSMUSG00000030826 | -0.591369703 | 1.42E-05 | 0.000298 | 1252.538 | 1078.64 | 1313.377 | 1130.037 | 1747.399 | 1409.009 | 2090.579 | 1947.876 |
| ENSMUSG00000030830 | -0.02072329 | 0.921 | 0.952855 | 70.4238 | 80.38804 | 60.45178 | 62.97152 | 61.43201 | 62.57295 | 73.28598 | 79.39415 |
| ENSMUSG00000030834 | 0.802951314 | 0.004887 | 0.025343 | 45.27245 | 40.79393 | 37.2648 | 53.48266 | 23.40267 | 27.93435 | 22.17865 | 28.27737 |
| ENSMUSG00000030835 | -0.421261614 | 0.000317 | 0.003238 | 1665.02 | 1617.359 | 2055.361 | 1690.742 | 2192.05 | 2063.79 | 2568.866 | 2590.642 |
| ENSMUSG00000030838 | -1.578939328 | 4.49E-06 | 0.000124 | 210.2654 | 125.9813 | 308.8838 | 237.2215 | 607.4943 | 348.6207 | 607.5022 | 1076.715 |
| ENSMUSG00000030839 | -0.17924823 | 0.318656 | 0.486848 | 219.3198 | 196.7707 | 279.0719 | 232.9083 | 271.0809 | 204.4795 | 255.5366 | 321.927 |
| ENSMUSG00000030842 | -0.299530945 | 0.006637 | 0.03176 | 1482.924 | 1689.349 | 1966.753 | 1607.93 | 2159.872 | 1824.672 | 2129.15 | 2191.496 |
| ENSMUSG00000030844 | 0.706597279 | 0.004084 | 0.022152 | 124.7507 | 203.9697 | 157.3403 | 160.448 | 94.5858 | 139.6718 | 86.78602 | 75.04379 |
| ENSMUSG00000030846 | 0.368357825 | 0.005827 | 0.02881 | 3561.432 | 3415.892 | 2842.89 | 3612.667 | 2546.991 | 3207.981 | 2428.08 | 2223.036 |
| ENSMUSG00000030847 | 0.383273818 | 9.21E-05 | 0.001252 | 789.7527 | 722.2926 | 833.0753 | 818.6297 | 535.3361 | 644.7249 | 652.8238 | 595.9999 |

| | | | | | | | | | | | |
|---|---|---|---|---|---|---|---|---|---|---|---|
| ENSMUSG00000030849 | 0.085725963 | 0.500357 | 0.656799 | 4760.649 | 3778.238 | 5374.412 | 4092.286 | 3719.075 | 4367.815 | 4618.945 | 4264.445 |
| ENSMUSG00000030850 | -0.085749063 | 0.392283 | 0.559719 | 1248.513 | 1174.625 | 1228.91 | 1193.871 | 1130.154 | 1227.994 | 1283.469 | 1503.051 |
| ENSMUSG00000030852 | -0.22730827 | 0.093111 | 0.209847 | 1490.973 | 1261.012 | 1931.973 | 1470.773 | 1625.511 | 1876.071 | 1896.757 | 1811.927 |
| ENSMUSG00000030854 | 1.812784793 | 2.58E-18 | 2.38E-15 | 226.3622 | 194.3711 | 183.8397 | 207.8923 | 54.60623 | 74.86406 | 62.6788 | 39.15328 |
| ENSMUSG00000030861 | 0.081129641 | 0.346882 | 0.515719 | 2421.573 | 2130.883 | 1983.315 | 2278.189 | 1959.974 | 2107.368 | 2236.187 | 2027.27 |
| ENSMUSG00000030862 | -0.532652426 | 0.371823 | 0.540897 | 107.6478 | 220.7672 | 674.9069 | 196.6782 | 456.3521 | 255.8787 | 459.0016 | 565.5474 |
| ENSMUSG00000030865 | -0.799870583 | 0.00471 | 0.024668 | 32.19374 | 21.59679 | 32.29616 | 35.36756 | 45.83023 | 43.57759 | 61.71451 | 61.99269 |
| ENSMUSG00000030867 | -0.029880606 | 0.750548 | 0.844651 | 1145.896 | 1195.022 | 1162.662 | 1087.768 | 1300.798 | 1008.989 | 1184.147 | 1189.825 |
| ENSMUSG00000030868 | 0.155689425 | 0.01135 | 0.046904 | 1908.485 | 1932.912 | 1898.849 | 2059.945 | 1830.284 | 1680.531 | 1703.899 | 1788 |
| ENSMUSG00000030869 | -0.542786773 | 3.27E-06 | 9.68E-05 | 1800.837 | 1730.143 | 1999.878 | 1835.663 | 2607.448 | 2181.114 | 2882.26 | 3062.657 |
| ENSMUSG00000030870 | 0.180246902 | 0.092744 | 0.209295 | 2710.31 | 2645.606 | 2435.462 | 2750.044 | 2175.473 | 2804.609 | 2192.794 | 2131.679 |
| ENSMUSG00000030871 | -0.127220976 | 0.145876 | 0.284387 | 379.2825 | 412.7386 | 416.5376 | 388.1806 | 445.6259 | 414.5458 | 447.4302 | 435.0364 |
| ENSMUSG00000030872 | -0.088830848 | 0.354091 | 0.522769 | 2454.773 | 2799.184 | 3136.868 | 2708.638 | 3013.094 | 2657.116 | 3096.333 | 3038.729 |
| ENSMUSG00000030873 | -0.307342471 | 0.246192 | 0.409107 | 1433.627 | 1057.043 | 2437.118 | 1504.415 | 1865.388 | 1308.445 | 2259.33 | 2528.649 |
| ENSMUSG00000030876 | -0.133381169 | 0.108405 | 0.233184 | 2675.099 | 3061.945 | 2942.263 | 3255.541 | 3166.186 | 3526.433 | 3304.619 | 3095.284 |
| ENSMUSG00000030878 | -1.080698237 | 1.78E-06 | 5.95E-05 | 352.119 | 314.3532 | 429.7873 | 423.5481 | 677.7024 | 525.1658 | 1064.575 | 949.467 |
| ENSMUSG00000030879 | -0.380627358 | 3.27E-05 | 0.000571 | 1275.677 | 1286.209 | 1385.422 | 1255.117 | 1739.599 | 1468.23 | 1696.185 | 1869.569 |
| ENSMUSG00000030880 | -0.278417688 | 0.000486 | 0.00448 | 1190.162 | 1145.83 | 1139.475 | 1075.691 | 1438.289 | 1233.581 | 1388.576 | 1456.284 |
| ENSMUSG00000030881 | 0.003627143 | 0.960653 | 0.977472 | 2637.875 | 2688.8 | 2677.269 | 2633.59 | 2865.852 | 2357.659 | 2719.295 | 2664.598 |
| ENSMUSG00000030882 | 0.249142783 | 0.196215 | 0.350492 | 99.59938 | 113.983 | 92.74794 | 100.0643 | 91.66046 | 102.7984 | 77.14313 | 69.60583 |
| ENSMUSG00000030884 | -0.63665159 | 1.39E-06 | 4.87E-05 | 4251.586 | 4031.4 | 4552.93 | 4405.418 | 6718.517 | 5091.874 | 7070.168 | 7926.364 |
| ENSMUSG00000030887 | 0.088653916 | 0.824722 | 0.893464 | 28.16952 | 49.19268 | 29.81184 | 33.64232 | 28.27823 | 56.98608 | 24.10723 | 22.83941 |
| ENSMUSG00000030888 | -0.182813854 | 0.061439 | 0.157577 | 767.6195 | 775.0847 | 662.4853 | 730.6421 | 864.9237 | 727.4105 | 864.0031 | 871.1605 |
| ENSMUSG00000030889 | 0.557194277 | 0.182325 | 0.332903 | 24.1453 | 27.5959 | 18.21835 | 28.46658 | 13.65156 | 27.93435 | 13.50005 | 11.9635 |
| ENSMUSG00000030890 | -0.388075389 | 0.029618 | 0.093727 | 733.4136 | 758.2872 | 924.9951 | 731.5047 | 1096.025 | 692.7719 | 1134.968 | 1196.35 |
| ENSMUSG00000030894 | -0.264608501 | 0.01973 | 0.069796 | 2260.604 | 1758.938 | 2326.152 | 2123.779 | 2583.07 | 2285.03 | 2462.795 | 2848.401 |
| ENSMUSG00000030895 | 2.315981253 | 0.004701 | 0.024642 | 144.8718 | 5.999108 | 18.21835 | 60.38364 | 7.80089 | 13.40849 | 18.32149 | 6.525547 |
| ENSMUSG00000030898 | -0.258561666 | 0.685394 | 0.79985 | 10.06054 | 8.398751 | 13.24971 | 8.626235 | 10.72622 | 10.05637 | 24.10723 | 3.262773 |
| ENSMUSG00000030905 | 0.394576211 | 0.056236 | 0.147826 | 1549.324 | 2158.479 | 1752.274 | 1813.235 | 1392.459 | 1994.513 | 1069.397 | 1076.715 |
| ENSMUSG00000030909 | -3.004471646 | 3.56E-08 | 2.42E-06 | 33.19979 | 8.398751 | 66.24853 | 42.26855 | 216.4747 | 125.1459 | 314.3583 | 556.8466 |
| ENSMUSG00000030917 | 0.342264645 | 0.001863 | 0.012474 | 1098.611 | 945.4594 | 1085.648 | 1093.807 | 954.634 | 746.4059 | 875.5746 | 754.7882 |
| ENSMUSG00000030921 | 0.31130913 | 0.423968 | 0.589124 | 135.8173 | 188.372 | 283.2125 | 77.63611 | 116.0382 | 87.15518 | 181.2864 | 167.489 |
| ENSMUSG00000030922 | -0.046560347 | 0.829713 | 0.896882 | 65.39353 | 71.98929 | 54.65504 | 74.18562 | 56.55646 | 79.33356 | 71.3574 | 67.43065 |
| ENSMUSG00000030924 | 0.411369666 | 0.00312 | 0.018303 | 310.8708 | 290.3568 | 312.1962 | 345.912 | 257.4294 | 270.4045 | 226.608 | 193.5912 |
| ENSMUSG00000030929 | 0.119001833 | 0.3797 | 0.547789 | 641.8627 | 640.7047 | 479.4737 | 662.4948 | 576.2908 | 619.0252 | 542.8948 | 492.6788 |
| ENSMUSG00000030930 | -0.388946452 | 0.042101 | 0.120915 | 723.3531 | 506.3247 | 763.5143 | 662.4948 | 881.5006 | 603.382 | 1014.432 | 979.9196 |
| ENSMUSG00000030934 | 0.134728076 | 0.030064 | 0.094648 | 3624.814 | 3954.612 | 3576.593 | 3863.691 | 3399.238 | 3616.94 | 3382.726 | 3279.087 |
| ENSMUSG00000030935 | 0.117722328 | 0.675838 | 0.792479 | 47.28455 | 35.99465 | 36.43669 | 37.95543 | 34.1289 | 36.87335 | 27.96439 | 46.76642 |
| ENSMUSG00000030942 | -0.03436814 | 0.514921 | 0.669147 | 2387.367 | 2349.251 | 2382.463 | 2371.352 | 2567.468 | 2328.608 | 2423.259 | 2398.138 |
| ENSMUSG00000030945 | -3.845412629 | 8.11E-08 | 4.80E-06 | 140.8476 | 23.99643 | 279.0719 | 137.1571 | 2227.154 | 824.6221 | 2090.579 | 3219.27 |
| ENSMUSG00000030946 | 0.207376694 | 0.114415 | 0.241901 | 315.9011 | 458.3318 | 397.4912 | 411.4714 | 346.1645 | 354.2076 | 332.6798 | 337.1532 |
| ENSMUSG00000030956 | 0.22520244 | 0.005523 | 0.02764 | 898.4065 | 971.8554 | 873.6525 | 888.5022 | 783.0144 | 753.1101 | 725.1455 | 843.9707 |
| ENSMUSG00000030960 | 0.133616815 | 0.074312 | 0.179906 | 676.0685 | 755.8876 | 720.4528 | 756.5208 | 667.9512 | 673.7766 | 665.3595 | 643.8539 |
| ENSMUSG00000030963 | -5.57917345 | 4.13E-05 | 0.000691 | 12.07265 | 1.199822 | 305.5713 | 37.95543 | 3635.215 | 665.955 | 5751.985 | 7019.313 |
| ENSMUSG00000030965 | -0.032105163 | 0.733827 | 0.833293 | 1297.81 | 1240.615 | 1031.821 | 1109.334 | 1214.014 | 1152.013 | 1159.076 | 1255.08 |
| ENSMUSG00000030966 | 0.049778769 | 0.746128 | 0.841259 | 206.2411 | 267.5602 | 290.6654 | 209.6175 | 244.7529 | 239.1181 | 238.1794 | 218.6058 |
| ENSMUSG00000030967 | 0.307601426 | 0.082479 | 0.192981 | 3610.729 | 3799.835 | 3224.647 | 3667.012 | 2495.31 | 4119.758 | 2606.474 | 2335.058 |
| ENSMUSG00000030970 | 0.010965545 | 0.86307 | 0.917928 | 2581.535 | 2486.03 | 2435.462 | 2379.116 | 2341.242 | 2642.59 | 2396.259 | 2427.503 |
| ENSMUSG00000030972 | -3.466329589 | 1.63E-07 | 8.43E-06 | 7.04238 | 3.599465 | 30.63995 | 11.21411 | 153.0925 | 39.10809 | 122.4647 | 274.073 |
| ENSMUSG00000030978 | -0.192347495 | 0.007008 | 0.032971 | 3066.454 | 3504.679 | 3046.604 | 3160.652 | 3795.133 | 3488.442 | 3515.798 | 3794.605 |
| ENSMUSG00000030979 | -0.236849985 | 0.209398 | 0.366526 | 296.786 | 341.9491 | 228.5574 | 282.9405 | 345.1894 | 251.4092 | 339.4298 | 416.5474 |
| ENSMUSG00000030980 | 0.031667773 | 0.692444 | 0.805328 | 3637.893 | 3687.052 | 3430.846 | 3261.579 | 3298.802 | 3827.006 | 3237.119 | 3348.693 |
| ENSMUSG00000030982 | -0.133897634 | 0.133438 | 0.266927 | 1586.548 | 1568.167 | 1643.792 | 1537.195 | 1810.782 | 1484.99 | 1880.364 | 1773.861 |
| ENSMUSG00000030983 | -0.11986312 | 0.177231 | 0.326281 | 2165.029 | 2478.831 | 2200.279 | 2258.348 | 2745.913 | 2483.923 | 2456.045 | 2200.197 |
| ENSMUSG00000030986 | -0.106795551 | 0.223504 | 0.383185 | 746.4923 | 674.2997 | 693.9534 | 666.808 | 789.8402 | 686.0677 | 719.3597 | 799.3794 |
| ENSMUSG00000030987 | 0.349200769 | 0.004541 | 0.023968 | 870.237 | 878.2694 | 776.764 | 948.0232 | 639.673 | 812.331 | 686.5739 | 588.3868 |
| ENSMUSG00000030990 | 0.088282667 | 0.420915 | 0.586339 | 999.012 | 983.8537 | 1006.978 | 921.2819 | 1049.22 | 943.0637 | 924.7533 | 760.2262 |
| ENSMUSG00000030994 | -0.414882638 | 0.339403 | 0.507907 | 12.07265 | 16.7975 | 24.8432 | 11.21411 | 19.50223 | 16.76061 | 20.25007 | 30.45255 |
| ENSMUSG00000030996 | 0.50445409 | 0.277425 | 0.443765 | 17.10292 | 15.59768 | 19.87456 | 21.56559 | 7.80089 | 7.821619 | 18.32149 | 18.48905 |
| ENSMUSG00000031004 | 0.150634613 | 0.111973 | 0.238496 | 7734.546 | 7504.884 | 7060.437 | 6856.132 | 6888.186 | 7445.064 | 6038.379 | 5892.569 |
| ENSMUSG00000031007 | -0.14268883 | 0.202192 | 0.357713 | 1987.963 | 1774.536 | 2146.452 | 2211.767 | 1970.7 | 2086.137 | 2398.187 | 2514.511 |
| ENSMUSG00000031010 | -0.138270366 | 0.125552 | 0.256106 | 2050.339 | 1996.503 | 1999.878 | 1843.426 | 2217.403 | 1859.31 | 2276.687 | 2327.445 |
| ENSMUSG00000031012 | 0.312266525 | 0.047608 | 0.131477 | 2145.914 | 2660.004 | 2038.799 | 2121.191 | 1799.08 | 2343.133 | 1548.648 | 1529.153 |
| ENSMUSG00000031015 | 0.115763358 | 0.123555 | 0.253229 | 1379.301 | 1474.581 | 1432.624 | 1388.824 | 1432.439 | 1320.736 | 1208.254 | 1274.657 |

| | | | | | | | | | | | |
|---|---|---|---|---|---|---|---|---|---|---|---|
| ENSMUSG00000031016 | 0.107350452 | 0.363467 | 0.532013 | 1231.411 | 1401.392 | 1046.727 | 1310.325 | 1129.179 | 1220.173 | 1263.219 | 1015.81 |
| ENSMUSG00000031021 | -0.508292673 | 0.000357 | 0.003535 | 657.9595 | 593.9117 | 797.4667 | 734.9552 | 1040.444 | 781.0445 | 1048.182 | 1094.117 |
| ENSMUSG00000031023 | -0.204499709 | 0.145225 | 0.283446 | 318.9192 | 321.5522 | 363.5388 | 295.0172 | 418.3228 | 298.3389 | 404.0372 | 375.2189 |
| ENSMUSG00000031024 | 0.123899898 | 0.255772 | 0.419652 | 1023.157 | 1043.845 | 1218.973 | 1174.893 | 911.7291 | 1040.275 | 973.9321 | 1172.423 |
| ENSMUSG00000031026 | 0.811969691 | 0.077322 | 0.184733 | 18.10898 | 15.59768 | 18.21835 | 18.97772 | 7.80089 | 16.76061 | 9.642892 | 6.525547 |
| ENSMUSG00000031027 | 1.438961248 | 0.000331 | 0.003341 | 25.15136 | 37.19447 | 28.15563 | 38.81806 | 11.70134 | 17.87799 | 10.60718 | 7.613138 |
| ENSMUSG00000031028 | 0.960554272 | 0.001697 | 0.011607 | 326.9677 | 123.5816 | 173.0743 | 288.9789 | 108.2374 | 135.2023 | 101.2504 | 125.073 |
| ENSMUSG00000031029 | -0.116799911 | 0.194526 | 0.34829 | 7257.676 | 9031.057 | 8889.725 | 7937.861 | 9503.435 | 8751.274 | 9238.854 | 8412.517 |
| ENSMUSG00000031059 | -0.498073257 | 4.47E-05 | 0.000729 | 3737.492 | 3869.424 | 4164.548 | 3889.569 | 5771.684 | 4322.003 | 5573.591 | 6451.59 |
| ENSMUSG00000031060 | -0.137332608 | 0.08076 | 0.190212 | 2086.557 | 2186.075 | 1989.112 | 1909.848 | 2234.955 | 2060.438 | 2250.651 | 2436.204 |
| ENSMUSG00000031065 | -0.075556936 | 0.258601 | 0.422514 | 3198.247 | 2967.159 | 3207.257 | 3332.315 | 3282.225 | 3177.812 | 3332.583 | 3601.014 |
| ENSMUSG00000031066 | 0.195337432 | 0.024729 | 0.082399 | 1097.605 | 1163.827 | 1003.665 | 1108.471 | 1054.095 | 946.4159 | 889.0746 | 926.6276 |
| ENSMUSG00000031068 | -0.110280168 | 0.119192 | 0.248201 | 3284.767 | 3875.424 | 3679.278 | 3602.316 | 4034.035 | 3697.391 | 3846.549 | 4007.773 |
| ENSMUSG00000031070 | 0.667590767 | 0.093695 | 0.210713 | 42.25428 | 109.1838 | 57.96746 | 62.10889 | 30.22845 | 64.8077 | 27.96439 | 47.85401 |
| ENSMUSG00000031072 | -0.108284078 | 0.158678 | 0.301634 | 537.233 | 529.1213 | 534.9569 | 504.6347 | 580.1912 | 539.6917 | 580.5021 | 567.7225 |
| ENSMUSG00000031073 | -1.073342496 | 0.046353 | 0.129228 | 14.08476 | 5.999108 | 7.45296 | 5.175741 | 12.67645 | 12.29112 | 25.07152 | 18.48905 |
| ENSMUSG00000031075 | -0.281903287 | 0.04104 | 0.118888 | 1074.466 | 1149.429 | 1576.715 | 1080.005 | 1455.841 | 1429.121 | 1451.255 | 1600.934 |
| ENSMUSG00000031077 | -0.349607026 | 0.022341 | 0.076273 | 364.1917 | 418.7377 | 436.4122 | 372.6533 | 540.2117 | 381.0246 | 515.8947 | 590.562 |
| ENSMUSG00000031078 | -0.055901462 | 0.496835 | 0.653524 | 5617.808 | 5158.033 | 5571.501 | 5418.138 | 6007.661 | 4914.211 | 5696.056 | 6007.853 |
| ENSMUSG00000031079 | 0.308933609 | 0.292774 | 0.460125 | 54.32694 | 73.18911 | 49.6864 | 39.68068 | 41.92979 | 56.98608 | 41.46443 | 33.71532 |
| ENSMUSG00000031090 | -0.244367951 | 0.276666 | 0.442803 | 262.5802 | 217.1677 | 210.3391 | 282.9405 | 242.8027 | 211.1837 | 289.2867 | 410.0218 |
| ENSMUSG00000031093 | 0.72013239 | 0.00014 | 0.001735 | 1005.048 | 1343.8 | 939.901 | 1187.833 | 653.3246 | 927.4205 | 586.2878 | 550.3211 |
| ENSMUSG00000031095 | 0.07919372 | 0.209664 | 0.366924 | 1461.797 | 1474.581 | 1368.032 | 1453.521 | 1383.683 | 1431.356 | 1326.862 | 1306.197 |
| ENSMUSG00000031097 | 0.10390894 | 0.705232 | 0.814617 | 73.44197 | 55.19179 | 53.82693 | 37.95543 | 58.50668 | 54.75133 | 43.39301 | 47.85401 |
| ENSMUSG00000031098 | -0.728775992 | 0.217449 | 0.375877 | 72.43591 | 16.7975 | 101.029 | 26.74133 | 128.7147 | 29.05173 | 108.0004 | 94.62042 |
| ENSMUSG00000031099 | 0.590966068 | 7.06E-08 | 4.25E-06 | 1220.344 | 1418.189 | 1284.393 | 1468.185 | 813.2428 | 1034.688 | 905.4675 | 826.5692 |
| ENSMUSG00000031101 | -0.090758658 | 0.827284 | 0.895137 | 25.15136 | 29.99554 | 37.2648 | 12.93935 | 34.1289 | 18.99536 | 25.07152 | 33.71532 |
| ENSMUSG00000031103 | 0.55100468 | 5.10E-08 | 3.31E-06 | 537.233 | 583.1133 | 599.5492 | 562.4305 | 347.1396 | 408.9589 | 374.1442 | 429.5985 |
| ENSMUSG00000031104 | 0.806947959 | 0.084026 | 0.195387 | 86.52067 | 27.5959 | 28.98373 | 40.5433 | 29.25334 | 36.87335 | 18.32149 | 20.66423 |
| ENSMUSG00000031105 | 0.236495449 | 0.051085 | 0.138101 | 273.6468 | 278.3586 | 236.8385 | 264.8254 | 224.2756 | 249.1744 | 213.1079 | 206.6423 |
| ENSMUSG00000031107 | 0.083455771 | 0.603185 | 0.740579 | 476.8698 | 526.7217 | 463.7397 | 431.3117 | 462.2028 | 575.4477 | 373.1799 | 380.6569 |
| ENSMUSG00000031109 | -0.399444317 | 0.001962 | 0.012949 | 269.6226 | 235.165 | 308.0557 | 244.9851 | 337.3885 | 367.6161 | 318.2154 | 376.3065 |
| ENSMUSG00000031111 | 1.685857034 | 7.00E-16 | 3.92E-13 | 237.4288 | 233.9652 | 259.1974 | 274.3143 | 58.50668 | 110.62 | 79.07171 | 65.25547 |
| ENSMUSG00000031112 | 0.453405505 | 0.069935 | 0.172234 | 558.3602 | 716.2935 | 458.7711 | 724.6037 | 406.6214 | 686.0677 | 374.1442 | 328.4525 |
| ENSMUSG00000031119 | -0.982687297 | 7.87E-06 | 0.000191 | 1108.672 | 793.082 | 1276.94 | 974.7645 | 1667.44 | 1488.342 | 2209.186 | 2846.226 |
| ENSMUSG00000031129 | -0.461580407 | 0.080537 | 0.189809 | 53.32088 | 93.58608 | 120.9036 | 85.39973 | 108.2374 | 112.8548 | 147.5362 | 118.5474 |
| ENSMUSG00000031133 | 0.133136612 | 0.564396 | 0.709411 | 279.6831 | 152.3773 | 163.137 | 172.5247 | 161.8685 | 150.8455 | 177.4292 | 209.9051 |
| ENSMUSG00000031134 | -0.006063991 | 0.934101 | 0.961183 | 3581.554 | 3823.831 | 3304.974 | 3741.198 | 3449.944 | 3909.692 | 3575.584 | 3574.912 |
| ENSMUSG00000031137 | 0.484720971 | 0.091888 | 0.208135 | 90.54489 | 100.785 | 180.5272 | 125.0804 | 77.03379 | 64.8077 | 102.2147 | 112.0219 |
| ENSMUSG00000031139 | 1.223789097 | 8.24E-06 | 0.000197 | 158.9566 | 194.3711 | 87.7793 | 189.7772 | 72.15824 | 63.69032 | 70.39311 | 63.08028 |
| ENSMUSG00000031142 | 0.468131441 | 0.0267 | 0.087156 | 69.41775 | 76.78858 | 81.15445 | 73.323 | 56.55646 | 61.45558 | 42.42872 | 57.64233 |
| ENSMUSG00000031143 | -0.177185587 | 0.604324 | 0.741075 | 25.15136 | 14.39786 | 25.67131 | 21.56559 | 27.30312 | 22.34748 | 22.17865 | 27.18978 |
| ENSMUSG00000031144 | 0.256821504 | 0.18648 | 0.33839 | 269.6226 | 232.7654 | 255.0568 | 404.5704 | 230.1263 | 268.1698 | 248.7866 | 227.3065 |
| ENSMUSG00000031145 | 0.07653164 | 0.617404 | 0.750131 | 390.3491 | 391.1418 | 332.0708 | 325.2091 | 391.9947 | 262.5829 | 369.3228 | 337.1532 |
| ENSMUSG00000031146 | -0.000341468 | 0.996855 | 0.998149 | 2006.072 | 2207.672 | 2489.289 | 2335.122 | 2413.4 | 2156.532 | 2283.437 | 2189.321 |
| ENSMUSG00000031147 | -0.181327239 | 0.5631 | 0.708471 | 35.2119 | 47.99286 | 49.6864 | 38.81806 | 42.9049 | 30.1691 | 54.00019 | 67.43065 |
| ENSMUSG00000031148 | 0.035246024 | 0.663746 | 0.783755 | 1270.647 | 1198.622 | 1118.772 | 1189.558 | 1204.262 | 1064.858 | 1138.826 | 1251.817 |
| ENSMUSG00000031149 | 0.275129697 | 0.026623 | 0.086996 | 435.6215 | 579.5138 | 554.0033 | 519.2993 | 480.7299 | 438.0106 | 396.3228 | 410.0218 |
| ENSMUSG00000031150 | -0.132921259 | 0.294043 | 0.461714 | 415.5004 | 343.149 | 491.8953 | 392.4937 | 468.0534 | 449.1844 | 432.9658 | 455.7007 |
| ENSMUSG00000031153 | 0.047803787 | 0.555232 | 0.702355 | 1491.979 | 1286.209 | 1315.861 | 1425.917 | 1284.222 | 1302.858 | 1305.648 | 1449.759 |
| ENSMUSG00000031154 | 0.075894762 | 0.112557 | 0.239278 | 2475.9 | 2523.225 | 2480.179 | 2421.384 | 2288.586 | 2428.054 | 2325.865 | 2350.284 |
| ENSMUSG00000031155 | 0.017539756 | 0.911088 | 0.947486 | 135.8173 | 106.7841 | 114.2787 | 135.4319 | 118.9636 | 129.6154 | 120.5361 | 118.5474 |
| ENSMUSG00000031156 | 0.049745751 | 0.642256 | 0.768658 | 603.6326 | 549.5183 | 544.8942 | 498.5964 | 506.0828 | 473.7666 | 574.7163 | 565.5474 |
| ENSMUSG00000031157 | -0.029126757 | 0.623972 | 0.754981 | 1475.882 | 1540.571 | 1526.201 | 1440.581 | 1578.705 | 1507.338 | 1459.934 | 1557.43 |
| ENSMUSG00000031158 | -0.228697279 | 0.041514 | 0.119829 | 948.7093 | 950.2587 | 944.0416 | 993.7423 | 1172.084 | 897.2514 | 1217.897 | 1207.226 |
| ENSMUSG00000031161 | -0.207367348 | 0.068136 | 0.168913 | 1793.795 | 1658.153 | 2019.752 | 1774.417 | 1943.397 | 1866.015 | 2074.186 | 2486.233 |
| ENSMUSG00000031165 | -0.194312182 | 0.308364 | 0.476744 | 74.44802 | 85.18733 | 79.49824 | 87.12497 | 89.71024 | 113.9722 | 91.60747 | 78.30656 |
| ENSMUSG00000031166 | 0.059623545 | 0.356445 | 0.524955 | 2173.077 | 1950.91 | 2066.126 | 2051.319 | 2007.754 | 1863.78 | 1957.507 | 2080.562 |
| ENSMUSG00000031167 | -0.178478412 | 0.010482 | 0.044148 | 15289.01 | 16874.29 | 17201.43 | 16669.34 | 19666.04 | 17132.7 | 19462.25 | 18467.3 |
| ENSMUSG00000031168 | -0.64579116 | 0.000776 | 0.006435 | 393.3673 | 392.3416 | 407.4285 | 412.334 | 596.7681 | 419.0153 | 665.3595 | 830.9196 |
| ENSMUSG00000031169 | 0.190239382 | 0.087956 | 0.201829 | 247.4894 | 301.1552 | 279.0719 | 278.6274 | 244.7529 | 250.2918 | 243.0009 | 230.5693 |
| ENSMUSG00000031170 | 0.970996254 | 0.025188 | 0.083461 | 17.10292 | 51.59233 | 44.71776 | 46.58167 | 22.42756 | 15.64324 | 27.96439 | 15.22628 |
| ENSMUSG00000031171 | -0.078467831 | 0.439034 | 0.602492 | 539.2451 | 531.5209 | 481.9581 | 582.2709 | 587.017 | 524.0485 | 610.395 | 530.7445 |
| ENSMUSG00000031174 | 0.2755723 | 0.069628 | 0.171547 | 832.007 | 872.2703 | 659.1729 | 789.3005 | 597.7432 | 831.3263 | 593.0378 | 581.8612 |

| | | | | | | | | | | | |
|---|---|---|---|---|---|---|---|---|---|---|---|
| ENSMUSG00000031176 | -0.076096967 | 0.312569 | 0.480916 | 1590.572 | 1515.375 | 1657.041 | 1658.825 | 1646.963 | 1584.436 | 1680.756 | 1861.956 |
| ENSMUSG00000031189 | -0.590787042 | 0.150203 | 0.290155 | 10.06054 | 21.59679 | 10.76539 | 18.11509 | 19.50223 | 27.93435 | 20.25007 | 22.83941 |
| ENSMUSG00000031196 | -0.287498406 | 0.278022 | 0.444257 | 100.6054 | 74.38894 | 155.684 | 85.39973 | 121.8889 | 103.9158 | 135.0005 | 149 |
| ENSMUSG00000031197 | 0.145252463 | 0.22755 | 0.38753 | 2230.423 | 2547.221 | 1996.565 | 2668.094 | 2132.568 | 2429.171 | 2075.15 | 1900.022 |
| ENSMUSG00000031198 | 0.103591381 | 0.181862 | 0.332454 | 1936.655 | 2174.077 | 2087.657 | 2346.336 | 2045.784 | 2002.334 | 2012.471 | 1892.408 |
| ENSMUSG00000031200 | 0.238758665 | 0.13669 | 0.271407 | 292.7618 | 263.9607 | 229.3855 | 223.4195 | 175.52 | 219.0053 | 250.7152 | 208.8175 |
| ENSMUSG00000031201 | 0.045986564 | 0.541253 | 0.690719 | 871.2431 | 1009.05 | 944.8697 | 926.4576 | 896.1273 | 930.7726 | 937.2891 | 866.8101 |
| ENSMUSG00000031202 | 0.744262294 | 0.001859 | 0.012452 | 122.7386 | 154.777 | 105.9976 | 150.0965 | 69.2329 | 111.7374 | 71.3574 | 66.34306 |
| ENSMUSG00000031207 | -0.188792018 | 0.176034 | 0.324519 | 3254.586 | 2627.609 | 3741.386 | 3472.06 | 3670.319 | 3068.309 | 3903.443 | 4288.372 |
| ENSMUSG00000031209 | 0.036128797 | 0.731806 | 0.831919 | 1644.899 | 1791.334 | 1999.049 | 1745.95 | 1762.026 | 1490.577 | 1932.435 | 1818.452 |
| ENSMUSG00000031210 | -0.481752855 | 0.659627 | 0.780834 | 4.024217 | 1.199822 | 38.92101 | 0.862623 | 13.65156 | 4.469496 | 20.25007 | 25.0146 |
| ENSMUSG00000031214 | 0.219676689 | 0.022397 | 0.076392 | 563.3904 | 567.5156 | 490.2391 | 571.9194 | 443.6756 | 502.8183 | 484.0732 | 451.3503 |
| ENSMUSG00000031216 | -0.697180562 | 0.00496 | 0.0256 | 515.0998 | 445.1338 | 717.1403 | 567.6063 | 772.2882 | 568.7434 | 1022.147 | 1279.007 |
| ENSMUSG00000031217 | -0.041854383 | 0.688403 | 0.802246 | 2394.409 | 2757.19 | 3048.261 | 2607.711 | 2611.348 | 2685.05 | 2652.759 | 3179.029 |
| ENSMUSG00000031220 | -0.038338181 | 0.924677 | 0.955253 | 14.08476 | 22.79661 | 14.07781 | 15.52722 | 20.47734 | 12.29112 | 19.28578 | 15.22628 |
| ENSMUSG00000031221 | -0.111513605 | 0.114345 | 0.241847 | 2504.069 | 2633.608 | 2571.271 | 2749.181 | 3024.795 | 2707.397 | 2934.332 | 2629.795 |
| ENSMUSG00000031224 | -0.427527568 | 0.220022 | 0.378834 | 15.09082 | 28.79572 | 36.43669 | 29.3292 | 28.27823 | 43.57759 | 39.53586 | 36.9781 |
| ENSMUSG00000031226 | 0.049900969 | 0.756963 | 0.848807 | 1204.247 | 998.2515 | 963.088 | 1163.679 | 914.6544 | 1391.131 | 911.2533 | 966.8685 |
| ENSMUSG00000031227 | 0.484873144 | 7.62E-06 | 0.000187 | 389.343 | 334.7502 | 361.8826 | 409.7462 | 258.4045 | 265.935 | 279.6439 | 266.4598 |
| ENSMUSG00000031229 | 0.461754051 | 0.004578 | 0.024118 | 7560.498 | 7945.218 | 6199.206 | 7230.51 | 4872.631 | 7132.199 | 4617.981 | 4387.342 |
| ENSMUSG00000031230 | 1.449924883 | 0.022457 | 0.076505 | 23.13925 | 25.19625 | 30.63995 | 18.97772 | 8.776002 | 21.23011 | 2.892867 | 3.262773 |
| ENSMUSG00000031231 | -0.720569266 | 9.49E-05 | 0.001281 | 2256.58 | 2330.053 | 3024.245 | 2308.38 | 4001.857 | 2778.909 | 4434.766 | 5131.255 |
| ENSMUSG00000031232 | -0.249880743 | 0.003814 | 0.021094 | 1692.183 | 1721.744 | 1822.663 | 1672.627 | 1777.628 | 2129.715 | 2108.9 | 2202.372 |
| ENSMUSG00000031239 | 0.472639885 | 0.040558 | 0.117861 | 2030.218 | 3519.077 | 2771.673 | 2707.775 | 1852.711 | 2944.281 | 1666.292 | 1484.562 |
| ENSMUSG00000031242 | 0.353729484 | 0.000312 | 0.0032 | 705.2441 | 905.8653 | 765.9986 | 763.4218 | 599.6935 | 633.5511 | 610.395 | 611.2262 |
| ENSMUSG00000031245 | 0.263481914 | 0.003668 | 0.020491 | 2124.787 | 2469.233 | 2002.362 | 2051.319 | 1682.067 | 1895.066 | 1873.614 | 1748.846 |
| ENSMUSG00000031246 | 0.351972759 | 0.001314 | 0.009604 | 5050.393 | 6079.496 | 5253.508 | 5912.421 | 4252.46 | 5120.925 | 4240.944 | 3854.423 |
| ENSMUSG00000031253 | -0.637640931 | 0.056022 | 0.147496 | 54.32694 | 67.19001 | 157.3403 | 80.22398 | 141.3911 | 112.8548 | 144.6434 | 162.0511 |
| ENSMUSG00000031255 | 0.469199837 | 0.064528 | 0.163321 | 287.7315 | 151.1775 | 179.6991 | 300.193 | 136.5156 | 179.8972 | 157.1791 | 191.416 |
| ENSMUSG00000031256 | -0.086160883 | 0.21705 | 0.375514 | 1462.803 | 1542.97 | 1422.687 | 1374.159 | 1510.447 | 1539.742 | 1463.791 | 1642.263 |
| ENSMUSG00000031257 | 0.208885794 | 0.683802 | 0.798548 | 20.12109 | 19.19714 | 11.59349 | 13.80198 | 11.70134 | 25.6996 | 7.714313 | 10.87591 |
| ENSMUSG00000031258 | -0.609644505 | 0.15432 | 0.295787 | 25.15136 | 16.7975 | 16.56213 | 11.21411 | 29.25334 | 33.52122 | 28.92867 | 14.13868 |
| ENSMUSG00000031262 | -0.02200658 | 0.894098 | 0.936094 | 404.4339 | 406.7395 | 291.4935 | 316.5828 | 404.6712 | 402.2547 | 309.5368 | 321.927 |
| ENSMUSG00000031264 | 0.314258231 | 0.43863 | 0.602154 | 27.16347 | 16.7975 | 29.81184 | 18.97772 | 25.35289 | 12.29112 | 15.42863 | 21.75182 |
| ENSMUSG00000031266 | 0.14719226 | 0.213097 | 0.370925 | 283.7073 | 307.1543 | 283.2125 | 289.8415 | 278.8818 | 235.7659 | 293.1439 | 240.3576 |
| ENSMUSG00000031273 | 0.630416917 | 0.000368 | 0.003615 | 545.2815 | 592.7118 | 624.3924 | 612.4627 | 310.0854 | 531.8701 | 329.7869 | 364.343 |
| ENSMUSG00000031274 | -0.029550793 | 0.759685 | 0.850154 | 1828.001 | 1996.503 | 2331.12 | 1991.798 | 1912.193 | 2069.377 | 2092.507 | 2245.876 |
| ENSMUSG00000031278 | 0.128510832 | 0.41053 | 0.576705 | 1997.018 | 1316.204 | 1283.565 | 1476.811 | 1284.222 | 1325.206 | 1593.97 | 1351.876 |
| ENSMUSG00000031283 | 1.217483551 | 1.25E-05 | 0.000274 | 151.9142 | 253.1623 | 155.684 | 227.7326 | 82.88446 | 119.559 | 81.00029 | 55.46715 |
| ENSMUSG00000031284 | 0.951141502 | 0.002043 | 0.013322 | 227.3683 | 171.5745 | 128.3565 | 197.5408 | 75.08357 | 155.315 | 73.28598 | 71.78101 |
| ENSMUSG00000031285 | 1.152461149 | 0.000333 | 0.003355 | 831.0009 | 1030.647 | 607.8303 | 953.199 | 365.6667 | 663.7202 | 311.4654 | 199.0292 |
| ENSMUSG00000031286 | -0.55802478 | 0.000662 | 0.005694 | 90.54489 | 97.18554 | 118.4192 | 118.1794 | 159.9183 | 173.193 | 142.7148 | 152.2628 |
| ENSMUSG00000031289 | 0.748086924 | 0.128023 | 0.259366 | 11.0666 | 23.99643 | 15.73403 | 23.29083 | 16.57689 | 10.05637 | 10.60718 | 6.525547 |
| ENSMUSG00000031290 | 0.740981268 | 6.25E-06 | 0.00016 | 1110.684 | 1340.201 | 948.1821 | 1176.618 | 640.6481 | 890.5472 | 620.0379 | 586.2116 |
| ENSMUSG00000031292 | 0.149532767 | 0.329657 | 0.497437 | 357.1493 | 296.3559 | 313.0243 | 273.4516 | 257.4294 | 316.2169 | 314.3583 | 229.4817 |
| ENSMUSG00000031293 | 0.050188823 | 0.675374 | 0.792365 | 290.7497 | 338.3497 | 310.54 | 285.5284 | 298.3841 | 334.0949 | 266.1438 | 283.8613 |
| ENSMUSG00000031294 | -2.812757041 | 0.000112 | 0.001463 | 7.04238 | 1.199822 | 7.45296 | 3.450494 | 52.65601 | 7.821619 | 25.07152 | 51.11678 |
| ENSMUSG00000031295 | 0.230219301 | 0.081142 | 0.190797 | 1001.024 | 859.0722 | 789.1856 | 970.4514 | 740.1095 | 916.2468 | 776.2528 | 653.6422 |
| ENSMUSG00000031297 | 0.863369996 | 4.63E-06 | 0.000126 | 127.7689 | 151.1775 | 118.4192 | 146.646 | 67.28268 | 63.69032 | 88.7146 | 78.30656 |
| ENSMUSG00000031298 | 0.30017262 | 0.287165 | 0.454071 | 165.999 | 266.3604 | 165.6213 | 160.448 | 142.3663 | 234.6486 | 127.2862 | 110.9343 |
| ENSMUSG00000031299 | -0.386590375 | 0.004435 | 0.023569 | 4324.022 | 4408.144 | 5146.683 | 4670.244 | 6113.948 | 4577.882 | 6561.023 | 6997.561 |
| ENSMUSG00000031302 | 1.719464556 | 5.07E-06 | 0.000136 | 200.2048 | 181.1731 | 231.0418 | 219.1064 | 42.9049 | 122.9112 | 56.89306 | 30.45255 |
| ENSMUSG00000031303 | -0.228712393 | 0.426426 | 0.591383 | 139.8416 | 109.1838 | 139.1219 | 127.6683 | 193.072 | 93.85942 | 107.0361 | 210.9927 |
| ENSMUSG00000031304 | -0.112818775 | 0.455903 | 0.617914 | 122.7386 | 142.7788 | 127.5284 | 122.4925 | 133.5902 | 131.8501 | 130.179 | 160.9635 |
| ENSMUSG00000031309 | -0.104712849 | 0.176779 | 0.325688 | 2999.048 | 2751.191 | 2779.126 | 2887.201 | 2982.865 | 3447.099 | 2890.939 | 2958.248 |
| ENSMUSG00000031310 | 0.2210651 | 0.000663 | 0.005697 | 1847.116 | 1644.955 | 1651.245 | 1744.225 | 1502.647 | 1511.807 | 1442.577 | 1453.022 |
| ENSMUSG00000031311 | -0.060702966 | 0.326266 | 0.494017 | 14443.92 | 15439.3 | 15111.29 | 15597.96 | 17143.43 | 14963.87 | 15591.59 | 15494.91 |
| ENSMUSG00000031312 | 0.569222558 | 0.195796 | 0.349952 | 35.2119 | 38.39429 | 35.60859 | 59.52102 | 14.62667 | 50.28183 | 30.85725 | 18.48905 |
| ENSMUSG00000031314 | 0.168043599 | 0.014051 | 0.054885 | 1571.457 | 1529.772 | 1440.077 | 1495.789 | 1271.545 | 1445.882 | 1329.755 | 1325.774 |
| ENSMUSG00000031320 | -0.052320114 | 0.566706 | 0.711144 | 46977.71 | 57661.02 | 51693.73 | 53741.44 | 59071.27 | 54732.34 | 55704.09 | 48320.58 |
| ENSMUSG00000031323 | -0.145452269 | 0.683897 | 0.798608 | 20.12109 | 34.79482 | 42.23344 | 24.15346 | 26.32801 | 43.57759 | 28.92867 | 35.89051 |
| ENSMUSG00000031327 | 0.221900139 | 0.143456 | 0.281124 | 424.5549 | 373.1445 | 403.2879 | 365.7524 | 300.3343 | 434.6585 | 307.6082 | 302.3503 |
| ENSMUSG00000031328 | 0.13651123 | 0.0678 | 0.1683 | 18938.97 | 20273.38 | 22539.41 | 20296.67 | 17469.12 | 18578.58 | 18678.28 | 19919.23 |
| ENSMUSG00000031333 | -0.255589467 | 0.0053 | 0.026792 | 703.232 | 669.5004 | 609.4865 | 614.1879 | 735.2339 | 738.5843 | 773.3599 | 849.4086 |

| | | | | | | | | | | | |
|---|---|---|---|---|---|---|---|---|---|---|---|
| ENSMUSG00000031337 | 0.102912268 | 0.493035 | 0.650007 | 258.556 | 196.7707 | 200.4018 | 238.0841 | 201.848 | 213.4185 | 234.3223 | 182.7153 |
| ENSMUSG00000031340 | 1.11548247 | 1.94E-10 | 3.17E-08 | 388.337 | 376.744 | 325.4459 | 412.334 | 193.072 | 211.1837 | 162.0006 | 127.2482 |
| ENSMUSG00000031342 | 0.532595262 | 2.94E-07 | 1.35E-05 | 954.7456 | 1017.449 | 1013.603 | 1057.576 | 648.449 | 825.7395 | 685.6096 | 637.3284 |
| ENSMUSG00000031343 | 0.43271823 | 0.144872 | 0.282968 | 152.9203 | 169.1748 | 164.7932 | 211.3428 | 96.53602 | 220.1227 | 97.39321 | 104.4087 |
| ENSMUSG00000031344 | 1.302015585 | 0.112134 | 0.238643 | 18.10898 | 21.59679 | 9.93728 | 25.01608 | 1.950223 | 22.34748 | 1.928578 | 4.350364 |
| ENSMUSG00000031347 | -0.046524406 | 0.629797 | 0.759014 | 1069.436 | 1337.801 | 1231.395 | 1248.216 | 1143.806 | 1338.614 | 1241.04 | 1323.598 |
| ENSMUSG00000031349 | -0.798905838 | 2.54E-05 | 0.000473 | 443.67 | 392.3416 | 477.8175 | 428.7239 | 718.657 | 521.8137 | 788.7885 | 1003.847 |
| ENSMUSG00000031351 | -0.367176878 | 0.019775 | 0.069889 | 243.4652 | 309.554 | 317.1648 | 276.0395 | 306.185 | 335.2122 | 439.7159 | 396.9707 |
| ENSMUSG00000031352 | -0.157617054 | 0.107148 | 0.231478 | 632.8082 | 572.3149 | 578.8465 | 615.9132 | 662.1006 | 620.1426 | 632.5737 | 763.4889 |
| ENSMUSG00000031353 | -0.280816929 | 0.009158 | 0.039986 | 6741.57 | 7339.308 | 8482.296 | 7480.671 | 9876.902 | 7763.515 | 8971.746 | 9889.466 |
| ENSMUSG00000031355 | 0.274358628 | 0.049483 | 0.135116 | 255.5378 | 326.3515 | 293.9779 | 255.3366 | 216.4747 | 270.4045 | 223.7151 | 224.0438 |
| ENSMUSG00000031357 | 0.01524524 | 0.823243 | 0.892753 | 1618.741 | 1647.355 | 1619.777 | 1766.653 | 1554.327 | 1596.728 | 1743.435 | 1689.029 |
| ENSMUSG00000031358 | -0.146726022 | 0.085645 | 0.198029 | 1160.987 | 1180.624 | 1064.117 | 1119.685 | 1290.072 | 1250.342 | 1353.862 | 1110.431 |
| ENSMUSG00000031360 | 0.196782051 | 0.030677 | 0.096043 | 1536.245 | 1464.982 | 1302.612 | 1543.233 | 1290.072 | 1148.661 | 1373.148 | 1286.62 |
| ENSMUSG00000031365 | 0.255354974 | 0.031635 | 0.09809 | 951.7274 | 968.256 | 852.1217 | 952.3363 | 781.0642 | 911.7773 | 785.8957 | 640.5912 |
| ENSMUSG00000031367 | 0.331691374 | 0.110317 | 0.23618 | 792.7708 | 673.0999 | 510.9418 | 759.9713 | 482.6801 | 762.0491 | 481.1803 | 449.1751 |
| ENSMUSG00000031370 | 0.08511011 | 0.344576 | 0.513276 | 1674.074 | 1631.757 | 1639.651 | 1784.768 | 1546.527 | 1828.024 | 1499.47 | 1473.686 |
| ENSMUSG00000031371 | -0.226140015 | 0.003196 | 0.018615 | 628.784 | 646.7038 | 619.4238 | 585.7214 | 766.4375 | 710.6499 | 727.074 | 692.7955 |
| ENSMUSG00000031373 | -0.332132598 | 0.041178 | 0.119094 | 308.8587 | 267.5602 | 366.0231 | 286.391 | 390.0445 | 331.8601 | 348.1084 | 479.6277 |
| ENSMUSG00000031375 | -0.31184245 | 0.116865 | 0.245132 | 3124.805 | 3501.079 | 5718.076 | 4230.306 | 4949.665 | 3985.673 | 5711.485 | 5929.547 |
| ENSMUSG00000031376 | 0.778658895 | 0.091864 | 0.208107 | 55.33299 | 13.19804 | 26.49941 | 33.64232 | 18.52711 | 16.76061 | 19.28578 | 20.66423 |
| ENSMUSG00000031377 | -0.604060542 | 0.341972 | 0.510758 | 27.16347 | 2.399643 | 30.63995 | 32.77969 | 30.22845 | 16.76061 | 45.32159 | 50.02919 |
| ENSMUSG00000031378 | 0.116370558 | 0.253458 | 0.417179 | 488.9424 | 451.1329 | 509.2856 | 467.5419 | 411.497 | 405.6068 | 466.716 | 486.1532 |
| ENSMUSG00000031379 | 0.434828335 | 0.002552 | 0.01566 | 187.1261 | 241.1641 | 202.8861 | 226.0074 | 151.1423 | 179.8972 | 148.5005 | 154.4379 |
| ENSMUSG00000031380 | 0.701673909 | 0.000461 | 0.004307 | 438.6397 | 481.1284 | 489.411 | 532.2387 | 247.6783 | 427.9543 | 229.5008 | 290.3868 |
| ENSMUSG00000031381 | -0.239511627 | 0.098377 | 0.218516 | 467.8153 | 344.3488 | 423.1625 | 452.0147 | 437.825 | 581.0345 | 529.3948 | 446.9999 |
| ENSMUSG00000031383 | -1.040405934 | 0.001666 | 0.011446 | 324.9556 | 239.9643 | 496.864 | 305.3687 | 544.1121 | 408.9589 | 658.6095 | 1202.876 |
| ENSMUSG00000031384 | -2.483460339 | 2.36E-06 | 7.36E-05 | 32.19374 | 10.79839 | 86.12309 | 40.5433 | 216.4747 | 115.0895 | 234.3223 | 389.3576 |
| ENSMUSG00000031385 | -0.131647333 | 0.814475 | 0.887294 | 5.030272 | 13.19804 | 14.90592 | 29.3292 | 12.67645 | 17.87799 | 23.14294 | 15.22628 |
| ENSMUSG00000031386 | 0.205578499 | 0.022168 | 0.075821 | 3081.544 | 3336.704 | 3300.833 | 3275.381 | 2698.133 | 3267.202 | 2564.045 | 2741.817 |
| ENSMUSG00000031387 | -0.151557051 | 0.189815 | 0.342466 | 687.1351 | 736.6904 | 753.577 | 733.23 | 904.9033 | 669.3071 | 767.5742 | 890.7371 |
| ENSMUSG00000031388 | -0.130813167 | 0.041824 | 0.120365 | 1068.43 | 1093.037 | 1086.476 | 1040.324 | 1240.342 | 1119.609 | 1184.147 | 1147.409 |
| ENSMUSG00000031389 | -0.206095556 | 0.393531 | 0.56085 | 187.1261 | 159.5763 | 288.1811 | 176.8378 | 252.5538 | 165.3714 | 236.2508 | 283.8613 |
| ENSMUSG00000031391 | -0.171058279 | 0.494077 | 0.650917 | 687.1351 | 391.1418 | 995.3842 | 795.3389 | 752.7859 | 679.3635 | 849.5388 | 951.6422 |
| ENSMUSG00000031392 | 0.170579708 | 0.03799 | 0.112489 | 2377.306 | 2428.439 | 2220.982 | 2292.853 | 2179.374 | 2261.565 | 1971.971 | 1865.219 |
| ENSMUSG00000031393 | 0.287413194 | 0.016382 | 0.061218 | 1816.934 | 1818.929 | 1681.056 | 1620.87 | 1361.255 | 1753.16 | 1316.255 | 1253.993 |
| ENSMUSG00000031394 | -0.017446225 | 0.965594 | 0.980631 | 17.10292 | 25.19625 | 15.73403 | 14.6646 | 12.67645 | 24.58223 | 17.3572 | 18.48905 |
| ENSMUSG00000031397 | -1.193686128 | NA | NA | 22.1332 | 10.79839 | 14.07781 | 5.175741 | 7.80089 | 99.44629 | 5.785735 | 6.525547 |
| ENSMUSG00000031398 | 0.458517317 | 0.00013 | 0.001639 | 910.4792 | 1040.245 | 847.1531 | 947.1606 | 754.7362 | 750.8754 | 641.2523 | 576.4233 |
| ENSMUSG00000031399 | -0.042298073 | 0.660334 | 0.781326 | 710.2744 | 766.686 | 798.2948 | 724.6037 | 714.7566 | 705.0631 | 826.3958 | 842.8831 |
| ENSMUSG00000031400 | -0.065673544 | 0.481184 | 0.640351 | 1275.677 | 1142.23 | 1130.366 | 1104.158 | 1291.047 | 1059.271 | 1256.469 | 1259.43 |
| ENSMUSG00000031402 | -0.079811923 | 0.30937 | 0.477801 | 1090.563 | 1102.636 | 1220.629 | 1256.842 | 1245.217 | 1165.421 | 1284.433 | 1245.292 |
| ENSMUSG00000031403 | -0.09647028 | 0.349814 | 0.518671 | 1996.012 | 2360.049 | 1870.693 | 2075.472 | 2355.869 | 2416.88 | 2108.9 | 1990.292 |
| ENSMUSG00000031409 | 2.208246456 | 2.17E-09 | 2.47E-07 | 68.4117 | 62.39072 | 46.37397 | 77.63611 | 8.776002 | 22.34748 | 13.50005 | 10.87591 |
| ENSMUSG00000031422 | -0.061697605 | 0.206285 | 0.362797 | 10041.43 | 11031.16 | 10637.86 | 10961.36 | 10995.36 | 11283.24 | 11279.29 | 10977.06 |
| ENSMUSG00000031425 | 0.726212692 | 0.011241 | 0.046601 | 403.4278 | 341.9491 | 247.6039 | 508.9479 | 186.2463 | 345.2686 | 216.0008 | 160.9635 |
| ENSMUSG00000031428 | 0.971020254 | 4.00E-15 | 1.85E-12 | 1108.672 | 1105.036 | 911.7454 | 1242.178 | 493.4063 | 612.321 | 585.3235 | 537.27 |
| ENSMUSG00000031429 | 0.030730182 | 0.675635 | 0.792479 | 971.8485 | 1048.644 | 944.0416 | 961.8252 | 1006.315 | 902.8383 | 988.3964 | 939.6787 |
| ENSMUSG00000031430 | 0.069314619 | 0.889145 | 0.933305 | 36.21796 | 8.398751 | 28.98373 | 26.74133 | 12.67645 | 27.93435 | 31.82154 | 23.927 |
| ENSMUSG00000031431 | 0.125754626 | 0.62296 | 0.754426 | 2755.583 | 1959.309 | 1696.79 | 3043.336 | 1678.167 | 2413.528 | 3081.868 | 1492.175 |
| ENSMUSG00000031432 | 0.063838017 | 0.38772 | 0.555313 | 947.7032 | 955.0579 | 890.2146 | 936.8091 | 926.3557 | 852.5564 | 944.0391 | 840.7079 |
| ENSMUSG00000031433 | 0.161378855 | 0.239845 | 0.401796 | 264.5923 | 229.1659 | 200.4018 | 239.8093 | 199.8978 | 189.9536 | 217.9294 | 226.2189 |
| ENSMUSG00000031434 | 0.089298287 | 0.432056 | 0.596508 | 355.1372 | 322.752 | 315.5086 | 339.011 | 274.0063 | 349.7381 | 312.4297 | 317.5766 |
| ENSMUSG00000031438 | -0.264745753 | 0.092079 | 0.208389 | 1392.379 | 1382.194 | 2081.86 | 1427.642 | 1772.752 | 1652.596 | 2048.15 | 2079.474 |
| ENSMUSG00000031441 | -0.260865047 | 0.072399 | 0.176687 | 3320.985 | 3160.33 | 3311.598 | 3302.985 | 3495.774 | 3218.037 | 3787.728 | 5191.072 |
| ENSMUSG00000031442 | 0.092751299 | 0.4695 | 0.629502 | 727.3773 | 518.3229 | 555.6596 | 647.8302 | 530.4606 | 567.626 | 610.395 | 589.4744 |
| ENSMUSG00000031444 | -2.530521442 | 3.11E-05 | 0.00055 | 23.13925 | 8.398751 | 18.21835 | 23.29083 | 136.5156 | 26.81698 | 52.07161 | 208.8175 |
| ENSMUSG00000031445 | -0.450925562 | 0.289209 | 0.456248 | 25.15136 | 17.99732 | 17.39024 | 32.77969 | 36.07912 | 17.87799 | 25.07152 | 48.9416 |
| ENSMUSG00000031446 | -0.092676843 | 0.156643 | 0.298843 | 3291.81 | 3085.941 | 2975.387 | 3087.329 | 3355.358 | 3066.075 | 3348.012 | 3493.343 |
| ENSMUSG00000031447 | -0.461901205 | 0.000418 | 0.003987 | 6139.95 | 6541.427 | 7817.327 | 6578.367 | 9461.505 | 7481.937 | 9303.462 | 11049.93 |
| ENSMUSG00000031450 | 0.107498581 | 0.380322 | 0.548377 | 307.8526 | 380.3434 | 342.008 | 379.5543 | 320.8116 | 372.0856 | 312.4297 | 303.4379 |
| ENSMUSG00000031451 | -0.842631186 | 0.04103 | 0.118883 | 1380.307 | 1702.547 | 3849.04 | 1746.813 | 3519.177 | 2369.95 | 4381.73 | 5294.393 |
| ENSMUSG00000031453 | -0.052200445 | 0.587685 | 0.72794 | 757.5589 | 790.6824 | 751.9208 | 817.7671 | 833.7202 | 689.4198 | 837.9673 | 870.0729 |
| ENSMUSG00000031458 | -0.413260005 | 0.013043 | 0.05215 | 311.8769 | 290.3568 | 380.9291 | 344.1868 | 454.4019 | 330.7427 | 456.1088 | 528.5693 |

| | | | | | | | | | | | |
|---|---|---|---|---|---|---|---|---|---|---|---|
| ENSMUSG00000031461 | -0.015376378 | 0.967942 | 0.981932 | 18.10898 | 20.39697 | 21.53077 | 29.3292 | 14.62667 | 31.28647 | 20.25007 | 25.0146 |
| ENSMUSG00000031465 | -1.436218068 | 5.31E-05 | 0.000829 | 149.9021 | 92.38626 | 216.1358 | 188.9145 | 441.7254 | 213.4185 | 403.0729 | 697.1459 |
| ENSMUSG00000031467 | -0.27586566 | 0.000201 | 0.002282 | 928.5882 | 1010.25 | 894.3552 | 906.6173 | 1065.797 | 1117.374 | 1179.326 | 1159.372 |
| ENSMUSG00000031478 | 0.312746458 | 0.115007 | 0.242689 | 267.6105 | 267.5602 | 221.1045 | 266.5507 | 158.9431 | 277.1088 | 217.9294 | 169.6642 |
| ENSMUSG00000031479 | -0.225092314 | 0.011637 | 0.047824 | 1789.771 | 1583.764 | 1833.428 | 1623.457 | 2115.016 | 1785.564 | 1963.293 | 2121.89 |
| ENSMUSG00000031480 | -0.300542106 | 0.115771 | 0.243773 | 311.8769 | 245.9634 | 423.1625 | 297.6051 | 334.4632 | 460.3581 | 357.7513 | 426.3357 |
| ENSMUSG00000031482 | -0.57291101 | 1.95E-05 | 0.000385 | 481.9 | 487.1275 | 536.6131 | 530.5134 | 789.8402 | 611.2036 | 720.324 | 909.2261 |
| ENSMUSG00000031483 | -0.190419335 | 0.020411 | 0.071549 | 2458.797 | 2576.017 | 2757.595 | 2461.927 | 2833.673 | 2638.12 | 3070.297 | 3159.452 |
| ENSMUSG00000031485 | -0.220326063 | 0.011297 | 0.046778 | 1751.541 | 1494.978 | 1548.559 | 1613.969 | 1878.064 | 1670.474 | 1907.364 | 2010.956 |
| ENSMUSG00000031486 | 0.017381328 | 0.878293 | 0.927015 | 2881.34 | 2783.586 | 3608.889 | 3059.726 | 2978.965 | 2671.641 | 3120.44 | 3418.299 |
| ENSMUSG00000031487 | -0.219184149 | 0.096888 | 0.216018 | 305.8405 | 377.9438 | 294.806 | 290.7041 | 406.6214 | 377.6724 | 365.4656 | 323.0146 |
| ENSMUSG00000031488 | 0.109146447 | 0.448906 | 0.611484 | 654.9414 | 703.0954 | 956.4632 | 655.5939 | 744.0099 | 650.3117 | 658.6095 | 702.5838 |
| ENSMUSG00000031490 | -0.180398108 | 0.05365 | 0.14313 | 1237.447 | 1209.42 | 1297.643 | 1287.897 | 1500.696 | 1210.116 | 1541.898 | 1449.759 |
| ENSMUSG00000031491 | -0.194359467 | 0.602729 | 0.740217 | 16.09687 | 20.39697 | 14.90592 | 28.46658 | 26.32801 | 18.99536 | 23.14294 | 22.83941 |
| ENSMUSG00000031492 | 0.17021399 | 0.69477 | 0.80727 | 11.0666 | 14.39786 | 13.24971 | 13.80198 | 9.751113 | 13.40849 | 10.60718 | 13.05109 |
| ENSMUSG00000031493 | 0.642936025 | 0.08508 | 0.197118 | 32.19374 | 40.79393 | 27.32752 | 37.09281 | 11.70134 | 20.11273 | 30.85725 | 25.0146 |
| ENSMUSG00000031497 | 0.646134956 | 0.012226 | 0.049679 | 428.5792 | 493.1267 | 423.1625 | 527.9256 | 218.4249 | 477.1187 | 294.1082 | 207.7299 |
| ENSMUSG00000031502 | -0.491136757 | 0.025424 | 0.08399 | 13941.9 | 13194.44 | 22367.16 | 15351.25 | 20042.44 | 15851.07 | 25559.45 | 29706.46 |
| ENSMUSG00000031503 | -0.52319561 | 0.022355 | 0.076305 | 6831.109 | 6291.864 | 11383.15 | 7211.532 | 9825.222 | 8060.737 | 13034.3 | 14665.08 |
| ENSMUSG00000031504 | -0.363317896 | 0.018565 | 0.067001 | 768.6255 | 627.5067 | 838.0439 | 758.2461 | 937.082 | 770.9881 | 1205.361 | 937.5035 |
| ENSMUSG00000031505 | -0.217152737 | 0.016307 | 0.061024 | 1331.01 | 1470.981 | 1445.874 | 1544.959 | 1839.06 | 1487.225 | 1672.077 | 1734.708 |
| ENSMUSG00000031506 | -0.410356503 | 0.356381 | 0.524955 | 24.1453 | 9.598572 | 14.07781 | 14.6646 | 21.45245 | 25.6996 | 12.53576 | 23.927 |
| ENSMUSG00000031508 | 0.122754196 | 0.296612 | 0.464315 | 3552.378 | 3558.671 | 2894.233 | 3634.233 | 3019.92 | 3728.677 | 2920.832 | 2857.102 |
| ENSMUSG00000031511 | -0.098315892 | 0.312652 | 0.480979 | 1879.31 | 1864.523 | 2278.949 | 1912.436 | 1996.053 | 1988.926 | 2191.829 | 2322.007 |
| ENSMUSG00000031512 | -0.568782744 | 0.439995 | 0.603263 | 7.04238 | 4.799286 | 23.18699 | 3.450494 | 9.751113 | 13.40849 | 8.678602 | 26.10219 |
| ENSMUSG00000031513 | 0.023549624 | 0.73662 | 0.834974 | 1297.81 | 1400.192 | 1264.519 | 1363.808 | 1379.783 | 1274.924 | 1341.326 | 1238.766 |
| ENSMUSG00000031516 | -0.028209751 | 0.599229 | 0.737488 | 1634.838 | 1643.756 | 1613.98 | 1709.72 | 1739.599 | 1652.596 | 1655.684 | 1684.679 |
| ENSMUSG00000031517 | -0.581547147 | 0.00249 | 0.015382 | 287.7315 | 401.9402 | 380.1009 | 375.2412 | 468.0534 | 434.6585 | 524.5733 | 735.2116 |
| ENSMUSG00000031519 | -0.492105304 | 0.256779 | 0.420395 | 20.12109 | 13.19804 | 16.56213 | 12.07673 | 14.62667 | 21.23011 | 33.75012 | 17.40146 |
| ENSMUSG00000031520 | 0.9076441 | 5.48E-07 | 2.27E-05 | 547.2936 | 589.1124 | 529.9882 | 597.7981 | 260.3547 | 426.8369 | 264.2152 | 256.6715 |
| ENSMUSG00000031521 | -0.03100662 | 0.756069 | 0.848334 | 528.1785 | 572.3149 | 592.0962 | 572.782 | 586.0419 | 512.8747 | 567.002 | 649.2919 |
| ENSMUSG00000031523 | 0.049263359 | 0.514929 | 0.669147 | 2038.266 | 2218.47 | 2288.059 | 2105.664 | 2106.24 | 1915.179 | 2083.829 | 2254.576 |
| ENSMUSG00000031527 | -0.039624225 | 0.589162 | 0.729378 | 1164.005 | 1129.032 | 1066.601 | 1149.014 | 1258.869 | 1106.2 | 1145.576 | 1120.219 |
| ENSMUSG00000031529 | 0.248750156 | 0.00127 | 0.009355 | 2516.142 | 2450.036 | 2464.445 | 2597.359 | 2035.057 | 2361.011 | 2107.936 | 1938.087 |
| ENSMUSG00000031530 | -0.094999406 | 0.357218 | 0.525702 | 537.233 | 461.9313 | 496.864 | 533.1013 | 577.2659 | 497.2315 | 588.2164 | 504.6423 |
| ENSMUSG00000031532 | -0.184823426 | 0.013114 | 0.052372 | 3412.536 | 3279.112 | 3357.144 | 3468.609 | 3878.993 | 3396.817 | 4099.193 | 3989.284 |
| ENSMUSG00000031533 | -0.205400852 | 0.025012 | 0.083101 | 642.8687 | 543.5192 | 649.2356 | 647.8302 | 718.657 | 668.1897 | 741.5384 | 739.5619 |
| ENSMUSG00000031534 | -0.044180912 | 0.706463 | 0.815414 | 462.785 | 556.7172 | 457.943 | 513.261 | 453.4268 | 576.565 | 488.8946 | 532.9196 |
| ENSMUSG00000031536 | 0.028001825 | 0.713682 | 0.820362 | 649.9111 | 667.1008 | 619.4238 | 641.7919 | 615.2952 | 667.0723 | 600.7521 | 644.9415 |
| ENSMUSG00000031537 | -0.021127318 | 0.721592 | 0.825853 | 1800.837 | 1742.141 | 1707.556 | 1764.065 | 1791.279 | 1666.005 | 1853.364 | 1804.314 |
| ENSMUSG00000031538 | -0.088489506 | 0.258504 | 0.422432 | 2472.882 | 3031.949 | 2684.722 | 2858.734 | 2962.388 | 3028.084 | 2925.653 | 2827.737 |
| ENSMUSG00000031539 | 0.030140015 | 0.852809 | 0.911639 | 191.1503 | 220.7672 | 271.619 | 208.7549 | 207.6987 | 196.6578 | 212.1436 | 258.8467 |
| ENSMUSG00000031540 | 0.232059318 | 0.002938 | 0.01751 | 1933.636 | 1882.52 | 1671.947 | 1885.695 | 1467.543 | 1672.709 | 1533.22 | 1604.197 |
| ENSMUSG00000031543 | 0.241065301 | 0.228659 | 0.3887 | 98.59333 | 73.18911 | 74.5296 | 80.22398 | 63.38224 | 78.21619 | 65.57166 | 69.60583 |
| ENSMUSG00000031545 | -0.238836627 | 0.006082 | 0.029755 | 2712.323 | 2712.796 | 2937.294 | 2731.929 | 3241.27 | 2852.656 | 3374.048 | 3624.941 |
| ENSMUSG00000031546 | -0.146241354 | 0.082508 | 0.193024 | 1019.133 | 1223.818 | 1136.162 | 1099.845 | 1293.973 | 1148.661 | 1289.255 | 1220.277 |
| ENSMUSG00000031548 | -0.042196889 | 0.799249 | 0.877144 | 3863.249 | 5678.755 | 6290.298 | 5042.034 | 4907.735 | 4723.14 | 5261.162 | 6602.765 |
| ENSMUSG00000031554 | 0.741145536 | 0.004955 | 0.025582 | 90.54489 | 89.98662 | 79.49824 | 113.0037 | 49.73068 | 75.98144 | 60.75022 | 36.9781 |
| ENSMUSG00000031555 | -0.210062824 | 0.077001 | 0.184275 | 1507.069 | 1191.423 | 1507.982 | 1440.581 | 1483.144 | 1459.291 | 1840.828 | 1752.109 |
| ENSMUSG00000031556 | -0.396467888 | 0.001323 | 0.00965 | 1305.859 | 1238.216 | 1484.795 | 1406.076 | 1686.943 | 1467.112 | 1980.65 | 2021.832 |
| ENSMUSG00000031557 | -0.060360628 | 0.448824 | 0.611428 | 560.3723 | 575.9143 | 526.6758 | 592.6223 | 575.3157 | 570.9782 | 604.6093 | 599.2627 |
| ENSMUSG00000031558 | 0.120944185 | 0.554189 | 0.70118 | 2810.916 | 2542.422 | 2529.038 | 2751.769 | 1991.177 | 3742.086 | 2047.186 | 2000.08 |
| ENSMUSG00000031561 | 0.461680529 | 1.00E-09 | 1.30E-07 | 3099.653 | 3445.887 | 3009.34 | 3206.372 | 2349.043 | 2496.214 | 2274.758 | 2143.642 |
| ENSMUSG00000031562 | 0.193714173 | 0.128561 | 0.26023 | 1103.642 | 1400.192 | 1035.133 | 1180.932 | 979.9869 | 1176.595 | 1063.611 | 903.7882 |
| ENSMUSG00000031563 | -0.308532383 | 0.010896 | 0.045495 | 2265.634 | 1953.309 | 2537.319 | 2089.274 | 2505.061 | 2393.415 | 2990.261 | 3069.182 |
| ENSMUSG00000031565 | 0.091487422 | 0.351087 | 0.51987 | 5816 | 6278.666 | 5944.149 | 6071.144 | 6146.127 | 6104.215 | 5701.842 | 4674.466 |
| ENSMUSG00000031568 | -0.197526116 | 0.066097 | 0.165668 | 988.9514 | 1031.847 | 1004.493 | 1004.956 | 1044.344 | 1044.745 | 1150.397 | 1382.328 |
| ENSMUSG00000031570 | 0.00059185 | 0.993847 | 0.996515 | 932.6124 | 981.454 | 945.6978 | 944.5727 | 950.7335 | 919.5989 | 887.146 | 1045.175 |
| ENSMUSG00000031574 | 3.859224174 | 0.003395 | 0.019358 | 14594.83 | 223.1668 | 192.9488 | 1741.637 | 157.968 | 214.5358 | 170.6792 | 611.2262 |
| ENSMUSG00000031575 | -0.057749155 | 0.341337 | 0.510057 | 2557.39 | 2567.618 | 2483.492 | 2805.252 | 2744.938 | 2777.792 | 2633.474 | 2685.262 |
| ENSMUSG00000031576 | 0.162880583 | 0.759582 | 0.850143 | 24.1453 | 13.19804 | 19.04645 | 12.07673 | 16.57689 | 4.469496 | 19.28578 | 20.66423 |
| ENSMUSG00000031577 | -0.001604345 | 0.984399 | 0.99122 | 741.4621 | 826.677 | 745.296 | 786.7126 | 790.8153 | 835.7958 | 746.3598 | 728.686 |
| ENSMUSG00000031578 | -0.062333473 | 0.494589 | 0.651408 | 778.6861 | 722.2926 | 682.3599 | 708.2139 | 813.2428 | 788.8661 | 680.7882 | 735.2116 |
| ENSMUSG00000031583 | 0.178801272 | 0.070241 | 0.172779 | 573.451 | 614.3086 | 498.5202 | 579.683 | 510.9583 | 520.6963 | 503.3589 | 463.3138 |

| | | | | | | | | | | | |
|---|---|---|---|---|---|---|---|---|---|---|---|
| ENSMUSG00000031584 | -0.724322656 | 5.28E-07 | 2.20E-05 | 1797.819 | 1570.566 | 1816.038 | 1466.46 | 2756.64 | 2196.757 | 2616.117 | 3419.386 |
| ENSMUSG00000031585 | -0.014080256 | 0.905411 | 0.943292 | 1115.714 | 1105.036 | 827.2785 | 1092.944 | 1060.921 | 1084.97 | 1083.861 | 948.3794 |
| ENSMUSG00000031586 | 0.046849042 | 0.65787 | 0.77956 | 2416.543 | 2322.854 | 2549.74 | 2980.364 | 2276.885 | 2780.027 | 2496.545 | 2392.7 |
| ENSMUSG00000031590 | 0.116735754 | 0.195096 | 0.348939 | 1504.051 | 1569.367 | 1446.702 | 1495.789 | 1419.762 | 1572.145 | 1312.398 | 1243.117 |
| ENSMUSG00000031591 | -0.03771095 | 0.662094 | 0.782505 | 2554.372 | 2457.235 | 2994.434 | 2741.417 | 2605.497 | 2643.707 | 2943.011 | 2845.138 |
| ENSMUSG00000031592 | -0.01651072 | 0.762771 | 0.852453 | 2053.357 | 2036.097 | 1904.645 | 1936.59 | 2035.057 | 1977.752 | 1961.364 | 2042.496 |
| ENSMUSG00000031594 | -0.181153222 | 0.626262 | 0.756432 | 73.44197 | 40.79393 | 69.56096 | 60.38364 | 72.15824 | 29.05173 | 86.78602 | 89.18247 |
| ENSMUSG00000031595 | 0.646564342 | 3.37E-08 | 2.34E-06 | 553.3299 | 687.4977 | 727.9057 | 672.8463 | 461.2276 | 393.3157 | 430.073 | 402.4087 |
| ENSMUSG00000031596 | 0.202473635 | 0.07418 | 0.179688 | 743.4742 | 686.2979 | 807.404 | 740.9936 | 627.9717 | 707.2978 | 545.7877 | 711.2846 |
| ENSMUSG00000031600 | 0.008881149 | 0.937185 | 0.962692 | 969.8364 | 853.0731 | 838.872 | 832.4317 | 747.9104 | 1001.167 | 833.1458 | 891.8247 |
| ENSMUSG00000031601 | 0.162795857 | 0.159077 | 0.302057 | 2394.409 | 2519.625 | 2163.015 | 2721.577 | 2119.892 | 2585.604 | 2119.508 | 1928.299 |
| ENSMUSG00000031603 | 0.523056317 | 0.431826 | 0.596362 | 157.9505 | 137.9795 | 58.79557 | 62.97152 | 49.73068 | 194.4231 | 27.0001 | 19.57664 |
| ENSMUSG00000031604 | -1.065594256 | 1.77E-08 | 1.37E-06 | 735.4257 | 435.5352 | 726.2495 | 518.4367 | 1194.511 | 1105.083 | 1525.505 | 1234.416 |
| ENSMUSG00000031605 | -0.248726763 | 0.07018 | 0.172653 | 841.0614 | 680.2988 | 719.6247 | 720.2906 | 772.2882 | 741.9364 | 987.4321 | 1016.898 |
| ENSMUSG00000031608 | 0.37993431 | 0.001293 | 0.009468 | 1261.592 | 1258.613 | 956.4632 | 1156.778 | 841.5211 | 1004.519 | 868.8245 | 843.9707 |
| ENSMUSG00000031609 | -0.168680276 | 0.167668 | 0.313587 | 549.3057 | 514.7234 | 524.1915 | 686.6483 | 677.7024 | 581.0345 | 632.5737 | 667.7809 |
| ENSMUSG00000031613 | -0.189587879 | 0.356455 | 0.524955 | 219.3198 | 367.1454 | 381.7572 | 337.2858 | 289.6081 | 360.9118 | 432.0015 | 406.7591 |
| ENSMUSG00000031616 | 0.374291043 | 0.008674 | 0.038451 | 1537.251 | 1646.155 | 1588.309 | 1584.639 | 1103.826 | 1616.84 | 1047.218 | 1137.62 |
| ENSMUSG00000031617 | 0.207075944 | 0.028904 | 0.09207 | 1028.188 | 1160.227 | 951.4945 | 1136.938 | 998.514 | 917.3641 | 921.8604 | 863.5473 |
| ENSMUSG00000031618 | -0.708549782 | 0.019835 | 0.070063 | 61.36932 | 71.98929 | 139.1219 | 96.61383 | 140.416 | 109.5027 | 152.3577 | 203.3795 |
| ENSMUSG00000031622 | -0.17880002 | 0.018078 | 0.065834 | 2948.745 | 2938.363 | 3072.276 | 2916.53 | 3527.953 | 2956.572 | 3418.405 | 3539.021 |
| ENSMUSG00000031626 | 0.711044936 | 0.000392 | 0.003799 | 2722.383 | 3195.125 | 2587.833 | 2535.25 | 1590.407 | 2479.453 | 1391.469 | 1283.357 |
| ENSMUSG00000031627 | 0.273182013 | 0.050204 | 0.136381 | 599.6084 | 434.3354 | 491.0672 | 445.1137 | 374.4427 | 452.5365 | 374.1442 | 430.6861 |
| ENSMUSG00000031628 | -0.035284627 | 0.652452 | 0.77555 | 1519.142 | 1708.546 | 1528.685 | 1707.132 | 1770.802 | 1689.47 | 1589.149 | 1571.569 |
| ENSMUSG00000031629 | -0.205103191 | 0.077662 | 0.185281 | 200.2048 | 214.7681 | 210.3391 | 200.1287 | 246.7032 | 215.6532 | 255.5366 | 231.6569 |
| ENSMUSG00000031631 | 0.042382781 | 0.624918 | 0.755551 | 861.1825 | 935.8608 | 840.5282 | 906.6173 | 823.9691 | 944.1811 | 883.2889 | 788.5035 |
| ENSMUSG00000031633 | -0.514832662 | 2.71E-05 | 0.000495 | 10064.57 | 11094.75 | 11321.87 | 11183.91 | 16148.82 | 12197.26 | 15743.95 | 18299.81 |
| ENSMUSG00000031634 | -0.069149449 | 0.499331 | 0.655685 | 956.7577 | 881.8688 | 859.5747 | 963.5504 | 1039.469 | 815.6831 | 956.5749 | 1028.861 |
| ENSMUSG00000031636 | 0.505530013 | 0.007598 | 0.034881 | 611.681 | 843.4745 | 876.1368 | 854.8599 | 485.6054 | 767.636 | 478.2874 | 514.4306 |
| ENSMUSG00000031637 | 0.370257336 | 0.481669 | 0.640559 | 12.07265 | 11.99822 | 26.49941 | 18.11509 | 9.751113 | 20.11273 | 6.750024 | 17.40146 |
| ENSMUSG00000031639 | 0.108391592 | 0.484164 | 0.642397 | 190.1443 | 235.165 | 271.619 | 196.6782 | 220.3752 | 203.3621 | 195.7507 | 209.9051 |
| ENSMUSG00000031641 | -0.054076313 | 0.493651 | 0.65051 | 642.8687 | 665.901 | 583.8152 | 610.7374 | 631.8721 | 651.4291 | 671.1453 | 639.5036 |
| ENSMUSG00000031642 | 0.005391317 | 0.975901 | 0.986862 | 1404.452 | 982.6538 | 1018.571 | 1117.097 | 864.9237 | 1312.915 | 1344.219 | 984.2699 |
| ENSMUSG00000031644 | 0.228458782 | 0.076915 | 0.18419 | 826.9767 | 757.0874 | 700.5782 | 778.949 | 573.3654 | 807.8615 | 636.4308 | 598.1751 |
| ENSMUSG00000031647 | -0.446956313 | 0.092348 | 0.20869 | 373.2462 | 347.9482 | 562.2844 | 422.6855 | 530.4606 | 331.8601 | 654.7523 | 810.2554 |
| ENSMUSG00000031652 | 0.09061601 | 0.304848 | 0.473394 | 1074.466 | 1114.634 | 948.1821 | 1013.583 | 946.8331 | 991.1108 | 900.6461 | 1057.139 |
| ENSMUSG00000031654 | -0.374505208 | 0.54121 | 0.690719 | 4.024217 | 13.19804 | 33.12427 | 29.3292 | 25.35289 | 35.75597 | 28.92867 | 14.13868 |
| ENSMUSG00000031657 | -0.194838997 | 0.041279 | 0.119337 | 1094.587 | 1106.235 | 972.1972 | 1038.599 | 1256.918 | 1073.797 | 1147.504 | 1338.825 |
| ENSMUSG00000031659 | -0.245735232 | 0.366276 | 0.534852 | 185.114 | 215.9679 | 395.0069 | 222.5569 | 289.6081 | 211.1837 | 362.5727 | 345.854 |
| ENSMUSG00000031660 | -0.127312243 | 0.044979 | 0.126449 | 2285.755 | 2438.037 | 2458.649 | 2345.473 | 2574.294 | 2641.472 | 2430.973 | 2762.481 |
| ENSMUSG00000031661 | -0.021872964 | 0.923812 | 0.954694 | 1533.227 | 1641.356 | 2736.064 | 1682.978 | 1948.272 | 1353.14 | 1873.614 | 2536.262 |
| ENSMUSG00000031662 | -0.18084474 | 0.437674 | 0.601289 | 70.4238 | 50.3925 | 57.96746 | 63.83414 | 50.70579 | 78.21619 | 73.28598 | 73.95619 |
| ENSMUSG00000031665 | 0.268163552 | 0.039037 | 0.11447 | 4307.925 | 4441.739 | 3364.597 | 3868.004 | 3441.168 | 3920.866 | 3024.011 | 2883.204 |
| ENSMUSG00000031666 | 0.135618032 | 0.100497 | 0.221597 | 1252.538 | 1063.042 | 1069.914 | 1201.635 | 1013.141 | 1038.041 | 1087.718 | 1037.562 |
| ENSMUSG00000031667 | -0.489694533 | 6.64E-05 | 0.000991 | 1291.774 | 1185.424 | 1296.815 | 1335.341 | 1602.108 | 1525.216 | 1895.792 | 2153.43 |
| ENSMUSG00000031668 | -0.071940468 | 0.560059 | 0.706118 | 845.0857 | 836.2756 | 860.4028 | 810.0035 | 807.3922 | 720.7063 | 1007.682 | 986.4451 |
| ENSMUSG00000031669 | -0.085644141 | 0.597171 | 0.736036 | 237.4288 | 333.5504 | 274.1033 | 238.9467 | 332.513 | 250.2918 | 262.2867 | 302.3503 |
| ENSMUSG00000031671 | -0.090946952 | 0.377506 | 0.545596 | 331.9979 | 328.7511 | 303.087 | 325.2091 | 377.3681 | 331.8601 | 317.2511 | 344.7664 |
| ENSMUSG00000031672 | -0.616695254 | 0.000173 | 0.002029 | 4029.248 | 4020.602 | 5097.824 | 4357.111 | 6696.089 | 4760.014 | 7155.026 | 8231.977 |
| ENSMUSG00000031673 | 0.155278319 | 0.033565 | 0.102561 | 9120.889 | 10790 | 9913.264 | 9385.344 | 9047.083 | 8317.733 | 9159.783 | 8677.889 |
| ENSMUSG00000031681 | 0.22317178 | 0.0664 | 0.166099 | 1248.513 | 1265.812 | 1200.755 | 1277.545 | 945.858 | 1334.145 | 970.0749 | 1028.861 |
| ENSMUSG00000031683 | -0.207608824 | 0.001958 | 0.012938 | 1412.5 | 1594.563 | 1517.091 | 1415.565 | 1684.017 | 1777.742 | 1726.078 | 1667.277 |
| ENSMUSG00000031684 | 0.001372323 | 0.986283 | 0.992283 | 631.8021 | 693.4968 | 657.5167 | 630.5778 | 617.2455 | 632.4337 | 688.5025 | 669.9561 |
| ENSMUSG00000031691 | 0.144806457 | 0.062252 | 0.159064 | 2356.179 | 2283.26 | 2221.81 | 2256.623 | 2202.776 | 2197.875 | 2003.793 | 1841.292 |
| ENSMUSG00000031696 | -0.153745055 | 0.005126 | 0.026178 | 2923.594 | 2859.175 | 2937.294 | 2923.431 | 3277.349 | 3024.732 | 3259.297 | 3392.197 |
| ENSMUSG00000031697 | -0.063163503 | 0.487991 | 0.645802 | 1147.908 | 1250.214 | 1180.052 | 1180.069 | 1287.147 | 1395.6 | 1179.326 | 1108.255 |
| ENSMUSG00000031698 | -1.378210607 | 0.000283 | 0.002972 | 38.23007 | 14.39786 | 25.67131 | 28.46658 | 65.33246 | 50.28183 | 55.92877 | 107.6715 |
| ENSMUSG00000031700 | 0.435095167 | 5.09E-05 | 0.000802 | 725.3652 | 693.4968 | 565.5968 | 707.3513 | 473.9041 | 521.8137 | 503.3589 | 490.5036 |
| ENSMUSG00000031701 | -0.206295978 | 0.00086 | 0.006949 | 4205.307 | 4414.143 | 4210.922 | 4298.453 | 4941.864 | 4526.482 | 5080.84 | 5208.474 |
| ENSMUSG00000031703 | -0.089480274 | 0.230185 | 0.390428 | 2093.599 | 2098.488 | 2306.277 | 2338.572 | 2208.627 | 2300.673 | 2367.33 | 2531.912 |
| ENSMUSG00000031706 | 0.138699865 | 0.144095 | 0.282148 | 693.1715 | 664.7011 | 690.6409 | 818.6297 | 629.9219 | 686.0677 | 647.038 | 644.9415 |
| ENSMUSG00000031708 | -0.146407612 | 0.067461 | 0.16764 | 4656.02 | 4452.538 | 4697.021 | 4741.841 | 5405.042 | 4510.839 | 5044.197 | 5569.554 |
| ENSMUSG00000031709 | 0.610039728 | 2.16E-08 | 1.62E-06 | 1016.115 | 1023.448 | 1011.946 | 991.1544 | 719.6321 | 748.6406 | 628.7165 | 551.4087 |
| ENSMUSG00000031711 | -0.041902456 | 0.615684 | 0.748949 | 1566.427 | 1472.181 | 1305.096 | 1356.044 | 1549.452 | 1443.647 | 1480.184 | 1388.854 |

| | | | | | | | | | | | |
|---|---|---|---|---|---|---|---|---|---|---|---|
| ENSMUSG00000031712 | -0.241055875 | 0.276689 | 0.442803 | 83.50251 | 79.18822 | 111.7944 | 105.2401 | 89.71024 | 98.32892 | 121.5004 | 141.3868 |
| ENSMUSG00000031714 | 0.312761346 | 0.002544 | 0.015622 | 2006.072 | 2061.293 | 1843.365 | 2208.316 | 1550.427 | 1915.179 | 1559.256 | 1512.839 |
| ENSMUSG00000031715 | -0.067770858 | 0.409792 | 0.575975 | 3459.821 | 3539.474 | 3013.48 | 3331.452 | 3276.374 | 3846.002 | 3389.476 | 3471.591 |
| ENSMUSG00000031722 | 0.227686672 | 0.688649 | 0.802482 | 280.6892 | 25.19625 | 58.79557 | 161.3106 | 116.0382 | 105.0332 | 119.5719 | 108.7591 |
| ENSMUSG00000031723 | -0.039707897 | 0.736182 | 0.834773 | 284.7134 | 355.1472 | 320.4773 | 292.4294 | 319.8365 | 345.2686 | 328.8226 | 291.4744 |
| ENSMUSG00000031725 | -3.785898288 | 0.000214 | 0.002389 | 0 | 2.399643 | 5.796746 | 0 | 23.40267 | 5.58687 | 34.71441 | 51.11678 |
| ENSMUSG00000031728 | 0.01118792 | 0.911055 | 0.947486 | 1260.586 | 1412.19 | 1228.91 | 1363.808 | 1306.649 | 1492.812 | 1269.005 | 1155.022 |
| ENSMUSG00000031729 | 0.091722616 | 0.127748 | 0.259119 | 4678.153 | 4306.159 | 4104.925 | 4571.042 | 4187.128 | 4194.622 | 4131.979 | 4057.802 |
| ENSMUSG00000031730 | -0.086365116 | 0.397997 | 0.565163 | 472.8455 | 556.7172 | 527.5039 | 453.74 | 564.5894 | 503.9357 | 513.0018 | 550.3211 |
| ENSMUSG00000031731 | -0.009797266 | 0.857341 | 0.914643 | 2503.063 | 2414.041 | 2468.586 | 2412.758 | 2430.952 | 2385.594 | 2428.08 | 2622.182 |
| ENSMUSG00000031732 | 0.116758101 | 0.287093 | 0.454071 | 685.123 | 692.297 | 568.9093 | 723.7411 | 615.2952 | 679.3635 | 597.8593 | 568.8101 |
| ENSMUSG00000031734 | -1.083319547 | 4.88E-05 | 0.000782 | 395.3794 | 220.7672 | 567.253 | 377.8291 | 720.6073 | 631.3164 | 961.3963 | 999.4962 |
| ENSMUSG00000031736 | -0.387346678 | 0.130403 | 0.262799 | 55.33299 | 50.3925 | 77.84202 | 54.34528 | 59.48179 | 96.09417 | 87.75031 | 69.60583 |
| ENSMUSG00000031737 | -0.638132272 | 0.095785 | 0.214305 | 114.6902 | 39.59411 | 96.06037 | 82.81186 | 87.76002 | 84.92043 | 195.7507 | 151.1752 |
| ENSMUSG00000031740 | 0.410920507 | 2.70E-05 | 0.000495 | 4568.493 | 5639.161 | 5578.954 | 5474.209 | 4049.637 | 4354.407 | 4029.764 | 3557.51 |
| ENSMUSG00000031748 | 0.814011375 | 1.23E-05 | 0.000269 | 450.7124 | 446.3336 | 375.9604 | 609.0122 | 250.6036 | 318.4516 | 218.8936 | 283.8613 |
| ENSMUSG00000031749 | 0.014647509 | 0.872145 | 0.9234 | 1047.303 | 1199.822 | 1170.943 | 1145.564 | 1152.582 | 987.7587 | 1212.111 | 1162.635 |
| ENSMUSG00000031750 | -0.334820467 | 0.121067 | 0.25028 | 73.44197 | 68.38983 | 94.40416 | 88.85022 | 101.4116 | 90.5073 | 131.1433 | 88.09488 |
| ENSMUSG00000031751 | -0.298101601 | 0.000886 | 0.007097 | 3332.052 | 2925.165 | 3278.474 | 3037.297 | 3702.498 | 3477.268 | 3950.693 | 4330.788 |
| ENSMUSG00000031753 | -0.173878108 | 0.054573 | 0.145045 | 1470.851 | 1520.174 | 1401.156 | 1401.763 | 1726.922 | 1407.891 | 1628.684 | 1768.423 |
| ENSMUSG00000031754 | 0.143367539 | 0.099909 | 0.2207 | 2540.287 | 2675.602 | 2399.025 | 2601.672 | 2492.385 | 2504.035 | 2184.115 | 2067.511 |
| ENSMUSG00000031755 | 0.11755264 | 0.194002 | 0.347453 | 781.7042 | 808.6797 | 727.9057 | 720.2906 | 726.4579 | 725.1758 | 726.1097 | 618.8393 |
| ENSMUSG00000031756 | 0.021933536 | 0.843255 | 0.905412 | 674.0564 | 715.0936 | 621.08 | 737.5431 | 762.537 | 696.1241 | 661.5024 | 584.0364 |
| ENSMUSG00000031758 | -0.148005396 | 0.461045 | 0.622234 | 59.35721 | 69.58965 | 57.96746 | 62.10889 | 73.13335 | 73.74669 | 58.82164 | 69.60583 |
| ENSMUSG00000031760 | 1.286792324 | 8.55E-06 | 0.000203 | 79.47829 | 81.58786 | 94.40416 | 79.36136 | 30.22845 | 54.75133 | 27.96439 | 25.0146 |
| ENSMUSG00000031762 | -0.0535945 | 0.803566 | 0.879998 | 949.7153 | 513.5236 | 817.3412 | 890.2274 | 905.8784 | 653.6638 | 707.7882 | 1025.598 |
| ENSMUSG00000031765 | -0.774479706 | 0.004557 | 0.024024 | 1146.902 | 517.1231 | 1046.727 | 1022.209 | 1731.798 | 973.2328 | 1620.97 | 2062.073 |
| ENSMUSG00000031766 | -2.81066857 | 3.23E-06 | 9.59E-05 | 160.9687 | 73.18911 | 341.1799 | 161.3106 | 992.6633 | 454.7713 | 1540.934 | 2184.97 |
| ENSMUSG00000031767 | 0.207418707 | 0.071317 | 0.174751 | 257.5499 | 280.7582 | 255.0568 | 256.1992 | 207.6987 | 215.6532 | 246.858 | 237.0949 |
| ENSMUSG00000031770 | -0.697961673 | 0.002707 | 0.016383 | 1146.902 | 855.4728 | 969.7129 | 1169.717 | 1238.391 | 1259.281 | 2563.081 | 1658.576 |
| ENSMUSG00000031774 | -0.092343728 | 0.088458 | 0.202604 | 1581.517 | 1599.362 | 1615.636 | 1498.377 | 1703.519 | 1667.122 | 1665.327 | 1672.715 |
| ENSMUSG00000031775 | -0.991076752 | 0.105052 | 0.228493 | 24.1453 | 11.99822 | 102.6852 | 20.70296 | 79.95913 | 45.81234 | 88.7146 | 104.4087 |
| ENSMUSG00000031776 | -0.193360785 | 0.031181 | 0.097042 | 2416.543 | 2735.593 | 3119.478 | 2795.763 | 3301.727 | 2999.032 | 3172.511 | 3184.467 |
| ENSMUSG00000031778 | -0.725092051 | 0.033891 | 0.10344 | 188.1322 | 167.975 | 414.8814 | 182.0136 | 318.8614 | 254.7613 | 441.6444 | 562.2846 |
| ENSMUSG00000031781 | -0.556584012 | 0.001047 | 0.008098 | 1133.823 | 905.8653 | 1146.1 | 984.2534 | 1624.535 | 1108.435 | 1473.434 | 1928.299 |
| ENSMUSG00000031782 | -0.779109017 | 4.47E-05 | 0.00073 | 1087.545 | 893.867 | 1095.585 | 894.5406 | 1736.673 | 1112.905 | 1803.221 | 2163.219 |
| ENSMUSG00000031783 | -0.179551329 | 0.025698 | 0.084646 | 1516.124 | 1436.186 | 1445.874 | 1554.448 | 1723.022 | 1488.342 | 1793.578 | 1735.795 |
| ENSMUSG00000031785 | -0.87810262 | 0.00415 | 0.022424 | 1438.658 | 1059.442 | 2699.628 | 1369.846 | 2585.02 | 1857.076 | 3433.834 | 4198.102 |
| ENSMUSG00000031787 | -0.138340085 | 0.094763 | 0.212638 | 805.8495 | 838.6753 | 883.5898 | 856.5851 | 966.3353 | 831.3263 | 985.5035 | 941.8539 |
| ENSMUSG00000031788 | -0.437409433 | 0.020892 | 0.072697 | 1066.418 | 992.2524 | 1324.971 | 1028.247 | 1543.601 | 1002.285 | 1525.505 | 1904.372 |
| ENSMUSG00000031789 | 0.266186502 | 0.069047 | 0.170616 | 237.4288 | 219.5673 | 228.5574 | 213.9306 | 182.3458 | 231.2964 | 166.822 | 168.5766 |
| ENSMUSG00000031790 | 0.126480581 | 0.339785 | 0.508231 | 530.1906 | 527.9215 | 719.6247 | 604.6991 | 568.4899 | 532.9874 | 489.8589 | 594.9123 |
| ENSMUSG00000031791 | -0.379692313 | 0.026982 | 0.087728 | 359.1614 | 409.1391 | 422.3344 | 411.4714 | 495.3565 | 375.4377 | 562.1806 | 651.4671 |
| ENSMUSG00000031792 | 0.127481229 | 0.115492 | 0.243459 | 852.128 | 889.0678 | 864.5433 | 850.5468 | 815.1931 | 739.7017 | 741.5384 | 866.8101 |
| ENSMUSG00000031796 | 0.233240782 | 0.000418 | 0.003992 | 2220.362 | 2099.688 | 2018.924 | 2194.514 | 1865.388 | 1902.888 | 1788.756 | 1702.08 |
| ENSMUSG00000031799 | 0.077182843 | 0.276513 | 0.44271 | 13113.92 | 15637.27 | 15286.85 | 14559.36 | 13794.9 | 13839.8 | 13454.73 | 14455.17 |
| ENSMUSG00000031802 | 1.048838402 | 0.113143 | 0.240074 | 75.45408 | 59.99108 | 22.35888 | 19.84034 | 12.67645 | 49.16446 | 16.39292 | 7.613138 |
| ENSMUSG00000031803 | -0.831462817 | 7.49E-05 | 0.00109 | 68.4117 | 55.19179 | 64.59232 | 57.79577 | 100.4365 | 86.03781 | 124.3933 | 127.2482 |
| ENSMUSG00000031805 | -0.071175249 | 0.710924 | 0.818885 | 256.5439 | 145.1784 | 226.0731 | 200.1287 | 222.3254 | 188.8362 | 217.9294 | 242.5328 |
| ENSMUSG00000031807 | -0.39806017 | 1.84E-05 | 0.000367 | 1866.231 | 1856.124 | 2047.908 | 1920.2 | 2771.266 | 2165.471 | 2653.724 | 2543.876 |
| ENSMUSG00000031808 | 0.365800917 | 3.38E-07 | 1.52E-05 | 2354.167 | 2054.094 | 2084.344 | 2127.23 | 1603.083 | 1772.155 | 1642.184 | 1673.803 |
| ENSMUSG00000031812 | -0.052291079 | 0.479239 | 0.638386 | 4569.499 | 4589.317 | 4860.986 | 5056.699 | 5312.406 | 4608.051 | 5173.411 | 4687.518 |
| ENSMUSG00000031813 | -0.190937994 | 0.067173 | 0.167374 | 1351.131 | 1327.003 | 1554.356 | 1400.901 | 1613.809 | 1362.079 | 1680.756 | 1776.036 |
| ENSMUSG00000031816 | 0.002951736 | 0.970682 | 0.983617 | 1054.345 | 1070.241 | 915.0578 | 1029.972 | 1048.245 | 1039.158 | 955.6106 | 1014.722 |
| ENSMUSG00000031818 | -0.535608819 | 6.81E-05 | 0.001009 | 10644.06 | 11014.36 | 12965.67 | 11963.73 | 17358.93 | 12845.33 | 17812.35 | 19516.82 |
| ENSMUSG00000031819 | -0.127834781 | 0.039995 | 0.116705 | 1766.631 | 1912.516 | 1847.506 | 1765.79 | 2056.51 | 1868.249 | 1997.043 | 2041.408 |
| ENSMUSG00000031820 | -0.221362791 | 0.008315 | 0.037339 | 1657.978 | 1676.151 | 1778.773 | 1743.362 | 2029.207 | 1737.517 | 2085.757 | 2141.467 |
| ENSMUSG00000031821 | -0.07058356 | 0.483677 | 0.641981 | 689.1472 | 893.867 | 762.6862 | 733.23 | 852.2473 | 783.2792 | 783.0028 | 810.2554 |
| ENSMUSG00000031822 | -0.165816655 | 0.236092 | 0.397076 | 349.1009 | 389.942 | 405.7722 | 301.9182 | 464.153 | 350.8555 | 399.2157 | 406.7591 |
| ENSMUSG00000031823 | -0.329197443 | 0.007953 | 0.036101 | 889.3521 | 849.4736 | 977.1658 | 905.7547 | 1172.084 | 900.6035 | 1173.54 | 1305.109 |
| ENSMUSG00000031824 | -0.313120314 | 0.039613 | 0.115854 | 399.4036 | 268.76 | 315.5086 | 313.995 | 403.6961 | 340.7991 | 430.073 | 438.2992 |
| ENSMUSG00000031825 | 0.4698941 | 0.012963 | 0.051885 | 1428.597 | 1535.772 | 1214.832 | 1313.776 | 967.3104 | 1334.145 | 987.4321 | 676.4817 |
| ENSMUSG00000031826 | -0.147532891 | 0.020255 | 0.071109 | 2081.526 | 2127.284 | 1964.269 | 2096.175 | 2188.15 | 2272.739 | 2238.115 | 2459.043 |
| ENSMUSG00000031827 | -0.227798547 | 0.057764 | 0.150663 | 2622.784 | 2813.581 | 3059.854 | 2842.344 | 3450.919 | 2763.266 | 3095.368 | 3969.707 |

| | | | | | | | | | | | |
|---|---|---|---|---|---|---|---|---|---|---|---|
| ENSMUSG00000031828 | -0.290423444 | 0.078814 | 0.187255 | 371.2341 | 320.3523 | 467.8802 | 420.0976 | 532.4108 | 374.3203 | 501.4304 | 526.3941 |
| ENSMUSG00000031832 | -0.080889882 | 0.469052 | 0.629175 | 354.1311 | 313.1534 | 330.4145 | 364.0271 | 406.6214 | 329.6254 | 350.037 | 354.5547 |
| ENSMUSG00000031833 | 0.049061236 | 0.610139 | 0.7452 | 904.4429 | 878.2694 | 863.7152 | 910.0678 | 994.6135 | 836.9132 | 831.2173 | 773.2773 |
| ENSMUSG00000031834 | -0.202158736 | 0.023377 | 0.078947 | 1372.258 | 1516.574 | 1577.543 | 1499.24 | 1831.259 | 1502.868 | 1724.149 | 1804.314 |
| ENSMUSG00000031835 | -0.038604295 | 0.417862 | 0.583496 | 6065.502 | 6264.268 | 6283.673 | 6122.039 | 6547.872 | 6126.562 | 6562.952 | 6164.466 |
| ENSMUSG00000031837 | 0.010652068 | 0.957715 | 0.975959 | 79.47829 | 92.38626 | 65.42042 | 82.81186 | 78.98402 | 84.92043 | 85.82174 | 66.34306 |
| ENSMUSG00000031838 | -0.335217086 | 0.012603 | 0.050785 | 823.9585 | 933.4612 | 1060.805 | 946.298 | 1278.371 | 930.7726 | 1232.362 | 1308.372 |
| ENSMUSG00000031839 | -0.014721971 | 0.824043 | 0.893455 | 3978.945 | 4121.387 | 4204.297 | 4204.427 | 4221.257 | 3814.715 | 4150.301 | 4492.839 |
| ENSMUSG00000031840 | 0.033060045 | 0.843082 | 0.905412 | 368.2159 | 237.5647 | 261.6817 | 337.2858 | 277.9067 | 328.508 | 283.501 | 289.2992 |
| ENSMUSG00000031841 | -0.32590098 | 0.227696 | 0.387636 | 89.53884 | 76.78858 | 84.46688 | 138.0198 | 91.66046 | 151.9629 | 101.2504 | 144.6496 |
| ENSMUSG00000031842 | -1.512936228 | 0.000187 | 0.00215 | 155.9384 | 62.39072 | 237.6666 | 168.2116 | 379.3183 | 214.5358 | 549.6448 | 641.6787 |
| ENSMUSG00000031843 | -0.159644208 | 0.07457 | 0.180328 | 819.9343 | 919.0633 | 1020.227 | 954.0616 | 1035.568 | 1100.613 | 1008.646 | 1007.109 |
| ENSMUSG00000031844 | -2.958488405 | 0.000157 | 0.001887 | 26.15741 | 5.999108 | 73.70149 | 33.64232 | 320.8116 | 75.98144 | 243.9652 | 448.0875 |
| ENSMUSG00000031847 | -0.231186763 | 0.43834 | 0.601907 | 34.20585 | 28.79572 | 29.81184 | 35.36756 | 26.32801 | 49.16446 | 36.64299 | 39.15328 |
| ENSMUSG00000031848 | -0.316607125 | 0.000293 | 0.003048 | 2594.614 | 2989.955 | 2926.529 | 2935.508 | 3818.536 | 3214.685 | 3394.298 | 3827.233 |
| ENSMUSG00000031849 | 1.036929207 | 0.093358 | 0.2102 | 10.06054 | 10.79839 | 17.39024 | 32.77969 | 7.80089 | 3.352122 | 10.60718 | 13.05109 |
| ENSMUSG00000031851 | -0.135205074 | 0.291021 | 0.458127 | 357.1493 | 377.9438 | 411.569 | 409.7462 | 462.2028 | 382.1419 | 374.1442 | 492.6788 |
| ENSMUSG00000031853 | -0.080504776 | 0.367566 | 0.53607 | 576.4691 | 521.9224 | 516.7385 | 525.3377 | 565.5646 | 548.6307 | 616.1808 | 530.7445 |
| ENSMUSG00000031858 | 0.068564923 | 0.340005 | 0.508478 | 2524.19 | 2300.058 | 2149.765 | 2287.678 | 2306.138 | 2233.631 | 2191.829 | 2096.876 |
| ENSMUSG00000031860 | 0.606116315 | 0.00108 | 0.008282 | 144.8718 | 115.1829 | 117.5911 | 125.943 | 72.15824 | 99.44629 | 73.28598 | 87.00729 |
| ENSMUSG00000031861 | -0.146087878 | 0.149971 | 0.289889 | 288.7376 | 310.7538 | 322.1335 | 299.3304 | 316.9112 | 353.0902 | 324.0012 | 358.9051 |
| ENSMUSG00000031862 | -0.059921534 | 0.333454 | 0.501623 | 1752.547 | 1731.342 | 1787.882 | 1777.004 | 1981.426 | 1795.62 | 1752.113 | 1819.54 |
| ENSMUSG00000031864 | -0.303960078 | 0.002082 | 0.013514 | 902.4308 | 975.4549 | 845.4969 | 868.6619 | 1062.871 | 1003.402 | 1097.361 | 1267.044 |
| ENSMUSG00000031865 | -0.212482929 | 0.049448 | 0.135042 | 2759.607 | 2877.172 | 3334.785 | 3014.006 | 3540.629 | 2919.699 | 3521.584 | 3907.715 |
| ENSMUSG00000031870 | 0.093749999 | 0.61245 | 0.746738 | 130.7871 | 139.1793 | 114.2787 | 142.3329 | 104.3369 | 157.5497 | 117.6433 | 114.1971 |
| ENSMUSG00000031871 | -0.331781173 | 0.037827 | 0.112177 | 1706.268 | 1474.581 | 2347.682 | 1814.097 | 2410.475 | 1908.475 | 2266.08 | 2660.248 |
| ENSMUSG00000031872 | -0.204940478 | 0.38066 | 0.548577 | 143.8658 | 81.58786 | 149.8873 | 113.0037 | 136.5156 | 113.9722 | 144.6434 | 169.6642 |
| ENSMUSG00000031875 | 0.13355451 | 0.04505 | 0.126612 | 2664.032 | 2724.795 | 3006.027 | 3000.205 | 2623.049 | 2657.116 | 2490.759 | 2623.27 |
| ENSMUSG00000031877 | 1.594122024 | 0.000389 | 0.003779 | 44.26639 | 91.18644 | 57.13936 | 56.93315 | 21.45245 | 37.99072 | 13.50005 | 9.78832 |
| ENSMUSG00000031878 | 0.070871551 | 0.45612 | 0.618021 | 1491.979 | 1631.757 | 1459.124 | 1677.803 | 1511.423 | 1685 | 1371.219 | 1393.204 |
| ENSMUSG00000031879 | -0.320046661 | 0.042633 | 0.122008 | 657.9595 | 625.107 | 587.1276 | 582.2709 | 869.7993 | 545.2786 | 764.6813 | 879.8612 |
| ENSMUSG00000031880 | 0.340616627 | 0.001059 | 0.008162 | 309.8647 | 308.3541 | 311.3681 | 288.9789 | 230.1263 | 240.2354 | 232.3937 | 259.9343 |
| ENSMUSG00000031881 | -0.74443623 | 0.05164 | 0.139072 | 6003.126 | 5074.045 | 11283.78 | 5797.693 | 10792.53 | 6832.743 | 13157.73 | 16393.26 |
| ENSMUSG00000031883 | -0.80363726 | 0.035692 | 0.10767 | 20.12109 | 35.99465 | 41.40533 | 18.97772 | 71.18313 | 36.87335 | 40.50014 | 54.37955 |
| ENSMUSG00000031885 | 0.154738091 | 0.101583 | 0.223524 | 3208.307 | 3495.08 | 2983.668 | 3412.539 | 3031.621 | 3311.897 | 2750.153 | 2672.211 |
| ENSMUSG00000031886 | -1.101427805 | 0.029885 | 0.094285 | 11.0666 | 14.39786 | 27.32752 | 12.07673 | 29.25334 | 15.64324 | 38.57157 | 56.55474 |
| ENSMUSG00000031887 | -0.136517788 | 0.410787 | 0.576758 | 262.5802 | 225.5664 | 293.9779 | 244.1224 | 314.961 | 212.3011 | 316.2868 | 284.9489 |
| ENSMUSG00000031889 | 0.024414176 | 0.789505 | 0.8709 | 677.0746 | 741.4897 | 628.5329 | 720.2906 | 742.0597 | 661.4855 | 656.6809 | 656.905 |
| ENSMUSG00000031891 | 0.58579039 | 0.001195 | 0.008947 | 990.9635 | 1154.228 | 1196.614 | 1092.944 | 568.4899 | 753.1101 | 620.0379 | 1014.722 |
| ENSMUSG00000031893 | 0.526245134 | 0.345497 | 0.514314 | 17.10292 | 10.79839 | 8.281066 | 14.6646 | 8.776002 | 5.58687 | 5.785735 | 15.22628 |
| ENSMUSG00000031896 | 0.499313464 | 0.289857 | 0.457118 | 13.07871 | 31.19536 | 13.24971 | 18.97772 | 17.552 | 14.52586 | 11.57147 | 9.78832 |
| ENSMUSG00000031897 | -0.239194062 | 0.192834 | 0.346067 | 415.5004 | 434.3354 | 606.1741 | 401.9825 | 549.9628 | 432.4238 | 539.0376 | 673.2189 |
| ENSMUSG00000031901 | -0.107315178 | 0.403457 | 0.570459 | 277.671 | 253.1623 | 263.3379 | 307.094 | 263.2801 | 292.752 | 290.251 | 342.5912 |
| ENSMUSG00000031902 | -0.033703692 | 0.560298 | 0.706246 | 1785.746 | 1841.726 | 1727.43 | 1768.378 | 1769.827 | 1916.297 | 1842.757 | 1759.722 |
| ENSMUSG00000031903 | 0.295967076 | 0.002323 | 0.014609 | 408.4581 | 401.9402 | 401.6317 | 391.6311 | 318.8614 | 364.264 | 319.1797 | 304.5255 |
| ENSMUSG00000031904 | -0.168384071 | 0.0931 | 0.209847 | 1073.46 | 1071.441 | 1209.864 | 1083.455 | 1294.948 | 1053.684 | 1336.505 | 1302.934 |
| ENSMUSG00000031906 | 1.820560737 | 0.000583 | 0.005151 | 273.6468 | 380.3434 | 192.1207 | 345.0494 | 56.55646 | 202.2447 | 55.92877 | 22.83941 |
| ENSMUSG00000031907 | 0.056317263 | 0.675149 | 0.792182 | 190.1443 | 242.3639 | 231.8699 | 228.5952 | 235.9769 | 213.4185 | 216.9651 | 191.416 |
| ENSMUSG00000031910 | -0.619369008 | 0.004555 | 0.024019 | 69.41775 | 53.99197 | 73.70149 | 56.93315 | 85.8098 | 91.62468 | 90.64318 | 123.9854 |
| ENSMUSG00000031913 | -0.075185878 | 0.350098 | 0.518822 | 1609.687 | 1623.359 | 1526.201 | 1656.237 | 1672.316 | 1509.572 | 1726.078 | 1848.905 |
| ENSMUSG00000031916 | -0.163226214 | 0.037351 | 0.11122 | 836.0312 | 787.0829 | 789.1856 | 834.1569 | 989.738 | 843.6174 | 902.5747 | 898.3502 |
| ENSMUSG00000031917 | -0.123033981 | 0.159198 | 0.302155 | 1156.963 | 1250.214 | 1267.003 | 1337.929 | 1484.119 | 1306.21 | 1243.933 | 1425.832 |
| ENSMUSG00000031918 | -0.123716551 | 0.042173 | 0.121051 | 2286.762 | 2270.062 | 2480.179 | 2367.901 | 2696.183 | 2490.627 | 2599.724 | 2463.394 |
| ENSMUSG00000031919 | 0.025625794 | 0.837217 | 0.901479 | 329.9858 | 303.5548 | 368.5075 | 354.5383 | 344.2143 | 284.9304 | 331.7155 | 374.1313 |
| ENSMUSG00000031921 | 0.217490685 | 0.029713 | 0.093902 | 1181.108 | 1295.807 | 1120.428 | 1292.21 | 1019.966 | 1205.647 | 956.5749 | 1023.423 |
| ENSMUSG00000031922 | 0.024905695 | 0.72561 | 0.828345 | 1644.899 | 1614.96 | 1609.011 | 1758.889 | 1612.834 | 1773.273 | 1560.22 | 1571.569 |
| ENSMUSG00000031924 | -0.27966491 | 0.065388 | 0.164684 | 5280.779 | 4318.158 | 5901.088 | 4390.754 | 6198.783 | 4796.887 | 5853.235 | 7298.824 |
| ENSMUSG00000031925 | 0.642847384 | 0.001097 | 0.008387 | 860.1765 | 1057.043 | 851.2936 | 940.2596 | 493.4063 | 867.0823 | 495.6446 | 519.8685 |
| ENSMUSG00000031928 | -0.053754726 | 0.492842 | 0.649892 | 858.1644 | 944.2595 | 869.512 | 890.2274 | 979.0118 | 958.707 | 849.5388 | 908.1386 |
| ENSMUSG00000031930 | -0.027903106 | 0.721444 | 0.825786 | 1934.643 | 2219.67 | 2007.33 | 1919.337 | 2124.768 | 2192.288 | 1935.328 | 1982.679 |
| ENSMUSG00000031931 | 0.076538928 | 0.345264 | 0.514053 | 734.4197 | 825.4772 | 693.9534 | 762.5592 | 731.3335 | 712.8847 | 707.7882 | 703.6714 |
| ENSMUSG00000031932 | -0.848037651 | 0.154664 | 0.296293 | 17.10292 | 3.599465 | 4.96864 | 15.52722 | 15.60178 | 13.40849 | 27.0001 | 18.48905 |
| ENSMUSG00000031934 | 0.037057285 | 0.685463 | 0.79988 | 741.4621 | 837.4754 | 744.4679 | 779.8116 | 785.9397 | 813.4483 | 750.217 | 672.1313 |
| ENSMUSG00000031935 | -0.122666047 | 0.112309 | 0.238882 | 1192.174 | 1071.441 | 995.3842 | 1113.647 | 1215.964 | 1154.247 | 1193.79 | 1194.175 |

| | | | | | | | | | | | |
|---|---|---|---|---|---|---|---|---|---|---|---|
| ENSMUSG00000031937 | 0.037573299 | 0.734922 | 0.834023 | 298.7981 | 321.5522 | 284.0406 | 268.2759 | 274.0063 | 306.1605 | 269.0367 | 291.4744 |
| ENSMUSG00000031938 | -1.227519834 | 2.63E-07 | 1.25E-05 | 1416.525 | 1650.954 | 1701.759 | 1658.825 | 4463.084 | 2167.706 | 3109.833 | 5311.795 |
| ENSMUSG00000031939 | 0.289913699 | 0.053414 | 0.142561 | 3644.935 | 3828.631 | 2968.762 | 3619.568 | 3051.123 | 3596.827 | 2506.188 | 2347.022 |
| ENSMUSG00000031948 | -0.126167107 | 0.065359 | 0.164657 | 4300.882 | 4832.881 | 4345.904 | 4499.444 | 5287.054 | 4642.689 | 4860.982 | 4824.554 |
| ENSMUSG00000031949 | 0.271715761 | 0.065554 | 0.164783 | 341.0524 | 315.5531 | 277.4157 | 362.3019 | 248.6534 | 301.691 | 297.0011 | 226.2189 |
| ENSMUSG00000031950 | -0.25388035 | 0.006057 | 0.029662 | 1278.695 | 1160.227 | 1228.91 | 1216.299 | 1393.434 | 1276.041 | 1566.97 | 1587.883 |
| ENSMUSG00000031951 | -0.373593812 | 0.001917 | 0.012726 | 424.5549 | 538.7199 | 430.6154 | 417.5098 | 653.3246 | 543.0438 | 557.3591 | 587.2992 |
| ENSMUSG00000031952 | 1.882127809 | 7.22E-16 | 3.92E-13 | 116.7023 | 127.1811 | 108.482 | 106.9653 | 37.05423 | 20.11273 | 31.82154 | 34.80291 |
| ENSMUSG00000031953 | -0.308672267 | 0.021447 | 0.074001 | 335.0161 | 285.5575 | 287.353 | 248.4356 | 344.2143 | 319.569 | 363.537 | 403.4963 |
| ENSMUSG00000031954 | 0.064878985 | 0.355447 | 0.524098 | 3471.894 | 4035 | 3685.903 | 3698.067 | 3520.152 | 3813.598 | 3459.87 | 3440.051 |
| ENSMUSG00000031955 | -0.543942521 | 0.002279 | 0.01441 | 1093.581 | 1007.85 | 1524.544 | 1162.816 | 1699.619 | 1344.201 | 1734.756 | 2206.722 |
| ENSMUSG00000031958 | -0.962274179 | 0.003998 | 0.021826 | 96.58122 | 63.59054 | 124.216 | 67.28463 | 164.7938 | 92.74205 | 182.2507 | 246.8832 |
| ENSMUSG00000031959 | 0.121623761 | 0.230179 | 0.390428 | 948.7093 | 881.8688 | 764.3424 | 868.6619 | 820.0686 | 702.8283 | 827.3601 | 829.832 |
| ENSMUSG00000031960 | -0.326681908 | 0.008641 | 0.038378 | 2474.894 | 2346.851 | 2608.536 | 2121.191 | 2947.761 | 2498.448 | 2928.546 | 3604.277 |
| ENSMUSG00000031962 | -0.08288217 | 0.858767 | 0.915304 | 9.054489 | 29.99554 | 28.98373 | 39.68068 | 36.07912 | 33.52122 | 24.10723 | 20.66423 |
| ENSMUSG00000031963 | -0.19723097 | 0.358662 | 0.526985 | 1170.041 | 971.8554 | 697.2658 | 877.2881 | 1015.091 | 1473.816 | 983.5749 | 787.4159 |
| ENSMUSG00000031965 | 3.442690713 | 0.002514 | 0.015492 | 36.21796 | 4.799286 | 0.828107 | 58.6584 | 3.900445 | 1.117374 | 0.964289 | 3.262773 |
| ENSMUSG00000031967 | -0.272749596 | 0.002958 | 0.017587 | 1544.293 | 1472.181 | 1417.719 | 1242.178 | 1757.151 | 1562.089 | 1742.471 | 1791.263 |
| ENSMUSG00000031969 | -0.23099723 | 0.00459 | 0.02416 | 1097.605 | 1097.837 | 990.4155 | 1093.807 | 1248.142 | 1134.135 | 1297.933 | 1338.825 |
| ENSMUSG00000031970 | -0.146739679 | 0.537213 | 0.687647 | 116.7023 | 77.9884 | 93.57605 | 136.2945 | 119.9387 | 134.0849 | 130.179 | 87.00729 |
| ENSMUSG00000031971 | -0.284188656 | 0.159015 | 0.302041 | 113.6841 | 122.3818 | 94.40416 | 115.5915 | 142.3663 | 172.0756 | 123.429 | 104.4087 |
| ENSMUSG00000031972 | 0.413861768 | 0.357383 | 0.52586 | 202.2169 | 110.3836 | 159.8246 | 153.547 | 51.6809 | 189.9536 | 177.4292 | 51.11678 |
| ENSMUSG00000031974 | -0.353782605 | 0.000317 | 0.003238 | 586.5297 | 569.9152 | 563.9406 | 604.6991 | 759.6117 | 669.3071 | 699.1096 | 843.9707 |
| ENSMUSG00000031976 | -0.09819573 | 0.315149 | 0.483544 | 448.7002 | 487.1275 | 463.7397 | 434.7622 | 540.2117 | 488.2925 | 442.6087 | 490.5036 |
| ENSMUSG00000031979 | -0.12872199 | 0.098207 | 0.218244 | 919.5337 | 799.0811 | 903.4643 | 822.9428 | 963.41 | 901.7209 | 961.3963 | 941.8539 |
| ENSMUSG00000031980 | -2.008803888 | 3.09E-05 | 0.000548 | 73.44197 | 55.19179 | 60.45178 | 33.64232 | 173.5698 | 60.3382 | 269.0367 | 392.6204 |
| ENSMUSG00000031982 | -0.047214569 | 0.759651 | 0.850154 | 184.1079 | 203.9697 | 180.5272 | 154.4096 | 172.5947 | 221.2401 | 182.2507 | 169.6642 |
| ENSMUSG00000031983 | 0.049087294 | 0.555117 | 0.702258 | 2341.088 | 2579.616 | 2899.201 | 2498.158 | 2521.638 | 2362.129 | 2489.795 | 2601.518 |
| ENSMUSG00000031984 | -0.238342211 | 0.000357 | 0.003535 | 1290.768 | 1288.608 | 1187.505 | 1199.047 | 1507.522 | 1387.779 | 1442.577 | 1513.927 |
| ENSMUSG00000031985 | -0.111230976 | 0.148122 | 0.287611 | 2487.972 | 2496.829 | 2212.701 | 2423.972 | 2666.929 | 2350.955 | 2618.045 | 2751.605 |
| ENSMUSG00000031986 | -0.083514167 | 0.462176 | 0.623214 | 702.2259 | 614.3086 | 629.361 | 583.1335 | 769.3628 | 689.4198 | 615.2165 | 604.7006 |
| ENSMUSG00000031987 | -0.003020283 | 0.97791 | 0.987865 | 1373.264 | 1722.944 | 1638.823 | 1788.219 | 1547.502 | 1790.033 | 1491.755 | 1709.693 |
| ENSMUSG00000031988 | -0.145928912 | 0.073016 | 0.177713 | 1555.36 | 1577.765 | 1638.823 | 1582.914 | 1705.47 | 1566.558 | 1830.221 | 1929.387 |
| ENSMUSG00000031990 | 0.202645327 | 0.015918 | 0.060128 | 1298.816 | 1238.216 | 1320.83 | 1423.329 | 1269.595 | 1117.374 | 1072.29 | 1132.182 |
| ENSMUSG00000031993 | -0.08311615 | 0.3083 | 0.476744 | 1129.799 | 1018.648 | 1182.536 | 1083.455 | 1124.303 | 1154.247 | 1126.29 | 1276.832 |
| ENSMUSG00000031994 | -0.358035697 | 0.3855 | 0.552948 | 18.10898 | 28.79572 | 28.98373 | 34.50494 | 24.37778 | 59.22083 | 35.6787 | 22.83941 |
| ENSMUSG00000031995 | -0.444611011 | 0.029753 | 0.093981 | 1260.586 | 1292.208 | 1941.082 | 1238.727 | 1792.255 | 1392.248 | 2263.187 | 2355.722 |
| ENSMUSG00000031996 | -0.03845036 | 0.709645 | 0.81832 | 9769.794 | 8666.311 | 10367.07 | 9424.162 | 9258.682 | 8471.93 | 10644.79 | 10886.79 |
| ENSMUSG00000031997 | -1.104623957 | 0.030914 | 0.096572 | 19.11503 | 8.398751 | 31.46805 | 11.21411 | 18.52711 | 36.87335 | 48.21446 | 48.9416 |
| ENSMUSG00000032000 | 0.293082884 | 0.047523 | 0.13134 | 459.7668 | 323.9518 | 339.5237 | 361.4392 | 267.1805 | 317.3342 | 331.7155 | 295.8248 |
| ENSMUSG00000032002 | -0.137013426 | 0.1289 | 0.260595 | 1918.546 | 1797.333 | 1859.099 | 1833.938 | 1862.463 | 2364.364 | 1952.686 | 1970.715 |
| ENSMUSG00000032006 | -0.037138505 | 0.697608 | 0.809242 | 489.9485 | 488.3274 | 508.4575 | 475.3055 | 446.601 | 493.8794 | 545.7877 | 527.4817 |
| ENSMUSG00000032009 | 0.636058201 | 7.36E-07 | 2.89E-05 | 3092.611 | 4007.404 | 3043.292 | 3299.535 | 2132.568 | 2580.017 | 1979.686 | 1956.576 |
| ENSMUSG00000032010 | -1.068793517 | 0.000126 | 0.001592 | 839.0493 | 453.5325 | 714.656 | 756.5208 | 895.1522 | 1415.713 | 2265.115 | 1223.54 |
| ENSMUSG00000032011 | -0.012749747 | 0.9598 | 0.977262 | 965.8122 | 871.0704 | 1768.836 | 991.1544 | 991.6882 | 862.6128 | 1267.076 | 1518.277 |
| ENSMUSG00000032012 | -0.204847587 | 0.282386 | 0.449014 | 255.5378 | 265.1606 | 349.461 | 244.9851 | 275.9565 | 273.7567 | 316.2868 | 420.8978 |
| ENSMUSG00000032013 | -0.7480003 | 0.150704 | 0.290819 | 45.27245 | 26.39607 | 149.0592 | 55.2079 | 169.6694 | 56.98608 | 135.0005 | 103.3212 |
| ENSMUSG00000032014 | -0.486678537 | 0.009731 | 0.041727 | 901.4247 | 1023.448 | 1317.518 | 1021.346 | 1414.887 | 1100.613 | 1480.184 | 1980.503 |
| ENSMUSG00000032015 | -1.496835728 | 0.021255 | 0.073599 | 1.006054 | 3.599465 | 13.24971 | 7.763611 | 18.52711 | 21.23011 | 20.25007 | 14.13868 |
| ENSMUSG00000032017 | 0.883431056 | 0.000364 | 0.003591 | 445.6821 | 604.7101 | 492.7234 | 446.839 | 274.9814 | 400.0199 | 237.2151 | 166.4014 |
| ENSMUSG00000032018 | -0.366560708 | 0.001102 | 0.008401 | 805.8495 | 776.2845 | 669.1102 | 678.0221 | 951.7086 | 855.9086 | 1079.04 | 884.2116 |
| ENSMUSG00000032020 | 0.638200893 | 0.001726 | 0.011754 | 898.4065 | 910.6645 | 704.7187 | 773.7733 | 478.7797 | 770.9881 | 431.0373 | 431.7737 |
| ENSMUSG00000032021 | 0.475534142 | 0.144167 | 0.282247 | 49.29666 | 35.99465 | 36.43669 | 47.44429 | 33.15378 | 31.28647 | 39.53586 | 17.40146 |
| ENSMUSG00000032023 | 0.809300451 | 0.080257 | 0.189366 | 63.38142 | 71.98929 | 39.74912 | 69.8725 | 33.15378 | 64.8077 | 29.89296 | 11.9635 |
| ENSMUSG00000032024 | 0.470516069 | 0.017609 | 0.064558 | 3886.388 | 4682.903 | 3755.464 | 4085.385 | 3107.68 | 4180.096 | 2470.509 | 2084.912 |
| ENSMUSG00000032026 | 0.017125754 | 0.874047 | 0.92474 | 1371.252 | 1484.179 | 1143.615 | 1372.434 | 1343.703 | 1381.074 | 1165.826 | 1414.956 |
| ENSMUSG00000032028 | 0.523826727 | 0.14377 | 0.281589 | 44.26639 | 25.19625 | 33.95237 | 21.56559 | 20.47734 | 22.34748 | 19.28578 | 25.0146 |
| ENSMUSG00000032030 | 0.138491179 | 0.189766 | 0.342466 | 1528.197 | 1566.967 | 1347.329 | 1491.476 | 1276.421 | 1593.375 | 1235.254 | 1285.533 |
| ENSMUSG00000032033 | -0.830889337 | 0.202779 | 0.358441 | 5.030272 | 4.799286 | 14.07781 | 8.626235 | 5.850668 | 26.81698 | 15.42863 | 10.87591 |
| ENSMUSG00000032034 | -0.748071984 | 0.102989 | 0.225502 | 79.47829 | 38.39429 | 176.3867 | 81.08661 | 135.5405 | 77.09881 | 205.3936 | 214.2554 |
| ENSMUSG00000032035 | -0.071164187 | 0.586631 | 0.726928 | 2049.333 | 2046.896 | 2357.62 | 2190.201 | 2168.648 | 1848.137 | 2261.258 | 2804.897 |
| ENSMUSG00000032036 | 1.023851885 | 6.79E-06 | 0.000172 | 128.775 | 169.1748 | 153.1997 | 146.646 | 69.2329 | 101.681 | 52.07161 | 71.78101 |
| ENSMUSG00000032038 | 0.28526069 | 0.028326 | 0.090824 | 435.6215 | 448.7333 | 451.3181 | 544.3154 | 414.4223 | 435.7759 | 355.8227 | 338.2408 |
| ENSMUSG00000032040 | -0.142383271 | 0.11034 | 0.23618 | 992.9756 | 1225.018 | 1041.758 | 1096.394 | 1278.371 | 1135.252 | 1204.397 | 1184.387 |

| | | | | | | | | | | | |
|---|---|---|---|---|---|---|---|---|---|---|---|
| ENSMUSG00000032041 | 0.06701624 | 0.615743 | 0.748972 | 283.7073 | 314.3532 | 308.8838 | 341.5989 | 293.5085 | 346.386 | 303.7511 | 249.0584 |
| ENSMUSG00000032042 | -0.118955138 | 0.22161 | 0.380751 | 3285.774 | 3288.711 | 3435.814 | 3453.082 | 3895.57 | 3039.258 | 3659.477 | 4026.262 |
| ENSMUSG00000032044 | -0.001720265 | 0.988092 | 0.993258 | 502.0211 | 538.7199 | 482.7862 | 527.9256 | 540.2117 | 443.5975 | 589.1807 | 477.4525 |
| ENSMUSG00000032046 | -0.271696839 | 0.034662 | 0.105306 | 1089.557 | 1199.822 | 1563.465 | 1210.261 | 1621.61 | 1355.375 | 1527.434 | 1610.722 |
| ENSMUSG00000032047 | -0.584293213 | 4.44E-05 | 0.000728 | 3156.999 | 3383.497 | 3762.917 | 3442.73 | 5099.832 | 3864.997 | 5368.198 | 6277.576 |
| ENSMUSG00000032050 | -0.231327612 | 0.00236 | 0.014765 | 6599.717 | 5804.737 | 6273.736 | 5792.517 | 6575.176 | 7084.152 | 7505.063 | 7564.196 |
| ENSMUSG00000032051 | 1.024336371 | 0.169326 | 0.315632 | 5489.033 | 609.5093 | 660.001 | 1435.405 | 1015.091 | 762.0491 | 961.3963 | 1289.883 |
| ENSMUSG00000032058 | -0.128576325 | 0.092627 | 0.209097 | 1368.234 | 1361.797 | 1295.159 | 1270.644 | 1550.427 | 1335.262 | 1382.791 | 1517.19 |
| ENSMUSG00000032059 | -0.277838573 | 0.002403 | 0.014969 | 645.8869 | 613.1088 | 690.6409 | 653.006 | 777.1637 | 719.5889 | 781.0742 | 880.9488 |
| ENSMUSG00000032060 | -1.745679108 | 1.18E-06 | 4.32E-05 | 255.5378 | 296.3559 | 667.4539 | 279.49 | 1102.851 | 644.7249 | 1530.327 | 1752.109 |
| ENSMUSG00000032062 | -0.608823866 | 0.00756 | 0.034739 | 159.9626 | 117.5825 | 220.2764 | 181.1509 | 239.8774 | 198.8926 | 282.5367 | 317.5766 |
| ENSMUSG00000032064 | 0.283298947 | 0.023017 | 0.078073 | 856.1523 | 825.4772 | 751.9208 | 901.4416 | 681.6028 | 823.5047 | 651.8595 | 584.0364 |
| ENSMUSG00000032067 | 0.124023708 | 0.03698 | 0.110528 | 1520.148 | 1457.783 | 1545.247 | 1539.783 | 1352.479 | 1410.126 | 1347.112 | 1459.547 |
| ENSMUSG00000032068 | -0.467177705 | 0.153988 | 0.295488 | 746.4923 | 543.5192 | 1804.444 | 711.6644 | 1369.056 | 1065.975 | 1238.147 | 1591.146 |
| ENSMUSG00000032076 | 0.480978976 | 0.00483 | 0.025102 | 3273.701 | 3637.859 | 2416.415 | 3502.251 | 2458.256 | 2829.191 | 2077.079 | 1827.153 |
| ENSMUSG00000032077 | -0.012440165 | 0.900034 | 0.93991 | 531.1967 | 637.1052 | 530.8164 | 599.5233 | 625.0463 | 569.8608 | 552.5377 | 567.7225 |
| ENSMUSG00000032078 | 0.00695199 | 0.948893 | 0.971043 | 1183.12 | 1336.601 | 1095.585 | 1240.453 | 1393.434 | 1074.914 | 1150.397 | 1209.401 |
| ENSMUSG00000032080 | 1.297748101 | 0.376562 | 0.544772 | 82.49646 | 400.7404 | 124.216 | 4.313117 | 183.3209 | 59.22083 | 2.892867 | 3.262773 |
| ENSMUSG00000032083 | 2.246141729 | 3.76E-06 | 0.000108 | 519.124 | 239.9643 | 101.8571 | 498.5964 | 72.15824 | 118.4417 | 55.92877 | 40.24087 |
| ENSMUSG00000032085 | -0.516145662 | 0.074913 | 0.180874 | 584.5176 | 754.6877 | 1719.977 | 1142.114 | 1459.742 | 1292.802 | 1440.648 | 1818.452 |
| ENSMUSG00000032086 | 0.121742416 | 0.141715 | 0.278593 | 1593.59 | 1710.946 | 1584.996 | 1576.876 | 1301.774 | 1543.094 | 1578.541 | 1518.277 |
| ENSMUSG00000032087 | -0.140247543 | 0.63477 | 0.762467 | 31.18769 | 33.595 | 57.13936 | 41.40593 | 45.83023 | 36.87335 | 45.32159 | 53.29196 |
| ENSMUSG00000032089 | 0.344183471 | 0.462947 | 0.623933 | 36.21796 | 17.99732 | 33.95237 | 42.26855 | 13.65156 | 16.76061 | 44.3573 | 28.27737 |
| ENSMUSG00000032091 | -0.567331181 | 0.356702 | 0.525276 | 8.048435 | 23.99643 | 11.59349 | 3.450494 | 18.52711 | 24.58223 | 12.53576 | 13.05109 |
| ENSMUSG00000032092 | -0.774719394 | 0.012633 | 0.050861 | 184.1079 | 151.1775 | 264.9941 | 173.3873 | 359.8161 | 160.9019 | 335.5726 | 468.7518 |
| ENSMUSG00000032096 | -0.139408635 | 0.017479 | 0.064197 | 5454.827 | 5657.159 | 5654.312 | 5751.111 | 6120.774 | 5755.594 | 6376.844 | 6548.386 |
| ENSMUSG00000032097 | 0.213810277 | 0.035194 | 0.106518 | 8714.443 | 9164.237 | 7753.562 | 8580.516 | 7225.575 | 8675.292 | 6828.132 | 6770.255 |
| ENSMUSG00000032098 | -3.424677111 | 0.000654 | 0.005641 | 0 | 0 | 2.48432 | 4.313117 | 13.65156 | 4.469496 | 24.10723 | 34.80291 |
| ENSMUSG00000032101 | -0.055211355 | 0.817202 | 0.888971 | 100.6054 | 89.98662 | 104.3414 | 124.2178 | 90.68535 | 151.9629 | 82.92887 | 112.0219 |
| ENSMUSG00000032103 | -0.430649259 | 0.099701 | 0.220467 | 30.18163 | 27.5959 | 30.63995 | 38.81806 | 39.00445 | 44.69496 | 45.32159 | 43.50364 |
| ENSMUSG00000032105 | -1.937798134 | 4.53E-06 | 0.000124 | 106.6418 | 39.59411 | 122.5598 | 50.89479 | 360.7912 | 147.4934 | 259.3938 | 460.051 |
| ENSMUSG00000032109 | 0.078585104 | 0.495299 | 0.652059 | 266.6044 | 269.9598 | 281.5563 | 244.1224 | 234.0267 | 279.3435 | 251.6795 | 241.4452 |
| ENSMUSG00000032112 | 0.026773782 | 0.683353 | 0.798175 | 1173.059 | 1099.037 | 1104.694 | 1206.81 | 1163.308 | 1105.083 | 1098.325 | 1134.358 |
| ENSMUSG00000032113 | 0.05580424 | 0.718493 | 0.824146 | 495.9848 | 662.3015 | 443.037 | 536.5518 | 476.8294 | 613.4384 | 488.8946 | 475.2773 |
| ENSMUSG00000032114 | -1.702396406 | 1.52E-12 | 4.24E-10 | 579.4873 | 373.1445 | 592.9243 | 598.6607 | 1766.902 | 1050.332 | 1815.756 | 2350.284 |
| ENSMUSG00000032115 | -0.324357883 | 0.008107 | 0.036639 | 2255.574 | 2044.496 | 2220.982 | 2034.929 | 2391.948 | 2241.452 | 2931.439 | 3148.576 |
| ENSMUSG00000032116 | -0.042926303 | 0.544176 | 0.692833 | 4359.234 | 4566.521 | 4316.92 | 4256.184 | 4467.96 | 4094.059 | 4554.338 | 4907.211 |
| ENSMUSG00000032118 | 2.189543507 | 9.94E-16 | 5.24E-13 | 394.3733 | 449.9331 | 327.9302 | 421.8229 | 107.2622 | 124.0285 | 71.3574 | 46.76642 |
| ENSMUSG00000032119 | 0.074195607 | 0.346887 | 0.515719 | 702.2259 | 680.2988 | 697.2658 | 679.7473 | 667.9512 | 632.4337 | 707.7882 | 611.2262 |
| ENSMUSG00000032120 | -0.665165131 | 0.002966 | 0.017615 | 319.9253 | 229.1659 | 364.3669 | 334.6979 | 506.0828 | 329.6254 | 493.7161 | 652.5547 |
| ENSMUSG00000032121 | -0.171435846 | 0.11623 | 0.244349 | 547.2936 | 427.1365 | 466.224 | 477.0308 | 531.4357 | 508.4052 | 525.5376 | 595.9999 |
| ENSMUSG00000032122 | 0.252267652 | 0.318276 | 0.486549 | 117.7084 | 62.39072 | 81.15445 | 68.14726 | 72.15824 | 74.86406 | 64.60737 | 65.25547 |
| ENSMUSG00000032123 | -0.190897129 | 0.034517 | 0.104985 | 868.2249 | 817.0785 | 867.0276 | 811.7287 | 1000.464 | 836.9132 | 1038.539 | 962.5181 |
| ENSMUSG00000032125 | -0.280019372 | 0.068475 | 0.169521 | 704.238 | 563.9161 | 928.3075 | 704.7634 | 821.0437 | 842.5001 | 973.9321 | 889.6495 |
| ENSMUSG00000032126 | -0.280226185 | 0.003395 | 0.019358 | 994.9878 | 955.0579 | 889.3865 | 916.1062 | 1242.292 | 985.524 | 1126.29 | 1202.876 |
| ENSMUSG00000032127 | -0.086604132 | 0.296595 | 0.464315 | 1697.214 | 1497.377 | 1634.682 | 1546.684 | 1732.773 | 1507.338 | 1805.149 | 1724.919 |
| ENSMUSG00000032128 | 0.157557493 | 0.620448 | 0.752486 | 54.32694 | 55.19179 | 69.56096 | 36.23019 | 41.92979 | 69.27719 | 35.6787 | 46.76642 |
| ENSMUSG00000032131 | -0.106257235 | 0.537422 | 0.687697 | 97.58727 | 89.98662 | 82.81066 | 90.57547 | 108.2374 | 101.681 | 86.78602 | 91.35765 |
| ENSMUSG00000032135 | -0.027063179 | 0.712726 | 0.819906 | 2964.842 | 2844.777 | 3140.18 | 3146.851 | 3060.874 | 2873.886 | 3021.118 | 3374.795 |
| ENSMUSG00000032171 | -0.299068338 | 0.010612 | 0.044574 | 1434.634 | 1521.374 | 1624.745 | 1550.997 | 2120.867 | 1502.868 | 1901.578 | 2018.569 |
| ENSMUSG00000032172 | -0.327860758 | 0.108445 | 0.233244 | 66.39959 | 68.38983 | 91.09173 | 86.26235 | 95.56091 | 117.3243 | 84.85745 | 96.79561 |
| ENSMUSG00000032174 | -1.139241204 | 0.043893 | 0.1244 | 7.04238 | 5.999108 | 14.90592 | 7.763611 | 17.552 | 10.05637 | 17.3572 | 34.80291 |
| ENSMUSG00000032175 | -0.141992584 | 0.185584 | 0.337199 | 543.2694 | 419.9375 | 468.7084 | 453.74 | 487.5557 | 550.8654 | 525.5376 | 518.7809 |
| ENSMUSG00000032177 | 0.713543861 | 2.27E-08 | 1.69E-06 | 423.5489 | 493.1267 | 375.1323 | 368.3402 | 246.7032 | 255.8787 | 252.6438 | 254.4963 |
| ENSMUSG00000032178 | 0.043693417 | 0.524085 | 0.677274 | 4448.772 | 4894.072 | 4556.243 | 4529.636 | 4781.946 | 4498.548 | 4499.373 | 4093.693 |
| ENSMUSG00000032179 | 0.384859454 | 0.254918 | 0.41887 | 28.16952 | 29.99554 | 21.53077 | 32.77969 | 16.57689 | 26.81698 | 23.14294 | 19.57664 |
| ENSMUSG00000032180 | -0.297693893 | 0.00029 | 0.003026 | 1065.412 | 1208.22 | 1131.194 | 1230.964 | 1453.891 | 1293.919 | 1462.827 | 1485.649 |
| ENSMUSG00000032181 | 1.098501792 | 0.033982 | 0.103683 | 123.7447 | 31.19536 | 40.57722 | 154.4096 | 27.30312 | 53.63396 | 40.50014 | 42.41605 |
| ENSMUSG00000032182 | -0.193449002 | 0.076722 | 0.183916 | 490.9545 | 472.7297 | 469.5365 | 466.6793 | 568.4899 | 448.067 | 567.9663 | 586.2116 |
| ENSMUSG00000032184 | 0.004039465 | 0.964117 | 0.979778 | 355.1372 | 347.9482 | 366.0231 | 346.7746 | 359.8161 | 367.6161 | 346.1798 | 339.3284 |
| ENSMUSG00000032185 | -0.073620762 | 0.444861 | 0.607797 | 1493.991 | 1384.594 | 1479.827 | 1417.29 | 1749.35 | 1331.91 | 1473.434 | 1522.628 |
| ENSMUSG00000032186 | 0.338817266 | 0.00061 | 0.005335 | 477.8758 | 454.7324 | 437.2403 | 509.8105 | 368.5921 | 385.4941 | 342.3227 | 391.5328 |
| ENSMUSG00000032187 | 0.006468274 | 0.932613 | 0.960099 | 4534.287 | 4608.515 | 4613.382 | 4123.34 | 4180.302 | 4813.648 | 4173.444 | 4633.138 |
| ENSMUSG00000032192 | -0.02032684 | 0.826173 | 0.894472 | 874.2612 | 819.4781 | 999.5247 | 962.6878 | 862.9735 | 929.6552 | 938.2534 | 983.1823 |

| | | | | | | | | | | | |
|---|---|---|---|---|---|---|---|---|---|---|---|
| ENSMUSG00000032193 | -0.936436627 | 1.50E-05 | 0.000312 | 799.8132 | 442.7341 | 754.4051 | 535.6892 | 1130.154 | 1115.139 | 1589.149 | 1014.722 |
| ENSMUSG00000032194 | 0.277081762 | 0.001414 | 0.010154 | 2184.144 | 2422.44 | 2384.947 | 2313.556 | 1948.272 | 2167.706 | 1812.864 | 1751.022 |
| ENSMUSG00000032198 | 0.275760447 | 1.64E-05 | 0.000334 | 2452.761 | 2244.866 | 2480.179 | 2492.119 | 1923.895 | 2000.1 | 2108.9 | 1958.752 |
| ENSMUSG00000032199 | -0.110387337 | 0.057694 | 0.15054 | 8182.24 | 8404.75 | 8223.927 | 8468.375 | 9658.478 | 8588.137 | 9131.818 | 8543.028 |
| ENSMUSG00000032202 | -0.391206482 | 0.082326 | 0.192817 | 323.9495 | 326.3515 | 295.6341 | 345.912 | 397.8454 | 258.1134 | 582.4307 | 454.6131 |
| ENSMUSG00000032207 | 0.763376301 | 0.189964 | 0.342514 | 22.1332 | 10.79839 | 9.93728 | 9.488858 | 9.751113 | 3.352122 | 7.714313 | 9.78832 |
| ENSMUSG00000032212 | 0.241366997 | 0.060738 | 0.156387 | 4324.022 | 4321.757 | 3679.278 | 3904.234 | 3021.87 | 4296.303 | 3348.976 | 3061.569 |
| ENSMUSG00000032215 | 0.013824214 | 0.887715 | 0.932533 | 1741.48 | 1788.934 | 1690.994 | 1871.03 | 1880.99 | 1966.578 | 1646.042 | 1531.328 |
| ENSMUSG00000032216 | 0.118002293 | 0.003116 | 0.018294 | 39712.99 | 42604.46 | 41260.41 | 41911.43 | 38630.98 | 37794.06 | 38472.24 | 37590.41 |
| ENSMUSG00000032217 | 0.198550315 | 0.018323 | 0.066361 | 1166.017 | 1099.037 | 1006.978 | 1017.033 | 925.3806 | 1012.341 | 899.6818 | 898.3502 |
| ENSMUSG00000032218 | 0.029382484 | 0.728195 | 0.829347 | 2428.615 | 2324.054 | 2293.027 | 2689.66 | 2592.821 | 2380.007 | 2381.794 | 2184.97 |
| ENSMUSG00000032220 | 0.217123305 | 0.047705 | 0.131627 | 1325.98 | 1175.825 | 1210.692 | 1118.823 | 980.962 | 940.829 | 1227.54 | 1004.934 |
| ENSMUSG00000032221 | -0.054573051 | 0.6663 | 0.785715 | 443.67 | 520.7225 | 419.8501 | 417.5098 | 419.2979 | 540.8091 | 433.9301 | 475.2773 |
| ENSMUSG00000032224 | -0.109060484 | 0.563351 | 0.708643 | 485.9243 | 471.5299 | 501.0045 | 443.3885 | 433.9245 | 364.264 | 596.895 | 655.8174 |
| ENSMUSG00000032226 | 0.321702172 | 0.294223 | 0.461789 | 110.666 | 171.5745 | 168.1056 | 120.7673 | 90.68535 | 187.7188 | 97.39321 | 81.56933 |
| ENSMUSG00000032228 | 0.212981497 | 0.002047 | 0.01334 | 6700.322 | 6904.973 | 6583.448 | 6816.451 | 6111.998 | 6285.229 | 5520.555 | 5380.313 |
| ENSMUSG00000032231 | -0.115608662 | 0.321048 | 0.489084 | 7541.383 | 7520.481 | 9582.85 | 7117.506 | 8841.334 | 7519.928 | 8692.103 | 9360.896 |
| ENSMUSG00000032232 | -0.403938189 | 0.019971 | 0.070341 | 3161.023 | 2701.998 | 3622.138 | 3038.16 | 3508.45 | 3101.83 | 5035.518 | 4925.7 |
| ENSMUSG00000032235 | 0.09855141 | 0.341537 | 0.510315 | 829.9948 | 759.487 | 698.0939 | 750.4824 | 660.1504 | 811.2136 | 707.7882 | 657.9926 |
| ENSMUSG00000032238 | 0.653356574 | 0.000344 | 0.003432 | 1152.938 | 1052.243 | 945.6978 | 1133.487 | 620.1708 | 962.0591 | 597.8593 | 544.8831 |
| ENSMUSG00000032239 | 0.124235204 | 0.078015 | 0.18574 | 2312.919 | 2334.853 | 2198.623 | 2404.132 | 2235.93 | 2228.044 | 2075.15 | 1946.788 |
| ENSMUSG00000032243 | 0.197727238 | 0.335517 | 0.503725 | 104.6297 | 111.5834 | 114.2787 | 107.8279 | 78.98402 | 125.1459 | 77.14313 | 102.2336 |
| ENSMUSG00000032244 | -0.006903368 | 0.915052 | 0.949629 | 2683.147 | 2865.174 | 2640.832 | 2474.867 | 2762.49 | 2712.984 | 2595.866 | 2637.408 |
| ENSMUSG00000032245 | -0.172345488 | 0.074663 | 0.180461 | 874.2612 | 877.0695 | 1083.163 | 957.5121 | 1108.702 | 1078.266 | 991.2893 | 1100.642 |
| ENSMUSG00000032246 | -1.718115738 | 2.99E-08 | 2.13E-06 | 335.0161 | 255.562 | 593.7525 | 347.6373 | 1184.76 | 758.697 | 1154.254 | 1947.876 |
| ENSMUSG00000032249 | -0.050323901 | 0.367581 | 0.53607 | 10635 | 11146.34 | 10577.41 | 10720.68 | 11347.37 | 10333.48 | 11250.36 | 11672.03 |
| ENSMUSG00000032251 | 0.096738764 | 0.311453 | 0.480077 | 521.1362 | 610.7092 | 578.8465 | 629.7152 | 574.3406 | 564.2739 | 511.0733 | 539.4452 |
| ENSMUSG00000032252 | 0.154481956 | 0.076801 | 0.18406 | 1213.302 | 1078.64 | 1046.727 | 1168.855 | 981.9371 | 1108.435 | 1017.325 | 942.9415 |
| ENSMUSG00000032253 | 0.510446156 | 0.0143 | 0.055669 | 3677.129 | 4082.993 | 2982.012 | 3612.667 | 2178.399 | 3797.955 | 2068.4 | 2032.708 |
| ENSMUSG00000032254 | 0.09954774 | 0.311079 | 0.479748 | 2200.241 | 2066.093 | 1984.143 | 2260.936 | 2011.655 | 2291.734 | 1803.221 | 1839.117 |
| ENSMUSG00000032258 | 0.146693098 | 0.230709 | 0.391167 | 452.7245 | 452.3327 | 412.3971 | 445.1137 | 369.5672 | 483.823 | 366.4299 | 373.0437 |
| ENSMUSG00000032261 | -0.006932059 | 0.964869 | 0.980218 | 787.7406 | 778.6842 | 918.3703 | 884.1891 | 873.6997 | 630.199 | 825.4315 | 1057.139 |
| ENSMUSG00000032262 | 0.09390959 | 0.626663 | 0.756669 | 276.6649 | 251.9625 | 247.6039 | 284.6658 | 268.1556 | 284.9304 | 279.6439 | 160.9635 |
| ENSMUSG00000032263 | -0.219984235 | 0.01687 | 0.062507 | 593.5721 | 621.5076 | 609.4865 | 701.3129 | 721.5824 | 682.7156 | 786.86 | 751.5254 |
| ENSMUSG00000032264 | -0.071717622 | 0.492537 | 0.649627 | 856.1523 | 963.4567 | 766.8267 | 923.0071 | 955.6091 | 817.9178 | 929.5748 | 981.0072 |
| ENSMUSG00000032265 | 0.378111894 | 0.118326 | 0.247126 | 302.8224 | 344.3488 | 260.0255 | 313.1323 | 181.3707 | 360.9118 | 221.7865 | 175.1022 |
| ENSMUSG00000032267 | 0.134311585 | 0.146703 | 0.285743 | 882.3097 | 872.2703 | 785.8732 | 877.2881 | 787.8899 | 854.7912 | 779.1456 | 690.6203 |
| ENSMUSG00000032269 | 1.575700292 | 1.63E-08 | 1.28E-06 | 82.49646 | 104.3845 | 75.3577 | 95.75121 | 30.22845 | 41.34284 | 29.89296 | 18.48905 |
| ENSMUSG00000032271 | 0.443604797 | 0.1899 | 0.342466 | 54.32694 | 58.79126 | 52.17072 | 93.16334 | 29.25334 | 43.57759 | 68.46453 | 48.9416 |
| ENSMUSG00000032278 | -1.795099026 | 0.00273 | 0.01649 | 371.2341 | 178.7734 | 939.0729 | 298.4677 | 1328.102 | 706.1804 | 2034.65 | 2137.116 |
| ENSMUSG00000032279 | -0.913890093 | 1.21E-05 | 0.000266 | 1615.723 | 1534.572 | 2093.454 | 1632.084 | 3095.978 | 2049.264 | 3483.012 | 4328.613 |
| ENSMUSG00000032280 | 0.255672646 | 0.025661 | 0.084587 | 2377.306 | 2324.054 | 2602.739 | 2279.914 | 1623.56 | 2276.091 | 1997.043 | 2133.854 |
| ENSMUSG00000032281 | 1.967748045 | 0.025444 | 0.084007 | 644.8808 | 41.99375 | 78.67013 | 156.9975 | 38.02934 | 43.57759 | 43.39301 | 110.9343 |
| ENSMUSG00000032285 | -0.144405907 | 0.481522 | 0.640521 | 124.7507 | 94.7859 | 129.1846 | 124.2178 | 115.0631 | 122.9112 | 172.6078 | 113.1095 |
| ENSMUSG00000032288 | -0.079370443 | 0.365547 | 0.534244 | 1913.515 | 1937.712 | 1953.504 | 1994.386 | 2215.453 | 1767.686 | 2210.151 | 2044.671 |
| ENSMUSG00000032289 | 0.209715285 | 0.163918 | 0.308862 | 1485.942 | 1754.139 | 2110.016 | 1561.349 | 1394.409 | 1246.989 | 1546.72 | 1789.087 |
| ENSMUSG00000032290 | -1.88E-05 | 0.999735 | 0.999735 | 2421.573 | 2388.845 | 2431.321 | 2503.333 | 2367.57 | 2330.842 | 2577.545 | 2469.919 |
| ENSMUSG00000032291 | 0.181735429 | 0.306367 | 0.474992 | 174.0474 | 152.3773 | 207.8548 | 151.8217 | 141.3911 | 177.6625 | 128.2505 | 159.8759 |
| ENSMUSG00000032292 | 0.248526202 | 0.251761 | 0.415385 | 76.46013 | 103.1847 | 105.1695 | 94.88858 | 78.98402 | 101.681 | 63.64308 | 76.13138 |
| ENSMUSG00000032293 | 0.221372259 | 0.04115 | 0.119091 | 3695.238 | 3583.867 | 3127.759 | 3583.338 | 2824.897 | 3586.771 | 2749.188 | 2839.7 |
| ENSMUSG00000032294 | -0.380322267 | 0.008675 | 0.038451 | 14719.58 | 15958.83 | 18427.86 | 15150.26 | 21183.32 | 15629.83 | 21897.08 | 24928.68 |
| ENSMUSG00000032295 | 0.00723975 | 0.906476 | 0.943913 | 1649.929 | 1665.352 | 1608.183 | 1670.039 | 1582.606 | 1565.441 | 1660.506 | 1751.022 |
| ENSMUSG00000032297 | 1.956401486 | 0.0286 | 0.091413 | 22.1332 | 9.598572 | 1.656213 | 53.48266 | 6.825779 | 3.352122 | 6.750024 | 5.437955 |
| ENSMUSG00000032298 | 0.227311567 | 0.089199 | 0.203896 | 361.1735 | 313.1534 | 272.4471 | 364.0271 | 291.5583 | 296.1041 | 272.8938 | 258.8467 |
| ENSMUSG00000032299 | -0.072312986 | 0.278072 | 0.444297 | 1237.447 | 1383.394 | 1334.08 | 1362.082 | 1455.841 | 1349.788 | 1359.648 | 1423.657 |
| ENSMUSG00000032300 | 0.058647553 | 0.598072 | 0.736699 | 573.451 | 461.9313 | 582.159 | 492.558 | 475.8543 | 536.3396 | 528.4305 | 488.3284 |
| ENSMUSG00000032301 | -0.128613485 | 0.018687 | 0.06728 | 3959.83 | 3889.821 | 4177.798 | 3920.624 | 4124.721 | 4394.632 | 4386.551 | 4535.255 |
| ENSMUSG00000032303 | 1.961205261 | 0.008625 | 0.038338 | 64.38748 | 7.198929 | 9.93728 | 67.28463 | 10.72622 | 11.17374 | 7.714313 | 8.700729 |
| ENSMUSG00000032305 | -0.047427425 | 0.629166 | 0.7586 | 531.1967 | 568.7154 | 492.7234 | 493.4206 | 549.9628 | 519.579 | 494.6803 | 588.3868 |
| ENSMUSG00000032306 | -0.315018949 | 0.016932 | 0.062663 | 642.8687 | 573.5147 | 571.3936 | 600.386 | 682.5779 | 612.321 | 783.0028 | 892.9123 |
| ENSMUSG00000032307 | -0.070721169 | 0.49083 | 0.648072 | 1187.144 | 1169.826 | 1201.583 | 1282.721 | 1170.134 | 1503.986 | 1249.719 | 1163.722 |
| ENSMUSG00000032308 | -0.033183526 | 0.76279 | 0.852453 | 350.1069 | 317.9527 | 328.7583 | 318.3081 | 373.4676 | 298.3389 | 347.1441 | 325.1897 |
| ENSMUSG00000032309 | -0.025841275 | 0.727699 | 0.829154 | 1369.24 | 1521.374 | 1363.064 | 1491.476 | 1489.97 | 1377.722 | 1425.219 | 1553.08 |
| ENSMUSG00000032311 | 0.302376772 | 0.224114 | 0.383875 | 100.6054 | 81.58786 | 78.67013 | 101.7896 | 56.55646 | 99.44629 | 81.00029 | 57.64233 |

| | | | | | | | | | | | |
|---|---|---|---|---|---|---|---|---|---|---|---|
| ENSMUSG00000032312 | -0.097229454 | 0.156267 | 0.29834 | 1532.221 | 1694.148 | 1696.79 | 1683.841 | 1848.811 | 1796.738 | 1651.827 | 1771.686 |
| ENSMUSG00000032313 | -0.209793397 | 0.607301 | 0.74321 | 10.06054 | 31.19536 | 20.70267 | 23.29083 | 28.27823 | 22.34748 | 20.25007 | 27.18978 |
| ENSMUSG00000032314 | -0.673398552 | 6.00E-05 | 0.000916 | 2568.457 | 2297.658 | 2755.939 | 2591.321 | 3987.23 | 2845.952 | 4240.944 | 5216.087 |
| ENSMUSG00000032316 | -0.038182648 | 0.618335 | 0.750937 | 1602.645 | 1479.38 | 1398.672 | 1494.064 | 1580.655 | 1396.718 | 1571.791 | 1582.445 |
| ENSMUSG00000032320 | 0.14196149 | 0.132918 | 0.266268 | 3584.572 | 3954.612 | 3859.805 | 4194.075 | 3650.817 | 3863.88 | 3585.227 | 3032.204 |
| ENSMUSG00000032322 | 0.262262892 | 0.351505 | 0.520274 | 55.33299 | 44.3934 | 77.84202 | 62.97152 | 43.88001 | 65.92507 | 53.0359 | 39.15328 |
| ENSMUSG00000032323 | 3.922666418 | 0.02682 | 0.087407 | 8176.204 | 23.99643 | 41.40533 | 942.8475 | 17.552 | 15.64324 | 22.17865 | 550.3211 |
| ENSMUSG00000032324 | 0.038220122 | 0.548791 | 0.697048 | 6614.807 | 7591.271 | 7154.841 | 7686.838 | 7183.645 | 7088.621 | 7057.632 | 6957.32 |
| ENSMUSG00000032327 | -0.359070517 | 0.016263 | 0.060915 | 317.9132 | 461.9313 | 409.9128 | 323.4838 | 541.1868 | 462.5929 | 471.5374 | 462.2262 |
| ENSMUSG00000032328 | -0.080157202 | 0.359161 | 0.52755 | 2842.104 | 2648.006 | 2692.175 | 2801.801 | 2585.02 | 2756.562 | 3004.725 | 3267.124 |
| ENSMUSG00000032329 | 0.289881858 | 0.000212 | 0.002375 | 2472.882 | 2388.845 | 2341.886 | 2582.695 | 1989.227 | 2222.457 | 1952.686 | 1842.379 |
| ENSMUSG00000032330 | -0.491226927 | 0.0005 | 0.004592 | 2393.403 | 2490.83 | 3054.057 | 2571.481 | 3635.215 | 2863.83 | 3909.228 | 4366.678 |
| ENSMUSG00000032332 | 0.632200007 | 9.15E-05 | 0.00125 | 5146.974 | 7368.104 | 5718.904 | 6165.17 | 3677.145 | 5092.991 | 3732.763 | 3238.846 |
| ENSMUSG00000032333 | 0.124565711 | 0.106777 | 0.231028 | 930.6003 | 845.8742 | 859.5747 | 885.9143 | 821.0437 | 867.0823 | 787.8242 | 755.8758 |
| ENSMUSG00000032334 | 0.16803401 | 0.13638 | 0.271085 | 2593.608 | 3214.322 | 3068.963 | 2855.284 | 2663.029 | 2972.215 | 2601.652 | 2204.547 |
| ENSMUSG00000032336 | -0.02531309 | 0.711045 | 0.818913 | 3418.573 | 3635.459 | 3805.978 | 3827.46 | 3489.923 | 3668.339 | 3857.157 | 3935.992 |
| ENSMUSG00000032338 | 0.197729303 | 0.613529 | 0.747313 | 40.24217 | 25.19625 | 68.73285 | 69.8725 | 32.17867 | 37.99072 | 48.21446 | 60.9051 |
| ENSMUSG00000032340 | 0.178428659 | 0.131794 | 0.264793 | 4167.077 | 5101.641 | 4802.19 | 4733.215 | 3744.427 | 5049.414 | 4062.55 | 3760.89 |
| ENSMUSG00000032342 | -0.294716985 | 0.021778 | 0.074836 | 430.5913 | 392.3416 | 339.5237 | 331.2474 | 431.9743 | 411.1937 | 492.7518 | 492.6788 |
| ENSMUSG00000032343 | -0.074176572 | 0.722234 | 0.826177 | 55.33299 | 75.58876 | 70.38906 | 58.6584 | 76.05868 | 68.15982 | 66.53595 | 61.99269 |
| ENSMUSG00000032344 | 0.420039216 | 0.06236 | 0.159231 | 153.9263 | 176.3738 | 148.2311 | 124.2178 | 93.61069 | 156.4324 | 89.67889 | 110.9343 |
| ENSMUSG00000032346 | 1.270025632 | 4.99E-07 | 2.11E-05 | 159.9626 | 105.5843 | 151.5435 | 169.9368 | 52.65601 | 75.98144 | 43.39301 | 72.8686 |
| ENSMUSG00000032348 | 0.336921657 | 4.33E-06 | 0.000121 | 3128.829 | 3300.709 | 3480.532 | 3478.098 | 2844.4 | 2448.167 | 2579.474 | 2728.766 |
| ENSMUSG00000032349 | 0.327491147 | 0.003951 | 0.021669 | 2763.631 | 2326.454 | 2361.76 | 2593.046 | 1980.451 | 2383.359 | 1875.542 | 1767.336 |
| ENSMUSG00000032350 | -0.537453231 | 0.028273 | 0.090718 | 2871.279 | 1916.115 | 3418.424 | 2678.446 | 3047.223 | 2847.069 | 5900.485 | 4004.51 |
| ENSMUSG00000032352 | -0.261918596 | 0.022554 | 0.076743 | 1130.805 | 981.454 | 1234.707 | 1231.826 | 1252.043 | 1243.637 | 1463.791 | 1535.679 |
| ENSMUSG00000032353 | 0.042722343 | 0.603147 | 0.740579 | 1227.386 | 1490.178 | 1268.659 | 1278.408 | 1319.326 | 1238.05 | 1271.897 | 1275.744 |
| ENSMUSG00000032356 | 0.315893709 | 0.380527 | 0.548577 | 36.21796 | 35.99465 | 34.78048 | 25.01608 | 26.32801 | 40.22547 | 23.14294 | 16.31387 |
| ENSMUSG00000032357 | -2.565311935 | 1.22E-06 | 4.42E-05 | 43.26034 | 17.99732 | 127.5284 | 53.48266 | 368.5921 | 147.4934 | 330.7512 | 593.8247 |
| ENSMUSG00000032358 | -0.644572958 | 0.046648 | 0.129698 | 28.16952 | 34.79482 | 40.57722 | 21.56559 | 58.50668 | 32.40385 | 54.00019 | 50.02919 |
| ENSMUSG00000032359 | -0.565147937 | 0.008195 | 0.036934 | 1688.159 | 1227.417 | 1835.912 | 1410.389 | 2351.968 | 1436.943 | 2362.508 | 2966.948 |
| ENSMUSG00000032363 | 0.189479608 | 0.07716 | 0.184511 | 659.9717 | 783.4835 | 753.577 | 663.3575 | 578.241 | 626.8469 | 594.0021 | 709.1094 |
| ENSMUSG00000032366 | -0.200047975 | 0.167937 | 0.313962 | 9042.417 | 8519.933 | 12713.92 | 9523.363 | 10555.58 | 10501.08 | 11154.9 | 13512.23 |
| ENSMUSG00000032369 | 0.359362335 | 2.06E-06 | 6.57E-05 | 2427.609 | 2190.874 | 2212.701 | 2489.531 | 1762.026 | 1855.958 | 1932.435 | 1716.219 |
| ENSMUSG00000032370 | -0.602013546 | 0.000486 | 0.004476 | 467.8153 | 406.7395 | 545.7223 | 492.558 | 693.3041 | 550.8654 | 729.0026 | 933.1532 |
| ENSMUSG00000032372 | -0.560445587 | 0.00058 | 0.005136 | 1157.969 | 896.2667 | 997.0404 | 1161.954 | 1421.712 | 1186.651 | 1691.363 | 1915.248 |
| ENSMUSG00000032373 | -1.268192431 | 0.011569 | 0.04764 | 744.4802 | 605.9099 | 1918.723 | 695.2745 | 2326.616 | 1169.891 | 2513.902 | 3540.109 |
| ENSMUSG00000032374 | 0.344751644 | 0.009325 | 0.040449 | 2567.451 | 2151.28 | 1674.432 | 2161.734 | 1600.158 | 1781.094 | 1782.971 | 1570.482 |
| ENSMUSG00000032375 | -0.325081508 | 0.01842 | 0.066595 | 360.1675 | 350.3479 | 452.9743 | 422.6855 | 509.0081 | 477.1187 | 429.1087 | 576.4233 |
| ENSMUSG00000032376 | 0.053736171 | 0.535811 | 0.686541 | 1795.807 | 1781.735 | 1798.648 | 1852.053 | 1751.3 | 1925.236 | 1783.935 | 1504.138 |
| ENSMUSG00000032377 | 0.878167058 | 0.000361 | 0.003568 | 512.0817 | 616.7083 | 411.569 | 682.3352 | 270.1058 | 415.6632 | 329.7869 | 193.5912 |
| ENSMUSG00000032380 | -0.223084659 | 0.204378 | 0.360404 | 125.7568 | 136.7797 | 98.54469 | 103.5148 | 143.3414 | 148.6108 | 121.5004 | 126.1606 |
| ENSMUSG00000032381 | -0.191513246 | 0.041562 | 0.119911 | 1549.324 | 1474.581 | 1653.729 | 1708.857 | 1792.255 | 1595.61 | 1934.364 | 1973.978 |
| ENSMUSG00000032382 | -0.205230479 | 0.036361 | 0.109243 | 2391.391 | 2385.245 | 2329.464 | 2552.503 | 2783.943 | 2326.373 | 2997.011 | 3026.766 |
| ENSMUSG00000032383 | -0.079794366 | 0.296311 | 0.464124 | 5393.457 | 6451.44 | 6241.44 | 6041.815 | 6754.596 | 5996.947 | 6367.201 | 6378.722 |
| ENSMUSG00000032384 | 0.371747723 | 0.066081 | 0.165659 | 963.8001 | 968.256 | 982.9626 | 941.1222 | 728.4081 | 1099.496 | 565.0735 | 588.3868 |
| ENSMUSG00000032386 | 0.163129526 | 0.048048 | 0.132413 | 1064.406 | 1142.23 | 1079.851 | 1231.826 | 989.738 | 1081.618 | 945.9677 | 1020.16 |
| ENSMUSG00000032387 | -0.360775678 | 0.000425 | 0.004035 | 568.4207 | 441.5343 | 496.0359 | 490.8328 | 668.9264 | 600.0299 | 664.3952 | 631.8904 |
| ENSMUSG00000032388 | 0.098045459 | 0.193258 | 0.346501 | 2150.944 | 2411.641 | 2331.948 | 2436.049 | 2216.428 | 2324.138 | 2181.222 | 1995.73 |
| ENSMUSG00000032392 | 0.038955252 | 0.736503 | 0.834945 | 320.9313 | 307.1543 | 278.2438 | 320.8959 | 323.737 | 315.0995 | 290.251 | 264.2846 |
| ENSMUSG00000032393 | -0.016995827 | 0.764118 | 0.853562 | 2631.838 | 2624.01 | 2558.849 | 2548.19 | 2749.814 | 2515.209 | 2705.795 | 2509.073 |
| ENSMUSG00000032394 | -0.474099271 | 0.186545 | 0.338399 | 96.58122 | 125.9813 | 73.70149 | 89.71284 | 118.9636 | 240.2354 | 92.57176 | 83.74451 |
| ENSMUSG00000032396 | -0.233740595 | 0.021284 | 0.073658 | 670.0322 | 602.3104 | 641.7826 | 604.6991 | 757.6615 | 681.5982 | 679.8239 | 843.9707 |
| ENSMUSG00000032397 | -0.156733863 | 0.139704 | 0.275791 | 1378.294 | 1712.145 | 1562.637 | 1547.547 | 1877.089 | 1892.832 | 1584.327 | 1556.343 |
| ENSMUSG00000032398 | -0.151300794 | 0.220451 | 0.379325 | 583.5115 | 619.1079 | 655.0323 | 621.0889 | 730.3584 | 556.4523 | 671.1453 | 795.0291 |
| ENSMUSG00000032399 | -0.167943889 | 0.021171 | 0.073392 | 51614.61 | 58853.65 | 59419.14 | 57901.88 | 68915.99 | 61246.63 | 64941.02 | 60806.13 |
| ENSMUSG00000032400 | 0.197074447 | 0.191075 | 0.343745 | 771.6437 | 829.0767 | 580.5027 | 734.0926 | 646.4988 | 764.2839 | 565.0735 | 565.5474 |
| ENSMUSG00000032402 | 0.123366855 | 0.01993 | 0.070247 | 2931.642 | 2874.772 | 3092.15 | 3116.659 | 2692.282 | 2797.905 | 2734.724 | 2813.598 |
| ENSMUSG00000032403 | -0.334718566 | 0.011755 | 0.048188 | 197.1867 | 157.1766 | 165.6213 | 172.5247 | 235.0018 | 200.01 | 211.1793 | 227.3065 |
| ENSMUSG00000032405 | 0.193575692 | 0.070912 | 0.174014 | 1506.063 | 1394.193 | 1447.53 | 1481.125 | 1144.781 | 1536.389 | 1189.933 | 1228.978 |
| ENSMUSG00000032407 | 0.228649667 | 0.126392 | 0.257329 | 4030.254 | 4057.796 | 3096.291 | 3503.114 | 2775.167 | 4032.603 | 2943.011 | 2783.146 |
| ENSMUSG00000032409 | 0.104361916 | 0.361005 | 0.529584 | 1020.139 | 934.661 | 977.9939 | 906.6173 | 868.8242 | 1079.383 | 807.11 | 817.8685 |
| ENSMUSG00000032410 | 0.460006873 | 0.000531 | 0.004803 | 1987.963 | 1817.73 | 1601.558 | 1858.954 | 1177.934 | 1650.362 | 1284.433 | 1170.248 |
| ENSMUSG00000032411 | 0.165012651 | 0.15515 | 0.296977 | 2274.689 | 2294.059 | 1955.988 | 2442.087 | 2122.817 | 2250.391 | 1950.757 | 1672.715 |

| | | | | | | | | | | | |
|---|---|---|---|---|---|---|---|---|---|---|---|
| ENSMUSG00000032412 | -0.148619596 | 0.077814 | 0.185448 | 3073.496 | 3449.487 | 3577.421 | 3217.586 | 3583.534 | 3355.474 | 3812.799 | 4011.036 |
| ENSMUSG00000032413 | -0.055008857 | 0.652446 | 0.77555 | 724.3591 | 634.7056 | 563.1125 | 633.1656 | 637.7228 | 779.9271 | 629.6808 | 606.8758 |
| ENSMUSG00000032415 | -0.086287725 | 0.625311 | 0.75573 | 232.3986 | 231.5656 | 212.8234 | 213.068 | 316.9112 | 212.3011 | 198.6436 | 215.343 |
| ENSMUSG00000032417 | 0.758533095 | 0.040824 | 0.11838 | 117.7084 | 43.19358 | 46.37397 | 66.42201 | 39.00445 | 43.57759 | 33.75012 | 45.67883 |
| ENSMUSG00000032418 | 0.183768395 | 0.71134 | 0.818995 | 1536.245 | 334.7502 | 528.332 | 566.7436 | 678.6775 | 567.626 | 516.859 | 848.321 |
| ENSMUSG00000032419 | 0.972948213 | 0.008328 | 0.037379 | 190.1443 | 328.7511 | 230.2136 | 195.8155 | 143.3414 | 200.01 | 59.78593 | 78.30656 |
| ENSMUSG00000032420 | -0.19746497 | 0.336325 | 0.504611 | 273.6468 | 206.3693 | 200.4018 | 299.3304 | 278.8818 | 301.691 | 206.3579 | 338.2408 |
| ENSMUSG00000032422 | -0.111869457 | 0.273725 | 0.43956 | 916.5155 | 856.6726 | 834.7315 | 791.0257 | 974.1362 | 788.8661 | 902.5747 | 1004.934 |
| ENSMUSG00000032423 | 0.120696967 | 0.189834 | 0.342466 | 5092.647 | 5727.948 | 4966.984 | 5303.409 | 4520.616 | 5589.105 | 4567.838 | 4720.145 |
| ENSMUSG00000032425 | 0.150686891 | 0.123157 | 0.252779 | 622.7476 | 503.925 | 527.5039 | 580.5456 | 511.9334 | 491.6446 | 520.7161 | 489.416 |
| ENSMUSG00000032431 | 0.048062536 | 0.581021 | 0.722352 | 2204.265 | 2761.989 | 2476.867 | 2417.934 | 2482.633 | 2331.96 | 2235.222 | 2485.146 |
| ENSMUSG00000032434 | -0.243886835 | 0.018507 | 0.066842 | 1846.11 | 1745.74 | 1971.722 | 1780.455 | 2347.093 | 1792.268 | 2258.365 | 2299.168 |
| ENSMUSG00000032435 | -0.117147616 | 0.208442 | 0.365615 | 1590.572 | 1426.588 | 1594.105 | 1627.771 | 1683.042 | 1467.112 | 1791.649 | 1827.153 |
| ENSMUSG00000032436 | 0.090252948 | 0.260428 | 0.424448 | 867.2189 | 880.669 | 920.8546 | 846.2336 | 884.426 | 776.575 | 776.2528 | 864.6349 |
| ENSMUSG00000032437 | -0.2578525 | 0.000746 | 0.006244 | 3411.53 | 3429.09 | 3495.438 | 3334.902 | 3739.552 | 3824.772 | 4428.98 | 4352.54 |
| ENSMUSG00000032440 | 0.118071297 | 0.083029 | 0.193881 | 2850.152 | 2952.761 | 3226.303 | 2874.261 | 2837.574 | 2676.111 | 2605.509 | 2851.664 |
| ENSMUSG00000032443 | 0.958926036 | 0.015789 | 0.059844 | 20.12109 | 21.59679 | 18.21835 | 22.42821 | 11.70134 | 11.17374 | 9.642892 | 9.78832 |
| ENSMUSG00000032449 | 0.291167592 | 0.009933 | 0.042377 | 2894.418 | 3193.925 | 2935.638 | 3216.723 | 2381.222 | 3041.492 | 2392.401 | 2189.321 |
| ENSMUSG00000032452 | 0.121323847 | 0.487849 | 0.645722 | 121.7326 | 119.9822 | 122.5598 | 142.3329 | 119.9387 | 137.437 | 116.679 | 92.44524 |
| ENSMUSG00000032454 | -1.69926738 | 0.000647 | 0.005588 | 18.10898 | 37.19447 | 17.39024 | 6.900988 | 72.15824 | 32.40385 | 67.50024 | 82.65692 |
| ENSMUSG00000032456 | -0.722680457 | 0.002231 | 0.014227 | 233.4046 | 124.7814 | 197.0894 | 173.3873 | 292.5334 | 217.8879 | 305.6797 | 388.27 |
| ENSMUSG00000032458 | -0.173367149 | 0.006109 | 0.02985 | 4648.977 | 4805.285 | 5072.153 | 4685.771 | 5386.515 | 5130.982 | 5315.162 | 5834.926 |
| ENSMUSG00000032459 | -0.351534182 | 0.003414 | 0.019436 | 496.9909 | 525.5218 | 534.1288 | 536.5518 | 752.7859 | 560.9218 | 615.2165 | 741.7371 |
| ENSMUSG00000032462 | -0.267134882 | 0.184362 | 0.335494 | 279.6831 | 171.5745 | 178.0429 | 171.6621 | 218.4249 | 223.4748 | 240.108 | 281.6861 |
| ENSMUSG00000032463 | -0.086389679 | 0.292009 | 0.459252 | 626.7719 | 616.7083 | 710.5155 | 685.7857 | 688.4286 | 700.5936 | 729.9669 | 688.4452 |
| ENSMUSG00000032468 | -0.037461157 | 0.742077 | 0.838997 | 935.6306 | 741.4897 | 746.9522 | 869.5245 | 773.2633 | 838.0306 | 823.5029 | 947.2918 |
| ENSMUSG00000032469 | -0.027157585 | 0.786679 | 0.868825 | 599.6084 | 587.9126 | 538.2693 | 620.2263 | 586.0419 | 605.6168 | 534.2162 | 665.6057 |
| ENSMUSG00000032470 | -0.364059635 | 0.056757 | 0.148796 | 557.3541 | 433.1356 | 539.0974 | 492.558 | 741.0846 | 429.0717 | 649.9309 | 783.0656 |
| ENSMUSG00000032475 | 0.101825547 | 0.41603 | 0.581855 | 877.2794 | 931.0615 | 729.5619 | 820.3549 | 782.0393 | 655.8986 | 789.7528 | 898.3502 |
| ENSMUSG00000032477 | 0.163654111 | 0.122269 | 0.251777 | 1157.969 | 1376.195 | 1061.633 | 1142.114 | 1019.966 | 985.524 | 1053.004 | 1166.985 |
| ENSMUSG00000032478 | 0.23528666 | 0.016953 | 0.062705 | 695.1836 | 805.0802 | 694.7815 | 669.3958 | 612.3699 | 559.8044 | 592.0735 | 665.6057 |
| ENSMUSG00000032479 | 0.309523331 | 0.001209 | 0.009011 | 4560.444 | 4958.862 | 5061.388 | 4997.178 | 3550.38 | 4625.929 | 3808.942 | 3815.27 |
| ENSMUSG00000032480 | 0.033323025 | 0.639855 | 0.766639 | 2546.324 | 2560.419 | 2382.463 | 2749.181 | 2578.194 | 2336.429 | 2614.188 | 2473.182 |
| ENSMUSG00000032481 | 0.121069178 | 0.256364 | 0.420089 | 3519.178 | 3688.251 | 3297.521 | 3214.998 | 3095.003 | 3763.316 | 2859.117 | 2897.343 |
| ENSMUSG00000032482 | -0.227462223 | 0.220056 | 0.378858 | 130.7871 | 98.38537 | 150.7154 | 104.3774 | 130.6649 | 149.7281 | 137.8934 | 151.1752 |
| ENSMUSG00000032484 | 0.747394564 | 0.214637 | 0.372761 | 125.7568 | 16.7975 | 22.35888 | 56.07053 | 38.02934 | 21.23011 | 38.57157 | 33.71532 |
| ENSMUSG00000032485 | -0.188960237 | 0.038472 | 0.11337 | 1411.494 | 1408.59 | 1405.297 | 1333.616 | 1607.959 | 1346.436 | 1677.863 | 1702.08 |
| ENSMUSG00000032487 | -1.021089518 | 0.028009 | 0.090126 | 39.23612 | 25.19625 | 37.2648 | 15.52722 | 35.10401 | 46.92971 | 109.929 | 45.67883 |
| ENSMUSG00000032489 | -0.115979186 | 0.739615 | 0.837239 | 42.25428 | 83.98751 | 45.54586 | 70.73513 | 45.83023 | 99.44629 | 52.07161 | 65.25547 |
| ENSMUSG00000032491 | 0.109224607 | 0.24676 | 0.409845 | 601.6205 | 645.504 | 622.7362 | 708.2139 | 656.2499 | 592.2083 | 570.8592 | 570.9853 |
| ENSMUSG00000032492 | -1.350933067 | 5.46E-06 | 0.000143 | 969.8364 | 901.066 | 1609.839 | 1023.071 | 2550.891 | 1620.192 | 2621.902 | 4698.393 |
| ENSMUSG00000032495 | 0.116020838 | 0.810459 | 0.88445 | 48.29061 | 20.39697 | 37.2648 | 31.05445 | 12.67645 | 27.93435 | 29.89296 | 56.55474 |
| ENSMUSG00000032496 | -0.003589263 | 0.994091 | 0.996626 | 40.24217 | 11.99822 | 22.35888 | 31.91707 | 39.97956 | 21.23011 | 14.46434 | 31.54014 |
| ENSMUSG00000032497 | -0.14819962 | 0.099712 | 0.220467 | 1336.04 | 1111.035 | 1204.895 | 1149.014 | 1254.968 | 1296.154 | 1307.576 | 1463.898 |
| ENSMUSG00000032498 | -0.117747816 | 0.233519 | 0.394086 | 734.4197 | 757.0874 | 635.9859 | 765.147 | 864.9237 | 785.514 | 734.7883 | 750.4379 |
| ENSMUSG00000032500 | -0.337979902 | 0.315881 | 0.484252 | 189.1382 | 133.1802 | 341.1799 | 177.7004 | 217.4498 | 164.254 | 303.7511 | 379.5693 |
| ENSMUSG00000032501 | 0.161109227 | 0.127971 | 0.259325 | 981.9091 | 909.4647 | 923.3389 | 915.2435 | 823.9691 | 948.6506 | 864.0031 | 699.3211 |
| ENSMUSG00000032502 | 0.313661327 | 0.216164 | 0.374544 | 82.49646 | 63.59054 | 72.04528 | 86.26235 | 60.4569 | 84.92043 | 45.32159 | 55.46715 |
| ENSMUSG00000032503 | 0.827950934 | 0.110907 | 0.236943 | 19.11503 | 28.79572 | 12.4216 | 15.52722 | 10.72622 | 15.64324 | 11.57147 | 4.350364 |
| ENSMUSG00000032504 | -0.115907954 | 0.115017 | 0.242689 | 2743.51 | 2646.806 | 2949.716 | 2735.379 | 2979.94 | 2726.393 | 3095.368 | 3202.956 |
| ENSMUSG00000032507 | -0.112419137 | 0.719443 | 0.824494 | 75.45408 | 61.1909 | 84.46688 | 79.36136 | 85.8098 | 39.10809 | 92.57176 | 107.6715 |
| ENSMUSG00000032508 | -0.083748828 | 0.539701 | 0.689371 | 557.3541 | 488.3274 | 631.0173 | 508.9479 | 587.9921 | 468.1797 | 631.6094 | 629.7152 |
| ENSMUSG00000032511 | 0.10245541 | 0.745162 | 0.840853 | 31.18769 | 41.99375 | 33.12427 | 54.34528 | 36.07912 | 45.81234 | 27.0001 | 41.32846 |
| ENSMUSG00000032512 | -0.162170966 | 0.038286 | 0.112999 | 1353.143 | 1222.618 | 1228.91 | 1211.123 | 1394.409 | 1280.511 | 1441.612 | 1494.35 |
| ENSMUSG00000032513 | -0.193225116 | 0.112016 | 0.238534 | 479.8879 | 567.5156 | 526.6758 | 496.0085 | 620.1708 | 484.9404 | 597.8593 | 661.2554 |
| ENSMUSG00000032514 | -0.55894636 | 0.225359 | 0.385123 | 10.06054 | 15.59768 | 15.73403 | 9.488858 | 13.65156 | 14.52586 | 17.3572 | 29.36496 |
| ENSMUSG00000032515 | -0.313397688 | 0.14915 | 0.289028 | 248.4954 | 154.777 | 303.087 | 231.1831 | 264.2552 | 248.057 | 351.9655 | 303.4379 |
| ENSMUSG00000032518 | -0.208304524 | 0.011528 | 0.047501 | 43471.61 | 51345.16 | 49602.76 | 47841.1 | 60980.54 | 51671.85 | 56706.95 | 52761.22 |
| ENSMUSG00000032519 | -0.282716057 | 0.027199 | 0.088176 | 532.2028 | 459.5316 | 490.2391 | 603.8364 | 666.001 | 619.0252 | 708.7525 | 544.8831 |
| ENSMUSG00000032525 | 0.192837641 | 0.036813 | 0.110225 | 2756.589 | 2324.054 | 2249.966 | 2370.489 | 2234.955 | 2262.683 | 2008.614 | 1980.503 |
| ENSMUSG00000032526 | -0.504543259 | 3.60E-07 | 1.61E-05 | 418.5186 | 416.3381 | 451.3181 | 443.3885 | 618.2206 | 545.2786 | 608.4665 | 684.0948 |
| ENSMUSG00000032527 | -0.468150237 | 0.000182 | 0.002108 | 3150.962 | 3070.343 | 3268.537 | 2886.338 | 4210.531 | 3337.596 | 4604.481 | 4967.028 |
| ENSMUSG00000032528 | -0.093076339 | 0.851118 | 0.910569 | 11.0666 | 8.398751 | 12.4216 | 7.763611 | 7.80089 | 10.05637 | 10.60718 | 14.13868 |
| ENSMUSG00000032531 | -0.124402338 | 0.361546 | 0.530099 | 2489.985 | 1745.74 | 1927.004 | 1886.558 | 2204.727 | 1840.315 | 2329.723 | 2399.226 |

| | | | | | | | | | | | |
|---|---|---|---|---|---|---|---|---|---|---|---|
| ENSMUSG00000032532 | -2.307072597 | 0.009987 | 0.042531 | 2.012109 | 4.799286 | 47.20208 | 2.58787 | 65.33246 | 24.58223 | 82.92887 | 109.8467 |
| ENSMUSG00000032534 | -0.014546036 | 0.87045 | 0.922376 | 1317.931 | 1469.781 | 1339.048 | 1434.543 | 1448.04 | 1548.68 | 1373.148 | 1246.379 |
| ENSMUSG00000032536 | -0.025152822 | 0.695315 | 0.807649 | 1468.839 | 1421.789 | 1511.295 | 1494.064 | 1496.796 | 1491.694 | 1599.756 | 1412.781 |
| ENSMUSG00000032537 | -0.398011161 | 0.005382 | 0.027036 | 196.1806 | 166.7752 | 224.4169 | 194.9529 | 274.0063 | 230.1791 | 286.3939 | 242.5328 |
| ENSMUSG00000032540 | -0.300939123 | 0.066102 | 0.165668 | 423.5489 | 382.7431 | 395.0069 | 444.2511 | 543.137 | 363.1466 | 512.0375 | 609.051 |
| ENSMUSG00000032547 | 0.171221969 | 0.099486 | 0.220192 | 3370.282 | 3930.615 | 3330.645 | 3662.699 | 3125.232 | 3714.151 | 2900.582 | 2953.897 |
| ENSMUSG00000032548 | -0.950726671 | 0.001027 | 0.007993 | 292.7618 | 203.9697 | 421.5063 | 271.7264 | 474.8792 | 345.2686 | 722.2526 | 760.2262 |
| ENSMUSG00000032549 | 0.359554805 | 0.041366 | 0.119548 | 710.2744 | 544.719 | 743.6398 | 872.1124 | 425.1485 | 600.0299 | 610.395 | 604.7006 |
| ENSMUSG00000032551 | 0.146881931 | 0.11091 | 0.236943 | 632.8082 | 595.1115 | 645.9232 | 736.6805 | 603.5939 | 591.0909 | 579.5378 | 587.2992 |
| ENSMUSG00000032553 | -0.174997244 | 0.067347 | 0.167583 | 942.6729 | 1019.848 | 1061.633 | 929.9081 | 1143.806 | 969.8807 | 1140.754 | 1208.314 |
| ENSMUSG00000032554 | 1.033201036 | 0.000786 | 0.006496 | 458.7608 | 190.7716 | 195.4332 | 395.0816 | 127.7396 | 179.8972 | 146.572 | 152.2628 |
| ENSMUSG00000032555 | -0.004170971 | 0.958686 | 0.976693 | 2733.45 | 2734.393 | 2546.428 | 2552.503 | 2718.61 | 2929.755 | 2547.652 | 2399.226 |
| ENSMUSG00000032556 | 0.606600655 | 0.325421 | 0.493262 | 16.09687 | 47.99286 | 28.15563 | 18.11509 | 15.60178 | 35.75597 | 4.821446 | 16.31387 |
| ENSMUSG00000032557 | 0.18736171 | 0.037023 | 0.110565 | 1844.098 | 1924.514 | 1724.946 | 1656.237 | 1478.269 | 1746.456 | 1452.219 | 1600.934 |
| ENSMUSG00000032558 | 0.392646603 | 0.003039 | 0.017943 | 782.7103 | 833.876 | 690.6409 | 856.5851 | 661.1255 | 690.5372 | 502.3947 | 555.759 |
| ENSMUSG00000032560 | -0.231928973 | 0.000297 | 0.00307 | 1626.79 | 1622.159 | 1593.277 | 1658.825 | 1998.003 | 1776.625 | 1860.114 | 1998.992 |
| ENSMUSG00000032561 | -0.60812413 | 0.005692 | 0.028318 | 53.32088 | 68.38983 | 57.96746 | 69.8725 | 76.05868 | 89.38993 | 96.42892 | 118.5474 |
| ENSMUSG00000032562 | -0.079296766 | 0.376939 | 0.545034 | 6461.887 | 6964.964 | 7660.814 | 7041.596 | 7614.644 | 6478.535 | 7611.134 | 8015.546 |
| ENSMUSG00000032563 | -0.165183687 | 0.005093 | 0.02606 | 1486.948 | 1647.355 | 1531.997 | 1569.112 | 1794.205 | 1764.334 | 1721.256 | 1707.518 |
| ENSMUSG00000032565 | 0.214289116 | 0.085692 | 0.198113 | 368.2159 | 353.9474 | 453.8024 | 381.2796 | 334.4632 | 370.9682 | 339.4298 | 300.1751 |
| ENSMUSG00000032567 | -0.27491336 | 0.040023 | 0.116765 | 250.5075 | 226.7663 | 204.5423 | 195.8155 | 250.6036 | 246.9397 | 264.2152 | 298 |
| ENSMUSG00000032570 | -0.217228721 | 0.02049 | 0.07174 | 1931.624 | 1869.322 | 2185.373 | 1860.679 | 2157.921 | 2072.729 | 2380.83 | 2513.423 |
| ENSMUSG00000032571 | 0.054042904 | 0.422101 | 0.587229 | 1071.448 | 1143.43 | 1125.397 | 1037.736 | 1038.494 | 1024.632 | 1104.111 | 1046.263 |
| ENSMUSG00000032572 | 0.129568618 | 0.724091 | 0.827323 | 52.31483 | 103.1847 | 125.8722 | 56.93315 | 62.40712 | 52.51658 | 107.0361 | 87.00729 |
| ENSMUSG00000032575 | -0.144638516 | 0.004607 | 0.024237 | 2448.736 | 2416.441 | 2506.679 | 2419.659 | 2717.635 | 2573.313 | 2795.474 | 2737.467 |
| ENSMUSG00000032577 | -0.588338095 | 0.016189 | 0.060702 | 289.7437 | 214.7681 | 376.7885 | 231.1831 | 387.1192 | 281.5783 | 469.6088 | 536.1824 |
| ENSMUSG00000032578 | -0.092030016 | 0.396412 | 0.563679 | 592.566 | 513.5236 | 609.4865 | 570.1941 | 550.9379 | 574.3303 | 693.3239 | 619.9269 |
| ENSMUSG00000032579 | 0.197766836 | 0.106055 | 0.23006 | 337.0282 | 352.7475 | 331.2427 | 322.6212 | 329.5876 | 313.9821 | 247.8223 | 279.5109 |
| ENSMUSG00000032580 | 0.342563495 | 0.000698 | 0.005931 | 7161.095 | 6359.054 | 5311.476 | 6257.471 | 4692.236 | 5337.696 | 4828.196 | 4926.788 |
| ENSMUSG00000032582 | 0.063480165 | 0.474513 | 0.634013 | 2472.882 | 2535.223 | 2095.938 | 2279.051 | 2236.905 | 2453.754 | 2101.186 | 2183.883 |
| ENSMUSG00000032583 | -0.444580984 | 0.00629 | 0.030455 | 228.3743 | 209.9688 | 221.9326 | 258.787 | 318.8614 | 236.8833 | 324.0012 | 371.9562 |
| ENSMUSG00000032584 | 0.856630515 | 0.00656 | 0.031503 | 178.0716 | 263.9607 | 216.1358 | 241.5346 | 113.1129 | 214.5358 | 93.53605 | 76.13138 |
| ENSMUSG00000032586 | -0.060683328 | 0.560527 | 0.706343 | 436.6276 | 429.5361 | 441.3808 | 433.037 | 527.5352 | 429.0717 | 426.2158 | 431.7737 |
| ENSMUSG00000032589 | 0.231161759 | 0.133673 | 0.267172 | 119.7205 | 109.1838 | 110.9663 | 118.1794 | 93.61069 | 106.1505 | 95.46463 | 95.70802 |
| ENSMUSG00000032590 | -0.292926589 | 0.013887 | 0.054426 | 1339.058 | 1383.394 | 1320.002 | 1342.242 | 1720.096 | 1317.384 | 1620.97 | 1937 |
| ENSMUSG00000032593 | 0.073987948 | 0.666073 | 0.785506 | 107.6478 | 136.7797 | 125.8722 | 147.5086 | 138.4658 | 129.6154 | 110.8933 | 113.1095 |
| ENSMUSG00000032594 | 0.219522106 | 0.016992 | 0.062834 | 4666.08 | 4817.283 | 4311.951 | 5005.804 | 4157.875 | 4385.693 | 4125.229 | 3477.029 |
| ENSMUSG00000032595 | 0.831087357 | 0.00198 | 0.013044 | 124.7507 | 187.1722 | 122.5598 | 126.8057 | 71.18313 | 117.3243 | 62.6788 | 64.16787 |
| ENSMUSG00000032596 | 0.511861319 | 0.014207 | 0.055412 | 419.5247 | 597.5111 | 727.0776 | 406.2957 | 419.2979 | 312.8647 | 390.5371 | 385.0072 |
| ENSMUSG00000032598 | -0.482866814 | 0.000189 | 0.002171 | 221.332 | 196.7707 | 260.8536 | 238.9467 | 316.9112 | 289.3999 | 341.3584 | 339.3284 |
| ENSMUSG00000032599 | 0.110107581 | 0.256927 | 0.42059 | 3823.007 | 3904.219 | 3367.91 | 3738.61 | 3263.698 | 3954.387 | 3447.334 | 3076.795 |
| ENSMUSG00000032601 | -0.265345831 | 0.000778 | 0.006448 | 1315.919 | 1337.801 | 1339.877 | 1352.594 | 1521.174 | 1470.464 | 1660.506 | 1773.861 |
| ENSMUSG00000032602 | -0.219044356 | 0.206881 | 0.363479 | 1196.199 | 1264.612 | 1273.628 | 1218.887 | 1394.409 | 1082.735 | 1295.04 | 1993.554 |
| ENSMUSG00000032604 | -0.114441175 | 0.140618 | 0.277122 | 4122.811 | 4098.59 | 4178.626 | 4377.814 | 4911.636 | 4027.016 | 4761.66 | 4461.299 |
| ENSMUSG00000032606 | 0.008243266 | 0.920273 | 0.95244 | 718.3228 | 694.6967 | 721.2809 | 732.3673 | 755.7113 | 650.3117 | 756.967 | 686.27 |
| ENSMUSG00000032607 | -0.390330218 | 0.013798 | 0.054192 | 629.79 | 646.7038 | 706.375 | 669.3958 | 898.0775 | 641.3727 | 837.9673 | 1099.555 |
| ENSMUSG00000032609 | 0.28200103 | 0.001199 | 0.008972 | 936.6366 | 937.0606 | 1099.726 | 1032.56 | 828.8446 | 819.0352 | 777.2171 | 874.4232 |
| ENSMUSG00000032611 | 0.501467568 | 0.30998 | 0.478464 | 11.0666 | 21.59679 | 28.98373 | 26.74133 | 15.60178 | 25.6996 | 8.678602 | 13.05109 |
| ENSMUSG00000032612 | -0.21044653 | 0.00516 | 0.026322 | 2795.825 | 2847.177 | 2886.78 | 2691.385 | 3386.562 | 2925.285 | 3157.083 | 3512.919 |
| ENSMUSG00000032615 | -0.388793024 | 0.000655 | 0.005649 | 441.6579 | 442.7341 | 528.332 | 491.6954 | 612.3699 | 556.4523 | 616.1808 | 712.3722 |
| ENSMUSG00000032621 | 0.405794113 | 0.016009 | 0.060323 | 5527.263 | 4974.46 | 4090.019 | 4880.724 | 3411.914 | 5005.836 | 3146.476 | 3134.438 |
| ENSMUSG00000032624 | 0.292985993 | 0.002639 | 0.016038 | 3604.693 | 3624.661 | 3118.65 | 3354.743 | 2785.893 | 3214.685 | 2654.688 | 2527.562 |
| ENSMUSG00000032625 | -0.256887721 | 0.167469 | 0.31347 | 265.5983 | 231.5656 | 342.008 | 283.8031 | 278.8818 | 280.4609 | 375.1085 | 410.0218 |
| ENSMUSG00000032633 | 0.070005452 | 0.631455 | 0.760118 | 1371.252 | 980.2542 | 1135.334 | 1289.622 | 1195.486 | 898.3688 | 1241.04 | 1215.927 |
| ENSMUSG00000032637 | 0.023678631 | 0.757316 | 0.848813 | 2591.596 | 2218.47 | 2560.506 | 2438.637 | 2542.115 | 2248.157 | 2391.437 | 2471.007 |
| ENSMUSG00000032640 | 0.119455674 | 0.302619 | 0.470882 | 1546.306 | 1635.357 | 1361.407 | 1578.601 | 1358.33 | 1696.174 | 1297.933 | 1282.27 |
| ENSMUSG00000032641 | -0.089933869 | 0.539132 | 0.688978 | 212.2775 | 194.3711 | 227.7293 | 192.365 | 261.3298 | 201.1273 | 224.6794 | 192.5036 |
| ENSMUSG00000032643 | -0.300861793 | 0.081881 | 0.192069 | 220.3259 | 266.3604 | 334.5551 | 290.7041 | 408.5716 | 280.4609 | 339.4298 | 342.5912 |
| ENSMUSG00000032648 | 0.003917179 | 0.976857 | 0.987311 | 848.1038 | 605.9099 | 721.2809 | 643.5171 | 693.3041 | 651.4291 | 657.6452 | 810.2554 |
| ENSMUSG00000032649 | -0.889032743 | 0.016038 | 0.060347 | 71.42986 | 37.19447 | 91.91984 | 72.46037 | 84.83468 | 84.92043 | 136.9291 | 201.2044 |
| ENSMUSG00000032652 | 0.063009228 | 0.625023 | 0.755628 | 460.7729 | 424.7368 | 366.8512 | 453.74 | 385.169 | 357.5597 | 467.6802 | 420.8978 |
| ENSMUSG00000032656 | 0.054133955 | 0.686527 | 0.800666 | 301.8163 | 340.7493 | 276.5876 | 256.1992 | 274.9814 | 320.6864 | 264.2152 | 269.7226 |
| ENSMUSG00000032657 | 0.262145071 | 0.026394 | 0.086387 | 920.5397 | 1135.031 | 966.4004 | 985.116 | 897.1024 | 940.829 | 786.86 | 714.5473 |
| ENSMUSG00000032661 | 0.938166819 | 0.277608 | 0.444018 | 3.018163 | 13.19804 | 36.43669 | 5.175741 | 5.850668 | 3.352122 | 14.46434 | 6.525547 |

| | | | | | | | | | | | |
|---|---|---|---|---|---|---|---|---|---|---|---|
| ENSMUSG00000032666 | 0.300710087 | 0.007049 | 0.033083 | 1493.991 | 1245.415 | 1217.317 | 1481.987 | 1041.419 | 1244.755 | 1122.433 | 1008.197 |
| ENSMUSG00000032667 | 0.193395563 | 0.020322 | 0.071278 | 1253.544 | 1100.236 | 1124.569 | 1225.788 | 1071.647 | 1076.031 | 1031.789 | 935.3283 |
| ENSMUSG00000032673 | -0.057201162 | 0.562993 | 0.708434 | 408.4581 | 411.5388 | 388.382 | 462.3662 | 464.153 | 413.4284 | 421.3944 | 439.3868 |
| ENSMUSG00000032679 | 0.100168582 | 0.488553 | 0.646314 | 1010.079 | 687.4977 | 828.9347 | 913.5183 | 666.001 | 843.6174 | 831.2173 | 871.1605 |
| ENSMUSG00000032680 | -0.572356916 | 0.029322 | 0.092982 | 42.25428 | 38.39429 | 57.13936 | 50.03216 | 67.28268 | 54.75133 | 65.57166 | 93.53283 |
| ENSMUSG00000032688 | 0.074099434 | 0.60959 | 0.744777 | 527.1725 | 531.5209 | 486.0986 | 495.1459 | 443.6756 | 623.4947 | 461.8945 | 408.9342 |
| ENSMUSG00000032690 | 1.608992814 | 0.006226 | 0.030285 | 66.39959 | 291.5566 | 231.0418 | 36.23019 | 39.00445 | 36.87335 | 81.96458 | 46.76642 |
| ENSMUSG00000032698 | -0.116612972 | 0.522927 | 0.676229 | 665.0019 | 851.8733 | 1167.63 | 833.2943 | 1010.215 | 788.8661 | 908.3604 | 1108.255 |
| ENSMUSG00000032702 | -0.134474287 | 0.09614 | 0.214762 | 1547.312 | 1472.181 | 1569.262 | 1589.815 | 1521.174 | 1642.54 | 1793.578 | 1828.241 |
| ENSMUSG00000032705 | 0.234442755 | 0.003736 | 0.020813 | 847.0978 | 821.8778 | 770.1392 | 847.0963 | 712.8064 | 740.819 | 699.1096 | 639.5036 |
| ENSMUSG00000032712 | 0.380840304 | 0.137118 | 0.272112 | 2555.378 | 2573.617 | 1973.378 | 2394.643 | 1411.961 | 3016.91 | 1655.684 | 1209.401 |
| ENSMUSG00000032714 | 0.170370692 | 0.009772 | 0.041855 | 1201.229 | 1215.419 | 1210.692 | 1173.168 | 1142.83 | 1024.632 | 1024.075 | 1072.365 |
| ENSMUSG00000032715 | 0.330955891 | 0.29031 | 0.457397 | 31.18769 | 27.5959 | 39.74912 | 33.64232 | 29.25334 | 29.05173 | 18.32149 | 29.36496 |
| ENSMUSG00000032717 | 0.426523441 | 0.00018 | 0.002091 | 713.2925 | 839.8751 | 856.2623 | 850.5468 | 568.4899 | 673.7766 | 532.2876 | 653.6422 |
| ENSMUSG00000032718 | -0.027739628 | 0.90384 | 0.942474 | 215.2956 | 219.5673 | 308.8838 | 163.0358 | 211.5992 | 182.132 | 243.0009 | 288.2116 |
| ENSMUSG00000032719 | 0.212197742 | 0.326225 | 0.493995 | 122.7386 | 142.7788 | 211.1672 | 160.448 | 150.1671 | 160.9019 | 115.7147 | 125.073 |
| ENSMUSG00000032724 | -0.484142244 | 0.024358 | 0.081456 | 189.1382 | 185.9723 | 141.6062 | 156.9975 | 229.1512 | 187.7188 | 196.715 | 327.3649 |
| ENSMUSG00000032725 | 0.984787343 | 0.000781 | 0.00647 | 57.3451 | 65.99018 | 44.71776 | 50.03216 | 36.07912 | 21.23011 | 28.92867 | 22.83941 |
| ENSMUSG00000032727 | 0.281719162 | 0.106602 | 0.230828 | 1204.247 | 1157.828 | 950.6664 | 1079.142 | 856.1477 | 1230.229 | 821.5744 | 704.759 |
| ENSMUSG00000032733 | 0.152368426 | 0.090186 | 0.20549 | 608.6629 | 514.7234 | 529.1601 | 583.1335 | 500.2321 | 491.6446 | 514.9304 | 505.7299 |
| ENSMUSG00000032735 | 0.741834148 | 0.00031 | 0.003189 | 251.5136 | 320.3523 | 240.979 | 267.4133 | 158.9431 | 222.3574 | 123.429 | 141.3868 |
| ENSMUSG00000032737 | 0.01173334 | 0.878713 | 0.927246 | 2445.718 | 2445.236 | 2459.477 | 2651.705 | 2341.242 | 2323.021 | 2512.938 | 2746.167 |
| ENSMUSG00000032740 | 0.584482498 | 0.016656 | 0.061928 | 2300.846 | 2775.187 | 1976.691 | 2058.22 | 1166.233 | 2462.693 | 1240.076 | 1207.226 |
| ENSMUSG00000032741 | 0.052840543 | 0.61292 | 0.746981 | 2216.338 | 2178.876 | 2023.065 | 2080.648 | 2394.873 | 1850.372 | 1820.578 | 2125.153 |
| ENSMUSG00000032743 | 0.242810909 | 0.003321 | 0.019101 | 1130.805 | 1065.442 | 1001.181 | 1084.318 | 963.41 | 922.951 | 915.1104 | 814.6057 |
| ENSMUSG00000032744 | 0.015448299 | 0.872863 | 0.923925 | 1353.143 | 1295.807 | 1458.296 | 1230.964 | 1333.952 | 1181.064 | 1288.29 | 1479.124 |
| ENSMUSG00000032745 | 0.156869885 | 0.110897 | 0.236943 | 3394.427 | 3825.031 | 3150.946 | 3598.003 | 2862.927 | 3547.663 | 3187.94 | 2928.883 |
| ENSMUSG00000032750 | 0.541669558 | 0.262237 | 0.426418 | 73.44197 | 43.19358 | 48.85829 | 39.68068 | 22.42756 | 74.86406 | 20.25007 | 23.927 |
| ENSMUSG00000032754 | 0.412909493 | 5.44E-05 | 0.000845 | 702.2259 | 747.4888 | 713.8279 | 787.5752 | 596.7681 | 573.2129 | 575.6806 | 469.8393 |
| ENSMUSG00000032757 | 0.205500851 | 0.018654 | 0.067217 | 1518.136 | 1426.588 | 1273.628 | 1495.789 | 1195.486 | 1335.262 | 1262.255 | 1161.547 |
| ENSMUSG00000032758 | -7.642415729 | 1.46E-07 | 7.71E-06 | 1.006054 | 0 | 50.5145 | 6.900988 | 3269.548 | 612.321 | 1371.219 | 6473.342 |
| ENSMUSG00000032763 | -0.172333489 | 0.086928 | 0.200218 | 879.2915 | 919.0633 | 1011.118 | 919.5566 | 1106.751 | 889.4298 | 1092.54 | 1113.693 |
| ENSMUSG00000032766 | -0.015065647 | 0.894702 | 0.936266 | 786.7345 | 806.2801 | 1035.133 | 894.5406 | 811.2926 | 889.4298 | 905.4675 | 958.1677 |
| ENSMUSG00000032773 | 0.828816021 | 0.038297 | 0.112999 | 45.27245 | 27.5959 | 39.74912 | 43.9938 | 34.1289 | 17.87799 | 25.07152 | 10.87591 |
| ENSMUSG00000032776 | -0.602376948 | 0.080301 | 0.189421 | 165.999 | 125.9813 | 318.8211 | 194.0903 | 210.624 | 197.7752 | 381.8585 | 433.9488 |
| ENSMUSG00000032777 | -0.095890607 | 0.270826 | 0.436005 | 1985.951 | 2170.477 | 2086.829 | 2067.709 | 2164.747 | 2023.564 | 2150.365 | 2542.788 |
| ENSMUSG00000032782 | 0.16534128 | 0.134322 | 0.268006 | 633.8142 | 681.4986 | 539.9255 | 684.0604 | 528.5103 | 611.2036 | 583.3949 | 539.4452 |
| ENSMUSG00000032783 | 0.043297299 | 0.736399 | 0.834878 | 444.676 | 447.5334 | 371.8199 | 461.5036 | 463.1779 | 376.5551 | 372.2156 | 461.1386 |
| ENSMUSG00000032786 | -0.004989498 | 0.986741 | 0.992387 | 2029.212 | 734.2908 | 936.5886 | 1010.132 | 1115.527 | 817.9178 | 1403.041 | 1389.941 |
| ENSMUSG00000032788 | -0.714683085 | 0.000196 | 0.002231 | 595.5842 | 647.9036 | 797.4667 | 553.8043 | 1063.846 | 806.7441 | 957.5391 | 1431.27 |
| ENSMUSG00000032796 | -0.606193848 | 0.000898 | 0.007179 | 2051.345 | 1346.2 | 2153.077 | 1868.442 | 2283.711 | 2424.702 | 3142.618 | 3446.576 |
| ENSMUSG00000032802 | -0.149908491 | 0.637525 | 0.764831 | 993.9817 | 311.9536 | 573.0498 | 584.8587 | 627.9717 | 452.5365 | 794.5743 | 859.197 |
| ENSMUSG00000032803 | -0.13683603 | 0.007866 | 0.035802 | 3650.971 | 3912.618 | 3570.796 | 3737.748 | 3960.902 | 4086.237 | 4123.3 | 4175.262 |
| ENSMUSG00000032806 | -0.266196332 | 0.047322 | 0.130902 | 299.8042 | 293.9563 | 285.6968 | 299.3304 | 353.9654 | 298.3389 | 334.6083 | 430.6861 |
| ENSMUSG00000032812 | 0.603385011 | 1.10E-08 | 9.47E-07 | 4632.88 | 5368.002 | 4986.858 | 5224.048 | 3226.643 | 3940.978 | 2923.725 | 3213.832 |
| ENSMUSG00000032815 | 0.092603221 | 0.419511 | 0.585046 | 961.788 | 889.0678 | 1078.195 | 832.4317 | 886.3762 | 774.3402 | 972.0035 | 895.0875 |
| ENSMUSG00000032816 | 0.423115987 | 0.00997 | 0.042487 | 605.6447 | 752.2881 | 774.2797 | 778.949 | 525.585 | 709.5326 | 448.3945 | 489.416 |
| ENSMUSG00000032826 | 0.563032443 | 0.000603 | 0.005288 | 812.8919 | 749.8885 | 641.7826 | 874.7002 | 488.5308 | 671.5418 | 498.5375 | 426.3357 |
| ENSMUSG00000032827 | 0.586179333 | 3.58E-05 | 0.000614 | 2808.904 | 2703.198 | 2259.903 | 2686.21 | 1693.768 | 2205.696 | 1634.47 | 1432.357 |
| ENSMUSG00000032834 | 0.026946781 | 0.70806 | 0.816595 | 877.2794 | 959.8572 | 934.9324 | 921.2819 | 949.7584 | 881.6082 | 856.2888 | 936.4159 |
| ENSMUSG00000032839 | -0.500033378 | 0.000864 | 0.006969 | 281.6952 | 193.1713 | 221.9326 | 231.1831 | 317.8863 | 287.1651 | 345.2155 | 363.2554 |
| ENSMUSG00000032840 | -0.222723662 | 0.105939 | 0.229961 | 531.1967 | 581.9134 | 637.6421 | 583.9961 | 686.4784 | 525.1658 | 748.2884 | 764.5765 |
| ENSMUSG00000032841 | 0.108329112 | 0.460231 | 0.621639 | 986.9393 | 1162.627 | 1030.165 | 1063.615 | 1097.975 | 714.002 | 1036.611 | 1085.416 |
| ENSMUSG00000032842 | 0.069650773 | 0.556063 | 0.702924 | 356.1432 | 349.1481 | 342.8361 | 321.7586 | 276.9316 | 365.3813 | 320.144 | 343.6788 |
| ENSMUSG00000032846 | 0.126969824 | 0.461194 | 0.622349 | 950.7214 | 785.8831 | 759.3738 | 887.6396 | 628.9468 | 1035.806 | 654.7523 | 780.8904 |
| ENSMUSG00000032849 | 0.04042558 | 0.64752 | 0.77184 | 940.6608 | 896.2667 | 927.4794 | 827.2559 | 908.8037 | 778.8097 | 927.6462 | 874.4232 |
| ENSMUSG00000032850 | -0.028925924 | 0.836544 | 0.901206 | 229.3804 | 313.1534 | 264.166 | 253.6113 | 251.5787 | 308.3953 | 269.0367 | 251.2335 |
| ENSMUSG00000032854 | -1.023684675 | 0.009654 | 0.041504 | 117.7084 | 125.9813 | 333.727 | 135.4319 | 330.5627 | 185.4841 | 415.6086 | 519.8685 |
| ENSMUSG00000032855 | 0.222956666 | 0.000686 | 0.005847 | 1709.286 | 1721.744 | 1659.526 | 1897.772 | 1493.871 | 1507.338 | 1514.898 | 1472.598 |
| ENSMUSG00000032860 | -1.822529546 | 1.43E-05 | 0.000299 | 39.23612 | 20.39697 | 54.65504 | 31.05445 | 168.6943 | 51.39921 | 147.5362 | 149 |
| ENSMUSG00000032867 | -0.283242687 | 0.00248 | 0.015333 | 2228.41 | 2320.455 | 2521.585 | 2080.648 | 2806.37 | 2495.096 | 2756.903 | 3077.883 |
| ENSMUSG00000032869 | -0.044120942 | 0.668486 | 0.787337 | 794.7829 | 901.066 | 999.5247 | 924.7324 | 969.2606 | 823.5047 | 983.5749 | 957.0802 |
| ENSMUSG00000032870 | -0.102012898 | 0.302459 | 0.470753 | 948.7093 | 889.0678 | 1036.79 | 975.6272 | 1039.469 | 955.3549 | 960.432 | 1181.124 |
| ENSMUSG00000032872 | -0.459718025 | 3.70E-06 | 0.000107 | 1551.336 | 1466.182 | 1315.033 | 1496.652 | 2199.851 | 1721.873 | 2072.257 | 2018.569 |

| | | | | | | | | | | | |
|---|---|---|---|---|---|---|---|---|---|---|---|
| ENSMUSG00000032875 | -0.090793158 | 0.351815 | 0.520605 | 2201.247 | 1926.913 | 2291.371 | 2168.635 | 2210.577 | 2019.095 | 2359.616 | 2560.189 |
| ENSMUSG00000032878 | -0.21844965 | 0.397735 | 0.564974 | 150.9082 | 103.1847 | 110.9663 | 155.2722 | 114.088 | 148.6108 | 217.9294 | 125.073 |
| ENSMUSG00000032883 | -0.053680331 | 0.589949 | 0.730011 | 2483.948 | 2534.023 | 1927.832 | 2384.291 | 2319.79 | 2521.913 | 2430.973 | 2406.839 |
| ENSMUSG00000032890 | 0.502245745 | 0.065495 | 0.164684 | 101.6115 | 56.39161 | 62.9361 | 89.71284 | 62.40712 | 62.57295 | 50.14304 | 44.59123 |
| ENSMUSG00000032897 | -0.301047948 | 0.000524 | 0.004747 | 1641.881 | 1557.368 | 1779.601 | 1641.573 | 2036.032 | 1795.62 | 2142.651 | 2184.97 |
| ENSMUSG00000032898 | -0.099608237 | 0.078152 | 0.186006 | 2350.143 | 2240.067 | 2322.011 | 2238.508 | 2517.737 | 2300.673 | 2522.58 | 2462.306 |
| ENSMUSG00000032899 | 0.573638233 | 0.026914 | 0.087576 | 188.1322 | 268.76 | 284.0406 | 242.3972 | 195.0223 | 225.7096 | 136.9291 | 103.3212 |
| ENSMUSG00000032902 | 0.287684442 | 0.034526 | 0.104995 | 2896.43 | 4165.78 | 3042.464 | 3346.979 | 3163.261 | 2790.083 | 2618.045 | 2444.905 |
| ENSMUSG00000032905 | 0.012513158 | 0.865553 | 0.91951 | 1292.78 | 1181.824 | 1197.442 | 1329.303 | 1310.55 | 1258.163 | 1212.111 | 1178.949 |
| ENSMUSG00000032908 | 0.044999041 | 0.699432 | 0.810259 | 1303.846 | 1028.247 | 1413.578 | 1295.66 | 1118.453 | 1190.003 | 1241.04 | 1342.087 |
| ENSMUSG00000032911 | -0.390625072 | 0.014937 | 0.057527 | 494.9787 | 404.3399 | 597.893 | 550.3538 | 576.2908 | 595.5604 | 726.1097 | 790.6787 |
| ENSMUSG00000032913 | 0.261453081 | 0.011792 | 0.048308 | 1106.66 | 1143.43 | 1041.758 | 1124.861 | 939.0322 | 1058.153 | 902.5747 | 784.1532 |
| ENSMUSG00000032915 | 0.011998819 | 0.984024 | 0.991004 | 12.07265 | 20.39697 | 7.45296 | 6.900988 | 10.72622 | 18.99536 | 10.60718 | 5.437955 |
| ENSMUSG00000032925 | 1.566782291 | 0.006024 | 0.02956 | 46.2785 | 32.39518 | 13.24971 | 41.40593 | 4.875557 | 20.11273 | 12.53576 | 7.613138 |
| ENSMUSG00000032932 | 0.192147139 | 0.003629 | 0.020376 | 1564.415 | 1512.975 | 1471.545 | 1590.678 | 1296.898 | 1439.178 | 1345.183 | 1294.233 |
| ENSMUSG00000032936 | 0.600132406 | 0.018803 | 0.067514 | 158.9566 | 244.7636 | 153.1997 | 240.672 | 156.9929 | 163.1366 | 110.8933 | 94.62042 |
| ENSMUSG00000032939 | -0.17613323 | 0.012013 | 0.049006 | 1376.282 | 1370.196 | 1437.593 | 1293.073 | 1509.472 | 1471.582 | 1543.827 | 1664.014 |
| ENSMUSG00000032940 | 0.815569394 | 0.002152 | 0.013845 | 89.53884 | 82.78769 | 68.73285 | 103.5148 | 59.48179 | 59.22083 | 44.3573 | 32.62773 |
| ENSMUSG00000032946 | 0.087898856 | 0.497716 | 0.654123 | 1040.26 | 785.8831 | 887.7303 | 942.8475 | 727.433 | 813.4483 | 959.4677 | 941.8539 |
| ENSMUSG00000032952 | 0.309654926 | 0.041072 | 0.118926 | 203.223 | 220.7672 | 183.8397 | 214.7932 | 140.416 | 196.6578 | 156.2148 | 170.7518 |
| ENSMUSG00000032959 | -0.222680461 | 0.007935 | 0.036028 | 7303.955 | 7384.902 | 7996.198 | 7156.325 | 9311.338 | 7617.139 | 8630.388 | 9263.013 |
| ENSMUSG00000032965 | 0.074020265 | 0.342441 | 0.511165 | 1047.303 | 1081.039 | 985.4469 | 1113.647 | 1088.224 | 984.4066 | 983.5749 | 957.0802 |
| ENSMUSG00000032966 | -0.084066312 | 0.175754 | 0.324274 | 6177.174 | 7075.348 | 7069.546 | 6873.384 | 7286.032 | 7303.157 | 6997.846 | 7241.181 |
| ENSMUSG00000032968 | 0.819958712 | 0.029224 | 0.092817 | 1187.144 | 343.149 | 322.9616 | 538.2771 | 352.9903 | 417.8979 | 298.9296 | 284.9489 |
| ENSMUSG00000032977 | -0.215223859 | 0.02571 | 0.084658 | 756.5529 | 760.6869 | 787.5294 | 784.9874 | 939.0322 | 759.8144 | 912.2175 | 975.5692 |
| ENSMUSG00000032978 | -3.469699302 | 2.58E-11 | 5.41E-09 | 11.0666 | 14.39786 | 14.90592 | 5.175741 | 154.0676 | 41.34284 | 104.1432 | 202.2919 |
| ENSMUSG00000032985 | 0.289679052 | 0.467745 | 0.628016 | 27.16347 | 53.99197 | 18.21835 | 37.95543 | 33.15378 | 30.1691 | 24.10723 | 23.927 |
| ENSMUSG00000032988 | 0.438991186 | 0.323971 | 0.491966 | 23.13925 | 47.99286 | 35.60859 | 26.74133 | 16.57689 | 13.40849 | 39.53586 | 28.27737 |
| ENSMUSG00000032997 | -0.085472522 | 0.375956 | 0.544226 | 1210.283 | 1343.8 | 1470.717 | 1195.596 | 1354.43 | 1390.013 | 1291.183 | 1505.226 |
| ENSMUSG00000032998 | 0.201209885 | 0.067608 | 0.167869 | 1753.553 | 1776.936 | 1719.149 | 1896.046 | 1524.099 | 1876.071 | 1373.148 | 1444.321 |
| ENSMUSG00000033004 | 0.192544206 | 0.113011 | 0.239961 | 3201.265 | 2833.978 | 2815.563 | 2907.904 | 2340.267 | 3169.99 | 2556.331 | 2224.124 |
| ENSMUSG00000033006 | 0.934330751 | 0.135721 | 0.270154 | 59.35721 | 13.19804 | 10.76539 | 45.71905 | 20.47734 | 17.87799 | 8.678602 | 20.66423 |
| ENSMUSG00000033007 | -0.009492182 | 0.977427 | 0.987546 | 21.12714 | 38.39429 | 30.63995 | 50.03216 | 39.00445 | 37.99072 | 32.78583 | 31.54014 |
| ENSMUSG00000033009 | 0.115140712 | 0.379606 | 0.547773 | 340.0464 | 326.3515 | 315.5086 | 297.6051 | 276.9316 | 263.7003 | 282.5367 | 357.8175 |
| ENSMUSG00000033014 | 0.142315711 | 0.043029 | 0.122781 | 2164.023 | 2104.487 | 1970.894 | 1970.232 | 1852.711 | 1960.992 | 1719.328 | 1904.372 |
| ENSMUSG00000033016 | 0.341759181 | 0.007639 | 0.035009 | 361.1735 | 387.5424 | 366.8512 | 337.2858 | 281.8072 | 340.7991 | 278.6796 | 244.708 |
| ENSMUSG00000033020 | -0.112048907 | 0.215416 | 0.373588 | 1631.82 | 1715.745 | 1590.793 | 1616.556 | 1880.015 | 1520.746 | 1755.006 | 1925.036 |
| ENSMUSG00000033021 | 0.042594286 | 0.57199 | 0.714829 | 995.9938 | 1021.048 | 982.1345 | 1069.653 | 937.082 | 943.0637 | 1002.861 | 1068.014 |
| ENSMUSG00000033022 | -0.476930622 | 0.000168 | 0.001983 | 732.4076 | 789.4826 | 839.7001 | 786.7126 | 1018.016 | 890.5472 | 1241.04 | 1232.241 |
| ENSMUSG00000033029 | -0.493113169 | 0.023835 | 0.080243 | 51.30877 | 61.1909 | 70.38906 | 52.62003 | 79.95913 | 78.21619 | 100.2861 | 72.8686 |
| ENSMUSG00000033031 | 0.181364223 | 0.219969 | 0.378778 | 1044.284 | 1048.644 | 781.7327 | 1101.57 | 863.9486 | 1062.623 | 797.4671 | 781.978 |
| ENSMUSG00000033032 | -0.471373758 | 0.000438 | 0.004131 | 1024.163 | 915.4638 | 1351.47 | 1065.34 | 1505.572 | 1336.379 | 1643.149 | 1559.606 |
| ENSMUSG00000033033 | 0.232609126 | 0.225777 | 0.385543 | 146.8839 | 189.5718 | 238.4947 | 163.0358 | 174.5449 | 166.4887 | 148.5005 | 139.2117 |
| ENSMUSG00000033036 | 0.077340378 | 0.807398 | 0.882415 | 26.15741 | 28.79572 | 36.43669 | 21.56559 | 24.37778 | 25.6996 | 29.89296 | 27.18978 |
| ENSMUSG00000033039 | -0.034983463 | 0.755936 | 0.848334 | 1028.188 | 807.4799 | 1087.304 | 888.5022 | 993.6384 | 908.4251 | 989.3607 | 1016.898 |
| ENSMUSG00000033047 | -0.280548523 | 0.003361 | 0.019239 | 5052.405 | 5468.787 | 5563.22 | 5482.835 | 7099.785 | 5462.842 | 6910.096 | 6722.401 |
| ENSMUSG00000033053 | 0.398166186 | 0.350037 | 0.518773 | 33.19979 | 46.79304 | 29.81184 | 43.9938 | 33.15378 | 34.6386 | 38.57157 | 9.78832 |
| ENSMUSG00000033054 | 0.053702168 | 0.570848 | 0.714034 | 981.9091 | 1005.45 | 889.3865 | 849.6841 | 921.4802 | 977.7023 | 862.0745 | 825.4816 |
| ENSMUSG00000033055 | -0.280305114 | 0.007255 | 0.03375 | 566.4086 | 634.7056 | 651.7199 | 594.3476 | 784.9646 | 631.3164 | 773.3599 | 780.8904 |
| ENSMUSG00000033059 | 0.015460914 | 0.877299 | 0.92652 | 3037.278 | 3607.863 | 3710.746 | 3188.256 | 3608.887 | 2943.163 | 3336.441 | 3509.656 |
| ENSMUSG00000033060 | -0.443064526 | 0.003138 | 0.01837 | 792.7708 | 743.8894 | 980.4783 | 793.6136 | 964.3851 | 950.8854 | 1268.04 | 1320.336 |
| ENSMUSG00000033061 | 3.514084796 | 0.000644 | 0.005577 | 62.37537 | 4.799286 | 2.48432 | 104.3774 | 3.900445 | 2.234748 | 5.785735 | 3.262773 |
| ENSMUSG00000033063 | -0.843168119 | 0.147499 | 0.286664 | 3.018163 | 9.598572 | 14.07781 | 13.80198 | 16.57689 | 10.05637 | 17.3572 | 29.36496 |
| ENSMUSG00000033065 | -0.131523777 | 0.113165 | 0.240093 | 1837.055 | 1846.525 | 1862.412 | 1877.069 | 2118.917 | 1747.573 | 2116.615 | 2146.905 |
| ENSMUSG00000033066 | 0.467700281 | 0.000934 | 0.00743 | 811.8859 | 911.8644 | 929.1356 | 966.1383 | 690.3788 | 792.2182 | 622.9308 | 512.2554 |
| ENSMUSG00000033068 | -0.276200901 | 0.002952 | 0.017557 | 383.3067 | 430.7359 | 422.3344 | 401.1199 | 539.2366 | 468.1797 | 483.1089 | 490.5036 |
| ENSMUSG00000033075 | 0.043713003 | 0.588597 | 0.728826 | 1308.877 | 1409.79 | 1226.426 | 1262.018 | 1299.823 | 1346.436 | 1160.04 | 1242.029 |
| ENSMUSG00000033082 | -0.236542132 | 0.226709 | 0.386669 | 225.3562 | 155.9768 | 188.8083 | 217.3811 | 183.3209 | 202.2447 | 282.5367 | 261.0219 |
| ENSMUSG00000033083 | 0.123605392 | 0.484416 | 0.642684 | 1427.591 | 1258.613 | 958.9475 | 1277.545 | 1070.672 | 1489.46 | 1034.682 | 923.3648 |
| ENSMUSG00000033088 | -0.000662063 | 0.991017 | 0.994824 | 1636.85 | 1610.16 | 1534.482 | 1652.787 | 1638.187 | 1529.685 | 1640.256 | 1625.949 |
| ENSMUSG00000033096 | -0.362246261 | 0.001136 | 0.008611 | 1277.689 | 1237.016 | 1399.5 | 1263.743 | 1683.042 | 1375.488 | 1684.613 | 1914.16 |
| ENSMUSG00000033099 | -0.167309525 | 0.052303 | 0.140368 | 849.1099 | 1067.841 | 932.4481 | 919.5566 | 1047.27 | 1063.74 | 1051.075 | 1065.839 |
| ENSMUSG00000033102 | 0.158384706 | 0.206149 | 0.362677 | 726.3712 | 710.2943 | 693.9534 | 765.147 | 564.5894 | 793.3356 | 658.6095 | 579.686 |
| ENSMUSG00000033105 | -0.781404348 | 1.42E-05 | 0.000298 | 549.3057 | 423.537 | 617.7675 | 517.5741 | 886.3762 | 657.016 | 1097.361 | 985.3575 |

| | | | | | | | | | | | |
|---|---|---|---|---|---|---|---|---|---|---|---|
| ENSMUSG00000033106 | -0.02506277 | 0.798726 | 0.876883 | 1207.265 | 1352.199 | 1190.817 | 1295.66 | 1390.509 | 1401.187 | 1207.29 | 1133.27 |
| ENSMUSG00000033107 | -0.118793553 | 0.704479 | 0.81399 | 54.32694 | 33.595 | 45.54586 | 31.05445 | 49.73068 | 31.28647 | 55.92877 | 41.32846 |
| ENSMUSG00000033111 | 0.062710467 | 0.565445 | 0.710019 | 607.6568 | 686.2979 | 539.9255 | 597.7981 | 602.6188 | 570.9782 | 521.6804 | 629.7152 |
| ENSMUSG00000033114 | 0.018634818 | 0.889821 | 0.933475 | 200.2048 | 199.1704 | 218.6202 | 193.2277 | 195.9974 | 173.193 | 201.5364 | 230.5693 |
| ENSMUSG00000033124 | -0.435896818 | 0.020442 | 0.07163 | 320.9313 | 307.1543 | 356.914 | 304.5061 | 494.3814 | 283.813 | 475.3946 | 490.5036 |
| ENSMUSG00000033126 | -0.006103505 | 0.9676 | 0.981804 | 132.7992 | 176.3738 | 177.2148 | 150.9591 | 154.0676 | 164.254 | 153.322 | 168.5766 |
| ENSMUSG00000033128 | -0.180003418 | 0.049058 | 0.134374 | 1151.932 | 1294.607 | 1305.096 | 1180.932 | 1331.027 | 1267.102 | 1443.541 | 1545.467 |
| ENSMUSG00000033147 | 0.103085244 | 0.468316 | 0.628589 | 291.7558 | 283.1579 | 308.8838 | 304.5061 | 255.4792 | 343.0338 | 275.7867 | 233.8321 |
| ENSMUSG00000033149 | -0.356726398 | 0.000263 | 0.002813 | 3368.27 | 3155.531 | 3463.97 | 3368.545 | 4076.94 | 3723.09 | 4346.051 | 4959.415 |
| ENSMUSG00000033152 | 0.117656523 | 0.300716 | 0.469148 | 1009.073 | 1382.194 | 1267.003 | 1329.303 | 1146.731 | 1231.346 | 1077.111 | 1141.971 |
| ENSMUSG00000033157 | 0.151168024 | 0.215413 | 0.373588 | 649.9111 | 681.4986 | 558.1439 | 582.2709 | 519.7343 | 660.3681 | 515.8947 | 528.5693 |
| ENSMUSG00000033159 | -0.1331283 | 0.079885 | 0.188801 | 1434.634 | 1348.599 | 1549.388 | 1362.082 | 1555.303 | 1490.577 | 1670.149 | 1531.328 |
| ENSMUSG00000033161 | -1.607511684 | 0.000401 | 0.003866 | 21725.74 | 12473.34 | 34326.68 | 19340.02 | 54802.23 | 27755.57 | 90391.5 | 94803.14 |
| ENSMUSG00000033166 | 0.000638644 | 0.994407 | 0.996743 | 1042.272 | 1100.236 | 1077.367 | 1140.388 | 1145.756 | 1213.468 | 1010.575 | 989.7079 |
| ENSMUSG00000033170 | -0.356758717 | 0.049134 | 0.134463 | 479.8879 | 383.9429 | 554.8314 | 458.9157 | 591.8926 | 432.4238 | 666.3238 | 715.6349 |
| ENSMUSG00000033174 | 0.190723013 | 0.096156 | 0.214771 | 1271.653 | 1243.015 | 1231.395 | 1110.196 | 861.0233 | 1145.308 | 1079.04 | 1170.248 |
| ENSMUSG00000033177 | 0.452015656 | 0.371195 | 0.540197 | 9.054489 | 10.79839 | 15.73403 | 18.11509 | 6.825779 | 13.40849 | 7.714313 | 11.9635 |
| ENSMUSG00000033184 | -0.07402253 | 0.308378 | 0.476744 | 3084.563 | 3235.919 | 3234.585 | 3442.73 | 3087.202 | 3467.212 | 3533.155 | 3597.751 |
| ENSMUSG00000033186 | 0.266996706 | 0.052889 | 0.141529 | 2444.712 | 2433.238 | 2060.329 | 2515.41 | 1888.791 | 2485.04 | 1834.078 | 1648.788 |
| ENSMUSG00000033187 | -0.839658256 | 0.036581 | 0.109743 | 16.09687 | 10.79839 | 11.59349 | 11.21411 | 21.45245 | 29.05173 | 22.17865 | 16.31387 |
| ENSMUSG00000033191 | -0.522584484 | 0.011659 | 0.047904 | 1484.936 | 1282.609 | 2288.887 | 1671.764 | 2283.711 | 1757.629 | 2721.224 | 2906.043 |
| ENSMUSG00000033192 | 0.053554338 | 0.649077 | 0.772713 | 359.1614 | 376.744 | 339.5237 | 360.5766 | 373.4676 | 387.7288 | 320.144 | 301.2627 |
| ENSMUSG00000033207 | 0.502261225 | 0.013872 | 0.054415 | 147.89 | 122.3818 | 127.5284 | 142.3329 | 66.30757 | 96.09417 | 111.8575 | 107.6715 |
| ENSMUSG00000033208 | 1.058249335 | 0.046072 | 0.12858 | 13.07871 | 16.7975 | 33.95237 | 35.36756 | 11.70134 | 20.11273 | 7.714313 | 8.700729 |
| ENSMUSG00000033209 | 0.249168274 | 0.085177 | 0.197242 | 1881.322 | 2048.095 | 1818.522 | 1909.848 | 1694.743 | 2036.973 | 1438.719 | 1272.482 |
| ENSMUSG00000033214 | 1.265719231 | 0.051616 | 0.139029 | 41.24823 | 26.39607 | 18.21835 | 31.91707 | 4.875557 | 31.28647 | 7.714313 | 5.437955 |
| ENSMUSG00000033216 | -0.213622938 | 0.051865 | 0.139494 | 570.4328 | 604.7101 | 703.0625 | 565.0184 | 699.1548 | 636.9032 | 772.3956 | 725.4233 |
| ENSMUSG00000033220 | -0.003790783 | 0.986554 | 0.992307 | 74.44802 | 68.38983 | 112.6225 | 82.81186 | 93.61069 | 80.45093 | 92.57176 | 73.95619 |
| ENSMUSG00000033222 | 0.02309417 | 0.80327 | 0.879779 | 863.1946 | 985.0535 | 810.7164 | 898.8537 | 947.8082 | 869.317 | 869.7888 | 809.1678 |
| ENSMUSG00000033227 | -0.703392814 | 0.042951 | 0.122653 | 46.2785 | 53.99197 | 125.0441 | 74.18562 | 112.1378 | 170.9582 | 96.42892 | 110.9343 |
| ENSMUSG00000033228 | -0.060429919 | 0.240853 | 0.403048 | 3566.463 | 3594.665 | 3534.359 | 3557.459 | 3546.48 | 3916.396 | 3774.228 | 3626.029 |
| ENSMUSG00000033233 | 0.148813822 | 0.211667 | 0.369168 | 487.9364 | 581.9134 | 459.5992 | 498.5964 | 503.1574 | 486.0577 | 410.7872 | 426.3357 |
| ENSMUSG00000033237 | 0.40202057 | 0.014677 | 0.056742 | 3604.693 | 3664.255 | 2964.622 | 3672.188 | 2420.226 | 3573.362 | 2392.401 | 2138.204 |
| ENSMUSG00000033253 | 0.183570662 | 0.058713 | 0.152279 | 960.7819 | 920.2631 | 793.3262 | 866.074 | 811.2926 | 839.1479 | 754.0741 | 710.197 |
| ENSMUSG00000033256 | 0.079956828 | 0.558813 | 0.705145 | 309.8647 | 243.5638 | 311.3681 | 358.8514 | 296.4338 | 293.8694 | 280.6081 | 290.3868 |
| ENSMUSG00000033257 | 0.210077755 | 0.107022 | 0.231369 | 1771.662 | 1720.544 | 1613.152 | 1720.934 | 1435.364 | 1859.31 | 1384.719 | 1222.452 |
| ENSMUSG00000033272 | -0.384132017 | 0.000968 | 0.007642 | 1927.6 | 2153.68 | 2488.46 | 2116.878 | 3019.92 | 2373.303 | 2839.832 | 3105.073 |
| ENSMUSG00000033276 | 0.101755139 | 0.489197 | 0.646795 | 203.223 | 172.7743 | 188.8083 | 238.0841 | 202.8232 | 193.3057 | 174.5363 | 179.4525 |
| ENSMUSG00000033278 | -0.230404598 | 0.1215 | 0.250741 | 939.6548 | 710.2943 | 1026.852 | 895.4032 | 1152.582 | 832.4437 | 1110.861 | 1097.379 |
| ENSMUSG00000033282 | 0.323948828 | 0.033127 | 0.101547 | 429.5852 | 461.9313 | 390.8663 | 475.3055 | 302.2845 | 449.1844 | 333.6441 | 319.7518 |
| ENSMUSG00000033285 | -0.011389073 | 0.899924 | 0.939848 | 1335.034 | 1293.408 | 1074.054 | 1190.42 | 1286.172 | 1179.947 | 1262.255 | 1197.438 |
| ENSMUSG00000033287 | -0.415065867 | 0.001696 | 0.011602 | 622.7476 | 707.8947 | 803.2634 | 665.9453 | 1007.29 | 762.0491 | 916.0747 | 1048.438 |
| ENSMUSG00000033294 | -0.153326862 | 0.045796 | 0.127985 | 588.5418 | 550.7181 | 554.8314 | 554.6669 | 651.3744 | 596.6778 | 609.4308 | 642.7663 |
| ENSMUSG00000033295 | -0.127322262 | 0.11332 | 0.240365 | 4086.593 | 3724.246 | 4558.727 | 4087.11 | 4309.992 | 4290.717 | 4660.41 | 4720.145 |
| ENSMUSG00000033306 | 0.237057199 | 0.057924 | 0.150867 | 3515.154 | 3172.328 | 3450.72 | 3247.777 | 2397.799 | 3557.719 | 2708.688 | 2696.138 |
| ENSMUSG00000033307 | -0.786281484 | 1.39E-06 | 4.86E-05 | 3011.121 | 3545.473 | 3976.568 | 3389.248 | 6068.118 | 4330.942 | 6138.665 | 7473.926 |
| ENSMUSG00000033308 | 0.763095105 | 9.00E-05 | 0.001237 | 585.5236 | 392.3416 | 364.3669 | 469.2672 | 228.176 | 332.9775 | 272.8938 | 233.8321 |
| ENSMUSG00000033313 | 0.047754215 | 0.892524 | 0.934962 | 27.16347 | 40.79393 | 51.34261 | 51.75741 | 57.53157 | 35.75597 | 26.03581 | 46.76642 |
| ENSMUSG00000033318 | -0.632488474 | 0.003194 | 0.018605 | 423.5489 | 382.7431 | 607.8303 | 506.36 | 706.9557 | 513.9921 | 794.5743 | 964.6933 |
| ENSMUSG00000033319 | -0.060591582 | 0.349022 | 0.517643 | 1078.49 | 1091.838 | 1114.632 | 1118.823 | 1083.349 | 1124.078 | 1191.861 | 1195.263 |
| ENSMUSG00000033323 | -0.116587053 | 0.196701 | 0.351025 | 809.8738 | 769.0856 | 815.685 | 727.1916 | 912.7042 | 753.1101 | 840.8602 | 876.5984 |
| ENSMUSG00000033326 | 0.022131398 | 0.809982 | 0.884139 | 3036.272 | 2825.58 | 3074.76 | 2792.312 | 2937.035 | 2458.223 | 3107.904 | 3046.343 |
| ENSMUSG00000033327 | 1.632360822 | 0.019059 | 0.068197 | 822.9525 | 1270.611 | 702.2344 | 963.5504 | 278.8818 | 730.7627 | 137.8934 | 65.25547 |
| ENSMUSG00000033335 | -0.403257063 | 0.011868 | 0.048577 | 1537.251 | 1319.804 | 1895.536 | 1636.397 | 2031.157 | 1613.488 | 2307.544 | 2500.372 |
| ENSMUSG00000033342 | 0.607357479 | 0.11936 | 0.248355 | 23.13925 | 20.39697 | 28.15563 | 39.68068 | 14.62667 | 25.6996 | 18.32149 | 15.22628 |
| ENSMUSG00000033350 | -0.522721704 | 0.016061 | 0.060359 | 628.784 | 346.7484 | 591.2681 | 455.4652 | 721.5824 | 530.7527 | 784.9314 | 870.0729 |
| ENSMUSG00000033352 | 0.064225285 | 0.289178 | 0.456248 | 1819.952 | 1841.726 | 1857.443 | 1919.337 | 1723.997 | 1885.01 | 1695.22 | 1815.19 |
| ENSMUSG00000033355 | 0.352855276 | 0.437509 | 0.601202 | 219.3198 | 461.9313 | 556.4877 | 87.9876 | 210.624 | 206.7142 | 310.5011 | 309.9635 |
| ENSMUSG00000033356 | 0.12847914 | 0.326524 | 0.494204 | 276.6649 | 320.3523 | 250.9163 | 308.8192 | 281.8072 | 252.5265 | 237.2151 | 284.9489 |
| ENSMUSG00000033361 | 0.59018574 | 0.000107 | 0.001418 | 1334.028 | 1337.801 | 1082.335 | 1385.373 | 815.1931 | 1081.618 | 834.1101 | 683.0072 |
| ENSMUSG00000033364 | 0.120520755 | 0.183789 | 0.334834 | 982.9151 | 1094.237 | 914.2297 | 933.3586 | 864.9237 | 985.524 | 867.8602 | 888.5619 |
| ENSMUSG00000033365 | -0.230043973 | 0.008762 | 0.038733 | 805.8495 | 849.4736 | 932.4481 | 914.3809 | 958.5344 | 983.2892 | 1111.825 | 1057.139 |
| ENSMUSG00000033373 | 0.033652687 | 0.735913 | 0.834773 | 553.3299 | 625.107 | 621.08 | 647.8302 | 584.0917 | 549.7481 | 671.1453 | 585.124 |
| ENSMUSG00000033377 | 0.68751184 | 0.021331 | 0.073719 | 235.4167 | 131.9804 | 104.3414 | 159.5853 | 99.46135 | 93.85942 | 69.42882 | 129.4233 |

| | | | | | | | | | | | |
|---|---|---|---|---|---|---|---|---|---|---|---|
| ENSMUSG00000033379 | -0.573445987 | 0.001863 | 0.012474 | 2130.823 | 1457.783 | 2152.249 | 1986.622 | 2880.479 | 2044.795 | 3011.475 | 3565.124 |
| ENSMUSG00000033382 | -0.020100831 | 0.800223 | 0.877926 | 1159.981 | 967.0562 | 1032.649 | 1067.065 | 1036.543 | 1088.322 | 1113.754 | 1049.525 |
| ENSMUSG00000033386 | 0.080649819 | 0.53053 | 0.682263 | 672.0443 | 601.1106 | 631.0173 | 539.1397 | 478.7797 | 613.4384 | 560.252 | 659.0802 |
| ENSMUSG00000033389 | -0.046326085 | 0.763356 | 0.852918 | 341.0524 | 329.9509 | 455.4586 | 417.5098 | 344.2143 | 369.8508 | 429.1087 | 454.6131 |
| ENSMUSG00000033392 | 0.261356512 | 0.023438 | 0.079105 | 1498.015 | 1186.623 | 1189.161 | 1402.626 | 1058.971 | 1258.163 | 1066.504 | 1020.16 |
| ENSMUSG00000033396 | -0.049784862 | 0.604313 | 0.741075 | 845.0857 | 845.8742 | 869.512 | 897.9911 | 898.0775 | 762.0491 | 943.0748 | 976.6568 |
| ENSMUSG00000033400 | -0.156536261 | 0.031115 | 0.096904 | 1300.828 | 1283.809 | 1240.504 | 1290.485 | 1496.796 | 1280.511 | 1463.791 | 1457.372 |
| ENSMUSG00000033405 | -0.176758917 | 0.331859 | 0.499738 | 142.8597 | 141.5789 | 164.7932 | 126.8057 | 198.9227 | 128.498 | 170.6792 | 152.2628 |
| ENSMUSG00000033411 | 0.059933677 | 0.530458 | 0.682263 | 1252.538 | 1364.197 | 1123.741 | 1198.184 | 1248.142 | 1248.107 | 1188.004 | 1048.438 |
| ENSMUSG00000033416 | -0.678650961 | 5.22E-05 | 0.000819 | 955.7516 | 922.6628 | 1030.165 | 1015.308 | 1590.407 | 1109.552 | 1552.506 | 2029.445 |
| ENSMUSG00000033417 | 0.019655804 | 0.750875 | 0.844855 | 1837.055 | 1784.135 | 1688.509 | 1800.295 | 1667.44 | 1842.55 | 1728.006 | 1776.036 |
| ENSMUSG00000033419 | -0.372560395 | 0.095484 | 0.213737 | 1287.75 | 1073.84 | 818.9975 | 1372.434 | 1999.953 | 1055.919 | 1593.006 | 1244.204 |
| ENSMUSG00000033420 | 0.429353109 | 0.024832 | 0.082652 | 1311.895 | 1832.127 | 1563.465 | 1662.275 | 1106.751 | 1667.122 | 1027.932 | 928.8028 |
| ENSMUSG00000033423 | 0.053134224 | 0.551522 | 0.699148 | 1802.849 | 2022.899 | 1972.55 | 2021.989 | 2057.485 | 2011.273 | 1766.578 | 1702.08 |
| ENSMUSG00000033427 | -1.83422307 | 4.57E-08 | 3.01E-06 | 62.37537 | 55.19179 | 70.38906 | 45.71905 | 239.8774 | 96.09417 | 200.5721 | 296.9124 |
| ENSMUSG00000033429 | -0.555930055 | 0.000164 | 0.001944 | 477.8758 | 465.5308 | 532.4726 | 479.6187 | 722.5575 | 537.4569 | 770.467 | 845.0583 |
| ENSMUSG00000033430 | -0.002975762 | 0.965843 | 0.980776 | 1034.224 | 1088.238 | 1130.366 | 1019.621 | 1097 | 1064.858 | 1101.218 | 1016.898 |
| ENSMUSG00000033434 | -0.047871203 | 0.622091 | 0.753664 | 424.5549 | 406.7395 | 473.677 | 491.6954 | 476.8294 | 464.8276 | 469.6088 | 450.2627 |
| ENSMUSG00000033436 | 0.208877469 | 0.002946 | 0.017545 | 3200.259 | 3598.265 | 3447.408 | 3443.593 | 2782.968 | 3203.512 | 2872.617 | 2986.525 |
| ENSMUSG00000033439 | 0.421206985 | 0.019151 | 0.068393 | 466.8092 | 374.3443 | 311.3681 | 357.9887 | 263.2801 | 343.0338 | 297.0011 | 224.0438 |
| ENSMUSG00000033444 | 0.035650678 | 0.622423 | 0.753967 | 2527.209 | 2504.028 | 2697.143 | 2490.394 | 2449.48 | 2289.5 | 2714.474 | 2515.598 |
| ENSMUSG00000033446 | 0.173877928 | 0.088415 | 0.202555 | 937.6427 | 791.8822 | 717.1403 | 800.5146 | 733.2837 | 748.6406 | 689.4668 | 705.8466 |
| ENSMUSG00000033450 | -0.159578639 | 0.731551 | 0.831681 | 41.24823 | 56.39161 | 38.09291 | 39.68068 | 19.50223 | 92.74205 | 52.07161 | 31.54014 |
| ENSMUSG00000033453 | -0.439474893 | 0.037243 | 0.110954 | 240.447 | 292.7565 | 464.5678 | 350.2251 | 373.4676 | 492.762 | 453.2159 | 512.2554 |
| ENSMUSG00000033454 | 0.391145343 | 0.000378 | 0.00369 | 1315.919 | 1388.194 | 1088.132 | 1330.165 | 959.5095 | 1084.97 | 869.7888 | 990.7955 |
| ENSMUSG00000033458 | -0.054028345 | 0.693952 | 0.806574 | 213.2835 | 231.5656 | 226.9012 | 235.4962 | 262.3049 | 189.9536 | 235.2866 | 253.4087 |
| ENSMUSG00000033460 | 0.359323295 | 0.002444 | 0.015178 | 1458.779 | 1433.787 | 1079.023 | 1324.99 | 979.9869 | 1149.778 | 1019.254 | 977.7444 |
| ENSMUSG00000033467 | -0.055119465 | 0.671954 | 0.789942 | 420.5307 | 478.7288 | 568.0811 | 498.5964 | 548.9877 | 551.9828 | 438.7516 | 505.7299 |
| ENSMUSG00000033475 | 0.145500173 | 0.421929 | 0.587229 | 107.6478 | 101.9848 | 99.3728 | 91.43809 | 103.3618 | 98.32892 | 75.21455 | 84.8321 |
| ENSMUSG00000033478 | 0.121283211 | 0.204254 | 0.360255 | 1423.567 | 1368.996 | 1411.922 | 1620.87 | 1302.749 | 1519.629 | 1287.326 | 1249.642 |
| ENSMUSG00000033486 | 0.197346983 | 0.450399 | 0.612885 | 78.47224 | 99.58519 | 76.18581 | 73.323 | 67.28268 | 99.44629 | 48.21446 | 70.69342 |
| ENSMUSG00000033487 | 0.262838397 | 0.018617 | 0.067094 | 3364.246 | 3099.139 | 2835.437 | 3384.072 | 2565.518 | 3094.009 | 2638.295 | 2273.065 |
| ENSMUSG00000033488 | -0.710611796 | 0.000328 | 0.003324 | 601.6205 | 537.52 | 587.9557 | 540.8649 | 970.2358 | 631.3164 | 830.253 | 1280.095 |
| ENSMUSG00000033491 | -0.551166205 | 0.153544 | 0.294877 | 393.3673 | 113.983 | 152.3716 | 192.365 | 345.1894 | 454.7713 | 204.4293 | 244.708 |
| ENSMUSG00000033499 | 0.12705427 | 0.226004 | 0.385786 | 3221.386 | 2856.775 | 2810.594 | 2759.533 | 2422.176 | 3154.347 | 2552.473 | 2538.438 |
| ENSMUSG00000033502 | 0.12556335 | 0.356396 | 0.524955 | 856.1523 | 805.0802 | 654.2042 | 828.1186 | 591.8926 | 754.2275 | 708.7525 | 826.5692 |
| ENSMUSG00000033508 | 0.211849508 | 0.557417 | 0.703981 | 25.15136 | 27.5959 | 45.54586 | 46.58167 | 27.30312 | 41.34284 | 35.6787 | 21.75182 |
| ENSMUSG00000033510 | 0.208928809 | 0.713291 | 0.820218 | 12.07265 | 5.999108 | 14.07781 | 24.15346 | 8.776002 | 17.87799 | 7.714313 | 15.22628 |
| ENSMUSG00000033526 | -0.29459182 | 0.069625 | 0.171547 | 463.7911 | 376.744 | 543.238 | 421.8229 | 520.7094 | 449.1844 | 588.2164 | 659.0802 |
| ENSMUSG00000033530 | -0.266443875 | 0.012535 | 0.0506 | 621.7416 | 500.3256 | 568.0811 | 589.1718 | 651.3744 | 621.26 | 741.5384 | 729.7736 |
| ENSMUSG00000033533 | -5.070626963 | 8.28E-09 | 7.38E-07 | 14.08476 | 2.399643 | 46.37397 | 18.97772 | 780.089 | 175.4277 | 488.8946 | 1322.511 |
| ENSMUSG00000033538 | 0.497151568 | 0.002961 | 0.017594 | 257.5499 | 266.3604 | 218.6202 | 248.4356 | 151.1423 | 189.9536 | 215.0365 | 144.6496 |
| ENSMUSG00000033540 | 0.393975145 | 0.003393 | 0.019358 | 445.6821 | 503.925 | 439.7246 | 487.3823 | 345.1894 | 433.5411 | 297.9654 | 352.3795 |
| ENSMUSG00000033542 | -0.218614073 | 0.002237 | 0.014252 | 1480.912 | 1512.975 | 1729.087 | 1534.607 | 1810.782 | 1794.503 | 1801.292 | 1880.445 |
| ENSMUSG00000033543 | 0.135561873 | 0.227661 | 0.387612 | 1702.244 | 1632.957 | 1527.029 | 1749.4 | 1449.015 | 1814.616 | 1427.148 | 1329.036 |
| ENSMUSG00000033544 | 0.839934018 | 0.088602 | 0.202855 | 63.38142 | 164.3756 | 45.54586 | 67.28463 | 31.20356 | 88.27255 | 35.6787 | 34.80291 |
| ENSMUSG00000033545 | 0.21070899 | 0.005718 | 0.028408 | 1586.548 | 1499.777 | 1618.12 | 1707.132 | 1350.529 | 1517.394 | 1333.612 | 1344.263 |
| ENSMUSG00000033554 | -0.006033114 | 0.958962 | 0.976798 | 410.4702 | 365.9456 | 329.5864 | 381.2796 | 360.7912 | 379.9072 | 337.5012 | 415.4598 |
| ENSMUSG00000033557 | -0.302623493 | 0.000305 | 0.003142 | 2112.714 | 1931.713 | 2106.703 | 2003.012 | 2468.007 | 2220.222 | 2710.617 | 2659.16 |
| ENSMUSG00000033565 | 0.389329191 | 0.000564 | 0.005023 | 3462.839 | 4015.803 | 3637.872 | 3608.354 | 2817.097 | 3376.705 | 2448.33 | 2600.43 |
| ENSMUSG00000033569 | 0.551161589 | 0.232581 | 0.393087 | 54.32694 | 34.79482 | 29.81184 | 73.323 | 19.50223 | 60.3382 | 27.0001 | 25.0146 |
| ENSMUSG00000033577 | -0.246695138 | 0.083211 | 0.194153 | 3516.16 | 2766.788 | 3889.617 | 3214.998 | 3427.516 | 3565.541 | 4103.05 | 4791.926 |
| ENSMUSG00000033578 | 1.252539096 | 0.000131 | 0.001649 | 1740.474 | 950.2587 | 689.8128 | 935.9465 | 331.5378 | 747.5233 | 339.4298 | 393.708 |
| ENSMUSG00000033579 | -1.100702493 | 0.006725 | 0.032041 | 36.21796 | 40.79393 | 73.70149 | 54.34528 | 132.6151 | 42.46022 | 113.7861 | 152.2628 |
| ENSMUSG00000033581 | 0.159817028 | 0.029574 | 0.093639 | 2062.411 | 1893.318 | 1934.457 | 2029.753 | 1739.599 | 1928.588 | 1789.721 | 1633.562 |
| ENSMUSG00000033585 | 0.435630018 | 2.00E-05 | 0.000392 | 2465.839 | 2272.462 | 2314.558 | 2650.842 | 1827.359 | 2071.612 | 1657.613 | 1620.511 |
| ENSMUSG00000033589 | 0.106497225 | 0.139514 | 0.275505 | 722.347 | 724.6922 | 692.2971 | 760.8339 | 690.3788 | 650.3117 | 668.2524 | 684.0948 |
| ENSMUSG00000033590 | -0.018899198 | 0.869103 | 0.921481 | 345.0766 | 363.5459 | 409.9128 | 329.5222 | 339.3387 | 351.9728 | 379.9299 | 396.9707 |
| ENSMUSG00000033594 | -0.560783 | 0.010632 | 0.044636 | 72.43591 | 65.99018 | 82.81066 | 53.48266 | 88.73513 | 88.27255 | 119.5719 | 108.7591 |
| ENSMUSG00000033595 | 0.596097561 | 0.001293 | 0.009468 | 271.6347 | 295.1561 | 266.6503 | 282.0779 | 232.0765 | 212.3011 | 137.8934 | 155.5255 |
| ENSMUSG00000033596 | -0.024369042 | 0.770409 | 0.857476 | 2095.611 | 2062.493 | 1762.211 | 1883.107 | 2022.381 | 2102.898 | 1959.436 | 1846.73 |
| ENSMUSG00000033597 | 0.379982928 | 0.073017 | 0.177713 | 253.5257 | 173.9741 | 197.0894 | 246.7103 | 162.8436 | 217.8879 | 165.8577 | 123.9854 |
| ENSMUSG00000033610 | -0.914731689 | 0.000479 | 0.004437 | 412.4823 | 309.554 | 351.1172 | 364.0271 | 651.3744 | 354.2076 | 749.2527 | 954.905 |
| ENSMUSG00000033615 | 1.06163193 | 0.017002 | 0.062847 | 49.29666 | 26.39607 | 34.78048 | 94.02596 | 25.35289 | 30.1691 | 21.21436 | 21.75182 |

| | | | | | | | | | | | |
|---|---|---|---|---|---|---|---|---|---|---|---|
| ENSMUSG00000033618 | -0.048461267 | 0.824613 | 0.893455 | 80.48435 | 68.38983 | 57.96746 | 79.36136 | 64.35735 | 78.21619 | 87.75031 | 65.25547 |
| ENSMUSG00000033623 | 0.199876611 | 0.019905 | 0.070188 | 2307.889 | 2258.064 | 1914.583 | 2186.751 | 1875.139 | 1977.752 | 1734.756 | 1956.576 |
| ENSMUSG00000033624 | -0.465838942 | 0.003119 | 0.018301 | 487.9364 | 401.9402 | 434.756 | 368.3402 | 624.0712 | 436.8933 | 627.7522 | 648.2043 |
| ENSMUSG00000033628 | 0.016691541 | 0.795243 | 0.874099 | 1037.242 | 1105.036 | 1038.446 | 994.6049 | 1035.568 | 1050.332 | 1031.789 | 1004.934 |
| ENSMUSG00000033629 | 0.080893286 | 0.569186 | 0.713058 | 2212.314 | 1650.954 | 1434.281 | 1732.148 | 1554.327 | 1800.09 | 1507.184 | 1784.737 |
| ENSMUSG00000033632 | 0.204482725 | 0.014454 | 0.056127 | 532.2028 | 533.9206 | 549.0347 | 590.8971 | 470.9788 | 500.5836 | 459.0016 | 487.2408 |
| ENSMUSG00000033633 | -1.361540016 | 6.59E-07 | 2.64E-05 | 853.1341 | 331.1507 | 829.7628 | 704.7634 | 1537.751 | 1343.084 | 1927.614 | 2182.795 |
| ENSMUSG00000033634 | -2.630723911 | 2.73E-08 | 1.97E-06 | 4.024217 | 13.19804 | 13.24971 | 12.07673 | 77.03379 | 32.40385 | 58.82164 | 95.70802 |
| ENSMUSG00000033644 | -1.555221709 | NA | NA | 11.0666 | 19.19714 | 19.87456 | 16.38985 | 7.80089 | 165.3714 | 9.642892 | 13.05109 |
| ENSMUSG00000033653 | -0.278121488 | 0.079802 | 0.188683 | 537.233 | 497.9259 | 521.7072 | 440.8006 | 630.897 | 450.3018 | 600.7521 | 739.5619 |
| ENSMUSG00000033658 | -0.017136751 | 0.822767 | 0.89248 | 738.4439 | 671.9001 | 745.296 | 705.626 | 756.6864 | 745.2885 | 696.2168 | 700.4087 |
| ENSMUSG00000033669 | -0.24354518 | 0.083339 | 0.194351 | 752.5287 | 677.8992 | 741.9835 | 646.9676 | 782.0393 | 655.8986 | 985.5035 | 913.5765 |
| ENSMUSG00000033671 | -0.001715037 | 0.983066 | 0.990742 | 1022.151 | 917.8635 | 927.4794 | 961.8252 | 898.0775 | 941.9464 | 945.0034 | 1050.613 |
| ENSMUSG00000033676 | 0.161016079 | 0.348313 | 0.516965 | 127.7689 | 130.7805 | 162.3089 | 152.6844 | 143.3414 | 144.1413 | 109.929 | 117.4598 |
| ENSMUSG00000033684 | 0.058889561 | 0.538563 | 0.688441 | 1805.868 | 1482.979 | 1878.146 | 1858.091 | 1658.664 | 1667.122 | 1809.006 | 1613.985 |
| ENSMUSG00000033685 | -0.06189811 | 0.679095 | 0.794865 | 3468.875 | 3508.278 | 4311.123 | 3929.25 | 4034.035 | 2982.271 | 4009.514 | 4860.445 |
| ENSMUSG00000033688 | 1.397218057 | 0.052209 | 0.140259 | 24.1453 | 8.398751 | 5.796746 | 25.8787 | 3.900445 | 4.469496 | 11.57147 | 4.350364 |
| ENSMUSG00000033697 | -0.122342082 | 0.184062 | 0.335148 | 592.566 | 541.1195 | 513.4261 | 495.1459 | 572.3903 | 565.3913 | 571.8235 | 619.9269 |
| ENSMUSG00000033701 | -0.081060537 | 0.309979 | 0.478464 | 1336.04 | 1461.383 | 1521.232 | 1574.288 | 1463.642 | 1598.962 | 1523.577 | 1652.051 |
| ENSMUSG00000033703 | -0.37723183 | 0.000989 | 0.00778 | 386.3249 | 429.5361 | 449.6619 | 452.0147 | 553.8632 | 493.8794 | 540.9662 | 643.8539 |
| ENSMUSG00000033705 | 0.2845854 | 0.000395 | 0.003816 | 777.68 | 735.4906 | 862.059 | 765.147 | 641.6232 | 644.7249 | 656.6809 | 638.416 |
| ENSMUSG00000033706 | -0.115035271 | 0.118235 | 0.247046 | 1010.079 | 1089.438 | 1084.82 | 1042.049 | 1233.516 | 1139.722 | 1064.575 | 1138.708 |
| ENSMUSG00000033707 | -0.472538643 | 0.013305 | 0.052892 | 296.786 | 279.5584 | 263.3379 | 219.969 | 380.2934 | 272.6393 | 338.4655 | 477.4525 |
| ENSMUSG00000033712 | 0.021018687 | 0.696345 | 0.808286 | 2878.322 | 3136.333 | 3081.385 | 3119.247 | 3071.601 | 3067.192 | 2992.189 | 2907.131 |
| ENSMUSG00000033713 | 0.237354673 | 0.243995 | 0.406566 | 431.5973 | 455.9322 | 327.9302 | 300.193 | 313.9858 | 415.6632 | 307.6082 | 246.8832 |
| ENSMUSG00000033715 | 1.229568433 | 0.009018 | 0.039559 | 167.005 | 247.1632 | 153.1997 | 212.2054 | 60.4569 | 177.6625 | 68.46453 | 26.10219 |
| ENSMUSG00000033717 | -0.334630956 | 0.151144 | 0.291424 | 238.4349 | 196.7707 | 303.9151 | 214.7932 | 217.4498 | 291.6346 | 424.2872 | 270.8102 |
| ENSMUSG00000033720 | 0.07013405 | 0.585298 | 0.725623 | 363.1856 | 388.7422 | 351.1172 | 295.8799 | 299.3592 | 356.4423 | 314.3583 | 361.0802 |
| ENSMUSG00000033721 | 0.230863294 | 0.031726 | 0.098292 | 1147.908 | 1154.228 | 1228.082 | 1136.938 | 866.874 | 898.3688 | 1124.361 | 1087.591 |
| ENSMUSG00000033722 | -0.585568201 | 0.000389 | 0.003779 | 500.009 | 477.529 | 556.4877 | 490.8328 | 705.9806 | 574.3303 | 775.2885 | 984.2699 |
| ENSMUSG00000033726 | -1.72827484 | 0.000153 | 0.00186 | 144.8718 | 45.59322 | 184.6678 | 135.4319 | 342.2641 | 159.7845 | 545.7877 | 647.1167 |
| ENSMUSG00000033728 | -0.140051306 | 0.128793 | 0.260464 | 840.0554 | 872.2703 | 749.4365 | 814.3166 | 957.5593 | 805.6267 | 887.146 | 954.905 |
| ENSMUSG00000033732 | -0.113765062 | 0.077777 | 0.185435 | 5246.573 | 5118.439 | 5001.764 | 5079.127 | 5822.39 | 4999.132 | 5675.806 | 5621.758 |
| ENSMUSG00000033735 | -0.449772247 | 0.000639 | 0.005551 | 1368.234 | 1283.809 | 1517.091 | 1279.271 | 1894.641 | 1490.577 | 1819.614 | 2238.262 |
| ENSMUSG00000033737 | 1.723451984 | 9.90E-20 | 1.22E-16 | 212.2775 | 243.5638 | 178.871 | 216.5185 | 68.25779 | 69.27719 | 71.3574 | 47.85401 |
| ENSMUSG00000033739 | -0.279170475 | 0.040695 | 0.118149 | 241.453 | 193.1713 | 192.9488 | 175.9752 | 245.728 | 253.6439 | 233.358 | 241.4452 |
| ENSMUSG00000033751 | -0.271700015 | 0.000683 | 0.005824 | 966.8182 | 1006.65 | 938.2448 | 954.9242 | 1160.382 | 1045.862 | 1200.54 | 1257.255 |
| ENSMUSG00000033752 | 0.247358474 | 0.12149 | 0.250741 | 315.9011 | 314.3532 | 307.2276 | 401.1199 | 310.0854 | 325.1559 | 267.1081 | 226.2189 |
| ENSMUSG00000033760 | 0.37118079 | 0.007709 | 0.035234 | 1430.609 | 1407.391 | 1204.067 | 1397.45 | 967.3104 | 1349.788 | 926.6819 | 962.5181 |
| ENSMUSG00000033762 | -0.327514922 | 0.007858 | 0.035773 | 187.1261 | 196.7707 | 173.0743 | 174.2499 | 249.6285 | 223.4748 | 210.215 | 231.6569 |
| ENSMUSG00000033763 | 0.225515237 | 0.0067 | 0.031964 | 1412.5 | 1560.968 | 1516.263 | 1306.012 | 1239.366 | 1309.562 | 1165.826 | 1240.941 |
| ENSMUSG00000033767 | 0.347192695 | 0.017107 | 0.063131 | 1431.615 | 1380.995 | 1290.19 | 1444.032 | 1022.892 | 1431.356 | 998.0393 | 909.2261 |
| ENSMUSG00000033768 | -0.004577005 | 0.982159 | 0.990044 | 514.0938 | 418.7377 | 438.0684 | 479.6187 | 629.9219 | 308.3953 | 483.1089 | 433.9488 |
| ENSMUSG00000033769 | -0.128065475 | 0.119652 | 0.248542 | 1227.386 | 1294.607 | 1320.83 | 1083.455 | 1361.255 | 1358.727 | 1307.576 | 1354.051 |
| ENSMUSG00000033770 | -2.40623725 | 2.34E-05 | 0.000445 | 262.5802 | 83.98751 | 345.3205 | 155.2722 | 936.1069 | 438.0106 | 1518.755 | 1600.934 |
| ENSMUSG00000033773 | 0.044827903 | 0.627367 | 0.757172 | 551.3178 | 583.1133 | 563.1125 | 597.7981 | 589.9423 | 528.5179 | 599.7879 | 504.6423 |
| ENSMUSG00000033777 | 0.528199253 | 0.028342 | 0.090843 | 50.30272 | 45.59322 | 48.03018 | 51.75741 | 35.10401 | 31.28647 | 36.64299 | 32.62773 |
| ENSMUSG00000033781 | -0.495514788 | 0.00318 | 0.018551 | 541.2572 | 556.7172 | 576.3622 | 626.2647 | 864.9237 | 586.6214 | 763.717 | 1028.861 |
| ENSMUSG00000033788 | -0.379965184 | 0.016227 | 0.060821 | 495.9848 | 430.7359 | 618.5957 | 439.0754 | 595.793 | 598.9125 | 603.645 | 787.4159 |
| ENSMUSG00000033790 | 0.191655683 | 0.120921 | 0.250136 | 813.898 | 898.6663 | 684.8442 | 759.1087 | 750.8357 | 756.4623 | 652.8238 | 600.3503 |
| ENSMUSG00000033792 | -0.039961716 | 0.815418 | 0.887554 | 637.8385 | 652.7029 | 526.6758 | 522.7498 | 606.5192 | 427.9543 | 706.824 | 661.2554 |
| ENSMUSG00000033793 | -0.349648322 | 0.040198 | 0.117074 | 1726.389 | 1127.832 | 1496.389 | 1260.293 | 1609.909 | 1461.525 | 1894.828 | 2184.97 |
| ENSMUSG00000033799 | 0.202624476 | 0.052274 | 0.140316 | 1270.647 | 1408.59 | 1108.835 | 1186.97 | 1123.328 | 1182.182 | 996.1107 | 1017.985 |
| ENSMUSG00000033805 | 0.956474117 | 0.048336 | 0.13299 | 55.33299 | 59.99108 | 41.40533 | 55.2079 | 20.47734 | 56.98608 | 22.17865 | 9.78832 |
| ENSMUSG00000033808 | 0.229034566 | 0.038963 | 0.114307 | 1843.092 | 1629.358 | 1594.933 | 1684.704 | 1362.23 | 1712.934 | 1427.148 | 1259.43 |
| ENSMUSG00000033809 | -0.3055463 | 0.026354 | 0.086328 | 244.4712 | 260.3613 | 294.806 | 217.3811 | 345.1894 | 284.9304 | 296.0368 | 330.6277 |
| ENSMUSG00000033813 | -0.123283667 | 0.028677 | 0.091535 | 2767.656 | 2991.155 | 2711.221 | 2830.268 | 3165.211 | 3142.056 | 2983.511 | 3011.54 |
| ENSMUSG00000033819 | -0.28389221 | 0.005194 | 0.026408 | 726.3712 | 845.8742 | 924.167 | 780.6743 | 1014.116 | 909.5425 | 1009.611 | 1057.139 |
| ENSMUSG00000033825 | 0.097163926 | 0.895473 | 0.936737 | 63.38142 | 100.785 | 71.21717 | 80.22398 | 55.58134 | 212.3011 | 15.42863 | 11.9635 |
| ENSMUSG00000033826 | -0.161869289 | 0.657537 | 0.779315 | 20.12109 | 14.39786 | 23.18699 | 25.01608 | 16.57689 | 29.05173 | 23.14294 | 25.0146 |
| ENSMUSG00000033831 | 2.176567342 | 0.080882 | 0.190379 | 892.3702 | 3.599465 | 46.37397 | 442.5259 | 50.70579 | 39.10809 | 109.929 | 106.5839 |
| ENSMUSG00000033845 | -0.051573503 | 0.45476 | 0.616955 | 1338.052 | 1279.01 | 1176.74 | 1291.347 | 1325.176 | 1252.576 | 1332.648 | 1356.226 |
| ENSMUSG00000033847 | 0.600453976 | 0.268687 | 0.43328 | 17.10292 | 27.5959 | 16.56213 | 24.15346 | 19.50223 | 11.17374 | 3.857157 | 21.75182 |
| ENSMUSG00000033849 | 0.191205663 | 0.653607 | 0.776402 | 18.10898 | 20.39697 | 15.73403 | 25.01608 | 10.72622 | 13.40849 | 19.28578 | 26.10219 |

| | | | | | | | | | | | |
|---|---|---|---|---|---|---|---|---|---|---|---|
| ENSMUSG00000033854 | 0.470275309 | 0.288425 | 0.455361 | 16.09687 | 17.99732 | 10.76539 | 19.84034 | 13.65156 | 11.17374 | 8.678602 | 13.05109 |
| ENSMUSG00000033855 | -0.133795157 | 0.434852 | 0.59894 | 470.8334 | 473.9295 | 499.3483 | 454.6026 | 562.6392 | 358.6771 | 638.3594 | 522.0437 |
| ENSMUSG00000033857 | -0.319102249 | 0.039777 | 0.116162 | 234.4107 | 317.9527 | 251.7444 | 207.0296 | 325.6872 | 288.2825 | 340.3941 | 302.3503 |
| ENSMUSG00000033860 | 3.576408778 | 0.008645 | 0.03838 | 618.7234 | 2.399643 | 7.45296 | 315.7202 | 23.40267 | 12.29112 | 17.3572 | 26.10219 |
| ENSMUSG00000033862 | 0.179524795 | 0.038047 | 0.112572 | 1240.465 | 1337.801 | 1109.663 | 1315.501 | 1122.353 | 1164.304 | 1089.647 | 1038.649 |
| ENSMUSG00000033863 | 0.077364092 | 0.869693 | 0.921839 | 2119.757 | 1090.638 | 1228.91 | 2313.556 | 768.3877 | 2554.317 | 2048.15 | 1029.949 |
| ENSMUSG00000033871 | -0.182175346 | 0.416227 | 0.582046 | 148.896 | 122.3818 | 143.2624 | 89.71284 | 129.6898 | 153.0803 | 114.7504 | 175.1022 |
| ENSMUSG00000033880 | 0.177957029 | 0.692951 | 0.805613 | 741.4621 | 1827.328 | 2182.061 | 661.6322 | 1226.69 | 886.0777 | 1512.97 | 1158.285 |
| ENSMUSG00000033882 | 2.082576482 | 2.43E-07 | 1.19E-05 | 131.7931 | 211.1686 | 110.1382 | 178.5631 | 22.42756 | 65.92507 | 42.42872 | 18.48905 |
| ENSMUSG00000033883 | 0.104637149 | 0.515816 | 0.669849 | 563.3904 | 518.3229 | 471.1927 | 473.5803 | 470.0037 | 606.7341 | 436.823 | 370.8686 |
| ENSMUSG00000033885 | -0.177215494 | 0.125357 | 0.2559 | 753.5347 | 682.6985 | 850.4655 | 725.4664 | 811.2926 | 738.5843 | 911.2533 | 947.2918 |
| ENSMUSG00000033898 | 0.963242258 | 0.047281 | 0.130807 | 23.13925 | 10.79839 | 15.73403 | 14.6646 | 8.776002 | 10.05637 | 6.750024 | 7.613138 |
| ENSMUSG00000033900 | 0.371736235 | 0.051327 | 0.138554 | 566.4086 | 566.3158 | 350.2891 | 529.6508 | 373.4676 | 480.4709 | 374.1442 | 326.2773 |
| ENSMUSG00000033902 | 0.283150402 | 0.003869 | 0.021314 | 925.57 | 859.0722 | 897.6676 | 957.5121 | 733.2837 | 867.0823 | 691.3953 | 702.5838 |
| ENSMUSG00000033904 | 0.266632223 | 0.007809 | 0.035595 | 1463.809 | 1306.606 | 1265.347 | 1408.664 | 1164.283 | 1260.398 | 1123.397 | 977.7444 |
| ENSMUSG00000033906 | 0.09442807 | 0.30062 | 0.469038 | 578.4813 | 574.7145 | 514.2542 | 576.2325 | 498.2819 | 573.2129 | 502.3947 | 527.4817 |
| ENSMUSG00000033909 | 0.113751484 | 0.136584 | 0.271378 | 1040.26 | 1003.051 | 904.2924 | 910.0678 | 880.5255 | 898.3688 | 888.1103 | 893.9999 |
| ENSMUSG00000033910 | 0.077380706 | 0.661515 | 0.782072 | 2200.241 | 3390.696 | 3632.904 | 2696.561 | 3063.8 | 2272.739 | 2697.117 | 3263.861 |
| ENSMUSG00000033916 | -0.157886362 | 0.034915 | 0.105882 | 2221.368 | 2255.664 | 2479.351 | 2281.639 | 2626.95 | 2340.899 | 2588.152 | 2752.693 |
| ENSMUSG00000033917 | -0.46432874 | 0.002065 | 0.013422 | 1363.204 | 1312.605 | 1782.085 | 1406.939 | 1963.874 | 1622.427 | 2047.186 | 2461.219 |
| ENSMUSG00000033918 | -0.101649893 | 0.113542 | 0.240638 | 1458.779 | 1404.991 | 1515.435 | 1382.785 | 1581.631 | 1536.389 | 1465.72 | 1602.022 |
| ENSMUSG00000033931 | -0.029895074 | 0.710627 | 0.818736 | 984.9272 | 1171.026 | 1086.476 | 992.8796 | 1066.772 | 1101.731 | 1054.932 | 1097.379 |
| ENSMUSG00000033933 | 0.194874262 | 0.054752 | 0.145418 | 1305.859 | 1313.805 | 1355.611 | 1331.891 | 1327.126 | 1233.581 | 1018.289 | 1058.226 |
| ENSMUSG00000033938 | -0.521397514 | 0.000285 | 0.002988 | 1457.773 | 1618.559 | 1743.164 | 1550.997 | 2241.781 | 1749.808 | 2333.58 | 2819.036 |
| ENSMUSG00000033940 | -0.017681706 | 0.756127 | 0.848334 | 3557.408 | 3589.866 | 3633.732 | 3742.061 | 3899.47 | 3480.62 | 3582.334 | 3740.226 |
| ENSMUSG00000033943 | 0.291052388 | 0.006442 | 0.031083 | 2632.844 | 2588.015 | 2259.075 | 2549.052 | 1892.691 | 2434.758 | 1993.186 | 1876.095 |
| ENSMUSG00000033948 | -0.424607013 | 0.049318 | 0.134807 | 246.4833 | 164.3756 | 266.6503 | 184.6014 | 234.0267 | 263.7003 | 284.4653 | 377.3941 |
| ENSMUSG00000033949 | 0.072479728 | 0.806048 | 0.881513 | 92.557 | 79.18822 | 67.90474 | 75.91087 | 45.83023 | 84.92043 | 108.0004 | 60.9051 |
| ENSMUSG00000033952 | 0.347392383 | 0.015371 | 0.058709 | 1524.172 | 1400.192 | 1072.398 | 1460.422 | 1196.462 | 1191.121 | 965.2535 | 935.3283 |
| ENSMUSG00000033953 | 0.132397971 | 0.133254 | 0.266682 | 2727.413 | 2620.41 | 2382.463 | 2648.254 | 2169.623 | 2630.299 | 2457.973 | 2209.985 |
| ENSMUSG00000033955 | -0.042220278 | 0.686277 | 0.800526 | 1492.985 | 1460.183 | 1553.528 | 1372.434 | 1350.529 | 1354.257 | 1666.292 | 1682.503 |
| ENSMUSG00000033960 | 0.11372043 | 0.307003 | 0.475531 | 1260.586 | 1003.051 | 965.5723 | 1200.772 | 966.3353 | 1063.74 | 1076.147 | 988.6203 |
| ENSMUSG00000033961 | -0.040388938 | 0.692117 | 0.805049 | 459.7668 | 412.7386 | 369.3356 | 388.1806 | 420.273 | 427.9543 | 417.5372 | 407.8467 |
| ENSMUSG00000033964 | -0.001777241 | 0.97991 | 0.988858 | 922.5518 | 855.4728 | 820.6537 | 870.3871 | 854.1975 | 845.8522 | 896.7889 | 874.4232 |
| ENSMUSG00000033965 | 0.662477025 | 3.94E-05 | 0.000666 | 907.461 | 1262.212 | 1252.097 | 1203.36 | 685.5032 | 935.2421 | 648.9666 | 653.6422 |
| ENSMUSG00000033966 | -0.4895733 | 0.276715 | 0.442803 | 48.29061 | 28.79572 | 49.6864 | 50.03216 | 38.02934 | 27.93435 | 110.8933 | 71.78101 |
| ENSMUSG00000033967 | -0.285245814 | 0.350492 | 0.519114 | 45.27245 | 25.19625 | 25.67131 | 34.50494 | 39.97956 | 39.10809 | 46.28588 | 33.71532 |
| ENSMUSG00000033970 | 0.00632826 | 0.927112 | 0.956765 | 1263.604 | 1402.591 | 1373.001 | 1359.495 | 1441.215 | 1315.149 | 1309.505 | 1307.284 |
| ENSMUSG00000033972 | 0.193484375 | 0.240877 | 0.403048 | 394.3733 | 433.1356 | 418.1938 | 488.2449 | 359.8161 | 499.4662 | 345.2155 | 313.2262 |
| ENSMUSG00000033981 | 0.910752931 | 0.010524 | 0.044254 | 117.7084 | 97.18554 | 60.45178 | 162.1732 | 71.18313 | 65.92507 | 61.71451 | 33.71532 |
| ENSMUSG00000033983 | 0.099523756 | 0.296314 | 0.464124 | 640.8566 | 706.6949 | 584.6433 | 650.4181 | 658.2001 | 569.8608 | 588.2164 | 589.4744 |
| ENSMUSG00000033985 | -0.037906622 | 0.781179 | 0.865089 | 320.9313 | 315.5531 | 333.727 | 293.292 | 285.7076 | 279.3435 | 378.0014 | 353.4671 |
| ENSMUSG00000033987 | 0.298812926 | 0.243478 | 0.406076 | 51.30877 | 64.79036 | 59.62368 | 69.00988 | 45.83023 | 68.15982 | 46.28588 | 39.15328 |
| ENSMUSG00000033991 | -0.154174582 | 0.133243 | 0.266682 | 543.2694 | 505.1249 | 459.5992 | 477.0308 | 610.4197 | 502.8183 | 534.2162 | 557.9342 |
| ENSMUSG00000033998 | -0.251993995 | 0.087251 | 0.20071 | 1152.938 | 1198.622 | 1477.342 | 1201.635 | 1262.769 | 1279.393 | 1794.542 | 1655.314 |
| ENSMUSG00000034006 | -0.146031148 | 0.251364 | 0.415004 | 549.3057 | 490.727 | 627.7048 | 619.3637 | 630.897 | 534.1048 | 702.9668 | 665.6057 |
| ENSMUSG00000034007 | 0.14587097 | 0.167287 | 0.313224 | 730.3955 | 797.8813 | 744.4679 | 735.8178 | 630.897 | 789.9835 | 694.2882 | 603.6131 |
| ENSMUSG00000034021 | 0.328630971 | 0.002353 | 0.014735 | 2960.818 | 3280.312 | 2750.97 | 2984.677 | 2211.552 | 2844.834 | 2189.901 | 2290.467 |
| ENSMUSG00000034022 | -0.019461722 | 0.783951 | 0.8671 | 2107.684 | 2361.249 | 2226.779 | 2195.377 | 2448.504 | 2179.997 | 2238.115 | 2140.379 |
| ENSMUSG00000034023 | 0.13724412 | 0.233535 | 0.394086 | 422.5428 | 388.7422 | 371.8199 | 341.5989 | 343.2392 | 388.8462 | 324.0012 | 329.5401 |
| ENSMUSG00000034024 | -0.021169339 | 0.696503 | 0.808418 | 9976.035 | 9798.942 | 9793.189 | 10447.23 | 10661.87 | 10017.26 | 10331.39 | 9595.816 |
| ENSMUSG00000034031 | 0.020265019 | 0.937065 | 0.962676 | 96.58122 | 49.19268 | 72.87338 | 75.04824 | 89.71024 | 64.8077 | 70.39311 | 65.25547 |
| ENSMUSG00000034032 | -0.050490062 | 0.662317 | 0.782719 | 767.6195 | 945.4594 | 794.1543 | 835.8822 | 947.8082 | 740.819 | 829.2887 | 940.7663 |
| ENSMUSG00000034035 | 0.185344496 | 0.309602 | 0.478081 | 105.6357 | 117.5825 | 112.6225 | 116.4542 | 93.61069 | 115.0895 | 78.10742 | 112.0219 |
| ENSMUSG00000034037 | 0.184553486 | 0.038107 | 0.11269 | 995.9938 | 941.8599 | 1024.368 | 1015.308 | 777.1637 | 859.2607 | 967.182 | 898.3502 |
| ENSMUSG00000034040 | 0.861164645 | 2.56E-05 | 0.000474 | 333.004 | 458.3318 | 433.0998 | 385.5927 | 186.2463 | 311.7474 | 178.3935 | 210.9927 |
| ENSMUSG00000034041 | -0.280870608 | 0.215674 | 0.37387 | 98.59333 | 86.38715 | 137.4657 | 88.85022 | 105.312 | 107.2679 | 139.8219 | 149 |
| ENSMUSG00000034042 | 0.136038545 | 0.169793 | 0.31628 | 2683.147 | 2570.018 | 2292.199 | 2555.091 | 2236.905 | 2671.641 | 2196.651 | 2086 |
| ENSMUSG00000034055 | 0.326938376 | 0.014614 | 0.056566 | 587.5357 | 722.2926 | 584.6433 | 674.5716 | 532.4108 | 597.7951 | 428.1444 | 489.416 |
| ENSMUSG00000034059 | 0.635563903 | 4.61E-05 | 0.000747 | 347.0888 | 334.7502 | 245.1196 | 319.1707 | 205.7485 | 226.8269 | 178.3935 | 190.3284 |
| ENSMUSG00000034063 | 0.168319353 | 0.698152 | 0.809552 | 25.15136 | 31.19536 | 21.53077 | 23.29083 | 36.07912 | 26.81698 | 12.53576 | 14.13868 |
| ENSMUSG00000034064 | 0.061219576 | 0.458937 | 0.620478 | 1441.676 | 1685.749 | 1623.089 | 1615.694 | 1575.78 | 1540.859 | 1602.649 | 1380.153 |
| ENSMUSG00000034066 | -0.695622376 | 0.000258 | 0.002771 | 164.9929 | 129.5807 | 177.2148 | 176.8378 | 291.5583 | 192.1883 | 260.3581 | 308.8759 |
| ENSMUSG00000034071 | -0.293755684 | 0.286881 | 0.453972 | 82.49646 | 51.59233 | 67.07664 | 55.2079 | 96.53602 | 50.28183 | 87.75031 | 79.39415 |

| | | | | | | | | | | | |
|---|---|---|---|---|---|---|---|---|---|---|---|
| ENSMUSG00000034075 | -0.302449826 | 0.00126 | 0.009306 | 1821.964 | 1779.335 | 1905.473 | 1700.231 | 2164.747 | 1917.414 | 2397.223 | 2407.927 |
| ENSMUSG00000034083 | 0.095806683 | 0.390916 | 0.558288 | 1018.127 | 975.4549 | 860.4028 | 993.7423 | 946.8331 | 1024.632 | 850.503 | 777.6276 |
| ENSMUSG00000034088 | -0.314347088 | 0.008346 | 0.037423 | 8016.241 | 7545.678 | 8705.885 | 7406.485 | 10063.15 | 7864.079 | 10107.68 | 11351.19 |
| ENSMUSG00000034098 | 0.752344133 | 0.050641 | 0.137247 | 21.12714 | 17.99732 | 19.04645 | 25.01608 | 11.70134 | 13.40849 | 12.53576 | 11.9635 |
| ENSMUSG00000034101 | 0.310936926 | 0.000464 | 0.00433 | 1639.869 | 1577.765 | 1655.385 | 1658.825 | 1209.138 | 1522.981 | 1270.933 | 1265.956 |
| ENSMUSG00000034105 | -0.075074846 | 0.416001 | 0.581855 | 448.7002 | 477.529 | 475.3332 | 493.4206 | 507.0579 | 462.5929 | 546.752 | 478.5401 |
| ENSMUSG00000034107 | 0.506277755 | 0.190474 | 0.343064 | 20.12109 | 15.59768 | 27.32752 | 19.84034 | 15.60178 | 16.76061 | 13.50005 | 13.05109 |
| ENSMUSG00000034108 | -0.296586735 | 0.025786 | 0.084832 | 1006.054 | 785.8831 | 992.0717 | 928.1829 | 1090.174 | 957.5896 | 1156.183 | 1358.401 |
| ENSMUSG00000034109 | -0.013752438 | 0.845285 | 0.906852 | 2765.643 | 2923.965 | 2653.254 | 2783.686 | 2552.841 | 2962.159 | 2803.189 | 2913.657 |
| ENSMUSG00000034110 | -0.175387699 | 0.357926 | 0.526282 | 144.8718 | 153.5772 | 122.5598 | 105.2401 | 146.2667 | 126.2633 | 179.3578 | 139.2117 |
| ENSMUSG00000034111 | 0.283619074 | 0.024876 | 0.082752 | 562.3844 | 471.5299 | 404.116 | 439.938 | 370.5423 | 392.1983 | 405.0014 | 373.0437 |
| ENSMUSG00000034112 | -0.114561673 | 0.585548 | 0.725829 | 130.7871 | 152.3773 | 201.2299 | 138.8824 | 182.3458 | 125.1459 | 196.715 | 170.7518 |
| ENSMUSG00000034116 | 0.445571193 | 0.101896 | 0.223827 | 62.37537 | 49.19268 | 71.21717 | 50.89479 | 46.80534 | 29.05173 | 51.10733 | 44.59123 |
| ENSMUSG00000034118 | -0.005399082 | 0.962398 | 0.978526 | 1119.738 | 1360.598 | 1608.183 | 1343.105 | 1394.409 | 1303.976 | 1371.219 | 1384.503 |
| ENSMUSG00000034120 | -0.028905835 | 0.623848 | 0.754981 | 8317.051 | 8827.087 | 7861.216 | 8741.826 | 8385.957 | 8813.847 | 8359.423 | 8868.218 |
| ENSMUSG00000034121 | 0.157293387 | 0.07564 | 0.181937 | 523.1483 | 577.1142 | 544.8942 | 578.8204 | 514.8588 | 531.8701 | 486.0017 | 461.1386 |
| ENSMUSG00000034126 | -0.070527367 | 0.39966 | 0.567034 | 1064.406 | 1031.847 | 1006.978 | 941.1222 | 994.6135 | 1006.754 | 1079.04 | 1164.81 |
| ENSMUSG00000034127 | -0.293022226 | 0.295092 | 0.462839 | 951.7274 | 1004.251 | 2256.591 | 1074.829 | 1573.83 | 1184.417 | 1775.256 | 1946.788 |
| ENSMUSG00000034135 | -0.096978836 | 0.273074 | 0.438705 | 1419.543 | 1166.227 | 1418.547 | 1268.057 | 1371.006 | 1352.023 | 1437.755 | 1482.387 |
| ENSMUSG00000034145 | -0.377900792 | 0.524102 | 0.677274 | 11.0666 | 5.999108 | 5.796746 | 14.6646 | 4.875557 | 15.64324 | 13.50005 | 15.22628 |
| ENSMUSG00000034152 | -0.242502462 | 9.80E-05 | 0.001316 | 1601.639 | 1546.57 | 1621.433 | 1560.486 | 1891.716 | 1741.986 | 1883.257 | 1972.89 |
| ENSMUSG00000034154 | -0.013448553 | 0.842387 | 0.904953 | 1232.417 | 1143.43 | 1136.162 | 1122.273 | 1129.179 | 1239.168 | 1161.004 | 1148.496 |
| ENSMUSG00000034156 | -0.27522177 | 0.120812 | 0.250023 | 225.3562 | 148.7779 | 167.2775 | 194.0903 | 229.1512 | 229.0617 | 250.7152 | 181.6277 |
| ENSMUSG00000034157 | -0.176551896 | 0.032849 | 0.100924 | 707.2562 | 740.2899 | 725.4214 | 736.6805 | 824.9442 | 731.88 | 853.3959 | 876.5984 |
| ENSMUSG00000034158 | 0.277560143 | 0.031173 | 0.097037 | 3280.743 | 3051.146 | 2953.028 | 3435.829 | 2357.819 | 3320.836 | 2477.259 | 2340.496 |
| ENSMUSG00000034159 | -2.33314632 | 2.33E-05 | 0.000442 | 11.0666 | 5.999108 | 20.70267 | 3.450494 | 67.28268 | 29.05173 | 46.28588 | 67.43065 |
| ENSMUSG00000034160 | 0.486165285 | 8.52E-05 | 0.00119 | 10735.61 | 10414.45 | 9247.467 | 11346.95 | 7512.258 | 9078.665 | 6847.417 | 6364.583 |
| ENSMUSG00000034161 | -0.154707227 | 0.305328 | 0.47378 | 606.6508 | 765.4861 | 564.7687 | 660.7696 | 904.9033 | 683.8329 | 657.6452 | 641.6787 |
| ENSMUSG00000034163 | 0.330466302 | 0.002607 | 0.015903 | 2315.937 | 1994.103 | 1853.303 | 2101.351 | 1581.631 | 1921.883 | 1559.256 | 1510.664 |
| ENSMUSG00000034164 | -0.206556423 | 0.143334 | 0.280975 | 1737.456 | 1583.764 | 2325.323 | 1721.796 | 2065.286 | 1859.31 | 2154.222 | 2427.503 |
| ENSMUSG00000034165 | -0.075522004 | 0.433081 | 0.597256 | 2909.509 | 3169.928 | 3324.848 | 3335.765 | 3084.277 | 3042.61 | 3496.513 | 3803.306 |
| ENSMUSG00000034168 | 0.187146206 | 0.039779 | 0.116162 | 2295.816 | 2188.474 | 2627.582 | 2268.7 | 1868.313 | 2011.273 | 2123.365 | 2240.438 |
| ENSMUSG00000034171 | -0.35664819 | 0.045239 | 0.126946 | 449.7063 | 364.7457 | 485.2705 | 359.714 | 581.1663 | 410.0763 | 481.1803 | 653.6422 |
| ENSMUSG00000034173 | 0.060417472 | 0.792788 | 0.873221 | 419.5247 | 443.934 | 289.8373 | 416.6471 | 449.5263 | 457.006 | 361.6084 | 236.0073 |
| ENSMUSG00000034175 | -0.06541821 | 0.414381 | 0.580277 | 506.0453 | 505.1249 | 563.1125 | 544.3154 | 558.7388 | 545.2786 | 554.4663 | 562.2846 |
| ENSMUSG00000034177 | 0.387110042 | 8.56E-05 | 0.001194 | 565.4025 | 521.9224 | 588.7838 | 484.7944 | 405.6463 | 429.0717 | 421.3944 | 396.9707 |
| ENSMUSG00000034187 | -0.23981482 | 0.084549 | 0.196221 | 996.9999 | 807.4799 | 871.9963 | 775.4985 | 927.3309 | 883.8429 | 1012.504 | 1252.905 |
| ENSMUSG00000034189 | -0.067956484 | 0.361677 | 0.530157 | 779.6921 | 842.2747 | 751.9208 | 856.5851 | 855.1726 | 852.5564 | 839.8959 | 836.3575 |
| ENSMUSG00000034190 | -0.012223827 | 0.877373 | 0.92652 | 1726.389 | 1878.921 | 1690.994 | 1882.244 | 1863.438 | 1626.897 | 1844.685 | 1901.109 |
| ENSMUSG00000034192 | -0.123662745 | 0.139316 | 0.275255 | 1230.404 | 1493.778 | 1470.717 | 1324.99 | 1466.567 | 1558.737 | 1514.898 | 1472.598 |
| ENSMUSG00000034194 | -0.285567074 | 1.65E-06 | 5.60E-05 | 1332.016 | 1318.604 | 1308.408 | 1408.664 | 1708.395 | 1624.662 | 1585.291 | 1625.949 |
| ENSMUSG00000034201 | -0.010810471 | 0.920772 | 0.952725 | 810.8798 | 711.4942 | 845.4969 | 823.8054 | 802.5166 | 710.6499 | 779.1456 | 926.6276 |
| ENSMUSG00000034203 | -0.635599902 | 0.000699 | 0.005942 | 592.566 | 512.3238 | 489.411 | 513.261 | 814.2179 | 545.2786 | 870.7531 | 1043 |
| ENSMUSG00000034205 | 0.036175142 | 0.81428 | 0.887216 | 1602.645 | 1702.547 | 2266.528 | 1907.261 | 1411.961 | 2102.898 | 1854.328 | 1928.299 |
| ENSMUSG00000034206 | 0.294892898 | 0.089828 | 0.204901 | 302.8224 | 260.3613 | 299.7746 | 309.6818 | 235.0018 | 312.8647 | 224.6794 | 184.8905 |
| ENSMUSG00000034209 | 0.152901684 | 0.498717 | 0.655065 | 51.30877 | 53.99197 | 62.9361 | 65.55939 | 56.55646 | 56.98608 | 54.96448 | 42.41605 |
| ENSMUSG00000034210 | 0.045219036 | 0.534233 | 0.685327 | 1094.587 | 1069.041 | 997.8685 | 1012.72 | 999.4891 | 1077.149 | 1014.432 | 951.6422 |
| ENSMUSG00000034211 | -0.164987075 | 0.011379 | 0.047004 | 1500.027 | 1517.774 | 1515.435 | 1620.87 | 1787.379 | 1663.77 | 1642.184 | 1808.664 |
| ENSMUSG00000034212 | -0.238096346 | 0.588006 | 0.728241 | 13.07871 | 8.398751 | 16.56213 | 12.93935 | 12.67645 | 17.87799 | 18.32149 | 11.9635 |
| ENSMUSG00000034216 | -0.09481412 | 0.36029 | 0.528788 | 686.1291 | 669.5004 | 672.4226 | 546.9033 | 630.897 | 699.4762 | 708.7525 | 709.1094 |
| ENSMUSG00000034218 | -0.027415081 | 0.820873 | 0.89144 | 590.5539 | 615.5084 | 542.4098 | 464.0914 | 602.6188 | 496.1141 | 578.5735 | 573.1605 |
| ENSMUSG00000034220 | -0.251468194 | 0.00218 | 0.013975 | 1844.098 | 1893.318 | 2088.485 | 1795.119 | 2142.32 | 2131.95 | 2363.473 | 2436.204 |
| ENSMUSG00000034226 | -0.717556951 | 0.082124 | 0.192468 | 29.17558 | 16.7975 | 43.88965 | 29.3292 | 50.70579 | 25.6996 | 48.21446 | 72.8686 |
| ENSMUSG00000034227 | 0.137407956 | 0.506289 | 0.66209 | 133.8052 | 117.5825 | 165.6213 | 119.042 | 92.63557 | 124.0285 | 123.429 | 149 |
| ENSMUSG00000034235 | 0.110733523 | 0.137838 | 0.273093 | 871.2431 | 907.0651 | 833.0753 | 898.8537 | 785.9397 | 765.4013 | 849.5388 | 847.2335 |
| ENSMUSG00000034243 | 0.146882197 | 0.084344 | 0.195879 | 3361.228 | 2964.759 | 2946.403 | 2906.179 | 2631.825 | 3078.366 | 2642.152 | 2648.284 |
| ENSMUSG00000034245 | -0.017429369 | 0.864359 | 0.91877 | 1155.956 | 1181.824 | 1235.535 | 1057.576 | 1330.052 | 1038.041 | 1209.219 | 1107.168 |
| ENSMUSG00000034247 | 0.083989398 | 0.424242 | 0.589372 | 549.3057 | 652.7029 | 588.7838 | 548.6285 | 586.0419 | 499.4662 | 522.6447 | 595.9999 |
| ENSMUSG00000034248 | 0.290667316 | 0.028982 | 0.092256 | 1286.744 | 916.6637 | 907.6049 | 1043.774 | 911.7291 | 889.4298 | 781.0742 | 814.6057 |
| ENSMUSG00000034252 | 0.139767339 | 0.072873 | 0.177504 | 4657.026 | 4638.51 | 4284.624 | 4808.263 | 3862.416 | 4640.455 | 4149.336 | 4039.313 |
| ENSMUSG00000034254 | -0.24455027 | 0.016826 | 0.062382 | 1365.216 | 1391.793 | 1503.014 | 1428.505 | 1605.033 | 1448.117 | 1755.971 | 1931.562 |
| ENSMUSG00000034255 | -0.367838595 | 0.057869 | 0.15083 | 598.6023 | 392.3416 | 657.5167 | 533.1013 | 666.001 | 535.2222 | 787.8242 | 828.7444 |
| ENSMUSG00000034258 | -1.103023951 | 5.73E-05 | 0.000878 | 108.6539 | 95.98572 | 173.0743 | 107.8279 | 283.7574 | 159.7845 | 268.0724 | 333.8905 |
| ENSMUSG00000034259 | -0.284330427 | 0.004743 | 0.024772 | 620.7355 | 730.6913 | 776.764 | 760.8339 | 853.2224 | 819.0352 | 945.0034 | 902.7006 |

| | | | | | | | | | | | |
|---|---|---|---|---|---|---|---|---|---|---|---|
| ENSMUSG00000034263 | -0.267409253 | 0.000592 | 0.005218 | 1341.07 | 1243.015 | 1355.611 | 1406.076 | 1687.918 | 1477.169 | 1567.934 | 1705.343 |
| ENSMUSG00000034265 | -0.444138818 | 0.11568 | 0.243665 | 187.1261 | 99.58519 | 203.7142 | 157.8601 | 168.6943 | 162.0192 | 257.4652 | 295.8248 |
| ENSMUSG00000034266 | 0.306355154 | 0.124553 | 0.25468 | 83.50251 | 125.9813 | 94.40416 | 101.7896 | 78.98402 | 71.51194 | 84.85745 | 91.35765 |
| ENSMUSG00000034269 | 0.207507916 | 0.122866 | 0.252404 | 5535.311 | 4916.869 | 4128.94 | 4768.583 | 3493.824 | 4923.15 | 4507.088 | 3832.671 |
| ENSMUSG00000034271 | 0.034831878 | 0.834178 | 0.899643 | 431.5973 | 470.33 | 447.1776 | 540.0023 | 436.8499 | 382.1419 | 607.5022 | 416.5474 |
| ENSMUSG00000034274 | -0.21041343 | 0.041578 | 0.11992 | 886.3339 | 903.4656 | 929.1356 | 849.6841 | 1024.842 | 893.8993 | 1012.504 | 1197.438 |
| ENSMUSG00000034275 | 0.81562406 | 0.006073 | 0.029723 | 374.2522 | 280.7582 | 242.6352 | 286.391 | 102.3867 | 279.3435 | 151.3934 | 140.2993 |
| ENSMUSG00000034278 | 0.033912886 | 0.739928 | 0.837336 | 433.6094 | 521.9224 | 506.8013 | 489.1075 | 435.8748 | 515.1095 | 467.6802 | 488.3284 |
| ENSMUSG00000034282 | -0.863123968 | 1.27E-05 | 0.000276 | 312.8829 | 319.1525 | 428.1311 | 301.0556 | 705.9806 | 417.8979 | 661.5024 | 691.7079 |
| ENSMUSG00000034285 | -0.128274176 | 0.050645 | 0.137247 | 1791.783 | 1997.703 | 1960.128 | 1948.666 | 2103.315 | 1998.982 | 2083.829 | 2227.387 |
| ENSMUSG00000034290 | 0.011427732 | 0.848943 | 0.909191 | 3721.395 | 3727.846 | 3666.856 | 3811.933 | 3964.803 | 3479.503 | 3780.978 | 3581.437 |
| ENSMUSG00000034292 | -0.172838354 | 0.199691 | 0.354614 | 484.9182 | 485.9277 | 439.7246 | 482.2065 | 571.4152 | 426.8369 | 504.3232 | 629.7152 |
| ENSMUSG00000034295 | -0.962307418 | 0.000242 | 0.002641 | 885.3278 | 583.1133 | 1311.721 | 872.975 | 1632.336 | 1155.365 | 2058.757 | 2275.241 |
| ENSMUSG00000034297 | 0.278256181 | 0.013272 | 0.052815 | 2328.01 | 2344.451 | 2206.076 | 2511.96 | 1684.992 | 2334.194 | 1809.006 | 1917.423 |
| ENSMUSG00000034300 | 0.195558291 | 0.025676 | 0.084619 | 1464.815 | 1551.369 | 1297.643 | 1552.722 | 1233.516 | 1336.379 | 1204.397 | 1347.525 |
| ENSMUSG00000034303 | 0.42887087 | 0.082202 | 0.192602 | 155.9384 | 178.7734 | 122.5598 | 152.6844 | 104.3369 | 162.0192 | 78.10742 | 108.7591 |
| ENSMUSG00000034308 | -0.312603908 | 0.149734 | 0.289644 | 128.775 | 98.38537 | 149.0592 | 106.1027 | 182.3458 | 110.62 | 151.3934 | 155.5255 |
| ENSMUSG00000034310 | 0.954350515 | 0.002475 | 0.015325 | 36.21796 | 51.59233 | 44.71776 | 43.13117 | 17.552 | 32.40385 | 19.28578 | 21.75182 |
| ENSMUSG00000034311 | -0.040680631 | 0.648345 | 0.77224 | 1003.036 | 1113.434 | 1036.79 | 1042.049 | 1074.573 | 1219.055 | 1036.611 | 984.2699 |
| ENSMUSG00000034312 | -0.144030577 | 0.213666 | 0.371775 | 913.4974 | 905.8653 | 1109.663 | 900.5789 | 1026.792 | 977.7023 | 997.075 | 1233.328 |
| ENSMUSG00000034317 | 0.36467142 | 0.004671 | 0.024538 | 1775.686 | 1752.939 | 1618.948 | 1928.826 | 1406.111 | 1688.352 | 1171.611 | 1231.153 |
| ENSMUSG00000034320 | -0.247517663 | 0.112158 | 0.238643 | 663.9959 | 574.7145 | 857.0904 | 548.6285 | 845.4215 | 712.8847 | 829.2887 | 753.7006 |
| ENSMUSG00000034321 | -0.044852726 | 0.619276 | 0.751581 | 581.4994 | 658.702 | 688.1566 | 677.1594 | 695.2544 | 706.1804 | 659.5738 | 628.6276 |
| ENSMUSG00000034324 | 0.016867905 | 0.966482 | 0.981263 | 136.8234 | 94.7859 | 169.7619 | 124.2178 | 74.10846 | 253.6439 | 115.7147 | 77.21897 |
| ENSMUSG00000034327 | 0.022273538 | 0.838088 | 0.902068 | 727.3773 | 687.4977 | 599.5492 | 721.1532 | 701.105 | 588.8561 | 664.3952 | 737.3868 |
| ENSMUSG00000034329 | 0.069299802 | 0.532387 | 0.683958 | 249.5015 | 289.157 | 264.9941 | 290.7041 | 247.6783 | 279.3435 | 258.4295 | 257.7591 |
| ENSMUSG00000034330 | -0.710936053 | 0.050932 | 0.137866 | 174.0474 | 141.5789 | 347.8048 | 142.3329 | 285.7076 | 177.6625 | 389.5728 | 467.6642 |
| ENSMUSG00000034333 | -0.432016879 | 0.000467 | 0.004346 | 375.2583 | 307.1543 | 319.6492 | 291.5667 | 398.8205 | 413.4284 | 445.5016 | 487.2408 |
| ENSMUSG00000034334 | 0.32008995 | 0.125297 | 0.255863 | 83.50251 | 104.3845 | 92.74794 | 112.1411 | 70.20801 | 91.62468 | 89.67889 | 63.08028 |
| ENSMUSG00000034336 | 1.61884036 | 0.010284 | 0.043496 | 163.9869 | 31.19536 | 24.8432 | 215.6559 | 41.92979 | 37.99072 | 33.75012 | 28.27737 |
| ENSMUSG00000034341 | -0.371415233 | 0.007656 | 0.035035 | 1486.948 | 1181.824 | 1481.483 | 1382.785 | 1804.931 | 1416.83 | 1793.578 | 2144.73 |
| ENSMUSG00000034342 | 0.237656457 | 0.002947 | 0.017545 | 1465.821 | 1458.983 | 1430.968 | 1424.191 | 1152.582 | 1358.727 | 1127.254 | 1265.956 |
| ENSMUSG00000034343 | -0.337690315 | 0.001448 | 0.010317 | 662.9898 | 598.7109 | 670.7664 | 645.2424 | 806.4171 | 731.88 | 770.467 | 951.6422 |
| ENSMUSG00000034345 | -0.126212035 | 0.043722 | 0.124058 | 1623.772 | 1709.746 | 1633.026 | 1716.621 | 1818.583 | 1763.216 | 1759.828 | 1952.226 |
| ENSMUSG00000034349 | 0.282633029 | 0.111303 | 0.23755 | 6001.114 | 6150.285 | 4489.994 | 5308.585 | 3924.823 | 6239.417 | 4069.3 | 3810.919 |
| ENSMUSG00000034353 | -0.065157465 | 0.786861 | 0.868975 | 73.44197 | 97.18554 | 110.9663 | 125.943 | 89.71024 | 128.498 | 123.429 | 85.9197 |
| ENSMUSG00000034354 | 0.023642593 | 0.710317 | 0.818711 | 2476.906 | 2420.04 | 2523.241 | 2252.31 | 2251.532 | 2453.754 | 2431.937 | 2378.562 |
| ENSMUSG00000034361 | 0.308223287 | 0.005861 | 0.028931 | 370.228 | 382.7431 | 463.7397 | 409.7462 | 341.289 | 307.2779 | 339.4298 | 327.3649 |
| ENSMUSG00000034371 | -0.900442851 | 0.000125 | 0.001587 | 499.003 | 542.3193 | 595.4087 | 571.0568 | 1034.593 | 602.2646 | 988.3964 | 1496.525 |
| ENSMUSG00000034377 | 0.154012036 | 0.17505 | 0.323397 | 1508.075 | 1430.187 | 1545.247 | 1527.706 | 1240.342 | 1659.301 | 1283.469 | 1222.452 |
| ENSMUSG00000034379 | -0.092991642 | 0.521529 | 0.674846 | 251.5136 | 277.1588 | 251.7444 | 224.2821 | 294.4836 | 274.874 | 287.3582 | 212.0803 |
| ENSMUSG00000034382 | -0.260911069 | 0.283798 | 0.450329 | 242.4591 | 292.7565 | 438.8965 | 251.8861 | 392.9699 | 256.996 | 356.787 | 463.3138 |
| ENSMUSG00000034387 | -0.319478067 | 0.267621 | 0.431862 | 43.26034 | 58.79126 | 75.3577 | 56.07053 | 52.65601 | 59.22083 | 87.75031 | 92.44524 |
| ENSMUSG00000034390 | -0.015073442 | 0.902038 | 0.94115 | 999.012 | 703.0954 | 867.0276 | 846.2336 | 809.3424 | 806.7441 | 923.789 | 913.5765 |
| ENSMUSG00000034394 | -0.109426227 | 0.763406 | 0.852922 | 16.09687 | 23.99643 | 20.70267 | 32.77969 | 28.27823 | 30.1691 | 22.17865 | 20.66423 |
| ENSMUSG00000034401 | -0.126653107 | 0.247946 | 0.411297 | 735.4257 | 644.3042 | 796.6386 | 796.2015 | 776.1886 | 785.514 | 925.7176 | 761.3138 |
| ENSMUSG00000034402 | 0.468130058 | 0.127653 | 0.259067 | 41.24823 | 38.39429 | 32.29616 | 43.9938 | 23.40267 | 36.87335 | 31.82154 | 20.66423 |
| ENSMUSG00000034403 | -0.117218537 | 0.23676 | 0.397964 | 3536.281 | 3576.668 | 3872.227 | 3589.376 | 3860.466 | 3371.118 | 4033.622 | 4543.956 |
| ENSMUSG00000034412 | -0.4733531 | 0.004188 | 0.02259 | 523.1483 | 536.3202 | 523.3634 | 521.0246 | 740.1095 | 506.1705 | 784.9314 | 888.5619 |
| ENSMUSG00000034413 | -0.018650732 | 0.890011 | 0.933475 | 522.1422 | 419.9375 | 608.6584 | 496.0085 | 492.4312 | 486.0577 | 515.8947 | 582.9488 |
| ENSMUSG00000034422 | 0.295608779 | 0.240665 | 0.402912 | 282.7013 | 351.5477 | 434.756 | 204.4418 | 260.3547 | 191.071 | 321.1083 | 264.2846 |
| ENSMUSG00000034424 | -0.593745651 | 1.74E-06 | 5.83E-05 | 1472.864 | 1392.993 | 1490.592 | 1424.191 | 2055.535 | 1769.921 | 2256.437 | 2642.846 |
| ENSMUSG00000034427 | -1.768137699 | 0.000179 | 0.002078 | 122.7386 | 33.595 | 188.8083 | 90.57547 | 328.6125 | 179.8972 | 361.6084 | 617.7517 |
| ENSMUSG00000034429 | -0.093826302 | 0.382838 | 0.550842 | 323.9495 | 358.7466 | 376.7885 | 303.6435 | 359.8161 | 373.2029 | 352.9298 | 368.6934 |
| ENSMUSG00000034430 | -0.218747873 | 0.004795 | 0.024968 | 587.5357 | 556.7172 | 548.2066 | 546.0407 | 663.0757 | 607.8515 | 670.181 | 661.2554 |
| ENSMUSG00000034432 | -0.076592774 | 0.171478 | 0.318725 | 2037.26 | 2164.478 | 2182.061 | 2264.387 | 2296.387 | 2239.218 | 2257.401 | 2329.62 |
| ENSMUSG00000034435 | -0.395931164 | 0.059527 | 0.153912 | 465.8032 | 585.5129 | 910.0892 | 531.3761 | 723.5326 | 740.819 | 889.0746 | 928.8028 |
| ENSMUSG00000034437 | 0.078750246 | 0.73778 | 0.835931 | 49.29666 | 63.59054 | 53.82693 | 56.07053 | 45.83023 | 43.57759 | 57.85735 | 63.08028 |
| ENSMUSG00000034442 | -0.091350324 | 0.374372 | 0.543108 | 355.1372 | 358.7466 | 332.0708 | 346.7746 | 410.5219 | 337.447 | 378.0014 | 354.5547 |
| ENSMUSG00000034445 | -0.332713973 | 0.054666 | 0.145231 | 210.2654 | 172.7743 | 183.0116 | 195.8155 | 246.7032 | 181.0146 | 236.2508 | 295.8248 |
| ENSMUSG00000034449 | -0.31476243 | 0.127681 | 0.259096 | 211.2714 | 123.5816 | 182.1835 | 166.4863 | 204.7734 | 175.4277 | 205.3936 | 266.4598 |
| ENSMUSG00000034450 | 0.346588756 | 0.166483 | 0.312079 | 210.2654 | 249.5629 | 212.8234 | 241.5346 | 183.3209 | 229.0617 | 209.2507 | 96.79561 |
| ENSMUSG00000034452 | 0.069482526 | 0.611411 | 0.746062 | 277.671 | 320.3523 | 305.5713 | 320.0333 | 305.2098 | 324.0385 | 232.3937 | 305.6131 |
| ENSMUSG00000034453 | 0.013141919 | 0.879895 | 0.927943 | 1146.902 | 1181.824 | 1035.133 | 1098.12 | 1134.054 | 977.7023 | 1167.754 | 1136.533 |

| | | | | | | | | | | | |
|---|---|---|---|---|---|---|---|---|---|---|---|
| ENSMUSG00000034457 | 0.974534493 | 0.001995 | 0.013106 | 117.7084 | 129.5807 | 85.29498 | 158.7227 | 67.28268 | 92.74205 | 52.07161 | 38.06569 |
| ENSMUSG00000034459 | 0.641359609 | 0.324525 | 0.492495 | 42.25428 | 145.1784 | 173.9024 | 14.6646 | 42.9049 | 39.10809 | 102.2147 | 56.55474 |
| ENSMUSG00000034460 | 0.08699219 | 0.55917 | 0.705403 | 345.0766 | 333.5504 | 305.5713 | 289.8415 | 297.4089 | 374.3203 | 263.2509 | 264.2846 |
| ENSMUSG00000034462 | 0.148591824 | 0.007321 | 0.033955 | 3900.473 | 3737.444 | 3552.577 | 3934.426 | 3327.08 | 3497.381 | 3408.762 | 3411.773 |
| ENSMUSG00000034463 | 0.430222169 | 0.002329 | 0.014615 | 1269.641 | 1524.973 | 1370.516 | 1381.06 | 1054.095 | 1297.271 | 903.5389 | 861.3721 |
| ENSMUSG00000034471 | -0.29640543 | 0.073157 | 0.177961 | 890.3581 | 783.4835 | 1011.118 | 826.3933 | 1087.249 | 783.2792 | 1111.825 | 1331.211 |
| ENSMUSG00000034472 | 0.227927278 | 0.319089 | 0.487145 | 149.9021 | 148.7779 | 192.9488 | 154.4096 | 116.0382 | 200.01 | 127.2862 | 109.8467 |
| ENSMUSG00000034473 | -0.08569848 | 0.245181 | 0.408105 | 821.9464 | 866.2711 | 867.0276 | 791.0257 | 895.1522 | 830.209 | 897.7532 | 925.54 |
| ENSMUSG00000034480 | 0.277905972 | 0.015735 | 0.059716 | 425.561 | 447.5334 | 431.4436 | 403.7078 | 318.8614 | 407.8415 | 364.5013 | 317.5766 |
| ENSMUSG00000034484 | 0.147507971 | 0.008671 | 0.038451 | 3416.561 | 3811.833 | 3656.091 | 3564.36 | 3351.458 | 3158.817 | 3304.619 | 3224.708 |
| ENSMUSG00000034485 | -0.069496512 | 0.300126 | 0.468307 | 2382.337 | 2284.46 | 2476.867 | 2531.8 | 2528.464 | 2429.171 | 2463.759 | 2737.467 |
| ENSMUSG00000034487 | 0.475323209 | 0.000183 | 0.002122 | 1838.061 | 2344.451 | 1816.038 | 2168.635 | 1447.065 | 1738.634 | 1319.148 | 1369.277 |
| ENSMUSG00000034488 | 0.082821375 | 0.679022 | 0.794865 | 91.55095 | 75.58876 | 102.6852 | 100.9269 | 77.03379 | 107.2679 | 79.07171 | 89.18247 |
| ENSMUSG00000034501 | 0.342868576 | 0.000164 | 0.001941 | 397.3915 | 393.5415 | 399.1474 | 408.0209 | 311.0605 | 317.3342 | 334.6083 | 296.9124 |
| ENSMUSG00000034509 | -0.1171807 | 0.271298 | 0.436499 | 379.2825 | 329.9509 | 356.0859 | 351.9504 | 396.8703 | 335.2122 | 406.93 | 398.0583 |
| ENSMUSG00000034518 | 0.052974306 | 0.540134 | 0.689686 | 1476.888 | 1499.777 | 1293.503 | 1275.82 | 1391.484 | 1387.779 | 1252.612 | 1309.46 |
| ENSMUSG00000034520 | 0.186362949 | 0.084377 | 0.195931 | 1933.636 | 2121.284 | 1902.161 | 2241.958 | 1923.895 | 2026.917 | 1698.113 | 1556.343 |
| ENSMUSG00000034522 | 0.318034603 | 2.35E-06 | 7.34E-05 | 1630.814 | 1606.561 | 1703.415 | 1550.134 | 1288.122 | 1333.027 | 1367.362 | 1218.102 |
| ENSMUSG00000034525 | 0.077467879 | 0.254545 | 0.418333 | 2752.565 | 2506.427 | 2481.836 | 2649.117 | 2589.896 | 2555.435 | 2335.508 | 2366.598 |
| ENSMUSG00000034532 | -0.032632722 | 0.855974 | 0.913721 | 182.0958 | 199.1704 | 162.3089 | 199.266 | 196.9725 | 221.2401 | 203.465 | 137.0365 |
| ENSMUSG00000034538 | 0.197254613 | 0.530353 | 0.682199 | 65.39353 | 82.78769 | 37.2648 | 46.58167 | 56.55646 | 41.34284 | 57.85735 | 44.59123 |
| ENSMUSG00000034543 | 0.203981532 | 0.028568 | 0.091332 | 2150.944 | 2415.241 | 2077.72 | 2309.243 | 1939.496 | 2204.579 | 1780.078 | 1847.817 |
| ENSMUSG00000034544 | 0.042929973 | 0.57256 | 0.715181 | 1645.905 | 1694.148 | 1547.731 | 1608.793 | 1487.045 | 1747.573 | 1517.791 | 1553.08 |
| ENSMUSG00000034551 | 0.52163388 | 0.114429 | 0.241901 | 98.59333 | 122.3818 | 106.8258 | 99.2017 | 48.75557 | 129.6154 | 69.42882 | 50.02919 |
| ENSMUSG00000034555 | 0.949861704 | 0.334433 | 0.502631 | 23.13925 | 19.19714 | 2.48432 | 12.07673 | 0.975111 | 21.23011 | 1.928578 | 5.437955 |
| ENSMUSG00000034557 | 0.018396977 | 0.783624 | 0.866969 | 1261.592 | 1265.812 | 1298.471 | 1298.248 | 1268.62 | 1357.61 | 1259.362 | 1175.686 |
| ENSMUSG00000034560 | 0.177099269 | 0.030784 | 0.09628 | 1439.664 | 1274.21 | 1165.146 | 1276.683 | 1150.631 | 1125.196 | 1139.79 | 1141.971 |
| ENSMUSG00000034563 | -0.2435009 | 0.136703 | 0.271407 | 211.2714 | 205.1695 | 183.8397 | 179.4257 | 220.3752 | 177.6625 | 256.5009 | 266.4598 |
| ENSMUSG00000034566 | -0.503691098 | 0.000272 | 0.00287 | 4220.398 | 4162.181 | 5175.666 | 4375.226 | 6163.679 | 5015.892 | 6548.488 | 7701.232 |
| ENSMUSG00000034570 | -0.267044134 | 0.256082 | 0.419812 | 76.46013 | 59.99108 | 88.60741 | 81.08661 | 79.95913 | 74.86406 | 92.57176 | 122.8978 |
| ENSMUSG00000034573 | -0.307116515 | 0.024895 | 0.082785 | 3520.184 | 2817.181 | 4207.61 | 3468.609 | 3936.524 | 3871.701 | 4740.446 | 4794.102 |
| ENSMUSG00000034574 | 0.060320373 | 0.605345 | 0.741849 | 1046.297 | 986.2533 | 1040.93 | 906.6173 | 921.4802 | 1143.074 | 925.7176 | 827.6568 |
| ENSMUSG00000034575 | 0.090442214 | 0.258092 | 0.421945 | 1011.085 | 877.0695 | 900.1519 | 936.8091 | 828.8446 | 909.5425 | 913.1818 | 848.321 |
| ENSMUSG00000034579 | 0.358887557 | 0.26691 | 0.431387 | 37.22401 | 28.79572 | 40.57722 | 43.13117 | 20.47734 | 41.34284 | 29.89296 | 26.10219 |
| ENSMUSG00000034584 | 0.187171198 | 0.432106 | 0.596508 | 179.0777 | 178.7734 | 221.9326 | 144.0581 | 192.0969 | 105.0332 | 142.7148 | 195.7664 |
| ENSMUSG00000034586 | 0.091771301 | 0.680104 | 0.795588 | 235.4167 | 119.9822 | 178.871 | 213.068 | 204.7734 | 177.6625 | 157.1791 | 163.1387 |
| ENSMUSG00000034587 | -0.017938886 | 0.878208 | 0.927008 | 212.2775 | 251.9625 | 213.6515 | 223.4195 | 237.9272 | 224.5922 | 216.0008 | 231.6569 |
| ENSMUSG00000034591 | 0.32183465 | 0.123774 | 0.253453 | 74.44802 | 65.99018 | 71.21717 | 62.97152 | 54.60623 | 62.57295 | 56.89306 | 45.67883 |
| ENSMUSG00000034593 | 0.146748453 | 0.016597 | 0.061758 | 1429.603 | 1425.388 | 1422.687 | 1483.712 | 1221.814 | 1354.257 | 1328.79 | 1301.847 |
| ENSMUSG00000034595 | -0.10282167 | 0.334187 | 0.502383 | 694.1775 | 662.3015 | 829.7628 | 691.824 | 821.0437 | 692.7719 | 809.0386 | 770.0145 |
| ENSMUSG00000034601 | 0.361411913 | 0.017258 | 0.063586 | 601.6205 | 574.7145 | 432.2717 | 461.5036 | 385.169 | 471.5319 | 398.2514 | 354.5547 |
| ENSMUSG00000034602 | 0.152393545 | 0.040431 | 0.117586 | 2769.668 | 2488.43 | 2779.126 | 2773.335 | 2375.371 | 2524.148 | 2589.116 | 2241.525 |
| ENSMUSG00000034607 | -0.797719519 | 0.170547 | 0.317363 | 8.048435 | 13.19804 | 57.13936 | 23.29083 | 59.48179 | 26.81698 | 47.25017 | 44.59123 |
| ENSMUSG00000034610 | 0.243043548 | 0.018742 | 0.067399 | 2368.252 | 2462.034 | 2108.359 | 2331.671 | 1944.372 | 2247.039 | 1958.471 | 1681.416 |
| ENSMUSG00000034612 | 0.023653121 | 0.863163 | 0.917928 | 1026.175 | 752.2881 | 1128.709 | 932.496 | 882.4757 | 873.7865 | 1026.004 | 998.4086 |
| ENSMUSG00000034613 | -0.266303959 | 0.047236 | 0.130749 | 1264.61 | 980.2542 | 1296.815 | 1090.356 | 1191.586 | 1307.328 | 1446.434 | 1629.211 |
| ENSMUSG00000034614 | 0.414856511 | 0.027487 | 0.088858 | 1148.914 | 1081.039 | 968.0567 | 1168.855 | 835.6704 | 1001.167 | 915.1104 | 523.1313 |
| ENSMUSG00000034616 | -0.002552175 | 0.984896 | 0.991442 | 630.7961 | 520.7225 | 711.3436 | 684.9231 | 662.1006 | 532.9874 | 666.3238 | 693.8831 |
| ENSMUSG00000034617 | -0.071657777 | 0.55072 | 0.698537 | 493.9727 | 589.1124 | 485.2705 | 471.8551 | 541.1868 | 465.945 | 529.3948 | 603.6131 |
| ENSMUSG00000034620 | -0.205888876 | 0.003357 | 0.019229 | 1124.769 | 1168.626 | 1180.88 | 1243.04 | 1360.28 | 1443.647 | 1369.291 | 1270.306 |
| ENSMUSG00000034621 | 0.205645477 | 0.022406 | 0.076393 | 1757.577 | 1626.958 | 1539.45 | 1669.176 | 1469.493 | 1592.258 | 1374.112 | 1281.182 |
| ENSMUSG00000034634 | 2.147491294 | 0.00219 | 0.014014 | 767.6195 | 63.59054 | 261.6817 | 345.0494 | 87.76002 | 121.7938 | 52.07161 | 63.08028 |
| ENSMUSG00000034636 | 0.095400038 | 0.084249 | 0.195708 | 1715.323 | 1703.747 | 1585.824 | 1628.633 | 1551.402 | 1552.033 | 1548.648 | 1551.992 |
| ENSMUSG00000034639 | 0.093495716 | 0.567172 | 0.711413 | 228.3743 | 280.7582 | 194.6051 | 197.5408 | 199.8978 | 226.8269 | 227.5722 | 187.0657 |
| ENSMUSG00000034640 | 0.440670753 | 0.023483 | 0.079219 | 820.9404 | 824.2774 | 640.9545 | 801.3772 | 464.153 | 806.7441 | 540.9662 | 463.3138 |
| ENSMUSG00000034645 | 0.802867027 | 0.152931 | 0.293979 | 22.1332 | 9.598572 | 9.93728 | 10.35148 | 7.80089 | 4.469496 | 8.678602 | 8.700729 |
| ENSMUSG00000034647 | 0.56005885 | 0.007507 | 0.034582 | 1736.45 | 1674.951 | 1366.376 | 1463.872 | 810.3175 | 1578.85 | 993.2178 | 851.5838 |
| ENSMUSG00000034648 | 0.581260235 | 0.000735 | 0.006171 | 1272.659 | 1356.998 | 1123.741 | 1324.127 | 891.2517 | 1112.905 | 769.5028 | 619.9269 |
| ENSMUSG00000034652 | 0.468038233 | 0.094499 | 0.212176 | 91.55095 | 113.983 | 92.74794 | 90.57547 | 105.312 | 56.98608 | 46.28588 | 71.78101 |
| ENSMUSG00000034653 | 0.007337278 | 0.950599 | 0.971873 | 415.5004 | 363.5459 | 366.8512 | 422.6855 | 399.7956 | 425.7195 | 404.0372 | 331.7153 |
| ENSMUSG00000034656 | 0.545042602 | 2.50E-06 | 7.74E-05 | 429.5852 | 441.5343 | 390.0382 | 404.5704 | 246.7032 | 322.9211 | 280.6081 | 291.4744 |
| ENSMUSG00000034659 | -0.256820034 | 0.06407 | 0.162494 | 1203.241 | 1376.195 | 1496.389 | 1200.772 | 1610.884 | 1279.393 | 1510.077 | 1904.372 |
| ENSMUSG00000034663 | -0.089361193 | 0.538301 | 0.6882 | 472.8455 | 364.7457 | 449.6619 | 404.5704 | 412.4721 | 557.5697 | 419.4658 | 413.2846 |
| ENSMUSG00000034664 | 0.237865606 | 0.183163 | 0.334038 | 164.9929 | 157.1766 | 238.4947 | 197.5408 | 146.2667 | 151.9629 | 178.3935 | 168.5766 |

| | | | | | | | | | | | |
|---|---|---|---|---|---|---|---|---|---|---|---|
| ENSMUSG00000034667 | -0.242817624 | 1.60E-06 | 5.46E-05 | 2099.635 | 2122.484 | 2150.593 | 2062.533 | 2585.02 | 2397.885 | 2505.223 | 2490.584 |
| ENSMUSG00000034673 | -0.108977923 | 0.123202 | 0.25284 | 2336.058 | 2727.194 | 2377.494 | 2391.192 | 2574.294 | 2735.332 | 2599.724 | 2688.525 |
| ENSMUSG00000034674 | -0.616403427 | 3.58E-05 | 0.000614 | 2062.411 | 1327.003 | 1406.953 | 1400.901 | 2509.937 | 2162.119 | 2353.83 | 2474.27 |
| ENSMUSG00000034675 | -0.024705295 | 0.740805 | 0.837968 | 3551.372 | 3603.064 | 4075.113 | 3933.563 | 3747.353 | 3686.217 | 3834.014 | 4163.299 |
| ENSMUSG00000034681 | -0.075719272 | 0.658264 | 0.779727 | 2227.404 | 2426.039 | 3368.738 | 3193.432 | 2711.785 | 3298.488 | 3432.869 | 2379.649 |
| ENSMUSG00000034684 | 0.255196926 | 8.50E-05 | 0.001189 | 2520.166 | 2721.195 | 2510.819 | 2545.602 | 2193.025 | 2306.26 | 2053.936 | 2072.949 |
| ENSMUSG00000034685 | 0.013862405 | 0.900158 | 0.939987 | 1890.376 | 1888.519 | 2455.336 | 1848.602 | 1930.72 | 1972.165 | 1958.471 | 2147.992 |
| ENSMUSG00000034686 | 0.218420612 | 0.318211 | 0.48653 | 96.58122 | 82.78769 | 97.71658 | 106.1027 | 65.33246 | 82.68568 | 108.9647 | 72.8686 |
| ENSMUSG00000034687 | 0.16506229 | 0.109722 | 0.235359 | 4220.398 | 4165.78 | 5118.527 | 4305.354 | 3426.541 | 4203.561 | 4215.872 | 4042.576 |
| ENSMUSG00000034701 | 0.059531583 | 0.796933 | 0.875383 | 48.29061 | 74.38894 | 81.98256 | 69.00988 | 62.40712 | 60.3382 | 68.46453 | 71.78101 |
| ENSMUSG00000034706 | 0.831567626 | 0.005202 | 0.026435 | 38.23007 | 40.79393 | 57.13936 | 66.42201 | 29.25334 | 27.93435 | 27.0001 | 30.45255 |
| ENSMUSG00000034707 | -0.331994631 | 0.005353 | 0.026934 | 2924.6 | 2795.584 | 3286.755 | 2887.201 | 3782.457 | 2988.976 | 3936.228 | 4265.532 |
| ENSMUSG00000034708 | -0.016133746 | 0.852292 | 0.911279 | 3012.127 | 3237.118 | 3741.386 | 3206.372 | 3346.582 | 3094.009 | 3498.441 | 3408.51 |
| ENSMUSG00000034709 | -0.101002254 | 0.170175 | 0.3168 | 1129.799 | 1184.224 | 1170.943 | 1069.653 | 1211.088 | 1134.135 | 1237.183 | 1299.671 |
| ENSMUSG00000034714 | -0.628779943 | 0.023337 | 0.078854 | 415.5004 | 199.1704 | 305.5713 | 238.9467 | 483.6552 | 281.5783 | 432.9658 | 594.9123 |
| ENSMUSG00000034723 | 0.326385929 | 0.000581 | 0.005142 | 1323.968 | 1392.993 | 1364.72 | 1402.626 | 1062.871 | 1280.511 | 1004.789 | 1027.774 |
| ENSMUSG00000034724 | 0.132616311 | 0.299247 | 0.467409 | 2659.002 | 2492.029 | 2403.165 | 2578.382 | 1836.135 | 2822.487 | 2266.08 | 2319.832 |
| ENSMUSG00000034729 | -0.038102881 | 0.541938 | 0.691162 | 1066.418 | 999.4513 | 988.7593 | 1023.934 | 1015.091 | 1058.153 | 1071.325 | 1043 |
| ENSMUSG00000034730 | -0.184477275 | 0.5949 | 0.734069 | 41.24823 | 25.19625 | 50.5145 | 35.36756 | 38.02934 | 65.92507 | 33.75012 | 36.9781 |
| ENSMUSG00000034731 | -0.400241745 | 0.026816 | 0.087407 | 86.52067 | 92.38626 | 96.88848 | 69.00988 | 119.9387 | 101.681 | 113.7861 | 118.5474 |
| ENSMUSG00000034738 | -0.504515377 | 0.006755 | 0.032129 | 186.1201 | 141.5789 | 143.2624 | 131.1188 | 201.848 | 174.3104 | 263.2509 | 213.1679 |
| ENSMUSG00000034739 | -2.724208979 | 6.01E-07 | 2.44E-05 | 18.10898 | 8.398751 | 34.78048 | 7.763611 | 105.312 | 44.69496 | 159.1077 | 151.1752 |
| ENSMUSG00000034744 | -0.114818553 | 0.250016 | 0.413794 | 699.2078 | 796.6815 | 774.2797 | 717.7027 | 773.2633 | 727.4105 | 817.7172 | 915.7517 |
| ENSMUSG00000034748 | 0.078051366 | 0.384384 | 0.552315 | 727.3773 | 725.892 | 798.2948 | 802.2398 | 788.865 | 749.758 | 671.1453 | 686.27 |
| ENSMUSG00000034751 | 0.225501256 | 0.127842 | 0.259223 | 668.0201 | 603.5102 | 598.7211 | 610.7374 | 444.6508 | 690.5372 | 495.6446 | 492.6788 |
| ENSMUSG00000034755 | 0.605800623 | 0.345733 | 0.514502 | 18.10898 | 53.99197 | 39.74912 | 30.19182 | 12.67645 | 58.10345 | 8.678602 | 14.13868 |
| ENSMUSG00000034757 | 0.077711243 | 0.358116 | 0.526477 | 1326.986 | 1106.235 | 1222.285 | 1180.932 | 1187.686 | 1051.449 | 1168.718 | 1175.686 |
| ENSMUSG00000034758 | 0.37912604 | 0.016778 | 0.062231 | 233.4046 | 239.9643 | 260.0255 | 236.3588 | 160.8934 | 240.2354 | 176.4649 | 169.6642 |
| ENSMUSG00000034761 | 0.230897185 | 0.001136 | 0.008611 | 2447.73 | 2226.869 | 2343.542 | 2580.107 | 2019.456 | 2173.293 | 2023.079 | 1967.452 |
| ENSMUSG00000034762 | -1.681436616 | 0.008416 | 0.037657 | 16.09687 | 8.398751 | 54.65504 | 6.900988 | 39.00445 | 56.98608 | 79.07171 | 103.3212 |
| ENSMUSG00000034764 | -0.403312629 | 0.071855 | 0.175813 | 227.3683 | 116.3827 | 218.6202 | 199.266 | 212.5743 | 241.3528 | 257.4652 | 299.0875 |
| ENSMUSG00000034765 | -0.245121653 | 0.214006 | 0.37205 | 317.9132 | 251.9625 | 178.871 | 238.0841 | 307.1601 | 321.8037 | 304.7154 | 233.8321 |
| ENSMUSG00000034768 | -0.86133626 | 0.142938 | 0.280407 | 18.10898 | 13.19804 | 4.96864 | 6.900988 | 30.22845 | 22.34748 | 9.642892 | 15.22628 |
| ENSMUSG00000034771 | 0.350331891 | 0.001679 | 0.011504 | 1206.259 | 1235.816 | 1083.163 | 1038.599 | 916.6046 | 1034.688 | 815.7886 | 811.343 |
| ENSMUSG00000034773 | -0.02813009 | 0.862676 | 0.917774 | 206.2411 | 231.5656 | 162.3089 | 170.7995 | 194.0472 | 210.0663 | 171.6435 | 207.7299 |
| ENSMUSG00000034780 | 0.081427274 | 0.762468 | 0.852338 | 117.7084 | 124.7814 | 118.4192 | 92.30071 | 97.51113 | 162.0192 | 71.3574 | 97.8832 |
| ENSMUSG00000034781 | -0.249344925 | 0.014317 | 0.055685 | 2042.29 | 2018.1 | 2340.229 | 2096.175 | 2672.78 | 2198.992 | 2394.33 | 2836.438 |
| ENSMUSG00000034785 | -1.679704803 | 0.050198 | 0.136381 | 0 | 3.599465 | 17.39024 | 4.313117 | 13.65156 | 17.87799 | 17.3572 | 33.71532 |
| ENSMUSG00000034786 | -0.286475024 | 0.081027 | 0.190588 | 198.1927 | 229.1659 | 231.8699 | 212.2054 | 252.5538 | 202.2447 | 313.394 | 293.6496 |
| ENSMUSG00000034789 | 0.013023832 | 0.860248 | 0.91625 | 1835.043 | 1876.521 | 1680.228 | 2013.363 | 1806.881 | 1889.48 | 1854.328 | 1786.912 |
| ENSMUSG00000034792 | 0.281485315 | 0.526792 | 0.67975 | 15.09082 | 14.39786 | 19.04645 | 11.21411 | 16.57689 | 8.938993 | 11.57147 | 11.9635 |
| ENSMUSG00000034793 | -0.187997495 | 0.031099 | 0.096889 | 778.6861 | 676.6993 | 721.2809 | 667.6706 | 820.0686 | 760.9318 | 791.6814 | 867.8977 |
| ENSMUSG00000034795 | 1.185360851 | 0.002121 | 0.013698 | 132.7992 | 206.3693 | 101.8571 | 135.4319 | 46.80534 | 118.4417 | 49.17875 | 39.15328 |
| ENSMUSG00000034796 | -0.143944882 | 0.656663 | 0.77873 | 153.9263 | 61.1909 | 105.1695 | 163.0358 | 132.6151 | 91.62468 | 163.9292 | 146.8248 |
| ENSMUSG00000034799 | 0.431609625 | 0.034781 | 0.10556 | 160.9687 | 149.9777 | 110.9663 | 188.0519 | 124.8142 | 117.3243 | 112.8218 | 96.79561 |
| ENSMUSG00000034800 | -0.154685118 | 0.325202 | 0.493053 | 141.8537 | 142.7788 | 120.0755 | 159.5853 | 153.0925 | 138.5544 | 175.5006 | 159.8759 |
| ENSMUSG00000034801 | 0.016736759 | 0.877115 | 0.926493 | 810.8798 | 680.2988 | 673.2507 | 748.7572 | 687.4535 | 822.3873 | 709.7168 | 661.2554 |
| ENSMUSG00000034807 | -0.159763542 | 0.06272 | 0.159863 | 2795.825 | 3078.742 | 2996.09 | 2945.859 | 3527.953 | 2861.595 | 3523.513 | 3283.437 |
| ENSMUSG00000034810 | 0.100654952 | 0.563986 | 0.709199 | 224.3501 | 177.5736 | 212.8234 | 278.6274 | 236.952 | 217.8879 | 199.6079 | 180.5401 |
| ENSMUSG00000034813 | 0.267432834 | 0.024307 | 0.081315 | 506.0453 | 518.3229 | 606.1741 | 481.3439 | 389.0694 | 422.3674 | 463.8231 | 480.7153 |
| ENSMUSG00000034818 | 0.440744558 | 0.114436 | 0.241901 | 54.32694 | 44.3934 | 34.78048 | 51.75741 | 32.17867 | 29.05173 | 33.75012 | 41.32846 |
| ENSMUSG00000034820 | 0.265991704 | 0.032733 | 0.100666 | 3279.737 | 2939.563 | 3029.214 | 3205.509 | 2583.07 | 3164.403 | 2512.938 | 2097.963 |
| ENSMUSG00000034825 | -1.312591051 | 0.011224 | 0.046557 | 27.16347 | 9.598572 | 52.17072 | 25.01608 | 76.05868 | 33.52122 | 62.6788 | 113.1095 |
| ENSMUSG00000034826 | 0.002803312 | 0.97573 | 0.986757 | 1227.386 | 1377.395 | 1126.225 | 1253.392 | 1330.052 | 1267.102 | 1249.719 | 1122.394 |
| ENSMUSG00000034829 | 0.079578155 | 0.793189 | 0.873246 | 44.26639 | 65.99018 | 45.54586 | 41.40593 | 50.70579 | 63.69032 | 36.64299 | 34.80291 |
| ENSMUSG00000034832 | 0.403102755 | 0.002507 | 0.015472 | 3107.702 | 2640.807 | 2668.16 | 2986.403 | 1934.621 | 2715.219 | 2117.579 | 1857.606 |
| ENSMUSG00000034837 | -0.011104772 | 0.912046 | 0.948028 | 1885.346 | 2178.876 | 2307.105 | 2022.852 | 2103.315 | 2323.021 | 1852.399 | 2182.795 |
| ENSMUSG00000034839 | -0.007864302 | 0.969892 | 0.983264 | 94.56911 | 127.1811 | 127.5284 | 87.12497 | 122.864 | 100.5637 | 121.5004 | 92.44524 |
| ENSMUSG00000034842 | -0.521107054 | 0.064153 | 0.162639 | 52.31483 | 33.595 | 39.74912 | 69.00988 | 63.38224 | 62.57295 | 72.32169 | 82.65692 |
| ENSMUSG00000034845 | -0.909922568 | 0.001554 | 0.01081 | 1289.762 | 881.8688 | 2019.752 | 1317.226 | 2462.156 | 1461.525 | 2874.546 | 3555.335 |
| ENSMUSG00000034848 | 0.242993373 | 0.013787 | 0.054173 | 639.8506 | 715.0936 | 661.6572 | 682.3352 | 532.4108 | 512.8747 | 630.6451 | 602.5255 |
| ENSMUSG00000034850 | 0.127865694 | 0.031784 | 0.098457 | 2147.926 | 2176.476 | 2309.589 | 2175.536 | 1975.576 | 2006.804 | 1944.007 | 2140.379 |
| ENSMUSG00000034853 | 0.550190395 | 0.002109 | 0.013635 | 225.3562 | 238.7645 | 177.2148 | 241.5346 | 132.6151 | 128.498 | 156.2148 | 184.8905 |
| ENSMUSG00000034854 | -0.404863611 | 0.090878 | 0.206558 | 212.2775 | 202.7698 | 268.3065 | 219.969 | 298.3841 | 177.6625 | 317.2511 | 403.4963 |

| | | | | | | | | | | | |
|---|---|---|---|---|---|---|---|---|---|---|---|
| ENSMUSG00000034855 | -0.973646137 | 0.140641 | 0.277138 | 13.07871 | 5.999108 | 34.78048 | 4.313117 | 23.40267 | 18.99536 | 42.42872 | 30.45255 |
| ENSMUSG00000034858 | 0.017499721 | 0.826233 | 0.894472 | 812.8919 | 749.8885 | 699.7501 | 747.8946 | 779.1139 | 736.3495 | 713.574 | 742.8247 |
| ENSMUSG00000034863 | 0.130565473 | 0.110186 | 0.23608 | 741.4621 | 754.6877 | 662.4853 | 705.626 | 653.3246 | 692.7719 | 646.0737 | 621.0145 |
| ENSMUSG00000034867 | 0.094558762 | 0.19591 | 0.350055 | 1038.248 | 1067.841 | 965.5723 | 1096.394 | 969.2606 | 984.4066 | 1017.325 | 929.8904 |
| ENSMUSG00000034868 | 0.022742219 | 0.758493 | 0.849288 | 6500.117 | 6612.216 | 7375.118 | 6702.585 | 6983.747 | 6063.989 | 6753.881 | 6964.933 |
| ENSMUSG00000034871 | -0.242012087 | 0.625184 | 0.755716 | 21.12714 | 16.7975 | 32.29616 | 18.11509 | 41.92979 | 15.64324 | 12.53576 | 34.80291 |
| ENSMUSG00000034875 | -0.967700199 | 1.18E-06 | 4.31E-05 | 554.3359 | 565.1159 | 732.0463 | 547.7659 | 1067.747 | 823.5047 | 1223.683 | 1579.182 |
| ENSMUSG00000034880 | -0.258909145 | 0.018415 | 0.066587 | 950.7214 | 913.0642 | 881.1055 | 993.7423 | 1105.776 | 944.1811 | 1128.218 | 1295.321 |
| ENSMUSG00000034881 | -0.177317154 | 0.477147 | 0.636625 | 131.7931 | 136.7797 | 178.0429 | 134.5693 | 169.6694 | 106.1505 | 150.4291 | 231.6569 |
| ENSMUSG00000034883 | -0.467615008 | 0.04637 | 0.129237 | 56.33904 | 98.38537 | 96.88848 | 94.02596 | 142.3663 | 115.0895 | 123.429 | 96.79561 |
| ENSMUSG00000034889 | -0.236732965 | 0.018088 | 0.065858 | 888.346 | 1006.65 | 1074.882 | 971.3141 | 1195.486 | 1024.632 | 1145.576 | 1279.007 |
| ENSMUSG00000034891 | 0.162614329 | 0.370374 | 0.539513 | 84.50857 | 100.785 | 89.43552 | 80.22398 | 71.18313 | 86.03781 | 82.92887 | 76.13138 |
| ENSMUSG00000034892 | -0.229897594 | 0.040158 | 0.117013 | 7947.829 | 9111.445 | 10241.19 | 8239.78 | 11166 | 8848.485 | 10756.65 | 10908.54 |
| ENSMUSG00000034893 | 0.060126484 | 0.353482 | 0.522236 | 1439.664 | 1319.804 | 1324.143 | 1325.852 | 1303.724 | 1346.436 | 1305.648 | 1232.241 |
| ENSMUSG00000034898 | -0.009978005 | 0.96068 | 0.977472 | 206.2411 | 133.1802 | 178.0429 | 169.0742 | 172.5947 | 203.3621 | 128.2505 | 189.2408 |
| ENSMUSG00000034902 | -0.25093258 | 0.030307 | 0.095156 | 1753.553 | 1715.745 | 1965.097 | 1785.631 | 2173.523 | 1748.69 | 2179.294 | 2491.671 |
| ENSMUSG00000034903 | -0.096827837 | 0.592821 | 0.732398 | 1988.969 | 1163.827 | 1426.828 | 1508.728 | 1343.703 | 1448.117 | 1822.507 | 1897.846 |
| ENSMUSG00000034906 | 0.132285395 | 0.084024 | 0.195387 | 1439.664 | 1584.964 | 1446.702 | 1463.872 | 1409.036 | 1432.474 | 1342.291 | 1227.89 |
| ENSMUSG00000034908 | -0.020906434 | 0.804845 | 0.880483 | 3137.884 | 2654.005 | 2747.658 | 2914.805 | 3122.306 | 2626.946 | 2952.653 | 2919.094 |
| ENSMUSG00000034910 | 0.478915915 | 0.020192 | 0.070946 | 1558.378 | 1773.336 | 1339.877 | 1502.69 | 1020.942 | 1635.836 | 947.8962 | 825.4816 |
| ENSMUSG00000034911 | -0.21001535 | 0.289938 | 0.457144 | 579.4873 | 410.339 | 751.9208 | 596.9355 | 708.9059 | 576.565 | 590.145 | 833.0948 |
| ENSMUSG00000034912 | 0.56697397 | 0.051003 | 0.137961 | 140.8476 | 145.1784 | 121.7317 | 138.8824 | 63.38224 | 154.1976 | 78.10742 | 73.95619 |
| ENSMUSG00000034917 | -0.04925352 | 0.800294 | 0.877926 | 523.1483 | 362.3461 | 574.706 | 493.4206 | 496.3317 | 366.4987 | 552.5377 | 607.9634 |
| ENSMUSG00000034918 | -2.985566145 | 5.46E-10 | 7.47E-08 | 33.19979 | 32.39518 | 98.54469 | 50.03216 | 450.5014 | 137.437 | 392.4657 | 723.2481 |
| ENSMUSG00000034919 | -0.410664836 | 0.014159 | 0.055272 | 109.6599 | 139.1793 | 120.0755 | 94.02596 | 163.8187 | 157.5497 | 146.572 | 144.6496 |
| ENSMUSG00000034926 | -1.254050183 | 5.67E-09 | 5.30E-07 | 961.788 | 777.4844 | 1316.69 | 965.2757 | 2571.369 | 1546.446 | 2556.331 | 2921.27 |
| ENSMUSG00000034928 | 0.025890131 | 0.706083 | 0.815104 | 2344.107 | 2072.092 | 2218.498 | 2368.764 | 2239.831 | 2095.076 | 2214.008 | 2298.08 |
| ENSMUSG00000034930 | -0.262194252 | 0.150165 | 0.290143 | 397.3915 | 394.7413 | 391.6944 | 378.6917 | 495.3565 | 329.6254 | 432.9658 | 615.5766 |
| ENSMUSG00000034931 | -0.087370295 | 0.193464 | 0.346682 | 1198.211 | 1162.627 | 1275.284 | 1189.558 | 1294.948 | 1198.942 | 1318.183 | 1317.073 |
| ENSMUSG00000034932 | -0.382654609 | 0.001788 | 0.012107 | 821.9464 | 787.0829 | 1041.758 | 821.2176 | 1157.457 | 978.8197 | 1165.826 | 1227.89 |
| ENSMUSG00000034936 | -0.633482587 | 0.060326 | 0.155584 | 271.6347 | 181.1731 | 262.5098 | 295.0172 | 230.1263 | 308.3953 | 718.3954 | 311.0511 |
| ENSMUSG00000034940 | 0.078657536 | 0.231204 | 0.39178 | 1098.611 | 1129.032 | 1010.29 | 1092.081 | 1019.966 | 1008.989 | 1047.218 | 1019.073 |
| ENSMUSG00000034947 | -1.161263911 | 0.001949 | 0.012893 | 70.4238 | 87.58697 | 120.0755 | 89.71284 | 257.4294 | 87.15518 | 159.1077 | 319.7518 |
| ENSMUSG00000034949 | 0.344490798 | 0.005983 | 0.029416 | 369.2219 | 374.3443 | 335.3832 | 359.714 | 270.1058 | 330.7427 | 291.2153 | 240.3576 |
| ENSMUSG00000034951 | -0.299004154 | 0.063919 | 0.16218 | 899.4126 | 770.2854 | 970.541 | 792.751 | 1044.344 | 773.2229 | 1130.147 | 1276.832 |
| ENSMUSG00000034957 | -0.163063734 | 0.311767 | 0.480258 | 180.0837 | 209.9688 | 236.0104 | 170.7995 | 225.2507 | 208.949 | 198.6436 | 259.9343 |
| ENSMUSG00000034958 | 1.870440041 | 2.92E-06 | 8.84E-05 | 59.35721 | 37.19447 | 49.6864 | 94.88858 | 13.65156 | 13.40849 | 16.39292 | 22.83941 |
| ENSMUSG00000034959 | 1.098508955 | 0.006752 | 0.032129 | 71.42986 | 31.19536 | 38.09291 | 57.79577 | 19.50223 | 17.87799 | 36.64299 | 18.48905 |
| ENSMUSG00000034968 | 0.372349014 | 0.14751 | 0.286664 | 76.46013 | 75.58876 | 67.90474 | 64.69676 | 60.4569 | 74.86406 | 45.32159 | 39.15328 |
| ENSMUSG00000034973 | -0.062063841 | 0.433423 | 0.597583 | 634.8203 | 685.0981 | 615.2832 | 632.303 | 688.4286 | 701.7109 | 655.7166 | 630.8028 |
| ENSMUSG00000034974 | -0.27284024 | 0.001108 | 0.008439 | 870.237 | 910.6645 | 920.8546 | 920.4193 | 1101.876 | 975.4676 | 1098.325 | 1200.701 |
| ENSMUSG00000034981 | 0.246350926 | 0.094917 | 0.212854 | 566.4086 | 823.0776 | 858.7466 | 664.2201 | 608.4695 | 563.1565 | 630.6451 | 652.5547 |
| ENSMUSG00000034993 | 0.046451679 | 0.726339 | 0.828716 | 2032.23 | 1818.929 | 2381.635 | 2133.268 | 2138.419 | 1654.831 | 1937.257 | 2373.124 |
| ENSMUSG00000034994 | -0.189340424 | 0.073864 | 0.179209 | 83032.68 | 93936.43 | 94470.4 | 93964.71 | 117534.1 | 86060.15 | 109801.7 | 103251.5 |
| ENSMUSG00000034997 | -0.898460263 | 0.063103 | 0.160705 | 6.036326 | 15.59768 | 16.56213 | 6.900988 | 17.552 | 17.87799 | 21.21436 | 27.18978 |
| ENSMUSG00000034998 | -0.05018752 | 0.616658 | 0.749494 | 544.2754 | 561.5165 | 458.7711 | 545.178 | 533.3859 | 588.8561 | 522.6447 | 537.27 |
| ENSMUSG00000035000 | -1.226885286 | 8.12E-07 | 3.13E-05 | 1248.513 | 1024.648 | 1845.85 | 1406.939 | 2711.785 | 2226.927 | 3158.047 | 4841.956 |
| ENSMUSG00000035004 | 0.058912554 | 0.797809 | 0.875981 | 87.52673 | 69.58965 | 75.3577 | 70.73513 | 56.55646 | 86.03781 | 87.75031 | 60.9051 |
| ENSMUSG00000035007 | -0.033837602 | 0.78078 | 0.864743 | 669.0261 | 571.1151 | 650.8918 | 659.907 | 706.9557 | 525.1658 | 691.3953 | 688.4452 |
| ENSMUSG00000035011 | 0.071050139 | 0.456035 | 0.617957 | 863.1946 | 693.4968 | 774.2797 | 717.7027 | 686.4784 | 729.6453 | 730.9312 | 756.9634 |
| ENSMUSG00000035021 | -0.339054034 | 0.001166 | 0.008785 | 567.4147 | 519.5227 | 510.9418 | 468.4046 | 624.0712 | 588.8561 | 711.6454 | 686.27 |
| ENSMUSG00000035024 | -0.005310457 | 0.96671 | 0.981333 | 1050.321 | 988.6529 | 793.3262 | 817.7671 | 1036.543 | 819.0352 | 909.3247 | 893.9999 |
| ENSMUSG00000035027 | -0.266651108 | 0.000296 | 0.003066 | 2548.336 | 2615.611 | 2799.829 | 2530.937 | 3134.983 | 2958.807 | 3066.44 | 3467.24 |
| ENSMUSG00000035032 | -0.390142684 | 0.45331 | 0.615723 | 5.030272 | 13.19804 | 11.59349 | 10.35148 | 13.65156 | 7.821619 | 12.53576 | 18.48905 |
| ENSMUSG00000035041 | 0.383290305 | 0.580325 | 0.721633 | 24.1453 | 16.7975 | 4.140533 | 24.15346 | 20.47734 | 20.11273 | 4.821446 | 7.613138 |
| ENSMUSG00000035045 | 0.410441542 | 0.061942 | 0.158493 | 210.2654 | 159.5763 | 165.6213 | 144.0581 | 102.3867 | 163.1366 | 144.6434 | 101.146 |
| ENSMUSG00000035047 | -0.035633436 | 0.77962 | 0.863998 | 559.3662 | 515.9233 | 524.1915 | 460.6409 | 565.5646 | 424.6022 | 540.9662 | 578.5985 |
| ENSMUSG00000035048 | -0.314387103 | 0.0006 | 0.005266 | 1293.786 | 1446.985 | 1346.501 | 1326.715 | 1703.519 | 1479.403 | 1661.47 | 1884.795 |
| ENSMUSG00000035049 | -0.41316619 | 0.013953 | 0.054594 | 429.5852 | 356.347 | 466.224 | 408.0209 | 543.137 | 415.6632 | 562.1806 | 691.7079 |
| ENSMUSG00000035051 | 0.279555048 | 0.00955 | 0.041173 | 1238.453 | 1150.629 | 1043.414 | 1129.174 | 949.7584 | 1088.322 | 892.9318 | 826.5692 |
| ENSMUSG00000035062 | 0.387994628 | 0.001056 | 0.008143 | 543.2694 | 661.1017 | 580.5027 | 644.3797 | 421.2481 | 509.5226 | 507.2161 | 417.635 |
| ENSMUSG00000035064 | 0.039952188 | 0.71854 | 0.824149 | 1184.126 | 1078.64 | 1204.895 | 1264.606 | 1248.142 | 1000.05 | 1299.862 | 1054.963 |
| ENSMUSG00000035067 | 0.62979045 | 0.000184 | 0.002122 | 309.8647 | 280.7582 | 324.6178 | 309.6818 | 151.1423 | 253.6439 | 200.5721 | 188.1533 |
| ENSMUSG00000035069 | -0.249447786 | 0.035627 | 0.107545 | 323.9495 | 327.5513 | 337.8675 | 320.8959 | 361.7663 | 355.325 | 377.0371 | 464.4014 |

| | | | | | | | | | | | |
|---|---|---|---|---|---|---|---|---|---|---|---|
| ENSMUSG00000035078 | -0.376149404 | 0.022193 | 0.075893 | 483.9121 | 488.3274 | 521.7072 | 425.2734 | 714.7566 | 439.128 | 654.7523 | 680.832 |
| ENSMUSG00000035085 | 0.39855554 | 0.18676 | 0.338689 | 48.29061 | 31.19536 | 57.13936 | 41.40593 | 33.15378 | 36.87335 | 26.03581 | 40.24087 |
| ENSMUSG00000035086 | -0.156072789 | 0.023764 | 0.080064 | 2664.032 | 2729.594 | 2861.108 | 2911.354 | 3073.551 | 2859.36 | 3306.548 | 3204.043 |
| ENSMUSG00000035093 | 0.103303248 | 0.276489 | 0.44271 | 2013.115 | 1739.741 | 1560.981 | 1778.73 | 1570.904 | 1759.864 | 1593.006 | 1678.153 |
| ENSMUSG00000035095 | -2.2891324 | 3.27E-18 | 2.87E-15 | 33.19979 | 32.39518 | 38.09291 | 40.5433 | 124.8142 | 147.4934 | 207.3222 | 228.3941 |
| ENSMUSG00000035104 | -1.062821572 | 7.09E-05 | 0.001042 | 442.6639 | 377.9438 | 585.4714 | 442.5259 | 810.3175 | 534.1048 | 1239.112 | 1280.095 |
| ENSMUSG00000035105 | -0.166055792 | 0.192966 | 0.346202 | 447.6942 | 475.1293 | 591.2681 | 510.6731 | 599.6935 | 516.2268 | 521.6804 | 637.3284 |
| ENSMUSG00000035107 | -0.04137364 | 0.623994 | 0.754981 | 721.341 | 701.8956 | 640.1264 | 644.3797 | 672.8268 | 692.7719 | 669.2167 | 749.3503 |
| ENSMUSG00000035109 | 0.183945677 | 0.439917 | 0.603203 | 99.59938 | 74.38894 | 66.24853 | 84.5371 | 84.83468 | 82.68568 | 62.6788 | 55.46715 |
| ENSMUSG00000035112 | -0.421204434 | 0.00253 | 0.015558 | 1295.798 | 1029.447 | 1423.515 | 1131.762 | 1480.219 | 1409.009 | 1782.971 | 1866.306 |
| ENSMUSG00000035121 | -0.191930812 | 0.482547 | 0.641034 | 70.4238 | 34.79482 | 56.31125 | 61.24627 | 65.33246 | 74.86406 | 65.57166 | 50.02919 |
| ENSMUSG00000035125 | 0.271685091 | 0.029098 | 0.092576 | 645.8869 | 645.504 | 503.4888 | 625.402 | 564.5894 | 494.9967 | 499.5018 | 442.6496 |
| ENSMUSG00000035126 | 0.029692426 | 0.849401 | 0.909599 | 118.7144 | 119.9822 | 134.9814 | 137.1571 | 144.3165 | 125.1459 | 116.679 | 115.2847 |
| ENSMUSG00000035131 | 0.294437972 | 0.836706 | 0.901267 | 9.054489 | 17.99732 | 7.45296 | 17.25247 | 0.975111 | 41.34284 | 0 | 0 |
| ENSMUSG00000035133 | -0.156943349 | 0.150043 | 0.289999 | 1706.268 | 1365.397 | 1364.72 | 1301.699 | 1508.497 | 1497.281 | 1708.72 | 1681.416 |
| ENSMUSG00000035139 | -0.063804984 | 0.396549 | 0.563802 | 831.0009 | 733.091 | 752.7489 | 796.2015 | 842.4962 | 783.2792 | 801.3243 | 827.6568 |
| ENSMUSG00000035142 | -0.229230583 | 0.011418 | 0.047144 | 326.9677 | 344.3488 | 332.0708 | 355.4009 | 413.4472 | 384.3767 | 408.8586 | 385.0072 |
| ENSMUSG00000035150 | -0.070138609 | 0.689982 | 0.803616 | 2699.244 | 1654.554 | 1728.259 | 2562.854 | 2247.632 | 2571.078 | 2069.365 | 2189.321 |
| ENSMUSG00000035151 | 0.200852494 | 0.057745 | 0.150636 | 603.6326 | 601.1106 | 504.3169 | 524.4751 | 487.5557 | 529.6353 | 447.4302 | 476.3649 |
| ENSMUSG00000035152 | 0.021997194 | 0.638629 | 0.765758 | 5062.466 | 5228.822 | 5059.732 | 4884.174 | 4794.622 | 5065.057 | 5073.125 | 4992.043 |
| ENSMUSG00000035158 | -0.630921915 | 0.000224 | 0.002479 | 284.7134 | 199.1704 | 275.7595 | 229.4578 | 308.1352 | 368.7335 | 448.3945 | 408.9342 |
| ENSMUSG00000035161 | 0.063869867 | 0.545298 | 0.693831 | 680.0927 | 628.7065 | 553.1752 | 670.2585 | 568.4899 | 577.6824 | 603.645 | 672.1313 |
| ENSMUSG00000035164 | -0.022307475 | 0.878152 | 0.927008 | 397.3915 | 266.3604 | 332.0708 | 335.5605 | 305.2098 | 330.7427 | 343.2869 | 375.2189 |
| ENSMUSG00000035165 | -1.204409857 | 0.005151 | 0.026292 | 14.08476 | 11.99822 | 17.39024 | 16.38985 | 21.45245 | 55.8687 | 24.10723 | 38.06569 |
| ENSMUSG00000035168 | -0.198451732 | 0.067864 | 0.168393 | 2398.434 | 2267.663 | 2861.108 | 2202.278 | 2596.721 | 2766.618 | 3130.083 | 2673.299 |
| ENSMUSG00000035171 | -0.057450966 | 0.521228 | 0.674694 | 1030.2 | 1067.841 | 1045.071 | 1193.871 | 1228.64 | 1136.369 | 1128.218 | 1020.16 |
| ENSMUSG00000035172 | -0.271311484 | 0.027644 | 0.089225 | 453.7305 | 485.9277 | 529.1601 | 452.0147 | 644.5486 | 482.7056 | 630.6451 | 559.0218 |
| ENSMUSG00000035173 | -0.005208125 | 0.961592 | 0.978074 | 1423.567 | 1096.637 | 1117.944 | 1092.081 | 1136.98 | 1255.928 | 1184.147 | 1170.248 |
| ENSMUSG00000035181 | -0.132041799 | 0.155802 | 0.297848 | 779.6921 | 795.4817 | 875.3087 | 775.4985 | 896.1273 | 806.7441 | 847.6102 | 986.4451 |
| ENSMUSG00000035183 | 0.109108139 | 0.575291 | 0.717501 | 76.46013 | 121.182 | 93.57605 | 94.02596 | 86.78491 | 90.5073 | 93.53605 | 84.8321 |
| ENSMUSG00000035184 | 0.121141559 | 0.313398 | 0.481484 | 261.5741 | 241.1641 | 238.4947 | 240.672 | 215.4996 | 232.4138 | 251.6795 | 202.2919 |
| ENSMUSG00000035187 | 0.857222551 | 0.051469 | 0.138734 | 29.17558 | 20.39697 | 56.31125 | 40.5433 | 19.50223 | 30.1691 | 12.53576 | 19.57664 |
| ENSMUSG00000035189 | 0.976348651 | 0.000884 | 0.007086 | 132.7992 | 173.9741 | 197.9175 | 183.7388 | 60.4569 | 136.3196 | 59.78593 | 94.62042 |
| ENSMUSG00000035198 | -0.401274687 | 0.004072 | 0.022119 | 630.7961 | 761.8867 | 802.4353 | 708.2139 | 1047.27 | 770.9881 | 904.5032 | 1111.518 |
| ENSMUSG00000035199 | 0.055706101 | 0.434241 | 0.598443 | 1800.837 | 1763.738 | 1866.552 | 2007.325 | 1712.295 | 1730.812 | 1821.542 | 1896.759 |
| ENSMUSG00000035200 | 0.09480514 | 0.836472 | 0.901206 | 48.29061 | 20.39697 | 45.54586 | 26.74133 | 21.45245 | 34.6386 | 20.25007 | 56.55474 |
| ENSMUSG00000035202 | -0.210185984 | 0.357888 | 0.526267 | 5056.429 | 4069.795 | 2797.344 | 3055.412 | 6147.102 | 3448.216 | 3899.585 | 3830.496 |
| ENSMUSG00000035203 | -0.201710719 | 0.056048 | 0.147522 | 2327.004 | 2471.632 | 2950.544 | 2368.764 | 2906.807 | 2578.899 | 2984.475 | 3168.153 |
| ENSMUSG00000035206 | -0.455623047 | 0.001454 | 0.010346 | 646.893 | 625.107 | 747.7803 | 611.6001 | 856.1477 | 732.9974 | 907.3961 | 1113.693 |
| ENSMUSG00000035208 | 0.699271179 | 0.198346 | 0.352939 | 14.08476 | 39.59411 | 57.96746 | 18.11509 | 12.67645 | 17.87799 | 30.85725 | 18.48905 |
| ENSMUSG00000035211 | -0.010512325 | 0.986318 | 0.992283 | 18.10898 | 10.79839 | 4.96864 | 12.07673 | 9.751113 | 10.05637 | 5.785735 | 20.66423 |
| ENSMUSG00000035212 | 0.057675732 | 0.42569 | 0.590702 | 2671.074 | 2777.587 | 3028.386 | 3058 | 2771.266 | 2600.13 | 2812.831 | 2902.781 |
| ENSMUSG00000035215 | -0.144276443 | 0.105357 | 0.229003 | 1513.106 | 1593.363 | 1590.793 | 1509.591 | 1952.173 | 1564.324 | 1597.827 | 1743.409 |
| ENSMUSG00000035226 | 0.566331012 | 0.4655 | 0.626323 | 283.7073 | 15.59768 | 14.90592 | 69.00988 | 81.90935 | 46.92971 | 64.60737 | 65.25547 |
| ENSMUSG00000035227 | -0.015723634 | 0.77147 | 0.858295 | 3906.509 | 4076.994 | 3709.918 | 4021.551 | 4002.832 | 4107.467 | 3915.978 | 3856.598 |
| ENSMUSG00000035228 | 0.37781371 | 0.00628 | 0.030431 | 263.5862 | 349.1481 | 310.54 | 276.9021 | 256.4543 | 238.0007 | 211.1793 | 216.4306 |
| ENSMUSG00000035232 | -0.202176166 | 0.020493 | 0.07174 | 643.8748 | 718.6931 | 747.7803 | 712.527 | 821.0437 | 738.5843 | 830.253 | 858.1094 |
| ENSMUSG00000035234 | 0.100719656 | 0.44505 | 0.607929 | 493.9727 | 496.7261 | 411.569 | 424.4108 | 414.4223 | 509.5226 | 396.3228 | 381.7445 |
| ENSMUSG00000035235 | -0.097579544 | 0.410652 | 0.576745 | 256.5439 | 287.9572 | 228.5574 | 252.7487 | 279.8569 | 275.9914 | 251.6795 | 287.124 |
| ENSMUSG00000035236 | 0.513548419 | 0.006904 | 0.032666 | 1597.614 | 1696.548 | 1360.579 | 1622.595 | 1037.518 | 1579.967 | 902.5747 | 877.686 |
| ENSMUSG00000035237 | 0.764345463 | 0.02613 | 0.085764 | 38.23007 | 32.39518 | 24.01509 | 35.36756 | 24.37778 | 18.99536 | 14.46434 | 18.48905 |
| ENSMUSG00000035239 | 0.108726533 | 0.669732 | 0.788252 | 160.9687 | 178.7734 | 146.5749 | 140.6076 | 117.0134 | 221.2401 | 140.7862 | 102.2336 |
| ENSMUSG00000035242 | -0.152455209 | 0.022862 | 0.077618 | 10941.85 | 11583.08 | 11793.89 | 12239.76 | 13755.9 | 11933.56 | 12803.83 | 13255.56 |
| ENSMUSG00000035245 | -0.323896854 | 0.045761 | 0.127925 | 787.7406 | 638.3051 | 1034.305 | 824.6681 | 1062.871 | 854.7912 | 1026.968 | 1171.336 |
| ENSMUSG00000035246 | 1.07431834 | 6.32E-10 | 8.52E-08 | 133.8052 | 158.3764 | 139.1219 | 137.1571 | 59.48179 | 74.86406 | 74.25027 | 60.9051 |
| ENSMUSG00000035247 | -0.065643471 | 0.305069 | 0.473524 | 3990.012 | 3885.022 | 3863.117 | 3789.505 | 4049.637 | 3701.86 | 4266.015 | 4229.642 |
| ENSMUSG00000035248 | 0.013928218 | 0.852374 | 0.911279 | 1683.129 | 1618.559 | 1759.727 | 1604.48 | 1720.096 | 1488.342 | 1721.256 | 1671.628 |
| ENSMUSG00000035258 | 0.671318326 | 0.080249 | 0.189366 | 398.3975 | 331.1507 | 293.9779 | 410.6088 | 168.6943 | 448.067 | 183.2149 | 101.146 |
| ENSMUSG00000035266 | 0.025397664 | 0.802039 | 0.879265 | 356.1432 | 355.1472 | 315.5086 | 311.4071 | 338.3636 | 326.2732 | 314.3583 | 332.8029 |
| ENSMUSG00000035268 | -0.292831186 | 0.000764 | 0.006351 | 1489.967 | 1630.557 | 1716.665 | 1698.506 | 1970.7 | 1795.62 | 2063.579 | 2178.445 |
| ENSMUSG00000035270 | 0.152800369 | 0.428574 | 0.59347 | 91.55095 | 111.5834 | 126.7003 | 99.2017 | 84.83468 | 87.15518 | 100.2861 | 114.1971 |
| ENSMUSG00000035273 | 0.507258026 | 0.007762 | 0.035442 | 182.0958 | 287.9572 | 245.1196 | 249.2982 | 196.9725 | 186.6015 | 157.1791 | 137.0365 |
| ENSMUSG00000035274 | 0.281631507 | 0.030123 | 0.09475 | 785.7285 | 889.0678 | 794.9824 | 779.8116 | 649.4241 | 817.9178 | 652.8238 | 552.4963 |
| ENSMUSG00000035275 | -0.307437975 | 0.006975 | 0.032896 | 743.4742 | 855.4728 | 653.3761 | 763.4218 | 989.738 | 873.7865 | 1019.254 | 842.8831 |

| | | | | | | | | | | | |
|---|---|---|---|---|---|---|---|---|---|---|---|
| ENSMUSG00000035278 | -0.164927295 | 0.012524 | 0.050582 | 1007.06 | 989.8528 | 937.4167 | 979.0777 | 1127.229 | 1031.336 | 1099.29 | 1125.657 |
| ENSMUSG00000035279 | 0.332514202 | 0.074027 | 0.179481 | 215.2956 | 293.9563 | 270.7909 | 205.3044 | 244.7529 | 191.071 | 175.5006 | 169.6642 |
| ENSMUSG00000035283 | 0.483333222 | 0.276265 | 0.442485 | 20.12109 | 15.59768 | 24.01509 | 17.25247 | 13.65156 | 18.99536 | 6.750024 | 16.31387 |
| ENSMUSG00000035284 | 0.130797295 | 0.247758 | 0.411023 | 402.4217 | 345.5486 | 401.6317 | 375.2412 | 365.6667 | 385.4941 | 333.6441 | 309.9635 |
| ENSMUSG00000035285 | 0.033767129 | 0.891644 | 0.934533 | 102.6175 | 125.9813 | 158.9965 | 119.042 | 110.1876 | 84.92043 | 135.0005 | 165.3138 |
| ENSMUSG00000035293 | 0.316223184 | 0.030067 | 0.094648 | 1670.05 | 1618.559 | 1242.16 | 1557.035 | 1210.113 | 1517.394 | 1103.147 | 1058.226 |
| ENSMUSG00000035295 | 0.550519275 | 0.227857 | 0.387732 | 25.15136 | 15.59768 | 14.07781 | 17.25247 | 15.60178 | 12.29112 | 14.46434 | 6.525547 |
| ENSMUSG00000035296 | 1.307897979 | 0.003299 | 0.019041 | 188.1322 | 201.57 | 102.6852 | 180.2883 | 64.35735 | 129.6154 | 54.96448 | 22.83941 |
| ENSMUSG00000035297 | -0.181662132 | 0.004497 | 0.023786 | 2383.343 | 2574.817 | 2443.743 | 2369.627 | 2978.965 | 2743.153 | 2642.152 | 2713.54 |
| ENSMUSG00000035298 | 0.429375923 | 0.104605 | 0.227851 | 51.30877 | 49.19268 | 58.79557 | 41.40593 | 35.10401 | 48.04709 | 33.75012 | 32.62773 |
| ENSMUSG00000035299 | 0.184840481 | 0.305055 | 0.473524 | 611.681 | 401.9402 | 379.2728 | 514.9862 | 488.5308 | 402.2547 | 363.537 | 424.1605 |
| ENSMUSG00000035305 | 0.743847523 | 0.001018 | 0.007941 | 1132.817 | 1398.992 | 1000.353 | 1273.232 | 703.0553 | 1071.562 | 596.895 | 498.1167 |
| ENSMUSG00000035310 | 0.029261402 | 0.802695 | 0.87942 | 564.3965 | 464.3309 | 429.7873 | 456.3278 | 430.9992 | 483.823 | 462.8588 | 498.1167 |
| ENSMUSG00000035311 | -0.032815683 | 0.6684 | 0.787337 | 1210.283 | 1010.25 | 1148.584 | 1113.647 | 1137.955 | 1150.895 | 1119.54 | 1182.212 |
| ENSMUSG00000035314 | 0.120776932 | 0.221716 | 0.380835 | 543.2694 | 637.1052 | 617.7675 | 639.204 | 581.1663 | 589.9735 | 570.8592 | 499.2043 |
| ENSMUSG00000035325 | -0.075622148 | 0.264962 | 0.429148 | 3546.342 | 3721.846 | 3844.071 | 3451.357 | 3802.934 | 3619.175 | 3786.764 | 4138.284 |
| ENSMUSG00000035329 | 0.064524921 | 0.650186 | 0.773634 | 524.1543 | 469.1302 | 426.4749 | 420.0976 | 408.5716 | 545.2786 | 404.0372 | 401.3211 |
| ENSMUSG00000035342 | -0.050117851 | 0.480083 | 0.639095 | 3086.575 | 3147.132 | 2939.779 | 2841.482 | 3152.535 | 2852.656 | 3100.19 | 3329.116 |
| ENSMUSG00000035349 | -1.761121671 | 0.008213 | 0.036999 | 17.10292 | 7.198929 | 56.31125 | 17.25247 | 89.71024 | 20.11273 | 82.92887 | 141.3868 |
| ENSMUSG00000035351 | -0.083671983 | 0.30429 | 0.472686 | 586.5297 | 593.9117 | 554.8314 | 647.8302 | 622.121 | 643.6075 | 632.5737 | 627.5401 |
| ENSMUSG00000035352 | 0.170413303 | 0.781467 | 0.865244 | 9.054489 | 25.19625 | 57.96746 | 12.93935 | 26.32801 | 31.28647 | 23.14294 | 13.05109 |
| ENSMUSG00000035354 | -0.396891069 | 0.003875 | 0.021338 | 757.5589 | 697.0963 | 1029.337 | 778.0864 | 1114.552 | 955.3549 | 1088.682 | 1140.883 |
| ENSMUSG00000035356 | 0.211287997 | 0.115865 | 0.243805 | 382.3007 | 310.7538 | 290.6654 | 353.6756 | 263.2801 | 325.1559 | 292.1796 | 275.1605 |
| ENSMUSG00000035357 | -0.061616974 | 0.750749 | 0.844816 | 747.4984 | 1034.246 | 1373.829 | 959.2373 | 1016.066 | 1046.98 | 905.4675 | 1327.949 |
| ENSMUSG00000035364 | 0.021920651 | 0.941837 | 0.966348 | 46.2785 | 62.39072 | 43.88965 | 38.81806 | 57.53157 | 58.10345 | 32.78583 | 39.15328 |
| ENSMUSG00000035365 | 0.191310543 | 0.132348 | 0.265502 | 311.8769 | 277.1588 | 271.619 | 313.995 | 233.0516 | 280.4609 | 233.358 | 283.8613 |
| ENSMUSG00000035367 | 0.191748912 | 0.202008 | 0.35757 | 398.3975 | 433.1356 | 395.0069 | 410.6088 | 279.8569 | 444.7149 | 372.2156 | 337.1532 |
| ENSMUSG00000035370 | 0.116950785 | 0.644547 | 0.770057 | 206.2411 | 141.5789 | 130.0127 | 144.0581 | 181.3707 | 92.74205 | 132.1076 | 166.4014 |
| ENSMUSG00000035372 | -0.265343595 | 0.007619 | 0.034968 | 437.6336 | 481.1284 | 448.8338 | 404.5704 | 576.2908 | 508.4052 | 493.7161 | 548.1459 |
| ENSMUSG00000035373 | 1.369864424 | 0.074185 | 0.179688 | 7.04238 | 20.39697 | 31.46805 | 4.313117 | 10.72622 | 4.469496 | 4.821446 | 4.350364 |
| ENSMUSG00000035376 | -0.20480535 | 0.016016 | 0.060326 | 870.237 | 847.074 | 785.8732 | 934.2212 | 1042.394 | 925.1858 | 1021.182 | 971.2188 |
| ENSMUSG00000035378 | -0.211670941 | 0.100963 | 0.222465 | 285.7194 | 319.1525 | 314.6805 | 242.3972 | 326.6623 | 317.3342 | 326.894 | 373.0437 |
| ENSMUSG00000035382 | -0.434532525 | 0.018572 | 0.06701 | 568.4207 | 512.3238 | 660.001 | 570.1941 | 857.1228 | 511.7573 | 861.1102 | 893.9999 |
| ENSMUSG00000035385 | 0.383647714 | 0.578673 | 0.720064 | 12.07265 | 14.39786 | 26.49941 | 3.450494 | 15.60178 | 6.704245 | 5.785735 | 15.22628 |
| ENSMUSG00000035390 | 0.299992804 | 0.006701 | 0.031964 | 436.6276 | 428.3363 | 396.6631 | 509.8105 | 368.5921 | 354.2076 | 370.287 | 345.854 |
| ENSMUSG00000035392 | -0.005024258 | 0.960575 | 0.977472 | 719.3289 | 693.4968 | 767.6548 | 615.9132 | 676.7272 | 757.5796 | 725.1455 | 647.1167 |
| ENSMUSG00000035394 | 0.888755201 | 0.019175 | 0.068437 | 126.7628 | 97.18554 | 123.3879 | 106.1027 | 37.05423 | 119.559 | 42.42872 | 46.76642 |
| ENSMUSG00000035397 | -0.31762269 | 0.05065 | 0.137247 | 326.9677 | 293.9563 | 335.3832 | 291.5667 | 448.5512 | 283.813 | 435.8587 | 386.0948 |
| ENSMUSG00000035399 | -0.116111081 | 0.189983 | 0.342514 | 597.5963 | 614.3086 | 636.814 | 670.2585 | 704.0304 | 621.26 | 669.2167 | 737.3868 |
| ENSMUSG00000035401 | 0.484974376 | 0.003659 | 0.020463 | 2149.938 | 2126.084 | 1544.419 | 2027.165 | 1248.142 | 1833.611 | 1180.29 | 1345.35 |
| ENSMUSG00000035403 | -0.756994029 | 0.000394 | 0.003815 | 103.6236 | 116.3827 | 114.2787 | 106.9653 | 120.9138 | 198.8926 | 204.4293 | 221.8686 |
| ENSMUSG00000035407 | -0.444173111 | 0.129222 | 0.261046 | 299.8042 | 423.537 | 735.3587 | 364.8897 | 595.793 | 436.8933 | 594.0021 | 855.9342 |
| ENSMUSG00000035413 | -0.029016843 | 0.707944 | 0.816562 | 1720.353 | 1894.518 | 1657.041 | 1831.35 | 1946.322 | 1819.085 | 1801.292 | 1675.978 |
| ENSMUSG00000035431 | 0.376513281 | 0.066281 | 0.165981 | 76.46013 | 64.79036 | 66.24853 | 66.42201 | 54.60623 | 48.04709 | 57.85735 | 50.02919 |
| ENSMUSG00000035437 | 0.175497663 | 0.127242 | 0.258405 | 2732.444 | 2751.191 | 2293.027 | 2359.275 | 2053.584 | 2673.876 | 2115.65 | 2130.591 |
| ENSMUSG00000035439 | -0.168646713 | 0.053186 | 0.142096 | 411.4762 | 423.537 | 385.0696 | 389.9058 | 472.929 | 458.1234 | 436.823 | 438.2992 |
| ENSMUSG00000035441 | -0.439705357 | 0.028897 | 0.092064 | 534.2149 | 358.7466 | 611.9708 | 421.8229 | 616.2703 | 511.7573 | 679.8239 | 808.0802 |
| ENSMUSG00000035443 | -0.015926731 | 0.825597 | 0.894046 | 1237.447 | 1372.596 | 1295.159 | 1232.689 | 1262.769 | 1393.365 | 1241.04 | 1296.409 |
| ENSMUSG00000035451 | -0.673808047 | 0.235375 | 0.396285 | 45.27245 | 33.595 | 204.5423 | 72.46037 | 216.4747 | 60.3382 | 111.8575 | 180.5401 |
| ENSMUSG00000035455 | -0.070003552 | 0.591944 | 0.731743 | 791.7648 | 926.2622 | 631.8454 | 688.3735 | 811.2926 | 835.7958 | 782.0385 | 754.7882 |
| ENSMUSG00000035456 | -0.151963584 | 0.712727 | 0.819906 | 16.09687 | 19.19714 | 14.07781 | 13.80198 | 12.67645 | 25.6996 | 16.39292 | 15.22628 |
| ENSMUSG00000035458 | 0.506558281 | 0.013613 | 0.05367 | 214.2896 | 277.1588 | 216.1358 | 233.771 | 189.1716 | 210.0663 | 119.5719 | 143.562 |
| ENSMUSG00000035459 | 0.736938736 | 0.168289 | 0.314398 | 78.47224 | 13.19804 | 40.57722 | 45.71905 | 16.57689 | 43.57759 | 22.17865 | 25.0146 |
| ENSMUSG00000035469 | 0.238665352 | 0.038368 | 0.113119 | 2693.208 | 2845.977 | 2411.447 | 2713.814 | 2418.276 | 2613.538 | 2104.079 | 1901.109 |
| ENSMUSG00000035472 | -0.538532725 | 0.061362 | 0.157418 | 65.39353 | 35.99465 | 40.57722 | 48.30692 | 69.2329 | 52.51658 | 63.64308 | 91.35765 |
| ENSMUSG00000035473 | -0.182714121 | 0.119839 | 0.248762 | 580.4934 | 610.7092 | 751.9208 | 679.7473 | 717.6819 | 739.7017 | 682.7167 | 841.7955 |
| ENSMUSG00000035476 | 0.104153449 | 0.217788 | 0.376323 | 683.1109 | 706.6949 | 719.6247 | 753.9329 | 632.8472 | 737.4669 | 651.8595 | 644.9415 |
| ENSMUSG00000035478 | -0.325310808 | 0.000269 | 0.002849 | 3798.861 | 4456.137 | 4517.322 | 4235.481 | 5611.766 | 4703.028 | 5407.734 | 5585.868 |
| ENSMUSG00000035486 | 0.031267046 | 0.897372 | 0.937817 | 41.24823 | 43.19358 | 45.54586 | 44.85642 | 35.10401 | 44.69496 | 49.17875 | 42.41605 |
| ENSMUSG00000035493 | 0.172139255 | 0.343729 | 0.51247 | 8885.472 | 12340.16 | 10834.95 | 10946.69 | 10397.61 | 12487.77 | 8373.887 | 6910.554 |
| ENSMUSG00000035495 | -0.097936486 | 0.366635 | 0.535165 | 873.2552 | 909.4647 | 756.8895 | 690.9614 | 886.3762 | 822.3873 | 899.6818 | 842.8831 |
| ENSMUSG00000035498 | -0.489500064 | 9.23E-06 | 0.000214 | 378.2764 | 379.1436 | 433.9279 | 364.8897 | 539.2366 | 506.1705 | 620.0379 | 519.8685 |
| ENSMUSG00000035504 | -0.618853967 | 0.000615 | 0.005376 | 194.1685 | 267.5602 | 267.4784 | 243.2598 | 420.273 | 284.9304 | 346.1798 | 441.562 |
| ENSMUSG00000035505 | -0.03743897 | 0.796654 | 0.875233 | 285.7194 | 207.5691 | 244.2915 | 266.5507 | 243.7778 | 286.0478 | 228.5365 | 275.1605 |

| | | | | | | | | | | | |
|---|---|---|---|---|---|---|---|---|---|---|---|
| ENSMUSG00000035506 | 0.17010098 | 0.571354 | 0.714306 | 38.23007 | 39.59411 | 37.2648 | 46.58167 | 33.15378 | 44.69496 | 23.14294 | 43.50364 |
| ENSMUSG00000035509 | 0.274452672 | 0.291494 | 0.458716 | 73.44197 | 59.99108 | 77.01392 | 55.2079 | 58.50668 | 70.39457 | 53.0359 | 38.06569 |
| ENSMUSG00000035513 | -0.84794309 | 0.000915 | 0.007299 | 78.47224 | 77.9884 | 73.70149 | 74.18562 | 89.71024 | 136.3196 | 125.3576 | 196.854 |
| ENSMUSG00000035517 | -0.203911119 | 0.202512 | 0.358141 | 696.1896 | 767.8858 | 910.0892 | 581.4082 | 862.9735 | 730.7627 | 809.0386 | 1001.671 |
| ENSMUSG00000035521 | -0.321898233 | 0.003769 | 0.020938 | 718.3228 | 631.1061 | 757.7176 | 705.626 | 893.202 | 751.9928 | 877.5031 | 996.2334 |
| ENSMUSG00000035529 | 0.1037586 | 0.119953 | 0.248895 | 1179.096 | 1147.029 | 1135.334 | 1199.909 | 1149.656 | 1108.435 | 1041.432 | 1038.649 |
| ENSMUSG00000035530 | -0.111169868 | 0.104307 | 0.22735 | 7702.352 | 7398.1 | 8788.696 | 8257.032 | 8571.228 | 8546.794 | 8809.746 | 8801.875 |
| ENSMUSG00000035539 | 0.234956804 | 0.501468 | 0.657789 | 40.24217 | 22.79661 | 25.67131 | 28.46658 | 19.50223 | 24.58223 | 22.17865 | 33.71532 |
| ENSMUSG00000035540 | -0.320982519 | 0.736215 | 0.834773 | 93.56306 | 0 | 8.281066 | 78.49874 | 45.83023 | 41.34284 | 75.21455 | 63.08028 |
| ENSMUSG00000035545 | 0.289804536 | 1.20E-05 | 0.000264 | 4734.492 | 4194.576 | 4500.76 | 4654.716 | 3878.018 | 3827.006 | 3558.227 | 3533.583 |
| ENSMUSG00000035547 | -0.05265107 | 0.786164 | 0.868451 | 755.5468 | 650.3033 | 1039.274 | 789.3005 | 841.5211 | 595.5604 | 919.9319 | 999.4962 |
| ENSMUSG00000035557 | -0.139924316 | 0.870979 | 0.922556 | 9.054489 | 21.59679 | 9.93728 | 8.626235 | 6.825779 | 40.22547 | 3.857157 | 3.262773 |
| ENSMUSG00000035559 | -0.529101142 | 0.007156 | 0.033419 | 500.009 | 323.9518 | 474.5051 | 371.7907 | 581.1663 | 442.4801 | 660.5381 | 727.5984 |
| ENSMUSG00000035561 | -0.389776968 | 0.068659 | 0.169839 | 187.1261 | 152.3773 | 195.4332 | 229.4578 | 253.5289 | 197.7752 | 212.1436 | 340.416 |
| ENSMUSG00000035566 | 0.108617556 | 0.492643 | 0.649676 | 644.8808 | 518.3229 | 655.0323 | 569.3315 | 457.3272 | 703.9457 | 491.7875 | 564.4598 |
| ENSMUSG00000035569 | 0.438512019 | 0.012307 | 0.049933 | 6620.844 | 6096.293 | 5214.587 | 5486.285 | 3771.731 | 6060.637 | 3885.121 | 3562.948 |
| ENSMUSG00000035572 | -0.012697301 | 0.846511 | 0.907743 | 992.9756 | 1009.05 | 1014.431 | 991.1544 | 1036.543 | 1000.05 | 1056.861 | 947.2918 |
| ENSMUSG00000035575 | -0.071471373 | 0.387246 | 0.554697 | 1422.561 | 1520.174 | 1315.861 | 1311.188 | 1494.846 | 1337.497 | 1471.505 | 1542.204 |
| ENSMUSG00000035578 | -0.38559304 | 0.072152 | 0.176275 | 84.50857 | 97.18554 | 90.26362 | 103.5148 | 126.7645 | 101.681 | 99.32178 | 163.1387 |
| ENSMUSG00000035580 | 0.388803014 | 0.373646 | 0.542437 | 37.22401 | 65.99018 | 36.43669 | 36.23019 | 20.47734 | 62.57295 | 25.07152 | 26.10219 |
| ENSMUSG00000035585 | -0.128017314 | 0.126106 | 0.256974 | 3176.114 | 3363.1 | 3537.672 | 3324.551 | 4017.459 | 3245.972 | 3565.941 | 3815.27 |
| ENSMUSG00000035594 | -0.346576861 | 0.494234 | 0.651076 | 12.07265 | 14.39786 | 13.24971 | 25.01608 | 10.72622 | 13.40849 | 27.0001 | 31.54014 |
| ENSMUSG00000035595 | -0.150602674 | 0.436044 | 0.600086 | 131.7931 | 94.7859 | 107.6539 | 87.12497 | 112.1378 | 131.8501 | 121.5004 | 102.2336 |
| ENSMUSG00000035596 | -0.159445631 | 0.056249 | 0.147841 | 1159.981 | 1090.638 | 1265.347 | 1175.756 | 1356.38 | 1176.595 | 1364.469 | 1345.35 |
| ENSMUSG00000035597 | 0.49662949 | 0.001121 | 0.008519 | 2655.983 | 2703.198 | 2025.549 | 2818.191 | 1827.359 | 2223.574 | 1710.649 | 1469.336 |
| ENSMUSG00000035601 | 0.227616378 | 0.060647 | 0.15626 | 441.6579 | 548.3184 | 489.411 | 579.683 | 482.6801 | 410.0763 | 452.2516 | 412.197 |
| ENSMUSG00000035606 | 0.27215657 | 0.534587 | 0.685591 | 24.1453 | 58.79126 | 26.49941 | 27.60395 | 32.17867 | 39.10809 | 15.42863 | 26.10219 |
| ENSMUSG00000035614 | 0.077078822 | 0.412783 | 0.578635 | 974.8667 | 863.8715 | 758.5457 | 850.5468 | 829.8197 | 795.5704 | 816.7529 | 823.3065 |
| ENSMUSG00000035615 | 0.268347571 | 0.258656 | 0.422517 | 58.35115 | 68.38983 | 56.31125 | 81.94923 | 53.63112 | 65.92507 | 54.96448 | 45.67883 |
| ENSMUSG00000035620 | 0.116807435 | 0.192204 | 0.34549 | 1065.412 | 1059.442 | 914.2297 | 1136.075 | 913.6793 | 959.8243 | 999.9679 | 975.5692 |
| ENSMUSG00000035621 | -0.064196923 | 0.522637 | 0.675983 | 2790.795 | 2249.665 | 2653.254 | 2512.822 | 2796.619 | 2375.537 | 2565.973 | 2935.408 |
| ENSMUSG00000035623 | 0.200685458 | 0.051872 | 0.139494 | 1888.364 | 1850.125 | 1572.574 | 1775.279 | 1449.015 | 1776.625 | 1408.826 | 1531.328 |
| ENSMUSG00000035629 | 0.098868631 | 0.286647 | 0.453796 | 765.6074 | 848.2738 | 883.5898 | 857.4478 | 772.2882 | 699.4762 | 810.9672 | 850.4962 |
| ENSMUSG00000035632 | 0.035812311 | 0.674019 | 0.791346 | 1673.068 | 1526.173 | 1428.484 | 1357.769 | 1488.995 | 1386.661 | 1457.041 | 1501.963 |
| ENSMUSG00000035637 | -1.04408482 | 0.000331 | 0.003341 | 920.5397 | 494.3265 | 658.3448 | 552.9417 | 1383.683 | 739.7017 | 1247.79 | 2044.671 |
| ENSMUSG00000035638 | -0.581332162 | 0.328372 | 0.496188 | 188.1322 | 133.1802 | 588.7838 | 165.6237 | 385.169 | 200.01 | 392.4657 | 632.978 |
| ENSMUSG00000035640 | 0.368384754 | 1.86E-06 | 6.10E-05 | 672.0443 | 623.9072 | 664.9696 | 647.8302 | 521.6845 | 488.2925 | 523.609 | 487.2408 |
| ENSMUSG00000035642 | -0.289687735 | 0.031326 | 0.09733 | 401.4157 | 315.5531 | 394.1788 | 416.6471 | 492.4312 | 408.9589 | 437.7873 | 531.832 |
| ENSMUSG00000035649 | 0.490568148 | 0.002613 | 0.015928 | 1569.445 | 1403.791 | 1170.943 | 1507.866 | 861.0233 | 1329.675 | 897.7532 | 935.3283 |
| ENSMUSG00000035653 | 0.333783272 | 0.562715 | 0.708229 | 25.15136 | 19.19714 | 91.91984 | 18.11509 | 20.47734 | 31.28647 | 36.64299 | 34.80291 |
| ENSMUSG00000035666 | 0.010620234 | 0.886425 | 0.932078 | 1875.285 | 1881.32 | 1798.648 | 1859.816 | 1959.974 | 1707.348 | 1724.149 | 1967.452 |
| ENSMUSG00000035671 | -0.152220791 | 0.26108 | 0.425173 | 1327.992 | 1023.448 | 1055.008 | 1001.506 | 1388.559 | 1014.576 | 1296.969 | 1196.35 |
| ENSMUSG00000035673 | -0.190334372 | 0.207058 | 0.363672 | 770.6376 | 529.1213 | 751.0927 | 572.782 | 794.7157 | 688.3024 | 823.5029 | 688.4452 |
| ENSMUSG00000035674 | -0.454651107 | 0.003115 | 0.018293 | 1612.705 | 1508.176 | 1831.772 | 1603.617 | 2183.274 | 1664.887 | 2342.258 | 2796.197 |
| ENSMUSG00000035678 | 0.195602644 | 0.324122 | 0.492102 | 76.46013 | 73.18911 | 81.15445 | 79.36136 | 72.15824 | 52.51658 | 74.25027 | 71.78101 |
| ENSMUSG00000035681 | 0.40339743 | 0.029611 | 0.093727 | 548.2996 | 433.1356 | 482.7862 | 536.5518 | 273.0312 | 486.0577 | 413.6801 | 341.5036 |
| ENSMUSG00000035683 | -0.015440335 | 0.8153 | 0.887529 | 864.2007 | 895.0669 | 880.2773 | 922.1445 | 931.2313 | 911.7773 | 899.6818 | 857.0218 |
| ENSMUSG00000035686 | -0.065757405 | 0.921855 | 0.953418 | 83.50251 | 13.19804 | 17.39024 | 31.91707 | 20.47734 | 33.52122 | 79.07171 | 19.57664 |
| ENSMUSG00000035692 | 0.317257466 | 0.718021 | 0.823903 | 56.33904 | 263.9607 | 510.1137 | 34.50494 | 114.088 | 82.68568 | 269.0367 | 228.3941 |
| ENSMUSG00000035694 | 0.501333189 | 0.105482 | 0.229195 | 93.56306 | 104.3845 | 118.4192 | 107.8279 | 78.98402 | 117.3243 | 63.64308 | 40.24087 |
| ENSMUSG00000035696 | 0.351734045 | 0.01833 | 0.06637 | 2558.396 | 2658.805 | 2355.963 | 2676.721 | 1901.467 | 2673.876 | 1714.506 | 1743.409 |
| ENSMUSG00000035697 | 0.511868133 | 0.000225 | 0.002482 | 336.0222 | 405.5397 | 368.5075 | 382.1422 | 245.728 | 319.569 | 226.608 | 255.5839 |
| ENSMUSG00000035699 | -2.966333336 | 5.18E-05 | 0.000814 | 5.030272 | 8.398751 | 19.04645 | 5.175741 | 117.0134 | 14.52586 | 30.85725 | 133.7737 |
| ENSMUSG00000035704 | -0.006337438 | 0.9587 | 0.976693 | 579.4873 | 532.7208 | 463.7397 | 540.0023 | 511.9334 | 623.4947 | 474.4303 | 515.5182 |
| ENSMUSG00000035711 | 0.645896342 | 0.015244 | 0.058358 | 66.39959 | 51.59233 | 49.6864 | 58.6584 | 29.25334 | 44.69496 | 41.46443 | 29.36496 |
| ENSMUSG00000035713 | 0.628802101 | 2.17E-05 | 0.000421 | 238.4349 | 195.5709 | 213.6515 | 228.5952 | 145.2916 | 166.4887 | 132.1076 | 123.9854 |
| ENSMUSG00000035722 | 0.21810376 | 0.136202 | 0.270789 | 448.7002 | 367.1454 | 427.303 | 425.2734 | 347.1396 | 282.6956 | 378.9656 | 426.3357 |
| ENSMUSG00000035725 | 0.138355774 | 0.219859 | 0.378695 | 754.5408 | 713.8938 | 732.8744 | 766.8723 | 556.7886 | 763.1665 | 658.6095 | 721.0729 |
| ENSMUSG00000035726 | -0.092139262 | 0.181682 | 0.3322 | 3046.333 | 3228.72 | 2826.328 | 2979.502 | 3461.645 | 3155.464 | 3089.582 | 3163.802 |
| ENSMUSG00000035735 | 0.101911028 | 0.512578 | 0.667795 | 192.1564 | 219.5673 | 193.777 | 243.2598 | 221.3503 | 202.2447 | 161.0363 | 206.6423 |
| ENSMUSG00000035745 | 0.601245782 | 0.204686 | 0.36081 | 10.06054 | 31.19536 | 20.70267 | 18.97772 | 9.751113 | 17.87799 | 12.53576 | 13.05109 |
| ENSMUSG00000035754 | -0.379828737 | 0.000892 | 0.007133 | 1031.206 | 1273.011 | 1224.77 | 1170.58 | 1606.008 | 1317.384 | 1450.291 | 1740.146 |
| ENSMUSG00000035757 | -0.285988295 | 0.017934 | 0.065471 | 413.4883 | 405.5397 | 415.7095 | 426.9986 | 557.7637 | 408.9589 | 505.2875 | 553.5839 |
| ENSMUSG00000035759 | 0.114695564 | 0.44859 | 0.61128 | 200.2048 | 239.9643 | 187.9802 | 165.6237 | 175.52 | 174.3104 | 191.8935 | 188.1533 |

| | | | | | | | | | | | |
|---|---|---|---|---|---|---|---|---|---|---|---|
| ENSMUSG00000035762 | -0.110685574 | 0.269241 | 0.434022 | 534.2149 | 476.3291 | 489.411 | 503.7721 | 493.4063 | 604.4994 | 559.2877 | 507.905 |
| ENSMUSG00000035764 | -0.041348806 | 0.664508 | 0.784504 | 837.0372 | 917.8635 | 738.6711 | 772.048 | 821.0437 | 862.6128 | 882.3246 | 788.5035 |
| ENSMUSG00000035765 | -0.036566877 | 0.640326 | 0.766904 | 790.7587 | 848.2738 | 814.8569 | 894.5406 | 890.2766 | 889.4298 | 855.3245 | 799.3794 |
| ENSMUSG00000035769 | -1.482700315 | 9.13E-07 | 3.46E-05 | 356.1432 | 245.9634 | 507.6294 | 339.011 | 997.5389 | 537.4569 | 1004.789 | 1512.839 |
| ENSMUSG00000035770 | 0.151146287 | 0.074982 | 0.181015 | 3703.286 | 3286.311 | 3054.057 | 3243.464 | 3198.365 | 3101.83 | 2866.832 | 2796.197 |
| ENSMUSG00000035772 | -0.299078662 | 8.05E-05 | 0.001142 | 855.1462 | 836.2756 | 907.6049 | 843.6458 | 1091.15 | 1095.027 | 966.2177 | 1087.591 |
| ENSMUSG00000035773 | -0.466317912 | 0.220087 | 0.378875 | 17.10292 | 25.19625 | 21.53077 | 31.05445 | 39.00445 | 17.87799 | 32.78583 | 41.32846 |
| ENSMUSG00000035775 | 0.283410783 | 0.238572 | 0.399881 | 221.332 | 172.7743 | 262.5098 | 200.9913 | 235.0018 | 158.6671 | 195.7507 | 115.2847 |
| ENSMUSG00000035776 | 0.022380724 | 0.726948 | 0.829129 | 1232.417 | 1346.2 | 1379.626 | 1362.945 | 1326.151 | 1328.558 | 1277.683 | 1309.46 |
| ENSMUSG00000035778 | 0.314965387 | 0.003873 | 0.02133 | 1009.073 | 1105.036 | 989.5874 | 1017.896 | 938.0571 | 846.9696 | 838.9316 | 686.27 |
| ENSMUSG00000035781 | -0.040090334 | 0.37663 | 0.544772 | 2415.537 | 2370.847 | 2494.257 | 2411.033 | 2510.912 | 2500.683 | 2459.902 | 2498.197 |
| ENSMUSG00000035783 | -0.22514746 | 0.312399 | 0.480769 | 4012.145 | 6886.976 | 9459.462 | 6974.311 | 7756.035 | 7348.969 | 7199.383 | 9646.933 |
| ENSMUSG00000035790 | -0.190040063 | 0.039167 | 0.114777 | 555.342 | 676.6993 | 611.1427 | 574.5072 | 666.001 | 669.3071 | 688.5025 | 730.8612 |
| ENSMUSG00000035798 | 0.260758182 | 0.013115 | 0.052372 | 1366.222 | 1345 | 1194.958 | 1378.472 | 1035.568 | 1300.623 | 1045.289 | 1029.949 |
| ENSMUSG00000035799 | 0.372774326 | 0.069127 | 0.170721 | 434.6155 | 680.2988 | 367.6793 | 550.3538 | 419.2979 | 453.6539 | 363.537 | 331.7153 |
| ENSMUSG00000035805 | 0.184269939 | 0.529999 | 0.681935 | 32.19374 | 32.39518 | 42.23344 | 29.3292 | 26.32801 | 36.87335 | 27.96439 | 29.36496 |
| ENSMUSG00000035818 | 0.042111155 | 0.92029 | 0.95244 | 136.8234 | 196.7707 | 116.763 | 133.7066 | 122.864 | 287.1651 | 89.67889 | 67.43065 |
| ENSMUSG00000035824 | 0.169014129 | 0.140382 | 0.276804 | 729.3894 | 811.0794 | 766.8267 | 801.3772 | 739.1344 | 787.7487 | 662.4667 | 575.3357 |
| ENSMUSG00000035828 | -1.25291117 | 2.82E-05 | 0.000514 | 1506.063 | 773.8849 | 1449.187 | 1757.164 | 2314.914 | 2458.223 | 5331.555 | 2973.474 |
| ENSMUSG00000035829 | 0.020933491 | 0.834855 | 0.900162 | 463.7911 | 453.5325 | 409.0847 | 483.9318 | 453.4268 | 463.7102 | 403.0729 | 464.4014 |
| ENSMUSG00000035834 | -0.259005679 | 0.036958 | 0.1105 | 195.1745 | 189.5718 | 209.511 | 221.6942 | 260.3547 | 254.7613 | 247.8223 | 215.343 |
| ENSMUSG00000035835 | 0.348407507 | 0.000324 | 0.003286 | 1890.376 | 1928.113 | 1812.725 | 2122.054 | 1693.768 | 1593.375 | 1458.005 | 1344.263 |
| ENSMUSG00000035840 | 0.112659187 | 0.318089 | 0.486423 | 607.6568 | 488.3274 | 469.5365 | 514.9862 | 474.8792 | 448.067 | 485.0374 | 515.5182 |
| ENSMUSG00000035842 | 0.270955137 | 0.002056 | 0.013383 | 623.7537 | 646.7038 | 615.2832 | 592.6223 | 564.5894 | 484.9404 | 516.859 | 483.978 |
| ENSMUSG00000035845 | -0.11484753 | 0.504051 | 0.65977 | 295.78 | 245.9634 | 307.2276 | 239.8093 | 325.6872 | 220.1227 | 294.1082 | 339.3284 |
| ENSMUSG00000035847 | 0.433052723 | 6.33E-05 | 0.000954 | 964.8061 | 907.0651 | 915.8859 | 1163.679 | 697.2046 | 788.8661 | 740.5741 | 702.5838 |
| ENSMUSG00000035849 | 0.869120565 | 0.099645 | 0.22043 | 19.11503 | 13.19804 | 13.24971 | 13.80198 | 5.850668 | 11.17374 | 3.857157 | 11.9635 |
| ENSMUSG00000035851 | 0.284615306 | 0.034802 | 0.105608 | 3805.904 | 3601.864 | 2995.262 | 3439.28 | 2469.957 | 3555.484 | 2768.474 | 2569.978 |
| ENSMUSG00000035852 | -1.340835531 | 9.25E-07 | 3.49E-05 | 167.005 | 125.9813 | 169.7619 | 156.1349 | 385.169 | 206.7142 | 429.1087 | 548.1459 |
| ENSMUSG00000035863 | -0.389922479 | 0.002809 | 0.01686 | 2003.054 | 2250.865 | 2730.268 | 2215.217 | 3017.969 | 2574.43 | 2919.868 | 3544.459 |
| ENSMUSG00000035864 | 0.408981474 | 0.041788 | 0.120336 | 319.9253 | 297.5557 | 263.3379 | 467.5419 | 272.0561 | 272.6393 | 213.1079 | 258.8467 |
| ENSMUSG00000035868 | 0.529298139 | 0.043513 | 0.123608 | 135.8173 | 149.9777 | 76.18581 | 93.16334 | 80.93424 | 83.80306 | 65.57166 | 83.74451 |
| ENSMUSG00000035873 | -0.21717187 | 0.061811 | 0.158289 | 765.6074 | 788.2827 | 846.325 | 895.4032 | 892.2268 | 815.6831 | 1058.79 | 1065.839 |
| ENSMUSG00000035875 | -0.309046733 | 0.486645 | 0.644622 | 17.10292 | 31.19536 | 18.21835 | 25.01608 | 32.17867 | 43.57759 | 24.10723 | 13.05109 |
| ENSMUSG00000035877 | -0.317870404 | 0.108479 | 0.233289 | 1534.233 | 950.2587 | 1276.112 | 1420.741 | 1195.486 | 1673.826 | 2102.15 | 1488.912 |
| ENSMUSG00000035878 | -1.465100017 | 1.33E-07 | 7.16E-06 | 134.8113 | 143.9786 | 173.9024 | 169.0742 | 395.8952 | 234.6486 | 441.6444 | 646.0291 |
| ENSMUSG00000035885 | -0.795455506 | 8.79E-05 | 0.001216 | 4522.214 | 4254.567 | 6021.991 | 4647.815 | 8421.061 | 5370.1 | 9047.925 | 10915.06 |
| ENSMUSG00000035890 | -0.334463527 | 0.000486 | 0.00448 | 1162.999 | 1219.019 | 1202.411 | 1229.238 | 1587.481 | 1269.337 | 1577.577 | 1633.562 |
| ENSMUSG00000035891 | 0.244047332 | 0.0024 | 0.014962 | 1061.387 | 935.8608 | 1004.493 | 1018.758 | 878.5753 | 819.0352 | 906.4318 | 790.6787 |
| ENSMUSG00000035898 | 0.361964424 | 0.017443 | 0.064118 | 869.231 | 791.8822 | 679.8755 | 791.8884 | 565.5646 | 788.8661 | 558.3234 | 525.3065 |
| ENSMUSG00000035900 | 0.309436171 | 4.85E-05 | 0.000778 | 1639.869 | 1673.751 | 1520.404 | 1544.959 | 1368.081 | 1331.91 | 1269.969 | 1173.511 |
| ENSMUSG00000035901 | 0.191190267 | 0.003878 | 0.021346 | 3827.031 | 3769.839 | 3492.126 | 4086.247 | 3357.308 | 3188.986 | 3396.226 | 3347.605 |
| ENSMUSG00000035910 | -0.794995267 | 0.001813 | 0.012232 | 1409.482 | 1012.649 | 2397.369 | 1387.099 | 2519.688 | 2033.621 | 2689.402 | 3530.321 |
| ENSMUSG00000035914 | 0.00035245 | 0.996327 | 0.997787 | 1177.084 | 1288.608 | 1405.297 | 1234.414 | 1304.699 | 1280.511 | 1279.612 | 1240.941 |
| ENSMUSG00000035916 | -0.260360214 | 0.542598 | 0.691778 | 17.10292 | 26.39607 | 11.59349 | 15.52722 | 20.47734 | 13.40849 | 21.21436 | 28.27737 |
| ENSMUSG00000035919 | -0.07501977 | 0.5279 | 0.680556 | 495.9848 | 457.132 | 491.8953 | 545.178 | 593.8428 | 481.5882 | 562.1806 | 458.9634 |
| ENSMUSG00000035929 | 0.237697808 | 0.019207 | 0.068496 | 849.1099 | 857.8724 | 1000.353 | 858.3104 | 687.4535 | 721.8237 | 840.8602 | 775.4524 |
| ENSMUSG00000035930 | 1.574808274 | 0.061722 | 0.158104 | 27.16347 | 53.99197 | 33.12427 | 52.62003 | 3.900445 | 43.57759 | 7.714313 | 1.087591 |
| ENSMUSG00000035933 | -0.223456565 | 0.101856 | 0.223792 | 595.5842 | 511.124 | 525.8477 | 458.0531 | 600.6686 | 497.2315 | 698.1454 | 642.7663 |
| ENSMUSG00000035934 | 0.074554014 | 0.602493 | 0.740217 | 300.8103 | 301.1552 | 323.7897 | 235.4962 | 252.5538 | 315.0995 | 270.9653 | 264.2846 |
| ENSMUSG00000035936 | -0.347645581 | 0.003643 | 0.020416 | 599.6084 | 644.3042 | 733.7025 | 615.9132 | 792.7655 | 718.4715 | 830.253 | 960.3429 |
| ENSMUSG00000035941 | 0.116668268 | 0.154723 | 0.296344 | 1674.074 | 1794.933 | 1570.09 | 1606.205 | 1429.513 | 1603.432 | 1445.469 | 1648.788 |
| ENSMUSG00000035944 | -0.165465237 | 0.073919 | 0.179271 | 651.9232 | 614.3086 | 554.8314 | 564.1558 | 666.001 | 626.8469 | 719.3597 | 657.9926 |
| ENSMUSG00000035948 | 0.372924231 | 0.106359 | 0.230448 | 228.3743 | 140.3791 | 132.4971 | 171.6621 | 139.4409 | 97.21155 | 148.5005 | 133.7737 |
| ENSMUSG00000035949 | 0.104836669 | 0.123543 | 0.253229 | 1839.067 | 1847.725 | 1904.645 | 2003.874 | 1851.736 | 1825.789 | 1755.006 | 1632.474 |
| ENSMUSG00000035953 | -0.104574449 | 0.183878 | 0.334911 | 1083.521 | 1113.434 | 1157.693 | 1213.711 | 1258.869 | 1187.769 | 1324.933 | 1140.883 |
| ENSMUSG00000035954 | 0.016561973 | 0.90834 | 0.945278 | 405.4399 | 353.9474 | 327.9302 | 389.9058 | 321.7867 | 435.7759 | 386.68 | 316.489 |
| ENSMUSG00000035958 | 0.084802142 | 0.348028 | 0.516667 | 1359.179 | 1327.003 | 1164.318 | 1308.6 | 1291.047 | 1306.21 | 1113.754 | 1151.759 |
| ENSMUSG00000035960 | -0.411098251 | 0.001527 | 0.010688 | 1384.331 | 1384.594 | 1436.765 | 1526.844 | 2182.299 | 1454.821 | 1928.578 | 2056.635 |
| ENSMUSG00000035964 | 0.302004588 | 0.272575 | 0.438057 | 91.55095 | 148.7779 | 178.871 | 166.4863 | 157.968 | 128.498 | 104.1432 | 84.8321 |
| ENSMUSG00000035967 | 0.351606507 | 0.001984 | 0.01306 | 3139.896 | 2682.801 | 2468.586 | 3064.039 | 2032.132 | 2536.439 | 2282.472 | 2049.022 |
| ENSMUSG00000035969 | -0.030099946 | 0.748001 | 0.842597 | 637.8385 | 577.1142 | 670.7664 | 624.5394 | 625.0463 | 631.3164 | 711.6454 | 597.0875 |
| ENSMUSG00000035983 | -0.188824024 | 0.517783 | 0.671803 | 29.17558 | 25.19625 | 28.15563 | 24.15346 | 26.32801 | 29.05173 | 34.71441 | 31.54014 |
| ENSMUSG00000035984 | -0.044219094 | 0.905769 | 0.943371 | 28.16952 | 37.19447 | 59.62368 | 49.16954 | 57.53157 | 43.57759 | 54.96448 | 23.927 |

| | | | | | | | | | | | |
|---|---|---|---|---|---|---|---|---|---|---|---|
| ENSMUSG00000035992 | 0.393118796 | 0.006716 | 0.03202 | 1185.132 | 1238.216 | 1048.383 | 1184.382 | 802.5166 | 1158.717 | 788.7885 | 796.1167 |
| ENSMUSG00000036002 | -0.052662235 | 0.712637 | 0.819906 | 592.566 | 429.5361 | 507.6294 | 502.9095 | 472.929 | 493.8794 | 637.3951 | 504.6423 |
| ENSMUSG00000036006 | 0.438481682 | 0.022523 | 0.076682 | 196.1806 | 205.1695 | 188.8083 | 216.5185 | 120.9138 | 186.6015 | 172.6078 | 115.2847 |
| ENSMUSG00000036009 | 0.080241917 | 0.617559 | 0.750167 | 283.7073 | 344.3488 | 279.9 | 366.615 | 339.3387 | 344.1512 | 256.5009 | 265.3722 |
| ENSMUSG00000036019 | 0.182565268 | 0.198656 | 0.353251 | 1110.684 | 1262.212 | 1162.662 | 1020.484 | 1029.718 | 1246.989 | 917.039 | 820.0437 |
| ENSMUSG00000036022 | 0.323917951 | 0.029628 | 0.093727 | 391.3551 | 334.7502 | 292.3216 | 325.2091 | 247.6783 | 325.1559 | 243.0009 | 257.7591 |
| ENSMUSG00000036023 | -0.011556623 | 0.890232 | 0.933635 | 1214.308 | 1310.205 | 1103.866 | 1323.264 | 1268.62 | 1254.811 | 1287.326 | 1176.774 |
| ENSMUSG00000036026 | -0.174157808 | 0.164136 | 0.309111 | 1310.889 | 1129.032 | 1388.735 | 1229.238 | 1437.314 | 1144.191 | 1512.97 | 1613.985 |
| ENSMUSG00000036030 | 0.162784715 | 0.384891 | 0.552504 | 402.4217 | 381.5432 | 277.4157 | 332.9727 | 319.8365 | 400.0199 | 268.0724 | 256.6715 |
| ENSMUSG00000036036 | 0.628056262 | 2.74E-07 | 1.28E-05 | 2009.091 | 2062.493 | 1755.586 | 2064.258 | 1251.068 | 1549.798 | 1219.826 | 1085.416 |
| ENSMUSG00000036040 | -0.560190393 | 0.102625 | 0.224947 | 50.30272 | 23.99643 | 36.43669 | 57.79577 | 51.6809 | 50.28183 | 87.75031 | 59.81751 |
| ENSMUSG00000036046 | -0.106670966 | 0.121369 | 0.250642 | 1183.12 | 1207.02 | 1310.893 | 1258.568 | 1358.33 | 1280.511 | 1295.04 | 1410.606 |
| ENSMUSG00000036052 | -0.228555899 | 0.067092 | 0.167299 | 362.1796 | 297.5557 | 263.3379 | 306.2313 | 374.4427 | 347.5033 | 370.287 | 345.854 |
| ENSMUSG00000036053 | 0.349086099 | 0.01924 | 0.068565 | 2346.119 | 2276.061 | 1956.816 | 2424.835 | 1664.515 | 2290.617 | 1673.042 | 1441.058 |
| ENSMUSG00000036054 | 0.041172539 | 0.702907 | 0.812784 | 939.6548 | 813.479 | 838.0439 | 790.1631 | 888.3264 | 903.9556 | 770.467 | 723.2481 |
| ENSMUSG00000036057 | 0.094061524 | 0.206053 | 0.362636 | 1052.333 | 976.6547 | 1004.493 | 923.0071 | 934.1566 | 892.7819 | 902.5747 | 976.6568 |
| ENSMUSG00000036061 | 0.794453401 | 5.74E-08 | 3.64E-06 | 1057.363 | 1353.399 | 906.7768 | 1218.887 | 617.2455 | 763.1665 | 622.9308 | 611.2262 |
| ENSMUSG00000036062 | 0.800609737 | 0.000424 | 0.004034 | 105.6357 | 74.38894 | 113.4506 | 115.5915 | 46.80534 | 61.45558 | 61.71451 | 66.34306 |
| ENSMUSG00000036073 | 0.00980444 | 0.928511 | 0.957673 | 1152.938 | 1015.049 | 1004.493 | 1216.299 | 1016.066 | 979.9371 | 1215.004 | 1148.496 |
| ENSMUSG00000036078 | -0.26274825 | 0.043794 | 0.124184 | 909.4731 | 915.4638 | 977.1658 | 920.4193 | 1162.333 | 848.0869 | 1186.076 | 1269.219 |
| ENSMUSG00000036083 | -4.009541891 | 7.91E-12 | 1.87E-09 | 13.07871 | 4.799286 | 34.78048 | 15.52722 | 338.3636 | 89.38993 | 177.4292 | 506.8174 |
| ENSMUSG00000036086 | 0.161171714 | 0.289056 | 0.456124 | 493.9727 | 577.1142 | 438.8965 | 544.3154 | 433.9245 | 566.5087 | 456.1088 | 379.5693 |
| ENSMUSG00000036087 | 0.050877019 | 0.547651 | 0.695961 | 1496.003 | 1390.593 | 1242.16 | 1363.808 | 1241.317 | 1357.61 | 1289.255 | 1412.781 |
| ENSMUSG00000036091 | -0.187628007 | 0.490197 | 0.647606 | 32.19374 | 41.99375 | 31.46805 | 37.09281 | 34.1289 | 43.57759 | 35.6787 | 48.9416 |
| ENSMUSG00000036093 | 0.023382022 | 0.754922 | 0.847857 | 2018.145 | 2188.474 | 1869.865 | 2032.341 | 1852.711 | 2072.729 | 2061.65 | 1987.029 |
| ENSMUSG00000036095 | -0.00131289 | 0.997923 | 0.998833 | 23.13925 | 19.19714 | 16.56213 | 50.03216 | 19.50223 | 45.81234 | 17.3572 | 27.18978 |
| ENSMUSG00000036097 | 0.174120145 | 0.182177 | 0.332766 | 1822.97 | 1865.722 | 1631.37 | 1830.487 | 1589.431 | 1977.752 | 1434.862 | 1335.562 |
| ENSMUSG00000036098 | -0.056978557 | 0.873457 | 0.924289 | 260.5681 | 421.1374 | 264.166 | 295.8799 | 306.185 | 591.0909 | 164.8934 | 229.4817 |
| ENSMUSG00000036099 | 0.263829832 | 0.003329 | 0.019112 | 1252.538 | 1264.612 | 1055.836 | 1185.245 | 937.082 | 1071.562 | 999.0036 | 952.7298 |
| ENSMUSG00000036103 | 0.056316085 | 0.66948 | 0.7881 | 1771.662 | 2723.595 | 2390.744 | 2284.227 | 2239.831 | 2362.129 | 2185.079 | 2030.533 |
| ENSMUSG00000036104 | 0.143779488 | 0.152957 | 0.293999 | 1486.948 | 1235.816 | 1305.924 | 1300.836 | 1111.627 | 1100.613 | 1323.005 | 1288.795 |
| ENSMUSG00000036106 | -0.577847386 | 0.000355 | 0.003518 | 566.4086 | 547.1186 | 711.3436 | 512.3984 | 920.5051 | 653.6638 | 942.1105 | 973.394 |
| ENSMUSG00000036109 | -0.003035664 | 0.98619 | 0.992283 | 160.9687 | 183.5727 | 211.1672 | 151.8217 | 213.5494 | 151.9629 | 176.4649 | 166.4014 |
| ENSMUSG00000036110 | -2.184053654 | 0.003095 | 0.018205 | 3.018163 | 2.399643 | 19.04645 | 2.58787 | 28.27823 | 16.76061 | 32.78583 | 47.85401 |
| ENSMUSG00000036111 | 0.263681376 | 0.26364 | 0.427964 | 131.7931 | 101.9848 | 110.1382 | 164.7611 | 124.8142 | 118.4417 | 76.17884 | 105.4963 |
| ENSMUSG00000036112 | -0.035775957 | 0.362443 | 0.531001 | 6711.389 | 7035.753 | 6921.315 | 7035.557 | 6958.394 | 7188.068 | 7136.704 | 7117.196 |
| ENSMUSG00000036114 | -0.183016669 | 0.091332 | 0.207258 | 522.1422 | 562.7163 | 554.8314 | 549.4912 | 629.9219 | 516.2268 | 684.6453 | 652.5547 |
| ENSMUSG00000036120 | 0.073723786 | 0.464088 | 0.624924 | 575.4631 | 495.5263 | 515.0823 | 558.1174 | 545.0872 | 462.5929 | 542.8948 | 486.1532 |
| ENSMUSG00000036123 | -3.338832408 | 2.16E-06 | 6.84E-05 | 15.09082 | 3.599465 | 50.5145 | 11.21411 | 190.1467 | 41.34284 | 290.251 | 299.0875 |
| ENSMUSG00000036136 | -0.672595942 | 0.005187 | 0.0264 | 122.7386 | 140.3791 | 181.3554 | 159.5853 | 185.2711 | 188.8362 | 245.8937 | 344.7664 |
| ENSMUSG00000036138 | -0.283051987 | 0.023751 | 0.080036 | 1898.425 | 1845.326 | 2041.283 | 1718.346 | 2327.591 | 1828.024 | 2273.794 | 2700.489 |
| ENSMUSG00000036139 | 0.303596809 | 0.201459 | 0.356928 | 1119.738 | 1478.18 | 1074.882 | 1160.229 | 968.2855 | 1464.877 | 868.8245 | 613.4014 |
| ENSMUSG00000036144 | -1.417695559 | 0.000123 | 0.001566 | 91.55095 | 160.7761 | 71.21717 | 99.2017 | 435.8748 | 136.3196 | 326.894 | 226.2189 |
| ENSMUSG00000036151 | 0.33936974 | 0.527336 | 0.680215 | 12.07265 | 23.99643 | 9.109173 | 16.38985 | 10.72622 | 20.11273 | 6.750024 | 10.87591 |
| ENSMUSG00000036155 | 0.225417824 | 0.020986 | 0.072929 | 1318.937 | 1172.226 | 1235.535 | 1262.018 | 901.0028 | 1125.196 | 1107.004 | 1136.533 |
| ENSMUSG00000036158 | 0.358354124 | 8.29E-05 | 0.001166 | 1391.373 | 1358.198 | 1641.307 | 1386.236 | 1087.249 | 1112.905 | 1086.754 | 1223.54 |
| ENSMUSG00000036160 | -0.003533965 | 0.969214 | 0.982853 | 1005.048 | 1095.437 | 968.8848 | 936.8091 | 1106.751 | 935.2421 | 1026.004 | 941.8539 |
| ENSMUSG00000036167 | 0.114582023 | 0.281504 | 0.447843 | 1180.102 | 1365.397 | 1157.693 | 1197.321 | 1116.502 | 1309.562 | 1002.861 | 1096.292 |
| ENSMUSG00000036168 | 0.663471351 | 0.017936 | 0.065471 | 42.25428 | 69.58965 | 66.24853 | 63.83414 | 36.07912 | 46.92971 | 41.46443 | 28.27737 |
| ENSMUSG00000036169 | -1.883796051 | 0.007398 | 0.034209 | 637.8385 | 230.3657 | 2045.423 | 588.3092 | 2484.584 | 1594.493 | 3625.727 | 5221.525 |
| ENSMUSG00000036172 | -0.098854176 | 0.894633 | 0.936266 | 11.0666 | 5.999108 | 28.98373 | 2.58787 | 19.50223 | 11.17374 | 7.714313 | 14.13868 |
| ENSMUSG00000036180 | -0.218169319 | 0.007312 | 0.033948 | 1934.643 | 1768.537 | 2033.002 | 1896.909 | 2322.715 | 1988.926 | 2222.687 | 2348.109 |
| ENSMUSG00000036181 | -0.611332227 | 0.001817 | 0.01224 | 690.1533 | 716.2935 | 1175.911 | 920.4193 | 1484.119 | 1020.163 | 1483.077 | 1367.102 |
| ENSMUSG00000036185 | 0.24681397 | 0.529246 | 0.681503 | 30.18163 | 55.19179 | 86.12309 | 54.34528 | 27.30312 | 65.92507 | 50.14304 | 47.85401 |
| ENSMUSG00000036186 | 0.120095802 | 0.235492 | 0.396374 | 421.5368 | 506.3247 | 525.8477 | 489.9701 | 463.1779 | 458.1234 | 432.9658 | 435.0364 |
| ENSMUSG00000036188 | 0.042939444 | 0.515286 | 0.66935 | 924.564 | 981.454 | 1002.837 | 1017.033 | 931.2313 | 945.2985 | 952.7177 | 984.2699 |
| ENSMUSG00000036192 | 0.144923816 | 0.585274 | 0.725623 | 42.25428 | 35.99465 | 59.62368 | 51.75741 | 48.75557 | 43.57759 | 43.39301 | 36.9781 |
| ENSMUSG00000036196 | 0.50840589 | 0.123954 | 0.253674 | 66.39959 | 68.38983 | 57.13936 | 62.97152 | 34.1289 | 77.09881 | 34.71441 | 33.71532 |
| ENSMUSG00000036197 | 0.131699785 | 0.190328 | 0.342936 | 1465.821 | 1334.202 | 1158.521 | 1248.216 | 1172.084 | 1314.032 | 1110.861 | 1153.934 |
| ENSMUSG00000036198 | 1.404131056 | 0.00025 | 0.002707 | 47.28455 | 32.39518 | 49.6864 | 64.69676 | 9.751113 | 24.58223 | 22.17865 | 17.40146 |
| ENSMUSG00000036199 | -0.437671061 | 0.004698 | 0.024641 | 1956.776 | 2022.899 | 2350.995 | 2065.121 | 2783.943 | 2110.72 | 2881.296 | 3596.664 |
| ENSMUSG00000036202 | 0.112018354 | 0.416439 | 0.582046 | 1554.354 | 1589.764 | 1245.472 | 1483.712 | 1337.853 | 1687.235 | 1193.79 | 1214.839 |
| ENSMUSG00000036206 | -0.170705653 | 0.133127 | 0.266493 | 1221.35 | 1195.022 | 1449.187 | 1312.913 | 1465.592 | 1219.055 | 1518.755 | 1628.124 |
| ENSMUSG00000036208 | -0.361447316 | 0.002363 | 0.014777 | 316.9071 | 307.1543 | 361.8826 | 330.3848 | 421.2481 | 476.0014 | 365.4656 | 432.8613 |

| | | | | | | | | | | | |
|---|---|---|---|---|---|---|---|---|---|---|---|
| ENSMUSG00000036214 | -0.218850433 | 0.184309 | 0.335432 | 133.8052 | 136.7797 | 124.216 | 104.3774 | 161.8685 | 140.7891 | 125.3576 | 151.1752 |
| ENSMUSG00000036218 | 0.311648241 | 0.198039 | 0.35246 | 110.666 | 187.1722 | 101.029 | 107.8279 | 87.76002 | 107.2679 | 108.9647 | 102.2336 |
| ENSMUSG00000036223 | 0.13163858 | 0.284509 | 0.451069 | 303.8284 | 382.7431 | 332.8989 | 338.1484 | 322.7618 | 345.2686 | 289.2867 | 280.5985 |
| ENSMUSG00000036225 | -0.317210262 | 0.00401 | 0.02187 | 646.893 | 661.1017 | 686.5004 | 571.0568 | 783.0144 | 697.2414 | 809.0386 | 905.9634 |
| ENSMUSG00000036241 | 0.174303793 | 0.111993 | 0.238512 | 4649.983 | 4949.264 | 4864.298 | 5009.255 | 3939.45 | 5303.057 | 4007.586 | 4008.861 |
| ENSMUSG00000036242 | -0.617329282 | 0.077045 | 0.184333 | 68.4117 | 141.5789 | 239.3228 | 141.4703 | 225.2507 | 149.7281 | 275.7867 | 256.6715 |
| ENSMUSG00000036246 | 0.336806702 | 0.003824 | 0.021132 | 388.337 | 341.9491 | 385.0696 | 398.5321 | 342.2641 | 290.5173 | 298.9296 | 267.5474 |
| ENSMUSG00000036249 | 0.23569119 | 0.053902 | 0.143655 | 420.5307 | 395.9411 | 429.7873 | 353.6756 | 369.5672 | 296.1041 | 368.3585 | 323.0146 |
| ENSMUSG00000036256 | -0.712862213 | 0.002238 | 0.014252 | 6570.541 | 4819.683 | 8555.998 | 5918.46 | 9650.677 | 6872.968 | 11260.97 | 14611.79 |
| ENSMUSG00000036257 | -0.065927924 | 0.465121 | 0.62595 | 2084.545 | 1968.907 | 1647.104 | 2036.654 | 1955.098 | 2091.724 | 2054.9 | 1994.642 |
| ENSMUSG00000036264 | 0.952954898 | 0.112158 | 0.238643 | 35.2119 | 41.99375 | 24.8432 | 24.15346 | 21.45245 | 32.40385 | 4.821446 | 6.525547 |
| ENSMUSG00000036270 | 0.123068809 | 0.127527 | 0.25887 | 1892.388 | 1737.342 | 1828.459 | 1943.491 | 1649.888 | 1843.667 | 1766.578 | 1538.941 |
| ENSMUSG00000036273 | -0.338009517 | 0.076635 | 0.18378 | 396.3854 | 317.9527 | 375.1323 | 360.5766 | 447.5761 | 316.2169 | 470.5731 | 599.2627 |
| ENSMUSG00000036275 | -0.032955037 | 0.565347 | 0.710019 | 5841.152 | 6300.263 | 6404.577 | 6196.225 | 5937.453 | 6427.136 | 6457.845 | 6494.006 |
| ENSMUSG00000036278 | -1.097581726 | 1.52E-05 | 0.000315 | 323.9495 | 241.1641 | 308.0557 | 273.4516 | 607.4943 | 347.5033 | 612.3236 | 887.4743 |
| ENSMUSG00000036281 | 0.206534036 | 0.023723 | 0.079969 | 827.9827 | 830.2765 | 727.9057 | 874.7002 | 715.7317 | 756.4623 | 659.5738 | 693.8831 |
| ENSMUSG00000036282 | 0.090814705 | 0.549442 | 0.69758 | 1151.932 | 1147.029 | 922.5108 | 968.7262 | 907.8286 | 1270.454 | 865.9317 | 889.6495 |
| ENSMUSG00000036285 | -0.176275111 | 0.157711 | 0.300269 | 571.4389 | 476.3291 | 467.8802 | 492.558 | 572.3903 | 468.1797 | 609.4308 | 617.7517 |
| ENSMUSG00000036291 | -0.112256763 | 0.374841 | 0.543403 | 500.009 | 451.1329 | 366.8512 | 426.9986 | 461.2276 | 430.189 | 524.5733 | 466.5766 |
| ENSMUSG00000036292 | 0.490660331 | 0.000626 | 0.005449 | 345.0766 | 265.1606 | 264.9941 | 295.0172 | 181.3707 | 226.8269 | 221.7865 | 203.3795 |
| ENSMUSG00000036295 | 0.207071438 | 0.267473 | 0.431699 | 287.7315 | 208.7689 | 187.9802 | 257.0618 | 235.0018 | 164.254 | 212.1436 | 203.3795 |
| ENSMUSG00000036298 | 0.625789154 | 7.13E-05 | 0.001046 | 638.8445 | 698.2961 | 529.1601 | 684.0604 | 441.7254 | 464.8276 | 435.8587 | 308.8759 |
| ENSMUSG00000036299 | -0.271170625 | 0.002035 | 0.013289 | 1530.209 | 1432.587 | 1548.559 | 1689.879 | 1804.931 | 1702.878 | 1937.257 | 2042.496 |
| ENSMUSG00000036304 | -1.886985546 | 0.000528 | 0.00478 | 12.07265 | 7.198929 | 14.90592 | 8.626235 | 44.85512 | 13.40849 | 33.75012 | 67.43065 |
| ENSMUSG00000036305 | 0.05544542 | 0.695523 | 0.807789 | 149.9021 | 157.1766 | 145.7468 | 139.745 | 150.1671 | 137.437 | 126.3219 | 155.5255 |
| ENSMUSG00000036306 | -0.11443344 | 0.639258 | 0.766313 | 100.6054 | 99.58519 | 109.3101 | 62.10889 | 124.8142 | 105.0332 | 85.82174 | 85.9197 |
| ENSMUSG00000036309 | 0.015235262 | 0.859963 | 0.91605 | 7852.254 | 8918.273 | 7679.033 | 8495.979 | 8130.478 | 9135.651 | 7768.314 | 7563.108 |
| ENSMUSG00000036315 | -0.180436281 | 0.029293 | 0.092968 | 645.8869 | 687.4977 | 708.8593 | 673.7089 | 831.7699 | 711.7673 | 745.3955 | 788.5035 |
| ENSMUSG00000036323 | -0.024961392 | 0.557433 | 0.703981 | 4140.92 | 4042.199 | 4144.674 | 4138.868 | 4309.992 | 4096.293 | 4239.979 | 4107.832 |
| ENSMUSG00000036327 | -0.201670542 | 0.026388 | 0.086387 | 579.4873 | 518.3229 | 556.4877 | 522.7498 | 666.9761 | 563.1565 | 633.538 | 639.5036 |
| ENSMUSG00000036330 | -2.827499857 | 3.07E-05 | 0.000546 | 81.4904 | 16.7975 | 39.74912 | 97.47645 | 556.7886 | 130.7328 | 336.5369 | 650.3795 |
| ENSMUSG00000036333 | -0.250332966 | 0.006007 | 0.029501 | 3331.046 | 3153.131 | 3555.89 | 3397.011 | 4013.558 | 3441.512 | 4111.729 | 4418.883 |
| ENSMUSG00000036334 | 0.570440993 | 2.70E-05 | 0.000494 | 429.5852 | 559.1168 | 504.3169 | 420.0976 | 308.1352 | 360.9118 | 329.7869 | 288.2116 |
| ENSMUSG00000036339 | -0.12518047 | 0.192248 | 0.34549 | 732.4076 | 718.6931 | 701.4063 | 676.2968 | 696.2295 | 730.7627 | 780.1099 | 877.686 |
| ENSMUSG00000036352 | -0.243671857 | 0.013323 | 0.052912 | 1165.011 | 1156.628 | 1098.898 | 1216.299 | 1285.197 | 1219.055 | 1410.755 | 1574.832 |
| ENSMUSG00000036353 | 0.434761034 | 0.15146 | 0.291852 | 36.21796 | 32.39518 | 41.40533 | 32.77969 | 24.37778 | 34.6386 | 20.25007 | 27.18978 |
| ENSMUSG00000036356 | -0.439934066 | 0.078234 | 0.186164 | 114.6902 | 88.78679 | 160.6527 | 194.9529 | 163.8187 | 193.3057 | 192.8578 | 212.0803 |
| ENSMUSG00000036362 | 1.031544688 | 0.052564 | 0.140925 | 12.07265 | 20.39697 | 11.59349 | 11.21411 | 8.776002 | 4.469496 | 5.785735 | 7.613138 |
| ENSMUSG00000036368 | 0.656868299 | 0.029617 | 0.093727 | 710.2744 | 305.9545 | 243.4633 | 351.9504 | 243.7778 | 261.4655 | 263.2509 | 253.4087 |
| ENSMUSG00000036371 | -0.167188191 | 6.23E-05 | 0.000944 | 17383.61 | 18528.84 | 17352.15 | 17546.62 | 20206.26 | 19456.84 | 19900.04 | 19937.72 |
| ENSMUSG00000036372 | -0.101615515 | 0.065484 | 0.164684 | 1592.584 | 1637.756 | 1514.607 | 1558.761 | 1737.648 | 1672.709 | 1684.613 | 1661.839 |
| ENSMUSG00000036376 | -0.141351382 | 0.096091 | 0.214678 | 646.893 | 694.6967 | 750.2646 | 735.8178 | 819.0935 | 724.0584 | 786.86 | 790.6787 |
| ENSMUSG00000036377 | 0.375409501 | 0.00014 | 0.001736 | 601.6205 | 569.9152 | 501.0045 | 577.9577 | 454.4019 | 405.6068 | 453.2159 | 418.7226 |
| ENSMUSG00000036381 | -0.255285687 | 0.523468 | 0.676726 | 105.6357 | 32.39518 | 93.57605 | 54.34528 | 53.63112 | 75.98144 | 96.42892 | 116.3722 |
| ENSMUSG00000036390 | -0.458909213 | 0.01902 | 0.068123 | 591.56 | 601.1106 | 757.7176 | 774.6359 | 696.2295 | 777.6924 | 1220.79 | 1052.788 |
| ENSMUSG00000036391 | 0.259163824 | 0.019446 | 0.069097 | 687.1351 | 719.8929 | 641.7826 | 634.8909 | 527.5352 | 660.3681 | 506.2518 | 548.1459 |
| ENSMUSG00000036395 | -0.633875939 | 0.035121 | 0.106348 | 804.8435 | 853.0731 | 1878.146 | 926.4576 | 1568.954 | 1099.496 | 1934.364 | 2324.182 |
| ENSMUSG00000036398 | -0.195188517 | 0.036135 | 0.108758 | 891.3642 | 937.0606 | 971.3691 | 1009.269 | 1126.254 | 953.1201 | 1089.647 | 1193.087 |
| ENSMUSG00000036402 | -0.343380773 | 0.010268 | 0.043476 | 3016.151 | 2879.572 | 3507.86 | 3287.458 | 3704.448 | 3228.094 | 4458.873 | 4712.532 |
| ENSMUSG00000036403 | -0.009449119 | 0.940617 | 0.965411 | 468.8213 | 409.1391 | 339.5237 | 419.235 | 443.6756 | 426.8369 | 384.7514 | 390.4452 |
| ENSMUSG00000036411 | 0.709365485 | 0.000463 | 0.00432 | 146.8839 | 207.5691 | 161.4808 | 173.3873 | 94.5858 | 138.5544 | 101.2504 | 87.00729 |
| ENSMUSG00000036412 | -0.734370029 | 0.072699 | 0.177172 | 46.2785 | 139.1793 | 152.3716 | 87.12497 | 167.7191 | 102.7984 | 150.4291 | 286.0365 |
| ENSMUSG00000036422 | -1.05331289 | 0.000601 | 0.005272 | 18.10898 | 38.39429 | 20.70267 | 28.46658 | 59.48179 | 56.98608 | 46.28588 | 54.37955 |
| ENSMUSG00000036427 | -0.390428398 | 0.002747 | 0.01657 | 9314.051 | 9418.599 | 10778.64 | 9710.553 | 13170.83 | 9725.624 | 13949.41 | 14566.11 |
| ENSMUSG00000036430 | -0.268367783 | 0.021438 | 0.073984 | 633.8142 | 631.1061 | 621.08 | 589.1718 | 832.7451 | 650.3117 | 659.5738 | 837.4451 |
| ENSMUSG00000036432 | -0.618721138 | 0.001569 | 0.010893 | 147.89 | 117.5825 | 138.2938 | 128.5309 | 178.4454 | 173.193 | 190.9293 | 276.2481 |
| ENSMUSG00000036435 | 0.070491456 | 0.312292 | 0.480745 | 1374.27 | 1280.21 | 1213.176 | 1322.402 | 1245.217 | 1296.154 | 1185.111 | 1215.927 |
| ENSMUSG00000036437 | 1.202671705 | 0.010943 | 0.04565 | 41.24823 | 29.99554 | 21.53077 | 56.93315 | 10.72622 | 22.34748 | 9.642892 | 22.83941 |
| ENSMUSG00000036438 | -0.026229294 | 0.557561 | 0.704095 | 12705.46 | 13276.03 | 13402.91 | 13928.78 | 13442.88 | 13487.82 | 13685.19 | 13679.72 |
| ENSMUSG00000036442 | -0.050973744 | 0.505077 | 0.660879 | 984.9272 | 1012.649 | 1002.009 | 1048.95 | 1102.851 | 940.829 | 1072.29 | 1076.715 |
| ENSMUSG00000036446 | 0.413925104 | 0.454793 | 0.616955 | 376.2643 | 937.0606 | 712.9998 | 704.7634 | 502.1823 | 1042.51 | 223.7151 | 281.6861 |
| ENSMUSG00000036450 | -0.123344105 | 0.099298 | 0.219926 | 1615.723 | 1610.16 | 1647.104 | 1475.949 | 1884.89 | 1640.305 | 1724.149 | 1662.927 |
| ENSMUSG00000036452 | 0.019309673 | 0.886408 | 0.932078 | 618.7234 | 434.3354 | 470.3646 | 497.7338 | 479.7548 | 554.2176 | 502.3947 | 458.9634 |
| ENSMUSG00000036459 | 0.107290219 | 0.072193 | 0.176314 | 1088.551 | 1131.432 | 1079.023 | 1085.18 | 980.962 | 1027.984 | 1038.539 | 1020.16 |

| | | | | | | | | | | | |
|---|---|---|---|---|---|---|---|---|---|---|---|
| ENSMUSG00000036461 | -0.03760016 | 0.749902 | 0.844092 | 1405.458 | 1174.625 | 1525.372 | 1372.434 | 1292.022 | 1249.224 | 1493.684 | 1591.146 |
| ENSMUSG00000036466 | 0.099192815 | 0.679987 | 0.795555 | 109.6599 | 83.98751 | 122.5598 | 100.0643 | 69.2329 | 94.9768 | 94.50034 | 131.5985 |
| ENSMUSG00000036469 | 0.312391167 | 0.209308 | 0.366507 | 52.31483 | 55.19179 | 57.13936 | 42.26855 | 46.80534 | 46.92971 | 34.71441 | 38.06569 |
| ENSMUSG00000036473 | 0.191948282 | 0.040789 | 0.118329 | 862.1886 | 1029.447 | 882.7617 | 832.4317 | 758.6366 | 765.4013 | 788.7885 | 840.7079 |
| ENSMUSG00000036478 | 0.239933831 | 0.020261 | 0.071112 | 4632.88 | 4222.172 | 4091.675 | 4494.268 | 3297.826 | 4338.764 | 3750.121 | 3383.496 |
| ENSMUSG00000036480 | -0.148624351 | 0.74292 | 0.839394 | 8.048435 | 21.59679 | 17.39024 | 25.8787 | 22.42756 | 13.40849 | 23.14294 | 21.75182 |
| ENSMUSG00000036492 | -0.971870429 | 0.030244 | 0.095011 | 8.048435 | 13.19804 | 11.59349 | 12.93935 | 30.22845 | 16.76061 | 14.46434 | 28.27737 |
| ENSMUSG00000036499 | 0.037245772 | 0.716682 | 0.822669 | 1052.333 | 962.2569 | 1030.993 | 948.0232 | 877.6002 | 1091.674 | 879.4317 | 1046.263 |
| ENSMUSG00000036501 | 0.090843945 | 0.386622 | 0.554128 | 1112.696 | 914.264 | 930.7919 | 1087.768 | 1046.294 | 891.6645 | 888.1103 | 973.394 |
| ENSMUSG00000036502 | 0.152568087 | 0.529203 | 0.681503 | 516.1059 | 341.9491 | 372.648 | 569.3315 | 323.737 | 589.9735 | 344.2512 | 363.2554 |
| ENSMUSG00000036503 | 0.088122525 | 0.213935 | 0.37205 | 1555.36 | 1502.177 | 1535.31 | 1710.582 | 1415.862 | 1557.619 | 1503.327 | 1457.372 |
| ENSMUSG00000036504 | -0.137383997 | 0.194853 | 0.348638 | 983.9212 | 1207.02 | 1231.395 | 1137.8 | 1305.674 | 1090.557 | 1295.04 | 1323.598 |
| ENSMUSG00000036510 | 0.585625485 | 0.177412 | 0.326485 | 34.20585 | 33.595 | 17.39024 | 22.42821 | 12.67645 | 27.93435 | 12.53576 | 18.48905 |
| ENSMUSG00000036513 | -0.220699726 | 0.035101 | 0.106323 | 733.4136 | 920.2631 | 855.4342 | 844.5084 | 1028.742 | 905.073 | 902.5747 | 1070.19 |
| ENSMUSG00000036523 | 0.511616372 | 0.018836 | 0.067595 | 802.8314 | 536.3202 | 522.5353 | 621.9515 | 395.8952 | 612.321 | 405.0014 | 329.5401 |
| ENSMUSG00000036528 | 0.137602956 | 0.209011 | 0.366165 | 697.1957 | 680.2988 | 646.7513 | 614.1879 | 698.1797 | 569.8608 | 532.2876 | 595.9999 |
| ENSMUSG00000036529 | -0.021112039 | 0.817109 | 0.888971 | 1444.694 | 1249.014 | 1366.376 | 1434.543 | 1434.389 | 1318.501 | 1278.647 | 1547.642 |
| ENSMUSG00000036533 | 0.880390248 | 2.70E-08 | 1.96E-06 | 839.0493 | 1195.022 | 804.0915 | 964.4131 | 584.0917 | 564.2739 | 451.2873 | 464.4014 |
| ENSMUSG00000036534 | 0.021607921 | 0.81371 | 0.886897 | 571.4389 | 500.3256 | 514.2542 | 487.3823 | 537.2863 | 490.5272 | 523.609 | 489.416 |
| ENSMUSG00000036537 | -0.217122149 | 0.117141 | 0.245399 | 147.89 | 149.9777 | 139.1219 | 151.8217 | 176.4951 | 147.4934 | 188.0364 | 170.7518 |
| ENSMUSG00000036545 | 0.847154739 | 1.64E-05 | 0.000334 | 1032.212 | 1551.369 | 1101.382 | 1501.828 | 592.8677 | 968.7633 | 667.2881 | 654.7298 |
| ENSMUSG00000036550 | -0.084426002 | 0.080981 | 0.190515 | 5684.207 | 5724.349 | 5482.066 | 5447.467 | 6139.301 | 5707.547 | 5801.164 | 6029.605 |
| ENSMUSG00000036552 | 0.086555817 | 0.604433 | 0.741075 | 464.7971 | 523.1222 | 392.5225 | 347.6373 | 451.4765 | 458.1234 | 343.2869 | 371.9562 |
| ENSMUSG00000036553 | -0.697257363 | 0.005323 | 0.026853 | 484.9182 | 359.9465 | 671.5945 | 399.3947 | 773.2633 | 509.5226 | 783.0028 | 1043 |
| ENSMUSG00000036555 | 0.071487206 | 0.379706 | 0.547789 | 807.8616 | 925.0624 | 846.325 | 935.0839 | 871.7495 | 852.5564 | 801.3243 | 817.8685 |
| ENSMUSG00000036560 | 0.683589975 | 0.21687 | 0.375381 | 13.07871 | 14.39786 | 17.39024 | 33.64232 | 13.65156 | 18.99536 | 4.821446 | 11.9635 |
| ENSMUSG00000036561 | 0.121286438 | 0.122311 | 0.251807 | 963.8001 | 903.4656 | 972.1972 | 928.1829 | 943.9077 | 806.7441 | 878.4674 | 834.1824 |
| ENSMUSG00000036564 | -0.160312487 | 0.436941 | 0.600728 | 554.3359 | 404.3399 | 611.9708 | 634.8909 | 563.6143 | 439.128 | 695.2525 | 768.9269 |
| ENSMUSG00000036565 | 0.028509369 | 0.733547 | 0.833185 | 2817.958 | 2560.419 | 2903.342 | 2527.487 | 2687.407 | 2402.354 | 2639.259 | 2870.153 |
| ENSMUSG00000036568 | 0.446399417 | 4.91E-06 | 0.000133 | 709.2683 | 739.0901 | 653.3761 | 676.2968 | 524.6099 | 568.7434 | 465.7517 | 478.5401 |
| ENSMUSG00000036570 | 0.805077361 | 3.56E-06 | 0.000104 | 478.8819 | 613.1088 | 619.4238 | 678.8847 | 333.4881 | 438.0106 | 271.9295 | 326.2773 |
| ENSMUSG00000036572 | 0.510224011 | 0.003309 | 0.019066 | 2235.453 | 2373.247 | 1743.993 | 2138.444 | 1307.624 | 2019.095 | 1424.255 | 1210.489 |
| ENSMUSG00000036578 | 0.808757086 | 0.002026 | 0.013257 | 56.33904 | 50.3925 | 76.18581 | 68.14726 | 30.22845 | 43.57759 | 32.78583 | 38.06569 |
| ENSMUSG00000036580 | 0.120031318 | 0.147707 | 0.286957 | 1765.625 | 1773.336 | 1596.59 | 1628.633 | 1667.44 | 1519.629 | 1630.613 | 1400.817 |
| ENSMUSG00000036585 | -0.705340845 | 2.62E-07 | 1.25E-05 | 750.5165 | 794.2819 | 649.2356 | 664.2201 | 1212.063 | 1387.779 | 944.0391 | 1113.693 |
| ENSMUSG00000036591 | 0.079328609 | 0.265699 | 0.429889 | 1707.274 | 1551.369 | 1611.496 | 1569.112 | 1483.144 | 1639.188 | 1545.756 | 1428.007 |
| ENSMUSG00000036594 | -1.370541835 | 6.98E-05 | 0.001029 | 31.18769 | 25.19625 | 50.5145 | 35.36756 | 127.7396 | 52.51658 | 98.35749 | 91.35765 |
| ENSMUSG00000036596 | 0.361595895 | 0.320047 | 0.488124 | 85.51462 | 245.9634 | 136.6376 | 125.943 | 92.63557 | 179.8972 | 83.89316 | 105.4963 |
| ENSMUSG00000036598 | 0.411051023 | 0.165698 | 0.311164 | 45.27245 | 53.99197 | 38.09291 | 42.26855 | 42.9049 | 34.6386 | 34.71441 | 21.75182 |
| ENSMUSG00000036599 | 0.373756707 | 0.015708 | 0.059687 | 255.5378 | 353.9474 | 252.5725 | 271.7264 | 224.2756 | 220.1227 | 237.2151 | 190.3284 |
| ENSMUSG00000036602 | -0.466224164 | 0.488174 | 0.645999 | 108.6539 | 238.7645 | 835.5596 | 293.292 | 461.2276 | 233.5312 | 560.252 | 785.2408 |
| ENSMUSG00000036606 | -0.368177141 | 0.005026 | 0.025823 | 5512.172 | 5852.729 | 7310.525 | 5586.35 | 7736.533 | 6421.549 | 8378.709 | 8780.123 |
| ENSMUSG00000036611 | 0.199762387 | 0.209008 | 0.366165 | 220.3259 | 179.9732 | 164.7932 | 163.0358 | 173.5698 | 157.5497 | 139.8219 | 162.0511 |
| ENSMUSG00000036613 | -0.304867417 | 0.006609 | 0.03165 | 467.8153 | 461.9313 | 467.0521 | 444.2511 | 591.8926 | 464.8276 | 605.5736 | 610.1386 |
| ENSMUSG00000036615 | -0.053968491 | 0.53729 | 0.687647 | 782.7103 | 807.4799 | 829.7628 | 875.5628 | 815.1931 | 781.0445 | 918.9676 | 907.051 |
| ENSMUSG00000036617 | 0.602257636 | 0.000614 | 0.00537 | 2907.497 | 3167.529 | 2670.644 | 2632.727 | 1655.739 | 2653.763 | 1585.291 | 1600.934 |
| ENSMUSG00000036620 | -0.315438134 | 0.007978 | 0.036149 | 1648.923 | 1556.169 | 1816.866 | 1542.371 | 1959.974 | 1682.765 | 2182.186 | 2344.846 |
| ENSMUSG00000036622 | 0.002006895 | 0.97832 | 0.988064 | 1624.778 | 1851.325 | 1674.432 | 1802.883 | 1861.487 | 1674.944 | 1678.827 | 1724.919 |
| ENSMUSG00000036632 | -0.109314103 | 0.196901 | 0.351214 | 874.2612 | 987.4531 | 963.088 | 1024.797 | 1125.278 | 972.1155 | 1019.254 | 1035.387 |
| ENSMUSG00000036634 | 0.855892056 | 0.064671 | 0.163614 | 16.09687 | 22.79661 | 53.82693 | 27.60395 | 14.62667 | 15.64324 | 19.28578 | 17.40146 |
| ENSMUSG00000036636 | 0.048995342 | 0.521908 | 0.675148 | 885.3278 | 843.4745 | 875.3087 | 779.8116 | 826.8944 | 820.1526 | 846.6459 | 775.4524 |
| ENSMUSG00000036639 | -0.108008958 | 0.322565 | 0.490822 | 497.9969 | 647.9036 | 580.5027 | 598.6607 | 676.7272 | 611.2036 | 575.6806 | 640.5912 |
| ENSMUSG00000036641 | 0.047164717 | 0.867028 | 0.920283 | 93.56306 | 97.18554 | 131.669 | 144.9207 | 85.8098 | 170.9582 | 87.75031 | 109.8467 |
| ENSMUSG00000036644 | -0.065438802 | 0.532703 | 0.684218 | 2015.127 | 1806.931 | 2075.235 | 1896.909 | 2035.057 | 1717.404 | 2091.543 | 2313.306 |
| ENSMUSG00000036646 | -0.124929764 | 0.044763 | 0.126015 | 2240.483 | 2411.641 | 2403.994 | 2304.067 | 2689.357 | 2384.476 | 2553.438 | 2576.503 |
| ENSMUSG00000036655 | -0.158197564 | 0.741956 | 0.838976 | 36.21796 | 19.19714 | 21.53077 | 33.64232 | 37.05423 | 12.29112 | 23.14294 | 51.11678 |
| ENSMUSG00000036661 | 0.102132334 | 0.429593 | 0.594488 | 640.8566 | 527.9215 | 664.1415 | 601.2486 | 520.7094 | 490.5272 | 657.6452 | 600.3503 |
| ENSMUSG00000036667 | 0.346709114 | 4.51E-09 | 4.48E-07 | 3237.483 | 3225.12 | 2984.496 | 3126.148 | 2321.74 | 2498.448 | 2545.723 | 2518.861 |
| ENSMUSG00000036672 | -0.072669178 | 0.455105 | 0.61714 | 642.8687 | 712.694 | 648.4075 | 694.4119 | 666.001 | 793.3356 | 734.7883 | 642.7663 |
| ENSMUSG00000036676 | 0.083359052 | 0.296238 | 0.464123 | 942.6729 | 920.2631 | 874.4806 | 953.199 | 799.5913 | 930.7726 | 846.6459 | 908.1386 |
| ENSMUSG00000036678 | -0.006632502 | 0.943331 | 0.967283 | 741.4621 | 844.6744 | 804.0915 | 672.8463 | 753.761 | 781.0445 | 751.1813 | 788.5035 |
| ENSMUSG00000036686 | 0.032804772 | 0.699948 | 0.810633 | 686.1291 | 807.4799 | 702.2344 | 747.8946 | 746.9353 | 676.0113 | 718.3954 | 731.9488 |
| ENSMUSG00000036687 | -0.370125285 | 0.070737 | 0.173675 | 161.9748 | 157.1766 | 223.5888 | 162.1732 | 229.1512 | 173.193 | 217.9294 | 292.562 |
| ENSMUSG00000036693 | -0.008756772 | 0.882529 | 0.929731 | 1128.793 | 1119.433 | 1151.068 | 1077.417 | 1131.129 | 1103.966 | 1112.79 | 1156.109 |

| | | | | | | | | | | | |
|---|---|---|---|---|---|---|---|---|---|---|---|
| ENSMUSG00000036698 | 0.302351526 | 0.012555 | 0.050659 | 2030.218 | 1794.933 | 1575.887 | 1829.624 | 1280.321 | 1703.995 | 1350.969 | 1529.153 |
| ENSMUSG00000036699 | 1.024779127 | 1.04E-09 | 1.34E-07 | 204.229 | 169.1748 | 197.0894 | 246.7103 | 104.3369 | 99.44629 | 105.1075 | 93.53283 |
| ENSMUSG00000036707 | -0.185936126 | 0.000738 | 0.006192 | 2830.031 | 2877.172 | 3088.838 | 2912.217 | 3277.349 | 3430.338 | 3212.047 | 3406.335 |
| ENSMUSG00000036712 | 0.122460666 | 0.29 | 0.457144 | 1091.569 | 928.6619 | 778.4202 | 881.6012 | 893.202 | 839.1479 | 834.1101 | 811.343 |
| ENSMUSG00000036718 | -0.11710604 | 0.466569 | 0.627229 | 168.0111 | 172.7743 | 212.8234 | 166.4863 | 206.7236 | 156.4324 | 207.3222 | 210.9927 |
| ENSMUSG00000036721 | 0.014794376 | 0.906973 | 0.944324 | 430.5913 | 463.1311 | 418.1938 | 460.6409 | 468.0534 | 515.1095 | 390.5371 | 380.6569 |
| ENSMUSG00000036733 | 0.045819222 | 0.927922 | 0.957316 | 21.12714 | 14.39786 | 8.281066 | 25.01608 | 13.65156 | 26.81698 | 14.46434 | 11.9635 |
| ENSMUSG00000036737 | 0.039924532 | 0.678306 | 0.794492 | 1843.092 | 1553.769 | 1523.716 | 1754.576 | 1488.995 | 1786.681 | 1597.827 | 1621.598 |
| ENSMUSG00000036743 | 0.14881827 | 0.83181 | 0.898139 | 19.11503 | 8.398751 | 18.21835 | 16.38985 | 7.80089 | 29.05173 | 17.3572 | 2.175182 |
| ENSMUSG00000036745 | -0.108587596 | 0.661743 | 0.782191 | 119.7205 | 104.3845 | 157.3403 | 114.7289 | 86.78491 | 172.0756 | 129.2147 | 149 |
| ENSMUSG00000036748 | -0.077875473 | 0.282845 | 0.449379 | 2329.016 | 2404.442 | 2312.074 | 2306.655 | 2548.941 | 2190.053 | 2526.438 | 2601.518 |
| ENSMUSG00000036751 | -0.701182793 | 7.77E-05 | 0.001118 | 3074.502 | 3047.547 | 3964.146 | 3144.263 | 5295.83 | 3726.443 | 5679.663 | 6810.495 |
| ENSMUSG00000036752 | -0.37323918 | 0.000817 | 0.006684 | 5431.687 | 5306.811 | 6368.968 | 5827.884 | 7770.662 | 6117.623 | 7505.063 | 8316.809 |
| ENSMUSG00000036764 | 0.051057519 | 0.784187 | 0.867163 | 310.8708 | 235.165 | 284.0406 | 275.1769 | 245.728 | 195.5405 | 291.2153 | 334.9781 |
| ENSMUSG00000036766 | 0.559255211 | 0.064357 | 0.162993 | 42.25428 | 65.99018 | 48.85829 | 87.9876 | 36.07912 | 43.57759 | 43.39301 | 43.50364 |
| ENSMUSG00000036768 | 0.147773129 | 0.203042 | 0.358768 | 1294.792 | 1079.839 | 1185.849 | 1167.13 | 1141.855 | 1230.229 | 968.1463 | 928.8028 |
| ENSMUSG00000036769 | 0.375178599 | 0.070256 | 0.172791 | 188.1322 | 183.5727 | 114.2787 | 191.5024 | 126.7645 | 125.1459 | 119.5719 | 150.0876 |
| ENSMUSG00000036775 | 0.107828075 | 0.15426 | 0.295702 | 774.6619 | 785.8831 | 798.2948 | 837.6074 | 768.3877 | 712.8847 | 783.0028 | 701.4963 |
| ENSMUSG00000036777 | 0.162369623 | 0.041155 | 0.119091 | 1429.603 | 1404.991 | 1197.442 | 1364.67 | 1222.79 | 1207.881 | 1158.111 | 1230.066 |
| ENSMUSG00000036779 | 0.0699741 | 0.540339 | 0.689873 | 1305.859 | 1299.407 | 1228.082 | 1400.901 | 1215.964 | 1511.807 | 1119.54 | 1140.883 |
| ENSMUSG00000036781 | -0.232182833 | 7.64E-05 | 0.001106 | 2205.271 | 2223.269 | 2330.292 | 2224.706 | 2776.142 | 2575.547 | 2701.938 | 2498.197 |
| ENSMUSG00000036782 | 0.20326538 | 0.138841 | 0.274647 | 820.9404 | 1009.05 | 1166.802 | 992.8796 | 760.5868 | 792.2182 | 937.2891 | 976.6568 |
| ENSMUSG00000036790 | 0.423047615 | 0.40427 | 0.571173 | 32.19374 | 17.99732 | 44.71776 | 45.71905 | 14.62667 | 52.51658 | 20.25007 | 18.48905 |
| ENSMUSG00000036792 | 0.295373509 | 0.018705 | 0.067309 | 720.3349 | 699.496 | 650.0637 | 727.1916 | 570.4401 | 688.3024 | 547.7162 | 473.1021 |
| ENSMUSG00000036805 | 0.763674722 | 0.166769 | 0.312509 | 10.06054 | 44.3934 | 25.67131 | 29.3292 | 28.27823 | 13.40849 | 12.53576 | 9.78832 |
| ENSMUSG00000036810 | 0.134333167 | 0.181232 | 0.331597 | 1397.41 | 1360.598 | 1191.645 | 1415.565 | 1275.446 | 1327.44 | 1060.718 | 1224.628 |
| ENSMUSG00000036813 | -2.09049446 | 1.61E-06 | 5.49E-05 | 20.12109 | 16.7975 | 37.2648 | 27.60395 | 80.93424 | 52.51658 | 118.6076 | 184.8905 |
| ENSMUSG00000036814 | -2.483347701 | 6.82E-05 | 0.001009 | 4.024217 | 0 | 10.76539 | 6.900988 | 33.15378 | 18.99536 | 34.71441 | 39.15328 |
| ENSMUSG00000036815 | 1.107225031 | 0.000186 | 0.002142 | 173.0413 | 226.7663 | 192.9488 | 256.1992 | 72.15824 | 166.4887 | 79.07171 | 77.21897 |
| ENSMUSG00000036817 | -0.118799375 | 0.124826 | 0.255099 | 1939.673 | 1803.332 | 1755.586 | 1940.903 | 2015.555 | 1825.789 | 2141.686 | 2093.613 |
| ENSMUSG00000036819 | -0.214798118 | 0.060291 | 0.15554 | 492.9666 | 454.7324 | 405.7722 | 441.6632 | 501.2072 | 459.2408 | 532.2876 | 588.3868 |
| ENSMUSG00000036820 | -0.336029311 | 0.014458 | 0.056131 | 486.9303 | 473.9295 | 650.0637 | 591.7597 | 707.9308 | 597.7951 | 725.1455 | 753.7006 |
| ENSMUSG00000036822 | -0.092126263 | 0.101476 | 0.223356 | 1579.505 | 1646.155 | 1644.62 | 1682.978 | 1683.042 | 1714.052 | 1788.756 | 1801.051 |
| ENSMUSG00000036825 | -0.147610876 | 0.115769 | 0.243773 | 1045.29 | 1198.622 | 1049.211 | 985.9787 | 1155.507 | 1106.2 | 1310.469 | 1161.547 |
| ENSMUSG00000036832 | -1.855016188 | 0.000598 | 0.005253 | 19.11503 | 20.39697 | 9.109173 | 18.97772 | 86.78491 | 16.76061 | 48.21446 | 91.35765 |
| ENSMUSG00000036833 | 0.091630175 | 0.445674 | 0.60843 | 606.6508 | 483.5281 | 456.2868 | 526.2003 | 506.0828 | 478.2361 | 517.8233 | 441.562 |
| ENSMUSG00000036834 | 0.052593499 | 0.596973 | 0.73584 | 573.451 | 548.3184 | 535.785 | 532.2387 | 472.929 | 601.1473 | 521.6804 | 516.6058 |
| ENSMUSG00000036835 | -0.252286701 | 0.001264 | 0.009324 | 1210.283 | 1180.624 | 1383.766 | 1330.165 | 1571.879 | 1478.286 | 1570.827 | 1464.985 |
| ENSMUSG00000036840 | 0.187792008 | 0.170339 | 0.31701 | 1271.653 | 1412.19 | 1277.769 | 1565.662 | 1084.324 | 1524.098 | 1158.111 | 1087.591 |
| ENSMUSG00000036845 | 0.172001175 | 0.126504 | 0.257501 | 411.4762 | 365.9456 | 431.4436 | 397.6694 | 385.169 | 359.7945 | 374.1442 | 307.7883 |
| ENSMUSG00000036850 | -0.481817769 | 0.000169 | 0.001991 | 892.3702 | 775.0847 | 879.4492 | 845.371 | 1230.59 | 922.951 | 1237.183 | 1347.525 |
| ENSMUSG00000036853 | -0.044490041 | 0.821563 | 0.891669 | 718.3228 | 502.7252 | 929.9637 | 605.5617 | 733.2837 | 570.9782 | 803.2529 | 737.3868 |
| ENSMUSG00000036854 | 0.287278021 | 0.046774 | 0.129912 | 266.6044 | 305.9545 | 245.9477 | 269.1385 | 237.9272 | 242.4702 | 178.3935 | 231.6569 |
| ENSMUSG00000036856 | 0.023288513 | 0.903142 | 0.94209 | 3244.525 | 3703.849 | 4831.174 | 3102.857 | 3101.829 | 4970.08 | 3148.404 | 3425.912 |
| ENSMUSG00000036860 | -0.397809781 | 1.58E-06 | 5.42E-05 | 571.4389 | 538.7199 | 568.9093 | 580.5456 | 724.5077 | 744.1711 | 698.1454 | 814.6057 |
| ENSMUSG00000036862 | 0.351567557 | 0.000545 | 0.004895 | 1647.917 | 1935.312 | 2131.546 | 2012.501 | 1543.601 | 1651.479 | 1446.434 | 1417.131 |
| ENSMUSG00000036863 | 0.065296821 | 0.512606 | 0.667795 | 548.2996 | 591.512 | 523.3634 | 653.006 | 534.361 | 564.2739 | 538.0734 | 577.5109 |
| ENSMUSG00000036864 | 0.621557329 | 8.61E-06 | 0.000204 | 452.7245 | 484.7279 | 370.9918 | 464.0914 | 329.5876 | 306.1605 | 261.3224 | 253.4087 |
| ENSMUSG00000036867 | -0.13380105 | 0.319555 | 0.487736 | 563.3904 | 614.3086 | 750.2646 | 633.1656 | 651.3744 | 610.0863 | 731.8955 | 818.9561 |
| ENSMUSG00000036873 | -0.149414787 | 0.190105 | 0.342667 | 577.4752 | 561.5165 | 548.2066 | 592.6223 | 631.8721 | 556.4523 | 588.2164 | 752.613 |
| ENSMUSG00000036875 | 0.003244469 | 0.974841 | 0.986277 | 452.7245 | 518.3229 | 451.3181 | 443.3885 | 504.1325 | 480.4709 | 420.4301 | 453.5255 |
| ENSMUSG00000036879 | 0.26266103 | 0.003063 | 0.018044 | 1359.179 | 1461.383 | 1357.267 | 1343.105 | 1035.568 | 1299.506 | 1107.004 | 1160.46 |
| ENSMUSG00000036880 | -0.111084665 | 0.380604 | 0.548577 | 3713.347 | 4204.175 | 4364.95 | 4118.165 | 4449.433 | 3614.705 | 4231.301 | 5418.379 |
| ENSMUSG00000036882 | 0.1235636 | 0.365112 | 0.53387 | 349.1009 | 315.5531 | 388.382 | 364.8897 | 369.5672 | 263.7003 | 328.8226 | 340.416 |
| ENSMUSG00000036885 | 0.218586049 | 0.125424 | 0.255959 | 584.5176 | 587.9126 | 468.7084 | 703.9008 | 481.705 | 547.5133 | 532.2876 | 453.5255 |
| ENSMUSG00000036887 | 0.545740649 | 8.49E-05 | 0.001188 | 558.3602 | 698.2961 | 742.8116 | 659.907 | 413.4472 | 487.1751 | 527.4662 | 393.708 |
| ENSMUSG00000036890 | -0.002427299 | 0.979813 | 0.988814 | 516.1059 | 553.1177 | 533.3007 | 611.6001 | 517.7841 | 602.2646 | 548.6805 | 551.4087 |
| ENSMUSG00000036892 | -1.979390226 | 0.004995 | 0.025736 | 114.6902 | 14.39786 | 170.59 | 79.36136 | 390.0445 | 176.5451 | 336.5369 | 593.8247 |
| ENSMUSG00000036893 | 0.089962586 | 0.177784 | 0.32704 | 2528.215 | 2608.412 | 2524.069 | 2477.455 | 2322.715 | 2616.89 | 2253.544 | 2332.883 |
| ENSMUSG00000036894 | 0.612996837 | 0.000722 | 0.006102 | 404.4339 | 555.5174 | 389.2101 | 499.459 | 279.8569 | 391.0809 | 279.6439 | 257.7591 |
| ENSMUSG00000036896 | 0.527662393 | 1.85E-06 | 6.10E-05 | 581.4994 | 719.8929 | 699.7501 | 616.7758 | 427.0988 | 426.8369 | 500.4661 | 460.051 |
| ENSMUSG00000036898 | 0.187500751 | 0.187614 | 0.339805 | 1056.357 | 1072.64 | 916.714 | 1048.088 | 803.4917 | 1155.365 | 864.0031 | 772.1897 |
| ENSMUSG00000036899 | -2.660520712 | 3.71E-09 | 3.78E-07 | 92.557 | 50.3925 | 71.21717 | 37.95543 | 332.513 | 131.8501 | 409.8229 | 719.9853 |
| ENSMUSG00000036902 | 0.774605157 | 0.000144 | 0.001767 | 866.2128 | 1521.374 | 1103.038 | 1111.059 | 598.7183 | 873.7865 | 532.2876 | 685.1824 |

| | | | | | | | | | | | |
|---|---|---|---|---|---|---|---|---|---|---|---|
| ENSMUSG00000036904 | -0.107290187 | 0.477673 | 0.637035 | 413.4883 | 527.9215 | 418.1938 | 351.0878 | 440.7503 | 400.0199 | 515.8947 | 481.8029 |
| ENSMUSG00000036905 | 0.434008114 | 0.00675 | 0.032129 | 595.5842 | 839.8751 | 863.7152 | 598.6607 | 538.2614 | 490.5272 | 603.645 | 511.1678 |
| ENSMUSG00000036907 | -0.236439603 | 0.661511 | 0.782072 | 6.036326 | 14.39786 | 13.24971 | 10.35148 | 6.825779 | 17.87799 | 17.3572 | 9.78832 |
| ENSMUSG00000036908 | 0.229147106 | 0.036643 | 0.109822 | 569.4268 | 587.9126 | 642.6107 | 576.2325 | 436.8499 | 487.1751 | 538.0734 | 566.635 |
| ENSMUSG00000036913 | 0.952915813 | 0.119957 | 0.248895 | 30.18163 | 19.19714 | 12.4216 | 71.59775 | 8.776002 | 18.99536 | 15.42863 | 26.10219 |
| ENSMUSG00000036915 | -1.314968978 | 0.067574 | 0.167851 | 16.09687 | 2.399643 | 7.45296 | 6.038364 | 11.70134 | 7.821619 | 20.25007 | 40.24087 |
| ENSMUSG00000036916 | 0.374361456 | 0.002754 | 0.01658 | 1179.096 | 1054.643 | 905.1205 | 1161.091 | 828.8446 | 965.4112 | 779.1456 | 743.9123 |
| ENSMUSG00000036918 | -0.654562824 | 0.008608 | 0.038281 | 207.2472 | 244.7636 | 317.9929 | 216.5185 | 400.7707 | 235.7659 | 389.5728 | 527.4817 |
| ENSMUSG00000036923 | 1.343355645 | 5.50E-08 | 3.55E-06 | 128.775 | 169.1748 | 105.9976 | 125.0804 | 70.20801 | 49.16446 | 42.42872 | 45.67883 |
| ENSMUSG00000036928 | 0.041805855 | 0.874476 | 0.925049 | 117.7084 | 176.3738 | 111.7944 | 153.547 | 128.7147 | 198.8926 | 100.2861 | 115.2847 |
| ENSMUSG00000036931 | 0.124740184 | 0.675914 | 0.792479 | 64.38748 | 38.39429 | 39.74912 | 35.36756 | 35.10401 | 46.92971 | 36.64299 | 44.59123 |
| ENSMUSG00000036932 | -0.459900534 | 0.003995 | 0.021823 | 1730.413 | 1327.003 | 1538.622 | 1417.29 | 2015.555 | 1530.803 | 2180.258 | 2544.963 |
| ENSMUSG00000036934 | 0.089448365 | 0.393488 | 0.56085 | 609.6689 | 608.3095 | 524.1915 | 649.5555 | 547.0374 | 615.6731 | 568.9306 | 515.5182 |
| ENSMUSG00000036940 | -0.271442658 | 8.91E-07 | 3.39E-05 | 2239.477 | 2277.261 | 2175.436 | 2258.348 | 2613.298 | 2699.576 | 2862.01 | 2624.357 |
| ENSMUSG00000036941 | 0.158476824 | 0.149015 | 0.288808 | 610.675 | 631.1061 | 507.6294 | 639.204 | 561.6641 | 568.7434 | 494.6803 | 513.343 |
| ENSMUSG00000036943 | 0.058766922 | 0.426138 | 0.591116 | 1162.999 | 1157.828 | 1189.989 | 1208.536 | 1130.154 | 1143.074 | 1034.682 | 1226.803 |
| ENSMUSG00000036944 | 1.20552832 | 0.038263 | 0.112988 | 53.32088 | 11.99822 | 17.39024 | 21.56559 | 7.80089 | 12.29112 | 16.39292 | 8.700729 |
| ENSMUSG00000036948 | 0.030326433 | 0.770915 | 0.857781 | 597.5963 | 536.3202 | 486.9267 | 526.2003 | 499.257 | 496.1141 | 522.6447 | 582.9488 |
| ENSMUSG00000036955 | -0.015364734 | 0.884074 | 0.930774 | 1140.866 | 949.0588 | 949.8383 | 1046.362 | 1076.523 | 887.195 | 1081.932 | 1083.241 |
| ENSMUSG00000036957 | -0.278900695 | 0.081661 | 0.191675 | 286.7255 | 255.562 | 370.9918 | 275.1769 | 319.8365 | 324.0385 | 405.9657 | 394.7956 |
| ENSMUSG00000036959 | 0.320274682 | 0.005644 | 0.028107 | 454.7366 | 441.5343 | 389.2101 | 355.4009 | 315.9361 | 345.2686 | 329.7869 | 320.8394 |
| ENSMUSG00000036960 | -0.615887829 | 0.418056 | 0.583693 | 1.006054 | 13.19804 | 12.4216 | 12.07673 | 9.751113 | 6.704245 | 12.53576 | 30.45255 |
| ENSMUSG00000036961 | 0.375098007 | 0.205921 | 0.362467 | 61.36932 | 85.18733 | 80.32634 | 56.07053 | 44.85512 | 36.87335 | 58.82164 | 77.21897 |
| ENSMUSG00000036964 | 0.614727488 | 0.211544 | 0.369057 | 19.11503 | 40.79393 | 29.81184 | 23.29083 | 16.57689 | 34.6386 | 9.642892 | 13.05109 |
| ENSMUSG00000036966 | -0.257150417 | 0.058845 | 0.152512 | 486.9303 | 415.1383 | 593.7525 | 439.0754 | 602.6188 | 545.2786 | 627.7522 | 539.4452 |
| ENSMUSG00000036968 | 0.222269578 | 0.00665 | 0.031813 | 1429.603 | 1629.358 | 1351.47 | 1519.08 | 1342.728 | 1262.633 | 1255.504 | 1217.014 |
| ENSMUSG00000036975 | -0.233000999 | 0.132753 | 0.266153 | 262.5802 | 230.3657 | 252.5725 | 238.9467 | 307.1601 | 219.0053 | 338.4655 | 291.4744 |
| ENSMUSG00000036977 | 0.070568421 | 0.428438 | 0.593459 | 495.9848 | 531.5209 | 544.8942 | 540.8649 | 469.0285 | 517.3442 | 542.8948 | 483.978 |
| ENSMUSG00000036980 | -0.238966868 | 0.096942 | 0.216109 | 641.8627 | 706.6949 | 742.8116 | 690.9614 | 876.6251 | 614.5558 | 827.3601 | 964.6933 |
| ENSMUSG00000036983 | -0.231066447 | 0.080394 | 0.189543 | 203.223 | 248.3631 | 193.777 | 228.5952 | 234.0267 | 251.4092 | 256.5009 | 281.6861 |
| ENSMUSG00000036985 | -0.487750215 | 0.006404 | 0.030924 | 487.9364 | 415.1383 | 611.9708 | 570.1941 | 696.2295 | 555.3349 | 783.0028 | 892.9123 |
| ENSMUSG00000036986 | 0.293950734 | 0.000269 | 0.002849 | 1320.949 | 1313.805 | 1258.722 | 1195.596 | 1030.693 | 1091.674 | 1092.54 | 933.1532 |
| ENSMUSG00000036989 | 0.030253998 | 0.794683 | 0.873721 | 1051.327 | 1053.443 | 1089.788 | 1283.584 | 1130.154 | 912.8946 | 1153.29 | 1189.825 |
| ENSMUSG00000036990 | 0.128586194 | 0.270092 | 0.435085 | 2597.632 | 2183.675 | 2159.702 | 2505.921 | 2131.593 | 2566.608 | 2031.757 | 1913.073 |
| ENSMUSG00000036992 | -0.208983333 | 0.014664 | 0.056722 | 622.7476 | 631.1061 | 647.5794 | 588.3092 | 743.0348 | 649.1943 | 717.4311 | 766.7517 |
| ENSMUSG00000036995 | 0.565510581 | 0.000761 | 0.006341 | 335.0161 | 485.9277 | 469.5365 | 464.9541 | 330.5627 | 340.7991 | 258.4295 | 256.6715 |
| ENSMUSG00000037001 | -0.161568761 | 0.178248 | 0.327598 | 323.9495 | 343.149 | 286.5249 | 333.8353 | 367.617 | 383.2593 | 307.6082 | 380.6569 |
| ENSMUSG00000037003 | 0.109598607 | 0.246888 | 0.409947 | 1913.515 | 1833.327 | 1596.59 | 1852.053 | 1690.843 | 1624.662 | 1846.614 | 1503.051 |
| ENSMUSG00000037005 | -1.802565049 | 5.89E-05 | 0.000901 | 65.39353 | 44.3934 | 141.6062 | 66.42201 | 286.6827 | 113.9722 | 236.2508 | 475.2773 |
| ENSMUSG00000037007 | 0.076945192 | 0.627647 | 0.757411 | 588.5418 | 667.1008 | 510.1137 | 546.0407 | 522.6597 | 679.3635 | 559.2877 | 428.5109 |
| ENSMUSG00000037010 | -0.65175627 | 0.082398 | 0.192898 | 75.45408 | 109.1838 | 213.6515 | 127.6683 | 232.0765 | 129.6154 | 146.572 | 319.7518 |
| ENSMUSG00000037012 | -0.352048523 | 0.09689 | 0.216018 | 2307.889 | 2181.276 | 3008.511 | 2382.566 | 2939.961 | 1955.405 | 3621.87 | 4094.78 |
| ENSMUSG00000037013 | 0.055997153 | 0.315369 | 0.483707 | 2844.116 | 3034.349 | 2805.625 | 2924.294 | 2833.673 | 2742.036 | 2907.332 | 2676.562 |
| ENSMUSG00000037016 | 0.024083384 | 0.691212 | 0.804404 | 4160.035 | 4435.74 | 4181.11 | 4383.853 | 4181.277 | 4537.656 | 4004.693 | 4153.51 |
| ENSMUSG00000037017 | 0.082642966 | 0.188193 | 0.340452 | 1331.01 | 1335.401 | 1278.597 | 1287.897 | 1207.188 | 1252.576 | 1176.433 | 1305.109 |
| ENSMUSG00000037020 | -0.276594688 | 0.127214 | 0.258402 | 123.7447 | 172.7743 | 149.8873 | 141.4703 | 196.9725 | 158.6671 | 146.572 | 208.8175 |
| ENSMUSG00000037022 | -0.010307037 | 0.921739 | 0.953353 | 322.9434 | 387.5424 | 322.9616 | 343.3242 | 364.6916 | 343.0338 | 330.7512 | 344.7664 |
| ENSMUSG00000037029 | 0.087974203 | 0.210799 | 0.36828 | 2716.347 | 2622.81 | 2470.242 | 2597.359 | 2568.443 | 2233.631 | 2564.045 | 2419.89 |
| ENSMUSG00000037031 | 0.122680897 | 0.089156 | 0.203824 | 2219.356 | 2136.882 | 2423.04 | 2243.684 | 1911.218 | 2090.607 | 2188.936 | 2101.226 |
| ENSMUSG00000037032 | 0.103182523 | 0.261712 | 0.425894 | 1141.872 | 1346.2 | 1204.895 | 1361.22 | 1227.665 | 1062.623 | 1195.719 | 1217.014 |
| ENSMUSG00000037035 | 0.69332052 | 0.01946 | 0.069136 | 99.59938 | 91.18644 | 61.27989 | 100.9269 | 32.17867 | 68.15982 | 63.64308 | 54.37955 |
| ENSMUSG00000037049 | -0.631458378 | 7.25E-05 | 0.001061 | 843.0736 | 939.4603 | 1073.226 | 896.2658 | 1486.07 | 1046.98 | 1568.898 | 1710.781 |
| ENSMUSG00000037058 | 0.108218946 | 0.229495 | 0.389727 | 5525.251 | 5671.556 | 4907.36 | 6047.853 | 5262.676 | 5493.011 | 5187.876 | 4605.948 |
| ENSMUSG00000037060 | 0.233022175 | 0.04891 | 0.13412 | 765.6074 | 1018.648 | 980.4783 | 926.4576 | 745.9602 | 880.4908 | 722.2526 | 792.8539 |
| ENSMUSG00000037062 | 0.045497183 | 0.645844 | 0.770604 | 3349.155 | 3402.694 | 3196.492 | 3621.293 | 2903.881 | 3827.006 | 3113.69 | 3306.277 |
| ENSMUSG00000037070 | 0.024622463 | 0.762682 | 0.852453 | 1944.703 | 2248.466 | 1919.551 | 2246.272 | 2126.718 | 1988.926 | 1992.221 | 2106.664 |
| ENSMUSG00000037071 | 0.42589957 | 0.027213 | 0.088204 | 602.6266 | 725.892 | 573.0498 | 684.0604 | 439.7752 | 640.2554 | 508.1804 | 336.0656 |
| ENSMUSG00000037072 | -0.025866527 | 0.663456 | 0.783613 | 6833.121 | 7588.871 | 7224.402 | 7696.327 | 7650.723 | 7658.482 | 7395.134 | 7168.313 |
| ENSMUSG00000037075 | -0.004720137 | 0.960587 | 0.977472 | 1214.308 | 1370.196 | 1323.314 | 1394 | 1318.35 | 1481.638 | 1355.791 | 1163.722 |
| ENSMUSG00000037085 | -0.36744714 | 0.001181 | 0.008874 | 297.7921 | 285.5575 | 240.979 | 264.8254 | 378.3432 | 336.3296 | 350.037 | 336.0656 |
| ENSMUSG00000037089 | -0.302981742 | 0.000706 | 0.005989 | 1416.525 | 1625.758 | 1674.432 | 1524.256 | 2009.704 | 1710.7 | 2032.722 | 1944.613 |
| ENSMUSG00000037095 | 0.716895422 | 0.325124 | 0.493053 | 74.44802 | 4.799286 | 17.39024 | 32.77969 | 21.45245 | 7.821619 | 21.21436 | 28.27737 |
| ENSMUSG00000037096 | 0.13820452 | 0.524027 | 0.677271 | 69.41775 | 82.78769 | 93.57605 | 103.5148 | 88.73513 | 61.45558 | 83.89316 | 83.74451 |
| ENSMUSG00000037098 | -0.791623369 | 9.00E-06 | 0.000211 | 951.7274 | 872.2703 | 1117.944 | 984.2534 | 1571.879 | 1273.806 | 1641.22 | 2312.219 |

| | | | | | | | | | | | |
|---|---|---|---|---|---|---|---|---|---|---|---|
| ENSMUSG00000037103 | -0.077512746 | 0.307252 | 0.475725 | 1066.418 | 1011.45 | 1134.506 | 961.8252 | 1085.299 | 1093.909 | 1094.468 | 1133.27 |
| ENSMUSG00000037104 | 0.215541635 | 0.038628 | 0.113622 | 1024.163 | 844.6744 | 818.9975 | 878.1507 | 705.9806 | 840.2653 | 779.1456 | 746.0875 |
| ENSMUSG00000037108 | 0.310063564 | 0.073318 | 0.178282 | 676.0685 | 926.2622 | 683.188 | 784.1248 | 687.4535 | 763.1665 | 520.7161 | 503.5547 |
| ENSMUSG00000037110 | 0.143769404 | 0.242585 | 0.405136 | 688.1412 | 586.7127 | 620.2519 | 585.7214 | 505.1077 | 500.5836 | 574.7163 | 665.6057 |
| ENSMUSG00000037111 | 0.542060059 | 0.000161 | 0.001918 | 1431.615 | 1583.764 | 1265.347 | 1299.974 | 885.4011 | 1223.525 | 835.0744 | 888.5619 |
| ENSMUSG00000037112 | -0.311968919 | 0.098103 | 0.218092 | 196.1806 | 172.7743 | 179.6991 | 138.0198 | 248.6534 | 157.5497 | 223.7151 | 220.781 |
| ENSMUSG00000037119 | -0.226098375 | 0.000465 | 0.004333 | 3163.035 | 2896.369 | 3165.852 | 2973.463 | 3481.147 | 3355.474 | 3737.585 | 3696.722 |
| ENSMUSG00000037126 | 0.232516837 | 0.372828 | 0.541805 | 248.4954 | 278.3586 | 289.8373 | 331.2474 | 212.5743 | 391.0809 | 209.2507 | 165.3138 |
| ENSMUSG00000037129 | -0.584548463 | 0.255152 | 0.41907 | 22.1332 | 9.598572 | 52.17072 | 23.29083 | 51.6809 | 24.58223 | 33.75012 | 52.20437 |
| ENSMUSG00000037134 | 0.106937696 | 0.319734 | 0.487813 | 744.4802 | 718.6931 | 602.8616 | 664.2201 | 602.6188 | 706.1804 | 642.2166 | 581.8612 |
| ENSMUSG00000037138 | 0.3960608 | 9.33E-06 | 0.000215 | 991.9696 | 1046.244 | 1075.711 | 1067.928 | 705.0055 | 882.7255 | 812.8958 | 779.8028 |
| ENSMUSG00000037139 | -0.225885293 | 0.61666 | 0.749494 | 11.0666 | 13.19804 | 30.63995 | 17.25247 | 16.57689 | 26.81698 | 18.32149 | 23.927 |
| ENSMUSG00000037143 | -0.170392235 | 0.747236 | 0.842198 | 28.16952 | 7.198929 | 9.109173 | 16.38985 | 15.60178 | 14.52586 | 23.14294 | 15.22628 |
| ENSMUSG00000037148 | -0.018212053 | 0.820214 | 0.89104 | 1160.987 | 1367.797 | 1173.427 | 1188.695 | 1200.362 | 1203.412 | 1240.076 | 1304.022 |
| ENSMUSG00000037149 | -0.115515335 | 0.020369 | 0.07143 | 4100.678 | 4334.955 | 4192.704 | 4318.293 | 4794.622 | 4418.097 | 4591.945 | 4550.481 |
| ENSMUSG00000037151 | 0.0807398 | 0.513459 | 0.668226 | 276.6649 | 304.7547 | 294.806 | 302.7808 | 287.6578 | 232.4138 | 312.4297 | 280.5985 |
| ENSMUSG00000037152 | -0.446064715 | 0.000224 | 0.002474 | 858.1644 | 773.8849 | 915.0578 | 784.1248 | 1078.473 | 944.1811 | 1248.754 | 1268.131 |
| ENSMUSG00000037161 | 1.612271491 | 0.006719 | 0.032025 | 1180.102 | 215.9679 | 229.3855 | 456.3278 | 160.8934 | 144.1413 | 147.5362 | 228.3941 |
| ENSMUSG00000037166 | 0.230383191 | 0.036813 | 0.110225 | 385.3188 | 340.7493 | 394.1788 | 349.3625 | 336.4134 | 290.5173 | 293.1439 | 333.8905 |
| ENSMUSG00000037169 | -0.163852342 | 0.347406 | 0.516242 | 1717.335 | 1686.949 | 1241.332 | 1239.59 | 2090.639 | 1424.652 | 1693.292 | 1380.153 |
| ENSMUSG00000037172 | 0.063159391 | 0.481611 | 0.640528 | 864.2007 | 770.2854 | 703.8906 | 810.8661 | 781.0642 | 762.0491 | 716.4668 | 753.7006 |
| ENSMUSG00000037174 | 0.094130335 | 0.093054 | 0.209824 | 2736.468 | 2777.587 | 2754.283 | 2732.791 | 2739.088 | 2515.209 | 2606.474 | 2441.642 |
| ENSMUSG00000037185 | -0.802715564 | 0.185609 | 0.337199 | 19.11503 | 8.398751 | 52.17072 | 17.25247 | 46.80534 | 18.99536 | 31.82154 | 72.8686 |
| ENSMUSG00000037188 | -1.077958667 | 0.042779 | 0.122331 | 71.42986 | 26.39607 | 122.5598 | 52.62003 | 195.0223 | 40.22547 | 144.6434 | 197.9416 |
| ENSMUSG00000037190 | -0.827026231 | 5.56E-05 | 0.000859 | 224.3501 | 281.9581 | 351.1172 | 276.9021 | 549.9628 | 339.6817 | 578.5735 | 544.8831 |
| ENSMUSG00000037196 | -0.525808445 | 0.030705 | 0.096115 | 47.28455 | 38.39429 | 70.38906 | 62.10889 | 78.0089 | 75.98144 | 75.21455 | 88.09488 |
| ENSMUSG00000037197 | -0.050996326 | 0.504096 | 0.659783 | 3563.445 | 3905.419 | 3731.448 | 3960.304 | 4263.187 | 4071.711 | 3762.656 | 3607.54 |
| ENSMUSG00000037204 | -0.285008735 | 0.015851 | 0.059945 | 834.0191 | 727.0918 | 843.8407 | 852.272 | 1000.464 | 808.9788 | 1062.647 | 1098.467 |
| ENSMUSG00000037206 | 0.568094178 | 1.95E-06 | 6.32E-05 | 2795.825 | 3688.251 | 3376.191 | 3315.062 | 2346.118 | 2535.322 | 2045.257 | 1959.839 |
| ENSMUSG00000037210 | 0.093027167 | 0.450172 | 0.612621 | 1261.592 | 1166.227 | 1165.146 | 1173.168 | 1058.971 | 1396.718 | 1028.897 | 985.3575 |
| ENSMUSG00000037211 | 0.239943391 | 0.060721 | 0.156364 | 3785.783 | 3727.846 | 3101.259 | 3886.981 | 3088.178 | 3658.283 | 3009.546 | 2523.211 |
| ENSMUSG00000037214 | 0.022164155 | 0.823316 | 0.89278 | 304.8345 | 333.5504 | 319.6492 | 301.9182 | 318.8614 | 326.2732 | 302.7868 | 291.4744 |
| ENSMUSG00000037216 | -0.070815754 | 0.65875 | 0.780202 | 181.0898 | 171.5745 | 136.6376 | 158.7227 | 157.968 | 178.7799 | 193.8221 | 147.9124 |
| ENSMUSG00000037217 | 0.350851342 | 0.257395 | 0.421143 | 36.21796 | 32.39518 | 33.12427 | 56.07053 | 29.25334 | 33.52122 | 27.0001 | 34.80291 |
| ENSMUSG00000037221 | -0.045981362 | 0.644402 | 0.770028 | 615.7053 | 577.1142 | 674.9069 | 708.2139 | 723.5326 | 640.2554 | 628.7165 | 671.0437 |
| ENSMUSG00000037224 | 0.287032955 | 0.354226 | 0.52284 | 34.20585 | 31.19536 | 37.2648 | 33.64232 | 19.50223 | 26.81698 | 27.96439 | 38.06569 |
| ENSMUSG00000037225 | 0.478306601 | 0.010765 | 0.045083 | 556.3481 | 679.099 | 558.1439 | 587.4466 | 430.0241 | 564.2739 | 418.5015 | 295.8248 |
| ENSMUSG00000037234 | 0.36144435 | 0.002533 | 0.015566 | 2540.287 | 2481.231 | 2260.731 | 2505.921 | 1623.56 | 2337.547 | 1877.471 | 1781.474 |
| ENSMUSG00000037235 | -0.056159497 | 0.456708 | 0.618506 | 4973.933 | 4666.106 | 4622.491 | 4796.187 | 4666.883 | 4640.455 | 5529.234 | 4976.817 |
| ENSMUSG00000037236 | 0.163946805 | 0.143266 | 0.28087 | 8005.175 | 9211.03 | 7974.667 | 8871.22 | 7326.011 | 9169.172 | 7189.74 | 6718.05 |
| ENSMUSG00000037239 | 0.823137129 | 2.00E-07 | 1.00E-05 | 834.0191 | 745.0892 | 634.3297 | 783.2621 | 385.169 | 546.3959 | 405.9657 | 356.7299 |
| ENSMUSG00000037242 | -0.203138909 | 0.00839 | 0.037585 | 4473.924 | 4114.188 | 4786.456 | 4883.312 | 5086.181 | 5017.01 | 5321.912 | 5601.094 |
| ENSMUSG00000037243 | 0.066180377 | 0.492054 | 0.649178 | 534.2149 | 560.3167 | 513.4261 | 503.7721 | 559.7139 | 486.0577 | 508.1804 | 458.9634 |
| ENSMUSG00000037251 | 0.197984585 | 0.035427 | 0.107046 | 525.1604 | 567.5156 | 475.3332 | 509.8105 | 463.1779 | 420.1327 | 457.0731 | 466.5766 |
| ENSMUSG00000037253 | 0.213220931 | 0.090333 | 0.205724 | 1685.141 | 1701.347 | 1410.266 | 1570.837 | 1471.443 | 1615.723 | 1237.183 | 1166.985 |
| ENSMUSG00000037254 | 0.866336111 | 0.023989 | 0.080571 | 284.7134 | 160.7761 | 173.0743 | 254.4739 | 116.0382 | 222.3574 | 69.42882 | 71.78101 |
| ENSMUSG00000037257 | -0.386425923 | 1.27E-05 | 0.000276 | 786.7345 | 771.4852 | 828.1066 | 752.2077 | 1009.24 | 915.1294 | 1052.039 | 1126.744 |
| ENSMUSG00000037259 | 0.314955921 | 0.077577 | 0.185199 | 268.6165 | 289.157 | 211.1672 | 269.1385 | 228.176 | 254.7613 | 174.5363 | 176.1898 |
| ENSMUSG00000037260 | -0.307236471 | 0.016941 | 0.062681 | 1107.666 | 1045.045 | 1163.49 | 1093.807 | 1365.156 | 1054.801 | 1458.005 | 1579.182 |
| ENSMUSG00000037262 | 0.22088073 | 0.075316 | 0.181454 | 1048.309 | 893.867 | 766.8267 | 789.3005 | 763.5122 | 816.8005 | 731.8955 | 687.3576 |
| ENSMUSG00000037266 | 0.449770565 | 4.21E-06 | 0.000118 | 12812.1 | 11033.56 | 9758.409 | 12308.77 | 8427.887 | 8795.969 | 8506.959 | 7883.948 |
| ENSMUSG00000037270 | 0.09247928 | 0.314241 | 0.482338 | 1769.65 | 1553.769 | 1652.073 | 1610.518 | 1579.68 | 1715.169 | 1529.363 | 1354.051 |
| ENSMUSG00000037275 | -0.128425561 | 0.116356 | 0.244531 | 989.9575 | 969.4558 | 861.2309 | 973.9019 | 1085.299 | 954.2375 | 1040.468 | 1063.664 |
| ENSMUSG00000037278 | -0.736493869 | 5.49E-07 | 2.27E-05 | 507.0514 | 543.5192 | 510.9418 | 431.3117 | 927.3309 | 629.0816 | 846.6459 | 913.5765 |
| ENSMUSG00000037279 | 0.99147898 | 4.21E-05 | 0.000701 | 104.6297 | 73.18911 | 101.8571 | 108.6906 | 47.78045 | 64.8077 | 40.50014 | 43.50364 |
| ENSMUSG00000037280 | 0.372275851 | 0.123094 | 0.252702 | 95.57516 | 131.9804 | 72.87338 | 120.7673 | 86.78491 | 72.62932 | 71.3574 | 93.53283 |
| ENSMUSG00000037286 | 0.207986887 | 0.10182 | 0.223792 | 2693.208 | 2880.772 | 2466.93 | 2671.545 | 2153.046 | 2888.412 | 2275.722 | 1956.576 |
| ENSMUSG00000037287 | 0.082006639 | 0.540355 | 0.689873 | 1718.341 | 1347.4 | 1292.674 | 1275.82 | 1198.412 | 1562.089 | 1336.505 | 1225.715 |
| ENSMUSG00000037295 | -0.142136486 | 0.247495 | 0.410697 | 1140.866 | 938.2604 | 1110.491 | 896.2658 | 1253.018 | 957.5896 | 1139.79 | 1158.285 |
| ENSMUSG00000037296 | -0.104503601 | 0.19239 | 0.345566 | 645.8869 | 638.3051 | 671.5945 | 701.3129 | 736.209 | 689.4198 | 758.8956 | 673.2189 |
| ENSMUSG00000037300 | 0.190086455 | 0.018019 | 0.065722 | 1406.464 | 1370.196 | 1212.348 | 1329.303 | 1173.059 | 1252.576 | 1141.718 | 1091.941 |
| ENSMUSG00000037306 | -0.291729728 | 0.054277 | 0.144444 | 448.7002 | 361.1463 | 532.4726 | 404.5704 | 516.809 | 454.7713 | 573.7521 | 595.9999 |
| ENSMUSG00000037313 | 0.10479023 | 0.252672 | 0.41633 | 2389.379 | 2398.443 | 2331.948 | 2541.289 | 2395.848 | 2499.566 | 2107.936 | 1981.591 |
| ENSMUSG00000037315 | -0.088211413 | 0.463782 | 0.624694 | 463.7911 | 392.3416 | 430.6154 | 411.4714 | 384.1939 | 438.0106 | 472.5017 | 512.2554 |

| | | | | | | | | | | | |
|---|---|---|---|---|---|---|---|---|---|---|---|
| ENSMUSG00000037316 | -0.063510704 | 0.455551 | 0.617528 | 567.4147 | 578.314 | 522.5353 | 548.6285 | 568.4899 | 624.6121 | 579.5378 | 541.6204 |
| ENSMUSG00000037318 | -0.080807608 | 0.837477 | 0.90162 | 18.10898 | 17.99732 | 33.12427 | 15.52722 | 21.45245 | 26.81698 | 18.32149 | 23.927 |
| ENSMUSG00000037321 | 0.043786556 | 0.883573 | 0.930352 | 189.1382 | 289.157 | 372.648 | 132.844 | 232.0765 | 204.4795 | 282.5367 | 234.9197 |
| ENSMUSG00000037325 | -0.397088541 | 0.000349 | 0.003475 | 336.0222 | 316.7529 | 309.7119 | 294.1546 | 414.4223 | 400.0199 | 372.2156 | 467.6642 |
| ENSMUSG00000037326 | 0.034856821 | 0.724149 | 0.827323 | 895.3884 | 743.8894 | 768.483 | 725.4664 | 786.9148 | 717.3542 | 735.7526 | 817.8685 |
| ENSMUSG00000037331 | 0.130630081 | 0.25236 | 0.416113 | 4146.956 | 4052.997 | 3629.591 | 3539.344 | 3346.582 | 4243.787 | 3160.94 | 3286.7 |
| ENSMUSG00000037335 | 3.82387937 | 0.003985 | 0.021805 | 83.50251 | 9.598572 | 0.828107 | 91.43809 | 0 | 0 | 8.678602 | 4.350364 |
| ENSMUSG00000037336 | 0.58831095 | 0.056074 | 0.147526 | 80.48435 | 37.19447 | 52.17072 | 54.34528 | 39.97956 | 42.46022 | 37.60728 | 29.36496 |
| ENSMUSG00000037337 | 0.051303266 | 0.673024 | 0.790437 | 219.3198 | 254.3622 | 233.5261 | 256.1992 | 249.6285 | 226.8269 | 243.0009 | 208.8175 |
| ENSMUSG00000037339 | -0.095140247 | 0.167646 | 0.313587 | 812.8919 | 776.2845 | 784.217 | 808.2782 | 841.5211 | 900.6035 | 810.9672 | 848.321 |
| ENSMUSG00000037341 | -0.691894676 | 0.076591 | 0.183719 | 29.17558 | 19.19714 | 32.29616 | 12.07673 | 34.1289 | 27.93435 | 43.39301 | 44.59123 |
| ENSMUSG00000037343 | 0.020779492 | 0.756788 | 0.848662 | 1461.797 | 1532.172 | 1390.391 | 1481.987 | 1349.554 | 1511.807 | 1473.434 | 1445.409 |
| ENSMUSG00000037344 | 0.017608838 | 0.836948 | 0.901367 | 488.9424 | 445.1338 | 493.5516 | 475.3055 | 442.7005 | 469.2971 | 488.8946 | 481.8029 |
| ENSMUSG00000037347 | -0.520930821 | 0.091672 | 0.207774 | 65.39353 | 58.79126 | 62.9361 | 86.26235 | 88.73513 | 54.75133 | 108.0004 | 141.3868 |
| ENSMUSG00000037348 | -0.103125411 | 0.576839 | 0.718897 | 145.8779 | 129.5807 | 204.5423 | 169.0742 | 195.0223 | 151.9629 | 166.822 | 185.9781 |
| ENSMUSG00000037349 | -0.425608306 | 0.016472 | 0.061452 | 293.7679 | 215.9679 | 330.4145 | 283.8031 | 407.5965 | 297.2215 | 442.6087 | 364.343 |
| ENSMUSG00000037351 | -0.335538736 | 0.000812 | 0.00665 | 2846.128 | 2624.01 | 2949.716 | 2668.094 | 3521.127 | 2954.337 | 3574.62 | 3942.518 |
| ENSMUSG00000037353 | -0.039590411 | 0.614659 | 0.748195 | 909.4731 | 914.264 | 874.4806 | 897.9911 | 977.0615 | 824.6221 | 936.3248 | 954.905 |
| ENSMUSG00000037355 | 0.25703338 | 0.030053 | 0.094636 | 1040.26 | 1025.847 | 857.9185 | 922.1445 | 741.0846 | 956.4722 | 778.1814 | 741.7371 |
| ENSMUSG00000037358 | 0.052421015 | 0.864314 | 0.91877 | 35.2119 | 64.79036 | 42.23344 | 47.44429 | 37.05423 | 34.6386 | 57.85735 | 52.20437 |
| ENSMUSG00000037361 | -0.150086796 | 0.071193 | 0.174541 | 1670.05 | 1793.733 | 2009.815 | 1807.196 | 1912.193 | 1971.048 | 2008.614 | 2191.496 |
| ENSMUSG00000037362 | 0.799487669 | 0.008512 | 0.037966 | 271.6347 | 370.7449 | 254.2287 | 466.6793 | 244.7529 | 274.874 | 139.8219 | 123.9854 |
| ENSMUSG00000037363 | 0.564029636 | 0.007087 | 0.033199 | 385.3188 | 404.3399 | 257.5412 | 345.912 | 182.3458 | 312.8647 | 222.7508 | 224.0438 |
| ENSMUSG00000037364 | 0.112413176 | 0.012041 | 0.049101 | 3992.024 | 4075.794 | 4017.145 | 3923.212 | 3585.484 | 3736.499 | 3664.299 | 3821.795 |
| ENSMUSG00000037366 | -1.114466517 | 9.91E-05 | 0.00133 | 347.0888 | 277.1588 | 510.1137 | 397.6694 | 812.2677 | 439.128 | 884.2532 | 1184.387 |
| ENSMUSG00000037369 | 0.300165834 | 0.279142 | 0.445469 | 1854.158 | 1072.64 | 940.7291 | 1723.522 | 914.6544 | 1746.456 | 916.0747 | 964.6933 |
| ENSMUSG00000037370 | 0.030762555 | 0.809302 | 0.883606 | 496.9909 | 568.7154 | 502.6607 | 651.2807 | 597.7432 | 477.1187 | 565.0735 | 531.832 |
| ENSMUSG00000037373 | -0.138054377 | 0.018698 | 0.067295 | 6543.378 | 7328.51 | 7480.287 | 7018.305 | 7912.053 | 7648.426 | 7756.742 | 7902.437 |
| ENSMUSG00000037375 | 0.018077354 | 0.843195 | 0.905412 | 386.3249 | 415.1383 | 380.1009 | 376.1038 | 359.8161 | 386.6114 | 401.1443 | 388.27 |
| ENSMUSG00000037376 | -0.040078542 | 0.656463 | 0.778543 | 1067.424 | 1177.025 | 1159.349 | 1054.126 | 1252.043 | 1198.942 | 1030.825 | 1100.642 |
| ENSMUSG00000037379 | 0.184569296 | 0.624918 | 0.755551 | 91.55095 | 265.1606 | 247.6039 | 149.2339 | 158.9431 | 254.7613 | 98.35749 | 151.1752 |
| ENSMUSG00000037386 | 0.797862606 | 0.001201 | 0.00898 | 111.672 | 89.98662 | 72.04528 | 113.0037 | 48.75557 | 62.57295 | 45.32159 | 66.34306 |
| ENSMUSG00000037395 | 0.02314546 | 0.793273 | 0.873286 | 1006.054 | 838.6753 | 890.2146 | 1016.17 | 893.202 | 945.2985 | 918.0033 | 938.5911 |
| ENSMUSG00000037400 | 0.131457233 | 0.037701 | 0.111957 | 1922.57 | 1809.331 | 1738.196 | 1885.695 | 1632.336 | 1758.747 | 1627.72 | 1697.73 |
| ENSMUSG00000037405 | 0.196211947 | 0.341241 | 0.509955 | 307.8526 | 227.9661 | 362.7107 | 257.0618 | 272.0561 | 179.8972 | 266.1438 | 291.4744 |
| ENSMUSG00000037406 | -0.490572671 | 0.181996 | 0.332533 | 28.16952 | 19.19714 | 46.37397 | 20.70296 | 46.80534 | 40.22547 | 43.39301 | 31.54014 |
| ENSMUSG00000037408 | -0.071501522 | 0.551509 | 0.699148 | 271.6347 | 307.1543 | 303.9151 | 244.9851 | 282.7823 | 309.5126 | 310.5011 | 280.5985 |
| ENSMUSG00000037410 | 0.305654479 | 0.000191 | 0.002184 | 1415.518 | 1679.75 | 1663.666 | 1625.183 | 1334.927 | 1350.905 | 1241.04 | 1238.766 |
| ENSMUSG00000037411 | -0.13378304 | 0.731279 | 0.831495 | 52.31483 | 20.39697 | 33.12427 | 25.8787 | 46.80534 | 24.58223 | 41.46443 | 31.54014 |
| ENSMUSG00000037415 | 0.148446403 | 0.089454 | 0.204227 | 1295.798 | 1150.629 | 1079.023 | 1193.871 | 1073.598 | 1030.219 | 1001.896 | 1151.759 |
| ENSMUSG00000037416 | 0.017202251 | 0.875362 | 0.925381 | 1189.156 | 1189.023 | 1091.445 | 1124.861 | 1019.966 | 1328.558 | 1003.825 | 1188.737 |
| ENSMUSG00000037418 | 0.83476558 | 0.0001 | 0.001337 | 152.9203 | 188.372 | 118.4192 | 130.2561 | 79.95913 | 97.21155 | 84.85745 | 67.43065 |
| ENSMUSG00000037419 | -0.566758274 | 0.003125 | 0.018318 | 452.7245 | 506.3247 | 785.8732 | 445.1137 | 786.9148 | 735.2322 | 829.2887 | 895.0875 |
| ENSMUSG00000037426 | 0.193278663 | 0.045768 | 0.127927 | 713.2925 | 591.512 | 584.6433 | 622.8142 | 576.2908 | 545.2786 | 551.5734 | 523.1313 |
| ENSMUSG00000037428 | 1.075701581 | 0.056439 | 0.148213 | 54.32694 | 20.39697 | 15.73403 | 68.14726 | 10.72622 | 29.05173 | 17.3572 | 18.48905 |
| ENSMUSG00000037432 | 0.292404392 | 0.062534 | 0.159511 | 168.0111 | 177.5736 | 186.324 | 223.4195 | 133.5902 | 169.8409 | 164.8934 | 150.0876 |
| ENSMUSG00000037434 | -0.405019238 | 0.000481 | 0.00445 | 565.4025 | 571.1151 | 621.08 | 609.8748 | 692.329 | 709.5326 | 810.9672 | 924.4524 |
| ENSMUSG00000037440 | -0.832896023 | 0.078321 | 0.186298 | 131.7931 | 86.38715 | 267.4784 | 170.7995 | 243.7778 | 92.74205 | 303.7511 | 530.7445 |
| ENSMUSG00000037443 | -1.033932572 | 0.000409 | 0.003928 | 774.6619 | 735.4906 | 660.8291 | 688.3735 | 643.5735 | 2225.809 | 1563.113 | 1422.569 |
| ENSMUSG00000037446 | 0.188758065 | 0.159185 | 0.302155 | 352.119 | 467.9304 | 399.1474 | 411.4714 | 383.2187 | 402.2547 | 321.1083 | 323.0146 |
| ENSMUSG00000037447 | 0.516071872 | 0.009466 | 0.040932 | 186.1201 | 175.1739 | 157.3403 | 155.2722 | 86.78491 | 112.8548 | 153.322 | 117.4598 |
| ENSMUSG00000037455 | 0.429255911 | 7.84E-06 | 0.000191 | 767.6195 | 753.4879 | 689.8128 | 819.4923 | 543.137 | 610.0863 | 515.8947 | 582.9488 |
| ENSMUSG00000037458 | -0.071528195 | 0.41325 | 0.579071 | 2139.878 | 2081.69 | 1922.864 | 1980.584 | 2010.68 | 2054.851 | 2029.829 | 2441.642 |
| ENSMUSG00000037461 | 0.040109553 | 0.61148 | 0.746072 | 2538.275 | 2895.169 | 2466.102 | 2464.515 | 2561.617 | 2674.994 | 2403.973 | 2434.029 |
| ENSMUSG00000037463 | 0.387897768 | 0.029848 | 0.094199 | 181.0898 | 123.5816 | 155.684 | 175.9752 | 136.5156 | 105.0332 | 120.5361 | 125.073 |
| ENSMUSG00000037465 | -0.303203867 | 0.237937 | 0.399097 | 576.4691 | 647.9036 | 739.4992 | 715.1149 | 896.1273 | 555.3349 | 572.7878 | 1282.27 |
| ENSMUSG00000037466 | -0.244889306 | 0.025103 | 0.083342 | 532.2028 | 592.7118 | 526.6758 | 550.3538 | 751.8108 | 577.6824 | 665.3595 | 610.1386 |
| ENSMUSG00000037470 | -0.208577103 | 0.025263 | 0.083583 | 1871.261 | 1854.924 | 1845.85 | 1513.904 | 1902.442 | 2012.391 | 2157.115 | 2113.189 |
| ENSMUSG00000037474 | -0.088725061 | 0.555837 | 0.702687 | 1102.636 | 1581.365 | 1249.613 | 1127.449 | 1411.961 | 1563.206 | 1281.54 | 1122.394 |
| ENSMUSG00000037475 | 0.513806049 | 0.017192 | 0.063383 | 4237.501 | 4396.146 | 3134.384 | 3816.246 | 2209.602 | 4139.871 | 2408.794 | 2156.693 |
| ENSMUSG00000037477 | -2.616953957 | 4.22E-05 | 0.000701 | 6.036326 | 1.199822 | 4.140533 | 3.450494 | 26.32801 | 10.05637 | 21.21436 | 34.80291 |
| ENSMUSG00000037486 | 0.128027144 | 0.210768 | 0.368274 | 1351.131 | 1315.004 | 1143.615 | 1287.897 | 1190.611 | 1327.44 | 1044.325 | 1101.73 |
| ENSMUSG00000037487 | -0.025481465 | 0.681347 | 0.796487 | 4354.203 | 3995.406 | 4395.59 | 4338.996 | 4193.954 | 4153.28 | 4464.659 | 4582.021 |
| ENSMUSG00000037490 | 0.083307379 | 0.726735 | 0.828973 | 138.8355 | 116.3827 | 141.6062 | 117.3168 | 104.3369 | 80.45093 | 140.7862 | 159.8759 |

| | | | | | | | | | | | |
|---|---|---|---|---|---|---|---|---|---|---|---|
| ENSMUSG00000037492 | 0.936044013 | 0.02806 | 0.090224 | 86.52067 | 89.98662 | 47.20208 | 61.24627 | 22.42756 | 71.51194 | 27.96439 | 27.18978 |
| ENSMUSG00000037493 | 0.132678225 | 0.344559 | 0.513276 | 423.5489 | 597.5111 | 485.2705 | 585.7214 | 520.7094 | 481.5882 | 481.1803 | 423.0729 |
| ENSMUSG00000037499 | -0.049724022 | 0.557617 | 0.704117 | 1342.077 | 1653.354 | 1524.544 | 1473.361 | 1629.411 | 1560.972 | 1453.184 | 1557.43 |
| ENSMUSG00000037503 | 0.163626111 | 0.055646 | 0.146914 | 6873.363 | 7395.7 | 7072.859 | 7210.67 | 6353.825 | 7300.922 | 6093.343 | 5743.569 |
| ENSMUSG00000037509 | -0.203214169 | 0.375624 | 0.544104 | 183.1019 | 133.1802 | 264.9941 | 165.6237 | 213.5494 | 181.0146 | 211.1793 | 256.6715 |
| ENSMUSG00000037514 | 0.070613216 | 0.479122 | 0.638368 | 1290.768 | 1560.968 | 1229.738 | 1369.846 | 1388.559 | 1225.759 | 1217.897 | 1354.051 |
| ENSMUSG00000037519 | -0.116396833 | 0.158879 | 0.301844 | 2046.315 | 1694.148 | 2020.58 | 1931.414 | 2033.107 | 2006.804 | 2076.115 | 2228.474 |
| ENSMUSG00000037523 | -0.344242264 | 0.086905 | 0.200189 | 839.0493 | 717.4933 | 804.9196 | 703.0381 | 928.306 | 613.4384 | 1104.111 | 1244.204 |
| ENSMUSG00000037525 | -0.04982998 | 0.740199 | 0.83754 | 233.4046 | 256.7618 | 244.2915 | 215.6559 | 267.1805 | 289.3999 | 222.7508 | 203.3795 |
| ENSMUSG00000037526 | 0.018619904 | 0.840768 | 0.904003 | 716.3107 | 779.884 | 718.7966 | 806.553 | 766.4375 | 778.8097 | 779.1456 | 656.905 |
| ENSMUSG00000037531 | -0.061094489 | 0.410473 | 0.576705 | 707.2562 | 697.0963 | 672.4226 | 645.2424 | 741.0846 | 698.3588 | 681.7524 | 715.6349 |
| ENSMUSG00000037533 | 0.166357343 | 0.128272 | 0.259781 | 1109.678 | 1058.243 | 923.3389 | 1053.263 | 911.7291 | 1061.505 | 915.1104 | 803.7298 |
| ENSMUSG00000037536 | -0.155279464 | 0.044908 | 0.126301 | 606.6508 | 652.7029 | 673.2507 | 671.9837 | 741.0846 | 690.5372 | 756.967 | 712.3722 |
| ENSMUSG00000037541 | -0.681598932 | 0.001612 | 0.011115 | 281.6952 | 219.5673 | 380.1009 | 197.5408 | 364.6916 | 420.1327 | 468.6445 | 479.6277 |
| ENSMUSG00000037542 | -1.465996482 | 0.026257 | 0.086104 | 34.20585 | 2.399643 | 22.35888 | 28.46658 | 60.4569 | 22.34748 | 49.17875 | 110.9343 |
| ENSMUSG00000037544 | 0.011823083 | 0.923348 | 0.954293 | 1175.071 | 1125.433 | 917.5421 | 1202.497 | 1273.495 | 988.8761 | 1065.54 | 1053.876 |
| ENSMUSG00000037548 | -0.768439048 | 1.12E-09 | 1.43E-07 | 130.7871 | 130.7805 | 128.3565 | 126.8057 | 221.3503 | 231.2964 | 226.608 | 200.1168 |
| ENSMUSG00000037552 | 0.004706058 | 0.957603 | 0.975954 | 1540.269 | 1303.006 | 1530.341 | 1498.377 | 1600.158 | 1414.596 | 1481.148 | 1359.489 |
| ENSMUSG00000037553 | 0.344916244 | 0.000776 | 0.006435 | 1208.271 | 1370.196 | 1356.439 | 1325.852 | 1216.939 | 986.6413 | 987.4321 | 949.467 |
| ENSMUSG00000037563 | -0.151577005 | 0.092418 | 0.208797 | 23118.12 | 27646.29 | 27873.24 | 27357.24 | 31855.91 | 26916.42 | 30740.57 | 28222.99 |
| ENSMUSG00000037568 | 0.069189372 | 0.570331 | 0.71379 | 713.2925 | 839.8751 | 815.685 | 736.6805 | 806.4171 | 845.8522 | 662.4667 | 644.9415 |
| ENSMUSG00000037570 | -0.267304037 | 2.96E-05 | 0.000531 | 2166.035 | 2278.461 | 2401.509 | 2151.383 | 2782.968 | 2574.43 | 2791.617 | 2678.737 |
| ENSMUSG00000037572 | -0.043480005 | 0.769636 | 0.857029 | 901.4247 | 1172.226 | 785.0451 | 805.6903 | 986.8126 | 1045.862 | 899.6818 | 839.6203 |
| ENSMUSG00000037573 | -0.172970164 | 0.112565 | 0.239278 | 1174.065 | 987.4531 | 953.1507 | 971.3141 | 1060.921 | 1090.557 | 1316.255 | 1136.533 |
| ENSMUSG00000037579 | 1.723686956 | 1.15E-14 | 4.71E-12 | 168.0111 | 167.975 | 143.2624 | 128.5309 | 59.48179 | 42.46022 | 47.25017 | 33.71532 |
| ENSMUSG00000037580 | 1.038502192 | 0.13746 | 0.272558 | 237.4288 | 27.5959 | 19.04645 | 106.1027 | 47.78045 | 24.58223 | 41.46443 | 76.13138 |
| ENSMUSG00000037593 | 0.283078385 | 0.482324 | 0.640923 | 15.09082 | 31.19536 | 24.8432 | 22.42821 | 12.67645 | 24.58223 | 23.14294 | 16.31387 |
| ENSMUSG00000037594 | -0.996215643 | 2.69E-07 | 1.26E-05 | 139.8416 | 175.1739 | 214.4796 | 160.448 | 424.1734 | 268.1698 | 325.9297 | 358.9051 |
| ENSMUSG00000037600 | -0.853913227 | 5.45E-06 | 0.000143 | 179.0777 | 177.5736 | 235.1823 | 149.2339 | 313.0107 | 288.2825 | 321.1083 | 418.7226 |
| ENSMUSG00000037601 | -0.517821676 | 1.16E-07 | 6.31E-06 | 1255.556 | 1390.593 | 1452.499 | 1239.59 | 2007.754 | 1710.7 | 1819.614 | 2104.489 |
| ENSMUSG00000037605 | 0.832888234 | 0.007342 | 0.034038 | 975.8727 | 1339.001 | 991.2436 | 1031.698 | 497.3068 | 1115.139 | 459.9659 | 363.2554 |
| ENSMUSG00000037606 | -0.104425491 | 0.4106 | 0.576745 | 1092.575 | 1132.632 | 1362.235 | 1262.881 | 1383.683 | 1046.98 | 1342.291 | 1443.233 |
| ENSMUSG00000037608 | 0.110176144 | 0.178645 | 0.328132 | 6297.9 | 6241.472 | 5882.041 | 5954.69 | 5537.657 | 6440.544 | 5197.519 | 5408.59 |
| ENSMUSG00000037610 | -0.33258604 | 0.082402 | 0.192898 | 109.6599 | 100.785 | 95.23226 | 92.30071 | 122.864 | 160.9019 | 111.8575 | 105.4963 |
| ENSMUSG00000037613 | -0.173724944 | 0.380834 | 0.548644 | 79.47829 | 75.58876 | 98.54469 | 63.83414 | 91.66046 | 88.27255 | 90.64318 | 88.09488 |
| ENSMUSG00000037617 | 0.402476358 | 0.012908 | 0.051766 | 209.2593 | 184.7725 | 159.8246 | 226.87 | 131.64 | 147.4934 | 156.2148 | 155.5255 |
| ENSMUSG00000037621 | -0.214300002 | 0.448988 | 0.611551 | 219.3198 | 251.9625 | 443.037 | 295.8799 | 281.8072 | 235.7659 | 412.7158 | 475.2773 |
| ENSMUSG00000037622 | 0.004820483 | 0.953018 | 0.973191 | 912.4913 | 896.2667 | 942.3853 | 843.6458 | 870.7744 | 833.5611 | 898.7175 | 979.9196 |
| ENSMUSG00000037624 | 0.365312315 | 0.019911 | 0.070196 | 203.223 | 172.7743 | 158.1684 | 215.6559 | 156.9929 | 135.2023 | 139.8219 | 150.0876 |
| ENSMUSG00000037625 | -0.154426007 | 0.755416 | 0.848205 | 403.4278 | 441.5343 | 1299.299 | 560.7053 | 566.5397 | 439.128 | 856.2888 | 1149.584 |
| ENSMUSG00000037627 | -0.278908537 | 0.51037 | 0.66605 | 9.054489 | 15.59768 | 17.39024 | 13.80198 | 13.65156 | 16.76061 | 22.17865 | 15.22628 |
| ENSMUSG00000037628 | 0.203627534 | 0.125348 | 0.2559 | 348.0948 | 363.5459 | 353.6015 | 388.1806 | 371.5174 | 311.7474 | 321.1083 | 256.6715 |
| ENSMUSG00000037636 | -0.029773063 | 0.943882 | 0.967579 | 32.19374 | 20.39697 | 22.35888 | 14.6646 | 19.50223 | 16.76061 | 19.28578 | 35.89051 |
| ENSMUSG00000037638 | -0.370242866 | 0.079596 | 0.18848 | 143.8658 | 157.1766 | 254.2287 | 179.4257 | 282.7823 | 213.4185 | 219.8579 | 236.0073 |
| ENSMUSG00000037640 | 0.266199523 | 0.038233 | 0.112948 | 898.4065 | 1052.243 | 718.7966 | 848.8215 | 750.8357 | 663.7202 | 766.6099 | 739.5619 |
| ENSMUSG00000037643 | -0.05576568 | 0.382961 | 0.550971 | 2195.211 | 2103.287 | 2233.404 | 2410.17 | 2263.233 | 2335.312 | 2294.044 | 2409.014 |
| ENSMUSG00000037646 | -0.187529888 | 0.01373 | 0.054032 | 1156.963 | 1010.25 | 1089.788 | 1134.35 | 1182.81 | 1222.407 | 1269.969 | 1330.124 |
| ENSMUSG00000037649 | -0.384684688 | 0.058368 | 0.151661 | 122.7386 | 99.58519 | 116.763 | 117.3168 | 189.1716 | 107.2679 | 150.4291 | 149 |
| ENSMUSG00000037652 | 0.350815995 | 0.003426 | 0.019484 | 1291.774 | 1334.202 | 1264.519 | 1518.217 | 923.4304 | 1271.572 | 1043.361 | 1004.934 |
| ENSMUSG00000037653 | -0.022752427 | 0.958883 | 0.976771 | 15.09082 | 21.59679 | 14.90592 | 9.488858 | 13.65156 | 16.76061 | 11.57147 | 19.57664 |
| ENSMUSG00000037656 | -0.084939778 | 0.292496 | 0.459782 | 2089.575 | 2189.674 | 1983.315 | 1987.485 | 2270.059 | 1962.109 | 2144.579 | 2368.773 |
| ENSMUSG00000037660 | -1.377692208 | 0.141697 | 0.278593 | 11.0666 | 39.59411 | 170.59 | 24.15346 | 139.4409 | 34.6386 | 212.1436 | 252.3211 |
| ENSMUSG00000037661 | -0.456038043 | 0.044391 | 0.12535 | 161.9748 | 147.578 | 164.7932 | 114.7289 | 162.8436 | 163.1366 | 201.5364 | 280.5985 |
| ENSMUSG00000037664 | -0.290322679 | 0.237953 | 0.399097 | 9448.863 | 7146.137 | 13666.24 | 9110.167 | 10743.78 | 7592.557 | 13621.55 | 16192.06 |
| ENSMUSG00000037669 | 0.006538661 | 0.924411 | 0.955046 | 862.1886 | 919.0633 | 929.1356 | 966.1383 | 890.2766 | 901.7209 | 931.5033 | 938.5911 |
| ENSMUSG00000037674 | 0.114860203 | 0.322944 | 0.491041 | 1499.021 | 1598.162 | 1450.015 | 1541.508 | 1368.081 | 1724.108 | 1292.147 | 1238.766 |
| ENSMUSG00000037679 | 0.28332414 | 0.006191 | 0.03015 | 614.6992 | 685.0981 | 616.1113 | 625.402 | 502.1823 | 602.2646 | 512.0375 | 470.9269 |
| ENSMUSG00000037681 | -0.113175737 | 0.707073 | 0.815914 | 37.22401 | 46.79304 | 38.92101 | 24.15346 | 42.9049 | 32.40385 | 45.32159 | 36.9781 |
| ENSMUSG00000037685 | 0.709170293 | 7.63E-06 | 0.000187 | 1765.625 | 1803.332 | 1453.327 | 1888.283 | 1047.27 | 1352.023 | 1007.682 | 820.0437 |
| ENSMUSG00000037686 | -0.923752669 | 0.000453 | 0.004247 | 35.2119 | 25.19625 | 27.32752 | 34.50494 | 50.70579 | 51.39921 | 69.42882 | 60.9051 |
| ENSMUSG00000037692 | 0.395381308 | 0.003334 | 0.019134 | 909.4731 | 799.0811 | 709.6874 | 764.2844 | 523.6348 | 636.9032 | 542.8948 | 716.7225 |
| ENSMUSG00000037697 | 0.203375744 | 0.048917 | 0.13412 | 580.4934 | 598.7109 | 599.5492 | 546.9033 | 509.0081 | 581.0345 | 463.8231 | 466.5766 |
| ENSMUSG00000037703 | -1.765374763 | 1.76E-07 | 8.99E-06 | 417.5126 | 379.1436 | 780.0764 | 407.1583 | 1688.893 | 743.0538 | 1859.15 | 2456.868 |
| ENSMUSG00000037705 | -0.548138339 | 0.333491 | 0.501623 | 10.06054 | 20.39697 | 7.45296 | 4.313117 | 14.62667 | 20.11273 | 11.57147 | 14.13868 |

| | | | | | | | | | | | |
|---|---|---|---|---|---|---|---|---|---|---|---|
| ENSMUSG00000037706 | 0.019424121 | 0.757227 | 0.848812 | 6850.224 | 7200.129 | 7728.719 | 7237.411 | 7046.154 | 6957.888 | 6968.918 | 7659.904 |
| ENSMUSG00000037709 | 0.004355626 | 0.977018 | 0.987393 | 607.6568 | 449.9331 | 472.8489 | 601.2486 | 448.5512 | 509.5226 | 554.4663 | 614.489 |
| ENSMUSG00000037710 | -1.32710203 | 1.52E-07 | 7.98E-06 | 512.0817 | 437.9349 | 713.8279 | 492.558 | 1340.778 | 777.6924 | 1424.255 | 1870.657 |
| ENSMUSG00000037712 | -0.196070002 | 0.067306 | 0.167502 | 3851.176 | 3612.663 | 4220.86 | 3968.931 | 4415.304 | 3781.194 | 4547.588 | 5191.072 |
| ENSMUSG00000037720 | -0.157032324 | 0.022077 | 0.075579 | 1723.371 | 1726.543 | 1599.902 | 1783.043 | 1766.902 | 1946.466 | 1916.043 | 1989.204 |
| ENSMUSG00000037722 | -0.187780451 | 0.060792 | 0.156439 | 1040.26 | 899.8662 | 845.4969 | 1002.369 | 1088.224 | 1168.773 | 1081.932 | 975.5692 |
| ENSMUSG00000037725 | 0.355919506 | 0.002721 | 0.016459 | 1352.137 | 1414.59 | 1221.457 | 1519.943 | 1168.183 | 1221.29 | 959.4677 | 954.905 |
| ENSMUSG00000037730 | -0.023340396 | 0.79338 | 0.873352 | 944.685 | 932.2613 | 787.5294 | 885.0517 | 961.4597 | 858.1433 | 911.2533 | 871.1605 |
| ENSMUSG00000037731 | 0.291362639 | 0.5238 | 0.677073 | 21.12714 | 67.19001 | 90.26362 | 30.19182 | 44.85512 | 42.46022 | 46.28588 | 36.9781 |
| ENSMUSG00000037735 | 0.893379206 | 0.019307 | 0.068749 | 34.20585 | 38.39429 | 28.15563 | 24.15346 | 14.62667 | 20.11273 | 22.17865 | 9.78832 |
| ENSMUSG00000037736 | 0.052098557 | 0.481235 | 0.640351 | 2092.593 | 2153.68 | 1831.772 | 1926.238 | 2003.854 | 1941.996 | 1885.185 | 1882.62 |
| ENSMUSG00000037738 | 0.619466003 | 0.087472 | 0.201069 | 41.24823 | 25.19625 | 39.74912 | 43.13117 | 21.45245 | 37.99072 | 21.21436 | 17.40146 |
| ENSMUSG00000037740 | -0.600817919 | 2.49E-08 | 1.81E-06 | 915.5095 | 957.4576 | 983.7907 | 956.6495 | 1499.721 | 1201.177 | 1427.148 | 1655.314 |
| ENSMUSG00000037742 | -0.17793403 | 0.047674 | 0.1316 | 269689 | 309771.1 | 306712.5 | 305151.3 | 373206.3 | 296306.4 | 349439.1 | 328746.2 |
| ENSMUSG00000037747 | -1.473430297 | 0.000177 | 0.002063 | 92.557 | 63.59054 | 220.2764 | 116.4542 | 252.5538 | 210.0663 | 427.1801 | 482.8904 |
| ENSMUSG00000037750 | 0.104167952 | 0.403545 | 0.570526 | 338.0343 | 331.1507 | 377.6166 | 294.1546 | 345.1894 | 278.2261 | 311.4654 | 312.1386 |
| ENSMUSG00000037752 | 0.039928734 | 0.790744 | 0.871818 | 139.8416 | 152.3773 | 140.7781 | 128.5309 | 155.0427 | 125.1459 | 139.8219 | 123.9854 |
| ENSMUSG00000037754 | -0.569640629 | 0.014266 | 0.055587 | 223.3441 | 227.9661 | 309.7119 | 289.8415 | 480.7299 | 234.6486 | 405.0014 | 440.4744 |
| ENSMUSG00000037759 | -1.533448564 | 0.009574 | 0.041244 | 7.04238 | 7.198929 | 9.109173 | 2.58787 | 13.65156 | 12.29112 | 16.39292 | 32.62773 |
| ENSMUSG00000037761 | -0.193605003 | 0.026905 | 0.08756 | 478.8819 | 449.9331 | 454.6305 | 417.5098 | 545.0872 | 497.2315 | 507.2161 | 507.905 |
| ENSMUSG00000037762 | -0.631524525 | 0.005357 | 0.026941 | 187.1261 | 250.7627 | 209.511 | 155.2722 | 305.2098 | 214.5358 | 312.4297 | 408.9342 |
| ENSMUSG00000037771 | 0.269809577 | 0.321386 | 0.489397 | 39.23612 | 44.3934 | 46.37397 | 38.81806 | 42.9049 | 37.99072 | 32.78583 | 26.10219 |
| ENSMUSG00000037772 | -0.199233665 | 0.001799 | 0.01216 | 1379.301 | 1377.395 | 1415.234 | 1469.91 | 1730.823 | 1550.915 | 1604.577 | 1592.233 |
| ENSMUSG00000037773 | 0.145656924 | 0.24098 | 0.403112 | 760.5771 | 895.0669 | 644.267 | 740.9936 | 771.313 | 673.7766 | 668.2524 | 630.8028 |
| ENSMUSG00000037784 | 0.09561715 | 0.211844 | 0.369406 | 1045.29 | 965.8563 | 924.9951 | 1043.774 | 924.4055 | 884.9603 | 937.2891 | 977.7444 |
| ENSMUSG00000037787 | -0.289724318 | 0.002407 | 0.014983 | 792.7708 | 859.0722 | 828.1066 | 879.876 | 1180.86 | 968.7633 | 954.6463 | 1001.671 |
| ENSMUSG00000037788 | 0.025952515 | 0.756331 | 0.848356 | 1314.913 | 1360.598 | 1522.888 | 1520.805 | 1517.273 | 1406.774 | 1345.183 | 1351.876 |
| ENSMUSG00000037791 | 0.114487562 | 0.107967 | 0.232567 | 1683.129 | 1550.169 | 1787.054 | 1787.356 | 1609.909 | 1596.728 | 1506.22 | 1582.445 |
| ENSMUSG00000037795 | 0.249058225 | 0.015009 | 0.057724 | 2002.048 | 2055.294 | 1772.976 | 1915.024 | 1575.78 | 1914.062 | 1540.934 | 1485.649 |
| ENSMUSG00000037798 | 3.476871436 | 0.002233 | 0.014238 | 419.5247 | 7.198929 | 19.04645 | 238.9467 | 8.776002 | 16.76061 | 25.07152 | 10.87591 |
| ENSMUSG00000037805 | -0.163786738 | 0.046836 | 0.130007 | 21393.75 | 24927.49 | 24897.85 | 25316.27 | 29496.14 | 25843.75 | 27396.42 | 25403.95 |
| ENSMUSG00000037808 | 0.266491301 | 0.071358 | 0.17483 | 1837.055 | 1776.936 | 1570.918 | 1901.222 | 1488.995 | 1895.066 | 1278.647 | 1228.978 |
| ENSMUSG00000037813 | -0.52063075 | 0.022241 | 0.075999 | 131.7931 | 86.38715 | 153.1997 | 105.2401 | 151.1423 | 150.8455 | 165.8577 | 218.6058 |
| ENSMUSG00000037815 | -0.231190055 | 0.038237 | 0.112948 | 9726.534 | 9395.802 | 11347.55 | 10154.8 | 12189.87 | 9751.324 | 12555.04 | 13191.39 |
| ENSMUSG00000037816 | 0.030229387 | 0.790758 | 0.871818 | 520.1301 | 455.9322 | 525.0196 | 417.5098 | 478.7797 | 508.4052 | 432.9658 | 460.051 |
| ENSMUSG00000037818 | 0.198401825 | 0.311223 | 0.479901 | 398.3975 | 287.9572 | 299.7746 | 343.3242 | 208.6738 | 372.0856 | 283.501 | 295.8248 |
| ENSMUSG00000037820 | 0.144174871 | 0.26217 | 0.426384 | 1328.998 | 1342.6 | 1482.311 | 1316.363 | 1077.498 | 1243.637 | 1092.54 | 1538.941 |
| ENSMUSG00000037822 | 0.349397242 | 0.001346 | 0.009765 | 2678.117 | 2661.204 | 2485.148 | 2919.118 | 2094.539 | 2482.805 | 2043.329 | 1813.014 |
| ENSMUSG00000037824 | -0.236408698 | 0.021101 | 0.073203 | 2697.232 | 2573.617 | 2918.248 | 2663.781 | 3143.759 | 2674.994 | 3422.262 | 3545.547 |
| ENSMUSG00000037826 | 0.237105009 | 0.166186 | 0.311827 | 504.0332 | 382.7431 | 389.2101 | 481.3439 | 366.6419 | 408.9589 | 435.8587 | 279.5109 |
| ENSMUSG00000037827 | -1.881009124 | 0.004651 | 0.02445 | 5.030272 | 0 | 5.796746 | 6.900988 | 14.62667 | 10.05637 | 23.14294 | 19.57664 |
| ENSMUSG00000037833 | 0.335781666 | 0.223379 | 0.383102 | 56.33904 | 80.38804 | 57.13936 | 71.59775 | 39.00445 | 67.04245 | 41.46443 | 63.08028 |
| ENSMUSG00000037843 | 0.662043778 | 0.397398 | 0.564661 | 48.29061 | 4.799286 | 13.24971 | 104.3774 | 40.95467 | 13.40849 | 22.17865 | 31.54014 |
| ENSMUSG00000037845 | -0.254249853 | 0.048205 | 0.132708 | 193.1624 | 206.3693 | 207.8548 | 235.4962 | 277.9067 | 236.8833 | 262.2867 | 228.3941 |
| ENSMUSG00000037846 | 0.338223597 | 0.014308 | 0.055672 | 212.2775 | 241.1641 | 193.777 | 208.7549 | 169.6694 | 188.8362 | 167.7863 | 149 |
| ENSMUSG00000037847 | -0.249322678 | 0.110653 | 0.23675 | 380.2885 | 299.9554 | 370.1637 | 335.5605 | 379.3183 | 330.7427 | 440.6801 | 498.1167 |
| ENSMUSG00000037851 | -0.265079674 | 0.006275 | 0.030414 | 2162.011 | 2206.472 | 2172.124 | 2114.29 | 2696.183 | 2149.828 | 2713.51 | 2838.613 |
| ENSMUSG00000037852 | 0.812893118 | 6.81E-08 | 4.12E-06 | 6931.715 | 6675.807 | 5126.808 | 7502.237 | 3853.64 | 4526.482 | 3402.976 | 3151.839 |
| ENSMUSG00000037855 | 0.253329152 | 0.110323 | 0.23618 | 150.9082 | 176.3738 | 175.5586 | 147.5086 | 151.1423 | 147.4934 | 133.0719 | 113.1095 |
| ENSMUSG00000037857 | 0.040926771 | 0.5871 | 0.727362 | 1040.26 | 1173.425 | 1088.132 | 1107.609 | 1106.751 | 1115.139 | 987.4321 | 1075.628 |
| ENSMUSG00000037860 | 0.412140761 | 0.011341 | 0.046878 | 152.9203 | 137.9795 | 161.4808 | 125.0804 | 111.1627 | 106.1505 | 119.5719 | 96.79561 |
| ENSMUSG00000037868 | 0.154116617 | 0.625652 | 0.755894 | 35.2119 | 26.39607 | 28.98373 | 38.81806 | 36.07912 | 21.23011 | 32.78583 | 26.10219 |
| ENSMUSG00000037872 | 0.952836157 | 0.05719 | 0.149716 | 35.2119 | 49.19268 | 17.39024 | 62.10889 | 13.65156 | 35.75597 | 21.21436 | 14.13868 |
| ENSMUSG00000037876 | 0.111320731 | 0.102791 | 0.22523 | 2249.538 | 2080.491 | 1933.629 | 2167.773 | 1899.517 | 1958.757 | 2012.471 | 1932.649 |
| ENSMUSG00000037885 | -0.107608688 | 0.137431 | 0.272529 | 740.456 | 757.0874 | 790.0137 | 755.6582 | 874.6748 | 779.9271 | 802.2886 | 822.2189 |
| ENSMUSG00000037887 | 0.264522241 | 0.308523 | 0.476852 | 190.1443 | 87.58697 | 118.4192 | 117.3168 | 103.3618 | 126.2633 | 107.0361 | 91.35765 |
| ENSMUSG00000037890 | 0.156044652 | 0.19707 | 0.351414 | 898.4065 | 921.4629 | 706.375 | 857.4478 | 813.2428 | 770.9881 | 799.3957 | 649.2919 |
| ENSMUSG00000037892 | 0.154391656 | 0.145567 | 0.283934 | 1098.611 | 1371.396 | 1262.035 | 1090.356 | 1154.532 | 1097.261 | 972.9678 | 1106.08 |
| ENSMUSG00000037894 | -0.353390063 | 0.000157 | 0.001888 | 13672.28 | 14689.42 | 14952.29 | 15302.08 | 20647.01 | 15856.66 | 18778.57 | 19602.74 |
| ENSMUSG00000037896 | 0.053481089 | 0.587012 | 0.727302 | 1567.433 | 1418.189 | 1245.472 | 1437.993 | 1276.421 | 1314.032 | 1353.862 | 1517.19 |
| ENSMUSG00000037902 | 0.215490524 | 0.010517 | 0.044254 | 1332.016 | 1353.399 | 1549.388 | 1267.194 | 1131.129 | 1200.06 | 1201.504 | 1208.314 |
| ENSMUSG00000037904 | -0.993288687 | 1.49E-05 | 0.00031 | 108.6539 | 86.38715 | 110.9663 | 81.94923 | 143.3414 | 162.0192 | 246.858 | 220.781 |
| ENSMUSG00000037905 | -0.122482427 | 0.079183 | 0.187744 | 1555.36 | 1662.953 | 1743.164 | 1682.116 | 1882.94 | 1695.057 | 1764.649 | 1891.321 |
| ENSMUSG00000037907 | 0.0134788 | 0.883271 | 0.930247 | 758.565 | 859.0722 | 758.5457 | 843.6458 | 854.1975 | 841.3827 | 735.7526 | 756.9634 |

| | | | | | | | | | | | |
|---|---|---|---|---|---|---|---|---|---|---|---|
| ENSMUSG00000037916 | -0.592191936 | 9.13E-06 | 0.000213 | 3015.145 | 3168.729 | 3397.722 | 3133.911 | 4850.204 | 3633.701 | 4968.982 | 5716.379 |
| ENSMUSG00000037921 | 0.713680663 | 0.085499 | 0.197791 | 69.41775 | 123.5816 | 173.0743 | 40.5433 | 59.48179 | 60.3382 | 70.39311 | 57.64233 |
| ENSMUSG00000037926 | 0.289621416 | 0.10137 | 0.223203 | 985.9333 | 849.4736 | 735.3587 | 780.6743 | 590.9175 | 917.3641 | 678.8596 | 554.6715 |
| ENSMUSG00000037933 | -0.120561059 | 0.254067 | 0.417641 | 1135.835 | 1052.243 | 1168.458 | 1150.74 | 1221.814 | 1019.045 | 1291.183 | 1369.277 |
| ENSMUSG00000037935 | -0.056582268 | 0.410411 | 0.576669 | 6000.108 | 7057.35 | 6661.29 | 6574.054 | 7139.765 | 6973.532 | 6520.523 | 6708.262 |
| ENSMUSG00000037936 | 1.042409462 | 0.100013 | 0.220847 | 6868.333 | 1013.849 | 1418.547 | 1698.506 | 1289.097 | 1143.074 | 1319.148 | 1588.971 |
| ENSMUSG00000037938 | 0.146283996 | 0.123438 | 0.253103 | 425.561 | 400.7404 | 430.6154 | 438.2127 | 423.1983 | 365.3813 | 375.1085 | 368.6934 |
| ENSMUSG00000037940 | 0.097878835 | 0.449237 | 0.611846 | 324.9556 | 302.355 | 308.8838 | 248.4356 | 247.6783 | 278.2261 | 283.501 | 296.9124 |
| ENSMUSG00000037946 | -0.846824392 | 0.096027 | 0.214632 | 57.3451 | 37.19447 | 175.5586 | 43.9938 | 175.52 | 68.15982 | 145.6077 | 177.2773 |
| ENSMUSG00000037949 | -0.077337576 | 0.48202 | 0.640751 | 1001.024 | 901.066 | 950.6664 | 932.496 | 1074.573 | 810.0962 | 1034.682 | 1073.452 |
| ENSMUSG00000037953 | -3.116941946 | 0.000795 | 0.006545 | 2.012109 | 0 | 14.90592 | 5.175741 | 24.37778 | 10.05637 | 61.71451 | 100.0584 |
| ENSMUSG00000037957 | -0.108741488 | 0.19093 | 0.343551 | 853.1341 | 752.2881 | 807.404 | 855.7225 | 814.2179 | 889.4298 | 882.3246 | 942.9415 |
| ENSMUSG00000037958 | 0.135983742 | 0.277855 | 0.444068 | 1760.595 | 1806.931 | 1650.417 | 1587.227 | 1578.705 | 1895.066 | 1414.612 | 1304.022 |
| ENSMUSG00000037960 | -0.14848625 | 0.254078 | 0.417641 | 372.2401 | 373.1445 | 441.3808 | 454.6026 | 464.153 | 384.3767 | 459.0016 | 514.4306 |
| ENSMUSG00000037962 | -0.371379999 | 0.08303 | 0.193881 | 108.6539 | 117.5825 | 149.0592 | 87.9876 | 185.2711 | 131.8501 | 141.7505 | 140.2993 |
| ENSMUSG00000037965 | 0.329141672 | 0.004586 | 0.02415 | 2772.686 | 2450.036 | 2206.904 | 2530.075 | 1887.815 | 2372.185 | 1809.006 | 1859.781 |
| ENSMUSG00000037966 | -0.806198065 | 2.32E-07 | 1.14E-05 | 634.8203 | 647.9036 | 805.7478 | 640.9293 | 1230.59 | 883.8429 | 1288.29 | 1371.452 |
| ENSMUSG00000037971 | 0.004436448 | 0.953048 | 0.973191 | 760.5771 | 704.2952 | 712.9998 | 721.1532 | 770.3379 | 714.002 | 683.681 | 722.1605 |
| ENSMUSG00000037972 | 0.041748792 | 0.598109 | 0.736699 | 1854.158 | 1962.908 | 2246.653 | 2160.872 | 2003.854 | 1973.283 | 1967.15 | 2051.197 |
| ENSMUSG00000037977 | -1.120996445 | 0.00022 | 0.002437 | 26.15741 | 34.79482 | 39.74912 | 33.64232 | 86.78491 | 45.81234 | 67.50024 | 92.44524 |
| ENSMUSG00000037979 | -0.235616298 | 0.358273 | 0.526574 | 115.6963 | 85.18733 | 81.98256 | 122.4925 | 88.73513 | 94.9768 | 151.3934 | 142.4744 |
| ENSMUSG00000037989 | 0.358750186 | 0.057877 | 0.15083 | 246.4833 | 229.1659 | 272.4471 | 238.0841 | 144.3165 | 258.1134 | 179.3578 | 189.2408 |
| ENSMUSG00000037990 | 0.574854056 | 0.011174 | 0.04641 | 171.0292 | 167.975 | 158.1684 | 162.1732 | 94.5858 | 162.0192 | 106.0718 | 80.48174 |
| ENSMUSG00000037991 | -0.146057896 | 0.498506 | 0.654881 | 103.6236 | 157.1766 | 117.5911 | 104.3774 | 129.6898 | 170.9582 | 114.7504 | 117.4598 |
| ENSMUSG00000037992 | -0.0585143 | 0.539439 | 0.689165 | 950.7214 | 994.6521 | 1078.195 | 918.694 | 1149.656 | 1035.806 | 957.5391 | 962.5181 |
| ENSMUSG00000037993 | -0.153289928 | 0.015713 | 0.059687 | 1704.256 | 1642.556 | 1736.54 | 1671.764 | 1898.542 | 1730.812 | 1941.114 | 1942.438 |
| ENSMUSG00000037994 | 0.759318661 | 0.004736 | 0.024759 | 207.2472 | 229.1659 | 112.6225 | 241.5346 | 119.9387 | 115.0895 | 141.7505 | 89.18247 |
| ENSMUSG00000037995 | 0.170938475 | 0.164402 | 0.309391 | 891.3642 | 1071.441 | 757.7176 | 867.7992 | 873.6997 | 773.2229 | 737.6812 | 798.2919 |
| ENSMUSG00000037996 | 0.360529607 | 0.435782 | 0.599895 | 26.15741 | 10.79839 | 24.01509 | 31.91707 | 14.62667 | 29.05173 | 15.42863 | 14.13868 |
| ENSMUSG00000037997 | 0.437531223 | 1.79E-05 | 0.000361 | 692.1654 | 703.0954 | 592.9243 | 690.9614 | 488.5308 | 551.9828 | 473.466 | 463.3138 |
| ENSMUSG00000037999 | -0.23908198 | 0.13313 | 0.266493 | 196.1806 | 170.3747 | 197.0894 | 179.4257 | 177.4703 | 201.1273 | 243.0009 | 256.6715 |
| ENSMUSG00000038000 | -0.153383136 | 0.098773 | 0.219219 | 718.3228 | 719.8929 | 703.0625 | 679.7473 | 794.7157 | 696.1241 | 879.4317 | 763.4889 |
| ENSMUSG00000038002 | 0.272535801 | 0.093288 | 0.210094 | 1390.367 | 1307.805 | 1036.79 | 1369.846 | 972.186 | 1373.253 | 1010.575 | 870.0729 |
| ENSMUSG00000038005 | 0.078409664 | 0.335988 | 0.504269 | 1400.428 | 1557.368 | 1410.266 | 1448.345 | 1355.405 | 1534.155 | 1289.255 | 1329.036 |
| ENSMUSG00000038007 | -0.50893678 | 0.122782 | 0.252347 | 1222.356 | 604.7101 | 1860.756 | 992.017 | 2110.141 | 901.7209 | 1815.756 | 1833.679 |
| ENSMUSG00000038009 | -0.232156549 | 0.425385 | 0.590383 | 212.2775 | 125.9813 | 207.0267 | 213.068 | 233.0516 | 151.9629 | 165.8577 | 341.5036 |
| ENSMUSG00000038010 | 0.367079017 | 0.018459 | 0.066696 | 163.9869 | 178.7734 | 144.9187 | 162.1732 | 113.1129 | 135.2023 | 112.8218 | 142.4744 |
| ENSMUSG00000038011 | -0.448432839 | 0.245428 | 0.408405 | 30.18163 | 14.39786 | 15.73403 | 18.97772 | 21.45245 | 34.6386 | 23.14294 | 29.36496 |
| ENSMUSG00000038013 | 0.200914938 | 0.202337 | 0.3579 | 813.898 | 788.2827 | 721.2809 | 722.8785 | 558.7388 | 887.195 | 612.3236 | 592.7371 |
| ENSMUSG00000038014 | 0.080167373 | 0.23758 | 0.398725 | 6128.883 | 6380.651 | 5855.542 | 6534.373 | 5486.951 | 6328.807 | 5806.949 | 5931.722 |
| ENSMUSG00000038020 | 0.042688026 | 0.756026 | 0.848334 | 183.1019 | 185.9723 | 187.1521 | 191.5024 | 205.7485 | 166.4887 | 194.7864 | 157.7007 |
| ENSMUSG00000038022 | 0.050439974 | 0.660943 | 0.781719 | 365.1977 | 400.7404 | 358.5702 | 323.4838 | 371.5174 | 358.6771 | 357.7513 | 306.7007 |
| ENSMUSG00000038023 | -0.219120762 | 0.017786 | 0.065068 | 1186.138 | 1223.818 | 1222.285 | 1136.938 | 1383.683 | 1289.45 | 1280.576 | 1597.671 |
| ENSMUSG00000038024 | 0.16162122 | 0.082375 | 0.192898 | 999.012 | 932.2613 | 794.1543 | 891.0901 | 770.3379 | 839.1479 | 801.3243 | 820.0437 |
| ENSMUSG00000038025 | 0.015951732 | 0.834035 | 0.899541 | 1135.835 | 1072.64 | 1236.363 | 1145.564 | 1139.905 | 1057.036 | 1159.076 | 1187.649 |
| ENSMUSG00000038026 | 0.003493786 | 0.992036 | 0.995436 | 23.13925 | 19.19714 | 26.49941 | 18.97772 | 25.35289 | 25.6996 | 16.39292 | 20.66423 |
| ENSMUSG00000038028 | -0.486029822 | 0.003447 | 0.019576 | 132.7992 | 141.5789 | 148.2311 | 146.646 | 216.4747 | 150.8455 | 201.5364 | 228.3941 |
| ENSMUSG00000038034 | 0.124662348 | 0.094644 | 0.212448 | 1094.587 | 949.0588 | 1061.633 | 1061.027 | 977.0615 | 955.3549 | 982.6107 | 909.2261 |
| ENSMUSG00000038037 | 0.251027896 | 0.233243 | 0.393874 | 108.6539 | 105.5843 | 110.9663 | 96.61383 | 91.66046 | 105.0332 | 60.75022 | 97.8832 |
| ENSMUSG00000038039 | 0.110381988 | 0.155605 | 0.29757 | 1193.18 | 1100.236 | 1112.147 | 1060.164 | 1077.498 | 1098.379 | 990.325 | 970.1312 |
| ENSMUSG00000038042 | 0.110021192 | 0.15883 | 0.301808 | 708.2623 | 808.6797 | 771.7954 | 792.751 | 711.8313 | 731.88 | 677.8953 | 734.124 |
| ENSMUSG00000038045 | -0.274948965 | 0.537107 | 0.687629 | 14.08476 | 13.19804 | 18.21835 | 9.488858 | 10.72622 | 22.34748 | 15.42863 | 18.48905 |
| ENSMUSG00000038046 | -0.415376186 | 0.006663 | 0.031853 | 257.5499 | 272.3595 | 227.7293 | 272.589 | 352.0152 | 261.4655 | 373.1799 | 385.0072 |
| ENSMUSG00000038047 | 0.154243207 | 0.234635 | 0.395501 | 962.794 | 1019.848 | 764.3424 | 867.7992 | 782.0393 | 959.8243 | 737.6812 | 766.7517 |
| ENSMUSG00000038055 | -0.015664881 | 0.877159 | 0.926493 | 402.4217 | 488.3274 | 467.0521 | 408.8835 | 446.601 | 438.0106 | 464.7874 | 433.9488 |
| ENSMUSG00000038056 | 0.362283788 | 0.06058 | 0.156133 | 2237.465 | 2042.096 | 1850.818 | 2021.127 | 1475.343 | 2312.964 | 1343.255 | 1210.489 |
| ENSMUSG00000038057 | -1.731359546 | 0.000891 | 0.007125 | 5.030272 | 5.999108 | 5.796746 | 4.313117 | 12.67645 | 23.46486 | 14.46434 | 19.57664 |
| ENSMUSG00000038058 | 0.034466932 | 0.863135 | 0.917928 | 150.9082 | 197.9706 | 267.4784 | 214.7932 | 178.4454 | 213.4185 | 187.0721 | 234.9197 |
| ENSMUSG00000038059 | 0.265992594 | 0.026006 | 0.085434 | 897.4005 | 779.884 | 743.6398 | 820.3549 | 790.8153 | 638.0206 | 681.7524 | 582.9488 |
| ENSMUSG00000038060 | -1.018756787 | 0.000293 | 0.003047 | 185.114 | 116.3827 | 208.6829 | 137.1571 | 253.5289 | 219.0053 | 387.6442 | 453.5255 |
| ENSMUSG00000038065 | -0.197431444 | 0.264879 | 0.429083 | 1289.762 | 976.6547 | 1026.024 | 1320.677 | 1242.292 | 1146.426 | 1776.221 | 1124.569 |
| ENSMUSG00000038068 | -0.087663061 | 0.714578 | 0.821054 | 199.1988 | 134.38 | 269.9628 | 165.6237 | 254.5041 | 174.3104 | 207.3222 | 182.7153 |
| ENSMUSG00000038069 | -0.016157039 | 0.889925 | 0.933475 | 652.9293 | 584.3131 | 597.893 | 727.1916 | 677.7024 | 728.5279 | 606.5379 | 580.7736 |
| ENSMUSG00000038070 | 0.239822148 | 0.005008 | 0.025762 | 1195.193 | 1273.011 | 1223.113 | 1236.139 | 1004.365 | 1192.238 | 984.5392 | 992.9707 |

| | | | | | | | | | | | |
|---|---|---|---|---|---|---|---|---|---|---|---|
| ENSMUSG00000038072 | -1.433434142 | 1.00E-07 | 5.66E-06 | 1369.24 | 1064.242 | 1653.729 | 1513.042 | 3958.952 | 1911.827 | 3780.978 | 5477.109 |
| ENSMUSG00000038074 | 0.155011321 | 0.103247 | 0.22588 | 707.2562 | 800.281 | 631.0173 | 698.725 | 616.2703 | 651.4291 | 623.8951 | 652.5547 |
| ENSMUSG00000038077 | 0.637811739 | 0.005163 | 0.026322 | 134.8113 | 95.98572 | 134.9814 | 161.3106 | 70.20801 | 90.5073 | 105.1075 | 73.95619 |
| ENSMUSG00000038079 | 0.21988992 | 0.031795 | 0.098473 | 862.1886 | 949.0588 | 862.8871 | 953.199 | 847.3717 | 743.0538 | 845.6816 | 675.3941 |
| ENSMUSG00000038080 | -0.398554556 | 0.000665 | 0.005713 | 434.6155 | 499.1258 | 505.145 | 503.7721 | 555.8134 | 730.7627 | 602.6807 | 674.3065 |
| ENSMUSG00000038084 | -0.133596135 | 0.036633 | 0.109822 | 1977.903 | 1931.713 | 1931.145 | 1883.107 | 1976.551 | 2080.551 | 2144.579 | 2271.978 |
| ENSMUSG00000038085 | -0.30966518 | 0.050469 | 0.136938 | 238.4349 | 271.1597 | 356.0859 | 318.3081 | 356.8907 | 317.3342 | 391.5014 | 404.5839 |
| ENSMUSG00000038086 | -0.086731555 | 0.800984 | 0.878562 | 30.18163 | 46.79304 | 27.32752 | 43.13117 | 28.27823 | 34.6386 | 35.6787 | 57.64233 |
| ENSMUSG00000038094 | -2.33412082 | 9.66E-06 | 0.000221 | 13.07871 | 4.799286 | 18.21835 | 6.900988 | 41.92979 | 30.1691 | 53.0359 | 94.62042 |
| ENSMUSG00000038095 | 0.172818193 | 0.140234 | 0.276573 | 3562.438 | 3322.306 | 3211.398 | 3155.477 | 2762.49 | 3631.466 | 2768.474 | 2593.905 |
| ENSMUSG00000038102 | -0.090481817 | 0.196587 | 0.350992 | 1565.421 | 1466.182 | 1476.514 | 1519.08 | 1663.54 | 1464.877 | 1595.899 | 1692.292 |
| ENSMUSG00000038112 | 0.869407616 | 0.006624 | 0.031706 | 2382.337 | 2622.81 | 1514.607 | 2566.305 | 1234.491 | 2130.832 | 980.6821 | 627.5401 |
| ENSMUSG00000038115 | -0.188025512 | 0.648615 | 0.772344 | 31.18769 | 26.39607 | 27.32752 | 24.15346 | 29.25334 | 54.75133 | 23.14294 | 17.40146 |
| ENSMUSG00000038116 | 0.104713875 | 0.241841 | 0.404258 | 1547.312 | 1494.978 | 1445.874 | 1500.102 | 1376.857 | 1590.023 | 1361.576 | 1240.941 |
| ENSMUSG00000038119 | 0.521175915 | 0.023143 | 0.078386 | 1308.877 | 1644.955 | 1493.904 | 1615.694 | 850.2971 | 1662.653 | 944.0391 | 768.9269 |
| ENSMUSG00000038121 | -0.326116922 | 0.000181 | 0.0021 | 1285.737 | 1120.633 | 1209.036 | 1205.085 | 1440.239 | 1393.365 | 1527.434 | 1684.679 |
| ENSMUSG00000038122 | -0.064664445 | 0.569832 | 0.713532 | 424.5549 | 424.7368 | 350.2891 | 347.6373 | 378.3432 | 423.4848 | 405.9657 | 406.7591 |
| ENSMUSG00000038126 | 0.156267586 | 0.388264 | 0.555834 | 892.3702 | 718.6931 | 696.4377 | 739.2683 | 617.2455 | 936.3595 | 647.038 | 534.0072 |
| ENSMUSG00000038127 | 0.095458592 | 0.262131 | 0.426384 | 3670.086 | 4145.383 | 3849.04 | 3748.099 | 3459.695 | 4107.467 | 3455.048 | 3403.073 |
| ENSMUSG00000038128 | 0.46419929 | 0.040124 | 0.116969 | 114.6902 | 151.1775 | 91.09173 | 118.1794 | 77.03379 | 103.9158 | 92.57176 | 69.60583 |
| ENSMUSG00000038132 | 1.202365325 | 0.002483 | 0.015349 | 257.5499 | 261.5611 | 214.4796 | 246.7103 | 101.4116 | 205.5968 | 79.07171 | 40.24087 |
| ENSMUSG00000038135 | 0.176191295 | 0.543315 | 0.692406 | 43.26034 | 46.79304 | 33.12427 | 30.19182 | 34.1289 | 35.75597 | 27.0001 | 38.06569 |
| ENSMUSG00000038141 | 0.048401047 | 0.609148 | 0.744335 | 640.8566 | 613.1088 | 683.188 | 559.8426 | 626.0215 | 602.2646 | 563.1449 | 624.2773 |
| ENSMUSG00000038143 | 0.411109602 | 0.001528 | 0.010693 | 1478.9 | 1652.154 | 1498.045 | 1487.163 | 993.6384 | 1450.352 | 1117.611 | 1038.649 |
| ENSMUSG00000038145 | -0.099690611 | 0.342466 | 0.511165 | 2130.823 | 1754.139 | 1767.18 | 1947.804 | 1890.741 | 1987.809 | 2346.116 | 1918.511 |
| ENSMUSG00000038146 | 0.123912059 | 0.23402 | 0.394761 | 1083.521 | 989.8528 | 1242.988 | 1067.065 | 902.9531 | 1014.576 | 996.1107 | 1113.693 |
| ENSMUSG00000038147 | 0.622912573 | 0.024416 | 0.081559 | 50.30272 | 58.79126 | 57.13936 | 40.5433 | 26.32801 | 32.40385 | 42.42872 | 32.62773 |
| ENSMUSG00000038148 | -2.457870068 | 2.55E-05 | 0.000474 | 19.11503 | 17.99732 | 99.3728 | 30.19182 | 151.1423 | 97.21155 | 286.3939 | 386.0948 |
| ENSMUSG00000038150 | 0.163095737 | 0.002181 | 0.013977 | 1580.511 | 1625.758 | 1623.917 | 1542.371 | 1420.737 | 1449.234 | 1398.219 | 1422.569 |
| ENSMUSG00000038151 | -0.233926003 | 0.405848 | 0.572524 | 36.21796 | 45.59322 | 71.21717 | 52.62003 | 69.2329 | 68.15982 | 56.89306 | 48.9416 |
| ENSMUSG00000038152 | -0.409700845 | 0.105439 | 0.229127 | 62.37537 | 41.99375 | 57.96746 | 51.75741 | 92.63557 | 65.92507 | 54.96448 | 71.78101 |
| ENSMUSG00000038155 | -0.080912159 | 0.88354 | 0.930352 | 7.04238 | 20.39697 | 9.93728 | 7.763611 | 16.57689 | 8.938993 | 12.53576 | 8.700729 |
| ENSMUSG00000038156 | 0.344157689 | 0.067814 | 0.168314 | 838.0433 | 485.9277 | 840.5282 | 843.6458 | 521.6845 | 556.4523 | 656.6809 | 637.3284 |
| ENSMUSG00000038160 | -0.172570832 | 0.039386 | 0.115345 | 763.5953 | 831.4763 | 823.9661 | 907.4799 | 898.0775 | 917.3641 | 931.5033 | 1004.934 |
| ENSMUSG00000038167 | -0.924780933 | 0.000575 | 0.005118 | 80.48435 | 79.18822 | 139.95 | 76.77349 | 232.0765 | 140.7891 | 152.3577 | 191.416 |
| ENSMUSG00000038168 | -0.983508641 | 0.001073 | 0.008244 | 449.7063 | 278.3586 | 599.5492 | 469.2672 | 775.2135 | 492.762 | 989.3607 | 1298.584 |
| ENSMUSG00000038170 | -0.082391244 | 0.468234 | 0.628535 | 1214.308 | 1115.834 | 1024.368 | 1130.899 | 1042.394 | 1067.092 | 1339.398 | 1298.584 |
| ENSMUSG00000038172 | 0.20848115 | 0.095461 | 0.213737 | 462.785 | 364.7457 | 376.7885 | 440.8006 | 356.8907 | 382.1419 | 372.2156 | 313.2262 |
| ENSMUSG00000038173 | -0.809785705 | 0.007487 | 0.034522 | 35.2119 | 31.19536 | 28.15563 | 38.81806 | 55.58134 | 43.57759 | 50.14304 | 84.8321 |
| ENSMUSG00000038174 | 0.234561192 | 0.153941 | 0.295429 | 603.6326 | 641.9045 | 520.051 | 579.683 | 417.3476 | 665.955 | 451.2873 | 458.9634 |
| ENSMUSG00000038175 | 0.500365293 | 0.011512 | 0.047468 | 1953.758 | 1803.332 | 1717.493 | 1906.398 | 1219.864 | 1822.437 | 1321.076 | 854.8466 |
| ENSMUSG00000038178 | -1.437725 | 1.42E-06 | 4.95E-05 | 277.671 | 231.5656 | 370.1637 | 256.1992 | 795.6908 | 372.0856 | 767.5742 | 1143.058 |
| ENSMUSG00000038180 | 0.211489472 | 0.393742 | 0.561064 | 97.58727 | 73.18911 | 62.9361 | 61.24627 | 74.10846 | 59.22083 | 51.10733 | 69.60583 |
| ENSMUSG00000038181 | -0.216992547 | 0.15705 | 0.299495 | 846.0917 | 861.4719 | 902.6362 | 847.9589 | 1101.876 | 705.0631 | 1058.79 | 1152.847 |
| ENSMUSG00000038187 | -0.008060745 | 0.908648 | 0.945375 | 586.5297 | 589.1124 | 608.6584 | 606.4243 | 599.6935 | 606.7341 | 593.0378 | 606.8758 |
| ENSMUSG00000038188 | 0.060410407 | 0.560472 | 0.706328 | 420.5307 | 386.3425 | 486.0986 | 450.2895 | 435.8748 | 406.7242 | 405.9657 | 427.4233 |
| ENSMUSG00000038193 | 2.377581594 | 0.056844 | 0.148915 | 204.229 | 26.39607 | 8.281066 | 263.9628 | 7.80089 | 75.98144 | 7.714313 | 5.437955 |
| ENSMUSG00000038195 | -0.833402933 | 0.001582 | 0.010966 | 46.2785 | 37.19447 | 52.99882 | 43.9938 | 73.13335 | 59.22083 | 81.96458 | 108.7591 |
| ENSMUSG00000038199 | -0.024494677 | 0.959415 | 0.977097 | 10.06054 | 14.39786 | 7.45296 | 17.25247 | 10.72622 | 13.40849 | 10.60718 | 15.22628 |
| ENSMUSG00000038201 | 2.282916102 | 0.000338 | 0.00339 | 13.07871 | 21.59679 | 9.93728 | 31.05445 | 2.925334 | 4.469496 | 3.857157 | 4.350364 |
| ENSMUSG00000038203 | 0.786654814 | 0.435538 | 0.599694 | 9.054489 | 9.598572 | 6.624853 | 34.50494 | 10.72622 | 21.23011 | 2.892867 | 0 |
| ENSMUSG00000038204 | -0.340493528 | 0.471702 | 0.631172 | 9.054489 | 15.59768 | 27.32752 | 23.29083 | 37.05423 | 16.76061 | 19.28578 | 22.83941 |
| ENSMUSG00000038205 | 0.195010266 | 0.097504 | 0.2171 | 568.4207 | 533.9206 | 549.8628 | 615.0506 | 474.8792 | 587.7388 | 434.8944 | 486.1532 |
| ENSMUSG00000038206 | -0.253209447 | 0.005789 | 0.028665 | 636.8324 | 583.1133 | 621.9081 | 635.7535 | 690.3788 | 708.4152 | 722.2526 | 835.27 |
| ENSMUSG00000038208 | -0.43589872 | 0.000625 | 0.005449 | 357.1493 | 386.3425 | 472.0208 | 411.4714 | 531.4357 | 482.7056 | 578.5735 | 611.2262 |
| ENSMUSG00000038210 | 0.107002682 | 0.110208 | 0.236096 | 1409.482 | 1512.975 | 1466.577 | 1529.431 | 1339.803 | 1384.427 | 1465.72 | 1304.022 |
| ENSMUSG00000038212 | -0.006519184 | 0.932201 | 0.959977 | 1465.821 | 1522.574 | 1397.016 | 1482.85 | 1318.35 | 1512.925 | 1504.291 | 1558.518 |
| ENSMUSG00000038213 | 0.013893082 | 0.943004 | 0.967108 | 210.2654 | 268.76 | 279.0719 | 182.8762 | 228.176 | 192.1883 | 222.7508 | 288.2116 |
| ENSMUSG00000038214 | -0.142423539 | 0.097794 | 0.217587 | 536.227 | 563.9161 | 556.4877 | 516.7115 | 561.6641 | 601.1473 | 582.4307 | 653.6422 |
| ENSMUSG00000038215 | 0.186058547 | 0.10506 | 0.228493 | 863.1946 | 815.8786 | 756.8895 | 905.7547 | 733.2837 | 855.9086 | 712.6097 | 636.2408 |
| ENSMUSG00000038216 | 0.170192516 | 0.891549 | 0.934507 | 513.0877 | 9.598572 | 14.07781 | 60.38364 | 216.4747 | 25.6996 | 160.072 | 128.3357 |
| ENSMUSG00000038217 | 0.083586708 | 0.61447 | 0.748118 | 152.9203 | 215.9679 | 204.5423 | 194.0903 | 172.5947 | 202.2447 | 153.322 | 196.854 |
| ENSMUSG00000038224 | -3.216198602 | 0.000114 | 0.001488 | 111.672 | 9.598572 | 168.9338 | 108.6906 | 987.7878 | 278.2261 | 770.467 | 1674.89 |
| ENSMUSG00000038225 | 0.343155925 | 0.027374 | 0.088571 | 450.7124 | 496.7261 | 365.195 | 373.516 | 332.513 | 395.5504 | 316.2868 | 282.7737 |

| | | | | | | | | | | | |
|---|---|---|---|---|---|---|---|---|---|---|---|
| ENSMUSG00000038227 | 0.377281103 | 0.002568 | 0.015735 | 4345.149 | 4285.763 | 3746.354 | 4931.619 | 3486.023 | 3888.462 | 3126.225 | 2825.562 |
| ENSMUSG00000038233 | 0.655667281 | 0.166329 | 0.311956 | 31.18769 | 86.38715 | 33.95237 | 61.24627 | 36.07912 | 56.98608 | 22.17865 | 19.57664 |
| ENSMUSG00000038235 | 0.027523396 | 0.636322 | 0.763786 | 3899.467 | 4049.398 | 4334.31 | 4174.235 | 4172.501 | 3877.288 | 3955.514 | 4144.81 |
| ENSMUSG00000038236 | 0.072430835 | 0.316341 | 0.484663 | 903.4368 | 897.4665 | 868.6839 | 931.6334 | 796.6659 | 901.7209 | 848.5745 | 879.8612 |
| ENSMUSG00000038239 | -0.3214956 | 0.128221 | 0.259706 | 90.54489 | 94.7859 | 100.2009 | 68.14726 | 114.088 | 88.27255 | 105.1075 | 133.7737 |
| ENSMUSG00000038240 | -0.253205542 | 0.013925 | 0.054528 | 252.5196 | 237.5647 | 247.6039 | 266.5507 | 284.7325 | 297.2215 | 314.3583 | 302.3503 |
| ENSMUSG00000038241 | 0.204662514 | 0.000371 | 0.00364 | 1485.942 | 1475.78 | 1455.811 | 1402.626 | 1217.914 | 1302.858 | 1254.54 | 1274.657 |
| ENSMUSG00000038242 | 0.988417521 | 0.023942 | 0.080488 | 22.1332 | 23.99643 | 28.15563 | 25.01608 | 10.72622 | 22.34748 | 7.714313 | 9.78832 |
| ENSMUSG00000038244 | -0.222592264 | 0.209013 | 0.366165 | 470.8334 | 333.5504 | 332.8989 | 495.1459 | 505.1077 | 392.1983 | 475.3946 | 532.9196 |
| ENSMUSG00000038248 | 0.598218561 | 0.01559 | 0.059351 | 402.4217 | 429.5361 | 280.7281 | 364.8897 | 240.8525 | 364.264 | 196.715 | 174.0146 |
| ENSMUSG00000038250 | 0.088966282 | 0.264138 | 0.428339 | 825.9706 | 890.2676 | 818.1694 | 827.2559 | 782.0393 | 852.5564 | 727.074 | 798.2919 |
| ENSMUSG00000038252 | 0.086882035 | 0.149909 | 0.289823 | 4316.979 | 4585.718 | 4221.688 | 4244.97 | 4304.141 | 4196.857 | 3975.764 | 3871.824 |
| ENSMUSG00000038253 | 0.528667136 | 0.039663 | 0.115972 | 1575.481 | 1876.521 | 1293.503 | 1769.241 | 999.4891 | 1853.724 | 857.2531 | 805.905 |
| ENSMUSG00000038256 | 0.397708472 | 0.002384 | 0.014882 | 2038.266 | 2080.491 | 1773.804 | 1911.574 | 1540.676 | 1809.029 | 1350.969 | 1222.452 |
| ENSMUSG00000038260 | -0.099394094 | 0.301202 | 0.469588 | 856.1523 | 881.8688 | 896.0114 | 861.7609 | 1039.469 | 808.9788 | 990.325 | 903.7882 |
| ENSMUSG00000038264 | -0.498047372 | 0.04068 | 0.118141 | 270.6286 | 178.7734 | 313.8524 | 263.9628 | 286.6827 | 328.508 | 527.4662 | 309.9635 |
| ENSMUSG00000038267 | -0.80463903 | 0.001065 | 0.008197 | 355.1372 | 293.9563 | 494.3797 | 287.2536 | 636.7477 | 395.5504 | 669.2167 | 799.3794 |
| ENSMUSG00000038268 | 0.108111321 | 0.407254 | 0.57363 | 294.7739 | 399.5406 | 303.9151 | 344.1868 | 310.0854 | 331.8601 | 310.5011 | 290.3868 |
| ENSMUSG00000038271 | 0.303053385 | 0.010773 | 0.045094 | 649.9111 | 665.901 | 669.9383 | 759.1087 | 635.7726 | 585.504 | 536.1448 | 467.6642 |
| ENSMUSG00000038274 | -0.173849161 | 0.038844 | 0.113995 | 16134.09 | 18017.72 | 17808.43 | 17215.38 | 21262.3 | 17060.07 | 20330.11 | 19378.7 |
| ENSMUSG00000038276 | -0.080717164 | 0.858759 | 0.915304 | 21.12714 | 14.39786 | 42.23344 | 37.09281 | 23.40267 | 46.92971 | 31.82154 | 20.66423 |
| ENSMUSG00000038279 | -0.176505576 | 0.179599 | 0.32949 | 900.4186 | 988.6529 | 899.3238 | 894.5406 | 1123.328 | 787.7487 | 1095.432 | 1152.847 |
| ENSMUSG00000038280 | -0.1490732 | 0.386961 | 0.554354 | 607.6568 | 431.9358 | 625.2205 | 543.4528 | 551.913 | 505.0531 | 621.0022 | 773.2773 |
| ENSMUSG00000038286 | -0.755447608 | 8.68E-05 | 0.001208 | 543.2694 | 530.3211 | 635.9859 | 597.7981 | 1024.842 | 638.0206 | 971.0392 | 1262.693 |
| ENSMUSG00000038290 | 0.157616872 | 0.059337 | 0.153551 | 1625.784 | 1892.119 | 1876.49 | 1793.394 | 1635.262 | 1750.925 | 1492.72 | 1566.131 |
| ENSMUSG00000038291 | -0.09272149 | 0.466877 | 0.627489 | 560.3723 | 453.5325 | 483.6143 | 524.4751 | 595.793 | 444.7149 | 584.3592 | 530.7445 |
| ENSMUSG00000038292 | 0.087768779 | 0.843115 | 0.905412 | 13.07871 | 35.99465 | 18.21835 | 22.42821 | 22.42756 | 17.87799 | 28.92867 | 14.13868 |
| ENSMUSG00000038295 | -1.012348671 | 0.027169 | 0.088122 | 9.054489 | 8.398751 | 7.45296 | 12.93935 | 22.42756 | 17.87799 | 12.53576 | 23.927 |
| ENSMUSG00000038296 | -0.898475876 | 0.002649 | 0.016086 | 655.9474 | 499.1258 | 1062.461 | 602.1112 | 1192.561 | 712.8847 | 1556.363 | 1796.7 |
| ENSMUSG00000038298 | -3.155569977 | 0.000116 | 0.001505 | 251.5136 | 35.99465 | 499.3483 | 319.1707 | 2699.108 | 673.7766 | 2007.65 | 4479.788 |
| ENSMUSG00000038299 | -0.205306876 | 0.005209 | 0.026462 | 1295.798 | 1377.395 | 1339.048 | 1318.089 | 1663.54 | 1573.263 | 1496.577 | 1409.518 |
| ENSMUSG00000038301 | 0.812201138 | 5.54E-08 | 3.55E-06 | 563.3904 | 637.1052 | 453.8024 | 602.1112 | 323.737 | 381.0246 | 289.2867 | 290.3868 |
| ENSMUSG00000038302 | -0.471521113 | 0.009706 | 0.041667 | 155.9384 | 128.3809 | 145.7468 | 133.7066 | 155.0427 | 170.9582 | 234.3223 | 221.8686 |
| ENSMUSG00000038304 | 0.684741748 | 0.206375 | 0.362903 | 17.10292 | 11.99822 | 14.90592 | 16.38985 | 2.925334 | 15.64324 | 9.642892 | 9.78832 |
| ENSMUSG00000038305 | 0.057690488 | 0.511989 | 0.667368 | 1252.538 | 1533.372 | 1379.626 | 1296.523 | 1367.106 | 1336.379 | 1323.005 | 1217.014 |
| ENSMUSG00000038312 | -0.01598076 | 0.856193 | 0.913827 | 846.0917 | 709.0945 | 746.9522 | 752.2077 | 740.1095 | 732.9974 | 800.36 | 815.6933 |
| ENSMUSG00000038319 | -0.139899552 | 0.377745 | 0.545813 | 338.0343 | 393.5415 | 443.8652 | 301.0556 | 450.5014 | 365.3813 | 361.6084 | 449.1751 |
| ENSMUSG00000038323 | -0.280952091 | 0.015747 | 0.059733 | 210.2654 | 213.5682 | 189.6364 | 220.8316 | 236.952 | 245.8223 | 261.3224 | 268.635 |
| ENSMUSG00000038324 | -0.083719279 | 0.211041 | 0.368438 | 3799.867 | 3804.634 | 3909.491 | 4006.024 | 4300.241 | 3705.213 | 4266.015 | 4175.262 |
| ENSMUSG00000038331 | -1.202080683 | 0.004079 | 0.022134 | 32.19374 | 22.79661 | 77.01392 | 44.85642 | 74.10846 | 67.04245 | 127.2862 | 141.3868 |
| ENSMUSG00000038332 | 0.592617757 | 0.000522 | 0.004735 | 2641.899 | 2496.829 | 1907.958 | 3088.192 | 1524.099 | 1962.109 | 1871.685 | 1362.752 |
| ENSMUSG00000038335 | -0.164376224 | 0.108059 | 0.232685 | 1192.174 | 1436.186 | 1262.035 | 1202.497 | 1622.585 | 1300.623 | 1375.076 | 1404.08 |
| ENSMUSG00000038342 | 0.231309226 | 0.031185 | 0.097042 | 1554.354 | 1384.594 | 1425.172 | 1546.684 | 1160.382 | 1378.84 | 1406.898 | 1089.766 |
| ENSMUSG00000038344 | 0.10859871 | 0.234431 | 0.395292 | 678.0806 | 679.099 | 619.4238 | 678.8847 | 565.5646 | 686.0677 | 614.2522 | 597.0875 |
| ENSMUSG00000038346 | 0.089256876 | 0.181579 | 0.3321 | 1798.825 | 1834.527 | 1784.57 | 1939.178 | 1761.051 | 1836.963 | 1618.077 | 1702.08 |
| ENSMUSG00000038347 | -0.020387703 | 0.95406 | 0.973813 | 79.47829 | 56.39161 | 77.01392 | 87.12497 | 53.63112 | 135.2023 | 65.57166 | 51.11678 |
| ENSMUSG00000038349 | 0.08352063 | 0.585192 | 0.725623 | 847.0978 | 644.3042 | 780.0764 | 784.9874 | 642.5984 | 910.6599 | 729.0026 | 604.7006 |
| ENSMUSG00000038351 | -0.070523983 | 0.488906 | 0.646689 | 507.0514 | 484.7279 | 516.7385 | 521.0246 | 500.2321 | 488.2925 | 530.359 | 614.489 |
| ENSMUSG00000038352 | 0.811151508 | 0.000957 | 0.007588 | 73.44197 | 82.78769 | 95.23226 | 75.04824 | 40.95467 | 60.3382 | 34.71441 | 51.11678 |
| ENSMUSG00000038354 | 0.850593817 | 1.82E-06 | 6.03E-05 | 269.6226 | 283.1579 | 232.698 | 276.0395 | 162.8436 | 184.3667 | 118.6076 | 122.8978 |
| ENSMUSG00000038357 | 0.035819343 | 0.957382 | 0.975954 | 29.17558 | 5.999108 | 4.96864 | 18.11509 | 22.42756 | 8.938993 | 14.46434 | 10.87591 |
| ENSMUSG00000038365 | 0.06448673 | 0.536103 | 0.686694 | 790.7587 | 861.4719 | 763.5143 | 710.8018 | 795.6908 | 655.8986 | 726.1097 | 808.0802 |
| ENSMUSG00000038366 | -0.039611795 | 0.771896 | 0.858471 | 3597.65 | 3631.86 | 4837.799 | 3711.869 | 4187.128 | 3348.77 | 4020.122 | 4664.678 |
| ENSMUSG00000038368 | -0.405178399 | 4.64E-05 | 0.00075 | 423.5489 | 447.5334 | 399.1474 | 367.4776 | 526.5601 | 508.4052 | 576.6449 | 551.4087 |
| ENSMUSG00000038369 | 0.132609088 | 0.1399 | 0.276031 | 1371.252 | 1310.205 | 1350.642 | 1322.402 | 1139.905 | 1413.478 | 1139.79 | 1194.175 |
| ENSMUSG00000038370 | 0.347067829 | 0.053208 | 0.14212 | 246.4833 | 178.7734 | 202.8861 | 198.4034 | 160.8934 | 205.5968 | 138.8576 | 145.7372 |
| ENSMUSG00000038371 | 0.035534442 | 0.663852 | 0.78383 | 2834.055 | 2348.051 | 2679.753 | 2771.609 | 2435.828 | 2681.698 | 2719.295 | 2542.788 |
| ENSMUSG00000038372 | -0.120701776 | 0.356228 | 0.524872 | 452.7245 | 655.1026 | 518.3948 | 573.6446 | 571.4152 | 632.4337 | 630.6451 | 554.6715 |
| ENSMUSG00000038374 | -0.03700164 | 0.526531 | 0.679534 | 3287.786 | 3361.9 | 3314.083 | 3437.555 | 3674.219 | 3488.442 | 3309.44 | 3276.912 |
| ENSMUSG00000038375 | -0.856100691 | 0.004406 | 0.023459 | 902.4308 | 416.3381 | 972.1972 | 703.0381 | 1172.084 | 810.0962 | 1497.541 | 1942.438 |
| ENSMUSG00000038379 | 0.086664038 | 0.456958 | 0.618708 | 764.6013 | 722.2926 | 604.5178 | 626.2647 | 652.3495 | 702.8283 | 635.4666 | 565.5474 |
| ENSMUSG00000038383 | -0.128275455 | 0.190126 | 0.342672 | 971.8485 | 950.2587 | 981.3064 | 907.4799 | 987.7878 | 933.0074 | 1038.539 | 1206.139 |
| ENSMUSG00000038384 | 0.468815363 | 0.003636 | 0.0204 | 783.7163 | 751.0883 | 684.8442 | 720.2906 | 538.2614 | 695.0067 | 481.1803 | 410.0218 |
| ENSMUSG00000038387 | -0.121992992 | 0.35033 | 0.519044 | 774.6619 | 671.9001 | 905.1205 | 919.5566 | 872.7246 | 778.8097 | 930.539 | 982.0948 |

| | | | | | | | | | | | |
|---|---|---|---|---|---|---|---|---|---|---|---|
| ENSMUSG00000038388 | 0.162614046 | 0.109695 | 0.235355 | 2298.834 | 2274.862 | 1897.192 | 2376.528 | 1907.318 | 2222.457 | 1941.114 | 1832.591 |
| ENSMUSG00000038390 | 0.488533317 | 1.42E-05 | 0.000298 | 653.9353 | 740.2899 | 643.4389 | 737.5431 | 483.6552 | 555.3349 | 513.9661 | 424.1605 |
| ENSMUSG00000038393 | 0.203930281 | 0.38339 | 0.551379 | 10084.69 | 12979.67 | 13445.97 | 16096.55 | 11853.45 | 6864.029 | 10806.79 | 16148.55 |
| ENSMUSG00000038398 | -0.102722442 | 0.206884 | 0.363479 | 1866.231 | 2102.087 | 1995.737 | 2143.619 | 2180.349 | 2388.946 | 2125.293 | 2012.044 |
| ENSMUSG00000038400 | 0.440081918 | 0.022071 | 0.075574 | 1078.49 | 1397.792 | 1045.071 | 1103.295 | 699.1548 | 1205.647 | 793.61 | 710.197 |
| ENSMUSG00000038406 | 0.102533924 | 0.057604 | 0.150422 | 1958.788 | 1940.111 | 1933.629 | 1911.574 | 1876.114 | 1735.282 | 1753.078 | 1846.73 |
| ENSMUSG00000038412 | -0.484960651 | 0.000118 | 0.001515 | 817.9222 | 683.8983 | 901.8081 | 904.0294 | 1139.905 | 991.1108 | 1218.862 | 1284.445 |
| ENSMUSG00000038415 | -1.157046443 | 0.121741 | 0.251128 | 158.9566 | 50.3925 | 578.8465 | 94.88858 | 470.0037 | 318.4516 | 603.645 | 578.5985 |
| ENSMUSG00000038416 | -0.072931105 | 0.330279 | 0.498145 | 3087.581 | 3521.476 | 2955.513 | 2997.617 | 3435.317 | 3311.897 | 3240.976 | 3217.094 |
| ENSMUSG00000038417 | -0.195398876 | 0.051522 | 0.138835 | 545.2815 | 507.5245 | 509.2856 | 582.2709 | 608.4695 | 541.9264 | 651.8595 | 653.6422 |
| ENSMUSG00000038418 | 0.281039658 | 0.544857 | 0.693414 | 2011.103 | 2008.501 | 1837.569 | 3797.269 | 2400.724 | 702.8283 | 2671.081 | 2170.832 |
| ENSMUSG00000038422 | -0.978751855 | 0.001209 | 0.009011 | 90.54489 | 75.58876 | 65.42042 | 50.89479 | 140.416 | 82.68568 | 134.0362 | 197.9416 |
| ENSMUSG00000038425 | 0.209116595 | 0.159113 | 0.302088 | 414.4944 | 481.1284 | 338.6956 | 429.5865 | 327.6374 | 405.6068 | 389.5728 | 314.3138 |
| ENSMUSG00000038429 | -0.194033957 | 0.028614 | 0.091413 | 2574.493 | 2522.025 | 2851.999 | 2673.27 | 3066.725 | 2635.885 | 3163.833 | 3286.7 |
| ENSMUSG00000038437 | 0.264880047 | 4.17E-05 | 0.000696 | 2629.826 | 2926.365 | 2592.802 | 2742.28 | 2279.81 | 2374.42 | 2186.044 | 2220.861 |
| ENSMUSG00000038446 | 0.190318846 | 0.062606 | 0.159615 | 1529.203 | 1440.986 | 1251.269 | 1299.974 | 1304.699 | 1282.745 | 1171.611 | 1076.715 |
| ENSMUSG00000038451 | -0.170801942 | 0.291407 | 0.458657 | 263.5862 | 243.5638 | 275.7595 | 250.1608 | 309.1103 | 215.6532 | 289.2867 | 349.1167 |
| ENSMUSG00000038453 | 0.589867033 | 0.001485 | 0.010486 | 573.451 | 418.7377 | 491.0672 | 390.7684 | 232.0765 | 369.8508 | 301.8225 | 342.5912 |
| ENSMUSG00000038456 | 0.394772522 | 2.41E-05 | 0.000454 | 1568.439 | 1856.124 | 1599.902 | 1696.78 | 1297.873 | 1381.074 | 1302.755 | 1127.832 |
| ENSMUSG00000038459 | -0.768683429 | 1.05E-06 | 3.89E-05 | 1121.751 | 1095.437 | 1420.203 | 1230.101 | 1736.673 | 1710.7 | 2309.473 | 2539.525 |
| ENSMUSG00000038462 | -0.726956365 | 3.46E-05 | 0.000598 | 3126.817 | 2838.778 | 3879.68 | 3353.018 | 5396.266 | 3771.138 | 6012.343 | 6668.021 |
| ENSMUSG00000038463 | -0.54599868 | 0.031306 | 0.09733 | 248.4954 | 248.3631 | 388.382 | 385.5927 | 430.9992 | 291.6346 | 565.0735 | 569.8977 |
| ENSMUSG00000038467 | -0.162230509 | 0.074911 | 0.180874 | 4639.923 | 4705.7 | 5290.773 | 4647.815 | 5309.481 | 4717.553 | 5628.556 | 5925.196 |
| ENSMUSG00000038473 | -0.204798871 | 0.250746 | 0.414335 | 142.8597 | 116.3827 | 130.0127 | 94.02596 | 136.5156 | 130.7328 | 135.0005 | 154.4379 |
| ENSMUSG00000038481 | 0.18314533 | 0.059318 | 0.15354 | 1704.256 | 1642.556 | 1440.906 | 1817.548 | 1481.194 | 1576.615 | 1426.184 | 1333.387 |
| ENSMUSG00000038482 | -0.074618398 | 0.303304 | 0.47171 | 2903.473 | 3379.897 | 2970.418 | 3177.042 | 3465.546 | 3219.155 | 3124.297 | 3276.912 |
| ENSMUSG00000038485 | -0.016475377 | 0.867655 | 0.920789 | 1038.248 | 974.2551 | 826.4504 | 891.9527 | 938.0571 | 1010.106 | 950.7891 | 871.1605 |
| ENSMUSG00000038486 | -0.060622839 | 0.707484 | 0.816185 | 554.3359 | 431.9358 | 653.3761 | 548.6285 | 534.361 | 468.1797 | 628.7165 | 653.6422 |
| ENSMUSG00000038489 | -0.389986318 | 0.004433 | 0.023569 | 581.4994 | 574.7145 | 708.8593 | 568.4689 | 784.9646 | 658.1333 | 799.3957 | 948.3794 |
| ENSMUSG00000038495 | 0.19475744 | 0.012668 | 0.050983 | 2023.175 | 2056.494 | 1872.349 | 1926.238 | 1613.809 | 1912.944 | 1707.756 | 1647.7 |
| ENSMUSG00000038496 | -0.31452414 | 0.144455 | 0.282595 | 70.4238 | 65.99018 | 78.67013 | 66.42201 | 100.4365 | 62.57295 | 94.50034 | 92.44524 |
| ENSMUSG00000038497 | -0.014425286 | 0.884196 | 0.930838 | 840.0554 | 842.2747 | 844.6688 | 959.2373 | 841.5211 | 798.9225 | 884.2532 | 998.4086 |
| ENSMUSG00000038500 | 0.040945799 | 0.68665 | 0.800759 | 694.1775 | 747.4888 | 693.9534 | 698.725 | 739.1344 | 768.7534 | 631.6094 | 614.489 |
| ENSMUSG00000038502 | -0.046016397 | 0.453846 | 0.616122 | 7109.786 | 7263.72 | 7443.022 | 7710.991 | 7621.47 | 8192.587 | 7511.813 | 7162.875 |
| ENSMUSG00000038503 | -0.053161151 | 0.384869 | 0.552504 | 2275.695 | 2631.209 | 2470.242 | 2479.18 | 2568.443 | 2548.73 | 2565.973 | 2539.525 |
| ENSMUSG00000038506 | -0.203615432 | 0.164026 | 0.308999 | 469.8274 | 537.52 | 409.9128 | 485.657 | 512.9085 | 463.7102 | 538.0734 | 674.3065 |
| ENSMUSG00000038507 | 0.302189582 | 0.085617 | 0.197989 | 657.9595 | 957.4576 | 1066.601 | 646.105 | 683.553 | 664.8376 | 683.681 | 666.6933 |
| ENSMUSG00000038508 | -0.009842901 | 0.970617 | 0.983617 | 68.4117 | 44.3934 | 78.67013 | 47.44429 | 62.40712 | 61.45558 | 67.50024 | 50.02919 |
| ENSMUSG00000038510 | 0.016867983 | 0.863338 | 0.917949 | 746.4923 | 862.6717 | 675.735 | 807.4156 | 796.6659 | 773.2229 | 730.9312 | 751.5254 |
| ENSMUSG00000038515 | -0.188207461 | 0.155986 | 0.298083 | 309.8647 | 283.1579 | 301.4308 | 347.6373 | 376.393 | 320.6864 | 312.4297 | 407.8467 |
| ENSMUSG00000038517 | 0.091899014 | 0.361682 | 0.530157 | 434.6155 | 508.7243 | 534.9569 | 486.5197 | 492.4312 | 461.4755 | 436.823 | 453.5255 |
| ENSMUSG00000038518 | 0.077981865 | 0.546281 | 0.69465 | 2567.451 | 2525.624 | 2344.37 | 2512.822 | 2106.24 | 3022.497 | 2160.972 | 2138.204 |
| ENSMUSG00000038520 | 0.092341173 | 0.286763 | 0.453937 | 1696.208 | 1598.162 | 1576.715 | 1630.358 | 1644.038 | 1603.432 | 1530.327 | 1319.248 |
| ENSMUSG00000038521 | 0.673080947 | 2.04E-05 | 0.000398 | 1410.488 | 1853.724 | 1495.561 | 1996.973 | 892.2268 | 1241.403 | 1168.718 | 935.3283 |
| ENSMUSG00000038522 | -2.827099368 | 3.76E-05 | 0.000641 | 55.33299 | 21.59679 | 143.2624 | 63.83414 | 461.2276 | 187.7188 | 434.8944 | 936.4159 |
| ENSMUSG00000038524 | 0.26658818 | 0.030881 | 0.096501 | 243.4652 | 226.7663 | 272.4471 | 259.6497 | 223.3005 | 194.4231 | 195.7507 | 221.8686 |
| ENSMUSG00000038525 | 0.235956623 | 0.028777 | 0.091745 | 3089.593 | 3553.871 | 3219.679 | 3275.381 | 2969.214 | 3190.103 | 2642.152 | 2353.547 |
| ENSMUSG00000038526 | 0.386011536 | 0.105959 | 0.229961 | 265.5983 | 165.5754 | 285.6968 | 258.787 | 212.5743 | 126.2633 | 179.3578 | 229.4817 |
| ENSMUSG00000038527 | -0.110033894 | 0.61495 | 0.7485 | 72.43591 | 79.18822 | 67.07664 | 69.00988 | 62.40712 | 70.39457 | 76.17884 | 101.146 |
| ENSMUSG00000038528 | -3.056681767 | 1.56E-12 | 4.29E-10 | 35.2119 | 13.19804 | 59.62368 | 23.29083 | 371.5174 | 149.7281 | 231.4294 | 349.1167 |
| ENSMUSG00000038530 | 1.482531221 | 0.069593 | 0.171539 | 440.6518 | 35.99465 | 71.21717 | 287.2536 | 71.18313 | 31.28647 | 80.036 | 116.3722 |
| ENSMUSG00000038533 | 0.184325864 | 0.05039 | 0.136764 | 1534.233 | 1534.572 | 1623.089 | 1568.25 | 1354.43 | 1610.136 | 1270.933 | 1276.832 |
| ENSMUSG00000038534 | 0.035887324 | 0.73128 | 0.831495 | 415.5004 | 418.7377 | 357.7421 | 375.2412 | 362.7414 | 367.6161 | 378.0014 | 417.635 |
| ENSMUSG00000038535 | 0.357392487 | 0.027266 | 0.08833 | 1580.511 | 1620.959 | 1270.316 | 1675.215 | 1213.038 | 1537.507 | 1109.897 | 937.5035 |
| ENSMUSG00000038538 | 0.410665516 | 0.048483 | 0.133274 | 2855.182 | 2597.614 | 2053.704 | 2322.182 | 1551.402 | 2763.266 | 1551.541 | 1528.065 |
| ENSMUSG00000038539 | -0.054046709 | 0.630259 | 0.759281 | 904.4429 | 1184.224 | 1188.333 | 1023.071 | 1099.926 | 1049.214 | 1095.432 | 1219.19 |
| ENSMUSG00000038540 | 0.835848605 | 0.030221 | 0.094971 | 38.23007 | 57.59143 | 43.06154 | 25.01608 | 24.37778 | 25.6996 | 27.96439 | 13.05109 |
| ENSMUSG00000038541 | 2.259871517 | 0.001048 | 0.008098 | 58.35115 | 17.99732 | 18.21835 | 21.56559 | 3.900445 | 5.58687 | 1.928578 | 13.05109 |
| ENSMUSG00000038542 | 0.024078553 | 0.7622 | 0.852142 | 981.9091 | 973.0553 | 844.6688 | 931.6334 | 913.6793 | 952.0027 | 929.5748 | 870.0729 |
| ENSMUSG00000038543 | -0.031012166 | 0.837083 | 0.901459 | 181.0898 | 208.7689 | 251.7444 | 228.5952 | 255.4792 | 215.6532 | 205.3936 | 214.2554 |
| ENSMUSG00000038544 | -0.08453263 | 0.265106 | 0.429306 | 823.9585 | 835.0758 | 891.0427 | 850.5468 | 969.2606 | 898.3688 | 838.9316 | 901.613 |
| ENSMUSG00000038545 | -0.041078338 | 0.503633 | 0.65955 | 3257.604 | 3240.718 | 3415.94 | 3267.618 | 3297.826 | 3167.756 | 3588.12 | 3509.656 |
| ENSMUSG00000038546 | 0.420854291 | 0.003968 | 0.021742 | 1693.189 | 1746.94 | 1520.404 | 1809.784 | 1089.199 | 1657.066 | 1183.183 | 1128.92 |
| ENSMUSG00000038550 | -0.160685357 | 0.603839 | 0.740916 | 81.4904 | 74.38894 | 40.57722 | 64.69676 | 71.18313 | 99.44629 | 71.3574 | 48.9416 |

| | | | | | | | | | | | |
|---|---|---|---|---|---|---|---|---|---|---|---|
| ENSMUSG00000038552 | -0.046687258 | 0.603578 | 0.740729 | 652.9293 | 739.0901 | 710.5155 | 699.5877 | 672.8268 | 703.9457 | 710.6811 | 806.9926 |
| ENSMUSG00000038555 | 0.009550052 | 0.946551 | 0.969279 | 229.3804 | 194.3711 | 229.3855 | 266.5507 | 208.6738 | 238.0007 | 216.0008 | 254.4963 |
| ENSMUSG00000038563 | -0.113447791 | 0.244284 | 0.406905 | 600.6145 | 542.3193 | 587.9557 | 618.501 | 653.3246 | 582.1519 | 599.7879 | 709.1094 |
| ENSMUSG00000038564 | 0.079535272 | 0.254285 | 0.417943 | 1223.362 | 1347.4 | 1233.879 | 1269.782 | 1174.034 | 1225.759 | 1266.112 | 1132.182 |
| ENSMUSG00000038567 | -3.738826048 | 2.18E-12 | 5.74E-10 | 24.1453 | 3.599465 | 39.74912 | 42.26855 | 619.1957 | 196.6578 | 267.1081 | 396.9707 |
| ENSMUSG00000038569 | 0.009275042 | 0.942346 | 0.966541 | 272.6407 | 281.9581 | 242.6352 | 282.0779 | 276.9316 | 263.7003 | 300.8582 | 228.3941 |
| ENSMUSG00000038570 | 0.406715959 | 0.119485 | 0.24843 | 69.41775 | 82.78769 | 53.82693 | 90.57547 | 58.50668 | 44.69496 | 69.42882 | 50.02919 |
| ENSMUSG00000038572 | 0.291525622 | 0.295837 | 0.463653 | 39.23612 | 45.59322 | 49.6864 | 37.09281 | 32.17867 | 41.34284 | 40.50014 | 26.10219 |
| ENSMUSG00000038576 | 0.100531493 | 0.437417 | 0.60116 | 888.346 | 902.2658 | 1106.35 | 948.0232 | 866.874 | 734.1148 | 1009.611 | 976.6568 |
| ENSMUSG00000038578 | -0.02817972 | 0.740375 | 0.837688 | 802.8314 | 711.4942 | 775.1078 | 693.5493 | 793.7406 | 717.3542 | 742.5027 | 788.5035 |
| ENSMUSG00000038580 | 0.823337445 | 1.68E-11 | 3.69E-09 | 331.9979 | 382.7431 | 407.4285 | 422.6855 | 215.4996 | 238.0007 | 210.215 | 210.9927 |
| ENSMUSG00000038582 | -0.349784129 | 0.055259 | 0.146299 | 510.0696 | 375.5441 | 462.0835 | 375.2412 | 534.361 | 400.0199 | 595.9307 | 665.6057 |
| ENSMUSG00000038583 | 1.037063941 | 0.063301 | 0.161055 | 50.30272 | 50.3925 | 27.32752 | 52.62003 | 11.70134 | 45.81234 | 24.10723 | 6.525547 |
| ENSMUSG00000038587 | 0.36131205 | 0.00052 | 0.004724 | 5832.097 | 6860.58 | 7777.577 | 6829.39 | 4776.095 | 5357.809 | 5749.092 | 5370.525 |
| ENSMUSG00000038593 | 0.256810457 | 0.013444 | 0.053245 | 954.7456 | 903.4656 | 971.3691 | 958.3747 | 810.3175 | 922.951 | 746.3598 | 692.7955 |
| ENSMUSG00000038594 | -0.099035526 | 0.63443 | 0.762319 | 98.59333 | 77.9884 | 94.40416 | 91.43809 | 117.0134 | 86.03781 | 109.929 | 75.04379 |
| ENSMUSG00000038600 | -2.335483164 | 0.000256 | 0.002755 | 266.6044 | 141.5789 | 658.3448 | 296.7425 | 1757.151 | 481.5882 | 1998.007 | 2647.197 |
| ENSMUSG00000038602 | 0.327796474 | 0.065704 | 0.165029 | 210.2654 | 153.5772 | 167.2775 | 164.7611 | 109.2125 | 153.0803 | 139.8219 | 153.3503 |
| ENSMUSG00000038604 | -0.524585886 | 0.000325 | 0.003302 | 865.2067 | 682.6985 | 969.7129 | 830.7064 | 1153.557 | 979.9371 | 1331.683 | 1355.138 |
| ENSMUSG00000038605 | -0.233148871 | 0.097898 | 0.217792 | 476.8698 | 355.1472 | 379.2728 | 381.2796 | 430.0241 | 406.7242 | 521.6804 | 513.343 |
| ENSMUSG00000038607 | 0.192378237 | 0.105684 | 0.229522 | 443.67 | 512.3238 | 558.972 | 546.0407 | 515.8339 | 427.9543 | 440.6801 | 419.8102 |
| ENSMUSG00000038608 | 0.593792375 | 0.006378 | 0.03084 | 450.7124 | 502.7252 | 346.1486 | 470.1298 | 294.4836 | 415.6632 | 227.5722 | 234.9197 |
| ENSMUSG00000038611 | 0.015641875 | 0.859726 | 0.915903 | 1615.723 | 1595.763 | 1628.058 | 1423.329 | 1535.8 | 1442.53 | 1477.291 | 1739.058 |
| ENSMUSG00000038612 | 0.129884671 | 0.103805 | 0.226596 | 6218.422 | 5260.018 | 5472.129 | 6172.071 | 4999.396 | 5180.146 | 5382.662 | 5572.817 |
| ENSMUSG00000038615 | -0.317836293 | 0.015572 | 0.059294 | 4085.587 | 3753.042 | 4553.758 | 3818.834 | 4593.749 | 4102.998 | 5774.164 | 5737.043 |
| ENSMUSG00000038618 | -0.65364696 | 0.00153 | 0.010695 | 416.5065 | 436.735 | 611.9708 | 456.3278 | 780.089 | 500.5836 | 812.8958 | 930.978 |
| ENSMUSG00000038619 | -0.037173205 | 0.505565 | 0.661418 | 2906.491 | 3097.939 | 2827.156 | 2944.997 | 3070.626 | 2973.332 | 3129.118 | 2902.781 |
| ENSMUSG00000038622 | 0.011333823 | 0.927999 | 0.957316 | 920.5397 | 671.9001 | 693.9534 | 765.147 | 709.881 | 814.5657 | 802.2886 | 701.4963 |
| ENSMUSG00000038623 | -0.546176613 | 0.02639 | 0.086387 | 212.2775 | 128.3809 | 216.9639 | 172.5247 | 242.8027 | 179.8972 | 340.3941 | 304.5255 |
| ENSMUSG00000038628 | -0.038093816 | 0.602264 | 0.740138 | 1312.901 | 1397.792 | 1192.474 | 1269.782 | 1338.828 | 1379.957 | 1297.933 | 1288.795 |
| ENSMUSG00000038630 | 0.355088539 | 0.162498 | 0.306841 | 69.41775 | 79.18822 | 56.31125 | 48.30692 | 40.95467 | 49.16446 | 49.17875 | 57.64233 |
| ENSMUSG00000038633 | -0.173254936 | 0.149882 | 0.289823 | 1644.899 | 1606.561 | 2085.173 | 1865.855 | 1961.924 | 1849.254 | 1942.078 | 2373.124 |
| ENSMUSG00000038637 | 0.16939648 | 0.106848 | 0.231093 | 429.5852 | 435.5352 | 455.4586 | 535.6892 | 403.6961 | 422.3674 | 408.8586 | 418.7226 |
| ENSMUSG00000038641 | -1.415674978 | 0.056364 | 0.148073 | 13.07871 | 3.599465 | 0.828107 | 10.35148 | 15.60178 | 10.05637 | 19.28578 | 29.36496 |
| ENSMUSG00000038642 | 0.356910519 | 0.043192 | 0.123138 | 341.0524 | 343.149 | 386.7258 | 315.7202 | 245.728 | 217.8879 | 362.5727 | 255.5839 |
| ENSMUSG00000038644 | -0.221619215 | 0.101463 | 0.223353 | 788.7466 | 1165.027 | 894.3552 | 853.9973 | 1089.199 | 972.1155 | 1083.861 | 1166.985 |
| ENSMUSG00000038646 | -0.093665996 | 0.200653 | 0.356032 | 1793.795 | 1856.124 | 1911.27 | 2128.092 | 1990.202 | 2090.607 | 2098.293 | 2030.533 |
| ENSMUSG00000038648 | 0.375584693 | 0.010679 | 0.044794 | 3828.037 | 3880.223 | 3178.273 | 3607.491 | 2431.928 | 3656.048 | 2657.581 | 2426.416 |
| ENSMUSG00000038650 | -0.09413123 | 0.381434 | 0.549379 | 2459.803 | 2578.416 | 3015.136 | 2442.95 | 2965.313 | 2393.415 | 2838.867 | 3007.189 |
| ENSMUSG00000038658 | 0.117720083 | 0.200838 | 0.356273 | 1498.015 | 1419.389 | 1193.302 | 1312.05 | 1259.844 | 1322.971 | 1194.754 | 1217.014 |
| ENSMUSG00000038664 | 0.006402522 | 0.952943 | 0.973191 | 1918.546 | 1806.931 | 1933.629 | 1682.978 | 1647.938 | 2176.645 | 1795.506 | 1691.204 |
| ENSMUSG00000038665 | 0.308131505 | 0.388028 | 0.555667 | 50.30272 | 47.99286 | 37.2648 | 43.9938 | 42.9049 | 54.75133 | 20.25007 | 27.18978 |
| ENSMUSG00000038668 | 0.965274717 | 1.61E-08 | 1.26E-06 | 1166.017 | 1690.549 | 1218.973 | 1477.674 | 680.6277 | 906.1904 | 673.0738 | 584.0364 |
| ENSMUSG00000038670 | 0.295805475 | 0.471539 | 0.631172 | 25.15136 | 51.59233 | 33.12427 | 44.85642 | 18.52711 | 48.04709 | 37.60728 | 21.75182 |
| ENSMUSG00000038671 | 0.042506659 | 0.497605 | 0.654039 | 1274.671 | 1222.618 | 1277.769 | 1302.561 | 1309.574 | 1221.29 | 1199.576 | 1201.788 |
| ENSMUSG00000038677 | -0.062347809 | 0.786511 | 0.868693 | 2358.191 | 2100.888 | 4130.596 | 2470.554 | 2122.817 | 2706.28 | 3138.761 | 3583.613 |
| ENSMUSG00000038679 | -0.115746742 | 0.513601 | 0.66829 | 655.9474 | 508.7243 | 802.4353 | 633.1656 | 600.6686 | 889.4298 | 618.1094 | 713.4598 |
| ENSMUSG00000038683 | -0.119269333 | 0.031307 | 0.09733 | 1344.089 | 1312.605 | 1319.174 | 1349.143 | 1449.991 | 1438.06 | 1396.291 | 1501.963 |
| ENSMUSG00000038685 | 0.015750085 | 0.858001 | 0.914917 | 1050.321 | 1100.236 | 955.6351 | 940.2596 | 1040.444 | 1048.097 | 995.1464 | 913.5765 |
| ENSMUSG00000038690 | -0.504261239 | 0.000295 | 0.003061 | 2806.892 | 3139.933 | 3567.483 | 2963.112 | 4499.164 | 3435.925 | 4480.087 | 5283.518 |
| ENSMUSG00000038692 | 0.657646258 | 0.001089 | 0.008336 | 2576.505 | 2261.664 | 1990.768 | 2236.783 | 1186.71 | 2143.124 | 1170.647 | 1247.467 |
| ENSMUSG00000038695 | -0.098614667 | 0.15402 | 0.295488 | 703.232 | 751.0883 | 751.0927 | 713.3896 | 762.537 | 762.0491 | 800.36 | 799.3794 |
| ENSMUSG00000038696 | -0.215969507 | 0.024227 | 0.081141 | 1676.087 | 1733.742 | 2018.924 | 1757.164 | 2156.946 | 1841.433 | 2091.543 | 2258.927 |
| ENSMUSG00000038697 | -0.287960869 | 0.000159 | 0.001898 | 1148.914 | 1078.64 | 1275.284 | 1128.312 | 1478.269 | 1420.182 | 1397.255 | 1363.839 |
| ENSMUSG00000038700 | 0.43023247 | 0.01154 | 0.047539 | 1029.194 | 1005.45 | 1032.649 | 1055.851 | 742.0597 | 1041.393 | 708.7525 | 568.8101 |
| ENSMUSG00000038702 | 0.645599632 | 4.08E-05 | 0.000685 | 633.8142 | 910.6645 | 761.8581 | 828.9812 | 469.0285 | 619.0252 | 426.2158 | 490.5036 |
| ENSMUSG00000038704 | -2.084019539 | 0.001383 | 0.009973 | 11.0666 | 7.198929 | 5.796746 | 7.763611 | 28.27823 | 7.821619 | 28.92867 | 69.60583 |
| ENSMUSG00000038705 | -0.06449502 | 0.5623 | 0.707802 | 876.2733 | 793.082 | 862.8871 | 860.8982 | 890.2766 | 1049.214 | 846.6459 | 764.5765 |
| ENSMUSG00000038708 | -0.124088471 | 0.266553 | 0.431024 | 3052.369 | 2579.616 | 2929.013 | 2421.384 | 2679.606 | 2856.008 | 2998.939 | 3435.7 |
| ENSMUSG00000038712 | -0.192466721 | 0.182195 | 0.332766 | 1836.049 | 1485.379 | 1729.915 | 1566.524 | 1852.711 | 1451.469 | 1974.864 | 2283.941 |
| ENSMUSG00000038717 | -0.541183715 | 2.85E-05 | 0.000518 | 2788.783 | 2795.584 | 3415.94 | 2910.492 | 4197.854 | 3472.799 | 4638.231 | 5025.758 |
| ENSMUSG00000038718 | 0.575032945 | 2.19E-05 | 0.000423 | 1357.167 | 1172.226 | 1135.334 | 1232.689 | 788.865 | 1038.041 | 748.2884 | 713.4598 |
| ENSMUSG00000038721 | 0.221468601 | 0.003167 | 0.018493 | 2079.514 | 2145.281 | 2183.717 | 2148.795 | 1747.399 | 2028.034 | 1874.578 | 1691.204 |
| ENSMUSG00000038722 | -0.12373641 | 0.020549 | 0.071883 | 1753.553 | 1844.126 | 1787.054 | 1861.541 | 1935.596 | 2042.56 | 1931.471 | 1987.029 |

| | | | | | | | | | | | |
|---|---|---|---|---|---|---|---|---|---|---|---|
| ENSMUSG00000038725 | 1.725423938 | 0.166655 | 0.312357 | 36.21796 | 209.9688 | 32.29616 | 84.5371 | 19.50223 | 74.86406 | 15.42863 | 0 |
| ENSMUSG00000038729 | -0.630250035 | 0.071867 | 0.175819 | 28.16952 | 35.99465 | 22.35888 | 15.52722 | 46.80534 | 29.05173 | 38.57157 | 41.32846 |
| ENSMUSG00000038732 | 0.23044665 | 0.022374 | 0.076356 | 763.5953 | 729.4915 | 858.7466 | 695.2745 | 622.121 | 717.3542 | 628.7165 | 631.8904 |
| ENSMUSG00000038733 | 0.197209688 | 0.171335 | 0.318509 | 5638.935 | 5108.84 | 4822.893 | 5493.186 | 3972.603 | 6048.346 | 4317.123 | 4036.051 |
| ENSMUSG00000038736 | -0.104002045 | 0.311455 | 0.480077 | 629.79 | 545.9188 | 496.864 | 528.7882 | 585.0668 | 565.3913 | 632.5737 | 579.686 |
| ENSMUSG00000038738 | 0.091391566 | 0.645637 | 0.770506 | 98.59333 | 109.1838 | 125.8722 | 98.33908 | 89.71024 | 91.62468 | 94.50034 | 130.5109 |
| ENSMUSG00000038740 | 0.260029662 | 0.020464 | 0.071685 | 1436.646 | 1793.733 | 1684.369 | 1374.159 | 1288.122 | 1430.239 | 1195.719 | 1336.649 |
| ENSMUSG00000038742 | -0.561047121 | 0.022149 | 0.075785 | 34.20585 | 43.19358 | 43.06154 | 47.44429 | 54.60623 | 75.98144 | 53.0359 | 65.25547 |
| ENSMUSG00000038745 | -1.637573944 | 0.000291 | 0.003035 | 29.17558 | 17.99732 | 72.87338 | 43.9938 | 137.4907 | 68.15982 | 112.8218 | 195.7664 |
| ENSMUSG00000038759 | -0.131381788 | 0.065416 | 0.164684 | 1841.079 | 2007.301 | 1817.694 | 1809.784 | 2079.912 | 2140.889 | 1877.471 | 2087.087 |
| ENSMUSG00000038762 | -0.165067895 | 0.008236 | 0.037056 | 3286.78 | 3438.689 | 3295.864 | 3085.604 | 3658.618 | 3489.559 | 3639.227 | 3903.364 |
| ENSMUSG00000038763 | -0.071440466 | 0.872108 | 0.9234 | 26.15741 | 14.39786 | 12.4216 | 22.42821 | 14.62667 | 13.40849 | 25.07152 | 26.10219 |
| ENSMUSG00000038764 | 0.168910694 | 0.111212 | 0.237397 | 1250.526 | 1085.838 | 973.8534 | 1036.873 | 879.5504 | 1050.332 | 941.1462 | 995.1458 |
| ENSMUSG00000038765 | 0.308043332 | 0.055371 | 0.146491 | 330.9919 | 343.149 | 282.3844 | 343.3242 | 275.9565 | 326.2732 | 215.0365 | 232.7445 |
| ENSMUSG00000038766 | 0.28291472 | 0.002248 | 0.014301 | 2472.882 | 2290.459 | 1948.535 | 2180.712 | 1765.927 | 1976.635 | 1755.006 | 1809.752 |
| ENSMUSG00000038773 | 0.11939729 | 0.037571 | 0.111678 | 2424.591 | 2260.464 | 2230.091 | 2304.93 | 2172.548 | 2192.288 | 2079.972 | 2042.496 |
| ENSMUSG00000038774 | -0.308180631 | 0.012187 | 0.049543 | 515.0998 | 403.14 | 515.0823 | 434.7622 | 545.0872 | 527.4006 | 602.6807 | 640.5912 |
| ENSMUSG00000038775 | -0.269110132 | 0.096651 | 0.215616 | 1074.466 | 989.8528 | 1236.363 | 1107.609 | 1305.674 | 962.0591 | 1397.255 | 1648.788 |
| ENSMUSG00000038776 | -0.315649796 | 0.043294 | 0.123188 | 452.7245 | 365.9456 | 404.116 | 439.0754 | 557.7637 | 386.6114 | 518.7876 | 605.7882 |
| ENSMUSG00000038777 | 0.359112035 | 0.001406 | 0.01011 | 459.7668 | 511.124 | 412.3971 | 489.9701 | 372.4925 | 392.1983 | 368.3585 | 325.1897 |
| ENSMUSG00000038780 | -0.058791492 | 0.40788 | 0.574162 | 1409.482 | 1282.609 | 1476.514 | 1339.654 | 1495.821 | 1384.427 | 1455.112 | 1405.168 |
| ENSMUSG00000038781 | -0.038971612 | 0.850826 | 0.910548 | 329.9858 | 272.3595 | 457.1149 | 343.3242 | 345.1894 | 264.8177 | 435.8587 | 396.9707 |
| ENSMUSG00000038782 | -1.410267916 | 0.001997 | 0.013119 | 22.1332 | 9.598572 | 28.98373 | 23.29083 | 39.97956 | 30.1691 | 69.42882 | 85.9197 |
| ENSMUSG00000038784 | 0.387951303 | 0.004716 | 0.024686 | 887.3399 | 935.8608 | 737.843 | 847.9589 | 610.4197 | 810.0962 | 578.5735 | 605.7882 |
| ENSMUSG00000038793 | -0.560566645 | 0.317802 | 0.486138 | 16.09687 | 4.799286 | 7.45296 | 15.52722 | 13.65156 | 10.05637 | 15.42863 | 26.10219 |
| ENSMUSG00000038797 | 0.007694388 | 0.938999 | 0.96418 | 451.7184 | 412.7386 | 427.303 | 388.1806 | 380.2934 | 411.1937 | 448.3945 | 430.6861 |
| ENSMUSG00000038803 | -0.270919505 | 0.016837 | 0.06241 | 1766.631 | 1827.328 | 2035.486 | 1880.519 | 2306.138 | 1890.597 | 2208.222 | 2658.073 |
| ENSMUSG00000038805 | 1.850745226 | NA | NA | 1175.071 | 1256.213 | 799.951 | 1080.867 | 25.35289 | 1112.905 | 27.0001 | 30.45255 |
| ENSMUSG00000038806 | 0.052411196 | 0.578344 | 0.719946 | 1272.659 | 1292.208 | 1235.535 | 1426.779 | 1431.463 | 1211.234 | 1242.969 | 1153.934 |
| ENSMUSG00000038807 | -0.693429091 | 0.005186 | 0.0264 | 527.1725 | 357.5468 | 607.8303 | 416.6471 | 636.7477 | 506.1705 | 991.2893 | 954.905 |
| ENSMUSG00000038811 | -0.49482703 | 0.049445 | 0.135042 | 133.8052 | 82.78769 | 155.684 | 138.0198 | 141.3911 | 146.376 | 203.465 | 230.5693 |
| ENSMUSG00000038812 | 0.000712048 | 0.98937 | 0.99393 | 1914.521 | 1953.309 | 1996.565 | 1940.04 | 2025.306 | 1945.348 | 1967.15 | 1861.956 |
| ENSMUSG00000038816 | -0.199690879 | 0.007044 | 0.033083 | 1363.204 | 1259.813 | 1437.593 | 1366.396 | 1521.174 | 1484.99 | 1682.685 | 1547.642 |
| ENSMUSG00000038822 | 0.269755635 | 0.045061 | 0.12662 | 936.6366 | 859.0722 | 749.4365 | 878.1507 | 708.9059 | 865.9649 | 666.3238 | 598.1751 |
| ENSMUSG00000038827 | -0.006095244 | 0.951435 | 0.972512 | 668.0201 | 662.3015 | 571.3936 | 667.6706 | 689.4037 | 602.2646 | 596.895 | 689.5327 |
| ENSMUSG00000038828 | -0.188787038 | 0.03215 | 0.099357 | 1609.687 | 1804.532 | 1815.21 | 1741.637 | 2056.51 | 1731.93 | 2022.114 | 2133.854 |
| ENSMUSG00000038831 | 0.201824415 | 0.101659 | 0.223572 | 536.227 | 417.5379 | 492.7234 | 458.9157 | 373.4676 | 472.6492 | 391.5014 | 421.9853 |
| ENSMUSG00000038836 | -0.101971835 | 0.388662 | 0.55606 | 355.1372 | 317.9527 | 304.7432 | 339.011 | 323.737 | 389.9636 | 380.8942 | 318.6642 |
| ENSMUSG00000038838 | -0.172157681 | 0.060912 | 0.156655 | 704.238 | 622.7074 | 693.1253 | 632.303 | 686.4784 | 715.1194 | 792.6457 | 796.1167 |
| ENSMUSG00000038843 | -1.438490993 | 1.39E-08 | 1.11E-06 | 427.5731 | 393.5415 | 673.2507 | 463.2288 | 1087.249 | 983.2892 | 1208.254 | 2031.62 |
| ENSMUSG00000038844 | -0.511734462 | 0.002641 | 0.016041 | 1007.06 | 844.6744 | 1156.037 | 883.3265 | 1300.798 | 1026.867 | 1640.256 | 1581.357 |
| ENSMUSG00000038845 | -0.596291304 | 0.000137 | 0.001706 | 3170.077 | 2919.166 | 3354.66 | 2919.981 | 4551.82 | 3375.587 | 4896.66 | 5868.642 |
| ENSMUSG00000038848 | 0.03228832 | 0.690364 | 0.803922 | 1981.927 | 2099.688 | 1950.191 | 1970.232 | 1933.646 | 2206.814 | 1807.078 | 1877.182 |
| ENSMUSG00000038855 | 0.365795415 | 8.69E-06 | 0.000205 | 2577.511 | 2654.005 | 2417.243 | 2696.561 | 1942.422 | 2236.983 | 2018.257 | 1830.416 |
| ENSMUSG00000038859 | 0.096429276 | 0.437513 | 0.601202 | 564.3965 | 604.7101 | 593.7525 | 604.6991 | 450.5014 | 629.0816 | 523.609 | 613.4014 |
| ENSMUSG00000038860 | 0.592171594 | 0.017076 | 0.063045 | 769.6316 | 709.0945 | 650.8918 | 747.8946 | 486.5805 | 747.5233 | 366.4299 | 308.8759 |
| ENSMUSG00000038861 | -0.032813522 | 0.606251 | 0.742598 | 1468.839 | 1390.593 | 1514.607 | 1406.076 | 1520.199 | 1395.6 | 1484.041 | 1515.014 |
| ENSMUSG00000038866 | -0.044667609 | 0.570602 | 0.713919 | 655.9474 | 671.9001 | 703.8906 | 740.131 | 734.2588 | 732.9974 | 722.2526 | 672.1313 |
| ENSMUSG00000038871 | -0.11202147 | 0.270075 | 0.435085 | 1187.144 | 1189.023 | 1397.016 | 1484.575 | 1414.887 | 1335.262 | 1404.969 | 1532.416 |
| ENSMUSG00000038872 | 0.945825299 | 0.002678 | 0.016234 | 1344.089 | 1227.417 | 931.62 | 1085.18 | 403.6961 | 1116.257 | 423.3229 | 439.3868 |
| ENSMUSG00000038876 | -0.068773623 | 0.402948 | 0.569914 | 679.0867 | 686.2979 | 666.6258 | 694.4119 | 748.8855 | 653.6638 | 766.6099 | 687.3576 |
| ENSMUSG00000038879 | 0.220539328 | 0.129461 | 0.261379 | 219.3198 | 181.1731 | 173.0743 | 197.5408 | 158.9431 | 186.6015 | 148.5005 | 168.5766 |
| ENSMUSG00000038880 | -0.66279892 | 0.000133 | 0.001664 | 637.8385 | 674.2997 | 801.6072 | 690.9614 | 1214.989 | 762.0491 | 1134.004 | 1330.124 |
| ENSMUSG00000038884 | 0.733880169 | 1.35E-10 | 2.38E-08 | 1004.042 | 1040.245 | 1161.834 | 877.2881 | 668.9264 | 550.8654 | 630.6451 | 603.6131 |
| ENSMUSG00000038886 | 0.198250285 | 0.019443 | 0.069097 | 2466.845 | 2514.826 | 2436.29 | 2504.196 | 2308.088 | 2309.612 | 2172.543 | 1856.518 |
| ENSMUSG00000038888 | -0.007415948 | 0.965746 | 0.980731 | 486.9303 | 688.6976 | 486.0986 | 590.8971 | 641.6232 | 440.2454 | 529.3948 | 650.3795 |
| ENSMUSG00000038893 | 0.199480966 | 0.077979 | 0.1857 | 540.2512 | 627.5067 | 518.3948 | 564.1558 | 416.3725 | 513.9921 | 519.7519 | 507.905 |
| ENSMUSG00000038894 | 0.258943696 | 0.074494 | 0.180215 | 1938.667 | 1589.764 | 1756.414 | 1856.366 | 1449.015 | 1471.582 | 1875.542 | 1171.336 |
| ENSMUSG00000038895 | 0.144080752 | 0.14048 | 0.276915 | 499.003 | 506.3247 | 520.8791 | 592.6223 | 478.7797 | 517.3442 | 470.5731 | 453.5255 |
| ENSMUSG00000038900 | -0.242822929 | 0.006038 | 0.0296 | 31202.78 | 36606.56 | 36359.68 | 35085.49 | 46154.94 | 37759.42 | 41563.76 | 39300.1 |
| ENSMUSG00000038902 | 0.189440038 | 0.038592 | 0.113578 | 2239.477 | 2366.048 | 2220.982 | 2227.294 | 2019.456 | 2271.622 | 1894.828 | 1753.197 |
| ENSMUSG00000038903 | -0.420552781 | 0.061473 | 0.15764 | 65.39353 | 65.99018 | 81.98256 | 100.9269 | 97.51113 | 87.15518 | 111.8575 | 126.1606 |
| ENSMUSG00000038909 | -0.095650399 | 0.157694 | 0.300269 | 2166.035 | 2277.261 | 2370.869 | 2302.342 | 2605.497 | 2381.124 | 2487.866 | 2266.54 |
| ENSMUSG00000038910 | 0.636792766 | 0.000163 | 0.001932 | 903.4368 | 919.0633 | 780.0764 | 945.4353 | 546.0623 | 775.4576 | 478.2874 | 482.8904 |

| | | | | | | | | | | | |
|---|---|---|---|---|---|---|---|---|---|---|---|
| ENSMUSG00000038914 | -0.023821175 | 0.675862 | 0.792479 | 2011.103 | 2120.085 | 1977.519 | 1998.699 | 2077.962 | 1992.278 | 2007.65 | 2159.956 |
| ENSMUSG00000038916 | 0.622239067 | 0.209829 | 0.367176 | 25.15136 | 16.7975 | 12.4216 | 44.85642 | 18.52711 | 12.29112 | 16.39292 | 17.40146 |
| ENSMUSG00000038917 | 1.11731494 | 0.003301 | 0.019044 | 40.24217 | 53.99197 | 33.12427 | 55.2079 | 22.42756 | 32.40385 | 17.3572 | 11.9635 |
| ENSMUSG00000038930 | -0.134254158 | 0.149921 | 0.289823 | 311.8769 | 327.5513 | 315.5086 | 299.3304 | 348.1147 | 351.9728 | 347.1441 | 327.3649 |
| ENSMUSG00000038932 | 2.610058434 | 3.32E-10 | 4.78E-08 | 144.8718 | 175.1739 | 131.669 | 166.4863 | 23.40267 | 51.39921 | 9.642892 | 17.40146 |
| ENSMUSG00000038936 | 0.058221345 | 0.439158 | 0.602565 | 1149.92 | 1238.216 | 1079.851 | 1223.2 | 1119.428 | 1138.604 | 1175.468 | 1068.014 |
| ENSMUSG00000038943 | -0.083760409 | 0.411884 | 0.577639 | 1619.748 | 1576.565 | 1268.659 | 1544.096 | 1754.225 | 1554.267 | 1509.113 | 1546.555 |
| ENSMUSG00000038949 | -0.021795322 | 0.79577 | 0.874471 | 746.4923 | 680.2988 | 680.7037 | 725.4664 | 695.2544 | 674.894 | 719.3597 | 787.4159 |
| ENSMUSG00000038954 | 0.126790344 | 0.246047 | 0.408918 | 417.5126 | 430.7359 | 484.4424 | 412.334 | 360.7912 | 419.0153 | 436.823 | 382.8321 |
| ENSMUSG00000038957 | -0.017571835 | 0.828925 | 0.896493 | 1310.889 | 1258.613 | 1211.52 | 1243.903 | 1343.703 | 1124.078 | 1342.291 | 1272.482 |
| ENSMUSG00000038963 | -1.844700665 | 9.48E-06 | 0.000218 | 65.39353 | 47.99286 | 139.1219 | 36.23019 | 227.2009 | 167.6061 | 258.4295 | 387.1824 |
| ENSMUSG00000038965 | -0.030980491 | 0.553313 | 0.700407 | 3437.688 | 3691.851 | 3719.855 | 3702.38 | 3842.914 | 3623.644 | 3652.727 | 3748.926 |
| ENSMUSG00000038967 | -0.670675721 | 1.45E-05 | 0.000303 | 1049.315 | 920.2631 | 1146.928 | 1003.231 | 1558.228 | 1217.938 | 1830.221 | 1953.314 |
| ENSMUSG00000038970 | -0.043330674 | 0.664666 | 0.784591 | 911.4852 | 734.2908 | 871.9963 | 791.0257 | 776.1886 | 849.2043 | 908.3604 | 878.7736 |
| ENSMUSG00000038975 | -0.017206079 | 0.829535 | 0.896882 | 1799.831 | 1720.544 | 1798.648 | 1859.816 | 1949.248 | 1612.371 | 1788.756 | 1915.248 |
| ENSMUSG00000038976 | 0.088484297 | 0.367047 | 0.535638 | 2659.002 | 2352.85 | 2943.091 | 2666.369 | 2353.919 | 2289.5 | 2582.366 | 2767.919 |
| ENSMUSG00000038980 | -0.539343204 | 0.319738 | 0.487813 | 23.13925 | 15.59768 | 14.90592 | 8.626235 | 21.45245 | 10.05637 | 17.3572 | 41.32846 |
| ENSMUSG00000038982 | -0.28500425 | 0.000615 | 0.005376 | 544.2754 | 573.5147 | 512.598 | 543.4528 | 678.6775 | 610.0863 | 661.5024 | 693.8831 |
| ENSMUSG00000038984 | -0.000962101 | 0.994627 | 0.996842 | 179.0777 | 220.7672 | 187.1521 | 175.1126 | 180.3956 | 197.7752 | 212.1436 | 169.6642 |
| ENSMUSG00000038987 | -0.651486655 | 0.121773 | 0.251139 | 29.17558 | 10.79839 | 24.8432 | 11.21411 | 25.35289 | 33.52122 | 30.85725 | 30.45255 |
| ENSMUSG00000038990 | 0.031814537 | 0.716585 | 0.822622 | 728.3834 | 623.9072 | 631.0173 | 698.725 | 637.7228 | 652.5465 | 694.2882 | 639.5036 |
| ENSMUSG00000038991 | -0.186322369 | 0.015442 | 0.058895 | 7022.259 | 7480.887 | 8086.461 | 7252.938 | 8183.134 | 7833.91 | 8792.389 | 9148.816 |
| ENSMUSG00000039000 | -0.161961587 | 0.022918 | 0.077773 | 1699.226 | 1712.145 | 1588.309 | 1510.454 | 1845.886 | 1797.855 | 1723.185 | 1911.985 |
| ENSMUSG00000039001 | -0.104949851 | 0.179793 | 0.329705 | 11880.5 | 13681.56 | 13450.11 | 12408.84 | 14982.59 | 13066.57 | 14227.12 | 13020.64 |
| ENSMUSG00000039004 | -0.092379198 | 0.610608 | 0.745576 | 317.9132 | 239.9643 | 366.8512 | 326.9343 | 258.4045 | 322.9211 | 390.5371 | 365.4306 |
| ENSMUSG00000039005 | 0.422229796 | 0.009256 | 0.040241 | 150.9082 | 146.3782 | 163.137 | 150.9591 | 110.1876 | 113.9722 | 95.46463 | 138.1241 |
| ENSMUSG00000039007 | -0.453231395 | 0.007162 | 0.033436 | 525.1604 | 473.9295 | 732.0463 | 540.0023 | 720.6073 | 650.3117 | 826.3958 | 915.7517 |
| ENSMUSG00000039016 | -0.482148635 | 1.36E-05 | 0.000292 | 1057.363 | 1039.045 | 1183.364 | 1076.554 | 1462.667 | 1282.745 | 1593.97 | 1747.759 |
| ENSMUSG00000039018 | -0.143557338 | 0.191856 | 0.345015 | 549.3057 | 567.5156 | 535.785 | 550.3538 | 628.9468 | 509.5226 | 691.3953 | 600.3503 |
| ENSMUSG00000039021 | 0.217786547 | 0.733393 | 0.833185 | 5.030272 | 11.99822 | 16.56213 | 13.80198 | 9.751113 | 20.11273 | 4.821446 | 6.525547 |
| ENSMUSG00000039031 | -0.087477906 | 0.339351 | 0.507871 | 1371.252 | 1257.413 | 1281.081 | 1411.252 | 1259.844 | 1403.422 | 1407.862 | 1585.708 |
| ENSMUSG00000039032 | 0.796209347 | 0.012434 | 0.050314 | 100.6054 | 101.9848 | 57.13936 | 99.2017 | 55.58134 | 73.74669 | 32.78583 | 44.59123 |
| ENSMUSG00000039033 | -0.028912159 | 0.819653 | 0.890713 | 416.5065 | 435.5352 | 374.3042 | 414.9219 | 402.721 | 481.5882 | 441.6444 | 346.9416 |
| ENSMUSG00000039037 | 1.016261069 | 0.000181 | 0.0021 | 363.1856 | 284.3577 | 206.1986 | 416.6471 | 161.8685 | 212.3011 | 147.5362 | 106.5839 |
| ENSMUSG00000039041 | -0.370180512 | 0.000107 | 0.001418 | 1694.196 | 1721.744 | 1838.397 | 1741.637 | 2361.72 | 1893.949 | 2336.473 | 2450.343 |
| ENSMUSG00000039043 | -0.164533196 | 0.153621 | 0.294937 | 882.3097 | 879.4692 | 916.714 | 802.2398 | 1041.419 | 805.6267 | 945.9677 | 1107.168 |
| ENSMUSG00000039046 | 0.164317629 | 0.056041 | 0.147522 | 1839.067 | 1682.15 | 1576.715 | 1675.215 | 1515.323 | 1678.296 | 1423.291 | 1426.92 |
| ENSMUSG00000039047 | -0.20308581 | 0.074086 | 0.179534 | 898.4065 | 951.4585 | 1170.115 | 1006.682 | 1053.12 | 1084.97 | 1246.826 | 1253.993 |
| ENSMUSG00000039048 | -0.454639804 | 0.000385 | 0.003751 | 833.013 | 829.0767 | 880.2773 | 778.949 | 1167.208 | 878.256 | 1266.112 | 1238.766 |
| ENSMUSG00000039050 | -0.188344938 | 0.028625 | 0.091433 | 1030.2 | 1027.047 | 1084.82 | 1077.417 | 1310.55 | 1053.684 | 1237.183 | 1206.139 |
| ENSMUSG00000039055 | -0.10150048 | 0.472959 | 0.632348 | 316.9071 | 363.5459 | 261.6817 | 307.9566 | 383.2187 | 299.4563 | 337.5012 | 316.489 |
| ENSMUSG00000039057 | 0.870341241 | 0.002029 | 0.013272 | 108.6539 | 79.18822 | 90.26362 | 125.0804 | 46.80534 | 73.74669 | 64.60737 | 35.89051 |
| ENSMUSG00000039058 | -0.144490785 | 0.313182 | 0.481393 | 270.6286 | 265.1606 | 246.7758 | 245.8477 | 291.5583 | 345.2686 | 261.3224 | 238.1824 |
| ENSMUSG00000039062 | -2.370201944 | 3.45E-07 | 1.54E-05 | 1033.218 | 1096.637 | 1714.181 | 1178.344 | 5110.558 | 2844.834 | 6095.272 | 11917.82 |
| ENSMUSG00000039063 | -0.854484483 | 0.000139 | 0.001728 | 159.9626 | 131.9804 | 127.5284 | 131.1188 | 235.9769 | 163.1366 | 266.1438 | 329.5401 |
| ENSMUSG00000039065 | -0.198004555 | 0.073507 | 0.178552 | 306.8466 | 311.9536 | 293.1497 | 263.9628 | 339.3387 | 305.0431 | 362.5727 | 338.2408 |
| ENSMUSG00000039067 | -0.032259956 | 0.574478 | 0.716825 | 3701.274 | 3834.63 | 3445.752 | 3661.837 | 3901.42 | 3736.499 | 3597.763 | 3732.613 |
| ENSMUSG00000039068 | -0.034933827 | 0.501464 | 0.657789 | 2752.565 | 2842.377 | 2753.455 | 2764.708 | 2952.637 | 2867.182 | 2873.582 | 2687.438 |
| ENSMUSG00000039069 | -0.018273829 | 0.869341 | 0.921582 | 376.2643 | 463.1311 | 453.8024 | 373.516 | 432.9494 | 422.3674 | 406.93 | 424.1605 |
| ENSMUSG00000039081 | 0.035717205 | 0.693432 | 0.806121 | 4526.239 | 4505.33 | 5160.761 | 4269.124 | 4027.21 | 4532.069 | 4545.659 | 4908.299 |
| ENSMUSG00000039084 | 0.007908578 | 0.991187 | 0.994852 | 12.07265 | 4.799286 | 14.07781 | 12.07673 | 5.850668 | 16.76061 | 18.32149 | 2.175182 |
| ENSMUSG00000039086 | 0.002935928 | 0.98187 | 0.989807 | 414.4944 | 383.9429 | 316.3367 | 361.4392 | 387.1192 | 403.372 | 360.6441 | 319.7518 |
| ENSMUSG00000039087 | 0.13588417 | 0.232305 | 0.392945 | 799.8132 | 627.5067 | 799.951 | 722.0159 | 633.8224 | 693.8893 | 619.0736 | 741.7371 |
| ENSMUSG00000039089 | 0.68163725 | 4.06E-06 | 0.000114 | 1464.815 | 1570.566 | 1167.63 | 1457.834 | 866.874 | 1088.322 | 854.3602 | 718.8977 |
| ENSMUSG00000039092 | 0.992888324 | 0.008178 | 0.036876 | 54.32694 | 63.59054 | 97.71658 | 62.10889 | 57.53157 | 20.11273 | 27.0001 | 34.80291 |
| ENSMUSG00000039096 | 0.230259826 | 0.13025 | 0.262661 | 479.8879 | 417.5379 | 378.4447 | 452.8773 | 419.2979 | 425.7195 | 304.7154 | 324.1021 |
| ENSMUSG00000039099 | 0.900284531 | 0.243612 | 0.406189 | 34.20585 | 5.999108 | 4.140533 | 24.15346 | 5.850668 | 8.938993 | 5.785735 | 16.31387 |
| ENSMUSG00000039100 | -0.045432482 | 0.629893 | 0.759023 | 2894.418 | 2376.846 | 2462.789 | 2621.513 | 2446.554 | 2561.021 | 2726.045 | 2954.985 |
| ENSMUSG00000039103 | 0.434023456 | 0.023065 | 0.078219 | 646.893 | 805.0802 | 808.2321 | 583.9961 | 425.1485 | 710.6499 | 449.3588 | 520.9561 |
| ENSMUSG00000039105 | -0.251404811 | 0.001447 | 0.010317 | 3170.077 | 3001.953 | 3251.975 | 3271.068 | 3596.211 | 3480.62 | 3822.442 | 4216.591 |
| ENSMUSG00000039108 | -0.015744421 | 0.840596 | 0.903871 | 1986.957 | 1985.705 | 1972.55 | 2166.91 | 2085.763 | 2260.448 | 1919.9 | 1938.087 |
| ENSMUSG00000039109 | 0.899869858 | 4.31E-06 | 0.00012 | 709.2683 | 799.0811 | 660.8291 | 778.0864 | 445.6259 | 429.0717 | 459.9659 | 243.6204 |
| ENSMUSG00000039110 | -0.246654785 | 0.304031 | 0.472601 | 93.56306 | 76.78858 | 121.7317 | 76.77349 | 119.9387 | 78.21619 | 119.5719 | 120.7226 |
| ENSMUSG00000039114 | 0.10652661 | 0.862105 | 0.917537 | 23.13925 | 25.19625 | 12.4216 | 54.34528 | 21.45245 | 55.8687 | 11.57147 | 18.48905 |

| | | | | | | | | | | | |
|---|---|---|---|---|---|---|---|---|---|---|---|
| ENSMUSG00000039115 | 0.276738828 | 0.004783 | 0.024944 | 3732.462 | 4770.49 | 4586.055 | 4012.924 | 3338.781 | 3821.419 | 3453.119 | 3503.131 |
| ENSMUSG00000039116 | 0.245442704 | 0.100241 | 0.221292 | 694.1775 | 641.9045 | 953.9788 | 722.8785 | 551.913 | 716.2368 | 645.1095 | 631.8904 |
| ENSMUSG00000039117 | 0.21787988 | 0.015062 | 0.057854 | 784.7224 | 724.6922 | 734.5306 | 835.0195 | 645.5237 | 722.941 | 662.4667 | 618.8393 |
| ENSMUSG00000039126 | 0.890889831 | 8.18E-05 | 0.001157 | 287.7315 | 208.7689 | 240.1509 | 257.0618 | 112.1378 | 197.7752 | 109.929 | 117.4598 |
| ENSMUSG00000039128 | -0.111183021 | 0.151814 | 0.292289 | 2294.81 | 2735.593 | 2264.872 | 2435.186 | 2698.133 | 2582.252 | 2576.581 | 2646.109 |
| ENSMUSG00000039130 | -0.071313701 | 0.487301 | 0.645251 | 971.8485 | 1091.838 | 1086.476 | 1128.312 | 1266.67 | 1162.069 | 1089.647 | 976.6568 |
| ENSMUSG00000039131 | -1.951632197 | 1.20E-05 | 0.000265 | 145.8779 | 63.59054 | 319.6492 | 138.8824 | 697.2046 | 290.5173 | 652.8238 | 948.3794 |
| ENSMUSG00000039137 | 0.291729921 | 0.013948 | 0.054589 | 675.0625 | 662.3015 | 578.0184 | 669.3958 | 602.6188 | 565.3913 | 472.5017 | 469.8393 |
| ENSMUSG00000039145 | 0.476893817 | 0.002105 | 0.013614 | 251.5136 | 278.3586 | 348.6329 | 260.5123 | 182.3458 | 196.6578 | 231.4294 | 208.8175 |
| ENSMUSG00000039148 | -0.088953247 | 0.089467 | 0.204231 | 2754.577 | 2695.999 | 2924.045 | 2842.344 | 2914.608 | 2939.811 | 2998.939 | 3084.408 |
| ENSMUSG00000039153 | 0.775958034 | 0.000164 | 0.001939 | 522.1422 | 622.7074 | 446.3495 | 555.5295 | 328.6125 | 398.9026 | 323.0369 | 202.2919 |
| ENSMUSG00000039154 | -0.751110407 | 0.006675 | 0.031889 | 340.0464 | 218.3675 | 379.2728 | 351.9504 | 470.9788 | 341.9165 | 521.6804 | 838.5327 |
| ENSMUSG00000039155 | -0.125269316 | 0.812107 | 0.885673 | 11.0666 | 9.598572 | 9.93728 | 7.763611 | 10.72622 | 6.704245 | 16.39292 | 7.613138 |
| ENSMUSG00000039156 | 0.260091181 | 0.001387 | 0.009988 | 1033.218 | 1103.836 | 1098.898 | 1186.107 | 1006.315 | 918.4815 | 875.5746 | 892.9123 |
| ENSMUSG00000039157 | -0.099424779 | 0.373757 | 0.542513 | 1077.484 | 956.2578 | 1170.943 | 991.1544 | 1011.19 | 1030.219 | 1184.147 | 1272.482 |
| ENSMUSG00000039158 | 0.30303807 | 0.004353 | 0.023267 | 274.6528 | 289.157 | 300.6027 | 272.589 | 223.3005 | 222.3574 | 243.9652 | 231.6569 |
| ENSMUSG00000039159 | 0.282254476 | 1.40E-05 | 0.000296 | 3468.875 | 3779.438 | 3439.127 | 3420.302 | 2723.486 | 3072.779 | 2892.867 | 2909.306 |
| ENSMUSG00000039163 | -0.0565121 | 0.52628 | 0.679434 | 626.7719 | 559.1168 | 567.253 | 627.1273 | 575.3157 | 667.0723 | 619.0736 | 616.6641 |
| ENSMUSG00000039164 | -0.16734486 | 0.300935 | 0.469291 | 148.896 | 172.7743 | 147.403 | 119.9047 | 148.2169 | 183.2494 | 165.8577 | 162.0511 |
| ENSMUSG00000039166 | 0.12104832 | 0.169216 | 0.315546 | 719.3289 | 776.2845 | 682.3599 | 747.8946 | 665.0259 | 709.5326 | 704.8954 | 607.9634 |
| ENSMUSG00000039167 | -0.504159246 | 0.01415 | 0.055247 | 536.227 | 391.1418 | 723.7652 | 431.3117 | 744.0099 | 591.0909 | 784.9314 | 836.3575 |
| ENSMUSG00000039168 | 0.077224269 | 0.330484 | 0.498318 | 2072.472 | 2330.053 | 2538.147 | 2176.399 | 2213.503 | 2111.837 | 2171.579 | 2145.817 |
| ENSMUSG00000039176 | 0.028423165 | 0.668913 | 0.78769 | 1761.601 | 1642.556 | 1689.338 | 1665.726 | 1752.275 | 1535.272 | 1701.97 | 1635.737 |
| ENSMUSG00000039178 | -0.016220319 | 0.892123 | 0.934822 | 551.3178 | 637.1052 | 558.1439 | 621.9515 | 709.881 | 538.5743 | 549.6448 | 594.9123 |
| ENSMUSG00000039179 | 0.008152588 | 0.985157 | 0.991442 | 26.15741 | 27.5959 | 9.109173 | 22.42821 | 22.42756 | 22.34748 | 15.42863 | 23.927 |
| ENSMUSG00000039182 | -0.10884436 | 0.445181 | 0.608026 | 326.9677 | 367.1454 | 288.1811 | 301.9182 | 358.841 | 278.2261 | 355.8227 | 388.27 |
| ENSMUSG00000039183 | -0.355058248 | 0.000176 | 0.002053 | 762.5892 | 757.0874 | 719.6247 | 687.5109 | 1024.842 | 817.9178 | 929.5748 | 966.8685 |
| ENSMUSG00000039187 | -0.051742401 | 0.633753 | 0.761891 | 279.6831 | 285.5575 | 262.5098 | 262.2375 | 312.0356 | 274.874 | 275.7867 | 264.2846 |
| ENSMUSG00000039191 | 0.148582852 | 0.092268 | 0.208585 | 1649.929 | 1828.528 | 1611.496 | 1726.11 | 1526.049 | 1720.756 | 1515.863 | 1384.503 |
| ENSMUSG00000039193 | -0.091969107 | 0.878015 | 0.927008 | 8.048435 | 3.599465 | 14.90592 | 21.56559 | 15.60178 | 12.29112 | 15.42863 | 8.700729 |
| ENSMUSG00000039194 | 0.557770861 | 0.003037 | 0.017935 | 113.6841 | 163.1757 | 122.5598 | 138.8824 | 78.0089 | 100.5637 | 87.75031 | 98.97079 |
| ENSMUSG00000039195 | -0.527642045 | 0.001476 | 0.010446 | 294.7739 | 235.165 | 329.5864 | 261.3749 | 364.6916 | 344.1512 | 410.7872 | 499.2043 |
| ENSMUSG00000039197 | -0.375440254 | 2.72E-08 | 1.97E-06 | 1214.308 | 1233.417 | 1174.255 | 1169.717 | 1517.273 | 1445.882 | 1630.613 | 1617.248 |
| ENSMUSG00000039199 | 0.30906432 | 0.006054 | 0.029652 | 1774.68 | 1677.351 | 1541.935 | 1753.714 | 1494.846 | 1536.389 | 1255.504 | 1159.372 |
| ENSMUSG00000039200 | -0.283144095 | 0.256336 | 0.42008 | 53.32088 | 45.59322 | 38.92101 | 40.5433 | 47.78045 | 67.04245 | 54.96448 | 46.76642 |
| ENSMUSG00000039201 | -0.133162587 | 0.320735 | 0.488689 | 226.3622 | 238.7645 | 242.6352 | 241.5346 | 265.2303 | 216.7706 | 253.6081 | 305.6131 |
| ENSMUSG00000039202 | 0.162943456 | 0.381713 | 0.549695 | 3231.447 | 1962.908 | 1994.909 | 1990.072 | 1958.024 | 1682.765 | 2233.294 | 2324.182 |
| ENSMUSG00000039205 | 0.079274386 | 0.218173 | 0.376811 | 1452.742 | 1421.789 | 1411.922 | 1294.798 | 1329.077 | 1333.027 | 1339.398 | 1277.92 |
| ENSMUSG00000039206 | 0.176013738 | 0.14782 | 0.287144 | 179.0777 | 195.5709 | 190.4645 | 178.5631 | 164.7938 | 170.9582 | 160.072 | 162.0511 |
| ENSMUSG00000039208 | -0.417278387 | 0.075197 | 0.181251 | 225.3562 | 267.5602 | 323.7897 | 266.5507 | 373.4676 | 229.0617 | 342.3227 | 502.4671 |
| ENSMUSG00000039210 | 0.049149635 | 0.581981 | 0.722845 | 759.571 | 626.3068 | 640.9545 | 680.6099 | 632.8472 | 663.7202 | 655.7166 | 665.6057 |
| ENSMUSG00000039217 | 0.589706968 | 0.113332 | 0.240365 | 38.23007 | 69.58965 | 43.88965 | 69.00988 | 29.25334 | 60.3382 | 27.96439 | 29.36496 |
| ENSMUSG00000039218 | 0.20962798 | 0.016341 | 0.061103 | 8757.703 | 7922.422 | 8706.713 | 8819.463 | 6953.519 | 8332.259 | 7616.92 | 6681.072 |
| ENSMUSG00000039219 | 0.499740788 | 0.005355 | 0.026935 | 3259.616 | 3225.12 | 2370.869 | 3075.253 | 1793.23 | 2909.642 | 1927.614 | 1807.576 |
| ENSMUSG00000039220 | -0.099859055 | 0.378386 | 0.546525 | 1294.792 | 1283.809 | 1251.269 | 1058.439 | 1258.869 | 1125.196 | 1384.719 | 1467.16 |
| ENSMUSG00000039221 | 0.118810494 | 0.299302 | 0.467416 | 5601.711 | 6218.675 | 5205.478 | 6196.225 | 5612.741 | 6200.309 | 5024.911 | 4547.218 |
| ENSMUSG00000039230 | -0.235910434 | 0.058598 | 0.152065 | 1243.483 | 1362.997 | 1445.874 | 1212.849 | 1573.83 | 1229.112 | 1676.899 | 1719.482 |
| ENSMUSG00000039231 | -0.022555846 | 0.796806 | 0.875296 | 1342.077 | 1362.997 | 1276.112 | 1331.891 | 1457.791 | 1438.06 | 1323.005 | 1175.686 |
| ENSMUSG00000039232 | 0.542991659 | 0.020966 | 0.072885 | 131.7931 | 115.1829 | 157.3403 | 122.4925 | 128.7147 | 74.86406 | 83.89316 | 73.95619 |
| ENSMUSG00000039233 | 0.257649581 | 0.01445 | 0.056123 | 1323.968 | 1366.597 | 1196.614 | 1319.814 | 1099.926 | 1234.698 | 1101.218 | 917.9269 |
| ENSMUSG00000039234 | 0.239687743 | 0.003797 | 0.021024 | 1630.814 | 1733.742 | 1898.849 | 1762.34 | 1432.439 | 1375.488 | 1529.363 | 1615.073 |
| ENSMUSG00000039236 | 0.272623455 | 0.419489 | 0.585046 | 55.33299 | 40.79393 | 46.37397 | 81.08661 | 49.73068 | 65.92507 | 36.64299 | 33.71532 |
| ENSMUSG00000039238 | -0.895445849 | 0.014225 | 0.05547 | 28.16952 | 13.19804 | 28.98373 | 18.97772 | 53.63112 | 29.05173 | 45.32159 | 39.15328 |
| ENSMUSG00000039239 | 0.507428907 | 0.000123 | 0.001572 | 798.8072 | 819.4781 | 935.7605 | 835.0195 | 544.1121 | 730.7627 | 607.5022 | 503.5547 |
| ENSMUSG00000039242 | 0.176977048 | 0.107736 | 0.232341 | 1401.434 | 1443.385 | 1127.053 | 1354.319 | 1157.457 | 1298.389 | 1207.29 | 1045.175 |
| ENSMUSG00000039244 | -0.293111584 | 0.016646 | 0.061903 | 549.3057 | 680.2988 | 687.3285 | 584.8587 | 821.0437 | 683.8329 | 704.8954 | 854.8466 |
| ENSMUSG00000039246 | -0.222957002 | 0.07403 | 0.179481 | 495.9848 | 477.529 | 467.8802 | 406.2957 | 636.7477 | 535.2222 | 483.1089 | 499.2043 |
| ENSMUSG00000039252 | -0.635081223 | 0.015049 | 0.057816 | 223.3441 | 255.562 | 486.9267 | 271.7264 | 368.5921 | 439.128 | 542.8948 | 574.2481 |
| ENSMUSG00000039253 | -0.118905193 | 0.270182 | 0.435158 | 312.8829 | 317.9527 | 329.5864 | 305.3687 | 329.5876 | 311.7474 | 345.2155 | 388.27 |
| ENSMUSG00000039254 | -0.28457415 | 0.00614 | 0.029934 | 493.9727 | 453.5325 | 476.1613 | 473.5803 | 589.9423 | 503.9357 | 564.1092 | 653.6422 |
| ENSMUSG00000039257 | 1.307343247 | 3.97E-06 | 0.000112 | 583.5115 | 854.2729 | 654.2042 | 835.8822 | 257.4294 | 496.1141 | 243.0009 | 187.0657 |
| ENSMUSG00000039262 | 0.218508489 | 0.003074 | 0.0181 | 4211.344 | 4277.364 | 4164.548 | 4227.718 | 3514.301 | 4096.293 | 3481.084 | 3418.299 |
| ENSMUSG00000039263 | 0.041581924 | 0.405475 | 0.572348 | 1786.753 | 1770.937 | 1748.133 | 1736.461 | 1748.375 | 1677.179 | 1679.792 | 1734.708 |
| ENSMUSG00000039270 | 0.251213167 | 0.002015 | 0.013208 | 1573.469 | 1557.368 | 1500.529 | 1588.952 | 1168.183 | 1388.896 | 1408.826 | 1260.518 |

| | | | | | | | | | | | |
|---|---|---|---|---|---|---|---|---|---|---|---|
| ENSMUSG00000039275 | -0.063940645 | 0.345164 | 0.513945 | 1376.282 | 1346.2 | 1469.061 | 1302.561 | 1411.961 | 1460.408 | 1377.969 | 1496.525 |
| ENSMUSG00000039278 | 1.606980411 | 0.006801 | 0.032317 | 156.9445 | 32.39518 | 38.92101 | 244.1224 | 50.70579 | 33.52122 | 31.82154 | 39.15328 |
| ENSMUSG00000039285 | 0.127823539 | 0.059325 | 0.15354 | 2027.2 | 1964.108 | 1967.581 | 2192.789 | 1800.055 | 1986.691 | 1820.578 | 1857.606 |
| ENSMUSG00000039286 | 0.139088557 | 0.12323 | 0.252844 | 2476.906 | 2139.282 | 2315.386 | 2137.581 | 1823.458 | 2221.34 | 2141.686 | 2051.197 |
| ENSMUSG00000039294 | 0.160395248 | 0.038182 | 0.112841 | 952.7335 | 923.8626 | 822.3099 | 860.8982 | 799.5913 | 798.9225 | 810.0029 | 772.1897 |
| ENSMUSG00000039298 | 0.058127511 | 0.406645 | 0.573149 | 1866.231 | 1698.947 | 1670.291 | 1715.758 | 1703.519 | 1774.39 | 1606.506 | 1592.233 |
| ENSMUSG00000039304 | -0.2850258 | 0.316405 | 0.484686 | 207.2472 | 177.5736 | 338.6956 | 177.7004 | 279.8569 | 186.6015 | 244.9294 | 388.27 |
| ENSMUSG00000039307 | 0.066845274 | 0.502496 | 0.658496 | 723.3531 | 820.6779 | 727.9057 | 732.3673 | 683.553 | 816.8005 | 652.8238 | 714.5473 |
| ENSMUSG00000039308 | 0.003343924 | 0.955929 | 0.975 | 1042.272 | 1095.437 | 1019.399 | 1033.423 | 1052.145 | 1044.745 | 1059.754 | 1019.073 |
| ENSMUSG00000039316 | -0.355577421 | 0.025237 | 0.083544 | 207.2472 | 266.3604 | 262.5098 | 192.365 | 279.8569 | 280.4609 | 278.6796 | 348.0291 |
| ENSMUSG00000039318 | -0.094574393 | 0.219141 | 0.377925 | 1188.15 | 1063.042 | 1121.256 | 1104.158 | 1182.81 | 1099.496 | 1283.469 | 1213.752 |
| ENSMUSG00000039323 | 0.164998407 | 0.04496 | 0.126415 | 3063.436 | 3016.351 | 3600.608 | 3140.812 | 2799.545 | 2723.041 | 3083.797 | 2832.087 |
| ENSMUSG00000039328 | 0.126453056 | 0.232554 | 0.393087 | 815.9101 | 1016.249 | 838.872 | 758.2461 | 767.4126 | 773.2229 | 794.5743 | 801.5546 |
| ENSMUSG00000039345 | -0.109795837 | 0.415024 | 0.580938 | 201.2109 | 249.5629 | 209.511 | 207.0296 | 253.5289 | 202.2447 | 245.8937 | 230.5693 |
| ENSMUSG00000039347 | 0.160668225 | 0.106155 | 0.230174 | 774.6619 | 859.0722 | 720.4528 | 875.5628 | 768.3877 | 757.5796 | 670.181 | 691.7079 |
| ENSMUSG00000039349 | -0.781623737 | 0.000933 | 0.007425 | 1165.011 | 893.867 | 1755.586 | 1307.737 | 2128.668 | 1473.816 | 2344.187 | 2862.54 |
| ENSMUSG00000039354 | 0.042298663 | 0.541886 | 0.691157 | 1149.92 | 1076.24 | 1005.321 | 1065.34 | 1024.842 | 1072.679 | 1047.218 | 1025.598 |
| ENSMUSG00000039356 | -0.231586333 | 0.002059 | 0.013396 | 863.1946 | 1010.25 | 942.3853 | 923.0071 | 1142.83 | 1050.332 | 1089.647 | 1102.817 |
| ENSMUSG00000039357 | 0.055877617 | 0.46061 | 0.62196 | 856.1523 | 934.661 | 844.6688 | 922.1445 | 886.3762 | 806.7441 | 838.9316 | 887.4743 |
| ENSMUSG00000039361 | 0.010350139 | 0.894601 | 0.936266 | 6292.87 | 5827.533 | 6348.265 | 6766.419 | 5615.666 | 6554.516 | 6391.309 | 6499.444 |
| ENSMUSG00000039364 | -3.139586349 | 5.04E-05 | 0.000798 | 5.030272 | 2.399643 | 0 | 2.58787 | 32.17867 | 11.17374 | 14.46434 | 28.27737 |
| ENSMUSG00000039367 | 0.028260971 | 0.527919 | 0.680556 | 3749.565 | 3751.842 | 3797.697 | 3742.923 | 3579.634 | 3616.94 | 3727.942 | 3827.233 |
| ENSMUSG00000039372 | 1.242210686 | 0.07901 | 0.187542 | 26.15741 | 4.799286 | 5.796746 | 21.56559 | 6.825779 | 5.58687 | 5.785735 | 6.525547 |
| ENSMUSG00000039375 | 0.255473835 | 0.268672 | 0.43328 | 61.36932 | 64.79036 | 67.90474 | 58.6584 | 53.63112 | 61.45558 | 38.57157 | 58.72992 |
| ENSMUSG00000039376 | 0.491275634 | 0.171821 | 0.319092 | 35.2119 | 29.99554 | 29.81184 | 26.74133 | 17.552 | 22.34748 | 14.46434 | 32.62773 |
| ENSMUSG00000039377 | -0.390442985 | 0.090547 | 0.206111 | 147.89 | 115.1829 | 134.9814 | 100.0643 | 119.9387 | 210.0663 | 184.1792 | 139.2117 |
| ENSMUSG00000039382 | 0.186906275 | 0.044159 | 0.124893 | 1205.253 | 1129.032 | 1017.743 | 1187.833 | 1046.294 | 949.768 | 1073.254 | 915.7517 |
| ENSMUSG00000039384 | 0.049710057 | 0.836386 | 0.901206 | 98.59333 | 116.3827 | 171.4181 | 114.7289 | 88.73513 | 136.3196 | 139.8219 | 120.7226 |
| ENSMUSG00000039385 | -0.829658807 | 0.003856 | 0.021278 | 322.9434 | 184.7725 | 339.5237 | 244.1224 | 399.7956 | 293.8694 | 549.6448 | 698.2335 |
| ENSMUSG00000039391 | 0.650387418 | 0.078527 | 0.186692 | 39.23612 | 32.39518 | 25.67131 | 29.3292 | 16.57689 | 27.93435 | 12.53576 | 23.927 |
| ENSMUSG00000039395 | 0.018719342 | 0.909436 | 0.946034 | 132.7992 | 117.5825 | 125.0441 | 143.1955 | 126.7645 | 106.1505 | 147.5362 | 131.5985 |
| ENSMUSG00000039396 | 0.504185292 | 0.010238 | 0.043407 | 952.7335 | 1040.245 | 722.9371 | 920.4193 | 647.4739 | 865.9649 | 585.3235 | 464.4014 |
| ENSMUSG00000039405 | 0.04374324 | 0.651946 | 0.775328 | 1707.274 | 1632.957 | 1544.419 | 1744.225 | 1447.065 | 1623.545 | 1847.578 | 1511.752 |
| ENSMUSG00000039410 | 0.114205251 | 0.352833 | 0.521562 | 843.0736 | 727.0918 | 966.4004 | 808.2782 | 661.1255 | 852.5564 | 805.1815 | 775.4524 |
| ENSMUSG00000039414 | -0.259843853 | 0.131427 | 0.264258 | 527.1725 | 387.5424 | 421.5063 | 366.615 | 450.5014 | 414.5458 | 570.8592 | 602.5255 |
| ENSMUSG00000039419 | 0.681218517 | 1.81E-06 | 5.99E-05 | 226.3622 | 256.7618 | 202.058 | 212.2054 | 137.4907 | 156.4324 | 135.9648 | 128.3357 |
| ENSMUSG00000039427 | -0.233865122 | 0.082827 | 0.19355 | 582.5055 | 559.1168 | 635.1578 | 554.6669 | 612.3699 | 567.626 | 759.8599 | 802.6422 |
| ENSMUSG00000039428 | -0.428935314 | 0.001026 | 0.00799 | 1030.2 | 835.0758 | 1095.585 | 974.7645 | 1161.358 | 1174.36 | 1494.648 | 1471.511 |
| ENSMUSG00000039431 | 0.573163973 | 0.000376 | 0.003675 | 367.2098 | 380.3434 | 276.5876 | 365.7524 | 223.3005 | 286.0478 | 210.215 | 214.2554 |
| ENSMUSG00000039438 | -3.997899626 | 2.83E-09 | 3.09E-07 | 82.49646 | 26.39607 | 188.8083 | 86.26235 | 1553.352 | 545.2786 | 1243.933 | 2802.722 |
| ENSMUSG00000039449 | 0.057725387 | 0.481571 | 0.640521 | 1579.505 | 1341.4 | 1360.579 | 1471.636 | 1384.658 | 1461.525 | 1392.434 | 1289.883 |
| ENSMUSG00000039450 | -0.98365184 | 1.93E-05 | 0.000383 | 470.8334 | 341.9491 | 475.3332 | 458.0531 | 966.3353 | 550.8654 | 748.2884 | 1189.825 |
| ENSMUSG00000039452 | 0.26211633 | 0.250852 | 0.414441 | 212.2775 | 295.1561 | 205.3704 | 235.4962 | 172.5947 | 287.1651 | 169.7149 | 160.9635 |
| ENSMUSG00000039456 | 0.191761479 | 0.051193 | 0.138272 | 1117.726 | 1099.037 | 1095.585 | 1154.19 | 949.7584 | 1097.261 | 1029.861 | 834.1824 |
| ENSMUSG00000039457 | -0.654826507 | 0.014592 | 0.056517 | 378.2764 | 442.7341 | 558.972 | 287.2536 | 744.985 | 394.4331 | 875.5746 | 609.051 |
| ENSMUSG00000039458 | -0.181401463 | 0.018116 | 0.065906 | 952.7335 | 884.2685 | 908.433 | 999.7806 | 1060.921 | 986.6413 | 1086.754 | 1114.781 |
| ENSMUSG00000039461 | -0.404617377 | 0.016491 | 0.061485 | 505.0393 | 577.1142 | 575.5341 | 514.1236 | 721.5824 | 531.8701 | 685.6096 | 935.3283 |
| ENSMUSG00000039463 | 0.103933877 | 0.208876 | 0.366165 | 1189.156 | 1232.217 | 1159.349 | 1186.97 | 1230.59 | 1115.139 | 998.0393 | 1090.854 |
| ENSMUSG00000039470 | 0.195711973 | 0.149558 | 0.289495 | 497.9969 | 499.1258 | 392.5225 | 457.1905 | 432.9494 | 459.2408 | 349.0727 | 369.781 |
| ENSMUSG00000039473 | 0.113131196 | 0.180613 | 0.330793 | 2588.578 | 2372.047 | 2378.322 | 2291.991 | 2011.655 | 2482.805 | 2191.829 | 2219.773 |
| ENSMUSG00000039474 | -0.260857107 | 0.134926 | 0.268977 | 506.0453 | 488.3274 | 696.4377 | 475.3055 | 629.9219 | 574.3303 | 573.7521 | 820.0437 |
| ENSMUSG00000039476 | 0.72332256 | 2.00E-07 | 1.00E-05 | 269.6226 | 308.3541 | 278.2438 | 242.3972 | 141.3911 | 176.5451 | 169.7149 | 177.2773 |
| ENSMUSG00000039477 | 0.220404842 | 0.00372 | 0.020734 | 2294.81 | 2078.091 | 2188.686 | 2206.591 | 1852.711 | 2021.33 | 1701.97 | 1954.401 |
| ENSMUSG00000039478 | 0.166916849 | 0.186391 | 0.33832 | 691.1593 | 626.3068 | 628.5329 | 709.0765 | 558.7388 | 729.6453 | 536.1448 | 542.708 |
| ENSMUSG00000039480 | -0.053926599 | 0.606256 | 0.742598 | 376.2643 | 430.7359 | 433.9279 | 431.3117 | 480.7299 | 394.4331 | 426.2158 | 433.9488 |
| ENSMUSG00000039481 | 0.077564938 | 0.529994 | 0.681935 | 261.5741 | 303.5548 | 250.0882 | 269.1385 | 260.3547 | 232.4138 | 286.3939 | 244.708 |
| ENSMUSG00000039483 | -0.224866244 | 0.017067 | 0.063045 | 713.2925 | 623.9072 | 632.6735 | 609.8748 | 789.8402 | 674.894 | 790.7171 | 756.9634 |
| ENSMUSG00000039485 | 0.386530648 | 0.000409 | 0.00393 | 324.9556 | 313.1534 | 310.54 | 343.3242 | 233.0516 | 254.7613 | 230.4651 | 271.8978 |
| ENSMUSG00000039488 | 0.830947955 | 0.042023 | 0.12073 | 45.27245 | 52.79215 | 32.29616 | 53.48266 | 31.20356 | 41.34284 | 13.50005 | 17.40146 |
| ENSMUSG00000039492 | -0.287527317 | 0.546785 | 0.6951 | 10.06054 | 20.39697 | 43.88965 | 17.25247 | 25.35289 | 32.40385 | 29.89296 | 25.0146 |
| ENSMUSG00000039496 | -0.646554975 | 0.145462 | 0.28379 | 9.054489 | 23.99643 | 19.04645 | 10.35148 | 17.552 | 24.58223 | 30.85725 | 23.927 |
| ENSMUSG00000039497 | 0.157455817 | 0.15471 | 0.296344 | 726.3712 | 950.2587 | 914.2297 | 906.6173 | 818.1184 | 798.9225 | 706.824 | 812.4305 |
| ENSMUSG00000039501 | -0.043561485 | 0.786109 | 0.868451 | 695.1836 | 693.4968 | 1027.68 | 664.2201 | 843.4713 | 716.2368 | 754.0741 | 863.5473 |
| ENSMUSG00000039509 | -0.047805866 | 0.484101 | 0.642397 | 1127.787 | 1285.009 | 1194.13 | 1176.618 | 1216.939 | 1290.567 | 1232.362 | 1201.788 |

| | | | | | | | | | | | |
|---|---|---|---|---|---|---|---|---|---|---|---|
| ENSMUSG00000039512 | -0.159218404 | 0.225017 | 0.384774 | 812.8919 | 848.2738 | 694.7815 | 647.8302 | 940.9824 | 730.7627 | 774.3242 | 903.7882 |
| ENSMUSG00000039515 | -0.215310421 | 0.00558 | 0.027863 | 4124.823 | 4349.353 | 4740.082 | 4426.984 | 5178.816 | 4592.408 | 5319.983 | 5391.189 |
| ENSMUSG00000039519 | 0.670823749 | 0.000118 | 0.001515 | 614.6992 | 699.496 | 671.5945 | 673.7089 | 351.0401 | 582.1519 | 381.8585 | 356.7299 |
| ENSMUSG00000039523 | -0.134495363 | 0.219766 | 0.378606 | 702.2259 | 628.7065 | 558.972 | 653.006 | 780.089 | 627.9642 | 695.2525 | 685.1824 |
| ENSMUSG00000039529 | -0.359900212 | 0.066987 | 0.167136 | 283.7073 | 278.3586 | 427.303 | 237.2215 | 403.6961 | 357.5597 | 363.537 | 451.3503 |
| ENSMUSG00000039530 | -0.087695481 | 0.168475 | 0.314619 | 2021.163 | 2188.474 | 2293.027 | 2213.492 | 2347.093 | 2316.317 | 2389.509 | 2211.073 |
| ENSMUSG00000039531 | 0.334088064 | 0.020193 | 0.070946 | 310.8708 | 355.1472 | 261.6817 | 335.5605 | 249.6285 | 286.0478 | 220.8222 | 244.708 |
| ENSMUSG00000039533 | 0.899631807 | 0.034676 | 0.10533 | 31.18769 | 25.19625 | 30.63995 | 40.5433 | 17.552 | 12.29112 | 9.642892 | 29.36496 |
| ENSMUSG00000039536 | 0.113760279 | 0.136453 | 0.271172 | 2866.249 | 2725.995 | 2687.206 | 2962.249 | 2469.957 | 2888.412 | 2565.973 | 2467.744 |
| ENSMUSG00000039539 | 0.620090126 | 0.320467 | 0.488405 | 10.06054 | 15.59768 | 13.24971 | 11.21411 | 5.850668 | 17.87799 | 5.785735 | 3.262773 |
| ENSMUSG00000039542 | 0.11085561 | 0.310763 | 0.479392 | 3982.969 | 3623.461 | 4549.618 | 4061.231 | 3646.916 | 4403.571 | 3585.227 | 3386.759 |
| ENSMUSG00000039543 | 0.350355851 | 0.279475 | 0.445876 | 40.24217 | 40.79393 | 49.6864 | 61.24627 | 27.30312 | 29.05173 | 39.53586 | 55.46715 |
| ENSMUSG00000039545 | 0.735372129 | 0.036054 | 0.10855 | 42.25428 | 93.58608 | 50.5145 | 45.71905 | 29.25334 | 44.69496 | 27.96439 | 36.9781 |
| ENSMUSG00000039546 | -1.949828824 | 1.54E-05 | 0.000318 | 50.30272 | 34.79482 | 153.1997 | 49.16954 | 297.4089 | 153.0803 | 364.5013 | 300.1751 |
| ENSMUSG00000039552 | 0.162890201 | 0.814932 | 0.887383 | 15.09082 | 1.199822 | 19.87456 | 13.80198 | 17.552 | 11.17374 | 7.714313 | 8.700729 |
| ENSMUSG00000039556 | 0.140861072 | 0.637389 | 0.764767 | 64.38748 | 80.38804 | 79.49824 | 40.5433 | 56.55646 | 50.28183 | 80.036 | 52.20437 |
| ENSMUSG00000039563 | 0.071155988 | 0.859813 | 0.915943 | 22.1332 | 13.19804 | 18.21835 | 17.25247 | 12.67645 | 21.23011 | 13.50005 | 20.66423 |
| ENSMUSG00000039568 | -0.199754903 | 0.098761 | 0.219219 | 680.0927 | 793.082 | 833.0753 | 711.6644 | 856.1477 | 725.1758 | 906.4318 | 977.7444 |
| ENSMUSG00000039577 | -0.196229937 | 0.2194 | 0.378197 | 124.7507 | 140.3791 | 128.3565 | 106.1027 | 156.9929 | 128.498 | 150.4291 | 133.7737 |
| ENSMUSG00000039578 | -0.530785644 | 0.01729 | 0.063668 | 202.2169 | 182.3729 | 304.7432 | 251.0234 | 287.6578 | 267.0524 | 365.4656 | 441.562 |
| ENSMUSG00000039579 | -0.227825796 | 0.31291 | 0.481091 | 57.3451 | 75.58876 | 86.9512 | 63.83414 | 70.20801 | 80.45093 | 101.2504 | 80.48174 |
| ENSMUSG00000039585 | 0.002815104 | 0.98024 | 0.988885 | 1635.844 | 1517.774 | 1615.636 | 1545.821 | 1410.986 | 1935.292 | 1471.505 | 1487.825 |
| ENSMUSG00000039599 | 0.058420376 | 0.355998 | 0.524617 | 1108.672 | 1175.825 | 1143.615 | 1166.267 | 1153.557 | 1125.196 | 1065.54 | 1066.927 |
| ENSMUSG00000039601 | -0.523178786 | 0.016457 | 0.061433 | 249.5015 | 231.5656 | 441.3808 | 307.094 | 439.7752 | 363.1466 | 460.9302 | 506.8174 |
| ENSMUSG00000039607 | 0.773483648 | 0.000219 | 0.00243 | 2822.989 | 3286.311 | 2587.833 | 3014.869 | 1512.398 | 2604.599 | 1412.684 | 1322.511 |
| ENSMUSG00000039611 | -0.455793208 | 0.011036 | 0.045976 | 433.6094 | 369.545 | 560.6282 | 470.9924 | 630.897 | 467.0624 | 689.4668 | 731.9488 |
| ENSMUSG00000039615 | -0.165665437 | 0.08833 | 0.20241 | 1798.825 | 1912.516 | 1953.504 | 1802.02 | 2085.763 | 1787.799 | 2130.115 | 2370.949 |
| ENSMUSG00000039616 | -0.925890899 | 0.122181 | 0.251727 | 27.16347 | 20.39697 | 34.78048 | 20.70296 | 53.63112 | 6.704245 | 53.0359 | 82.65692 |
| ENSMUSG00000039617 | 0.138216121 | 0.666053 | 0.785506 | 49.29666 | 21.59679 | 49.6864 | 46.58167 | 41.92979 | 35.75597 | 40.50014 | 34.80291 |
| ENSMUSG00000039620 | -0.108864641 | 0.568241 | 0.712463 | 220.3259 | 170.3747 | 260.8536 | 282.9405 | 250.6036 | 240.2354 | 301.8225 | 217.5182 |
| ENSMUSG00000039621 | 0.085763615 | 0.371715 | 0.540783 | 616.7113 | 663.5013 | 687.3285 | 688.3735 | 591.8926 | 681.5982 | 566.0377 | 666.6933 |
| ENSMUSG00000039623 | 0.064857413 | 0.527796 | 0.680556 | 389.343 | 386.3425 | 327.9302 | 366.615 | 350.065 | 338.5644 | 341.3584 | 373.0437 |
| ENSMUSG00000039628 | -0.496348707 | 0.026801 | 0.087377 | 419.5247 | 507.5245 | 287.353 | 345.0494 | 672.8268 | 379.9072 | 537.1091 | 605.7882 |
| ENSMUSG00000039629 | 0.225060541 | 0.383275 | 0.551257 | 101.6115 | 77.9884 | 57.96746 | 107.8279 | 76.05868 | 84.92043 | 59.78593 | 75.04379 |
| ENSMUSG00000039630 | 0.032759302 | 0.574643 | 0.716935 | 18959.09 | 19522.3 | 18142.16 | 19446.98 | 17882.57 | 20028.93 | 18017.74 | 18433.58 |
| ENSMUSG00000039632 | 0.207915547 | 0.386939 | 0.554354 | 65.39353 | 45.59322 | 60.45178 | 68.14726 | 45.83023 | 56.98608 | 45.32159 | 60.9051 |
| ENSMUSG00000039633 | -0.169284942 | 0.206842 | 0.363479 | 355.1372 | 267.5602 | 293.1497 | 289.8415 | 333.4881 | 387.7288 | 305.6797 | 330.6277 |
| ENSMUSG00000039634 | 0.119833501 | 0.41106 | 0.576931 | 319.9253 | 269.9598 | 239.3228 | 299.3304 | 236.952 | 231.2964 | 293.1439 | 276.2481 |
| ENSMUSG00000039637 | -0.015377986 | 0.851179 | 0.910582 | 674.0564 | 706.6949 | 742.8116 | 698.725 | 664.0508 | 724.0584 | 695.2525 | 772.1897 |
| ENSMUSG00000039639 | -1.580036643 | 0.007826 | 0.035655 | 40.24217 | 19.19714 | 75.3577 | 23.29083 | 87.76002 | 36.87335 | 96.42892 | 253.4087 |
| ENSMUSG00000039640 | -0.840803818 | 8.12E-07 | 3.13E-05 | 1475.882 | 1464.982 | 1970.066 | 1559.623 | 2868.777 | 2134.185 | 2930.475 | 3658.656 |
| ENSMUSG00000039646 | 0.031425983 | 0.679989 | 0.795555 | 1130.805 | 1201.021 | 1153.553 | 1180.069 | 1147.706 | 1069.327 | 1255.504 | 1088.679 |
| ENSMUSG00000039648 | -0.334771813 | 0.029163 | 0.092704 | 493.9727 | 413.9384 | 446.3495 | 455.4652 | 554.8383 | 425.7195 | 638.3594 | 663.4306 |
| ENSMUSG00000039652 | -0.438777667 | 0.025798 | 0.084857 | 92.557 | 64.79036 | 91.09173 | 93.16334 | 96.53602 | 127.3806 | 112.8218 | 129.4233 |
| ENSMUSG00000039656 | -0.091480538 | 0.170315 | 0.316996 | 1896.412 | 1887.319 | 1957.644 | 2011.638 | 2129.643 | 1882.775 | 2138.793 | 2109.927 |
| ENSMUSG00000039660 | -0.13710946 | 0.117579 | 0.246092 | 1140.866 | 1232.217 | 1063.289 | 1170.58 | 1376.857 | 1153.13 | 1314.326 | 1215.927 |
| ENSMUSG00000039661 | 1.260939404 | 2.59E-05 | 0.000479 | 170.0232 | 95.98572 | 104.3414 | 200.9913 | 57.53157 | 51.39921 | 76.17884 | 53.29196 |
| ENSMUSG00000039662 | -0.196984463 | 0.039422 | 0.115432 | 1377.288 | 1451.784 | 1392.047 | 1371.571 | 1611.859 | 1365.431 | 1623.863 | 1807.576 |
| ENSMUSG00000039670 | -0.351411419 | 0.034068 | 0.103877 | 142.8597 | 166.7752 | 164.7932 | 131.9814 | 171.6196 | 166.4887 | 208.2865 | 226.2189 |
| ENSMUSG00000039671 | 0.173021181 | 0.09585 | 0.214399 | 2153.962 | 2112.886 | 2225.951 | 2103.939 | 1706.445 | 1870.484 | 1764.649 | 2286.116 |
| ENSMUSG00000039672 | 0.375094599 | 0.43524 | 0.599372 | 13.07871 | 9.598572 | 19.87456 | 12.07673 | 9.751113 | 14.52586 | 8.678602 | 9.78832 |
| ENSMUSG00000039676 | -0.172803452 | 0.710862 | 0.818885 | 16.09687 | 17.99732 | 44.71776 | 14.6646 | 23.40267 | 33.52122 | 21.21436 | 28.27737 |
| ENSMUSG00000039678 | -0.42436455 | 0.002895 | 0.017287 | 1165.011 | 1167.426 | 1207.379 | 1132.625 | 1616.735 | 1178.83 | 1542.863 | 1931.562 |
| ENSMUSG00000039680 | -0.922266279 | 1.13E-05 | 0.000252 | 959.7759 | 916.6637 | 1373.001 | 1035.148 | 1779.578 | 1401.187 | 2675.902 | 2265.452 |
| ENSMUSG00000039682 | -0.777056684 | 2.18E-05 | 0.000422 | 1485.942 | 1566.967 | 1630.542 | 1489.751 | 2939.961 | 1755.395 | 2457.009 | 3425.912 |
| ENSMUSG00000039683 | 0.423519737 | 0.006623 | 0.031706 | 536.227 | 662.3015 | 695.6096 | 688.3735 | 387.1192 | 600.0299 | 461.8945 | 478.5401 |
| ENSMUSG00000039684 | -0.583974122 | 0.251141 | 0.414808 | 7.04238 | 9.598572 | 9.93728 | 9.488858 | 14.62667 | 14.52586 | 6.750024 | 18.48905 |
| ENSMUSG00000039686 | 0.095711685 | 0.251288 | 0.415 | 1256.562 | 1327.003 | 1193.302 | 1277.545 | 1228.64 | 1259.281 | 1193.79 | 1045.175 |
| ENSMUSG00000039693 | -0.105086782 | 0.274089 | 0.439954 | 833.013 | 932.2613 | 988.7593 | 931.6334 | 1065.797 | 1011.224 | 1017.325 | 870.0729 |
| ENSMUSG00000039697 | -0.340648096 | 0.001367 | 0.009889 | 674.0564 | 721.0927 | 711.3436 | 649.5555 | 779.1139 | 807.8615 | 891.0032 | 1011.46 |
| ENSMUSG00000039699 | -0.199942201 | 0.712164 | 0.819637 | 7.04238 | 10.79839 | 11.59349 | 8.626235 | 16.57689 | 5.58687 | 12.53576 | 8.700729 |
| ENSMUSG00000039701 | -0.075680966 | 0.679727 | 0.795399 | 730.3955 | 508.7243 | 617.7675 | 670.2585 | 535.3361 | 662.6028 | 884.2532 | 581.8612 |
| ENSMUSG00000039703 | -0.167668175 | 0.045926 | 0.128272 | 1745.504 | 1576.565 | 1760.555 | 1823.586 | 2011.655 | 1738.634 | 1933.4 | 2077.299 |
| ENSMUSG00000039704 | -0.496558401 | 0.018243 | 0.066201 | 160.9687 | 110.3836 | 155.684 | 109.5532 | 201.848 | 159.7845 | 163.9292 | 232.7445 |

| | | | | | | | | | | | |
|---|---|---|---|---|---|---|---|---|---|---|---|
| ENSMUSG00000039706 | 0.432300483 | 7.50E-08 | 4.48E-06 | 2162.011 | 2199.273 | 2327.808 | 2466.241 | 1589.431 | 1629.131 | 1707.756 | 1861.956 |
| ENSMUSG00000039710 | -5.842807431 | 1.29E-06 | 4.58E-05 | 0 | 0 | 10.76539 | 3.450494 | 175.52 | 52.51658 | 246.858 | 361.0802 |
| ENSMUSG00000039713 | -0.227615243 | 0.044753 | 0.126005 | 753.5347 | 800.281 | 992.0717 | 747.0319 | 1009.24 | 903.9556 | 970.0749 | 974.4816 |
| ENSMUSG00000039714 | 0.098912254 | 0.665684 | 0.785381 | 55.33299 | 82.78769 | 63.76421 | 84.5371 | 57.53157 | 68.15982 | 65.57166 | 76.13138 |
| ENSMUSG00000039715 | -0.140030005 | 0.174631 | 0.322816 | 791.7648 | 926.2622 | 947.354 | 881.6012 | 1045.319 | 846.9696 | 1043.361 | 971.2188 |
| ENSMUSG00000039716 | 0.739709802 | 0.066291 | 0.165983 | 68.4117 | 50.3925 | 49.6864 | 54.34528 | 20.47734 | 63.69032 | 27.0001 | 22.83941 |
| ENSMUSG00000039720 | 0.828858315 | 0.229785 | 0.390041 | 21.12714 | 15.59768 | 14.90592 | 6.900988 | 2.925334 | 18.99536 | 5.785735 | 5.437955 |
| ENSMUSG00000039725 | -0.165929132 | 0.073125 | 0.177906 | 551.3178 | 577.1142 | 597.0649 | 558.98 | 719.6321 | 604.4994 | 598.8236 | 639.5036 |
| ENSMUSG00000039728 | 2.498750346 | 0.002842 | 0.017029 | 158.9566 | 13.19804 | 8.281066 | 40.5433 | 13.65156 | 11.17374 | 7.714313 | 6.525547 |
| ENSMUSG00000039735 | 0.085259868 | 0.175545 | 0.323954 | 4467.887 | 4315.758 | 4171.173 | 4781.522 | 4008.683 | 4281.778 | 4258.301 | 4171.999 |
| ENSMUSG00000039737 | -0.007073348 | 0.944068 | 0.967635 | 606.6508 | 575.9143 | 587.1276 | 503.7721 | 611.3948 | 589.9735 | 516.859 | 565.5474 |
| ENSMUSG00000039738 | -0.240253946 | 0.075383 | 0.181533 | 727.3773 | 595.1115 | 635.1578 | 607.2869 | 751.8108 | 623.4947 | 911.2533 | 741.7371 |
| ENSMUSG00000039740 | 0.106813824 | 0.258205 | 0.422055 | 1648.923 | 1368.996 | 1416.062 | 1579.464 | 1395.384 | 1414.596 | 1512.005 | 1262.693 |
| ENSMUSG00000039741 | 0.022022919 | 0.845338 | 0.906857 | 1184.126 | 1027.047 | 1026.852 | 979.9403 | 1051.17 | 1208.999 | 918.9676 | 975.5692 |
| ENSMUSG00000039745 | 0.389816881 | 0.067247 | 0.16744 | 315.9011 | 170.3747 | 230.2136 | 220.8316 | 166.744 | 148.6108 | 202.5007 | 197.9416 |
| ENSMUSG00000039747 | -0.281252854 | 0.108236 | 0.232904 | 246.4833 | 226.7663 | 269.1347 | 229.4578 | 381.2685 | 233.5312 | 304.7154 | 261.0219 |
| ENSMUSG00000039748 | -0.197509626 | 0.069008 | 0.170565 | 452.7245 | 446.3336 | 428.9592 | 381.2796 | 535.3361 | 432.4238 | 503.3589 | 485.0656 |
| ENSMUSG00000039753 | 0.191475468 | 0.012152 | 0.049446 | 2077.502 | 2079.291 | 1849.162 | 2151.383 | 1694.743 | 1917.414 | 1773.328 | 1757.547 |
| ENSMUSG00000039754 | -0.043224899 | 0.757123 | 0.848808 | 310.8708 | 313.1534 | 225.245 | 270.0012 | 304.2347 | 291.6346 | 268.0724 | 286.0365 |
| ENSMUSG00000039756 | -0.171896562 | 0.000876 | 0.007033 | 1611.699 | 1607.761 | 1562.637 | 1643.298 | 1843.935 | 1833.611 | 1760.792 | 1799.963 |
| ENSMUSG00000039759 | 0.251840738 | 0.002447 | 0.015181 | 922.5518 | 896.2667 | 818.1694 | 945.4353 | 802.5166 | 724.0584 | 762.7527 | 716.7225 |
| ENSMUSG00000039763 | -0.023264807 | 0.904533 | 0.942794 | 193.1624 | 146.3782 | 140.7781 | 179.4257 | 186.2463 | 155.315 | 198.6436 | 129.4233 |
| ENSMUSG00000039765 | 0.253143816 | 0.007535 | 0.034672 | 1167.023 | 976.6547 | 980.4783 | 1076.554 | 856.1477 | 956.4722 | 814.8243 | 899.4378 |
| ENSMUSG00000039768 | -0.305628268 | 0.015762 | 0.05977 | 1886.352 | 1646.155 | 1774.633 | 1750.263 | 2301.263 | 1729.695 | 2106.972 | 2585.204 |
| ENSMUSG00000039770 | 0.166668346 | 0.115851 | 0.243805 | 1377.288 | 1252.614 | 1402.813 | 1379.335 | 1174.034 | 1410.126 | 1201.504 | 1038.649 |
| ENSMUSG00000039771 | -0.244581078 | 0.000985 | 0.007755 | 943.679 | 1058.243 | 1035.961 | 943.7101 | 1244.242 | 1136.369 | 1165.826 | 1166.985 |
| ENSMUSG00000039774 | -0.02273974 | 0.858208 | 0.915078 | 312.8829 | 327.5513 | 399.1474 | 325.2091 | 314.961 | 329.6254 | 384.7514 | 358.9051 |
| ENSMUSG00000039781 | 0.03522323 | 0.620659 | 0.752618 | 918.5276 | 890.2676 | 839.7001 | 903.1668 | 832.7451 | 901.7209 | 893.8961 | 836.3575 |
| ENSMUSG00000039782 | -0.154778298 | 0.668726 | 0.78757 | 373.2462 | 130.7805 | 495.2078 | 382.1422 | 247.6783 | 336.3296 | 495.6446 | 460.051 |
| ENSMUSG00000039783 | -0.705391332 | 0.077617 | 0.185221 | 11.0666 | 14.39786 | 9.109173 | 13.80198 | 18.52711 | 16.76061 | 21.21436 | 21.75182 |
| ENSMUSG00000039787 | 0.284941464 | 0.056061 | 0.147526 | 323.9495 | 469.1302 | 454.6305 | 395.9442 | 383.2187 | 326.2732 | 331.7155 | 306.7007 |
| ENSMUSG00000039789 | -0.020766371 | 0.88728 | 0.932394 | 202.2169 | 245.9634 | 226.9012 | 204.4418 | 216.4747 | 259.2308 | 229.5008 | 185.9781 |
| ENSMUSG00000039795 | 0.047830712 | 0.543111 | 0.692194 | 1102.636 | 1046.244 | 975.5096 | 1078.279 | 958.5344 | 976.585 | 1087.718 | 1040.825 |
| ENSMUSG00000039798 | 0.054223221 | 0.875521 | 0.925381 | 34.20585 | 28.79572 | 30.63995 | 29.3292 | 35.10401 | 32.40385 | 15.42863 | 35.89051 |
| ENSMUSG00000039801 | 0.314566658 | 0.003373 | 0.019279 | 1192.174 | 1183.024 | 1141.131 | 1156.778 | 899.0526 | 1112.905 | 941.1462 | 804.8174 |
| ENSMUSG00000039804 | -0.053957012 | 0.458142 | 0.619812 | 1938.667 | 2006.102 | 1826.803 | 1748.538 | 2060.41 | 1976.635 | 1836.971 | 1927.211 |
| ENSMUSG00000039809 | 0.619576457 | 0.104316 | 0.22735 | 38.23007 | 32.39518 | 28.15563 | 22.42821 | 12.67645 | 15.64324 | 23.14294 | 27.18978 |
| ENSMUSG00000039810 | -0.043468445 | 0.747902 | 0.842597 | 453.7305 | 387.5424 | 402.4598 | 415.7845 | 464.153 | 334.0949 | 472.5017 | 438.2992 |
| ENSMUSG00000039813 | -0.543784152 | 0.055075 | 0.146002 | 128.775 | 113.983 | 240.979 | 148.3712 | 267.1805 | 154.1976 | 221.7865 | 280.5985 |
| ENSMUSG00000039814 | 0.108906884 | 0.481214 | 0.640351 | 197.1867 | 177.5736 | 143.2624 | 171.6621 | 174.5449 | 141.9065 | 161.0363 | 159.8759 |
| ENSMUSG00000039824 | 0.702180033 | 1.82E-05 | 0.000365 | 245.4773 | 179.9732 | 231.0418 | 213.068 | 135.5405 | 153.0803 | 135.9648 | 110.9343 |
| ENSMUSG00000039826 | -0.242682539 | 0.059413 | 0.153683 | 780.6982 | 896.2667 | 824.7942 | 759.9713 | 1097.975 | 758.697 | 1013.468 | 985.3575 |
| ENSMUSG00000039828 | 0.178439697 | 0.072168 | 0.176275 | 911.4852 | 957.4576 | 898.4957 | 913.5183 | 826.8944 | 937.4769 | 772.3956 | 715.6349 |
| ENSMUSG00000039831 | -0.1344469 | 0.084137 | 0.195572 | 2015.127 | 2044.496 | 2217.67 | 1977.996 | 2167.672 | 2296.204 | 2116.615 | 2485.146 |
| ENSMUSG00000039834 | -0.128079988 | 0.26173 | 0.425894 | 417.5126 | 453.5325 | 463.7397 | 426.9986 | 510.9583 | 397.7852 | 497.5732 | 517.6934 |
| ENSMUSG00000039835 | 0.16631266 | 0.011368 | 0.046969 | 907.461 | 885.4683 | 844.6688 | 874.7002 | 757.6615 | 781.0445 | 795.5386 | 793.9415 |
| ENSMUSG00000039838 | 1.865465914 | 0.009796 | 0.04193 | 23.13925 | 21.59679 | 5.796746 | 35.36756 | 8.776002 | 7.821619 | 5.785735 | 1.087591 |
| ENSMUSG00000039840 | -0.085133517 | 0.488705 | 0.646469 | 620.7355 | 502.7252 | 469.5365 | 472.7177 | 508.033 | 527.4006 | 557.3591 | 597.0875 |
| ENSMUSG00000039841 | 0.004290445 | 0.976324 | 0.986966 | 336.0222 | 297.5557 | 258.3693 | 332.11 | 293.5085 | 354.2076 | 311.4654 | 261.0219 |
| ENSMUSG00000039842 | -0.305967993 | 0.032653 | 0.100492 | 340.0464 | 321.5522 | 308.0557 | 241.5346 | 413.4472 | 394.4331 | 351.9655 | 334.9781 |
| ENSMUSG00000039844 | -0.196355374 | 0.003894 | 0.02143 | 2279.719 | 2460.834 | 2644.973 | 2360.138 | 2854.151 | 2660.468 | 2835.01 | 2817.949 |
| ENSMUSG00000039849 | -0.076729587 | 0.252948 | 0.41662 | 2079.514 | 2067.293 | 2269.84 | 2047.868 | 2391.948 | 2147.593 | 2200.508 | 2188.233 |
| ENSMUSG00000039850 | -0.041599782 | 0.553252 | 0.700378 | 919.5337 | 1036.646 | 1009.462 | 990.2918 | 1034.593 | 968.7633 | 1020.218 | 1046.263 |
| ENSMUSG00000039852 | 0.521892981 | 0.000732 | 0.006159 | 3902.485 | 4255.767 | 3584.046 | 4005.161 | 2541.14 | 3701.86 | 2394.33 | 2330.708 |
| ENSMUSG00000039853 | -0.390421165 | 0.291552 | 0.458729 | 23.13925 | 35.99465 | 53.82693 | 24.15346 | 42.9049 | 32.40385 | 57.85735 | 46.76642 |
| ENSMUSG00000039860 | 0.20204137 | 0.470171 | 0.630173 | 51.30877 | 39.59411 | 57.13936 | 60.38364 | 37.05423 | 34.6386 | 55.92877 | 54.37955 |
| ENSMUSG00000039865 | 0.150425695 | 0.302579 | 0.47086 | 449.7063 | 391.1418 | 548.2066 | 398.5321 | 363.7165 | 391.0809 | 396.3228 | 462.2262 |
| ENSMUSG00000039873 | -0.280962541 | 0.102456 | 0.224629 | 114.6902 | 119.9822 | 103.5133 | 99.2017 | 151.1423 | 113.9722 | 118.6076 | 145.7372 |
| ENSMUSG00000039878 | -2.658234437 | 3.41E-06 | 0.0001 | 116.7023 | 35.99465 | 181.3554 | 123.3552 | 752.7859 | 321.8037 | 609.4308 | 1208.314 |
| ENSMUSG00000039879 | 0.294992227 | 0.021582 | 0.07437 | 959.7759 | 832.6761 | 952.3226 | 1149.877 | 707.9308 | 911.7773 | 801.3243 | 756.9634 |
| ENSMUSG00000039883 | 0.160853761 | 0.24254 | 0.405097 | 2246.519 | 2478.831 | 2206.904 | 2875.124 | 1965.824 | 2564.374 | 2396.259 | 1846.73 |
| ENSMUSG00000039886 | -0.365011567 | 0.000406 | 0.003904 | 1043.278 | 952.6583 | 1036.79 | 1241.315 | 1444.14 | 1251.459 | 1383.755 | 1429.095 |
| ENSMUSG00000039887 | 0.093826891 | 0.29555 | 0.463282 | 458.7608 | 515.9233 | 457.1149 | 464.0914 | 446.601 | 441.3628 | 418.5015 | 467.6642 |
| ENSMUSG00000039891 | 0.917127788 | 0.000358 | 0.003544 | 73.44197 | 74.38894 | 55.48314 | 59.52102 | 36.07912 | 43.57759 | 30.85725 | 28.27737 |

| | | | | | | | | | | | |
|---|---|---|---|---|---|---|---|---|---|---|---|
| ENSMUSG00000039899 | -0.010667495 | 0.93426 | 0.961185 | 461.7789 | 437.9349 | 536.6131 | 521.0246 | 499.257 | 499.4662 | 411.7515 | 565.5474 |
| ENSMUSG00000039901 | -0.066533095 | 0.399439 | 0.566906 | 495.9848 | 485.9277 | 532.4726 | 489.9701 | 515.8339 | 524.0485 | 537.1091 | 524.2189 |
| ENSMUSG00000039903 | -0.167864184 | 0.645435 | 0.770415 | 38.23007 | 23.99643 | 27.32752 | 34.50494 | 37.05423 | 32.40385 | 50.14304 | 19.57664 |
| ENSMUSG00000039904 | 0.092762485 | 0.748331 | 0.842865 | 30.18163 | 43.19358 | 44.71776 | 57.79577 | 39.97956 | 39.10809 | 36.64299 | 50.02919 |
| ENSMUSG00000039908 | 0.075599149 | 0.57486 | 0.717108 | 380.2885 | 278.3586 | 298.9465 | 354.5383 | 300.3343 | 292.752 | 337.5012 | 315.4014 |
| ENSMUSG00000039910 | -0.133449572 | 0.374637 | 0.543364 | 2335.052 | 1740.941 | 1918.723 | 1588.952 | 1834.184 | 1864.897 | 2112.758 | 2506.897 |
| ENSMUSG00000039911 | 0.40666832 | 0.048852 | 0.133988 | 690.1533 | 673.0999 | 457.943 | 602.1112 | 377.3681 | 450.3018 | 628.7165 | 369.781 |
| ENSMUSG00000039913 | 1.132977701 | 0.001376 | 0.009945 | 49.29666 | 83.98751 | 58.79557 | 37.95543 | 17.552 | 29.05173 | 32.78583 | 25.0146 |
| ENSMUSG00000039914 | -0.098598704 | 0.362927 | 0.531351 | 371.2341 | 339.5495 | 302.2589 | 319.1707 | 382.2436 | 351.9728 | 341.3584 | 348.0291 |
| ENSMUSG00000039917 | -0.065954467 | 0.443863 | 0.606906 | 917.5216 | 998.2515 | 904.2924 | 828.1186 | 947.8082 | 1017.928 | 894.8603 | 954.905 |
| ENSMUSG00000039929 | 0.089387378 | 0.384487 | 0.552315 | 413.4883 | 388.7422 | 349.461 | 395.0816 | 388.0943 | 341.9165 | 374.1442 | 346.9416 |
| ENSMUSG00000039934 | 0.814668677 | 0.000424 | 0.004034 | 173.0413 | 189.5718 | 148.2311 | 235.4962 | 97.51113 | 140.7891 | 108.0004 | 78.30656 |
| ENSMUSG00000039936 | -0.009050888 | 0.94562 | 0.968715 | 239.4409 | 211.1686 | 226.9012 | 233.771 | 259.3796 | 195.5405 | 243.9652 | 217.5182 |
| ENSMUSG00000039942 | -0.43319801 | 0.155287 | 0.297177 | 223.3441 | 85.18733 | 103.5133 | 107.8279 | 157.968 | 178.7799 | 206.3579 | 158.7883 |
| ENSMUSG00000039943 | 0.051907358 | 0.647895 | 0.771952 | 969.8364 | 1003.051 | 833.9034 | 1058.439 | 903.9282 | 1073.797 | 906.4318 | 843.9707 |
| ENSMUSG00000039952 | -0.314636047 | 0.032308 | 0.099696 | 5226.452 | 5149.634 | 7380.914 | 5643.283 | 6585.902 | 6345.567 | 7589.92 | 8585.444 |
| ENSMUSG00000039953 | 0.123552723 | 0.05745 | 0.150164 | 4643.947 | 4853.278 | 5145.855 | 4718.551 | 4156.9 | 4723.14 | 4512.873 | 4382.992 |
| ENSMUSG00000039954 | -1.541633626 | 0.003797 | 0.021024 | 39.23612 | 17.99732 | 53.82693 | 54.34528 | 94.5858 | 32.40385 | 145.6077 | 210.9927 |
| ENSMUSG00000039956 | 4.226006561 | 0.005417 | 0.02719 | 1075.472 | 4.799286 | 9.93728 | 110.4158 | 4.875557 | 10.05637 | 5.785735 | 43.50364 |
| ENSMUSG00000039958 | 0.324210833 | 0.071036 | 0.174224 | 638.8445 | 627.5067 | 481.13 | 675.4342 | 449.5263 | 629.0816 | 480.216 | 376.3065 |
| ENSMUSG00000039959 | 0.292483472 | 9.93E-05 | 0.001331 | 3127.823 | 2793.185 | 2692.175 | 2919.981 | 2271.034 | 2524.148 | 2238.115 | 2385.087 |
| ENSMUSG00000039960 | 0.10547961 | 0.235218 | 0.396138 | 1080.502 | 1143.43 | 986.275 | 985.9787 | 986.8126 | 1063.74 | 921.8604 | 924.4524 |
| ENSMUSG00000039963 | 0.270933434 | 0.077797 | 0.185446 | 217.3077 | 248.3631 | 184.6678 | 181.1509 | 176.4951 | 185.4841 | 167.7863 | 156.6131 |
| ENSMUSG00000039967 | 0.241181704 | 0.195 | 0.348834 | 1956.776 | 1928.113 | 1620.605 | 1788.219 | 1378.807 | 2228.044 | 1301.79 | 1262.693 |
| ENSMUSG00000039968 | 0.434170331 | 0.011738 | 0.048154 | 1625.784 | 1610.16 | 1368.86 | 1637.259 | 1003.39 | 1606.784 | 1058.79 | 951.6422 |
| ENSMUSG00000039976 | -0.003112147 | 0.976057 | 0.986896 | 1615.723 | 1680.95 | 2030.517 | 1836.525 | 1834.184 | 1573.263 | 1866.864 | 1907.635 |
| ENSMUSG00000039977 | 0.208106475 | 0.638197 | 0.765487 | 16.09687 | 16.7975 | 11.59349 | 12.93935 | 16.57689 | 8.938993 | 10.60718 | 13.05109 |
| ENSMUSG00000039982 | 0.423990631 | 0.000414 | 0.003965 | 1057.363 | 1135.031 | 1039.274 | 1095.532 | 749.8606 | 998.9324 | 716.4668 | 761.3138 |
| ENSMUSG00000039983 | -0.091796374 | 0.400035 | 0.567446 | 642.8687 | 617.9081 | 612.7989 | 590.8971 | 679.6526 | 538.5743 | 700.0739 | 705.8466 |
| ENSMUSG00000039985 | 0.029805304 | 0.779239 | 0.863813 | 1868.243 | 1803.332 | 1579.199 | 1885.695 | 1889.766 | 1893.949 | 1718.363 | 1486.737 |
| ENSMUSG00000039987 | 0.110008725 | 0.479796 | 0.638851 | 495.9848 | 455.9322 | 332.8989 | 485.657 | 436.8499 | 453.6539 | 378.9656 | 369.781 |
| ENSMUSG00000039988 | -0.147345239 | 0.086468 | 0.199496 | 1203.241 | 1189.023 | 1213.176 | 1185.245 | 1277.396 | 1219.055 | 1292.147 | 1518.277 |
| ENSMUSG00000039989 | 0.267261271 | 0.122329 | 0.251807 | 657.9595 | 391.1418 | 481.9581 | 430.4491 | 372.4925 | 427.9543 | 395.3586 | 435.0364 |
| ENSMUSG00000039990 | -0.075001819 | 0.451201 | 0.613705 | 958.7698 | 812.2792 | 787.5294 | 740.131 | 885.4011 | 834.6784 | 858.2174 | 893.9999 |
| ENSMUSG00000039994 | -0.098159224 | 0.389999 | 0.557408 | 1004.042 | 1197.422 | 968.8848 | 935.9465 | 1247.167 | 1070.444 | 987.4321 | 1085.416 |
| ENSMUSG00000039997 | 0.727328663 | 0.007377 | 0.034135 | 239.4409 | 416.3381 | 336.2113 | 194.9529 | 204.7734 | 224.5922 | 146.572 | 140.2993 |
| ENSMUSG00000040003 | 0.096948475 | 0.596277 | 0.735213 | 279.6831 | 277.1588 | 206.1986 | 250.1608 | 193.072 | 301.691 | 243.0009 | 208.8175 |
| ENSMUSG00000040006 | -0.042955468 | 0.658983 | 0.780378 | 1468.839 | 1275.41 | 1441.734 | 1584.639 | 1360.28 | 1409.009 | 1584.327 | 1595.496 |
| ENSMUSG00000040007 | -0.094384555 | 0.561897 | 0.70744 | 408.4581 | 321.5522 | 483.6143 | 409.7462 | 417.3476 | 354.2076 | 461.8945 | 502.4671 |
| ENSMUSG00000040009 | 0.115883396 | 0.340889 | 0.509511 | 221.332 | 266.3604 | 240.979 | 253.6113 | 242.8027 | 227.9443 | 203.465 | 231.6569 |
| ENSMUSG00000040010 | -0.044733732 | 0.572721 | 0.715262 | 808.8677 | 746.289 | 819.8256 | 722.8785 | 785.9397 | 806.7441 | 765.6456 | 839.6203 |
| ENSMUSG00000040013 | -4.947914067 | NA | NA | 0 | 0 | 0.828107 | 3.450494 | 0 | 131.8501 | 1.928578 | 0 |
| ENSMUSG00000040016 | -1.337603606 | 0.000142 | 0.001748 | 157.9505 | 194.3711 | 288.1811 | 197.5408 | 437.825 | 220.1227 | 709.7168 | 751.5254 |
| ENSMUSG00000040018 | -0.164124876 | 0.158341 | 0.301192 | 975.8727 | 724.6922 | 915.8859 | 889.3648 | 958.5344 | 879.3734 | 1029.861 | 1063.664 |
| ENSMUSG00000040021 | 0.132180229 | 0.065484 | 0.164684 | 1800.837 | 1728.943 | 1675.26 | 1809.784 | 1494.846 | 1729.695 | 1637.363 | 1540.029 |
| ENSMUSG00000040022 | 0.235805782 | 0.007856 | 0.035773 | 1425.579 | 1440.986 | 1416.89 | 1469.048 | 1105.776 | 1371.018 | 1281.54 | 1127.832 |
| ENSMUSG00000040025 | -0.017630832 | 0.825928 | 0.894299 | 1649.929 | 1784.135 | 1526.201 | 1622.595 | 1646.963 | 1583.319 | 1608.434 | 1820.627 |
| ENSMUSG00000040026 | 1.855795974 | 0.004999 | 0.02575 | 32.19374 | 8.398751 | 11.59349 | 14.6646 | 6.825779 | 4.469496 | 3.857157 | 3.262773 |
| ENSMUSG00000040028 | 0.030998891 | 0.833083 | 0.898985 | 2058.387 | 2050.495 | 1798.648 | 1890.008 | 1647.938 | 2526.383 | 1749.221 | 1708.606 |
| ENSMUSG00000040029 | 0.050523054 | 0.41451 | 0.580308 | 1371.252 | 1301.806 | 1313.377 | 1386.236 | 1284.222 | 1368.783 | 1250.683 | 1287.708 |
| ENSMUSG00000040033 | 0.251868843 | 0.179805 | 0.329705 | 492.9666 | 579.5138 | 707.2031 | 377.8291 | 433.9245 | 422.3674 | 489.8589 | 465.489 |
| ENSMUSG00000040034 | 0.007702164 | 0.943816 | 0.967564 | 471.8395 | 526.7217 | 427.303 | 442.5259 | 493.4063 | 469.2971 | 413.6801 | 478.5401 |
| ENSMUSG00000040035 | 0.869692237 | 0.065223 | 0.164448 | 117.7084 | 29.99554 | 38.92101 | 123.3552 | 39.97956 | 41.34284 | 47.25017 | 41.32846 |
| ENSMUSG00000040037 | 0.671978467 | 0.000157 | 0.001885 | 1015.109 | 1432.587 | 1329.111 | 1223.2 | 787.8899 | 1031.336 | 714.5383 | 604.7006 |
| ENSMUSG00000040040 | 0.115198552 | 0.306782 | 0.475352 | 822.9525 | 847.074 | 772.6235 | 861.7609 | 798.6162 | 859.2607 | 766.6099 | 625.3649 |
| ENSMUSG00000040043 | 0.14963564 | 0.02203 | 0.075461 | 3050.357 | 2987.556 | 2965.45 | 3185.669 | 2703.984 | 2994.563 | 2619.974 | 2673.299 |
| ENSMUSG00000040044 | 0.116443951 | 0.126808 | 0.257937 | 1680.111 | 1680.95 | 1811.897 | 1842.564 | 1547.502 | 1783.329 | 1593.97 | 1551.992 |
| ENSMUSG00000040046 | -0.801159662 | 0.025807 | 0.084869 | 14.08476 | 17.99732 | 12.4216 | 21.56559 | 34.1289 | 25.6996 | 30.85725 | 23.927 |
| ENSMUSG00000040048 | -0.652292562 | 1.08E-05 | 0.000243 | 1848.122 | 1726.543 | 2119.953 | 1922.788 | 2937.035 | 2243.687 | 3133.94 | 3659.744 |
| ENSMUSG00000040054 | 0.115752972 | 0.166416 | 0.312068 | 2120.763 | 1727.743 | 1859.099 | 1974.545 | 1684.992 | 1869.367 | 1741.506 | 1797.788 |
| ENSMUSG00000040055 | 0.249570093 | 0.53814 | 0.688091 | 35.2119 | 25.19625 | 72.87338 | 47.44429 | 55.58134 | 26.81698 | 36.64299 | 33.71532 |
| ENSMUSG00000040061 | -0.226146327 | 0.501621 | 0.657943 | 72.43591 | 44.3934 | 126.7003 | 76.77349 | 105.312 | 63.69032 | 96.42892 | 110.9343 |
| ENSMUSG00000040078 | -0.076282346 | 0.590012 | 0.730039 | 286.7255 | 387.5424 | 286.5249 | 351.9504 | 368.5921 | 346.386 | 304.7154 | 362.1678 |
| ENSMUSG00000040084 | -0.154228854 | 0.081336 | 0.19111 | 2045.309 | 2154.879 | 1854.131 | 2084.961 | 2521.638 | 2048.147 | 2214.972 | 2267.627 |

| | | | | | | | | | | | |
|---|---|---|---|---|---|---|---|---|---|---|---|
| ENSMUSG00000040093 | 0.086453469 | 0.334358 | 0.50258 | 2000.036 | 2134.483 | 2439.602 | 2166.91 | 1915.119 | 2058.203 | 2258.365 | 2003.343 |
| ENSMUSG00000040097 | -0.064746564 | 0.534863 | 0.685754 | 1173.059 | 1180.624 | 1131.194 | 1019.621 | 1208.163 | 991.1108 | 1284.433 | 1223.54 |
| ENSMUSG00000040102 | 0.191593578 | 0.009788 | 0.041905 | 658.9656 | 661.1017 | 665.7977 | 662.4948 | 584.0917 | 615.6731 | 571.8235 | 548.1459 |
| ENSMUSG00000040105 | -0.371632606 | 0.033988 | 0.103683 | 572.4449 | 423.537 | 580.5027 | 553.8043 | 568.4899 | 563.1565 | 826.3958 | 800.467 |
| ENSMUSG00000040111 | 0.195820452 | 0.449575 | 0.61217 | 1821.964 | 771.4852 | 891.0427 | 1080.867 | 799.5913 | 933.0074 | 1009.611 | 1244.204 |
| ENSMUSG00000040112 | -0.345542815 | 6.97E-06 | 0.000175 | 951.7274 | 1027.047 | 1055.008 | 991.1544 | 1272.52 | 1217.938 | 1227.54 | 1397.555 |
| ENSMUSG00000040118 | 0.636918182 | 0.00551 | 0.027591 | 2446.724 | 2216.07 | 1743.993 | 2304.067 | 1327.126 | 2148.71 | 1137.861 | 988.6203 |
| ENSMUSG00000040121 | 0.729314536 | 0.043934 | 0.124457 | 86.52067 | 101.9848 | 52.17072 | 107.8279 | 36.07912 | 86.03781 | 50.14304 | 38.06569 |
| ENSMUSG00000040123 | 0.301340189 | 0.009474 | 0.04095 | 2574.493 | 2524.425 | 2037.142 | 2549.915 | 1950.223 | 2291.734 | 1818.649 | 1798.876 |
| ENSMUSG00000040124 | 0.080654821 | 0.432628 | 0.596978 | 506.0453 | 604.7101 | 530.8164 | 535.6892 | 557.7637 | 536.3396 | 499.5018 | 462.2262 |
| ENSMUSG00000040125 | 0.03406074 | 0.94357 | 0.96742 | 11.0666 | 8.398751 | 14.90592 | 8.626235 | 10.72622 | 10.05637 | 9.642892 | 11.9635 |
| ENSMUSG00000040128 | 0.219765503 | 0.018843 | 0.067607 | 2628.82 | 2532.823 | 2317.87 | 2737.104 | 2151.096 | 2439.228 | 2239.079 | 1943.525 |
| ENSMUSG00000040133 | 0.278939939 | 0.261991 | 0.426243 | 76.46013 | 76.78858 | 79.49824 | 117.3168 | 61.43201 | 60.3382 | 75.21455 | 92.44524 |
| ENSMUSG00000040136 | 0.151150423 | 0.620857 | 0.752761 | 27.16347 | 44.3934 | 34.78048 | 48.30692 | 38.02934 | 27.93435 | 31.82154 | 41.32846 |
| ENSMUSG00000040138 | 0.483175546 | 0.475116 | 0.634588 | 19.11503 | 9.598572 | 62.9361 | 10.35148 | 14.62667 | 11.17374 | 27.96439 | 19.57664 |
| ENSMUSG00000040139 | -0.228152064 | 0.062181 | 0.158943 | 361.1735 | 341.9491 | 337.0394 | 357.9887 | 437.825 | 470.4145 | 381.8585 | 348.0291 |
| ENSMUSG00000040140 | 0.522068653 | 0.268159 | 0.432618 | 16.09687 | 11.99822 | 13.24971 | 20.70296 | 10.72622 | 15.64324 | 10.60718 | 6.525547 |
| ENSMUSG00000040146 | -0.442583363 | 0.017653 | 0.064683 | 746.4923 | 565.1159 | 1020.227 | 671.9837 | 964.3851 | 864.8476 | 1106.04 | 1150.671 |
| ENSMUSG00000040147 | -0.03072389 | 0.876116 | 0.925815 | 79.47829 | 74.38894 | 96.88848 | 100.0643 | 86.78491 | 88.27255 | 79.07171 | 106.5839 |
| ENSMUSG00000040151 | 0.177935448 | 0.037097 | 0.110672 | 2654.977 | 2801.583 | 2481.836 | 2697.424 | 2366.595 | 2643.707 | 2241.008 | 2149.08 |
| ENSMUSG00000040152 | 0.052402958 | 0.753043 | 0.846365 | 2204.265 | 1746.94 | 2407.306 | 1927.964 | 1895.616 | 1482.755 | 2184.115 | 2429.678 |
| ENSMUSG00000040158 | -0.380777832 | 0.00947 | 0.04094 | 918.5276 | 1055.843 | 1185.849 | 1096.394 | 1410.986 | 1061.505 | 1415.576 | 1655.314 |
| ENSMUSG00000040164 | 0.075156941 | 0.821546 | 0.891669 | 24.1453 | 34.79482 | 24.8432 | 25.8787 | 27.30312 | 22.34748 | 33.75012 | 19.57664 |
| ENSMUSG00000040167 | 0.137370492 | 0.418719 | 0.584292 | 322.9434 | 379.1436 | 235.1823 | 285.5284 | 286.6827 | 310.63 | 261.3224 | 250.146 |
| ENSMUSG00000040170 | 0.595736328 | 0.002124 | 0.013708 | 1176.078 | 1385.794 | 1023.54 | 1728.697 | 998.514 | 1048.097 | 700.0739 | 770.0145 |
| ENSMUSG00000040174 | -0.169921014 | 0.007624 | 0.034974 | 848.1038 | 874.6699 | 837.2158 | 904.892 | 956.5842 | 981.0545 | 972.0035 | 988.6203 |
| ENSMUSG00000040177 | -0.288837726 | 0.019627 | 0.06958 | 354.1311 | 295.1561 | 300.6027 | 290.7041 | 425.1485 | 383.2593 | 374.1442 | 331.7153 |
| ENSMUSG00000040181 | -0.439849787 | 0.042822 | 0.122436 | 726.3712 | 613.1088 | 813.2007 | 974.7645 | 1246.192 | 734.1148 | 902.5747 | 1361.664 |
| ENSMUSG00000040183 | 0.299214564 | 0.028504 | 0.091143 | 1242.477 | 1166.227 | 981.3064 | 1217.162 | 1115.527 | 995.5803 | 838.9316 | 792.8539 |
| ENSMUSG00000040187 | 0.558422795 | 0.005349 | 0.026934 | 129.781 | 125.9813 | 129.1846 | 120.7673 | 83.85957 | 116.2069 | 72.32169 | 71.78101 |
| ENSMUSG00000040188 | -0.265547975 | 0.030936 | 0.09662 | 1415.518 | 1408.59 | 1646.276 | 1201.635 | 1535.8 | 1588.906 | 1742.471 | 1952.226 |
| ENSMUSG00000040189 | 0.853402258 | 1.11E-11 | 2.58E-09 | 572.4449 | 604.7101 | 510.9418 | 532.2387 | 302.2845 | 362.0292 | 261.3224 | 303.4379 |
| ENSMUSG00000040195 | 0.273673468 | 0.008451 | 0.037774 | 665.0019 | 634.7056 | 547.3785 | 608.1496 | 553.8632 | 492.762 | 520.7161 | 460.051 |
| ENSMUSG00000040204 | -0.05350076 | 0.686326 | 0.800533 | 2705.28 | 3543.073 | 2544.772 | 2700.874 | 3144.734 | 3391.23 | 2710.617 | 2677.649 |
| ENSMUSG00000040209 | 0.171098609 | 0.2164 | 0.37475 | 3174.101 | 3226.32 | 2965.45 | 3057.138 | 2541.14 | 3619.175 | 2486.902 | 2387.262 |
| ENSMUSG00000040212 | 0.490353961 | 0.000185 | 0.002131 | 1059.375 | 1494.978 | 1366.376 | 1230.964 | 1031.668 | 934.1247 | 804.2172 | 896.1751 |
| ENSMUSG00000040213 | -0.444162222 | 0.000417 | 0.003987 | 440.6518 | 410.339 | 442.2089 | 500.3216 | 631.8721 | 500.5836 | 686.5739 | 622.1021 |
| ENSMUSG00000040219 | -0.331512045 | 0.140954 | 0.277628 | 236.4228 | 159.5763 | 242.6352 | 158.7227 | 217.4498 | 206.7142 | 247.8223 | 332.8029 |
| ENSMUSG00000040220 | -0.179636994 | 0.043528 | 0.123629 | 877.2794 | 903.4656 | 956.4632 | 994.6049 | 1081.398 | 934.1247 | 1082.897 | 1130.007 |
| ENSMUSG00000040225 | 0.378443719 | 0.001604 | 0.011078 | 7512.208 | 7068.149 | 6494.84 | 7072.65 | 5025.724 | 6725.475 | 5178.233 | 4724.496 |
| ENSMUSG00000040229 | 0.70597182 | 0.072442 | 0.176733 | 19.11503 | 21.59679 | 25.67131 | 24.15346 | 16.57689 | 17.87799 | 12.53576 | 8.700729 |
| ENSMUSG00000040231 | -0.20403634 | 0.608847 | 0.744259 | 17.10292 | 17.99732 | 10.76539 | 12.93935 | 16.57689 | 15.64324 | 18.32149 | 16.31387 |
| ENSMUSG00000040234 | 0.567117112 | 7.67E-18 | 6.44E-15 | 1833.031 | 2073.292 | 1913.754 | 1959.881 | 1284.222 | 1381.074 | 1296.005 | 1288.795 |
| ENSMUSG00000040236 | -0.214326562 | 0.058916 | 0.152633 | 1091.569 | 1192.623 | 1148.584 | 1132.625 | 1449.015 | 1064.858 | 1321.076 | 1459.547 |
| ENSMUSG00000040242 | -0.054223641 | 0.473568 | 0.632933 | 2501.051 | 2252.065 | 2078.548 | 2323.908 | 2470.932 | 2356.542 | 2269.937 | 2406.839 |
| ENSMUSG00000040247 | 0.303880863 | 0.109073 | 0.23428 | 90.54489 | 119.9822 | 112.6225 | 110.4158 | 101.4116 | 75.98144 | 95.46463 | 77.21897 |
| ENSMUSG00000040249 | 0.491249757 | 2.45E-06 | 7.62E-05 | 3447.748 | 3747.043 | 4030.395 | 4138.868 | 2560.642 | 3202.394 | 2646.009 | 2524.299 |
| ENSMUSG00000040250 | 0.110336042 | 0.221223 | 0.380229 | 1605.663 | 1638.956 | 1367.204 | 1539.783 | 1412.936 | 1547.563 | 1433.898 | 1300.759 |
| ENSMUSG00000040253 | 0.764239815 | 0.012567 | 0.050698 | 307.8526 | 693.4968 | 486.0986 | 261.3749 | 202.8232 | 302.8084 | 306.644 | 216.4306 |
| ENSMUSG00000040254 | 0.640911458 | 0.00027 | 0.002858 | 1317.931 | 1794.933 | 1329.111 | 1726.972 | 903.9282 | 1293.919 | 964.2892 | 793.9415 |
| ENSMUSG00000040258 | 0.348128998 | 0.587797 | 0.72803 | 16.09687 | 3.599465 | 11.59349 | 20.70296 | 16.57689 | 10.05637 | 4.821446 | 9.78832 |
| ENSMUSG00000040260 | 0.300628533 | 3.91E-05 | 0.000662 | 1598.62 | 1694.148 | 1830.944 | 1620.87 | 1316.4 | 1451.469 | 1346.148 | 1364.927 |
| ENSMUSG00000040263 | -0.265530027 | 0.032084 | 0.099221 | 715.3046 | 860.272 | 782.5608 | 594.3476 | 928.306 | 907.3078 | 833.1458 | 876.5984 |
| ENSMUSG00000040265 | -0.224552437 | 0.180598 | 0.330793 | 243.4652 | 211.1686 | 309.7119 | 246.7103 | 265.2303 | 254.7613 | 346.1798 | 317.5766 |
| ENSMUSG00000040268 | 0.135328575 | 0.133449 | 0.266927 | 3248.55 | 2825.58 | 3250.319 | 3220.174 | 2481.658 | 2964.393 | 2998.939 | 2981.087 |
| ENSMUSG00000040269 | -0.427283715 | 0.00078 | 0.006465 | 508.0574 | 497.9259 | 602.0335 | 522.7498 | 719.6321 | 584.3867 | 754.0741 | 809.1678 |
| ENSMUSG00000040270 | 0.41123521 | 0.083098 | 0.193963 | 976.8788 | 914.264 | 764.3424 | 890.2274 | 610.4197 | 1027.984 | 600.7521 | 427.4233 |
| ENSMUSG00000040272 | 0.291888129 | 0.023228 | 0.078575 | 735.4257 | 785.8831 | 738.6711 | 706.4886 | 626.9966 | 690.5372 | 635.4666 | 468.7518 |
| ENSMUSG00000040274 | 0.393182758 | 0.004175 | 0.022536 | 1166.017 | 1232.217 | 1389.563 | 1262.881 | 759.6117 | 1166.539 | 1001.896 | 920.1021 |
| ENSMUSG00000040276 | 0.268373073 | 0.131811 | 0.264798 | 105.6357 | 104.3845 | 130.8408 | 131.1188 | 84.83468 | 108.3853 | 95.46463 | 105.4963 |
| ENSMUSG00000040280 | -0.697775867 | 0.00531 | 0.026818 | 133.8052 | 104.3845 | 202.058 | 206.167 | 233.0516 | 219.0053 | 270.9653 | 329.5401 |
| ENSMUSG00000040282 | 0.26565033 | 0.024015 | 0.080612 | 386.3249 | 442.7341 | 365.195 | 419.235 | 345.1894 | 369.8508 | 332.6798 | 292.562 |
| ENSMUSG00000040283 | -0.371598235 | 0.135047 | 0.26916 | 203.223 | 100.785 | 136.6376 | 144.0581 | 166.744 | 157.5497 | 183.2149 | 250.146 |
| ENSMUSG00000040287 | 0.462716444 | 0.099186 | 0.219818 | 56.33904 | 59.99108 | 70.38906 | 56.93315 | 43.88001 | 49.16446 | 27.0001 | 57.64233 |

| | | | | | | | | | | | |
|---|---|---|---|---|---|---|---|---|---|---|---|
| ENSMUSG00000040289 | -0.227458654 | 0.115503 | 0.243459 | 806.8556 | 643.1043 | 660.8291 | 771.1854 | 869.7993 | 655.8986 | 977.7892 | 870.0729 |
| ENSMUSG00000040296 | 0.42563037 | 0.015722 | 0.059687 | 795.789 | 1055.843 | 1040.93 | 652.1434 | 578.241 | 611.2036 | 759.8599 | 688.4452 |
| ENSMUSG00000040297 | 0.084910009 | 0.250267 | 0.413857 | 1604.657 | 1466.182 | 1416.89 | 1563.936 | 1331.027 | 1490.577 | 1416.541 | 1469.336 |
| ENSMUSG00000040302 | 0.081063685 | 0.601329 | 0.739186 | 455.7426 | 475.1293 | 414.0533 | 422.6855 | 405.6463 | 541.9264 | 346.1798 | 377.3941 |
| ENSMUSG00000040310 | -0.139337713 | 0.780698 | 0.864704 | 12.07265 | 16.7975 | 13.24971 | 14.6646 | 8.776002 | 14.52586 | 27.96439 | 10.87591 |
| ENSMUSG00000040312 | 0.209378496 | 0.088057 | 0.201935 | 665.0019 | 795.4817 | 722.9371 | 680.6099 | 660.1504 | 654.7812 | 661.5024 | 498.1167 |
| ENSMUSG00000040321 | 0.328754173 | 0.038631 | 0.113622 | 458.7608 | 526.7217 | 430.6154 | 510.6731 | 416.3725 | 473.7666 | 333.6441 | 309.9635 |
| ENSMUSG00000040322 | 0.284635484 | 0.007106 | 0.033261 | 1357.167 | 1169.826 | 1289.362 | 1434.543 | 1126.254 | 1148.661 | 1119.54 | 917.9269 |
| ENSMUSG00000040323 | -0.967353121 | 0.004479 | 0.02373 | 14.08476 | 16.7975 | 18.21835 | 19.84034 | 39.97956 | 23.46486 | 35.6787 | 35.89051 |
| ENSMUSG00000040325 | 0.113045057 | 0.172289 | 0.319734 | 1114.708 | 1058.243 | 927.4794 | 1016.17 | 985.8375 | 960.9417 | 902.5747 | 953.8174 |
| ENSMUSG00000040327 | -0.061490415 | 0.566804 | 0.711182 | 513.0877 | 434.3354 | 522.5353 | 445.9763 | 494.3814 | 461.4755 | 497.5732 | 548.1459 |
| ENSMUSG00000040329 | 0.348297921 | 0.388322 | 0.555851 | 19.11503 | 25.19625 | 15.73403 | 23.29083 | 13.65156 | 22.34748 | 18.32149 | 10.87591 |
| ENSMUSG00000040331 | -0.202173228 | 0.001103 | 0.008412 | 2260.604 | 2303.657 | 2037.142 | 2273.013 | 2515.787 | 2625.829 | 2513.902 | 2549.314 |
| ENSMUSG00000040339 | 0.27454808 | 0.047375 | 0.131009 | 884.3218 | 944.2595 | 925.8232 | 968.7262 | 696.2295 | 997.8151 | 705.8597 | 679.7444 |
| ENSMUSG00000040345 | 0.011873701 | 0.955573 | 0.974791 | 76.46013 | 101.9848 | 110.9663 | 82.81186 | 83.85957 | 92.74205 | 112.8218 | 79.39415 |
| ENSMUSG00000040350 | -0.930056241 | 3.98E-05 | 0.00067 | 179.0777 | 117.5825 | 183.0116 | 173.3873 | 294.4836 | 221.2401 | 317.2511 | 414.3722 |
| ENSMUSG00000040351 | -0.374683712 | 7.32E-05 | 0.001068 | 975.8727 | 985.0535 | 953.9788 | 976.4898 | 1165.258 | 1165.421 | 1248.754 | 1466.073 |
| ENSMUSG00000040354 | -0.312353347 | 0.00138 | 0.009965 | 1460.791 | 1457.783 | 1444.218 | 1355.182 | 1728.872 | 1543.094 | 1781.042 | 2045.759 |
| ENSMUSG00000040356 | 0.019094979 | 0.807822 | 0.882512 | 1823.977 | 1572.966 | 1666.979 | 1762.34 | 1539.701 | 1730.812 | 1723.185 | 1746.671 |
| ENSMUSG00000040359 | -0.013472871 | 0.831457 | 0.897939 | 1307.871 | 1281.409 | 1296.815 | 1430.23 | 1348.579 | 1358.727 | 1323.969 | 1338.825 |
| ENSMUSG00000040363 | 0.208035652 | 0.013928 | 0.054528 | 1920.558 | 1882.52 | 1936.113 | 1989.21 | 1636.237 | 1915.179 | 1607.47 | 1534.591 |
| ENSMUSG00000040365 | -0.049289821 | 0.519723 | 0.673278 | 1899.431 | 1794.933 | 2006.502 | 1832.212 | 1938.521 | 1771.038 | 1996.079 | 2091.438 |
| ENSMUSG00000040370 | -0.112369412 | 0.385047 | 0.552598 | 880.2976 | 617.9081 | 674.0788 | 747.8946 | 773.2633 | 716.2368 | 864.0031 | 803.7298 |
| ENSMUSG00000040372 | -0.585167519 | 0.028178 | 0.090506 | 27.16347 | 32.39518 | 37.2648 | 28.46658 | 50.70579 | 52.51658 | 38.57157 | 46.76642 |
| ENSMUSG00000040373 | 0.95942509 | 0.080342 | 0.189494 | 37.22401 | 10.79839 | 14.07781 | 42.26855 | 13.65156 | 17.87799 | 11.57147 | 10.87591 |
| ENSMUSG00000040374 | -0.064920498 | 0.210643 | 0.368171 | 1435.64 | 1473.381 | 1420.203 | 1430.23 | 1469.493 | 1505.103 | 1523.577 | 1523.715 |
| ENSMUSG00000040380 | 0.749866648 | 0.022389 | 0.07638 | 57.3451 | 63.59054 | 67.07664 | 37.95543 | 19.50223 | 40.22547 | 35.6787 | 39.15328 |
| ENSMUSG00000040383 | -0.018846728 | 0.817193 | 0.888971 | 2119.757 | 1965.308 | 1912.098 | 1902.085 | 2012.63 | 1774.39 | 2095.4 | 2117.54 |
| ENSMUSG00000040385 | -0.383370433 | 8.84E-05 | 0.001222 | 7668.146 | 8444.344 | 8613.965 | 8356.234 | 11410.75 | 9052.965 | 10758.57 | 11929.79 |
| ENSMUSG00000040387 | 0.538657231 | 0.019033 | 0.068142 | 90.54489 | 83.98751 | 99.3728 | 102.6522 | 61.43201 | 88.27255 | 62.6788 | 47.85401 |
| ENSMUSG00000040389 | -0.096992553 | 0.421491 | 0.586832 | 316.9071 | 245.9634 | 308.8838 | 275.1769 | 301.3094 | 321.8037 | 288.3225 | 318.6642 |
| ENSMUSG00000040390 | -0.245278622 | 0.024194 | 0.081113 | 636.8324 | 616.7083 | 571.3936 | 648.6929 | 788.865 | 614.5558 | 732.8598 | 793.9415 |
| ENSMUSG00000040396 | -0.057909604 | 0.389198 | 0.556608 | 1313.907 | 1473.381 | 1326.627 | 1290.485 | 1405.135 | 1400.07 | 1438.719 | 1374.715 |
| ENSMUSG00000040405 | 0.595653997 | 0.168009 | 0.314065 | 24.1453 | 13.19804 | 19.04645 | 13.80198 | 10.72622 | 12.29112 | 10.60718 | 13.05109 |
| ENSMUSG00000040407 | 0.139877484 | 0.168889 | 0.315147 | 2688.177 | 2390.044 | 2468.586 | 2433.461 | 2045.784 | 2680.58 | 2133.008 | 2201.284 |
| ENSMUSG00000040410 | -0.022960344 | 0.802328 | 0.879311 | 552.3238 | 471.5299 | 568.0811 | 555.5295 | 551.913 | 543.0438 | 525.5376 | 566.635 |
| ENSMUSG00000040412 | 1.011353397 | 0.02775 | 0.089474 | 159.9626 | 67.19001 | 49.6864 | 169.0742 | 34.1289 | 53.63396 | 87.75031 | 45.67883 |
| ENSMUSG00000040414 | -0.087207414 | 0.352692 | 0.521546 | 995.9938 | 916.6637 | 900.98 | 871.2497 | 1041.419 | 937.4769 | 1059.754 | 872.2481 |
| ENSMUSG00000040415 | 0.323091171 | 3.32E-05 | 0.000578 | 2823.995 | 2717.596 | 2771.673 | 3092.505 | 2467.032 | 2280.561 | 2285.365 | 2084.912 |
| ENSMUSG00000040420 | 0.242165433 | 0.602328 | 0.740165 | 13.07871 | 13.19804 | 11.59349 | 18.11509 | 16.57689 | 10.05637 | 7.714313 | 13.05109 |
| ENSMUSG00000040423 | 0.340513082 | 0.013746 | 0.054066 | 1466.827 | 1285.009 | 1305.096 | 1493.201 | 1007.29 | 1400.07 | 1032.754 | 945.1167 |
| ENSMUSG00000040424 | 0.093750017 | 0.628264 | 0.757958 | 89.53884 | 116.3827 | 110.9663 | 106.9653 | 99.46135 | 116.2069 | 105.1075 | 76.13138 |
| ENSMUSG00000040428 | 0.618679065 | 4.68E-09 | 4.58E-07 | 493.9727 | 616.7083 | 544.0661 | 526.2003 | 361.7663 | 374.3203 | 353.8941 | 328.4525 |
| ENSMUSG00000040429 | 0.34877043 | 0.128296 | 0.259802 | 198.1927 | 172.7743 | 129.1846 | 171.6621 | 95.56091 | 173.193 | 116.679 | 142.4744 |
| ENSMUSG00000040430 | 0.237127381 | 0.074327 | 0.179906 | 1524.172 | 1353.399 | 1660.354 | 1585.502 | 1054.095 | 1576.615 | 1261.29 | 1307.284 |
| ENSMUSG00000040432 | -1.203243184 | 0.020886 | 0.072688 | 9.054489 | 8.398751 | 8.281066 | 4.313117 | 25.35289 | 11.17374 | 13.50005 | 18.48905 |
| ENSMUSG00000040433 | -0.096274785 | 0.32693 | 0.494615 | 927.5821 | 723.4924 | 844.6688 | 806.553 | 862.9735 | 845.8522 | 871.7174 | 952.7298 |
| ENSMUSG00000040434 | -0.555196659 | 0.001188 | 0.008902 | 576.4691 | 579.5138 | 642.6107 | 494.2833 | 783.0144 | 614.5558 | 942.1105 | 1028.861 |
| ENSMUSG00000040435 | -0.100372281 | 0.54833 | 0.696723 | 609.6689 | 407.9393 | 545.7223 | 543.4528 | 565.5646 | 435.7759 | 618.1094 | 640.5912 |
| ENSMUSG00000040441 | 0.770725005 | 0.003776 | 0.020951 | 121.7326 | 93.58608 | 96.06037 | 130.2561 | 69.2329 | 39.10809 | 64.60737 | 85.9197 |
| ENSMUSG00000040446 | 0.019478741 | 0.777882 | 0.86305 | 972.8546 | 934.661 | 883.5898 | 983.3908 | 956.5842 | 937.4769 | 924.7533 | 903.7882 |
| ENSMUSG00000040447 | -0.519988845 | 0.003223 | 0.018717 | 358.1554 | 279.5584 | 476.9894 | 370.9281 | 525.585 | 452.5365 | 540.0019 | 616.6641 |
| ENSMUSG00000040451 | -0.376785999 | 0.012806 | 0.051414 | 1173.059 | 962.2569 | 1269.487 | 1242.178 | 1316.4 | 1234.698 | 1769.471 | 1716.219 |
| ENSMUSG00000040455 | 0.383230009 | 0.007045 | 0.033083 | 821.9464 | 613.1088 | 554.0033 | 709.0765 | 521.6845 | 562.0392 | 467.6802 | 517.6934 |
| ENSMUSG00000040459 | 0.355534905 | 0.012675 | 0.050998 | 9695.346 | 9437.796 | 8007.791 | 9864.962 | 7160.242 | 9225.041 | 6606.345 | 5931.722 |
| ENSMUSG00000040462 | -0.12108416 | 0.274822 | 0.4409 | 3282.755 | 3067.944 | 3605.576 | 3384.072 | 3471.396 | 3021.38 | 3921.764 | 4095.868 |
| ENSMUSG00000040463 | -0.151672104 | 0.00153 | 0.010695 | 4702.298 | 4852.078 | 5010.873 | 4665.93 | 5391.39 | 5157.799 | 5389.412 | 5423.817 |
| ENSMUSG00000040464 | -0.003603504 | 0.972202 | 0.984619 | 966.8182 | 898.6663 | 818.9975 | 937.6717 | 847.3717 | 1040.275 | 832.1815 | 912.4889 |
| ENSMUSG00000040466 | 0.342482122 | 0.000102 | 0.001359 | 607.6568 | 633.5058 | 663.3134 | 669.3958 | 551.913 | 512.8747 | 468.6445 | 498.1167 |
| ENSMUSG00000040471 | -2.141901614 | 0.001916 | 0.012726 | 17.10292 | 2.399643 | 24.01509 | 5.175741 | 78.0089 | 16.76061 | 48.21446 | 73.95619 |
| ENSMUSG00000040472 | -0.410249766 | 0.001482 | 0.010473 | 607.6568 | 753.4879 | 712.1717 | 631.4404 | 937.082 | 725.1758 | 926.6819 | 1002.759 |
| ENSMUSG00000040473 | 0.124324372 | 0.59917 | 0.737465 | 219.3198 | 199.1704 | 146.5749 | 241.5346 | 177.4703 | 246.9397 | 186.1078 | 129.4233 |
| ENSMUSG00000040479 | -0.248871031 | 0.003398 | 0.019369 | 1676.087 | 1737.342 | 1961.785 | 1791.669 | 2065.286 | 1972.165 | 2136.865 | 2345.934 |
| ENSMUSG00000040481 | 0.444443047 | 0.008164 | 0.036829 | 4376.336 | 4000.205 | 3263.568 | 3503.977 | 2517.737 | 3761.081 | 2513.902 | 2336.146 |

| | | | | | | | | | | | |
|---|---|---|---|---|---|---|---|---|---|---|---|
| ENSMUSG00000040482 | -0.158565421 | 0.024608 | 0.082043 | 561.3783 | 568.7154 | 581.3309 | 584.8587 | 654.2997 | 631.3164 | 630.6451 | 648.2043 |
| ENSMUSG00000040483 | 0.338968875 | 0.38498 | 0.552546 | 164.9929 | 415.1383 | 439.7246 | 117.3168 | 206.7236 | 187.7188 | 259.3938 | 244.708 |
| ENSMUSG00000040488 | 0.507088194 | 9.14E-06 | 0.000213 | 2696.226 | 3720.647 | 3574.936 | 3440.142 | 2346.118 | 2586.721 | 2237.151 | 2281.766 |
| ENSMUSG00000040489 | 0.537965763 | 0.19509 | 0.348939 | 33.19979 | 39.59411 | 27.32752 | 19.84034 | 30.22845 | 12.29112 | 15.42863 | 23.927 |
| ENSMUSG00000040490 | 0.246561278 | 0.597314 | 0.736063 | 20.12109 | 53.99197 | 55.48314 | 37.09281 | 23.40267 | 51.39921 | 18.32149 | 47.85401 |
| ENSMUSG00000040502 | -0.334349566 | 0.013995 | 0.054744 | 360.1675 | 374.3443 | 333.727 | 339.8737 | 403.6961 | 372.0856 | 464.7874 | 532.9196 |
| ENSMUSG00000040505 | 0.367750643 | 0.085913 | 0.198475 | 161.9748 | 251.9625 | 188.8083 | 184.6014 | 197.9476 | 117.3243 | 151.3934 | 141.3868 |
| ENSMUSG00000040506 | 0.075350233 | 0.266975 | 0.431387 | 1505.057 | 1450.584 | 1610.667 | 1419.878 | 1380.758 | 1479.403 | 1395.326 | 1430.182 |
| ENSMUSG00000040511 | 0.601795356 | 3.00E-07 | 1.37E-05 | 524.1543 | 457.132 | 443.037 | 412.334 | 320.8116 | 321.8037 | 278.6796 | 288.2116 |
| ENSMUSG00000040520 | 0.338865879 | 0.002009 | 0.013178 | 999.012 | 1190.223 | 1069.914 | 1021.346 | 775.2135 | 993.3456 | 793.61 | 822.2189 |
| ENSMUSG00000040521 | -0.425273937 | 0.000506 | 0.004629 | 650.9172 | 657.5022 | 663.3134 | 566.7436 | 900.0277 | 686.0677 | 879.4317 | 940.7663 |
| ENSMUSG00000040522 | 0.710800596 | 0.23248 | 0.393025 | 26.15741 | 9.598572 | 14.07781 | 16.38985 | 6.825779 | 5.58687 | 20.25007 | 7.613138 |
| ENSMUSG00000040524 | 0.402554639 | 3.36E-09 | 3.52E-07 | 1215.314 | 1263.412 | 1344.017 | 1321.539 | 970.2358 | 1021.28 | 935.3605 | 969.0437 |
| ENSMUSG00000040525 | -0.686266575 | 0.010366 | 0.043758 | 137.8294 | 95.98572 | 216.1358 | 106.9653 | 238.9023 | 182.132 | 254.5723 | 222.9562 |
| ENSMUSG00000040528 | 0.50469605 | 0.114751 | 0.242317 | 56.33904 | 52.79215 | 29.81184 | 44.85642 | 30.22845 | 34.6386 | 24.10723 | 40.24087 |
| ENSMUSG00000040532 | -0.010611877 | 0.8932 | 0.935419 | 953.7395 | 917.8635 | 984.6188 | 985.116 | 1018.991 | 893.8993 | 923.789 | 1035.387 |
| ENSMUSG00000040536 | 1.376919154 | 0.000125 | 0.001589 | 35.2119 | 53.99197 | 29.81184 | 50.89479 | 18.52711 | 17.87799 | 12.53576 | 16.31387 |
| ENSMUSG00000040537 | 0.406730708 | 0.005864 | 0.028934 | 771.6437 | 749.8885 | 811.5445 | 826.3933 | 448.5512 | 736.3495 | 603.645 | 597.0875 |
| ENSMUSG00000040540 | -0.499126289 | 0.164393 | 0.309391 | 23.13925 | 13.19804 | 22.35888 | 18.97772 | 30.22845 | 18.99536 | 32.78583 | 28.27737 |
| ENSMUSG00000040543 | -0.3471115 | 0.237695 | 0.398881 | 52.31483 | 74.38894 | 94.40416 | 54.34528 | 68.25779 | 70.39457 | 92.57176 | 119.635 |
| ENSMUSG00000040548 | -0.178795321 | 0.356973 | 0.525467 | 554.3359 | 399.5406 | 395.0069 | 456.3278 | 431.9743 | 386.6114 | 622.9308 | 601.4379 |
| ENSMUSG00000040549 | -2.91E-05 | 0.999619 | 0.999733 | 2663.026 | 2658.805 | 2383.291 | 2639.628 | 2574.294 | 2674.994 | 2543.795 | 2548.226 |
| ENSMUSG00000040550 | -0.302260632 | 0.005877 | 0.028969 | 1115.714 | 1135.031 | 1151.896 | 1157.641 | 1410.011 | 1153.13 | 1433.898 | 1625.949 |
| ENSMUSG00000040552 | 0.613761482 | 0.05251 | 0.14082 | 77.46619 | 141.5789 | 125.8722 | 56.07053 | 67.28268 | 65.92507 | 70.39311 | 57.64233 |
| ENSMUSG00000040554 | 0.06342898 | 0.728953 | 0.829749 | 136.8234 | 157.1766 | 166.4494 | 174.2499 | 159.9183 | 183.2494 | 114.7504 | 151.1752 |
| ENSMUSG00000040557 | 0.007745133 | 0.934372 | 0.961247 | 877.2794 | 862.6717 | 805.7478 | 925.595 | 944.8829 | 860.3781 | 884.2532 | 761.3138 |
| ENSMUSG00000040560 | -0.076679071 | 0.544264 | 0.692898 | 547.2936 | 427.1365 | 476.1613 | 425.2734 | 485.6054 | 430.189 | 520.7161 | 541.6204 |
| ENSMUSG00000040562 | -0.131579738 | 0.185816 | 0.337442 | 2157.987 | 2082.89 | 2257.419 | 2074.61 | 2282.736 | 2381.124 | 2708.688 | 2018.569 |
| ENSMUSG00000040563 | -0.033819157 | 0.844756 | 0.906563 | 281.6952 | 181.1731 | 221.9326 | 280.3526 | 261.3298 | 216.7706 | 252.6438 | 258.8467 |
| ENSMUSG00000040564 | -0.431900397 | 0.111188 | 0.237374 | 91.55095 | 67.19001 | 114.2787 | 69.8725 | 105.312 | 89.38993 | 162.0006 | 106.5839 |
| ENSMUSG00000040565 | 0.023556943 | 0.861396 | 0.917047 | 1480.912 | 1046.244 | 1306.752 | 1289.622 | 1493.871 | 1157.6 | 1147.504 | 1243.117 |
| ENSMUSG00000040569 | -0.522749197 | 0.028639 | 0.091448 | 256.5439 | 230.3657 | 221.9326 | 263.1002 | 244.7529 | 536.3396 | 315.3226 | 301.2627 |
| ENSMUSG00000040570 | -0.050321275 | 0.804303 | 0.880231 | 140.8476 | 109.1838 | 97.71658 | 148.3712 | 133.5902 | 151.9629 | 115.7147 | 113.1095 |
| ENSMUSG00000040584 | 1.573589633 | 0.002584 | 0.015783 | 224.3501 | 117.5825 | 61.27989 | 170.7995 | 21.45245 | 101.681 | 38.57157 | 31.54014 |
| ENSMUSG00000040586 | 0.14329861 | 0.259453 | 0.423495 | 944.685 | 837.4754 | 933.2762 | 892.8153 | 776.1886 | 1014.576 | 778.1814 | 700.4087 |
| ENSMUSG00000040591 | 0.255939253 | 0.011423 | 0.047152 | 866.2128 | 1057.043 | 914.2297 | 1034.286 | 851.2722 | 862.6128 | 791.6814 | 735.2116 |
| ENSMUSG00000040592 | 0.100874316 | 0.474293 | 0.633764 | 142.8597 | 143.9786 | 167.2775 | 152.6844 | 132.6151 | 139.6718 | 138.8576 | 156.6131 |
| ENSMUSG00000040594 | 0.203907569 | 0.231489 | 0.392032 | 422.5428 | 423.537 | 372.648 | 414.9219 | 397.8454 | 433.5411 | 330.7512 | 255.5839 |
| ENSMUSG00000040596 | 0.13365177 | 0.118284 | 0.24712 | 1741.48 | 1892.119 | 1747.305 | 1794.257 | 1596.257 | 1835.846 | 1643.149 | 1463.898 |
| ENSMUSG00000040599 | -0.138038079 | 0.241993 | 0.404403 | 686.1291 | 730.6913 | 555.6596 | 582.2709 | 764.4873 | 691.6546 | 702.0025 | 647.1167 |
| ENSMUSG00000040605 | -0.116263668 | 0.518962 | 0.672618 | 113.6841 | 137.9795 | 170.59 | 127.6683 | 170.6445 | 139.6718 | 151.3934 | 134.8613 |
| ENSMUSG00000040606 | -0.22220746 | 0.092714 | 0.209262 | 1012.091 | 881.8688 | 1118.772 | 863.4861 | 1138.93 | 927.4205 | 1184.147 | 1272.482 |
| ENSMUSG00000040612 | -0.408428802 | 0.100551 | 0.221689 | 280.6892 | 233.9652 | 350.2891 | 238.0841 | 218.4249 | 445.8323 | 345.2155 | 456.7883 |
| ENSMUSG00000040613 | -0.44902745 | 0.173227 | 0.320991 | 57.3451 | 82.78769 | 97.71658 | 46.58167 | 142.3663 | 67.04245 | 95.46463 | 82.65692 |
| ENSMUSG00000040616 | -0.363767236 | 0.075241 | 0.181332 | 601.6205 | 563.9161 | 817.3412 | 632.303 | 887.3513 | 547.5133 | 891.0032 | 1040.825 |
| ENSMUSG00000040618 | 0.105105881 | 0.319528 | 0.487736 | 1754.559 | 1797.333 | 1512.951 | 1677.803 | 1553.352 | 1724.108 | 1661.47 | 1326.861 |
| ENSMUSG00000040620 | -0.107030162 | 0.116532 | 0.244705 | 1167.023 | 1141.03 | 1139.475 | 1091.219 | 1274.47 | 1280.511 | 1180.29 | 1151.759 |
| ENSMUSG00000040621 | -0.063203295 | 0.56209 | 0.707587 | 280.6892 | 343.149 | 298.1184 | 328.6596 | 345.1894 | 325.1559 | 323.0369 | 311.0511 |
| ENSMUSG00000040624 | 0.066222844 | 0.478145 | 0.637481 | 1021.145 | 862.6717 | 1060.805 | 971.3141 | 865.8988 | 1004.519 | 943.0748 | 932.0656 |
| ENSMUSG00000040629 | -5.239484736 | NA | NA | 3.018163 | 0 | 0 | 2.58787 | 0 | 212.3011 | 0 | 0 |
| ENSMUSG00000040631 | -0.396269495 | 0.02135 | 0.073761 | 598.6023 | 649.1034 | 872.8244 | 665.0827 | 929.2811 | 687.1851 | 967.182 | 1084.328 |
| ENSMUSG00000040632 | -0.063789118 | 0.710551 | 0.818711 | 221.332 | 253.1623 | 207.0267 | 226.0074 | 250.6036 | 281.5783 | 173.572 | 242.5328 |
| ENSMUSG00000040640 | -0.02501994 | 0.862212 | 0.917545 | 161.9748 | 141.5789 | 168.1056 | 171.6621 | 142.3663 | 162.0192 | 178.3935 | 174.0146 |
| ENSMUSG00000040648 | -0.043264439 | 0.469418 | 0.629438 | 1504.051 | 1544.17 | 1542.763 | 1521.668 | 1485.095 | 1651.479 | 1548.648 | 1616.16 |
| ENSMUSG00000040649 | 0.853469436 | 7.09E-07 | 2.82E-05 | 1474.876 | 1481.78 | 1445.874 | 1567.387 | 832.7451 | 1138.604 | 658.6095 | 675.3941 |
| ENSMUSG00000040651 | 0.490045632 | 0.000472 | 0.004373 | 1547.312 | 1625.758 | 1459.952 | 1523.393 | 927.3309 | 1426.887 | 1005.754 | 1024.511 |
| ENSMUSG00000040652 | 0.076671164 | 0.147233 | 0.286457 | 5169.107 | 5477.185 | 5638.578 | 5738.171 | 5167.115 | 5364.513 | 5137.733 | 5219.35 |
| ENSMUSG00000040653 | 0.12811076 | 0.618759 | 0.751194 | 96.58122 | 115.1829 | 94.40416 | 123.3552 | 81.90935 | 147.4934 | 75.21455 | 89.18247 |
| ENSMUSG00000040658 | -0.520008177 | 1.34E-06 | 4.73E-05 | 417.5126 | 488.3274 | 388.382 | 406.2957 | 666.001 | 565.3913 | 602.6807 | 597.0875 |
| ENSMUSG00000040659 | -0.137334661 | 0.375154 | 0.54362 | 1009.073 | 955.0579 | 1411.922 | 1120.548 | 1320.301 | 987.7587 | 1322.04 | 1318.16 |
| ENSMUSG00000040661 | -0.018312669 | 0.802244 | 0.879281 | 1317.931 | 1189.023 | 1199.098 | 1159.366 | 1274.47 | 1272.689 | 1161.004 | 1219.19 |
| ENSMUSG00000040663 | 0.120146959 | 0.433083 | 0.597256 | 363.1856 | 350.3479 | 411.569 | 410.6088 | 323.737 | 455.8886 | 323.0369 | 313.2262 |
| ENSMUSG00000040666 | -0.114727783 | 0.624299 | 0.7551 | 83.50251 | 136.7797 | 100.2009 | 116.4542 | 140.416 | 134.0849 | 85.82174 | 112.0219 |
| ENSMUSG00000040667 | -0.107949876 | 0.110671 | 0.236762 | 2449.742 | 2254.465 | 2294.683 | 2374.802 | 2609.398 | 2301.791 | 2566.938 | 2623.27 |

| | | | | | | | | | | | |
|---|---|---|---|---|---|---|---|---|---|---|---|
| ENSMUSG00000040669 | 0.052431884 | 0.64355 | 0.769509 | 1914.521 | 1788.934 | 1903.817 | 2042.692 | 1867.338 | 2172.175 | 1806.114 | 1533.503 |
| ENSMUSG00000040675 | -0.134268185 | 0.165741 | 0.311164 | 652.9293 | 759.487 | 730.39 | 693.5493 | 800.5664 | 681.5982 | 837.003 | 790.6787 |
| ENSMUSG00000040680 | -0.43303306 | 0.351405 | 0.520215 | 9.054489 | 10.79839 | 13.24971 | 15.52722 | 22.42756 | 14.52586 | 9.642892 | 19.57664 |
| ENSMUSG00000040681 | -0.098065959 | 0.165748 | 0.311164 | 11226.56 | 13565.18 | 12604.61 | 12921.24 | 13673.99 | 13774.99 | 13174.12 | 13231.63 |
| ENSMUSG00000040687 | -0.014934598 | 0.879705 | 0.927815 | 480.894 | 463.1311 | 487.7548 | 451.1521 | 428.0739 | 452.5365 | 520.7161 | 501.3795 |
| ENSMUSG00000040688 | -0.399522706 | 1.41E-05 | 0.000297 | 1110.684 | 1059.442 | 1132.022 | 1118.823 | 1537.751 | 1243.637 | 1478.255 | 1572.657 |
| ENSMUSG00000040690 | 0.246033303 | 0.082454 | 0.192948 | 1758.583 | 2589.215 | 2623.442 | 2230.744 | 1677.191 | 2028.034 | 1981.614 | 2072.949 |
| ENSMUSG00000040693 | 0.617967836 | 0.013861 | 0.054385 | 1389.361 | 1622.159 | 1019.399 | 1269.782 | 785.9397 | 1301.741 | 836.0387 | 529.6569 |
| ENSMUSG00000040694 | -0.108579965 | 0.751018 | 0.844912 | 28.16952 | 20.39697 | 24.01509 | 43.9938 | 33.15378 | 26.81698 | 32.78583 | 33.71532 |
| ENSMUSG00000040697 | -0.235616664 | 0.012475 | 0.050457 | 568.4207 | 587.9126 | 584.6433 | 599.5233 | 717.6819 | 596.6778 | 692.3596 | 748.2627 |
| ENSMUSG00000040699 | 0.012550904 | 0.892532 | 0.934962 | 1629.808 | 2025.299 | 2015.612 | 1924.513 | 1899.517 | 2023.564 | 1824.435 | 1782.562 |
| ENSMUSG00000040701 | 0.017024142 | 0.895303 | 0.936666 | 402.4217 | 322.752 | 408.2566 | 370.9281 | 344.2143 | 329.6254 | 415.6086 | 399.1459 |
| ENSMUSG00000040703 | 0.037083865 | 0.829628 | 0.896882 | 638.8445 | 873.4701 | 928.3075 | 703.9008 | 686.4784 | 640.2554 | 773.3599 | 964.6933 |
| ENSMUSG00000040706 | -1.88532556 | 0.009236 | 0.040213 | 13.07871 | 0 | 4.96864 | 3.450494 | 15.60178 | 18.99536 | 17.3572 | 28.27737 |
| ENSMUSG00000040710 | 0.790492256 | 0.003319 | 0.019096 | 178.0716 | 257.9616 | 154.8559 | 170.7995 | 91.66046 | 166.4887 | 90.64318 | 91.35765 |
| ENSMUSG00000040711 | 0.453454912 | 3.40E-07 | 1.53E-05 | 1482.924 | 1502.177 | 1650.417 | 1682.116 | 1110.652 | 1298.389 | 1085.79 | 1123.482 |
| ENSMUSG00000040712 | 0.116837602 | 0.255996 | 0.419812 | 729.3894 | 565.1159 | 608.6584 | 685.7857 | 604.569 | 585.504 | 608.4665 | 590.562 |
| ENSMUSG00000040713 | -0.296644824 | 0.004783 | 0.024944 | 739.45 | 897.4665 | 884.4179 | 726.329 | 1051.17 | 1008.989 | 910.289 | 1016.898 |
| ENSMUSG00000040714 | -0.217102689 | 0.14371 | 0.281501 | 130.7871 | 141.5789 | 139.1219 | 168.2116 | 188.1965 | 159.7845 | 159.1077 | 167.489 |
| ENSMUSG00000040717 | 0.37439262 | 2.80E-06 | 8.57E-05 | 2413.524 | 2558.02 | 2277.293 | 2352.374 | 1921.944 | 1997.865 | 1818.649 | 1665.102 |
| ENSMUSG00000040720 | -0.046835807 | 0.457699 | 0.61944 | 2114.726 | 1989.304 | 1973.378 | 1942.628 | 1927.795 | 2111.837 | 2117.579 | 2127.328 |
| ENSMUSG00000040721 | 0.766521476 | 3.17E-05 | 0.000559 | 345.0766 | 425.9366 | 257.5412 | 312.2697 | 194.0472 | 232.4138 | 175.5006 | 184.8905 |
| ENSMUSG00000040722 | -0.122295758 | 0.48733 | 0.645251 | 788.7466 | 736.6904 | 1205.723 | 772.048 | 1036.543 | 835.7958 | 892.9318 | 1050.613 |
| ENSMUSG00000040723 | 0.149313692 | 0.123608 | 0.253282 | 2837.073 | 2537.623 | 2588.661 | 2944.997 | 2262.258 | 2610.186 | 2731.831 | 2232.825 |
| ENSMUSG00000040724 | 1.135989858 | 0.000557 | 0.004971 | 192.1564 | 212.3684 | 171.4181 | 213.068 | 69.2329 | 165.3714 | 58.82164 | 66.34306 |
| ENSMUSG00000040725 | -0.092298699 | 0.195156 | 0.348978 | 5579.577 | 5302.011 | 5894.463 | 5560.471 | 6052.516 | 5355.574 | 6182.058 | 6224.284 |
| ENSMUSG00000040726 | -0.257049991 | 0.596293 | 0.735213 | 28.16952 | 20.39697 | 17.39024 | 9.488858 | 34.1289 | 18.99536 | 12.53576 | 23.927 |
| ENSMUSG00000040728 | -0.092135843 | 0.490279 | 0.647606 | 984.9272 | 853.0731 | 1052.524 | 853.1346 | 897.1024 | 1215.703 | 995.1464 | 885.2991 |
| ENSMUSG00000040729 | 0.127101859 | 0.256197 | 0.419963 | 336.0222 | 311.9536 | 273.2752 | 295.0172 | 285.7076 | 290.5173 | 269.0367 | 266.4598 |
| ENSMUSG00000040731 | -0.254437022 | 0.000734 | 0.006166 | 10306.02 | 10516.44 | 11727.65 | 10625.8 | 13089.89 | 11586.05 | 13407.48 | 13421.96 |
| ENSMUSG00000040732 | -0.205025738 | 0.365628 | 0.534244 | 429.5852 | 421.1374 | 699.7501 | 582.2709 | 567.5148 | 429.0717 | 696.2168 | 767.8393 |
| ENSMUSG00000040734 | 0.040205871 | 0.844952 | 0.906601 | 104.6297 | 106.7841 | 160.6527 | 139.745 | 144.3165 | 112.8548 | 109.929 | 132.6861 |
| ENSMUSG00000040738 | 0.019423735 | 0.838632 | 0.90239 | 1233.423 | 1096.637 | 1003.665 | 1178.344 | 1124.303 | 1208.999 | 1094.468 | 1023.423 |
| ENSMUSG00000040740 | -2.726239084 | 5.37E-07 | 2.23E-05 | 16.09687 | 11.99822 | 18.21835 | 13.80198 | 130.6649 | 25.6996 | 66.53595 | 176.1898 |
| ENSMUSG00000040746 | 0.224630773 | 0.003163 | 0.018476 | 2253.562 | 2247.266 | 2206.904 | 2340.298 | 1867.338 | 2136.419 | 1977.757 | 1762.985 |
| ENSMUSG00000040747 | 0.528557113 | 0.018608 | 0.067083 | 91.55095 | 107.9839 | 119.2474 | 79.36136 | 73.13335 | 62.57295 | 82.92887 | 56.55474 |
| ENSMUSG00000040749 | -0.137588552 | 0.251967 | 0.415613 | 376.2643 | 374.3443 | 405.7722 | 335.5605 | 462.2028 | 433.5411 | 370.287 | 375.2189 |
| ENSMUSG00000040751 | 0.489677878 | 0.15685 | 0.299207 | 32.19374 | 47.99286 | 45.54586 | 62.10889 | 32.17867 | 50.28183 | 27.96439 | 23.927 |
| ENSMUSG00000040752 | 0.693375747 | 0.214404 | 0.372602 | 42.25428 | 92.38626 | 48.85829 | 53.48266 | 35.10401 | 79.33356 | 21.21436 | 10.87591 |
| ENSMUSG00000040759 | 0.028076117 | 0.957591 | 0.975954 | 14.08476 | 11.99822 | 17.39024 | 40.5433 | 14.62667 | 31.28647 | 23.14294 | 14.13868 |
| ENSMUSG00000040760 | 0.276990798 | 0.012331 | 0.049983 | 2074.484 | 2024.099 | 1652.073 | 1880.519 | 1527.999 | 1826.907 | 1489.827 | 1451.934 |
| ENSMUSG00000040761 | 0.22459838 | 0.03002 | 0.094613 | 806.8556 | 622.7074 | 717.9685 | 721.1532 | 595.793 | 667.0723 | 604.6093 | 590.562 |
| ENSMUSG00000040767 | -0.159543571 | 0.129255 | 0.261057 | 585.5236 | 571.1151 | 564.7687 | 546.9033 | 707.9308 | 539.6917 | 667.2881 | 615.5766 |
| ENSMUSG00000040771 | -0.050418453 | 0.44695 | 0.609585 | 1172.053 | 1207.02 | 1121.256 | 1189.558 | 1255.943 | 1264.867 | 1147.504 | 1186.562 |
| ENSMUSG00000040774 | -0.085917101 | 0.351529 | 0.520274 | 1004.042 | 1149.429 | 989.5874 | 1081.73 | 1023.867 | 1206.764 | 1062.647 | 1189.825 |
| ENSMUSG00000040782 | -0.020332379 | 0.757097 | 0.848808 | 2224.386 | 2358.849 | 2204.42 | 2332.534 | 2195.951 | 2507.387 | 2286.33 | 2260.014 |
| ENSMUSG00000040785 | 0.271059159 | 0.022219 | 0.075941 | 8903.581 | 9257.823 | 7712.985 | 8708.184 | 6726.318 | 8801.556 | 6760.631 | 6370.021 |
| ENSMUSG00000040794 | 0.279404328 | 0.034047 | 0.103828 | 305.8405 | 407.9393 | 384.2415 | 366.615 | 311.0605 | 308.3953 | 262.2867 | 325.1897 |
| ENSMUSG00000040795 | -0.049687822 | 0.664353 | 0.784372 | 341.0524 | 319.1525 | 309.7119 | 304.5061 | 331.5378 | 286.0478 | 368.3585 | 330.6277 |
| ENSMUSG00000040797 | 0.179135474 | 0.424599 | 0.589675 | 64.38748 | 70.78947 | 83.63877 | 97.47645 | 72.15824 | 80.45093 | 71.3574 | 56.55474 |
| ENSMUSG00000040808 | -2.441163717 | 4.65E-15 | 2.09E-12 | 1246.501 | 853.0731 | 1661.182 | 1080.005 | 4698.086 | 3428.104 | 8626.531 | 9542.524 |
| ENSMUSG00000040809 | 0.252559408 | 0.722212 | 0.826177 | 40.24217 | 3.599465 | 12.4216 | 21.56559 | 18.52711 | 6.704245 | 27.0001 | 13.05109 |
| ENSMUSG00000040811 | 0.204820776 | 0.043254 | 0.123157 | 1199.217 | 1055.843 | 1040.102 | 1074.829 | 917.5797 | 823.5047 | 1048.182 | 1000.584 |
| ENSMUSG00000040812 | 0.238584395 | 0.388258 | 0.555834 | 63.38142 | 44.3934 | 43.88965 | 61.24627 | 53.63112 | 39.10809 | 34.71441 | 53.29196 |
| ENSMUSG00000040813 | -0.383866747 | 0.016082 | 0.060413 | 1064.406 | 974.2551 | 1342.361 | 1129.174 | 1494.846 | 1116.257 | 1489.827 | 1786.912 |
| ENSMUSG00000040818 | 0.058268926 | 0.564115 | 0.709266 | 1355.155 | 1421.789 | 1107.179 | 1374.159 | 1182.81 | 1358.727 | 1248.754 | 1257.255 |
| ENSMUSG00000040820 | -0.209000642 | 0.020183 | 0.070946 | 354.1311 | 359.9465 | 382.5853 | 351.0878 | 406.6214 | 401.1373 | 426.2158 | 440.4744 |
| ENSMUSG00000040822 | -0.140942811 | 0.15265 | 0.293622 | 1010.079 | 1046.244 | 1136.162 | 1171.443 | 1268.62 | 1050.332 | 1181.254 | 1313.81 |
| ENSMUSG00000040824 | -0.206718163 | 0.021835 | 0.074979 | 3493.021 | 3973.809 | 3772.026 | 3608.354 | 4765.369 | 3742.086 | 4266.015 | 4356.89 |
| ENSMUSG00000040828 | 0.225067851 | 0.225307 | 0.385096 | 88.53278 | 103.1847 | 89.43552 | 75.04824 | 76.05868 | 72.62932 | 74.25027 | 80.48174 |
| ENSMUSG00000040829 | 0.575828298 | 0.01624 | 0.060854 | 74.44802 | 74.38894 | 85.29498 | 54.34528 | 47.78045 | 40.22547 | 54.00019 | 51.11678 |
| ENSMUSG00000040836 | 0.234469408 | 0.002186 | 0.013994 | 706.2502 | 746.289 | 715.4841 | 672.8463 | 603.5939 | 635.7859 | 595.9307 | 577.5109 |
| ENSMUSG00000040840 | 0.195259651 | 0.310656 | 0.479267 | 240.447 | 251.9625 | 190.4645 | 232.9083 | 270.1058 | 182.132 | 170.6792 | 175.1022 |
| ENSMUSG00000040841 | 0.112271752 | 0.305086 | 0.473524 | 843.0736 | 1095.437 | 1003.665 | 993.7423 | 912.7042 | 1000.05 | 917.039 | 810.2554 |

| | | | | | | | | | | | |
|---|---|---|---|---|---|---|---|---|---|---|---|
| ENSMUSG00000040842 | -0.166358011 | 0.045333 | 0.127133 | 3000.054 | 2994.755 | 3064.823 | 2949.31 | 3558.181 | 2892.882 | 3433.834 | 3590.138 |
| ENSMUSG00000040843 | 0.01599112 | 0.819275 | 0.890648 | 1446.706 | 1378.595 | 1458.296 | 1563.936 | 1383.683 | 1491.694 | 1400.148 | 1513.927 |
| ENSMUSG00000040848 | 0.192874911 | 0.003954 | 0.021671 | 2754.577 | 2943.162 | 2855.312 | 2963.112 | 2523.588 | 2678.346 | 2566.938 | 2305.693 |
| ENSMUSG00000040850 | -0.082893801 | 0.21299 | 0.370828 | 2816.952 | 2731.994 | 2646.629 | 2584.42 | 2660.104 | 3071.661 | 2836.939 | 2848.401 |
| ENSMUSG00000040852 | 0.173086386 | 0.017502 | 0.064258 | 1228.392 | 1312.605 | 1196.614 | 1329.303 | 1113.577 | 1110.67 | 1198.611 | 1068.014 |
| ENSMUSG00000040855 | 0.307579831 | 0.018594 | 0.067064 | 378.2764 | 419.9375 | 468.7084 | 408.0209 | 338.3636 | 402.2547 | 304.7154 | 309.9635 |
| ENSMUSG00000040856 | 1.03812962 | 0.069224 | 0.170848 | 2569.463 | 4468.135 | 3132.727 | 4120.752 | 1526.049 | 3860.528 | 1019.254 | 553.5839 |
| ENSMUSG00000040857 | -0.284827815 | 0.010192 | 0.043241 | 1526.184 | 1482.979 | 1586.652 | 1481.987 | 1844.911 | 1498.399 | 1966.186 | 2094.7 |
| ENSMUSG00000040859 | 0.03766565 | 0.669213 | 0.787855 | 1090.563 | 1097.837 | 1051.695 | 1027.385 | 974.1362 | 966.5286 | 1180.29 | 1033.212 |
| ENSMUSG00000040860 | -0.113874036 | 0.21925 | 0.378035 | 850.1159 | 1039.045 | 973.0253 | 905.7547 | 962.4349 | 1007.871 | 994.1821 | 1111.518 |
| ENSMUSG00000040865 | 0.355991098 | 0.004097 | 0.022205 | 1411.494 | 1350.999 | 1224.77 | 1383.648 | 992.6633 | 1302.858 | 927.6462 | 974.4816 |
| ENSMUSG00000040867 | 0.347409385 | 0.007029 | 0.033062 | 262.5802 | 268.76 | 277.4157 | 247.5729 | 216.4747 | 231.2964 | 209.2507 | 172.927 |
| ENSMUSG00000040875 | -0.48461841 | 0.028348 | 0.090844 | 221.332 | 218.3675 | 269.9628 | 207.0296 | 303.2596 | 202.2447 | 394.3943 | 382.8321 |
| ENSMUSG00000040877 | -0.032788392 | 0.787533 | 0.869461 | 181.0898 | 190.7716 | 200.4018 | 201.8539 | 192.0969 | 216.7706 | 186.1078 | 199.0292 |
| ENSMUSG00000040883 | -0.21252725 | 0.295731 | 0.463526 | 720.3349 | 387.5424 | 501.8326 | 440.8006 | 677.7024 | 492.762 | 564.1092 | 641.6787 |
| ENSMUSG00000040888 | -0.252465294 | 0.010693 | 0.044832 | 466.8092 | 598.7109 | 551.519 | 550.3538 | 651.3744 | 613.4384 | 645.1095 | 669.9561 |
| ENSMUSG00000040891 | 0.913655516 | 0.074875 | 0.18083 | 9.054489 | 19.19714 | 15.73403 | 21.56559 | 7.80089 | 5.58687 | 12.53576 | 8.700729 |
| ENSMUSG00000040896 | -0.175666162 | 0.638575 | 0.765743 | 301.8163 | 260.3613 | 852.9498 | 337.2858 | 470.9788 | 312.8647 | 529.3948 | 667.7809 |
| ENSMUSG00000040904 | -0.690522288 | 0.120306 | 0.249311 | 21.12714 | 10.79839 | 40.57722 | 20.70296 | 38.02934 | 24.58223 | 50.14304 | 39.15328 |
| ENSMUSG00000040907 | 0.200455388 | 0.043238 | 0.123138 | 620.7355 | 645.504 | 616.9394 | 709.0765 | 546.0623 | 613.4384 | 506.2518 | 592.7371 |
| ENSMUSG00000040913 | -0.101112811 | 0.451608 | 0.614001 | 699.2078 | 754.6877 | 695.6096 | 779.8116 | 831.7699 | 601.1473 | 806.1457 | 901.613 |
| ENSMUSG00000040918 | -0.104153767 | 0.317325 | 0.485577 | 351.113 | 337.1499 | 387.5539 | 346.7746 | 358.841 | 379.9072 | 423.3229 | 368.6934 |
| ENSMUSG00000040928 | 0.391969951 | 0.026339 | 0.086314 | 2440.688 | 2430.838 | 1688.509 | 2131.543 | 1678.167 | 2128.598 | 1539.006 | 1276.832 |
| ENSMUSG00000040929 | 0.503987501 | 0.013362 | 0.053047 | 707.2562 | 703.0954 | 590.44 | 719.428 | 409.5467 | 684.9503 | 479.2517 | 344.7664 |
| ENSMUSG00000040936 | 0.357071804 | 0.001272 | 0.009359 | 265.5983 | 313.1534 | 284.8687 | 295.0172 | 226.2258 | 235.7659 | 230.4651 | 210.9927 |
| ENSMUSG00000040938 | 0.119283446 | 0.315859 | 0.484252 | 187.1261 | 183.5727 | 197.0894 | 203.5791 | 182.3458 | 179.8972 | 178.3935 | 170.7518 |
| ENSMUSG00000040940 | 0.198170703 | 0.000158 | 0.001895 | 2439.682 | 2418.84 | 2269.84 | 2466.241 | 2064.311 | 2096.194 | 2087.686 | 2113.189 |
| ENSMUSG00000040943 | 0.178386551 | 0.048741 | 0.133803 | 1378.294 | 1270.611 | 1392.875 | 1317.226 | 1188.661 | 1344.201 | 1140.754 | 1064.752 |
| ENSMUSG00000040945 | 0.070602885 | 0.357668 | 0.526139 | 5689.237 | 6642.212 | 5881.213 | 5844.274 | 5736.58 | 6173.492 | 5652.663 | 5341.16 |
| ENSMUSG00000040952 | -0.138099912 | 0.105127 | 0.228558 | 21086.9 | 25305.44 | 24445.71 | 24048.22 | 27889.16 | 24239.2 | 27656.78 | 24629.59 |
| ENSMUSG00000040957 | -0.361396875 | 0.079583 | 0.18848 | 359.1614 | 229.1659 | 328.7583 | 313.995 | 319.8365 | 315.0995 | 494.6803 | 453.5255 |
| ENSMUSG00000040964 | -0.220648999 | 0.201138 | 0.356627 | 961.788 | 974.2551 | 1377.969 | 1009.269 | 1179.885 | 1001.167 | 1277.683 | 1581.357 |
| ENSMUSG00000040966 | -5.174584012 | 1.37E-09 | 1.71E-07 | 9.054489 | 4.799286 | 45.54586 | 18.11509 | 883.4508 | 151.9629 | 500.4661 | 1282.27 |
| ENSMUSG00000040969 | 0.016248656 | 0.914069 | 0.949091 | 409.4641 | 356.347 | 417.3657 | 353.6756 | 296.4338 | 433.5411 | 363.537 | 428.5109 |
| ENSMUSG00000040972 | 0.585812135 | 0.073702 | 0.178909 | 55.33299 | 46.79304 | 60.45178 | 61.24627 | 50.70579 | 18.99536 | 33.75012 | 45.67883 |
| ENSMUSG00000040978 | -2.421682467 | 2.77E-08 | 1.99E-06 | 26.15741 | 15.59768 | 35.60859 | 31.91707 | 154.0676 | 60.3382 | 126.3219 | 249.0584 |
| ENSMUSG00000040987 | 0.44965898 | 0.002182 | 0.013977 | 175.0535 | 208.7689 | 195.4332 | 173.3873 | 121.8889 | 148.6108 | 150.4291 | 129.4233 |
| ENSMUSG00000040990 | 0.236664645 | 0.043471 | 0.123545 | 1089.557 | 1329.402 | 1353.126 | 1277.545 | 1112.602 | 1231.346 | 1011.539 | 930.978 |
| ENSMUSG00000040997 | 0.082156975 | 0.390212 | 0.557613 | 2775.704 | 2273.662 | 2468.586 | 2512.822 | 2475.808 | 2060.438 | 2497.509 | 2441.642 |
| ENSMUSG00000040998 | -0.5501498 | 0.015087 | 0.057912 | 6603.741 | 6181.481 | 10881.32 | 6664.629 | 9719.91 | 7909.891 | 12224.29 | 14560.67 |
| ENSMUSG00000041000 | 0.191007584 | 0.132774 | 0.266153 | 475.8637 | 545.9188 | 549.8628 | 426.9986 | 493.4063 | 416.7805 | 394.3943 | 444.8248 |
| ENSMUSG00000041012 | -0.069789374 | 0.560875 | 0.706583 | 408.4581 | 345.5486 | 350.2891 | 402.8452 | 361.7663 | 359.7945 | 428.1444 | 432.8613 |
| ENSMUSG00000041014 | 1.166380406 | 0.004771 | 0.024904 | 29.17558 | 49.19268 | 30.63995 | 47.44429 | 20.47734 | 26.81698 | 11.57147 | 10.87591 |
| ENSMUSG00000041020 | 0.438387853 | 0.262393 | 0.426597 | 201.2109 | 119.9822 | 104.3414 | 166.4863 | 83.85957 | 208.949 | 76.17884 | 68.51824 |
| ENSMUSG00000041025 | -0.051942417 | 0.65041 | 0.7738 | 631.8021 | 608.3095 | 726.2495 | 623.6768 | 742.0597 | 621.26 | 592.0735 | 731.9488 |
| ENSMUSG00000041028 | -0.414406507 | 0.005771 | 0.028599 | 4821.012 | 3723.046 | 4525.603 | 4124.203 | 5470.374 | 4352.172 | 6163.736 | 6930.13 |
| ENSMUSG00000041035 | -0.220818325 | 0.021243 | 0.073573 | 697.1957 | 658.702 | 765.1705 | 693.5493 | 895.1522 | 741.9364 | 782.0385 | 863.5473 |
| ENSMUSG00000041037 | 0.267197684 | 0.001585 | 0.010978 | 1288.756 | 1210.62 | 1172.599 | 1060.164 | 933.1815 | 976.585 | 989.3607 | 1031.036 |
| ENSMUSG00000041040 | 0.068867421 | 0.564462 | 0.709411 | 1752.547 | 1552.569 | 1441.734 | 1900.36 | 1388.559 | 1715.169 | 1725.113 | 1509.576 |
| ENSMUSG00000041044 | -0.086287934 | 0.727245 | 0.829154 | 46.2785 | 40.79393 | 62.108 | 53.48266 | 50.70579 | 65.92507 | 47.25017 | 53.29196 |
| ENSMUSG00000041046 | -0.38913146 | 0.054847 | 0.145543 | 338.0343 | 244.7636 | 376.7885 | 331.2474 | 360.7912 | 334.0949 | 560.252 | 437.2116 |
| ENSMUSG00000041052 | -5.026631181 | 8.38E-09 | 7.40E-07 | 1.006054 | 4.799286 | 19.87456 | 6.038364 | 209.6489 | 79.33356 | 267.1081 | 490.5036 |
| ENSMUSG00000041057 | -0.087442252 | 0.163227 | 0.307935 | 2355.173 | 2524.425 | 2233.404 | 2364.451 | 2649.377 | 2496.214 | 2451.223 | 2465.569 |
| ENSMUSG00000041058 | 0.33863131 | 0.000968 | 0.007642 | 2702.262 | 2915.566 | 2425.524 | 2969.15 | 1927.795 | 2442.58 | 2214.972 | 2122.978 |
| ENSMUSG00000041064 | 0.36700081 | 0.012278 | 0.049834 | 477.8758 | 451.1329 | 423.9906 | 517.5741 | 412.4721 | 419.0153 | 299.8939 | 319.7518 |
| ENSMUSG00000041073 | 0.523542635 | 0.180791 | 0.331052 | 51.30877 | 31.19536 | 67.07664 | 92.30071 | 34.1289 | 29.05173 | 47.25017 | 58.72992 |
| ENSMUSG00000041075 | 0.398712975 | 0.001789 | 0.012109 | 2653.971 | 3083.541 | 2724.471 | 2429.148 | 1888.791 | 2415.763 | 1732.828 | 2224.124 |
| ENSMUSG00000041078 | 0.094586017 | 0.661498 | 0.782072 | 129.781 | 123.5816 | 111.7944 | 137.1571 | 86.78491 | 156.4324 | 102.2147 | 126.1606 |
| ENSMUSG00000041079 | -0.292858027 | 0.021283 | 0.073658 | 279.6831 | 310.7538 | 329.5864 | 263.9628 | 376.393 | 309.5126 | 375.1085 | 388.27 |
| ENSMUSG00000041084 | -0.081246918 | 0.132801 | 0.26616 | 3125.811 | 3444.688 | 3185.726 | 3360.781 | 3551.355 | 3406.874 | 3436.727 | 3477.029 |
| ENSMUSG00000041096 | 0.22313784 | 0.018368 | 0.066469 | 1242.477 | 1277.81 | 1309.237 | 1398.313 | 1039.469 | 1263.75 | 1161.968 | 1015.81 |
| ENSMUSG00000041112 | -0.13458609 | 0.213914 | 0.37205 | 250.5075 | 261.5611 | 303.087 | 281.2153 | 291.5583 | 301.691 | 304.7154 | 308.8759 |
| ENSMUSG00000041115 | 0.02967821 | 0.833718 | 0.89941 | 380.2885 | 284.3577 | 361.0545 | 338.1484 | 330.5627 | 319.569 | 295.0725 | 393.708 |
| ENSMUSG00000041117 | 0.082331499 | 0.354337 | 0.522963 | 1952.752 | 2098.488 | 2068.61 | 1875.343 | 1853.687 | 1652.596 | 2006.686 | 2035.971 |

| | | | | | | | | | | | |
|---|---|---|---|---|---|---|---|---|---|---|---|
| ENSMUSG00000041119 | -0.110845495 | 0.134022 | 0.267618 | 990.9635 | 1037.846 | 940.7291 | 1065.34 | 1127.229 | 1022.397 | 1089.647 | 1114.781 |
| ENSMUSG00000041120 | 0.748243287 | 0.001076 | 0.008262 | 1767.638 | 1941.311 | 1545.247 | 2246.272 | 1043.369 | 1679.413 | 989.3607 | 753.7006 |
| ENSMUSG00000041124 | 0.076546302 | 0.279096 | 0.445456 | 1518.136 | 1731.342 | 1543.591 | 1616.556 | 1515.323 | 1438.06 | 1588.184 | 1531.328 |
| ENSMUSG00000041126 | 0.016966715 | 0.82424 | 0.893455 | 4521.208 | 4799.286 | 4541.337 | 4788.423 | 5024.749 | 4770.07 | 4510.945 | 4124.145 |
| ENSMUSG00000041130 | -0.188305778 | 0.111818 | 0.238386 | 1190.162 | 1169.826 | 1210.692 | 1163.679 | 1413.911 | 1118.491 | 1252.612 | 1609.635 |
| ENSMUSG00000041132 | 0.706699354 | 0.001339 | 0.009733 | 559.3662 | 361.1463 | 317.9929 | 479.6187 | 255.4792 | 291.6346 | 309.5368 | 195.7664 |
| ENSMUSG00000041133 | -0.102998467 | 0.027146 | 0.088078 | 4107.72 | 4266.565 | 4264.749 | 4068.995 | 4438.707 | 4337.646 | 4567.838 | 4597.248 |
| ENSMUSG00000041134 | -0.386759677 | 0.142275 | 0.279433 | 131.7931 | 59.99108 | 86.9512 | 78.49874 | 125.7894 | 103.9158 | 102.2147 | 135.9489 |
| ENSMUSG00000041135 | 0.288336694 | 0.12612 | 0.256974 | 261.5741 | 293.9563 | 218.6202 | 251.8861 | 196.9725 | 264.8177 | 222.7508 | 154.4379 |
| ENSMUSG00000041141 | 0.906872683 | 0.009921 | 0.042337 | 73.44197 | 76.78858 | 52.17072 | 62.97152 | 29.25334 | 61.45558 | 26.03581 | 25.0146 |
| ENSMUSG00000041143 | -0.458797176 | 0.006105 | 0.029848 | 380.2885 | 310.7538 | 436.4122 | 343.3242 | 548.9877 | 377.6724 | 542.8948 | 553.5839 |
| ENSMUSG00000041144 | 0.36356072 | 0.044052 | 0.124678 | 300.8103 | 273.5593 | 318.8211 | 261.3749 | 216.4747 | 301.691 | 196.715 | 183.8029 |
| ENSMUSG00000041147 | 0.32042526 | 0.00157 | 0.010894 | 703.232 | 730.6913 | 622.7362 | 677.1594 | 575.3157 | 601.1473 | 515.8947 | 494.8539 |
| ENSMUSG00000041153 | -0.012238272 | 0.931287 | 0.95941 | 289.7437 | 361.1463 | 274.1033 | 295.0172 | 316.9112 | 349.7381 | 299.8939 | 261.0219 |
| ENSMUSG00000041161 | -0.488020949 | 0.01299 | 0.051973 | 165.999 | 151.1775 | 154.8559 | 158.7227 | 242.8027 | 145.2586 | 238.1794 | 257.7591 |
| ENSMUSG00000041164 | -0.062312907 | 0.436431 | 0.600273 | 1393.385 | 1506.976 | 1604.871 | 1393.137 | 1498.746 | 1470.464 | 1525.505 | 1665.102 |
| ENSMUSG00000041168 | -0.201664075 | 0.165057 | 0.310466 | 3562.438 | 2378.046 | 2565.474 | 2742.28 | 3402.163 | 2749.858 | 3150.333 | 3633.642 |
| ENSMUSG00000041180 | 0.37020303 | 0.244543 | 0.407283 | 228.3743 | 123.5816 | 99.3728 | 172.5247 | 89.71024 | 138.5544 | 163.9292 | 90.27006 |
| ENSMUSG00000041187 | 0.178067719 | 0.039585 | 0.115854 | 583.5115 | 620.3077 | 667.4539 | 609.0122 | 548.9877 | 517.3442 | 539.0376 | 588.3868 |
| ENSMUSG00000041189 | 0.578322155 | 0.000137 | 0.001703 | 560.3723 | 494.3265 | 586.2995 | 496.0085 | 312.0356 | 431.3064 | 396.3228 | 292.562 |
| ENSMUSG00000041193 | 0.594640565 | 0.142825 | 0.280273 | 71.42986 | 52.79215 | 36.43669 | 60.38364 | 19.50223 | 63.69032 | 29.89296 | 33.71532 |
| ENSMUSG00000041199 | -0.153307446 | 0.136449 | 0.271172 | 359.1614 | 347.9482 | 313.0243 | 333.8353 | 412.4721 | 350.8555 | 369.3228 | 369.781 |
| ENSMUSG00000041203 | -0.099752163 | 0.092101 | 0.208414 | 3500.063 | 3563.47 | 3749.667 | 3704.968 | 3990.155 | 3608.001 | 4000.836 | 3959.919 |
| ENSMUSG00000041205 | -0.024831468 | 0.946435 | 0.969279 | 22.1332 | 22.79661 | 20.70267 | 13.80198 | 16.57689 | 18.99536 | 25.07152 | 19.57664 |
| ENSMUSG00000041215 | 0.151509715 | 0.051128 | 0.138178 | 2460.809 | 2439.237 | 2196.139 | 2297.166 | 2115.992 | 2316.317 | 2065.507 | 1956.576 |
| ENSMUSG00000041219 | 0.078486141 | 0.367778 | 0.536239 | 1670.05 | 1439.786 | 1378.798 | 1481.987 | 1481.194 | 1483.873 | 1323.969 | 1364.927 |
| ENSMUSG00000041220 | -0.149522807 | 0.128564 | 0.26023 | 2402.458 | 2334.853 | 2302.965 | 2317.007 | 2820.022 | 2178.879 | 2513.902 | 2864.715 |
| ENSMUSG00000041225 | 0.266301881 | 0.014473 | 0.056174 | 1401.434 | 1428.987 | 1218.145 | 1457.834 | 1124.303 | 1328.558 | 1116.647 | 1008.197 |
| ENSMUSG00000041229 | 0.119190537 | 0.364213 | 0.532935 | 658.9656 | 505.1249 | 506.8013 | 558.98 | 485.6054 | 486.0577 | 599.7879 | 480.7153 |
| ENSMUSG00000041231 | -0.211794653 | 0.006245 | 0.030339 | 1068.43 | 1153.028 | 1177.568 | 1237.865 | 1273.495 | 1327.44 | 1328.79 | 1444.321 |
| ENSMUSG00000041235 | 0.50659915 | 0.010513 | 0.044246 | 1302.84 | 1101.436 | 915.8859 | 1077.417 | 727.433 | 1096.144 | 675.0024 | 597.0875 |
| ENSMUSG00000041236 | -0.068096021 | 0.322781 | 0.490873 | 1732.426 | 1766.137 | 1755.586 | 1709.72 | 1881.965 | 1644.775 | 1918.935 | 1851.08 |
| ENSMUSG00000041237 | -2.585125729 | 2.25E-06 | 7.05E-05 | 23.13925 | 7.198929 | 51.34261 | 23.29083 | 205.7485 | 70.39457 | 106.0718 | 253.4087 |
| ENSMUSG00000041238 | -0.028883148 | 0.736089 | 0.834773 | 1084.527 | 1033.046 | 1138.647 | 1003.231 | 991.6882 | 1057.036 | 1114.718 | 1184.387 |
| ENSMUSG00000041241 | -0.406347135 | 0.005827 | 0.02881 | 747.4984 | 754.6877 | 814.0288 | 614.1879 | 1062.871 | 745.2885 | 986.4678 | 1087.591 |
| ENSMUSG00000041245 | 0.00157269 | 0.992973 | 0.995995 | 179.0777 | 182.3729 | 146.5749 | 145.7834 | 141.3911 | 166.4887 | 201.5364 | 141.3868 |
| ENSMUSG00000041248 | -2.838261867 | 0.001668 | 0.011454 | 64.38748 | 44.3934 | 539.0974 | 105.2401 | 1073.598 | 403.372 | 1806.114 | 2106.664 |
| ENSMUSG00000041258 | 0.042760191 | 0.568805 | 0.712804 | 1038.248 | 1043.845 | 991.2436 | 1113.647 | 954.634 | 1086.088 | 1016.361 | 1009.285 |
| ENSMUSG00000041261 | -0.281366842 | 0.023844 | 0.080261 | 296.786 | 295.1561 | 305.5713 | 302.7808 | 321.7867 | 367.6161 | 336.5369 | 435.0364 |
| ENSMUSG00000041263 | -0.000884401 | 0.995855 | 0.99765 | 247.4894 | 256.7618 | 390.0382 | 320.8959 | 274.9814 | 315.0995 | 318.2154 | 311.0511 |
| ENSMUSG00000041264 | 0.031539215 | 0.727439 | 0.829154 | 1399.422 | 1331.802 | 1368.86 | 1301.699 | 1361.255 | 1495.047 | 1242.004 | 1187.649 |
| ENSMUSG00000041268 | -0.121578365 | 0.252656 | 0.41633 | 861.1825 | 699.496 | 716.3122 | 827.2559 | 805.4419 | 773.2229 | 896.7889 | 902.7006 |
| ENSMUSG00000041272 | 0.479843198 | 0.002052 | 0.013363 | 246.4833 | 229.1659 | 187.1521 | 206.167 | 143.3414 | 177.6625 | 163.9292 | 137.0365 |
| ENSMUSG00000041278 | -0.149469678 | 0.070978 | 0.174105 | 1032.212 | 1084.639 | 957.2913 | 1092.081 | 1214.989 | 1057.036 | 1134.968 | 1210.489 |
| ENSMUSG00000041287 | 0.478081592 | 0.383017 | 0.550971 | 12.07265 | 15.59768 | 12.4216 | 15.52722 | 6.825779 | 20.11273 | 6.750024 | 6.525547 |
| ENSMUSG00000041293 | -0.390600755 | 0.227435 | 0.387405 | 58.35115 | 61.1909 | 152.3716 | 92.30071 | 104.3369 | 109.5027 | 131.1433 | 134.8613 |
| ENSMUSG00000041297 | 0.070021297 | 0.494795 | 0.651583 | 1938.667 | 1730.143 | 1719.977 | 1921.925 | 1582.606 | 2035.856 | 1622.899 | 1726.007 |
| ENSMUSG00000041298 | 0.42186932 | 0.000254 | 0.002745 | 471.8395 | 508.7243 | 395.835 | 463.2288 | 333.4881 | 313.9821 | 362.5727 | 359.9926 |
| ENSMUSG00000041301 | -0.654958699 | 0.000211 | 0.002369 | 259.562 | 305.9545 | 176.3867 | 220.8316 | 382.2436 | 410.0763 | 345.2155 | 373.0437 |
| ENSMUSG00000041303 | -0.00878809 | 0.930266 | 0.958826 | 855.1462 | 883.0686 | 709.6874 | 865.2114 | 900.0277 | 841.3827 | 806.1457 | 781.978 |
| ENSMUSG00000041308 | 0.283066026 | 0.015604 | 0.059391 | 1543.287 | 1378.595 | 1355.611 | 1509.591 | 1231.566 | 1410.126 | 1096.397 | 1019.073 |
| ENSMUSG00000041313 | 0.075900394 | 0.529233 | 0.681503 | 1319.943 | 950.2587 | 1196.614 | 1002.369 | 1055.07 | 994.4629 | 1083.861 | 1108.255 |
| ENSMUSG00000041319 | -0.319679986 | 0.005032 | 0.025846 | 548.2996 | 579.5138 | 612.7989 | 571.9194 | 736.209 | 591.0909 | 756.0027 | 802.6422 |
| ENSMUSG00000041323 | -2.513968935 | 0.000608 | 0.005327 | 8.048435 | 4.799286 | 77.01392 | 18.11509 | 111.1627 | 62.57295 | 216.0008 | 231.6569 |
| ENSMUSG00000041324 | 1.131558592 | 0.008766 | 0.038736 | 56.33904 | 57.59143 | 28.15563 | 52.62003 | 19.50223 | 39.10809 | 18.32149 | 11.9635 |
| ENSMUSG00000041328 | 0.138185445 | 0.058317 | 0.151607 | 2508.094 | 2266.463 | 2097.594 | 2303.205 | 2131.593 | 2139.771 | 2026.936 | 2037.058 |
| ENSMUSG00000041329 | 0.101919203 | 0.289891 | 0.457128 | 620.7355 | 632.3059 | 759.3738 | 701.3129 | 664.0508 | 605.6168 | 642.2166 | 619.9269 |
| ENSMUSG00000041341 | -0.105561851 | 0.112732 | 0.239506 | 1014.103 | 965.8563 | 915.0578 | 982.5282 | 1027.767 | 1044.745 | 1080.968 | 1015.81 |
| ENSMUSG00000041343 | -0.338125274 | 0.036224 | 0.108956 | 164.9929 | 136.7797 | 139.1219 | 136.2945 | 210.624 | 147.4934 | 183.2149 | 187.0657 |
| ENSMUSG00000041346 | -0.15056434 | 0.165079 | 0.310476 | 344.0706 | 377.9438 | 356.914 | 300.193 | 406.6214 | 367.6161 | 379.9299 | 373.0437 |
| ENSMUSG00000041351 | -0.082645278 | 0.691214 | 0.804404 | 700.2138 | 537.52 | 933.2762 | 546.9033 | 668.9264 | 573.2129 | 756.967 | 880.9488 |
| ENSMUSG00000041353 | 0.24045446 | 0.035655 | 0.107577 | 587.5357 | 531.5209 | 556.4877 | 676.2968 | 470.9788 | 560.9218 | 489.8589 | 472.0145 |
| ENSMUSG00000041354 | 0.080148535 | 0.21389 | 0.37205 | 1293.786 | 1277.81 | 1255.41 | 1371.571 | 1286.172 | 1198.942 | 1188.004 | 1245.292 |
| ENSMUSG00000041355 | -0.232460581 | 0.000103 | 0.00137 | 4661.05 | 5117.239 | 5013.358 | 5186.092 | 5890.647 | 5601.396 | 5807.914 | 6170.992 |

| | | | | | | | | | | | |
|---|---|---|---|---|---|---|---|---|---|---|---|
| ENSMUSG00000041360 | -0.103732803 | 0.077254 | 0.184614 | 2094.605 | 2138.082 | 1946.051 | 2037.517 | 2200.826 | 2105.133 | 2253.544 | 2262.189 |
| ENSMUSG00000041361 | -0.046160483 | 0.824174 | 0.893455 | 297.7921 | 160.7761 | 266.6503 | 271.7264 | 226.2258 | 282.6956 | 292.1796 | 230.5693 |
| ENSMUSG00000041362 | -0.002300379 | 0.98578 | 0.992016 | 196.1806 | 200.3702 | 196.2613 | 176.8378 | 188.1965 | 202.2447 | 170.6792 | 209.9051 |
| ENSMUSG00000041372 | -0.315264918 | 0.114455 | 0.241913 | 220.3259 | 165.5754 | 248.432 | 217.3811 | 246.7032 | 194.4231 | 310.5011 | 309.9635 |
| ENSMUSG00000041375 | 0.05091927 | 0.455104 | 0.61714 | 1677.093 | 1534.572 | 1523.716 | 1580.326 | 1576.755 | 1558.737 | 1414.612 | 1547.642 |
| ENSMUSG00000041378 | -0.425148739 | 0.010463 | 0.044093 | 1429.603 | 1132.632 | 1835.912 | 1575.15 | 1797.13 | 1752.043 | 2043.329 | 2432.941 |
| ENSMUSG00000041390 | 0.674669698 | 1.63E-10 | 2.79E-08 | 1133.823 | 1343.8 | 1028.508 | 1157.641 | 732.3086 | 789.9835 | 721.2883 | 675.3941 |
| ENSMUSG00000041396 | -0.004709721 | 0.974052 | 0.985749 | 201.2109 | 193.1713 | 159.8246 | 169.0742 | 157.968 | 194.4231 | 189.965 | 181.6277 |
| ENSMUSG00000041406 | -0.097571783 | 0.410886 | 0.576811 | 443.67 | 521.9224 | 451.3181 | 425.2734 | 523.6348 | 550.8654 | 451.2873 | 442.6496 |
| ENSMUSG00000041408 | 0.063549096 | 0.431335 | 0.596045 | 3340.1 | 2992.355 | 2941.435 | 3201.196 | 2861.952 | 3306.31 | 2788.724 | 2983.262 |
| ENSMUSG00000041415 | -0.151539545 | 0.052458 | 0.140714 | 2200.241 | 2002.502 | 2006.502 | 2051.319 | 2243.731 | 2559.904 | 2217.865 | 2155.606 |
| ENSMUSG00000041417 | 0.654926327 | 0.002302 | 0.014516 | 6131.901 | 4817.283 | 4109.065 | 6225.554 | 2457.28 | 4639.337 | 3704.799 | 2716.803 |
| ENSMUSG00000041420 | -0.148806671 | 0.235847 | 0.396755 | 219.3198 | 244.7636 | 250.0882 | 260.5123 | 308.1352 | 249.1744 | 250.7152 | 272.9854 |
| ENSMUSG00000041423 | 0.666763338 | 5.17E-07 | 2.17E-05 | 309.8647 | 325.1516 | 332.0708 | 313.1323 | 241.8276 | 173.193 | 187.0721 | 203.3795 |
| ENSMUSG00000041426 | -0.315663334 | 0.005935 | 0.029204 | 570.4328 | 549.5183 | 502.6607 | 487.3823 | 619.1957 | 569.8608 | 693.3239 | 740.6495 |
| ENSMUSG00000041429 | -0.080694328 | 0.55579 | 0.702687 | 281.6952 | 313.1534 | 227.7293 | 264.8254 | 291.5583 | 269.2872 | 267.1081 | 318.6642 |
| ENSMUSG00000041431 | -0.047392687 | 0.618466 | 0.751043 | 2565.439 | 2334.853 | 2125.75 | 2623.238 | 2764.441 | 2432.523 | 2417.473 | 2355.722 |
| ENSMUSG00000041438 | -0.285194851 | 0.001365 | 0.009873 | 1145.896 | 1132.632 | 1138.647 | 1151.602 | 1578.705 | 1234.698 | 1402.076 | 1349.701 |
| ENSMUSG00000041439 | -0.072142443 | 0.579457 | 0.72065 | 477.8758 | 544.719 | 563.9406 | 560.7053 | 549.9628 | 467.0624 | 569.8949 | 671.0437 |
| ENSMUSG00000041440 | 0.003247723 | 0.975991 | 0.986896 | 280.6892 | 277.1588 | 297.2903 | 271.7264 | 265.2303 | 283.813 | 266.1438 | 311.0511 |
| ENSMUSG00000041444 | -0.006520468 | 0.967429 | 0.981738 | 1180.102 | 1238.216 | 1326.627 | 1350.006 | 985.8375 | 1629.131 | 1104.111 | 1401.905 |
| ENSMUSG00000041445 | -0.502634952 | 0.023355 | 0.078886 | 508.0574 | 379.1436 | 677.3912 | 432.1744 | 638.6979 | 529.6353 | 745.3955 | 917.9269 |
| ENSMUSG00000041453 | -0.085859426 | 0.252058 | 0.415689 | 28929.09 | 34773.23 | 32895.71 | 31934.32 | 36050.84 | 33826.27 | 34468.52 | 32064.36 |
| ENSMUSG00000041459 | 0.032559943 | 0.700805 | 0.811268 | 11667.21 | 11604.67 | 9691.332 | 11114.9 | 10368.36 | 11721.25 | 10844.4 | 10157.01 |
| ENSMUSG00000041460 | 0.144077547 | 0.535335 | 0.686216 | 71.42986 | 87.58697 | 103.5133 | 79.36136 | 101.4116 | 71.51194 | 62.6788 | 73.95619 |
| ENSMUSG00000041468 | 0.3546655 | 0.477425 | 0.636797 | 37.22401 | 70.78947 | 33.95237 | 33.64232 | 40.95467 | 56.98608 | 27.96439 | 10.87591 |
| ENSMUSG00000041471 | -0.090463394 | 0.420015 | 0.585395 | 237.4288 | 233.9652 | 213.6515 | 237.2215 | 225.2507 | 253.6439 | 255.5366 | 246.8832 |
| ENSMUSG00000041476 | 0.297993247 | 0.184138 | 0.335253 | 146.8839 | 113.983 | 120.9036 | 109.5532 | 77.03379 | 137.437 | 93.53605 | 92.44524 |
| ENSMUSG00000041477 | 0.263318294 | 0.003199 | 0.01862 | 583.5115 | 578.314 | 578.8465 | 589.1718 | 480.7299 | 534.1048 | 491.7875 | 435.0364 |
| ENSMUSG00000041479 | -0.331859855 | 0.310023 | 0.47849 | 31.18769 | 37.19447 | 70.38906 | 47.44429 | 59.48179 | 74.86406 | 44.3573 | 57.64233 |
| ENSMUSG00000041481 | 0.508935281 | 1.53E-08 | 1.21E-06 | 1197.205 | 1117.034 | 1152.724 | 1310.325 | 843.4713 | 911.7773 | 843.753 | 760.2262 |
| ENSMUSG00000041482 | 0.579340666 | 0.001008 | 0.007893 | 1771.662 | 2342.052 | 1759.727 | 1849.465 | 1409.036 | 1653.714 | 1127.254 | 977.7444 |
| ENSMUSG00000041483 | 0.240516979 | 0.035588 | 0.107475 | 1081.508 | 1091.838 | 942.3853 | 1016.17 | 828.8446 | 1048.097 | 838.9316 | 780.8904 |
| ENSMUSG00000041488 | -0.391475412 | 6.59E-08 | 4.03E-06 | 1052.333 | 1024.648 | 1133.678 | 1083.455 | 1427.563 | 1327.44 | 1372.183 | 1510.664 |
| ENSMUSG00000041491 | -0.031912076 | 0.696041 | 0.808238 | 541.2572 | 583.1133 | 543.238 | 565.0184 | 588.9672 | 593.3256 | 527.4662 | 572.0729 |
| ENSMUSG00000041498 | 0.250557171 | 0.102346 | 0.22451 | 474.8577 | 410.339 | 326.274 | 362.3019 | 318.8614 | 385.4941 | 319.1797 | 298 |
| ENSMUSG00000041506 | -0.312396931 | 0.004436 | 0.023569 | 773.6558 | 784.6833 | 855.4342 | 788.4379 | 1071.647 | 811.2136 | 1058.79 | 1034.299 |
| ENSMUSG00000041515 | 0.117627475 | 0.544897 | 0.693416 | 433.6094 | 290.3568 | 350.2891 | 288.9789 | 299.3592 | 256.996 | 296.0368 | 404.5839 |
| ENSMUSG00000041523 | -0.750303154 | 0.2644 | 0.428689 | 182.0958 | 104.3845 | 398.3193 | 144.9207 | 626.9966 | 81.56831 | 320.144 | 367.6058 |
| ENSMUSG00000041528 | -0.078423461 | 0.41523 | 0.581139 | 973.8606 | 922.6628 | 992.8999 | 920.4193 | 1027.767 | 865.9649 | 1005.754 | 1123.482 |
| ENSMUSG00000041530 | 0.239791551 | 0.000341 | 0.003409 | 2166.035 | 2300.058 | 2279.778 | 2361.001 | 1904.392 | 2079.433 | 1924.721 | 1805.401 |
| ENSMUSG00000041534 | 0.065105901 | 0.550528 | 0.698381 | 800.8193 | 1011.45 | 1043.414 | 948.0232 | 920.5051 | 978.8197 | 824.4672 | 913.5765 |
| ENSMUSG00000041540 | 0.610838888 | 0.00101 | 0.007894 | 406.446 | 309.554 | 286.5249 | 340.7363 | 219.4 | 283.813 | 200.5721 | 176.1898 |
| ENSMUSG00000041544 | -1.462153683 | 0.003371 | 0.019279 | 3.018163 | 7.198929 | 9.93728 | 12.07673 | 29.25334 | 16.76061 | 22.17865 | 21.75182 |
| ENSMUSG00000041548 | 0.229170672 | 0.39263 | 0.560025 | 256.5439 | 153.5772 | 182.1835 | 285.5284 | 132.6151 | 259.2308 | 189.965 | 168.5766 |
| ENSMUSG00000041552 | 1.10372495 | 0.012143 | 0.049419 | 46.2785 | 57.59143 | 19.87456 | 64.69676 | 23.40267 | 27.93435 | 13.50005 | 22.83941 |
| ENSMUSG00000041556 | -0.179528189 | 0.353072 | 0.521776 | 81.4904 | 91.18644 | 89.43552 | 107.8279 | 124.8142 | 87.15518 | 109.929 | 96.79561 |
| ENSMUSG00000041559 | 0.646132792 | 0.002631 | 0.016007 | 352.119 | 731.8911 | 492.7234 | 507.2226 | 294.4836 | 379.9072 | 309.5368 | 346.9416 |
| ENSMUSG00000041560 | -0.098336402 | 0.237245 | 0.398307 | 8155.077 | 8845.084 | 8462.422 | 8978.185 | 10152.86 | 8537.855 | 9754.749 | 8421.218 |
| ENSMUSG00000041565 | -0.082543465 | 0.830486 | 0.897601 | 20.12109 | 15.59768 | 14.07781 | 19.84034 | 22.42756 | 15.64324 | 21.21436 | 14.13868 |
| ENSMUSG00000041570 | 0.135458377 | 0.211611 | 0.369104 | 1708.28 | 1380.995 | 1286.05 | 1384.511 | 1222.79 | 1395.6 | 1261.29 | 1363.839 |
| ENSMUSG00000041571 | 0.170123976 | 0.019548 | 0.069352 | 5168.101 | 5894.723 | 5949.118 | 5865.84 | 5474.275 | 4867.282 | 5044.197 | 4946.364 |
| ENSMUSG00000041577 | 0.848135971 | 0.00403 | 0.021947 | 110.666 | 243.5638 | 197.9175 | 185.4641 | 109.2125 | 141.9065 | 86.78602 | 71.78101 |
| ENSMUSG00000041578 | -0.023960986 | 0.889205 | 0.933305 | 115.6963 | 148.7779 | 141.6062 | 150.0965 | 131.64 | 162.0192 | 118.6076 | 154.4379 |
| ENSMUSG00000041592 | -0.125615817 | 0.574726 | 0.71699 | 293.7679 | 229.1659 | 411.569 | 268.2759 | 263.2801 | 289.3999 | 334.6083 | 427.4233 |
| ENSMUSG00000041594 | -0.188676382 | 0.079782 | 0.188683 | 536.227 | 664.7011 | 669.1102 | 653.8686 | 668.9264 | 686.0677 | 731.8955 | 790.6787 |
| ENSMUSG00000041596 | 0.465695993 | 0.248778 | 0.412196 | 65.39353 | 116.3827 | 117.5911 | 125.943 | 80.93424 | 131.8501 | 32.78583 | 63.08028 |
| ENSMUSG00000041598 | -0.036585415 | 0.766914 | 0.855443 | 2874.297 | 2398.443 | 2861.108 | 2758.67 | 2601.597 | 2542.026 | 3455.048 | 2574.328 |
| ENSMUSG00000041605 | -0.293421551 | 0.187929 | 0.340174 | 336.0222 | 232.7654 | 393.3507 | 264.8254 | 283.7574 | 321.8037 | 445.5016 | 454.6131 |
| ENSMUSG00000041607 | -0.158242769 | 0.326969 | 0.49463 | 401.4157 | 362.3461 | 587.1276 | 478.756 | 549.9628 | 471.5319 | 522.6447 | 501.3795 |
| ENSMUSG00000041608 | -0.338057029 | 0.478506 | 0.637833 | 69.41775 | 16.7975 | 42.23344 | 31.05445 | 39.00445 | 81.56831 | 30.85725 | 51.11678 |
| ENSMUSG00000041609 | -0.922288469 | 3.48E-06 | 0.000102 | 308.8587 | 369.545 | 375.1323 | 295.8799 | 597.7432 | 429.0717 | 757.9313 | 771.1021 |
| ENSMUSG00000041617 | -0.283292792 | 0.202089 | 0.357578 | 107.6478 | 99.58519 | 157.3403 | 136.2945 | 182.3458 | 151.9629 | 163.9292 | 113.1095 |
| ENSMUSG00000041623 | -0.189584384 | 0.17211 | 0.319434 | 464.7971 | 476.3291 | 423.9906 | 465.8167 | 531.4357 | 398.9026 | 556.3948 | 599.2627 |

| | | | | | | | | | | | |
|---|---|---|---|---|---|---|---|---|---|---|---|
| ENSMUSG00000041624 | 0.439535838 | 0.104444 | 0.227536 | 66.39959 | 53.99197 | 68.73285 | 57.79577 | 39.00445 | 64.8077 | 45.32159 | 33.71532 |
| ENSMUSG00000041625 | -0.562492642 | 0.003098 | 0.018214 | 343.0645 | 349.1481 | 476.9894 | 325.2091 | 529.4854 | 404.4894 | 650.8952 | 623.1897 |
| ENSMUSG00000041629 | -0.28127186 | 0.001275 | 0.009376 | 1158.975 | 1225.018 | 1258.722 | 1185.245 | 1556.278 | 1271.572 | 1485.005 | 1553.08 |
| ENSMUSG00000041632 | -0.276255955 | 0.001263 | 0.009321 | 930.6003 | 913.0642 | 881.9336 | 834.1569 | 1089.199 | 977.7023 | 1056.861 | 1184.387 |
| ENSMUSG00000041633 | 0.387661877 | 0.000108 | 0.001425 | 881.3036 | 937.0606 | 765.9986 | 824.6681 | 585.0668 | 700.5936 | 666.3238 | 651.4671 |
| ENSMUSG00000041638 | -0.201771448 | 0.014202 | 0.055412 | 2109.696 | 2166.878 | 2309.589 | 1934.864 | 2400.724 | 2327.49 | 2392.401 | 2679.824 |
| ENSMUSG00000041642 | 0.292694599 | 0.012939 | 0.051854 | 262.5802 | 261.5611 | 239.3228 | 246.7103 | 228.176 | 195.5405 | 208.2865 | 190.3284 |
| ENSMUSG00000041644 | -2.999824438 | 1.57E-07 | 8.19E-06 | 21.12714 | 7.198929 | 89.43552 | 29.3292 | 376.393 | 153.0803 | 328.8226 | 327.3649 |
| ENSMUSG00000041645 | -0.117698824 | 0.113623 | 0.240733 | 2143.902 | 2255.664 | 2394.884 | 2056.494 | 2398.774 | 2238.1 | 2506.188 | 2459.043 |
| ENSMUSG00000041649 | 0.868654654 | 4.60E-06 | 0.000126 | 130.7871 | 155.9768 | 118.4192 | 160.448 | 74.10846 | 84.92043 | 84.85745 | 65.25547 |
| ENSMUSG00000041650 | -1.339496529 | 4.88E-06 | 0.000132 | 325.9616 | 296.3559 | 431.4436 | 276.0395 | 904.9033 | 393.3157 | 896.7889 | 1171.336 |
| ENSMUSG00000041653 | -0.489400984 | 0.196289 | 0.350562 | 18.10898 | 10.79839 | 17.39024 | 20.70296 | 19.50223 | 29.05173 | 27.0001 | 19.57664 |
| ENSMUSG00000041654 | -0.381311425 | 0.047264 | 0.130781 | 346.0827 | 417.5379 | 403.2879 | 314.8576 | 475.8543 | 337.447 | 497.5732 | 617.7517 |
| ENSMUSG00000041658 | 0.709761542 | 1.34E-08 | 1.09E-06 | 400.4096 | 424.7368 | 344.4924 | 413.1967 | 231.1014 | 271.5219 | 248.7866 | 215.343 |
| ENSMUSG00000041660 | 1.132809919 | 0.005257 | 0.026624 | 18.10898 | 28.79572 | 33.95237 | 31.91707 | 11.70134 | 16.76061 | 14.46434 | 8.700729 |
| ENSMUSG00000041669 | -1.010936487 | 0.038521 | 0.11341 | 8.048435 | 13.19804 | 18.21835 | 6.038364 | 23.40267 | 15.64324 | 21.21436 | 31.54014 |
| ENSMUSG00000041670 | 0.572957168 | 0.092406 | 0.208796 | 88.53278 | 115.1829 | 81.98256 | 164.7611 | 85.8098 | 107.2679 | 67.50024 | 42.41605 |
| ENSMUSG00000041671 | -0.335227023 | 0.000889 | 0.007112 | 372.2401 | 351.5477 | 391.6944 | 371.7907 | 488.5308 | 429.0717 | 444.5373 | 516.6058 |
| ENSMUSG00000041673 | 0.812334239 | 0.144304 | 0.282425 | 49.29666 | 95.98572 | 75.3577 | 48.30692 | 19.50223 | 91.62468 | 17.3572 | 25.0146 |
| ENSMUSG00000041674 | 0.668642787 | 0.123792 | 0.253462 | 19.11503 | 31.19536 | 19.87456 | 29.3292 | 14.62667 | 25.6996 | 12.53576 | 9.78832 |
| ENSMUSG00000041679 | -0.134703649 | 0.394378 | 0.561649 | 140.8476 | 172.7743 | 195.4332 | 167.349 | 159.9183 | 198.8926 | 203.465 | 181.6277 |
| ENSMUSG00000041684 | 0.086057803 | 0.303816 | 0.472347 | 699.2078 | 754.6877 | 642.6107 | 711.6644 | 653.3246 | 700.5936 | 637.3951 | 651.4671 |
| ENSMUSG00000041685 | 0.395599251 | 0.002007 | 0.013175 | 2140.884 | 2082.89 | 1872.349 | 2221.255 | 1396.359 | 1982.222 | 1504.291 | 1441.058 |
| ENSMUSG00000041688 | 0.136223485 | 0.265587 | 0.429745 | 1813.916 | 2178.876 | 1911.27 | 1990.935 | 1802.981 | 2183.349 | 1566.97 | 1630.299 |
| ENSMUSG00000041695 | -0.366983139 | 0.391783 | 0.559211 | 77.46619 | 23.99643 | 35.60859 | 42.26855 | 40.95467 | 91.62468 | 56.89306 | 42.41605 |
| ENSMUSG00000041696 | -0.322800592 | 0.114121 | 0.241462 | 146.8839 | 136.7797 | 223.5888 | 206.167 | 188.1965 | 215.6532 | 233.358 | 258.8467 |
| ENSMUSG00000041697 | -0.860042905 | 8.48E-06 | 0.000202 | 4343.137 | 3738.644 | 4947.937 | 3922.349 | 7685.827 | 4983.488 | 8088.458 | 10013.45 |
| ENSMUSG00000041700 | -0.232717337 | 0.587334 | 0.727603 | 13.07871 | 32.39518 | 31.46805 | 31.91707 | 42.9049 | 25.6996 | 40.50014 | 18.48905 |
| ENSMUSG00000041702 | -0.040496823 | 0.758249 | 0.849269 | 607.6568 | 592.7118 | 548.2066 | 500.3216 | 622.121 | 468.1797 | 563.1449 | 656.905 |
| ENSMUSG00000041712 | -0.123356978 | 0.125818 | 0.2565 | 1406.464 | 1575.366 | 1416.89 | 1504.415 | 1747.399 | 1462.643 | 1606.506 | 1608.547 |
| ENSMUSG00000041716 | -0.007383027 | 0.947356 | 0.969686 | 353.1251 | 403.14 | 361.8826 | 331.2474 | 383.2187 | 360.9118 | 385.7157 | 323.0146 |
| ENSMUSG00000041718 | 0.112807082 | 0.639448 | 0.766468 | 673.0504 | 772.6851 | 529.9882 | 646.105 | 509.0081 | 954.2375 | 486.0017 | 475.2773 |
| ENSMUSG00000041720 | -0.234463318 | 0.001886 | 0.012582 | 1665.02 | 1653.354 | 1813.554 | 1623.457 | 1919.019 | 1839.198 | 2098.293 | 2092.525 |
| ENSMUSG00000041729 | 0.133457182 | 0.096384 | 0.215176 | 636.8324 | 656.3024 | 634.3297 | 613.3253 | 529.4854 | 588.8561 | 596.895 | 600.3503 |
| ENSMUSG00000041731 | 0.740871131 | 1.55E-07 | 8.10E-06 | 985.9333 | 1273.011 | 1115.46 | 1259.43 | 678.6775 | 858.1433 | 620.0379 | 616.6641 |
| ENSMUSG00000041733 | -0.371382234 | 0.00064 | 0.005551 | 632.8082 | 550.7181 | 625.2205 | 583.1335 | 756.6864 | 678.2461 | 769.5028 | 891.8247 |
| ENSMUSG00000041734 | 0.314781601 | 0.001405 | 0.010107 | 2301.852 | 2644.407 | 2504.194 | 2366.176 | 1858.562 | 2308.495 | 1829.257 | 1896.759 |
| ENSMUSG00000041736 | -0.127496401 | 0.576955 | 0.718897 | 1008.066 | 607.1097 | 833.0753 | 697.8624 | 819.0935 | 557.5697 | 929.5748 | 1131.095 |
| ENSMUSG00000041737 | -1.152644128 | 0.04292 | 0.122598 | 205.2351 | 129.5807 | 536.6131 | 150.0965 | 657.225 | 274.874 | 505.2875 | 835.27 |
| ENSMUSG00000041740 | -0.144987666 | 0.025749 | 0.084756 | 4520.202 | 4910.87 | 5209.619 | 4913.503 | 5315.332 | 5212.55 | 5380.734 | 5716.379 |
| ENSMUSG00000041741 | 0.557791266 | 4.67E-08 | 3.07E-06 | 1846.11 | 2145.281 | 1946.051 | 2153.108 | 1253.018 | 1575.497 | 1400.148 | 1268.131 |
| ENSMUSG00000041747 | -0.03033329 | 0.745211 | 0.840853 | 1147.908 | 1114.634 | 968.8848 | 969.5888 | 1092.125 | 1154.247 | 1031.789 | 1008.197 |
| ENSMUSG00000041750 | 0.671080296 | 0.046598 | 0.129599 | 39.23612 | 34.79482 | 40.57722 | 34.50494 | 29.25334 | 30.1691 | 21.21436 | 13.05109 |
| ENSMUSG00000041757 | -0.384558726 | 0.158473 | 0.301338 | 495.9848 | 353.9474 | 793.3262 | 534.8266 | 738.1593 | 451.4191 | 706.824 | 949.467 |
| ENSMUSG00000041762 | -0.241390501 | 0.235741 | 0.396684 | 235.4167 | 179.9732 | 167.2775 | 282.9405 | 215.4996 | 245.8223 | 264.2152 | 299.0875 |
| ENSMUSG00000041763 | 0.069881215 | 0.390737 | 0.558171 | 1871.261 | 1798.532 | 1621.433 | 1846.877 | 1632.336 | 1855.958 | 1712.578 | 1598.759 |
| ENSMUSG00000041765 | -0.296090377 | 0.001227 | 0.009108 | 984.9272 | 1058.243 | 1070.742 | 1090.356 | 1274.47 | 1154.247 | 1273.826 | 1460.635 |
| ENSMUSG00000041769 | 0.077241801 | 0.376897 | 0.545034 | 1567.433 | 1607.761 | 1470.717 | 1626.908 | 1381.733 | 1609.019 | 1599.756 | 1354.051 |
| ENSMUSG00000041771 | 0.06192904 | 0.860658 | 0.916562 | 19.11503 | 23.99643 | 21.53077 | 33.64232 | 22.42756 | 26.81698 | 18.32149 | 27.18978 |
| ENSMUSG00000041773 | 0.693759849 | 3.20E-05 | 0.000562 | 4495.051 | 4342.154 | 3249.49 | 3741.198 | 2443.629 | 3188.986 | 1979.686 | 2173.007 |
| ENSMUSG00000041774 | 0.07342032 | 0.674176 | 0.791463 | 183.1019 | 183.5727 | 188.8083 | 175.1126 | 218.4249 | 130.7328 | 177.4292 | 166.4014 |
| ENSMUSG00000041775 | 0.425650394 | 0.000541 | 0.004873 | 618.7234 | 527.9215 | 504.3169 | 600.386 | 354.9405 | 446.9496 | 455.1445 | 419.8102 |
| ENSMUSG00000041777 | 0.335235452 | 0.004825 | 0.025084 | 1769.65 | 1794.933 | 1628.886 | 1846.014 | 1276.421 | 1692.822 | 1413.648 | 1197.438 |
| ENSMUSG00000041779 | 0.19424801 | 0.08194 | 0.192183 | 539.2451 | 580.7136 | 587.9557 | 508.9479 | 425.1485 | 553.1002 | 466.716 | 493.7664 |
| ENSMUSG00000041781 | -0.164240841 | 0.027327 | 0.088451 | 1364.21 | 1404.991 | 1336.564 | 1184.382 | 1476.319 | 1440.295 | 1509.113 | 1496.525 |
| ENSMUSG00000041782 | -0.792526706 | 0.006681 | 0.031901 | 353.1251 | 285.5575 | 744.4679 | 362.3019 | 736.209 | 557.5697 | 796.5028 | 936.4159 |
| ENSMUSG00000041789 | -0.170007864 | 0.706896 | 0.815762 | 16.09687 | 37.19447 | 50.5145 | 25.8787 | 39.00445 | 20.11273 | 54.00019 | 32.62773 |
| ENSMUSG00000041794 | 0.265053391 | 0.516158 | 0.670199 | 40.24217 | 25.19625 | 38.09291 | 79.36136 | 36.07912 | 53.63396 | 29.89296 | 33.71532 |
| ENSMUSG00000041797 | 0.83729459 | 8.00E-05 | 0.001136 | 179.0777 | 251.9625 | 173.0743 | 237.2215 | 109.2125 | 147.4934 | 123.429 | 90.27006 |
| ENSMUSG00000041798 | -0.386794694 | 0.201416 | 0.356888 | 23.13925 | 31.19536 | 31.46805 | 20.70296 | 34.1289 | 33.52122 | 40.50014 | 30.45255 |
| ENSMUSG00000041801 | -0.281757052 | 0.067929 | 0.16853 | 920.5397 | 1269.411 | 1223.942 | 1129.174 | 1580.655 | 1051.449 | 1399.184 | 1490 |
| ENSMUSG00000041809 | 0.464250359 | 0.110472 | 0.23639 | 97.58727 | 97.18554 | 74.5296 | 87.9876 | 72.15824 | 86.03781 | 66.53595 | 33.71532 |
| ENSMUSG00000041815 | -0.060147741 | 0.29537 | 0.463117 | 3259.616 | 3169.928 | 3209.741 | 2951.898 | 3435.317 | 3203.512 | 3264.119 | 3220.357 |
| ENSMUSG00000041817 | 0.168948526 | 0.257518 | 0.421268 | 388.337 | 375.5441 | 349.461 | 372.6533 | 296.4338 | 427.9543 | 305.6797 | 292.562 |

| | | | | | | | | | | | |
|---|---|---|---|---|---|---|---|---|---|---|---|
| ENSMUSG00000041827 | 0.435634545 | 0.312427 | 0.480769 | 48.29061 | 111.5834 | 172.2462 | 59.52102 | 64.35735 | 46.92971 | 103.1789 | 75.04379 |
| ENSMUSG00000041828 | 1.217468518 | 0.000534 | 0.004826 | 210.2654 | 215.9679 | 147.403 | 422.6855 | 93.61069 | 159.7845 | 98.35749 | 77.21897 |
| ENSMUSG00000041831 | -0.516907233 | 0.408866 | 0.57507 | 8.048435 | 11.99822 | 14.07781 | 12.93935 | 5.850668 | 13.40849 | 11.57147 | 36.9781 |
| ENSMUSG00000041836 | 0.483218303 | 0.002289 | 0.014455 | 161.9748 | 171.5745 | 135.8095 | 163.0358 | 117.0134 | 119.559 | 118.6076 | 95.70802 |
| ENSMUSG00000041837 | -0.135059743 | 0.279976 | 0.446314 | 621.7416 | 569.9152 | 518.3948 | 524.4751 | 613.345 | 722.941 | 592.0735 | 524.2189 |
| ENSMUSG00000041840 | -0.005850095 | 0.956981 | 0.975803 | 696.1896 | 806.2801 | 646.7513 | 704.7634 | 741.0846 | 792.2182 | 691.3953 | 637.3284 |
| ENSMUSG00000041841 | -0.138031939 | 0.107056 | 0.231384 | 12147.1 | 14802.2 | 15184.16 | 14217.76 | 16498.88 | 15124.78 | 15505.77 | 14879.33 |
| ENSMUSG00000041842 | -0.199525199 | 0.419806 | 0.58528 | 158.9566 | 100.785 | 120.9036 | 92.30071 | 153.0925 | 93.85942 | 136.9291 | 158.7883 |
| ENSMUSG00000041845 | -0.128873144 | 0.189494 | 0.342068 | 606.6508 | 572.3149 | 629.361 | 635.7535 | 668.9264 | 578.7998 | 702.0025 | 724.3357 |
| ENSMUSG00000041846 | 0.129317467 | 0.16803 | 0.314073 | 2073.478 | 2135.682 | 1876.49 | 2254.898 | 1726.922 | 2134.185 | 1881.328 | 1883.708 |
| ENSMUSG00000041849 | 0.140631841 | 0.419839 | 0.585282 | 110.666 | 128.3809 | 135.8095 | 142.3329 | 116.0382 | 119.559 | 136.9291 | 96.79561 |
| ENSMUSG00000041852 | 0.045170082 | 0.544167 | 0.692833 | 1559.384 | 1602.962 | 1663.666 | 1504.415 | 1528.975 | 1667.122 | 1521.648 | 1418.219 |
| ENSMUSG00000041859 | -0.129480819 | 0.256572 | 0.42028 | 2297.828 | 2999.554 | 2661.535 | 2158.284 | 2823.922 | 2597.895 | 2741.474 | 2899.518 |
| ENSMUSG00000041870 | -0.251609599 | 0.037773 | 0.112085 | 2154.968 | 2177.676 | 2552.225 | 2210.904 | 2657.178 | 2215.753 | 2782.939 | 3174.678 |
| ENSMUSG00000041878 | 0.099742154 | 0.789858 | 0.87119 | 18.10898 | 29.99554 | 24.8432 | 23.29083 | 19.50223 | 26.81698 | 28.92867 | 14.13868 |
| ENSMUSG00000041879 | 0.133672741 | 0.073201 | 0.178021 | 3576.523 | 3947.413 | 4032.051 | 3564.36 | 3476.272 | 3706.33 | 3259.297 | 3341.08 |
| ENSMUSG00000041881 | -0.251281581 | 0.00106 | 0.008165 | 1816.934 | 1937.712 | 2043.767 | 1833.938 | 2302.238 | 2076.081 | 2268.008 | 2438.379 |
| ENSMUSG00000041886 | 0.076218268 | 0.709836 | 0.818489 | 97.58727 | 124.7814 | 120.9036 | 95.75121 | 109.2125 | 90.5073 | 129.2147 | 85.9197 |
| ENSMUSG00000041889 | -0.479312748 | 0.005441 | 0.027286 | 346.0827 | 350.3479 | 519.2229 | 375.2412 | 485.6054 | 546.3959 | 517.8233 | 672.1313 |
| ENSMUSG00000041890 | 0.187515258 | 0.022307 | 0.076182 | 1454.755 | 1331.802 | 1289.362 | 1376.747 | 1182.81 | 1329.675 | 1146.54 | 1130.007 |
| ENSMUSG00000041891 | -0.08589217 | 0.165813 | 0.311254 | 3860.231 | 3899.42 | 3739.73 | 3587.651 | 3806.835 | 3850.471 | 4239.015 | 4111.094 |
| ENSMUSG00000041895 | 0.089255165 | 0.177502 | 0.326585 | 1438.658 | 1464.982 | 1416.062 | 1514.767 | 1431.463 | 1419.065 | 1360.612 | 1272.482 |
| ENSMUSG00000041907 | 0.633698779 | 0.000127 | 0.001609 | 270.6286 | 359.9465 | 262.5098 | 330.3848 | 222.3254 | 208.949 | 195.7507 | 159.8759 |
| ENSMUSG00000041912 | 0.390576081 | 0.003227 | 0.018734 | 388.337 | 344.3488 | 312.1962 | 401.1199 | 277.9067 | 306.1605 | 278.6796 | 240.3576 |
| ENSMUSG00000041915 | 0.050464531 | 0.555522 | 0.702577 | 1422.561 | 1425.388 | 1318.346 | 1495.789 | 1301.774 | 1500.633 | 1248.754 | 1418.219 |
| ENSMUSG00000041920 | -0.238484412 | 0.043238 | 0.123138 | 342.0585 | 377.9438 | 372.648 | 341.5989 | 456.3521 | 435.7759 | 351.9655 | 448.0875 |
| ENSMUSG00000041921 | 0.246974823 | 0.001284 | 0.009421 | 1461.797 | 1470.981 | 1326.627 | 1463.009 | 1185.735 | 1301.741 | 1216.933 | 1115.868 |
| ENSMUSG00000041923 | 1.278529822 | 0.001745 | 0.011852 | 60.36326 | 52.79215 | 48.85829 | 58.6584 | 15.60178 | 43.57759 | 20.25007 | 11.9635 |
| ENSMUSG00000041926 | -0.072043592 | 0.409973 | 0.576141 | 1051.327 | 1059.442 | 1085.648 | 948.8858 | 1099.926 | 1068.21 | 993.2178 | 1196.35 |
| ENSMUSG00000041930 | -0.317704542 | 0.316036 | 0.484327 | 154.9324 | 100.785 | 166.4494 | 155.2722 | 166.744 | 98.32892 | 168.7506 | 287.124 |
| ENSMUSG00000041935 | 0.055299084 | 0.629379 | 0.758785 | 3402.476 | 2753.59 | 2752.626 | 3235.701 | 2842.449 | 3262.732 | 3073.19 | 2510.16 |
| ENSMUSG00000041936 | -0.34889742 | 0.037762 | 0.112084 | 6357.257 | 6018.305 | 9158.031 | 6509.357 | 8309.899 | 7084.152 | 9984.25 | 10339.73 |
| ENSMUSG00000041939 | -0.527958003 | 3.55E-05 | 0.000612 | 577.4752 | 530.3211 | 568.9093 | 545.178 | 811.2926 | 624.6121 | 884.2532 | 883.124 |
| ENSMUSG00000041945 | -0.692937865 | 0.068151 | 0.168921 | 55.33299 | 55.19179 | 86.12309 | 41.40593 | 109.2125 | 41.34284 | 105.1075 | 129.4233 |
| ENSMUSG00000041949 | 0.085679313 | 0.485854 | 0.644184 | 404.4339 | 399.5406 | 336.2113 | 388.1806 | 375.4179 | 307.2779 | 353.8941 | 401.3211 |
| ENSMUSG00000041954 | 0.132978983 | 0.602712 | 0.740217 | 68.4117 | 65.99018 | 54.65504 | 45.71905 | 61.43201 | 40.22547 | 60.75022 | 50.02919 |
| ENSMUSG00000041957 | -0.672529654 | 5.01E-06 | 0.000135 | 335.0161 | 326.3515 | 450.49 | 343.3242 | 579.2161 | 486.0577 | 637.3951 | 619.9269 |
| ENSMUSG00000041958 | -0.265322271 | 0.004632 | 0.024358 | 1049.315 | 1060.642 | 1107.179 | 1084.318 | 1274.47 | 1111.787 | 1345.183 | 1438.883 |
| ENSMUSG00000041959 | 0.18320185 | 0.020866 | 0.072634 | 4301.888 | 4637.31 | 5129.292 | 4938.52 | 3934.574 | 4137.636 | 4500.338 | 4170.912 |
| ENSMUSG00000041961 | 0.301423229 | 0.020903 | 0.072722 | 960.7819 | 945.4594 | 1002.837 | 977.3524 | 674.777 | 992.2282 | 783.9671 | 704.759 |
| ENSMUSG00000041966 | 0.006846072 | 0.942017 | 0.966348 | 727.3773 | 601.1106 | 651.7199 | 653.8686 | 608.4695 | 679.3635 | 636.4308 | 700.4087 |
| ENSMUSG00000041974 | -0.141221368 | 0.242486 | 0.405043 | 494.9787 | 399.5406 | 388.382 | 468.4046 | 454.4019 | 463.7102 | 540.0019 | 473.1021 |
| ENSMUSG00000041975 | -0.074361286 | 0.628655 | 0.758208 | 499.003 | 400.7404 | 334.5551 | 383.0048 | 403.6961 | 378.7898 | 493.7161 | 424.1605 |
| ENSMUSG00000041977 | -0.053495823 | 0.444429 | 0.607449 | 1097.605 | 998.2515 | 1040.93 | 999.7806 | 1016.066 | 1061.505 | 1097.361 | 1119.131 |
| ENSMUSG00000041986 | -0.169964899 | 0.456319 | 0.618115 | 53.32088 | 49.19268 | 45.54586 | 59.52102 | 47.78045 | 65.92507 | 62.6788 | 57.64233 |
| ENSMUSG00000041992 | -0.269034537 | 0.120831 | 0.250033 | 317.9132 | 280.7582 | 414.8814 | 332.11 | 426.1236 | 307.2779 | 446.4659 | 443.7372 |
| ENSMUSG00000041995 | -0.067872809 | 0.395633 | 0.562932 | 1168.029 | 1198.622 | 1286.05 | 1184.382 | 1242.292 | 1179.947 | 1401.112 | 1246.379 |
| ENSMUSG00000041997 | 0.028536775 | 0.716884 | 0.822761 | 2317.949 | 2314.456 | 2326.152 | 2452.439 | 2189.125 | 2602.364 | 2281.508 | 2156.693 |
| ENSMUSG00000042010 | -0.101784229 | 0.607127 | 0.743047 | 343.0645 | 256.7618 | 329.5864 | 328.6596 | 333.4881 | 229.0617 | 366.4299 | 421.9853 |
| ENSMUSG00000042015 | -0.005256609 | 0.932135 | 0.959964 | 1019.133 | 982.6538 | 1024.368 | 1003.231 | 1044.344 | 975.4676 | 991.2893 | 1034.299 |
| ENSMUSG00000042029 | 0.079408017 | 0.50429 | 0.65999 | 928.5882 | 1016.249 | 788.3575 | 860.8982 | 846.3966 | 983.2892 | 764.6813 | 804.8174 |
| ENSMUSG00000042032 | -0.051359394 | 0.274991 | 0.441133 | 2306.883 | 2363.648 | 2250.794 | 2384.291 | 2388.048 | 2428.054 | 2414.58 | 2410.102 |
| ENSMUSG00000042035 | 0.420024208 | 0.001188 | 0.008902 | 1735.444 | 1734.942 | 1557.669 | 1808.059 | 1129.179 | 1598.962 | 1265.147 | 1116.956 |
| ENSMUSG00000042042 | 0.066794268 | 0.39552 | 0.562859 | 848.1038 | 861.4719 | 920.8546 | 918.694 | 804.4668 | 926.3031 | 828.3244 | 834.1824 |
| ENSMUSG00000042043 | -0.161498117 | 7.88E-05 | 0.00113 | 3762.643 | 3666.655 | 3743.87 | 3704.968 | 4192.004 | 4166.688 | 4235.158 | 4046.926 |
| ENSMUSG00000042046 | -0.118173967 | 0.210188 | 0.367699 | 1171.047 | 1145.83 | 1373.829 | 1219.75 | 1416.837 | 1185.534 | 1350.969 | 1379.065 |
| ENSMUSG00000042050 | 0.136718505 | 0.27115 | 0.436327 | 783.7163 | 874.6699 | 638.4702 | 698.725 | 694.2793 | 686.0677 | 735.7526 | 603.6131 |
| ENSMUSG00000042055 | 0.076495242 | 0.193366 | 0.346575 | 2172.071 | 2022.899 | 1963.441 | 2041.83 | 1883.915 | 2012.391 | 1921.828 | 1958.752 |
| ENSMUSG00000042063 | 0.077914676 | 0.40232 | 0.569418 | 1021.145 | 1100.236 | 1055.836 | 1090.356 | 1057.996 | 1130.783 | 960.432 | 893.9999 |
| ENSMUSG00000042064 | -0.152821475 | 0.552784 | 0.700117 | 80.48435 | 69.58965 | 95.23226 | 80.22398 | 81.90935 | 60.3382 | 95.46463 | 125.073 |
| ENSMUSG00000042066 | -0.054469379 | 0.70094 | 0.811323 | 409.4641 | 382.7431 | 372.648 | 443.3885 | 424.1734 | 322.9211 | 457.0731 | 465.489 |
| ENSMUSG00000042073 | -0.371067369 | 0.006996 | 0.032932 | 717.3168 | 746.289 | 805.7478 | 804.8277 | 958.5344 | 776.575 | 1044.325 | 1197.438 |
| ENSMUSG00000042078 | 0.954945641 | 0.024019 | 0.080612 | 45.27245 | 27.5959 | 24.8432 | 61.24627 | 18.52711 | 30.1691 | 16.39292 | 17.40146 |
| ENSMUSG00000042079 | -0.224055234 | 1.30E-06 | 4.63E-05 | 7176.186 | 7353.706 | 7551.504 | 7530.703 | 8809.156 | 8276.39 | 8897.496 | 8605.021 |

| | | | | | | | | | | | |
|---|---|---|---|---|---|---|---|---|---|---|---|
| ENSMUSG00000042082 | -0.079499431 | 0.686108 | 0.80043 | 462.785 | 476.3291 | 491.8953 | 435.6249 | 599.6935 | 317.3342 | 462.8588 | 591.6496 |
| ENSMUSG00000042087 | -0.229028124 | 0.201299 | 0.356789 | 109.6599 | 115.1829 | 106.8258 | 120.7673 | 107.2622 | 118.4417 | 154.2863 | 150.0876 |
| ENSMUSG00000042096 | -2.514046737 | 7.92E-05 | 0.001131 | 180.0837 | 34.79482 | 273.2752 | 131.1188 | 867.8491 | 444.7149 | 763.717 | 1464.985 |
| ENSMUSG00000042097 | 0.033596026 | 0.818543 | 0.890168 | 349.1009 | 325.1516 | 273.2752 | 310.5445 | 285.7076 | 262.5829 | 308.5725 | 370.8686 |
| ENSMUSG00000042099 | -0.198136891 | 0.104695 | 0.227968 | 1125.775 | 975.4549 | 1249.613 | 1061.027 | 1216.939 | 1064.858 | 1396.291 | 1385.591 |
| ENSMUSG00000042102 | -3.058555929 | 1.38E-07 | 7.36E-06 | 16.09687 | 4.799286 | 28.98373 | 33.64232 | 195.0223 | 69.27719 | 106.0718 | 332.8029 |
| ENSMUSG00000042104 | 0.385818736 | 0.025241 | 0.083544 | 880.2976 | 807.4799 | 640.1264 | 769.4602 | 623.0961 | 760.9318 | 513.0018 | 473.1021 |
| ENSMUSG00000042105 | 0.189089039 | 0.022083 | 0.075581 | 1759.589 | 1748.14 | 1702.587 | 1889.145 | 1443.165 | 1737.517 | 1476.327 | 1573.744 |
| ENSMUSG00000042106 | -0.138608726 | 0.513508 | 0.668226 | 308.8587 | 488.3274 | 529.1601 | 401.9825 | 589.9423 | 360.9118 | 427.1801 | 524.2189 |
| ENSMUSG00000042109 | -0.100247535 | 0.470748 | 0.630616 | 534.2149 | 604.7101 | 806.5759 | 709.0765 | 737.1841 | 662.6028 | 709.7168 | 739.5619 |
| ENSMUSG00000042111 | -0.070286428 | 0.470776 | 0.630616 | 930.6003 | 820.6779 | 912.5735 | 838.47 | 1046.294 | 836.9132 | 898.7175 | 895.0875 |
| ENSMUSG00000042115 | -1.04068572 | 0.201154 | 0.356627 | 591.56 | 68.38983 | 162.3089 | 156.9975 | 587.9921 | 98.32892 | 670.181 | 657.9926 |
| ENSMUSG00000042116 | -1.09981623 | 1.20E-06 | 4.38E-05 | 193.1624 | 269.9598 | 251.7444 | 263.9628 | 487.5557 | 351.9728 | 519.7519 | 738.4743 |
| ENSMUSG00000042118 | -2.666749086 | 0.002451 | 0.015197 | 42.25428 | 5.999108 | 68.73285 | 52.62003 | 264.2552 | 43.57759 | 207.3222 | 564.4598 |
| ENSMUSG00000042121 | 0.201912723 | 0.158631 | 0.301575 | 743.4742 | 812.2792 | 808.2321 | 808.2782 | 666.9761 | 893.8993 | 603.645 | 594.9123 |
| ENSMUSG00000042129 | 0.27939499 | 0.157347 | 0.299783 | 327.9737 | 278.3586 | 216.9639 | 270.8638 | 236.952 | 214.5358 | 280.6081 | 167.489 |
| ENSMUSG00000042133 | 0.208696186 | 0.189865 | 0.342466 | 4815.982 | 5146.035 | 3960.006 | 4176.823 | 3569.882 | 5214.785 | 3357.655 | 3518.357 |
| ENSMUSG00000042138 | 0.492266435 | 0.001524 | 0.010673 | 1075.472 | 1275.41 | 913.4016 | 1211.986 | 776.1886 | 985.524 | 762.7527 | 656.905 |
| ENSMUSG00000042148 | -0.376075028 | 0.004397 | 0.023435 | 472.8455 | 464.3309 | 578.0184 | 497.7338 | 613.345 | 577.6824 | 637.3951 | 787.4159 |
| ENSMUSG00000042155 | -0.086204443 | 0.36551 | 0.534244 | 1043.278 | 1094.237 | 1011.118 | 1067.065 | 1210.113 | 1233.581 | 1029.861 | 1000.584 |
| ENSMUSG00000042156 | 0.349163291 | 0.0047 | 0.024642 | 1364.21 | 1384.594 | 1269.487 | 1504.415 | 1071.647 | 1319.619 | 1025.039 | 920.1021 |
| ENSMUSG00000042165 | -0.170618974 | 0.632201 | 0.760868 | 31.18769 | 20.39697 | 26.49941 | 14.6646 | 22.42756 | 26.81698 | 27.96439 | 27.18978 |
| ENSMUSG00000042167 | 0.083161589 | 0.427751 | 0.592774 | 1133.823 | 1180.624 | 1138.647 | 1270.644 | 1166.233 | 1230.229 | 1134.968 | 927.7152 |
| ENSMUSG00000042178 | -0.082973702 | 0.445608 | 0.608384 | 551.3178 | 563.9161 | 704.7187 | 558.98 | 644.5486 | 620.1426 | 602.6807 | 655.8174 |
| ENSMUSG00000042182 | 1.039070263 | 0.001507 | 0.010598 | 44.26639 | 58.79126 | 30.63995 | 51.75741 | 24.37778 | 18.99536 | 20.25007 | 26.10219 |
| ENSMUSG00000042185 | 0.126085675 | 0.177435 | 0.326495 | 1588.56 | 1384.594 | 1348.158 | 1482.85 | 1347.604 | 1390.013 | 1411.719 | 1168.073 |
| ENSMUSG00000042190 | -0.402318826 | 0.169257 | 0.315546 | 82.49646 | 105.5843 | 136.6376 | 73.323 | 132.6151 | 92.74205 | 111.8575 | 189.2408 |
| ENSMUSG00000042195 | -0.618554061 | 0.001239 | 0.009173 | 430.5913 | 353.9474 | 570.5655 | 472.7177 | 665.0259 | 538.5743 | 721.2883 | 885.2991 |
| ENSMUSG00000042197 | 0.209606145 | 0.039517 | 0.115692 | 1774.68 | 1787.734 | 1459.952 | 1748.538 | 1476.319 | 1641.423 | 1373.148 | 1362.752 |
| ENSMUSG00000042198 | -0.100380234 | 0.292853 | 0.460188 | 1006.054 | 1083.439 | 944.8697 | 1201.635 | 1117.478 | 1201.177 | 1135.933 | 1086.503 |
| ENSMUSG00000042202 | 0.268235421 | 0.05637 | 0.148073 | 1399.422 | 1340.201 | 1199.098 | 1209.398 | 946.8331 | 1358.727 | 1059.754 | 909.2261 |
| ENSMUSG00000042203 | -0.038560003 | 0.680492 | 0.795992 | 738.4439 | 643.1043 | 703.0625 | 734.0926 | 735.2339 | 644.7249 | 732.8598 | 784.1532 |
| ENSMUSG00000042207 | 0.226508324 | 0.016852 | 0.062455 | 3768.68 | 3798.635 | 3484.673 | 3614.392 | 3040.397 | 3667.222 | 3018.225 | 2809.248 |
| ENSMUSG00000042208 | 0.048091701 | 0.656215 | 0.778448 | 681.0988 | 729.4915 | 606.1741 | 770.3228 | 673.8019 | 745.2885 | 625.8237 | 650.3795 |
| ENSMUSG00000042210 | 0.046749174 | 0.670212 | 0.788576 | 710.2744 | 935.8608 | 795.8105 | 863.4861 | 877.6002 | 800.0399 | 761.7884 | 758.051 |
| ENSMUSG00000042211 | -0.1110393 | 0.101099 | 0.222739 | 1739.468 | 1560.968 | 1633.026 | 1608.793 | 1814.682 | 1641.423 | 1778.149 | 1831.503 |
| ENSMUSG00000042213 | 0.4922099 | 0.008514 | 0.037966 | 260.5681 | 253.1623 | 226.0731 | 290.7041 | 156.9929 | 242.4702 | 147.5362 | 187.0657 |
| ENSMUSG00000042215 | -0.182973877 | 0.142015 | 0.279081 | 806.8556 | 779.884 | 913.4016 | 878.1507 | 957.5593 | 796.6877 | 939.2176 | 1144.146 |
| ENSMUSG00000042216 | 0.018673688 | 0.925459 | 0.955704 | 212.2775 | 145.1784 | 218.6202 | 233.771 | 228.176 | 155.315 | 223.7151 | 193.5912 |
| ENSMUSG00000042225 | 0.485081264 | 0.004893 | 0.025347 | 755.5468 | 662.3015 | 489.411 | 678.0221 | 401.7459 | 576.565 | 417.5372 | 451.3503 |
| ENSMUSG00000042228 | -0.147621593 | 0.282633 | 0.449214 | 619.7295 | 472.7297 | 648.4075 | 535.6892 | 604.569 | 558.687 | 629.6808 | 731.9488 |
| ENSMUSG00000042229 | -0.226051785 | 0.076895 | 0.18419 | 784.7224 | 751.0883 | 687.3285 | 760.8339 | 926.3557 | 680.4808 | 894.8603 | 986.4451 |
| ENSMUSG00000042240 | 3.161050619 | NA | NA | 4.024217 | 5.999108 | 63.76421 | 1.725247 | 0.975111 | 4.469496 | 0.964289 | 2.175182 |
| ENSMUSG00000042246 | 0.458732038 | 0.022705 | 0.077159 | 107.6478 | 76.78858 | 107.6539 | 121.6299 | 71.18313 | 81.56831 | 77.14313 | 72.8686 |
| ENSMUSG00000042249 | 0.463901883 | 0.000142 | 0.00175 | 773.6558 | 599.9108 | 718.7966 | 765.147 | 483.6552 | 592.2083 | 524.5733 | 474.1897 |
| ENSMUSG00000042251 | -0.208786805 | 0.416537 | 0.582046 | 145.8779 | 118.7823 | 175.5586 | 106.1027 | 107.2622 | 192.1883 | 137.8934 | 195.7664 |
| ENSMUSG00000042254 | 1.046386929 | 0.001903 | 0.012672 | 42.25428 | 77.9884 | 59.62368 | 92.30071 | 41.92979 | 24.58223 | 37.60728 | 27.18978 |
| ENSMUSG00000042256 | 0.28855181 | 0.498858 | 0.655203 | 14.08476 | 17.99732 | 18.21835 | 12.93935 | 8.776002 | 12.29112 | 15.42863 | 15.22628 |
| ENSMUSG00000042258 | 2.559912468 | 1.63E-05 | 0.000333 | 127.7689 | 47.99286 | 22.35888 | 174.2499 | 17.552 | 20.11273 | 12.53576 | 13.05109 |
| ENSMUSG00000042271 | 0.054902019 | 0.503125 | 0.65912 | 635.8264 | 595.1115 | 568.9093 | 592.6223 | 587.9921 | 608.9689 | 548.6805 | 556.8466 |
| ENSMUSG00000042272 | -0.026657073 | 0.739158 | 0.836877 | 2300.846 | 2272.462 | 2388.26 | 2401.544 | 2164.747 | 2464.927 | 2268.008 | 2645.022 |
| ENSMUSG00000042275 | -0.272482424 | 0.0055 | 0.027553 | 562.3844 | 567.5156 | 562.2844 | 529.6508 | 640.6481 | 593.3256 | 701.0382 | 747.1751 |
| ENSMUSG00000042282 | -0.031362685 | 0.907647 | 0.944866 | 35.2119 | 51.59233 | 35.60859 | 38.81806 | 39.97956 | 50.28183 | 35.6787 | 38.06569 |
| ENSMUSG00000042284 | -1.03930864 | 5.45E-05 | 0.000845 | 396.3854 | 373.1445 | 554.8314 | 355.4009 | 884.426 | 493.8794 | 864.0031 | 1211.576 |
| ENSMUSG00000042286 | 0.239062452 | 0.046141 | 0.128754 | 1033.218 | 969.4558 | 1294.331 | 1188.695 | 889.3015 | 1045.862 | 874.6103 | 996.2334 |
| ENSMUSG00000042289 | 0.857531632 | 0.0192 | 0.068496 | 24.1453 | 40.79393 | 48.85829 | 47.44429 | 28.27823 | 14.52586 | 21.21436 | 25.0146 |
| ENSMUSG00000042292 | 0.043770352 | 0.656842 | 0.778842 | 559.3662 | 593.9117 | 615.2832 | 625.402 | 531.4357 | 661.4855 | 566.0377 | 566.635 |
| ENSMUSG00000042293 | -0.446410262 | 0.000105 | 0.001397 | 195.1745 | 201.57 | 221.1045 | 226.87 | 273.0312 | 297.2215 | 275.7867 | 308.8759 |
| ENSMUSG00000042298 | 0.119220793 | 0.329419 | 0.497241 | 938.6487 | 885.4683 | 871.9963 | 939.397 | 770.3379 | 1038.041 | 740.5741 | 800.467 |
| ENSMUSG00000042302 | 0.242936797 | 0.049039 | 0.134366 | 1016.115 | 1015.049 | 951.4945 | 968.7262 | 759.6117 | 998.9324 | 885.2175 | 694.9707 |
| ENSMUSG00000042303 | -0.438676482 | 0.011268 | 0.04669 | 421.5368 | 315.5531 | 514.2542 | 411.4714 | 537.2863 | 461.4755 | 619.0736 | 639.5036 |
| ENSMUSG00000042305 | -0.13576629 | 0.130428 | 0.26282 | 1953.758 | 1834.527 | 2046.251 | 2108.252 | 2198.876 | 2461.575 | 1998.971 | 2072.949 |
| ENSMUSG00000042306 | -0.439743313 | 0.312402 | 0.480769 | 248.4954 | 131.9804 | 598.7211 | 213.068 | 527.5352 | 196.6578 | 438.7516 | 455.7007 |
| ENSMUSG00000042308 | 0.112098588 | 0.147971 | 0.287378 | 636.8324 | 669.5004 | 693.1253 | 632.303 | 585.0668 | 639.138 | 597.8593 | 614.489 |

| | | | | | | | | | | | |
|---|---|---|---|---|---|---|---|---|---|---|---|
| ENSMUSG00000042312 | -0.190697811 | 0.227496 | 0.387474 | 407.452 | 398.3407 | 444.6933 | 424.4108 | 496.3317 | 340.7991 | 546.752 | 527.4817 |
| ENSMUSG00000042320 | 0.39551388 | 0.119395 | 0.2484 | 89.53884 | 64.79036 | 54.65504 | 85.39973 | 44.85512 | 58.10345 | 55.92877 | 65.25547 |
| ENSMUSG00000042323 | 0.314980723 | 0.07554 | 0.181836 | 4541.329 | 4615.713 | 3619.654 | 4010.337 | 2903.881 | 4749.957 | 2940.118 | 2900.605 |
| ENSMUSG00000042328 | -0.08735355 | 0.35714 | 0.525643 | 582.5055 | 548.3184 | 541.5817 | 604.6991 | 623.0961 | 557.5697 | 666.3238 | 570.9853 |
| ENSMUSG00000042331 | -0.505965552 | 0.006015 | 0.029526 | 520.1301 | 533.9206 | 737.0149 | 464.0914 | 787.8899 | 611.2036 | 864.9674 | 939.6787 |
| ENSMUSG00000042333 | 0.944073987 | 0.00504 | 0.025865 | 50.30272 | 51.59233 | 48.85829 | 30.19182 | 32.17867 | 22.34748 | 17.3572 | 21.75182 |
| ENSMUSG00000042340 | 0.253995894 | 0.106278 | 0.230327 | 250.5075 | 341.9491 | 241.8071 | 251.0234 | 205.7485 | 235.7659 | 213.1079 | 253.4087 |
| ENSMUSG00000042348 | -0.083173288 | 0.267098 | 0.431449 | 1028.188 | 1109.835 | 1165.146 | 1184.382 | 1215.964 | 1190.003 | 1226.576 | 1123.482 |
| ENSMUSG00000042349 | 0.498015557 | 4.99E-06 | 0.000134 | 404.4339 | 362.3461 | 373.4761 | 337.2858 | 272.0561 | 277.1088 | 247.8223 | 249.0584 |
| ENSMUSG00000042350 | -0.122171751 | 0.055185 | 0.14622 | 1115.714 | 1060.642 | 1101.382 | 1060.164 | 1242.292 | 1124.078 | 1178.361 | 1175.686 |
| ENSMUSG00000042353 | -1.205490391 | 0.010279 | 0.043496 | 12.07265 | 4.799286 | 18.21835 | 15.52722 | 30.22845 | 33.52122 | 37.60728 | 17.40146 |
| ENSMUSG00000042354 | -0.103536166 | 0.250332 | 0.413915 | 1711.298 | 1709.746 | 1443.39 | 1663.138 | 1904.392 | 1671.592 | 1626.756 | 1806.489 |
| ENSMUSG00000042357 | 0.847769328 | 0.021762 | 0.074822 | 82.49646 | 169.1748 | 82.81066 | 105.2401 | 66.30757 | 91.62468 | 31.82154 | 54.37955 |
| ENSMUSG00000042359 | -0.395497356 | 0.032121 | 0.099285 | 522.1422 | 365.9456 | 499.3483 | 438.2127 | 607.4943 | 445.8323 | 585.3235 | 764.5765 |
| ENSMUSG00000042363 | 0.138999099 | 0.192326 | 0.345491 | 1413.506 | 1416.989 | 1175.911 | 1508.728 | 1302.749 | 1273.806 | 1328.79 | 1100.642 |
| ENSMUSG00000042364 | -0.038403961 | 0.679098 | 0.794865 | 2016.133 | 2136.882 | 2455.336 | 2313.556 | 2201.801 | 2240.335 | 2168.686 | 2556.927 |
| ENSMUSG00000042367 | 0.119623706 | 0.774003 | 0.859972 | 27.16347 | 61.1909 | 52.99882 | 26.74133 | 57.53157 | 30.1691 | 25.07152 | 41.32846 |
| ENSMUSG00000042369 | -0.205337876 | 0.006239 | 0.030325 | 1143.884 | 1226.218 | 1324.143 | 1225.788 | 1507.522 | 1355.375 | 1375.076 | 1436.708 |
| ENSMUSG00000042371 | -4.917652246 | 8.55E-17 | 6.07E-14 | 6.036326 | 4.799286 | 15.73403 | 6.038364 | 261.3298 | 63.69032 | 197.6793 | 478.5401 |
| ENSMUSG00000042377 | -0.328228759 | 0.083482 | 0.194528 | 103.6236 | 85.18733 | 120.9036 | 91.43809 | 118.9636 | 106.1505 | 140.7862 | 139.2117 |
| ENSMUSG00000042379 | -0.637539004 | 0.019107 | 0.068286 | 123.7447 | 56.39161 | 112.6225 | 106.1027 | 187.2214 | 122.9112 | 132.1076 | 180.5401 |
| ENSMUSG00000042380 | -0.418443183 | 0.000918 | 0.007323 | 459.7668 | 455.9322 | 574.706 | 474.4429 | 607.4943 | 582.1519 | 701.0382 | 738.4743 |
| ENSMUSG00000042388 | 0.121069253 | 0.677567 | 0.793929 | 50.30272 | 28.79572 | 43.88965 | 39.68068 | 30.22845 | 46.92971 | 37.60728 | 35.89051 |
| ENSMUSG00000042389 | -0.104643586 | 0.251562 | 0.415205 | 452.7245 | 503.925 | 458.7711 | 452.0147 | 542.1619 | 507.2878 | 485.0374 | 469.8393 |
| ENSMUSG00000042390 | 0.317910006 | 0.052793 | 0.141354 | 3007.096 | 2854.375 | 2686.378 | 2775.922 | 2015.555 | 3156.582 | 2041.4 | 1871.744 |
| ENSMUSG00000042396 | 0.075645888 | 0.282189 | 0.448817 | 2250.544 | 2117.685 | 2132.375 | 2362.726 | 2108.191 | 2238.1 | 2106.972 | 1959.839 |
| ENSMUSG00000042401 | -0.319715769 | 0.312708 | 0.481026 | 126.7628 | 177.5736 | 165.6213 | 123.3552 | 181.3707 | 308.3953 | 139.8219 | 110.9343 |
| ENSMUSG00000042404 | 0.270653738 | 0.002531 | 0.015562 | 412.4823 | 398.3407 | 404.9441 | 395.9442 | 350.065 | 320.6864 | 344.2512 | 319.7518 |
| ENSMUSG00000042406 | -0.074967834 | 0.262168 | 0.426384 | 4536.299 | 4597.716 | 5111.074 | 4973.887 | 5273.402 | 4753.309 | 5054.804 | 5167.145 |
| ENSMUSG00000042408 | 0.433900316 | 3.95E-07 | 1.74E-05 | 1084.527 | 983.8537 | 933.2762 | 1111.059 | 776.1886 | 778.8097 | 729.0026 | 761.3138 |
| ENSMUSG00000042410 | -0.458061772 | 6.79E-05 | 0.001007 | 1768.644 | 1817.73 | 1816.866 | 1806.334 | 2421.201 | 2045.912 | 2452.187 | 2984.35 |
| ENSMUSG00000042417 | -1.286083699 | 0.171919 | 0.319177 | 12.07265 | 4.799286 | 16.56213 | 22.42821 | 16.57689 | 109.5027 | 6.750024 | 4.350364 |
| ENSMUSG00000042419 | 0.070580287 | 0.578073 | 0.719775 | 509.0635 | 635.9054 | 681.5318 | 691.824 | 641.6232 | 570.9782 | 642.2166 | 543.7955 |
| ENSMUSG00000042423 | 0.268297043 | 0.0011 | 0.008398 | 1105.654 | 959.8572 | 987.9312 | 1080.005 | 846.3966 | 922.951 | 841.8244 | 823.3065 |
| ENSMUSG00000042426 | -0.3944989 | 2.08E-06 | 6.61E-05 | 647.899 | 712.694 | 646.7513 | 614.1879 | 823.9691 | 840.2653 | 849.5388 | 927.7152 |
| ENSMUSG00000042428 | -0.629822305 | 6.36E-05 | 0.000956 | 554.3359 | 543.5192 | 631.0173 | 534.8266 | 980.962 | 652.5465 | 821.5744 | 1048.438 |
| ENSMUSG00000042429 | 0.566269596 | 0.000329 | 0.003329 | 1711.298 | 1500.977 | 1316.69 | 1384.511 | 919.53 | 1278.276 | 1002.861 | 792.8539 |
| ENSMUSG00000042436 | 0.463514645 | 0.007414 | 0.034276 | 3934.679 | 6341.057 | 5313.132 | 5305.997 | 4046.712 | 4680.68 | 3291.119 | 3134.438 |
| ENSMUSG00000042439 | 0.324488233 | 0.00048 | 0.004444 | 2218.35 | 2219.67 | 1949.363 | 2097.038 | 1712.295 | 1921.883 | 1542.863 | 1597.671 |
| ENSMUSG00000042444 | 0.194825365 | 0.038091 | 0.112661 | 980.903 | 885.4683 | 939.0729 | 939.397 | 716.7068 | 911.7773 | 837.003 | 809.1678 |
| ENSMUSG00000042446 | 0.113299167 | 0.198625 | 0.353231 | 2205.271 | 2367.248 | 2053.704 | 2142.757 | 1912.193 | 2278.326 | 1873.614 | 2041.408 |
| ENSMUSG00000042447 | 0.088132928 | 0.316515 | 0.48478 | 1277.689 | 1345 | 1356.439 | 1319.814 | 1270.57 | 1392.248 | 1228.504 | 1094.117 |
| ENSMUSG00000042448 | -0.296487034 | 0.372459 | 0.541481 | 50.30272 | 33.595 | 35.60859 | 44.85642 | 59.48179 | 60.3382 | 55.92877 | 26.10219 |
| ENSMUSG00000042453 | 0.609182867 | 0.001214 | 0.009039 | 3619.784 | 2620.41 | 2174.608 | 3328.001 | 2143.295 | 2097.311 | 2036.579 | 1420.394 |
| ENSMUSG00000042460 | -0.025398376 | 0.82454 | 0.893455 | 813.898 | 817.0785 | 741.9835 | 744.4441 | 653.3246 | 895.0167 | 785.8957 | 838.5327 |
| ENSMUSG00000042462 | -0.441653877 | 8.46E-05 | 0.001185 | 821.9464 | 1076.24 | 1023.54 | 1009.269 | 1454.866 | 1182.182 | 1384.719 | 1314.898 |
| ENSMUSG00000042472 | 0.018349312 | 0.757997 | 0.849141 | 1275.677 | 1323.403 | 1241.332 | 1231.826 | 1230.59 | 1299.506 | 1228.504 | 1246.379 |
| ENSMUSG00000042473 | 0.269815079 | 0.065425 | 0.164684 | 470.8334 | 457.132 | 370.9918 | 408.8835 | 282.7823 | 349.7381 | 375.1085 | 407.8467 |
| ENSMUSG00000042476 | 0.771202988 | 0.056756 | 0.148796 | 27.16347 | 25.19625 | 15.73403 | 31.05445 | 11.70134 | 18.99536 | 15.42863 | 11.9635 |
| ENSMUSG00000042485 | 0.052616391 | 0.817801 | 0.889465 | 88.53278 | 83.98751 | 100.2009 | 144.0581 | 85.8098 | 101.681 | 113.7861 | 102.2336 |
| ENSMUSG00000042487 | -0.153662377 | 0.135208 | 0.269381 | 1004.042 | 1174.625 | 1049.211 | 915.2435 | 1188.661 | 1025.749 | 1186.076 | 1202.876 |
| ENSMUSG00000042489 | -0.081664488 | 0.543558 | 0.692476 | 545.2815 | 718.6931 | 665.7977 | 527.9256 | 621.1459 | 584.3867 | 665.3595 | 727.5984 |
| ENSMUSG00000042492 | -0.031167774 | 0.725812 | 0.828514 | 1202.235 | 1222.618 | 1202.411 | 1200.772 | 1193.536 | 1106.2 | 1221.754 | 1411.693 |
| ENSMUSG00000042496 | 0.348737092 | 0.031377 | 0.097437 | 478.8819 | 482.3283 | 393.3507 | 404.5704 | 281.8072 | 429.0717 | 367.3942 | 302.3503 |
| ENSMUSG00000042498 | 0.060266474 | 0.812797 | 0.886216 | 36.21796 | 47.99286 | 48.85829 | 59.52102 | 43.88001 | 43.57759 | 53.0359 | 44.59123 |
| ENSMUSG00000042499 | -0.2021448 | 0.012447 | 0.050354 | 1392.379 | 1397.792 | 1425.172 | 1339.654 | 1572.855 | 1415.713 | 1717.399 | 1682.503 |
| ENSMUSG00000042500 | 0.030548442 | 0.795657 | 0.874398 | 232.3986 | 274.7591 | 245.1196 | 233.771 | 249.6285 | 244.7049 | 250.7152 | 217.5182 |
| ENSMUSG00000042501 | -1.470983916 | 0.193014 | 0.346255 | 1.006054 | 2.399643 | 31.46805 | 0 | 33.15378 | 14.52586 | 17.3572 | 32.62773 |
| ENSMUSG00000042502 | -0.120935127 | 0.058006 | 0.15106 | 2127.805 | 2200.473 | 2222.638 | 2069.434 | 2497.26 | 2195.64 | 2367.33 | 2310.043 |
| ENSMUSG00000042505 | -0.051391656 | 0.652472 | 0.77555 | 237.4288 | 217.1677 | 265.8222 | 236.3588 | 240.8525 | 250.2918 | 245.8937 | 257.7591 |
| ENSMUSG00000042506 | 0.078475598 | 0.132195 | 0.265309 | 4555.414 | 4714.099 | 4657.272 | 4735.803 | 4439.682 | 4690.736 | 4270.837 | 4275.321 |
| ENSMUSG00000042507 | 0.115956397 | 0.233442 | 0.394002 | 743.4742 | 620.3077 | 696.4377 | 756.5208 | 627.9717 | 711.7673 | 639.3237 | 624.2773 |
| ENSMUSG00000042508 | 0.213260277 | 0.055205 | 0.14622 | 1504.051 | 1746.94 | 1450.015 | 1626.045 | 1448.04 | 1555.385 | 1222.719 | 1230.066 |
| ENSMUSG00000042510 | -1.023778539 | 0.033115 | 0.10154 | 148.896 | 86.38715 | 447.1776 | 123.3552 | 488.5308 | 200.01 | 395.3586 | 556.8466 |

| | | | | | | | | | | | |
|---|---|---|---|---|---|---|---|---|---|---|---|
| ENSMUSG00000042514 | 0.3125255 | 0.05112 | 0.138178 | 661.9838 | 449.9331 | 538.2693 | 565.881 | 394.9201 | 395.5504 | 448.3945 | 547.0583 |
| ENSMUSG00000042515 | 0.958141878 | 1.23E-06 | 4.42E-05 | 703.232 | 807.4799 | 539.9255 | 724.6037 | 342.2641 | 478.2361 | 343.2869 | 264.2846 |
| ENSMUSG00000042520 | -0.039045986 | 0.607484 | 0.743335 | 4664.068 | 4168.18 | 4380.684 | 4153.532 | 4533.292 | 4008.021 | 4596.766 | 4703.831 |
| ENSMUSG00000042523 | 0.315594319 | 0.004295 | 0.023021 | 602.6266 | 596.3113 | 558.972 | 604.6991 | 463.1779 | 563.1565 | 446.4659 | 426.3357 |
| ENSMUSG00000042524 | 0.141594517 | 0.196647 | 0.351004 | 2244.507 | 2181.276 | 2252.45 | 2470.554 | 1874.164 | 2469.397 | 2106.972 | 1845.642 |
| ENSMUSG00000042535 | -0.228018655 | 0.009448 | 0.040883 | 1343.083 | 1424.188 | 1440.906 | 1226.651 | 1639.162 | 1426.887 | 1682.685 | 1612.898 |
| ENSMUSG00000042541 | -0.166951504 | 0.007681 | 0.035142 | 3296.84 | 3325.905 | 3512 | 3204.646 | 3887.769 | 3565.541 | 3591.013 | 3932.729 |
| ENSMUSG00000042548 | -0.000908049 | 0.989707 | 0.994138 | 2791.801 | 2691.2 | 2608.536 | 2768.159 | 2812.221 | 2935.342 | 2603.581 | 2515.598 |
| ENSMUSG00000042549 | -0.665855191 | 0.028366 | 0.090844 | 30.18163 | 21.59679 | 36.43669 | 29.3292 | 43.88001 | 33.52122 | 55.92877 | 54.37955 |
| ENSMUSG00000042557 | -0.033122628 | 0.619526 | 0.751691 | 1906.473 | 2024.099 | 1863.24 | 1906.398 | 2079.912 | 2049.264 | 1867.828 | 1878.27 |
| ENSMUSG00000042558 | -0.163984146 | 0.024444 | 0.081609 | 1174.065 | 1294.607 | 1191.645 | 1217.162 | 1476.319 | 1310.68 | 1301.79 | 1371.452 |
| ENSMUSG00000042564 | 0.312870589 | 0.122273 | 0.251777 | 226.3622 | 179.9732 | 150.7154 | 175.9752 | 150.1671 | 188.8362 | 130.179 | 120.7226 |
| ENSMUSG00000042569 | -0.121328259 | 0.31179 | 0.480258 | 504.0332 | 530.3211 | 520.8791 | 514.1236 | 542.1619 | 512.8747 | 509.1447 | 687.3576 |
| ENSMUSG00000042570 | 0.114838261 | 0.089297 | 0.203993 | 873.2552 | 866.2711 | 815.685 | 842.7832 | 783.9895 | 802.2746 | 799.3957 | 749.3503 |
| ENSMUSG00000042572 | -0.13084291 | 0.151915 | 0.292392 | 2533.245 | 2200.473 | 2524.897 | 2477.455 | 2689.357 | 2324.138 | 2893.832 | 2754.868 |
| ENSMUSG00000042579 | -0.04937445 | 0.701917 | 0.812145 | 347.0888 | 332.3506 | 323.7897 | 348.4999 | 347.1396 | 283.813 | 361.6084 | 405.6715 |
| ENSMUSG00000042581 | 1.382511024 | 0.000151 | 0.00184 | 150.9082 | 113.983 | 103.5133 | 185.4641 | 35.10401 | 96.09417 | 38.57157 | 43.50364 |
| ENSMUSG00000042589 | 0.538857713 | 1.08E-05 | 0.000242 | 717.3168 | 586.7127 | 786.7013 | 628.8525 | 466.1032 | 448.067 | 442.6087 | 517.6934 |
| ENSMUSG00000042590 | -0.139960439 | 0.031665 | 0.098155 | 1388.355 | 1347.4 | 1264.519 | 1367.258 | 1425.613 | 1516.277 | 1429.077 | 1543.292 |
| ENSMUSG00000042594 | -0.256394065 | 0.005821 | 0.028795 | 1143.884 | 1073.84 | 1274.456 | 1133.487 | 1448.04 | 1213.468 | 1424.255 | 1442.146 |
| ENSMUSG00000042595 | 0.076989833 | 0.620973 | 0.752803 | 756.5529 | 647.9036 | 679.0474 | 655.5939 | 532.4108 | 852.5564 | 623.8951 | 589.4744 |
| ENSMUSG00000042599 | 0.458087154 | 0.009167 | 0.040005 | 1686.147 | 1370.196 | 1135.334 | 1422.466 | 883.4508 | 1296.154 | 1093.504 | 813.5181 |
| ENSMUSG00000042605 | 0.151041398 | 0.085073 | 0.197118 | 1567.433 | 1371.396 | 1436.765 | 1415.565 | 1149.656 | 1397.835 | 1323.005 | 1347.525 |
| ENSMUSG00000042606 | -0.197376789 | 0.040573 | 0.117886 | 1898.425 | 2252.065 | 1923.692 | 1787.356 | 2412.425 | 2090.607 | 2142.651 | 2362.248 |
| ENSMUSG00000042607 | 0.817770044 | 1.04E-06 | 3.86E-05 | 921.5458 | 1021.048 | 751.9208 | 932.496 | 595.793 | 608.9689 | 457.0731 | 394.7956 |
| ENSMUSG00000042608 | -0.196114469 | 0.077001 | 0.184275 | 959.7759 | 898.6663 | 1014.431 | 896.2658 | 1021.917 | 915.1294 | 1175.468 | 1206.139 |
| ENSMUSG00000042613 | -0.205763927 | 0.097067 | 0.216361 | 1965.83 | 1768.537 | 2395.712 | 2020.264 | 2105.265 | 2120.776 | 2569.831 | 2608.043 |
| ENSMUSG00000042616 | -0.109515169 | 0.283248 | 0.449727 | 353.1251 | 411.5388 | 413.2252 | 392.4937 | 465.1281 | 408.9589 | 403.0729 | 416.5474 |
| ENSMUSG00000042622 | -0.412588772 | 0.065275 | 0.164512 | 314.895 | 200.3702 | 283.2125 | 274.3143 | 313.9858 | 264.8177 | 497.5732 | 352.3795 |
| ENSMUSG00000042625 | 0.114519141 | 0.113766 | 0.240897 | 3452.779 | 3291.11 | 2919.076 | 3057.138 | 2937.035 | 3047.079 | 2928.546 | 2832.087 |
| ENSMUSG00000042626 | 0.163538197 | 0.01539 | 0.058748 | 3810.934 | 4110.589 | 3945.928 | 3884.394 | 3870.217 | 3397.935 | 3377.905 | 3413.948 |
| ENSMUSG00000042628 | 0.105149396 | 0.286473 | 0.45356 | 1507.069 | 1253.814 | 1271.144 | 1363.808 | 1233.516 | 1122.961 | 1387.612 | 1271.394 |
| ENSMUSG00000042632 | -0.021400271 | 0.831449 | 0.897939 | 387.3309 | 341.9491 | 371.8199 | 356.2635 | 384.1939 | 331.8601 | 375.1085 | 388.27 |
| ENSMUSG00000042642 | -0.143464211 | 0.173697 | 0.321637 | 675.0625 | 649.1034 | 604.5178 | 654.7312 | 719.6321 | 598.9125 | 755.0384 | 777.6276 |
| ENSMUSG00000042644 | 0.097428353 | 0.505179 | 0.660965 | 1187.144 | 1060.642 | 1325.799 | 1017.896 | 990.7131 | 902.8383 | 1060.718 | 1338.825 |
| ENSMUSG00000042647 | -0.37836867 | 0.016269 | 0.060915 | 151.9142 | 130.7805 | 134.1533 | 153.547 | 165.7689 | 162.0192 | 216.0008 | 197.9416 |
| ENSMUSG00000042650 | -0.148643409 | 0.054136 | 0.144176 | 2509.1 | 2128.483 | 2438.774 | 2299.754 | 2543.09 | 2438.11 | 2725.081 | 2690.7 |
| ENSMUSG00000042655 | -0.751027757 | 0.177872 | 0.32709 | 5.030272 | 9.598572 | 4.96864 | 12.93935 | 11.70134 | 17.87799 | 16.39292 | 8.700729 |
| ENSMUSG00000042659 | 0.131780395 | 0.646657 | 0.77119 | 780.6982 | 1175.825 | 1295.159 | 1227.513 | 1264.719 | 680.4808 | 630.6451 | 1512.839 |
| ENSMUSG00000042660 | -0.181727926 | 0.094353 | 0.211925 | 611.681 | 626.3068 | 534.9569 | 573.6446 | 663.0757 | 584.3867 | 652.8238 | 758.051 |
| ENSMUSG00000042662 | -1.443841536 | 7.16E-05 | 0.001051 | 81.4904 | 59.99108 | 91.91984 | 66.42201 | 152.1174 | 92.74205 | 265.1795 | 306.7007 |
| ENSMUSG00000042670 | 0.025300143 | 0.731492 | 0.831665 | 945.6911 | 987.4531 | 919.1984 | 974.7645 | 948.7833 | 969.8807 | 979.7178 | 859.197 |
| ENSMUSG00000042671 | -0.152732052 | 0.500097 | 0.65655 | 46.2785 | 62.39072 | 48.03018 | 50.89479 | 58.50668 | 56.98608 | 49.17875 | 65.25547 |
| ENSMUSG00000042672 | -1.5954937 | 9.42E-05 | 0.001274 | 31.18769 | 28.79572 | 45.54586 | 33.64232 | 107.2622 | 42.46022 | 99.32178 | 172.927 |
| ENSMUSG00000042675 | 0.290645953 | 0.012605 | 0.050785 | 2786.771 | 2656.405 | 2572.927 | 3154.614 | 2524.563 | 2236.983 | 2483.045 | 1886.971 |
| ENSMUSG00000042677 | -0.134758447 | 0.609928 | 0.745057 | 47.28455 | 33.595 | 52.17072 | 43.9938 | 51.6809 | 36.87335 | 49.17875 | 57.64233 |
| ENSMUSG00000042678 | -0.185269161 | 0.521776 | 0.675051 | 63.38142 | 61.1909 | 63.76421 | 89.71284 | 78.98402 | 112.8548 | 75.21455 | 50.02919 |
| ENSMUSG00000042680 | -0.304360014 | 0.006545 | 0.031463 | 489.9485 | 411.5388 | 439.7246 | 425.2734 | 530.4606 | 482.7056 | 617.1451 | 550.3211 |
| ENSMUSG00000042682 | -0.055284042 | 0.312163 | 0.480616 | 2119.757 | 2242.466 | 2186.202 | 2341.16 | 2312.964 | 2355.425 | 2287.294 | 2282.854 |
| ENSMUSG00000042684 | -0.431490744 | 0.085958 | 0.198529 | 358.1554 | 265.1606 | 434.756 | 322.6212 | 613.345 | 298.3389 | 381.8585 | 569.8977 |
| ENSMUSG00000042686 | 0.403412808 | 0.109788 | 0.235472 | 162.9808 | 119.9822 | 91.91984 | 150.0965 | 73.13335 | 120.6764 | 111.8575 | 91.35765 |
| ENSMUSG00000042688 | -0.051669384 | 0.496972 | 0.653568 | 1904.461 | 1836.927 | 1883.943 | 1916.749 | 2187.175 | 1928.588 | 1814.792 | 1886.971 |
| ENSMUSG00000042694 | -0.039384721 | 0.690973 | 0.804327 | 589.5479 | 699.496 | 712.1717 | 622.8142 | 701.105 | 614.5558 | 675.0024 | 704.759 |
| ENSMUSG00000042699 | -0.165819268 | 0.013949 | 0.054589 | 5391.445 | 5579.17 | 5940.009 | 5539.768 | 6183.181 | 5799.172 | 6501.238 | 6702.824 |
| ENSMUSG00000042700 | 0.082543641 | 0.336053 | 0.504284 | 2228.41 | 2168.078 | 1917.895 | 1981.446 | 1943.397 | 2164.354 | 1895.792 | 1828.241 |
| ENSMUSG00000042705 | -0.023100377 | 0.743238 | 0.839486 | 933.6184 | 872.2703 | 961.4318 | 946.298 | 919.53 | 956.4722 | 994.1821 | 907.051 |
| ENSMUSG00000042708 | -0.652600767 | 0.233261 | 0.393874 | 38.23007 | 32.39518 | 28.15563 | 18.97772 | 37.05423 | 101.681 | 28.92867 | 17.40146 |
| ENSMUSG00000042709 | -0.202023547 | 0.028316 | 0.090822 | 763.5953 | 800.281 | 843.8407 | 872.1124 | 1039.469 | 844.7348 | 941.1462 | 948.3794 |
| ENSMUSG00000042712 | 0.090005668 | 0.232401 | 0.392999 | 8768.77 | 8119.192 | 8904.631 | 8511.506 | 7286.032 | 8724.457 | 8424.994 | 7798.028 |
| ENSMUSG00000042719 | -0.190024552 | 0.19634 | 0.350591 | 770.6376 | 772.6851 | 671.5945 | 756.5208 | 983.8873 | 623.4947 | 872.6817 | 907.051 |
| ENSMUSG00000042724 | -0.579260734 | 0.009547 | 0.041169 | 206.2411 | 163.1757 | 179.6991 | 123.3552 | 223.3005 | 185.4841 | 288.3225 | 306.7007 |
| ENSMUSG00000042726 | 0.023892674 | 0.748994 | 0.843439 | 1341.07 | 1450.584 | 1520.404 | 1582.051 | 1432.439 | 1507.338 | 1463.791 | 1397.555 |
| ENSMUSG00000042729 | -0.153309877 | 0.487365 | 0.645252 | 120.7265 | 106.7841 | 115.1068 | 91.43809 | 142.3663 | 139.6718 | 118.6076 | 81.56933 |
| ENSMUSG00000042734 | -0.500970675 | 0.03936 | 0.115288 | 43.26034 | 49.19268 | 70.38906 | 57.79577 | 73.13335 | 63.69032 | 86.78602 | 90.27006 |

| | | | | | | | | | | | |
|---|---|---|---|---|---|---|---|---|---|---|---|
| ENSMUSG00000042737 | -0.103223585 | 0.324799 | 0.492725 | 579.4873 | 628.7065 | 751.9208 | 679.7473 | 671.8517 | 695.0067 | 719.3597 | 753.7006 |
| ENSMUSG00000042742 | 0.181528429 | 0.271411 | 0.436604 | 937.6427 | 812.2792 | 785.8732 | 835.8822 | 683.553 | 997.8151 | 702.9668 | 589.4744 |
| ENSMUSG00000042743 | 0.382112349 | 0.017606 | 0.064558 | 410.4702 | 375.5441 | 290.6654 | 361.4392 | 339.3387 | 259.2308 | 251.6795 | 251.2335 |
| ENSMUSG00000042744 | 0.140021639 | 0.192307 | 0.34549 | 975.8727 | 811.0794 | 891.0427 | 819.4923 | 811.2926 | 890.5472 | 702.0025 | 772.1897 |
| ENSMUSG00000042745 | -0.087744268 | 0.619599 | 0.75173 | 1929.612 | 1889.719 | 2307.933 | 2183.3 | 2041.883 | 1552.033 | 2372.151 | 2866.89 |
| ENSMUSG00000042747 | -0.078946785 | 0.366032 | 0.534678 | 1537.251 | 1688.149 | 1854.959 | 1644.16 | 1938.521 | 1649.244 | 1775.256 | 1740.146 |
| ENSMUSG00000042750 | 0.120789559 | 0.241087 | 0.403217 | 1319.943 | 1334.202 | 1170.115 | 1355.182 | 1120.403 | 1373.253 | 1203.433 | 1065.839 |
| ENSMUSG00000042751 | 0.154390267 | 0.648272 | 0.772203 | 59.35721 | 39.59411 | 101.8571 | 108.6906 | 72.15824 | 81.56831 | 58.82164 | 67.43065 |
| ENSMUSG00000042757 | 0.291985601 | 0.275604 | 0.441808 | 107.6478 | 107.9839 | 117.5911 | 130.2561 | 83.85957 | 144.1413 | 61.71451 | 90.27006 |
| ENSMUSG00000042759 | -0.444783843 | 0.092219 | 0.208526 | 36.21796 | 32.39518 | 51.34261 | 31.91707 | 50.70579 | 48.04709 | 56.89306 | 52.20437 |
| ENSMUSG00000042761 | 1.649306571 | 0.001627 | 0.011205 | 15.09082 | 31.19536 | 25.67131 | 24.15346 | 6.825779 | 6.704245 | 13.50005 | 3.262773 |
| ENSMUSG00000042763 | -0.142835497 | 0.604809 | 0.741291 | 56.33904 | 43.19358 | 52.17072 | 47.44429 | 57.53157 | 33.52122 | 65.57166 | 63.08028 |
| ENSMUSG00000042766 | 0.880975307 | 1.36E-06 | 4.80E-05 | 273.6468 | 274.7591 | 227.7293 | 250.1608 | 170.6445 | 127.3806 | 103.1789 | 155.5255 |
| ENSMUSG00000042770 | -0.566886712 | 0.004536 | 0.023944 | 711.2804 | 785.8831 | 825.6223 | 866.9366 | 1085.299 | 783.2792 | 1242.004 | 1615.073 |
| ENSMUSG00000042772 | -0.023551327 | 0.698046 | 0.809552 | 2012.109 | 1941.311 | 1926.176 | 1922.788 | 2021.406 | 2017.978 | 1836.971 | 2055.547 |
| ENSMUSG00000042784 | -0.633389888 | 0.068082 | 0.168841 | 323.9495 | 247.1632 | 800.7791 | 350.2251 | 604.569 | 469.2971 | 694.2882 | 905.9634 |
| ENSMUSG00000042787 | -0.020915979 | 0.875349 | 0.925381 | 349.1009 | 395.9411 | 291.4935 | 301.9182 | 312.0356 | 328.508 | 349.0727 | 364.343 |
| ENSMUSG00000042788 | 1.256287998 | 0.001474 | 0.010444 | 59.35721 | 53.99197 | 43.06154 | 52.62003 | 10.72622 | 34.6386 | 14.46434 | 28.27737 |
| ENSMUSG00000042790 | 0.092284822 | 0.347491 | 0.516273 | 769.6316 | 689.8974 | 634.3297 | 619.3637 | 657.225 | 602.2646 | 628.7165 | 653.6422 |
| ENSMUSG00000042793 | 0.566622308 | 0.244015 | 0.406566 | 91.55095 | 207.5691 | 121.7317 | 142.3329 | 63.38224 | 213.4185 | 56.89306 | 46.76642 |
| ENSMUSG00000042797 | -1.269100385 | 1.29E-06 | 4.58E-05 | 211.2714 | 169.1748 | 287.353 | 221.6942 | 549.9628 | 325.1559 | 505.2875 | 766.7517 |
| ENSMUSG00000042804 | -0.168878305 | 0.316959 | 0.485098 | 399.4036 | 508.7243 | 648.4075 | 503.7721 | 556.7886 | 507.2878 | 560.252 | 693.8831 |
| ENSMUSG00000042807 | -0.504569524 | 0.014298 | 0.055669 | 170.0232 | 104.3845 | 175.5586 | 153.547 | 205.7485 | 187.7188 | 267.1081 | 197.9416 |
| ENSMUSG00000042810 | 0.158870586 | 0.019464 | 0.069137 | 1507.069 | 1400.192 | 1376.313 | 1451.795 | 1334.927 | 1341.966 | 1239.112 | 1221.365 |
| ENSMUSG00000042812 | 1.068257174 | 0.081999 | 0.192296 | 58.35115 | 103.1847 | 74.5296 | 98.33908 | 55.58134 | 82.68568 | 11.57147 | 9.78832 |
| ENSMUSG00000042814 | -0.020327373 | 0.886994 | 0.932224 | 265.5983 | 202.7698 | 242.6352 | 295.8799 | 272.0561 | 246.9397 | 244.9294 | 259.9343 |
| ENSMUSG00000042821 | -0.333922956 | 0.127113 | 0.258285 | 181.0898 | 159.5763 | 200.4018 | 190.6398 | 251.5787 | 141.9065 | 248.7866 | 280.5985 |
| ENSMUSG00000042826 | -0.039791546 | 0.819136 | 0.890602 | 183.1019 | 220.7672 | 278.2438 | 227.7326 | 195.0223 | 256.996 | 263.2509 | 221.8686 |
| ENSMUSG00000042828 | 0.50879813 | 0.193478 | 0.346682 | 25.15136 | 21.59679 | 14.07781 | 23.29083 | 16.57689 | 15.64324 | 12.53576 | 14.13868 |
| ENSMUSG00000042831 | -0.309598394 | 0.001533 | 0.0107 | 724.3591 | 761.8867 | 726.2495 | 741.8562 | 918.5549 | 776.575 | 962.3606 | 1001.671 |
| ENSMUSG00000042834 | 0.267490883 | 0.00612 | 0.029877 | 13269.86 | 17421.41 | 16572.07 | 16344.13 | 12858.79 | 14295.68 | 12888.69 | 12799.86 |
| ENSMUSG00000042842 | -0.346963873 | 0.180884 | 0.331139 | 199.1988 | 205.1695 | 327.1021 | 196.6782 | 298.3841 | 181.0146 | 362.5727 | 339.3284 |
| ENSMUSG00000042851 | 0.445527038 | 0.00144 | 0.010278 | 556.3481 | 494.3265 | 443.8652 | 470.1298 | 338.3636 | 443.5975 | 344.2512 | 316.489 |
| ENSMUSG00000042854 | -0.258723932 | 0.095265 | 0.213504 | 161.9748 | 207.5691 | 192.1207 | 190.6398 | 216.4747 | 187.7188 | 227.5722 | 267.5474 |
| ENSMUSG00000042857 | 0.038441472 | 0.852611 | 0.91148 | 135.8173 | 172.7743 | 153.1997 | 119.9047 | 110.1876 | 179.8972 | 148.5005 | 127.2482 |
| ENSMUSG00000042870 | -0.652768194 | 0.001033 | 0.00802 | 329.9858 | 265.1606 | 429.7873 | 267.4133 | 478.7797 | 415.6632 | 507.2161 | 632.978 |
| ENSMUSG00000042873 | 0.426551372 | 0.135026 | 0.269148 | 56.33904 | 76.78858 | 52.99882 | 104.3774 | 52.65601 | 55.8687 | 60.75022 | 46.76642 |
| ENSMUSG00000042901 | 0.199283703 | 0.036424 | 0.109395 | 1560.39 | 1874.121 | 1609.011 | 1742.499 | 1509.472 | 1638.07 | 1404.969 | 1356.226 |
| ENSMUSG00000042903 | 0.208104544 | 0.01975 | 0.06984 | 2477.912 | 2369.648 | 2370.869 | 2718.989 | 2299.312 | 1967.696 | 2326.83 | 2007.693 |
| ENSMUSG00000042918 | 0.333617256 | 0.149425 | 0.289398 | 153.9263 | 194.3711 | 163.9651 | 213.068 | 142.3663 | 204.4795 | 120.5361 | 108.7591 |
| ENSMUSG00000042942 | 0.148178023 | 0.318313 | 0.486564 | 276.6649 | 188.372 | 225.245 | 232.0457 | 190.1467 | 212.3011 | 215.0365 | 216.4306 |
| ENSMUSG00000042961 | 0.592424537 | 3.27E-07 | 1.48E-05 | 1308.877 | 1563.367 | 1488.108 | 1551.86 | 818.1184 | 1115.139 | 1030.825 | 958.1677 |
| ENSMUSG00000042978 | 0.030660948 | 0.594823 | 0.734023 | 1673.068 | 1707.346 | 1784.57 | 1794.257 | 1696.694 | 1743.104 | 1634.47 | 1744.496 |
| ENSMUSG00000042985 | 0.571528895 | 0.112158 | 0.238643 | 588.5418 | 1563.367 | 798.2948 | 765.147 | 668.9264 | 1013.458 | 474.4303 | 342.5912 |
| ENSMUSG00000042988 | 0.793673199 | 0.001049 | 0.008103 | 250.5075 | 370.7449 | 398.3193 | 271.7264 | 167.7191 | 261.4655 | 135.0005 | 181.6277 |
| ENSMUSG00000042992 | -0.358407757 | 0.006097 | 0.029818 | 207.2472 | 217.1677 | 198.7456 | 206.167 | 258.4045 | 245.8223 | 243.0009 | 315.4014 |
| ENSMUSG00000042997 | -0.223985204 | 0.023128 | 0.078349 | 704.238 | 614.3086 | 768.483 | 638.3414 | 753.761 | 775.4576 | 830.253 | 827.6568 |
| ENSMUSG00000043003 | 0.139466542 | 0.396292 | 0.563567 | 342.0585 | 385.1427 | 331.2427 | 348.4999 | 385.169 | 356.4423 | 296.0368 | 238.1824 |
| ENSMUSG00000043004 | 0.565053083 | 2.87E-05 | 0.00052 | 3172.089 | 3599.465 | 3366.253 | 3722.22 | 2135.494 | 3011.323 | 2047.186 | 2176.27 |
| ENSMUSG00000043008 | -0.211534414 | 0.096852 | 0.215986 | 334.01 | 345.5486 | 289.8373 | 298.4677 | 389.0694 | 343.0338 | 412.7158 | 318.6642 |
| ENSMUSG00000043015 | 0.311543922 | 0.079294 | 0.187911 | 652.9293 | 814.6788 | 587.1276 | 615.9132 | 509.0081 | 716.2368 | 457.0731 | 468.7518 |
| ENSMUSG00000043017 | 0.159853048 | 0.538337 | 0.6882 | 58.35115 | 81.58786 | 70.38906 | 43.9938 | 66.30757 | 53.63396 | 52.07161 | 54.37955 |
| ENSMUSG00000043019 | 0.080856899 | 0.580473 | 0.721769 | 1710.292 | 1280.21 | 1228.91 | 1315.501 | 1179.885 | 1182.182 | 1276.719 | 1594.409 |
| ENSMUSG00000043024 | 0.433675703 | 0.342225 | 0.511012 | 14.08476 | 11.99822 | 17.39024 | 12.07673 | 10.72622 | 13.40849 | 9.642892 | 7.613138 |
| ENSMUSG00000043029 | 0.197464386 | 0.588203 | 0.728387 | 23.13925 | 26.39607 | 23.18699 | 15.52722 | 16.57689 | 16.76061 | 19.28578 | 23.927 |
| ENSMUSG00000043051 | -0.205712052 | 0.547216 | 0.695504 | 24.1453 | 46.79304 | 47.20208 | 41.40593 | 54.60623 | 26.81698 | 51.10733 | 51.11678 |
| ENSMUSG00000043059 | 0.07388862 | 0.421225 | 0.586621 | 670.0322 | 712.694 | 708.8593 | 659.907 | 698.1797 | 582.1519 | 630.6451 | 701.4963 |
| ENSMUSG00000043061 | 0.466828261 | 0.003586 | 0.020184 | 1765.625 | 2186.075 | 1639.651 | 1874.481 | 1318.35 | 1768.803 | 1183.183 | 1131.095 |
| ENSMUSG00000043065 | 0.109372403 | 0.474932 | 0.63448 | 629.79 | 676.6993 | 435.5841 | 644.3797 | 497.3068 | 588.8561 | 575.6806 | 548.1459 |
| ENSMUSG00000043067 | 0.181181858 | 0.031488 | 0.097721 | 1700.232 | 1904.117 | 1763.039 | 1737.324 | 1518.248 | 1769.921 | 1499.47 | 1478.036 |
| ENSMUSG00000043068 | 0.215415536 | 0.116418 | 0.244577 | 147.89 | 152.3773 | 159.8246 | 175.9752 | 130.6649 | 135.2023 | 140.7862 | 142.4744 |
| ENSMUSG00000043079 | -0.836295979 | 0.00345 | 0.019588 | 468.8213 | 255.562 | 534.9569 | 521.8872 | 597.7432 | 557.5697 | 867.8602 | 1160.46 |
| ENSMUSG00000043085 | -0.885137493 | 0.003405 | 0.019397 | 63.38142 | 58.79126 | 96.06037 | 60.38364 | 124.8142 | 86.03781 | 111.8575 | 193.5912 |
| ENSMUSG00000043088 | 0.102409841 | 0.529534 | 0.681668 | 254.5318 | 241.1641 | 328.7583 | 227.7326 | 231.1014 | 255.8787 | 216.0008 | 279.5109 |

| | | | | | | | | | | | |
|---|---|---|---|---|---|---|---|---|---|---|---|
| ENSMUSG00000043090 | 0.007426644 | 0.947985 | 0.970222 | 982.9151 | 890.2676 | 777.5921 | 785.85 | 872.7246 | 956.4722 | 792.6457 | 795.0291 |
| ENSMUSG00000043091 | -0.196180122 | 0.084235 | 0.195701 | 2037.26 | 1899.317 | 2344.37 | 2051.319 | 2325.64 | 2021.33 | 2466.652 | 2735.292 |
| ENSMUSG00000043099 | 0.400680734 | 0.011729 | 0.048149 | 2038.266 | 3022.35 | 3043.292 | 2720.714 | 2250.557 | 2244.805 | 1610.363 | 2094.7 |
| ENSMUSG00000043102 | 0.165290252 | 0.794314 | 0.873694 | 11.0666 | 10.79839 | 7.45296 | 14.6646 | 3.900445 | 20.11273 | 5.785735 | 9.78832 |
| ENSMUSG00000043110 | 1.011705738 | NA | NA | 84.50857 | 237.5647 | 85.29498 | 103.5148 | 26.32801 | 187.7188 | 31.82154 | 7.613138 |
| ENSMUSG00000043131 | 0.076915433 | 0.080364 | 0.189523 | 3350.161 | 3269.514 | 3160.055 | 3263.305 | 3054.049 | 3109.652 | 3080.904 | 3119.211 |
| ENSMUSG00000043140 | -0.202649321 | 0.084555 | 0.196221 | 647.899 | 712.694 | 573.8779 | 634.0283 | 806.4171 | 626.8469 | 715.5026 | 802.6422 |
| ENSMUSG00000043144 | -1.96346104 | 0.012116 | 0.049352 | 7.04238 | 9.598572 | 48.85829 | 7.763611 | 69.2329 | 13.40849 | 54.96448 | 150.0876 |
| ENSMUSG00000043153 | -0.783613307 | 0.0008 | 0.006574 | 146.8839 | 128.3809 | 208.6829 | 151.8217 | 250.6036 | 193.3057 | 293.1439 | 359.9926 |
| ENSMUSG00000043154 | 0.120657921 | 0.218301 | 0.376926 | 2290.786 | 2177.676 | 2406.478 | 2222.981 | 1839.06 | 2373.303 | 2216.901 | 1941.35 |
| ENSMUSG00000043155 | -1.315807526 | 0.008877 | 0.039101 | 15.09082 | 13.19804 | 13.24971 | 13.80198 | 36.07912 | 11.17374 | 32.78583 | 57.64233 |
| ENSMUSG00000043157 | 0.81949766 | 0.016766 | 0.06223 | 33.19979 | 37.19447 | 33.12427 | 27.60395 | 13.65156 | 16.76061 | 26.03581 | 17.40146 |
| ENSMUSG00000043162 | -0.363940365 | 0.004527 | 0.023914 | 603.6326 | 755.8876 | 763.5143 | 665.9453 | 860.0482 | 753.1101 | 988.3964 | 986.4451 |
| ENSMUSG00000043165 | -0.017364021 | 0.935469 | 0.961785 | 116.7023 | 145.1784 | 146.5749 | 150.9591 | 117.9885 | 155.315 | 185.1435 | 107.6715 |
| ENSMUSG00000043168 | 0.767229909 | 7.58E-07 | 2.97E-05 | 922.5518 | 641.9045 | 607.8303 | 753.0703 | 419.2979 | 395.5504 | 414.6443 | 489.416 |
| ENSMUSG00000043183 | -0.03023682 | 0.829723 | 0.896882 | 457.7547 | 424.7368 | 361.8826 | 401.1199 | 486.5805 | 454.7713 | 385.7157 | 351.2919 |
| ENSMUSG00000043190 | 0.048860994 | 0.766058 | 0.854851 | 321.9374 | 488.3274 | 360.2264 | 376.1038 | 389.0694 | 404.4894 | 396.3228 | 302.3503 |
| ENSMUSG00000043192 | -0.343667406 | 0.235574 | 0.39644 | 29.17558 | 23.99643 | 31.46805 | 22.42821 | 36.07912 | 32.40385 | 30.85725 | 36.9781 |
| ENSMUSG00000043207 | -0.256857745 | 0.000267 | 0.002835 | 1219.338 | 1175.825 | 1165.974 | 1182.657 | 1336.878 | 1401.187 | 1381.826 | 1549.817 |
| ENSMUSG00000043219 | 0.250352558 | 0.107934 | 0.232523 | 259.562 | 233.9652 | 200.4018 | 256.1992 | 234.0267 | 202.2447 | 193.8221 | 167.489 |
| ENSMUSG00000043230 | -0.479932684 | 0.236686 | 0.397876 | 22.1332 | 13.19804 | 19.04645 | 9.488858 | 21.45245 | 26.81698 | 19.28578 | 21.75182 |
| ENSMUSG00000043241 | 0.310169612 | 0.016682 | 0.061985 | 2169.053 | 2285.66 | 1777.117 | 1904.673 | 1505.572 | 1990.043 | 1595.899 | 1469.336 |
| ENSMUSG00000043243 | -0.104120325 | 0.611888 | 0.746201 | 66.39959 | 55.19179 | 61.27989 | 60.38364 | 73.13335 | 70.39457 | 61.71451 | 56.55474 |
| ENSMUSG00000043251 | -0.235652686 | 0.244214 | 0.406826 | 275.6589 | 190.7716 | 351.9453 | 233.771 | 320.8116 | 279.3435 | 280.6081 | 361.0802 |
| ENSMUSG00000043252 | 0.144460821 | 0.028137 | 0.090419 | 1384.331 | 1510.575 | 1446.702 | 1532.882 | 1260.819 | 1362.079 | 1324.933 | 1368.19 |
| ENSMUSG00000043257 | -0.165774863 | 0.129818 | 0.26205 | 431.5973 | 455.9322 | 502.6607 | 428.7239 | 493.4063 | 490.5272 | 474.4303 | 584.0364 |
| ENSMUSG00000043259 | 0.002180384 | 0.990348 | 0.994551 | 201.2109 | 169.1748 | 150.7154 | 176.8378 | 190.1467 | 135.2023 | 162.9649 | 207.7299 |
| ENSMUSG00000043262 | 0.167706361 | 0.150581 | 0.290642 | 720.3349 | 664.7011 | 559.8001 | 733.23 | 642.5984 | 622.3774 | 560.252 | 557.9342 |
| ENSMUSG00000043263 | 0.892647064 | 0.193099 | 0.34636 | 13.07871 | 31.19536 | 50.5145 | 4.313117 | 12.67645 | 13.40849 | 16.39292 | 10.87591 |
| ENSMUSG00000043279 | 0.147100203 | 0.182189 | 0.332766 | 937.6427 | 917.8635 | 1058.32 | 807.4156 | 779.1139 | 917.3641 | 875.5746 | 789.5911 |
| ENSMUSG00000043284 | -0.307021032 | 0.009937 | 0.04238 | 1326.986 | 1207.02 | 1280.253 | 1268.057 | 1767.877 | 1233.581 | 1645.077 | 1640.087 |
| ENSMUSG00000043289 | 1.527678196 | 0.042867 | 0.122527 | 8.048435 | 20.39697 | 19.87456 | 18.97772 | 1.950223 | 15.64324 | 3.857157 | 2.175182 |
| ENSMUSG00000043290 | -0.178996297 | 0.596963 | 0.73584 | 34.20585 | 27.5959 | 32.29616 | 20.70296 | 33.15378 | 27.93435 | 44.3573 | 23.927 |
| ENSMUSG00000043300 | 0.283145942 | 0.009439 | 0.040855 | 584.5176 | 584.3131 | 516.7385 | 638.3414 | 431.9743 | 531.8701 | 482.1446 | 464.4014 |
| ENSMUSG00000043311 | -0.307171355 | 0.113063 | 0.24001 | 228.3743 | 247.1632 | 255.0568 | 177.7004 | 283.7574 | 203.3621 | 296.0368 | 339.3284 |
| ENSMUSG00000043313 | -0.119162064 | 0.552384 | 0.69976 | 114.6902 | 139.1793 | 134.1533 | 133.7066 | 133.5902 | 168.7235 | 101.2504 | 164.2263 |
| ENSMUSG00000043323 | -0.002352484 | 0.973429 | 0.985497 | 1234.429 | 1090.638 | 1126.225 | 1152.465 | 1204.262 | 1101.731 | 1131.111 | 1174.598 |
| ENSMUSG00000043333 | -0.082872524 | 0.581274 | 0.722532 | 137.8294 | 158.3764 | 146.5749 | 132.844 | 156.0178 | 131.8501 | 172.6078 | 146.8248 |
| ENSMUSG00000043336 | -0.037880671 | 0.753807 | 0.846914 | 1344.089 | 1283.809 | 1456.64 | 1362.945 | 1403.185 | 1136.369 | 1661.47 | 1391.029 |
| ENSMUSG00000043340 | -0.339906502 | 0.48647 | 0.644509 | 6.036326 | 9.598572 | 19.04645 | 18.97772 | 16.57689 | 17.87799 | 20.25007 | 14.13868 |
| ENSMUSG00000043342 | -0.0939356 | 0.325712 | 0.49356 | 2862.225 | 3025.95 | 3642.841 | 3304.711 | 3151.56 | 3506.32 | 3649.834 | 3396.547 |
| ENSMUSG00000043366 | 0.629416188 | 0.263094 | 0.427359 | 28.16952 | 7.198929 | 19.87456 | 35.36756 | 11.70134 | 8.938993 | 14.46434 | 23.927 |
| ENSMUSG00000043372 | 0.071122671 | 0.597306 | 0.736063 | 197.1867 | 208.7689 | 173.9024 | 172.5247 | 169.6694 | 195.5405 | 168.7506 | 180.5401 |
| ENSMUSG00000043384 | 0.364091629 | 7.10E-06 | 0.000177 | 3542.317 | 3474.683 | 3066.479 | 3557.459 | 2537.24 | 2917.464 | 2583.331 | 2560.189 |
| ENSMUSG00000043388 | 0.996626142 | 0.015002 | 0.057719 | 73.44197 | 41.99375 | 43.06154 | 109.5532 | 34.1289 | 48.04709 | 30.85725 | 21.75182 |
| ENSMUSG00000043391 | 0.435374643 | 1.22E-06 | 4.42E-05 | 1124.769 | 1335.401 | 1303.44 | 1327.578 | 926.3557 | 1003.402 | 866.896 | 970.1312 |
| ENSMUSG00000043398 | -0.366790209 | 0.32336 | 0.491365 | 48.29061 | 37.19447 | 49.6864 | 70.73513 | 83.85957 | 29.05173 | 79.07171 | 73.95619 |
| ENSMUSG00000043410 | 0.814246723 | 0.018309 | 0.066321 | 186.1201 | 214.7681 | 117.5911 | 152.6844 | 74.10846 | 169.8409 | 70.39311 | 67.43065 |
| ENSMUSG00000043411 | 0.228322662 | 0.007334 | 0.03401 | 3370.282 | 3169.928 | 2880.155 | 3269.343 | 2665.954 | 3024.732 | 2544.759 | 2597.168 |
| ENSMUSG00000043415 | 0.665380024 | 1.81E-07 | 9.22E-06 | 354.1311 | 375.5441 | 409.9128 | 445.9763 | 247.6783 | 280.4609 | 220.8222 | 253.4087 |
| ENSMUSG00000043418 | 0.132707426 | 0.665602 | 0.785344 | 29.17558 | 43.19358 | 30.63995 | 39.68068 | 22.42756 | 36.87335 | 33.75012 | 36.9781 |
| ENSMUSG00000043419 | -0.081994431 | 0.552245 | 0.699721 | 322.9434 | 381.5432 | 328.7583 | 371.7907 | 417.3476 | 411.1937 | 304.7154 | 353.4671 |
| ENSMUSG00000043421 | -0.12183937 | 0.366581 | 0.535128 | 266.6044 | 280.7582 | 314.6805 | 263.1002 | 358.841 | 265.935 | 288.3225 | 311.0511 |
| ENSMUSG00000043424 | -0.005486595 | 0.983166 | 0.990789 | 42.25428 | 74.38894 | 43.88965 | 52.62003 | 57.53157 | 50.28183 | 51.10733 | 53.29196 |
| ENSMUSG00000043430 | 1.136123149 | 0.05673 | 0.148796 | 26.15741 | 9.598572 | 28.98373 | 11.21411 | 9.751113 | 3.352122 | 10.60718 | 10.87591 |
| ENSMUSG00000043432 | -0.220557231 | 0.160471 | 0.303969 | 130.7871 | 110.3836 | 114.2787 | 111.2784 | 139.4409 | 122.9112 | 154.2863 | 126.1606 |
| ENSMUSG00000043439 | -0.170623939 | 0.267385 | 0.431699 | 302.8224 | 284.3577 | 345.3205 | 363.1645 | 399.7956 | 338.5644 | 425.2515 | 295.8248 |
| ENSMUSG00000043445 | -0.68013026 | 0.000134 | 0.001675 | 595.5842 | 476.3291 | 641.7826 | 495.1459 | 900.0277 | 627.9642 | 956.5749 | 1056.051 |
| ENSMUSG00000043448 | -0.123622309 | 0.804451 | 0.880289 | 8.048435 | 15.59768 | 35.60859 | 30.19182 | 28.27823 | 24.58223 | 17.3572 | 28.27737 |
| ENSMUSG00000043456 | 0.241789416 | 0.501761 | 0.657987 | 123.7447 | 128.3809 | 275.7595 | 112.1411 | 89.71024 | 101.681 | 165.8577 | 184.8905 |
| ENSMUSG00000043458 | 0.735933599 | 0.114647 | 0.242208 | 15.09082 | 26.39607 | 12.4216 | 27.60395 | 10.72622 | 16.76061 | 11.57147 | 9.78832 |
| ENSMUSG00000043460 | -0.797748366 | 0.156381 | 0.298465 | 4.024217 | 10.79839 | 19.87456 | 6.900988 | 18.52711 | 12.29112 | 20.25007 | 21.75182 |
| ENSMUSG00000043461 | 0.121632298 | 0.746824 | 0.841887 | 240.447 | 147.578 | 465.3959 | 208.7549 | 391.0196 | 172.0756 | 180.3221 | 233.8321 |
| ENSMUSG00000043463 | 0.226503912 | 0.692901 | 0.805613 | 15.09082 | 9.598572 | 4.96864 | 20.70296 | 14.62667 | 10.05637 | 9.642892 | 8.700729 |

| | | | | | | | | | | | |
|---|---|---|---|---|---|---|---|---|---|---|---|
| ENSMUSG00000043467 | 0.109747721 | 0.357412 | 0.52586 | 459.7668 | 439.1347 | 459.5992 | 376.1038 | 383.2187 | 462.5929 | 375.1085 | 387.1824 |
| ENSMUSG00000043483 | 0.03175233 | 0.84426 | 0.906226 | 199.1988 | 261.5611 | 216.9639 | 247.5729 | 206.7236 | 279.3435 | 219.8579 | 199.0292 |
| ENSMUSG00000043487 | -0.200938615 | 0.160689 | 0.304203 | 178.0716 | 158.3764 | 178.0429 | 139.745 | 197.9476 | 174.3104 | 200.5721 | 178.3649 |
| ENSMUSG00000043496 | 0.346497779 | 0.003417 | 0.01944 | 5422.633 | 7401.699 | 6321.766 | 6791.435 | 5573.736 | 5522.063 | 4554.338 | 4748.423 |
| ENSMUSG00000043498 | 0.82893124 | 0.051502 | 0.138802 | 107.6478 | 128.3809 | 128.3565 | 143.1955 | 41.92979 | 149.7281 | 58.82164 | 35.89051 |
| ENSMUSG00000043501 | 0.666775343 | 0.24523 | 0.408111 | 14.08476 | 23.99643 | 29.81184 | 15.52722 | 9.751113 | 27.93435 | 8.678602 | 6.525547 |
| ENSMUSG00000043505 | -0.584500574 | 0.045695 | 0.127839 | 201.2109 | 103.1847 | 209.511 | 183.7388 | 244.7529 | 159.7845 | 282.5367 | 361.0802 |
| ENSMUSG00000043510 | -0.110366169 | 0.208868 | 0.366165 | 531.1967 | 603.5102 | 561.4563 | 614.1879 | 663.0757 | 631.3164 | 602.6807 | 595.9999 |
| ENSMUSG00000043518 | 0.255578075 | 0.231374 | 0.391909 | 146.8839 | 124.7814 | 131.669 | 151.8217 | 105.312 | 139.6718 | 139.8219 | 80.48174 |
| ENSMUSG00000043531 | 0.300309117 | 0.075612 | 0.181937 | 295.78 | 250.7627 | 398.3193 | 304.5061 | 288.6329 | 230.1791 | 240.108 | 257.7591 |
| ENSMUSG00000043535 | 0.119889312 | 0.042708 | 0.122167 | 2042.29 | 2076.891 | 1949.363 | 1943.491 | 1752.275 | 1858.193 | 1925.685 | 1832.591 |
| ENSMUSG00000043541 | 0.090161456 | 0.7441 | 0.839946 | 34.20585 | 34.79482 | 38.92101 | 39.68068 | 26.32801 | 41.34284 | 32.78583 | 39.15328 |
| ENSMUSG00000043542 | 0.376131844 | 0.002309 | 0.014546 | 969.8364 | 1000.651 | 805.7478 | 1054.989 | 696.2295 | 849.2043 | 751.1813 | 654.7298 |
| ENSMUSG00000043556 | 0.230696471 | 0.222035 | 0.381233 | 152.9203 | 131.9804 | 114.2787 | 112.1411 | 117.9885 | 96.09417 | 94.50034 | 126.1606 |
| ENSMUSG00000043557 | 0.565458622 | 0.003909 | 0.021475 | 140.8476 | 127.1811 | 118.4192 | 123.3552 | 78.0089 | 107.2679 | 66.53595 | 93.53283 |
| ENSMUSG00000043572 | -0.216551182 | 0.140483 | 0.276915 | 287.7315 | 255.562 | 313.0243 | 216.5185 | 300.3343 | 320.6864 | 285.4296 | 341.5036 |
| ENSMUSG00000043587 | -0.099451711 | 0.566581 | 0.711059 | 359.1614 | 338.3497 | 467.8802 | 354.5383 | 389.0694 | 310.63 | 458.0374 | 472.0145 |
| ENSMUSG00000043592 | -1.183548421 | 0.002624 | 0.015975 | 37.22401 | 15.59768 | 57.96746 | 31.91707 | 76.05868 | 68.15982 | 65.57166 | 117.4598 |
| ENSMUSG00000043602 | -0.090398884 | 0.642969 | 0.769112 | 169.0171 | 155.9768 | 183.8397 | 127.6683 | 189.1716 | 194.4231 | 167.7863 | 126.1606 |
| ENSMUSG00000043614 | 0.13949569 | 0.076458 | 0.183476 | 711.2804 | 655.1026 | 717.1403 | 704.7634 | 623.0961 | 683.8329 | 617.1451 | 611.2262 |
| ENSMUSG00000043621 | -0.146163742 | 0.566581 | 0.711059 | 50.30272 | 101.9848 | 67.07664 | 61.24627 | 71.18313 | 74.86406 | 81.00029 | 81.56933 |
| ENSMUSG00000043629 | 0.101728937 | 0.68957 | 0.803352 | 142.8597 | 106.7841 | 195.4332 | 112.1411 | 92.63557 | 144.1413 | 141.7505 | 142.4744 |
| ENSMUSG00000043631 | 0.34563637 | 0.045372 | 0.127206 | 213.2835 | 259.1615 | 218.6202 | 320.8959 | 199.8978 | 173.193 | 212.1436 | 210.9927 |
| ENSMUSG00000043633 | -0.133103056 | 0.515775 | 0.669844 | 119.7205 | 165.5754 | 125.0441 | 111.2784 | 145.2916 | 177.6625 | 133.0719 | 114.1971 |
| ENSMUSG00000043635 | 0.007730924 | 0.970479 | 0.983563 | 87.52673 | 81.58786 | 78.67013 | 61.24627 | 66.30757 | 70.39457 | 83.89316 | 85.9197 |
| ENSMUSG00000043639 | 0.556708279 | 0.000261 | 0.002796 | 467.8153 | 471.5299 | 386.7258 | 394.2189 | 341.289 | 229.0617 | 313.394 | 282.7737 |
| ENSMUSG00000043644 | -0.004611421 | 0.979667 | 0.988775 | 84.50857 | 82.78769 | 76.18581 | 82.81186 | 74.10846 | 73.74669 | 88.7146 | 90.27006 |
| ENSMUSG00000043648 | -0.231992346 | 0.354126 | 0.522769 | 56.33904 | 62.39072 | 51.34261 | 59.52102 | 79.95913 | 83.80306 | 50.14304 | 55.46715 |
| ENSMUSG00000043659 | -1.063875685 | 0.337649 | 0.506104 | 0 | 5.999108 | 55.48314 | 2.58787 | 27.30312 | 11.17374 | 38.57157 | 57.64233 |
| ENSMUSG00000043661 | 0.395517106 | 0.527892 | 0.680556 | 21.12714 | 22.79661 | 18.21835 | 7.763611 | 4.875557 | 27.93435 | 10.60718 | 9.78832 |
| ENSMUSG00000043664 | 0.071800886 | 0.60165 | 0.739531 | 216.3017 | 261.5611 | 275.7595 | 211.3428 | 217.4498 | 230.1791 | 245.8937 | 224.0438 |
| ENSMUSG00000043668 | 0.425672166 | 0.00294 | 0.017518 | 1212.295 | 1037.846 | 1090.616 | 1151.602 | 669.9015 | 1024.632 | 906.4318 | 744.9999 |
| ENSMUSG00000043670 | 0.409922657 | 0.260094 | 0.424166 | 25.15136 | 26.39607 | 21.53077 | 34.50494 | 20.47734 | 27.93435 | 14.46434 | 18.48905 |
| ENSMUSG00000043671 | 0.23039924 | 0.152869 | 0.293922 | 758.565 | 735.4906 | 611.1427 | 728.0542 | 619.1957 | 768.7534 | 561.2163 | 465.489 |
| ENSMUSG00000043673 | -0.189757371 | 0.427471 | 0.592509 | 170.0232 | 211.1686 | 297.2903 | 228.5952 | 260.3547 | 172.0756 | 274.8224 | 328.4525 |
| ENSMUSG00000043681 | -0.470309284 | 0.36271 | 0.531242 | 50.30272 | 22.79661 | 64.59232 | 34.50494 | 110.1876 | 20.11273 | 60.75022 | 47.85401 |
| ENSMUSG00000043683 | -0.396335984 | 0.004057 | 0.022062 | 915.5095 | 983.8537 | 1056.664 | 911.793 | 1348.579 | 968.7633 | 1288.29 | 1484.562 |
| ENSMUSG00000043687 | 0.375036109 | 0.356752 | 0.525308 | 40.24217 | 29.99554 | 24.01509 | 14.6646 | 23.40267 | 16.76061 | 19.28578 | 23.927 |
| ENSMUSG00000043702 | -0.316123293 | 0.009154 | 0.039984 | 469.8274 | 543.5192 | 570.5655 | 541.7276 | 712.8064 | 569.8608 | 616.1808 | 748.2627 |
| ENSMUSG00000043705 | -0.691630378 | 0.195501 | 0.349493 | 14.08476 | 4.799286 | 27.32752 | 12.93935 | 31.20356 | 14.52586 | 26.03581 | 25.0146 |
| ENSMUSG00000043716 | -0.059448542 | 0.519842 | 0.673341 | 38715.99 | 47948.47 | 43475.6 | 45392.11 | 47530.83 | 49376.76 | 45112.34 | 40894.51 |
| ENSMUSG00000043719 | 1.236880901 | 0.13913 | 0.275033 | 233.4046 | 719.8929 | 424.8187 | 370.9281 | 152.1174 | 488.2925 | 71.3574 | 30.45255 |
| ENSMUSG00000043733 | -0.223529027 | 0.015149 | 0.058087 | 2946.733 | 2861.574 | 3351.348 | 2916.53 | 3559.156 | 3076.131 | 3695.156 | 3771.766 |
| ENSMUSG00000043740 | 1.060103936 | 0.097945 | 0.217858 | 22.1332 | 7.198929 | 18.21835 | 9.488858 | 6.825779 | 2.234748 | 10.60718 | 7.613138 |
| ENSMUSG00000043753 | 0.030114211 | NA | NA | 32.19374 | 33.595 | 24.01509 | 22.42821 | 11.70134 | 88.27255 | 5.785735 | 4.350364 |
| ENSMUSG00000043760 | -1.042613573 | 0.000992 | 0.00779 | 432.6034 | 254.3622 | 548.2066 | 293.292 | 745.9602 | 419.0153 | 843.753 | 1141.971 |
| ENSMUSG00000043773 | 1.004069451 | 1.05E-06 | 3.88E-05 | 281.6952 | 424.7368 | 346.1486 | 328.6596 | 124.8142 | 212.3011 | 155.2506 | 196.854 |
| ENSMUSG00000043782 | -0.840443623 | 0.042107 | 0.120915 | 40.24217 | 14.39786 | 53.82693 | 37.09281 | 71.18313 | 37.99072 | 67.50024 | 85.9197 |
| ENSMUSG00000043789 | -0.521688668 | 0.178416 | 0.327822 | 32.19374 | 31.19536 | 44.71776 | 34.50494 | 45.83023 | 29.05173 | 41.46443 | 89.18247 |
| ENSMUSG00000043794 | 1.217064514 | 0.021684 | 0.074648 | 53.32088 | 23.99643 | 19.04645 | 24.15346 | 8.776002 | 20.11273 | 16.39292 | 6.525547 |
| ENSMUSG00000043795 | -0.065346463 | 0.826212 | 0.894472 | 185.114 | 177.5736 | 131.669 | 116.4542 | 135.5405 | 259.2308 | 115.7147 | 128.3357 |
| ENSMUSG00000043801 | -0.238917643 | 0.070433 | 0.173067 | 165.999 | 177.5736 | 171.4181 | 178.5631 | 231.1014 | 191.071 | 212.1436 | 182.7153 |
| ENSMUSG00000043807 | 0.081908771 | 0.842108 | 0.904759 | 33.19979 | 14.39786 | 33.95237 | 32.77969 | 28.27823 | 40.22547 | 23.14294 | 17.40146 |
| ENSMUSG00000043811 | -1.038235706 | 0.017643 | 0.064671 | 7.04238 | 20.39697 | 22.35888 | 33.64232 | 42.9049 | 52.51658 | 33.75012 | 43.50364 |
| ENSMUSG00000043822 | 0.365968363 | 0.103862 | 0.226662 | 227.3683 | 212.3684 | 295.6341 | 272.589 | 208.6738 | 268.1698 | 168.7506 | 138.1241 |
| ENSMUSG00000043831 | -0.001113158 | 0.989194 | 0.993825 | 586.5297 | 614.3086 | 542.4098 | 549.4912 | 575.3157 | 548.6307 | 593.0378 | 572.0729 |
| ENSMUSG00000043832 | 0.258049553 | 0.536858 | 0.687405 | 19.11503 | 23.99643 | 24.8432 | 12.93935 | 21.45245 | 12.29112 | 20.25007 | 13.05109 |
| ENSMUSG00000043833 | 0.270015057 | 0.161863 | 0.30586 | 140.8476 | 122.3818 | 91.91984 | 125.943 | 88.73513 | 100.5637 | 111.8575 | 96.79561 |
| ENSMUSG00000043843 | 0.191848922 | 0.641823 | 0.768381 | 21.12714 | 16.7975 | 30.63995 | 28.46658 | 11.70134 | 26.81698 | 19.28578 | 28.27737 |
| ENSMUSG00000043850 | 1.010307136 | 0.000158 | 0.001891 | 210.2654 | 203.9697 | 130.0127 | 171.6621 | 123.8391 | 56.98608 | 81.96458 | 91.35765 |
| ENSMUSG00000043857 | 0.515453897 | 0.433088 | 0.597256 | 13.07871 | 9.598572 | 4.140533 | 28.46658 | 10.72622 | 7.821619 | 12.53576 | 7.613138 |
| ENSMUSG00000043866 | -0.315975148 | 0.007775 | 0.035492 | 1226.38 | 1444.585 | 1522.888 | 1214.574 | 1607.959 | 1500.633 | 1675.935 | 1947.876 |
| ENSMUSG00000043872 | 0.095886942 | 0.559672 | 0.705754 | 230.3864 | 239.9643 | 179.6991 | 219.1064 | 236.952 | 163.1366 | 200.5721 | 209.9051 |
| ENSMUSG00000043881 | 0.287898566 | 0.009152 | 0.039984 | 600.6145 | 650.3033 | 511.7699 | 599.5233 | 457.3272 | 539.6917 | 460.9302 | 475.2773 |

| | | | | | | | | | | | |
|---|---|---|---|---|---|---|---|---|---|---|---|
| ENSMUSG00000043885 | 0.079013635 | 0.296556 | 0.464315 | 1197.205 | 1126.632 | 1175.083 | 1294.798 | 1139.905 | 1182.182 | 1161.968 | 1057.139 |
| ENSMUSG00000043889 | -0.189211428 | 0.537595 | 0.687763 | 37.22401 | 63.59054 | 37.2648 | 42.26855 | 47.78045 | 34.6386 | 62.6788 | 58.72992 |
| ENSMUSG00000043895 | 0.016514744 | 0.89638 | 0.93742 | 741.4621 | 929.8617 | 789.1856 | 711.6644 | 850.2971 | 724.0584 | 686.5739 | 872.2481 |
| ENSMUSG00000043903 | 0.156622052 | 0.760014 | 0.850471 | 22.1332 | 23.99643 | 56.31125 | 43.13117 | 19.50223 | 41.34284 | 15.42863 | 55.46715 |
| ENSMUSG00000043909 | 0.110109245 | 0.126173 | 0.257025 | 2272.677 | 2277.261 | 2215.185 | 2232.47 | 2098.44 | 2307.378 | 1939.186 | 1992.467 |
| ENSMUSG00000043913 | 0.826879681 | 0.041068 | 0.118926 | 48.29061 | 38.39429 | 38.92101 | 41.40593 | 26.32801 | 37.99072 | 9.642892 | 20.66423 |
| ENSMUSG00000043923 | 0.223146586 | 0.020397 | 0.071514 | 518.118 | 514.7234 | 509.2856 | 555.5295 | 478.7797 | 481.5882 | 435.8587 | 401.3211 |
| ENSMUSG00000043924 | -0.217914362 | 0.493604 | 0.65051 | 59.35721 | 31.19536 | 39.74912 | 31.05445 | 41.92979 | 41.34284 | 61.71451 | 42.41605 |
| ENSMUSG00000043929 | 0.145650956 | 0.412772 | 0.578635 | 365.1977 | 362.3461 | 265.8222 | 328.6596 | 281.8072 | 386.6114 | 276.751 | 249.0584 |
| ENSMUSG00000043940 | 0.089033466 | 0.318267 | 0.486549 | 2599.644 | 2184.875 | 2534.006 | 2309.243 | 2126.718 | 2480.57 | 2317.187 | 2130.591 |
| ENSMUSG00000043943 | -0.02403324 | 0.90254 | 0.941515 | 287.7315 | 230.3657 | 409.0847 | 282.9405 | 312.0356 | 325.1559 | 343.2869 | 252.3211 |
| ENSMUSG00000043953 | -0.030382126 | 0.914042 | 0.949091 | 53.32088 | 45.59322 | 49.6864 | 40.5433 | 31.20356 | 48.04709 | 61.71451 | 52.20437 |
| ENSMUSG00000043962 | 0.00686239 | 0.857004 | 0.914481 | 7536.353 | 7434.094 | 7499.334 | 7493.61 | 7298.708 | 7544.51 | 7660.313 | 7318.4 |
| ENSMUSG00000043964 | 0.08022758 | 0.483655 | 0.641981 | 585.5236 | 641.9045 | 662.4853 | 634.8909 | 593.8428 | 524.0485 | 563.1449 | 708.0218 |
| ENSMUSG00000043969 | 0.123430312 | 0.415417 | 0.581269 | 2319.961 | 1623.359 | 1635.511 | 1831.35 | 1574.805 | 2060.438 | 1684.613 | 1483.474 |
| ENSMUSG00000043972 | 0.850767441 | 0.121013 | 0.250271 | 12.07265 | 21.59679 | 12.4216 | 13.80198 | 3.900445 | 5.58687 | 12.53576 | 10.87591 |
| ENSMUSG00000043987 | 0.129956362 | 0.133283 | 0.26671 | 1011.085 | 847.074 | 872.8244 | 903.1668 | 774.2384 | 844.7348 | 837.9673 | 865.7225 |
| ENSMUSG00000043991 | 0.346707574 | 0.11138 | 0.237673 | 731.4015 | 833.876 | 567.253 | 598.6607 | 454.4019 | 785.514 | 457.0731 | 450.2627 |
| ENSMUSG00000043993 | 0.129824054 | 0.51686 | 0.670922 | 587.5357 | 545.9188 | 598.7211 | 646.9676 | 531.4357 | 784.3966 | 451.2873 | 408.9342 |
| ENSMUSG00000043998 | -0.359249899 | 0.009387 | 0.040662 | 1302.84 | 1345 | 1555.184 | 1397.45 | 1733.748 | 1415.713 | 1836.007 | 2200.197 |
| ENSMUSG00000043999 | 0.147218517 | 0.764352 | 0.853773 | 20.12109 | 13.19804 | 17.39024 | 19.84034 | 13.65156 | 30.1691 | 9.642892 | 10.87591 |
| ENSMUSG00000044005 | 0.319057257 | 0.492571 | 0.649627 | 32.19374 | 29.99554 | 18.21835 | 36.23019 | 27.30312 | 7.821619 | 32.78583 | 25.0146 |
| ENSMUSG00000044006 | 2.152166243 | 1.34E-12 | 3.88E-10 | 102.6175 | 140.3791 | 139.1219 | 186.3267 | 44.85512 | 35.75597 | 20.25007 | 27.18978 |
| ENSMUSG00000044017 | 1.454518551 | 7.30E-11 | 1.39E-08 | 429.5852 | 616.7083 | 395.0069 | 496.8711 | 194.0472 | 233.5312 | 155.2506 | 123.9854 |
| ENSMUSG00000044018 | -0.387162079 | 2.44E-05 | 0.000457 | 1560.39 | 1463.782 | 1551.872 | 1522.53 | 1933.646 | 1724.108 | 2118.543 | 2200.197 |
| ENSMUSG00000044022 | 0.505472791 | 0.039348 | 0.11527 | 49.29666 | 55.19179 | 77.01392 | 60.38364 | 45.83023 | 43.57759 | 40.50014 | 41.32846 |
| ENSMUSG00000044024 | -0.307255326 | 0.530162 | 0.682096 | 20.12109 | 13.19804 | 10.76539 | 7.763611 | 11.70134 | 15.64324 | 23.14294 | 13.05109 |
| ENSMUSG00000044026 | -0.250909844 | 0.240748 | 0.403015 | 843.0736 | 719.8929 | 1397.016 | 791.8884 | 1111.627 | 912.8946 | 1277.683 | 1164.81 |
| ENSMUSG00000044030 | -0.09455439 | 0.267073 | 0.431449 | 1101.63 | 1126.632 | 1017.743 | 1038.599 | 1118.453 | 1023.515 | 1201.504 | 1226.803 |
| ENSMUSG00000044033 | -0.019699919 | 0.919365 | 0.952207 | 535.2209 | 431.9358 | 602.0335 | 485.657 | 498.2819 | 386.6114 | 491.7875 | 708.0218 |
| ENSMUSG00000044034 | -1.418178318 | 0.000744 | 0.00623 | 33.19979 | 29.99554 | 67.90474 | 25.8787 | 98.48624 | 56.98608 | 97.39321 | 168.5766 |
| ENSMUSG00000044037 | -0.283187189 | 0.164999 | 0.310389 | 461.7789 | 350.3479 | 656.6886 | 462.3662 | 604.569 | 462.5929 | 592.0735 | 693.8831 |
| ENSMUSG00000044042 | 0.338313347 | 0.002335 | 0.014636 | 1006.054 | 944.2595 | 949.0102 | 858.3104 | 708.9059 | 874.9039 | 733.8241 | 654.7298 |
| ENSMUSG00000044043 | 0.20430995 | 0.466563 | 0.627229 | 61.36932 | 101.9848 | 72.04528 | 77.63611 | 70.20801 | 89.38993 | 43.39301 | 68.51824 |
| ENSMUSG00000044052 | -0.607667566 | 0.261018 | 0.425111 | 15.09082 | 8.398751 | 15.73403 | 6.900988 | 20.47734 | 12.29112 | 9.642892 | 28.27737 |
| ENSMUSG00000044060 | 0.482147198 | 0.096048 | 0.214635 | 93.56306 | 91.18644 | 65.42042 | 98.33908 | 42.9049 | 93.85942 | 59.78593 | 53.29196 |
| ENSMUSG00000044066 | -0.022533029 | 0.678888 | 0.794865 | 1321.955 | 1397.792 | 1327.455 | 1362.945 | 1358.33 | 1392.248 | 1385.684 | 1356.226 |
| ENSMUSG00000044068 | 0.121901786 | 0.327569 | 0.495339 | 474.8577 | 412.7386 | 429.7873 | 459.7783 | 386.1441 | 454.7713 | 451.2873 | 341.5036 |
| ENSMUSG00000044071 | 1.948823242 | 1.27E-09 | 1.60E-07 | 342.0585 | 374.3443 | 369.3356 | 409.7462 | 63.38224 | 179.8972 | 82.92887 | 61.99269 |
| ENSMUSG00000044072 | 0.019649158 | 0.919497 | 0.952207 | 69.41775 | 82.78769 | 77.84202 | 75.04824 | 67.28268 | 69.27719 | 89.67889 | 73.95619 |
| ENSMUSG00000044080 | -0.405734333 | 0.065217 | 0.164448 | 1041.266 | 1072.64 | 1593.277 | 1077.417 | 1587.481 | 1022.397 | 1661.47 | 2068.598 |
| ENSMUSG00000044081 | 0.285644433 | 0.443117 | 0.60632 | 18.10898 | 38.39429 | 38.92101 | 30.19182 | 23.40267 | 29.05173 | 18.32149 | 32.62773 |
| ENSMUSG00000044092 | -0.266782846 | 0.621068 | 0.75282 | 19.11503 | 5.999108 | 13.24971 | 9.488858 | 11.70134 | 8.938993 | 14.46434 | 22.83941 |
| ENSMUSG00000044098 | 0.174781552 | 0.157537 | 0.300113 | 717.3168 | 725.892 | 537.4412 | 619.3637 | 621.1459 | 587.7388 | 555.4306 | 535.0948 |
| ENSMUSG00000044117 | 0.169127469 | 0.518779 | 0.672487 | 72.43591 | 44.3934 | 53.82693 | 74.18562 | 44.85512 | 65.92507 | 55.92877 | 52.20437 |
| ENSMUSG00000044122 | 0.399304524 | 0.087145 | 0.200618 | 69.41775 | 67.19001 | 82.81066 | 84.5371 | 50.70579 | 54.75133 | 49.17875 | 77.21897 |
| ENSMUSG00000044134 | -0.065170476 | 0.526177 | 0.679432 | 653.9353 | 603.5102 | 622.7362 | 641.7919 | 703.0553 | 551.9828 | 686.5739 | 696.0583 |
| ENSMUSG00000044139 | -2.017936068 | 4.60E-05 | 0.000747 | 12.07265 | 17.99732 | 14.90592 | 5.175741 | 43.88001 | 34.6386 | 34.71441 | 88.09488 |
| ENSMUSG00000044141 | -0.693868192 | 0.144623 | 0.28269 | 6.036326 | 14.39786 | 9.109173 | 15.52722 | 20.47734 | 17.87799 | 22.17865 | 11.9635 |
| ENSMUSG00000044145 | -0.060140215 | 0.747357 | 0.842283 | 76.46013 | 69.58965 | 67.90474 | 69.00988 | 73.13335 | 79.33356 | 65.57166 | 77.21897 |
| ENSMUSG00000044147 | -0.26364131 | 0.001643 | 0.011307 | 2496.021 | 2495.629 | 2524.069 | 2546.465 | 2825.873 | 2706.28 | 3212.047 | 3335.642 |
| ENSMUSG00000044148 | 0.13832218 | 0.156027 | 0.298129 | 1183.12 | 1335.401 | 1203.239 | 1282.721 | 1154.532 | 1296.154 | 1016.361 | 1079.978 |
| ENSMUSG00000044149 | 0.069543445 | 0.681438 | 0.796493 | 143.8658 | 194.3711 | 151.5435 | 172.5247 | 137.4907 | 188.8362 | 156.2148 | 147.9124 |
| ENSMUSG00000044150 | 0.444053268 | 0.008914 | 0.039209 | 661.9838 | 725.892 | 539.9255 | 703.9008 | 487.5557 | 626.8469 | 400.18 | 419.8102 |
| ENSMUSG00000044155 | -0.103950849 | 0.189372 | 0.341947 | 1582.524 | 1805.731 | 1554.356 | 1764.065 | 1827.359 | 1851.489 | 1847.578 | 1677.065 |
| ENSMUSG00000044156 | -1.895422583 | 0.011791 | 0.048308 | 16.09687 | 7.198929 | 114.2787 | 11.21411 | 80.93424 | 87.15518 | 187.0721 | 201.2044 |
| ENSMUSG00000044164 | 0.154932207 | 0.632313 | 0.760923 | 43.26034 | 57.59143 | 80.32634 | 53.48266 | 39.00445 | 78.21619 | 41.46443 | 53.29196 |
| ENSMUSG00000044167 | 0.395784907 | 0.040411 | 0.117563 | 1472.864 | 1578.965 | 1414.406 | 1763.202 | 897.1024 | 1687.235 | 1170.647 | 981.0072 |
| ENSMUSG00000044177 | 0.738862233 | 0.000246 | 0.002673 | 170.0232 | 257.9616 | 236.0104 | 200.9913 | 122.864 | 167.6061 | 112.8218 | 115.2847 |
| ENSMUSG00000044197 | -0.688334952 | 0.000205 | 0.002321 | 259.562 | 171.5745 | 291.4935 | 198.4034 | 417.3476 | 329.6254 | 368.3585 | 371.9562 |
| ENSMUSG00000044199 | -0.387457134 | 0.447453 | 0.610046 | 11.0666 | 10.79839 | 26.49941 | 15.52722 | 28.27823 | 8.938993 | 24.10723 | 22.83941 |
| ENSMUSG00000044201 | 0.106104666 | 0.497578 | 0.654039 | 600.6145 | 511.124 | 451.3181 | 645.2424 | 535.3361 | 602.2646 | 485.0374 | 429.5985 |
| ENSMUSG00000044211 | -0.193837585 | 0.70615 | 0.815104 | 19.11503 | 22.79661 | 4.96864 | 16.38985 | 17.552 | 23.46486 | 16.39292 | 14.13868 |
| ENSMUSG00000044216 | 0.468515327 | 0.384671 | 0.552404 | 15.09082 | 14.39786 | 16.56213 | 12.07673 | 5.850668 | 11.17374 | 5.785735 | 19.57664 |

| | | | | | | | | | | | |
|---|---|---|---|---|---|---|---|---|---|---|---|
| ENSMUSG00000044217 | 0.623079413 | 0.125484 | 0.256017 | 44.26639 | 55.19179 | 97.71658 | 50.03216 | 30.22845 | 63.69032 | 44.3573 | 22.83941 |
| ENSMUSG00000044221 | -0.23129945 | 0.003339 | 0.019157 | 2636.868 | 2590.415 | 2541.459 | 2660.331 | 2817.097 | 2876.121 | 3125.261 | 3424.824 |
| ENSMUSG00000044224 | -0.195727157 | 0.060996 | 0.156745 | 978.8909 | 818.2783 | 779.2483 | 897.1284 | 889.3015 | 1050.332 | 1048.182 | 990.7955 |
| ENSMUSG00000044229 | 0.041615863 | 0.697067 | 0.808775 | 427.5731 | 500.3256 | 487.7548 | 455.4652 | 504.1325 | 403.372 | 459.9659 | 448.0875 |
| ENSMUSG00000044231 | 0.037719362 | 0.851442 | 0.910624 | 96.58122 | 81.58786 | 97.71658 | 80.22398 | 72.15824 | 88.27255 | 80.036 | 107.6715 |
| ENSMUSG00000044244 | 0.164900993 | 0.311496 | 0.480082 | 154.9324 | 164.3756 | 205.3704 | 198.4034 | 172.5947 | 174.3104 | 164.8934 | 134.8613 |
| ENSMUSG00000044249 | -2.422913863 | 0.001156 | 0.008736 | 2.012109 | 3.599465 | 31.46805 | 11.21411 | 57.53157 | 27.93435 | 58.82164 | 118.5474 |
| ENSMUSG00000044250 | 0.301576966 | 0.032737 | 0.100666 | 273.6468 | 211.1686 | 242.6352 | 236.3588 | 194.0472 | 217.8879 | 203.465 | 167.489 |
| ENSMUSG00000044252 | -0.231169159 | 0.046278 | 0.129097 | 1058.369 | 1171.026 | 1214.832 | 1169.717 | 1293.973 | 1118.491 | 1458.97 | 1544.379 |
| ENSMUSG00000044254 | -0.848294579 | 0.00035 | 0.003478 | 119.7205 | 88.78679 | 143.2624 | 124.2178 | 202.8232 | 144.1413 | 263.2509 | 249.0584 |
| ENSMUSG00000044258 | -0.501600238 | 0.006678 | 0.031897 | 178.0716 | 119.9822 | 213.6515 | 186.3267 | 238.9023 | 227.9443 | 259.3938 | 266.4598 |
| ENSMUSG00000044268 | 0.905510708 | 0.186934 | 0.338872 | 5.030272 | 29.99554 | 7.45296 | 12.93935 | 5.850668 | 10.05637 | 5.785735 | 7.613138 |
| ENSMUSG00000044279 | -0.909954863 | 0.00013 | 0.001638 | 387.3309 | 293.9563 | 523.3634 | 332.9727 | 690.3788 | 481.5882 | 778.1814 | 941.8539 |
| ENSMUSG00000044285 | -0.012610093 | 0.897744 | 0.938049 | 399.4036 | 455.9322 | 465.3959 | 478.756 | 482.6801 | 427.9543 | 458.0374 | 446.9999 |
| ENSMUSG00000044288 | 0.837767665 | 0.019643 | 0.069614 | 129.781 | 58.79126 | 52.99882 | 77.63611 | 41.92979 | 56.98608 | 49.17875 | 30.45255 |
| ENSMUSG00000044296 | -0.071282928 | 0.840255 | 0.903715 | 40.24217 | 37.19447 | 26.49941 | 23.29083 | 37.05423 | 41.34284 | 34.71441 | 19.57664 |
| ENSMUSG00000044308 | -0.109886419 | 0.0739 | 0.179265 | 2943.715 | 2639.607 | 2802.313 | 2634.452 | 2868.777 | 2961.041 | 2982.546 | 3082.233 |
| ENSMUSG00000044313 | -0.910301612 | 0.001046 | 0.008096 | 40.24217 | 45.59322 | 52.99882 | 33.64232 | 74.10846 | 59.22083 | 81.00029 | 109.8467 |
| ENSMUSG00000044317 | 0.193849085 | 0.183446 | 0.334355 | 214.2896 | 226.7663 | 203.7142 | 163.0358 | 172.5947 | 173.193 | 185.1435 | 172.927 |
| ENSMUSG00000044320 | -0.067103863 | 0.754451 | 0.847512 | 69.41775 | 97.18554 | 83.63877 | 85.39973 | 95.56091 | 63.69032 | 96.42892 | 94.62042 |
| ENSMUSG00000044328 | 0.048736518 | 0.638303 | 0.765516 | 572.4449 | 682.6985 | 608.6584 | 596.9355 | 670.8766 | 572.0955 | 587.2521 | 544.8831 |
| ENSMUSG00000044337 | 0.056872823 | 0.548816 | 0.697048 | 1863.213 | 1926.913 | 1638.823 | 1769.241 | 1869.288 | 1655.948 | 1536.113 | 1855.43 |
| ENSMUSG00000044338 | -0.105741326 | 0.477708 | 0.637036 | 1554.354 | 1480.58 | 2275.637 | 1659.688 | 1718.146 | 2000.1 | 1744.399 | 2042.496 |
| ENSMUSG00000044339 | -0.003418729 | 0.985113 | 0.991442 | 118.7144 | 193.1713 | 155.684 | 150.0965 | 170.6445 | 144.1413 | 162.9649 | 139.2117 |
| ENSMUSG00000044340 | -0.044418297 | 0.628595 | 0.758208 | 530.1906 | 481.1284 | 545.7223 | 554.6669 | 518.7592 | 596.6778 | 531.3233 | 536.1824 |
| ENSMUSG00000044345 | -0.127409864 | 0.102706 | 0.22507 | 1333.022 | 1339.001 | 1568.434 | 1350.006 | 1564.079 | 1493.929 | 1494.648 | 1558.518 |
| ENSMUSG00000044348 | 0.27998579 | 0.051964 | 0.1397 | 422.5428 | 380.3434 | 362.7107 | 483.9318 | 319.8365 | 393.3157 | 353.8941 | 292.562 |
| ENSMUSG00000044349 | 0.100638154 | 0.724873 | 0.82773 | 235.4167 | 152.3773 | 185.4959 | 400.2573 | 217.4498 | 210.0663 | 219.8579 | 262.1095 |
| ENSMUSG00000044350 | 0.818363412 | 1.81E-09 | 2.17E-07 | 447.6942 | 392.3416 | 427.303 | 413.1967 | 215.4996 | 297.2215 | 215.0365 | 227.3065 |
| ENSMUSG00000044352 | -0.508671601 | 0.284678 | 0.45126 | 31.18769 | 10.79839 | 53.82693 | 30.19182 | 67.28268 | 30.1691 | 38.57157 | 44.59123 |
| ENSMUSG00000044362 | 0.278669548 | 0.375829 | 0.544226 | 28.16952 | 46.79304 | 38.09291 | 29.3292 | 33.15378 | 26.81698 | 23.14294 | 33.71532 |
| ENSMUSG00000044365 | 0.775787713 | 0.000714 | 0.006047 | 642.8687 | 711.4942 | 507.6294 | 620.2263 | 319.8365 | 551.9828 | 309.5368 | 268.635 |
| ENSMUSG00000044367 | -0.551222222 | 0.00102 | 0.00796 | 316.9071 | 279.5584 | 328.7583 | 302.7808 | 446.601 | 345.2686 | 427.1801 | 581.8612 |
| ENSMUSG00000044375 | 0.236341739 | 0.291474 | 0.458716 | 53.32088 | 74.38894 | 81.15445 | 68.14726 | 50.70579 | 59.22083 | 63.64308 | 61.99269 |
| ENSMUSG00000044390 | -0.479797104 | 0.035383 | 0.106949 | 42.25428 | 58.79126 | 58.79557 | 70.73513 | 85.8098 | 78.21619 | 69.42882 | 89.18247 |
| ENSMUSG00000044393 | -0.071394934 | 0.541356 | 0.690721 | 544.2754 | 461.9313 | 640.1264 | 515.8488 | 548.9877 | 563.1565 | 594.0021 | 569.8977 |
| ENSMUSG00000044408 | 0.04792744 | 0.608884 | 0.744259 | 2444.712 | 2885.571 | 2509.163 | 2781.098 | 2475.808 | 2912.994 | 2460.866 | 2423.153 |
| ENSMUSG00000044424 | -0.107912599 | 0.462514 | 0.623578 | 135.8173 | 141.5789 | 130.0127 | 161.3106 | 137.4907 | 157.5497 | 164.8934 | 153.3503 |
| ENSMUSG00000044433 | -0.627531576 | 0.000737 | 0.006182 | 372.2401 | 309.554 | 452.9743 | 346.7746 | 643.5735 | 405.6068 | 589.1807 | 652.5547 |
| ENSMUSG00000044442 | -0.035268892 | 0.694693 | 0.807231 | 1337.046 | 1402.591 | 1311.721 | 1374.159 | 1369.056 | 1595.61 | 1259.362 | 1336.649 |
| ENSMUSG00000044444 | -2.373969863 | 0.000117 | 0.001515 | 17.10292 | 2.399643 | 28.15563 | 7.763611 | 74.10846 | 43.57759 | 48.21446 | 125.073 |
| ENSMUSG00000044447 | -0.364270868 | 0.034138 | 0.104071 | 607.6568 | 521.9224 | 407.4285 | 461.5036 | 810.3175 | 563.1565 | 635.4666 | 560.1094 |
| ENSMUSG00000044452 | -0.290318067 | 0.000646 | 0.005588 | 640.8566 | 651.5031 | 586.2995 | 625.402 | 824.9442 | 706.1804 | 783.9671 | 741.7371 |
| ENSMUSG00000044456 | -0.006989902 | 0.945888 | 0.968881 | 323.9495 | 344.3488 | 378.4447 | 331.2474 | 359.8161 | 360.9118 | 322.0726 | 343.6788 |
| ENSMUSG00000044461 | -0.143241135 | 0.365627 | 0.534244 | 1392.379 | 1478.18 | 1015.259 | 1452.658 | 1655.739 | 1500.633 | 1584.327 | 1151.759 |
| ENSMUSG00000044465 | 0.122443205 | 0.128454 | 0.260065 | 1018.127 | 1097.837 | 933.2762 | 973.0393 | 914.6544 | 969.8807 | 874.6103 | 932.0656 |
| ENSMUSG00000044468 | 0.642673732 | 0.002621 | 0.015962 | 748.5044 | 628.7065 | 436.4122 | 671.9837 | 320.8116 | 508.4052 | 440.6801 | 321.927 |
| ENSMUSG00000044469 | 0.108971163 | 0.375052 | 0.543582 | 200.2048 | 189.5718 | 183.8397 | 196.6782 | 162.8436 | 178.7799 | 190.9293 | 181.6277 |
| ENSMUSG00000044475 | -0.239876099 | 0.013232 | 0.052723 | 871.2431 | 879.4692 | 867.8558 | 1029.11 | 1102.851 | 978.8197 | 1052.039 | 1175.686 |
| ENSMUSG00000044477 | -0.212968254 | 0.008904 | 0.039182 | 2491.997 | 2652.805 | 3109.54 | 2706.05 | 3202.266 | 3100.713 | 3146.476 | 3259.51 |
| ENSMUSG00000044496 | -0.26854095 | 0.020635 | 0.072021 | 1422.561 | 1315.004 | 1333.252 | 1213.711 | 1538.726 | 1324.088 | 1645.077 | 1856.518 |
| ENSMUSG00000044499 | 1.417870638 | 0.010276 | 0.043496 | 33.19979 | 19.19714 | 12.4216 | 36.23019 | 5.850668 | 7.821619 | 16.39292 | 7.613138 |
| ENSMUSG00000044501 | 0.061053429 | 0.741121 | 0.838275 | 138.8355 | 158.3764 | 139.1219 | 192.365 | 145.2916 | 160.9019 | 174.5363 | 121.8102 |
| ENSMUSG00000044502 | -0.21432492 | 0.002764 | 0.016632 | 1985.951 | 2080.491 | 2153.077 | 2130.68 | 2381.222 | 2210.166 | 2511.009 | 2586.292 |
| ENSMUSG00000044505 | 0.083796673 | 0.841306 | 0.904265 | 25.15136 | 14.39786 | 14.07781 | 16.38985 | 11.70134 | 15.64324 | 21.21436 | 17.40146 |
| ENSMUSG00000044522 | -0.572787587 | 0.340239 | 0.508746 | 10.06054 | 2.399643 | 23.18699 | 13.80198 | 13.65156 | 17.87799 | 17.3572 | 26.10219 |
| ENSMUSG00000044528 | 0.541753354 | 0.001449 | 0.010319 | 287.7315 | 359.9465 | 322.9616 | 375.2412 | 224.2756 | 284.9304 | 238.1794 | 177.2773 |
| ENSMUSG00000044533 | -0.242157413 | 0.012181 | 0.049536 | 40892.09 | 49958.17 | 50081.4 | 50400.5 | 61254.54 | 50970.14 | 59069.46 | 55004.92 |
| ENSMUSG00000044534 | 0.206011703 | 0.70218 | 0.812223 | 31.18769 | 11.99822 | 14.07781 | 15.52722 | 17.552 | 5.58687 | 21.21436 | 18.48905 |
| ENSMUSG00000044548 | 0.990606181 | 9.19E-10 | 1.20E-07 | 1320.949 | 2208.871 | 1814.382 | 1848.602 | 960.4846 | 1023.515 | 818.6815 | 816.7809 |
| ENSMUSG00000044550 | 0.969348414 | 3.58E-25 | 8.25E-22 | 688.1412 | 669.5004 | 607.8303 | 664.2201 | 365.6667 | 326.2732 | 334.6083 | 314.3138 |
| ENSMUSG00000044551 | -0.059303859 | 0.727349 | 0.829154 | 170.0232 | 242.3639 | 221.1045 | 212.2054 | 246.7032 | 207.8316 | 179.3578 | 246.8832 |
| ENSMUSG00000044556 | 0.187989421 | 0.758543 | 0.849288 | 15.09082 | 5.999108 | 10.76539 | 12.93935 | 3.900445 | 16.76061 | 12.53576 | 6.525547 |
| ENSMUSG00000044562 | -0.161448448 | 0.159548 | 0.302632 | 667.014 | 586.7127 | 763.5143 | 688.3735 | 736.209 | 697.2414 | 733.8241 | 863.5473 |

| | | | | | | | | | | | |
|---|---|---|---|---|---|---|---|---|---|---|---|
| ENSMUSG00000044566 | -0.19606221 | 0.403649 | 0.570586 | 58.35115 | 104.3845 | 71.21717 | 87.12497 | 84.83468 | 94.9768 | 107.0361 | 79.39415 |
| ENSMUSG00000044573 | -0.019497511 | 0.815846 | 0.887863 | 2361.21 | 2910.767 | 2486.804 | 2551.64 | 2636.701 | 2771.088 | 2517.759 | 2521.036 |
| ENSMUSG00000044576 | 1.821351562 | 2.62E-09 | 2.89E-07 | 59.35721 | 75.58876 | 62.108 | 51.75741 | 14.62667 | 21.23011 | 21.21436 | 13.05109 |
| ENSMUSG00000044583 | -0.082575443 | 0.772718 | 0.858991 | 31.18769 | 40.79393 | 39.74912 | 31.05445 | 32.17867 | 46.92971 | 41.46443 | 30.45255 |
| ENSMUSG00000044595 | 0.444686349 | 0.296098 | 0.463983 | 58.35115 | 32.39518 | 32.29616 | 37.09281 | 18.52711 | 49.16446 | 32.78583 | 17.40146 |
| ENSMUSG00000044600 | -0.224895127 | 0.001204 | 0.008985 | 2383.343 | 2535.223 | 2651.597 | 2459.34 | 3124.257 | 2747.623 | 2826.332 | 3023.503 |
| ENSMUSG00000044609 | 0.295419723 | 0.473845 | 0.633257 | 19.11503 | 17.99732 | 12.4216 | 15.52722 | 15.60178 | 13.40849 | 10.60718 | 13.05109 |
| ENSMUSG00000044617 | -0.011998048 | 0.901504 | 0.940806 | 531.1967 | 477.529 | 535.785 | 592.6223 | 560.689 | 546.3959 | 547.7162 | 503.5547 |
| ENSMUSG00000044626 | -0.324707908 | 0.440983 | 0.604305 | 41.24823 | 21.59679 | 47.20208 | 23.29083 | 60.4569 | 21.23011 | 45.32159 | 40.24087 |
| ENSMUSG00000044627 | -0.187593101 | 0.007102 | 0.03326 | 4242.531 | 4288.162 | 4653.131 | 4288.101 | 5214.895 | 4525.365 | 4967.053 | 5192.16 |
| ENSMUSG00000044628 | -0.684176112 | 7.76E-05 | 0.001117 | 143.8658 | 141.5789 | 158.9965 | 124.2178 | 230.1263 | 175.4277 | 238.1794 | 269.7226 |
| ENSMUSG00000044629 | 0.57214225 | 9.10E-05 | 0.001245 | 770.6376 | 831.4763 | 674.0788 | 811.7287 | 512.9085 | 654.7812 | 476.3588 | 432.8613 |
| ENSMUSG00000044636 | -0.149022048 | 0.315325 | 0.48368 | 400.4096 | 562.7163 | 436.4122 | 477.0308 | 554.8383 | 598.9125 | 482.1446 | 442.6496 |
| ENSMUSG00000044641 | -0.723501484 | 0.000413 | 0.003957 | 726.3712 | 789.4826 | 939.901 | 747.8946 | 1195.486 | 860.3781 | 1505.255 | 1729.27 |
| ENSMUSG00000044645 | 0.78838114 | 0.056719 | 0.148796 | 25.15136 | 15.59768 | 19.04645 | 23.29083 | 16.57689 | 10.05637 | 10.60718 | 10.87591 |
| ENSMUSG00000044646 | 0.655086061 | 0.000145 | 0.001771 | 665.0019 | 919.0633 | 812.3726 | 907.4799 | 490.481 | 692.7719 | 450.323 | 465.489 |
| ENSMUSG00000044647 | 0.995634534 | 2.96E-09 | 3.18E-07 | 210.2654 | 193.1713 | 176.3867 | 212.2054 | 91.66046 | 124.0285 | 86.78602 | 95.70802 |
| ENSMUSG00000044667 | -0.142503439 | 0.689787 | 0.803554 | 89.53884 | 34.79482 | 86.9512 | 80.22398 | 46.80534 | 89.38993 | 102.2147 | 84.8321 |
| ENSMUSG00000044674 | 0.369219392 | 9.24E-06 | 0.000214 | 3987.999 | 5057.248 | 4574.461 | 4593.47 | 3376.81 | 3580.067 | 3537.977 | 3604.277 |
| ENSMUSG00000044676 | 0.629818459 | 0.000801 | 0.006575 | 539.2451 | 616.7083 | 353.6015 | 447.7016 | 298.3841 | 363.1466 | 276.751 | 325.1897 |
| ENSMUSG00000044681 | -4.101112944 | NA | NA | 3.018163 | 0 | 2.48432 | 4.313117 | 0.975111 | 167.6061 | 0 | 1.087591 |
| ENSMUSG00000044694 | 0.055855073 | 0.831618 | 0.897984 | 73.44197 | 55.19179 | 68.73285 | 74.18562 | 51.6809 | 60.3382 | 55.92877 | 94.62042 |
| ENSMUSG00000044700 | -0.018538261 | 0.777896 | 0.86305 | 1143.884 | 1205.821 | 1220.629 | 1204.222 | 1223.765 | 1124.078 | 1238.147 | 1249.642 |
| ENSMUSG00000044702 | 0.001209225 | 0.993869 | 0.996515 | 247.4894 | 244.7636 | 196.2613 | 250.1608 | 283.7574 | 198.8926 | 224.6794 | 228.3941 |
| ENSMUSG00000044707 | -0.292332199 | 0.027281 | 0.088347 | 273.6468 | 273.5593 | 247.6039 | 217.3811 | 296.4338 | 301.691 | 353.8941 | 283.8613 |
| ENSMUSG00000044708 | -1.41130867 | 3.10E-05 | 0.000549 | 184.1079 | 91.18644 | 221.1045 | 154.4096 | 306.185 | 258.1134 | 616.1808 | 553.5839 |
| ENSMUSG00000044709 | -0.131380193 | 0.180511 | 0.330704 | 911.4852 | 953.8581 | 1098.069 | 1054.989 | 1202.312 | 991.1108 | 1119.54 | 1090.854 |
| ENSMUSG00000044712 | 0.266983661 | 0.016645 | 0.061903 | 388.337 | 429.5361 | 403.2879 | 445.9763 | 349.0898 | 395.5504 | 323.0369 | 318.6642 |
| ENSMUSG00000044715 | -0.2213725 | 0.021654 | 0.074576 | 405.4399 | 453.5325 | 411.569 | 414.0593 | 518.7592 | 450.3018 | 467.6802 | 524.2189 |
| ENSMUSG00000044716 | -0.165132289 | 0.617474 | 0.750138 | 31.18769 | 25.19625 | 43.88965 | 27.60395 | 29.25334 | 33.52122 | 31.82154 | 50.02919 |
| ENSMUSG00000044724 | 0.465697534 | 0.206747 | 0.363402 | 19.11503 | 27.5959 | 19.04645 | 23.29083 | 17.552 | 15.64324 | 12.53576 | 18.48905 |
| ENSMUSG00000044730 | 0.201437919 | 0.03125 | 0.09721 | 600.6145 | 585.5129 | 579.6746 | 580.5456 | 553.8632 | 494.9967 | 539.0376 | 450.2627 |
| ENSMUSG00000044734 | 0.382194156 | 0.067117 | 0.167299 | 347.0888 | 356.347 | 284.0406 | 270.0012 | 204.7734 | 322.9211 | 177.4292 | 259.9343 |
| ENSMUSG00000044748 | -2.935445253 | 6.95E-16 | 3.92E-13 | 128.775 | 79.18822 | 153.1997 | 94.02596 | 722.5575 | 357.5597 | 1067.468 | 1339.912 |
| ENSMUSG00000044749 | 0.37482033 | 0.133308 | 0.266732 | 93.56306 | 107.9839 | 80.32634 | 124.2178 | 77.03379 | 93.85942 | 89.67889 | 52.20437 |
| ENSMUSG00000044751 | -0.391149617 | 0.477537 | 0.6369 | 9.054489 | 8.398751 | 8.281066 | 9.488858 | 13.65156 | 4.469496 | 10.60718 | 17.40146 |
| ENSMUSG00000044757 | -0.49269659 | 0.337761 | 0.506148 | 9.054489 | 11.99822 | 10.76539 | 12.93935 | 20.47734 | 5.58687 | 20.25007 | 16.31387 |
| ENSMUSG00000044763 | -0.226885109 | 0.036955 | 0.1105 | 611.681 | 508.7243 | 467.0521 | 535.6892 | 615.2952 | 650.3117 | 641.2523 | 576.4233 |
| ENSMUSG00000044768 | -0.378913807 | 5.78E-05 | 0.000886 | 518.118 | 491.9268 | 513.4261 | 519.2993 | 582.1415 | 661.4855 | 705.8597 | 709.1094 |
| ENSMUSG00000044770 | -0.2462469 | 0.509962 | 0.665714 | 55.33299 | 52.79215 | 77.01392 | 65.55939 | 72.15824 | 33.52122 | 69.42882 | 122.8978 |
| ENSMUSG00000044783 | 0.374932319 | 0.003047 | 0.017963 | 7899.539 | 7872.029 | 6033.585 | 7378.881 | 5780.46 | 6533.286 | 5445.341 | 4744.072 |
| ENSMUSG00000044786 | -0.214576414 | 0.472569 | 0.631964 | 562.3844 | 277.1588 | 392.5225 | 610.7374 | 425.1485 | 417.8979 | 837.003 | 458.9634 |
| ENSMUSG00000044788 | -0.93146334 | 0.032819 | 0.100851 | 120.7265 | 52.79215 | 132.4971 | 81.94923 | 153.0925 | 67.04245 | 229.5008 | 291.4744 |
| ENSMUSG00000044791 | 0.090330855 | 0.087739 | 0.201482 | 2905.485 | 2818.381 | 2738.549 | 2683.622 | 2539.19 | 2697.341 | 2644.081 | 2586.292 |
| ENSMUSG00000044792 | -0.226817433 | 0.010161 | 0.043136 | 2346.119 | 2486.03 | 2548.084 | 2699.149 | 2981.89 | 2572.195 | 3056.797 | 3185.554 |
| ENSMUSG00000044795 | -0.073131772 | 0.5088 | 0.664543 | 495.9848 | 482.3283 | 496.0359 | 439.0754 | 578.241 | 443.5975 | 500.4661 | 488.3284 |
| ENSMUSG00000044807 | 0.688318026 | 7.51E-06 | 0.000185 | 1738.462 | 1916.115 | 1377.141 | 1810.647 | 1013.141 | 1347.553 | 957.5391 | 927.7152 |
| ENSMUSG00000044811 | -0.005170056 | 0.990242 | 0.994499 | 24.1453 | 15.59768 | 33.95237 | 17.25247 | 20.47734 | 31.28647 | 14.46434 | 26.10219 |
| ENSMUSG00000044813 | -0.135426949 | 0.375951 | 0.544226 | 746.4923 | 683.8983 | 1007.806 | 762.5592 | 728.4081 | 855.9086 | 968.1463 | 966.8685 |
| ENSMUSG00000044816 | 0.042928811 | 0.935258 | 0.961682 | 49.29666 | 8.398751 | 42.23344 | 46.58167 | 39.97956 | 24.58223 | 24.10723 | 54.37955 |
| ENSMUSG00000044819 | -2.899681883 | 1.16E-10 | 2.08E-08 | 31.18769 | 14.39786 | 46.37397 | 38.81806 | 141.3911 | 121.7938 | 348.1084 | 371.9562 |
| ENSMUSG00000044820 | 0.778793789 | 0.14461 | 0.28269 | 15.09082 | 33.595 | 23.18699 | 18.11509 | 16.57689 | 21.23011 | 4.821446 | 9.78832 |
| ENSMUSG00000044827 | 0.748899686 | 0.022633 | 0.076955 | 39.23612 | 52.79215 | 66.24853 | 33.64232 | 35.10401 | 29.05173 | 26.03581 | 23.927 |
| ENSMUSG00000044835 | 0.605270786 | 2.94E-06 | 8.88E-05 | 275.6589 | 296.3559 | 286.5249 | 357.1261 | 201.848 | 202.2447 | 206.3579 | 189.2408 |
| ENSMUSG00000044847 | 0.01821427 | 0.882171 | 0.929407 | 471.8395 | 374.3443 | 381.7572 | 401.9825 | 388.0943 | 459.2408 | 395.3586 | 367.6058 |
| ENSMUSG00000044854 | -0.181619312 | 0.610032 | 0.745119 | 30.18163 | 26.39607 | 32.29616 | 17.25247 | 28.27823 | 20.11273 | 34.71441 | 36.9781 |
| ENSMUSG00000044857 | -0.196525952 | 0.055518 | 0.146735 | 1265.616 | 1325.803 | 1309.237 | 1263.743 | 1461.692 | 1260.398 | 1473.434 | 1721.657 |
| ENSMUSG00000044863 | -0.067815835 | 0.74718 | 0.842186 | 108.6539 | 121.182 | 82.81066 | 111.2784 | 133.5902 | 121.7938 | 89.67889 | 97.8832 |
| ENSMUSG00000044864 | 0.01430594 | 0.877197 | 0.926493 | 1598.62 | 1586.164 | 1970.894 | 1818.41 | 1667.44 | 1828.024 | 1717.399 | 1698.817 |
| ENSMUSG00000044867 | -0.472115068 | 0.295015 | 0.462756 | 10.06054 | 17.99732 | 19.87456 | 17.25247 | 24.37778 | 35.75597 | 14.46434 | 16.31387 |
| ENSMUSG00000044876 | 0.017146093 | 0.809682 | 0.883969 | 1043.278 | 985.0535 | 984.6188 | 988.5665 | 965.3602 | 922.951 | 1014.432 | 1050.613 |
| ENSMUSG00000044881 | -0.386133525 | 0.199347 | 0.354242 | 148.896 | 189.5718 | 138.2938 | 192.365 | 219.4 | 135.2023 | 163.9292 | 355.6423 |
| ENSMUSG00000044894 | -0.796643599 | 9.04E-05 | 0.001241 | 1926.594 | 1808.131 | 2255.762 | 1711.445 | 3211.042 | 2128.598 | 3552.441 | 4487.401 |
| ENSMUSG00000044906 | 0.398813349 | 0.014666 | 0.056722 | 998.0059 | 1037.846 | 833.0753 | 983.3908 | 741.0846 | 929.6552 | 707.7882 | 542.708 |

| | | | | | | | | | | | |
|---|---|---|---|---|---|---|---|---|---|---|---|
| ENSMUSG00000044912 | 1.906498578 | 2.22E-10 | 3.56E-08 | 112.6781 | 118.7823 | 145.7468 | 169.9368 | 38.02934 | 55.8687 | 30.85725 | 21.75182 |
| ENSMUSG00000044916 | 0.580974655 | 0.039695 | 0.116029 | 56.33904 | 45.59322 | 51.34261 | 40.5433 | 35.10401 | 41.34284 | 24.10723 | 29.36496 |
| ENSMUSG00000044921 | -0.06996782 | 0.794436 | 0.873694 | 276.6649 | 395.9411 | 510.9418 | 332.9727 | 461.2276 | 230.1791 | 390.5371 | 510.0802 |
| ENSMUSG00000044927 | 0.072343518 | 0.546704 | 0.695045 | 1898.425 | 2136.882 | 1744.821 | 1800.295 | 1675.241 | 2155.415 | 1804.185 | 1572.657 |
| ENSMUSG00000044933 | -0.465148734 | 0.254021 | 0.41762 | 16.09687 | 26.39607 | 24.8432 | 12.93935 | 17.552 | 37.99072 | 28.92867 | 26.10219 |
| ENSMUSG00000044934 | -0.017736981 | 0.91362 | 0.948837 | 574.457 | 686.2979 | 546.5504 | 550.3538 | 577.2659 | 767.636 | 572.7878 | 467.6642 |
| ENSMUSG00000044937 | 0.103351778 | 0.649157 | 0.772758 | 133.8052 | 100.785 | 90.26362 | 133.7066 | 92.63557 | 140.7891 | 96.42892 | 97.8832 |
| ENSMUSG00000044938 | -0.24526909 | 0.61538 | 0.748718 | 14.08476 | 10.79839 | 11.59349 | 12.07673 | 11.70134 | 24.58223 | 8.678602 | 13.05109 |
| ENSMUSG00000044948 | -0.131876615 | 0.476157 | 0.635644 | 570.4328 | 407.9393 | 510.1137 | 504.6347 | 504.1325 | 448.067 | 743.4669 | 488.3284 |
| ENSMUSG00000044949 | -0.141484715 | 0.213021 | 0.370828 | 531.1967 | 544.719 | 554.8314 | 594.3476 | 696.2295 | 547.5133 | 662.4667 | 547.0583 |
| ENSMUSG00000044950 | 0.201598578 | 0.081875 | 0.192069 | 1204.247 | 1223.818 | 924.167 | 1066.203 | 906.8535 | 1052.566 | 985.5035 | 893.9999 |
| ENSMUSG00000044952 | -0.189845204 | 0.544105 | 0.692833 | 82.49646 | 45.59322 | 48.85829 | 64.69676 | 68.25779 | 41.34284 | 80.036 | 85.9197 |
| ENSMUSG00000044966 | -0.369311438 | 0.18266 | 0.333317 | 47.28455 | 40.79393 | 33.12427 | 28.46658 | 49.73068 | 46.92971 | 55.92877 | 39.15328 |
| ENSMUSG00000044968 | 0.1878113 | 0.070274 | 0.172791 | 624.7598 | 663.5013 | 657.5167 | 653.8686 | 562.6392 | 663.7202 | 555.4306 | 501.3795 |
| ENSMUSG00000044976 | -1.456052848 | 0.005261 | 0.02664 | 29.17558 | 19.19714 | 64.59232 | 41.40593 | 84.83468 | 36.87335 | 105.1075 | 199.0292 |
| ENSMUSG00000044982 | -0.278045221 | 0.000856 | 0.006926 | 411.4762 | 428.3363 | 433.9279 | 432.1744 | 491.4561 | 545.2786 | 502.3947 | 531.832 |
| ENSMUSG00000044986 | -0.921289878 | 0.001353 | 0.009799 | 700.2138 | 350.3479 | 552.3471 | 550.3538 | 1011.19 | 583.2693 | 944.0391 | 1541.117 |
| ENSMUSG00000044991 | 0.068962694 | 0.652847 | 0.775799 | 174.0474 | 197.9706 | 173.0743 | 198.4034 | 171.6196 | 185.4841 | 145.6077 | 206.6423 |
| ENSMUSG00000045004 | 0.544873043 | 0.280612 | 0.446983 | 11.0666 | 19.19714 | 19.87456 | 24.15346 | 12.67645 | 22.34748 | 8.678602 | 7.613138 |
| ENSMUSG00000045005 | -0.26197133 | 0.095592 | 0.213927 | 292.7618 | 195.5709 | 239.3228 | 233.771 | 261.3298 | 264.8177 | 299.8939 | 328.4525 |
| ENSMUSG00000045007 | -0.213241563 | 0.266765 | 0.431236 | 79.47829 | 93.58608 | 119.2474 | 98.33908 | 127.7396 | 97.21155 | 114.7504 | 114.1971 |
| ENSMUSG00000045008 | 0.115617918 | 0.723996 | 0.827323 | 39.23612 | 39.59411 | 27.32752 | 41.40593 | 40.95467 | 43.57759 | 22.17865 | 29.36496 |
| ENSMUSG00000045009 | 0.376934694 | 0.225844 | 0.385585 | 41.24823 | 52.79215 | 50.5145 | 41.40593 | 39.00445 | 51.39921 | 28.92867 | 23.927 |
| ENSMUSG00000045010 | -2.201982785 | 0.068611 | 0.169744 | 2.012109 | 3.599465 | 4.96864 | 7.763611 | 3.900445 | 71.51194 | 9.642892 | 0 |
| ENSMUSG00000045027 | -0.078565402 | 0.862395 | 0.917656 | 47.28455 | 19.19714 | 27.32752 | 24.15346 | 51.6809 | 18.99536 | 31.82154 | 21.75182 |
| ENSMUSG00000045031 | 0.168794183 | 0.519064 | 0.672647 | 77.46619 | 75.58876 | 87.7793 | 122.4925 | 62.40712 | 109.5027 | 80.036 | 72.8686 |
| ENSMUSG00000045034 | -1.189604193 | 0.026701 | 0.087156 | 8.048435 | 4.799286 | 14.90592 | 10.35148 | 28.27823 | 11.17374 | 17.3572 | 31.54014 |
| ENSMUSG00000045038 | 0.557220404 | 5.91E-06 | 0.000153 | 871.2431 | 878.2694 | 834.7315 | 917.8314 | 504.1325 | 725.1758 | 600.7521 | 551.4087 |
| ENSMUSG00000045039 | 0.192444016 | 0.01861 | 0.067083 | 2259.598 | 2434.438 | 2045.423 | 2225.569 | 1836.135 | 2107.368 | 1883.257 | 2016.394 |
| ENSMUSG00000045045 | -0.32614353 | 0.000213 | 0.002383 | 1261.592 | 1247.814 | 1479.827 | 1422.466 | 1690.843 | 1563.206 | 1795.506 | 1740.146 |
| ENSMUSG00000045053 | 0.627978161 | 0.174302 | 0.322434 | 15.09082 | 10.79839 | 14.90592 | 16.38985 | 9.751113 | 12.29112 | 7.714313 | 7.613138 |
| ENSMUSG00000045055 | -0.000549131 | 0.99865 | 0.9993 | 23.13925 | 27.5959 | 24.8432 | 24.15346 | 26.32801 | 31.28647 | 18.32149 | 23.927 |
| ENSMUSG00000045062 | -0.02209965 | 0.897471 | 0.937817 | 112.6781 | 121.182 | 96.06037 | 113.8663 | 97.51113 | 125.1459 | 122.4647 | 104.4087 |
| ENSMUSG00000045064 | -0.193255152 | 0.172996 | 0.32066 | 145.8779 | 184.7725 | 157.3403 | 169.0742 | 187.2214 | 186.6015 | 167.7863 | 208.8175 |
| ENSMUSG00000045071 | 0.236106155 | 0.026623 | 0.086996 | 1041.266 | 1171.026 | 1123.741 | 1221.475 | 941.9575 | 1134.135 | 924.7533 | 870.0729 |
| ENSMUSG00000045078 | 0.093539262 | 0.109168 | 0.234444 | 1441.676 | 1486.579 | 1473.202 | 1416.428 | 1308.599 | 1426.887 | 1344.219 | 1373.628 |
| ENSMUSG00000045092 | -0.53500441 | 0.010332 | 0.043643 | 490.9545 | 290.3568 | 483.6143 | 501.1842 | 506.0828 | 622.3774 | 813.8601 | 619.9269 |
| ENSMUSG00000045094 | -0.772159071 | 0.077656 | 0.185281 | 73.44197 | 26.39607 | 64.59232 | 50.89479 | 69.2329 | 43.57759 | 152.3577 | 103.3212 |
| ENSMUSG00000045095 | 0.32526593 | 3.18E-05 | 0.000559 | 1514.112 | 1500.977 | 1454.983 | 1532.019 | 1192.561 | 1320.736 | 1198.611 | 1078.89 |
| ENSMUSG00000045098 | 0.021568919 | 0.779705 | 0.863998 | 2531.233 | 2421.24 | 2260.731 | 2097.9 | 2251.532 | 2316.317 | 2376.973 | 2223.036 |
| ENSMUSG00000045100 | -0.031908503 | 0.812551 | 0.886 | 349.1009 | 325.1516 | 399.1474 | 356.2635 | 417.3476 | 298.3389 | 378.0014 | 368.6934 |
| ENSMUSG00000045102 | -0.507047291 | 0.407633 | 0.573902 | 15.09082 | 19.19714 | 5.796746 | 23.29083 | 15.60178 | 46.92971 | 14.46434 | 13.05109 |
| ENSMUSG00000045103 | 0.053924385 | 0.688213 | 0.802075 | 504.0332 | 425.9366 | 462.0835 | 530.5134 | 437.825 | 564.2739 | 413.6801 | 439.3868 |
| ENSMUSG00000045104 | -0.811410448 | 0.07644 | 0.183455 | 16.09687 | 19.19714 | 19.04645 | 13.80198 | 15.60178 | 20.11273 | 33.75012 | 50.02919 |
| ENSMUSG00000045106 | -0.199581183 | 0.367107 | 0.535684 | 111.672 | 149.9777 | 124.216 | 115.5915 | 134.5654 | 200.01 | 113.7861 | 127.2482 |
| ENSMUSG00000045107 | -0.251912674 | 0.141309 | 0.278097 | 259.562 | 207.5691 | 301.4308 | 271.7264 | 313.0107 | 240.2354 | 354.8584 | 332.8029 |
| ENSMUSG00000045114 | -0.289940164 | 0.324892 | 0.492826 | 33.19979 | 26.39607 | 29.81184 | 22.42821 | 33.15378 | 41.34284 | 30.85725 | 31.54014 |
| ENSMUSG00000045128 | -0.147595106 | 0.100269 | 0.221306 | 33085.1 | 40234.82 | 35564.7 | 37803.61 | 44800.51 | 39601.97 | 41217.58 | 36864.99 |
| ENSMUSG00000045136 | -0.0140404 | 0.935043 | 0.96165 | 3522.196 | 2987.556 | 3718.199 | 4037.078 | 3565.982 | 2518.561 | 3867.764 | 4453.686 |
| ENSMUSG00000045140 | -0.144158445 | 0.442203 | 0.605393 | 212.2775 | 194.3711 | 138.2938 | 163.8985 | 189.1716 | 164.254 | 193.8221 | 233.8321 |
| ENSMUSG00000045160 | -0.738842774 | 3.93E-06 | 0.000111 | 477.8758 | 407.9393 | 525.8477 | 429.5865 | 783.9895 | 604.4994 | 724.1812 | 962.5181 |
| ENSMUSG00000045165 | -0.216603147 | 0.280549 | 0.44694 | 60.36326 | 89.98662 | 93.57605 | 76.77349 | 94.5858 | 87.15518 | 99.32178 | 91.35765 |
| ENSMUSG00000045176 | 0.037830237 | 0.73814 | 0.836081 | 412.4823 | 339.5495 | 406.6004 | 398.5321 | 352.0152 | 359.7945 | 423.3229 | 383.9197 |
| ENSMUSG00000045180 | -0.143032869 | 0.42365 | 0.588949 | 761.5831 | 505.1249 | 791.6699 | 778.949 | 626.9966 | 769.8708 | 942.1105 | 797.2043 |
| ENSMUSG00000045193 | 0.172923129 | 0.064521 | 0.163321 | 6923.666 | 7707.654 | 7667.439 | 7670.448 | 7032.503 | 7390.312 | 6421.202 | 5740.306 |
| ENSMUSG00000045201 | -0.314847492 | 0.362406 | 0.531001 | 26.15741 | 27.5959 | 33.95237 | 46.58167 | 43.88001 | 34.6386 | 29.89296 | 59.81751 |
| ENSMUSG00000045205 | 0.196641605 | 0.080012 | 0.188964 | 1249.52 | 1103.836 | 1042.586 | 1213.711 | 950.7335 | 1190.003 | 931.5033 | 951.6422 |
| ENSMUSG00000045210 | 0.19794117 | 0.044162 | 0.124893 | 1244.489 | 1213.02 | 1023.54 | 1128.312 | 986.8126 | 1088.322 | 1038.539 | 901.613 |
| ENSMUSG00000045211 | -0.504274685 | 0.000197 | 0.002239 | 184.1079 | 167.975 | 183.0116 | 175.1126 | 239.8774 | 214.5358 | 284.4653 | 268.635 |
| ENSMUSG00000045215 | 0.628133894 | 0.006208 | 0.030206 | 848.1038 | 1091.838 | 709.6874 | 934.2212 | 603.5939 | 839.1479 | 457.0731 | 418.7226 |
| ENSMUSG00000045216 | -0.120830509 | 0.29786 | 0.465794 | 2596.626 | 2392.444 | 3051.573 | 2853.559 | 2707.884 | 2621.36 | 3341.262 | 3179.029 |
| ENSMUSG00000045231 | 0.413505178 | 0.373301 | 0.54215 | 30.18163 | 27.5959 | 43.88965 | 28.46658 | 39.97956 | 32.40385 | 12.53576 | 13.05109 |
| ENSMUSG00000045237 | -0.096638244 | 0.418626 | 0.584254 | 502.0211 | 536.3202 | 582.159 | 521.8872 | 540.2117 | 511.7573 | 684.6453 | 553.5839 |
| ENSMUSG00000045246 | 0.577273459 | 0.20904 | 0.366178 | 48.29061 | 58.79126 | 28.98373 | 16.38985 | 23.40267 | 25.6996 | 34.71441 | 17.40146 |

| | | | | | | | | | | | |
|---|---|---|---|---|---|---|---|---|---|---|---|
| ENSMUSG00000045248 | -0.150458392 | 0.199111 | 0.353889 | 310.8708 | 311.9536 | 397.4912 | 354.5383 | 364.6916 | 404.4894 | 380.8942 | 380.6569 |
| ENSMUSG00000045251 | 0.117727272 | 0.394025 | 0.561424 | 399.4036 | 404.3399 | 396.6631 | 517.5741 | 427.0988 | 426.8369 | 394.3943 | 336.0656 |
| ENSMUSG00000045252 | -0.066664574 | 0.553087 | 0.700314 | 624.7598 | 611.909 | 659.1729 | 583.1335 | 673.8019 | 538.5743 | 654.7523 | 728.686 |
| ENSMUSG00000045257 | -0.378484413 | 0.012604 | 0.050785 | 195.1745 | 214.7681 | 216.1358 | 162.1732 | 249.6285 | 222.3574 | 256.5009 | 294.7372 |
| ENSMUSG00000045259 | -0.34068075 | 0.079042 | 0.187578 | 71.42986 | 94.7859 | 97.71658 | 75.04824 | 118.9636 | 96.09417 | 97.39321 | 116.3722 |
| ENSMUSG00000045268 | -0.345122217 | 0.074544 | 0.180288 | 293.7679 | 226.7663 | 216.9639 | 203.5791 | 312.0356 | 220.1227 | 366.4299 | 294.7372 |
| ENSMUSG00000045273 | 0.158296517 | 0.310521 | 0.479098 | 657.9595 | 725.892 | 500.1764 | 651.2807 | 521.6845 | 683.8329 | 580.5021 | 483.978 |
| ENSMUSG00000045275 | -0.087524528 | 0.541482 | 0.690786 | 144.8718 | 165.5754 | 136.6376 | 163.0358 | 152.1174 | 181.0146 | 161.0363 | 153.3503 |
| ENSMUSG00000045281 | -0.234001555 | 0.450102 | 0.612571 | 27.16347 | 32.39518 | 38.09291 | 29.3292 | 42.9049 | 23.46486 | 41.46443 | 41.32846 |
| ENSMUSG00000045282 | 0.268721153 | 0.178023 | 0.327284 | 95.57516 | 104.3845 | 96.88848 | 80.22398 | 72.15824 | 82.68568 | 64.60737 | 93.53283 |
| ENSMUSG00000045284 | 0.494850735 | 0.015248 | 0.05836 | 269.6226 | 307.1543 | 259.1974 | 253.6113 | 169.6694 | 272.6393 | 185.1435 | 145.7372 |
| ENSMUSG00000045287 | 0.661599728 | 9.26E-06 | 0.000214 | 360.1675 | 326.3515 | 292.3216 | 312.2697 | 181.3707 | 242.4702 | 173.572 | 219.6934 |
| ENSMUSG00000045291 | -1.732613505 | 0.010544 | 0.044329 | 22.1332 | 1.199822 | 43.88965 | 18.11509 | 68.25779 | 36.87335 | 64.60737 | 116.3722 |
| ENSMUSG00000045294 | -0.917268764 | 6.72E-09 | 6.08E-07 | 1031.206 | 970.6556 | 1223.942 | 974.7645 | 2032.132 | 1834.728 | 2521.616 | 1545.467 |
| ENSMUSG00000045302 | -0.1076573 | 0.257842 | 0.421574 | 2320.967 | 1952.11 | 1930.317 | 2096.175 | 2268.109 | 1972.165 | 2397.223 | 2303.518 |
| ENSMUSG00000045312 | 0.370077288 | 0.000844 | 0.006845 | 4251.586 | 4739.295 | 4486.682 | 4283.788 | 3128.157 | 3990.143 | 2930.475 | 3695.635 |
| ENSMUSG00000045314 | -0.520787093 | 0.073355 | 0.178347 | 107.6478 | 82.78769 | 170.59 | 74.18562 | 160.8934 | 120.6764 | 155.2506 | 189.2408 |
| ENSMUSG00000045316 | -0.698749389 | 1.59E-05 | 0.000327 | 744.4802 | 736.6904 | 728.7338 | 753.9329 | 1229.615 | 841.3827 | 1246.826 | 1492.175 |
| ENSMUSG00000045318 | 0.252100684 | 0.468023 | 0.628343 | 31.18769 | 27.5959 | 43.06154 | 42.26855 | 17.552 | 34.6386 | 30.85725 | 39.15328 |
| ENSMUSG00000045319 | 0.032820555 | 0.834374 | 0.89975 | 530.1906 | 392.3416 | 361.0545 | 376.9665 | 384.1939 | 467.0624 | 406.93 | 364.343 |
| ENSMUSG00000045327 | 0.298440702 | 0.373606 | 0.542422 | 74.44802 | 79.18822 | 54.65504 | 50.03216 | 40.95467 | 86.03781 | 40.50014 | 42.41605 |
| ENSMUSG00000045328 | 0.452530918 | 0.006172 | 0.030069 | 2421.573 | 2529.224 | 1753.93 | 2141.032 | 1528.975 | 2092.842 | 1379.898 | 1461.722 |
| ENSMUSG00000045333 | 0.351994241 | 0.017032 | 0.062944 | 599.6084 | 701.8956 | 762.6862 | 781.5369 | 550.9379 | 686.0677 | 533.2519 | 461.1386 |
| ENSMUSG00000045345 | 0.40584629 | 0.184274 | 0.3354 | 29.17558 | 34.79482 | 29.81184 | 31.05445 | 28.27823 | 24.58223 | 18.32149 | 22.83941 |
| ENSMUSG00000045348 | 0.301594685 | 0.073389 | 0.178403 | 309.8647 | 242.3639 | 334.5551 | 345.0494 | 249.6285 | 244.7049 | 300.8582 | 205.5547 |
| ENSMUSG00000045349 | -0.218164286 | 0.409838 | 0.575996 | 42.25428 | 38.39429 | 41.40533 | 49.16954 | 65.33246 | 43.57759 | 39.53586 | 51.11678 |
| ENSMUSG00000045362 | -0.750969534 | 0.086019 | 0.19862 | 9.054489 | 9.598572 | 9.109173 | 14.6646 | 22.42756 | 13.40849 | 18.32149 | 17.40146 |
| ENSMUSG00000045374 | -0.084744519 | 0.341633 | 0.510375 | 491.9606 | 471.5299 | 540.7536 | 539.1397 | 518.7592 | 527.4006 | 564.1092 | 561.197 |
| ENSMUSG00000045377 | -0.386877838 | 0.01332 | 0.052912 | 366.2038 | 290.3568 | 430.6154 | 340.7363 | 439.7752 | 397.7852 | 495.6446 | 537.27 |
| ENSMUSG00000045382 | 0.583914403 | 0.028409 | 0.090916 | 2027.2 | 2050.495 | 1324.143 | 1337.929 | 1153.557 | 1734.165 | 757.9313 | 850.4962 |
| ENSMUSG00000045392 | 0.875712507 | 0.00583 | 0.02881 | 793.7769 | 819.4781 | 573.8779 | 1050.675 | 274.0063 | 746.4059 | 461.8945 | 282.7737 |
| ENSMUSG00000045394 | 0.028141667 | 0.654433 | 0.777033 | 8056.483 | 8105.994 | 8729.072 | 8458.023 | 7613.669 | 8435.057 | 8656.424 | 8004.67 |
| ENSMUSG00000045409 | 0.209723095 | 0.01224 | 0.049724 | 1194.187 | 1253.814 | 1069.086 | 1262.018 | 1047.27 | 1097.261 | 994.1821 | 991.8831 |
| ENSMUSG00000045410 | 0.178711866 | 0.011632 | 0.047812 | 776.674 | 766.686 | 735.3587 | 807.4156 | 682.5779 | 696.1241 | 688.5025 | 659.0802 |
| ENSMUSG00000045411 | 0.113658214 | 0.197166 | 0.35155 | 4394.445 | 4340.954 | 4228.312 | 4257.047 | 4338.27 | 4370.05 | 3622.834 | 3584.7 |
| ENSMUSG00000045414 | -0.03689297 | 0.821615 | 0.891669 | 845.0857 | 980.2542 | 1336.564 | 950.6111 | 992.6633 | 897.2514 | 1111.825 | 1219.19 |
| ENSMUSG00000045427 | 0.010776846 | 0.819199 | 0.890618 | 3658.014 | 3726.646 | 3584.046 | 3825.735 | 3751.253 | 3620.292 | 3726.013 | 3583.613 |
| ENSMUSG00000045435 | 0.012961315 | 0.893111 | 0.93538 | 443.67 | 412.7386 | 441.3808 | 493.4206 | 450.5014 | 464.8276 | 449.3588 | 413.2846 |
| ENSMUSG00000045438 | -0.243379263 | 0.00045 | 0.004226 | 922.5518 | 980.2542 | 986.275 | 1038.599 | 1220.839 | 1162.069 | 1123.397 | 1145.233 |
| ENSMUSG00000045440 | 1.923036721 | 0.02654 | 0.086801 | 26.15741 | 16.7975 | 18.21835 | 44.85642 | 2.925334 | 22.34748 | 1.928578 | 1.087591 |
| ENSMUSG00000045441 | 0.384747048 | 0.066644 | 0.166573 | 118.7144 | 113.983 | 131.669 | 99.2017 | 76.05868 | 108.3853 | 101.2504 | 69.60583 |
| ENSMUSG00000045455 | -0.405726673 | 0.424996 | 0.589975 | 10.06054 | 5.999108 | 9.109173 | 10.35148 | 15.60178 | 14.52586 | 8.678602 | 8.700729 |
| ENSMUSG00000045464 | 0.068133251 | 0.645406 | 0.770415 | 187.1261 | 175.1739 | 137.4657 | 163.8985 | 155.0427 | 159.7845 | 164.8934 | 151.1752 |
| ENSMUSG00000045466 | 0.172987616 | 0.230832 | 0.391243 | 462.785 | 509.9242 | 484.4424 | 470.1298 | 468.0534 | 491.6446 | 429.1087 | 319.7518 |
| ENSMUSG00000045467 | -0.232897834 | 0.373288 | 0.54215 | 47.28455 | 32.39518 | 33.95237 | 34.50494 | 42.9049 | 37.99072 | 47.25017 | 45.67883 |
| ENSMUSG00000045482 | 0.309812857 | 0.004069 | 0.022106 | 3553.384 | 3545.473 | 3264.396 | 3537.619 | 2639.626 | 3412.46 | 2553.438 | 2610.219 |
| ENSMUSG00000045498 | 0.284078889 | 0.366149 | 0.534751 | 29.17558 | 37.19447 | 41.40533 | 38.81806 | 25.35289 | 43.57759 | 25.07152 | 27.18978 |
| ENSMUSG00000045503 | -0.228468389 | 0.011114 | 0.046242 | 773.6558 | 856.6726 | 888.5584 | 921.2819 | 1063.846 | 915.1294 | 1046.254 | 1006.022 |
| ENSMUSG00000045506 | 0.032080763 | 0.911476 | 0.947783 | 62.37537 | 58.79126 | 90.26362 | 72.46037 | 73.13335 | 100.5637 | 51.10733 | 54.37955 |
| ENSMUSG00000045515 | -0.093918659 | 0.539321 | 0.689165 | 1841.079 | 1404.991 | 1696.79 | 1543.233 | 1306.649 | 1658.183 | 1970.043 | 1989.204 |
| ENSMUSG00000045519 | 0.148014548 | 0.319636 | 0.487789 | 418.5186 | 368.3452 | 322.1335 | 390.7684 | 345.1894 | 401.1373 | 332.6798 | 274.073 |
| ENSMUSG00000045532 | 0.721258722 | 0.00044 | 0.004145 | 177.0656 | 142.7788 | 115.9349 | 135.4319 | 69.2329 | 88.27255 | 100.2861 | 88.09488 |
| ENSMUSG00000045534 | -0.305560096 | 0.337712 | 0.506116 | 140.8476 | 81.58786 | 124.216 | 169.9368 | 112.1378 | 163.1366 | 247.8223 | 116.3722 |
| ENSMUSG00000045538 | -0.127689435 | 0.281493 | 0.447843 | 367.2098 | 404.3399 | 394.1788 | 470.1298 | 407.5965 | 459.2408 | 428.1444 | 494.8539 |
| ENSMUSG00000045545 | 0.29216136 | 0.520514 | 0.673959 | 83.50251 | 220.7672 | 167.2775 | 154.4096 | 199.8978 | 63.69032 | 189.0007 | 57.64233 |
| ENSMUSG00000045555 | -0.264805727 | 0.410759 | 0.576758 | 69.41775 | 44.3934 | 72.04528 | 51.75741 | 47.78045 | 55.8687 | 75.21455 | 107.6715 |
| ENSMUSG00000045569 | 2.070613027 | 0.034399 | 0.104695 | 841.0614 | 37.19447 | 46.37397 | 176.8378 | 54.60623 | 88.27255 | 44.3573 | 75.04379 |
| ENSMUSG00000045573 | 0.748868815 | 3.54E-10 | 5.07E-08 | 9974.023 | 10966.37 | 12954.07 | 12087.94 | 6031.063 | 6188.018 | 7712.385 | 7432.597 |
| ENSMUSG00000045576 | -0.068033999 | 0.545939 | 0.694455 | 724.3591 | 662.3015 | 600.3773 | 621.0889 | 628.9468 | 630.199 | 682.7167 | 790.6787 |
| ENSMUSG00000045589 | 0.128337866 | 0.740671 | 0.83792 | 20.12109 | 16.7975 | 34.78048 | 18.97772 | 16.57689 | 22.34748 | 23.14294 | 21.75182 |
| ENSMUSG00000045594 | -0.688653142 | 0.001741 | 0.011834 | 1237.447 | 1084.639 | 1476.514 | 1200.772 | 2182.299 | 1215.703 | 1922.793 | 2738.554 |
| ENSMUSG00000045598 | 0.000614532 | 0.993232 | 0.996146 | 783.7163 | 881.8688 | 792.498 | 800.5146 | 818.1184 | 810.0962 | 834.1101 | 789.5911 |
| ENSMUSG00000045613 | 0.177073135 | 0.794195 | 0.873694 | 11.0666 | 14.39786 | 6.624853 | 17.25247 | 5.850668 | 25.6996 | 5.785735 | 6.525547 |
| ENSMUSG00000045624 | 0.250634808 | 0.100463 | 0.22155 | 1875.285 | 2004.902 | 1552.7 | 1603.617 | 1351.504 | 1917.414 | 1339.398 | 1305.109 |

| | | | | | | | | | | | |
|---|---|---|---|---|---|---|---|---|---|---|---|
| ENSMUSG00000045625 | -1.209445241 | 0.00238 | 0.014867 | 13.07871 | 22.79661 | 19.87456 | 25.01608 | 41.92979 | 32.40385 | 36.64299 | 76.13138 |
| ENSMUSG00000045629 | -0.838555885 | 0.000774 | 0.006424 | 230.3864 | 225.5664 | 330.4145 | 286.391 | 499.257 | 287.1651 | 469.6088 | 664.5182 |
| ENSMUSG00000045636 | 0.021346446 | 0.865901 | 0.919615 | 1203.241 | 1033.046 | 1314.205 | 1306.875 | 1107.726 | 1031.336 | 1385.684 | 1263.781 |
| ENSMUSG00000045639 | -0.137091937 | 0.045716 | 0.127857 | 1229.398 | 1175.825 | 1100.554 | 1106.746 | 1295.923 | 1289.45 | 1274.79 | 1207.226 |
| ENSMUSG00000045657 | 0.394407749 | 0.234848 | 0.395723 | 33.19979 | 34.79482 | 28.15563 | 22.42821 | 23.40267 | 16.76061 | 22.17865 | 27.18978 |
| ENSMUSG00000045658 | 0.386302748 | 0.070269 | 0.172791 | 442.6639 | 874.6699 | 748.6084 | 676.2968 | 586.0419 | 607.8515 | 478.2874 | 425.2481 |
| ENSMUSG00000045659 | -0.172574461 | 0.214599 | 0.372744 | 605.6447 | 591.512 | 700.5782 | 596.9355 | 604.569 | 600.0299 | 783.0028 | 825.4816 |
| ENSMUSG00000045662 | 0.703206261 | 0.389441 | 0.556741 | 3.018163 | 25.19625 | 16.56213 | 14.6646 | 1.950223 | 20.11273 | 4.821446 | 9.78832 |
| ENSMUSG00000045664 | -0.160493666 | 0.299817 | 0.467943 | 340.0464 | 331.1507 | 404.9441 | 376.1038 | 418.3228 | 321.8037 | 385.7157 | 499.2043 |
| ENSMUSG00000045665 | -0.297211293 | 0.04867 | 0.133669 | 893.3763 | 977.8546 | 1211.52 | 966.1383 | 1325.176 | 956.4722 | 1261.29 | 1433.445 |
| ENSMUSG00000045667 | -0.421981664 | 0.00232 | 0.014605 | 1089.557 | 1196.222 | 1474.03 | 1202.497 | 1521.174 | 1458.173 | 1647.97 | 2024.007 |
| ENSMUSG00000045671 | 0.471702201 | 0.001527 | 0.010688 | 1860.195 | 1856.124 | 1483.139 | 1760.615 | 1222.79 | 1593.375 | 1190.897 | 1011.46 |
| ENSMUSG00000045672 | 0.489745759 | 0.000414 | 0.003961 | 1474.876 | 1744.541 | 1448.359 | 1366.396 | 968.2855 | 1352.023 | 1010.575 | 965.7809 |
| ENSMUSG00000045679 | 0.405280095 | 0.036409 | 0.109369 | 155.9384 | 184.7725 | 166.4494 | 200.9913 | 128.7147 | 166.4887 | 141.7505 | 97.8832 |
| ENSMUSG00000045680 | 0.058056262 | 0.635342 | 0.76294 | 4111.744 | 4475.334 | 5290.773 | 4121.615 | 4130.571 | 3662.752 | 4677.767 | 4819.116 |
| ENSMUSG00000045689 | 0.378478566 | 0.122717 | 0.25224 | 51.30877 | 58.79126 | 53.82693 | 54.34528 | 31.20356 | 41.34284 | 44.3573 | 51.11678 |
| ENSMUSG00000045690 | 0.139543465 | 0.207259 | 0.36392 | 308.8587 | 307.1543 | 334.5551 | 309.6818 | 312.0356 | 305.0431 | 268.0724 | 259.9343 |
| ENSMUSG00000045691 | -0.126356276 | 0.172439 | 0.319892 | 708.2623 | 692.297 | 732.0463 | 778.0864 | 841.5211 | 693.8893 | 822.5387 | 820.0437 |
| ENSMUSG00000045709 | 0.604824472 | 0.037166 | 0.110761 | 50.30272 | 47.99286 | 38.92101 | 43.13117 | 27.30312 | 41.34284 | 26.03581 | 23.927 |
| ENSMUSG00000045725 | -0.926187098 | 0.000109 | 0.001427 | 98.59333 | 98.38537 | 114.2787 | 92.30071 | 152.1174 | 134.0849 | 245.8937 | 234.9197 |
| ENSMUSG00000045730 | 0.322856663 | 0.515694 | 0.669825 | 38.23007 | 21.59679 | 14.90592 | 36.23019 | 10.72622 | 27.93435 | 33.75012 | 16.31387 |
| ENSMUSG00000045733 | 0.678249945 | 0.085881 | 0.198449 | 21.12714 | 17.99732 | 20.70267 | 24.15346 | 15.60178 | 16.76061 | 10.60718 | 9.78832 |
| ENSMUSG00000045751 | 0.016979619 | 0.870441 | 0.922376 | 572.4449 | 651.5031 | 584.6433 | 627.1273 | 608.4695 | 684.9503 | 574.7163 | 538.3576 |
| ENSMUSG00000045752 | -0.121023881 | 0.110758 | 0.236865 | 1115.714 | 1189.023 | 1248.785 | 1172.305 | 1240.342 | 1207.881 | 1301.79 | 1391.029 |
| ENSMUSG00000045757 | -0.080152471 | 0.517094 | 0.671078 | 310.8708 | 310.7538 | 260.0255 | 260.5123 | 324.7121 | 315.0995 | 292.1796 | 271.8978 |
| ENSMUSG00000045761 | 1.745762408 | 0.001387 | 0.009988 | 42.25428 | 55.19179 | 44.71776 | 31.91707 | 2.925334 | 14.52586 | 8.678602 | 26.10219 |
| ENSMUSG00000045763 | 0.695717274 | 2.04E-09 | 2.35E-07 | 2668.056 | 3029.549 | 2798.172 | 2875.124 | 1417.812 | 2002.334 | 1702.935 | 1898.934 |
| ENSMUSG00000045767 | 0.083719627 | 0.122475 | 0.252024 | 3826.025 | 4136.985 | 3797.697 | 4047.429 | 3778.556 | 3811.363 | 3736.621 | 3585.788 |
| ENSMUSG00000045775 | -1.618666938 | 6.24E-06 | 0.00016 | 172.0353 | 172.7743 | 390.8663 | 203.5791 | 645.5237 | 357.5597 | 764.6813 | 1120.219 |
| ENSMUSG00000045776 | 0.32477619 | 0.148051 | 0.287504 | 61.36932 | 61.1909 | 62.108 | 55.2079 | 58.50668 | 41.34284 | 44.3573 | 46.76642 |
| ENSMUSG00000045777 | -0.494351894 | 0.378544 | 0.546711 | 21.12714 | 3.599465 | 39.74912 | 18.97772 | 31.20356 | 26.81698 | 34.71441 | 26.10219 |
| ENSMUSG00000045790 | 0.061170866 | 0.634231 | 0.762269 | 230.3864 | 230.3657 | 220.2764 | 271.7264 | 206.7236 | 224.5922 | 248.7866 | 233.8321 |
| ENSMUSG00000045795 | 0.046348198 | 0.65109 | 0.774459 | 339.0403 | 329.9509 | 366.8512 | 332.9727 | 315.9361 | 344.1512 | 309.5368 | 358.9051 |
| ENSMUSG00000045799 | -0.068055142 | 0.469378 | 0.629429 | 849.1099 | 1025.847 | 938.2448 | 927.3203 | 1027.767 | 1004.519 | 870.7531 | 1016.898 |
| ENSMUSG00000045813 | 0.414141094 | 0.254787 | 0.418694 | 22.1332 | 29.99554 | 19.04645 | 18.97772 | 16.57689 | 15.64324 | 18.32149 | 16.31387 |
| ENSMUSG00000045817 | 0.075420216 | 0.421307 | 0.586625 | 1844.098 | 1638.956 | 1498.873 | 1474.224 | 1500.696 | 1641.423 | 1482.112 | 1500.876 |
| ENSMUSG00000045822 | -0.046873992 | 0.808355 | 0.88299 | 127.7689 | 103.1847 | 114.2787 | 115.5915 | 138.4658 | 86.03781 | 129.2147 | 121.8102 |
| ENSMUSG00000045826 | 0.543971701 | 0.307174 | 0.475645 | 12.07265 | 9.598572 | 19.87456 | 20.70296 | 15.60178 | 5.58687 | 7.714313 | 14.13868 |
| ENSMUSG00000045827 | -0.298878402 | 0.187076 | 0.339035 | 389.343 | 488.3274 | 659.1729 | 385.5927 | 588.9672 | 407.8415 | 651.8595 | 716.7225 |
| ENSMUSG00000045838 | -0.078054926 | 0.762792 | 0.852453 | 125.7568 | 98.38537 | 151.5435 | 140.6076 | 128.7147 | 83.80306 | 174.5363 | 158.7883 |
| ENSMUSG00000045854 | -0.033620202 | 0.660572 | 0.781458 | 762.5892 | 853.0731 | 803.2634 | 790.1631 | 783.9895 | 796.6877 | 877.5031 | 823.3065 |
| ENSMUSG00000045867 | 0.090112585 | 0.616028 | 0.749122 | 208.2533 | 229.1659 | 243.4633 | 292.4294 | 228.176 | 274.874 | 175.5006 | 238.1824 |
| ENSMUSG00000045871 | 0.764450172 | 0.019127 | 0.068334 | 84.50857 | 173.9741 | 105.9976 | 117.3168 | 49.73068 | 81.56831 | 50.14304 | 102.2336 |
| ENSMUSG00000045875 | 0.676642117 | 0.001622 | 0.011176 | 208.2533 | 176.3738 | 151.5435 | 195.8155 | 109.2125 | 157.5497 | 89.67889 | 102.2336 |
| ENSMUSG00000045876 | 0.181538587 | 0.388663 | 0.55606 | 60.36326 | 57.59143 | 67.07664 | 66.42201 | 51.6809 | 65.92507 | 52.07161 | 53.29196 |
| ENSMUSG00000045877 | -0.229216743 | 0.411704 | 0.577518 | 59.35721 | 40.79393 | 53.82693 | 40.5433 | 49.73068 | 79.33356 | 53.0359 | 46.76642 |
| ENSMUSG00000045886 | -0.005642403 | 0.99059 | 0.994686 | 42.25428 | 13.19804 | 20.70267 | 17.25247 | 14.62667 | 33.52122 | 23.14294 | 22.83941 |
| ENSMUSG00000045896 | -0.044093078 | 0.651404 | 0.774783 | 2043.296 | 2072.092 | 1838.397 | 2095.312 | 2257.383 | 2093.959 | 2182.186 | 1761.898 |
| ENSMUSG00000045912 | -0.699341726 | 0.020563 | 0.071916 | 19.11503 | 21.59679 | 18.21835 | 26.74133 | 33.15378 | 33.52122 | 36.64299 | 35.89051 |
| ENSMUSG00000045915 | -0.238094579 | 0.408276 | 0.574587 | 45.27245 | 52.79215 | 71.21717 | 50.03216 | 84.83468 | 63.69032 | 42.42872 | 68.51824 |
| ENSMUSG00000045917 | -0.096146718 | 0.458883 | 0.62045 | 212.2775 | 225.5664 | 250.9163 | 198.4034 | 236.952 | 221.2401 | 228.5365 | 262.1095 |
| ENSMUSG00000045930 | -0.761261796 | 0.00344 | 0.019546 | 538.2391 | 321.5522 | 776.764 | 533.9639 | 822.9939 | 670.4245 | 1037.575 | 1151.759 |
| ENSMUSG00000045932 | 0.497143577 | 1.95E-05 | 0.000385 | 872.2491 | 1071.441 | 809.8883 | 879.0133 | 615.2952 | 588.8561 | 709.7168 | 655.8174 |
| ENSMUSG00000045934 | -0.089316363 | 0.444551 | 0.60748 | 631.8021 | 533.9206 | 554.0033 | 550.3538 | 645.5237 | 500.5836 | 613.2879 | 654.7298 |
| ENSMUSG00000045948 | -0.265043079 | 0.003501 | 0.019796 | 675.0625 | 728.2917 | 753.577 | 646.9676 | 846.3966 | 774.3402 | 834.1101 | 913.5765 |
| ENSMUSG00000045954 | 0.249710608 | 0.02457 | 0.081941 | 881.3036 | 1053.443 | 984.6188 | 963.5504 | 693.3041 | 867.0823 | 792.6457 | 913.5765 |
| ENSMUSG00000045962 | -0.013525225 | 0.878096 | 0.927008 | 4662.056 | 4116.588 | 4456.87 | 4575.355 | 3890.694 | 4699.675 | 4538.909 | 4853.919 |
| ENSMUSG00000045967 | 0.425096785 | 0.340798 | 0.5095 | 16.09687 | 14.39786 | 14.90592 | 17.25247 | 6.825779 | 16.76061 | 12.53576 | 10.87591 |
| ENSMUSG00000045968 | -0.386782245 | 0.18607 | 0.337837 | 25.15136 | 26.39607 | 28.98373 | 43.13117 | 39.00445 | 37.99072 | 44.3573 | 41.32846 |
| ENSMUSG00000045969 | -0.095110567 | 0.207373 | 0.364047 | 1906.473 | 2012.101 | 1880.63 | 2223.843 | 2152.071 | 2077.198 | 2232.329 | 2107.752 |
| ENSMUSG00000045973 | -0.088504811 | 0.099198 | 0.219818 | 2884.358 | 2880.772 | 2728.611 | 2904.453 | 3017.969 | 3057.136 | 2890.939 | 3152.927 |
| ENSMUSG00000045975 | -0.086069209 | 0.471333 | 0.630997 | 527.1725 | 517.1231 | 587.9557 | 508.0852 | 507.0579 | 503.9357 | 617.1451 | 644.9415 |
| ENSMUSG00000045980 | -0.270613795 | 0.160318 | 0.303749 | 206.2411 | 166.7752 | 209.511 | 179.4257 | 206.7236 | 174.3104 | 240.108 | 299.0875 |
| ENSMUSG00000045983 | -0.199428232 | 0.030957 | 0.09664 | 9310.027 | 9350.209 | 10391.08 | 8831.539 | 10610.19 | 9606.065 | 11097.04 | 12186.46 |

| | | | | | | | | | | | |
|---|---|---|---|---|---|---|---|---|---|---|---|
| ENSMUSG00000045994 | 0.488602608 | 0.208517 | 0.365712 | 35.2119 | 28.79572 | 27.32752 | 40.5433 | 14.62667 | 39.10809 | 20.25007 | 20.66423 |
| ENSMUSG00000045996 | -0.162788958 | 0.047238 | 0.130749 | 627.7779 | 622.7074 | 693.9534 | 640.0666 | 706.9557 | 767.636 | 744.4312 | 677.5692 |
| ENSMUSG00000046010 | -0.201814115 | 0.029974 | 0.094508 | 556.3481 | 500.3256 | 494.3797 | 488.2449 | 592.8677 | 544.1612 | 568.9306 | 638.416 |
| ENSMUSG00000046020 | 0.332222706 | 0.003325 | 0.019108 | 1578.499 | 1874.121 | 1609.839 | 1712.308 | 1368.081 | 1585.554 | 1192.826 | 1234.416 |
| ENSMUSG00000046027 | -0.26008494 | 0.0032 | 0.01862 | 833.013 | 993.4522 | 873.6525 | 811.7287 | 1089.199 | 1008.989 | 1019.254 | 1082.153 |
| ENSMUSG00000046032 | -0.217487732 | 0.022003 | 0.075423 | 1049.315 | 1019.848 | 1247.129 | 1004.956 | 1271.545 | 1225.759 | 1195.719 | 1335.562 |
| ENSMUSG00000046034 | -0.352935989 | 2.18E-05 | 0.000422 | 701.2199 | 703.0954 | 650.8918 | 652.1434 | 813.2428 | 820.1526 | 881.3603 | 939.6787 |
| ENSMUSG00000046049 | -0.061971529 | 0.724159 | 0.827323 | 90.54489 | 110.3836 | 106.8258 | 104.3774 | 102.3867 | 113.9722 | 90.64318 | 123.9854 |
| ENSMUSG00000046056 | 0.696676002 | 0.007908 | 0.035924 | 277.671 | 169.1748 | 140.7781 | 212.2054 | 131.64 | 158.6671 | 92.57176 | 110.9343 |
| ENSMUSG00000046058 | -0.048753466 | 0.721155 | 0.825522 | 577.4752 | 401.9402 | 483.6143 | 484.7944 | 463.1779 | 551.9828 | 459.9659 | 542.708 |
| ENSMUSG00000046062 | 0.113746855 | 0.168052 | 0.314084 | 2171.065 | 2049.295 | 1924.52 | 2051.319 | 1895.616 | 2118.541 | 1786.828 | 1773.861 |
| ENSMUSG00000046070 | -3.283491251 | 1.57E-09 | 1.91E-07 | 8.048435 | 8.398751 | 4.96864 | 3.450494 | 59.48179 | 23.46486 | 58.82164 | 95.70802 |
| ENSMUSG00000046079 | -0.026355384 | 0.781636 | 0.865364 | 1681.117 | 1464.982 | 1608.183 | 1559.623 | 1567.004 | 1429.121 | 1605.541 | 1830.416 |
| ENSMUSG00000046080 | -0.283089878 | 0.338803 | 0.507282 | 54.32694 | 34.79482 | 32.29616 | 35.36756 | 37.05423 | 58.10345 | 53.0359 | 42.41605 |
| ENSMUSG00000046082 | -2.816001533 | 9.00E-05 | 0.001237 | 171.0292 | 32.39518 | 390.8663 | 219.969 | 1563.103 | 630.199 | 1284.433 | 2261.102 |
| ENSMUSG00000046085 | -0.44944384 | 0.171659 | 0.318855 | 27.16347 | 23.99643 | 35.60859 | 25.01608 | 52.65601 | 33.52122 | 39.53586 | 27.18978 |
| ENSMUSG00000046093 | -0.177059243 | 0.479174 | 0.638386 | 86.52067 | 47.99286 | 71.21717 | 68.14726 | 64.35735 | 69.27719 | 96.42892 | 80.48174 |
| ENSMUSG00000046096 | 0.050473984 | 0.57458 | 0.716905 | 1095.593 | 928.6619 | 945.6978 | 979.0777 | 878.5753 | 1012.341 | 1000.932 | 922.2772 |
| ENSMUSG00000046101 | -1.210862742 | 0.000673 | 0.005762 | 45.27245 | 46.79304 | 40.57722 | 50.89479 | 70.20801 | 75.98144 | 89.67889 | 189.2408 |
| ENSMUSG00000046110 | -0.21477791 | 0.757513 | 0.848813 | 4.024217 | 23.99643 | 3.312427 | 14.6646 | 9.751113 | 15.64324 | 16.39292 | 10.87591 |
| ENSMUSG00000046111 | 0.068156715 | 0.533083 | 0.684566 | 1111.69 | 1153.028 | 937.4167 | 1011.857 | 1057.021 | 1095.027 | 999.0036 | 864.6349 |
| ENSMUSG00000046138 | 0.32971489 | 0.02824 | 0.090627 | 1743.492 | 1700.147 | 1478.998 | 1574.288 | 1214.989 | 1705.113 | 1169.683 | 1079.978 |
| ENSMUSG00000046139 | 0.039113251 | 0.602701 | 0.740217 | 911.4852 | 814.6788 | 854.606 | 825.5307 | 804.4668 | 801.1572 | 878.4674 | 830.9196 |
| ENSMUSG00000046152 | 0.420759221 | 8.81E-07 | 3.36E-05 | 887.3399 | 970.6556 | 838.0439 | 887.6396 | 682.5779 | 711.7673 | 667.2881 | 612.3138 |
| ENSMUSG00000046157 | -0.169214015 | 0.339815 | 0.508235 | 192.1564 | 239.9643 | 213.6515 | 162.1732 | 214.5245 | 184.3667 | 262.2867 | 244.708 |
| ENSMUSG00000046159 | -0.225865049 | 0.203806 | 0.35981 | 83.50251 | 97.18554 | 110.1382 | 107.8279 | 109.2125 | 137.437 | 113.7861 | 107.6715 |
| ENSMUSG00000046167 | 0.880838423 | 8.06E-06 | 0.000194 | 400.4096 | 427.1365 | 434.756 | 522.7498 | 203.7983 | 344.1512 | 203.465 | 219.6934 |
| ENSMUSG00000046168 | 0.32705774 | 0.514948 | 0.669147 | 15.09082 | 15.59768 | 9.109173 | 8.626235 | 10.72622 | 11.17374 | 9.642892 | 6.525547 |
| ENSMUSG00000046169 | 0.921517631 | 1.33E-05 | 0.000286 | 519.124 | 595.1115 | 352.7734 | 476.1682 | 274.0063 | 274.874 | 297.9654 | 177.2773 |
| ENSMUSG00000046178 | 2.029364739 | 0.010865 | 0.045396 | 11.0666 | 16.7975 | 11.59349 | 28.46658 | 0 | 7.821619 | 6.750024 | 2.175182 |
| ENSMUSG00000046179 | -0.067369996 | 0.56131 | 0.706846 | 445.6821 | 455.9322 | 411.569 | 405.433 | 500.2321 | 490.5272 | 409.8229 | 398.0583 |
| ENSMUSG00000046182 | 0.712242046 | 0.012944 | 0.051857 | 219.3198 | 229.1659 | 204.5423 | 230.3205 | 158.9431 | 193.3057 | 120.5361 | 66.34306 |
| ENSMUSG00000046185 | -0.136702724 | 0.227409 | 0.387396 | 449.7063 | 496.7261 | 431.4436 | 395.0816 | 530.4606 | 434.6585 | 471.5374 | 507.905 |
| ENSMUSG00000046186 | 0.12707956 | 0.153384 | 0.294696 | 450.7124 | 449.9331 | 477.8175 | 433.8996 | 393.945 | 413.4284 | 444.5373 | 407.8467 |
| ENSMUSG00000046191 | 0.500188007 | 0.003372 | 0.019279 | 198.1927 | 173.9741 | 167.2775 | 168.2116 | 114.088 | 159.7845 | 111.8575 | 115.2847 |
| ENSMUSG00000046201 | 0.057280132 | 0.42466 | 0.589675 | 1646.911 | 1808.131 | 1881.458 | 1783.043 | 1642.087 | 1826.907 | 1648.934 | 1728.182 |
| ENSMUSG00000046203 | -0.034994135 | 0.943488 | 0.96739 | 29.17558 | 15.59768 | 81.98256 | 37.09281 | 56.55646 | 40.22547 | 29.89296 | 42.41605 |
| ENSMUSG00000046204 | 0.413956153 | 0.121734 | 0.251128 | 47.28455 | 47.99286 | 41.40533 | 53.48266 | 42.9049 | 37.99072 | 26.03581 | 35.89051 |
| ENSMUSG00000046207 | -0.331979197 | 0.067087 | 0.167299 | 129.781 | 85.18733 | 131.669 | 103.5148 | 134.5654 | 146.376 | 143.6791 | 144.6496 |
| ENSMUSG00000046213 | 2.560957697 | NA | NA | 70.4238 | 0 | 1.656213 | 3.450494 | 0.975111 | 6.704245 | 1.928578 | 3.262773 |
| ENSMUSG00000046219 | 0.182909617 | 0.673846 | 0.791227 | 22.1332 | 20.39697 | 16.56213 | 19.84034 | 10.72622 | 25.6996 | 22.17865 | 10.87591 |
| ENSMUSG00000046223 | 0.383732879 | 0.067438 | 0.16764 | 325.9616 | 275.959 | 379.2728 | 330.3848 | 240.8525 | 164.254 | 314.3583 | 286.0365 |
| ENSMUSG00000046229 | -0.338475409 | 0.002959 | 0.017587 | 1247.507 | 1235.816 | 1477.342 | 1350.868 | 1729.847 | 1452.586 | 1588.184 | 1948.963 |
| ENSMUSG00000046230 | -0.259385301 | 0.061811 | 0.158289 | 375.2583 | 313.1534 | 405.7722 | 320.0333 | 357.8658 | 432.4238 | 428.1444 | 477.4525 |
| ENSMUSG00000046269 | 0.129321587 | 0.42765 | 0.59268 | 226.3622 | 164.3756 | 181.3554 | 207.0296 | 199.8978 | 158.6671 | 194.7864 | 158.7883 |
| ENSMUSG00000046275 | -0.064964425 | 0.831375 | 0.897939 | 37.22401 | 33.595 | 28.15563 | 38.81806 | 29.25334 | 26.81698 | 45.32159 | 42.41605 |
| ENSMUSG00000046280 | -0.427114863 | 0.025318 | 0.083681 | 370.228 | 295.1561 | 536.6131 | 372.6533 | 524.6099 | 462.5929 | 513.0018 | 621.0145 |
| ENSMUSG00000046287 | 2.105632144 | 2.08E-10 | 3.37E-08 | 67.40564 | 51.59233 | 71.21717 | 82.81186 | 16.57689 | 17.87799 | 20.25007 | 8.700729 |
| ENSMUSG00000046295 | 0.292674265 | 0.0282 | 0.090562 | 626.7719 | 550.7181 | 486.0986 | 608.1496 | 490.481 | 532.9874 | 404.0372 | 427.4233 |
| ENSMUSG00000046311 | 0.00270723 | 0.967654 | 0.981805 | 1301.834 | 1253.814 | 1279.425 | 1229.238 | 1264.719 | 1358.727 | 1196.683 | 1236.591 |
| ENSMUSG00000046312 | 0.204927979 | 0.088237 | 0.202222 | 566.4086 | 757.0874 | 593.7525 | 604.6991 | 502.1823 | 576.565 | 578.5735 | 527.4817 |
| ENSMUSG00000046314 | 0.277814103 | 0.056758 | 0.148796 | 1739.468 | 1728.943 | 1417.719 | 1779.592 | 1438.289 | 1703.995 | 1234.29 | 1121.306 |
| ENSMUSG00000046318 | 0.069459773 | 0.724266 | 0.827323 | 101.6115 | 123.5816 | 121.7317 | 87.12497 | 90.68535 | 98.32892 | 123.429 | 100.0584 |
| ENSMUSG00000046321 | 0.017166799 | 0.969112 | 0.982852 | 15.09082 | 19.19714 | 29.81184 | 21.56559 | 9.751113 | 26.81698 | 27.0001 | 21.75182 |
| ENSMUSG00000046323 | -4.589847547 | NA | NA | 2.012109 | 0 | 2.48432 | 2.58787 | 0.975111 | 170.9582 | 0 | 0 |
| ENSMUSG00000046324 | -0.365453921 | 0.00705 | 0.033083 | 1942.691 | 1505.776 | 2017.268 | 1871.893 | 2387.072 | 1888.362 | 2574.652 | 2605.868 |
| ENSMUSG00000046329 | 0.031056194 | 0.697373 | 0.80906 | 1270.647 | 1112.235 | 1255.41 | 1193.008 | 1130.154 | 1124.078 | 1197.647 | 1280.095 |
| ENSMUSG00000046330 | -0.107547352 | 0.093804 | 0.210846 | 17125.06 | 19258.34 | 19168.18 | 18829.35 | 21223.3 | 19691.48 | 20233.68 | 18987.17 |
| ENSMUSG00000046337 | -0.568800666 | 0.223685 | 0.383425 | 6.036326 | 11.99822 | 17.39024 | 9.488858 | 16.57689 | 18.99536 | 17.3572 | 14.13868 |
| ENSMUSG00000046338 | -0.820741129 | 0.252605 | 0.416293 | 32.19374 | 8.398751 | 17.39024 | 21.56559 | 20.47734 | 92.74205 | 10.60718 | 17.40146 |
| ENSMUSG00000046341 | 0.839596975 | 0.000796 | 0.006549 | 846.0917 | 921.4629 | 756.8895 | 959.2373 | 772.2882 | 478.2361 | 371.2513 | 324.1021 |
| ENSMUSG00000046351 | 0.248207936 | 0.021965 | 0.075351 | 1063.399 | 1093.037 | 860.4028 | 1021.346 | 765.4624 | 921.8336 | 822.5387 | 888.5619 |
| ENSMUSG00000046352 | -2.162344655 | 3.32E-12 | 8.40E-10 | 29.17558 | 32.39518 | 33.95237 | 35.36756 | 163.8187 | 88.27255 | 121.5004 | 213.1679 |
| ENSMUSG00000046364 | -0.164361983 | 0.046403 | 0.129251 | 23424.97 | 27157.96 | 28330.36 | 26693.88 | 32035.33 | 27896.36 | 30196.72 | 28221.9 |

| | | | | | | | | | | | |
|---|---|---|---|---|---|---|---|---|---|---|---|
| ENSMUSG00000046378 | 0.708377113 | 0.372178 | 0.541244 | 16.09687 | 7.198929 | 22.35888 | 20.70296 | 13.65156 | 22.34748 | 3.857157 | 1.087591 |
| ENSMUSG00000046380 | -0.134326111 | 0.480339 | 0.63939 | 241.453 | 364.7457 | 346.9767 | 285.5284 | 371.5174 | 248.057 | 375.1085 | 363.2554 |
| ENSMUSG00000046387 | 0.421536916 | 0.00553 | 0.027658 | 278.6771 | 361.1463 | 312.1962 | 306.2313 | 189.1716 | 263.7003 | 224.6794 | 262.1095 |
| ENSMUSG00000046402 | 0.09314322 | 0.461518 | 0.622645 | 4664.068 | 6345.856 | 5361.162 | 5826.159 | 5769.734 | 5752.242 | 4762.624 | 4523.291 |
| ENSMUSG00000046404 | 0.20697148 | 0.184612 | 0.33585 | 393.3673 | 431.9358 | 409.0847 | 485.657 | 316.9112 | 478.2361 | 361.6084 | 334.9781 |
| ENSMUSG00000046408 | 0.162055945 | 0.629595 | 0.758916 | 44.26639 | 22.79661 | 33.12427 | 35.36756 | 30.22845 | 39.10809 | 21.21436 | 31.54014 |
| ENSMUSG00000046410 | 0.113088129 | 0.44327 | 0.606369 | 117.7084 | 140.3791 | 126.7003 | 120.7673 | 120.9138 | 112.8548 | 112.8218 | 119.635 |
| ENSMUSG00000046413 | -0.605334609 | 0.232814 | 0.393409 | 27.16347 | 9.598572 | 31.46805 | 12.07673 | 19.50223 | 41.34284 | 21.21436 | 41.32846 |
| ENSMUSG00000046415 | 0.474587143 | 0.154845 | 0.296485 | 35.2119 | 51.59233 | 25.67131 | 32.77969 | 22.42756 | 31.28647 | 26.03581 | 23.927 |
| ENSMUSG00000046417 | 0.21193027 | 0.083486 | 0.194528 | 682.1049 | 682.6985 | 503.4888 | 621.9515 | 557.7637 | 498.3488 | 531.3233 | 559.0218 |
| ENSMUSG00000046432 | -0.185172794 | 0.00647 | 0.031202 | 4488.008 | 4440.539 | 4866.783 | 4592.607 | 5187.592 | 4782.361 | 5416.412 | 5522.788 |
| ENSMUSG00000046434 | -0.011964985 | 0.891569 | 0.934507 | 48892.23 | 52257.03 | 47697.29 | 54410.84 | 57380.42 | 51776.88 | 50320.47 | 45470.01 |
| ENSMUSG00000046441 | 0.002782674 | 0.983818 | 0.990959 | 274.6528 | 315.5531 | 241.8071 | 259.6497 | 313.9858 | 261.4655 | 263.2509 | 246.8832 |
| ENSMUSG00000046442 | 0.266279038 | 0.129097 | 0.260851 | 451.7184 | 299.9554 | 298.9465 | 413.1967 | 344.2143 | 300.5736 | 303.7511 | 268.635 |
| ENSMUSG00000046447 | 0.151986404 | 0.25602 | 0.419812 | 3815.964 | 4098.59 | 3283.443 | 3882.668 | 3118.406 | 4276.191 | 3231.333 | 2946.284 |
| ENSMUSG00000046449 | 0.336297454 | 0.156512 | 0.298685 | 223.3441 | 157.1766 | 168.9338 | 194.9529 | 90.68535 | 184.3667 | 160.072 | 155.5255 |
| ENSMUSG00000046456 | 0.939228539 | 0.103857 | 0.226662 | 28.16952 | 16.7975 | 15.73403 | 31.91707 | 12.67645 | 23.46486 | 3.857157 | 8.700729 |
| ENSMUSG00000046463 | 0.182833975 | 0.577244 | 0.71911 | 32.19374 | 76.78858 | 61.27989 | 43.9938 | 58.50668 | 46.92971 | 44.3573 | 38.06569 |
| ENSMUSG00000046470 | -0.366922163 | 0.094914 | 0.212854 | 455.7426 | 292.7565 | 506.8013 | 438.2127 | 591.8926 | 414.5458 | 465.7517 | 714.5473 |
| ENSMUSG00000046480 | -1.807726344 | 0.002243 | 0.014277 | 19.11503 | 10.79839 | 71.21717 | 38.81806 | 71.18313 | 65.92507 | 113.7861 | 242.5328 |
| ENSMUSG00000046491 | 0.28610674 | 0.107457 | 0.231929 | 406.446 | 621.5076 | 518.3948 | 524.4751 | 427.0988 | 522.9311 | 330.7512 | 417.635 |
| ENSMUSG00000046500 | 0.118404238 | 0.844182 | 0.906196 | 16.09687 | 10.79839 | 9.109173 | 12.07673 | 8.776002 | 23.46486 | 5.785735 | 6.525547 |
| ENSMUSG00000046516 | -0.233533376 | 0.141629 | 0.278489 | 589.5479 | 393.5415 | 438.0684 | 389.9058 | 549.9628 | 521.8137 | 470.5731 | 587.2992 |
| ENSMUSG00000046519 | 0.169075603 | 0.065891 | 0.165361 | 1411.494 | 1391.793 | 1207.379 | 1505.278 | 1199.387 | 1281.628 | 1281.54 | 1141.971 |
| ENSMUSG00000046532 | 0.497663335 | 0.331601 | 0.499409 | 273.6468 | 199.1704 | 151.5435 | 217.3811 | 67.28268 | 366.4987 | 69.42882 | 93.53283 |
| ENSMUSG00000046541 | -0.518119547 | 0.010145 | 0.043081 | 137.8294 | 100.785 | 120.0755 | 103.5148 | 194.0472 | 132.9675 | 192.8578 | 141.3868 |
| ENSMUSG00000046546 | -0.128102386 | 0.400654 | 0.567964 | 799.8132 | 734.2908 | 1119.6 | 876.4255 | 874.6748 | 874.9039 | 1010.575 | 1101.73 |
| ENSMUSG00000046550 | 0.404694484 | 0.013883 | 0.054426 | 220.3259 | 220.7672 | 202.058 | 232.0457 | 206.7236 | 130.7328 | 158.1434 | 164.2263 |
| ENSMUSG00000046556 | 0.121575807 | 0.280492 | 0.446927 | 853.1341 | 1019.848 | 917.5421 | 893.6779 | 870.7744 | 949.768 | 709.7168 | 855.9342 |
| ENSMUSG00000046561 | 0.696892079 | 0.016242 | 0.060854 | 85.51462 | 137.9795 | 78.67013 | 123.3552 | 92.63557 | 54.75133 | 51.10733 | 63.08028 |
| ENSMUSG00000046562 | -0.278204286 | 0.007032 | 0.033064 | 1832.025 | 1714.545 | 1835.084 | 1790.806 | 2260.308 | 1778.86 | 2257.401 | 2401.401 |
| ENSMUSG00000046567 | 0.004753122 | 0.974159 | 0.985749 | 270.6286 | 219.5673 | 189.6364 | 201.8539 | 224.2756 | 215.6532 | 224.6794 | 212.0803 |
| ENSMUSG00000046572 | 0.092587817 | 0.358311 | 0.526574 | 816.9161 | 752.2881 | 748.6084 | 766.8723 | 780.089 | 750.8754 | 610.395 | 752.613 |
| ENSMUSG00000046573 | -0.169723631 | 0.05912 | 0.153118 | 307.8526 | 314.3532 | 315.5086 | 329.5222 | 352.9903 | 365.3813 | 354.8584 | 353.4671 |
| ENSMUSG00000046574 | 0.287280262 | 0.00941 | 0.040738 | 976.8788 | 913.0642 | 928.3075 | 915.2435 | 765.4624 | 916.2468 | 687.5382 | 691.7079 |
| ENSMUSG00000046580 | -0.782156055 | 0.000935 | 0.007436 | 36.21796 | 39.59411 | 46.37397 | 36.23019 | 79.95913 | 59.22083 | 60.75022 | 72.8686 |
| ENSMUSG00000046589 | -0.188625883 | 0.420164 | 0.585558 | 115.6963 | 81.58786 | 130.0127 | 88.85022 | 143.3414 | 89.38993 | 110.8933 | 131.5985 |
| ENSMUSG00000046591 | -0.034852016 | 0.81412 | 0.887187 | 453.7305 | 499.1258 | 411.569 | 434.7622 | 453.4268 | 583.2693 | 410.7872 | 394.7956 |
| ENSMUSG00000046593 | -0.01372804 | 0.98157 | 0.989771 | 10.06054 | 23.99643 | 14.07781 | 24.15346 | 18.52711 | 32.40385 | 4.821446 | 17.40146 |
| ENSMUSG00000046598 | -2.180880061 | 0.000255 | 0.002751 | 266.6044 | 175.1739 | 636.814 | 257.9244 | 1306.649 | 442.4801 | 1952.686 | 2361.16 |
| ENSMUSG00000046603 | -0.165437855 | 0.291617 | 0.458791 | 190.1443 | 153.5772 | 175.5586 | 211.3428 | 174.5449 | 205.5968 | 209.2507 | 232.7445 |
| ENSMUSG00000046605 | 0.242892454 | 0.184457 | 0.335635 | 463.7911 | 365.9456 | 351.1172 | 423.5481 | 342.2641 | 449.1844 | 282.5367 | 282.7737 |
| ENSMUSG00000046610 | -0.963101364 | 0.000437 | 0.004124 | 45.27245 | 51.59233 | 45.54586 | 37.95543 | 115.0631 | 55.8687 | 90.64318 | 88.09488 |
| ENSMUSG00000046613 | 0.986984369 | 0.000106 | 0.0014 | 61.36932 | 47.99286 | 57.13936 | 61.24627 | 29.25334 | 29.05173 | 32.78583 | 23.927 |
| ENSMUSG00000046618 | 0.762465611 | 0.000226 | 0.002491 | 512.0817 | 369.545 | 310.54 | 567.6063 | 256.4543 | 278.2261 | 277.7153 | 225.1314 |
| ENSMUSG00000046623 | -0.464069373 | 0.394496 | 0.561749 | 32.19374 | 32.39518 | 35.60859 | 14.6646 | 48.75557 | 8.938993 | 40.50014 | 59.81751 |
| ENSMUSG00000046636 | -0.127792368 | 0.648766 | 0.772442 | 33.19979 | 57.59143 | 44.71776 | 58.6584 | 41.92979 | 49.16446 | 57.85735 | 63.08028 |
| ENSMUSG00000046637 | -0.009352968 | 0.986802 | 0.992394 | 12.07265 | 10.79839 | 12.4216 | 13.80198 | 24.37778 | 6.704245 | 11.57147 | 6.525547 |
| ENSMUSG00000046658 | 0.075238948 | 0.370538 | 0.539654 | 621.7416 | 611.909 | 659.1729 | 673.7089 | 631.8721 | 574.3303 | 648.9666 | 581.8612 |
| ENSMUSG00000046667 | 0.576747187 | 0.011625 | 0.047793 | 205.2351 | 307.1543 | 172.2462 | 215.6559 | 124.8142 | 189.9536 | 128.2505 | 159.8759 |
| ENSMUSG00000046668 | -0.045202198 | 0.523495 | 0.676726 | 4660.044 | 5102.841 | 4822.893 | 4785.835 | 4975.993 | 5493.011 | 4641.124 | 4877.846 |
| ENSMUSG00000046671 | -0.313282014 | 4.14E-06 | 0.000116 | 1927.6 | 1838.127 | 2042.111 | 1876.206 | 2367.57 | 2220.222 | 2453.152 | 2510.16 |
| ENSMUSG00000046675 | -0.53784913 | 0.000357 | 0.003535 | 322.9434 | 262.7609 | 353.6015 | 319.1707 | 405.6463 | 416.7805 | 445.5016 | 563.3722 |
| ENSMUSG00000046679 | 0.090347457 | 0.510421 | 0.66605 | 385.3188 | 537.52 | 484.4424 | 512.3984 | 505.1077 | 416.7805 | 470.5731 | 408.9342 |
| ENSMUSG00000046687 | -2.885678473 | 4.85E-05 | 0.000778 | 2.012109 | 2.399643 | 9.109173 | 15.52722 | 98.48624 | 17.87799 | 39.53586 | 63.08028 |
| ENSMUSG00000046688 | -0.308462892 | 0.065411 | 0.164684 | 145.8779 | 113.983 | 164.7932 | 144.9207 | 156.9929 | 169.8409 | 176.4649 | 205.5547 |
| ENSMUSG00000046691 | -0.256754018 | 0.000155 | 0.001874 | 3248.55 | 3595.865 | 3396.065 | 3198.608 | 3996.006 | 3725.325 | 4191.765 | 4137.196 |
| ENSMUSG00000046694 | 0.111373084 | 0.632387 | 0.760935 | 193.1624 | 167.975 | 167.2775 | 248.4356 | 190.1467 | 167.6061 | 237.2151 | 123.9854 |
| ENSMUSG00000046699 | 0.267616795 | 0.646672 | 0.77119 | 72.43591 | 91.18644 | 48.85829 | 55.2079 | 41.92979 | 137.437 | 20.25007 | 22.83941 |
| ENSMUSG00000046707 | 0.060249945 | 0.512027 | 0.66737 | 1610.693 | 1588.564 | 1682.713 | 1701.956 | 1577.73 | 1778.86 | 1592.041 | 1368.19 |
| ENSMUSG00000046709 | 1.08049168 | 0.002638 | 0.016038 | 159.9626 | 82.78769 | 59.62368 | 130.2561 | 42.9049 | 71.51194 | 50.14304 | 40.24087 |
| ENSMUSG00000046711 | -0.329959921 | 0.046789 | 0.129933 | 1775.686 | 1834.527 | 2667.331 | 2444.675 | 2645.477 | 2365.481 | 3349.941 | 2605.868 |
| ENSMUSG00000046714 | -0.385039118 | 0.007479 | 0.03451 | 918.5276 | 926.2622 | 1165.974 | 1098.12 | 1213.038 | 1102.848 | 1487.898 | 1565.044 |
| ENSMUSG00000046718 | 0.18000771 | 0.73921 | 0.836882 | 678.0806 | 1770.937 | 2605.223 | 569.3315 | 1056.046 | 907.3078 | 1676.899 | 1323.598 |

| | | | | | | | | | | | |
|---|---|---|---|---|---|---|---|---|---|---|---|
| ENSMUSG00000046719 | 0.391633236 | 0.211049 | 0.368438 | 86.52067 | 147.578 | 99.3728 | 131.9814 | 64.35735 | 143.0239 | 69.42882 | 78.30656 |
| ENSMUSG00000046721 | -0.232177727 | 0.070109 | 0.172594 | 241.453 | 293.9563 | 244.2915 | 267.4133 | 344.2143 | 268.1698 | 298.9296 | 315.4014 |
| ENSMUSG00000046722 | -0.314329051 | 0.016016 | 0.060326 | 2026.193 | 1860.923 | 2346.026 | 2006.462 | 2495.31 | 2044.795 | 2756.903 | 2950.635 |
| ENSMUSG00000046727 | -1.257101363 | 0.019169 | 0.068437 | 342.0585 | 261.5611 | 1026.852 | 381.2796 | 1097.975 | 601.1473 | 1405.934 | 1705.343 |
| ENSMUSG00000046731 | -0.350568777 | 0.029142 | 0.092653 | 223.3441 | 221.967 | 207.8548 | 176.8378 | 312.0356 | 208.949 | 269.0367 | 265.3722 |
| ENSMUSG00000046733 | -0.101385071 | 0.629634 | 0.758916 | 259.562 | 261.5611 | 380.1009 | 284.6658 | 355.9156 | 219.0053 | 385.7157 | 312.1386 |
| ENSMUSG00000046743 | 1.141209431 | 4.27E-07 | 1.85E-05 | 5342.149 | 6344.656 | 5493.659 | 5966.767 | 2421.201 | 4147.693 | 1953.65 | 1972.89 |
| ENSMUSG00000046747 | 0.024595627 | 0.954893 | 0.97432 | 19.11503 | 15.59768 | 9.109173 | 17.25247 | 16.57689 | 16.76061 | 15.42863 | 10.87591 |
| ENSMUSG00000046750 | 0.87723341 | 0.025113 | 0.083346 | 23.13925 | 25.19625 | 30.63995 | 31.05445 | 7.80089 | 18.99536 | 18.32149 | 15.22628 |
| ENSMUSG00000046753 | -0.007199525 | 0.932598 | 0.960099 | 543.2694 | 537.52 | 484.4424 | 550.3538 | 558.7388 | 518.4616 | 526.5019 | 519.8685 |
| ENSMUSG00000046756 | -0.388109381 | 0.000124 | 0.001575 | 1298.816 | 1388.194 | 1389.563 | 1251.667 | 1771.777 | 1499.516 | 1708.72 | 1991.379 |
| ENSMUSG00000046761 | -0.491150518 | 0.01921 | 0.068496 | 221.332 | 182.3729 | 316.3367 | 203.5791 | 317.8863 | 263.7003 | 318.2154 | 401.3211 |
| ENSMUSG00000046768 | 0.086905772 | 0.450509 | 0.612944 | 1097.605 | 1520.174 | 1486.451 | 1356.044 | 1271.545 | 1369.901 | 1256.469 | 1243.117 |
| ENSMUSG00000046774 | -1.754232904 | NA | NA | 3.018163 | 7.198929 | 3.312427 | 7.763611 | 0.975111 | 67.04245 | 3.857157 | 0 |
| ENSMUSG00000046782 | 0.450065432 | 0.381873 | 0.549831 | 29.17558 | 8.398751 | 38.09291 | 18.97772 | 21.45245 | 20.11273 | 17.3572 | 10.87591 |
| ENSMUSG00000046785 | 0.104613542 | 0.171906 | 0.319177 | 2255.574 | 2030.098 | 2001.534 | 2119.466 | 2007.754 | 2112.954 | 1873.614 | 1824.978 |
| ENSMUSG00000046791 | -0.232705208 | 0.093566 | 0.210593 | 286.7255 | 349.1481 | 366.8512 | 368.3402 | 430.0241 | 332.9775 | 445.5016 | 402.4087 |
| ENSMUSG00000046792 | -0.351350918 | 0.011939 | 0.048816 | 902.4308 | 891.4674 | 1127.053 | 954.0616 | 1362.23 | 956.4722 | 1275.755 | 1350.788 |
| ENSMUSG00000046793 | 0.835405491 | 0.063498 | 0.161378 | 18.10898 | 15.59768 | 13.24971 | 25.01608 | 8.776002 | 12.29112 | 8.678602 | 10.87591 |
| ENSMUSG00000046794 | 0.804163666 | 0.001868 | 0.012502 | 818.9282 | 859.0722 | 501.8326 | 668.5332 | 403.6961 | 608.9689 | 345.2155 | 272.9854 |
| ENSMUSG00000046798 | -0.433958824 | 5.40E-05 | 0.00084 | 871.2431 | 827.8769 | 847.1531 | 838.47 | 1147.706 | 960.9417 | 1135.933 | 1327.949 |
| ENSMUSG00000046805 | 0.059667167 | 0.804199 | 0.880231 | 198.1927 | 256.7618 | 276.5876 | 156.1349 | 232.0765 | 143.0239 | 236.2508 | 239.27 |
| ENSMUSG00000046806 | -0.127631463 | 0.424663 | 0.589675 | 182.0958 | 149.9777 | 181.3554 | 134.5693 | 185.2711 | 154.1976 | 180.3221 | 188.1533 |
| ENSMUSG00000046807 | 0.070203094 | 0.743833 | 0.839833 | 268.6165 | 147.578 | 219.4483 | 189.7772 | 179.4205 | 168.7235 | 195.7507 | 243.6204 |
| ENSMUSG00000046808 | -0.141381791 | 0.33265 | 0.500725 | 177.0656 | 169.1748 | 135.8095 | 182.0136 | 172.5947 | 181.0146 | 194.7864 | 182.7153 |
| ENSMUSG00000046811 | -0.150102188 | 0.832434 | 0.898707 | 8.048435 | 2.399643 | 21.53077 | 13.80198 | 23.40267 | 6.704245 | 12.53576 | 8.700729 |
| ENSMUSG00000046814 | -0.988753014 | 0.007156 | 0.033419 | 47.28455 | 25.19625 | 65.42042 | 52.62003 | 94.5858 | 55.8687 | 88.7146 | 141.3868 |
| ENSMUSG00000046818 | 0.268787988 | 0.144223 | 0.282297 | 199.1988 | 255.562 | 289.8373 | 256.1992 | 160.8934 | 194.4231 | 239.1437 | 237.0949 |
| ENSMUSG00000046822 | -0.026590946 | 0.816738 | 0.88878 | 733.4136 | 749.8885 | 780.9046 | 756.5208 | 713.7815 | 668.1897 | 771.4313 | 924.4524 |
| ENSMUSG00000046826 | -0.607883143 | 0.020727 | 0.072244 | 43.26034 | 68.38983 | 57.13936 | 56.07053 | 77.03379 | 63.69032 | 84.85745 | 116.3722 |
| ENSMUSG00000046834 | 0.502231418 | 0.27393 | 0.439753 | 18.10898 | 28.79572 | 29.81184 | 60.38364 | 35.10401 | 16.76061 | 21.21436 | 23.927 |
| ENSMUSG00000046836 | 0.017311373 | 0.808934 | 0.883359 | 1817.94 | 1612.56 | 1663.666 | 1720.071 | 1768.852 | 1574.38 | 1745.363 | 1643.35 |
| ENSMUSG00000046840 | -3.816258746 | 0.00039 | 0.00379 | 0 | 1.199822 | 4.140533 | 1.725247 | 25.35289 | 1.117374 | 26.03581 | 50.02919 |
| ENSMUSG00000046841 | 0.251172755 | 0.01511 | 0.057966 | 3233.459 | 3861.026 | 3722.339 | 3428.928 | 2689.357 | 3466.094 | 2816.689 | 2998.489 |
| ENSMUSG00000046844 | 0.832089399 | 0.000327 | 0.003313 | 389.343 | 268.76 | 250.0882 | 452.0147 | 215.4996 | 221.2401 | 175.5006 | 152.2628 |
| ENSMUSG00000046861 | -0.411796895 | 0.001603 | 0.011078 | 727.3773 | 578.314 | 623.5643 | 659.0443 | 861.9984 | 703.9457 | 876.5389 | 1001.671 |
| ENSMUSG00000046862 | 0.121204896 | 0.26516 | 0.429317 | 646.893 | 687.4977 | 628.5329 | 722.0159 | 588.9672 | 690.5372 | 654.7523 | 534.0072 |
| ENSMUSG00000046865 | -0.319941288 | 0.003122 | 0.018309 | 3977.939 | 4584.518 | 4447.761 | 4093.148 | 5796.062 | 4364.463 | 5405.805 | 5780.547 |
| ENSMUSG00000046873 | 0.115888362 | 0.305651 | 0.474161 | 987.9454 | 812.2792 | 746.1241 | 879.0133 | 816.1682 | 846.9696 | 711.6454 | 786.3284 |
| ENSMUSG00000046876 | 0.287287241 | 0.044543 | 0.125719 | 1104.648 | 878.2694 | 1116.288 | 1212.849 | 799.5913 | 1082.735 | 864.0031 | 790.6787 |
| ENSMUSG00000046879 | 0.283349836 | 0.087809 | 0.201617 | 745.4863 | 970.6556 | 988.7593 | 670.2585 | 704.0304 | 556.4523 | 779.1456 | 731.9488 |
| ENSMUSG00000046897 | 0.096660483 | 0.195849 | 0.350014 | 4930.672 | 4738.095 | 4515.665 | 5004.942 | 4602.525 | 4898.568 | 4302.658 | 4142.634 |
| ENSMUSG00000046908 | 0.257557966 | 0.602608 | 0.740217 | 31.18769 | 14.39786 | 16.56213 | 9.488858 | 11.70134 | 12.29112 | 18.32149 | 17.40146 |
| ENSMUSG00000046909 | -0.320155226 | 0.01552 | 0.059131 | 306.8466 | 359.9465 | 380.9291 | 344.1868 | 484.6303 | 355.325 | 442.6087 | 454.6131 |
| ENSMUSG00000046916 | -0.38665559 | 0.066852 | 0.166901 | 176.0595 | 160.7761 | 266.6503 | 161.3106 | 251.5787 | 224.5922 | 221.7865 | 304.5255 |
| ENSMUSG00000046947 | 0.057131671 | 0.574441 | 0.716825 | 458.7608 | 517.1231 | 419.022 | 473.5803 | 430.0241 | 446.9496 | 483.1089 | 431.7737 |
| ENSMUSG00000046949 | 0.087892301 | 0.277848 | 0.444068 | 527.1725 | 551.9179 | 500.1764 | 528.7882 | 500.2321 | 511.7573 | 470.5731 | 499.2043 |
| ENSMUSG00000046959 | -2.012515508 | 7.77E-07 | 3.03E-05 | 85.51462 | 47.99286 | 104.3414 | 62.97152 | 322.7618 | 120.6764 | 256.5009 | 516.6058 |
| ENSMUSG00000046961 | 1.187980137 | 1.14E-07 | 6.24E-06 | 163.9869 | 143.9786 | 102.6852 | 170.7995 | 72.15824 | 56.98608 | 65.57166 | 59.81751 |
| ENSMUSG00000046962 | 0.041438606 | 0.685618 | 0.800011 | 496.9909 | 454.7324 | 439.7246 | 502.0469 | 497.3068 | 408.9589 | 474.4303 | 457.8758 |
| ENSMUSG00000046982 | 0.277341303 | 0.177298 | 0.326339 | 2174.083 | 2615.611 | 2008.987 | 2378.253 | 1655.739 | 2838.13 | 1594.934 | 1483.474 |
| ENSMUSG00000046985 | -0.081151216 | 0.351119 | 0.519875 | 1478.9 | 1416.989 | 1366.376 | 1550.134 | 1461.692 | 1657.066 | 1638.327 | 1392.117 |
| ENSMUSG00000046994 | -0.268412158 | 0.043241 | 0.123138 | 339.0403 | 273.5593 | 255.8849 | 263.9628 | 310.0854 | 336.3296 | 340.3941 | 376.3065 |
| ENSMUSG00000046997 | -0.080861408 | 0.456706 | 0.618506 | 962.794 | 934.661 | 1149.412 | 1064.477 | 1038.494 | 1001.167 | 1063.611 | 1249.642 |
| ENSMUSG00000046999 | -0.752792302 | 0.000198 | 0.002246 | 110.666 | 85.18733 | 112.6225 | 108.6906 | 177.4703 | 163.1366 | 137.8934 | 227.3065 |
| ENSMUSG00000047003 | -0.257415296 | 0.008714 | 0.038558 | 489.9485 | 567.5156 | 597.0649 | 546.0407 | 715.7317 | 619.0252 | 632.5737 | 663.4306 |
| ENSMUSG00000047013 | 0.045637468 | 0.90377 | 0.942474 | 25.15136 | 25.19625 | 34.78048 | 27.60395 | 28.27823 | 34.6386 | 13.50005 | 33.71532 |
| ENSMUSG00000047022 | -0.124429681 | 0.364763 | 0.533444 | 335.0161 | 239.9643 | 274.9314 | 279.49 | 326.6623 | 288.2825 | 333.6441 | 282.7737 |
| ENSMUSG00000047030 | -0.250647375 | 0.033134 | 0.101547 | 443.67 | 401.9402 | 483.6143 | 526.2003 | 510.9583 | 518.4616 | 583.3949 | 599.2627 |
| ENSMUSG00000047033 | 0.338656263 | 0.225878 | 0.385608 | 41.24823 | 41.99375 | 47.20208 | 31.91707 | 28.27823 | 39.10809 | 31.82154 | 29.36496 |
| ENSMUSG00000047034 | 0.053393392 | 0.887593 | 0.932533 | 16.09687 | 33.595 | 19.04645 | 22.42821 | 23.40267 | 23.46486 | 16.39292 | 23.927 |
| ENSMUSG00000047036 | 0.24295158 | 0.017847 | 0.065262 | 4430.663 | 4369.75 | 3707.433 | 4403.693 | 3489.923 | 4148.81 | 3336.441 | 3314.978 |
| ENSMUSG00000047037 | 0.228110009 | 0.086224 | 0.199042 | 190.1443 | 200.3702 | 169.7619 | 200.1287 | 153.0925 | 162.0192 | 155.2506 | 178.3649 |
| ENSMUSG00000047040 | -0.978352333 | 0.000427 | 0.004053 | 281.6952 | 214.7681 | 409.0847 | 258.787 | 490.481 | 358.6771 | 621.0022 | 826.5692 |

| | | | | | | | | | | | |
|---|---|---|---|---|---|---|---|---|---|---|---|
| ENSMUSG00000047044 | 0.074223886 | 0.610286 | 0.7453 | 605.6447 | 542.3193 | 413.2252 | 428.7239 | 445.6259 | 479.3535 | 497.5732 | 464.4014 |
| ENSMUSG00000047045 | 0.12368466 | 0.081364 | 0.191131 | 3255.592 | 3039.148 | 3493.782 | 3364.232 | 2939.961 | 2857.126 | 3055.832 | 3224.708 |
| ENSMUSG00000047067 | -0.378156415 | 0.002183 | 0.013977 | 248.4954 | 235.165 | 234.3542 | 238.9467 | 308.1352 | 260.3482 | 342.3227 | 331.7153 |
| ENSMUSG00000047084 | -0.014391632 | 0.851822 | 0.9109 | 862.1886 | 921.4629 | 822.3099 | 885.9143 | 841.5211 | 949.768 | 854.3602 | 879.8612 |
| ENSMUSG00000047085 | 1.412181961 | 8.09E-11 | 1.52E-08 | 367.2098 | 361.1463 | 353.6015 | 421.8229 | 88.73513 | 189.9536 | 131.1433 | 156.6131 |
| ENSMUSG00000047090 | 0.15866817 | 0.114311 | 0.241803 | 505.0393 | 545.9188 | 571.3936 | 537.4144 | 491.4561 | 422.3674 | 528.4305 | 491.5912 |
| ENSMUSG00000047098 | -0.052381299 | 0.641871 | 0.768381 | 674.0564 | 682.6985 | 669.9383 | 570.1941 | 694.2793 | 570.9782 | 735.7526 | 688.4452 |
| ENSMUSG00000047108 | 0.53575413 | 0.393215 | 0.560681 | 23.13925 | 28.79572 | 8.281066 | 18.97772 | 6.825779 | 26.81698 | 5.785735 | 15.22628 |
| ENSMUSG00000047109 | -2.46302389 | NA | NA | 7.04238 | 2.399643 | 2.48432 | 2.58787 | 4.875557 | 3.352122 | 62.6788 | 8.700729 |
| ENSMUSG00000047115 | 0.285546015 | 0.112964 | 0.239914 | 221.332 | 195.5709 | 231.8699 | 187.1893 | 171.6196 | 185.4841 | 203.465 | 125.073 |
| ENSMUSG00000047117 | 0.798352141 | 1.40E-06 | 4.89E-05 | 149.9021 | 152.3773 | 130.8408 | 165.6237 | 81.90935 | 84.92043 | 97.39321 | 79.39415 |
| ENSMUSG00000047123 | -0.250353876 | 0.12837 | 0.259923 | 359.1614 | 295.1561 | 402.4598 | 296.7425 | 382.2436 | 327.3906 | 422.3587 | 479.6277 |
| ENSMUSG00000047126 | -0.184941732 | 0.018116 | 0.065906 | 6587.644 | 6326.659 | 7288.995 | 6649.965 | 7226.55 | 7118.79 | 7884.992 | 8300.495 |
| ENSMUSG00000047139 | -0.145404349 | 0.020599 | 0.071986 | 13304.06 | 14165.09 | 14804.89 | 15158.88 | 15773.4 | 15418.65 | 16799.85 | 15534.06 |
| ENSMUSG00000047141 | 0.200160298 | 0.124593 | 0.254706 | 814.904 | 703.0954 | 698.0939 | 729.7795 | 618.2206 | 784.3966 | 622.9308 | 539.4452 |
| ENSMUSG00000047143 | -0.463205513 | 0.231823 | 0.392418 | 36.21796 | 21.59679 | 18.21835 | 28.46658 | 37.05423 | 52.51658 | 22.17865 | 32.62773 |
| ENSMUSG00000047146 | 0.601863 | 1.05E-05 | 0.000238 | 952.7335 | 1076.24 | 857.0904 | 871.2497 | 621.1459 | 755.3449 | 536.1448 | 562.2846 |
| ENSMUSG00000047153 | -0.013034499 | 0.914546 | 0.949313 | 760.5771 | 773.8849 | 632.6735 | 641.7919 | 815.1931 | 625.7295 | 708.7525 | 679.7444 |
| ENSMUSG00000047155 | -0.2098173 | 0.504044 | 0.65977 | 54.32694 | 33.595 | 25.67131 | 50.03216 | 55.58134 | 45.81234 | 46.28588 | 41.32846 |
| ENSMUSG00000047161 | -0.413636448 | 0.073002 | 0.177713 | 71.42986 | 68.38983 | 53.82693 | 56.93315 | 61.43201 | 94.9768 | 81.00029 | 95.70802 |
| ENSMUSG00000047180 | 0.346115583 | 0.000825 | 0.006722 | 739.45 | 680.2988 | 804.0915 | 730.6421 | 569.465 | 565.3913 | 528.4305 | 664.5182 |
| ENSMUSG00000047181 | 0.292850576 | 3.06E-05 | 0.000545 | 1464.815 | 1731.342 | 1608.183 | 1622.595 | 1320.301 | 1298.389 | 1313.362 | 1311.635 |
| ENSMUSG00000047182 | 0.419716019 | 0.383472 | 0.55141 | 11.0666 | 33.595 | 27.32752 | 14.6646 | 10.72622 | 18.99536 | 16.39292 | 18.48905 |
| ENSMUSG00000047187 | -0.054834551 | 0.229133 | 0.389221 | 5791.855 | 5821.534 | 5795.09 | 5956.415 | 6012.536 | 5793.585 | 6221.594 | 6241.685 |
| ENSMUSG00000047193 | 0.254941838 | 0.026842 | 0.087449 | 1286.744 | 1210.62 | 1111.319 | 1212.849 | 1025.817 | 1167.656 | 1015.396 | 830.9196 |
| ENSMUSG00000047205 | 0.024500938 | 0.838302 | 0.902245 | 341.0524 | 333.5504 | 353.6015 | 362.3019 | 292.5334 | 325.1559 | 359.6799 | 391.5328 |
| ENSMUSG00000047213 | 0.06652368 | 0.343781 | 0.512506 | 3787.795 | 3856.226 | 3778.651 | 4093.148 | 3408.014 | 4020.312 | 3759.763 | 3631.467 |
| ENSMUSG00000047215 | -0.092442436 | 0.193126 | 0.34636 | 24051.74 | 26745.22 | 26450.55 | 26896.6 | 29907.64 | 26942.12 | 28467.74 | 25716.09 |
| ENSMUSG00000047216 | 0.925923564 | 0.334373 | 0.50258 | 29.17558 | 1.199822 | 1.656213 | 31.91707 | 9.751113 | 6.704245 | 9.642892 | 7.613138 |
| ENSMUSG00000047220 | 0.288582443 | 0.362591 | 0.531109 | 53.32088 | 131.9804 | 123.3879 | 69.8725 | 84.83468 | 67.04245 | 79.07171 | 78.30656 |
| ENSMUSG00000047221 | -0.164060034 | 0.25061 | 0.41419 | 244.4712 | 311.9536 | 245.9477 | 251.8861 | 268.1556 | 348.6207 | 288.3225 | 274.073 |
| ENSMUSG00000047227 | 0.332307738 | 0.264873 | 0.429083 | 106.6418 | 119.9822 | 94.40416 | 134.5693 | 87.76002 | 141.9065 | 78.10742 | 54.37955 |
| ENSMUSG00000047228 | 1.28922013 | 0.000571 | 0.005078 | 53.32088 | 64.79036 | 47.20208 | 56.93315 | 30.22845 | 32.40385 | 18.32149 | 9.78832 |
| ENSMUSG00000047230 | -3.099248609 | 4.91E-06 | 0.000133 | 269.6226 | 62.39072 | 577.1903 | 329.5222 | 2871.703 | 1093.909 | 2568.866 | 4087.167 |
| ENSMUSG00000047238 | 0.280915671 | 0.00014 | 0.001734 | 1369.24 | 1466.182 | 1551.872 | 1535.47 | 1173.059 | 1298.389 | 1233.326 | 1173.511 |
| ENSMUSG00000047242 | -0.108005807 | 0.59246 | 0.732205 | 174.0474 | 159.5763 | 141.6062 | 170.7995 | 165.7689 | 239.1181 | 139.8219 | 152.2628 |
| ENSMUSG00000047246 | 0.182197607 | 0.570718 | 0.713976 | 22.1332 | 34.79482 | 33.12427 | 31.05445 | 34.1289 | 22.34748 | 27.0001 | 22.83941 |
| ENSMUSG00000047248 | 0.09958014 | 0.195599 | 0.349634 | 1270.647 | 1360.598 | 1160.177 | 1191.283 | 1149.656 | 1216.82 | 1157.147 | 1121.306 |
| ENSMUSG00000047250 | -0.046608715 | 0.814973 | 0.887383 | 894.3823 | 1046.244 | 1751.446 | 1163.679 | 1199.387 | 1140.839 | 1440.648 | 1236.591 |
| ENSMUSG00000047259 | -0.058199395 | 0.920002 | 0.9524 | 23.13925 | 7.198929 | 9.109173 | 10.35148 | 14.62667 | 11.17374 | 19.28578 | 6.525547 |
| ENSMUSG00000047260 | -0.406906203 | 4.98E-05 | 0.000792 | 1211.289 | 1125.433 | 1303.44 | 1195.596 | 1674.266 | 1366.549 | 1603.613 | 1769.511 |
| ENSMUSG00000047261 | 1.869615794 | 2.28E-05 | 0.000436 | 581.4994 | 170.3747 | 209.511 | 835.8822 | 107.2622 | 124.0285 | 120.5361 | 140.2993 |
| ENSMUSG00000047264 | -0.034343926 | 0.6563 | 0.778499 | 1134.829 | 1193.822 | 1146.1 | 1317.226 | 1273.495 | 1166.539 | 1254.54 | 1212.664 |
| ENSMUSG00000047281 | -1.164851614 | 0.003023 | 0.017883 | 137.8294 | 111.5834 | 331.2427 | 121.6299 | 438.8001 | 193.3057 | 452.2516 | 492.6788 |
| ENSMUSG00000047284 | -0.081860059 | 0.491569 | 0.648724 | 1342.077 | 1306.606 | 1214.832 | 1241.315 | 1482.169 | 1074.914 | 1332.648 | 1510.664 |
| ENSMUSG00000047298 | 0.154895221 | 0.434861 | 0.59894 | 68.4117 | 85.18733 | 85.29498 | 94.02596 | 78.98402 | 84.92043 | 69.42882 | 66.34306 |
| ENSMUSG00000047307 | 0.02908714 | 0.949807 | 0.971602 | 20.12109 | 26.39607 | 18.21835 | 24.15346 | 23.40267 | 23.46486 | 7.714313 | 32.62773 |
| ENSMUSG00000047330 | -0.592550234 | 0.001215 | 0.009039 | 85.51462 | 79.18822 | 86.12309 | 70.73513 | 106.2871 | 107.2679 | 133.0719 | 138.1241 |
| ENSMUSG00000047342 | 0.363486205 | 0.004327 | 0.02316 | 472.8455 | 543.5192 | 426.4749 | 462.3662 | 364.6916 | 426.8369 | 366.4299 | 320.8394 |
| ENSMUSG00000047343 | 0.509794579 | 0.284212 | 0.450792 | 12.07265 | 11.99822 | 10.76539 | 13.80198 | 8.776002 | 10.05637 | 6.750024 | 8.700729 |
| ENSMUSG00000047344 | 1.197162494 | 5.68E-09 | 5.30E-07 | 135.8173 | 109.1838 | 104.3414 | 110.4158 | 52.65601 | 44.69496 | 58.82164 | 43.50364 |
| ENSMUSG00000047361 | 0.81254279 | 0.027699 | 0.089348 | 132.7992 | 61.1909 | 110.9663 | 92.30071 | 52.65601 | 93.85942 | 32.78583 | 47.85401 |
| ENSMUSG00000047363 | 0.402027323 | 0.268302 | 0.432776 | 29.17558 | 25.19625 | 28.98373 | 25.8787 | 27.30312 | 20.11273 | 11.57147 | 23.927 |
| ENSMUSG00000047368 | 0.01700012 | 0.903868 | 0.942474 | 1165.011 | 1233.417 | 1002.837 | 1284.446 | 996.5638 | 1393.365 | 1242.004 | 998.4086 |
| ENSMUSG00000047369 | 0.27133332 | 0.42192 | 0.587229 | 59.35721 | 40.79393 | 60.45178 | 63.83414 | 33.15378 | 75.98144 | 34.71441 | 43.50364 |
| ENSMUSG00000047370 | -0.019899354 | 0.96922 | 0.982853 | 13.07871 | 11.99822 | 19.87456 | 12.93935 | 27.30312 | 10.05637 | 11.57147 | 9.78832 |
| ENSMUSG00000047371 | -0.252304028 | 0.038662 | 0.113652 | 900.4186 | 977.8546 | 1014.431 | 941.9849 | 1127.229 | 909.5425 | 1230.433 | 1299.671 |
| ENSMUSG00000047379 | -0.267565482 | 0.022305 | 0.076182 | 889.3521 | 910.6645 | 1017.743 | 986.8413 | 1167.208 | 925.1858 | 1214.04 | 1274.657 |
| ENSMUSG00000047384 | 1.318481794 | 0.019826 | 0.070043 | 18.10898 | 25.19625 | 19.04645 | 34.50494 | 2.925334 | 18.99536 | 9.642892 | 7.613138 |
| ENSMUSG00000047388 | -0.211098474 | 0.001871 | 0.012515 | 872.2491 | 903.4656 | 952.3226 | 854.8599 | 1024.842 | 1035.806 | 1018.289 | 1070.19 |
| ENSMUSG00000047407 | 0.399928243 | 0.002571 | 0.015735 | 1963.818 | 1919.714 | 1435.937 | 1647.611 | 1404.16 | 1488.342 | 1230.433 | 1155.022 |
| ENSMUSG00000047409 | -0.032609163 | 0.705525 | 0.814739 | 2379.319 | 2402.043 | 2143.968 | 2338.572 | 2253.482 | 2624.712 | 2456.045 | 2139.292 |
| ENSMUSG00000047412 | 0.295210958 | 0.071941 | 0.175931 | 1550.33 | 1568.167 | 1319.174 | 1638.122 | 1145.756 | 1670.474 | 1133.04 | 1002.759 |
| ENSMUSG00000047414 | 0.51573411 | 0.000242 | 0.002642 | 1484.936 | 2014.5 | 1891.396 | 1673.49 | 1004.365 | 1401.187 | 1182.219 | 1354.051 |

| | | | | | | | | | | | |
|---|---|---|---|---|---|---|---|---|---|---|---|
| ENSMUSG00000047415 | -0.126302811 | 0.768257 | 0.856029 | 21.12714 | 32.39518 | 12.4216 | 15.52722 | 21.45245 | 15.64324 | 22.17865 | 28.27737 |
| ENSMUSG00000047417 | -0.132160231 | 0.091796 | 0.208003 | 1181.108 | 1077.44 | 1280.253 | 1180.932 | 1372.957 | 1273.806 | 1238.147 | 1293.146 |
| ENSMUSG00000047419 | 0.455217917 | 0.147403 | 0.286613 | 47.28455 | 38.39429 | 28.98373 | 38.81806 | 33.15378 | 33.52122 | 22.17865 | 22.83941 |
| ENSMUSG00000047420 | 1.384016749 | 0.083016 | 0.193881 | 50.30272 | 76.78858 | 19.04645 | 52.62003 | 5.850668 | 58.10345 | 4.821446 | 7.613138 |
| ENSMUSG00000047423 | 0.031578237 | 0.695196 | 0.807562 | 1174.065 | 1295.807 | 1387.907 | 1274.958 | 1216.939 | 1185.534 | 1286.362 | 1334.474 |
| ENSMUSG00000047428 | -0.883378172 | 0.189891 | 0.342466 | 13.07871 | 8.398751 | 14.07781 | 8.626235 | 31.20356 | 3.352122 | 33.75012 | 13.05109 |
| ENSMUSG00000047434 | -0.314012547 | 0.063801 | 0.161968 | 770.6376 | 797.8813 | 1164.318 | 783.2621 | 1053.12 | 908.4251 | 1124.361 | 1287.708 |
| ENSMUSG00000047443 | 0.615382917 | 0.037775 | 0.112085 | 49.29666 | 41.99375 | 38.09291 | 34.50494 | 19.50223 | 31.28647 | 28.92867 | 27.18978 |
| ENSMUSG00000047446 | 0.251796329 | 0.036328 | 0.109187 | 1501.033 | 1385.794 | 1835.912 | 1601.892 | 1127.229 | 1398.952 | 1340.362 | 1449.759 |
| ENSMUSG00000047454 | -0.267966255 | 0.028394 | 0.090901 | 579.4873 | 629.9063 | 539.9255 | 579.683 | 646.4988 | 615.6731 | 696.2168 | 843.9707 |
| ENSMUSG00000047459 | -0.042920129 | 0.353505 | 0.522236 | 3463.845 | 3561.07 | 3624.623 | 3486.724 | 3780.507 | 3574.48 | 3581.37 | 3626.029 |
| ENSMUSG00000047466 | -0.166458282 | 0.225365 | 0.385123 | 343.0645 | 308.3541 | 373.4761 | 349.3625 | 388.0943 | 308.3953 | 406.93 | 440.4744 |
| ENSMUSG00000047473 | 0.027830624 | 0.837682 | 0.901788 | 617.7174 | 526.7217 | 428.9592 | 504.6347 | 448.5512 | 554.2176 | 510.109 | 524.2189 |
| ENSMUSG00000047485 | 0.552939191 | 0.167519 | 0.313498 | 23.13925 | 16.7975 | 28.15563 | 28.46658 | 12.67645 | 13.40849 | 15.42863 | 25.0146 |
| ENSMUSG00000047496 | -0.082910992 | 0.687806 | 0.801753 | 172.0353 | 172.7743 | 257.5412 | 193.2277 | 175.52 | 179.8972 | 221.7865 | 267.5474 |
| ENSMUSG00000047497 | 0.120560522 | 0.425797 | 0.590777 | 347.0888 | 356.347 | 372.648 | 358.8514 | 415.3974 | 273.7567 | 343.2869 | 286.0365 |
| ENSMUSG00000047501 | -0.262910863 | 0.217375 | 0.375825 | 1906.473 | 1656.954 | 2989.465 | 1864.129 | 2628.9 | 1829.141 | 2522.58 | 3121.386 |
| ENSMUSG00000047502 | -0.947800895 | 0.0032 | 0.01862 | 32.19374 | 19.19714 | 25.67131 | 23.29083 | 49.73068 | 46.92971 | 33.75012 | 64.16787 |
| ENSMUSG00000047507 | 0.102191242 | 0.758334 | 0.849288 | 42.25428 | 55.19179 | 89.43552 | 48.30692 | 62.40712 | 35.75597 | 54.00019 | 67.43065 |
| ENSMUSG00000047514 | 0.08380064 | 0.373733 | 0.542513 | 1841.079 | 1720.544 | 1739.024 | 2036.654 | 1707.42 | 1531.92 | 1827.328 | 1857.606 |
| ENSMUSG00000047515 | -0.199255457 | 0.583144 | 0.723773 | 26.15741 | 56.39161 | 28.15563 | 31.91707 | 27.30312 | 53.63396 | 41.46443 | 40.24087 |
| ENSMUSG00000047534 | 0.255990153 | 0.11929 | 0.248293 | 1139.86 | 1065.442 | 790.0137 | 1086.906 | 857.1228 | 1073.797 | 762.7527 | 724.3357 |
| ENSMUSG00000047539 | -0.076693552 | 0.55389 | 0.700898 | 1232.417 | 1093.037 | 950.6664 | 969.5888 | 1177.934 | 1238.05 | 1139.79 | 919.0145 |
| ENSMUSG00000047547 | -0.574419116 | 0.000504 | 0.004619 | 889.3521 | 855.4728 | 984.6188 | 818.6297 | 1306.649 | 944.1811 | 1372.183 | 1660.752 |
| ENSMUSG00000047554 | -0.089020891 | 0.257655 | 0.42138 | 954.7456 | 923.8626 | 855.4342 | 974.7645 | 1030.693 | 912.8946 | 997.075 | 1001.671 |
| ENSMUSG00000047557 | -0.031279388 | 0.785156 | 0.867819 | 381.2946 | 437.9349 | 423.1625 | 449.4268 | 497.3068 | 433.5411 | 395.3586 | 402.4087 |
| ENSMUSG00000047562 | -0.594848713 | 0.489804 | 0.647274 | 9.054489 | 11.99822 | 13.24971 | 5.175741 | 5.850668 | 44.69496 | 3.857157 | 5.437955 |
| ENSMUSG00000047583 | -0.50150665 | 0.004313 | 0.023105 | 338.0343 | 380.3434 | 301.4308 | 297.6051 | 425.1485 | 423.4848 | 392.4657 | 622.1021 |
| ENSMUSG00000047586 | -3.442456611 | 0.000267 | 0.002839 | 15.09082 | 1.199822 | 66.24853 | 9.488858 | 270.1058 | 135.2023 | 288.3225 | 312.1386 |
| ENSMUSG00000047592 | -0.087675236 | 0.875064 | 0.925241 | 29.17558 | 7.198929 | 26.49941 | 10.35148 | 19.50223 | 11.17374 | 19.28578 | 28.27737 |
| ENSMUSG00000047603 | 0.112072543 | 0.411568 | 0.577415 | 341.0524 | 291.5566 | 272.4471 | 297.6051 | 307.1601 | 230.1791 | 296.0368 | 277.3357 |
| ENSMUSG00000047604 | -0.286895612 | 0.010842 | 0.045334 | 681.0988 | 653.9027 | 721.2809 | 848.8215 | 940.9824 | 823.5047 | 953.682 | 828.7444 |
| ENSMUSG00000047613 | -0.179927837 | 0.057608 | 0.150422 | 680.0927 | 765.4861 | 731.2182 | 709.9391 | 848.3468 | 707.2978 | 873.646 | 837.4451 |
| ENSMUSG00000047617 | -0.094035345 | 0.367559 | 0.53607 | 413.4883 | 507.5245 | 491.0672 | 439.938 | 511.9334 | 502.8183 | 452.2516 | 508.9926 |
| ENSMUSG00000047632 | 0.163454595 | 0.358453 | 0.526721 | 510.0696 | 368.3452 | 566.4249 | 627.9899 | 541.1868 | 469.2971 | 432.9658 | 410.0218 |
| ENSMUSG00000047635 | -0.119062705 | 0.189096 | 0.341613 | 330.9919 | 333.5504 | 343.6643 | 336.4232 | 354.9405 | 376.5551 | 349.0727 | 381.7445 |
| ENSMUSG00000047638 | -2.192345088 | 0.002658 | 0.016124 | 118.7144 | 8.398751 | 137.4657 | 59.52102 | 365.6667 | 221.2401 | 447.4302 | 449.1751 |
| ENSMUSG00000047641 | -0.309550968 | 0.632023 | 0.760703 | 6.036326 | 5.999108 | 20.70267 | 6.038364 | 16.57689 | 6.704245 | 15.42863 | 9.78832 |
| ENSMUSG00000047648 | 0.128445581 | 0.30777 | 0.476267 | 893.3763 | 794.2819 | 660.001 | 776.3611 | 763.5122 | 778.8097 | 700.0739 | 613.4014 |
| ENSMUSG00000047649 | 0.025654748 | 0.79889 | 0.876929 | 631.8021 | 650.3033 | 563.1125 | 625.402 | 643.5735 | 633.5511 | 526.5019 | 622.1021 |
| ENSMUSG00000047654 | -0.847848927 | 1.46E-05 | 0.000304 | 48.29061 | 45.59322 | 57.96746 | 61.24627 | 94.5858 | 105.0332 | 94.50034 | 92.44524 |
| ENSMUSG00000047656 | -0.060442953 | 0.584059 | 0.724617 | 307.8526 | 297.5557 | 302.2589 | 272.589 | 343.2392 | 294.9868 | 305.6797 | 284.9489 |
| ENSMUSG00000047658 | 0.284343334 | 0.221683 | 0.380835 | 87.52673 | 101.9848 | 88.60741 | 58.6584 | 78.0089 | 71.51194 | 62.6788 | 63.08028 |
| ENSMUSG00000047669 | -0.014842465 | 0.914694 | 0.949364 | 281.6952 | 271.1597 | 239.3228 | 232.9083 | 223.3005 | 273.7567 | 294.1082 | 242.5328 |
| ENSMUSG00000047675 | -0.028311714 | 0.696237 | 0.808286 | 36523.8 | 43116.79 | 39051.02 | 39835.95 | 42529.48 | 41091.43 | 40562.82 | 37480.56 |
| ENSMUSG00000047676 | -0.150010523 | 0.366351 | 0.534907 | 212.2775 | 315.5531 | 279.9 | 249.2982 | 348.1147 | 297.2215 | 259.3938 | 266.4598 |
| ENSMUSG00000047686 | 1.81358642 | 0.009729 | 0.041727 | 128.775 | 220.7672 | 129.1846 | 199.266 | 46.80534 | 121.7938 | 21.21436 | 3.262773 |
| ENSMUSG00000047692 | 0.450550073 | 0.499656 | 0.656019 | 18.10898 | 11.99822 | 8.281066 | 13.80198 | 3.900445 | 22.34748 | 6.750024 | 5.437955 |
| ENSMUSG00000047694 | -0.003606824 | 0.970905 | 0.983692 | 1170.041 | 1034.246 | 997.0404 | 1083.455 | 1127.229 | 1163.186 | 931.5033 | 1074.54 |
| ENSMUSG00000047710 | 0.128750151 | 0.110072 | 0.23589 | 1289.762 | 1478.18 | 1382.938 | 1404.351 | 1175.984 | 1360.962 | 1323.969 | 1219.19 |
| ENSMUSG00000047712 | 0.758996737 | 1.34E-05 | 0.000288 | 588.5418 | 557.917 | 461.2554 | 590.0345 | 298.3841 | 431.3064 | 310.5011 | 258.8467 |
| ENSMUSG00000047714 | 0.047725651 | 0.58634 | 0.726713 | 2804.88 | 2897.569 | 3024.245 | 3245.19 | 2639.626 | 3082.835 | 3142.618 | 2720.065 |
| ENSMUSG00000047719 | -0.604141732 | 9.55E-05 | 0.001288 | 410.4702 | 344.3488 | 392.5225 | 414.9219 | 596.7681 | 452.5365 | 601.7164 | 725.4233 |
| ENSMUSG00000047721 | -0.243106089 | 0.016571 | 0.061674 | 952.7335 | 1010.25 | 1030.993 | 1081.73 | 1299.823 | 1027.984 | 1179.326 | 1317.073 |
| ENSMUSG00000047728 | -2.496457718 | 6.18E-08 | 3.85E-06 | 42.25428 | 21.59679 | 99.3728 | 37.95543 | 248.6534 | 158.6671 | 251.6795 | 482.8904 |
| ENSMUSG00000047730 | -0.273543899 | 0.604111 | 0.741013 | 21.12714 | 20.39697 | 64.59232 | 13.80198 | 39.00445 | 49.16446 | 33.75012 | 23.927 |
| ENSMUSG00000047731 | -0.101385869 | 0.453339 | 0.615723 | 2758.601 | 2117.685 | 2774.157 | 2442.087 | 2640.601 | 2236.983 | 3217.833 | 2733.116 |
| ENSMUSG00000047735 | -0.486281291 | 0.064032 | 0.162442 | 393.3673 | 448.7333 | 630.1891 | 366.615 | 622.121 | 392.1983 | 641.2523 | 921.1897 |
| ENSMUSG00000047746 | -2.474973024 | 1.06E-05 | 0.000238 | 13.07871 | 3.599465 | 22.35888 | 14.6646 | 83.85957 | 29.05173 | 61.71451 | 128.3357 |
| ENSMUSG00000047747 | 0.339153628 | 0.003373 | 0.019279 | 904.4429 | 917.8635 | 795.8105 | 841.0579 | 626.0215 | 811.2136 | 688.5025 | 607.9634 |
| ENSMUSG00000047749 | 0.299924705 | 0.010419 | 0.043953 | 995.9938 | 998.2515 | 913.4016 | 1039.461 | 845.4215 | 943.0637 | 724.1812 | 693.8831 |
| ENSMUSG00000047757 | -0.078877974 | 0.710502 | 0.818711 | 172.0353 | 169.1748 | 161.4808 | 155.2722 | 175.52 | 244.7049 | 135.9648 | 139.2117 |
| ENSMUSG00000047759 | 0.367242706 | 0.187161 | 0.339116 | 263.5862 | 337.1499 | 182.1835 | 210.4801 | 184.296 | 279.3435 | 180.3221 | 125.073 |
| ENSMUSG00000047766 | 0.374203623 | 5.36E-05 | 0.000835 | 790.7587 | 875.8697 | 756.0614 | 845.371 | 633.8224 | 680.4808 | 638.3594 | 566.635 |

| | | | | | | | | | | | |
|---|---|---|---|---|---|---|---|---|---|---|---|
| ENSMUSG00000047767 | -0.207270403 | 0.182037 | 0.332574 | 344.0706 | 268.76 | 248.432 | 275.1769 | 330.5627 | 336.3296 | 270.001 | 375.2189 |
| ENSMUSG00000047777 | 0.160305255 | 0.135075 | 0.269187 | 908.4671 | 981.454 | 878.6211 | 1051.538 | 920.5051 | 940.829 | 795.5386 | 761.3138 |
| ENSMUSG00000047786 | 1.377404263 | 2.07E-09 | 2.38E-07 | 460.7729 | 632.3059 | 568.9093 | 541.7276 | 188.1965 | 325.1559 | 163.9292 | 171.8394 |
| ENSMUSG00000047787 | 0.732856338 | 0.00469 | 0.024612 | 333.004 | 368.3452 | 374.3042 | 319.1707 | 149.192 | 344.1512 | 169.7149 | 177.2773 |
| ENSMUSG00000047789 | 0.392114129 | 0.011169 | 0.046405 | 993.9817 | 740.2899 | 640.9545 | 873.8376 | 625.0463 | 703.9457 | 591.1093 | 555.759 |
| ENSMUSG00000047793 | 0.201041046 | 0.205471 | 0.361918 | 656.9535 | 529.1213 | 823.9661 | 649.5555 | 688.4286 | 576.565 | 523.609 | 527.4817 |
| ENSMUSG00000047797 | -1.355588262 | 1.25E-05 | 0.000272 | 227.3683 | 167.975 | 308.8838 | 238.0841 | 664.0508 | 290.5173 | 577.6092 | 882.0364 |
| ENSMUSG00000047798 | 0.021633675 | 0.970953 | 0.983692 | 57.3451 | 4.799286 | 25.67131 | 40.5433 | 21.45245 | 30.1691 | 40.50014 | 34.80291 |
| ENSMUSG00000047804 | -0.068314592 | 0.38527 | 0.552791 | 582.5055 | 547.1186 | 568.9093 | 541.7276 | 567.5148 | 568.7434 | 589.1807 | 624.2773 |
| ENSMUSG00000047810 | 0.094287001 | 0.848904 | 0.909191 | 39.23612 | 45.59322 | 141.6062 | 31.91707 | 58.50668 | 40.22547 | 64.60737 | 79.39415 |
| ENSMUSG00000047821 | 0.040461423 | 0.699538 | 0.810259 | 555.342 | 683.8983 | 699.7501 | 604.6991 | 635.7726 | 645.8422 | 618.1094 | 573.1605 |
| ENSMUSG00000047822 | -1.433299522 | 0.004966 | 0.025622 | 15.09082 | 15.59768 | 13.24971 | 8.626235 | 38.02934 | 11.17374 | 39.53586 | 52.20437 |
| ENSMUSG00000047824 | 0.134261905 | 0.084599 | 0.196299 | 1479.906 | 1679.75 | 1528.685 | 1658.825 | 1432.439 | 1569.911 | 1400.148 | 1380.153 |
| ENSMUSG00000047832 | -0.082819738 | 0.346776 | 0.515638 | 1373.264 | 1521.374 | 1503.014 | 1622.595 | 1760.076 | 1619.075 | 1513.934 | 1483.474 |
| ENSMUSG00000047842 | 0.286882895 | 0.171959 | 0.319219 | 90.54489 | 134.38 | 88.60741 | 108.6906 | 81.90935 | 102.7984 | 81.96458 | 78.30656 |
| ENSMUSG00000047843 | -0.125542073 | 0.163676 | 0.308553 | 1610.693 | 1846.525 | 2027.205 | 1977.133 | 2097.464 | 2116.307 | 1977.757 | 1952.226 |
| ENSMUSG00000047844 | 0.536453319 | 2.75E-11 | 5.70E-09 | 1356.161 | 1339.001 | 1460.78 | 1457.834 | 932.2064 | 1058.153 | 990.325 | 892.9123 |
| ENSMUSG00000047861 | -1.277355455 | 0.160493 | 0.303969 | 49.29666 | 20.39697 | 382.5853 | 52.62003 | 222.3254 | 179.8972 | 377.0371 | 445.9123 |
| ENSMUSG00000047866 | -0.103820489 | 0.300807 | 0.469207 | 1745.504 | 1589.764 | 1624.745 | 1849.465 | 1803.956 | 1602.314 | 1830.221 | 2082.737 |
| ENSMUSG00000047867 | -0.371197492 | 0.029623 | 0.093727 | 1099.617 | 776.2845 | 1386.251 | 1040.324 | 1331.027 | 1222.407 | 1523.577 | 1492.175 |
| ENSMUSG00000047875 | -0.027314209 | 0.909341 | 0.94599 | 100.6054 | 104.3845 | 89.43552 | 123.3552 | 106.2871 | 135.2023 | 115.7147 | 68.51824 |
| ENSMUSG00000047878 | 0.309732154 | 0.068337 | 0.169337 | 707.2562 | 719.8929 | 823.9661 | 586.584 | 494.3814 | 470.4145 | 594.0021 | 730.8612 |
| ENSMUSG00000047879 | -0.088595854 | 0.232084 | 0.392678 | 1607.675 | 1599.362 | 1491.42 | 1577.738 | 1701.569 | 1800.09 | 1519.72 | 1652.051 |
| ENSMUSG00000047880 | 0.911379629 | 0.116878 | 0.245132 | 51.30877 | 44.3934 | 30.63995 | 42.26855 | 11.70134 | 55.8687 | 11.57147 | 10.87591 |
| ENSMUSG00000047881 | 0.067902479 | 0.397073 | 0.564329 | 1250.526 | 1386.994 | 1276.112 | 1231.826 | 1106.751 | 1287.215 | 1272.862 | 1239.854 |
| ENSMUSG00000047888 | 0.549153385 | 0.011958 | 0.048861 | 2774.698 | 2897.569 | 2290.543 | 2594.771 | 1426.588 | 2782.261 | 1652.792 | 1354.051 |
| ENSMUSG00000047897 | -2.959875007 | 0.000225 | 0.002482 | 2.012109 | 20.39697 | 7.45296 | 10.35148 | 117.0134 | 5.58687 | 91.60747 | 95.70802 |
| ENSMUSG00000047898 | 0.04411892 | 0.949953 | 0.971623 | 8.048435 | 14.39786 | 48.03018 | 12.93935 | 5.850668 | 16.76061 | 33.75012 | 25.0146 |
| ENSMUSG00000047905 | 0.305523048 | 0.421245 | 0.586621 | 40.24217 | 25.19625 | 23.18699 | 21.56559 | 27.30312 | 16.76061 | 27.0001 | 17.40146 |
| ENSMUSG00000047907 | 0.706351262 | 0.005663 | 0.028185 | 1477.894 | 1655.754 | 1171.771 | 1370.709 | 774.2384 | 1425.769 | 672.1095 | 606.8758 |
| ENSMUSG00000047909 | 0.272605024 | 0.03699 | 0.110538 | 2771.68 | 2615.611 | 2359.276 | 2655.155 | 2480.683 | 2453.754 | 1917.007 | 1758.635 |
| ENSMUSG00000047910 | 0.278276457 | 0.217589 | 0.37606 | 110.666 | 140.3791 | 130.0127 | 146.646 | 115.0631 | 147.4934 | 92.57176 | 80.48174 |
| ENSMUSG00000047911 | 0.56391057 | 0.132708 | 0.266107 | 46.2785 | 44.3934 | 29.81184 | 38.81806 | 16.57689 | 43.57759 | 27.0001 | 20.66423 |
| ENSMUSG00000047921 | -0.419643961 | 0.00966 | 0.041508 | 741.4621 | 809.8795 | 932.4481 | 733.23 | 1140.88 | 775.4576 | 1118.575 | 1268.131 |
| ENSMUSG00000047945 | 0.025071224 | 0.743301 | 0.839506 | 10765.79 | 11777.45 | 12327.2 | 11716.15 | 11474.13 | 12624.09 | 10661.18 | 11028.17 |
| ENSMUSG00000047963 | 0.165828036 | 0.179183 | 0.328916 | 751.5226 | 995.8519 | 1017.743 | 1065.34 | 808.3673 | 877.1387 | 902.5747 | 827.6568 |
| ENSMUSG00000047965 | 0.080080886 | 0.801859 | 0.879143 | 40.24217 | 27.5959 | 39.74912 | 46.58167 | 37.05423 | 49.16446 | 36.64299 | 23.927 |
| ENSMUSG00000047976 | 0.610316064 | 0.191122 | 0.343795 | 14.08476 | 11.99822 | 18.21835 | 15.52722 | 6.825779 | 14.52586 | 9.642892 | 8.700729 |
| ENSMUSG00000047986 | -0.675392924 | 0.005899 | 0.029057 | 465.8032 | 299.9554 | 486.9267 | 304.5061 | 554.8383 | 458.1234 | 588.2164 | 887.4743 |
| ENSMUSG00000047988 | 0.809981042 | 0.050103 | 0.136267 | 35.2119 | 44.3934 | 33.95237 | 24.15346 | 28.27823 | 24.58223 | 15.42863 | 9.78832 |
| ENSMUSG00000047989 | -0.052974302 | 0.539378 | 0.689165 | 807.8616 | 889.0678 | 957.2913 | 928.1829 | 975.1113 | 897.2514 | 874.6103 | 972.3064 |
| ENSMUSG00000047990 | 1.021263222 | 0.021301 | 0.073704 | 131.7931 | 129.5807 | 75.3577 | 94.88858 | 41.92979 | 108.3853 | 38.57157 | 23.927 |
| ENSMUSG00000047996 | 0.642258478 | 0.000548 | 0.004911 | 473.8516 | 435.5352 | 396.6631 | 458.0531 | 236.952 | 382.1419 | 217.9294 | 294.7372 |
| ENSMUSG00000048000 | 0.011316143 | 0.915357 | 0.949794 | 2695.22 | 2433.238 | 2417.243 | 2569.755 | 2135.494 | 2400.12 | 2577.545 | 2924.532 |
| ENSMUSG00000048001 | -1.094353429 | 0.076866 | 0.184144 | 9.054489 | 1.199822 | 10.76539 | 6.038364 | 17.552 | 12.29112 | 9.642892 | 19.57664 |
| ENSMUSG00000048003 | 0.106051251 | 0.821901 | 0.891822 | 13.07871 | 21.59679 | 11.59349 | 12.07673 | 14.62667 | 7.821619 | 12.53576 | 18.48905 |
| ENSMUSG00000048004 | 1.419673979 | 0.00016 | 0.001911 | 112.6781 | 77.9884 | 41.40533 | 111.2784 | 35.10401 | 41.34284 | 27.96439 | 23.927 |
| ENSMUSG00000048007 | -0.454194994 | 7.71E-05 | 0.001113 | 250.5075 | 287.9572 | 236.8385 | 265.688 | 340.3138 | 327.3906 | 385.7157 | 368.6934 |
| ENSMUSG00000048012 | -0.044794045 | 0.8603 | 0.91625 | 54.32694 | 46.79304 | 57.96746 | 57.79577 | 77.03379 | 46.92971 | 52.07161 | 47.85401 |
| ENSMUSG00000048015 | 0.142783642 | 0.640901 | 0.767484 | 26.15741 | 29.99554 | 39.74912 | 36.23019 | 27.30312 | 32.40385 | 24.10723 | 36.9781 |
| ENSMUSG00000048022 | -0.502793945 | 0.002382 | 0.014871 | 902.4308 | 844.6744 | 1009.462 | 905.7547 | 1226.69 | 1024.632 | 1188.004 | 1752.109 |
| ENSMUSG00000048027 | -0.03949056 | 0.730438 | 0.830979 | 366.2038 | 327.5513 | 375.1323 | 338.1484 | 337.3885 | 406.7242 | 326.894 | 378.4817 |
| ENSMUSG00000048029 | -1.335479141 | 0.003651 | 0.020442 | 14.08476 | 5.999108 | 4.96864 | 11.21411 | 22.42756 | 20.11273 | 25.07152 | 23.927 |
| ENSMUSG00000048038 | 0.554347146 | 0.496136 | 0.652768 | 20.12109 | 15.59768 | 6.624853 | 34.50494 | 14.62667 | 29.05173 | 7.714313 | 1.087591 |
| ENSMUSG00000048039 | 0.118105048 | 0.348301 | 0.516965 | 1549.324 | 1744.541 | 1446.702 | 1661.413 | 1480.219 | 1793.385 | 1369.291 | 1255.08 |
| ENSMUSG00000048040 | -0.316443668 | 0.030383 | 0.095285 | 471.8395 | 638.3051 | 592.9243 | 540.0023 | 692.329 | 614.5558 | 630.6451 | 854.8466 |
| ENSMUSG00000048047 | 0.157189193 | 0.196046 | 0.35023 | 1040.26 | 1109.835 | 868.6839 | 1223.2 | 995.5886 | 995.5803 | 856.2888 | 955.9926 |
| ENSMUSG00000048058 | 0.035529248 | 0.643072 | 0.769186 | 983.9212 | 1101.436 | 1151.896 | 1123.998 | 1022.892 | 1091.674 | 1092.54 | 1050.613 |
| ENSMUSG00000048070 | 3.147749901 | 0.006928 | 0.032741 | 60.36326 | 0 | 8.281066 | 85.39973 | 7.80089 | 1.117374 | 1.928578 | 6.525547 |
| ENSMUSG00000048076 | -0.2146738 | 0.00893 | 0.039261 | 11151.11 | 11505.09 | 12388.48 | 11591.93 | 13482.86 | 11970.43 | 13671.69 | 14995.71 |
| ENSMUSG00000048078 | 0.408711462 | 0.020708 | 0.072206 | 2662.02 | 2501.628 | 2289.715 | 2568.03 | 1573.83 | 2678.346 | 1757.899 | 1540.029 |
| ENSMUSG00000048087 | -0.421064749 | 0.328975 | 0.496849 | 26.15741 | 10.79839 | 19.87456 | 24.15346 | 39.00445 | 32.40385 | 19.28578 | 18.48905 |
| ENSMUSG00000048096 | 0.325619807 | 0.276412 | 0.442681 | 109.6599 | 141.5789 | 153.1997 | 241.5346 | 133.5902 | 154.1976 | 80.036 | 149 |
| ENSMUSG00000048100 | -0.027112809 | 0.78373 | 0.866969 | 785.7285 | 710.2943 | 619.4238 | 670.2585 | 692.329 | 755.3449 | 681.7524 | 706.9342 |

| | | | | | | | | | | | |
|---|---|---|---|---|---|---|---|---|---|---|---|
| ENSMUSG00000048106 | 0.120806192 | 0.201921 | 0.357508 | 465.8032 | 455.9322 | 433.9279 | 457.1905 | 383.2187 | 449.1844 | 439.7159 | 394.7956 |
| ENSMUSG00000048108 | -2.038508286 | 0.001481 | 0.010472 | 186.1201 | 80.38804 | 443.8652 | 191.5024 | 604.569 | 357.5597 | 1304.683 | 1441.058 |
| ENSMUSG00000048109 | -0.066157333 | 0.627493 | 0.757276 | 439.6458 | 417.5379 | 357.7421 | 357.9887 | 361.7663 | 489.4099 | 385.7157 | 408.9342 |
| ENSMUSG00000048118 | 0.41914975 | 0.015337 | 0.058604 | 1416.525 | 1416.989 | 1031.821 | 1170.58 | 835.6704 | 1230.229 | 931.5033 | 767.8393 |
| ENSMUSG00000048120 | -0.554247593 | 0.015935 | 0.060156 | 439.6458 | 269.9598 | 448.8338 | 405.433 | 597.7432 | 370.9682 | 720.324 | 609.051 |
| ENSMUSG00000048126 | 0.614355253 | 0.001097 | 0.008386 | 2368.252 | 3322.306 | 3020.933 | 2780.236 | 1671.341 | 2655.998 | 1530.327 | 1649.876 |
| ENSMUSG00000048138 | -0.866079624 | 0.33808 | 0.506544 | 22.1332 | 35.99465 | 270.7909 | 25.01608 | 115.0631 | 89.38993 | 204.4293 | 237.0949 |
| ENSMUSG00000048142 | 0.666198603 | 0.014851 | 0.057258 | 118.7144 | 62.39072 | 124.216 | 106.9653 | 60.4569 | 56.98608 | 58.82164 | 84.8321 |
| ENSMUSG00000048148 | -0.153492369 | 0.756301 | 0.848356 | 12.07265 | 19.19714 | 22.35888 | 12.07673 | 15.60178 | 32.40385 | 14.46434 | 10.87591 |
| ENSMUSG00000048153 | -0.114781363 | 0.884286 | 0.930838 | 1.006054 | 8.398751 | 9.93728 | 25.01608 | 8.776002 | 20.11273 | 7.714313 | 11.9635 |
| ENSMUSG00000048154 | 0.396215402 | 0.005589 | 0.027892 | 1855.164 | 1638.956 | 1499.701 | 1507.866 | 1206.213 | 1568.793 | 1060.718 | 1104.993 |
| ENSMUSG00000048163 | 0.14098502 | 0.734862 | 0.834007 | 30.18163 | 50.3925 | 45.54586 | 18.11509 | 38.02934 | 21.23011 | 43.39301 | 27.18978 |
| ENSMUSG00000048170 | 0.024994927 | 0.808833 | 0.883359 | 1935.649 | 1960.508 | 1607.355 | 1702.819 | 1659.639 | 2014.625 | 1725.113 | 1680.328 |
| ENSMUSG00000048174 | 0.406619781 | 0.017585 | 0.064496 | 199.1988 | 183.5727 | 184.6678 | 225.1447 | 142.3663 | 185.4841 | 151.3934 | 119.635 |
| ENSMUSG00000048175 | -0.096753927 | 0.25141 | 0.415028 | 898.4065 | 861.4719 | 954.8069 | 855.7225 | 910.754 | 893.8993 | 966.2177 | 1049.525 |
| ENSMUSG00000048186 | -0.209715967 | 0.192462 | 0.345566 | 296.786 | 214.7681 | 291.4935 | 220.8316 | 279.8569 | 259.2308 | 321.1083 | 325.1897 |
| ENSMUSG00000048191 | -0.138537342 | 0.716727 | 0.822669 | 18.10898 | 20.39697 | 19.87456 | 15.52722 | 17.552 | 13.40849 | 26.03581 | 23.927 |
| ENSMUSG00000048200 | -0.133990188 | 0.383228 | 0.551232 | 210.2654 | 187.1722 | 255.0568 | 225.1447 | 258.4045 | 197.7752 | 245.8937 | 263.197 |
| ENSMUSG00000048216 | 1.832930824 | 1.73E-06 | 5.82E-05 | 82.49646 | 59.99108 | 69.56096 | 106.9653 | 9.751113 | 34.6386 | 25.07152 | 20.66423 |
| ENSMUSG00000048218 | 0.115989443 | 0.165355 | 0.310736 | 726.3712 | 766.686 | 795.8105 | 821.2176 | 688.4286 | 782.1619 | 719.3597 | 683.0072 |
| ENSMUSG00000048222 | 0.133171037 | 0.21797 | 0.376565 | 641.8627 | 699.496 | 639.2983 | 691.824 | 571.4152 | 710.6499 | 614.2522 | 540.5328 |
| ENSMUSG00000048232 | -0.052637809 | 0.599297 | 0.737523 | 732.4076 | 704.2952 | 775.9359 | 708.2139 | 730.3584 | 673.7766 | 866.896 | 758.051 |
| ENSMUSG00000048234 | -0.530135746 | 0.000536 | 0.004837 | 1185.132 | 1173.425 | 1137.819 | 1036.873 | 1617.71 | 1185.534 | 1753.078 | 1988.117 |
| ENSMUSG00000048240 | 0.182912789 | 0.248459 | 0.411889 | 201.2109 | 241.1641 | 190.4645 | 223.4195 | 210.624 | 212.3011 | 175.5006 | 154.4379 |
| ENSMUSG00000048249 | 0.140648111 | 0.473226 | 0.632568 | 1117.726 | 971.8554 | 1006.978 | 1150.74 | 813.2428 | 1292.802 | 1081.932 | 665.6057 |
| ENSMUSG00000048251 | 0.299970785 | 0.10142 | 0.223287 | 149.9021 | 147.578 | 133.3252 | 107.8279 | 92.63557 | 117.3243 | 104.1432 | 122.8978 |
| ENSMUSG00000048264 | 0.181157347 | 0.148816 | 0.288612 | 815.9101 | 680.2988 | 702.2344 | 646.105 | 587.9921 | 748.6406 | 570.8592 | 602.5255 |
| ENSMUSG00000048271 | 0.247791156 | 0.035761 | 0.107826 | 2705.28 | 2559.219 | 2827.156 | 3095.956 | 2288.586 | 2824.722 | 2300.794 | 2010.956 |
| ENSMUSG00000048277 | -0.473590626 | 0.000603 | 0.005288 | 930.6003 | 1051.044 | 1095.585 | 926.4576 | 1400.26 | 1093.909 | 1388.576 | 1675.978 |
| ENSMUSG00000048279 | 0.664521548 | 0.001215 | 0.00904 | 666.008 | 665.901 | 502.6607 | 625.402 | 464.153 | 479.3535 | 351.0013 | 256.6715 |
| ENSMUSG00000048280 | 0.298285021 | 0.01747 | 0.064188 | 389.343 | 431.9358 | 330.4145 | 367.4776 | 289.6081 | 343.0338 | 315.3226 | 284.9489 |
| ENSMUSG00000048281 | 0.185670641 | 0.416508 | 0.582046 | 87.52673 | 115.1829 | 75.3577 | 89.71284 | 82.88446 | 71.51194 | 65.57166 | 102.2336 |
| ENSMUSG00000048285 | 0.188221777 | 0.029982 | 0.09451 | 948.7093 | 923.8626 | 923.3389 | 984.2534 | 803.4917 | 783.2792 | 940.1819 | 789.5911 |
| ENSMUSG00000048304 | 1.116690151 | 0.032363 | 0.099797 | 20.12109 | 14.39786 | 10.76539 | 17.25247 | 10.72622 | 7.821619 | 5.785735 | 4.350364 |
| ENSMUSG00000048307 | -0.019243516 | 0.769448 | 0.856923 | 1754.559 | 1725.343 | 1768.836 | 1870.168 | 1852.711 | 1678.296 | 1896.757 | 1786.912 |
| ENSMUSG00000048310 | -0.109232642 | 0.282579 | 0.449168 | 1384.331 | 1692.948 | 1623.917 | 1531.157 | 1749.35 | 1476.051 | 1850.471 | 1643.35 |
| ENSMUSG00000048327 | 0.053138956 | 0.593494 | 0.732922 | 1288.756 | 1438.586 | 1215.661 | 1385.373 | 1393.434 | 1390.013 | 1185.111 | 1164.81 |
| ENSMUSG00000048329 | -0.824248762 | 0.000363 | 0.003587 | 48.29061 | 51.59233 | 68.73285 | 40.5433 | 79.95913 | 101.681 | 97.39321 | 92.44524 |
| ENSMUSG00000048330 | 0.791689844 | 0.037981 | 0.112482 | 254.5318 | 92.38626 | 71.21717 | 123.3552 | 75.08357 | 78.21619 | 71.3574 | 88.09488 |
| ENSMUSG00000048332 | 0.103090901 | 0.390232 | 0.557613 | 3370.282 | 4447.738 | 4947.937 | 4239.794 | 4207.605 | 3728.677 | 4047.122 | 3850.072 |
| ENSMUSG00000048347 | 0.408786 | 0.232734 | 0.39331 | 81.4904 | 97.18554 | 57.13936 | 86.26235 | 42.9049 | 94.9768 | 67.50024 | 36.9781 |
| ENSMUSG00000048349 | -0.417447741 | 0.458493 | 0.62015 | 18.10898 | 16.7975 | 20.70267 | 8.626235 | 13.65156 | 44.69496 | 12.53576 | 15.22628 |
| ENSMUSG00000048351 | -0.379523885 | 0.00154 | 0.010734 | 460.7729 | 447.5334 | 438.8965 | 405.433 | 619.1957 | 470.4145 | 557.3591 | 630.8028 |
| ENSMUSG00000048355 | 0.023773248 | 0.863516 | 0.918066 | 403.4278 | 441.5343 | 352.7734 | 405.433 | 349.0898 | 470.4145 | 347.1441 | 410.0218 |
| ENSMUSG00000048368 | 1.485888349 | 0.000863 | 0.006961 | 177.0656 | 119.9822 | 63.76421 | 190.6398 | 31.20356 | 89.38993 | 47.25017 | 29.36496 |
| ENSMUSG00000048371 | -0.222546063 | 0.024482 | 0.081706 | 457.7547 | 427.1365 | 427.303 | 434.7622 | 507.0579 | 455.8886 | 569.8949 | 503.5547 |
| ENSMUSG00000048373 | -1.786963815 | 2.07E-06 | 6.59E-05 | 79.47829 | 64.79036 | 163.9651 | 86.26235 | 288.6329 | 172.0756 | 523.609 | 380.6569 |
| ENSMUSG00000048376 | -0.051910254 | 0.747968 | 0.842597 | 6250.616 | 4680.504 | 4373.231 | 5379.32 | 5346.535 | 4887.394 | 6817.524 | 4388.43 |
| ENSMUSG00000048379 | 0.143072142 | 0.283202 | 0.449692 | 984.9272 | 941.8599 | 819.8256 | 934.2212 | 730.3584 | 1042.51 | 798.4314 | 762.4014 |
| ENSMUSG00000048385 | 0.065345815 | 0.845724 | 0.907112 | 21.12714 | 17.99732 | 24.01509 | 21.56559 | 22.42756 | 18.99536 | 19.28578 | 20.66423 |
| ENSMUSG00000048387 | 0.345003298 | 0.098316 | 0.218405 | 827.9827 | 1167.426 | 674.9069 | 948.0232 | 783.9895 | 896.134 | 637.3951 | 529.6569 |
| ENSMUSG00000048388 | 0.507422099 | 0.00194 | 0.012845 | 361.1735 | 394.7413 | 344.4924 | 433.037 | 280.8321 | 340.7991 | 230.4651 | 227.3065 |
| ENSMUSG00000048402 | 0.699583388 | 0.000114 | 0.001489 | 465.8032 | 525.5218 | 642.6107 | 509.8105 | 311.0605 | 443.5975 | 276.751 | 290.3868 |
| ENSMUSG00000048406 | -0.135699021 | 0.337388 | 0.505794 | 183.1019 | 189.5718 | 224.4169 | 210.4801 | 198.9227 | 227.9443 | 255.5366 | 206.6423 |
| ENSMUSG00000048410 | -0.058151605 | 0.617244 | 0.750007 | 514.0938 | 481.1284 | 549.0347 | 480.4813 | 563.6143 | 449.1844 | 503.3589 | 592.7371 |
| ENSMUSG00000048416 | 0.48535647 | 0.05527 | 0.146309 | 118.7144 | 148.7779 | 88.60741 | 121.6299 | 77.03379 | 105.0332 | 99.32178 | 58.72992 |
| ENSMUSG00000048424 | -2.178508572 | 0.001705 | 0.011648 | 45.27245 | 43.19358 | 309.7119 | 76.77349 | 494.3814 | 379.9072 | 645.1095 | 634.0656 |
| ENSMUSG00000048429 | -0.268200485 | 0.000547 | 0.004911 | 1205.253 | 1065.442 | 1254.582 | 1196.459 | 1477.294 | 1331.91 | 1425.219 | 1457.372 |
| ENSMUSG00000048439 | -0.05310085 | 0.699498 | 0.810259 | 269.6226 | 344.3488 | 317.1648 | 274.3143 | 298.3841 | 366.4987 | 289.2867 | 295.8248 |
| ENSMUSG00000048440 | 0.293928319 | 0.000276 | 0.00291 | 776.674 | 772.6851 | 812.3726 | 721.1532 | 645.5237 | 663.7202 | 617.1451 | 588.3868 |
| ENSMUSG00000048442 | -1.41380008 | 0.002558 | 0.015683 | 22.1332 | 13.19804 | 48.03018 | 30.19182 | 69.2329 | 35.75597 | 83.89316 | 116.3722 |
| ENSMUSG00000048445 | 0.14933627 | 0.438175 | 0.601843 | 466.8092 | 371.9447 | 272.4471 | 307.094 | 367.617 | 325.1559 | 316.2868 | 267.5474 |
| ENSMUSG00000048450 | -1.008393887 | 0.063426 | 0.161238 | 15.09082 | 5.999108 | 22.35888 | 13.80198 | 10.72622 | 39.10809 | 28.92867 | 38.06569 |
| ENSMUSG00000048458 | 0.420269658 | 0.038485 | 0.113374 | 106.6418 | 155.9768 | 101.029 | 113.8663 | 85.8098 | 101.681 | 81.00029 | 87.00729 |

| | | | | | | | | | | | |
|---|---|---|---|---|---|---|---|---|---|---|---|
| ENSMUSG00000048478 | -0.005371903 | 0.971771 | 0.984304 | 169.0171 | 177.5736 | 162.3089 | 180.2883 | 171.6196 | 207.8316 | 168.7506 | 143.562 |
| ENSMUSG00000048481 | 0.126449554 | 0.399122 | 0.566587 | 144.8718 | 125.9813 | 134.9814 | 130.2561 | 120.9138 | 128.498 | 108.0004 | 134.8613 |
| ENSMUSG00000048482 | -0.579804346 | 0.104008 | 0.226926 | 27.16347 | 20.39697 | 24.01509 | 18.11509 | 28.27823 | 22.34748 | 36.64299 | 46.76642 |
| ENSMUSG00000048485 | -0.858221508 | 0.013457 | 0.053275 | 70.4238 | 62.39072 | 89.43552 | 75.04824 | 136.5156 | 56.98608 | 150.4291 | 195.7664 |
| ENSMUSG00000048486 | -0.431226206 | 0.031033 | 0.096765 | 371.2341 | 241.1641 | 284.0406 | 288.9789 | 358.841 | 298.3389 | 442.6087 | 499.2043 |
| ENSMUSG00000048489 | -1.013450877 | 0.005035 | 0.025854 | 369.2219 | 88.78679 | 250.0882 | 265.688 | 526.5601 | 356.4423 | 661.5024 | 423.0729 |
| ENSMUSG00000048490 | 0.223952635 | 0.007508 | 0.034582 | 1712.305 | 1593.363 | 1715.009 | 1817.548 | 1300.798 | 1550.915 | 1538.041 | 1469.336 |
| ENSMUSG00000048495 | 0.077742402 | 0.569209 | 0.713058 | 320.9313 | 351.5477 | 297.2903 | 331.2474 | 285.7076 | 378.7898 | 286.3939 | 281.6861 |
| ENSMUSG00000048497 | 0.17196587 | 0.027937 | 0.089952 | 1051.327 | 1142.23 | 966.4004 | 1039.461 | 938.0571 | 918.4815 | 972.0035 | 892.9123 |
| ENSMUSG00000048502 | 0.63056325 | 0.029103 | 0.092576 | 65.39353 | 39.59411 | 67.07664 | 75.04824 | 38.02934 | 44.69496 | 47.25017 | 30.45255 |
| ENSMUSG00000048503 | 0.192672953 | 0.059585 | 0.154042 | 464.7971 | 434.3354 | 459.5992 | 514.1236 | 446.601 | 413.4284 | 406.93 | 373.0437 |
| ENSMUSG00000048520 | 0.488460501 | 0.090238 | 0.205554 | 32.19374 | 38.39429 | 35.60859 | 48.30692 | 27.30312 | 30.1691 | 28.92867 | 23.927 |
| ENSMUSG00000048534 | 0.359243294 | 0.634866 | 0.762533 | 23.13925 | 2.399643 | 13.24971 | 20.70296 | 7.80089 | 3.352122 | 21.21436 | 14.13868 |
| ENSMUSG00000048537 | -0.032452863 | 0.773476 | 0.859536 | 1612.705 | 1568.167 | 1849.99 | 1657.962 | 1770.802 | 1422.417 | 1695.22 | 1954.401 |
| ENSMUSG00000048546 | 0.015313344 | 0.882654 | 0.929757 | 838.0433 | 712.694 | 699.7501 | 830.7064 | 762.537 | 688.3024 | 827.3601 | 770.0145 |
| ENSMUSG00000048550 | 0.044525459 | 0.728038 | 0.829271 | 193.1624 | 212.3684 | 171.4181 | 195.8155 | 176.4951 | 182.132 | 194.7864 | 193.5912 |
| ENSMUSG00000048572 | -1.872178389 | 2.35E-21 | 4.34E-18 | 536.227 | 561.5165 | 642.6107 | 797.0641 | 3169.112 | 1902.888 | 2349.973 | 1871.744 |
| ENSMUSG00000048574 | -0.378226112 | 0.475063 | 0.634588 | 17.10292 | 8.398751 | 10.76539 | 6.900988 | 16.57689 | 12.29112 | 19.28578 | 7.613138 |
| ENSMUSG00000048578 | -0.053198254 | 0.455948 | 0.617925 | 9747.661 | 10296.87 | 9329.449 | 8805.661 | 10137.26 | 9197.106 | 9949.536 | 10322.33 |
| ENSMUSG00000048581 | 0.413042504 | 0.005763 | 0.028566 | 220.3259 | 253.1623 | 255.8849 | 208.7549 | 200.8729 | 188.8362 | 162.9649 | 151.1752 |
| ENSMUSG00000048582 | -1.228192135 | 0.059679 | 0.154234 | 21.12714 | 3.599465 | 27.32752 | 31.05445 | 37.05423 | 17.87799 | 42.42872 | 98.97079 |
| ENSMUSG00000048583 | 0.454604958 | 0.01869 | 0.06728 | 18245.8 | 23713.27 | 20770.57 | 21139.45 | 13890.46 | 22398.88 | 11768.18 | 13143.54 |
| ENSMUSG00000048603 | -0.247737636 | 0.267224 | 0.431561 | 92.557 | 92.38626 | 81.98256 | 91.43809 | 108.2374 | 118.4417 | 128.2505 | 69.60583 |
| ENSMUSG00000048612 | -0.355859851 | 0.033132 | 0.101547 | 1810.898 | 1710.946 | 2370.869 | 1660.55 | 2440.704 | 1839.198 | 2438.687 | 2949.547 |
| ENSMUSG00000048616 | -0.254648837 | 0.402682 | 0.569693 | 114.6902 | 79.18822 | 96.06037 | 120.7673 | 128.7147 | 61.45558 | 163.9292 | 135.9489 |
| ENSMUSG00000048617 | 0.116329109 | 0.5718 | 0.714738 | 73.44197 | 88.78679 | 86.12309 | 78.49874 | 83.85957 | 83.80306 | 56.89306 | 77.21897 |
| ENSMUSG00000048636 | -0.952321801 | 0.01817 | 0.066019 | 48.29061 | 23.99643 | 68.73285 | 74.18562 | 92.63557 | 60.3382 | 106.0718 | 159.8759 |
| ENSMUSG00000048644 | 0.240923521 | 0.016006 | 0.060322 | 1726.389 | 1774.536 | 1716.665 | 2012.501 | 1686.943 | 1622.427 | 1462.827 | 1346.438 |
| ENSMUSG00000048647 | 0.145245319 | 0.752504 | 0.846016 | 28.16952 | 34.79482 | 28.98373 | 37.95543 | 23.40267 | 58.10345 | 22.17865 | 14.13868 |
| ENSMUSG00000048661 | 0.001676762 | 0.989144 | 0.993825 | 973.8606 | 735.4906 | 792.498 | 918.694 | 799.5913 | 975.4676 | 854.3602 | 789.5911 |
| ENSMUSG00000048668 | 0.087728354 | 0.264899 | 0.429083 | 1199.217 | 1136.231 | 1088.132 | 1175.756 | 1186.71 | 1069.327 | 1062.647 | 1007.109 |
| ENSMUSG00000048677 | 0.260646358 | 0.005254 | 0.026624 | 666.008 | 679.099 | 688.1566 | 739.2683 | 547.0374 | 645.8422 | 533.2519 | 591.6496 |
| ENSMUSG00000048696 | -0.034480899 | 0.744464 | 0.840254 | 749.5105 | 670.7002 | 694.7815 | 732.3673 | 842.4962 | 667.0723 | 745.3955 | 660.1678 |
| ENSMUSG00000048699 | 1.457662744 | 0.012323 | 0.049967 | 10.06054 | 23.99643 | 13.24971 | 24.15346 | 8.776002 | 8.938993 | 2.892867 | 5.437955 |
| ENSMUSG00000048701 | 0.245519291 | 0.042247 | 0.121185 | 2421.573 | 2364.848 | 2090.141 | 2388.604 | 1968.75 | 2380.007 | 1773.328 | 1693.379 |
| ENSMUSG00000048706 | 0.31854816 | 0.004686 | 0.024598 | 968.8303 | 1172.226 | 1040.102 | 1303.424 | 961.4597 | 860.3781 | 857.2531 | 916.8393 |
| ENSMUSG00000048707 | -1.175263588 | 1.60E-10 | 2.76E-08 | 212.2775 | 196.7707 | 269.1347 | 198.4034 | 507.0579 | 368.7335 | 497.5732 | 609.051 |
| ENSMUSG00000048709 | 0.070905735 | 0.880077 | 0.927943 | 7.04238 | 17.99732 | 15.73403 | 14.6646 | 13.65156 | 16.76061 | 12.53576 | 9.78832 |
| ENSMUSG00000048720 | 0.391042548 | 1.83E-05 | 0.000366 | 624.7598 | 549.5183 | 532.4726 | 559.8426 | 424.1734 | 457.006 | 425.2515 | 421.9853 |
| ENSMUSG00000048721 | -1.27395539 | 0.012076 | 0.049211 | 13.07871 | 8.398751 | 24.01509 | 12.07673 | 35.10401 | 15.64324 | 55.92877 | 33.71532 |
| ENSMUSG00000048728 | 0.043498926 | 0.781669 | 0.865364 | 113.6841 | 155.9768 | 133.3252 | 132.844 | 133.5902 | 132.9675 | 118.6076 | 133.7737 |
| ENSMUSG00000048731 | 0.003539185 | 0.983652 | 0.990959 | 235.4167 | 208.7689 | 210.3391 | 271.7264 | 258.4045 | 172.0756 | 249.7509 | 243.6204 |
| ENSMUSG00000048732 | -0.531949155 | 0.025897 | 0.085107 | 109.6599 | 155.9768 | 152.3716 | 120.7673 | 242.8027 | 127.3806 | 179.3578 | 228.3941 |
| ENSMUSG00000048747 | 0.17858053 | 0.767286 | 0.855545 | 8.048435 | 27.5959 | 9.93728 | 10.35148 | 5.850668 | 16.76061 | 16.39292 | 9.78832 |
| ENSMUSG00000048752 | 2.653897226 | 1.87E-07 | 9.48E-06 | 29.17558 | 23.99643 | 53.82693 | 29.3292 | 4.875557 | 6.704245 | 3.857157 | 6.525547 |
| ENSMUSG00000048755 | -0.384066106 | 0.000311 | 0.003198 | 719.3289 | 787.0829 | 734.5306 | 796.2015 | 1089.199 | 836.9132 | 964.2892 | 1071.277 |
| ENSMUSG00000048756 | 0.170071885 | 0.033064 | 0.101436 | 2647.935 | 2769.188 | 2410.618 | 2769.884 | 2235.93 | 2377.772 | 2566.938 | 2235 |
| ENSMUSG00000048758 | -0.269690325 | 0.003702 | 0.02065 | 20497.35 | 24715.12 | 24785.23 | 23508.22 | 30600.94 | 25286.18 | 29520.75 | 27315.94 |
| ENSMUSG00000048763 | 0.387368278 | 0.000875 | 0.007031 | 1839.067 | 1677.351 | 1764.695 | 1784.768 | 1415.862 | 1601.197 | 1267.076 | 1119.131 |
| ENSMUSG00000048772 | -0.643507042 | 0.005331 | 0.026873 | 220.3259 | 141.5789 | 152.3716 | 121.6299 | 232.0765 | 194.4231 | 249.7509 | 316.489 |
| ENSMUSG00000048776 | 1.114214822 | 0.000296 | 0.003066 | 106.6418 | 134.38 | 64.59232 | 96.61383 | 36.07912 | 65.92507 | 44.3573 | 39.15328 |
| ENSMUSG00000048779 | 0.271991153 | 0.131716 | 0.264722 | 90.54489 | 82.78769 | 106.8258 | 92.30071 | 71.18313 | 75.98144 | 80.036 | 82.65692 |
| ENSMUSG00000048782 | -1.537837257 | 0.023176 | 0.078469 | 10.06054 | 7.198929 | 3.312427 | 18.97772 | 23.40267 | 15.64324 | 15.42863 | 60.9051 |
| ENSMUSG00000048787 | 0.348702331 | 0.013282 | 0.052818 | 329.9858 | 353.9474 | 261.6817 | 301.9182 | 248.6534 | 264.8177 | 211.1793 | 253.4087 |
| ENSMUSG00000048794 | -0.029988981 | 0.938684 | 0.963964 | 20.12109 | 13.19804 | 21.53077 | 18.11509 | 21.45245 | 12.29112 | 18.32149 | 22.83941 |
| ENSMUSG00000048796 | 0.018752281 | 0.864772 | 0.91903 | 297.7921 | 317.9527 | 346.9767 | 352.813 | 351.0401 | 301.691 | 333.6441 | 313.2262 |
| ENSMUSG00000048799 | 0.153523271 | 0.055817 | 0.147195 | 898.4065 | 976.6547 | 921.6827 | 872.975 | 797.641 | 905.073 | 799.3957 | 796.1167 |
| ENSMUSG00000048807 | -0.160925976 | 0.326083 | 0.493883 | 222.338 | 188.372 | 261.6817 | 203.5791 | 240.8525 | 198.8926 | 276.751 | 264.2846 |
| ENSMUSG00000048814 | 0.024243683 | 0.872031 | 0.9234 | 274.6528 | 224.3666 | 195.4332 | 209.6175 | 195.9974 | 234.6486 | 224.6794 | 232.7445 |
| ENSMUSG00000048826 | -0.058906436 | 0.662594 | 0.782896 | 1339.058 | 1293.408 | 1663.666 | 1290.485 | 1524.099 | 1188.886 | 1430.041 | 1678.153 |
| ENSMUSG00000048827 | 0.580458212 | 0.051457 | 0.138728 | 61.36932 | 38.39429 | 44.71776 | 52.62003 | 24.37778 | 43.57759 | 29.89296 | 34.80291 |
| ENSMUSG00000048832 | -0.155691865 | 0.253289 | 0.417012 | 532.2028 | 493.1267 | 616.1113 | 416.6471 | 587.017 | 524.0485 | 616.1808 | 565.5474 |
| ENSMUSG00000048833 | -0.118099711 | 0.233325 | 0.393913 | 1307.871 | 1227.417 | 1387.907 | 1340.517 | 1395.384 | 1271.572 | 1391.469 | 1657.489 |
| ENSMUSG00000048834 | -1.231657111 | 0.008719 | 0.03857 | 13.07871 | 17.99732 | 19.87456 | 14.6646 | 19.50223 | 70.39457 | 28.92867 | 35.89051 |

| | | | | | | | | | | | |
|---|---|---|---|---|---|---|---|---|---|---|---|
| ENSMUSG00000048856 | -0.063579802 | 0.690696 | 0.804105 | 185.114 | 153.5772 | 148.2311 | 144.0581 | 168.6943 | 188.8362 | 162.0006 | 139.2117 |
| ENSMUSG00000048865 | 0.3045493 | 0.233416 | 0.393993 | 66.39959 | 64.79036 | 70.38906 | 46.58167 | 46.80534 | 37.99072 | 56.89306 | 58.72992 |
| ENSMUSG00000048874 | 0.123472683 | 0.09864 | 0.218994 | 2807.898 | 2404.442 | 2427.181 | 2398.956 | 2208.627 | 2321.903 | 2375.044 | 2308.956 |
| ENSMUSG00000048878 | 0.063035338 | 0.529979 | 0.681935 | 2737.474 | 2483.631 | 2683.065 | 2725.028 | 2392.923 | 2206.814 | 2649.867 | 2926.708 |
| ENSMUSG00000048895 | 0.184180963 | 0.475259 | 0.634734 | 81.4904 | 71.98929 | 72.04528 | 128.5309 | 78.0089 | 79.33356 | 89.67889 | 65.25547 |
| ENSMUSG00000048897 | -0.048552594 | 0.633695 | 0.761871 | 850.1159 | 793.082 | 958.1194 | 828.1186 | 900.0277 | 798.9225 | 860.1459 | 990.7955 |
| ENSMUSG00000048899 | -0.52242592 | 0.054178 | 0.144266 | 48.29061 | 47.99286 | 55.48314 | 68.14726 | 60.4569 | 65.92507 | 80.036 | 110.9343 |
| ENSMUSG00000048905 | 1.041526138 | 0.012065 | 0.049182 | 33.19979 | 44.3934 | 56.31125 | 24.15346 | 20.47734 | 10.05637 | 23.14294 | 22.83941 |
| ENSMUSG00000048911 | -0.108267811 | 0.287214 | 0.454092 | 346.0827 | 417.5379 | 393.3507 | 405.433 | 454.4019 | 414.5458 | 403.0729 | 411.1094 |
| ENSMUSG00000048915 | 0.642584881 | 0.02054 | 0.071879 | 883.3157 | 1113.434 | 839.7001 | 975.6272 | 513.8837 | 1039.158 | 513.9661 | 375.2189 |
| ENSMUSG00000048920 | -0.255686318 | 0.084219 | 0.195701 | 907.461 | 771.4852 | 872.8244 | 815.1792 | 1001.439 | 754.2275 | 1044.325 | 1220.277 |
| ENSMUSG00000048921 | -0.20461432 | 0.319688 | 0.487813 | 216.3017 | 160.7761 | 139.95 | 126.8057 | 195.0223 | 212.3011 | 175.5006 | 157.7007 |
| ENSMUSG00000048922 | 0.18476343 | 0.097361 | 0.216806 | 996.9999 | 921.4629 | 809.8883 | 984.2534 | 903.9282 | 869.317 | 729.9669 | 762.4014 |
| ENSMUSG00000048924 | 0.570980064 | 2.39E-06 | 7.45E-05 | 588.5418 | 611.909 | 497.6921 | 519.2993 | 374.4427 | 426.8369 | 355.8227 | 333.8905 |
| ENSMUSG00000048930 | 0.097709341 | 0.265142 | 0.429317 | 955.7516 | 999.4513 | 1048.383 | 939.397 | 861.0233 | 859.2607 | 945.9677 | 1019.073 |
| ENSMUSG00000048949 | -0.172866664 | 0.596311 | 0.735213 | 38.23007 | 20.39697 | 41.40533 | 25.8787 | 36.07912 | 41.34284 | 29.89296 | 35.89051 |
| ENSMUSG00000048960 | -0.243717055 | 0.144777 | 0.282897 | 424.5549 | 298.7556 | 475.3332 | 355.4009 | 406.6214 | 454.7713 | 448.3945 | 534.0072 |
| ENSMUSG00000048965 | -0.026737845 | 0.893424 | 0.935548 | 80.48435 | 92.38626 | 122.5598 | 104.3774 | 98.48624 | 87.15518 | 109.929 | 113.1095 |
| ENSMUSG00000048967 | 0.745439366 | 0.001419 | 0.010174 | 85.51462 | 79.18822 | 93.57605 | 105.2401 | 41.92979 | 45.81234 | 66.53595 | 63.08028 |
| ENSMUSG00000048970 | -0.180867227 | 0.174699 | 0.32291 | 458.7608 | 405.5397 | 483.6143 | 439.938 | 507.0579 | 407.8415 | 590.145 | 522.0437 |
| ENSMUSG00000048978 | 0.20489349 | 0.489647 | 0.647159 | 34.20585 | 40.79393 | 28.98373 | 39.68068 | 39.97956 | 26.81698 | 30.85725 | 26.10219 |
| ENSMUSG00000048988 | -0.372699207 | 0.471156 | 0.630989 | 34.20585 | 23.99643 | 85.29498 | 29.3292 | 56.55646 | 24.58223 | 50.14304 | 93.53283 |
| ENSMUSG00000048997 | 0.178605343 | 0.062735 | 0.159878 | 647.899 | 631.1061 | 675.735 | 615.0506 | 580.1912 | 630.199 | 502.3947 | 560.1094 |
| ENSMUSG00000049001 | -0.13844647 | 0.518794 | 0.672487 | 278.6771 | 182.3729 | 246.7758 | 252.7487 | 175.52 | 286.0478 | 310.5011 | 287.124 |
| ENSMUSG00000049036 | -0.12798967 | 0.375999 | 0.544226 | 134.8113 | 154.777 | 173.9024 | 142.3329 | 179.4205 | 159.7845 | 159.1077 | 164.2263 |
| ENSMUSG00000049037 | 0.328799824 | 0.185258 | 0.336793 | 51.30877 | 67.19001 | 53.82693 | 54.34528 | 44.85512 | 45.81234 | 54.96448 | 33.71532 |
| ENSMUSG00000049038 | 0.226644359 | 0.041182 | 0.119094 | 285.7194 | 326.3515 | 289.8373 | 312.2697 | 260.3547 | 263.7003 | 277.7153 | 233.8321 |
| ENSMUSG00000049044 | 1.005705068 | 0.003201 | 0.01862 | 1164.005 | 406.7395 | 395.835 | 462.3662 | 265.2303 | 382.1419 | 297.0011 | 265.3722 |
| ENSMUSG00000049047 | 0.26561939 | 0.000273 | 0.002877 | 1168.029 | 1203.421 | 1071.57 | 1197.321 | 993.6384 | 970.9981 | 984.5392 | 907.051 |
| ENSMUSG00000049076 | 0.214046764 | 0.026358 | 0.086328 | 1743.492 | 1551.369 | 1565.122 | 1813.235 | 1350.529 | 1545.328 | 1553.47 | 1305.109 |
| ENSMUSG00000049086 | 0.030329431 | 0.874551 | 0.925069 | 404.4339 | 404.3399 | 349.461 | 468.4046 | 347.1396 | 544.1612 | 394.3943 | 307.7883 |
| ENSMUSG00000049090 | -0.052067667 | 0.477213 | 0.636625 | 1418.537 | 1508.176 | 1314.205 | 1400.038 | 1514.348 | 1354.257 | 1485.97 | 1486.737 |
| ENSMUSG00000049091 | -0.989521172 | 1.23E-06 | 4.42E-05 | 1035.23 | 705.4951 | 1093.929 | 947.1606 | 1824.433 | 1292.802 | 1937.257 | 2457.956 |
| ENSMUSG00000049092 | 0.381285894 | 0.226623 | 0.386557 | 96.58122 | 127.1811 | 103.5133 | 124.2178 | 63.38224 | 148.6108 | 74.25027 | 60.9051 |
| ENSMUSG00000049097 | 0.522044627 | 0.060765 | 0.156433 | 47.28455 | 39.59411 | 47.20208 | 39.68068 | 35.10401 | 21.23011 | 31.82154 | 32.62773 |
| ENSMUSG00000049100 | 0.290262485 | 0.402906 | 0.569898 | 82.49646 | 52.79215 | 141.6062 | 112.1411 | 56.55646 | 67.04245 | 110.8933 | 84.8321 |
| ENSMUSG00000049103 | -0.348906467 | 0.382141 | 0.550015 | 25.15136 | 11.99822 | 30.63995 | 23.29083 | 39.00445 | 21.23011 | 23.14294 | 33.71532 |
| ENSMUSG00000049106 | 0.122255584 | 0.036352 | 0.109234 | 1757.577 | 1773.336 | 1842.537 | 1764.928 | 1608.934 | 1743.104 | 1583.363 | 1627.036 |
| ENSMUSG00000049107 | 0.064690632 | 0.745994 | 0.841259 | 330.9919 | 423.537 | 239.3228 | 283.8031 | 334.4632 | 341.9165 | 294.1082 | 247.9708 |
| ENSMUSG00000049112 | -0.29360104 | 0.224402 | 0.384119 | 440.6518 | 224.3666 | 374.3042 | 346.7746 | 407.5965 | 312.8647 | 573.7521 | 405.6715 |
| ENSMUSG00000049115 | -0.384890859 | 0.272718 | 0.438211 | 699.2078 | 212.3684 | 314.6805 | 422.6855 | 446.601 | 353.0902 | 759.8599 | 593.8247 |
| ENSMUSG00000049119 | 0.309284052 | 0.075604 | 0.181937 | 393.3673 | 441.5343 | 521.7072 | 419.235 | 302.2845 | 476.0014 | 323.0369 | 333.8905 |
| ENSMUSG00000049122 | 0.455416248 | 0.175391 | 0.323752 | 37.22401 | 39.59411 | 34.78048 | 35.36756 | 31.20356 | 37.99072 | 16.39292 | 21.75182 |
| ENSMUSG00000049124 | -0.095923118 | 0.221721 | 0.380835 | 788.7466 | 784.6833 | 892.6989 | 794.4762 | 913.6793 | 865.9649 | 873.646 | 834.1824 |
| ENSMUSG00000049128 | 0.226932321 | 0.655358 | 0.777782 | 133.8052 | 33.595 | 164.7932 | 43.9938 | 97.51113 | 79.33356 | 99.32178 | 45.67883 |
| ENSMUSG00000049130 | 0.586715887 | 0.021322 | 0.073719 | 62.37537 | 89.98662 | 88.60741 | 61.24627 | 61.43201 | 43.57759 | 42.42872 | 53.29196 |
| ENSMUSG00000049134 | -0.968955782 | 0.003265 | 0.018892 | 20.12109 | 28.79572 | 24.8432 | 18.97772 | 58.50668 | 39.10809 | 31.82154 | 51.11678 |
| ENSMUSG00000049152 | -7.179486936 | 1.39E-09 | 1.72E-07 | 0 | 0 | 4.96864 | 1.725247 | 231.1014 | 32.40385 | 293.1439 | 460.051 |
| ENSMUSG00000049164 | 0.195710856 | 0.25844 | 0.422364 | 762.5892 | 830.2765 | 823.138 | 694.4119 | 572.3903 | 929.6552 | 560.252 | 654.7298 |
| ENSMUSG00000049176 | 0.709943131 | 1.64E-05 | 0.000335 | 191.1503 | 145.1784 | 180.5272 | 168.2116 | 98.48624 | 122.9112 | 104.1432 | 94.62042 |
| ENSMUSG00000049184 | 0.506053278 | 0.002575 | 0.015739 | 447.6942 | 458.3318 | 404.9441 | 460.6409 | 252.5538 | 405.6068 | 329.7869 | 259.9343 |
| ENSMUSG00000049191 | 0.100468432 | 0.382412 | 0.550315 | 464.7971 | 537.52 | 457.1149 | 546.0407 | 476.8294 | 494.9967 | 405.0014 | 493.7664 |
| ENSMUSG00000049225 | 0.619900722 | 0.011296 | 0.046778 | 1294.792 | 1003.051 | 719.6247 | 1438.856 | 690.3788 | 878.256 | 843.753 | 487.2408 |
| ENSMUSG00000049230 | -0.045676865 | 0.815493 | 0.887583 | 149.9021 | 167.975 | 139.1219 | 169.0742 | 175.52 | 116.2069 | 152.3577 | 201.2044 |
| ENSMUSG00000049232 | 0.083349499 | 0.511797 | 0.667212 | 579.4873 | 560.3167 | 621.08 | 726.329 | 556.7886 | 674.894 | 519.7519 | 600.3503 |
| ENSMUSG00000049233 | 0.524468498 | 0.399272 | 0.566756 | 17.10292 | 3.599465 | 11.59349 | 13.80198 | 5.850668 | 13.40849 | 6.750024 | 6.525547 |
| ENSMUSG00000049241 | 0.187781682 | 0.28813 | 0.455012 | 152.9203 | 149.9777 | 172.2462 | 119.042 | 150.1671 | 113.9722 | 125.3576 | 131.5985 |
| ENSMUSG00000049252 | 1.035743753 | 0.098789 | 0.219221 | 12.07265 | 28.79572 | 13.24971 | 12.93935 | 1.950223 | 13.40849 | 9.642892 | 7.613138 |
| ENSMUSG00000049265 | 3.048695187 | 0.00536 | 0.026945 | 2000.036 | 63.59054 | 72.87338 | 477.0308 | 55.58134 | 69.27719 | 45.32159 | 145.7372 |
| ENSMUSG00000049281 | 2.842650727 | 8.14E-06 | 0.000195 | 284.7134 | 82.78769 | 36.43669 | 272.589 | 19.50223 | 45.81234 | 18.32149 | 10.87591 |
| ENSMUSG00000049285 | -0.313628968 | 0.043218 | 0.123138 | 130.7871 | 159.5763 | 159.8246 | 128.5309 | 182.3458 | 166.4887 | 204.4293 | 164.2263 |
| ENSMUSG00000049287 | -0.608848956 | 0.008659 | 0.038435 | 237.4288 | 202.7698 | 275.7595 | 156.9975 | 328.6125 | 235.7659 | 335.5726 | 431.7737 |
| ENSMUSG00000049288 | 0.427503879 | 5.62E-08 | 3.58E-06 | 3047.339 | 3340.303 | 3308.286 | 3357.331 | 2499.21 | 2664.937 | 2323.937 | 2218.686 |
| ENSMUSG00000049295 | 0.077510574 | 0.163793 | 0.30871 | 2510.106 | 2535.223 | 2412.275 | 2335.984 | 2387.072 | 2361.011 | 2287.294 | 2240.438 |

| | | | | | | | | | | | |
|---|---|---|---|---|---|---|---|---|---|---|---|
| ENSMUSG00000049299 | -0.040399644 | 0.634391 | 0.762319 | 1075.472 | 1085.838 | 1171.771 | 1224.063 | 1190.611 | 1067.092 | 1163.897 | 1268.131 |
| ENSMUSG00000049300 | -0.030807365 | 0.792743 | 0.873221 | 526.1664 | 586.7127 | 577.1903 | 709.0765 | 631.8721 | 621.26 | 643.1809 | 555.759 |
| ENSMUSG00000049303 | 0.534812863 | 0.012478 | 0.050458 | 242.4591 | 191.9714 | 162.3089 | 153.547 | 133.5902 | 156.4324 | 99.32178 | 128.3357 |
| ENSMUSG00000049305 | -0.195978225 | 0.013206 | 0.052628 | 1369.24 | 1475.78 | 1514.607 | 1430.23 | 1731.798 | 1474.934 | 1697.149 | 1727.095 |
| ENSMUSG00000049307 | -0.53469952 | 0.002031 | 0.013279 | 122.7386 | 123.5816 | 123.3879 | 126.8057 | 150.1671 | 184.3667 | 225.6437 | 158.7883 |
| ENSMUSG00000049313 | -0.019433618 | 0.880073 | 0.927943 | 1569.445 | 1451.784 | 1902.989 | 1575.15 | 1489.97 | 1432.474 | 1755.971 | 1911.985 |
| ENSMUSG00000049321 | 0.22779404 | 0.149513 | 0.289488 | 402.4217 | 357.5468 | 366.8512 | 384.7301 | 326.6623 | 410.0763 | 304.7154 | 250.146 |
| ENSMUSG00000049323 | -0.188911189 | 0.044892 | 0.126281 | 675.0625 | 641.9045 | 725.4214 | 610.7374 | 739.1344 | 697.2414 | 822.5387 | 765.6641 |
| ENSMUSG00000049327 | -0.162570617 | 0.006989 | 0.032922 | 1684.135 | 1620.959 | 1562.637 | 1618.282 | 1893.666 | 1722.991 | 1852.399 | 1785.825 |
| ENSMUSG00000049336 | 0.985864916 | 1.30E-05 | 0.000281 | 207.2472 | 202.7698 | 221.9326 | 153.547 | 79.95913 | 136.3196 | 79.07171 | 102.2336 |
| ENSMUSG00000049339 | -0.203582819 | 0.026593 | 0.086943 | 1708.28 | 1839.326 | 2067.782 | 1843.426 | 2154.996 | 1909.592 | 2300.794 | 2225.211 |
| ENSMUSG00000049353 | 0.233732359 | 0.392916 | 0.56032 | 46.2785 | 37.19447 | 42.23344 | 43.9938 | 29.25334 | 30.1691 | 38.57157 | 46.76642 |
| ENSMUSG00000049354 | -0.005038301 | 0.930346 | 0.958826 | 3564.451 | 3621.061 | 3687.559 | 3541.069 | 3701.523 | 3376.705 | 3564.977 | 3820.707 |
| ENSMUSG00000049357 | 0.56807476 | 0.014984 | 0.057674 | 109.6599 | 130.7805 | 104.3414 | 127.6683 | 70.20801 | 111.7374 | 78.10742 | 58.72992 |
| ENSMUSG00000049382 | -0.648226962 | 0.091217 | 0.207073 | 3112.732 | 2480.031 | 6060.084 | 3001.067 | 6195.857 | 3528.667 | 5624.699 | 7618.576 |
| ENSMUSG00000049396 | -0.421441225 | 0.215542 | 0.373736 | 25.15136 | 33.595 | 28.98373 | 31.91707 | 50.70579 | 29.05173 | 27.0001 | 53.29196 |
| ENSMUSG00000049401 | -0.054201383 | 0.608549 | 0.744128 | 2416.543 | 2907.168 | 3199.804 | 2502.471 | 2850.25 | 2676.111 | 2917.939 | 3003.927 |
| ENSMUSG00000049404 | -1.040311459 | 0.020726 | 0.072244 | 23.13925 | 27.5959 | 38.92101 | 14.6646 | 32.17867 | 32.40385 | 77.14313 | 72.8686 |
| ENSMUSG00000049409 | 0.345451444 | 0.097551 | 0.217178 | 157.9505 | 244.7636 | 159.8246 | 200.1287 | 154.0676 | 186.6015 | 129.2147 | 129.4233 |
| ENSMUSG00000049411 | 0.008714793 | 0.934848 | 0.961523 | 350.1069 | 333.5504 | 358.5702 | 335.5605 | 316.9112 | 315.0995 | 357.7513 | 380.6569 |
| ENSMUSG00000049420 | 0.289101122 | 0.372241 | 0.541259 | 171.0292 | 68.38983 | 83.63877 | 94.02596 | 75.08357 | 64.8077 | 100.2861 | 101.146 |
| ENSMUSG00000049421 | 0.26338242 | 0.000508 | 0.004643 | 2392.397 | 2567.618 | 2239.2 | 2278.189 | 1965.824 | 2134.185 | 1933.4 | 1858.693 |
| ENSMUSG00000049422 | -1.426944157 | 0.000758 | 0.006321 | 983.9212 | 533.9206 | 1434.281 | 736.6805 | 2194 | 1296.154 | 2691.331 | 3739.138 |
| ENSMUSG00000049436 | -0.3837335 | 0.255506 | 0.419389 | 746.4923 | 645.504 | 1283.565 | 762.5592 | 1774.703 | 501.701 | 1190.897 | 1019.073 |
| ENSMUSG00000049439 | -0.145864165 | 0.095131 | 0.213231 | 1034.224 | 855.4728 | 930.7919 | 962.6878 | 1055.07 | 969.8807 | 1116.647 | 1045.175 |
| ENSMUSG00000049470 | 0.182980497 | 0.001347 | 0.009765 | 4160.035 | 3798.635 | 4050.27 | 3824.01 | 3374.86 | 3639.287 | 3499.405 | 3436.788 |
| ENSMUSG00000049482 | -0.236296103 | 0.004257 | 0.022873 | 580.4934 | 518.3229 | 581.3309 | 552.9417 | 649.4241 | 619.0252 | 671.1453 | 693.8831 |
| ENSMUSG00000049488 | 0.197754769 | 0.083626 | 0.1948 | 488.9424 | 549.5183 | 443.037 | 508.0852 | 475.8543 | 459.2408 | 390.5371 | 406.7591 |
| ENSMUSG00000049489 | -0.312903733 | 0.005502 | 0.027557 | 546.2875 | 475.1293 | 587.1276 | 596.9355 | 710.8561 | 595.5604 | 710.6811 | 726.5108 |
| ENSMUSG00000049493 | -0.887670728 | 5.30E-05 | 0.000828 | 338.0343 | 263.9607 | 426.4749 | 314.8576 | 585.0668 | 431.3064 | 642.2166 | 829.832 |
| ENSMUSG00000049502 | 0.431852296 | 0.032291 | 0.09966 | 309.8647 | 454.7324 | 327.1021 | 240.672 | 244.7529 | 274.874 | 251.6795 | 214.2554 |
| ENSMUSG00000049504 | 0.155500058 | 0.227198 | 0.387145 | 832.007 | 748.6886 | 748.6084 | 651.2807 | 679.6526 | 796.6877 | 603.645 | 595.9999 |
| ENSMUSG00000049511 | 0.016636552 | 0.969564 | 0.983052 | 17.10292 | 21.59679 | 19.87456 | 11.21411 | 17.552 | 17.87799 | 23.14294 | 9.78832 |
| ENSMUSG00000049515 | 0.153864129 | 0.696168 | 0.808283 | 22.1332 | 11.99822 | 23.18699 | 24.15346 | 15.60178 | 15.64324 | 24.10723 | 18.48905 |
| ENSMUSG00000049516 | -0.149218279 | 0.137696 | 0.272879 | 804.8435 | 748.6886 | 664.1415 | 660.7696 | 822.9939 | 755.3449 | 753.1098 | 857.0218 |
| ENSMUSG00000049517 | -0.123732931 | 0.068209 | 0.169044 | 23729.8 | 27051.18 | 26117.66 | 26502.38 | 30011 | 27176.77 | 28747.39 | 26722.11 |
| ENSMUSG00000049521 | -0.452375313 | 0.014304 | 0.055671 | 716.3107 | 674.2997 | 1051.695 | 732.3673 | 1024.842 | 836.9132 | 1201.504 | 1283.357 |
| ENSMUSG00000049532 | 0.209964754 | 0.036731 | 0.110051 | 1922.57 | 2248.466 | 1909.614 | 2084.098 | 1873.189 | 1956.522 | 1566.97 | 1660.752 |
| ENSMUSG00000049536 | 0.143892259 | 0.354653 | 0.523262 | 433.6094 | 587.9126 | 418.1938 | 384.7301 | 439.7752 | 381.0246 | 380.8942 | 445.9123 |
| ENSMUSG00000049538 | 0.025028979 | 0.862177 | 0.917545 | 1227.386 | 995.8519 | 1344.845 | 1065.34 | 899.0526 | 1286.098 | 1174.504 | 1197.438 |
| ENSMUSG00000049550 | -0.077653191 | 0.303569 | 0.472003 | 1419.543 | 1303.006 | 1210.692 | 1367.258 | 1369.056 | 1411.243 | 1464.755 | 1346.438 |
| ENSMUSG00000049551 | 0.331293653 | 0.320324 | 0.488405 | 33.19979 | 23.99643 | 36.43669 | 37.09281 | 33.15378 | 16.76061 | 26.03581 | 28.27737 |
| ENSMUSG00000049553 | -0.206634975 | 0.007317 | 0.033955 | 1364.21 | 1358.198 | 1270.316 | 1222.337 | 1379.783 | 1584.436 | 1488.862 | 1562.868 |
| ENSMUSG00000049555 | -0.256483374 | 0.491048 | 0.648215 | 58.35115 | 33.595 | 96.88848 | 48.30692 | 56.55646 | 54.75133 | 71.3574 | 102.2336 |
| ENSMUSG00000049556 | 0.209152081 | 0.203947 | 0.359886 | 223.3441 | 189.5718 | 256.7131 | 283.8031 | 231.1014 | 197.7752 | 183.2149 | 215.343 |
| ENSMUSG00000049562 | -0.513198641 | 0.044825 | 0.126151 | 118.7144 | 106.7841 | 175.5586 | 93.16334 | 181.3707 | 122.9112 | 207.3222 | 194.6788 |
| ENSMUSG00000049571 | 0.679693706 | 0.155386 | 0.297305 | 16.09687 | 16.7975 | 14.07781 | 23.29083 | 5.850668 | 15.64324 | 8.678602 | 14.13868 |
| ENSMUSG00000049577 | -0.232017343 | 0.104296 | 0.22735 | 522.1422 | 491.9268 | 721.2809 | 527.063 | 719.6321 | 631.3164 | 617.1451 | 692.7955 |
| ENSMUSG00000049580 | 0.20232651 | 0.089802 | 0.204868 | 514.0938 | 447.5334 | 453.8024 | 403.7078 | 397.8454 | 446.9496 | 376.0728 | 359.9926 |
| ENSMUSG00000049588 | 0.331561926 | 0.09995 | 0.220735 | 88.53278 | 100.785 | 89.43552 | 92.30071 | 87.76002 | 68.15982 | 81.00029 | 56.55474 |
| ENSMUSG00000049598 | -1.195023976 | 0.02132 | 0.073719 | 33.19979 | 4.799286 | 28.98373 | 18.11509 | 72.15824 | 41.34284 | 51.10733 | 31.54014 |
| ENSMUSG00000049600 | -0.083523935 | 0.406985 | 0.573383 | 344.0706 | 346.7484 | 359.3983 | 346.7746 | 380.2934 | 405.6068 | 364.5013 | 330.6277 |
| ENSMUSG00000049606 | 0.141244594 | 0.175215 | 0.323572 | 1785.746 | 1725.343 | 1438.421 | 1423.329 | 1361.255 | 1562.089 | 1445.469 | 1406.255 |
| ENSMUSG00000049624 | -0.294331875 | 0.015217 | 0.058302 | 617.7174 | 704.2952 | 688.1566 | 654.7312 | 857.1228 | 668.1897 | 797.4671 | 944.0291 |
| ENSMUSG00000049625 | 0.045629325 | 0.873802 | 0.924601 | 35.2119 | 43.19358 | 54.65504 | 40.5433 | 34.1289 | 34.6386 | 54.00019 | 45.67883 |
| ENSMUSG00000049630 | 0.002012089 | 0.996151 | 0.997718 | 19.11503 | 27.5959 | 59.62368 | 39.68068 | 27.30312 | 36.87335 | 31.82154 | 51.11678 |
| ENSMUSG00000049643 | 0.097883908 | 0.354748 | 0.523305 | 451.7184 | 555.5174 | 491.0672 | 502.9095 | 512.9085 | 439.128 | 478.2874 | 436.124 |
| ENSMUSG00000049649 | -0.500751257 | 0.268714 | 0.433286 | 11.0666 | 15.59768 | 10.76539 | 7.763611 | 10.72622 | 16.76061 | 18.32149 | 17.40146 |
| ENSMUSG00000049657 | -0.037098203 | 0.640334 | 0.766904 | 843.0736 | 865.0713 | 877.793 | 809.1408 | 835.6704 | 813.4483 | 945.9677 | 886.3867 |
| ENSMUSG00000049658 | 0.059284491 | 0.578491 | 0.719984 | 907.461 | 853.0731 | 706.375 | 822.9428 | 770.3379 | 877.1387 | 751.1813 | 756.9634 |
| ENSMUSG00000049659 | -0.180529151 | 0.086414 | 0.199408 | 1270.647 | 1103.836 | 1217.317 | 1098.12 | 1155.507 | 1263.75 | 1476.327 | 1420.394 |
| ENSMUSG00000049670 | -0.076109133 | 0.434693 | 0.598843 | 808.8677 | 787.0829 | 801.6072 | 831.569 | 915.6295 | 718.4715 | 861.1102 | 908.1386 |
| ENSMUSG00000049672 | -0.037597356 | 0.745314 | 0.840853 | 761.5831 | 794.2819 | 683.188 | 684.9231 | 811.2926 | 791.1009 | 778.1814 | 616.6641 |
| ENSMUSG00000049676 | 2.165058752 | 6.02E-31 | 2.78E-27 | 253.5257 | 224.3666 | 183.8397 | 200.1287 | 49.73068 | 46.92971 | 48.21446 | 46.76642 |

| | | | | | | | | | | | |
|---|---|---|---|---|---|---|---|---|---|---|---|
| ENSMUSG00000049680 | -0.275233159 | 0.074619 | 0.180423 | 709.2683 | 757.0874 | 818.1694 | 749.6198 | 841.5211 | 695.0067 | 998.0393 | 1137.62 |
| ENSMUSG00000049686 | -0.057109903 | 0.595511 | 0.734635 | 371.2341 | 388.7422 | 400.8036 | 383.8675 | 391.0196 | 368.7335 | 382.8228 | 465.489 |
| ENSMUSG00000049687 | 0.512218646 | 3.77E-05 | 0.000643 | 319.9253 | 345.5486 | 345.3205 | 380.417 | 234.0267 | 280.4609 | 247.8223 | 214.2554 |
| ENSMUSG00000049690 | -0.166296679 | 0.468722 | 0.62887 | 239.4409 | 122.3818 | 198.7456 | 169.0742 | 183.3209 | 167.6061 | 235.2866 | 233.8321 |
| ENSMUSG00000049717 | -0.202469344 | 0.188794 | 0.341288 | 190.1443 | 163.1757 | 152.3716 | 138.0198 | 164.7938 | 201.1273 | 180.3221 | 193.5912 |
| ENSMUSG00000049721 | -1.707820514 | 4.43E-05 | 0.000728 | 84.50857 | 73.18911 | 197.9175 | 97.47645 | 308.1352 | 150.8455 | 494.6803 | 529.6569 |
| ENSMUSG00000049728 | -0.088995079 | 0.439786 | 0.603068 | 315.9011 | 311.9536 | 263.3379 | 308.8192 | 293.5085 | 348.6207 | 324.0012 | 308.8759 |
| ENSMUSG00000049739 | -0.126562878 | 0.172894 | 0.320567 | 1037.242 | 1051.044 | 995.3842 | 883.3265 | 1155.507 | 979.9371 | 1059.754 | 1131.095 |
| ENSMUSG00000049744 | -0.496819828 | 0.210728 | 0.36826 | 15.09082 | 16.7975 | 29.81184 | 15.52722 | 24.37778 | 27.93435 | 36.64299 | 20.66423 |
| ENSMUSG00000049751 | -0.285986967 | 0.003756 | 0.020898 | 4100.678 | 4349.353 | 4601.789 | 4349.348 | 5730.729 | 4402.454 | 5400.019 | 5682.663 |
| ENSMUSG00000049755 | -0.284408134 | 0.065189 | 0.164406 | 617.7174 | 580.7136 | 477.8175 | 541.7276 | 638.6979 | 526.2832 | 719.3597 | 814.6057 |
| ENSMUSG00000049758 | 0.908102452 | 0.064884 | 0.163997 | 16.09687 | 11.99822 | 21.53077 | 20.70296 | 11.70134 | 8.938993 | 12.53576 | 4.350364 |
| ENSMUSG00000049760 | -0.573947213 | 0.000144 | 0.001766 | 1915.527 | 1887.319 | 2143.14 | 1952.117 | 2966.289 | 2111.837 | 3124.297 | 3555.335 |
| ENSMUSG00000049764 | -0.019943737 | 0.868005 | 0.921001 | 483.9121 | 445.1338 | 554.0033 | 585.7214 | 476.8294 | 570.9782 | 517.8233 | 537.27 |
| ENSMUSG00000049775 | 0.007749977 | 0.9601 | 0.977364 | 14913.75 | 15867.64 | 24221.29 | 16660.71 | 16324.34 | 17120.41 | 18501.82 | 19336.28 |
| ENSMUSG00000049791 | -0.433984307 | 0.000284 | 0.002979 | 3319.979 | 2572.417 | 3097.119 | 2966.562 | 3674.219 | 3555.484 | 4678.731 | 4245.956 |
| ENSMUSG00000049792 | -0.054819753 | 0.568794 | 0.712804 | 1150.926 | 1330.602 | 1215.661 | 1250.804 | 1416.837 | 1137.487 | 1220.79 | 1361.664 |
| ENSMUSG00000049799 | -2.19785419 | 4.32E-09 | 4.33E-07 | 27.16347 | 51.59233 | 41.40533 | 43.9938 | 276.9316 | 94.9768 | 134.0362 | 245.7956 |
| ENSMUSG00000049800 | 0.222178629 | 0.046692 | 0.129778 | 902.4308 | 1027.047 | 956.4632 | 979.9403 | 793.7406 | 988.8761 | 729.0026 | 803.7298 |
| ENSMUSG00000049804 | 0.457586656 | 4.37E-08 | 2.90E-06 | 2771.68 | 2903.568 | 2794.032 | 2900.14 | 1971.675 | 2372.185 | 1908.328 | 2029.445 |
| ENSMUSG00000049807 | 0.019596114 | 0.866049 | 0.919627 | 843.0736 | 832.6761 | 1059.148 | 938.5344 | 926.3557 | 801.1572 | 886.1817 | 1013.635 |
| ENSMUSG00000049811 | 0.293547309 | 0.237267 | 0.398307 | 225.3562 | 124.7814 | 109.3101 | 150.0965 | 129.6898 | 119.559 | 110.8933 | 137.0365 |
| ENSMUSG00000049823 | -0.123695551 | 0.168183 | 0.314296 | 1082.514 | 1186.623 | 1335.736 | 1196.459 | 1391.484 | 1349.788 | 1268.04 | 1224.628 |
| ENSMUSG00000049858 | -0.314153167 | 0.184248 | 0.335386 | 219.3198 | 268.76 | 269.9628 | 255.3366 | 268.1556 | 194.4231 | 411.7515 | 385.0072 |
| ENSMUSG00000049866 | 0.012248629 | 0.915365 | 0.949794 | 3125.811 | 2345.651 | 2748.486 | 2870.811 | 2370.496 | 2724.158 | 3014.368 | 2890.817 |
| ENSMUSG00000049871 | 0.057744369 | 0.719467 | 0.824494 | 133.8052 | 118.7823 | 139.95 | 112.1411 | 131.64 | 122.9112 | 106.0718 | 125.073 |
| ENSMUSG00000049872 | -0.190579845 | 0.623122 | 0.754516 | 16.09687 | 43.19358 | 38.92101 | 34.50494 | 28.27823 | 41.34284 | 29.89296 | 52.20437 |
| ENSMUSG00000049878 | 0.027628204 | 0.825172 | 0.893879 | 1367.228 | 1337.801 | 1411.094 | 1444.894 | 1371.982 | 1643.657 | 1354.826 | 1086.503 |
| ENSMUSG00000049881 | -0.155523064 | 0.05545 | 0.146659 | 978.8909 | 1049.844 | 1102.21 | 1065.34 | 1262.769 | 1065.975 | 1189.933 | 1155.022 |
| ENSMUSG00000049882 | -0.307119329 | 0.005388 | 0.027059 | 231.3925 | 215.9679 | 225.245 | 223.4195 | 269.1307 | 255.8787 | 302.7868 | 280.5985 |
| ENSMUSG00000049891 | 0.461783119 | 0.430518 | 0.595268 | 17.10292 | 9.598572 | 14.07781 | 10.35148 | 7.80089 | 3.352122 | 8.678602 | 17.40146 |
| ENSMUSG00000049892 | -0.561954154 | 0.11205 | 0.238579 | 144.8718 | 70.78947 | 125.8722 | 147.5086 | 94.5858 | 201.1273 | 279.6439 | 147.9124 |
| ENSMUSG00000049897 | 0.600973751 | 0.264029 | 0.428237 | 19.11503 | 33.595 | 14.90592 | 25.8787 | 5.850668 | 26.81698 | 10.60718 | 18.48905 |
| ENSMUSG00000049904 | -0.178198141 | 0.167654 | 0.313587 | 239.4409 | 239.9643 | 255.8849 | 267.4133 | 308.1352 | 319.569 | 246.858 | 262.1095 |
| ENSMUSG00000049907 | 0.155843422 | 0.053863 | 0.143573 | 1733.432 | 1535.772 | 1577.543 | 1795.119 | 1539.701 | 1384.427 | 1550.577 | 1487.825 |
| ENSMUSG00000049908 | -0.320374386 | 0.621212 | 0.752895 | 17.10292 | 4.799286 | 9.109173 | 18.97772 | 13.65156 | 31.28647 | 11.57147 | 6.525547 |
| ENSMUSG00000049916 | -0.066538332 | 0.667099 | 0.786407 | 418.5186 | 389.942 | 338.6956 | 389.9058 | 514.8588 | 385.4941 | 352.9298 | 354.5547 |
| ENSMUSG00000049922 | -0.240514178 | 0.137491 | 0.272589 | 355.1372 | 335.95 | 344.4924 | 297.6051 | 379.3183 | 302.8084 | 396.3228 | 495.9415 |
| ENSMUSG00000049928 | 0.275526802 | 0.546133 | 0.694558 | 10.06054 | 13.19804 | 23.18699 | 26.74133 | 13.65156 | 16.76061 | 13.50005 | 17.40146 |
| ENSMUSG00000049929 | 0.689975839 | 0.00721 | 0.033611 | 824.9646 | 914.264 | 551.519 | 799.652 | 398.8205 | 750.8754 | 423.3229 | 342.5912 |
| ENSMUSG00000049932 | -0.077743704 | 0.367298 | 0.535835 | 2763.631 | 3094.34 | 3212.226 | 3252.091 | 3619.613 | 3130.882 | 3199.511 | 3055.043 |
| ENSMUSG00000049939 | -0.416848221 | 0.136681 | 0.271407 | 62.37537 | 75.58876 | 114.2787 | 62.97152 | 100.4365 | 100.5637 | 81.00029 | 140.2993 |
| ENSMUSG00000049940 | -0.136076136 | 0.02094 | 0.072808 | 1655.965 | 1736.142 | 1785.398 | 1738.186 | 1914.143 | 1819.085 | 1872.65 | 1995.73 |
| ENSMUSG00000049946 | -0.663713993 | 0.187653 | 0.339841 | 10.06054 | 8.398751 | 14.07781 | 20.70296 | 20.47734 | 10.05637 | 23.14294 | 31.54014 |
| ENSMUSG00000049950 | -0.41084649 | 0.008707 | 0.038535 | 180.0837 | 203.9697 | 138.2938 | 175.1126 | 257.4294 | 229.0617 | 226.608 | 209.9051 |
| ENSMUSG00000049957 | -0.061657858 | 0.605649 | 0.742173 | 588.5418 | 718.6931 | 594.5806 | 626.2647 | 681.6028 | 610.0863 | 585.3235 | 759.1386 |
| ENSMUSG00000049960 | -0.308507319 | 0.019333 | 0.068816 | 757.5589 | 716.2935 | 900.1519 | 800.5146 | 1011.19 | 794.453 | 986.4678 | 1141.971 |
| ENSMUSG00000049969 | -0.376305371 | 0.000879 | 0.007046 | 733.4136 | 842.2747 | 943.2135 | 930.7707 | 1004.365 | 1119.609 | 1111.825 | 1246.379 |
| ENSMUSG00000049971 | 0.480724672 | 0.018413 | 0.066587 | 108.6539 | 127.1811 | 82.81066 | 109.5532 | 78.98402 | 68.15982 | 71.3574 | 87.00729 |
| ENSMUSG00000049988 | 0.269760257 | 0.340855 | 0.509511 | 50.30272 | 47.99286 | 65.42042 | 35.36756 | 48.75557 | 36.87335 | 42.42872 | 36.9781 |
| ENSMUSG00000049999 | 0.084947482 | 0.746041 | 0.841259 | 98.59333 | 51.59233 | 67.90474 | 99.2017 | 82.88446 | 81.56831 | 69.42882 | 66.34306 |
| ENSMUSG00000050002 | -0.018311841 | 0.874084 | 0.92474 | 409.4641 | 368.3452 | 374.3042 | 351.9504 | 369.5672 | 328.508 | 402.1086 | 421.9853 |
| ENSMUSG00000050010 | 0.492972191 | 7.32E-05 | 0.001068 | 6856.26 | 6483.836 | 5776.872 | 7026.068 | 4494.288 | 5630.448 | 4583.266 | 3868.561 |
| ENSMUSG00000050017 | -0.085690716 | 0.147437 | 0.286613 | 1674.074 | 1725.343 | 1735.712 | 1720.071 | 1904.392 | 1738.634 | 1751.149 | 1880.445 |
| ENSMUSG00000050022 | 0.188116926 | 0.348853 | 0.517452 | 97.58727 | 95.98572 | 112.6225 | 86.26235 | 83.85957 | 70.39457 | 82.92887 | 107.6715 |
| ENSMUSG00000050029 | 0.188302883 | 0.151071 | 0.291313 | 1476.888 | 1593.363 | 1409.437 | 1500.965 | 1234.491 | 1670.474 | 1204.397 | 1139.795 |
| ENSMUSG00000050035 | 0.121859038 | 0.757852 | 0.849068 | 28.16952 | 26.39607 | 21.53077 | 24.15346 | 23.40267 | 33.52122 | 24.10723 | 10.87591 |
| ENSMUSG00000050043 | -0.60537124 | 9.22E-05 | 0.001253 | 1556.366 | 1575.366 | 1927.832 | 1525.981 | 2512.862 | 1906.24 | 2471.473 | 3130.087 |
| ENSMUSG00000050050 | 0.44519703 | 0.035057 | 0.106225 | 100.6054 | 77.9884 | 101.029 | 75.91087 | 58.50668 | 64.8077 | 77.14313 | 60.9051 |
| ENSMUSG00000050052 | -1.08299075 | 0.005925 | 0.02916 | 182.0958 | 129.5807 | 452.9743 | 156.1349 | 467.0783 | 280.4609 | 615.2165 | 590.562 |
| ENSMUSG00000050064 | -0.068580399 | 0.571181 | 0.714257 | 840.0554 | 748.6886 | 794.9824 | 731.5047 | 677.7024 | 962.0591 | 799.3957 | 829.832 |
| ENSMUSG00000050069 | -1.28317478 | 0.14222 | 0.279384 | 24.1453 | 31.19536 | 216.9639 | 24.15346 | 124.8142 | 88.27255 | 327.8583 | 181.6277 |
| ENSMUSG00000050071 | 0.348060346 | 0.000255 | 0.002751 | 1413.506 | 1315.004 | 1598.246 | 1555.31 | 1111.627 | 1119.609 | 1283.469 | 1109.343 |
| ENSMUSG00000050074 | -0.643749949 | 0.064944 | 0.164102 | 409.4641 | 303.5548 | 1121.256 | 443.3885 | 913.6793 | 719.5889 | 978.7535 | 949.467 |

| | | | | | | | | | | | |
|---|---|---|---|---|---|---|---|---|---|---|---|
| ENSMUSG00000050075 | 1.361683394 | 0.110128 | 0.235983 | 44.26639 | 33.595 | 19.04645 | 25.8787 | 5.850668 | 36.87335 | 0.964289 | 4.350364 |
| ENSMUSG00000050079 | -0.170289453 | 0.109651 | 0.235317 | 853.1341 | 694.6967 | 679.0474 | 759.9713 | 806.4171 | 835.7958 | 787.8242 | 932.0656 |
| ENSMUSG00000050088 | 0.009846915 | 0.912172 | 0.948044 | 928.5882 | 940.6601 | 914.2297 | 1085.18 | 984.8624 | 928.5379 | 905.4675 | 1025.598 |
| ENSMUSG00000050097 | -0.713255243 | 0.097963 | 0.217858 | 7.04238 | 22.79661 | 17.39024 | 15.52722 | 31.20356 | 25.6996 | 18.32149 | 27.18978 |
| ENSMUSG00000050100 | -2.972412918 | 0.000492 | 0.004524 | 4.024217 | 1.199822 | 12.4216 | 1.725247 | 28.27823 | 6.704245 | 46.28588 | 73.95619 |
| ENSMUSG00000050103 | -1.073468149 | 0.028205 | 0.090562 | 6.036326 | 8.398751 | 14.07781 | 5.175741 | 15.60178 | 20.11273 | 17.3572 | 18.48905 |
| ENSMUSG00000050105 | -0.11596941 | 0.404276 | 0.571173 | 213.2835 | 241.1641 | 259.1974 | 198.4034 | 252.5538 | 229.0617 | 230.4651 | 276.2481 |
| ENSMUSG00000050106 | -0.006352328 | 0.979501 | 0.988724 | 74.44802 | 91.18644 | 111.7944 | 102.6522 | 108.2374 | 61.45558 | 99.32178 | 113.1095 |
| ENSMUSG00000050107 | -0.244479994 | 0.101767 | 0.22373 | 202.2169 | 167.975 | 198.7456 | 160.448 | 249.6285 | 206.7142 | 192.8578 | 215.343 |
| ENSMUSG00000050108 | 0.529042485 | 0.298477 | 0.466483 | 25.15136 | 22.79661 | 9.109173 | 14.6646 | 9.751113 | 12.29112 | 18.32149 | 8.700729 |
| ENSMUSG00000050132 | -0.157314007 | 0.582529 | 0.723204 | 40.24217 | 28.79572 | 32.29616 | 47.44429 | 36.07912 | 37.99072 | 51.10733 | 41.32846 |
| ENSMUSG00000050144 | -0.293551312 | 0.03695 | 0.1105 | 763.5953 | 671.9001 | 740.3273 | 695.2745 | 987.7878 | 677.1287 | 835.0744 | 1019.073 |
| ENSMUSG00000050147 | -0.53712465 | 0.137093 | 0.272092 | 24.1453 | 41.99375 | 80.32634 | 50.89479 | 68.25779 | 60.3382 | 71.3574 | 88.09488 |
| ENSMUSG00000050148 | 0.21652397 | 0.02516 | 0.083399 | 1363.204 | 1430.187 | 1363.064 | 1299.974 | 1088.224 | 1372.135 | 1150.397 | 1085.416 |
| ENSMUSG00000050164 | -0.317429886 | 0.50073 | 0.657055 | 34.20585 | 56.39161 | 31.46805 | 53.48266 | 29.25334 | 70.39457 | 93.53605 | 25.0146 |
| ENSMUSG00000050174 | -0.134346892 | 0.438484 | 0.601998 | 113.6841 | 93.58608 | 117.5911 | 101.7896 | 122.864 | 102.7984 | 108.0004 | 135.9489 |
| ENSMUSG00000050188 | -0.004590425 | 0.966872 | 0.981442 | 341.0524 | 340.7493 | 308.0557 | 308.8192 | 333.4881 | 291.6346 | 315.3226 | 359.9926 |
| ENSMUSG00000050192 | -0.297662981 | 0.002136 | 0.013757 | 380.2885 | 369.545 | 450.49 | 412.334 | 497.3068 | 502.8183 | 504.3232 | 482.8904 |
| ENSMUSG00000050195 | 0.145560754 | 0.74959 | 0.84382 | 33.19979 | 26.39607 | 14.90592 | 25.8787 | 24.37778 | 36.87335 | 17.3572 | 11.9635 |
| ENSMUSG00000050199 | -0.290967261 | 0.06572 | 0.165046 | 4224.422 | 2805.183 | 3863.946 | 4452.862 | 3876.067 | 4623.694 | 5264.055 | 5015.97 |
| ENSMUSG00000050212 | 0.037000256 | 0.668113 | 0.787099 | 2358.191 | 2741.592 | 2864.421 | 2713.814 | 2806.37 | 2409.059 | 2639.259 | 2552.576 |
| ENSMUSG00000050213 | -0.069990834 | 0.397247 | 0.564489 | 747.4984 | 721.0927 | 688.9847 | 647.8302 | 782.0393 | 699.4762 | 736.7169 | 722.1605 |
| ENSMUSG00000050222 | 0.178249336 | 0.307116 | 0.475594 | 154.9324 | 213.5682 | 216.1358 | 202.7165 | 156.9929 | 213.4185 | 169.7149 | 156.6131 |
| ENSMUSG00000050229 | 0.048059309 | 0.62433 | 0.7551 | 1214.308 | 1198.622 | 1199.927 | 1195.596 | 1101.876 | 1376.605 | 1113.754 | 1060.401 |
| ENSMUSG00000050234 | -0.791170397 | 0.012751 | 0.051239 | 146.8839 | 109.1838 | 293.9779 | 178.5631 | 312.0356 | 225.7096 | 281.5724 | 444.8248 |
| ENSMUSG00000050240 | 0.233597358 | 0.24096 | 0.403112 | 221.332 | 254.3622 | 231.8699 | 229.4578 | 165.7689 | 279.3435 | 158.1434 | 194.6788 |
| ENSMUSG00000050244 | -0.212972202 | 0.004527 | 0.023914 | 943.679 | 956.2578 | 892.6989 | 866.9366 | 1110.652 | 1041.393 | 988.3964 | 1097.379 |
| ENSMUSG00000050248 | -0.086441768 | 0.31569 | 0.48408 | 1175.071 | 1368.996 | 1393.703 | 1301.699 | 1432.439 | 1261.515 | 1447.398 | 1420.394 |
| ENSMUSG00000050270 | -0.16692166 | 0.360608 | 0.529213 | 91.55095 | 115.1829 | 84.46688 | 87.12497 | 111.1627 | 117.3243 | 99.32178 | 94.62042 |
| ENSMUSG00000050271 | -0.49627005 | 0.004855 | 0.025204 | 400.4096 | 302.355 | 441.3808 | 381.2796 | 659.1752 | 417.8979 | 499.5018 | 577.5109 |
| ENSMUSG00000050272 | 0.258159575 | 0.366947 | 0.535534 | 64.38748 | 45.59322 | 33.95237 | 53.48266 | 39.97956 | 48.04709 | 35.6787 | 41.32846 |
| ENSMUSG00000050276 | 3.031545628 | 0.000151 | 0.001834 | 34.20585 | 13.19804 | 19.04645 | 60.38364 | 5.850668 | 7.821619 | 1.928578 | 0 |
| ENSMUSG00000050288 | 0.659411052 | 1.39E-05 | 0.000294 | 3957.818 | 4955.263 | 4128.112 | 4225.13 | 2654.253 | 3563.306 | 2218.829 | 2496.022 |
| ENSMUSG00000050295 | 0.162736874 | 0.409653 | 0.575883 | 1893.394 | 2062.493 | 1722.462 | 2085.824 | 1320.301 | 2548.73 | 1601.684 | 1466.073 |
| ENSMUSG00000050299 | -0.092151931 | 0.446275 | 0.608934 | 262.5802 | 311.9536 | 330.4145 | 284.6658 | 320.8116 | 282.6956 | 334.6083 | 329.5401 |
| ENSMUSG00000050310 | 0.20628403 | 0.17152 | 0.318725 | 1503.045 | 1460.183 | 1302.612 | 1472.498 | 1231.566 | 1625.779 | 1110.861 | 1006.022 |
| ENSMUSG00000050312 | -0.214536292 | 0.029196 | 0.092758 | 280.6892 | 314.3532 | 334.5551 | 314.8576 | 357.8658 | 364.264 | 362.5727 | 361.0802 |
| ENSMUSG00000050315 | -0.248093249 | 0.362908 | 0.531351 | 169.0171 | 255.562 | 395.835 | 302.7808 | 256.4543 | 347.5033 | 286.3939 | 445.9123 |
| ENSMUSG00000050323 | -0.79200341 | 3.89E-09 | 3.95E-07 | 173.0413 | 175.1739 | 196.2613 | 178.5631 | 313.0107 | 278.2261 | 294.1082 | 368.6934 |
| ENSMUSG00000050332 | 0.338082727 | 0.044642 | 0.125859 | 1012.091 | 967.0562 | 862.8871 | 914.3809 | 682.5779 | 1023.515 | 635.4666 | 630.8028 |
| ENSMUSG00000050334 | 0.86423171 | 0.006986 | 0.032918 | 291.7558 | 267.5602 | 209.511 | 253.6113 | 123.8391 | 251.4092 | 94.50034 | 92.44524 |
| ENSMUSG00000050335 | -0.548596464 | 0.107784 | 0.23239 | 555.342 | 428.3363 | 1006.978 | 472.7177 | 836.6455 | 460.3581 | 927.6462 | 1380.153 |
| ENSMUSG00000050345 | 0.912950786 | 0.327184 | 0.494879 | 49.29666 | 5.999108 | 4.140533 | 4.313117 | 5.850668 | 10.05637 | 3.857157 | 14.13868 |
| ENSMUSG00000050357 | 0.327407665 | 0.224786 | 0.384523 | 60.36326 | 55.19179 | 52.17072 | 64.69676 | 43.88001 | 64.8077 | 32.78583 | 44.59123 |
| ENSMUSG00000050359 | -0.901775747 | 0.169658 | 0.316093 | 693.1715 | 283.1579 | 1545.247 | 511.5357 | 2375.371 | 470.4145 | 1447.398 | 1374.715 |
| ENSMUSG00000050368 | 0.088020927 | 0.385632 | 0.553051 | 1519.142 | 1642.556 | 1676.916 | 1816.685 | 1393.434 | 1796.738 | 1529.363 | 1545.467 |
| ENSMUSG00000050370 | -1.015195204 | 0.11685 | 0.245132 | 10.06054 | 1.199822 | 19.87456 | 8.626235 | 11.70134 | 21.23011 | 25.07152 | 23.927 |
| ENSMUSG00000050373 | 0.079668995 | 0.380654 | 0.548577 | 719.3289 | 649.1034 | 655.8605 | 769.4602 | 684.5281 | 686.0677 | 652.8238 | 622.1021 |
| ENSMUSG00000050377 | 0.28928539 | 0.354697 | 0.523285 | 55.33299 | 49.19268 | 35.60859 | 81.94923 | 50.70579 | 37.99072 | 46.28588 | 46.76642 |
| ENSMUSG00000050379 | 0.241264701 | 0.053824 | 0.14351 | 1455.761 | 1618.559 | 1312.549 | 1528.569 | 1307.624 | 1488.342 | 1125.325 | 1082.153 |
| ENSMUSG00000050382 | 0.009229445 | 0.934442 | 0.961265 | 563.3904 | 560.3167 | 518.3948 | 427.8613 | 526.5601 | 512.8747 | 498.5375 | 515.5182 |
| ENSMUSG00000050390 | -0.275513532 | 0.191001 | 0.343646 | 866.2128 | 764.2863 | 1298.471 | 861.7609 | 1127.229 | 857.0259 | 1099.29 | 1507.401 |
| ENSMUSG00000050394 | 0.604812449 | 1.65E-05 | 0.000337 | 503.0272 | 646.7038 | 568.0811 | 603.8364 | 359.8161 | 471.5319 | 351.0013 | 344.7664 |
| ENSMUSG00000050395 | -0.992611101 | 0.130493 | 0.262922 | 6.036326 | 15.59768 | 16.56213 | 5.175741 | 19.50223 | 5.58687 | 26.03581 | 34.80291 |
| ENSMUSG00000050410 | -0.107185553 | 0.404476 | 0.571375 | 679.0867 | 952.6583 | 908.433 | 752.2077 | 955.6091 | 911.7773 | 859.1816 | 817.8685 |
| ENSMUSG00000050428 | 0.065078041 | 0.531215 | 0.682784 | 405.4399 | 382.7431 | 394.1788 | 346.7746 | 341.289 | 349.7381 | 372.2156 | 398.0583 |
| ENSMUSG00000050447 | 0.162438551 | 0.49199 | 0.64914 | 301.8163 | 220.7672 | 489.411 | 285.5284 | 267.1805 | 267.0524 | 307.6082 | 319.7518 |
| ENSMUSG00000050471 | 0.069351326 | 0.405603 | 0.572441 | 551.3178 | 575.9143 | 594.5806 | 588.3092 | 581.1663 | 577.6824 | 526.5019 | 517.6934 |
| ENSMUSG00000050473 | 2.563481718 | 0.000272 | 0.002874 | 61.36932 | 8.398751 | 15.73403 | 30.19182 | 3.900445 | 4.469496 | 5.785735 | 5.437955 |
| ENSMUSG00000050490 | -0.182225484 | 0.673045 | 0.790437 | 12.07265 | 25.19625 | 18.21835 | 12.07673 | 15.60178 | 26.81698 | 16.39292 | 17.40146 |
| ENSMUSG00000050493 | -0.710488505 | 0.010949 | 0.045658 | 369.2219 | 154.777 | 300.6027 | 297.6051 | 371.5174 | 335.2122 | 582.4307 | 549.2335 |
| ENSMUSG00000050503 | 0.702378084 | 0.054212 | 0.144314 | 117.7084 | 214.7681 | 149.0592 | 190.6398 | 99.46135 | 167.6061 | 45.32159 | 101.146 |
| ENSMUSG00000050505 | 0.696259958 | 0.236065 | 0.397076 | 19.11503 | 49.19268 | 23.18699 | 39.68068 | 9.751113 | 45.81234 | 12.53576 | 13.05109 |
| ENSMUSG00000050520 | -0.379460439 | 0.043299 | 0.123188 | 2614.735 | 2367.248 | 3942.616 | 2654.292 | 3435.317 | 2972.215 | 4512.873 | 4144.81 |

| | | | | | | | | | | | |
|---|---|---|---|---|---|---|---|---|---|---|---|
| ENSMUSG00000050530 | -0.160033599 | 0.05144 | 0.138718 | 2188.168 | 2073.292 | 2507.507 | 2208.316 | 2418.276 | 2486.157 | 2428.08 | 2704.839 |
| ENSMUSG00000050534 | -0.818096796 | 0.033111 | 0.10154 | 16.09687 | 28.79572 | 15.73403 | 16.38985 | 48.75557 | 27.93435 | 29.89296 | 27.18978 |
| ENSMUSG00000050538 | 0.407482359 | 0.054863 | 0.145564 | 127.7689 | 131.9804 | 115.9349 | 140.6076 | 123.8391 | 92.74205 | 104.1432 | 67.43065 |
| ENSMUSG00000050541 | -0.358170922 | 0.41116 | 0.576974 | 37.22401 | 23.99643 | 77.01392 | 69.00988 | 55.58134 | 48.04709 | 54.00019 | 109.8467 |
| ENSMUSG00000050545 | -0.074322012 | 0.785798 | 0.868372 | 90.54489 | 65.99018 | 57.13936 | 48.30692 | 63.38224 | 62.57295 | 88.7146 | 59.81751 |
| ENSMUSG00000050549 | -0.474788879 | 0.004547 | 0.023986 | 226.3622 | 206.3693 | 304.7432 | 201.8539 | 297.4089 | 337.447 | 365.4656 | 307.7883 |
| ENSMUSG00000050552 | -0.228652309 | 0.051067 | 0.138074 | 969.8364 | 868.6708 | 1024.368 | 930.7707 | 1180.86 | 914.012 | 1092.54 | 1259.43 |
| ENSMUSG00000050555 | 0.375364661 | 0.003863 | 0.021297 | 387.3309 | 411.5388 | 322.1335 | 368.3402 | 267.1805 | 331.8601 | 284.4653 | 263.197 |
| ENSMUSG00000050556 | 0.135871293 | 0.388784 | 0.556189 | 542.2633 | 773.8849 | 765.1705 | 665.9453 | 604.569 | 734.1148 | 510.109 | 652.5547 |
| ENSMUSG00000050565 | -0.024055874 | 0.702249 | 0.812227 | 3332.052 | 3025.95 | 3164.195 | 3311.612 | 3166.186 | 3101.83 | 3367.298 | 3417.211 |
| ENSMUSG00000050567 | 0.128788373 | 0.056113 | 0.147609 | 1155.956 | 1131.432 | 1241.332 | 1207.673 | 1083.349 | 1023.515 | 1107.004 | 1121.306 |
| ENSMUSG00000050587 | -0.003557372 | 0.988217 | 0.993293 | 64.38748 | 70.78947 | 72.87338 | 81.08661 | 48.75557 | 74.86406 | 90.64318 | 76.13138 |
| ENSMUSG00000050592 | -0.081702333 | 0.723233 | 0.826859 | 45.27245 | 52.79215 | 43.06154 | 44.85642 | 48.75557 | 42.46022 | 49.17875 | 55.46715 |
| ENSMUSG00000050605 | 0.197255584 | 0.070794 | 0.173776 | 773.6558 | 939.4603 | 731.2182 | 784.9874 | 711.8313 | 741.9364 | 730.9312 | 627.5401 |
| ENSMUSG00000050608 | -0.315849596 | 5.38E-05 | 0.000839 | 2615.741 | 2555.62 | 2588.661 | 2637.04 | 3140.834 | 2909.642 | 3284.369 | 3607.54 |
| ENSMUSG00000050619 | -0.022491743 | 0.85555 | 0.913423 | 689.1472 | 634.7056 | 756.8895 | 698.725 | 675.7521 | 588.8561 | 732.8598 | 827.6568 |
| ENSMUSG00000050621 | -0.10138178 | 0.422072 | 0.587229 | 225.3562 | 266.3604 | 250.0882 | 247.5729 | 294.4836 | 230.1791 | 254.5723 | 280.5985 |
| ENSMUSG00000050623 | 0.466693095 | 0.159876 | 0.303067 | 73.44197 | 53.99197 | 113.4506 | 71.59775 | 59.48179 | 35.75597 | 80.036 | 51.11678 |
| ENSMUSG00000050627 | -0.358183923 | 0.007975 | 0.036144 | 906.455 | 715.0936 | 938.2448 | 950.6111 | 1029.718 | 967.646 | 1265.147 | 1240.941 |
| ENSMUSG00000050628 | -0.424129324 | 0.000123 | 0.001567 | 1619.748 | 1446.985 | 1742.336 | 1531.157 | 2221.304 | 1766.568 | 2165.793 | 2355.722 |
| ENSMUSG00000050635 | 0.543200504 | NA | NA | 28.16952 | 14.39786 | 1132.022 | 62.97152 | 230.1263 | 170.9582 | 141.7505 | 306.7007 |
| ENSMUSG00000050640 | -0.001088226 | 0.988953 | 0.993799 | 1300.828 | 1097.837 | 1136.99 | 1193.008 | 1220.839 | 1143.074 | 1220.79 | 1147.409 |
| ENSMUSG00000050645 | -4.028964731 | 3.44E-07 | 1.54E-05 | 3.018163 | 2.399643 | 17.39024 | 1.725247 | 134.5654 | 17.87799 | 98.35749 | 157.7007 |
| ENSMUSG00000050663 | 1.435948278 | 0.015263 | 0.058395 | 18.10898 | 33.595 | 23.18699 | 34.50494 | 2.925334 | 17.87799 | 4.821446 | 15.22628 |
| ENSMUSG00000050666 | -0.194338742 | 0.326398 | 0.494094 | 736.4318 | 592.7118 | 1057.492 | 865.2114 | 841.5211 | 740.819 | 1097.361 | 1044.087 |
| ENSMUSG00000050668 | -0.059564677 | 0.574027 | 0.716553 | 574.457 | 543.5192 | 461.2554 | 514.1236 | 496.3317 | 563.1565 | 535.1805 | 585.124 |
| ENSMUSG00000050671 | 0.182561498 | 0.682375 | 0.797284 | 26.15741 | 21.59679 | 16.56213 | 27.60395 | 34.1289 | 13.40849 | 12.53576 | 20.66423 |
| ENSMUSG00000050675 | 0.579866636 | 0.130162 | 0.262572 | 64.38748 | 69.58965 | 72.04528 | 87.12497 | 40.95467 | 89.38993 | 42.42872 | 23.927 |
| ENSMUSG00000050677 | -1.105566405 | 0.010935 | 0.04564 | 16.09687 | 17.99732 | 7.45296 | 9.488858 | 22.42756 | 21.23011 | 35.6787 | 28.27737 |
| ENSMUSG00000050697 | 0.120656174 | 0.105018 | 0.228481 | 1238.453 | 1107.435 | 1096.413 | 1204.222 | 1026.792 | 1112.905 | 1095.432 | 1039.737 |
| ENSMUSG00000050700 | 0.411031 | 0.245068 | 0.407954 | 173.0413 | 265.1606 | 279.9 | 277.7648 | 176.4951 | 341.9165 | 121.5004 | 109.8467 |
| ENSMUSG00000050702 | 1.22113185 | 0.027298 | 0.088387 | 42.25428 | 52.79215 | 18.21835 | 21.56559 | 8.776002 | 26.81698 | 14.46434 | 7.613138 |
| ENSMUSG00000050705 | -0.366761225 | 0.005986 | 0.02942 | 520.1301 | 448.7333 | 489.411 | 480.4813 | 600.6686 | 505.0531 | 649.9309 | 744.9999 |
| ENSMUSG00000050708 | -0.493658961 | 0.000725 | 0.006116 | 20797.16 | 19811.45 | 26021.59 | 21820.92 | 33878.29 | 23893.93 | 29971.07 | 36799.73 |
| ENSMUSG00000050711 | 1.761967144 | 0.015529 | 0.059143 | 860.1765 | 113.983 | 81.98256 | 376.1038 | 97.51113 | 94.9768 | 108.0004 | 121.8102 |
| ENSMUSG00000050714 | 0.31423713 | 0.078375 | 0.18638 | 490.9545 | 479.9286 | 373.4761 | 437.3501 | 287.6578 | 477.1187 | 336.5369 | 331.7153 |
| ENSMUSG00000050721 | 0.163286307 | 0.146896 | 0.285922 | 527.1725 | 574.7145 | 512.598 | 444.2511 | 500.2321 | 436.8933 | 429.1087 | 468.7518 |
| ENSMUSG00000050730 | -0.24188011 | 0.003093 | 0.018196 | 709.2683 | 752.2881 | 753.577 | 668.5332 | 789.8402 | 864.8476 | 855.3245 | 899.4378 |
| ENSMUSG00000050732 | -0.384978678 | 0.009114 | 0.039867 | 1525.178 | 1484.179 | 1935.285 | 1678.665 | 2116.967 | 1682.765 | 2251.615 | 2600.43 |
| ENSMUSG00000050737 | -1.117283682 | 0.00049 | 0.004507 | 378.2764 | 303.5548 | 731.2182 | 395.9442 | 782.0393 | 576.565 | 1258.397 | 1310.547 |
| ENSMUSG00000050751 | -0.156804435 | 0.662579 | 0.782896 | 33.19979 | 32.39518 | 36.43669 | 43.9938 | 51.6809 | 55.8687 | 21.21436 | 34.80291 |
| ENSMUSG00000050761 | 0.131127025 | 0.769326 | 0.856839 | 30.18163 | 23.99643 | 25.67131 | 81.08661 | 34.1289 | 35.75597 | 37.60728 | 40.24087 |
| ENSMUSG00000050762 | -0.866741297 | 0.205357 | 0.36182 | 10.06054 | 5.999108 | 8.281066 | 6.038364 | 13.65156 | 2.234748 | 24.10723 | 15.22628 |
| ENSMUSG00000050777 | -1.611157178 | 2.70E-06 | 8.31E-05 | 313.889 | 211.1686 | 454.6305 | 268.2759 | 1145.756 | 410.0763 | 887.146 | 1372.54 |
| ENSMUSG00000050786 | -0.247190248 | 0.072463 | 0.176738 | 187.1261 | 205.1695 | 208.6829 | 232.0457 | 244.7529 | 210.0663 | 258.4295 | 276.2481 |
| ENSMUSG00000050796 | -0.253523113 | 0.033316 | 0.101953 | 360.1675 | 374.3443 | 410.7409 | 306.2313 | 452.4516 | 400.0199 | 431.0373 | 445.9123 |
| ENSMUSG00000050812 | 0.010667919 | 0.83906 | 0.902746 | 3156.999 | 3106.338 | 3095.463 | 3108.895 | 2917.533 | 3237.033 | 3148.404 | 3074.62 |
| ENSMUSG00000050821 | 0.355174314 | 0.127049 | 0.25821 | 135.8173 | 107.9839 | 120.9036 | 193.2277 | 120.9138 | 108.3853 | 120.5361 | 87.00729 |
| ENSMUSG00000050822 | 0.557901017 | 0.026214 | 0.08598 | 177.0656 | 89.98662 | 101.029 | 131.9814 | 92.63557 | 89.38993 | 83.89316 | 73.95619 |
| ENSMUSG00000050830 | -0.83135725 | 0.01501 | 0.057724 | 25.15136 | 35.99465 | 52.99882 | 21.56559 | 60.4569 | 53.63396 | 53.0359 | 75.04379 |
| ENSMUSG00000050840 | -0.393821317 | 0.424716 | 0.589675 | 12.07265 | 13.19804 | 24.01509 | 6.900988 | 24.37778 | 18.99536 | 14.46434 | 16.31387 |
| ENSMUSG00000050846 | -0.595505217 | 9.82E-07 | 3.67E-05 | 471.8395 | 527.9215 | 562.2844 | 534.8266 | 942.9326 | 703.9457 | 728.0383 | 793.9415 |
| ENSMUSG00000050854 | -1.486821926 | 4.36E-05 | 0.000719 | 58.35115 | 57.59143 | 110.1382 | 62.10889 | 180.3956 | 94.9768 | 260.3581 | 274.073 |
| ENSMUSG00000050855 | 0.442491935 | 0.0006 | 0.005266 | 275.6589 | 291.5566 | 226.9012 | 257.0618 | 195.9974 | 198.8926 | 197.6793 | 178.3649 |
| ENSMUSG00000050856 | -0.730186935 | 0.000124 | 0.001581 | 1048.309 | 1070.241 | 1264.519 | 944.5727 | 1861.487 | 1195.59 | 1802.256 | 2319.832 |
| ENSMUSG00000050866 | -3.175850256 | 9.10E-06 | 0.000213 | 80.48435 | 16.7975 | 216.1358 | 67.28463 | 864.9237 | 420.1327 | 829.2887 | 1332.299 |
| ENSMUSG00000050875 | 0.612354023 | 3.44E-06 | 0.000101 | 1028.188 | 1107.435 | 850.4655 | 1117.097 | 673.8019 | 782.1619 | 651.8595 | 575.3357 |
| ENSMUSG00000050890 | -0.070187492 | 0.555811 | 0.702687 | 468.8213 | 439.1347 | 438.8965 | 385.5927 | 449.5263 | 518.4616 | 394.3943 | 456.7883 |
| ENSMUSG00000050891 | 0.034171639 | 0.799617 | 0.877444 | 334.01 | 358.7466 | 310.54 | 369.2029 | 322.7618 | 379.9072 | 361.6084 | 275.1605 |
| ENSMUSG00000050896 | 1.814263203 | 3.80E-20 | 5.39E-17 | 157.9505 | 139.1793 | 147.403 | 165.6237 | 40.95467 | 36.87335 | 54.96448 | 40.24087 |
| ENSMUSG00000050908 | 0.207010917 | 0.132071 | 0.265176 | 197.1867 | 262.7609 | 243.4633 | 230.3205 | 214.5245 | 192.1883 | 186.1078 | 215.343 |
| ENSMUSG00000050910 | -0.209435152 | 0.105 | 0.22847 | 485.9243 | 383.9429 | 420.6782 | 380.417 | 505.1077 | 476.0014 | 414.6443 | 537.27 |
| ENSMUSG00000050912 | -0.240762733 | 0.014274 | 0.055602 | 2532.239 | 2310.856 | 2786.579 | 2547.327 | 3113.53 | 2573.313 | 3238.083 | 3102.897 |
| ENSMUSG00000050914 | -0.29655856 | 0.302282 | 0.4706 | 98.59333 | 57.59143 | 95.23226 | 69.00988 | 115.0631 | 67.04245 | 82.92887 | 129.4233 |

| | | | | | | | | | | | |
|---|---|---|---|---|---|---|---|---|---|---|---|
| ENSMUSG00000050919 | -1.038327075 | 0.053177 | 0.142096 | 32.19374 | 8.398751 | 38.09291 | 36.23019 | 49.73068 | 22.34748 | 93.53605 | 71.78101 |
| ENSMUSG00000050926 | 0.823048069 | 0.009181 | 0.040038 | 38.23007 | 52.79215 | 42.23344 | 50.89479 | 25.35289 | 37.99072 | 19.28578 | 21.75182 |
| ENSMUSG00000050930 | -0.070102472 | 0.606526 | 0.742606 | 173.0413 | 179.9732 | 192.1207 | 162.1732 | 192.0969 | 207.8316 | 175.5006 | 167.489 |
| ENSMUSG00000050931 | -0.516577143 | 0.006119 | 0.029877 | 680.0927 | 478.7288 | 689.8128 | 646.105 | 729.3833 | 765.4013 | 1199.576 | 876.5984 |
| ENSMUSG00000050944 | 0.653177856 | 0.113474 | 0.240582 | 32.19374 | 22.79661 | 17.39024 | 26.74133 | 10.72622 | 18.99536 | 21.21436 | 11.9635 |
| ENSMUSG00000050945 | 0.041654424 | 0.834971 | 0.900183 | 125.7568 | 143.9786 | 86.12309 | 131.1188 | 112.1378 | 122.9112 | 127.2862 | 108.7591 |
| ENSMUSG00000050947 | 0.303127719 | 0.000403 | 0.003882 | 574.457 | 619.1079 | 569.7374 | 593.485 | 506.0828 | 498.3488 | 458.0374 | 445.9123 |
| ENSMUSG00000050953 | 0.497978708 | 0.006483 | 0.031237 | 4229.453 | 4029.001 | 2407.306 | 3459.983 | 2551.866 | 2949.868 | 2435.794 | 2063.16 |
| ENSMUSG00000050954 | 0.126150445 | 0.559847 | 0.705919 | 132.7992 | 106.7841 | 130.8408 | 94.02596 | 89.71024 | 128.498 | 86.78602 | 121.8102 |
| ENSMUSG00000050957 | -0.771402735 | 0.037391 | 0.111305 | 22.1332 | 29.99554 | 37.2648 | 31.05445 | 52.65601 | 24.58223 | 66.53595 | 61.99269 |
| ENSMUSG00000050965 | 0.045435829 | 0.645761 | 0.770604 | 967.8243 | 992.2524 | 1205.723 | 1040.324 | 946.8331 | 998.9324 | 1034.682 | 1099.555 |
| ENSMUSG00000050967 | 1.162965063 | 0.027567 | 0.089052 | 38.23007 | 10.79839 | 19.87456 | 24.15346 | 5.850668 | 12.29112 | 9.642892 | 14.13868 |
| ENSMUSG00000050973 | -0.286973524 | 0.237219 | 0.3983 | 91.55095 | 73.18911 | 99.3728 | 53.48266 | 92.63557 | 89.38993 | 90.64318 | 115.2847 |
| ENSMUSG00000050989 | -0.023079082 | 0.813522 | 0.886798 | 1183.12 | 1329.402 | 1489.764 | 1420.741 | 1284.222 | 1489.46 | 1421.362 | 1319.248 |
| ENSMUSG00000051000 | -0.439973688 | 0.003768 | 0.020938 | 264.5923 | 254.3622 | 303.087 | 212.2054 | 378.3432 | 301.691 | 390.5371 | 331.7153 |
| ENSMUSG00000051007 | 0.15563015 | 0.157841 | 0.300382 | 1279.701 | 1514.175 | 1325.799 | 1396.587 | 1306.649 | 1337.497 | 1286.362 | 1019.073 |
| ENSMUSG00000051022 | -0.039255552 | 0.85549 | 0.913423 | 157.9505 | 128.3809 | 140.7781 | 169.9368 | 213.5494 | 126.2633 | 129.2147 | 144.6496 |
| ENSMUSG00000051034 | 0.018262561 | 0.916564 | 0.95061 | 150.9082 | 200.3702 | 154.8559 | 147.5086 | 181.3707 | 179.8972 | 141.7505 | 140.2993 |
| ENSMUSG00000051037 | 0.14555738 | 0.619174 | 0.751531 | 73.44197 | 86.38715 | 79.49824 | 48.30692 | 63.38224 | 91.62468 | 57.85735 | 46.76642 |
| ENSMUSG00000051041 | 0.261441565 | 0.020426 | 0.071587 | 852.128 | 1066.641 | 1058.32 | 1029.972 | 907.8286 | 765.4013 | 904.5032 | 763.4889 |
| ENSMUSG00000051043 | -0.658509971 | 0.000731 | 0.006153 | 1985.951 | 1778.136 | 2896.717 | 2165.185 | 3170.087 | 2606.834 | 3706.728 | 4451.51 |
| ENSMUSG00000051048 | 0.176821618 | 0.418178 | 0.583777 | 412.4823 | 439.1347 | 353.6015 | 498.5964 | 387.1192 | 535.2222 | 276.751 | 308.8759 |
| ENSMUSG00000051065 | 0.31022356 | 0.026184 | 0.085926 | 944.685 | 797.8813 | 675.735 | 938.5344 | 691.3539 | 682.7156 | 750.217 | 581.8612 |
| ENSMUSG00000051067 | 0.074129659 | 0.806497 | 0.881775 | 42.25428 | 43.19358 | 34.78048 | 41.40593 | 49.73068 | 32.40385 | 45.32159 | 25.0146 |
| ENSMUSG00000051074 | -0.449573666 | 0.043836 | 0.124276 | 52.31483 | 39.59411 | 56.31125 | 49.16954 | 66.30757 | 79.33356 | 60.75022 | 65.25547 |
| ENSMUSG00000051076 | -0.603324732 | 0.20203 | 0.35757 | 27.16347 | 25.19625 | 72.04528 | 15.52722 | 57.53157 | 64.8077 | 41.46443 | 50.02919 |
| ENSMUSG00000051098 | 0.402971137 | 0.051746 | 0.139277 | 234.4107 | 211.1686 | 190.4645 | 236.3588 | 139.4409 | 234.6486 | 148.5005 | 138.1241 |
| ENSMUSG00000051107 | -0.072470775 | 0.889698 | 0.933475 | 20.12109 | 19.19714 | 15.73403 | 5.175741 | 15.60178 | 10.05637 | 16.39292 | 20.66423 |
| ENSMUSG00000051111 | 1.881366183 | 0.000177 | 0.002061 | 140.8476 | 56.39161 | 51.34261 | 199.266 | 18.52711 | 49.16446 | 26.03581 | 28.27737 |
| ENSMUSG00000051113 | -0.046289936 | 0.692097 | 0.805049 | 202.2169 | 209.9688 | 198.7456 | 200.1287 | 207.6987 | 230.1791 | 203.465 | 195.7664 |
| ENSMUSG00000051124 | -0.404542071 | 0.063389 | 0.161165 | 140.8476 | 94.7859 | 143.2624 | 144.9207 | 163.8187 | 139.6718 | 162.9649 | 229.4817 |
| ENSMUSG00000051146 | 0.138590265 | 0.739261 | 0.836889 | 96.58122 | 27.5959 | 34.78048 | 52.62003 | 41.92979 | 51.39921 | 59.78593 | 39.15328 |
| ENSMUSG00000051147 | -0.273643792 | 0.132877 | 0.266242 | 111.672 | 80.38804 | 99.3728 | 83.67448 | 100.4365 | 118.4417 | 123.429 | 112.0219 |
| ENSMUSG00000051149 | -0.430021766 | 0.10486 | 0.228252 | 86.52067 | 79.18822 | 82.81066 | 63.83414 | 109.2125 | 62.57295 | 136.9291 | 110.9343 |
| ENSMUSG00000051154 | -0.052524777 | 0.395372 | 0.562692 | 2227.404 | 2296.458 | 2265.7 | 2441.224 | 2487.509 | 2478.336 | 2323.937 | 2285.029 |
| ENSMUSG00000051159 | -1.632765622 | 5.91E-07 | 2.41E-05 | 553.3299 | 595.1115 | 481.9581 | 580.5456 | 2593.796 | 630.199 | 1955.578 | 1674.89 |
| ENSMUSG00000051166 | 1.066244 | 5.88E-07 | 2.41E-05 | 745.4863 | 575.9143 | 472.0208 | 753.0703 | 301.3094 | 410.0763 | 248.7866 | 256.6715 |
| ENSMUSG00000051169 | -0.412230204 | 0.085217 | 0.19731 | 72.43591 | 67.19001 | 72.04528 | 68.14726 | 92.63557 | 98.32892 | 61.71451 | 120.7226 |
| ENSMUSG00000051177 | -0.378535385 | 0.14823 | 0.287688 | 465.8032 | 249.5629 | 508.4575 | 329.5222 | 399.7956 | 385.4941 | 573.7521 | 662.343 |
| ENSMUSG00000051184 | -0.180616335 | 0.409729 | 0.575929 | 123.7447 | 68.38983 | 122.5598 | 113.0037 | 108.2374 | 121.7938 | 121.5004 | 135.9489 |
| ENSMUSG00000051185 | -0.23339039 | 0.009542 | 0.041155 | 638.8445 | 617.9081 | 589.6119 | 626.2647 | 677.7024 | 681.5982 | 735.7526 | 811.343 |
| ENSMUSG00000051212 | -0.718070064 | 0.187829 | 0.340127 | 7.04238 | 11.99822 | 16.56213 | 12.07673 | 19.50223 | 26.81698 | 6.750024 | 26.10219 |
| ENSMUSG00000051220 | -0.048478227 | 0.64389 | 0.769677 | 412.4823 | 448.7333 | 380.1009 | 361.4392 | 420.273 | 395.5504 | 401.1443 | 436.124 |
| ENSMUSG00000051223 | -0.035282422 | 0.432487 | 0.596842 | 8294.918 | 8797.091 | 8891.381 | 8778.057 | 9121.191 | 8832.842 | 8677.638 | 8992.203 |
| ENSMUSG00000051224 | 0.06413812 | 0.687221 | 0.801255 | 378.2764 | 365.9456 | 322.9616 | 425.2734 | 405.6463 | 416.7805 | 309.5368 | 295.8248 |
| ENSMUSG00000051228 | -0.183698314 | 0.773911 | 0.859922 | 28.16952 | 15.59768 | 45.54586 | 7.763611 | 26.32801 | 18.99536 | 53.0359 | 11.9635 |
| ENSMUSG00000051232 | -0.136567102 | 0.090899 | 0.206579 | 630.7961 | 562.7163 | 563.9406 | 579.683 | 660.1504 | 641.3727 | 659.5738 | 606.8758 |
| ENSMUSG00000051234 | -0.13217852 | 0.0327 | 0.100617 | 2460.809 | 2779.986 | 2778.298 | 2723.302 | 2916.558 | 2924.168 | 2893.832 | 3039.817 |
| ENSMUSG00000051235 | 0.02718175 | 0.841142 | 0.904194 | 226.3622 | 229.1659 | 194.6051 | 228.5952 | 206.7236 | 234.6486 | 186.1078 | 234.9197 |
| ENSMUSG00000051236 | 0.045398772 | 0.612957 | 0.746981 | 1050.321 | 895.0669 | 1055.008 | 1061.027 | 947.8082 | 1067.092 | 997.075 | 928.8028 |
| ENSMUSG00000051238 | -0.290955575 | 0.002904 | 0.017327 | 398.3975 | 363.5459 | 423.1625 | 400.2573 | 515.8339 | 449.1844 | 466.716 | 511.1678 |
| ENSMUSG00000051242 | 0.33420227 | 0.066459 | 0.166202 | 127.7689 | 128.3809 | 154.0278 | 125.0804 | 112.1378 | 126.2633 | 103.1789 | 83.74451 |
| ENSMUSG00000051243 | 0.87898067 | 0.029307 | 0.092968 | 92.557 | 41.99375 | 49.6864 | 129.3935 | 31.20356 | 52.51658 | 48.21446 | 39.15328 |
| ENSMUSG00000051246 | -0.015253269 | 0.962393 | 0.978526 | 22.1332 | 26.39607 | 28.15563 | 32.77969 | 27.30312 | 29.05173 | 20.25007 | 34.80291 |
| ENSMUSG00000051255 | -0.030481999 | 0.821597 | 0.891669 | 307.8526 | 386.3425 | 320.4773 | 360.5766 | 327.6374 | 415.6632 | 346.1798 | 314.3138 |
| ENSMUSG00000051256 | -0.392691979 | 0.00736 | 0.034102 | 717.3168 | 693.4968 | 712.9998 | 700.4503 | 913.6793 | 703.9457 | 928.6105 | 1161.547 |
| ENSMUSG00000051278 | 0.156220235 | 0.449572 | 0.61217 | 493.9727 | 563.9161 | 463.7397 | 468.4046 | 388.0943 | 655.8986 | 399.2157 | 342.5912 |
| ENSMUSG00000051285 | 0.420859405 | 0.000483 | 0.004461 | 2295.816 | 2462.034 | 1881.458 | 2340.298 | 1534.825 | 1977.752 | 1625.792 | 1568.306 |
| ENSMUSG00000051306 | 0.19566675 | 0.078714 | 0.18709 | 1116.72 | 1100.236 | 941.5572 | 1105.021 | 945.858 | 1051.449 | 934.3962 | 789.5911 |
| ENSMUSG00000051316 | -0.079059289 | 0.459095 | 0.620555 | 679.0867 | 695.8965 | 810.7164 | 715.1149 | 766.4375 | 876.0213 | 698.1454 | 727.5984 |
| ENSMUSG00000051319 | -0.250322996 | 0.107144 | 0.231478 | 400.4096 | 333.5504 | 332.8989 | 343.3242 | 421.2481 | 316.2169 | 447.4302 | 491.5912 |
| ENSMUSG00000051323 | 0.734326957 | 0.019646 | 0.069614 | 441.6579 | 541.1195 | 513.4261 | 526.2003 | 195.9974 | 565.3913 | 233.358 | 221.8686 |
| ENSMUSG00000051329 | 0.141574404 | 0.084475 | 0.196085 | 957.7637 | 1049.844 | 863.7152 | 974.7645 | 896.1273 | 852.5564 | 866.896 | 865.7225 |
| ENSMUSG00000051331 | 0.777504257 | 0.000725 | 0.006116 | 473.8516 | 345.5486 | 297.2903 | 319.1707 | 224.2756 | 283.813 | 169.7149 | 159.8759 |

| | | | | | | | | | | | |
|---|---|---|---|---|---|---|---|---|---|---|---|
| ENSMUSG00000051335 | 0.219478581 | 0.182948 | 0.333743 | 102.6175 | 112.7832 | 102.6852 | 125.943 | 102.3867 | 94.9768 | 92.57176 | 91.35765 |
| ENSMUSG00000051339 | 0.37503979 | 2.04E-05 | 0.000399 | 1030.2 | 1006.65 | 1141.131 | 1176.618 | 804.4668 | 915.1294 | 827.3601 | 815.6933 |
| ENSMUSG00000051341 | 0.499963794 | 0.041908 | 0.120494 | 305.8405 | 163.1757 | 168.9338 | 166.4863 | 125.7894 | 130.7328 | 162.9649 | 149 |
| ENSMUSG00000051343 | -0.152772794 | 0.362386 | 0.531001 | 1015.109 | 1031.847 | 1530.341 | 1098.982 | 1474.368 | 1057.036 | 1228.504 | 1441.058 |
| ENSMUSG00000051344 | 0.342294869 | 0.063295 | 0.161055 | 423.5489 | 367.1454 | 322.9616 | 379.5543 | 205.7485 | 357.5597 | 291.2153 | 324.1021 |
| ENSMUSG00000051346 | -0.412011503 | 0.010836 | 0.045318 | 301.8163 | 302.355 | 382.5853 | 310.5445 | 436.8499 | 328.508 | 453.2159 | 508.9926 |
| ENSMUSG00000051351 | 0.025126382 | 0.802851 | 0.879477 | 789.7527 | 909.4647 | 793.3262 | 830.7064 | 706.9557 | 859.2607 | 885.2175 | 812.4305 |
| ENSMUSG00000051355 | -0.164090448 | 0.067296 | 0.167501 | 1263.604 | 1436.186 | 1427.656 | 1505.278 | 1705.47 | 1421.3 | 1621.934 | 1561.781 |
| ENSMUSG00000051359 | -0.025314183 | 0.786246 | 0.868451 | 914.5034 | 812.2792 | 939.0729 | 948.0232 | 901.978 | 883.8429 | 864.9674 | 1032.124 |
| ENSMUSG00000051367 | 0.727390123 | 0.226344 | 0.386224 | 24.1453 | 28.79572 | 20.70267 | 34.50494 | 26.32801 | 27.93435 | 6.750024 | 4.350364 |
| ENSMUSG00000051373 | -0.336260204 | 0.174046 | 0.322088 | 61.36932 | 80.38804 | 57.13936 | 63.83414 | 92.63557 | 56.98608 | 78.10742 | 102.2336 |
| ENSMUSG00000051375 | -0.116599715 | 0.497141 | 0.653697 | 318.9192 | 287.9572 | 415.7095 | 346.7746 | 346.1645 | 293.8694 | 420.4301 | 426.3357 |
| ENSMUSG00000051378 | 0.043243875 | 0.691059 | 0.804376 | 602.6266 | 602.3104 | 566.4249 | 672.8463 | 667.9512 | 524.0485 | 567.002 | 612.3138 |
| ENSMUSG00000051379 | 0.116809326 | 0.486134 | 0.644361 | 537.233 | 585.5129 | 521.7072 | 524.4751 | 499.257 | 665.955 | 389.5728 | 445.9123 |
| ENSMUSG00000051390 | -0.156562701 | 0.109079 | 0.23428 | 1366.222 | 1277.81 | 1618.948 | 1486.3 | 1659.639 | 1450.352 | 1658.577 | 1644.438 |
| ENSMUSG00000051391 | 0.001203862 | 0.98746 | 0.992894 | 3189.192 | 2902.368 | 3025.902 | 3070.077 | 3037.472 | 2787.848 | 2976.761 | 3376.97 |
| ENSMUSG00000051396 | -0.07649647 | 0.672564 | 0.790252 | 196.1806 | 188.372 | 144.0906 | 133.7066 | 164.7938 | 187.7188 | 190.9293 | 152.2628 |
| ENSMUSG00000051397 | -0.624422135 | 0.156216 | 0.298311 | 1311.895 | 848.2738 | 2814.734 | 1284.446 | 2631.825 | 1409.009 | 2540.902 | 3069.182 |
| ENSMUSG00000051403 | -0.142604376 | 0.126523 | 0.257511 | 1303.846 | 1319.804 | 1363.892 | 1217.162 | 1552.377 | 1251.459 | 1392.434 | 1547.642 |
| ENSMUSG00000051412 | 0.197767986 | 0.036641 | 0.109822 | 872.2491 | 939.4603 | 822.3099 | 1009.269 | 739.1344 | 838.0306 | 821.5744 | 777.6276 |
| ENSMUSG00000051413 | 0.297346238 | 0.004153 | 0.022429 | 817.9222 | 729.4915 | 682.3599 | 690.0988 | 611.3948 | 657.016 | 545.7877 | 561.197 |
| ENSMUSG00000051427 | 0.13603517 | 0.234232 | 0.395045 | 907.461 | 951.4585 | 788.3575 | 889.3648 | 840.5459 | 915.1294 | 774.3242 | 686.27 |
| ENSMUSG00000051435 | 0.276776186 | 0.519733 | 0.673278 | 23.13925 | 11.99822 | 23.18699 | 15.52722 | 11.70134 | 16.76061 | 12.53576 | 20.66423 |
| ENSMUSG00000051439 | -0.045031944 | 0.849612 | 0.909644 | 64.38748 | 58.79126 | 89.43552 | 87.12497 | 83.85957 | 69.27719 | 64.60737 | 93.53283 |
| ENSMUSG00000051444 | 0.217147067 | 0.138956 | 0.274785 | 319.9253 | 365.9456 | 283.2125 | 316.5828 | 307.1601 | 311.7474 | 255.5366 | 229.4817 |
| ENSMUSG00000051451 | 0.135341757 | 0.112981 | 0.239923 | 3568.475 | 3141.133 | 2900.029 | 3378.034 | 3097.929 | 3070.544 | 2877.439 | 2777.708 |
| ENSMUSG00000051455 | 2.1551098 | 2.63E-13 | 9.14E-11 | 70.4238 | 97.18554 | 101.029 | 87.9876 | 14.62667 | 29.05173 | 16.39292 | 20.66423 |
| ENSMUSG00000051457 | -0.054539888 | 0.862939 | 0.917928 | 35.2119 | 23.99643 | 26.49941 | 43.9938 | 28.27823 | 32.40385 | 35.6787 | 39.15328 |
| ENSMUSG00000051469 | -0.121182553 | 0.030526 | 0.095668 | 1837.055 | 1706.146 | 1711.696 | 1789.081 | 1903.417 | 1982.222 | 1913.15 | 1864.131 |
| ENSMUSG00000051483 | -0.990228929 | 4.47E-06 | 0.000124 | 1711.298 | 1724.144 | 2375.01 | 1871.893 | 3987.23 | 2377.772 | 3711.549 | 5186.722 |
| ENSMUSG00000051486 | -0.326718191 | 0.339743 | 0.508231 | 26.15741 | 50.3925 | 36.43669 | 43.9938 | 38.02934 | 72.62932 | 50.14304 | 35.89051 |
| ENSMUSG00000051495 | 0.176416871 | 0.212959 | 0.370828 | 2654.977 | 2358.849 | 2373.354 | 2608.573 | 1811.757 | 2810.196 | 2282.472 | 1942.438 |
| ENSMUSG00000051497 | -1.481039677 | 5.25E-06 | 0.00014 | 629.79 | 353.9474 | 973.0253 | 602.9738 | 1681.092 | 931.89 | 2105.043 | 2431.854 |
| ENSMUSG00000051498 | 0.30215437 | 0.376746 | 0.54488 | 40.24217 | 38.39429 | 44.71776 | 30.19182 | 22.42756 | 41.34284 | 21.21436 | 40.24087 |
| ENSMUSG00000051499 | -0.075943086 | 0.705323 | 0.814655 | 60.36326 | 71.98929 | 57.13936 | 57.79577 | 65.33246 | 69.27719 | 61.71451 | 63.08028 |
| ENSMUSG00000051502 | -0.975176838 | 0.003668 | 0.020491 | 49.29666 | 47.99286 | 103.5133 | 59.52102 | 96.53602 | 91.62468 | 163.9292 | 162.0511 |
| ENSMUSG00000051506 | 0.156559209 | 0.687106 | 0.80119 | 31.18769 | 28.79572 | 47.20208 | 17.25247 | 34.1289 | 21.23011 | 27.0001 | 29.36496 |
| ENSMUSG00000051510 | -0.045715826 | 0.619023 | 0.751427 | 1728.401 | 1442.185 | 1531.169 | 1682.116 | 1479.244 | 1701.761 | 1654.72 | 1757.547 |
| ENSMUSG00000051515 | 0.123443291 | 0.706638 | 0.815556 | 32.19374 | 45.59322 | 66.24853 | 71.59775 | 48.75557 | 60.3382 | 54.00019 | 35.89051 |
| ENSMUSG00000051517 | 0.205342597 | 0.066874 | 0.166901 | 466.8092 | 416.3381 | 366.8512 | 434.7622 | 393.945 | 367.6161 | 349.0727 | 349.1167 |
| ENSMUSG00000051518 | -0.241952458 | 0.002839 | 0.017017 | 631.8021 | 647.9036 | 725.4214 | 672.8463 | 841.5211 | 781.0445 | 762.7527 | 785.2408 |
| ENSMUSG00000051527 | 0.551480208 | 0.037204 | 0.110857 | 298.7981 | 287.9572 | 215.3077 | 240.672 | 150.1671 | 267.0524 | 184.1792 | 109.8467 |
| ENSMUSG00000051537 | 0.607023123 | 0.056138 | 0.147645 | 31.18769 | 38.39429 | 24.8432 | 31.91707 | 21.45245 | 20.11273 | 22.17865 | 18.48905 |
| ENSMUSG00000051550 | 0.146070651 | 0.083033 | 0.193881 | 698.2017 | 772.6851 | 708.0312 | 752.2077 | 669.9015 | 620.1426 | 637.3951 | 719.9853 |
| ENSMUSG00000051557 | -0.26807571 | 0.028887 | 0.092055 | 351.113 | 313.1534 | 356.914 | 317.4454 | 463.1779 | 386.6114 | 349.0727 | 414.3722 |
| ENSMUSG00000051579 | 0.006891009 | 0.934622 | 0.961343 | 2317.949 | 2695.999 | 2736.892 | 2742.28 | 2532.364 | 2856.008 | 2541.866 | 2515.598 |
| ENSMUSG00000051586 | 0.258824161 | 0.038661 | 0.113652 | 1114.708 | 826.677 | 944.8697 | 937.6717 | 717.6819 | 928.5379 | 790.7171 | 761.3138 |
| ENSMUSG00000051615 | -0.268629267 | 0.018384 | 0.066516 | 786.7345 | 754.6877 | 860.4028 | 833.2943 | 872.7246 | 946.4159 | 913.1818 | 1169.16 |
| ENSMUSG00000051617 | 1.320744314 | 0.026831 | 0.087429 | 24.1453 | 7.198929 | 16.56213 | 11.21411 | 6.825779 | 7.821619 | 4.821446 | 4.350364 |
| ENSMUSG00000051627 | -0.218615568 | 0.661899 | 0.782325 | 14.08476 | 20.39697 | 9.93728 | 12.07673 | 13.65156 | 27.93435 | 9.642892 | 14.13868 |
| ENSMUSG00000051650 | -1.418415635 | 0.000339 | 0.003394 | 72.43591 | 43.19358 | 92.74794 | 58.6584 | 160.8934 | 71.51194 | 218.8936 | 264.2846 |
| ENSMUSG00000051652 | -0.087612566 | 0.73793 | 0.835997 | 126.7628 | 131.9804 | 207.0267 | 187.1893 | 174.5449 | 124.0285 | 152.3577 | 244.708 |
| ENSMUSG00000051669 | -0.799370389 | 0.015036 | 0.057778 | 129.781 | 87.58697 | 162.3089 | 81.94923 | 195.0223 | 116.2069 | 189.965 | 303.4379 |
| ENSMUSG00000051671 | -0.538775133 | 0.002074 | 0.013477 | 352.119 | 308.3541 | 352.7734 | 365.7524 | 536.3112 | 357.5597 | 483.1089 | 627.5401 |
| ENSMUSG00000051674 | 0.015353889 | 0.884948 | 0.931269 | 790.7587 | 866.2711 | 719.6247 | 775.4985 | 763.5122 | 698.3588 | 760.8242 | 892.9123 |
| ENSMUSG00000051675 | 0.071451689 | 0.406751 | 0.573149 | 1098.611 | 1019.848 | 942.3853 | 966.1383 | 987.7878 | 869.317 | 1001.896 | 969.0437 |
| ENSMUSG00000051678 | 0.550495093 | 0.021014 | 0.072999 | 71.42986 | 86.38715 | 86.9512 | 59.52102 | 51.6809 | 45.81234 | 63.64308 | 45.67883 |
| ENSMUSG00000051695 | -0.210638341 | 0.020704 | 0.072206 | 6322.046 | 6577.422 | 6913.862 | 6660.316 | 7439.124 | 6647.258 | 7867.635 | 8682.24 |
| ENSMUSG00000051703 | 0.021524797 | 0.939939 | 0.964823 | 38.23007 | 32.39518 | 27.32752 | 41.40593 | 37.05423 | 39.10809 | 31.82154 | 29.36496 |
| ENSMUSG00000051705 | 0.023077006 | 0.912071 | 0.948028 | 223.3441 | 175.1739 | 131.669 | 134.5693 | 144.3165 | 159.7845 | 165.8577 | 182.7153 |
| ENSMUSG00000051721 | 0.152240303 | 0.386826 | 0.55432 | 139.8416 | 135.5798 | 117.5911 | 123.3552 | 95.56091 | 105.0332 | 127.2862 | 135.9489 |
| ENSMUSG00000051726 | -0.843888949 | 0.104325 | 0.22735 | 24.1453 | 10.79839 | 64.59232 | 28.46658 | 38.02934 | 43.57759 | 85.82174 | 64.16787 |
| ENSMUSG00000051727 | 0.204564126 | 0.700157 | 0.810722 | 95.57516 | 153.5772 | 101.029 | 89.71284 | 56.55646 | 232.4138 | 55.92877 | 36.9781 |
| ENSMUSG00000051730 | 0.072417095 | 0.519368 | 0.672947 | 390.3491 | 343.149 | 399.9755 | 405.433 | 409.5467 | 348.6207 | 332.6798 | 375.2189 |

| | | | | | | | | | | | |
|---|---|---|---|---|---|---|---|---|---|---|---|
| ENSMUSG00000051735 | -0.196204438 | 0.447014 | 0.609627 | 63.38142 | 68.38983 | 77.01392 | 71.59775 | 77.03379 | 65.92507 | 61.71451 | 117.4598 |
| ENSMUSG00000051736 | 0.070863164 | 0.724356 | 0.827323 | 179.0777 | 221.967 | 144.9187 | 194.9529 | 174.5449 | 219.0053 | 175.5006 | 134.8613 |
| ENSMUSG00000051747 | 0.788306592 | 0.137982 | 0.273269 | 56.33904 | 43.19358 | 81.15445 | 31.05445 | 9.751113 | 60.3382 | 27.0001 | 26.10219 |
| ENSMUSG00000051768 | 0.110321385 | 0.094105 | 0.211419 | 1924.582 | 2117.685 | 2027.205 | 2055.632 | 1928.77 | 1997.865 | 1804.185 | 1795.613 |
| ENSMUSG00000051786 | 0.044676847 | 0.627125 | 0.757029 | 786.7345 | 719.8929 | 703.0625 | 789.3005 | 783.0144 | 695.0067 | 664.3952 | 765.6641 |
| ENSMUSG00000051790 | 0.599067107 | 2.12E-06 | 6.73E-05 | 2112.714 | 2092.489 | 2063.642 | 2160.009 | 1275.446 | 1754.277 | 1361.576 | 1174.598 |
| ENSMUSG00000051811 | 0.495568046 | 1.38E-05 | 0.000294 | 949.7153 | 1077.44 | 982.9626 | 970.4514 | 807.3922 | 753.1101 | 668.2524 | 592.7371 |
| ENSMUSG00000051817 | -0.06101328 | 0.428474 | 0.593464 | 2473.888 | 2556.82 | 2490.117 | 2593.909 | 2960.438 | 2638.12 | 2484.009 | 2467.744 |
| ENSMUSG00000051839 | 0.893360643 | 0.194112 | 0.347616 | 101.6115 | 15.59768 | 15.73403 | 31.05445 | 23.40267 | 37.99072 | 9.642892 | 17.40146 |
| ENSMUSG00000051844 | 1.381539428 | 0.005085 | 0.026035 | 21.12714 | 16.7975 | 14.90592 | 17.25247 | 8.776002 | 8.938993 | 4.821446 | 4.350364 |
| ENSMUSG00000051851 | -0.210536748 | 0.330724 | 0.498558 | 394.3733 | 371.9447 | 467.0521 | 468.4046 | 424.1734 | 332.9775 | 530.359 | 683.0072 |
| ENSMUSG00000051853 | -0.148823757 | 0.062343 | 0.15921 | 3936.691 | 3694.251 | 3888.789 | 3852.477 | 4453.333 | 3753.26 | 4192.729 | 4644.014 |
| ENSMUSG00000051855 | -0.067576854 | 0.550498 | 0.698381 | 21018.49 | 23474.51 | 20041.84 | 24934.13 | 27465.96 | 22838.01 | 23020.48 | 20432.57 |
| ENSMUSG00000051860 | 0.022223311 | 0.942106 | 0.966348 | 32.19374 | 35.99465 | 38.09291 | 22.42821 | 31.20356 | 34.6386 | 26.03581 | 34.80291 |
| ENSMUSG00000051864 | -0.260567122 | 0.082602 | 0.193195 | 637.8385 | 551.9179 | 816.5131 | 678.0221 | 836.6455 | 652.5465 | 873.646 | 855.9342 |
| ENSMUSG00000051890 | 0.344605838 | 0.071261 | 0.174639 | 139.8416 | 158.3764 | 115.9349 | 144.9207 | 83.85957 | 115.0895 | 127.2862 | 113.1095 |
| ENSMUSG00000051910 | -0.428121396 | 0.073607 | 0.178749 | 348.0948 | 260.3613 | 531.6445 | 333.8353 | 396.8703 | 413.4284 | 568.9306 | 606.8758 |
| ENSMUSG00000051920 | -0.211680171 | 0.813427 | 0.886746 | 10.06054 | 22.79661 | 8.281066 | 5.175741 | 9.751113 | 35.75597 | 6.750024 | 1.087591 |
| ENSMUSG00000051934 | -0.088167862 | 0.182687 | 0.333335 | 1699.226 | 1780.535 | 1666.979 | 1737.324 | 1883.915 | 1910.71 | 1842.757 | 1677.065 |
| ENSMUSG00000051950 | 0.336995846 | 0.002056 | 0.013383 | 1399.422 | 1384.594 | 1253.753 | 1186.97 | 1014.116 | 1201.177 | 1011.539 | 908.1386 |
| ENSMUSG00000051951 | 0.096763718 | 0.843255 | 0.905412 | 22.1332 | 25.19625 | 26.49941 | 18.97772 | 16.57689 | 36.87335 | 7.714313 | 26.10219 |
| ENSMUSG00000051977 | -0.087375748 | 0.655513 | 0.777916 | 86.52067 | 107.9839 | 98.54469 | 93.16334 | 108.2374 | 92.74205 | 82.92887 | 126.1606 |
| ENSMUSG00000051978 | -0.295279901 | 0.029862 | 0.094227 | 233.4046 | 199.1704 | 229.3855 | 206.167 | 281.8072 | 219.0053 | 278.6796 | 286.0365 |
| ENSMUSG00000051980 | -2.640358309 | 0.009136 | 0.039948 | 11.0666 | 2.399643 | 81.98256 | 18.97772 | 142.3663 | 41.34284 | 241.0723 | 291.4744 |
| ENSMUSG00000051984 | 0.008011892 | 0.975619 | 0.986739 | 87.52673 | 61.1909 | 81.15445 | 112.1411 | 93.61069 | 60.3382 | 104.1432 | 82.65692 |
| ENSMUSG00000051985 | -1.249180622 | 0.007806 | 0.035589 | 13.07871 | 21.59679 | 24.01509 | 9.488858 | 63.38224 | 23.46486 | 28.92867 | 45.67883 |
| ENSMUSG00000051989 | 0.072518467 | 0.402193 | 0.569282 | 780.6982 | 788.2827 | 693.9534 | 768.5975 | 726.4579 | 665.955 | 707.7882 | 779.8028 |
| ENSMUSG00000051998 | -0.376159117 | 0.455283 | 0.617301 | 29.17558 | 10.79839 | 25.67131 | 9.488858 | 19.50223 | 21.23011 | 19.28578 | 38.06569 |
| ENSMUSG00000052031 | -0.178131261 | 0.015872 | 0.059989 | 935.6306 | 921.4629 | 850.4655 | 875.5628 | 1075.548 | 1030.219 | 963.3249 | 981.0072 |
| ENSMUSG00000052033 | -0.123347093 | 0.177328 | 0.326362 | 527.1725 | 581.9134 | 536.6131 | 617.6384 | 572.3903 | 644.7249 | 625.8237 | 623.1897 |
| ENSMUSG00000052040 | -0.02426422 | 0.878458 | 0.927104 | 1252.538 | 1017.449 | 1127.053 | 1382.785 | 1014.116 | 1319.619 | 1497.541 | 1031.036 |
| ENSMUSG00000052056 | 0.461274203 | 5.54E-06 | 0.000145 | 1610.693 | 1380.995 | 1328.283 | 1364.67 | 1113.577 | 1108.435 | 935.3605 | 971.2188 |
| ENSMUSG00000052062 | 0.356579249 | 0.038676 | 0.113664 | 788.7466 | 857.8724 | 643.4389 | 706.4886 | 519.7343 | 776.575 | 567.9663 | 475.2773 |
| ENSMUSG00000052085 | -0.915864348 | 0.006271 | 0.030411 | 290.7497 | 185.9723 | 547.3785 | 309.6818 | 570.4401 | 365.3813 | 747.3241 | 836.3575 |
| ENSMUSG00000052087 | -0.577815873 | 0.348369 | 0.517007 | 24.1453 | 9.598572 | 25.67131 | 13.80198 | 30.22845 | 5.58687 | 26.03581 | 47.85401 |
| ENSMUSG00000052102 | -0.681326736 | 0.000422 | 0.004017 | 768.6255 | 563.9161 | 808.2321 | 685.7857 | 1072.622 | 822.3873 | 1125.325 | 1515.014 |
| ENSMUSG00000052105 | -0.041504413 | 0.658235 | 0.779727 | 614.6992 | 506.3247 | 521.7072 | 564.1558 | 549.9628 | 592.2083 | 552.5377 | 578.5985 |
| ENSMUSG00000052117 | 0.674995553 | 2.85E-06 | 8.66E-05 | 391.3551 | 333.5504 | 336.2113 | 357.1261 | 199.8978 | 277.1088 | 198.6436 | 214.2554 |
| ENSMUSG00000052125 | 0.214080519 | 0.482012 | 0.640751 | 70.4238 | 71.98929 | 48.03018 | 84.5371 | 37.05423 | 81.56831 | 57.85735 | 60.9051 |
| ENSMUSG00000052131 | 3.411395445 | 0.013588 | 0.053608 | 14259.81 | 37.19447 | 389.2101 | 1431.092 | 218.4249 | 241.3528 | 288.3225 | 766.7517 |
| ENSMUSG00000052133 | -0.011837181 | 0.94708 | 0.969541 | 258.556 | 158.3764 | 209.511 | 175.1126 | 203.7983 | 216.7706 | 201.5364 | 187.0657 |
| ENSMUSG00000052135 | 0.615101519 | 0.054383 | 0.144646 | 87.52673 | 100.785 | 84.46688 | 80.22398 | 29.25334 | 84.92043 | 67.50024 | 48.9416 |
| ENSMUSG00000052137 | 0.525110096 | 0.000401 | 0.003866 | 452.7245 | 482.3283 | 361.0545 | 442.5259 | 255.4792 | 362.0292 | 291.2153 | 299.0875 |
| ENSMUSG00000052139 | -0.04152733 | 0.563931 | 0.709179 | 1448.718 | 1468.582 | 1560.153 | 1402.626 | 1420.737 | 1474.934 | 1621.934 | 1534.591 |
| ENSMUSG00000052142 | 0.296264808 | 0.513837 | 0.668361 | 13.07871 | 13.19804 | 14.90592 | 16.38985 | 13.65156 | 11.17374 | 15.42863 | 6.525547 |
| ENSMUSG00000052143 | -0.416180158 | 0.276951 | 0.443045 | 20.12109 | 22.79661 | 30.63995 | 13.80198 | 33.15378 | 18.99536 | 33.75012 | 30.45255 |
| ENSMUSG00000052144 | 0.123936655 | 0.214528 | 0.372744 | 2525.196 | 2415.241 | 2202.764 | 2268.7 | 2071.136 | 2527.5 | 2078.043 | 1959.839 |
| ENSMUSG00000052146 | -0.130694516 | 0.133582 | 0.267103 | 19225.7 | 22248.29 | 23261.52 | 21596.64 | 25684.43 | 21674.82 | 24556.59 | 22601.23 |
| ENSMUSG00000052151 | -0.699160901 | 0.002476 | 0.015326 | 696.1896 | 619.1079 | 1090.616 | 667.6706 | 1216.939 | 839.1479 | 1359.648 | 1577.007 |
| ENSMUSG00000052154 | 0.092689309 | 0.825516 | 0.894011 | 17.10292 | 23.99643 | 14.90592 | 11.21411 | 12.67645 | 16.76061 | 14.46434 | 18.48905 |
| ENSMUSG00000052155 | 0.123483286 | 0.304143 | 0.472636 | 1113.702 | 1175.825 | 1142.787 | 1261.156 | 962.4349 | 1312.915 | 1080.968 | 953.8174 |
| ENSMUSG00000052160 | 0.203993785 | 0.527451 | 0.680315 | 55.33299 | 134.38 | 147.403 | 94.88858 | 83.85957 | 80.45093 | 123.429 | 87.00729 |
| ENSMUSG00000052180 | 1.123845327 | 0.121509 | 0.250741 | 15.09082 | 27.5959 | 7.45296 | 36.23019 | 12.67645 | 11.17374 | 14.46434 | 1.087591 |
| ENSMUSG00000052188 | 0.172939595 | 0.495647 | 0.652332 | 161.9748 | 172.7743 | 109.3101 | 216.5185 | 181.3707 | 106.1505 | 158.1434 | 139.2117 |
| ENSMUSG00000052212 | 0.617653718 | 0.244993 | 0.407865 | 16.09687 | 7.198929 | 10.76539 | 14.6646 | 6.825779 | 10.05637 | 9.642892 | 5.437955 |
| ENSMUSG00000052214 | -0.449080167 | 5.69E-05 | 0.000874 | 787.7406 | 837.4754 | 992.8999 | 845.371 | 1176.959 | 1020.163 | 1264.183 | 1269.219 |
| ENSMUSG00000052221 | 0.255839562 | 0.198464 | 0.353079 | 170.0232 | 193.1713 | 192.1207 | 169.9368 | 168.6943 | 197.7752 | 120.5361 | 120.7226 |
| ENSMUSG00000052229 | 0.556116374 | 0.282316 | 0.448943 | 10.06054 | 11.99822 | 14.07781 | 11.21411 | 7.80089 | 12.29112 | 4.821446 | 7.613138 |
| ENSMUSG00000052234 | -2.840750146 | 4.50E-06 | 0.000124 | 12.07265 | 7.198929 | 6.624853 | 11.21411 | 35.10401 | 32.40385 | 159.1077 | 39.15328 |
| ENSMUSG00000052241 | -0.290259079 | 0.193356 | 0.346575 | 60.36326 | 53.99197 | 49.6864 | 62.10889 | 86.78491 | 62.57295 | 63.64308 | 63.08028 |
| ENSMUSG00000052248 | -0.025954098 | 0.892513 | 0.934962 | 79.47829 | 75.58876 | 94.40416 | 87.12497 | 75.08357 | 78.21619 | 101.2504 | 89.18247 |
| ENSMUSG00000052253 | -0.115579493 | 0.155888 | 0.297926 | 1657.978 | 1586.164 | 1743.164 | 1797.707 | 1920.969 | 1638.07 | 1870.721 | 1923.949 |
| ENSMUSG00000052270 | 0.001547658 | 0.997327 | 0.998409 | 14.08476 | 11.99822 | 9.109173 | 10.35148 | 13.65156 | 8.938993 | 11.57147 | 10.87591 |
| ENSMUSG00000052271 | 0.134283353 | 0.639987 | 0.766639 | 34.20585 | 41.99375 | 33.12427 | 33.64232 | 29.25334 | 24.58223 | 38.57157 | 36.9781 |

| | | | | | | | | | | | |
|---|---|---|---|---|---|---|---|---|---|---|---|
| ENSMUSG00000052291 | 0.200146646 | 0.218748 | 0.377522 | 186.1201 | 212.3684 | 181.3554 | 251.8861 | 158.9431 | 192.1883 | 172.6078 | 201.2044 |
| ENSMUSG00000052293 | -0.126460375 | 0.583025 | 0.723674 | 101.6115 | 88.78679 | 72.87338 | 94.88858 | 94.5858 | 130.7328 | 91.60747 | 73.95619 |
| ENSMUSG00000052296 | -0.346563065 | 0.002165 | 0.013919 | 1840.073 | 1734.942 | 2040.455 | 1820.998 | 2259.333 | 1990.043 | 2464.723 | 2743.992 |
| ENSMUSG00000052298 | 0.020650933 | 0.728019 | 0.829271 | 2313.925 | 2480.031 | 2280.606 | 2350.649 | 2279.81 | 2266.035 | 2464.723 | 2274.153 |
| ENSMUSG00000052299 | 0.037364548 | 0.674408 | 0.791543 | 1518.136 | 1394.193 | 1250.441 | 1386.236 | 1261.794 | 1471.582 | 1367.362 | 1305.109 |
| ENSMUSG00000052301 | -0.822098221 | 0.001014 | 0.007919 | 156.9445 | 73.18911 | 125.8722 | 103.5148 | 195.0223 | 157.5497 | 235.2866 | 226.2189 |
| ENSMUSG00000052302 | -0.194799758 | 0.33713 | 0.50549 | 74.44802 | 55.19179 | 85.29498 | 81.94923 | 79.95913 | 89.38993 | 81.96458 | 91.35765 |
| ENSMUSG00000052305 | 0.430298152 | 0.364503 | 0.533191 | 7772.776 | 4464.536 | 5639.406 | 15291.73 | 6322.622 | 8428.353 | 3955.514 | 5908.882 |
| ENSMUSG00000052310 | -0.199712769 | 0.059172 | 0.153209 | 2608.699 | 2595.214 | 3074.76 | 2824.229 | 3314.403 | 2686.167 | 3218.797 | 3534.671 |
| ENSMUSG00000052316 | 0.154588234 | 0.592196 | 0.731956 | 30.18163 | 40.79393 | 49.6864 | 37.09281 | 33.15378 | 39.10809 | 28.92867 | 41.32846 |
| ENSMUSG00000052331 | 0.215026738 | 0.160718 | 0.304207 | 455.7426 | 453.5325 | 370.1637 | 507.2226 | 352.0152 | 478.2361 | 362.5727 | 346.9416 |
| ENSMUSG00000052336 | -0.280414109 | 0.526221 | 0.679434 | 45.27245 | 211.1686 | 228.5574 | 87.9876 | 167.7191 | 210.0663 | 132.1076 | 185.9781 |
| ENSMUSG00000052337 | -0.504823939 | 0.000542 | 0.004878 | 3753.589 | 3228.72 | 3927.71 | 3481.548 | 5041.325 | 3802.424 | 5433.769 | 6144.89 |
| ENSMUSG00000052353 | 0.978101 | 4.68E-10 | 6.49E-08 | 278.6771 | 277.1588 | 210.3391 | 279.49 | 132.6151 | 116.2069 | 133.0719 | 147.9124 |
| ENSMUSG00000052364 | 0.346390894 | 0.017714 | 0.064893 | 260.5681 | 310.7538 | 236.8385 | 308.8192 | 245.728 | 223.4748 | 215.0365 | 192.5036 |
| ENSMUSG00000052369 | -0.040078618 | 0.616303 | 0.749357 | 835.0251 | 941.8599 | 838.0439 | 864.3487 | 905.8784 | 922.951 | 923.789 | 820.0437 |
| ENSMUSG00000052371 | 0.255964477 | 0.065884 | 0.165361 | 539.2451 | 476.3291 | 418.1938 | 441.6632 | 349.0898 | 458.1234 | 416.5729 | 345.854 |
| ENSMUSG00000052372 | 1.750502293 | 0.015946 | 0.060178 | 54.32694 | 69.58965 | 51.34261 | 46.58167 | 0.975111 | 43.57759 | 8.678602 | 13.05109 |
| ENSMUSG00000052373 | 0.359894542 | 0.04564 | 0.127741 | 395.3794 | 446.3336 | 427.303 | 420.0976 | 321.7867 | 440.2454 | 316.2868 | 238.1824 |
| ENSMUSG00000052374 | 0.097863692 | 0.80255 | 0.87942 | 29.17558 | 15.59768 | 26.49941 | 17.25247 | 15.60178 | 27.93435 | 20.25007 | 19.57664 |
| ENSMUSG00000052384 | -0.319009115 | 0.221573 | 0.380724 | 226.3622 | 194.3711 | 386.7258 | 191.5024 | 284.7325 | 239.1181 | 350.037 | 374.1313 |
| ENSMUSG00000052387 | 0.730093527 | 2.41E-05 | 0.000454 | 162.9808 | 143.9786 | 160.6527 | 195.8155 | 97.51113 | 84.92043 | 104.1432 | 114.1971 |
| ENSMUSG00000052392 | -2.411064524 | 3.15E-08 | 2.23E-06 | 50.30272 | 19.19714 | 48.03018 | 37.09281 | 189.1716 | 81.56831 | 208.2865 | 346.9416 |
| ENSMUSG00000052395 | -0.059798498 | 0.425532 | 0.590542 | 777.68 | 796.6815 | 779.2483 | 756.5208 | 811.2926 | 877.1387 | 794.5743 | 758.051 |
| ENSMUSG00000052396 | -2.376894552 | 4.15E-05 | 0.000694 | 21.12714 | 15.59768 | 48.85829 | 32.77969 | 206.7236 | 49.16446 | 66.53595 | 295.8248 |
| ENSMUSG00000052397 | -0.692192331 | 0.000141 | 0.001742 | 5687.225 | 5353.604 | 7002.47 | 5954.69 | 8909.592 | 6613.737 | 11148.15 | 12104.89 |
| ENSMUSG00000052406 | -0.082507567 | 0.317823 | 0.486138 | 1146.902 | 1144.63 | 981.3064 | 1129.174 | 1214.014 | 1106.2 | 1126.29 | 1210.489 |
| ENSMUSG00000052407 | 0.59350326 | 0.001001 | 0.007842 | 173.0413 | 181.1731 | 139.95 | 148.3712 | 110.1876 | 128.498 | 89.67889 | 96.79561 |
| ENSMUSG00000052415 | 1.211888583 | 5.65E-16 | 3.48E-13 | 2214.326 | 1912.516 | 2773.329 | 1978.858 | 870.7744 | 914.012 | 891.9675 | 1158.285 |
| ENSMUSG00000052419 | 0.00305824 | 0.961752 | 0.978182 | 976.8788 | 1025.847 | 1049.211 | 1068.791 | 1006.315 | 1015.693 | 1035.647 | 1057.139 |
| ENSMUSG00000052423 | -0.076990041 | 0.471058 | 0.630903 | 956.7577 | 829.0767 | 969.7129 | 1006.682 | 948.7833 | 869.317 | 1073.254 | 1079.978 |
| ENSMUSG00000052428 | -0.285725101 | 0.008337 | 0.037408 | 1462.803 | 1427.788 | 1651.245 | 1400.901 | 1718.146 | 1581.084 | 1852.399 | 2094.7 |
| ENSMUSG00000052430 | 0.46083761 | 0.001486 | 0.010486 | 1028.188 | 1101.436 | 1077.367 | 1022.209 | 679.6526 | 1012.341 | 717.4311 | 664.5182 |
| ENSMUSG00000052446 | 0.006654782 | 0.953652 | 0.973604 | 554.3359 | 598.7109 | 565.5968 | 514.9862 | 506.0828 | 608.9689 | 489.8589 | 618.8393 |
| ENSMUSG00000052456 | -0.421361497 | 0.001251 | 0.009255 | 1423.567 | 1430.187 | 1688.509 | 1455.246 | 2105.265 | 1576.615 | 2016.329 | 2335.058 |
| ENSMUSG00000052459 | -0.922488513 | 0.000168 | 0.00198 | 2442.7 | 1962.908 | 3654.435 | 2449.851 | 4372.399 | 3256.028 | 5315.162 | 6980.16 |
| ENSMUSG00000052485 | -0.486969637 | 0.048834 | 0.133961 | 215.2956 | 157.1766 | 320.4773 | 232.0457 | 345.1894 | 236.8833 | 301.8225 | 415.4598 |
| ENSMUSG00000052488 | 0.00796671 | 0.894063 | 0.936094 | 1600.632 | 1574.166 | 1697.619 | 1558.761 | 1651.839 | 1622.427 | 1580.47 | 1543.292 |
| ENSMUSG00000052496 | -0.524748344 | 0.11784 | 0.246482 | 33.19979 | 25.19625 | 26.49941 | 18.11509 | 25.35289 | 39.10809 | 45.32159 | 38.06569 |
| ENSMUSG00000052504 | -0.238427875 | 0.421109 | 0.586533 | 140.8476 | 188.372 | 358.5702 | 222.5569 | 191.1218 | 275.9914 | 263.2509 | 345.854 |
| ENSMUSG00000052512 | 0.722254774 | 0.000292 | 0.00304 | 2613.729 | 2446.436 | 1960.957 | 2208.316 | 1268.62 | 2049.264 | 1116.647 | 1160.46 |
| ENSMUSG00000052516 | 0.283265543 | 0.072527 | 0.176847 | 3158.005 | 3273.113 | 2355.963 | 3449.631 | 2910.707 | 2812.431 | 2305.615 | 2025.095 |
| ENSMUSG00000052520 | -5.110139959 | 2.39E-09 | 2.69E-07 | 20.12109 | 9.598572 | 101.8571 | 32.77969 | 1768.852 | 379.9072 | 887.146 | 2659.16 |
| ENSMUSG00000052525 | 0.321674301 | 0.546252 | 0.69465 | 28.16952 | 28.79572 | 35.60859 | 42.26855 | 18.52711 | 61.45558 | 14.46434 | 14.13868 |
| ENSMUSG00000052533 | -0.159189021 | 0.050318 | 0.13665 | 1102.636 | 1259.813 | 1122.084 | 1109.334 | 1241.317 | 1210.116 | 1279.612 | 1394.292 |
| ENSMUSG00000052534 | 0.665201157 | 0.001024 | 0.00798 | 13049.53 | 16940.28 | 14398.29 | 15604 | 7857.447 | 14267.75 | 7826.171 | 7880.685 |
| ENSMUSG00000052539 | 0.089127429 | 0.290068 | 0.457172 | 950.7214 | 980.2542 | 933.2762 | 835.8822 | 822.9939 | 926.3031 | 888.1103 | 838.5327 |
| ENSMUSG00000052544 | 0.214213739 | 0.003754 | 0.020893 | 759.571 | 787.0829 | 744.4679 | 787.5752 | 627.9717 | 706.1804 | 666.3238 | 653.6422 |
| ENSMUSG00000052557 | 0.548089417 | 0.001085 | 0.008307 | 732.4076 | 700.6958 | 611.1427 | 640.9293 | 429.049 | 616.7905 | 424.2872 | 366.5182 |
| ENSMUSG00000052560 | 0.338913519 | 0.006413 | 0.030958 | 452.7245 | 503.925 | 515.0823 | 459.7783 | 362.7414 | 453.6539 | 382.8228 | 328.4525 |
| ENSMUSG00000052562 | -3.233211126 | 1.11E-12 | 3.30E-10 | 7.04238 | 11.99822 | 7.45296 | 4.313117 | 69.2329 | 60.3382 | 44.3573 | 109.8467 |
| ENSMUSG00000052563 | 0.416738147 | 0.001466 | 0.0104 | 334.01 | 287.9572 | 313.0243 | 352.813 | 251.5787 | 267.0524 | 244.9294 | 202.2919 |
| ENSMUSG00000052566 | -0.27416799 | 0.004925 | 0.025461 | 743.4742 | 761.8867 | 828.1066 | 685.7857 | 904.9033 | 813.4483 | 965.2535 | 966.8685 |
| ENSMUSG00000052572 | 0.858798033 | 0.003453 | 0.019598 | 202.2169 | 98.38537 | 97.71658 | 140.6076 | 61.43201 | 96.09417 | 69.42882 | 70.69342 |
| ENSMUSG00000052581 | 1.061644579 | 0.009274 | 0.04031 | 26.15741 | 34.79482 | 16.56213 | 29.3292 | 11.70134 | 15.64324 | 10.60718 | 13.05109 |
| ENSMUSG00000052584 | 0.015272787 | 0.927944 | 0.957316 | 175.0535 | 142.7788 | 184.6678 | 182.8762 | 182.3458 | 202.2447 | 152.3577 | 143.562 |
| ENSMUSG00000052593 | -0.089023332 | 0.262941 | 0.427261 | 1621.76 | 1668.952 | 1693.478 | 1667.451 | 1669.391 | 1606.784 | 1872.65 | 1926.124 |
| ENSMUSG00000052595 | -2.676927263 | 1.26E-05 | 0.000274 | 7.04238 | 3.599465 | 9.109173 | 4.313117 | 35.10401 | 15.64324 | 28.92867 | 76.13138 |
| ENSMUSG00000052605 | -0.83856926 | 0.00798 | 0.036152 | 67.40564 | 55.19179 | 74.5296 | 42.26855 | 111.1627 | 58.10345 | 119.5719 | 139.2117 |
| ENSMUSG00000052609 | -0.378169175 | 0.028753 | 0.091698 | 1161.993 | 892.6672 | 1392.047 | 1074.829 | 1387.583 | 1130.783 | 1690.399 | 1670.54 |
| ENSMUSG00000052613 | 0.911930851 | 0.000449 | 0.004215 | 109.6599 | 135.5798 | 120.9036 | 133.7066 | 58.50668 | 102.7984 | 54.96448 | 50.02919 |
| ENSMUSG00000052616 | -0.231738222 | 0.533331 | 0.684646 | 22.1332 | 19.19714 | 36.43669 | 43.13117 | 24.37778 | 41.34284 | 36.64299 | 41.32846 |
| ENSMUSG00000052629 | 1.324779613 | 0.000678 | 0.005799 | 32.19374 | 29.99554 | 29.81184 | 39.68068 | 8.776002 | 17.87799 | 17.3572 | 8.700729 |
| ENSMUSG00000052632 | 0.091699143 | 0.347374 | 0.516236 | 817.9222 | 741.4897 | 790.0137 | 917.8314 | 756.6864 | 772.1055 | 830.253 | 709.1094 |

| | | | | | | | | | | | |
|---|---|---|---|---|---|---|---|---|---|---|---|
| ENSMUSG00000052656 | -0.186602635 | 0.023923 | 0.080468 | 925.57 | 883.0686 | 941.5572 | 920.4193 | 965.3602 | 1107.318 | 1129.183 | 977.7444 |
| ENSMUSG00000052658 | -0.294000473 | 0.346359 | 0.515184 | 36.21796 | 22.79661 | 37.2648 | 24.15346 | 32.17867 | 31.28647 | 42.42872 | 42.41605 |
| ENSMUSG00000052673 | -0.856832035 | 0.046178 | 0.128838 | 34.20585 | 25.19625 | 25.67131 | 28.46658 | 61.43201 | 16.76061 | 60.75022 | 66.34306 |
| ENSMUSG00000052675 | 0.233343185 | 0.236904 | 0.398024 | 202.2169 | 183.5727 | 143.2624 | 186.3267 | 130.6649 | 198.8926 | 149.4648 | 129.4233 |
| ENSMUSG00000052676 | 0.793842774 | 0.00068 | 0.005809 | 461.7789 | 553.1177 | 330.4145 | 413.1967 | 196.9725 | 359.7945 | 252.6438 | 204.4671 |
| ENSMUSG00000052681 | -0.021799386 | 0.674033 | 0.791346 | 3966.872 | 4165.78 | 4067.66 | 4295.865 | 4303.166 | 4028.134 | 4134.872 | 4280.759 |
| ENSMUSG00000052684 | -0.080745657 | 0.603297 | 0.740618 | 1565.421 | 1181.824 | 1707.556 | 1484.575 | 1213.038 | 1596.728 | 1811.899 | 1662.927 |
| ENSMUSG00000052688 | 0.817721257 | 0.00878 | 0.038786 | 114.6902 | 155.9768 | 111.7944 | 113.8663 | 70.20801 | 113.9722 | 54.96448 | 42.41605 |
| ENSMUSG00000052698 | -0.194777126 | 0.128668 | 0.260297 | 1088.551 | 1157.828 | 1437.593 | 1175.756 | 1264.719 | 1251.459 | 1416.541 | 1632.474 |
| ENSMUSG00000052707 | 0.397802104 | 0.022548 | 0.076743 | 4961.86 | 4837.68 | 3683.418 | 4452 | 3153.51 | 4649.394 | 3088.618 | 2721.153 |
| ENSMUSG00000052712 | -0.28488686 | 0.011602 | 0.047743 | 1052.333 | 1139.83 | 1332.424 | 1246.491 | 1449.015 | 1250.342 | 1503.327 | 1612.898 |
| ENSMUSG00000052713 | 0.208268018 | 0.012882 | 0.051672 | 1210.283 | 1189.023 | 1113.803 | 1314.638 | 993.6384 | 1046.98 | 1007.682 | 1132.182 |
| ENSMUSG00000052724 | 0.077749618 | 0.755805 | 0.848334 | 54.32694 | 55.19179 | 43.88965 | 46.58167 | 36.07912 | 49.16446 | 46.28588 | 57.64233 |
| ENSMUSG00000052726 | 0.884107115 | 6.88E-08 | 4.15E-06 | 326.9677 | 326.3515 | 346.9767 | 371.7907 | 144.3165 | 236.8833 | 190.9293 | 172.927 |
| ENSMUSG00000052727 | 0.809418172 | 0.003113 | 0.018288 | 1752.547 | 1350.999 | 1180.88 | 2342.885 | 817.1433 | 1481.638 | 735.7526 | 748.2627 |
| ENSMUSG00000052738 | -0.738977675 | 3.71E-06 | 0.000107 | 2081.526 | 2016.9 | 2388.26 | 1969.369 | 3685.921 | 2512.974 | 3566.906 | 4348.189 |
| ENSMUSG00000052748 | 0.281751514 | 0.1611 | 0.304774 | 558.3602 | 404.3399 | 349.461 | 401.9825 | 269.1307 | 450.3018 | 355.8227 | 334.9781 |
| ENSMUSG00000052749 | 0.081789431 | 0.905634 | 0.943335 | 23.13925 | 4.799286 | 11.59349 | 6.900988 | 7.80089 | 13.40849 | 18.32149 | 4.350364 |
| ENSMUSG00000052751 | -0.066051162 | 0.458051 | 0.619779 | 1094.587 | 962.2569 | 900.1519 | 899.7163 | 1030.693 | 960.9417 | 1024.075 | 1017.985 |
| ENSMUSG00000052752 | -0.133658806 | 0.123106 | 0.252702 | 2166.035 | 2081.69 | 2456.992 | 2354.962 | 2322.715 | 2344.251 | 2588.152 | 2689.613 |
| ENSMUSG00000052760 | 0.938452259 | 0.000602 | 0.00528 | 101.6115 | 86.38715 | 108.482 | 131.1188 | 38.02934 | 71.51194 | 44.3573 | 70.69342 |
| ENSMUSG00000052763 | -0.059028025 | 0.461229 | 0.622349 | 875.2673 | 947.859 | 1024.368 | 979.0777 | 973.1611 | 946.4159 | 1052.039 | 1016.898 |
| ENSMUSG00000052769 | -0.66466438 | 0.146757 | 0.285743 | 6.036326 | 11.99822 | 15.73403 | 12.93935 | 12.67645 | 22.34748 | 21.21436 | 18.48905 |
| ENSMUSG00000052776 | 0.081507271 | 0.886467 | 0.932078 | 41.24823 | 137.9795 | 144.9187 | 19.84034 | 68.25779 | 53.63396 | 124.3933 | 78.30656 |
| ENSMUSG00000052783 | 0.396292062 | 0.126789 | 0.257937 | 77.46619 | 110.3836 | 140.7781 | 128.5309 | 60.4569 | 100.5637 | 88.7146 | 98.97079 |
| ENSMUSG00000052794 | 0.231923802 | 0.025544 | 0.084292 | 455.7426 | 507.5245 | 433.9279 | 482.2065 | 384.1939 | 440.2454 | 409.8229 | 364.343 |
| ENSMUSG00000052798 | -0.027093756 | 0.762966 | 0.852533 | 1204.247 | 1322.203 | 1073.226 | 1200.772 | 1320.301 | 1150.895 | 1183.183 | 1231.153 |
| ENSMUSG00000052812 | -0.137886949 | 0.406259 | 0.572907 | 500.009 | 365.9456 | 332.0708 | 326.0717 | 466.1032 | 408.9589 | 431.0373 | 368.6934 |
| ENSMUSG00000052821 | 1.192835954 | 2.17E-07 | 1.07E-05 | 105.6357 | 109.1838 | 95.23226 | 102.6522 | 44.85512 | 62.57295 | 38.57157 | 34.80291 |
| ENSMUSG00000052825 | 0.099181115 | 0.648489 | 0.772265 | 58.35115 | 63.59054 | 57.96746 | 55.2079 | 56.55646 | 64.8077 | 52.07161 | 45.67883 |
| ENSMUSG00000052833 | -0.137550661 | 0.027924 | 0.089945 | 3328.028 | 3666.655 | 3685.903 | 3519.504 | 4153.974 | 3762.199 | 3806.049 | 3896.839 |
| ENSMUSG00000052837 | -0.322213172 | 0.29405 | 0.461714 | 486.9303 | 202.7698 | 374.3042 | 389.9058 | 463.1779 | 246.9397 | 576.6449 | 531.832 |
| ENSMUSG00000052852 | -0.281441965 | 0.099246 | 0.219875 | 244.4712 | 171.5745 | 250.9163 | 262.2375 | 234.0267 | 296.1041 | 310.5011 | 292.562 |
| ENSMUSG00000052854 | 1.114146432 | 0.00017 | 0.002 | 443.67 | 777.4844 | 550.6909 | 679.7473 | 279.8569 | 463.7102 | 204.4293 | 184.8905 |
| ENSMUSG00000052861 | -0.046172908 | 0.93046 | 0.958826 | 23.13925 | 9.598572 | 20.70267 | 8.626235 | 17.552 | 24.58223 | 12.53576 | 9.78832 |
| ENSMUSG00000052889 | 0.57338483 | 0.057238 | 0.149798 | 83.50251 | 88.78679 | 91.91984 | 119.042 | 51.6809 | 105.0332 | 54.00019 | 47.85401 |
| ENSMUSG00000052906 | -0.068905333 | 0.439039 | 0.602492 | 914.5034 | 1041.445 | 992.8999 | 998.0554 | 1054.095 | 957.5896 | 975.8606 | 1151.759 |
| ENSMUSG00000052911 | 0.350605334 | 2.50E-05 | 0.000467 | 3584.572 | 4102.19 | 4055.238 | 4283.788 | 3044.298 | 3452.686 | 3102.118 | 2971.299 |
| ENSMUSG00000052912 | 0.872969322 | 0.016 | 0.060315 | 39.23612 | 27.5959 | 29.81184 | 25.8787 | 23.40267 | 15.64324 | 13.50005 | 14.13868 |
| ENSMUSG00000052914 | -0.328070893 | 0.041133 | 0.119064 | 696.1896 | 733.091 | 759.3738 | 727.1916 | 753.761 | 719.5889 | 1090.611 | 1096.292 |
| ENSMUSG00000052915 | 0.209929844 | 0.011949 | 0.048832 | 4462.857 | 4216.173 | 3716.543 | 4203.564 | 3440.193 | 3929.805 | 3388.512 | 3592.313 |
| ENSMUSG00000052917 | 0.369999557 | 0.000178 | 0.002074 | 1142.878 | 1006.65 | 891.8708 | 1051.538 | 744.985 | 789.9835 | 790.7171 | 840.7079 |
| ENSMUSG00000052920 | 0.018298647 | 0.924401 | 0.955046 | 117.7084 | 98.38537 | 115.9349 | 95.75121 | 104.3369 | 80.45093 | 115.7147 | 121.8102 |
| ENSMUSG00000052921 | -0.184409382 | 0.145892 | 0.284388 | 570.4328 | 554.3175 | 737.843 | 561.5679 | 657.225 | 668.1897 | 656.6809 | 776.54 |
| ENSMUSG00000052926 | -0.254918091 | 0.013678 | 0.053858 | 1253.544 | 1053.443 | 1014.431 | 1098.982 | 1433.414 | 1211.234 | 1405.934 | 1221.365 |
| ENSMUSG00000052928 | 0.200932731 | 0.10542 | 0.229113 | 486.9303 | 433.1356 | 569.7374 | 572.782 | 487.5557 | 462.5929 | 428.1444 | 419.8102 |
| ENSMUSG00000052934 | -0.122519245 | 0.177094 | 0.326095 | 704.238 | 755.8876 | 660.8291 | 726.329 | 851.2722 | 721.8237 | 729.9669 | 792.8539 |
| ENSMUSG00000052942 | -0.055631215 | 0.743774 | 0.839833 | 639.8506 | 524.322 | 858.7466 | 622.8142 | 568.4899 | 778.8097 | 723.2169 | 683.0072 |
| ENSMUSG00000052949 | -0.050330286 | 0.605107 | 0.741607 | 618.7234 | 713.8938 | 640.1264 | 609.0122 | 648.449 | 747.5233 | 650.8952 | 624.2773 |
| ENSMUSG00000052951 | 0.603643889 | 0.001959 | 0.012938 | 320.9313 | 359.9465 | 260.0255 | 332.9727 | 193.072 | 275.9914 | 212.1436 | 156.6131 |
| ENSMUSG00000052955 | 0.127284952 | 0.793691 | 0.873486 | 17.10292 | 16.7975 | 10.76539 | 17.25247 | 14.62667 | 18.99536 | 5.785735 | 17.40146 |
| ENSMUSG00000052957 | 0.349757722 | 0.044092 | 0.12477 | 6725.473 | 8398.751 | 6021.991 | 8094.859 | 5710.252 | 7588.087 | 5109.768 | 4537.43 |
| ENSMUSG00000052962 | -0.182439372 | 0.01793 | 0.065471 | 619.7295 | 705.4951 | 663.3134 | 634.0283 | 776.1886 | 737.4669 | 729.9669 | 728.686 |
| ENSMUSG00000052981 | -0.296011572 | 0.352088 | 0.520893 | 42.25428 | 26.39607 | 37.2648 | 38.81806 | 28.27823 | 42.46022 | 58.82164 | 48.9416 |
| ENSMUSG00000052997 | -0.114778656 | 0.061692 | 0.158048 | 3644.935 | 4030.201 | 3603.92 | 3522.954 | 4074.99 | 4048.246 | 3890.907 | 4005.598 |
| ENSMUSG00000053007 | 1.011174814 | 0.004102 | 0.02223 | 238.4349 | 226.7663 | 181.3554 | 239.8093 | 98.48624 | 206.7142 | 65.57166 | 69.60583 |
| ENSMUSG00000053012 | -0.015311634 | 0.863219 | 0.917928 | 1549.324 | 1732.542 | 1541.106 | 1615.694 | 1441.215 | 1577.732 | 1751.149 | 1734.708 |
| ENSMUSG00000053024 | 0.463604344 | 0.220125 | 0.378905 | 20.12109 | 21.59679 | 21.53077 | 20.70296 | 12.67645 | 18.99536 | 18.32149 | 10.87591 |
| ENSMUSG00000053025 | 0.656997061 | 0.002279 | 0.01441 | 248.4954 | 301.1552 | 276.5876 | 289.8415 | 159.9183 | 260.3482 | 152.3577 | 135.9489 |
| ENSMUSG00000053038 | -0.364749614 | 0.170792 | 0.317692 | 41.24823 | 44.3934 | 27.32752 | 43.13117 | 54.60623 | 42.46022 | 48.21446 | 54.37955 |
| ENSMUSG00000053040 | -0.694517639 | 4.95E-05 | 0.000788 | 172.0353 | 145.1784 | 164.7932 | 199.266 | 313.9858 | 215.6532 | 268.0724 | 306.7007 |
| ENSMUSG00000053046 | 0.689672026 | 0.009522 | 0.041098 | 80.48435 | 53.99197 | 57.96746 | 71.59775 | 34.1289 | 54.75133 | 38.57157 | 36.9781 |
| ENSMUSG00000053062 | 0.154856669 | 0.33932 | 0.507866 | 341.0524 | 471.5299 | 342.008 | 440.8006 | 378.3432 | 416.7805 | 328.8226 | 307.7883 |
| ENSMUSG00000053070 | 0.141609503 | 0.422585 | 0.587698 | 268.6165 | 280.7582 | 252.5725 | 326.9343 | 278.8818 | 303.9258 | 254.5723 | 185.9781 |

| | | | | | | | | | | | |
|---|---|---|---|---|---|---|---|---|---|---|---|
| ENSMUSG00000053080 | 0.363343258 | 1.16E-06 | 4.27E-05 | 3690.207 | 3983.407 | 3600.608 | 3942.189 | 3109.63 | 3138.704 | 2702.903 | 2876.678 |
| ENSMUSG00000053091 | -0.192262446 | 0.093779 | 0.210839 | 399.4036 | 452.3327 | 399.1474 | 369.2029 | 512.9085 | 403.372 | 466.716 | 463.3138 |
| ENSMUSG00000053093 | 1.019322561 | 0.017773 | 0.065041 | 48.29061 | 71.98929 | 89.43552 | 40.5433 | 39.00445 | 48.04709 | 20.25007 | 16.31387 |
| ENSMUSG00000053094 | -0.033040156 | 0.518579 | 0.672477 | 1737.456 | 1734.942 | 1806.929 | 1770.966 | 1842.96 | 1813.498 | 1814.792 | 1744.496 |
| ENSMUSG00000053110 | 0.276600887 | 7.24E-05 | 0.001061 | 5584.608 | 5954.714 | 5591.376 | 5781.303 | 4497.213 | 5237.132 | 4489.73 | 4691.868 |
| ENSMUSG00000053111 | 0.269213269 | 0.534381 | 0.685422 | 17.10292 | 29.99554 | 19.04645 | 15.52722 | 15.60178 | 23.46486 | 18.32149 | 9.78832 |
| ENSMUSG00000053113 | 0.29255146 | 0.03288 | 0.100987 | 455.7426 | 490.727 | 476.9894 | 452.0147 | 412.4721 | 452.5365 | 363.537 | 302.3503 |
| ENSMUSG00000053117 | 0.905759563 | 0.000123 | 0.001567 | 302.8224 | 265.1606 | 214.4796 | 270.8638 | 135.5405 | 207.8316 | 113.7861 | 105.4963 |
| ENSMUSG00000053119 | -0.167205764 | 0.02258 | 0.076817 | 2601.657 | 2531.623 | 2740.205 | 2457.614 | 2839.524 | 2654.881 | 3019.189 | 3087.671 |
| ENSMUSG00000053128 | 0.252135894 | 0.067375 | 0.167607 | 575.4631 | 569.9152 | 588.7838 | 687.5109 | 493.4063 | 630.199 | 471.5374 | 440.4744 |
| ENSMUSG00000053134 | -0.023970949 | 0.728723 | 0.829692 | 743.4742 | 806.2801 | 771.7954 | 762.5592 | 799.5913 | 750.8754 | 806.1457 | 775.4524 |
| ENSMUSG00000053137 | -0.381996512 | 0.00077 | 0.006395 | 317.9132 | 304.7547 | 348.6329 | 353.6756 | 423.1983 | 383.2593 | 484.0732 | 438.2992 |
| ENSMUSG00000053141 | 0.709063561 | 1.36E-05 | 0.000291 | 268.6165 | 184.7725 | 227.7293 | 232.9083 | 130.6649 | 137.437 | 161.0363 | 130.5109 |
| ENSMUSG00000053153 | 1.196233828 | 0.000103 | 0.00137 | 49.29666 | 38.39429 | 39.74912 | 56.93315 | 21.45245 | 23.46486 | 16.39292 | 19.57664 |
| ENSMUSG00000053158 | -0.045005927 | 0.757467 | 0.848813 | 331.9979 | 301.1552 | 424.8187 | 339.011 | 361.7663 | 307.2779 | 379.9299 | 394.7956 |
| ENSMUSG00000053161 | -0.330228103 | 0.495515 | 0.652245 | 10.06054 | 14.39786 | 14.90592 | 17.25247 | 12.67645 | 21.23011 | 9.642892 | 28.27737 |
| ENSMUSG00000053164 | -0.22850821 | 0.615412 | 0.748718 | 15.09082 | 16.7975 | 21.53077 | 24.15346 | 16.57689 | 41.34284 | 15.42863 | 18.48905 |
| ENSMUSG00000053166 | 1.527429495 | 1.55E-06 | 5.34E-05 | 403.4278 | 587.9126 | 434.756 | 467.5419 | 137.4907 | 298.3389 | 103.1789 | 118.5474 |
| ENSMUSG00000053173 | -0.290365926 | 0.216225 | 0.374591 | 78.47224 | 61.1909 | 45.54586 | 69.8725 | 81.90935 | 87.15518 | 68.46453 | 73.95619 |
| ENSMUSG00000053175 | 0.119178611 | 0.662826 | 0.78302 | 59.35721 | 51.59233 | 69.56096 | 37.09281 | 48.75557 | 45.81234 | 46.28588 | 59.81751 |
| ENSMUSG00000053178 | 0.450920057 | 0.070365 | 0.172946 | 83.50251 | 105.5843 | 90.26362 | 108.6906 | 52.65601 | 100.5637 | 71.3574 | 59.81751 |
| ENSMUSG00000053182 | 1.059918509 | 0.000134 | 0.001675 | 168.0111 | 129.5807 | 94.40416 | 125.943 | 53.63112 | 89.38993 | 59.78593 | 45.67883 |
| ENSMUSG00000053192 | 0.410326167 | 0.015308 | 0.058531 | 564.3965 | 555.5174 | 386.7258 | 655.5939 | 455.377 | 377.6724 | 391.5014 | 401.3211 |
| ENSMUSG00000053198 | -0.032146135 | 0.820539 | 0.891235 | 338.0343 | 445.1338 | 487.7548 | 426.9986 | 442.7005 | 386.6114 | 419.4658 | 488.3284 |
| ENSMUSG00000053199 | 0.75604657 | 0.03311 | 0.10154 | 135.8173 | 262.7609 | 219.4483 | 174.2499 | 100.4365 | 204.4795 | 64.60737 | 100.0584 |
| ENSMUSG00000053205 | -0.20783607 | 0.230204 | 0.390428 | 161.9748 | 159.5763 | 150.7154 | 134.5693 | 202.8232 | 144.1413 | 151.3934 | 201.2044 |
| ENSMUSG00000053214 | 1.449263203 | 0.007966 | 0.036131 | 57.3451 | 15.59768 | 37.2648 | 26.74133 | 4.875557 | 18.99536 | 12.53576 | 14.13868 |
| ENSMUSG00000053216 | 0.188378829 | 0.68623 | 0.800522 | 9.054489 | 19.19714 | 28.15563 | 32.77969 | 20.47734 | 14.52586 | 24.10723 | 19.57664 |
| ENSMUSG00000053219 | -0.110534433 | 0.629401 | 0.758785 | 113.6841 | 62.39072 | 75.3577 | 74.18562 | 89.71024 | 83.80306 | 84.85745 | 93.53283 |
| ENSMUSG00000053226 | 0.304954382 | 0.110842 | 0.236908 | 176.0595 | 167.975 | 155.684 | 150.9591 | 125.7894 | 111.7374 | 111.8575 | 177.2773 |
| ENSMUSG00000053253 | -0.294810087 | 0.003346 | 0.019176 | 1412.5 | 1431.387 | 1572.574 | 1502.69 | 1781.528 | 1546.446 | 1888.078 | 2046.846 |
| ENSMUSG00000053263 | 0.728743772 | 0.065128 | 0.164345 | 36.21796 | 51.59233 | 53.82693 | 50.03216 | 25.35289 | 51.39921 | 24.10723 | 15.22628 |
| ENSMUSG00000053279 | 1.085402865 | 0.000186 | 0.002139 | 764.6013 | 932.2613 | 644.267 | 835.0195 | 301.3094 | 648.077 | 317.2511 | 230.5693 |
| ENSMUSG00000053286 | 0.089046304 | 0.598532 | 0.736956 | 766.6134 | 757.0874 | 509.2856 | 610.7374 | 500.2321 | 606.7341 | 677.8953 | 698.2335 |
| ENSMUSG00000053289 | 0.050982771 | 0.544105 | 0.692833 | 1288.756 | 1286.209 | 1227.254 | 1204.222 | 1094.075 | 1352.023 | 1194.754 | 1192 |
| ENSMUSG00000053291 | -0.135090431 | 0.369223 | 0.53809 | 268.6165 | 284.3577 | 280.7281 | 314.8576 | 377.3681 | 255.8787 | 286.3939 | 341.5036 |
| ENSMUSG00000053293 | 0.148471648 | 0.096027 | 0.214632 | 1050.321 | 1123.033 | 1061.633 | 1221.475 | 986.8126 | 1107.318 | 931.5033 | 997.321 |
| ENSMUSG00000053297 | -1.167730822 | 0.000155 | 0.001873 | 98.59333 | 81.58786 | 123.3879 | 111.2784 | 241.8276 | 120.6764 | 217.9294 | 353.4671 |
| ENSMUSG00000053303 | -4.571905625 | 1.75E-05 | 0.000354 | 2.012109 | 0 | 1.656213 | 0 | 12.67645 | 5.58687 | 37.60728 | 33.71532 |
| ENSMUSG00000053310 | 0.03189377 | 0.895429 | 0.936737 | 153.9263 | 130.7805 | 161.4808 | 119.042 | 110.1876 | 201.1273 | 107.0361 | 135.9489 |
| ENSMUSG00000053317 | -0.165075179 | 0.122879 | 0.252404 | 1854.158 | 1937.712 | 2204.42 | 1908.123 | 2230.08 | 1888.362 | 2224.615 | 2521.036 |
| ENSMUSG00000053329 | -0.732709431 | 2.42E-06 | 7.52E-05 | 1314.913 | 1333.002 | 1489.764 | 1326.715 | 2328.566 | 1614.606 | 2373.116 | 2764.657 |
| ENSMUSG00000053332 | 0.174026665 | 0.020012 | 0.070419 | 9072.598 | 9808.541 | 8370.502 | 8992.85 | 8038.818 | 8312.146 | 8421.137 | 7347.765 |
| ENSMUSG00000053333 | 0.048283129 | 0.497144 | 0.653697 | 784.7224 | 803.8804 | 818.9975 | 769.4602 | 813.2428 | 750.8754 | 772.3956 | 734.124 |
| ENSMUSG00000053334 | 0.105297316 | 0.490425 | 0.647723 | 134.8113 | 163.1757 | 158.9965 | 136.2945 | 130.6649 | 150.8455 | 123.429 | 146.8248 |
| ENSMUSG00000053347 | 0.209288482 | 0.192611 | 0.3458 | 519.124 | 548.3184 | 452.9743 | 532.2387 | 381.2685 | 577.6824 | 444.5373 | 371.9562 |
| ENSMUSG00000053358 | -0.579035801 | 0.116895 | 0.245132 | 34.20585 | 17.99732 | 30.63995 | 25.8787 | 34.1289 | 32.40385 | 63.64308 | 32.62773 |
| ENSMUSG00000053368 | 0.047573201 | 0.92052 | 0.952564 | 26.15741 | 57.59143 | 33.12427 | 52.62003 | 32.17867 | 80.45093 | 22.17865 | 29.36496 |
| ENSMUSG00000053390 | 0.127424589 | 0.378319 | 0.546471 | 438.6397 | 455.9322 | 419.8501 | 408.8835 | 378.3432 | 500.5836 | 371.2513 | 327.3649 |
| ENSMUSG00000053395 | 2.573273409 | 3.30E-07 | 1.49E-05 | 74.44802 | 39.59411 | 70.38906 | 37.95543 | 10.72622 | 5.58687 | 3.857157 | 17.40146 |
| ENSMUSG00000053398 | -0.228765017 | 0.051187 | 0.138272 | 4910.551 | 5762.743 | 5897.775 | 5732.996 | 6645.384 | 5323.17 | 6642.988 | 7525.043 |
| ENSMUSG00000053399 | 0.355363162 | 0.019806 | 0.069986 | 254.5318 | 314.3532 | 364.3669 | 257.9244 | 238.9023 | 243.5876 | 220.8222 | 228.3941 |
| ENSMUSG00000053411 | 0.323449117 | 0.013748 | 0.054066 | 538.2391 | 410.339 | 475.3332 | 548.6285 | 433.9245 | 397.7852 | 401.1443 | 344.7664 |
| ENSMUSG00000053414 | 0.279238404 | 0.068932 | 0.1704 | 1614.717 | 1798.532 | 1460.78 | 1645.886 | 1299.823 | 1772.155 | 1162.933 | 1137.62 |
| ENSMUSG00000053436 | -0.301448204 | 0.000414 | 0.003962 | 1818.946 | 2018.1 | 1951.847 | 1882.244 | 2336.367 | 2126.363 | 2334.544 | 2654.81 |
| ENSMUSG00000053441 | 0.801624255 | 0.014856 | 0.057265 | 507.0514 | 563.9161 | 395.835 | 504.6347 | 246.7032 | 522.9311 | 198.6436 | 163.1387 |
| ENSMUSG00000053442 | -0.343208159 | 0.564977 | 0.709818 | 4.024217 | 13.19804 | 14.07781 | 10.35148 | 22.42756 | 11.17374 | 12.53576 | 6.525547 |
| ENSMUSG00000053453 | 0.006789634 | 0.938964 | 0.96418 | 1656.972 | 1876.521 | 1646.276 | 1884.832 | 1821.508 | 1912.944 | 1664.363 | 1631.387 |
| ENSMUSG00000053460 | 0.38712725 | 0.000548 | 0.004911 | 795.789 | 896.2667 | 686.5004 | 749.6198 | 654.2997 | 607.8515 | 548.6805 | 577.5109 |
| ENSMUSG00000053461 | 0.715198643 | 0.047053 | 0.130406 | 27.16347 | 26.39607 | 22.35888 | 22.42821 | 17.552 | 15.64324 | 15.42863 | 10.87591 |
| ENSMUSG00000053470 | 0.162656412 | 0.035065 | 0.106232 | 2411.512 | 2188.474 | 2138.171 | 2310.968 | 2046.759 | 2145.358 | 2064.543 | 1827.153 |
| ENSMUSG00000053475 | 0.612062878 | 5.42E-06 | 0.000143 | 214.2896 | 212.3684 | 199.5737 | 217.3811 | 134.5654 | 150.8455 | 147.5362 | 118.5474 |
| ENSMUSG00000053477 | 0.811589815 | 0.037916 | 0.112359 | 8167.149 | 9195.432 | 6502.293 | 7767.925 | 3373.885 | 8496.513 | 3322.94 | 2829.912 |
| ENSMUSG00000053483 | 0.053928411 | 0.432875 | 0.597141 | 1266.622 | 1333.002 | 1194.13 | 1282.721 | 1239.366 | 1241.403 | 1261.29 | 1143.058 |

| | | | | | | | | | | | |
|---|---|---|---|---|---|---|---|---|---|---|---|
| ENSMUSG00000053510 | -0.12179819 | 0.01869 | 0.06728 | 3113.738 | 3010.352 | 3092.15 | 3208.097 | 3362.184 | 3223.624 | 3499.405 | 3435.7 |
| ENSMUSG00000053519 | -0.469931321 | 0.311794 | 0.480258 | 32.19374 | 17.99732 | 51.34261 | 26.74133 | 32.17867 | 23.46486 | 49.17875 | 73.95619 |
| ENSMUSG00000053522 | 0.928960952 | 0.046397 | 0.129251 | 124.7507 | 304.7547 | 152.3716 | 209.6175 | 67.28268 | 224.5922 | 67.50024 | 56.55474 |
| ENSMUSG00000053536 | 0.010671548 | 0.865971 | 0.919627 | 1745.504 | 1925.714 | 1797.819 | 1855.503 | 1770.802 | 1862.663 | 1737.649 | 1898.934 |
| ENSMUSG00000053550 | 0.246541319 | 0.085233 | 0.197322 | 241.453 | 214.7681 | 273.2752 | 238.9467 | 218.4249 | 179.8972 | 187.0721 | 232.7445 |
| ENSMUSG00000053552 | -0.07360544 | 0.419726 | 0.585258 | 640.8566 | 614.3086 | 612.7989 | 640.9293 | 749.8606 | 627.9642 | 631.6094 | 629.7152 |
| ENSMUSG00000053553 | -0.285334271 | 0.017652 | 0.064683 | 598.6023 | 651.5031 | 663.3134 | 613.3253 | 893.202 | 627.9642 | 774.3242 | 781.978 |
| ENSMUSG00000053559 | -1.104039671 | 1.40E-05 | 0.000296 | 439.6458 | 302.355 | 686.5004 | 445.9763 | 975.1113 | 706.1804 | 1078.075 | 1274.657 |
| ENSMUSG00000053560 | -0.100297068 | 0.527299 | 0.680214 | 1285.737 | 1301.806 | 1427.656 | 1773.554 | 1552.377 | 1178.83 | 1685.577 | 1790.175 |
| ENSMUSG00000053565 | -0.058154798 | 0.472123 | 0.63146 | 4466.881 | 5264.817 | 5181.463 | 5113.632 | 5649.795 | 4903.038 | 5295.876 | 4999.656 |
| ENSMUSG00000053574 | 0.047206969 | 0.857601 | 0.914643 | 102.6175 | 75.58876 | 57.96746 | 87.9876 | 87.76002 | 93.85942 | 76.17884 | 55.46715 |
| ENSMUSG00000053580 | -0.200246592 | 0.257313 | 0.421045 | 916.5155 | 736.6904 | 1175.083 | 998.918 | 886.3762 | 1371.018 | 1119.54 | 1024.511 |
| ENSMUSG00000053581 | -0.000777525 | 0.994445 | 0.996743 | 679.0867 | 801.4808 | 796.6386 | 694.4119 | 840.5459 | 747.5233 | 733.8241 | 649.2919 |
| ENSMUSG00000053600 | -0.054813285 | 0.733706 | 0.833261 | 266.6044 | 265.1606 | 255.0568 | 292.4294 | 269.1307 | 329.6254 | 312.4297 | 209.9051 |
| ENSMUSG00000053604 | -0.014896084 | 0.880259 | 0.928028 | 901.4247 | 950.2587 | 751.0927 | 904.0294 | 946.8331 | 855.9086 | 838.9316 | 897.2626 |
| ENSMUSG00000053613 | 0.378871054 | 0.361379 | 0.529959 | 25.15136 | 31.19536 | 54.65504 | 25.01608 | 16.57689 | 24.58223 | 26.03581 | 38.06569 |
| ENSMUSG00000053617 | 0.361336278 | 1.93E-06 | 6.28E-05 | 2704.274 | 2879.572 | 2909.139 | 3048.511 | 2096.489 | 2467.162 | 2164.829 | 2260.014 |
| ENSMUSG00000053626 | -0.067506404 | 0.825433 | 0.893982 | 40.24217 | 33.595 | 34.78048 | 41.40593 | 25.35289 | 52.51658 | 35.6787 | 44.59123 |
| ENSMUSG00000053641 | 0.213631762 | 0.154147 | 0.295564 | 490.9545 | 338.3497 | 386.7258 | 448.5642 | 330.5627 | 414.5458 | 331.7155 | 361.0802 |
| ENSMUSG00000053644 | -0.119372968 | 0.169971 | 0.316547 | 2709.304 | 2727.194 | 2637.52 | 2503.333 | 3091.103 | 2511.857 | 2783.903 | 3099.635 |
| ENSMUSG00000053646 | -0.026346054 | 0.752761 | 0.846203 | 2760.613 | 2232.868 | 2572.099 | 2489.531 | 2504.086 | 2430.289 | 2585.259 | 2724.416 |
| ENSMUSG00000053647 | -0.578083468 | 0.033727 | 0.102988 | 98.59333 | 89.98662 | 147.403 | 100.0643 | 138.4658 | 120.6764 | 152.3577 | 241.4452 |
| ENSMUSG00000053675 | 0.342727968 | 0.59214 | 0.731936 | 5.030272 | 27.5959 | 52.17072 | 28.46658 | 10.72622 | 20.11273 | 24.10723 | 34.80291 |
| ENSMUSG00000053684 | 0.09086242 | 0.46562 | 0.626438 | 226.3622 | 196.7707 | 216.1358 | 189.7772 | 197.9476 | 189.9536 | 206.3579 | 183.8029 |
| ENSMUSG00000053687 | 0.718827366 | 0.106268 | 0.230327 | 22.1332 | 14.39786 | 17.39024 | 20.70296 | 8.776002 | 7.821619 | 11.57147 | 17.40146 |
| ENSMUSG00000053693 | 0.8503762 | 0.001544 | 0.01075 | 54.32694 | 41.99375 | 51.34261 | 56.93315 | 22.42756 | 31.28647 | 27.96439 | 32.62773 |
| ENSMUSG00000053702 | 0.292468473 | 0.058711 | 0.152279 | 303.8284 | 268.76 | 309.7119 | 248.4356 | 183.3209 | 240.2354 | 232.3937 | 268.635 |
| ENSMUSG00000053714 | 0.425573096 | 0.25578 | 0.419652 | 38.23007 | 58.79126 | 38.09291 | 49.16954 | 24.37778 | 52.51658 | 40.50014 | 19.57664 |
| ENSMUSG00000053716 | -0.070464576 | 0.354446 | 0.523068 | 885.3278 | 1016.249 | 945.6978 | 942.8475 | 1031.668 | 1031.336 | 980.6821 | 933.1532 |
| ENSMUSG00000053730 | -0.188961849 | 0.145371 | 0.283642 | 444.676 | 509.9242 | 485.2705 | 497.7338 | 554.8383 | 455.8886 | 535.1805 | 662.343 |
| ENSMUSG00000053740 | 0.222372057 | 0.633678 | 0.761871 | 13.07871 | 19.19714 | 15.73403 | 7.763611 | 12.67645 | 11.17374 | 12.53576 | 10.87591 |
| ENSMUSG00000053746 | -0.765414921 | 0.002149 | 0.013833 | 74.44802 | 73.18911 | 101.8571 | 87.9876 | 159.9183 | 88.27255 | 158.1434 | 168.5766 |
| ENSMUSG00000053749 | -0.49819305 | 0.235466 | 0.396374 | 13.07871 | 16.7975 | 20.70267 | 15.52722 | 11.70134 | 27.93435 | 26.03581 | 28.27737 |
| ENSMUSG00000053754 | 0.271535625 | 0.002365 | 0.014782 | 4396.458 | 4871.275 | 4250.671 | 4576.218 | 3636.19 | 4292.951 | 3592.941 | 3467.24 |
| ENSMUSG00000053768 | -0.619918093 | 4.13E-05 | 0.000691 | 1853.152 | 1805.731 | 2229.263 | 1864.129 | 3028.696 | 2177.762 | 3203.369 | 3505.306 |
| ENSMUSG00000053769 | 0.120781479 | 0.423446 | 0.588709 | 122.7386 | 128.3809 | 124.216 | 115.5915 | 104.3369 | 105.0332 | 118.6076 | 122.8978 |
| ENSMUSG00000053773 | 0.039502861 | 0.939472 | 0.964398 | 7.04238 | 13.19804 | 7.45296 | 13.80198 | 12.67645 | 11.17374 | 8.678602 | 7.613138 |
| ENSMUSG00000053774 | 0.342209491 | 0.018534 | 0.066927 | 3409.518 | 3244.317 | 2958.825 | 3462.571 | 2277.86 | 3354.357 | 2545.723 | 2137.116 |
| ENSMUSG00000053799 | -0.781177353 | 1.39E-05 | 0.000295 | 510.0696 | 397.1409 | 594.5806 | 452.8773 | 864.9237 | 636.9032 | 818.6815 | 1041.912 |
| ENSMUSG00000053801 | -0.275529864 | 0.021632 | 0.074527 | 748.5044 | 879.4692 | 811.5445 | 819.4923 | 1097 | 821.27 | 922.8247 | 1101.73 |
| ENSMUSG00000053819 | 0.26500628 | 0.040533 | 0.117827 | 1713.311 | 1354.599 | 1463.264 | 1476.811 | 1048.245 | 1495.047 | 1280.576 | 1177.861 |
| ENSMUSG00000053821 | -0.249457884 | 0.370642 | 0.53969 | 49.29666 | 32.39518 | 55.48314 | 46.58167 | 63.38224 | 45.81234 | 66.53595 | 43.50364 |
| ENSMUSG00000053825 | 0.880953873 | 0.007067 | 0.033143 | 96.58122 | 74.38894 | 70.38906 | 148.3712 | 41.92979 | 74.86406 | 52.07161 | 43.50364 |
| ENSMUSG00000053835 | 0.384685231 | 0.235553 | 0.39644 | 44.26639 | 55.19179 | 55.48314 | 25.01608 | 34.1289 | 36.87335 | 33.75012 | 32.62773 |
| ENSMUSG00000053838 | -0.048908137 | 0.352293 | 0.52107 | 2069.454 | 1968.907 | 2101.735 | 1992.66 | 2173.523 | 2095.076 | 2072.257 | 2075.124 |
| ENSMUSG00000053841 | -0.038882702 | 0.541562 | 0.690839 | 2693.208 | 2639.607 | 2673.956 | 2626.689 | 2810.271 | 2474.984 | 2802.224 | 2834.262 |
| ENSMUSG00000053846 | -0.592592203 | 0.017539 | 0.064352 | 322.9434 | 213.5682 | 346.9767 | 249.2982 | 395.8952 | 278.2261 | 598.8236 | 436.124 |
| ENSMUSG00000053862 | -3.788001849 | 6.97E-11 | 1.34E-08 | 10.06054 | 2.399643 | 20.70267 | 8.626235 | 158.9431 | 49.16446 | 133.0719 | 246.8832 |
| ENSMUSG00000053870 | 0.112060245 | 0.427901 | 0.592924 | 390.3491 | 368.3452 | 275.7595 | 347.6373 | 302.2845 | 351.9728 | 328.8226 | 293.6496 |
| ENSMUSG00000053877 | -0.222255907 | 0.199622 | 0.354566 | 119.7205 | 109.1838 | 111.7944 | 121.6299 | 143.3414 | 159.7845 | 130.179 | 106.5839 |
| ENSMUSG00000053886 | -0.731153003 | 0.002052 | 0.013363 | 311.8769 | 278.3586 | 469.5365 | 282.9405 | 490.481 | 397.7852 | 605.5736 | 737.3868 |
| ENSMUSG00000053897 | -0.398232274 | 0.063228 | 0.160933 | 744.4802 | 904.6654 | 918.3703 | 1022.209 | 1004.365 | 782.1619 | 1338.433 | 1606.372 |
| ENSMUSG00000053898 | -0.433105959 | 0.023086 | 0.078235 | 2025.187 | 2004.902 | 2350.167 | 1930.551 | 2945.811 | 1898.419 | 2564.045 | 3813.094 |
| ENSMUSG00000053907 | 0.069560859 | 0.298678 | 0.466718 | 7132.925 | 7218.126 | 6762.319 | 7774.826 | 7163.168 | 7058.452 | 6430.844 | 6876.838 |
| ENSMUSG00000053916 | -0.480989142 | 0.11131 | 0.23755 | 46.2785 | 49.19268 | 44.71776 | 29.3292 | 71.18313 | 36.87335 | 60.75022 | 66.34306 |
| ENSMUSG00000053925 | 0.332731902 | 0.352165 | 0.520922 | 84.50857 | 71.98929 | 56.31125 | 55.2079 | 55.58134 | 87.15518 | 39.53586 | 30.45255 |
| ENSMUSG00000053929 | -0.157529422 | 0.046022 | 0.128481 | 2348.131 | 2294.059 | 2459.477 | 2339.435 | 2606.473 | 2334.194 | 2726.045 | 2864.715 |
| ENSMUSG00000053930 | 0.45794776 | 0.325728 | 0.49356 | 18.10898 | 11.99822 | 19.87456 | 11.21411 | 8.776002 | 14.52586 | 8.678602 | 13.05109 |
| ENSMUSG00000053931 | -0.104937292 | 0.077419 | 0.184893 | 8933.763 | 8267.97 | 9219.311 | 9313.746 | 9904.206 | 9172.524 | 9762.464 | 9597.991 |
| ENSMUSG00000053950 | 0.054363712 | 0.461793 | 0.622924 | 1003.036 | 1076.24 | 1079.851 | 1002.369 | 1030.693 | 978.8197 | 1059.754 | 935.3283 |
| ENSMUSG00000053957 | -0.027791851 | 0.955618 | 0.974791 | 9.054489 | 15.59768 | 12.4216 | 12.07673 | 19.50223 | 8.938993 | 13.50005 | 7.613138 |
| ENSMUSG00000053963 | -0.737995398 | 0.055702 | 0.146946 | 19.11503 | 34.79482 | 39.74912 | 27.60395 | 47.78045 | 81.56831 | 37.60728 | 35.89051 |
| ENSMUSG00000053964 | 0.436964842 | 0.012979 | 0.051938 | 245.4773 | 259.1615 | 197.0894 | 236.3588 | 147.2418 | 195.5405 | 143.6791 | 206.6423 |
| ENSMUSG00000053965 | 0.491156945 | 0.001112 | 0.008467 | 944.685 | 856.6726 | 893.5271 | 967.0009 | 559.7139 | 853.6738 | 642.2166 | 551.4087 |

| | | | | | | | | | | | |
|---|---|---|---|---|---|---|---|---|---|---|---|
| ENSMUSG00000053977 | -1.489192981 | 0.02806 | 0.090224 | 6.036326 | 7.198929 | 7.45296 | 4.313117 | 5.850668 | 10.05637 | 18.32149 | 35.89051 |
| ENSMUSG00000053985 | 0.374346274 | 0.120087 | 0.249025 | 105.6357 | 177.5736 | 103.5133 | 131.9814 | 106.2871 | 94.9768 | 78.10742 | 119.635 |
| ENSMUSG00000054000 | -0.156630954 | 0.287067 | 0.454071 | 228.3743 | 217.1677 | 208.6829 | 226.0074 | 224.2756 | 200.01 | 277.7153 | 278.4233 |
| ENSMUSG00000054003 | -1.165886903 | NA | NA | 38.23007 | 45.59322 | 19.87456 | 25.01608 | 32.17867 | 201.1273 | 28.92867 | 26.10219 |
| ENSMUSG00000054008 | -0.225179888 | 0.073667 | 0.178862 | 1994 | 1750.54 | 2087.657 | 1733.011 | 2254.457 | 1768.803 | 2293.08 | 2527.562 |
| ENSMUSG00000054013 | 0.336075026 | 0.514364 | 0.66878 | 25.15136 | 8.398751 | 9.109173 | 20.70296 | 12.67645 | 11.17374 | 14.46434 | 11.9635 |
| ENSMUSG00000054021 | -0.115368297 | 0.389125 | 0.556608 | 242.4591 | 238.7645 | 236.0104 | 223.4195 | 218.4249 | 255.8787 | 243.0009 | 302.3503 |
| ENSMUSG00000054027 | 0.653679265 | 9.69E-06 | 0.000221 | 611.681 | 409.1391 | 499.3483 | 448.5642 | 309.1103 | 284.9304 | 334.6083 | 323.0146 |
| ENSMUSG00000054034 | 0.297737203 | 0.166259 | 0.311869 | 125.7568 | 125.9813 | 186.324 | 109.5532 | 99.46135 | 106.1505 | 113.7861 | 127.2482 |
| ENSMUSG00000054046 | 1.412585609 | 0.000122 | 0.00156 | 52.31483 | 33.595 | 49.6864 | 41.40593 | 14.62667 | 26.81698 | 11.57147 | 14.13868 |
| ENSMUSG00000054051 | 0.224540685 | 0.00906 | 0.039699 | 782.7103 | 710.2943 | 645.9232 | 730.6421 | 610.4197 | 617.9079 | 608.4665 | 617.7517 |
| ENSMUSG00000054052 | -5.313492612 | 8.64E-09 | 7.57E-07 | 7.04238 | 1.199822 | 12.4216 | 6.038364 | 389.0694 | 32.40385 | 155.2506 | 498.1167 |
| ENSMUSG00000054057 | 0.046758109 | 0.698494 | 0.809762 | 295.78 | 392.3416 | 365.195 | 321.7586 | 344.2143 | 320.6864 | 323.0369 | 341.5036 |
| ENSMUSG00000054065 | -0.85504723 | 0.00108 | 0.008282 | 173.0413 | 159.5763 | 252.5725 | 132.844 | 341.289 | 203.3621 | 362.5727 | 392.6204 |
| ENSMUSG00000054072 | 0.188926442 | 0.595914 | 0.735026 | 171.0292 | 367.1454 | 462.9116 | 150.9591 | 222.3254 | 211.1837 | 337.5012 | 239.27 |
| ENSMUSG00000054074 | 0.053995609 | 0.615309 | 0.748691 | 808.8677 | 951.4585 | 771.7954 | 848.8215 | 839.5708 | 912.8946 | 740.5741 | 761.3138 |
| ENSMUSG00000054079 | -0.282422428 | 0.003185 | 0.018573 | 739.45 | 745.0892 | 650.8918 | 770.3228 | 927.3309 | 821.27 | 826.3958 | 957.0802 |
| ENSMUSG00000054083 | 0.758537232 | 0.155472 | 0.297421 | 25.15136 | 31.19536 | 13.24971 | 34.50494 | 21.45245 | 18.99536 | 16.39292 | 4.350364 |
| ENSMUSG00000054091 | -0.101253476 | 0.39145 | 0.558919 | 1096.599 | 991.0526 | 1103.866 | 1127.449 | 1126.254 | 940.829 | 1256.469 | 1310.547 |
| ENSMUSG00000054099 | 0.074953541 | 0.588143 | 0.728361 | 423.5489 | 431.9358 | 380.9291 | 470.1298 | 377.3681 | 498.3488 | 390.5371 | 354.5547 |
| ENSMUSG00000054115 | -0.024584065 | 0.821179 | 0.891615 | 943.679 | 1025.847 | 786.7013 | 844.5084 | 962.4349 | 891.6645 | 838.9316 | 964.6933 |
| ENSMUSG00000054135 | 1.243780501 | 0.029317 | 0.092982 | 43.26034 | 20.39697 | 14.07781 | 31.05445 | 4.875557 | 20.11273 | 13.50005 | 7.613138 |
| ENSMUSG00000054146 | 0.805993555 | 0.0019 | 0.012662 | 135.8173 | 232.7654 | 178.0429 | 150.0965 | 79.95913 | 143.0239 | 90.64318 | 84.8321 |
| ENSMUSG00000054150 | -0.089039612 | 0.694311 | 0.806889 | 155.9384 | 93.58608 | 99.3728 | 114.7289 | 140.416 | 107.2679 | 103.1789 | 142.4744 |
| ENSMUSG00000054161 | -0.528054267 | 0.349828 | 0.518671 | 10.06054 | 3.599465 | 11.59349 | 12.07673 | 20.47734 | 15.64324 | 7.714313 | 10.87591 |
| ENSMUSG00000054162 | 0.721415763 | 0.003265 | 0.018892 | 160.9687 | 98.38537 | 111.7944 | 150.9591 | 67.28268 | 97.21155 | 89.67889 | 63.08028 |
| ENSMUSG00000054178 | -0.017174165 | 0.914538 | 0.949313 | 131.7931 | 106.7841 | 121.7317 | 117.3168 | 132.6151 | 105.0332 | 116.679 | 129.4233 |
| ENSMUSG00000054181 | 0.252745788 | 0.391345 | 0.558814 | 38.23007 | 35.99465 | 35.60859 | 32.77969 | 26.32801 | 40.22547 | 23.14294 | 30.45255 |
| ENSMUSG00000054191 | -0.429311057 | 0.335937 | 0.504269 | 34.20585 | 23.99643 | 18.21835 | 14.6646 | 26.32801 | 18.99536 | 50.14304 | 26.10219 |
| ENSMUSG00000054196 | 0.355029806 | 0.077853 | 0.185497 | 405.4399 | 477.529 | 391.6944 | 483.0692 | 306.185 | 493.8794 | 301.8225 | 272.9854 |
| ENSMUSG00000054199 | 0.183698641 | 0.028459 | 0.091029 | 1006.054 | 1011.45 | 896.8395 | 869.5245 | 794.7157 | 868.1997 | 835.0744 | 829.832 |
| ENSMUSG00000054200 | -0.288564149 | 0.60763 | 0.743464 | 50.30272 | 49.19268 | 45.54586 | 43.13117 | 28.27823 | 27.93435 | 143.6791 | 29.36496 |
| ENSMUSG00000054203 | 2.391721521 | 5.26E-05 | 0.000824 | 18.10898 | 21.59679 | 13.24971 | 18.11509 | 2.925334 | 5.58687 | 2.892867 | 2.175182 |
| ENSMUSG00000054206 | -0.005726491 | 0.987701 | 0.993028 | 29.17558 | 33.595 | 30.63995 | 29.3292 | 29.25334 | 51.39921 | 22.17865 | 20.66423 |
| ENSMUSG00000054226 | -0.228048055 | 0.007564 | 0.034746 | 1080.502 | 1066.641 | 997.8685 | 1075.691 | 1295.923 | 1146.426 | 1143.647 | 1356.226 |
| ENSMUSG00000054237 | -0.008508281 | 0.944313 | 0.967726 | 639.8506 | 725.892 | 630.1891 | 678.8847 | 581.1663 | 808.9788 | 638.3594 | 662.343 |
| ENSMUSG00000054252 | -1.090116463 | 0.000584 | 0.005158 | 258.556 | 178.7734 | 453.8024 | 251.8861 | 579.2161 | 353.0902 | 661.5024 | 842.8831 |
| ENSMUSG00000054256 | 0.480781209 | 0.00172 | 0.011727 | 838.0433 | 841.0749 | 751.0927 | 771.1854 | 592.8677 | 738.5843 | 500.4661 | 462.2262 |
| ENSMUSG00000054263 | 0.148723864 | 0.299141 | 0.467283 | 1389.361 | 1186.623 | 1622.261 | 1624.32 | 1118.453 | 1535.272 | 1395.326 | 1207.226 |
| ENSMUSG00000054277 | -0.115110048 | 0.263536 | 0.427853 | 961.788 | 829.0767 | 921.6827 | 913.5183 | 1006.315 | 835.7958 | 1085.79 | 999.4962 |
| ENSMUSG00000054280 | 0.31950898 | 0.000678 | 0.005799 | 989.9575 | 815.8786 | 969.7129 | 898.8537 | 735.2339 | 677.1287 | 751.1813 | 783.0656 |
| ENSMUSG00000054293 | 0.038363376 | 0.925514 | 0.955704 | 25.15136 | 16.7975 | 21.53077 | 14.6646 | 24.37778 | 12.29112 | 15.42863 | 23.927 |
| ENSMUSG00000054302 | -0.037628142 | 0.558372 | 0.704826 | 1061.387 | 1093.037 | 1091.445 | 1144.701 | 1098.95 | 1092.792 | 1129.183 | 1187.649 |
| ENSMUSG00000054304 | 0.006696933 | 0.991282 | 0.994893 | 38.23007 | 29.99554 | 13.24971 | 15.52722 | 9.751113 | 50.28183 | 12.53576 | 23.927 |
| ENSMUSG00000054309 | -0.129292802 | 0.021992 | 0.075415 | 2233.441 | 2376.846 | 2284.746 | 2244.546 | 2521.638 | 2348.72 | 2558.259 | 2562.365 |
| ENSMUSG00000054312 | -0.328507492 | 0.00026 | 0.002786 | 1189.156 | 1252.614 | 1279.425 | 1297.386 | 1682.067 | 1404.539 | 1486.934 | 1729.27 |
| ENSMUSG00000054320 | 0.402293461 | 0.512789 | 0.66798 | 7.04238 | 22.79661 | 16.56213 | 26.74133 | 11.70134 | 26.81698 | 11.57147 | 5.437955 |
| ENSMUSG00000054321 | -0.744712045 | 8.69E-08 | 5.08E-06 | 133.8052 | 154.777 | 159.8246 | 127.6683 | 215.4996 | 248.057 | 256.5009 | 244.708 |
| ENSMUSG00000054342 | 0.556289253 | 0.004473 | 0.023717 | 492.9666 | 772.6851 | 476.1613 | 529.6508 | 450.5014 | 425.7195 | 360.6441 | 305.6131 |
| ENSMUSG00000054364 | -0.259890102 | 0.042752 | 0.122272 | 2448.736 | 2237.667 | 2756.767 | 2652.567 | 2777.117 | 2487.275 | 3349.941 | 3477.029 |
| ENSMUSG00000054381 | -0.507929221 | 0.007134 | 0.033334 | 229.3804 | 196.7707 | 168.1056 | 142.3329 | 291.5583 | 294.9868 | 223.7151 | 234.9197 |
| ENSMUSG00000054383 | 0.212583119 | 0.68905 | 0.802898 | 29.17558 | 26.39607 | 7.45296 | 18.97772 | 24.37778 | 14.52586 | 21.21436 | 9.78832 |
| ENSMUSG00000054387 | 0.369988298 | 0.000577 | 0.00512 | 4312.955 | 3856.226 | 3701.637 | 4163.884 | 2940.936 | 3716.386 | 2886.117 | 2865.803 |
| ENSMUSG00000054400 | 0.370286012 | 0.025264 | 0.083583 | 534.2149 | 472.7297 | 342.8361 | 412.334 | 331.5378 | 372.0856 | 369.3228 | 288.2116 |
| ENSMUSG00000054404 | -0.003150277 | 0.982869 | 0.990598 | 995.9938 | 1028.247 | 1299.299 | 837.6074 | 901.0028 | 1069.327 | 1141.718 | 1059.314 |
| ENSMUSG00000054405 | -0.064171498 | 0.231213 | 0.39178 | 3609.723 | 3897.02 | 3550.093 | 3758.451 | 3949.201 | 3919.748 | 3733.728 | 3881.613 |
| ENSMUSG00000054408 | -0.322995383 | 0.00014 | 0.001734 | 2111.708 | 2228.069 | 2136.515 | 2193.652 | 2521.638 | 2452.636 | 3009.546 | 2859.277 |
| ENSMUSG00000054414 | 0.074690877 | 0.234316 | 0.395152 | 993.9817 | 1096.637 | 1063.289 | 1047.225 | 1008.265 | 993.3456 | 986.4678 | 999.4962 |
| ENSMUSG00000054418 | -0.101107273 | 0.808268 | 0.882948 | 19.11503 | 14.39786 | 16.56213 | 12.07673 | 15.60178 | 15.64324 | 23.14294 | 11.9635 |
| ENSMUSG00000054423 | 0.854072342 | 0.002748 | 0.016571 | 106.6418 | 85.18733 | 78.67013 | 163.0358 | 58.50668 | 71.51194 | 54.00019 | 56.55474 |
| ENSMUSG00000054426 | 0.17432524 | 0.394071 | 0.561441 | 130.7871 | 115.1829 | 109.3101 | 119.9047 | 81.90935 | 130.7328 | 120.5361 | 88.09488 |
| ENSMUSG00000054428 | -0.365016019 | 0.004486 | 0.023747 | 3940.715 | 3599.465 | 4167.861 | 3853.339 | 5115.434 | 3953.27 | 4919.803 | 6054.62 |
| ENSMUSG00000054434 | -0.354350159 | 0.009019 | 0.039559 | 234.4107 | 194.3711 | 202.058 | 200.9913 | 235.0018 | 251.4092 | 270.9653 | 306.7007 |
| ENSMUSG00000054435 | -0.474935741 | 0.006348 | 0.030712 | 730.3955 | 536.3202 | 733.7025 | 720.2906 | 871.7495 | 727.4105 | 985.5035 | 1199.613 |

| | | | | | | | | | | | |
|---|---|---|---|---|---|---|---|---|---|---|---|
| ENSMUSG00000054446 | 0.291797516 | 0.576956 | 0.718897 | 12.07265 | 23.99643 | 14.07781 | 11.21411 | 16.57689 | 17.87799 | 8.678602 | 6.525547 |
| ENSMUSG00000054452 | -0.24702532 | 0.027657 | 0.089254 | 6938.757 | 7213.327 | 8443.375 | 6927.729 | 8526.373 | 7433.89 | 9095.175 | 9982.999 |
| ENSMUSG00000054453 | 0.505025093 | 0.226403 | 0.386289 | 21.12714 | 35.99465 | 22.35888 | 36.23019 | 26.32801 | 11.17374 | 27.0001 | 16.31387 |
| ENSMUSG00000054455 | -0.137840477 | 0.088682 | 0.202915 | 2854.176 | 2640.807 | 2908.31 | 2913.08 | 2967.264 | 2834.778 | 3226.512 | 3425.912 |
| ENSMUSG00000054457 | 1.476849842 | 0.000266 | 0.00283 | 21.12714 | 23.99643 | 28.98373 | 25.8787 | 10.72622 | 8.938993 | 8.678602 | 7.613138 |
| ENSMUSG00000054459 | 1.128301874 | 0.023322 | 0.07882 | 72.43591 | 25.19625 | 21.53077 | 75.04824 | 17.552 | 27.93435 | 25.07152 | 18.48905 |
| ENSMUSG00000054469 | -0.188545787 | 0.062166 | 0.158933 | 1097.605 | 962.2569 | 934.9324 | 1008.407 | 1102.851 | 1062.623 | 1080.004 | 1317.073 |
| ENSMUSG00000054474 | -1.376484302 | 5.91E-08 | 3.74E-06 | 372.2401 | 377.9438 | 477.8175 | 404.5704 | 1115.527 | 607.8515 | 932.4676 | 1584.62 |
| ENSMUSG00000054477 | 0.418152183 | 0.135555 | 0.269999 | 113.6841 | 59.99108 | 65.42042 | 106.9653 | 61.43201 | 69.27719 | 74.25027 | 54.37955 |
| ENSMUSG00000054484 | -0.115250293 | 0.439357 | 0.602708 | 200.2048 | 188.372 | 192.9488 | 254.4739 | 226.2258 | 202.2447 | 228.5365 | 250.146 |
| ENSMUSG00000054499 | -0.270863408 | 0.037053 | 0.110607 | 568.4207 | 532.7208 | 529.1601 | 558.1174 | 626.9966 | 525.1658 | 743.4669 | 743.9123 |
| ENSMUSG00000054509 | 0.193386381 | 0.04332 | 0.123211 | 322.9434 | 314.3532 | 320.4773 | 315.7202 | 289.6081 | 272.6393 | 285.4296 | 265.3722 |
| ENSMUSG00000054514 | 0.287553426 | 0.271145 | 0.436327 | 40.24217 | 47.99286 | 37.2648 | 35.36756 | 33.15378 | 32.40385 | 33.75012 | 31.54014 |
| ENSMUSG00000054517 | 0.332975963 | 0.083262 | 0.194221 | 231.3925 | 289.157 | 197.0894 | 191.5024 | 159.9183 | 157.5497 | 179.3578 | 222.9562 |
| ENSMUSG00000054519 | -0.077170893 | 0.608452 | 0.744118 | 460.7729 | 467.9304 | 428.9592 | 410.6088 | 471.9539 | 591.0909 | 422.3587 | 379.5693 |
| ENSMUSG00000054520 | 0.196243579 | 0.117794 | 0.246459 | 246.4833 | 286.7573 | 270.7909 | 238.0841 | 240.8525 | 244.7049 | 203.465 | 219.6934 |
| ENSMUSG00000054545 | -0.747362702 | 0.005343 | 0.026915 | 54.32694 | 34.79482 | 33.95237 | 33.64232 | 63.38224 | 52.51658 | 71.3574 | 75.04379 |
| ENSMUSG00000054555 | -0.406928608 | 0.192461 | 0.345566 | 201.2109 | 262.7609 | 570.5655 | 237.2215 | 345.1894 | 396.6678 | 449.3588 | 497.0291 |
| ENSMUSG00000054556 | 0.373864091 | 0.406757 | 0.573149 | 27.16347 | 31.19536 | 25.67131 | 47.44429 | 16.57689 | 46.92971 | 22.17865 | 16.31387 |
| ENSMUSG00000054580 | 0.663346831 | 1.72E-06 | 5.78E-05 | 521.1362 | 433.1356 | 511.7699 | 572.782 | 313.0107 | 387.7288 | 299.8939 | 289.2992 |
| ENSMUSG00000054582 | 0.466704469 | 0.122213 | 0.251736 | 82.49646 | 106.7841 | 60.45178 | 59.52102 | 51.6809 | 75.98144 | 55.92877 | 39.15328 |
| ENSMUSG00000054604 | 0.164064454 | 0.136613 | 0.271398 | 3273.701 | 3610.263 | 3074.76 | 3485.862 | 2873.653 | 3582.301 | 2905.403 | 2637.408 |
| ENSMUSG00000054611 | 0.212176974 | 0.025572 | 0.084355 | 3041.302 | 3043.947 | 2497.57 | 2643.078 | 2378.296 | 2647.059 | 2255.472 | 2406.839 |
| ENSMUSG00000054612 | 0.076050805 | 0.565254 | 0.710014 | 173.0413 | 196.7707 | 186.324 | 164.7611 | 181.3707 | 177.6625 | 167.7863 | 155.5255 |
| ENSMUSG00000054619 | -0.545064208 | 0.002023 | 0.013247 | 808.8677 | 715.0936 | 758.5457 | 775.4985 | 1183.785 | 741.9364 | 1149.433 | 1386.679 |
| ENSMUSG00000054622 | 0.157238274 | 0.8264 | 0.894548 | 7.04238 | 15.59768 | 4.140533 | 25.8787 | 13.65156 | 17.87799 | 3.857157 | 11.9635 |
| ENSMUSG00000054626 | -0.412873618 | 0.559398 | 0.705627 | 10.06054 | 5.999108 | 12.4216 | 12.07673 | 8.776002 | 33.52122 | 4.821446 | 7.613138 |
| ENSMUSG00000054630 | -2.32018316 | 4.17E-05 | 0.000696 | 20.12109 | 1.199822 | 19.04645 | 11.21411 | 46.80534 | 49.16446 | 60.75022 | 104.4087 |
| ENSMUSG00000054640 | 0.043598486 | 0.811238 | 0.885092 | 744.4802 | 757.0874 | 843.0126 | 637.4788 | 739.1344 | 487.1751 | 790.7171 | 875.5108 |
| ENSMUSG00000054641 | 0.515705092 | 0.006561 | 0.031503 | 178.0716 | 140.3791 | 178.871 | 151.8217 | 119.9387 | 83.80306 | 115.7147 | 134.8613 |
| ENSMUSG00000054648 | -0.023110317 | 0.793002 | 0.873246 | 605.6447 | 625.107 | 553.1752 | 571.0568 | 630.897 | 608.9689 | 603.645 | 544.8831 |
| ENSMUSG00000054659 | -0.632288157 | 0.043212 | 0.123138 | 39.23612 | 38.39429 | 38.09291 | 27.60395 | 81.90935 | 41.34284 | 52.07161 | 45.67883 |
| ENSMUSG00000054662 | -0.953971849 | 0.000505 | 0.00462 | 97.58727 | 95.98572 | 128.3565 | 75.04824 | 157.968 | 136.3196 | 193.8221 | 281.6861 |
| ENSMUSG00000054667 | 1.619376224 | 0.041874 | 0.120472 | 45.27245 | 31.19536 | 28.98373 | 25.01608 | 0.975111 | 27.93435 | 11.57147 | 2.175182 |
| ENSMUSG00000054675 | 0.481172317 | 1.97E-06 | 6.36E-05 | 891.3642 | 1054.643 | 1074.054 | 1071.378 | 705.0055 | 756.4623 | 668.2524 | 803.7298 |
| ENSMUSG00000054676 | 0.098499845 | 0.467698 | 0.627998 | 553.3299 | 597.5111 | 539.9255 | 478.756 | 561.6641 | 398.9026 | 518.7876 | 543.7955 |
| ENSMUSG00000054679 | 0.490423575 | 0.058757 | 0.152371 | 115.6963 | 121.182 | 120.0755 | 143.1955 | 56.55646 | 124.0285 | 102.2147 | 73.95619 |
| ENSMUSG00000054690 | -0.7744876 | 0.000255 | 0.002751 | 1017.121 | 671.9001 | 1315.861 | 895.4032 | 1514.348 | 1308.445 | 1821.542 | 2031.62 |
| ENSMUSG00000054693 | 0.11442387 | 0.225676 | 0.385513 | 4698.274 | 5496.382 | 4989.342 | 5153.313 | 4381.175 | 5444.964 | 4508.052 | 4452.598 |
| ENSMUSG00000054702 | -0.123059023 | 0.373152 | 0.542019 | 284.7134 | 266.3604 | 339.5237 | 344.1868 | 287.6578 | 353.0902 | 366.4299 | 341.5036 |
| ENSMUSG00000054708 | 0.109859569 | 0.304563 | 0.47303 | 1106.66 | 1019.848 | 1066.601 | 1256.842 | 1106.751 | 1130.783 | 927.6462 | 961.4305 |
| ENSMUSG00000054715 | -0.110053302 | 0.666706 | 0.785994 | 314.895 | 340.7493 | 185.4959 | 241.5346 | 289.6081 | 191.071 | 303.7511 | 381.7445 |
| ENSMUSG00000054716 | -0.024725662 | 0.780198 | 0.864358 | 1168.029 | 1287.409 | 1124.569 | 1305.149 | 1314.45 | 1185.534 | 1318.183 | 1147.409 |
| ENSMUSG00000054717 | -0.070353476 | 0.285778 | 0.452692 | 16142.14 | 16364.37 | 15202.38 | 16756.46 | 17930.35 | 17504.78 | 16689.92 | 15559.08 |
| ENSMUSG00000054720 | 0.152789744 | 0.172473 | 0.319892 | 675.0625 | 531.5209 | 625.2205 | 647.8302 | 507.0579 | 598.9125 | 598.8236 | 528.5693 |
| ENSMUSG00000054723 | -0.097078565 | 0.316006 | 0.484322 | 574.457 | 632.3059 | 694.7815 | 617.6384 | 721.5824 | 702.8283 | 622.9308 | 649.2919 |
| ENSMUSG00000054728 | 0.481363456 | 0.004613 | 0.024264 | 1283.725 | 1237.016 | 1353.954 | 1106.746 | 812.2677 | 1214.586 | 699.1096 | 843.9707 |
| ENSMUSG00000054733 | -0.811512722 | 6.67E-08 | 4.06E-06 | 459.7668 | 460.7315 | 526.6758 | 480.4813 | 918.5549 | 657.016 | 778.1814 | 1031.036 |
| ENSMUSG00000054737 | 0.502084731 | 0.001117 | 0.008499 | 422.5428 | 428.3363 | 317.1648 | 427.8613 | 234.0267 | 322.9211 | 293.1439 | 276.2481 |
| ENSMUSG00000054752 | 0.485613315 | 0.052006 | 0.139774 | 405.4399 | 398.3407 | 402.4598 | 494.2833 | 273.0312 | 483.823 | 247.8223 | 210.9927 |
| ENSMUSG00000054763 | -2.393492649 | 0.000524 | 0.004749 | 8.048435 | 1.199822 | 28.98373 | 23.29083 | 61.43201 | 33.52122 | 129.2147 | 103.3212 |
| ENSMUSG00000054766 | -0.062868113 | 0.402144 | 0.569261 | 11178.27 | 12961.67 | 11394.75 | 12156.09 | 12450.22 | 13519.11 | 11928.26 | 11914.56 |
| ENSMUSG00000054770 | 0.139441136 | 0.3018 | 0.470203 | 806.8556 | 769.0856 | 747.7803 | 812.5913 | 723.5326 | 890.5472 | 638.3594 | 595.9999 |
| ENSMUSG00000054792 | -0.033283917 | 0.71935 | 0.824462 | 460.7729 | 495.5263 | 500.1764 | 466.6793 | 478.7797 | 453.6539 | 493.7161 | 541.6204 |
| ENSMUSG00000054793 | 0.239851536 | 0.005712 | 0.028384 | 1405.458 | 1533.372 | 1575.059 | 1532.882 | 1191.586 | 1182.182 | 1331.683 | 1416.044 |
| ENSMUSG00000054808 | -0.35728355 | 0.006365 | 0.030785 | 8077.61 | 7924.821 | 9263.201 | 7746.359 | 10502.92 | 8309.911 | 10791.36 | 12685.66 |
| ENSMUSG00000054814 | -0.0901409 | 0.312859 | 0.481091 | 1103.642 | 991.0526 | 989.5874 | 1123.998 | 1139.905 | 997.8151 | 1171.611 | 1170.248 |
| ENSMUSG00000054823 | 0.091277815 | 0.282745 | 0.449315 | 2641.899 | 2331.253 | 2237.544 | 2417.934 | 2107.216 | 2474.984 | 2155.186 | 2302.43 |
| ENSMUSG00000054836 | -0.076113783 | 0.464594 | 0.625468 | 375.2583 | 363.5459 | 444.6933 | 368.3402 | 405.6463 | 417.8979 | 397.2871 | 418.7226 |
| ENSMUSG00000054843 | -0.111186026 | 0.448831 | 0.611428 | 1972.873 | 2432.038 | 2906.654 | 2156.559 | 2388.048 | 2176.645 | 2705.795 | 2957.16 |
| ENSMUSG00000054850 | -0.27877088 | 0.129072 | 0.260829 | 211.2714 | 172.7743 | 250.9163 | 174.2499 | 236.952 | 197.7752 | 260.3581 | 288.2116 |
| ENSMUSG00000054855 | 0.189866833 | 0.640163 | 0.7668 | 43.26034 | 28.79572 | 31.46805 | 14.6646 | 31.20356 | 21.23011 | 29.89296 | 20.66423 |
| ENSMUSG00000054863 | 0.128849408 | 0.236945 | 0.398057 | 580.4934 | 703.0954 | 638.4702 | 598.6607 | 528.5103 | 638.0206 | 607.5022 | 529.6569 |
| ENSMUSG00000054871 | 0.675833506 | 0.001343 | 0.009748 | 542.2633 | 621.5076 | 650.8918 | 828.1186 | 354.9405 | 576.565 | 409.8229 | 314.3138 |

| | | | | | | | | | | | |
|---|---|---|---|---|---|---|---|---|---|---|---|
| ENSMUSG00000054874 | -0.028515747 | 0.654136 | 0.776731 | 1696.208 | 1808.131 | 1828.459 | 1705.407 | 1881.965 | 1733.047 | 1720.292 | 1842.379 |
| ENSMUSG00000054889 | -0.598758862 | 0.016914 | 0.062633 | 974.8667 | 575.9143 | 1346.501 | 847.0963 | 1276.421 | 1059.271 | 1794.542 | 1543.292 |
| ENSMUSG00000054892 | -0.892086883 | 0.165667 | 0.311139 | 7.04238 | 3.599465 | 18.21835 | 4.313117 | 17.552 | 8.938993 | 15.42863 | 20.66423 |
| ENSMUSG00000054893 | 0.297903376 | 0.027664 | 0.08926 | 254.5318 | 241.1641 | 231.0418 | 208.7549 | 211.5992 | 198.8926 | 166.822 | 182.7153 |
| ENSMUSG00000054894 | -0.214864705 | 0.062962 | 0.160412 | 230.3864 | 217.1677 | 238.4947 | 226.87 | 253.5289 | 287.1651 | 240.108 | 281.6861 |
| ENSMUSG00000054901 | -0.230784201 | 0.420625 | 0.585984 | 25.15136 | 27.5959 | 36.43669 | 36.23019 | 33.15378 | 42.46022 | 32.78583 | 40.24087 |
| ENSMUSG00000054905 | 0.168414475 | 0.762848 | 0.852453 | 41.24823 | 10.79839 | 15.73403 | 20.70296 | 22.42756 | 12.29112 | 11.57147 | 32.62773 |
| ENSMUSG00000054909 | -0.175268258 | 0.727475 | 0.829154 | 15.09082 | 19.19714 | 7.45296 | 11.21411 | 12.67645 | 21.23011 | 16.39292 | 8.700729 |
| ENSMUSG00000054920 | 0.072308046 | 0.565032 | 0.709838 | 1539.263 | 1422.988 | 1694.306 | 1659.688 | 1426.588 | 1219.055 | 1626.756 | 1736.883 |
| ENSMUSG00000054931 | 0.054350229 | 0.777745 | 0.862987 | 152.9203 | 155.9768 | 151.5435 | 106.1027 | 122.864 | 150.8455 | 154.2863 | 116.3722 |
| ENSMUSG00000054932 | -1.180626494 | 0.127195 | 0.258402 | 1145.896 | 91.18644 | 589.6119 | 1195.596 | 1626.486 | 646.9596 | 1995.114 | 2583.029 |
| ENSMUSG00000054934 | 0.428239746 | 0.05158 | 0.138974 | 125.7568 | 93.58608 | 114.2787 | 119.9047 | 70.20801 | 105.0332 | 97.39321 | 65.25547 |
| ENSMUSG00000054939 | 0.248358381 | 0.281455 | 0.447843 | 150.9082 | 111.5834 | 89.43552 | 93.16334 | 94.5858 | 75.98144 | 97.39321 | 105.4963 |
| ENSMUSG00000054942 | 0.116718833 | 0.137756 | 0.272969 | 1271.653 | 1214.219 | 1144.443 | 1140.388 | 1055.07 | 1136.369 | 1176.433 | 1028.861 |
| ENSMUSG00000054945 | 0.356412889 | 0.179307 | 0.329085 | 89.53884 | 79.18822 | 66.24853 | 78.49874 | 49.73068 | 86.03781 | 65.57166 | 43.50364 |
| ENSMUSG00000054951 | 0.014004116 | 0.950091 | 0.971623 | 104.6297 | 95.98572 | 131.669 | 86.26235 | 94.5858 | 109.5027 | 129.2147 | 81.56933 |
| ENSMUSG00000054966 | 0.993570875 | 0.000354 | 0.003512 | 299.8042 | 527.9215 | 394.1788 | 383.8675 | 238.9023 | 288.2825 | 147.5362 | 131.5985 |
| ENSMUSG00000054967 | -0.036806249 | 0.820369 | 0.891148 | 247.4894 | 267.5602 | 218.6202 | 217.3811 | 302.2845 | 206.7142 | 216.9651 | 246.8832 |
| ENSMUSG00000054976 | 1.060753217 | 0.105228 | 0.22875 | 25.15136 | 8.398751 | 11.59349 | 36.23019 | 5.850668 | 14.52586 | 14.46434 | 4.350364 |
| ENSMUSG00000054978 | -0.130906822 | 0.703516 | 0.813207 | 33.19979 | 44.3934 | 43.88965 | 36.23019 | 29.25334 | 29.05173 | 50.14304 | 64.16787 |
| ENSMUSG00000054988 | 1.216870369 | 6.32E-05 | 0.000953 | 102.6175 | 87.58697 | 53.82693 | 90.57547 | 38.02934 | 46.92971 | 33.75012 | 25.0146 |
| ENSMUSG00000054999 | 0.020314577 | 0.951583 | 0.972579 | 57.3451 | 106.7841 | 73.70149 | 70.73513 | 80.93424 | 117.3243 | 50.14304 | 55.46715 |
| ENSMUSG00000055003 | -0.299140095 | 0.603942 | 0.740916 | 17.10292 | 7.198929 | 18.21835 | 5.175741 | 11.70134 | 10.05637 | 14.46434 | 22.83941 |
| ENSMUSG00000055013 | 0.121663293 | 0.166079 | 0.311658 | 1858.182 | 1811.731 | 1940.254 | 1935.727 | 1534.825 | 1948.7 | 1679.792 | 1777.124 |
| ENSMUSG00000055022 | -0.023159397 | 0.907571 | 0.94484 | 147.89 | 164.3756 | 139.95 | 181.1509 | 195.0223 | 129.6154 | 188.0364 | 129.4233 |
| ENSMUSG00000055024 | 0.221146535 | 0.012525 | 0.050582 | 2035.248 | 1746.94 | 1720.806 | 1892.596 | 1470.468 | 1739.751 | 1540.934 | 1595.496 |
| ENSMUSG00000055026 | -0.129131172 | 0.692849 | 0.805613 | 35.2119 | 37.19447 | 30.63995 | 33.64232 | 22.42756 | 53.63396 | 36.64299 | 36.9781 |
| ENSMUSG00000055027 | -0.596357566 | 0.251352 | 0.415004 | 7.04238 | 10.79839 | 7.45296 | 8.626235 | 9.751113 | 20.11273 | 12.53576 | 8.700729 |
| ENSMUSG00000055041 | -0.13380678 | 0.332702 | 0.500763 | 456.7487 | 451.1329 | 523.3634 | 423.5481 | 513.8837 | 430.189 | 475.3946 | 616.6641 |
| ENSMUSG00000055044 | -0.29044437 | 0.18663 | 0.33852 | 281.6952 | 287.9572 | 472.8489 | 331.2474 | 411.497 | 301.691 | 468.6445 | 500.2919 |
| ENSMUSG00000055053 | 0.390137876 | 0.00082 | 0.006696 | 1721.359 | 1907.716 | 1812.725 | 1591.54 | 1283.246 | 1567.676 | 1131.111 | 1385.591 |
| ENSMUSG00000055065 | 0.32828618 | 0.001135 | 0.008607 | 17878.59 | 16783.1 | 15823.46 | 17391.35 | 12599.41 | 16109.18 | 13038.15 | 12316.97 |
| ENSMUSG00000055067 | 0.092407071 | 0.277693 | 0.444018 | 623.7537 | 687.4977 | 664.9696 | 715.9775 | 679.6526 | 600.0299 | 615.2165 | 629.7152 |
| ENSMUSG00000055093 | -0.123746976 | 0.611035 | 0.7459 | 103.6236 | 161.9759 | 221.1045 | 139.745 | 159.9183 | 167.6061 | 203.465 | 152.2628 |
| ENSMUSG00000055114 | -2.090302239 | 7.88E-07 | 3.05E-05 | 30.18163 | 11.99822 | 43.06154 | 42.26855 | 83.85957 | 97.21155 | 164.8934 | 202.2919 |
| ENSMUSG00000055116 | -0.12813492 | 0.491549 | 0.648724 | 462.785 | 386.3425 | 371.8199 | 364.8897 | 420.273 | 301.691 | 542.8948 | 466.5766 |
| ENSMUSG00000055128 | -0.007594264 | 0.927982 | 0.957316 | 426.567 | 423.537 | 446.3495 | 451.1521 | 440.7503 | 460.3581 | 419.4658 | 439.3868 |
| ENSMUSG00000055134 | 0.020617662 | 0.949121 | 0.971183 | 40.24217 | 35.99465 | 21.53077 | 36.23019 | 24.37778 | 36.87335 | 36.64299 | 33.71532 |
| ENSMUSG00000055137 | -1.797811454 | 1.14E-05 | 0.000254 | 66.39959 | 64.79036 | 107.6539 | 65.55939 | 284.7325 | 93.85942 | 233.358 | 448.0875 |
| ENSMUSG00000055148 | -0.268641979 | 0.10033 | 0.221414 | 578.4813 | 430.7359 | 541.5817 | 535.6892 | 505.1077 | 546.3959 | 739.6098 | 724.3357 |
| ENSMUSG00000055150 | -0.415182657 | 0.119162 | 0.248167 | 39.23612 | 39.59411 | 28.98373 | 33.64232 | 48.75557 | 48.04709 | 36.64299 | 54.37955 |
| ENSMUSG00000055172 | 0.652477919 | 6.11E-07 | 2.47E-05 | 623.7537 | 867.471 | 751.9208 | 790.1631 | 422.2232 | 535.2222 | 474.4303 | 498.1167 |
| ENSMUSG00000055184 | 0.029945579 | 0.920246 | 0.95244 | 61.36932 | 65.99018 | 41.40533 | 78.49874 | 77.03379 | 42.46022 | 69.42882 | 52.20437 |
| ENSMUSG00000055188 | -0.454375706 | 0.122712 | 0.25224 | 37.22401 | 33.595 | 36.43669 | 31.05445 | 57.53157 | 59.22083 | 32.78583 | 40.24087 |
| ENSMUSG00000055198 | 0.437289887 | 0.265263 | 0.429407 | 25.15136 | 32.39518 | 20.70267 | 19.84034 | 24.37778 | 18.99536 | 17.3572 | 10.87591 |
| ENSMUSG00000055200 | 0.179209428 | 0.114976 | 0.24268 | 411.4762 | 373.1445 | 407.4285 | 379.5543 | 400.7707 | 313.9821 | 324.0012 | 349.1167 |
| ENSMUSG00000055202 | 0.88299963 | 0.000805 | 0.006603 | 249.5015 | 309.554 | 237.6666 | 323.4838 | 150.1671 | 229.0617 | 135.9648 | 92.44524 |
| ENSMUSG00000055204 | 0.050175489 | 0.611349 | 0.746052 | 4165.065 | 3906.619 | 3644.497 | 3662.699 | 3289.05 | 4285.13 | 3536.048 | 3743.489 |
| ENSMUSG00000055210 | -0.042028334 | 0.805574 | 0.881048 | 718.3228 | 778.6842 | 574.706 | 677.1594 | 729.3833 | 906.1904 | 647.038 | 545.9707 |
| ENSMUSG00000055214 | 1.022225378 | 0.000391 | 0.003799 | 53.32088 | 74.38894 | 63.76421 | 81.08661 | 32.17867 | 31.28647 | 46.28588 | 23.927 |
| ENSMUSG00000055216 | 0.578854871 | 0.384194 | 0.552197 | 27.16347 | 27.5959 | 20.70267 | 19.84034 | 3.900445 | 39.10809 | 14.46434 | 6.525547 |
| ENSMUSG00000055228 | 0.165395109 | 0.256441 | 0.42014 | 207.2472 | 161.9759 | 169.7619 | 167.349 | 172.5947 | 145.2586 | 160.072 | 151.1752 |
| ENSMUSG00000055235 | -0.101617795 | 0.47027 | 0.63026 | 615.7053 | 659.9018 | 884.4179 | 710.8018 | 746.9353 | 680.4808 | 777.2171 | 878.7736 |
| ENSMUSG00000055239 | -0.102238926 | 0.092154 | 0.20843 | 2216.338 | 2006.102 | 2213.529 | 2089.274 | 2270.059 | 2205.696 | 2373.116 | 2305.693 |
| ENSMUSG00000055240 | 0.18866793 | 0.157716 | 0.300269 | 1065.412 | 1165.027 | 929.9637 | 938.5344 | 904.9033 | 1087.205 | 821.5744 | 780.8904 |
| ENSMUSG00000055254 | 0.90720516 | 0.001781 | 0.012076 | 662.9898 | 1214.219 | 881.1055 | 1006.682 | 407.5965 | 848.0869 | 393.43 | 358.9051 |
| ENSMUSG00000055296 | 0.217885547 | 0.091449 | 0.207421 | 1747.516 | 1434.987 | 1574.231 | 1568.25 | 1198.412 | 1688.352 | 1333.612 | 1220.277 |
| ENSMUSG00000055301 | 1.199250799 | 0.139033 | 0.274909 | 183.1019 | 8.398751 | 22.35888 | 25.8787 | 17.552 | 21.23011 | 22.17865 | 43.50364 |
| ENSMUSG00000055302 | -0.068263336 | 0.188539 | 0.340946 | 10638.02 | 11487.09 | 10555.88 | 11113.18 | 11804.7 | 11634.1 | 10972.65 | 11500.19 |
| ENSMUSG00000055305 | 0.081498657 | 0.510657 | 0.666244 | 349.1009 | 391.1418 | 307.2276 | 326.9343 | 345.1894 | 283.813 | 321.1083 | 344.7664 |
| ENSMUSG00000055312 | -3.628685621 | 5.37E-15 | 2.36E-12 | 14.08476 | 16.7975 | 10.76539 | 15.52722 | 214.5245 | 54.75133 | 166.822 | 267.5474 |
| ENSMUSG00000055313 | 0.020338898 | 0.912292 | 0.948044 | 125.7568 | 133.1802 | 99.3728 | 95.75121 | 101.4116 | 121.7938 | 103.1789 | 119.635 |
| ENSMUSG00000055319 | -0.030225279 | 0.707214 | 0.815963 | 1412.5 | 1595.763 | 1365.548 | 1568.25 | 1560.178 | 1413.478 | 1540.934 | 1548.73 |
| ENSMUSG00000055320 | 0.415763379 | 0.043171 | 0.123111 | 3983.975 | 3507.078 | 3592.327 | 3610.942 | 2176.448 | 4176.744 | 2563.081 | 2100.138 |

| | | | | | | | | | | | |
|---|---|---|---|---|---|---|---|---|---|---|---|
| ENSMUSG00000055322 | -0.337588618 | 0.009639 | 0.041458 | 3965.866 | 3229.92 | 3776.994 | 3930.113 | 3993.081 | 4128.697 | 5539.841 | 5172.583 |
| ENSMUSG00000055323 | 0.586045571 | 0.173334 | 0.321126 | 50.30272 | 50.3925 | 24.8432 | 37.09281 | 23.40267 | 48.04709 | 18.32149 | 18.48905 |
| ENSMUSG00000055333 | 0.04758006 | 0.889209 | 0.933305 | 44.26639 | 45.59322 | 67.90474 | 32.77969 | 63.38224 | 49.16446 | 41.46443 | 30.45255 |
| ENSMUSG00000055334 | -0.038165571 | 0.727951 | 0.829271 | 274.6528 | 280.7582 | 310.54 | 278.6274 | 299.3592 | 312.8647 | 264.2152 | 301.2627 |
| ENSMUSG00000055341 | -0.168058152 | 0.736938 | 0.835233 | 21.12714 | 9.598572 | 10.76539 | 13.80198 | 20.47734 | 14.52586 | 7.714313 | 19.57664 |
| ENSMUSG00000055368 | 3.98928519 | 0.001687 | 0.011549 | 334.01 | 3.599465 | 10.76539 | 356.2635 | 17.552 | 7.821619 | 13.50005 | 5.437955 |
| ENSMUSG00000055370 | -0.638004476 | 0.332973 | 0.501088 | 6.036326 | 3.599465 | 11.59349 | 20.70296 | 14.62667 | 6.704245 | 24.10723 | 20.66423 |
| ENSMUSG00000055371 | 0.038658518 | 0.493181 | 0.650107 | 1182.114 | 1246.615 | 1230.566 | 1234.414 | 1210.113 | 1163.186 | 1186.076 | 1203.963 |
| ENSMUSG00000055373 | -2.001327643 | 0.000394 | 0.00381 | 183.1019 | 61.1909 | 340.3518 | 169.9368 | 490.481 | 585.504 | 640.288 | 1308.372 |
| ENSMUSG00000055401 | -0.198402749 | 0.080553 | 0.189822 | 421.5368 | 411.5388 | 397.4912 | 400.2573 | 523.6348 | 403.372 | 435.8587 | 506.8174 |
| ENSMUSG00000055407 | 0.947851046 | 2.50E-06 | 7.74E-05 | 147.89 | 135.5798 | 150.7154 | 213.068 | 78.98402 | 90.5073 | 88.7146 | 78.30656 |
| ENSMUSG00000055409 | 0.788838251 | 0.01365 | 0.053783 | 857.1583 | 1179.425 | 796.6386 | 960.9626 | 533.3859 | 984.4066 | 357.7513 | 320.8394 |
| ENSMUSG00000055413 | -0.885347936 | 0.104733 | 0.228023 | 5.030272 | 9.598572 | 26.49941 | 8.626235 | 20.47734 | 23.46486 | 23.14294 | 26.10219 |
| ENSMUSG00000055415 | -0.410731679 | 0.278119 | 0.444297 | 110.666 | 62.39072 | 200.4018 | 74.18562 | 168.6943 | 93.85942 | 161.0363 | 172.927 |
| ENSMUSG00000055421 | 0.569145112 | 0.007968 | 0.036131 | 317.9132 | 213.5682 | 256.7131 | 313.995 | 156.0178 | 239.1181 | 206.3579 | 142.4744 |
| ENSMUSG00000055430 | 0.709369688 | 0.063832 | 0.162024 | 113.6841 | 44.3934 | 62.9361 | 148.3712 | 48.75557 | 52.51658 | 67.50024 | 57.64233 |
| ENSMUSG00000055435 | -0.279824298 | 0.163295 | 0.307999 | 2780.734 | 2037.297 | 2379.15 | 2333.397 | 1820.533 | 3708.565 | 3193.726 | 2849.489 |
| ENSMUSG00000055436 | 0.30222965 | 0.02285 | 0.077594 | 9443.832 | 9140.24 | 8027.666 | 9205.918 | 6647.334 | 9317.783 | 6702.774 | 6380.897 |
| ENSMUSG00000055447 | 0.60817665 | 0.001342 | 0.009747 | 3066.454 | 2898.769 | 2437.118 | 2828.542 | 1616.735 | 2575.547 | 1855.292 | 1320.336 |
| ENSMUSG00000055452 | 0.120547959 | 0.819474 | 0.890707 | 9.054489 | 14.39786 | 15.73403 | 12.07673 | 10.72622 | 4.469496 | 14.46434 | 17.40146 |
| ENSMUSG00000055480 | 0.20918457 | 0.510826 | 0.666363 | 48.29061 | 118.7823 | 72.87338 | 57.79577 | 59.48179 | 79.33356 | 57.85735 | 59.81751 |
| ENSMUSG00000055485 | -0.174554254 | 0.108237 | 0.232904 | 545.2815 | 472.7297 | 524.1915 | 449.4268 | 559.7139 | 614.5558 | 573.7521 | 500.2919 |
| ENSMUSG00000055489 | -0.340540571 | 0.513516 | 0.668226 | 86.52067 | 46.79304 | 355.2577 | 138.0198 | 190.1467 | 117.3243 | 236.2508 | 251.2335 |
| ENSMUSG00000055491 | -0.105243278 | 0.384606 | 0.552354 | 656.9535 | 646.7038 | 676.5631 | 561.5679 | 780.089 | 658.1333 | 577.6092 | 717.8101 |
| ENSMUSG00000055493 | 0.091846826 | 0.446091 | 0.608774 | 291.7558 | 284.3577 | 295.6341 | 249.2982 | 261.3298 | 232.4138 | 270.001 | 287.124 |
| ENSMUSG00000055494 | -0.056974492 | 0.88993 | 0.933475 | 12.07265 | 22.79661 | 19.87456 | 28.46658 | 28.27823 | 24.58223 | 15.42863 | 18.48905 |
| ENSMUSG00000055497 | 0.523524717 | 0.311813 | 0.480258 | 19.11503 | 22.79661 | 11.59349 | 23.29083 | 6.825779 | 23.46486 | 14.46434 | 8.700729 |
| ENSMUSG00000055523 | -1.805989445 | 0.000772 | 0.006413 | 6.036326 | 13.19804 | 42.23344 | 12.07673 | 83.85957 | 67.04245 | 57.85735 | 51.11678 |
| ENSMUSG00000055531 | 0.298888276 | 0.027915 | 0.089945 | 4516.178 | 4758.492 | 4046.957 | 4338.996 | 3312.453 | 4632.633 | 3312.333 | 3098.547 |
| ENSMUSG00000055538 | 0.488503354 | 2.25E-06 | 7.05E-05 | 1793.795 | 2273.662 | 1989.112 | 2153.971 | 1445.115 | 1576.615 | 1530.327 | 1298.584 |
| ENSMUSG00000055541 | 0.545220752 | 0.226942 | 0.386817 | 22.1332 | 55.19179 | 43.88965 | 18.97772 | 21.45245 | 34.6386 | 21.21436 | 18.48905 |
| ENSMUSG00000055553 | -0.14052653 | 0.117505 | 0.245966 | 1331.01 | 1250.214 | 1348.986 | 1390.549 | 1470.468 | 1270.454 | 1548.648 | 1577.007 |
| ENSMUSG00000055555 | -0.72866033 | 0.208955 | 0.366165 | 11.0666 | 3.599465 | 8.281066 | 12.07673 | 7.80089 | 24.58223 | 12.53576 | 14.13868 |
| ENSMUSG00000055560 | 0.287042633 | 0.406373 | 0.572907 | 45.27245 | 40.79393 | 20.70267 | 50.89479 | 30.22845 | 29.05173 | 34.71441 | 34.80291 |
| ENSMUSG00000055567 | 0.337909689 | 0.265853 | 0.430063 | 49.29666 | 40.79393 | 61.27989 | 73.323 | 49.73068 | 58.10345 | 39.53586 | 31.54014 |
| ENSMUSG00000055612 | -0.05726157 | 0.564814 | 0.70971 | 1964.824 | 2291.659 | 1815.21 | 1952.98 | 2295.412 | 2109.602 | 2048.15 | 1890.233 |
| ENSMUSG00000055629 | 0.034902256 | 0.700125 | 0.810722 | 921.5458 | 919.0633 | 818.9975 | 939.397 | 959.5095 | 892.7819 | 864.0031 | 793.9415 |
| ENSMUSG00000055632 | -0.202131714 | 0.655651 | 0.77803 | 13.07871 | 52.79215 | 62.108 | 47.44429 | 51.6809 | 69.27719 | 35.6787 | 45.67883 |
| ENSMUSG00000055633 | 0.014712801 | 0.880926 | 0.928519 | 513.0877 | 537.52 | 578.8465 | 501.1842 | 561.6641 | 508.4052 | 559.2877 | 478.5401 |
| ENSMUSG00000055639 | 0.035287753 | 0.772105 | 0.858587 | 2259.598 | 1962.908 | 2080.204 | 2160.872 | 1715.221 | 2505.153 | 2076.115 | 1965.277 |
| ENSMUSG00000055652 | -0.33844222 | 0.001219 | 0.009053 | 413.4883 | 431.9358 | 399.1474 | 383.8675 | 537.2863 | 441.3628 | 530.359 | 545.9707 |
| ENSMUSG00000055653 | 0.311467751 | 0.001029 | 0.007993 | 31295.33 | 35587.91 | 29400.27 | 33929.57 | 27385.03 | 28189.11 | 26532.42 | 22818.75 |
| ENSMUSG00000055660 | -0.164374954 | 0.307782 | 0.476267 | 202.2169 | 142.7788 | 152.3716 | 157.8601 | 172.5947 | 166.4887 | 201.5364 | 193.5912 |
| ENSMUSG00000055670 | 0.126754494 | 0.224774 | 0.384523 | 1295.798 | 1141.03 | 1177.568 | 1143.839 | 1059.946 | 1269.337 | 1069.397 | 960.3429 |
| ENSMUSG00000055675 | -0.815404684 | 4.99E-05 | 0.000794 | 561.3783 | 385.1427 | 641.7826 | 533.9639 | 773.2633 | 737.4669 | 1128.218 | 1099.555 |
| ENSMUSG00000055676 | 0.050946076 | 0.848538 | 0.909127 | 44.26639 | 37.19447 | 58.79557 | 55.2079 | 54.60623 | 44.69496 | 37.60728 | 53.29196 |
| ENSMUSG00000055681 | -0.156897571 | 0.0543 | 0.144444 | 3202.271 | 3161.53 | 3242.866 | 3177.042 | 3608.887 | 3096.244 | 3632.477 | 3914.24 |
| ENSMUSG00000055692 | 0.220377176 | 0.126484 | 0.257489 | 414.4944 | 319.1525 | 375.9604 | 414.9219 | 322.7618 | 377.6724 | 271.9295 | 339.3284 |
| ENSMUSG00000055707 | 0.013076995 | 0.90066 | 0.940351 | 622.7476 | 760.6869 | 689.8128 | 590.0345 | 651.3744 | 624.6121 | 661.5024 | 698.2335 |
| ENSMUSG00000055717 | -0.035110066 | 0.844794 | 0.906563 | 221.332 | 212.3684 | 194.6051 | 213.9306 | 160.8934 | 268.1698 | 195.7507 | 239.27 |
| ENSMUSG00000055720 | -0.080368718 | 0.229892 | 0.390187 | 1084.527 | 1151.829 | 1199.098 | 1163.679 | 1273.495 | 1174.36 | 1237.183 | 1177.861 |
| ENSMUSG00000055723 | 0.102784114 | 0.125755 | 0.256401 | 2172.071 | 2252.065 | 2196.139 | 2274.738 | 1989.227 | 2142.006 | 2222.687 | 1928.299 |
| ENSMUSG00000055725 | -0.702169327 | 0.007186 | 0.033518 | 59.35721 | 65.99018 | 50.5145 | 57.79577 | 126.7645 | 62.57295 | 86.78602 | 102.2336 |
| ENSMUSG00000055730 | 0.487993387 | 0.260497 | 0.424525 | 40.24217 | 46.79304 | 35.60859 | 50.89479 | 18.52711 | 59.22083 | 28.92867 | 17.40146 |
| ENSMUSG00000055733 | 0.639820346 | 0.01844 | 0.06664 | 92.557 | 87.58697 | 63.76421 | 97.47645 | 37.05423 | 73.74669 | 51.10733 | 57.64233 |
| ENSMUSG00000055737 | 0.067694303 | 0.515246 | 0.669344 | 2897.437 | 3294.71 | 2661.535 | 3378.896 | 2841.474 | 3241.502 | 2676.867 | 2911.481 |
| ENSMUSG00000055745 | -0.09167135 | 0.299336 | 0.46743 | 642.8687 | 691.0972 | 721.2809 | 605.5617 | 734.2588 | 673.7766 | 694.2882 | 731.9488 |
| ENSMUSG00000055748 | -0.740672367 | 0.478763 | 0.638074 | 2.012109 | 0 | 28.15563 | 2.58787 | 15.60178 | 12.29112 | 13.50005 | 14.13868 |
| ENSMUSG00000055760 | -0.191291657 | 0.114625 | 0.242189 | 294.7739 | 353.9474 | 267.4784 | 301.9182 | 346.1645 | 318.4516 | 352.9298 | 368.6934 |
| ENSMUSG00000055761 | 0.47931533 | 0.09263 | 0.209097 | 88.53278 | 64.79036 | 57.96746 | 62.97152 | 45.83023 | 67.04245 | 51.10733 | 32.62773 |
| ENSMUSG00000055762 | -0.073116939 | 0.274274 | 0.440173 | 4245.549 | 4826.882 | 4695.365 | 4554.652 | 5093.981 | 4675.093 | 4927.518 | 4574.408 |
| ENSMUSG00000055775 | 1.925425605 | 0.015825 | 0.059901 | 59.35721 | 46.79304 | 169.7619 | 41.40593 | 1.950223 | 35.75597 | 40.50014 | 5.437955 |
| ENSMUSG00000055782 | 0.045701751 | 0.817025 | 0.888935 | 268.6165 | 291.5566 | 204.5423 | 302.7808 | 266.2054 | 217.8879 | 333.6441 | 214.2554 |
| ENSMUSG00000055799 | 0.150148312 | 0.067845 | 0.168368 | 1152.938 | 1163.827 | 1127.053 | 1149.014 | 1022.892 | 1154.247 | 1031.789 | 929.8904 |

| | | | | | | | | | | | |
|---|---|---|---|---|---|---|---|---|---|---|---|
| ENSMUSG00000055805 | -0.538222392 | 0.042569 | 0.121868 | 142.8597 | 87.58697 | 106.8258 | 140.6076 | 187.2214 | 103.9158 | 195.7507 | 207.7299 |
| ENSMUSG00000055809 | 0.304169984 | 0.043994 | 0.124553 | 259.562 | 274.7591 | 236.8385 | 288.9789 | 262.3049 | 214.5358 | 195.7507 | 184.8905 |
| ENSMUSG00000055817 | -0.019926481 | 0.751956 | 0.845503 | 1264.61 | 1282.609 | 1217.317 | 1337.929 | 1322.251 | 1232.464 | 1310.469 | 1306.197 |
| ENSMUSG00000055818 | -0.286288937 | 0.543099 | 0.692194 | 27.16347 | 19.19714 | 18.21835 | 7.763611 | 28.27823 | 17.87799 | 15.42863 | 26.10219 |
| ENSMUSG00000055827 | -0.593880733 | 0.625219 | 0.755716 | 40.24217 | 7.198929 | 740.3273 | 54.34528 | 507.0579 | 244.7049 | 170.6792 | 349.1167 |
| ENSMUSG00000055833 | -0.090773705 | 0.733442 | 0.833185 | 40.24217 | 43.19358 | 39.74912 | 37.95543 | 37.05423 | 58.10345 | 38.57157 | 38.06569 |
| ENSMUSG00000055835 | 0.166464938 | 0.153215 | 0.294463 | 724.3591 | 747.4888 | 704.7187 | 759.9713 | 643.5735 | 769.8708 | 661.5024 | 541.6204 |
| ENSMUSG00000055839 | -0.129291537 | 0.120666 | 0.249778 | 2855.182 | 3270.714 | 3431.674 | 3113.208 | 3515.276 | 3175.577 | 3461.798 | 3706.51 |
| ENSMUSG00000055850 | -0.172057384 | 0.33488 | 0.503097 | 2388.373 | 1486.579 | 1620.605 | 1677.803 | 2067.236 | 1556.502 | 2092.507 | 2365.511 |
| ENSMUSG00000055862 | 0.269190204 | 0.062557 | 0.159535 | 394.3733 | 453.5325 | 332.8989 | 396.8068 | 314.961 | 386.6114 | 324.0012 | 281.6861 |
| ENSMUSG00000055865 | -0.317054856 | 0.285935 | 0.452863 | 27.16347 | 29.99554 | 37.2648 | 30.19182 | 33.15378 | 30.1691 | 42.42872 | 50.02919 |
| ENSMUSG00000055866 | 0.030915448 | 0.917825 | 0.951436 | 555.342 | 325.1516 | 313.8524 | 470.9924 | 330.5627 | 541.9264 | 530.359 | 227.3065 |
| ENSMUSG00000055884 | -0.207139055 | 0.079016 | 0.187542 | 529.1846 | 478.7288 | 419.8501 | 510.6731 | 632.8472 | 564.2739 | 506.2518 | 532.9196 |
| ENSMUSG00000055897 | 0.643440605 | 0.001826 | 0.012286 | 3269.677 | 2957.56 | 2273.153 | 2862.185 | 1557.253 | 2640.355 | 1727.042 | 1349.701 |
| ENSMUSG00000055900 | -0.075009134 | 0.43137 | 0.596045 | 349.1009 | 340.7493 | 316.3367 | 332.9727 | 368.5921 | 355.325 | 355.8227 | 328.4525 |
| ENSMUSG00000055912 | -0.750616998 | 0.000983 | 0.007746 | 303.8284 | 259.1615 | 351.1172 | 366.615 | 575.3157 | 351.9728 | 486.0017 | 743.9123 |
| ENSMUSG00000055917 | -0.050611602 | 0.619794 | 0.751867 | 1036.236 | 971.8554 | 971.3691 | 1094.669 | 1122.353 | 1086.088 | 1123.397 | 887.4743 |
| ENSMUSG00000055923 | 0.223481291 | 0.055483 | 0.146683 | 456.7487 | 493.1267 | 398.3193 | 420.0976 | 398.8205 | 411.1937 | 366.4299 | 334.9781 |
| ENSMUSG00000055926 | 0.399873093 | 0.316739 | 0.484882 | 23.13925 | 19.19714 | 32.29616 | 17.25247 | 15.60178 | 12.29112 | 20.25007 | 21.75182 |
| ENSMUSG00000055932 | -0.016543549 | 0.781913 | 0.865498 | 2485.96 | 2644.407 | 2435.462 | 2375.665 | 2528.464 | 2422.467 | 2618.045 | 2478.62 |
| ENSMUSG00000055943 | -0.259739456 | 0.027951 | 0.089981 | 1820.958 | 1527.373 | 1835.084 | 1739.049 | 2082.838 | 1686.118 | 2194.722 | 2326.357 |
| ENSMUSG00000055945 | 0.620894848 | 0.093508 | 0.210486 | 26.15741 | 29.99554 | 17.39024 | 24.15346 | 13.65156 | 14.52586 | 16.39292 | 18.48905 |
| ENSMUSG00000055963 | 0.361651065 | 0.025157 | 0.083399 | 259.562 | 367.1454 | 324.6178 | 326.9343 | 247.6783 | 301.691 | 234.3223 | 210.9927 |
| ENSMUSG00000055976 | -0.716013774 | 0.223957 | 0.383749 | 49.29666 | 28.79572 | 75.3577 | 34.50494 | 145.2916 | 13.40849 | 74.25027 | 76.13138 |
| ENSMUSG00000055978 | 0.096371982 | 0.815565 | 0.887608 | 19.11503 | 20.39697 | 38.92101 | 31.05445 | 24.37778 | 40.22547 | 18.32149 | 20.66423 |
| ENSMUSG00000055980 | 0.186934379 | 0.06536 | 0.164657 | 410.4702 | 454.7324 | 409.0847 | 399.3947 | 359.8161 | 349.7381 | 407.8943 | 349.1167 |
| ENSMUSG00000055991 | 0.107747914 | 0.214581 | 0.372744 | 809.8738 | 784.6833 | 753.577 | 741.8562 | 739.1344 | 755.3449 | 737.6812 | 632.978 |
| ENSMUSG00000055994 | -0.324182832 | 0.401906 | 0.569051 | 16.09687 | 27.5959 | 23.18699 | 25.01608 | 25.35289 | 18.99536 | 26.03581 | 44.59123 |
| ENSMUSG00000056004 | 0.59556805 | 0.020538 | 0.071879 | 141.8537 | 141.5789 | 118.4192 | 129.3935 | 90.68535 | 127.3806 | 79.07171 | 54.37955 |
| ENSMUSG00000056008 | -0.883268252 | 0.133023 | 0.266392 | 12.07265 | 5.999108 | 22.35888 | 10.35148 | 20.47734 | 8.938993 | 25.07152 | 40.24087 |
| ENSMUSG00000056014 | -0.099968423 | 0.495256 | 0.652059 | 251.5136 | 313.1534 | 238.4947 | 258.787 | 274.9814 | 336.3296 | 247.8223 | 277.3357 |
| ENSMUSG00000056019 | 0.108691254 | 0.412314 | 0.578199 | 308.8587 | 274.7591 | 295.6341 | 260.5123 | 241.8276 | 315.0995 | 244.9294 | 256.6715 |
| ENSMUSG00000056025 | 0.176409087 | 0.564454 | 0.709411 | 1700.232 | 567.5156 | 855.4342 | 866.074 | 747.9104 | 719.5889 | 1212.111 | 850.4962 |
| ENSMUSG00000056031 | 0.008916596 | 0.979059 | 0.988557 | 31.18769 | 46.79304 | 21.53077 | 32.77969 | 39.00445 | 30.1691 | 35.6787 | 25.0146 |
| ENSMUSG00000056035 | 3.397254334 | 0.004121 | 0.022315 | 83.50251 | 1.199822 | 1.656213 | 75.91087 | 0.975111 | 2.234748 | 6.750024 | 5.437955 |
| ENSMUSG00000056043 | -0.056089092 | 0.712579 | 0.819906 | 127.7689 | 151.1775 | 146.5749 | 138.0198 | 162.8436 | 158.6671 | 128.2505 | 135.9489 |
| ENSMUSG00000056050 | 0.09197041 | 0.120344 | 0.249335 | 1976.897 | 1992.904 | 2049.564 | 2018.539 | 1803.956 | 2022.447 | 1845.649 | 1873.919 |
| ENSMUSG00000056054 | 0.29250603 | 0.467502 | 0.62788 | 272.6407 | 67.19001 | 119.2474 | 179.4257 | 147.2418 | 77.09881 | 151.3934 | 145.7372 |
| ENSMUSG00000056055 | 0.200387254 | 0.064166 | 0.16265 | 832.007 | 1099.037 | 1005.321 | 963.5504 | 790.8153 | 900.6035 | 804.2172 | 898.3502 |
| ENSMUSG00000056069 | -0.842117151 | 0.001737 | 0.011811 | 263.5862 | 214.7681 | 427.303 | 280.3526 | 510.9583 | 336.3296 | 722.2526 | 559.0218 |
| ENSMUSG00000056071 | 0.267150577 | 0.570204 | 0.713761 | 415.5004 | 83.98751 | 146.5749 | 171.6621 | 203.7983 | 75.98144 | 183.2149 | 216.4306 |
| ENSMUSG00000056073 | 0.882823592 | 0.00658 | 0.031552 | 300.8103 | 285.5575 | 178.871 | 305.3687 | 160.8934 | 231.2964 | 108.0004 | 80.48174 |
| ENSMUSG00000056076 | -0.338237545 | 9.29E-05 | 0.001259 | 5749.601 | 6051.9 | 6426.107 | 5730.408 | 8118.777 | 6609.268 | 7491.563 | 8067.751 |
| ENSMUSG00000056091 | 0.570074127 | 0.009895 | 0.042285 | 1285.737 | 693.4968 | 592.0962 | 857.4478 | 601.6437 | 569.8608 | 560.252 | 577.5109 |
| ENSMUSG00000056116 | 0.102078158 | 0.491056 | 0.648215 | 615.7053 | 866.2711 | 877.793 | 634.8909 | 670.8766 | 716.2368 | 757.9313 | 643.8539 |
| ENSMUSG00000056121 | -0.358656794 | 0.001339 | 0.009733 | 955.7516 | 820.6779 | 877.793 | 787.5752 | 1184.76 | 922.951 | 1162.933 | 1140.883 |
| ENSMUSG00000056124 | 0.29376292 | 0.040804 | 0.118353 | 1721.359 | 1712.145 | 1699.275 | 1602.754 | 1071.647 | 1735.282 | 1435.827 | 1252.905 |
| ENSMUSG00000056130 | 0.467244178 | 0.117305 | 0.245573 | 34.20585 | 53.99197 | 42.23344 | 38.81806 | 35.10401 | 36.87335 | 26.03581 | 23.927 |
| ENSMUSG00000056131 | -0.315967716 | 0.013736 | 0.054041 | 494.9787 | 501.5254 | 433.0998 | 453.74 | 519.7343 | 506.1705 | 648.0023 | 667.7809 |
| ENSMUSG00000056144 | 0.4927372 | 0.192742 | 0.345969 | 136.8234 | 209.9688 | 260.8536 | 62.97152 | 132.6151 | 125.1459 | 119.5719 | 98.97079 |
| ENSMUSG00000056145 | 0.655067773 | 0.030904 | 0.096556 | 98.59333 | 141.5789 | 83.63877 | 80.22398 | 50.70579 | 97.21155 | 54.96448 | 53.29196 |
| ENSMUSG00000056153 | 0.080512987 | 0.360124 | 0.528671 | 1291.774 | 1424.188 | 1286.05 | 1324.99 | 1179.885 | 1392.248 | 1315.29 | 1148.496 |
| ENSMUSG00000056155 | 0.105907243 | 0.678671 | 0.794819 | 59.35721 | 89.98662 | 48.85829 | 61.24627 | 53.63112 | 61.45558 | 55.92877 | 68.51824 |
| ENSMUSG00000056158 | -0.107971268 | 0.801836 | 0.879143 | 11.0666 | 15.59768 | 18.21835 | 25.8787 | 14.62667 | 26.81698 | 19.28578 | 16.31387 |
| ENSMUSG00000056162 | -0.847508536 | 0.000725 | 0.006116 | 102.6175 | 74.38894 | 95.23226 | 68.14726 | 144.3165 | 105.0332 | 160.072 | 203.3795 |
| ENSMUSG00000056167 | 0.044040976 | 0.586399 | 0.726738 | 2919.57 | 2985.156 | 2549.74 | 2919.981 | 2760.54 | 2896.234 | 2859.117 | 2512.335 |
| ENSMUSG00000056174 | 0.486220495 | 0.011945 | 0.048827 | 955.7516 | 1550.169 | 1367.204 | 1161.954 | 974.1362 | 1140.839 | 738.6455 | 740.6495 |
| ENSMUSG00000056185 | 0.77532852 | 1.50E-08 | 1.19E-06 | 355.1372 | 370.7449 | 310.54 | 394.2189 | 195.9974 | 222.3574 | 182.2507 | 236.0073 |
| ENSMUSG00000056201 | -0.214936579 | 0.011241 | 0.046601 | 17619.03 | 19374.72 | 21072.83 | 18849.19 | 23393.9 | 20028.93 | 22019.54 | 23831.3 |
| ENSMUSG00000056204 | -0.551594667 | 7.73E-05 | 0.001114 | 1198.211 | 1065.442 | 1141.959 | 1028.247 | 1615.759 | 1257.046 | 1644.113 | 1981.591 |
| ENSMUSG00000056209 | -0.1286706 | 0.217051 | 0.375514 | 1709.286 | 1868.122 | 1989.94 | 1834.8 | 2317.84 | 1762.099 | 1901.578 | 2111.014 |
| ENSMUSG00000056211 | 0.232349619 | 0.080481 | 0.189701 | 1684.135 | 1800.932 | 1526.201 | 1626.908 | 1213.038 | 1788.916 | 1339.398 | 1309.46 |
| ENSMUSG00000056214 | 0.06481856 | 0.564354 | 0.709411 | 1057.363 | 1021.048 | 1048.383 | 1064.477 | 1140.88 | 1107.318 | 903.5389 | 855.9342 |
| ENSMUSG00000056215 | -0.623524558 | 0.04981 | 0.135749 | 26.15741 | 23.99643 | 36.43669 | 24.15346 | 58.50668 | 41.34284 | 36.64299 | 34.80291 |

| | | | | | | | | | | | |
|---|---|---|---|---|---|---|---|---|---|---|---|
| ENSMUSG00000056216 | 0.029264391 | 0.802367 | 0.879311 | 1169.035 | 1229.817 | 915.0578 | 1020.484 | 979.0118 | 1011.224 | 1149.433 | 1102.817 |
| ENSMUSG00000056220 | 0.037576981 | 0.751504 | 0.84515 | 1238.453 | 1451.784 | 1630.542 | 1331.891 | 1425.613 | 1168.773 | 1479.22 | 1433.445 |
| ENSMUSG00000056222 | 1.102332392 | 0.005079 | 0.026022 | 132.7992 | 51.59233 | 62.9361 | 122.4925 | 41.92979 | 51.39921 | 54.00019 | 25.0146 |
| ENSMUSG00000056228 | 0.093953489 | 0.415336 | 0.581203 | 583.5115 | 493.1267 | 563.9406 | 537.4144 | 496.3317 | 452.5365 | 496.6089 | 597.0875 |
| ENSMUSG00000056234 | -0.592636571 | 0.000627 | 0.005456 | 1208.271 | 1144.63 | 1183.364 | 1174.893 | 1856.612 | 1182.182 | 2237.151 | 1827.153 |
| ENSMUSG00000056258 | 1.515657025 | 0.031639 | 0.09809 | 36.21796 | 11.99822 | 10.76539 | 73.323 | 7.80089 | 20.11273 | 6.750024 | 11.9635 |
| ENSMUSG00000056260 | 0.343274246 | 0.036146 | 0.108774 | 792.7708 | 817.0785 | 708.0312 | 751.3451 | 523.6348 | 794.453 | 625.8237 | 475.2773 |
| ENSMUSG00000056267 | 0.292311041 | 0.041955 | 0.120575 | 565.4025 | 517.1231 | 470.3646 | 548.6285 | 457.3272 | 517.3442 | 393.43 | 348.0291 |
| ENSMUSG00000056268 | 0.162018993 | 0.361627 | 0.530157 | 746.4923 | 671.9001 | 722.109 | 720.2906 | 526.5601 | 901.7209 | 583.3949 | 547.0583 |
| ENSMUSG00000056293 | -0.713448235 | NA | NA | 45.27245 | 5.999108 | 897.6676 | 53.48266 | 623.0961 | 335.2122 | 266.1438 | 419.8102 |
| ENSMUSG00000056296 | 0.862885285 | 0.000504 | 0.004618 | 77.46619 | 106.7841 | 89.43552 | 119.042 | 50.70579 | 72.62932 | 45.32159 | 47.85401 |
| ENSMUSG00000056305 | -0.397997734 | 1.52E-07 | 7.98E-06 | 983.9212 | 988.6529 | 1016.915 | 1008.407 | 1427.563 | 1208.999 | 1270.933 | 1360.576 |
| ENSMUSG00000056310 | -0.157779226 | 0.046468 | 0.129353 | 833.013 | 850.6735 | 801.6072 | 819.4923 | 924.4055 | 833.5611 | 929.5748 | 996.2334 |
| ENSMUSG00000056313 | -0.173608764 | 0.315451 | 0.483792 | 1512.1 | 1029.447 | 1068.258 | 1286.172 | 1112.602 | 1357.61 | 1750.185 | 1301.847 |
| ENSMUSG00000056342 | -0.025633434 | 0.773517 | 0.859536 | 2630.832 | 2422.44 | 2393.228 | 2118.603 | 2270.059 | 2642.59 | 2384.687 | 2438.379 |
| ENSMUSG00000056366 | -2.092426007 | 0.012066 | 0.049182 | 2.012109 | 3.599465 | 11.59349 | 1.725247 | 28.27823 | 4.469496 | 15.42863 | 33.71532 |
| ENSMUSG00000056367 | 0.003977173 | 0.983599 | 0.990959 | 189.1382 | 171.5745 | 125.8722 | 175.1126 | 197.9476 | 176.5451 | 142.7148 | 141.3868 |
| ENSMUSG00000056383 | 0.31866466 | 0.044297 | 0.125159 | 366.2038 | 370.7449 | 289.8373 | 272.589 | 214.5245 | 278.2261 | 269.0367 | 278.4233 |
| ENSMUSG00000056394 | -0.091382225 | 0.386053 | 0.553527 | 1766.631 | 2393.644 | 2049.564 | 1869.305 | 2203.752 | 2161.002 | 2038.507 | 2200.197 |
| ENSMUSG00000056411 | -0.172358908 | 0.665353 | 0.785151 | 17.10292 | 15.59768 | 29.81184 | 14.6646 | 25.35289 | 18.99536 | 18.32149 | 25.0146 |
| ENSMUSG00000056412 | 0.196567895 | 0.578562 | 0.719984 | 19.11503 | 19.19714 | 22.35888 | 19.84034 | 15.60178 | 20.11273 | 15.42863 | 19.57664 |
| ENSMUSG00000056413 | -0.763571688 | 0.003377 | 0.019298 | 124.7507 | 93.58608 | 166.4494 | 129.3935 | 233.0516 | 137.437 | 219.8579 | 284.9489 |
| ENSMUSG00000056418 | 0.335875678 | 0.036881 | 0.110409 | 116.7023 | 104.3845 | 102.6852 | 113.8663 | 83.85957 | 91.62468 | 87.75031 | 83.74451 |
| ENSMUSG00000056427 | -0.320860367 | 0.17114 | 0.318195 | 220.3259 | 142.7788 | 236.8385 | 330.3848 | 257.4294 | 298.3389 | 292.1796 | 317.5766 |
| ENSMUSG00000056429 | -0.073574677 | 0.336503 | 0.504754 | 2151.95 | 1955.709 | 2099.25 | 2016.814 | 2084.788 | 1974.4 | 2300.794 | 2294.817 |
| ENSMUSG00000056445 | 0.240005576 | 0.460014 | 0.621478 | 111.672 | 107.9839 | 107.6539 | 138.0198 | 89.71024 | 167.6061 | 81.96458 | 55.46715 |
| ENSMUSG00000056458 | -0.50471454 | 0.00399 | 0.02181 | 130.7871 | 123.5816 | 184.6678 | 121.6299 | 200.8729 | 194.4231 | 210.215 | 192.5036 |
| ENSMUSG00000056459 | 0.233618686 | 0.082033 | 0.192351 | 294.7739 | 401.9402 | 376.7885 | 320.0333 | 282.7823 | 293.8694 | 323.0369 | 283.8613 |
| ENSMUSG00000056468 | 0.615046635 | 0.011313 | 0.046782 | 142.8597 | 185.9723 | 104.3414 | 169.0742 | 74.10846 | 121.7938 | 102.2147 | 94.62042 |
| ENSMUSG00000056476 | 0.646512851 | 0.000103 | 0.00137 | 281.6952 | 290.3568 | 265.8222 | 256.1992 | 219.4 | 176.5451 | 168.7506 | 132.6861 |
| ENSMUSG00000056481 | -0.006220859 | 0.97465 | 0.986138 | 560.3723 | 916.6637 | 1094.757 | 696.9998 | 793.7406 | 833.5611 | 774.3242 | 882.0364 |
| ENSMUSG00000056486 | 0.169215857 | 0.06742 | 0.16764 | 502.0211 | 537.52 | 517.5666 | 550.3538 | 417.3476 | 488.2925 | 489.8589 | 479.6277 |
| ENSMUSG00000056487 | -4.687453614 | 5.04E-07 | 2.12E-05 | 0 | 0 | 3.312427 | 0.862623 | 30.22845 | 10.05637 | 30.85725 | 46.76642 |
| ENSMUSG00000056492 | -0.633405139 | 0.012129 | 0.049394 | 1511.094 | 1064.242 | 2510.819 | 1362.945 | 2371.471 | 1842.55 | 2894.796 | 2898.43 |
| ENSMUSG00000056493 | 0.165276741 | 0.096637 | 0.215612 | 1030.2 | 862.6717 | 952.3226 | 878.1507 | 774.2384 | 766.5186 | 876.5389 | 903.7882 |
| ENSMUSG00000056498 | 0.477200902 | 0.123964 | 0.253674 | 38.23007 | 49.19268 | 45.54586 | 51.75741 | 44.85512 | 20.11273 | 36.64299 | 30.45255 |
| ENSMUSG00000056501 | 0.182681905 | 0.568822 | 0.712804 | 583.5115 | 436.735 | 236.8385 | 282.0779 | 206.7236 | 330.7427 | 471.5374 | 345.854 |
| ENSMUSG00000056508 | 0.257119874 | 0.507327 | 0.663071 | 39.23612 | 44.3934 | 42.23344 | 46.58167 | 16.57689 | 56.98608 | 27.96439 | 43.50364 |
| ENSMUSG00000056515 | 0.213955765 | 0.008898 | 0.039168 | 1467.833 | 1751.739 | 1586.652 | 1494.064 | 1440.239 | 1348.671 | 1296.969 | 1342.087 |
| ENSMUSG00000056529 | 0.429116827 | 0.384723 | 0.552435 | 14.08476 | 55.19179 | 25.67131 | 26.74133 | 32.17867 | 20.11273 | 13.50005 | 23.927 |
| ENSMUSG00000056531 | 0.327559622 | 0.098969 | 0.219531 | 266.6044 | 218.3675 | 197.0894 | 201.8539 | 138.4658 | 231.2964 | 180.3221 | 154.4379 |
| ENSMUSG00000056536 | 0.041548445 | 0.573225 | 0.715794 | 998.0059 | 977.8546 | 1007.806 | 919.5566 | 948.7833 | 987.7587 | 878.4674 | 978.832 |
| ENSMUSG00000056537 | 0.156025478 | 0.35036 | 0.519044 | 2323.986 | 2375.647 | 2151.421 | 2198.827 | 1815.657 | 2825.839 | 1855.292 | 1625.949 |
| ENSMUSG00000056553 | -0.082482493 | 0.806679 | 0.88179 | 38.23007 | 17.99732 | 34.78048 | 40.5433 | 39.97956 | 41.34284 | 29.89296 | 29.36496 |
| ENSMUSG00000056569 | 0.107946035 | 0.821848 | 0.891817 | 48.29061 | 25.19625 | 23.18699 | 65.55939 | 31.20356 | 67.04245 | 27.96439 | 25.0146 |
| ENSMUSG00000056579 | 0.264003502 | 0.01158 | 0.047664 | 4213.356 | 4519.728 | 4205.954 | 4634.013 | 3447.994 | 4401.337 | 3462.762 | 3323.678 |
| ENSMUSG00000056592 | -0.039761254 | 0.873009 | 0.924027 | 179.0777 | 287.9572 | 147.403 | 139.745 | 189.1716 | 195.5405 | 220.8222 | 166.4014 |
| ENSMUSG00000056596 | -0.047210882 | 0.803863 | 0.880139 | 155.9384 | 127.1811 | 114.2787 | 162.1732 | 177.4703 | 127.3806 | 136.9291 | 135.9489 |
| ENSMUSG00000056598 | -0.453984028 | 0.042986 | 0.122715 | 62.37537 | 52.79215 | 70.38906 | 73.323 | 101.4116 | 93.85942 | 95.46463 | 65.25547 |
| ENSMUSG00000056602 | 0.578063526 | 0.000837 | 0.006802 | 956.7577 | 937.0606 | 635.1578 | 857.4478 | 496.3317 | 646.9596 | 653.7881 | 469.8393 |
| ENSMUSG00000056608 | 0.226628383 | 0.115603 | 0.243585 | 1169.035 | 951.4585 | 967.2285 | 1045.5 | 761.5619 | 1119.609 | 781.0742 | 872.2481 |
| ENSMUSG00000056612 | -0.039219635 | 0.72079 | 0.82531 | 3381.349 | 3708.648 | 3609.717 | 3497.938 | 3864.366 | 2965.511 | 3591.977 | 4165.474 |
| ENSMUSG00000056629 | -0.288538525 | 0.000203 | 0.002298 | 1211.289 | 1138.631 | 1162.662 | 1228.376 | 1518.248 | 1309.562 | 1420.398 | 1543.292 |
| ENSMUSG00000056643 | -1.925204624 | 8.95E-06 | 0.000211 | 16.09687 | 9.598572 | 15.73403 | 9.488858 | 33.15378 | 31.28647 | 55.92877 | 73.95619 |
| ENSMUSG00000056648 | 0.357985108 | 3.63E-08 | 2.45E-06 | 3982.969 | 3781.837 | 3891.273 | 3729.984 | 3020.895 | 3215.803 | 3009.546 | 2759.219 |
| ENSMUSG00000056656 | 0.916645054 | 0.09943 | 0.220139 | 19.11503 | 8.398751 | 14.07781 | 12.07673 | 8.776002 | 3.352122 | 9.642892 | 6.525547 |
| ENSMUSG00000056665 | -1.219733565 | 0.00155 | 0.010786 | 56.33904 | 55.19179 | 87.7793 | 49.16954 | 167.7191 | 54.75133 | 160.072 | 196.854 |
| ENSMUSG00000056666 | -0.99703062 | 2.38E-05 | 0.00045 | 404.4339 | 412.7386 | 511.7699 | 341.5989 | 822.9939 | 501.701 | 867.8602 | 1141.971 |
| ENSMUSG00000056671 | -0.713023758 | 1.96E-05 | 0.000387 | 144.8718 | 171.5745 | 198.7456 | 162.1732 | 264.2552 | 231.2964 | 334.6083 | 280.5985 |
| ENSMUSG00000056673 | -0.445904848 | 0.788818 | 0.870406 | 4.024217 | 1153.028 | 1123.741 | 9.488858 | 1082.374 | 7.821619 | 1037.575 | 991.8831 |
| ENSMUSG00000056679 | 0.53164709 | 9.45E-05 | 0.001278 | 443.67 | 465.5308 | 421.5063 | 583.1335 | 314.961 | 368.7335 | 324.9654 | 316.489 |
| ENSMUSG00000056687 | -0.125999737 | 0.611253 | 0.746018 | 89.53884 | 62.39072 | 103.5133 | 98.33908 | 67.28268 | 101.681 | 104.1432 | 115.2847 |
| ENSMUSG00000056692 | -0.229690857 | 0.018158 | 0.066009 | 2932.648 | 2716.396 | 3025.902 | 2601.672 | 3212.992 | 2893.999 | 3399.119 | 3717.386 |
| ENSMUSG00000056698 | -0.184973418 | 0.099518 | 0.220229 | 331.9979 | 301.1552 | 334.5551 | 331.2474 | 355.9156 | 322.9211 | 392.4657 | 406.7591 |

| | | | | | | | | | | | |
|---|---|---|---|---|---|---|---|---|---|---|---|
| ENSMUSG00000056708 | -0.257711659 | 0.109276 | 0.234647 | 1016.115 | 1037.846 | 1326.627 | 996.3301 | 1393.434 | 981.0545 | 1298.898 | 1560.693 |
| ENSMUSG00000056718 | 0.345295011 | 0.212278 | 0.370023 | 50.30272 | 35.99465 | 57.96746 | 56.07053 | 41.92979 | 46.92971 | 40.50014 | 29.36496 |
| ENSMUSG00000056724 | -0.601903267 | 0.003459 | 0.019605 | 471.8395 | 285.5575 | 514.2542 | 375.2412 | 631.8721 | 491.6446 | 620.0379 | 759.1386 |
| ENSMUSG00000056735 | 0.320040862 | 0.131552 | 0.26445 | 101.6115 | 110.3836 | 75.3577 | 84.5371 | 72.15824 | 75.98144 | 85.82174 | 61.99269 |
| ENSMUSG00000056737 | 0.003573857 | 0.978966 | 0.988554 | 1925.588 | 1768.537 | 2463.617 | 2083.236 | 2178.399 | 1678.296 | 2267.044 | 2099.051 |
| ENSMUSG00000056749 | -0.075932176 | 0.741205 | 0.838318 | 536.227 | 356.347 | 287.353 | 510.6731 | 421.2481 | 395.5504 | 579.5378 | 385.0072 |
| ENSMUSG00000056752 | -0.32458695 | 0.495279 | 0.652059 | 7.04238 | 7.198929 | 14.07781 | 12.93935 | 12.67645 | 12.29112 | 14.46434 | 13.05109 |
| ENSMUSG00000056753 | -0.34496215 | 0.065564 | 0.164783 | 79.47829 | 73.18911 | 79.49824 | 86.26235 | 97.51113 | 81.56831 | 113.7861 | 112.0219 |
| ENSMUSG00000056755 | 0.900424141 | 2.99E-05 | 0.000534 | 87.52673 | 121.182 | 96.88848 | 112.1411 | 55.58134 | 69.27719 | 52.07161 | 46.76642 |
| ENSMUSG00000056758 | 0.364747795 | 0.056541 | 0.148438 | 1899.431 | 2081.69 | 1534.482 | 2005.6 | 1497.771 | 2005.687 | 1258.397 | 1078.89 |
| ENSMUSG00000056763 | 0.271888149 | 0.078319 | 0.186298 | 1579.505 | 1425.388 | 1379.626 | 1488.888 | 1109.677 | 1638.07 | 1083.861 | 1034.299 |
| ENSMUSG00000056770 | -0.078123863 | 0.246615 | 0.409679 | 3033.254 | 2844.777 | 2747.658 | 3204.646 | 3152.535 | 3164.403 | 3149.368 | 3023.503 |
| ENSMUSG00000056820 | -0.06100744 | 0.197891 | 0.352298 | 3008.103 | 3082.342 | 2912.451 | 3044.198 | 3144.734 | 3035.905 | 3177.333 | 3204.043 |
| ENSMUSG00000056824 | -0.127798298 | 0.745291 | 0.840853 | 29.17558 | 22.79661 | 43.88965 | 23.29083 | 37.05423 | 16.76061 | 37.60728 | 39.15328 |
| ENSMUSG00000056832 | 0.07650154 | 0.56473 | 0.709652 | 415.5004 | 428.3363 | 395.0069 | 376.9665 | 387.1192 | 464.8276 | 326.894 | 353.4671 |
| ENSMUSG00000056836 | -0.333515927 | 0.497787 | 0.65413 | 6.036326 | 9.598572 | 13.24971 | 8.626235 | 12.67645 | 12.29112 | 9.642892 | 13.05109 |
| ENSMUSG00000056851 | 0.028837348 | 0.642866 | 0.76908 | 9048.453 | 9137.841 | 9392.385 | 9902.918 | 9633.125 | 9591.539 | 8811.674 | 8707.254 |
| ENSMUSG00000056870 | 0.678340407 | 2.54E-05 | 0.000473 | 982.9151 | 1059.442 | 941.5572 | 1099.845 | 557.7637 | 862.6128 | 538.0734 | 594.9123 |
| ENSMUSG00000056880 | -0.106706835 | 0.756117 | 0.848334 | 21.12714 | 32.39518 | 19.04645 | 19.84034 | 25.35289 | 25.6996 | 20.25007 | 27.18978 |
| ENSMUSG00000056888 | -0.250521805 | 0.570313 | 0.71379 | 23.13925 | 17.99732 | 16.56213 | 21.56559 | 23.40267 | 39.10809 | 11.57147 | 20.66423 |
| ENSMUSG00000056895 | -0.285545617 | 0.084388 | 0.195932 | 355.1372 | 501.5254 | 452.9743 | 478.756 | 665.0259 | 440.2454 | 503.3589 | 569.8977 |
| ENSMUSG00000056899 | -0.438305264 | 0.058074 | 0.151172 | 66.39959 | 38.39429 | 57.96746 | 56.07053 | 84.83468 | 69.27719 | 75.21455 | 68.51824 |
| ENSMUSG00000056900 | -0.021289862 | 0.920044 | 0.9524 | 156.9445 | 124.7814 | 104.3414 | 137.1571 | 118.9636 | 143.0239 | 104.1432 | 165.3138 |
| ENSMUSG00000056904 | 0.161270325 | 0.675174 | 0.792182 | 15.09082 | 27.5959 | 24.8432 | 19.84034 | 22.42756 | 18.99536 | 23.14294 | 13.05109 |
| ENSMUSG00000056917 | -0.126871177 | 0.07934 | 0.187995 | 1045.29 | 1028.247 | 1136.99 | 1025.659 | 1093.1 | 1146.426 | 1168.718 | 1221.365 |
| ENSMUSG00000056919 | 0.170177878 | 0.064977 | 0.164142 | 435.6215 | 457.132 | 427.303 | 458.0531 | 358.841 | 411.1937 | 396.3228 | 414.3722 |
| ENSMUSG00000056938 | -0.081063403 | 0.442426 | 0.605653 | 337.0282 | 334.7502 | 331.2427 | 335.5605 | 360.7912 | 354.2076 | 310.5011 | 391.5328 |
| ENSMUSG00000056941 | -0.075962572 | 0.374314 | 0.543066 | 1233.423 | 1202.221 | 1199.098 | 1217.162 | 1316.4 | 1108.435 | 1296.005 | 1392.117 |
| ENSMUSG00000056947 | 0.724335608 | 0.063326 | 0.161073 | 47.28455 | 23.99643 | 28.15563 | 58.6584 | 18.52711 | 26.81698 | 27.96439 | 22.83941 |
| ENSMUSG00000056952 | -0.556204019 | 1.13E-08 | 9.52E-07 | 786.7345 | 772.6851 | 944.0416 | 822.9428 | 1268.62 | 1105.083 | 1222.719 | 1299.671 |
| ENSMUSG00000056962 | -0.218155686 | 0.025058 | 0.083207 | 1246.501 | 1111.035 | 1074.054 | 1143.839 | 1315.425 | 1152.013 | 1418.469 | 1434.533 |
| ENSMUSG00000056966 | 1.414467431 | 0.157677 | 0.300269 | 27.16347 | 3.599465 | 10.76539 | 39.68068 | 7.80089 | 18.99536 | 3.857157 | 0 |
| ENSMUSG00000056972 | 2.331402018 | 0.000812 | 0.00665 | 43.26034 | 17.99732 | 5.796746 | 37.09281 | 3.900445 | 5.58687 | 5.785735 | 5.437955 |
| ENSMUSG00000056973 | -0.49112721 | 0.17297 | 0.32066 | 33.19979 | 19.19714 | 42.23344 | 34.50494 | 30.22845 | 64.8077 | 49.17875 | 39.15328 |
| ENSMUSG00000056999 | 0.062514475 | 0.61517 | 0.748571 | 2745.522 | 2001.302 | 2810.594 | 2486.944 | 2693.257 | 2217.988 | 2419.402 | 2290.467 |
| ENSMUSG00000057036 | -0.205660858 | 0.079812 | 0.188683 | 393.3673 | 470.33 | 476.1613 | 481.3439 | 589.9423 | 494.9967 | 544.8234 | 469.8393 |
| ENSMUSG00000057054 | -0.088888579 | 0.7398 | 0.837336 | 66.39959 | 40.79393 | 56.31125 | 62.97152 | 58.50668 | 70.39457 | 70.39311 | 42.41605 |
| ENSMUSG00000057058 | 0.942617796 | 2.08E-15 | 1.04E-12 | 329.9858 | 299.9554 | 289.8373 | 339.8737 | 166.744 | 170.9582 | 156.2148 | 162.0511 |
| ENSMUSG00000057060 | -0.16278687 | 0.618544 | 0.751043 | 48.29061 | 32.39518 | 51.34261 | 62.10889 | 55.58134 | 40.22547 | 42.42872 | 80.48174 |
| ENSMUSG00000057069 | 0.270337382 | 0.233281 | 0.393874 | 205.2351 | 131.9804 | 114.2787 | 165.6237 | 119.9387 | 151.9629 | 132.1076 | 107.6715 |
| ENSMUSG00000057092 | -0.252592566 | 0.559463 | 0.705627 | 51.30877 | 49.19268 | 195.4332 | 74.18562 | 121.8889 | 101.681 | 98.35749 | 120.7226 |
| ENSMUSG00000057093 | 0.312845289 | 0.179417 | 0.329222 | 144.8718 | 148.7779 | 82.81066 | 107.8279 | 107.2622 | 92.74205 | 85.82174 | 102.2336 |
| ENSMUSG00000057098 | 0.809547477 | 0.001831 | 0.012303 | 1544.293 | 2006.102 | 1385.422 | 1691.605 | 828.8446 | 1563.206 | 733.8241 | 655.8174 |
| ENSMUSG00000057101 | -0.038607328 | 0.712894 | 0.819914 | 711.2804 | 814.6788 | 660.001 | 784.9874 | 824.9442 | 679.3635 | 767.5742 | 775.4524 |
| ENSMUSG00000057103 | -3.1100892 | 2.69E-15 | 1.31E-12 | 94.56911 | 69.58965 | 115.1068 | 100.9269 | 1002.414 | 308.3953 | 560.252 | 1417.131 |
| ENSMUSG00000057110 | 0.039918121 | 0.70787 | 0.816528 | 1723.371 | 1518.974 | 1440.906 | 1657.962 | 1804.931 | 1458.173 | 1474.398 | 1429.095 |
| ENSMUSG00000057113 | -0.027808227 | 0.764832 | 0.853994 | 29843.6 | 33794.17 | 29704.18 | 33691.49 | 34902.16 | 34867.66 | 31368.33 | 28365.46 |
| ENSMUSG00000057123 | -1.089738258 | 0.000157 | 0.001887 | 77.46619 | 51.59233 | 84.46688 | 69.00988 | 145.2916 | 116.2069 | 112.8218 | 229.4817 |
| ENSMUSG00000057130 | -0.182568596 | 0.032749 | 0.100686 | 593.5721 | 609.5093 | 626.0486 | 653.006 | 756.6864 | 727.4105 | 691.3953 | 642.7663 |
| ENSMUSG00000057132 | 0.230561776 | 0.121826 | 0.251178 | 204.229 | 191.9714 | 245.1196 | 221.6942 | 179.4205 | 214.5358 | 185.1435 | 158.7883 |
| ENSMUSG00000057133 | 0.205164894 | 0.077208 | 0.184574 | 3603.687 | 3544.273 | 3155.086 | 3307.298 | 2809.296 | 3590.123 | 2867.796 | 2538.438 |
| ENSMUSG00000057134 | -0.135337777 | 0.044335 | 0.125209 | 1102.636 | 1031.847 | 1100.554 | 1088.631 | 1149.656 | 1128.548 | 1221.754 | 1251.817 |
| ENSMUSG00000057137 | -0.141959982 | 0.537371 | 0.687679 | 74.44802 | 86.38715 | 85.29498 | 59.52102 | 90.68535 | 68.15982 | 73.28598 | 104.4087 |
| ENSMUSG00000057143 | 0.651220318 | 0.000757 | 0.006312 | 532.2028 | 839.8751 | 683.188 | 630.5778 | 341.289 | 408.9589 | 395.3586 | 564.4598 |
| ENSMUSG00000057147 | 0.024959099 | 0.814631 | 0.887294 | 764.6013 | 928.6619 | 722.9371 | 827.2559 | 725.4828 | 834.6784 | 782.0385 | 842.8831 |
| ENSMUSG00000057156 | -0.364579459 | 0.025266 | 0.083583 | 453.7305 | 406.7395 | 375.1323 | 295.8799 | 503.1574 | 400.0199 | 520.7161 | 544.8831 |
| ENSMUSG00000057173 | 0.704209725 | 0.253654 | 0.417294 | 22.1332 | 5.999108 | 14.90592 | 12.93935 | 2.925334 | 12.29112 | 8.678602 | 10.87591 |
| ENSMUSG00000057176 | -0.279734784 | 0.081602 | 0.19156 | 146.8839 | 146.3782 | 136.6376 | 196.6782 | 192.0969 | 179.8972 | 203.465 | 185.9781 |
| ENSMUSG00000057177 | -0.119400076 | 0.192128 | 0.34547 | 1651.941 | 1754.139 | 1675.26 | 1687.292 | 1722.047 | 1623.545 | 1949.793 | 2055.547 |
| ENSMUSG00000057181 | 0.238459031 | 0.052768 | 0.141306 | 1032.212 | 1141.03 | 1071.57 | 1031.698 | 920.5051 | 1107.318 | 783.0028 | 814.6057 |
| ENSMUSG00000057182 | 2.049990887 | 7.33E-06 | 0.000181 | 138.8355 | 57.59143 | 63.76421 | 144.9207 | 27.30312 | 35.75597 | 25.07152 | 9.78832 |
| ENSMUSG00000057191 | 0.542688861 | 0.040688 | 0.118146 | 115.6963 | 99.58519 | 86.12309 | 100.0643 | 65.33246 | 82.68568 | 86.78602 | 40.24087 |
| ENSMUSG00000057193 | 0.22079019 | 0.015023 | 0.057753 | 3125.811 | 3009.152 | 3323.192 | 3095.956 | 2748.839 | 2297.321 | 2775.224 | 2951.722 |
| ENSMUSG00000057219 | -0.09962808 | 0.572418 | 0.715077 | 99.59938 | 112.7832 | 101.8571 | 124.2178 | 108.2374 | 99.44629 | 129.2147 | 132.6861 |

| | | | | | | | | | | | |
|---|---|---|---|---|---|---|---|---|---|---|---|
| ENSMUSG00000057228 | -3.872240707 | 1.95E-06 | 6.32E-05 | 111.672 | 35.99465 | 496.0359 | 180.2883 | 2428.027 | 979.9371 | 3806.049 | 4861.532 |
| ENSMUSG00000057229 | -0.381461887 | 6.01E-06 | 0.000155 | 881.3036 | 950.2587 | 959.7756 | 916.1062 | 1214.989 | 1079.383 | 1319.148 | 1213.752 |
| ENSMUSG00000057230 | 0.155995308 | 0.126721 | 0.257848 | 1975.891 | 2057.694 | 1989.112 | 2026.303 | 1634.287 | 2168.823 | 1690.399 | 1732.533 |
| ENSMUSG00000057234 | -0.18108201 | 0.273931 | 0.439753 | 126.7628 | 112.7832 | 130.0127 | 107.8279 | 150.1671 | 110.62 | 139.8219 | 140.2993 |
| ENSMUSG00000057236 | -0.130259254 | 0.028473 | 0.09106 | 5765.698 | 6647.011 | 6389.671 | 6235.905 | 6992.523 | 6862.912 | 6843.56 | 6701.736 |
| ENSMUSG00000057246 | 0.253292172 | 0.560693 | 0.706503 | 51.30877 | 43.19358 | 24.8432 | 57.79577 | 33.15378 | 61.45558 | 36.64299 | 17.40146 |
| ENSMUSG00000057262 | -1.409645798 | 0.072279 | 0.176499 | 0 | 11.99822 | 2.48432 | 9.488858 | 16.57689 | 13.40849 | 15.42863 | 17.40146 |
| ENSMUSG00000057265 | -0.697076844 | 0.006311 | 0.030548 | 53.32088 | 45.59322 | 44.71776 | 46.58167 | 54.60623 | 86.03781 | 69.42882 | 98.97079 |
| ENSMUSG00000057278 | -0.126929764 | 0.06496 | 0.164122 | 2200.241 | 2320.455 | 2138.999 | 2144.482 | 2599.647 | 2247.039 | 2344.187 | 2416.627 |
| ENSMUSG00000057280 | 0.420947038 | 0.307431 | 0.475922 | 36.21796 | 28.79572 | 24.8432 | 31.91707 | 35.10401 | 23.46486 | 22.17865 | 9.78832 |
| ENSMUSG00000057286 | 0.040319413 | 0.639755 | 0.766629 | 795.789 | 851.8733 | 821.4818 | 742.7188 | 850.2971 | 726.2932 | 758.8956 | 784.1532 |
| ENSMUSG00000057315 | -0.580042865 | 0.009171 | 0.040005 | 1879.31 | 1644.955 | 2619.301 | 2059.945 | 2654.253 | 2104.015 | 3161.904 | 4346.014 |
| ENSMUSG00000057322 | -0.147677646 | 0.068378 | 0.169381 | 6136.932 | 6996.159 | 7610.3 | 6521.434 | 7780.413 | 7278.575 | 7780.849 | 7364.079 |
| ENSMUSG00000057329 | 0.257078451 | 0.224573 | 0.38434 | 1860.195 | 1925.714 | 1452.499 | 1765.79 | 1330.052 | 2194.523 | 1247.79 | 1088.679 |
| ENSMUSG00000057335 | 0.281338687 | 0.050256 | 0.136502 | 977.8848 | 1023.448 | 792.498 | 877.2881 | 744.0099 | 936.3595 | 641.2523 | 698.2335 |
| ENSMUSG00000057337 | -0.080264354 | 0.687381 | 0.801308 | 183.1019 | 142.7788 | 215.3077 | 164.7611 | 146.2667 | 172.0756 | 199.6079 | 230.5693 |
| ENSMUSG00000057342 | -0.28522767 | 0.045192 | 0.126893 | 628.784 | 515.9233 | 622.7362 | 540.0023 | 731.3335 | 540.8091 | 783.0028 | 756.9634 |
| ENSMUSG00000057359 | 0.074685857 | 0.699195 | 0.810259 | 90.54489 | 83.98751 | 107.6539 | 96.61383 | 85.8098 | 109.5027 | 75.21455 | 91.35765 |
| ENSMUSG00000057363 | -0.182308239 | 0.004767 | 0.024888 | 1270.647 | 1233.417 | 1281.909 | 1149.014 | 1354.43 | 1432.474 | 1408.826 | 1404.08 |
| ENSMUSG00000057367 | 0.235361402 | 0.001352 | 0.00979 | 1626.79 | 1638.956 | 1480.655 | 1623.457 | 1274.47 | 1439.178 | 1399.184 | 1296.409 |
| ENSMUSG00000057375 | -0.141217573 | 0.19044 | 0.343038 | 683.1109 | 700.6958 | 695.6096 | 796.2015 | 780.089 | 678.2461 | 846.6459 | 866.8101 |
| ENSMUSG00000057378 | 0.178993037 | 0.555328 | 0.702428 | 95.57516 | 61.1909 | 77.84202 | 73.323 | 47.78045 | 106.1505 | 54.00019 | 65.25547 |
| ENSMUSG00000057388 | -0.302379977 | 0.005629 | 0.028049 | 1371.252 | 1355.798 | 1455.811 | 1328.44 | 1726.922 | 1395.6 | 1726.078 | 1947.876 |
| ENSMUSG00000057396 | -0.027340779 | 0.875525 | 0.925381 | 115.6963 | 101.9848 | 96.88848 | 88.85022 | 110.1876 | 93.85942 | 111.8575 | 93.53283 |
| ENSMUSG00000057406 | -0.008273153 | 0.916029 | 0.950215 | 3153.98 | 3481.882 | 3107.884 | 3165.828 | 3266.623 | 3583.419 | 3130.083 | 3000.664 |
| ENSMUSG00000057409 | 0.235948898 | 0.083863 | 0.19518 | 299.8042 | 326.3515 | 277.4157 | 257.0618 | 242.8027 | 268.1698 | 264.2152 | 207.7299 |
| ENSMUSG00000057411 | -0.071532416 | 0.218711 | 0.377493 | 1416.525 | 1384.594 | 1346.501 | 1414.703 | 1492.895 | 1405.657 | 1512.005 | 1431.27 |
| ENSMUSG00000057421 | -0.095654371 | 0.085511 | 0.197794 | 2184.144 | 2121.284 | 2035.486 | 2153.971 | 2310.039 | 2209.049 | 2197.615 | 2358.985 |
| ENSMUSG00000057425 | -2.926605714 | 0.010727 | 0.044955 | 19.11503 | 0 | 53.82693 | 18.97772 | 174.5449 | 39.10809 | 124.3933 | 363.2554 |
| ENSMUSG00000057440 | 0.247968488 | 0.045192 | 0.126893 | 594.5781 | 535.1204 | 497.6921 | 622.8142 | 401.7459 | 472.6492 | 520.7161 | 500.2919 |
| ENSMUSG00000057454 | 0.856498532 | 0.161247 | 0.304978 | 97.58727 | 13.19804 | 31.46805 | 28.46658 | 33.15378 | 18.99536 | 28.92867 | 13.05109 |
| ENSMUSG00000057455 | 0.847007059 | 0.015028 | 0.057758 | 80.48435 | 43.19358 | 38.92101 | 85.39973 | 38.02934 | 37.99072 | 33.75012 | 28.27737 |
| ENSMUSG00000057457 | 0.542249175 | 0.054954 | 0.145724 | 116.7023 | 208.7689 | 112.6225 | 140.6076 | 95.56091 | 134.0849 | 99.32178 | 67.43065 |
| ENSMUSG00000057465 | -0.488351243 | 0.394136 | 0.561451 | 8.048435 | 5.999108 | 14.07781 | 6.900988 | 14.62667 | 18.99536 | 9.642892 | 6.525547 |
| ENSMUSG00000057469 | -0.113115765 | 0.237932 | 0.399097 | 902.4308 | 875.8697 | 771.7954 | 855.7225 | 906.8535 | 814.5657 | 961.3963 | 997.321 |
| ENSMUSG00000057497 | -0.091514956 | 0.404165 | 0.571145 | 1118.732 | 1171.026 | 928.3075 | 1092.081 | 1287.147 | 1059.271 | 1052.039 | 1189.825 |
| ENSMUSG00000057506 | -0.06306351 | 0.39513 | 0.562477 | 940.6608 | 875.8697 | 918.3703 | 970.4514 | 1043.369 | 930.7726 | 958.5034 | 939.6787 |
| ENSMUSG00000057522 | 0.232352296 | 0.00582 | 0.028795 | 4444.748 | 4508.929 | 3849.04 | 4346.76 | 3786.357 | 3755.494 | 3801.228 | 3250.81 |
| ENSMUSG00000057530 | 0.177815567 | 0.007656 | 0.035035 | 2746.528 | 2524.425 | 2919.904 | 2680.171 | 2399.749 | 2299.556 | 2415.544 | 2500.372 |
| ENSMUSG00000057531 | -0.428374794 | 0.002316 | 0.014586 | 1003.036 | 1054.643 | 1307.58 | 1101.57 | 1605.033 | 1174.36 | 1520.684 | 1712.956 |
| ENSMUSG00000057541 | -0.298358855 | 7.75E-06 | 0.000189 | 858.1644 | 818.2783 | 832.2472 | 803.1025 | 1067.747 | 996.6977 | 979.7178 | 1027.774 |
| ENSMUSG00000057551 | 0.192521467 | 0.07349 | 0.178552 | 1403.446 | 1340.201 | 1213.176 | 1457.834 | 1136.98 | 1386.661 | 1139.79 | 1075.628 |
| ENSMUSG00000057554 | 0.183961151 | 0.015845 | 0.059937 | 1186.138 | 1105.036 | 1242.988 | 1166.267 | 1062.871 | 1033.571 | 952.7177 | 1093.029 |
| ENSMUSG00000057561 | -0.150249746 | 0.087215 | 0.200669 | 2355.173 | 2583.216 | 2212.701 | 2541.289 | 2906.807 | 2401.237 | 2768.474 | 2674.386 |
| ENSMUSG00000057572 | 0.05504313 | 0.626567 | 0.756629 | 941.6669 | 982.6538 | 907.6049 | 908.3425 | 1070.672 | 791.1009 | 815.7886 | 920.1021 |
| ENSMUSG00000057594 | -0.038688718 | 0.720284 | 0.825124 | 392.3612 | 418.7377 | 405.7722 | 375.2412 | 470.9788 | 383.2593 | 392.4657 | 386.0948 |
| ENSMUSG00000057596 | 0.623666387 | 0.227091 | 0.387035 | 85.51462 | 43.19358 | 67.07664 | 12.07673 | 38.02934 | 29.05173 | 31.82154 | 35.89051 |
| ENSMUSG00000057604 | -0.842284325 | 5.58E-09 | 5.28E-07 | 296.786 | 382.7431 | 373.4761 | 360.5766 | 726.4579 | 581.0345 | 527.4662 | 699.3211 |
| ENSMUSG00000057605 | 0.210678332 | 0.460234 | 0.621639 | 46.2785 | 38.39429 | 42.23344 | 29.3292 | 29.25334 | 37.99072 | 29.89296 | 38.06569 |
| ENSMUSG00000057606 | 0.660795439 | 0.1118 | 0.238376 | 109.6599 | 136.7797 | 80.32634 | 94.88858 | 66.30757 | 126.2633 | 31.82154 | 42.41605 |
| ENSMUSG00000057614 | -1.164585669 | 3.21E-05 | 0.000563 | 727.3773 | 602.3104 | 1320.002 | 837.6074 | 1783.479 | 1173.243 | 2382.759 | 2481.883 |
| ENSMUSG00000057637 | 0.030825429 | 0.710318 | 0.818711 | 991.9696 | 964.6565 | 1041.758 | 943.7101 | 922.4553 | 886.0777 | 1004.789 | 1046.263 |
| ENSMUSG00000057649 | 0.127410832 | 0.109676 | 0.235343 | 1965.83 | 1986.904 | 1884.771 | 1903.81 | 1747.399 | 1996.748 | 1631.577 | 1711.868 |
| ENSMUSG00000057657 | 0.026249574 | 0.905458 | 0.943292 | 53.32088 | 67.19001 | 61.27989 | 53.48266 | 55.58134 | 55.8687 | 68.46453 | 50.02919 |
| ENSMUSG00000057666 | -0.593246449 | 3.54E-05 | 0.00061 | 23661.39 | 24024.03 | 28516.68 | 24046.49 | 39409.12 | 28139.95 | 38454.89 | 45236.18 |
| ENSMUSG00000057667 | -0.28517217 | 0.038638 | 0.113625 | 234.4107 | 278.3586 | 337.8675 | 302.7808 | 365.6667 | 368.7335 | 351.9655 | 321.927 |
| ENSMUSG00000057672 | -0.419985801 | 0.001448 | 0.010317 | 1080.502 | 1036.646 | 1162.662 | 1044.637 | 1497.771 | 1099.496 | 1540.934 | 1647.7 |
| ENSMUSG00000057691 | -0.048832302 | 0.529295 | 0.681503 | 1039.254 | 1204.621 | 1072.398 | 1067.065 | 1151.606 | 1157.6 | 1057.825 | 1163.722 |
| ENSMUSG00000057706 | 0.132658171 | 0.224439 | 0.384146 | 754.5408 | 995.8519 | 881.1055 | 842.7832 | 799.5913 | 854.7912 | 718.3954 | 795.0291 |
| ENSMUSG00000057716 | 1.11382548 | 0.042174 | 0.121051 | 48.29061 | 17.99732 | 26.49941 | 60.38364 | 20.47734 | 31.28647 | 9.642892 | 9.78832 |
| ENSMUSG00000057719 | 0.900433856 | 0.041729 | 0.120224 | 24.1453 | 39.59411 | 26.49941 | 22.42821 | 9.751113 | 22.34748 | 19.28578 | 8.700729 |
| ENSMUSG00000057722 | -0.662010267 | 0.02666 | 0.08707 | 254.5318 | 182.3729 | 465.3959 | 276.0395 | 411.497 | 317.3342 | 519.7519 | 618.8393 |
| ENSMUSG00000057729 | 0.212564964 | 0.166824 | 0.312579 | 138.8355 | 115.1829 | 129.1846 | 151.8217 | 116.0382 | 115.0895 | 115.7147 | 116.3722 |
| ENSMUSG00000057738 | -0.19543694 | 0.016346 | 0.061108 | 8078.616 | 7422.096 | 8836.726 | 8167.319 | 9045.132 | 8500.982 | 9813.571 | 9865.539 |

| | | | | | | | | | | | |
|---|---|---|---|---|---|---|---|---|---|---|---|
| ENSMUSG00000057751 | 0.429268887 | 0.092495 | 0.208895 | 535.2209 | 779.884 | 592.9243 | 686.6483 | 388.0943 | 779.9271 | 389.5728 | 369.781 |
| ENSMUSG00000057762 | -0.263300012 | 0.650086 | 0.773564 | 12.07265 | 3.599465 | 18.21835 | 18.97772 | 25.35289 | 10.05637 | 12.53576 | 16.31387 |
| ENSMUSG00000057766 | -0.839472128 | 0.000374 | 0.003658 | 128.775 | 97.18554 | 123.3879 | 100.9269 | 176.4951 | 136.3196 | 231.4294 | 262.1095 |
| ENSMUSG00000057777 | 1.288847538 | 0.038992 | 0.114373 | 23.13925 | 10.79839 | 7.45296 | 31.91707 | 6.825779 | 11.17374 | 6.750024 | 5.437955 |
| ENSMUSG00000057778 | -0.20967992 | 0.344368 | 0.51309 | 134.8113 | 99.58519 | 115.9349 | 114.7289 | 156.0178 | 87.15518 | 147.5362 | 146.8248 |
| ENSMUSG00000057788 | -0.261621122 | 0.006746 | 0.03212 | 1006.054 | 1106.235 | 1074.054 | 1015.308 | 1360.28 | 1070.444 | 1270.933 | 1332.299 |
| ENSMUSG00000057789 | -0.215261766 | 0.076295 | 0.183168 | 1125.775 | 1073.84 | 1267.003 | 1036.011 | 1318.35 | 1067.092 | 1364.469 | 1478.036 |
| ENSMUSG00000057835 | 0.121388632 | 0.509163 | 0.664889 | 128.775 | 107.9839 | 110.1382 | 122.4925 | 103.3618 | 131.8501 | 110.8933 | 85.9197 |
| ENSMUSG00000057836 | 0.418638183 | 0.207288 | 0.363936 | 39.23612 | 62.39072 | 108.482 | 64.69676 | 57.53157 | 46.92971 | 54.96448 | 46.76642 |
| ENSMUSG00000057841 | -0.237752089 | 0.00858 | 0.038183 | 24451.15 | 29092.07 | 28038.03 | 25581.96 | 34901.18 | 28916.52 | 32704.83 | 29834.8 |
| ENSMUSG00000057842 | 0.349006436 | 0.036448 | 0.109416 | 223.3441 | 190.7716 | 154.0278 | 210.4801 | 170.6445 | 156.4324 | 138.8576 | 144.6496 |
| ENSMUSG00000057858 | -0.082115194 | 0.25026 | 0.413857 | 916.5155 | 1009.05 | 939.901 | 895.4032 | 1040.444 | 1002.285 | 971.0392 | 962.5181 |
| ENSMUSG00000057863 | -0.33262789 | 0.000269 | 0.002853 | 11926.77 | 13411.61 | 14007.42 | 12954.02 | 17907.92 | 14238.7 | 16827.81 | 16885.94 |
| ENSMUSG00000057880 | -0.623913091 | 0.000213 | 0.002383 | 1197.205 | 981.454 | 1002.837 | 1080.867 | 1864.413 | 1140.839 | 1608.434 | 1954.401 |
| ENSMUSG00000057886 | 0.166807232 | 0.670995 | 0.789151 | 21.12714 | 52.79215 | 25.67131 | 37.95543 | 25.35289 | 43.57759 | 27.0001 | 26.10219 |
| ENSMUSG00000057894 | 0.221006508 | 0.123755 | 0.253443 | 980.903 | 860.272 | 712.9998 | 823.8054 | 676.7272 | 890.5472 | 686.5739 | 643.8539 |
| ENSMUSG00000057895 | -0.090219624 | 0.474218 | 0.63371 | 447.6942 | 519.5227 | 432.2717 | 441.6632 | 538.2614 | 549.7481 | 438.7516 | 430.6861 |
| ENSMUSG00000057897 | 0.204041415 | 0.061897 | 0.158433 | 333.004 | 278.3586 | 319.6492 | 333.8353 | 292.5334 | 264.8177 | 273.8581 | 268.635 |
| ENSMUSG00000057914 | 0.20832957 | 0.262813 | 0.427166 | 239.4409 | 259.1615 | 244.2915 | 233.771 | 175.52 | 292.752 | 182.2507 | 195.7664 |
| ENSMUSG00000057933 | -3.420592995 | 2.91E-05 | 0.000526 | 68.4117 | 26.39607 | 321.3054 | 102.6522 | 1669.391 | 460.3581 | 1079.04 | 2352.46 |
| ENSMUSG00000057948 | -0.061649297 | 0.841088 | 0.904189 | 22.1332 | 28.79572 | 32.29616 | 36.23019 | 32.17867 | 29.05173 | 37.60728 | 26.10219 |
| ENSMUSG00000057963 | -0.082941468 | 0.402714 | 0.569693 | 775.6679 | 689.8974 | 629.361 | 720.2906 | 811.2926 | 687.1851 | 725.1455 | 755.8758 |
| ENSMUSG00000057967 | 0.921479032 | 0.045862 | 0.128111 | 61.36932 | 136.7797 | 77.01392 | 87.9876 | 37.05423 | 97.21155 | 34.71441 | 22.83941 |
| ENSMUSG00000057969 | -0.857989538 | 6.98E-06 | 0.000175 | 618.7234 | 585.5129 | 804.0915 | 679.7473 | 1195.486 | 839.1479 | 1270.933 | 1569.394 |
| ENSMUSG00000057982 | -0.012917671 | 0.907723 | 0.944892 | 447.6942 | 467.9304 | 358.5702 | 419.235 | 421.2481 | 446.9496 | 412.7158 | 424.1605 |
| ENSMUSG00000058006 | -0.038377806 | 0.710043 | 0.818574 | 1254.55 | 1021.048 | 1075.711 | 1042.912 | 1139.905 | 1263.75 | 1042.397 | 1068.014 |
| ENSMUSG00000058013 | 0.039752782 | 0.498618 | 0.654982 | 7147.01 | 7170.133 | 7380.086 | 7141.66 | 6886.236 | 6513.174 | 7173.347 | 7482.627 |
| ENSMUSG00000058022 | -0.90423501 | 0.031094 | 0.096889 | 19.11503 | 17.99732 | 33.12427 | 18.11509 | 51.6809 | 21.23011 | 35.6787 | 57.64233 |
| ENSMUSG00000058046 | -0.175813475 | 0.622966 | 0.754426 | 18.10898 | 19.19714 | 20.70267 | 27.60395 | 30.22845 | 18.99536 | 27.96439 | 19.57664 |
| ENSMUSG00000058056 | -0.259203482 | 0.424558 | 0.589675 | 932.6124 | 915.4638 | 2553.053 | 1101.57 | 1390.509 | 1249.224 | 1792.614 | 2155.606 |
| ENSMUSG00000058064 | 0.871960581 | 0.000446 | 0.004196 | 1461.797 | 1808.131 | 1548.559 | 1691.605 | 1437.314 | 845.8522 | 675.9667 | 597.0875 |
| ENSMUSG00000058070 | -0.247268274 | 0.029788 | 0.094056 | 1281.713 | 1199.822 | 1544.419 | 1299.111 | 1598.207 | 1376.605 | 1574.684 | 1774.949 |
| ENSMUSG00000058076 | -0.473006084 | 8.47E-05 | 0.001187 | 3138.89 | 3082.342 | 3657.747 | 3144.263 | 4382.15 | 3740.968 | 4594.838 | 5360.736 |
| ENSMUSG00000058093 | 0.13831618 | 0.298613 | 0.466655 | 343.0645 | 343.149 | 309.7119 | 299.3304 | 295.4587 | 348.6207 | 281.5724 | 250.146 |
| ENSMUSG00000058099 | 0.024148023 | 0.925721 | 0.95581 | 60.36326 | 41.99375 | 68.73285 | 52.62003 | 60.4569 | 41.34284 | 54.96448 | 64.16787 |
| ENSMUSG00000058126 | -0.241073843 | 0.320414 | 0.488405 | 55.33299 | 58.79126 | 62.9361 | 51.75741 | 65.33246 | 49.16446 | 87.75031 | 67.43065 |
| ENSMUSG00000058135 | -0.455884743 | 0.013643 | 0.05377 | 1480.912 | 1316.204 | 1946.879 | 1595.853 | 2649.377 | 1510.69 | 2363.473 | 2174.095 |
| ENSMUSG00000058145 | 0.03363814 | 0.904823 | 0.942937 | 254.5318 | 447.5334 | 612.7989 | 275.1769 | 380.2934 | 414.5458 | 304.7154 | 454.6131 |
| ENSMUSG00000058152 | 0.222768022 | 0.372565 | 0.541549 | 76.46013 | 50.3925 | 73.70149 | 52.62003 | 58.50668 | 46.92971 | 62.6788 | 48.9416 |
| ENSMUSG00000058153 | 0.426709653 | 0.234126 | 0.394903 | 45.27245 | 51.59233 | 36.43669 | 75.04824 | 22.42756 | 52.51658 | 42.42872 | 38.06569 |
| ENSMUSG00000058159 | 0.138878862 | 0.739663 | 0.837241 | 23.13925 | 21.59679 | 23.18699 | 20.70296 | 12.67645 | 27.93435 | 27.96439 | 11.9635 |
| ENSMUSG00000058173 | -0.047402224 | 0.656888 | 0.778847 | 782.7103 | 921.4629 | 944.0416 | 783.2621 | 870.7744 | 817.9178 | 864.9674 | 991.8831 |
| ENSMUSG00000058174 | 0.251540544 | 0.036504 | 0.109564 | 313.889 | 326.3515 | 276.5876 | 364.0271 | 274.0063 | 272.6393 | 261.3224 | 267.5474 |
| ENSMUSG00000058192 | 0.349544942 | 0.000576 | 0.00512 | 528.1785 | 541.1195 | 522.5353 | 506.36 | 381.2685 | 471.5319 | 415.6086 | 378.4817 |
| ENSMUSG00000058230 | 0.050576052 | 0.331292 | 0.499089 | 2288.774 | 2181.276 | 2355.135 | 2208.316 | 2151.096 | 2213.518 | 2154.222 | 2208.897 |
| ENSMUSG00000058239 | -0.13040145 | 0.016409 | 0.061304 | 1865.225 | 1764.937 | 1874.005 | 1823.586 | 2076.987 | 1934.175 | 2029.829 | 1981.591 |
| ENSMUSG00000058240 | 0.145104585 | 0.102303 | 0.22448 | 1230.404 | 1208.22 | 1015.259 | 1268.057 | 1062.871 | 1073.797 | 1076.147 | 1054.963 |
| ENSMUSG00000058246 | 1.044448472 | 0.021374 | 0.073804 | 19.11503 | 25.19625 | 28.98373 | 13.80198 | 8.776002 | 7.821619 | 13.50005 | 11.9635 |
| ENSMUSG00000058248 | 0.017712776 | 0.922484 | 0.953784 | 122.7386 | 93.58608 | 136.6376 | 120.7673 | 130.6649 | 118.4417 | 121.5004 | 98.97079 |
| ENSMUSG00000058254 | 0.656197365 | 4.84E-16 | 3.08E-13 | 2577.511 | 2866.374 | 2769.189 | 3026.083 | 1794.205 | 1946.466 | 1714.506 | 1678.153 |
| ENSMUSG00000058258 | -1.366729234 | 2.35E-11 | 4.98E-09 | 181.0898 | 169.1748 | 266.6503 | 184.6014 | 527.5352 | 375.4377 | 613.2879 | 554.6715 |
| ENSMUSG00000058267 | -0.083548468 | 0.391861 | 0.559248 | 1060.381 | 1184.224 | 1134.506 | 1125.724 | 1365.156 | 1200.06 | 1052.039 | 1155.022 |
| ENSMUSG00000058290 | 0.154578411 | 0.111954 | 0.238489 | 706.2502 | 692.297 | 655.0323 | 696.9998 | 636.7477 | 698.3588 | 582.4307 | 553.5839 |
| ENSMUSG00000058291 | 0.112990569 | 0.208921 | 0.366165 | 1410.488 | 1592.163 | 1560.981 | 1523.393 | 1400.26 | 1565.441 | 1418.469 | 1244.204 |
| ENSMUSG00000058297 | 0.55622991 | 6.11E-05 | 0.000928 | 7934.751 | 9405.401 | 7250.902 | 8023.261 | 7005.2 | 5320.935 | 5105.911 | 4745.16 |
| ENSMUSG00000058298 | 0.10701207 | 0.367604 | 0.53607 | 526.1664 | 601.1106 | 570.5655 | 524.4751 | 495.3565 | 616.7905 | 487.9303 | 463.3138 |
| ENSMUSG00000058301 | 0.119377847 | 0.159657 | 0.302776 | 1580.511 | 1668.952 | 1500.529 | 1348.281 | 1366.131 | 1461.525 | 1357.719 | 1424.744 |
| ENSMUSG00000058317 | 0.232528549 | 0.002134 | 0.013753 | 759.571 | 808.6797 | 751.9208 | 797.0641 | 673.8019 | 703.9457 | 648.9666 | 625.3649 |
| ENSMUSG00000058318 | 0.371391507 | 0.002575 | 0.015739 | 2378.312 | 2232.868 | 2058.673 | 2298.892 | 1639.162 | 2147.593 | 1661.47 | 1485.649 |
| ENSMUSG00000058325 | -0.069133378 | 0.444898 | 0.607797 | 2010.097 | 1988.104 | 2100.907 | 1795.119 | 2207.652 | 1835.846 | 2217.865 | 2017.481 |
| ENSMUSG00000058331 | 0.270119966 | 0.095586 | 0.213927 | 173.0413 | 178.7734 | 198.7456 | 172.5247 | 132.6151 | 188.8362 | 143.6791 | 135.9489 |
| ENSMUSG00000058351 | -0.682950937 | 7.34E-05 | 0.001069 | 360.1675 | 332.3506 | 367.6793 | 266.5507 | 493.4063 | 415.6632 | 570.8592 | 649.2919 |
| ENSMUSG00000058355 | -0.194660972 | 0.001534 | 0.010701 | 2601.657 | 2676.802 | 2659.05 | 2726.753 | 3179.838 | 2825.839 | 3000.868 | 3197.518 |
| ENSMUSG00000058385 | -0.4361119 | 0.438979 | 0.602492 | 5.030272 | 23.99643 | 33.12427 | 25.8787 | 32.17867 | 15.64324 | 45.32159 | 26.10219 |

| | | | | | | | | | | | |
|---|---|---|---|---|---|---|---|---|---|---|---|
| ENSMUSG00000058388 | 0.116488283 | 0.140038 | 0.276218 | 1090.563 | 1280.21 | 1143.615 | 1252.529 | 1149.656 | 1097.261 | 1074.218 | 1073.452 |
| ENSMUSG00000058392 | -0.152218007 | 0.01824 | 0.066201 | 1018.127 | 1003.051 | 968.8848 | 974.7645 | 1146.731 | 1126.313 | 1084.825 | 1045.175 |
| ENSMUSG00000058396 | -0.712927647 | 0.023288 | 0.078718 | 327.9737 | 115.1829 | 331.2427 | 250.1608 | 334.4632 | 332.9775 | 477.3231 | 537.27 |
| ENSMUSG00000058402 | 0.392996092 | 0.040347 | 0.117435 | 310.8708 | 308.3541 | 217.792 | 288.1162 | 191.1218 | 282.6956 | 187.0721 | 195.7664 |
| ENSMUSG00000058407 | 0.011423737 | 0.870824 | 0.922508 | 2227.404 | 2337.252 | 2362.588 | 2322.182 | 2227.154 | 2529.735 | 2274.758 | 2146.905 |
| ENSMUSG00000058420 | -0.635734838 | 0.050429 | 0.13685 | 181.0898 | 81.58786 | 191.2926 | 163.0358 | 194.0472 | 150.8455 | 270.001 | 345.854 |
| ENSMUSG00000058440 | 0.091298108 | 0.133435 | 0.266927 | 1441.676 | 1366.597 | 1308.408 | 1426.779 | 1297.873 | 1322.971 | 1290.219 | 1292.058 |
| ENSMUSG00000058441 | 0.04472354 | 0.911934 | 0.948028 | 38.23007 | 22.79661 | 13.24971 | 28.46658 | 19.50223 | 24.58223 | 27.0001 | 28.27737 |
| ENSMUSG00000058443 | -0.086298336 | 0.76455 | 0.853838 | 40.24217 | 58.79126 | 67.07664 | 56.07053 | 41.92979 | 77.09881 | 67.50024 | 50.02919 |
| ENSMUSG00000058444 | 0.04830559 | 0.530871 | 0.682549 | 1009.073 | 913.0642 | 972.1972 | 982.5282 | 860.0482 | 1000.05 | 946.932 | 946.2042 |
| ENSMUSG00000058446 | -0.270056801 | 0.086393 | 0.199383 | 854.1401 | 591.512 | 818.1694 | 772.048 | 817.1433 | 781.0445 | 958.5034 | 1107.168 |
| ENSMUSG00000058454 | 0.024240277 | 0.946492 | 0.969279 | 1275.677 | 345.5486 | 452.1462 | 472.7177 | 617.2455 | 496.1141 | 635.4666 | 754.7882 |
| ENSMUSG00000058486 | -0.1391941 | 0.130745 | 0.263258 | 640.8566 | 616.7083 | 596.2368 | 611.6001 | 701.105 | 589.9735 | 695.2525 | 726.5108 |
| ENSMUSG00000058488 | -2.30897626 | 4.35E-05 | 0.000719 | 315.9011 | 191.9714 | 582.9871 | 412.334 | 1722.047 | 584.3867 | 1886.15 | 3259.51 |
| ENSMUSG00000058492 | -0.336685605 | 0.343191 | 0.511957 | 20.12109 | 15.59768 | 25.67131 | 17.25247 | 27.30312 | 20.11273 | 22.17865 | 30.45255 |
| ENSMUSG00000058498 | 0.377350918 | 0.152608 | 0.293572 | 42.25428 | 56.39161 | 65.42042 | 59.52102 | 47.78045 | 44.69496 | 48.21446 | 31.54014 |
| ENSMUSG00000058503 | 0.230470589 | 0.166868 | 0.31263 | 2401.452 | 2493.229 | 1878.146 | 2197.102 | 1694.743 | 2585.604 | 1701.97 | 1662.927 |
| ENSMUSG00000058542 | -1.077778659 | 0.030125 | 0.09475 | 8.048435 | 4.799286 | 13.24971 | 6.038364 | 17.552 | 13.40849 | 17.3572 | 20.66423 |
| ENSMUSG00000058546 | -0.201734751 | 0.029191 | 0.092758 | 21828.36 | 26199.3 | 26654.27 | 26113.34 | 31555.58 | 26128.68 | 29622 | 28615.61 |
| ENSMUSG00000058558 | -0.074694745 | 0.347926 | 0.516564 | 34091.16 | 40745.94 | 37967.86 | 38793.04 | 42621.14 | 38777.35 | 41353.54 | 36897.62 |
| ENSMUSG00000058569 | -0.158909449 | 0.063201 | 0.16091 | 5670.122 | 5732.747 | 6089.068 | 5547.532 | 6487.416 | 5546.645 | 6708.56 | 6979.072 |
| ENSMUSG00000058571 | 0.660508145 | 0.000582 | 0.00515 | 5510.16 | 6635.013 | 4952.078 | 6411.88 | 3811.71 | 5146.625 | 3202.404 | 2712.452 |
| ENSMUSG00000058586 | -0.162888218 | 0.120159 | 0.249148 | 580.4934 | 655.1026 | 662.4853 | 595.2102 | 706.9557 | 597.7951 | 728.0383 | 756.9634 |
| ENSMUSG00000058587 | 0.005990604 | 0.944094 | 0.967635 | 3240.501 | 3167.529 | 3462.314 | 3499.664 | 3155.46 | 3061.605 | 3330.655 | 3770.678 |
| ENSMUSG00000058589 | 0.736064504 | 0.000523 | 0.004747 | 88.53278 | 82.78769 | 101.029 | 79.36136 | 46.80534 | 59.22083 | 45.32159 | 60.9051 |
| ENSMUSG00000058594 | -0.139508144 | 0.124151 | 0.254028 | 2747.534 | 2725.995 | 2687.206 | 2951.898 | 3116.456 | 2605.716 | 3205.297 | 3312.802 |
| ENSMUSG00000058600 | -0.144603433 | 0.088098 | 0.20198 | 17045.58 | 19983.03 | 19863.79 | 18985.48 | 22718.14 | 18952.9 | 21793.9 | 20409.73 |
| ENSMUSG00000058603 | -0.038245829 | 0.814188 | 0.887208 | 172.0353 | 217.1677 | 214.4796 | 176.8378 | 242.8027 | 193.3057 | 189.965 | 174.0146 |
| ENSMUSG00000058620 | -2.256166946 | 2.14E-11 | 4.59E-09 | 30.18163 | 27.5959 | 57.13936 | 50.89479 | 216.4747 | 115.0895 | 192.8578 | 274.073 |
| ENSMUSG00000058624 | 0.96730643 | 0.015235 | 0.058346 | 146.8839 | 79.18822 | 44.71776 | 60.38364 | 35.10401 | 54.75133 | 43.39301 | 35.89051 |
| ENSMUSG00000058625 | 0.035619603 | 0.932788 | 0.960099 | 17.10292 | 17.99732 | 14.90592 | 12.07673 | 13.65156 | 13.40849 | 11.57147 | 21.75182 |
| ENSMUSG00000058626 | 0.606201223 | 0.525781 | 0.679063 | 1.006054 | 5.999108 | 33.95237 | 5.175741 | 4.875557 | 6.704245 | 13.50005 | 5.437955 |
| ENSMUSG00000058638 | 0.160513557 | 0.071566 | 0.175176 | 939.6548 | 981.454 | 938.2448 | 865.2114 | 826.8944 | 746.4059 | 913.1818 | 841.7955 |
| ENSMUSG00000058655 | 0.053796085 | 0.384077 | 0.552152 | 17384.62 | 17474.2 | 17086.32 | 18634.39 | 18105.87 | 16627.64 | 17366.85 | 15894.06 |
| ENSMUSG00000058672 | -0.515510139 | 0.051836 | 0.139437 | 1195.193 | 937.0606 | 1647.932 | 1422.466 | 1711.32 | 1029.102 | 2139.758 | 2559.102 |
| ENSMUSG00000058690 | 0.117938953 | 0.157103 | 0.299565 | 1496.003 | 1583.764 | 1578.371 | 1538.058 | 1320.301 | 1457.056 | 1331.683 | 1603.109 |
| ENSMUSG00000058704 | 0.115138262 | 0.215756 | 0.373966 | 1334.028 | 1416.989 | 1192.474 | 1293.073 | 1205.238 | 1355.375 | 1150.397 | 1121.306 |
| ENSMUSG00000058706 | 0.210143492 | 0.028612 | 0.091413 | 984.9272 | 1000.651 | 922.5108 | 1086.906 | 879.5504 | 965.4112 | 819.6458 | 789.5911 |
| ENSMUSG00000058709 | -0.440216152 | 1.62E-05 | 0.000332 | 1656.972 | 1910.116 | 1686.025 | 1573.425 | 2316.864 | 1985.574 | 2468.58 | 2485.146 |
| ENSMUSG00000058715 | 0.55592738 | 0.002753 | 0.016576 | 179.0777 | 179.9732 | 215.3077 | 226.0074 | 107.2622 | 158.6671 | 159.1077 | 120.7226 |
| ENSMUSG00000058729 | -0.020284095 | 0.879416 | 0.927685 | 511.0756 | 481.1284 | 376.7885 | 451.1521 | 407.5965 | 515.1095 | 435.8587 | 486.1532 |
| ENSMUSG00000058740 | 0.248456474 | 0.508197 | 0.66388 | 60.36326 | 43.19358 | 45.54586 | 86.26235 | 47.78045 | 80.45093 | 32.78583 | 38.06569 |
| ENSMUSG00000058743 | 0.030813539 | 0.879458 | 0.927685 | 79.47829 | 105.5843 | 99.3728 | 81.08661 | 94.5858 | 88.27255 | 71.3574 | 103.3212 |
| ENSMUSG00000058748 | 0.148587815 | 0.247582 | 0.410774 | 249.5015 | 309.554 | 293.1497 | 296.7425 | 267.1805 | 292.752 | 243.9652 | 232.7445 |
| ENSMUSG00000058756 | 0.249020186 | 0.003478 | 0.019693 | 2780.734 | 3301.909 | 3463.97 | 3394.423 | 2779.067 | 2628.064 | 2801.26 | 2682 |
| ENSMUSG00000058761 | 0.137911064 | 0.428818 | 0.593719 | 1569.445 | 1386.994 | 1218.973 | 1219.75 | 1061.896 | 1605.667 | 1291.183 | 944.0291 |
| ENSMUSG00000058793 | 0.062528959 | 0.372714 | 0.541681 | 2460.809 | 2314.456 | 2377.494 | 2537.838 | 2222.279 | 2312.964 | 2533.188 | 2212.16 |
| ENSMUSG00000058794 | 0.289822134 | 0.326047 | 0.493883 | 58.35115 | 46.79304 | 46.37397 | 42.26855 | 56.55646 | 39.10809 | 30.85725 | 31.54014 |
| ENSMUSG00000058799 | 0.153751936 | 0.155797 | 0.297848 | 13174.28 | 14765 | 12467.15 | 14566.26 | 12873.42 | 14239.82 | 11652.47 | 10648.6 |
| ENSMUSG00000058806 | -0.426461792 | 0.015959 | 0.060209 | 355.1372 | 307.1543 | 264.166 | 313.1323 | 405.6463 | 302.8084 | 497.5732 | 457.8758 |
| ENSMUSG00000058809 | -0.22362196 | 0.455978 | 0.617925 | 75.45408 | 40.79393 | 43.88965 | 41.40593 | 69.2329 | 43.57759 | 57.85735 | 64.16787 |
| ENSMUSG00000058812 | -1.220546011 | 0.006065 | 0.029689 | 13.07871 | 5.999108 | 10.76539 | 9.488858 | 23.40267 | 14.52586 | 25.07152 | 29.36496 |
| ENSMUSG00000058818 | 0.64022417 | 0.05682 | 0.148896 | 56.33904 | 38.39429 | 49.6864 | 32.77969 | 24.37778 | 20.11273 | 39.53586 | 29.36496 |
| ENSMUSG00000058831 | 0.529401738 | 0.02075 | 0.072312 | 77.46619 | 101.9848 | 110.1382 | 106.1027 | 83.85957 | 78.21619 | 55.92877 | 56.55474 |
| ENSMUSG00000058833 | -0.225703078 | 0.002999 | 0.017756 | 923.5579 | 910.6645 | 938.2448 | 908.3425 | 1051.17 | 1058.153 | 1008.646 | 1188.737 |
| ENSMUSG00000058835 | 0.332579946 | 0.003481 | 0.019699 | 2471.876 | 2648.006 | 2350.995 | 2940.683 | 1880.015 | 2440.345 | 2016.329 | 1932.649 |
| ENSMUSG00000058838 | -0.037671099 | 0.849778 | 0.909716 | 62.37537 | 71.98929 | 58.79557 | 66.42201 | 71.18313 | 64.8077 | 70.39311 | 58.72992 |
| ENSMUSG00000058881 | 0.284691651 | 0.000119 | 0.001533 | 2735.462 | 2729.594 | 2388.26 | 2745.731 | 2199.851 | 2080.551 | 2303.687 | 2112.102 |
| ENSMUSG00000058883 | -0.361621598 | 0.5312 | 0.682784 | 16.09687 | 7.198929 | 9.109173 | 11.21411 | 9.751113 | 5.58687 | 22.17865 | 18.48905 |
| ENSMUSG00000058886 | 0.061409194 | 0.357814 | 0.5262 | 1836.049 | 1907.716 | 1820.178 | 1817.548 | 1873.189 | 1838.08 | 1727.042 | 1632.474 |
| ENSMUSG00000058897 | 0.925065267 | 8.68E-06 | 0.000205 | 232.3986 | 271.1597 | 172.2462 | 234.6336 | 111.1627 | 149.7281 | 125.3576 | 92.44524 |
| ENSMUSG00000058900 | 0.258030078 | 0.105959 | 0.229961 | 125.7568 | 153.5772 | 128.3565 | 136.2945 | 130.6649 | 102.7984 | 111.8575 | 107.6715 |
| ENSMUSG00000058905 | 0.109903891 | 0.790514 | 0.871705 | 11.0666 | 27.5959 | 20.70267 | 18.97772 | 22.42756 | 18.99536 | 14.46434 | 16.31387 |
| ENSMUSG00000058914 | 0.817532212 | 0.000317 | 0.003241 | 1033.218 | 1384.594 | 987.9312 | 1191.283 | 580.1912 | 958.707 | 635.4666 | 433.9488 |

| | | | | | | | | | | | |
|---|---|---|---|---|---|---|---|---|---|---|---|
| ENSMUSG00000058921 | -2.632876974 | 0.001816 | 0.01224 | 3.018163 | 1.199822 | 11.59349 | 1.725247 | 19.50223 | 8.938993 | 22.17865 | 60.9051 |
| ENSMUSG00000058922 | -0.182400111 | 0.059719 | 0.154279 | 538.2391 | 550.7181 | 545.7223 | 629.7152 | 708.9059 | 622.3774 | 606.5379 | 632.978 |
| ENSMUSG00000058925 | 0.993719508 | 0.035968 | 0.108345 | 16.09687 | 17.99732 | 21.53077 | 24.15346 | 12.67645 | 14.52586 | 8.678602 | 4.350364 |
| ENSMUSG00000058927 | -0.856696831 | 0.027274 | 0.088341 | 17.10292 | 19.19714 | 25.67131 | 18.97772 | 33.15378 | 20.11273 | 40.50014 | 53.29196 |
| ENSMUSG00000058952 | -1.322179197 | 3.85E-07 | 1.70E-05 | 208.2533 | 173.9741 | 165.6213 | 153.547 | 535.3361 | 260.3482 | 358.7156 | 598.1751 |
| ENSMUSG00000058966 | 0.260091987 | 0.16528 | 0.310695 | 146.8839 | 184.7725 | 151.5435 | 168.2116 | 122.864 | 170.9582 | 107.0361 | 143.562 |
| ENSMUSG00000058975 | 0.122002968 | 0.706084 | 0.815104 | 30.18163 | 37.19447 | 34.78048 | 41.40593 | 27.30312 | 25.6996 | 49.17875 | 29.36496 |
| ENSMUSG00000058979 | -0.118060516 | 0.233261 | 0.393874 | 516.1059 | 511.124 | 464.5678 | 501.1842 | 548.0126 | 490.5272 | 516.859 | 605.7882 |
| ENSMUSG00000058997 | -0.178491988 | 0.121427 | 0.250705 | 1381.313 | 1358.198 | 1462.436 | 1372.434 | 1494.846 | 1349.788 | 1546.72 | 1918.511 |
| ENSMUSG00000059003 | 0.521853095 | 0.374125 | 0.54288 | 18.10898 | 28.79572 | 11.59349 | 10.35148 | 5.850668 | 22.34748 | 8.678602 | 10.87591 |
| ENSMUSG00000059005 | -0.124042793 | 0.016921 | 0.062635 | 26982.38 | 29897.15 | 29086.42 | 27980.92 | 32117.24 | 30108.76 | 30940.18 | 31006.13 |
| ENSMUSG00000059013 | -0.426059624 | 0.032265 | 0.099617 | 491.9606 | 386.3425 | 654.2042 | 519.2993 | 690.3788 | 497.2315 | 797.4671 | 774.3649 |
| ENSMUSG00000059022 | 0.625729235 | 1.37E-11 | 3.16E-09 | 1078.49 | 969.4558 | 1212.348 | 1081.73 | 659.1752 | 729.6453 | 712.6097 | 716.7225 |
| ENSMUSG00000059027 | 0.817379734 | 0.130012 | 0.262383 | 15.09082 | 14.39786 | 13.24971 | 14.6646 | 14.62667 | 3.352122 | 7.714313 | 6.525547 |
| ENSMUSG00000059040 | -0.423687674 | 0.315293 | 0.483672 | 27.16347 | 25.19625 | 40.57722 | 28.46658 | 53.63112 | 15.64324 | 39.53586 | 54.37955 |
| ENSMUSG00000059049 | 0.104489268 | 0.471887 | 0.631304 | 2789.789 | 2739.193 | 4137.221 | 3272.794 | 2763.465 | 2816.9 | 3360.548 | 3097.459 |
| ENSMUSG00000059058 | -0.334985008 | 0.428563 | 0.59347 | 28.16952 | 14.39786 | 19.04645 | 18.97772 | 24.37778 | 22.34748 | 39.53586 | 15.22628 |
| ENSMUSG00000059060 | -0.178766674 | 0.556858 | 0.703587 | 31.18769 | 41.99375 | 48.03018 | 69.8725 | 62.40712 | 45.81234 | 58.82164 | 50.02919 |
| ENSMUSG00000059070 | -0.176037629 | 0.043665 | 0.123943 | 24697.63 | 28846.11 | 27580.09 | 28269.03 | 34103.54 | 27532.1 | 31542.86 | 30407.96 |
| ENSMUSG00000059089 | 0.410420845 | 0.350449 | 0.519114 | 23.13925 | 38.39429 | 67.07664 | 28.46658 | 25.35289 | 18.99536 | 42.42872 | 31.54014 |
| ENSMUSG00000059119 | -0.141235169 | 0.069219 | 0.170848 | 5045.363 | 5856.329 | 5427.411 | 4929.031 | 6143.201 | 5447.199 | 5907.235 | 5940.423 |
| ENSMUSG00000059142 | 0.442621713 | 0.010285 | 0.043496 | 1307.871 | 1273.011 | 1297.643 | 1428.505 | 834.6953 | 1331.91 | 991.2893 | 748.2627 |
| ENSMUSG00000059146 | 1.085230526 | 4.19E-08 | 2.79E-06 | 587.5357 | 657.5022 | 541.5817 | 696.9998 | 302.2845 | 406.7242 | 229.5008 | 232.7445 |
| ENSMUSG00000059149 | -0.124449883 | 0.443934 | 0.606952 | 603.6326 | 532.7208 | 639.2983 | 524.4751 | 594.8179 | 462.5929 | 711.6454 | 738.4743 |
| ENSMUSG00000059173 | -0.039097428 | 0.715082 | 0.821407 | 560.3723 | 659.9018 | 668.2821 | 577.0951 | 577.2659 | 597.7951 | 668.2524 | 689.5327 |
| ENSMUSG00000059182 | 0.067442466 | 0.406399 | 0.572907 | 928.5882 | 841.0749 | 872.8244 | 923.0071 | 840.5459 | 895.0167 | 892.9318 | 775.4524 |
| ENSMUSG00000059183 | 0.007626374 | 0.924156 | 0.954942 | 516.1059 | 539.9197 | 515.9104 | 541.7276 | 530.4606 | 524.0485 | 494.6803 | 553.5839 |
| ENSMUSG00000059187 | 0.298224625 | 0.348426 | 0.51705 | 159.9626 | 105.5843 | 65.42042 | 139.745 | 94.5858 | 112.8548 | 111.8575 | 63.08028 |
| ENSMUSG00000059195 | -0.431664816 | 0.031025 | 0.096765 | 248.4954 | 244.7636 | 256.7131 | 228.5952 | 370.5423 | 206.7142 | 357.7513 | 383.9197 |
| ENSMUSG00000059203 | 1.491838403 | 0.019412 | 0.069032 | 14.08476 | 32.39518 | 14.07781 | 6.900988 | 4.875557 | 6.704245 | 7.714313 | 4.350364 |
| ENSMUSG00000059208 | -0.042109881 | 0.390778 | 0.558176 | 12528.39 | 13232.83 | 12301.52 | 12339.83 | 13231.29 | 12373.8 | 13176.05 | 13104.39 |
| ENSMUSG00000059213 | -2.653455874 | 0.001419 | 0.010174 | 38.23007 | 3.599465 | 87.7793 | 32.77969 | 142.3663 | 134.0849 | 303.7511 | 444.8248 |
| ENSMUSG00000059248 | -0.028599595 | 0.808437 | 0.883027 | 3823.007 | 4793.287 | 5213.759 | 4081.072 | 4777.07 | 4404.689 | 4128.122 | 4960.503 |
| ENSMUSG00000059263 | 0.037336526 | 0.54645 | 0.694817 | 4166.071 | 3768.639 | 3769.541 | 4204.427 | 3765.88 | 3895.166 | 3936.228 | 3907.715 |
| ENSMUSG00000059273 | 0.093960861 | 0.2586 | 0.422514 | 1603.651 | 1689.349 | 1633.026 | 1673.49 | 1376.857 | 1699.526 | 1488.862 | 1620.511 |
| ENSMUSG00000059277 | 0.13145254 | 0.460903 | 0.622087 | 162.9808 | 166.7752 | 134.1533 | 169.0742 | 181.3707 | 141.9065 | 129.2147 | 123.9854 |
| ENSMUSG00000059278 | -0.403573768 | 4.49E-06 | 0.000124 | 654.9414 | 659.9018 | 727.0776 | 642.6545 | 891.2517 | 807.8615 | 890.0389 | 963.6057 |
| ENSMUSG00000059288 | 0.149793395 | 0.269313 | 0.4341 | 628.784 | 620.3077 | 599.5492 | 633.1656 | 550.9379 | 708.4152 | 492.7518 | 486.1532 |
| ENSMUSG00000059291 | -0.1331057 | 0.05313 | 0.142028 | 26096.04 | 29946.35 | 28763.46 | 29054.02 | 33370.26 | 30677.51 | 31497.54 | 29316.02 |
| ENSMUSG00000059316 | -0.525091554 | 0.000823 | 0.006715 | 730.3955 | 770.2854 | 896.0114 | 845.371 | 1093.1 | 884.9603 | 1237.183 | 1451.934 |
| ENSMUSG00000059323 | -0.296042904 | 0.004053 | 0.022045 | 1577.493 | 1572.966 | 1821.006 | 1644.16 | 2096.489 | 1717.404 | 2061.65 | 2249.138 |
| ENSMUSG00000059325 | -0.808913443 | 0.016899 | 0.062603 | 256.5439 | 137.9795 | 468.7084 | 288.9789 | 595.793 | 307.2779 | 584.3592 | 534.0072 |
| ENSMUSG00000059326 | 0.112215269 | 0.120369 | 0.249357 | 893.3763 | 967.0562 | 901.8081 | 954.0616 | 903.9282 | 810.0962 | 876.5389 | 843.9707 |
| ENSMUSG00000059327 | 0.38128548 | 0.056812 | 0.148894 | 303.8284 | 296.3559 | 323.7897 | 351.0878 | 206.7236 | 349.7381 | 230.4651 | 193.5912 |
| ENSMUSG00000059336 | -1.265651492 | 0.015705 | 0.059687 | 81.4904 | 19.19714 | 171.4181 | 40.5433 | 227.2009 | 191.071 | 144.6434 | 191.416 |
| ENSMUSG00000059355 | -0.072029987 | 0.316825 | 0.484973 | 931.6063 | 1028.247 | 1008.634 | 1045.5 | 1074.573 | 1059.271 | 994.1821 | 1093.029 |
| ENSMUSG00000059361 | 0.764206169 | 0.075007 | 0.18103 | 23.13925 | 16.7975 | 34.78048 | 37.95543 | 23.40267 | 13.40849 | 11.57147 | 18.48905 |
| ENSMUSG00000059363 | -0.235133027 | 0.003573 | 0.020122 | 556.3481 | 626.3068 | 595.4087 | 577.0951 | 731.3335 | 696.1241 | 651.8595 | 690.6203 |
| ENSMUSG00000059395 | -0.0514865 | 0.88355 | 0.930352 | 45.27245 | 38.39429 | 28.15563 | 31.91707 | 34.1289 | 56.98608 | 24.10723 | 33.71532 |
| ENSMUSG00000059401 | 0.707139494 | 1.38E-06 | 4.83E-05 | 301.8163 | 320.3523 | 326.274 | 330.3848 | 149.192 | 206.7142 | 223.7151 | 204.4671 |
| ENSMUSG00000059409 | -0.174709373 | 0.038344 | 0.113112 | 2042.29 | 2176.476 | 2273.153 | 2087.549 | 2659.129 | 2177.762 | 2321.044 | 2525.387 |
| ENSMUSG00000059412 | -3.037393265 | 0.000243 | 0.002648 | 669.0261 | 183.5727 | 2863.593 | 838.47 | 9012.954 | 3364.413 | 10712.29 | 14309.44 |
| ENSMUSG00000059422 | 0.121010738 | 0.78323 | 0.866677 | 13.07871 | 13.19804 | 13.24971 | 19.84034 | 11.70134 | 18.99536 | 14.46434 | 9.78832 |
| ENSMUSG00000059423 | 0.272397336 | 0.080025 | 0.188964 | 418.5186 | 401.9402 | 303.9151 | 320.8959 | 303.2596 | 339.6817 | 297.9654 | 253.4087 |
| ENSMUSG00000059430 | 0.771467567 | 0.198514 | 0.353135 | 389.343 | 803.8804 | 591.2681 | 892.8153 | 464.153 | 774.3402 | 131.1433 | 199.0292 |
| ENSMUSG00000059434 | 0.408956838 | 0.389652 | 0.556999 | 23.13925 | 11.99822 | 14.90592 | 12.07673 | 7.80089 | 12.29112 | 10.60718 | 16.31387 |
| ENSMUSG00000059436 | -0.006258688 | 0.897404 | 0.937817 | 2025.187 | 1982.105 | 1967.581 | 2061.67 | 1999.953 | 2063.79 | 1997.043 | 2013.131 |
| ENSMUSG00000059439 | -0.171084013 | 0.201552 | 0.357026 | 791.7648 | 575.9143 | 826.4504 | 649.5555 | 747.9104 | 764.2839 | 845.6816 | 847.2335 |
| ENSMUSG00000059447 | -0.612294156 | 4.25E-05 | 0.000703 | 1857.176 | 1527.373 | 1917.895 | 1728.697 | 2723.486 | 2009.039 | 2769.438 | 3248.635 |
| ENSMUSG00000059456 | -0.1189475 | 0.466632 | 0.627252 | 227.3683 | 218.3675 | 294.806 | 265.688 | 248.6534 | 230.1791 | 320.144 | 295.8248 |
| ENSMUSG00000059461 | -0.424624649 | 0.159349 | 0.302349 | 33.19979 | 46.79304 | 25.67131 | 24.15346 | 43.88001 | 42.46022 | 46.28588 | 39.15328 |
| ENSMUSG00000059474 | 0.166677447 | 0.243101 | 0.405667 | 4390.421 | 4522.127 | 3805.978 | 4328.645 | 3629.364 | 4910.859 | 3542.798 | 3103.985 |
| ENSMUSG00000059475 | -0.015368039 | 0.873355 | 0.924258 | 799.8132 | 897.4665 | 709.6874 | 797.0641 | 802.5166 | 779.9271 | 779.1456 | 872.2481 |
| ENSMUSG00000059479 | 0.454195028 | 0.008936 | 0.039268 | 333.004 | 427.1365 | 321.3054 | 373.516 | 230.1263 | 343.0338 | 263.2509 | 225.1314 |

| | | | | | | | | | | | |
|---|---|---|---|---|---|---|---|---|---|---|---|
| ENSMUSG00000059481 | 3.009092296 | 0.003291 | 0.01901 | 97.58727 | 4.799286 | 2.48432 | 78.49874 | 4.875557 | 2.234748 | 4.821446 | 10.87591 |
| ENSMUSG00000059482 | 0.026565407 | 0.833658 | 0.899397 | 593.5721 | 596.3113 | 577.1903 | 662.4948 | 590.9175 | 724.0584 | 510.109 | 562.2846 |
| ENSMUSG00000059486 | 0.039031159 | 0.659383 | 0.780701 | 1269.641 | 1093.037 | 1059.148 | 1170.58 | 1118.453 | 1211.234 | 1086.754 | 1053.876 |
| ENSMUSG00000059493 | 0.546171318 | 0.028021 | 0.090144 | 1151.932 | 1458.983 | 1056.664 | 1361.22 | 686.4784 | 1402.304 | 702.0025 | 653.6422 |
| ENSMUSG00000059495 | 0.206569599 | 0.055479 | 0.146683 | 4755.619 | 4537.725 | 4252.328 | 4724.589 | 3429.466 | 4732.079 | 3921.764 | 3751.102 |
| ENSMUSG00000059498 | 0.435081624 | 0.001414 | 0.010154 | 278.6771 | 227.9661 | 277.4157 | 281.2153 | 193.072 | 176.5451 | 226.608 | 192.5036 |
| ENSMUSG00000059518 | -0.340249543 | 2.39E-05 | 0.000451 | 706.2502 | 703.0954 | 741.1554 | 701.3129 | 884.426 | 884.9603 | 846.6459 | 997.321 |
| ENSMUSG00000059534 | -0.8103801 | 8.77E-05 | 0.001216 | 1646.911 | 1594.563 | 2204.42 | 1687.292 | 3023.82 | 1998.982 | 3374.048 | 4114.357 |
| ENSMUSG00000059540 | 0.203995846 | 0.118086 | 0.246817 | 497.9969 | 520.7225 | 439.7246 | 523.6125 | 415.3974 | 521.8137 | 405.9657 | 377.3941 |
| ENSMUSG00000059552 | 0.042487374 | 0.546018 | 0.694507 | 2896.43 | 3135.134 | 2834.609 | 2705.187 | 2904.857 | 2842.6 | 2862.01 | 2618.919 |
| ENSMUSG00000059554 | 0.310007652 | 0.009524 | 0.041098 | 374.2522 | 374.3443 | 346.1486 | 389.9058 | 307.1601 | 345.2686 | 286.3939 | 258.8467 |
| ENSMUSG00000059555 | -0.007479867 | 0.937128 | 0.962687 | 573.451 | 533.9206 | 506.8013 | 474.4429 | 528.5103 | 493.8794 | 521.6804 | 552.4963 |
| ENSMUSG00000059586 | -0.009183961 | 0.914071 | 0.949091 | 1044.284 | 1077.44 | 1028.508 | 1189.558 | 1122.353 | 1166.539 | 1080.004 | 999.4962 |
| ENSMUSG00000059588 | -0.28086143 | 0.054289 | 0.144444 | 886.3339 | 868.6708 | 1256.238 | 946.298 | 1286.172 | 1011.224 | 1239.112 | 1274.657 |
| ENSMUSG00000059602 | -0.44632796 | 0.119715 | 0.248615 | 65.39353 | 53.99197 | 69.56096 | 66.42201 | 78.0089 | 50.28183 | 107.0361 | 113.1095 |
| ENSMUSG00000059625 | -3.384075285 | NA | NA | 3.018163 | 3.599465 | 0 | 4.313117 | 0 | 113.9722 | 0 | 0 |
| ENSMUSG00000059647 | 0.198302941 | 0.698596 | 0.809829 | 15.09082 | 17.99732 | 10.76539 | 12.07673 | 7.80089 | 18.99536 | 6.750024 | 15.22628 |
| ENSMUSG00000059657 | 0.158969135 | 0.779627 | 0.863998 | 57.3451 | 9.598572 | 19.87456 | 25.01608 | 34.1289 | 14.52586 | 31.82154 | 19.57664 |
| ENSMUSG00000059659 | 0.36293769 | 0.242049 | 0.404445 | 54.32694 | 85.18733 | 58.79557 | 69.00988 | 35.10401 | 78.21619 | 52.07161 | 42.41605 |
| ENSMUSG00000059668 | -1.785413391 | 0.098777 | 0.219219 | 3.018163 | 0 | 43.88965 | 2.58787 | 56.55646 | 11.17374 | 42.42872 | 61.99269 |
| ENSMUSG00000059669 | 0.103113817 | 0.155566 | 0.297525 | 734.4197 | 679.099 | 688.1566 | 737.5431 | 666.9761 | 673.7766 | 664.3952 | 639.5036 |
| ENSMUSG00000059674 | 0.282124666 | 0.058656 | 0.152173 | 311.8769 | 379.1436 | 334.5551 | 292.4294 | 244.7529 | 322.9211 | 233.358 | 282.7737 |
| ENSMUSG00000059689 | 0.18246005 | 0.02715 | 0.088078 | 816.9161 | 908.2649 | 847.1531 | 959.2373 | 764.4873 | 745.2885 | 811.9315 | 789.5911 |
| ENSMUSG00000059708 | 0.088804261 | 0.532262 | 0.683893 | 383.3067 | 350.3479 | 328.7583 | 328.6596 | 341.289 | 395.5504 | 278.6796 | 292.562 |
| ENSMUSG00000059713 | -0.015544964 | 0.870772 | 0.922508 | 609.6689 | 641.9045 | 727.0776 | 623.6768 | 600.6686 | 667.0723 | 695.2525 | 669.9561 |
| ENSMUSG00000059714 | -0.083183253 | 0.466982 | 0.627585 | 1041.266 | 949.0588 | 1122.084 | 1151.602 | 1018.016 | 1005.637 | 1245.862 | 1250.73 |
| ENSMUSG00000059734 | -0.734999854 | 2.08E-06 | 6.61E-05 | 1733.432 | 1716.945 | 1978.347 | 1730.423 | 3157.41 | 2133.067 | 2990.261 | 3635.817 |
| ENSMUSG00000059742 | 0.516337101 | 0.01943 | 0.06908 | 195.1745 | 220.7672 | 285.6968 | 212.2054 | 108.2374 | 203.3621 | 158.1434 | 170.7518 |
| ENSMUSG00000059743 | -0.780531597 | 2.99E-06 | 9.01E-05 | 730.3955 | 549.5183 | 818.9975 | 630.5778 | 1168.183 | 919.5989 | 1412.684 | 1190.912 |
| ENSMUSG00000059751 | 0.086308976 | 0.722701 | 0.826557 | 66.39959 | 113.983 | 74.5296 | 80.22398 | 71.18313 | 86.03781 | 92.57176 | 64.16787 |
| ENSMUSG00000059772 | 0.254059166 | 0.06915 | 0.170734 | 611.681 | 735.4906 | 615.2832 | 595.2102 | 570.4401 | 640.2554 | 481.1803 | 451.3503 |
| ENSMUSG00000059775 | -0.318370782 | 0.080273 | 0.18938 | 124.7507 | 169.1748 | 132.4971 | 127.6683 | 195.0223 | 134.0849 | 170.6792 | 188.1533 |
| ENSMUSG00000059776 | -0.251223035 | 0.599026 | 0.737435 | 10.06054 | 13.19804 | 11.59349 | 7.763611 | 14.62667 | 14.52586 | 13.50005 | 7.613138 |
| ENSMUSG00000059791 | -0.363915206 | 0.001851 | 0.012413 | 401.4157 | 487.1275 | 486.9267 | 455.4652 | 641.6232 | 512.8747 | 636.4308 | 563.3722 |
| ENSMUSG00000059796 | -0.236112034 | 0.000595 | 0.005238 | 18459.09 | 19332.72 | 19841.43 | 19252.89 | 24210.06 | 20358.56 | 22649.22 | 23338.62 |
| ENSMUSG00000059810 | -0.110433697 | 0.262688 | 0.427035 | 776.674 | 838.6753 | 868.6839 | 784.1248 | 918.5549 | 770.9881 | 858.2174 | 979.9196 |
| ENSMUSG00000059811 | -0.180738758 | 0.055917 | 0.147283 | 1464.815 | 1339.001 | 1147.756 | 1353.456 | 1475.343 | 1407.891 | 1597.827 | 1528.065 |
| ENSMUSG00000059820 | 0.022838604 | 0.770566 | 0.857548 | 1071.448 | 1060.642 | 955.6351 | 1095.532 | 1071.647 | 1064.858 | 1017.325 | 961.4305 |
| ENSMUSG00000059824 | -0.006915312 | 0.962886 | 0.978958 | 336.0222 | 329.9509 | 369.3356 | 309.6818 | 366.6419 | 406.7242 | 283.501 | 295.8248 |
| ENSMUSG00000059834 | -0.075898412 | 0.405844 | 0.572524 | 760.5771 | 742.6895 | 688.9847 | 766.8723 | 818.1184 | 786.6314 | 826.3958 | 685.1824 |
| ENSMUSG00000059839 | 0.542698554 | 0.000169 | 0.00199 | 475.8637 | 494.3265 | 336.2113 | 409.7462 | 303.2596 | 298.3389 | 293.1439 | 280.5985 |
| ENSMUSG00000059842 | 0.076699964 | 0.73992 | 0.837336 | 170.0232 | 139.1793 | 181.3554 | 133.7066 | 195.9974 | 154.1976 | 142.7148 | 98.97079 |
| ENSMUSG00000059851 | -0.082221201 | 0.239781 | 0.401724 | 1512.1 | 1570.566 | 1401.985 | 1412.115 | 1499.721 | 1509.572 | 1586.256 | 1641.175 |
| ENSMUSG00000059852 | -0.095547898 | 0.849852 | 0.909743 | 17.10292 | 19.19714 | 29.81184 | 29.3292 | 10.72622 | 15.64324 | 33.75012 | 42.41605 |
| ENSMUSG00000059857 | 1.455278819 | 5.44E-05 | 0.000845 | 815.9101 | 1163.827 | 704.7187 | 1101.57 | 302.2845 | 670.4245 | 207.3222 | 201.2044 |
| ENSMUSG00000059866 | 0.113847449 | 0.313338 | 0.481484 | 801.8253 | 609.5093 | 679.8755 | 720.2906 | 705.9806 | 665.955 | 642.2166 | 585.124 |
| ENSMUSG00000059878 | 0.036788328 | 0.646386 | 0.771051 | 2900.455 | 3070.343 | 2866.905 | 3240.014 | 3162.286 | 3082.835 | 2853.332 | 2674.386 |
| ENSMUSG00000059883 | 0.122189136 | 0.536695 | 0.687291 | 524.1543 | 538.7199 | 732.0463 | 471.8551 | 523.6348 | 383.2593 | 512.0375 | 664.5182 |
| ENSMUSG00000059890 | 0.13795612 | 0.036721 | 0.110036 | 2366.24 | 2309.656 | 2218.498 | 2368.764 | 2019.456 | 2262.683 | 2149.401 | 1987.029 |
| ENSMUSG00000059895 | -0.218594481 | 0.013886 | 0.054426 | 1407.47 | 1678.55 | 1754.758 | 1697.643 | 1962.899 | 1855.958 | 1809.971 | 1981.591 |
| ENSMUSG00000059897 | -0.073133873 | 0.639615 | 0.766541 | 333.004 | 377.9438 | 304.7432 | 297.6051 | 364.6916 | 377.6724 | 375.1085 | 261.0219 |
| ENSMUSG00000059900 | 0.135663128 | 0.739159 | 0.836877 | 37.22401 | 29.99554 | 65.42042 | 46.58167 | 62.40712 | 26.81698 | 49.17875 | 25.0146 |
| ENSMUSG00000059901 | 0.121530253 | 0.564249 | 0.709377 | 136.8234 | 104.3845 | 163.9651 | 156.1349 | 96.53602 | 139.6718 | 139.8219 | 142.4744 |
| ENSMUSG00000059920 | 0.037886884 | 0.711777 | 0.819301 | 888.346 | 962.2569 | 1008.634 | 970.4514 | 908.8037 | 1092.792 | 876.5389 | 854.8466 |
| ENSMUSG00000059921 | 0.125955719 | 0.388631 | 0.55606 | 1693.189 | 1992.904 | 1792.023 | 1890.008 | 1660.615 | 2197.875 | 1453.184 | 1441.058 |
| ENSMUSG00000059923 | -0.109193253 | 0.110737 | 0.236865 | 1890.376 | 2008.501 | 2162.186 | 1983.171 | 2116.967 | 2070.494 | 2276.687 | 2214.335 |
| ENSMUSG00000059939 | 0.259151495 | 0.017796 | 0.065089 | 859.1704 | 838.6753 | 726.2495 | 751.3451 | 646.4988 | 731.88 | 698.1454 | 574.2481 |
| ENSMUSG00000059970 | 0.957054479 | 6.49E-13 | 2.03E-10 | 444.676 | 476.3291 | 398.3193 | 523.6125 | 244.7529 | 210.0663 | 227.5722 | 266.4598 |
| ENSMUSG00000059974 | 0.473845271 | 0.05015 | 0.136353 | 493.9727 | 736.6904 | 653.3761 | 649.5555 | 325.6872 | 702.8283 | 378.0014 | 418.7226 |
| ENSMUSG00000059975 | 0.184767349 | 0.188313 | 0.340615 | 375.2583 | 431.9358 | 303.9151 | 360.5766 | 301.3094 | 366.4987 | 301.8225 | 323.0146 |
| ENSMUSG00000059981 | -0.053447251 | 0.50357 | 0.659515 | 2497.027 | 2244.866 | 2565.474 | 2333.397 | 2446.554 | 2289.5 | 2569.831 | 2701.576 |
| ENSMUSG00000059991 | -0.035561864 | 0.875092 | 0.925241 | 115.6963 | 176.3738 | 162.3089 | 173.3873 | 188.1965 | 144.1413 | 194.7864 | 115.2847 |
| ENSMUSG00000059995 | 0.080260706 | 0.256618 | 0.42028 | 1826.995 | 1698.947 | 1763.867 | 1833.938 | 1682.067 | 1535.272 | 1742.471 | 1779.299 |
| ENSMUSG00000060002 | -0.978963672 | 1.21E-07 | 6.61E-06 | 1705.262 | 1708.546 | 2216.841 | 1740.774 | 3212.992 | 2664.937 | 3750.121 | 4903.948 |

| | | | | | | | | | | | |
|---|---|---|---|---|---|---|---|---|---|---|---|
| ENSMUSG00000060012 | -0.556876202 | 0.032316 | 0.099704 | 273.6468 | 221.967 | 284.8687 | 202.7165 | 374.4427 | 210.0663 | 342.3227 | 519.8685 |
| ENSMUSG00000060019 | -0.103464906 | 0.56428 | 0.709377 | 88.53278 | 101.9848 | 108.482 | 112.1411 | 129.6898 | 94.9768 | 114.7504 | 102.2336 |
| ENSMUSG00000060029 | 0.1966511 | 0.675689 | 0.792479 | 49.29666 | 53.99197 | 28.15563 | 50.89479 | 25.35289 | 81.56831 | 24.10723 | 28.27737 |
| ENSMUSG00000060032 | -0.302987439 | 0.048334 | 0.13299 | 1786.753 | 1367.797 | 1536.966 | 1404.351 | 1963.874 | 1420.182 | 1877.471 | 2258.927 |
| ENSMUSG00000060034 | 0.637714149 | 0.292456 | 0.459782 | 14.08476 | 10.79839 | 9.93728 | 15.52722 | 5.850668 | 16.76061 | 6.750024 | 3.262773 |
| ENSMUSG00000060036 | -0.136603607 | 0.131746 | 0.264754 | 46475.69 | 55201.39 | 50353.02 | 50315.97 | 62347.64 | 50209.21 | 57102.31 | 52777.53 |
| ENSMUSG00000060038 | -0.10460355 | 0.203909 | 0.359853 | 617.7174 | 610.7092 | 611.9708 | 669.3958 | 721.5824 | 645.8422 | 691.3953 | 639.5036 |
| ENSMUSG00000060044 | 0.215387099 | 0.271725 | 0.436965 | 335.0161 | 297.5557 | 488.5829 | 319.1707 | 246.7032 | 311.7474 | 356.787 | 327.3649 |
| ENSMUSG00000060063 | 0.740842254 | 6.26E-10 | 8.49E-08 | 359.1614 | 340.7493 | 308.8838 | 321.7586 | 186.2463 | 187.7188 | 224.6794 | 195.7664 |
| ENSMUSG00000060073 | -0.088042351 | 0.124566 | 0.25468 | 3809.928 | 4040.999 | 4284.624 | 4114.714 | 4223.207 | 4423.684 | 4210.086 | 4422.145 |
| ENSMUSG00000060090 | 0.238597527 | 0.005363 | 0.026954 | 708.2623 | 765.4861 | 688.1566 | 772.9107 | 577.2659 | 619.0252 | 665.3595 | 624.2773 |
| ENSMUSG00000060098 | -0.231253801 | 0.024406 | 0.081555 | 929.5942 | 1043.845 | 856.2623 | 928.1829 | 1180.86 | 984.4066 | 1041.432 | 1199.613 |
| ENSMUSG00000060121 | 0.049795813 | 0.612821 | 0.746981 | 475.8637 | 488.3274 | 431.4436 | 470.9924 | 492.4312 | 424.6022 | 465.7517 | 416.5474 |
| ENSMUSG00000060126 | -0.149130269 | 0.064364 | 0.162993 | 61526.26 | 75394.39 | 73126.78 | 71444.2 | 81807.94 | 74033.86 | 80378.32 | 75923.65 |
| ENSMUSG00000060131 | 0.531405161 | 0.332893 | 0.501009 | 23.13925 | 28.79572 | 5.796746 | 16.38985 | 9.751113 | 14.52586 | 14.46434 | 11.9635 |
| ENSMUSG00000060143 | -0.303392517 | 0.032276 | 0.099628 | 3384.367 | 4278.564 | 5276.695 | 4344.172 | 5814.589 | 4548.83 | 5589.02 | 5378.138 |
| ENSMUSG00000060147 | 0.321042252 | 4.14E-05 | 0.000692 | 2879.328 | 2908.367 | 2691.347 | 2963.112 | 2468.007 | 2391.181 | 2245.829 | 2052.284 |
| ENSMUSG00000060149 | 0.11787115 | 0.568787 | 0.712804 | 189.1382 | 281.9581 | 281.5563 | 255.3366 | 227.2009 | 300.5736 | 228.5365 | 172.927 |
| ENSMUSG00000060152 | -0.556708352 | 0.000532 | 0.004815 | 489.9485 | 469.1302 | 543.238 | 458.0531 | 688.4286 | 528.5179 | 810.9672 | 855.9342 |
| ENSMUSG00000060166 | 0.007673507 | 0.926783 | 0.956532 | 1067.424 | 1054.643 | 1147.756 | 1021.346 | 1075.548 | 956.4722 | 1092.54 | 1144.146 |
| ENSMUSG00000060176 | 0.566994661 | 0.004044 | 0.022004 | 141.8537 | 99.58519 | 143.2624 | 159.5853 | 83.85957 | 103.9158 | 94.50034 | 87.00729 |
| ENSMUSG00000060181 | -0.038331991 | 0.702591 | 0.812521 | 502.0211 | 506.3247 | 585.4714 | 590.8971 | 547.0374 | 606.7341 | 527.4662 | 567.7225 |
| ENSMUSG00000060198 | 0.277600653 | 0.210347 | 0.367817 | 56.33904 | 64.79036 | 59.62368 | 50.03216 | 51.6809 | 49.16446 | 45.32159 | 43.50364 |
| ENSMUSG00000060204 | -0.243486297 | 0.516735 | 0.670807 | 13.07871 | 15.59768 | 18.21835 | 18.97772 | 22.42756 | 16.76061 | 16.39292 | 22.83941 |
| ENSMUSG00000060206 | 0.479597243 | 0.005224 | 0.026524 | 1474.876 | 1339.001 | 1171.771 | 1306.012 | 878.5753 | 1308.445 | 788.7885 | 820.0437 |
| ENSMUSG00000060212 | 0.293168689 | 0.377541 | 0.545603 | 40.24217 | 69.58965 | 30.63995 | 64.69676 | 48.75557 | 41.34284 | 40.50014 | 35.89051 |
| ENSMUSG00000060216 | 0.01000872 | 0.915818 | 0.950114 | 532.2028 | 623.9072 | 635.9859 | 576.2325 | 550.9379 | 593.3256 | 626.788 | 580.7736 |
| ENSMUSG00000060224 | -0.866046136 | 0.00037 | 0.003629 | 185.114 | 182.3729 | 231.8699 | 170.7995 | 352.9903 | 244.7049 | 287.3582 | 519.8685 |
| ENSMUSG00000060227 | 0.205701788 | 0.01825 | 0.066208 | 2006.072 | 2194.474 | 1880.63 | 2007.325 | 1621.61 | 1956.522 | 1760.792 | 1672.715 |
| ENSMUSG00000060240 | -0.230829152 | 0.511637 | 0.667063 | 58.35115 | 35.99465 | 55.48314 | 86.26235 | 67.28268 | 42.46022 | 68.46453 | 100.0584 |
| ENSMUSG00000060244 | 0.132719515 | 0.422058 | 0.587229 | 125.7568 | 134.38 | 135.8095 | 118.1794 | 104.3369 | 125.1459 | 103.1789 | 137.0365 |
| ENSMUSG00000060260 | -0.525060554 | 0.01089 | 0.04548 | 192.1564 | 165.5754 | 240.979 | 183.7388 | 255.4792 | 203.3621 | 332.6798 | 336.0656 |
| ENSMUSG00000060261 | 0.067344284 | 0.22764 | 0.387612 | 8454.881 | 9256.623 | 9131.532 | 8958.345 | 8086.598 | 8892.063 | 8505.03 | 8686.59 |
| ENSMUSG00000060268 | -0.301316805 | 0.624038 | 0.754981 | 10.06054 | 8.398751 | 12.4216 | 6.038364 | 17.552 | 16.76061 | 4.821446 | 6.525547 |
| ENSMUSG00000060275 | 0.386342103 | 0.3192 | 0.487276 | 34.20585 | 87.58697 | 65.42042 | 60.38364 | 39.00445 | 72.62932 | 27.96439 | 50.02919 |
| ENSMUSG00000060279 | -0.227974187 | 0.052727 | 0.141217 | 1542.281 | 1569.367 | 1865.724 | 1490.613 | 2065.286 | 1604.549 | 1820.578 | 2086 |
| ENSMUSG00000060284 | 0.107915449 | 0.778083 | 0.863206 | 35.2119 | 31.19536 | 52.17072 | 27.60395 | 44.85512 | 18.99536 | 43.39301 | 28.27737 |
| ENSMUSG00000060288 | -0.135315211 | 0.089131 | 0.203791 | 722.347 | 808.6797 | 763.5143 | 710.8018 | 861.0233 | 842.5001 | 773.3599 | 821.1313 |
| ENSMUSG00000060301 | 0.281704549 | 0.046702 | 0.129778 | 1335.034 | 1403.791 | 1070.742 | 1003.231 | 980.962 | 1148.661 | 946.932 | 879.8612 |
| ENSMUSG00000060314 | 0.146464963 | 0.630266 | 0.759281 | 86.52067 | 58.79126 | 74.5296 | 55.2079 | 52.65601 | 94.9768 | 56.89306 | 44.59123 |
| ENSMUSG00000060317 | -4.490562195 | 0.000742 | 0.006219 | 0 | 0 | 14.90592 | 20.70296 | 253.5289 | 84.92043 | 181.2864 | 287.124 |
| ENSMUSG00000060336 | 0.602670851 | 0.007909 | 0.035924 | 307.8526 | 290.3568 | 233.5261 | 326.9343 | 195.0223 | 272.6393 | 143.6791 | 152.2628 |
| ENSMUSG00000060371 | 2.587254458 | 3.89E-06 | 0.000111 | 161.9748 | 39.59411 | 37.2648 | 94.88858 | 12.67645 | 17.87799 | 6.750024 | 18.48905 |
| ENSMUSG00000060373 | -0.05697439 | 0.279921 | 0.446314 | 7232.525 | 8001.61 | 7703.876 | 7996.52 | 8034.917 | 8230.578 | 7852.207 | 8063.4 |
| ENSMUSG00000060376 | -0.525909148 | 8.16E-05 | 0.001155 | 712.2865 | 641.9045 | 755.2332 | 798.7894 | 1174.034 | 822.3873 | 1120.504 | 1072.365 |
| ENSMUSG00000060377 | 0.420545184 | 0.270375 | 0.435432 | 20.12109 | 17.99732 | 20.70267 | 19.84034 | 10.72622 | 13.40849 | 16.39292 | 18.48905 |
| ENSMUSG00000060380 | -0.631834726 | 0.132213 | 0.265317 | 19.11503 | 10.79839 | 28.15563 | 14.6646 | 39.00445 | 29.05173 | 25.07152 | 20.66423 |
| ENSMUSG00000060397 | 0.140614635 | 0.295279 | 0.463053 | 195.1745 | 239.9643 | 184.6678 | 206.167 | 195.0223 | 175.4277 | 192.8578 | 182.7153 |
| ENSMUSG00000060402 | -0.584547652 | 0.01803 | 0.065724 | 199.1988 | 118.7823 | 206.1986 | 160.448 | 219.4 | 191.071 | 267.1081 | 351.2919 |
| ENSMUSG00000060419 | -0.067192382 | 0.584926 | 0.7254 | 537.233 | 673.0999 | 712.9998 | 633.1656 | 716.7068 | 702.8283 | 684.6453 | 574.2481 |
| ENSMUSG00000060424 | -0.520604364 | 0.007966 | 0.036131 | 213.2835 | 153.5772 | 226.0731 | 213.9306 | 250.6036 | 231.2964 | 363.537 | 314.3138 |
| ENSMUSG00000060427 | -0.050072659 | 0.649293 | 0.77287 | 652.9293 | 689.8974 | 570.5655 | 646.9676 | 667.9512 | 757.5796 | 594.0021 | 629.7152 |
| ENSMUSG00000060429 | -0.185485196 | 0.453751 | 0.616039 | 142.8597 | 113.983 | 115.9349 | 104.3774 | 134.5654 | 88.27255 | 130.179 | 189.2408 |
| ENSMUSG00000060438 | -0.050020829 | 0.833179 | 0.898985 | 50.30272 | 55.19179 | 41.40533 | 43.13117 | 49.73068 | 41.34284 | 54.00019 | 50.02919 |
| ENSMUSG00000060441 | 0.93905307 | 0.04018 | 0.117057 | 43.26034 | 55.19179 | 46.37397 | 20.70296 | 25.35289 | 33.52122 | 16.39292 | 10.87591 |
| ENSMUSG00000060445 | -1.073740558 | 0.230754 | 0.391182 | 3.018163 | 8.398751 | 8.281066 | 14.6646 | 8.776002 | 52.51658 | 2.892867 | 8.700729 |
| ENSMUSG00000060450 | -0.178662437 | 0.00682 | 0.032358 | 1879.31 | 1926.913 | 2110.844 | 2042.692 | 2194.976 | 2163.236 | 2302.723 | 2353.547 |
| ENSMUSG00000060459 | -3.095046969 | 0.00012 | 0.001543 | 120.7265 | 63.59054 | 568.9093 | 141.4703 | 1377.832 | 586.6214 | 2700.01 | 2985.438 |
| ENSMUSG00000060461 | 1.455624048 | 0.043718 | 0.124058 | 19.11503 | 7.198929 | 22.35888 | 8.626235 | 1.950223 | 11.17374 | 4.821446 | 3.262773 |
| ENSMUSG00000060467 | -0.333389728 | 0.375692 | 0.544124 | 24.1453 | 10.79839 | 25.67131 | 17.25247 | 26.32801 | 26.81698 | 22.17865 | 23.927 |
| ENSMUSG00000060470 | -0.501936987 | 0.108114 | 0.232775 | 103.6236 | 45.59322 | 120.0755 | 69.8725 | 105.312 | 97.21155 | 143.6791 | 135.9489 |
| ENSMUSG00000060475 | -0.061408444 | 0.23017 | 0.390428 | 2885.364 | 2885.571 | 2745.173 | 2875.124 | 3109.63 | 2886.177 | 2966.153 | 2919.094 |
| ENSMUSG00000060477 | -0.032610331 | 0.825788 | 0.8942 | 333.004 | 328.7511 | 409.9128 | 327.7969 | 437.825 | 336.3296 | 311.4654 | 346.9416 |
| ENSMUSG00000060487 | 0.197499063 | 0.416339 | 0.582046 | 70.4238 | 119.9822 | 90.26362 | 88.85022 | 90.68535 | 87.15518 | 57.85735 | 85.9197 |

| | | | | | | | | | | | |
|---|---|---|---|---|---|---|---|---|---|---|---|
| ENSMUSG00000060510 | 0.304076127 | 0.022459 | 0.076505 | 3555.396 | 3785.437 | 3099.603 | 3583.338 | 2830.748 | 3549.898 | 2579.474 | 2398.138 |
| ENSMUSG00000060512 | -0.351367432 | 0.002329 | 0.014615 | 796.795 | 845.8742 | 954.8069 | 860.8982 | 1008.265 | 967.646 | 1165.826 | 1272.482 |
| ENSMUSG00000060519 | 0.215892531 | 0.057211 | 0.14975 | 541.2572 | 607.1097 | 637.6421 | 559.8426 | 541.1868 | 445.8323 | 562.1806 | 468.7518 |
| ENSMUSG00000060538 | -0.336562226 | 0.007995 | 0.03621 | 341.0524 | 370.7449 | 452.9743 | 351.0878 | 510.9583 | 429.0717 | 474.4303 | 501.3795 |
| ENSMUSG00000060548 | 0.527985268 | 0.002266 | 0.014359 | 1443.688 | 1588.564 | 1224.77 | 1321.539 | 913.6793 | 1320.736 | 792.6457 | 841.7955 |
| ENSMUSG00000060550 | -0.512670838 | 0.013849 | 0.054358 | 95.57516 | 105.5843 | 170.59 | 129.3935 | 164.7938 | 175.4277 | 180.3221 | 197.9416 |
| ENSMUSG00000060568 | 0.210123084 | 0.27646 | 0.44271 | 104.6297 | 74.38894 | 80.32634 | 96.61383 | 81.90935 | 78.21619 | 75.21455 | 72.8686 |
| ENSMUSG00000060572 | 0.471674764 | 0.000144 | 0.001766 | 5122.829 | 6649.411 | 5892.807 | 6134.978 | 4219.307 | 5118.691 | 4172.479 | 3652.131 |
| ENSMUSG00000060579 | -0.428271849 | 0.031365 | 0.09742 | 70.4238 | 57.59143 | 61.27989 | 68.14726 | 97.51113 | 72.62932 | 94.50034 | 81.56933 |
| ENSMUSG00000060586 | -1.376591223 | 1.17E-08 | 9.73E-07 | 65.39353 | 98.38537 | 96.88848 | 113.0037 | 314.961 | 170.9582 | 249.7509 | 234.9197 |
| ENSMUSG00000060591 | 0.086478423 | 0.278381 | 0.444638 | 3717.371 | 4265.366 | 4468.463 | 4278.613 | 4246.61 | 3787.898 | 3803.156 | 3919.678 |
| ENSMUSG00000060594 | 0.0969919 | 0.618191 | 0.750811 | 174.0474 | 230.3657 | 202.8861 | 210.4801 | 193.072 | 241.3528 | 193.8221 | 135.9489 |
| ENSMUSG00000060600 | 0.519446502 | 0.011505 | 0.047468 | 591.56 | 799.0811 | 544.8942 | 634.0283 | 436.8499 | 608.9689 | 426.2158 | 319.7518 |
| ENSMUSG00000060601 | -0.071487011 | 0.464797 | 0.625696 | 1795.807 | 1755.339 | 1989.94 | 1918.475 | 2122.817 | 1660.418 | 1981.614 | 2075.124 |
| ENSMUSG00000060636 | -0.089113693 | 0.211135 | 0.368553 | 18533.53 | 20687.32 | 20744.07 | 19996.48 | 22713.27 | 19748.47 | 22154.54 | 20436.92 |
| ENSMUSG00000060639 | -0.193349215 | 0.179365 | 0.329159 | 266.6044 | 340.7493 | 270.7909 | 324.3464 | 400.7707 | 299.4563 | 323.0369 | 349.1167 |
| ENSMUSG00000060657 | 0.475521176 | 0.002282 | 0.014425 | 2351.149 | 1990.504 | 1560.153 | 1783.905 | 1274.47 | 1703.995 | 1316.255 | 1232.241 |
| ENSMUSG00000060671 | 0.02781748 | 0.730178 | 0.830734 | 1607.675 | 1568.167 | 1512.123 | 1666.589 | 1659.639 | 1379.957 | 1575.648 | 1616.16 |
| ENSMUSG00000060675 | -0.776667705 | 0.000237 | 0.002595 | 304.8345 | 299.9554 | 333.727 | 309.6818 | 565.5646 | 339.6817 | 508.1804 | 725.4233 |
| ENSMUSG00000060679 | -0.052837465 | 0.388112 | 0.555703 | 1717.335 | 1588.564 | 1714.181 | 1730.423 | 1773.727 | 1834.728 | 1690.399 | 1709.693 |
| ENSMUSG00000060680 | -0.75092854 | 0.084053 | 0.1954 | 15.09082 | 15.59768 | 18.21835 | 6.038364 | 25.35289 | 23.46486 | 18.32149 | 25.0146 |
| ENSMUSG00000060681 | -0.070986337 | 0.262887 | 0.427212 | 808.8677 | 832.6761 | 827.2785 | 828.1186 | 897.1024 | 857.0259 | 840.8602 | 867.8977 |
| ENSMUSG00000060703 | 0.261410096 | 0.032448 | 0.099966 | 648.9051 | 700.6958 | 620.2519 | 770.3228 | 644.5486 | 610.0863 | 537.1091 | 493.7664 |
| ENSMUSG00000060708 | -0.122622623 | 0.354059 | 0.522769 | 423.5489 | 367.1454 | 392.5225 | 484.7944 | 472.929 | 381.0246 | 487.9303 | 475.2773 |
| ENSMUSG00000060716 | -1.002987508 | 0.000736 | 0.006174 | 215.2956 | 165.5754 | 293.9779 | 223.4195 | 512.9085 | 214.5358 | 517.8233 | 556.8466 |
| ENSMUSG00000060733 | -0.619638788 | 9.10E-06 | 0.000213 | 716.3107 | 619.1079 | 741.1554 | 671.1211 | 1057.021 | 811.2136 | 1127.254 | 1227.89 |
| ENSMUSG00000060739 | 0.110669106 | 0.161544 | 0.305415 | 6384.421 | 7058.55 | 6673.711 | 6962.234 | 6411.357 | 6900.902 | 6132.879 | 5633.722 |
| ENSMUSG00000060743 | -0.125171815 | 0.096026 | 0.214632 | 11566.61 | 13724.76 | 13072.49 | 13288.71 | 15139.58 | 13726.94 | 14008.23 | 13456.76 |
| ENSMUSG00000060771 | 0.274409985 | 0.089281 | 0.203986 | 245.4773 | 230.3657 | 237.6666 | 216.5185 | 196.9725 | 243.5876 | 164.8934 | 164.2263 |
| ENSMUSG00000060780 | 0.222429327 | 0.218822 | 0.377614 | 179.0777 | 196.7707 | 199.5737 | 189.7772 | 154.0676 | 217.8879 | 155.2506 | 129.4233 |
| ENSMUSG00000060791 | -0.133167322 | 0.463198 | 0.624014 | 133.8052 | 106.7841 | 159.8246 | 131.1188 | 121.8889 | 155.315 | 144.6434 | 164.2263 |
| ENSMUSG00000060798 | 0.162200484 | 0.378667 | 0.54683 | 324.9556 | 271.1597 | 271.619 | 351.0878 | 291.5583 | 348.6207 | 229.5008 | 220.781 |
| ENSMUSG00000060802 | 0.417823402 | 0.000429 | 0.004068 | 3402.476 | 4315.758 | 4466.807 | 3355.605 | 2835.624 | 2967.746 | 3130.083 | 2698.313 |
| ENSMUSG00000060803 | -0.064176487 | 0.466455 | 0.627229 | 1364.21 | 1647.355 | 1627.23 | 1504.415 | 1581.631 | 1508.455 | 1594.934 | 1736.883 |
| ENSMUSG00000060807 | -5.350673139 | 0.001465 | 0.0104 | 6.036326 | 0 | 388.382 | 22.42821 | 3957.977 | 551.9828 | 5103.983 | 7404.32 |
| ENSMUSG00000060860 | -0.110257128 | 0.060041 | 0.155025 | 2381.331 | 2494.429 | 2608.536 | 2682.759 | 2693.257 | 2723.041 | 2776.189 | 2787.496 |
| ENSMUSG00000060862 | -0.091842767 | 0.449382 | 0.611998 | 371.2341 | 327.5513 | 323.7897 | 276.9021 | 339.3387 | 373.2029 | 340.3941 | 330.6277 |
| ENSMUSG00000060882 | 0.693801859 | 2.18E-05 | 0.000422 | 212.2775 | 303.5548 | 291.4935 | 271.7264 | 156.9929 | 166.4887 | 149.4648 | 194.6788 |
| ENSMUSG00000060890 | 0.274061243 | 0.387916 | 0.555551 | 31.18769 | 28.79572 | 24.8432 | 29.3292 | 25.35289 | 23.46486 | 17.3572 | 28.27737 |
| ENSMUSG00000060904 | -0.006305372 | 0.920173 | 0.95244 | 3829.043 | 4396.146 | 4011.349 | 4290.689 | 4132.522 | 4259.43 | 4150.301 | 4054.54 |
| ENSMUSG00000060923 | 0.184259162 | 0.185148 | 0.336625 | 241.453 | 197.9706 | 217.792 | 179.4257 | 185.2711 | 182.132 | 176.4649 | 192.5036 |
| ENSMUSG00000060924 | 0.805337566 | 0.132257 | 0.265376 | 19.11503 | 9.598572 | 13.24971 | 35.36756 | 9.751113 | 11.17374 | 9.642892 | 14.13868 |
| ENSMUSG00000060935 | 0.443185156 | 0.012324 | 0.049967 | 1578.499 | 1866.922 | 1388.735 | 1824.449 | 1057.996 | 1677.179 | 1128.218 | 1034.299 |
| ENSMUSG00000060938 | -0.199909612 | 0.015527 | 0.059143 | 21295.15 | 25371.43 | 26075.42 | 24503.68 | 29999.3 | 26936.54 | 28083.96 | 26678.61 |
| ENSMUSG00000060950 | -0.275302286 | 0.050986 | 0.137935 | 321.9374 | 388.7422 | 358.5702 | 291.5667 | 449.5263 | 375.4377 | 359.6799 | 460.051 |
| ENSMUSG00000060961 | -0.687153588 | 4.46E-05 | 0.000729 | 1175.071 | 1306.606 | 1347.329 | 1469.048 | 2350.993 | 1588.906 | 1872.65 | 2718.978 |
| ENSMUSG00000060962 | 1.631870058 | 6.42E-06 | 0.000163 | 152.9203 | 261.5611 | 152.3716 | 215.6559 | 77.03379 | 65.92507 | 84.85745 | 23.927 |
| ENSMUSG00000060969 | -2.348342126 | 0.005215 | 0.026488 | 183.1019 | 23.99643 | 450.49 | 184.6014 | 631.8721 | 404.4894 | 1668.22 | 1586.795 |
| ENSMUSG00000060981 | -0.311946305 | 0.553379 | 0.700443 | 7.04238 | 23.99643 | 26.49941 | 10.35148 | 12.67645 | 22.34748 | 23.14294 | 26.10219 |
| ENSMUSG00000060985 | -2.058616549 | 6.83E-06 | 0.000172 | 23.13925 | 15.59768 | 36.43669 | 16.38985 | 39.00445 | 91.62468 | 101.2504 | 152.2628 |
| ENSMUSG00000060988 | 0.181348736 | 0.643906 | 0.769677 | 17.10292 | 39.59411 | 21.53077 | 22.42821 | 26.32801 | 23.46486 | 16.39292 | 21.75182 |
| ENSMUSG00000060989 | -0.486498847 | 0.065985 | 0.165533 | 37.22401 | 51.59233 | 36.43669 | 39.68068 | 74.10846 | 52.51658 | 46.28588 | 56.55474 |
| ENSMUSG00000060992 | -0.146191721 | 0.05832 | 0.151607 | 3060.417 | 3479.482 | 3505.375 | 3243.464 | 3785.382 | 3376.705 | 3621.87 | 3920.766 |
| ENSMUSG00000061013 | 1.374245746 | 3.95E-05 | 0.000667 | 650.9172 | 470.33 | 217.792 | 567.6063 | 177.4703 | 240.2354 | 204.4293 | 113.1095 |
| ENSMUSG00000061024 | -0.328711532 | 0.000278 | 0.002922 | 1159.981 | 1178.225 | 1174.255 | 1208.536 | 1614.784 | 1269.337 | 1485.005 | 1558.518 |
| ENSMUSG00000061028 | 0.260591931 | 0.001096 | 0.008382 | 1446.706 | 1536.971 | 1349.814 | 1560.486 | 1322.251 | 1233.581 | 1198.611 | 1162.635 |
| ENSMUSG00000061032 | -0.172588319 | 0.008505 | 0.03796 | 6337.136 | 6362.654 | 6687.789 | 6474.852 | 7312.36 | 6612.62 | 7442.384 | 7782.802 |
| ENSMUSG00000061039 | 0.724466851 | 0.115625 | 0.243604 | 37.22401 | 25.19625 | 37.2648 | 29.3292 | 17.552 | 36.87335 | 15.42863 | 8.700729 |
| ENSMUSG00000061046 | -0.158379576 | 0.081378 | 0.191131 | 943.679 | 1132.632 | 1122.913 | 948.0232 | 1118.453 | 1163.186 | 1153.29 | 1192 |
| ENSMUSG00000061048 | 0.365503275 | 4.68E-05 | 0.000754 | 1732.426 | 1868.122 | 1738.196 | 1649.336 | 1411.961 | 1462.643 | 1372.183 | 1174.598 |
| ENSMUSG00000061062 | 0.725150019 | 6.09E-05 | 0.000928 | 803.8374 | 877.0695 | 834.7315 | 886.777 | 727.433 | 474.884 | 405.9657 | 449.1751 |
| ENSMUSG00000061068 | 0.264027653 | 0.79529 | 0.874099 | 7.04238 | 46.79304 | 10.76539 | 17.25247 | 12.67645 | 51.39921 | 2.892867 | 1.087591 |
| ENSMUSG00000061079 | -0.153579457 | 0.214581 | 0.372744 | 665.0019 | 515.9233 | 612.7989 | 577.9577 | 729.3833 | 699.4762 | 653.7881 | 556.8466 |
| ENSMUSG00000061080 | 1.251108616 | 4.73E-09 | 4.59E-07 | 568.4207 | 595.1115 | 458.7711 | 686.6483 | 200.8729 | 350.8555 | 208.2865 | 210.9927 |

| | | | | | | | | | | | |
|---|---|---|---|---|---|---|---|---|---|---|---|
| ENSMUSG00000061082 | -0.398593199 | 0.475875 | 0.635418 | 4.024217 | 9.598572 | 15.73403 | 12.07673 | 9.751113 | 21.23011 | 13.50005 | 10.87591 |
| ENSMUSG00000061086 | 0.776912378 | 0.017261 | 0.063586 | 53.32088 | 98.38537 | 77.01392 | 70.73513 | 39.00445 | 68.15982 | 30.85725 | 36.9781 |
| ENSMUSG00000061099 | -0.038824623 | 0.899327 | 0.939331 | 51.30877 | 25.19625 | 42.23344 | 31.91707 | 39.00445 | 32.40385 | 41.46443 | 42.41605 |
| ENSMUSG00000061104 | -0.142537374 | 0.657258 | 0.779135 | 28.16952 | 28.79572 | 51.34261 | 34.50494 | 33.15378 | 40.22547 | 34.71441 | 51.11678 |
| ENSMUSG00000061111 | -0.0869032 | 0.201056 | 0.356523 | 2384.349 | 2538.822 | 2678.097 | 2568.03 | 2767.366 | 2486.157 | 2771.367 | 2776.62 |
| ENSMUSG00000061118 | -0.295158396 | 0.001778 | 0.012062 | 445.6821 | 490.727 | 514.2542 | 555.5295 | 603.5939 | 601.1473 | 632.5737 | 627.5401 |
| ENSMUSG00000061119 | 0.106127922 | 0.318711 | 0.486891 | 846.0917 | 1025.847 | 1051.695 | 1102.433 | 877.6002 | 906.1904 | 1019.254 | 938.5911 |
| ENSMUSG00000061126 | -1.088318538 | 0.164133 | 0.309111 | 4.024217 | 7.198929 | 8.281066 | 11.21411 | 34.1289 | 2.234748 | 20.25007 | 8.700729 |
| ENSMUSG00000061130 | -0.039298157 | 0.612329 | 0.74664 | 3052.369 | 2943.162 | 2620.957 | 3159.79 | 3057.949 | 2939.811 | 3178.297 | 2922.357 |
| ENSMUSG00000061132 | 0.444679923 | 0.003153 | 0.018437 | 702.2259 | 519.5227 | 744.4679 | 563.2931 | 398.8205 | 464.8276 | 477.3231 | 519.8685 |
| ENSMUSG00000061136 | 0.090427561 | 0.404189 | 0.571145 | 3725.419 | 3906.619 | 3375.363 | 3649.76 | 3170.087 | 4151.045 | 3152.261 | 3293.226 |
| ENSMUSG00000061143 | 0.308296583 | 4.94E-05 | 0.000787 | 1086.539 | 1037.846 | 1046.727 | 1090.356 | 885.4011 | 934.1247 | 816.7529 | 806.9926 |
| ENSMUSG00000061144 | -0.346930393 | 0.322567 | 0.490822 | 20.12109 | 25.19625 | 32.29616 | 19.84034 | 28.27823 | 21.23011 | 38.57157 | 35.89051 |
| ENSMUSG00000061167 | 0.339683877 | 0.232385 | 0.392999 | 59.35721 | 77.9884 | 48.03018 | 56.93315 | 44.85512 | 56.98608 | 56.89306 | 31.54014 |
| ENSMUSG00000061171 | 0.685854883 | 0.243946 | 0.406525 | 42.25428 | 9.598572 | 15.73403 | 34.50494 | 23.40267 | 8.938993 | 21.21436 | 9.78832 |
| ENSMUSG00000061175 | -0.288767284 | 0.045248 | 0.126946 | 555.342 | 410.339 | 534.1288 | 469.2672 | 488.5308 | 654.7812 | 661.5024 | 603.6131 |
| ENSMUSG00000061186 | 1.829898636 | 1.18E-08 | 9.75E-07 | 433.6094 | 343.149 | 332.0708 | 375.2412 | 99.46135 | 183.2494 | 81.96458 | 53.29196 |
| ENSMUSG00000061207 | 0.0233276 | 0.772593 | 0.858991 | 700.2138 | 691.0972 | 737.0149 | 715.1149 | 724.5077 | 736.3495 | 633.538 | 706.9342 |
| ENSMUSG00000061232 | -0.135233834 | 0.577463 | 0.719335 | 1017.121 | 969.4558 | 1404.469 | 852.272 | 1012.166 | 717.3542 | 1470.541 | 1460.635 |
| ENSMUSG00000061244 | -0.025859003 | 0.774379 | 0.8603 | 1491.979 | 1479.38 | 1361.407 | 1410.389 | 1368.081 | 1667.122 | 1346.148 | 1466.073 |
| ENSMUSG00000061273 | -0.208267257 | 0.006652 | 0.031813 | 891.3642 | 985.0535 | 1006.978 | 921.2819 | 1040.444 | 1119.609 | 1072.29 | 1164.81 |
| ENSMUSG00000061286 | -0.257493847 | 0.009636 | 0.041456 | 723.3531 | 789.4826 | 809.0602 | 678.8847 | 990.7131 | 874.9039 | 906.4318 | 812.4305 |
| ENSMUSG00000061288 | 0.126812189 | 0.203907 | 0.359853 | 626.7719 | 545.9188 | 602.0335 | 516.7115 | 496.3317 | 522.9311 | 513.9661 | 566.635 |
| ENSMUSG00000061306 | -0.343128 | 0.003105 | 0.018253 | 3055.387 | 3231.119 | 3361.285 | 3143.4 | 4071.09 | 3226.976 | 4242.872 | 4684.255 |
| ENSMUSG00000061313 | 0.28431867 | 0.027448 | 0.088765 | 1237.447 | 1173.425 | 1098.069 | 1255.98 | 1034.593 | 1177.712 | 894.8603 | 805.905 |
| ENSMUSG00000061315 | -0.092999319 | 0.139614 | 0.275644 | 19772.99 | 21834.35 | 21711.3 | 22418.72 | 24139.86 | 22857 | 22773.62 | 21675.69 |
| ENSMUSG00000061322 | -0.568753873 | 0.256313 | 0.42008 | 14.08476 | 9.598572 | 4.96864 | 8.626235 | 15.60178 | 12.29112 | 11.57147 | 15.22628 |
| ENSMUSG00000061330 | -0.048468975 | 0.792042 | 0.872764 | 90.54489 | 101.9848 | 74.5296 | 91.43809 | 81.90935 | 101.681 | 94.50034 | 91.35765 |
| ENSMUSG00000061353 | -0.199505663 | 0.148671 | 0.288403 | 5178.162 | 4759.692 | 6133.786 | 5707.117 | 4962.341 | 5853.923 | 7087.525 | 7108.495 |
| ENSMUSG00000061360 | -0.165231567 | 0.020186 | 0.070946 | 1554.354 | 1726.543 | 1561.809 | 1684.704 | 1950.223 | 1850.372 | 1779.114 | 1735.795 |
| ENSMUSG00000061371 | 0.220295023 | 0.039602 | 0.115854 | 302.8224 | 315.5531 | 290.6654 | 308.8192 | 261.3298 | 286.0478 | 237.2151 | 261.0219 |
| ENSMUSG00000061374 | -0.006195088 | 0.928446 | 0.957659 | 1488.96 | 1582.565 | 1437.593 | 1484.575 | 1615.759 | 1425.769 | 1463.791 | 1508.489 |
| ENSMUSG00000061393 | -0.024206525 | 0.891461 | 0.9345 | 352.119 | 292.7565 | 244.2915 | 219.969 | 306.185 | 305.0431 | 243.0009 | 271.8978 |
| ENSMUSG00000061410 | 0.087360155 | 0.220545 | 0.379345 | 2082.533 | 1988.104 | 2170.467 | 2321.32 | 2015.555 | 2111.837 | 1925.685 | 2013.131 |
| ENSMUSG00000061411 | 0.017150056 | 0.895606 | 0.93677 | 768.6255 | 752.2881 | 852.1217 | 818.6297 | 800.5664 | 958.707 | 643.1809 | 754.7882 |
| ENSMUSG00000061414 | 0.791450635 | 0.000152 | 0.001847 | 181.0898 | 257.9616 | 230.2136 | 165.6237 | 132.6151 | 141.9065 | 95.46463 | 112.0219 |
| ENSMUSG00000061436 | 0.030401734 | 0.874824 | 0.925152 | 765.6074 | 466.7306 | 539.9255 | 452.8773 | 499.257 | 655.8986 | 548.6805 | 475.2773 |
| ENSMUSG00000061451 | 0.599985067 | 0.160144 | 0.303486 | 59.35721 | 213.5682 | 93.57605 | 135.4319 | 59.48179 | 137.437 | 75.21455 | 58.72992 |
| ENSMUSG00000061455 | 0.176417139 | 0.079635 | 0.18855 | 1061.387 | 902.2658 | 919.1984 | 1006.682 | 930.2562 | 922.951 | 822.5387 | 766.7517 |
| ENSMUSG00000061458 | -0.083906251 | 0.505596 | 0.661418 | 526.1664 | 536.3202 | 481.9581 | 477.0308 | 505.1077 | 632.4337 | 449.3588 | 555.759 |
| ENSMUSG00000061461 | -0.235819279 | 0.073832 | 0.179154 | 641.8627 | 634.7056 | 839.7001 | 708.2139 | 816.1682 | 707.2978 | 872.6817 | 933.1532 |
| ENSMUSG00000061462 | -0.039496941 | 0.914276 | 0.94925 | 65.39353 | 58.79126 | 63.76421 | 39.68068 | 45.83023 | 102.7984 | 44.3573 | 41.32846 |
| ENSMUSG00000061474 | -0.501144963 | 0.000428 | 0.004063 | 412.4823 | 376.744 | 423.1625 | 402.8452 | 612.3699 | 443.5975 | 553.502 | 677.5692 |
| ENSMUSG00000061477 | -0.110521376 | 0.19085 | 0.343486 | 43344.85 | 52668.57 | 49112.52 | 49637.94 | 57406.75 | 50945.55 | 53221.05 | 48693.63 |
| ENSMUSG00000061479 | -0.139027329 | 0.068041 | 0.168786 | 2612.723 | 2805.183 | 2965.45 | 2737.104 | 3157.41 | 2762.149 | 3081.868 | 3244.284 |
| ENSMUSG00000061482 | -0.172068643 | 0.722745 | 0.826557 | 14.08476 | 16.7975 | 28.15563 | 30.19182 | 36.07912 | 35.75597 | 16.39292 | 13.05109 |
| ENSMUSG00000061510 | 0.657059629 | 0.090785 | 0.206447 | 32.19374 | 33.595 | 27.32752 | 18.11509 | 18.52711 | 22.34748 | 18.32149 | 10.87591 |
| ENSMUSG00000061518 | -0.692549642 | 0.000108 | 0.001422 | 3819.988 | 3635.459 | 4753.332 | 4037.941 | 6554.698 | 4448.266 | 6954.453 | 8301.583 |
| ENSMUSG00000061527 | 1.123299304 | 0.001975 | 0.013015 | 231.3925 | 93.58608 | 197.0894 | 141.4703 | 115.0631 | 74.86406 | 73.28598 | 41.32846 |
| ENSMUSG00000061531 | 0.679256547 | 0.243449 | 0.406063 | 13.07871 | 4.799286 | 17.39024 | 13.80198 | 4.875557 | 6.704245 | 8.678602 | 10.87591 |
| ENSMUSG00000061533 | 0.0329886 | 0.718778 | 0.824217 | 416.5065 | 397.1409 | 375.1323 | 403.7078 | 402.721 | 407.8415 | 372.2156 | 373.0437 |
| ENSMUSG00000061535 | -0.181061397 | 0.122546 | 0.252114 | 255.5378 | 230.3657 | 266.6503 | 260.5123 | 252.5538 | 306.1605 | 303.7511 | 289.2992 |
| ENSMUSG00000061536 | -0.060287084 | 0.533371 | 0.68465 | 316.9071 | 353.9474 | 325.4459 | 351.0878 | 370.5423 | 355.325 | 338.4655 | 339.3284 |
| ENSMUSG00000061544 | 0.260272703 | 0.075059 | 0.181045 | 563.3904 | 553.1177 | 458.7711 | 525.3377 | 434.8996 | 549.7481 | 381.8585 | 387.1824 |
| ENSMUSG00000061559 | -0.22520018 | 0.003792 | 0.021015 | 1374.27 | 1486.579 | 1392.875 | 1581.189 | 1766.902 | 1584.436 | 1681.72 | 1786.912 |
| ENSMUSG00000061576 | 0.298012313 | 0.183262 | 0.334151 | 259.562 | 389.942 | 558.972 | 339.011 | 345.1894 | 289.3999 | 300.8582 | 324.1021 |
| ENSMUSG00000061578 | 0.267480367 | 0.115143 | 0.242838 | 155.9384 | 119.9822 | 143.2624 | 135.4319 | 95.56091 | 131.8501 | 119.5719 | 115.2847 |
| ENSMUSG00000061589 | -0.053407351 | 0.418798 | 0.584292 | 1570.451 | 1374.995 | 1477.342 | 1438.856 | 1491.92 | 1498.399 | 1512.005 | 1583.533 |
| ENSMUSG00000061601 | 1.032003459 | 5.11E-05 | 0.000804 | 527.1725 | 500.3256 | 348.6329 | 578.8204 | 202.8232 | 363.1466 | 220.8222 | 169.6642 |
| ENSMUSG00000061603 | 0.63369466 | 0.116812 | 0.245126 | 93.56306 | 56.39161 | 49.6864 | 131.9814 | 46.80534 | 80.45093 | 31.82154 | 55.46715 |
| ENSMUSG00000061607 | 0.302117962 | 0.010564 | 0.044402 | 1154.95 | 1313.805 | 1159.349 | 1136.938 | 894.1771 | 1178.83 | 908.3604 | 883.124 |
| ENSMUSG00000061613 | -0.140121977 | 0.008559 | 0.038099 | 3391.409 | 3588.666 | 3580.733 | 3657.524 | 4063.289 | 4000.199 | 3764.585 | 3842.459 |
| ENSMUSG00000061615 | -0.644444636 | 0.217358 | 0.375825 | 5.030272 | 10.79839 | 10.76539 | 5.175741 | 11.70134 | 12.29112 | 14.46434 | 10.87591 |
| ENSMUSG00000061650 | -0.338727263 | 0.000809 | 0.006633 | 875.2673 | 911.8644 | 949.8383 | 970.4514 | 1289.097 | 997.8151 | 1253.576 | 1147.409 |

| | | | | | | | | | | | |
|---|---|---|---|---|---|---|---|---|---|---|---|
| ENSMUSG00000061654 | 0.155128755 | 0.644622 | 0.770092 | 36.21796 | 32.39518 | 26.49941 | 21.56559 | 31.20356 | 21.23011 | 21.21436 | 30.45255 |
| ENSMUSG00000061665 | 0.12532279 | 0.16649 | 0.312079 | 4135.889 | 4140.584 | 4322.717 | 4177.686 | 3394.362 | 4448.266 | 3720.228 | 3821.795 |
| ENSMUSG00000061666 | 0.226761334 | 0.092324 | 0.208661 | 699.2078 | 677.8992 | 635.9859 | 708.2139 | 542.1619 | 726.2932 | 573.7521 | 483.978 |
| ENSMUSG00000061684 | 0.670107865 | 0.165988 | 0.311518 | 20.12109 | 14.39786 | 13.24971 | 11.21411 | 11.70134 | 5.58687 | 9.642892 | 9.78832 |
| ENSMUSG00000061689 | 0.228166207 | 0.007212 | 0.033611 | 1416.525 | 1602.962 | 1524.544 | 1600.167 | 1275.446 | 1468.23 | 1236.219 | 1267.044 |
| ENSMUSG00000061702 | 1.001777446 | 0.057164 | 0.149688 | 35.2119 | 39.59411 | 21.53077 | 28.46658 | 10.72622 | 33.52122 | 10.60718 | 7.613138 |
| ENSMUSG00000061718 | 0.332075533 | 0.065094 | 0.164345 | 486.9303 | 669.5004 | 502.6607 | 636.6161 | 466.1032 | 546.3959 | 477.3231 | 332.8029 |
| ENSMUSG00000061723 | 0.42035454 | 0.161328 | 0.305068 | 286.7255 | 167.975 | 202.8861 | 164.7611 | 117.0134 | 246.9397 | 135.9648 | 115.2847 |
| ENSMUSG00000061724 | -0.073021033 | 0.827817 | 0.895602 | 22.1332 | 37.19447 | 21.53077 | 34.50494 | 34.1289 | 34.6386 | 27.96439 | 23.927 |
| ENSMUSG00000061731 | -0.08612184 | 0.485082 | 0.643429 | 2293.804 | 2673.202 | 2404.822 | 2378.253 | 2503.111 | 3149.878 | 2554.402 | 2141.467 |
| ENSMUSG00000061740 | 0.392871406 | 0.050032 | 0.136154 | 1517.13 | 1294.607 | 1127.881 | 1790.806 | 979.0118 | 1382.192 | 1214.04 | 789.5911 |
| ENSMUSG00000061742 | -2.675125951 | 0.003258 | 0.018873 | 37.22401 | 10.79839 | 173.9024 | 46.58167 | 366.6419 | 121.7938 | 317.2511 | 912.4889 |
| ENSMUSG00000061751 | 0.05043945 | 0.486045 | 0.644336 | 969.8364 | 905.8653 | 982.1345 | 998.0554 | 870.7744 | 977.7023 | 946.932 | 933.1532 |
| ENSMUSG00000061755 | 0.44887022 | 0.006257 | 0.030366 | 3502.075 | 3279.112 | 2791.547 | 2779.373 | 2015.555 | 3033.671 | 2144.579 | 1855.43 |
| ENSMUSG00000061758 | 0.123168705 | 0.374828 | 0.543403 | 805.8495 | 548.3184 | 733.7025 | 669.3958 | 555.8134 | 672.6592 | 683.681 | 622.1021 |
| ENSMUSG00000061759 | 0.182145687 | 0.038502 | 0.113404 | 743.4742 | 627.5067 | 732.0463 | 690.0988 | 628.9468 | 583.2693 | 607.5022 | 644.9415 |
| ENSMUSG00000061778 | -0.155410164 | 0.182302 | 0.332894 | 614.6992 | 583.1133 | 756.0614 | 661.6322 | 731.3335 | 642.4901 | 742.5027 | 800.467 |
| ENSMUSG00000061787 | -0.187604258 | 0.014913 | 0.057462 | 22479.28 | 26211.3 | 25401.34 | 25023.84 | 30754.04 | 26258.29 | 28423.39 | 27441.01 |
| ENSMUSG00000061808 | -3.780920791 | 1.27E-06 | 4.53E-05 | 90.54489 | 14.39786 | 115.9349 | 104.3774 | 1415.862 | 244.7049 | 868.8245 | 1947.876 |
| ENSMUSG00000061815 | -0.410229493 | 0.203851 | 0.35982 | 29.17558 | 21.59679 | 33.95237 | 26.74133 | 28.27823 | 31.28647 | 50.14304 | 39.15328 |
| ENSMUSG00000061816 | 0.99243501 | 0.102811 | 0.225242 | 136.8234 | 47.99286 | 79.49824 | 75.04824 | 16.57689 | 89.38993 | 54.00019 | 10.87591 |
| ENSMUSG00000061838 | -0.448147332 | 5.69E-05 | 0.000874 | 3113.738 | 3208.323 | 3537.672 | 3169.279 | 4678.584 | 3555.484 | 4739.481 | 4801.715 |
| ENSMUSG00000061848 | 0.197430644 | 0.356841 | 0.525355 | 68.4117 | 99.58519 | 76.18581 | 100.9269 | 73.13335 | 73.74669 | 85.82174 | 67.43065 |
| ENSMUSG00000061859 | -0.515234209 | 1.61E-05 | 0.000331 | 926.5761 | 854.2729 | 1078.195 | 940.2596 | 1236.441 | 1191.121 | 1521.648 | 1484.562 |
| ENSMUSG00000061878 | -0.252744371 | 0.345099 | 0.513889 | 252.5196 | 160.7761 | 322.1335 | 303.6435 | 261.3298 | 233.5312 | 442.6087 | 302.3503 |
| ENSMUSG00000061882 | 0.181823515 | 0.327454 | 0.495205 | 200.2048 | 173.9741 | 192.9488 | 228.5952 | 135.5405 | 220.1227 | 160.072 | 188.1533 |
| ENSMUSG00000061887 | 0.305003932 | 0.045222 | 0.126938 | 3874.315 | 4194.576 | 3815.915 | 4088.835 | 2877.553 | 4390.163 | 2891.903 | 2771.182 |
| ENSMUSG00000061894 | 0.209578151 | 0.107861 | 0.232476 | 248.4954 | 226.7663 | 233.5261 | 192.365 | 193.072 | 189.9536 | 189.0007 | 206.6423 |
| ENSMUSG00000061897 | -0.153592677 | 0.863743 | 0.918221 | 88.53278 | 67.19001 | 10.76539 | 8.626235 | 79.95913 | 10.05637 | 9.642892 | 94.62042 |
| ENSMUSG00000061898 | 0.329297038 | 0.025646 | 0.084552 | 1140.866 | 1041.445 | 910.9173 | 1031.698 | 902.9531 | 993.3456 | 725.1455 | 661.2554 |
| ENSMUSG00000061904 | -0.501650339 | 0.000512 | 0.004669 | 13666.24 | 10788.8 | 13608.28 | 12135.39 | 17902.07 | 13357.09 | 19129.57 | 20685.98 |
| ENSMUSG00000061906 | -1.961924582 | 0.040544 | 0.117839 | 4.024217 | 0 | 11.59349 | 0.862623 | 16.57689 | 5.58687 | 16.39292 | 27.18978 |
| ENSMUSG00000061911 | 0.80765971 | 0.100587 | 0.221743 | 30.18163 | 15.59768 | 15.73403 | 43.13117 | 9.751113 | 18.99536 | 18.32149 | 13.05109 |
| ENSMUSG00000061947 | -3.448548076 | 0.00194 | 0.012845 | 13.07871 | 2.399643 | 69.56096 | 5.175741 | 170.6445 | 39.10809 | 354.8584 | 424.1605 |
| ENSMUSG00000061950 | -0.181865031 | 0.005257 | 0.026624 | 1929.612 | 1878.921 | 2036.314 | 1929.689 | 2274.935 | 2026.917 | 2254.508 | 2264.365 |
| ENSMUSG00000061959 | -1.278075784 | 0.019686 | 0.069694 | 10.06054 | 11.99822 | 36.43669 | 11.21411 | 43.88001 | 23.46486 | 33.75012 | 69.60583 |
| ENSMUSG00000061979 | -0.285675967 | 0.001034 | 0.008024 | 784.7224 | 823.0776 | 833.9034 | 681.4726 | 950.7335 | 945.2985 | 936.3248 | 972.3064 |
| ENSMUSG00000061981 | 0.065217258 | 0.422521 | 0.587689 | 1261.592 | 1180.624 | 1344.017 | 1287.897 | 1200.362 | 1116.257 | 1210.183 | 1326.861 |
| ENSMUSG00000061983 | -0.165808333 | 0.031548 | 0.09787 | 29313.41 | 35085.18 | 34551.09 | 32507.1 | 39336.97 | 36395.11 | 36736.52 | 34996.51 |
| ENSMUSG00000061991 | -0.655097324 | 0.309974 | 0.478464 | 10.06054 | 4.799286 | 4.96864 | 13.80198 | 11.70134 | 4.469496 | 16.39292 | 20.66423 |
| ENSMUSG00000062006 | -0.134202655 | 0.066355 | 0.166054 | 17820.24 | 20240.99 | 20418.63 | 19536.7 | 23034.08 | 20777.57 | 21822.83 | 19985.57 |
| ENSMUSG00000062012 | -0.092859806 | 0.326993 | 0.49463 | 618.7234 | 634.7056 | 626.8767 | 615.9132 | 755.7113 | 600.0299 | 634.5023 | 669.9561 |
| ENSMUSG00000062014 | 0.04878252 | 0.455856 | 0.617896 | 2873.291 | 2542.422 | 2541.459 | 2806.977 | 2548.941 | 2621.36 | 2581.402 | 2656.985 |
| ENSMUSG00000062031 | 0.382576569 | 7.82E-07 | 3.04E-05 | 1279.701 | 1217.819 | 1139.475 | 1247.354 | 980.962 | 890.5472 | 987.4321 | 884.2116 |
| ENSMUSG00000062040 | 0.020787037 | 0.833882 | 0.899486 | 508.0574 | 555.5174 | 446.3495 | 485.657 | 508.033 | 483.823 | 505.2875 | 464.4014 |
| ENSMUSG00000062044 | 0.523769547 | 5.10E-07 | 2.14E-05 | 341.0524 | 365.9456 | 377.6166 | 373.516 | 271.0809 | 259.2308 | 230.4651 | 254.4963 |
| ENSMUSG00000062054 | -0.763247887 | 1.37E-05 | 0.000292 | 759.571 | 718.6931 | 818.9975 | 749.6198 | 1342.728 | 891.6645 | 1264.183 | 1673.803 |
| ENSMUSG00000062070 | -0.577716297 | 9.13E-05 | 0.001248 | 6195.283 | 6501.833 | 7503.474 | 6579.229 | 10173.34 | 7232.763 | 10870.43 | 11692.69 |
| ENSMUSG00000062074 | 0.592640642 | 0.084959 | 0.196936 | 35.2119 | 45.59322 | 31.46805 | 45.71905 | 33.15378 | 27.93435 | 28.92867 | 14.13868 |
| ENSMUSG00000062075 | -0.182910879 | 0.044043 | 0.124672 | 981.9091 | 1105.036 | 967.2285 | 918.694 | 1183.785 | 1016.81 | 1127.254 | 1175.686 |
| ENSMUSG00000062078 | 0.224183797 | 0.064818 | 0.163904 | 5158.041 | 5177.23 | 4610.898 | 5315.486 | 3846.814 | 5401.386 | 4166.693 | 3931.642 |
| ENSMUSG00000062081 | -0.595790232 | 0.206095 | 0.362636 | 8.048435 | 10.79839 | 24.8432 | 16.38985 | 27.30312 | 14.52586 | 26.03581 | 23.927 |
| ENSMUSG00000062093 | -0.014499567 | 0.961805 | 0.978182 | 37.22401 | 43.19358 | 46.37397 | 38.81806 | 39.00445 | 62.57295 | 34.71441 | 31.54014 |
| ENSMUSG00000062098 | -0.084177868 | 0.465037 | 0.625928 | 1307.871 | 1075.04 | 1171.771 | 1429.367 | 1165.258 | 1393.365 | 1414.612 | 1313.81 |
| ENSMUSG00000062101 | -0.007933755 | 0.972242 | 0.984619 | 135.8173 | 167.975 | 126.7003 | 153.547 | 157.968 | 177.6625 | 159.1077 | 91.35765 |
| ENSMUSG00000062110 | 0.060028818 | 0.560944 | 0.706583 | 394.3733 | 416.3381 | 397.4912 | 375.2412 | 351.0401 | 360.9118 | 427.1801 | 377.3941 |
| ENSMUSG00000062115 | 0.163500562 | 0.042479 | 0.121682 | 2145.914 | 1930.513 | 2032.174 | 1818.41 | 1903.417 | 1796.738 | 1675.935 | 1700.992 |
| ENSMUSG00000062116 | 0.025916173 | 0.869116 | 0.921481 | 311.8769 | 255.562 | 207.0267 | 224.2821 | 250.6036 | 235.7659 | 260.3581 | 231.6569 |
| ENSMUSG00000062127 | -0.068432478 | 0.380498 | 0.548577 | 1101.63 | 991.0526 | 1002.837 | 1060.164 | 1046.294 | 1019.045 | 1160.04 | 1132.182 |
| ENSMUSG00000062151 | -0.1281421 | 0.590632 | 0.730709 | 50.30272 | 59.99108 | 69.56096 | 47.44429 | 54.60623 | 75.98144 | 60.75022 | 57.64233 |
| ENSMUSG00000062157 | 0.182063159 | 0.319848 | 0.487903 | 221.332 | 175.1739 | 237.6666 | 187.1893 | 196.9725 | 221.2401 | 156.2148 | 151.1752 |
| ENSMUSG00000062169 | -0.152782199 | 0.07682 | 0.184081 | 1387.349 | 1219.019 | 1264.519 | 1358.632 | 1376.857 | 1334.145 | 1547.684 | 1556.343 |
| ENSMUSG00000062175 | -0.073160671 | 0.568143 | 0.712423 | 378.2764 | 375.5441 | 399.1474 | 399.3947 | 483.6552 | 336.3296 | 417.5372 | 394.7956 |
| ENSMUSG00000062184 | 0.321072113 | 0.065664 | 0.164952 | 953.7395 | 1287.409 | 960.6037 | 1049.813 | 827.8695 | 1126.313 | 768.5385 | 679.7444 |

| | | | | | | | | | | | |
|---|---|---|---|---|---|---|---|---|---|---|---|
| ENSMUSG00000062190 | 0.004091223 | 0.952762 | 0.973168 | 1108.672 | 1141.03 | 1116.288 | 1244.766 | 1193.536 | 1137.487 | 1150.397 | 1116.956 |
| ENSMUSG00000062198 | 0.341006402 | 0.02062 | 0.072021 | 495.9848 | 485.9277 | 386.7258 | 401.1199 | 322.7618 | 411.1937 | 363.537 | 298 |
| ENSMUSG00000062202 | -0.088737959 | 0.352784 | 0.521562 | 884.3218 | 812.2792 | 1016.915 | 929.0455 | 908.8037 | 1000.05 | 1033.718 | 936.4159 |
| ENSMUSG00000062203 | -0.393862406 | 6.82E-06 | 0.000172 | 2135.853 | 2187.275 | 2184.545 | 2282.502 | 2850.25 | 2523.031 | 2933.368 | 3243.197 |
| ENSMUSG00000062209 | -0.729824408 | 0.000166 | 0.001967 | 304.8345 | 242.3639 | 407.4285 | 320.0333 | 417.3476 | 481.5882 | 601.7164 | 617.7517 |
| ENSMUSG00000062210 | -0.477521244 | 0.005008 | 0.025762 | 632.8082 | 722.2926 | 905.1205 | 728.9169 | 1128.204 | 760.9318 | 1054.932 | 1219.19 |
| ENSMUSG00000062232 | -0.093702271 | 0.341053 | 0.509716 | 793.7769 | 695.8965 | 800.7791 | 863.4861 | 771.313 | 911.7773 | 868.8245 | 818.9561 |
| ENSMUSG00000062234 | -0.285380101 | 0.007625 | 0.034974 | 2014.121 | 1804.532 | 2251.622 | 1971.957 | 2472.882 | 2091.724 | 2534.152 | 2705.927 |
| ENSMUSG00000062248 | -0.023204228 | 0.71676 | 0.822669 | 1523.166 | 1491.378 | 1428.484 | 1626.908 | 1566.029 | 1518.511 | 1552.506 | 1531.328 |
| ENSMUSG00000062257 | 0.320163651 | 0.287936 | 0.454754 | 76.46013 | 85.18733 | 144.9187 | 75.04824 | 57.53157 | 84.92043 | 65.57166 | 98.97079 |
| ENSMUSG00000062270 | -0.050469949 | 0.394667 | 0.561948 | 10421.72 | 10849.99 | 10375.35 | 11070.91 | 11455.61 | 10234.03 | 11531.93 | 11012.95 |
| ENSMUSG00000062284 | 1.027558248 | 0.098098 | 0.218092 | 24.1453 | 10.79839 | 5.796746 | 13.80198 | 6.825779 | 6.704245 | 8.678602 | 4.350364 |
| ENSMUSG00000062296 | 1.395552985 | 2.76E-06 | 8.50E-05 | 449.7063 | 151.1775 | 275.7595 | 306.2313 | 106.2871 | 120.6764 | 99.32178 | 123.9854 |
| ENSMUSG00000062300 | -0.366100391 | 0.00211 | 0.013636 | 2118.75 | 1956.909 | 2458.649 | 2123.779 | 2930.209 | 2261.565 | 2934.332 | 3035.467 |
| ENSMUSG00000062309 | 0.189293483 | 0.47048 | 0.630404 | 90.54489 | 106.7841 | 121.7317 | 100.0643 | 91.66046 | 136.3196 | 65.57166 | 75.04379 |
| ENSMUSG00000062312 | -0.317063766 | 0.000481 | 0.00445 | 1643.893 | 1502.177 | 1890.567 | 1587.227 | 2094.539 | 1918.531 | 2090.579 | 2153.43 |
| ENSMUSG00000062319 | 2.190801102 | 0.010026 | 0.042676 | 27.16347 | 65.99018 | 9.109173 | 34.50494 | 3.900445 | 21.23011 | 3.857157 | 1.087591 |
| ENSMUSG00000062328 | -0.022394549 | 0.780154 | 0.864358 | 28310.37 | 33020.29 | 31928.48 | 32916.85 | 33850.01 | 32983.77 | 32433.87 | 28880.98 |
| ENSMUSG00000062329 | -0.426305232 | 0.308838 | 0.477139 | 29.17558 | 27.5959 | 20.70267 | 40.5433 | 58.50668 | 22.34748 | 51.10733 | 26.10219 |
| ENSMUSG00000062352 | -0.337493238 | 0.000683 | 0.005824 | 551.3178 | 577.1142 | 592.0962 | 602.1112 | 729.3833 | 631.3164 | 798.4314 | 775.4524 |
| ENSMUSG00000062353 | -0.032681628 | 0.848831 | 0.909191 | 126.7628 | 135.5798 | 179.6991 | 166.4863 | 173.5698 | 140.7891 | 162.9649 | 146.8248 |
| ENSMUSG00000062373 | 0.032266898 | 0.764455 | 0.853836 | 1148.914 | 1331.802 | 1087.304 | 1183.519 | 1029.718 | 1307.328 | 1215.004 | 1091.941 |
| ENSMUSG00000062376 | -0.031603101 | 0.781339 | 0.865154 | 607.6568 | 676.6993 | 626.0486 | 604.6991 | 610.4197 | 751.9928 | 653.7881 | 553.5839 |
| ENSMUSG00000062380 | 2.989050039 | 0.002798 | 0.016808 | 609.6689 | 51.59233 | 31.46805 | 925.595 | 57.53157 | 53.63396 | 49.17875 | 43.50364 |
| ENSMUSG00000062382 | -0.410779017 | 0.041362 | 0.119548 | 117.7084 | 88.78679 | 142.4343 | 123.3552 | 161.8685 | 140.7891 | 135.9648 | 192.5036 |
| ENSMUSG00000062393 | 2.386785565 | 1.94E-06 | 6.31E-05 | 141.8537 | 35.99465 | 43.88965 | 129.3935 | 17.552 | 16.76061 | 14.46434 | 18.48905 |
| ENSMUSG00000062397 | -0.151038722 | 0.015223 | 0.058313 | 3549.36 | 4104.589 | 3829.165 | 3856.79 | 4142.273 | 4180.096 | 4332.551 | 4374.291 |
| ENSMUSG00000062410 | -3.190583575 | 5.36E-06 | 0.000142 | 2.012109 | 3.599465 | 5.796746 | 4.313117 | 18.52711 | 13.40849 | 39.53586 | 73.95619 |
| ENSMUSG00000062421 | -0.166373965 | 0.112584 | 0.239278 | 1530.209 | 1780.535 | 1974.206 | 1641.573 | 2081.863 | 1727.46 | 1958.471 | 2005.518 |
| ENSMUSG00000062444 | 0.589955446 | 0.197361 | 0.351659 | 91.55095 | 39.59411 | 37.2648 | 86.26235 | 23.40267 | 73.74669 | 35.6787 | 36.9781 |
| ENSMUSG00000062456 | 0.122679303 | 0.611879 | 0.746201 | 239.4409 | 273.5593 | 179.6991 | 227.7326 | 190.1467 | 306.1605 | 201.5364 | 146.8248 |
| ENSMUSG00000062470 | 0.417360244 | 0.039227 | 0.114934 | 133.8052 | 93.58608 | 104.3414 | 100.9269 | 79.95913 | 98.32892 | 71.3574 | 75.04379 |
| ENSMUSG00000062480 | -2.017730679 | 1.41E-07 | 7.53E-06 | 54.32694 | 27.5959 | 42.23344 | 29.3292 | 219.4 | 90.5073 | 99.32178 | 213.1679 |
| ENSMUSG00000062488 | -0.206271559 | 0.722055 | 0.826091 | 46.2785 | 67.19001 | 205.3704 | 29.3292 | 86.78491 | 45.81234 | 145.6077 | 123.9854 |
| ENSMUSG00000062510 | 0.058856594 | 0.678926 | 0.794865 | 534.2149 | 681.4986 | 486.0986 | 507.2226 | 532.4108 | 601.1473 | 524.5733 | 458.9634 |
| ENSMUSG00000062515 | -0.372138855 | 0.017305 | 0.063687 | 5137.92 | 3333.104 | 4690.396 | 4815.164 | 5486.951 | 4795.77 | 6583.202 | 6403.736 |
| ENSMUSG00000062519 | 0.226270959 | 0.074816 | 0.180734 | 657.9595 | 665.901 | 609.4865 | 690.0988 | 513.8837 | 690.5372 | 550.6091 | 488.3284 |
| ENSMUSG00000062526 | -0.425267947 | 0.023076 | 0.078231 | 393.3673 | 332.3506 | 415.7095 | 373.516 | 485.6054 | 346.386 | 602.6807 | 600.3503 |
| ENSMUSG00000062542 | 0.000795213 | 0.996042 | 0.997718 | 157.9505 | 143.9786 | 145.7468 | 188.0519 | 163.8187 | 139.6718 | 151.3934 | 181.6277 |
| ENSMUSG00000062545 | -1.200449014 | 0.019895 | 0.070181 | 22.1332 | 34.79482 | 76.18581 | 25.8787 | 68.25779 | 41.34284 | 90.64318 | 166.4014 |
| ENSMUSG00000062554 | -0.220440437 | 0.666526 | 0.785864 | 11.0666 | 7.198929 | 9.93728 | 12.07673 | 18.52711 | 10.05637 | 7.714313 | 10.87591 |
| ENSMUSG00000062563 | -0.208015558 | 0.284541 | 0.451081 | 516.1059 | 482.3283 | 677.3912 | 530.5134 | 553.8632 | 476.0014 | 679.8239 | 840.7079 |
| ENSMUSG00000062580 | -0.343823368 | 0.000211 | 0.002369 | 1455.761 | 1392.993 | 1535.31 | 1364.67 | 1615.759 | 1779.977 | 1848.542 | 2054.46 |
| ENSMUSG00000062585 | 0.221460642 | 0.543929 | 0.692808 | 77.46619 | 29.99554 | 96.06037 | 69.8725 | 51.6809 | 43.57759 | 67.50024 | 72.8686 |
| ENSMUSG00000062588 | -0.175486606 | 0.610804 | 0.745716 | 30.18163 | 19.19714 | 19.04645 | 23.29083 | 26.32801 | 24.58223 | 20.25007 | 32.62773 |
| ENSMUSG00000062590 | 0.06892275 | 0.500404 | 0.656807 | 547.2936 | 569.9152 | 597.893 | 611.6001 | 564.5894 | 616.7905 | 482.1446 | 557.9342 |
| ENSMUSG00000062591 | 0.882018429 | 7.88E-14 | 3.03E-11 | 832.007 | 775.0847 | 893.5271 | 945.4353 | 391.0196 | 467.0624 | 507.2161 | 506.8174 |
| ENSMUSG00000062593 | 0.508135428 | 0.026983 | 0.087728 | 96.58122 | 75.58876 | 75.3577 | 82.81186 | 71.18313 | 65.92507 | 46.28588 | 48.9416 |
| ENSMUSG00000062604 | 0.241180326 | 0.02793 | 0.089945 | 2700.25 | 2608.412 | 2468.586 | 2544.739 | 2096.489 | 2595.66 | 2200.508 | 1840.204 |
| ENSMUSG00000062609 | -2.64689859 | 3.04E-06 | 9.16E-05 | 196.1806 | 69.58965 | 308.8838 | 234.6336 | 1547.502 | 486.0577 | 1121.468 | 1918.511 |
| ENSMUSG00000062611 | 0.075854456 | 0.583616 | 0.724151 | 246.4833 | 319.1525 | 275.7595 | 308.8192 | 281.8072 | 303.9258 | 271.9295 | 232.7445 |
| ENSMUSG00000062619 | -0.416484635 | 0.007253 | 0.03375 | 415.5004 | 369.545 | 515.0823 | 524.4751 | 626.0215 | 508.4052 | 603.645 | 701.4963 |
| ENSMUSG00000062627 | 0.259440753 | 0.027898 | 0.089919 | 1513.106 | 1666.552 | 1383.766 | 1450.933 | 1193.536 | 1520.746 | 1181.254 | 1127.832 |
| ENSMUSG00000062646 | -0.209407469 | 0.05791 | 0.150867 | 345.0766 | 340.7493 | 327.1021 | 333.8353 | 374.4427 | 336.3296 | 419.4658 | 425.2481 |
| ENSMUSG00000062647 | -0.16183025 | 0.043641 | 0.123912 | 25751.97 | 29287.64 | 30309.53 | 30131.44 | 34710.06 | 29621.59 | 32738.58 | 32118.74 |
| ENSMUSG00000062661 | -0.254616908 | 0.026071 | 0.0856 | 818.9282 | 718.6931 | 960.6037 | 968.7262 | 990.7131 | 1024.632 | 1026.004 | 1101.73 |
| ENSMUSG00000062683 | -0.292972143 | 0.02244 | 0.076468 | 4170.095 | 4285.763 | 5226.181 | 4488.23 | 5940.378 | 4394.632 | 5659.413 | 6268.875 |
| ENSMUSG00000062691 | -0.186360416 | 0.026931 | 0.087614 | 520.1301 | 520.7225 | 539.0974 | 576.2325 | 657.225 | 607.8515 | 611.3593 | 578.5985 |
| ENSMUSG00000062694 | -0.996676557 | 0.182431 | 0.33303 | 11.0666 | 3.599465 | 36.43669 | 7.763611 | 21.45245 | 10.05637 | 32.78583 | 54.37955 |
| ENSMUSG00000062704 | 0.812441652 | 0.097688 | 0.21743 | 10.06054 | 16.7975 | 17.39024 | 12.07673 | 7.80089 | 10.05637 | 7.714313 | 6.525547 |
| ENSMUSG00000062713 | -2.419322509 | 0.010474 | 0.044126 | 68.4117 | 5.999108 | 218.6202 | 54.34528 | 269.1307 | 187.7188 | 802.2886 | 601.4379 |
| ENSMUSG00000062727 | -0.548903106 | 0.191788 | 0.344927 | 12.07265 | 17.99732 | 14.90592 | 13.80198 | 16.57689 | 23.46486 | 14.46434 | 31.54014 |
| ENSMUSG00000062729 | 0.109260927 | 0.32111 | 0.489099 | 1412.5 | 1309.005 | 1155.209 | 1375.884 | 1031.668 | 1233.581 | 1263.219 | 1341 |
| ENSMUSG00000062743 | 0.522225446 | 0.021433 | 0.073978 | 128.775 | 196.7707 | 127.5284 | 142.3329 | 92.63557 | 137.437 | 93.53605 | 90.27006 |

| | | | | | | | | | | | |
|---|---|---|---|---|---|---|---|---|---|---|---|
| ENSMUSG00000062753 | -0.011571697 | 0.926351 | 0.956313 | 218.3138 | 207.5691 | 255.0568 | 257.9244 | 239.8774 | 251.4092 | 230.4651 | 228.3941 |
| ENSMUSG00000062758 | 0.00443525 | 0.992857 | 0.995933 | 24.1453 | 11.99822 | 14.07781 | 16.38985 | 22.42756 | 23.46486 | 7.714313 | 13.05109 |
| ENSMUSG00000062760 | -0.537797413 | 0.015948 | 0.060178 | 74.44802 | 124.7814 | 143.2624 | 113.0037 | 159.9183 | 154.1976 | 151.3934 | 196.854 |
| ENSMUSG00000062761 | 0.337420201 | 0.000726 | 0.006118 | 2017.139 | 1949.71 | 1733.227 | 2032.341 | 1491.92 | 1718.521 | 1569.863 | 1338.825 |
| ENSMUSG00000062762 | -0.148375721 | 0.013118 | 0.052372 | 3278.731 | 3381.097 | 3603.092 | 3447.906 | 3913.122 | 3580.067 | 3757.835 | 3947.956 |
| ENSMUSG00000062785 | -0.389095619 | 0.328447 | 0.496233 | 51.30877 | 23.99643 | 48.85829 | 35.36756 | 42.9049 | 26.81698 | 72.32169 | 67.43065 |
| ENSMUSG00000062794 | -0.288437798 | 0.267412 | 0.431699 | 88.53278 | 69.58965 | 44.71776 | 55.2079 | 80.93424 | 88.27255 | 69.42882 | 75.04379 |
| ENSMUSG00000062797 | -0.005021373 | 0.957108 | 0.975878 | 805.8495 | 902.2658 | 730.39 | 812.5913 | 748.8855 | 858.1433 | 827.3601 | 824.394 |
| ENSMUSG00000062822 | 0.342387356 | 0.00051 | 0.004655 | 2158.993 | 2002.502 | 1771.32 | 2029.753 | 1548.477 | 1741.986 | 1599.756 | 1388.854 |
| ENSMUSG00000062825 | -0.341326782 | 0.006478 | 0.03123 | 23898.82 | 24998.28 | 29791.14 | 25469.82 | 33532.13 | 26166.67 | 34625.7 | 37637.18 |
| ENSMUSG00000062846 | -0.367713702 | 0.253645 | 0.417294 | 19.11503 | 33.595 | 33.95237 | 34.50494 | 50.70579 | 39.10809 | 30.85725 | 35.89051 |
| ENSMUSG00000062859 | 0.497885299 | 0.005199 | 0.026428 | 297.7921 | 255.562 | 330.4145 | 317.4454 | 185.2711 | 167.6061 | 231.4294 | 267.5474 |
| ENSMUSG00000062861 | -0.044250028 | 0.667606 | 0.786753 | 395.3794 | 437.9349 | 416.5376 | 387.3179 | 441.7254 | 431.3064 | 443.573 | 368.6934 |
| ENSMUSG00000062866 | 0.441017987 | 3.91E-06 | 0.000111 | 2791.801 | 2661.204 | 2394.884 | 2782.823 | 2001.904 | 2212.401 | 1781.042 | 1835.854 |
| ENSMUSG00000062867 | -0.118783564 | 0.214427 | 0.372607 | 3477.93 | 4470.535 | 3718.199 | 3834.361 | 4549.869 | 3909.692 | 4149.336 | 4216.591 |
| ENSMUSG00000062901 | 0.165702492 | 0.117845 | 0.246482 | 3445.736 | 3142.333 | 3113.681 | 3340.078 | 3016.019 | 3197.925 | 3050.047 | 2362.248 |
| ENSMUSG00000062906 | 0.194550639 | 0.049858 | 0.135799 | 420.5307 | 454.7324 | 390.0382 | 421.8229 | 352.0152 | 400.0199 | 349.0727 | 371.9562 |
| ENSMUSG00000062908 | -0.826791657 | 0.000243 | 0.002648 | 3097.641 | 2055.294 | 2558.021 | 2502.471 | 4160.8 | 2825.839 | 4737.553 | 6392.86 |
| ENSMUSG00000062929 | 0.148349636 | 0.255482 | 0.419388 | 2947.739 | 3121.936 | 2736.892 | 3345.254 | 2720.561 | 3411.343 | 2457.973 | 2375.299 |
| ENSMUSG00000062931 | 0.088064056 | 0.515055 | 0.669227 | 326.9677 | 337.1499 | 323.7897 | 370.9281 | 346.1645 | 357.5597 | 320.144 | 254.4963 |
| ENSMUSG00000062933 | -0.238503281 | 0.004119 | 0.022314 | 1396.403 | 1619.759 | 1575.059 | 1578.601 | 1988.252 | 1698.409 | 1747.292 | 1843.467 |
| ENSMUSG00000062937 | 0.138334805 | 0.105712 | 0.229531 | 1296.804 | 1548.97 | 1411.922 | 1306.012 | 1318.35 | 1317.384 | 1214.04 | 1201.788 |
| ENSMUSG00000062939 | 0.274907685 | 0.47168 | 0.631172 | 34.20585 | 25.19625 | 38.92101 | 30.19182 | 20.47734 | 40.22547 | 30.85725 | 15.22628 |
| ENSMUSG00000062944 | 0.093805704 | 0.563725 | 0.709065 | 177.0656 | 220.7672 | 170.59 | 151.8217 | 174.5449 | 184.3667 | 149.4648 | 164.2263 |
| ENSMUSG00000062949 | 0.312272202 | 0.064434 | 0.163145 | 1041.266 | 1078.64 | 844.6688 | 1150.74 | 731.3335 | 1108.435 | 745.3955 | 729.7736 |
| ENSMUSG00000062960 | -0.11850752 | 0.343917 | 0.512667 | 1759.589 | 1378.595 | 1856.615 | 1663.138 | 1658.664 | 1668.24 | 1778.149 | 2127.328 |
| ENSMUSG00000062961 | -0.030732439 | 0.916276 | 0.950418 | 34.20585 | 43.19358 | 33.12427 | 30.19182 | 33.15378 | 36.87335 | 45.32159 | 27.18978 |
| ENSMUSG00000062963 | -0.188317836 | 0.005238 | 0.026573 | 2173.077 | 2330.053 | 2331.12 | 2329.946 | 2625.975 | 2429.171 | 2815.724 | 2568.89 |
| ENSMUSG00000062980 | 0.514944074 | 0.004709 | 0.024668 | 3637.893 | 3858.626 | 2562.99 | 3421.165 | 2498.235 | 3006.854 | 2164.829 | 1762.985 |
| ENSMUSG00000062981 | -0.388689126 | 0.000433 | 0.004094 | 1289.762 | 1413.39 | 1491.42 | 1422.466 | 1721.071 | 1575.497 | 1926.65 | 2131.679 |
| ENSMUSG00000062991 | -0.162809895 | 0.530478 | 0.682263 | 47.28455 | 62.39072 | 91.91984 | 69.8725 | 78.98402 | 60.3382 | 81.96458 | 83.74451 |
| ENSMUSG00000062995 | 0.298850088 | 0.000239 | 0.002614 | 603.6326 | 681.4986 | 645.0951 | 682.3352 | 548.0126 | 535.2222 | 528.4305 | 511.1678 |
| ENSMUSG00000062997 | -0.147011173 | 0.068382 | 0.169381 | 15017.37 | 17741.76 | 18008.83 | 17381.86 | 20265.74 | 17750.6 | 19055.32 | 18387.9 |
| ENSMUSG00000063011 | 0.800366019 | 0.002982 | 0.017679 | 500.009 | 975.4549 | 615.2832 | 802.2398 | 497.3068 | 450.3018 | 476.3588 | 236.0073 |
| ENSMUSG00000063015 | 0.116583719 | 0.153521 | 0.294867 | 7288.864 | 7292.515 | 6898.956 | 8309.652 | 6597.603 | 7393.664 | 7020.989 | 6466.817 |
| ENSMUSG00000063018 | -0.157223016 | 0.456778 | 0.61851 | 232.3986 | 170.3747 | 201.2299 | 211.3428 | 257.4294 | 145.2586 | 248.7866 | 257.7591 |
| ENSMUSG00000063019 | -0.179064453 | 0.020624 | 0.072021 | 1146.902 | 1307.805 | 1364.72 | 1297.386 | 1509.472 | 1404.539 | 1474.398 | 1405.168 |
| ENSMUSG00000063021 | -0.47179573 | 0.401784 | 0.568965 | 8.048435 | 3.599465 | 11.59349 | 10.35148 | 6.825779 | 12.29112 | 16.39292 | 11.9635 |
| ENSMUSG00000063047 | 0.187912508 | 0.126863 | 0.25798 | 390.3491 | 425.9366 | 325.4459 | 357.9887 | 350.065 | 319.569 | 298.9296 | 344.7664 |
| ENSMUSG00000063049 | -0.188048892 | 0.050334 | 0.136674 | 506.0453 | 551.9179 | 563.9406 | 591.7597 | 620.1708 | 589.9735 | 611.3593 | 702.5838 |
| ENSMUSG00000063052 | -0.027571402 | 0.712456 | 0.819871 | 1028.188 | 1079.839 | 945.6978 | 1013.583 | 1042.394 | 1091.674 | 1032.754 | 974.4816 |
| ENSMUSG00000063060 | -0.484929328 | 0.047058 | 0.130406 | 116.7023 | 79.18822 | 168.1056 | 107.8279 | 168.6943 | 146.376 | 150.4291 | 197.9416 |
| ENSMUSG00000063063 | 1.41284326 | 0.004362 | 0.023299 | 80.48435 | 27.5959 | 35.60859 | 107.8279 | 21.45245 | 35.75597 | 20.25007 | 17.40146 |
| ENSMUSG00000063065 | -0.073191578 | 0.297252 | 0.465079 | 1949.733 | 2090.089 | 2110.016 | 2189.338 | 2305.163 | 2047.029 | 2135.9 | 2286.116 |
| ENSMUSG00000063077 | 0.165653876 | 0.112378 | 0.239001 | 2096.617 | 1721.744 | 1705.072 | 2055.632 | 1562.128 | 1782.212 | 1597.827 | 1817.365 |
| ENSMUSG00000063087 | 0.319900583 | 0.205591 | 0.361991 | 121.7326 | 151.1775 | 144.9187 | 132.844 | 78.0089 | 167.6061 | 97.39321 | 98.97079 |
| ENSMUSG00000063108 | 0.016999952 | 0.897082 | 0.937817 | 826.9767 | 899.8662 | 871.9963 | 904.0294 | 897.1024 | 1059.271 | 801.3243 | 704.759 |
| ENSMUSG00000063130 | -1.350778963 | 0.011185 | 0.046441 | 18.10898 | 13.19804 | 56.31125 | 15.52722 | 58.50668 | 40.22547 | 57.85735 | 108.7591 |
| ENSMUSG00000063142 | 0.63124223 | 9.21E-05 | 0.001252 | 132.7992 | 133.1802 | 134.1533 | 125.943 | 80.93424 | 98.32892 | 75.21455 | 85.9197 |
| ENSMUSG00000063145 | 0.216205522 | 0.201303 | 0.356789 | 314.895 | 292.7565 | 235.1823 | 301.0556 | 250.6036 | 296.1041 | 246.858 | 190.3284 |
| ENSMUSG00000063146 | -0.09319792 | 0.44111 | 0.60439 | 866.2128 | 863.8715 | 1085.648 | 929.0455 | 1002.414 | 848.0869 | 1021.182 | 1125.657 |
| ENSMUSG00000063160 | 0.191707767 | 0.00626 | 0.030366 | 807.8616 | 853.0731 | 891.8708 | 884.1891 | 758.6366 | 774.3402 | 739.6098 | 739.5619 |
| ENSMUSG00000063171 | -0.110788507 | 0.400376 | 0.567711 | 1437.652 | 1466.182 | 2041.283 | 1727.835 | 1676.216 | 1745.338 | 1761.756 | 2027.27 |
| ENSMUSG00000063172 | 0.201308432 | 0.136625 | 0.271398 | 359.1614 | 427.1365 | 334.5551 | 391.6311 | 373.4676 | 341.9165 | 320.144 | 277.3357 |
| ENSMUSG00000063179 | -0.33141775 | 0.000548 | 0.004914 | 441.6579 | 440.3345 | 505.145 | 520.162 | 583.1166 | 615.6731 | 576.6449 | 630.8028 |
| ENSMUSG00000063200 | -0.027203616 | 0.581448 | 0.722532 | 2561.414 | 2771.588 | 2681.409 | 2633.59 | 2692.282 | 2771.088 | 2674.938 | 2710.277 |
| ENSMUSG00000063229 | -0.683322893 | 0.000163 | 0.001932 | 6746.601 | 6475.437 | 8344.831 | 6799.198 | 11619.43 | 7524.397 | 12188.62 | 14220.25 |
| ENSMUSG00000063234 | 0.077066535 | 0.892195 | 0.934845 | 5.030272 | 13.19804 | 18.21835 | 12.07673 | 12.67645 | 5.58687 | 11.57147 | 16.31387 |
| ENSMUSG00000063235 | -0.409491649 | 0.000328 | 0.003319 | 693.1715 | 760.6869 | 836.3877 | 772.9107 | 1154.532 | 860.3781 | 1052.039 | 1001.671 |
| ENSMUSG00000063236 | -0.023783896 | 0.796209 | 0.874797 | 1148.914 | 1187.823 | 1214.004 | 1220.612 | 1245.217 | 1372.135 | 1165.826 | 1069.102 |
| ENSMUSG00000063239 | -0.243005145 | 0.738894 | 0.83673 | 183.1019 | 121.182 | 22.35888 | 17.25247 | 121.8889 | 37.99072 | 54.96448 | 191.416 |
| ENSMUSG00000063245 | 0.650336393 | 0.03521 | 0.106548 | 39.23612 | 35.99465 | 49.6864 | 37.95543 | 18.52711 | 27.93435 | 33.75012 | 23.927 |
| ENSMUSG00000063253 | 0.084945402 | 0.302489 | 0.470759 | 981.9091 | 1025.847 | 914.2297 | 1094.669 | 930.2562 | 959.8243 | 908.3604 | 988.6203 |
| ENSMUSG00000063260 | -0.284325942 | 0.589433 | 0.729568 | 9.054489 | 14.39786 | 5.796746 | 14.6646 | 13.65156 | 11.17374 | 19.28578 | 8.700729 |

| | | | | | | | | | | | |
|---|---|---|---|---|---|---|---|---|---|---|---|
| ENSMUSG00000063268 | -0.323477744 | 0.087449 | 0.201066 | 155.9384 | 167.975 | 202.058 | 131.1188 | 176.4951 | 183.2494 | 215.0365 | 247.9708 |
| ENSMUSG00000063273 | 0.189754238 | 0.185708 | 0.337279 | 3671.092 | 3531.075 | 2831.297 | 3160.652 | 2748.839 | 3682.865 | 2576.581 | 2559.102 |
| ENSMUSG00000063275 | -0.107064571 | 0.250975 | 0.414608 | 387.3309 | 406.7395 | 455.4586 | 428.7239 | 465.1281 | 438.0106 | 438.7516 | 468.7518 |
| ENSMUSG00000063281 | -0.35102207 | 0.016503 | 0.061505 | 610.675 | 649.1034 | 497.6921 | 486.5197 | 788.865 | 592.2083 | 795.5386 | 679.7444 |
| ENSMUSG00000063286 | -0.088478248 | 0.702169 | 0.812223 | 51.30877 | 79.18822 | 65.42042 | 56.93315 | 73.13335 | 64.8077 | 54.96448 | 75.04379 |
| ENSMUSG00000063296 | -0.391073797 | 0.15463 | 0.296258 | 207.2472 | 205.1695 | 398.3193 | 236.3588 | 289.6081 | 250.2918 | 364.5013 | 470.9269 |
| ENSMUSG00000063297 | 1.408798341 | 1.35E-05 | 0.000289 | 302.8224 | 556.7172 | 358.5702 | 450.2895 | 131.64 | 261.4655 | 149.4648 | 85.9197 |
| ENSMUSG00000063316 | -0.069765589 | 0.369523 | 0.5384 | 18803.16 | 21654.38 | 20869.12 | 21486.23 | 23793.69 | 20275.87 | 21934.69 | 20908.94 |
| ENSMUSG00000063317 | 0.01679894 | 0.897403 | 0.937817 | 434.6155 | 335.95 | 359.3983 | 410.6088 | 392.9699 | 431.3064 | 358.7156 | 341.5036 |
| ENSMUSG00000063320 | -0.353892844 | 0.022085 | 0.075581 | 318.9192 | 297.5557 | 209.511 | 237.2215 | 339.3387 | 351.9728 | 331.7155 | 331.7153 |
| ENSMUSG00000063328 | -0.071606749 | 0.820411 | 0.891148 | 32.19374 | 39.59411 | 37.2648 | 27.60395 | 49.73068 | 35.75597 | 26.03581 | 31.54014 |
| ENSMUSG00000063334 | 0.096218708 | 0.187862 | 0.340153 | 1563.408 | 1571.766 | 1392.047 | 1561.349 | 1432.439 | 1512.925 | 1360.612 | 1387.766 |
| ENSMUSG00000063354 | 0.214075874 | 0.585303 | 0.725623 | 22.1332 | 26.39607 | 28.98373 | 18.97772 | 22.42756 | 31.28647 | 14.46434 | 15.22628 |
| ENSMUSG00000063358 | -0.11850368 | 0.004067 | 0.022103 | 4493.039 | 4352.953 | 4414.636 | 4491.681 | 4936.989 | 4689.619 | 4839.767 | 4806.065 |
| ENSMUSG00000063362 | 0.162785254 | 0.06949 | 0.171344 | 984.9272 | 1013.849 | 975.5096 | 890.2274 | 820.0686 | 972.1155 | 835.0744 | 824.394 |
| ENSMUSG00000063364 | 0.067382051 | 0.838685 | 0.902394 | 50.30272 | 39.59411 | 28.98373 | 39.68068 | 50.70579 | 41.34284 | 24.10723 | 34.80291 |
| ENSMUSG00000063382 | -0.038703188 | 0.793897 | 0.873661 | 543.2694 | 505.1249 | 549.0347 | 445.1137 | 482.6801 | 421.25 | 565.0735 | 628.6276 |
| ENSMUSG00000063383 | 0.401401121 | 0.058646 | 0.152167 | 99.59938 | 103.1847 | 134.1533 | 101.7896 | 74.10846 | 108.3853 | 77.14313 | 73.95619 |
| ENSMUSG00000063406 | 0.098671025 | 0.419967 | 0.585395 | 1046.297 | 927.462 | 868.6839 | 875.5628 | 757.6615 | 1039.158 | 825.4315 | 850.4962 |
| ENSMUSG00000063410 | -0.284067999 | 0.000216 | 0.002405 | 1540.269 | 1540.571 | 1677.744 | 1494.064 | 1923.895 | 1720.756 | 1996.079 | 1972.89 |
| ENSMUSG00000063412 | -0.031662694 | 0.919222 | 0.952207 | 29.17558 | 58.79126 | 34.78048 | 33.64232 | 33.15378 | 45.81234 | 40.50014 | 39.15328 |
| ENSMUSG00000063415 | 0.43735911 | 0.234946 | 0.395797 | 920.5397 | 793.082 | 915.8859 | 1265.469 | 533.3859 | 1450.352 | 529.3948 | 364.343 |
| ENSMUSG00000063428 | 1.723714683 | 6.48E-16 | 3.86E-13 | 182.0958 | 196.7707 | 179.6991 | 218.2437 | 38.02934 | 71.51194 | 67.50024 | 58.72992 |
| ENSMUSG00000063430 | 0.08598543 | 0.802238 | 0.879281 | 57.3451 | 61.1909 | 110.1382 | 74.18562 | 45.83023 | 55.8687 | 77.14313 | 107.6715 |
| ENSMUSG00000063434 | 0.704150051 | 0.034448 | 0.104826 | 86.52067 | 151.1775 | 120.0755 | 120.7673 | 46.80534 | 122.9112 | 54.96448 | 69.60583 |
| ENSMUSG00000063439 | -0.017462702 | 0.876412 | 0.925983 | 428.5792 | 403.14 | 461.2554 | 432.1744 | 457.3272 | 368.7335 | 441.6444 | 479.6277 |
| ENSMUSG00000063445 | -0.024491035 | 0.834931 | 0.900183 | 932.6124 | 1309.005 | 1069.914 | 1052.401 | 1214.989 | 1081.618 | 1074.218 | 1063.664 |
| ENSMUSG00000063446 | -0.063692599 | 0.794033 | 0.873694 | 167.005 | 161.9759 | 302.2589 | 179.4257 | 187.2214 | 175.4277 | 235.2866 | 251.2335 |
| ENSMUSG00000063450 | 0.133427417 | 0.199645 | 0.354566 | 2693.208 | 2321.655 | 2763.392 | 2538.701 | 2046.759 | 2220.222 | 2610.331 | 2530.824 |
| ENSMUSG00000063455 | 0.23280512 | 0.111724 | 0.238257 | 1150.926 | 1017.449 | 1036.79 | 1108.471 | 809.3424 | 1182.182 | 924.7533 | 755.8758 |
| ENSMUSG00000063457 | -0.202317032 | 0.020107 | 0.070711 | 23449.11 | 28320.59 | 28421.45 | 26742.19 | 33269.82 | 28189.11 | 30907.4 | 30663.54 |
| ENSMUSG00000063458 | -0.650904645 | 0.297554 | 0.465485 | 3.018163 | 26.39607 | 11.59349 | 10.35148 | 11.70134 | 24.58223 | 16.39292 | 27.18978 |
| ENSMUSG00000063480 | -0.194559147 | 0.000551 | 0.004928 | 2305.877 | 2433.238 | 2553.881 | 2393.78 | 2821.972 | 2664.937 | 2791.617 | 2808.16 |
| ENSMUSG00000063488 | 0.182045351 | 0.1691 | 0.315435 | 515.0998 | 489.5272 | 381.7572 | 474.4429 | 387.1192 | 446.9496 | 439.7159 | 364.343 |
| ENSMUSG00000063506 | -0.198428435 | 0.581449 | 0.722532 | 60.36326 | 50.3925 | 83.63877 | 31.91707 | 64.35735 | 40.22547 | 69.42882 | 85.9197 |
| ENSMUSG00000063511 | 0.121392512 | 0.012498 | 0.050512 | 10940.84 | 10300.47 | 10083.85 | 10385.99 | 9326.94 | 9806.075 | 9383.498 | 9829.648 |
| ENSMUSG00000063524 | -0.640147216 | 0.000253 | 0.002732 | 12013.3 | 11711.46 | 14260.82 | 11725.64 | 19691.4 | 12970.48 | 20453.54 | 24359.87 |
| ENSMUSG00000063531 | -0.448408969 | 0.272697 | 0.438211 | 38.23007 | 59.99108 | 125.0441 | 36.23019 | 86.78491 | 70.39457 | 94.50034 | 103.3212 |
| ENSMUSG00000063535 | 0.561214987 | 0.04762 | 0.131488 | 63.38142 | 98.38537 | 67.07664 | 50.03216 | 51.6809 | 35.75597 | 51.10733 | 48.9416 |
| ENSMUSG00000063550 | -0.000857752 | 0.990102 | 0.994413 | 1761.601 | 1650.954 | 1647.104 | 1582.051 | 1742.524 | 1726.343 | 1613.256 | 1561.781 |
| ENSMUSG00000063556 | -0.29756552 | 0.38449 | 0.552315 | 20.12109 | 22.79661 | 14.90592 | 18.97772 | 25.35289 | 20.11273 | 25.07152 | 22.83941 |
| ENSMUSG00000063558 | 0.940971024 | 0.000121 | 0.001551 | 109.6599 | 109.1838 | 74.5296 | 89.71284 | 54.60623 | 44.69496 | 60.75022 | 38.06569 |
| ENSMUSG00000063564 | -0.335448048 | 0.553247 | 0.700378 | 313.889 | 221.967 | 1112.975 | 276.9021 | 572.3903 | 405.6068 | 715.5026 | 737.3868 |
| ENSMUSG00000063568 | 0.8179673 | 2.37E-05 | 0.00045 | 178.0716 | 154.777 | 151.5435 | 194.9529 | 91.66046 | 126.2633 | 85.82174 | 82.65692 |
| ENSMUSG00000063576 | -0.101652455 | 0.214001 | 0.37205 | 2780.734 | 3005.553 | 2774.157 | 3036.435 | 3271.498 | 2728.628 | 3186.011 | 3254.073 |
| ENSMUSG00000063590 | -3.156670398 | 3.22E-06 | 9.58E-05 | 1.006054 | 5.999108 | 3.312427 | 7.763611 | 42.9049 | 20.11273 | 22.17865 | 76.13138 |
| ENSMUSG00000063594 | 0.366922193 | 0.122669 | 0.252221 | 145.8779 | 196.7707 | 156.5122 | 138.8824 | 133.5902 | 168.7235 | 87.75031 | 104.4087 |
| ENSMUSG00000063605 | 0.016416173 | 0.846214 | 0.907479 | 713.2925 | 788.2827 | 837.2158 | 751.3451 | 744.985 | 721.8237 | 801.3243 | 787.4159 |
| ENSMUSG00000063611 | 1.061967366 | 9.86E-05 | 0.001323 | 141.8537 | 116.3827 | 93.57605 | 131.9814 | 33.15378 | 69.27719 | 66.53595 | 63.08028 |
| ENSMUSG00000063623 | -0.036663407 | 0.882038 | 0.929373 | 63.38142 | 68.38983 | 69.56096 | 60.38364 | 45.83023 | 62.57295 | 73.28598 | 87.00729 |
| ENSMUSG00000063626 | 0.965276719 | 0.00652 | 0.031379 | 33.19979 | 74.38894 | 56.31125 | 37.09281 | 23.40267 | 31.28647 | 25.07152 | 22.83941 |
| ENSMUSG00000063628 | 0.040584822 | 0.903562 | 0.942315 | 28.16952 | 17.99732 | 24.8432 | 23.29083 | 24.37778 | 26.81698 | 18.32149 | 22.83941 |
| ENSMUSG00000063632 | 0.457262378 | 0.056503 | 0.14836 | 4179.15 | 3772.239 | 2586.177 | 3317.65 | 2231.055 | 3928.687 | 2038.507 | 1893.496 |
| ENSMUSG00000063646 | -0.072595885 | 0.881311 | 0.928766 | 26.15741 | 10.79839 | 22.35888 | 39.68068 | 40.95467 | 22.34748 | 15.42863 | 26.10219 |
| ENSMUSG00000063652 | 0.265407877 | 0.032941 | 0.101158 | 353.1251 | 369.545 | 372.648 | 416.6471 | 316.9112 | 363.1466 | 310.5011 | 268.635 |
| ENSMUSG00000063659 | -0.007790745 | 0.935446 | 0.961785 | 804.8435 | 862.6717 | 895.1833 | 913.5183 | 781.0642 | 984.4066 | 882.3246 | 850.4962 |
| ENSMUSG00000063663 | 0.128441238 | 0.25296 | 0.41662 | 542.2633 | 458.3318 | 434.756 | 465.8167 | 397.8454 | 469.2971 | 413.6801 | 458.9634 |
| ENSMUSG00000063681 | 0.472607133 | 0.072161 | 0.176275 | 161.9748 | 167.975 | 147.403 | 160.448 | 100.4365 | 183.2494 | 81.00029 | 95.70802 |
| ENSMUSG00000063683 | -2.12184324 | 8.47E-06 | 0.000202 | 74.44802 | 22.79661 | 178.0429 | 78.49874 | 370.5423 | 251.4092 | 352.9298 | 570.9853 |
| ENSMUSG00000063684 | -0.61369548 | 0.050631 | 0.137247 | 29.17558 | 22.79661 | 26.49941 | 18.97772 | 41.92979 | 26.81698 | 37.60728 | 42.41605 |
| ENSMUSG00000063687 | 0.366424595 | 0.154889 | 0.296538 | 77.46619 | 134.38 | 75.3577 | 74.18562 | 65.33246 | 73.74669 | 75.21455 | 64.16787 |
| ENSMUSG00000063694 | -0.936412481 | 2.95E-05 | 0.000529 | 1901.443 | 1706.146 | 2384.947 | 1808.921 | 3430.442 | 2244.805 | 4209.122 | 5047.51 |
| ENSMUSG00000063696 | -0.175087774 | 0.280669 | 0.447016 | 161.9748 | 233.9652 | 179.6991 | 201.8539 | 247.6783 | 196.6578 | 231.4294 | 199.0292 |
| ENSMUSG00000063698 | 0.214210931 | 0.181224 | 0.331597 | 198.1927 | 238.7645 | 177.2148 | 239.8093 | 163.8187 | 206.7142 | 170.6792 | 194.6788 |

| | | | | | | | | | | | |
|---|---|---|---|---|---|---|---|---|---|---|---|
| ENSMUSG00000063704 | -0.328067096 | 0.174498 | 0.322656 | 204.229 | 101.9848 | 181.3554 | 165.6237 | 164.7938 | 191.071 | 204.4293 | 262.1095 |
| ENSMUSG00000063714 | -0.089242075 | 0.62278 | 0.75435 | 151.9142 | 171.5745 | 215.3077 | 223.4195 | 223.3005 | 224.5922 | 198.6436 | 166.4014 |
| ENSMUSG00000063727 | 0.527632364 | 0.107226 | 0.231621 | 236.4228 | 415.1383 | 166.4494 | 203.5791 | 247.6783 | 177.6625 | 171.6435 | 109.8467 |
| ENSMUSG00000063730 | -3.863385664 | 3.31E-06 | 9.75E-05 | 2.012109 | 0 | 9.109173 | 0.862623 | 29.25334 | 18.99536 | 67.50024 | 66.34306 |
| ENSMUSG00000063754 | -0.437030096 | 0.121176 | 0.250411 | 32.19374 | 33.595 | 27.32752 | 26.74133 | 38.02934 | 40.22547 | 50.14304 | 32.62773 |
| ENSMUSG00000063760 | 0.133866478 | 0.258736 | 0.422587 | 691.1593 | 647.9036 | 742.8116 | 692.6867 | 552.8881 | 762.0491 | 610.395 | 606.8758 |
| ENSMUSG00000063765 | 0.217239808 | 0.135216 | 0.269381 | 176.0595 | 195.5709 | 194.6051 | 227.7326 | 193.072 | 172.0756 | 164.8934 | 153.3503 |
| ENSMUSG00000063785 | 0.058414875 | 0.36607 | 0.534678 | 1066.418 | 1101.436 | 1020.227 | 1095.532 | 993.6384 | 1070.444 | 1037.575 | 1010.372 |
| ENSMUSG00000063787 | -0.163170948 | 0.093157 | 0.209901 | 881.3036 | 1112.235 | 1014.431 | 957.5121 | 1105.776 | 1035.806 | 1088.682 | 1207.226 |
| ENSMUSG00000063796 | -1.854957495 | 2.94E-06 | 8.88E-05 | 148.896 | 91.18644 | 279.0719 | 163.8985 | 772.2882 | 309.5126 | 421.3944 | 972.3064 |
| ENSMUSG00000063800 | -0.048951044 | 0.409316 | 0.575555 | 1499.021 | 1581.365 | 1445.046 | 1548.409 | 1586.506 | 1521.864 | 1616.149 | 1553.08 |
| ENSMUSG00000063801 | 0.005264859 | 0.93424 | 0.961185 | 1642.887 | 1713.345 | 1757.242 | 1624.32 | 1742.524 | 1622.427 | 1602.649 | 1745.584 |
| ENSMUSG00000063802 | -0.265227879 | 0.012252 | 0.049763 | 1070.442 | 1058.243 | 1002.837 | 961.8252 | 1126.254 | 1102.848 | 1248.754 | 1439.971 |
| ENSMUSG00000063804 | 0.169786023 | 0.633021 | 0.761309 | 30.18163 | 22.79661 | 26.49941 | 34.50494 | 29.25334 | 34.6386 | 16.39292 | 21.75182 |
| ENSMUSG00000063808 | -0.031592839 | 0.724196 | 0.827323 | 813.898 | 712.694 | 699.7501 | 660.7696 | 738.1593 | 763.1665 | 718.3954 | 729.7736 |
| ENSMUSG00000063810 | 0.134545957 | 0.344195 | 0.512916 | 446.6881 | 455.9322 | 370.1637 | 396.8068 | 376.393 | 464.8276 | 330.7512 | 348.0291 |
| ENSMUSG00000063838 | 0.074252816 | 0.603995 | 0.740933 | 415.5004 | 424.7368 | 593.7525 | 446.839 | 493.4063 | 420.1327 | 435.8587 | 439.3868 |
| ENSMUSG00000063849 | 0.091677535 | 0.413457 | 0.579273 | 577.4752 | 514.7234 | 549.0347 | 546.9033 | 486.5805 | 592.2083 | 532.2876 | 443.7372 |
| ENSMUSG00000063851 | -0.859738019 | 0.151598 | 0.291964 | 18.10898 | 11.99822 | 60.45178 | 13.80198 | 53.63112 | 37.99072 | 23.14294 | 76.13138 |
| ENSMUSG00000063856 | -0.605160386 | 0.000691 | 0.005886 | 3917.576 | 3269.514 | 4586.055 | 3731.709 | 6346.024 | 4168.923 | 5589.02 | 7483.714 |
| ENSMUSG00000063870 | -0.062484587 | 0.318596 | 0.486795 | 11024.34 | 11369.51 | 11303.66 | 10706.88 | 10573.13 | 12054.23 | 11607.15 | 12136.43 |
| ENSMUSG00000063873 | 0.790075432 | 0.008667 | 0.038451 | 471.8395 | 746.289 | 546.5504 | 476.1682 | 274.0063 | 557.5697 | 254.5723 | 209.9051 |
| ENSMUSG00000063875 | 1.301275722 | 0.013495 | 0.053377 | 13.07871 | 22.79661 | 12.4216 | 16.38985 | 4.875557 | 10.05637 | 4.821446 | 6.525547 |
| ENSMUSG00000063882 | -0.383055502 | 0.00233 | 0.014615 | 4069.49 | 4039.799 | 4947.937 | 4172.51 | 5701.476 | 4451.618 | 6039.343 | 6278.663 |
| ENSMUSG00000063884 | -0.04888579 | 0.434553 | 0.598739 | 1543.287 | 1524.973 | 1471.545 | 1524.256 | 1617.71 | 1482.755 | 1519.72 | 1650.963 |
| ENSMUSG00000063887 | 0.905740351 | 0.018603 | 0.067083 | 87.52673 | 134.38 | 88.60741 | 123.3552 | 43.88001 | 110.62 | 40.50014 | 36.9781 |
| ENSMUSG00000063888 | -0.16023518 | 0.031381 | 0.097437 | 3061.423 | 3238.318 | 3106.228 | 3218.448 | 3719.075 | 3145.408 | 3484.941 | 3755.452 |
| ENSMUSG00000063889 | 0.171714206 | 0.287071 | 0.454071 | 542.2633 | 419.9375 | 352.7734 | 452.8773 | 329.5876 | 395.5504 | 445.5016 | 398.0583 |
| ENSMUSG00000063894 | 0.111960267 | 0.192406 | 0.345566 | 1098.611 | 940.6601 | 991.2436 | 1005.819 | 886.3762 | 1025.749 | 893.8961 | 932.0656 |
| ENSMUSG00000063895 | -0.176834998 | 0.029668 | 0.093812 | 870.237 | 837.4754 | 916.714 | 853.1346 | 1038.494 | 892.7819 | 960.432 | 1040.825 |
| ENSMUSG00000063897 | -0.165438806 | 0.225656 | 0.385513 | 1079.496 | 922.6628 | 891.8708 | 1071.378 | 1266.67 | 979.9371 | 937.2891 | 1263.781 |
| ENSMUSG00000063902 | -0.092405414 | 0.247379 | 0.410579 | 697.1957 | 713.8938 | 740.3273 | 768.5975 | 804.4668 | 731.88 | 748.2884 | 830.9196 |
| ENSMUSG00000063903 | -3.170319749 | 7.21E-05 | 0.001057 | 21.12714 | 20.39697 | 151.5435 | 26.74133 | 488.5308 | 203.3621 | 492.7518 | 799.3794 |
| ENSMUSG00000063904 | -0.143711538 | 0.149838 | 0.289784 | 1807.88 | 1684.549 | 1861.584 | 1704.544 | 2098.44 | 1659.301 | 1885.185 | 2155.606 |
| ENSMUSG00000063919 | 0.447554304 | 0.27909 | 0.445456 | 57.3451 | 35.99465 | 24.01509 | 55.2079 | 30.22845 | 48.04709 | 29.89296 | 18.48905 |
| ENSMUSG00000063931 | -0.585406256 | 0.000255 | 0.002751 | 1369.24 | 1438.586 | 1497.217 | 1327.578 | 2241.781 | 1507.338 | 2048.15 | 2653.722 |
| ENSMUSG00000063935 | -0.374427069 | 0.471939 | 0.631304 | 17.10292 | 8.398751 | 32.29616 | 10.35148 | 18.52711 | 15.64324 | 28.92867 | 26.10219 |
| ENSMUSG00000063952 | 0.003714324 | 0.963432 | 0.97919 | 1206.259 | 1028.247 | 1029.337 | 1068.791 | 1049.22 | 1068.21 | 1062.647 | 1141.971 |
| ENSMUSG00000063953 | -0.246403847 | 0.609941 | 0.745057 | 15.09082 | 16.7975 | 13.24971 | 17.25247 | 18.52711 | 32.40385 | 14.46434 | 8.700729 |
| ENSMUSG00000063972 | 0.129692982 | 0.652137 | 0.775356 | 164.9929 | 97.18554 | 160.6527 | 127.6683 | 92.63557 | 185.4841 | 135.0005 | 91.35765 |
| ENSMUSG00000064023 | 0.051965974 | 0.694061 | 0.80665 | 393.3673 | 543.5192 | 428.9592 | 411.4714 | 448.5512 | 407.8415 | 459.9659 | 393.708 |
| ENSMUSG00000064030 | -0.016691975 | 0.784733 | 0.867455 | 960.7819 | 1018.648 | 976.3377 | 950.6111 | 976.0864 | 1007.871 | 976.8249 | 988.6203 |
| ENSMUSG00000064032 | 0.155714872 | 0.700205 | 0.810727 | 14.08476 | 19.19714 | 22.35888 | 25.8787 | 22.42756 | 22.34748 | 11.57147 | 17.40146 |
| ENSMUSG00000064036 | -1.50948181 | 0.002029 | 0.013272 | 4.024217 | 14.39786 | 9.93728 | 6.900988 | 19.50223 | 34.6386 | 20.25007 | 25.0146 |
| ENSMUSG00000064037 | -0.003325421 | 0.963153 | 0.978992 | 1022.151 | 1034.246 | 1030.165 | 968.7262 | 1078.473 | 1051.449 | 974.8963 | 958.1677 |
| ENSMUSG00000064043 | 0.660985786 | 0.004287 | 0.022996 | 1267.628 | 1037.846 | 1019.399 | 948.8858 | 572.3903 | 1025.749 | 652.8238 | 452.4379 |
| ENSMUSG00000064061 | 0.425710097 | 0.003407 | 0.019401 | 1344.089 | 1599.362 | 1118.772 | 1365.533 | 874.6748 | 1219.055 | 922.8247 | 1023.423 |
| ENSMUSG00000064065 | 0.497083636 | 0.005657 | 0.028163 | 278.6771 | 281.9581 | 283.2125 | 238.9467 | 155.0427 | 252.5265 | 163.9292 | 196.854 |
| ENSMUSG00000064068 | -0.194040166 | 0.033676 | 0.10285 | 1057.363 | 1131.432 | 1127.881 | 1130.899 | 1405.135 | 1099.496 | 1298.898 | 1282.27 |
| ENSMUSG00000064080 | 0.842547679 | 0.000617 | 0.005391 | 1381.313 | 2372.047 | 1609.011 | 2134.131 | 1075.548 | 1479.403 | 957.5391 | 667.7809 |
| ENSMUSG00000064090 | 0.029235916 | 0.83671 | 0.901267 | 183.1019 | 235.165 | 221.9326 | 182.8762 | 183.3209 | 204.4795 | 202.5007 | 215.343 |
| ENSMUSG00000064105 | -0.910370152 | 0.000155 | 0.00187 | 266.6044 | 170.3747 | 204.5423 | 223.4195 | 397.8454 | 253.6439 | 429.1087 | 545.9707 |
| ENSMUSG00000064109 | 0.160968386 | 0.826303 | 0.894495 | 7.04238 | 26.39607 | 6.624853 | 6.038364 | 2.925334 | 10.05637 | 13.50005 | 14.13868 |
| ENSMUSG00000064115 | 0.258088368 | 0.384453 | 0.552315 | 46.2785 | 45.59322 | 46.37397 | 63.83414 | 40.95467 | 56.98608 | 44.3573 | 27.18978 |
| ENSMUSG00000064120 | -0.279208906 | 0.011607 | 0.047752 | 684.117 | 638.3051 | 645.9232 | 628.8525 | 830.7948 | 642.4901 | 856.2888 | 820.0437 |
| ENSMUSG00000064125 | 1.561127346 | 2.16E-13 | 7.67E-11 | 340.0464 | 415.1383 | 280.7281 | 337.2858 | 129.6898 | 148.6108 | 106.0718 | 80.48174 |
| ENSMUSG00000064127 | -0.075566071 | 0.333904 | 0.502081 | 1186.138 | 1301.806 | 1180.88 | 1228.376 | 1251.068 | 1404.539 | 1193.79 | 1310.547 |
| ENSMUSG00000064128 | -0.001819225 | 0.986471 | 0.992305 | 275.6589 | 261.5611 | 243.4633 | 282.0779 | 268.1556 | 264.8177 | 275.7867 | 254.4963 |
| ENSMUSG00000064138 | 0.169226728 | 0.02742 | 0.088689 | 981.9091 | 1037.846 | 973.8534 | 996.3301 | 890.2766 | 969.8807 | 864.9674 | 822.2189 |
| ENSMUSG00000064141 | -0.100267942 | 0.562832 | 0.708328 | 194.1685 | 159.5763 | 153.1997 | 150.9591 | 196.9725 | 202.2447 | 163.9292 | 141.3868 |
| ENSMUSG00000064145 | -0.112909707 | 0.088191 | 0.202142 | 1789.771 | 1817.73 | 1840.881 | 1820.136 | 2123.792 | 1990.043 | 1922.793 | 1822.803 |
| ENSMUSG00000064147 | 0.666173057 | 0.214813 | 0.372961 | 19.11503 | 7.198929 | 19.87456 | 25.8787 | 11.70134 | 10.05637 | 17.3572 | 6.525547 |
| ENSMUSG00000064158 | 0.275098828 | 0.607025 | 0.742971 | 18.10898 | 11.99822 | 19.04645 | 16.38985 | 7.80089 | 5.58687 | 20.25007 | 20.66423 |
| ENSMUSG00000064165 | -0.325759953 | 0.397741 | 0.564974 | 23.13925 | 33.595 | 14.90592 | 29.3292 | 37.05423 | 35.75597 | 19.28578 | 33.71532 |

| | | | | | | | | | | | |
|---|---|---|---|---|---|---|---|---|---|---|---|
| ENSMUSG00000064179 | 0.699982002 | 0.002169 | 0.013927 | 400.4096 | 653.9027 | 557.3158 | 421.8229 | 398.8205 | 353.0902 | 220.8222 | 278.4233 |
| ENSMUSG00000064181 | -0.476661921 | 9.07E-05 | 0.001242 | 2526.202 | 2168.078 | 2779.954 | 2623.238 | 3348.532 | 2946.515 | 3749.156 | 4011.036 |
| ENSMUSG00000064193 | -0.705791288 | 0.015782 | 0.059834 | 26.15741 | 33.595 | 36.43669 | 30.19182 | 56.55646 | 33.52122 | 54.96448 | 60.9051 |
| ENSMUSG00000064202 | -0.049777362 | 0.868178 | 0.921132 | 69.41775 | 44.3934 | 67.07664 | 81.08661 | 45.83023 | 58.10345 | 76.17884 | 92.44524 |
| ENSMUSG00000064210 | 0.297887081 | 0.001427 | 0.010218 | 3851.176 | 3682.252 | 4017.145 | 4004.298 | 2742.988 | 3612.47 | 3179.261 | 3122.474 |
| ENSMUSG00000064215 | 0.715472946 | 5.01E-05 | 0.000794 | 937.6427 | 730.6913 | 581.3309 | 577.9577 | 443.6756 | 481.5882 | 403.0729 | 392.6204 |
| ENSMUSG00000064225 | -1.997359981 | 9.45E-06 | 0.000217 | 46.2785 | 15.59768 | 61.27989 | 38.81806 | 175.52 | 79.33356 | 128.2505 | 267.5474 |
| ENSMUSG00000064246 | 0.598144214 | 0.083466 | 0.194528 | 56.33904 | 53.99197 | 49.6864 | 62.10889 | 23.40267 | 41.34284 | 57.85735 | 23.927 |
| ENSMUSG00000064247 | -0.032109915 | 0.858384 | 0.915187 | 139.8416 | 93.58608 | 137.4657 | 119.042 | 118.9636 | 139.6718 | 117.6433 | 127.2482 |
| ENSMUSG00000064254 | 0.27555558 | 0.002919 | 0.017406 | 718.3228 | 811.0794 | 824.7942 | 703.0381 | 663.0757 | 586.6214 | 644.1452 | 629.7152 |
| ENSMUSG00000064262 | -0.001090198 | 0.995753 | 0.997645 | 251.5136 | 199.1704 | 273.2752 | 269.1385 | 200.8729 | 193.3057 | 281.5724 | 319.7518 |
| ENSMUSG00000064264 | -0.304027707 | 0.001306 | 0.009561 | 510.0696 | 592.7118 | 592.9243 | 534.8266 | 641.6232 | 698.3588 | 668.2524 | 746.0875 |
| ENSMUSG00000064267 | 0.150060762 | 0.371391 | 0.540396 | 327.9737 | 421.1374 | 363.5388 | 347.6373 | 378.3432 | 368.7335 | 326.894 | 240.3576 |
| ENSMUSG00000064272 | -0.63528786 | 0.217433 | 0.375877 | 6.036326 | 5.999108 | 8.281066 | 11.21411 | 15.60178 | 13.40849 | 8.678602 | 11.9635 |
| ENSMUSG00000064281 | 0.63345016 | 0.203467 | 0.359416 | 20.12109 | 14.39786 | 13.24971 | 8.626235 | 6.825779 | 8.938993 | 10.60718 | 9.78832 |
| ENSMUSG00000064284 | -0.412066075 | 0.001265 | 0.009324 | 200.2048 | 260.3613 | 237.6666 | 235.4962 | 310.0854 | 274.874 | 336.5369 | 318.6642 |
| ENSMUSG00000064289 | 0.115971839 | 0.259584 | 0.423619 | 644.8808 | 514.7234 | 562.2844 | 558.98 | 560.689 | 546.3959 | 509.1447 | 489.416 |
| ENSMUSG00000064293 | 0.829284198 | 0.011063 | 0.046079 | 200.2048 | 305.9545 | 225.245 | 228.5952 | 162.8436 | 213.4185 | 86.78602 | 77.21897 |
| ENSMUSG00000064294 | 1.522217548 | 1.05E-07 | 5.87E-06 | 1543.287 | 1823.729 | 1065.773 | 1759.752 | 555.8134 | 836.9132 | 469.6088 | 293.6496 |
| ENSMUSG00000064302 | 0.107601292 | 0.159452 | 0.302482 | 1685.141 | 1564.567 | 1546.075 | 1518.217 | 1427.563 | 1544.211 | 1545.756 | 1341 |
| ENSMUSG00000064307 | -0.394543191 | 0.077931 | 0.185652 | 64.38748 | 46.79304 | 56.31125 | 44.85642 | 72.15824 | 60.3382 | 68.46453 | 78.30656 |
| ENSMUSG00000064317 | -0.090795498 | 0.743011 | 0.839394 | 28.16952 | 40.79393 | 33.95237 | 43.9938 | 39.00445 | 44.69496 | 32.78583 | 40.24087 |
| ENSMUSG00000064325 | 0.732737718 | 0.000941 | 0.007478 | 276.6649 | 223.1668 | 194.6051 | 336.4232 | 126.7645 | 196.6578 | 154.2863 | 143.562 |
| ENSMUSG00000064326 | -0.208501621 | 0.003617 | 0.020338 | 1516.124 | 1616.16 | 1681.056 | 1633.809 | 1945.347 | 1766.568 | 1972.936 | 1764.073 |
| ENSMUSG00000064329 | 0.932738194 | 0.001919 | 0.012734 | 86.52067 | 123.5816 | 94.40416 | 122.4925 | 60.4569 | 84.92043 | 40.50014 | 38.06569 |
| ENSMUSG00000064330 | 0.100489749 | 0.729522 | 0.830192 | 24.1453 | 41.99375 | 37.2648 | 39.68068 | 30.22845 | 34.6386 | 32.78583 | 35.89051 |
| ENSMUSG00000064336 | 0.060191973 | 0.837212 | 0.901479 | 38.23007 | 44.3934 | 40.57722 | 33.64232 | 45.83023 | 41.34284 | 38.57157 | 23.927 |
| ENSMUSG00000064338 | 0.234455295 | 0.468116 | 0.628422 | 66.39959 | 110.3836 | 91.09173 | 116.4542 | 52.65601 | 100.5637 | 115.7147 | 57.64233 |
| ENSMUSG00000064340 | -0.726542261 | 1.50E-06 | 5.18E-05 | 588.5418 | 761.8867 | 691.469 | 746.1693 | 1023.867 | 933.0074 | 1305.648 | 1349.701 |
| ENSMUSG00000064341 | -0.472792106 | 3.21E-10 | 4.66E-08 | 102019.9 | 113606.3 | 115107.7 | 112273 | 153005.7 | 138261.6 | 162154.9 | 161378.9 |
| ENSMUSG00000064342 | -1.14461806 | 0.042219 | 0.121125 | 12.07265 | 2.399643 | 6.624853 | 5.175741 | 15.60178 | 11.17374 | 15.42863 | 16.31387 |
| ENSMUSG00000064343 | 0.011285405 | 0.981376 | 0.989687 | 11.0666 | 19.19714 | 21.53077 | 25.01608 | 30.22845 | 8.938993 | 16.39292 | 20.66423 |
| ENSMUSG00000064345 | -0.555082514 | 1.42E-07 | 7.53E-06 | 39944.38 | 39482.53 | 42269.05 | 37962.33 | 58673.42 | 48581.19 | 59192.89 | 68131.06 |
| ENSMUSG00000064346 | -0.782226434 | 0.043261 | 0.123158 | 16.09687 | 35.99465 | 28.98373 | 20.70296 | 27.30312 | 36.87335 | 58.82164 | 51.11678 |
| ENSMUSG00000064347 | -0.626242191 | 6.88E-06 | 0.000173 | 215.2956 | 233.9652 | 231.0418 | 234.6336 | 332.513 | 294.9868 | 360.6441 | 424.1605 |
| ENSMUSG00000064348 | -0.051006278 | 0.647992 | 0.772019 | 1406.464 | 1562.168 | 1924.52 | 1770.103 | 1812.732 | 1622.427 | 1674.006 | 1797.788 |
| ENSMUSG00000064349 | 0.013321909 | 0.898729 | 0.938865 | 353.1251 | 405.5397 | 425.6468 | 417.5098 | 366.6419 | 424.6022 | 400.18 | 398.0583 |
| ENSMUSG00000064350 | -0.605727783 | 0.069043 | 0.170616 | 49.29666 | 29.99554 | 45.54586 | 31.05445 | 61.43201 | 33.52122 | 67.50024 | 75.04379 |
| ENSMUSG00000064351 | -0.398249939 | 8.89E-05 | 0.001228 | 223732.4 | 226610.3 | 241774 | 245347.4 | 305951.9 | 255200.4 | 330386.7 | 343953.9 |
| ENSMUSG00000064354 | -0.369297889 | 9.39E-06 | 0.000216 | 541.2572 | 538.7199 | 618.5957 | 573.6446 | 772.2882 | 728.5279 | 734.7883 | 703.6714 |
| ENSMUSG00000064356 | -0.388417518 | 0.004189 | 0.02259 | 148.896 | 184.7725 | 163.137 | 174.2499 | 240.8525 | 203.3621 | 229.5008 | 202.2919 |
| ENSMUSG00000064357 | -0.283021775 | 0.00104 | 0.008056 | 503.0272 | 533.9206 | 584.6433 | 587.4466 | 659.1752 | 690.5372 | 652.8238 | 690.6203 |
| ENSMUSG00000064358 | -0.176394003 | 0.021038 | 0.073067 | 679.0867 | 782.2836 | 755.2332 | 764.2844 | 811.2926 | 857.0259 | 836.0387 | 864.6349 |
| ENSMUSG00000064359 | 0.003531136 | 0.994919 | 0.997071 | 30.18163 | 14.39786 | 34.78048 | 19.84034 | 6.825779 | 32.40385 | 39.53586 | 20.66423 |
| ENSMUSG00000064360 | 0.16266834 | 0.393416 | 0.560816 | 586.5297 | 658.702 | 582.9871 | 615.0506 | 378.3432 | 734.1148 | 582.4307 | 488.3284 |
| ENSMUSG00000064361 | -0.119633767 | 0.820699 | 0.891357 | 13.07871 | 15.59768 | 5.796746 | 12.07673 | 6.825779 | 13.40849 | 12.53576 | 17.40146 |
| ENSMUSG00000064363 | -0.228154289 | 9.21E-06 | 0.000214 | 41431.33 | 42528.87 | 42655.77 | 40479.47 | 46466 | 49181.22 | 48280.99 | 51797.61 |
| ENSMUSG00000064364 | -0.440932714 | 0.407147 | 0.573567 | 25.15136 | 20.39697 | 28.15563 | 14.6646 | 7.80089 | 27.93435 | 46.28588 | 38.06569 |
| ENSMUSG00000064365 | -0.160786311 | 0.83424 | 0.899657 | 26.15741 | 5.999108 | 15.73403 | 4.313117 | 5.850668 | 5.58687 | 24.10723 | 22.83941 |
| ENSMUSG00000064366 | 0.039830677 | 0.899526 | 0.939485 | 124.7507 | 89.98662 | 70.38906 | 89.71284 | 46.80534 | 113.9722 | 107.0361 | 96.79561 |
| ENSMUSG00000064367 | -0.486849222 | 0.001159 | 0.008752 | 56025.15 | 54765.85 | 59003.43 | 45543.93 | 77568.15 | 55781.55 | 77994.6 | 90425.59 |
| ENSMUSG00000064368 | -0.226058882 | 0.078867 | 0.187333 | 22296.18 | 22171.5 | 20927.91 | 17434.48 | 20603.13 | 28632.71 | 21981.94 | 25662.8 |
| ENSMUSG00000064369 | -0.642666743 | 0.002344 | 0.014688 | 645.8869 | 465.5308 | 617.7675 | 398.5321 | 779.1139 | 614.5558 | 847.6102 | 1081.066 |
| ENSMUSG00000064370 | -0.37100357 | 1.90E-07 | 9.58E-06 | 115542.3 | 122898.9 | 124480.2 | 121736.9 | 154803.8 | 140489.7 | 163852 | 167639.1 |
| ENSMUSG00000064371 | -0.534049695 | 0.283957 | 0.450503 | 17.10292 | 8.398751 | 15.73403 | 17.25247 | 13.65156 | 11.17374 | 28.92867 | 31.54014 |
| ENSMUSG00000064372 | -0.757607016 | 2.01E-06 | 6.48E-05 | 1155.956 | 829.0767 | 896.0114 | 1074.829 | 1705.47 | 1280.511 | 1733.792 | 1969.627 |
| ENSMUSG00000064373 | 0.682065721 | 1.47E-11 | 3.35E-09 | 8330.13 | 8866.681 | 7579.66 | 9807.167 | 5296.805 | 5895.266 | 5502.234 | 4860.445 |
| ENSMUSG00000064390 | -0.360366751 | 0.420487 | 0.585947 | 12.07265 | 17.99732 | 18.21835 | 30.19182 | 36.07912 | 27.93435 | 14.46434 | 22.83941 |
| ENSMUSG00000064427 | -0.231363689 | 0.698181 | 0.809552 | 10.06054 | 11.99822 | 10.76539 | 16.38985 | 24.37778 | 11.17374 | 3.857157 | 18.48905 |
| ENSMUSG00000064437 | 0.007218945 | 0.980464 | 0.988984 | 28.16952 | 43.19358 | 38.09291 | 25.8787 | 34.1289 | 30.1691 | 34.71441 | 34.80291 |
| ENSMUSG00000064493 | 0.156086658 | 0.566834 | 0.711182 | 59.35721 | 39.59411 | 55.48314 | 81.94923 | 54.60623 | 48.04709 | 52.07161 | 58.72992 |
| ENSMUSG00000064602 | 0.209642572 | 0.691286 | 0.804437 | 27.16347 | 28.79572 | 19.04645 | 38.81806 | 28.27823 | 42.46022 | 21.21436 | 6.525547 |
| ENSMUSG00000064612 | 0.453543494 | 0.347744 | 0.516479 | 21.12714 | 15.59768 | 19.04645 | 28.46658 | 24.37778 | 10.05637 | 19.28578 | 7.613138 |
| ENSMUSG00000064647 | -0.063996659 | 0.882801 | 0.929859 | 19.11503 | 33.595 | 24.8432 | 56.93315 | 36.07912 | 42.46022 | 20.25007 | 42.41605 |

| | | | | | | | | | | | |
|---|---|---|---|---|---|---|---|---|---|---|---|
| ENSMUSG00000064672 | 0.121663826 | 0.768269 | 0.856029 | 31.18769 | 19.19714 | 18.21835 | 31.05445 | 25.35289 | 31.28647 | 23.14294 | 11.9635 |
| ENSMUSG00000064778 | -0.307515187 | 0.62351 | 0.754837 | 10.06054 | 3.599465 | 14.07781 | 13.80198 | 8.776002 | 5.58687 | 20.25007 | 17.40146 |
| ENSMUSG00000064844 | 0.463662446 | 0.16694 | 0.312673 | 46.2785 | 28.79572 | 38.92101 | 27.60395 | 18.52711 | 32.40385 | 25.07152 | 27.18978 |
| ENSMUSG00000064871 | 0.382154887 | 0.197587 | 0.351893 | 40.24217 | 39.59411 | 37.2648 | 57.79577 | 36.07912 | 42.46022 | 27.96439 | 28.27737 |
| ENSMUSG00000064917 | 0.1261309 | 0.762076 | 0.852086 | 12.07265 | 14.39786 | 14.90592 | 18.11509 | 16.57689 | 14.52586 | 10.60718 | 13.05109 |
| ENSMUSG00000064918 | -0.16731377 | 0.657061 | 0.778951 | 23.13925 | 15.59768 | 19.87456 | 20.70296 | 31.20356 | 16.76061 | 17.3572 | 23.927 |
| ENSMUSG00000064952 | 0.806432257 | 0.335397 | 0.503669 | 16.09687 | 16.7975 | 11.59349 | 12.07673 | 1.950223 | 24.58223 | 3.857157 | 2.175182 |
| ENSMUSG00000064984 | 0.376672466 | 0.374821 | 0.543403 | 17.10292 | 22.79661 | 19.87456 | 26.74133 | 17.552 | 20.11273 | 21.21436 | 7.613138 |
| ENSMUSG00000065037 | 0.154357221 | 0.724325 | 0.827323 | 75.45408 | 45.59322 | 32.29616 | 50.89479 | 77.03379 | 51.39921 | 31.82154 | 22.83941 |
| ENSMUSG00000065126 | 0.095052812 | 0.510337 | 0.66605 | 232.3986 | 245.9634 | 181.3554 | 197.5408 | 208.6738 | 183.2494 | 201.5364 | 205.5547 |
| ENSMUSG00000065147 | 0.308042074 | 0.272465 | 0.437956 | 64.38748 | 93.58608 | 64.59232 | 104.3774 | 72.15824 | 70.39457 | 43.39301 | 78.30656 |
| ENSMUSG00000065162 | 0.887700742 | 0.109712 | 0.235359 | 14.08476 | 10.79839 | 14.07781 | 14.6646 | 5.850668 | 13.40849 | 6.750024 | 3.262773 |
| ENSMUSG00000065191 | 0.210334109 | 0.600616 | 0.738618 | 18.10898 | 32.39518 | 21.53077 | 27.60395 | 14.62667 | 32.40385 | 16.39292 | 22.83941 |
| ENSMUSG00000065208 | 0.140274777 | 0.791466 | 0.87239 | 7.04238 | 8.398751 | 10.76539 | 18.97772 | 7.80089 | 12.29112 | 10.60718 | 10.87591 |
| ENSMUSG00000065219 | 0.1090174 | 0.833937 | 0.899488 | 14.08476 | 8.398751 | 18.21835 | 20.70296 | 10.72622 | 13.40849 | 8.678602 | 25.0146 |
| ENSMUSG00000065258 | -0.63008982 | 0.130282 | 0.262661 | 18.10898 | 26.39607 | 20.70267 | 22.42821 | 26.32801 | 22.34748 | 27.0001 | 59.81751 |
| ENSMUSG00000065304 | -0.47298614 | 0.368685 | 0.537391 | 5.030272 | 8.398751 | 10.76539 | 9.488858 | 13.65156 | 6.704245 | 13.50005 | 13.05109 |
| ENSMUSG00000065336 | 0.415145767 | 0.29847 | 0.466483 | 21.12714 | 16.7975 | 22.35888 | 19.84034 | 10.72622 | 22.34748 | 13.50005 | 14.13868 |
| ENSMUSG00000065406 | 0.459439657 | 0.227658 | 0.387612 | 17.10292 | 20.39697 | 23.18699 | 29.3292 | 16.57689 | 17.87799 | 19.28578 | 11.9635 |
| ENSMUSG00000065418 | 0.465471155 | 0.40211 | 0.569261 | 8.048435 | 15.59768 | 11.59349 | 12.93935 | 6.825779 | 8.938993 | 14.46434 | 4.350364 |
| ENSMUSG00000065431 | 0.794754725 | 0.016363 | 0.061157 | 36.21796 | 34.79482 | 42.23344 | 47.44429 | 23.40267 | 32.40385 | 23.14294 | 14.13868 |
| ENSMUSG00000065434 | 0.262580802 | 0.214026 | 0.37205 | 80.48435 | 61.1909 | 72.04528 | 92.30071 | 66.30757 | 64.8077 | 57.85735 | 67.43065 |
| ENSMUSG00000065445 | -0.773880244 | 0.107338 | 0.231782 | 9.054489 | 15.59768 | 5.796746 | 7.763611 | 16.57689 | 16.76061 | 17.3572 | 13.05109 |
| ENSMUSG00000065475 | 0.213871327 | 0.619762 | 0.751867 | 12.07265 | 37.19447 | 24.8432 | 29.3292 | 14.62667 | 23.46486 | 27.96439 | 22.83941 |
| ENSMUSG00000065503 | 0.127919481 | 0.66296 | 0.783128 | 44.26639 | 53.99197 | 50.5145 | 38.81806 | 26.32801 | 52.51658 | 48.21446 | 44.59123 |
| ENSMUSG00000065519 | 0.269358368 | 0.430124 | 0.594901 | 40.24217 | 22.79661 | 33.95237 | 37.09281 | 29.25334 | 30.1691 | 34.71441 | 17.40146 |
| ENSMUSG00000065530 | 1.606620184 | 0.014508 | 0.056278 | 28.16952 | 64.79036 | 34.78048 | 38.81806 | 3.900445 | 35.75597 | 7.714313 | 7.613138 |
| ENSMUSG00000065546 | 0.11656128 | 0.529272 | 0.681503 | 118.7144 | 98.38537 | 98.54469 | 118.1794 | 83.85957 | 119.559 | 104.1432 | 93.53283 |
| ENSMUSG00000065564 | 0.654792603 | 0.162015 | 0.306116 | 13.07871 | 21.59679 | 16.56213 | 16.38985 | 5.850668 | 15.64324 | 9.642892 | 11.9635 |
| ENSMUSG00000065591 | 0.118052591 | 0.749923 | 0.844092 | 20.12109 | 17.99732 | 27.32752 | 25.01608 | 24.37778 | 25.6996 | 13.50005 | 20.66423 |
| ENSMUSG00000065592 | -0.132137281 | 0.721827 | 0.82602 | 16.09687 | 46.79304 | 38.09291 | 30.19182 | 35.10401 | 39.10809 | 27.96439 | 41.32846 |
| ENSMUSG00000065599 | 0.176130992 | 0.648434 | 0.772265 | 19.11503 | 21.59679 | 33.95237 | 37.09281 | 28.27823 | 15.64324 | 24.10723 | 31.54014 |
| ENSMUSG00000065608 | 0.760311035 | 0.122603 | 0.252118 | 15.09082 | 22.79661 | 18.21835 | 25.01608 | 18.52711 | 11.17374 | 13.50005 | 4.350364 |
| ENSMUSG00000065637 | -0.390766561 | 0.317929 | 0.486219 | 13.07871 | 44.3934 | 29.81184 | 25.01608 | 40.95467 | 30.1691 | 33.75012 | 41.32846 |
| ENSMUSG00000065669 | 0.609997402 | 0.235223 | 0.396138 | 18.10898 | 19.19714 | 20.70267 | 17.25247 | 4.875557 | 23.46486 | 11.57147 | 9.78832 |
| ENSMUSG00000065676 | 0.47864918 | 0.373013 | 0.541945 | 11.0666 | 11.99822 | 33.12427 | 14.6646 | 18.52711 | 10.05637 | 9.642892 | 13.05109 |
| ENSMUSG00000065734 | 0.07833887 | 0.804369 | 0.880251 | 53.32088 | 49.19268 | 28.98373 | 52.62003 | 52.65601 | 49.16446 | 42.42872 | 29.36496 |
| ENSMUSG00000065738 | -0.596822195 | 0.005886 | 0.028998 | 50.30272 | 35.99465 | 54.65504 | 47.44429 | 70.20801 | 73.74669 | 69.42882 | 73.95619 |
| ENSMUSG00000065767 | -0.048359692 | 0.88831 | 0.93282 | 27.16347 | 26.39607 | 23.18699 | 26.74133 | 31.20356 | 35.75597 | 17.3572 | 22.83941 |
| ENSMUSG00000065818 | 0.291274661 | 0.675879 | 0.792479 | 12.07265 | 19.19714 | 4.140533 | 27.60395 | 23.40267 | 10.05637 | 4.821446 | 13.05109 |
| ENSMUSG00000065847 | 1.096597978 | 0.236338 | 0.397364 | 21.12714 | 31.19536 | 20.70267 | 21.56559 | 0.975111 | 34.6386 | 7.714313 | 1.087591 |
| ENSMUSG00000065878 | 1.343109404 | 0.019597 | 0.069487 | 13.07871 | 16.7975 | 9.93728 | 23.29083 | 4.875557 | 10.05637 | 6.750024 | 3.262773 |
| ENSMUSG00000065947 | -0.265324189 | 0.048472 | 0.133263 | 168.0111 | 158.3764 | 159.8246 | 142.3329 | 182.3458 | 169.8409 | 197.6793 | 204.4671 |
| ENSMUSG00000065952 | -0.403456854 | 0.090008 | 0.205164 | 58.35115 | 51.59233 | 39.74912 | 59.52102 | 65.33246 | 56.98608 | 73.28598 | 80.48174 |
| ENSMUSG00000065954 | 0.293602662 | 0.024763 | 0.082467 | 2237.465 | 1521.374 | 1701.759 | 1892.596 | 1488.02 | 1549.798 | 1620.006 | 1342.087 |
| ENSMUSG00000065979 | -0.388151978 | 0.002552 | 0.01566 | 1041.266 | 1019.848 | 1218.145 | 1149.014 | 1425.613 | 1160.952 | 1525.505 | 1685.766 |
| ENSMUSG00000065990 | -0.233816118 | 0.020933 | 0.0728 | 1491.979 | 1380.995 | 1633.854 | 1450.07 | 1778.603 | 1497.281 | 1795.506 | 1935.912 |
| ENSMUSG00000066000 | 0.250229126 | 0.32176 | 0.489886 | 103.6236 | 115.1829 | 74.5296 | 72.46037 | 59.48179 | 91.62468 | 67.50024 | 88.09488 |
| ENSMUSG00000066026 | 0.522184345 | 0.009584 | 0.041281 | 4009.127 | 4843.68 | 3742.214 | 4308.804 | 2464.106 | 4410.276 | 2464.723 | 2431.854 |
| ENSMUSG00000066036 | 0.025558172 | 0.775408 | 0.861066 | 2842.104 | 2731.994 | 3326.504 | 2903.591 | 2641.577 | 2847.069 | 3020.154 | 3093.109 |
| ENSMUSG00000066037 | 0.097458632 | 0.250247 | 0.413857 | 6734.528 | 7388.501 | 6832.708 | 7174.44 | 6714.616 | 7393.664 | 6290.058 | 5893.656 |
| ENSMUSG00000066042 | -0.099563889 | 0.570343 | 0.71379 | 258.556 | 224.3666 | 178.871 | 185.4641 | 243.7778 | 235.7659 | 243.9652 | 181.6277 |
| ENSMUSG00000066043 | 0.145983167 | 0.064188 | 0.162682 | 1377.288 | 1160.227 | 1379.626 | 1284.446 | 1160.382 | 1158.717 | 1169.683 | 1217.014 |
| ENSMUSG00000066057 | 0.07810855 | 0.552277 | 0.699721 | 263.5862 | 297.5557 | 226.9012 | 238.9467 | 229.1512 | 261.4655 | 229.5008 | 250.146 |
| ENSMUSG00000066058 | -1.226423849 | 0.000237 | 0.002598 | 554.3359 | 418.7377 | 980.4783 | 533.9639 | 1231.566 | 728.5279 | 1716.435 | 2146.905 |
| ENSMUSG00000066068 | 0.020969657 | 0.957604 | 0.975954 | 17.10292 | 16.7975 | 21.53077 | 33.64232 | 27.30312 | 24.58223 | 20.25007 | 16.31387 |
| ENSMUSG00000066072 | -4.751643353 | 2.72E-07 | 1.28E-05 | 11.0666 | 1.199822 | 20.70267 | 6.900988 | 246.7032 | 53.63396 | 166.822 | 616.6641 |
| ENSMUSG00000066097 | -3.038831453 | 3.99E-10 | 5.67E-08 | 33.19979 | 13.19804 | 41.40533 | 34.50494 | 235.9769 | 81.56831 | 237.2151 | 456.7883 |
| ENSMUSG00000066113 | 0.670171715 | 0.004845 | 0.025159 | 412.4823 | 818.2783 | 567.253 | 546.9033 | 314.961 | 508.4052 | 335.5726 | 314.3138 |
| ENSMUSG00000066116 | -0.348715497 | 0.377339 | 0.545463 | 10.06054 | 22.79661 | 19.87456 | 14.6646 | 23.40267 | 22.34748 | 22.17865 | 17.40146 |
| ENSMUSG00000066129 | 0.091424768 | 0.847092 | 0.908105 | 26.15741 | 15.59768 | 44.71776 | 15.52722 | 33.15378 | 25.6996 | 22.17865 | 15.22628 |
| ENSMUSG00000066141 | -0.243458073 | 0.362896 | 0.531351 | 44.26639 | 63.59054 | 41.40533 | 45.71905 | 61.43201 | 49.16446 | 72.32169 | 45.67883 |
| ENSMUSG00000066148 | -0.112568386 | 0.200334 | 0.35555 | 1112.696 | 1057.043 | 1012.774 | 1009.269 | 1188.661 | 1055.919 | 1242.004 | 1040.825 |
| ENSMUSG00000066149 | 0.100420066 | 0.197551 | 0.351863 | 2262.616 | 2667.203 | 2300.48 | 2392.918 | 2388.048 | 2267.152 | 2176.401 | 2139.292 |

| | | | | | | | | | | | |
|---|---|---|---|---|---|---|---|---|---|---|---|
| ENSMUSG00000066150 | -0.46112672 | 0.000236 | 0.002591 | 1535.239 | 1327.003 | 1494.732 | 1517.355 | 2117.942 | 1581.084 | 2079.007 | 2310.043 |
| ENSMUSG00000066151 | -0.11489617 | 0.303546 | 0.472003 | 987.9454 | 929.8617 | 1084.82 | 919.5566 | 1031.668 | 916.2468 | 1074.218 | 1226.803 |
| ENSMUSG00000066152 | -0.319528507 | 0.009787 | 0.041905 | 506.0453 | 483.5281 | 501.0045 | 487.3823 | 584.0917 | 519.579 | 743.4669 | 619.9269 |
| ENSMUSG00000066170 | 0.025283034 | 0.953716 | 0.973604 | 94.56911 | 55.19179 | 54.65504 | 103.5148 | 38.02934 | 84.92043 | 134.0362 | 45.67883 |
| ENSMUSG00000066175 | -0.283623254 | 0.204144 | 0.360115 | 73.44197 | 83.98751 | 109.3101 | 67.28463 | 95.56091 | 86.03781 | 107.0361 | 118.5474 |
| ENSMUSG00000066189 | 1.063672831 | 0.056671 | 0.148729 | 24.1453 | 9.598572 | 10.76539 | 31.05445 | 8.776002 | 10.05637 | 6.750024 | 10.87591 |
| ENSMUSG00000066191 | -0.336787978 | 0.025139 | 0.083374 | 317.9132 | 326.3515 | 423.1625 | 360.5766 | 434.8996 | 383.2593 | 444.5373 | 543.7955 |
| ENSMUSG00000066196 | 0.275938352 | 0.483353 | 0.641736 | 47.28455 | 41.99375 | 23.18699 | 33.64232 | 35.10401 | 43.57759 | 24.10723 | 17.40146 |
| ENSMUSG00000066232 | 0.004965791 | 0.937523 | 0.962923 | 4041.32 | 4079.393 | 3673.481 | 3851.614 | 3745.403 | 3823.654 | 3861.978 | 4156.773 |
| ENSMUSG00000066233 | -0.17048009 | 0.200893 | 0.356336 | 214.2896 | 293.9563 | 261.6817 | 278.6274 | 313.9858 | 301.691 | 269.0367 | 294.7372 |
| ENSMUSG00000066235 | -0.181855721 | 0.228573 | 0.388662 | 258.556 | 242.3639 | 211.1672 | 202.7165 | 233.0516 | 233.5312 | 259.3938 | 309.9635 |
| ENSMUSG00000066245 | -0.312040211 | 0.41337 | 0.579196 | 21.12714 | 15.59768 | 22.35888 | 18.11509 | 36.07912 | 18.99536 | 20.25007 | 20.66423 |
| ENSMUSG00000066258 | 0.465437296 | 0.21199 | 0.369591 | 297.7921 | 376.744 | 459.5992 | 121.6299 | 251.5787 | 269.2872 | 255.5366 | 132.6861 |
| ENSMUSG00000066278 | -0.19550084 | 0.326779 | 0.494469 | 432.6034 | 483.5281 | 568.0811 | 425.2734 | 574.3406 | 353.0902 | 608.4665 | 650.3795 |
| ENSMUSG00000066306 | 0.107053685 | 0.049512 | 0.135176 | 4171.101 | 4247.368 | 4394.762 | 4476.153 | 3828.287 | 3943.213 | 4104.015 | 4181.788 |
| ENSMUSG00000066315 | 0.056664713 | 0.709918 | 0.818517 | 128.775 | 159.5763 | 134.1533 | 146.646 | 118.9636 | 140.7891 | 137.8934 | 149 |
| ENSMUSG00000066324 | 0.18119126 | 0.061905 | 0.158433 | 3967.878 | 4099.79 | 3842.415 | 4167.334 | 3236.394 | 4203.561 | 3405.869 | 3335.642 |
| ENSMUSG00000066357 | 0.014992189 | 0.863548 | 0.918066 | 3562.438 | 4043.399 | 4063.519 | 3654.936 | 4024.284 | 3405.756 | 3654.656 | 4079.554 |
| ENSMUSG00000066361 | 2.128486888 | 0.008861 | 0.039057 | 16.09687 | 34.79482 | 12.4216 | 24.15346 | 5.850668 | 12.29112 | 1.928578 | 0 |
| ENSMUSG00000066362 | -0.514506939 | 0.029563 | 0.093619 | 34.20585 | 31.19536 | 38.09291 | 38.81806 | 48.75557 | 53.63396 | 51.10733 | 51.11678 |
| ENSMUSG00000066363 | 0.705113526 | 0.23688 | 0.398021 | 17.10292 | 5.999108 | 9.109173 | 16.38985 | 5.850668 | 11.17374 | 8.678602 | 4.350364 |
| ENSMUSG00000066366 | -1.007216454 | 0.186295 | 0.338213 | 57.3451 | 1.199822 | 39.74912 | 73.323 | 39.97956 | 46.92971 | 117.6433 | 141.3868 |
| ENSMUSG00000066392 | -0.408809166 | 0.388663 | 0.55606 | 72.43591 | 105.5843 | 351.1172 | 88.85022 | 177.4703 | 138.5544 | 264.2152 | 241.4452 |
| ENSMUSG00000066406 | 0.642022364 | 0.000292 | 0.003043 | 3375.312 | 3061.945 | 2682.237 | 3118.384 | 1824.433 | 2745.388 | 1739.578 | 1533.503 |
| ENSMUSG00000066407 | -0.294070197 | 0.518605 | 0.672477 | 15.09082 | 15.59768 | 19.87456 | 28.46658 | 25.35289 | 11.17374 | 36.64299 | 23.927 |
| ENSMUSG00000066415 | 0.161021508 | 0.106838 | 0.231093 | 2065.43 | 2236.467 | 1787.054 | 2122.916 | 1692.793 | 2052.616 | 1752.113 | 1845.642 |
| ENSMUSG00000066438 | -0.82284911 | 0.02031 | 0.071247 | 90.54489 | 52.79215 | 142.4343 | 67.28463 | 150.1671 | 94.9768 | 163.9292 | 217.5182 |
| ENSMUSG00000066440 | 0.197230042 | 0.024282 | 0.081245 | 699.2078 | 673.0999 | 648.4075 | 654.7312 | 630.897 | 612.321 | 549.6448 | 539.4452 |
| ENSMUSG00000066441 | -0.317325715 | 0.008689 | 0.038475 | 837.0372 | 701.8956 | 743.6398 | 790.1631 | 970.2358 | 782.1619 | 1091.575 | 984.2699 |
| ENSMUSG00000066442 | -0.147672041 | 0.233277 | 0.393874 | 208.2533 | 243.5638 | 233.5261 | 233.771 | 290.5832 | 248.057 | 238.1794 | 240.3576 |
| ENSMUSG00000066456 | 0.608868533 | 2.35E-10 | 3.68E-08 | 2312.919 | 2594.014 | 2323.667 | 2495.57 | 1536.775 | 1858.193 | 1470.541 | 1512.839 |
| ENSMUSG00000066475 | 0.195041923 | 0.730875 | 0.83127 | 11.0666 | 13.19804 | 14.07781 | 9.488858 | 13.65156 | 14.52586 | 2.892867 | 10.87591 |
| ENSMUSG00000066487 | -0.367660727 | 0.287228 | 0.454092 | 19.11503 | 32.39518 | 43.88965 | 46.58167 | 48.75557 | 42.46022 | 56.89306 | 35.89051 |
| ENSMUSG00000066538 | -0.473468226 | 0.005577 | 0.027859 | 122.7386 | 115.1829 | 117.5911 | 108.6906 | 187.2214 | 148.6108 | 130.179 | 178.3649 |
| ENSMUSG00000066551 | -0.068566356 | 0.285672 | 0.452563 | 20401.78 | 22265.09 | 20116.37 | 22103.86 | 23381.22 | 22918.46 | 21712.9 | 21002.47 |
| ENSMUSG00000066553 | -0.454666929 | 0.375618 | 0.544104 | 12.07265 | 14.39786 | 9.109173 | 13.80198 | 20.47734 | 5.58687 | 19.28578 | 21.75182 |
| ENSMUSG00000066554 | -0.109958321 | 0.644693 | 0.770114 | 117.7084 | 113.983 | 152.3716 | 180.2883 | 143.3414 | 107.2679 | 170.6792 | 189.2408 |
| ENSMUSG00000066568 | -0.043707278 | 0.362249 | 0.530898 | 4051.381 | 4148.983 | 4210.094 | 4194.075 | 4196.879 | 4489.609 | 4285.301 | 4146.985 |
| ENSMUSG00000066571 | -0.285986809 | 0.066445 | 0.166189 | 950.7214 | 820.6779 | 915.0578 | 830.7064 | 1124.303 | 760.9318 | 1243.933 | 1158.285 |
| ENSMUSG00000066595 | -0.392571081 | 1.21E-05 | 0.000266 | 384.3128 | 367.1454 | 391.6944 | 344.1868 | 479.7548 | 513.9921 | 482.1446 | 477.4525 |
| ENSMUSG00000066607 | 0.14508939 | 0.185074 | 0.336559 | 357.1493 | 377.9438 | 414.8814 | 362.3019 | 328.6125 | 385.4941 | 335.5726 | 319.7518 |
| ENSMUSG00000066613 | 0.231630808 | 0.04989 | 0.135867 | 846.0917 | 973.0553 | 799.1229 | 885.0517 | 750.8357 | 880.4908 | 667.2881 | 684.0948 |
| ENSMUSG00000066621 | 0.094243768 | 0.308715 | 0.47703 | 1392.379 | 1099.037 | 1201.583 | 1184.382 | 1141.855 | 1190.003 | 1164.861 | 1073.452 |
| ENSMUSG00000066637 | -0.274912423 | 0.084454 | 0.19606 | 244.4712 | 211.1686 | 208.6829 | 212.2054 | 223.3005 | 227.9443 | 319.1797 | 289.2992 |
| ENSMUSG00000066640 | 0.186317018 | 0.074125 | 0.179604 | 665.0019 | 669.5004 | 724.5933 | 678.8847 | 604.569 | 691.6546 | 523.609 | 589.4744 |
| ENSMUSG00000066643 | 0.051915783 | 0.493407 | 0.650359 | 866.2128 | 881.8688 | 788.3575 | 911.793 | 809.3424 | 821.27 | 847.6102 | 846.1459 |
| ENSMUSG00000066647 | -0.14714465 | 0.379285 | 0.547395 | 245.4773 | 217.1677 | 193.777 | 165.6237 | 232.0765 | 262.5829 | 216.0008 | 197.9416 |
| ENSMUSG00000066677 | 0.329286088 | 0.70584 | 0.815002 | 3.018163 | 16.7975 | 28.98373 | 5.175741 | 2.925334 | 4.469496 | 19.28578 | 16.31387 |
| ENSMUSG00000066687 | -0.196161352 | 0.77818 | 0.863236 | 379.2825 | 140.3791 | 83.63877 | 350.2251 | 141.3911 | 223.4748 | 596.895 | 130.5109 |
| ENSMUSG00000066705 | 0.143888266 | 0.323407 | 0.49138 | 6146.992 | 7704.054 | 6123.849 | 7210.67 | 7581.49 | 6488.591 | 5575.52 | 4957.24 |
| ENSMUSG00000066720 | -0.051181351 | 0.755575 | 0.848333 | 531.1967 | 724.6922 | 497.6921 | 466.6793 | 666.001 | 548.6307 | 578.5735 | 502.4671 |
| ENSMUSG00000066721 | 0.288212243 | 0.115556 | 0.243516 | 90.54489 | 134.38 | 129.1846 | 114.7289 | 101.4116 | 94.9768 | 92.57176 | 94.62042 |
| ENSMUSG00000066724 | 0.125111077 | 0.670271 | 0.788584 | 43.26034 | 57.59143 | 50.5145 | 29.3292 | 35.10401 | 48.04709 | 39.53586 | 42.41605 |
| ENSMUSG00000066735 | 0.05971467 | 0.542312 | 0.691509 | 1282.719 | 1322.203 | 1210.692 | 1306.875 | 1229.615 | 1421.3 | 1180.29 | 1083.241 |
| ENSMUSG00000066797 | -0.843563069 | 0.076499 | 0.183536 | 6.036326 | 8.398751 | 9.109173 | 9.488858 | 19.50223 | 11.17374 | 13.50005 | 15.22628 |
| ENSMUSG00000066798 | 0.264640892 | 0.055296 | 0.146335 | 532.2028 | 634.7056 | 490.2391 | 565.0184 | 508.033 | 528.5179 | 416.5729 | 394.7956 |
| ENSMUSG00000066800 | 0.376247929 | 0.022155 | 0.075792 | 1371.252 | 1280.21 | 1002.009 | 1400.038 | 864.9237 | 825.7395 | 1252.612 | 948.3794 |
| ENSMUSG00000066809 | -0.261963102 | 0.500605 | 0.656937 | 55.33299 | 117.5825 | 230.2136 | 95.75121 | 174.5449 | 109.5027 | 156.2148 | 158.7883 |
| ENSMUSG00000066829 | -0.211899193 | 0.04333 | 0.123219 | 617.7174 | 607.1097 | 667.4539 | 540.0023 | 752.7859 | 725.1758 | 717.4311 | 621.0145 |
| ENSMUSG00000066838 | 0.160626288 | 0.139257 | 0.275174 | 253.5257 | 226.7663 | 234.3542 | 246.7103 | 212.5743 | 216.7706 | 213.1079 | 218.6058 |
| ENSMUSG00000066839 | -0.758619504 | 2.28E-05 | 0.000435 | 547.2936 | 475.1293 | 584.6433 | 470.1298 | 861.0233 | 634.6685 | 874.6103 | 1145.233 |
| ENSMUSG00000066842 | 0.507798805 | 3.83E-06 | 0.00011 | 3641.917 | 4364.951 | 3853.18 | 4244.97 | 2951.662 | 3206.864 | 2781.01 | 2386.175 |
| ENSMUSG00000066861 | 1.500924936 | 0.084863 | 0.196764 | 7.04238 | 41.99375 | 48.85829 | 1.725247 | 9.751113 | 11.17374 | 7.714313 | 6.525547 |
| ENSMUSG00000066877 | -0.011054025 | 0.866519 | 0.919848 | 1742.486 | 1708.546 | 1861.584 | 1777.867 | 1740.574 | 1680.531 | 1845.649 | 1881.533 |

| | | | | | | | | | | | |
|---|---|---|---|---|---|---|---|---|---|---|---|
| ENSMUSG00000066878 | -0.323972943 | 0.5183 | 0.672184 | 15.09082 | 15.59768 | 14.07781 | 12.07673 | 31.20356 | 7.821619 | 17.3572 | 14.13868 |
| ENSMUSG00000066880 | -0.080936198 | 0.355698 | 0.524342 | 546.2875 | 536.3202 | 522.5353 | 520.162 | 525.585 | 623.4947 | 542.8948 | 556.8466 |
| ENSMUSG00000066892 | 0.066006509 | 0.530282 | 0.682155 | 624.7598 | 615.5084 | 722.109 | 697.8624 | 678.6775 | 663.7202 | 649.9309 | 551.4087 |
| ENSMUSG00000066894 | 0.13764928 | 0.072458 | 0.176738 | 1061.387 | 905.8653 | 1021.884 | 957.5121 | 877.6002 | 918.4815 | 910.289 | 884.2116 |
| ENSMUSG00000066900 | -0.247044136 | 0.003775 | 0.020951 | 2372.276 | 2446.436 | 2328.636 | 2327.358 | 2746.889 | 2463.81 | 2898.653 | 3132.262 |
| ENSMUSG00000066902 | -0.289717047 | 0.517793 | 0.671803 | 14.08476 | 15.59768 | 13.24971 | 11.21411 | 25.35289 | 15.64324 | 10.60718 | 14.13868 |
| ENSMUSG00000066952 | -0.386352773 | 0.126233 | 0.257062 | 61.36932 | 39.59411 | 70.38906 | 49.16954 | 68.25779 | 60.3382 | 83.89316 | 77.21897 |
| ENSMUSG00000066975 | 0.453206821 | 0.301927 | 0.470321 | 28.16952 | 14.39786 | 14.90592 | 17.25247 | 11.70134 | 10.05637 | 16.39292 | 16.31387 |
| ENSMUSG00000066979 | -0.115316434 | 0.102821 | 0.225242 | 3319.979 | 3185.526 | 3174.961 | 3276.244 | 3855.59 | 3209.098 | 3463.727 | 3504.218 |
| ENSMUSG00000067017 | 0.246475149 | 0.389384 | 0.556741 | 40.24217 | 44.3934 | 31.46805 | 48.30692 | 28.27823 | 30.1691 | 39.53586 | 40.24087 |
| ENSMUSG00000067038 | -0.215980134 | 0.220646 | 0.379484 | 115.6963 | 131.9804 | 149.8873 | 110.4158 | 154.0676 | 168.7235 | 144.6434 | 122.8978 |
| ENSMUSG00000067058 | -0.305613478 | 0.570168 | 0.713761 | 11.0666 | 28.79572 | 13.24971 | 12.07673 | 33.15378 | 10.05637 | 14.46434 | 21.75182 |
| ENSMUSG00000067071 | -0.048053452 | 0.523171 | 0.676427 | 1270.647 | 1135.031 | 1089.788 | 1180.932 | 1249.118 | 1236.933 | 1196.683 | 1150.671 |
| ENSMUSG00000067121 | 0.08803792 | 0.819768 | 0.890713 | 20.12109 | 19.19714 | 19.04645 | 19.84034 | 19.50223 | 15.64324 | 26.03581 | 11.9635 |
| ENSMUSG00000067144 | -5.674314347 | 1.68E-05 | 0.00034 | 0 | 1.199822 | 0 | 0.862623 | 15.60178 | 4.469496 | 15.42863 | 63.08028 |
| ENSMUSG00000067148 | -0.134007372 | 0.115849 | 0.243805 | 2237.465 | 2078.091 | 2057.845 | 2220.393 | 2594.771 | 2087.255 | 2442.544 | 2303.518 |
| ENSMUSG00000067149 | 1.049304532 | 0.014383 | 0.055907 | 20.12109 | 27.5959 | 19.87456 | 18.11509 | 13.65156 | 11.17374 | 9.642892 | 6.525547 |
| ENSMUSG00000067150 | -0.217634575 | 0.106399 | 0.230508 | 1243.483 | 1103.836 | 1170.115 | 1071.378 | 1404.16 | 1005.637 | 1433.898 | 1491.087 |
| ENSMUSG00000067158 | -0.100757528 | 0.196948 | 0.351263 | 1302.84 | 1312.605 | 1236.363 | 1336.204 | 1261.794 | 1520.746 | 1397.255 | 1384.503 |
| ENSMUSG00000067194 | -0.134624466 | 0.005304 | 0.026803 | 3254.586 | 3480.682 | 3463.142 | 3545.383 | 3790.258 | 3727.56 | 3769.406 | 3802.218 |
| ENSMUSG00000067199 | -0.372477013 | 0.005003 | 0.025762 | 316.9071 | 283.1579 | 365.195 | 326.9343 | 393.945 | 365.3813 | 481.1803 | 435.0364 |
| ENSMUSG00000067203 | -0.613071456 | 0.106355 | 0.230448 | 16.09687 | 9.598572 | 23.18699 | 16.38985 | 21.45245 | 26.81698 | 26.03581 | 27.18978 |
| ENSMUSG00000067206 | -1.87011629 | 0.00242 | 0.01505 | 7.04238 | 3.599465 | 18.21835 | 10.35148 | 37.05423 | 14.52586 | 27.96439 | 66.34306 |
| ENSMUSG00000067212 | -0.164895922 | 0.576634 | 0.718739 | 301.8163 | 410.339 | 574.706 | 281.2153 | 488.5308 | 245.8223 | 570.8592 | 452.4379 |
| ENSMUSG00000067215 | 1.540256968 | 4.20E-05 | 0.000699 | 46.2785 | 47.99286 | 28.98373 | 36.23019 | 9.751113 | 12.29112 | 19.28578 | 13.05109 |
| ENSMUSG00000067219 | -0.611688466 | 0.14824 | 0.287688 | 176.0595 | 79.18822 | 377.6166 | 142.3329 | 404.6712 | 210.0663 | 352.9298 | 218.6058 |
| ENSMUSG00000067220 | 0.213908059 | 0.177261 | 0.326304 | 141.8537 | 175.1739 | 177.2148 | 150.0965 | 133.5902 | 149.7281 | 119.5719 | 153.3503 |
| ENSMUSG00000067235 | -0.613601621 | 0.283715 | 0.450275 | 17.10292 | 7.198929 | 9.109173 | 29.3292 | 25.35289 | 11.17374 | 23.14294 | 36.9781 |
| ENSMUSG00000067242 | 0.554859172 | 0.072971 | 0.177672 | 117.7084 | 77.9884 | 92.74794 | 146.646 | 58.50668 | 115.0895 | 60.75022 | 63.08028 |
| ENSMUSG00000067274 | -0.227270305 | 0.020005 | 0.070419 | 57410.49 | 70526.71 | 66696.54 | 67233.74 | 86306.13 | 69024.67 | 78313.78 | 72899.06 |
| ENSMUSG00000067276 | 0.072487341 | 0.677223 | 0.793627 | 7345.203 | 8552.328 | 6272.908 | 8142.303 | 9605.821 | 6274.056 | 7357.526 | 5586.955 |
| ENSMUSG00000067279 | 0.274773291 | 0.265308 | 0.429407 | 656.9535 | 446.3336 | 663.3134 | 326.9343 | 564.5894 | 367.6161 | 383.7871 | 414.3722 |
| ENSMUSG00000067288 | -0.068251723 | 0.327884 | 0.495612 | 14339.29 | 15073.36 | 16090.94 | 15182.17 | 17064.45 | 14986.22 | 16528.88 | 15045.74 |
| ENSMUSG00000067297 | -0.101227468 | 0.867161 | 0.92037 | 14.08476 | 11.99822 | 9.93728 | 5.175741 | 3.900445 | 11.17374 | 11.57147 | 17.40146 |
| ENSMUSG00000067321 | -0.600723428 | 0.046688 | 0.129778 | 32.19374 | 40.79393 | 28.98373 | 29.3292 | 33.15378 | 64.8077 | 48.21446 | 52.20437 |
| ENSMUSG00000067336 | 0.439118339 | 0.004397 | 0.023435 | 2786.771 | 3031.949 | 2678.925 | 2909.629 | 1776.653 | 2851.539 | 1924.721 | 1861.956 |
| ENSMUSG00000067344 | 0.124621934 | 0.589104 | 0.72937 | 64.38748 | 65.99018 | 52.99882 | 70.73513 | 59.48179 | 72.62932 | 53.0359 | 47.85401 |
| ENSMUSG00000067365 | 0.02447871 | 0.721958 | 0.826067 | 1114.708 | 1205.821 | 1108.835 | 1181.794 | 1094.075 | 1182.182 | 1170.647 | 1084.328 |
| ENSMUSG00000067367 | -0.087577627 | 0.43758 | 0.601249 | 1856.17 | 1986.904 | 1556.84 | 1797.707 | 2011.655 | 2161.002 | 1773.328 | 1698.817 |
| ENSMUSG00000067369 | 0.03274325 | 0.713761 | 0.820362 | 616.7113 | 724.6922 | 651.7199 | 624.5394 | 664.0508 | 601.1473 | 621.0022 | 668.8685 |
| ENSMUSG00000067370 | -0.32990585 | 0.150442 | 0.290485 | 90.54489 | 113.983 | 100.2009 | 75.04824 | 95.56091 | 99.44629 | 153.322 | 127.2482 |
| ENSMUSG00000067377 | 0.142710427 | 0.109476 | 0.234988 | 1896.412 | 2261.664 | 2204.42 | 2131.543 | 1997.028 | 1954.287 | 2028.864 | 1711.868 |
| ENSMUSG00000067389 | 0.703600951 | 0.153247 | 0.294464 | 30.18163 | 13.19804 | 26.49941 | 14.6646 | 11.70134 | 21.23011 | 9.642892 | 9.78832 |
| ENSMUSG00000067399 | 0.516337409 | 0.483008 | 0.641437 | 21.12714 | 8.398751 | 3.312427 | 18.11509 | 10.72622 | 14.52586 | 2.892867 | 7.613138 |
| ENSMUSG00000067424 | -0.156299903 | 0.444882 | 0.607797 | 85.51462 | 129.5807 | 128.3565 | 124.2178 | 117.0134 | 158.6671 | 134.0362 | 112.0219 |
| ENSMUSG00000067430 | 0.140709041 | 0.429397 | 0.594341 | 454.7366 | 479.9286 | 443.037 | 394.2189 | 403.6961 | 538.5743 | 354.8584 | 309.9635 |
| ENSMUSG00000067438 | 1.691022998 | 0.047093 | 0.130462 | 55.33299 | 5.999108 | 4.140533 | 43.9938 | 3.900445 | 7.821619 | 13.50005 | 8.700729 |
| ENSMUSG00000067547 | -0.627728777 | 0.046755 | 0.129878 | 39.23612 | 34.79482 | 28.15563 | 37.09281 | 55.58134 | 31.28647 | 55.92877 | 71.78101 |
| ENSMUSG00000067567 | 0.381287175 | 0.000189 | 0.002171 | 579.4873 | 556.7172 | 505.145 | 619.3637 | 451.4765 | 461.4755 | 411.7515 | 411.1094 |
| ENSMUSG00000067575 | 0.209383144 | 0.723998 | 0.827323 | 17.10292 | 5.999108 | 37.2648 | 17.25247 | 9.751113 | 17.87799 | 26.03581 | 14.13868 |
| ENSMUSG00000067578 | -2.614909031 | 3.72E-12 | 9.16E-10 | 22.1332 | 7.198929 | 9.93728 | 18.11509 | 74.10846 | 100.5637 | 111.8575 | 67.43065 |
| ENSMUSG00000067586 | -0.182499167 | 0.398809 | 0.566186 | 927.5821 | 1274.21 | 1619.777 | 1040.324 | 1549.452 | 1005.637 | 1221.754 | 1741.233 |
| ENSMUSG00000067608 | -0.532084954 | 0.111131 | 0.237288 | 15.09082 | 26.39607 | 24.8432 | 17.25247 | 34.1289 | 27.93435 | 29.89296 | 28.27737 |
| ENSMUSG00000067629 | 0.536654348 | 8.66E-09 | 7.57E-07 | 816.9161 | 829.0767 | 732.0463 | 852.272 | 570.4401 | 607.8515 | 519.7519 | 528.5693 |
| ENSMUSG00000067642 | 1.78844328 | 0.01618 | 0.06069 | 23.13925 | 17.99732 | 21.53077 | 18.11509 | 0 | 14.52586 | 4.821446 | 4.350364 |
| ENSMUSG00000067653 | 0.351793333 | 0.067119 | 0.167299 | 105.6357 | 124.7814 | 82.81066 | 117.3168 | 87.76002 | 81.56831 | 83.89316 | 82.65692 |
| ENSMUSG00000067656 | -2.741472047 | 1.71E-06 | 5.78E-05 | 6.036326 | 4.799286 | 18.21835 | 13.80198 | 44.85512 | 32.40385 | 83.89316 | 130.5109 |
| ENSMUSG00000067704 | 0.489573077 | 0.218431 | 0.377115 | 25.15136 | 32.39518 | 25.67131 | 46.58167 | 16.57689 | 35.75597 | 23.14294 | 17.40146 |
| ENSMUSG00000067713 | -0.123886519 | 0.068767 | 0.170016 | 1107.666 | 1208.22 | 1270.316 | 1211.123 | 1344.678 | 1300.623 | 1269.969 | 1314.898 |
| ENSMUSG00000067719 | -0.411044091 | 0.107798 | 0.232393 | 51.30877 | 52.79215 | 73.70149 | 54.34528 | 61.43201 | 63.69032 | 85.82174 | 98.97079 |
| ENSMUSG00000067722 | -0.159273011 | 0.047132 | 0.130553 | 829.9948 | 866.2711 | 756.0614 | 871.2497 | 984.8624 | 890.5472 | 915.1104 | 916.8393 |
| ENSMUSG00000067736 | -0.570738652 | 0.059169 | 0.153209 | 56.33904 | 41.99375 | 81.98256 | 43.9938 | 86.78491 | 59.22083 | 81.00029 | 107.6715 |
| ENSMUSG00000067768 | 0.236613005 | 0.753456 | 0.846727 | 3.018163 | 13.19804 | 27.32752 | 16.38985 | 4.875557 | 20.11273 | 5.785735 | 20.66423 |
| ENSMUSG00000067780 | 0.084507179 | 0.822656 | 0.89248 | 245.4773 | 226.7663 | 266.6503 | 439.938 | 136.5156 | 507.2878 | 259.3938 | 209.9051 |

| | | | | | | | | | | | |
|---|---|---|---|---|---|---|---|---|---|---|---|
| ENSMUSG00000067786 | 0.847979425 | 4.72E-11 | 9.27E-09 | 2249.538 | 1983.305 | 1896.364 | 2765.571 | 1287.147 | 1270.454 | 1226.576 | 1158.285 |
| ENSMUSG00000067787 | 0.013911255 | 0.864448 | 0.918772 | 941.6669 | 975.4549 | 918.3703 | 904.892 | 872.7246 | 863.7302 | 948.8605 | 1016.898 |
| ENSMUSG00000067795 | 0.720616161 | 0.127976 | 0.259325 | 15.09082 | 25.19625 | 21.53077 | 24.15346 | 11.70134 | 23.46486 | 9.642892 | 7.613138 |
| ENSMUSG00000067818 | -0.136778367 | 0.44374 | 0.606822 | 3037.278 | 4574.92 | 5362.819 | 3747.236 | 4495.263 | 4222.557 | 4118.479 | 5549.977 |
| ENSMUSG00000067825 | -0.181771992 | 0.106158 | 0.230174 | 395.3794 | 425.9366 | 462.9116 | 394.2189 | 470.9788 | 415.6632 | 489.8589 | 527.4817 |
| ENSMUSG00000067847 | -0.411233356 | 0.002801 | 0.01682 | 933.6184 | 955.0579 | 1143.615 | 853.1346 | 1212.063 | 1112.905 | 1280.576 | 1562.868 |
| ENSMUSG00000067851 | 0.096388938 | 0.283994 | 0.450523 | 2907.497 | 2583.216 | 2579.552 | 2634.452 | 2190.1 | 2736.449 | 2469.545 | 2618.919 |
| ENSMUSG00000067869 | 0.018627866 | 0.969732 | 0.983156 | 31.18769 | 11.99822 | 21.53077 | 18.11509 | 29.25334 | 8.938993 | 26.03581 | 17.40146 |
| ENSMUSG00000067870 | -0.101010591 | 0.453352 | 0.615723 | 253.5257 | 333.5504 | 361.0545 | 315.7202 | 353.9654 | 340.7991 | 349.0727 | 312.1386 |
| ENSMUSG00000067873 | 0.057086272 | 0.461749 | 0.622911 | 3227.422 | 3490.281 | 3127.759 | 3154.614 | 3156.435 | 3443.747 | 2994.118 | 2898.43 |
| ENSMUSG00000067878 | 0.262652821 | 0.120448 | 0.249438 | 130.7871 | 155.9768 | 143.2624 | 138.0198 | 115.0631 | 144.1413 | 98.35749 | 116.3722 |
| ENSMUSG00000067889 | -0.418395073 | 0.192189 | 0.34549 | 579.4873 | 395.9411 | 1244.644 | 569.3315 | 754.7362 | 730.7627 | 1138.826 | 1106.08 |
| ENSMUSG00000067916 | 0.385172986 | 0.138314 | 0.273779 | 258.556 | 295.1561 | 213.6515 | 225.1447 | 127.7396 | 288.2825 | 194.7864 | 149 |
| ENSMUSG00000067924 | -0.207531493 | 0.058401 | 0.151704 | 467.8153 | 428.3363 | 496.864 | 423.5481 | 475.8543 | 579.9172 | 490.8232 | 554.6715 |
| ENSMUSG00000067925 | -0.084685317 | 0.433079 | 0.597256 | 373.2462 | 437.9349 | 485.2705 | 451.1521 | 464.153 | 474.884 | 439.7159 | 477.4525 |
| ENSMUSG00000067928 | 0.446192925 | 0.035113 | 0.106342 | 681.0988 | 820.6779 | 616.1113 | 617.6384 | 517.7841 | 714.002 | 389.5728 | 386.0948 |
| ENSMUSG00000067929 | 0.295381429 | 0.535762 | 0.686525 | 26.15741 | 33.595 | 23.18699 | 22.42821 | 12.67645 | 42.46022 | 13.50005 | 17.40146 |
| ENSMUSG00000067931 | 0.054564657 | 0.731115 | 0.831493 | 450.7124 | 409.1391 | 401.6317 | 370.0655 | 369.5672 | 513.9921 | 343.2869 | 344.7664 |
| ENSMUSG00000067942 | 0.32253167 | 0.014845 | 0.057247 | 821.9464 | 925.0624 | 894.3552 | 798.7894 | 636.7477 | 865.9649 | 625.8237 | 623.1897 |
| ENSMUSG00000067995 | -0.105107401 | 0.190729 | 0.343357 | 768.6255 | 730.6913 | 678.2193 | 785.85 | 836.6455 | 787.7487 | 773.3599 | 788.5035 |
| ENSMUSG00000068011 | -0.437559623 | 0.166959 | 0.312673 | 55.33299 | 50.3925 | 131.669 | 93.16334 | 111.1627 | 101.681 | 115.7147 | 121.8102 |
| ENSMUSG00000068014 | -0.197061544 | 0.715987 | 0.822105 | 6.036326 | 11.99822 | 9.109173 | 12.07673 | 8.776002 | 18.99536 | 8.678602 | 8.700729 |
| ENSMUSG00000068015 | 0.021355763 | 0.861884 | 0.917409 | 922.5518 | 813.479 | 989.5874 | 850.5468 | 773.2633 | 1045.862 | 900.6461 | 806.9926 |
| ENSMUSG00000068036 | 0.067210184 | 0.418998 | 0.584463 | 3815.964 | 4025.401 | 3815.915 | 3759.313 | 3458.72 | 4228.144 | 3526.405 | 3502.043 |
| ENSMUSG00000068037 | 0.860167634 | 0.093331 | 0.210166 | 50.30272 | 41.99375 | 39.74912 | 36.23019 | 22.42756 | 44.69496 | 20.25007 | 5.437955 |
| ENSMUSG00000068039 | -0.069890022 | 0.27472 | 0.440813 | 8290.894 | 9301.017 | 9064.455 | 9410.36 | 9742.337 | 9730.094 | 9478.962 | 8905.196 |
| ENSMUSG00000068040 | -0.011092795 | 0.879481 | 0.927685 | 1835.043 | 2115.285 | 1965.925 | 1841.701 | 1888.791 | 2006.804 | 1879.4 | 2040.321 |
| ENSMUSG00000068079 | -0.307838212 | 0.053857 | 0.143573 | 123.7447 | 165.5754 | 141.6062 | 125.0804 | 160.8934 | 159.7845 | 168.7506 | 196.854 |
| ENSMUSG00000068101 | -0.244943822 | 0.043508 | 0.123608 | 386.3249 | 520.7225 | 425.6468 | 457.1905 | 589.9423 | 510.64 | 520.7161 | 495.9415 |
| ENSMUSG00000068114 | -0.18829955 | 0.145009 | 0.283145 | 272.6407 | 281.9581 | 255.0568 | 251.0234 | 340.3138 | 318.4516 | 296.0368 | 251.2335 |
| ENSMUSG00000068115 | 0.216203835 | 0.037486 | 0.111481 | 870.237 | 719.8929 | 717.1403 | 805.6903 | 646.4988 | 749.758 | 641.2523 | 643.8539 |
| ENSMUSG00000068122 | 0.295719944 | 0.237947 | 0.399097 | 2435.658 | 2922.765 | 2108.359 | 3148.576 | 1926.82 | 3297.371 | 2125.293 | 1298.584 |
| ENSMUSG00000068130 | 0.29634595 | 0.289997 | 0.457144 | 132.7992 | 100.785 | 80.32634 | 86.26235 | 56.55646 | 116.2069 | 78.10742 | 75.04379 |
| ENSMUSG00000068134 | 0.138641221 | 0.465332 | 0.626188 | 379.2825 | 392.3416 | 347.8048 | 330.3848 | 293.5085 | 446.9496 | 333.6441 | 242.5328 |
| ENSMUSG00000068141 | 0.080578199 | 0.72704 | 0.829154 | 42.25428 | 53.99197 | 47.20208 | 49.16954 | 48.75557 | 46.92971 | 40.50014 | 45.67883 |
| ENSMUSG00000068154 | 1.39885047 | 0.057803 | 0.150724 | 15.09082 | 7.198929 | 5.796746 | 36.23019 | 7.80089 | 3.352122 | 6.750024 | 6.525547 |
| ENSMUSG00000068184 | -0.187574998 | 0.049094 | 0.134427 | 546.2875 | 571.1151 | 612.7989 | 562.4305 | 607.4943 | 622.3774 | 647.038 | 736.2992 |
| ENSMUSG00000068196 | 0.487924902 | 0.019301 | 0.068742 | 550.3117 | 693.4968 | 433.0998 | 631.4404 | 467.0783 | 438.0106 | 463.8231 | 275.1605 |
| ENSMUSG00000068205 | -0.826996845 | 6.34E-07 | 2.56E-05 | 226.3622 | 230.3657 | 202.8861 | 230.3205 | 405.6463 | 318.4516 | 349.0727 | 504.6423 |
| ENSMUSG00000068206 | 0.069394126 | 0.359085 | 0.527481 | 1269.641 | 1298.207 | 1188.333 | 1339.654 | 1202.312 | 1126.313 | 1228.504 | 1297.496 |
| ENSMUSG00000068220 | 0.379262888 | 0.016679 | 0.061985 | 4604.711 | 7120.941 | 6315.141 | 5758.874 | 4448.458 | 5526.532 | 3743.371 | 4579.846 |
| ENSMUSG00000068226 | -0.09572167 | 0.82812 | 0.895832 | 21.12714 | 11.99822 | 14.90592 | 13.80198 | 11.70134 | 12.29112 | 18.32149 | 23.927 |
| ENSMUSG00000068227 | -1.875004813 | 0.008153 | 0.036789 | 10.06054 | 3.599465 | 17.39024 | 15.52722 | 48.75557 | 5.58687 | 71.3574 | 46.76642 |
| ENSMUSG00000068240 | -0.379518975 | 0.027519 | 0.088933 | 224.3501 | 274.7591 | 269.9628 | 239.8093 | 357.8658 | 233.5312 | 351.9655 | 367.6058 |
| ENSMUSG00000068245 | 0.634432585 | 0.223728 | 0.383462 | 25.15136 | 55.19179 | 44.71776 | 19.84034 | 25.35289 | 7.821619 | 36.64299 | 22.83941 |
| ENSMUSG00000068250 | 0.510507845 | 0.021688 | 0.074648 | 658.9656 | 695.8965 | 561.4563 | 726.329 | 391.0196 | 702.8283 | 421.3944 | 340.416 |
| ENSMUSG00000068262 | -0.079152726 | 0.806484 | 0.881775 | 20.12109 | 20.39697 | 23.18699 | 25.8787 | 22.42756 | 27.93435 | 23.14294 | 21.75182 |
| ENSMUSG00000068263 | -1.024737415 | 0.006561 | 0.031503 | 30.18163 | 29.99554 | 56.31125 | 44.85642 | 93.61069 | 42.46022 | 73.28598 | 120.7226 |
| ENSMUSG00000068264 | -0.364585464 | 0.001283 | 0.009418 | 261.5741 | 280.7582 | 306.3995 | 287.2536 | 365.6667 | 320.6864 | 385.7157 | 391.5328 |
| ENSMUSG00000068267 | -0.192400938 | 0.10379 | 0.226596 | 4617.789 | 5654.759 | 6573.51 | 5543.219 | 6361.626 | 5723.19 | 6495.452 | 7005.174 |
| ENSMUSG00000068270 | -0.592680742 | 0.000435 | 0.004115 | 731.4015 | 601.1106 | 886.9022 | 621.9515 | 976.0864 | 870.4344 | 1177.397 | 1263.781 |
| ENSMUSG00000068284 | 0.068598407 | 0.381791 | 0.549764 | 1185.132 | 1000.651 | 1038.446 | 1075.691 | 1029.718 | 1070.444 | 998.0393 | 1003.847 |
| ENSMUSG00000068290 | -0.278255931 | 0.002456 | 0.015219 | 1879.31 | 1910.116 | 2104.219 | 2040.105 | 2528.464 | 2083.903 | 2383.723 | 2627.62 |
| ENSMUSG00000068299 | -0.180945072 | 0.361556 | 0.530099 | 271.6347 | 232.7654 | 214.4796 | 198.4034 | 268.1556 | 187.7188 | 247.8223 | 334.9781 |
| ENSMUSG00000068303 | 0.37993006 | 0.054548 | 0.145001 | 86.52067 | 109.1838 | 110.1382 | 87.12497 | 84.83468 | 79.33356 | 62.6788 | 75.04379 |
| ENSMUSG00000068323 | 0.442772235 | 0.012367 | 0.050099 | 174.0474 | 148.7779 | 193.777 | 184.6014 | 114.088 | 164.254 | 123.429 | 116.3722 |
| ENSMUSG00000068327 | 3.882013675 | 0.004776 | 0.024921 | 30.18163 | 1.199822 | 0 | 58.6584 | 1.950223 | 1.117374 | 1.928578 | 1.087591 |
| ENSMUSG00000068328 | -0.240646659 | 0.005589 | 0.027892 | 1581.517 | 1616.16 | 1626.401 | 1651.924 | 1986.302 | 1645.892 | 1930.507 | 2088.175 |
| ENSMUSG00000068329 | -0.082681422 | 0.342298 | 0.511039 | 1007.06 | 1067.841 | 1078.195 | 1167.13 | 1272.52 | 1077.149 | 1099.29 | 1126.744 |
| ENSMUSG00000068335 | 0.059656439 | 0.716046 | 0.822105 | 436.6276 | 383.9429 | 385.8977 | 444.2511 | 391.0196 | 417.8979 | 484.0732 | 290.3868 |
| ENSMUSG00000068348 | -2.198325014 | 0.004897 | 0.025356 | 4.024217 | 4.799286 | 6.624853 | 0.862623 | 15.60178 | 5.58687 | 15.42863 | 38.06569 |
| ENSMUSG00000068373 | 0.016872669 | 0.960635 | 0.977472 | 66.39959 | 67.19001 | 43.06154 | 85.39973 | 46.80534 | 103.9158 | 51.10733 | 57.64233 |
| ENSMUSG00000068391 | -0.129005871 | 0.279986 | 0.446314 | 1091.569 | 1372.596 | 1503.014 | 1185.245 | 1503.622 | 1271.572 | 1424.255 | 1434.533 |
| ENSMUSG00000068394 | -0.050865376 | 0.792042 | 0.872764 | 613.6932 | 674.2997 | 509.2856 | 570.1941 | 625.0463 | 833.5611 | 545.7877 | 446.9999 |

| | | | | | | | | | | | |
|---|---|---|---|---|---|---|---|---|---|---|---|
| ENSMUSG00000068396 | -0.322129427 | 0.09154 | 0.207539 | 75.45408 | 80.38804 | 77.84202 | 71.59775 | 116.0382 | 96.09417 | 83.89316 | 84.8321 |
| ENSMUSG00000068407 | 1.295371182 | 2.03E-08 | 1.54E-06 | 156.9445 | 265.1606 | 213.6515 | 188.9145 | 63.38224 | 102.7984 | 93.53605 | 76.13138 |
| ENSMUSG00000068417 | 0.069317229 | 0.821526 | 0.891669 | 27.16347 | 29.99554 | 44.71776 | 34.50494 | 29.25334 | 40.22547 | 27.0001 | 34.80291 |
| ENSMUSG00000068457 | -0.42716449 | 0.797339 | 0.875673 | 1.006054 | 657.5022 | 554.0033 | 11.21411 | 537.2863 | 4.469496 | 530.359 | 573.1605 |
| ENSMUSG00000068463 | 1.027270904 | 0.147426 | 0.286613 | 19.11503 | 14.39786 | 5.796746 | 20.70296 | 8.776002 | 14.52586 | 1.928578 | 4.350364 |
| ENSMUSG00000068466 | -0.006011707 | 0.985034 | 0.991442 | 39.23612 | 51.59233 | 30.63995 | 45.71905 | 45.83023 | 58.10345 | 30.85725 | 32.62773 |
| ENSMUSG00000068479 | 0.091354053 | 0.479422 | 0.638512 | 632.8082 | 807.4799 | 701.4063 | 652.1434 | 587.017 | 779.9271 | 605.5736 | 649.2919 |
| ENSMUSG00000068522 | 0.394233526 | 0.06618 | 0.165818 | 97.58727 | 104.3845 | 110.1382 | 134.5693 | 70.20801 | 111.7374 | 80.036 | 79.39415 |
| ENSMUSG00000068523 | -0.231144908 | 0.013119 | 0.052372 | 1838.061 | 2033.698 | 2394.884 | 2118.603 | 2468.007 | 2327.49 | 2508.116 | 2542.788 |
| ENSMUSG00000068551 | -0.000180041 | 0.998411 | 0.999169 | 824.9646 | 763.0865 | 872.8244 | 756.5208 | 822.0188 | 805.6267 | 730.9312 | 862.4597 |
| ENSMUSG00000068566 | 0.24768593 | 1.38E-05 | 0.000294 | 2869.267 | 2848.376 | 3113.681 | 2970.013 | 2449.48 | 2417.998 | 2596.831 | 2478.62 |
| ENSMUSG00000068580 | -0.113933563 | 0.211202 | 0.3686 | 597.5963 | 685.0981 | 664.1415 | 643.5171 | 716.7068 | 629.0816 | 753.1098 | 701.4963 |
| ENSMUSG00000068587 | -1.360304079 | 1.30E-05 | 0.000281 | 46.2785 | 27.5959 | 54.65504 | 38.81806 | 74.10846 | 88.27255 | 124.3933 | 145.7372 |
| ENSMUSG00000068601 | 0.540865433 | 0.313354 | 0.481484 | 20.12109 | 23.99643 | 24.01509 | 21.56559 | 18.52711 | 12.29112 | 3.857157 | 27.18978 |
| ENSMUSG00000068614 | 0.757934884 | 0.07231 | 0.176516 | 70.4238 | 79.18822 | 70.38906 | 106.1027 | 62.40712 | 81.56831 | 23.14294 | 26.10219 |
| ENSMUSG00000068615 | 1.420047021 | 0.001531 | 0.010697 | 56.33904 | 25.19625 | 23.18699 | 48.30692 | 10.72622 | 20.11273 | 12.53576 | 14.13868 |
| ENSMUSG00000068617 | 0.183399587 | 0.539433 | 0.689165 | 35.2119 | 40.79393 | 62.9361 | 37.95543 | 32.17867 | 42.46022 | 37.60728 | 44.59123 |
| ENSMUSG00000068663 | -0.380481931 | 0.001731 | 0.011775 | 439.6458 | 469.1302 | 469.5365 | 427.8613 | 549.9628 | 492.762 | 638.3594 | 668.8685 |
| ENSMUSG00000068686 | 0.301280544 | 0.28791 | 0.454754 | 54.32694 | 64.79036 | 33.12427 | 58.6584 | 41.92979 | 37.99072 | 43.39301 | 46.76642 |
| ENSMUSG00000068696 | -0.160325183 | 0.815204 | 0.887529 | 18.10898 | 21.59679 | 14.07781 | 18.11509 | 13.65156 | 52.51658 | 6.750024 | 7.613138 |
| ENSMUSG00000068697 | 0.229677589 | 0.422773 | 0.587871 | 49.29666 | 33.595 | 42.23344 | 42.26855 | 27.30312 | 31.28647 | 45.32159 | 39.15328 |
| ENSMUSG00000068699 | 0.70643144 | 4.03E-07 | 1.76E-05 | 1490.973 | 1600.562 | 1962.613 | 1635.534 | 982.9122 | 1276.041 | 929.5748 | 913.5765 |
| ENSMUSG00000068706 | -0.687416637 | 0.08432 | 0.195848 | 29.17558 | 43.19358 | 32.29616 | 29.3292 | 44.85512 | 23.46486 | 71.3574 | 75.04379 |
| ENSMUSG00000068732 | 0.088965073 | 0.289376 | 0.456473 | 1419.543 | 1324.603 | 1263.691 | 1473.361 | 1281.296 | 1196.708 | 1397.255 | 1276.832 |
| ENSMUSG00000068735 | -0.183700352 | 0.06986 | 0.172073 | 1021.145 | 856.6726 | 1101.382 | 967.0009 | 1210.113 | 1061.505 | 1057.825 | 1157.197 |
| ENSMUSG00000068739 | -0.261637628 | 0.01328 | 0.052818 | 3338.088 | 3222.721 | 3431.674 | 3431.516 | 4148.124 | 3238.15 | 4349.908 | 4356.89 |
| ENSMUSG00000068740 | -0.128311136 | 0.22689 | 0.386817 | 908.4671 | 784.6833 | 859.5747 | 720.2906 | 829.8197 | 958.707 | 952.7177 | 836.3575 |
| ENSMUSG00000068742 | -0.120422464 | 0.26952 | 0.434282 | 823.9585 | 680.2988 | 756.8895 | 700.4503 | 822.9939 | 857.0259 | 852.4316 | 687.3576 |
| ENSMUSG00000068744 | 0.434069968 | 0.004125 | 0.022334 | 1129.799 | 1245.415 | 1064.117 | 1200.772 | 1084.324 | 861.4954 | 834.1101 | 652.5547 |
| ENSMUSG00000068747 | 0.063676913 | 0.307113 | 0.475594 | 2741.498 | 2649.206 | 2827.984 | 2681.034 | 2420.226 | 2578.899 | 2726.045 | 2707.014 |
| ENSMUSG00000068748 | 1.428052825 | 0.000194 | 0.002211 | 401.4157 | 381.5432 | 230.2136 | 386.4553 | 87.76002 | 261.4655 | 87.75031 | 83.74451 |
| ENSMUSG00000068749 | -0.015266679 | 0.841003 | 0.90415 | 3192.21 | 3391.895 | 3065.651 | 3704.105 | 3398.263 | 3496.264 | 3265.083 | 3336.729 |
| ENSMUSG00000068758 | 0.618744413 | 8.75E-09 | 7.60E-07 | 317.9132 | 298.7556 | 340.3518 | 348.4999 | 213.5494 | 212.3011 | 211.1793 | 215.343 |
| ENSMUSG00000068762 | 0.098011831 | 0.684092 | 0.798636 | 149.9021 | 111.5834 | 110.9663 | 159.5853 | 102.3867 | 174.3104 | 102.2147 | 119.635 |
| ENSMUSG00000068794 | 0.771882182 | 0.216877 | 0.375381 | 57.3451 | 11.99822 | 11.59349 | 57.79577 | 18.52711 | 31.28647 | 15.42863 | 16.31387 |
| ENSMUSG00000068798 | 0.112610214 | 0.072308 | 0.176516 | 2231.429 | 2380.446 | 2294.683 | 2448.988 | 2201.801 | 2281.678 | 2066.472 | 2104.489 |
| ENSMUSG00000068823 | 0.033670232 | 0.589519 | 0.729625 | 10424.74 | 11079.15 | 9927.342 | 10797.46 | 10131.41 | 11058.65 | 10164.57 | 9897.079 |
| ENSMUSG00000068854 | 0.098511391 | 0.366867 | 0.535461 | 469.8274 | 527.9215 | 529.1601 | 607.2869 | 507.0579 | 491.6446 | 534.2162 | 461.1386 |
| ENSMUSG00000068856 | -0.211787224 | 0.061583 | 0.157858 | 923.5579 | 946.6592 | 1155.209 | 989.4291 | 1239.366 | 991.1108 | 1178.361 | 1243.117 |
| ENSMUSG00000068860 | 0.26701111 | 0.357337 | 0.525834 | 39.23612 | 39.59411 | 38.09291 | 28.46658 | 27.30312 | 24.58223 | 35.6787 | 32.62773 |
| ENSMUSG00000068874 | -1.776143 | 1.80E-10 | 3.00E-08 | 781.7042 | 539.9197 | 977.9939 | 703.0381 | 2774.192 | 1254.811 | 3188.904 | 3070.27 |
| ENSMUSG00000068876 | -0.701667892 | 0.00331 | 0.019066 | 468.8213 | 448.7333 | 775.1078 | 460.6409 | 826.8944 | 592.2083 | 966.2177 | 1119.131 |
| ENSMUSG00000068877 | -0.736306049 | 0.116884 | 0.245132 | 12.07265 | 10.79839 | 11.59349 | 11.21411 | 24.37778 | 10.05637 | 14.46434 | 27.18978 |
| ENSMUSG00000068882 | 0.085275956 | 0.330881 | 0.498714 | 7076.586 | 7629.665 | 6797.927 | 7145.11 | 6417.208 | 7799.271 | 6361.416 | 6426.576 |
| ENSMUSG00000068917 | 0.142319691 | 0.079588 | 0.18848 | 2106.678 | 2132.083 | 1754.758 | 2003.874 | 1764.951 | 1788.916 | 1805.149 | 1882.62 |
| ENSMUSG00000068921 | -0.207465959 | 0.000192 | 0.002192 | 2156.981 | 2118.885 | 2196.967 | 2112.565 | 2550.891 | 2337.547 | 2454.116 | 2571.065 |
| ENSMUSG00000068922 | -0.252910705 | 0.05768 | 0.15054 | 967.8243 | 1070.241 | 1218.973 | 1159.366 | 1486.07 | 1041.393 | 1311.433 | 1424.744 |
| ENSMUSG00000068923 | 0.868093978 | 2.44E-05 | 0.000457 | 1382.319 | 1123.033 | 1026.852 | 1400.901 | 594.8179 | 984.4066 | 577.6092 | 547.0583 |
| ENSMUSG00000068959 | 0.186046906 | 0.257189 | 0.420954 | 178.0716 | 219.5673 | 154.8559 | 218.2437 | 152.1174 | 172.0756 | 182.2507 | 169.6642 |
| ENSMUSG00000068962 | 0.552248439 | 0.003608 | 0.020294 | 81.4904 | 94.7859 | 80.32634 | 80.22398 | 58.50668 | 53.63396 | 57.85735 | 58.72992 |
| ENSMUSG00000068966 | 0.368386776 | 0.00386 | 0.021292 | 605.6447 | 585.5129 | 529.1601 | 532.2387 | 420.273 | 520.6963 | 437.7873 | 365.4306 |
| ENSMUSG00000069011 | -0.108101763 | 0.768313 | 0.856029 | 18.10898 | 14.39786 | 20.70267 | 20.70296 | 18.52711 | 22.34748 | 24.10723 | 15.22628 |
| ENSMUSG00000069014 | -0.275415687 | 0.446816 | 0.609447 | 19.11503 | 16.7975 | 14.07781 | 19.84034 | 18.52711 | 17.87799 | 25.07152 | 22.83941 |
| ENSMUSG00000069020 | -0.196722348 | 0.046999 | 0.130283 | 1233.423 | 1421.789 | 1402.813 | 1408.664 | 1758.126 | 1387.779 | 1631.577 | 1485.649 |
| ENSMUSG00000069041 | -0.905469868 | NA | NA | 22.1332 | 19.19714 | 11.59349 | 12.93935 | 0.975111 | 119.559 | 2.892867 | 0 |
| ENSMUSG00000069045 | -0.202826359 | 0.903452 | 0.942315 | 9.054489 | 2669.603 | 2360.104 | 25.8787 | 2050.659 | 10.05637 | 1804.185 | 1964.19 |
| ENSMUSG00000069049 | -0.345607871 | 0.838586 | 0.90239 | 4.024217 | 2436.838 | 2334.433 | 16.38985 | 1860.512 | 22.34748 | 2091.543 | 2114.277 |
| ENSMUSG00000069072 | 1.227200148 | 0.01099 | 0.045808 | 55.33299 | 23.99643 | 22.35888 | 65.55939 | 20.47734 | 24.58223 | 12.53576 | 14.13868 |
| ENSMUSG00000069089 | -0.006828026 | 0.936603 | 0.962308 | 2167.041 | 2194.474 | 1823.491 | 1941.765 | 2190.1 | 2029.151 | 1969.078 | 1970.715 |
| ENSMUSG00000069094 | 0.008029389 | 0.941455 | 0.966057 | 1524.172 | 1554.969 | 1288.534 | 1496.652 | 1357.355 | 1699.526 | 1454.148 | 1319.248 |
| ENSMUSG00000069114 | 0.252631414 | 0.110773 | 0.236865 | 1005.048 | 1000.651 | 899.3238 | 1046.362 | 655.2748 | 1099.496 | 788.7885 | 774.3649 |
| ENSMUSG00000069135 | 0.15086034 | 0.065016 | 0.164218 | 1104.648 | 1070.241 | 1122.913 | 1243.903 | 1018.991 | 1107.318 | 994.1821 | 974.4816 |
| ENSMUSG00000069136 | 0.696618756 | 0.146756 | 0.285743 | 24.1453 | 10.79839 | 16.56213 | 14.6646 | 13.65156 | 6.704245 | 10.60718 | 9.78832 |
| ENSMUSG00000069170 | 0.595860078 | 0.002887 | 0.017251 | 358.1554 | 355.1472 | 295.6341 | 343.3242 | 223.3005 | 310.63 | 196.715 | 164.2263 |

| | | | | | | | | | | | |
|---|---|---|---|---|---|---|---|---|---|---|---|
| ENSMUSG00000069171 | 0.521125843 | 0.006923 | 0.032734 | 1416.525 | 1780.535 | 1241.332 | 1417.29 | 831.7699 | 1441.413 | 914.1461 | 892.9123 |
| ENSMUSG00000069184 | 0.070023514 | 0.743377 | 0.83954 | 79.47829 | 112.7832 | 72.04528 | 105.2401 | 95.56091 | 77.09881 | 90.64318 | 87.00729 |
| ENSMUSG00000069188 | -0.349734212 | 0.378686 | 0.54683 | 24.1453 | 11.99822 | 13.24971 | 15.52722 | 22.42756 | 21.23011 | 16.39292 | 22.83941 |
| ENSMUSG00000069206 | 0.333602991 | 0.049198 | 0.134578 | 393.3673 | 479.9286 | 311.3681 | 335.5605 | 279.8569 | 338.5644 | 323.0369 | 262.1095 |
| ENSMUSG00000069208 | 0.188507099 | 0.208038 | 0.36501 | 362.1796 | 361.1463 | 281.5563 | 276.9021 | 306.185 | 299.4563 | 275.7867 | 240.3576 |
| ENSMUSG00000069227 | 0.351686919 | 0.283905 | 0.45046 | 44.26639 | 22.79661 | 38.92101 | 49.16954 | 26.32801 | 36.87335 | 27.0001 | 32.62773 |
| ENSMUSG00000069237 | 0.102836395 | 0.213253 | 0.371127 | 2008.084 | 2204.072 | 1929.488 | 2160.872 | 1787.379 | 2131.95 | 1901.578 | 1909.81 |
| ENSMUSG00000069255 | -0.007466747 | 0.941343 | 0.965995 | 336.0222 | 387.5424 | 343.6643 | 379.5543 | 380.2934 | 375.4377 | 337.5012 | 359.9926 |
| ENSMUSG00000069265 | 0.483712619 | 0.217033 | 0.375514 | 18.10898 | 35.99465 | 31.46805 | 33.64232 | 20.47734 | 18.99536 | 14.46434 | 31.54014 |
| ENSMUSG00000069266 | -0.457458311 | 0.400995 | 0.56824 | 6.036326 | 11.99822 | 8.281066 | 9.488858 | 8.776002 | 20.11273 | 12.53576 | 7.613138 |
| ENSMUSG00000069270 | -0.45077311 | 0.456576 | 0.618419 | 9.054489 | 10.79839 | 5.796746 | 8.626235 | 20.47734 | 5.58687 | 13.50005 | 6.525547 |
| ENSMUSG00000069272 | -0.507685456 | 0.138021 | 0.273316 | 26.15741 | 35.99465 | 30.63995 | 31.91707 | 60.4569 | 44.69496 | 24.10723 | 47.85401 |
| ENSMUSG00000069273 | -0.994030498 | 0.08717 | 0.200649 | 3.018163 | 26.39607 | 11.59349 | 14.6646 | 39.00445 | 20.11273 | 30.85725 | 19.57664 |
| ENSMUSG00000069300 | -0.613842792 | 0.113439 | 0.240537 | 13.07871 | 15.59768 | 16.56213 | 13.80198 | 21.45245 | 14.52586 | 27.0001 | 27.18978 |
| ENSMUSG00000069301 | -0.434677568 | 0.107904 | 0.232505 | 58.35115 | 53.99197 | 43.06154 | 37.95543 | 78.0089 | 53.63396 | 51.10733 | 77.21897 |
| ENSMUSG00000069310 | -0.674808718 | 0.076165 | 0.182915 | 28.16952 | 65.99018 | 80.32634 | 49.16954 | 122.864 | 53.63396 | 76.17884 | 104.4087 |
| ENSMUSG00000069372 | 1.223913652 | 2.67E-12 | 6.94E-10 | 321.9374 | 370.7449 | 264.9941 | 377.8291 | 139.4409 | 174.3104 | 135.0005 | 122.8978 |
| ENSMUSG00000069378 | 0.91517161 | 0.201506 | 0.356978 | 19.11503 | 8.398751 | 23.18699 | 75.04824 | 21.45245 | 23.46486 | 4.821446 | 17.40146 |
| ENSMUSG00000069456 | -3.667469728 | 8.75E-05 | 0.001214 | 15.09082 | 2.399643 | 60.45178 | 13.80198 | 336.4134 | 67.04245 | 277.7153 | 490.5036 |
| ENSMUSG00000069495 | 0.195894586 | 0.058814 | 0.152476 | 1849.128 | 1893.318 | 1785.398 | 2104.801 | 1496.796 | 1878.306 | 1772.363 | 1517.19 |
| ENSMUSG00000069516 | -0.02778351 | 0.887017 | 0.932224 | 302.8224 | 579.5138 | 554.8314 | 490.8328 | 520.7094 | 502.8183 | 486.966 | 454.6131 |
| ENSMUSG00000069520 | -0.209613597 | 0.018102 | 0.065882 | 929.5942 | 1012.649 | 981.3064 | 976.4898 | 1080.423 | 1026.867 | 1126.29 | 1275.744 |
| ENSMUSG00000069539 | -0.107696903 | 0.156571 | 0.298768 | 1227.386 | 1168.626 | 1277.769 | 1215.437 | 1233.516 | 1267.102 | 1441.612 | 1327.949 |
| ENSMUSG00000069540 | 0.170795087 | 0.757405 | 0.848813 | 17.10292 | 8.398751 | 14.07781 | 29.3292 | 18.52711 | 8.938993 | 24.10723 | 9.78832 |
| ENSMUSG00000069565 | -0.0161982 | 0.79565 | 0.874398 | 3505.093 | 3781.837 | 3268.537 | 3504.839 | 3571.833 | 3539.841 | 3445.405 | 3655.394 |
| ENSMUSG00000069581 | 0.696920299 | 0.035939 | 0.108273 | 29.17558 | 31.19536 | 24.8432 | 29.3292 | 16.57689 | 17.87799 | 14.46434 | 21.75182 |
| ENSMUSG00000069601 | 0.228812853 | 0.030851 | 0.096442 | 4205.307 | 4753.693 | 3812.603 | 3798.131 | 3190.564 | 4000.199 | 3464.691 | 3481.379 |
| ENSMUSG00000069631 | 0.161351825 | 0.084018 | 0.195387 | 691.1593 | 794.2819 | 726.2495 | 715.9775 | 620.1708 | 657.016 | 610.395 | 729.7736 |
| ENSMUSG00000069633 | -0.063643852 | 0.69806 | 0.809552 | 110.666 | 122.3818 | 139.1219 | 139.745 | 133.5902 | 117.3243 | 131.1433 | 154.4379 |
| ENSMUSG00000069662 | 0.49337731 | 0.011719 | 0.04814 | 8486.068 | 10636.42 | 8879.787 | 9289.592 | 5292.904 | 9842.948 | 5604.449 | 5751.182 |
| ENSMUSG00000069670 | 0.150462849 | 0.751337 | 0.845087 | 16.09687 | 23.99643 | 15.73403 | 25.8787 | 17.552 | 29.05173 | 7.714313 | 19.57664 |
| ENSMUSG00000069678 | 0.067259609 | 0.537514 | 0.687719 | 690.1533 | 590.3122 | 650.0637 | 646.9676 | 651.3744 | 626.8469 | 665.3595 | 516.6058 |
| ENSMUSG00000069682 | -0.106860931 | 0.424439 | 0.589601 | 464.7971 | 592.7118 | 543.238 | 525.3377 | 691.3539 | 543.0438 | 553.502 | 499.2043 |
| ENSMUSG00000069706 | -0.581710106 | 0.250765 | 0.414335 | 9.054489 | 5.999108 | 12.4216 | 12.07673 | 16.57689 | 10.05637 | 22.17865 | 10.87591 |
| ENSMUSG00000069713 | -0.480117385 | 0.152712 | 0.29368 | 20.12109 | 33.595 | 51.34261 | 37.09281 | 45.83023 | 52.51658 | 60.75022 | 40.24087 |
| ENSMUSG00000069727 | 0.281949092 | 0.052237 | 0.140292 | 231.3925 | 308.3541 | 248.432 | 251.0234 | 247.6783 | 198.8926 | 206.3579 | 199.0292 |
| ENSMUSG00000069729 | 0.08198388 | 0.36318 | 0.531677 | 2262.616 | 2242.466 | 2389.916 | 2063.395 | 1974.6 | 2411.293 | 2076.115 | 2003.343 |
| ENSMUSG00000069733 | 0.25939382 | 0.625478 | 0.755733 | 33.19979 | 10.79839 | 36.43669 | 21.56559 | 10.72622 | 17.87799 | 19.28578 | 38.06569 |
| ENSMUSG00000069743 | 0.085903662 | 0.602367 | 0.740165 | 192.1564 | 171.5745 | 155.684 | 162.1732 | 166.744 | 191.071 | 154.2863 | 129.4233 |
| ENSMUSG00000069744 | -0.344742977 | 0.0003 | 0.003096 | 2426.603 | 2566.418 | 2770.845 | 2505.921 | 3233.469 | 2842.6 | 3267.012 | 3699.985 |
| ENSMUSG00000069755 | -0.067747419 | 0.65697 | 0.778894 | 314.895 | 265.1606 | 286.5249 | 284.6658 | 299.3592 | 381.0246 | 270.001 | 257.7591 |
| ENSMUSG00000069763 | -0.62994425 | 0.001282 | 0.009415 | 592.566 | 593.9117 | 506.8013 | 615.9132 | 1082.374 | 600.0299 | 1101.218 | 787.4159 |
| ENSMUSG00000069769 | 0.407956973 | 0.005775 | 0.028615 | 5300.9 | 5196.427 | 4709.442 | 5356.892 | 3495.774 | 5175.677 | 3552.441 | 3275.824 |
| ENSMUSG00000069793 | 0.02837536 | 0.890901 | 0.934072 | 461.7789 | 772.6851 | 571.3936 | 396.8068 | 523.6348 | 549.7481 | 468.6445 | 615.5766 |
| ENSMUSG00000069804 | 0.734259834 | 0.221035 | 0.380046 | 62.37537 | 53.99197 | 29.81184 | 21.56559 | 19.50223 | 52.51658 | 22.17865 | 6.525547 |
| ENSMUSG00000069805 | -3.738094259 | 4.71E-07 | 2.00E-05 | 188.1322 | 59.99108 | 535.785 | 309.6818 | 4715.638 | 1217.938 | 2555.366 | 6112.262 |
| ENSMUSG00000069806 | 0.464157515 | 0.000203 | 0.002299 | 941.6669 | 896.2667 | 987.1031 | 1078.279 | 838.5957 | 587.7388 | 714.5383 | 688.4452 |
| ENSMUSG00000069808 | -0.466845382 | 0.000662 | 0.005694 | 233.4046 | 211.1686 | 265.8222 | 207.0296 | 327.6374 | 283.813 | 300.8582 | 357.8175 |
| ENSMUSG00000069814 | -0.055990172 | 0.880015 | 0.927943 | 25.15136 | 22.79661 | 35.60859 | 37.09281 | 41.92979 | 36.87335 | 29.89296 | 17.40146 |
| ENSMUSG00000069833 | 0.509638139 | 1.96E-07 | 9.87E-06 | 6129.889 | 5653.559 | 5850.573 | 6763.831 | 4124.721 | 4911.977 | 4239.979 | 3863.124 |
| ENSMUSG00000069835 | -0.026696634 | 0.804844 | 0.880483 | 354.1311 | 400.7404 | 390.8663 | 411.4714 | 374.4427 | 429.0717 | 359.6799 | 425.2481 |
| ENSMUSG00000069844 | -0.069727413 | 0.584432 | 0.724884 | 391.3551 | 393.5415 | 375.1323 | 371.7907 | 431.9743 | 470.4145 | 364.5013 | 340.416 |
| ENSMUSG00000069862 | 0.409946549 | 0.262362 | 0.426584 | 37.22401 | 28.79572 | 37.2648 | 21.56559 | 26.32801 | 31.28647 | 21.21436 | 15.22628 |
| ENSMUSG00000069874 | 0.427421915 | 0.168658 | 0.314896 | 129.781 | 271.1597 | 203.7142 | 101.7896 | 114.088 | 116.2069 | 161.0363 | 132.6861 |
| ENSMUSG00000069892 | 1.097422585 | 0.124185 | 0.25407 | 19.11503 | 17.99732 | 22.35888 | 6.038364 | 0.975111 | 13.40849 | 7.714313 | 8.700729 |
| ENSMUSG00000069895 | 0.099579576 | 0.1109 | 0.236943 | 1560.39 | 1482.979 | 1639.651 | 1529.431 | 1476.319 | 1404.539 | 1503.327 | 1416.044 |
| ENSMUSG00000069899 | -0.117000436 | 0.56696 | 0.711292 | 93.56306 | 65.99018 | 99.3728 | 95.75121 | 113.1129 | 96.09417 | 94.50034 | 82.65692 |
| ENSMUSG00000069910 | -0.083155862 | 0.536603 | 0.687269 | 419.5247 | 418.7377 | 333.727 | 354.5383 | 423.1983 | 340.7991 | 399.2157 | 450.2627 |
| ENSMUSG00000069911 | -0.752062916 | 0.011935 | 0.048807 | 29.17558 | 32.39518 | 24.8432 | 36.23019 | 71.18313 | 41.34284 | 44.3573 | 48.9416 |
| ENSMUSG00000069917 | 0.709149162 | 0.041822 | 0.120365 | 1501.033 | 676.6993 | 940.7291 | 2036.654 | 744.0099 | 1204.529 | 468.6445 | 737.3868 |
| ENSMUSG00000069919 | 0.401463048 | 0.350172 | 0.51889 | 7633.94 | 4741.695 | 5411.677 | 13368.08 | 6505.943 | 8195.939 | 3838.835 | 5047.51 |
| ENSMUSG00000069920 | -0.197879848 | 0.49093 | 0.648158 | 114.6902 | 191.9714 | 256.7131 | 153.547 | 160.8934 | 287.1651 | 166.822 | 208.8175 |
| ENSMUSG00000069972 | -0.182848284 | 0.296751 | 0.464453 | 97.58727 | 110.3836 | 131.669 | 100.9269 | 140.416 | 130.7328 | 117.6433 | 112.0219 |
| ENSMUSG00000070000 | 0.106655851 | 0.459036 | 0.620528 | 174.0474 | 176.3738 | 158.1684 | 166.4863 | 182.3458 | 138.5544 | 158.1434 | 145.7372 |

| | | | | | | | | | | | |
|---|---|---|---|---|---|---|---|---|---|---|---|
| ENSMUSG00000070002 | 0.00493758 | 0.951936 | 0.972676 | 797.8011 | 788.2827 | 735.3587 | 753.9329 | 805.4419 | 748.6406 | 808.0743 | 698.2335 |
| ENSMUSG00000070003 | -0.035921477 | 0.705364 | 0.814655 | 2338.07 | 2769.188 | 2640.004 | 2285.09 | 2783.943 | 2317.434 | 2545.723 | 2634.146 |
| ENSMUSG00000070023 | -3.681372842 | 5.68E-06 | 0.000148 | 2.012109 | 2.399643 | 1.656213 | 1.725247 | 26.32801 | 8.938993 | 13.50005 | 50.02919 |
| ENSMUSG00000070031 | 0.181522874 | 0.659619 | 0.780834 | 33.19979 | 73.18911 | 46.37397 | 34.50494 | 65.33246 | 23.46486 | 32.78583 | 42.41605 |
| ENSMUSG00000070034 | 0.261425746 | 0.475378 | 0.6348 | 56.33904 | 59.99108 | 105.1695 | 32.77969 | 50.70579 | 44.69496 | 63.64308 | 53.29196 |
| ENSMUSG00000070044 | 0.266477837 | 5.72E-05 | 0.000877 | 3062.429 | 3384.697 | 3564.171 | 3263.305 | 2656.203 | 2833.661 | 2777.153 | 2771.182 |
| ENSMUSG00000070047 | 0.4523951 | 4.02E-05 | 0.000676 | 5059.447 | 4699.701 | 5323.069 | 5022.194 | 3076.476 | 4076.181 | 3413.584 | 4130.671 |
| ENSMUSG00000070056 | -0.031797705 | 0.751356 | 0.845087 | 1193.18 | 1054.643 | 1096.413 | 1180.069 | 1010.215 | 1098.379 | 1255.504 | 1262.693 |
| ENSMUSG00000070063 | 0.256032701 | 0.525224 | 0.678439 | 42.25428 | 20.39697 | 20.70267 | 20.70296 | 17.552 | 27.93435 | 23.14294 | 18.48905 |
| ENSMUSG00000070167 | 0.450339211 | 0.115981 | 0.243937 | 100.6054 | 77.9884 | 67.07664 | 49.16954 | 60.4569 | 59.22083 | 54.96448 | 40.24087 |
| ENSMUSG00000070282 | 0.462577689 | 0.000844 | 0.006843 | 531.1967 | 533.9206 | 476.1613 | 585.7214 | 441.7254 | 425.7195 | 364.5013 | 311.0511 |
| ENSMUSG00000070283 | -0.395324234 | 0.05096 | 0.137885 | 318.9192 | 333.5504 | 305.5713 | 299.3304 | 442.7005 | 261.4655 | 417.5372 | 530.7445 |
| ENSMUSG00000070284 | -0.307763228 | 0.081195 | 0.190896 | 280.6892 | 392.3416 | 385.8977 | 342.4615 | 448.5512 | 322.9211 | 443.573 | 518.7809 |
| ENSMUSG00000070287 | 0.297994193 | 0.248748 | 0.412183 | 98.59333 | 57.59143 | 91.91984 | 82.81186 | 48.75557 | 73.74669 | 65.57166 | 82.65692 |
| ENSMUSG00000070291 | 1.319646571 | 4.62E-05 | 0.000749 | 78.47224 | 53.99197 | 50.5145 | 43.13117 | 20.47734 | 27.93435 | 17.3572 | 25.0146 |
| ENSMUSG00000070304 | 0.189717466 | 0.28307 | 0.449522 | 158.9566 | 124.7814 | 137.4657 | 139.745 | 103.3618 | 121.7938 | 115.7147 | 152.2628 |
| ENSMUSG00000070305 | -0.295950938 | 0.076747 | 0.183953 | 115.6963 | 111.5834 | 149.0592 | 105.2401 | 155.0427 | 139.6718 | 143.6791 | 154.4379 |
| ENSMUSG00000070315 | 0.720749975 | 5.39E-07 | 2.24E-05 | 284.7134 | 287.9572 | 249.2601 | 298.4677 | 176.4951 | 177.6625 | 136.9291 | 189.2408 |
| ENSMUSG00000070319 | -0.186258 | 0.005071 | 0.025988 | 3156.999 | 3153.131 | 3351.348 | 3259.854 | 3855.59 | 3339.831 | 3689.37 | 3818.532 |
| ENSMUSG00000070327 | 0.210002537 | 0.422591 | 0.587698 | 908.4671 | 1385.794 | 1816.038 | 705.626 | 920.5051 | 966.5286 | 1198.611 | 1077.803 |
| ENSMUSG00000070336 | -0.21062568 | 0.676521 | 0.792904 | 17.10292 | 22.79661 | 14.90592 | 10.35148 | 9.751113 | 29.05173 | 24.10723 | 11.9635 |
| ENSMUSG00000070337 | 0.527846982 | 0.06941 | 0.171192 | 67.40564 | 69.58965 | 65.42042 | 119.042 | 52.65601 | 61.45558 | 42.42872 | 67.43065 |
| ENSMUSG00000070343 | 0.093168116 | 0.50298 | 0.658977 | 358.1554 | 345.5486 | 336.2113 | 340.7363 | 374.4427 | 346.386 | 321.1083 | 251.2335 |
| ENSMUSG00000070348 | -0.443420506 | 0.00116 | 0.008753 | 4936.709 | 3917.417 | 4994.311 | 4500.307 | 5716.102 | 5053.883 | 7143.454 | 7039.977 |
| ENSMUSG00000070366 | 0.010092867 | 0.974213 | 0.985749 | 45.27245 | 28.79572 | 25.67131 | 31.05445 | 27.30312 | 35.75597 | 29.89296 | 36.9781 |
| ENSMUSG00000070371 | -0.163553713 | 0.231967 | 0.392539 | 196.1806 | 223.1668 | 181.3554 | 206.167 | 230.1263 | 192.1883 | 249.7509 | 228.3941 |
| ENSMUSG00000070372 | 0.007955213 | 0.879949 | 0.927943 | 2576.505 | 2648.006 | 2524.897 | 2689.66 | 2612.323 | 2525.265 | 2537.045 | 2705.927 |
| ENSMUSG00000070394 | -0.312581832 | 0.003328 | 0.019112 | 915.5095 | 892.6672 | 983.7907 | 985.116 | 1217.914 | 984.4066 | 1165.826 | 1324.686 |
| ENSMUSG00000070407 | 0.271138107 | 0.093759 | 0.210821 | 1786.753 | 1967.707 | 1460.78 | 1736.461 | 1363.206 | 1889.48 | 1342.291 | 1164.81 |
| ENSMUSG00000070420 | -0.126453184 | 0.375942 | 0.544226 | 321.9374 | 284.3577 | 327.1021 | 248.4356 | 374.4427 | 287.1651 | 323.0369 | 304.5255 |
| ENSMUSG00000070423 | -0.381750489 | 0.376541 | 0.544772 | 13.07871 | 19.19714 | 25.67131 | 10.35148 | 19.50223 | 18.99536 | 20.25007 | 30.45255 |
| ENSMUSG00000070424 | 0.952223925 | 0.000365 | 0.003592 | 119.7205 | 87.58697 | 91.91984 | 95.75121 | 73.13335 | 39.10809 | 53.0359 | 38.06569 |
| ENSMUSG00000070425 | 0.790788389 | 0.14421 | 0.282297 | 36.21796 | 53.99197 | 35.60859 | 43.13117 | 19.50223 | 55.8687 | 10.60718 | 11.9635 |
| ENSMUSG00000070426 | -0.194001273 | 0.120903 | 0.250136 | 674.0564 | 705.4951 | 751.9208 | 623.6768 | 774.2384 | 651.4291 | 803.2529 | 922.2772 |
| ENSMUSG00000070427 | 0.381047464 | 0.055527 | 0.146736 | 129.781 | 105.5843 | 154.8559 | 148.3712 | 93.61069 | 127.3806 | 89.67889 | 105.4963 |
| ENSMUSG00000070436 | -0.10727917 | 0.287512 | 0.454425 | 15701.49 | 19729.87 | 20132.1 | 17733.81 | 20003.43 | 18003.13 | 19353.28 | 21595.21 |
| ENSMUSG00000070461 | 0.328653652 | 0.048775 | 0.133839 | 122.7386 | 140.3791 | 125.0441 | 156.9975 | 120.9138 | 105.0332 | 111.8575 | 95.70802 |
| ENSMUSG00000070462 | -0.158611446 | 0.219623 | 0.37843 | 1007.06 | 926.2622 | 1055.008 | 1006.682 | 1139.905 | 878.256 | 1117.611 | 1324.686 |
| ENSMUSG00000070469 | -0.062606026 | 0.818971 | 0.890502 | 53.32088 | 85.18733 | 54.65504 | 68.14726 | 89.71024 | 49.16446 | 61.71451 | 70.69342 |
| ENSMUSG00000070473 | 0.31345555 | 2.78E-07 | 1.30E-05 | 1353.143 | 1414.59 | 1407.781 | 1455.246 | 1127.229 | 1188.886 | 1097.361 | 1120.219 |
| ENSMUSG00000070476 | 0.14084514 | 0.297174 | 0.465036 | 237.4288 | 217.1677 | 260.0255 | 217.3811 | 193.072 | 236.8833 | 192.8578 | 225.1314 |
| ENSMUSG00000070493 | -0.50382294 | 5.33E-05 | 0.000831 | 6725.473 | 6551.026 | 7455.444 | 6713.799 | 10029.02 | 7706.529 | 9542.606 | 11640.49 |
| ENSMUSG00000070498 | -0.482286301 | 0.058452 | 0.151791 | 73.44197 | 35.99465 | 69.56096 | 62.10889 | 87.76002 | 81.56831 | 95.46463 | 73.95619 |
| ENSMUSG00000070509 | 0.50813693 | 0.002101 | 0.013594 | 406.446 | 598.7109 | 450.49 | 425.2734 | 339.3387 | 383.2593 | 324.0012 | 274.073 |
| ENSMUSG00000070520 | -0.034552613 | 0.647552 | 0.77184 | 708.2623 | 745.0892 | 811.5445 | 779.8116 | 785.9397 | 762.0491 | 770.467 | 803.7298 |
| ENSMUSG00000070526 | -0.144924951 | 0.20739 | 0.364047 | 309.8647 | 347.9482 | 366.0231 | 345.912 | 427.0988 | 359.7945 | 386.68 | 340.416 |
| ENSMUSG00000070527 | 0.04534247 | 0.787187 | 0.869179 | 175.0535 | 179.9732 | 133.3252 | 180.2883 | 188.1965 | 162.0192 | 141.7505 | 154.4379 |
| ENSMUSG00000070532 | 1.627200583 | 0.003174 | 0.018523 | 20.12109 | 14.39786 | 10.76539 | 18.97772 | 3.900445 | 5.58687 | 3.857157 | 7.613138 |
| ENSMUSG00000070544 | 0.303206411 | 0.052105 | 0.139998 | 7188.258 | 6981.762 | 5308.992 | 6557.664 | 4609.351 | 6895.316 | 4962.232 | 4634.226 |
| ENSMUSG00000070565 | -0.027375047 | 0.831417 | 0.897939 | 524.1543 | 404.3399 | 427.303 | 529.6508 | 437.825 | 525.1658 | 497.5732 | 463.3138 |
| ENSMUSG00000070570 | 0.030583611 | 0.796806 | 0.875296 | 287.7315 | 307.1543 | 326.274 | 326.9343 | 287.6578 | 354.2076 | 280.6081 | 302.3503 |
| ENSMUSG00000070572 | -0.037664453 | 0.926985 | 0.956688 | 16.09687 | 22.79661 | 12.4216 | 13.80198 | 13.65156 | 17.87799 | 20.25007 | 14.13868 |
| ENSMUSG00000070576 | 0.552602394 | 0.003848 | 0.021239 | 344.0706 | 430.7359 | 306.3995 | 356.2635 | 198.9227 | 330.7427 | 220.8222 | 229.4817 |
| ENSMUSG00000070583 | -0.252646508 | 0.116479 | 0.24465 | 134.8113 | 119.9822 | 107.6539 | 133.7066 | 146.2667 | 127.3806 | 149.4648 | 167.489 |
| ENSMUSG00000070604 | 0.121191569 | 0.515083 | 0.669227 | 174.0474 | 128.3809 | 163.9651 | 152.6844 | 137.4907 | 108.3853 | 163.9292 | 159.8759 |
| ENSMUSG00000070605 | 0.31893877 | 0.252414 | 0.416144 | 187.1261 | 200.3702 | 165.6213 | 140.6076 | 108.2374 | 220.1227 | 133.0719 | 94.62042 |
| ENSMUSG00000070610 | -0.219907196 | 0.255668 | 0.419581 | 124.7507 | 105.5843 | 90.26362 | 83.67448 | 125.7894 | 119.559 | 124.3933 | 98.97079 |
| ENSMUSG00000070639 | -0.129511943 | 0.413886 | 0.57983 | 852.128 | 551.9179 | 864.5433 | 678.8847 | 777.1637 | 755.3449 | 777.2171 | 917.9269 |
| ENSMUSG00000070643 | -0.101424406 | 0.182646 | 0.333317 | 1158.975 | 1099.037 | 1105.522 | 1207.673 | 1186.71 | 1188.886 | 1343.255 | 1185.474 |
| ENSMUSG00000070644 | -0.531012641 | 0.108023 | 0.232662 | 144.8718 | 58.79126 | 113.4506 | 107.8279 | 106.2871 | 121.7938 | 231.4294 | 155.5255 |
| ENSMUSG00000070645 | -2.047304592 | 0.04413 | 0.12486 | 2745.522 | 97.18554 | 698.0939 | 369.2029 | 2255.432 | 1287.215 | 3787.728 | 8831.24 |
| ENSMUSG00000070661 | -0.888243609 | 0.046004 | 0.12845 | 98.59333 | 58.79126 | 226.0731 | 97.47645 | 170.6445 | 113.9722 | 361.6084 | 245.7956 |
| ENSMUSG00000070683 | 0.197318314 | 0.502114 | 0.658262 | 30.18163 | 26.39607 | 28.98373 | 31.05445 | 24.37778 | 25.6996 | 28.92867 | 22.83941 |
| ENSMUSG00000070690 | 0.108347426 | 0.809835 | 0.884031 | 13.07871 | 16.7975 | 26.49941 | 25.8787 | 15.60178 | 14.52586 | 15.42863 | 31.54014 |

| | | | | | | | | | | | |
|---|---|---|---|---|---|---|---|---|---|---|---|
| ENSMUSG00000070695 | 0.722709161 | 0.391669 | 0.559104 | 11.0666 | 10.79839 | 9.93728 | 23.29083 | 2.925334 | 24.58223 | 1.928578 | 4.350364 |
| ENSMUSG00000070697 | -0.060656042 | 0.312409 | 0.480769 | 1858.182 | 1814.13 | 1899.677 | 2002.149 | 2050.659 | 1893.949 | 1988.364 | 1969.627 |
| ENSMUSG00000070699 | -0.297215608 | 0.035624 | 0.107545 | 457.7547 | 447.5334 | 455.4586 | 359.714 | 579.2161 | 422.3674 | 575.6806 | 534.0072 |
| ENSMUSG00000070705 | -0.06586169 | 0.552676 | 0.700082 | 312.8829 | 363.5459 | 312.1962 | 370.0655 | 352.0152 | 384.3767 | 351.0013 | 333.8905 |
| ENSMUSG00000070709 | 0.359532539 | 0.008625 | 0.038338 | 285.7194 | 291.5566 | 250.0882 | 266.5507 | 218.4249 | 249.1744 | 203.465 | 180.5401 |
| ENSMUSG00000070713 | 0.06164595 | 0.736062 | 0.834773 | 3396.44 | 4094.991 | 3680.934 | 3814.521 | 5164.189 | 2820.252 | 3134.904 | 3238.846 |
| ENSMUSG00000070720 | -0.14706599 | 0.41819 | 0.583777 | 212.2775 | 231.5656 | 335.3832 | 269.1385 | 312.0356 | 234.6486 | 300.8582 | 315.4014 |
| ENSMUSG00000070729 | -0.208073174 | 0.294437 | 0.461968 | 57.3451 | 62.39072 | 56.31125 | 69.8725 | 73.13335 | 64.8077 | 76.17884 | 69.60583 |
| ENSMUSG00000070730 | -0.218982088 | 0.084231 | 0.195701 | 790.7587 | 782.2836 | 744.4679 | 798.7894 | 905.8784 | 716.2368 | 930.539 | 1073.452 |
| ENSMUSG00000070732 | 0.320545789 | 0.569864 | 0.713532 | 16.09687 | 22.79661 | 14.90592 | 15.52722 | 5.850668 | 25.6996 | 17.3572 | 6.525547 |
| ENSMUSG00000070733 | 0.170234968 | 0.194833 | 0.348638 | 2080.52 | 1779.335 | 1724.118 | 1894.321 | 1692.793 | 2031.386 | 1547.684 | 1374.715 |
| ENSMUSG00000070737 | -0.096404434 | 0.538992 | 0.688884 | 209.2593 | 191.9714 | 181.3554 | 211.3428 | 163.8187 | 226.8269 | 219.8579 | 239.27 |
| ENSMUSG00000070738 | -0.040926702 | 0.445471 | 0.608333 | 1914.521 | 1922.114 | 1821.835 | 1934.864 | 1891.716 | 1962.109 | 2018.257 | 1937 |
| ENSMUSG00000070780 | -0.848785309 | 0.000278 | 0.002922 | 783.7163 | 434.3354 | 875.3087 | 600.386 | 966.3353 | 1001.167 | 1386.648 | 1500.876 |
| ENSMUSG00000070802 | 0.139177868 | 0.252758 | 0.41636 | 396.3854 | 493.1267 | 481.9581 | 523.6125 | 390.0445 | 473.7666 | 419.4658 | 439.3868 |
| ENSMUSG00000070803 | -0.268039735 | 0.344967 | 0.513776 | 337.0282 | 314.3532 | 248.432 | 328.6596 | 216.4747 | 420.1327 | 567.9663 | 274.073 |
| ENSMUSG00000070806 | -0.139596629 | 0.681238 | 0.796487 | 36.21796 | 40.79393 | 27.32752 | 20.70296 | 39.97956 | 31.28647 | 25.07152 | 40.24087 |
| ENSMUSG00000070808 | 0.363940761 | 0.005742 | 0.028494 | 414.4944 | 431.9358 | 385.8977 | 403.7078 | 302.2845 | 388.8462 | 271.9295 | 308.8759 |
| ENSMUSG00000070814 | 0.130525157 | 0.294097 | 0.461749 | 385.3188 | 406.7395 | 493.5516 | 391.6311 | 408.5716 | 354.2076 | 357.7513 | 413.2846 |
| ENSMUSG00000070822 | 0.352165146 | 0.022329 | 0.076244 | 415.5004 | 368.3452 | 370.9918 | 495.1459 | 361.7663 | 331.8601 | 340.3941 | 258.8467 |
| ENSMUSG00000070858 | 0.077518661 | 0.528501 | 0.68098 | 373.2462 | 345.5486 | 413.2252 | 417.5098 | 370.5423 | 397.7852 | 391.5014 | 311.0511 |
| ENSMUSG00000070866 | 0.928898404 | 0.101137 | 0.222793 | 15.09082 | 14.39786 | 13.24971 | 23.29083 | 5.850668 | 16.76061 | 3.857157 | 8.700729 |
| ENSMUSG00000070867 | -0.641736618 | 0.004269 | 0.022917 | 511.0756 | 316.7529 | 539.0974 | 369.2029 | 464.153 | 806.7441 | 680.7882 | 760.2262 |
| ENSMUSG00000070871 | -0.160385891 | 0.101732 | 0.223706 | 738.4439 | 729.4915 | 749.4365 | 845.371 | 878.5753 | 953.1201 | 817.7172 | 776.54 |
| ENSMUSG00000070880 | 0.40486781 | 0.048302 | 0.132936 | 91.55095 | 82.78769 | 84.46688 | 87.9876 | 58.50668 | 84.92043 | 62.6788 | 56.55474 |
| ENSMUSG00000070883 | 0.654136338 | 0.000458 | 0.004282 | 201.2109 | 211.1686 | 144.0906 | 171.6621 | 105.312 | 129.6154 | 127.2862 | 98.97079 |
| ENSMUSG00000070891 | 0.635887566 | 0.421991 | 0.587229 | 32.19374 | 20.39697 | 12.4216 | 12.93935 | 1.950223 | 32.40385 | 11.57147 | 4.350364 |
| ENSMUSG00000070923 | 0.147242709 | 0.060707 | 0.156363 | 2454.773 | 2553.22 | 2196.967 | 2291.991 | 2090.639 | 2337.547 | 2074.186 | 2069.686 |
| ENSMUSG00000070934 | -0.095404462 | 0.269445 | 0.434199 | 2142.896 | 2297.658 | 2466.102 | 2437.774 | 2567.468 | 2202.344 | 2551.509 | 2663.511 |
| ENSMUSG00000070939 | -0.086891636 | 0.202216 | 0.35772 | 952.7335 | 997.0517 | 1055.836 | 1024.797 | 1096.025 | 1021.28 | 1059.754 | 1106.08 |
| ENSMUSG00000070942 | 0.303783717 | 0.001598 | 0.011054 | 566.4086 | 566.3158 | 597.0649 | 575.3699 | 520.7094 | 413.4284 | 475.3946 | 456.7883 |
| ENSMUSG00000070953 | 0.079708162 | 0.431136 | 0.595899 | 752.5287 | 665.901 | 655.0323 | 797.0641 | 682.5779 | 632.4337 | 736.7169 | 664.5182 |
| ENSMUSG00000070972 | 0.150618369 | 0.253021 | 0.416674 | 377.2704 | 440.3345 | 341.1799 | 368.3402 | 303.2596 | 370.9682 | 317.2511 | 382.8321 |
| ENSMUSG00000070985 | -3.037333041 | 1.11E-05 | 0.000247 | 1.006054 | 4.799286 | 7.45296 | 1.725247 | 28.27823 | 17.87799 | 21.21436 | 56.55474 |
| ENSMUSG00000070997 | -0.123894516 | 0.707248 | 0.815963 | 45.27245 | 27.5959 | 32.29616 | 37.95543 | 27.30312 | 56.98608 | 36.64299 | 35.89051 |
| ENSMUSG00000071001 | 0.018601478 | 0.944076 | 0.967635 | 33.19979 | 52.79215 | 50.5145 | 52.62003 | 38.02934 | 49.16446 | 54.00019 | 45.67883 |
| ENSMUSG00000071014 | -0.682407397 | 4.25E-05 | 0.000703 | 1031.206 | 995.8519 | 1180.052 | 1080.867 | 1823.458 | 1173.243 | 1830.221 | 2055.547 |
| ENSMUSG00000071035 | -0.973614751 | 0.055858 | 0.147209 | 7.04238 | 7.198929 | 9.109173 | 7.763611 | 12.67645 | 8.938993 | 22.17865 | 17.40146 |
| ENSMUSG00000071037 | -0.280178266 | 0.082452 | 0.192948 | 155.9384 | 133.1802 | 122.5598 | 119.042 | 139.4409 | 168.7235 | 178.3935 | 156.6131 |
| ENSMUSG00000071041 | -0.0572854 | 0.779644 | 0.863998 | 90.54489 | 93.58608 | 69.56096 | 63.83414 | 83.85957 | 87.15518 | 80.036 | 77.21897 |
| ENSMUSG00000071042 | -0.27489171 | 0.056425 | 0.148197 | 425.561 | 359.9465 | 541.5817 | 469.2672 | 491.4561 | 518.4616 | 584.3592 | 584.0364 |
| ENSMUSG00000071054 | 0.190879827 | 0.00453 | 0.023921 | 5355.227 | 4987.658 | 4690.396 | 4875.548 | 4333.395 | 4670.624 | 4219.729 | 4216.591 |
| ENSMUSG00000071064 | 0.335167387 | 0.047489 | 0.131285 | 1475.882 | 1514.175 | 1307.58 | 1268.919 | 1093.1 | 1495.047 | 896.7889 | 927.7152 |
| ENSMUSG00000071068 | 1.027060294 | 0.009943 | 0.04239 | 36.21796 | 29.99554 | 50.5145 | 31.91707 | 12.67645 | 30.1691 | 13.50005 | 17.40146 |
| ENSMUSG00000071072 | 0.013496478 | 0.881119 | 0.928616 | 3669.08 | 4176.579 | 3668.512 | 3770.527 | 3584.509 | 4330.942 | 3521.584 | 3704.335 |
| ENSMUSG00000071073 | -0.21695411 | 0.534064 | 0.685259 | 28.16952 | 26.39607 | 28.15563 | 39.68068 | 36.07912 | 52.51658 | 23.14294 | 31.54014 |
| ENSMUSG00000071074 | -0.144498534 | 0.152168 | 0.292817 | 1901.443 | 1640.156 | 1936.941 | 1893.459 | 2118.917 | 1728.578 | 2127.222 | 2176.27 |
| ENSMUSG00000071076 | -0.24821909 | 0.004331 | 0.02316 | 2669.062 | 2410.441 | 3047.432 | 2800.938 | 3253.946 | 3044.844 | 3378.869 | 3308.452 |
| ENSMUSG00000071078 | 0.21605997 | 0.343602 | 0.512322 | 68.4117 | 89.98662 | 88.60741 | 89.71284 | 52.65601 | 88.27255 | 69.42882 | 80.48174 |
| ENSMUSG00000071103 | 0.127009426 | 0.553603 | 0.700643 | 265.5983 | 224.3666 | 207.8548 | 208.7549 | 203.7983 | 246.9397 | 245.8937 | 132.6861 |
| ENSMUSG00000071112 | 0.212476762 | 0.576082 | 0.718293 | 18.10898 | 22.79661 | 26.49941 | 25.01608 | 22.42756 | 25.6996 | 11.57147 | 20.66423 |
| ENSMUSG00000071141 | -0.171682754 | 0.595344 | 0.734568 | 35.2119 | 26.39607 | 36.43669 | 37.09281 | 54.60623 | 27.93435 | 29.89296 | 40.24087 |
| ENSMUSG00000071172 | -0.035722965 | 0.565083 | 0.709854 | 11987.14 | 13351.61 | 12164.06 | 13433.64 | 13401.93 | 13418.55 | 12691.97 | 12698.71 |
| ENSMUSG00000071176 | 0.124824014 | 0.132186 | 0.265309 | 1279.701 | 1385.794 | 1164.318 | 1351.731 | 1240.342 | 1227.994 | 1161.968 | 1118.044 |
| ENSMUSG00000071177 | 1.867551158 | 0.183229 | 0.334125 | 52.31483 | 0 | 0 | 45.71905 | 2.925334 | 2.234748 | 8.678602 | 13.05109 |
| ENSMUSG00000071178 | -0.98722648 | 0.183564 | 0.334538 | 119.7205 | 3.599465 | 36.43669 | 94.02596 | 77.03379 | 51.39921 | 161.0363 | 214.2554 |
| ENSMUSG00000071180 | 0.07074667 | 0.392067 | 0.559455 | 1835.043 | 1972.507 | 1909.614 | 2233.332 | 1841.01 | 1991.161 | 1923.757 | 1816.277 |
| ENSMUSG00000071181 | 0.818728781 | 0.007722 | 0.035288 | 50.30272 | 67.19001 | 41.40533 | 52.62003 | 34.1289 | 36.87335 | 19.28578 | 29.36496 |
| ENSMUSG00000071192 | 0.136830384 | 0.617579 | 0.750167 | 60.36326 | 39.59411 | 62.108 | 71.59775 | 39.00445 | 62.57295 | 54.00019 | 58.72992 |
| ENSMUSG00000071202 | 0.274350475 | 0.423821 | 0.588965 | 31.18769 | 43.19358 | 46.37397 | 58.6584 | 27.30312 | 55.8687 | 36.64299 | 29.36496 |
| ENSMUSG00000071203 | 0.107294721 | 0.740167 | 0.83754 | 132.7992 | 53.99197 | 79.49824 | 93.16334 | 54.60623 | 97.21155 | 103.1789 | 79.39415 |
| ENSMUSG00000071226 | 0.568996679 | 0.007644 | 0.03501 | 1011.085 | 898.6663 | 642.6107 | 943.7101 | 492.4312 | 842.5001 | 529.3948 | 492.6788 |
| ENSMUSG00000071252 | -0.086352264 | 0.535669 | 0.686453 | 181.0898 | 183.5727 | 154.0278 | 156.1349 | 169.6694 | 163.1366 | 194.7864 | 185.9781 |
| ENSMUSG00000071253 | -0.524471984 | 0.000108 | 0.001424 | 588.5418 | 507.5245 | 656.6886 | 615.9132 | 840.5459 | 689.4198 | 908.3604 | 972.3064 |

| | | | | | | | | | | | |
|---|---|---|---|---|---|---|---|---|---|---|---|
| ENSMUSG00000071256 | -0.192223307 | 0.139193 | 0.275107 | 299.8042 | 277.1588 | 301.4308 | 263.1002 | 317.8863 | 272.6393 | 371.2513 | 341.5036 |
| ENSMUSG00000071266 | 0.269408697 | 0.079799 | 0.188683 | 489.9485 | 562.7163 | 366.8512 | 474.4429 | 354.9405 | 446.9496 | 367.3942 | 400.2335 |
| ENSMUSG00000071267 | 0.263607364 | 0.059685 | 0.154234 | 543.2694 | 495.5263 | 463.7397 | 482.2065 | 371.5174 | 520.6963 | 397.2871 | 364.343 |
| ENSMUSG00000071281 | 0.379070499 | 0.019888 | 0.07017 | 512.0817 | 566.3158 | 371.8199 | 406.2957 | 351.0401 | 310.63 | 391.5014 | 370.8686 |
| ENSMUSG00000071291 | 0.12232906 | 0.446521 | 0.609181 | 221.332 | 195.5709 | 159.8246 | 207.8923 | 195.9974 | 197.7752 | 171.6435 | 154.4379 |
| ENSMUSG00000071302 | 0.149488837 | 0.107239 | 0.231621 | 589.5479 | 633.5058 | 642.6107 | 562.4305 | 573.3654 | 569.8608 | 550.6091 | 493.7664 |
| ENSMUSG00000071317 | 0.207201393 | 0.509817 | 0.665571 | 59.35721 | 128.3809 | 76.18581 | 80.22398 | 75.08357 | 105.0332 | 61.71451 | 55.46715 |
| ENSMUSG00000071324 | 0.63570767 | 0.070325 | 0.17287 | 58.35115 | 43.19358 | 34.78048 | 51.75741 | 26.32801 | 43.57759 | 33.75012 | 17.40146 |
| ENSMUSG00000071335 | -2.908701708 | 2.05E-07 | 1.02E-05 | 9.054489 | 3.599465 | 23.18699 | 11.21411 | 93.61069 | 33.52122 | 94.50034 | 138.1241 |
| ENSMUSG00000071337 | 0.430508419 | 0.009093 | 0.039805 | 5583.602 | 5897.123 | 4292.905 | 5571.685 | 3845.839 | 5209.198 | 3593.906 | 3188.817 |
| ENSMUSG00000071347 | 0.669316521 | 0.173914 | 0.321876 | 29.17558 | 41.99375 | 23.18699 | 49.16954 | 24.37778 | 41.34284 | 10.60718 | 14.13868 |
| ENSMUSG00000071350 | -0.116165351 | 0.442781 | 0.606004 | 186.1201 | 214.7681 | 159.8246 | 207.8923 | 221.3503 | 188.8362 | 229.5008 | 190.3284 |
| ENSMUSG00000071359 | 0.178927536 | 0.051436 | 0.138718 | 1696.208 | 1829.728 | 1635.511 | 1814.097 | 1552.377 | 1735.282 | 1506.22 | 1367.102 |
| ENSMUSG00000071369 | -0.036013576 | 0.783664 | 0.866969 | 621.7416 | 543.5192 | 682.3599 | 523.6125 | 581.1663 | 521.8137 | 652.8238 | 676.4817 |
| ENSMUSG00000071379 | -0.013043546 | 0.888397 | 0.932825 | 1146.902 | 1118.234 | 1315.033 | 1193.008 | 1137.955 | 1122.961 | 1343.255 | 1214.839 |
| ENSMUSG00000071392 | 0.968359914 | 0.058283 | 0.151589 | 17.10292 | 14.39786 | 22.35888 | 11.21411 | 6.825779 | 13.40849 | 5.785735 | 7.613138 |
| ENSMUSG00000071398 | 0.596743228 | 0.242062 | 0.404445 | 13.07871 | 13.19804 | 20.70267 | 18.11509 | 15.60178 | 6.704245 | 5.785735 | 15.22628 |
| ENSMUSG00000071415 | 0.004604249 | 0.956945 | 0.975803 | 31962.35 | 38316.3 | 36994.84 | 36229.32 | 37795.31 | 36959.38 | 36394.2 | 31894.7 |
| ENSMUSG00000071419 | 0.090639399 | 0.8076 | 0.882437 | 20.12109 | 44.3934 | 33.95237 | 37.95543 | 46.80534 | 27.93435 | 29.89296 | 22.83941 |
| ENSMUSG00000071424 | 0.291531541 | 0.550002 | 0.697988 | 28.16952 | 14.39786 | 19.04645 | 31.05445 | 14.62667 | 33.52122 | 18.32149 | 9.78832 |
| ENSMUSG00000071451 | -0.048765257 | 0.672759 | 0.790252 | 471.8395 | 610.7092 | 618.5957 | 516.7115 | 611.3948 | 563.1565 | 557.3591 | 561.197 |
| ENSMUSG00000071454 | -0.0765549 | 0.236306 | 0.397348 | 1227.386 | 1374.995 | 1327.455 | 1263.743 | 1414.887 | 1347.553 | 1338.433 | 1372.54 |
| ENSMUSG00000071456 | 0.001856594 | 0.992092 | 0.995436 | 149.9021 | 111.5834 | 110.1382 | 114.7289 | 115.0631 | 113.9722 | 108.9647 | 147.9124 |
| ENSMUSG00000071477 | 0.039702319 | 0.560744 | 0.706519 | 1155.956 | 1262.212 | 1226.426 | 1255.117 | 1161.358 | 1235.816 | 1241.04 | 1127.832 |
| ENSMUSG00000071497 | 0.143272505 | 0.65585 | 0.778183 | 57.3451 | 79.18822 | 75.3577 | 56.93315 | 39.00445 | 97.21155 | 51.10733 | 56.55474 |
| ENSMUSG00000071506 | -0.720297282 | 0.006796 | 0.032303 | 48.29061 | 31.19536 | 53.82693 | 41.40593 | 77.03379 | 52.51658 | 76.17884 | 83.74451 |
| ENSMUSG00000071516 | -0.291792427 | 0.380063 | 0.548089 | 32.19374 | 68.38983 | 33.95237 | 35.36756 | 40.95467 | 52.51658 | 64.60737 | 47.85401 |
| ENSMUSG00000071528 | -0.612878368 | 1.39E-05 | 0.000294 | 2132.835 | 2151.28 | 2832.125 | 2153.108 | 3492.849 | 2852.656 | 3783.871 | 4049.102 |
| ENSMUSG00000071531 | 0.193793649 | 0.584661 | 0.72512 | 31.18769 | 47.99286 | 50.5145 | 30.19182 | 43.88001 | 29.05173 | 22.17865 | 44.59123 |
| ENSMUSG00000071532 | -0.20464494 | 0.380226 | 0.548281 | 61.36932 | 51.59233 | 78.67013 | 67.28463 | 93.61069 | 73.74669 | 70.39311 | 61.99269 |
| ENSMUSG00000071533 | 0.060404981 | 0.418912 | 0.584387 | 2710.31 | 2865.174 | 2591.146 | 2988.99 | 2628.9 | 2854.891 | 2717.367 | 2496.022 |
| ENSMUSG00000071537 | -0.744823492 | 0.065115 | 0.164345 | 31.18769 | 21.59679 | 64.59232 | 23.29083 | 45.83023 | 68.15982 | 71.3574 | 52.20437 |
| ENSMUSG00000071540 | 0.834799342 | 0.057922 | 0.150867 | 31.18769 | 29.99554 | 13.24971 | 20.70296 | 13.65156 | 16.76061 | 10.60718 | 11.9635 |
| ENSMUSG00000071547 | -0.248729869 | 0.007125 | 0.033325 | 3064.442 | 3636.659 | 3091.322 | 3122.697 | 4220.282 | 3434.808 | 3824.371 | 3858.773 |
| ENSMUSG00000071550 | 0.747965887 | 0.014267 | 0.055587 | 57.3451 | 50.3925 | 53.82693 | 62.10889 | 35.10401 | 44.69496 | 18.32149 | 35.89051 |
| ENSMUSG00000071551 | -0.985534133 | 0.012274 | 0.04983 | 175.0535 | 128.3809 | 410.7409 | 165.6237 | 292.5334 | 280.4609 | 524.5733 | 647.1167 |
| ENSMUSG00000071552 | 0.659236842 | 0.288854 | 0.455844 | 14.08476 | 4.799286 | 17.39024 | 18.97772 | 8.776002 | 15.64324 | 6.750024 | 4.350364 |
| ENSMUSG00000071553 | 0.204423675 | 0.263264 | 0.427486 | 410.4702 | 524.322 | 352.7734 | 486.5197 | 448.5512 | 420.1327 | 381.8585 | 287.124 |
| ENSMUSG00000071561 | 0.320623345 | 0.56886 | 0.712804 | 147.89 | 21.59679 | 33.12427 | 80.22398 | 75.08357 | 31.28647 | 55.92877 | 64.16787 |
| ENSMUSG00000071562 | -0.082867519 | 0.865357 | 0.919408 | 113.6841 | 29.99554 | 31.46805 | 80.22398 | 92.63557 | 34.6386 | 71.3574 | 71.78101 |
| ENSMUSG00000071567 | 1.52820265 | 2.38E-07 | 1.17E-05 | 98.59333 | 104.3845 | 109.3101 | 137.1571 | 32.17867 | 62.57295 | 35.6787 | 26.10219 |
| ENSMUSG00000071568 | 0.886657971 | 0.290206 | 0.457312 | 16.09687 | 43.19358 | 11.59349 | 14.6646 | 18.52711 | 23.46486 | 1.928578 | 2.175182 |
| ENSMUSG00000071573 | -0.044615418 | 0.832776 | 0.898936 | 101.6115 | 151.1775 | 177.2148 | 128.5309 | 129.6898 | 137.437 | 136.9291 | 172.927 |
| ENSMUSG00000071604 | -0.355736111 | 0.062166 | 0.158933 | 292.7618 | 243.5638 | 219.4483 | 298.4677 | 427.0988 | 249.1744 | 343.2869 | 328.4525 |
| ENSMUSG00000071632 | -0.290604119 | 0.00849 | 0.037901 | 607.6568 | 627.5067 | 547.3785 | 629.7152 | 666.001 | 663.7202 | 824.4672 | 793.9415 |
| ENSMUSG00000071636 | 0.040043155 | 0.944341 | 0.967726 | 15.09082 | 4.799286 | 14.90592 | 17.25247 | 10.72622 | 5.58687 | 17.3572 | 17.40146 |
| ENSMUSG00000071637 | 0.012289276 | 0.981863 | 0.989807 | 988.9514 | 505.1249 | 585.4714 | 1320.677 | 381.2685 | 851.4391 | 1645.077 | 493.7664 |
| ENSMUSG00000071644 | -0.086682734 | 0.345793 | 0.514549 | 19544.62 | 24820.71 | 22134.46 | 21870.09 | 25869.7 | 22286.03 | 23316.51 | 22365.22 |
| ENSMUSG00000071645 | 0.00124881 | 0.989721 | 0.994138 | 982.9151 | 961.057 | 881.1055 | 916.1062 | 934.1566 | 1025.749 | 972.0035 | 803.7298 |
| ENSMUSG00000071646 | -0.058420419 | 0.360232 | 0.528787 | 4255.61 | 4172.979 | 4757.473 | 4306.216 | 4716.613 | 4351.055 | 4577.481 | 4574.408 |
| ENSMUSG00000071647 | 0.110807712 | 0.045211 | 0.126927 | 1499.021 | 1505.776 | 1474.858 | 1478.537 | 1428.538 | 1390.013 | 1320.112 | 1377.978 |
| ENSMUSG00000071648 | 0.035900397 | 0.737837 | 0.835944 | 436.6276 | 535.1204 | 539.0974 | 496.8711 | 484.6303 | 511.7573 | 445.5016 | 517.6934 |
| ENSMUSG00000071649 | -0.00540257 | 0.960683 | 0.977472 | 1147.908 | 1103.836 | 1233.051 | 1129.174 | 1158.432 | 981.0545 | 1136.897 | 1356.226 |
| ENSMUSG00000071650 | -0.057697975 | 0.383682 | 0.55167 | 5243.555 | 5377.6 | 5486.206 | 5211.971 | 5507.429 | 5033.77 | 5744.271 | 5902.357 |
| ENSMUSG00000071652 | -0.066646455 | 0.395769 | 0.563082 | 941.6669 | 937.0606 | 872.8244 | 876.4255 | 1035.568 | 895.0167 | 946.932 | 916.8393 |
| ENSMUSG00000071653 | -0.109482534 | 0.427238 | 0.59233 | 235.4167 | 314.3532 | 289.8373 | 261.3749 | 320.8116 | 298.3389 | 255.5366 | 312.1386 |
| ENSMUSG00000071654 | -0.091755576 | 0.280624 | 0.446983 | 893.3763 | 905.8653 | 949.0102 | 1016.17 | 1109.677 | 968.7633 | 968.1463 | 966.8685 |
| ENSMUSG00000071655 | -0.241443294 | 0.000867 | 0.006982 | 3778.74 | 3858.626 | 4278.827 | 3958.579 | 4737.091 | 4276.191 | 4833.982 | 4922.437 |
| ENSMUSG00000071656 | 1.478725872 | 1.89E-05 | 0.000375 | 264.5923 | 343.149 | 207.0267 | 276.0395 | 121.8889 | 154.1976 | 68.46453 | 46.76642 |
| ENSMUSG00000071657 | 0.066774351 | 0.386308 | 0.553849 | 1001.024 | 1066.641 | 967.2285 | 1023.071 | 904.9033 | 943.0637 | 972.9678 | 1051.701 |
| ENSMUSG00000071658 | 1.700257742 | 1.21E-08 | 9.91E-07 | 196.1806 | 145.1784 | 108.482 | 255.3366 | 47.78045 | 65.92507 | 47.25017 | 56.55474 |
| ENSMUSG00000071659 | -0.090014434 | 0.081852 | 0.192069 | 7126.889 | 7306.913 | 7203.7 | 7139.072 | 7280.181 | 7413.777 | 8020.957 | 7912.225 |
| ENSMUSG00000071660 | 0.170259533 | 0.07909 | 0.187628 | 1402.44 | 1455.384 | 1154.381 | 1287.034 | 1237.416 | 1097.261 | 1225.612 | 1143.058 |
| ENSMUSG00000071661 | -0.221759398 | 0.436328 | 0.600244 | 50.30272 | 65.99018 | 53.82693 | 36.23019 | 76.05868 | 43.57759 | 63.64308 | 55.46715 |

| | | | | | | | | | | | |
|---|---|---|---|---|---|---|---|---|---|---|---|
| ENSMUSG00000071662 | -0.095570185 | 0.274746 | 0.440816 | 1351.131 | 1493.778 | 1310.065 | 1315.501 | 1561.153 | 1419.065 | 1313.362 | 1547.642 |
| ENSMUSG00000071669 | -0.601785901 | 0.010945 | 0.045651 | 184.1079 | 161.9759 | 177.2148 | 133.7066 | 275.9565 | 151.9629 | 249.7509 | 318.6642 |
| ENSMUSG00000071671 | 0.779507933 | 0.222604 | 0.381926 | 26.15741 | 11.99822 | 8.281066 | 6.900988 | 11.70134 | 6.704245 | 4.821446 | 7.613138 |
| ENSMUSG00000071679 | 1.668656087 | 0.000209 | 0.002351 | 39.23612 | 46.79304 | 30.63995 | 30.19182 | 5.850668 | 21.23011 | 10.60718 | 8.700729 |
| ENSMUSG00000071691 | 0.20282705 | 0.458578 | 0.62022 | 47.28455 | 59.99108 | 45.54586 | 37.09281 | 37.05423 | 46.92971 | 47.25017 | 32.62773 |
| ENSMUSG00000071708 | 0.12139406 | 0.18854 | 0.340946 | 1585.542 | 1718.144 | 1421.859 | 1748.538 | 1503.622 | 1607.901 | 1412.684 | 1425.832 |
| ENSMUSG00000071711 | -0.289023904 | 0.002331 | 0.014615 | 1577.493 | 1644.955 | 1770.492 | 1783.905 | 2136.469 | 1772.155 | 2127.222 | 2245.876 |
| ENSMUSG00000071713 | 0.28181179 | 0.376051 | 0.544259 | 131.7931 | 46.79304 | 111.7944 | 105.2401 | 74.10846 | 69.27719 | 81.00029 | 102.2336 |
| ENSMUSG00000071714 | 0.147730426 | 0.710561 | 0.818711 | 48.29061 | 13.19804 | 31.46805 | 35.36756 | 29.25334 | 26.81698 | 28.92867 | 31.54014 |
| ENSMUSG00000071715 | 0.53940768 | 0.102168 | 0.224277 | 38.23007 | 27.5959 | 33.12427 | 29.3292 | 17.552 | 25.6996 | 17.3572 | 28.27737 |
| ENSMUSG00000071722 | 0.352783753 | 0.008936 | 0.039268 | 204.229 | 279.5584 | 243.4633 | 245.8477 | 182.3458 | 197.7752 | 193.8221 | 187.0657 |
| ENSMUSG00000071723 | 0.753552334 | 1.14E-07 | 6.24E-06 | 484.9182 | 382.7431 | 344.4924 | 427.8613 | 241.8276 | 269.2872 | 222.7508 | 239.27 |
| ENSMUSG00000071724 | 0.202597352 | 0.403733 | 0.570586 | 58.35115 | 75.58876 | 77.84202 | 66.42201 | 61.43201 | 69.27719 | 69.42882 | 41.32846 |
| ENSMUSG00000071748 | 0.716665941 | 0.042881 | 0.122528 | 27.16347 | 22.79661 | 27.32752 | 25.8787 | 18.52711 | 18.99536 | 13.50005 | 11.9635 |
| ENSMUSG00000071757 | 0.239759182 | 0.00208 | 0.013509 | 752.5287 | 825.4772 | 754.4051 | 766.0097 | 699.1548 | 655.8986 | 620.0379 | 646.0291 |
| ENSMUSG00000071793 | -0.481682604 | 0.001042 | 0.008069 | 472.8455 | 475.1293 | 516.7385 | 489.1075 | 828.8446 | 519.579 | 659.5738 | 719.9853 |
| ENSMUSG00000071796 | -0.256399848 | 0.065436 | 0.164684 | 334.01 | 385.1427 | 438.0684 | 375.2412 | 518.7592 | 502.8183 | 397.2871 | 413.2846 |
| ENSMUSG00000071847 | -1.140345347 | 0.130911 | 0.263472 | 524.1543 | 587.9126 | 3603.92 | 696.9998 | 2627.925 | 1163.186 | 3814.728 | 4327.525 |
| ENSMUSG00000071855 | 0.273462452 | 0.025128 | 0.083355 | 877.2794 | 886.6681 | 660.8291 | 777.2238 | 702.0801 | 695.0067 | 655.7166 | 592.7371 |
| ENSMUSG00000071856 | 0.593628217 | 0.008637 | 0.038373 | 430.5913 | 494.3265 | 457.943 | 400.2573 | 240.8525 | 455.8886 | 242.0366 | 243.6204 |
| ENSMUSG00000071862 | -0.006807826 | 0.988756 | 0.99371 | 24.1453 | 20.39697 | 14.07781 | 18.11509 | 10.72622 | 35.75597 | 14.46434 | 16.31387 |
| ENSMUSG00000071866 | -0.205610121 | 0.008226 | 0.037019 | 21337.41 | 25159.06 | 25903.18 | 24173.3 | 29256.26 | 26277.29 | 27701.13 | 28130.54 |
| ENSMUSG00000071984 | 0.953882823 | 0.055198 | 0.14622 | 221.332 | 418.7377 | 264.166 | 295.0172 | 119.9387 | 362.0292 | 81.96458 | 55.46715 |
| ENSMUSG00000072066 | 0.267271373 | 0.214094 | 0.372133 | 126.7628 | 212.3684 | 138.2938 | 147.5086 | 121.8889 | 117.3243 | 159.1077 | 118.5474 |
| ENSMUSG00000072082 | -0.030491026 | 0.802174 | 0.879281 | 687.1351 | 712.694 | 551.519 | 698.725 | 710.8561 | 615.6731 | 623.8951 | 753.7006 |
| ENSMUSG00000072115 | 0.8112035 | 5.24E-06 | 0.00014 | 194.1685 | 184.7725 | 217.792 | 222.5569 | 101.4116 | 153.0803 | 110.8933 | 103.3212 |
| ENSMUSG00000072214 | -0.173721936 | 0.267542 | 0.431773 | 489.9485 | 364.7457 | 486.9267 | 427.8613 | 558.7388 | 385.4941 | 509.1447 | 543.7955 |
| ENSMUSG00000072235 | 0.134414795 | 0.197801 | 0.352206 | 7590.68 | 6735.798 | 7606.988 | 9104.128 | 7353.314 | 6505.352 | 6897.56 | 7523.955 |
| ENSMUSG00000072244 | 0.296917754 | 0.379081 | 0.547178 | 97.58727 | 85.18733 | 115.1068 | 55.2079 | 46.80534 | 111.7374 | 67.50024 | 61.99269 |
| ENSMUSG00000072258 | 0.191452552 | 0.181918 | 0.332463 | 537.233 | 622.7074 | 437.2403 | 527.9256 | 457.3272 | 525.1658 | 484.0732 | 391.5328 |
| ENSMUSG00000072294 | -0.133591761 | 0.64114 | 0.767671 | 55.33299 | 67.19001 | 54.65504 | 34.50494 | 50.70579 | 49.16446 | 75.21455 | 55.46715 |
| ENSMUSG00000072421 | 0.797154078 | 0.003324 | 0.019108 | 270.6286 | 268.76 | 231.0418 | 357.9887 | 244.7529 | 173.193 | 122.4647 | 108.7591 |
| ENSMUSG00000072437 | -0.086533827 | 0.805177 | 0.880717 | 54.32694 | 47.99286 | 41.40533 | 67.28463 | 51.6809 | 92.74205 | 44.3573 | 35.89051 |
| ENSMUSG00000072494 | -0.225790914 | 0.197505 | 0.351815 | 88.53278 | 127.1811 | 111.7944 | 106.1027 | 116.0382 | 116.2069 | 140.7862 | 132.6861 |
| ENSMUSG00000072501 | 0.307247655 | 0.030762 | 0.096237 | 2396.421 | 2228.069 | 1766.351 | 2176.399 | 1675.241 | 2155.415 | 1566.97 | 1525.89 |
| ENSMUSG00000072568 | -0.017818429 | 0.897023 | 0.937817 | 896.3944 | 922.6628 | 1106.35 | 1031.698 | 822.0188 | 1224.642 | 974.8963 | 988.6203 |
| ENSMUSG00000072571 | -0.263965116 | 0.445077 | 0.607929 | 24.1453 | 26.39607 | 32.29616 | 26.74133 | 27.30312 | 29.05173 | 50.14304 | 25.0146 |
| ENSMUSG00000072572 | 0.910815906 | 0.005121 | 0.026168 | 40.24217 | 35.99465 | 35.60859 | 44.85642 | 27.30312 | 24.58223 | 17.3572 | 14.13868 |
| ENSMUSG00000072573 | 0.392802628 | 0.190612 | 0.343246 | 45.27245 | 55.19179 | 43.06154 | 52.62003 | 40.95467 | 32.40385 | 51.10733 | 23.927 |
| ENSMUSG00000072575 | -1.045825317 | 0.055674 | 0.146914 | 19.11503 | 15.59768 | 33.95237 | 11.21411 | 50.70579 | 14.52586 | 31.82154 | 68.51824 |
| ENSMUSG00000072582 | -0.038495427 | 0.607741 | 0.743551 | 1082.514 | 1105.036 | 1013.603 | 1096.394 | 1179.885 | 1133.017 | 1073.254 | 1024.511 |
| ENSMUSG00000072591 | 1.86236247 | 4.20E-10 | 5.87E-08 | 92.557 | 133.1802 | 97.71658 | 102.6522 | 26.32801 | 43.57759 | 17.3572 | 30.45255 |
| ENSMUSG00000072612 | -0.411183705 | 0.347155 | 0.515993 | 12.07265 | 21.59679 | 11.59349 | 17.25247 | 19.50223 | 12.29112 | 21.21436 | 29.36496 |
| ENSMUSG00000072620 | -0.157688965 | 0.654513 | 0.777079 | 157.9505 | 200.3702 | 295.6341 | 118.1794 | 173.5698 | 121.7938 | 321.1083 | 244.708 |
| ENSMUSG00000072621 | 0.452848151 | 0.334814 | 0.503039 | 18.10898 | 35.99465 | 31.46805 | 14.6646 | 20.47734 | 26.81698 | 10.60718 | 15.22628 |
| ENSMUSG00000072623 | 0.501639818 | 1.78E-05 | 0.000358 | 514.0938 | 514.7234 | 479.4737 | 520.162 | 354.9405 | 427.9543 | 330.7512 | 319.7518 |
| ENSMUSG00000072640 | -0.034796 | 0.830876 | 0.897813 | 370.228 | 298.7556 | 289.0092 | 295.0172 | 241.8276 | 341.9165 | 361.6084 | 338.2408 |
| ENSMUSG00000072647 | 0.105369911 | 0.578853 | 0.720218 | 126.7628 | 148.7779 | 110.1382 | 106.9653 | 118.9636 | 122.9112 | 90.64318 | 123.9854 |
| ENSMUSG00000072653 | 0.21831192 | 0.084813 | 0.196695 | 509.0635 | 590.3122 | 506.8013 | 494.2833 | 483.6552 | 518.4616 | 416.5729 | 385.0072 |
| ENSMUSG00000072660 | -0.004418597 | 0.990958 | 0.994824 | 17.10292 | 27.5959 | 17.39024 | 32.77969 | 20.47734 | 30.1691 | 27.0001 | 17.40146 |
| ENSMUSG00000072662 | 0.218016017 | 0.629045 | 0.758529 | 28.16952 | 51.59233 | 115.9349 | 55.2079 | 44.85512 | 33.52122 | 69.42882 | 68.51824 |
| ENSMUSG00000072663 | 0.347869472 | 0.473295 | 0.632614 | 15.09082 | 21.59679 | 32.29616 | 10.35148 | 14.62667 | 21.23011 | 11.57147 | 15.22628 |
| ENSMUSG00000072664 | -4.178381539 | 0.000245 | 0.002668 | 0 | 0 | 3.312427 | 1.725247 | 13.65156 | 2.234748 | 29.89296 | 51.11678 |
| ENSMUSG00000072683 | 0.694923468 | 0.304819 | 0.473388 | 23.13925 | 4.799286 | 15.73403 | 6.900988 | 5.850668 | 7.821619 | 4.821446 | 13.05109 |
| ENSMUSG00000072688 | -0.114504423 | 0.765055 | 0.854012 | 44.26639 | 39.59411 | 44.71776 | 31.05445 | 31.20356 | 22.34748 | 65.57166 | 53.29196 |
| ENSMUSG00000072692 | 0.061383966 | 0.652125 | 0.775356 | 224.3501 | 291.5566 | 270.7909 | 316.5828 | 274.9814 | 273.7567 | 254.5723 | 254.4963 |
| ENSMUSG00000072694 | 0.058379259 | 0.55365 | 0.700643 | 606.6508 | 563.9161 | 621.9081 | 628.8525 | 596.7681 | 642.4901 | 579.5378 | 508.9926 |
| ENSMUSG00000072704 | -0.013488263 | 0.867748 | 0.920834 | 1699.226 | 1908.916 | 1766.351 | 1878.794 | 1656.714 | 1934.175 | 1786.828 | 1944.613 |
| ENSMUSG00000072720 | 0.317683317 | 0.597815 | 0.736533 | 14.08476 | 13.19804 | 17.39024 | 7.763611 | 10.72622 | 17.87799 | 2.892867 | 10.87591 |
| ENSMUSG00000072723 | 1.10459438 | 0.02715 | 0.088078 | 17.10292 | 9.598572 | 19.87456 | 17.25247 | 6.825779 | 10.05637 | 7.714313 | 5.437955 |
| ENSMUSG00000072761 | -0.289651767 | 0.19684 | 0.351173 | 52.31483 | 71.98929 | 73.70149 | 56.07053 | 66.30757 | 72.62932 | 78.10742 | 93.53283 |
| ENSMUSG00000072762 | -0.190147432 | 0.286795 | 0.453937 | 121.7326 | 111.5834 | 110.9663 | 115.5915 | 104.3369 | 150.8455 | 119.5719 | 151.1752 |
| ENSMUSG00000072763 | 0.371434868 | 0.0788 | 0.187244 | 92.557 | 121.182 | 89.43552 | 88.85022 | 72.15824 | 94.9768 | 65.57166 | 69.60583 |
| ENSMUSG00000072768 | -0.161471693 | 0.785929 | 0.868441 | 14.08476 | 7.198929 | 12.4216 | 6.900988 | 7.80089 | 5.58687 | 11.57147 | 20.66423 |

| | | | | | | | | | | | |
|---|---|---|---|---|---|---|---|---|---|---|---|
| ENSMUSG00000072770 | 0.8667703 | 5.03E-07 | 2.12E-05 | 314.895 | 344.3488 | 224.4169 | 283.8031 | 163.8187 | 177.6625 | 160.072 | 137.0365 |
| ENSMUSG00000072772 | -0.088128228 | 0.401827 | 0.568982 | 4017.175 | 4325.357 | 4036.192 | 4371.776 | 5017.923 | 3651.579 | 4665.231 | 4467.824 |
| ENSMUSG00000072774 | 0.157270629 | 0.501865 | 0.658076 | 108.6539 | 143.9786 | 138.2938 | 144.9207 | 117.9885 | 160.9019 | 122.4647 | 79.39415 |
| ENSMUSG00000072809 | -0.24741951 | 0.374668 | 0.543366 | 44.26639 | 29.99554 | 31.46805 | 39.68068 | 33.15378 | 50.28183 | 46.28588 | 43.50364 |
| ENSMUSG00000072812 | 1.59992123 | 0.001192 | 0.008927 | 165.999 | 237.5647 | 185.4959 | 238.0841 | 62.40712 | 145.2586 | 49.17875 | 16.31387 |
| ENSMUSG00000072825 | -0.183743933 | 0.079102 | 0.187628 | 1168.029 | 1030.647 | 1238.848 | 1080.005 | 1240.342 | 1117.374 | 1377.969 | 1397.555 |
| ENSMUSG00000072844 | 1.162437806 | 0.001319 | 0.009624 | 309.8647 | 236.3648 | 146.5749 | 140.6076 | 78.0089 | 157.5497 | 69.42882 | 67.43065 |
| ENSMUSG00000072847 | 0.490817652 | 0.001489 | 0.010503 | 245.4773 | 196.7707 | 205.3704 | 221.6942 | 153.0925 | 178.7799 | 127.2862 | 160.9635 |
| ENSMUSG00000072849 | 2.780199243 | 0.017727 | 0.064916 | 345.0766 | 10.79839 | 7.45296 | 366.615 | 17.552 | 36.87335 | 19.28578 | 32.62773 |
| ENSMUSG00000072872 | 0.004151794 | 0.968317 | 0.982153 | 782.7103 | 805.0802 | 877.793 | 979.9403 | 858.098 | 770.9881 | 907.3961 | 901.613 |
| ENSMUSG00000072875 | 0.452594531 | 0.002218 | 0.01416 | 533.2088 | 697.0963 | 559.8001 | 569.3315 | 407.5965 | 530.7527 | 420.4301 | 364.343 |
| ENSMUSG00000072884 | -0.499371262 | 0.284497 | 0.451069 | 6.036326 | 21.59679 | 14.07781 | 11.21411 | 21.45245 | 15.64324 | 18.32149 | 18.48905 |
| ENSMUSG00000072889 | 0.106719088 | 0.102269 | 0.224432 | 1682.123 | 1588.564 | 1649.588 | 1723.522 | 1520.199 | 1620.192 | 1589.149 | 1443.233 |
| ENSMUSG00000072893 | 0.533716569 | 1.50E-05 | 0.000312 | 869.231 | 831.4763 | 696.4377 | 979.9403 | 592.8677 | 631.3164 | 567.002 | 541.6204 |
| ENSMUSG00000072915 | 0.341893985 | 0.031822 | 0.098541 | 423.5489 | 417.5379 | 292.3216 | 351.0878 | 320.8116 | 320.6864 | 262.2867 | 265.3722 |
| ENSMUSG00000072940 | 0.203099947 | 0.262709 | 0.427035 | 93.56306 | 112.7832 | 110.1382 | 100.0643 | 104.3369 | 83.80306 | 96.42892 | 76.13138 |
| ENSMUSG00000072941 | -3.127979378 | 6.28E-06 | 0.00016 | 184.1079 | 34.79482 | 244.2915 | 127.6683 | 1405.135 | 398.9026 | 1158.111 | 2207.81 |
| ENSMUSG00000072944 | 2.32853931 | 1.59E-11 | 3.55E-09 | 194.1685 | 112.7832 | 85.29498 | 115.5915 | 22.42756 | 36.87335 | 19.28578 | 22.83941 |
| ENSMUSG00000072946 | -0.504867065 | 0.003572 | 0.020122 | 968.8303 | 803.8804 | 920.0265 | 931.6334 | 1264.719 | 899.4861 | 1328.79 | 1650.963 |
| ENSMUSG00000072949 | -0.057310074 | 0.727327 | 0.829154 | 1901.443 | 2282.061 | 2216.013 | 2197.965 | 2353.919 | 1671.592 | 2007.65 | 2912.569 |
| ENSMUSG00000072955 | 0.080694594 | 0.667559 | 0.786748 | 107.6478 | 87.58697 | 107.6539 | 117.3168 | 86.78491 | 121.7938 | 97.39321 | 93.53283 |
| ENSMUSG00000072962 | -0.543595684 | 0.293725 | 0.461282 | 13.07871 | 33.595 | 20.70267 | 12.07673 | 32.17867 | 11.17374 | 28.92867 | 42.41605 |
| ENSMUSG00000072964 | -0.073704762 | 0.441664 | 0.605014 | 1294.792 | 1464.982 | 1260.378 | 1406.939 | 1399.285 | 1277.159 | 1428.112 | 1604.197 |
| ENSMUSG00000072966 | 0.672066375 | 1.13E-08 | 9.52E-07 | 1041.266 | 1060.642 | 859.5747 | 1087.768 | 609.4446 | 689.4198 | 688.5025 | 552.4963 |
| ENSMUSG00000072969 | 0.270544992 | 0.079201 | 0.187764 | 601.6205 | 545.9188 | 598.7211 | 685.7857 | 546.0623 | 619.0252 | 449.3588 | 403.4963 |
| ENSMUSG00000072972 | 0.557116693 | 0.380768 | 0.548634 | 6.036326 | 25.19625 | 8.281066 | 11.21411 | 4.875557 | 8.938993 | 12.53576 | 7.613138 |
| ENSMUSG00000072974 | 0.873995768 | 0.087917 | 0.20179 | 11.0666 | 23.99643 | 13.24971 | 16.38985 | 5.850668 | 10.05637 | 12.53576 | 6.525547 |
| ENSMUSG00000072980 | -0.14193128 | 0.36606 | 0.534678 | 250.5075 | 261.5611 | 254.2287 | 335.5605 | 332.513 | 347.5033 | 271.9295 | 265.3722 |
| ENSMUSG00000073000 | 0.274670456 | 0.159137 | 0.302102 | 213.2835 | 206.3693 | 263.3379 | 183.7388 | 167.7191 | 157.5497 | 155.2506 | 237.0949 |
| ENSMUSG00000073002 | -0.187540051 | 0.235301 | 0.396223 | 378.2764 | 262.7609 | 388.382 | 334.6979 | 363.7165 | 340.7991 | 405.0014 | 446.9999 |
| ENSMUSG00000073016 | -0.386687004 | 0.311098 | 0.479748 | 104.6297 | 49.19268 | 60.45178 | 55.2079 | 94.5858 | 39.10809 | 104.1432 | 114.1971 |
| ENSMUSG00000073062 | 0.185128929 | 0.360908 | 0.529527 | 250.5075 | 200.3702 | 166.4494 | 226.87 | 194.0472 | 238.0007 | 156.2148 | 154.4379 |
| ENSMUSG00000073067 | -0.380918621 | 0.367251 | 0.535809 | 15.09082 | 20.39697 | 12.4216 | 9.488858 | 13.65156 | 20.11273 | 18.32149 | 21.75182 |
| ENSMUSG00000073073 | 0.412659547 | 0.263001 | 0.427321 | 50.30272 | 40.79393 | 28.98373 | 56.07053 | 36.07912 | 46.92971 | 30.85725 | 18.48905 |
| ENSMUSG00000073077 | 0.763279143 | 0.13566 | 0.27012 | 15.09082 | 22.79661 | 10.76539 | 13.80198 | 8.776002 | 14.52586 | 6.750024 | 6.525547 |
| ENSMUSG00000073096 | -0.407902616 | 0.001795 | 0.012147 | 315.9011 | 327.5513 | 357.7421 | 309.6818 | 427.0988 | 357.5597 | 501.4304 | 452.4379 |
| ENSMUSG00000073102 | 0.877193757 | 0.01537 | 0.058709 | 25.15136 | 23.99643 | 41.40533 | 35.36756 | 15.60178 | 20.11273 | 19.28578 | 14.13868 |
| ENSMUSG00000073125 | 0.295023404 | 0.500331 | 0.656799 | 77.46619 | 245.9634 | 392.5225 | 188.0519 | 153.0925 | 155.315 | 297.9654 | 130.5109 |
| ENSMUSG00000073131 | 0.184063414 | 0.073179 | 0.177991 | 1222.356 | 1107.435 | 1127.881 | 1178.344 | 916.6046 | 1202.295 | 966.2177 | 998.4086 |
| ENSMUSG00000073139 | -0.371012775 | 0.071261 | 0.174639 | 795.789 | 705.4951 | 1055.008 | 734.9552 | 885.4011 | 781.0445 | 1354.826 | 1236.591 |
| ENSMUSG00000073144 | -0.246879915 | 0.519855 | 0.673341 | 19.11503 | 11.99822 | 27.32752 | 21.56559 | 21.45245 | 20.11273 | 23.14294 | 31.54014 |
| ENSMUSG00000073147 | 0.035392904 | 0.645976 | 0.770635 | 889.3521 | 837.4754 | 913.4016 | 934.2212 | 833.7202 | 850.3217 | 943.0748 | 863.5473 |
| ENSMUSG00000073155 | 0.103363032 | 0.125428 | 0.255959 | 1505.057 | 1511.775 | 1533.653 | 1549.272 | 1497.771 | 1461.525 | 1424.255 | 1294.233 |
| ENSMUSG00000073158 | -0.162334214 | 0.601994 | 0.739904 | 33.19979 | 35.99465 | 30.63995 | 51.75741 | 34.1289 | 33.52122 | 51.10733 | 51.11678 |
| ENSMUSG00000073164 | 0.28304923 | 0.065105 | 0.164345 | 377.2704 | 460.7315 | 307.2276 | 345.0494 | 283.7574 | 329.6254 | 329.7869 | 278.4233 |
| ENSMUSG00000073176 | 0.3134043 | 0.023494 | 0.079242 | 309.8647 | 297.5557 | 240.979 | 245.8477 | 213.5494 | 224.5922 | 201.5364 | 239.27 |
| ENSMUSG00000073179 | 0.824935926 | 0.043797 | 0.124184 | 68.4117 | 86.38715 | 69.56096 | 66.42201 | 21.45245 | 79.33356 | 37.60728 | 26.10219 |
| ENSMUSG00000073197 | 0.238147779 | 0.599512 | 0.737591 | 43.26034 | 41.99375 | 33.95237 | 49.16954 | 33.15378 | 70.39457 | 22.17865 | 17.40146 |
| ENSMUSG00000073198 | 0.020842119 | 0.951105 | 0.972282 | 25.15136 | 33.595 | 24.8432 | 31.91707 | 18.52711 | 39.10809 | 27.0001 | 29.36496 |
| ENSMUSG00000073207 | -0.546120625 | 0.099257 | 0.219875 | 196.1806 | 87.58697 | 125.0441 | 112.1411 | 150.1671 | 121.7938 | 196.715 | 292.562 |
| ENSMUSG00000073209 | 1.141097131 | 3.96E-06 | 0.000112 | 277.671 | 422.3372 | 255.8849 | 350.2251 | 161.8685 | 203.3621 | 114.7504 | 112.0219 |
| ENSMUSG00000073233 | 0.14555969 | 0.722069 | 0.826091 | 14.08476 | 15.59768 | 12.4216 | 16.38985 | 11.70134 | 13.40849 | 12.53576 | 15.22628 |
| ENSMUSG00000073236 | 0.519747968 | 0.000555 | 0.004961 | 337.0282 | 446.3336 | 334.5551 | 377.8291 | 219.4 | 302.8084 | 260.3581 | 259.9343 |
| ENSMUSG00000073293 | 0.436857028 | 0.002201 | 0.014069 | 280.6892 | 274.7591 | 264.9941 | 307.9566 | 177.4703 | 253.6439 | 198.6436 | 205.5547 |
| ENSMUSG00000073295 | 0.210784427 | 0.129967 | 0.262322 | 286.7255 | 347.9482 | 301.4308 | 345.0494 | 238.9023 | 328.508 | 266.1438 | 274.073 |
| ENSMUSG00000073374 | 0.395405252 | 0.031111 | 0.096904 | 120.7265 | 123.5816 | 118.4192 | 121.6299 | 87.76002 | 108.3853 | 102.2147 | 69.60583 |
| ENSMUSG00000073400 | 0.757960203 | 0.282471 | 0.449073 | 49.29666 | 8.398751 | 9.109173 | 14.6646 | 4.875557 | 15.64324 | 12.53576 | 15.22628 |
| ENSMUSG00000073408 | -0.423949372 | 0.410711 | 0.576758 | 17.10292 | 21.59679 | 9.93728 | 15.52722 | 10.72622 | 35.75597 | 26.03581 | 13.05109 |
| ENSMUSG00000073409 | -1.04859875 | 0.004018 | 0.021898 | 37.22401 | 49.19268 | 94.40416 | 45.71905 | 84.83468 | 89.38993 | 123.429 | 172.927 |
| ENSMUSG00000073411 | -0.443561441 | 0.055544 | 0.146761 | 1058.369 | 1118.234 | 1728.259 | 1029.972 | 1575.78 | 1093.909 | 2101.186 | 1941.35 |
| ENSMUSG00000073412 | 0.475679421 | 0.210657 | 0.368171 | 20.12109 | 45.59322 | 38.09291 | 30.19182 | 32.17867 | 17.87799 | 26.03581 | 19.57664 |
| ENSMUSG00000073413 | -0.26869704 | 0.14143 | 0.278217 | 79.47829 | 88.78679 | 90.26362 | 75.04824 | 84.83468 | 105.0332 | 113.7861 | 97.8832 |
| ENSMUSG00000073414 | 0.495242958 | 0.245564 | 0.408521 | 22.1332 | 16.7975 | 24.8432 | 31.91707 | 22.42756 | 15.64324 | 21.21436 | 8.700729 |
| ENSMUSG00000073415 | 0.251575361 | 0.411296 | 0.577077 | 43.26034 | 47.99286 | 35.60859 | 30.19182 | 30.22845 | 41.34284 | 24.10723 | 35.89051 |

| | | | | | | | | | | | |
|---|---|---|---|---|---|---|---|---|---|---|---|
| ENSMUSG00000073418 | 2.09900742 | 0.000954 | 0.00757 | 437.6336 | 58.79126 | 56.31125 | 95.75121 | 25.35289 | 53.63396 | 27.96439 | 44.59123 |
| ENSMUSG00000073421 | -0.806044853 | 9.32E-05 | 0.001262 | 311.8769 | 310.7538 | 369.3356 | 395.0816 | 694.2793 | 377.6724 | 651.8595 | 702.5838 |
| ENSMUSG00000073422 | -0.183343118 | 0.478604 | 0.637909 | 51.30877 | 49.19268 | 36.43669 | 33.64232 | 41.92979 | 49.16446 | 50.14304 | 51.11678 |
| ENSMUSG00000073423 | 0.142256886 | 0.14789 | 0.287251 | 841.0614 | 937.0606 | 870.3401 | 934.2212 | 907.8286 | 851.4391 | 755.0384 | 730.8612 |
| ENSMUSG00000073427 | 0.351197028 | 0.007725 | 0.03529 | 204.229 | 259.1615 | 218.6202 | 215.6559 | 180.3956 | 178.7799 | 181.2864 | 160.9635 |
| ENSMUSG00000073433 | -0.516874157 | 0.121365 | 0.250642 | 92.557 | 55.19179 | 151.5435 | 113.8663 | 115.0631 | 108.3853 | 166.822 | 203.3795 |
| ENSMUSG00000073434 | 0.074047827 | 0.389317 | 0.556692 | 789.7527 | 867.471 | 756.0614 | 746.1693 | 789.8402 | 769.8708 | 741.5384 | 694.9707 |
| ENSMUSG00000073435 | 0.12930498 | 0.714119 | 0.820631 | 28.16952 | 29.99554 | 20.70267 | 33.64232 | 37.05423 | 21.23011 | 21.21436 | 22.83941 |
| ENSMUSG00000073436 | 0.067232114 | 0.609131 | 0.744335 | 471.8395 | 500.3256 | 618.5957 | 644.3797 | 487.5557 | 562.0392 | 561.2163 | 526.3941 |
| ENSMUSG00000073437 | 0.270223519 | 0.187957 | 0.340191 | 293.7679 | 292.7565 | 226.9012 | 184.6014 | 209.6489 | 231.2964 | 157.1791 | 228.3941 |
| ENSMUSG00000073448 | -0.359030507 | 0.427495 | 0.592509 | 11.0666 | 14.39786 | 16.56213 | 19.84034 | 14.62667 | 30.1691 | 23.14294 | 11.9635 |
| ENSMUSG00000073460 | 0.440412553 | 0.194715 | 0.348526 | 48.29061 | 77.9884 | 47.20208 | 48.30692 | 26.32801 | 61.45558 | 40.50014 | 34.80291 |
| ENSMUSG00000073468 | 0.034255525 | 0.642836 | 0.76908 | 1024.163 | 994.6521 | 959.7756 | 1092.081 | 954.634 | 967.646 | 1037.575 | 1015.81 |
| ENSMUSG00000073471 | -0.410686321 | 0.000468 | 0.00435 | 464.7971 | 467.9304 | 523.3634 | 494.2833 | 652.3495 | 539.6917 | 660.5381 | 741.7371 |
| ENSMUSG00000073478 | -0.094934086 | 0.665952 | 0.785506 | 61.36932 | 61.1909 | 81.15445 | 59.52102 | 67.28268 | 58.10345 | 78.10742 | 78.30656 |
| ENSMUSG00000073481 | -0.192108095 | 0.028075 | 0.090256 | 2409.5 | 2303.657 | 2630.895 | 2436.911 | 2759.565 | 2451.519 | 2977.725 | 2987.613 |
| ENSMUSG00000073482 | -0.306902588 | 0.455477 | 0.617474 | 14.08476 | 16.7975 | 9.93728 | 18.11509 | 16.57689 | 16.76061 | 15.42863 | 23.927 |
| ENSMUSG00000073485 | 0.047449411 | 0.885748 | 0.931739 | 46.2785 | 53.99197 | 37.2648 | 73.323 | 35.10401 | 67.04245 | 60.75022 | 41.32846 |
| ENSMUSG00000073489 | 1.279605977 | 0.0044 | 0.023438 | 443.67 | 563.9161 | 380.9291 | 459.7783 | 241.8276 | 363.1466 | 94.50034 | 61.99269 |
| ENSMUSG00000073490 | 0.710699645 | 0.001943 | 0.012857 | 104.6297 | 113.983 | 91.09173 | 77.63611 | 54.60623 | 64.8077 | 70.39311 | 45.67883 |
| ENSMUSG00000073514 | 0.274261454 | 0.447281 | 0.609946 | 41.24823 | 55.19179 | 43.06154 | 25.8787 | 38.02934 | 34.6386 | 43.39301 | 19.57664 |
| ENSMUSG00000073530 | -0.715059801 | 0.003145 | 0.018406 | 250.5075 | 253.1623 | 374.3042 | 295.0172 | 280.8321 | 515.1095 | 551.5734 | 580.7736 |
| ENSMUSG00000073538 | -3.176100406 | NA | NA | 4.024217 | 2.399643 | 1.656213 | 0.862623 | 2.925334 | 68.15982 | 5.785735 | 3.262773 |
| ENSMUSG00000073542 | -0.303341433 | 0.018542 | 0.066941 | 236.4228 | 197.9706 | 249.2601 | 204.4418 | 272.0561 | 298.3389 | 276.751 | 251.2335 |
| ENSMUSG00000073555 | 1.073685404 | 0.245471 | 0.40844 | 6.036326 | 39.59411 | 38.09291 | 2.58787 | 5.850668 | 14.52586 | 18.32149 | 2.175182 |
| ENSMUSG00000073557 | 0.379544919 | 0.0366 | 0.109781 | 757.5589 | 962.2569 | 890.2146 | 965.2757 | 649.4241 | 952.0027 | 592.0735 | 555.759 |
| ENSMUSG00000073563 | 0.017482837 | 0.782163 | 0.865651 | 1930.618 | 1943.711 | 1806.101 | 2027.165 | 1863.438 | 1996.748 | 1848.542 | 1906.547 |
| ENSMUSG00000073565 | 0.513272951 | 0.072509 | 0.176828 | 86.52067 | 75.58876 | 63.76421 | 83.67448 | 44.85512 | 72.62932 | 65.57166 | 33.71532 |
| ENSMUSG00000073590 | 0.26057753 | 0.429513 | 0.594457 | 32.19374 | 32.39518 | 35.60859 | 32.77969 | 36.07912 | 33.52122 | 25.07152 | 16.31387 |
| ENSMUSG00000073591 | 0.472231168 | 0.000213 | 0.002383 | 295.78 | 279.5584 | 260.0255 | 262.2375 | 189.1716 | 231.2964 | 188.0364 | 182.7153 |
| ENSMUSG00000073599 | -0.490001365 | 0.013367 | 0.053054 | 321.9374 | 293.9563 | 446.3495 | 334.6979 | 485.6054 | 357.5597 | 502.3947 | 618.8393 |
| ENSMUSG00000073600 | 0.211663224 | 0.252423 | 0.416144 | 241.453 | 182.3729 | 218.6202 | 213.068 | 159.9183 | 144.1413 | 220.8222 | 214.2554 |
| ENSMUSG00000073607 | 0.129889672 | 0.743988 | 0.839922 | 18.10898 | 55.19179 | 31.46805 | 25.8787 | 27.30312 | 31.28647 | 34.71441 | 25.0146 |
| ENSMUSG00000073609 | 0.099124412 | 0.363744 | 0.532291 | 480.894 | 465.5308 | 395.0069 | 463.2288 | 424.1734 | 374.3203 | 439.7159 | 443.7372 |
| ENSMUSG00000073616 | -0.265641417 | 0.000206 | 0.002326 | 1226.38 | 1205.821 | 1317.518 | 1271.507 | 1486.07 | 1398.952 | 1548.648 | 1606.372 |
| ENSMUSG00000073627 | -0.189010371 | 0.479409 | 0.638512 | 76.46013 | 67.19001 | 61.27989 | 37.95543 | 74.10846 | 62.57295 | 60.75022 | 78.30656 |
| ENSMUSG00000073633 | -0.489659306 | 0.026774 | 0.087334 | 85.51462 | 64.79036 | 88.60741 | 81.08661 | 111.1627 | 81.56831 | 117.6433 | 140.2993 |
| ENSMUSG00000073639 | 0.143469369 | 0.007268 | 0.033798 | 3832.061 | 4039.799 | 3649.466 | 3974.106 | 3531.853 | 3442.63 | 3569.798 | 3478.116 |
| ENSMUSG00000073640 | -0.367563732 | 0.128593 | 0.260261 | 30.18163 | 45.59322 | 43.88965 | 44.85642 | 55.58134 | 51.39921 | 52.07161 | 53.29196 |
| ENSMUSG00000073643 | -0.324032484 | 0.02861 | 0.091413 | 1190.162 | 1100.236 | 1051.695 | 1082.592 | 1579.68 | 1252.576 | 1647.97 | 1056.051 |
| ENSMUSG00000073647 | -0.55000706 | 0.005394 | 0.027083 | 190.1443 | 148.7779 | 178.871 | 138.8824 | 319.8365 | 203.3621 | 216.0008 | 221.8686 |
| ENSMUSG00000073650 | 0.668120364 | 0.019573 | 0.06943 | 49.29666 | 39.59411 | 74.5296 | 54.34528 | 33.15378 | 31.28647 | 37.60728 | 35.89051 |
| ENSMUSG00000073652 | 0.079128228 | 0.903985 | 0.942489 | 2.012109 | 26.39607 | 22.35888 | 13.80198 | 11.70134 | 18.99536 | 18.32149 | 11.9635 |
| ENSMUSG00000073664 | 0.066207234 | 0.638881 | 0.76591 | 839.0493 | 681.4986 | 799.951 | 784.9874 | 589.9423 | 900.6035 | 725.1455 | 753.7006 |
| ENSMUSG00000073676 | -0.178444849 | 0.169169 | 0.315531 | 5664.086 | 4060.196 | 4213.407 | 4345.035 | 5356.286 | 4499.665 | 5050.947 | 5782.722 |
| ENSMUSG00000073678 | 0.072306398 | 0.512308 | 0.667549 | 783.7163 | 861.4719 | 791.6699 | 858.3104 | 682.5779 | 920.7163 | 788.7885 | 742.8247 |
| ENSMUSG00000073680 | -1.483510643 | 0.008869 | 0.039084 | 6.036326 | 3.599465 | 15.73403 | 10.35148 | 20.47734 | 36.87335 | 13.50005 | 31.54014 |
| ENSMUSG00000073682 | 0.527016186 | 0.260122 | 0.424174 | 35.2119 | 33.595 | 20.70267 | 23.29083 | 13.65156 | 36.87335 | 11.57147 | 16.31387 |
| ENSMUSG00000073684 | -0.097554346 | 0.308342 | 0.476744 | 576.4691 | 680.2988 | 707.2031 | 646.9676 | 692.329 | 680.4808 | 666.3238 | 755.8758 |
| ENSMUSG00000073700 | -0.158886252 | 0.226845 | 0.386817 | 398.3975 | 381.5432 | 486.0986 | 376.1038 | 487.5557 | 391.0809 | 495.6446 | 460.051 |
| ENSMUSG00000073702 | -0.118510639 | 0.201796 | 0.357355 | 22199.6 | 28200.61 | 25619.13 | 25010.91 | 29980.77 | 25392.33 | 27917.14 | 26384.96 |
| ENSMUSG00000073705 | -0.160026967 | 0.23807 | 0.399185 | 267.6105 | 380.3434 | 308.8838 | 301.9182 | 375.4179 | 365.3813 | 319.1797 | 343.6788 |
| ENSMUSG00000073716 | 0.449354678 | 0.189442 | 0.342042 | 39.23612 | 39.59411 | 36.43669 | 31.91707 | 18.52711 | 42.46022 | 22.17865 | 25.0146 |
| ENSMUSG00000073725 | -0.01324063 | 0.859074 | 0.915425 | 1169.035 | 1160.227 | 1185.021 | 1183.519 | 1075.548 | 1160.952 | 1237.183 | 1269.219 |
| ENSMUSG00000073728 | 0.275671566 | 0.401275 | 0.56855 | 53.32088 | 37.19447 | 35.60859 | 52.62003 | 34.1289 | 55.8687 | 28.92867 | 29.36496 |
| ENSMUSG00000073733 | 0.311982494 | 0.315615 | 0.484004 | 63.38142 | 38.39429 | 75.3577 | 49.16954 | 61.43201 | 34.6386 | 45.32159 | 41.32846 |
| ENSMUSG00000073755 | 0.422869001 | 0.007488 | 0.034522 | 347.0888 | 484.7279 | 527.5039 | 454.6026 | 339.3387 | 393.3157 | 310.5011 | 311.0511 |
| ENSMUSG00000073758 | 0.280339347 | 0.024332 | 0.081381 | 435.6215 | 469.1302 | 472.0208 | 516.7115 | 431.9743 | 430.189 | 371.2513 | 326.2773 |
| ENSMUSG00000073771 | 0.309394991 | 0.066014 | 0.165559 | 145.8779 | 154.777 | 133.3252 | 174.2499 | 117.9885 | 147.4934 | 112.8218 | 113.1095 |
| ENSMUSG00000073775 | -0.169602339 | 0.123344 | 0.252965 | 457.7547 | 442.7341 | 457.943 | 519.2993 | 560.689 | 574.3303 | 458.0374 | 522.0437 |
| ENSMUSG00000073779 | 0.141390654 | 0.752936 | 0.846348 | 35.2119 | 20.39697 | 22.35888 | 39.68068 | 24.37778 | 46.92971 | 15.42863 | 20.66423 |
| ENSMUSG00000073787 | -0.239530609 | 0.263126 | 0.42736 | 61.36932 | 77.9884 | 71.21717 | 56.93315 | 87.76002 | 62.57295 | 78.10742 | 85.9197 |
| ENSMUSG00000073791 | -0.188865251 | 0.322032 | 0.490219 | 171.0292 | 201.57 | 145.7468 | 164.7611 | 140.416 | 224.5922 | 202.5007 | 209.9051 |
| ENSMUSG00000073792 | 0.288647118 | 0.086563 | 0.199626 | 631.8021 | 706.6949 | 582.159 | 684.9231 | 540.2117 | 705.0631 | 464.7874 | 423.0729 |

| | | | | | | | | | | | |
|---|---|---|---|---|---|---|---|---|---|---|---|
| ENSMUSG00000073795 | -0.187084912 | 0.777325 | 0.862728 | 10.06054 | 8.398751 | 6.624853 | 16.38985 | 12.67645 | 23.46486 | 3.857157 | 7.613138 |
| ENSMUSG00000073802 | -0.400557509 | 0.217655 | 0.376128 | 26.15741 | 32.39518 | 55.48314 | 40.5433 | 63.38224 | 37.99072 | 56.89306 | 46.76642 |
| ENSMUSG00000073805 | 0.571247034 | 0.271988 | 0.437265 | 218.3138 | 171.5745 | 129.1846 | 94.88858 | 62.40712 | 246.9397 | 53.0359 | 51.11678 |
| ENSMUSG00000073821 | 0.813132246 | 0.051361 | 0.138584 | 53.32088 | 52.79215 | 24.01509 | 49.16954 | 19.50223 | 42.46022 | 17.3572 | 22.83941 |
| ENSMUSG00000073838 | -0.579132317 | 0.000118 | 0.001515 | 1854.158 | 1778.136 | 2036.314 | 1751.126 | 2806.37 | 2016.86 | 2886.117 | 3375.883 |
| ENSMUSG00000073856 | 0.063601539 | 0.793604 | 0.873442 | 117.7084 | 69.58965 | 76.18581 | 80.22398 | 80.93424 | 87.15518 | 64.60737 | 96.79561 |
| ENSMUSG00000073867 | 0.340033466 | 0.290618 | 0.457766 | 52.31483 | 58.79126 | 68.73285 | 60.38364 | 26.32801 | 67.04245 | 57.85735 | 39.15328 |
| ENSMUSG00000073888 | 0.276928346 | 0.050788 | 0.13752 | 426.567 | 502.7252 | 428.1311 | 443.3885 | 396.8703 | 443.5975 | 341.3584 | 303.4379 |
| ENSMUSG00000073889 | 0.317002324 | 0.000205 | 0.002322 | 2652.965 | 3133.934 | 2748.486 | 2931.195 | 2310.039 | 2536.439 | 2159.043 | 2198.022 |
| ENSMUSG00000073910 | 0.267958855 | 0.019492 | 0.069213 | 356.1432 | 308.3541 | 346.1486 | 389.9058 | 282.7823 | 318.4516 | 284.4653 | 280.5985 |
| ENSMUSG00000073940 | 0.367921986 | 0.202569 | 0.358173 | 1636.85 | 769.0856 | 1131.194 | 2103.939 | 1204.262 | 1412.361 | 827.3601 | 928.8028 |
| ENSMUSG00000073968 | 0.056186633 | 0.550775 | 0.698537 | 415.5004 | 446.3336 | 401.6317 | 470.1298 | 422.2232 | 425.7195 | 397.2871 | 421.9853 |
| ENSMUSG00000073982 | 0.011236698 | 0.895184 | 0.936594 | 625.7658 | 685.0981 | 703.8906 | 690.0988 | 679.6526 | 608.9689 | 700.0739 | 694.9707 |
| ENSMUSG00000073985 | 0.107627329 | 0.596259 | 0.735213 | 91.55095 | 103.1847 | 100.2009 | 102.6522 | 81.90935 | 122.9112 | 77.14313 | 88.09488 |
| ENSMUSG00000073987 | 0.262505977 | 0.014588 | 0.056515 | 836.0312 | 805.0802 | 858.7466 | 960.9626 | 687.4535 | 769.8708 | 797.4671 | 631.8904 |
| ENSMUSG00000073988 | 0.436210936 | 0.130134 | 0.262545 | 228.3743 | 232.7654 | 180.5272 | 200.1287 | 137.4907 | 261.4655 | 114.7504 | 108.7591 |
| ENSMUSG00000074006 | -0.248686932 | 0.533291 | 0.684646 | 14.08476 | 15.59768 | 15.73403 | 19.84034 | 11.70134 | 23.46486 | 22.17865 | 20.66423 |
| ENSMUSG00000074024 | 0.684063407 | 0.046705 | 0.129778 | 279.6831 | 345.5486 | 193.777 | 220.8316 | 99.46135 | 287.1651 | 148.5005 | 112.0219 |
| ENSMUSG00000074028 | -4.309560554 | 3.26E-05 | 0.00057 | 3.018163 | 3.599465 | 43.88965 | 6.038364 | 190.1467 | 67.04245 | 272.8938 | 599.2627 |
| ENSMUSG00000074030 | -0.251978814 | 0.1134 | 0.240481 | 288.7376 | 302.355 | 268.3065 | 259.6497 | 371.5174 | 265.935 | 293.1439 | 400.2335 |
| ENSMUSG00000074039 | 0.099889176 | 0.689983 | 0.803616 | 47.28455 | 59.99108 | 43.06154 | 41.40593 | 38.02934 | 48.04709 | 49.17875 | 42.41605 |
| ENSMUSG00000074063 | -0.583472202 | 0.313745 | 0.481736 | 102.6175 | 10.79839 | 57.96746 | 56.07053 | 92.63557 | 42.46022 | 60.75022 | 145.7372 |
| ENSMUSG00000074064 | -0.492952281 | 0.00027 | 0.002855 | 427.5731 | 367.1454 | 369.3356 | 364.8897 | 527.5352 | 440.2454 | 544.8234 | 638.416 |
| ENSMUSG00000074071 | 0.869452487 | 0.019206 | 0.068496 | 41.24823 | 25.19625 | 48.85829 | 38.81806 | 18.52711 | 15.64324 | 31.82154 | 18.48905 |
| ENSMUSG00000074088 | -0.240293805 | 0.002619 | 0.015951 | 1853.152 | 2165.678 | 1943.566 | 2006.462 | 2524.563 | 2183.349 | 2272.83 | 2427.503 |
| ENSMUSG00000074093 | 0.05136487 | 0.712777 | 0.819906 | 439.6458 | 561.5165 | 389.2101 | 510.6731 | 425.1485 | 465.945 | 505.2875 | 435.0364 |
| ENSMUSG00000074102 | 0.096461296 | 0.353872 | 0.522675 | 1826.995 | 2058.894 | 1980.831 | 2217.805 | 1848.811 | 2201.227 | 1762.721 | 1751.022 |
| ENSMUSG00000074115 | -0.927399487 | 0.023813 | 0.080199 | 27.16347 | 47.99286 | 22.35888 | 23.29083 | 85.8098 | 49.16446 | 61.71451 | 30.45255 |
| ENSMUSG00000074121 | 0.120302935 | 0.545542 | 0.694093 | 99.59938 | 79.18822 | 101.029 | 105.2401 | 111.1627 | 79.33356 | 82.92887 | 81.56933 |
| ENSMUSG00000074129 | -0.083371181 | 0.354767 | 0.523305 | 21689.53 | 25954.54 | 24558.33 | 25356.82 | 28621.47 | 24435.85 | 26634.63 | 23668.16 |
| ENSMUSG00000074136 | 0.086662667 | 0.748653 | 0.843126 | 72.43591 | 71.98929 | 80.32634 | 49.16954 | 61.43201 | 89.38993 | 53.0359 | 54.37955 |
| ENSMUSG00000074151 | 0.260265692 | 0.699435 | 0.810259 | 3.018163 | 9.598572 | 22.35888 | 8.626235 | 9.751113 | 7.821619 | 12.53576 | 6.525547 |
| ENSMUSG00000074158 | 0.01967012 | 0.890718 | 0.933986 | 154.9324 | 177.5736 | 148.2311 | 183.7388 | 160.8934 | 154.1976 | 158.1434 | 181.6277 |
| ENSMUSG00000074165 | 0.343550939 | 0.013759 | 0.054098 | 964.8061 | 946.6592 | 845.4969 | 962.6878 | 689.4037 | 938.5942 | 679.8239 | 624.2773 |
| ENSMUSG00000074166 | 0.577357796 | 0.000144 | 0.001769 | 398.3975 | 385.1427 | 272.4471 | 379.5543 | 250.6036 | 253.6439 | 233.358 | 222.9562 |
| ENSMUSG00000074170 | 0.015011737 | 0.905802 | 0.943371 | 308.8587 | 369.545 | 324.6178 | 275.1769 | 343.2392 | 318.4516 | 310.5011 | 289.2992 |
| ENSMUSG00000074171 | -0.188726855 | 0.390564 | 0.558001 | 51.30877 | 64.79036 | 71.21717 | 53.48266 | 72.15824 | 56.98608 | 74.25027 | 70.69342 |
| ENSMUSG00000074178 | 0.149612252 | 0.471299 | 0.630997 | 147.89 | 142.7788 | 96.88848 | 104.3774 | 108.2374 | 118.4417 | 93.53605 | 121.8102 |
| ENSMUSG00000074179 | -2.910321179 | 1.30E-05 | 0.000281 | 36.21796 | 40.79393 | 119.2474 | 39.68068 | 589.9423 | 119.559 | 318.2154 | 749.3503 |
| ENSMUSG00000074182 | -0.016248202 | 0.854239 | 0.912586 | 870.237 | 807.4799 | 722.109 | 818.6297 | 862.9735 | 762.0491 | 822.5387 | 803.7298 |
| ENSMUSG00000074183 | -2.17932283 | 0.005702 | 0.028351 | 10.06054 | 2.399643 | 2.48432 | 1.725247 | 30.22845 | 7.821619 | 9.642892 | 27.18978 |
| ENSMUSG00000074194 | -0.022914429 | 0.878906 | 0.927396 | 142.8597 | 149.9777 | 141.6062 | 125.0804 | 136.5156 | 163.1366 | 140.7862 | 127.2482 |
| ENSMUSG00000074199 | 1.729629529 | 0.001211 | 0.009018 | 53.32088 | 55.19179 | 38.09291 | 37.09281 | 15.60178 | 27.93435 | 8.678602 | 3.262773 |
| ENSMUSG00000074203 | 0.933772033 | 0.019539 | 0.069334 | 40.24217 | 39.59411 | 38.92101 | 29.3292 | 27.30312 | 26.81698 | 13.50005 | 9.78832 |
| ENSMUSG00000074206 | -0.684071116 | 0.14461 | 0.28269 | 12.07265 | 13.19804 | 22.35888 | 15.52722 | 41.92979 | 14.52586 | 18.32149 | 27.18978 |
| ENSMUSG00000074207 | 0.662960668 | 0.345066 | 0.513882 | 9030.344 | 910.6645 | 2126.578 | 2211.767 | 2403.649 | 1426.887 | 2235.222 | 2952.81 |
| ENSMUSG00000074211 | -0.555641687 | 0.000138 | 0.001716 | 213.2835 | 194.3711 | 240.979 | 222.5569 | 274.0063 | 290.5173 | 347.1441 | 371.9562 |
| ENSMUSG00000074212 | 0.099441515 | 0.292908 | 0.460235 | 745.4863 | 778.6842 | 799.951 | 778.949 | 660.1504 | 832.4437 | 694.2882 | 712.3722 |
| ENSMUSG00000074213 | -0.204210004 | 0.7153 | 0.821607 | 13.07871 | 5.999108 | 14.07781 | 12.93935 | 12.67645 | 5.58687 | 12.53576 | 22.83941 |
| ENSMUSG00000074215 | 0.003191224 | 0.991184 | 0.994852 | 95.57516 | 137.9795 | 111.7944 | 94.88858 | 98.48624 | 172.0756 | 94.50034 | 73.95619 |
| ENSMUSG00000074217 | -1.177786437 | 0.021867 | 0.075029 | 48.29061 | 19.19714 | 65.42042 | 35.36756 | 84.83468 | 31.28647 | 96.42892 | 169.6642 |
| ENSMUSG00000074218 | -1.061718975 | 0.024278 | 0.081245 | 36.21796 | 41.99375 | 59.62368 | 31.91707 | 111.1627 | 23.46486 | 92.57176 | 127.2482 |
| ENSMUSG00000074219 | -0.537693821 | 0.101959 | 0.223912 | 18.10898 | 16.7975 | 21.53077 | 16.38985 | 28.27823 | 23.46486 | 26.03581 | 28.27737 |
| ENSMUSG00000074220 | -0.020198695 | 0.916621 | 0.950616 | 72.43591 | 59.99108 | 73.70149 | 68.14726 | 68.25779 | 68.15982 | 77.14313 | 65.25547 |
| ENSMUSG00000074221 | -0.040605677 | 0.654939 | 0.777385 | 783.7163 | 775.0847 | 705.5469 | 695.2745 | 789.8402 | 698.3588 | 729.0026 | 823.3065 |
| ENSMUSG00000074227 | -0.577416228 | 0.001124 | 0.008534 | 4410.542 | 3657.056 | 6031.101 | 4086.247 | 6264.115 | 5412.56 | 7631.384 | 7830.656 |
| ENSMUSG00000074228 | 0.155073714 | 0.78019 | 0.864358 | 13.07871 | 14.39786 | 12.4216 | 6.900988 | 4.875557 | 10.05637 | 9.642892 | 17.40146 |
| ENSMUSG00000074238 | -0.368146479 | 5.02E-06 | 0.000135 | 1623.772 | 1506.976 | 1744.821 | 1643.298 | 2207.652 | 1891.714 | 2136.865 | 2181.708 |
| ENSMUSG00000074247 | -0.159786258 | 0.081326 | 0.19111 | 1249.52 | 1201.021 | 1331.595 | 1268.057 | 1356.38 | 1238.05 | 1519.72 | 1529.153 |
| ENSMUSG00000074252 | 0.326489203 | 0.237181 | 0.398273 | 47.28455 | 50.3925 | 65.42042 | 56.93315 | 29.25334 | 53.63396 | 51.10733 | 42.41605 |
| ENSMUSG00000074259 | -0.724172079 | 0.000418 | 0.003987 | 97.58727 | 71.98929 | 86.12309 | 81.94923 | 128.7147 | 112.8548 | 137.8934 | 179.4525 |
| ENSMUSG00000074261 | -4.143633507 | 6.57E-08 | 4.03E-06 | 3.018163 | 0 | 15.73403 | 4.313117 | 149.192 | 31.28647 | 75.21455 | 164.2263 |
| ENSMUSG00000074264 | 0.187506242 | 0.619313 | 0.751581 | 85.51462 | 59.99108 | 47.20208 | 97.47645 | 50.70579 | 108.3853 | 54.96448 | 41.32846 |
| ENSMUSG00000074269 | -0.118951687 | 0.726066 | 0.828686 | 25.15136 | 26.39607 | 26.49941 | 19.84034 | 34.1289 | 20.11273 | 21.21436 | 30.45255 |

| | | | | | | | | | | | |
|---|---|---|---|---|---|---|---|---|---|---|---|
| ENSMUSG00000074272 | -0.311972137 | 0.024224 | 0.081141 | 211.2714 | 153.5772 | 172.2462 | 190.6398 | 211.5992 | 231.2964 | 226.608 | 236.0073 |
| ENSMUSG00000074277 | 0.245808502 | 0.37087 | 0.539895 | 113.6841 | 104.3845 | 132.4971 | 113.0037 | 125.7894 | 52.51658 | 118.6076 | 93.53283 |
| ENSMUSG00000074281 | 1.215057003 | 0.00153 | 0.010695 | 50.30272 | 61.1909 | 33.12427 | 45.71905 | 28.27823 | 26.81698 | 13.50005 | 13.05109 |
| ENSMUSG00000074282 | 0.039884284 | 0.811916 | 0.885545 | 228.3743 | 238.7645 | 211.1672 | 173.3873 | 185.2711 | 175.4277 | 238.1794 | 227.3065 |
| ENSMUSG00000074283 | -0.119362225 | 0.609023 | 0.744318 | 65.39353 | 77.9884 | 68.73285 | 70.73513 | 92.63557 | 51.39921 | 76.17884 | 85.9197 |
| ENSMUSG00000074284 | -0.486903069 | 0.158521 | 0.301397 | 22.1332 | 29.99554 | 33.95237 | 40.5433 | 35.10401 | 30.1691 | 53.0359 | 59.81751 |
| ENSMUSG00000074305 | 0.375959463 | 0.00351 | 0.019817 | 1715.323 | 1311.405 | 1589.137 | 1808.059 | 1058.971 | 1392.248 | 1267.076 | 1235.503 |
| ENSMUSG00000074336 | 4.131162518 | 0.002461 | 0.015244 | 97.58727 | 2.399643 | 4.140533 | 25.8787 | 0 | 0 | 0.964289 | 6.525547 |
| ENSMUSG00000074340 | -0.098448637 | 0.870207 | 0.922225 | 137.8294 | 109.1838 | 61.27989 | 97.47645 | 50.70579 | 291.6346 | 43.39301 | 48.9416 |
| ENSMUSG00000074342 | 0.267804511 | 0.570927 | 0.714063 | 14.08476 | 10.79839 | 21.53077 | 10.35148 | 11.70134 | 10.05637 | 10.60718 | 15.22628 |
| ENSMUSG00000074345 | 0.657125135 | 0.15074 | 0.290859 | 23.13925 | 17.99732 | 12.4216 | 25.01608 | 13.65156 | 11.17374 | 7.714313 | 17.40146 |
| ENSMUSG00000074346 | -0.424640553 | 0.568348 | 0.712549 | 6.036326 | 3.599465 | 33.95237 | 7.763611 | 12.67645 | 10.05637 | 23.14294 | 23.927 |
| ENSMUSG00000074355 | 0.253495859 | 0.370559 | 0.539654 | 77.46619 | 65.99018 | 74.5296 | 75.04824 | 39.00445 | 86.03781 | 49.17875 | 72.8686 |
| ENSMUSG00000074357 | 0.16405137 | 0.598251 | 0.736776 | 44.26639 | 47.99286 | 37.2648 | 33.64232 | 25.35289 | 51.39921 | 31.82154 | 36.9781 |
| ENSMUSG00000074358 | -0.017570474 | 0.850157 | 0.910016 | 591.56 | 566.3158 | 517.5666 | 562.4305 | 508.033 | 573.2129 | 599.7879 | 582.9488 |
| ENSMUSG00000074364 | 0.11036729 | 0.308373 | 0.476744 | 1432.621 | 1706.146 | 1880.63 | 1700.231 | 1462.667 | 1639.188 | 1425.219 | 1700.992 |
| ENSMUSG00000074384 | 0.078251536 | 0.644517 | 0.770057 | 126.7628 | 123.5816 | 115.1068 | 110.4158 | 126.7645 | 108.3853 | 122.4647 | 91.35765 |
| ENSMUSG00000074398 | 0.089185747 | 0.761744 | 0.851959 | 62.37537 | 33.595 | 45.54586 | 62.97152 | 61.43201 | 45.81234 | 45.32159 | 40.24087 |
| ENSMUSG00000074405 | 0.019214624 | 0.794361 | 0.873694 | 1007.06 | 949.0588 | 1073.226 | 987.7039 | 988.7629 | 940.829 | 987.4321 | 1050.613 |
| ENSMUSG00000074406 | 0.021226955 | 0.885136 | 0.931361 | 428.5792 | 512.3238 | 660.001 | 540.0023 | 549.9628 | 548.6307 | 548.6805 | 464.4014 |
| ENSMUSG00000074415 | 1.274344149 | 0.001856 | 0.012441 | 3906.509 | 5045.25 | 3573.28 | 4470.115 | 1488.02 | 3337.596 | 1194.754 | 1006.022 |
| ENSMUSG00000074445 | 1.559275716 | 0.010642 | 0.04467 | 38.23007 | 28.79572 | 107.6539 | 28.46658 | 26.32801 | 25.6996 | 10.60718 | 6.525547 |
| ENSMUSG00000074452 | -0.178558402 | 0.728119 | 0.829311 | 11.0666 | 14.39786 | 11.59349 | 7.763611 | 17.552 | 6.704245 | 9.642892 | 16.31387 |
| ENSMUSG00000074457 | 0.090290769 | 0.169733 | 0.3162 | 2686.165 | 2943.162 | 3141.008 | 2950.172 | 2726.411 | 2682.815 | 2830.189 | 2772.27 |
| ENSMUSG00000074466 | 0.059985438 | 0.808924 | 0.883359 | 97.58727 | 86.38715 | 81.98256 | 62.97152 | 59.48179 | 72.62932 | 104.1432 | 78.30656 |
| ENSMUSG00000074476 | -0.18231647 | 0.042304 | 0.121311 | 1013.097 | 1255.013 | 1107.179 | 1081.73 | 1335.902 | 1306.21 | 1209.219 | 1201.788 |
| ENSMUSG00000074479 | 0.285417182 | 0.540969 | 0.690512 | 10.06054 | 11.99822 | 12.4216 | 20.70296 | 10.72622 | 13.40849 | 9.642892 | 11.9635 |
| ENSMUSG00000074480 | 0.41074189 | 0.013433 | 0.053214 | 3755.601 | 3784.237 | 3066.479 | 3477.235 | 2525.538 | 3601.297 | 2118.543 | 2349.197 |
| ENSMUSG00000074482 | 1.311004798 | 0.048496 | 0.133289 | 15.09082 | 29.99554 | 9.109173 | 10.35148 | 11.70134 | 6.704245 | 2.892867 | 4.350364 |
| ENSMUSG00000074489 | 0.227801313 | 0.798843 | 0.876929 | 6.036326 | 2.399643 | 45.54586 | 4.313117 | 8.776002 | 11.17374 | 18.32149 | 11.9635 |
| ENSMUSG00000074500 | 0.40528941 | 0.006932 | 0.032746 | 174.0474 | 177.5736 | 156.5122 | 169.0742 | 131.64 | 118.4417 | 111.8575 | 149 |
| ENSMUSG00000074505 | 0.207556023 | 0.224053 | 0.383871 | 610.675 | 723.4924 | 671.5945 | 619.3637 | 553.8632 | 763.1665 | 442.6087 | 514.4306 |
| ENSMUSG00000074506 | -0.136415314 | 0.769585 | 0.857024 | 10.06054 | 15.59768 | 14.07781 | 12.07673 | 15.60178 | 20.11273 | 13.50005 | 7.613138 |
| ENSMUSG00000074513 | -0.06189635 | 0.433704 | 0.597888 | 627.7779 | 638.3051 | 631.8454 | 587.4466 | 679.6526 | 608.9689 | 670.181 | 631.8904 |
| ENSMUSG00000074516 | -0.092636211 | 0.570948 | 0.714063 | 86.52067 | 94.7859 | 94.40416 | 99.2017 | 110.1876 | 99.44629 | 93.53605 | 96.79561 |
| ENSMUSG00000074519 | -0.238529292 | 0.365585 | 0.534244 | 83.50251 | 112.7832 | 123.3879 | 100.0643 | 178.4454 | 91.62468 | 95.46463 | 129.4233 |
| ENSMUSG00000074521 | -0.319754109 | 0.385436 | 0.5529 | 17.10292 | 34.79482 | 17.39024 | 31.05445 | 33.15378 | 36.87335 | 24.10723 | 30.45255 |
| ENSMUSG00000074527 | -0.161363823 | 0.803034 | 0.879572 | 8.048435 | 25.19625 | 5.796746 | 7.763611 | 10.72622 | 13.40849 | 6.750024 | 20.66423 |
| ENSMUSG00000074529 | 0.226972533 | 0.719327 | 0.824462 | 31.18769 | 11.99822 | 7.45296 | 12.93935 | 15.60178 | 23.46486 | 8.678602 | 6.525547 |
| ENSMUSG00000074575 | 0.10324322 | 0.822746 | 0.89248 | 13.07871 | 21.59679 | 24.01509 | 17.25247 | 20.47734 | 27.93435 | 10.60718 | 11.9635 |
| ENSMUSG00000074576 | -0.343530566 | 0.04262 | 0.12199 | 226.3622 | 175.1739 | 174.7305 | 189.7772 | 249.6285 | 188.8362 | 285.4296 | 246.8832 |
| ENSMUSG00000074577 | -0.010055206 | 0.950021 | 0.971623 | 186.1201 | 167.975 | 232.698 | 216.5185 | 192.0969 | 200.01 | 182.2507 | 238.1824 |
| ENSMUSG00000074578 | 0.270140056 | 0.01565 | 0.059528 | 3063.436 | 3280.312 | 2879.327 | 3152.026 | 2765.416 | 2881.708 | 2530.295 | 2082.737 |
| ENSMUSG00000074579 | 0.517082808 | 0.011723 | 0.048144 | 135.8173 | 172.7743 | 134.9814 | 142.3329 | 95.56091 | 137.437 | 82.92887 | 93.53283 |
| ENSMUSG00000074580 | -0.141366503 | 0.57217 | 0.714912 | 58.35115 | 64.79036 | 61.27989 | 95.75121 | 61.43201 | 92.74205 | 81.00029 | 75.04379 |
| ENSMUSG00000074582 | -0.577467835 | 0.002177 | 0.013964 | 894.3823 | 621.5076 | 987.9312 | 752.2077 | 1151.606 | 907.3078 | 1327.826 | 1474.774 |
| ENSMUSG00000074604 | 1.524328916 | 1.67E-05 | 0.000339 | 66.39959 | 107.9839 | 62.108 | 81.08661 | 35.10401 | 35.75597 | 26.03581 | 13.05109 |
| ENSMUSG00000074607 | 1.428659349 | 2.08E-05 | 0.000406 | 116.7023 | 63.59054 | 61.27989 | 90.57547 | 39.97956 | 35.75597 | 27.96439 | 19.57664 |
| ENSMUSG00000074622 | -0.647187312 | 0.020571 | 0.071931 | 540.2512 | 308.3541 | 534.1288 | 395.9442 | 433.9245 | 544.1612 | 802.2886 | 1007.109 |
| ENSMUSG00000074625 | -1.334981399 | 0.112586 | 0.239278 | 16.09687 | 2.399643 | 36.43669 | 6.900988 | 76.05868 | 5.58687 | 40.50014 | 34.80291 |
| ENSMUSG00000074627 | -0.446960724 | 0.298957 | 0.467074 | 9.054489 | 9.598572 | 14.90592 | 12.93935 | 17.552 | 18.99536 | 13.50005 | 14.13868 |
| ENSMUSG00000074628 | -1.826232478 | 0.006984 | 0.032918 | 7.04238 | 7.198929 | 27.32752 | 6.900988 | 40.95467 | 10.05637 | 62.6788 | 59.81751 |
| ENSMUSG00000074629 | 0.249098535 | 0.443308 | 0.606369 | 69.41775 | 56.39161 | 44.71776 | 47.44429 | 43.88001 | 70.39457 | 38.57157 | 30.45255 |
| ENSMUSG00000074634 | -1.059197604 | 0.01444 | 0.056095 | 70.4238 | 20.39697 | 60.45178 | 30.19182 | 87.76002 | 55.8687 | 95.46463 | 140.2993 |
| ENSMUSG00000074635 | 0.072228021 | 0.833078 | 0.898985 | 31.18769 | 37.19447 | 43.88965 | 30.19182 | 25.35289 | 24.58223 | 34.71441 | 51.11678 |
| ENSMUSG00000074639 | -2.733147837 | 1.08E-05 | 0.000243 | 4.024217 | 1.199822 | 15.73403 | 6.038364 | 35.10401 | 33.52122 | 40.50014 | 76.13138 |
| ENSMUSG00000074643 | -0.285095248 | 0.416364 | 0.582046 | 64.38748 | 49.19268 | 35.60859 | 45.71905 | 62.40712 | 60.3382 | 30.85725 | 83.74451 |
| ENSMUSG00000074646 | 0.712565183 | 0.070168 | 0.172649 | 40.24217 | 51.59233 | 27.32752 | 40.5433 | 32.17867 | 15.64324 | 15.42863 | 33.71532 |
| ENSMUSG00000074649 | -0.03063231 | 0.85151 | 0.910624 | 433.6094 | 352.7475 | 406.6004 | 386.4553 | 436.8499 | 297.2215 | 393.43 | 486.1532 |
| ENSMUSG00000074650 | 0.955189261 | 0.093245 | 0.21005 | 21.12714 | 22.79661 | 19.04645 | 15.52722 | 16.57689 | 15.64324 | 3.857157 | 4.350364 |
| ENSMUSG00000074652 | -0.488621934 | 0.113552 | 0.240638 | 21.12714 | 33.595 | 28.15563 | 24.15346 | 33.15378 | 37.99072 | 30.85725 | 47.85401 |
| ENSMUSG00000074653 | -0.652620998 | 0.292479 | 0.459782 | 11.0666 | 3.599465 | 24.01509 | 5.175741 | 20.47734 | 17.87799 | 16.39292 | 15.22628 |
| ENSMUSG00000074656 | 0.148323062 | 0.258815 | 0.422679 | 7238.561 | 7138.938 | 6202.519 | 6875.109 | 5639.069 | 7935.591 | 5599.627 | 5598.919 |
| ENSMUSG00000074657 | 0.995860764 | 0.004798 | 0.02497 | 315.9011 | 128.3809 | 120.9036 | 322.6212 | 115.0631 | 130.7328 | 105.1075 | 94.62042 |

| | | | | | | | | | | | |
|---|---|---|---|---|---|---|---|---|---|---|---|
| ENSMUSG00000074671 | 0.292639322 | 0.009126 | 0.039911 | 369.2219 | 465.5308 | 434.756 | 420.0976 | 350.065 | 327.3906 | 324.9654 | 376.3065 |
| ENSMUSG00000074673 | -0.833017615 | 0.075548 | 0.181836 | 18.10898 | 10.79839 | 14.90592 | 12.93935 | 41.92979 | 17.87799 | 27.0001 | 14.13868 |
| ENSMUSG00000074676 | -0.115521689 | 0.778222 | 0.863236 | 46.2785 | 46.79304 | 64.59232 | 56.93315 | 63.38224 | 18.99536 | 66.53595 | 83.74451 |
| ENSMUSG00000074678 | 0.029891682 | 0.907895 | 0.945018 | 141.8537 | 123.5816 | 117.5911 | 116.4542 | 170.6445 | 75.98144 | 137.8934 | 103.3212 |
| ENSMUSG00000074682 | -0.012050448 | 0.854156 | 0.91255 | 1108.672 | 1156.628 | 1191.645 | 1218.887 | 1159.407 | 1242.52 | 1148.468 | 1169.16 |
| ENSMUSG00000074698 | 0.054460462 | 0.329255 | 0.497034 | 3915.564 | 3883.822 | 3850.696 | 3817.972 | 3506.5 | 3966.678 | 3765.549 | 3657.569 |
| ENSMUSG00000074715 | -0.987647989 | 0.003764 | 0.02093 | 34.20585 | 28.79572 | 43.06154 | 25.8787 | 35.10401 | 78.21619 | 72.32169 | 77.21897 |
| ENSMUSG00000074733 | 0.54618728 | 0.016672 | 0.061975 | 1405.458 | 1589.764 | 1229.738 | 1419.878 | 753.761 | 1524.098 | 837.9673 | 750.4379 |
| ENSMUSG00000074736 | -0.270781218 | 0.400908 | 0.568172 | 37.22401 | 35.99465 | 79.49824 | 41.40593 | 62.40712 | 62.57295 | 50.14304 | 60.9051 |
| ENSMUSG00000074738 | -0.013360008 | 0.915159 | 0.949686 | 450.7124 | 399.5406 | 378.4447 | 374.3786 | 464.153 | 393.3157 | 348.1084 | 411.1094 |
| ENSMUSG00000074743 | 0.216058409 | 0.242782 | 0.405354 | 543.2694 | 677.8992 | 443.8652 | 458.0531 | 341.289 | 477.1187 | 491.7875 | 515.5182 |
| ENSMUSG00000074746 | 0.048396377 | 0.61715 | 0.749991 | 1667.032 | 1835.727 | 1654.557 | 1850.327 | 1590.407 | 1968.813 | 1626.756 | 1591.146 |
| ENSMUSG00000074748 | 0.235499616 | 0.084973 | 0.196945 | 7035.338 | 7779.643 | 6798.755 | 7272.779 | 5385.54 | 8002.633 | 5539.841 | 5608.707 |
| ENSMUSG00000074749 | 0.104828649 | 0.45302 | 0.615499 | 1040.26 | 1349.799 | 1047.555 | 1215.437 | 1131.129 | 1217.938 | 1119.54 | 855.9342 |
| ENSMUSG00000074754 | -0.16236314 | 0.264864 | 0.429083 | 259.562 | 209.9688 | 274.1033 | 264.8254 | 301.3094 | 244.7049 | 261.3224 | 324.1021 |
| ENSMUSG00000074766 | 0.65120683 | 7.99E-05 | 0.001136 | 567.4147 | 664.7011 | 403.2879 | 506.36 | 358.841 | 367.6161 | 324.9654 | 309.9635 |
| ENSMUSG00000074768 | 1.406621153 | 0.159313 | 0.302311 | 30.18163 | 0 | 5.796746 | 46.58167 | 7.80089 | 11.17374 | 5.785735 | 6.525547 |
| ENSMUSG00000074771 | 1.016122368 | 0.132123 | 0.265252 | 8.048435 | 28.79572 | 9.93728 | 16.38985 | 10.72622 | 12.29112 | 5.785735 | 2.175182 |
| ENSMUSG00000074781 | 0.008825475 | 0.869944 | 0.922052 | 2788.783 | 2803.983 | 2832.125 | 2975.188 | 2757.615 | 2928.638 | 2737.617 | 2912.569 |
| ENSMUSG00000074782 | 0.626221332 | 0.04399 | 0.124553 | 84.50857 | 63.59054 | 55.48314 | 65.55939 | 35.10401 | 64.8077 | 46.28588 | 28.27737 |
| ENSMUSG00000074785 | 1.009110062 | 8.97E-08 | 5.17E-06 | 363.1856 | 423.537 | 352.7734 | 457.1905 | 152.1174 | 265.935 | 187.0721 | 189.2408 |
| ENSMUSG00000074793 | 0.178706428 | 0.106857 | 0.231093 | 1044.284 | 1386.994 | 1345.673 | 1309.462 | 1163.308 | 1148.661 | 1015.396 | 1166.985 |
| ENSMUSG00000074794 | -0.104259067 | 0.527743 | 0.680556 | 2682.141 | 2213.671 | 2412.275 | 3145.988 | 2156.946 | 3085.07 | 3415.512 | 2581.941 |
| ENSMUSG00000074796 | -1.221065319 | 0.010043 | 0.042724 | 83.50251 | 73.18911 | 202.058 | 56.07053 | 274.0063 | 89.38993 | 234.3223 | 370.8686 |
| ENSMUSG00000074797 | -0.269950431 | 0.027173 | 0.088122 | 1402.44 | 1596.962 | 1750.617 | 1534.607 | 2037.008 | 1534.155 | 1895.792 | 2111.014 |
| ENSMUSG00000074800 | -0.126030251 | 0.215918 | 0.374211 | 396.3854 | 465.5308 | 467.0521 | 430.4491 | 514.8588 | 457.006 | 500.4661 | 445.9123 |
| ENSMUSG00000074802 | 0.11756496 | 0.377385 | 0.545463 | 505.0393 | 490.727 | 456.2868 | 520.162 | 466.1032 | 555.3349 | 399.2157 | 398.0583 |
| ENSMUSG00000074807 | -0.051920598 | 0.713404 | 0.820246 | 273.6468 | 313.1534 | 364.3669 | 308.8192 | 316.9112 | 387.7288 | 294.1082 | 309.9635 |
| ENSMUSG00000074811 | -0.447517208 | 0.004794 | 0.024968 | 128.775 | 109.1838 | 127.5284 | 125.943 | 171.6196 | 144.1413 | 158.1434 | 197.9416 |
| ENSMUSG00000074813 | -0.07225321 | 0.870016 | 0.922075 | 22.1332 | 10.79839 | 17.39024 | 14.6646 | 14.62667 | 15.64324 | 12.53576 | 26.10219 |
| ENSMUSG00000074817 | 0.072533001 | 0.89482 | 0.936319 | 11.0666 | 16.7975 | 23.18699 | 6.038364 | 11.70134 | 20.11273 | 9.642892 | 13.05109 |
| ENSMUSG00000074818 | 0.830207294 | 3.70E-05 | 0.000632 | 292.7618 | 224.3666 | 208.6829 | 222.5569 | 157.968 | 153.0803 | 97.39321 | 125.073 |
| ENSMUSG00000074824 | 0.913935635 | 0.132402 | 0.265582 | 22.1332 | 9.598572 | 12.4216 | 12.93935 | 8.776002 | 4.469496 | 13.50005 | 3.262773 |
| ENSMUSG00000074825 | 0.432882713 | 2.45E-07 | 1.19E-05 | 717.3168 | 805.0802 | 703.8906 | 691.824 | 541.1868 | 551.9828 | 545.7877 | 518.7809 |
| ENSMUSG00000074826 | -0.43686859 | 0.108714 | 0.233632 | 40.24217 | 41.99375 | 47.20208 | 30.19182 | 50.70579 | 43.57759 | 52.07161 | 69.60583 |
| ENSMUSG00000074829 | 0.082918803 | 0.518306 | 0.672184 | 338.0343 | 406.7395 | 297.2903 | 386.4553 | 342.2641 | 318.4516 | 351.9655 | 332.8029 |
| ENSMUSG00000074862 | -0.056651186 | 0.83808 | 0.902068 | 61.36932 | 43.19358 | 41.40533 | 55.2079 | 65.33246 | 60.3382 | 43.39301 | 40.24087 |
| ENSMUSG00000074863 | -0.045943133 | 0.929643 | 0.958519 | 6.036326 | 10.79839 | 21.53077 | 17.25247 | 16.57689 | 17.87799 | 9.642892 | 14.13868 |
| ENSMUSG00000074865 | 0.197036278 | 0.507087 | 0.662804 | 80.48435 | 85.18733 | 64.59232 | 93.16334 | 56.55646 | 109.5027 | 48.21446 | 68.51824 |
| ENSMUSG00000074867 | 0.371841257 | 0.029527 | 0.09352 | 137.8294 | 178.7734 | 139.1219 | 157.8601 | 134.5654 | 117.3243 | 122.4647 | 97.8832 |
| ENSMUSG00000074873 | 0.557754613 | 0.013111 | 0.052372 | 377.2704 | 399.5406 | 375.9604 | 439.0754 | 201.848 | 405.6068 | 266.1438 | 208.8175 |
| ENSMUSG00000074874 | 0.574071514 | 0.038284 | 0.112999 | 100.6054 | 50.3925 | 82.81066 | 91.43809 | 48.75557 | 63.69032 | 44.3573 | 63.08028 |
| ENSMUSG00000074876 | -0.412046403 | 0.053399 | 0.142543 | 75.45408 | 69.58965 | 79.49824 | 70.73513 | 128.7147 | 81.56831 | 97.39321 | 84.8321 |
| ENSMUSG00000074884 | -0.384652439 | 0.00019 | 0.002172 | 3967.878 | 4596.516 | 4617.523 | 4132.829 | 5950.129 | 4863.929 | 5441.484 | 6348.269 |
| ENSMUSG00000074886 | -0.241554602 | 0.001837 | 0.012338 | 1869.249 | 1761.338 | 1928.66 | 1883.97 | 2350.993 | 1966.578 | 2295.008 | 2188.233 |
| ENSMUSG00000074890 | -0.726448711 | 3.57E-06 | 0.000104 | 269.6226 | 271.1597 | 328.7583 | 251.0234 | 476.8294 | 362.0292 | 473.466 | 542.708 |
| ENSMUSG00000074892 | -1.012304913 | 8.51E-05 | 0.00119 | 119.7205 | 164.3756 | 141.6062 | 102.6522 | 278.8818 | 175.4277 | 235.2866 | 374.1313 |
| ENSMUSG00000074896 | -0.400040075 | 0.553175 | 0.700377 | 83.50251 | 230.3657 | 467.8802 | 71.59775 | 200.8729 | 169.8409 | 419.4658 | 336.0656 |
| ENSMUSG00000074899 | -0.087311328 | 0.887425 | 0.932494 | 5.030272 | 4.799286 | 18.21835 | 12.93935 | 14.62667 | 13.40849 | 8.678602 | 7.613138 |
| ENSMUSG00000074909 | 0.17482001 | 0.044746 | 0.126005 | 722.347 | 778.6842 | 767.6548 | 853.9973 | 681.6028 | 747.5233 | 675.0024 | 664.5182 |
| ENSMUSG00000074912 | -0.167967183 | 0.66608 | 0.785506 | 29.17558 | 14.39786 | 24.8432 | 23.29083 | 18.52711 | 21.23011 | 26.03581 | 38.06569 |
| ENSMUSG00000074913 | 0.903432819 | 0.076997 | 0.184275 | 28.16952 | 35.99465 | 21.53077 | 33.64232 | 13.65156 | 31.28647 | 13.50005 | 5.437955 |
| ENSMUSG00000074916 | -0.107038742 | 0.229559 | 0.38976 | 551.3178 | 569.9152 | 653.3761 | 576.2325 | 607.4943 | 659.2507 | 618.1094 | 651.4671 |
| ENSMUSG00000074918 | -0.164635036 | 0.216315 | 0.374654 | 335.0161 | 239.9643 | 330.4145 | 338.1484 | 357.8658 | 350.8555 | 340.3941 | 349.1167 |
| ENSMUSG00000074919 | -4.596490607 | 3.81E-05 | 0.000647 | 0 | 0 | 3.312427 | 0 | 21.45245 | 6.704245 | 19.28578 | 40.24087 |
| ENSMUSG00000074922 | -0.225800426 | 0.037065 | 0.11062 | 441.6579 | 472.7297 | 412.3971 | 420.9603 | 441.7254 | 515.1095 | 563.1449 | 520.9561 |
| ENSMUSG00000074923 | -2.106875726 | 0.00022 | 0.002437 | 34.20585 | 11.99822 | 94.40416 | 23.29083 | 154.0676 | 77.09881 | 244.9294 | 233.8321 |
| ENSMUSG00000074925 | -0.6898973 | 0.008756 | 0.038718 | 51.30877 | 71.98929 | 52.99882 | 40.5433 | 84.83468 | 62.57295 | 96.42892 | 103.3212 |
| ENSMUSG00000074934 | -1.798000154 | 6.79E-05 | 0.001007 | 17.10292 | 8.398751 | 7.45296 | 6.900988 | 25.35289 | 30.1691 | 48.21446 | 33.71532 |
| ENSMUSG00000074968 | 1.19786417 | 0.000792 | 0.00653 | 39.23612 | 86.38715 | 75.3577 | 86.26235 | 22.42756 | 45.81234 | 27.0001 | 30.45255 |
| ENSMUSG00000074971 | 1.03518668 | 2.43E-05 | 0.000457 | 3679.141 | 3891.021 | 2376.666 | 4535.674 | 1748.375 | 2175.527 | 2081.9 | 1060.401 |
| ENSMUSG00000074987 | 0.288028567 | 0.122527 | 0.252103 | 729.3894 | 757.0874 | 534.9569 | 723.7411 | 612.3699 | 696.1241 | 535.1805 | 403.4963 |
| ENSMUSG00000074994 | 0.326749197 | 0.007287 | 0.03386 | 2081.526 | 2176.476 | 1952.675 | 2047.006 | 1551.402 | 2057.086 | 1537.077 | 1438.883 |
| ENSMUSG00000075000 | -0.197939858 | 0.051819 | 0.139412 | 389.343 | 328.7511 | 370.9918 | 347.6373 | 421.2481 | 443.5975 | 386.68 | 399.1459 |

| | | | | | | | | | | | |
|---|---|---|---|---|---|---|---|---|---|---|---|
| ENSMUSG00000075012 | -0.380091692 | 0.005609 | 0.027963 | 406.446 | 439.1347 | 540.7536 | 466.6793 | 570.4401 | 509.5226 | 668.2524 | 665.6057 |
| ENSMUSG00000075014 | -0.367466393 | 0.737682 | 0.835871 | 26.15741 | 19.19714 | 3.312427 | 0 | 7.80089 | 40.22547 | 5.785735 | 8.700729 |
| ENSMUSG00000075020 | 1.073049409 | 0.052367 | 0.140498 | 23.13925 | 14.39786 | 15.73403 | 25.8787 | 2.925334 | 16.76061 | 9.642892 | 8.700729 |
| ENSMUSG00000075025 | -3.29289393 | 0.002738 | 0.016518 | 10.06054 | 0 | 34.78048 | 7.763611 | 94.5858 | 30.1691 | 157.1791 | 237.0949 |
| ENSMUSG00000075027 | 0.572445896 | 0.30683 | 0.475352 | 15.09082 | 14.39786 | 17.39024 | 12.93935 | 15.60178 | 5.58687 | 3.857157 | 15.22628 |
| ENSMUSG00000075028 | 0.295549674 | 0.003248 | 0.018835 | 1236.441 | 1007.85 | 1098.069 | 1054.989 | 796.6659 | 967.646 | 891.0032 | 929.8904 |
| ENSMUSG00000075033 | -1.374320017 | 0.000238 | 0.002599 | 234.4107 | 111.5834 | 333.727 | 213.068 | 517.7841 | 262.5829 | 779.1456 | 758.051 |
| ENSMUSG00000075040 | 0.124145877 | 0.12522 | 0.255761 | 841.0614 | 813.479 | 811.5445 | 847.9589 | 779.1139 | 806.7441 | 777.2171 | 677.5692 |
| ENSMUSG00000075044 | -4.036426996 | 3.01E-19 | 3.47E-16 | 10.06054 | 4.799286 | 19.04645 | 10.35148 | 174.5449 | 92.74205 | 178.3935 | 295.8248 |
| ENSMUSG00000075053 | -0.17961322 | 0.552853 | 0.700117 | 24.1453 | 25.19625 | 33.12427 | 40.5433 | 32.17867 | 40.22547 | 36.64299 | 31.54014 |
| ENSMUSG00000075054 | -0.230334583 | 0.061856 | 0.158368 | 536.227 | 539.9197 | 619.4238 | 561.5679 | 720.6073 | 530.7527 | 683.681 | 713.4598 |
| ENSMUSG00000075224 | 0.734264665 | 0.078332 | 0.186301 | 21.12714 | 15.59768 | 19.87456 | 17.25247 | 9.751113 | 10.05637 | 9.642892 | 15.22628 |
| ENSMUSG00000075225 | -0.302737639 | 0.450018 | 0.612502 | 15.09082 | 13.19804 | 17.39024 | 18.11509 | 15.60178 | 22.34748 | 27.0001 | 14.13868 |
| ENSMUSG00000075227 | -0.374426966 | 0.044747 | 0.126005 | 413.4883 | 471.5299 | 556.4877 | 458.0531 | 636.7477 | 426.8369 | 622.9308 | 776.54 |
| ENSMUSG00000075229 | -0.034829943 | 0.634682 | 0.762411 | 819.9343 | 773.8849 | 806.5759 | 805.6903 | 774.2384 | 879.3734 | 798.4314 | 836.3575 |
| ENSMUSG00000075232 | -0.139913312 | 0.060519 | 0.156029 | 1415.518 | 1505.776 | 1250.441 | 1369.846 | 1518.248 | 1507.338 | 1545.756 | 1526.978 |
| ENSMUSG00000075254 | 0.418214913 | 9.72E-05 | 0.001308 | 1924.582 | 2195.673 | 2090.141 | 2141.032 | 1527.024 | 1868.249 | 1412.684 | 1443.233 |
| ENSMUSG00000075256 | -0.278862654 | 0.386877 | 0.55432 | 27.16347 | 19.19714 | 23.18699 | 18.97772 | 26.32801 | 25.6996 | 24.10723 | 31.54014 |
| ENSMUSG00000075266 | -0.013473713 | 0.921945 | 0.953458 | 619.7295 | 604.7101 | 584.6433 | 656.4565 | 650.3992 | 757.5796 | 584.3592 | 497.0291 |
| ENSMUSG00000075267 | -1.068308573 | 0.002279 | 0.01441 | 11.0666 | 19.19714 | 14.07781 | 13.80198 | 25.35289 | 30.1691 | 32.78583 | 32.62773 |
| ENSMUSG00000075268 | -0.06999056 | 0.857585 | 0.914643 | 29.17558 | 21.59679 | 22.35888 | 22.42821 | 39.97956 | 16.76061 | 19.28578 | 23.927 |
| ENSMUSG00000075270 | -0.937632445 | 0.018589 | 0.067061 | 24.1453 | 28.79572 | 33.12427 | 18.97772 | 46.80534 | 23.46486 | 58.82164 | 71.78101 |
| ENSMUSG00000075271 | -0.066988314 | 0.643941 | 0.769677 | 214.2896 | 268.76 | 217.792 | 197.5408 | 260.3547 | 227.9443 | 208.2865 | 241.4452 |
| ENSMUSG00000075273 | 0.108940563 | 0.239639 | 0.401559 | 394.3733 | 413.9384 | 368.5075 | 386.4553 | 343.2392 | 374.3203 | 375.1085 | 354.5547 |
| ENSMUSG00000075277 | 0.668706601 | 0.001684 | 0.011531 | 362.1796 | 399.5406 | 298.9465 | 308.8192 | 200.8729 | 303.9258 | 198.6436 | 157.7007 |
| ENSMUSG00000075279 | 0.056346841 | 0.847743 | 0.908531 | 30.18163 | 28.79572 | 27.32752 | 31.91707 | 32.17867 | 30.1691 | 22.17865 | 29.36496 |
| ENSMUSG00000075284 | 0.325327105 | 4.55E-05 | 0.00074 | 721.341 | 787.0829 | 816.5131 | 800.5146 | 662.1006 | 621.26 | 610.395 | 601.4379 |
| ENSMUSG00000075289 | -0.123219704 | 0.528821 | 0.681273 | 92.557 | 88.78679 | 72.04528 | 110.4158 | 100.4365 | 106.1505 | 87.75031 | 102.2336 |
| ENSMUSG00000075296 | -1.246094891 | 0.036887 | 0.110409 | 21.12714 | 31.19536 | 98.54469 | 35.36756 | 187.2214 | 26.81698 | 114.7504 | 114.1971 |
| ENSMUSG00000075297 | 0.211995129 | 0.356837 | 0.525355 | 43.26034 | 50.3925 | 48.85829 | 50.89479 | 44.85512 | 40.22547 | 40.50014 | 41.32846 |
| ENSMUSG00000075302 | 0.529147317 | 0.007683 | 0.035143 | 136.8234 | 166.7752 | 169.7619 | 139.745 | 86.78491 | 129.6154 | 121.5004 | 87.00729 |
| ENSMUSG00000075304 | 1.213160885 | 5.24E-13 | 1.70E-10 | 382.3007 | 471.5299 | 414.8814 | 376.9665 | 172.5947 | 206.7142 | 131.1433 | 200.1168 |
| ENSMUSG00000075307 | 0.59464006 | 0.222258 | 0.381505 | 15.09082 | 14.39786 | 10.76539 | 11.21411 | 8.776002 | 5.58687 | 10.60718 | 8.700729 |
| ENSMUSG00000075312 | -0.948260436 | 0.063616 | 0.161622 | 8.048435 | 5.999108 | 11.59349 | 5.175741 | 14.62667 | 12.29112 | 12.53576 | 20.66423 |
| ENSMUSG00000075316 | 1.456756039 | 0.001146 | 0.008674 | 249.5015 | 326.3515 | 159.8246 | 295.0172 | 102.3867 | 185.4841 | 44.3573 | 43.50364 |
| ENSMUSG00000075318 | 0.845451213 | 0.006715 | 0.03202 | 123.7447 | 158.3764 | 103.5133 | 144.0581 | 64.35735 | 122.9112 | 49.17875 | 58.72992 |
| ENSMUSG00000075324 | 0.724920127 | 0.002996 | 0.017744 | 667.014 | 767.8858 | 657.5167 | 641.7919 | 325.6872 | 667.0723 | 297.0011 | 365.4306 |
| ENSMUSG00000075327 | -0.010445974 | 0.916409 | 0.950503 | 561.3783 | 547.1186 | 562.2844 | 617.6384 | 507.0579 | 620.1426 | 567.002 | 614.489 |
| ENSMUSG00000075330 | -0.225543729 | 0.712862 | 0.819914 | 2.012109 | 16.7975 | 15.73403 | 12.93935 | 19.50223 | 12.29112 | 10.60718 | 13.05109 |
| ENSMUSG00000075334 | 0.496937326 | 0.006121 | 0.029877 | 4246.555 | 5144.835 | 4044.473 | 4399.38 | 3473.346 | 4039.307 | 2986.404 | 2138.204 |
| ENSMUSG00000075376 | 0.330399513 | 0.035887 | 0.108137 | 1718.341 | 1530.972 | 1401.156 | 1645.023 | 1098.95 | 1663.77 | 1205.361 | 1039.737 |
| ENSMUSG00000075389 | 0.106707463 | 0.320373 | 0.488405 | 1038.248 | 1111.035 | 1177.568 | 1366.396 | 1081.398 | 1188.886 | 1078.075 | 1013.635 |
| ENSMUSG00000075391 | -0.814606179 | 0.113031 | 0.239974 | 13.07871 | 15.59768 | 5.796746 | 11.21411 | 25.35289 | 11.17374 | 14.46434 | 28.27737 |
| ENSMUSG00000075394 | 0.904737188 | 0.002397 | 0.014943 | 1086.539 | 1408.59 | 933.2762 | 1281.859 | 573.3654 | 1099.496 | 455.1445 | 388.27 |
| ENSMUSG00000075408 | -0.072272699 | 0.636473 | 0.763916 | 327.9737 | 374.3443 | 423.9906 | 341.5989 | 434.8996 | 358.6771 | 312.4297 | 438.2992 |
| ENSMUSG00000075410 | -0.134762671 | 0.615866 | 0.748989 | 58.35115 | 58.79126 | 59.62368 | 58.6584 | 45.83023 | 94.9768 | 59.78593 | 58.72992 |
| ENSMUSG00000075415 | 0.187716003 | 0.010576 | 0.044441 | 1422.561 | 1406.191 | 1489.764 | 1475.949 | 1284.222 | 1376.605 | 1161.004 | 1270.306 |
| ENSMUSG00000075416 | -0.13343632 | 0.776285 | 0.861886 | 12.07265 | 15.59768 | 8.281066 | 12.07673 | 15.60178 | 8.938993 | 16.39292 | 10.87591 |
| ENSMUSG00000075419 | -0.427426497 | 0.050191 | 0.136381 | 432.6034 | 376.744 | 470.3646 | 370.0655 | 612.3699 | 329.6254 | 587.2521 | 689.5327 |
| ENSMUSG00000075420 | -0.305574263 | 0.374181 | 0.542916 | 88.53278 | 50.3925 | 153.1997 | 75.04824 | 98.48624 | 112.8548 | 93.53605 | 151.1752 |
| ENSMUSG00000075463 | -0.087705359 | 0.811119 | 0.885014 | 17.10292 | 35.99465 | 31.46805 | 24.15346 | 29.25334 | 18.99536 | 30.85725 | 35.89051 |
| ENSMUSG00000075467 | -0.269071623 | 0.009883 | 0.042258 | 1086.539 | 1173.425 | 1170.115 | 1061.89 | 1484.119 | 1138.604 | 1323.005 | 1464.985 |
| ENSMUSG00000075470 | -0.120761925 | 0.282932 | 0.449379 | 1149.92 | 1099.037 | 1163.49 | 1098.982 | 1319.326 | 975.4676 | 1299.862 | 1309.46 |
| ENSMUSG00000075478 | 0.496251928 | 0.127704 | 0.259113 | 70.4238 | 95.98572 | 58.79557 | 128.5309 | 46.80534 | 87.15518 | 52.07161 | 65.25547 |
| ENSMUSG00000075486 | -0.167704662 | 0.026705 | 0.087156 | 729.3894 | 692.297 | 761.03 | 779.8116 | 801.5415 | 825.7395 | 825.4315 | 880.9488 |
| ENSMUSG00000075502 | -0.018301419 | 0.95931 | 0.97709 | 52.31483 | 41.99375 | 24.01509 | 35.36756 | 52.65601 | 37.99072 | 25.07152 | 39.15328 |
| ENSMUSG00000075511 | -0.037834728 | 0.883486 | 0.930352 | 62.37537 | 79.18822 | 71.21717 | 62.10889 | 78.98402 | 80.45093 | 43.39301 | 79.39415 |
| ENSMUSG00000075514 | 0.169657831 | 0.257276 | 0.421045 | 177.0656 | 225.5664 | 200.4018 | 224.2821 | 185.2711 | 212.3011 | 160.072 | 178.3649 |
| ENSMUSG00000075520 | -0.639797521 | 0.156997 | 0.299425 | 18.10898 | 8.398751 | 28.98373 | 18.97772 | 19.50223 | 23.46486 | 37.60728 | 36.9781 |
| ENSMUSG00000075528 | -0.540211631 | 0.001475 | 0.010445 | 149.9021 | 157.1766 | 183.8397 | 174.2499 | 253.5289 | 181.0146 | 256.5009 | 277.3357 |
| ENSMUSG00000075534 | 0.268419398 | 0.452427 | 0.614874 | 46.2785 | 45.59322 | 78.67013 | 43.9938 | 40.95467 | 24.58223 | 52.07161 | 60.9051 |
| ENSMUSG00000075553 | -0.97830562 | 0.112431 | 0.239086 | 3.018163 | 14.39786 | 10.76539 | 4.313117 | 17.552 | 15.64324 | 21.21436 | 8.700729 |
| ENSMUSG00000075569 | 0.382981271 | 0.380656 | 0.548577 | 29.17558 | 70.78947 | 33.95237 | 37.95543 | 26.32801 | 56.98608 | 23.14294 | 25.0146 |
| ENSMUSG00000075576 | -0.067573076 | 0.807568 | 0.882437 | 70.4238 | 57.59143 | 51.34261 | 79.36136 | 88.73513 | 78.21619 | 51.10733 | 53.29196 |

| | | | | | | | | | | | |
|---|---|---|---|---|---|---|---|---|---|---|---|
| ENSMUSG00000075585 | -0.384674093 | 0.05591 | 0.147283 | 292.7618 | 248.3631 | 361.0545 | 324.3464 | 368.5921 | 297.2215 | 404.0372 | 534.0072 |
| ENSMUSG00000075588 | 0.068860938 | 0.522991 | 0.676265 | 768.6255 | 748.6886 | 664.1415 | 758.2461 | 791.7904 | 608.9689 | 724.1812 | 674.3065 |
| ENSMUSG00000075589 | 0.329785417 | 0.344871 | 0.513675 | 58.35115 | 50.3925 | 47.20208 | 91.43809 | 51.6809 | 69.27719 | 28.92867 | 47.85401 |
| ENSMUSG00000075590 | 0.157440405 | 0.352851 | 0.521562 | 1027.181 | 976.6547 | 804.0915 | 1001.506 | 851.2722 | 1117.374 | 797.4671 | 649.2919 |
| ENSMUSG00000075592 | -0.217210668 | 0.389923 | 0.557343 | 376.2643 | 274.7591 | 350.2891 | 246.7103 | 409.5467 | 206.7142 | 373.1799 | 461.1386 |
| ENSMUSG00000075595 | 0.206404653 | 0.061953 | 0.158498 | 1899.431 | 1865.722 | 1747.305 | 1845.152 | 1406.111 | 1936.409 | 1494.648 | 1541.117 |
| ENSMUSG00000075596 | 1.51252135 | 0.062003 | 0.158604 | 37.22401 | 37.19447 | 13.24971 | 26.74133 | 3.900445 | 30.1691 | 1.928578 | 4.350364 |
| ENSMUSG00000075600 | 0.105455569 | 0.460485 | 0.621841 | 210.2654 | 201.57 | 165.6213 | 175.9752 | 172.5947 | 157.5497 | 176.4649 | 191.416 |
| ENSMUSG00000075602 | -1.144026437 | 0.02137 | 0.073804 | 1748.522 | 1079.839 | 3784.447 | 1756.301 | 3902.395 | 2528.618 | 5174.376 | 6892.065 |
| ENSMUSG00000075604 | 4.195146597 | NA | NA | 8817.06 | 7.198929 | 4.140533 | 762.5592 | 0 | 1.117374 | 4.821446 | 517.6934 |
| ENSMUSG00000075700 | -0.343493908 | 0.000721 | 0.006099 | 3161.023 | 2939.563 | 2977.871 | 2900.14 | 3589.385 | 3340.949 | 3771.335 | 4497.189 |
| ENSMUSG00000075701 | -0.160665441 | 0.019209 | 0.068496 | 1521.154 | 1431.387 | 1488.108 | 1616.556 | 1651.839 | 1660.418 | 1807.078 | 1654.226 |
| ENSMUSG00000075702 | 0.494420548 | 3.16E-05 | 0.000559 | 1235.435 | 1734.942 | 1543.591 | 1420.741 | 1075.548 | 1131.9 | 961.3963 | 1043 |
| ENSMUSG00000075703 | 0.003501177 | 0.962956 | 0.978975 | 904.4429 | 909.4647 | 832.2472 | 878.1507 | 822.0188 | 939.7116 | 869.7888 | 883.124 |
| ENSMUSG00000075704 | -0.378515656 | 0.014011 | 0.054785 | 500.009 | 358.7466 | 395.0069 | 382.1422 | 602.6188 | 431.3064 | 514.9304 | 577.5109 |
| ENSMUSG00000075705 | -0.319946171 | 0.001913 | 0.012725 | 1017.121 | 907.0651 | 999.5247 | 1010.995 | 1289.097 | 1051.449 | 1191.861 | 1381.241 |
| ENSMUSG00000075706 | -0.592608298 | 0.00011 | 0.001438 | 3486.984 | 3061.945 | 3771.198 | 3470.334 | 5423.569 | 3789.016 | 5187.876 | 6397.211 |
| ENSMUSG00000075707 | 0.436260849 | 0.322718 | 0.490873 | 27.16347 | 20.39697 | 14.07781 | 25.01608 | 10.72622 | 20.11273 | 22.17865 | 10.87591 |
| ENSMUSG00000076122 | -0.104501932 | 0.826795 | 0.89487 | 20.12109 | 11.99822 | 11.59349 | 12.93935 | 8.776002 | 17.87799 | 22.17865 | 11.9635 |
| ENSMUSG00000076137 | 0.523198793 | 0.198028 | 0.35246 | 85.51462 | 69.58965 | 79.49824 | 56.93315 | 31.20356 | 98.32892 | 43.39301 | 30.45255 |
| ENSMUSG00000076258 | -0.283970288 | 0.273704 | 0.43956 | 6871.351 | 5108.84 | 3538.5 | 3801.582 | 8973.949 | 4806.943 | 4937.161 | 4802.802 |
| ENSMUSG00000076281 | 0.143677474 | 0.659749 | 0.780834 | 615.7053 | 563.9161 | 260.0255 | 337.2858 | 632.8472 | 373.2029 | 303.7511 | 296.9124 |
| ENSMUSG00000076431 | 0.419477575 | 0.004801 | 0.02498 | 3501.069 | 3935.415 | 3006.855 | 3007.105 | 2281.76 | 3220.272 | 2252.579 | 2301.343 |
| ENSMUSG00000076432 | -0.173588813 | 0.015151 | 0.058087 | 6385.427 | 7532.48 | 7442.194 | 6821.627 | 7888.65 | 7729.994 | 7876.314 | 8288.532 |
| ENSMUSG00000076434 | -0.556656001 | 0.150931 | 0.291135 | 13.07871 | 15.59768 | 13.24971 | 11.21411 | 20.47734 | 22.34748 | 15.42863 | 19.57664 |
| ENSMUSG00000076435 | 0.306919622 | 0.000399 | 0.003854 | 1208.271 | 1180.624 | 1138.647 | 1169.717 | 1078.473 | 907.3078 | 937.2891 | 871.1605 |
| ENSMUSG00000076437 | -0.130967511 | 0.118958 | 0.247866 | 2352.155 | 2669.603 | 2734.408 | 2683.622 | 3130.107 | 2609.069 | 2825.367 | 2866.89 |
| ENSMUSG00000076439 | 1.113399655 | 0.048458 | 0.133246 | 30.18163 | 16.7975 | 14.07781 | 15.52722 | 3.900445 | 14.52586 | 11.57147 | 5.437955 |
| ENSMUSG00000076441 | -3.085141731 | 1.81E-05 | 0.000363 | 516.1059 | 139.1793 | 1046.727 | 640.0666 | 7672.176 | 1477.169 | 3443.477 | 7287.948 |
| ENSMUSG00000076609 | -0.071278851 | 0.868668 | 0.921387 | 13.07871 | 11.99822 | 27.32752 | 20.70296 | 21.45245 | 23.46486 | 13.50005 | 19.57664 |
| ENSMUSG00000076617 | -0.192945745 | 0.670453 | 0.788748 | 24.1453 | 20.39697 | 45.54586 | 12.07673 | 25.35289 | 29.05173 | 25.07152 | 38.06569 |
| ENSMUSG00000076928 | -0.448548529 | 0.323607 | 0.491571 | 8.048435 | 9.598572 | 21.53077 | 13.80198 | 19.50223 | 17.87799 | 15.42863 | 20.66423 |
| ENSMUSG00000077167 | -0.075430851 | 0.859208 | 0.91551 | 25.15136 | 13.19804 | 22.35888 | 24.15346 | 17.552 | 25.6996 | 13.50005 | 33.71532 |
| ENSMUSG00000077192 | 0.146391056 | 0.427475 | 0.592509 | 133.8052 | 159.5763 | 128.3565 | 147.5086 | 156.0178 | 140.7891 | 110.8933 | 105.4963 |
| ENSMUSG00000077212 | -0.142126449 | 0.592583 | 0.732287 | 42.25428 | 39.59411 | 52.99882 | 51.75741 | 68.25779 | 49.16446 | 40.50014 | 48.9416 |
| ENSMUSG00000077222 | 0.000956423 | 0.997685 | 0.998659 | 35.2119 | 21.59679 | 39.74912 | 43.13117 | 34.1289 | 46.92971 | 32.78583 | 27.18978 |
| ENSMUSG00000077394 | -0.223715162 | 0.50322 | 0.659151 | 37.22401 | 25.19625 | 19.87456 | 25.8787 | 39.00445 | 32.40385 | 25.07152 | 29.36496 |
| ENSMUSG00000077426 | -0.282687817 | 0.501338 | 0.657712 | 13.07871 | 14.39786 | 9.93728 | 11.21411 | 16.57689 | 13.40849 | 15.42863 | 13.05109 |
| ENSMUSG00000077440 | -0.024205871 | 0.887812 | 0.932581 | 105.6357 | 133.1802 | 135.8095 | 121.6299 | 130.6649 | 137.437 | 134.0362 | 102.2336 |
| ENSMUSG00000077450 | 0.052417308 | 0.424193 | 0.589348 | 3586.584 | 3855.027 | 3968.287 | 3907.684 | 3615.713 | 3694.039 | 3494.584 | 3970.795 |
| ENSMUSG00000077457 | 0.697486128 | 0.279809 | 0.446185 | 14.08476 | 9.598572 | 15.73403 | 27.60395 | 13.65156 | 18.99536 | 5.785735 | 3.262773 |
| ENSMUSG00000077493 | 0.860640438 | 0.097637 | 0.217343 | 10.06054 | 14.39786 | 12.4216 | 18.97772 | 11.70134 | 6.704245 | 5.785735 | 6.525547 |
| ENSMUSG00000077506 | -0.226078726 | 0.608486 | 0.744118 | 15.09082 | 14.39786 | 7.45296 | 12.07673 | 14.62667 | 13.40849 | 14.46434 | 14.13868 |
| ENSMUSG00000077575 | -0.067720782 | 0.85329 | 0.911961 | 37.22401 | 22.79661 | 20.70267 | 35.36756 | 19.50223 | 39.10809 | 29.89296 | 33.71532 |
| ENSMUSG00000077704 | -0.466465346 | 0.260987 | 0.425111 | 29.17558 | 28.79572 | 11.59349 | 18.97772 | 42.9049 | 23.46486 | 25.07152 | 29.36496 |
| ENSMUSG00000077711 | -0.501792654 | 0.143005 | 0.280478 | 22.1332 | 38.39429 | 30.63995 | 37.09281 | 62.40712 | 52.51658 | 33.75012 | 32.62773 |
| ENSMUSG00000077734 | -0.510788664 | 0.376941 | 0.545034 | 17.10292 | 7.198929 | 19.04645 | 16.38985 | 32.17867 | 26.81698 | 21.21436 | 5.437955 |
| ENSMUSG00000077737 | 0.274488793 | 0.345571 | 0.514314 | 68.4117 | 49.19268 | 45.54586 | 44.85642 | 52.65601 | 51.39921 | 31.82154 | 35.89051 |
| ENSMUSG00000077797 | -0.483916295 | 0.266803 | 0.431258 | 22.1332 | 8.398751 | 17.39024 | 17.25247 | 29.25334 | 29.05173 | 17.3572 | 16.31387 |
| ENSMUSG00000078117 | 0.265529969 | 0.46718 | 0.627759 | 28.16952 | 31.19536 | 30.63995 | 31.05445 | 15.60178 | 37.99072 | 28.92867 | 18.48905 |
| ENSMUSG00000078122 | 0.644928572 | 0.147097 | 0.286223 | 29.17558 | 17.99732 | 24.8432 | 17.25247 | 10.72622 | 11.17374 | 11.57147 | 23.927 |
| ENSMUSG00000078125 | 0.892824011 | 0.09379 | 0.210839 | 10.06054 | 16.7975 | 32.29616 | 22.42821 | 13.65156 | 14.52586 | 10.60718 | 5.437955 |
| ENSMUSG00000078126 | -0.020930302 | 0.885426 | 0.931559 | 110.666 | 135.5798 | 123.3879 | 124.2178 | 124.8142 | 125.1459 | 127.2862 | 122.8978 |
| ENSMUSG00000078134 | -0.056422759 | 0.914455 | 0.949313 | 8.048435 | 9.598572 | 16.56213 | 14.6646 | 19.50223 | 7.821619 | 8.678602 | 15.22628 |
| ENSMUSG00000078137 | -1.909883874 | 0.000218 | 0.002419 | 5.030272 | 4.799286 | 11.59349 | 8.626235 | 35.10401 | 13.40849 | 33.75012 | 32.62773 |
| ENSMUSG00000078139 | -0.68004757 | 0.184045 | 0.335148 | 7.04238 | 5.999108 | 6.624853 | 12.93935 | 14.62667 | 13.40849 | 9.642892 | 15.22628 |
| ENSMUSG00000078141 | -0.459862622 | 0.456149 | 0.618021 | 6.036326 | 15.59768 | 8.281066 | 18.97772 | 19.50223 | 4.469496 | 21.21436 | 21.75182 |
| ENSMUSG00000078153 | -0.163858181 | 0.476534 | 0.635976 | 67.40564 | 73.18911 | 81.15445 | 71.59775 | 98.48624 | 64.8077 | 65.57166 | 100.0584 |
| ENSMUSG00000078154 | -0.093351233 | 0.697073 | 0.808775 | 54.32694 | 76.78858 | 52.99882 | 57.79577 | 54.60623 | 79.33356 | 55.92877 | 67.43065 |
| ENSMUSG00000078160 | 0.569744204 | 0.243707 | 0.406273 | 43.26034 | 45.59322 | 24.01509 | 18.97772 | 19.50223 | 39.10809 | 11.57147 | 18.48905 |
| ENSMUSG00000078183 | 0.160252465 | 0.759533 | 0.850138 | 13.07871 | 13.19804 | 9.109173 | 16.38985 | 14.62667 | 15.64324 | 11.57147 | 4.350364 |
| ENSMUSG00000078185 | -0.158689382 | 0.150148 | 0.29014 | 368.2159 | 343.149 | 350.2891 | 342.4615 | 419.2979 | 430.189 | 340.3941 | 378.4817 |
| ENSMUSG00000078190 | 0.827460291 | 0.000312 | 0.003201 | 2168.047 | 2284.46 | 1755.586 | 2660.331 | 975.1113 | 1931.94 | 1080.004 | 1011.46 |
| ENSMUSG00000078193 | -0.040180057 | 0.820121 | 0.89103 | 186.1201 | 238.7645 | 317.1648 | 234.6336 | 264.2552 | 232.4138 | 264.2152 | 244.708 |

| | | | | | | | | | | | |
|---|---|---|---|---|---|---|---|---|---|---|---|
| ENSMUSG00000078201 | -0.278431401 | 0.017996 | 0.065652 | 451.7184 | 435.5352 | 522.5353 | 458.9157 | 578.241 | 474.884 | 601.7164 | 613.4014 |
| ENSMUSG00000078202 | -0.902224668 | 0.001509 | 0.010607 | 183.1019 | 118.7823 | 203.7142 | 128.5309 | 301.3094 | 165.3714 | 342.3227 | 377.3941 |
| ENSMUSG00000078234 | -0.388910287 | 0.144434 | 0.28259 | 952.7335 | 769.0856 | 1392.875 | 741.8562 | 1221.814 | 882.7255 | 1056.861 | 1890.233 |
| ENSMUSG00000078238 | 0.512054786 | 0.067125 | 0.167299 | 77.46619 | 57.59143 | 62.9361 | 60.38364 | 55.58134 | 43.57759 | 27.96439 | 54.37955 |
| ENSMUSG00000078240 | 0.61663057 | 0.125403 | 0.255959 | 46.2785 | 34.79482 | 19.87456 | 25.8787 | 20.47734 | 25.6996 | 13.50005 | 22.83941 |
| ENSMUSG00000078247 | 0.383828804 | 0.133669 | 0.267172 | 360.1675 | 376.744 | 381.7572 | 402.8452 | 280.8321 | 467.0624 | 226.608 | 192.5036 |
| ENSMUSG00000078249 | -0.379597668 | 0.756059 | 0.848334 | 288.7376 | 304.7547 | 27.32752 | 27.60395 | 465.1281 | 30.1691 | 18.32149 | 329.5401 |
| ENSMUSG00000078302 | 0.639620029 | 0.001216 | 0.00904 | 1208.271 | 1201.021 | 859.5747 | 1240.453 | 626.9966 | 954.2375 | 781.0742 | 531.832 |
| ENSMUSG00000078307 | -0.039639802 | 0.930447 | 0.958826 | 14.08476 | 9.598572 | 14.07781 | 11.21411 | 11.70134 | 16.76061 | 13.50005 | 8.700729 |
| ENSMUSG00000078317 | -0.204098542 | 0.242771 | 0.405354 | 155.9384 | 178.7734 | 179.6991 | 159.5853 | 163.8187 | 169.8409 | 192.8578 | 250.146 |
| ENSMUSG00000078348 | -0.192575772 | 0.027411 | 0.088676 | 1547.312 | 1678.55 | 1876.49 | 1597.579 | 2019.456 | 1757.629 | 1948.828 | 1931.562 |
| ENSMUSG00000078349 | -1.176532742 | 0.000274 | 0.002891 | 59.35721 | 38.39429 | 55.48314 | 31.05445 | 84.83468 | 69.27719 | 122.4647 | 140.2993 |
| ENSMUSG00000078350 | -1.091015768 | 3.97E-08 | 2.67E-06 | 329.9858 | 317.9527 | 441.3808 | 386.4553 | 684.5281 | 567.626 | 897.7532 | 997.321 |
| ENSMUSG00000078365 | 0.596557186 | 0.296799 | 0.46449 | 16.09687 | 20.39697 | 9.109173 | 12.93935 | 3.900445 | 14.52586 | 13.50005 | 6.525547 |
| ENSMUSG00000078377 | -0.056142433 | 0.797702 | 0.875947 | 70.4238 | 87.58697 | 77.01392 | 82.81186 | 95.56091 | 80.45093 | 94.50034 | 58.72992 |
| ENSMUSG00000078427 | 0.046622265 | 0.486512 | 0.644509 | 1598.62 | 1864.523 | 1745.649 | 1734.736 | 1674.266 | 1729.695 | 1654.72 | 1661.839 |
| ENSMUSG00000078429 | 0.149287666 | 0.028452 | 0.091024 | 11737.64 | 12371.36 | 12267.57 | 12524.43 | 11049.96 | 11803.94 | 11309.18 | 9930.794 |
| ENSMUSG00000078435 | 0.020502604 | 0.909085 | 0.945776 | 92.557 | 112.7832 | 90.26362 | 98.33908 | 99.46135 | 111.7374 | 82.92887 | 93.53283 |
| ENSMUSG00000078439 | -1.527791596 | 5.63E-05 | 0.000866 | 101.6115 | 69.58965 | 192.1207 | 128.5309 | 308.1352 | 185.4841 | 342.3227 | 586.2116 |
| ENSMUSG00000078440 | -0.411259132 | 0.000468 | 0.004349 | 882.3097 | 940.6601 | 1078.195 | 973.9019 | 1259.844 | 1076.031 | 1358.683 | 1460.635 |
| ENSMUSG00000078444 | -0.169083654 | 0.590914 | 0.73091 | 36.21796 | 26.39607 | 27.32752 | 23.29083 | 25.35289 | 27.93435 | 34.71441 | 39.15328 |
| ENSMUSG00000078451 | -0.837211298 | 0.002214 | 0.014142 | 50.30272 | 23.99643 | 37.2648 | 35.36756 | 57.53157 | 64.8077 | 76.17884 | 65.25547 |
| ENSMUSG00000078452 | 0.445618672 | 0.047844 | 0.131989 | 61.36932 | 87.58697 | 76.18581 | 88.85022 | 58.50668 | 45.81234 | 62.6788 | 63.08028 |
| ENSMUSG00000078453 | 0.111175378 | 0.137864 | 0.273093 | 1331.01 | 1480.58 | 1413.578 | 1382.785 | 1215.964 | 1251.459 | 1317.219 | 1406.255 |
| ENSMUSG00000078484 | 0.091797964 | 0.256053 | 0.419812 | 1074.466 | 1013.849 | 924.167 | 1055.851 | 980.962 | 983.2892 | 889.0746 | 963.6057 |
| ENSMUSG00000078485 | 0.136759212 | 0.521468 | 0.674815 | 144.8718 | 172.7743 | 154.0278 | 145.7834 | 134.5654 | 197.7752 | 119.5719 | 109.8467 |
| ENSMUSG00000078486 | 0.076746008 | 0.732343 | 0.832274 | 93.56306 | 76.78858 | 73.70149 | 71.59775 | 65.33246 | 99.44629 | 68.46453 | 66.34306 |
| ENSMUSG00000078490 | 0.352597861 | 0.210924 | 0.368359 | 43.26034 | 74.38894 | 58.79557 | 65.55939 | 42.9049 | 61.45558 | 50.14304 | 34.80291 |
| ENSMUSG00000078495 | 0.036432969 | 0.777296 | 0.862728 | 314.895 | 403.14 | 329.5864 | 326.9343 | 300.3343 | 379.9072 | 329.7869 | 328.4525 |
| ENSMUSG00000078496 | -0.226517052 | 0.56326 | 0.708577 | 17.10292 | 22.79661 | 15.73403 | 19.84034 | 32.17867 | 15.64324 | 22.17865 | 17.40146 |
| ENSMUSG00000078502 | -0.026288981 | 0.931686 | 0.95966 | 109.6599 | 118.7823 | 77.01392 | 78.49874 | 111.1627 | 143.0239 | 67.50024 | 68.51824 |
| ENSMUSG00000078515 | 0.056174399 | 0.443298 | 0.606369 | 1821.964 | 1665.352 | 1594.933 | 1671.764 | 1645.988 | 1566.558 | 1546.72 | 1735.795 |
| ENSMUSG00000078517 | -0.053917934 | 0.575485 | 0.717646 | 801.8253 | 913.0642 | 761.8581 | 756.5208 | 909.7789 | 776.575 | 846.6459 | 816.7809 |
| ENSMUSG00000078519 | -0.102349856 | 0.676105 | 0.792619 | 60.36326 | 75.58876 | 55.48314 | 48.30692 | 53.63112 | 79.33356 | 66.53595 | 56.55474 |
| ENSMUSG00000078521 | -0.88048701 | 0.005632 | 0.028056 | 55.33299 | 88.78679 | 48.85829 | 34.50494 | 78.0089 | 94.9768 | 129.2147 | 113.1095 |
| ENSMUSG00000078532 | -0.287862109 | 0.060773 | 0.156433 | 329.9858 | 338.3497 | 349.461 | 367.4776 | 470.0037 | 317.3342 | 406.93 | 497.0291 |
| ENSMUSG00000078546 | 0.400842319 | 0.00967 | 0.041541 | 247.4894 | 292.7565 | 256.7131 | 276.9021 | 200.8729 | 254.7613 | 187.0721 | 170.7518 |
| ENSMUSG00000078552 | -0.568449935 | 0.15008 | 0.290039 | 10.06054 | 21.59679 | 19.04645 | 14.6646 | 20.47734 | 21.23011 | 31.82154 | 22.83941 |
| ENSMUSG00000078566 | -0.171089648 | 0.346095 | 0.514916 | 1135.835 | 791.8822 | 832.2472 | 1243.903 | 1103.826 | 888.3124 | 1266.112 | 1250.73 |
| ENSMUSG00000078570 | -0.532967548 | 6.90E-05 | 0.001019 | 433.6094 | 469.1302 | 486.0986 | 483.0692 | 665.0259 | 532.9874 | 723.2169 | 787.4159 |
| ENSMUSG00000078572 | -0.617058324 | 1.23E-06 | 4.42E-05 | 323.9495 | 326.3515 | 414.8814 | 370.9281 | 539.2366 | 505.0531 | 529.3948 | 634.0656 |
| ENSMUSG00000078578 | -0.100081179 | 0.073411 | 0.178414 | 8972.999 | 9349.009 | 9144.782 | 9965.889 | 9856.425 | 10484.32 | 9706.535 | 10078.71 |
| ENSMUSG00000078580 | -0.081668442 | 0.493154 | 0.650107 | 262.5802 | 281.9581 | 261.6817 | 219.1064 | 281.8072 | 260.3482 | 270.001 | 269.7226 |
| ENSMUSG00000078584 | -0.043348921 | 0.600898 | 0.738753 | 840.0554 | 735.4906 | 843.8407 | 822.0802 | 815.1931 | 783.2792 | 858.2174 | 887.4743 |
| ENSMUSG00000078588 | 0.997420544 | 0.024539 | 0.081868 | 43.26034 | 70.78947 | 38.09291 | 36.23019 | 14.62667 | 41.34284 | 13.50005 | 25.0146 |
| ENSMUSG00000078591 | 1.293392479 | 0.015209 | 0.058284 | 13.07871 | 17.99732 | 9.93728 | 17.25247 | 7.80089 | 5.58687 | 5.785735 | 4.350364 |
| ENSMUSG00000078607 | 1.107288742 | 2.39E-07 | 1.17E-05 | 204.229 | 226.7663 | 183.8397 | 242.3972 | 116.0382 | 126.2633 | 84.85745 | 70.69342 |
| ENSMUSG00000078612 | -0.430833824 | 0.159054 | 0.302057 | 60.36326 | 28.79572 | 53.82693 | 43.13117 | 49.73068 | 67.04245 | 81.96458 | 53.29196 |
| ENSMUSG00000078619 | 0.130297119 | 0.037641 | 0.111832 | 2598.638 | 2937.163 | 2787.407 | 2793.175 | 2671.805 | 2543.143 | 2468.58 | 2469.919 |
| ENSMUSG00000078622 | -0.027744163 | 0.751347 | 0.845087 | 3164.041 | 2850.776 | 2889.264 | 2865.635 | 2879.504 | 2652.646 | 3213.011 | 3251.897 |
| ENSMUSG00000078630 | -0.55691125 | 0.23187 | 0.392425 | 13.07871 | 8.398751 | 8.281066 | 20.70296 | 16.57689 | 23.46486 | 15.42863 | 19.57664 |
| ENSMUSG00000078640 | 0.603689031 | 0.006963 | 0.032856 | 84.50857 | 116.3827 | 123.3879 | 114.7289 | 66.30757 | 94.9768 | 67.50024 | 60.9051 |
| ENSMUSG00000078650 | -3.253300152 | 5.91E-06 | 0.000153 | 140.8476 | 11.99822 | 78.67013 | 160.448 | 1343.703 | 496.1141 | 810.0029 | 1091.941 |
| ENSMUSG00000078651 | 0.293662896 | 0.12177 | 0.251139 | 214.2896 | 129.5807 | 175.5586 | 162.1732 | 140.416 | 159.7845 | 132.1076 | 125.073 |
| ENSMUSG00000078652 | -0.339852038 | 0.00027 | 0.002858 | 1778.704 | 1811.731 | 1855.787 | 1920.2 | 2383.172 | 2061.555 | 2211.115 | 2668.949 |
| ENSMUSG00000078653 | 0.027984142 | 0.918738 | 0.952073 | 63.38142 | 57.59143 | 43.06154 | 69.00988 | 68.25779 | 54.75133 | 65.57166 | 39.15328 |
| ENSMUSG00000078656 | -0.100210596 | 0.68324 | 0.798094 | 52.31483 | 103.1847 | 94.40416 | 87.12497 | 102.3867 | 89.38993 | 90.64318 | 78.30656 |
| ENSMUSG00000078670 | -0.130620844 | 0.438293 | 0.601907 | 633.8142 | 689.8974 | 963.088 | 649.5555 | 851.2722 | 654.7812 | 795.5386 | 914.6641 |
| ENSMUSG00000078671 | 0.305448952 | 0.000513 | 0.004674 | 4274.725 | 3672.654 | 4006.38 | 4069.858 | 3185.689 | 3425.869 | 3496.513 | 2859.277 |
| ENSMUSG00000078676 | 0.073948166 | 0.26314 | 0.42736 | 1701.238 | 1676.151 | 1577.543 | 1600.167 | 1563.103 | 1655.948 | 1461.862 | 1545.467 |
| ENSMUSG00000078681 | -0.249720597 | 0.005739 | 0.028486 | 670.0322 | 706.6949 | 785.8732 | 708.2139 | 848.3468 | 773.2229 | 902.5747 | 890.7371 |
| ENSMUSG00000078684 | 0.114566134 | 0.279542 | 0.445915 | 304.8345 | 273.5593 | 272.4471 | 270.0012 | 256.4543 | 245.8223 | 259.3938 | 272.9854 |
| ENSMUSG00000078695 | -0.456230027 | 0.002517 | 0.015498 | 394.3733 | 341.9491 | 493.5516 | 412.334 | 571.4152 | 479.3535 | 541.9305 | 664.5182 |
| ENSMUSG00000078700 | -0.009415071 | 0.959529 | 0.977106 | 119.7205 | 94.7859 | 92.74794 | 94.88858 | 102.3867 | 86.03781 | 97.39321 | 118.5474 |

| | | | | | | | | | | | |
|---|---|---|---|---|---|---|---|---|---|---|---|
| ENSMUSG00000078706 | -0.067111718 | 0.561849 | 0.707428 | 1313.907 | 1220.219 | 1238.848 | 1199.909 | 1343.703 | 1053.684 | 1509.113 | 1300.759 |
| ENSMUSG00000078713 | -0.562398015 | 3.80E-06 | 0.000109 | 1122.757 | 1157.828 | 1320.002 | 1261.156 | 1802.981 | 1486.108 | 1763.685 | 2129.503 |
| ENSMUSG00000078716 | 0.139724612 | 0.390744 | 0.558171 | 427.5731 | 404.3399 | 438.8965 | 464.9541 | 402.721 | 494.9967 | 388.6085 | 290.3868 |
| ENSMUSG00000078762 | 0.262029987 | 0.107913 | 0.232505 | 232.3986 | 285.5575 | 215.3077 | 239.8093 | 203.7983 | 248.057 | 171.6435 | 187.0657 |
| ENSMUSG00000078763 | 1.067815482 | 0.076907 | 0.18419 | 11.0666 | 16.7975 | 22.35888 | 9.488858 | 4.875557 | 4.469496 | 13.50005 | 5.437955 |
| ENSMUSG00000078765 | 0.213258803 | 0.137618 | 0.272753 | 168.0111 | 140.3791 | 180.5272 | 177.7004 | 139.4409 | 155.315 | 142.7148 | 140.2993 |
| ENSMUSG00000078768 | 0.155345181 | 0.343474 | 0.512173 | 313.889 | 322.752 | 296.4622 | 302.7808 | 290.5832 | 353.0902 | 251.6795 | 214.2554 |
| ENSMUSG00000078771 | 0.34155019 | 0.121667 | 0.251033 | 61.36932 | 70.78947 | 75.3577 | 54.34528 | 53.63112 | 55.8687 | 49.17875 | 47.85401 |
| ENSMUSG00000078772 | 1.030123802 | 0.159673 | 0.302776 | 13.07871 | 19.19714 | 10.76539 | 19.84034 | 2.925334 | 20.11273 | 5.785735 | 2.175182 |
| ENSMUSG00000078773 | 0.294902604 | 0.03073 | 0.096177 | 316.9071 | 350.3479 | 275.7595 | 294.1546 | 285.7076 | 260.3482 | 243.0009 | 216.4306 |
| ENSMUSG00000078776 | -0.009680995 | 0.987977 | 0.993251 | 4.024217 | 17.99732 | 10.76539 | 11.21411 | 3.900445 | 12.29112 | 10.60718 | 17.40146 |
| ENSMUSG00000078779 | 0.098113061 | 0.598557 | 0.736956 | 363.1856 | 345.5486 | 250.9163 | 289.8415 | 364.6916 | 226.8269 | 281.5724 | 291.4744 |
| ENSMUSG00000078784 | 0.191868777 | 0.0763 | 0.183168 | 1245.495 | 1341.4 | 1228.082 | 1432.818 | 1131.129 | 1350.905 | 1079.04 | 1034.299 |
| ENSMUSG00000078786 | 0.268813788 | 0.058137 | 0.151252 | 318.9192 | 296.3559 | 305.5713 | 294.1546 | 235.0018 | 317.3342 | 222.7508 | 234.9197 |
| ENSMUSG00000078787 | -0.237468452 | 0.629561 | 0.758916 | 19.11503 | 8.398751 | 26.49941 | 15.52722 | 11.70134 | 27.93435 | 16.39292 | 27.18978 |
| ENSMUSG00000078789 | -0.219468556 | 0.217056 | 0.375514 | 390.3491 | 376.744 | 408.2566 | 358.8514 | 537.2863 | 302.8084 | 449.3588 | 495.9415 |
| ENSMUSG00000078794 | 0.678032878 | 9.14E-07 | 3.46E-05 | 1287.75 | 1756.539 | 1485.623 | 1532.882 | 1040.444 | 1092.792 | 823.5029 | 832.0072 |
| ENSMUSG00000078812 | -0.448029889 | 3.42E-08 | 2.35E-06 | 9794.945 | 10520.04 | 11065.16 | 10231.58 | 14582.79 | 12648.67 | 13971.59 | 15563.43 |
| ENSMUSG00000078813 | -0.27335875 | 0.037107 | 0.110672 | 360.1675 | 275.959 | 343.6643 | 301.0556 | 399.7956 | 344.1512 | 372.2156 | 433.9488 |
| ENSMUSG00000078816 | 0.429734288 | 0.099849 | 0.220685 | 43.26034 | 49.19268 | 53.82693 | 63.83414 | 34.1289 | 49.16446 | 35.6787 | 38.06569 |
| ENSMUSG00000078851 | -0.137330208 | 0.248713 | 0.412161 | 287.7315 | 365.9456 | 361.0545 | 365.7524 | 385.169 | 406.7242 | 378.9656 | 348.0291 |
| ENSMUSG00000078853 | 0.213670815 | 0.381574 | 0.549538 | 133.8052 | 236.3648 | 217.792 | 131.9814 | 127.7396 | 147.4934 | 179.3578 | 165.3138 |
| ENSMUSG00000078861 | -0.069242399 | 0.645063 | 0.770202 | 210.2654 | 207.5691 | 168.1056 | 199.266 | 202.8232 | 233.5312 | 171.6435 | 215.343 |
| ENSMUSG00000078862 | -0.221175082 | 0.083845 | 0.195162 | 227.3683 | 190.7716 | 219.4483 | 189.7772 | 239.8774 | 231.2964 | 267.1081 | 226.2189 |
| ENSMUSG00000078864 | 0.110399594 | 0.643689 | 0.769625 | 66.39959 | 100.785 | 61.27989 | 79.36136 | 84.83468 | 71.51194 | 59.78593 | 67.43065 |
| ENSMUSG00000078866 | 0.512601378 | 0.007302 | 0.033914 | 335.0161 | 405.5397 | 303.087 | 404.5704 | 248.6534 | 336.3296 | 196.715 | 233.8321 |
| ENSMUSG00000078867 | 0.244624774 | 0.330381 | 0.498218 | 51.30877 | 85.18733 | 64.59232 | 55.2079 | 64.35735 | 52.51658 | 49.17875 | 48.9416 |
| ENSMUSG00000078870 | 0.050580497 | 0.770104 | 0.85724 | 232.3986 | 231.5656 | 189.6364 | 255.3366 | 231.1014 | 261.4655 | 216.0008 | 168.5766 |
| ENSMUSG00000078877 | -0.159837332 | 0.35778 | 0.5262 | 161.9748 | 130.7805 | 110.9663 | 113.8663 | 151.1423 | 132.9675 | 149.4648 | 142.4744 |
| ENSMUSG00000078878 | 0.069781832 | 0.60328 | 0.740618 | 206.2411 | 218.3675 | 206.1986 | 196.6782 | 208.6738 | 211.1837 | 162.9649 | 205.5547 |
| ENSMUSG00000078897 | -0.653121751 | 0.070487 | 0.173155 | 13.07871 | 28.79572 | 40.57722 | 30.19182 | 40.95467 | 44.69496 | 40.50014 | 52.20437 |
| ENSMUSG00000078902 | 0.425784112 | 0.235822 | 0.396749 | 53.32088 | 58.79126 | 28.98373 | 47.44429 | 19.50223 | 42.46022 | 35.6787 | 42.41605 |
| ENSMUSG00000078903 | 0.32343376 | 0.542384 | 0.691553 | 17.10292 | 9.598572 | 11.59349 | 6.900988 | 7.80089 | 6.704245 | 10.60718 | 10.87591 |
| ENSMUSG00000078908 | 0.05018389 | 0.582004 | 0.722845 | 637.8385 | 614.3086 | 618.5957 | 532.2387 | 561.6641 | 559.8044 | 595.9307 | 601.4379 |
| ENSMUSG00000078919 | 0.32899068 | 0.078749 | 0.187147 | 536.227 | 497.9259 | 517.5666 | 457.1905 | 339.3387 | 556.4523 | 395.3586 | 308.8759 |
| ENSMUSG00000078920 | 0.182251591 | 0.624038 | 0.754981 | 50.30272 | 73.18911 | 78.67013 | 26.74133 | 37.05423 | 49.16446 | 60.75022 | 54.37955 |
| ENSMUSG00000078921 | 0.473835548 | 0.467416 | 0.62788 | 9.054489 | 21.59679 | 11.59349 | 8.626235 | 9.751113 | 5.58687 | 17.3572 | 3.262773 |
| ENSMUSG00000078922 | 0.361980138 | 0.188172 | 0.340447 | 63.38142 | 77.9884 | 56.31125 | 68.14726 | 39.00445 | 71.51194 | 56.89306 | 39.15328 |
| ENSMUSG00000078923 | -0.376468654 | 0.012751 | 0.051239 | 179.0777 | 166.7752 | 221.9326 | 206.167 | 279.8569 | 217.8879 | 238.1794 | 271.8978 |
| ENSMUSG00000078931 | -0.390837699 | 0.25606 | 0.419812 | 29.17558 | 19.19714 | 27.32752 | 16.38985 | 35.10401 | 32.40385 | 23.14294 | 30.45255 |
| ENSMUSG00000078937 | -2.324285216 | 5.55E-08 | 3.55E-06 | 26.15741 | 20.39697 | 25.67131 | 20.70296 | 109.2125 | 40.22547 | 145.6077 | 170.7518 |
| ENSMUSG00000078941 | -0.335763019 | 0.012721 | 0.05116 | 1063.399 | 1135.031 | 1246.3 | 1205.085 | 1722.047 | 1133.017 | 1457.041 | 1556.343 |
| ENSMUSG00000078942 | -0.267432867 | 0.470369 | 0.6303 | 41.24823 | 21.59679 | 30.63995 | 17.25247 | 31.20356 | 26.81698 | 42.42872 | 32.62773 |
| ENSMUSG00000078945 | -0.902405152 | 0.011143 | 0.046329 | 39.23612 | 20.39697 | 17.39024 | 20.70296 | 47.78045 | 33.52122 | 47.25017 | 53.29196 |
| ENSMUSG00000078949 | -2.840374991 | 8.75E-05 | 0.001214 | 11.0666 | 2.399643 | 52.99882 | 18.97772 | 98.48624 | 49.16446 | 194.7864 | 275.1605 |
| ENSMUSG00000078954 | -0.442717565 | 0.023396 | 0.078998 | 148.896 | 128.3809 | 166.4494 | 131.9814 | 176.4951 | 147.4934 | 236.2508 | 222.9562 |
| ENSMUSG00000078956 | 0.542476822 | 0.299016 | 0.467127 | 19.11503 | 19.19714 | 7.45296 | 12.07673 | 6.825779 | 11.17374 | 12.53576 | 8.700729 |
| ENSMUSG00000078963 | -0.664820726 | 0.03498 | 0.106043 | 39.23612 | 26.39607 | 48.85829 | 37.95543 | 48.75557 | 43.57759 | 69.42882 | 81.56933 |
| ENSMUSG00000078970 | 0.114422521 | 0.276649 | 0.442803 | 545.2815 | 621.5076 | 544.0661 | 605.5617 | 576.2908 | 464.8276 | 529.3948 | 566.635 |
| ENSMUSG00000078974 | -0.115741712 | 0.145301 | 0.283565 | 1564.415 | 1822.529 | 1883.943 | 1657.1 | 1946.322 | 1873.836 | 1862.042 | 1823.89 |
| ENSMUSG00000078994 | 0.278784912 | 0.182263 | 0.332856 | 203.223 | 172.7743 | 142.4343 | 151.8217 | 138.4658 | 97.21155 | 163.9292 | 151.1752 |
| ENSMUSG00000078995 | 0.792627247 | 0.031512 | 0.097778 | 64.38748 | 56.39161 | 43.06154 | 32.77969 | 33.15378 | 15.64324 | 37.60728 | 26.10219 |
| ENSMUSG00000079000 | -1.737244531 | 0.00257 | 0.015735 | 9.054489 | 3.599465 | 11.59349 | 4.313117 | 21.45245 | 12.29112 | 38.57157 | 23.927 |
| ENSMUSG00000079003 | 0.137196577 | 0.087657 | 0.201344 | 1011.085 | 1003.051 | 1102.21 | 1039.461 | 884.426 | 966.5286 | 901.6104 | 1031.036 |
| ENSMUSG00000079015 | 1.787961045 | 0.116379 | 0.244551 | 43.26034 | 1.199822 | 0.828107 | 31.91707 | 6.825779 | 6.704245 | 7.714313 | 1.087591 |
| ENSMUSG00000079017 | 0.842223718 | 0.32337 | 0.491365 | 99.59938 | 953.8581 | 819.8256 | 102.6522 | 177.4703 | 206.7142 | 511.0733 | 206.6423 |
| ENSMUSG00000079018 | -0.428571118 | 0.102886 | 0.225315 | 534.2149 | 303.5548 | 533.3007 | 483.0692 | 547.0374 | 458.1234 | 958.5034 | 532.9196 |
| ENSMUSG00000079020 | -0.213323432 | 0.063621 | 0.161622 | 894.3823 | 785.8831 | 1058.32 | 951.4737 | 1006.315 | 1046.98 | 1026.004 | 1205.051 |
| ENSMUSG00000079022 | -0.03996818 | 0.925507 | 0.955704 | 19.11503 | 15.59768 | 19.04645 | 12.07673 | 9.751113 | 20.11273 | 16.39292 | 21.75182 |
| ENSMUSG00000079036 | 0.174396509 | 0.308427 | 0.476744 | 1430.609 | 1368.996 | 1133.678 | 1369.846 | 1217.914 | 1554.267 | 1022.147 | 904.8758 |
| ENSMUSG00000079037 | -0.254872753 | 0.008441 | 0.03774 | 1490.973 | 1245.415 | 1349.814 | 1576.013 | 1635.262 | 1648.127 | 1844.685 | 1631.387 |
| ENSMUSG00000079038 | 0.588319866 | 0.051147 | 0.13821 | 138.8355 | 148.7779 | 116.763 | 117.3168 | 85.8098 | 139.6718 | 70.39311 | 51.11678 |
| ENSMUSG00000079042 | 0.4010706 | 0.116523 | 0.244705 | 436.6276 | 278.3586 | 737.843 | 526.2003 | 369.5672 | 364.264 | 406.93 | 359.9926 |
| ENSMUSG00000079043 | -0.080248426 | 0.671517 | 0.789547 | 165.999 | 167.975 | 116.763 | 135.4319 | 177.4703 | 168.7235 | 142.7148 | 128.3357 |

| | | | | | | | | | | | |
|---|---|---|---|---|---|---|---|---|---|---|---|
| ENSMUSG00000079051 | 0.918730512 | 0.192159 | 0.34549 | 15.09082 | 21.59679 | 7.45296 | 12.93935 | 4.875557 | 13.40849 | 10.60718 | 1.087591 |
| ENSMUSG00000079055 | 0.273903413 | 0.347789 | 0.516479 | 42.25428 | 27.5959 | 47.20208 | 48.30692 | 31.20356 | 40.22547 | 29.89296 | 36.9781 |
| ENSMUSG00000079056 | -0.182217492 | 0.325168 | 0.493053 | 94.56911 | 146.3782 | 153.1997 | 120.7673 | 147.2418 | 145.2586 | 141.7505 | 150.0876 |
| ENSMUSG00000079057 | 0.064546748 | 0.727797 | 0.829201 | 210.2654 | 200.3702 | 239.3228 | 179.4257 | 175.52 | 153.0803 | 228.5365 | 236.0073 |
| ENSMUSG00000079065 | 0.367006331 | 0.088893 | 0.203323 | 789.7527 | 794.2819 | 568.0811 | 582.2709 | 455.377 | 750.8754 | 513.0018 | 400.2335 |
| ENSMUSG00000079067 | -0.060875859 | 0.831479 | 0.897939 | 49.29666 | 43.19358 | 41.40533 | 62.97152 | 49.73068 | 71.51194 | 40.50014 | 44.59123 |
| ENSMUSG00000079083 | 0.202546802 | 0.236981 | 0.398082 | 451.7184 | 489.5272 | 300.6027 | 354.5383 | 322.7618 | 374.3203 | 358.7156 | 328.4525 |
| ENSMUSG00000079084 | 0.415490855 | 4.52E-06 | 0.000124 | 890.3581 | 892.6672 | 849.6374 | 996.3301 | 656.2499 | 744.1711 | 686.5739 | 635.1532 |
| ENSMUSG00000079103 | -2.822612692 | 2.86E-05 | 0.000519 | 5.030272 | 3.599465 | 5.796746 | 3.450494 | 19.50223 | 10.05637 | 35.6787 | 61.99269 |
| ENSMUSG00000079104 | -0.127937088 | 0.296236 | 0.464123 | 757.5589 | 776.2845 | 751.9208 | 780.6743 | 925.3806 | 657.016 | 837.003 | 929.8904 |
| ENSMUSG00000079105 | -0.269077065 | 0.612956 | 0.746981 | 35.2119 | 14.39786 | 14.90592 | 13.80198 | 10.72622 | 32.40385 | 33.75012 | 17.40146 |
| ENSMUSG00000079108 | -0.177245941 | 0.184735 | 0.336008 | 713.2925 | 808.6797 | 720.4528 | 707.3513 | 965.3602 | 651.4291 | 813.8601 | 901.613 |
| ENSMUSG00000079109 | -0.157665599 | 0.143928 | 0.28187 | 519.124 | 595.1115 | 464.5678 | 528.7882 | 607.4943 | 625.7295 | 582.4307 | 530.7445 |
| ENSMUSG00000079110 | 0.171685298 | 0.725848 | 0.828514 | 12.07265 | 33.595 | 34.78048 | 12.07673 | 21.45245 | 20.11273 | 15.42863 | 25.0146 |
| ENSMUSG00000079111 | 0.018866282 | 0.843197 | 0.905412 | 2467.851 | 3019.951 | 2712.049 | 2581.832 | 2536.265 | 2982.271 | 2462.795 | 2659.16 |
| ENSMUSG00000079139 | 0.051513604 | 0.537252 | 0.687647 | 999.012 | 1015.049 | 922.5108 | 952.3363 | 998.514 | 991.1108 | 911.2533 | 848.321 |
| ENSMUSG00000079144 | 0.065705504 | 0.550832 | 0.698561 | 333.004 | 326.3515 | 302.2589 | 342.4615 | 332.513 | 336.3296 | 295.0725 | 281.6861 |
| ENSMUSG00000079157 | 0.797037134 | 0.048642 | 0.133631 | 95.57516 | 59.99108 | 31.46805 | 102.6522 | 39.97956 | 55.8687 | 40.50014 | 30.45255 |
| ENSMUSG00000079164 | -0.601428858 | 0.035708 | 0.1077 | 21.12714 | 27.5959 | 28.98373 | 33.64232 | 39.97956 | 40.22547 | 51.10733 | 38.06569 |
| ENSMUSG00000079165 | 1.117479264 | 0.018301 | 0.066318 | 37.22401 | 20.39697 | 24.8432 | 59.52102 | 16.57689 | 23.46486 | 9.642892 | 16.31387 |
| ENSMUSG00000079173 | -0.013091075 | 0.970284 | 0.983446 | 35.2119 | 46.79304 | 64.59232 | 39.68068 | 42.9049 | 70.39457 | 28.92867 | 46.76642 |
| ENSMUSG00000079177 | 0.765620395 | 0.232902 | 0.393519 | 18.10898 | 20.39697 | 19.87456 | 7.763611 | 1.950223 | 16.76061 | 11.57147 | 8.700729 |
| ENSMUSG00000079179 | 0.154440518 | 0.20936 | 0.366525 | 1043.278 | 904.6654 | 854.606 | 1076.554 | 953.6589 | 937.4769 | 861.1102 | 733.0364 |
| ENSMUSG00000079184 | 0.296998476 | 0.091968 | 0.208232 | 1812.91 | 2025.299 | 1439.249 | 1648.473 | 1236.441 | 1919.649 | 1199.576 | 1281.182 |
| ENSMUSG00000079197 | -0.081808115 | 0.359916 | 0.528449 | 1425.579 | 1608.961 | 1731.571 | 1412.115 | 1659.639 | 1583.319 | 1585.291 | 1710.781 |
| ENSMUSG00000079215 | -0.187470725 | 0.001271 | 0.009359 | 4102.69 | 4459.737 | 4098.3 | 4294.14 | 4980.869 | 4538.774 | 4843.624 | 4937.664 |
| ENSMUSG00000079225 | -0.162833308 | 0.35715 | 0.525643 | 75.45408 | 92.38626 | 74.5296 | 87.12497 | 99.46135 | 86.03781 | 91.60747 | 90.27006 |
| ENSMUSG00000079227 | 0.740530745 | 0.193861 | 0.347234 | 17.10292 | 23.99643 | 25.67131 | 10.35148 | 21.45245 | 6.704245 | 5.785735 | 11.9635 |
| ENSMUSG00000079235 | 0.055893559 | 0.89861 | 0.938795 | 23.13925 | 13.19804 | 24.8432 | 10.35148 | 20.47734 | 16.76061 | 14.46434 | 17.40146 |
| ENSMUSG00000079242 | 0.238277172 | 0.431407 | 0.596051 | 39.23612 | 37.19447 | 29.81184 | 37.95543 | 29.25334 | 26.81698 | 41.46443 | 23.927 |
| ENSMUSG00000079259 | 0.737673844 | 0.009981 | 0.042515 | 194.1685 | 172.7743 | 202.8861 | 258.787 | 105.312 | 204.4795 | 100.2861 | 88.09488 |
| ENSMUSG00000079260 | -0.209559059 | 0.181931 | 0.332463 | 133.8052 | 106.7841 | 125.0441 | 107.8279 | 139.4409 | 128.498 | 129.2147 | 151.1752 |
| ENSMUSG00000079261 | -1.020952105 | 0.062314 | 0.159178 | 11.0666 | 13.19804 | 9.93728 | 8.626235 | 26.32801 | 14.52586 | 8.678602 | 36.9781 |
| ENSMUSG00000079262 | -4.638955917 | 3.34E-05 | 0.000581 | 11.0666 | 0 | 30.63995 | 6.900988 | 254.5041 | 54.75133 | 284.4653 | 625.3649 |
| ENSMUSG00000079277 | 0.375598533 | 0.011896 | 0.048671 | 449.7063 | 359.9465 | 481.9581 | 352.813 | 329.5876 | 337.447 | 332.6798 | 268.635 |
| ENSMUSG00000079283 | -0.178821846 | 0.090009 | 0.205164 | 353.1251 | 361.1463 | 367.6793 | 386.4553 | 376.393 | 461.4755 | 436.823 | 389.3576 |
| ENSMUSG00000079293 | -0.331398376 | 0.527283 | 0.680214 | 7.04238 | 8.398751 | 11.59349 | 10.35148 | 7.80089 | 7.821619 | 15.42863 | 16.31387 |
| ENSMUSG00000079297 | -0.291188665 | 0.367767 | 0.536239 | 30.18163 | 25.19625 | 24.01509 | 29.3292 | 30.22845 | 48.04709 | 27.0001 | 28.27737 |
| ENSMUSG00000079316 | 0.050474795 | 0.623311 | 0.754646 | 1319.943 | 1237.016 | 1169.287 | 1365.533 | 1143.806 | 1116.257 | 1415.576 | 1239.854 |
| ENSMUSG00000079317 | -0.10611983 | 0.396035 | 0.563287 | 230.3864 | 250.7627 | 247.6039 | 264.8254 | 229.1512 | 273.7567 | 269.0367 | 299.0875 |
| ENSMUSG00000079323 | 0.466097027 | 0.069081 | 0.170632 | 60.36326 | 56.39161 | 61.27989 | 58.6584 | 39.00445 | 60.3382 | 33.75012 | 39.15328 |
| ENSMUSG00000079330 | 0.626958028 | 0.059672 | 0.154234 | 47.28455 | 86.38715 | 57.96746 | 61.24627 | 47.78045 | 55.8687 | 35.6787 | 23.927 |
| ENSMUSG00000079334 | 0.073043342 | 0.543931 | 0.692808 | 372.2401 | 377.9438 | 403.2879 | 411.4714 | 361.7663 | 310.63 | 398.2514 | 417.635 |
| ENSMUSG00000079339 | -0.450589132 | 0.376482 | 0.544755 | 12.07265 | 11.99822 | 25.67131 | 11.21411 | 12.67645 | 14.52586 | 23.14294 | 33.71532 |
| ENSMUSG00000079343 | 0.47425371 | 0.12307 | 0.252685 | 55.33299 | 63.59054 | 36.43669 | 38.81806 | 25.35289 | 36.87335 | 38.57157 | 38.06569 |
| ENSMUSG00000079355 | 1.116166395 | 6.39E-05 | 0.000959 | 153.9263 | 253.1623 | 183.8397 | 167.349 | 76.05868 | 127.3806 | 90.64318 | 55.46715 |
| ENSMUSG00000079357 | -0.106935594 | 0.813971 | 0.88713 | 11.0666 | 16.7975 | 11.59349 | 15.52722 | 15.60178 | 21.23011 | 13.50005 | 8.700729 |
| ENSMUSG00000079363 | 0.039527275 | 0.936583 | 0.962308 | 14.08476 | 57.59143 | 45.54586 | 18.11509 | 22.42756 | 33.52122 | 41.46443 | 33.71532 |
| ENSMUSG00000079410 | 0.639702012 | 0.02441 | 0.081555 | 38.23007 | 43.19358 | 45.54586 | 34.50494 | 21.45245 | 30.1691 | 25.07152 | 27.18978 |
| ENSMUSG00000079414 | 0.806170533 | 0.004027 | 0.021935 | 90.54489 | 86.38715 | 95.23226 | 136.2945 | 71.18313 | 73.74669 | 51.10733 | 38.06569 |
| ENSMUSG00000079415 | -0.943753571 | 0.068738 | 0.16999 | 10.06054 | 10.79839 | 24.8432 | 10.35148 | 20.47734 | 16.76061 | 46.28588 | 25.0146 |
| ENSMUSG00000079418 | -0.135880063 | 0.165156 | 0.310557 | 457.7547 | 397.1409 | 409.9128 | 424.4108 | 437.825 | 434.6585 | 496.6089 | 487.2408 |
| ENSMUSG00000079419 | 0.138575896 | 0.620013 | 0.752034 | 119.7205 | 81.58786 | 105.9976 | 78.49874 | 81.90935 | 71.51194 | 130.179 | 66.34306 |
| ENSMUSG00000079426 | -0.157977448 | 0.03103 | 0.096765 | 3108.708 | 3425.49 | 3744.698 | 3366.819 | 3897.52 | 3651.579 | 3758.799 | 3919.678 |
| ENSMUSG00000079427 | -0.094786575 | 0.430978 | 0.595726 | 281.6952 | 335.95 | 317.9929 | 337.2858 | 372.4925 | 302.8084 | 362.5727 | 319.7518 |
| ENSMUSG00000079429 | 0.521526784 | 0.005021 | 0.025805 | 495.9848 | 477.529 | 422.3344 | 505.4974 | 292.5334 | 467.0624 | 276.751 | 289.2992 |
| ENSMUSG00000079434 | 0.004325292 | 0.989098 | 0.993825 | 40.24217 | 22.79661 | 35.60859 | 48.30692 | 43.88001 | 39.10809 | 29.89296 | 34.80291 |
| ENSMUSG00000079435 | -0.089974807 | 0.253038 | 0.416674 | 16540.54 | 19848.65 | 18813.75 | 17575.95 | 20507.57 | 19054.58 | 19864.36 | 18031.17 |
| ENSMUSG00000079442 | -0.292351905 | 0.001916 | 0.012726 | 377.2704 | 437.9349 | 450.49 | 408.0209 | 522.6597 | 487.1751 | 510.109 | 529.6569 |
| ENSMUSG00000079444 | -0.114813607 | 0.732069 | 0.832065 | 23.13925 | 55.19179 | 62.108 | 66.42201 | 61.43201 | 55.8687 | 54.00019 | 53.29196 |
| ENSMUSG00000079445 | -0.657602675 | 0.001513 | 0.010621 | 109.6599 | 106.7841 | 172.2462 | 107.8279 | 183.3209 | 177.6625 | 195.7507 | 229.4817 |
| ENSMUSG00000079450 | 0.408585287 | 0.084626 | 0.196338 | 158.9566 | 242.3639 | 176.3867 | 198.4034 | 128.7147 | 204.4795 | 145.6077 | 105.4963 |
| ENSMUSG00000079465 | -0.58809528 | 2.48E-10 | 3.85E-08 | 588.5418 | 553.1177 | 571.3936 | 565.0184 | 760.5868 | 834.6784 | 947.8962 | 882.0364 |
| ENSMUSG00000079467 | -0.07954477 | 0.787033 | 0.869061 | 30.18163 | 26.39607 | 24.8432 | 30.19182 | 31.20356 | 33.52122 | 24.10723 | 29.36496 |

| | | | | | | | | | | | |
|---|---|---|---|---|---|---|---|---|---|---|---|
| ENSMUSG00000079469 | 0.267904944 | 0.003361 | 0.019239 | 437.6336 | 434.3354 | 427.303 | 426.136 | 364.6916 | 381.0246 | 325.9297 | 362.1678 |
| ENSMUSG00000079470 | -0.029226061 | 0.857561 | 0.914643 | 362.1796 | 325.1516 | 301.4308 | 325.2091 | 308.1352 | 441.3628 | 303.7511 | 288.2116 |
| ENSMUSG00000079471 | -2.800639693 | 0.000508 | 0.004643 | 3.018163 | 1.199822 | 26.49941 | 6.038364 | 40.95467 | 24.58223 | 72.32169 | 122.8978 |
| ENSMUSG00000079477 | -0.223930861 | 0.024373 | 0.081492 | 5301.906 | 4832.881 | 5680.811 | 5366.381 | 6035.939 | 5311.996 | 6427.952 | 6966.021 |
| ENSMUSG00000079478 | -0.148591135 | 0.06679 | 0.166804 | 856.1523 | 945.4594 | 809.0602 | 915.2435 | 1014.116 | 938.5942 | 1011.539 | 938.5911 |
| ENSMUSG00000079480 | -0.037951 | 0.654778 | 0.777293 | 657.9595 | 677.8992 | 688.9847 | 631.4404 | 737.1841 | 620.1426 | 676.931 | 690.6203 |
| ENSMUSG00000079481 | 0.216650306 | 0.217842 | 0.37638 | 477.8758 | 526.7217 | 427.303 | 441.6632 | 385.169 | 546.3959 | 333.6441 | 346.9416 |
| ENSMUSG00000079484 | -0.197618503 | 0.171733 | 0.31896 | 133.8052 | 159.5763 | 162.3089 | 138.0198 | 181.3707 | 166.4887 | 152.3577 | 180.5401 |
| ENSMUSG00000079487 | 0.201289156 | 0.011392 | 0.047047 | 1799.831 | 1736.142 | 1697.619 | 1864.129 | 1682.067 | 1534.155 | 1567.934 | 1387.766 |
| ENSMUSG00000079491 | -0.410417767 | 0.349348 | 0.518043 | 13.07871 | 14.39786 | 20.70267 | 10.35148 | 15.60178 | 13.40849 | 26.03581 | 22.83941 |
| ENSMUSG00000079497 | -0.911877783 | 0.064792 | 0.163876 | 16.09687 | 15.59768 | 41.40533 | 31.91707 | 49.73068 | 22.34748 | 43.39301 | 83.74451 |
| ENSMUSG00000079499 | -0.283421527 | 0.371289 | 0.540291 | 64.38748 | 44.3934 | 53.82693 | 37.95543 | 63.38224 | 42.46022 | 49.17875 | 89.18247 |
| ENSMUSG00000079502 | -0.15978027 | 0.789548 | 0.8709 | 13.07871 | 15.59768 | 7.45296 | 7.763611 | 9.751113 | 23.46486 | 5.785735 | 9.78832 |
| ENSMUSG00000079505 | 0.036245751 | 0.931861 | 0.959787 | 19.11503 | 22.79661 | 21.53077 | 20.70296 | 11.70134 | 15.64324 | 33.75012 | 20.66423 |
| ENSMUSG00000079508 | -0.512215573 | 0.000697 | 0.005924 | 512.0817 | 419.9375 | 536.6131 | 428.7239 | 583.1166 | 578.7998 | 770.467 | 775.4524 |
| ENSMUSG00000079509 | 0.249461092 | 0.008344 | 0.037423 | 1744.498 | 1809.331 | 1562.637 | 1776.142 | 1340.778 | 1645.892 | 1350.005 | 1461.722 |
| ENSMUSG00000079511 | -0.038267194 | 0.846925 | 0.908032 | 108.6539 | 117.5825 | 91.91984 | 125.0804 | 120.9138 | 139.6718 | 95.46463 | 98.97079 |
| ENSMUSG00000079523 | -0.01608919 | 0.854981 | 0.913062 | 10847.28 | 12960.47 | 13745.74 | 12259.61 | 12544.81 | 12154.8 | 12136.54 | 13537.25 |
| ENSMUSG00000079547 | -0.752649905 | 0.001272 | 0.009359 | 79.47829 | 94.7859 | 115.1068 | 96.61383 | 188.1965 | 109.5027 | 153.322 | 200.1168 |
| ENSMUSG00000079550 | 0.375175163 | 0.150358 | 0.290364 | 81.4904 | 94.7859 | 105.1695 | 100.0643 | 62.40712 | 110.62 | 54.96448 | 67.43065 |
| ENSMUSG00000079553 | -0.020167941 | 0.774816 | 0.860668 | 1615.723 | 1552.569 | 1613.152 | 1572.563 | 1761.051 | 1609.019 | 1578.541 | 1494.35 |
| ENSMUSG00000079554 | 0.589534647 | 0.099329 | 0.219967 | 27.16347 | 33.595 | 19.87456 | 29.3292 | 13.65156 | 20.11273 | 17.3572 | 21.75182 |
| ENSMUSG00000079555 | 0.070207319 | 0.521298 | 0.674712 | 501.0151 | 517.1231 | 439.7246 | 472.7177 | 456.3521 | 490.5272 | 491.7875 | 396.9707 |
| ENSMUSG00000079557 | 0.140425088 | 0.078158 | 0.186006 | 940.6608 | 968.256 | 1080.679 | 1042.912 | 923.4304 | 901.7209 | 867.8602 | 970.1312 |
| ENSMUSG00000079559 | -0.048632201 | 0.848816 | 0.909191 | 54.32694 | 67.19001 | 96.06037 | 62.97152 | 61.43201 | 89.38993 | 73.28598 | 67.43065 |
| ENSMUSG00000079560 | 0.337667565 | 0.049384 | 0.134948 | 389.343 | 436.735 | 314.6805 | 346.7746 | 288.6329 | 374.3203 | 267.1081 | 245.7956 |
| ENSMUSG00000079562 | -0.221479202 | 0.017558 | 0.064409 | 2899.449 | 2969.558 | 3119.478 | 3045.061 | 3490.898 | 2994.563 | 3591.013 | 3954.481 |
| ENSMUSG00000079564 | 0.329016477 | 0.668425 | 0.787337 | 6.036326 | 15.59768 | 30.63995 | 14.6646 | 2.925334 | 32.40385 | 8.678602 | 9.78832 |
| ENSMUSG00000079588 | -0.209599262 | 0.713772 | 0.820362 | 7.04238 | 9.598572 | 8.281066 | 13.80198 | 4.875557 | 11.17374 | 10.60718 | 18.48905 |
| ENSMUSG00000079593 | 0.179277834 | 0.293601 | 0.461155 | 121.7326 | 93.58608 | 135.8095 | 120.7673 | 103.3618 | 105.0332 | 105.1075 | 105.4963 |
| ENSMUSG00000079597 | 0.101119941 | 0.835457 | 0.900443 | 123.7447 | 23.99643 | 34.78048 | 84.5371 | 78.98402 | 50.28183 | 66.53595 | 53.29196 |
| ENSMUSG00000079598 | -0.664361518 | 0.066335 | 0.166045 | 50.30272 | 22.79661 | 26.49941 | 37.09281 | 37.05423 | 78.21619 | 51.10733 | 51.11678 |
| ENSMUSG00000079602 | 0.182982345 | 0.302122 | 0.470457 | 199.1988 | 184.7725 | 226.9012 | 188.9145 | 137.4907 | 189.9536 | 216.9651 | 160.9635 |
| ENSMUSG00000079604 | 0.193634036 | 0.69951 | 0.810259 | 10.06054 | 9.598572 | 19.04645 | 13.80198 | 13.65156 | 15.64324 | 10.60718 | 6.525547 |
| ENSMUSG00000079605 | 0.307920151 | 0.551354 | 0.69903 | 24.1453 | 15.59768 | 14.07781 | 9.488858 | 19.50223 | 6.704245 | 13.50005 | 10.87591 |
| ENSMUSG00000079608 | 0.003148358 | 0.988037 | 0.993257 | 71.42986 | 59.99108 | 68.73285 | 86.26235 | 79.95913 | 60.3382 | 72.32169 | 73.95619 |
| ENSMUSG00000079610 | 0.054617372 | 0.630264 | 0.759281 | 322.9434 | 345.5486 | 355.2577 | 332.11 | 308.1352 | 362.0292 | 288.3225 | 349.1167 |
| ENSMUSG00000079614 | -0.054372044 | 0.477228 | 0.636625 | 2324.992 | 2403.243 | 2170.467 | 2308.38 | 2335.392 | 2651.529 | 2350.937 | 2220.861 |
| ENSMUSG00000079620 | -0.314351454 | 0.505938 | 0.661724 | 14.08476 | 16.7975 | 32.29616 | 12.07673 | 16.57689 | 26.81698 | 17.3572 | 33.71532 |
| ENSMUSG00000079641 | -0.073855068 | 0.360906 | 0.529527 | 13394.61 | 15643.27 | 16257.39 | 15047.6 | 16902.58 | 15476.75 | 16270.45 | 14861.93 |
| ENSMUSG00000079645 | 0.342625812 | 0.405737 | 0.572499 | 30.18163 | 33.595 | 16.56213 | 25.8787 | 16.57689 | 23.46486 | 29.89296 | 13.05109 |
| ENSMUSG00000079652 | 0.667009109 | 0.150486 | 0.290519 | 16.09687 | 20.39697 | 13.24971 | 18.11509 | 9.751113 | 16.76061 | 9.642892 | 6.525547 |
| ENSMUSG00000079654 | 0.333028687 | 0.253681 | 0.417294 | 53.32088 | 35.99465 | 44.71776 | 67.28463 | 46.80534 | 43.57759 | 38.57157 | 31.54014 |
| ENSMUSG00000079657 | 0.543460968 | 0.131129 | 0.263802 | 46.2785 | 41.99375 | 54.65504 | 54.34528 | 46.80534 | 18.99536 | 46.28588 | 22.83941 |
| ENSMUSG00000079658 | 0.041345818 | 0.720378 | 0.825128 | 1892.388 | 2166.878 | 1879.802 | 2003.012 | 1874.164 | 2344.251 | 1724.149 | 1774.949 |
| ENSMUSG00000079659 | 0.150155138 | 0.217156 | 0.375651 | 481.9 | 509.9242 | 396.6631 | 439.938 | 401.7459 | 461.4755 | 370.287 | 412.197 |
| ENSMUSG00000079669 | 0.68334021 | 7.38E-05 | 0.001074 | 248.4954 | 307.1543 | 241.8071 | 279.49 | 161.8685 | 208.949 | 168.7506 | 130.5109 |
| ENSMUSG00000079671 | -0.215767956 | 0.437672 | 0.601289 | 43.26034 | 29.99554 | 49.6864 | 35.36756 | 41.92979 | 40.22547 | 45.32159 | 57.64233 |
| ENSMUSG00000079677 | 0.094433672 | 0.853804 | 0.912279 | 31.18769 | 13.19804 | 13.24971 | 28.46658 | 16.57689 | 35.75597 | 12.53576 | 16.31387 |
| ENSMUSG00000079685 | -1.2239439 | 0.001188 | 0.008902 | 16.09687 | 15.59768 | 16.56213 | 8.626235 | 25.35289 | 29.05173 | 37.60728 | 40.24087 |
| ENSMUSG00000079697 | 0.930712502 | 0.040616 | 0.117993 | 56.33904 | 69.58965 | 31.46805 | 59.52102 | 16.57689 | 54.75133 | 24.10723 | 18.48905 |
| ENSMUSG00000079737 | -0.311288407 | 0.193637 | 0.3469 | 49.29666 | 91.18644 | 70.38906 | 90.57547 | 100.4365 | 87.15518 | 84.85745 | 101.146 |
| ENSMUSG00000079834 | 0.022757446 | 0.858433 | 0.915187 | 338.0343 | 309.554 | 364.3669 | 389.0432 | 398.8205 | 328.508 | 310.5011 | 343.6788 |
| ENSMUSG00000079845 | 0.677528378 | 0.513757 | 0.668338 | 6.036326 | 2.399643 | 34.78048 | 25.8787 | 1.950223 | 33.52122 | 3.857157 | 4.350364 |
| ENSMUSG00000079884 | -0.12201794 | 0.644354 | 0.77002 | 39.23612 | 61.1909 | 42.23344 | 56.07053 | 65.33246 | 42.46022 | 54.96448 | 52.20437 |
| ENSMUSG00000079906 | -0.274811606 | 0.396624 | 0.563821 | 26.15741 | 19.19714 | 28.15563 | 30.19182 | 33.15378 | 41.34284 | 25.07152 | 27.18978 |
| ENSMUSG00000079941 | -1.00204387 | 0.003159 | 0.018463 | 15.09082 | 16.7975 | 21.53077 | 14.6646 | 40.95467 | 26.81698 | 30.85725 | 38.06569 |
| ENSMUSG00000079942 | -0.187072742 | 0.513226 | 0.668037 | 41.24823 | 33.595 | 41.40533 | 25.8787 | 37.05423 | 48.04709 | 34.71441 | 42.41605 |
| ENSMUSG00000080021 | -0.025191882 | 0.961397 | 0.977983 | 14.08476 | 20.39697 | 6.624853 | 15.52722 | 18.52711 | 18.99536 | 8.678602 | 10.87591 |
| ENSMUSG00000080059 | 0.556632725 | 0.143594 | 0.281335 | 97.58727 | 87.58697 | 59.62368 | 71.59775 | 36.07912 | 92.74205 | 56.89306 | 29.36496 |
| ENSMUSG00000080076 | 0.001288388 | 0.997967 | 0.998833 | 8.048435 | 7.198929 | 16.56213 | 13.80198 | 14.62667 | 12.29112 | 10.60718 | 8.700729 |
| ENSMUSG00000080115 | -2.076249156 | 2.94E-10 | 4.41E-08 | 12.07265 | 27.5959 | 14.90592 | 13.80198 | 60.4569 | 68.15982 | 66.53595 | 88.09488 |
| ENSMUSG00000080242 | -1.220352349 | 0.343179 | 0.511957 | 343.0645 | 319.1525 | 31.46805 | 38.81806 | 666.9761 | 24.58223 | 34.71441 | 979.9196 |
| ENSMUSG00000080268 | -0.320193809 | 0.002178 | 0.013964 | 820.9404 | 716.2935 | 807.404 | 790.1631 | 1028.742 | 833.5611 | 970.0749 | 1083.241 |

| | | | | | | | | | | | |
|---|---|---|---|---|---|---|---|---|---|---|---|
| ENSMUSG00000080316 | 0.432054201 | 0.001263 | 0.00932 | 1419.543 | 1361.797 | 1170.115 | 1212.849 | 949.7584 | 1177.712 | 894.8603 | 804.8174 |
| ENSMUSG00000080364 | -0.111695577 | 0.739155 | 0.836877 | 43.26034 | 64.79036 | 28.98373 | 53.48266 | 45.83023 | 72.62932 | 46.28588 | 40.24087 |
| ENSMUSG00000080486 | 0.236392508 | 0.429203 | 0.594163 | 76.46013 | 86.38715 | 47.20208 | 55.2079 | 74.10846 | 56.98608 | 40.50014 | 52.20437 |
| ENSMUSG00000080715 | 0.717501157 | 0.1605 | 0.303969 | 39.23612 | 22.79661 | 24.8432 | 34.50494 | 12.67645 | 34.6386 | 20.25007 | 6.525547 |
| ENSMUSG00000080717 | -0.076089672 | 0.743209 | 0.839486 | 95.57516 | 113.983 | 149.0592 | 80.22398 | 104.3369 | 107.2679 | 116.679 | 134.8613 |
| ENSMUSG00000080727 | 0.131807779 | 0.476282 | 0.635732 | 80.48435 | 109.1838 | 108.482 | 119.042 | 96.53602 | 99.44629 | 87.75031 | 97.8832 |
| ENSMUSG00000080746 | -0.396861747 | 0.136586 | 0.271378 | 32.19374 | 32.39518 | 36.43669 | 29.3292 | 48.75557 | 33.52122 | 42.42872 | 46.76642 |
| ENSMUSG00000080772 | 1.139237898 | 0.080014 | 0.188964 | 16.09687 | 23.99643 | 10.76539 | 14.6646 | 5.850668 | 8.938993 | 13.50005 | 1.087591 |
| ENSMUSG00000080776 | -0.06441419 | 0.742333 | 0.839079 | 132.7992 | 172.7743 | 135.8095 | 138.8824 | 159.9183 | 192.1883 | 135.0005 | 118.5474 |
| ENSMUSG00000080783 | -0.052142512 | 0.914052 | 0.949091 | 10.06054 | 8.398751 | 25.67131 | 17.25247 | 16.57689 | 14.52586 | 19.28578 | 14.13868 |
| ENSMUSG00000080797 | -0.381769997 | 0.391037 | 0.558417 | 17.10292 | 17.99732 | 8.281066 | 11.21411 | 22.42756 | 15.64324 | 13.50005 | 18.48905 |
| ENSMUSG00000080848 | 0.051479539 | 0.728586 | 0.829664 | 203.223 | 255.562 | 266.6503 | 225.1447 | 224.2756 | 269.2872 | 219.8579 | 204.4671 |
| ENSMUSG00000080850 | 0.914559889 | 0.052858 | 0.141465 | 19.11503 | 19.19714 | 25.67131 | 30.19182 | 11.70134 | 22.34748 | 6.750024 | 9.78832 |
| ENSMUSG00000080875 | 0.051201667 | 0.814643 | 0.887294 | 52.31483 | 62.39072 | 68.73285 | 67.28463 | 68.25779 | 49.16446 | 62.6788 | 61.99269 |
| ENSMUSG00000080885 | 0.991295165 | 0.065463 | 0.164684 | 28.16952 | 25.19625 | 15.73403 | 18.11509 | 6.825779 | 20.11273 | 12.53576 | 4.350364 |
| ENSMUSG00000080893 | 2.066561877 | 0.000129 | 0.001623 | 18.10898 | 93.58608 | 109.3101 | 96.61383 | 18.52711 | 10.05637 | 22.17865 | 25.0146 |
| ENSMUSG00000080896 | -0.439215712 | 0.399636 | 0.567034 | 5.030272 | 15.59768 | 14.07781 | 7.763611 | 10.72622 | 18.99536 | 12.53576 | 15.22628 |
| ENSMUSG00000080902 | 0.110083243 | 0.80268 | 0.87942 | 14.08476 | 11.99822 | 11.59349 | 15.52722 | 15.60178 | 13.40849 | 11.57147 | 8.700729 |
| ENSMUSG00000080921 | -0.100195189 | 0.422109 | 0.587229 | 243.4652 | 244.7636 | 257.5412 | 213.068 | 272.0561 | 223.4748 | 262.2867 | 268.635 |
| ENSMUSG00000080942 | 0.590185365 | 0.099923 | 0.2207 | 24.1453 | 31.19536 | 34.78048 | 31.05445 | 11.70134 | 22.34748 | 26.03581 | 20.66423 |
| ENSMUSG00000081004 | -0.156488927 | 0.764874 | 0.853994 | 20.12109 | 4.799286 | 10.76539 | 15.52722 | 11.70134 | 14.52586 | 19.28578 | 11.9635 |
| ENSMUSG00000081044 | -3.036668775 | 1.86E-05 | 0.000371 | 3.018163 | 1.199822 | 10.76539 | 7.763611 | 22.42756 | 103.9158 | 38.57157 | 27.18978 |
| ENSMUSG00000081046 | 0.740942553 | 0.297913 | 0.465838 | 15.09082 | 15.59768 | 16.56213 | 7.763611 | 0.975111 | 10.05637 | 5.785735 | 16.31387 |
| ENSMUSG00000081049 | -0.205828079 | 0.222237 | 0.381505 | 167.005 | 147.578 | 226.0731 | 160.448 | 195.0223 | 191.071 | 212.1436 | 213.1679 |
| ENSMUSG00000081051 | -0.244480192 | 0.022658 | 0.077026 | 334.01 | 383.9429 | 332.8989 | 390.7684 | 438.8001 | 414.5458 | 457.0731 | 394.7956 |
| ENSMUSG00000081087 | -0.002987607 | 0.991031 | 0.994824 | 33.19979 | 49.19268 | 47.20208 | 53.48266 | 47.78045 | 41.34284 | 54.96448 | 39.15328 |
| ENSMUSG00000081094 | -0.111124099 | 0.397961 | 0.565157 | 366.2038 | 489.5272 | 481.13 | 466.6793 | 523.6348 | 516.2268 | 487.9303 | 419.8102 |
| ENSMUSG00000081111 | 0.444612975 | 0.429863 | 0.594675 | 18.10898 | 7.198929 | 7.45296 | 14.6646 | 12.67645 | 7.821619 | 6.750024 | 7.613138 |
| ENSMUSG00000081113 | -0.260531693 | 0.502468 | 0.658496 | 19.11503 | 15.59768 | 16.56213 | 12.07673 | 18.52711 | 24.58223 | 15.42863 | 17.40146 |
| ENSMUSG00000081123 | 0.237331937 | 0.665835 | 0.785418 | 10.06054 | 51.59233 | 15.73403 | 21.56559 | 22.42756 | 27.93435 | 15.42863 | 17.40146 |
| ENSMUSG00000081137 | 0.302955086 | 0.418066 | 0.583693 | 52.31483 | 67.19001 | 58.79557 | 75.91087 | 41.92979 | 93.85942 | 31.82154 | 39.15328 |
| ENSMUSG00000081176 | 0.118817749 | 0.722918 | 0.826653 | 23.13925 | 29.99554 | 19.04645 | 31.05445 | 22.42756 | 22.34748 | 27.96439 | 21.75182 |
| ENSMUSG00000081208 | 0.415991008 | 0.35556 | 0.524181 | 18.10898 | 16.7975 | 19.87456 | 25.01608 | 8.776002 | 22.34748 | 19.28578 | 9.78832 |
| ENSMUSG00000081214 | -0.002550047 | 0.992506 | 0.995642 | 62.37537 | 58.79126 | 100.2009 | 77.63611 | 51.6809 | 90.5073 | 81.00029 | 78.30656 |
| ENSMUSG00000081219 | -1.294391532 | 0.023281 | 0.07871 | 4.024217 | 16.7975 | 16.56213 | 4.313117 | 19.50223 | 21.23011 | 34.71441 | 26.10219 |
| ENSMUSG00000081248 | 1.080004513 | 0.050122 | 0.136298 | 22.1332 | 15.59768 | 14.07781 | 16.38985 | 2.925334 | 13.40849 | 10.60718 | 5.437955 |
| ENSMUSG00000081249 | 0.379299591 | 0.435839 | 0.599929 | 8.048435 | 21.59679 | 17.39024 | 21.56559 | 16.57689 | 16.76061 | 9.642892 | 9.78832 |
| ENSMUSG00000081265 | 1.365081307 | 0.046958 | 0.130236 | 18.10898 | 10.79839 | 8.281066 | 21.56559 | 5.850668 | 11.17374 | 4.821446 | 1.087591 |
| ENSMUSG00000081272 | -0.254594261 | 0.137849 | 0.273093 | 109.6599 | 157.1766 | 120.9036 | 131.1188 | 155.0427 | 130.7328 | 160.072 | 170.7518 |
| ENSMUSG00000081274 | 0.393706332 | 0.36252 | 0.531048 | 18.10898 | 15.59768 | 18.21835 | 24.15346 | 16.57689 | 21.23011 | 8.678602 | 11.9635 |
| ENSMUSG00000081281 | -0.300515362 | 0.534369 | 0.685422 | 11.0666 | 8.398751 | 11.59349 | 12.07673 | 20.47734 | 12.29112 | 8.678602 | 11.9635 |
| ENSMUSG00000081303 | 0.699564247 | 0.172362 | 0.319838 | 20.12109 | 11.99822 | 19.87456 | 11.21411 | 14.62667 | 11.17374 | 7.714313 | 5.437955 |
| ENSMUSG00000081344 | -0.1003085 | 0.462083 | 0.623179 | 213.2835 | 213.5682 | 254.2287 | 188.0519 | 226.2258 | 233.5312 | 251.6795 | 220.781 |
| ENSMUSG00000081378 | 0.449540261 | 0.305365 | 0.473796 | 34.20585 | 38.39429 | 12.4216 | 25.01608 | 22.42756 | 23.46486 | 14.46434 | 19.57664 |
| ENSMUSG00000081382 | 0.323511504 | 0.533725 | 0.685009 | 11.0666 | 11.99822 | 16.56213 | 8.626235 | 7.80089 | 13.40849 | 5.785735 | 11.9635 |
| ENSMUSG00000081400 | 0.178950258 | 0.452871 | 0.615342 | 67.40564 | 91.18644 | 52.99882 | 75.04824 | 64.35735 | 70.39457 | 63.64308 | 53.29196 |
| ENSMUSG00000081406 | -0.075870508 | 0.459645 | 0.621026 | 429.5852 | 541.1195 | 505.145 | 493.4206 | 553.8632 | 511.7573 | 522.6447 | 485.0656 |
| ENSMUSG00000081434 | -0.074630219 | 0.87859 | 0.927169 | 15.09082 | 16.7975 | 6.624853 | 10.35148 | 14.62667 | 12.29112 | 9.642892 | 14.13868 |
| ENSMUSG00000081453 | -0.140330797 | 0.635934 | 0.763518 | 32.19374 | 28.79572 | 31.46805 | 47.44429 | 44.85512 | 30.1691 | 40.50014 | 39.15328 |
| ENSMUSG00000081476 | -0.206749104 | 0.311682 | 0.480258 | 71.42986 | 91.18644 | 62.9361 | 69.8725 | 90.68535 | 79.33356 | 74.25027 | 94.62042 |
| ENSMUSG00000081485 | -0.276665116 | 0.199546 | 0.354519 | 69.41775 | 53.99197 | 73.70149 | 66.42201 | 64.35735 | 77.09881 | 83.89316 | 95.70802 |
| ENSMUSG00000081487 | 0.63928514 | 0.310918 | 0.479591 | 89.53884 | 56.39161 | 39.74912 | 43.9938 | 18.52711 | 90.5073 | 29.89296 | 8.700729 |
| ENSMUSG00000081534 | -0.33308255 | 0.007108 | 0.033261 | 1798.825 | 1702.547 | 2104.219 | 1915.887 | 2458.256 | 1892.832 | 2462.795 | 2663.511 |
| ENSMUSG00000081559 | -0.061022846 | 0.894523 | 0.936266 | 19.11503 | 11.99822 | 14.90592 | 31.05445 | 20.47734 | 30.1691 | 16.39292 | 14.13868 |
| ENSMUSG00000081564 | 0.769269515 | 0.129118 | 0.260865 | 60.36326 | 69.58965 | 44.71776 | 31.05445 | 24.37778 | 63.69032 | 16.39292 | 16.31387 |
| ENSMUSG00000081603 | -0.074926451 | 0.719127 | 0.824361 | 90.54489 | 92.38626 | 72.04528 | 94.88858 | 109.2125 | 101.681 | 73.28598 | 83.74451 |
| ENSMUSG00000081604 | 0.085144447 | 0.668086 | 0.787099 | 66.39959 | 81.58786 | 73.70149 | 70.73513 | 60.4569 | 80.45093 | 70.39311 | 64.16787 |
| ENSMUSG00000081648 | 1.140109378 | 0.041459 | 0.119727 | 51.30877 | 37.19447 | 17.39024 | 26.74133 | 6.825779 | 30.1691 | 13.50005 | 9.78832 |
| ENSMUSG00000081683 | 0.101751692 | 0.706126 | 0.815104 | 779.6921 | 962.2569 | 1675.26 | 800.5146 | 933.1815 | 708.4152 | 964.2892 | 1325.774 |
| ENSMUSG00000081684 | 0.385581422 | 0.415791 | 0.581704 | 21.12714 | 19.19714 | 14.90592 | 14.6646 | 14.62667 | 20.11273 | 5.785735 | 13.05109 |
| ENSMUSG00000081730 | 1.004110597 | 0.075044 | 0.181045 | 19.11503 | 40.79393 | 24.8432 | 21.56559 | 7.80089 | 29.05173 | 8.678602 | 7.613138 |
| ENSMUSG00000081738 | -0.376586359 | 0.346479 | 0.51528 | 13.07871 | 19.19714 | 15.73403 | 16.38985 | 17.552 | 31.28647 | 16.39292 | 18.48905 |
| ENSMUSG00000081752 | -0.236816892 | 0.544848 | 0.693414 | 12.07265 | 15.59768 | 15.73403 | 20.70296 | 23.40267 | 15.64324 | 20.25007 | 16.31387 |
| ENSMUSG00000081769 | 0.023809146 | 0.957474 | 0.975954 | 49.29666 | 40.79393 | 24.01509 | 20.70296 | 15.60178 | 48.04709 | 33.75012 | 34.80291 |

| | | | | | | | | | | | |
|---|---|---|---|---|---|---|---|---|---|---|---|
| ENSMUSG00000081788 | 0.034405734 | 0.918748 | 0.952073 | 35.2119 | 38.39429 | 18.21835 | 28.46658 | 33.15378 | 27.93435 | 28.92867 | 26.10219 |
| ENSMUSG00000081792 | 0.060117587 | 0.864803 | 0.91903 | 30.18163 | 44.3934 | 24.8432 | 26.74133 | 22.42756 | 42.46022 | 25.07152 | 30.45255 |
| ENSMUSG00000081824 | -0.596894856 | 0.171513 | 0.318725 | 11.0666 | 7.198929 | 15.73403 | 18.97772 | 24.37778 | 17.87799 | 16.39292 | 22.83941 |
| ENSMUSG00000081855 | 0.172399773 | 0.45149 | 0.613917 | 80.48435 | 67.19001 | 89.43552 | 87.9876 | 69.2329 | 97.21155 | 60.75022 | 63.08028 |
| ENSMUSG00000081865 | 0.171917716 | 0.73612 | 0.834773 | 12.07265 | 20.39697 | 13.24971 | 9.488858 | 13.65156 | 6.704245 | 9.642892 | 18.48905 |
| ENSMUSG00000081887 | 0.545088522 | 0.385781 | 0.553222 | 11.0666 | 15.59768 | 21.53077 | 5.175741 | 5.850668 | 7.821619 | 6.750024 | 16.31387 |
| ENSMUSG00000081895 | -0.268858316 | 0.065971 | 0.165519 | 175.0535 | 197.9706 | 231.0418 | 227.7326 | 276.9316 | 248.057 | 261.3224 | 217.5182 |
| ENSMUSG00000081926 | 0.114376922 | 0.729272 | 0.83001 | 23.13925 | 35.99465 | 30.63995 | 24.15346 | 30.22845 | 26.81698 | 28.92867 | 18.48905 |
| ENSMUSG00000081944 | 0.751860209 | 0.012941 | 0.051854 | 45.27245 | 44.3934 | 42.23344 | 66.42201 | 23.40267 | 37.99072 | 30.85725 | 26.10219 |
| ENSMUSG00000081953 | -0.217830568 | 0.607022 | 0.742971 | 12.07265 | 15.59768 | 19.04645 | 14.6646 | 18.52711 | 10.05637 | 23.14294 | 19.57664 |
| ENSMUSG00000081968 | 0.812346058 | 0.324724 | 0.492692 | 26.15741 | 16.7975 | 14.90592 | 18.11509 | 3.900445 | 33.52122 | 2.892867 | 3.262773 |
| ENSMUSG00000081984 | -0.518514875 | 0.014736 | 0.056929 | 97.58727 | 118.7823 | 135.8095 | 73.323 | 155.0427 | 145.2586 | 145.6077 | 163.1387 |
| ENSMUSG00000081992 | -0.255037815 | 0.453266 | 0.615723 | 22.1332 | 29.99554 | 17.39024 | 23.29083 | 31.20356 | 20.11273 | 30.85725 | 27.18978 |
| ENSMUSG00000081996 | 0.39231533 | 0.225557 | 0.385381 | 43.26034 | 51.59233 | 54.65504 | 35.36756 | 36.07912 | 48.04709 | 21.21436 | 35.89051 |
| ENSMUSG00000081998 | 0.455570404 | 0.224238 | 0.383974 | 46.2785 | 49.19268 | 37.2648 | 31.05445 | 19.50223 | 31.28647 | 47.25017 | 20.66423 |
| ENSMUSG00000081999 | -0.34739183 | 0.174794 | 0.322987 | 36.21796 | 46.79304 | 48.85829 | 39.68068 | 53.63112 | 69.27719 | 52.07161 | 43.50364 |
| ENSMUSG00000082016 | -0.609861173 | 0.202939 | 0.358657 | 16.09687 | 14.39786 | 31.46805 | 14.6646 | 38.02934 | 12.29112 | 36.64299 | 30.45255 |
| ENSMUSG00000082029 | 0.420892983 | 0.139376 | 0.27532 | 42.25428 | 44.3934 | 32.29616 | 39.68068 | 22.42756 | 30.1691 | 31.82154 | 33.71532 |
| ENSMUSG00000082035 | 0.490664601 | 0.146763 | 0.285743 | 31.18769 | 37.19447 | 31.46805 | 30.19182 | 24.37778 | 33.52122 | 18.32149 | 16.31387 |
| ENSMUSG00000082044 | -0.161621793 | 0.164498 | 0.30951 | 223.3441 | 261.5611 | 252.5725 | 281.2153 | 277.9067 | 287.1651 | 301.8225 | 272.9854 |
| ENSMUSG00000082045 | -0.239786375 | 0.16788 | 0.313889 | 151.9142 | 170.3747 | 154.0278 | 180.2883 | 241.8276 | 154.1976 | 180.3221 | 197.9416 |
| ENSMUSG00000082063 | 0.048078492 | 0.890028 | 0.933475 | 19.11503 | 23.99643 | 24.8432 | 17.25247 | 20.47734 | 17.87799 | 24.10723 | 19.57664 |
| ENSMUSG00000082064 | 0.268756447 | 0.490302 | 0.647606 | 20.12109 | 16.7975 | 14.90592 | 15.52722 | 14.62667 | 14.52586 | 13.50005 | 13.05109 |
| ENSMUSG00000082078 | 1.099213651 | 0.045189 | 0.126893 | 52.31483 | 47.99286 | 33.95237 | 36.23019 | 12.67645 | 46.92971 | 11.57147 | 8.700729 |
| ENSMUSG00000082088 | -0.859086462 | 0.119883 | 0.248827 | 11.0666 | 3.599465 | 17.39024 | 6.900988 | 11.70134 | 22.34748 | 17.3572 | 20.66423 |
| ENSMUSG00000082100 | 0.024350562 | 0.972824 | 0.98497 | 37.22401 | 44.3934 | 19.87456 | 37.09281 | 53.63112 | 68.15982 | 6.750024 | 7.613138 |
| ENSMUSG00000082107 | -0.158102392 | 0.733876 | 0.833297 | 14.08476 | 7.198929 | 17.39024 | 12.07673 | 18.52711 | 12.29112 | 15.42863 | 10.87591 |
| ENSMUSG00000082127 | 0.189422738 | 0.408613 | 0.574884 | 462.785 | 583.1133 | 390.0382 | 523.6125 | 435.8748 | 615.6731 | 368.3585 | 298 |
| ENSMUSG00000082132 | 1.102770127 | 0.155817 | 0.297848 | 21.12714 | 23.99643 | 7.45296 | 16.38985 | 1.950223 | 21.23011 | 5.785735 | 3.262773 |
| ENSMUSG00000082144 | 0.261614211 | 0.699302 | 0.810259 | 8.048435 | 21.59679 | 9.93728 | 18.11509 | 5.850668 | 27.93435 | 5.785735 | 8.700729 |
| ENSMUSG00000082152 | -1.699251457 | 0.008188 | 0.036914 | 2.012109 | 7.198929 | 19.87456 | 15.52722 | 42.9049 | 15.64324 | 32.78583 | 55.46715 |
| ENSMUSG00000082163 | 0.067664978 | 0.712701 | 0.819906 | 134.8113 | 165.5754 | 149.0592 | 186.3267 | 157.968 | 176.5451 | 117.6433 | 155.5255 |
| ENSMUSG00000082171 | -0.807216922 | 0.106578 | 0.230815 | 10.06054 | 8.398751 | 4.96864 | 8.626235 | 18.52711 | 13.40849 | 10.60718 | 13.05109 |
| ENSMUSG00000082193 | -0.325333814 | 0.302945 | 0.471231 | 20.12109 | 29.99554 | 24.8432 | 28.46658 | 39.00445 | 23.46486 | 32.78583 | 33.71532 |
| ENSMUSG00000082226 | 0.999477218 | 0.003927 | 0.021557 | 28.16952 | 33.595 | 37.2648 | 49.16954 | 21.45245 | 22.34748 | 14.46434 | 16.31387 |
| ENSMUSG00000082229 | 0.701774533 | 0.014654 | 0.0567 | 48.29061 | 45.59322 | 59.62368 | 43.9938 | 27.30312 | 22.34748 | 31.82154 | 40.24087 |
| ENSMUSG00000082264 | 0.328778807 | 0.529629 | 0.681737 | 7.04238 | 15.59768 | 11.59349 | 18.11509 | 7.80089 | 11.17374 | 7.714313 | 15.22628 |
| ENSMUSG00000082284 | 0.097388253 | 0.732687 | 0.832561 | 64.38748 | 92.38626 | 80.32634 | 99.2017 | 62.40712 | 120.6764 | 64.60737 | 67.43065 |
| ENSMUSG00000082286 | 0.582408658 | 0.003047 | 0.017963 | 224.3501 | 268.76 | 274.1033 | 322.6212 | 165.7689 | 230.1791 | 196.715 | 135.9489 |
| ENSMUSG00000082307 | -0.068811913 | 0.91323 | 0.948699 | 15.09082 | 17.99732 | 3.312427 | 13.80198 | 7.80089 | 8.938993 | 22.17865 | 13.05109 |
| ENSMUSG00000082319 | -0.16380672 | 0.642132 | 0.768609 | 19.11503 | 19.19714 | 23.18699 | 21.56559 | 19.50223 | 22.34748 | 19.28578 | 32.62773 |
| ENSMUSG00000082361 | -0.403803031 | 0.045836 | 0.128058 | 313.889 | 284.3577 | 496.864 | 350.2251 | 470.0037 | 449.1844 | 591.1093 | 404.5839 |
| ENSMUSG00000082383 | 0.628027975 | 0.090004 | 0.205164 | 46.2785 | 34.79482 | 31.46805 | 34.50494 | 23.40267 | 36.87335 | 13.50005 | 21.75182 |
| ENSMUSG00000082399 | 0.195241385 | 0.753452 | 0.846727 | 21.12714 | 14.39786 | 14.90592 | 5.175741 | 6.825779 | 23.46486 | 9.642892 | 8.700729 |
| ENSMUSG00000082419 | 1.07885643 | 0.085564 | 0.19789 | 17.10292 | 21.59679 | 14.90592 | 11.21411 | 7.80089 | 15.64324 | 1.928578 | 5.437955 |
| ENSMUSG00000082424 | -0.211110138 | 0.572284 | 0.715006 | 44.26639 | 39.59411 | 39.74912 | 44.85642 | 66.30757 | 18.99536 | 55.92877 | 53.29196 |
| ENSMUSG00000082454 | 0.071620171 | 0.779128 | 0.863784 | 44.26639 | 59.99108 | 37.2648 | 56.07053 | 41.92979 | 50.28183 | 48.21446 | 46.76642 |
| ENSMUSG00000082465 | 0.062241426 | 0.85872 | 0.915304 | 16.09687 | 29.99554 | 24.01509 | 25.01608 | 18.52711 | 21.23011 | 26.03581 | 25.0146 |
| ENSMUSG00000082475 | -0.302820985 | 0.489196 | 0.646795 | 15.09082 | 10.79839 | 10.76539 | 10.35148 | 17.552 | 14.52586 | 10.60718 | 15.22628 |
| ENSMUSG00000082536 | -0.215002484 | 0.023068 | 0.078219 | 968.8303 | 1107.435 | 901.8081 | 1020.484 | 1264.719 | 1064.858 | 1159.076 | 1146.321 |
| ENSMUSG00000082575 | 0.13809016 | 0.635145 | 0.762769 | 36.21796 | 55.19179 | 37.2648 | 63.83414 | 39.97956 | 40.22547 | 42.42872 | 52.20437 |
| ENSMUSG00000082585 | 0.077825147 | 0.722792 | 0.826559 | 90.54489 | 137.9795 | 92.74794 | 86.26235 | 93.61069 | 110.62 | 99.32178 | 80.48174 |
| ENSMUSG00000082588 | -0.006957011 | 0.990648 | 0.994691 | 9.054489 | 19.19714 | 5.796746 | 18.11509 | 21.45245 | 12.29112 | 11.57147 | 6.525547 |
| ENSMUSG00000082674 | 0.232937119 | 0.306739 | 0.475352 | 103.6236 | 91.18644 | 95.23226 | 75.04824 | 79.95913 | 72.62932 | 99.32178 | 57.64233 |
| ENSMUSG00000082691 | -0.580640872 | 0.079748 | 0.188683 | 28.16952 | 32.39518 | 28.98373 | 37.95543 | 44.85512 | 29.05173 | 47.25017 | 69.60583 |
| ENSMUSG00000082762 | -0.082751293 | 0.829686 | 0.896882 | 13.07871 | 21.59679 | 15.73403 | 15.52722 | 15.60178 | 16.76061 | 16.39292 | 20.66423 |
| ENSMUSG00000082809 | -0.351389844 | 0.258972 | 0.422872 | 30.18163 | 39.59411 | 23.18699 | 22.42821 | 39.97956 | 40.22547 | 34.71441 | 30.45255 |
| ENSMUSG00000082820 | 0.603551928 | 0.263167 | 0.427366 | 17.10292 | 25.19625 | 14.90592 | 18.97772 | 2.925334 | 14.52586 | 17.3572 | 15.22628 |
| ENSMUSG00000082860 | 0.105968326 | 0.737676 | 0.835871 | 37.22401 | 35.99465 | 28.15563 | 25.8787 | 28.27823 | 35.75597 | 32.78583 | 20.66423 |
| ENSMUSG00000082895 | 0.438291368 | 0.199629 | 0.354566 | 23.13925 | 21.59679 | 28.98373 | 25.01608 | 16.57689 | 21.23011 | 20.25007 | 15.22628 |
| ENSMUSG00000082896 | -0.090871297 | 0.821301 | 0.891669 | 18.10898 | 14.39786 | 19.04645 | 10.35148 | 17.552 | 16.76061 | 16.39292 | 15.22628 |
| ENSMUSG00000082901 | 0.131430934 | 0.703537 | 0.813207 | 58.35115 | 50.3925 | 26.49941 | 42.26855 | 49.73068 | 27.93435 | 48.21446 | 34.80291 |
| ENSMUSG00000082908 | 0.77661447 | 0.486525 | 0.644509 | 567.4147 | 569.9152 | 510.9418 | 536.5518 | 615.2952 | 624.6121 | 20.25007 | 15.22628 |
| ENSMUSG00000082922 | 0.636808574 | 0.104033 | 0.226954 | 41.24823 | 29.99554 | 33.12427 | 37.09281 | 38.02934 | 22.34748 | 17.3572 | 13.05109 |

| | | | | | | | | | | | |
|---|---|---|---|---|---|---|---|---|---|---|---|
| ENSMUSG00000082925 | -0.625524698 | 0.277765 | 0.444068 | 6.036326 | 8.398751 | 7.45296 | 10.35148 | 15.60178 | 20.11273 | 7.714313 | 6.525547 |
| ENSMUSG00000082935 | 0.031515575 | 0.933183 | 0.960345 | 23.13925 | 19.19714 | 16.56213 | 16.38985 | 22.42756 | 17.87799 | 13.50005 | 19.57664 |
| ENSMUSG00000082938 | 0.178314987 | 0.646835 | 0.771238 | 19.11503 | 22.79661 | 14.07781 | 18.11509 | 15.60178 | 18.99536 | 18.32149 | 11.9635 |
| ENSMUSG00000082969 | -0.012953087 | 0.981714 | 0.989807 | 12.07265 | 15.59768 | 28.98373 | 12.07673 | 8.776002 | 33.52122 | 12.53576 | 15.22628 |
| ENSMUSG00000082985 | 0.885789479 | 0.139783 | 0.275902 | 28.16952 | 16.7975 | 16.56213 | 18.97772 | 3.900445 | 18.99536 | 16.39292 | 4.350364 |
| ENSMUSG00000083012 | 0.040137988 | 0.65519 | 0.777633 | 702.2259 | 707.8947 | 806.5759 | 828.1186 | 775.2135 | 706.1804 | 748.2884 | 735.2116 |
| ENSMUSG00000083022 | 0.08240593 | 0.78949 | 0.8709 | 54.32694 | 46.79304 | 40.57722 | 52.62003 | 39.00445 | 71.51194 | 37.60728 | 35.89051 |
| ENSMUSG00000083061 | -0.239489343 | 0.079684 | 0.188618 | 128.775 | 149.9777 | 158.1684 | 141.4703 | 179.4205 | 175.4277 | 160.072 | 168.5766 |
| ENSMUSG00000083064 | 0.128377827 | 0.75852 | 0.849288 | 19.11503 | 14.39786 | 12.4216 | 17.25247 | 15.60178 | 17.87799 | 14.46434 | 9.78832 |
| ENSMUSG00000083087 | 0.785919413 | 0.099101 | 0.219744 | 14.08476 | 22.79661 | 13.24971 | 18.97772 | 13.65156 | 5.58687 | 11.57147 | 8.700729 |
| ENSMUSG00000083097 | -0.344009091 | 0.246016 | 0.408918 | 37.22401 | 31.19536 | 28.98373 | 30.19182 | 54.60623 | 36.87335 | 30.85725 | 39.15328 |
| ENSMUSG00000083104 | 0.396316316 | 0.446683 | 0.609311 | 18.10898 | 13.19804 | 12.4216 | 13.80198 | 13.65156 | 17.87799 | 5.785735 | 6.525547 |
| ENSMUSG00000083139 | -0.279406495 | 0.721882 | 0.826031 | 6.036326 | 10.79839 | 10.76539 | 12.07673 | 4.875557 | 33.52122 | 5.785735 | 4.350364 |
| ENSMUSG00000083161 | -0.133241729 | 0.801017 | 0.878562 | 28.16952 | 9.598572 | 10.76539 | 8.626235 | 18.52711 | 14.52586 | 17.3572 | 11.9635 |
| ENSMUSG00000083179 | 0.078315514 | 0.875728 | 0.925524 | 36.21796 | 14.39786 | 14.90592 | 12.93935 | 12.67645 | 27.93435 | 16.39292 | 17.40146 |
| ENSMUSG00000083240 | -0.168037201 | 0.760479 | 0.850871 | 12.07265 | 4.799286 | 13.24971 | 7.763611 | 14.62667 | 6.704245 | 9.642892 | 11.9635 |
| ENSMUSG00000083246 | 0.168024304 | 0.743553 | 0.839647 | 17.10292 | 27.5959 | 20.70267 | 18.11509 | 23.40267 | 32.40385 | 8.678602 | 9.78832 |
| ENSMUSG00000083261 | -0.094501255 | 0.811941 | 0.885545 | 19.11503 | 31.19536 | 11.59349 | 26.74133 | 22.42756 | 22.34748 | 26.03581 | 22.83941 |
| ENSMUSG00000083282 | -0.030154539 | 0.831577 | 0.897984 | 2015.127 | 1340.201 | 1315.861 | 1580.326 | 1617.71 | 1520.746 | 1637.363 | 1607.46 |
| ENSMUSG00000083306 | 1.2322468 | 0.108145 | 0.232788 | 43.26034 | 77.9884 | 34.78048 | 18.11509 | 12.67645 | 51.39921 | 5.785735 | 4.350364 |
| ENSMUSG00000083307 | 0.587372674 | 0.002986 | 0.017691 | 146.8839 | 151.1775 | 213.6515 | 172.5247 | 124.8142 | 131.8501 | 90.64318 | 109.8467 |
| ENSMUSG00000083311 | 0.468530718 | 0.310978 | 0.479643 | 20.12109 | 25.19625 | 14.90592 | 12.07673 | 17.552 | 13.40849 | 7.714313 | 13.05109 |
| ENSMUSG00000083327 | -0.391505071 | 0.008113 | 0.036656 | 105.6357 | 127.1811 | 129.1846 | 124.2178 | 173.5698 | 143.0239 | 161.0363 | 159.8759 |
| ENSMUSG00000083332 | 0.780337714 | 0.198287 | 0.352867 | 9.054489 | 16.7975 | 17.39024 | 21.56559 | 7.80089 | 12.29112 | 15.42863 | 2.175182 |
| ENSMUSG00000083337 | 0.152569203 | 0.555455 | 0.702541 | 51.30877 | 47.99286 | 46.37397 | 47.44429 | 39.00445 | 58.10345 | 33.75012 | 43.50364 |
| ENSMUSG00000083340 | 1.394889929 | 0.048699 | 0.133728 | 39.23612 | 57.59143 | 28.15563 | 34.50494 | 6.825779 | 40.22547 | 11.57147 | 2.175182 |
| ENSMUSG00000083364 | -0.678593518 | 0.034837 | 0.105698 | 26.15741 | 28.79572 | 22.35888 | 18.97772 | 47.78045 | 37.99072 | 27.0001 | 40.24087 |
| ENSMUSG00000083367 | -0.165109834 | 0.63613 | 0.763654 | 32.19374 | 17.99732 | 21.53077 | 25.8787 | 32.17867 | 31.28647 | 19.28578 | 27.18978 |
| ENSMUSG00000083396 | 0.141532802 | 0.549613 | 0.697734 | 89.53884 | 88.78679 | 55.48314 | 69.00988 | 72.15824 | 78.21619 | 60.75022 | 61.99269 |
| ENSMUSG00000083397 | 0.679548675 | 0.095395 | 0.213692 | 21.12714 | 19.19714 | 28.98373 | 37.95543 | 17.552 | 14.52586 | 23.14294 | 11.9635 |
| ENSMUSG00000083411 | -0.030207251 | 0.946591 | 0.969279 | 12.07265 | 17.99732 | 13.24971 | 8.626235 | 14.62667 | 10.05637 | 13.50005 | 14.13868 |
| ENSMUSG00000083421 | 2.176475393 | 0.013431 | 0.053214 | 63.38142 | 91.18644 | 29.81184 | 62.10889 | 4.875557 | 45.81234 | 1.928578 | 2.175182 |
| ENSMUSG00000083424 | -0.382605648 | 0.625396 | 0.755732 | 8.048435 | 2.399643 | 21.53077 | 4.313117 | 15.60178 | 12.29112 | 3.857157 | 16.31387 |
| ENSMUSG00000083477 | 0.007030303 | 0.989183 | 0.993825 | 10.06054 | 9.598572 | 8.281066 | 12.93935 | 6.825779 | 15.64324 | 7.714313 | 10.87591 |
| ENSMUSG00000083496 | -0.202531981 | 0.512993 | 0.668016 | 21.12714 | 33.595 | 26.49941 | 24.15346 | 36.07912 | 29.05173 | 27.0001 | 28.27737 |
| ENSMUSG00000083538 | 0.65332992 | 0.27574 | 0.441988 | 24.1453 | 27.5959 | 19.87456 | 25.01608 | 8.776002 | 36.87335 | 10.60718 | 5.437955 |
| ENSMUSG00000083563 | -0.341946082 | 0.208795 | 0.36613 | 39.23612 | 77.9884 | 52.99882 | 61.24627 | 81.90935 | 55.8687 | 70.39311 | 83.74451 |
| ENSMUSG00000083567 | -0.029601846 | 0.91357 | 0.948837 | 39.23612 | 64.79036 | 62.9361 | 45.71905 | 40.95467 | 58.10345 | 54.96448 | 63.08028 |
| ENSMUSG00000083596 | 0.051718617 | 0.88366 | 0.930391 | 20.12109 | 37.19447 | 43.06154 | 23.29083 | 35.10401 | 27.93435 | 28.92867 | 27.18978 |
| ENSMUSG00000083610 | 0.159974316 | 0.778365 | 0.86326 | 16.09687 | 4.799286 | 9.109173 | 12.07673 | 12.67645 | 5.58687 | 8.678602 | 10.87591 |
| ENSMUSG00000083619 | 0.077790046 | 0.758153 | 0.849263 | 44.26639 | 43.19358 | 47.20208 | 61.24627 | 56.55646 | 45.81234 | 41.46443 | 42.41605 |
| ENSMUSG00000083621 | -0.124437014 | 0.323058 | 0.491133 | 172.0353 | 185.9723 | 158.1684 | 178.5631 | 197.9476 | 187.7188 | 176.4649 | 193.5912 |
| ENSMUSG00000083626 | -0.856630149 | 0.147632 | 0.286841 | 2.012109 | 9.598572 | 9.109173 | 10.35148 | 20.47734 | 10.05637 | 10.60718 | 15.22628 |
| ENSMUSG00000083674 | 0.58992776 | 0.138543 | 0.274145 | 29.17558 | 83.98751 | 35.60859 | 48.30692 | 23.40267 | 31.28647 | 39.53586 | 35.89051 |
| ENSMUSG00000083716 | -0.295444893 | 0.006754 | 0.032129 | 230.3864 | 241.1641 | 242.6352 | 277.7648 | 308.1352 | 310.63 | 293.1439 | 307.7883 |
| ENSMUSG00000083718 | 0.633420202 | 0.000456 | 0.004272 | 183.1019 | 207.5691 | 150.7154 | 158.7227 | 113.1129 | 135.2023 | 106.0718 | 95.70802 |
| ENSMUSG00000083734 | 0.860677526 | 0.040851 | 0.118432 | 21.12714 | 26.39607 | 13.24971 | 22.42821 | 10.72622 | 12.29112 | 11.57147 | 10.87591 |
| ENSMUSG00000083757 | 0.226970084 | 0.567656 | 0.711923 | 15.09082 | 37.19447 | 32.29616 | 20.70296 | 27.30312 | 23.46486 | 20.25007 | 18.48905 |
| ENSMUSG00000083773 | -0.60577902 | 0.175421 | 0.323757 | 12.07265 | 7.198929 | 10.76539 | 10.35148 | 13.65156 | 18.99536 | 17.3572 | 11.9635 |
| ENSMUSG00000083783 | 0.942136792 | 0.077263 | 0.184614 | 19.11503 | 16.7975 | 9.109173 | 11.21411 | 8.776002 | 8.938993 | 4.821446 | 6.525547 |
| ENSMUSG00000083798 | -0.885836565 | 0.06315 | 0.160802 | 6.036326 | 8.398751 | 8.281066 | 12.07673 | 14.62667 | 16.76061 | 21.21436 | 11.9635 |
| ENSMUSG00000083811 | 0.094069899 | 0.824664 | 0.893455 | 34.20585 | 20.39697 | 20.70267 | 17.25247 | 30.22845 | 12.29112 | 25.07152 | 18.48905 |
| ENSMUSG00000083816 | -0.416375569 | 0.074297 | 0.179904 | 44.26639 | 45.59322 | 38.09291 | 40.5433 | 55.58134 | 48.04709 | 54.00019 | 66.34306 |
| ENSMUSG00000083827 | 1.04892684 | 0.075081 | 0.181049 | 88.53278 | 92.38626 | 62.108 | 69.00988 | 21.45245 | 98.32892 | 18.32149 | 13.05109 |
| ENSMUSG00000083833 | -0.072407887 | 0.810206 | 0.884279 | 25.15136 | 31.19536 | 34.78048 | 41.40593 | 31.20356 | 27.93435 | 40.50014 | 40.24087 |
| ENSMUSG00000083844 | -0.261369491 | 0.007285 | 0.03386 | 437.6336 | 459.5316 | 458.7711 | 394.2189 | 510.9583 | 483.823 | 544.8234 | 555.759 |
| ENSMUSG00000083863 | -0.814030286 | 0.034632 | 0.105247 | 21.12714 | 9.598572 | 16.56213 | 16.38985 | 33.15378 | 18.99536 | 28.92867 | 31.54014 |
| ENSMUSG00000083879 | 0.081593294 | 0.880907 | 0.928519 | 13.07871 | 7.198929 | 13.24971 | 9.488858 | 14.62667 | 11.17374 | 10.60718 | 4.350364 |
| ENSMUSG00000083897 | 1.314894795 | 0.085133 | 0.197167 | 25.15136 | 26.39607 | 2.48432 | 17.25247 | 5.850668 | 12.29112 | 2.892867 | 7.613138 |
| ENSMUSG00000083899 | 0.048238262 | 0.841587 | 0.904462 | 113.6841 | 122.3818 | 76.18581 | 88.85022 | 91.66046 | 128.498 | 83.89316 | 82.65692 |
| ENSMUSG00000083906 | 0.404804843 | 0.479568 | 0.638593 | 22.1332 | 20.39697 | 9.109173 | 13.80198 | 9.751113 | 23.46486 | 10.60718 | 5.437955 |
| ENSMUSG00000083907 | -0.198594368 | 0.58995 | 0.730011 | 14.08476 | 29.99554 | 25.67131 | 29.3292 | 25.35289 | 22.34748 | 27.96439 | 38.06569 |
| ENSMUSG00000083964 | 0.258722039 | 0.679848 | 0.795491 | 24.1453 | 9.598572 | 22.35888 | 8.626235 | 16.57689 | 24.58223 | 6.750024 | 6.525547 |
| ENSMUSG00000083992 | -0.275016885 | 0.116589 | 0.244797 | 148.896 | 124.7814 | 134.9814 | 117.3168 | 187.2214 | 136.3196 | 174.5363 | 137.0365 |

| | | | | | | | | | | | |
|---|---|---|---|---|---|---|---|---|---|---|---|
| ENSMUSG00000084007 | 0.590807802 | 0.223434 | 0.383136 | 22.1332 | 21.59679 | 20.70267 | 22.42821 | 22.42756 | 20.11273 | 9.642892 | 5.437955 |
| ENSMUSG00000084013 | -0.212029273 | 0.63352 | 0.761759 | 21.12714 | 17.99732 | 9.109173 | 14.6646 | 23.40267 | 21.23011 | 14.46434 | 13.05109 |
| ENSMUSG00000084025 | 2.368890969 | 0.013786 | 0.054173 | 32.19374 | 28.79572 | 8.281066 | 7.763611 | 0 | 10.05637 | 3.857157 | 1.087591 |
| ENSMUSG00000084033 | 0.152176557 | 0.767625 | 0.855731 | 9.054489 | 15.59768 | 8.281066 | 12.07673 | 11.70134 | 12.29112 | 10.60718 | 5.437955 |
| ENSMUSG00000084067 | -0.418117631 | 0.373114 | 0.542019 | 13.07871 | 4.799286 | 10.76539 | 14.6646 | 14.62667 | 14.52586 | 15.42863 | 14.13868 |
| ENSMUSG00000084085 | 1.104553546 | 0.010674 | 0.044781 | 27.16347 | 35.99465 | 20.70267 | 39.68068 | 13.65156 | 23.46486 | 9.642892 | 10.87591 |
| ENSMUSG00000084106 | -0.026050536 | 0.86864 | 0.921387 | 261.5741 | 359.9465 | 294.806 | 353.6756 | 358.841 | 362.0292 | 294.1082 | 277.3357 |
| ENSMUSG00000084111 | -0.015411496 | 0.924748 | 0.955253 | 116.7023 | 131.9804 | 132.4971 | 121.6299 | 143.3414 | 108.3853 | 139.8219 | 115.2847 |
| ENSMUSG00000084128 | -1.622346723 | 3.03E-05 | 0.00054 | 217.3077 | 128.3809 | 427.303 | 156.9975 | 723.5326 | 355.325 | 824.4672 | 963.6057 |
| ENSMUSG00000084131 | -0.266579053 | 0.372247 | 0.541259 | 27.16347 | 34.79482 | 33.12427 | 41.40593 | 40.95467 | 27.93435 | 45.32159 | 50.02919 |
| ENSMUSG00000084146 | 0.050828992 | 0.935989 | 0.962106 | 9.054489 | 19.19714 | 8.281066 | 6.900988 | 9.751113 | 16.76061 | 11.57147 | 3.262773 |
| ENSMUSG00000084159 | -0.258968949 | 0.070837 | 0.173853 | 135.8173 | 121.182 | 111.7944 | 115.5915 | 143.3414 | 140.7891 | 150.4291 | 143.562 |
| ENSMUSG00000084166 | -0.059553848 | 0.787754 | 0.869649 | 56.33904 | 81.58786 | 65.42042 | 64.69676 | 68.25779 | 61.45558 | 84.85745 | 63.08028 |
| ENSMUSG00000084221 | 0.120917673 | 0.824628 | 0.893455 | 8.048435 | 9.598572 | 10.76539 | 14.6646 | 5.850668 | 15.64324 | 6.750024 | 11.9635 |
| ENSMUSG00000084235 | 0.098062092 | 0.7516 | 0.845207 | 32.19374 | 32.39518 | 33.12427 | 38.81806 | 29.25334 | 40.22547 | 37.60728 | 20.66423 |
| ENSMUSG00000084260 | -0.322006474 | 0.400797 | 0.56809 | 21.12714 | 16.7975 | 12.4216 | 14.6646 | 16.57689 | 24.58223 | 21.21436 | 18.48905 |
| ENSMUSG00000084274 | 0.437538412 | 0.086509 | 0.19954 | 64.38748 | 64.79036 | 65.42042 | 50.89479 | 37.05423 | 60.3382 | 37.60728 | 46.76642 |
| ENSMUSG00000084284 | 0.082008572 | 0.828497 | 0.896134 | 15.09082 | 19.19714 | 20.70267 | 20.70296 | 21.45245 | 15.64324 | 21.21436 | 13.05109 |
| ENSMUSG00000084304 | -0.448795687 | 0.354829 | 0.523354 | 8.048435 | 7.198929 | 9.93728 | 12.93935 | 16.57689 | 11.17374 | 9.642892 | 15.22628 |
| ENSMUSG00000084319 | 0.038516287 | 0.748214 | 0.842785 | 1239.459 | 1534.572 | 1259.55 | 1461.284 | 1384.658 | 1501.751 | 1340.362 | 1121.306 |
| ENSMUSG00000084349 | -0.047027317 | 0.654759 | 0.777293 | 575.4631 | 650.3033 | 556.4877 | 602.9738 | 701.105 | 565.3913 | 619.0736 | 574.2481 |
| ENSMUSG00000084350 | 0.409620581 | 0.045635 | 0.127741 | 145.8779 | 189.5718 | 164.7932 | 155.2722 | 105.312 | 167.6061 | 120.5361 | 100.0584 |
| ENSMUSG00000084381 | 0.246341392 | 0.678735 | 0.794843 | 8.048435 | 10.79839 | 22.35888 | 12.07673 | 16.57689 | 15.64324 | 8.678602 | 4.350364 |
| ENSMUSG00000084383 | 1.961722996 | 0.021569 | 0.074347 | 62.37537 | 26.39607 | 8.281066 | 24.15346 | 3.900445 | 22.34748 | 2.892867 | 2.175182 |
| ENSMUSG00000084391 | 1.176426364 | 0.140979 | 0.277628 | 74.44802 | 92.38626 | 53.82693 | 29.3292 | 13.65156 | 80.45093 | 14.46434 | 2.175182 |
| ENSMUSG00000084399 | 1.165018922 | 0.087184 | 0.200658 | 19.11503 | 29.99554 | 18.21835 | 12.07673 | 4.875557 | 20.11273 | 1.928578 | 8.700729 |
| ENSMUSG00000084407 | 0.209710885 | 0.430467 | 0.595242 | 60.36326 | 107.9839 | 74.5296 | 54.34528 | 58.50668 | 69.27719 | 63.64308 | 64.16787 |
| ENSMUSG00000084411 | 1.675480533 | 0.013429 | 0.053214 | 92.557 | 57.59143 | 42.23344 | 72.46037 | 8.776002 | 59.22083 | 7.714313 | 7.613138 |
| ENSMUSG00000084416 | -0.153761028 | 0.20693 | 0.363517 | 355.1372 | 411.5388 | 476.1613 | 475.3055 | 488.5308 | 484.9404 | 492.7518 | 448.0875 |
| ENSMUSG00000084433 | 0.423061518 | 0.405215 | 0.572112 | 10.06054 | 9.598572 | 19.04645 | 25.8787 | 8.776002 | 15.64324 | 12.53576 | 11.9635 |
| ENSMUSG00000084519 | 0.473530923 | 0.403133 | 0.570045 | 11.0666 | 21.59679 | 13.24971 | 6.038364 | 9.751113 | 10.05637 | 11.57147 | 5.437955 |
| ENSMUSG00000084592 | 0.767332389 | 0.077023 | 0.184304 | 21.12714 | 26.39607 | 28.15563 | 25.8787 | 9.751113 | 26.81698 | 10.60718 | 13.05109 |
| ENSMUSG00000084603 | -0.50908716 | 0.331946 | 0.499788 | 11.0666 | 5.999108 | 8.281066 | 7.763611 | 16.57689 | 13.40849 | 9.642892 | 7.613138 |
| ENSMUSG00000084611 | 1.307616947 | 0.204834 | 0.360967 | 16.09687 | 25.19625 | 15.73403 | 18.11509 | 0.975111 | 25.6996 | 3.857157 | 0 |
| ENSMUSG00000084771 | -0.883499343 | 0.032801 | 0.100827 | 10.06054 | 13.19804 | 9.93728 | 11.21411 | 21.45245 | 14.52586 | 19.28578 | 26.10219 |
| ENSMUSG00000084772 | -0.112398652 | 0.85095 | 0.910548 | 8.048435 | 5.999108 | 14.90592 | 21.56559 | 7.80089 | 23.46486 | 14.46434 | 9.78832 |
| ENSMUSG00000084786 | -0.225451456 | 0.025234 | 0.083544 | 2300.846 | 2406.842 | 2375.838 | 2388.604 | 2974.089 | 2255.978 | 2895.76 | 2946.284 |
| ENSMUSG00000084790 | 0.222705048 | 0.436056 | 0.600086 | 60.36326 | 55.19179 | 52.99882 | 39.68068 | 36.07912 | 59.22083 | 49.17875 | 33.71532 |
| ENSMUSG00000084792 | 0.893736502 | 0.03067 | 0.096038 | 74.44802 | 69.58965 | 72.04528 | 98.33908 | 24.37778 | 83.80306 | 26.03581 | 35.89051 |
| ENSMUSG00000084797 | 0.098043638 | 0.821565 | 0.891669 | 29.17558 | 13.19804 | 31.46805 | 12.93935 | 17.552 | 20.11273 | 22.17865 | 21.75182 |
| ENSMUSG00000084799 | 0.434442662 | 0.008198 | 0.036938 | 179.0777 | 195.5709 | 162.3089 | 167.349 | 118.9636 | 149.7281 | 144.6434 | 106.5839 |
| ENSMUSG00000084800 | 0.110494381 | 0.86513 | 0.91922 | 12.07265 | 17.99732 | 15.73403 | 9.488858 | 3.900445 | 29.05173 | 8.678602 | 9.78832 |
| ENSMUSG00000084803 | -0.343837341 | 0.487162 | 0.645214 | 14.08476 | 19.19714 | 28.98373 | 19.84034 | 11.70134 | 17.87799 | 43.39301 | 31.54014 |
| ENSMUSG00000084808 | 0.216681156 | 0.139807 | 0.275907 | 149.9021 | 147.578 | 139.1219 | 126.8057 | 130.6649 | 124.0285 | 114.7504 | 114.1971 |
| ENSMUSG00000084817 | 0.209897072 | 0.330088 | 0.497965 | 80.48435 | 92.38626 | 78.67013 | 102.6522 | 73.13335 | 97.21155 | 75.21455 | 60.9051 |
| ENSMUSG00000084822 | 0.529776604 | 0.362244 | 0.530898 | 12.07265 | 9.598572 | 17.39024 | 10.35148 | 10.72622 | 13.40849 | 2.892867 | 7.613138 |
| ENSMUSG00000084826 | 0.153433675 | 0.747934 | 0.842597 | 15.09082 | 16.7975 | 9.109173 | 10.35148 | 12.67645 | 11.17374 | 7.714313 | 14.13868 |
| ENSMUSG00000084830 | -0.81870713 | 0.004157 | 0.022448 | 37.22401 | 20.39697 | 33.12427 | 25.8787 | 45.83023 | 61.45558 | 52.07161 | 47.85401 |
| ENSMUSG00000084835 | -0.1847899 | 0.496975 | 0.653568 | 34.20585 | 29.99554 | 41.40533 | 42.26855 | 48.75557 | 37.99072 | 34.71441 | 47.85401 |
| ENSMUSG00000084838 | -0.02935402 | 0.90835 | 0.945278 | 43.26034 | 37.19447 | 43.06154 | 45.71905 | 40.95467 | 50.28183 | 33.75012 | 48.9416 |
| ENSMUSG00000084844 | 0.188365415 | 0.205403 | 0.361868 | 306.8466 | 278.3586 | 302.2589 | 347.6373 | 278.8818 | 238.0007 | 238.1794 | 330.6277 |
| ENSMUSG00000084845 | 0.601192308 | 0.005162 | 0.026322 | 103.6236 | 113.983 | 94.40416 | 134.5693 | 55.58134 | 78.21619 | 75.21455 | 85.9197 |
| ENSMUSG00000084846 | 0.519476096 | 0.016689 | 0.061987 | 117.7084 | 110.3836 | 105.1695 | 114.7289 | 72.15824 | 109.5027 | 68.46453 | 63.08028 |
| ENSMUSG00000084854 | -3.353048086 | 4.09E-09 | 4.13E-07 | 7.04238 | 4.799286 | 13.24971 | 7.763611 | 118.9636 | 26.81698 | 58.82164 | 135.9489 |
| ENSMUSG00000084862 | -0.466047019 | 0.348282 | 0.516965 | 15.09082 | 13.19804 | 13.24971 | 9.488858 | 25.35289 | 6.704245 | 18.32149 | 19.57664 |
| ENSMUSG00000084866 | -0.006586656 | 0.973641 | 0.985571 | 80.48435 | 76.78858 | 81.98256 | 88.85022 | 75.08357 | 65.92507 | 97.39321 | 91.35765 |
| ENSMUSG00000084880 | 0.080271479 | 0.768123 | 0.856029 | 47.28455 | 44.3934 | 48.85829 | 47.44429 | 41.92979 | 36.87335 | 62.6788 | 35.89051 |
| ENSMUSG00000084881 | 0.36774856 | 0.407388 | 0.573711 | 25.15136 | 32.39518 | 12.4216 | 24.15346 | 20.47734 | 24.58223 | 16.39292 | 10.87591 |
| ENSMUSG00000084883 | -0.636324786 | 0.006201 | 0.030184 | 150.9082 | 143.9786 | 192.9488 | 148.3712 | 276.9316 | 148.6108 | 285.4296 | 278.4233 |
| ENSMUSG00000084885 | 0.878876783 | 0.000458 | 0.004286 | 62.37537 | 58.79126 | 56.31125 | 73.323 | 37.05423 | 31.28647 | 40.50014 | 27.18978 |
| ENSMUSG00000084897 | 0.247816502 | 0.718432 | 0.824127 | 316.9071 | 53.99197 | 33.12427 | 106.9653 | 90.68535 | 236.8833 | 54.00019 | 48.9416 |
| ENSMUSG00000084899 | -0.299800869 | 0.348648 | 0.517296 | 26.15741 | 20.39697 | 32.29616 | 23.29083 | 24.37778 | 34.6386 | 29.89296 | 38.06569 |
| ENSMUSG00000084904 | 0.744036143 | 0.088892 | 0.203323 | 23.13925 | 33.595 | 26.49941 | 25.01608 | 18.52711 | 26.81698 | 10.60718 | 8.700729 |
| ENSMUSG00000084910 | 0.198975862 | 0.124547 | 0.25468 | 534.2149 | 537.52 | 457.943 | 566.7436 | 442.7005 | 492.762 | 512.0375 | 377.3941 |

| | | | | | | | | | | | |
|---|---|---|---|---|---|---|---|---|---|---|---|
| ENSMUSG00000084911 | 0.256185989 | 0.214822 | 0.372961 | 226.3622 | 196.7707 | 149.0592 | 200.9913 | 145.2916 | 206.7142 | 168.7506 | 126.1606 |
| ENSMUSG00000084915 | 0.434613793 | 0.23185 | 0.392425 | 31.18769 | 17.99732 | 25.67131 | 24.15346 | 13.65156 | 18.99536 | 18.32149 | 22.83941 |
| ENSMUSG00000084917 | 0.811939773 | 0.087089 | 0.200513 | 25.15136 | 38.39429 | 19.04645 | 20.70296 | 10.72622 | 25.6996 | 13.50005 | 8.700729 |
| ENSMUSG00000084925 | 0.113462043 | 0.705507 | 0.814739 | 47.28455 | 89.98662 | 70.38906 | 56.93315 | 53.63112 | 79.33356 | 68.46453 | 42.41605 |
| ENSMUSG00000084938 | 1.152583919 | 0.018252 | 0.066208 | 16.09687 | 33.595 | 18.21835 | 18.97772 | 13.65156 | 5.58687 | 11.57147 | 7.613138 |
| ENSMUSG00000084941 | 0.309457637 | 0.483323 | 0.641736 | 122.7386 | 88.78679 | 62.9361 | 98.33908 | 63.38224 | 153.0803 | 43.39301 | 41.32846 |
| ENSMUSG00000084946 | 0.709818725 | 0.006929 | 0.032741 | 58.35115 | 52.79215 | 44.71776 | 43.9938 | 35.10401 | 31.28647 | 27.0001 | 28.27737 |
| ENSMUSG00000084947 | 0.922580879 | 0.260731 | 0.424793 | 20.12109 | 27.5959 | 13.24971 | 18.11509 | 0 | 24.58223 | 9.642892 | 7.613138 |
| ENSMUSG00000084950 | 0.361008332 | 0.145794 | 0.284282 | 178.0716 | 176.3738 | 137.4657 | 201.8539 | 134.5654 | 192.1883 | 124.3933 | 89.18247 |
| ENSMUSG00000084957 | 0.029233555 | 0.659357 | 0.780701 | 857.1583 | 832.6761 | 803.2634 | 822.9428 | 825.9193 | 776.575 | 817.7172 | 826.5692 |
| ENSMUSG00000084965 | 5.157604895 | 0.003271 | 0.018918 | 324.9556 | 0 | 1.656213 | 34.50494 | 3.900445 | 1.117374 | 2.892867 | 2.175182 |
| ENSMUSG00000084968 | 0.270106684 | 0.462312 | 0.623351 | 32.19374 | 63.59054 | 67.07664 | 28.46658 | 40.95467 | 42.46022 | 44.3573 | 30.45255 |
| ENSMUSG00000084970 | -0.820875962 | 0.116979 | 0.245141 | 14.08476 | 4.799286 | 14.90592 | 12.07673 | 24.37778 | 16.76061 | 10.60718 | 30.45255 |
| ENSMUSG00000084974 | 0.548075938 | 0.268559 | 0.433148 | 18.10898 | 17.99732 | 14.90592 | 28.46658 | 7.80089 | 24.58223 | 11.57147 | 10.87591 |
| ENSMUSG00000084978 | 0.979712581 | 0.122855 | 0.252404 | 23.13925 | 34.79482 | 11.59349 | 18.97772 | 5.850668 | 25.6996 | 5.785735 | 7.613138 |
| ENSMUSG00000084981 | -1.287450038 | 0.088863 | 0.203305 | 9.054489 | 0 | 7.45296 | 11.21411 | 14.62667 | 7.821619 | 29.89296 | 16.31387 |
| ENSMUSG00000084984 | 0.247936837 | 0.718694 | 0.824171 | 22.1332 | 5.999108 | 22.35888 | 16.38985 | 25.35289 | 2.234748 | 12.53576 | 16.31387 |
| ENSMUSG00000084989 | 0.730369178 | 0.029082 | 0.092542 | 33.19979 | 31.19536 | 39.74912 | 44.85642 | 15.60178 | 31.28647 | 24.10723 | 19.57664 |
| ENSMUSG00000084998 | 0.186110447 | 0.468528 | 0.628746 | 80.48435 | 56.39161 | 68.73285 | 79.36136 | 82.88446 | 68.15982 | 52.07161 | 47.85401 |
| ENSMUSG00000085001 | 1.001937184 | 0.125051 | 0.255444 | 20.12109 | 7.198929 | 12.4216 | 12.93935 | 3.900445 | 13.40849 | 2.892867 | 6.525547 |
| ENSMUSG00000085006 | -0.496806308 | 0.17026 | 0.316925 | 23.13925 | 19.19714 | 15.73403 | 18.97772 | 23.40267 | 37.99072 | 20.25007 | 27.18978 |
| ENSMUSG00000085015 | -0.042866182 | 0.91814 | 0.951643 | 30.18163 | 22.79661 | 44.71776 | 18.11509 | 22.42756 | 45.81234 | 27.96439 | 23.927 |
| ENSMUSG00000085016 | 0.344246343 | 0.328004 | 0.495712 | 35.2119 | 39.59411 | 18.21835 | 36.23019 | 21.45245 | 25.6996 | 28.92867 | 25.0146 |
| ENSMUSG00000085024 | 0.82571977 | 6.01E-09 | 5.55E-07 | 418.5186 | 443.934 | 378.4447 | 433.037 | 213.5494 | 294.9868 | 207.3222 | 229.4817 |
| ENSMUSG00000085028 | 0.57128443 | 0.008852 | 0.039036 | 204.229 | 172.7743 | 159.8246 | 138.8824 | 112.1378 | 153.0803 | 89.67889 | 100.0584 |
| ENSMUSG00000085030 | 0.164372551 | 0.611098 | 0.745927 | 23.13925 | 28.79572 | 33.12427 | 21.56559 | 22.42756 | 26.81698 | 22.17865 | 23.927 |
| ENSMUSG00000085036 | 0.265409013 | 0.719705 | 0.824664 | 25.15136 | 13.19804 | 21.53077 | 16.38985 | 4.875557 | 44.69496 | 6.750024 | 7.613138 |
| ENSMUSG00000085037 | -0.052127568 | 0.634476 | 0.762319 | 306.8466 | 277.1588 | 284.8687 | 301.9182 | 294.4836 | 343.0338 | 292.1796 | 286.0365 |
| ENSMUSG00000085039 | 0.032286581 | 0.901598 | 0.940849 | 49.29666 | 49.19268 | 43.88965 | 42.26855 | 50.70579 | 51.39921 | 47.25017 | 30.45255 |
| ENSMUSG00000085042 | 0.63509396 | 0.000451 | 0.004226 | 158.9566 | 171.5745 | 151.5435 | 138.0198 | 124.8142 | 88.27255 | 84.85745 | 100.0584 |
| ENSMUSG00000085054 | 0.532675299 | 0.05282 | 0.141404 | 44.26639 | 35.99465 | 39.74912 | 41.40593 | 24.37778 | 34.6386 | 27.0001 | 26.10219 |
| ENSMUSG00000085057 | 0.069462352 | 0.875014 | 0.925241 | 32.19374 | 11.99822 | 24.01509 | 14.6646 | 26.32801 | 18.99536 | 15.42863 | 18.48905 |
| ENSMUSG00000085063 | 0.652083064 | 0.288727 | 0.455682 | 21.12714 | 29.99554 | 10.76539 | 9.488858 | 10.72622 | 21.23011 | 7.714313 | 5.437955 |
| ENSMUSG00000085069 | -0.33579982 | 0.108977 | 0.234115 | 349.1009 | 303.5548 | 378.4447 | 320.0333 | 365.6667 | 287.1651 | 544.8234 | 507.905 |
| ENSMUSG00000085071 | -0.10035815 | 0.680938 | 0.796362 | 77.46619 | 82.78769 | 58.79557 | 66.42201 | 62.40712 | 59.22083 | 92.57176 | 90.27006 |
| ENSMUSG00000085072 | -0.335214653 | 0.409186 | 0.575474 | 32.19374 | 23.99643 | 20.70267 | 16.38985 | 20.47734 | 46.92971 | 24.10723 | 26.10219 |
| ENSMUSG00000085078 | 0.742792099 | 0.318899 | 0.487005 | 32.19374 | 33.595 | 13.24971 | 18.11509 | 7.80089 | 40.22547 | 5.785735 | 4.350364 |
| ENSMUSG00000085083 | 0.250334686 | 0.51906 | 0.672647 | 20.12109 | 23.99643 | 19.87456 | 14.6646 | 17.552 | 11.17374 | 19.28578 | 17.40146 |
| ENSMUSG00000085084 | 0.705663829 | 0.030743 | 0.0962 | 94.56911 | 106.7841 | 87.7793 | 107.8279 | 52.65601 | 106.1505 | 39.53586 | 45.67883 |
| ENSMUSG00000085085 | -0.45683962 | 0.440717 | 0.603986 | 7.04238 | 16.7975 | 13.24971 | 12.93935 | 30.22845 | 6.704245 | 9.642892 | 21.75182 |
| ENSMUSG00000085088 | 0.394730085 | 0.400556 | 0.56788 | 21.12714 | 44.3934 | 25.67131 | 36.23019 | 15.60178 | 45.81234 | 19.28578 | 16.31387 |
| ENSMUSG00000085095 | -0.021538241 | 0.967513 | 0.981769 | 14.08476 | 10.79839 | 9.93728 | 19.84034 | 10.72622 | 24.58223 | 7.714313 | 13.05109 |
| ENSMUSG00000085105 | 0.101387486 | 0.727111 | 0.829154 | 32.19374 | 29.99554 | 44.71776 | 44.85642 | 37.05423 | 31.28647 | 30.85725 | 43.50364 |
| ENSMUSG00000085111 | 0.058913397 | 0.792204 | 0.872891 | 88.53278 | 110.3836 | 96.88848 | 72.46037 | 81.90935 | 110.62 | 70.39311 | 90.27006 |
| ENSMUSG00000085112 | 0.866588842 | 0.066071 | 0.165658 | 17.10292 | 14.39786 | 21.53077 | 13.80198 | 12.67645 | 6.704245 | 10.60718 | 6.525547 |
| ENSMUSG00000085125 | 1.707162809 | 0.017088 | 0.063075 | 25.15136 | 21.59679 | 14.07781 | 13.80198 | 0 | 4.469496 | 9.642892 | 8.700729 |
| ENSMUSG00000085133 | 0.218210784 | 0.283715 | 0.450275 | 444.676 | 307.1543 | 395.835 | 408.0209 | 274.0063 | 469.2971 | 304.7154 | 291.4744 |
| ENSMUSG00000085139 | 1.37992628 | 0.025107 | 0.083342 | 56.33904 | 87.58697 | 36.43669 | 89.71284 | 16.57689 | 65.92507 | 9.642892 | 11.9635 |
| ENSMUSG00000085148 | -0.861351785 | 0.001462 | 0.010395 | 279.6831 | 135.5798 | 277.4157 | 221.6942 | 361.7663 | 291.6346 | 449.3588 | 561.197 |
| ENSMUSG00000085151 | 0.541249277 | 0.151049 | 0.291301 | 38.23007 | 55.19179 | 30.63995 | 42.26855 | 33.15378 | 41.34284 | 24.10723 | 15.22628 |
| ENSMUSG00000085154 | -0.457699932 | 0.159583 | 0.302668 | 38.23007 | 28.79572 | 39.74912 | 37.09281 | 40.95467 | 32.40385 | 51.10733 | 73.95619 |
| ENSMUSG00000085156 | -0.04193617 | 0.686102 | 0.80043 | 437.6336 | 393.5415 | 402.4598 | 427.8613 | 457.3272 | 457.006 | 421.3944 | 375.2189 |
| ENSMUSG00000085162 | 0.856043572 | 0.220861 | 0.379783 | 20.12109 | 19.19714 | 16.56213 | 30.19182 | 1.950223 | 27.93435 | 12.53576 | 5.437955 |
| ENSMUSG00000085169 | 0.149696446 | 0.623724 | 0.754981 | 44.26639 | 51.59233 | 63.76421 | 64.69676 | 42.9049 | 63.69032 | 64.60737 | 31.54014 |
| ENSMUSG00000085170 | 0.71016292 | 0.362892 | 0.531351 | 10.06054 | 10.79839 | 14.07781 | 17.25247 | 4.875557 | 22.34748 | 2.892867 | 2.175182 |
| ENSMUSG00000085171 | 0.151738276 | 0.69059 | 0.804033 | 21.12714 | 20.39697 | 22.35888 | 17.25247 | 22.42756 | 21.23011 | 18.32149 | 10.87591 |
| ENSMUSG00000085172 | -0.207513832 | 0.578875 | 0.720218 | 19.11503 | 21.59679 | 27.32752 | 37.09281 | 19.50223 | 37.99072 | 36.64299 | 28.27737 |
| ENSMUSG00000085174 | 0.054531291 | 0.921206 | 0.953014 | 27.16347 | 7.198929 | 12.4216 | 24.15346 | 10.72622 | 13.40849 | 16.39292 | 28.27737 |
| ENSMUSG00000085175 | 0.161032187 | 0.405648 | 0.57246 | 93.56306 | 128.3809 | 96.88848 | 93.16334 | 81.90935 | 90.5073 | 107.0361 | 87.00729 |
| ENSMUSG00000085178 | 0.362205772 | 0.435721 | 0.599856 | 43.26034 | 34.79482 | 22.35888 | 34.50494 | 16.57689 | 48.04709 | 13.50005 | 27.18978 |
| ENSMUSG00000085180 | -0.322178458 | 0.010308 | 0.043553 | 423.5489 | 326.3515 | 335.3832 | 338.1484 | 398.8205 | 486.0577 | 458.0374 | 437.2116 |
| ENSMUSG00000085181 | 0.180102719 | 0.804972 | 0.880546 | 17.10292 | 9.598572 | 14.07781 | 14.6646 | 1.950223 | 30.1691 | 9.642892 | 7.613138 |
| ENSMUSG00000085184 | 0.190559892 | 0.639471 | 0.766468 | 23.13925 | 31.19536 | 21.53077 | 15.52722 | 15.60178 | 13.40849 | 23.14294 | 27.18978 |
| ENSMUSG00000085189 | 0.705220317 | 0.234956 | 0.395797 | 7.04238 | 22.79661 | 14.07781 | 12.93935 | 6.825779 | 11.17374 | 3.857157 | 13.05109 |

| | | | | | | | | | | | |
|---|---|---|---|---|---|---|---|---|---|---|---|
| ENSMUSG00000085192 | 0.926365365 | 0.134643 | 0.268587 | 46.2785 | 29.99554 | 55.48314 | 38.81806 | 11.70134 | 54.75133 | 18.32149 | 5.437955 |
| ENSMUSG00000085196 | 0.342185953 | 0.603929 | 0.740916 | 13.07871 | 15.59768 | 9.109173 | 9.488858 | 1.950223 | 11.17374 | 17.3572 | 6.525547 |
| ENSMUSG00000085201 | 0.744461581 | 0.095385 | 0.213692 | 43.26034 | 44.3934 | 25.67131 | 16.38985 | 28.27823 | 15.64324 | 16.39292 | 16.31387 |
| ENSMUSG00000085203 | -2.341030283 | 0.000144 | 0.001766 | 10.06054 | 8.398751 | 3.312427 | 3.450494 | 24.37778 | 12.29112 | 49.17875 | 39.15328 |
| ENSMUSG00000085204 | -0.00430222 | 0.991772 | 0.995331 | 32.19374 | 26.39607 | 39.74912 | 29.3292 | 21.45245 | 17.87799 | 33.75012 | 55.46715 |
| ENSMUSG00000085207 | -0.438720943 | 0.095464 | 0.213737 | 37.22401 | 26.39607 | 36.43669 | 36.23019 | 39.00445 | 43.57759 | 50.14304 | 53.29196 |
| ENSMUSG00000085208 | 0.12329665 | 0.052353 | 0.14048 | 1372.258 | 1510.575 | 1375.485 | 1425.917 | 1278.371 | 1293.919 | 1280.576 | 1362.752 |
| ENSMUSG00000085211 | 0.098082589 | 0.855611 | 0.913423 | 12.07265 | 7.198929 | 10.76539 | 14.6646 | 11.70134 | 4.469496 | 10.60718 | 15.22628 |
| ENSMUSG00000085213 | -0.115411233 | 0.757146 | 0.848808 | 26.15741 | 20.39697 | 18.21835 | 24.15346 | 31.20356 | 27.93435 | 14.46434 | 22.83941 |
| ENSMUSG00000085218 | 0.074827441 | 0.748466 | 0.842966 | 57.3451 | 61.1909 | 45.54586 | 68.14726 | 54.60623 | 50.28183 | 63.64308 | 51.11678 |
| ENSMUSG00000085227 | -0.399185143 | 0.087442 | 0.201066 | 38.23007 | 55.19179 | 48.85829 | 44.85642 | 57.53157 | 56.98608 | 57.85735 | 73.95619 |
| ENSMUSG00000085229 | 1.597961286 | 0.001798 | 0.012158 | 36.21796 | 27.5959 | 22.35888 | 31.05445 | 6.825779 | 20.11273 | 7.714313 | 4.350364 |
| ENSMUSG00000085232 | 0.419335493 | 0.617112 | 0.749991 | 13.07871 | 22.79661 | 9.93728 | 7.763611 | 4.875557 | 29.05173 | 2.892867 | 3.262773 |
| ENSMUSG00000085235 | -0.337289411 | 0.099907 | 0.2207 | 251.5136 | 166.7752 | 231.8699 | 189.7772 | 254.5041 | 195.5405 | 281.5724 | 330.6277 |
| ENSMUSG00000085236 | 0.138764954 | 0.76832 | 0.856029 | 23.13925 | 9.598572 | 17.39024 | 15.52722 | 11.70134 | 10.05637 | 15.42863 | 22.83941 |
| ENSMUSG00000085241 | -0.044754063 | 0.745235 | 0.840853 | 185.114 | 231.5656 | 235.1823 | 198.4034 | 222.3254 | 195.5405 | 237.2151 | 220.781 |
| ENSMUSG00000085251 | -2.816941244 | 0.00068 | 0.005806 | 4.024217 | 0 | 5.796746 | 0.862623 | 21.45245 | 6.704245 | 21.21436 | 28.27737 |
| ENSMUSG00000085255 | 0.533831971 | 0.297591 | 0.465488 | 7.04238 | 14.39786 | 19.87456 | 12.07673 | 9.751113 | 7.821619 | 9.642892 | 9.78832 |
| ENSMUSG00000085260 | -0.640945922 | 0.237452 | 0.398582 | 10.06054 | 7.198929 | 5.796746 | 12.93935 | 13.65156 | 13.40849 | 7.714313 | 21.75182 |
| ENSMUSG00000085269 | 0.147531252 | 0.583018 | 0.723674 | 77.46619 | 61.1909 | 42.23344 | 48.30692 | 49.73068 | 50.28183 | 47.25017 | 58.72992 |
| ENSMUSG00000085271 | -0.383209337 | 0.449787 | 0.612329 | 5.030272 | 17.99732 | 10.76539 | 23.29083 | 17.552 | 21.23011 | 18.32149 | 17.40146 |
| ENSMUSG00000085280 | 0.471600383 | 0.378933 | 0.5471 | 19.11503 | 23.99643 | 28.15563 | 12.07673 | 5.850668 | 26.81698 | 12.53576 | 15.22628 |
| ENSMUSG00000085282 | 0.190029045 | 0.417237 | 0.5828 | 105.6357 | 189.5718 | 115.1068 | 128.5309 | 113.1129 | 138.5544 | 95.46463 | 123.9854 |
| ENSMUSG00000085286 | 0.331588076 | 0.409283 | 0.575555 | 19.11503 | 33.595 | 22.35888 | 21.56559 | 25.35289 | 23.46486 | 15.42863 | 11.9635 |
| ENSMUSG00000085287 | 0.108271627 | 0.552163 | 0.699721 | 181.0898 | 183.5727 | 202.058 | 232.0457 | 156.0178 | 233.5312 | 197.6793 | 155.5255 |
| ENSMUSG00000085311 | 0.408346533 | 0.571276 | 0.714304 | 25.15136 | 8.398751 | 14.90592 | 34.50494 | 19.50223 | 32.40385 | 7.714313 | 3.262773 |
| ENSMUSG00000085315 | 0.47077866 | 0.073397 | 0.178403 | 57.3451 | 39.59411 | 49.6864 | 43.13117 | 30.22845 | 36.87335 | 38.57157 | 31.54014 |
| ENSMUSG00000085316 | 0.903715324 | 0.138572 | 0.274173 | 27.16347 | 9.598572 | 14.07781 | 11.21411 | 3.900445 | 5.58687 | 9.642892 | 14.13868 |
| ENSMUSG00000085323 | 0.19510683 | 0.425712 | 0.590702 | 110.666 | 88.78679 | 60.45178 | 87.9876 | 66.30757 | 92.74205 | 69.42882 | 75.04379 |
| ENSMUSG00000085327 | -0.003433958 | 0.993134 | 0.996103 | 14.08476 | 31.19536 | 22.35888 | 23.29083 | 21.45245 | 29.05173 | 26.03581 | 14.13868 |
| ENSMUSG00000085334 | 0.731124742 | 0.01379 | 0.054173 | 2437.67 | 2380.446 | 1824.319 | 2565.442 | 1026.792 | 2520.796 | 1104.111 | 896.1751 |
| ENSMUSG00000085335 | 0.356579622 | 0.513978 | 0.66845 | 41.24823 | 31.19536 | 23.18699 | 33.64232 | 14.62667 | 58.10345 | 14.46434 | 14.13868 |
| ENSMUSG00000085337 | 0.393502925 | 0.098544 | 0.218807 | 100.6054 | 113.983 | 112.6225 | 150.0965 | 78.0089 | 124.0285 | 91.60747 | 70.69342 |
| ENSMUSG00000085338 | -0.149436199 | 0.801761 | 0.879143 | 19.11503 | 5.999108 | 6.624853 | 9.488858 | 7.80089 | 8.938993 | 10.60718 | 18.48905 |
| ENSMUSG00000085342 | 0.088173158 | 0.771571 | 0.858355 | 56.33904 | 53.99197 | 46.37397 | 48.30692 | 65.33246 | 58.10345 | 39.53586 | 29.36496 |
| ENSMUSG00000085347 | 0.229245498 | 0.419991 | 0.585395 | 45.27245 | 45.59322 | 53.82693 | 49.16954 | 29.25334 | 58.10345 | 35.6787 | 43.50364 |
| ENSMUSG00000085355 | 0.132394219 | 0.417595 | 0.583211 | 150.9082 | 201.57 | 175.5586 | 159.5853 | 182.3458 | 159.7845 | 148.5005 | 134.8613 |
| ENSMUSG00000085360 | -0.453952344 | 0.22869 | 0.3887 | 28.16952 | 40.79393 | 58.79557 | 45.71905 | 49.73068 | 31.28647 | 81.00029 | 76.13138 |
| ENSMUSG00000085363 | 0.757211278 | 6.71E-07 | 2.68E-05 | 406.446 | 443.934 | 360.2264 | 383.8675 | 274.0063 | 207.8316 | 266.1438 | 192.5036 |
| ENSMUSG00000085376 | 0.015145348 | 0.965403 | 0.980598 | 44.26639 | 28.79572 | 24.01509 | 39.68068 | 21.45245 | 42.46022 | 33.75012 | 38.06569 |
| ENSMUSG00000085379 | 0.119646801 | 0.545823 | 0.694403 | 105.6357 | 95.98572 | 81.98256 | 90.57547 | 95.56091 | 94.9768 | 66.53595 | 87.00729 |
| ENSMUSG00000085385 | 0.113396487 | 0.436264 | 0.600201 | 1954.764 | 1691.748 | 1410.266 | 1649.336 | 1678.167 | 1813.498 | 1460.898 | 1245.292 |
| ENSMUSG00000085391 | 0.735766309 | 0.329787 | 0.497592 | 17.10292 | 28.79572 | 29.81184 | 20.70296 | 8.776002 | 41.34284 | 4.821446 | 3.262773 |
| ENSMUSG00000085395 | -0.070648137 | 0.875542 | 0.925381 | 19.11503 | 25.19625 | 46.37397 | 16.38985 | 21.45245 | 32.40385 | 20.25007 | 39.15328 |
| ENSMUSG00000085396 | 0.227953622 | 0.009504 | 0.04105 | 4637.911 | 4170.58 | 3987.333 | 4417.495 | 4035.011 | 3781.194 | 3608.37 | 3271.474 |
| ENSMUSG00000085399 | 0.538732732 | 0.007475 | 0.034499 | 1268.635 | 1481.78 | 915.8859 | 1224.925 | 903.9282 | 1100.613 | 733.8241 | 627.5401 |
| ENSMUSG00000085403 | 0.250467325 | 0.713348 | 0.820233 | 19.11503 | 8.398751 | 21.53077 | 25.01608 | 10.72622 | 24.58223 | 25.07152 | 2.175182 |
| ENSMUSG00000085407 | 0.789178406 | 0.267382 | 0.431699 | 15.09082 | 19.19714 | 19.87456 | 14.6646 | 9.751113 | 14.52586 | 15.42863 | 0 |
| ENSMUSG00000085408 | 0.000189145 | 0.999585 | 0.999733 | 32.19374 | 32.39518 | 24.8432 | 24.15346 | 21.45245 | 42.46022 | 19.28578 | 30.45255 |
| ENSMUSG00000085412 | 1.378512585 | 0.002516 | 0.015496 | 68.4117 | 63.59054 | 48.85829 | 79.36136 | 22.42756 | 12.29112 | 14.46434 | 51.11678 |
| ENSMUSG00000085415 | 0.097539192 | 0.870525 | 0.922402 | 16.09687 | 8.398751 | 9.109173 | 11.21411 | 7.80089 | 21.23011 | 7.714313 | 5.437955 |
| ENSMUSG00000085419 | 0.561733073 | 0.266957 | 0.431387 | 20.12109 | 16.7975 | 9.93728 | 10.35148 | 5.850668 | 11.17374 | 10.60718 | 10.87591 |
| ENSMUSG00000085421 | -0.347883921 | 0.575187 | 0.71742 | 14.08476 | 4.799286 | 19.04645 | 6.038364 | 7.80089 | 10.05637 | 21.21436 | 17.40146 |
| ENSMUSG00000085433 | -0.325803767 | 0.114123 | 0.241462 | 80.48435 | 94.7859 | 92.74794 | 104.3774 | 86.78491 | 110.62 | 134.0362 | 135.9489 |
| ENSMUSG00000085436 | -0.058108144 | 0.734751 | 0.833942 | 113.6841 | 115.1829 | 137.4657 | 127.6683 | 118.9636 | 158.6671 | 122.4647 | 116.3722 |
| ENSMUSG00000085438 | -0.043974573 | 0.31985 | 0.487903 | 3697.25 | 3736.244 | 3561.687 | 3633.37 | 3719.075 | 3790.133 | 3697.085 | 3871.824 |
| ENSMUSG00000085440 | 0.381185184 | 0.624123 | 0.754985 | 21.12714 | 7.198929 | 13.24971 | 5.175741 | 14.62667 | 13.40849 | 6.750024 | 1.087591 |
| ENSMUSG00000085442 | 0.453062947 | 0.156302 | 0.298378 | 35.2119 | 37.19447 | 24.01509 | 27.60395 | 19.50223 | 23.46486 | 24.10723 | 22.83941 |
| ENSMUSG00000085445 | 0.477888197 | 0.150326 | 0.290364 | 33.19979 | 47.99286 | 27.32752 | 40.5433 | 18.52711 | 29.05173 | 32.78583 | 26.10219 |
| ENSMUSG00000085448 | 0.695907672 | 0.189838 | 0.342466 | 11.0666 | 21.59679 | 19.87456 | 21.56559 | 14.62667 | 18.99536 | 5.785735 | 6.525547 |
| ENSMUSG00000085457 | 0.325568675 | 0.168555 | 0.314737 | 95.57516 | 70.78947 | 65.42042 | 56.93315 | 62.40712 | 54.75133 | 56.89306 | 55.46715 |
| ENSMUSG00000085465 | -0.524288906 | 0.263736 | 0.428017 | 21.12714 | 9.598572 | 11.59349 | 12.07673 | 19.50223 | 11.17374 | 19.28578 | 28.27737 |
| ENSMUSG00000085492 | -0.13332345 | 0.460741 | 0.62196 | 153.9263 | 173.9741 | 142.4343 | 163.8985 | 174.5449 | 150.8455 | 145.6077 | 224.0438 |
| ENSMUSG00000085497 | 0.334592665 | 0.582256 | 0.722932 | 6.036326 | 17.99732 | 15.73403 | 15.52722 | 13.65156 | 18.99536 | 4.821446 | 6.525547 |

| | | | | | | | | | | | |
|---|---|---|---|---|---|---|---|---|---|---|---|
| ENSMUSG00000085498 | -0.129292613 | 0.724742 | 0.827662 | 14.08476 | 20.39697 | 17.39024 | 15.52722 | 17.552 | 17.87799 | 17.3572 | 20.66423 |
| ENSMUSG00000085501 | -0.01295191 | 0.954132 | 0.973813 | 52.31483 | 61.1909 | 68.73285 | 57.79577 | 54.60623 | 75.98144 | 54.96448 | 57.64233 |
| ENSMUSG00000085511 | 0.495119585 | 0.304401 | 0.472818 | 13.07871 | 16.7975 | 24.01509 | 24.15346 | 21.45245 | 16.76061 | 7.714313 | 9.78832 |
| ENSMUSG00000085517 | 0.978997939 | 7.26E-05 | 0.001062 | 73.44197 | 97.18554 | 112.6225 | 116.4542 | 65.33246 | 43.57759 | 48.21446 | 45.67883 |
| ENSMUSG00000085521 | 0.606503575 | 0.085046 | 0.197087 | 94.56911 | 62.39072 | 90.26362 | 66.42201 | 28.27823 | 84.92043 | 50.14304 | 43.50364 |
| ENSMUSG00000085525 | 0.31078497 | 0.495333 | 0.652059 | 24.1453 | 21.59679 | 28.15563 | 12.07673 | 12.67645 | 24.58223 | 21.21436 | 10.87591 |
| ENSMUSG00000085541 | -1.52269888 | 0.001089 | 0.008336 | 56.33904 | 49.19268 | 154.0278 | 67.28463 | 278.8818 | 115.0895 | 145.6077 | 402.4087 |
| ENSMUSG00000085558 | 0.520257114 | 0.106043 | 0.23006 | 50.30272 | 70.78947 | 53.82693 | 47.44429 | 31.20356 | 39.10809 | 26.03581 | 58.72992 |
| ENSMUSG00000085562 | -0.328903992 | 0.154423 | 0.295923 | 150.9082 | 235.165 | 142.4343 | 156.1349 | 276.9316 | 182.132 | 227.5722 | 169.6642 |
| ENSMUSG00000085566 | 0.512398732 | 0.001401 | 0.010083 | 162.9808 | 189.5718 | 158.1684 | 177.7004 | 133.5902 | 96.09417 | 129.2147 | 121.8102 |
| ENSMUSG00000085568 | 0.733216415 | 0.069062 | 0.170626 | 26.15741 | 21.59679 | 33.95237 | 31.05445 | 18.52711 | 12.29112 | 26.03581 | 10.87591 |
| ENSMUSG00000085573 | 0.165568421 | 0.723852 | 0.827323 | 20.12109 | 39.59411 | 19.04645 | 14.6646 | 23.40267 | 15.64324 | 29.89296 | 13.05109 |
| ENSMUSG00000085582 | 0.649138021 | 0.015791 | 0.059844 | 160.9687 | 243.5638 | 107.6539 | 174.2499 | 134.5654 | 98.32892 | 113.7861 | 89.18247 |
| ENSMUSG00000085584 | 0.587600168 | 0.144744 | 0.282892 | 17.10292 | 25.19625 | 15.73403 | 24.15346 | 16.57689 | 12.29112 | 14.46434 | 10.87591 |
| ENSMUSG00000085586 | -0.107362806 | 0.68398 | 0.798636 | 84.50857 | 49.19268 | 94.40416 | 69.8725 | 73.13335 | 63.69032 | 97.39321 | 88.09488 |
| ENSMUSG00000085589 | 0.13464074 | 0.79462 | 0.873721 | 19.11503 | 10.79839 | 17.39024 | 6.038364 | 9.751113 | 11.17374 | 12.53576 | 15.22628 |
| ENSMUSG00000085596 | 0.042168695 | 0.922519 | 0.953784 | 21.12714 | 17.99732 | 12.4216 | 12.93935 | 11.70134 | 21.23011 | 17.3572 | 11.9635 |
| ENSMUSG00000085601 | -0.024397244 | 0.923373 | 0.954293 | 92.557 | 149.9777 | 118.4192 | 97.47645 | 117.0134 | 159.7845 | 87.75031 | 101.146 |
| ENSMUSG00000085603 | 1.423262073 | 6.33E-05 | 0.000954 | 68.4117 | 63.59054 | 39.74912 | 57.79577 | 20.47734 | 26.81698 | 27.0001 | 10.87591 |
| ENSMUSG00000085606 | 0.160346624 | 0.716426 | 0.822491 | 23.13925 | 23.99643 | 19.04645 | 12.07673 | 26.32801 | 15.64324 | 16.39292 | 10.87591 |
| ENSMUSG00000085612 | -0.095331445 | 0.844854 | 0.906563 | 10.06054 | 11.99822 | 11.59349 | 19.84034 | 8.776002 | 16.76061 | 11.57147 | 20.66423 |
| ENSMUSG00000085614 | 0.513920668 | 0.103102 | 0.225724 | 67.40564 | 64.79036 | 41.40533 | 50.03216 | 39.97956 | 52.51658 | 39.53586 | 23.927 |
| ENSMUSG00000085615 | -0.201715201 | 0.326499 | 0.494204 | 131.7931 | 123.5816 | 122.5598 | 103.5148 | 117.0134 | 111.7374 | 182.2507 | 141.3868 |
| ENSMUSG00000085622 | 0.199625566 | 0.218968 | 0.377759 | 233.4046 | 224.3666 | 167.2775 | 188.0519 | 166.744 | 204.4795 | 167.7863 | 167.489 |
| ENSMUSG00000085628 | -0.349757557 | 0.357594 | 0.526087 | 13.07871 | 28.79572 | 17.39024 | 18.97772 | 21.45245 | 30.1691 | 20.25007 | 27.18978 |
| ENSMUSG00000085633 | -0.70372104 | 0.186909 | 0.338866 | 8.048435 | 7.198929 | 7.45296 | 15.52722 | 10.72622 | 25.6996 | 12.53576 | 14.13868 |
| ENSMUSG00000085636 | -0.056205417 | 0.922503 | 0.953784 | 16.09687 | 17.99732 | 14.07781 | 12.07673 | 22.42756 | 3.352122 | 12.53576 | 23.927 |
| ENSMUSG00000085637 | 0.162692824 | 0.742189 | 0.839029 | 26.15741 | 10.79839 | 14.90592 | 14.6646 | 7.80089 | 22.34748 | 14.46434 | 15.22628 |
| ENSMUSG00000085642 | 0.210401109 | 0.467349 | 0.62788 | 44.26639 | 51.59233 | 52.17072 | 44.85642 | 41.92979 | 49.16446 | 25.07152 | 51.11678 |
| ENSMUSG00000085645 | 0.245005505 | 0.041433 | 0.11967 | 492.9666 | 471.5299 | 473.677 | 410.6088 | 342.2641 | 363.1466 | 415.6086 | 438.2992 |
| ENSMUSG00000085651 | -1.880206986 | 0.021187 | 0.073432 | 15.09082 | 1.199822 | 40.57722 | 10.35148 | 47.78045 | 12.29112 | 83.89316 | 105.4963 |
| ENSMUSG00000085655 | -0.141417301 | 0.604311 | 0.741075 | 36.21796 | 52.79215 | 40.57722 | 36.23019 | 37.05423 | 56.98608 | 40.50014 | 47.85401 |
| ENSMUSG00000085658 | 0.772127352 | 0.148149 | 0.287623 | 23.13925 | 33.595 | 32.29616 | 27.60395 | 10.72622 | 36.87335 | 14.46434 | 6.525547 |
| ENSMUSG00000085663 | 1.102609509 | 0.00812 | 0.03667 | 46.2785 | 40.79393 | 44.71776 | 56.07053 | 11.70134 | 39.10809 | 24.10723 | 13.05109 |
| ENSMUSG00000085665 | -0.300918076 | 0.570724 | 0.713976 | 25.15136 | 17.99732 | 14.07781 | 8.626235 | 13.65156 | 16.76061 | 13.50005 | 36.9781 |
| ENSMUSG00000085666 | 5.695206587 | 8.15E-08 | 4.80E-06 | 4.024217 | 255.562 | 310.54 | 281.2153 | 2.925334 | 3.352122 | 5.785735 | 4.350364 |
| ENSMUSG00000085667 | 0.569025548 | 0.023864 | 0.080312 | 119.7205 | 105.5843 | 97.71658 | 131.9814 | 90.68535 | 94.9768 | 74.25027 | 46.76642 |
| ENSMUSG00000085682 | 0.222236086 | 0.652242 | 0.77543 | 15.09082 | 45.59322 | 26.49941 | 16.38985 | 29.25334 | 21.23011 | 11.57147 | 26.10219 |
| ENSMUSG00000085683 | -0.46515662 | 0.456291 | 0.618115 | 10.06054 | 9.598572 | 23.18699 | 5.175741 | 20.47734 | 6.704245 | 14.46434 | 25.0146 |
| ENSMUSG00000085687 | 0.220217198 | 0.31454 | 0.482757 | 138.8355 | 153.5772 | 142.4343 | 119.042 | 109.2125 | 167.6061 | 104.1432 | 94.62042 |
| ENSMUSG00000085689 | -2.619825151 | 1.85E-08 | 1.42E-06 | 8.048435 | 3.599465 | 8.281066 | 4.313117 | 33.15378 | 25.6996 | 45.32159 | 46.76642 |
| ENSMUSG00000085696 | 0.092608117 | 0.639976 | 0.766639 | 155.9384 | 128.3809 | 132.4971 | 144.0581 | 133.5902 | 174.3104 | 106.0718 | 113.1095 |
| ENSMUSG00000085702 | 0.521012262 | 0.028742 | 0.09168 | 77.46619 | 80.38804 | 77.84202 | 57.79577 | 46.80534 | 65.92507 | 48.21446 | 43.50364 |
| ENSMUSG00000085704 | -0.092376828 | 0.57809 | 0.719775 | 128.775 | 147.578 | 108.482 | 116.4542 | 145.2916 | 127.3806 | 139.8219 | 118.5474 |
| ENSMUSG00000085705 | 0.493111625 | 0.416572 | 0.582046 | 8.048435 | 14.39786 | 18.21835 | 12.07673 | 13.65156 | 8.938993 | 12.53576 | 2.175182 |
| ENSMUSG00000085711 | -0.564345315 | 0.099202 | 0.219818 | 20.12109 | 15.59768 | 20.70267 | 15.52722 | 27.30312 | 21.23011 | 31.82154 | 26.10219 |
| ENSMUSG00000085730 | 0.409858839 | 0.377137 | 0.545276 | 11.0666 | 13.19804 | 24.8432 | 18.97772 | 14.62667 | 16.76061 | 9.642892 | 10.87591 |
| ENSMUSG00000085733 | -0.60222732 | 0.216411 | 0.37475 | 19.11503 | 13.19804 | 14.90592 | 9.488858 | 32.17867 | 10.05637 | 17.3572 | 26.10219 |
| ENSMUSG00000085735 | 0.287988665 | 0.604429 | 0.741075 | 13.07871 | 4.799286 | 18.21835 | 20.70296 | 16.57689 | 12.29112 | 10.60718 | 7.613138 |
| ENSMUSG00000085741 | 0.294623651 | 0.199561 | 0.354519 | 211.2714 | 199.1704 | 145.7468 | 195.8155 | 146.2667 | 215.6532 | 137.8934 | 113.1095 |
| ENSMUSG00000085743 | 0.893545975 | 0.004487 | 0.023747 | 267.6105 | 447.5334 | 338.6956 | 361.4392 | 178.4454 | 327.3906 | 138.8576 | 117.4598 |
| ENSMUSG00000085761 | 0.524988538 | 0.13591 | 0.270344 | 25.15136 | 27.5959 | 30.63995 | 30.19182 | 12.67645 | 23.46486 | 17.3572 | 26.10219 |
| ENSMUSG00000085766 | 0.000207912 | 0.999496 | 0.999733 | 31.18769 | 39.59411 | 45.54586 | 33.64232 | 39.00445 | 45.81234 | 21.21436 | 44.59123 |
| ENSMUSG00000085767 | 0.346066035 | 0.011286 | 0.046753 | 227.3683 | 221.967 | 194.6051 | 201.8539 | 186.2463 | 173.193 | 154.2863 | 150.0876 |
| ENSMUSG00000085771 | 0.411730312 | 0.654051 | 0.77673 | 24.1453 | 7.198929 | 13.24971 | 10.35148 | 0.975111 | 30.1691 | 2.892867 | 7.613138 |
| ENSMUSG00000085772 | -2.198496132 | 0.001704 | 0.011646 | 10.06054 | 0 | 9.93728 | 5.175741 | 14.62667 | 22.34748 | 30.85725 | 50.02919 |
| ENSMUSG00000085776 | 0.07236819 | 0.833149 | 0.898985 | 21.12714 | 25.19625 | 19.04645 | 18.97772 | 16.57689 | 21.23011 | 20.25007 | 21.75182 |
| ENSMUSG00000085779 | 0.967268032 | 0.20432 | 0.360337 | 19.11503 | 19.19714 | 23.18699 | 17.25247 | 8.776002 | 26.81698 | 3.857157 | 1.087591 |
| ENSMUSG00000085783 | -0.548630805 | 0.118416 | 0.247227 | 17.10292 | 35.99465 | 20.70267 | 17.25247 | 32.17867 | 32.40385 | 29.89296 | 36.9781 |
| ENSMUSG00000085784 | 1.317040812 | 0.062187 | 0.158943 | 20.12109 | 56.39161 | 19.87456 | 33.64232 | 6.825779 | 33.52122 | 8.678602 | 3.262773 |
| ENSMUSG00000085787 | 0.212070798 | 0.689563 | 0.803352 | 8.048435 | 15.59768 | 11.59349 | 8.626235 | 7.80089 | 11.17374 | 5.785735 | 13.05109 |
| ENSMUSG00000085788 | -0.191334067 | 0.669156 | 0.787855 | 23.13925 | 26.39607 | 19.04645 | 24.15346 | 39.97956 | 35.75597 | 12.53576 | 17.40146 |
| ENSMUSG00000085793 | -0.159666089 | 0.275875 | 0.442091 | 334.01 | 304.7547 | 313.0243 | 291.5667 | 354.9405 | 373.2029 | 395.3586 | 264.2846 |
| ENSMUSG00000085794 | -0.428432596 | 0.253489 | 0.417192 | 14.08476 | 20.39697 | 24.01509 | 19.84034 | 17.552 | 34.6386 | 29.89296 | 23.927 |

| | | | | | | | | | | | |
|---|---|---|---|---|---|---|---|---|---|---|---|
| ENSMUSG00000085795 | 0.18874467 | 0.32237 | 0.490613 | 795.789 | 871.0704 | 925.8232 | 709.0765 | 834.6953 | 920.7163 | 510.109 | 631.8904 |
| ENSMUSG00000085811 | 1.837888625 | 0.029794 | 0.094061 | 28.16952 | 19.19714 | 11.59349 | 7.763611 | 1.950223 | 8.938993 | 7.714313 | 0 |
| ENSMUSG00000085819 | 0.438082611 | 0.333268 | 0.50145 | 19.11503 | 10.79839 | 14.07781 | 19.84034 | 13.65156 | 14.52586 | 11.57147 | 7.613138 |
| ENSMUSG00000085826 | -2.309167876 | 1.22E-05 | 0.000268 | 9.054489 | 5.999108 | 30.63995 | 8.626235 | 50.70579 | 49.16446 | 77.14313 | 96.79561 |
| ENSMUSG00000085828 | 0.272096223 | 0.385617 | 0.553051 | 89.53884 | 134.38 | 112.6225 | 103.5148 | 92.63557 | 147.4934 | 64.60737 | 59.81751 |
| ENSMUSG00000085829 | -0.042799448 | 0.844053 | 0.90611 | 187.1261 | 229.1659 | 161.4808 | 189.7772 | 178.4454 | 255.8787 | 217.9294 | 137.0365 |
| ENSMUSG00000085833 | 0.11506149 | 0.740543 | 0.837826 | 25.15136 | 28.79572 | 24.01509 | 21.56559 | 31.20356 | 23.46486 | 20.25007 | 16.31387 |
| ENSMUSG00000085837 | 0.286710087 | 0.347689 | 0.516454 | 33.19979 | 31.19536 | 28.98373 | 36.23019 | 30.22845 | 32.40385 | 24.10723 | 19.57664 |
| ENSMUSG00000085852 | 0.368704967 | 0.512964 | 0.668016 | 8.048435 | 11.99822 | 11.59349 | 17.25247 | 6.825779 | 12.29112 | 14.46434 | 4.350364 |
| ENSMUSG00000085862 | -0.040760803 | 0.935541 | 0.961806 | 12.07265 | 7.198929 | 9.93728 | 18.11509 | 14.62667 | 12.29112 | 14.46434 | 7.613138 |
| ENSMUSG00000085870 | 0.524039036 | 0.104313 | 0.22735 | 259.562 | 273.5593 | 149.8873 | 227.7326 | 137.4907 | 268.1698 | 125.3576 | 102.2336 |
| ENSMUSG00000085873 | -0.04179683 | 0.931675 | 0.95966 | 18.10898 | 9.598572 | 32.29616 | 22.42821 | 34.1289 | 18.99536 | 18.32149 | 14.13868 |
| ENSMUSG00000085875 | -0.748176122 | 0.104106 | 0.227059 | 12.07265 | 14.39786 | 8.281066 | 16.38985 | 20.47734 | 13.40849 | 32.78583 | 18.48905 |
| ENSMUSG00000085881 | -0.118012871 | 0.837416 | 0.901608 | 12.07265 | 5.999108 | 9.109173 | 12.93935 | 9.751113 | 20.11273 | 7.714313 | 6.525547 |
| ENSMUSG00000085882 | -0.140319301 | 0.30869 | 0.47703 | 415.5004 | 359.9465 | 451.3181 | 444.2511 | 449.5263 | 374.3203 | 502.3947 | 517.6934 |
| ENSMUSG00000085890 | 1.112823499 | 0.04873 | 0.133795 | 16.09687 | 37.19447 | 29.81184 | 25.01608 | 13.65156 | 23.46486 | 9.642892 | 3.262773 |
| ENSMUSG00000085891 | 1.049540412 | 0.006274 | 0.030414 | 54.32694 | 85.18733 | 66.24853 | 87.9876 | 39.97956 | 49.16446 | 39.53586 | 13.05109 |
| ENSMUSG00000085894 | 0.226118074 | 0.424022 | 0.589155 | 64.38748 | 86.38715 | 62.108 | 80.22398 | 54.60623 | 94.9768 | 51.10733 | 50.02919 |
| ENSMUSG00000085900 | 1.015668273 | 0.066342 | 0.166045 | 14.08476 | 17.99732 | 9.93728 | 14.6646 | 8.776002 | 3.352122 | 4.821446 | 10.87591 |
| ENSMUSG00000085912 | 1.941300587 | 0.000259 | 0.002781 | 22.1332 | 44.3934 | 14.90592 | 26.74133 | 8.776002 | 8.938993 | 4.821446 | 5.437955 |
| ENSMUSG00000085913 | 0.321680361 | 0.248145 | 0.411541 | 86.52067 | 67.19001 | 59.62368 | 64.69676 | 45.83023 | 73.74669 | 64.60737 | 38.06569 |
| ENSMUSG00000085923 | 0.649795428 | 0.2816 | 0.44792 | 16.09687 | 9.598572 | 16.56213 | 6.900988 | 7.80089 | 11.17374 | 2.892867 | 9.78832 |
| ENSMUSG00000085925 | 1.212244465 | 0.004478 | 0.02373 | 87.52673 | 106.7841 | 59.62368 | 88.85022 | 39.00445 | 69.27719 | 20.25007 | 19.57664 |
| ENSMUSG00000085929 | -0.00602158 | 0.985009 | 0.991442 | 46.2785 | 29.99554 | 24.8432 | 38.81806 | 42.9049 | 34.6386 | 27.0001 | 35.89051 |
| ENSMUSG00000085933 | -0.039510094 | 0.942957 | 0.967108 | 7.04238 | 8.398751 | 18.21835 | 7.763611 | 9.751113 | 7.821619 | 14.46434 | 10.87591 |
| ENSMUSG00000085936 | 0.216616351 | 0.187921 | 0.340174 | 102.6175 | 112.7832 | 97.71658 | 117.3168 | 99.46135 | 92.74205 | 91.60747 | 85.9197 |
| ENSMUSG00000085939 | -0.600386388 | 0.033189 | 0.101653 | 71.42986 | 47.99286 | 48.85829 | 68.14726 | 94.5858 | 56.98608 | 89.67889 | 117.4598 |
| ENSMUSG00000085940 | -0.642003099 | 0.429665 | 0.594534 | 16.09687 | 1.199822 | 9.93728 | 7.763611 | 13.65156 | 17.87799 | 21.21436 | 2.175182 |
| ENSMUSG00000085945 | 0.176205571 | 0.466539 | 0.627229 | 102.6175 | 124.7814 | 158.1684 | 125.0804 | 75.08357 | 145.2586 | 116.679 | 116.3722 |
| ENSMUSG00000085956 | -0.060559324 | 0.854789 | 0.913001 | 32.19374 | 35.99465 | 35.60859 | 41.40593 | 22.42756 | 52.51658 | 32.78583 | 44.59123 |
| ENSMUSG00000085957 | -0.163279431 | 0.699401 | 0.810259 | 23.13925 | 9.598572 | 19.04645 | 17.25247 | 13.65156 | 20.11273 | 18.32149 | 26.10219 |
| ENSMUSG00000085962 | 0.308787725 | 0.334762 | 0.503039 | 45.27245 | 41.99375 | 27.32752 | 43.9938 | 42.9049 | 26.81698 | 25.07152 | 32.62773 |
| ENSMUSG00000085971 | 0.144261359 | 0.719239 | 0.824438 | 24.1453 | 43.19358 | 30.63995 | 25.01608 | 42.9049 | 29.05173 | 22.17865 | 16.31387 |
| ENSMUSG00000085982 | 1.756786875 | 0.000157 | 0.001889 | 36.21796 | 47.99286 | 25.67131 | 21.56559 | 5.850668 | 12.29112 | 7.714313 | 13.05109 |
| ENSMUSG00000085995 | -0.433823314 | 0.266587 | 0.431024 | 18.10898 | 35.99465 | 16.56213 | 35.36756 | 48.75557 | 32.40385 | 32.78583 | 28.27737 |
| ENSMUSG00000085996 | 0.636512005 | 0.000423 | 0.004024 | 111.672 | 122.3818 | 152.3716 | 140.6076 | 78.0089 | 86.03781 | 95.46463 | 80.48174 |
| ENSMUSG00000085998 | -0.751714762 | 0.130564 | 0.263031 | 9.054489 | 19.19714 | 10.76539 | 11.21411 | 11.70134 | 14.52586 | 27.0001 | 30.45255 |
| ENSMUSG00000086003 | -1.153883739 | 0.010356 | 0.043728 | 6.036326 | 11.99822 | 9.109173 | 8.626235 | 20.47734 | 13.40849 | 23.14294 | 21.75182 |
| ENSMUSG00000086010 | -3.325501248 | 4.72E-11 | 9.27E-09 | 3.018163 | 11.99822 | 24.01509 | 10.35148 | 94.5858 | 90.5073 | 108.9647 | 206.6423 |
| ENSMUSG00000086013 | 0.104207128 | 0.581611 | 0.722577 | 102.6175 | 106.7841 | 93.57605 | 119.9047 | 78.0089 | 113.9722 | 105.1075 | 96.79561 |
| ENSMUSG00000086015 | 0.360507051 | 0.320288 | 0.488405 | 41.24823 | 29.99554 | 25.67131 | 33.64232 | 26.32801 | 13.40849 | 28.92867 | 32.62773 |
| ENSMUSG00000086034 | -0.485829052 | 0.345386 | 0.514192 | 13.07871 | 14.39786 | 9.109173 | 12.07673 | 7.80089 | 15.64324 | 28.92867 | 15.22628 |
| ENSMUSG00000086040 | 0.279643624 | 0.273716 | 0.43956 | 175.0535 | 151.1775 | 177.2148 | 232.9083 | 87.76002 | 168.7235 | 167.7863 | 183.8029 |
| ENSMUSG00000086043 | -0.099188131 | 0.814845 | 0.887383 | 40.24217 | 35.99465 | 48.85829 | 30.19182 | 63.38224 | 16.76061 | 32.78583 | 53.29196 |
| ENSMUSG00000086044 | 0.905764532 | 0.067204 | 0.167403 | 18.10898 | 15.59768 | 11.59349 | 18.11509 | 6.825779 | 6.704245 | 13.50005 | 6.525547 |
| ENSMUSG00000086054 | -0.293631875 | 0.25599 | 0.419812 | 77.46619 | 62.39072 | 75.3577 | 43.13117 | 61.43201 | 89.38993 | 79.07171 | 87.00729 |
| ENSMUSG00000086058 | -0.3549139 | 0.497558 | 0.654039 | 16.09687 | 3.599465 | 11.59349 | 9.488858 | 12.67645 | 15.64324 | 12.53576 | 11.9635 |
| ENSMUSG00000086061 | -1.139600802 | 0.017969 | 0.065568 | 33.19979 | 9.598572 | 23.18699 | 21.56559 | 54.60623 | 18.99536 | 55.92877 | 64.16787 |
| ENSMUSG00000086067 | 2.329235733 | 7.94E-05 | 0.001132 | 25.15136 | 23.99643 | 20.70267 | 14.6646 | 1.950223 | 5.58687 | 1.928578 | 7.613138 |
| ENSMUSG00000086075 | -0.536550113 | 0.185315 | 0.336864 | 18.10898 | 10.79839 | 27.32752 | 33.64232 | 27.30312 | 40.22547 | 27.96439 | 36.9781 |
| ENSMUSG00000086077 | -0.079364751 | 0.756511 | 0.848507 | 65.39353 | 47.99286 | 86.9512 | 66.42201 | 85.8098 | 55.8687 | 66.53595 | 75.04379 |
| ENSMUSG00000086096 | 1.311443132 | 0.195387 | 0.349322 | 18.10898 | 2.399643 | 1.656213 | 43.13117 | 1.950223 | 11.17374 | 5.785735 | 7.613138 |
| ENSMUSG00000086103 | 0.78144557 | 0.039764 | 0.116156 | 66.39959 | 59.99108 | 47.20208 | 49.16954 | 40.95467 | 49.16446 | 24.10723 | 15.22628 |
| ENSMUSG00000086105 | 1.565032518 | 0.008821 | 0.038937 | 35.2119 | 37.19447 | 39.74912 | 30.19182 | 2.925334 | 27.93435 | 5.785735 | 11.9635 |
| ENSMUSG00000086108 | -0.688417194 | 0.043014 | 0.122757 | 22.1332 | 17.99732 | 30.63995 | 21.56559 | 30.22845 | 29.05173 | 49.17875 | 41.32846 |
| ENSMUSG00000086111 | -0.134343632 | 0.794321 | 0.873694 | 10.06054 | 11.99822 | 23.18699 | 8.626235 | 12.67645 | 20.11273 | 9.642892 | 17.40146 |
| ENSMUSG00000086119 | -0.054691085 | 0.610986 | 0.74589 | 467.8153 | 601.1106 | 505.145 | 529.6508 | 546.0623 | 498.3488 | 577.6092 | 559.0218 |
| ENSMUSG00000086140 | -0.714094121 | 0.124398 | 0.254449 | 12.07265 | 5.999108 | 8.281066 | 11.21411 | 13.65156 | 20.11273 | 15.42863 | 13.05109 |
| ENSMUSG00000086158 | -0.145758512 | 0.276074 | 0.442294 | 150.9082 | 181.1731 | 167.2775 | 187.1893 | 193.072 | 204.4795 | 176.4649 | 185.9781 |
| ENSMUSG00000086181 | 0.416050459 | 0.385346 | 0.552857 | 19.11503 | 13.19804 | 9.93728 | 10.35148 | 9.751113 | 8.938993 | 10.60718 | 9.78832 |
| ENSMUSG00000086199 | 0.510447391 | 0.187913 | 0.340174 | 44.26639 | 44.3934 | 34.78048 | 26.74133 | 14.62667 | 39.10809 | 30.85725 | 20.66423 |
| ENSMUSG00000086212 | -1.058579155 | 0.053667 | 0.143131 | 9.054489 | 5.999108 | 15.73403 | 8.626235 | 10.72622 | 32.40385 | 26.03581 | 14.13868 |
| ENSMUSG00000086213 | -1.930064362 | 7.15E-06 | 0.000178 | 27.16347 | 11.99822 | 44.71776 | 24.15346 | 78.98402 | 60.3382 | 144.6434 | 131.5985 |
| ENSMUSG00000086228 | 0.121755855 | 0.747537 | 0.842434 | 25.15136 | 26.39607 | 34.78048 | 24.15346 | 33.15378 | 13.40849 | 31.82154 | 22.83941 |

| | | | | | | | | | | | |
|---|---|---|---|---|---|---|---|---|---|---|---|
| ENSMUSG00000086237 | -0.546691781 | 0.320469 | 0.488405 | 7.04238 | 4.799286 | 12.4216 | 10.35148 | 12.67645 | 16.76061 | 15.42863 | 6.525547 |
| ENSMUSG00000086240 | 0.681360773 | 0.028694 | 0.091574 | 91.55095 | 106.7841 | 77.01392 | 98.33908 | 59.48179 | 91.62468 | 48.21446 | 33.71532 |
| ENSMUSG00000086243 | -1.758459506 | 0.02133 | 0.073719 | 12.07265 | 7.198929 | 29.81184 | 4.313117 | 43.88001 | 7.821619 | 36.64299 | 93.53283 |
| ENSMUSG00000086245 | -0.173222499 | 0.563856 | 0.709134 | 38.23007 | 40.79393 | 51.34261 | 32.77969 | 41.92979 | 48.04709 | 32.78583 | 61.99269 |
| ENSMUSG00000086247 | 0.545685742 | 0.107067 | 0.231384 | 43.26034 | 73.18911 | 62.108 | 81.94923 | 34.1289 | 59.22083 | 56.89306 | 28.27737 |
| ENSMUSG00000086265 | -1.285173973 | 0.093894 | 0.210995 | 3.018163 | 5.999108 | 29.81184 | 6.038364 | 34.1289 | 12.29112 | 16.39292 | 47.85401 |
| ENSMUSG00000086266 | 0.443613419 | 0.009918 | 0.042337 | 269.6226 | 334.7502 | 297.2903 | 293.292 | 194.0472 | 292.752 | 197.6793 | 194.6788 |
| ENSMUSG00000086283 | 0.956160205 | 0.000246 | 0.002673 | 73.44197 | 124.7814 | 86.12309 | 75.04824 | 39.97956 | 55.8687 | 45.32159 | 43.50364 |
| ENSMUSG00000086287 | -0.285738852 | 0.596307 | 0.735213 | 14.08476 | 14.39786 | 4.140533 | 10.35148 | 16.57689 | 14.52586 | 9.642892 | 10.87591 |
| ENSMUSG00000086288 | 0.275299701 | 0.143412 | 0.281067 | 109.6599 | 134.38 | 100.2009 | 127.6683 | 96.53602 | 117.3243 | 90.64318 | 84.8321 |
| ENSMUSG00000086290 | 0.34737647 | 0.000458 | 0.004282 | 862.1886 | 837.4754 | 732.8744 | 803.9651 | 674.777 | 681.5982 | 622.9308 | 562.2846 |
| ENSMUSG00000086296 | -0.005242078 | 0.992224 | 0.995514 | 7.04238 | 39.59411 | 22.35888 | 21.56559 | 15.60178 | 17.87799 | 22.17865 | 34.80291 |
| ENSMUSG00000086308 | 0.573630378 | 0.024382 | 0.081507 | 239.4409 | 279.5584 | 250.0882 | 276.0395 | 195.9974 | 249.1744 | 158.1434 | 98.97079 |
| ENSMUSG00000086316 | 0.154938555 | 0.121104 | 0.250318 | 779.6921 | 842.2747 | 854.606 | 957.5121 | 755.7113 | 854.7912 | 770.467 | 705.8466 |
| ENSMUSG00000086322 | -0.086714875 | 0.731172 | 0.831495 | 43.26034 | 32.39518 | 51.34261 | 48.30692 | 51.6809 | 44.69496 | 42.42872 | 48.9416 |
| ENSMUSG00000086324 | -0.552678611 | 0.442908 | 0.606134 | 13.07871 | 10.79839 | 1.656213 | 12.07673 | 25.35289 | 5.58687 | 10.60718 | 13.05109 |
| ENSMUSG00000086328 | 0.15041937 | 0.353997 | 0.522769 | 320.9313 | 331.1507 | 289.8373 | 311.4071 | 363.7165 | 282.6956 | 251.6795 | 229.4817 |
| ENSMUSG00000086330 | -0.093701225 | 0.862415 | 0.917656 | 34.20585 | 10.79839 | 13.24971 | 15.52722 | 9.751113 | 31.28647 | 17.3572 | 20.66423 |
| ENSMUSG00000086331 | -1.121342914 | 0.108059 | 0.232685 | 15.09082 | 7.198929 | 2.48432 | 6.038364 | 7.80089 | 16.76061 | 30.85725 | 10.87591 |
| ENSMUSG00000086341 | 0.21320834 | 0.77505 | 0.860825 | 21.12714 | 4.799286 | 7.45296 | 12.93935 | 19.50223 | 4.469496 | 12.53576 | 3.262773 |
| ENSMUSG00000086342 | -0.690597961 | 0.09949 | 0.220192 | 16.09687 | 13.19804 | 7.45296 | 11.21411 | 16.57689 | 20.11273 | 19.28578 | 20.66423 |
| ENSMUSG00000086350 | -0.115169956 | 0.718631 | 0.824151 | 35.2119 | 27.5959 | 26.49941 | 30.19182 | 29.25334 | 44.69496 | 32.78583 | 22.83941 |
| ENSMUSG00000086358 | 0.316650732 | 0.53604 | 0.68669 | 11.0666 | 9.598572 | 20.70267 | 24.15346 | 7.80089 | 15.64324 | 11.57147 | 18.48905 |
| ENSMUSG00000086368 | 1.343810526 | 0.017952 | 0.065518 | 21.12714 | 19.19714 | 19.04645 | 6.900988 | 4.875557 | 8.938993 | 5.785735 | 6.525547 |
| ENSMUSG00000086370 | 0.397998006 | 0.077722 | 0.185351 | 458.7608 | 355.1472 | 362.7107 | 478.756 | 251.5787 | 468.1797 | 274.8224 | 263.197 |
| ENSMUSG00000086382 | 0.255315052 | 0.242882 | 0.405447 | 57.3451 | 79.18822 | 72.87338 | 65.55939 | 54.60623 | 60.3382 | 49.17875 | 66.34306 |
| ENSMUSG00000086386 | 0.446149874 | 0.066489 | 0.166232 | 66.39959 | 136.7797 | 101.8571 | 108.6906 | 77.03379 | 72.62932 | 78.10742 | 75.04379 |
| ENSMUSG00000086389 | -0.272324936 | 0.531161 | 0.682784 | 33.19979 | 17.99732 | 13.24971 | 23.29083 | 17.552 | 25.6996 | 39.53586 | 22.83941 |
| ENSMUSG00000086390 | -1.515038694 | 9.25E-05 | 0.001255 | 65.39353 | 43.19358 | 137.4657 | 75.04824 | 219.4 | 120.6764 | 258.4295 | 323.0146 |
| ENSMUSG00000086392 | -0.587913 | 0.186469 | 0.33839 | 9.054489 | 13.19804 | 9.93728 | 12.93935 | 24.37778 | 13.40849 | 15.42863 | 14.13868 |
| ENSMUSG00000086395 | 0.184830484 | 0.457413 | 0.619143 | 49.29666 | 46.79304 | 55.48314 | 67.28463 | 50.70579 | 52.51658 | 53.0359 | 36.9781 |
| ENSMUSG00000086401 | 0.157736131 | 0.576628 | 0.718739 | 40.24217 | 47.99286 | 35.60859 | 48.30692 | 31.20356 | 51.39921 | 32.78583 | 39.15328 |
| ENSMUSG00000086421 | 0.191643473 | 0.514561 | 0.668973 | 46.2785 | 49.19268 | 34.78048 | 37.95543 | 37.05423 | 49.16446 | 27.0001 | 33.71532 |
| ENSMUSG00000086427 | 0.370921795 | 0.000116 | 0.001506 | 1410.488 | 1635.357 | 1411.094 | 1625.183 | 1244.242 | 1248.107 | 1164.861 | 1044.087 |
| ENSMUSG00000086429 | 0.173128562 | 0.549922 | 0.697988 | 289.7437 | 259.1615 | 372.648 | 263.9628 | 373.4676 | 338.5644 | 163.9292 | 176.1898 |
| ENSMUSG00000086436 | -0.014064328 | 0.97682 | 0.987311 | 15.09082 | 15.59768 | 18.21835 | 8.626235 | 16.57689 | 8.938993 | 21.21436 | 10.87591 |
| ENSMUSG00000086442 | 0.793734774 | 0.114738 | 0.242317 | 39.23612 | 29.99554 | 23.18699 | 25.01608 | 15.60178 | 31.28647 | 15.42863 | 5.437955 |
| ENSMUSG00000086446 | 0.955079105 | 0.104645 | 0.227912 | 75.45408 | 53.99197 | 42.23344 | 65.55939 | 18.52711 | 65.92507 | 5.785735 | 32.62773 |
| ENSMUSG00000086448 | 0.140241135 | 0.574166 | 0.716582 | 60.36326 | 58.79126 | 80.32634 | 65.55939 | 59.48179 | 41.34284 | 69.42882 | 70.69342 |
| ENSMUSG00000086451 | 0.881238878 | 0.004348 | 0.023246 | 57.3451 | 103.1847 | 105.9976 | 75.91087 | 40.95467 | 67.04245 | 43.39301 | 34.80291 |
| ENSMUSG00000086453 | 0.441546054 | 0.31293 | 0.481091 | 23.13925 | 21.59679 | 33.12427 | 30.19182 | 26.32801 | 30.1691 | 10.60718 | 13.05109 |
| ENSMUSG00000086454 | 0.500596146 | 0.18885 | 0.341288 | 27.16347 | 29.99554 | 32.29616 | 36.23019 | 29.25334 | 24.58223 | 25.07152 | 9.78832 |
| ENSMUSG00000086465 | -0.513601003 | 0.309216 | 0.477644 | 15.09082 | 9.598572 | 8.281066 | 6.900988 | 9.751113 | 12.29112 | 20.25007 | 14.13868 |
| ENSMUSG00000086468 | 0.174712888 | 0.569633 | 0.713384 | 27.16347 | 37.19447 | 29.81184 | 29.3292 | 24.37778 | 27.93435 | 22.17865 | 34.80291 |
| ENSMUSG00000086472 | -0.152725749 | 0.636672 | 0.764056 | 37.22401 | 27.5959 | 19.87456 | 27.60395 | 27.30312 | 30.1691 | 28.92867 | 38.06569 |
| ENSMUSG00000086477 | 0.269446503 | 0.214607 | 0.372744 | 137.8294 | 135.5798 | 88.60741 | 99.2017 | 98.48624 | 110.62 | 82.92887 | 89.18247 |
| ENSMUSG00000086495 | 0.054698153 | 0.865131 | 0.91922 | 35.2119 | 23.99643 | 25.67131 | 33.64232 | 23.40267 | 34.6386 | 23.14294 | 33.71532 |
| ENSMUSG00000086496 | 0.735261326 | 0.034301 | 0.104482 | 33.19979 | 32.39518 | 44.71776 | 43.13117 | 23.40267 | 34.6386 | 15.42863 | 19.57664 |
| ENSMUSG00000086502 | 0.177887779 | 0.321251 | 0.489274 | 136.8234 | 103.1847 | 119.2474 | 116.4542 | 93.61069 | 121.7938 | 92.57176 | 114.1971 |
| ENSMUSG00000086503 | 1.119856268 | 0.589227 | 0.729391 | 25857.61 | 95.98572 | 100.2009 | 25110.97 | 53.63112 | 23243.62 | 99.32178 | 146.8248 |
| ENSMUSG00000086506 | -0.468604212 | 0.34453 | 0.513276 | 11.0666 | 4.799286 | 15.73403 | 18.97772 | 21.45245 | 21.23011 | 14.46434 | 14.13868 |
| ENSMUSG00000086515 | 0.43444559 | 0.131777 | 0.264789 | 42.25428 | 46.79304 | 35.60859 | 49.16954 | 35.10401 | 33.52122 | 37.60728 | 21.75182 |
| ENSMUSG00000086533 | 0.235402048 | 0.362274 | 0.530898 | 62.37537 | 44.3934 | 73.70149 | 76.77349 | 46.80534 | 67.04245 | 50.14304 | 56.55474 |
| ENSMUSG00000086536 | 0.38942497 | 0.487861 | 0.645722 | 30.18163 | 22.79661 | 23.18699 | 22.42821 | 2.925334 | 21.23011 | 24.10723 | 27.18978 |
| ENSMUSG00000086537 | 0.113470135 | 0.695094 | 0.807493 | 38.23007 | 53.99197 | 45.54586 | 35.36756 | 49.73068 | 42.46022 | 38.57157 | 28.27737 |
| ENSMUSG00000086539 | -0.341823506 | 0.365488 | 0.534244 | 19.11503 | 15.59768 | 41.40533 | 24.15346 | 34.1289 | 30.1691 | 32.78583 | 31.54014 |
| ENSMUSG00000086541 | 0.028586076 | 0.921329 | 0.953074 | 39.23612 | 44.3934 | 32.29616 | 43.13117 | 41.92979 | 51.39921 | 30.85725 | 31.54014 |
| ENSMUSG00000086544 | 0.893216301 | 0.003903 | 0.021455 | 101.6115 | 62.39072 | 49.6864 | 79.36136 | 39.00445 | 46.92971 | 43.39301 | 28.27737 |
| ENSMUSG00000086549 | -0.136032375 | 0.512226 | 0.667488 | 99.59938 | 68.38983 | 106.8258 | 100.0643 | 84.83468 | 106.1505 | 105.1075 | 118.5474 |
| ENSMUSG00000086555 | -0.411797382 | 0.40269 | 0.569693 | 9.054489 | 9.598572 | 14.90592 | 9.488858 | 21.45245 | 15.64324 | 9.642892 | 10.87591 |
| ENSMUSG00000086560 | 0.971776783 | 0.099263 | 0.219875 | 28.16952 | 14.39786 | 11.59349 | 6.900988 | 8.776002 | 6.704245 | 6.750024 | 8.700729 |
| ENSMUSG00000086567 | 0.408055082 | 0.020703 | 0.072206 | 365.1977 | 358.7466 | 363.5388 | 328.6596 | 261.3298 | 359.7945 | 240.108 | 206.6423 |
| ENSMUSG00000086583 | 0.118113728 | 0.392425 | 0.559836 | 945.6911 | 1197.422 | 988.7593 | 1161.091 | 1038.494 | 1149.778 | 971.0392 | 795.0291 |
| ENSMUSG00000086584 | -0.145014848 | 0.841301 | 0.904265 | 25.15136 | 3.599465 | 3.312427 | 18.11509 | 8.776002 | 13.40849 | 19.28578 | 14.13868 |

| | | | | | | | | | | | |
|---|---|---|---|---|---|---|---|---|---|---|---|
| ENSMUSG00000086587 | 2.207679531 | 1.72E-06 | 5.78E-05 | 88.53278 | 110.3836 | 93.57605 | 115.5915 | 19.50223 | 48.04709 | 12.53576 | 8.700729 |
| ENSMUSG00000086591 | -0.018455413 | 0.96153 | 0.978064 | 19.11503 | 21.59679 | 15.73403 | 25.8787 | 18.52711 | 23.46486 | 27.0001 | 14.13868 |
| ENSMUSG00000086596 | 1.036256091 | 0.000227 | 0.002501 | 50.30272 | 40.79393 | 52.17072 | 56.07053 | 30.22845 | 20.11273 | 24.10723 | 22.83941 |
| ENSMUSG00000086598 | 0.693625193 | 0.104448 | 0.227536 | 16.09687 | 32.39518 | 35.60859 | 36.23019 | 18.52711 | 17.87799 | 27.0001 | 10.87591 |
| ENSMUSG00000086600 | -0.739710781 | 0.098219 | 0.218244 | 9.054489 | 10.79839 | 13.24971 | 23.29083 | 25.35289 | 15.64324 | 28.92867 | 25.0146 |
| ENSMUSG00000086602 | -0.159146786 | 0.721827 | 0.82602 | 20.12109 | 9.598572 | 14.07781 | 10.35148 | 15.60178 | 11.17374 | 17.3572 | 16.31387 |
| ENSMUSG00000086603 | -0.269890796 | 0.653893 | 0.776692 | 18.10898 | 15.59768 | 8.281066 | 4.313117 | 9.751113 | 11.17374 | 23.14294 | 10.87591 |
| ENSMUSG00000086604 | -0.106786601 | 0.725102 | 0.827919 | 33.19979 | 68.38983 | 41.40533 | 37.09281 | 56.55646 | 46.92971 | 47.25017 | 41.32846 |
| ENSMUSG00000086606 | -0.52989916 | 0.358958 | 0.527337 | 12.07265 | 15.59768 | 7.45296 | 7.763611 | 20.47734 | 4.469496 | 16.39292 | 19.57664 |
| ENSMUSG00000086607 | -0.603968239 | 0.111866 | 0.238407 | 24.1453 | 11.99822 | 28.15563 | 15.52722 | 33.15378 | 23.46486 | 29.89296 | 35.89051 |
| ENSMUSG00000086610 | 0.120906559 | 0.728788 | 0.829716 | 27.16347 | 33.595 | 52.99882 | 25.01608 | 27.30312 | 29.05173 | 33.75012 | 38.06569 |
| ENSMUSG00000086629 | 0.413340742 | 0.014811 | 0.057151 | 236.4228 | 256.7618 | 215.3077 | 242.3972 | 153.0925 | 231.2964 | 177.4292 | 152.2628 |
| ENSMUSG00000086638 | -0.65142716 | 0.160679 | 0.304203 | 8.048435 | 14.39786 | 15.73403 | 7.763611 | 15.60178 | 20.11273 | 23.14294 | 13.05109 |
| ENSMUSG00000086644 | -0.273534583 | 0.547211 | 0.695504 | 26.15741 | 14.39786 | 55.48314 | 21.56559 | 25.35289 | 43.57759 | 36.64299 | 38.06569 |
| ENSMUSG00000086645 | -1.78131373 | 0.002569 | 0.015735 | 5.030272 | 3.599465 | 9.93728 | 6.900988 | 21.45245 | 8.938993 | 35.6787 | 22.83941 |
| ENSMUSG00000086657 | -0.005178844 | 0.986302 | 0.992283 | 34.20585 | 44.3934 | 35.60859 | 26.74133 | 43.88001 | 32.40385 | 36.64299 | 27.18978 |
| ENSMUSG00000086677 | 0.493414522 | 0.261004 | 0.425111 | 12.07265 | 13.19804 | 16.56213 | 18.97772 | 9.751113 | 13.40849 | 9.642892 | 10.87591 |
| ENSMUSG00000086681 | -0.10772033 | 0.830767 | 0.897748 | 12.07265 | 11.99822 | 10.76539 | 12.07673 | 6.825779 | 15.64324 | 19.28578 | 8.700729 |
| ENSMUSG00000086682 | 0.427426825 | 0.089677 | 0.204634 | 141.8537 | 118.7823 | 81.98256 | 113.0037 | 78.0089 | 109.5027 | 88.7146 | 61.99269 |
| ENSMUSG00000086688 | -0.323128523 | 0.401626 | 0.568792 | 22.1332 | 16.7975 | 23.18699 | 17.25247 | 35.10401 | 16.76061 | 19.28578 | 28.27737 |
| ENSMUSG00000086693 | 0.376966536 | 0.530217 | 0.682119 | 21.12714 | 13.19804 | 15.73403 | 4.313117 | 7.80089 | 10.05637 | 7.714313 | 16.31387 |
| ENSMUSG00000086695 | 0.224889661 | 0.463624 | 0.624527 | 42.25428 | 31.19536 | 43.88965 | 45.71905 | 27.30312 | 50.28183 | 33.75012 | 29.36496 |
| ENSMUSG00000086697 | 0.407616494 | 0.270113 | 0.435085 | 20.12109 | 29.99554 | 29.81184 | 23.29083 | 22.42756 | 25.6996 | 13.50005 | 16.31387 |
| ENSMUSG00000086704 | 1.443828347 | 0.077385 | 0.184836 | 51.30877 | 57.59143 | 33.95237 | 35.36756 | 4.875557 | 51.39921 | 1.928578 | 7.613138 |
| ENSMUSG00000086712 | -0.025908664 | 0.967992 | 0.981932 | 5.030272 | 23.99643 | 18.21835 | 20.70296 | 21.45245 | 31.28647 | 6.750024 | 9.78832 |
| ENSMUSG00000086714 | -0.223766617 | 0.560964 | 0.706583 | 29.17558 | 20.39697 | 33.12427 | 18.97772 | 24.37778 | 43.57759 | 31.82154 | 19.57664 |
| ENSMUSG00000086715 | 0.264035907 | 0.555722 | 0.702687 | 13.07871 | 9.598572 | 17.39024 | 19.84034 | 15.60178 | 13.40849 | 10.60718 | 10.87591 |
| ENSMUSG00000086716 | -0.666953962 | 0.29461 | 0.4622 | 3.018163 | 4.799286 | 15.73403 | 6.900988 | 12.67645 | 7.821619 | 13.50005 | 15.22628 |
| ENSMUSG00000086718 | -0.364656895 | 0.437388 | 0.60116 | 9.054489 | 10.79839 | 9.109173 | 16.38985 | 20.47734 | 13.40849 | 11.57147 | 13.05109 |
| ENSMUSG00000086725 | 0.107113127 | 0.488264 | 0.646025 | 197.1867 | 163.1757 | 168.1056 | 181.1509 | 162.8436 | 145.2586 | 152.3577 | 199.0292 |
| ENSMUSG00000086728 | -0.074678185 | 0.764528 | 0.853838 | 46.2785 | 35.99465 | 42.23344 | 46.58167 | 38.02934 | 52.51658 | 49.17875 | 41.32846 |
| ENSMUSG00000086740 | -0.020417799 | 0.943323 | 0.967283 | 35.2119 | 34.79482 | 31.46805 | 31.91707 | 32.17867 | 24.58223 | 35.6787 | 42.41605 |
| ENSMUSG00000086741 | 0.918546748 | 0.096616 | 0.215606 | 32.19374 | 9.598572 | 28.98373 | 12.93935 | 14.62667 | 13.40849 | 6.750024 | 9.78832 |
| ENSMUSG00000086742 | 0.482038664 | 0.319083 | 0.487145 | 64.38748 | 31.19536 | 59.62368 | 46.58167 | 10.72622 | 59.22083 | 31.82154 | 43.50364 |
| ENSMUSG00000086743 | 0.276065143 | 0.52715 | 0.680117 | 26.15741 | 32.39518 | 26.49941 | 18.97772 | 38.02934 | 14.52586 | 17.3572 | 15.22628 |
| ENSMUSG00000086746 | 0.089045037 | 0.715028 | 0.821396 | 55.33299 | 88.78679 | 85.29498 | 81.94923 | 57.53157 | 67.04245 | 78.10742 | 90.27006 |
| ENSMUSG00000086748 | 4.340419945 | 1.91E-18 | 1.86E-15 | 165.999 | 185.9723 | 127.5284 | 185.4641 | 1.950223 | 21.23011 | 4.821446 | 5.437955 |
| ENSMUSG00000086753 | 0.312734619 | 0.220279 | 0.379065 | 163.9869 | 148.7779 | 153.1997 | 177.7004 | 99.46135 | 204.4795 | 108.9647 | 106.5839 |
| ENSMUSG00000086765 | 0.358906033 | 0.410502 | 0.576705 | 88.53278 | 69.58965 | 43.06154 | 88.85022 | 36.07912 | 68.15982 | 95.46463 | 26.10219 |
| ENSMUSG00000086770 | 1.086378728 | 0.010856 | 0.045379 | 28.16952 | 20.39697 | 20.70267 | 26.74133 | 16.57689 | 7.821619 | 7.714313 | 13.05109 |
| ENSMUSG00000086774 | -0.430701946 | 0.449791 | 0.612329 | 5.030272 | 8.398751 | 16.56213 | 9.488858 | 10.72622 | 20.11273 | 15.42863 | 7.613138 |
| ENSMUSG00000086779 | 0.183706139 | 0.720849 | 0.82531 | 19.11503 | 11.99822 | 19.87456 | 18.11509 | 10.72622 | 29.05173 | 7.714313 | 14.13868 |
| ENSMUSG00000086782 | 0.524301654 | 0.006166 | 0.030052 | 238.4349 | 200.3702 | 154.0278 | 217.3811 | 159.9183 | 148.6108 | 141.7505 | 112.0219 |
| ENSMUSG00000086784 | -0.328341336 | 0.026688 | 0.087148 | 660.9777 | 676.6993 | 572.2217 | 618.501 | 952.6837 | 604.4994 | 763.717 | 850.4962 |
| ENSMUSG00000086786 | 0.174225108 | 0.710519 | 0.818711 | 11.0666 | 13.19804 | 14.07781 | 9.488858 | 9.751113 | 13.40849 | 11.57147 | 7.613138 |
| ENSMUSG00000086794 | 0.091404256 | 0.726216 | 0.828695 | 48.29061 | 49.19268 | 35.60859 | 43.9938 | 43.88001 | 40.22547 | 48.21446 | 32.62773 |
| ENSMUSG00000086796 | -0.633289757 | 0.03677 | 0.110148 | 26.15741 | 16.7975 | 22.35888 | 29.3292 | 36.07912 | 41.34284 | 32.78583 | 38.06569 |
| ENSMUSG00000086804 | -0.377862439 | 0.689944 | 0.803616 | 11.0666 | 8.398751 | 0 | 20.70296 | 7.80089 | 22.34748 | 4.821446 | 17.40146 |
| ENSMUSG00000086807 | -1.417701851 | 0.020009 | 0.070419 | 10.06054 | 3.599465 | 14.90592 | 14.6646 | 34.1289 | 13.40849 | 16.39292 | 53.29196 |
| ENSMUSG00000086810 | 0.123991511 | 0.75145 | 0.845141 | 39.23612 | 20.39697 | 33.95237 | 50.89479 | 22.42756 | 25.6996 | 38.57157 | 46.76642 |
| ENSMUSG00000086825 | 1.421450315 | 4.44E-05 | 0.000728 | 170.0232 | 243.5638 | 151.5435 | 229.4578 | 67.28268 | 132.9675 | 51.10733 | 45.67883 |
| ENSMUSG00000086826 | 0.291826174 | 0.198571 | 0.353192 | 125.7568 | 121.182 | 91.09173 | 112.1411 | 95.56091 | 118.4417 | 65.57166 | 88.09488 |
| ENSMUSG00000086837 | 0.52216666 | 0.089421 | 0.204176 | 53.32088 | 46.79304 | 48.03018 | 65.55939 | 37.05423 | 50.28183 | 22.17865 | 40.24087 |
| ENSMUSG00000086841 | 0.113446163 | 0.324186 | 0.492159 | 2999.048 | 3006.753 | 2826.328 | 3003.655 | 3237.37 | 2774.44 | 2713.51 | 2213.248 |
| ENSMUSG00000086843 | 0.308659059 | 0.090607 | 0.206196 | 254.5318 | 314.3532 | 205.3704 | 292.4294 | 240.8525 | 240.2354 | 204.4293 | 174.0146 |
| ENSMUSG00000086844 | -0.909453811 | 0.057819 | 0.150744 | 14.08476 | 9.598572 | 35.60859 | 16.38985 | 24.37778 | 29.05173 | 49.17875 | 41.32846 |
| ENSMUSG00000086847 | -0.064244272 | NA | NA | 10.06054 | 5.999108 | 110.1382 | 12.93935 | 36.07912 | 26.81698 | 46.28588 | 36.9781 |
| ENSMUSG00000086851 | -0.203097089 | 0.638726 | 0.765824 | 23.13925 | 10.79839 | 14.07781 | 15.52722 | 19.50223 | 24.58223 | 17.3572 | 11.9635 |
| ENSMUSG00000086852 | -0.447373346 | NA | NA | 2.012109 | 1.199822 | 38.92101 | 0.862623 | 15.60178 | 20.11273 | 11.57147 | 11.9635 |
| ENSMUSG00000086859 | 0.268078267 | 4.54E-05 | 0.000739 | 1411.494 | 1339.001 | 1397.016 | 1518.217 | 1171.109 | 1153.13 | 1211.147 | 1172.423 |
| ENSMUSG00000086873 | 0.139283861 | 0.78214 | 0.865651 | 19.11503 | 15.59768 | 14.90592 | 18.97772 | 7.80089 | 25.6996 | 8.678602 | 20.66423 |
| ENSMUSG00000086877 | 0.60864053 | 0.003193 | 0.018605 | 187.1261 | 141.5789 | 144.9187 | 194.9529 | 93.61069 | 140.7891 | 93.53605 | 112.0219 |
| ENSMUSG00000086881 | -0.224259402 | 0.503684 | 0.659571 | 63.38142 | 39.59411 | 58.79557 | 47.44429 | 36.07912 | 78.21619 | 82.92887 | 47.85401 |
| ENSMUSG00000086889 | 0.354784901 | 0.212977 | 0.370828 | 82.49646 | 113.983 | 71.21717 | 66.42201 | 62.40712 | 92.74205 | 52.07161 | 53.29196 |

| | | | | | | | | | | | |
|---|---|---|---|---|---|---|---|---|---|---|---|
| ENSMUSG00000086890 | -0.099445454 | 0.854875 | 0.913001 | 17.10292 | 10.79839 | 9.93728 | 10.35148 | 14.62667 | 18.99536 | 13.50005 | 4.350364 |
| ENSMUSG00000086894 | 0.275122165 | 0.341591 | 0.510353 | 86.52067 | 71.98929 | 62.108 | 63.83414 | 38.02934 | 86.03781 | 56.89306 | 54.37955 |
| ENSMUSG00000086903 | 0.707384574 | 0.082263 | 0.19272 | 76.46013 | 76.78858 | 59.62368 | 75.91087 | 43.88001 | 82.68568 | 29.89296 | 20.66423 |
| ENSMUSG00000086904 | 0.274902126 | 0.464262 | 0.625067 | 88.53278 | 52.79215 | 37.2648 | 80.22398 | 48.75557 | 83.80306 | 41.46443 | 40.24087 |
| ENSMUSG00000086914 | -0.185291118 | 0.46738 | 0.62788 | 48.29061 | 57.59143 | 56.31125 | 50.03216 | 54.60623 | 42.46022 | 77.14313 | 66.34306 |
| ENSMUSG00000086915 | 0.104884679 | 0.753581 | 0.846815 | 37.22401 | 28.79572 | 26.49941 | 22.42821 | 30.22845 | 23.46486 | 20.25007 | 32.62773 |
| ENSMUSG00000086916 | 0.684502183 | 0.175089 | 0.323435 | 32.19374 | 25.19625 | 28.98373 | 28.46658 | 14.62667 | 37.99072 | 10.60718 | 8.700729 |
| ENSMUSG00000086918 | 0.440088135 | 0.306828 | 0.475352 | 33.19979 | 27.5959 | 32.29616 | 49.16954 | 22.42756 | 44.69496 | 12.53576 | 26.10219 |
| ENSMUSG00000086920 | -0.016338676 | 0.977433 | 0.987546 | 20.12109 | 14.39786 | 7.45296 | 6.900988 | 5.850668 | 16.76061 | 14.46434 | 11.9635 |
| ENSMUSG00000086922 | -0.159770855 | 0.521116 | 0.674643 | 47.28455 | 46.79304 | 33.95237 | 38.81806 | 41.92979 | 52.51658 | 45.32159 | 45.67883 |
| ENSMUSG00000086924 | -1.14478528 | 0.007704 | 0.035228 | 14.08476 | 7.198929 | 22.35888 | 14.6646 | 27.30312 | 23.46486 | 35.6787 | 44.59123 |
| ENSMUSG00000086930 | 0.205781372 | 0.711568 | 0.819194 | 18.10898 | 7.198929 | 9.109173 | 15.52722 | 4.875557 | 11.17374 | 14.46434 | 13.05109 |
| ENSMUSG00000086938 | -0.220401777 | 0.341938 | 0.510749 | 125.7568 | 130.7805 | 99.3728 | 82.81186 | 127.7396 | 94.9768 | 125.3576 | 160.9635 |
| ENSMUSG00000086942 | -0.083417725 | 0.833295 | 0.899058 | 19.11503 | 20.39697 | 19.87456 | 14.6646 | 20.47734 | 24.58223 | 22.17865 | 10.87591 |
| ENSMUSG00000086961 | -0.397640057 | 0.429288 | 0.594236 | 7.04238 | 10.79839 | 16.56213 | 10.35148 | 13.65156 | 15.64324 | 21.21436 | 8.700729 |
| ENSMUSG00000086965 | -0.340300209 | 0.303001 | 0.471278 | 79.47829 | 79.18822 | 57.13936 | 44.85642 | 117.0134 | 46.92971 | 76.17884 | 88.09488 |
| ENSMUSG00000086968 | 0.070644785 | 0.55075 | 0.698537 | 439.6458 | 407.9393 | 373.4761 | 412.334 | 413.4472 | 329.6254 | 379.9299 | 430.6861 |
| ENSMUSG00000087006 | -0.643035429 | 0.033286 | 0.101896 | 195.1745 | 135.5798 | 341.1799 | 229.4578 | 278.8818 | 252.5265 | 405.0014 | 474.1897 |
| ENSMUSG00000087013 | -0.010784326 | 0.960382 | 0.977472 | 105.6357 | 113.983 | 168.1056 | 101.7896 | 131.64 | 105.0332 | 120.5361 | 137.0365 |
| ENSMUSG00000087014 | -0.003685439 | 0.994142 | 0.996626 | 11.0666 | 9.598572 | 11.59349 | 9.488858 | 14.62667 | 5.58687 | 10.60718 | 10.87591 |
| ENSMUSG00000087020 | -0.465746736 | 0.381954 | 0.549871 | 15.09082 | 8.398751 | 6.624853 | 8.626235 | 11.70134 | 14.52586 | 7.714313 | 19.57664 |
| ENSMUSG00000087022 | -2.270349491 | 1.16E-05 | 0.000257 | 9.054489 | 4.799286 | 14.07781 | 12.07673 | 21.45245 | 41.34284 | 52.07161 | 81.56933 |
| ENSMUSG00000087024 | 1.619095443 | 0.005596 | 0.027922 | 37.22401 | 25.19625 | 21.53077 | 23.29083 | 2.925334 | 3.352122 | 15.42863 | 13.05109 |
| ENSMUSG00000087026 | 0.299670601 | 0.482659 | 0.64109 | 13.07871 | 20.39697 | 24.8432 | 26.74133 | 16.57689 | 11.17374 | 25.07152 | 16.31387 |
| ENSMUSG00000087028 | -0.152318345 | 0.832792 | 0.898936 | 10.06054 | 5.999108 | 19.04645 | 6.900988 | 23.40267 | 5.58687 | 3.857157 | 14.13868 |
| ENSMUSG00000087029 | 0.52970212 | 0.19283 | 0.346067 | 132.7992 | 177.5736 | 129.1846 | 166.4863 | 92.63557 | 184.3667 | 108.0004 | 34.80291 |
| ENSMUSG00000087042 | 0.715958902 | 0.006796 | 0.032303 | 66.39959 | 53.99197 | 46.37397 | 43.9938 | 34.1289 | 31.28647 | 31.82154 | 30.45255 |
| ENSMUSG00000087050 | -0.418620147 | 0.313929 | 0.481979 | 14.08476 | 13.19804 | 18.21835 | 18.97772 | 17.552 | 30.1691 | 13.50005 | 26.10219 |
| ENSMUSG00000087060 | 0.391123124 | 0.035529 | 0.10732 | 184.1079 | 260.3613 | 195.4332 | 164.7611 | 153.0925 | 162.0192 | 167.7863 | 128.3357 |
| ENSMUSG00000087063 | 1.149756203 | 0.058418 | 0.151725 | 12.07265 | 23.99643 | 19.87456 | 16.38985 | 6.825779 | 17.87799 | 4.821446 | 3.262773 |
| ENSMUSG00000087075 | -1.311112639 | 0.002289 | 0.014455 | 21.12714 | 39.59411 | 52.99882 | 22.42821 | 98.48624 | 36.87335 | 101.2504 | 101.146 |
| ENSMUSG00000087088 | 0.416296519 | 0.019009 | 0.068097 | 96.58122 | 110.3836 | 96.06037 | 95.75121 | 66.30757 | 71.51194 | 83.89316 | 76.13138 |
| ENSMUSG00000087095 | -0.00443547 | 0.964471 | 0.979975 | 403.4278 | 363.5459 | 389.2101 | 395.0816 | 374.4427 | 425.7195 | 402.1086 | 355.6423 |
| ENSMUSG00000087107 | -0.075887075 | 0.726743 | 0.828973 | 84.50857 | 68.38983 | 97.71658 | 113.8663 | 91.66046 | 116.2069 | 88.7146 | 90.27006 |
| ENSMUSG00000087120 | 0.305781796 | 0.543501 | 0.692451 | 23.13925 | 27.5959 | 20.70267 | 20.70296 | 38.02934 | 13.40849 | 10.60718 | 11.9635 |
| ENSMUSG00000087128 | 0.342533837 | 0.274234 | 0.440147 | 33.19979 | 40.79393 | 32.29616 | 40.5433 | 23.40267 | 41.34284 | 23.14294 | 28.27737 |
| ENSMUSG00000087132 | -0.754716689 | 0.100684 | 0.221931 | 20.12109 | 15.59768 | 21.53077 | 13.80198 | 20.47734 | 14.52586 | 41.46443 | 43.50364 |
| ENSMUSG00000087135 | 0.858919504 | 0.010867 | 0.045396 | 66.39959 | 37.19447 | 38.09291 | 59.52102 | 23.40267 | 26.81698 | 23.14294 | 38.06569 |
| ENSMUSG00000087138 | 0.292475714 | 0.044854 | 0.126192 | 543.2694 | 471.5299 | 401.6317 | 426.136 | 367.617 | 420.1327 | 405.0014 | 309.9635 |
| ENSMUSG00000087141 | -0.088593815 | 0.632664 | 0.761035 | 77.46619 | 80.38804 | 92.74794 | 77.63611 | 78.0089 | 79.33356 | 95.46463 | 96.79561 |
| ENSMUSG00000087142 | 0.236604026 | 0.422126 | 0.587229 | 60.36326 | 57.59143 | 38.92101 | 43.13117 | 49.73068 | 51.39921 | 29.89296 | 38.06569 |
| ENSMUSG00000087143 | 0.391519926 | 0.003593 | 0.020216 | 322.9434 | 357.5468 | 338.6956 | 380.417 | 278.8818 | 312.8647 | 251.6795 | 224.0438 |
| ENSMUSG00000087150 | 0.125466992 | 0.514377 | 0.66878 | 302.8224 | 311.9536 | 366.8512 | 380.417 | 214.5245 | 306.1605 | 386.68 | 342.5912 |
| ENSMUSG00000087153 | -0.145991688 | 0.434368 | 0.598574 | 117.7084 | 86.38715 | 105.1695 | 106.9653 | 124.8142 | 91.62468 | 130.179 | 114.1971 |
| ENSMUSG00000087159 | -0.325437536 | 0.520198 | 0.673644 | 31.18769 | 27.5959 | 18.21835 | 20.70296 | 10.72622 | 55.8687 | 31.82154 | 23.927 |
| ENSMUSG00000087165 | -0.344651739 | 0.097332 | 0.216795 | 111.672 | 104.3845 | 91.91984 | 126.8057 | 135.5405 | 99.44629 | 161.0363 | 155.5255 |
| ENSMUSG00000087166 | 0.322000974 | 0.21568 | 0.37387 | 65.39353 | 50.3925 | 82.81066 | 56.07053 | 61.43201 | 41.34284 | 47.25017 | 54.37955 |
| ENSMUSG00000087168 | 0.451679949 | 0.229926 | 0.390207 | 21.12714 | 19.19714 | 22.35888 | 19.84034 | 16.57689 | 12.29112 | 19.28578 | 11.9635 |
| ENSMUSG00000087174 | 0.392452192 | 0.113514 | 0.240613 | 1065.412 | 830.2765 | 601.2054 | 1056.714 | 655.2748 | 982.1718 | 587.2521 | 482.8904 |
| ENSMUSG00000087177 | 0.394418174 | 0.011104 | 0.04622 | 595.5842 | 649.1034 | 513.4261 | 657.3191 | 401.7459 | 589.9735 | 432.9658 | 413.2846 |
| ENSMUSG00000087178 | 0.372646033 | 0.118495 | 0.247362 | 124.7507 | 94.7859 | 83.63877 | 74.18562 | 71.18313 | 71.51194 | 87.75031 | 59.81751 |
| ENSMUSG00000087179 | -0.012471002 | 0.949274 | 0.971271 | 319.9253 | 291.5566 | 327.1021 | 255.3366 | 350.065 | 203.3621 | 290.251 | 359.9926 |
| ENSMUSG00000087187 | 0.657705258 | 0.110779 | 0.236865 | 14.08476 | 23.99643 | 28.15563 | 25.8787 | 12.67645 | 15.64324 | 19.28578 | 10.87591 |
| ENSMUSG00000087204 | 1.52084835 | 0.003212 | 0.018674 | 43.26034 | 92.38626 | 57.13936 | 70.73513 | 21.45245 | 49.16446 | 10.60718 | 10.87591 |
| ENSMUSG00000087211 | -0.698681514 | 2.45E-07 | 1.19E-05 | 362.1796 | 307.1543 | 274.9314 | 273.4516 | 549.9628 | 464.8276 | 516.859 | 440.4744 |
| ENSMUSG00000087213 | -0.107325948 | 0.713069 | 0.820065 | 37.22401 | 45.59322 | 49.6864 | 44.85642 | 44.85512 | 44.69496 | 33.75012 | 68.51824 |
| ENSMUSG00000087221 | -0.074302605 | 0.583533 | 0.724151 | 149.9021 | 139.1793 | 138.2938 | 134.5693 | 149.192 | 143.0239 | 159.1077 | 139.2117 |
| ENSMUSG00000087222 | -0.444933302 | 0.044732 | 0.126005 | 39.23612 | 45.59322 | 48.85829 | 40.5433 | 62.40712 | 55.8687 | 55.92877 | 63.08028 |
| ENSMUSG00000087223 | -2.124546514 | 0.030325 | 0.095184 | 1.006054 | 0 | 5.796746 | 8.626235 | 14.62667 | 3.352122 | 16.39292 | 34.80291 |
| ENSMUSG00000087231 | 0.43128951 | 0.010074 | 0.042828 | 225.3562 | 205.1695 | 196.2613 | 209.6175 | 156.0178 | 197.7752 | 140.7862 | 126.1606 |
| ENSMUSG00000087249 | 0.036649667 | 0.918039 | 0.951605 | 67.40564 | 25.19625 | 33.12427 | 42.26855 | 45.83023 | 46.92971 | 37.60728 | 33.71532 |
| ENSMUSG00000087253 | -0.430283577 | 0.192293 | 0.34549 | 30.18163 | 19.19714 | 21.53077 | 33.64232 | 41.92979 | 25.6996 | 36.64299 | 36.9781 |
| ENSMUSG00000087259 | 0.174493367 | 0.320879 | 0.488867 | 460.7729 | 284.3577 | 412.3971 | 491.6954 | 337.3885 | 387.7288 | 387.6442 | 351.2919 |
| ENSMUSG00000087260 | -0.155251605 | 0.118042 | 0.246781 | 1126.781 | 1100.236 | 1096.413 | 1232.689 | 1384.658 | 1083.853 | 1255.504 | 1349.701 |

| | | | | | | | | | | | |
|---|---|---|---|---|---|---|---|---|---|---|---|
| ENSMUSG00000087263 | 0.214681963 | 0.549454 | 0.69758 | 94.56911 | 37.19447 | 39.74912 | 58.6584 | 37.05423 | 58.10345 | 49.17875 | 54.37955 |
| ENSMUSG00000087264 | 1.163994789 | 0.001882 | 0.012567 | 29.17558 | 51.59233 | 43.88965 | 30.19182 | 15.60178 | 11.17374 | 21.21436 | 20.66423 |
| ENSMUSG00000087266 | 0.094204197 | 0.879462 | 0.927685 | 10.06054 | 14.39786 | 6.624853 | 14.6646 | 3.900445 | 17.87799 | 14.46434 | 6.525547 |
| ENSMUSG00000087268 | -0.380388352 | 0.552946 | 0.700183 | 7.04238 | 2.399643 | 10.76539 | 23.29083 | 15.60178 | 11.17374 | 14.46434 | 16.31387 |
| ENSMUSG00000087269 | 0.142194977 | 0.411055 | 0.576931 | 152.9203 | 139.1793 | 147.403 | 166.4863 | 118.9636 | 175.4277 | 128.2505 | 128.3357 |
| ENSMUSG00000087281 | -0.101070372 | 0.795094 | 0.873988 | 22.1332 | 13.19804 | 35.60859 | 23.29083 | 26.32801 | 31.28647 | 24.10723 | 20.66423 |
| ENSMUSG00000087291 | -0.33354429 | 0.45627 | 0.618115 | 12.07265 | 20.39697 | 9.109173 | 12.07673 | 17.552 | 12.29112 | 21.21436 | 15.22628 |
| ENSMUSG00000087298 | 0.337236612 | 0.465098 | 0.62595 | 18.10898 | 25.19625 | 17.39024 | 12.93935 | 21.45245 | 13.40849 | 7.714313 | 15.22628 |
| ENSMUSG00000087299 | -0.213265485 | 0.491708 | 0.648814 | 31.18769 | 29.99554 | 31.46805 | 25.01608 | 30.22845 | 43.57759 | 38.57157 | 23.927 |
| ENSMUSG00000087305 | 0.012280346 | 0.946807 | 0.969393 | 76.46013 | 106.7841 | 89.43552 | 91.43809 | 89.71024 | 98.32892 | 80.036 | 92.44524 |
| ENSMUSG00000087307 | -0.227880994 | 0.599401 | 0.737591 | 16.09687 | 15.59768 | 21.53077 | 33.64232 | 31.20356 | 35.75597 | 18.32149 | 17.40146 |
| ENSMUSG00000087314 | 0.270393587 | 0.542813 | 0.692005 | 21.12714 | 31.19536 | 14.07781 | 18.11509 | 23.40267 | 21.23011 | 10.60718 | 14.13868 |
| ENSMUSG00000087326 | 0.476023854 | 0.239956 | 0.401945 | 22.1332 | 31.19536 | 16.56213 | 21.56559 | 19.50223 | 21.23011 | 12.53576 | 11.9635 |
| ENSMUSG00000087327 | 0.667237907 | 0.235316 | 0.396223 | 16.09687 | 8.398751 | 8.281066 | 22.42821 | 10.72622 | 10.05637 | 8.678602 | 5.437955 |
| ENSMUSG00000087331 | -0.414347853 | 0.038276 | 0.112999 | 60.36326 | 56.39161 | 62.9361 | 62.10889 | 93.61069 | 67.04245 | 78.10742 | 83.74451 |
| ENSMUSG00000087333 | 0.403950685 | 3.56E-05 | 0.000612 | 766.6134 | 691.0972 | 783.3889 | 683.1978 | 504.1325 | 608.9689 | 563.1449 | 536.1824 |
| ENSMUSG00000087334 | 0.370808476 | 0.479933 | 0.638987 | 16.09687 | 8.398751 | 10.76539 | 10.35148 | 9.751113 | 11.17374 | 4.821446 | 9.78832 |
| ENSMUSG00000087336 | -0.176165975 | 0.511603 | 0.667063 | 69.41775 | 35.99465 | 56.31125 | 43.9938 | 52.65601 | 63.69032 | 63.64308 | 53.29196 |
| ENSMUSG00000087338 | 0.472938472 | 0.337825 | 0.506202 | 16.09687 | 17.99732 | 19.87456 | 17.25247 | 5.850668 | 23.46486 | 11.57147 | 10.87591 |
| ENSMUSG00000087341 | 0.760054391 | 0.042874 | 0.122528 | 37.22401 | 35.99465 | 36.43669 | 26.74133 | 12.67645 | 32.40385 | 17.3572 | 18.48905 |
| ENSMUSG00000087347 | 0.084210034 | 0.878155 | 0.927008 | 24.1453 | 4.799286 | 14.90592 | 10.35148 | 11.70134 | 13.40849 | 15.42863 | 10.87591 |
| ENSMUSG00000087354 | 0.255030411 | 0.52855 | 0.68098 | 60.36326 | 29.99554 | 33.95237 | 48.30692 | 31.20356 | 60.3382 | 21.21436 | 32.62773 |
| ENSMUSG00000087356 | -0.476716634 | 0.27655 | 0.44271 | 16.09687 | 10.79839 | 11.59349 | 17.25247 | 15.60178 | 15.64324 | 16.39292 | 30.45255 |
| ENSMUSG00000087361 | 0.282444889 | 0.593328 | 0.732826 | 15.09082 | 13.19804 | 5.796746 | 19.84034 | 8.776002 | 10.05637 | 14.46434 | 10.87591 |
| ENSMUSG00000087364 | -0.846369721 | 0.181395 | 0.331863 | 13.07871 | 16.7975 | 45.54586 | 18.97772 | 68.25779 | 7.821619 | 45.32159 | 48.9416 |
| ENSMUSG00000087365 | -0.090860639 | 0.498227 | 0.654655 | 346.0827 | 291.5566 | 313.8524 | 265.688 | 310.0854 | 281.5783 | 347.1441 | 356.7299 |
| ENSMUSG00000087366 | 0.505344155 | 0.039723 | 0.116065 | 136.8234 | 104.3845 | 107.6539 | 94.88858 | 72.15824 | 110.62 | 59.78593 | 70.69342 |
| ENSMUSG00000087367 | 0.599203916 | 0.194975 | 0.348823 | 10.06054 | 20.39697 | 13.24971 | 20.70296 | 10.72622 | 12.29112 | 9.642892 | 9.78832 |
| ENSMUSG00000087368 | 0.398806098 | 0.035154 | 0.10643 | 196.1806 | 227.9661 | 135.8095 | 187.1893 | 145.2916 | 148.6108 | 146.572 | 123.9854 |
| ENSMUSG00000087370 | 0.249637842 | 0.110365 | 0.236189 | 670.0322 | 545.9188 | 539.9255 | 639.204 | 432.9494 | 650.3117 | 459.9659 | 473.1021 |
| ENSMUSG00000087371 | 0.732374625 | 0.062249 | 0.159064 | 20.12109 | 32.39518 | 29.81184 | 24.15346 | 13.65156 | 21.23011 | 19.28578 | 9.78832 |
| ENSMUSG00000087373 | 0.535143088 | 0.231616 | 0.392139 | 28.16952 | 28.79572 | 42.23344 | 33.64232 | 13.65156 | 42.46022 | 23.14294 | 13.05109 |
| ENSMUSG00000087377 | -0.172016031 | 0.603572 | 0.740729 | 32.19374 | 38.39429 | 27.32752 | 25.8787 | 28.27823 | 51.39921 | 28.92867 | 30.45255 |
| ENSMUSG00000087380 | -1.895099681 | 0.000431 | 0.004079 | 11.0666 | 3.599465 | 8.281066 | 6.900988 | 24.37778 | 45.81234 | 27.96439 | 14.13868 |
| ENSMUSG00000087381 | 0.46173211 | 0.012597 | 0.050782 | 136.8234 | 105.5843 | 127.5284 | 111.2784 | 91.66046 | 99.44629 | 71.3574 | 88.09488 |
| ENSMUSG00000087382 | 1.317410995 | 0.034206 | 0.104245 | 30.18163 | 20.39697 | 9.109173 | 20.70296 | 7.80089 | 14.52586 | 7.714313 | 2.175182 |
| ENSMUSG00000087385 | -0.145725632 | 0.31187 | 0.480297 | 147.89 | 122.3818 | 134.9814 | 131.9814 | 145.2916 | 141.9065 | 165.8577 | 141.3868 |
| ENSMUSG00000087390 | 0.494147168 | 0.341937 | 0.510749 | 20.12109 | 31.19536 | 14.07781 | 19.84034 | 5.850668 | 24.58223 | 11.57147 | 18.48905 |
| ENSMUSG00000087396 | 0.361219819 | 0.021089 | 0.073174 | 256.5439 | 287.9572 | 284.0406 | 337.2858 | 212.5743 | 272.6393 | 235.2866 | 188.1533 |
| ENSMUSG00000087397 | -0.62269175 | 0.167552 | 0.313498 | 25.15136 | 14.39786 | 16.56213 | 13.80198 | 29.25334 | 42.46022 | 14.46434 | 21.75182 |
| ENSMUSG00000087400 | 0.095248256 | 0.711204 | 0.818994 | 59.35721 | 75.58876 | 115.9349 | 108.6906 | 91.66046 | 77.09881 | 78.10742 | 91.35765 |
| ENSMUSG00000087403 | 0.308292088 | 0.225711 | 0.385537 | 110.666 | 97.18554 | 77.84202 | 58.6584 | 64.35735 | 81.56831 | 60.75022 | 70.69342 |
| ENSMUSG00000087408 | 0.302330969 | 0.162589 | 0.306979 | 65.39353 | 58.79126 | 66.24853 | 55.2079 | 53.63112 | 54.75133 | 44.3573 | 46.76642 |
| ENSMUSG00000087411 | 0.872657854 | 0.10035 | 0.221433 | 22.1332 | 10.79839 | 20.70267 | 14.6646 | 6.825779 | 16.76061 | 8.678602 | 5.437955 |
| ENSMUSG00000087412 | -0.042254312 | 0.749096 | 0.843451 | 217.3077 | 253.1623 | 217.792 | 258.787 | 270.1058 | 224.5922 | 222.7508 | 256.6715 |
| ENSMUSG00000087413 | 1.249225399 | 0.001027 | 0.007993 | 23.13925 | 29.99554 | 24.8432 | 34.50494 | 14.62667 | 11.17374 | 11.57147 | 9.78832 |
| ENSMUSG00000087424 | 0.429305043 | 0.1753 | 0.323695 | 25.15136 | 39.59411 | 39.74912 | 41.40593 | 21.45245 | 32.40385 | 30.85725 | 23.927 |
| ENSMUSG00000087433 | 0.004765799 | 0.988234 | 0.993293 | 32.19374 | 52.79215 | 59.62368 | 37.95543 | 57.53157 | 51.39921 | 42.42872 | 30.45255 |
| ENSMUSG00000087436 | 0.139649503 | 0.759218 | 0.84986 | 24.1453 | 15.59768 | 17.39024 | 12.07673 | 12.67645 | 24.58223 | 10.60718 | 15.22628 |
| ENSMUSG00000087439 | -0.469622741 | 0.351611 | 0.520353 | 13.07871 | 5.999108 | 6.624853 | 11.21411 | 11.70134 | 10.05637 | 16.39292 | 13.05109 |
| ENSMUSG00000087440 | -0.072806553 | 0.788605 | 0.870224 | 44.26639 | 69.58965 | 66.24853 | 44.85642 | 53.63112 | 68.15982 | 68.46453 | 45.67883 |
| ENSMUSG00000087443 | -0.409303697 | 0.189307 | 0.341864 | 25.15136 | 25.19625 | 21.53077 | 17.25247 | 28.27823 | 30.1691 | 31.82154 | 27.18978 |
| ENSMUSG00000087444 | -2.260423572 | 3.74E-05 | 0.000638 | 6.036326 | 3.599465 | 9.109173 | 6.038364 | 24.37778 | 20.11273 | 22.17865 | 54.37955 |
| ENSMUSG00000087445 | 0.499381605 | 0.074321 | 0.179906 | 102.6175 | 100.785 | 86.12309 | 102.6522 | 70.20801 | 100.5637 | 67.50024 | 39.15328 |
| ENSMUSG00000087478 | -0.15285341 | 0.625356 | 0.755732 | 35.2119 | 28.79572 | 30.63995 | 31.91707 | 26.32801 | 48.04709 | 39.53586 | 27.18978 |
| ENSMUSG00000087479 | 1.100501194 | 0.000375 | 0.003665 | 97.58727 | 106.7841 | 70.38906 | 140.6076 | 47.78045 | 61.45558 | 54.96448 | 29.36496 |
| ENSMUSG00000087480 | 0.80421036 | 0.010124 | 0.043004 | 43.26034 | 58.79126 | 30.63995 | 43.9938 | 24.37778 | 27.93435 | 22.17865 | 26.10219 |
| ENSMUSG00000087481 | -0.352170109 | 0.49585 | 0.652554 | 20.12109 | 7.198929 | 9.93728 | 13.80198 | 12.67645 | 10.05637 | 25.07152 | 17.40146 |
| ENSMUSG00000087484 | 0.13124546 | 0.608735 | 0.744176 | 70.4238 | 67.19001 | 79.49824 | 81.94923 | 90.68535 | 78.21619 | 47.25017 | 57.64233 |
| ENSMUSG00000087497 | 0.103888316 | 0.487066 | 0.645134 | 563.3904 | 719.8929 | 667.4539 | 703.9008 | 607.4943 | 774.3402 | 576.6449 | 512.2554 |
| ENSMUSG00000087501 | 0.534407264 | 0.235791 | 0.396732 | 25.15136 | 43.19358 | 24.01509 | 29.3292 | 18.52711 | 37.99072 | 15.42863 | 11.9635 |
| ENSMUSG00000087502 | 0.174602195 | 0.594466 | 0.733828 | 85.51462 | 37.19447 | 41.40533 | 43.13117 | 48.75557 | 45.81234 | 44.3573 | 44.59123 |
| ENSMUSG00000087507 | 1.362934633 | 0.019065 | 0.068204 | 78.47224 | 73.18911 | 52.17072 | 54.34528 | 31.20356 | 51.39921 | 13.50005 | 4.350364 |
| ENSMUSG00000087516 | 0.373373703 | 0.135815 | 0.270228 | 510.0696 | 284.3577 | 640.9545 | 358.8514 | 287.6578 | 322.9211 | 404.0372 | 371.9562 |

| | | | | | | | | | | | |
|---|---|---|---|---|---|---|---|---|---|---|---|
| ENSMUSG00000087518 | -2.199271257 | 0.019519 | 0.069291 | 0 | 0 | 13.24971 | 7.763611 | 19.50223 | 24.58223 | 20.25007 | 34.80291 |
| ENSMUSG00000087519 | -0.304870921 | 0.44331 | 0.606369 | 21.12714 | 13.19804 | 19.04645 | 13.80198 | 15.60178 | 29.05173 | 18.32149 | 20.66423 |
| ENSMUSG00000087523 | 0.227189873 | 0.287143 | 0.454071 | 130.7871 | 118.7823 | 96.88848 | 131.9814 | 96.53602 | 130.7328 | 104.1432 | 77.21897 |
| ENSMUSG00000087526 | 1.717195527 | 0.018145 | 0.066 | 7.04238 | 22.79661 | 9.109173 | 23.29083 | 4.875557 | 5.58687 | 0.964289 | 7.613138 |
| ENSMUSG00000087528 | -0.17316171 | 0.396432 | 0.563679 | 85.51462 | 100.785 | 86.9512 | 80.22398 | 108.2374 | 122.9112 | 82.92887 | 83.74451 |
| ENSMUSG00000087535 | -0.382660728 | 0.216765 | 0.375292 | 26.15741 | 27.5959 | 23.18699 | 25.8787 | 23.40267 | 33.52122 | 35.6787 | 41.32846 |
| ENSMUSG00000087543 | 0.228801111 | 0.505732 | 0.661501 | 102.6175 | 64.79036 | 60.45178 | 37.09281 | 69.2329 | 41.34284 | 55.92877 | 58.72992 |
| ENSMUSG00000087556 | 0.474284015 | 0.269338 | 0.434103 | 49.29666 | 58.79126 | 38.92101 | 52.62003 | 22.42756 | 71.51194 | 25.07152 | 25.0146 |
| ENSMUSG00000087574 | 0.169345523 | 0.425169 | 0.590172 | 139.8416 | 129.5807 | 104.3414 | 130.2561 | 77.03379 | 130.7328 | 117.6433 | 122.8978 |
| ENSMUSG00000087578 | 0.189948003 | 0.574024 | 0.716553 | 24.1453 | 17.99732 | 30.63995 | 27.60395 | 18.52711 | 24.58223 | 24.10723 | 21.75182 |
| ENSMUSG00000087589 | 0.262362809 | 0.032627 | 0.100446 | 222.338 | 263.9607 | 250.9163 | 250.1608 | 223.3005 | 205.5968 | 209.2507 | 183.8029 |
| ENSMUSG00000087590 | 0.113336267 | 0.394908 | 0.562234 | 661.9838 | 749.8885 | 633.5016 | 782.3995 | 699.1548 | 763.1665 | 597.8593 | 553.5839 |
| ENSMUSG00000087594 | 0.269769365 | 0.278974 | 0.445354 | 78.47224 | 73.18911 | 112.6225 | 68.14726 | 79.95913 | 61.45558 | 56.89306 | 78.30656 |
| ENSMUSG00000087595 | -0.273160583 | 0.275522 | 0.441755 | 77.46619 | 65.99018 | 89.43552 | 106.1027 | 140.416 | 89.38993 | 99.32178 | 81.56933 |
| ENSMUSG00000087598 | 0.138334884 | 0.268777 | 0.43335 | 500.009 | 622.7074 | 517.5666 | 493.4206 | 552.8881 | 480.4709 | 478.2874 | 423.0729 |
| ENSMUSG00000087610 | 0.468851553 | 0.325407 | 0.493262 | 26.15741 | 20.39697 | 11.59349 | 12.07673 | 11.70134 | 8.938993 | 16.39292 | 13.05109 |
| ENSMUSG00000087611 | 0.24186932 | 0.490293 | 0.647606 | 21.12714 | 21.59679 | 19.87456 | 26.74133 | 14.62667 | 21.23011 | 18.32149 | 21.75182 |
| ENSMUSG00000087612 | -0.79115124 | 0.136031 | 0.270537 | 10.06054 | 17.99732 | 7.45296 | 10.35148 | 20.47734 | 33.52122 | 12.53576 | 11.9635 |
| ENSMUSG00000087613 | -1.309753695 | 0.003457 | 0.019605 | 58.35115 | 33.595 | 81.98256 | 70.73513 | 81.90935 | 84.92043 | 149.4648 | 292.562 |
| ENSMUSG00000087620 | 1.714075635 | 3.49E-05 | 0.000603 | 25.15136 | 27.5959 | 23.18699 | 25.8787 | 6.825779 | 8.938993 | 8.678602 | 6.525547 |
| ENSMUSG00000087623 | 0.712086438 | 0.139328 | 0.275255 | 25.15136 | 25.19625 | 21.53077 | 10.35148 | 14.62667 | 15.64324 | 6.750024 | 13.05109 |
| ENSMUSG00000087624 | 0.408690886 | 0.245671 | 0.408625 | 33.19979 | 28.79572 | 30.63995 | 25.8787 | 19.50223 | 23.46486 | 31.82154 | 14.13868 |
| ENSMUSG00000087626 | 0.371261478 | 0.15234 | 0.293087 | 43.26034 | 53.99197 | 43.88965 | 46.58167 | 42.9049 | 37.99072 | 27.96439 | 35.89051 |
| ENSMUSG00000087627 | -0.051938172 | 0.95006 | 0.971623 | 6.036326 | 1.199822 | 24.01509 | 12.93935 | 23.40267 | 6.704245 | 9.642892 | 6.525547 |
| ENSMUSG00000087648 | -0.431190537 | 0.416333 | 0.582046 | 12.07265 | 15.59768 | 24.8432 | 13.80198 | 29.25334 | 10.05637 | 13.50005 | 36.9781 |
| ENSMUSG00000087651 | -0.011738603 | 0.939373 | 0.964398 | 251.5136 | 199.1704 | 278.2438 | 241.5346 | 234.0267 | 208.949 | 280.6081 | 256.6715 |
| ENSMUSG00000087658 | 0.214574594 | 0.180419 | 0.330568 | 146.8839 | 125.9813 | 135.8095 | 153.547 | 140.416 | 125.1459 | 110.8933 | 108.7591 |
| ENSMUSG00000087662 | -0.278825227 | NA | NA | 2.012109 | 0 | 52.99882 | 4.313117 | 17.552 | 24.58223 | 8.678602 | 21.75182 |
| ENSMUSG00000087672 | 0.597204866 | 0.088004 | 0.201891 | 98.59333 | 86.38715 | 57.13936 | 44.85642 | 42.9049 | 69.27719 | 33.75012 | 43.50364 |
| ENSMUSG00000087674 | 1.224194569 | 0.0056 | 0.027933 | 31.18769 | 19.19714 | 18.21835 | 18.11509 | 9.751113 | 8.938993 | 10.60718 | 7.613138 |
| ENSMUSG00000087679 | -0.211685916 | 0.007129 | 0.033333 | 1157.969 | 1270.611 | 1253.753 | 1147.289 | 1426.588 | 1300.623 | 1351.933 | 1511.752 |
| ENSMUSG00000087687 | -0.097013696 | 0.377851 | 0.545879 | 356.1432 | 316.7529 | 337.8675 | 321.7586 | 352.9903 | 315.0995 | 396.3228 | 359.9926 |
| ENSMUSG00000087691 | 0.943587167 | 0.007255 | 0.03375 | 30.18163 | 49.19268 | 33.12427 | 36.23019 | 19.50223 | 25.6996 | 12.53576 | 19.57664 |
| ENSMUSG00000087700 | -0.942049848 | 0.018883 | 0.067699 | 15.09082 | 8.398751 | 14.07781 | 15.52722 | 21.45245 | 18.99536 | 28.92867 | 33.71532 |
| ENSMUSG00000087701 | 0.250291313 | 0.544359 | 0.69297 | 28.16952 | 26.39607 | 28.15563 | 17.25247 | 26.32801 | 30.1691 | 14.46434 | 13.05109 |
| ENSMUSG00000087795 | 0.074118727 | 0.866131 | 0.919627 | 12.07265 | 15.59768 | 19.87456 | 18.97772 | 24.37778 | 11.17374 | 12.53576 | 15.22628 |
| ENSMUSG00000088088 | -1.151183575 | 0.095069 | 0.213129 | 11.0666 | 9.598572 | 3.312427 | 6.038364 | 23.40267 | 24.58223 | 3.857157 | 14.13868 |
| ENSMUSG00000088139 | 0.158086185 | 0.672163 | 0.790055 | 19.11503 | 39.59411 | 43.06154 | 28.46658 | 21.45245 | 37.99072 | 25.07152 | 32.62773 |
| ENSMUSG00000088185 | -0.98252398 | 0.049712 | 0.135582 | 17.10292 | 5.999108 | 13.24971 | 6.900988 | 22.42756 | 12.29112 | 26.03581 | 25.0146 |
| ENSMUSG00000088378 | 0.547755647 | 0.286834 | 0.453937 | 21.12714 | 13.19804 | 9.93728 | 15.52722 | 7.80089 | 13.40849 | 5.785735 | 14.13868 |
| ENSMUSG00000088549 | 0.618468012 | 0.430661 | 0.595421 | 13.07871 | 19.19714 | 18.21835 | 11.21411 | 3.900445 | 25.6996 | 9.642892 | 1.087591 |
| ENSMUSG00000088609 | -0.41438754 | 0.412526 | 0.578395 | 29.17558 | 17.99732 | 9.93728 | 23.29083 | 40.95467 | 33.52122 | 19.28578 | 13.05109 |
| ENSMUSG00000088632 | 0.505416781 | 0.30632 | 0.474959 | 21.12714 | 11.99822 | 13.24971 | 24.15346 | 15.60178 | 15.64324 | 5.785735 | 13.05109 |
| ENSMUSG00000088984 | 0.211941086 | 0.636274 | 0.763777 | 25.15136 | 29.99554 | 27.32752 | 37.95543 | 14.62667 | 44.69496 | 30.85725 | 14.13868 |
| ENSMUSG00000089235 | -0.645823749 | 0.128777 | 0.260461 | 25.15136 | 14.39786 | 27.32752 | 10.35148 | 37.05423 | 21.23011 | 27.0001 | 35.89051 |
| ENSMUSG00000089255 | 0.396389809 | 0.476182 | 0.635644 | 6.036326 | 22.79661 | 9.109173 | 12.93935 | 8.776002 | 8.938993 | 10.60718 | 9.78832 |
| ENSMUSG00000089317 | 0.324867095 | 0.564097 | 0.709266 | 8.048435 | 8.398751 | 15.73403 | 16.38985 | 8.776002 | 14.52586 | 11.57147 | 4.350364 |
| ENSMUSG00000089396 | 0.746988769 | 0.22027 | 0.379065 | 4.024217 | 15.59768 | 17.39024 | 12.93935 | 7.80089 | 8.938993 | 8.678602 | 4.350364 |
| ENSMUSG00000089417 | -0.392409647 | 0.192856 | 0.346072 | 44.26639 | 51.59233 | 37.2648 | 32.77969 | 56.55646 | 35.75597 | 51.10733 | 72.8686 |
| ENSMUSG00000089542 | 0.272143059 | 0.486102 | 0.644361 | 28.16952 | 28.79572 | 24.01509 | 30.19182 | 38.02934 | 20.11273 | 19.28578 | 14.13868 |
| ENSMUSG00000089635 | 0.863919448 | 0.177883 | 0.32709 | 105.6357 | 14.39786 | 17.39024 | 35.36756 | 19.50223 | 17.87799 | 35.6787 | 21.75182 |
| ENSMUSG00000089636 | 0.515865527 | 0.146198 | 0.284947 | 32.19374 | 26.39607 | 24.01509 | 30.19182 | 22.42756 | 18.99536 | 25.07152 | 11.9635 |
| ENSMUSG00000089639 | 0.929430814 | 0.253929 | 0.417545 | 30.18163 | 35.99465 | 16.56213 | 19.84034 | 10.72622 | 36.87335 | 0.964289 | 5.437955 |
| ENSMUSG00000089651 | 0.97756364 | 0.046991 | 0.13028 | 32.19374 | 58.79126 | 40.57722 | 44.85642 | 18.52711 | 46.92971 | 13.50005 | 10.87591 |
| ENSMUSG00000089661 | 0.905676002 | 0.000233 | 0.002554 | 172.0353 | 219.5673 | 141.6062 | 202.7165 | 97.51113 | 128.498 | 103.1789 | 63.08028 |
| ENSMUSG00000089662 | -0.972704723 | 0.11722 | 0.245482 | 11.0666 | 2.399643 | 4.96864 | 10.35148 | 10.72622 | 22.34748 | 15.42863 | 8.700729 |
| ENSMUSG00000089663 | 0.098192447 | 0.766907 | 0.855443 | 31.18769 | 21.59679 | 27.32752 | 25.01608 | 24.37778 | 23.46486 | 31.82154 | 18.48905 |
| ENSMUSG00000089665 | 0.510302322 | 0.253898 | 0.41753 | 30.18163 | 32.39518 | 28.98373 | 26.74133 | 14.62667 | 18.99536 | 10.60718 | 39.15328 |
| ENSMUSG00000089669 | 0.735233916 | 0.033473 | 0.102307 | 35.2119 | 55.19179 | 38.92101 | 25.8787 | 28.27823 | 20.11273 | 22.17865 | 21.75182 |
| ENSMUSG00000089678 | -3.205778529 | 0.000194 | 0.002211 | 38.23007 | 5.999108 | 99.3728 | 26.74133 | 417.3476 | 115.0895 | 292.1796 | 751.5254 |
| ENSMUSG00000089682 | 0.082231439 | 0.306424 | 0.475001 | 1299.822 | 1119.433 | 1236.363 | 1166.267 | 1140.88 | 1051.449 | 1162.933 | 1200.701 |
| ENSMUSG00000089684 | 1.542076219 | 0.001471 | 0.010426 | 35.2119 | 22.79661 | 16.56213 | 33.64232 | 12.67645 | 12.29112 | 6.750024 | 5.437955 |
| ENSMUSG00000089687 | -0.222982003 | 0.534734 | 0.685637 | 26.15741 | 16.7975 | 18.21835 | 18.97772 | 30.22845 | 18.99536 | 20.25007 | 23.927 |
| ENSMUSG00000089704 | -1.165229474 | 0.007075 | 0.033165 | 72.43591 | 59.99108 | 81.98256 | 32.77969 | 138.4658 | 49.16446 | 162.0006 | 204.4671 |

| | | | | | | | | | | | |
|---|---|---|---|---|---|---|---|---|---|---|---|
| ENSMUSG00000089706 | 0.04575191 | 0.886789 | 0.932224 | 29.17558 | 23.99643 | 38.09291 | 39.68068 | 28.27823 | 36.87335 | 24.10723 | 39.15328 |
| ENSMUSG00000089712 | -3.385593343 | 3.32E-09 | 3.50E-07 | 18.10898 | 3.599465 | 12.4216 | 12.93935 | 93.61069 | 40.22547 | 238.1794 | 123.9854 |
| ENSMUSG00000089715 | 0.558302261 | 9.28E-06 | 0.000214 | 4437.706 | 4703.3 | 4222.516 | 4399.38 | 2839.524 | 3835.945 | 2702.903 | 2685.262 |
| ENSMUSG00000089726 | 0.146555702 | 0.447723 | 0.610316 | 520.1301 | 383.9429 | 445.5214 | 637.4788 | 463.1779 | 559.8044 | 373.1799 | 401.3211 |
| ENSMUSG00000089736 | 0.625777557 | 0.006951 | 0.032819 | 155.9384 | 135.5798 | 182.1835 | 196.6782 | 106.2871 | 146.376 | 108.0004 | 75.04379 |
| ENSMUSG00000089762 | -0.11432838 | 0.385393 | 0.552881 | 730.3955 | 839.8751 | 905.1205 | 699.5877 | 849.3219 | 756.4623 | 810.9672 | 1020.16 |
| ENSMUSG00000089764 | 0.440671556 | 0.23549 | 0.396374 | 65.39353 | 76.78858 | 77.84202 | 91.43809 | 52.65601 | 105.0332 | 36.64299 | 35.89051 |
| ENSMUSG00000089768 | 0.008908784 | 0.962053 | 0.97838 | 72.43591 | 77.9884 | 76.18581 | 84.5371 | 71.18313 | 84.92043 | 84.85745 | 68.51824 |
| ENSMUSG00000089769 | 0.492885048 | 0.089283 | 0.203986 | 63.38142 | 85.18733 | 50.5145 | 49.16954 | 56.55646 | 43.57759 | 34.71441 | 40.24087 |
| ENSMUSG00000089774 | -1.097304592 | 6.10E-05 | 0.000928 | 914.5034 | 725.892 | 1039.274 | 905.7547 | 1089.199 | 1790.033 | 3134.904 | 1658.576 |
| ENSMUSG00000089782 | -0.222365256 | 0.227907 | 0.38778 | 131.7931 | 105.5843 | 104.3414 | 147.5086 | 155.0427 | 157.5497 | 122.4647 | 137.0365 |
| ENSMUSG00000089789 | -0.618910434 | 0.16695 | 0.312673 | 10.06054 | 10.79839 | 9.109173 | 9.488858 | 11.70134 | 18.99536 | 12.53576 | 17.40146 |
| ENSMUSG00000089804 | -0.237785303 | 0.082087 | 0.192406 | 389.343 | 338.3497 | 470.3646 | 338.1484 | 476.8294 | 422.3674 | 478.2874 | 436.124 |
| ENSMUSG00000089809 | 0.1736711 | 0.431577 | 0.596141 | 566.4086 | 367.1454 | 510.1137 | 485.657 | 367.617 | 432.4238 | 599.7879 | 311.0511 |
| ENSMUSG00000089810 | 0.559120941 | 0.00448 | 0.02373 | 590.5539 | 571.1151 | 533.3007 | 613.3253 | 318.8614 | 572.0955 | 353.8941 | 323.0146 |
| ENSMUSG00000089812 | 0.508084877 | 0.096572 | 0.215545 | 93.56306 | 106.7841 | 76.18581 | 142.3329 | 90.68535 | 92.74205 | 67.50024 | 43.50364 |
| ENSMUSG00000089817 | 0.531991497 | 0.007283 | 0.033856 | 113.6841 | 106.7841 | 103.5133 | 98.33908 | 63.38224 | 79.33356 | 59.78593 | 90.27006 |
| ENSMUSG00000089818 | 0.802336513 | 0.049313 | 0.134807 | 25.15136 | 26.39607 | 24.8432 | 29.3292 | 9.751113 | 25.6996 | 11.57147 | 14.13868 |
| ENSMUSG00000089824 | 0.131341143 | 0.565725 | 0.710273 | 903.4368 | 1156.628 | 1058.32 | 1072.241 | 789.8402 | 1507.338 | 804.2172 | 725.4233 |
| ENSMUSG00000089827 | -0.25541768 | 0.702702 | 0.812599 | 6.036326 | 13.19804 | 10.76539 | 6.900988 | 3.900445 | 6.704245 | 22.17865 | 10.87591 |
| ENSMUSG00000089829 | 1.302538957 | 0.007473 | 0.034499 | 32.19374 | 41.99375 | 16.56213 | 34.50494 | 18.52711 | 16.76061 | 8.678602 | 6.525547 |
| ENSMUSG00000089832 | -0.28931833 | 0.068377 | 0.169381 | 479.8879 | 593.9117 | 645.9232 | 455.4652 | 677.7024 | 541.9264 | 662.4667 | 775.4524 |
| ENSMUSG00000089842 | -0.064430931 | 0.898095 | 0.93831 | 13.07871 | 10.79839 | 10.76539 | 6.900988 | 11.70134 | 6.704245 | 10.60718 | 14.13868 |
| ENSMUSG00000089844 | 0.516416812 | 0.347511 | 0.516273 | 8.048435 | 19.19714 | 16.56213 | 16.38985 | 4.875557 | 16.76061 | 7.714313 | 13.05109 |
| ENSMUSG00000089847 | -0.074528024 | 0.451488 | 0.613917 | 445.6821 | 493.1267 | 550.6909 | 494.2833 | 542.1619 | 486.0577 | 519.7519 | 542.708 |
| ENSMUSG00000089854 | 1.384737553 | 0.007133 | 0.033334 | 20.12109 | 20.39697 | 14.07781 | 18.11509 | 8.776002 | 8.938993 | 7.714313 | 2.175182 |
| ENSMUSG00000089857 | 0.086213328 | 0.609737 | 0.744907 | 179.0777 | 205.1695 | 144.9187 | 197.5408 | 196.9725 | 166.4887 | 147.5362 | 171.8394 |
| ENSMUSG00000089862 | -0.169854184 | 0.33359 | 0.501731 | 258.556 | 272.3595 | 253.4006 | 293.292 | 286.6827 | 232.4138 | 294.1082 | 399.1459 |
| ENSMUSG00000089872 | 0.12045147 | 0.103259 | 0.22588 | 755.5468 | 691.0972 | 757.7176 | 760.8339 | 700.1299 | 703.9457 | 668.2524 | 659.0802 |
| ENSMUSG00000089875 | 0.516767702 | 0.051057 | 0.138067 | 77.46619 | 70.78947 | 73.70149 | 103.5148 | 38.02934 | 72.62932 | 54.96448 | 63.08028 |
| ENSMUSG00000089876 | -0.237326188 | 0.323612 | 0.491571 | 152.9203 | 128.3809 | 211.1672 | 162.1732 | 172.5947 | 145.2586 | 185.1435 | 270.8102 |
| ENSMUSG00000089877 | 1.167397643 | 0.126951 | 0.258098 | 15.09082 | 43.19358 | 23.18699 | 27.60395 | 6.825779 | 33.52122 | 1.928578 | 6.525547 |
| ENSMUSG00000089886 | 0.691096273 | 0.19682 | 0.351171 | 20.12109 | 15.59768 | 14.07781 | 11.21411 | 16.57689 | 4.469496 | 8.678602 | 7.613138 |
| ENSMUSG00000089889 | -0.051805337 | 0.819577 | 0.890713 | 105.6357 | 112.7832 | 106.8258 | 105.2401 | 113.1129 | 146.376 | 114.7504 | 71.78101 |
| ENSMUSG00000089901 | 0.33791469 | 0.077938 | 0.185652 | 686.1291 | 795.4817 | 588.7838 | 740.9936 | 578.241 | 748.6406 | 486.966 | 410.0218 |
| ENSMUSG00000089911 | -0.304824753 | 1.98E-05 | 0.000389 | 1738.462 | 1546.57 | 1580.027 | 1544.959 | 2011.655 | 1852.606 | 1969.078 | 2083.825 |
| ENSMUSG00000089917 | 0.026471644 | 0.858937 | 0.915379 | 669.0261 | 559.1168 | 626.8767 | 629.7152 | 548.9877 | 479.3535 | 688.5025 | 723.2481 |
| ENSMUSG00000089924 | 1.56179416 | 0.00438 | 0.023367 | 14.08476 | 20.39697 | 14.07781 | 16.38985 | 3.900445 | 10.05637 | 3.857157 | 4.350364 |
| ENSMUSG00000089940 | 0.673024907 | 0.118324 | 0.247126 | 120.7265 | 170.3747 | 70.38906 | 81.08661 | 55.58134 | 131.8501 | 45.32159 | 44.59123 |
| ENSMUSG00000089960 | -1.09411286 | 0.155356 | 0.297279 | 12.07265 | 0 | 19.04645 | 12.07673 | 12.67645 | 14.52586 | 26.03581 | 40.24087 |
| ENSMUSG00000089968 | 0.218890595 | 0.57932 | 0.720529 | 32.19374 | 39.59411 | 17.39024 | 31.91707 | 32.17867 | 33.52122 | 20.25007 | 17.40146 |
| ENSMUSG00000089972 | 1.824922648 | 0.00215 | 0.013838 | 23.13925 | 23.99643 | 14.90592 | 19.84034 | 1.950223 | 12.29112 | 5.785735 | 3.262773 |
| ENSMUSG00000089979 | 0.232486692 | 0.594044 | 0.733356 | 24.1453 | 10.79839 | 22.35888 | 22.42821 | 16.57689 | 14.52586 | 25.07152 | 11.9635 |
| ENSMUSG00000089984 | -0.141828806 | 0.699173 | 0.810259 | 18.10898 | 27.5959 | 19.87456 | 31.91707 | 36.07912 | 18.99536 | 27.0001 | 25.0146 |
| ENSMUSG00000089991 | 0.358531982 | 0.471304 | 0.630997 | 23.13925 | 31.19536 | 10.76539 | 18.97772 | 22.42756 | 21.23011 | 12.53576 | 8.700729 |
| ENSMUSG00000089993 | 0.170303319 | 0.690855 | 0.80424 | 16.09687 | 10.79839 | 13.24971 | 19.84034 | 10.72622 | 14.52586 | 15.42863 | 13.05109 |
| ENSMUSG00000089996 | 0.206709348 | 0.490206 | 0.647606 | 41.24823 | 46.79304 | 52.17072 | 81.08661 | 56.55646 | 39.10809 | 53.0359 | 43.50364 |
| ENSMUSG00000089998 | -0.331533448 | 0.381313 | 0.549249 | 23.13925 | 32.39518 | 32.29616 | 13.80198 | 28.27823 | 41.34284 | 32.78583 | 25.0146 |
| ENSMUSG00000089999 | 0.007750511 | 0.984586 | 0.9913 | 19.11503 | 22.79661 | 17.39024 | 18.97772 | 24.37778 | 10.05637 | 24.10723 | 18.48905 |
| ENSMUSG00000090000 | 0.031032828 | 0.580936 | 0.722295 | 1773.674 | 1911.316 | 1842.537 | 1899.497 | 1785.429 | 1889.48 | 1774.292 | 1820.627 |
| ENSMUSG00000090002 | -1.100106848 | 0.083094 | 0.193963 | 3.018163 | 13.19804 | 14.07781 | 6.900988 | 21.45245 | 8.938993 | 14.46434 | 34.80291 |
| ENSMUSG00000090015 | 0.047641443 | 0.735586 | 0.834534 | 336.0222 | 445.1338 | 361.8826 | 357.1261 | 417.3476 | 376.5551 | 311.4654 | 343.6788 |
| ENSMUSG00000090019 | -0.451770726 | 0.00712 | 0.033308 | 252.5196 | 254.3622 | 307.2276 | 270.8638 | 353.9654 | 289.3999 | 370.287 | 472.0145 |
| ENSMUSG00000090026 | 0.606614032 | 0.208539 | 0.365717 | 22.1332 | 17.99732 | 15.73403 | 12.93935 | 7.80089 | 18.99536 | 7.714313 | 10.87591 |
| ENSMUSG00000090031 | 0.1372877 | 0.471655 | 0.631172 | 147.89 | 112.7832 | 108.482 | 138.0198 | 122.864 | 134.0849 | 110.8933 | 93.53283 |
| ENSMUSG00000090035 | 0.181465051 | 0.158887 | 0.301844 | 777.68 | 851.8733 | 1009.462 | 870.3871 | 649.4241 | 859.2607 | 736.7169 | 852.6714 |
| ENSMUSG00000090061 | 2.190933457 | 1.98E-09 | 2.31E-07 | 277.671 | 363.5459 | 205.3704 | 376.1038 | 47.78045 | 126.2633 | 54.00019 | 40.24087 |
| ENSMUSG00000090066 | 0.340880175 | 0.329543 | 0.497331 | 41.24823 | 41.99375 | 33.95237 | 25.8787 | 29.25334 | 40.22547 | 18.32149 | 25.0146 |
| ENSMUSG00000090069 | -0.22846458 | 0.573342 | 0.715844 | 29.17558 | 15.59768 | 26.49941 | 19.84034 | 26.32801 | 36.87335 | 29.89296 | 14.13868 |
| ENSMUSG00000090070 | 0.265445013 | 0.558927 | 0.705241 | 18.10898 | 14.39786 | 14.90592 | 23.29083 | 17.552 | 11.17374 | 8.678602 | 21.75182 |
| ENSMUSG00000090071 | 0.338027052 | 0.243318 | 0.405881 | 42.25428 | 32.39518 | 42.23344 | 59.52102 | 32.17867 | 40.22547 | 36.64299 | 31.54014 |
| ENSMUSG00000090077 | 0.269942907 | 0.352572 | 0.521441 | 37.22401 | 67.19001 | 47.20208 | 42.26855 | 35.10401 | 50.28183 | 35.6787 | 39.15328 |
| ENSMUSG00000090083 | 0.111839593 | 0.524701 | 0.677859 | 377.2704 | 446.3336 | 342.008 | 424.4108 | 351.0401 | 488.2925 | 304.7154 | 327.3649 |
| ENSMUSG00000090084 | 0.412266015 | 0.001402 | 0.010091 | 618.7234 | 547.1186 | 501.8326 | 556.3922 | 360.7912 | 416.7805 | 496.6089 | 395.8832 |

| | | | | | | | | | | | |
|---|---|---|---|---|---|---|---|---|---|---|---|
| ENSMUSG00000090086 | 0.700893861 | 6.87E-05 | 0.001017 | 667.014 | 643.1043 | 492.7234 | 626.2647 | 301.3094 | 486.0577 | 325.9297 | 381.7445 |
| ENSMUSG00000090093 | -0.295004646 | 0.313489 | 0.481504 | 54.32694 | 51.59233 | 37.2648 | 42.26855 | 59.48179 | 42.46022 | 45.32159 | 79.39415 |
| ENSMUSG00000090100 | 0.556617346 | 0.000564 | 0.005023 | 670.0322 | 785.8831 | 612.7989 | 711.6644 | 436.8499 | 621.26 | 395.3586 | 437.2116 |
| ENSMUSG00000090101 | 0.397354101 | 0.433394 | 0.597583 | 15.09082 | 10.79839 | 13.24971 | 15.52722 | 13.65156 | 4.469496 | 14.46434 | 8.700729 |
| ENSMUSG00000090110 | 0.671142853 | 0.003123 | 0.018309 | 251.5136 | 299.9554 | 216.1358 | 264.8254 | 136.5156 | 240.2354 | 135.9648 | 135.9489 |
| ENSMUSG00000090112 | 0.269649006 | 0.018052 | 0.065766 | 1227.386 | 1233.417 | 1027.68 | 1290.485 | 979.0118 | 1152.013 | 914.1461 | 919.0145 |
| ENSMUSG00000090113 | 0.657365326 | 0.10338 | 0.226011 | 35.2119 | 38.39429 | 70.38906 | 42.26855 | 14.62667 | 41.34284 | 35.6787 | 27.18978 |
| ENSMUSG00000090115 | 0.224145372 | 0.155479 | 0.297421 | 462.785 | 501.5254 | 369.3356 | 389.9058 | 378.3432 | 436.8933 | 299.8939 | 358.9051 |
| ENSMUSG00000090122 | -0.520330912 | 0.037527 | 0.111572 | 95.57516 | 141.5789 | 220.2764 | 133.7066 | 208.6738 | 222.3574 | 184.1792 | 234.9197 |
| ENSMUSG00000090124 | -0.258854574 | 0.130894 | 0.263472 | 87.52673 | 99.58519 | 101.8571 | 112.1411 | 121.8889 | 100.5637 | 126.3219 | 131.5985 |
| ENSMUSG00000090125 | -0.056244733 | 0.880663 | 0.928348 | 36.21796 | 32.39518 | 38.92101 | 18.11509 | 33.15378 | 45.81234 | 29.89296 | 21.75182 |
| ENSMUSG00000090136 | -0.178288861 | 0.555705 | 0.702687 | 27.16347 | 34.79482 | 31.46805 | 31.05445 | 33.15378 | 46.92971 | 35.6787 | 25.0146 |
| ENSMUSG00000090137 | -0.298076811 | 0.004991 | 0.025723 | 6665.11 | 7653.662 | 7756.047 | 7532.428 | 10168.46 | 7524.397 | 9242.712 | 9465.305 |
| ENSMUSG00000090150 | -0.401927023 | 0.000831 | 0.006763 | 574.457 | 599.9108 | 630.1891 | 554.6669 | 738.1593 | 649.1943 | 833.1458 | 896.1751 |
| ENSMUSG00000090157 | 0.575258907 | 0.143255 | 0.28087 | 27.16347 | 23.99643 | 23.18699 | 25.01608 | 16.57689 | 17.87799 | 8.678602 | 23.927 |
| ENSMUSG00000090165 | -4.94433169 | 2.12E-16 | 1.45E-13 | 5.030272 | 1.199822 | 7.45296 | 9.488858 | 206.7236 | 53.63396 | 144.6434 | 328.4525 |
| ENSMUSG00000090171 | -3.638456587 | 1.58E-06 | 5.40E-05 | 50.30272 | 17.99732 | 103.5133 | 55.2079 | 1049.22 | 145.2586 | 435.8587 | 1202.876 |
| ENSMUSG00000090173 | 0.117492974 | 0.704101 | 0.813757 | 38.23007 | 28.79572 | 33.12427 | 33.64232 | 22.42756 | 42.46022 | 30.85725 | 28.27737 |
| ENSMUSG00000090175 | -4.778302061 | 2.31E-10 | 3.65E-08 | 3.018163 | 2.399643 | 1.656213 | 1.725247 | 47.78045 | 13.40849 | 81.00029 | 95.70802 |
| ENSMUSG00000090191 | -0.653509857 | 0.006013 | 0.029523 | 90.54489 | 93.58608 | 56.31125 | 100.9269 | 103.3618 | 135.2023 | 141.7505 | 155.5255 |
| ENSMUSG00000090194 | 0.81673533 | 0.108163 | 0.232799 | 19.11503 | 16.7975 | 13.24971 | 18.97772 | 5.850668 | 17.87799 | 8.678602 | 6.525547 |
| ENSMUSG00000090200 | 2.426269251 | 0.018858 | 0.067648 | 54.32694 | 22.79661 | 16.56213 | 12.07673 | 0.975111 | 16.76061 | 0 | 2.175182 |
| ENSMUSG00000090207 | -0.014144115 | 0.95453 | 0.974112 | 45.27245 | 50.3925 | 54.65504 | 56.07053 | 56.55646 | 43.57759 | 65.57166 | 42.41605 |
| ENSMUSG00000090210 | 0.548625987 | 0.007551 | 0.03472 | 122.7386 | 147.578 | 120.9036 | 154.4096 | 95.56091 | 116.2069 | 92.57176 | 68.51824 |
| ENSMUSG00000090213 | -0.551227355 | 0.001256 | 0.009284 | 967.8243 | 893.867 | 1059.148 | 980.8029 | 1315.425 | 1003.402 | 1688.47 | 1710.781 |
| ENSMUSG00000090215 | 0.887035393 | 0.221829 | 0.380951 | 12.07265 | 8.398751 | 9.109173 | 30.19182 | 4.875557 | 4.469496 | 4.821446 | 18.48905 |
| ENSMUSG00000090222 | 0.713836579 | 0.001147 | 0.008679 | 74.44802 | 92.38626 | 101.8571 | 81.94923 | 41.92979 | 56.98608 | 59.78593 | 55.46715 |
| ENSMUSG00000090223 | -0.251841789 | 0.105049 | 0.228493 | 358.1554 | 262.7609 | 330.4145 | 409.7462 | 415.3974 | 359.7945 | 396.3228 | 452.4379 |
| ENSMUSG00000090226 | 1.52291224 | 0.118369 | 0.247186 | 34.20585 | 26.39607 | 9.109173 | 31.91707 | 1.950223 | 26.81698 | 6.750024 | 0 |
| ENSMUSG00000090231 | -1.127702664 | 0.002616 | 0.015938 | 40.24217 | 39.59411 | 62.9361 | 31.91707 | 115.0631 | 45.81234 | 87.75031 | 133.7737 |
| ENSMUSG00000090235 | 0.389775891 | 0.122981 | 0.252586 | 67.40564 | 55.19179 | 60.45178 | 49.16954 | 47.78045 | 51.39921 | 46.28588 | 31.54014 |
| ENSMUSG00000090236 | -2.093216104 | 0.000547 | 0.004911 | 10.06054 | 9.598572 | 36.43669 | 14.6646 | 39.00445 | 36.87335 | 71.3574 | 157.7007 |
| ENSMUSG00000090247 | -0.270149596 | 0.063639 | 0.161646 | 649.9111 | 857.8724 | 997.8685 | 908.3425 | 1003.39 | 917.3641 | 1033.718 | 1164.81 |
| ENSMUSG00000090248 | 0.463041844 | 0.245205 | 0.408107 | 26.15741 | 26.39607 | 20.70267 | 20.70296 | 11.70134 | 26.81698 | 13.50005 | 16.31387 |
| ENSMUSG00000090257 | -0.772244897 | 0.006992 | 0.03293 | 74.44802 | 45.59322 | 109.3101 | 75.04824 | 95.56091 | 146.376 | 128.2505 | 153.3503 |
| ENSMUSG00000090258 | -0.030851369 | 0.725231 | 0.828015 | 896.3944 | 745.0892 | 867.0276 | 851.4094 | 829.8197 | 812.331 | 897.7532 | 896.1751 |
| ENSMUSG00000090264 | -0.716241205 | 0.119504 | 0.24843 | 26.15741 | 13.19804 | 36.43669 | 49.16954 | 38.02934 | 30.1691 | 72.32169 | 66.34306 |
| ENSMUSG00000090266 | 0.134109401 | 0.133699 | 0.267196 | 616.7113 | 653.9027 | 589.6119 | 619.3637 | 594.8179 | 606.7341 | 537.1091 | 518.7809 |
| ENSMUSG00000090272 | 0.095254435 | 0.811373 | 0.885135 | 104.6297 | 163.1757 | 138.2938 | 75.91087 | 59.48179 | 65.92507 | 162.0006 | 163.1387 |
| ENSMUSG00000090273 | 0.325752429 | 0.371069 | 0.540056 | 27.16347 | 25.19625 | 19.87456 | 28.46658 | 16.57689 | 22.34748 | 27.0001 | 14.13868 |
| ENSMUSG00000090290 | 0.04207906 | 0.726133 | 0.828686 | 623.7537 | 532.7208 | 486.9267 | 489.1075 | 577.2659 | 472.6492 | 516.859 | 501.3795 |
| ENSMUSG00000090291 | -0.56809603 | 0.117285 | 0.245573 | 18.10898 | 21.59679 | 24.8432 | 35.36756 | 23.40267 | 36.87335 | 44.3573 | 44.59123 |
| ENSMUSG00000090307 | 0.722579637 | 0.190712 | 0.343357 | 63.38142 | 16.7975 | 44.71776 | 21.56559 | 30.22845 | 16.76061 | 31.82154 | 9.78832 |
| ENSMUSG00000090327 | 0.112542783 | 0.875956 | 0.925713 | 11.0666 | 14.39786 | 12.4216 | 8.626235 | 16.57689 | 17.87799 | 0.964289 | 7.613138 |
| ENSMUSG00000090329 | 1.395979957 | 0.011107 | 0.04622 | 17.10292 | 26.39607 | 9.109173 | 18.97772 | 6.825779 | 5.58687 | 4.821446 | 9.78832 |
| ENSMUSG00000090330 | 0.241109171 | 0.385995 | 0.553486 | 157.9505 | 140.3791 | 140.7781 | 163.0358 | 108.2374 | 208.949 | 105.1075 | 88.09488 |
| ENSMUSG00000090363 | 0.487754084 | 0.324347 | 0.492323 | 17.10292 | 19.19714 | 10.76539 | 12.07673 | 7.80089 | 16.76061 | 7.714313 | 9.78832 |
| ENSMUSG00000090370 | -0.750940873 | 0.034565 | 0.105077 | 17.10292 | 25.19625 | 20.70267 | 18.97772 | 20.47734 | 41.34284 | 35.6787 | 40.24087 |
| ENSMUSG00000090381 | -0.162554897 | 0.781781 | 0.865437 | 17.10292 | 8.398751 | 6.624853 | 15.52722 | 14.62667 | 22.34748 | 5.785735 | 10.87591 |
| ENSMUSG00000090386 | 0.991124215 | 0.091546 | 0.207539 | 587.5357 | 771.4852 | 442.2089 | 436.4875 | 195.9974 | 646.9596 | 160.072 | 122.8978 |
| ENSMUSG00000090394 | -0.085543153 | 0.489938 | 0.647404 | 563.3904 | 638.3051 | 671.5945 | 637.4788 | 772.2882 | 709.5326 | 617.1451 | 565.5474 |
| ENSMUSG00000090397 | 0.15334991 | 0.630634 | 0.759477 | 46.2785 | 25.19625 | 34.78048 | 43.9938 | 26.32801 | 43.57759 | 35.6787 | 30.45255 |
| ENSMUSG00000090399 | -0.577671113 | 0.162035 | 0.306124 | 15.09082 | 21.59679 | 14.90592 | 11.21411 | 31.20356 | 26.81698 | 18.32149 | 16.31387 |
| ENSMUSG00000090459 | 0.747669505 | 0.162136 | 0.306251 | 29.17558 | 13.19804 | 21.53077 | 21.56559 | 6.825779 | 23.46486 | 14.46434 | 6.525547 |
| ENSMUSG00000090486 | -1.35532944 | 0.030167 | 0.094851 | 28.16952 | 15.59768 | 15.73403 | 18.97772 | 47.78045 | 11.17374 | 29.89296 | 112.0219 |
| ENSMUSG00000090509 | -0.671420626 | 0.098153 | 0.218177 | 16.09687 | 17.99732 | 32.29616 | 33.64232 | 28.27823 | 31.28647 | 40.50014 | 60.9051 |
| ENSMUSG00000090516 | 0.543277172 | 0.09183 | 0.208054 | 46.2785 | 39.59411 | 38.92101 | 43.13117 | 22.42756 | 34.6386 | 39.53586 | 18.48905 |
| ENSMUSG00000090523 | 0.243929377 | 0.013478 | 0.053323 | 649.9111 | 788.2827 | 712.9998 | 752.2077 | 598.7183 | 677.1287 | 606.5379 | 568.8101 |
| ENSMUSG00000090551 | -0.150347775 | 0.80078 | 0.878406 | 20.12109 | 10.79839 | 8.281066 | 13.80198 | 17.552 | 22.34748 | 3.857157 | 15.22628 |
| ENSMUSG00000090553 | 0.054268219 | 0.384798 | 0.5525 | 2612.723 | 2705.598 | 2811.422 | 2755.219 | 2578.194 | 2823.604 | 2598.759 | 2486.233 |
| ENSMUSG00000090555 | -1.392252205 | 0.105695 | 0.229522 | 9.054489 | 0 | 22.35888 | 6.038364 | 14.62667 | 11.17374 | 26.03581 | 47.85401 |
| ENSMUSG00000090564 | 0.147830856 | 0.66799 | 0.787054 | 45.27245 | 50.3925 | 33.95237 | 37.95543 | 38.02934 | 22.34748 | 56.89306 | 32.62773 |
| ENSMUSG00000090576 | 0.068820612 | 0.848844 | 0.909191 | 49.29666 | 37.19447 | 48.03018 | 22.42821 | 38.02934 | 48.04709 | 39.53586 | 23.927 |
| ENSMUSG00000090582 | 0.297437977 | 0.602636 | 0.740217 | 17.10292 | 25.19625 | 12.4216 | 9.488858 | 24.37778 | 7.821619 | 11.57147 | 7.613138 |

| | | | | | | | | | | | |
|---|---|---|---|---|---|---|---|---|---|---|---|
| ENSMUSG00000090592 | 1.527167886 | 0.204155 | 0.360115 | 39.23612 | 38.39429 | 3.312427 | 17.25247 | 2.925334 | 30.1691 | 0 | 1.087591 |
| ENSMUSG00000090602 | -0.03978348 | 0.913582 | 0.948837 | 21.12714 | 16.7975 | 14.90592 | 20.70296 | 19.50223 | 22.34748 | 15.42863 | 18.48905 |
| ENSMUSG00000090622 | 0.128294914 | 0.440089 | 0.603304 | 171.0292 | 128.3809 | 131.669 | 137.1571 | 115.0631 | 132.9675 | 145.6077 | 126.1606 |
| ENSMUSG00000090625 | 0.35301958 | 0.164071 | 0.309052 | 152.9203 | 137.9795 | 142.4343 | 128.5309 | 92.63557 | 170.9582 | 95.46463 | 81.56933 |
| ENSMUSG00000090626 | 0.0042996 | 0.967771 | 0.981869 | 384.3128 | 368.3452 | 324.6178 | 362.3019 | 360.7912 | 340.7991 | 338.4655 | 393.708 |
| ENSMUSG00000090638 | -1.148483806 | 0.162604 | 0.306979 | 7.04238 | 7.198929 | 84.46688 | 11.21411 | 63.38224 | 16.76061 | 71.3574 | 93.53283 |
| ENSMUSG00000090641 | 0.381380624 | 0.170884 | 0.317832 | 145.8779 | 103.1847 | 80.32634 | 78.49874 | 66.30757 | 103.9158 | 76.17884 | 66.34306 |
| ENSMUSG00000090659 | 0.042792392 | 0.905921 | 0.943442 | 51.30877 | 74.38894 | 42.23344 | 31.91707 | 35.10401 | 34.6386 | 59.78593 | 63.08028 |
| ENSMUSG00000090673 | -0.654212849 | 0.011994 | 0.048962 | 45.27245 | 49.19268 | 82.81066 | 46.58167 | 84.83468 | 99.44629 | 80.036 | 90.27006 |
| ENSMUSG00000090674 | -0.191699826 | 0.405291 | 0.572175 | 61.36932 | 69.58965 | 45.54586 | 72.46037 | 78.98402 | 62.57295 | 66.53595 | 75.04379 |
| ENSMUSG00000090691 | 0.026862347 | 0.967376 | 0.981738 | 8.048435 | 20.39697 | 14.90592 | 4.313117 | 9.751113 | 18.99536 | 4.821446 | 13.05109 |
| ENSMUSG00000090698 | -0.481345665 | 0.074516 | 0.180244 | 190.1443 | 77.9884 | 154.8559 | 152.6844 | 176.4951 | 164.254 | 234.3223 | 230.5693 |
| ENSMUSG00000090733 | -0.156184058 | 0.054084 | 0.144058 | 14168.26 | 15637.27 | 17075.56 | 15264.99 | 18217.03 | 15849.95 | 18141.17 | 17043.64 |
| ENSMUSG00000090761 | 0.311249366 | 0.167362 | 0.313301 | 257.5499 | 238.7645 | 223.5888 | 263.1002 | 163.8187 | 296.1041 | 184.1792 | 149 |
| ENSMUSG00000090778 | 0.754451359 | 0.246784 | 0.409849 | 92.557 | 97.18554 | 38.09291 | 72.46037 | 14.62667 | 118.4417 | 29.89296 | 15.22628 |
| ENSMUSG00000090779 | 0.643599723 | 0.061859 | 0.158368 | 57.3451 | 89.98662 | 56.31125 | 62.10889 | 29.25334 | 70.39457 | 35.6787 | 34.80291 |
| ENSMUSG00000090799 | 0.415965757 | 0.348562 | 0.51721 | 19.11503 | 13.19804 | 25.67131 | 15.52722 | 11.70134 | 8.938993 | 17.3572 | 17.40146 |
| ENSMUSG00000090812 | 0.074797471 | 0.859079 | 0.915425 | 19.11503 | 22.79661 | 24.8432 | 9.488858 | 20.47734 | 17.87799 | 17.3572 | 16.31387 |
| ENSMUSG00000090817 | -3.279211627 | 3.17E-05 | 0.000559 | 53.32088 | 3.599465 | 82.81066 | 44.85642 | 423.1983 | 204.4795 | 440.6801 | 728.686 |
| ENSMUSG00000090841 | -0.320297997 | 0.011826 | 0.048438 | 9844.242 | 11443.9 | 13147.85 | 10683.59 | 14291.23 | 11896.68 | 13582.01 | 16567.28 |
| ENSMUSG00000090862 | -0.134259172 | 0.082823 | 0.19355 | 13621.98 | 15688.87 | 14816.48 | 14162.55 | 17165.86 | 15308.03 | 16749.7 | 14745.56 |
| ENSMUSG00000090863 | -1.850074792 | 0.037688 | 0.111936 | 7.04238 | 8.398751 | 33.95237 | 0.862623 | 26.32801 | 11.17374 | 50.14304 | 94.62042 |
| ENSMUSG00000090873 | 0.084522988 | 0.769855 | 0.857118 | 30.18163 | 27.5959 | 38.09291 | 37.09281 | 34.1289 | 26.81698 | 36.64299 | 28.27737 |
| ENSMUSG00000090877 | -1.37665303 | 0.019143 | 0.068376 | 197.1867 | 105.5843 | 559.8001 | 168.2116 | 554.8383 | 372.0856 | 751.1813 | 1000.584 |
| ENSMUSG00000090891 | 0.524476961 | 0.24041 | 0.402559 | 21.12714 | 11.99822 | 16.56213 | 23.29083 | 8.776002 | 17.87799 | 10.60718 | 14.13868 |
| ENSMUSG00000090919 | 0.373703609 | 0.015076 | 0.057883 | 319.9253 | 363.5459 | 311.3681 | 385.5927 | 270.1058 | 329.6254 | 232.3937 | 233.8321 |
| ENSMUSG00000090925 | 0.690426085 | 0.049305 | 0.134807 | 41.24823 | 43.19358 | 38.92101 | 41.40593 | 26.32801 | 41.34284 | 17.3572 | 17.40146 |
| ENSMUSG00000090935 | 0.042395671 | 0.564553 | 0.709478 | 1229.398 | 1105.036 | 1143.615 | 1155.053 | 1042.394 | 1124.078 | 1192.826 | 1140.883 |
| ENSMUSG00000090936 | -0.118209424 | 0.775286 | 0.860983 | 24.1453 | 16.7975 | 9.93728 | 19.84034 | 16.57689 | 16.76061 | 20.25007 | 22.83941 |
| ENSMUSG00000090946 | 0.074040459 | 0.609054 | 0.744318 | 622.7476 | 525.5218 | 461.2554 | 560.7053 | 479.7548 | 612.321 | 534.2162 | 435.0364 |
| ENSMUSG00000090952 | -0.237723084 | 0.043866 | 0.124342 | 263.5862 | 213.5682 | 240.979 | 222.5569 | 262.3049 | 288.2825 | 268.0724 | 292.562 |
| ENSMUSG00000090958 | -0.254125114 | 0.189163 | 0.341637 | 480.894 | 445.1338 | 770.1392 | 550.3538 | 696.2295 | 549.7481 | 648.0023 | 788.5035 |
| ENSMUSG00000090963 | 0.020374587 | 0.951018 | 0.972247 | 28.16952 | 26.39607 | 32.29616 | 18.11509 | 27.30312 | 27.93435 | 21.21436 | 27.18978 |
| ENSMUSG00000091002 | -0.301958735 | 0.571836 | 0.714738 | 7.04238 | 43.19358 | 51.34261 | 32.77969 | 25.35289 | 43.57759 | 40.50014 | 56.55474 |
| ENSMUSG00000091007 | 0.011270917 | 0.976263 | 0.986958 | 27.16347 | 29.99554 | 29.81184 | 22.42821 | 32.17867 | 12.29112 | 30.85725 | 32.62773 |
| ENSMUSG00000091021 | 0.250911979 | 0.108364 | 0.233123 | 208.2533 | 257.9616 | 179.6991 | 214.7932 | 190.1467 | 170.9582 | 162.0006 | 197.9416 |
| ENSMUSG00000091050 | 0.103617948 | 0.613485 | 0.747309 | 82.49646 | 69.58965 | 64.59232 | 69.8725 | 61.43201 | 78.21619 | 58.82164 | 68.51824 |
| ENSMUSG00000091086 | -0.166962805 | 0.320314 | 0.488405 | 99.59938 | 112.7832 | 106.8258 | 106.1027 | 127.7396 | 131.8501 | 121.5004 | 95.70802 |
| ENSMUSG00000091102 | -0.815150757 | 0.126202 | 0.257055 | 16.09687 | 9.598572 | 9.93728 | 3.450494 | 12.67645 | 15.64324 | 19.28578 | 20.66423 |
| ENSMUSG00000091119 | 0.505422665 | 0.14638 | 0.285218 | 31.18769 | 57.59143 | 43.88965 | 37.95543 | 17.552 | 39.10809 | 30.85725 | 32.62773 |
| ENSMUSG00000091133 | -0.391274721 | 0.242357 | 0.404864 | 20.12109 | 22.79661 | 24.01509 | 20.70296 | 28.27823 | 30.1691 | 36.64299 | 19.57664 |
| ENSMUSG00000091144 | -0.358696088 | 0.198881 | 0.353549 | 28.16952 | 32.39518 | 23.18699 | 30.19182 | 38.02934 | 37.99072 | 37.60728 | 31.54014 |
| ENSMUSG00000091155 | -0.299805407 | 0.538185 | 0.688101 | 17.10292 | 14.39786 | 14.07781 | 34.50494 | 15.60178 | 40.22547 | 25.07152 | 18.48905 |
| ENSMUSG00000091183 | 0.421900302 | 0.061495 | 0.157675 | 124.7507 | 203.9697 | 118.4192 | 140.6076 | 89.71024 | 117.3243 | 123.429 | 106.5839 |
| ENSMUSG00000091243 | 0.22075967 | 0.367988 | 0.536461 | 115.6963 | 106.7841 | 91.09173 | 142.3329 | 125.7894 | 67.04245 | 93.53605 | 104.4087 |
| ENSMUSG00000091264 | 0.044133211 | 0.616443 | 0.749428 | 861.1825 | 844.6744 | 789.1856 | 779.8116 | 720.6073 | 869.317 | 812.8958 | 772.1897 |
| ENSMUSG00000091272 | -0.805629275 | 0.072778 | 0.177343 | 14.08476 | 10.79839 | 18.21835 | 24.15346 | 19.50223 | 41.34284 | 19.28578 | 39.15328 |
| ENSMUSG00000091277 | 0.658190356 | 0.118215 | 0.247035 | 127.7689 | 117.5825 | 121.7317 | 144.0581 | 129.6898 | 119.559 | 35.6787 | 39.15328 |
| ENSMUSG00000091313 | 0.677481355 | 0.227858 | 0.387732 | 42.25428 | 29.99554 | 21.53077 | 39.68068 | 16.57689 | 46.92971 | 11.57147 | 8.700729 |
| ENSMUSG00000091318 | 1.743147388 | NA | NA | 63.38142 | 101.9848 | 48.03018 | 46.58167 | 3.900445 | 68.15982 | 5.785735 | 0 |
| ENSMUSG00000091329 | -0.421733769 | 0.437904 | 0.60156 | 10.06054 | 3.599465 | 15.73403 | 7.763611 | 11.70134 | 12.29112 | 11.57147 | 15.22628 |
| ENSMUSG00000091337 | 0.357857236 | 0.002632 | 0.016007 | 4057.417 | 4553.323 | 3911.976 | 4730.627 | 3270.523 | 4016.96 | 3305.583 | 2870.153 |
| ENSMUSG00000091345 | 2.547160453 | 0.002044 | 0.013323 | 432.6034 | 1767.337 | 1089.788 | 916.9688 | 104.3369 | 493.8794 | 77.14313 | 44.59123 |
| ENSMUSG00000091366 | 0.009633771 | 0.987346 | 0.992833 | 6.036326 | 5.999108 | 13.24971 | 16.38985 | 9.751113 | 4.469496 | 13.50005 | 14.13868 |
| ENSMUSG00000091387 | -0.012243922 | 0.932137 | 0.959964 | 296.786 | 255.562 | 255.0568 | 272.589 | 296.4338 | 258.1134 | 312.4297 | 220.781 |
| ENSMUSG00000091402 | 0.047185367 | 0.915931 | 0.950167 | 17.10292 | 9.598572 | 12.4216 | 18.11509 | 15.60178 | 17.87799 | 12.53576 | 9.78832 |
| ENSMUSG00000091421 | -0.222791993 | 0.365224 | 0.533949 | 59.35721 | 45.59322 | 38.09291 | 58.6584 | 56.55646 | 52.51658 | 58.82164 | 67.43065 |
| ENSMUSG00000091423 | -0.067774442 | 0.855605 | 0.913423 | 20.12109 | 11.99822 | 24.8432 | 27.60395 | 20.47734 | 21.23011 | 23.14294 | 25.0146 |
| ENSMUSG00000091449 | 0.031093841 | 0.919483 | 0.952207 | 28.16952 | 35.99465 | 24.01509 | 28.46658 | 31.20356 | 22.34748 | 27.0001 | 32.62773 |
| ENSMUSG00000091457 | 1.713511417 | 0.001101 | 0.008401 | 21.12714 | 33.595 | 19.87456 | 17.25247 | 7.80089 | 10.05637 | 7.714313 | 2.175182 |
| ENSMUSG00000091460 | -0.205856319 | 0.630536 | 0.759418 | 13.07871 | 10.79839 | 14.07781 | 12.07673 | 16.57689 | 17.87799 | 10.60718 | 13.05109 |
| ENSMUSG00000091472 | 0.795305173 | 0.038467 | 0.11337 | 43.26034 | 34.79482 | 25.67131 | 30.19182 | 18.52711 | 30.1691 | 15.42863 | 13.05109 |
| ENSMUSG00000091474 | 0.323383225 | 0.024172 | 0.081052 | 652.9293 | 544.719 | 470.3646 | 616.7758 | 471.9539 | 517.3442 | 451.2873 | 385.0072 |
| ENSMUSG00000091475 | 0.13546158 | 0.436637 | 0.6004 | 173.0413 | 193.1713 | 146.5749 | 187.1893 | 184.296 | 176.5451 | 145.6077 | 129.4233 |

| | | | | | | | | | | | |
|---|---|---|---|---|---|---|---|---|---|---|---|
| ENSMUSG00000091476 | -0.675626684 | 0.086515 | 0.19954 | 23.13925 | 14.39786 | 16.56213 | 23.29083 | 30.22845 | 16.76061 | 36.64299 | 40.24087 |
| ENSMUSG00000091478 | -0.548125859 | 0.116972 | 0.245141 | 35.2119 | 27.5959 | 21.53077 | 34.50494 | 46.80534 | 24.58223 | 44.3573 | 57.64233 |
| ENSMUSG00000091492 | 0.473494668 | 0.248451 | 0.411889 | 22.1332 | 15.59768 | 24.8432 | 20.70296 | 8.776002 | 17.87799 | 14.46434 | 19.57664 |
| ENSMUSG00000091498 | -0.870917913 | 0.000841 | 0.006826 | 84.50857 | 100.785 | 109.3101 | 85.39973 | 154.0676 | 108.3853 | 190.9293 | 241.4452 |
| ENSMUSG00000091509 | 0.776225624 | 0.016449 | 0.061416 | 714.2986 | 802.6806 | 510.1137 | 607.2869 | 276.9316 | 716.2368 | 276.751 | 268.635 |
| ENSMUSG00000091512 | -0.079392711 | 0.249803 | 0.413494 | 1336.04 | 1246.615 | 1298.471 | 1350.006 | 1380.758 | 1308.445 | 1350.969 | 1490 |
| ENSMUSG00000091514 | 0.303248449 | 0.62434 | 0.7551 | 14.08476 | 17.99732 | 7.45296 | 6.900988 | 15.60178 | 8.938993 | 3.857157 | 8.700729 |
| ENSMUSG00000091515 | 0.077001915 | 0.726276 | 0.828695 | 79.47829 | 80.38804 | 62.9361 | 58.6584 | 63.38224 | 58.10345 | 64.60737 | 79.39415 |
| ENSMUSG00000091534 | 0.111927158 | 0.835235 | 0.900414 | 16.09687 | 5.999108 | 12.4216 | 7.763611 | 6.825779 | 10.05637 | 10.60718 | 11.9635 |
| ENSMUSG00000091537 | -0.244708786 | 0.001176 | 0.008841 | 3175.108 | 3177.127 | 3135.212 | 2963.974 | 3700.547 | 3299.606 | 3735.656 | 4014.299 |
| ENSMUSG00000091549 | 0.341696427 | 0.077235 | 0.184596 | 419.5247 | 622.7074 | 606.1741 | 351.0878 | 383.2187 | 406.7242 | 386.68 | 400.2335 |
| ENSMUSG00000091562 | -0.138565881 | 0.696943 | 0.808775 | 102.6175 | 65.99018 | 53.82693 | 51.75741 | 101.4116 | 39.10809 | 73.28598 | 87.00729 |
| ENSMUSG00000091575 | -0.022839504 | 0.969576 | 0.983052 | 24.1453 | 8.398751 | 24.01509 | 8.626235 | 7.80089 | 27.93435 | 11.57147 | 19.57664 |
| ENSMUSG00000091577 | 0.853190343 | 0.255465 | 0.419388 | 5.030272 | 17.99732 | 9.93728 | 37.95543 | 6.825779 | 16.76061 | 12.53576 | 3.262773 |
| ENSMUSG00000091586 | 0.566884993 | 9.32E-06 | 0.000215 | 474.8577 | 490.727 | 385.0696 | 449.4268 | 328.6125 | 300.5736 | 323.0369 | 259.9343 |
| ENSMUSG00000091604 | -0.538378875 | 0.165206 | 0.310588 | 22.1332 | 10.79839 | 16.56213 | 23.29083 | 34.1289 | 18.99536 | 26.03581 | 27.18978 |
| ENSMUSG00000091625 | 0.163352253 | 0.241335 | 0.403522 | 782.7103 | 833.876 | 683.188 | 871.2497 | 682.5779 | 851.4391 | 718.3954 | 578.5985 |
| ENSMUSG00000091639 | -0.625983583 | 0.148159 | 0.287623 | 11.0666 | 14.39786 | 19.87456 | 13.80198 | 33.15378 | 13.40849 | 24.10723 | 20.66423 |
| ENSMUSG00000091648 | -0.251606089 | 0.489611 | 0.647157 | 17.10292 | 26.39607 | 14.07781 | 23.29083 | 25.35289 | 21.23011 | 27.96439 | 20.66423 |
| ENSMUSG00000091649 | 0.805034618 | 0.274345 | 0.440249 | 10.06054 | 39.59411 | 43.88965 | 11.21411 | 14.62667 | 17.87799 | 25.07152 | 2.175182 |
| ENSMUSG00000091655 | -0.813174507 | 0.204614 | 0.360718 | 6.036326 | 9.598572 | 3.312427 | 13.80198 | 16.57689 | 13.40849 | 5.785735 | 21.75182 |
| ENSMUSG00000091705 | -1.280529012 | 0.00012 | 0.001537 | 18.10898 | 23.99643 | 17.39024 | 14.6646 | 58.50668 | 41.34284 | 33.75012 | 44.59123 |
| ENSMUSG00000091721 | -1.313168467 | 0.115804 | 0.243805 | 6.036326 | 0 | 14.90592 | 5.175741 | 8.776002 | 8.938993 | 20.25007 | 28.27737 |
| ENSMUSG00000091735 | -0.387049168 | 0.105588 | 0.229366 | 49.29666 | 34.79482 | 51.34261 | 50.03216 | 67.28268 | 67.04245 | 59.78593 | 50.02919 |
| ENSMUSG00000091754 | 0.391870392 | 0.18361 | 0.334588 | 106.6418 | 113.983 | 68.73285 | 72.46037 | 66.30757 | 97.21155 | 64.60737 | 46.76642 |
| ENSMUSG00000091764 | -0.027382069 | 0.872014 | 0.9234 | 123.7447 | 183.5727 | 154.8559 | 163.0358 | 176.4951 | 165.3714 | 137.8934 | 156.6131 |
| ENSMUSG00000091784 | -0.066152845 | 0.836367 | 0.901206 | 38.23007 | 27.5959 | 33.12427 | 31.05445 | 24.37778 | 49.16446 | 31.82154 | 31.54014 |
| ENSMUSG00000091803 | -0.074445614 | 0.407408 | 0.573711 | 478.8819 | 559.1168 | 501.8326 | 467.5419 | 531.4357 | 522.9311 | 515.8947 | 539.4452 |
| ENSMUSG00000091811 | 0.011874339 | 0.911759 | 0.948024 | 859.1704 | 967.0562 | 873.6525 | 888.5022 | 962.4349 | 981.0545 | 864.0031 | 749.3503 |
| ENSMUSG00000091831 | 0.343716569 | 0.460676 | 0.62196 | 42.25428 | 34.79482 | 34.78048 | 25.01608 | 11.70134 | 48.04709 | 30.85725 | 17.40146 |
| ENSMUSG00000091845 | -0.11357064 | 0.71355 | 0.820362 | 25.15136 | 25.19625 | 33.12427 | 44.85642 | 39.00445 | 34.6386 | 34.71441 | 31.54014 |
| ENSMUSG00000091890 | -0.360882467 | 0.498494 | 0.654881 | 17.10292 | 9.598572 | 31.46805 | 12.93935 | 31.20356 | 10.05637 | 29.89296 | 20.66423 |
| ENSMUSG00000091896 | 0.053356313 | 0.392414 | 0.559836 | 3117.762 | 3507.078 | 3460.658 | 3406.5 | 3199.34 | 3428.104 | 3195.654 | 3180.116 |
| ENSMUSG00000091898 | 0.203428862 | 0.463928 | 0.624845 | 46.2785 | 62.39072 | 53.82693 | 37.95543 | 44.85512 | 33.52122 | 40.50014 | 54.37955 |
| ENSMUSG00000091905 | 0.164827738 | 0.703005 | 0.812828 | 20.12109 | 16.7975 | 16.56213 | 13.80198 | 21.45245 | 14.52586 | 8.678602 | 15.22628 |
| ENSMUSG00000091931 | 0.056643803 | 0.686415 | 0.800586 | 225.3562 | 230.3657 | 198.7456 | 256.1992 | 209.6489 | 251.4092 | 196.715 | 218.6058 |
| ENSMUSG00000091952 | 0.021710525 | 0.952523 | 0.972978 | 34.20585 | 39.59411 | 37.2648 | 36.23019 | 21.45245 | 59.22083 | 27.0001 | 38.06569 |
| ENSMUSG00000091955 | 0.013549387 | 0.956949 | 0.975803 | 51.30877 | 67.19001 | 75.3577 | 57.79577 | 72.15824 | 73.74669 | 57.85735 | 45.67883 |
| ENSMUSG00000091956 | -0.362736491 | 0.572031 | 0.714829 | 12.07265 | 11.99822 | 7.45296 | 11.21411 | 3.900445 | 27.93435 | 14.46434 | 8.700729 |
| ENSMUSG00000091957 | -0.070969631 | 0.674887 | 0.791945 | 134.8113 | 183.5727 | 153.1997 | 175.1126 | 204.7734 | 155.315 | 161.0363 | 156.6131 |
| ENSMUSG00000091968 | -0.733855383 | 0.137974 | 0.273269 | 13.07871 | 8.398751 | 14.90592 | 5.175741 | 14.62667 | 14.52586 | 16.39292 | 23.927 |
| ENSMUSG00000091971 | -0.963441237 | 0.003557 | 0.020054 | 98.59333 | 106.7841 | 217.792 | 103.5148 | 214.5245 | 165.3714 | 289.2867 | 359.9926 |
| ENSMUSG00000091972 | -3.0083745 | 0.002468 | 0.015282 | 3.018163 | 0 | 10.76539 | 2.58787 | 31.20356 | 2.234748 | 57.85735 | 43.50364 |
| ENSMUSG00000091985 | 0.435869975 | 0.068752 | 0.170001 | 203.223 | 221.967 | 195.4332 | 171.6621 | 122.864 | 223.4748 | 124.3933 | 115.2847 |
| ENSMUSG00000091993 | -0.137497455 | 0.644552 | 0.770057 | 32.19374 | 46.79304 | 43.88965 | 39.68068 | 54.60623 | 55.8687 | 36.64299 | 31.54014 |
| ENSMUSG00000091994 | -0.361353367 | 0.003782 | 0.02097 | 164.9929 | 163.1757 | 178.871 | 168.2116 | 202.8232 | 208.949 | 215.0365 | 242.5328 |
| ENSMUSG00000092014 | 1.296272918 | 0.006956 | 0.032832 | 15.09082 | 22.79661 | 13.24971 | 22.42821 | 8.776002 | 5.58687 | 7.714313 | 7.613138 |
| ENSMUSG00000092029 | -0.201659066 | 0.591644 | 0.731496 | 25.15136 | 17.99732 | 17.39024 | 17.25247 | 22.42756 | 31.28647 | 17.3572 | 18.48905 |
| ENSMUSG00000092035 | 1.02058843 | 0.000376 | 0.003675 | 2013.115 | 1061.842 | 934.1043 | 1589.815 | 724.5077 | 1008.989 | 543.8591 | 482.8904 |
| ENSMUSG00000092060 | 0.688103406 | 0.000786 | 0.006497 | 579.4873 | 548.3184 | 470.3646 | 489.1075 | 273.0312 | 469.2971 | 306.644 | 246.8832 |
| ENSMUSG00000092071 | -1.417170758 | 0.054298 | 0.144444 | 1.006054 | 4.799286 | 14.90592 | 14.6646 | 14.62667 | 12.29112 | 39.53586 | 29.36496 |
| ENSMUSG00000092072 | 0.074926162 | 0.848851 | 0.909191 | 12.07265 | 26.39607 | 19.04645 | 27.60395 | 16.57689 | 22.34748 | 21.21436 | 20.66423 |
| ENSMUSG00000092074 | -0.414913573 | 0.000213 | 0.002383 | 192.1564 | 220.7672 | 194.6051 | 206.167 | 274.0063 | 281.5783 | 253.6081 | 274.073 |
| ENSMUSG00000092083 | 0.047024759 | 0.884285 | 0.930838 | 51.30877 | 67.19001 | 125.8722 | 59.52102 | 59.48179 | 75.98144 | 76.17884 | 83.74451 |
| ENSMUSG00000092085 | 0.069859711 | 0.924301 | 0.955039 | 17.10292 | 9.598572 | 8.281066 | 9.488858 | 16.57689 | 17.87799 | 6.750024 | 1.087591 |
| ENSMUSG00000092086 | -0.042816452 | 0.894718 | 0.936266 | 34.20585 | 29.99554 | 40.57722 | 33.64232 | 51.6809 | 32.40385 | 24.10723 | 34.80291 |
| ENSMUSG00000092118 | -0.018380989 | 0.90993 | 0.946495 | 190.1443 | 205.1695 | 136.6376 | 163.0358 | 180.3956 | 169.8409 | 179.3578 | 170.7518 |
| ENSMUSG00000092124 | -0.542763149 | 0.14902 | 0.288808 | 25.15136 | 21.59679 | 33.12427 | 14.6646 | 35.10401 | 23.46486 | 33.75012 | 45.67883 |
| ENSMUSG00000092164 | -0.517095582 | 0.21479 | 0.372961 | 147.89 | 34.79482 | 134.1533 | 106.1027 | 118.9636 | 97.21155 | 204.4293 | 185.9781 |
| ENSMUSG00000092167 | 0.434362424 | 0.408275 | 0.574587 | 27.16347 | 39.59411 | 21.53077 | 13.80198 | 13.65156 | 33.52122 | 19.28578 | 8.700729 |
| ENSMUSG00000092183 | 0.00771476 | 0.983934 | 0.991004 | 22.1332 | 23.99643 | 25.67131 | 16.38985 | 17.552 | 14.52586 | 26.03581 | 29.36496 |
| ENSMUSG00000092187 | 0.375509387 | 0.226859 | 0.386817 | 77.46619 | 98.38537 | 77.84202 | 99.2017 | 48.75557 | 112.8548 | 59.78593 | 51.11678 |
| ENSMUSG00000092190 | 1.021102057 | 0.013023 | 0.05208 | 34.20585 | 37.19447 | 17.39024 | 27.60395 | 15.60178 | 12.29112 | 10.60718 | 18.48905 |
| ENSMUSG00000092192 | -0.745700837 | 0.025775 | 0.084815 | 15.09082 | 17.99732 | 15.73403 | 18.97772 | 26.32801 | 33.52122 | 29.89296 | 23.927 |

| | | | | | | | | | | | |
|---|---|---|---|---|---|---|---|---|---|---|---|
| ENSMUSG00000092203 | 0.120688838 | 0.283794 | 0.450329 | 1814.922 | 1715.745 | 1574.231 | 1676.077 | 1788.354 | 1630.249 | 1530.327 | 1285.533 |
| ENSMUSG00000092232 | -0.051451022 | 0.919092 | 0.952207 | 29.17558 | 9.598572 | 35.60859 | 22.42821 | 43.88001 | 18.99536 | 17.3572 | 20.66423 |
| ENSMUSG00000092233 | -0.888414127 | 0.145055 | 0.283203 | 2.012109 | 5.999108 | 9.93728 | 12.07673 | 10.72622 | 14.52586 | 20.25007 | 10.87591 |
| ENSMUSG00000092241 | -0.527234804 | 0.145112 | 0.283257 | 13.07871 | 23.99643 | 19.87456 | 24.15346 | 36.07912 | 32.40385 | 21.21436 | 27.18978 |
| ENSMUSG00000092260 | 0.108856463 | 0.602671 | 0.740217 | 155.9384 | 193.1713 | 123.3879 | 136.2945 | 124.8142 | 177.6625 | 141.7505 | 118.5474 |
| ENSMUSG00000092274 | -0.809273754 | 0.001544 | 0.01075 | 3955.806 | 2428.439 | 5001.764 | 3309.024 | 5846.767 | 3986.791 | 7938.993 | 7980.743 |
| ENSMUSG00000092281 | 0.279689181 | 0.41921 | 0.584714 | 26.15741 | 35.99465 | 34.78048 | 31.91707 | 26.32801 | 40.22547 | 19.28578 | 20.66423 |
| ENSMUSG00000092286 | 0.818479257 | 0.047521 | 0.13134 | 93.56306 | 83.98751 | 56.31125 | 72.46037 | 38.02934 | 83.80306 | 26.03581 | 26.10219 |
| ENSMUSG00000092300 | 0.286250022 | 0.270722 | 0.435914 | 54.32694 | 61.1909 | 52.99882 | 60.38364 | 41.92979 | 37.99072 | 65.57166 | 41.32846 |
| ENSMUSG00000092323 | 0.534711776 | 0.460207 | 0.621639 | 51.30877 | 20.39697 | 28.98373 | 43.13117 | 6.825779 | 70.39457 | 10.60718 | 11.9635 |
| ENSMUSG00000092335 | 0.166977079 | 0.603588 | 0.740729 | 31.18769 | 27.5959 | 20.70267 | 28.46658 | 23.40267 | 25.6996 | 27.0001 | 19.57664 |
| ENSMUSG00000092341 | 0.393687832 | 0.071136 | 0.174447 | 51440.57 | 46278.32 | 37798.93 | 39102.72 | 23845.37 | 50462.85 | 30493.72 | 28116.4 |
| ENSMUSG00000092349 | -0.143128261 | 0.671155 | 0.789222 | 24.1453 | 21.59679 | 29.81184 | 20.70296 | 35.10401 | 24.58223 | 26.03581 | 20.66423 |
| ENSMUSG00000092354 | 0.491220818 | 0.250574 | 0.414167 | 32.19374 | 29.99554 | 17.39024 | 15.52722 | 12.67645 | 13.40849 | 20.25007 | 20.66423 |
| ENSMUSG00000092369 | 0.528940716 | 0.121063 | 0.25028 | 34.20585 | 29.99554 | 23.18699 | 36.23019 | 23.40267 | 15.64324 | 19.28578 | 27.18978 |
| ENSMUSG00000092373 | 0.472617078 | 0.331211 | 0.499088 | 12.07265 | 11.99822 | 17.39024 | 21.56559 | 5.850668 | 13.40849 | 11.57147 | 15.22628 |
| ENSMUSG00000092402 | -0.079795308 | 0.814065 | 0.88718 | 30.18163 | 52.79215 | 42.23344 | 37.95543 | 58.50668 | 30.1691 | 52.07161 | 30.45255 |
| ENSMUSG00000092412 | 0.223602551 | 0.242967 | 0.405517 | 119.7205 | 136.7797 | 135.8095 | 131.1188 | 115.0631 | 140.7891 | 109.929 | 82.65692 |
| ENSMUSG00000092416 | 0.031927487 | 0.790348 | 0.87162 | 530.1906 | 597.5111 | 462.0835 | 517.5741 | 497.3068 | 593.3256 | 492.7518 | 475.2773 |
| ENSMUSG00000092417 | -0.117661684 | 0.306142 | 0.474723 | 346.0827 | 381.5432 | 361.8826 | 392.4937 | 466.1032 | 400.0199 | 371.2513 | 369.781 |
| ENSMUSG00000092454 | 0.234326699 | 0.575391 | 0.717577 | 14.08476 | 20.39697 | 28.15563 | 25.01608 | 19.50223 | 21.23011 | 10.60718 | 23.927 |
| ENSMUSG00000092469 | 1.226351877 | 0.029936 | 0.094427 | 108.6539 | 110.3836 | 72.87338 | 63.83414 | 12.67645 | 93.85942 | 24.10723 | 21.75182 |
| ENSMUSG00000092470 | -1.6966224 | 4.54E-05 | 0.000739 | 19.11503 | 19.19714 | 38.09291 | 30.19182 | 49.73068 | 56.98608 | 109.929 | 131.5985 |
| ENSMUSG00000092471 | 5.322460876 | 0.055148 | 0.146153 | 993.9817 | 2.399643 | 0.828107 | 89.71284 | 0 | 0 | 0 | 27.18978 |
| ENSMUSG00000092486 | 0.26544961 | 0.048834 | 0.133961 | 939.6548 | 1221.418 | 1038.446 | 1145.564 | 938.0571 | 1063.74 | 864.0031 | 748.2627 |
| ENSMUSG00000092526 | 1.188008904 | 0.016036 | 0.060347 | 88.53278 | 87.58697 | 45.54586 | 56.93315 | 15.60178 | 65.92507 | 19.28578 | 21.75182 |
| ENSMUSG00000092540 | 1.1766553 | 0.038368 | 0.113119 | 22.1332 | 26.39607 | 21.53077 | 25.8787 | 5.850668 | 24.58223 | 4.821446 | 7.613138 |
| ENSMUSG00000092558 | 0.051857325 | 0.513765 | 0.668338 | 938.6487 | 970.6556 | 833.0753 | 884.1891 | 895.1522 | 889.4298 | 825.4315 | 884.2116 |
| ENSMUSG00000092564 | -0.304180718 | 0.19468 | 0.348497 | 117.7084 | 67.19001 | 90.26362 | 109.5532 | 98.48624 | 102.7984 | 135.9648 | 139.2117 |
| ENSMUSG00000092569 | -0.227514432 | 0.251303 | 0.415 | 66.39959 | 55.19179 | 68.73285 | 57.79577 | 73.13335 | 71.51194 | 80.036 | 66.34306 |
| ENSMUSG00000092572 | 1.546791358 | 0.03803 | 0.112563 | 13.07871 | 25.19625 | 23.18699 | 43.13117 | 7.80089 | 22.34748 | 4.821446 | 1.087591 |
| ENSMUSG00000092593 | -0.049141399 | 0.901473 | 0.940806 | 21.12714 | 14.39786 | 16.56213 | 14.6646 | 21.45245 | 20.11273 | 14.46434 | 13.05109 |
| ENSMUSG00000092595 | 0.257547832 | 0.211244 | 0.368638 | 70.4238 | 65.99018 | 60.45178 | 73.323 | 54.60623 | 50.28183 | 56.89306 | 64.16787 |
| ENSMUSG00000092607 | -0.030052643 | 0.670895 | 0.789117 | 845.0857 | 865.0713 | 828.1066 | 936.8091 | 880.5255 | 898.3688 | 886.1817 | 884.2116 |
| ENSMUSG00000092625 | 0.596147221 | 0.483506 | 0.641846 | 15.09082 | 29.99554 | 4.96864 | 3.450494 | 4.875557 | 20.11273 | 5.785735 | 4.350364 |
| ENSMUSG00000092626 | -1.344712797 | 0.005293 | 0.026772 | 5.030272 | 7.198929 | 9.93728 | 10.35148 | 24.37778 | 14.52586 | 16.39292 | 28.27737 |
| ENSMUSG00000092680 | -0.234825343 | 0.437047 | 0.60083 | 34.20585 | 44.3934 | 29.81184 | 33.64232 | 50.70579 | 26.81698 | 42.42872 | 45.67883 |
| ENSMUSG00000092695 | 1.038624943 | 0.279609 | 0.445982 | 23.13925 | 22.79661 | 9.109173 | 23.29083 | 0 | 29.05173 | 3.857157 | 5.437955 |
| ENSMUSG00000092713 | -0.719435593 | 0.157799 | 0.300364 | 9.054489 | 7.198929 | 9.109173 | 6.900988 | 18.52711 | 13.40849 | 13.50005 | 7.613138 |
| ENSMUSG00000092741 | 0.034457309 | 0.907104 | 0.944408 | 65.39353 | 82.78769 | 79.49824 | 87.9876 | 70.20801 | 79.33356 | 45.32159 | 114.1971 |
| ENSMUSG00000092812 | -1.242128203 | 0.019363 | 0.068895 | 4.024217 | 5.999108 | 8.281066 | 8.626235 | 15.60178 | 24.58223 | 12.53576 | 11.9635 |
| ENSMUSG00000092819 | -0.022029467 | 0.951413 | 0.972512 | 18.10898 | 29.99554 | 27.32752 | 18.11509 | 30.22845 | 18.99536 | 21.21436 | 23.927 |
| ENSMUSG00000092827 | -0.054432192 | 0.853358 | 0.911961 | 25.15136 | 26.39607 | 35.60859 | 32.77969 | 33.15378 | 25.6996 | 34.71441 | 31.54014 |
| ENSMUSG00000092837 | 0.209952878 | 0.621264 | 0.752909 | 52.31483 | 21.59679 | 27.32752 | 36.23019 | 46.80534 | 22.34748 | 19.28578 | 30.45255 |
| ENSMUSG00000092887 | -0.179234871 | 0.661386 | 0.782069 | 18.10898 | 20.39697 | 27.32752 | 25.8787 | 26.32801 | 42.46022 | 16.39292 | 19.57664 |
| ENSMUSG00000092889 | -0.023003108 | 0.965296 | 0.980544 | 17.10292 | 10.79839 | 11.59349 | 15.52722 | 25.35289 | 12.29112 | 11.57147 | 6.525547 |
| ENSMUSG00000092981 | 0.520428389 | 0.049042 | 0.134366 | 220.3259 | 190.7716 | 180.5272 | 211.3428 | 89.71024 | 213.4185 | 147.5362 | 109.8467 |
| ENSMUSG00000093044 | 0.869114016 | 0.075084 | 0.181049 | 11.0666 | 21.59679 | 19.87456 | 18.11509 | 7.80089 | 14.52586 | 5.785735 | 10.87591 |
| ENSMUSG00000093080 | 0.629461535 | 0.266043 | 0.430333 | 31.18769 | 16.7975 | 11.59349 | 8.626235 | 10.72622 | 15.64324 | 6.750024 | 10.87591 |
| ENSMUSG00000093178 | 0.295367723 | 0.559512 | 0.705641 | 16.09687 | 21.59679 | 14.90592 | 7.763611 | 15.60178 | 15.64324 | 7.714313 | 9.78832 |
| ENSMUSG00000093402 | -0.679200932 | 0.153433 | 0.294729 | 10.06054 | 20.39697 | 14.07781 | 15.52722 | 34.1289 | 13.40849 | 15.42863 | 32.62773 |
| ENSMUSG00000093424 | 0.264264157 | 0.373125 | 0.542019 | 38.23007 | 52.79215 | 29.81184 | 39.68068 | 33.15378 | 33.52122 | 27.0001 | 39.15328 |
| ENSMUSG00000093436 | -0.35227823 | 0.359319 | 0.5277 | 21.12714 | 14.39786 | 29.81184 | 14.6646 | 26.32801 | 20.11273 | 26.03581 | 30.45255 |
| ENSMUSG00000093445 | -0.13333601 | 0.797122 | 0.875505 | 22.1332 | 33.595 | 14.90592 | 6.038364 | 17.552 | 22.34748 | 20.25007 | 22.83941 |
| ENSMUSG00000093460 | 1.338732563 | 0.019074 | 0.068209 | 67.40564 | 85.18733 | 62.9361 | 78.49874 | 19.50223 | 73.74669 | 12.53576 | 10.87591 |
| ENSMUSG00000093483 | -0.073845688 | 0.657332 | 0.779173 | 139.8416 | 141.5789 | 122.5598 | 119.042 | 133.5902 | 111.7374 | 146.572 | 156.6131 |
| ENSMUSG00000093514 | 0.636546276 | 0.066768 | 0.166804 | 58.35115 | 34.79482 | 31.46805 | 35.36756 | 21.45245 | 24.58223 | 34.71441 | 21.75182 |
| ENSMUSG00000093553 | -0.462709277 | 0.055198 | 0.14622 | 38.23007 | 37.19447 | 48.85829 | 36.23019 | 59.48179 | 53.63396 | 60.75022 | 47.85401 |
| ENSMUSG00000093561 | 1.305587656 | 0.029257 | 0.092906 | 44.26639 | 44.3934 | 24.01509 | 29.3292 | 14.62667 | 29.05173 | 11.57147 | 2.175182 |
| ENSMUSG00000093565 | 0.416273368 | 0.004714 | 0.024683 | 200.2048 | 225.5664 | 183.0116 | 174.2499 | 156.0178 | 147.4934 | 129.2147 | 152.2628 |
| ENSMUSG00000093577 | -0.372368185 | 0.116791 | 0.245109 | 81.4904 | 65.99018 | 108.482 | 98.33908 | 99.46135 | 136.3196 | 135.0005 | 90.27006 |
| ENSMUSG00000093579 | 0.019757492 | 0.962408 | 0.978526 | 14.08476 | 23.99643 | 20.70267 | 19.84034 | 16.57689 | 29.05173 | 11.57147 | 20.66423 |
| ENSMUSG00000093593 | -0.167212564 | 0.718628 | 0.824151 | 19.11503 | 16.7975 | 29.81184 | 11.21411 | 21.45245 | 27.93435 | 11.57147 | 26.10219 |
| ENSMUSG00000093594 | -0.01061782 | 0.963163 | 0.978992 | 90.54489 | 131.9804 | 134.1533 | 149.2339 | 110.1876 | 170.9582 | 116.679 | 113.1095 |

| | | | | | | | | | | | |
|---|---|---|---|---|---|---|---|---|---|---|---|
| ENSMUSG00000093599 | -1.02155728 | 0.083694 | 0.194883 | 6.036326 | 2.399643 | 11.59349 | 14.6646 | 26.32801 | 18.99536 | 12.53576 | 14.13868 |
| ENSMUSG00000093606 | 0.24022632 | 0.183365 | 0.334274 | 81.4904 | 85.18733 | 94.40416 | 99.2017 | 74.10846 | 75.98144 | 84.85745 | 70.69342 |
| ENSMUSG00000093619 | 1.205782592 | 0.015116 | 0.057976 | 14.08476 | 23.99643 | 15.73403 | 27.60395 | 4.875557 | 12.29112 | 10.60718 | 7.613138 |
| ENSMUSG00000093622 | 1.141442093 | 0.00855 | 0.038069 | 38.23007 | 53.99197 | 33.95237 | 37.09281 | 12.67645 | 33.52122 | 9.642892 | 18.48905 |
| ENSMUSG00000093629 | 0.871086268 | 0.002022 | 0.013243 | 67.40564 | 79.18822 | 50.5145 | 74.18562 | 27.30312 | 48.04709 | 31.82154 | 41.32846 |
| ENSMUSG00000093637 | 0.195021036 | 0.724309 | 0.827323 | 11.0666 | 13.19804 | 16.56213 | 11.21411 | 18.52711 | 7.821619 | 14.46434 | 4.350364 |
| ENSMUSG00000093650 | 0.352337943 | 0.546999 | 0.695324 | 19.11503 | 23.99643 | 24.01509 | 12.93935 | 3.900445 | 29.05173 | 11.57147 | 18.48905 |
| ENSMUSG00000093661 | 0.279730487 | 0.003745 | 0.020854 | 856.1523 | 837.4754 | 833.9034 | 885.0517 | 681.6028 | 789.9835 | 729.0026 | 611.2262 |
| ENSMUSG00000093668 | 0.825187266 | 0.181847 | 0.332454 | 33.19979 | 39.59411 | 24.01509 | 24.15346 | 6.825779 | 42.46022 | 11.57147 | 7.613138 |
| ENSMUSG00000093672 | 0.521674915 | 0.400677 | 0.567964 | 18.10898 | 7.198929 | 8.281066 | 18.11509 | 14.62667 | 11.17374 | 3.857157 | 6.525547 |
| ENSMUSG00000093674 | -0.18880907 | 0.024215 | 0.081141 | 21328.35 | 24190.8 | 24766.19 | 23039.81 | 28367.94 | 23508.43 | 26763.85 | 27732.49 |
| ENSMUSG00000093677 | 0.515037216 | 0.142483 | 0.279723 | 41.24823 | 34.79482 | 33.95237 | 27.60395 | 15.60178 | 33.52122 | 19.28578 | 28.27737 |
| ENSMUSG00000093686 | -0.699384899 | 0.177549 | 0.326639 | 8.048435 | 13.19804 | 5.796746 | 9.488858 | 9.751113 | 12.29112 | 22.17865 | 14.13868 |
| ENSMUSG00000093726 | 0.340330741 | 0.248514 | 0.411905 | 91.55095 | 99.58519 | 55.48314 | 93.16334 | 48.75557 | 91.62468 | 74.25027 | 53.29196 |
| ENSMUSG00000093734 | 0.225214413 | 0.261447 | 0.425621 | 92.557 | 87.58697 | 68.73285 | 81.94923 | 68.25779 | 60.3382 | 79.07171 | 73.95619 |
| ENSMUSG00000093739 | 0.341524442 | 0.503505 | 0.659477 | 17.10292 | 28.79572 | 18.21835 | 16.38985 | 7.80089 | 30.1691 | 12.53576 | 13.05109 |
| ENSMUSG00000093749 | 1.171237011 | 0.106644 | 0.23085 | 24.1453 | 34.79482 | 16.56213 | 25.8787 | 6.825779 | 31.28647 | 3.857157 | 3.262773 |
| ENSMUSG00000093760 | 1.544546982 | 0.00614 | 0.029934 | 15.09082 | 17.99732 | 21.53077 | 31.91707 | 11.70134 | 10.05637 | 5.785735 | 2.175182 |
| ENSMUSG00000093772 | 0.747951397 | 0.006234 | 0.030311 | 84.50857 | 110.3836 | 89.43552 | 94.02596 | 53.63112 | 86.03781 | 42.42872 | 43.50364 |
| ENSMUSG00000093798 | 0.01660783 | 0.937309 | 0.962765 | 54.32694 | 58.79126 | 60.45178 | 71.59775 | 56.55646 | 61.45558 | 56.89306 | 68.51824 |
| ENSMUSG00000093803 | 0.178207921 | 0.01657 | 0.061674 | 1356.161 | 1439.786 | 1346.501 | 1470.773 | 1286.172 | 1306.21 | 1136.897 | 1232.241 |
| ENSMUSG00000093826 | -0.44394053 | 0.255272 | 0.419218 | 15.09082 | 15.59768 | 20.70267 | 12.07673 | 23.40267 | 15.64324 | 20.25007 | 27.18978 |
| ENSMUSG00000093846 | -0.188774688 | 0.644279 | 0.769981 | 13.07871 | 11.99822 | 13.24971 | 16.38985 | 15.60178 | 15.64324 | 19.28578 | 11.9635 |
| ENSMUSG00000093865 | 0.076206493 | 0.872167 | 0.9234 | 15.09082 | 10.79839 | 9.93728 | 13.80198 | 10.72622 | 10.05637 | 17.3572 | 8.700729 |
| ENSMUSG00000093904 | -0.289328167 | 0.000853 | 0.006913 | 4465.875 | 4355.352 | 4867.611 | 4903.152 | 6229.986 | 5019.244 | 5610.234 | 5864.291 |
| ENSMUSG00000093908 | -0.302950075 | 0.495545 | 0.652245 | 11.0666 | 8.398751 | 23.18699 | 16.38985 | 20.47734 | 18.99536 | 19.28578 | 15.22628 |
| ENSMUSG00000093910 | 0.224024594 | 0.24572 | 0.40867 | 75.45408 | 68.38983 | 77.01392 | 68.14726 | 55.58134 | 62.57295 | 65.57166 | 64.16787 |
| ENSMUSG00000093930 | -1.254239488 | 1.09E-13 | 4.07E-11 | 2469.863 | 2015.7 | 2804.797 | 2271.288 | 5098.857 | 4267.252 | 6638.167 | 6808.32 |
| ENSMUSG00000093954 | -0.107349493 | 0.786009 | 0.868441 | 41.24823 | 63.59054 | 33.95237 | 35.36756 | 70.20801 | 22.34748 | 42.42872 | 51.11678 |
| ENSMUSG00000093979 | 0.253175835 | 0.323366 | 0.491365 | 124.7507 | 128.3809 | 145.7468 | 113.8663 | 89.71024 | 162.0192 | 104.1432 | 75.04379 |
| ENSMUSG00000093985 | 1.40799692 | 0.018241 | 0.066201 | 39.23612 | 32.39518 | 33.12427 | 28.46658 | 4.875557 | 26.81698 | 15.42863 | 3.262773 |
| ENSMUSG00000093989 | -0.232906398 | 0.278784 | 0.445128 | 78.47224 | 77.9884 | 82.81066 | 71.59775 | 100.4365 | 71.51194 | 78.10742 | 115.2847 |
| ENSMUSG00000094090 | -0.262218455 | 0.395877 | 0.563149 | 19.11503 | 32.39518 | 24.01509 | 24.15346 | 30.22845 | 29.05173 | 27.96439 | 31.54014 |
| ENSMUSG00000094114 | -0.701640622 | 0.001118 | 0.008503 | 112.6781 | 118.7823 | 118.4192 | 106.1027 | 121.8889 | 222.3574 | 216.9651 | 180.5401 |
| ENSMUSG00000094158 | 0.86330824 | 0.173438 | 0.321255 | 15.09082 | 22.79661 | 13.24971 | 5.175741 | 7.80089 | 7.821619 | 11.57147 | 3.262773 |
| ENSMUSG00000094248 | -0.232404973 | 0.325183 | 0.493053 | 110.666 | 89.98662 | 93.57605 | 87.12497 | 152.1174 | 82.68568 | 119.5719 | 92.44524 |
| ENSMUSG00000094281 | 0.172794539 | 0.70418 | 0.813797 | 16.09687 | 14.39786 | 11.59349 | 13.80198 | 9.751113 | 7.821619 | 15.42863 | 16.31387 |
| ENSMUSG00000094344 | 0.252075607 | 0.481536 | 0.640521 | 17.10292 | 27.5959 | 19.87456 | 20.70296 | 19.50223 | 17.87799 | 17.3572 | 16.31387 |
| ENSMUSG00000094365 | -0.518430477 | 0.122317 | 0.251807 | 18.10898 | 26.39607 | 19.04645 | 23.29083 | 38.02934 | 35.75597 | 26.03581 | 23.927 |
| ENSMUSG00000094392 | 0.134225831 | 0.659947 | 0.780969 | 34.20585 | 28.79572 | 40.57722 | 29.3292 | 23.40267 | 30.1691 | 38.57157 | 29.36496 |
| ENSMUSG00000094410 | -0.063730482 | 0.644743 | 0.770114 | 164.9929 | 157.1766 | 161.4808 | 153.547 | 176.4951 | 150.8455 | 186.1078 | 151.1752 |
| ENSMUSG00000094441 | -0.169314337 | 0.228478 | 0.388572 | 243.4652 | 278.3586 | 269.9628 | 265.688 | 342.2641 | 328.508 | 249.7509 | 268.635 |
| ENSMUSG00000094483 | 0.309471992 | 0.057042 | 0.149392 | 5653.019 | 6087.894 | 5337.975 | 5887.405 | 3853.64 | 6430.488 | 4091.479 | 4157.861 |
| ENSMUSG00000094518 | 0.103982863 | 0.520116 | 0.673585 | 105.6357 | 130.7805 | 114.2787 | 124.2178 | 110.1876 | 125.1459 | 105.1075 | 101.146 |
| ENSMUSG00000094526 | -0.03596217 | 0.932902 | 0.960162 | 14.08476 | 16.7975 | 38.09291 | 22.42821 | 29.25334 | 18.99536 | 20.25007 | 26.10219 |
| ENSMUSG00000094530 | -0.503632821 | 0.302857 | 0.471133 | 14.08476 | 7.198929 | 12.4216 | 17.25247 | 18.52711 | 17.87799 | 9.642892 | 27.18978 |
| ENSMUSG00000094551 | 0.355282268 | 0.168755 | 0.315007 | 70.4238 | 58.79126 | 44.71776 | 81.08661 | 50.70579 | 53.63396 | 47.25017 | 47.85401 |
| ENSMUSG00000094568 | 0.189392658 | 0.681428 | 0.796493 | 20.12109 | 25.19625 | 24.8432 | 16.38985 | 30.22845 | 8.938993 | 13.50005 | 22.83941 |
| ENSMUSG00000094595 | 0.176513801 | 0.446655 | 0.609311 | 69.41775 | 100.785 | 94.40416 | 86.26235 | 66.30757 | 100.5637 | 62.6788 | 81.56933 |
| ENSMUSG00000094626 | -0.799839305 | 0.017002 | 0.062847 | 20.12109 | 16.7975 | 24.01509 | 13.80198 | 39.00445 | 30.1691 | 31.82154 | 29.36496 |
| ENSMUSG00000094627 | 0.678150237 | 0.018827 | 0.067576 | 180.0837 | 236.3648 | 164.7932 | 219.969 | 188.1965 | 143.0239 | 81.96458 | 87.00729 |
| ENSMUSG00000094634 | 0.521130432 | 0.043207 | 0.123138 | 63.38142 | 73.18911 | 55.48314 | 62.97152 | 35.10401 | 59.22083 | 47.25017 | 35.89051 |
| ENSMUSG00000094638 | 0.052393511 | 0.900657 | 0.940351 | 18.10898 | 27.5959 | 15.73403 | 17.25247 | 15.60178 | 29.05173 | 13.50005 | 17.40146 |
| ENSMUSG00000094685 | -0.463725057 | 0.441708 | 0.605029 | 8.048435 | 13.19804 | 8.281066 | 14.6646 | 22.42756 | 3.352122 | 17.3572 | 17.40146 |
| ENSMUSG00000094724 | -0.757848014 | 0.000395 | 0.00382 | 721.341 | 585.5129 | 919.1984 | 769.4602 | 1331.027 | 820.1526 | 1274.79 | 1642.263 |
| ENSMUSG00000094777 | 0.051232223 | 0.829528 | 0.896882 | 58.35115 | 49.19268 | 67.90474 | 51.75741 | 61.43201 | 42.46022 | 54.96448 | 60.9051 |
| ENSMUSG00000094786 | 0.0218055 | 0.941307 | 0.965995 | 41.24823 | 85.18733 | 53.82693 | 54.34528 | 62.40712 | 43.57759 | 53.0359 | 70.69342 |
| ENSMUSG00000094840 | 0.693273532 | 0.25867 | 0.422517 | 24.1453 | 47.99286 | 30.63995 | 19.84034 | 9.751113 | 45.81234 | 10.60718 | 9.78832 |
| ENSMUSG00000094870 | 0.001257839 | 0.986369 | 0.992283 | 1085.533 | 1053.443 | 989.5874 | 1146.427 | 1069.697 | 1108.435 | 1065.54 | 1027.774 |
| ENSMUSG00000094910 | 0.369266582 | 1.31E-05 | 0.000284 | 877.2794 | 963.4567 | 1012.774 | 929.9081 | 695.2544 | 798.9225 | 729.9669 | 706.9342 |
| ENSMUSG00000094936 | 0.489605521 | 0.014754 | 0.056968 | 100.6054 | 115.1829 | 118.4192 | 110.4158 | 72.15824 | 96.09417 | 88.7146 | 59.81751 |
| ENSMUSG00000094942 | 0.435334145 | 0.103597 | 0.226379 | 48.29061 | 92.38626 | 85.29498 | 78.49874 | 46.80534 | 51.39921 | 62.6788 | 64.16787 |
| ENSMUSG00000094955 | -0.22546057 | 0.677848 | 0.794142 | 7.04238 | 10.79839 | 20.70267 | 9.488858 | 13.65156 | 7.821619 | 14.46434 | 20.66423 |
| ENSMUSG00000094958 | 0.239691072 | 0.520306 | 0.673736 | 19.11503 | 19.19714 | 20.70267 | 16.38985 | 14.62667 | 16.76061 | 19.28578 | 13.05109 |

| | | | | | | | | | | | |
|---|---|---|---|---|---|---|---|---|---|---|---|
| ENSMUSG00000094974 | -0.111534261 | 0.827611 | 0.895439 | 16.09687 | 8.398751 | 8.281066 | 22.42821 | 9.751113 | 17.87799 | 18.32149 | 14.13868 |
| ENSMUSG00000094989 | 0.160636654 | 0.641429 | 0.767968 | 18.10898 | 40.79393 | 38.09291 | 30.19182 | 31.20356 | 31.28647 | 25.07152 | 26.10219 |
| ENSMUSG00000095026 | -1.513637776 | 0.071502 | 0.175064 | 10.06054 | 0 | 11.59349 | 6.038364 | 8.776002 | 8.938993 | 21.21436 | 41.32846 |
| ENSMUSG00000095040 | 0.70593099 | 0.17432 | 0.322435 | 24.1453 | 14.39786 | 10.76539 | 15.52722 | 6.825779 | 16.76061 | 8.678602 | 7.613138 |
| ENSMUSG00000095041 | 0.80113706 | 4.37E-07 | 1.89E-05 | 3153.98 | 2776.387 | 2228.435 | 2619.788 | 1742.524 | 1583.319 | 1144.611 | 1715.131 |
| ENSMUSG00000095079 | -0.013887598 | 0.98094 | 0.98941 | 7.04238 | 4.799286 | 21.53077 | 20.70296 | 16.57689 | 14.52586 | 13.50005 | 10.87591 |
| ENSMUSG00000095098 | -0.256416422 | 0.037603 | 0.11175 | 1045.29 | 1306.606 | 1445.874 | 1143.839 | 1539.701 | 1325.206 | 1423.291 | 1615.073 |
| ENSMUSG00000095105 | 0.794923194 | 0.004736 | 0.024759 | 153.9263 | 217.1677 | 273.2752 | 133.7066 | 85.8098 | 102.7984 | 108.0004 | 152.2628 |
| ENSMUSG00000095115 | 0.429576781 | 3.20E-06 | 9.56E-05 | 1381.313 | 1545.37 | 1551.872 | 1649.336 | 1099.926 | 1252.576 | 1034.682 | 1165.898 |
| ENSMUSG00000095123 | 0.560555247 | 0.000242 | 0.002642 | 1135.835 | 1078.64 | 1009.462 | 1377.61 | 679.6526 | 979.9371 | 771.4313 | 690.6203 |
| ENSMUSG00000095134 | -0.373295465 | 0.423694 | 0.588965 | 18.10898 | 10.79839 | 18.21835 | 12.07673 | 21.45245 | 30.1691 | 11.57147 | 14.13868 |
| ENSMUSG00000095159 | -0.332494266 | 0.510857 | 0.666363 | 13.07871 | 9.598572 | 6.624853 | 11.21411 | 9.751113 | 10.05637 | 11.57147 | 19.57664 |
| ENSMUSG00000095193 | -0.320253367 | 0.532845 | 0.684308 | 27.16347 | 23.99643 | 16.56213 | 8.626235 | 34.1289 | 13.40849 | 15.42863 | 31.54014 |
| ENSMUSG00000095224 | -0.205810708 | 0.718121 | 0.823924 | 17.10292 | 22.79661 | 10.76539 | 18.97772 | 37.05423 | 5.58687 | 17.3572 | 19.57664 |
| ENSMUSG00000095253 | 0.181016169 | 0.295497 | 0.463271 | 871.2431 | 861.4719 | 760.2019 | 740.131 | 686.4784 | 969.8807 | 637.3951 | 557.9342 |
| ENSMUSG00000095315 | -0.369103837 | 0.178423 | 0.327822 | 28.16952 | 32.39518 | 33.95237 | 27.60395 | 46.80534 | 41.34284 | 35.6787 | 33.71532 |
| ENSMUSG00000095325 | -0.137003726 | 0.119495 | 0.24843 | 393.3673 | 352.7475 | 389.2101 | 370.0655 | 418.3228 | 425.7195 | 405.9657 | 407.8467 |
| ENSMUSG00000095332 | 0.456093591 | 0.187098 | 0.339035 | 23.13925 | 27.5959 | 30.63995 | 37.09281 | 27.30312 | 20.11273 | 15.42863 | 23.927 |
| ENSMUSG00000095362 | -0.033386165 | 0.73012 | 0.830719 | 313.889 | 328.7511 | 323.7897 | 330.3848 | 346.1645 | 320.6864 | 350.037 | 308.8759 |
| ENSMUSG00000095385 | -0.663605449 | 0.092905 | 0.209563 | 13.07871 | 11.99822 | 26.49941 | 20.70296 | 20.47734 | 31.28647 | 30.85725 | 33.71532 |
| ENSMUSG00000095403 | -0.570151081 | 0.004877 | 0.025305 | 174.0474 | 127.1811 | 179.6991 | 135.4319 | 264.2552 | 164.254 | 245.8937 | 241.4452 |
| ENSMUSG00000095407 | 0.103112288 | 0.672757 | 0.790252 | 90.54489 | 83.98751 | 64.59232 | 100.0643 | 82.88446 | 77.09881 | 98.35749 | 56.55474 |
| ENSMUSG00000095427 | 0.044604341 | 0.893104 | 0.93538 | 28.16952 | 40.79393 | 36.43669 | 31.05445 | 44.85512 | 26.81698 | 22.17865 | 38.06569 |
| ENSMUSG00000095432 | 0.019270902 | 0.901256 | 0.940707 | 329.9858 | 431.9358 | 304.7432 | 332.9727 | 313.0107 | 413.4284 | 333.6441 | 318.6642 |
| ENSMUSG00000095440 | -0.188204302 | 0.255827 | 0.419692 | 177.0656 | 209.9688 | 174.7305 | 154.4096 | 194.0472 | 187.7188 | 182.2507 | 250.146 |
| ENSMUSG00000095478 | -0.545245723 | 0.110219 | 0.236096 | 23.13925 | 25.19625 | 14.90592 | 18.11509 | 35.10401 | 24.58223 | 26.03581 | 31.54014 |
| ENSMUSG00000095526 | 0.162707854 | 0.714139 | 0.820631 | 28.16952 | 59.99108 | 28.98373 | 28.46658 | 23.40267 | 56.98608 | 23.14294 | 26.10219 |
| ENSMUSG00000095567 | -0.150581309 | 0.018767 | 0.067448 | 2407.488 | 2612.011 | 2443.743 | 2497.295 | 2942.886 | 2662.702 | 2618.045 | 2828.824 |
| ENSMUSG00000095588 | -0.193302099 | 0.614315 | 0.748023 | 14.08476 | 14.39786 | 22.35888 | 15.52722 | 21.45245 | 21.23011 | 15.42863 | 18.48905 |
| ENSMUSG00000095597 | -0.038543211 | 0.766134 | 0.854869 | 281.6952 | 346.7484 | 283.2125 | 295.8799 | 310.0854 | 354.2076 | 301.8225 | 271.8978 |
| ENSMUSG00000095620 | 0.693903656 | 0.245892 | 0.408808 | 50.30272 | 5.999108 | 20.70267 | 45.71905 | 20.47734 | 13.40849 | 18.32149 | 23.927 |
| ENSMUSG00000095677 | -0.12168177 | 0.226943 | 0.386817 | 487.9364 | 527.9215 | 553.1752 | 602.9738 | 643.5735 | 589.9735 | 573.7521 | 557.9342 |
| ENSMUSG00000095681 | 0.134902463 | 0.810425 | 0.88445 | 8.048435 | 8.398751 | 19.87456 | 11.21411 | 6.825779 | 17.87799 | 10.60718 | 8.700729 |
| ENSMUSG00000095687 | -0.692378719 | 0.000534 | 0.004823 | 721.341 | 724.6922 | 1006.978 | 791.8884 | 1363.206 | 884.9603 | 1291.183 | 1706.43 |
| ENSMUSG00000095690 | -0.313997374 | 0.396597 | 0.563821 | 24.1453 | 27.5959 | 19.87456 | 30.19182 | 30.22845 | 21.23011 | 26.03581 | 48.9416 |
| ENSMUSG00000095742 | -0.432849399 | 0.125929 | 0.256641 | 211.2714 | 244.7636 | 374.3042 | 131.9814 | 301.3094 | 299.4563 | 388.6085 | 309.9635 |
| ENSMUSG00000095762 | 0.256501809 | 0.48153 | 0.640521 | 16.09687 | 26.39607 | 30.63995 | 28.46658 | 26.32801 | 22.34748 | 20.25007 | 16.31387 |
| ENSMUSG00000095789 | -0.058319742 | 0.809214 | 0.883562 | 50.30272 | 58.79126 | 60.45178 | 63.83414 | 52.65601 | 68.15982 | 47.25017 | 76.13138 |
| ENSMUSG00000095836 | 0.408288757 | 0.431733 | 0.596279 | 20.12109 | 11.99822 | 8.281066 | 9.488858 | 8.776002 | 10.05637 | 7.714313 | 10.87591 |
| ENSMUSG00000095847 | -0.192463108 | 0.581248 | 0.722532 | 22.1332 | 17.99732 | 17.39024 | 18.97772 | 21.45245 | 16.76061 | 26.03581 | 22.83941 |
| ENSMUSG00000095868 | 0.127700991 | 0.702338 | 0.812279 | 33.19979 | 46.79304 | 25.67131 | 57.79577 | 33.15378 | 33.52122 | 43.39301 | 39.15328 |
| ENSMUSG00000095930 | 0.081435167 | 0.600692 | 0.738618 | 141.8537 | 116.3827 | 135.8095 | 117.3168 | 111.1627 | 116.2069 | 128.2505 | 128.3357 |
| ENSMUSG00000095990 | -0.349926181 | 0.060801 | 0.15644 | 68.4117 | 92.38626 | 91.09173 | 80.22398 | 112.1378 | 117.3243 | 102.2147 | 91.35765 |
| ENSMUSG00000096006 | 0.36302641 | 0.260699 | 0.424778 | 46.2785 | 47.99286 | 56.31125 | 76.77349 | 39.97956 | 67.04245 | 31.82154 | 39.15328 |
| ENSMUSG00000096054 | 0.072523954 | 0.639988 | 0.766639 | 626.7719 | 637.1052 | 729.5619 | 725.4664 | 478.7797 | 626.8469 | 726.1097 | 755.8758 |
| ENSMUSG00000096056 | 0.564689094 | 0.366372 | 0.534907 | 10.06054 | 16.7975 | 15.73403 | 42.26855 | 9.751113 | 27.93435 | 12.53576 | 7.613138 |
| ENSMUSG00000096140 | -0.123670787 | 0.792919 | 0.873246 | 14.08476 | 34.79482 | 13.24971 | 18.97772 | 17.552 | 33.52122 | 21.21436 | 15.22628 |
| ENSMUSG00000096145 | 0.055958307 | 0.571375 | 0.714306 | 912.4913 | 1123.033 | 1021.055 | 987.7039 | 1092.125 | 945.2985 | 919.9319 | 929.8904 |
| ENSMUSG00000096146 | 0.264046017 | 0.45049 | 0.612944 | 22.1332 | 33.595 | 38.09291 | 31.91707 | 20.47734 | 18.99536 | 33.75012 | 31.54014 |
| ENSMUSG00000096188 | 0.210641614 | 0.000119 | 0.001531 | 2857.194 | 2687.6 | 2741.033 | 2762.12 | 2412.425 | 2410.176 | 2254.508 | 2473.182 |
| ENSMUSG00000096199 | -0.186361279 | 0.149536 | 0.289495 | 349.1009 | 370.7449 | 440.5527 | 320.0333 | 447.5761 | 394.4331 | 444.5373 | 398.0583 |
| ENSMUSG00000096210 | 0.00496746 | 0.954338 | 0.973969 | 19800.16 | 20560.14 | 23318.65 | 23553.07 | 22949.24 | 20301.57 | 23060.01 | 20623.99 |
| ENSMUSG00000096215 | -0.885624675 | 5.00E-05 | 0.000794 | 45.27245 | 63.59054 | 81.15445 | 77.63611 | 134.5654 | 113.9722 | 124.3933 | 123.9854 |
| ENSMUSG00000096221 | 0.263158895 | 0.305033 | 0.473524 | 82.49646 | 128.3809 | 96.06037 | 81.08661 | 86.78491 | 96.09417 | 85.82174 | 53.29196 |
| ENSMUSG00000096225 | -3.461415111 | NA | NA | 1.006054 | 1.199822 | 4.96864 | 0.862623 | 0 | 89.38993 | 0 | 0 |
| ENSMUSG00000096252 | 0.677855923 | 0.256043 | 0.419812 | 20.12109 | 10.79839 | 4.96864 | 13.80198 | 8.776002 | 5.58687 | 6.750024 | 9.78832 |
| ENSMUSG00000096255 | -0.04586214 | 0.856782 | 0.914297 | 471.8395 | 364.7457 | 264.9941 | 313.1323 | 453.4268 | 261.4655 | 277.7153 | 466.5766 |
| ENSMUSG00000096279 | -0.162348194 | 0.797431 | 0.875722 | 12.07265 | 19.19714 | 22.35888 | 3.450494 | 18.52711 | 17.87799 | 6.750024 | 20.66423 |
| ENSMUSG00000096299 | 0.773502841 | 0.105073 | 0.228494 | 19.11503 | 31.19536 | 16.56213 | 26.74133 | 6.825779 | 14.52586 | 22.17865 | 10.87591 |
| ENSMUSG00000096351 | 0.461422562 | 0.046369 | 0.129237 | 70.4238 | 62.39072 | 51.34261 | 69.8725 | 41.92979 | 53.63396 | 43.39301 | 45.67883 |
| ENSMUSG00000096361 | -0.492348255 | 0.186523 | 0.338393 | 18.10898 | 22.79661 | 29.81184 | 14.6646 | 31.20356 | 27.93435 | 22.17865 | 39.15328 |
| ENSMUSG00000096370 | 0.601539633 | 0.046634 | 0.129679 | 152.9203 | 147.578 | 98.54469 | 177.7004 | 71.18313 | 151.9629 | 78.10742 | 79.39415 |
| ENSMUSG00000096401 | -0.192176891 | 0.048132 | 0.132565 | 348.0948 | 357.5468 | 349.461 | 365.7524 | 394.9201 | 425.7195 | 368.3585 | 436.124 |
| ENSMUSG00000096403 | -0.309490447 | 0.806285 | 0.88172 | 251.5136 | 232.7654 | 24.8432 | 18.11509 | 296.4338 | 11.17374 | 20.25007 | 325.1897 |

| | | | | | | | | | | | |
|---|---|---|---|---|---|---|---|---|---|---|---|
| ENSMUSG00000096433 | 0.239741497 | 0.189866 | 0.342466 | 197.1867 | 245.9634 | 226.0731 | 157.8601 | 154.0676 | 202.2447 | 183.2149 | 159.8759 |
| ENSMUSG00000096458 | -0.247101708 | 0.576181 | 0.718319 | 23.13925 | 21.59679 | 20.70267 | 9.488858 | 19.50223 | 13.40849 | 26.03581 | 29.36496 |
| ENSMUSG00000096472 | -0.203148739 | 0.042094 | 0.120915 | 516.1059 | 500.3256 | 464.5678 | 521.0246 | 622.121 | 512.8747 | 614.2522 | 552.4963 |
| ENSMUSG00000096474 | 0.018647897 | 0.964429 | 0.979975 | 14.08476 | 26.39607 | 44.71776 | 24.15346 | 21.45245 | 32.40385 | 30.85725 | 23.927 |
| ENSMUSG00000096544 | -0.256000988 | 0.069377 | 0.171134 | 210.2654 | 218.3675 | 267.4784 | 251.8861 | 306.185 | 248.057 | 266.1438 | 314.3138 |
| ENSMUSG00000096606 | 1.529647847 | 2.53E-07 | 1.21E-05 | 88.53278 | 92.38626 | 62.9361 | 89.71284 | 39.00445 | 33.52122 | 24.10723 | 18.48905 |
| ENSMUSG00000096617 | -0.079341101 | 0.841465 | 0.904383 | 19.11503 | 27.5959 | 21.53077 | 12.93935 | 21.45245 | 14.52586 | 27.0001 | 21.75182 |
| ENSMUSG00000096681 | 0.313218352 | 0.54128 | 0.690719 | 20.12109 | 34.79482 | 10.76539 | 17.25247 | 20.47734 | 8.938993 | 23.14294 | 13.05109 |
| ENSMUSG00000096687 | 0.238643424 | 0.043038 | 0.122788 | 486.9303 | 469.1302 | 424.8187 | 484.7944 | 383.2187 | 462.5929 | 389.5728 | 345.854 |
| ENSMUSG00000096696 | -0.111204822 | 0.727698 | 0.829154 | 44.26639 | 92.38626 | 69.56096 | 67.28463 | 82.88446 | 100.5637 | 64.60737 | 46.76642 |
| ENSMUSG00000096712 | -0.050398107 | 0.936318 | 0.962298 | 7.04238 | 28.79572 | 14.07781 | 8.626235 | 11.70134 | 15.64324 | 6.750024 | 26.10219 |
| ENSMUSG00000096727 | 0.398577153 | 0.375624 | 0.544104 | 40.24217 | 33.595 | 85.29498 | 37.95543 | 31.20356 | 23.46486 | 64.60737 | 30.45255 |
| ENSMUSG00000096740 | 0.168533298 | 0.287942 | 0.454754 | 251.5136 | 343.149 | 250.9163 | 260.5123 | 223.3005 | 269.2872 | 216.0008 | 274.073 |
| ENSMUSG00000096768 | 0.400890109 | 0.130393 | 0.262799 | 3027.218 | 1694.148 | 1495.561 | 1994.386 | 1244.242 | 2344.251 | 1203.433 | 1428.007 |
| ENSMUSG00000096780 | 0.433277263 | 0.033476 | 0.102307 | 476.8698 | 467.9304 | 288.1811 | 311.4071 | 289.6081 | 273.7567 | 246.858 | 331.7153 |
| ENSMUSG00000096789 | -0.217182318 | 0.591681 | 0.731496 | 15.09082 | 14.39786 | 11.59349 | 17.25247 | 20.47734 | 16.76061 | 18.32149 | 11.9635 |
| ENSMUSG00000096795 | -0.366938565 | 0.06907 | 0.170626 | 106.6418 | 70.78947 | 90.26362 | 74.18562 | 103.3618 | 125.1459 | 97.39321 | 116.3722 |
| ENSMUSG00000096841 | 0.124168991 | 0.803638 | 0.880025 | 17.10292 | 11.99822 | 9.109173 | 11.21411 | 17.552 | 8.938993 | 8.678602 | 9.78832 |
| ENSMUSG00000096842 | -0.362054141 | 0.0099 | 0.042286 | 2973.897 | 3395.495 | 4368.262 | 3196.883 | 4872.631 | 3759.964 | 4487.802 | 4790.839 |
| ENSMUSG00000096847 | 0.439743154 | 0.00679 | 0.032292 | 141.8537 | 143.9786 | 185.4959 | 175.1126 | 126.7645 | 109.5027 | 127.2862 | 114.1971 |
| ENSMUSG00000096856 | 0.233100587 | 0.682244 | 0.797182 | 11.0666 | 16.7975 | 4.96864 | 12.93935 | 6.825779 | 14.52586 | 9.642892 | 7.613138 |
| ENSMUSG00000096862 | 1.478834726 | 0.000163 | 0.001932 | 28.16952 | 47.99286 | 33.95237 | 25.8787 | 14.62667 | 13.40849 | 10.60718 | 9.78832 |
| ENSMUSG00000096883 | -0.531457582 | 0.05166 | 0.139108 | 34.20585 | 50.3925 | 52.17072 | 35.36756 | 74.10846 | 70.39457 | 54.96448 | 48.9416 |
| ENSMUSG00000096910 | -0.149232564 | 0.414078 | 0.579966 | 231.3925 | 181.1731 | 245.1196 | 248.4356 | 315.9361 | 263.7003 | 204.4293 | 222.9562 |
| ENSMUSG00000096914 | -0.281387173 | 0.434148 | 0.598404 | 20.12109 | 28.79572 | 33.12427 | 35.36756 | 36.07912 | 29.05173 | 25.07152 | 53.29196 |
| ENSMUSG00000096916 | -0.263455121 | 0.097175 | 0.216524 | 169.0171 | 169.1748 | 123.3879 | 155.2722 | 203.7983 | 191.071 | 166.822 | 176.1898 |
| ENSMUSG00000096917 | 0.092396194 | 0.71479 | 0.821225 | 50.30272 | 43.19358 | 45.54586 | 62.97152 | 51.6809 | 55.8687 | 41.46443 | 41.32846 |
| ENSMUSG00000096918 | -0.924807964 | 0.02838 | 0.090872 | 13.07871 | 14.39786 | 9.93728 | 8.626235 | 20.47734 | 14.52586 | 25.07152 | 26.10219 |
| ENSMUSG00000096919 | -0.137603438 | 0.75601 | 0.848334 | 18.10898 | 13.19804 | 18.21835 | 8.626235 | 16.57689 | 12.29112 | 14.46434 | 20.66423 |
| ENSMUSG00000096924 | 0.248599964 | 0.592832 | 0.732398 | 28.16952 | 28.79572 | 12.4216 | 24.15346 | 12.67645 | 32.40385 | 19.28578 | 14.13868 |
| ENSMUSG00000096926 | 0.339345548 | 0.431567 | 0.596141 | 40.24217 | 37.19447 | 36.43669 | 25.8787 | 21.45245 | 52.51658 | 18.32149 | 18.48905 |
| ENSMUSG00000096929 | 0.328879292 | 0.232444 | 0.393014 | 61.36932 | 40.79393 | 49.6864 | 44.85642 | 29.25334 | 45.81234 | 36.64299 | 45.67883 |
| ENSMUSG00000096932 | 0.084769284 | 0.869583 | 0.921775 | 17.10292 | 11.99822 | 13.24971 | 23.29083 | 16.57689 | 16.76061 | 23.14294 | 5.437955 |
| ENSMUSG00000096935 | 0.085586698 | 0.669863 | 0.788305 | 69.41775 | 82.78769 | 79.49824 | 87.12497 | 86.78491 | 79.33356 | 60.75022 | 73.95619 |
| ENSMUSG00000096938 | -0.895292593 | 0.07448 | 0.180205 | 19.11503 | 15.59768 | 40.57722 | 11.21411 | 33.15378 | 64.8077 | 24.10723 | 40.24087 |
| ENSMUSG00000096942 | -0.128716432 | 0.35048 | 0.519114 | 175.0535 | 202.7698 | 161.4808 | 160.448 | 204.7734 | 183.2494 | 195.7507 | 177.2773 |
| ENSMUSG00000096943 | 0.75664414 | 0.026905 | 0.08756 | 34.20585 | 32.39518 | 22.35888 | 37.09281 | 16.57689 | 22.34748 | 19.28578 | 16.31387 |
| ENSMUSG00000096944 | 0.612172603 | 0.150575 | 0.290642 | 33.19979 | 33.595 | 32.29616 | 49.16954 | 14.62667 | 43.57759 | 24.10723 | 15.22628 |
| ENSMUSG00000096948 | -0.246554725 | 0.66733 | 0.786628 | 10.06054 | 19.19714 | 15.73403 | 14.6646 | 3.900445 | 20.11273 | 26.03581 | 20.66423 |
| ENSMUSG00000096950 | 0.053088482 | 0.863178 | 0.917928 | 29.17558 | 25.19625 | 28.98373 | 28.46658 | 26.32801 | 30.1691 | 31.82154 | 19.57664 |
| ENSMUSG00000096954 | -0.076535181 | 0.832591 | 0.898824 | 34.20585 | 22.79661 | 20.70267 | 20.70296 | 34.1289 | 23.46486 | 27.0001 | 18.48905 |
| ENSMUSG00000096956 | 0.137351187 | 0.381897 | 0.549831 | 674.0564 | 922.6628 | 1112.975 | 957.5121 | 788.865 | 820.1526 | 780.1099 | 947.2918 |
| ENSMUSG00000096957 | -0.117571837 | 0.593827 | 0.733138 | 87.52673 | 75.58876 | 80.32634 | 69.8725 | 72.15824 | 73.74669 | 112.8218 | 80.48174 |
| ENSMUSG00000096959 | 0.383715135 | 0.476626 | 0.636053 | 17.10292 | 14.39786 | 11.59349 | 21.56559 | 6.825779 | 22.34748 | 6.750024 | 14.13868 |
| ENSMUSG00000096960 | 0.124477183 | 0.866129 | 0.919627 | 17.10292 | 5.999108 | 9.93728 | 13.80198 | 3.900445 | 24.58223 | 11.57147 | 3.262773 |
| ENSMUSG00000096967 | -0.117150739 | 0.831041 | 0.897887 | 12.07265 | 23.99643 | 13.24971 | 10.35148 | 5.850668 | 25.6996 | 15.42863 | 17.40146 |
| ENSMUSG00000096968 | 0.52129127 | 0.312182 | 0.480616 | 13.07871 | 29.99554 | 13.24971 | 20.70296 | 10.72622 | 23.46486 | 10.60718 | 8.700729 |
| ENSMUSG00000096971 | -0.126600197 | 0.598311 | 0.7368 | 153.9263 | 127.1811 | 115.1068 | 94.02596 | 103.3618 | 126.2633 | 122.4647 | 182.7153 |
| ENSMUSG00000096974 | 0.193886605 | 0.575741 | 0.717916 | 39.23612 | 34.79482 | 19.87456 | 24.15346 | 27.30312 | 24.58223 | 22.17865 | 28.27737 |
| ENSMUSG00000096975 | -0.303784926 | 0.33385 | 0.502081 | 37.22401 | 20.39697 | 33.12427 | 34.50494 | 28.27823 | 48.04709 | 39.53586 | 40.24087 |
| ENSMUSG00000096976 | 0.376556556 | 0.403108 | 0.570045 | 35.2119 | 37.19447 | 28.15563 | 46.58167 | 26.32801 | 50.28183 | 26.03581 | 10.87591 |
| ENSMUSG00000096977 | 0.811188055 | 0.172478 | 0.319892 | 14.08476 | 23.99643 | 9.109173 | 15.52722 | 3.900445 | 15.64324 | 10.60718 | 5.437955 |
| ENSMUSG00000096980 | 0.151215911 | 0.664628 | 0.784591 | 25.15136 | 23.99643 | 19.87456 | 28.46658 | 27.30312 | 25.6996 | 17.3572 | 17.40146 |
| ENSMUSG00000096981 | 0.134850986 | 0.44809 | 0.610689 | 95.57516 | 100.785 | 109.3101 | 83.67448 | 86.78491 | 99.44629 | 84.85745 | 83.74451 |
| ENSMUSG00000096983 | 0.182642878 | 0.625908 | 0.756104 | 46.2785 | 34.79482 | 31.46805 | 51.75741 | 40.95467 | 54.75133 | 20.25007 | 29.36496 |
| ENSMUSG00000096988 | -0.011274772 | 0.972047 | 0.98453 | 21.12714 | 33.595 | 28.15563 | 41.40593 | 29.25334 | 35.75597 | 27.0001 | 33.71532 |
| ENSMUSG00000096990 | 0.508673904 | 0.458809 | 0.62042 | 16.09687 | 19.19714 | 15.73403 | 25.8787 | 7.80089 | 34.6386 | 8.678602 | 3.262773 |
| ENSMUSG00000096991 | 0.989460024 | 0.094839 | 0.212782 | 72.43591 | 55.19179 | 18.21835 | 29.3292 | 14.62667 | 43.57759 | 21.21436 | 8.700729 |
| ENSMUSG00000096995 | -0.381589423 | 0.1542 | 0.295618 | 251.5136 | 133.1802 | 158.9965 | 235.4962 | 163.8187 | 258.1134 | 300.8582 | 293.6496 |
| ENSMUSG00000096996 | 0.26368433 | 0.240096 | 0.402143 | 160.9687 | 142.7788 | 98.54469 | 114.7289 | 116.0382 | 98.32892 | 128.2505 | 85.9197 |
| ENSMUSG00000097000 | -0.0435912 | 0.872617 | 0.923718 | 47.28455 | 34.79482 | 31.46805 | 44.85642 | 43.88001 | 35.75597 | 46.28588 | 36.9781 |
| ENSMUSG00000097002 | 0.775815033 | 0.17063 | 0.317421 | 13.07871 | 14.39786 | 20.70267 | 13.80198 | 2.925334 | 6.704245 | 11.57147 | 15.22628 |
| ENSMUSG00000097003 | 0.779219786 | 0.007361 | 0.034102 | 176.0595 | 235.165 | 156.5122 | 181.1509 | 87.76002 | 179.8972 | 91.60747 | 77.21897 |
| ENSMUSG00000097004 | 0.088400504 | 0.786219 | 0.868451 | 41.24823 | 49.19268 | 38.09291 | 56.07053 | 44.85512 | 23.46486 | 57.85735 | 46.76642 |

| | | | | | | | | | | | |
|---|---|---|---|---|---|---|---|---|---|---|---|
| ENSMUSG00000097006 | 0.112720996 | 0.277897 | 0.444096 | 926.5761 | 947.859 | 900.1519 | 819.4923 | 866.874 | 941.9464 | 746.3598 | 767.8393 |
| ENSMUSG00000097011 | -0.091257121 | 0.853232 | 0.911961 | 12.07265 | 14.39786 | 19.04645 | 11.21411 | 24.37778 | 16.76061 | 10.60718 | 8.700729 |
| ENSMUSG00000097013 | -0.023403108 | 0.969358 | 0.982939 | 33.19979 | 11.99822 | 9.93728 | 5.175741 | 13.65156 | 17.87799 | 16.39292 | 13.05109 |
| ENSMUSG00000097017 | 0.586292053 | 0.324672 | 0.492653 | 13.07871 | 15.59768 | 10.76539 | 9.488858 | 7.80089 | 3.352122 | 15.42863 | 5.437955 |
| ENSMUSG00000097021 | 0.79858266 | 0.088659 | 0.202912 | 21.12714 | 28.79572 | 24.01509 | 9.488858 | 11.70134 | 13.40849 | 10.60718 | 11.9635 |
| ENSMUSG00000097022 | -0.82462141 | 0.162723 | 0.307141 | 17.10292 | 5.999108 | 4.140533 | 5.175741 | 12.67645 | 13.40849 | 13.50005 | 17.40146 |
| ENSMUSG00000097025 | -0.093404191 | 0.803917 | 0.880139 | 24.1453 | 19.19714 | 26.49941 | 17.25247 | 27.30312 | 20.11273 | 15.42863 | 30.45255 |
| ENSMUSG00000097027 | -0.147941397 | 0.715894 | 0.822105 | 20.12109 | 44.3934 | 15.73403 | 21.56559 | 24.37778 | 25.6996 | 32.78583 | 28.27737 |
| ENSMUSG00000097032 | -0.074710426 | 0.76051 | 0.850871 | 46.2785 | 39.59411 | 43.88965 | 50.89479 | 39.00445 | 51.39921 | 55.92877 | 44.59123 |
| ENSMUSG00000097033 | 0.162639141 | 0.330692 | 0.498551 | 213.2835 | 220.7672 | 216.1358 | 244.1224 | 172.5947 | 261.4655 | 188.0364 | 178.3649 |
| ENSMUSG00000097039 | 0.310141641 | 0.115431 | 0.243363 | 139.8416 | 149.9777 | 120.0755 | 135.4319 | 120.9138 | 121.7938 | 119.5719 | 76.13138 |
| ENSMUSG00000097042 | 0.27589697 | 0.018154 | 0.066009 | 420.5307 | 446.3336 | 417.3657 | 477.8934 | 345.1894 | 429.0717 | 344.2512 | 338.2408 |
| ENSMUSG00000097045 | 0.392404709 | 0.279299 | 0.44568 | 52.31483 | 34.79482 | 23.18699 | 56.93315 | 27.30312 | 39.10809 | 31.82154 | 29.36496 |
| ENSMUSG00000097047 | -0.730792886 | 0.048396 | 0.133114 | 21.12714 | 11.99822 | 21.53077 | 35.36756 | 44.85512 | 33.52122 | 34.71441 | 38.06569 |
| ENSMUSG00000097048 | 0.385551578 | 0.014595 | 0.056517 | 833.013 | 841.0749 | 651.7199 | 917.8314 | 544.1121 | 712.8847 | 709.7168 | 515.5182 |
| ENSMUSG00000097050 | 0.106260783 | 0.75457 | 0.847514 | 21.12714 | 35.99465 | 33.95237 | 24.15346 | 24.37778 | 22.34748 | 34.71441 | 25.0146 |
| ENSMUSG00000097051 | 1.086821335 | 0.061013 | 0.156753 | 81.4904 | 68.38983 | 47.20208 | 85.39973 | 14.62667 | 82.68568 | 24.10723 | 11.9635 |
| ENSMUSG00000097052 | 0.005949994 | 0.973834 | 0.985583 | 108.6539 | 112.7832 | 88.60741 | 108.6906 | 113.1129 | 96.09417 | 117.6433 | 88.09488 |
| ENSMUSG00000097055 | -0.08098344 | 0.780665 | 0.864704 | 51.30877 | 43.19358 | 30.63995 | 35.36756 | 31.20356 | 44.69496 | 45.32159 | 47.85401 |
| ENSMUSG00000097056 | 0.097538579 | 0.785987 | 0.868441 | 32.19374 | 37.19447 | 53.82693 | 52.62003 | 56.55646 | 21.23011 | 44.3573 | 42.41605 |
| ENSMUSG00000097057 | -0.410846618 | 0.03825 | 0.112968 | 71.42986 | 62.39072 | 70.38906 | 62.97152 | 97.51113 | 102.7984 | 75.21455 | 80.48174 |
| ENSMUSG00000097058 | -0.786922212 | 0.083488 | 0.194528 | 9.054489 | 13.19804 | 10.76539 | 9.488858 | 27.30312 | 14.52586 | 13.50005 | 17.40146 |
| ENSMUSG00000097059 | -0.080894805 | 0.476027 | 0.635575 | 681.0988 | 743.8894 | 735.3587 | 789.3005 | 742.0597 | 659.2507 | 841.8244 | 876.5984 |
| ENSMUSG00000097061 | 0.354433042 | 0.044743 | 0.126005 | 308.8587 | 296.3559 | 255.0568 | 369.2029 | 274.0063 | 236.8833 | 266.1438 | 183.8029 |
| ENSMUSG00000097062 | -0.42031719 | 0.02318 | 0.078469 | 161.9748 | 112.7832 | 169.7619 | 125.943 | 187.2214 | 172.0756 | 180.3221 | 226.2189 |
| ENSMUSG00000097063 | 0.423755425 | 0.054629 | 0.145174 | 98.59333 | 112.7832 | 108.482 | 100.9269 | 61.43201 | 63.69032 | 104.1432 | 83.74451 |
| ENSMUSG00000097065 | -0.029720034 | 0.960219 | 0.977431 | 14.08476 | 13.19804 | 14.07781 | 5.175741 | 9.751113 | 16.76061 | 16.39292 | 4.350364 |
| ENSMUSG00000097068 | 1.290586316 | 0.00579 | 0.028665 | 17.10292 | 16.7975 | 18.21835 | 18.11509 | 9.751113 | 4.469496 | 6.750024 | 7.613138 |
| ENSMUSG00000097073 | -0.608172721 | 0.022204 | 0.075903 | 31.18769 | 28.79572 | 30.63995 | 31.05445 | 54.60623 | 35.75597 | 49.17875 | 45.67883 |
| ENSMUSG00000097075 | 0.248736419 | 0.596345 | 0.735213 | 34.20585 | 7.198929 | 25.67131 | 24.15346 | 20.47734 | 21.23011 | 18.32149 | 17.40146 |
| ENSMUSG00000097077 | -0.146786328 | 0.660495 | 0.781448 | 40.24217 | 40.79393 | 22.35888 | 27.60395 | 34.1289 | 25.6996 | 43.39301 | 40.24087 |
| ENSMUSG00000097078 | -0.10219503 | 0.823016 | 0.892656 | 19.11503 | 29.99554 | 27.32752 | 28.46658 | 16.57689 | 17.87799 | 25.07152 | 53.29196 |
| ENSMUSG00000097080 | -0.063250652 | 0.597242 | 0.736063 | 280.6892 | 291.5566 | 361.8826 | 320.8959 | 307.1601 | 341.9165 | 326.894 | 339.3284 |
| ENSMUSG00000097081 | -0.567459083 | 0.166244 | 0.311869 | 18.10898 | 29.99554 | 55.48314 | 37.95543 | 61.43201 | 32.40385 | 45.32159 | 71.78101 |
| ENSMUSG00000097084 | -1.243625529 | 0.123831 | 0.253513 | 91.55095 | 68.38983 | 577.1903 | 155.2722 | 448.5512 | 162.0192 | 648.0023 | 855.9342 |
| ENSMUSG00000097088 | 0.231927603 | 0.527905 | 0.680556 | 19.11503 | 20.39697 | 24.8432 | 15.52722 | 16.57689 | 15.64324 | 17.3572 | 18.48905 |
| ENSMUSG00000097090 | 0.080724857 | 0.847512 | 0.908344 | 32.19374 | 27.5959 | 13.24971 | 18.11509 | 16.57689 | 17.87799 | 27.96439 | 22.83941 |
| ENSMUSG00000097093 | -0.451967971 | 0.116864 | 0.245132 | 25.15136 | 25.19625 | 20.70267 | 25.8787 | 32.17867 | 32.40385 | 31.82154 | 35.89051 |
| ENSMUSG00000097098 | 1.059803766 | 0.046585 | 0.129599 | 42.25428 | 35.99465 | 10.76539 | 44.85642 | 10.72622 | 23.46486 | 10.60718 | 19.57664 |
| ENSMUSG00000097099 | 0.250172671 | 0.103766 | 0.226585 | 185.114 | 176.3738 | 149.8873 | 176.8378 | 147.2418 | 119.559 | 149.4648 | 160.9635 |
| ENSMUSG00000097100 | -0.610537221 | 0.263651 | 0.427964 | 24.1453 | 31.19536 | 6.624853 | 7.763611 | 24.37778 | 25.6996 | 23.14294 | 31.54014 |
| ENSMUSG00000097101 | -1.026038268 | 0.007087 | 0.033199 | 30.18163 | 26.39607 | 38.92101 | 37.95543 | 67.28268 | 32.40385 | 65.57166 | 107.6715 |
| ENSMUSG00000097103 | 0.53398257 | 0.112632 | 0.239321 | 91.55095 | 69.58965 | 57.13936 | 82.81186 | 45.83023 | 69.27719 | 66.53595 | 26.10219 |
| ENSMUSG00000097113 | -0.56882693 | 0.159023 | 0.302041 | 19.11503 | 13.19804 | 14.90592 | 10.35148 | 19.50223 | 21.23011 | 27.96439 | 16.31387 |
| ENSMUSG00000097119 | 0.316994203 | 0.029826 | 0.094146 | 1123.763 | 1045.045 | 780.0764 | 976.4898 | 778.1388 | 929.6552 | 768.5385 | 673.2189 |
| ENSMUSG00000097121 | -0.002501874 | 0.990871 | 0.994824 | 72.43591 | 61.1909 | 91.09173 | 62.97152 | 66.30757 | 82.68568 | 73.28598 | 67.43065 |
| ENSMUSG00000097123 | 0.182193126 | 0.56108 | 0.706653 | 80.48435 | 98.38537 | 73.70149 | 73.323 | 45.83023 | 116.2069 | 56.89306 | 68.51824 |
| ENSMUSG00000097124 | 1.186814896 | 0.021271 | 0.07364 | 478.8819 | 98.38537 | 77.84202 | 158.7227 | 67.28268 | 118.4417 | 84.85745 | 87.00729 |
| ENSMUSG00000097125 | 0.138673508 | 0.716026 | 0.822105 | 46.2785 | 23.99643 | 27.32752 | 31.91707 | 39.97956 | 31.28647 | 29.89296 | 16.31387 |
| ENSMUSG00000097128 | 0.103411953 | 0.838446 | 0.902295 | 13.07871 | 11.99822 | 21.53077 | 12.07673 | 5.850668 | 13.40849 | 17.3572 | 18.48905 |
| ENSMUSG00000097129 | -0.715566396 | 0.113726 | 0.240848 | 11.0666 | 16.7975 | 14.90592 | 7.763611 | 15.60178 | 14.52586 | 26.03581 | 26.10219 |
| ENSMUSG00000097131 | 0.506016457 | 0.004564 | 0.024053 | 207.2472 | 215.9679 | 166.4494 | 159.5853 | 116.0382 | 158.6671 | 120.5361 | 131.5985 |
| ENSMUSG00000097133 | 0.165689896 | 0.658134 | 0.779672 | 23.13925 | 23.99643 | 19.04645 | 24.15346 | 14.62667 | 29.05173 | 15.42863 | 21.75182 |
| ENSMUSG00000097134 | -0.416968791 | 0.343994 | 0.512741 | 11.0666 | 9.598572 | 16.56213 | 14.6646 | 18.52711 | 11.17374 | 16.39292 | 23.927 |
| ENSMUSG00000097136 | 1.12019579 | 0.025173 | 0.083426 | 70.4238 | 52.79215 | 37.2648 | 43.9938 | 11.70134 | 44.69496 | 27.96439 | 9.78832 |
| ENSMUSG00000097140 | 0.40830869 | 0.206385 | 0.362903 | 34.20585 | 28.79572 | 39.74912 | 39.68068 | 23.40267 | 39.10809 | 25.07152 | 20.66423 |
| ENSMUSG00000097141 | -0.207829442 | 0.144471 | 0.282595 | 128.775 | 122.3818 | 131.669 | 126.8057 | 148.2169 | 128.498 | 153.322 | 158.7883 |
| ENSMUSG00000097145 | -0.03661499 | 0.930431 | 0.958826 | 45.27245 | 43.19358 | 24.8432 | 22.42821 | 17.552 | 49.16446 | 31.82154 | 40.24087 |
| ENSMUSG00000097146 | 1.656892306 | 0.001159 | 0.008752 | 27.16347 | 37.19447 | 21.53077 | 44.85642 | 9.751113 | 17.87799 | 10.60718 | 3.262773 |
| ENSMUSG00000097150 | 0.314420382 | 0.593691 | 0.733018 | 4.024217 | 15.59768 | 12.4216 | 15.52722 | 6.825779 | 6.704245 | 10.60718 | 14.13868 |
| ENSMUSG00000097151 | 0.867217398 | 0.128907 | 0.260595 | 17.10292 | 10.79839 | 14.90592 | 8.626235 | 6.825779 | 10.05637 | 2.892867 | 8.700729 |
| ENSMUSG00000097152 | -0.728209362 | 0.286365 | 0.453452 | 17.10292 | 9.598572 | 3.312427 | 2.58787 | 16.57689 | 8.938993 | 14.46434 | 13.05109 |
| ENSMUSG00000097154 | 1.07883599 | 0.127839 | 0.259223 | 5.030272 | 32.39518 | 9.93728 | 12.07673 | 10.72622 | 5.58687 | 3.857157 | 7.613138 |
| ENSMUSG00000097158 | -0.11606408 | 0.77185 | 0.858471 | 13.07871 | 14.39786 | 23.18699 | 21.56559 | 26.32801 | 15.64324 | 17.3572 | 19.57664 |

| | | | | | | | | | | | |
|---|---|---|---|---|---|---|---|---|---|---|---|
| ENSMUSG00000097162 | -0.250943258 | 0.11901 | 0.247933 | 111.672 | 121.182 | 119.2474 | 91.43809 | 121.8889 | 131.8501 | 136.9291 | 135.9489 |
| ENSMUSG00000097164 | -0.650478627 | 5.06E-08 | 3.30E-06 | 343.0645 | 248.3631 | 280.7281 | 293.292 | 458.3023 | 479.3535 | 458.0374 | 436.124 |
| ENSMUSG00000097165 | -0.015110197 | 0.977886 | 0.987865 | 6.036326 | 27.5959 | 24.01509 | 23.29083 | 14.62667 | 12.29112 | 31.82154 | 22.83941 |
| ENSMUSG00000097166 | 0.351794624 | 0.483854 | 0.64217 | 7.04238 | 11.99822 | 14.90592 | 15.52722 | 11.70134 | 6.704245 | 9.642892 | 10.87591 |
| ENSMUSG00000097167 | 0.273413741 | 0.159235 | 0.302194 | 108.6539 | 184.7725 | 139.1219 | 140.6076 | 113.1129 | 112.8548 | 130.179 | 116.3722 |
| ENSMUSG00000097170 | -0.016822888 | 0.971317 | 0.983952 | 10.06054 | 14.39786 | 11.59349 | 9.488858 | 9.751113 | 14.52586 | 8.678602 | 13.05109 |
| ENSMUSG00000097174 | -0.002266024 | 0.990553 | 0.994686 | 112.6781 | 127.1811 | 99.3728 | 116.4542 | 84.83468 | 124.0285 | 128.2505 | 118.5474 |
| ENSMUSG00000097176 | 0.936368761 | 0.083035 | 0.193881 | 47.28455 | 21.59679 | 11.59349 | 19.84034 | 9.751113 | 15.64324 | 9.642892 | 17.40146 |
| ENSMUSG00000097177 | -0.029965314 | 0.870929 | 0.922556 | 95.57516 | 99.58519 | 86.12309 | 91.43809 | 79.95913 | 103.9158 | 85.82174 | 110.9343 |
| ENSMUSG00000097178 | 0.611173733 | 0.075635 | 0.181937 | 65.39353 | 80.38804 | 50.5145 | 65.55939 | 29.25334 | 63.69032 | 52.07161 | 26.10219 |
| ENSMUSG00000097180 | 0.262833552 | 0.223766 | 0.383492 | 172.0353 | 220.7672 | 116.763 | 141.4703 | 138.4658 | 125.1459 | 135.9648 | 140.2993 |
| ENSMUSG00000097183 | 0.175290568 | 0.331283 | 0.499089 | 499.003 | 331.1507 | 428.9592 | 511.5357 | 394.9201 | 306.1605 | 459.0016 | 408.9342 |
| ENSMUSG00000097184 | -0.431013726 | 0.299298 | 0.467416 | 11.0666 | 13.19804 | 19.04645 | 12.07673 | 24.37778 | 15.64324 | 15.42863 | 19.57664 |
| ENSMUSG00000097185 | 0.356320261 | 0.537308 | 0.687647 | 13.07871 | 11.99822 | 23.18699 | 6.900988 | 12.67645 | 5.58687 | 8.678602 | 16.31387 |
| ENSMUSG00000097187 | -0.124073363 | 0.260242 | 0.424259 | 210.2654 | 223.1668 | 224.4169 | 200.9913 | 233.0516 | 231.2964 | 241.0723 | 229.4817 |
| ENSMUSG00000097193 | -0.128177373 | 0.678944 | 0.794865 | 29.17558 | 27.5959 | 28.98373 | 23.29083 | 31.20356 | 29.05173 | 36.64299 | 21.75182 |
| ENSMUSG00000097195 | 0.216174117 | 0.076133 | 0.182871 | 1448.718 | 1720.544 | 1413.578 | 1527.706 | 1454.866 | 1479.403 | 1219.826 | 1103.905 |
| ENSMUSG00000097197 | -0.092213586 | 0.834774 | 0.900128 | 13.07871 | 19.19714 | 28.98373 | 23.29083 | 31.20356 | 29.05173 | 17.3572 | 13.05109 |
| ENSMUSG00000097201 | 1.522578149 | 5.27E-05 | 0.000825 | 361.1735 | 404.3399 | 394.1788 | 567.6063 | 125.7894 | 296.1041 | 106.0718 | 73.95619 |
| ENSMUSG00000097204 | 0.254523906 | 0.052262 | 0.140316 | 168.0111 | 164.3756 | 178.0429 | 188.0519 | 142.3663 | 158.6671 | 141.7505 | 144.6496 |
| ENSMUSG00000097207 | -0.508024894 | 0.247586 | 0.410774 | 14.08476 | 16.7975 | 18.21835 | 14.6646 | 20.47734 | 18.99536 | 13.50005 | 38.06569 |
| ENSMUSG00000097208 | -0.577474126 | 0.351987 | 0.520786 | 12.07265 | 13.19804 | 7.45296 | 2.58787 | 9.751113 | 8.938993 | 12.53576 | 20.66423 |
| ENSMUSG00000097209 | -1.319878977 | 0.003325 | 0.019108 | 12.07265 | 8.398751 | 28.15563 | 13.80198 | 47.78045 | 24.58223 | 44.3573 | 41.32846 |
| ENSMUSG00000097216 | 0.759772314 | 0.014982 | 0.057674 | 41.24823 | 34.79482 | 54.65504 | 49.16954 | 26.32801 | 36.87335 | 21.21436 | 22.83941 |
| ENSMUSG00000097217 | 0.054204689 | 0.857213 | 0.914599 | 41.24823 | 43.19358 | 41.40533 | 31.91707 | 25.35289 | 35.75597 | 50.14304 | 40.24087 |
| ENSMUSG00000097221 | -0.874482669 | 0.025001 | 0.083078 | 74.44802 | 63.59054 | 81.15445 | 45.71905 | 123.8391 | 44.69496 | 150.4291 | 166.4014 |
| ENSMUSG00000097222 | 0.507468391 | 0.206084 | 0.362636 | 66.39959 | 35.99465 | 56.31125 | 33.64232 | 26.32801 | 55.8687 | 21.21436 | 32.62773 |
| ENSMUSG00000097231 | 0.174530611 | 0.611625 | 0.746175 | 35.2119 | 21.59679 | 34.78048 | 40.5433 | 37.05423 | 34.6386 | 20.25007 | 26.10219 |
| ENSMUSG00000097233 | -0.098065089 | 0.768641 | 0.856335 | 23.13925 | 17.99732 | 22.35888 | 25.8787 | 18.52711 | 27.93435 | 26.03581 | 23.927 |
| ENSMUSG00000097234 | -0.054287624 | 0.930924 | 0.959196 | 27.16347 | 32.39518 | 7.45296 | 16.38985 | 12.67645 | 45.81234 | 12.53576 | 15.22628 |
| ENSMUSG00000097236 | 0.245258009 | 0.606011 | 0.742468 | 11.0666 | 21.59679 | 14.90592 | 38.81806 | 19.50223 | 16.76061 | 18.32149 | 18.48905 |
| ENSMUSG00000097239 | -0.212472805 | 0.599938 | 0.737967 | 21.12714 | 14.39786 | 18.21835 | 18.11509 | 28.27823 | 12.29112 | 25.07152 | 17.40146 |
| ENSMUSG00000097240 | 0.950874109 | 0.08747 | 0.201069 | 18.10898 | 34.79482 | 17.39024 | 22.42821 | 16.57689 | 20.11273 | 6.750024 | 4.350364 |
| ENSMUSG00000097242 | 0.269402429 | 0.21941 | 0.378197 | 206.2411 | 188.372 | 152.3716 | 162.1732 | 141.3911 | 203.3621 | 109.929 | 133.7737 |
| ENSMUSG00000097243 | 0.171439973 | 0.579154 | 0.720371 | 32.19374 | 32.39518 | 28.98373 | 25.8787 | 19.50223 | 32.40385 | 27.96439 | 26.10219 |
| ENSMUSG00000097246 | 0.980380054 | 0.038522 | 0.11341 | 46.2785 | 52.79215 | 19.04645 | 31.05445 | 15.60178 | 32.40385 | 15.42863 | 11.9635 |
| ENSMUSG00000097251 | -0.115772533 | 0.805359 | 0.880864 | 14.08476 | 17.99732 | 17.39024 | 12.93935 | 6.825779 | 21.23011 | 22.17865 | 17.40146 |
| ENSMUSG00000097253 | 0.128663268 | 0.610301 | 0.7453 | 60.36326 | 70.78947 | 72.04528 | 56.07053 | 80.93424 | 45.81234 | 53.0359 | 56.55474 |
| ENSMUSG00000097254 | 0.39854577 | 0.444216 | 0.607248 | 39.23612 | 13.19804 | 20.70267 | 14.6646 | 11.70134 | 10.05637 | 24.10723 | 20.66423 |
| ENSMUSG00000097257 | 0.056176324 | 0.878237 | 0.927008 | 20.12109 | 26.39607 | 16.56213 | 26.74133 | 20.47734 | 29.05173 | 19.28578 | 17.40146 |
| ENSMUSG00000097258 | 0.198372035 | 0.51111 | 0.66663 | 91.55095 | 91.18644 | 101.8571 | 128.5309 | 61.43201 | 141.9065 | 92.57176 | 65.25547 |
| ENSMUSG00000097261 | 0.849665982 | 0.077592 | 0.185201 | 23.13925 | 8.398751 | 20.70267 | 19.84034 | 7.80089 | 12.29112 | 10.60718 | 9.78832 |
| ENSMUSG00000097263 | 0.869164658 | 0.056366 | 0.148073 | 160.9687 | 189.5718 | 125.8722 | 152.6844 | 52.65601 | 194.4231 | 52.07161 | 45.67883 |
| ENSMUSG00000097267 | -0.181546071 | 0.727471 | 0.829154 | 14.08476 | 7.198929 | 13.24971 | 9.488858 | 10.72622 | 18.99536 | 6.750024 | 14.13868 |
| ENSMUSG00000097271 | -0.453171027 | 0.432044 | 0.596508 | 10.06054 | 4.799286 | 7.45296 | 11.21411 | 5.850668 | 8.938993 | 17.3572 | 14.13868 |
| ENSMUSG00000097275 | 0.502843456 | 0.036208 | 0.108926 | 113.6841 | 142.7788 | 158.1684 | 115.5915 | 71.18313 | 131.8501 | 82.92887 | 89.18247 |
| ENSMUSG00000097276 | 0.042854602 | 0.886177 | 0.932038 | 29.17558 | 21.59679 | 30.63995 | 28.46658 | 24.37778 | 27.93435 | 27.96439 | 27.18978 |
| ENSMUSG00000097277 | 0.436944525 | 0.015323 | 0.058574 | 373.2462 | 373.1445 | 308.0557 | 453.74 | 267.1805 | 357.5597 | 230.4651 | 259.9343 |
| ENSMUSG00000097282 | 1.467981005 | 0.000558 | 0.004977 | 27.16347 | 43.19358 | 25.67131 | 38.81806 | 17.552 | 14.52586 | 6.750024 | 9.78832 |
| ENSMUSG00000097284 | 0.30649805 | 0.176505 | 0.325238 | 73.44197 | 85.18733 | 74.5296 | 63.83414 | 73.13335 | 65.92507 | 49.17875 | 51.11678 |
| ENSMUSG00000097287 | 0.156505131 | 0.28084 | 0.447174 | 222.338 | 193.1713 | 232.698 | 188.9145 | 212.5743 | 162.0192 | 185.1435 | 191.416 |
| ENSMUSG00000097289 | 0.365592268 | 0.503848 | 0.659691 | 13.07871 | 8.398751 | 13.24971 | 10.35148 | 12.67645 | 7.821619 | 3.857157 | 10.87591 |
| ENSMUSG00000097290 | 0.291476181 | 0.015918 | 0.060128 | 416.5065 | 460.7315 | 443.8652 | 522.7498 | 322.7618 | 388.8462 | 397.2871 | 399.1459 |
| ENSMUSG00000097292 | 0.139054959 | 0.773148 | 0.859281 | 26.15741 | 22.79661 | 12.4216 | 13.80198 | 16.57689 | 27.93435 | 12.53576 | 10.87591 |
| ENSMUSG00000097296 | 0.657537539 | 0.001611 | 0.011111 | 99.59938 | 76.78858 | 77.84202 | 80.22398 | 48.75557 | 61.45558 | 52.07161 | 50.02919 |
| ENSMUSG00000097303 | 0.290727887 | 0.541347 | 0.690721 | 22.1332 | 23.99643 | 8.281066 | 14.6646 | 13.65156 | 11.17374 | 13.50005 | 17.40146 |
| ENSMUSG00000097305 | 1.254771006 | 0.009805 | 0.041957 | 67.40564 | 32.39518 | 42.23344 | 38.81806 | 15.60178 | 37.99072 | 9.642892 | 13.05109 |
| ENSMUSG00000097309 | 0.12427467 | 0.680762 | 0.796207 | 96.58122 | 86.38715 | 50.5145 | 59.52102 | 56.55646 | 91.62468 | 67.50024 | 52.20437 |
| ENSMUSG00000097310 | -0.648708579 | 0.352798 | 0.521562 | 2.012109 | 9.598572 | 18.21835 | 8.626235 | 19.50223 | 6.704245 | 10.60718 | 23.927 |
| ENSMUSG00000097316 | 0.007477482 | 0.976145 | 0.986896 | 73.44197 | 99.58519 | 66.24853 | 75.91087 | 78.98402 | 60.3382 | 67.50024 | 105.4963 |
| ENSMUSG00000097317 | 0.35700149 | 0.521161 | 0.674654 | 10.06054 | 17.99732 | 15.73403 | 39.68068 | 7.80089 | 16.76061 | 19.28578 | 21.75182 |
| ENSMUSG00000097318 | -0.277541938 | 0.290567 | 0.457764 | 47.28455 | 58.79126 | 49.6864 | 34.50494 | 62.40712 | 43.57759 | 58.82164 | 64.16787 |
| ENSMUSG00000097320 | -0.151797121 | 0.181536 | 0.332054 | 264.5923 | 249.5629 | 311.3681 | 282.9405 | 311.0605 | 289.3999 | 322.0726 | 312.1386 |
| ENSMUSG00000097321 | 0.150235606 | 0.634481 | 0.762319 | 44.26639 | 26.39607 | 47.20208 | 44.85642 | 30.22845 | 51.39921 | 32.78583 | 33.71532 |

| | | | | | | | | | | | |
|---|---|---|---|---|---|---|---|---|---|---|---|
| ENSMUSG00000097322 | -0.015251449 | 0.96669 | 0.981333 | 36.21796 | 39.59411 | 32.29616 | 38.81806 | 25.35289 | 59.22083 | 23.14294 | 41.32846 |
| ENSMUSG00000097324 | -0.150007854 | 0.662746 | 0.782975 | 31.18769 | 61.1909 | 67.90474 | 51.75741 | 79.95913 | 53.63396 | 36.64299 | 65.25547 |
| ENSMUSG00000097325 | -0.085504924 | 0.794709 | 0.873721 | 18.10898 | 32.39518 | 27.32752 | 24.15346 | 23.40267 | 32.40385 | 25.07152 | 27.18978 |
| ENSMUSG00000097326 | -0.190208231 | 0.614514 | 0.748118 | 18.10898 | 29.99554 | 29.81184 | 25.01608 | 28.27823 | 16.76061 | 41.46443 | 30.45255 |
| ENSMUSG00000097327 | 0.387955698 | 0.069454 | 0.171277 | 143.8658 | 112.7832 | 146.5749 | 124.2178 | 119.9387 | 75.98144 | 85.82174 | 121.8102 |
| ENSMUSG00000097328 | 0.037544869 | 0.812448 | 0.88594 | 208.2533 | 275.959 | 245.9477 | 247.5729 | 224.2756 | 293.8694 | 205.3936 | 229.4817 |
| ENSMUSG00000097330 | 0.910795416 | 0.116317 | 0.244477 | 43.26034 | 52.79215 | 24.01509 | 24.15346 | 19.50223 | 39.10809 | 12.53576 | 5.437955 |
| ENSMUSG00000097331 | -0.273605161 | 0.313607 | 0.481605 | 69.41775 | 61.1909 | 50.5145 | 77.63611 | 105.312 | 80.45093 | 73.28598 | 53.29196 |
| ENSMUSG00000097333 | 0.069263546 | 0.566428 | 0.710963 | 365.1977 | 363.5459 | 322.1335 | 315.7202 | 327.6374 | 364.264 | 327.8583 | 280.5985 |
| ENSMUSG00000097334 | -0.714462701 | 0.111907 | 0.238466 | 15.09082 | 14.39786 | 14.07781 | 8.626235 | 25.35289 | 11.17374 | 20.25007 | 28.27737 |
| ENSMUSG00000097335 | 0.661394184 | 0.207431 | 0.364083 | 24.1453 | 35.99465 | 14.07781 | 34.50494 | 10.72622 | 33.52122 | 11.57147 | 13.05109 |
| ENSMUSG00000097336 | 1.944827773 | 0.022486 | 0.076582 | 36.21796 | 95.98572 | 29.81184 | 63.83414 | 21.45245 | 32.40385 | 4.821446 | 0 |
| ENSMUSG00000097337 | 0.595417645 | 0.157583 | 0.300171 | 41.24823 | 37.19447 | 36.43669 | 28.46658 | 19.50223 | 40.22547 | 10.60718 | 25.0146 |
| ENSMUSG00000097340 | 0.38581865 | 0.203511 | 0.35946 | 163.9869 | 154.777 | 101.029 | 107.8279 | 69.2329 | 155.315 | 78.10742 | 101.146 |
| ENSMUSG00000097343 | 0.014274415 | 0.947751 | 0.970036 | 91.55095 | 118.7823 | 111.7944 | 70.73513 | 85.8098 | 100.5637 | 108.9647 | 92.44524 |
| ENSMUSG00000097345 | -0.21301444 | 0.481386 | 0.640461 | 26.15741 | 25.19625 | 21.53077 | 25.01608 | 33.15378 | 25.6996 | 27.96439 | 26.10219 |
| ENSMUSG00000097347 | 0.44292827 | 0.006848 | 0.032457 | 255.5378 | 245.9634 | 260.0255 | 299.3304 | 217.4498 | 231.2964 | 180.3221 | 152.2628 |
| ENSMUSG00000097350 | -0.350008852 | 0.034383 | 0.104663 | 117.7084 | 101.9848 | 110.1382 | 131.9814 | 170.6445 | 132.9675 | 135.0005 | 151.1752 |
| ENSMUSG00000097353 | 0.730747968 | 8.92E-05 | 0.001229 | 163.9869 | 154.777 | 111.7944 | 163.8985 | 84.83468 | 90.5073 | 87.75031 | 94.62042 |
| ENSMUSG00000097354 | 0.278871092 | 0.324489 | 0.492495 | 72.43591 | 92.38626 | 112.6225 | 94.02596 | 43.88001 | 98.32892 | 78.10742 | 87.00729 |
| ENSMUSG00000097357 | -0.268708206 | 0.534069 | 0.685259 | 34.20585 | 23.99643 | 19.04645 | 18.97772 | 20.47734 | 16.76061 | 45.32159 | 32.62773 |
| ENSMUSG00000097358 | 0.606149797 | 0.049584 | 0.135313 | 122.7386 | 117.5825 | 88.60741 | 98.33908 | 69.2329 | 109.5027 | 62.6788 | 39.15328 |
| ENSMUSG00000097360 | 0.099741602 | 0.750348 | 0.844519 | 26.15741 | 34.79482 | 28.98373 | 28.46658 | 20.47734 | 35.75597 | 26.03581 | 28.27737 |
| ENSMUSG00000097364 | 0.896918735 | NA | NA | 328.9798 | 325.1516 | 181.3554 | 307.9566 | 78.0089 | 423.4848 | 61.71451 | 51.11678 |
| ENSMUSG00000097365 | 0.641351104 | 0.13142 | 0.264258 | 34.20585 | 46.79304 | 36.43669 | 43.13117 | 32.17867 | 42.46022 | 16.39292 | 11.9635 |
| ENSMUSG00000097372 | 1.093146495 | 0.120273 | 0.2493 | 14.08476 | 26.39607 | 6.624853 | 18.11509 | 2.925334 | 15.64324 | 8.678602 | 3.262773 |
| ENSMUSG00000097375 | 0.162645417 | 0.333327 | 0.501498 | 304.8345 | 299.9554 | 250.0882 | 316.5828 | 235.9769 | 334.0949 | 262.2867 | 214.2554 |
| ENSMUSG00000097378 | 0.421393028 | 0.048022 | 0.132382 | 91.55095 | 85.18733 | 73.70149 | 77.63611 | 59.48179 | 71.51194 | 66.53595 | 46.76642 |
| ENSMUSG00000097379 | 0.47248224 | 0.192236 | 0.34549 | 95.57516 | 69.58965 | 83.63877 | 87.12497 | 60.4569 | 102.7984 | 51.10733 | 28.27737 |
| ENSMUSG00000097380 | 0.569516737 | 0.229005 | 0.389075 | 29.17558 | 52.79215 | 45.54586 | 25.01608 | 14.62667 | 46.92971 | 26.03581 | 15.22628 |
| ENSMUSG00000097381 | 1.117160193 | 0.005735 | 0.028477 | 26.15741 | 27.5959 | 19.04645 | 25.8787 | 8.776002 | 12.29112 | 15.42863 | 8.700729 |
| ENSMUSG00000097383 | -0.953535322 | 0.063313 | 0.161061 | 15.09082 | 14.39786 | 24.01509 | 15.52722 | 23.40267 | 12.29112 | 55.92877 | 42.41605 |
| ENSMUSG00000097384 | -0.92954561 | 0.102486 | 0.224668 | 7.04238 | 3.599465 | 8.281066 | 13.80198 | 18.52711 | 17.87799 | 19.28578 | 7.613138 |
| ENSMUSG00000097385 | 0.916273394 | 0.012861 | 0.051599 | 92.557 | 94.7859 | 57.13936 | 72.46037 | 32.17867 | 70.39457 | 42.42872 | 22.83941 |
| ENSMUSG00000097391 | 1.141555982 | 0.000318 | 0.003241 | 285.7194 | 273.5593 | 245.9477 | 328.6596 | 113.1129 | 221.2401 | 113.7861 | 66.34306 |
| ENSMUSG00000097392 | 0.294694628 | 0.101749 | 0.223717 | 694.1775 | 663.5013 | 626.0486 | 694.4119 | 458.3023 | 772.1055 | 499.5018 | 454.6131 |
| ENSMUSG00000097397 | -0.074539451 | 0.719807 | 0.82468 | 69.41775 | 92.38626 | 73.70149 | 61.24627 | 82.88446 | 72.62932 | 70.39311 | 84.8321 |
| ENSMUSG00000097402 | 0.341284246 | 0.543413 | 0.692451 | 23.13925 | 16.7975 | 11.59349 | 19.84034 | 6.825779 | 25.6996 | 17.3572 | 6.525547 |
| ENSMUSG00000097403 | 0.063004367 | 0.769688 | 0.857036 | 104.6297 | 94.7859 | 110.9663 | 106.1027 | 79.95913 | 135.2023 | 102.2147 | 82.65692 |
| ENSMUSG00000097404 | -0.588155951 | 0.281556 | 0.447887 | 5.030272 | 8.398751 | 12.4216 | 15.52722 | 10.72622 | 10.05637 | 19.28578 | 22.83941 |
| ENSMUSG00000097405 | -0.041379746 | 0.913523 | 0.948837 | 52.31483 | 38.39429 | 58.79557 | 29.3292 | 33.15378 | 27.93435 | 57.85735 | 65.25547 |
| ENSMUSG00000097407 | -0.03076386 | 0.966398 | 0.981231 | 19.11503 | 3.599465 | 12.4216 | 8.626235 | 3.900445 | 5.58687 | 21.21436 | 14.13868 |
| ENSMUSG00000097410 | 0.459692851 | 0.306719 | 0.475352 | 63.38142 | 79.18822 | 48.85829 | 46.58167 | 35.10401 | 87.15518 | 27.96439 | 22.83941 |
| ENSMUSG00000097411 | -0.202653943 | 0.581518 | 0.722532 | 25.15136 | 29.99554 | 18.21835 | 19.84034 | 24.37778 | 30.1691 | 17.3572 | 34.80291 |
| ENSMUSG00000097412 | -0.007139181 | 0.964561 | 0.980014 | 132.7992 | 139.1793 | 106.8258 | 120.7673 | 137.4907 | 116.2069 | 117.6433 | 128.3357 |
| ENSMUSG00000097413 | -0.507025012 | 0.27578 | 0.442015 | 13.07871 | 5.999108 | 13.24971 | 14.6646 | 12.67645 | 18.99536 | 13.50005 | 22.83941 |
| ENSMUSG00000097414 | 0.164687898 | 0.382996 | 0.550971 | 139.8416 | 122.3818 | 110.9663 | 161.3106 | 129.6898 | 108.3853 | 135.0005 | 103.3212 |
| ENSMUSG00000097415 | 0.737777129 | 0.001454 | 0.010346 | 1668.038 | 2196.873 | 1536.966 | 1902.947 | 996.5638 | 1677.179 | 949.8248 | 756.9634 |
| ENSMUSG00000097416 | 0.267158099 | 0.670742 | 0.788987 | 5.030272 | 16.7975 | 16.56213 | 8.626235 | 5.850668 | 16.76061 | 5.785735 | 10.87591 |
| ENSMUSG00000097417 | -0.232001864 | 0.407504 | 0.573764 | 36.21796 | 41.99375 | 29.81184 | 31.05445 | 40.95467 | 31.28647 | 49.17875 | 40.24087 |
| ENSMUSG00000097418 | 1.052134554 | 0.004065 | 0.0221 | 37.22401 | 52.79215 | 42.23344 | 46.58167 | 17.552 | 34.6386 | 22.17865 | 11.9635 |
| ENSMUSG00000097421 | -2.282489686 | 0.000189 | 0.002171 | 9.054489 | 1.199822 | 19.87456 | 10.35148 | 70.20801 | 25.6996 | 37.60728 | 67.43065 |
| ENSMUSG00000097422 | 1.081058196 | 0.040457 | 0.117623 | 11.0666 | 25.19625 | 28.15563 | 12.07673 | 7.80089 | 11.17374 | 10.60718 | 6.525547 |
| ENSMUSG00000097430 | 0.332138946 | 0.145069 | 0.283203 | 61.36932 | 87.58697 | 85.29498 | 69.00988 | 49.73068 | 67.04245 | 68.46453 | 55.46715 |
| ENSMUSG00000097431 | 0.226653909 | 0.041771 | 0.120306 | 306.8466 | 335.95 | 357.7421 | 335.5605 | 298.3841 | 283.813 | 254.5723 | 306.7007 |
| ENSMUSG00000097439 | 0.208857426 | 0.471253 | 0.630997 | 35.2119 | 46.79304 | 46.37397 | 57.79577 | 42.9049 | 45.81234 | 45.32159 | 27.18978 |
| ENSMUSG00000097440 | -0.17960596 | 0.659242 | 0.780635 | 18.10898 | 21.59679 | 23.18699 | 18.11509 | 19.50223 | 17.87799 | 37.60728 | 16.31387 |
| ENSMUSG00000097443 | 0.645151804 | 0.237885 | 0.399097 | 25.15136 | 39.59411 | 18.21835 | 15.52722 | 8.776002 | 27.93435 | 18.32149 | 7.613138 |
| ENSMUSG00000097445 | -0.263947239 | 0.253535 | 0.417231 | 54.32694 | 49.19268 | 60.45178 | 45.71905 | 49.73068 | 70.39457 | 70.39311 | 61.99269 |
| ENSMUSG00000097451 | 1.458432286 | 0.00026 | 0.002785 | 6121.841 | 5047.649 | 4285.452 | 6668.942 | 1845.886 | 3662.752 | 1405.934 | 1136.533 |
| ENSMUSG00000097457 | 0.721351903 | 0.11856 | 0.247389 | 18.10898 | 21.59679 | 19.87456 | 28.46658 | 8.776002 | 23.46486 | 8.678602 | 13.05109 |
| ENSMUSG00000097462 | 0.354683322 | 0.083608 | 0.194784 | 152.9203 | 155.9768 | 105.1695 | 178.5631 | 120.9138 | 116.2069 | 126.3219 | 98.97079 |
| ENSMUSG00000097463 | 0.315338441 | 0.401556 | 0.568792 | 19.11503 | 19.19714 | 16.56213 | 17.25247 | 15.60178 | 13.40849 | 13.50005 | 15.22628 |
| ENSMUSG00000097464 | 1.381533849 | 0.011932 | 0.048807 | 133.8052 | 122.3818 | 65.42042 | 119.042 | 20.47734 | 105.0332 | 21.21436 | 22.83941 |

| | | | | | | | | | | | |
|---|---|---|---|---|---|---|---|---|---|---|---|
| ENSMUSG00000097467 | -0.0574219 | 0.792374 | 0.872973 | 68.4117 | 76.78858 | 55.48314 | 85.39973 | 82.88446 | 70.39457 | 78.10742 | 65.25547 |
| ENSMUSG00000097470 | 0.476587567 | 0.102647 | 0.224967 | 73.44197 | 71.98929 | 56.31125 | 62.97152 | 58.50668 | 43.57759 | 59.78593 | 27.18978 |
| ENSMUSG00000097472 | 0.532777888 | 0.107633 | 0.232214 | 91.55095 | 88.78679 | 63.76421 | 102.6522 | 66.30757 | 93.85942 | 39.53586 | 40.24087 |
| ENSMUSG00000097476 | 0.098661844 | 0.891305 | 0.93439 | 9.054489 | 22.79661 | 9.93728 | 7.763611 | 2.925334 | 25.6996 | 7.714313 | 9.78832 |
| ENSMUSG00000097479 | -0.136632446 | 0.699508 | 0.810259 | 36.21796 | 22.79661 | 42.23344 | 62.10889 | 40.95467 | 50.28183 | 34.71441 | 55.46715 |
| ENSMUSG00000097481 | -0.619808373 | 0.128025 | 0.259366 | 24.1453 | 22.79661 | 11.59349 | 15.52722 | 29.25334 | 16.76061 | 34.71441 | 31.54014 |
| ENSMUSG00000097487 | 0.31251332 | 0.072883 | 0.177504 | 187.1261 | 173.9741 | 130.0127 | 179.4257 | 136.5156 | 154.1976 | 122.4647 | 126.1606 |
| ENSMUSG00000097488 | 0.231096259 | 0.291896 | 0.459113 | 125.7568 | 118.7823 | 126.7003 | 148.3712 | 78.98402 | 149.7281 | 108.9647 | 106.5839 |
| ENSMUSG00000097493 | 0.096113722 | 0.551696 | 0.69932 | 255.5378 | 315.5531 | 344.4924 | 301.9182 | 240.8525 | 354.2076 | 264.2152 | 281.6861 |
| ENSMUSG00000097494 | 0.631722151 | 0.060546 | 0.156067 | 33.19979 | 35.99465 | 42.23344 | 52.62003 | 25.35289 | 35.75597 | 28.92867 | 16.31387 |
| ENSMUSG00000097503 | -0.711629442 | 0.063236 | 0.160933 | 20.12109 | 21.59679 | 14.90592 | 16.38985 | 25.35289 | 18.99536 | 42.42872 | 31.54014 |
| ENSMUSG00000097504 | 0.429415037 | 0.111956 | 0.238489 | 537.233 | 441.5343 | 272.4471 | 376.1038 | 274.0063 | 455.8886 | 240.108 | 238.1824 |
| ENSMUSG00000097507 | 1.147262916 | 0.074074 | 0.179534 | 56.33904 | 37.19447 | 37.2648 | 40.5433 | 17.552 | 48.04709 | 7.714313 | 4.350364 |
| ENSMUSG00000097509 | -0.334423667 | 0.512956 | 0.668016 | 10.06054 | 20.39697 | 7.45296 | 9.488858 | 15.60178 | 17.87799 | 16.39292 | 8.700729 |
| ENSMUSG00000097514 | -0.338239705 | 0.583635 | 0.724151 | 15.09082 | 13.19804 | 8.281066 | 9.488858 | 6.825779 | 23.46486 | 22.17865 | 5.437955 |
| ENSMUSG00000097515 | 0.650121528 | 0.256696 | 0.420325 | 17.10292 | 23.99643 | 10.76539 | 6.900988 | 7.80089 | 5.58687 | 12.53576 | 10.87591 |
| ENSMUSG00000097518 | -0.458471317 | 0.027124 | 0.088055 | 64.38748 | 69.58965 | 64.59232 | 77.63611 | 98.48624 | 105.0332 | 105.1075 | 70.69342 |
| ENSMUSG00000097519 | -0.778672648 | 0.090509 | 0.206049 | 22.1332 | 10.79839 | 14.07781 | 17.25247 | 21.45245 | 41.34284 | 33.75012 | 14.13868 |
| ENSMUSG00000097523 | -0.533378195 | 0.207158 | 0.363779 | 9.054489 | 16.7975 | 14.90592 | 11.21411 | 14.62667 | 24.58223 | 16.39292 | 19.57664 |
| ENSMUSG00000097526 | -0.375359974 | 0.526615 | 0.67957 | 5.030272 | 17.99732 | 8.281066 | 14.6646 | 8.776002 | 11.17374 | 26.03581 | 13.05109 |
| ENSMUSG00000097527 | -0.106187277 | 0.836886 | 0.901352 | 9.054489 | 14.39786 | 8.281066 | 10.35148 | 5.850668 | 14.52586 | 10.60718 | 14.13868 |
| ENSMUSG00000097530 | -0.073140732 | 0.821001 | 0.891475 | 28.16952 | 32.39518 | 24.8432 | 28.46658 | 38.02934 | 33.52122 | 27.96439 | 19.57664 |
| ENSMUSG00000097532 | 0.533007464 | 0.147252 | 0.286465 | 31.18769 | 40.79393 | 38.09291 | 27.60395 | 13.65156 | 35.75597 | 23.14294 | 22.83941 |
| ENSMUSG00000097533 | 0.055279085 | 0.918354 | 0.951771 | 10.06054 | 13.19804 | 12.4216 | 18.97772 | 20.47734 | 12.29112 | 4.821446 | 15.22628 |
| ENSMUSG00000097534 | 0.239071088 | 0.431601 | 0.596141 | 27.16347 | 39.59411 | 41.40533 | 53.48266 | 35.10401 | 32.40385 | 29.89296 | 40.24087 |
| ENSMUSG00000097536 | 0.443039712 | 0.12617 | 0.257025 | 123.7447 | 97.18554 | 84.46688 | 55.2079 | 72.15824 | 79.33356 | 59.78593 | 53.29196 |
| ENSMUSG00000097537 | 0.326173502 | 0.045244 | 0.126946 | 170.0232 | 172.7743 | 153.1997 | 148.3712 | 121.8889 | 158.6671 | 114.7504 | 118.5474 |
| ENSMUSG00000097545 | 0.462163655 | 0.014249 | 0.055538 | 136.8234 | 152.3773 | 137.4657 | 108.6906 | 93.61069 | 118.4417 | 84.85745 | 91.35765 |
| ENSMUSG00000097554 | -0.855064043 | 0.065053 | 0.164288 | 10.06054 | 16.7975 | 11.59349 | 6.900988 | 24.37778 | 24.58223 | 12.53576 | 19.57664 |
| ENSMUSG00000097557 | 0.617208652 | 0.178533 | 0.327959 | 10.06054 | 21.59679 | 14.90592 | 15.52722 | 9.751113 | 8.938993 | 10.60718 | 10.87591 |
| ENSMUSG00000097558 | -0.303158421 | 0.442051 | 0.605319 | 17.10292 | 13.19804 | 18.21835 | 12.07673 | 16.57689 | 15.64324 | 24.10723 | 18.48905 |
| ENSMUSG00000097561 | -0.074475028 | 0.869155 | 0.921481 | 19.11503 | 15.59768 | 15.73403 | 10.35148 | 14.62667 | 20.11273 | 20.25007 | 8.700729 |
| ENSMUSG00000097564 | -0.292385907 | 0.503191 | 0.659151 | 13.07871 | 13.19804 | 12.4216 | 8.626235 | 18.52711 | 13.40849 | 12.53576 | 13.05109 |
| ENSMUSG00000097566 | -0.004004828 | 0.992513 | 0.995642 | 26.15741 | 21.59679 | 26.49941 | 13.80198 | 12.67645 | 18.99536 | 24.10723 | 32.62773 |
| ENSMUSG00000097567 | 0.162374299 | 0.638073 | 0.765389 | 30.18163 | 65.99018 | 65.42042 | 61.24627 | 32.17867 | 46.92971 | 54.00019 | 66.34306 |
| ENSMUSG00000097571 | 0.072635034 | 0.73568 | 0.834576 | 181.0898 | 121.182 | 131.669 | 142.3329 | 112.1378 | 182.132 | 124.3933 | 130.5109 |
| ENSMUSG00000097572 | 0.675235826 | 0.058096 | 0.151187 | 76.46013 | 101.9848 | 91.91984 | 36.23019 | 47.78045 | 55.8687 | 54.00019 | 33.71532 |
| ENSMUSG00000097573 | -0.47727367 | 0.026311 | 0.086252 | 50.30272 | 59.99108 | 58.79557 | 63.83414 | 70.20801 | 82.68568 | 72.32169 | 100.0584 |
| ENSMUSG00000097574 | 0.264721823 | 0.187534 | 0.339692 | 281.6952 | 286.7573 | 322.9616 | 300.193 | 222.3254 | 354.2076 | 228.5365 | 188.1533 |
| ENSMUSG00000097576 | 0.220147188 | 0.681039 | 0.796429 | 7.04238 | 19.19714 | 10.76539 | 20.70296 | 7.80089 | 10.05637 | 17.3572 | 14.13868 |
| ENSMUSG00000097577 | 0.248829184 | 0.234877 | 0.395735 | 67.40564 | 92.38626 | 71.21717 | 84.5371 | 72.15824 | 55.8687 | 63.64308 | 72.8686 |
| ENSMUSG00000097578 | 0.003750455 | 0.987675 | 0.993028 | 78.47224 | 53.99197 | 78.67013 | 54.34528 | 57.53157 | 62.57295 | 81.00029 | 64.16787 |
| ENSMUSG00000097579 | 0.453295579 | 0.420507 | 0.585947 | 15.09082 | 11.99822 | 17.39024 | 5.175741 | 5.850668 | 11.17374 | 10.60718 | 8.700729 |
| ENSMUSG00000097583 | 1.020313686 | 0.01495 | 0.057567 | 115.6963 | 86.38715 | 55.48314 | 76.77349 | 31.20356 | 78.21619 | 32.78583 | 22.83941 |
| ENSMUSG00000097585 | 0.541946291 | 0.099675 | 0.220467 | 60.36326 | 46.79304 | 58.79557 | 38.81806 | 37.05423 | 42.46022 | 19.28578 | 42.41605 |
| ENSMUSG00000097589 | 0.23623763 | 0.324068 | 0.49206 | 549.3057 | 463.1311 | 329.5864 | 327.7969 | 329.5876 | 489.4099 | 335.5726 | 262.1095 |
| ENSMUSG00000097595 | 0.405757251 | 0.29114 | 0.458275 | 30.18163 | 16.7975 | 25.67131 | 19.84034 | 21.45245 | 20.11273 | 15.42863 | 13.05109 |
| ENSMUSG00000097601 | -0.080529683 | 0.842251 | 0.904859 | 29.17558 | 67.19001 | 49.6864 | 56.07053 | 25.35289 | 80.45093 | 47.25017 | 60.9051 |
| ENSMUSG00000097604 | -0.282265837 | 0.592706 | 0.732341 | 12.07265 | 4.799286 | 10.76539 | 8.626235 | 11.70134 | 10.05637 | 7.714313 | 15.22628 |
| ENSMUSG00000097610 | 1.085026338 | 0.006815 | 0.032357 | 26.15741 | 35.99465 | 23.18699 | 25.8787 | 15.60178 | 7.821619 | 17.3572 | 10.87591 |
| ENSMUSG00000097616 | 0.053749754 | 0.82905 | 0.896575 | 37.22401 | 47.99286 | 48.85829 | 49.16954 | 41.92979 | 53.63396 | 43.39301 | 38.06569 |
| ENSMUSG00000097617 | -0.044070277 | 0.749125 | 0.843451 | 173.0413 | 170.3747 | 164.7932 | 178.5631 | 180.3956 | 205.5968 | 163.9292 | 158.7883 |
| ENSMUSG00000097618 | 0.356895845 | 0.087374 | 0.200969 | 87.52673 | 92.38626 | 83.63877 | 89.71284 | 76.05868 | 83.80306 | 64.60737 | 51.11678 |
| ENSMUSG00000097619 | 1.538291582 | 0.003111 | 0.018278 | 17.10292 | 23.99643 | 16.56213 | 22.42821 | 11.70134 | 4.469496 | 7.714313 | 3.262773 |
| ENSMUSG00000097620 | -0.584452234 | 0.393507 | 0.56085 | 5.030272 | 11.99822 | 10.76539 | 7.763611 | 26.32801 | 3.352122 | 14.46434 | 8.700729 |
| ENSMUSG00000097621 | 1.871227876 | 0.000461 | 0.004303 | 23.13925 | 17.99732 | 21.53077 | 12.93935 | 2.925334 | 4.469496 | 6.750024 | 6.525547 |
| ENSMUSG00000097622 | -0.438587544 | 0.111078 | 0.237221 | 42.25428 | 52.79215 | 72.04528 | 38.81806 | 68.25779 | 58.10345 | 68.46453 | 84.8321 |
| ENSMUSG00000097626 | -0.145657113 | 0.633441 | 0.761714 | 43.26034 | 43.19358 | 24.01509 | 41.40593 | 41.92979 | 44.69496 | 31.82154 | 48.9416 |
| ENSMUSG00000097633 | 1.194909529 | 0.096068 | 0.214652 | 34.20585 | 23.99643 | 25.67131 | 10.35148 | 12.67645 | 22.34748 | 1.928578 | 4.350364 |
| ENSMUSG00000097635 | -0.039881319 | 0.934252 | 0.961185 | 13.07871 | 21.59679 | 14.90592 | 11.21411 | 9.751113 | 21.23011 | 21.21436 | 9.78832 |
| ENSMUSG00000097636 | 1.097873317 | 0.018493 | 0.066804 | 43.26034 | 35.99465 | 21.53077 | 23.29083 | 11.70134 | 20.11273 | 19.28578 | 6.525547 |
| ENSMUSG00000097640 | 0.312888721 | 0.028889 | 0.092055 | 499.003 | 529.1213 | 514.2542 | 587.4466 | 399.7956 | 507.2878 | 468.6445 | 339.3284 |
| ENSMUSG00000097643 | -0.087164897 | 0.734114 | 0.833414 | 48.29061 | 52.79215 | 51.34261 | 37.09281 | 58.50668 | 39.10809 | 46.28588 | 56.55474 |
| ENSMUSG00000097649 | 0.88508277 | 0.002219 | 0.01416 | 92.557 | 59.99108 | 64.59232 | 70.73513 | 41.92979 | 51.39921 | 25.07152 | 38.06569 |

| | | | | | | | | | | | |
|---|---|---|---|---|---|---|---|---|---|---|---|
| ENSMUSG00000097651 | -3.07821116 | 0.000995 | 0.007805 | 1.006054 | 0 | 9.93728 | 1.725247 | 29.25334 | 6.704245 | 32.78583 | 42.41605 |
| ENSMUSG00000097658 | 0.200252988 | 0.679005 | 0.794865 | 24.1453 | 13.19804 | 24.8432 | 42.26855 | 32.17867 | 21.23011 | 27.0001 | 10.87591 |
| ENSMUSG00000097660 | 0.931930673 | 0.003158 | 0.018463 | 49.29666 | 34.79482 | 34.78048 | 45.71905 | 20.47734 | 22.34748 | 27.0001 | 16.31387 |
| ENSMUSG00000097662 | 0.278616389 | 0.652651 | 0.775616 | 11.0666 | 15.59768 | 12.4216 | 6.900988 | 5.850668 | 18.99536 | 8.678602 | 4.350364 |
| ENSMUSG00000097664 | 0.261584188 | 0.548846 | 0.697048 | 61.36932 | 33.595 | 49.6864 | 57.79577 | 43.88001 | 65.92507 | 45.32159 | 14.13868 |
| ENSMUSG00000097666 | -0.100924427 | 0.787006 | 0.869061 | 34.20585 | 52.79215 | 25.67131 | 25.01608 | 41.92979 | 49.16446 | 27.0001 | 28.27737 |
| ENSMUSG00000097667 | 1.00940759 | 0.077091 | 0.184394 | 14.08476 | 21.59679 | 22.35888 | 18.97772 | 2.925334 | 14.52586 | 15.42863 | 5.437955 |
| ENSMUSG00000097674 | 0.569220038 | 0.088191 | 0.202142 | 42.25428 | 22.79661 | 40.57722 | 35.36756 | 29.25334 | 24.58223 | 23.14294 | 18.48905 |
| ENSMUSG00000097675 | -0.776099927 | 0.088566 | 0.202802 | 13.07871 | 8.398751 | 29.81184 | 16.38985 | 22.42756 | 32.40385 | 25.07152 | 38.06569 |
| ENSMUSG00000097680 | 0.232874267 | 0.436489 | 0.600286 | 36.21796 | 39.59411 | 38.92101 | 28.46658 | 23.40267 | 39.10809 | 27.96439 | 31.54014 |
| ENSMUSG00000097684 | 0.443648211 | 0.390888 | 0.558288 | 30.18163 | 52.79215 | 49.6864 | 43.13117 | 21.45245 | 72.62932 | 19.28578 | 16.31387 |
| ENSMUSG00000097692 | -0.276663321 | 0.323974 | 0.491966 | 47.28455 | 53.99197 | 39.74912 | 28.46658 | 41.92979 | 50.28183 | 54.96448 | 56.55474 |
| ENSMUSG00000097697 | 0.332607963 | 0.594511 | 0.733835 | 7.04238 | 11.99822 | 36.43669 | 28.46658 | 9.751113 | 24.58223 | 23.14294 | 9.78832 |
| ENSMUSG00000097699 | 0.396646485 | 0.17613 | 0.324612 | 44.26639 | 47.99286 | 52.99882 | 68.14726 | 32.17867 | 51.39921 | 30.85725 | 48.9416 |
| ENSMUSG00000097702 | 0.67356147 | 0.347061 | 0.515896 | 6.036326 | 19.19714 | 10.76539 | 17.25247 | 11.70134 | 15.64324 | 3.857157 | 2.175182 |
| ENSMUSG00000097703 | 0.491381856 | 0.432101 | 0.596508 | 12.07265 | 17.99732 | 9.109173 | 10.35148 | 3.900445 | 17.87799 | 4.821446 | 8.700729 |
| ENSMUSG00000097706 | -0.844563699 | 0.079879 | 0.188801 | 9.054489 | 4.799286 | 8.281066 | 9.488858 | 16.57689 | 14.52586 | 15.42863 | 10.87591 |
| ENSMUSG00000097707 | 1.226567448 | 0.180537 | 0.330718 | 59.35721 | 25.19625 | 18.21835 | 8.626235 | 4.875557 | 36.87335 | 3.857157 | 2.175182 |
| ENSMUSG00000097709 | 0.276555235 | 0.101847 | 0.223792 | 161.9748 | 214.7681 | 169.7619 | 148.3712 | 122.864 | 148.6108 | 151.3934 | 149 |
| ENSMUSG00000097714 | -1.014640827 | 0.028367 | 0.090844 | 9.054489 | 8.398751 | 7.45296 | 7.763611 | 19.50223 | 17.87799 | 17.3572 | 10.87591 |
| ENSMUSG00000097715 | -0.395688103 | 0.026545 | 0.086801 | 219.3198 | 182.3729 | 211.9953 | 219.1064 | 345.1894 | 212.3011 | 243.0009 | 295.8248 |
| ENSMUSG00000097718 | 0.045457047 | 0.886959 | 0.932224 | 28.16952 | 26.39607 | 21.53077 | 23.29083 | 21.45245 | 25.6996 | 27.96439 | 20.66423 |
| ENSMUSG00000097722 | 0.940659613 | 0.018429 | 0.066615 | 35.2119 | 57.59143 | 38.09291 | 44.85642 | 20.47734 | 35.75597 | 10.60718 | 25.0146 |
| ENSMUSG00000097723 | 0.238779367 | 0.669546 | 0.7881 | 17.10292 | 13.19804 | 5.796746 | 12.93935 | 13.65156 | 6.704245 | 6.750024 | 14.13868 |
| ENSMUSG00000097727 | -0.030333675 | 0.953714 | 0.973604 | 12.07265 | 7.198929 | 7.45296 | 14.6646 | 11.70134 | 14.52586 | 7.714313 | 8.700729 |
| ENSMUSG00000097728 | 0.334955335 | 0.410887 | 0.576811 | 23.13925 | 34.79482 | 18.21835 | 14.6646 | 17.552 | 14.52586 | 19.28578 | 19.57664 |
| ENSMUSG00000097729 | 0.226677584 | 0.163959 | 0.308905 | 690.1533 | 640.7047 | 635.1578 | 735.8178 | 525.585 | 754.2275 | 584.3592 | 445.9123 |
| ENSMUSG00000097730 | 0.421243387 | 0.067296 | 0.167501 | 111.672 | 111.5834 | 84.46688 | 101.7896 | 67.28268 | 103.9158 | 59.78593 | 75.04379 |
| ENSMUSG00000097732 | 0.36729179 | 0.371062 | 0.540056 | 23.13925 | 34.79482 | 39.74912 | 28.46658 | 15.60178 | 40.22547 | 26.03581 | 16.31387 |
| ENSMUSG00000097734 | 0.567677236 | 0.204397 | 0.360404 | 16.09687 | 31.19536 | 17.39024 | 25.01608 | 19.50223 | 17.87799 | 7.714313 | 15.22628 |
| ENSMUSG00000097737 | 0.484270738 | 0.146773 | 0.285743 | 58.35115 | 88.78679 | 57.13936 | 76.77349 | 39.97956 | 82.68568 | 42.42872 | 35.89051 |
| ENSMUSG00000097743 | 0.482681019 | 0.037102 | 0.110672 | 392.3612 | 397.1409 | 272.4471 | 343.3242 | 221.3503 | 370.9682 | 221.7865 | 191.416 |
| ENSMUSG00000097745 | -0.010725896 | 0.984389 | 0.99122 | 11.0666 | 11.99822 | 7.45296 | 19.84034 | 13.65156 | 20.11273 | 10.60718 | 6.525547 |
| ENSMUSG00000097746 | -0.188336438 | 0.670222 | 0.788576 | 10.06054 | 19.19714 | 11.59349 | 14.6646 | 15.60178 | 10.05637 | 17.3572 | 19.57664 |
| ENSMUSG00000097747 | -0.083432932 | 0.825201 | 0.893879 | 23.13925 | 41.99375 | 36.43669 | 34.50494 | 18.52711 | 51.39921 | 38.57157 | 35.89051 |
| ENSMUSG00000097748 | -0.212719287 | 0.704443 | 0.81399 | 12.07265 | 10.79839 | 11.59349 | 12.07673 | 11.70134 | 24.58223 | 13.50005 | 4.350364 |
| ENSMUSG00000097750 | 0.095180112 | 0.534688 | 0.685626 | 395.3794 | 417.5379 | 313.0243 | 304.5061 | 282.7823 | 338.5644 | 343.2869 | 371.9562 |
| ENSMUSG00000097752 | 0.451875651 | 0.210309 | 0.367806 | 36.21796 | 39.59411 | 40.57722 | 24.15346 | 27.30312 | 36.87335 | 22.17865 | 16.31387 |
| ENSMUSG00000097755 | 0.556460303 | 0.130292 | 0.262661 | 33.19979 | 49.19268 | 24.8432 | 35.36756 | 34.1289 | 22.34748 | 23.14294 | 16.31387 |
| ENSMUSG00000097756 | 1.678561937 | 0.008949 | 0.039297 | 25.15136 | 77.9884 | 31.46805 | 50.89479 | 5.850668 | 34.6386 | 5.785735 | 11.9635 |
| ENSMUSG00000097757 | -0.272307124 | 0.460786 | 0.621975 | 28.16952 | 23.99643 | 20.70267 | 18.11509 | 16.57689 | 36.87335 | 27.0001 | 29.36496 |
| ENSMUSG00000097760 | 0.399609443 | 0.19623 | 0.350492 | 36.21796 | 37.19447 | 28.15563 | 29.3292 | 30.22845 | 18.99536 | 24.10723 | 25.0146 |
| ENSMUSG00000097762 | -1.40629789 | 0.001812 | 0.012229 | 10.06054 | 13.19804 | 33.12427 | 11.21411 | 46.80534 | 40.22547 | 36.64299 | 57.64233 |
| ENSMUSG00000097767 | 3.861724126 | 5.73E-47 | 1.06E-42 | 2567.451 | 2366.048 | 1733.227 | 2360.138 | 145.2916 | 245.8223 | 147.5362 | 82.65692 |
| ENSMUSG00000097768 | -0.858555051 | 0.000932 | 0.007425 | 58.35115 | 37.19447 | 71.21717 | 47.44429 | 77.03379 | 98.32892 | 116.679 | 98.97079 |
| ENSMUSG00000097769 | 0.146979276 | 0.160036 | 0.303341 | 532.2028 | 526.7217 | 529.9882 | 576.2325 | 539.2366 | 521.8137 | 469.6088 | 425.2481 |
| ENSMUSG00000097772 | -0.019414282 | 0.853312 | 0.911961 | 400.4096 | 374.3443 | 361.0545 | 424.4108 | 417.3476 | 422.3674 | 366.4299 | 376.3065 |
| ENSMUSG00000097773 | 0.166198583 | 0.697013 | 0.808775 | 45.27245 | 23.99643 | 29.81184 | 22.42821 | 39.97956 | 33.52122 | 17.3572 | 17.40146 |
| ENSMUSG00000097777 | 0.295116605 | 0.100805 | 0.22217 | 96.58122 | 88.78679 | 81.15445 | 88.85022 | 69.2329 | 80.45093 | 72.32169 | 67.43065 |
| ENSMUSG00000097781 | -0.00035142 | 0.999094 | 0.999528 | 48.29061 | 46.79304 | 72.87338 | 46.58167 | 49.73068 | 41.34284 | 79.07171 | 44.59123 |
| ENSMUSG00000097784 | -0.06506233 | 0.772895 | 0.859063 | 73.44197 | 79.18822 | 60.45178 | 75.91087 | 60.4569 | 96.09417 | 79.07171 | 66.34306 |
| ENSMUSG00000097785 | 0.17380552 | 0.477131 | 0.636625 | 46.2785 | 67.19001 | 53.82693 | 61.24627 | 53.63112 | 44.69496 | 60.75022 | 42.41605 |
| ENSMUSG00000097786 | 0.583020678 | 0.260413 | 0.424448 | 10.06054 | 9.598572 | 14.07781 | 13.80198 | 9.751113 | 5.58687 | 5.785735 | 10.87591 |
| ENSMUSG00000097787 | -0.121206869 | 0.558092 | 0.704573 | 126.7628 | 107.9839 | 158.9965 | 115.5915 | 109.2125 | 128.498 | 148.5005 | 169.6642 |
| ENSMUSG00000097788 | -0.172090965 | 0.698289 | 0.809625 | 25.15136 | 20.39697 | 9.109173 | 19.84034 | 18.52711 | 22.34748 | 14.46434 | 28.27737 |
| ENSMUSG00000097789 | -0.258297506 | 0.572589 | 0.715181 | 32.19374 | 32.39518 | 71.21717 | 31.91707 | 41.92979 | 33.52122 | 33.75012 | 92.44524 |
| ENSMUSG00000097791 | 0.674272705 | 0.037529 | 0.111572 | 28.16952 | 40.79393 | 32.29616 | 45.71905 | 26.32801 | 25.6996 | 17.3572 | 22.83941 |
| ENSMUSG00000097792 | 0.077180562 | 0.72034 | 0.825128 | 85.51462 | 93.58608 | 78.67013 | 110.4158 | 84.83468 | 111.7374 | 80.036 | 72.8686 |
| ENSMUSG00000097793 | 0.019462856 | 0.927336 | 0.956943 | 65.39353 | 50.3925 | 59.62368 | 64.69676 | 68.25779 | 53.63396 | 57.85735 | 57.64233 |
| ENSMUSG00000097796 | 1.541061297 | 5.17E-06 | 0.000138 | 90.54489 | 73.18911 | 60.45178 | 75.04824 | 25.35289 | 41.34284 | 22.17865 | 14.13868 |
| ENSMUSG00000097797 | 0.538615721 | 0.038797 | 0.113874 | 117.7084 | 117.5825 | 73.70149 | 129.3935 | 78.98402 | 96.09417 | 54.96448 | 71.78101 |
| ENSMUSG00000097806 | 0.151082156 | 0.615878 | 0.748989 | 37.22401 | 43.19358 | 35.60859 | 37.09281 | 46.80534 | 34.6386 | 32.78583 | 22.83941 |
| ENSMUSG00000097814 | 0.531963678 | 0.004365 | 0.023305 | 256.5439 | 323.9518 | 193.777 | 279.49 | 170.6445 | 206.7142 | 189.965 | 159.8759 |
| ENSMUSG00000097815 | 0.056936619 | 0.569878 | 0.713532 | 981.9091 | 1047.444 | 886.0741 | 1061.89 | 930.2562 | 928.5379 | 1078.075 | 883.124 |

| | | | | | | | | | | | |
|---|---|---|---|---|---|---|---|---|---|---|---|
| ENSMUSG00000097816 | -0.722371654 | 0.225789 | 0.385543 | 7.04238 | 19.19714 | 3.312427 | 7.763611 | 17.552 | 17.87799 | 11.57147 | 13.05109 |
| ENSMUSG00000097819 | 0.382258288 | 0.282465 | 0.449073 | 162.9808 | 237.5647 | 242.6352 | 260.5123 | 130.6649 | 329.6254 | 108.9647 | 125.073 |
| ENSMUSG00000097820 | 0.328227887 | 0.160631 | 0.304186 | 202.2169 | 249.5629 | 216.9639 | 217.3811 | 145.2916 | 265.935 | 131.1433 | 164.2263 |
| ENSMUSG00000097823 | 0.128698675 | 0.683032 | 0.797952 | 39.23612 | 50.3925 | 33.12427 | 50.03216 | 26.32801 | 36.87335 | 54.00019 | 40.24087 |
| ENSMUSG00000097825 | 0.654389958 | 0.297827 | 0.465782 | 22.1332 | 13.19804 | 21.53077 | 33.64232 | 11.70134 | 24.58223 | 19.28578 | 2.175182 |
| ENSMUSG00000097827 | 0.019655084 | 0.95716 | 0.975878 | 20.12109 | 21.59679 | 19.87456 | 31.91707 | 22.42756 | 15.64324 | 26.03581 | 28.27737 |
| ENSMUSG00000097828 | 0.080539516 | 0.784694 | 0.867455 | 89.53884 | 63.59054 | 62.108 | 92.30071 | 54.60623 | 110.62 | 64.60737 | 61.99269 |
| ENSMUSG00000097831 | 0.092576479 | 0.868494 | 0.92134 | 5.030272 | 14.39786 | 11.59349 | 12.93935 | 8.776002 | 13.40849 | 13.50005 | 5.437955 |
| ENSMUSG00000097832 | -0.924367814 | 0.083042 | 0.193881 | 14.08476 | 7.198929 | 7.45296 | 6.900988 | 10.72622 | 12.29112 | 17.3572 | 27.18978 |
| ENSMUSG00000097834 | -0.365709629 | 0.57857 | 0.719984 | 6.036326 | 8.398751 | 24.8432 | 3.450494 | 14.62667 | 14.52586 | 14.46434 | 11.9635 |
| ENSMUSG00000097835 | 0.021258325 | 0.943068 | 0.967121 | 32.19374 | 40.79393 | 36.43669 | 27.60395 | 26.32801 | 30.1691 | 35.6787 | 42.41605 |
| ENSMUSG00000097838 | 1.0326474 | 0.053016 | 0.141745 | 16.09687 | 15.59768 | 14.90592 | 8.626235 | 3.900445 | 6.704245 | 8.678602 | 7.613138 |
| ENSMUSG00000097840 | 1.334792085 | 0.020756 | 0.072317 | 35.2119 | 29.99554 | 25.67131 | 39.68068 | 20.47734 | 14.52586 | 15.42863 | 1.087591 |
| ENSMUSG00000097842 | 0.697431189 | 0.003265 | 0.018892 | 189.1382 | 100.785 | 125.0441 | 166.4863 | 93.61069 | 99.44629 | 76.17884 | 90.27006 |
| ENSMUSG00000097850 | 0.966938747 | 0.159869 | 0.303067 | 25.15136 | 10.79839 | 14.07781 | 12.07673 | 5.850668 | 18.99536 | 2.892867 | 4.350364 |
| ENSMUSG00000097855 | 0.104603926 | 0.6366 | 0.764019 | 70.4238 | 82.78769 | 72.87338 | 85.39973 | 82.88446 | 88.27255 | 58.82164 | 59.81751 |
| ENSMUSG00000097857 | 1.156078486 | 0.006817 | 0.032358 | 21.12714 | 19.19714 | 23.18699 | 18.97772 | 6.825779 | 10.05637 | 12.53576 | 7.613138 |
| ENSMUSG00000097858 | -0.802498191 | 0.086805 | 0.200059 | 16.09687 | 7.198929 | 14.90592 | 24.15346 | 17.552 | 39.10809 | 31.82154 | 21.75182 |
| ENSMUSG00000097863 | 0.45558285 | 0.172453 | 0.319892 | 41.24823 | 34.79482 | 26.49941 | 37.09281 | 20.47734 | 35.75597 | 26.03581 | 19.57664 |
| ENSMUSG00000097867 | 0.026236353 | 0.895564 | 0.93677 | 79.47829 | 69.58965 | 70.38906 | 69.8725 | 59.48179 | 82.68568 | 66.53595 | 76.13138 |
| ENSMUSG00000097868 | 0.300034921 | 0.263813 | 0.428038 | 55.33299 | 41.99375 | 51.34261 | 51.75741 | 30.22845 | 46.92971 | 35.6787 | 51.11678 |
| ENSMUSG00000097874 | 0.158400539 | 0.747971 | 0.842597 | 19.11503 | 15.59768 | 15.73403 | 10.35148 | 12.67645 | 23.46486 | 8.678602 | 9.78832 |
| ENSMUSG00000097875 | -0.130864579 | 0.708253 | 0.816766 | 26.15741 | 34.79482 | 30.63995 | 39.68068 | 23.40267 | 54.75133 | 35.6787 | 30.45255 |
| ENSMUSG00000097876 | -0.307108102 | 0.421799 | 0.587172 | 56.33904 | 25.19625 | 41.40533 | 44.85642 | 52.65601 | 79.33356 | 48.21446 | 28.27737 |
| ENSMUSG00000097877 | -0.06984412 | 0.896957 | 0.937817 | 17.10292 | 13.19804 | 12.4216 | 14.6646 | 10.72622 | 26.81698 | 17.3572 | 5.437955 |
| ENSMUSG00000097882 | -0.016419424 | 0.949135 | 0.971183 | 35.2119 | 51.59233 | 63.76421 | 55.2079 | 51.6809 | 46.92971 | 56.89306 | 53.29196 |
| ENSMUSG00000097885 | 0.551047309 | 0.045521 | 0.127544 | 110.666 | 135.5798 | 109.3101 | 108.6906 | 83.85957 | 112.8548 | 75.21455 | 44.59123 |
| ENSMUSG00000097887 | -0.21378326 | 0.672521 | 0.790252 | 10.06054 | 7.198929 | 10.76539 | 15.52722 | 11.70134 | 13.40849 | 7.714313 | 18.48905 |
| ENSMUSG00000097888 | 0.693835742 | 0.17944 | 0.329231 | 10.06054 | 23.99643 | 9.109173 | 20.70296 | 9.751113 | 10.05637 | 10.60718 | 8.700729 |
| ENSMUSG00000097889 | -0.226370638 | 0.596084 | 0.735186 | 12.07265 | 16.7975 | 15.73403 | 29.3292 | 24.37778 | 20.11273 | 27.0001 | 15.22628 |
| ENSMUSG00000097890 | -0.150827657 | 0.793564 | 0.873442 | 17.10292 | 10.79839 | 12.4216 | 6.038364 | 11.70134 | 23.46486 | 7.714313 | 8.700729 |
| ENSMUSG00000097893 | -1.079618109 | 0.034382 | 0.104663 | 6.036326 | 7.198929 | 5.796746 | 10.35148 | 14.62667 | 13.40849 | 11.57147 | 22.83941 |
| ENSMUSG00000097906 | -0.20818887 | 0.26658 | 0.431024 | 94.56911 | 113.983 | 118.4192 | 85.39973 | 137.4907 | 106.1505 | 108.9647 | 122.8978 |
| ENSMUSG00000097908 | 0.217197119 | 0.171888 | 0.319177 | 132.7992 | 171.5745 | 169.7619 | 157.8601 | 115.0631 | 143.0239 | 145.6077 | 140.2993 |
| ENSMUSG00000097912 | 0.293385756 | 0.63267 | 0.761035 | 11.0666 | 9.598572 | 9.93728 | 15.52722 | 6.825779 | 20.11273 | 6.750024 | 4.350364 |
| ENSMUSG00000097915 | 0.216440863 | 0.470334 | 0.630299 | 48.29061 | 35.99465 | 32.29616 | 37.09281 | 35.10401 | 24.58223 | 29.89296 | 42.41605 |
| ENSMUSG00000097919 | 0.396742457 | 0.412548 | 0.578395 | 12.07265 | 20.39697 | 19.87456 | 9.488858 | 11.70134 | 12.29112 | 7.714313 | 15.22628 |
| ENSMUSG00000097924 | -0.062489338 | 0.788217 | 0.86993 | 79.47829 | 81.58786 | 83.63877 | 53.48266 | 78.98402 | 91.62468 | 78.10742 | 61.99269 |
| ENSMUSG00000097928 | -0.104502847 | 0.722577 | 0.826518 | 61.36932 | 31.19536 | 46.37397 | 66.42201 | 53.63112 | 44.69496 | 63.64308 | 59.81751 |
| ENSMUSG00000097934 | 0.941123004 | 0.08082 | 0.190281 | 19.11503 | 14.39786 | 12.4216 | 29.3292 | 10.72622 | 15.64324 | 8.678602 | 4.350364 |
| ENSMUSG00000097944 | 0.278295314 | 0.22997 | 0.390215 | 149.9021 | 95.98572 | 84.46688 | 101.7896 | 78.0089 | 92.74205 | 100.2861 | 84.8321 |
| ENSMUSG00000097960 | 0.067329903 | 0.858674 | 0.915304 | 16.09687 | 26.39607 | 18.21835 | 16.38985 | 19.50223 | 14.52586 | 21.21436 | 17.40146 |
| ENSMUSG00000097970 | -1.07998655 | 0.00086 | 0.006949 | 27.16347 | 16.7975 | 19.04645 | 20.70296 | 40.95467 | 34.6386 | 58.82164 | 42.41605 |
| ENSMUSG00000097971 | -0.510356843 | 0.002691 | 0.016303 | 1567.433 | 1461.383 | 2273.153 | 1880.519 | 2629.875 | 2143.124 | 2325.865 | 3136.613 |
| ENSMUSG00000097974 | 0.575537943 | 0.128807 | 0.260464 | 39.23612 | 22.79661 | 24.8432 | 31.91707 | 22.42756 | 27.93435 | 14.46434 | 15.22628 |
| ENSMUSG00000097990 | 0.093925845 | 0.819335 | 0.890662 | 18.10898 | 16.7975 | 21.53077 | 25.01608 | 13.65156 | 14.52586 | 18.32149 | 30.45255 |
| ENSMUSG00000097993 | 0.312769342 | 0.28681 | 0.453937 | 90.54489 | 70.78947 | 82.81066 | 65.55939 | 48.75557 | 81.56831 | 39.53586 | 80.48174 |
| ENSMUSG00000097994 | 0.148975101 | 0.526416 | 0.679534 | 112.6781 | 151.1775 | 142.4343 | 100.9269 | 86.78491 | 154.1976 | 109.929 | 106.5839 |
| ENSMUSG00000097997 | -0.72313561 | 0.212328 | 0.370075 | 10.06054 | 7.198929 | 10.76539 | 5.175741 | 8.776002 | 7.821619 | 14.46434 | 23.927 |
| ENSMUSG00000098021 | 0.057428551 | 0.963793 | 0.979503 | 0 | 41.99375 | 0.828107 | 25.8787 | 16.57689 | 1.117374 | 24.10723 | 23.927 |
| ENSMUSG00000098022 | 0.170041797 | 0.75752 | 0.848813 | 28.16952 | 8.398751 | 13.24971 | 13.80198 | 22.42756 | 14.52586 | 7.714313 | 11.9635 |
| ENSMUSG00000098024 | 0.188749457 | 0.621672 | 0.753254 | 113.6841 | 61.1909 | 52.99882 | 63.83414 | 52.65601 | 107.2679 | 55.92877 | 40.24087 |
| ENSMUSG00000098041 | 0.6437785 | 0.134299 | 0.268006 | 139.8416 | 127.1811 | 111.7944 | 100.0643 | 38.02934 | 160.9019 | 47.25017 | 60.9051 |
| ENSMUSG00000098050 | 0.178434784 | 0.565288 | 0.710014 | 25.15136 | 32.39518 | 36.43669 | 24.15346 | 24.37778 | 25.6996 | 25.07152 | 29.36496 |
| ENSMUSG00000098066 | 0.312501692 | 0.244555 | 0.407283 | 40.24217 | 40.79393 | 45.54586 | 36.23019 | 38.02934 | 32.40385 | 26.03581 | 34.80291 |
| ENSMUSG00000098068 | 0.452799276 | 0.607822 | 0.743601 | 22.1332 | 14.39786 | 7.45296 | 8.626235 | 0 | 21.23011 | 9.642892 | 7.613138 |
| ENSMUSG00000098076 | -0.30464561 | 0.416545 | 0.582046 | 15.09082 | 21.59679 | 27.32752 | 15.52722 | 26.32801 | 17.87799 | 28.92867 | 25.0146 |
| ENSMUSG00000098087 | -0.42243665 | 0.015682 | 0.059628 | 131.7931 | 103.1847 | 102.6852 | 128.5309 | 130.6649 | 151.9629 | 165.8577 | 177.2773 |
| ENSMUSG00000098090 | 0.411722353 | 0.019265 | 0.06864 | 495.9848 | 479.9286 | 354.4296 | 469.2672 | 363.7165 | 410.0763 | 319.1797 | 258.8467 |
| ENSMUSG00000098098 | 0.385486617 | 0.016045 | 0.060359 | 288.7376 | 349.1481 | 303.087 | 357.9887 | 260.3547 | 307.2779 | 215.0365 | 212.0803 |
| ENSMUSG00000098112 | -0.058703911 | 0.862666 | 0.917774 | 44.26639 | 50.3925 | 53.82693 | 23.29083 | 59.48179 | 39.10809 | 41.46443 | 38.06569 |
| ENSMUSG00000098120 | 0.071968495 | 0.691656 | 0.804766 | 127.7689 | 164.3756 | 151.5435 | 110.4158 | 127.7396 | 131.8501 | 117.6433 | 149 |
| ENSMUSG00000098123 | -0.987897893 | 0.046814 | 0.129965 | 8.048435 | 4.799286 | 6.624853 | 7.763611 | 12.67645 | 11.17374 | 16.39292 | 14.13868 |
| ENSMUSG00000098132 | 0.412300074 | 0.000158 | 0.001891 | 1199.217 | 1205.821 | 1045.899 | 1209.398 | 791.7904 | 1025.749 | 878.4674 | 805.905 |

| | | | | | | | | | | | |
|---|---|---|---|---|---|---|---|---|---|---|---|
| ENSMUSG00000098134 | -0.164981276 | 0.089944 | 0.205116 | 496.9909 | 425.9366 | 494.3797 | 542.5902 | 552.8881 | 537.4569 | 549.6448 | 562.2846 |
| ENSMUSG00000098146 | 0.885677924 | 0.121358 | 0.250642 | 54.32694 | 43.19358 | 24.8432 | 19.84034 | 8.776002 | 39.10809 | 19.28578 | 9.78832 |
| ENSMUSG00000098158 | -0.862204206 | 0.120344 | 0.249335 | 13.07871 | 8.398751 | 9.109173 | 7.763611 | 23.40267 | 6.704245 | 26.03581 | 13.05109 |
| ENSMUSG00000098176 | -0.336838653 | 0.007571 | 0.034766 | 147.89 | 160.7761 | 173.9024 | 161.3106 | 207.6987 | 187.7188 | 204.4293 | 214.2554 |
| ENSMUSG00000098178 | 0.159217731 | 0.451383 | 0.613862 | 52138.77 | 40515.57 | 29089.73 | 38587.74 | 48992.52 | 35526.91 | 32403.97 | 26653.59 |
| ENSMUSG00000098183 | 0.312588952 | 0.454113 | 0.616212 | 102.6175 | 50.3925 | 27.32752 | 45.71905 | 43.88001 | 54.75133 | 49.17875 | 33.71532 |
| ENSMUSG00000098184 | -0.805613967 | 0.21799 | 0.376565 | 4.024217 | 10.79839 | 8.281066 | 7.763611 | 28.27823 | 7.821619 | 8.678602 | 8.700729 |
| ENSMUSG00000098188 | 0.160532282 | 0.155831 | 0.297848 | 560.3723 | 608.3095 | 717.1403 | 691.824 | 519.7343 | 579.9172 | 597.8593 | 612.3138 |
| ENSMUSG00000098196 | 0.52227721 | 0.091217 | 0.207073 | 67.40564 | 43.19358 | 38.92101 | 43.13117 | 29.25334 | 25.6996 | 38.57157 | 40.24087 |
| ENSMUSG00000098202 | 2.066986402 | 0.007708 | 0.035234 | 19.11503 | 43.19358 | 21.53077 | 37.09281 | 5.850668 | 20.11273 | 0.964289 | 2.175182 |
| ENSMUSG00000098206 | 0.803886765 | 0.037826 | 0.112177 | 123.7447 | 196.7707 | 110.1382 | 109.5532 | 73.13335 | 140.7891 | 44.3573 | 51.11678 |
| ENSMUSG00000098221 | -0.065864068 | 0.884778 | 0.931197 | 10.06054 | 17.99732 | 27.32752 | 26.74133 | 17.552 | 27.93435 | 27.0001 | 14.13868 |
| ENSMUSG00000098233 | 0.089204495 | 0.802162 | 0.879281 | 25.15136 | 20.39697 | 31.46805 | 22.42821 | 15.60178 | 22.34748 | 28.92867 | 27.18978 |
| ENSMUSG00000098234 | 0.175922562 | 0.141303 | 0.278097 | 1291.774 | 1219.019 | 1271.144 | 1361.22 | 1141.855 | 1372.135 | 1095.432 | 945.1167 |
| ENSMUSG00000098235 | 0.718368735 | 0.327821 | 0.495558 | 25.15136 | 20.39697 | 12.4216 | 16.38985 | 3.900445 | 31.28647 | 4.821446 | 5.437955 |
| ENSMUSG00000098243 | 0.761040715 | 0.034875 | 0.105796 | 21.12714 | 26.39607 | 28.98373 | 21.56559 | 13.65156 | 14.52586 | 13.50005 | 16.31387 |
| ENSMUSG00000098259 | 0.47361631 | 0.225149 | 0.384861 | 48.29061 | 32.39518 | 44.71776 | 52.62003 | 28.27823 | 56.98608 | 23.14294 | 20.66423 |
| ENSMUSG00000098269 | 0.235722669 | 0.569587 | 0.713376 | 42.25428 | 17.99732 | 25.67131 | 19.84034 | 24.37778 | 18.99536 | 29.89296 | 16.31387 |
| ENSMUSG00000098274 | 0.131527455 | 0.241012 | 0.403128 | 5632.898 | 6659.01 | 5859.683 | 6049.579 | 5573.736 | 6455.07 | 5454.02 | 4608.123 |
| ENSMUSG00000098293 | 0.411367592 | 0.37026 | 0.539389 | 19.11503 | 22.79661 | 22.35888 | 31.05445 | 25.35289 | 7.821619 | 25.07152 | 13.05109 |
| ENSMUSG00000098294 | 1.835714185 | 0.002309 | 0.014546 | 12.07265 | 31.19536 | 17.39024 | 10.35148 | 4.875557 | 5.58687 | 3.857157 | 5.437955 |
| ENSMUSG00000098318 | 0.18908949 | 0.161106 | 0.304774 | 640.8566 | 573.5147 | 465.3959 | 577.9577 | 577.2659 | 476.0014 | 474.4303 | 450.2627 |
| ENSMUSG00000098332 | -0.170748197 | 0.126906 | 0.258034 | 324.9556 | 261.5611 | 308.8838 | 287.2536 | 312.0356 | 343.0338 | 325.9297 | 353.4671 |
| ENSMUSG00000098374 | 0.530181036 | 0.001917 | 0.012726 | 209.2593 | 183.5727 | 174.7305 | 233.771 | 122.864 | 162.0192 | 121.5004 | 150.0876 |
| ENSMUSG00000098404 | -0.423470918 | 0.404997 | 0.572028 | 8.048435 | 14.39786 | 10.76539 | 6.900988 | 10.72622 | 11.17374 | 20.25007 | 10.87591 |
| ENSMUSG00000098424 | 0.286659772 | 0.361232 | 0.529809 | 65.39353 | 67.19001 | 65.42042 | 69.8725 | 27.30312 | 75.98144 | 64.60737 | 52.20437 |
| ENSMUSG00000098447 | 0.162500182 | 0.729447 | 0.830159 | 17.10292 | 7.198929 | 16.56213 | 15.52722 | 9.751113 | 14.52586 | 10.60718 | 16.31387 |
| ENSMUSG00000098449 | -0.100927397 | 0.817551 | 0.889298 | 11.0666 | 13.19804 | 19.87456 | 18.11509 | 18.52711 | 11.17374 | 23.14294 | 14.13868 |
| ENSMUSG00000098488 | 0.330397923 | 0.248303 | 0.411704 | 62.37537 | 53.99197 | 43.06154 | 68.14726 | 44.85512 | 60.3382 | 44.3573 | 31.54014 |
| ENSMUSG00000098526 | 1.34297388 | 0.032705 | 0.100617 | 17.10292 | 26.39607 | 6.624853 | 21.56559 | 6.825779 | 12.29112 | 5.785735 | 3.262773 |
| ENSMUSG00000098557 | 0.549205639 | 0.053278 | 0.142281 | 810.8798 | 1227.417 | 1013.603 | 1184.382 | 499.257 | 1246.989 | 612.3236 | 537.27 |
| ENSMUSG00000098620 | 0.605688642 | 0.240861 | 0.403048 | 63.38142 | 59.99108 | 30.63995 | 40.5433 | 15.60178 | 67.04245 | 28.92867 | 16.31387 |
| ENSMUSG00000098645 | 1.30875155 | 0.085953 | 0.198529 | 25.15136 | 21.59679 | 7.45296 | 14.6646 | 4.875557 | 17.87799 | 1.928578 | 3.262773 |
| ENSMUSG00000098661 | 0.174607771 | 0.732198 | 0.83216 | 13.07871 | 14.39786 | 17.39024 | 10.35148 | 19.50223 | 11.17374 | 12.53576 | 5.437955 |
| ENSMUSG00000098662 | 1.007604232 | 0.058082 | 0.151172 | 35.2119 | 34.79482 | 17.39024 | 34.50494 | 15.60178 | 21.23011 | 20.25007 | 3.262773 |
| ENSMUSG00000098679 | -0.566564795 | 0.351085 | 0.51987 | 23.13925 | 13.19804 | 4.140533 | 7.763611 | 21.45245 | 24.58223 | 10.60718 | 14.13868 |
| ENSMUSG00000098683 | 0.58053369 | 0.09262 | 0.209097 | 39.23612 | 22.79661 | 24.01509 | 29.3292 | 19.50223 | 17.87799 | 21.21436 | 18.48905 |
| ENSMUSG00000098702 | 0.148095521 | 0.217378 | 0.375825 | 384.3128 | 352.7475 | 342.8361 | 354.5383 | 302.2845 | 364.264 | 348.1084 | 279.5109 |
| ENSMUSG00000098708 | -0.256350345 | 0.678276 | 0.794492 | 33.19979 | 20.39697 | 19.04645 | 31.91707 | 18.52711 | 18.99536 | 76.17884 | 10.87591 |
| ENSMUSG00000098749 | 0.108825058 | 0.787971 | 0.869836 | 16.09687 | 13.19804 | 27.32752 | 17.25247 | 14.62667 | 20.11273 | 18.32149 | 16.31387 |
| ENSMUSG00000098766 | 0.882727167 | 0.086844 | 0.2001 | 18.10898 | 20.39697 | 7.45296 | 17.25247 | 8.776002 | 8.938993 | 9.642892 | 6.525547 |
| ENSMUSG00000098789 | -0.017503455 | 0.950594 | 0.971873 | 80.48435 | 51.59233 | 80.32634 | 93.16334 | 86.78491 | 48.04709 | 82.92887 | 92.44524 |
| ENSMUSG00000098813 | 0.367134728 | 0.471668 | 0.631172 | 18.10898 | 31.19536 | 14.90592 | 15.52722 | 27.30312 | 8.938993 | 10.60718 | 14.13868 |
| ENSMUSG00000098891 | 1.971496312 | 8.31E-05 | 0.001167 | 25.15136 | 33.595 | 24.01509 | 43.9938 | 8.776002 | 14.52586 | 3.857157 | 5.437955 |
| ENSMUSG00000098905 | 0.570506589 | 0.002225 | 0.014197 | 150.9082 | 182.3729 | 115.1068 | 139.745 | 95.56091 | 101.681 | 102.2147 | 94.62042 |
| ENSMUSG00000098912 | 0.571433203 | 0.000944 | 0.007498 | 324.9556 | 293.9563 | 243.4633 | 302.7808 | 167.7191 | 252.5265 | 188.0364 | 176.1898 |
| ENSMUSG00000098919 | 0.149741109 | 0.772649 | 0.858991 | 15.09082 | 13.19804 | 19.04645 | 13.80198 | 12.67645 | 17.87799 | 20.25007 | 4.350364 |
| ENSMUSG00000098923 | -0.03437402 | 0.66572 | 0.785381 | 580.4934 | 615.5084 | 588.7838 | 553.8043 | 634.7975 | 585.504 | 577.6092 | 593.8247 |
| ENSMUSG00000098973 | -0.049209042 | 0.904676 | 0.94289 | 349.1009 | 175.1739 | 101.8571 | 144.0581 | 324.7121 | 118.4417 | 193.8221 | 158.7883 |
| ENSMUSG00000098979 | 2.252537281 | 2.72E-10 | 4.14E-08 | 276.6649 | 359.9465 | 178.0429 | 318.3081 | 57.53157 | 105.0332 | 38.57157 | 36.9781 |
| ENSMUSG00000099032 | 1.369570025 | 7.74E-21 | 1.30E-17 | 810.8798 | 809.8795 | 691.469 | 822.0802 | 264.2552 | 386.6114 | 262.2867 | 301.2627 |
| ENSMUSG00000099034 | 0.304358652 | 0.037449 | 0.111406 | 133.8052 | 135.5798 | 124.216 | 138.8824 | 100.4365 | 113.9722 | 107.0361 | 109.8467 |
| ENSMUSG00000099041 | -0.089503749 | 0.778313 | 0.863254 | 21.12714 | 31.19536 | 32.29616 | 22.42821 | 23.40267 | 29.05173 | 29.89296 | 31.54014 |
| ENSMUSG00000099083 | 0.419127122 | 2.93E-05 | 0.000527 | 1529.203 | 1533.372 | 1284.393 | 1583.777 | 1081.398 | 1244.755 | 1044.325 | 1064.752 |
| ENSMUSG00000099094 | 0.201360743 | 0.305121 | 0.473538 | 101.6115 | 99.58519 | 137.4657 | 134.5693 | 95.56091 | 92.74205 | 122.4647 | 102.2336 |
| ENSMUSG00000099098 | 0.642458279 | 0.25039 | 0.413922 | 24.1453 | 7.198929 | 19.87456 | 15.52722 | 5.850668 | 16.76061 | 7.714313 | 13.05109 |
| ENSMUSG00000099136 | 1.573455475 | 0.004892 | 0.025347 | 40.24217 | 31.19536 | 24.8432 | 23.29083 | 3.900445 | 22.34748 | 7.714313 | 6.525547 |
| ENSMUSG00000099137 | 1.088035418 | 0.000959 | 0.007598 | 113.6841 | 105.5843 | 74.5296 | 89.71284 | 25.35289 | 72.62932 | 41.46443 | 41.32846 |
| ENSMUSG00000099270 | 0.695379903 | 0.076235 | 0.183058 | 49.29666 | 35.99465 | 38.92101 | 44.85642 | 32.17867 | 40.22547 | 20.25007 | 11.9635 |
| ENSMUSG00000099307 | 2.162518003 | 2.17E-07 | 1.07E-05 | 28.16952 | 35.99465 | 35.60859 | 29.3292 | 8.776002 | 7.821619 | 7.714313 | 4.350364 |
| ENSMUSG00000099314 | 0.082000088 | 0.889853 | 0.933475 | 6.036326 | 5.999108 | 15.73403 | 12.07673 | 11.70134 | 13.40849 | 7.714313 | 5.437955 |
| ENSMUSG00000099338 | -0.26221561 | 0.623861 | 0.754981 | 25.15136 | 7.198929 | 19.87456 | 13.80198 | 22.42756 | 7.821619 | 23.14294 | 26.10219 |
| ENSMUSG00000099354 | 0.531327838 | 0.329052 | 0.496849 | 20.12109 | 32.39518 | 19.04645 | 44.85642 | 14.62667 | 37.99072 | 7.714313 | 20.66423 |
| ENSMUSG00000099377 | -0.458086737 | 0.355409 | 0.524084 | 6.036326 | 8.398751 | 11.59349 | 11.21411 | 16.57689 | 13.40849 | 8.678602 | 13.05109 |

| | | | | | | | | | | | |
|---|---|---|---|---|---|---|---|---|---|---|---|
| ENSMUSG00000099397 | -0.288230411 | 0.444033 | 0.607042 | 25.15136 | 16.7975 | 25.67131 | 14.6646 | 18.52711 | 26.81698 | 23.14294 | 32.62773 |
| ENSMUSG00000099413 | -2.781754549 | 3.49E-05 | 0.000603 | 4.024217 | 1.199822 | 13.24971 | 7.763611 | 33.15378 | 16.76061 | 64.60737 | 70.69342 |
| ENSMUSG00000099465 | 0.301570477 | 0.315956 | 0.484317 | 43.26034 | 41.99375 | 28.98373 | 43.9938 | 39.97956 | 34.6386 | 25.07152 | 28.27737 |
| ENSMUSG00000099470 | 0.660352476 | 0.12268 | 0.252221 | 19.11503 | 26.39607 | 24.8432 | 18.11509 | 8.776002 | 10.05637 | 16.39292 | 20.66423 |
| ENSMUSG00000099471 | -0.090818797 | 0.761582 | 0.851916 | 27.16347 | 22.79661 | 24.01509 | 27.60395 | 25.35289 | 25.6996 | 27.0001 | 30.45255 |
| ENSMUSG00000099478 | 0.678261724 | 2.93E-05 | 0.000527 | 240.447 | 253.1623 | 209.511 | 251.0234 | 146.2667 | 184.3667 | 121.5004 | 144.6496 |
| ENSMUSG00000099481 | 0.233000455 | 0.091234 | 0.207085 | 565.4025 | 638.3051 | 529.9882 | 515.8488 | 498.2819 | 572.0955 | 440.6801 | 401.3211 |
| ENSMUSG00000099492 | -0.428928769 | 0.189125 | 0.341613 | 21.12714 | 26.39607 | 17.39024 | 23.29083 | 23.40267 | 27.93435 | 30.85725 | 35.89051 |
| ENSMUSG00000099521 | -0.151266199 | 0.51798 | 0.671951 | 70.4238 | 76.78858 | 74.5296 | 106.1027 | 79.95913 | 72.62932 | 101.2504 | 110.9343 |
| ENSMUSG00000099597 | 0.699370077 | 0.021113 | 0.073217 | 890.3581 | 998.2515 | 903.4643 | 1191.283 | 443.6756 | 1119.609 | 492.7518 | 398.0583 |
| ENSMUSG00000099632 | -0.021500663 | 0.926553 | 0.956348 | 76.46013 | 79.18822 | 91.91984 | 61.24627 | 80.93424 | 84.92043 | 90.64318 | 56.55474 |
| ENSMUSG00000099635 | 1.766287596 | 0.210994 | 0.368412 | 15.09082 | 23.99643 | 11.59349 | 17.25247 | 0 | 20.11273 | 0 | 0 |
| ENSMUSG00000099647 | 0.508306454 | 0.324289 | 0.492275 | 9.054489 | 26.39607 | 10.76539 | 27.60395 | 11.70134 | 14.52586 | 13.50005 | 11.9635 |
| ENSMUSG00000099655 | -0.325300315 | 0.210847 | 0.368293 | 33.19979 | 35.99465 | 34.78048 | 31.05445 | 34.1289 | 46.92971 | 41.46443 | 46.76642 |
| ENSMUSG00000099681 | -0.436314928 | 0.006048 | 0.02964 | 199.1988 | 226.7663 | 274.1033 | 181.1509 | 295.4587 | 282.6956 | 283.501 | 331.7153 |
| ENSMUSG00000099689 | 0.505893153 | 0.000398 | 0.00385 | 1051.327 | 1111.035 | 887.7303 | 1057.576 | 788.865 | 845.8522 | 693.3239 | 563.3722 |
| ENSMUSG00000099696 | -0.230684178 | 0.632422 | 0.760935 | 30.18163 | 23.99643 | 28.15563 | 12.07673 | 21.45245 | 20.11273 | 50.14304 | 18.48905 |
| ENSMUSG00000099703 | -0.079636106 | 0.836463 | 0.901206 | 32.19374 | 17.99732 | 28.15563 | 20.70296 | 21.45245 | 16.76061 | 30.85725 | 35.89051 |
| ENSMUSG00000099708 | 0.040425496 | 0.872275 | 0.923462 | 42.25428 | 44.3934 | 43.06154 | 34.50494 | 41.92979 | 41.34284 | 33.75012 | 42.41605 |
| ENSMUSG00000099757 | 1.131334639 | 0.008664 | 0.038447 | 17.10292 | 37.19447 | 24.8432 | 38.81806 | 11.70134 | 17.87799 | 14.46434 | 9.78832 |
| ENSMUSG00000099758 | 0.857799409 | 0.171149 | 0.318195 | 29.17558 | 27.5959 | 12.4216 | 16.38985 | 13.65156 | 23.46486 | 6.750024 | 3.262773 |
| ENSMUSG00000099759 | 0.250173062 | 0.656382 | 0.778543 | 17.10292 | 31.19536 | 82.81066 | 55.2079 | 33.15378 | 74.86406 | 20.25007 | 29.36496 |
| ENSMUSG00000099760 | 0.435441476 | 0.186376 | 0.33832 | 47.28455 | 39.59411 | 52.99882 | 62.97152 | 23.40267 | 44.69496 | 52.07161 | 30.45255 |
| ENSMUSG00000099764 | -0.212517568 | 0.166922 | 0.312673 | 154.9324 | 232.7654 | 201.2299 | 193.2277 | 248.6534 | 213.4185 | 225.6437 | 216.4306 |
| ENSMUSG00000099773 | 0.214381053 | 0.761997 | 0.852086 | 29.17558 | 29.99554 | 24.8432 | 13.80198 | 1.950223 | 41.34284 | 27.96439 | 13.05109 |
| ENSMUSG00000099825 | 0.558888061 | 0.13352 | 0.267011 | 21.12714 | 27.5959 | 17.39024 | 24.15346 | 12.67645 | 14.52586 | 17.3572 | 16.31387 |
| ENSMUSG00000099843 | 0.104972347 | 0.685156 | 0.79974 | 75.45408 | 79.18822 | 61.27989 | 62.10889 | 78.0089 | 58.10345 | 43.39301 | 78.30656 |
| ENSMUSG00000099853 | 0.449991975 | 0.114436 | 0.241901 | 51.30877 | 33.595 | 42.23344 | 60.38364 | 39.00445 | 31.28647 | 32.78583 | 34.80291 |
| ENSMUSG00000099858 | -1.226963098 | 0.050765 | 0.1375 | 12.07265 | 8.398751 | 5.796746 | 8.626235 | 36.07912 | 5.58687 | 14.46434 | 25.0146 |
| ENSMUSG00000099875 | -0.371286129 | 0.112871 | 0.239757 | 49.29666 | 71.98929 | 68.73285 | 50.03216 | 78.0089 | 61.45558 | 82.92887 | 87.00729 |
| ENSMUSG00000099876 | 0.293223683 | 0.874695 | 0.925069 | 0 | 197.9706 | 147.403 | 0.862623 | 66.30757 | 1.117374 | 121.5004 | 93.53283 |
| ENSMUSG00000099881 | 0.060825931 | 0.567262 | 0.711477 | 572.4449 | 610.7092 | 648.4075 | 714.2523 | 546.0623 | 613.4384 | 628.7165 | 655.8174 |
| ENSMUSG00000099913 | 0.069907604 | 0.91323 | 0.948699 | 23.13925 | 21.59679 | 7.45296 | 24.15346 | 7.80089 | 40.22547 | 9.642892 | 15.22628 |
| ENSMUSG00000099923 | 0.311377478 | 0.420593 | 0.585984 | 22.1332 | 23.99643 | 21.53077 | 18.97772 | 22.42756 | 16.76061 | 20.25007 | 9.78832 |
| ENSMUSG00000099966 | 0.002396845 | 0.980069 | 0.988885 | 539.2451 | 476.3291 | 518.3948 | 496.8711 | 475.8543 | 560.9218 | 476.3588 | 517.6934 |
| ENSMUSG00000099998 | -0.470676479 | 0.401332 | 0.568587 | 11.0666 | 13.19804 | 9.93728 | 4.313117 | 18.52711 | 16.76061 | 7.714313 | 9.78832 |
| ENSMUSG00000100005 | 0.707776434 | 0.000392 | 0.003804 | 1932.63 | 1878.921 | 1524.544 | 1554.448 | 985.8375 | 1517.394 | 940.1819 | 775.4524 |
| ENSMUSG00000100017 | -0.119830125 | 0.406295 | 0.572907 | 365.1977 | 364.7457 | 402.4598 | 352.813 | 375.4179 | 513.9921 | 356.787 | 369.781 |
| ENSMUSG00000100053 | 1.122062899 | 0.115519 | 0.243464 | 38.23007 | 43.19358 | 9.109173 | 25.01608 | 2.925334 | 29.05173 | 13.50005 | 7.613138 |
| ENSMUSG00000100071 | 0.369625304 | 0.467566 | 0.62788 | 13.07871 | 16.7975 | 16.56213 | 22.42821 | 9.751113 | 25.6996 | 9.642892 | 8.700729 |
| ENSMUSG00000100075 | 0.060154907 | 0.821343 | 0.891669 | 81.4904 | 67.19001 | 122.5598 | 144.9207 | 108.2374 | 91.62468 | 101.2504 | 100.0584 |
| ENSMUSG00000100078 | -0.076550678 | 0.79039 | 0.87162 | 26.15741 | 47.99286 | 33.12427 | 42.26855 | 41.92979 | 39.10809 | 34.71441 | 41.32846 |
| ENSMUSG00000100084 | -0.453236556 | 0.058886 | 0.152598 | 53.32088 | 41.99375 | 59.62368 | 44.85642 | 63.38224 | 82.68568 | 73.28598 | 55.46715 |
| ENSMUSG00000100131 | -0.478511355 | 9.01E-05 | 0.001237 | 174.0474 | 178.7734 | 179.6991 | 161.3106 | 242.8027 | 261.4655 | 217.9294 | 244.708 |
| ENSMUSG00000100147 | -0.223889415 | 0.724892 | 0.82773 | 7.04238 | 4.799286 | 33.12427 | 18.11509 | 18.52711 | 11.17374 | 23.14294 | 21.75182 |
| ENSMUSG00000100151 | 5.327651127 | NA | NA | 292.7618 | 0 | 0 | 10.35148 | 0 | 3.352122 | 0.964289 | 3.262773 |
| ENSMUSG00000100153 | 0.270078961 | 0.097289 | 0.216725 | 127.7689 | 143.9786 | 129.1846 | 108.6906 | 111.1627 | 103.9158 | 109.929 | 95.70802 |
| ENSMUSG00000100158 | 1.445117766 | 0.00295 | 0.01755 | 257.5499 | 392.3416 | 155.684 | 189.7772 | 78.0089 | 196.6578 | 54.00019 | 36.9781 |
| ENSMUSG00000100164 | -0.439512376 | 0.001259 | 0.009302 | 162.9808 | 190.7716 | 189.6364 | 199.266 | 290.5832 | 230.1791 | 236.2508 | 250.146 |
| ENSMUSG00000100183 | 0.394637487 | 0.251752 | 0.415385 | 93.56306 | 105.5843 | 48.85829 | 84.5371 | 61.43201 | 40.22547 | 95.46463 | 54.37955 |
| ENSMUSG00000100199 | -0.376802721 | 0.195353 | 0.349297 | 32.19374 | 27.5959 | 32.29616 | 25.8787 | 32.17867 | 31.28647 | 47.25017 | 42.41605 |
| ENSMUSG00000100201 | -0.111890309 | 0.766301 | 0.854966 | 15.09082 | 31.19536 | 19.87456 | 21.56559 | 24.37778 | 30.1691 | 22.17865 | 17.40146 |
| ENSMUSG00000100209 | -0.062834269 | 0.750988 | 0.844912 | 66.39959 | 68.38983 | 90.26362 | 72.46037 | 86.78491 | 73.74669 | 75.21455 | 76.13138 |
| ENSMUSG00000100213 | 0.743334554 | 0.115817 | 0.243805 | 128.775 | 117.5825 | 78.67013 | 65.55939 | 36.07912 | 122.9112 | 47.25017 | 27.18978 |
| ENSMUSG00000100215 | 0.057018123 | 0.711275 | 0.818994 | 204.229 | 185.9723 | 157.3403 | 201.8539 | 162.8436 | 211.1837 | 173.572 | 172.927 |
| ENSMUSG00000100228 | -0.199467785 | 0.648179 | 0.772142 | 22.1332 | 26.39607 | 17.39024 | 12.07673 | 18.52711 | 35.75597 | 17.3572 | 17.40146 |
| ENSMUSG00000100235 | 0.084192138 | 0.724105 | 0.827323 | 41.24823 | 50.3925 | 43.06154 | 49.16954 | 43.88001 | 36.87335 | 44.3573 | 47.85401 |
| ENSMUSG00000100252 | 0.266907619 | 0.160451 | 0.303969 | 93.56306 | 129.5807 | 101.8571 | 96.61383 | 79.95913 | 101.681 | 81.96458 | 85.9197 |
| ENSMUSG00000100280 | 0.113734194 | 0.677613 | 0.793933 | 49.29666 | 46.79304 | 52.17072 | 36.23019 | 31.20356 | 51.39921 | 40.50014 | 47.85401 |
| ENSMUSG00000100303 | 0.237878683 | 0.526118 | 0.679403 | 53.32088 | 27.5959 | 52.17072 | 57.79577 | 20.47734 | 45.81234 | 53.0359 | 43.50364 |
| ENSMUSG00000100320 | -0.17235473 | 0.693805 | 0.806504 | 16.09687 | 11.99822 | 23.18699 | 12.07673 | 19.50223 | 11.17374 | 18.32149 | 22.83941 |
| ENSMUSG00000100345 | 0.187024757 | 0.405724 | 0.572499 | 221.332 | 313.1534 | 197.0894 | 296.7425 | 255.4792 | 278.2261 | 205.3936 | 163.1387 |
| ENSMUSG00000100347 | -0.308967139 | 0.598734 | 0.737125 | 10.06054 | 2.399643 | 10.76539 | 12.07673 | 6.825779 | 13.40849 | 13.50005 | 10.87591 |
| ENSMUSG00000100394 | -0.291086435 | 0.701264 | 0.811596 | 19.11503 | 10.79839 | 10.76539 | 13.80198 | 5.850668 | 46.92971 | 7.714313 | 6.525547 |

| | | | | | | | | | | | |
|---|---|---|---|---|---|---|---|---|---|---|---|
| ENSMUSG00000100455 | 0.130434816 | 0.559043 | 0.70529 | 100.6054 | 65.99018 | 78.67013 | 97.47645 | 80.93424 | 93.85942 | 75.21455 | 64.16787 |
| ENSMUSG00000100480 | 0.792125577 | 0.210355 | 0.367817 | 10.06054 | 10.79839 | 19.87456 | 12.07673 | 3.900445 | 12.29112 | 2.892867 | 11.9635 |
| ENSMUSG00000100490 | 0.543057431 | 0.336733 | 0.505017 | 19.11503 | 23.99643 | 19.04645 | 13.80198 | 6.825779 | 27.93435 | 6.750024 | 10.87591 |
| ENSMUSG00000100505 | 0.569609187 | 0.180019 | 0.329967 | 15.09082 | 14.39786 | 14.90592 | 18.11509 | 11.70134 | 11.17374 | 10.60718 | 8.700729 |
| ENSMUSG00000100510 | 2.180960863 | 0.129316 | 0.261122 | 111.672 | 5.999108 | 4.140533 | 150.0965 | 4.875557 | 50.28183 | 3.857157 | 1.087591 |
| ENSMUSG00000100514 | -0.263763433 | 0.471979 | 0.631312 | 42.25428 | 35.99465 | 28.98373 | 28.46658 | 53.63112 | 31.28647 | 23.14294 | 54.37955 |
| ENSMUSG00000100555 | 1.500528138 | 0.081868 | 0.192069 | 38.23007 | 64.79036 | 30.63995 | 29.3292 | 2.925334 | 46.92971 | 5.785735 | 2.175182 |
| ENSMUSG00000100594 | -0.4426926 | 0.212826 | 0.370803 | 26.15741 | 22.79661 | 14.07781 | 32.77969 | 30.22845 | 27.93435 | 32.78583 | 39.15328 |
| ENSMUSG00000100599 | -0.124792601 | 0.653001 | 0.775933 | 32.19374 | 43.19358 | 43.06154 | 52.62003 | 36.07912 | 56.98608 | 43.39301 | 51.11678 |
| ENSMUSG00000100600 | 1.606060573 | 0.00804 | 0.036377 | 61.36932 | 100.785 | 45.54586 | 79.36136 | 14.62667 | 60.3382 | 7.714313 | 11.9635 |
| ENSMUSG00000100622 | 1.176568047 | 0.077152 | 0.184511 | 63.38142 | 74.38894 | 32.29616 | 53.48266 | 13.65156 | 63.69032 | 17.3572 | 4.350364 |
| ENSMUSG00000100629 | 0.640435598 | 0.0044 | 0.023438 | 119.7205 | 91.18644 | 77.84202 | 94.88858 | 56.55646 | 52.51658 | 74.25027 | 61.99269 |
| ENSMUSG00000100632 | 0.14552267 | 0.718058 | 0.823903 | 16.09687 | 14.39786 | 19.87456 | 17.25247 | 14.62667 | 13.40849 | 21.21436 | 11.9635 |
| ENSMUSG00000100642 | 0.092487276 | 0.706057 | 0.815104 | 175.0535 | 122.3818 | 92.74794 | 98.33908 | 123.8391 | 121.7938 | 117.6433 | 93.53283 |
| ENSMUSG00000100680 | -0.176690226 | 0.379105 | 0.547178 | 97.58727 | 81.58786 | 96.06037 | 99.2017 | 119.9387 | 112.8548 | 114.7504 | 76.13138 |
| ENSMUSG00000100691 | -0.169172573 | 0.059803 | 0.154475 | 504.0332 | 507.5245 | 529.1601 | 533.1013 | 625.0463 | 570.9782 | 611.3593 | 524.2189 |
| ENSMUSG00000100706 | 0.659613773 | 0.012847 | 0.051568 | 89.53884 | 100.785 | 65.42042 | 82.81186 | 63.38224 | 64.8077 | 35.6787 | 50.02919 |
| ENSMUSG00000100707 | 0.593048419 | 0.001414 | 0.010154 | 115.6963 | 129.5807 | 108.482 | 107.8279 | 73.13335 | 93.85942 | 73.28598 | 65.25547 |
| ENSMUSG00000100720 | 0.559297039 | 0.190768 | 0.343394 | 13.07871 | 19.19714 | 17.39024 | 21.56559 | 15.60178 | 13.40849 | 10.60718 | 8.700729 |
| ENSMUSG00000100747 | -0.150662038 | 0.645406 | 0.770415 | 24.1453 | 23.99643 | 30.63995 | 25.01608 | 35.10401 | 18.99536 | 32.78583 | 28.27737 |
| ENSMUSG00000100755 | 0.051522879 | 0.815143 | 0.887515 | 185.114 | 197.9706 | 185.4959 | 187.1893 | 160.8934 | 241.3528 | 208.2865 | 118.5474 |
| ENSMUSG00000100801 | -0.243612275 | 0.457311 | 0.619095 | 27.16347 | 28.79572 | 26.49941 | 17.25247 | 34.1289 | 23.46486 | 27.0001 | 32.62773 |
| ENSMUSG00000100819 | 0.202412989 | 0.733709 | 0.833261 | 10.06054 | 21.59679 | 9.93728 | 13.80198 | 3.900445 | 17.87799 | 17.3572 | 8.700729 |
| ENSMUSG00000100826 | 0.941460836 | 6.27E-07 | 2.53E-05 | 597.5963 | 537.52 | 577.1903 | 684.0604 | 248.6534 | 438.0106 | 293.1439 | 269.7226 |
| ENSMUSG00000100860 | 1.771500185 | 0.000467 | 0.004345 | 38.23007 | 82.78769 | 43.06154 | 79.36136 | 22.42756 | 27.93435 | 4.821446 | 16.31387 |
| ENSMUSG00000100862 | -0.313973323 | 2.65E-05 | 0.000487 | 709.2683 | 764.2863 | 726.2495 | 691.824 | 883.4508 | 947.5332 | 914.1461 | 846.1459 |
| ENSMUSG00000100863 | -0.008999882 | 0.975588 | 0.986739 | 42.25428 | 69.58965 | 39.74912 | 40.5433 | 53.63112 | 44.69496 | 55.92877 | 36.9781 |
| ENSMUSG00000100876 | 0.074882602 | 0.727711 | 0.829154 | 109.6599 | 154.777 | 92.74794 | 118.1794 | 112.1378 | 126.2633 | 121.5004 | 89.18247 |
| ENSMUSG00000100927 | -1.066784153 | 0.034647 | 0.105275 | 8.048435 | 7.198929 | 5.796746 | 6.038364 | 18.52711 | 10.05637 | 14.46434 | 13.05109 |
| ENSMUSG00000100954 | -0.712027176 | 0.000118 | 0.001515 | 106.6418 | 88.78679 | 120.9036 | 87.12497 | 155.0427 | 149.7281 | 158.1434 | 200.1168 |
| ENSMUSG00000100967 | 0.360813609 | 0.280758 | 0.447081 | 76.46013 | 86.38715 | 50.5145 | 43.13117 | 50.70579 | 68.15982 | 47.25017 | 32.62773 |
| ENSMUSG00000100975 | 1.2807365 | 3.21E-05 | 0.000563 | 109.6599 | 68.38983 | 92.74794 | 56.93315 | 25.35289 | 43.57759 | 33.75012 | 32.62773 |
| ENSMUSG00000100980 | 0.057065566 | 0.919672 | 0.952281 | 11.0666 | 11.99822 | 19.87456 | 21.56559 | 7.80089 | 31.28647 | 9.642892 | 14.13868 |
| ENSMUSG00000101009 | 0.749477419 | 0.007881 | 0.035842 | 57.3451 | 76.78858 | 39.74912 | 63.83414 | 33.15378 | 36.87335 | 34.71441 | 35.89051 |
| ENSMUSG00000101013 | 0.101917197 | 0.762844 | 0.852453 | 22.1332 | 17.99732 | 25.67131 | 32.77969 | 22.42756 | 22.34748 | 25.07152 | 22.83941 |
| ENSMUSG00000101059 | 0.726373455 | 0.650928 | 0.774317 | 0 | 33.595 | 20.70267 | 0 | 20.47734 | 0 | 6.750024 | 5.437955 |
| ENSMUSG00000101111 | -0.274217567 | 0.008854 | 0.039036 | 542.2633 | 689.8974 | 630.1891 | 653.8686 | 775.2135 | 717.3542 | 718.3954 | 830.9196 |
| ENSMUSG00000101135 | 1.490207891 | 0.021763 | 0.074822 | 26.15741 | 32.39518 | 19.04645 | 12.07673 | 4.875557 | 18.99536 | 3.857157 | 4.350364 |
| ENSMUSG00000101162 | 0.63134558 | 0.149365 | 0.289355 | 25.15136 | 20.39697 | 14.07781 | 19.84034 | 11.70134 | 17.87799 | 7.714313 | 14.13868 |
| ENSMUSG00000101166 | -0.444424934 | 0.425313 | 0.590327 | 12.07265 | 8.398751 | 4.140533 | 13.80198 | 10.72622 | 14.52586 | 8.678602 | 18.48905 |
| ENSMUSG00000101172 | -0.523484173 | 0.130654 | 0.263105 | 18.10898 | 16.7975 | 15.73403 | 23.29083 | 23.40267 | 33.52122 | 27.0001 | 22.83941 |
| ENSMUSG00000101174 | 0.55316828 | 2.76E-05 | 0.000504 | 1306.865 | 1269.411 | 1080.679 | 1105.021 | 885.4011 | 946.4159 | 689.4668 | 723.2481 |
| ENSMUSG00000101188 | -0.282996568 | 0.016921 | 0.062635 | 261.5741 | 251.9625 | 260.0255 | 254.4739 | 359.8161 | 274.874 | 310.5011 | 304.5255 |
| ENSMUSG00000101191 | -0.372752161 | 0.431236 | 0.595993 | 15.09082 | 11.99822 | 20.70267 | 20.70296 | 30.22845 | 24.58223 | 8.678602 | 26.10219 |
| ENSMUSG00000101210 | 0.276286647 | 0.667712 | 0.786827 | 17.10292 | 14.39786 | 11.59349 | 6.900988 | 10.72622 | 14.52586 | 1.928578 | 14.13868 |
| ENSMUSG00000101225 | -0.23662846 | 0.063884 | 0.162113 | 156.9445 | 194.3711 | 176.3867 | 170.7995 | 210.624 | 195.5405 | 196.715 | 218.6058 |
| ENSMUSG00000101236 | -1.793489585 | 0.004299 | 0.023037 | 9.054489 | 4.799286 | 4.140533 | 3.450494 | 7.80089 | 30.1691 | 15.42863 | 20.66423 |
| ENSMUSG00000101249 | -0.255538109 | 0.022554 | 0.076743 | 244.4712 | 286.7573 | 289.0092 | 275.1769 | 296.4338 | 325.1559 | 346.1798 | 340.416 |
| ENSMUSG00000101304 | -0.058361377 | 0.86617 | 0.919627 | 21.12714 | 22.79661 | 37.2648 | 23.29083 | 22.42756 | 24.58223 | 28.92867 | 33.71532 |
| ENSMUSG00000101320 | -0.02579055 | 0.907958 | 0.945031 | 102.6175 | 107.9839 | 135.8095 | 163.8985 | 126.7645 | 99.44629 | 155.2506 | 139.2117 |
| ENSMUSG00000101335 | 0.319254534 | 0.192296 | 0.34549 | 153.9263 | 169.1748 | 138.2938 | 181.1509 | 118.9636 | 192.1883 | 92.57176 | 112.0219 |
| ENSMUSG00000101337 | 0.537738304 | 0.118393 | 0.247207 | 43.26034 | 43.19358 | 42.23344 | 39.68068 | 17.552 | 46.92971 | 27.0001 | 25.0146 |
| ENSMUSG00000101355 | 0.182131399 | 0.629066 | 0.758529 | 17.10292 | 41.99375 | 23.18699 | 30.19182 | 26.32801 | 27.93435 | 19.28578 | 25.0146 |
| ENSMUSG00000101356 | 1.128856172 | 0.000653 | 0.005639 | 208.2533 | 219.5673 | 136.6376 | 279.49 | 92.63557 | 159.7845 | 66.53595 | 67.43065 |
| ENSMUSG00000101389 | 1.119808431 | 0.000371 | 0.003637 | 178.0716 | 116.3827 | 143.2624 | 164.7611 | 74.10846 | 63.69032 | 104.1432 | 34.80291 |
| ENSMUSG00000101431 | -0.012212721 | 0.970026 | 0.983292 | 25.15136 | 45.59322 | 35.60859 | 33.64232 | 32.17867 | 26.81698 | 32.78583 | 48.9416 |
| ENSMUSG00000101438 | -0.017171144 | 0.966695 | 0.981333 | 13.07871 | 23.99643 | 16.56213 | 16.38985 | 15.60178 | 13.40849 | 16.39292 | 25.0146 |
| ENSMUSG00000101450 | 0.360267534 | 0.483401 | 0.641753 | 26.15741 | 7.198929 | 14.07781 | 16.38985 | 11.70134 | 16.76061 | 8.678602 | 13.05109 |
| ENSMUSG00000101451 | 0.002927905 | 0.995231 | 0.997284 | 25.15136 | 21.59679 | 24.8432 | 13.80198 | 14.62667 | 41.34284 | 15.42863 | 14.13868 |
| ENSMUSG00000101462 | 0.702040152 | 0.106015 | 0.230052 | 42.25428 | 53.99197 | 52.17072 | 44.85642 | 17.552 | 59.22083 | 25.07152 | 17.40146 |
| ENSMUSG00000101493 | 0.646863468 | 0.017299 | 0.063687 | 112.6781 | 107.9839 | 72.87338 | 79.36136 | 76.05868 | 69.27719 | 46.28588 | 45.67883 |
| ENSMUSG00000101517 | -2.031230085 | 0.000173 | 0.002024 | 3.018163 | 4.799286 | 9.109173 | 4.313117 | 22.42756 | 13.40849 | 25.07152 | 27.18978 |
| ENSMUSG00000101523 | 0.174311391 | 0.589115 | 0.72937 | 26.15741 | 28.79572 | 24.8432 | 34.50494 | 22.42756 | 21.23011 | 24.10723 | 33.71532 |
| ENSMUSG00000101556 | 1.392869285 | 0.005869 | 0.028941 | 18.10898 | 11.99822 | 19.87456 | 14.6646 | 5.850668 | 7.821619 | 6.750024 | 4.350364 |

| | | | | | | | | | | | |
|---|---|---|---|---|---|---|---|---|---|---|---|
| ENSMUSG00000101559 | 0.899223903 | 0.202721 | 0.358389 | 55.33299 | 61.1909 | 40.57722 | 50.89479 | 12.67645 | 81.56831 | 6.750024 | 10.87591 |
| ENSMUSG00000101586 | -0.079991572 | 0.836555 | 0.901206 | 30.18163 | 25.19625 | 18.21835 | 25.8787 | 29.25334 | 39.10809 | 19.28578 | 17.40146 |
| ENSMUSG00000101587 | 1.343130671 | 0.050917 | 0.13785 | 7.04238 | 15.59768 | 33.95237 | 28.46658 | 7.80089 | 11.17374 | 1.928578 | 13.05109 |
| ENSMUSG00000101588 | 0.776125567 | 0.044224 | 0.125039 | 66.39959 | 69.58965 | 33.95237 | 50.03216 | 26.32801 | 51.39921 | 30.85725 | 19.57664 |
| ENSMUSG00000101599 | 0.853877463 | 0.001243 | 0.009201 | 469.8274 | 500.3256 | 374.3042 | 414.9219 | 170.6445 | 396.6678 | 222.7508 | 183.8029 |
| ENSMUSG00000101603 | -0.008134991 | 0.983974 | 0.991004 | 22.1332 | 10.79839 | 21.53077 | 15.52722 | 21.45245 | 14.52586 | 16.39292 | 18.48905 |
| ENSMUSG00000101609 | 0.962698913 | 0.032075 | 0.099208 | 3451.772 | 4124.986 | 2361.76 | 2749.181 | 1163.308 | 3212.451 | 1203.433 | 930.978 |
| ENSMUSG00000101655 | 0.391446871 | 0.441608 | 0.604982 | 15.09082 | 15.59768 | 23.18699 | 17.25247 | 15.60178 | 4.469496 | 12.53576 | 21.75182 |
| ENSMUSG00000101678 | 0.781931237 | 0.085133 | 0.197167 | 30.18163 | 19.19714 | 43.88965 | 44.85642 | 16.57689 | 32.40385 | 10.60718 | 21.75182 |
| ENSMUSG00000101767 | 0.327413601 | 0.453438 | 0.615795 | 26.15741 | 13.19804 | 19.87456 | 30.19182 | 20.47734 | 24.58223 | 10.60718 | 16.31387 |
| ENSMUSG00000101792 | 1.355384098 | 0.018049 | 0.065766 | 10.06054 | 21.59679 | 11.59349 | 22.42821 | 9.751113 | 3.352122 | 5.785735 | 6.525547 |
| ENSMUSG00000101795 | -0.157274684 | 0.532675 | 0.684218 | 56.33904 | 47.99286 | 54.65504 | 36.23019 | 55.58134 | 44.69496 | 54.00019 | 63.08028 |
| ENSMUSG00000101828 | 0.452902649 | 0.255363 | 0.419287 | 32.19374 | 32.39518 | 14.90592 | 25.01608 | 14.62667 | 24.58223 | 19.28578 | 17.40146 |
| ENSMUSG00000101845 | 0.009830548 | 0.982792 | 0.990592 | 21.12714 | 15.59768 | 14.90592 | 14.6646 | 7.80089 | 24.58223 | 17.3572 | 16.31387 |
| ENSMUSG00000101849 | -0.373372896 | 0.384217 | 0.552197 | 10.06054 | 8.398751 | 14.90592 | 12.93935 | 15.60178 | 13.40849 | 15.42863 | 16.31387 |
| ENSMUSG00000101856 | -0.097544466 | 0.437977 | 0.601615 | 168.0111 | 175.1739 | 166.4494 | 182.8762 | 167.7191 | 197.7752 | 189.0007 | 187.0657 |
| ENSMUSG00000101872 | 0.188207283 | 0.514866 | 0.669147 | 48.29061 | 38.39429 | 28.98373 | 46.58167 | 41.92979 | 32.40385 | 31.82154 | 35.89051 |
| ENSMUSG00000101892 | -0.02825571 | 0.775558 | 0.861181 | 555.342 | 529.1213 | 603.6897 | 484.7944 | 563.6143 | 582.1519 | 522.6447 | 549.2335 |
| ENSMUSG00000101939 | -0.166459064 | 0.438739 | 0.602213 | 70.4238 | 56.39161 | 52.99882 | 62.97152 | 69.2329 | 73.74669 | 55.92877 | 73.95619 |
| ENSMUSG00000101970 | 0.467393522 | 0.002033 | 0.013285 | 1467.833 | 1547.77 | 1537.794 | 1907.261 | 958.5344 | 1467.112 | 1199.576 | 1049.525 |
| ENSMUSG00000102014 | 0.305029333 | 0.500435 | 0.656807 | 14.08476 | 26.39607 | 14.07781 | 24.15346 | 13.65156 | 24.58223 | 14.46434 | 10.87591 |
| ENSMUSG00000102018 | 0.010716925 | 0.963029 | 0.978992 | 111.672 | 87.58697 | 85.29498 | 89.71284 | 89.71024 | 96.09417 | 65.57166 | 120.7226 |
| ENSMUSG00000102059 | -0.022799786 | 0.950054 | 0.971623 | 36.21796 | 32.39518 | 34.78048 | 26.74133 | 19.50223 | 50.28183 | 26.03581 | 36.9781 |
| ENSMUSG00000102062 | -0.226903096 | 0.689133 | 0.802945 | 16.09687 | 7.198929 | 13.24971 | 5.175741 | 17.552 | 10.05637 | 13.50005 | 7.613138 |
| ENSMUSG00000102070 | -0.302065913 | 2.18E-05 | 0.000422 | 569.4268 | 580.7136 | 593.7525 | 571.9194 | 722.5575 | 682.7156 | 712.6097 | 737.3868 |
| ENSMUSG00000102091 | 1.517916791 | 0.00533 | 0.026873 | 46.2785 | 50.3925 | 19.87456 | 62.10889 | 5.850668 | 30.1691 | 12.53576 | 14.13868 |
| ENSMUSG00000102101 | -0.517879131 | 0.016523 | 0.061545 | 81.4904 | 76.78858 | 105.1695 | 105.2401 | 109.2125 | 110.62 | 149.4648 | 160.9635 |
| ENSMUSG00000102112 | -1.355593033 | 0.007068 | 0.033143 | 12.07265 | 16.7975 | 21.53077 | 6.900988 | 26.32801 | 18.99536 | 52.07161 | 48.9416 |
| ENSMUSG00000102145 | -0.876261612 | 0.048656 | 0.133649 | 14.08476 | 8.398751 | 12.4216 | 9.488858 | 24.37778 | 18.99536 | 12.53576 | 26.10219 |
| ENSMUSG00000102151 | 0.739651813 | 0.092312 | 0.208659 | 21.12714 | 13.19804 | 18.21835 | 23.29083 | 10.72622 | 16.76061 | 8.678602 | 9.78832 |
| ENSMUSG00000102153 | 1.124891819 | 0.156118 | 0.298241 | 49.29666 | 20.39697 | 31.46805 | 15.52722 | 4.875557 | 31.28647 | 16.39292 | 1.087591 |
| ENSMUSG00000102171 | 0.430478041 | 0.064821 | 0.163904 | 146.8839 | 139.1793 | 130.8408 | 130.2561 | 124.8142 | 130.7328 | 73.28598 | 77.21897 |
| ENSMUSG00000102189 | -0.002212435 | 0.996219 | 0.997733 | 37.22401 | 25.19625 | 12.4216 | 12.93935 | 27.30312 | 17.87799 | 22.17865 | 19.57664 |
| ENSMUSG00000102202 | 0.814408726 | 0.043757 | 0.124128 | 19.11503 | 15.59768 | 24.8432 | 25.01608 | 12.67645 | 14.52586 | 11.57147 | 9.78832 |
| ENSMUSG00000102241 | 0.582811846 | 0.323289 | 0.491365 | 19.11503 | 13.19804 | 15.73403 | 6.900988 | 4.875557 | 16.76061 | 8.678602 | 6.525547 |
| ENSMUSG00000102243 | 0.817835507 | 0.142299 | 0.27945 | 22.1332 | 25.19625 | 14.07781 | 9.488858 | 5.850668 | 17.87799 | 7.714313 | 8.700729 |
| ENSMUSG00000102246 | 0.635401982 | 0.311684 | 0.480258 | 10.06054 | 21.59679 | 34.78048 | 22.42821 | 6.825779 | 27.93435 | 17.3572 | 5.437955 |
| ENSMUSG00000102252 | 0.265119684 | 0.060784 | 0.156439 | 637.8385 | 523.1222 | 525.8477 | 737.5431 | 519.7343 | 444.7149 | 528.4305 | 525.3065 |
| ENSMUSG00000102275 | 1.133001532 | 0.015804 | 0.059879 | 80.48435 | 69.58965 | 57.13936 | 79.36136 | 25.35289 | 70.39457 | 21.21436 | 14.13868 |
| ENSMUSG00000102277 | 0.887510776 | 0.21295 | 0.370828 | 76.46013 | 101.9848 | 33.95237 | 46.58167 | 20.47734 | 93.85942 | 20.25007 | 5.437955 |
| ENSMUSG00000102282 | 1.026884437 | 0.012134 | 0.049404 | 37.22401 | 38.39429 | 28.98373 | 40.5433 | 11.70134 | 32.40385 | 14.46434 | 13.05109 |
| ENSMUSG00000102297 | 1.244583143 | 0.017902 | 0.065414 | 124.7507 | 176.3738 | 81.15445 | 101.7896 | 45.83023 | 110.62 | 33.75012 | 14.13868 |
| ENSMUSG00000102302 | 0.335143153 | 0.279494 | 0.445876 | 66.39959 | 45.59322 | 45.54586 | 50.89479 | 45.83023 | 58.10345 | 27.96439 | 33.71532 |
| ENSMUSG00000102305 | 0.560446291 | 0.287823 | 0.454754 | 36.21796 | 19.19714 | 23.18699 | 27.60395 | 16.57689 | 35.75597 | 13.50005 | 6.525547 |
| ENSMUSG00000102319 | 1.08326703 | 0.071497 | 0.175064 | 64.38748 | 83.98751 | 43.88965 | 65.55939 | 17.552 | 78.21619 | 16.39292 | 9.78832 |
| ENSMUSG00000102323 | 0.336671904 | 0.346985 | 0.515824 | 23.13925 | 16.7975 | 22.35888 | 24.15346 | 20.47734 | 14.52586 | 17.3572 | 16.31387 |
| ENSMUSG00000102326 | -0.545615386 | 0.54515 | 0.69369 | 16.09687 | 17.99732 | 7.45296 | 5.175741 | 8.776002 | 50.28183 | 6.750024 | 2.175182 |
| ENSMUSG00000102331 | 0.929465165 | 0.209378 | 0.366525 | 33.19979 | 85.18733 | 31.46805 | 65.55939 | 11.70134 | 81.56831 | 13.50005 | 6.525547 |
| ENSMUSG00000102336 | 0.045982342 | 0.887174 | 0.932336 | 31.18769 | 50.3925 | 33.95237 | 35.36756 | 41.92979 | 39.10809 | 22.17865 | 42.41605 |
| ENSMUSG00000102344 | 1.107229547 | 0.06746 | 0.16764 | 57.3451 | 58.79126 | 33.12427 | 37.95543 | 15.60178 | 52.51658 | 13.50005 | 5.437955 |
| ENSMUSG00000102365 | 0.979542449 | 0.101818 | 0.223792 | 10.06054 | 9.598572 | 14.07781 | 21.56559 | 4.875557 | 13.40849 | 4.821446 | 5.437955 |
| ENSMUSG00000102386 | 1.589202085 | 0.105864 | 0.229835 | 15.09082 | 20.39697 | 3.312427 | 36.23019 | 1.950223 | 18.99536 | 0.964289 | 3.262773 |
| ENSMUSG00000102389 | -1.184991331 | 0.203772 | 0.35981 | 6.036326 | 0 | 15.73403 | 2.58787 | 7.80089 | 7.821619 | 12.53576 | 28.27737 |
| ENSMUSG00000102411 | 0.085689473 | 0.792653 | 0.873177 | 122.7386 | 55.19179 | 57.96746 | 62.10889 | 76.05868 | 50.28183 | 80.036 | 73.95619 |
| ENSMUSG00000102425 | 0.361326444 | 0.41283 | 0.57864 | 12.07265 | 20.39697 | 19.04645 | 18.97772 | 17.552 | 17.87799 | 8.678602 | 10.87591 |
| ENSMUSG00000102426 | 0.785035741 | 0.009624 | 0.041433 | 229.3804 | 279.5584 | 186.324 | 243.2598 | 100.4365 | 235.7659 | 101.2504 | 107.6715 |
| ENSMUSG00000102428 | 0.986098508 | 0.000378 | 0.00369 | 172.0353 | 215.9679 | 158.9965 | 143.1955 | 70.20801 | 136.3196 | 81.00029 | 60.9051 |
| ENSMUSG00000102440 | 0.011058488 | 0.978881 | 0.988554 | 27.16347 | 46.79304 | 24.01509 | 38.81806 | 57.53157 | 22.34748 | 30.85725 | 23.927 |
| ENSMUSG00000102448 | 0.550485278 | 0.415854 | 0.581748 | 20.12109 | 22.79661 | 15.73403 | 16.38985 | 6.825779 | 33.52122 | 6.750024 | 4.350364 |
| ENSMUSG00000102452 | 0.623743046 | 0.145807 | 0.284282 | 23.13925 | 22.79661 | 35.60859 | 21.56559 | 10.72622 | 23.46486 | 22.17865 | 10.87591 |
| ENSMUSG00000102460 | 1.192990512 | 0.105126 | 0.228558 | 503.0272 | 519.5227 | 259.1974 | 351.9504 | 116.0382 | 476.0014 | 60.75022 | 61.99269 |
| ENSMUSG00000102478 | 0.598550389 | 0.259882 | 0.423918 | 7.04238 | 19.19714 | 12.4216 | 12.07673 | 6.825779 | 8.938993 | 6.750024 | 10.87591 |
| ENSMUSG00000102496 | 0.250333051 | 0.577842 | 0.719661 | 48.29061 | 20.39697 | 22.35888 | 15.52722 | 19.50223 | 29.05173 | 19.28578 | 21.75182 |
| ENSMUSG00000102533 | 0.823978802 | 0.116978 | 0.245141 | 23.13925 | 26.39607 | 14.07781 | 11.21411 | 9.751113 | 14.52586 | 4.821446 | 13.05109 |

| | | | | | | | | | | | |
|---|---|---|---|---|---|---|---|---|---|---|---|
| ENSMUSG00000102543 | 0.213825965 | 0.63233 | 0.760923 | 45.27245 | 71.98929 | 54.65504 | 34.50494 | 47.78045 | 78.21619 | 19.28578 | 32.62773 |
| ENSMUSG00000102545 | -0.250788038 | 0.665339 | 0.785151 | 19.11503 | 20.39697 | 15.73403 | 9.488858 | 16.57689 | 4.469496 | 26.03581 | 29.36496 |
| ENSMUSG00000102553 | 0.350682303 | 0.284853 | 0.451421 | 43.26034 | 23.99643 | 28.98373 | 37.95543 | 24.37778 | 33.52122 | 25.07152 | 22.83941 |
| ENSMUSG00000102555 | 0.175986419 | 0.669224 | 0.787855 | 37.22401 | 33.595 | 21.53077 | 18.11509 | 34.1289 | 23.46486 | 14.46434 | 25.0146 |
| ENSMUSG00000102559 | 0.500317726 | 0.228096 | 0.388065 | 16.09687 | 22.79661 | 21.53077 | 15.52722 | 18.52711 | 13.40849 | 10.60718 | 10.87591 |
| ENSMUSG00000102562 | 1.220820364 | 0.005099 | 0.026084 | 66.39959 | 73.18911 | 46.37397 | 43.13117 | 29.25334 | 42.46022 | 13.50005 | 13.05109 |
| ENSMUSG00000102570 | 1.035426426 | 0.032411 | 0.099911 | 40.24217 | 31.19536 | 29.81184 | 26.74133 | 10.72622 | 27.93435 | 5.785735 | 18.48905 |
| ENSMUSG00000102571 | 1.067128248 | 0.09735 | 0.216806 | 35.2119 | 23.99643 | 16.56213 | 25.01608 | 5.850668 | 30.1691 | 4.821446 | 7.613138 |
| ENSMUSG00000102573 | -0.002671749 | 0.99592 | 0.99765 | 13.07871 | 8.398751 | 5.796746 | 13.80198 | 11.70134 | 11.17374 | 10.60718 | 7.613138 |
| ENSMUSG00000102577 | 1.40025404 | 0.000129 | 0.001621 | 37.22401 | 28.79572 | 25.67131 | 31.05445 | 12.67645 | 12.29112 | 10.60718 | 10.87591 |
| ENSMUSG00000102588 | 0.889055642 | 0.363433 | 0.532005 | 50.30272 | 62.39072 | 19.04645 | 48.30692 | 9.751113 | 82.68568 | 3.857157 | 1.087591 |
| ENSMUSG00000102593 | 1.192601329 | 0.031893 | 0.098719 | 53.32088 | 56.39161 | 31.46805 | 37.09281 | 22.42756 | 40.22547 | 6.750024 | 8.700729 |
| ENSMUSG00000102602 | 0.158326359 | 0.484649 | 0.642947 | 247.4894 | 225.5664 | 225.245 | 213.068 | 162.8436 | 301.691 | 201.5364 | 151.1752 |
| ENSMUSG00000102615 | -1.04530759 | 0.0161 | 0.060456 | 23.13925 | 8.398751 | 26.49941 | 17.25247 | 34.1289 | 32.40385 | 29.89296 | 60.9051 |
| ENSMUSG00000102630 | 1.296262053 | 0.085419 | 0.197648 | 24.1453 | 10.79839 | 24.01509 | 17.25247 | 9.751113 | 15.64324 | 5.785735 | 0 |
| ENSMUSG00000102644 | 0.213919266 | 0.082526 | 0.193042 | 478.8819 | 430.7359 | 349.461 | 392.4937 | 350.065 | 364.264 | 365.4656 | 341.5036 |
| ENSMUSG00000102649 | 0.846561202 | 0.165326 | 0.310719 | 104.6297 | 100.785 | 62.9361 | 77.63611 | 28.27823 | 128.498 | 16.39292 | 19.57664 |
| ENSMUSG00000102664 | 0.113365062 | 0.824653 | 0.893455 | 23.13925 | 11.99822 | 12.4216 | 11.21411 | 13.65156 | 22.34748 | 8.678602 | 9.78832 |
| ENSMUSG00000102670 | 1.501565468 | 0.075843 | 0.182356 | 34.20585 | 29.99554 | 12.4216 | 21.56559 | 0.975111 | 20.11273 | 12.53576 | 1.087591 |
| ENSMUSG00000102691 | 0.850447609 | 0.108544 | 0.233384 | 15.09082 | 19.19714 | 21.53077 | 49.16954 | 13.65156 | 16.76061 | 7.714313 | 20.66423 |
| ENSMUSG00000102700 | 0.769855103 | 0.160146 | 0.303486 | 36.21796 | 34.79482 | 19.04645 | 21.56559 | 9.751113 | 34.6386 | 13.50005 | 7.613138 |
| ENSMUSG00000102704 | 0.128365684 | 0.751116 | 0.844971 | 70.4238 | 55.19179 | 54.65504 | 75.91087 | 43.88001 | 110.62 | 51.10733 | 29.36496 |
| ENSMUSG00000102719 | 0.165470402 | 0.777233 | 0.862728 | 40.24217 | 38.39429 | 32.29616 | 16.38985 | 14.62667 | 64.8077 | 12.53576 | 21.75182 |
| ENSMUSG00000102721 | 1.191858784 | 0.037925 | 0.112369 | 15.09082 | 20.39697 | 9.93728 | 14.6646 | 3.900445 | 12.29112 | 5.785735 | 4.350364 |
| ENSMUSG00000102732 | 1.397495548 | 0.091294 | 0.207196 | 6.036326 | 23.99643 | 16.56213 | 12.07673 | 1.950223 | 14.52586 | 4.821446 | 1.087591 |
| ENSMUSG00000102742 | 0.148616544 | 0.648492 | 0.772265 | 35.2119 | 40.79393 | 51.34261 | 27.60395 | 36.07912 | 46.92971 | 29.89296 | 27.18978 |
| ENSMUSG00000102748 | -0.028080204 | 0.944204 | 0.967693 | 40.24217 | 41.99375 | 50.5145 | 23.29083 | 53.63112 | 26.81698 | 54.00019 | 23.927 |
| ENSMUSG00000102752 | 0.197683835 | 0.169044 | 0.315362 | 598.6023 | 572.3149 | 635.1578 | 515.8488 | 441.7254 | 638.0206 | 492.7518 | 453.5255 |
| ENSMUSG00000102758 | 0.452148445 | 0.025123 | 0.083355 | 255.5378 | 297.5557 | 244.2915 | 301.9182 | 170.6445 | 281.5783 | 191.8935 | 159.8759 |
| ENSMUSG00000102764 | 0.68422444 | 0.369478 | 0.538378 | 23.13925 | 10.79839 | 4.96864 | 11.21411 | 6.825779 | 14.52586 | 8.678602 | 1.087591 |
| ENSMUSG00000102776 | 0.618657721 | 0.190886 | 0.343506 | 54.32694 | 68.38983 | 39.74912 | 37.95543 | 25.35289 | 65.92507 | 15.42863 | 23.927 |
| ENSMUSG00000102779 | 0.100731913 | 0.904296 | 0.9426 | 10.06054 | 22.79661 | 5.796746 | 12.07673 | 6.825779 | 32.40385 | 5.785735 | 2.175182 |
| ENSMUSG00000102780 | 0.039640571 | 0.912051 | 0.948028 | 45.27245 | 31.19536 | 24.8432 | 43.9938 | 35.10401 | 52.51658 | 27.96439 | 26.10219 |
| ENSMUSG00000102786 | 2.026092265 | 0.011972 | 0.048893 | 29.17558 | 13.19804 | 6.624853 | 15.52722 | 2.925334 | 10.05637 | 1.928578 | 1.087591 |
| ENSMUSG00000102805 | -0.72047239 | 0.288713 | 0.455682 | 6.036326 | 8.398751 | 17.39024 | 8.626235 | 24.37778 | 3.352122 | 26.03581 | 13.05109 |
| ENSMUSG00000102813 | 0.240128506 | 0.225062 | 0.384784 | 91.55095 | 73.18911 | 90.26362 | 76.77349 | 76.05868 | 63.69032 | 79.07171 | 61.99269 |
| ENSMUSG00000102824 | -0.134868428 | 0.557294 | 0.703981 | 54.32694 | 67.19001 | 52.99882 | 54.34528 | 61.43201 | 53.63396 | 55.92877 | 79.39415 |
| ENSMUSG00000102840 | -0.120434304 | 0.59948 | 0.737591 | 44.26639 | 55.19179 | 52.17072 | 45.71905 | 57.53157 | 59.22083 | 44.3573 | 53.29196 |
| ENSMUSG00000102854 | 0.922190816 | 0.187073 | 0.339035 | 34.20585 | 40.79393 | 14.90592 | 18.11509 | 7.80089 | 36.87335 | 4.821446 | 7.613138 |
| ENSMUSG00000102861 | 0.380519734 | 0.182367 | 0.332947 | 112.6781 | 128.3809 | 133.3252 | 134.5693 | 75.08357 | 162.0192 | 81.96458 | 72.8686 |
| ENSMUSG00000102865 | 1.222833449 | 0.223469 | 0.383161 | 31.18769 | 46.79304 | 13.24971 | 15.52722 | 0.975111 | 37.99072 | 5.785735 | 1.087591 |
| ENSMUSG00000102869 | -0.34206545 | 0.058323 | 0.151607 | 1065.412 | 1012.649 | 1101.382 | 976.4898 | 1553.352 | 840.2653 | 1421.362 | 1451.934 |
| ENSMUSG00000102882 | 0.559287324 | 0.131375 | 0.264232 | 38.23007 | 26.39607 | 19.87456 | 31.91707 | 13.65156 | 23.46486 | 20.25007 | 21.75182 |
| ENSMUSG00000102899 | 1.03885939 | 0.015384 | 0.058734 | 22.1332 | 22.79661 | 21.53077 | 14.6646 | 10.72622 | 12.29112 | 7.714313 | 8.700729 |
| ENSMUSG00000102912 | -0.048132745 | 0.929067 | 0.958086 | 18.10898 | 8.398751 | 24.01509 | 6.900988 | 18.52711 | 17.87799 | 12.53576 | 10.87591 |
| ENSMUSG00000102918 | -0.082613645 | 0.652511 | 0.77555 | 353.1251 | 430.7359 | 551.519 | 401.1199 | 458.3023 | 350.8555 | 488.8946 | 541.6204 |
| ENSMUSG00000102960 | 0.380827385 | 0.539843 | 0.689457 | 7.04238 | 17.99732 | 9.109173 | 12.93935 | 4.875557 | 16.76061 | 4.821446 | 9.78832 |
| ENSMUSG00000102964 | 0.045202596 | 0.925447 | 0.955704 | 18.10898 | 32.39518 | 23.18699 | 17.25247 | 12.67645 | 32.40385 | 11.57147 | 31.54014 |
| ENSMUSG00000102976 | 0.249393516 | 0.075736 | 0.182146 | 644.8808 | 668.3006 | 481.13 | 603.8364 | 483.6552 | 591.0909 | 455.1445 | 486.1532 |
| ENSMUSG00000102997 | 0.240584816 | 0.271604 | 0.436865 | 73.44197 | 63.59054 | 79.49824 | 69.8725 | 58.50668 | 77.09881 | 57.85735 | 50.02919 |
| ENSMUSG00000103002 | 1.365747027 | 0.054773 | 0.145442 | 20.12109 | 17.99732 | 4.140533 | 19.84034 | 2.925334 | 11.17374 | 6.750024 | 3.262773 |
| ENSMUSG00000103009 | -0.272878182 | 0.23198 | 0.392539 | 84.50857 | 111.5834 | 133.3252 | 126.8057 | 149.192 | 129.6154 | 102.2147 | 171.8394 |
| ENSMUSG00000103013 | 1.162532949 | 0.023228 | 0.078575 | 30.18163 | 17.99732 | 13.24971 | 20.70296 | 7.80089 | 14.52586 | 4.821446 | 9.78832 |
| ENSMUSG00000103015 | 0.071062375 | 0.885378 | 0.931559 | 14.08476 | 10.79839 | 12.4216 | 23.29083 | 15.60178 | 22.34748 | 11.57147 | 8.700729 |
| ENSMUSG00000103030 | 0.335409527 | 0.52922 | 0.681503 | 9.054489 | 16.7975 | 14.07781 | 11.21411 | 6.825779 | 15.64324 | 12.53576 | 5.437955 |
| ENSMUSG00000103034 | 0.224834309 | 0.321105 | 0.489099 | 80.48435 | 92.38626 | 65.42042 | 91.43809 | 80.93424 | 71.51194 | 76.17884 | 52.20437 |
| ENSMUSG00000103037 | 0.110635182 | 0.73866 | 0.836517 | 26.15741 | 34.79482 | 35.60859 | 22.42821 | 24.37778 | 21.23011 | 34.71441 | 29.36496 |
| ENSMUSG00000103038 | 0.764214326 | 0.416927 | 0.582497 | 50.30272 | 98.38537 | 25.67131 | 19.84034 | 12.67645 | 92.74205 | 6.750024 | 2.175182 |
| ENSMUSG00000103039 | -0.054217514 | 0.842974 | 0.905412 | 46.2785 | 39.59411 | 43.88965 | 47.44429 | 60.4569 | 51.39921 | 37.60728 | 34.80291 |
| ENSMUSG00000103041 | 0.153565891 | 0.20894 | 0.366165 | 336.0222 | 380.3434 | 317.9929 | 357.1261 | 319.8365 | 358.6771 | 285.4296 | 286.0365 |
| ENSMUSG00000103046 | -0.085198981 | 0.744277 | 0.840094 | 107.6478 | 61.1909 | 57.13936 | 81.94923 | 96.53602 | 78.21619 | 81.96458 | 69.60583 |
| ENSMUSG00000103047 | 1.069531337 | 0.019172 | 0.068437 | 55.33299 | 63.59054 | 47.20208 | 48.30692 | 22.42756 | 52.51658 | 14.46434 | 13.05109 |
| ENSMUSG00000103081 | 0.680779188 | 0.166242 | 0.311869 | 69.41775 | 41.99375 | 43.88965 | 29.3292 | 23.40267 | 55.8687 | 24.10723 | 11.9635 |
| ENSMUSG00000103085 | 1.393869922 | 0.076666 | 0.183808 | 43.26034 | 20.39697 | 14.90592 | 18.97772 | 3.900445 | 24.58223 | 7.714313 | 1.087591 |

| | | | | | | | | | | | |
|---|---|---|---|---|---|---|---|---|---|---|---|
| ENSMUSG00000103088 | 0.166537639 | 0.39032 | 0.557694 | 160.9687 | 147.578 | 195.4332 | 125.943 | 157.968 | 115.0895 | 134.0362 | 154.4379 |
| ENSMUSG00000103098 | 0.847454324 | 0.088629 | 0.20287 | 20.12109 | 10.79839 | 26.49941 | 13.80198 | 11.70134 | 8.938993 | 12.53576 | 6.525547 |
| ENSMUSG00000103103 | -0.316022813 | 0.270893 | 0.436075 | 27.16347 | 34.79482 | 43.88965 | 31.05445 | 44.85512 | 45.81234 | 32.78583 | 47.85401 |
| ENSMUSG00000103108 | 0.090618172 | 0.791451 | 0.87239 | 26.15741 | 22.79661 | 37.2648 | 20.70296 | 30.22845 | 23.46486 | 22.17865 | 25.0146 |
| ENSMUSG00000103115 | 0.844123751 | 0.010267 | 0.043476 | 66.39959 | 50.3925 | 37.2648 | 49.16954 | 26.32801 | 40.22547 | 27.0001 | 19.57664 |
| ENSMUSG00000103119 | 0.130067819 | 0.698136 | 0.809552 | 25.15136 | 28.79572 | 27.32752 | 27.60395 | 28.27823 | 22.34748 | 16.39292 | 32.62773 |
| ENSMUSG00000103133 | 0.124998274 | 0.573968 | 0.716553 | 62.37537 | 58.79126 | 54.65504 | 59.52102 | 58.50668 | 62.57295 | 52.07161 | 42.41605 |
| ENSMUSG00000103138 | 0.336013279 | 0.52985 | 0.681904 | 34.20585 | 9.598572 | 20.70267 | 18.11509 | 24.37778 | 20.11273 | 13.50005 | 7.613138 |
| ENSMUSG00000103144 | 0.29568868 | 0.502841 | 0.658841 | 17.10292 | 55.19179 | 47.20208 | 34.50494 | 43.88001 | 26.81698 | 35.6787 | 18.48905 |
| ENSMUSG00000103149 | 0.331304698 | 0.53813 | 0.688091 | 23.13925 | 23.99643 | 6.624853 | 13.80198 | 18.52711 | 10.05637 | 14.46434 | 9.78832 |
| ENSMUSG00000103160 | 0.359119399 | 0.137511 | 0.2726 | 73.44197 | 64.79036 | 71.21717 | 67.28463 | 42.9049 | 74.86406 | 52.07161 | 46.76642 |
| ENSMUSG00000103183 | 0.761096124 | 2.43E-05 | 0.000457 | 366.2038 | 403.14 | 319.6492 | 366.615 | 195.9974 | 292.752 | 190.9293 | 179.4525 |
| ENSMUSG00000103251 | 1.39039885 | 0.156219 | 0.298311 | 19.11503 | 16.7975 | 14.07781 | 12.07673 | 3.900445 | 18.99536 | 0.964289 | 0 |
| ENSMUSG00000103283 | 0.534730809 | 0.027074 | 0.087924 | 97.58727 | 100.785 | 82.81066 | 87.9876 | 49.73068 | 90.5073 | 53.0359 | 61.99269 |
| ENSMUSG00000103284 | 1.304869223 | 0.027304 | 0.088391 | 135.8173 | 136.7797 | 53.82693 | 86.26235 | 20.47734 | 106.1505 | 22.17865 | 18.48905 |
| ENSMUSG00000103285 | 0.735150288 | 0.237038 | 0.398088 | 91.55095 | 74.38894 | 23.18699 | 37.95543 | 27.30312 | 75.98144 | 22.17865 | 10.87591 |
| ENSMUSG00000103286 | 0.12831609 | 0.77825 | 0.863236 | 14.08476 | 28.79572 | 15.73403 | 18.97772 | 23.40267 | 16.76061 | 21.21436 | 8.700729 |
| ENSMUSG00000103303 | 0.783798622 | 0.28403 | 0.450542 | 17.10292 | 14.39786 | 9.93728 | 25.8787 | 7.80089 | 24.58223 | 4.821446 | 2.175182 |
| ENSMUSG00000103309 | 0.121622625 | 0.618791 | 0.751194 | 44.26639 | 51.59233 | 56.31125 | 61.24627 | 51.6809 | 59.22083 | 39.53586 | 46.76642 |
| ENSMUSG00000103332 | -0.012563714 | 0.973811 | 0.985583 | 24.1453 | 29.99554 | 42.23344 | 19.84034 | 34.1289 | 17.87799 | 35.6787 | 29.36496 |
| ENSMUSG00000103334 | 0.437568112 | 0.116697 | 0.244967 | 184.1079 | 277.1588 | 163.137 | 226.0074 | 171.6196 | 227.9443 | 123.429 | 104.4087 |
| ENSMUSG00000103364 | 1.200375964 | 0.045252 | 0.126946 | 106.6418 | 99.58519 | 60.45178 | 84.5371 | 15.60178 | 101.681 | 16.39292 | 19.57664 |
| ENSMUSG00000103388 | 0.289532128 | 0.228697 | 0.3887 | 83.50251 | 56.39161 | 69.56096 | 68.14726 | 62.40712 | 70.39457 | 47.25017 | 47.85401 |
| ENSMUSG00000103400 | -0.029906894 | 0.951605 | 0.972579 | 7.04238 | 19.19714 | 23.18699 | 20.70296 | 17.552 | 10.05637 | 20.25007 | 23.927 |
| ENSMUSG00000103409 | 0.056490913 | 0.912779 | 0.948443 | 11.0666 | 9.598572 | 9.109173 | 12.07673 | 6.825779 | 6.704245 | 11.57147 | 15.22628 |
| ENSMUSG00000103421 | -0.577717826 | 0.019937 | 0.070247 | 47.28455 | 50.3925 | 63.76421 | 56.07053 | 70.20801 | 61.45558 | 89.67889 | 104.4087 |
| ENSMUSG00000103436 | -0.211586648 | 0.440473 | 0.603785 | 36.21796 | 28.79572 | 27.32752 | 29.3292 | 34.1289 | 36.87335 | 31.82154 | 38.06569 |
| ENSMUSG00000103445 | 0.379530282 | 0.317921 | 0.486219 | 29.17558 | 35.99465 | 19.87456 | 18.11509 | 16.57689 | 21.23011 | 23.14294 | 17.40146 |
| ENSMUSG00000103469 | -0.502568583 | 0.195871 | 0.350019 | 23.13925 | 9.598572 | 22.35888 | 29.3292 | 28.27823 | 25.6996 | 37.60728 | 29.36496 |
| ENSMUSG00000103472 | -0.424524635 | 0.110348 | 0.23618 | 87.52673 | 43.19358 | 61.27989 | 67.28463 | 105.312 | 69.27719 | 94.50034 | 79.39415 |
| ENSMUSG00000103482 | 2.113056958 | 0.049078 | 0.134408 | 31.18769 | 31.19536 | 15.73403 | 25.01608 | 3.900445 | 20.11273 | 0 | 0 |
| ENSMUSG00000103486 | -0.322771921 | 0.290904 | 0.458022 | 22.1332 | 25.19625 | 33.12427 | 21.56559 | 28.27823 | 32.40385 | 33.75012 | 33.71532 |
| ENSMUSG00000103509 | 0.393031593 | 0.311271 | 0.47993 | 35.2119 | 21.59679 | 21.53077 | 18.11509 | 14.62667 | 15.64324 | 21.21436 | 21.75182 |
| ENSMUSG00000103520 | 0.232924751 | 0.327779 | 0.495558 | 108.6539 | 152.3773 | 148.2311 | 125.943 | 97.51113 | 158.6671 | 115.7147 | 83.74451 |
| ENSMUSG00000103529 | 0.621129459 | 0.05144 | 0.138718 | 39.23612 | 45.59322 | 37.2648 | 49.16954 | 18.52711 | 40.22547 | 27.96439 | 25.0146 |
| ENSMUSG00000103532 | 0.016639399 | 0.978954 | 0.988554 | 9.054489 | 25.19625 | 17.39024 | 12.07673 | 6.825779 | 18.99536 | 6.750024 | 30.45255 |
| ENSMUSG00000103546 | 1.246494411 | 0.001447 | 0.010317 | 68.4117 | 59.99108 | 52.99882 | 64.69676 | 11.70134 | 45.81234 | 26.03581 | 20.66423 |
| ENSMUSG00000103558 | 0.469928697 | 0.28716 | 0.454071 | 27.16347 | 31.19536 | 29.81184 | 25.01608 | 11.70134 | 37.99072 | 19.28578 | 13.05109 |
| ENSMUSG00000103562 | 0.770066871 | 0.116252 | 0.244368 | 9.054489 | 22.79661 | 18.21835 | 18.97772 | 6.825779 | 13.40849 | 11.57147 | 8.700729 |
| ENSMUSG00000103567 | -0.176420667 | 0.517126 | 0.671078 | 43.26034 | 37.19447 | 46.37397 | 27.60395 | 39.00445 | 42.46022 | 45.32159 | 47.85401 |
| ENSMUSG00000103574 | 0.707662365 | 0.013153 | 0.052465 | 48.29061 | 45.59322 | 43.06154 | 60.38364 | 23.40267 | 37.99072 | 33.75012 | 26.10219 |
| ENSMUSG00000103585 | 0.442248952 | 0.115651 | 0.243632 | 70.4238 | 77.9884 | 51.34261 | 51.75741 | 38.02934 | 63.69032 | 40.50014 | 42.41605 |
| ENSMUSG00000103590 | -0.034355717 | 0.959354 | 0.97709 | 8.048435 | 21.59679 | 10.76539 | 17.25247 | 19.50223 | 24.58223 | 12.53576 | 2.175182 |
| ENSMUSG00000103591 | 0.842450123 | 0.256606 | 0.42028 | 15.09082 | 20.39697 | 11.59349 | 10.35148 | 4.875557 | 21.23011 | 3.857157 | 2.175182 |
| ENSMUSG00000103593 | 0.228965236 | 0.653202 | 0.776071 | 22.1332 | 9.598572 | 13.24971 | 12.07673 | 11.70134 | 8.938993 | 8.678602 | 19.57664 |
| ENSMUSG00000103621 | 1.512295043 | 0.009722 | 0.041725 | 35.2119 | 41.99375 | 26.49941 | 43.13117 | 6.825779 | 29.05173 | 12.53576 | 3.262773 |
| ENSMUSG00000103622 | -0.049361198 | 0.804085 | 0.880201 | 92.557 | 142.7788 | 113.4506 | 100.9269 | 101.4116 | 127.3806 | 105.1075 | 130.5109 |
| ENSMUSG00000103646 | 1.30968553 | 0.051263 | 0.138441 | 23.13925 | 17.99732 | 14.90592 | 22.42821 | 1.950223 | 18.99536 | 7.714313 | 3.262773 |
| ENSMUSG00000103649 | 0.652518353 | 0.011225 | 0.046557 | 117.7084 | 105.5843 | 78.67013 | 113.0037 | 69.2329 | 88.27255 | 61.71451 | 44.59123 |
| ENSMUSG00000103653 | 0.696795082 | 0.261901 | 0.426134 | 5.030272 | 19.19714 | 28.98373 | 22.42821 | 13.65156 | 18.99536 | 6.750024 | 7.613138 |
| ENSMUSG00000103657 | 1.33970802 | 0.053622 | 0.143075 | 28.16952 | 23.99643 | 9.93728 | 16.38985 | 2.925334 | 18.99536 | 3.857157 | 5.437955 |
| ENSMUSG00000103662 | 0.492437717 | 0.158348 | 0.301192 | 48.29061 | 74.38894 | 62.9361 | 74.18562 | 46.80534 | 75.98144 | 31.82154 | 30.45255 |
| ENSMUSG00000103668 | 0.184591226 | 0.696743 | 0.808595 | 13.07871 | 14.39786 | 19.87456 | 17.25247 | 5.850668 | 15.64324 | 17.3572 | 18.48905 |
| ENSMUSG00000103672 | 0.681377814 | 0.052533 | 0.140861 | 67.40564 | 43.19358 | 52.17072 | 35.36756 | 20.47734 | 43.57759 | 34.71441 | 25.0146 |
| ENSMUSG00000103677 | -0.111627421 | 0.671498 | 0.789547 | 40.24217 | 53.99197 | 43.88965 | 48.30692 | 66.30757 | 39.10809 | 49.17875 | 45.67883 |
| ENSMUSG00000103686 | 0.497877064 | 0.202034 | 0.35757 | 41.24823 | 44.3934 | 19.87456 | 37.95543 | 33.15378 | 21.23011 | 29.89296 | 16.31387 |
| ENSMUSG00000103693 | 1.210046803 | 0.044839 | 0.126171 | 16.09687 | 46.79304 | 14.90592 | 12.07673 | 6.825779 | 14.52586 | 10.60718 | 6.525547 |
| ENSMUSG00000103697 | 0.468267475 | 0.173801 | 0.321743 | 37.22401 | 33.595 | 40.57722 | 32.77969 | 16.57689 | 40.22547 | 23.14294 | 25.0146 |
| ENSMUSG00000103711 | 0.260639716 | 0.432837 | 0.597133 | 62.37537 | 25.19625 | 37.2648 | 44.85642 | 33.15378 | 31.28647 | 39.53586 | 38.06569 |
| ENSMUSG00000103715 | -0.108082803 | 0.578458 | 0.719984 | 115.6963 | 85.18733 | 78.67013 | 90.57547 | 86.78491 | 99.44629 | 108.9647 | 103.3212 |
| ENSMUSG00000103726 | -0.424852383 | 0.369619 | 0.538498 | 20.12109 | 13.19804 | 9.93728 | 12.93935 | 15.60178 | 10.05637 | 22.17865 | 27.18978 |
| ENSMUSG00000103731 | 0.695005966 | 0.138411 | 0.273942 | 15.09082 | 14.39786 | 19.04645 | 18.11509 | 11.70134 | 15.64324 | 8.678602 | 5.437955 |
| ENSMUSG00000103738 | 1.573620065 | 0.013785 | 0.054173 | 19.11503 | 25.19625 | 17.39024 | 6.900988 | 1.950223 | 5.58687 | 6.750024 | 8.700729 |
| ENSMUSG00000103739 | 0.330545016 | 0.513923 | 0.668426 | 16.09687 | 7.198929 | 12.4216 | 17.25247 | 8.776002 | 15.64324 | 10.60718 | 7.613138 |

| | | | | | | | | | | | |
|---|---|---|---|---|---|---|---|---|---|---|---|
| ENSMUSG00000103746 | 0.022162456 | 0.949609 | 0.971453 | 45.27245 | 32.39518 | 42.23344 | 37.95543 | 18.52711 | 46.92971 | 48.21446 | 42.41605 |
| ENSMUSG00000103749 | 0.152372263 | 0.608025 | 0.74375 | 56.33904 | 37.19447 | 40.57722 | 38.81806 | 50.70579 | 27.93435 | 40.50014 | 35.89051 |
| ENSMUSG00000103751 | 1.247004951 | 0.136828 | 0.271626 | 10.06054 | 22.79661 | 10.76539 | 13.80198 | 2.925334 | 14.52586 | 6.750024 | 0 |
| ENSMUSG00000103755 | -0.028336908 | 0.953003 | 0.973191 | 22.1332 | 11.99822 | 9.109173 | 18.11509 | 11.70134 | 12.29112 | 23.14294 | 15.22628 |
| ENSMUSG00000103756 | -0.201888337 | 0.623785 | 0.754981 | 22.1332 | 17.99732 | 19.04645 | 9.488858 | 17.552 | 24.58223 | 20.25007 | 16.31387 |
| ENSMUSG00000103793 | 0.297948284 | 0.2287 | 0.3887 | 68.4117 | 75.58876 | 45.54586 | 68.14726 | 56.55646 | 58.10345 | 48.21446 | 45.67883 |
| ENSMUSG00000103804 | 0.223962745 | 0.728853 | 0.829737 | 14.08476 | 16.7975 | 8.281066 | 8.626235 | 9.751113 | 10.05637 | 18.32149 | 2.175182 |
| ENSMUSG00000103818 | 1.025784995 | 0.174527 | 0.322656 | 27.16347 | 20.39697 | 10.76539 | 22.42821 | 3.900445 | 27.93435 | 4.821446 | 3.262773 |
| ENSMUSG00000103821 | 0.428220995 | 0.255663 | 0.419581 | 41.24823 | 23.99643 | 22.35888 | 25.8787 | 17.552 | 20.11273 | 28.92867 | 17.40146 |
| ENSMUSG00000103822 | 0.804427642 | 0.248151 | 0.411541 | 18.10898 | 11.99822 | 21.53077 | 12.07673 | 8.776002 | 20.11273 | 6.750024 | 1.087591 |
| ENSMUSG00000103828 | 1.361512789 | 0.070718 | 0.173673 | 21.12714 | 22.79661 | 15.73403 | 17.25247 | 1.950223 | 21.23011 | 4.821446 | 2.175182 |
| ENSMUSG00000103835 | 0.462211297 | 0.470706 | 0.630616 | 25.15136 | 10.79839 | 5.796746 | 16.38985 | 6.825779 | 20.11273 | 6.750024 | 8.700729 |
| ENSMUSG00000103839 | 0.828064246 | 0.006565 | 0.031504 | 146.8839 | 122.3818 | 77.84202 | 106.9653 | 62.40712 | 96.09417 | 47.25017 | 50.02919 |
| ENSMUSG00000103851 | 1.082392169 | 0.050783 | 0.13752 | 69.41775 | 133.1802 | 84.46688 | 77.63611 | 26.32801 | 106.1505 | 18.32149 | 21.75182 |
| ENSMUSG00000103857 | 0.801096813 | 0.119615 | 0.248522 | 26.15741 | 46.79304 | 32.29616 | 51.75741 | 14.62667 | 48.04709 | 11.57147 | 16.31387 |
| ENSMUSG00000103864 | -0.226766557 | 0.681548 | 0.79652 | 26.15741 | 19.19714 | 13.24971 | 5.175741 | 15.60178 | 16.76061 | 13.50005 | 28.27737 |
| ENSMUSG00000103867 | 1.069706353 | 0.115347 | 0.243212 | 25.15136 | 22.79661 | 15.73403 | 7.763611 | 6.825779 | 18.99536 | 2.892867 | 5.437955 |
| ENSMUSG00000103882 | 1.267897754 | 0.082435 | 0.192948 | 42.25428 | 17.99732 | 13.24971 | 15.52722 | 4.875557 | 22.34748 | 7.714313 | 2.175182 |
| ENSMUSG00000103891 | 0.000234143 | 0.999525 | 0.999733 | 117.7084 | 98.38537 | 48.85829 | 74.18562 | 86.78491 | 128.498 | 83.89316 | 39.15328 |
| ENSMUSG00000103897 | -0.059062118 | 0.89256 | 0.934962 | 18.10898 | 29.99554 | 28.98373 | 12.07673 | 20.47734 | 14.52586 | 27.96439 | 29.36496 |
| ENSMUSG00000103898 | 0.046989415 | 0.892058 | 0.934807 | 78.47224 | 62.39072 | 42.23344 | 45.71905 | 68.25779 | 75.98144 | 38.57157 | 38.06569 |
| ENSMUSG00000103901 | 0.333614148 | 0.401631 | 0.568792 | 41.24823 | 47.99286 | 20.70267 | 25.8787 | 35.10401 | 29.05173 | 25.07152 | 17.40146 |
| ENSMUSG00000103906 | -0.289446402 | 0.206166 | 0.362677 | 73.44197 | 65.99018 | 59.62368 | 56.93315 | 83.85957 | 54.75133 | 85.82174 | 87.00729 |
| ENSMUSG00000103916 | 1.026316002 | 0.144844 | 0.282942 | 110.666 | 159.5763 | 62.9361 | 79.36136 | 14.62667 | 145.2586 | 30.85725 | 11.9635 |
| ENSMUSG00000103928 | -0.067048236 | 0.858459 | 0.915187 | 79.47829 | 35.99465 | 32.29616 | 44.85642 | 51.6809 | 71.51194 | 39.53586 | 39.15328 |
| ENSMUSG00000103931 | 1.311344239 | 0.169594 | 0.316036 | 20.12109 | 11.99822 | 12.4216 | 16.38985 | 0 | 17.87799 | 5.785735 | 1.087591 |
| ENSMUSG00000103932 | 1.697796219 | 0.006836 | 0.032416 | 131.7931 | 208.7689 | 71.21717 | 99.2017 | 27.30312 | 102.7984 | 13.50005 | 14.13868 |
| ENSMUSG00000103945 | 0.348193392 | 0.415638 | 0.581533 | 44.26639 | 37.19447 | 14.90592 | 20.70296 | 23.40267 | 25.6996 | 19.28578 | 22.83941 |
| ENSMUSG00000103957 | 0.043341899 | 0.948173 | 0.970361 | 16.09687 | 11.99822 | 6.624853 | 6.900988 | 9.751113 | 2.234748 | 10.60718 | 17.40146 |
| ENSMUSG00000103959 | 2.043844335 | 0.060334 | 0.155584 | 44.26639 | 29.99554 | 18.21835 | 13.80198 | 0 | 20.11273 | 5.785735 | 0 |
| ENSMUSG00000103966 | 0.600030017 | 0.127405 | 0.258678 | 26.15741 | 16.7975 | 21.53077 | 31.91707 | 17.552 | 12.29112 | 13.50005 | 20.66423 |
| ENSMUSG00000103970 | 1.388162017 | 0.041718 | 0.120224 | 33.19979 | 16.7975 | 17.39024 | 18.97772 | 0.975111 | 17.87799 | 5.785735 | 8.700729 |
| ENSMUSG00000103973 | 0.812261007 | 0.123286 | 0.252903 | 31.18769 | 38.39429 | 26.49941 | 17.25247 | 9.751113 | 33.52122 | 11.57147 | 9.78832 |
| ENSMUSG00000103976 | 0.887795745 | 0.218884 | 0.377684 | 18.10898 | 22.79661 | 12.4216 | 18.97772 | 2.925334 | 23.46486 | 10.60718 | 2.175182 |
| ENSMUSG00000103983 | 0.832201565 | 0.123879 | 0.253569 | 229.3804 | 190.7716 | 112.6225 | 113.8663 | 78.0089 | 208.949 | 49.17875 | 27.18978 |
| ENSMUSG00000103984 | 0.870778444 | 0.325943 | 0.493811 | 14.08476 | 16.7975 | 19.87456 | 12.93935 | 1.950223 | 26.81698 | 0.964289 | 5.437955 |
| ENSMUSG00000104002 | 0.172647574 | 0.657512 | 0.779315 | 37.22401 | 19.19714 | 19.87456 | 19.84034 | 26.32801 | 20.11273 | 22.17865 | 16.31387 |
| ENSMUSG00000104010 | 0.497370615 | 0.368844 | 0.537581 | 27.16347 | 23.99643 | 8.281066 | 8.626235 | 9.751113 | 13.40849 | 12.53576 | 11.9635 |
| ENSMUSG00000104011 | -0.688208273 | 0.087023 | 0.200411 | 34.20585 | 8.398751 | 32.29616 | 31.05445 | 42.9049 | 41.34284 | 50.14304 | 38.06569 |
| ENSMUSG00000104018 | -0.207629801 | 0.683631 | 0.798398 | 8.048435 | 16.7975 | 6.624853 | 19.84034 | 16.57689 | 15.64324 | 11.57147 | 15.22628 |
| ENSMUSG00000104028 | -0.030210652 | 0.952274 | 0.972832 | 14.08476 | 17.99732 | 9.93728 | 23.29083 | 19.50223 | 6.704245 | 17.3572 | 22.83941 |
| ENSMUSG00000104043 | 1.274495686 | 0.084736 | 0.196541 | 12.07265 | 19.19714 | 17.39024 | 27.60395 | 0.975111 | 17.87799 | 9.642892 | 3.262773 |
| ENSMUSG00000104044 | 0.897837237 | 0.275993 | 0.442202 | 13.07871 | 16.7975 | 11.59349 | 19.84034 | 1.950223 | 22.34748 | 7.714313 | 1.087591 |
| ENSMUSG00000104060 | 1.288147663 | 0.011745 | 0.048169 | 15.09082 | 14.39786 | 14.90592 | 13.80198 | 5.850668 | 8.938993 | 4.821446 | 4.350364 |
| ENSMUSG00000104063 | 0.372443754 | 0.1395 | 0.275505 | 79.47829 | 119.9822 | 85.29498 | 75.04824 | 78.0089 | 68.15982 | 81.00029 | 48.9416 |
| ENSMUSG00000104090 | -0.548799805 | 0.131609 | 0.264536 | 27.16347 | 20.39697 | 17.39024 | 15.52722 | 38.02934 | 30.1691 | 27.96439 | 20.66423 |
| ENSMUSG00000104091 | 1.376040329 | 0.001715 | 0.011699 | 24.1453 | 25.19625 | 25.67131 | 34.50494 | 11.70134 | 15.64324 | 10.60718 | 4.350364 |
| ENSMUSG00000104094 | 0.215501437 | 0.53776 | 0.687843 | 32.19374 | 23.99643 | 32.29616 | 25.8787 | 18.52711 | 36.87335 | 23.14294 | 20.66423 |
| ENSMUSG00000104097 | 1.10225198 | 0.012318 | 0.049966 | 17.10292 | 21.59679 | 17.39024 | 24.15346 | 5.850668 | 12.29112 | 9.642892 | 9.78832 |
| ENSMUSG00000104107 | 0.507682177 | 0.28986 | 0.457118 | 34.20585 | 28.79572 | 18.21835 | 47.44429 | 27.30312 | 36.87335 | 13.50005 | 13.05109 |
| ENSMUSG00000104113 | -0.455944969 | 0.322818 | 0.49089 | 8.048435 | 13.19804 | 14.90592 | 10.35148 | 16.57689 | 10.05637 | 15.42863 | 21.75182 |
| ENSMUSG00000104126 | 0.078174664 | 0.879587 | 0.927743 | 8.048435 | 22.79661 | 28.15563 | 31.05445 | 10.72622 | 22.34748 | 30.85725 | 21.75182 |
| ENSMUSG00000104145 | 1.198127447 | NA | NA | 120.7265 | 248.3631 | 104.3414 | 127.6683 | 23.40267 | 194.4231 | 27.0001 | 17.40146 |
| ENSMUSG00000104156 | 0.539173985 | 0.00021 | 0.002368 | 179.0777 | 167.975 | 197.0894 | 191.5024 | 120.9138 | 147.4934 | 124.3933 | 115.2847 |
| ENSMUSG00000104164 | 0.731519456 | 0.142917 | 0.280395 | 21.12714 | 19.19714 | 15.73403 | 15.52722 | 9.751113 | 18.99536 | 4.821446 | 9.78832 |
| ENSMUSG00000104168 | 0.145648844 | 0.767235 | 0.855545 | 36.21796 | 21.59679 | 18.21835 | 18.97772 | 12.67645 | 12.29112 | 37.60728 | 22.83941 |
| ENSMUSG00000104184 | 0.915236762 | 0.00142 | 0.010174 | 111.672 | 166.7752 | 154.0278 | 125.943 | 73.13335 | 115.0895 | 54.96448 | 53.29196 |
| ENSMUSG00000104195 | -0.377643486 | 0.059875 | 0.15464 | 74.44802 | 61.1909 | 75.3577 | 63.83414 | 98.48624 | 94.9768 | 71.3574 | 93.53283 |
| ENSMUSG00000104197 | 0.892230707 | 0.144579 | 0.28269 | 21.12714 | 23.99643 | 15.73403 | 21.56559 | 9.751113 | 25.6996 | 4.821446 | 4.350364 |
| ENSMUSG00000104204 | 0.221783856 | 0.489689 | 0.647168 | 62.37537 | 52.79215 | 54.65504 | 75.04824 | 64.35735 | 74.86406 | 37.60728 | 33.71532 |
| ENSMUSG00000104212 | 0.56854612 | 0.185701 | 0.337279 | 15.09082 | 16.7975 | 15.73403 | 15.52722 | 9.751113 | 14.52586 | 8.678602 | 9.78832 |
| ENSMUSG00000104231 | 0.396932151 | 0.282912 | 0.449379 | 44.26639 | 32.39518 | 49.6864 | 25.8787 | 39.00445 | 31.28647 | 17.3572 | 28.27737 |
| ENSMUSG00000104241 | 0.293856387 | 0.490218 | 0.647606 | 29.17558 | 16.7975 | 14.07781 | 20.70296 | 22.42756 | 12.29112 | 13.50005 | 17.40146 |
| ENSMUSG00000104244 | 1.086129976 | 0.016081 | 0.060413 | 95.57516 | 157.1766 | 94.40416 | 119.9047 | 36.07912 | 108.3853 | 54.96448 | 20.66423 |

| | | | | | | | | | | | |
|---|---|---|---|---|---|---|---|---|---|---|---|
| ENSMUSG00000104245 | 0.410188712 | 0.172455 | 0.319892 | 45.27245 | 40.79393 | 52.17072 | 62.97152 | 38.02934 | 52.51658 | 35.6787 | 26.10219 |
| ENSMUSG00000104253 | -0.126403351 | 0.698354 | 0.80965 | 33.19979 | 31.19536 | 33.95237 | 33.64232 | 33.15378 | 54.75133 | 32.78583 | 23.927 |
| ENSMUSG00000104263 | 0.949667729 | 0.006591 | 0.031598 | 65.39353 | 68.38983 | 68.73285 | 58.6584 | 14.62667 | 43.57759 | 30.85725 | 46.76642 |
| ENSMUSG00000104272 | 0.485809089 | 0.392745 | 0.560119 | 28.16952 | 34.79482 | 10.76539 | 16.38985 | 5.850668 | 25.6996 | 18.32149 | 14.13868 |
| ENSMUSG00000104273 | -0.232323024 | 0.527811 | 0.680556 | 20.12109 | 31.19536 | 22.35888 | 25.8787 | 30.22845 | 43.57759 | 21.21436 | 21.75182 |
| ENSMUSG00000104277 | 1.046189979 | 0.083658 | 0.194849 | 43.26034 | 59.99108 | 34.78048 | 58.6584 | 12.67645 | 61.45558 | 10.60718 | 10.87591 |
| ENSMUSG00000104279 | 0.906930288 | 0.003279 | 0.018947 | 60.36326 | 95.98572 | 84.46688 | 56.93315 | 33.15378 | 45.81234 | 52.07161 | 27.18978 |
| ENSMUSG00000104292 | 0.998702838 | 0.192433 | 0.345566 | 42.25428 | 38.39429 | 19.87456 | 19.84034 | 5.850668 | 43.57759 | 7.714313 | 3.262773 |
| ENSMUSG00000104293 | 1.264372284 | 0.018191 | 0.066063 | 64.38748 | 49.19268 | 29.81184 | 34.50494 | 16.57689 | 39.10809 | 7.714313 | 10.87591 |
| ENSMUSG00000104336 | -1.290766816 | 0.070971 | 0.174105 | 1.006054 | 9.598572 | 9.93728 | 3.450494 | 16.57689 | 6.704245 | 14.46434 | 20.66423 |
| ENSMUSG00000104339 | 0.981974931 | 0.031015 | 0.096765 | 149.9021 | 98.38537 | 60.45178 | 75.91087 | 30.22845 | 97.21155 | 32.78583 | 34.80291 |
| ENSMUSG00000104342 | 0.479113214 | 0.085325 | 0.197461 | 112.6781 | 135.5798 | 104.3414 | 120.7673 | 65.33246 | 136.3196 | 65.57166 | 72.8686 |
| ENSMUSG00000104344 | -0.252724311 | 0.141409 | 0.278205 | 92.557 | 100.785 | 80.32634 | 79.36136 | 110.1876 | 105.0332 | 107.0361 | 95.70802 |
| ENSMUSG00000104346 | 0.169287978 | 0.572065 | 0.714829 | 75.45408 | 119.9822 | 91.09173 | 55.2079 | 81.90935 | 50.28183 | 81.00029 | 89.18247 |
| ENSMUSG00000104347 | 0.142903251 | 0.753024 | 0.846365 | 16.09687 | 11.99822 | 25.67131 | 25.01608 | 16.57689 | 26.81698 | 10.60718 | 18.48905 |
| ENSMUSG00000104361 | -0.634155534 | 0.222466 | 0.381724 | 12.07265 | 7.198929 | 10.76539 | 6.900988 | 10.72622 | 16.76061 | 21.21436 | 8.700729 |
| ENSMUSG00000104375 | -7.182620914 | 1.65E-10 | 2.80E-08 | 0 | 0 | 0.828107 | 0 | 24.37778 | 33.52122 | 60.75022 | 51.11678 |
| ENSMUSG00000104376 | -1.036254065 | 0.010251 | 0.04344 | 16.09687 | 14.39786 | 19.04645 | 7.763611 | 23.40267 | 24.58223 | 35.6787 | 33.71532 |
| ENSMUSG00000104378 | 1.04697365 | 0.225084 | 0.384785 | 20.12109 | 16.7975 | 9.109173 | 9.488858 | 2.925334 | 20.11273 | 2.892867 | 1.087591 |
| ENSMUSG00000104379 | 0.491772745 | 0.224402 | 0.384119 | 21.12714 | 22.79661 | 14.90592 | 22.42821 | 11.70134 | 13.40849 | 20.25007 | 11.9635 |
| ENSMUSG00000104383 | 1.32902576 | 0.018966 | 0.067955 | 28.16952 | 25.19625 | 25.67131 | 30.19182 | 5.850668 | 25.6996 | 4.821446 | 7.613138 |
| ENSMUSG00000104388 | 1.187705802 | 0.122065 | 0.251544 | 99.59938 | 128.3809 | 33.12427 | 54.34528 | 18.52711 | 98.32892 | 17.3572 | 4.350364 |
| ENSMUSG00000104392 | 1.136225692 | 0.007635 | 0.035002 | 75.45408 | 55.19179 | 50.5145 | 65.55939 | 13.65156 | 46.92971 | 36.64299 | 15.22628 |
| ENSMUSG00000104394 | 0.228116706 | 0.529075 | 0.681503 | 157.9505 | 173.9741 | 190.4645 | 166.4863 | 105.312 | 289.3999 | 96.42892 | 97.8832 |
| ENSMUSG00000104399 | 0.532739994 | 0.099 | 0.219557 | 86.52067 | 94.7859 | 72.04528 | 101.7896 | 56.55646 | 102.7984 | 46.28588 | 40.24087 |
| ENSMUSG00000104402 | 0.89006029 | 0.126865 | 0.25798 | 31.18769 | 89.98662 | 36.43669 | 39.68068 | 13.65156 | 60.3382 | 18.32149 | 14.13868 |
| ENSMUSG00000104420 | 0.818621921 | 0.090862 | 0.206558 | 14.08476 | 20.39697 | 29.81184 | 18.11509 | 6.825779 | 18.99536 | 11.57147 | 9.78832 |
| ENSMUSG00000104432 | 0.66319952 | 0.114064 | 0.241446 | 101.6115 | 103.1847 | 63.76421 | 64.69676 | 36.07912 | 103.9158 | 34.71441 | 35.89051 |
| ENSMUSG00000104435 | 0.178664964 | 0.453623 | 0.61591 | 271.6347 | 303.5548 | 236.0104 | 278.6274 | 232.0765 | 362.0292 | 203.465 | 165.3138 |
| ENSMUSG00000104436 | 0.30659211 | 0.723765 | 0.827323 | 21.12714 | 13.19804 | 9.93728 | 1.725247 | 1.950223 | 20.11273 | 9.642892 | 5.437955 |
| ENSMUSG00000104443 | 0.122628709 | 0.804275 | 0.880231 | 14.08476 | 16.7975 | 14.90592 | 22.42821 | 5.850668 | 17.87799 | 15.42863 | 23.927 |
| ENSMUSG00000104444 | -2.264110225 | 0.001229 | 0.009119 | 19.11503 | 7.198929 | 67.90474 | 9.488858 | 118.9636 | 43.57759 | 96.42892 | 242.5328 |
| ENSMUSG00000104445 | -2.559638657 | 0.000116 | 0.001507 | 13.07871 | 5.999108 | 43.06154 | 6.038364 | 75.08357 | 45.81234 | 92.57176 | 192.5036 |
| ENSMUSG00000104449 | 1.255228051 | 0.047353 | 0.130969 | 35.2119 | 20.39697 | 35.60859 | 35.36756 | 9.751113 | 33.52122 | 4.821446 | 5.437955 |
| ENSMUSG00000104453 | 0.750836904 | 0.010827 | 0.045291 | 155.9384 | 136.7797 | 120.9036 | 201.8539 | 69.2329 | 146.376 | 77.14313 | 73.95619 |
| ENSMUSG00000104467 | 0.444802087 | 0.181687 | 0.3322 | 109.6599 | 129.5807 | 126.7003 | 129.3935 | 75.08357 | 162.0192 | 75.21455 | 52.20437 |
| ENSMUSG00000104476 | 1.109588539 | NA | NA | 77.46619 | 129.5807 | 90.26362 | 86.26235 | 28.27823 | 129.6154 | 12.53576 | 7.613138 |
| ENSMUSG00000104496 | -0.179904395 | 0.671009 | 0.789151 | 20.12109 | 23.99643 | 16.56213 | 10.35148 | 14.62667 | 24.58223 | 24.10723 | 16.31387 |
| ENSMUSG00000104515 | 0.679450813 | 0.393626 | 0.560943 | 24.1453 | 34.79482 | 6.624853 | 9.488858 | 4.875557 | 27.93435 | 10.60718 | 3.262773 |
| ENSMUSG00000104517 | 0.432064 | 0.275162 | 0.441292 | 22.1332 | 32.39518 | 31.46805 | 30.19182 | 18.52711 | 34.6386 | 21.21436 | 11.9635 |
| ENSMUSG00000104524 | 1.57215712 | 0.015808 | 0.059881 | 55.33299 | 73.18911 | 43.88965 | 37.95543 | 6.825779 | 42.46022 | 18.32149 | 3.262773 |
| ENSMUSG00000104528 | -0.039143889 | 0.911987 | 0.948028 | 26.15741 | 16.7975 | 22.35888 | 28.46658 | 17.552 | 22.34748 | 29.89296 | 27.18978 |
| ENSMUSG00000104546 | 0.132968618 | 0.606391 | 0.742598 | 84.50857 | 63.59054 | 57.13936 | 71.59775 | 66.30757 | 83.80306 | 47.25017 | 55.46715 |
| ENSMUSG00000104548 | 0.522825298 | 0.106026 | 0.230052 | 84.50857 | 86.38715 | 102.6852 | 116.4542 | 69.2329 | 107.2679 | 59.78593 | 35.89051 |
| ENSMUSG00000104569 | 0.50417276 | 0.27192 | 0.437195 | 25.15136 | 17.99732 | 13.24971 | 38.81806 | 12.67645 | 21.23011 | 18.32149 | 15.22628 |
| ENSMUSG00000104597 | 1.238218197 | 0.001133 | 0.0086 | 61.36932 | 57.59143 | 68.73285 | 98.33908 | 28.27823 | 52.51658 | 17.3572 | 23.927 |
| ENSMUSG00000104605 | -0.436898724 | 0.125879 | 0.256596 | 27.16347 | 45.59322 | 38.92101 | 33.64232 | 62.40712 | 45.81234 | 48.21446 | 39.15328 |
| ENSMUSG00000104610 | -0.013533149 | 0.979275 | 0.988704 | 15.09082 | 38.39429 | 11.59349 | 28.46658 | 16.57689 | 31.28647 | 32.78583 | 13.05109 |
| ENSMUSG00000104621 | 0.98947135 | 0.067583 | 0.167851 | 28.16952 | 19.19714 | 27.32752 | 25.01608 | 6.825779 | 26.81698 | 11.57147 | 5.437955 |
| ENSMUSG00000104626 | 1.199725057 | 0.112258 | 0.238801 | 45.27245 | 76.78858 | 49.6864 | 30.19182 | 4.875557 | 65.92507 | 8.678602 | 8.700729 |
| ENSMUSG00000104649 | 0.275915945 | 0.515146 | 0.669261 | 20.12109 | 15.59768 | 14.07781 | 12.93935 | 13.65156 | 10.05637 | 11.57147 | 16.31387 |
| ENSMUSG00000104671 | 0.491277045 | 0.228563 | 0.388662 | 39.23612 | 45.59322 | 41.40533 | 34.50494 | 20.47734 | 53.63396 | 23.14294 | 17.40146 |
| ENSMUSG00000104682 | 0.685431121 | 0.063375 | 0.161154 | 113.6841 | 119.9822 | 103.5133 | 106.9653 | 50.70579 | 134.0849 | 46.28588 | 45.67883 |
| ENSMUSG00000104684 | 0.564357448 | 0.287474 | 0.454404 | 20.12109 | 11.99822 | 33.95237 | 10.35148 | 15.60178 | 10.05637 | 16.39292 | 9.78832 |
| ENSMUSG00000104686 | -0.230713544 | 0.504481 | 0.660194 | 20.12109 | 20.39697 | 25.67131 | 30.19182 | 19.50223 | 26.81698 | 31.82154 | 35.89051 |
| ENSMUSG00000104693 | -1.772145159 | 0.002514 | 0.015492 | 4.024217 | 3.599465 | 14.07781 | 10.35148 | 21.45245 | 15.64324 | 45.32159 | 29.36496 |
| ENSMUSG00000104699 | -0.007323598 | 0.989804 | 0.994168 | 12.07265 | 4.799286 | 17.39024 | 15.52722 | 18.52711 | 14.52586 | 5.785735 | 11.9635 |
| ENSMUSG00000104713 | 1.690110796 | 0.007999 | 0.03622 | 32.19374 | 103.1847 | 126.7003 | 18.11509 | 8.776002 | 24.58223 | 32.78583 | 20.66423 |
| ENSMUSG00000104719 | 0.477619954 | 0.270796 | 0.435995 | 28.16952 | 34.79482 | 18.21835 | 35.36756 | 11.70134 | 33.52122 | 22.17865 | 16.31387 |
| ENSMUSG00000104720 | 0.650993093 | 0.108668 | 0.233561 | 28.16952 | 31.19536 | 23.18699 | 21.56559 | 13.65156 | 22.34748 | 21.21436 | 8.700729 |
| ENSMUSG00000104724 | 0.392933417 | 0.273799 | 0.439641 | 42.25428 | 29.99554 | 30.63995 | 37.09281 | 15.60178 | 37.99072 | 22.17865 | 31.54014 |
| ENSMUSG00000104728 | 0.170676555 | 0.611362 | 0.746052 | 44.26639 | 38.39429 | 63.76421 | 52.62003 | 39.97956 | 69.27719 | 29.89296 | 39.15328 |
| ENSMUSG00000104735 | 1.264044596 | 0.100383 | 0.221451 | 18.10898 | 32.39518 | 23.18699 | 6.038364 | 5.850668 | 20.11273 | 1.928578 | 5.437955 |
| ENSMUSG00000104737 | -1.471797779 | 0.00831 | 0.037333 | 28.16952 | 4.799286 | 10.76539 | 12.93935 | 36.07912 | 18.99536 | 57.85735 | 44.59123 |

| | | | | | | | | | | | |
|---|---|---|---|---|---|---|---|---|---|---|---|
| ENSMUSG00000104761 | -0.077544659 | 0.861885 | 0.917409 | 23.13925 | 10.79839 | 18.21835 | 23.29083 | 16.57689 | 32.40385 | 17.3572 | 14.13868 |
| ENSMUSG00000104784 | -0.322244782 | 0.278178 | 0.444354 | 49.29666 | 76.78858 | 38.92101 | 40.5433 | 53.63112 | 53.63396 | 73.28598 | 73.95619 |
| ENSMUSG00000104786 | 0.063926194 | 0.896892 | 0.937817 | 13.07871 | 19.19714 | 6.624853 | 13.80198 | 8.776002 | 13.40849 | 13.50005 | 14.13868 |
| ENSMUSG00000104795 | 0.493494454 | 0.086474 | 0.199496 | 85.51462 | 83.98751 | 56.31125 | 69.8725 | 41.92979 | 78.21619 | 46.28588 | 43.50364 |
| ENSMUSG00000104802 | -0.226840475 | 0.578914 | 0.720218 | 17.10292 | 45.59322 | 23.18699 | 19.84034 | 40.95467 | 26.81698 | 25.07152 | 29.36496 |
| ENSMUSG00000104814 | 0.603805717 | 0.283371 | 0.449884 | 102.6175 | 57.59143 | 24.01509 | 37.95543 | 18.52711 | 73.74669 | 28.92867 | 25.0146 |
| ENSMUSG00000104822 | 0.50862957 | 0.275244 | 0.441385 | 25.15136 | 22.79661 | 13.24971 | 11.21411 | 13.65156 | 13.40849 | 14.46434 | 8.700729 |
| ENSMUSG00000104830 | 0.773325532 | 0.082664 | 0.193291 | 21.12714 | 21.59679 | 14.90592 | 15.52722 | 10.72622 | 15.64324 | 8.678602 | 7.613138 |
| ENSMUSG00000104850 | 0.595288221 | 0.060855 | 0.156535 | 66.39959 | 47.99286 | 33.12427 | 50.89479 | 36.07912 | 39.10809 | 31.82154 | 23.927 |
| ENSMUSG00000104851 | 1.432647703 | 0.009502 | 0.04105 | 152.9203 | 134.38 | 90.26362 | 122.4925 | 25.35289 | 110.62 | 37.60728 | 11.9635 |
| ENSMUSG00000104852 | 0.826735492 | 0.178914 | 0.32856 | 90.54489 | 86.38715 | 58.79557 | 48.30692 | 21.45245 | 106.1505 | 15.42863 | 17.40146 |
| ENSMUSG00000104868 | 1.876500867 | 0.000105 | 0.00139 | 44.26639 | 53.99197 | 26.49941 | 28.46658 | 9.751113 | 17.87799 | 9.642892 | 4.350364 |
| ENSMUSG00000104871 | 1.81282216 | 0.000379 | 0.003694 | 75.45408 | 104.3845 | 62.108 | 66.42201 | 13.65156 | 50.28183 | 15.42863 | 8.700729 |
| ENSMUSG00000104891 | 0.607456452 | 0.264832 | 0.429083 | 79.47829 | 103.1847 | 67.90474 | 69.8725 | 29.25334 | 126.2633 | 37.60728 | 17.40146 |
| ENSMUSG00000104894 | 1.47034399 | 0.006544 | 0.031463 | 18.10898 | 31.19536 | 15.73403 | 19.84034 | 5.850668 | 10.05637 | 2.892867 | 11.9635 |
| ENSMUSG00000104897 | -0.290080399 | 0.608008 | 0.74375 | 11.0666 | 8.398751 | 4.140533 | 13.80198 | 8.776002 | 16.76061 | 11.57147 | 8.700729 |
| ENSMUSG00000104912 | 0.613778 | 0.495996 | 0.652653 | 10.06054 | 14.39786 | 12.4216 | 13.80198 | 7.80089 | 23.46486 | 0.964289 | 1.087591 |
| ENSMUSG00000104913 | -0.328524449 | 0.174444 | 0.322632 | 58.35115 | 67.19001 | 46.37397 | 56.07053 | 81.90935 | 51.39921 | 80.036 | 70.69342 |
| ENSMUSG00000104918 | 0.950483117 | 0.00034 | 0.003407 | 238.4349 | 219.5673 | 168.1056 | 244.1224 | 103.3618 | 172.0756 | 101.2504 | 73.95619 |
| ENSMUSG00000104943 | 0.960680337 | 0.036548 | 0.109661 | 44.26639 | 26.39607 | 18.21835 | 27.60395 | 8.776002 | 22.34748 | 11.57147 | 17.40146 |
| ENSMUSG00000104950 | 0.33455433 | 0.210217 | 0.367715 | 85.51462 | 81.58786 | 115.9349 | 113.8663 | 58.50668 | 109.5027 | 61.71451 | 87.00729 |
| ENSMUSG00000104951 | 0.59718242 | 0.447999 | 0.61061 | 10.06054 | 28.79572 | 10.76539 | 6.038364 | 3.900445 | 21.23011 | 2.892867 | 8.700729 |
| ENSMUSG00000104960 | 0.013652367 | 0.829915 | 0.897037 | 1772.668 | 1967.707 | 1842.537 | 1797.707 | 1912.193 | 1854.841 | 1763.685 | 1776.036 |
| ENSMUSG00000104965 | 0.006398525 | 0.988477 | 0.993483 | 26.15741 | 25.19625 | 29.81184 | 35.36756 | 9.751113 | 36.87335 | 29.89296 | 40.24087 |
| ENSMUSG00000104969 | 0.596306513 | 0.197351 | 0.351659 | 19.11503 | 19.19714 | 19.87456 | 18.11509 | 9.751113 | 17.87799 | 17.3572 | 5.437955 |
| ENSMUSG00000104970 | -0.211691568 | 0.682909 | 0.797858 | 8.048435 | 13.19804 | 14.90592 | 6.900988 | 7.80089 | 11.17374 | 13.50005 | 17.40146 |
| ENSMUSG00000104973 | 1.189371367 | 0.010281 | 0.043496 | 68.4117 | 80.38804 | 67.07664 | 66.42201 | 21.45245 | 68.15982 | 18.32149 | 16.31387 |
| ENSMUSG00000105006 | -0.361127672 | 0.232922 | 0.393519 | 34.20585 | 31.19536 | 38.92101 | 26.74133 | 30.22845 | 46.92971 | 37.60728 | 54.37955 |
| ENSMUSG00000105021 | 0.578014907 | 0.198039 | 0.35246 | 15.09082 | 27.5959 | 20.70267 | 21.56559 | 13.65156 | 20.11273 | 16.39292 | 6.525547 |
| ENSMUSG00000105031 | 0.549676463 | 0.335969 | 0.504269 | 9.054489 | 19.19714 | 9.109173 | 15.52722 | 4.875557 | 7.821619 | 15.42863 | 7.613138 |
| ENSMUSG00000105033 | 1.340120476 | 0.053788 | 0.143436 | 66.39959 | 74.38894 | 41.40533 | 48.30692 | 2.925334 | 53.63396 | 27.0001 | 7.613138 |
| ENSMUSG00000105041 | 1.141189162 | 0.115945 | 0.243888 | 53.32088 | 49.19268 | 32.29616 | 18.11509 | 15.60178 | 44.69496 | 4.821446 | 4.350364 |
| ENSMUSG00000105044 | 0.308280516 | 0.612312 | 0.74664 | 25.15136 | 7.198929 | 4.96864 | 18.97772 | 10.72622 | 11.17374 | 10.60718 | 13.05109 |
| ENSMUSG00000105048 | -0.045491449 | 0.936458 | 0.962308 | 13.07871 | 22.79661 | 9.109173 | 12.93935 | 10.72622 | 27.93435 | 6.750024 | 14.13868 |
| ENSMUSG00000105076 | 0.050926365 | 0.918911 | 0.952188 | 12.07265 | 15.59768 | 13.24971 | 18.11509 | 14.62667 | 7.821619 | 9.642892 | 25.0146 |
| ENSMUSG00000105095 | 0.150548368 | 0.365693 | 0.534297 | 134.8113 | 134.38 | 108.482 | 145.7834 | 114.088 | 131.8501 | 118.6076 | 106.5839 |
| ENSMUSG00000105100 | -0.009692991 | 0.98162 | 0.989771 | 23.13925 | 13.19804 | 27.32752 | 15.52722 | 14.62667 | 15.64324 | 26.03581 | 23.927 |
| ENSMUSG00000105107 | 1.275731857 | 2.66E-07 | 1.26E-05 | 71.42986 | 74.38894 | 77.01392 | 72.46037 | 32.17867 | 40.22547 | 27.0001 | 22.83941 |
| ENSMUSG00000105112 | -0.907069967 | 0.061158 | 0.157008 | 7.04238 | 11.99822 | 4.96864 | 12.07673 | 18.52711 | 13.40849 | 15.42863 | 19.57664 |
| ENSMUSG00000105135 | 0.183676663 | 0.543033 | 0.692189 | 67.40564 | 69.58965 | 35.60859 | 71.59775 | 46.80534 | 69.27719 | 45.32159 | 53.29196 |
| ENSMUSG00000105136 | 0.958704209 | 0.175931 | 0.324376 | 70.4238 | 51.59233 | 41.40533 | 55.2079 | 5.850668 | 81.56831 | 13.50005 | 11.9635 |
| ENSMUSG00000105139 | 0.715315008 | 0.013886 | 0.054426 | 53.32088 | 55.19179 | 52.99882 | 55.2079 | 27.30312 | 50.28183 | 29.89296 | 25.0146 |
| ENSMUSG00000105140 | -0.458972321 | 0.256944 | 0.42059 | 15.09082 | 10.79839 | 14.07781 | 12.07673 | 22.42756 | 13.40849 | 18.32149 | 17.40146 |
| ENSMUSG00000105156 | 0.500878449 | 0.371886 | 0.540904 | 17.10292 | 21.59679 | 15.73403 | 43.13117 | 12.67645 | 33.52122 | 14.46434 | 8.700729 |
| ENSMUSG00000105161 | 0.489466537 | 0.03272 | 0.100646 | 145.8779 | 139.1793 | 169.7619 | 167.349 | 106.2871 | 160.9019 | 92.57176 | 84.8321 |
| ENSMUSG00000105168 | -4.452351792 | 0.000377 | 0.003684 | 0 | 0 | 2.48432 | 0.862623 | 39.97956 | 1.117374 | 12.53576 | 25.0146 |
| ENSMUSG00000105176 | 0.481498636 | 0.299474 | 0.467605 | 85.51462 | 81.58786 | 58.79557 | 85.39973 | 38.02934 | 116.2069 | 47.25017 | 21.75182 |
| ENSMUSG00000105201 | 0.681704691 | 0.106609 | 0.230828 | 64.38748 | 81.58786 | 57.13936 | 87.12497 | 18.52711 | 82.68568 | 45.32159 | 34.80291 |
| ENSMUSG00000105224 | 1.330849747 | 0.024523 | 0.08183 | 134.8113 | 173.9741 | 87.7793 | 109.5532 | 24.37778 | 134.0849 | 20.25007 | 22.83941 |
| ENSMUSG00000105271 | 2.181750458 | 0.000386 | 0.003751 | 35.2119 | 47.99286 | 25.67131 | 25.8787 | 1.950223 | 7.821619 | 16.39292 | 3.262773 |
| ENSMUSG00000105272 | -0.070113574 | 0.726105 | 0.828686 | 79.47829 | 69.58965 | 85.29498 | 77.63611 | 64.35735 | 88.27255 | 81.96458 | 94.62042 |
| ENSMUSG00000105282 | 1.039601643 | 0.052996 | 0.141745 | 114.6902 | 106.7841 | 58.79557 | 87.9876 | 25.35289 | 108.3853 | 25.07152 | 20.66423 |
| ENSMUSG00000105302 | 1.545111415 | 0.000317 | 0.003241 | 34.20585 | 41.99375 | 32.29616 | 27.60395 | 12.67645 | 17.87799 | 11.57147 | 4.350364 |
| ENSMUSG00000105304 | 1.35442358 | 0.011219 | 0.046557 | 47.28455 | 50.3925 | 22.35888 | 28.46658 | 13.65156 | 29.05173 | 6.750024 | 8.700729 |
| ENSMUSG00000105340 | 0.312122622 | 0.250403 | 0.413922 | 64.38748 | 45.59322 | 45.54586 | 53.48266 | 39.97956 | 42.46022 | 54.00019 | 31.54014 |
| ENSMUSG00000105345 | 0.115327562 | 0.770807 | 0.857712 | 90.54489 | 49.19268 | 50.5145 | 23.29083 | 48.75557 | 52.51658 | 54.00019 | 41.32846 |
| ENSMUSG00000105349 | -0.13274097 | 0.755048 | 0.847926 | 45.27245 | 25.19625 | 24.01509 | 28.46658 | 34.1289 | 56.98608 | 22.17865 | 21.75182 |
| ENSMUSG00000105359 | 0.18089422 | 0.745991 | 0.841259 | 19.11503 | 9.598572 | 7.45296 | 10.35148 | 16.57689 | 7.821619 | 8.678602 | 7.613138 |
| ENSMUSG00000105366 | 1.163638806 | 0.000965 | 0.007629 | 91.55095 | 134.38 | 110.1382 | 127.6683 | 45.83023 | 92.74205 | 29.89296 | 39.15328 |
| ENSMUSG00000105368 | -0.402453037 | 0.316427 | 0.484686 | 17.10292 | 19.19714 | 9.93728 | 16.38985 | 14.62667 | 24.58223 | 21.21436 | 21.75182 |
| ENSMUSG00000105378 | 1.179198456 | 0.007795 | 0.035549 | 26.15741 | 22.79661 | 22.35888 | 30.19182 | 7.80089 | 20.11273 | 8.678602 | 8.700729 |
| ENSMUSG00000105383 | 0.076266802 | 0.646628 | 0.77119 | 152.9203 | 143.9786 | 170.59 | 125.0804 | 163.8187 | 132.9675 | 128.2505 | 137.0365 |
| ENSMUSG00000105388 | -0.129194661 | 0.405009 | 0.572028 | 204.229 | 259.1615 | 262.5098 | 229.4578 | 306.185 | 230.1791 | 279.6439 | 227.3065 |
| ENSMUSG00000105402 | -1.764631977 | 0.007532 | 0.034672 | 10.06054 | 5.999108 | 9.109173 | 8.626235 | 40.95467 | 4.469496 | 24.10723 | 45.67883 |

| | | | | | | | | | | | |
|---|---|---|---|---|---|---|---|---|---|---|---|
| ENSMUSG00000105403 | 1.163066058 | 0.000467 | 0.004346 | 77.46619 | 45.59322 | 56.31125 | 81.08661 | 23.40267 | 37.99072 | 35.6787 | 19.57664 |
| ENSMUSG00000105408 | 0.360747529 | 0.097261 | 0.216688 | 94.56911 | 61.1909 | 77.84202 | 87.9876 | 58.50668 | 72.62932 | 60.75022 | 59.81751 |
| ENSMUSG00000105419 | -0.259814698 | 0.46975 | 0.629746 | 23.13925 | 35.99465 | 23.18699 | 39.68068 | 25.35289 | 52.51658 | 35.6787 | 32.62773 |
| ENSMUSG00000105429 | 0.355572271 | 0.46271 | 0.62375 | 33.19979 | 34.79482 | 22.35888 | 31.91707 | 22.42756 | 46.92971 | 15.42863 | 10.87591 |
| ENSMUSG00000105434 | 1.082948179 | 0.018172 | 0.066019 | 79.47829 | 119.9822 | 67.07664 | 75.04824 | 20.47734 | 84.92043 | 28.92867 | 27.18978 |
| ENSMUSG00000105442 | -0.372730511 | 0.587639 | 0.727933 | 15.09082 | 2.399643 | 9.93728 | 11.21411 | 3.900445 | 21.23011 | 12.53576 | 13.05109 |
| ENSMUSG00000105444 | -2.453342035 | 0.000127 | 0.001607 | 3.018163 | 1.199822 | 8.281066 | 6.038364 | 27.30312 | 17.87799 | 15.42863 | 44.59123 |
| ENSMUSG00000105449 | 0.492800935 | 0.070414 | 0.173043 | 118.7144 | 93.58608 | 95.23226 | 100.0643 | 63.38224 | 110.62 | 49.17875 | 67.43065 |
| ENSMUSG00000105471 | 0.222571106 | 0.642518 | 0.768773 | 27.16347 | 9.598572 | 14.07781 | 22.42821 | 18.52711 | 21.23011 | 12.53576 | 10.87591 |
| ENSMUSG00000105504 | 0.324027902 | 0.533919 | 0.685162 | 17.10292 | 15.59768 | 35.60859 | 25.8787 | 19.50223 | 13.40849 | 33.75012 | 8.700729 |
| ENSMUSG00000105509 | -0.045798767 | 0.798478 | 0.876664 | 116.7023 | 106.7841 | 103.5133 | 91.43809 | 93.61069 | 126.2633 | 113.7861 | 97.8832 |
| ENSMUSG00000105519 | 0.653095614 | 0.179067 | 0.32881 | 54.32694 | 40.79393 | 50.5145 | 40.5433 | 13.65156 | 63.69032 | 23.14294 | 18.48905 |
| ENSMUSG00000105542 | 0.485854616 | 0.44996 | 0.612494 | 30.18163 | 17.99732 | 28.15563 | 25.01608 | 5.850668 | 45.81234 | 12.53576 | 8.700729 |
| ENSMUSG00000105550 | -0.254633152 | 0.612701 | 0.746981 | 15.09082 | 7.198929 | 18.21835 | 7.763611 | 9.751113 | 17.87799 | 15.42863 | 15.22628 |
| ENSMUSG00000105552 | 0.419024781 | 0.259188 | 0.423101 | 25.15136 | 37.19447 | 29.81184 | 27.60395 | 20.47734 | 34.6386 | 20.25007 | 14.13868 |
| ENSMUSG00000105556 | -0.259889757 | 0.430022 | 0.594816 | 60.36326 | 50.3925 | 33.12427 | 38.81806 | 56.55646 | 72.62932 | 55.92877 | 32.62773 |
| ENSMUSG00000105565 | -0.095710865 | 0.770738 | 0.857687 | 33.19979 | 52.79215 | 48.03018 | 27.60395 | 33.15378 | 55.8687 | 35.6787 | 47.85401 |
| ENSMUSG00000105568 | 2.092182739 | 0.004384 | 0.023381 | 54.32694 | 43.19358 | 20.70267 | 25.8787 | 2.925334 | 21.23011 | 8.678602 | 1.087591 |
| ENSMUSG00000105572 | 0.324439245 | 0.267723 | 0.43199 | 214.2896 | 154.777 | 244.2915 | 213.9306 | 158.9431 | 261.4655 | 143.6791 | 97.8832 |
| ENSMUSG00000105601 | -0.143895345 | 0.836863 | 0.901352 | 6.036326 | 9.598572 | 11.59349 | 11.21411 | 11.70134 | 12.29112 | 17.3572 | 1.087591 |
| ENSMUSG00000105607 | 0.650850675 | 0.299722 | 0.467835 | 39.23612 | 8.398751 | 14.90592 | 10.35148 | 14.62667 | 15.64324 | 9.642892 | 6.525547 |
| ENSMUSG00000105609 | -1.157027247 | 0.032878 | 0.100987 | 7.04238 | 8.398751 | 5.796746 | 6.900988 | 15.60178 | 6.704245 | 21.21436 | 18.48905 |
| ENSMUSG00000105613 | 0.092664008 | 0.8136 | 0.88683 | 22.1332 | 22.79661 | 30.63995 | 31.91707 | 12.67645 | 35.75597 | 25.07152 | 28.27737 |
| ENSMUSG00000105632 | 0.183510923 | 0.744815 | 0.840598 | 7.04238 | 7.198929 | 26.49941 | 14.6646 | 12.67645 | 11.17374 | 11.57147 | 14.13868 |
| ENSMUSG00000105636 | 0.445906934 | 0.585415 | 0.725713 | 18.10898 | 9.598572 | 13.24971 | 8.626235 | 1.950223 | 25.6996 | 5.785735 | 3.262773 |
| ENSMUSG00000105655 | 0.865730595 | 0.055905 | 0.147283 | 101.6115 | 98.38537 | 74.5296 | 57.79577 | 30.22845 | 96.09417 | 27.0001 | 29.36496 |
| ENSMUSG00000105677 | 0.686206722 | 0.009163 | 0.04 | 85.51462 | 82.78769 | 63.76421 | 103.5148 | 39.97956 | 69.27719 | 54.00019 | 45.67883 |
| ENSMUSG00000105679 | 0.347476754 | 0.487796 | 0.645722 | 49.29666 | 44.3934 | 18.21835 | 17.25247 | 17.552 | 39.10809 | 27.96439 | 16.31387 |
| ENSMUSG00000105691 | 1.149503442 | 0.049842 | 0.135799 | 40.24217 | 39.59411 | 38.92101 | 35.36756 | 3.900445 | 40.22547 | 10.60718 | 15.22628 |
| ENSMUSG00000105692 | 0.354866251 | 0.017383 | 0.063922 | 203.223 | 163.1757 | 164.7932 | 207.8923 | 144.3165 | 153.0803 | 135.9648 | 145.7372 |
| ENSMUSG00000105699 | 0.128330037 | 0.803982 | 0.88014 | 15.09082 | 21.59679 | 8.281066 | 9.488858 | 10.72622 | 17.87799 | 8.678602 | 11.9635 |
| ENSMUSG00000105703 | 0.183153439 | 0.539817 | 0.689457 | 2495.015 | 1451.784 | 2230.919 | 3103.719 | 1301.774 | 1885.01 | 3368.262 | 1620.511 |
| ENSMUSG00000105708 | 0.58978596 | 0.127206 | 0.258402 | 24.1453 | 51.59233 | 32.29616 | 43.13117 | 21.45245 | 33.52122 | 29.89296 | 15.22628 |
| ENSMUSG00000105710 | -0.517730443 | 0.135817 | 0.270228 | 22.1332 | 21.59679 | 22.35888 | 18.11509 | 28.27823 | 43.57759 | 26.03581 | 22.83941 |
| ENSMUSG00000105719 | 0.132903864 | 0.771954 | 0.858471 | 10.06054 | 21.59679 | 26.49941 | 17.25247 | 12.67645 | 21.23011 | 12.53576 | 22.83941 |
| ENSMUSG00000105726 | 1.693682838 | 0.089975 | 0.205161 | 18.10898 | 15.59768 | 17.39024 | 12.07673 | 0 | 16.76061 | 1.928578 | 1.087591 |
| ENSMUSG00000105741 | 0.72417356 | 0.263934 | 0.428197 | 28.16952 | 9.598572 | 11.59349 | 18.97772 | 2.925334 | 15.64324 | 16.39292 | 6.525547 |
| ENSMUSG00000105742 | 0.247013903 | 0.560174 | 0.706138 | 31.18769 | 15.59768 | 14.90592 | 17.25247 | 12.67645 | 20.11273 | 16.39292 | 17.40146 |
| ENSMUSG00000105746 | 1.032804822 | 0.119629 | 0.248522 | 26.15741 | 3.599465 | 24.01509 | 20.70296 | 6.825779 | 16.76061 | 6.750024 | 6.525547 |
| ENSMUSG00000105748 | 1.297144401 | 0.033984 | 0.103683 | 43.26034 | 25.19625 | 15.73403 | 22.42821 | 7.80089 | 24.58223 | 5.785735 | 5.437955 |
| ENSMUSG00000105802 | 0.33977656 | 0.429784 | 0.59461 | 19.11503 | 11.99822 | 19.87456 | 19.84034 | 20.47734 | 13.40849 | 12.53576 | 9.78832 |
| ENSMUSG00000105813 | -0.764382417 | 0.008386 | 0.037578 | 25.15136 | 29.99554 | 48.03018 | 34.50494 | 53.63112 | 71.51194 | 60.75022 | 50.02919 |
| ENSMUSG00000105814 | 0.219747785 | 0.489049 | 0.646694 | 103.6236 | 105.5843 | 81.98256 | 84.5371 | 72.15824 | 136.3196 | 58.82164 | 55.46715 |
| ENSMUSG00000105837 | -0.228531452 | 0.653147 | 0.776057 | 11.0666 | 10.79839 | 13.24971 | 7.763611 | 8.776002 | 20.11273 | 8.678602 | 13.05109 |
| ENSMUSG00000105851 | -0.322562105 | 0.228701 | 0.3887 | 48.29061 | 49.19268 | 37.2648 | 29.3292 | 57.53157 | 46.92971 | 53.0359 | 45.67883 |
| ENSMUSG00000105852 | 0.407259507 | 0.329557 | 0.497331 | 33.19979 | 28.79572 | 24.8432 | 29.3292 | 24.37778 | 36.87335 | 13.50005 | 13.05109 |
| ENSMUSG00000105861 | 0.875132549 | 0.030209 | 0.09495 | 55.33299 | 34.79482 | 33.95237 | 50.89479 | 14.62667 | 41.34284 | 18.32149 | 21.75182 |
| ENSMUSG00000105868 | 0.863609424 | 0.076996 | 0.184275 | 39.23612 | 23.99643 | 26.49941 | 33.64232 | 7.80089 | 31.28647 | 19.28578 | 9.78832 |
| ENSMUSG00000105873 | 0.595478681 | 0.297233 | 0.465079 | 14.08476 | 13.19804 | 16.56213 | 22.42821 | 4.875557 | 22.34748 | 10.60718 | 6.525547 |
| ENSMUSG00000105879 | -0.392187715 | 0.08154 | 0.191438 | 97.58727 | 124.7814 | 63.76421 | 113.8663 | 133.5902 | 124.0285 | 132.1076 | 132.6861 |
| ENSMUSG00000105881 | 0.260899973 | 0.039094 | 0.1146 | 306.8466 | 296.3559 | 260.8536 | 271.7264 | 201.848 | 255.8787 | 246.858 | 242.5328 |
| ENSMUSG00000105890 | 1.932392029 | 0.019884 | 0.070169 | 19.11503 | 19.19714 | 7.45296 | 19.84034 | 1.950223 | 10.05637 | 0 | 5.437955 |
| ENSMUSG00000105895 | 0.403752713 | 0.427923 | 0.592924 | 15.09082 | 17.99732 | 14.90592 | 12.93935 | 3.900445 | 16.76061 | 13.50005 | 11.9635 |
| ENSMUSG00000105911 | 0.092271065 | 0.783868 | 0.86707 | 33.19979 | 22.79661 | 23.18699 | 34.50494 | 21.45245 | 35.75597 | 25.07152 | 25.0146 |
| ENSMUSG00000105936 | 0.931138891 | 0.083717 | 0.194889 | 23.13925 | 35.99465 | 31.46805 | 31.91707 | 10.72622 | 35.75597 | 11.57147 | 6.525547 |
| ENSMUSG00000105939 | 1.454889075 | 0.224222 | 0.383974 | 13.07871 | 26.39607 | 4.140533 | 19.84034 | 0.975111 | 21.23011 | 0 | 1.087591 |
| ENSMUSG00000105945 | 0.389130787 | 0.380761 | 0.548634 | 15.09082 | 31.19536 | 21.53077 | 25.8787 | 12.67645 | 23.46486 | 10.60718 | 25.0146 |
| ENSMUSG00000105957 | 0.017091653 | 0.925475 | 0.955704 | 125.7568 | 124.7814 | 154.8559 | 137.1571 | 116.0382 | 155.315 | 111.8575 | 155.5255 |
| ENSMUSG00000105959 | 1.974280441 | 0.003914 | 0.021492 | 19.11503 | 23.99643 | 10.76539 | 25.01608 | 3.900445 | 12.29112 | 1.928578 | 2.175182 |
| ENSMUSG00000105961 | 1.218936059 | 0.091113 | 0.206901 | 20.12109 | 2.399643 | 24.8432 | 15.52722 | 4.875557 | 7.821619 | 3.857157 | 10.87591 |
| ENSMUSG00000105962 | 1.669380476 | 0.044307 | 0.125167 | 37.22401 | 43.19358 | 15.73403 | 37.09281 | 6.825779 | 31.28647 | 2.892867 | 1.087591 |
| ENSMUSG00000105981 | 0.804071169 | 0.072599 | 0.176999 | 26.15741 | 32.39518 | 27.32752 | 19.84034 | 24.37778 | 15.64324 | 13.50005 | 6.525547 |
| ENSMUSG00000105985 | 1.007665473 | 0.057375 | 0.150029 | 146.8839 | 131.9804 | 86.9512 | 106.1027 | 32.17867 | 144.1413 | 33.75012 | 25.0146 |
| ENSMUSG00000105987 | -0.20543 | 0.017499 | 0.064257 | 1598.62 | 1982.105 | 1860.756 | 1723.522 | 2078.937 | 1940.879 | 2067.436 | 2170.832 |

| | | | | | | | | | | | |
|---|---|---|---|---|---|---|---|---|---|---|---|
| ENSMUSG00000105990 | 0.864691519 | 0.123676 | 0.253358 | 20.12109 | 29.99554 | 18.21835 | 19.84034 | 9.751113 | 25.6996 | 8.678602 | 4.350364 |
| ENSMUSG00000106018 | 1.648785514 | 0.024547 | 0.081878 | 18.10898 | 26.39607 | 7.45296 | 21.56559 | 3.900445 | 14.52586 | 1.928578 | 3.262773 |
| ENSMUSG00000106019 | -0.394142519 | 0.013528 | 0.053434 | 150.9082 | 155.9768 | 202.8861 | 178.5631 | 228.176 | 188.8362 | 253.6081 | 236.0073 |
| ENSMUSG00000106022 | 0.183165169 | 0.629839 | 0.759014 | 25.15136 | 31.19536 | 34.78048 | 22.42821 | 34.1289 | 31.28647 | 19.28578 | 15.22628 |
| ENSMUSG00000106024 | 1.679925308 | 0.042968 | 0.122682 | 125.7568 | 116.3827 | 40.57722 | 45.71905 | 9.751113 | 78.21619 | 12.53576 | 2.175182 |
| ENSMUSG00000106025 | 0.796610638 | 0.100458 | 0.22155 | 45.27245 | 37.19447 | 27.32752 | 27.60395 | 11.70134 | 40.22547 | 15.42863 | 11.9635 |
| ENSMUSG00000106032 | 1.291061019 | 0.037607 | 0.11175 | 17.10292 | 15.59768 | 9.109173 | 19.84034 | 4.875557 | 13.40849 | 3.857157 | 3.262773 |
| ENSMUSG00000106035 | -0.088678226 | 0.858878 | 0.91537 | 19.11503 | 20.39697 | 8.281066 | 13.80198 | 12.67645 | 8.938993 | 20.25007 | 22.83941 |
| ENSMUSG00000106037 | 0.080924119 | 0.615831 | 0.748989 | 123.7447 | 134.38 | 129.1846 | 127.6683 | 138.4658 | 132.9675 | 113.7861 | 101.146 |
| ENSMUSG00000106056 | 0.225057425 | 0.412504 | 0.578395 | 64.38748 | 77.9884 | 62.9361 | 58.6584 | 60.4569 | 68.15982 | 63.64308 | 32.62773 |
| ENSMUSG00000106057 | 0.940289995 | 0.218235 | 0.376883 | 21.12714 | 32.39518 | 8.281066 | 13.80198 | 4.875557 | 24.58223 | 7.714313 | 2.175182 |
| ENSMUSG00000106062 | 0.677218527 | 0.001721 | 0.01173 | 90.54489 | 95.98572 | 81.15445 | 87.12497 | 57.53157 | 63.69032 | 60.75022 | 39.15328 |
| ENSMUSG00000106063 | 1.139552135 | 0.065611 | 0.164856 | 17.10292 | 34.79482 | 20.70267 | 20.70296 | 4.875557 | 24.58223 | 8.678602 | 4.350364 |
| ENSMUSG00000106078 | -3.3523187 | 5.94E-08 | 3.74E-06 | 3.018163 | 2.399643 | 9.93728 | 6.038364 | 40.95467 | 23.46486 | 56.89306 | 103.3212 |
| ENSMUSG00000106086 | -0.53860231 | 0.309509 | 0.477977 | 16.09687 | 14.39786 | 21.53077 | 10.35148 | 40.95467 | 10.05637 | 24.10723 | 15.22628 |
| ENSMUSG00000106087 | 0.058811196 | 0.887684 | 0.932533 | 48.29061 | 73.18911 | 43.06154 | 50.89479 | 37.05423 | 99.44629 | 37.60728 | 32.62773 |
| ENSMUSG00000106099 | -0.17299141 | 0.515725 | 0.669825 | 74.44802 | 70.78947 | 105.1695 | 92.30071 | 113.1129 | 89.38993 | 122.4647 | 61.99269 |
| ENSMUSG00000106107 | -0.755895832 | 0.139564 | 0.275575 | 4.024217 | 8.398751 | 10.76539 | 17.25247 | 19.50223 | 16.76061 | 14.46434 | 18.48905 |
| ENSMUSG00000106111 | -1.791282704 | 0.00441 | 0.023471 | 6.036326 | 1.199822 | 9.109173 | 6.038364 | 17.552 | 10.05637 | 19.28578 | 32.62773 |
| ENSMUSG00000106121 | 1.305569678 | 0.033455 | 0.102277 | 9.054489 | 16.7975 | 14.90592 | 17.25247 | 2.925334 | 3.352122 | 11.57147 | 5.437955 |
| ENSMUSG00000106139 | 0.717261731 | 0.130126 | 0.262545 | 14.08476 | 31.19536 | 18.21835 | 31.05445 | 12.67645 | 23.46486 | 10.60718 | 10.87591 |
| ENSMUSG00000106164 | -0.0312956 | 0.947109 | 0.969541 | 7.04238 | 11.99822 | 14.07781 | 13.80198 | 11.70134 | 10.05637 | 15.42863 | 10.87591 |
| ENSMUSG00000106166 | 0.93215099 | 0.119922 | 0.248879 | 20.12109 | 13.19804 | 9.109173 | 25.8787 | 5.850668 | 14.52586 | 3.857157 | 11.9635 |
| ENSMUSG00000106186 | 1.32380994 | 0.051162 | 0.138229 | 17.10292 | 28.79572 | 14.90592 | 18.11509 | 4.875557 | 17.87799 | 7.714313 | 1.087591 |
| ENSMUSG00000106205 | 0.137496873 | 0.518198 | 0.672184 | 87.52673 | 63.59054 | 99.3728 | 103.5148 | 85.8098 | 74.86406 | 87.75031 | 75.04379 |
| ENSMUSG00000106214 | 0.674683948 | 0.508908 | 0.66462 | 34.20585 | 28.79572 | 25.67131 | 17.25247 | 5.850668 | 55.8687 | 4.821446 | 0 |
| ENSMUSG00000106229 | 0.673632151 | 0.260177 | 0.424206 | 10.06054 | 15.59768 | 19.04645 | 10.35148 | 3.900445 | 16.76061 | 8.678602 | 5.437955 |
| ENSMUSG00000106237 | -0.000736228 | 0.998579 | 0.999282 | 30.18163 | 16.7975 | 13.24971 | 25.01608 | 20.47734 | 14.52586 | 25.07152 | 25.0146 |
| ENSMUSG00000106241 | -0.465288937 | 0.33303 | 0.501134 | 10.06054 | 13.19804 | 11.59349 | 5.175741 | 14.62667 | 12.29112 | 12.53576 | 15.22628 |
| ENSMUSG00000106251 | 0.475835372 | 0.205681 | 0.362081 | 37.22401 | 47.99286 | 31.46805 | 38.81806 | 28.27823 | 45.81234 | 20.25007 | 17.40146 |
| ENSMUSG00000106262 | 0.616984655 | 0.03043 | 0.095415 | 42.25428 | 43.19358 | 43.88965 | 45.71905 | 39.00445 | 27.93435 | 23.14294 | 23.927 |
| ENSMUSG00000106288 | 0.452290825 | 0.534127 | 0.685286 | 17.10292 | 29.99554 | 7.45296 | 7.763611 | 3.900445 | 22.34748 | 13.50005 | 5.437955 |
| ENSMUSG00000106290 | -1.409995667 | 0.001102 | 0.008401 | 11.0666 | 9.598572 | 6.624853 | 10.35148 | 20.47734 | 20.11273 | 34.71441 | 23.927 |
| ENSMUSG00000106291 | 0.425067775 | 0.290052 | 0.457172 | 36.21796 | 45.59322 | 24.01509 | 29.3292 | 20.47734 | 41.34284 | 22.17865 | 16.31387 |
| ENSMUSG00000106317 | 0.182715455 | 0.680676 | 0.796156 | 48.29061 | 52.79215 | 43.88965 | 50.03216 | 29.25334 | 88.27255 | 22.17865 | 32.62773 |
| ENSMUSG00000106320 | 1.479282554 | 0.009884 | 0.042258 | 67.40564 | 61.1909 | 35.60859 | 43.13117 | 13.65156 | 44.69496 | 8.678602 | 7.613138 |
| ENSMUSG00000106332 | 1.198985842 | 0.162756 | 0.307141 | 35.2119 | 69.58965 | 24.8432 | 42.26855 | 11.70134 | 55.8687 | 0.964289 | 6.525547 |
| ENSMUSG00000106375 | 0.90356072 | 0.062446 | 0.15938 | 36.21796 | 35.99465 | 45.54586 | 26.74133 | 10.72622 | 39.10809 | 10.60718 | 17.40146 |
| ENSMUSG00000106379 | -0.145458341 | 0.647446 | 0.77184 | 24.1453 | 29.99554 | 28.15563 | 23.29083 | 23.40267 | 23.46486 | 34.71441 | 34.80291 |
| ENSMUSG00000106383 | 0.564446736 | 0.047651 | 0.131554 | 181.0898 | 154.777 | 108.482 | 138.0198 | 89.71024 | 153.0803 | 77.14313 | 73.95619 |
| ENSMUSG00000106386 | 0.942534788 | 0.067413 | 0.16764 | 238.4349 | 307.1543 | 183.0116 | 208.7549 | 80.93424 | 300.5736 | 51.10733 | 55.46715 |
| ENSMUSG00000106391 | 0.318467155 | 0.443386 | 0.606427 | 16.09687 | 19.19714 | 20.70267 | 17.25247 | 18.52711 | 18.99536 | 12.53576 | 8.700729 |
| ENSMUSG00000106396 | 0.727167624 | 0.198585 | 0.353192 | 12.07265 | 13.19804 | 16.56213 | 17.25247 | 6.825779 | 15.64324 | 2.892867 | 10.87591 |
| ENSMUSG00000106416 | 0.64114279 | 0.094623 | 0.212427 | 32.19374 | 41.99375 | 19.04645 | 25.01608 | 21.45245 | 18.99536 | 20.25007 | 14.13868 |
| ENSMUSG00000106417 | -0.462116506 | 0.451728 | 0.61406 | 10.06054 | 4.799286 | 7.45296 | 11.21411 | 4.875557 | 7.821619 | 15.42863 | 18.48905 |
| ENSMUSG00000106427 | 0.690347001 | 0.080733 | 0.190197 | 50.30272 | 49.19268 | 31.46805 | 32.77969 | 22.42756 | 42.46022 | 21.21436 | 15.22628 |
| ENSMUSG00000106430 | 0.429012825 | 0.215267 | 0.37347 | 20.12109 | 31.19536 | 35.60859 | 39.68068 | 20.47734 | 29.05173 | 20.25007 | 25.0146 |
| ENSMUSG00000106446 | 1.701673484 | 0.011153 | 0.046362 | 47.28455 | 41.99375 | 18.21835 | 33.64232 | 8.776002 | 26.81698 | 5.785735 | 2.175182 |
| ENSMUSG00000106457 | 0.431172895 | 0.330865 | 0.498714 | 24.1453 | 35.99465 | 22.35888 | 25.8787 | 14.62667 | 34.6386 | 10.60718 | 20.66423 |
| ENSMUSG00000106461 | 0.605123698 | 0.057585 | 0.150408 | 49.29666 | 59.99108 | 51.34261 | 42.26855 | 31.20356 | 44.69496 | 38.57157 | 18.48905 |
| ENSMUSG00000106464 | -0.115036703 | 0.760176 | 0.850601 | 34.20585 | 29.99554 | 42.23344 | 21.56559 | 22.42756 | 26.81698 | 49.17875 | 40.24087 |
| ENSMUSG00000106466 | -1.345044346 | 0.133802 | 0.267352 | 2.012109 | 1.199822 | 19.87456 | 5.175741 | 8.776002 | 7.821619 | 34.71441 | 21.75182 |
| ENSMUSG00000106472 | 1.607209239 | 0.065897 | 0.165361 | 26.15741 | 21.59679 | 13.24971 | 10.35148 | 2.925334 | 16.76061 | 3.857157 | 0 |
| ENSMUSG00000106498 | 0.052518836 | 0.919579 | 0.952239 | 9.054489 | 7.198929 | 12.4216 | 19.84034 | 15.60178 | 10.05637 | 8.678602 | 13.05109 |
| ENSMUSG00000106508 | -0.424273324 | 0.210478 | 0.367996 | 14.08476 | 26.39607 | 22.35888 | 19.84034 | 24.37778 | 25.6996 | 28.92867 | 31.54014 |
| ENSMUSG00000106526 | -1.653644432 | 0.004217 | 0.022693 | 11.0666 | 5.999108 | 9.93728 | 6.900988 | 22.42756 | 12.29112 | 19.28578 | 53.29196 |
| ENSMUSG00000106528 | 0.60010402 | 0.367871 | 0.536332 | 83.50251 | 110.3836 | 42.23344 | 63.83414 | 36.07912 | 130.7328 | 20.25007 | 10.87591 |
| ENSMUSG00000106535 | 0.118905087 | 0.749531 | 0.843806 | 21.12714 | 33.595 | 39.74912 | 18.97772 | 28.27823 | 23.46486 | 21.21436 | 31.54014 |
| ENSMUSG00000106543 | 0.777208934 | 0.241957 | 0.404379 | 17.10292 | 21.59679 | 11.59349 | 9.488858 | 3.900445 | 18.99536 | 8.678602 | 3.262773 |
| ENSMUSG00000106547 | 0.301071103 | 0.454023 | 0.616181 | 97.58727 | 61.1909 | 88.60741 | 83.67448 | 39.97956 | 131.8501 | 51.10733 | 46.76642 |
| ENSMUSG00000106590 | -0.283349939 | 0.652084 | 0.775356 | 5.030272 | 14.39786 | 14.90592 | 6.038364 | 5.850668 | 20.11273 | 14.46434 | 8.700729 |
| ENSMUSG00000106617 | 0.853528282 | 0.000146 | 0.001784 | 87.52673 | 69.58965 | 85.29498 | 81.94923 | 35.10401 | 53.63396 | 49.17875 | 42.41605 |
| ENSMUSG00000106631 | 1.285292315 | 0.033165 | 0.10161 | 25.15136 | 22.79661 | 6.624853 | 19.84034 | 3.900445 | 12.29112 | 6.750024 | 7.613138 |
| ENSMUSG00000106636 | 1.230698114 | 0.13205 | 0.265176 | 87.52673 | 49.19268 | 19.87456 | 44.85642 | 1.950223 | 52.51658 | 26.03581 | 5.437955 |

| | | | | | | | | | | | |
|---|---|---|---|---|---|---|---|---|---|---|---|
| ENSMUSG00000106641 | -0.08913126 | 0.733551 | 0.833185 | 69.41775 | 43.19358 | 69.56096 | 53.48266 | 46.80534 | 74.86406 | 68.46453 | 61.99269 |
| ENSMUSG00000106662 | 0.008073623 | 0.979498 | 0.988724 | 21.12714 | 25.19625 | 24.8432 | 22.42821 | 21.45245 | 24.58223 | 23.14294 | 23.927 |
| ENSMUSG00000106665 | 0.347289842 | 0.481885 | 0.640708 | 20.12109 | 23.99643 | 23.18699 | 25.8787 | 9.751113 | 35.75597 | 18.32149 | 9.78832 |
| ENSMUSG00000106678 | 0.311564781 | 0.384876 | 0.552504 | 35.2119 | 33.595 | 38.09291 | 24.15346 | 20.47734 | 40.22547 | 19.28578 | 26.10219 |
| ENSMUSG00000106706 | 0.836447282 | 0.200148 | 0.355255 | 70.4238 | 43.19358 | 42.23344 | 30.19182 | 14.62667 | 69.27719 | 8.678602 | 11.9635 |
| ENSMUSG00000106717 | 0.482475648 | 0.379421 | 0.547549 | 45.27245 | 31.19536 | 31.46805 | 37.09281 | 12.67645 | 54.75133 | 27.96439 | 8.700729 |
| ENSMUSG00000106732 | 1.341686946 | 0.010458 | 0.044087 | 25.15136 | 34.79482 | 16.56213 | 18.97772 | 4.875557 | 15.64324 | 11.57147 | 5.437955 |
| ENSMUSG00000106734 | 0.436196193 | 0.043725 | 0.124058 | 214.2896 | 256.7618 | 312.1962 | 172.5247 | 142.3663 | 198.8926 | 195.7507 | 169.6642 |
| ENSMUSG00000106746 | 0.411180067 | 0.644766 | 0.770114 | 6.036326 | 10.79839 | 11.59349 | 19.84034 | 4.875557 | 26.81698 | 3.857157 | 1.087591 |
| ENSMUSG00000106749 | 0.934886095 | 0.32635 | 0.494063 | 15.09082 | 19.19714 | 10.76539 | 11.21411 | 2.925334 | 22.34748 | 0 | 4.350364 |
| ENSMUSG00000106751 | 0.738092504 | 0.347506 | 0.516273 | 57.3451 | 39.59411 | 16.56213 | 25.01608 | 5.850668 | 60.3382 | 11.57147 | 5.437955 |
| ENSMUSG00000106767 | 0.206441207 | 0.522055 | 0.675291 | 60.36326 | 34.79482 | 48.85829 | 49.16954 | 31.20356 | 63.69032 | 36.64299 | 36.9781 |
| ENSMUSG00000106772 | 1.000541723 | 0.075993 | 0.18262 | 38.23007 | 29.99554 | 21.53077 | 25.01608 | 5.850668 | 32.40385 | 9.642892 | 9.78832 |
| ENSMUSG00000106825 | 0.276098817 | 0.345579 | 0.514314 | 31.18769 | 59.99108 | 43.06154 | 53.48266 | 39.00445 | 42.46022 | 41.46443 | 31.54014 |
| ENSMUSG00000106826 | 1.000247917 | 0.122058 | 0.251544 | 47.28455 | 52.79215 | 35.60859 | 68.14726 | 9.751113 | 69.27719 | 13.50005 | 9.78832 |
| ENSMUSG00000106831 | -0.459164532 | 0.14068 | 0.277185 | 23.13925 | 31.19536 | 21.53077 | 35.36756 | 42.9049 | 31.28647 | 33.75012 | 44.59123 |
| ENSMUSG00000106838 | 0.669839004 | 0.04008 | 0.116897 | 60.36326 | 49.19268 | 38.92101 | 43.13117 | 17.552 | 35.75597 | 37.60728 | 29.36496 |
| ENSMUSG00000106847 | 0.487895679 | 3.21E-05 | 0.000563 | 1028.188 | 789.4826 | 832.2472 | 796.2015 | 549.9628 | 625.7295 | 648.9666 | 632.978 |
| ENSMUSG00000106858 | 0.270977365 | 0.636004 | 0.763553 | 18.10898 | 4.799286 | 19.04645 | 10.35148 | 10.72622 | 15.64324 | 6.750024 | 10.87591 |
| ENSMUSG00000106864 | 0.412302395 | 0.008174 | 0.036868 | 2509.1 | 2511.226 | 2111.672 | 2584.42 | 1707.42 | 2432.523 | 1648.934 | 1512.839 |
| ENSMUSG00000106870 | 0.185458477 | 0.506036 | 0.661805 | 80.48435 | 64.79036 | 52.99882 | 66.42201 | 63.38224 | 80.45093 | 44.3573 | 44.59123 |
| ENSMUSG00000106874 | -0.058815486 | 0.86109 | 0.916827 | 35.2119 | 37.19447 | 29.81184 | 24.15346 | 38.02934 | 18.99536 | 38.57157 | 34.80291 |
| ENSMUSG00000106887 | 1.35808927 | 0.04329 | 0.123188 | 26.15741 | 27.5959 | 16.56213 | 20.70296 | 20.47734 | 7.821619 | 5.785735 | 1.087591 |
| ENSMUSG00000106891 | 0.242761412 | 0.356918 | 0.525428 | 53.32088 | 46.79304 | 58.79557 | 60.38364 | 39.00445 | 60.3382 | 52.07161 | 34.80291 |
| ENSMUSG00000106904 | -0.031684743 | 0.943766 | 0.967564 | 9.054489 | 17.99732 | 15.73403 | 16.38985 | 12.67645 | 12.29112 | 22.17865 | 13.05109 |
| ENSMUSG00000106915 | -0.315810524 | 0.583645 | 0.724151 | 12.07265 | 5.999108 | 15.73403 | 8.626235 | 14.62667 | 15.64324 | 4.821446 | 18.48905 |
| ENSMUSG00000106918 | -0.040139931 | 0.642334 | 0.768671 | 1334.028 | 1317.404 | 1513.779 | 1422.466 | 1458.767 | 1536.389 | 1470.541 | 1283.357 |
| ENSMUSG00000106926 | 0.105467645 | 0.698141 | 0.809552 | 30.18163 | 41.99375 | 33.95237 | 39.68068 | 38.02934 | 34.6386 | 29.89296 | 32.62773 |
| ENSMUSG00000106927 | 0.335193104 | 0.557376 | 0.703981 | 11.0666 | 14.39786 | 12.4216 | 23.29083 | 4.875557 | 17.87799 | 18.32149 | 7.613138 |
| ENSMUSG00000106928 | 0.389016929 | 0.078245 | 0.186166 | 65.39353 | 79.18822 | 73.70149 | 84.5371 | 51.6809 | 60.3382 | 48.21446 | 71.78101 |
| ENSMUSG00000106943 | 0.085835398 | 0.614308 | 0.748023 | 119.7205 | 116.3827 | 107.6539 | 131.9814 | 126.7645 | 119.559 | 93.53605 | 108.7591 |
| ENSMUSG00000106944 | 0.014354965 | 0.977068 | 0.987393 | 20.12109 | 28.79572 | 17.39024 | 14.6646 | 15.60178 | 37.99072 | 15.42863 | 10.87591 |
| ENSMUSG00000106947 | 0.790502677 | 0.00306 | 0.018035 | 76.46013 | 105.5843 | 61.27989 | 84.5371 | 49.73068 | 56.98608 | 48.21446 | 33.71532 |
| ENSMUSG00000106951 | -0.380602505 | 0.080459 | 0.189674 | 94.56911 | 101.9848 | 108.482 | 65.55939 | 120.9138 | 97.21155 | 137.8934 | 125.073 |
| ENSMUSG00000106978 | 0.130571449 | 0.430834 | 0.595571 | 170.0232 | 167.975 | 166.4494 | 142.3329 | 114.088 | 157.5497 | 157.1791 | 162.0511 |
| ENSMUSG00000106981 | 0.04726554 | 0.848529 | 0.909127 | 61.36932 | 67.19001 | 67.90474 | 50.89479 | 57.53157 | 49.16446 | 52.07161 | 80.48174 |
| ENSMUSG00000106990 | 0.461812819 | 0.08077 | 0.190212 | 133.8052 | 87.58697 | 72.87338 | 121.6299 | 64.35735 | 97.21155 | 72.32169 | 68.51824 |
| ENSMUSG00000106991 | 0.997977308 | 0.186485 | 0.33839 | 25.15136 | 13.19804 | 19.87456 | 18.97772 | 0.975111 | 24.58223 | 4.821446 | 8.700729 |
| ENSMUSG00000106992 | 0.769134042 | 0.299608 | 0.467712 | 16.09687 | 21.59679 | 10.76539 | 9.488858 | 1.950223 | 20.11273 | 8.678602 | 3.262773 |
| ENSMUSG00000107000 | 0.411520446 | 0.024904 | 0.0828 | 172.0353 | 131.9804 | 119.2474 | 129.3935 | 89.71024 | 108.3853 | 104.1432 | 113.1095 |
| ENSMUSG00000107002 | -0.532821299 | 0.000142 | 0.001748 | 1092.575 | 1202.221 | 1268.659 | 1233.552 | 1786.404 | 1310.68 | 1770.435 | 2072.949 |
| ENSMUSG00000107009 | 2.363473155 | 0.003026 | 0.017883 | 22.1332 | 26.39607 | 24.01509 | 15.52722 | 0.975111 | 13.40849 | 1.928578 | 1.087591 |
| ENSMUSG00000107015 | 1.289061537 | 0.022728 | 0.077221 | 17.10292 | 29.99554 | 14.07781 | 21.56559 | 4.875557 | 7.821619 | 16.39292 | 4.350364 |
| ENSMUSG00000107019 | 0.127694278 | 0.606444 | 0.742598 | 50.30272 | 58.79126 | 71.21717 | 62.97152 | 41.92979 | 65.92507 | 64.60737 | 51.11678 |
| ENSMUSG00000107023 | 0.735647142 | 0.009998 | 0.042568 | 386.3249 | 465.5308 | 429.7873 | 464.0914 | 213.5494 | 457.006 | 202.5007 | 176.1898 |
| ENSMUSG00000107035 | -0.517896772 | 0.322744 | 0.490873 | 6.036326 | 7.198929 | 9.93728 | 12.07673 | 9.751113 | 12.29112 | 9.642892 | 19.57664 |
| ENSMUSG00000107041 | 0.849705051 | 0.225537 | 0.385381 | 29.17558 | 19.19714 | 18.21835 | 19.84034 | 5.850668 | 32.40385 | 6.750024 | 3.262773 |
| ENSMUSG00000107061 | 0.598361109 | 0.265568 | 0.429745 | 19.11503 | 7.198929 | 26.49941 | 15.52722 | 12.67645 | 16.76061 | 8.678602 | 7.613138 |
| ENSMUSG00000107068 | 0.480782887 | 0.071518 | 0.175081 | 107.6478 | 119.9822 | 110.9663 | 117.3168 | 75.08357 | 128.498 | 60.75022 | 63.08028 |
| ENSMUSG00000107086 | 0.437904638 | 0.123518 | 0.25321 | 59.35721 | 53.99197 | 40.57722 | 44.85642 | 31.20356 | 32.40385 | 32.78583 | 50.02919 |
| ENSMUSG00000107091 | 0.809747252 | 0.042688 | 0.122149 | 126.7628 | 151.1775 | 81.15445 | 94.88858 | 45.83023 | 119.559 | 59.78593 | 33.71532 |
| ENSMUSG00000107092 | -0.335051584 | 0.403566 | 0.570526 | 13.07871 | 19.19714 | 18.21835 | 17.25247 | 12.67645 | 24.58223 | 20.25007 | 28.27737 |
| ENSMUSG00000107096 | 0.36681032 | 0.106693 | 0.230901 | 102.6175 | 77.9884 | 69.56096 | 93.16334 | 50.70579 | 72.62932 | 70.39311 | 72.8686 |
| ENSMUSG00000107102 | 0.716823919 | 0.24387 | 0.406473 | 19.11503 | 27.5959 | 28.98373 | 22.42821 | 8.776002 | 36.87335 | 5.785735 | 8.700729 |
| ENSMUSG00000107120 | 0.497076647 | 0.318798 | 0.486971 | 31.18769 | 40.79393 | 26.49941 | 31.05445 | 21.45245 | 46.92971 | 11.57147 | 11.9635 |
| ENSMUSG00000107121 | -0.175753918 | 0.571173 | 0.714257 | 43.26034 | 28.79572 | 57.13936 | 35.36756 | 37.05423 | 41.34284 | 50.14304 | 58.72992 |
| ENSMUSG00000107134 | 0.088255743 | 0.844421 | 0.906347 | 24.1453 | 16.7975 | 11.59349 | 29.3292 | 16.57689 | 29.05173 | 15.42863 | 16.31387 |
| ENSMUSG00000107136 | 1.828669295 | 0.03806 | 0.112588 | 24.1453 | 19.19714 | 13.24971 | 15.52722 | 2.925334 | 15.64324 | 1.928578 | 0 |
| ENSMUSG00000107143 | 0.349144275 | 0.185888 | 0.337539 | 67.40564 | 71.98929 | 59.62368 | 45.71905 | 46.80534 | 34.6386 | 54.00019 | 55.46715 |
| ENSMUSG00000107168 | 0.690453821 | 0.163509 | 0.308308 | 46.2785 | 28.79572 | 19.04645 | 12.93935 | 19.50223 | 20.11273 | 15.42863 | 10.87591 |
| ENSMUSG00000107171 | 0.012890668 | 0.980008 | 0.988885 | 19.11503 | 9.598572 | 24.01509 | 8.626235 | 15.60178 | 22.34748 | 12.53576 | 10.87591 |
| ENSMUSG00000107173 | 0.735653931 | 0.294182 | 0.461764 | 10.06054 | 15.59768 | 9.109173 | 16.38985 | 1.950223 | 17.87799 | 6.750024 | 4.350364 |
| ENSMUSG00000107176 | -0.049017716 | 0.660028 | 0.781014 | 399.4036 | 484.7279 | 423.1625 | 462.3662 | 511.9334 | 463.7102 | 431.0373 | 421.9853 |
| ENSMUSG00000107182 | 0.26423933 | 0.190559 | 0.343185 | 83.50251 | 98.38537 | 69.56096 | 72.46037 | 70.20801 | 62.57295 | 67.50024 | 67.43065 |

| | | | | | | | | | | | |
|---|---|---|---|---|---|---|---|---|---|---|---|
| ENSMUSG00000107191 | 0.231374701 | 0.475339 | 0.634795 | 21.12714 | 22.79661 | 26.49941 | 28.46658 | 20.47734 | 23.46486 | 21.21436 | 19.57664 |
| ENSMUSG00000107195 | -0.248675287 | 0.559678 | 0.705754 | 28.16952 | 19.19714 | 19.87456 | 11.21411 | 32.17867 | 23.46486 | 15.42863 | 21.75182 |
| ENSMUSG00000107200 | 0.665968623 | 0.066476 | 0.166221 | 59.35721 | 52.79215 | 53.82693 | 66.42201 | 35.10401 | 63.69032 | 21.21436 | 27.18978 |
| ENSMUSG00000107201 | 0.391468423 | 0.148292 | 0.287728 | 113.6841 | 95.98572 | 81.15445 | 105.2401 | 56.55646 | 115.0895 | 64.60737 | 66.34306 |
| ENSMUSG00000107215 | 0.154850387 | 0.806576 | 0.881777 | 17.10292 | 13.19804 | 10.76539 | 5.175741 | 4.875557 | 5.58687 | 14.46434 | 16.31387 |
| ENSMUSG00000107216 | 0.590369626 | 0.358212 | 0.526533 | 32.19374 | 5.999108 | 17.39024 | 15.52722 | 8.776002 | 17.87799 | 16.39292 | 4.350364 |
| ENSMUSG00000107227 | 0.567288909 | 0.330172 | 0.498051 | 15.09082 | 13.19804 | 5.796746 | 14.6646 | 10.72622 | 3.352122 | 8.678602 | 9.78832 |
| ENSMUSG00000107230 | 0.266865506 | 0.613096 | 0.747032 | 6.036326 | 19.19714 | 12.4216 | 20.70296 | 8.776002 | 10.05637 | 15.42863 | 14.13868 |
| ENSMUSG00000107276 | 0.686850155 | 0.243284 | 0.405862 | 20.12109 | 21.59679 | 33.95237 | 10.35148 | 5.850668 | 25.6996 | 13.50005 | 8.700729 |
| ENSMUSG00000107277 | 0.04356878 | 0.853916 | 0.912346 | 156.9445 | 107.9839 | 125.0441 | 112.1411 | 110.1876 | 172.0756 | 108.9647 | 96.79561 |
| ENSMUSG00000107278 | -0.085992806 | 0.844582 | 0.906467 | 25.15136 | 39.59411 | 21.53077 | 32.77969 | 25.35289 | 53.63396 | 31.82154 | 15.22628 |
| ENSMUSG00000107283 | 0.140940886 | 0.028366 | 0.090844 | 1319.943 | 1362.997 | 1294.331 | 1345.693 | 1193.536 | 1289.45 | 1199.576 | 1144.146 |
| ENSMUSG00000107300 | 1.339542122 | 0.033041 | 0.10138 | 12.07265 | 17.99732 | 10.76539 | 17.25247 | 8.776002 | 7.821619 | 0.964289 | 5.437955 |
| ENSMUSG00000107304 | -0.364130571 | 0.441771 | 0.60507 | 29.17558 | 16.7975 | 26.49941 | 15.52722 | 13.65156 | 36.87335 | 43.39301 | 19.57664 |
| ENSMUSG00000107317 | -0.664239613 | 0.051721 | 0.13923 | 20.12109 | 14.39786 | 16.56213 | 20.70296 | 36.07912 | 26.81698 | 24.10723 | 27.18978 |
| ENSMUSG00000107319 | 0.279436341 | 0.561476 | 0.707007 | 14.08476 | 7.198929 | 15.73403 | 14.6646 | 13.65156 | 7.821619 | 10.60718 | 10.87591 |
| ENSMUSG00000107320 | 0.286605465 | 0.509186 | 0.664889 | 26.15741 | 22.79661 | 14.90592 | 18.11509 | 24.37778 | 18.99536 | 13.50005 | 9.78832 |
| ENSMUSG00000107331 | 0.502418142 | 0.070158 | 0.172649 | 129.781 | 145.1784 | 139.95 | 159.5853 | 81.90935 | 166.4887 | 81.96458 | 76.13138 |
| ENSMUSG00000107335 | 0.387620085 | 0.394098 | 0.561441 | 27.16347 | 14.39786 | 24.8432 | 23.29083 | 21.45245 | 10.05637 | 26.03581 | 10.87591 |
| ENSMUSG00000107336 | 0.62094376 | 0.067107 | 0.167299 | 51.30877 | 67.19001 | 52.17072 | 75.91087 | 31.20356 | 64.8077 | 38.57157 | 26.10219 |
| ENSMUSG00000107352 | 0.351563338 | 0.248182 | 0.411541 | 146.8839 | 131.9804 | 142.4343 | 144.0581 | 81.90935 | 189.9536 | 100.2861 | 71.78101 |
| ENSMUSG00000107355 | 0.311271371 | 0.322141 | 0.490346 | 54.32694 | 41.99375 | 43.88965 | 51.75741 | 29.25334 | 59.22083 | 38.57157 | 28.27737 |
| ENSMUSG00000107362 | 1.140320063 | 0.013615 | 0.05367 | 53.32088 | 67.19001 | 32.29616 | 35.36756 | 12.67645 | 40.22547 | 18.32149 | 14.13868 |
| ENSMUSG00000107374 | -0.157578716 | 0.595518 | 0.734635 | 36.21796 | 32.39518 | 24.8432 | 36.23019 | 29.25334 | 39.10809 | 44.3573 | 31.54014 |
| ENSMUSG00000107383 | 0.029699372 | 0.838369 | 0.902265 | 262.5802 | 370.7449 | 284.8687 | 302.7808 | 342.2641 | 291.6346 | 279.6439 | 279.5109 |
| ENSMUSG00000107390 | 0.027233016 | 0.965035 | 0.980332 | 16.09687 | 13.19804 | 15.73403 | 25.01608 | 3.900445 | 35.75597 | 14.46434 | 15.22628 |
| ENSMUSG00000107402 | 0.41202958 | 0.08706 | 0.200472 | 65.39353 | 63.59054 | 58.79557 | 52.62003 | 39.97956 | 49.16446 | 54.96448 | 35.89051 |
| ENSMUSG00000107411 | 1.264826518 | 0.058306 | 0.151607 | 18.10898 | 28.79572 | 7.45296 | 12.07673 | 5.850668 | 13.40849 | 2.892867 | 5.437955 |
| ENSMUSG00000107428 | 1.000688743 | 0.045649 | 0.127749 | 21.12714 | 21.59679 | 29.81184 | 30.19182 | 15.60178 | 20.11273 | 12.53576 | 3.262773 |
| ENSMUSG00000107432 | 0.237413569 | 0.647814 | 0.771952 | 25.15136 | 22.79661 | 19.87456 | 10.35148 | 30.22845 | 11.17374 | 14.46434 | 9.78832 |
| ENSMUSG00000107436 | 1.599776129 | 0.062864 | 0.160185 | 31.18769 | 21.59679 | 12.4216 | 29.3292 | 13.65156 | 15.64324 | 1.928578 | 0 |
| ENSMUSG00000107451 | -0.754507322 | 0.092466 | 0.208881 | 16.09687 | 7.198929 | 9.109173 | 9.488858 | 18.52711 | 15.64324 | 21.21436 | 15.22628 |
| ENSMUSG00000107468 | 1.015043815 | 0.09361 | 0.210617 | 34.20585 | 55.19179 | 36.43669 | 39.68068 | 10.72622 | 51.39921 | 13.50005 | 6.525547 |
| ENSMUSG00000107470 | -0.082542121 | 0.83796 | 0.902035 | 20.12109 | 19.19714 | 12.4216 | 12.93935 | 20.47734 | 13.40849 | 17.3572 | 16.31387 |
| ENSMUSG00000107476 | 0.030429565 | 0.77936 | 0.863844 | 463.7911 | 491.9268 | 438.8965 | 408.8835 | 457.3272 | 403.372 | 486.0017 | 414.3722 |
| ENSMUSG00000107481 | -0.32509434 | 0.590596 | 0.730709 | 9.054489 | 19.19714 | 10.76539 | 12.07673 | 9.751113 | 34.6386 | 8.678602 | 10.87591 |
| ENSMUSG00000107491 | 0.850942166 | 0.174845 | 0.323049 | 48.29061 | 39.59411 | 19.04645 | 27.60395 | 9.751113 | 45.81234 | 11.57147 | 7.613138 |
| ENSMUSG00000107499 | 0.124956007 | 0.678866 | 0.794865 | 48.29061 | 52.79215 | 45.54586 | 53.48266 | 55.58134 | 25.6996 | 58.82164 | 42.41605 |
| ENSMUSG00000107502 | -1.213478537 | 0.083142 | 0.194017 | 20.12109 | 2.399643 | 21.53077 | 10.35148 | 53.63112 | 7.821619 | 31.82154 | 33.71532 |
| ENSMUSG00000107505 | 0.639051512 | 0.148904 | 0.288674 | 47.28455 | 63.59054 | 22.35888 | 39.68068 | 15.60178 | 39.10809 | 20.25007 | 35.89051 |
| ENSMUSG00000107509 | 0.209996611 | 0.629802 | 0.759014 | 22.1332 | 16.7975 | 14.07781 | 26.74133 | 25.35289 | 16.76061 | 10.60718 | 16.31387 |
| ENSMUSG00000107524 | 0.437949657 | 0.3636 | 0.532123 | 15.09082 | 21.59679 | 9.93728 | 16.38985 | 15.60178 | 11.17374 | 12.53576 | 6.525547 |
| ENSMUSG00000107529 | 1.15927114 | 0.013146 | 0.052448 | 55.33299 | 55.19179 | 36.43669 | 37.09281 | 14.62667 | 42.46022 | 12.53576 | 13.05109 |
| ENSMUSG00000107531 | 1.146989912 | 0.079737 | 0.188683 | 29.17558 | 37.19447 | 14.07781 | 11.21411 | 4.875557 | 22.34748 | 8.678602 | 5.437955 |
| ENSMUSG00000107556 | 0.908469202 | 0.344056 | 0.512751 | 15.09082 | 15.59768 | 9.93728 | 12.93935 | 0 | 17.87799 | 9.642892 | 1.087591 |
| ENSMUSG00000107560 | 0.919703741 | 0.020631 | 0.072021 | 56.33904 | 25.19625 | 43.06154 | 54.34528 | 24.37778 | 31.28647 | 12.53576 | 27.18978 |
| ENSMUSG00000107561 | -0.179191048 | 0.730804 | 0.831241 | 24.1453 | 31.19536 | 14.90592 | 19.84034 | 15.60178 | 52.51658 | 17.3572 | 16.31387 |
| ENSMUSG00000107567 | 0.275804498 | 0.536028 | 0.68669 | 21.12714 | 14.39786 | 23.18699 | 12.07673 | 17.552 | 17.87799 | 14.46434 | 8.700729 |
| ENSMUSG00000107577 | 0.407999511 | 0.117006 | 0.245171 | 161.9748 | 213.5682 | 208.6829 | 228.5952 | 118.9636 | 240.2354 | 136.9291 | 117.4598 |
| ENSMUSG00000107597 | -0.014748414 | 0.980218 | 0.988885 | 23.13925 | 13.19804 | 6.624853 | 6.900988 | 14.62667 | 17.87799 | 7.714313 | 9.78832 |
| ENSMUSG00000107608 | 0.27881625 | 0.229578 | 0.38976 | 60.36326 | 57.59143 | 79.49824 | 56.07053 | 53.63112 | 60.3382 | 49.17875 | 46.76642 |
| ENSMUSG00000107610 | 0.832367203 | 0.006814 | 0.032357 | 59.35721 | 80.38804 | 63.76421 | 55.2079 | 26.32801 | 50.28183 | 43.39301 | 25.0146 |
| ENSMUSG00000107624 | 0.707583184 | 0.179651 | 0.329508 | 57.3451 | 39.59411 | 21.53077 | 24.15346 | 21.45245 | 41.34284 | 12.53576 | 11.9635 |
| ENSMUSG00000107647 | -0.080144854 | 0.888193 | 0.93282 | 14.08476 | 14.39786 | 12.4216 | 8.626235 | 23.40267 | 5.58687 | 7.714313 | 15.22628 |
| ENSMUSG00000107655 | 0.768217986 | 0.024096 | 0.080842 | 293.7679 | 209.9688 | 232.698 | 359.714 | 117.9885 | 272.6393 | 167.7863 | 85.9197 |
| ENSMUSG00000107667 | 1.15008814 | 0.000631 | 0.005483 | 49.29666 | 45.59322 | 39.74912 | 31.05445 | 12.67645 | 20.11273 | 23.14294 | 18.48905 |
| ENSMUSG00000107689 | -1.469773989 | 0.029459 | 0.093354 | 11.0666 | 0 | 11.59349 | 13.80198 | 32.17867 | 14.52586 | 21.21436 | 34.80291 |
| ENSMUSG00000107726 | 0.593912711 | 0.250166 | 0.413857 | 41.24823 | 27.5959 | 18.21835 | 24.15346 | 13.65156 | 36.87335 | 13.50005 | 9.78832 |
| ENSMUSG00000107728 | 0.316981443 | 0.255054 | 0.418946 | 36.21796 | 52.79215 | 62.108 | 49.16954 | 39.97956 | 32.40385 | 49.17875 | 39.15328 |
| ENSMUSG00000107741 | 0.39735588 | 0.025207 | 0.083492 | 153.9263 | 177.5736 | 156.5122 | 131.9814 | 101.4116 | 103.9158 | 139.8219 | 123.9854 |
| ENSMUSG00000107742 | -0.206391022 | 0.571904 | 0.714774 | 29.17558 | 32.39518 | 26.49941 | 22.42821 | 34.1289 | 15.64324 | 37.60728 | 39.15328 |
| ENSMUSG00000107747 | 0.428963465 | 0.326087 | 0.493883 | 17.10292 | 22.79661 | 11.59349 | 14.6646 | 12.67645 | 14.52586 | 11.57147 | 9.78832 |
| ENSMUSG00000107749 | -0.147877427 | 0.708059 | 0.816595 | 21.12714 | 19.19714 | 26.49941 | 26.74133 | 15.60178 | 18.99536 | 33.75012 | 35.89051 |
| ENSMUSG00000107750 | 0.171110974 | 0.801762 | 0.879143 | 15.09082 | 17.99732 | 8.281066 | 7.763611 | 10.72622 | 23.46486 | 3.857157 | 5.437955 |

| | | | | | | | | | | | |
|---|---|---|---|---|---|---|---|---|---|---|---|
| ENSMUSG00000107756 | -0.063200452 | 0.894344 | 0.936192 | 17.10292 | 20.39697 | 26.49941 | 16.38985 | 10.72622 | 33.52122 | 27.0001 | 13.05109 |
| ENSMUSG00000107802 | -0.426415599 | 0.215144 | 0.373325 | 24.1453 | 15.59768 | 20.70267 | 21.56559 | 27.30312 | 26.81698 | 35.6787 | 20.66423 |
| ENSMUSG00000107814 | 0.369846818 | 0.457871 | 0.619627 | 22.1332 | 9.598572 | 9.93728 | 20.70296 | 10.72622 | 14.52586 | 14.46434 | 8.700729 |
| ENSMUSG00000107838 | -0.233452769 | 0.394928 | 0.562234 | 49.29666 | 41.99375 | 41.40533 | 62.10889 | 45.83023 | 72.62932 | 64.60737 | 46.76642 |
| ENSMUSG00000107865 | 2.296032446 | 0.0132 | 0.052619 | 22.1332 | 19.19714 | 16.56213 | 25.01608 | 5.850668 | 11.17374 | 0 | 0 |
| ENSMUSG00000107868 | -0.013036902 | 0.952017 | 0.972676 | 103.6236 | 107.9839 | 83.63877 | 106.1027 | 90.68535 | 129.6154 | 108.0004 | 76.13138 |
| ENSMUSG00000107877 | 0.536268484 | 0.136657 | 0.271402 | 34.20585 | 34.79482 | 19.04645 | 31.05445 | 24.37778 | 23.46486 | 16.39292 | 17.40146 |
| ENSMUSG00000107879 | 0.115648526 | 0.702202 | 0.812223 | 29.17558 | 23.99643 | 24.8432 | 29.3292 | 23.40267 | 26.81698 | 23.14294 | 26.10219 |
| ENSMUSG00000107881 | 0.366452327 | 0.027923 | 0.089945 | 301.8163 | 199.1704 | 264.166 | 244.9851 | 169.6694 | 198.8926 | 223.7151 | 192.5036 |
| ENSMUSG00000107884 | 1.007568586 | 0.119319 | 0.248319 | 23.13925 | 34.79482 | 16.56213 | 18.11509 | 6.825779 | 27.93435 | 3.857157 | 7.613138 |
| ENSMUSG00000107894 | 1.271581924 | 0.032558 | 0.100248 | 46.2785 | 56.39161 | 30.63995 | 34.50494 | 5.850668 | 42.46022 | 10.60718 | 10.87591 |
| ENSMUSG00000107899 | -0.125710577 | 0.887919 | 0.932608 | 7.04238 | 11.99822 | 1.656213 | 20.70296 | 0.975111 | 15.64324 | 14.46434 | 14.13868 |
| ENSMUSG00000107909 | 0.354536439 | 0.34793 | 0.516564 | 20.12109 | 16.7975 | 20.70267 | 28.46658 | 21.45245 | 14.52586 | 16.39292 | 15.22628 |
| ENSMUSG00000107932 | 1.089047541 | 0.15664 | 0.298843 | 31.18769 | 31.19536 | 20.70267 | 26.74133 | 2.925334 | 37.99072 | 7.714313 | 3.262773 |
| ENSMUSG00000107937 | -0.125049514 | 0.819731 | 0.890713 | 18.10898 | 10.79839 | 19.04645 | 6.900988 | 9.751113 | 21.23011 | 8.678602 | 20.66423 |
| ENSMUSG00000107944 | -0.007064911 | 0.978197 | 0.988047 | 63.38142 | 50.3925 | 70.38906 | 53.48266 | 77.03379 | 42.46022 | 61.71451 | 57.64233 |
| ENSMUSG00000107946 | 0.238818592 | 0.487212 | 0.645234 | 23.13925 | 22.79661 | 41.40533 | 28.46658 | 24.37778 | 27.93435 | 26.03581 | 20.66423 |
| ENSMUSG00000107961 | -0.711465723 | 0.111491 | 0.237827 | 8.048435 | 11.99822 | 12.4216 | 10.35148 | 14.62667 | 22.34748 | 21.21436 | 11.9635 |
| ENSMUSG00000107962 | 0.605529862 | 0.064092 | 0.162528 | 321.9374 | 212.3684 | 121.7317 | 191.5024 | 122.864 | 206.7142 | 128.2505 | 98.97079 |
| ENSMUSG00000107990 | 0.54100849 | 0.06547 | 0.164684 | 57.3451 | 47.99286 | 43.06154 | 63.83414 | 39.97956 | 49.16446 | 29.89296 | 27.18978 |
| ENSMUSG00000107994 | -0.189588374 | 0.379791 | 0.547868 | 74.44802 | 57.59143 | 58.79557 | 51.75741 | 78.0089 | 63.69032 | 63.64308 | 70.69342 |
| ENSMUSG00000107997 | -0.134033587 | 0.769772 | 0.857077 | 16.09687 | 13.19804 | 35.60859 | 33.64232 | 18.52711 | 40.22547 | 27.96439 | 22.83941 |
| ENSMUSG00000108010 | -0.476072495 | 0.333933 | 0.502083 | 15.09082 | 10.79839 | 41.40533 | 15.52722 | 20.47734 | 36.87335 | 28.92867 | 30.45255 |
| ENSMUSG00000108030 | 0.688222992 | 0.201704 | 0.35726 | 14.08476 | 13.19804 | 19.04645 | 12.07673 | 3.900445 | 14.52586 | 11.57147 | 6.525547 |
| ENSMUSG00000108034 | 0.635730769 | 0.061189 | 0.157046 | 46.2785 | 40.79393 | 41.40533 | 26.74133 | 28.27823 | 27.93435 | 27.96439 | 15.22628 |
| ENSMUSG00000108035 | 1.673947942 | 0.005318 | 0.026853 | 86.52067 | 73.18911 | 24.01509 | 50.03216 | 9.751113 | 42.46022 | 12.53576 | 8.700729 |
| ENSMUSG00000108037 | 1.012563139 | 0.001436 | 0.010261 | 96.58122 | 87.58697 | 49.6864 | 78.49874 | 54.60623 | 31.28647 | 38.57157 | 29.36496 |
| ENSMUSG00000108042 | 0.614853129 | 0.400916 | 0.568172 | 28.16952 | 14.39786 | 10.76539 | 8.626235 | 1.950223 | 22.34748 | 8.678602 | 7.613138 |
| ENSMUSG00000108049 | 0.351776515 | 0.118705 | 0.247522 | 87.52673 | 99.58519 | 89.43552 | 99.2017 | 63.38224 | 100.5637 | 74.25027 | 56.55474 |
| ENSMUSG00000108064 | 0.271947791 | 0.620951 | 0.752803 | 20.12109 | 11.99822 | 19.04645 | 6.038364 | 11.70134 | 15.64324 | 13.50005 | 6.525547 |
| ENSMUSG00000108080 | 0.585107629 | 0.127971 | 0.259325 | 17.10292 | 22.79661 | 24.01509 | 30.19182 | 19.50223 | 15.64324 | 11.57147 | 16.31387 |
| ENSMUSG00000108084 | -0.451633426 | 0.120983 | 0.250238 | 115.6963 | 110.3836 | 219.4483 | 106.1027 | 188.1965 | 129.6154 | 214.0722 | 224.0438 |
| ENSMUSG00000108101 | -0.209178162 | 0.486007 | 0.644332 | 27.16347 | 40.79393 | 46.37397 | 29.3292 | 36.07912 | 51.39921 | 36.64299 | 42.41605 |
| ENSMUSG00000108120 | -0.221546582 | 0.672113 | 0.790047 | 9.054489 | 4.799286 | 14.90592 | 8.626235 | 11.70134 | 12.29112 | 10.60718 | 9.78832 |
| ENSMUSG00000108122 | -0.143742619 | 0.831188 | 0.897939 | 24.1453 | 5.999108 | 8.281066 | 18.97772 | 6.825779 | 29.05173 | 20.25007 | 7.613138 |
| ENSMUSG00000108137 | 0.007212303 | 0.984242 | 0.99117 | 26.15741 | 25.19625 | 24.8432 | 29.3292 | 15.60178 | 39.10809 | 27.96439 | 22.83941 |
| ENSMUSG00000108148 | 1.491665307 | 0.071395 | 0.174895 | 27.16347 | 34.79482 | 19.04645 | 50.03216 | 3.900445 | 36.87335 | 2.892867 | 3.262773 |
| ENSMUSG00000108169 | 1.231184029 | 0.144982 | 0.283122 | 16.09687 | 28.79572 | 13.24971 | 25.8787 | 2.925334 | 27.93435 | 1.928578 | 3.262773 |
| ENSMUSG00000108171 | 1.183753335 | 0.083684 | 0.194883 | 27.16347 | 15.59768 | 11.59349 | 12.07673 | 2.925334 | 14.52586 | 9.642892 | 2.175182 |
| ENSMUSG00000108176 | 0.523892582 | 0.173719 | 0.321644 | 27.16347 | 46.79304 | 47.20208 | 62.10889 | 27.30312 | 49.16446 | 20.25007 | 31.54014 |
| ENSMUSG00000108181 | 0.562508941 | 0.059978 | 0.154884 | 71.42986 | 53.99197 | 37.2648 | 58.6584 | 32.17867 | 42.46022 | 44.3573 | 30.45255 |
| ENSMUSG00000108190 | -0.251225133 | 0.445833 | 0.608601 | 70.4238 | 59.99108 | 69.56096 | 30.19182 | 71.18313 | 45.81234 | 80.036 | 76.13138 |
| ENSMUSG00000108199 | 0.251543974 | 0.644899 | 0.770145 | 31.18769 | 15.59768 | 12.4216 | 19.84034 | 14.62667 | 31.28647 | 7.714313 | 13.05109 |
| ENSMUSG00000108207 | -0.475804782 | 0.074188 | 0.179688 | 47.28455 | 46.79304 | 82.81066 | 45.71905 | 79.95913 | 69.27719 | 76.17884 | 85.9197 |
| ENSMUSG00000108218 | 0.590746644 | 0.011846 | 0.048508 | 490.9545 | 519.5227 | 346.9767 | 440.8006 | 342.2641 | 406.7242 | 241.0723 | 203.3795 |
| ENSMUSG00000108219 | 0.844410114 | 0.275964 | 0.442194 | 27.16347 | 20.39697 | 11.59349 | 4.313117 | 1.950223 | 17.87799 | 5.785735 | 9.78832 |
| ENSMUSG00000108231 | -0.176937091 | 0.659723 | 0.780834 | 14.08476 | 22.79661 | 19.04645 | 14.6646 | 27.30312 | 21.23011 | 14.46434 | 16.31387 |
| ENSMUSG00000108236 | -0.620505402 | 0.076409 | 0.183406 | 29.17558 | 21.59679 | 36.43669 | 26.74133 | 30.22845 | 32.40385 | 55.92877 | 57.64233 |
| ENSMUSG00000108238 | 0.217044563 | 0.771848 | 0.858471 | 36.21796 | 39.59411 | 12.4216 | 29.3292 | 2.925334 | 59.22083 | 27.0001 | 11.9635 |
| ENSMUSG00000108255 | 0.525793258 | 0.011311 | 0.046782 | 200.2048 | 278.3586 | 226.9012 | 283.8031 | 205.7485 | 203.3621 | 150.4291 | 127.2482 |
| ENSMUSG00000108256 | 1.359426291 | 0.003462 | 0.019613 | 48.29061 | 28.79572 | 28.98373 | 28.46658 | 10.72622 | 24.58223 | 7.714313 | 9.78832 |
| ENSMUSG00000108290 | -1.321673604 | 1.11E-05 | 0.000248 | 172.0353 | 89.98662 | 230.2136 | 150.0965 | 282.7823 | 343.0338 | 416.5729 | 567.7225 |
| ENSMUSG00000108291 | 0.790649769 | 0.171509 | 0.318725 | 30.18163 | 32.39518 | 22.35888 | 23.29083 | 8.776002 | 36.87335 | 10.60718 | 6.525547 |
| ENSMUSG00000108298 | -0.379620011 | 0.464073 | 0.624924 | 18.10898 | 5.999108 | 22.35888 | 9.488858 | 20.47734 | 11.17374 | 22.17865 | 19.57664 |
| ENSMUSG00000108314 | 0.539683346 | 0.210874 | 0.368306 | 62.37537 | 43.19358 | 23.18699 | 21.56559 | 26.32801 | 18.99536 | 27.96439 | 29.36496 |
| ENSMUSG00000108318 | -1.485631051 | 0.01438 | 0.055907 | 23.13925 | 2.399643 | 34.78048 | 19.84034 | 39.97956 | 31.28647 | 86.78602 | 68.51824 |
| ENSMUSG00000108321 | 0.733216276 | 0.275097 | 0.441265 | 16.09687 | 31.19536 | 9.93728 | 15.52722 | 9.751113 | 23.46486 | 2.892867 | 7.613138 |
| ENSMUSG00000108348 | 1.18632166 | 6.45E-05 | 0.000965 | 78.47224 | 71.98929 | 68.73285 | 92.30071 | 25.35289 | 48.04709 | 23.14294 | 41.32846 |
| ENSMUSG00000108350 | 0.427438122 | 0.009505 | 0.04105 | 282.7013 | 271.1597 | 322.9616 | 309.6818 | 235.0018 | 275.9914 | 196.715 | 176.1898 |
| ENSMUSG00000108353 | -1.079518116 | 0.065342 | 0.164657 | 8.048435 | 3.599465 | 5.796746 | 8.626235 | 11.70134 | 20.11273 | 17.3572 | 6.525547 |
| ENSMUSG00000108358 | 0.877212791 | 0.001933 | 0.012811 | 53.32088 | 80.38804 | 65.42042 | 53.48266 | 28.27823 | 46.92971 | 31.82154 | 30.45255 |
| ENSMUSG00000108366 | 0.08476724 | 0.846557 | 0.907743 | 14.08476 | 37.19447 | 19.87456 | 18.11509 | 17.552 | 30.1691 | 17.3572 | 18.48905 |
| ENSMUSG00000108368 | 0.118396525 | 0.792613 | 0.873177 | 33.19979 | 33.595 | 15.73403 | 15.52722 | 26.32801 | 27.93435 | 12.53576 | 22.83941 |
| ENSMUSG00000108402 | 0.71982569 | 0.003025 | 0.017883 | 98.59333 | 104.3845 | 79.49824 | 75.04824 | 47.78045 | 72.62932 | 44.3573 | 52.20437 |

| | | | | | | | | | | | |
|---|---|---|---|---|---|---|---|---|---|---|---|
| ENSMUSG00000108414 | 0.268739202 | 0.047462 | 0.13123 | 2883.352 | 3251.516 | 2822.187 | 3307.298 | 2783.943 | 2992.328 | 2420.366 | 1982.679 |
| ENSMUSG00000108425 | -0.30302863 | 0.537846 | 0.687906 | 20.12109 | 13.19804 | 8.281066 | 9.488858 | 15.60178 | 22.34748 | 10.60718 | 14.13868 |
| ENSMUSG00000108434 | 0.440576498 | 0.364677 | 0.533361 | 18.10898 | 16.7975 | 57.13936 | 37.95543 | 21.45245 | 21.23011 | 19.28578 | 34.80291 |
| ENSMUSG00000108435 | -0.785307893 | 0.001142 | 0.008647 | 58.35115 | 37.19447 | 43.06154 | 42.26855 | 88.73513 | 87.15518 | 64.60737 | 71.78101 |
| ENSMUSG00000108446 | 0.712668576 | 0.189605 | 0.342235 | 54.32694 | 53.99197 | 18.21835 | 27.60395 | 25.35289 | 37.99072 | 7.714313 | 22.83941 |
| ENSMUSG00000108452 | -0.399003094 | 0.022676 | 0.077074 | 272.6407 | 287.9572 | 438.0684 | 357.1261 | 429.049 | 392.1983 | 503.3589 | 466.5766 |
| ENSMUSG00000108456 | 0.47985497 | 0.17964 | 0.329508 | 38.23007 | 57.59143 | 48.03018 | 55.2079 | 43.88001 | 51.39921 | 28.92867 | 18.48905 |
| ENSMUSG00000108461 | -0.327667729 | 0.111455 | 0.237806 | 102.6175 | 64.79036 | 97.71658 | 108.6906 | 112.1378 | 122.9112 | 132.1076 | 104.4087 |
| ENSMUSG00000108465 | 0.288601041 | 0.485861 | 0.644184 | 62.37537 | 49.19268 | 39.74912 | 34.50494 | 44.85512 | 56.98608 | 34.71441 | 15.22628 |
| ENSMUSG00000108477 | -0.458204995 | 0.110346 | 0.23618 | 23.13925 | 25.19625 | 23.18699 | 30.19182 | 36.07912 | 33.52122 | 38.57157 | 31.54014 |
| ENSMUSG00000108478 | 1.48805589 | 0.092529 | 0.208946 | 34.20585 | 38.39429 | 22.35888 | 8.626235 | 0.975111 | 27.93435 | 3.857157 | 4.350364 |
| ENSMUSG00000108486 | 0.731200486 | 0.099111 | 0.219744 | 47.28455 | 53.99197 | 42.23344 | 74.18562 | 17.552 | 63.69032 | 21.21436 | 29.36496 |
| ENSMUSG00000108494 | 1.130967853 | 0.032432 | 0.099944 | 59.35721 | 79.18822 | 31.46805 | 45.71905 | 17.552 | 53.63396 | 15.42863 | 11.9635 |
| ENSMUSG00000108500 | 0.915867125 | 0.102891 | 0.225315 | 21.12714 | 19.19714 | 14.90592 | 12.07673 | 3.900445 | 16.76061 | 9.642892 | 5.437955 |
| ENSMUSG00000108528 | 0.222849877 | 0.598177 | 0.736733 | 17.10292 | 28.79572 | 14.90592 | 17.25247 | 15.60178 | 22.34748 | 17.3572 | 10.87591 |
| ENSMUSG00000108551 | 0.224784265 | 0.639941 | 0.766639 | 34.20585 | 28.79572 | 19.04645 | 25.01608 | 11.70134 | 43.57759 | 21.21436 | 15.22628 |
| ENSMUSG00000108555 | 0.371937991 | 0.072426 | 0.176717 | 100.6054 | 81.58786 | 81.98256 | 78.49874 | 75.08357 | 73.74669 | 54.00019 | 61.99269 |
| ENSMUSG00000108557 | 1.297463591 | 0.142021 | 0.279081 | 38.23007 | 47.99286 | 11.59349 | 19.84034 | 5.850668 | 35.75597 | 0.964289 | 5.437955 |
| ENSMUSG00000108563 | 0.41416848 | 0.067445 | 0.16764 | 80.48435 | 58.79126 | 66.24853 | 91.43809 | 61.43201 | 53.63396 | 53.0359 | 55.46715 |
| ENSMUSG00000108573 | 0.06651807 | 0.822271 | 0.892171 | 43.26034 | 43.19358 | 30.63995 | 44.85642 | 44.85512 | 35.75597 | 27.0001 | 46.76642 |
| ENSMUSG00000108591 | 0.045964195 | 0.747126 | 0.842177 | 240.447 | 227.9661 | 178.0429 | 192.365 | 195.0223 | 201.1273 | 199.6079 | 214.2554 |
| ENSMUSG00000108592 | 0.350834087 | 0.226557 | 0.386481 | 35.2119 | 35.99465 | 39.74912 | 35.36756 | 31.20356 | 36.87335 | 23.14294 | 23.927 |
| ENSMUSG00000108601 | 1.585882774 | 0.001916 | 0.012726 | 98.59333 | 143.9786 | 83.63877 | 98.33908 | 45.83023 | 64.8077 | 23.14294 | 7.613138 |
| ENSMUSG00000108607 | 0.740156776 | 0.050058 | 0.136196 | 26.15741 | 28.79572 | 32.29616 | 25.8787 | 16.57689 | 20.11273 | 22.17865 | 8.700729 |
| ENSMUSG00000108617 | 0.313576773 | 0.40312 | 0.570045 | 20.12109 | 39.59411 | 22.35888 | 21.56559 | 22.42756 | 23.46486 | 20.25007 | 16.31387 |
| ENSMUSG00000108621 | 0.277255868 | 0.016507 | 0.061509 | 1446.706 | 1396.592 | 1173.427 | 1321.539 | 1108.702 | 1292.802 | 992.2536 | 1010.372 |
| ENSMUSG00000108624 | -0.489362288 | 0.38863 | 0.55606 | 5.030272 | 16.7975 | 14.07781 | 7.763611 | 11.70134 | 23.46486 | 8.678602 | 17.40146 |
| ENSMUSG00000108633 | 1.102501904 | 0.023518 | 0.079306 | 165.999 | 173.9741 | 121.7317 | 135.4319 | 32.17867 | 159.7845 | 54.96448 | 31.54014 |
| ENSMUSG00000108635 | 0.768746473 | 0.414411 | 0.580277 | 19.11503 | 22.79661 | 10.76539 | 6.038364 | 7.80089 | 22.34748 | 0 | 4.350364 |
| ENSMUSG00000108648 | 0.856152232 | 0.279728 | 0.446133 | 15.09082 | 17.99732 | 11.59349 | 18.97772 | 5.850668 | 24.58223 | 3.857157 | 1.087591 |
| ENSMUSG00000108655 | 0.367630571 | 0.118327 | 0.247126 | 89.53884 | 79.18822 | 64.59232 | 96.61383 | 52.65601 | 69.27719 | 78.10742 | 55.46715 |
| ENSMUSG00000108658 | 1.257187089 | 5.95E-09 | 5.52E-07 | 128.775 | 160.7761 | 179.6991 | 153.547 | 60.4569 | 78.21619 | 75.21455 | 46.76642 |
| ENSMUSG00000108660 | 0.109804538 | 0.830578 | 0.897636 | 13.07871 | 15.59768 | 9.93728 | 6.900988 | 6.825779 | 13.40849 | 9.642892 | 11.9635 |
| ENSMUSG00000108678 | 0.945719322 | 0.184502 | 0.335684 | 14.08476 | 38.39429 | 19.04645 | 29.3292 | 1.950223 | 31.28647 | 10.60718 | 8.700729 |
| ENSMUSG00000108681 | -0.093430411 | 0.779733 | 0.863998 | 31.18769 | 21.59679 | 33.95237 | 36.23019 | 43.88001 | 33.52122 | 22.17865 | 32.62773 |
| ENSMUSG00000108688 | -0.067311597 | 0.896484 | 0.937476 | 8.048435 | 10.79839 | 13.24971 | 12.93935 | 14.62667 | 16.76061 | 10.60718 | 5.437955 |
| ENSMUSG00000108701 | 0.011374295 | 0.97072 | 0.983617 | 39.23612 | 55.19179 | 50.5145 | 71.59775 | 58.50668 | 67.04245 | 56.89306 | 32.62773 |
| ENSMUSG00000108723 | 0.107141806 | 0.695644 | 0.807828 | 27.16347 | 35.99465 | 35.60859 | 34.50494 | 31.20356 | 30.1691 | 30.85725 | 31.54014 |
| ENSMUSG00000108732 | 0.370569937 | 0.295323 | 0.463083 | 25.15136 | 23.99643 | 24.01509 | 31.05445 | 24.37778 | 22.34748 | 12.53576 | 21.75182 |
| ENSMUSG00000108737 | 1.064329643 | 0.025529 | 0.084257 | 34.20585 | 39.59411 | 25.67131 | 40.5433 | 10.72622 | 32.40385 | 16.39292 | 7.613138 |
| ENSMUSG00000108738 | 0.290918546 | 0.331611 | 0.499409 | 61.36932 | 79.18822 | 39.74912 | 44.85642 | 43.88001 | 46.92971 | 37.60728 | 54.37955 |
| ENSMUSG00000108753 | 0.576864877 | 0.052463 | 0.140714 | 40.24217 | 44.3934 | 48.85829 | 42.26855 | 23.40267 | 40.22547 | 32.78583 | 21.75182 |
| ENSMUSG00000108763 | -1.466111558 | 0.016516 | 0.06153 | 12.07265 | 2.399643 | 13.24971 | 9.488858 | 35.10401 | 13.40849 | 14.46434 | 41.32846 |
| ENSMUSG00000108801 | 0.924627451 | 0.260947 | 0.425107 | 36.21796 | 5.999108 | 5.796746 | 5.175741 | 7.80089 | 4.469496 | 4.821446 | 10.87591 |
| ENSMUSG00000108806 | -0.27620912 | 0.546107 | 0.694558 | 13.07871 | 7.198929 | 19.87456 | 18.97772 | 15.60178 | 17.87799 | 14.46434 | 25.0146 |
| ENSMUSG00000108811 | 0.487835256 | 0.365109 | 0.53387 | 9.054489 | 21.59679 | 11.59349 | 7.763611 | 8.776002 | 10.05637 | 8.678602 | 7.613138 |
| ENSMUSG00000108825 | 0.091650066 | 0.751738 | 0.84531 | 83.50251 | 59.99108 | 51.34261 | 62.97152 | 58.50668 | 87.15518 | 44.3573 | 52.20437 |
| ENSMUSG00000108841 | 0.343371169 | 0.316872 | 0.485004 | 27.16347 | 32.39518 | 33.95237 | 27.60395 | 17.552 | 34.6386 | 25.07152 | 18.48905 |
| ENSMUSG00000108842 | 0.209884582 | 0.645996 | 0.770635 | 31.18769 | 34.79482 | 16.56213 | 17.25247 | 18.52711 | 29.05173 | 27.0001 | 10.87591 |
| ENSMUSG00000108866 | 0.775631674 | NA | NA | 2.012109 | 0 | 0.828107 | 207.0296 | 0.975111 | 0 | 0.964289 | 120.7226 |
| ENSMUSG00000108869 | 0.705104739 | 0.213138 | 0.370961 | 15.09082 | 11.99822 | 37.2648 | 34.50494 | 8.776002 | 20.11273 | 8.678602 | 23.927 |
| ENSMUSG00000108878 | 0.304634119 | 0.460724 | 0.62196 | 19.11503 | 44.3934 | 32.29616 | 24.15346 | 27.30312 | 34.6386 | 16.39292 | 18.48905 |
| ENSMUSG00000108883 | 0.634340306 | 0.217346 | 0.375825 | 22.1332 | 17.99732 | 9.109173 | 19.84034 | 7.80089 | 14.52586 | 15.42863 | 6.525547 |
| ENSMUSG00000108884 | -2.855008524 | 1.32E-08 | 1.08E-06 | 29.17558 | 4.799286 | 45.54586 | 12.93935 | 114.088 | 178.7799 | 191.8935 | 191.416 |
| ENSMUSG00000108923 | 0.373885788 | 0.28538 | 0.452179 | 37.22401 | 22.79661 | 24.8432 | 33.64232 | 17.552 | 18.99536 | 28.92867 | 26.10219 |
| ENSMUSG00000108929 | 0.688890698 | 0.280381 | 0.446867 | 14.08476 | 21.59679 | 9.109173 | 9.488858 | 1.950223 | 7.821619 | 10.60718 | 13.05109 |
| ENSMUSG00000108934 | -0.495409989 | 0.294343 | 0.461894 | 6.036326 | 13.19804 | 14.07781 | 22.42821 | 16.57689 | 18.99536 | 25.07152 | 18.48905 |
| ENSMUSG00000108940 | 0.298285448 | 0.478976 | 0.638266 | 14.08476 | 29.99554 | 25.67131 | 18.11509 | 23.40267 | 14.52586 | 12.53576 | 20.66423 |
| ENSMUSG00000108954 | 0.558953109 | 0.236103 | 0.397076 | 75.45408 | 57.59143 | 54.65504 | 50.89479 | 33.15378 | 86.03781 | 19.28578 | 23.927 |
| ENSMUSG00000108957 | 0.150729657 | 0.611178 | 0.745976 | 34.20585 | 41.99375 | 33.95237 | 27.60395 | 30.22845 | 39.10809 | 26.03581 | 28.27737 |
| ENSMUSG00000108984 | 1.126554813 | 0.115253 | 0.243042 | 25.15136 | 31.19536 | 23.18699 | 31.05445 | 5.850668 | 35.75597 | 2.892867 | 6.525547 |
| ENSMUSG00000108989 | 1.101913825 | 0.179091 | 0.32882 | 9.054489 | 10.79839 | 12.4216 | 23.29083 | 8.776002 | 13.40849 | 3.857157 | 0 |
| ENSMUSG00000109005 | 0.462365722 | 0.13979 | 0.275902 | 119.7205 | 77.9884 | 164.7932 | 154.4096 | 97.51113 | 137.437 | 73.28598 | 68.51824 |
| ENSMUSG00000109008 | -0.382340837 | 0.261698 | 0.425894 | 33.19979 | 61.1909 | 78.67013 | 45.71905 | 62.40712 | 49.16446 | 77.14313 | 96.79561 |

| | | | | | | | | | | | |
|---|---|---|---|---|---|---|---|---|---|---|---|
| ENSMUSG00000109015 | 0.323759454 | 0.193129 | 0.34636 | 70.4238 | 80.38804 | 47.20208 | 69.00988 | 46.80534 | 62.57295 | 53.0359 | 50.02919 |
| ENSMUSG00000109021 | 0.233886177 | 0.542077 | 0.691258 | 39.23612 | 26.39607 | 31.46805 | 18.11509 | 17.552 | 25.6996 | 21.21436 | 33.71532 |
| ENSMUSG00000109028 | 1.368354719 | 0.010607 | 0.044562 | 25.15136 | 14.39786 | 26.49941 | 25.01608 | 9.751113 | 16.76061 | 4.821446 | 4.350364 |
| ENSMUSG00000109032 | -0.129334961 | 0.771931 | 0.858471 | 32.19374 | 44.3934 | 38.09291 | 55.2079 | 34.1289 | 92.74205 | 27.0001 | 32.62773 |
| ENSMUSG00000109038 | 0.322727393 | 0.499514 | 0.655879 | 13.07871 | 10.79839 | 10.76539 | 14.6646 | 6.825779 | 13.40849 | 8.678602 | 10.87591 |
| ENSMUSG00000109051 | 2.299939147 | 0.005166 | 0.026328 | 20.12109 | 35.99465 | 19.87456 | 7.763611 | 3.900445 | 11.17374 | 0.964289 | 1.087591 |
| ENSMUSG00000109089 | 0.824327735 | 0.278406 | 0.44464 | 13.07871 | 3.599465 | 22.35888 | 17.25247 | 12.67645 | 13.40849 | 3.857157 | 2.175182 |
| ENSMUSG00000109093 | -3.324654173 | 1.91E-13 | 6.93E-11 | 35.2119 | 8.398751 | 72.87338 | 31.05445 | 276.9316 | 327.3906 | 385.7157 | 502.4671 |
| ENSMUSG00000109095 | 2.672562341 | 0.001991 | 0.013087 | 21.12714 | 19.19714 | 4.140533 | 34.50494 | 3.900445 | 5.58687 | 2.892867 | 0 |
| ENSMUSG00000109097 | 0.920432492 | 0.00079 | 0.00652 | 86.52067 | 127.1811 | 89.43552 | 123.3552 | 48.75557 | 80.45093 | 55.92877 | 40.24087 |
| ENSMUSG00000109105 | -2.436330601 | NA | NA | 5.030272 | 3.599465 | 3.312427 | 5.175741 | 2.925334 | 86.03781 | 2.892867 | 1.087591 |
| ENSMUSG00000109108 | 0.14634132 | 0.669175 | 0.787855 | 36.21796 | 19.19714 | 36.43669 | 24.15346 | 21.45245 | 27.93435 | 27.96439 | 28.27737 |
| ENSMUSG00000109111 | 1.01506831 | 0.04548 | 0.127448 | 21.12714 | 35.99465 | 23.18699 | 28.46658 | 24.37778 | 4.469496 | 13.50005 | 10.87591 |
| ENSMUSG00000109115 | 0.301450821 | 0.463209 | 0.624014 | 17.10292 | 20.39697 | 12.4216 | 15.52722 | 12.67645 | 11.17374 | 13.50005 | 15.22628 |
| ENSMUSG00000109118 | 0.057986199 | 0.816986 | 0.888935 | 114.6902 | 123.5816 | 120.9036 | 89.71284 | 91.66046 | 158.6671 | 98.35749 | 82.65692 |
| ENSMUSG00000109130 | -1.047540412 | 0.240876 | 0.403048 | 3.018163 | 2.399643 | 41.40533 | 10.35148 | 32.17867 | 10.05637 | 17.3572 | 59.81751 |
| ENSMUSG00000109143 | 1.057037204 | 0.033013 | 0.101329 | 22.1332 | 17.99732 | 23.18699 | 18.97772 | 11.70134 | 11.17374 | 2.892867 | 14.13868 |
| ENSMUSG00000109147 | -0.134776023 | 0.411066 | 0.576931 | 100.6054 | 100.785 | 92.74794 | 82.81186 | 101.4116 | 105.0332 | 108.0004 | 97.8832 |
| ENSMUSG00000109157 | 0.786639846 | 0.067688 | 0.168045 | 108.6539 | 112.7832 | 105.9976 | 119.9047 | 45.83023 | 139.6718 | 38.57157 | 35.89051 |
| ENSMUSG00000109165 | 0.097599425 | 0.866583 | 0.919863 | 23.13925 | 16.7975 | 14.90592 | 6.900988 | 15.60178 | 25.6996 | 8.678602 | 7.613138 |
| ENSMUSG00000109179 | 0.251564617 | 0.257833 | 0.421574 | 108.6539 | 113.983 | 66.24853 | 101.7896 | 77.03379 | 81.56831 | 79.07171 | 89.18247 |
| ENSMUSG00000109198 | -0.026654644 | 0.942068 | 0.966348 | 40.24217 | 38.39429 | 36.43669 | 53.48266 | 23.40267 | 69.27719 | 38.57157 | 41.32846 |
| ENSMUSG00000109216 | 0.489054064 | 0.1838 | 0.334834 | 20.12109 | 23.99643 | 19.04645 | 26.74133 | 19.50223 | 15.64324 | 12.53576 | 16.31387 |
| ENSMUSG00000109222 | -0.141336348 | 0.618609 | 0.751072 | 68.4117 | 53.99197 | 38.92101 | 63.83414 | 43.88001 | 77.09881 | 67.50024 | 59.81751 |
| ENSMUSG00000109224 | -0.199676051 | 0.46071 | 0.62196 | 55.33299 | 37.19447 | 46.37397 | 37.09281 | 48.75557 | 51.39921 | 62.6788 | 39.15328 |
| ENSMUSG00000109231 | 0.498728861 | 0.117221 | 0.245482 | 44.26639 | 33.595 | 29.81184 | 43.9938 | 32.17867 | 32.40385 | 22.17865 | 20.66423 |
| ENSMUSG00000109243 | 0.674717288 | 0.175401 | 0.323752 | 42.25428 | 38.39429 | 19.87456 | 35.36756 | 21.45245 | 35.75597 | 21.21436 | 6.525547 |
| ENSMUSG00000109244 | 0.222255689 | 0.51919 | 0.672764 | 18.10898 | 21.59679 | 32.29616 | 29.3292 | 23.40267 | 21.23011 | 21.21436 | 21.75182 |
| ENSMUSG00000109265 | 0.818485237 | 0.164266 | 0.309253 | 18.10898 | 5.999108 | 15.73403 | 11.21411 | 5.850668 | 12.29112 | 5.785735 | 5.437955 |
| ENSMUSG00000109274 | -0.293662949 | 0.129252 | 0.261057 | 365.1977 | 304.7547 | 445.5214 | 343.3242 | 435.8748 | 316.2169 | 492.7518 | 544.8831 |
| ENSMUSG00000109279 | -0.064230555 | 0.885564 | 0.931652 | 15.09082 | 28.79572 | 14.90592 | 12.93935 | 11.70134 | 24.58223 | 17.3572 | 20.66423 |
| ENSMUSG00000109282 | -1.666833527 | 0.171579 | 0.31877 | 5.030272 | 0 | 23.18699 | 2.58787 | 11.70134 | 2.234748 | 7.714313 | 77.21897 |
| ENSMUSG00000109293 | -0.278626744 | 0.493942 | 0.650785 | 17.10292 | 29.99554 | 30.63995 | 18.97772 | 28.27823 | 42.46022 | 16.39292 | 30.45255 |
| ENSMUSG00000109304 | -0.374842552 | 0.497318 | 0.653879 | 15.09082 | 3.599465 | 14.07781 | 10.35148 | 11.70134 | 17.87799 | 18.32149 | 8.700729 |
| ENSMUSG00000109311 | -2.444018244 | 5.04E-05 | 0.000798 | 30.18163 | 1.199822 | 45.54586 | 22.42821 | 128.7147 | 91.62468 | 129.2147 | 195.7664 |
| ENSMUSG00000109321 | 1.189499471 | 0.001484 | 0.010486 | 115.6963 | 105.5843 | 58.79557 | 96.61383 | 28.27823 | 70.39457 | 40.50014 | 26.10219 |
| ENSMUSG00000109324 | -0.34157266 | 6.42E-06 | 0.000163 | 6676.177 | 7513.282 | 7452.132 | 7133.896 | 9726.735 | 8192.587 | 9254.283 | 9285.853 |
| ENSMUSG00000109325 | -0.552991012 | 0.371028 | 0.540056 | 12.07265 | 3.599465 | 15.73403 | 7.763611 | 20.47734 | 5.58687 | 13.50005 | 18.48905 |
| ENSMUSG00000109336 | 0.166683112 | 0.010984 | 0.045793 | 1279.701 | 1174.625 | 1207.379 | 1199.909 | 1043.369 | 1144.191 | 1080.968 | 1064.752 |
| ENSMUSG00000109361 | 1.144120719 | 0.033341 | 0.101995 | 21.12714 | 22.79661 | 31.46805 | 54.34528 | 8.776002 | 23.46486 | 20.25007 | 6.525547 |
| ENSMUSG00000109378 | 0.526111496 | 0.000336 | 0.003379 | 232.3986 | 262.7609 | 209.511 | 276.9021 | 168.6943 | 186.6015 | 175.5006 | 150.0876 |
| ENSMUSG00000109379 | -0.603902275 | 0.318499 | 0.486689 | 10.06054 | 8.398751 | 11.59349 | 6.038364 | 4.875557 | 24.58223 | 10.60718 | 15.22628 |
| ENSMUSG00000109383 | 0.287340325 | 0.548807 | 0.697048 | 20.12109 | 22.79661 | 16.56213 | 14.6646 | 24.37778 | 16.76061 | 12.53576 | 6.525547 |
| ENSMUSG00000109390 | 0.193360796 | 0.672334 | 0.790206 | 27.16347 | 25.19625 | 30.63995 | 8.626235 | 22.42756 | 16.76061 | 23.14294 | 17.40146 |
| ENSMUSG00000109394 | 1.140380458 | 3.41E-09 | 3.56E-07 | 152.9203 | 160.7761 | 152.3716 | 163.0358 | 57.53157 | 96.09417 | 68.46453 | 64.16787 |
| ENSMUSG00000109398 | 0.546112493 | 0.046425 | 0.129272 | 118.7144 | 97.18554 | 101.8571 | 100.9269 | 72.15824 | 109.5027 | 48.21446 | 57.64233 |
| ENSMUSG00000109408 | -0.074971456 | 0.794681 | 0.873721 | 75.45408 | 40.79393 | 59.62368 | 50.03216 | 49.73068 | 80.45093 | 49.17875 | 59.81751 |
| ENSMUSG00000109416 | 0.703148939 | 0.346428 | 0.515245 | 24.1453 | 26.39607 | 16.56213 | 19.84034 | 5.850668 | 36.87335 | 8.678602 | 2.175182 |
| ENSMUSG00000109420 | -0.369701457 | 0.423799 | 0.588965 | 13.07871 | 10.79839 | 10.76539 | 7.763611 | 11.70134 | 10.05637 | 16.39292 | 16.31387 |
| ENSMUSG00000109429 | 0.848429748 | 0.25494 | 0.41887 | 32.19374 | 16.7975 | 8.281066 | 10.35148 | 3.900445 | 22.34748 | 3.857157 | 7.613138 |
| ENSMUSG00000109455 | -1.017621485 | 0.167603 | 0.313561 | 2.012109 | 8.398751 | 22.35888 | 17.25247 | 21.45245 | 6.704245 | 33.75012 | 40.24087 |
| ENSMUSG00000109485 | 0.028756135 | 0.954082 | 0.973813 | 7.04238 | 15.59768 | 9.109173 | 12.93935 | 12.67645 | 12.29112 | 7.714313 | 10.87591 |
| ENSMUSG00000109498 | 0.471595912 | 0.436157 | 0.600142 | 9.054489 | 8.398751 | 15.73403 | 18.97772 | 13.65156 | 3.352122 | 6.750024 | 14.13868 |
| ENSMUSG00000109509 | 0.02653065 | 0.897303 | 0.937817 | 104.6297 | 116.3827 | 103.5133 | 88.85022 | 97.51113 | 132.9675 | 84.85745 | 90.27006 |
| ENSMUSG00000109511 | -0.276118096 | 0.008637 | 0.038373 | 1540.269 | 1871.722 | 1946.879 | 1636.397 | 2153.046 | 1869.367 | 2212.079 | 2235 |
| ENSMUSG00000109517 | 0.790307747 | 0.106982 | 0.231311 | 23.13925 | 27.5959 | 13.24971 | 34.50494 | 15.60178 | 21.23011 | 13.50005 | 6.525547 |
| ENSMUSG00000109524 | 0.833202393 | 0.282886 | 0.449379 | 21.12714 | 9.598572 | 7.45296 | 17.25247 | 2.925334 | 18.99536 | 1.928578 | 7.613138 |
| ENSMUSG00000109536 | 0.569344457 | 0.041371 | 0.119548 | 146.8839 | 101.9848 | 120.9036 | 232.9083 | 88.73513 | 93.85942 | 106.0718 | 118.5474 |
| ENSMUSG00000109539 | 0.215156459 | 0.48228 | 0.640923 | 37.22401 | 28.79572 | 27.32752 | 27.60395 | 24.37778 | 26.81698 | 30.85725 | 21.75182 |
| ENSMUSG00000109554 | 1.147046579 | 0.001169 | 0.008803 | 59.35721 | 77.9884 | 45.54586 | 77.63611 | 18.52711 | 41.34284 | 21.21436 | 36.9781 |
| ENSMUSG00000109555 | 0.677975026 | 0.190706 | 0.343357 | 52.31483 | 37.19447 | 37.2648 | 30.19182 | 9.751113 | 51.39921 | 23.14294 | 14.13868 |
| ENSMUSG00000109564 | 1.4658541 | 0.255014 | 0.418922 | 65.39353 | 225.5664 | 43.06154 | 79.36136 | 16.57689 | 112.8548 | 20.25007 | 0 |
| ENSMUSG00000109587 | 0.224614296 | 0.477851 | 0.637135 | 42.25428 | 52.79215 | 43.06154 | 43.13117 | 26.32801 | 59.22083 | 31.82154 | 38.06569 |
| ENSMUSG00000109600 | 0.930129675 | 0.10664 | 0.23085 | 22.1332 | 15.59768 | 14.07781 | 8.626235 | 12.67645 | 6.704245 | 8.678602 | 3.262773 |

| | | | | | | | | | | | |
|---|---|---|---|---|---|---|---|---|---|---|---|
| ENSMUSG00000109628 | -2.247085174 | 0.004198 | 0.02262 | 10.06054 | 0 | 29.81184 | 18.11509 | 55.58134 | 25.6996 | 65.57166 | 131.5985 |
| ENSMUSG00000109640 | 0.352510893 | 0.672415 | 0.79025 | 12.07265 | 20.39697 | 18.21835 | 12.07673 | 8.776002 | 34.6386 | 4.821446 | 1.087591 |
| ENSMUSG00000109642 | 1.547705779 | 0.001511 | 0.010619 | 42.25428 | 44.3934 | 19.87456 | 29.3292 | 7.80089 | 15.64324 | 5.785735 | 17.40146 |
| ENSMUSG00000109644 | -2.200703676 | 7.54E-09 | 6.79E-07 | 183.1019 | 57.59143 | 213.6515 | 166.4863 | 708.9059 | 382.1419 | 716.4668 | 1052.788 |
| ENSMUSG00000109695 | 0.12236712 | 0.66646 | 0.785854 | 34.20585 | 37.19447 | 43.88965 | 34.50494 | 37.05423 | 41.34284 | 25.07152 | 34.80291 |
| ENSMUSG00000109714 | 0.386929686 | 0.32956 | 0.497331 | 32.19374 | 45.59322 | 29.81184 | 21.56559 | 36.07912 | 14.52586 | 23.14294 | 23.927 |
| ENSMUSG00000109715 | -0.818517386 | 0.030354 | 0.095243 | 15.09082 | 14.39786 | 26.49941 | 17.25247 | 26.32801 | 26.81698 | 31.82154 | 45.67883 |
| ENSMUSG00000109718 | -0.287991235 | 0.640466 | 0.767014 | 5.030272 | 5.999108 | 14.90592 | 17.25247 | 12.67645 | 17.87799 | 5.785735 | 17.40146 |
| ENSMUSG00000109724 | 0.47616464 | 0.096826 | 0.215953 | 45.27245 | 55.19179 | 48.03018 | 31.91707 | 32.17867 | 36.87335 | 32.78583 | 27.18978 |
| ENSMUSG00000109727 | 0.446168034 | 0.14621 | 0.284947 | 72.43591 | 45.59322 | 49.6864 | 40.5433 | 33.15378 | 52.51658 | 33.75012 | 33.71532 |
| ENSMUSG00000109751 | 1.18844411 | 0.249231 | 0.412686 | 19.11503 | 38.39429 | 17.39024 | 12.07673 | 0 | 31.28647 | 4.821446 | 2.175182 |
| ENSMUSG00000109771 | -0.131574387 | 0.648123 | 0.772125 | 47.28455 | 55.19179 | 33.95237 | 36.23019 | 57.53157 | 52.51658 | 40.50014 | 36.9781 |
| ENSMUSG00000109780 | 0.13332427 | 0.678654 | 0.794819 | 35.2119 | 35.99465 | 38.92101 | 55.2079 | 27.30312 | 45.81234 | 29.89296 | 48.9416 |
| ENSMUSG00000109783 | -0.665111335 | 0.116646 | 0.244888 | 19.11503 | 15.59768 | 15.73403 | 12.93935 | 27.30312 | 36.87335 | 23.14294 | 13.05109 |
| ENSMUSG00000109787 | 1.802462654 | 0.000148 | 0.001814 | 48.29061 | 63.59054 | 42.23344 | 66.42201 | 7.80089 | 33.52122 | 12.53576 | 9.78832 |
| ENSMUSG00000109799 | 0.310054104 | 0.482953 | 0.641435 | 23.13925 | 16.7975 | 14.07781 | 17.25247 | 14.62667 | 20.11273 | 7.714313 | 15.22628 |
| ENSMUSG00000109807 | 0.117001415 | 0.745664 | 0.841093 | 24.1453 | 38.39429 | 21.53077 | 35.36756 | 20.47734 | 34.6386 | 32.78583 | 21.75182 |
| ENSMUSG00000109810 | 0.149082309 | 0.6057 | 0.742186 | 35.2119 | 63.59054 | 42.23344 | 48.30692 | 46.80534 | 50.28183 | 32.78583 | 40.24087 |
| ENSMUSG00000109812 | 1.566106732 | 0.013389 | 0.053106 | 74.44802 | 77.9884 | 55.48314 | 50.89479 | 6.825779 | 56.98608 | 17.3572 | 6.525547 |
| ENSMUSG00000109814 | 0.09712141 | 0.786049 | 0.868441 | 66.39959 | 40.79393 | 39.74912 | 38.81806 | 34.1289 | 34.6386 | 69.42882 | 34.80291 |
| ENSMUSG00000109827 | 1.52873156 | 2.21E-05 | 0.000425 | 50.30272 | 70.78947 | 35.60859 | 66.42201 | 23.40267 | 21.23011 | 20.25007 | 11.9635 |
| ENSMUSG00000109829 | 0.127193581 | 0.684094 | 0.798636 | 30.18163 | 22.79661 | 28.15563 | 35.36756 | 32.17867 | 27.93435 | 22.17865 | 25.0146 |
| ENSMUSG00000109836 | -0.130765254 | 0.643247 | 0.76925 | 30.18163 | 39.59411 | 35.60859 | 34.50494 | 39.97956 | 26.81698 | 45.32159 | 40.24087 |
| ENSMUSG00000109844 | 0.415909351 | 0.477199 | 0.636625 | 9.054489 | 14.39786 | 12.4216 | 18.11509 | 17.552 | 7.821619 | 11.57147 | 3.262773 |
| ENSMUSG00000109857 | 0.055445987 | 0.897358 | 0.937817 | 10.06054 | 19.19714 | 17.39024 | 14.6646 | 18.52711 | 14.52586 | 10.60718 | 15.22628 |
| ENSMUSG00000109865 | -0.064835565 | 0.298103 | 0.466056 | 1356.161 | 1341.4 | 1414.406 | 1461.284 | 1453.891 | 1509.572 | 1393.398 | 1479.124 |
| ENSMUSG00000109866 | 0.668007504 | 0.0577 | 0.15054 | 132.7992 | 213.5682 | 149.0592 | 163.8985 | 86.78491 | 183.2494 | 54.00019 | 91.35765 |
| ENSMUSG00000109873 | 1.116581285 | 0.061028 | 0.156753 | 46.2785 | 33.595 | 24.8432 | 33.64232 | 6.825779 | 39.10809 | 10.60718 | 7.613138 |
| ENSMUSG00000109901 | -0.076810544 | 0.414473 | 0.5803 | 1180.102 | 1117.034 | 1150.24 | 1155.053 | 1228.64 | 1031.336 | 1251.647 | 1342.087 |
| ENSMUSG00000109904 | 0.033316717 | 0.893997 | 0.936094 | 63.38142 | 70.78947 | 67.90474 | 87.9876 | 83.85957 | 81.56831 | 71.3574 | 46.76642 |
| ENSMUSG00000109917 | 0.146557661 | 0.620605 | 0.752604 | 59.35721 | 58.79126 | 49.6864 | 60.38364 | 41.92979 | 73.74669 | 57.85735 | 32.62773 |
| ENSMUSG00000109933 | 0.100800273 | 0.767162 | 0.855545 | 27.16347 | 26.39607 | 34.78048 | 25.8787 | 37.05423 | 27.93435 | 23.14294 | 18.48905 |
| ENSMUSG00000109936 | -0.031980561 | 0.866279 | 0.919646 | 172.0353 | 195.5709 | 123.3879 | 180.2883 | 202.8232 | 168.7235 | 149.4648 | 163.1387 |
| ENSMUSG00000109946 | 0.118740106 | 0.41179 | 0.577594 | 216.3017 | 241.1641 | 264.9941 | 263.1002 | 240.8525 | 200.01 | 263.2509 | 203.3795 |
| ENSMUSG00000109953 | 0.135116989 | 0.769264 | 0.856822 | 16.09687 | 9.598572 | 10.76539 | 15.52722 | 14.62667 | 13.40849 | 8.678602 | 10.87591 |
| ENSMUSG00000109956 | 0.012018705 | 0.975743 | 0.986757 | 18.10898 | 41.99375 | 19.04645 | 27.60395 | 28.27823 | 31.28647 | 18.32149 | 27.18978 |
| ENSMUSG00000109975 | -0.2402284 | 0.300894 | 0.469266 | 57.3451 | 37.19447 | 62.108 | 61.24627 | 65.33246 | 61.45558 | 68.46453 | 64.16787 |
| ENSMUSG00000110010 | 0.188124666 | 0.424819 | 0.589775 | 62.37537 | 65.99018 | 55.48314 | 80.22398 | 51.6809 | 63.69032 | 68.46453 | 47.85401 |
| ENSMUSG00000110020 | 0.93894378 | 0.044623 | 0.125859 | 13.07871 | 29.99554 | 29.81184 | 16.38985 | 14.62667 | 12.29112 | 8.678602 | 10.87591 |
| ENSMUSG00000110030 | 0.534490674 | 0.208352 | 0.365492 | 24.1453 | 17.99732 | 21.53077 | 14.6646 | 9.751113 | 20.11273 | 12.53576 | 11.9635 |
| ENSMUSG00000110057 | -0.181938306 | 0.489034 | 0.646694 | 47.28455 | 61.1909 | 67.07664 | 48.30692 | 84.83468 | 63.69032 | 54.00019 | 51.11678 |
| ENSMUSG00000110060 | -1.082624883 | 0.073081 | 0.177846 | 17.10292 | 9.598572 | 19.04645 | 4.313117 | 20.47734 | 11.17374 | 28.92867 | 45.67883 |
| ENSMUSG00000110062 | -1.28359079 | 0.033804 | 0.103207 | 16.09687 | 4.799286 | 13.24971 | 22.42821 | 33.15378 | 13.40849 | 25.07152 | 67.43065 |
| ENSMUSG00000110071 | 1.420135778 | 0.013007 | 0.052027 | 8.048435 | 19.19714 | 15.73403 | 20.70296 | 2.925334 | 6.704245 | 7.714313 | 6.525547 |
| ENSMUSG00000110079 | 1.367833033 | 0.006924 | 0.032734 | 70.4238 | 91.18644 | 50.5145 | 40.5433 | 13.65156 | 50.28183 | 23.14294 | 10.87591 |
| ENSMUSG00000110084 | 0.615359216 | 0.267963 | 0.432339 | 13.07871 | 20.39697 | 13.24971 | 16.38985 | 2.925334 | 15.64324 | 8.678602 | 14.13868 |
| ENSMUSG00000110101 | 0.028530431 | 0.932733 | 0.960099 | 35.2119 | 32.39518 | 28.98373 | 57.79577 | 50.70579 | 34.6386 | 34.71441 | 31.54014 |
| ENSMUSG00000110105 | 0.06789516 | 0.83532 | 0.900426 | 33.19979 | 17.99732 | 34.78048 | 31.91707 | 24.37778 | 31.28647 | 31.82154 | 26.10219 |
| ENSMUSG00000110120 | 0.11552627 | 0.8394 | 0.9029 | 26.15741 | 32.39518 | 9.109173 | 12.93935 | 22.42756 | 29.05173 | 13.50005 | 8.700729 |
| ENSMUSG00000110124 | 0.064322311 | 0.880142 | 0.927958 | 27.16347 | 10.79839 | 19.04645 | 17.25247 | 14.62667 | 23.46486 | 19.28578 | 14.13868 |
| ENSMUSG00000110125 | 0.642321414 | 0.043983 | 0.124553 | 49.29666 | 61.1909 | 45.54586 | 50.89479 | 40.95467 | 44.69496 | 27.0001 | 19.57664 |
| ENSMUSG00000110129 | -0.177138377 | 0.384967 | 0.552546 | 75.45408 | 63.59054 | 74.5296 | 76.77349 | 78.98402 | 98.32892 | 86.78602 | 65.25547 |
| ENSMUSG00000110138 | 1.436529391 | 0.01648 | 0.061458 | 29.17558 | 17.99732 | 7.45296 | 18.11509 | 4.875557 | 7.821619 | 9.642892 | 4.350364 |
| ENSMUSG00000110140 | -0.504493383 | 0.279948 | 0.446314 | 16.09687 | 8.398751 | 8.281066 | 10.35148 | 14.62667 | 13.40849 | 12.53576 | 20.66423 |
| ENSMUSG00000110148 | -0.094737589 | 0.755075 | 0.847926 | 30.18163 | 28.79572 | 25.67131 | 30.19182 | 26.32801 | 41.34284 | 27.0001 | 28.27737 |
| ENSMUSG00000110156 | -0.32242825 | 0.206963 | 0.363539 | 38.23007 | 43.19358 | 34.78048 | 36.23019 | 43.88001 | 48.04709 | 39.53586 | 58.72992 |
| ENSMUSG00000110159 | 0.800512276 | 0.11883 | 0.247643 | 126.7628 | 147.578 | 66.24853 | 111.2784 | 38.02934 | 146.376 | 50.14304 | 25.0146 |
| ENSMUSG00000110161 | 0.042535994 | 0.922815 | 0.953984 | 20.12109 | 34.79482 | 13.24971 | 18.97772 | 17.552 | 29.05173 | 14.46434 | 22.83941 |
| ENSMUSG00000110165 | -0.901003477 | 0.091002 | 0.206763 | 6.036326 | 13.19804 | 9.109173 | 4.313117 | 9.751113 | 16.76061 | 18.32149 | 15.22628 |
| ENSMUSG00000110170 | 0.098276333 | 0.779046 | 0.863756 | 74.44802 | 45.59322 | 45.54586 | 43.13117 | 62.40712 | 65.92507 | 31.82154 | 34.80291 |
| ENSMUSG00000110185 | 0.826858387 | 0.003824 | 0.021132 | 273.6468 | 343.149 | 203.7142 | 285.5284 | 111.1627 | 253.6439 | 126.3219 | 132.6861 |
| ENSMUSG00000110195 | -0.666009803 | 0.02423 | 0.081141 | 68.4117 | 53.99197 | 56.31125 | 49.16954 | 106.2871 | 54.75133 | 76.17884 | 123.9854 |
| ENSMUSG00000110197 | 0.024163831 | 0.95986 | 0.977262 | 22.1332 | 8.398751 | 11.59349 | 22.42821 | 17.552 | 11.17374 | 14.46434 | 20.66423 |
| ENSMUSG00000110206 | 0.443863641 | 0.095828 | 0.214375 | 95.57516 | 110.3836 | 83.63877 | 115.5915 | 45.83023 | 99.44629 | 69.42882 | 83.74451 |

| | | | | | | | | | | | |
|---|---|---|---|---|---|---|---|---|---|---|---|
| ENSMUSG00000110218 | 0.334709564 | 0.153241 | 0.294464 | 84.50857 | 52.79215 | 90.26362 | 70.73513 | 58.50668 | 64.8077 | 54.96448 | 59.81751 |
| ENSMUSG00000110234 | 0.277818701 | 0.520989 | 0.674527 | 19.11503 | 19.19714 | 25.67131 | 20.70296 | 23.40267 | 7.821619 | 23.14294 | 15.22628 |
| ENSMUSG00000110256 | -0.30804615 | 0.536117 | 0.686694 | 13.07871 | 8.398751 | 9.93728 | 7.763611 | 14.62667 | 15.64324 | 10.60718 | 7.613138 |
| ENSMUSG00000110269 | 0.64427129 | 0.287464 | 0.454404 | 14.08476 | 25.19625 | 28.15563 | 26.74133 | 13.65156 | 33.52122 | 4.821446 | 8.700729 |
| ENSMUSG00000110275 | 0.086894935 | 0.612791 | 0.746981 | 135.8173 | 154.777 | 144.0906 | 141.4703 | 153.0925 | 106.1505 | 156.2148 | 125.073 |
| ENSMUSG00000110277 | 0.092698334 | 0.627733 | 0.757466 | 141.8537 | 221.967 | 155.684 | 135.4319 | 156.9929 | 143.0239 | 161.0363 | 150.0876 |
| ENSMUSG00000110279 | 0.349336854 | 0.174778 | 0.322987 | 40.24217 | 49.19268 | 52.99882 | 55.2079 | 40.95467 | 30.1691 | 38.57157 | 45.67883 |
| ENSMUSG00000110282 | 0.971772304 | 0.177829 | 0.327072 | 49.29666 | 44.3934 | 28.98373 | 31.91707 | 4.875557 | 56.98608 | 9.642892 | 7.613138 |
| ENSMUSG00000110289 | -0.093809587 | 0.799712 | 0.877497 | 19.11503 | 38.39429 | 43.06154 | 45.71905 | 31.20356 | 46.92971 | 50.14304 | 28.27737 |
| ENSMUSG00000110317 | 0.196521471 | 0.484162 | 0.642397 | 147.89 | 110.3836 | 134.9814 | 154.4096 | 88.73513 | 192.1883 | 107.0361 | 91.35765 |
| ENSMUSG00000110331 | 0.591126286 | 0.0018 | 0.012165 | 94.56911 | 88.78679 | 114.2787 | 106.9653 | 76.05868 | 65.92507 | 67.50024 | 59.81751 |
| ENSMUSG00000110340 | -0.485344275 | 0.266577 | 0.431024 | 14.08476 | 11.99822 | 9.109173 | 23.29083 | 20.47734 | 15.64324 | 23.14294 | 22.83941 |
| ENSMUSG00000110344 | -0.108944956 | 0.338576 | 0.507122 | 201.2109 | 218.3675 | 231.0418 | 238.9467 | 235.0018 | 250.2918 | 239.1437 | 237.0949 |
| ENSMUSG00000110350 | -0.418807229 | 0.123404 | 0.25306 | 62.37537 | 73.18911 | 75.3577 | 75.04824 | 99.46135 | 55.8687 | 96.42892 | 130.5109 |
| ENSMUSG00000110353 | -1.323105961 | 0.164627 | 0.30972 | 0 | 15.59768 | 7.45296 | 2.58787 | 7.80089 | 6.704245 | 18.32149 | 30.45255 |
| ENSMUSG00000110357 | -0.244630076 | 0.227764 | 0.387716 | 88.53278 | 76.78858 | 68.73285 | 80.22398 | 89.71024 | 78.21619 | 87.75031 | 116.3722 |
| ENSMUSG00000110360 | 0.312622321 | 0.386424 | 0.553973 | 38.23007 | 32.39518 | 31.46805 | 29.3292 | 16.57689 | 41.34284 | 21.21436 | 27.18978 |
| ENSMUSG00000110386 | -1.036730829 | 0.051403 | 0.138676 | 8.048435 | 9.598572 | 16.56213 | 15.52722 | 10.72622 | 18.99536 | 39.53586 | 33.71532 |
| ENSMUSG00000110393 | 0.388247802 | 0.36066 | 0.529247 | 79.47829 | 88.78679 | 79.49824 | 104.3774 | 37.05423 | 134.0849 | 63.64308 | 34.80291 |
| ENSMUSG00000110404 | 0.140627264 | 0.732986 | 0.83285 | 25.15136 | 21.59679 | 16.56213 | 23.29083 | 13.65156 | 21.23011 | 13.50005 | 30.45255 |
| ENSMUSG00000110409 | -0.268751147 | 0.360279 | 0.528788 | 39.23612 | 31.19536 | 55.48314 | 39.68068 | 37.05423 | 61.45558 | 55.92877 | 46.76642 |
| ENSMUSG00000110411 | 0.773569827 | 0.063851 | 0.16205 | 32.19374 | 31.19536 | 28.98373 | 16.38985 | 16.57689 | 23.46486 | 11.57147 | 11.9635 |
| ENSMUSG00000110419 | 0.563515202 | 0.07957 | 0.18848 | 48.29061 | 29.99554 | 34.78048 | 31.91707 | 18.52711 | 23.46486 | 27.96439 | 28.27737 |
| ENSMUSG00000110424 | 0.646058505 | 0.000746 | 0.006244 | 152.9203 | 177.5736 | 136.6376 | 169.0742 | 91.66046 | 124.0285 | 110.8933 | 79.39415 |
| ENSMUSG00000110427 | 0.246403691 | 0.696305 | 0.808286 | 6.036326 | 9.598572 | 18.21835 | 10.35148 | 2.925334 | 10.05637 | 10.60718 | 14.13868 |
| ENSMUSG00000110433 | 0.190280666 | 0.728897 | 0.829737 | 22.1332 | 7.198929 | 12.4216 | 21.56559 | 15.60178 | 21.23011 | 12.53576 | 6.525547 |
| ENSMUSG00000110444 | 0.029342907 | 0.817903 | 0.889524 | 262.5802 | 257.9616 | 251.7444 | 207.0296 | 248.6534 | 254.7613 | 229.5008 | 225.1314 |
| ENSMUSG00000110458 | -0.773443473 | 0.003456 | 0.019605 | 56.33904 | 65.99018 | 67.90474 | 71.59775 | 156.0178 | 72.62932 | 101.2504 | 117.4598 |
| ENSMUSG00000110476 | -0.943995979 | 0.173482 | 0.321304 | 19.11503 | 1.199822 | 10.76539 | 5.175741 | 15.60178 | 12.29112 | 27.0001 | 15.22628 |
| ENSMUSG00000110480 | 1.349139314 | 0.102399 | 0.224557 | 48.29061 | 32.39518 | 12.4216 | 12.93935 | 2.925334 | 27.93435 | 8.678602 | 2.175182 |
| ENSMUSG00000110497 | 0.844757586 | 0.001465 | 0.0104 | 371.2341 | 411.5388 | 269.1347 | 323.4838 | 203.7983 | 289.3999 | 139.8219 | 132.6861 |
| ENSMUSG00000110504 | 0.290647444 | 0.707532 | 0.816189 | 10.06054 | 9.598572 | 13.24971 | 20.70296 | 5.850668 | 30.1691 | 2.892867 | 5.437955 |
| ENSMUSG00000110521 | 1.141380179 | 0.028033 | 0.090168 | 18.10898 | 20.39697 | 14.07781 | 9.488858 | 4.875557 | 6.704245 | 8.678602 | 7.613138 |
| ENSMUSG00000110528 | -0.074641948 | 0.794566 | 0.873721 | 38.23007 | 70.78947 | 52.17072 | 70.73513 | 66.30757 | 51.39921 | 50.14304 | 76.13138 |
| ENSMUSG00000110529 | 0.173649428 | 0.660723 | 0.781535 | 44.26639 | 44.3934 | 24.8432 | 37.95543 | 17.552 | 52.51658 | 34.71441 | 29.36496 |
| ENSMUSG00000110540 | 0.053482604 | 0.915595 | 0.949979 | 7.04238 | 21.59679 | 26.49941 | 17.25247 | 11.70134 | 18.99536 | 14.46434 | 25.0146 |
| ENSMUSG00000110545 | -0.363336182 | 0.267469 | 0.431699 | 17.10292 | 22.79661 | 23.18699 | 29.3292 | 35.10401 | 31.28647 | 28.92867 | 23.927 |
| ENSMUSG00000110547 | 0.418561194 | 0.448661 | 0.611295 | 12.07265 | 22.79661 | 9.93728 | 23.29083 | 6.825779 | 18.99536 | 17.3572 | 7.613138 |
| ENSMUSG00000110557 | 0.008627483 | 0.983254 | 0.990824 | 59.35721 | 70.78947 | 47.20208 | 86.26235 | 69.2329 | 117.3243 | 36.64299 | 39.15328 |
| ENSMUSG00000110575 | -0.30332156 | 0.44345 | 0.606471 | 15.09082 | 13.19804 | 27.32752 | 15.52722 | 24.37778 | 25.6996 | 20.25007 | 18.48905 |
| ENSMUSG00000110588 | 0.125091175 | 0.661352 | 0.782069 | 64.38748 | 41.99375 | 74.5296 | 78.49874 | 55.58134 | 42.46022 | 71.3574 | 69.60583 |
| ENSMUSG00000110598 | 0.189602873 | 0.783707 | 0.866969 | 23.13925 | 20.39697 | 6.624853 | 4.313117 | 11.70134 | 20.11273 | 5.785735 | 9.78832 |
| ENSMUSG00000110601 | 0.991403048 | 0.121264 | 0.250508 | 13.07871 | 13.19804 | 9.93728 | 18.11509 | 11.70134 | 5.58687 | 8.678602 | 1.087591 |
| ENSMUSG00000110605 | -0.012294952 | 0.956084 | 0.97505 | 78.47224 | 58.79126 | 65.42042 | 63.83414 | 69.2329 | 50.28183 | 77.14313 | 71.78101 |
| ENSMUSG00000110611 | -0.307086417 | 0.410633 | 0.576745 | 15.09082 | 32.39518 | 19.87456 | 18.11509 | 19.50223 | 27.93435 | 27.96439 | 29.36496 |
| ENSMUSG00000110615 | 0.527924419 | 0.200022 | 0.355064 | 22.1332 | 15.59768 | 20.70267 | 26.74133 | 13.65156 | 14.52586 | 21.21436 | 9.78832 |
| ENSMUSG00000110622 | -1.104581973 | 0.066255 | 0.165939 | 16.09687 | 9.598572 | 9.93728 | 6.038364 | 17.552 | 6.704245 | 27.96439 | 36.9781 |
| ENSMUSG00000110630 | -0.373200049 | 0.267131 | 0.431449 | 29.17558 | 16.7975 | 28.98373 | 24.15346 | 22.42756 | 35.75597 | 37.60728 | 33.71532 |
| ENSMUSG00000110631 | 0.678813722 | 0.004604 | 0.024228 | 2938.685 | 2403.243 | 1710.868 | 2361.863 | 1419.762 | 1725.226 | 1872.65 | 862.4597 |
| ENSMUSG00000110639 | 0.945166635 | 0.229972 | 0.390215 | 10.06054 | 26.39607 | 12.4216 | 12.93935 | 0.975111 | 20.11273 | 6.750024 | 4.350364 |
| ENSMUSG00000110647 | -0.567838677 | 0.379997 | 0.548037 | 12.07265 | 11.99822 | 23.18699 | 4.313117 | 22.42756 | 5.58687 | 20.25007 | 28.27737 |
| ENSMUSG00000110649 | -0.470033943 | 0.231331 | 0.391871 | 44.26639 | 35.99465 | 67.90474 | 24.15346 | 68.25779 | 39.10809 | 48.21446 | 83.74451 |
| ENSMUSG00000110652 | 0.208973604 | 0.566727 | 0.711144 | 17.10292 | 27.5959 | 20.70267 | 22.42821 | 13.65156 | 22.34748 | 20.25007 | 19.57664 |
| ENSMUSG00000110656 | -0.316800374 | 0.038348 | 0.113112 | 152.9203 | 190.7716 | 202.8861 | 151.8217 | 197.9476 | 239.1181 | 219.8579 | 213.1679 |
| ENSMUSG00000110661 | 1.457828868 | 2.17E-05 | 0.000422 | 35.2119 | 41.99375 | 50.5145 | 43.13117 | 13.65156 | 16.76061 | 10.60718 | 21.75182 |
| ENSMUSG00000110669 | -1.018761611 | 0.012581 | 0.050741 | 20.12109 | 19.19714 | 24.01509 | 18.97772 | 46.80534 | 17.87799 | 42.42872 | 59.81751 |
| ENSMUSG00000110679 | -0.149523852 | 0.643941 | 0.769677 | 23.13925 | 26.39607 | 25.67131 | 25.01608 | 21.45245 | 23.46486 | 36.64299 | 29.36496 |
| ENSMUSG00000110686 | 0.047018814 | 0.913498 | 0.948837 | 20.12109 | 27.5959 | 24.01509 | 19.84034 | 27.30312 | 34.6386 | 12.53576 | 14.13868 |
| ENSMUSG00000110697 | -0.537563389 | 0.38343 | 0.551394 | 7.04238 | 16.7975 | 4.96864 | 23.29083 | 19.50223 | 10.05637 | 15.42863 | 30.45255 |
| ENSMUSG00000110702 | 0.189991293 | 0.633835 | 0.761939 | 20.12109 | 29.99554 | 59.62368 | 28.46658 | 33.15378 | 29.05173 | 26.03581 | 33.71532 |
| ENSMUSG00000110744 | -0.189400136 | 0.718168 | 0.823926 | 10.06054 | 10.79839 | 15.73403 | 14.6646 | 10.72622 | 6.704245 | 16.39292 | 25.0146 |
| ENSMUSG00000110750 | -2.21294019 | 0.000256 | 0.002754 | 8.048435 | 4.799286 | 4.96864 | 2.58787 | 23.40267 | 8.938993 | 30.85725 | 30.45255 |
| ENSMUSG00000110758 | 1.206371741 | 0.103947 | 0.22682 | 28.16952 | 55.19179 | 15.73403 | 30.19182 | 4.875557 | 39.10809 | 6.750024 | 5.437955 |
| ENSMUSG00000110759 | 1.523375494 | 0.076647 | 0.183785 | 19.11503 | 21.59679 | 15.73403 | 5.175741 | 0 | 12.29112 | 3.857157 | 5.437955 |

| | | | | | | | | | | | |
|---|---|---|---|---|---|---|---|---|---|---|---|
| ENSMUSG00000110768 | -0.229263074 | 0.714993 | 0.821396 | 8.048435 | 9.598572 | 12.4216 | 12.07673 | 15.60178 | 12.29112 | 19.28578 | 2.175182 |
| ENSMUSG00000110791 | 1.370236365 | 0.040398 | 0.117544 | 24.1453 | 40.79393 | 20.70267 | 18.97772 | 6.825779 | 25.6996 | 4.821446 | 3.262773 |
| ENSMUSG00000110795 | 0.515407892 | 0.349948 | 0.518737 | 26.15741 | 14.39786 | 14.07781 | 18.97772 | 12.67645 | 20.11273 | 15.42863 | 3.262773 |
| ENSMUSG00000110815 | -0.23299694 | 0.702002 | 0.812145 | 10.06054 | 14.39786 | 43.88965 | 11.21411 | 14.62667 | 14.52586 | 36.64299 | 28.27737 |
| ENSMUSG00000110824 | 0.220729225 | 0.608487 | 0.744118 | 14.08476 | 20.39697 | 18.21835 | 18.97772 | 14.62667 | 24.58223 | 10.60718 | 11.9635 |
| ENSMUSG00000110830 | 0.167309842 | 0.593277 | 0.732826 | 31.18769 | 22.79661 | 28.15563 | 26.74133 | 23.40267 | 30.1691 | 22.17865 | 21.75182 |
| ENSMUSG00000110834 | -0.238152651 | 0.287112 | 0.454071 | 42.25428 | 50.3925 | 49.6864 | 42.26855 | 54.60623 | 48.04709 | 56.89306 | 57.64233 |
| ENSMUSG00000110841 | -0.391642509 | 0.003075 | 0.0181 | 209.2593 | 208.7689 | 212.8234 | 219.969 | 306.185 | 226.8269 | 282.5367 | 300.1751 |
| ENSMUSG00000110863 | 0.511007466 | 0.048961 | 0.134207 | 49.29666 | 46.79304 | 40.57722 | 40.5433 | 34.1289 | 32.40385 | 27.0001 | 30.45255 |
| ENSMUSG00000110874 | -1.635070216 | 4.46E-08 | 2.95E-06 | 73.44197 | 29.99554 | 72.87338 | 64.69676 | 154.0676 | 150.8455 | 200.5721 | 247.9708 |
| ENSMUSG00000110899 | 0.40235311 | 0.347374 | 0.516236 | 16.09687 | 19.19714 | 16.56213 | 24.15346 | 13.65156 | 7.821619 | 16.39292 | 19.57664 |
| ENSMUSG00000110928 | 0.980348579 | 0.208029 | 0.36501 | 62.37537 | 21.59679 | 10.76539 | 24.15346 | 6.825779 | 29.05173 | 22.17865 | 2.175182 |
| ENSMUSG00000110945 | 0.211520719 | 0.389411 | 0.556741 | 208.2533 | 227.9661 | 148.2311 | 175.9752 | 144.3165 | 240.2354 | 147.5362 | 123.9854 |
| ENSMUSG00000110949 | -0.492600896 | 0.009069 | 0.039721 | 65.39353 | 59.99108 | 71.21717 | 73.323 | 94.5858 | 88.27255 | 110.8933 | 87.00729 |
| ENSMUSG00000111014 | -2.207221821 | 0.018256 | 0.066209 | 4.024217 | 1.199822 | 15.73403 | 3.450494 | 8.776002 | 16.76061 | 79.07171 | 9.78832 |
| ENSMUSG00000111055 | -0.377109223 | 0.467546 | 0.62788 | 10.06054 | 19.19714 | 23.18699 | 5.175741 | 17.552 | 20.11273 | 16.39292 | 20.66423 |
| ENSMUSG00000111063 | 0.577916398 | 1.02E-06 | 3.81E-05 | 366.2038 | 321.5522 | 302.2589 | 319.1707 | 222.3254 | 239.1181 | 203.465 | 212.0803 |
| ENSMUSG00000111067 | 0.491811741 | 0.226076 | 0.385874 | 19.11503 | 23.99643 | 20.70267 | 17.25247 | 21.45245 | 11.17374 | 12.53576 | 11.9635 |
| ENSMUSG00000111080 | 0.238790033 | 0.106249 | 0.230318 | 400.4096 | 457.132 | 385.0696 | 439.938 | 312.0356 | 448.067 | 355.8227 | 309.9635 |
| ENSMUSG00000111082 | -0.877622167 | 0.027033 | 0.087854 | 9.054489 | 15.59768 | 23.18699 | 13.80198 | 25.35289 | 26.81698 | 35.6787 | 26.10219 |
| ENSMUSG00000111088 | -0.158891049 | 0.585102 | 0.725569 | 52.31483 | 38.39429 | 65.42042 | 31.91707 | 55.58134 | 56.98608 | 48.21446 | 50.02919 |
| ENSMUSG00000111090 | 1.034559304 | 0.104231 | 0.22728 | 17.10292 | 15.59768 | 7.45296 | 32.77969 | 4.875557 | 16.76061 | 7.714313 | 6.525547 |
| ENSMUSG00000111118 | -1.333312324 | 0.039028 | 0.114462 | 10.06054 | 2.399643 | 9.109173 | 3.450494 | 7.80089 | 14.52586 | 25.07152 | 16.31387 |
| ENSMUSG00000111128 | 0.365805555 | 0.009399 | 0.0407 | 401.4157 | 401.9402 | 362.7107 | 336.4232 | 264.2552 | 345.2686 | 308.5725 | 246.8832 |
| ENSMUSG00000111147 | 0.713181573 | 0.338637 | 0.507173 | 10.06054 | 5.999108 | 21.53077 | 15.52722 | 2.925334 | 18.99536 | 7.714313 | 3.262773 |
| ENSMUSG00000111163 | 0.843517621 | 0.040104 | 0.116929 | 56.33904 | 70.78947 | 88.60741 | 76.77349 | 36.07912 | 78.21619 | 21.21436 | 28.27737 |
| ENSMUSG00000111186 | -1.905281335 | 0.05446 | 0.144809 | 4.024217 | 0 | 27.32752 | 3.450494 | 25.35289 | 7.821619 | 42.42872 | 56.55474 |
| ENSMUSG00000111189 | 0.608817979 | 0.347826 | 0.516492 | 9.054489 | 29.99554 | 9.93728 | 7.763611 | 7.80089 | 15.64324 | 5.785735 | 7.613138 |
| ENSMUSG00000111212 | 1.687182971 | 0.003865 | 0.021297 | 123.7447 | 134.38 | 46.37397 | 38.81806 | 13.65156 | 58.10345 | 15.42863 | 19.57664 |
| ENSMUSG00000111236 | 1.148094115 | 0.000194 | 0.002211 | 50.30272 | 51.59233 | 54.65504 | 37.95543 | 16.57689 | 24.58223 | 27.96439 | 18.48905 |
| ENSMUSG00000111246 | -0.274183887 | 0.493663 | 0.65051 | 18.10898 | 19.19714 | 14.07781 | 24.15346 | 20.47734 | 17.87799 | 18.32149 | 34.80291 |
| ENSMUSG00000111253 | 0.020061941 | 0.978192 | 0.988047 | 7.04238 | 4.799286 | 18.21835 | 25.01608 | 2.925334 | 17.87799 | 12.53576 | 21.75182 |
| ENSMUSG00000111264 | 0.590909975 | 0.058546 | 0.151973 | 51.30877 | 44.3934 | 35.60859 | 56.93315 | 30.22845 | 41.34284 | 32.78583 | 20.66423 |
| ENSMUSG00000111283 | -0.282542356 | 0.604358 | 0.741075 | 29.17558 | 21.59679 | 103.5133 | 25.01608 | 38.02934 | 39.10809 | 71.3574 | 70.69342 |
| ENSMUSG00000111291 | 0.411882211 | 0.040826 | 0.11838 | 223.3441 | 221.967 | 149.0592 | 200.9913 | 142.3663 | 187.7188 | 120.5361 | 146.8248 |
| ENSMUSG00000111293 | 0.307194387 | 0.560081 | 0.706118 | 10.06054 | 16.7975 | 19.87456 | 7.763611 | 9.751113 | 8.938993 | 16.39292 | 8.700729 |
| ENSMUSG00000111312 | -0.378598782 | 0.421757 | 0.587157 | 20.12109 | 19.19714 | 38.92101 | 18.11509 | 26.32801 | 20.11273 | 23.14294 | 56.55474 |
| ENSMUSG00000111325 | 0.565195753 | 0.17989 | 0.329821 | 22.1332 | 32.39518 | 19.04645 | 18.11509 | 16.57689 | 20.11273 | 8.678602 | 16.31387 |
| ENSMUSG00000111329 | -0.53655659 | 0.142749 | 0.280155 | 21.12714 | 21.59679 | 40.57722 | 16.38985 | 39.97956 | 39.10809 | 32.78583 | 33.71532 |
| ENSMUSG00000111334 | 0.132536464 | 0.831178 | 0.897939 | 20.12109 | 14.39786 | 28.98373 | 4.313117 | 7.80089 | 13.40849 | 21.21436 | 19.57664 |
| ENSMUSG00000111343 | 0.629752228 | 0.01242 | 0.05027 | 45.27245 | 50.3925 | 64.59232 | 58.6584 | 37.05423 | 36.87335 | 37.60728 | 30.45255 |
| ENSMUSG00000111345 | 0.40375964 | 0.335231 | 0.503502 | 20.12109 | 22.79661 | 13.24971 | 22.42821 | 14.62667 | 21.23011 | 12.53576 | 10.87591 |
| ENSMUSG00000111348 | -0.211687852 | 0.484744 | 0.643027 | 30.18163 | 37.19447 | 31.46805 | 23.29083 | 39.00445 | 31.28647 | 27.96439 | 42.41605 |
| ENSMUSG00000111356 | -0.763598045 | 0.02244 | 0.076468 | 20.12109 | 13.19804 | 23.18699 | 19.84034 | 30.22845 | 26.81698 | 33.75012 | 40.24087 |
| ENSMUSG00000111361 | 0.829693773 | 0.189487 | 0.342068 | 42.25428 | 62.39072 | 20.70267 | 26.74133 | 14.62667 | 51.39921 | 8.678602 | 10.87591 |
| ENSMUSG00000111365 | 0.469391854 | 0.409595 | 0.575873 | 6.036326 | 13.19804 | 16.56213 | 10.35148 | 4.875557 | 11.17374 | 6.750024 | 10.87591 |
| ENSMUSG00000111368 | -1.343758668 | 0.008387 | 0.037578 | 22.1332 | 5.999108 | 23.18699 | 19.84034 | 40.95467 | 18.99536 | 64.60737 | 57.64233 |
| ENSMUSG00000111375 | -0.472894888 | 0.438732 | 0.602213 | 16.09687 | 5.999108 | 8.281066 | 14.6646 | 15.60178 | 4.469496 | 25.07152 | 17.40146 |
| ENSMUSG00000111390 | 0.377711714 | 0.077692 | 0.185305 | 287.7315 | 176.3738 | 194.6051 | 229.4578 | 197.9476 | 200.01 | 153.322 | 132.6861 |
| ENSMUSG00000111394 | 0.158479008 | 0.118217 | 0.247035 | 474.8577 | 429.5361 | 544.0661 | 494.2833 | 448.5512 | 429.0717 | 443.573 | 423.0729 |
| ENSMUSG00000111434 | 0.855162808 | 0.263499 | 0.427829 | 34.20585 | 15.59768 | 10.76539 | 11.21411 | 12.67645 | 20.11273 | 5.785735 | 1.087591 |
| ENSMUSG00000111447 | 0.276928782 | 0.609353 | 0.744537 | 20.12109 | 5.999108 | 12.4216 | 14.6646 | 14.62667 | 6.704245 | 8.678602 | 14.13868 |
| ENSMUSG00000111465 | -0.289521183 | 0.540072 | 0.689654 | 15.09082 | 7.198929 | 13.24971 | 20.70296 | 18.52711 | 16.76061 | 23.14294 | 10.87591 |
| ENSMUSG00000111481 | 0.984897284 | 0.128954 | 0.260619 | 11.0666 | 25.19625 | 4.140533 | 18.97772 | 8.776002 | 6.704245 | 8.678602 | 5.437955 |
| ENSMUSG00000111496 | 0.470908524 | 0.108634 | 0.23354 | 64.38748 | 47.99286 | 60.45178 | 52.62003 | 30.22845 | 58.10345 | 30.85725 | 44.59123 |
| ENSMUSG00000111497 | 0.219817007 | 0.633173 | 0.761442 | 19.11503 | 31.19536 | 9.93728 | 17.25247 | 20.47734 | 14.52586 | 17.3572 | 13.05109 |
| ENSMUSG00000111514 | 0.042265514 | 0.922319 | 0.953784 | 25.15136 | 27.5959 | 22.35888 | 35.36756 | 18.52711 | 49.16446 | 17.3572 | 22.83941 |
| ENSMUSG00000111521 | -0.13682774 | 0.503788 | 0.659661 | 77.46619 | 104.3845 | 107.6539 | 100.9269 | 101.4116 | 96.09417 | 95.46463 | 137.0365 |
| ENSMUSG00000111531 | 0.296369806 | 0.68796 | 0.801831 | 8.048435 | 10.79839 | 26.49941 | 2.58787 | 8.776002 | 5.58687 | 9.642892 | 15.22628 |
| ENSMUSG00000111547 | -0.128344134 | 0.711783 | 0.819301 | 66.39959 | 38.39429 | 36.43669 | 25.8787 | 39.00445 | 44.69496 | 56.89306 | 41.32846 |
| ENSMUSG00000111583 | -0.23149861 | 0.521315 | 0.674712 | 22.1332 | 15.59768 | 18.21835 | 18.11509 | 22.42756 | 24.58223 | 15.42863 | 25.0146 |
| ENSMUSG00000111593 | 1.115152051 | 0.209368 | 0.366525 | 28.16952 | 35.99465 | 14.07781 | 25.8787 | 3.900445 | 37.99072 | 0.964289 | 5.437955 |
| ENSMUSG00000111594 | -0.544341129 | 0.050735 | 0.137438 | 91.55095 | 79.18822 | 98.54469 | 71.59775 | 188.1965 | 82.68568 | 113.7861 | 112.0219 |
| ENSMUSG00000111656 | -0.010392169 | 0.97951 | 0.988724 | 14.08476 | 25.19625 | 14.90592 | 18.11509 | 13.65156 | 20.11273 | 23.14294 | 15.22628 |

| | | | | | | | | | | | |
|---|---|---|---|---|---|---|---|---|---|---|---|
| ENSMUSG00000111662 | 0.101691184 | 0.767847 | 0.855915 | 32.19374 | 31.19536 | 22.35888 | 21.56559 | 28.27823 | 31.28647 | 20.25007 | 19.57664 |
| ENSMUSG00000111664 | 0.796009239 | 0.054777 | 0.145442 | 42.25428 | 29.99554 | 21.53077 | 47.44429 | 20.47734 | 26.81698 | 23.14294 | 10.87591 |
| ENSMUSG00000111681 | 0.554553462 | 0.055642 | 0.146914 | 58.35115 | 73.18911 | 50.5145 | 81.94923 | 52.65601 | 48.04709 | 50.14304 | 28.27737 |
| ENSMUSG00000111686 | -0.339013555 | 0.552731 | 0.700103 | 10.06054 | 19.19714 | 7.45296 | 6.900988 | 14.62667 | 12.29112 | 6.750024 | 20.66423 |
| ENSMUSG00000111722 | 0.296469889 | 0.518747 | 0.672487 | 28.16952 | 17.99732 | 14.90592 | 15.52722 | 16.57689 | 23.46486 | 13.50005 | 8.700729 |
| ENSMUSG00000111740 | 0.192659837 | 0.432497 | 0.596842 | 93.56306 | 101.9848 | 69.56096 | 74.18562 | 97.51113 | 71.51194 | 69.42882 | 56.55474 |
| ENSMUSG00000111745 | 1.709832367 | 0.053184 | 0.142096 | 22.1332 | 27.5959 | 11.59349 | 18.97772 | 4.875557 | 17.87799 | 1.928578 | 0 |
| ENSMUSG00000111761 | 0.130903532 | 0.756763 | 0.848662 | 22.1332 | 23.99643 | 29.81184 | 31.05445 | 18.52711 | 12.29112 | 27.0001 | 40.24087 |
| ENSMUSG00000111793 | 0.237980629 | 0.219682 | 0.378497 | 117.7084 | 85.18733 | 81.98256 | 103.5148 | 76.05868 | 83.80306 | 84.85745 | 84.8321 |
| ENSMUSG00000111828 | -0.328387477 | 0.466317 | 0.627229 | 22.1332 | 14.39786 | 9.109173 | 12.93935 | 16.57689 | 12.29112 | 21.21436 | 22.83941 |
| ENSMUSG00000111847 | 0.465042029 | 0.513166 | 0.668037 | 27.16347 | 7.198929 | 16.56213 | 18.11509 | 4.875557 | 31.28647 | 6.750024 | 7.613138 |
| ENSMUSG00000111857 | -0.119641876 | 0.779165 | 0.863784 | 15.09082 | 40.79393 | 30.63995 | 18.97772 | 22.42756 | 40.22547 | 21.21436 | 30.45255 |
| ENSMUSG00000111877 | -0.002759819 | 0.989407 | 0.99393 | 86.52067 | 104.3845 | 91.09173 | 118.1794 | 125.7894 | 90.5073 | 101.2504 | 82.65692 |
