## Supplementary material for "Lysine-specific demethylase 1a is obligatory for gene regulation during kidney development": Suppl Table-2

Supplementary Table-2

| ID | log2FoldChange | pvalue | padj | KKO-1 | KKO-2 | KKO-3 | KKO-4 | KWT-1 | KWT-2 | KWT-3 | KWT-4 |
|---|---|---|---|---|---|---|---|---|---|---|---|
| ENSMUSG00000000049 | 2.874240897 | 0.008411 | 0.03765 | 117.7084 | 1.199822 | 3.312427 | 115.5915 | 4.875557 | 4.469496 | 15.42863 | 7.613138 |
| ENSMUSG00000000154 | -1.335082452 | 0.001083 | 0.008295 | 78.47224 | 76.78858 | 136.6376 | 64.69676 | 239.8774 | 90.5073 | 190.9293 | 379.5693 |
| ENSMUSG00000000159 | -1.090125269 | 2.70E-10 | 4.14E-08 | 404.4339 | 392.3416 | 484.4424 | 359.714 | 942.9326 | 621.26 | 913.1818 | 1016.898 |
| ENSMUSG00000000202 | 1.245229166 | 9.48E-05 | 0.001281 | 54.32694 | 103.1847 | 62.9361 | 56.07053 | 31.20356 | 35.75597 | 23.14294 | 26.10219 |
| ENSMUSG00000000214 | 3.826258112 | 0.004683 | 0.024588 | 610.675 | 8.398751 | 4.140533 | 444.2511 | 8.776002 | 22.34748 | 11.57147 | 32.62773 |
| ENSMUSG00000000223 | 1.067523913 | 0.001226 | 0.009106 | 91.55095 | 109.1838 | 96.88848 | 87.12497 | 35.10401 | 81.56831 | 33.75012 | 33.71532 |
| ENSMUSG00000000385 | -1.535046527 | 0.001534 | 0.010701 | 908.4671 | 554.3175 | 2042.111 | 952.3363 | 3530.878 | 1622.427 | 3598.727 | 4167.649 |
| ENSMUSG00000000560 | 1.130104087 | 0.001341 | 0.009746 | 216.3017 | 263.9607 | 175.5586 | 197.5408 | 98.48624 | 168.7235 | 78.10742 | 44.59123 |
| ENSMUSG00000000673 | -3.119130414 | 1.80E-06 | 5.98E-05 | 10.06054 | 5.999108 | 6.624853 | 12.07673 | 77.03379 | 20.11273 | 36.64299 | 169.6642 |
| ENSMUSG00000000739 | 1.1428241 | 8.01E-08 | 4.77E-06 | 339.0403 | 439.1347 | 336.2113 | 474.4429 | 142.3663 | 239.1181 | 194.7864 | 143.562 |
| ENSMUSG00000000805 | -2.427067873 | 5.31E-06 | 0.000141 | 146.8839 | 56.39161 | 278.2438 | 196.6782 | 865.8988 | 497.2315 | 841.8244 | 1447.584 |
| ENSMUSG00000000861 | 1.515011403 | 1.82E-10 | 3.00E-08 | 297.7921 | 329.9509 | 227.7293 | 265.688 | 79.95913 | 145.2586 | 75.21455 | 92.44524 |
| ENSMUSG00000000889 | 3.693565762 | 0.002777 | 0.016703 | 444.676 | 11.99822 | 8.281066 | 556.3922 | 18.52711 | 10.05637 | 31.82154 | 18.48905 |
| ENSMUSG00000001076 | 1.268371504 | 4.48E-07 | 1.92E-05 | 179.0777 | 272.3595 | 198.7456 | 220.8316 | 115.0631 | 82.68568 | 107.0361 | 55.46715 |
| ENSMUSG00000001095 | -2.568567286 | 1.66E-07 | 8.51E-06 | 9.054489 | 10.79839 | 24.01509 | 13.80198 | 102.3867 | 35.75597 | 69.42882 | 138.1241 |
| ENSMUSG00000001249 | -1.718618889 | 1.24E-07 | 6.69E-06 | 418.5186 | 315.5531 | 731.2182 | 458.0531 | 1624.535 | 751.9928 | 1705.828 | 2252.401 |
| ENSMUSG00000001260 | 1.036964455 | 0.009726 | 0.041725 | 173.0413 | 290.3568 | 132.4971 | 205.3044 | 96.53602 | 177.6625 | 69.42882 | 46.76642 |
| ENSMUSG00000001493 | -1.45138218 | 2.97E-09 | 3.18E-07 | 159.9626 | 206.3693 | 192.9488 | 200.9913 | 756.6864 | 319.569 | 485.0374 | 516.6058 |
| ENSMUSG00000001504 | -1.848453805 | 0.006141 | 0.029934 | 113.6841 | 27.5959 | 219.4483 | 104.3774 | 241.8276 | 198.8926 | 595.9307 | 640.5912 |
| ENSMUSG00000001642 | -1.185603713 | 3.00E-05 | 0.000535 | 1831.019 | 1677.351 | 4328.513 | 1770.103 | 6270.941 | 4631.516 | 5394.234 | 5559.766 |
| ENSMUSG00000001661 | 1.010336101 | 4.37E-05 | 0.000721 | 1097.605 | 1129.032 | 787.5294 | 1071.378 | 418.3228 | 813.4483 | 401.1443 | 395.8832 |
| ENSMUSG00000001670 | 2.988764174 | 0.004445 | 0.023594 | 82.49646 | 8.398751 | 0.828107 | 68.14726 | 4.875557 | 3.352122 | 9.642892 | 2.175182 |
| ENSMUSG00000001827 | -1.67406505 | 0.00042 | 0.004005 | 333.004 | 313.1534 | 823.138 | 456.3278 | 1646.963 | 746.4059 | 1378.934 | 2375.299 |
| ENSMUSG00000001864 | -1.233598652 | 1.05E-05 | 0.000236 | 1395.397 | 1166.227 | 2216.841 | 1418.153 | 3621.563 | 1926.353 | 3914.05 | 5112.766 |
| ENSMUSG00000002032 | -1.225891688 | 0.000174 | 0.002036 | 186.1201 | 166.7752 | 247.6039 | 255.3366 | 497.3068 | 224.5922 | 495.6446 | 786.3284 |
| ENSMUSG00000002076 | -1.692100809 | 3.88E-06 | 0.000111 | 20.12109 | 5.999108 | 16.56213 | 17.25247 | 39.00445 | 49.16446 | 51.10733 | 57.64233 |
| ENSMUSG00000002204 | -2.948546453 | 3.03E-08 | 2.15E-06 | 396.3854 | 397.1409 | 775.1078 | 420.9603 | 3322.204 | 1191.121 | 4097.265 | 6751.765 |
| ENSMUSG00000002769 | -1.259737685 | 0.000167 | 0.001967 | 53.32088 | 33.595 | 39.74912 | 60.38364 | 134.5654 | 58.10345 | 113.7861 | 142.4744 |
| ENSMUSG00000002831 | 1.01935656 | 0.005085 | 0.026035 | 28.16952 | 28.79572 | 29.81184 | 41.40593 | 18.52711 | 12.29112 | 20.25007 | 11.9635 |
| ENSMUSG00000002847 | -2.438215362 | 0.001814 | 0.012234 | 199.1988 | 31.19536 | 355.2577 | 93.16334 | 828.8446 | 289.3999 | 1191.861 | 1371.452 |
| ENSMUSG00000002908 | -1.483810538 | 1.83E-05 | 0.000365 | 85.51462 | 61.1909 | 100.2009 | 70.73513 | 221.3503 | 98.32892 | 235.2866 | 334.9781 |
| ENSMUSG00000003038 | -1.055901554 | 3.01E-09 | 3.21E-07 | 4549.378 | 4956.463 | 4748.363 | 4908.328 | 6324.572 | 12207.31 | 10783.65 | 10524.62 |
| ENSMUSG00000003410 | 1.882440649 | 0.007633 | 0.034999 | 82.49646 | 25.19625 | 20.70267 | 251.8861 | 32.17867 | 31.28647 | 20.25007 | 19.57664 |
| ENSMUSG00000003974 | 1.682097861 | 0.00031 | 0.003189 | 27.16347 | 21.59679 | 24.01509 | 16.38985 | 5.850668 | 5.58687 | 9.642892 | 6.525547 |
| ENSMUSG00000004031 | 1.596443018 | 0.002171 | 0.013939 | 33.19979 | 16.7975 | 29.81184 | 64.69676 | 9.751113 | 18.99536 | 8.678602 | 10.87591 |
| ENSMUSG00000004098 | 1.379957252 | 0.000627 | 0.005456 | 445.6821 | 470.33 | 311.3681 | 367.4776 | 163.8187 | 289.3999 | 103.1789 | 56.55474 |
| ENSMUSG00000004341 | -3.257660957 | 4.44E-07 | 1.91E-05 | 48.29061 | 33.595 | 110.9663 | 37.95543 | 442.7005 | 144.1413 | 857.2531 | 767.8393 |
| ENSMUSG00000004655 | -1.993536476 | 0.000652 | 0.005628 | 884.3218 | 932.2613 | 2998.574 | 1168.855 | 6283.617 | 2153.18 | 7025.811 | 8369.013 |
| ENSMUSG00000004668 | -3.058995119 | 0.003404 | 0.019397 | 16.09687 | 9.598572 | 224.4169 | 31.05445 | 527.5352 | 160.9019 | 885.2175 | 773.2773 |
| ENSMUSG00000004814 | 1.788560466 | 0.000902 | 0.007205 | 20.12109 | 19.19714 | 27.32752 | 23.29083 | 1.950223 | 12.29112 | 6.750024 | 5.437955 |
| ENSMUSG00000004872 | 2.817048786 | 0.002123 | 0.013707 | 36.21796 | 53.99197 | 53.82693 | 39.68068 | 2.925334 | 22.34748 | 0 | 1.087591 |
| ENSMUSG00000004885 | 1.082293785 | 1.10E-10 | 2.00E-08 | 483.9121 | 729.4915 | 639.2983 | 590.0345 | 281.8072 | 354.2076 | 243.9652 | 274.073 |
| ENSMUSG00000004952 | 1.084677924 | 4.00E-16 | 2.64E-13 | 1173.059 | 1202.221 | 1086.476 | 1221.475 | 567.5148 | 679.3635 | 517.8233 | 443.7372 |
| ENSMUSG00000005125 | -1.136274736 | 6.49E-08 | 4.02E-06 | 3672.098 | 2484.83 | 3699.152 | 3913.723 | 8975.9 | 4749.957 | 8140.529 | 8404.904 |
| ENSMUSG00000005364 | -2.494827935 | 0.006599 | 0.031619 | 4.024217 | 5.999108 | 9.109173 | 0 | 27.30312 | 3.352122 | 45.32159 | 31.54014 |
| ENSMUSG00000005950 | -1.569345178 | 0.006202 | 0.030184 | 9.054489 | 14.39786 | 6.624853 | 8.626235 | 35.10401 | 7.821619 | 27.96439 | 42.41605 |
| ENSMUSG00000006014 | 1.586974992 | 5.39E-06 | 0.000143 | 97.58727 | 87.58697 | 57.96746 | 100.9269 | 19.50223 | 45.81234 | 29.89296 | 19.57664 |
| ENSMUSG00000006216 | -2.861607153 | 0.00012 | 0.001538 | 176.0595 | 83.98751 | 642.6107 | 156.9975 | 1367.106 | 725.1758 | 2425.187 | 3188.817 |
| ENSMUSG00000006310 | -1.12500474 | 7.67E-05 | 0.001108 | 18.10898 | 19.19714 | 19.87456 | 22.42821 | 45.83023 | 39.10809 | 42.42872 | 46.76642 |
| ENSMUSG00000006342 | -1.557815914 | 1.33E-06 | 4.72E-05 | 195.1745 | 154.777 | 323.7897 | 244.9851 | 571.4152 | 405.6068 | 606.5379 | 1125.657 |
| ENSMUSG00000006345 | -2.890728516 | 2.61E-05 | 0.000482 | 518.118 | 161.9759 | 1337.392 | 565.0184 | 4682.485 | 1748.69 | 4379.801 | 8347.262 |
| ENSMUSG00000006378 | -1.044095857 | 1.50E-11 | 3.37E-09 | 396.3854 | 371.9447 | 503.4888 | 450.2895 | 937.082 | 695.0067 | 884.2532 | 1039.737 |
| ENSMUSG00000006457 | 1.16603983 | 2.72E-12 | 6.98E-10 | 517.1119 | 611.909 | 431.4436 | 533.9639 | 287.6578 | 205.5968 | 242.0366 | 195.7664 |
| ENSMUSG00000006469 | -3.852462647 | 5.41E-05 | 0.000842 | 1.006054 | 1.199822 | 4.96864 | 0 | 49.73068 | 12.29112 | 8.678602 | 35.89051 |
| ENSMUSG00000006519 | -1.199217546 | 8.76E-07 | 3.35E-05 | 901.4247 | 973.0553 | 1369.688 | 1029.972 | 2678.631 | 1465.995 | 2177.365 | 3494.43 |
| ENSMUSG00000006538 | -3.672258131 | 9.94E-06 | 0.000226 | 15.09082 | 9.598572 | 87.7793 | 16.38985 | 377.3681 | 122.9112 | 614.2522 | 535.0948 |
| ENSMUSG00000006586 | 1.284872196 | 0.007912 | 0.035933 | 124.7507 | 195.5709 | 95.23226 | 134.5693 | 37.05423 | 127.3806 | 27.96439 | 33.71532 |
| ENSMUSG00000006649 | -1.92680269 | 1.16E-05 | 0.000258 | 90.54489 | 45.59322 | 194.6051 | 87.9876 | 246.7032 | 244.7049 | 455.1445 | 650.3795 |
| ENSMUSG00000006724 | -3.970560773 | 1.24E-12 | 3.62E-10 | 55.33299 | 26.39607 | 130.0127 | 105.2401 | 1611.859 | 604.4994 | 1074.218 | 1693.379 |
| ENSMUSG00000006958 | 1.502001172 | 2.08E-08 | 1.56E-06 | 583.5115 | 826.677 | 609.4865 | 684.0604 | 229.1512 | 378.7898 | 204.4293 | 142.4744 |
| ENSMUSG00000007021 | 1.302463351 | 0.001983 | 0.013056 | 74.44802 | 44.3934 | 57.96746 | 107.8279 | 26.32801 | 14.52586 | 27.0001 | 47.85401 |
| ENSMUSG00000007034 | -1.370332491 | 2.21E-05 | 0.000425 | 159.9626 | 94.7859 | 196.2613 | 142.3329 | 405.6463 | 208.949 | 330.7512 | 591.6496 |

| | | | | | | | | | | | |
|---|---|---|---|---|---|---|---|---|---|---|---|
| ENSMUSG00000007038 | -1.023860878 | 1.54E-05 | 0.000318 | 813.898 | 595.1115 | 998.6966 | 870.3871 | 1665.49 | 994.4629 | 1836.971 | 2171.919 |
| ENSMUSG00000007877 | -1.861138297 | 2.78E-05 | 0.000507 | 22.1332 | 8.398751 | 7.45296 | 12.07673 | 61.43201 | 30.1691 | 48.21446 | 41.32846 |
| ENSMUSG00000007908 | 1.078411305 | 4.82E-08 | 3.16E-06 | 167.005 | 171.5745 | 138.2938 | 196.6782 | 73.13335 | 96.09417 | 64.60737 | 85.9197 |
| ENSMUSG00000007946 | 3.483160398 | 0.002265 | 0.014359 | 82.49646 | 8.398751 | 5.796746 | 118.1794 | 0.975111 | 14.52586 | 3.857157 | 0 |
| ENSMUSG00000008658 | 1.302031086 | 0.00148 | 0.010472 | 623.7537 | 773.8849 | 467.0521 | 691.824 | 193.072 | 559.8044 | 148.5005 | 135.9489 |
| ENSMUSG00000008845 | 1.19771114 | 1.23E-06 | 4.42E-05 | 427.5731 | 303.5548 | 240.979 | 327.7969 | 116.0382 | 177.6625 | 171.6435 | 101.146 |
| ENSMUSG00000009145 | -1.021159934 | 0.00288 | 0.017225 | 108.6539 | 41.99375 | 74.5296 | 105.2401 | 157.968 | 110.62 | 157.1791 | 246.8832 |
| ENSMUSG00000009356 | -5.537024401 | 2.24E-10 | 3.57E-08 | 0 | 0 | 3.312427 | 1.725247 | 35.10401 | 25.6996 | 126.3219 | 67.43065 |
| ENSMUSG00000009378 | -2.493031591 | 0.000333 | 0.003357 | 133.8052 | 32.39518 | 345.3205 | 127.6683 | 902.9531 | 404.4894 | 1020.218 | 1274.657 |
| ENSMUSG00000009772 | -1.136624507 | 1.52E-10 | 2.64E-08 | 110.666 | 93.58608 | 134.1533 | 109.5532 | 231.1014 | 203.3621 | 285.4296 | 268.635 |
| ENSMUSG00000010044 | -1.98536623 | 1.13E-08 | 9.52E-07 | 13.07871 | 25.19625 | 27.32752 | 25.8787 | 89.71024 | 55.8687 | 92.57176 | 125.073 |
| ENSMUSG00000010064 | -1.086796083 | 0.000177 | 0.002063 | 293.7679 | 256.7618 | 342.8361 | 324.3464 | 617.2455 | 332.9775 | 604.6093 | 1033.212 |
| ENSMUSG00000010122 | -2.90255208 | 1.10E-05 | 0.000246 | 98.59333 | 35.99465 | 278.2438 | 122.4925 | 1127.229 | 414.5458 | 858.2174 | 1608.547 |
| ENSMUSG00000010476 | 1.022481864 | 0.000263 | 0.002808 | 312.8829 | 513.5236 | 327.1021 | 446.839 | 198.9227 | 303.9258 | 152.3577 | 132.6861 |
| ENSMUSG00000010505 | 1.853997916 | 0.00137 | 0.009903 | 42.25428 | 16.7975 | 15.73403 | 67.28463 | 9.751113 | 11.17374 | 7.714313 | 10.87591 |
| ENSMUSG00000010651 | -2.429228994 | 4.02E-08 | 2.69E-06 | 91.55095 | 55.19179 | 200.4018 | 100.0643 | 620.1708 | 244.7049 | 516.859 | 1033.212 |
| ENSMUSG00000011034 | -2.442223861 | 1.55E-08 | 1.22E-06 | 59.35721 | 35.99465 | 110.1382 | 52.62003 | 325.6872 | 151.9629 | 318.2154 | 612.3138 |
| ENSMUSG00000011751 | 1.609701475 | 3.06E-17 | 2.35E-14 | 152.9203 | 193.1713 | 159.8246 | 178.5631 | 57.53157 | 67.04245 | 55.92877 | 43.50364 |
| ENSMUSG00000012520 | 4.735000042 | 0.008486 | 0.037894 | 112.6781 | 3.599465 | 5.796746 | 194.0903 | 10.72622 | 0 | 0 | 1.087591 |
| ENSMUSG00000013483 | -1.215782467 | 0.007839 | 0.035703 | 10.06054 | 22.79661 | 9.93728 | 13.80198 | 34.1289 | 23.46486 | 22.17865 | 50.02919 |
| ENSMUSG00000014747 | 1.031186079 | 0.002256 | 0.014333 | 33.19979 | 44.3934 | 51.34261 | 55.2079 | 26.32801 | 25.6996 | 25.07152 | 13.05109 |
| ENSMUSG00000015337 | -1.320942895 | 2.18E-06 | 6.88E-05 | 143.8658 | 145.1784 | 219.4483 | 144.0581 | 340.3138 | 251.4092 | 434.8944 | 605.7882 |
| ENSMUSG00000015401 | -3.221421816 | 2.92E-07 | 1.35E-05 | 160.9687 | 81.58786 | 523.3634 | 176.8378 | 2451.43 | 1103.966 | 1915.078 | 3330.204 |
| ENSMUSG00000015405 | -1.301904198 | 0.004586 | 0.02415 | 844.0796 | 319.1525 | 880.2773 | 615.0506 | 1350.529 | 833.5611 | 1698.113 | 2674.386 |
| ENSMUSG00000015962 | -3.782088492 | 6.25E-06 | 0.00016 | 3.018163 | 1.199822 | 12.4216 | 0 | 37.05423 | 27.93435 | 101.2504 | 68.51824 |
| ENSMUSG00000016150 | 1.010314642 | 0.006548 | 0.031473 | 38.23007 | 31.19536 | 20.70267 | 34.50494 | 14.62667 | 14.52586 | 19.28578 | 13.05109 |
| ENSMUSG00000016194 | -1.559891389 | 5.07E-05 | 0.0008 | 42.25428 | 27.5959 | 27.32752 | 25.8787 | 118.9636 | 39.10809 | 89.67889 | 114.1971 |
| ENSMUSG00000016319 | -1.285689171 | 2.48E-07 | 1.20E-05 | 4903.509 | 5234.821 | 7362.696 | 4583.119 | 13273.22 | 7408.19 | 15104.63 | 18056.19 |
| ENSMUSG00000016756 | -1.560195727 | 0.000411 | 0.003947 | 19.11503 | 37.19447 | 24.01509 | 24.15346 | 48.75557 | 42.46022 | 75.21455 | 140.2993 |
| ENSMUSG00000016995 | 1.929493761 | 1.30E-05 | 0.000281 | 33.19979 | 73.18911 | 53.82693 | 73.323 | 16.57689 | 25.6996 | 11.57147 | 7.613138 |
| ENSMUSG00000017002 | 2.368217345 | 3.67E-06 | 0.000107 | 178.0716 | 387.5424 | 219.4483 | 396.8068 | 78.98402 | 71.51194 | 69.42882 | 8.700729 |
| ENSMUSG00000017417 | 1.335890433 | 3.26E-09 | 3.46E-07 | 149.9021 | 166.7752 | 147.403 | 106.9653 | 42.9049 | 56.98608 | 59.78593 | 66.34306 |
| ENSMUSG00000017453 | -1.659787862 | 1.50E-06 | 5.18E-05 | 113.6841 | 65.99018 | 173.0743 | 129.3935 | 360.7912 | 201.1273 | 405.9657 | 560.1094 |
| ENSMUSG00000017868 | -2.559350758 | 0.000658 | 0.005671 | 79.47829 | 15.59768 | 159.8246 | 50.89479 | 480.7299 | 141.9065 | 435.8587 | 747.1751 |
| ENSMUSG00000017950 | -2.26204967 | 0.000792 | 0.006528 | 705.2441 | 79.18822 | 843.0126 | 656.4565 | 2614.273 | 1533.037 | 2914.082 | 3896.839 |
| ENSMUSG00000018102 | -1.209706855 | 2.87E-08 | 2.05E-06 | 369.2219 | 359.9465 | 470.3646 | 347.6373 | 999.4891 | 574.3303 | 819.6458 | 1186.562 |
| ENSMUSG00000018126 | -1.879179527 | 2.50E-06 | 7.74E-05 | 164.9929 | 89.98662 | 346.9767 | 179.4257 | 576.2908 | 377.6724 | 878.4674 | 1047.35 |
| ENSMUSG00000018339 | -2.534035817 | 1.87E-07 | 9.48E-06 | 3185.168 | 1402.591 | 4210.922 | 3283.145 | 23916.56 | 6982.471 | 16035.16 | 23047.14 |
| ENSMUSG00000018381 | -1.070204867 | 0.001384 | 0.009976 | 46.2785 | 26.39607 | 52.99882 | 27.60395 | 82.88446 | 51.39921 | 87.75031 | 101.146 |
| ENSMUSG00000018459 | -2.898627159 | 8.10E-06 | 0.000195 | 17.10292 | 9.598572 | 57.96746 | 30.19182 | 317.8863 | 52.51658 | 101.2504 | 390.4452 |
| ENSMUSG00000018648 | -1.29564148 | 2.98E-05 | 0.000533 | 112.6781 | 63.59054 | 178.0429 | 105.2401 | 242.8027 | 221.2401 | 273.8581 | 394.7956 |
| ENSMUSG00000018796 | -1.666557315 | 3.57E-08 | 2.42E-06 | 639.8506 | 482.3283 | 949.8383 | 714.2523 | 1993.128 | 1070.444 | 2774.26 | 3011.54 |
| ENSMUSG00000018924 | -1.980865676 | 8.71E-11 | 1.62E-08 | 24.1453 | 11.99822 | 17.39024 | 21.56559 | 66.30757 | 97.21155 | 68.46453 | 67.43065 |
| ENSMUSG00000018927 | 1.497839986 | 2.97E-10 | 4.42E-08 | 334.01 | 296.3559 | 186.324 | 262.2375 | 113.1129 | 106.1505 | 94.50034 | 67.43065 |
| ENSMUSG00000019122 | 1.705873676 | 9.12E-08 | 5.25E-06 | 428.5792 | 243.5638 | 212.8234 | 351.9504 | 89.71024 | 128.498 | 110.8933 | 50.02919 |
| ENSMUSG00000019232 | -3.604779915 | 6.84E-11 | 1.33E-08 | 20.12109 | 3.599465 | 13.24971 | 16.38985 | 161.8685 | 44.69496 | 200.5721 | 247.9708 |
| ENSMUSG00000019762 | -2.400344786 | 1.82E-09 | 2.17E-07 | 21.12714 | 10.79839 | 31.46805 | 25.8787 | 119.9387 | 60.3382 | 129.2147 | 167.489 |
| ENSMUSG00000019767 | -1.610796836 | 0.010858 | 0.045379 | 13.07871 | 3.599465 | 10.76539 | 9.488858 | 17.552 | 20.11273 | 14.46434 | 61.99269 |
| ENSMUSG00000019775 | 1.163688735 | 0.003314 | 0.019073 | 98.59333 | 131.9804 | 86.9512 | 131.1188 | 38.02934 | 90.5073 | 50.14304 | 21.75182 |
| ENSMUSG00000019817 | 1.1387554 | 4.95E-06 | 0.000134 | 10512.26 | 11387.51 | 8303.425 | 12150.91 | 4282.689 | 7751.224 | 4050.014 | 3151.839 |
| ENSMUSG00000019888 | 1.985029152 | 0.00167 | 0.011463 | 111.672 | 14.39786 | 24.8432 | 37.09281 | 10.72622 | 12.29112 | 12.53576 | 11.9635 |
| ENSMUSG00000019987 | 1.533319166 | 0.003129 | 0.018333 | 148.896 | 33.595 | 34.78048 | 90.57547 | 29.25334 | 26.81698 | 32.78583 | 17.40146 |
| ENSMUSG00000020017 | 3.682129046 | 9.04E-06 | 0.000212 | 12.07265 | 22.79661 | 15.73403 | 54.34528 | 2.925334 | 0 | 1.928578 | 3.262773 |
| ENSMUSG00000020051 | -4.371501582 | 3.06E-07 | 1.39E-05 | 33.19979 | 4.799286 | 33.95237 | 42.26855 | 1150.631 | 135.2023 | 234.3223 | 851.5838 |
| ENSMUSG00000020062 | -4.876815617 | 3.08E-07 | 1.40E-05 | 6.036326 | 7.198929 | 73.70149 | 13.80198 | 628.9468 | 205.5968 | 648.9666 | 1490 |
| ENSMUSG00000020159 | -1.808603259 | 3.58E-05 | 0.000614 | 39.23612 | 39.59411 | 23.18699 | 23.29083 | 72.15824 | 210.0663 | 63.64308 | 91.35765 |
| ENSMUSG00000020258 | -1.265166811 | 1.42E-05 | 0.000298 | 84.50857 | 75.58876 | 110.9663 | 99.2017 | 199.8978 | 139.6718 | 201.5364 | 351.2919 |
| ENSMUSG00000020321 | -1.1230685 | 4.63E-07 | 1.98E-05 | 5155.023 | 4426.142 | 6289.47 | 4979.063 | 10855.91 | 6752.292 | 12210.79 | 15596.06 |
| ENSMUSG00000020432 | -1.470661687 | 6.38E-07 | 2.57E-05 | 1632.826 | 1135.031 | 1994.909 | 1546.684 | 5249.999 | 2045.912 | 3941.05 | 6252.561 |
| ENSMUSG00000020469 | 2.392814541 | 1.10E-08 | 9.47E-07 | 33.19979 | 65.99018 | 49.6864 | 77.63611 | 8.776002 | 10.05637 | 15.42863 | 8.700729 |
| ENSMUSG00000020475 | -1.82864481 | 2.05E-07 | 1.02E-05 | 126.7628 | 85.18733 | 150.7154 | 118.1794 | 318.8614 | 198.8926 | 511.0733 | 681.9196 |
| ENSMUSG00000020486 | -1.107751129 | 8.54E-06 | 0.000203 | 924.564 | 675.4995 | 1392.047 | 935.9465 | 2076.987 | 1345.318 | 2383.723 | 2663.511 |
| ENSMUSG00000020566 | -1.561837548 | 0.007788 | 0.035527 | 41.24823 | 33.595 | 217.792 | 43.9938 | 168.6943 | 134.0849 | 314.3583 | 379.5693 |
| ENSMUSG00000020607 | 1.04107406 | 0.000116 | 0.001503 | 817.9222 | 1243.015 | 804.0915 | 935.0839 | 440.7503 | 717.3542 | 416.5729 | 271.8978 |

| | | | | | | | | | | | |
|---|---|---|---|---|---|---|---|---|---|---|---|
| ENSMUSG00000020609 | 1.734187592 | 6.24E-05 | 0.000944 | 59.35721 | 31.19536 | 25.67131 | 48.30692 | 15.60178 | 8.938993 | 9.642892 | 15.22628 |
| ENSMUSG00000020620 | 1.090761413 | 5.07E-05 | 0.0008 | 276.6649 | 243.5638 | 154.0278 | 247.5729 | 91.66046 | 158.6671 | 103.1789 | 79.39415 |
| ENSMUSG00000020651 | -1.649642411 | 0.000861 | 0.006955 | 31.18769 | 27.5959 | 43.88965 | 39.68068 | 102.3867 | 29.05173 | 110.8933 | 205.5547 |
| ENSMUSG00000020681 | -2.449873437 | 0.000145 | 0.001776 | 179.0777 | 111.5834 | 509.2856 | 242.3972 | 1413.911 | 489.4099 | 1252.612 | 2542.788 |
| ENSMUSG00000020704 | -1.023430711 | 6.21E-09 | 5.65E-07 | 280.6892 | 227.9661 | 222.7607 | 266.5507 | 539.2366 | 446.9496 | 642.2166 | 399.1459 |
| ENSMUSG00000020723 | 1.133639439 | 6.71E-08 | 4.07E-06 | 353.1251 | 303.5548 | 325.4459 | 385.5927 | 118.9636 | 224.5922 | 148.5005 | 132.6861 |
| ENSMUSG00000020774 | -2.481020269 | 0.000196 | 0.00223 | 96.58122 | 17.99732 | 174.7305 | 103.5148 | 481.705 | 261.4655 | 571.8235 | 882.0364 |
| ENSMUSG00000020782 | -1.068767259 | 1.46E-06 | 5.06E-05 | 751.5226 | 641.9045 | 1048.383 | 660.7696 | 1525.074 | 1084.97 | 1873.614 | 2027.27 |
| ENSMUSG00000020811 | -1.182433202 | 1.93E-06 | 6.28E-05 | 247.4894 | 262.7609 | 438.0684 | 281.2153 | 636.7477 | 487.1751 | 701.0382 | 969.0437 |
| ENSMUSG00000020839 | -1.730609818 | 0.007033 | 0.033064 | 21.12714 | 16.7975 | 134.1533 | 28.46658 | 149.192 | 77.09881 | 147.5362 | 294.7372 |
| ENSMUSG00000020848 | 1.191005608 | 5.89E-14 | 2.31E-11 | 325.9616 | 296.3559 | 278.2438 | 394.2189 | 145.2916 | 147.4934 | 150.4291 | 123.9854 |
| ENSMUSG00000020887 | -1.723158226 | 0.004949 | 0.025556 | 5.030272 | 5.999108 | 11.59349 | 7.763611 | 12.67645 | 11.17374 | 40.50014 | 36.9781 |
| ENSMUSG00000020902 | 1.301761483 | 8.30E-09 | 7.38E-07 | 5648.995 | 8017.207 | 6619.056 | 7033.832 | 2581.12 | 4269.486 | 2247.758 | 1983.766 |
| ENSMUSG00000020904 | -2.226759968 | 2.24E-05 | 0.00043 | 30.18163 | 9.598572 | 52.17072 | 24.15346 | 129.6898 | 56.98608 | 243.0009 | 117.4598 |
| ENSMUSG00000020963 | -1.350650666 | 0.00338 | 0.019307 | 57.3451 | 11.99822 | 27.32752 | 32.77969 | 66.30757 | 52.51658 | 127.2862 | 84.8321 |
| ENSMUSG00000021048 | -1.008424864 | 5.00E-07 | 2.11E-05 | 1453.749 | 1409.79 | 1734.055 | 1456.108 | 2961.413 | 1899.536 | 3332.583 | 3986.021 |
| ENSMUSG00000021091 | 2.438610958 | 9.67E-06 | 0.000221 | 268.6165 | 41.99375 | 80.32634 | 191.5024 | 26.32801 | 15.64324 | 42.42872 | 22.83941 |
| ENSMUSG00000021098 | 1.713430075 | 8.29E-05 | 0.001166 | 269.6226 | 289.157 | 208.6829 | 301.9182 | 64.35735 | 178.7799 | 44.3573 | 39.15328 |
| ENSMUSG00000021118 | -1.046031369 | 0.000759 | 0.006322 | 40.24217 | 37.19447 | 67.07664 | 33.64232 | 100.4365 | 62.57295 | 114.7504 | 91.35765 |
| ENSMUSG00000021194 | 3.036095014 | 0.0057 | 0.028348 | 1591.578 | 64.79036 | 44.71776 | 1147.289 | 74.10846 | 45.81234 | 76.17884 | 151.1752 |
| ENSMUSG00000021207 | -2.83811215 | 4.81E-07 | 2.04E-05 | 6.036326 | 3.599465 | 9.109173 | 8.626235 | 33.15378 | 21.23011 | 57.85735 | 87.00729 |
| ENSMUSG00000021228 | -3.045649416 | 1.40E-05 | 0.000296 | 13.07871 | 7.198929 | 36.43669 | 6.900988 | 148.2169 | 34.6386 | 61.71451 | 284.9489 |
| ENSMUSG00000021236 | -1.439738138 | 8.12E-07 | 3.13E-05 | 868.2249 | 705.4951 | 1110.491 | 775.4985 | 2090.639 | 1187.769 | 2372.151 | 3736.963 |
| ENSMUSG00000021238 | -1.06674792 | 4.96E-09 | 4.74E-07 | 1885.346 | 1382.194 | 1538.622 | 1907.261 | 3684.946 | 2414.645 | 3687.442 | 4277.496 |
| ENSMUSG00000021255 | -2.603649274 | 7.50E-06 | 0.000185 | 55.33299 | 71.98929 | 207.0267 | 69.00988 | 527.5352 | 259.2308 | 1003.825 | 664.5182 |
| ENSMUSG00000021268 | 1.658559114 | 0.000624 | 0.005443 | 8713.437 | 10857.19 | 7024 | 10075.44 | 2831.723 | 5572.345 | 1811.899 | 1399.73 |
| ENSMUSG00000021278 | -2.47375294 | 0.000385 | 0.003746 | 141.8537 | 35.99465 | 336.2113 | 147.5086 | 948.7833 | 385.4941 | 777.2171 | 1567.219 |
| ENSMUSG00000021281 | -1.201068579 | 0.000112 | 0.001469 | 391.3551 | 250.7627 | 565.5968 | 324.3464 | 987.7878 | 440.2454 | 968.1463 | 1128.92 |
| ENSMUSG00000021322 | -1.435435224 | 0.000269 | 0.002849 | 27.16347 | 29.99554 | 32.29616 | 27.60395 | 100.4365 | 35.75597 | 60.75022 | 119.635 |
| ENSMUSG00000021335 | -3.610677438 | 1.12E-08 | 9.52E-07 | 52.31483 | 17.99732 | 119.2474 | 56.07053 | 894.1771 | 313.9821 | 600.7521 | 1200.701 |
| ENSMUSG00000021336 | -1.009108673 | 0.000973 | 0.007672 | 65.39353 | 38.39429 | 76.18581 | 42.26855 | 144.3165 | 82.68568 | 92.57176 | 129.4233 |
| ENSMUSG00000021364 | -1.236996503 | 1.65E-05 | 0.000336 | 207.2472 | 137.9795 | 251.7444 | 203.5791 | 493.4063 | 263.7003 | 445.5016 | 687.3576 |
| ENSMUSG00000021376 | -1.192754244 | 1.52E-05 | 0.000315 | 122.7386 | 145.1784 | 142.4343 | 132.844 | 322.7618 | 162.0192 | 310.5011 | 445.9123 |
| ENSMUSG00000021384 | -2.278960502 | 1.88E-09 | 2.20E-07 | 102.6175 | 47.99286 | 153.1997 | 112.1411 | 415.3974 | 268.1698 | 537.1091 | 804.8174 |
| ENSMUSG00000021388 | 1.118140677 | 7.18E-07 | 2.84E-05 | 1882.328 | 2223.269 | 1411.922 | 2299.754 | 947.8082 | 1239.168 | 805.1815 | 609.051 |
| ENSMUSG00000021390 | 1.274779245 | 1.75E-08 | 1.35E-06 | 3975.927 | 4632.511 | 2990.293 | 4987.689 | 1525.074 | 2507.387 | 1568.898 | 1253.993 |
| ENSMUSG00000021416 | -2.701804457 | 1.05E-07 | 5.87E-06 | 13.07871 | 20.39697 | 17.39024 | 19.84034 | 122.864 | 32.40385 | 94.50034 | 209.9051 |
| ENSMUSG00000021456 | -2.757720064 | 2.95E-07 | 1.35E-05 | 16.09687 | 13.19804 | 64.59232 | 18.11509 | 186.2463 | 93.85942 | 162.0006 | 321.927 |
| ENSMUSG00000021469 | -1.294015603 | 0.000876 | 0.007032 | 19.11503 | 11.99822 | 27.32752 | 16.38985 | 57.53157 | 43.57759 | 27.96439 | 56.55474 |
| ENSMUSG00000021490 | -3.04464571 | 8.27E-05 | 0.001166 | 535.2209 | 105.5843 | 1284.393 | 628.8525 | 6439.635 | 1949.818 | 3478.191 | 9211.897 |
| ENSMUSG00000021534 | 1.911850251 | 0.000579 | 0.005135 | 12.07265 | 16.7975 | 16.56213 | 16.38985 | 4.875557 | 3.352122 | 4.821446 | 3.262773 |
| ENSMUSG00000021565 | -2.762476486 | 5.62E-09 | 5.29E-07 | 31.18769 | 14.39786 | 51.34261 | 21.56559 | 284.7325 | 72.62932 | 190.9293 | 261.0219 |
| ENSMUSG00000021612 | -3.691801868 | 7.18E-06 | 0.000179 | 106.6418 | 17.99732 | 364.3669 | 153.547 | 1953.148 | 835.7958 | 2301.758 | 3220.357 |
| ENSMUSG00000021647 | 1.923403309 | 0.003663 | 0.02048 | 62.37537 | 19.19714 | 10.76539 | 61.24627 | 12.67645 | 4.469496 | 14.46434 | 8.700729 |
| ENSMUSG00000021700 | 1.795172055 | 0.000511 | 0.004657 | 121.7326 | 39.59411 | 44.71776 | 197.5408 | 35.10401 | 25.6996 | 24.10723 | 31.54014 |
| ENSMUSG00000021751 | -6.148751376 | 4.19E-11 | 8.40E-09 | 0 | 2.399643 | 3.312427 | 0 | 119.9387 | 15.64324 | 82.92887 | 185.9781 |
| ENSMUSG00000021765 | 1.284222812 | 0.011319 | 0.046796 | 126.7628 | 266.3604 | 166.4494 | 172.5247 | 75.08357 | 147.4934 | 61.71451 | 16.31387 |
| ENSMUSG00000021822 | -2.429339789 | 0.009252 | 0.040236 | 419.5247 | 206.3693 | 3204.773 | 536.5518 | 4974.043 | 1752.043 | 6910.096 | 9890.553 |
| ENSMUSG00000021876 | 1.039620679 | 4.18E-12 | 1.01E-09 | 2857.194 | 3467.484 | 2913.279 | 3397.011 | 1633.311 | 1866.015 | 1492.72 | 1153.934 |
| ENSMUSG00000021903 | 1.713988677 | 4.71E-07 | 2.00E-05 | 613.6932 | 341.9491 | 237.6666 | 585.7214 | 127.7396 | 179.8972 | 155.2506 | 79.39415 |
| ENSMUSG00000021904 | -1.33109632 | 8.19E-06 | 0.000196 | 371.2341 | 226.7663 | 555.6596 | 327.7969 | 646.4988 | 689.4198 | 1116.647 | 1279.007 |
| ENSMUSG00000021922 | 3.645006374 | 5.45E-06 | 0.000143 | 60.36326 | 19.19714 | 13.24971 | 38.81806 | 0 | 3.352122 | 1.928578 | 5.437955 |
| ENSMUSG00000021947 | -1.374064737 | 3.75E-07 | 1.66E-05 | 466.8092 | 435.5352 | 626.8767 | 494.2833 | 1515.323 | 695.0067 | 1137.861 | 1898.934 |
| ENSMUSG00000021950 | 1.301099415 | 1.97E-05 | 0.000389 | 265.5983 | 334.7502 | 235.1823 | 345.912 | 187.2214 | 73.74669 | 137.8934 | 79.39415 |
| ENSMUSG00000021991 | 1.055022453 | 8.90E-05 | 0.001228 | 283.7073 | 434.3354 | 382.5853 | 368.3402 | 175.52 | 268.1698 | 159.1077 | 104.4087 |
| ENSMUSG00000021999 | -1.973204701 | 0.000151 | 0.001836 | 47.28455 | 11.99822 | 76.18581 | 69.00988 | 147.2418 | 92.74205 | 226.608 | 340.416 |
| ENSMUSG00000022025 | 1.16589502 | 5.00E-05 | 0.000794 | 91.55095 | 100.785 | 97.71658 | 79.36136 | 29.25334 | 46.92971 | 59.78593 | 28.27737 |
| ENSMUSG00000022040 | -1.005135463 | 0.00033 | 0.003338 | 138.8355 | 104.3845 | 214.4796 | 133.7066 | 280.8321 | 202.2447 | 287.3582 | 419.8102 |
| ENSMUSG00000022044 | 2.68129849 | 0.000217 | 0.002418 | 84.50857 | 17.99732 | 10.76539 | 105.2401 | 10.72622 | 7.821619 | 5.785735 | 9.78832 |
| ENSMUSG00000022053 | 1.536601284 | 0.005677 | 0.028246 | 55.33299 | 142.7788 | 43.88965 | 87.9876 | 28.27823 | 53.63396 | 24.10723 | 7.613138 |
| ENSMUSG00000022099 | -1.287046801 | 0.000813 | 0.00665 | 395.3794 | 195.5709 | 590.44 | 316.5828 | 792.7655 | 405.6068 | 1022.147 | 1437.795 |
| ENSMUSG00000022103 | 1.306774131 | 2.76E-10 | 4.18E-08 | 282.7013 | 488.3274 | 378.4447 | 383.8675 | 183.3209 | 173.193 | 140.7862 | 121.8102 |
| ENSMUSG00000022123 | -1.504916161 | 1.78E-06 | 5.95E-05 | 68.4117 | 40.79393 | 115.9349 | 73.323 | 181.3707 | 157.5497 | 247.8223 | 265.3722 |
| ENSMUSG00000022132 | -2.356029569 | 0.002477 | 0.015326 | 123.7447 | 99.58519 | 845.4969 | 199.266 | 1317.375 | 724.0584 | 2124.329 | 2329.62 |

| | | | | | | | | | | | |
|---|---|---|---|---|---|---|---|---|---|---|---|
| ENSMUSG00000022208 | 1.782806275 | 8.38E-18 | 6.72E-15 | 149.9021 | 172.7743 | 144.0906 | 169.9368 | 42.9049 | 55.8687 | 33.75012 | 53.29196 |
| ENSMUSG00000022215 | -2.420550806 | 0.001798 | 0.012158 | 4.024217 | 11.99822 | 24.01509 | 2.58787 | 78.0089 | 14.52586 | 29.89296 | 106.5839 |
| ENSMUSG00000022218 | -1.550549858 | 1.87E-06 | 6.11E-05 | 41.24823 | 43.19358 | 57.13936 | 56.93315 | 134.5654 | 90.5073 | 121.5004 | 237.0949 |
| ENSMUSG00000022219 | -1.897865831 | 7.94E-05 | 0.001132 | 64.38748 | 25.19625 | 95.23226 | 52.62003 | 202.8232 | 82.68568 | 203.465 | 399.1459 |
| ENSMUSG00000022237 | -1.314140885 | 8.43E-08 | 4.95E-06 | 150.9082 | 163.1757 | 223.5888 | 177.7004 | 420.273 | 297.2215 | 414.6443 | 649.2919 |
| ENSMUSG00000022244 | -1.147290803 | 7.25E-06 | 0.00018 | 288.7376 | 319.1525 | 335.3832 | 288.1162 | 648.449 | 394.4331 | 638.3594 | 1046.263 |
| ENSMUSG00000022270 | -1.266684644 | 5.99E-08 | 3.75E-06 | 577.4752 | 350.3479 | 658.3448 | 587.4466 | 1240.342 | 887.195 | 1686.542 | 1420.394 |
| ENSMUSG00000022323 | -1.417347786 | 1.88E-08 | 1.44E-06 | 863.1946 | 686.2979 | 1153.553 | 983.3908 | 2526.513 | 1499.516 | 2206.294 | 3618.416 |
| ENSMUSG00000022330 | -1.42592984 | 1.38E-05 | 0.000294 | 428.5792 | 116.3827 | 313.0243 | 306.2313 | 542.1619 | 812.331 | 857.2531 | 920.1021 |
| ENSMUSG00000022351 | -1.061273975 | 1.60E-12 | 4.33E-10 | 889.3521 | 892.6672 | 924.167 | 817.7671 | 1766.902 | 1347.553 | 2165.793 | 2071.861 |
| ENSMUSG00000022366 | -6.283077507 | 8.78E-09 | 7.60E-07 | 0 | 3.599465 | 0 | 0 | 48.75557 | 20.11273 | 38.57157 | 130.5109 |
| ENSMUSG00000022376 | 1.378351721 | 0.001178 | 0.008849 | 196.1806 | 239.9643 | 194.6051 | 213.9306 | 64.35735 | 173.193 | 53.0359 | 34.80291 |
| ENSMUSG00000022383 | -1.500917255 | 0.000748 | 0.006253 | 52.31483 | 16.7975 | 54.65504 | 18.97772 | 96.53602 | 59.22083 | 130.179 | 119.635 |
| ENSMUSG00000022429 | 1.601642035 | 2.51E-09 | 2.79E-07 | 227.3683 | 266.3604 | 222.7607 | 216.5185 | 68.25779 | 119.559 | 75.21455 | 44.59123 |
| ENSMUSG00000022445 | -3.863399547 | 4.62E-09 | 4.56E-07 | 67.40564 | 21.59679 | 163.137 | 78.49874 | 1295.923 | 476.0014 | 1086.754 | 1964.19 |
| ENSMUSG00000022512 | 1.017838586 | 2.33E-05 | 0.000443 | 6673.159 | 6888.175 | 4421.261 | 7740.321 | 3095.978 | 4508.604 | 3111.761 | 1987.029 |
| ENSMUSG00000022523 | 1.01071381 | 5.53E-06 | 0.000144 | 335.0161 | 338.3497 | 279.0719 | 331.2474 | 135.5405 | 240.2354 | 136.9291 | 125.073 |
| ENSMUSG00000022575 | 1.002502513 | 3.45E-09 | 3.58E-07 | 530.1906 | 758.2872 | 613.627 | 634.0283 | 306.185 | 406.7242 | 266.1438 | 287.124 |
| ENSMUSG00000022580 | -1.391387438 | 7.30E-06 | 0.000181 | 173.0413 | 182.3729 | 271.619 | 203.5791 | 366.6419 | 328.508 | 609.4308 | 876.5984 |
| ENSMUSG00000022589 | 10.00473679 | 9.26E-05 | 0.001256 | 1670.05 | 0 | 1.656213 | 439.938 | 0 | 0 | 0.964289 | 1.087591 |
| ENSMUSG00000022613 | -5.937722768 | 2.44E-09 | 2.73E-07 | 4.024217 | 3.599465 | 26.49941 | 18.97772 | 1627.461 | 177.6625 | 183.2149 | 1289.883 |
| ENSMUSG00000022658 | 1.545818987 | 0.000126 | 0.0016 | 184.1079 | 93.58608 | 93.57605 | 207.0296 | 52.65601 | 48.04709 | 74.25027 | 22.83941 |
| ENSMUSG00000022679 | -1.557217989 | 7.81E-10 | 1.04E-07 | 201.2109 | 158.3764 | 174.7305 | 168.2116 | 477.8045 | 287.1651 | 676.931 | 625.3649 |
| ENSMUSG00000022742 | -1.020684549 | 2.03E-05 | 0.000397 | 518.118 | 423.537 | 591.2681 | 479.6187 | 1100.901 | 597.7951 | 951.7534 | 1434.533 |
| ENSMUSG00000022763 | -1.141427363 | 1.92E-05 | 0.00038 | 32.19374 | 25.19625 | 33.12427 | 28.46658 | 78.98402 | 71.51194 | 49.17875 | 64.16787 |
| ENSMUSG00000022780 | 1.141599418 | 0.000409 | 0.003927 | 111.672 | 169.1748 | 147.403 | 90.57547 | 51.6809 | 87.15518 | 34.71441 | 61.99269 |
| ENSMUSG00000022790 | -1.298935786 | 6.99E-05 | 0.00103 | 279.6831 | 89.98662 | 229.3855 | 196.6782 | 571.4152 | 320.6864 | 436.823 | 631.8904 |
| ENSMUSG00000022821 | -3.49123003 | 2.86E-07 | 1.33E-05 | 75.45408 | 10.79839 | 63.76421 | 41.40593 | 620.1708 | 183.2494 | 435.8587 | 917.9269 |
| ENSMUSG00000022829 | -1.511696289 | 0.002873 | 0.017193 | 9.054489 | 3.599465 | 11.59349 | 6.900988 | 18.52711 | 18.99536 | 18.32149 | 34.80291 |
| ENSMUSG00000022838 | -1.149652862 | 0.000518 | 0.004709 | 89.53884 | 38.39429 | 102.6852 | 77.63611 | 151.1423 | 115.0895 | 179.3578 | 241.4452 |
| ENSMUSG00000022853 | -1.949806543 | 1.70E-07 | 8.72E-06 | 170.0232 | 111.5834 | 207.0267 | 158.7227 | 767.4126 | 253.6439 | 464.7874 | 1017.985 |
| ENSMUSG00000022860 | 1.871222642 | 7.90E-05 | 0.001131 | 86.52067 | 209.9688 | 122.5598 | 204.4418 | 35.10401 | 82.68568 | 37.60728 | 15.22628 |
| ENSMUSG00000022868 | 3.439510106 | 0.00744 | 0.034372 | 990.9635 | 16.7975 | 15.73403 | 916.1062 | 12.67645 | 41.34284 | 70.39311 | 54.37955 |
| ENSMUSG00000022935 | 1.265022336 | 0.000217 | 0.002413 | 152.9203 | 124.7814 | 157.3403 | 210.4801 | 53.63112 | 120.6764 | 45.32159 | 50.02919 |
| ENSMUSG00000023019 | -1.97574196 | 6.57E-08 | 4.03E-06 | 260.5681 | 167.975 | 334.5551 | 244.1224 | 1024.842 | 427.9543 | 772.3956 | 1740.146 |
| ENSMUSG00000023046 | 1.38809093 | 0.000964 | 0.007629 | 2826.007 | 3606.664 | 2178.749 | 3216.723 | 1033.618 | 1997.865 | 944.0391 | 543.7955 |
| ENSMUSG00000023073 | -4.577577047 | 3.35E-08 | 2.33E-06 | 2.012109 | 4.799286 | 2.48432 | 0.862623 | 31.20356 | 10.05637 | 91.60747 | 102.2336 |
| ENSMUSG00000023078 | 1.229665392 | 0.00014 | 0.001734 | 50.30272 | 85.18733 | 89.43552 | 56.07053 | 21.45245 | 35.75597 | 26.03581 | 36.9781 |
| ENSMUSG00000023120 | -5.4525277 | 2.11E-05 | 0.000411 | 0 | 0 | 10.76539 | 1.725247 | 114.088 | 30.1691 | 212.1436 | 204.4671 |
| ENSMUSG00000023122 | -3.438552607 | 1.92E-06 | 6.28E-05 | 13.07871 | 2.399643 | 34.78048 | 3.450494 | 148.2169 | 46.92971 | 143.6791 | 250.146 |
| ENSMUSG00000023153 | -2.469702097 | 1.22E-05 | 0.000268 | 11.0666 | 1.199822 | 14.90592 | 6.038364 | 34.1289 | 34.6386 | 45.32159 | 73.95619 |
| ENSMUSG00000023484 | 3.549787245 | 0.000353 | 0.003511 | 637.8385 | 75.58876 | 58.79557 | 1006.682 | 29.25334 | 69.27719 | 41.46443 | 11.9635 |
| ENSMUSG00000023829 | -3.163770175 | 8.10E-06 | 0.000195 | 60.36326 | 10.79839 | 107.6539 | 47.44429 | 612.3699 | 203.3621 | 372.2156 | 845.0583 |
| ENSMUSG00000023914 | -3.657771973 | 3.28E-05 | 0.000573 | 169.0171 | 64.79036 | 857.0904 | 257.9244 | 4263.187 | 1259.281 | 3851.371 | 7656.641 |
| ENSMUSG00000023931 | 1.072475961 | 0.01052 | 0.044254 | 26.15741 | 47.99286 | 24.8432 | 50.03216 | 18.52711 | 25.6996 | 12.53576 | 14.13868 |
| ENSMUSG00000023959 | -1.272839763 | 4.46E-06 | 0.000124 | 275.6589 | 175.1739 | 341.1799 | 254.4739 | 534.361 | 393.3157 | 694.2882 | 910.3137 |
| ENSMUSG00000024011 | 1.336768093 | 0.00018 | 0.002091 | 348.0948 | 601.1106 | 332.0708 | 475.3055 | 202.8232 | 290.5173 | 111.8575 | 90.27006 |
| ENSMUSG00000024064 | -1.483637194 | 2.30E-05 | 0.000438 | 168.0111 | 87.58697 | 206.1986 | 217.3811 | 449.5263 | 235.7659 | 512.0375 | 705.8466 |
| ENSMUSG00000024066 | 1.49932047 | 2.50E-07 | 1.21E-05 | 870.237 | 634.7056 | 343.6643 | 721.1532 | 263.2801 | 274.874 | 224.6794 | 145.7372 |
| ENSMUSG00000024076 | 1.222536486 | 9.85E-08 | 5.59E-06 | 150.9082 | 135.5798 | 102.6852 | 162.1732 | 60.4569 | 69.27719 | 61.71451 | 44.59123 |
| ENSMUSG00000024114 | 1.039774439 | 0.000104 | 0.001383 | 111.672 | 152.3773 | 101.8571 | 112.1411 | 55.58134 | 34.6386 | 69.42882 | 71.78101 |
| ENSMUSG00000024131 | -4.233535956 | 1.35E-08 | 1.09E-06 | 66.39959 | 15.59768 | 103.5133 | 56.07053 | 1446.09 | 279.3435 | 729.0026 | 2099.051 |
| ENSMUSG00000024164 | 1.834888712 | 6.03E-06 | 0.000155 | 528.1785 | 1391.793 | 507.6294 | 846.2336 | 213.5494 | 353.0902 | 259.3938 | 91.35765 |
| ENSMUSG00000024245 | -1.126471319 | 0.000225 | 0.002482 | 269.6226 | 197.9706 | 490.2391 | 285.5284 | 556.7886 | 439.128 | 810.9672 | 911.4013 |
| ENSMUSG00000024268 | 1.117413524 | 0.003045 | 0.01796 | 1025.169 | 827.8769 | 661.6572 | 964.4131 | 289.6081 | 824.6221 | 284.4653 | 205.5547 |
| ENSMUSG00000024313 | -5.228780073 | 3.48E-08 | 2.39E-06 | 7.04238 | 7.198929 | 63.76421 | 12.07673 | 776.1886 | 175.4277 | 687.5382 | 1755.372 |
| ENSMUSG00000024330 | -1.049741062 | 4.80E-05 | 0.000771 | 145.8779 | 213.5682 | 149.8873 | 153.547 | 511.9334 | 238.0007 | 341.3584 | 277.3357 |
| ENSMUSG00000024347 | 1.286324004 | 0.00049 | 0.004507 | 106.6418 | 80.38804 | 86.9512 | 138.8824 | 23.40267 | 72.62932 | 42.42872 | 31.54014 |
| ENSMUSG00000024354 | -2.629234026 | 1.80E-05 | 0.000363 | 151.9142 | 53.99197 | 332.0708 | 144.0581 | 1004.365 | 453.6539 | 1006.718 | 1759.722 |
| ENSMUSG00000024366 | 1.863140234 | 0.000364 | 0.003587 | 46.2785 | 23.99643 | 28.15563 | 95.75121 | 12.67645 | 15.64324 | 15.42863 | 9.78832 |
| ENSMUSG00000024386 | -2.232133332 | 0.000177 | 0.002063 | 21.12714 | 4.799286 | 38.92101 | 22.42821 | 131.64 | 33.52122 | 80.036 | 168.5766 |
| ENSMUSG00000024388 | -1.875016164 | 6.60E-08 | 4.03E-06 | 51.30877 | 26.39607 | 64.59232 | 41.40593 | 147.2418 | 106.1505 | 166.822 | 257.7591 |
| ENSMUSG00000024391 | -2.463233726 | 5.34E-10 | 7.35E-08 | 63.38142 | 26.39607 | 28.98373 | 37.95543 | 235.9769 | 102.7984 | 192.8578 | 332.8029 |
| ENSMUSG00000024459 | 1.122664977 | 0.000623 | 0.005433 | 111.672 | 113.983 | 80.32634 | 125.943 | 41.92979 | 83.80306 | 41.46443 | 31.54014 |

| | | | | | | | | | | | |
|---|---|---|---|---|---|---|---|---|---|---|---|
| ENSMUSG00000024481 | 1.951193469 | 0.003952 | 0.021671 | 60.36326 | 107.9839 | 37.2648 | 96.61383 | 19.50223 | 40.22547 | 17.3572 | 1.087591 |
| ENSMUSG00000024503 | -2.621590653 | 0.000306 | 0.003155 | 102.6175 | 27.5959 | 313.8524 | 99.2017 | 784.9646 | 368.7335 | 959.4677 | 1234.416 |
| ENSMUSG00000024552 | -1.56598976 | 0.000823 | 0.006715 | 88.53278 | 99.58519 | 384.2415 | 108.6906 | 378.3432 | 311.7474 | 743.4669 | 586.2116 |
| ENSMUSG00000024565 | -1.224547991 | 0.000522 | 0.004735 | 72.43591 | 25.19625 | 57.96746 | 47.44429 | 92.63557 | 83.80306 | 136.9291 | 163.1387 |
| ENSMUSG00000024593 | 1.426343265 | 3.20E-08 | 2.26E-06 | 526.1664 | 441.5343 | 382.5853 | 433.8996 | 134.5654 | 269.2872 | 119.5719 | 141.3868 |
| ENSMUSG00000024610 | -1.066020768 | 0.000695 | 0.005913 | 43.26034 | 47.99286 | 95.23226 | 68.14726 | 181.3707 | 93.85942 | 132.1076 | 128.3357 |
| ENSMUSG00000024650 | -4.559778162 | 1.03E-06 | 3.82E-05 | 60.36326 | 9.598572 | 227.7293 | 56.07053 | 2274.935 | 611.2036 | 1394.362 | 4074.116 |
| ENSMUSG00000024664 | -1.365913954 | 1.01E-05 | 0.000229 | 400.4096 | 281.9581 | 464.5678 | 401.9825 | 1027.767 | 496.1141 | 851.4673 | 1619.423 |
| ENSMUSG00000024678 | 2.103231877 | 0.002266 | 0.014359 | 72.43591 | 61.1909 | 33.95237 | 83.67448 | 13.65156 | 37.99072 | 3.857157 | 3.262773 |
| ENSMUSG00000024679 | 1.113360627 | 0.000133 | 0.001673 | 271.6347 | 145.1784 | 223.5888 | 181.1509 | 91.66046 | 84.92043 | 140.7862 | 61.99269 |
| ENSMUSG00000024680 | -2.330851848 | 8.58E-07 | 3.29E-05 | 22.1332 | 28.79572 | 4.96864 | 12.93935 | 62.40712 | 63.69032 | 104.1432 | 109.8467 |
| ENSMUSG00000024694 | -2.737531005 | 5.27E-05 | 0.000824 | 316.9071 | 34.79482 | 410.7409 | 339.011 | 1937.546 | 1055.919 | 2195.686 | 2161.043 |
| ENSMUSG00000024757 | -4.972972177 | 4.41E-05 | 0.000725 | 14.08476 | 0 | 78.67013 | 22.42821 | 980.962 | 203.3621 | 743.4669 | 1699.905 |
| ENSMUSG00000024810 | 1.184438856 | 8.45E-08 | 4.95E-06 | 6305.949 | 6181.481 | 4281.311 | 6681.019 | 2542.115 | 3657.165 | 2399.151 | 1719.482 |
| ENSMUSG00000024866 | -1.058892735 | 3.90E-05 | 0.00066 | 442.6639 | 429.5361 | 553.1752 | 478.756 | 1182.81 | 551.9828 | 828.3244 | 1405.168 |
| ENSMUSG00000024892 | -2.037307432 | 2.85E-06 | 8.66E-05 | 753.5347 | 427.1365 | 565.5968 | 508.9479 | 2866.827 | 991.1108 | 1683.649 | 3716.299 |
| ENSMUSG00000024899 | -1.091619428 | 8.81E-08 | 5.11E-06 | 554.3359 | 554.3175 | 438.8965 | 565.881 | 1094.075 | 763.1665 | 1087.718 | 1557.43 |
| ENSMUSG00000024935 | -2.629325979 | 8.99E-11 | 1.66E-08 | 48.29061 | 25.19625 | 103.5133 | 49.16954 | 271.0809 | 221.2401 | 390.5371 | 525.3065 |
| ENSMUSG00000024955 | -1.075674739 | 0.000135 | 0.001689 | 537.233 | 386.3425 | 788.3575 | 513.261 | 1119.428 | 651.4291 | 1327.826 | 1594.409 |
| ENSMUSG00000024972 | -1.350193183 | 0.000631 | 0.005483 | 30.18163 | 27.5959 | 50.5145 | 29.3292 | 55.58134 | 71.51194 | 71.3574 | 154.4379 |
| ENSMUSG00000025056 | 1.493748171 | 0.009868 | 0.042218 | 141.8537 | 34.79482 | 20.70267 | 72.46037 | 18.52711 | 35.75597 | 23.14294 | 18.48905 |
| ENSMUSG00000025059 | -1.588480945 | 2.81E-09 | 3.08E-07 | 588.5418 | 353.9474 | 734.5306 | 612.4627 | 1533.85 | 1125.196 | 1716.435 | 2515.598 |
| ENSMUSG00000025176 | -1.804755162 | 5.59E-07 | 2.29E-05 | 334.01 | 166.7752 | 438.0684 | 332.9727 | 1129.179 | 513.9921 | 1004.789 | 1799.963 |
| ENSMUSG00000025194 | -3.111950633 | 3.41E-13 | 1.14E-10 | 48.29061 | 21.59679 | 50.5145 | 30.19182 | 447.5761 | 130.7328 | 238.1794 | 490.5036 |
| ENSMUSG00000025196 | -1.29065985 | 4.94E-05 | 0.000787 | 577.4752 | 250.7627 | 755.2332 | 556.3922 | 1195.486 | 845.8522 | 1313.362 | 1884.795 |
| ENSMUSG00000025226 | -1.097455621 | 9.65E-06 | 0.000221 | 175.0535 | 133.1802 | 164.7932 | 164.7611 | 391.9947 | 210.0663 | 294.1082 | 469.8393 |
| ENSMUSG00000025243 | -4.050333588 | 3.12E-11 | 6.39E-09 | 10.06054 | 7.198929 | 35.60859 | 6.900988 | 285.7076 | 81.56831 | 176.4649 | 457.8758 |
| ENSMUSG00000025255 | 1.245736482 | 0.002753 | 0.016576 | 114.6902 | 113.983 | 51.34261 | 89.71284 | 36.07912 | 69.27719 | 29.89296 | 20.66423 |
| ENSMUSG00000025347 | -4.33948442 | 1.42E-06 | 4.95E-05 | 5.030272 | 2.399643 | 31.46805 | 12.07673 | 258.4045 | 71.51194 | 162.9649 | 549.2335 |
| ENSMUSG00000025400 | -1.716211352 | 0.001921 | 0.012745 | 5.030272 | 4.799286 | 8.281066 | 6.038364 | 34.1289 | 13.40849 | 18.32149 | 14.13868 |
| ENSMUSG00000025418 | -2.450628883 | 0.001517 | 0.01064 | 87.52673 | 44.3934 | 316.3367 | 65.55939 | 638.6979 | 230.1791 | 608.4665 | 1334.474 |
| ENSMUSG00000025427 | 1.263683315 | 8.40E-10 | 1.11E-07 | 190.1443 | 202.7698 | 146.5749 | 168.2116 | 70.20801 | 96.09417 | 69.42882 | 58.72992 |
| ENSMUSG00000025467 | -1.342591806 | 0.000563 | 0.005013 | 11.0666 | 14.39786 | 13.24971 | 7.763611 | 32.17867 | 35.75597 | 24.10723 | 25.0146 |
| ENSMUSG00000025475 | 1.092632264 | 0.001504 | 0.010583 | 79.47829 | 67.19001 | 57.13936 | 67.28463 | 29.25334 | 53.63396 | 26.03581 | 18.48905 |
| ENSMUSG00000025479 | -3.424689237 | 1.23E-07 | 6.68E-06 | 14.08476 | 2.399643 | 7.45296 | 25.01608 | 156.0178 | 40.22547 | 93.53605 | 241.4452 |
| ENSMUSG00000025494 | -1.877582383 | 3.27E-08 | 2.29E-06 | 155.9384 | 101.9848 | 251.7444 | 138.0198 | 609.4446 | 324.0385 | 532.2876 | 919.0145 |
| ENSMUSG00000025497 | -2.314805506 | 4.11E-05 | 0.000689 | 117.7084 | 89.98662 | 269.1347 | 100.0643 | 674.777 | 272.6393 | 648.9666 | 1276.832 |
| ENSMUSG00000025504 | -1.419302796 | 1.33E-05 | 0.000286 | 458.7608 | 367.1454 | 767.6548 | 373.516 | 1260.819 | 629.0816 | 1572.756 | 1801.051 |
| ENSMUSG00000025608 | -1.136143995 | 2.00E-05 | 0.000392 | 1864.219 | 1397.792 | 3039.151 | 2085.824 | 3096.954 | 3794.602 | 5057.697 | 6489.656 |
| ENSMUSG00000025701 | 1.65192938 | 0.000223 | 0.002465 | 29.17558 | 35.99465 | 25.67131 | 24.15346 | 10.72622 | 13.40849 | 3.857157 | 8.700729 |
| ENSMUSG00000025732 | -1.469895001 | 8.06E-06 | 0.000194 | 347.0888 | 154.777 | 278.2438 | 166.4863 | 595.793 | 378.7898 | 653.7881 | 995.1458 |
| ENSMUSG00000025762 | -1.003961148 | 2.31E-08 | 1.71E-06 | 409.4641 | 335.95 | 402.4598 | 381.2796 | 623.0961 | 632.4337 | 777.2171 | 1036.474 |
| ENSMUSG00000025792 | -1.217626256 | 1.06E-05 | 0.000238 | 893.3763 | 782.2836 | 1283.565 | 873.8376 | 2289.561 | 1202.295 | 2049.114 | 3375.883 |
| ENSMUSG00000025911 | -1.679507065 | 2.36E-08 | 1.74E-06 | 243.4652 | 179.9732 | 294.806 | 235.4962 | 881.5006 | 358.6771 | 729.0026 | 1088.679 |
| ENSMUSG00000025937 | -1.639609521 | 1.82E-09 | 2.17E-07 | 503.0272 | 393.5415 | 703.0625 | 523.6125 | 1618.685 | 939.7116 | 1585.291 | 2476.445 |
| ENSMUSG00000025977 | 3.223396968 | 5.56E-08 | 3.55E-06 | 28.16952 | 17.99732 | 33.95237 | 25.8787 | 1.950223 | 3.352122 | 1.928578 | 4.350364 |
| ENSMUSG00000025991 | 2.616224666 | 0.000498 | 0.004571 | 126.7628 | 17.99732 | 10.76539 | 148.3712 | 12.67645 | 13.40849 | 11.57147 | 11.9635 |
| ENSMUSG00000026058 | 1.309505671 | 0.00123 | 0.009119 | 148.896 | 179.9732 | 157.3403 | 191.5024 | 47.78045 | 143.0239 | 46.28588 | 36.9781 |
| ENSMUSG00000026107 | -1.429503907 | 1.87E-05 | 0.000372 | 303.8284 | 232.7654 | 450.49 | 235.4962 | 822.0188 | 357.5597 | 1071.325 | 1044.087 |
| ENSMUSG00000026167 | -2.445566701 | 0.000133 | 0.001669 | 3.018163 | 2.399643 | 11.59349 | 1.725247 | 30.22845 | 18.99536 | 23.14294 | 32.62773 |
| ENSMUSG00000026175 | -2.319144863 | 9.71E-05 | 0.001308 | 112.6781 | 27.5959 | 212.8234 | 109.5532 | 619.1957 | 319.569 | 540.0019 | 834.1824 |
| ENSMUSG00000026202 | -1.537182333 | 6.24E-06 | 0.00016 | 527.1725 | 419.9375 | 908.433 | 569.3315 | 1485.095 | 804.5093 | 2008.614 | 2742.905 |
| ENSMUSG00000026204 | 1.616973253 | 0.002396 | 0.014942 | 177.0656 | 40.79393 | 50.5145 | 200.9913 | 50.70579 | 40.22547 | 23.14294 | 39.15328 |
| ENSMUSG00000026205 | -1.702891512 | 7.21E-07 | 2.85E-05 | 48.29061 | 35.99465 | 66.24853 | 46.58167 | 155.0427 | 79.33356 | 179.3578 | 230.5693 |
| ENSMUSG00000026247 | 3.47232237 | 2.62E-14 | 1.05E-11 | 191.1503 | 95.98572 | 62.108 | 159.5853 | 8.776002 | 18.99536 | 9.642892 | 8.700729 |
| ENSMUSG00000026295 | -3.781394294 | 1.99E-06 | 6.43E-05 | 115.6963 | 28.79572 | 411.569 | 138.0198 | 3226.643 | 744.1711 | 1914.114 | 3667.357 |
| ENSMUSG00000026348 | -5.6311638 | 2.01E-20 | 3.09E-17 | 3.018163 | 3.599465 | 5.796746 | 5.175741 | 307.1601 | 77.09881 | 105.1075 | 394.7956 |
| ENSMUSG00000026368 | -3.498695397 | 2.65E-05 | 0.000487 | 8.048435 | 0 | 2.48432 | 7.763611 | 65.33246 | 11.17374 | 29.89296 | 103.3212 |
| ENSMUSG00000026394 | -2.060048061 | 0.005861 | 0.028931 | 98.59333 | 38.39429 | 279.9 | 56.07053 | 450.5014 | 156.4324 | 583.3949 | 784.1532 |
| ENSMUSG00000026437 | -1.235844007 | 0.000176 | 0.002056 | 192.1564 | 158.3764 | 244.2915 | 152.6844 | 533.3859 | 183.2494 | 445.5016 | 599.2627 |
| ENSMUSG00000026535 | 2.138205585 | 0.008938 | 0.039268 | 45.27245 | 93.58608 | 76.18581 | 31.05445 | 28.27823 | 24.58223 | 1.928578 | 1.087591 |
| ENSMUSG00000026542 | -2.117172233 | 0.011081 | 0.046135 | 8.048435 | 0 | 3.312427 | 17.25247 | 18.52711 | 15.64324 | 30.85725 | 60.9051 |
| ENSMUSG00000026576 | -1.726853917 | 0.001014 | 0.007919 | 5303.919 | 2560.419 | 11402.2 | 5633.794 | 15996.7 | 10829.59 | 26881.49 | 28716.76 |
| ENSMUSG00000026587 | 2.85634451 | 8.62E-28 | 2.65E-24 | 405.4399 | 311.9536 | 284.8687 | 452.8773 | 36.07912 | 75.98144 | 47.25017 | 42.41605 |

| | | | | | | | | | | | |
|---|---|---|---|---|---|---|---|---|---|---|---|
| ENSMUSG00000026614 | 1.316142293 | 0.002327 | 0.014615 | 62.37537 | 76.78858 | 55.48314 | 44.85642 | 12.67645 | 46.92971 | 17.3572 | 19.57664 |
| ENSMUSG00000026617 | -1.535396696 | 1.13E-07 | 6.24E-06 | 603.6326 | 505.1249 | 808.2321 | 556.3922 | 1687.918 | 849.2043 | 2122.4 | 2512.335 |
| ENSMUSG00000026649 | -1.337510853 | 9.22E-05 | 0.001253 | 22.1332 | 19.19714 | 19.87456 | 18.11509 | 45.83023 | 30.1691 | 58.82164 | 65.25547 |
| ENSMUSG00000026726 | -3.124019044 | 3.18E-06 | 9.51E-05 | 72.43591 | 51.59233 | 303.9151 | 144.9207 | 1235.466 | 524.0485 | 1069.397 | 2171.919 |
| ENSMUSG00000026764 | 1.027960553 | 6.44E-05 | 0.000965 | 1412.5 | 1129.032 | 946.5259 | 1364.67 | 529.4854 | 947.5332 | 513.0018 | 390.4452 |
| ENSMUSG00000026834 | 2.393758305 | 0.00579 | 0.028665 | 112.6781 | 14.39786 | 8.281066 | 39.68068 | 12.67645 | 7.821619 | 11.57147 | 1.087591 |
| ENSMUSG00000026866 | -1.972272197 | 4.65E-05 | 0.000751 | 23.13925 | 11.99822 | 46.37397 | 24.15346 | 109.2125 | 48.04709 | 84.85745 | 176.1898 |
| ENSMUSG00000026938 | 1.479339163 | 0.00099 | 0.007785 | 118.7144 | 341.9491 | 156.5122 | 171.6621 | 73.13335 | 129.6154 | 45.32159 | 34.80291 |
| ENSMUSG00000026956 | -1.192911422 | 5.06E-05 | 0.000799 | 380.2885 | 279.5584 | 664.9696 | 360.5766 | 1109.677 | 568.7434 | 971.0392 | 1207.226 |
| ENSMUSG00000026971 | -1.661881406 | 0.001615 | 0.011128 | 65.39353 | 43.19358 | 253.4006 | 73.323 | 274.0063 | 186.6015 | 320.144 | 600.3503 |
| ENSMUSG00000026981 | 1.159544463 | 0.009734 | 0.041731 | 55.33299 | 69.58965 | 72.87338 | 67.28463 | 55.58134 | 16.76061 | 33.75012 | 11.9635 |
| ENSMUSG00000026994 | -1.110557889 | 0.000722 | 0.006102 | 264.5923 | 214.7681 | 539.9255 | 265.688 | 574.3406 | 423.4848 | 974.8963 | 804.8174 |
| ENSMUSG00000027070 | -2.241107802 | 7.91E-06 | 0.000192 | 1172.053 | 447.5334 | 2086.001 | 1198.184 | 5810.688 | 2947.633 | 5414.484 | 9015.043 |
| ENSMUSG00000027199 | -3.002615965 | 1.86E-09 | 2.20E-07 | 205.2351 | 153.5772 | 469.5365 | 313.995 | 2908.757 | 1078.266 | 1644.113 | 3531.408 |
| ENSMUSG00000027227 | -1.142881087 | 2.26E-05 | 0.000433 | 1100.623 | 861.4719 | 1184.192 | 1178.344 | 2807.345 | 1234.698 | 2033.686 | 3475.941 |
| ENSMUSG00000027238 | 1.130964371 | 0.001471 | 0.010426 | 386.3249 | 130.7805 | 147.403 | 179.4257 | 101.4116 | 118.4417 | 73.28598 | 92.44524 |
| ENSMUSG00000027249 | 2.723884424 | 0.009283 | 0.040329 | 63.38142 | 4.799286 | 4.140533 | 59.52102 | 1.950223 | 1.117374 | 13.50005 | 3.262773 |
| ENSMUSG00000027270 | -1.42194908 | 1.13E-06 | 4.15E-05 | 27.16347 | 40.79393 | 43.88965 | 40.5433 | 135.5405 | 65.92507 | 108.9647 | 97.8832 |
| ENSMUSG00000027318 | 1.215480718 | 0.004127 | 0.022339 | 271.6347 | 473.9295 | 221.1045 | 339.011 | 122.864 | 270.4045 | 115.7147 | 53.29196 |
| ENSMUSG00000027339 | 1.071096548 | 6.30E-05 | 0.00095 | 1196.199 | 1364.197 | 876.9649 | 1242.178 | 493.4063 | 901.7209 | 482.1446 | 350.2043 |
| ENSMUSG00000027350 | 2.306723415 | 0.006567 | 0.031504 | 1306.865 | 122.3818 | 123.3879 | 1159.366 | 117.9885 | 120.6764 | 105.1075 | 204.4671 |
| ENSMUSG00000027359 | -3.949421069 | 5.19E-07 | 2.17E-05 | 125.7568 | 28.79572 | 423.9906 | 170.7995 | 2991.641 | 1130.783 | 2665.295 | 4798.452 |
| ENSMUSG00000027360 | -2.02097751 | 0.000867 | 0.006982 | 517.1119 | 165.5754 | 1163.49 | 693.5493 | 1994.103 | 1582.202 | 2419.402 | 4315.561 |
| ENSMUSG00000027375 | -1.086662009 | 0.010353 | 0.043725 | 1749.529 | 1481.78 | 3863.946 | 1962.468 | 4602.525 | 2667.172 | 5406.769 | 6562.525 |
| ENSMUSG00000027400 | -2.97412148 | 5.47E-06 | 0.000143 | 2.012109 | 4.799286 | 2.48432 | 6.038364 | 49.73068 | 11.17374 | 32.78583 | 26.10219 |
| ENSMUSG00000027419 | -1.144682399 | 0.000813 | 0.00665 | 37.22401 | 19.19714 | 38.92101 | 34.50494 | 55.58134 | 49.16446 | 88.7146 | 95.70802 |
| ENSMUSG00000027452 | -1.769342405 | 3.73E-06 | 0.000107 | 306.8466 | 166.7752 | 511.7699 | 276.0395 | 1050.195 | 441.3628 | 1324.933 | 1487.825 |
| ENSMUSG00000027489 | 1.130753899 | 0.002727 | 0.016482 | 24.1453 | 39.59411 | 34.78048 | 36.23019 | 12.67645 | 22.34748 | 10.60718 | 16.31387 |
| ENSMUSG00000027500 | 1.933319499 | 0.002689 | 0.0163 | 1583.53 | 1313.805 | 892.6989 | 2166.91 | 228.176 | 948.6506 | 226.608 | 156.6131 |
| ENSMUSG00000027513 | -3.860321624 | 1.10E-13 | 4.07E-11 | 263.5862 | 101.9848 | 246.7758 | 288.9789 | 4061.339 | 1080.501 | 3388.512 | 4568.97 |
| ENSMUSG00000027520 | 1.061992503 | 2.06E-06 | 6.57E-05 | 402.4217 | 365.9456 | 221.1045 | 443.3885 | 176.4951 | 189.9536 | 172.6078 | 146.8248 |
| ENSMUSG00000027562 | -1.194662199 | 7.09E-06 | 0.000177 | 978.8909 | 867.471 | 1546.903 | 1026.522 | 2453.38 | 1488.342 | 2453.152 | 3724.999 |
| ENSMUSG00000027581 | 2.458883764 | 0.00092 | 0.007332 | 206.2411 | 34.79482 | 26.49941 | 314.8576 | 27.30312 | 32.40385 | 38.57157 | 7.613138 |
| ENSMUSG00000027610 | -1.458421354 | 2.67E-07 | 1.26E-05 | 673.0504 | 469.1302 | 751.0927 | 579.683 | 1552.377 | 902.8383 | 1679.792 | 2663.511 |
| ENSMUSG00000027612 | 1.602478155 | 0.012115 | 0.049352 | 22.1332 | 23.99643 | 5.796746 | 24.15346 | 2.925334 | 8.938993 | 7.714313 | 5.437955 |
| ENSMUSG00000027690 | -2.365064103 | 0.000406 | 0.003904 | 101.6115 | 16.7975 | 180.5272 | 135.4319 | 566.5397 | 357.5597 | 413.6801 | 903.7882 |
| ENSMUSG00000027709 | -1.194343334 | 2.84E-06 | 8.64E-05 | 949.7153 | 848.2738 | 958.1194 | 853.9973 | 2163.772 | 1051.449 | 2041.4 | 3005.014 |
| ENSMUSG00000027761 | -2.224328388 | 0.000158 | 0.001895 | 20.12109 | 10.79839 | 100.2009 | 44.85642 | 180.3956 | 102.7984 | 200.5721 | 343.6788 |
| ENSMUSG00000027762 | -5.116213779 | 4.84E-20 | 6.38E-17 | 3.018163 | 3.599465 | 6.624853 | 5.175741 | 209.6489 | 65.92507 | 104.1432 | 270.8102 |
| ENSMUSG00000027833 | 3.654136851 | 0.002727 | 0.016482 | 18.10898 | 5.999108 | 2.48432 | 50.89479 | 0 | 0 | 2.892867 | 3.262773 |
| ENSMUSG00000027840 | 1.06620754 | 0.001677 | 0.011496 | 210.2654 | 266.3604 | 154.8559 | 215.6559 | 77.03379 | 183.2494 | 77.14313 | 67.43065 |
| ENSMUSG00000027870 | -2.333951908 | 0.000626 | 0.005449 | 472.8455 | 56.39161 | 439.7246 | 269.1385 | 1461.692 | 688.3024 | 1488.862 | 2605.868 |
| ENSMUSG00000027895 | 1.074702826 | 0.000287 | 0.003003 | 144.8718 | 103.1847 | 78.67013 | 121.6299 | 44.85512 | 77.09881 | 53.0359 | 38.06569 |
| ENSMUSG00000028023 | -3.374290411 | 1.23E-06 | 4.42E-05 | 5.030272 | 8.398751 | 39.74912 | 6.038364 | 114.088 | 46.92971 | 176.4649 | 282.7737 |
| ENSMUSG00000028024 | -1.193570204 | 0.004381 | 0.023367 | 1554.354 | 795.4817 | 2146.452 | 1437.993 | 2656.203 | 1909.592 | 3600.656 | 5408.59 |
| ENSMUSG00000028137 | 1.243281001 | 0.001275 | 0.009377 | 158.9566 | 64.79036 | 81.15445 | 197.5408 | 45.83023 | 64.8077 | 61.71451 | 40.24087 |
| ENSMUSG00000028172 | 1.942226275 | 1.17E-08 | 9.73E-07 | 232.3986 | 467.9304 | 279.0719 | 372.6533 | 92.63557 | 144.1413 | 50.14304 | 65.25547 |
| ENSMUSG00000028176 | 1.513727494 | 0.002573 | 0.015735 | 65.39353 | 57.59143 | 33.12427 | 66.42201 | 10.72622 | 41.34284 | 16.39292 | 9.78832 |
| ENSMUSG00000028179 | -1.188040851 | 6.01E-07 | 2.44E-05 | 470.8334 | 353.9474 | 578.8465 | 602.9738 | 1079.448 | 727.4105 | 1272.862 | 1496.525 |
| ENSMUSG00000028197 | 1.154947292 | 1.39E-05 | 0.000294 | 614.6992 | 658.702 | 469.5365 | 821.2176 | 309.1103 | 417.8979 | 250.7152 | 174.0146 |
| ENSMUSG00000028199 | -1.003841333 | 2.47E-07 | 1.20E-05 | 921.5458 | 970.6556 | 1212.348 | 1016.17 | 2002.879 | 1378.84 | 2140.722 | 2742.905 |
| ENSMUSG00000028238 | -1.015297224 | 0.009938 | 0.04238 | 185.114 | 131.9804 | 443.037 | 170.7995 | 366.6419 | 275.9914 | 597.8593 | 643.8539 |
| ENSMUSG00000028270 | 1.120092563 | 0.00199 | 0.013086 | 108.6539 | 280.7582 | 188.8083 | 119.042 | 53.63112 | 118.4417 | 80.036 | 68.51824 |
| ENSMUSG00000028307 | -3.063767691 | 3.50E-05 | 0.000604 | 1417.531 | 223.1668 | 2804.797 | 1874.481 | 15074.25 | 5379.039 | 11604.26 | 20792.57 |
| ENSMUSG00000028327 | 1.201172437 | 1.12E-06 | 4.15E-05 | 250.5075 | 134.38 | 177.2148 | 162.1732 | 66.30757 | 96.09417 | 66.53595 | 87.00729 |
| ENSMUSG00000028341 | 2.83029428 | 0.002088 | 0.013543 | 184.1079 | 7.198929 | 7.45296 | 53.48266 | 9.751113 | 5.58687 | 13.50005 | 6.525547 |
| ENSMUSG00000028427 | -1.204203717 | 0.009659 | 0.041508 | 30.18163 | 38.39429 | 46.37397 | 21.56559 | 78.98402 | 31.28647 | 61.71451 | 142.4744 |
| ENSMUSG00000028523 | 1.944708517 | 0.001235 | 0.009155 | 17.10292 | 25.19625 | 12.4216 | 18.11509 | 0.975111 | 6.704245 | 4.821446 | 6.525547 |
| ENSMUSG00000028527 | -1.776205685 | 4.86E-09 | 4.69E-07 | 275.6589 | 285.5575 | 349.461 | 320.8959 | 1029.718 | 562.0392 | 817.7172 | 1810.839 |
| ENSMUSG00000028536 | -2.09854898 | 8.32E-09 | 7.38E-07 | 72.43591 | 74.38894 | 110.1382 | 69.8725 | 337.3885 | 143.0239 | 357.7513 | 563.3722 |
| ENSMUSG00000028544 | -1.652801749 | 4.55E-05 | 0.00074 | 44.26639 | 20.39697 | 63.76421 | 36.23019 | 124.8142 | 78.21619 | 106.0718 | 212.0803 |
| ENSMUSG00000028571 | -4.791852713 | 6.04E-35 | 3.71E-31 | 9.054489 | 5.999108 | 15.73403 | 6.900988 | 290.5832 | 151.9629 | 256.5009 | 365.4306 |
| ENSMUSG00000028644 | 1.458942459 | 0.001383 | 0.009973 | 80.48435 | 25.19625 | 32.29616 | 35.36756 | 16.57689 | 15.64324 | 13.50005 | 17.40146 |
| ENSMUSG00000028661 | -1.572085602 | 0.006508 | 0.03133 | 10.06054 | 10.79839 | 32.29616 | 16.38985 | 40.95467 | 16.76061 | 61.71451 | 89.18247 |

| | | | | | | | | | | | |
|---|---|---|---|---|---|---|---|---|---|---|---|
| ENSMUSG00000028699 | -1.455859072 | 0.000272 | 0.002873 | 47.28455 | 28.79572 | 88.60741 | 31.05445 | 132.6151 | 82.68568 | 141.7505 | 182.7153 |
| ENSMUSG00000028712 | -4.550915923 | 1.20E-08 | 9.91E-07 | 18.10898 | 5.999108 | 62.9361 | 31.05445 | 676.7272 | 198.8926 | 547.7162 | 1358.401 |
| ENSMUSG00000028715 | -2.440454421 | 0.002272 | 0.014388 | 21.12714 | 0 | 5.796746 | 12.93935 | 74.10846 | 20.11273 | 29.89296 | 93.53283 |
| ENSMUSG00000028716 | -1.70893299 | 0.005006 | 0.025762 | 899.4126 | 513.5236 | 2466.102 | 949.7485 | 4156.9 | 1633.601 | 4023.979 | 5974.138 |
| ENSMUSG00000028737 | -1.304137114 | 9.89E-08 | 5.60E-06 | 717.3168 | 569.9152 | 780.0764 | 718.5654 | 1771.777 | 964.2938 | 1669.185 | 2476.445 |
| ENSMUSG00000028755 | -1.115561313 | 1.84E-06 | 6.06E-05 | 109.6599 | 121.182 | 149.8873 | 135.4319 | 318.8614 | 191.071 | 237.2151 | 373.0437 |
| ENSMUSG00000028773 | -2.655731342 | 0.000366 | 0.003599 | 246.4833 | 113.983 | 739.4992 | 213.068 | 1864.413 | 568.7434 | 2356.723 | 3487.905 |
| ENSMUSG00000028776 | -1.208813419 | 0.000159 | 0.001898 | 1255.556 | 686.2979 | 1970.894 | 1146.427 | 2638.651 | 1631.366 | 3873.55 | 3554.248 |
| ENSMUSG00000028845 | 1.257637144 | 4.22E-06 | 0.000118 | 109.6599 | 146.3782 | 139.1219 | 164.7611 | 44.85512 | 84.92043 | 64.60737 | 40.24087 |
| ENSMUSG00000028970 | 2.409277944 | 0.002602 | 0.015881 | 848.1038 | 141.5789 | 66.24853 | 184.6014 | 47.78045 | 93.85942 | 46.28588 | 45.67883 |
| ENSMUSG00000029032 | -1.001127846 | 7.61E-07 | 2.98E-05 | 487.9364 | 460.7315 | 689.8128 | 512.3984 | 988.7629 | 754.2275 | 1227.54 | 1337.737 |
| ENSMUSG00000029053 | -1.123616355 | 4.65E-06 | 0.000126 | 218.3138 | 196.7707 | 313.8524 | 269.1385 | 545.0872 | 329.6254 | 593.0378 | 710.197 |
| ENSMUSG00000029082 | -2.109104783 | 6.70E-07 | 2.67E-05 | 38.23007 | 26.39607 | 71.21717 | 30.19182 | 186.2463 | 73.74669 | 232.3937 | 227.3065 |
| ENSMUSG00000029151 | 2.927377171 | 4.72E-29 | 1.74E-25 | 286.7255 | 202.7698 | 213.6515 | 248.4356 | 27.30312 | 45.81234 | 33.75012 | 18.48905 |
| ENSMUSG00000029154 | -1.985956642 | 6.28E-05 | 0.000949 | 35.2119 | 9.598572 | 47.20208 | 25.8787 | 76.05868 | 60.3382 | 152.3577 | 181.6277 |
| ENSMUSG00000029161 | -2.198293983 | 8.78E-08 | 5.11E-06 | 122.7386 | 70.78947 | 181.3554 | 132.844 | 659.1752 | 220.1227 | 465.7517 | 989.7079 |
| ENSMUSG00000029168 | 1.172004869 | 3.62E-09 | 3.71E-07 | 379.2825 | 321.5522 | 260.8536 | 467.5419 | 165.7689 | 139.6718 | 168.7506 | 159.8759 |
| ENSMUSG00000029188 | -2.867176583 | 3.06E-07 | 1.39E-05 | 2.012109 | 8.398751 | 24.01509 | 10.35148 | 69.2329 | 51.39921 | 91.60747 | 119.635 |
| ENSMUSG00000029193 | -2.869818902 | 0.000204 | 0.002312 | 5.030272 | 1.199822 | 38.92101 | 12.07673 | 72.15824 | 48.04709 | 91.60747 | 212.0803 |
| ENSMUSG00000029206 | 1.236128888 | 3.42E-05 | 0.000593 | 113.6841 | 99.58519 | 96.88848 | 95.75121 | 23.40267 | 65.92507 | 43.39301 | 40.24087 |
| ENSMUSG00000029269 | -1.669342357 | 7.65E-07 | 2.98E-05 | 18.10898 | 14.39786 | 26.49941 | 22.42821 | 57.53157 | 45.81234 | 68.46453 | 90.27006 |
| ENSMUSG00000029272 | 2.505406315 | 4.24E-07 | 1.84E-05 | 66.39959 | 140.3791 | 60.45178 | 123.3552 | 16.57689 | 32.40385 | 14.46434 | 5.437955 |
| ENSMUSG00000029273 | -2.864320882 | 3.21E-06 | 9.56E-05 | 438.6397 | 81.58786 | 647.5794 | 523.6125 | 3008.218 | 1648.127 | 3447.334 | 4218.766 |
| ENSMUSG00000029304 | -2.506112048 | 0.001029 | 0.007993 | 799.8132 | 382.7431 | 3035.011 | 802.2398 | 8444.464 | 2699.576 | 7180.097 | 10196.17 |
| ENSMUSG00000029352 | 1.196450267 | 0.001056 | 0.008143 | 42.25428 | 55.19179 | 52.17072 | 66.42201 | 16.57689 | 39.10809 | 15.42863 | 23.927 |
| ENSMUSG00000029368 | 2.989708023 | 0.01069 | 0.044829 | 2127.805 | 40.79393 | 56.31125 | 1977.133 | 83.85957 | 131.8501 | 143.6791 | 169.6642 |
| ENSMUSG00000029369 | -1.126771517 | 0.000132 | 0.001655 | 135.8173 | 146.3782 | 239.3228 | 163.0358 | 355.9156 | 211.1837 | 381.8585 | 548.1459 |
| ENSMUSG00000029370 | -1.411976013 | 0.000264 | 0.002818 | 143.8658 | 75.58876 | 250.9163 | 160.448 | 325.6872 | 217.8879 | 536.1448 | 602.5255 |
| ENSMUSG00000029373 | 1.154920441 | 1.29E-18 | 1.33E-15 | 541.2572 | 527.9215 | 505.145 | 505.4974 | 210.624 | 292.752 | 221.7865 | 209.9051 |
| ENSMUSG00000029442 | 1.119962521 | 0.000308 | 0.003171 | 236.4228 | 303.5548 | 212.8234 | 266.5507 | 100.4365 | 204.4795 | 91.60747 | 72.8686 |
| ENSMUSG00000029452 | -1.659184745 | 6.38E-05 | 0.000959 | 75.45408 | 38.39429 | 149.8873 | 71.59775 | 239.8774 | 140.7891 | 284.4653 | 398.0583 |
| ENSMUSG00000029468 | 1.059594237 | 4.03E-10 | 5.67E-08 | 168.0111 | 157.1766 | 165.6213 | 184.6014 | 64.35735 | 89.38993 | 90.64318 | 80.48174 |
| ENSMUSG00000029499 | -1.715113853 | 2.60E-07 | 1.24E-05 | 118.7144 | 49.19268 | 116.763 | 125.943 | 317.8863 | 207.8316 | 327.8583 | 499.2043 |
| ENSMUSG00000029553 | -1.924467955 | 3.82E-07 | 1.69E-05 | 20.12109 | 20.39697 | 42.23344 | 28.46658 | 89.71024 | 64.8077 | 106.0718 | 165.3138 |
| ENSMUSG00000029556 | -1.601848662 | 1.71E-06 | 5.78E-05 | 126.7628 | 57.59143 | 93.57605 | 67.28463 | 222.3254 | 167.6061 | 244.9294 | 414.3722 |
| ENSMUSG00000029563 | 1.218192309 | 0.003718 | 0.020731 | 371.2341 | 607.1097 | 373.4761 | 589.1718 | 151.1423 | 433.5411 | 162.9649 | 87.00729 |
| ENSMUSG00000029618 | -1.62801185 | 5.87E-06 | 0.000152 | 36.21796 | 59.99108 | 34.78048 | 46.58167 | 210.624 | 146.376 | 119.5719 | 69.60583 |
| ENSMUSG00000029700 | -3.211759482 | 1.12E-08 | 9.52E-07 | 5.030272 | 5.999108 | 16.56213 | 18.97772 | 172.5947 | 36.87335 | 91.60747 | 137.0365 |
| ENSMUSG00000029819 | 2.446354074 | 2.68E-07 | 1.26E-05 | 822.9525 | 148.7779 | 206.1986 | 669.3958 | 103.3618 | 75.98144 | 87.75031 | 71.78101 |
| ENSMUSG00000029829 | -2.220610276 | 0.000118 | 0.001515 | 161.9748 | 76.78858 | 392.5225 | 136.2945 | 700.1299 | 460.3581 | 1089.647 | 1331.211 |
| ENSMUSG00000030004 | -5.380759626 | 5.21E-24 | 1.07E-20 | 8.048435 | 1.199822 | 8.281066 | 5.175741 | 290.5832 | 87.15518 | 218.8936 | 373.0437 |
| ENSMUSG00000030043 | 1.912931415 | 8.31E-10 | 1.10E-07 | 61.36932 | 67.19001 | 57.13936 | 56.93315 | 14.62667 | 24.58223 | 13.50005 | 11.9635 |
| ENSMUSG00000030077 | 1.195972264 | 0.009451 | 0.040887 | 38.23007 | 20.39697 | 20.70267 | 55.2079 | 11.70134 | 17.87799 | 16.39292 | 13.05109 |
| ENSMUSG00000030088 | -2.06857743 | 0.000797 | 0.006558 | 458.7608 | 91.18644 | 407.4285 | 269.1385 | 1173.059 | 442.4801 | 1617.113 | 1914.16 |
| ENSMUSG00000030108 | -2.570414224 | 7.44E-06 | 0.000184 | 131.7931 | 53.99197 | 219.4483 | 110.4158 | 692.329 | 291.6346 | 738.6455 | 1344.263 |
| ENSMUSG00000030109 | -3.373332539 | 1.37E-12 | 3.88E-10 | 25.15136 | 4.799286 | 33.95237 | 8.626235 | 210.624 | 126.2633 | 198.6436 | 225.1314 |
| ENSMUSG00000030116 | 1.167913664 | 1.31E-06 | 4.64E-05 | 1191.168 | 1762.538 | 1416.89 | 1649.336 | 723.5326 | 992.2282 | 508.1804 | 455.7007 |
| ENSMUSG00000030160 | -2.1323115 | 6.02E-13 | 1.91E-10 | 516.1059 | 277.1588 | 593.7525 | 428.7239 | 1553.352 | 1168.773 | 2448.33 | 2796.197 |
| ENSMUSG00000030209 | 1.75399373 | 5.63E-05 | 0.000866 | 45.27245 | 46.79304 | 48.03018 | 40.5433 | 10.72622 | 26.81698 | 6.750024 | 9.78832 |
| ENSMUSG00000030223 | -1.445003849 | 0.000819 | 0.006695 | 120.7265 | 62.39072 | 266.6503 | 147.5086 | 266.2054 | 233.5312 | 509.1447 | 621.0145 |
| ENSMUSG00000030226 | 1.508075603 | 0.00045 | 0.004225 | 44.26639 | 46.79304 | 23.18699 | 52.62003 | 9.751113 | 23.46486 | 11.57147 | 14.13868 |
| ENSMUSG00000030237 | 1.192736343 | 0.001313 | 0.009599 | 528.1785 | 551.9179 | 401.6317 | 527.9256 | 163.8187 | 446.9496 | 150.4291 | 118.5474 |
| ENSMUSG00000030306 | 1.138671599 | 1.61E-05 | 0.000331 | 2258.592 | 1940.111 | 1348.158 | 2245.409 | 723.5326 | 1339.732 | 925.7176 | 550.3211 |
| ENSMUSG00000030329 | 2.045833463 | 1.28E-10 | 2.26E-08 | 215.2956 | 249.5629 | 240.1509 | 210.4801 | 71.18313 | 84.92043 | 38.57157 | 27.18978 |
| ENSMUSG00000030340 | -2.223041826 | 0.001066 | 0.008203 | 305.8405 | 75.58876 | 494.3797 | 169.9368 | 879.5504 | 449.1844 | 1942.078 | 1613.985 |
| ENSMUSG00000030364 | -2.585756858 | 0.00072 | 0.006093 | 8.048435 | 34.79482 | 8.281066 | 0.862623 | 115.0631 | 32.40385 | 55.92877 | 103.3212 |
| ENSMUSG00000030468 | -4.124328764 | 5.32E-09 | 5.06E-07 | 5.030272 | 5.999108 | 7.45296 | 12.93935 | 250.6036 | 12.29112 | 138.8576 | 150.0876 |
| ENSMUSG00000030492 | -2.223035781 | 8.86E-08 | 5.13E-06 | 64.38748 | 25.19625 | 101.029 | 70.73513 | 334.4632 | 159.7845 | 254.5723 | 477.4525 |
| ENSMUSG00000030543 | 1.103111331 | 0.00992 | 0.042337 | 25.15136 | 16.7975 | 29.81184 | 31.91707 | 11.70134 | 17.87799 | 11.57147 | 7.613138 |
| ENSMUSG00000030584 | 1.321155163 | 0.001207 | 0.009002 | 64.38748 | 52.79215 | 31.46805 | 86.26235 | 28.27823 | 23.46486 | 28.92867 | 13.05109 |
| ENSMUSG00000030621 | -1.160659653 | 1.13E-05 | 0.000252 | 316.9071 | 166.7752 | 339.5237 | 263.9628 | 494.3814 | 427.9543 | 714.5383 | 797.2043 |
| ENSMUSG00000030630 | -1.652129095 | 1.83E-07 | 9.33E-06 | 294.7739 | 183.5727 | 381.7572 | 318.3081 | 1010.215 | 435.7759 | 946.932 | 1314.898 |
| ENSMUSG00000030650 | -2.493651911 | 0.00014 | 0.001735 | 4.024217 | 2.399643 | 19.87456 | 5.175741 | 36.07912 | 32.40385 | 33.75012 | 79.39415 |
| ENSMUSG00000030674 | -1.973134879 | 1.10E-07 | 6.14E-06 | 129.781 | 86.38715 | 217.792 | 139.745 | 448.5512 | 306.1605 | 504.3232 | 998.4086 |

| | | | | | | | | | | | |
|---|---|---|---|---|---|---|---|---|---|---|---|
| ENSMUSG00000030699 | 1.342301035 | 0.000175 | 0.002044 | 40.24217 | 31.19536 | 43.06154 | 55.2079 | 11.70134 | 22.34748 | 19.28578 | 14.13868 |
| ENSMUSG00000030730 | 1.272268705 | 7.68E-06 | 0.000188 | 133.8052 | 182.3729 | 133.3252 | 150.9591 | 58.50668 | 99.44629 | 45.32159 | 45.67883 |
| ENSMUSG00000030747 | -1.696218784 | 9.33E-10 | 1.21E-07 | 240.447 | 197.9706 | 332.8989 | 221.6942 | 913.6793 | 426.8369 | 789.7528 | 1090.854 |
| ENSMUSG00000030769 | -3.234341524 | 4.63E-06 | 0.000126 | 2.012109 | 7.198929 | 19.04645 | 9.488858 | 134.5654 | 24.58223 | 42.42872 | 157.7007 |
| ENSMUSG00000030781 | -2.882209946 | 1.55E-06 | 5.33E-05 | 21.12714 | 7.198929 | 51.34261 | 28.46658 | 333.4881 | 62.57295 | 114.7504 | 292.562 |
| ENSMUSG00000030787 | 1.152302911 | 1.82E-05 | 0.000365 | 1072.454 | 1201.021 | 799.1229 | 1144.701 | 365.6667 | 708.4152 | 544.8234 | 278.4233 |
| ENSMUSG00000030800 | -1.341099451 | 4.64E-06 | 0.000126 | 798.8072 | 622.7074 | 1195.786 | 806.553 | 1936.571 | 1143.074 | 2355.758 | 3242.109 |
| ENSMUSG00000030838 | -1.578939328 | 4.49E-06 | 0.000124 | 210.2654 | 125.9813 | 308.8838 | 237.2215 | 607.4943 | 348.6207 | 607.5022 | 1076.715 |
| ENSMUSG00000030854 | 1.812784793 | 2.58E-18 | 2.38E-15 | 226.3622 | 194.3711 | 183.8397 | 207.8923 | 54.60623 | 74.86406 | 62.6788 | 39.15328 |
| ENSMUSG00000030878 | -1.080698237 | 1.78E-06 | 5.95E-05 | 352.119 | 314.3532 | 429.7873 | 423.5481 | 677.7024 | 525.1658 | 1064.575 | 949.467 |
| ENSMUSG00000030895 | 2.315981253 | 0.004701 | 0.024642 | 144.8718 | 5.999108 | 18.21835 | 60.38364 | 7.80089 | 13.40849 | 18.32149 | 6.525547 |
| ENSMUSG00000030909 | -3.004471646 | 3.56E-08 | 2.42E-06 | 33.19979 | 8.398751 | 66.24853 | 42.26855 | 216.4747 | 125.1459 | 314.3583 | 556.8466 |
| ENSMUSG00000030945 | -3.845412629 | 8.11E-08 | 4.80E-06 | 140.8476 | 23.99643 | 279.0719 | 137.1571 | 2227.154 | 824.6221 | 2090.579 | 3219.27 |
| ENSMUSG00000030963 | -5.57917345 | 4.13E-05 | 0.000691 | 12.07265 | 1.199822 | 305.5713 | 37.95543 | 3635.215 | 665.955 | 5751.985 | 7019.313 |
| ENSMUSG00000030972 | -3.466329589 | 1.63E-07 | 8.43E-06 | 7.04238 | 3.599465 | 30.63995 | 11.21411 | 153.0925 | 39.10809 | 122.4647 | 274.073 |
| ENSMUSG00000031027 | 1.438961248 | 0.000331 | 0.003341 | 25.15136 | 37.19447 | 28.15563 | 38.81806 | 11.70134 | 17.87799 | 10.60718 | 7.613138 |
| ENSMUSG00000031111 | 1.685857034 | 7.00E-16 | 3.92E-13 | 237.4288 | 233.9652 | 259.1974 | 274.3143 | 58.50668 | 110.62 | 79.07171 | 65.25547 |
| ENSMUSG00000031139 | 1.223789097 | 8.24E-06 | 0.000197 | 158.9566 | 194.3711 | 87.7793 | 189.7772 | 72.15824 | 63.69032 | 70.39311 | 63.08028 |
| ENSMUSG00000031283 | 1.217483551 | 1.25E-05 | 0.000274 | 151.9142 | 253.1623 | 155.684 | 227.7326 | 82.88446 | 119.559 | 81.00029 | 55.46715 |
| ENSMUSG00000031285 | 1.152461149 | 0.000333 | 0.003355 | 831.0009 | 1030.647 | 607.8303 | 953.199 | 365.6667 | 663.7202 | 311.4654 | 199.0292 |
| ENSMUSG00000031294 | -2.812757041 | 0.000112 | 0.001463 | 7.04238 | 1.199822 | 7.45296 | 3.450494 | 52.65601 | 7.821619 | 25.07152 | 51.11678 |
| ENSMUSG00000031302 | 1.719464556 | 5.07E-06 | 0.000136 | 200.2048 | 181.1731 | 231.0418 | 219.1064 | 42.9049 | 122.9112 | 56.89306 | 30.45255 |
| ENSMUSG00000031340 | 1.11548247 | 1.94E-10 | 3.17E-08 | 388.337 | 376.744 | 325.4459 | 412.334 | 193.072 | 211.1837 | 162.0006 | 127.2482 |
| ENSMUSG00000031383 | -1.040405934 | 0.001666 | 0.011446 | 324.9556 | 239.9643 | 496.864 | 305.3687 | 544.1121 | 408.9589 | 658.6095 | 1202.876 |
| ENSMUSG00000031384 | -2.483460339 | 2.36E-06 | 7.36E-05 | 32.19374 | 10.79839 | 86.12309 | 40.5433 | 216.4747 | 115.0895 | 234.3223 | 389.3576 |
| ENSMUSG00000031409 | 2.208246456 | 2.17E-09 | 2.47E-07 | 68.4117 | 62.39072 | 46.37397 | 77.63611 | 8.776002 | 22.34748 | 13.50005 | 10.87591 |
| ENSMUSG00000031444 | -2.530521442 | 3.11E-05 | 0.00055 | 23.13925 | 8.398751 | 18.21835 | 23.29083 | 136.5156 | 26.81698 | 52.07161 | 208.8175 |
| ENSMUSG00000031465 | -1.436218068 | 5.31E-05 | 0.000829 | 149.9021 | 92.38626 | 216.1358 | 188.9145 | 441.7254 | 213.4185 | 403.0729 | 697.1459 |
| ENSMUSG00000031574 | 3.859224174 | 0.003395 | 0.019358 | 14594.83 | 223.1668 | 192.9488 | 1741.637 | 157.968 | 214.5358 | 170.6792 | 611.2262 |
| ENSMUSG00000031604 | -1.065594256 | 1.77E-08 | 1.37E-06 | 735.4257 | 435.5352 | 726.2495 | 518.4367 | 1194.511 | 1105.083 | 1525.505 | 1234.416 |
| ENSMUSG00000031698 | -1.378210607 | 0.000283 | 0.002972 | 38.23007 | 14.39786 | 25.67131 | 28.46658 | 65.33246 | 50.28183 | 55.92877 | 107.6715 |
| ENSMUSG00000031725 | -3.785898288 | 0.000214 | 0.002389 | 0 | 2.399643 | 5.796746 | 0 | 23.40267 | 5.58687 | 34.71441 | 51.11678 |
| ENSMUSG00000031734 | -1.083319547 | 4.88E-05 | 0.000782 | 395.3794 | 220.7672 | 567.253 | 377.8291 | 720.6073 | 631.3164 | 961.3963 | 999.4962 |
| ENSMUSG00000031760 | 1.286792324 | 8.55E-06 | 0.000203 | 79.47829 | 81.58786 | 94.40416 | 79.36136 | 30.22845 | 54.75133 | 27.96439 | 25.0146 |
| ENSMUSG00000031766 | -2.81066857 | 3.23E-06 | 9.59E-05 | 160.9687 | 73.18911 | 341.1799 | 161.3106 | 992.6633 | 454.7713 | 1540.934 | 2184.97 |
| ENSMUSG00000031842 | -1.512936228 | 0.000187 | 0.00215 | 155.9384 | 62.39072 | 237.6666 | 168.2116 | 379.3183 | 214.5358 | 549.6448 | 641.6787 |
| ENSMUSG00000031844 | -2.958488405 | 0.000157 | 0.001887 | 26.15741 | 5.999108 | 73.70149 | 33.64232 | 320.8116 | 75.98144 | 243.9652 | 448.0875 |
| ENSMUSG00000031877 | 1.594122024 | 0.000389 | 0.003779 | 44.26639 | 91.18644 | 57.13936 | 56.93315 | 21.45245 | 37.99072 | 13.50005 | 9.78832 |
| ENSMUSG00000031906 | 1.820560737 | 0.000583 | 0.005151 | 273.6468 | 380.3434 | 192.1207 | 345.0494 | 56.55646 | 202.2447 | 55.92877 | 22.83941 |
| ENSMUSG00000031938 | -1.227519834 | 2.63E-07 | 1.25E-05 | 1416.525 | 1650.954 | 1701.759 | 1658.825 | 4463.084 | 2167.706 | 3109.833 | 5311.795 |
| ENSMUSG00000031952 | 1.882127809 | 7.22E-16 | 3.92E-13 | 116.7023 | 127.1811 | 108.482 | 106.9653 | 37.05423 | 20.11273 | 31.82154 | 34.80291 |
| ENSMUSG00000031965 | 3.442690713 | 0.002514 | 0.015492 | 36.21796 | 4.799286 | 0.828107 | 58.6584 | 3.900445 | 1.117374 | 0.964289 | 3.262773 |
| ENSMUSG00000031980 | -2.008803888 | 3.09E-05 | 0.000548 | 73.44197 | 55.19179 | 60.45178 | 33.64232 | 173.5698 | 60.3382 | 269.0367 | 392.6204 |
| ENSMUSG00000032010 | -1.068793517 | 0.000126 | 0.001592 | 839.0493 | 453.5325 | 714.656 | 756.5208 | 895.1522 | 1415.713 | 2265.115 | 1223.54 |
| ENSMUSG00000032036 | 1.023851885 | 6.79E-06 | 0.000172 | 128.775 | 169.1748 | 153.1997 | 146.646 | 69.2329 | 101.681 | 52.07161 | 71.78101 |
| ENSMUSG00000032060 | -1.745679108 | 1.18E-06 | 4.32E-05 | 255.5378 | 296.3559 | 667.4539 | 279.49 | 1102.851 | 644.7249 | 1530.327 | 1752.109 |
| ENSMUSG00000032083 | 2.246141729 | 3.76E-06 | 0.000108 | 519.124 | 239.9643 | 101.8571 | 498.5964 | 72.15824 | 118.4417 | 55.92877 | 40.24087 |
| ENSMUSG00000032098 | -3.424677111 | 0.000654 | 0.005641 | 0 | 0 | 2.48432 | 4.313117 | 13.65156 | 4.469496 | 24.10723 | 34.80291 |
| ENSMUSG00000032105 | -1.937798134 | 4.53E-06 | 0.000124 | 106.6418 | 39.59411 | 122.5598 | 50.89479 | 360.7912 | 147.4934 | 259.3938 | 460.051 |
| ENSMUSG00000032114 | -1.702396406 | 1.52E-12 | 4.24E-10 | 579.4873 | 373.1445 | 592.9243 | 598.6607 | 1766.902 | 1050.332 | 1815.756 | 2350.284 |
| ENSMUSG00000032118 | 2.189543507 | 9.94E-16 | 5.24E-13 | 394.3733 | 449.9331 | 327.9302 | 421.8229 | 107.2622 | 124.0285 | 71.3574 | 46.76642 |
| ENSMUSG00000032246 | -1.718115738 | 2.99E-08 | 2.13E-06 | 335.0161 | 255.562 | 593.7525 | 347.6373 | 1184.76 | 758.697 | 1154.254 | 1947.876 |
| ENSMUSG00000032269 | 1.575700292 | 1.63E-08 | 1.28E-06 | 82.49646 | 104.3845 | 75.3577 | 95.75121 | 30.22845 | 41.34284 | 29.89296 | 18.48905 |
| ENSMUSG00000032278 | -1.795099026 | 0.00273 | 0.01649 | 371.2341 | 178.7734 | 939.0729 | 298.4677 | 1328.102 | 706.1804 | 2034.65 | 2137.116 |
| ENSMUSG00000032303 | 1.961205261 | 0.008625 | 0.038338 | 64.38748 | 7.198929 | 9.93728 | 67.28463 | 10.72622 | 11.17374 | 7.714313 | 8.700729 |
| ENSMUSG00000032346 | 1.270025632 | 4.99E-07 | 2.11E-05 | 159.9626 | 105.5843 | 151.5435 | 169.9368 | 52.65601 | 75.98144 | 43.39301 | 72.8686 |
| ENSMUSG00000032357 | -2.565311935 | 1.22E-06 | 4.42E-05 | 43.26034 | 17.99732 | 127.5284 | 53.48266 | 368.5921 | 147.4934 | 330.7512 | 593.8247 |
| ENSMUSG00000032373 | -1.268192431 | 0.011569 | 0.04764 | 744.4802 | 605.9099 | 1918.723 | 695.2745 | 2326.616 | 1169.891 | 2513.902 | 3540.109 |
| ENSMUSG00000032454 | -1.69926738 | 0.000647 | 0.005588 | 18.10898 | 37.19447 | 17.39024 | 6.900988 | 72.15824 | 32.40385 | 67.50024 | 82.65692 |
| ENSMUSG00000032492 | -1.350933067 | 5.46E-06 | 0.000143 | 969.8364 | 901.066 | 1609.839 | 1023.071 | 2550.891 | 1620.192 | 2621.902 | 4698.393 |
| ENSMUSG00000032532 | -2.307072597 | 0.009987 | 0.042531 | 2.012109 | 4.799286 | 47.20208 | 2.58787 | 65.33246 | 24.58223 | 82.92887 | 109.8467 |
| ENSMUSG00000032554 | 1.033201036 | 0.000786 | 0.006496 | 458.7608 | 190.7716 | 195.4332 | 395.0816 | 127.7396 | 179.8972 | 146.572 | 152.2628 |
| ENSMUSG00000032690 | 1.608992814 | 0.006226 | 0.030285 | 66.39959 | 291.5566 | 231.0418 | 36.23019 | 39.00445 | 36.87335 | 81.96458 | 46.76642 |
| ENSMUSG00000032758 | -7.642415729 | 1.46E-07 | 7.71E-06 | 1.006054 | 0 | 50.5145 | 6.900988 | 3269.548 | 612.321 | 1371.219 | 6473.342 |

| | | | | | | | | | | | |
|---|---|---|---|---|---|---|---|---|---|---|---|
| ENSMUSG00000032854 | -1.023684675 | 0.009654 | 0.041504 | 117.7084 | 125.9813 | 333.727 | 135.4319 | 330.5627 | 185.4841 | 415.6086 | 519.8685 |
| ENSMUSG00000032860 | -1.822529546 | 1.43E-05 | 0.000299 | 39.23612 | 20.39697 | 54.65504 | 31.05445 | 168.6943 | 51.39921 | 147.5362 | 149 |
| ENSMUSG00000032925 | 1.566782291 | 0.006024 | 0.02956 | 46.2785 | 32.39518 | 13.24971 | 41.40593 | 4.875557 | 20.11273 | 12.53576 | 7.613138 |
| ENSMUSG00000032978 | -3.469699302 | 2.58E-11 | 5.41E-09 | 11.0666 | 14.39786 | 14.90592 | 5.175741 | 154.0676 | 41.34284 | 104.1432 | 202.2919 |
| ENSMUSG00000033061 | 3.514084796 | 0.000644 | 0.005577 | 62.37537 | 4.799286 | 2.48432 | 104.3774 | 3.900445 | 2.234748 | 5.785735 | 3.262773 |
| ENSMUSG00000033161 | -1.607511684 | 0.000401 | 0.003866 | 21725.74 | 12473.34 | 34326.68 | 19340.02 | 54802.23 | 27755.57 | 90391.5 | 94803.14 |
| ENSMUSG00000033427 | -1.83422307 | 4.57E-08 | 3.01E-06 | 62.37537 | 55.19179 | 70.38906 | 45.71905 | 239.8774 | 96.09417 | 200.5721 | 296.9124 |
| ENSMUSG00000033533 | -5.070626963 | 8.28E-09 | 7.38E-07 | 14.08476 | 2.399643 | 46.37397 | 18.97772 | 780.089 | 175.4277 | 488.8946 | 1322.511 |
| ENSMUSG00000033578 | 1.252539096 | 0.000131 | 0.001649 | 1740.474 | 950.2587 | 689.8128 | 935.9465 | 331.5378 | 747.5233 | 339.4298 | 393.708 |
| ENSMUSG00000033579 | -1.100702493 | 0.006725 | 0.032041 | 36.21796 | 40.79393 | 73.70149 | 54.34528 | 132.6151 | 42.46022 | 113.7861 | 152.2628 |
| ENSMUSG00000033633 | -1.361540016 | 6.59E-07 | 2.64E-05 | 853.1341 | 331.1507 | 829.7628 | 704.7634 | 1537.751 | 1343.084 | 1927.614 | 2182.795 |
| ENSMUSG00000033634 | -2.630723911 | 2.73E-08 | 1.97E-06 | 4.024217 | 13.19804 | 13.24971 | 12.07673 | 77.03379 | 32.40385 | 58.82164 | 95.70802 |
| ENSMUSG00000033715 | 1.229568433 | 0.009018 | 0.039559 | 167.005 | 247.1632 | 153.1997 | 212.2054 | 60.4569 | 177.6625 | 68.46453 | 26.10219 |
| ENSMUSG00000033726 | -1.72827484 | 0.000153 | 0.00186 | 144.8718 | 45.59322 | 184.6678 | 135.4319 | 342.2641 | 159.7845 | 545.7877 | 647.1167 |
| ENSMUSG00000033737 | 1.723451984 | 9.90E-20 | 1.22E-16 | 212.2775 | 243.5638 | 178.871 | 216.5185 | 68.25779 | 69.27719 | 71.3574 | 47.85401 |
| ENSMUSG00000033770 | -2.40623725 | 2.34E-05 | 0.000445 | 262.5802 | 83.98751 | 345.3205 | 155.2722 | 936.1069 | 438.0106 | 1518.755 | 1600.934 |
| ENSMUSG00000033860 | 3.576408778 | 0.008645 | 0.03838 | 618.7234 | 2.399643 | 7.45296 | 315.7202 | 23.40267 | 12.29112 | 17.3572 | 26.10219 |
| ENSMUSG00000033882 | 2.082576482 | 2.43E-07 | 1.19E-05 | 131.7931 | 211.1686 | 110.1382 | 178.5631 | 22.42756 | 65.92507 | 42.42872 | 18.48905 |
| ENSMUSG00000034159 | -2.33314632 | 2.33E-05 | 0.000442 | 11.0666 | 5.999108 | 20.70267 | 3.450494 | 67.28268 | 29.05173 | 46.28588 | 67.43065 |
| ENSMUSG00000034258 | -1.103023951 | 5.73E-05 | 0.000878 | 108.6539 | 95.98572 | 173.0743 | 107.8279 | 283.7574 | 159.7845 | 268.0724 | 333.8905 |
| ENSMUSG00000034336 | 1.61884036 | 0.010284 | 0.043496 | 163.9869 | 31.19536 | 24.8432 | 215.6559 | 41.92979 | 37.99072 | 33.75012 | 28.27737 |
| ENSMUSG00000034427 | -1.768137699 | 0.000179 | 0.002078 | 122.7386 | 33.595 | 188.8083 | 90.57547 | 328.6125 | 179.8972 | 361.6084 | 617.7517 |
| ENSMUSG00000034634 | 2.147491294 | 0.00219 | 0.014014 | 767.6195 | 63.59054 | 261.6817 | 345.0494 | 87.76002 | 121.7938 | 52.07161 | 63.08028 |
| ENSMUSG00000034739 | -2.724208979 | 6.01E-07 | 2.44E-05 | 18.10898 | 8.398751 | 34.78048 | 7.763611 | 105.312 | 44.69496 | 159.1077 | 151.1752 |
| ENSMUSG00000034762 | -1.681436616 | 0.008416 | 0.037657 | 16.09687 | 8.398751 | 54.65504 | 6.900988 | 39.00445 | 56.98608 | 79.07171 | 103.3212 |
| ENSMUSG00000034795 | 1.185360851 | 0.002121 | 0.013698 | 132.7992 | 206.3693 | 101.8571 | 135.4319 | 46.80534 | 118.4417 | 49.17875 | 39.15328 |
| ENSMUSG00000034825 | -1.312591051 | 0.011224 | 0.046557 | 27.16347 | 9.598572 | 52.17072 | 25.01608 | 76.05868 | 33.52122 | 62.6788 | 113.1095 |
| ENSMUSG00000034918 | -2.985566145 | 5.46E-10 | 7.47E-08 | 33.19979 | 32.39518 | 98.54469 | 50.03216 | 450.5014 | 137.437 | 392.4657 | 723.2481 |
| ENSMUSG00000034926 | -1.254050183 | 5.67E-09 | 5.30E-07 | 961.788 | 777.4844 | 1316.69 | 965.2757 | 2571.369 | 1546.446 | 2556.331 | 2921.27 |
| ENSMUSG00000034947 | -1.161263911 | 0.001949 | 0.012893 | 70.4238 | 87.58697 | 120.0755 | 89.71284 | 257.4294 | 87.15518 | 159.1077 | 319.7518 |
| ENSMUSG00000034958 | 1.870440041 | 2.92E-06 | 8.84E-05 | 59.35721 | 37.19447 | 49.6864 | 94.88858 | 13.65156 | 13.40849 | 16.39292 | 22.83941 |
| ENSMUSG00000034959 | 1.098508955 | 0.006752 | 0.032129 | 71.42986 | 31.19536 | 38.09291 | 57.79577 | 19.50223 | 17.87799 | 36.64299 | 18.48905 |
| ENSMUSG00000035000 | -1.226885286 | 8.12E-07 | 3.13E-05 | 1248.513 | 1024.648 | 1845.85 | 1406.939 | 2711.785 | 2226.927 | 3158.047 | 4841.956 |
| ENSMUSG00000035095 | -2.2891324 | 3.27E-18 | 2.87E-15 | 33.19979 | 32.39518 | 38.09291 | 40.5433 | 124.8142 | 147.4934 | 207.3222 | 228.3941 |
| ENSMUSG00000035104 | -1.062821572 | 7.09E-05 | 0.001042 | 442.6639 | 377.9438 | 585.4714 | 442.5259 | 810.3175 | 534.1048 | 1239.112 | 1280.095 |
| ENSMUSG00000035165 | -1.204409857 | 0.005151 | 0.026292 | 14.08476 | 11.99822 | 17.39024 | 16.38985 | 21.45245 | 55.8687 | 24.10723 | 38.06569 |
| ENSMUSG00000035246 | 1.07431834 | 6.32E-10 | 8.52E-08 | 133.8052 | 158.3764 | 139.1219 | 137.1571 | 59.48179 | 74.86406 | 74.25027 | 60.9051 |
| ENSMUSG00000035296 | 1.307897979 | 0.003299 | 0.019041 | 188.1322 | 201.57 | 102.6852 | 180.2883 | 64.35735 | 129.6154 | 54.96448 | 22.83941 |
| ENSMUSG00000035349 | -1.761121671 | 0.008213 | 0.036999 | 17.10292 | 7.198929 | 56.31125 | 17.25247 | 89.71024 | 20.11273 | 82.92887 | 141.3868 |
| ENSMUSG00000035637 | -1.04408482 | 0.000331 | 0.003341 | 920.5397 | 494.3265 | 658.3448 | 552.9417 | 1383.683 | 739.7017 | 1247.79 | 2044.671 |
| ENSMUSG00000035699 | -2.966333336 | 5.18E-05 | 0.000814 | 5.030272 | 8.398751 | 19.04645 | 5.175741 | 117.0134 | 14.52586 | 30.85725 | 133.7737 |
| ENSMUSG00000035769 | -1.482700315 | 9.13E-07 | 3.46E-05 | 356.1432 | 245.9634 | 507.6294 | 339.011 | 997.5389 | 537.4569 | 1004.789 | 1512.839 |
| ENSMUSG00000035828 | -1.25291117 | 2.82E-05 | 0.000514 | 1506.063 | 773.8849 | 1449.187 | 1757.164 | 2314.914 | 2458.223 | 5331.555 | 2973.474 |
| ENSMUSG00000035852 | -1.340835531 | 9.25E-07 | 3.49E-05 | 167.005 | 125.9813 | 169.7619 | 156.1349 | 385.169 | 206.7142 | 429.1087 | 548.1459 |
| ENSMUSG00000035878 | -1.465100017 | 1.33E-07 | 7.16E-06 | 134.8113 | 143.9786 | 173.9024 | 169.0742 | 395.8952 | 234.6486 | 441.6444 | 646.0291 |
| ENSMUSG00000036083 | -4.009541891 | 7.91E-12 | 1.87E-09 | 13.07871 | 4.799286 | 34.78048 | 15.52722 | 338.3636 | 89.38993 | 177.4292 | 506.8174 |
| ENSMUSG00000036110 | -2.184053654 | 0.003095 | 0.018205 | 3.018163 | 2.399643 | 19.04645 | 2.58787 | 28.27823 | 16.76061 | 32.78583 | 47.85401 |
| ENSMUSG00000036123 | -3.338832408 | 2.16E-06 | 6.84E-05 | 15.09082 | 3.599465 | 50.5145 | 11.21411 | 190.1467 | 41.34284 | 290.251 | 299.0875 |
| ENSMUSG00000036144 | -1.417695559 | 0.000123 | 0.001566 | 91.55095 | 160.7761 | 71.21717 | 99.2017 | 435.8748 | 136.3196 | 326.894 | 226.2189 |
| ENSMUSG00000036169 | -1.883796051 | 0.007398 | 0.034209 | 637.8385 | 230.3657 | 2045.423 | 588.3092 | 2484.584 | 1594.493 | 3625.727 | 5221.525 |
| ENSMUSG00000036198 | 1.404131056 | 0.00025 | 0.002707 | 47.28455 | 32.39518 | 49.6864 | 64.69676 | 9.751113 | 24.58223 | 22.17865 | 17.40146 |
| ENSMUSG00000036278 | -1.097581726 | 1.52E-05 | 0.000315 | 323.9495 | 241.1641 | 308.0557 | 273.4516 | 607.4943 | 347.5033 | 612.3236 | 887.4743 |
| ENSMUSG00000036304 | -1.886985546 | 0.000528 | 0.00478 | 12.07265 | 7.198929 | 14.90592 | 8.626235 | 44.85512 | 13.40849 | 33.75012 | 67.43065 |
| ENSMUSG00000036330 | -2.827499857 | 3.07E-05 | 0.000546 | 81.4904 | 16.7975 | 39.74912 | 97.47645 | 556.7886 | 130.7328 | 336.5369 | 650.3795 |
| ENSMUSG00000036422 | -1.05331289 | 0.000601 | 0.005272 | 18.10898 | 38.39429 | 20.70267 | 28.46658 | 59.48179 | 56.98608 | 46.28588 | 54.37955 |
| ENSMUSG00000036437 | 1.202671705 | 0.010943 | 0.04565 | 41.24823 | 29.99554 | 21.53077 | 56.93315 | 10.72622 | 22.34748 | 9.642892 | 22.83941 |
| ENSMUSG00000036594 | -1.370541835 | 6.98E-05 | 0.001029 | 31.18769 | 25.19625 | 50.5145 | 35.36756 | 127.7396 | 52.51658 | 98.35749 | 91.35765 |
| ENSMUSG00000036699 | 1.024779127 | 1.04E-09 | 1.34E-07 | 204.229 | 169.1748 | 197.0894 | 246.7103 | 104.3369 | 99.44629 | 105.1075 | 93.53283 |
| ENSMUSG00000036813 | -2.09049446 | 1.61E-06 | 5.49E-05 | 20.12109 | 16.7975 | 37.2648 | 27.60395 | 80.93424 | 52.51658 | 118.6076 | 184.8905 |
| ENSMUSG00000036814 | -2.483347701 | 6.82E-05 | 0.001009 | 4.024217 | 0 | 10.76539 | 6.900988 | 33.15378 | 18.99536 | 34.71441 | 39.15328 |
| ENSMUSG00000036815 | 1.107225031 | 0.000186 | 0.002142 | 173.0413 | 226.7663 | 192.9488 | 256.1992 | 72.15824 | 166.4887 | 79.07171 | 77.21897 |
| ENSMUSG00000036832 | -1.855016188 | 0.000598 | 0.005253 | 19.11503 | 20.39697 | 9.109173 | 18.97772 | 86.78491 | 16.76061 | 48.21446 | 91.35765 |
| ENSMUSG00000036892 | -1.979390226 | 0.004995 | 0.025736 | 114.6902 | 14.39786 | 170.59 | 79.36136 | 390.0445 | 176.5451 | 336.5369 | 593.8247 |
| ENSMUSG00000036899 | -2.660520712 | 3.71E-09 | 3.78E-07 | 92.557 | 50.3925 | 71.21717 | 37.95543 | 332.513 | 131.8501 | 409.8229 | 719.9853 |

| | | | | | | | | | | | |
|---|---|---|---|---|---|---|---|---|---|---|---|
| ENSMUSG00000036923 | 1.343355645 | 5.50E-08 | 3.55E-06 | 128.775 | 169.1748 | 105.9976 | 125.0804 | 70.20801 | 49.16446 | 42.42872 | 45.67883 |
| ENSMUSG00000037005 | -1.802565049 | 5.89E-05 | 0.000901 | 65.39353 | 44.3934 | 141.6062 | 66.42201 | 286.6827 | 113.9722 | 236.2508 | 475.2773 |
| ENSMUSG00000037161 | 1.612271491 | 0.006719 | 0.032025 | 1180.102 | 215.9679 | 229.3855 | 456.3278 | 160.8934 | 144.1413 | 147.5362 | 228.3941 |
| ENSMUSG00000037335 | 3.82387937 | 0.003985 | 0.021805 | 83.50251 | 9.598572 | 0.828107 | 91.43809 | 0 | 0 | 8.678602 | 4.350364 |
| ENSMUSG00000037366 | -1.114466517 | 9.91E-05 | 0.00133 | 347.0888 | 277.1588 | 510.1137 | 397.6694 | 812.2677 | 439.128 | 884.2532 | 1184.387 |
| ENSMUSG00000037443 | -1.033932572 | 0.000409 | 0.003928 | 774.6619 | 735.4906 | 660.8291 | 688.3735 | 643.5735 | 2225.809 | 1563.113 | 1422.569 |
| ENSMUSG00000037477 | -2.616953957 | 4.22E-05 | 0.000701 | 6.036326 | 1.199822 | 4.140533 | 3.450494 | 26.32801 | 10.05637 | 21.21436 | 34.80291 |
| ENSMUSG00000037579 | 1.723686956 | 1.15E-14 | 4.71E-12 | 168.0111 | 167.975 | 143.2624 | 128.5309 | 59.48179 | 42.46022 | 47.25017 | 33.71532 |
| ENSMUSG00000037703 | -1.765374763 | 1.76E-07 | 8.99E-06 | 417.5126 | 379.1436 | 780.0764 | 407.1583 | 1688.893 | 743.0538 | 1859.15 | 2456.868 |
| ENSMUSG00000037710 | -1.32710203 | 1.52E-07 | 7.98E-06 | 512.0817 | 437.9349 | 713.8279 | 492.558 | 1340.778 | 777.6924 | 1424.255 | 1870.657 |
| ENSMUSG00000037747 | -1.473430297 | 0.000177 | 0.002063 | 92.557 | 63.59054 | 220.2764 | 116.4542 | 252.5538 | 210.0663 | 427.1801 | 482.8904 |
| ENSMUSG00000037759 | -1.533448564 | 0.009574 | 0.041244 | 7.04238 | 7.198929 | 9.109173 | 2.58787 | 13.65156 | 12.29112 | 16.39292 | 32.62773 |
| ENSMUSG00000037798 | 3.476871436 | 0.002233 | 0.014238 | 419.5247 | 7.198929 | 19.04645 | 238.9467 | 8.776002 | 16.76061 | 25.07152 | 10.87591 |
| ENSMUSG00000037827 | -1.881009124 | 0.004651 | 0.02445 | 5.030272 | 0 | 5.796746 | 6.900988 | 14.62667 | 10.05637 | 23.14294 | 19.57664 |
| ENSMUSG00000037953 | -3.116941946 | 0.000795 | 0.006545 | 2.012109 | 0 | 14.90592 | 5.175741 | 24.37778 | 10.05637 | 61.71451 | 100.0584 |
| ENSMUSG00000037977 | -1.120996445 | 0.00022 | 0.002437 | 26.15741 | 34.79482 | 39.74912 | 33.64232 | 86.78491 | 45.81234 | 67.50024 | 92.44524 |
| ENSMUSG00000038057 | -1.731359546 | 0.000891 | 0.007125 | 5.030272 | 5.999108 | 5.796746 | 4.313117 | 12.67645 | 23.46486 | 14.46434 | 19.57664 |
| ENSMUSG00000038060 | -1.018756787 | 0.000293 | 0.003047 | 185.114 | 116.3827 | 208.6829 | 137.1571 | 253.5289 | 219.0053 | 387.6442 | 453.5255 |
| ENSMUSG00000038072 | -1.433434142 | 1.00E-07 | 5.66E-06 | 1369.24 | 1064.242 | 1653.729 | 1513.042 | 3958.952 | 1911.827 | 3780.978 | 5477.109 |
| ENSMUSG00000038094 | -2.33412082 | 9.66E-06 | 0.000221 | 13.07871 | 4.799286 | 18.21835 | 6.900988 | 41.92979 | 30.1691 | 53.0359 | 94.62042 |
| ENSMUSG00000038132 | 1.202365325 | 0.002483 | 0.015349 | 257.5499 | 261.5611 | 214.4796 | 246.7103 | 101.4116 | 205.5968 | 79.07171 | 40.24087 |
| ENSMUSG00000038148 | -2.457870068 | 2.55E-05 | 0.000474 | 19.11503 | 17.99732 | 99.3728 | 30.19182 | 151.1423 | 97.21155 | 286.3939 | 386.0948 |
| ENSMUSG00000038178 | -1.437725 | 1.42E-06 | 4.95E-05 | 277.671 | 231.5656 | 370.1637 | 256.1992 | 795.6908 | 372.0856 | 767.5742 | 1143.058 |
| ENSMUSG00000038201 | 2.282916102 | 0.000338 | 0.00339 | 13.07871 | 21.59679 | 9.93728 | 31.05445 | 2.925334 | 4.469496 | 3.857157 | 4.350364 |
| ENSMUSG00000038224 | -3.216198602 | 0.000114 | 0.001488 | 111.672 | 9.598572 | 168.9338 | 108.6906 | 987.7878 | 278.2261 | 770.467 | 1674.89 |
| ENSMUSG00000038298 | -3.155569977 | 0.000116 | 0.001505 | 251.5136 | 35.99465 | 499.3483 | 319.1707 | 2699.108 | 673.7766 | 2007.65 | 4479.788 |
| ENSMUSG00000038331 | -1.202080683 | 0.004079 | 0.022134 | 32.19374 | 22.79661 | 77.01392 | 44.85642 | 74.10846 | 67.04245 | 127.2862 | 141.3868 |
| ENSMUSG00000038522 | -2.827099368 | 3.76E-05 | 0.000641 | 55.33299 | 21.59679 | 143.2624 | 63.83414 | 461.2276 | 187.7188 | 434.8944 | 936.4159 |
| ENSMUSG00000038528 | -3.056681767 | 1.56E-12 | 4.29E-10 | 35.2119 | 13.19804 | 59.62368 | 23.29083 | 371.5174 | 149.7281 | 231.4294 | 349.1167 |
| ENSMUSG00000038541 | 2.259871517 | 0.001048 | 0.008098 | 58.35115 | 17.99732 | 18.21835 | 21.56559 | 3.900445 | 5.58687 | 1.928578 | 13.05109 |
| ENSMUSG00000038567 | -3.738826048 | 2.18E-12 | 5.74E-10 | 24.1453 | 3.599465 | 39.74912 | 42.26855 | 619.1957 | 196.6578 | 267.1081 | 396.9707 |
| ENSMUSG00000038600 | -2.335483164 | 0.000256 | 0.002755 | 266.6044 | 141.5789 | 658.3448 | 296.7425 | 1757.151 | 481.5882 | 1998.007 | 2647.197 |
| ENSMUSG00000038704 | -2.084019539 | 0.001383 | 0.009973 | 11.0666 | 7.198929 | 5.796746 | 7.763611 | 28.27823 | 7.821619 | 28.92867 | 69.60583 |
| ENSMUSG00000038745 | -1.637573944 | 0.000291 | 0.003035 | 29.17558 | 17.99732 | 72.87338 | 43.9938 | 137.4907 | 68.15982 | 112.8218 | 195.7664 |
| ENSMUSG00000038782 | -1.410267916 | 0.001997 | 0.013119 | 22.1332 | 9.598572 | 28.98373 | 23.29083 | 39.97956 | 30.1691 | 69.42882 | 85.9197 |
| ENSMUSG00000038843 | -1.438490993 | 1.39E-08 | 1.11E-06 | 427.5731 | 393.5415 | 673.2507 | 463.2288 | 1087.249 | 983.2892 | 1208.254 | 2031.62 |
| ENSMUSG00000038917 | 1.11731494 | 0.003301 | 0.019044 | 40.24217 | 53.99197 | 33.12427 | 55.2079 | 22.42756 | 32.40385 | 17.3572 | 11.9635 |
| ENSMUSG00000038932 | 2.610058434 | 3.32E-10 | 4.78E-08 | 144.8718 | 175.1739 | 131.669 | 166.4863 | 23.40267 | 51.39921 | 9.642892 | 17.40146 |
| ENSMUSG00000038963 | -1.844700665 | 9.48E-06 | 0.000218 | 65.39353 | 47.99286 | 139.1219 | 36.23019 | 227.2009 | 167.6061 | 258.4295 | 387.1824 |
| ENSMUSG00000039037 | 1.016261069 | 0.000181 | 0.0021 | 363.1856 | 284.3577 | 206.1986 | 416.6471 | 161.8685 | 212.3011 | 147.5362 | 106.5839 |
| ENSMUSG00000039062 | -2.370201944 | 3.45E-07 | 1.54E-05 | 1033.218 | 1096.637 | 1714.181 | 1178.344 | 5110.558 | 2844.834 | 6095.272 | 11917.82 |
| ENSMUSG00000039131 | -1.951632197 | 1.20E-05 | 0.000265 | 145.8779 | 63.59054 | 319.6492 | 138.8824 | 697.2046 | 290.5173 | 652.8238 | 948.3794 |
| ENSMUSG00000039257 | 1.307343247 | 3.97E-06 | 0.000112 | 583.5115 | 854.2729 | 654.2042 | 835.8822 | 257.4294 | 496.1141 | 243.0009 | 187.0657 |
| ENSMUSG00000039278 | 1.606980411 | 0.006801 | 0.032317 | 156.9445 | 32.39518 | 38.92101 | 244.1224 | 50.70579 | 33.52122 | 31.82154 | 39.15328 |
| ENSMUSG00000039364 | -3.139586349 | 5.04E-05 | 0.000798 | 5.030272 | 2.399643 | 0 | 2.58787 | 32.17867 | 11.17374 | 14.46434 | 28.27737 |
| ENSMUSG00000039438 | -3.997899626 | 2.83E-09 | 3.09E-07 | 82.49646 | 26.39607 | 188.8083 | 86.26235 | 1553.352 | 545.2786 | 1243.933 | 2802.722 |
| ENSMUSG00000039546 | -1.949828824 | 1.54E-05 | 0.000318 | 50.30272 | 34.79482 | 153.1997 | 49.16954 | 297.4089 | 153.0803 | 364.5013 | 300.1751 |
| ENSMUSG00000039639 | -1.580036643 | 0.007826 | 0.035655 | 40.24217 | 19.19714 | 75.3577 | 23.29083 | 87.76002 | 36.87335 | 96.42892 | 253.4087 |
| ENSMUSG00000039661 | 1.260939404 | 2.59E-05 | 0.000479 | 170.0232 | 95.98572 | 104.3414 | 200.9913 | 57.53157 | 51.39921 | 76.17884 | 53.29196 |
| ENSMUSG00000039710 | -5.842807431 | 1.29E-06 | 4.58E-05 | 0 | 0 | 10.76539 | 3.450494 | 175.52 | 52.51658 | 246.858 | 361.0802 |
| ENSMUSG00000039728 | 2.498750346 | 0.002842 | 0.017029 | 158.9566 | 13.19804 | 8.281066 | 40.5433 | 13.65156 | 11.17374 | 7.714313 | 6.525547 |
| ENSMUSG00000039838 | 1.865465914 | 0.009796 | 0.04193 | 23.13925 | 21.59679 | 5.796746 | 35.36756 | 8.776002 | 7.821619 | 5.785735 | 1.087591 |
| ENSMUSG00000039878 | -2.658234437 | 3.41E-06 | 0.0001 | 116.7023 | 35.99465 | 181.3554 | 123.3552 | 752.7859 | 321.8037 | 609.4308 | 1208.314 |
| ENSMUSG00000039913 | 1.132977701 | 0.001376 | 0.009945 | 49.29666 | 83.98751 | 58.79557 | 37.95543 | 17.552 | 29.05173 | 32.78583 | 25.0146 |
| ENSMUSG00000039954 | -1.541633626 | 0.003797 | 0.021024 | 39.23612 | 17.99732 | 53.82693 | 54.34528 | 94.5858 | 32.40385 | 145.6077 | 210.9927 |
| ENSMUSG00000039956 | 4.226006561 | 0.005417 | 0.02719 | 1075.472 | 4.799286 | 9.93728 | 110.4158 | 4.875557 | 10.05637 | 5.785735 | 43.50364 |
| ENSMUSG00000040016 | -1.337603606 | 0.000142 | 0.001748 | 157.9505 | 194.3711 | 288.1811 | 197.5408 | 437.825 | 220.1227 | 709.7168 | 751.5254 |
| ENSMUSG00000040026 | 1.855795974 | 0.004999 | 0.02575 | 32.19374 | 8.398751 | 11.59349 | 14.6646 | 6.825779 | 4.469496 | 3.857157 | 3.262773 |
| ENSMUSG00000040471 | -2.141901614 | 0.001916 | 0.012726 | 17.10292 | 2.399643 | 24.01509 | 5.175741 | 78.0089 | 16.76061 | 48.21446 | 73.95619 |
| ENSMUSG00000040536 | 1.376919154 | 0.000125 | 0.001589 | 35.2119 | 53.99197 | 29.81184 | 50.89479 | 18.52711 | 17.87799 | 12.53576 | 16.31387 |
| ENSMUSG00000040584 | 1.573589633 | 0.002584 | 0.015783 | 224.3501 | 117.5825 | 61.27989 | 170.7995 | 21.45245 | 101.681 | 38.57157 | 31.54014 |
| ENSMUSG00000040706 | -1.88532556 | 0.009236 | 0.040213 | 13.07871 | 0 | 4.96864 | 3.450494 | 15.60178 | 18.99536 | 17.3572 | 28.27737 |
| ENSMUSG00000040724 | 1.135989858 | 0.000557 | 0.004971 | 192.1564 | 212.3684 | 171.4181 | 213.068 | 69.2329 | 165.3714 | 58.82164 | 66.34306 |
| ENSMUSG00000040740 | -2.726239084 | 5.37E-07 | 2.23E-05 | 16.09687 | 11.99822 | 18.21835 | 13.80198 | 130.6649 | 25.6996 | 66.53595 | 176.1898 |

| | | | | | | | | | | | |
|---|---|---|---|---|---|---|---|---|---|---|---|
| ENSMUSG00000040808 | -2.441163717 | 4.65E-15 | 2.09E-12 | 1246.501 | 853.0731 | 1661.182 | 1080.005 | 4698.086 | 3428.104 | 8626.531 | 9542.524 |
| ENSMUSG00000040966 | -5.174584012 | 1.37E-09 | 1.71E-07 | 9.054489 | 4.799286 | 45.54586 | 18.11509 | 883.4508 | 151.9629 | 500.4661 | 1282.27 |
| ENSMUSG00000040978 | -2.421682467 | 2.77E-08 | 1.99E-06 | 26.15741 | 15.59768 | 35.60859 | 31.91707 | 154.0676 | 60.3382 | 126.3219 | 249.0584 |
| ENSMUSG00000041014 | 1.166380406 | 0.004771 | 0.024904 | 29.17558 | 49.19268 | 30.63995 | 47.44429 | 20.47734 | 26.81698 | 11.57147 | 10.87591 |
| ENSMUSG00000041052 | -5.026631181 | 8.38E-09 | 7.40E-07 | 1.006054 | 4.799286 | 19.87456 | 6.038364 | 209.6489 | 79.33356 | 267.1081 | 490.5036 |
| ENSMUSG00000041237 | -2.585125729 | 2.25E-06 | 7.05E-05 | 23.13925 | 7.198929 | 51.34261 | 23.29083 | 205.7485 | 70.39457 | 106.0718 | 253.4087 |
| ENSMUSG00000041248 | -2.838261867 | 0.001668 | 0.011454 | 64.38748 | 44.3934 | 539.0974 | 105.2401 | 1073.598 | 403.372 | 1806.114 | 2106.664 |
| ENSMUSG00000041323 | -2.513968935 | 0.000608 | 0.005327 | 8.048435 | 4.799286 | 77.01392 | 18.11509 | 111.1627 | 62.57295 | 216.0008 | 231.6569 |
| ENSMUSG00000041324 | 1.131558592 | 0.008766 | 0.038736 | 56.33904 | 57.59143 | 28.15563 | 52.62003 | 19.50223 | 39.10809 | 18.32149 | 11.9635 |
| ENSMUSG00000041544 | -1.462153683 | 0.003371 | 0.019279 | 3.018163 | 7.198929 | 9.93728 | 12.07673 | 29.25334 | 16.76061 | 22.17865 | 21.75182 |
| ENSMUSG00000041552 | 1.10372495 | 0.012143 | 0.049419 | 46.2785 | 57.59143 | 19.87456 | 64.69676 | 23.40267 | 27.93435 | 13.50005 | 22.83941 |
| ENSMUSG00000041644 | -2.999824438 | 1.57E-07 | 8.19E-06 | 21.12714 | 7.198929 | 89.43552 | 29.3292 | 376.393 | 153.0803 | 328.8226 | 327.3649 |
| ENSMUSG00000041650 | -1.339496529 | 4.88E-06 | 0.000132 | 325.9616 | 296.3559 | 431.4436 | 276.0395 | 904.9033 | 393.3157 | 896.7889 | 1171.336 |
| ENSMUSG00000041660 | 1.132809919 | 0.005257 | 0.026624 | 18.10898 | 28.79572 | 33.95237 | 31.91707 | 11.70134 | 16.76061 | 14.46434 | 8.700729 |
| ENSMUSG00000041828 | 1.217468518 | 0.000534 | 0.004826 | 210.2654 | 215.9679 | 147.403 | 422.6855 | 93.61069 | 159.7845 | 98.35749 | 77.21897 |
| ENSMUSG00000041923 | 1.278529822 | 0.001745 | 0.011852 | 60.36326 | 52.79215 | 48.85829 | 58.6584 | 15.60178 | 43.57759 | 20.25007 | 11.9635 |
| ENSMUSG00000042096 | -2.514046737 | 7.92E-05 | 0.001131 | 180.0837 | 34.79482 | 273.2752 | 131.1188 | 867.8491 | 444.7149 | 763.717 | 1464.985 |
| ENSMUSG00000042102 | -3.058555929 | 1.38E-07 | 7.36E-06 | 16.09687 | 4.799286 | 28.98373 | 33.64232 | 195.0223 | 69.27719 | 106.0718 | 332.8029 |
| ENSMUSG00000042116 | -1.09981623 | 1.20E-06 | 4.38E-05 | 193.1624 | 269.9598 | 251.7444 | 263.9628 | 487.5557 | 351.9728 | 519.7519 | 738.4743 |
| ENSMUSG00000042118 | -2.666749086 | 0.002451 | 0.015197 | 42.25428 | 5.999108 | 68.73285 | 52.62003 | 264.2552 | 43.57759 | 207.3222 | 564.4598 |
| ENSMUSG00000042182 | 1.039070263 | 0.001507 | 0.010598 | 44.26639 | 58.79126 | 30.63995 | 51.75741 | 24.37778 | 18.99536 | 20.25007 | 26.10219 |
| ENSMUSG00000042254 | 1.046386929 | 0.001903 | 0.012672 | 42.25428 | 77.9884 | 59.62368 | 92.30071 | 41.92979 | 24.58223 | 37.60728 | 27.18978 |
| ENSMUSG00000042258 | 2.559912468 | 1.63E-05 | 0.000333 | 127.7689 | 47.99286 | 22.35888 | 174.2499 | 17.552 | 20.11273 | 12.53576 | 13.05109 |
| ENSMUSG00000042284 | -1.03930864 | 5.45E-05 | 0.000845 | 396.3854 | 373.1445 | 554.8314 | 355.4009 | 884.426 | 493.8794 | 864.0031 | 1211.576 |
| ENSMUSG00000042353 | -1.205490391 | 0.010279 | 0.043496 | 12.07265 | 4.799286 | 18.21835 | 15.52722 | 30.22845 | 33.52122 | 37.60728 | 17.40146 |
| ENSMUSG00000042371 | -4.917652246 | 8.55E-17 | 6.07E-14 | 6.036326 | 4.799286 | 15.73403 | 6.038364 | 261.3298 | 63.69032 | 197.6793 | 478.5401 |
| ENSMUSG00000042581 | 1.382511024 | 0.000151 | 0.00184 | 150.9082 | 113.983 | 103.5133 | 185.4641 | 35.10401 | 96.09417 | 38.57157 | 43.50364 |
| ENSMUSG00000042662 | -1.443841536 | 7.16E-05 | 0.001051 | 81.4904 | 59.99108 | 91.91984 | 66.42201 | 152.1174 | 92.74205 | 265.1795 | 306.7007 |
| ENSMUSG00000042672 | -1.5954937 | 9.42E-05 | 0.001274 | 31.18769 | 28.79572 | 45.54586 | 33.64232 | 107.2622 | 42.46022 | 99.32178 | 172.927 |
| ENSMUSG00000042761 | 1.649306571 | 0.001627 | 0.011205 | 15.09082 | 31.19536 | 25.67131 | 24.15346 | 6.825779 | 6.704245 | 13.50005 | 3.262773 |
| ENSMUSG00000042788 | 1.256287998 | 0.001474 | 0.010444 | 59.35721 | 53.99197 | 43.06154 | 52.62003 | 10.72622 | 34.6386 | 14.46434 | 28.27737 |
| ENSMUSG00000042797 | -1.269100385 | 1.29E-06 | 4.58E-05 | 211.2714 | 169.1748 | 287.353 | 221.6942 | 549.9628 | 325.1559 | 505.2875 | 766.7517 |
| ENSMUSG00000043144 | -1.96346104 | 0.012116 | 0.049352 | 7.04238 | 9.598572 | 48.85829 | 7.763611 | 69.2329 | 13.40849 | 54.96448 | 150.0876 |
| ENSMUSG00000043155 | -1.315807526 | 0.008877 | 0.039101 | 15.09082 | 13.19804 | 13.24971 | 13.80198 | 36.07912 | 11.17374 | 32.78583 | 57.64233 |
| ENSMUSG00000043592 | -1.183548421 | 0.002624 | 0.015975 | 37.22401 | 15.59768 | 57.96746 | 31.91707 | 76.05868 | 68.15982 | 65.57166 | 117.4598 |
| ENSMUSG00000043760 | -1.042613573 | 0.000992 | 0.00779 | 432.6034 | 254.3622 | 548.2066 | 293.292 | 745.9602 | 419.0153 | 843.753 | 1141.971 |
| ENSMUSG00000043773 | 1.004069451 | 1.05E-06 | 3.88E-05 | 281.6952 | 424.7368 | 346.1486 | 328.6596 | 124.8142 | 212.3011 | 155.2506 | 196.854 |
| ENSMUSG00000043850 | 1.010307136 | 0.000158 | 0.001891 | 210.2654 | 203.9697 | 130.0127 | 171.6621 | 123.8391 | 56.98608 | 81.96458 | 91.35765 |
| ENSMUSG00000044006 | 2.152166243 | 1.34E-12 | 3.88E-10 | 102.6175 | 140.3791 | 139.1219 | 186.3267 | 44.85512 | 35.75597 | 20.25007 | 27.18978 |
| ENSMUSG00000044017 | 1.454518551 | 7.30E-11 | 1.39E-08 | 429.5852 | 616.7083 | 395.0069 | 496.8711 | 194.0472 | 233.5312 | 155.2506 | 123.9854 |
| ENSMUSG00000044034 | -1.418178318 | 0.000744 | 0.00623 | 33.19979 | 29.99554 | 67.90474 | 25.8787 | 98.48624 | 56.98608 | 97.39321 | 168.5766 |
| ENSMUSG00000044071 | 1.948823242 | 1.27E-09 | 1.60E-07 | 342.0585 | 374.3443 | 369.3356 | 409.7462 | 63.38224 | 179.8972 | 82.92887 | 61.99269 |
| ENSMUSG00000044139 | -2.017936068 | 4.60E-05 | 0.000747 | 12.07265 | 17.99732 | 14.90592 | 5.175741 | 43.88001 | 34.6386 | 34.71441 | 88.09488 |
| ENSMUSG00000044156 | -1.895422583 | 0.011791 | 0.048308 | 16.09687 | 7.198929 | 114.2787 | 11.21411 | 80.93424 | 87.15518 | 187.0721 | 201.2044 |
| ENSMUSG00000044249 | -2.422913863 | 0.001156 | 0.008736 | 2.012109 | 3.599465 | 31.46805 | 11.21411 | 57.53157 | 27.93435 | 58.82164 | 118.5474 |
| ENSMUSG00000044444 | -2.373969863 | 0.000117 | 0.001515 | 17.10292 | 2.399643 | 28.15563 | 7.763611 | 74.10846 | 43.57759 | 48.21446 | 125.073 |
| ENSMUSG00000044499 | 1.417870638 | 0.010276 | 0.043496 | 33.19979 | 19.19714 | 12.4216 | 36.23019 | 5.850668 | 7.821619 | 16.39292 | 7.613138 |
| ENSMUSG00000044576 | 1.821351562 | 2.62E-09 | 2.89E-07 | 59.35721 | 75.58876 | 62.108 | 51.75741 | 14.62667 | 21.23011 | 21.21436 | 13.05109 |
| ENSMUSG00000044708 | -1.41130867 | 3.10E-05 | 0.000549 | 184.1079 | 91.18644 | 221.1045 | 154.4096 | 306.185 | 258.1134 | 616.1808 | 553.5839 |
| ENSMUSG00000044748 | -2.935445253 | 6.95E-16 | 3.92E-13 | 128.775 | 79.18822 | 153.1997 | 94.02596 | 722.5575 | 357.5597 | 1067.468 | 1339.912 |
| ENSMUSG00000044819 | -2.899681883 | 1.16E-10 | 2.08E-08 | 31.18769 | 14.39786 | 46.37397 | 38.81806 | 141.3911 | 121.7938 | 348.1084 | 371.9562 |
| ENSMUSG00000044912 | 1.906498578 | 2.22E-10 | 3.56E-08 | 112.6781 | 118.7823 | 145.7468 | 169.9368 | 38.02934 | 55.8687 | 30.85725 | 21.75182 |
| ENSMUSG00000044976 | -1.456052848 | 0.005261 | 0.02664 | 29.17558 | 19.19714 | 64.59232 | 41.40593 | 84.83468 | 36.87335 | 105.1075 | 199.0292 |
| ENSMUSG00000045291 | -1.732613505 | 0.010544 | 0.044329 | 22.1332 | 1.199822 | 43.88965 | 18.11509 | 68.25779 | 36.87335 | 64.60737 | 116.3722 |
| ENSMUSG00000045625 | -1.209445241 | 0.00238 | 0.014867 | 13.07871 | 22.79661 | 19.87456 | 25.01608 | 41.92979 | 32.40385 | 36.64299 | 76.13138 |
| ENSMUSG00000045761 | 1.745762408 | 0.001387 | 0.009988 | 42.25428 | 55.19179 | 44.71776 | 31.91707 | 2.925334 | 14.52586 | 8.678602 | 26.10219 |
| ENSMUSG00000045775 | -1.618666938 | 6.24E-06 | 0.00016 | 172.0353 | 172.7743 | 390.8663 | 203.5791 | 645.5237 | 357.5597 | 764.6813 | 1120.219 |
| ENSMUSG00000046070 | -3.283491251 | 1.57E-09 | 1.91E-07 | 8.048435 | 8.398751 | 4.96864 | 3.450494 | 59.48179 | 23.46486 | 58.82164 | 95.70802 |
| ENSMUSG00000046082 | -2.816001533 | 9.00E-05 | 0.001237 | 171.0292 | 32.39518 | 390.8663 | 219.969 | 1563.103 | 630.199 | 1284.433 | 2261.102 |
| ENSMUSG00000046101 | -1.210862742 | 0.000673 | 0.005762 | 45.27245 | 46.79304 | 40.57722 | 50.89479 | 70.20801 | 75.98144 | 89.67889 | 189.2408 |
| ENSMUSG00000046178 | 2.029364739 | 0.010865 | 0.045396 | 11.0666 | 16.7975 | 11.59349 | 28.46658 | 0 | 7.821619 | 6.750024 | 2.175182 |
| ENSMUSG00000046287 | 2.105632144 | 2.08E-10 | 3.37E-08 | 67.40564 | 51.59233 | 71.21717 | 82.81186 | 16.57689 | 17.87799 | 20.25007 | 8.700729 |
| ENSMUSG00000046352 | -2.162344655 | 3.32E-12 | 8.40E-10 | 29.17558 | 32.39518 | 33.95237 | 35.36756 | 163.8187 | 88.27255 | 121.5004 | 213.1679 |
| ENSMUSG00000046480 | -1.807726344 | 0.002243 | 0.014277 | 19.11503 | 10.79839 | 71.21717 | 38.81806 | 71.18313 | 65.92507 | 113.7861 | 242.5528 |

| | | | | | | | | | | | |
|---|---|---|---|---|---|---|---|---|---|---|---|
| ENSMUSG00000046598 | -2.180880061 | 0.000255 | 0.002751 | 266.6044 | 175.1739 | 636.814 | 257.9244 | 1306.649 | 442.4801 | 1952.686 | 2361.16 |
| ENSMUSG00000046687 | -2.885678473 | 4.85E-05 | 0.000778 | 2.012109 | 2.399643 | 9.109173 | 15.52722 | 98.48624 | 17.87799 | 39.53586 | 63.08028 |
| ENSMUSG00000046709 | 1.08049168 | 0.002638 | 0.016038 | 159.9626 | 82.78769 | 59.62368 | 130.2561 | 42.9049 | 71.51194 | 50.14304 | 40.24087 |
| ENSMUSG00000046743 | 1.141209431 | 4.27E-07 | 1.85E-05 | 5342.149 | 6344.656 | 5493.659 | 5966.767 | 2421.201 | 4147.693 | 1953.65 | 1972.89 |
| ENSMUSG00000046840 | -3.816258746 | 0.00039 | 0.00379 | 0 | 1.199822 | 4.140533 | 1.725247 | 25.35289 | 1.117374 | 26.03581 | 50.02919 |
| ENSMUSG00000046959 | -2.012515508 | 7.77E-07 | 3.03E-05 | 85.51462 | 47.99286 | 104.3414 | 62.97152 | 322.7618 | 120.6764 | 256.5009 | 516.6058 |
| ENSMUSG00000046961 | 1.187980137 | 1.14E-07 | 6.24E-06 | 163.9869 | 143.9786 | 102.6852 | 170.7995 | 72.15824 | 56.98608 | 65.57166 | 59.81751 |
| ENSMUSG00000047085 | 1.412181961 | 8.09E-11 | 1.52E-08 | 367.2098 | 361.1463 | 353.6015 | 421.8229 | 88.73513 | 189.9536 | 131.1433 | 156.6131 |
| ENSMUSG00000047228 | 1.28922013 | 0.000571 | 0.005078 | 53.32088 | 64.79036 | 47.20208 | 56.93315 | 30.22845 | 32.40385 | 18.32149 | 9.78832 |
| ENSMUSG00000047230 | -3.099248609 | 4.91E-06 | 0.000133 | 269.6226 | 62.39072 | 577.1903 | 329.5222 | 2871.703 | 1093.909 | 2568.866 | 4087.167 |
| ENSMUSG00000047261 | 1.869615794 | 2.28E-05 | 0.000436 | 581.4994 | 170.3747 | 209.511 | 835.8822 | 107.2622 | 124.0285 | 120.5361 | 140.2993 |
| ENSMUSG00000047281 | -1.164851614 | 0.003023 | 0.017883 | 137.8294 | 111.5834 | 331.2427 | 121.6299 | 438.8001 | 193.3057 | 452.2516 | 492.6788 |
| ENSMUSG00000047344 | 1.197162494 | 5.68E-09 | 5.30E-07 | 135.8173 | 109.1838 | 104.3414 | 110.4158 | 52.65601 | 44.69496 | 58.82164 | 43.50364 |
| ENSMUSG00000047586 | -3.442456611 | 0.000267 | 0.002839 | 15.09082 | 1.199822 | 66.24853 | 9.488858 | 270.1058 | 135.2023 | 288.3225 | 312.1386 |
| ENSMUSG00000047638 | -2.192345088 | 0.002658 | 0.016124 | 118.7144 | 8.398751 | 137.4657 | 59.52102 | 365.6667 | 221.2401 | 447.4302 | 449.1751 |
| ENSMUSG00000047686 | 1.81358642 | 0.009729 | 0.041727 | 128.775 | 220.7672 | 129.1846 | 199.266 | 46.80534 | 121.7938 | 21.21436 | 3.262773 |
| ENSMUSG00000047728 | -2.496457718 | 6.18E-08 | 3.85E-06 | 42.25428 | 21.59679 | 99.3728 | 37.95543 | 248.6534 | 158.6671 | 251.6795 | 482.8904 |
| ENSMUSG00000047746 | -2.474973024 | 1.06E-05 | 0.000238 | 13.07871 | 3.599465 | 22.35888 | 14.6646 | 83.85957 | 29.05173 | 61.71451 | 128.3357 |
| ENSMUSG00000047786 | 1.377404263 | 2.07E-09 | 2.38E-07 | 460.7729 | 632.3059 | 568.9093 | 541.7276 | 188.1965 | 325.1559 | 163.9292 | 171.8394 |
| ENSMUSG00000047797 | -1.355588262 | 1.25E-05 | 0.000272 | 227.3683 | 167.975 | 308.8838 | 238.0841 | 664.0508 | 290.5173 | 577.6092 | 882.0364 |
| ENSMUSG00000047822 | -1.433299522 | 0.004966 | 0.025622 | 15.09082 | 15.59768 | 13.24971 | 8.626235 | 38.02934 | 11.17374 | 39.53586 | 52.20437 |
| ENSMUSG00000047897 | -2.959875007 | 0.000225 | 0.002482 | 2.012109 | 20.39697 | 7.45296 | 10.35148 | 117.0134 | 5.58687 | 91.60747 | 95.70802 |
| ENSMUSG00000048004 | 1.419673979 | 0.00016 | 0.001911 | 112.6781 | 77.9884 | 41.40533 | 111.2784 | 35.10401 | 41.34284 | 27.96439 | 23.927 |
| ENSMUSG00000048029 | -1.335479141 | 0.003651 | 0.020442 | 14.08476 | 5.999108 | 4.96864 | 11.21411 | 22.42756 | 20.11273 | 25.07152 | 23.927 |
| ENSMUSG00000048070 | 3.147749901 | 0.006928 | 0.032741 | 60.36326 | 0 | 8.281066 | 85.39973 | 7.80089 | 1.117374 | 1.928578 | 6.525547 |
| ENSMUSG00000048108 | -2.038508286 | 0.001481 | 0.010472 | 186.1201 | 80.38804 | 443.8652 | 191.5024 | 604.569 | 357.5597 | 1304.683 | 1441.058 |
| ENSMUSG00000048216 | 1.832930824 | 1.73E-06 | 5.82E-05 | 82.49646 | 59.99108 | 69.56096 | 106.9653 | 9.751113 | 34.6386 | 25.07152 | 20.66423 |
| ENSMUSG00000048368 | 1.485888349 | 0.000863 | 0.006961 | 177.0656 | 119.9822 | 63.76421 | 190.6398 | 31.20356 | 89.38993 | 47.25017 | 29.36496 |
| ENSMUSG00000048373 | -1.786963815 | 2.07E-06 | 6.59E-05 | 79.47829 | 64.79036 | 163.9651 | 86.26235 | 288.6329 | 172.0756 | 523.609 | 380.6569 |
| ENSMUSG00000048424 | -2.178508572 | 0.001705 | 0.011648 | 45.27245 | 43.19358 | 309.7119 | 76.77349 | 494.3814 | 379.9072 | 645.1095 | 634.0656 |
| ENSMUSG00000048442 | -1.41380008 | 0.002558 | 0.015683 | 22.1332 | 13.19804 | 48.03018 | 30.19182 | 69.2329 | 35.75597 | 83.89316 | 116.3722 |
| ENSMUSG00000048489 | -1.013450877 | 0.005035 | 0.025854 | 369.2219 | 88.78679 | 250.0882 | 265.688 | 526.5601 | 356.4423 | 661.5024 | 423.0729 |
| ENSMUSG00000048572 | -1.872178389 | 2.35E-21 | 4.34E-18 | 536.227 | 561.5165 | 642.6107 | 797.0641 | 3169.112 | 1902.888 | 2349.973 | 1871.744 |
| ENSMUSG00000048699 | 1.457662744 | 0.012323 | 0.049967 | 10.06054 | 23.99643 | 13.24971 | 24.15346 | 8.776002 | 8.938993 | 2.892867 | 5.437955 |
| ENSMUSG00000048707 | -1.175263588 | 1.60E-10 | 2.76E-08 | 212.2775 | 196.7707 | 269.1347 | 198.4034 | 507.0579 | 368.7335 | 497.5732 | 609.051 |
| ENSMUSG00000048721 | -1.27395539 | 0.012076 | 0.049211 | 13.07871 | 8.398751 | 24.01509 | 12.07673 | 35.10401 | 15.64324 | 55.92877 | 33.71532 |
| ENSMUSG00000048752 | 2.653897226 | 1.87E-07 | 9.48E-06 | 29.17558 | 23.99643 | 53.82693 | 29.3292 | 4.875557 | 6.704245 | 3.857157 | 6.525547 |
| ENSMUSG00000048776 | 1.114214822 | 0.000296 | 0.003066 | 106.6418 | 134.38 | 64.59232 | 96.61383 | 36.07912 | 65.92507 | 44.3573 | 39.15328 |
| ENSMUSG00000048834 | -1.231657111 | 0.008719 | 0.03857 | 13.07871 | 17.99732 | 19.87456 | 14.6646 | 19.50223 | 70.39457 | 28.92867 | 35.89051 |
| ENSMUSG00000048905 | 1.041526138 | 0.012065 | 0.049182 | 33.19979 | 44.3934 | 56.31125 | 24.15346 | 20.47734 | 10.05637 | 23.14294 | 22.83941 |
| ENSMUSG00000049044 | 1.005705068 | 0.003201 | 0.01862 | 1164.005 | 406.7395 | 395.835 | 462.3662 | 265.2303 | 382.1419 | 297.0011 | 265.3722 |
| ENSMUSG00000049152 | -7.179486936 | 1.39E-09 | 1.72E-07 | 0 | 0 | 4.96864 | 1.725247 | 231.1014 | 32.40385 | 293.1439 | 460.051 |
| ENSMUSG00000049265 | 3.048695187 | 0.00536 | 0.026945 | 2000.036 | 63.59054 | 72.87338 | 477.0308 | 55.58134 | 69.27719 | 45.32159 | 145.7372 |
| ENSMUSG00000049281 | 2.842650727 | 8.14E-06 | 0.000195 | 284.7134 | 82.78769 | 36.43669 | 272.589 | 19.50223 | 45.81234 | 18.32149 | 10.87591 |
| ENSMUSG00000049422 | -1.426944157 | 0.000758 | 0.006321 | 983.9212 | 533.9206 | 1434.281 | 736.6805 | 2194 | 1296.154 | 2691.331 | 3739.138 |
| ENSMUSG00000049676 | 2.165058752 | 6.02E-31 | 2.78E-27 | 253.5257 | 224.3666 | 183.8397 | 200.1287 | 49.73068 | 46.92971 | 48.21446 | 46.76642 |
| ENSMUSG00000049721 | -1.707820514 | 4.43E-05 | 0.000728 | 84.50857 | 73.18911 | 197.9175 | 97.47645 | 308.1352 | 150.8455 | 494.6803 | 529.6569 |
| ENSMUSG00000049799 | -2.19785419 | 4.32E-09 | 4.33E-07 | 27.16347 | 51.59233 | 41.40533 | 43.9938 | 276.9316 | 94.9768 | 134.0362 | 245.7956 |
| ENSMUSG00000050052 | -1.08299075 | 0.005925 | 0.02916 | 182.0958 | 129.5807 | 452.9743 | 156.1349 | 467.0783 | 280.4609 | 615.2165 | 590.562 |
| ENSMUSG00000050100 | -2.972412918 | 0.000492 | 0.004524 | 4.024217 | 1.199822 | 12.4216 | 1.725247 | 28.27823 | 6.704245 | 46.28588 | 73.95619 |
| ENSMUSG00000050276 | 3.031545628 | 0.000151 | 0.001834 | 34.20585 | 13.19804 | 19.04645 | 60.38364 | 5.850668 | 7.821619 | 1.928578 | 0 |
| ENSMUSG00000050473 | 2.563481718 | 0.000272 | 0.002874 | 61.36932 | 8.398751 | 15.73403 | 30.19182 | 3.900445 | 4.469496 | 5.785735 | 5.437955 |
| ENSMUSG00000050645 | -4.028964731 | 3.44E-07 | 1.54E-05 | 3.018163 | 2.399643 | 17.39024 | 1.725247 | 134.5654 | 17.87799 | 98.35749 | 157.7007 |
| ENSMUSG00000050677 | -1.105566405 | 0.010935 | 0.04564 | 16.09687 | 17.99732 | 7.45296 | 9.488858 | 22.42756 | 21.23011 | 35.6787 | 28.27737 |
| ENSMUSG00000050737 | -1.117283682 | 0.00049 | 0.004507 | 378.2764 | 303.5548 | 731.2182 | 395.9442 | 782.0393 | 576.565 | 1258.397 | 1310.547 |
| ENSMUSG00000050777 | -1.611157178 | 2.70E-06 | 8.31E-05 | 313.889 | 211.1686 | 454.6305 | 268.2759 | 1145.756 | 410.0763 | 887.146 | 1372.54 |
| ENSMUSG00000050854 | -1.486821926 | 4.36E-05 | 0.000719 | 58.35115 | 57.59143 | 110.1382 | 62.10889 | 180.3956 | 94.9768 | 260.3581 | 274.073 |
| ENSMUSG00000050866 | -3.175850256 | 9.10E-06 | 0.000213 | 80.48435 | 16.7975 | 216.1358 | 67.28463 | 864.9237 | 420.1327 | 829.2887 | 1332.299 |
| ENSMUSG00000050896 | 1.814263203 | 3.80E-20 | 5.39E-17 | 157.9505 | 139.1793 | 147.403 | 165.6237 | 40.95467 | 36.87335 | 54.96448 | 40.24087 |
| ENSMUSG00000051111 | 1.881366183 | 0.000177 | 0.002061 | 140.8476 | 56.39161 | 51.34261 | 199.266 | 18.52711 | 49.16446 | 26.03581 | 28.27737 |
| ENSMUSG00000051159 | -1.632765622 | 5.91E-07 | 2.41E-05 | 553.3299 | 595.1115 | 481.9581 | 580.5456 | 2593.796 | 630.199 | 1955.578 | 1674.89 |
| ENSMUSG00000051166 | 1.066244 | 5.88E-07 | 2.41E-05 | 745.4863 | 575.9143 | 472.0208 | 753.0703 | 301.3094 | 410.0763 | 248.7866 | 256.6715 |
| ENSMUSG00000051455 | 2.1551098 | 2.63E-13 | 9.14E-11 | 70.4238 | 97.18554 | 101.029 | 87.9876 | 14.62667 | 29.05173 | 16.39292 | 20.66423 |
| ENSMUSG00000051497 | -1.481039677 | 5.25E-06 | 0.00014 | 629.79 | 353.9474 | 973.0253 | 602.9738 | 1681.092 | 931.89 | 2105.043 | 2431.854 |

| | | | | | | | | | | | |
|---|---|---|---|---|---|---|---|---|---|---|---|
| ENSMUSG00000051650 | -1.418415635 | 0.000339 | 0.003394 | 72.43591 | 43.19358 | 92.74794 | 58.6584 | 160.8934 | 71.51194 | 218.8936 | 264.2846 |
| ENSMUSG00000051844 | 1.381539428 | 0.005085 | 0.026035 | 21.12714 | 16.7975 | 14.90592 | 17.25247 | 8.776002 | 8.938993 | 4.821446 | 4.350364 |
| ENSMUSG00000051980 | -2.640358309 | 0.009136 | 0.039948 | 11.0666 | 2.399643 | 81.98256 | 18.97772 | 142.3663 | 41.34284 | 241.0723 | 291.4744 |
| ENSMUSG00000051985 | -1.249180622 | 0.007806 | 0.035589 | 13.07871 | 21.59679 | 24.01509 | 9.488858 | 63.38224 | 23.46486 | 28.92867 | 45.67883 |
| ENSMUSG00000052234 | -2.840750146 | 4.50E-06 | 0.000124 | 12.07265 | 7.198929 | 6.624853 | 11.21411 | 35.10401 | 32.40385 | 159.1077 | 39.15328 |
| ENSMUSG00000052392 | -2.411064524 | 3.15E-08 | 2.23E-06 | 50.30272 | 19.19714 | 48.03018 | 37.09281 | 189.1716 | 81.56831 | 208.2865 | 346.9416 |
| ENSMUSG00000052396 | -2.376894552 | 4.15E-05 | 0.000694 | 21.12714 | 15.59768 | 48.85829 | 32.77969 | 206.7236 | 49.16446 | 66.53595 | 295.8248 |
| ENSMUSG00000052415 | 1.211888583 | 5.65E-16 | 3.48E-13 | 2214.326 | 1912.516 | 2773.329 | 1978.858 | 870.7744 | 914.012 | 891.9675 | 1158.285 |
| ENSMUSG00000052520 | -5.110139959 | 2.39E-09 | 2.69E-07 | 20.12109 | 9.598572 | 101.8571 | 32.77969 | 1768.852 | 379.9072 | 887.146 | 2659.16 |
| ENSMUSG00000052562 | -3.233211126 | 1.11E-12 | 3.30E-10 | 7.04238 | 11.99822 | 7.45296 | 4.313117 | 69.2329 | 60.3382 | 44.3573 | 109.8467 |
| ENSMUSG00000052581 | 1.061644579 | 0.009274 | 0.04031 | 26.15741 | 34.79482 | 16.56213 | 29.3292 | 11.70134 | 15.64324 | 10.60718 | 13.05109 |
| ENSMUSG00000052595 | -2.676927263 | 1.26E-05 | 0.000274 | 7.04238 | 3.599465 | 9.109173 | 4.313117 | 35.10401 | 15.64324 | 28.92867 | 76.13138 |
| ENSMUSG00000052629 | 1.324779613 | 0.000678 | 0.005799 | 32.19374 | 29.99554 | 29.81184 | 39.68068 | 8.776002 | 17.87799 | 17.3572 | 8.700729 |
| ENSMUSG00000052821 | 1.192835954 | 2.17E-07 | 1.07E-05 | 105.6357 | 109.1838 | 95.23226 | 102.6522 | 44.85512 | 62.57295 | 38.57157 | 34.80291 |
| ENSMUSG00000052854 | 1.114146432 | 0.00017 | 0.002 | 443.67 | 777.4844 | 550.6909 | 679.7473 | 279.8569 | 463.7102 | 204.4293 | 184.8905 |
| ENSMUSG00000053007 | 1.011174814 | 0.004102 | 0.02223 | 238.4349 | 226.7663 | 181.3554 | 239.8093 | 98.48624 | 206.7142 | 65.57166 | 69.60583 |
| ENSMUSG00000053153 | 1.196233828 | 0.000103 | 0.00137 | 49.29666 | 38.39429 | 39.74912 | 56.93315 | 21.45245 | 23.46486 | 16.39292 | 19.57664 |
| ENSMUSG00000053166 | 1.527429495 | 1.55E-06 | 5.34E-05 | 403.4278 | 587.9126 | 434.756 | 467.5419 | 137.4907 | 298.3389 | 103.1789 | 118.5474 |
| ENSMUSG00000053182 | 1.059918509 | 0.000134 | 0.001675 | 168.0111 | 129.5807 | 94.40416 | 125.943 | 53.63112 | 89.38993 | 59.78593 | 45.67883 |
| ENSMUSG00000053214 | 1.449263203 | 0.007966 | 0.036131 | 57.3451 | 15.59768 | 37.2648 | 26.74133 | 4.875557 | 18.99536 | 12.53576 | 14.13868 |
| ENSMUSG00000053279 | 1.085402865 | 0.000186 | 0.002139 | 764.6013 | 932.2613 | 644.267 | 835.0195 | 301.3094 | 648.077 | 317.2511 | 230.5693 |
| ENSMUSG00000053297 | -1.167730822 | 0.000155 | 0.001873 | 98.59333 | 81.58786 | 123.3879 | 111.2784 | 241.8276 | 120.6764 | 217.9294 | 353.4671 |
| ENSMUSG00000053303 | -4.571905625 | 1.75E-05 | 0.000354 | 2.012109 | 0 | 1.656213 | 0 | 12.67645 | 5.58687 | 37.60728 | 33.71532 |
| ENSMUSG00000053395 | 2.573273409 | 3.30E-07 | 1.49E-05 | 74.44802 | 39.59411 | 70.38906 | 37.95543 | 10.72622 | 5.58687 | 3.857157 | 17.40146 |
| ENSMUSG00000053559 | -1.104039671 | 1.40E-05 | 0.000296 | 439.6458 | 302.355 | 686.5004 | 445.9763 | 975.1113 | 706.1804 | 1078.075 | 1274.657 |
| ENSMUSG00000053862 | -3.788001849 | 6.97E-11 | 1.34E-08 | 10.06054 | 2.399643 | 20.70267 | 8.626235 | 158.9431 | 49.16446 | 133.0719 | 246.8832 |
| ENSMUSG00000054046 | 1.412585609 | 0.000122 | 0.00156 | 52.31483 | 33.595 | 49.6864 | 41.40593 | 14.62667 | 26.81698 | 11.57147 | 14.13868 |
| ENSMUSG00000054052 | -5.313492612 | 8.64E-09 | 7.57E-07 | 7.04238 | 1.199822 | 12.4216 | 6.038364 | 389.0694 | 32.40385 | 155.2506 | 498.1167 |
| ENSMUSG00000054203 | 2.391721521 | 5.26E-05 | 0.000824 | 18.10898 | 21.59679 | 13.24971 | 18.11509 | 2.925334 | 5.58687 | 2.892867 | 2.175182 |
| ENSMUSG00000054252 | -1.090116463 | 0.000584 | 0.005158 | 258.556 | 178.7734 | 453.8024 | 251.8861 | 579.2161 | 353.0902 | 661.5024 | 842.8831 |
| ENSMUSG00000054457 | 1.476849842 | 0.000266 | 0.00283 | 21.12714 | 23.99643 | 28.98373 | 25.8787 | 10.72622 | 8.938993 | 8.678602 | 7.613138 |
| ENSMUSG00000054474 | -1.376484302 | 5.91E-08 | 3.74E-06 | 372.2401 | 377.9438 | 477.8175 | 404.5704 | 1115.527 | 607.8515 | 932.4676 | 1584.62 |
| ENSMUSG00000054630 | -2.32018316 | 4.17E-05 | 0.000696 | 20.12109 | 1.199822 | 19.04645 | 11.21411 | 46.80534 | 49.16446 | 60.75022 | 104.4087 |
| ENSMUSG00000054763 | -2.393492649 | 0.000524 | 0.004749 | 8.048435 | 1.199822 | 28.98373 | 23.29083 | 61.43201 | 33.52122 | 129.2147 | 103.3212 |
| ENSMUSG00000054988 | 1.216870369 | 6.32E-05 | 0.000953 | 102.6175 | 87.58697 | 53.82693 | 90.57547 | 38.02934 | 46.92971 | 33.75012 | 25.0146 |
| ENSMUSG00000055114 | -2.090302239 | 7.88E-07 | 3.05E-05 | 30.18163 | 11.99822 | 43.06154 | 42.26855 | 83.85957 | 97.21155 | 164.8934 | 202.2919 |
| ENSMUSG00000055137 | -1.797811454 | 1.14E-05 | 0.000254 | 66.39959 | 64.79036 | 107.6539 | 65.55939 | 284.7325 | 93.85942 | 233.358 | 448.0875 |
| ENSMUSG00000055214 | 1.022225378 | 0.000391 | 0.003799 | 53.32088 | 74.38894 | 63.76421 | 81.08661 | 32.17867 | 31.28647 | 46.28588 | 23.927 |
| ENSMUSG00000055312 | -3.628685621 | 5.37E-15 | 2.36E-12 | 14.08476 | 16.7975 | 10.76539 | 15.52722 | 214.5245 | 54.75133 | 166.822 | 267.5474 |
| ENSMUSG00000055368 | 3.98928519 | 0.001687 | 0.011549 | 334.01 | 3.599465 | 10.76539 | 356.2635 | 17.552 | 7.821619 | 13.50005 | 5.437955 |
| ENSMUSG00000055373 | -2.001327643 | 0.000394 | 0.00381 | 183.1019 | 61.1909 | 340.3518 | 169.9368 | 490.481 | 585.504 | 640.288 | 1308.372 |
| ENSMUSG00000055523 | -1.805989445 | 0.000772 | 0.006413 | 6.036326 | 13.19804 | 42.23344 | 12.07673 | 83.85957 | 67.04245 | 57.85735 | 51.11678 |
| ENSMUSG00000056035 | 3.397254334 | 0.004121 | 0.022315 | 83.50251 | 1.199822 | 1.656213 | 75.91087 | 0.975111 | 2.234748 | 6.750024 | 5.437955 |
| ENSMUSG00000056222 | 1.102332392 | 0.005079 | 0.026022 | 132.7992 | 51.59233 | 62.9361 | 122.4925 | 41.92979 | 51.39921 | 54.00019 | 25.0146 |
| ENSMUSG00000056366 | -2.092426007 | 0.012066 | 0.049182 | 2.012109 | 3.599465 | 11.59349 | 1.725247 | 28.27823 | 4.469496 | 15.42863 | 33.71532 |
| ENSMUSG00000056487 | -4.687453614 | 5.04E-07 | 2.12E-05 | 0 | 0 | 3.312427 | 0.862623 | 30.22845 | 10.05637 | 30.85725 | 46.76642 |
| ENSMUSG00000056643 | -1.925204624 | 8.95E-06 | 0.000211 | 16.09687 | 9.598572 | 15.73403 | 9.488858 | 33.15378 | 31.28647 | 55.92877 | 73.95619 |
| ENSMUSG00000056665 | -1.219733565 | 0.00155 | 0.010786 | 56.33904 | 55.19179 | 87.7793 | 49.16954 | 167.7191 | 54.75133 | 160.072 | 196.854 |
| ENSMUSG00000056972 | 2.331402018 | 0.000812 | 0.00665 | 43.26034 | 17.99732 | 5.796746 | 37.09281 | 3.900445 | 5.58687 | 5.785735 | 5.437955 |
| ENSMUSG00000057103 | -3.1100892 | 2.69E-15 | 1.31E-12 | 94.56911 | 69.58965 | 115.1068 | 100.9269 | 1002.414 | 308.3953 | 560.252 | 1417.131 |
| ENSMUSG00000057123 | -1.089738258 | 0.000157 | 0.001887 | 77.46619 | 51.59233 | 84.46688 | 69.00988 | 145.2916 | 116.2069 | 112.8218 | 229.4817 |
| ENSMUSG00000057182 | 2.049990887 | 7.33E-06 | 0.000181 | 138.8355 | 57.59143 | 63.76421 | 144.9207 | 27.30312 | 35.75597 | 25.07152 | 9.78832 |
| ENSMUSG00000057228 | -3.872240707 | 1.95E-06 | 6.32E-05 | 111.672 | 35.99465 | 496.0359 | 180.2883 | 2428.027 | 979.9371 | 3806.049 | 4861.532 |
| ENSMUSG00000057425 | -2.926605714 | 0.010727 | 0.044955 | 19.11503 | 0 | 53.82693 | 18.97772 | 174.5449 | 39.10809 | 124.3933 | 363.2554 |
| ENSMUSG00000057614 | -1.164585669 | 3.21E-05 | 0.000563 | 727.3773 | 602.3104 | 1320.002 | 837.6074 | 1783.479 | 1173.243 | 2382.759 | 2481.883 |
| ENSMUSG00000057933 | -3.420592995 | 2.91E-05 | 0.000526 | 68.4117 | 26.39607 | 321.3054 | 102.6522 | 1669.391 | 460.3581 | 1079.04 | 2352.46 |
| ENSMUSG00000058258 | -1.366729234 | 2.35E-11 | 4.98E-09 | 181.0898 | 169.1748 | 266.6503 | 184.6014 | 527.5352 | 375.4377 | 613.2879 | 554.6715 |
| ENSMUSG00000058488 | -2.30897626 | 4.35E-05 | 0.000719 | 315.9011 | 191.9714 | 582.9871 | 412.334 | 1722.047 | 584.3867 | 1886.15 | 3259.51 |
| ENSMUSG00000058620 | -2.256166946 | 2.14E-11 | 4.59E-09 | 30.18163 | 27.5959 | 57.13936 | 50.89479 | 216.4747 | 115.0895 | 192.8578 | 274.073 |
| ENSMUSG00000058812 | -1.220546011 | 0.006065 | 0.029689 | 13.07871 | 5.999108 | 10.76539 | 9.488858 | 23.40267 | 14.52586 | 25.07152 | 29.36496 |
| ENSMUSG00000058921 | -2.632876974 | 0.001816 | 0.01224 | 3.018163 | 1.199822 | 11.59349 | 1.725247 | 19.50223 | 8.938993 | 22.17865 | 60.9051 |
| ENSMUSG00000058952 | -1.322179197 | 3.85E-07 | 1.70E-05 | 208.2533 | 173.9741 | 165.6213 | 153.547 | 535.3361 | 260.3482 | 358.7156 | 598.1751 |
| ENSMUSG00000059146 | 1.085230526 | 4.19E-08 | 2.79E-06 | 587.5357 | 657.5022 | 541.5817 | 696.9998 | 302.2845 | 406.7242 | 229.5008 | 232.7445 |
| ENSMUSG00000059213 | -2.653455874 | 0.001419 | 0.010174 | 38.23007 | 3.599465 | 87.7793 | 32.77969 | 142.3663 | 134.0849 | 303.7511 | 444.8248 |

| | | | | | | | | | | | |
|---|---|---|---|---|---|---|---|---|---|---|---|
| ENSMUSG00000059412 | -3.037393265 | 0.000243 | 0.002648 | 669.0261 | 183.5727 | 2863.593 | 838.47 | 9012.954 | 3364.413 | 10712.29 | 14309.44 |
| ENSMUSG00000059481 | 3.009092296 | 0.003291 | 0.01901 | 97.58727 | 4.799286 | 2.48432 | 78.49874 | 4.875557 | 2.234748 | 4.821446 | 10.87591 |
| ENSMUSG00000059857 | 1.455278819 | 5.44E-05 | 0.000845 | 815.9101 | 1163.827 | 704.7187 | 1101.57 | 302.2845 | 670.4245 | 207.3222 | 201.2044 |
| ENSMUSG00000060317 | -4.490562195 | 0.000742 | 0.006219 | 0 | 0 | 14.90592 | 20.70296 | 253.5289 | 84.92043 | 181.2864 | 287.124 |
| ENSMUSG00000060371 | 2.587254458 | 3.89E-06 | 0.000111 | 161.9748 | 39.59411 | 37.2648 | 94.88858 | 12.67645 | 17.87799 | 6.750024 | 18.48905 |
| ENSMUSG00000060459 | -3.095046969 | 0.00012 | 0.001543 | 120.7265 | 63.59054 | 568.9093 | 141.4703 | 1377.832 | 586.6214 | 2700.01 | 2985.438 |
| ENSMUSG00000060586 | -1.376591223 | 1.17E-08 | 9.73E-07 | 65.39353 | 98.38537 | 96.88848 | 113.0037 | 314.961 | 170.9582 | 249.7509 | 234.9197 |
| ENSMUSG00000060716 | -1.002987508 | 0.000736 | 0.006174 | 215.2956 | 165.5754 | 293.9779 | 223.4195 | 512.9085 | 214.5358 | 517.8233 | 556.8466 |
| ENSMUSG00000060807 | -5.350673139 | 0.001465 | 0.0104 | 6.036326 | 0 | 388.382 | 22.42821 | 3957.977 | 551.9828 | 5103.983 | 7404.32 |
| ENSMUSG00000060962 | 1.631870058 | 6.42E-06 | 0.000163 | 152.9203 | 261.5611 | 152.3716 | 215.6559 | 77.03379 | 65.92507 | 84.85745 | 23.927 |
| ENSMUSG00000060969 | -2.348342126 | 0.005215 | 0.026488 | 183.1019 | 23.99643 | 450.49 | 184.6014 | 631.8721 | 404.4894 | 1668.22 | 1586.795 |
| ENSMUSG00000060985 | -2.058616549 | 6.83E-06 | 0.000172 | 23.13925 | 15.59768 | 36.43669 | 16.38985 | 39.00445 | 91.62468 | 101.2504 | 152.2628 |
| ENSMUSG00000061013 | 1.374245746 | 3.95E-05 | 0.000667 | 650.9172 | 470.33 | 217.792 | 567.6063 | 177.4703 | 240.2354 | 204.4293 | 113.1095 |
| ENSMUSG00000061080 | 1.251108616 | 4.73E-09 | 4.59E-07 | 568.4207 | 595.1115 | 458.7711 | 686.6483 | 200.8729 | 350.8555 | 208.2865 | 210.9927 |
| ENSMUSG00000061186 | 1.829898636 | 1.18E-08 | 9.75E-07 | 433.6094 | 343.149 | 332.0708 | 375.2412 | 99.46135 | 183.2494 | 81.96458 | 53.29196 |
| ENSMUSG00000061527 | 1.123299304 | 0.001975 | 0.013015 | 231.3925 | 93.58608 | 197.0894 | 141.4703 | 115.0631 | 74.86406 | 73.28598 | 41.32846 |
| ENSMUSG00000061601 | 1.032003459 | 5.11E-05 | 0.000804 | 527.1725 | 500.3256 | 348.6329 | 578.8204 | 202.8232 | 363.1466 | 220.8222 | 169.6642 |
| ENSMUSG00000061742 | -2.675125951 | 0.003258 | 0.018873 | 37.22401 | 10.79839 | 173.9024 | 46.58167 | 366.6419 | 121.7938 | 317.2511 | 912.4889 |
| ENSMUSG00000061808 | -3.780920791 | 1.27E-06 | 4.53E-05 | 90.54489 | 14.39786 | 115.9349 | 104.3774 | 1415.862 | 244.7049 | 868.8245 | 1947.876 |
| ENSMUSG00000061947 | -3.448548076 | 0.00194 | 0.012845 | 13.07871 | 2.399643 | 69.56096 | 5.175741 | 170.6445 | 39.10809 | 354.8584 | 424.1605 |
| ENSMUSG00000062296 | 1.395552985 | 2.76E-06 | 8.50E-05 | 449.7063 | 151.1775 | 275.7595 | 306.2313 | 106.2871 | 120.6764 | 99.32178 | 123.9854 |
| ENSMUSG00000062319 | 2.190801102 | 0.010026 | 0.042676 | 27.16347 | 65.99018 | 9.109173 | 34.50494 | 3.900445 | 21.23011 | 3.857157 | 1.087591 |
| ENSMUSG00000062380 | 2.989050039 | 0.002798 | 0.016808 | 609.6689 | 51.59233 | 31.46805 | 925.595 | 57.53157 | 53.63396 | 49.17875 | 43.50364 |
| ENSMUSG00000062393 | 2.386785565 | 1.94E-06 | 6.31E-05 | 141.8537 | 35.99465 | 43.88965 | 129.3935 | 17.552 | 16.76061 | 14.46434 | 18.48905 |
| ENSMUSG00000062410 | -3.190583575 | 5.36E-06 | 0.000142 | 2.012109 | 3.599465 | 5.796746 | 4.313117 | 18.52711 | 13.40849 | 39.53586 | 73.95619 |
| ENSMUSG00000062480 | -2.017730679 | 1.41E-07 | 7.53E-06 | 54.32694 | 27.5959 | 42.23344 | 29.3292 | 219.4 | 90.5073 | 99.32178 | 213.1679 |
| ENSMUSG00000062609 | -2.64689859 | 3.04E-06 | 9.16E-05 | 196.1806 | 69.58965 | 308.8838 | 234.6336 | 1547.502 | 486.0577 | 1121.468 | 1918.511 |
| ENSMUSG00000062713 | -2.419322509 | 0.010474 | 0.044126 | 68.4117 | 5.999108 | 218.6202 | 54.34528 | 269.1307 | 187.7188 | 802.2886 | 601.4379 |
| ENSMUSG00000063063 | 1.41284326 | 0.004362 | 0.023299 | 80.48435 | 27.5959 | 35.60859 | 107.8279 | 21.45245 | 35.75597 | 20.25007 | 17.40146 |
| ENSMUSG00000063130 | -1.350778963 | 0.011185 | 0.046441 | 18.10898 | 13.19804 | 56.31125 | 15.52722 | 58.50668 | 40.22547 | 57.85735 | 108.7591 |
| ENSMUSG00000063297 | 1.408798341 | 1.35E-05 | 0.000289 | 302.8224 | 556.7172 | 358.5702 | 450.2895 | 131.64 | 261.4655 | 149.4648 | 85.9197 |
| ENSMUSG00000063428 | 1.723714683 | 6.48E-16 | 3.86E-13 | 182.0958 | 196.7707 | 179.6991 | 218.2437 | 38.02934 | 71.51194 | 67.50024 | 58.72992 |
| ENSMUSG00000063590 | -3.156670398 | 3.22E-06 | 9.58E-05 | 1.006054 | 5.999108 | 3.312427 | 7.763611 | 42.9049 | 20.11273 | 22.17865 | 76.13138 |
| ENSMUSG00000063611 | 1.061967366 | 9.86E-05 | 0.001323 | 141.8537 | 116.3827 | 93.57605 | 131.9814 | 33.15378 | 69.27719 | 66.53595 | 63.08028 |
| ENSMUSG00000063683 | -2.12184324 | 8.47E-06 | 0.000202 | 74.44802 | 22.79661 | 178.0429 | 78.49874 | 370.5423 | 251.4092 | 352.9298 | 570.9853 |
| ENSMUSG00000063730 | -3.863385664 | 3.31E-06 | 9.75E-05 | 2.012109 | 0 | 9.109173 | 0.862623 | 29.25334 | 18.99536 | 67.50024 | 66.34306 |
| ENSMUSG00000063796 | -1.854957495 | 2.94E-06 | 8.88E-05 | 148.896 | 91.18644 | 279.0719 | 163.8985 | 772.2882 | 309.5126 | 421.3944 | 972.3064 |
| ENSMUSG00000063903 | -3.170319749 | 7.21E-05 | 0.001057 | 21.12714 | 20.39697 | 151.5435 | 26.74133 | 488.5308 | 203.3621 | 492.7518 | 799.3794 |
| ENSMUSG00000064036 | -1.50948181 | 0.002029 | 0.013272 | 4.024217 | 14.39786 | 9.93728 | 6.900988 | 19.50223 | 34.6386 | 20.25007 | 25.0146 |
| ENSMUSG00000064125 | 1.561127346 | 2.16E-13 | 7.67E-11 | 340.0464 | 415.1383 | 280.7281 | 337.2858 | 129.6898 | 148.6108 | 106.0718 | 80.48174 |
| ENSMUSG00000064225 | -1.997359981 | 9.45E-06 | 0.000217 | 46.2785 | 15.59768 | 61.27989 | 38.81806 | 175.52 | 79.33356 | 128.2505 | 267.5474 |
| ENSMUSG00000064294 | 1.522217548 | 1.05E-07 | 5.87E-06 | 1543.287 | 1823.729 | 1065.773 | 1759.752 | 555.8134 | 836.9132 | 469.6088 | 293.6496 |
| ENSMUSG00000066058 | -1.226423849 | 0.000237 | 0.002598 | 554.3359 | 418.7377 | 980.4783 | 533.9639 | 1231.566 | 728.5279 | 1716.435 | 2146.905 |
| ENSMUSG00000066072 | -4.751643353 | 2.72E-07 | 1.28E-05 | 11.0666 | 1.199822 | 20.70267 | 6.900988 | 246.7032 | 53.63396 | 166.822 | 616.6641 |
| ENSMUSG00000066097 | -3.038831453 | 3.99E-10 | 5.67E-08 | 33.19979 | 13.19804 | 41.40533 | 34.50494 | 235.9769 | 81.56831 | 237.2151 | 456.7883 |
| ENSMUSG00000066361 | 2.128486888 | 0.008861 | 0.039057 | 16.09687 | 34.79482 | 12.4216 | 24.15346 | 5.850668 | 12.29112 | 1.928578 | 0 |
| ENSMUSG00000067144 | -5.674314347 | 1.68E-05 | 0.00034 | 0 | 1.199822 | 0 | 0.862623 | 15.60178 | 4.469496 | 15.42863 | 63.08028 |
| ENSMUSG00000067206 | -1.87011629 | 0.00242 | 0.01505 | 7.04238 | 3.599465 | 18.21835 | 10.35148 | 37.05423 | 14.52586 | 27.96439 | 66.34306 |
| ENSMUSG00000067215 | 1.540256968 | 4.20E-05 | 0.000699 | 46.2785 | 47.99286 | 28.98373 | 36.23019 | 9.751113 | 12.29112 | 19.28578 | 13.05109 |
| ENSMUSG00000067578 | -2.614909031 | 3.72E-12 | 9.16E-10 | 22.1332 | 7.198929 | 9.93728 | 18.11509 | 74.10846 | 100.5637 | 111.8575 | 67.43065 |
| ENSMUSG00000067656 | -2.741472047 | 1.71E-06 | 5.78E-05 | 6.036326 | 4.799286 | 18.21835 | 13.80198 | 44.85512 | 32.40385 | 83.89316 | 130.5109 |
| ENSMUSG00000068227 | -1.875004813 | 0.008153 | 0.036789 | 10.06054 | 3.599465 | 17.39024 | 15.52722 | 48.75557 | 5.58687 | 71.3574 | 46.76642 |
| ENSMUSG00000068263 | -1.024737415 | 0.006561 | 0.031503 | 30.18163 | 29.99554 | 56.31125 | 44.85642 | 93.61069 | 42.46022 | 73.28598 | 120.7226 |
| ENSMUSG00000068327 | 3.882013675 | 0.004776 | 0.024921 | 30.18163 | 1.199822 | 0 | 58.6584 | 1.950223 | 1.117374 | 1.928578 | 1.087591 |
| ENSMUSG00000068348 | -2.198325014 | 0.004897 | 0.025356 | 4.024217 | 4.799286 | 6.624853 | 0.862623 | 15.60178 | 5.58687 | 15.42863 | 38.06569 |
| ENSMUSG00000068407 | 1.295371182 | 2.03E-08 | 1.54E-06 | 156.9445 | 265.1606 | 213.6515 | 188.9145 | 63.38224 | 102.7984 | 93.53605 | 76.13138 |
| ENSMUSG00000068587 | -1.360304079 | 1.30E-05 | 0.000281 | 46.2785 | 27.5959 | 54.65504 | 38.81806 | 74.10846 | 88.27255 | 124.3933 | 145.7372 |
| ENSMUSG00000068615 | 1.420047021 | 0.001531 | 0.010697 | 56.33904 | 25.19625 | 23.18699 | 48.30692 | 10.72622 | 20.11273 | 12.53576 | 14.13868 |
| ENSMUSG00000068748 | 1.428052825 | 0.000194 | 0.002211 | 401.4157 | 381.5432 | 230.2136 | 386.4553 | 87.76002 | 261.4655 | 87.75031 | 83.74451 |
| ENSMUSG00000068874 | -1.776143 | 1.80E-10 | 3.00E-08 | 781.7042 | 539.9197 | 977.9939 | 703.0381 | 2774.192 | 1254.811 | 3188.904 | 3070.27 |
| ENSMUSG00000069072 | 1.227200148 | 0.01099 | 0.045808 | 55.33299 | 23.99643 | 22.35888 | 65.55939 | 20.47734 | 24.58223 | 12.53576 | 14.13868 |
| ENSMUSG00000069372 | 1.223913652 | 2.67E-12 | 6.94E-10 | 321.9374 | 370.7449 | 264.9941 | 377.8291 | 139.4409 | 174.3104 | 135.0005 | 122.8978 |
| ENSMUSG00000069456 | -3.667469728 | 8.75E-05 | 0.001214 | 15.09082 | 2.399643 | 60.45178 | 13.80198 | 336.4134 | 67.04245 | 277.7153 | 490.5036 |
| ENSMUSG00000069805 | -3.738094259 | 4.71E-07 | 2.00E-05 | 188.1322 | 59.99108 | 535.785 | 309.6818 | 4715.638 | 1217.938 | 2555.366 | 6112.262 |

| | | | | | | | | | | | |
|---|---|---|---|---|---|---|---|---|---|---|---|
| ENSMUSG00000070023 | -3.681372842 | 5.68E-06 | 0.000148 | 2.012109 | 2.399643 | 1.656213 | 1.725247 | 26.32801 | 8.938993 | 13.50005 | 50.02919 |
| ENSMUSG00000070291 | 1.319646571 | 4.62E-05 | 0.000749 | 78.47224 | 53.99197 | 50.5145 | 43.13117 | 20.47734 | 27.93435 | 17.3572 | 25.0146 |
| ENSMUSG00000070532 | 1.627200583 | 0.003174 | 0.018523 | 20.12109 | 14.39786 | 10.76539 | 18.97772 | 3.900445 | 5.58687 | 3.857157 | 7.613138 |
| ENSMUSG00000070985 | -3.037333041 | 1.11E-05 | 0.000247 | 1.006054 | 4.799286 | 7.45296 | 1.725247 | 28.27823 | 17.87799 | 21.21436 | 56.55474 |
| ENSMUSG00000071068 | 1.027060294 | 0.009943 | 0.04239 | 36.21796 | 29.99554 | 50.5145 | 31.91707 | 12.67645 | 30.1691 | 13.50005 | 17.40146 |
| ENSMUSG00000071335 | -2.908701708 | 2.05E-07 | 1.02E-05 | 9.054489 | 3.599465 | 23.18699 | 11.21411 | 93.61069 | 33.52122 | 94.50034 | 138.1241 |
| ENSMUSG00000071567 | 1.52820265 | 2.38E-07 | 1.17E-05 | 98.59333 | 104.3845 | 109.3101 | 137.1571 | 32.17867 | 62.57295 | 35.6787 | 26.10219 |
| ENSMUSG00000071656 | 1.478725872 | 1.89E-05 | 0.000375 | 264.5923 | 343.149 | 207.0267 | 276.0395 | 121.8889 | 154.1976 | 68.46453 | 46.76642 |
| ENSMUSG00000071658 | 1.700257742 | 1.21E-08 | 9.91E-07 | 196.1806 | 145.1784 | 108.482 | 255.3366 | 47.78045 | 65.92507 | 47.25017 | 56.55474 |
| ENSMUSG00000071679 | 1.668656087 | 0.000209 | 0.002351 | 39.23612 | 46.79304 | 30.63995 | 30.19182 | 5.850668 | 21.23011 | 10.60718 | 8.700729 |
| ENSMUSG00000072591 | 1.86236247 | 4.20E-10 | 5.87E-08 | 92.557 | 133.1802 | 97.71658 | 102.6522 | 26.32801 | 43.57759 | 17.3572 | 30.45255 |
| ENSMUSG00000072664 | -4.178381539 | 0.000245 | 0.002668 | 0 | 0 | 3.312427 | 1.725247 | 13.65156 | 2.234748 | 29.89296 | 51.11678 |
| ENSMUSG00000072812 | 1.59992123 | 0.001192 | 0.008927 | 165.999 | 237.5647 | 185.4959 | 238.0841 | 62.40712 | 145.2586 | 49.17875 | 16.31387 |
| ENSMUSG00000072844 | 1.162437806 | 0.001319 | 0.009624 | 309.8647 | 236.3648 | 146.5749 | 140.6076 | 78.0089 | 157.5497 | 69.42882 | 67.43065 |
| ENSMUSG00000072941 | -3.127979378 | 6.28E-06 | 0.00016 | 184.1079 | 34.79482 | 244.2915 | 127.6683 | 1405.135 | 398.9026 | 1158.111 | 2207.81 |
| ENSMUSG00000072944 | 2.32853931 | 1.59E-11 | 3.55E-09 | 194.1685 | 112.7832 | 85.29498 | 115.5915 | 22.42756 | 36.87335 | 19.28578 | 22.83941 |
| ENSMUSG00000073209 | 1.141097131 | 3.96E-06 | 0.000112 | 277.671 | 422.3372 | 255.8849 | 350.2251 | 161.8685 | 203.3621 | 114.7504 | 112.0219 |
| ENSMUSG00000073409 | -1.04859875 | 0.004018 | 0.021898 | 37.22401 | 49.19268 | 94.40416 | 45.71905 | 84.83468 | 89.38993 | 123.429 | 172.927 |
| ENSMUSG00000073418 | 2.09900742 | 0.000954 | 0.00757 | 437.6336 | 58.79126 | 56.31125 | 95.75121 | 25.35289 | 53.63396 | 27.96439 | 44.59123 |
| ENSMUSG00000073489 | 1.279605977 | 0.0044 | 0.023438 | 443.67 | 563.9161 | 380.9291 | 459.7783 | 241.8276 | 363.1466 | 94.50034 | 61.99269 |
| ENSMUSG00000073680 | -1.483510643 | 0.008869 | 0.039084 | 6.036326 | 3.599465 | 15.73403 | 10.35148 | 20.47734 | 36.87335 | 13.50005 | 31.54014 |
| ENSMUSG00000074028 | -4.309560554 | 3.26E-05 | 0.00057 | 3.018163 | 3.599465 | 43.88965 | 6.038364 | 190.1467 | 67.04245 | 272.8938 | 599.2627 |
| ENSMUSG00000074179 | -2.910321179 | 1.30E-05 | 0.000281 | 36.21796 | 40.79393 | 119.2474 | 39.68068 | 589.9423 | 119.559 | 318.2154 | 749.3503 |
| ENSMUSG00000074183 | -2.17932283 | 0.005702 | 0.028351 | 10.06054 | 2.399643 | 2.48432 | 1.725247 | 30.22845 | 7.821619 | 9.642892 | 27.18978 |
| ENSMUSG00000074199 | 1.729629529 | 0.001211 | 0.009018 | 53.32088 | 55.19179 | 38.09291 | 37.09281 | 15.60178 | 27.93435 | 8.678602 | 3.262773 |
| ENSMUSG00000074261 | -4.143633507 | 6.57E-08 | 4.03E-06 | 3.018163 | 0 | 15.73403 | 4.313117 | 149.192 | 31.28647 | 75.21455 | 164.2263 |
| ENSMUSG00000074281 | 1.215057003 | 0.00153 | 0.010695 | 50.30272 | 61.1909 | 33.12427 | 45.71905 | 28.27823 | 26.81698 | 13.50005 | 13.05109 |
| ENSMUSG00000074336 | 4.131162518 | 0.002461 | 0.015244 | 97.58727 | 2.399643 | 4.140533 | 25.8787 | 0 | 0 | 0.964289 | 6.525547 |
| ENSMUSG00000074415 | 1.274344149 | 0.001856 | 0.012441 | 3906.509 | 5045.25 | 3573.28 | 4470.115 | 1488.02 | 3337.596 | 1194.754 | 1006.022 |
| ENSMUSG00000074445 | 1.559275716 | 0.010642 | 0.04467 | 38.23007 | 28.79572 | 107.6539 | 28.46658 | 26.32801 | 25.6996 | 10.60718 | 6.525547 |
| ENSMUSG00000074604 | 1.524328916 | 1.67E-05 | 0.000339 | 66.39959 | 107.9839 | 62.108 | 81.08661 | 35.10401 | 35.75597 | 26.03581 | 13.05109 |
| ENSMUSG00000074607 | 1.428659349 | 2.08E-05 | 0.000406 | 116.7023 | 63.59054 | 61.27989 | 90.57547 | 39.97956 | 35.75597 | 27.96439 | 19.57664 |
| ENSMUSG00000074628 | -1.826232478 | 0.006984 | 0.032918 | 7.04238 | 7.198929 | 27.32752 | 6.900988 | 40.95467 | 10.05637 | 62.6788 | 59.81751 |
| ENSMUSG00000074639 | -2.733147837 | 1.08E-05 | 0.000243 | 4.024217 | 1.199822 | 15.73403 | 6.038364 | 35.10401 | 33.52122 | 40.50014 | 76.13138 |
| ENSMUSG00000074785 | 1.009110062 | 8.97E-08 | 5.17E-06 | 363.1856 | 423.537 | 352.7734 | 457.1905 | 152.1174 | 265.935 | 187.0721 | 189.2408 |
| ENSMUSG00000074796 | -1.221065319 | 0.010043 | 0.042724 | 83.50251 | 73.18911 | 202.058 | 56.07053 | 274.0063 | 89.38993 | 234.3223 | 370.8686 |
| ENSMUSG00000074892 | -1.012304913 | 8.51E-05 | 0.00119 | 119.7205 | 164.3756 | 141.6062 | 102.6522 | 278.8818 | 175.4277 | 235.2866 | 374.1313 |
| ENSMUSG00000074919 | -4.596490607 | 3.81E-05 | 0.000647 | 0 | 0 | 3.312427 | 0 | 21.45245 | 6.704245 | 19.28578 | 40.24087 |
| ENSMUSG00000074923 | -2.106875726 | 0.00022 | 0.002437 | 34.20585 | 11.99822 | 94.40416 | 23.29083 | 154.0676 | 77.09881 | 244.9294 | 233.8321 |
| ENSMUSG00000074934 | -1.798000154 | 6.79E-05 | 0.001007 | 17.10292 | 8.398751 | 7.45296 | 6.900988 | 25.35289 | 30.1691 | 48.21446 | 33.71532 |
| ENSMUSG00000074968 | 1.19786417 | 0.000792 | 0.00653 | 39.23612 | 86.38715 | 75.3577 | 86.26235 | 22.42756 | 45.81234 | 27.0001 | 30.45255 |
| ENSMUSG00000074971 | 1.03518668 | 2.43E-05 | 0.000457 | 3679.141 | 3891.021 | 2376.666 | 4535.674 | 1748.375 | 2175.527 | 2081.9 | 1060.401 |
| ENSMUSG00000075025 | -3.29289393 | 0.002738 | 0.016518 | 10.06054 | 0 | 34.78048 | 7.763611 | 94.5858 | 30.1691 | 157.1791 | 237.0949 |
| ENSMUSG00000075033 | -1.374320017 | 0.000238 | 0.002599 | 234.4107 | 111.5834 | 333.727 | 213.068 | 517.7841 | 262.5829 | 779.1456 | 758.051 |
| ENSMUSG00000075044 | -4.036426996 | 3.01E-19 | 3.47E-16 | 10.06054 | 4.799286 | 19.04645 | 10.35148 | 174.5449 | 92.74205 | 178.3935 | 295.8248 |
| ENSMUSG00000075267 | -1.068308573 | 0.002279 | 0.01441 | 11.0666 | 19.19714 | 14.07781 | 13.80198 | 25.35289 | 30.1691 | 32.78583 | 32.62773 |
| ENSMUSG00000075304 | 1.213160885 | 5.24E-13 | 1.70E-10 | 382.3007 | 471.5299 | 414.8814 | 376.9665 | 172.5947 | 206.7142 | 131.1433 | 200.1168 |
| ENSMUSG00000075316 | 1.456756039 | 0.001146 | 0.008674 | 249.5015 | 326.3515 | 159.8246 | 295.0172 | 102.3867 | 185.4841 | 44.3573 | 43.50364 |
| ENSMUSG00000076441 | -3.085141731 | 1.81E-05 | 0.000363 | 516.1059 | 139.1793 | 1046.727 | 640.0666 | 7672.176 | 1477.169 | 3443.477 | 7287.948 |
| ENSMUSG00000078137 | -1.909883874 | 0.000218 | 0.002419 | 5.030272 | 4.799286 | 11.59349 | 8.626235 | 35.10401 | 13.40849 | 33.75012 | 32.62773 |
| ENSMUSG00000078349 | -1.176532742 | 0.000274 | 0.002891 | 59.35721 | 38.39429 | 55.48314 | 31.05445 | 84.83468 | 69.27719 | 122.4647 | 140.2993 |
| ENSMUSG00000078350 | -1.091015768 | 3.97E-08 | 2.67E-06 | 329.9858 | 317.9527 | 441.3808 | 386.4553 | 684.5281 | 567.626 | 897.7532 | 997.321 |
| ENSMUSG00000078439 | -1.527791596 | 5.63E-05 | 0.000866 | 101.6115 | 69.58965 | 192.1207 | 128.5309 | 308.1352 | 185.4841 | 342.3227 | 586.2116 |
| ENSMUSG00000078607 | 1.107288742 | 2.39E-07 | 1.17E-05 | 204.229 | 226.7663 | 183.8397 | 242.3972 | 116.0382 | 126.2633 | 84.85745 | 70.69342 |
| ENSMUSG00000078650 | -3.253300152 | 5.91E-06 | 0.000153 | 140.8476 | 11.99822 | 78.67013 | 160.448 | 1343.703 | 496.1141 | 810.0029 | 1091.941 |
| ENSMUSG00000078937 | -2.324285216 | 5.55E-08 | 3.55E-06 | 26.15741 | 20.39697 | 25.67131 | 20.70296 | 109.2125 | 40.22547 | 145.6077 | 170.7518 |
| ENSMUSG00000078949 | -2.840374991 | 8.75E-05 | 0.001214 | 11.0666 | 2.399643 | 52.99882 | 18.97772 | 98.48624 | 49.16446 | 194.7864 | 275.1605 |
| ENSMUSG00000079000 | -1.737244531 | 0.00257 | 0.015735 | 9.054489 | 3.599465 | 11.59349 | 4.313117 | 21.45245 | 12.29112 | 38.57157 | 23.927 |
| ENSMUSG00000079103 | -2.822612692 | 2.86E-05 | 0.000519 | 5.030272 | 3.599465 | 5.796746 | 3.450494 | 19.50223 | 10.05637 | 35.6787 | 61.99269 |
| ENSMUSG00000079262 | -4.638955917 | 3.34E-05 | 0.000581 | 11.0666 | 0 | 30.63995 | 6.900988 | 254.5041 | 54.75133 | 284.4653 | 625.3649 |
| ENSMUSG00000079355 | 1.116166395 | 6.39E-05 | 0.000959 | 153.9263 | 253.1623 | 183.8397 | 167.349 | 76.05868 | 127.3806 | 90.64318 | 55.46715 |
| ENSMUSG00000079471 | -2.800639693 | 0.000508 | 0.004643 | 3.018163 | 1.199822 | 26.49941 | 6.038364 | 40.95467 | 24.58223 | 72.32169 | 122.8978 |
| ENSMUSG00000079685 | -1.2239439 | 0.001188 | 0.008902 | 16.09687 | 15.59768 | 16.56213 | 8.626235 | 25.35289 | 29.05173 | 37.60728 | 40.24087 |
| ENSMUSG00000079941 | -1.00204387 | 0.003159 | 0.018463 | 15.09082 | 16.7975 | 21.53077 | 14.6646 | 40.95467 | 26.81698 | 30.85725 | 38.06569 |

| | | | | | | | | | | | |
|---|---|---|---|---|---|---|---|---|---|---|---|
| ENSMUSG00000080115 | -2.076249156 | 2.94E-10 | 4.41E-08 | 12.07265 | 27.5959 | 14.90592 | 13.80198 | 60.4569 | 68.15982 | 66.53595 | 88.09488 |
| ENSMUSG00000080893 | 2.066561877 | 0.000129 | 0.001623 | 18.10898 | 93.58608 | 109.3101 | 96.61383 | 18.52711 | 10.05637 | 22.17865 | 25.0146 |
| ENSMUSG00000081044 | -3.036668775 | 1.86E-05 | 0.000371 | 3.018163 | 1.199822 | 10.76539 | 7.763611 | 22.42756 | 103.9158 | 38.57157 | 27.18978 |
| ENSMUSG00000082152 | -1.699251457 | 0.008188 | 0.036914 | 2.012109 | 7.198929 | 19.87456 | 15.52722 | 42.9049 | 15.64324 | 32.78583 | 55.46715 |
| ENSMUSG00000084085 | 1.104553546 | 0.010674 | 0.044781 | 27.16347 | 35.99465 | 20.70267 | 39.68068 | 13.65156 | 23.46486 | 9.642892 | 10.87591 |
| ENSMUSG00000084128 | -1.622346723 | 3.03E-05 | 0.00054 | 217.3077 | 128.3809 | 427.303 | 156.9975 | 723.5326 | 355.325 | 824.4672 | 963.6057 |
| ENSMUSG00000084854 | -3.353048086 | 4.09E-09 | 4.13E-07 | 7.04238 | 4.799286 | 13.24971 | 7.763611 | 118.9636 | 26.81698 | 58.82164 | 135.9489 |
| ENSMUSG00000084965 | 5.157604895 | 0.003271 | 0.018918 | 324.9556 | 0 | 1.656213 | 34.50494 | 3.900445 | 1.117374 | 2.892867 | 2.175182 |
| ENSMUSG00000085203 | -2.341030283 | 0.000144 | 0.001766 | 10.06054 | 8.398751 | 3.312427 | 3.450494 | 24.37778 | 12.29112 | 49.17875 | 39.15328 |
| ENSMUSG00000085229 | 1.597961286 | 0.001798 | 0.012158 | 36.21796 | 27.5959 | 22.35888 | 31.05445 | 6.825779 | 20.11273 | 7.714313 | 4.350364 |
| ENSMUSG00000085251 | -2.816941244 | 0.00068 | 0.005806 | 4.024217 | 0 | 5.796746 | 0.862623 | 21.45245 | 6.704245 | 21.21436 | 28.27737 |
| ENSMUSG00000085412 | 1.378512585 | 0.002516 | 0.015496 | 68.4117 | 63.59054 | 48.85829 | 79.36136 | 22.42756 | 12.29112 | 14.46434 | 51.11678 |
| ENSMUSG00000085541 | -1.52269888 | 0.001089 | 0.008336 | 56.33904 | 49.19268 | 154.0278 | 67.28463 | 278.8818 | 115.0895 | 145.6077 | 402.4087 |
| ENSMUSG00000085603 | 1.423262073 | 6.33E-05 | 0.000954 | 68.4117 | 63.59054 | 39.74912 | 57.79577 | 20.47734 | 26.81698 | 27.0001 | 10.87591 |
| ENSMUSG00000085663 | 1.102609509 | 0.00812 | 0.03667 | 46.2785 | 40.79393 | 44.71776 | 56.07053 | 11.70134 | 39.10809 | 24.10723 | 13.05109 |
| ENSMUSG00000085666 | 5.695206587 | 8.15E-08 | 4.80E-06 | 4.024217 | 255.562 | 310.54 | 281.2153 | 2.925334 | 3.352122 | 5.785735 | 4.350364 |
| ENSMUSG00000085689 | -2.619825151 | 1.85E-08 | 1.42E-06 | 8.048435 | 3.599465 | 8.281066 | 4.313117 | 33.15378 | 25.6996 | 45.32159 | 46.76642 |
| ENSMUSG00000085772 | -2.198496132 | 0.001704 | 0.011646 | 10.06054 | 0 | 9.93728 | 5.175741 | 14.62667 | 22.34748 | 30.85725 | 50.02919 |
| ENSMUSG00000085826 | -2.309167876 | 1.22E-05 | 0.000268 | 9.054489 | 5.999108 | 30.63995 | 8.626235 | 50.70579 | 49.16446 | 77.14313 | 96.79561 |
| ENSMUSG00000085891 | 1.049540412 | 0.006274 | 0.030414 | 54.32694 | 85.18733 | 66.24853 | 87.9876 | 39.97956 | 49.16446 | 39.53586 | 13.05109 |
| ENSMUSG00000085912 | 1.941300587 | 0.000259 | 0.002781 | 22.1332 | 44.3934 | 14.90592 | 26.74133 | 8.776002 | 8.938993 | 4.821446 | 5.437955 |
| ENSMUSG00000085925 | 1.212244465 | 0.004478 | 0.02373 | 87.52673 | 106.7841 | 59.62368 | 88.85022 | 39.00445 | 69.27719 | 20.25007 | 19.57664 |
| ENSMUSG00000085982 | 1.756786875 | 0.000157 | 0.001889 | 36.21796 | 47.99286 | 25.67131 | 21.56559 | 5.850668 | 12.29112 | 7.714313 | 13.05109 |
| ENSMUSG00000086003 | -1.153883739 | 0.010356 | 0.043728 | 6.036326 | 11.99822 | 9.109173 | 8.626235 | 20.47734 | 13.40849 | 23.14294 | 21.75182 |
| ENSMUSG00000086010 | -3.325501248 | 4.72E-11 | 9.27E-09 | 3.018163 | 11.99822 | 24.01509 | 10.35148 | 94.5858 | 90.5073 | 108.9647 | 206.6423 |
| ENSMUSG00000086067 | 2.329235733 | 7.94E-05 | 0.001132 | 25.15136 | 23.99643 | 20.70267 | 14.6646 | 1.950223 | 5.58687 | 1.928578 | 7.613138 |
| ENSMUSG00000086105 | 1.565032518 | 0.008821 | 0.038937 | 35.2119 | 37.19447 | 39.74912 | 30.19182 | 2.925334 | 27.93435 | 5.785735 | 11.9635 |
| ENSMUSG00000086213 | -1.930064362 | 7.15E-06 | 0.000178 | 27.16347 | 11.99822 | 44.71776 | 24.15346 | 78.98402 | 60.3382 | 144.6434 | 131.5985 |
| ENSMUSG00000086390 | -1.515038694 | 9.25E-05 | 0.001255 | 65.39353 | 43.19358 | 137.4657 | 75.04824 | 219.4 | 120.6764 | 258.4295 | 323.0146 |
| ENSMUSG00000086587 | 2.207679531 | 1.72E-06 | 5.78E-05 | 88.53278 | 110.3836 | 93.57605 | 115.5915 | 19.50223 | 48.04709 | 12.53576 | 8.700729 |
| ENSMUSG00000086596 | 1.036256091 | 0.000227 | 0.002501 | 50.30272 | 40.79393 | 52.17072 | 56.07053 | 30.22845 | 20.11273 | 24.10723 | 22.83941 |
| ENSMUSG00000086645 | -1.78131373 | 0.002569 | 0.015735 | 5.030272 | 3.599465 | 9.93728 | 6.900988 | 21.45245 | 8.938993 | 35.6787 | 22.83941 |
| ENSMUSG00000086748 | 4.340419945 | 1.91E-18 | 1.86E-15 | 165.999 | 185.9723 | 127.5284 | 185.4641 | 1.950223 | 21.23011 | 4.821446 | 5.437955 |
| ENSMUSG00000086770 | 1.086378728 | 0.010856 | 0.045379 | 28.16952 | 20.39697 | 20.70267 | 26.74133 | 16.57689 | 7.821619 | 7.714313 | 13.05109 |
| ENSMUSG00000086825 | 1.421450315 | 4.44E-05 | 0.000728 | 170.0232 | 243.5638 | 151.5435 | 229.4578 | 67.28268 | 132.9675 | 51.10733 | 45.67883 |
| ENSMUSG00000086924 | -1.14478528 | 0.007704 | 0.035228 | 14.08476 | 7.198929 | 22.35888 | 14.6646 | 27.30312 | 23.46486 | 35.6787 | 44.59123 |
| ENSMUSG00000087022 | -2.270349491 | 1.16E-05 | 0.000257 | 9.054489 | 4.799286 | 14.07781 | 12.07673 | 21.45245 | 41.34284 | 52.07161 | 81.56933 |
| ENSMUSG00000087024 | 1.619095443 | 0.005596 | 0.027922 | 37.22401 | 25.19625 | 21.53077 | 23.29083 | 2.925334 | 3.352122 | 15.42863 | 13.05109 |
| ENSMUSG00000087075 | -1.311112639 | 0.002289 | 0.014455 | 21.12714 | 39.59411 | 52.99882 | 22.42821 | 98.48624 | 36.87335 | 101.2504 | 101.146 |
| ENSMUSG00000087204 | 1.52084835 | 0.003212 | 0.018674 | 43.26034 | 92.38626 | 57.13936 | 70.73513 | 21.45245 | 49.16446 | 10.60718 | 10.87591 |
| ENSMUSG00000087264 | 1.163994789 | 0.001882 | 0.012567 | 29.17558 | 51.59233 | 43.88965 | 30.19182 | 15.60178 | 11.17374 | 21.21436 | 20.66423 |
| ENSMUSG00000087380 | -1.895099681 | 0.000431 | 0.004079 | 11.0666 | 3.599465 | 8.281066 | 6.900988 | 24.37778 | 45.81234 | 27.96439 | 14.13868 |
| ENSMUSG00000087413 | 1.249225399 | 0.001027 | 0.007993 | 23.13925 | 29.99554 | 24.8432 | 34.50494 | 14.62667 | 11.17374 | 11.57147 | 9.78832 |
| ENSMUSG00000087444 | -2.260423572 | 3.74E-05 | 0.000638 | 6.036326 | 3.599465 | 9.109173 | 6.038364 | 24.37778 | 20.11273 | 22.17865 | 54.37955 |
| ENSMUSG00000087479 | 1.100501194 | 0.000375 | 0.003665 | 97.58727 | 106.7841 | 70.38906 | 140.6076 | 47.78045 | 61.45558 | 54.96448 | 29.36496 |
| ENSMUSG00000087613 | -1.309753695 | 0.003457 | 0.019605 | 58.35115 | 33.595 | 81.98256 | 70.73513 | 81.90935 | 84.92043 | 149.4648 | 292.562 |
| ENSMUSG00000087620 | 1.714075635 | 3.49E-05 | 0.000603 | 25.15136 | 27.5959 | 23.18699 | 25.8787 | 6.825779 | 8.938993 | 8.678602 | 6.525547 |
| ENSMUSG00000087674 | 1.224194569 | 0.0056 | 0.027933 | 31.18769 | 19.19714 | 18.21835 | 18.11509 | 9.751113 | 8.938993 | 10.60718 | 7.613138 |
| ENSMUSG00000089678 | -3.205778529 | 0.000194 | 0.002211 | 38.23007 | 5.999108 | 99.3728 | 26.74133 | 417.3476 | 115.0895 | 292.1796 | 751.5254 |
| ENSMUSG00000089684 | 1.542076219 | 0.001471 | 0.010426 | 35.2119 | 22.79661 | 16.56213 | 33.64232 | 12.67645 | 12.29112 | 6.750024 | 5.437955 |
| ENSMUSG00000089704 | -1.165229474 | 0.007075 | 0.033165 | 72.43591 | 59.99108 | 81.98256 | 32.77969 | 138.4658 | 49.16446 | 162.0006 | 204.4671 |
| ENSMUSG00000089712 | -3.385593343 | 3.32E-09 | 3.50E-07 | 18.10898 | 3.599465 | 12.4216 | 12.93935 | 93.61069 | 40.22547 | 238.1794 | 123.9854 |
| ENSMUSG00000089774 | -1.097304592 | 6.10E-05 | 0.000928 | 914.5034 | 725.892 | 1039.274 | 905.7547 | 1089.199 | 1790.033 | 3134.904 | 1658.576 |
| ENSMUSG00000089829 | 1.302538957 | 0.007473 | 0.034499 | 32.19374 | 41.99375 | 16.56213 | 34.50494 | 18.52711 | 16.76061 | 8.678602 | 6.525547 |
| ENSMUSG00000089854 | 1.384737553 | 0.007133 | 0.033334 | 20.12109 | 20.39697 | 14.07781 | 18.11509 | 8.776002 | 8.938993 | 7.714313 | 2.175182 |
| ENSMUSG00000089924 | 1.56179416 | 0.00438 | 0.023367 | 14.08476 | 20.39697 | 14.07781 | 16.38985 | 3.900445 | 10.05637 | 3.857157 | 4.350364 |
| ENSMUSG00000089972 | 1.824922648 | 0.00215 | 0.013838 | 23.13925 | 23.99643 | 14.90592 | 19.84034 | 1.950223 | 12.29112 | 5.785735 | 3.262773 |
| ENSMUSG00000090061 | 2.190933457 | 1.98E-09 | 2.31E-07 | 277.671 | 363.5459 | 205.3704 | 376.1038 | 47.78045 | 126.2633 | 54.00019 | 40.24087 |
| ENSMUSG00000090165 | -4.94433169 | 2.12E-16 | 1.45E-13 | 5.030272 | 1.199822 | 7.45296 | 9.488858 | 206.7236 | 53.63396 | 144.6434 | 328.4525 |
| ENSMUSG00000090171 | -3.638456587 | 1.58E-06 | 5.40E-05 | 50.30272 | 17.99732 | 103.5133 | 55.2079 | 1049.22 | 145.2586 | 435.8587 | 1202.876 |
| ENSMUSG00000090175 | -4.778302061 | 2.31E-10 | 3.65E-08 | 3.018163 | 2.399643 | 1.656213 | 1.725247 | 47.78045 | 13.40849 | 81.00029 | 95.70802 |
| ENSMUSG00000090231 | -1.127702664 | 0.002616 | 0.015938 | 40.24217 | 39.59411 | 62.9361 | 31.91707 | 115.0631 | 45.81234 | 87.75031 | 133.7737 |
| ENSMUSG00000090236 | -2.093216104 | 0.000547 | 0.004911 | 10.06054 | 9.598572 | 36.43669 | 14.6646 | 39.00445 | 36.87335 | 71.3574 | 157.7007 |
| ENSMUSG00000090329 | 1.395979957 | 0.011107 | 0.04622 | 17.10292 | 26.39607 | 9.109173 | 18.97772 | 6.825779 | 5.58687 | 4.821446 | 9.78832 |

| | | | | | | | | | | | |
|---|---|---|---|---|---|---|---|---|---|---|---|
| ENSMUSG00000090817 | -3.279211627 | 3.17E-05 | 0.000559 | 53.32088 | 3.599465 | 82.81066 | 44.85642 | 423.1983 | 204.4795 | 440.6801 | 728.686 |
| ENSMUSG00000091345 | 2.547160453 | 0.002044 | 0.013323 | 432.6034 | 1767.337 | 1089.788 | 916.9688 | 104.3369 | 493.8794 | 77.14313 | 44.59123 |
| ENSMUSG00000091457 | 1.713511417 | 0.001101 | 0.008401 | 21.12714 | 33.595 | 19.87456 | 17.25247 | 7.80089 | 10.05637 | 7.714313 | 2.175182 |
| ENSMUSG00000091705 | -1.280529012 | 0.00012 | 0.001537 | 18.10898 | 23.99643 | 17.39024 | 14.6646 | 58.50668 | 41.34284 | 33.75012 | 44.59123 |
| ENSMUSG00000091972 | -3.0083745 | 0.002468 | 0.015282 | 3.018163 | 0 | 10.76539 | 2.58787 | 31.20356 | 2.234748 | 57.85735 | 43.50364 |
| ENSMUSG00000092014 | 1.296272918 | 0.006956 | 0.032832 | 15.09082 | 22.79661 | 13.24971 | 22.42821 | 8.776002 | 5.58687 | 7.714313 | 7.613138 |
| ENSMUSG00000092035 | 1.02058843 | 0.000376 | 0.003675 | 2013.115 | 1061.842 | 934.1043 | 1589.815 | 724.5077 | 1008.989 | 543.8591 | 482.8904 |
| ENSMUSG00000092470 | -1.6966224 | 4.54E-05 | 0.000739 | 19.11503 | 19.19714 | 38.09291 | 30.19182 | 49.73068 | 56.98608 | 109.929 | 131.5985 |
| ENSMUSG00000092626 | -1.344712797 | 0.005293 | 0.026772 | 5.030272 | 7.198929 | 9.93728 | 10.35148 | 24.37778 | 14.52586 | 16.39292 | 28.27737 |
| ENSMUSG00000093622 | 1.141442093 | 0.00855 | 0.038069 | 38.23007 | 53.99197 | 33.95237 | 37.09281 | 12.67645 | 33.52122 | 9.642892 | 18.48905 |
| ENSMUSG00000093760 | 1.544546982 | 0.00614 | 0.029934 | 15.09082 | 17.99732 | 21.53077 | 31.91707 | 11.70134 | 10.05637 | 5.785735 | 2.175182 |
| ENSMUSG00000093930 | -1.254239488 | 1.09E-13 | 4.07E-11 | 2469.863 | 2015.7 | 2804.797 | 2271.288 | 5098.857 | 4267.252 | 6638.167 | 6808.32 |
| ENSMUSG00000096606 | 1.529647847 | 2.53E-07 | 1.21E-05 | 88.53278 | 92.38626 | 62.9361 | 89.71284 | 39.00445 | 33.52122 | 24.10723 | 18.48905 |
| ENSMUSG00000096862 | 1.478834726 | 0.000163 | 0.001932 | 28.16952 | 47.99286 | 33.95237 | 25.8787 | 14.62667 | 13.40849 | 10.60718 | 9.78832 |
| ENSMUSG00000097068 | 1.290586316 | 0.00579 | 0.028665 | 17.10292 | 16.7975 | 18.21835 | 18.11509 | 9.751113 | 4.469496 | 6.750024 | 7.613138 |
| ENSMUSG00000097101 | -1.026038268 | 0.007087 | 0.033199 | 30.18163 | 26.39607 | 38.92101 | 37.95543 | 67.28268 | 32.40385 | 65.57166 | 107.6715 |
| ENSMUSG00000097146 | 1.656892306 | 0.001159 | 0.008752 | 27.16347 | 37.19447 | 21.53077 | 44.85642 | 9.751113 | 17.87799 | 10.60718 | 3.262773 |
| ENSMUSG00000097201 | 1.522578149 | 5.27E-05 | 0.000825 | 361.1735 | 404.3399 | 394.1788 | 567.6063 | 125.7894 | 296.1041 | 106.0718 | 73.95619 |
| ENSMUSG00000097209 | -1.319878977 | 0.003325 | 0.019108 | 12.07265 | 8.398751 | 28.15563 | 13.80198 | 47.78045 | 24.58223 | 44.3573 | 41.32846 |
| ENSMUSG00000097282 | 1.467981005 | 0.000558 | 0.004977 | 27.16347 | 43.19358 | 25.67131 | 38.81806 | 17.552 | 14.52586 | 6.750024 | 9.78832 |
| ENSMUSG00000097305 | 1.254771006 | 0.009805 | 0.041957 | 67.40564 | 32.39518 | 42.23344 | 38.81806 | 15.60178 | 37.99072 | 9.642892 | 13.05109 |
| ENSMUSG00000097381 | 1.117160193 | 0.005735 | 0.028477 | 26.15741 | 27.5959 | 19.04645 | 25.8787 | 8.776002 | 12.29112 | 15.42863 | 8.700729 |
| ENSMUSG00000097391 | 1.141555982 | 0.000318 | 0.003241 | 285.7194 | 273.5593 | 245.9477 | 328.6596 | 113.1129 | 221.2401 | 113.7861 | 66.34306 |
| ENSMUSG00000097418 | 1.052134554 | 0.004065 | 0.0221 | 37.22401 | 52.79215 | 42.23344 | 46.58167 | 17.552 | 34.6386 | 22.17865 | 11.9635 |
| ENSMUSG00000097421 | -2.282489686 | 0.000189 | 0.002171 | 9.054489 | 1.199822 | 19.87456 | 10.35148 | 70.20801 | 25.6996 | 37.60728 | 67.43065 |
| ENSMUSG00000097451 | 1.458432286 | 0.00026 | 0.002785 | 6121.841 | 5047.649 | 4285.452 | 6668.942 | 1845.886 | 3662.752 | 1405.934 | 1136.533 |
| ENSMUSG00000097464 | 1.381533849 | 0.011932 | 0.048807 | 133.8052 | 122.3818 | 65.42042 | 119.042 | 20.47734 | 105.0332 | 21.21436 | 22.83941 |
| ENSMUSG00000097610 | 1.085026338 | 0.006815 | 0.032357 | 26.15741 | 35.99465 | 23.18699 | 25.8787 | 15.60178 | 7.821619 | 17.3572 | 10.87591 |
| ENSMUSG00000097619 | 1.538291582 | 0.003111 | 0.018278 | 17.10292 | 23.99643 | 16.56213 | 22.42821 | 11.70134 | 4.469496 | 7.714313 | 3.262773 |
| ENSMUSG00000097621 | 1.871227876 | 0.000461 | 0.004303 | 23.13925 | 17.99732 | 21.53077 | 12.93935 | 2.925334 | 4.469496 | 6.750024 | 6.525547 |
| ENSMUSG00000097651 | -3.07821116 | 0.000995 | 0.007805 | 1.006054 | 0 | 9.93728 | 1.725247 | 29.25334 | 6.704245 | 32.78583 | 42.41605 |
| ENSMUSG00000097756 | 1.678561937 | 0.008949 | 0.039297 | 25.15136 | 77.9884 | 31.46805 | 50.89479 | 5.850668 | 34.6386 | 5.785735 | 11.9635 |
| ENSMUSG00000097762 | -1.40629789 | 0.001812 | 0.012229 | 10.06054 | 13.19804 | 33.12427 | 11.21411 | 46.80534 | 40.22547 | 36.64299 | 57.64233 |
| ENSMUSG00000097767 | 3.861724126 | 5.73E-47 | 1.06E-42 | 2567.451 | 2366.048 | 1733.227 | 2360.138 | 145.2916 | 245.8223 | 147.5362 | 82.65692 |
| ENSMUSG00000097796 | 1.541061297 | 5.17E-06 | 0.000138 | 90.54489 | 73.18911 | 60.45178 | 75.04824 | 25.35289 | 41.34284 | 22.17865 | 14.13868 |
| ENSMUSG00000097857 | 1.156078486 | 0.006817 | 0.032358 | 21.12714 | 19.19714 | 23.18699 | 18.97772 | 6.825779 | 10.05637 | 12.53576 | 7.613138 |
| ENSMUSG00000097970 | -1.07998655 | 0.00086 | 0.006949 | 27.16347 | 16.7975 | 19.04645 | 20.70296 | 40.95467 | 34.6386 | 58.82164 | 42.41605 |
| ENSMUSG00000098202 | 2.066986402 | 0.007708 | 0.035234 | 19.11503 | 43.19358 | 21.53077 | 37.09281 | 5.850668 | 20.11273 | 0.964289 | 2.175182 |
| ENSMUSG00000098294 | 1.835714185 | 0.002309 | 0.014546 | 12.07265 | 31.19536 | 17.39024 | 10.35148 | 4.875557 | 5.58687 | 3.857157 | 5.437955 |
| ENSMUSG00000098891 | 1.971496312 | 8.31E-05 | 0.001167 | 25.15136 | 33.595 | 24.01509 | 43.9938 | 8.776002 | 14.52586 | 3.857157 | 5.437955 |
| ENSMUSG00000098979 | 2.252537281 | 2.72E-10 | 4.14E-08 | 276.6649 | 359.9465 | 178.0429 | 318.3081 | 57.53157 | 105.0332 | 38.57157 | 36.9781 |
| ENSMUSG00000099032 | 1.369570025 | 7.74E-21 | 1.30E-17 | 810.8798 | 809.8795 | 691.469 | 822.0802 | 264.2552 | 386.6114 | 262.2867 | 301.2627 |
| ENSMUSG00000099136 | 1.573455475 | 0.004892 | 0.025347 | 40.24217 | 31.19536 | 24.8432 | 23.29083 | 3.900445 | 22.34748 | 7.714313 | 6.525547 |
| ENSMUSG00000099137 | 1.088035418 | 0.000959 | 0.007598 | 113.6841 | 105.5843 | 74.5296 | 89.71284 | 25.35289 | 72.62932 | 41.46443 | 41.32846 |
| ENSMUSG00000099307 | 2.162518003 | 2.17E-07 | 1.07E-05 | 28.16952 | 35.99465 | 35.60859 | 29.3292 | 8.776002 | 7.821619 | 7.714313 | 4.350364 |
| ENSMUSG00000099413 | -2.781754549 | 3.49E-05 | 0.000603 | 4.024217 | 1.199822 | 13.24971 | 7.763611 | 33.15378 | 16.76061 | 64.60737 | 70.69342 |
| ENSMUSG00000099757 | 1.131334639 | 0.008664 | 0.038447 | 17.10292 | 37.19447 | 24.8432 | 38.81806 | 11.70134 | 17.87799 | 14.46434 | 9.78832 |
| ENSMUSG00000100158 | 1.445117766 | 0.00295 | 0.01755 | 257.5499 | 392.3416 | 155.684 | 189.7772 | 78.0089 | 196.6578 | 54.00019 | 36.9781 |
| ENSMUSG00000100600 | 1.606060573 | 0.00804 | 0.036377 | 61.36932 | 100.785 | 45.54586 | 79.36136 | 14.62667 | 60.3382 | 7.714313 | 11.9635 |
| ENSMUSG00000100860 | 1.771500185 | 0.000467 | 0.004345 | 38.23007 | 82.78769 | 43.06154 | 79.36136 | 22.42756 | 27.93435 | 4.821446 | 16.31387 |
| ENSMUSG00000100975 | 1.2807365 | 3.21E-05 | 0.000563 | 109.6599 | 68.38983 | 92.74794 | 56.93315 | 25.35289 | 43.57759 | 33.75012 | 32.62773 |
| ENSMUSG00000101236 | -1.793489585 | 0.004299 | 0.023037 | 9.054489 | 4.799286 | 4.140533 | 3.450494 | 7.80089 | 30.1691 | 15.42863 | 20.66423 |
| ENSMUSG00000101356 | 1.128856172 | 0.000653 | 0.005639 | 208.2533 | 219.5673 | 136.6376 | 279.49 | 92.63557 | 159.7845 | 66.53595 | 67.43065 |
| ENSMUSG00000101389 | 1.119808431 | 0.000371 | 0.003637 | 178.0716 | 116.3827 | 143.2624 | 164.7611 | 74.10846 | 63.69032 | 104.1432 | 34.80291 |
| ENSMUSG00000101517 | -2.031230085 | 0.000173 | 0.002024 | 3.018163 | 4.799286 | 9.109173 | 4.313117 | 22.42756 | 13.40849 | 25.07152 | 27.18978 |
| ENSMUSG00000101556 | 1.392869285 | 0.005869 | 0.028941 | 18.10898 | 11.99822 | 19.87456 | 14.6646 | 5.850668 | 7.821619 | 6.750024 | 4.350364 |
| ENSMUSG00000102091 | 1.517916791 | 0.00533 | 0.026873 | 46.2785 | 50.3925 | 19.87456 | 62.10889 | 5.850668 | 30.1691 | 12.53576 | 14.13868 |
| ENSMUSG00000102112 | -1.355593033 | 0.007068 | 0.033143 | 12.07265 | 16.7975 | 21.53077 | 6.900988 | 26.32801 | 18.99536 | 52.07161 | 48.9416 |
| ENSMUSG00000102282 | 1.026884437 | 0.012134 | 0.049404 | 37.22401 | 38.39429 | 28.98373 | 40.5433 | 11.70134 | 32.40385 | 14.46434 | 13.05109 |
| ENSMUSG00000102562 | 1.220820364 | 0.005099 | 0.026084 | 66.39959 | 73.18911 | 46.37397 | 43.13117 | 29.25334 | 42.46022 | 13.50005 | 13.05109 |
| ENSMUSG00000102577 | 1.40025404 | 0.000129 | 0.001621 | 37.22401 | 28.79572 | 25.67131 | 31.05445 | 12.67645 | 12.29112 | 10.60718 | 10.87591 |
| ENSMUSG00000102786 | 2.026092265 | 0.011972 | 0.048893 | 29.17558 | 13.19804 | 6.624853 | 15.52722 | 2.925334 | 10.05637 | 1.928578 | 1.087591 |
| ENSMUSG00000103546 | 1.246494411 | 0.001447 | 0.010317 | 68.4117 | 59.99108 | 52.99882 | 64.69676 | 11.70134 | 45.81234 | 26.03581 | 20.66423 |
| ENSMUSG00000103621 | 1.512295043 | 0.009722 | 0.041725 | 35.2119 | 41.99375 | 26.49941 | 43.13117 | 6.825779 | 29.05173 | 12.53576 | 3.262773 |

| | | | | | | | | | | | |
|---|---|---|---|---|---|---|---|---|---|---|---|
| ENSMUSG00000103932 | 1.697796219 | 0.006836 | 0.032416 | 131.7931 | 208.7689 | 71.21717 | 99.2017 | 27.30312 | 102.7984 | 13.50005 | 14.13868 |
| ENSMUSG00000104060 | 1.288147663 | 0.011745 | 0.048169 | 15.09082 | 14.39786 | 14.90592 | 13.80198 | 5.850668 | 8.938993 | 4.821446 | 4.350364 |
| ENSMUSG00000104091 | 1.376040329 | 0.001715 | 0.011699 | 24.1453 | 25.19625 | 25.67131 | 34.50494 | 11.70134 | 15.64324 | 10.60718 | 4.350364 |
| ENSMUSG00000104097 | 1.10225198 | 0.012318 | 0.049966 | 17.10292 | 21.59679 | 17.39024 | 24.15346 | 5.850668 | 12.29112 | 9.642892 | 9.78832 |
| ENSMUSG00000104375 | -7.182620914 | 1.65E-10 | 2.80E-08 | 0 | 0 | 0.828107 | 0 | 24.37778 | 33.52122 | 60.75022 | 51.11678 |
| ENSMUSG00000104376 | -1.036254065 | 0.010251 | 0.04344 | 16.09687 | 14.39786 | 19.04645 | 7.763611 | 23.40267 | 24.58223 | 35.6787 | 33.71532 |
| ENSMUSG00000104392 | 1.136225692 | 0.007635 | 0.035002 | 75.45408 | 55.19179 | 50.5145 | 65.55939 | 13.65156 | 46.92971 | 36.64299 | 15.22628 |
| ENSMUSG00000104444 | -2.264110225 | 0.001229 | 0.009119 | 19.11503 | 7.198929 | 67.90474 | 9.488858 | 118.9636 | 43.57759 | 96.42892 | 242.5328 |
| ENSMUSG00000104445 | -2.559638657 | 0.000116 | 0.001507 | 13.07871 | 5.999108 | 43.06154 | 6.038364 | 75.08357 | 45.81234 | 92.57176 | 192.5036 |
| ENSMUSG00000104597 | 1.238218197 | 0.001133 | 0.0086 | 61.36932 | 57.59143 | 68.73285 | 98.33908 | 28.27823 | 52.51658 | 17.3572 | 23.927 |
| ENSMUSG00000104693 | -1.772145159 | 0.002514 | 0.015492 | 4.024217 | 3.599465 | 14.07781 | 10.35148 | 21.45245 | 15.64324 | 45.32159 | 29.36496 |
| ENSMUSG00000104713 | 1.690110796 | 0.007999 | 0.03622 | 32.19374 | 103.1847 | 126.7003 | 18.11509 | 8.776002 | 24.58223 | 32.78583 | 20.66423 |
| ENSMUSG00000104737 | -1.471797779 | 0.00831 | 0.037333 | 28.16952 | 4.799286 | 10.76539 | 12.93935 | 36.07912 | 18.99536 | 57.85735 | 44.59123 |
| ENSMUSG00000104851 | 1.432647703 | 0.009502 | 0.04105 | 152.9203 | 134.38 | 90.26362 | 122.4925 | 25.35289 | 110.62 | 37.60728 | 11.9635 |
| ENSMUSG00000104868 | 1.876500867 | 0.000105 | 0.00139 | 44.26639 | 53.99197 | 26.49941 | 28.46658 | 9.751113 | 17.87799 | 9.642892 | 4.350364 |
| ENSMUSG00000104871 | 1.81282216 | 0.000379 | 0.003694 | 75.45408 | 104.3845 | 62.108 | 66.42201 | 13.65156 | 50.28183 | 15.42863 | 8.700729 |
| ENSMUSG00000104894 | 1.47034399 | 0.006544 | 0.031463 | 18.10898 | 31.19536 | 15.73403 | 19.84034 | 5.850668 | 10.05637 | 2.892867 | 11.9635 |
| ENSMUSG00000104973 | 1.189371367 | 0.010281 | 0.043496 | 68.4117 | 80.38804 | 67.07664 | 66.42201 | 21.45245 | 68.15982 | 18.32149 | 16.31387 |
| ENSMUSG00000105107 | 1.275731857 | 2.66E-07 | 1.26E-05 | 71.42986 | 74.38894 | 77.01392 | 72.46037 | 32.17867 | 40.22547 | 27.0001 | 22.83941 |
| ENSMUSG00000105168 | -4.452351792 | 0.000377 | 0.003684 | 0 | 0 | 2.48432 | 0.862623 | 39.97956 | 1.117374 | 12.53576 | 25.0146 |
| ENSMUSG00000105271 | 2.181750458 | 0.000386 | 0.003751 | 35.2119 | 47.99286 | 25.67131 | 25.8787 | 1.950223 | 7.821619 | 16.39292 | 3.262773 |
| ENSMUSG00000105302 | 1.545111415 | 0.000317 | 0.003241 | 34.20585 | 41.99375 | 32.29616 | 27.60395 | 12.67645 | 17.87799 | 11.57147 | 4.350364 |
| ENSMUSG00000105304 | 1.35442358 | 0.011219 | 0.046557 | 47.28455 | 50.3925 | 22.35888 | 28.46658 | 13.65156 | 29.05173 | 6.750024 | 8.700729 |
| ENSMUSG00000105366 | 1.163638806 | 0.000965 | 0.007629 | 91.55095 | 134.38 | 110.1382 | 127.6683 | 45.83023 | 92.74205 | 29.89296 | 39.15328 |
| ENSMUSG00000105378 | 1.179198456 | 0.007795 | 0.035549 | 26.15741 | 22.79661 | 22.35888 | 30.19182 | 7.80089 | 20.11273 | 8.678602 | 8.700729 |
| ENSMUSG00000105402 | -1.764631977 | 0.007532 | 0.034672 | 10.06054 | 5.999108 | 9.109173 | 8.626235 | 40.95467 | 4.469496 | 24.10723 | 45.67883 |
| ENSMUSG00000105403 | 1.163066058 | 0.000467 | 0.004346 | 77.46619 | 45.59322 | 56.31125 | 81.08661 | 23.40267 | 37.99072 | 35.6787 | 19.57664 |
| ENSMUSG00000105444 | -2.453342035 | 0.000127 | 0.001607 | 3.018163 | 1.199822 | 8.281066 | 6.038364 | 27.30312 | 17.87799 | 15.42863 | 44.59123 |
| ENSMUSG00000105568 | 2.092182739 | 0.004384 | 0.023381 | 54.32694 | 43.19358 | 20.70267 | 25.8787 | 2.925334 | 21.23011 | 8.678602 | 1.087591 |
| ENSMUSG00000105959 | 1.974280441 | 0.003914 | 0.021492 | 19.11503 | 23.99643 | 10.76539 | 25.01608 | 3.900445 | 12.29112 | 1.928578 | 2.175182 |
| ENSMUSG00000106078 | -3.3523187 | 5.94E-08 | 3.74E-06 | 3.018163 | 2.399643 | 9.93728 | 6.038364 | 40.95467 | 23.46486 | 56.89306 | 103.3212 |
| ENSMUSG00000106111 | -1.791282704 | 0.00441 | 0.023471 | 6.036326 | 1.199822 | 9.109173 | 6.038364 | 17.552 | 10.05637 | 19.28578 | 32.62773 |
| ENSMUSG00000106290 | -1.409995667 | 0.001102 | 0.008401 | 11.0666 | 9.598572 | 6.624853 | 10.35148 | 20.47734 | 20.11273 | 34.71441 | 23.927 |
| ENSMUSG00000106320 | 1.479282554 | 0.009884 | 0.042258 | 67.40564 | 61.1909 | 35.60859 | 43.13117 | 13.65156 | 44.69496 | 8.678602 | 7.613138 |
| ENSMUSG00000106446 | 1.701673484 | 0.011153 | 0.046362 | 47.28455 | 41.99375 | 18.21835 | 33.64232 | 8.776002 | 26.81698 | 5.785735 | 2.175182 |
| ENSMUSG00000106526 | -1.653644432 | 0.004217 | 0.022693 | 11.0666 | 5.999108 | 9.93728 | 6.900988 | 22.42756 | 12.29112 | 19.28578 | 53.29196 |
| ENSMUSG00000106732 | 1.341686946 | 0.010458 | 0.044087 | 25.15136 | 34.79482 | 16.56213 | 18.97772 | 4.875557 | 15.64324 | 11.57147 | 5.437955 |
| ENSMUSG00000107009 | 2.363473155 | 0.003026 | 0.017883 | 22.1332 | 26.39607 | 24.01509 | 15.52722 | 0.975111 | 13.40849 | 1.928578 | 1.087591 |
| ENSMUSG00000107667 | 1.15008814 | 0.000631 | 0.005483 | 49.29666 | 45.59322 | 39.74912 | 31.05445 | 12.67645 | 20.11273 | 23.14294 | 18.48905 |
| ENSMUSG00000108035 | 1.673947942 | 0.005318 | 0.026853 | 86.52067 | 73.18911 | 24.01509 | 50.03216 | 9.751113 | 42.46022 | 12.53576 | 8.700729 |
| ENSMUSG00000108037 | 1.012563139 | 0.001436 | 0.010261 | 96.58122 | 87.58697 | 49.6864 | 78.49874 | 54.60623 | 31.28647 | 38.57157 | 29.36496 |
| ENSMUSG00000108256 | 1.359426291 | 0.003462 | 0.019613 | 48.29061 | 28.79572 | 28.98373 | 28.46658 | 10.72622 | 24.58223 | 7.714313 | 9.78832 |
| ENSMUSG00000108290 | -1.321673604 | 1.11E-05 | 0.000248 | 172.0353 | 89.98662 | 230.2136 | 150.0965 | 282.7823 | 343.0338 | 416.5729 | 567.7225 |
| ENSMUSG00000108348 | 1.18632166 | 6.45E-05 | 0.000965 | 78.47224 | 71.98929 | 68.73285 | 92.30071 | 25.35289 | 48.04709 | 23.14294 | 41.32846 |
| ENSMUSG00000108601 | 1.585882774 | 0.001916 | 0.012726 | 98.59333 | 143.9786 | 83.63877 | 98.33908 | 45.83023 | 64.8077 | 23.14294 | 7.613138 |
| ENSMUSG00000108658 | 1.257187089 | 5.95E-09 | 5.52E-07 | 128.775 | 160.7761 | 179.6991 | 153.547 | 60.4569 | 78.21619 | 75.21455 | 46.76642 |
| ENSMUSG00000108884 | -2.855008524 | 1.32E-08 | 1.08E-06 | 29.17558 | 4.799286 | 45.54586 | 12.93935 | 114.088 | 178.7799 | 191.8935 | 191.416 |
| ENSMUSG00000109028 | 1.368354719 | 0.010607 | 0.044562 | 25.15136 | 14.39786 | 26.49941 | 25.01608 | 9.751113 | 16.76061 | 4.821446 | 4.350364 |
| ENSMUSG00000109051 | 2.299939147 | 0.005166 | 0.026328 | 20.12109 | 35.99465 | 19.87456 | 7.763611 | 3.900445 | 11.17374 | 0.964289 | 1.087591 |
| ENSMUSG00000109093 | -3.324654173 | 1.91E-13 | 6.93E-11 | 35.2119 | 8.398751 | 72.87338 | 31.05445 | 276.9316 | 327.3906 | 385.7157 | 502.4671 |
| ENSMUSG00000109095 | 2.672562341 | 0.001991 | 0.013087 | 21.12714 | 19.19714 | 4.140533 | 34.50494 | 3.900445 | 5.58687 | 2.892867 | 0 |
| ENSMUSG00000109311 | -2.444018244 | 5.04E-05 | 0.000798 | 30.18163 | 1.199822 | 45.54586 | 22.42821 | 128.7147 | 91.62468 | 129.2147 | 195.7664 |
| ENSMUSG00000109321 | 1.189499471 | 0.001484 | 0.010486 | 115.6963 | 105.5843 | 58.79557 | 96.61383 | 28.27823 | 70.39457 | 40.50014 | 26.10219 |
| ENSMUSG00000109394 | 1.140380458 | 3.41E-09 | 3.56E-07 | 152.9203 | 160.7761 | 152.3716 | 163.0358 | 57.53157 | 96.09417 | 68.46453 | 64.16787 |
| ENSMUSG00000109554 | 1.147046579 | 0.001169 | 0.008803 | 59.35721 | 77.9884 | 45.54586 | 77.63611 | 18.52711 | 41.34284 | 21.21436 | 36.9781 |
| ENSMUSG00000109628 | -2.247085174 | 0.004198 | 0.02262 | 10.06054 | 0 | 29.81184 | 18.11509 | 55.58134 | 25.6996 | 65.57166 | 131.5985 |
| ENSMUSG00000109642 | 1.547705779 | 0.001511 | 0.010619 | 42.25428 | 44.3934 | 19.87456 | 29.3292 | 7.80089 | 15.64324 | 5.785735 | 17.40146 |
| ENSMUSG00000109644 | -2.200703676 | 7.54E-09 | 6.79E-07 | 183.1019 | 57.59143 | 213.6515 | 166.4863 | 708.9059 | 382.1419 | 716.4668 | 1052.788 |
| ENSMUSG00000109787 | 1.802462654 | 0.000148 | 0.001814 | 48.29061 | 63.59054 | 42.23344 | 66.42201 | 7.80089 | 33.52122 | 12.53576 | 9.78832 |
| ENSMUSG00000109827 | 1.52873156 | 2.21E-05 | 0.000425 | 50.30272 | 70.78947 | 35.60859 | 66.42201 | 23.40267 | 21.23011 | 20.25007 | 11.9635 |
| ENSMUSG00000110079 | 1.367833033 | 0.006924 | 0.032734 | 70.4238 | 91.18644 | 50.5145 | 40.5433 | 13.65156 | 50.28183 | 23.14294 | 10.87591 |
| ENSMUSG00000110661 | 1.457828868 | 2.17E-05 | 0.000422 | 35.2119 | 41.99375 | 50.5145 | 43.13117 | 13.65156 | 16.76061 | 10.60718 | 21.75182 |
| ENSMUSG00000110750 | -2.21294019 | 0.000256 | 0.002754 | 8.048435 | 4.799286 | 4.96864 | 2.58787 | 23.40267 | 8.938993 | 30.85725 | 30.45255 |
| ENSMUSG00000110874 | -1.635070216 | 4.46E-08 | 2.95E-06 | 73.44197 | 29.99554 | 72.87338 | 64.69676 | 154.0676 | 150.8455 | 200.5721 | 247.9708 |

| | | | | | | | | | | | |
|---|---|---|---|---|---|---|---|---|---|---|---|
| ENSMUSG00000111212 | 1.687182971 | 0.003865 | 0.021297 | 123.7447 | 134.38 | 46.37397 | 38.81806 | 13.65156 | 58.10345 | 15.42863 | 19.57664 |
| ENSMUSG00000111236 | 1.148094115 | 0.000194 | 0.002211 | 50.30272 | 51.59233 | 54.65504 | 37.95543 | 16.57689 | 24.58223 | 27.96439 | 18.48905 |
| ENSMUSG00000111368 | -1.343758668 | 0.008387 | 0.037578 | 22.1332 | 5.999108 | 23.18699 | 19.84034 | 40.95467 | 18.99536 | 64.60737 | 57.64233 |
