## Supplementary material for "Lysine-specific demethylase 1a is obligatory for gene regulation during kidney development": Suppl Table-3

**Supplementary Table-3.Potential HNF4a target genes down regulated in Kdm1a knockout kidneys**

Proc
Iyd
Xylb
Entpd8
Qprt
Glyctk
Cideb
Anks4b
Cfb
Aspdh
Aadat
Ephx2
Cyp2d26
Haao
Ces2e
Pklr
Hnf1a
Sord
Tmem25
Pafah2
Slc23a1
G6pc
Aqp11
Pck1
Slc6a12
Itga1
Mettl7b
Hmgcs1
Hpdl
Acox2
Entpd5
Aldh8a1
Pipox
4931406C07Rik
Aqp4
Sgk2
Rdh16f2
Slc47a1
Dppa3
Pdzk1
Mfsd4b1
Slc22a7
Agxt2
Them7
F10
Kynu

Upb1
Acy3
2610528J11Rik
F13b
Spp2
Cct6b
Slc26a1
Slc22a4
Cth
Gpd1
Ces1e
Slc22a18
Sult1b1
Ehhadh
Etnppl
Treh
Acsl1
Rdh16
Eaf2
Acot4
Tmem37
Cpb2
Susd2
Hsd3b3
Aldh1l1
Cyp2e1
Gss
Osr2
Fah
A1cf
Eci3
Fbp1
Slc25a34
Slc10a5
Rbp2
Bpnt1
Bdh1
Hsd17b2
Slc6a13
Pcx
Hgd
Gcat
H2-Q6
Usp2
Slc25a10
Fgfr3
Aadac

Ak4
Ppara
Gk
Slc10a1
Grhpr
Fbxl15
Kng2
Atp6v1b1
Amacr
Ggt6
Igfals
Agmat
Keg1
Cpn1
Tlr12
Serpinf2
Cyp2j5
Dqx1
Tpmt
Hykk
Cdhr5
Cgref1
Cldn14
Slc51b
Erich4
Agmo
Rassf6
Ugt3a2
Prodh2
Selenbp1
Acnat2
Cdkl1
P2ry2
Rarres1
Acat3
Tmem106a
Ttr
Gm11992
Gimd1
B3gnt2
Acmsd
Clec2h
Hpn
Abhd15
Sult1d1
Galnt2
Pxmp2

Nabp1
Dpp4
Gipc2
Cryz
Slc22a30
Cyp4a31
Abcc2
Ttc36
Ugt2b37
Cmah
Glyat
Slc6a19
Slc22a1
Serpina6
Slc25a5
Nat8f2
Ugt3a1
Serpina1a
Slc16a12
Slc3a1
Cdk18
Acsm1
Dbil5
Arhgef16
Pcca
Atp1b1
Angptl8
Cfap126
Cfi
Tgm1
Calml4
Aldh6a1
Tldc2
Idi1
Slc22a27
Acsm5
Enpep
Slc6a20a
Ccdc96
Cyp24a1
Ndrg1
Ranbp3l
Susd3
Rhpn1
Ankrd33b
Slc17a1
Serpina10

Pdzd3
Amdhd1
Fbp2
Hsd11b1
Ugt2b5
Cda
Slco4a1
Gal3st1
Tmem125
Esrp2
Foxq1
Ass1
Guca2b
Chst13
Slc5a8
Slc18a1
Tmem267
Flvcr2
Aifm3
Nat8f1
Igsf5
Klhdc8a
Sugct
Krt4
Pim3
Prkcz
Acnat1
Nuak2
Dgat2
Tcn2
Mep1a
Ace
Car2
Cisd1
Slc22a28
Cyp4a10
Cdhr2
P2rx5
Dhcr24
Cnpy1
Macrod1
Pkhd1
H2-Q2
Pde4c
Llgl2
Stxbp5l
Slc37a4

Gsta1
Pah
Slc17a4
Vil1
Clrn3
Myo15b
Papss2
Slc17a2
Mpv17l
Nr1h4
Ccno
Larp1b
Cpox
Myo7b
Gjb1
Pla1a
Slc4a9
Paqr9
Lactb2
Slc9a3
Cryl1
Slc16a5
Elovl2
Slc27a2
Slc2a2
Mep1b
Mdh1
Apcs
Aspa
Amn
Defb1
Mccc1
Gm10639
Tmprss2
Tprn
Mcrip2
Aoc1
Fitm1
Cystm1
Gsta2
Slc22a26
Mthfd1
Dio1
Acot3
Smim1
6430571L13Rik
Aldh4a1

Slc38a3
Gnmt
Acaa1b
Msmo1
Ces1f
Tmem116
Irx3
Gjb2
Sgk1
Vwa1
Hoga1
Sult1c2
Dao
Dusp9
Hnf4a
Cox7a1
Slc5a11
Sqle
Slc5a3
Nat8
Zmynd10
Cyp4a14
Tdrp
Aldob
Eva1a
Mro
Spink1
Hmgn2
Ugt1a1
Esrra
Phyhipl
Emx1
Cfap52
Galnt11
Eps8l2
Akr1d1
Gnai1
Hspa1b
Xpnpep2
Slc23a3
Tmem163
Smim24
Slc5a12
Ggt1
Slc5a1
Aqp1
Fads3

Cpt1b
Baiap2l2
Neu1
Apom
Slc7a9
A4gnt
Cep85
Tnfaip2
Rtn4r
Nlrp6
Lzts3
Dmgdh
Bhmt2
Dazl
Tinagl1
Corin
Atp1a1
Smagp
Miox
Mogat2
Slc39a5
Dusp14
Angptl4
Ccdc170
Npb
Kcnn1
Galnt3
Fabp3
Them6
Ihh
Plek2
Slc10a2
Adra1d
Zdhhc23
AA986860
Hsd3b2
Slc17a3
Grem2
Rida
Tbx10
Chchd10
Mal
Lrrc19
Calml3
Eno4
Slc44a4
Slc34a3

Adra2b
Agt
Anpep
Ugt8a
Napsa
Dmtn
Podxl
Insrr
Tuba4a
Nrip3
Slc13a2
Akr1b3
C4bp
Trpv5
Mgam
Prap1
Bst1
Kl
Ush1c
Tmigd1
Tdrd5
Lpar3
Sod3
Endog
Adhfe1
Aif1l
Slc51a
Igsf11
Slc43a2
Pigz
Plekhh1
Wscd1
Dcst1
Ddn
Thnsl2
Glis1
Atp6v0a4
Gucy2g
Prima1
Cdr2
Nxpe3
Tmem178
Fa2h
Msx1
Ripply2
Cnnm1
Mfsd4b5

Scube2
Gatm
Slc6a20b
Lgals12
Itgb6
Ankrd34b
Tmem252
Sall3
Irx1
Ch25h
Afm
Pitx2
Stk31
Lrp2
Esrrb
Apcdd1
Aoah
Acss1
Clic5
Galnt14
Stk32a
Pth1r
Ak7
Slc22a2
Sycp1
Sigirr
Pcsk2
Wnt7b
Plau
Irx2
Fam167a
Hnf4g
Pak6
Cldn10
Satb2
Cited1
Slc1a1
Mcmdc2
Casr
Spp1
Asic2
Gcnt1
Cck
Me3
Slc26a4
Piwil2
Tmem45b

Ugt1a2
Paqr5
Afp
Grem1
Scin
Dusp15
Slc13a3
Wdr72
Kcnj10
Abca13
Tmem174
Ocm
Slc34a2
Ugt2b38
Bsnd
Slc5a9
Nphs2
Ptpro
Gng13
Krt7
Slc34a1
Ace2
Cldn2
Rasgrp1
Foxl1
Ptgs2
Mettl7a2
Sycp2
Npsr1
Slco1a6
Tmem52
Slc4a11
Dlec1
Fut9
Ptger3
Slc22a6
Umod
Adora2b
Rdh19
Sycp3
Car12
Cwh43
Taf7l
S100g
Smim5
Aldh3b2
Tshr

Slc5a10
Fxyd2
Cubn
Gja3
Sucnr1
Sectm1b
Gcm1
Mael
Efcab6
Mylk3
Cyba
Hao2
Sim2
Sema3g
Nobox
Clec18a
Cbln4
Trpc6
Ckmt1
Slc13a1
Hes5
Prss8
Mov10l1
Ugt1a9
Meox2
Gdf7
Pou2f3
Epha8
Folr1
Insc
Hsd3b6
Cyp2j13
Cckar
Ankrd63
Eddm3b
Kcne3
Tmem72
Scn4b
Rhbg
Pfn3
Slc14a2
Kcnj16
Ly6a
Slc22a8
Wnt10a
Lbhd2
Car4

Scel
Gpx3
Atp6v1c2
Cryab
Pdzk1ip1
Hmx2
Foxi1
Hsf2bp
B3galt5
Atp12a
Slc22a12
Ptger2
Atp6v0d2
Msx2
Spns3
Tmem213
Pcdh8
Slc22a29
Sohlh1
Upk1a
Fndc9
Sostdc1
Tac2
Car15
Vstm2a
Asb9
Defb29
Slc12a3
Siglecg
Fkbp6
Nts
Kcnj15
Zfp366
Pik3c2g
Aqp2
Atp13a4
Cyp4f39
