## Supplementary material for "Lysine-specific demethylase 1a is obligatory for gene regulation during kidney development": Suppl Table-4

**Supplementary Table-4**

| ID | log2FoldChange | pvalue | padj | KKO-1 | KKO-2 | KKO-3 | KKO-4 | KWT-1 | KWT-2 | KWT-3 | KWT-4 |
|---|---|---|---|---|---|---|---|---|---|---|---|
| ENSMUSG00000024935 | -2.629325979 | 8.99E-11 | 1.66E-08 | 48.29061 | 25.19625 | 103.5133 | 49.16954 | 271.0809 | 221.2401 | 390.5371 | 525.3065 |
| ENSMUSG00000005089 | 0.073189052 | 0.775181 | 0.860918 | 60.36326 | 65.99018 | 81.98256 | 79.36136 | 56.55646 | 97.21155 | 64.60737 | 56.55474 |
| ENSMUSG00000005360 | 0.849322123 | 0.002528 | 0.015553 | 1241.471 | 1456.583 | 938.2448 | 1343.967 | 580.1912 | 1179.947 | 547.7162 | 456.7883 |
| ENSMUSG00000020142 | -0.140719609 | 0.231315 | 0.391871 | 355.1372 | 284.3577 | 335.3832 | 319.1707 | 393.945 | 364.264 | 348.1084 | 321.927 |
| ENSMUSG00000001918 | 0.089454044 | 0.317598 | 0.485914 | 2416.543 | 2300.058 | 2379.978 | 2339.435 | 2274.935 | 1873.836 | 2377.937 | 2340.496 |
| ENSMUSG00000005357 | 0.174863798 | 0.613047 | 0.747022 | 31.18769 | 33.595 | 24.01509 | 17.25247 | 21.45245 | 24.58223 | 23.14294 | 23.927 |
| ENSMUSG00000008932 | 0.159832738 | 0.731917 | 0.831995 | 19.11503 | 26.39607 | 9.93728 | 37.95543 | 21.45245 | 23.46486 | 22.17865 | 16.31387 |
| ENSMUSG00000028645 | -0.927202588 | 9.17E-05 | 0.001252 | 1324.974 | 892.6672 | 1589.137 | 1157.641 | 2124.768 | 1588.906 | 3354.762 | 2374.211 |
| ENSMUSG00000027690 | -2.365064103 | 0.000406 | 0.003904 | 101.6115 | 16.7975 | 180.5272 | 135.4319 | 566.5397 | 357.5597 | 413.6801 | 903.7882 |
| ENSMUSG00000003153 | 0.523154285 | 0.000436 | 0.004116 | 393.3673 | 323.9518 | 370.1637 | 453.74 | 286.6827 | 305.0431 | 248.7866 | 233.8321 |
| ENSMUSG00000018566 | -0.236037337 | 0.054802 | 0.145465 | 497.9969 | 479.9286 | 479.4737 | 525.3377 | 649.4241 | 496.1141 | 660.5381 | 527.4817 |
| ENSMUSG00000085028 | 0.57128443 | 0.008852 | 0.039036 | 204.229 | 172.7743 | 159.8246 | 138.8824 | 112.1378 | 153.0803 | 89.67889 | 100.0584 |
| ENSMUSG00000028976 | -0.482626019 | 0.17005 | 0.316662 | 100.6054 | 103.1847 | 100.2009 | 106.1027 | 109.2125 | 74.86406 | 129.2147 | 259.9343 |
| ENSMUSG00000026791 | -0.07161116 | 0.630502 | 0.759418 | 608.6629 | 555.5174 | 811.5445 | 664.2201 | 612.3699 | 630.199 | 787.8242 | 747.1751 |
| ENSMUSG00000005107 | -0.478126972 | 0.063684 | 0.161739 | 203.223 | 165.5754 | 209.511 | 201.8539 | 213.5494 | 168.7235 | 381.8585 | 323.0146 |
| ENSMUSG00000027661 | -0.262985269 | 0.144408 | 0.28259 | 183.1019 | 153.5772 | 172.2462 | 135.4319 | 211.5992 | 150.8455 | 225.6437 | 183.8029 |
| ENSMUSG00000037490 | 0.083307379 | 0.726735 | 0.828973 | 138.8355 | 116.3827 | 141.6062 | 117.3168 | 104.3369 | 80.45093 | 140.7862 | 159.8759 |
| ENSMUSG00000036298 | 0.625789154 | 7.13E-05 | 0.001046 | 638.8445 | 698.2961 | 529.1601 | 684.0604 | 441.7254 | 464.8276 | 435.8587 | 308.8759 |
| ENSMUSG00000024131 | -4.233535956 | 1.35E-08 | 1.09E-06 | 66.39959 | 15.59768 | 103.5133 | 56.07053 | 1446.09 | 279.3435 | 729.0026 | 2099.051 |
| ENSMUSG00000010095 | -0.429953573 | 0.007365 | 0.03411 | 1890.376 | 1706.146 | 2308.761 | 1810.647 | 2429.002 | 1991.161 | 2758.831 | 3218.182 |
| ENSMUSG00000006574 | -0.189456979 | 0.755316 | 0.848145 | 349.1009 | 23.99643 | 102.6852 | 111.2784 | 148.2169 | 98.32892 | 152.3577 | 270.8102 |
| ENSMUSG00000029141 | -0.041997982 | 0.675475 | 0.792434 | 891.3642 | 935.8608 | 779.2483 | 742.7188 | 842.4962 | 815.6831 | 879.4317 | 904.8758 |
| ENSMUSG00000028962 | -0.210627883 | 0.113729 | 0.240848 | 1421.555 | 1478.18 | 1465.749 | 1357.769 | 1508.497 | 1326.323 | 1767.542 | 2019.657 |
| ENSMUSG00000006576 | 0.42567303 | 4.63E-05 | 0.000749 | 1741.48 | 1564.567 | 1310.893 | 1681.253 | 1198.412 | 1201.177 | 1104.111 | 1184.387 |
| ENSMUSG00000060961 | -0.687153588 | 4.46E-05 | 0.000729 | 1175.071 | 1306.606 | 1347.329 | 1469.048 | 2350.993 | 1588.906 | 1872.65 | 2718.978 |
| ENSMUSG00000068323 | 0.442772235 | 0.012367 | 0.050099 | 174.0474 | 148.7779 | 193.777 | 184.6014 | 114.088 | 164.254 | 123.429 | 116.3722 |
| ENSMUSG00000021733 | 0.277600489 | 0.214017 | 0.37205 | 1487.954 | 1330.602 | 1216.489 | 1454.383 | 978.0366 | 1743.104 | 1024.075 | 784.1532 |
| ENSMUSG00000023032 | 0.122683986 | 0.291818 | 0.459069 | 657.9595 | 571.1151 | 549.8628 | 508.9479 | 517.7841 | 505.0531 | 486.0017 | 591.6496 |
| ENSMUSG00000024485 | -1.675977914 | 0.048155 | 0.132611 | 148.896 | 69.58965 | 772.6235 | 121.6299 | 779.1139 | 339.6817 | 1079.04 | 1359.489 |
| ENSMUSG00000026904 | 0.060443166 | 0.8314 | 0.897939 | 45.27245 | 55.19179 | 33.12427 | 39.68068 | 34.1289 | 52.51658 | 40.50014 | 38.06569 |
| ENSMUSG00000074796 | -1.221065319 | 0.010043 | 0.042724 | 83.50251 | 73.18911 | 202.058 | 56.07053 | 274.0063 | 89.38993 | 234.3223 | 370.8686 |
| ENSMUSG00000011034 | -2.442223861 | 1.55E-08 | 1.22E-06 | 59.35721 | 35.99465 | 110.1382 | 52.62003 | 325.6872 | 151.9629 | 318.2154 | 612.3138 |
| ENSMUSG00000030781 | -2.882209946 | 1.55E-06 | 5.33E-05 | 21.12714 | 7.198929 | 51.34261 | 28.46658 | 333.4881 | 62.57295 | 114.7504 | 292.562 |
| ENSMUSG00000089774 | -1.097304592 | 6.10E-05 | 0.000928 | 914.5034 | 725.892 | 1039.274 | 905.7547 | 1089.199 | 1790.033 | 3134.904 | 1658.576 |
| ENSMUSG00000006641 | -0.763781821 | 0.002098 | 0.013581 | 298.7981 | 239.9643 | 275.7595 | 251.0234 | 506.0828 | 271.5219 | 383.7871 | 648.2043 |
| ENSMUSG00000023945 | 0.540108428 | 0.366498 | 0.535049 | 15.09082 | 19.19714 | 10.76539 | 10.35148 | 4.875557 | 7.821619 | 5.785735 | 19.57664 |
| ENSMUSG00000020062 | -4.876815617 | 3.08E-07 | 1.40E-05 | 6.036326 | 7.198929 | 73.70149 | 13.80198 | 628.9468 | 205.5968 | 648.9666 | 1490 |
| ENSMUSG00000028544 | -1.652801749 | 4.55E-05 | 0.00074 | 44.26639 | 20.39697 | 63.76421 | 36.23019 | 124.8142 | 78.21619 | 106.0718 | 212.0803 |
| ENSMUSG00000042371 | -4.917652246 | 8.55E-17 | 6.07E-14 | 6.036326 | 4.799286 | 15.73403 | 6.038364 | 261.3298 | 63.69032 | 197.6793 | 478.5401 |
| ENSMUSG00000030769 | -3.234341524 | 4.63E-06 | 0.000126 | 2.012109 | 7.198929 | 19.04645 | 9.488858 | 134.5654 | 24.58223 | 42.42872 | 157.7007 |
| ENSMUSG00000041644 | -2.999824438 | 1.57E-07 | 8.19E-06 | 21.12714 | 7.198929 | 89.43552 | 29.3292 | 376.393 | 153.0803 | 328.8226 | 327.3649 |
| ENSMUSG00000030310 | 0.437625082 | 0.016689 | 0.061987 | 116.7023 | 103.1847 | 99.3728 | 93.16334 | 73.13335 | 89.38993 | 72.32169 | 69.60583 |
| ENSMUSG00000055368 | 3.98928519 | 0.001687 | 0.011549 | 334.01 | 3.599465 | 10.76539 | 356.2635 | 17.552 | 7.821619 | 13.50005 | 5.437955 |
| ENSMUSG00000020838 | -0.281912994 | 0.445912 | 0.608619 | 61.36932 | 44.3934 | 53.82693 | 49.16954 | 78.98402 | 34.6386 | 41.46443 | 98.97079 |
| ENSMUSG00000039728 | 2.498750346 | 0.002842 | 0.017029 | 158.9566 | 13.19804 | 8.281066 | 40.5433 | 13.65156 | 11.17374 | 7.714313 | 6.525547 |
| ENSMUSG00000030096 | 0.130375827 | 0.211268 | 0.368646 | 5621.832 | 4356.552 | 4731.801 | 4843.631 | 4126.671 | 4551.065 | 5007.554 | 4179.613 |
| ENSMUSG00000019558 | -0.327204521 | 0.018048 | 0.065766 | 1440.67 | 1148.229 | 1478.998 | 1266.331 | 1628.436 | 1338.614 | 1777.185 | 1950.051 |
| ENSMUSG00000028542 | -0.0079559 | 0.946459 | 0.969279 | 325.9616 | 310.7538 | 346.9767 | 330.3848 | 379.3183 | 298.3389 | 343.2869 | 300.1751 |
| ENSMUSG00000030307 | -0.105024344 | 0.690014 | 0.803616 | 35.2119 | 39.59411 | 61.27989 | 47.44429 | 48.75557 | 45.81234 | 53.0359 | 51.11678 |
| ENSMUSG00000030109 | -3.373332539 | 1.37E-12 | 3.88E-10 | 25.15136 | 4.799286 | 33.95237 | 8.626235 | 210.624 | 126.2633 | 198.6436 | 225.1314 |
| ENSMUSG00000030108 | -2.570414224 | 7.44E-06 | 0.000184 | 131.7931 | 53.99197 | 219.4483 | 110.4158 | 692.329 | 291.6346 | 738.6455 | 1344.263 |
| ENSMUSG00000019894 | -0.33041047 | 0.240966 | 0.403112 | 137.8294 | 140.3791 | 93.57605 | 147.5086 | 133.5902 | 110.62 | 159.1077 | 249.0584 |
| ENSMUSG00000027894 | -0.889202686 | 9.36E-07 | 3.53E-05 | 162.9808 | 170.3747 | 236.0104 | 191.5024 | 351.0401 | 275.9914 | 363.537 | 421.9853 |
| ENSMUSG00000021612 | -3.691801868 | 7.18E-06 | 0.000179 | 106.6418 | 17.99732 | 364.3669 | 153.547 | 1953.148 | 835.7958 | 2301.758 | 3220.357 |
| ENSMUSG00000021565 | -2.762476486 | 5.62E-09 | 5.29E-07 | 31.18769 | 14.39786 | 51.34261 | 21.56559 | 284.7325 | 72.62932 | 190.9293 | 261.0219 |
| ENSMUSG00000036814 | -2.483347701 | 6.82E-05 | 0.001009 | 4.024217 | 0 | 10.76539 | 6.900988 | 33.15378 | 18.99536 | 34.71441 | 39.15328 |
| ENSMUSG00000025243 | -4.050333588 | 3.12E-11 | 6.39E-09 | 10.06054 | 7.198929 | 35.60859 | 6.900988 | 285.7076 | 81.56831 | 176.4649 | 457.8758 |
| ENSMUSG00000041313 | 0.075900394 | 0.529233 | 0.681503 | 1319.943 | 950.2587 | 1196.614 | 1002.369 | 1055.07 | 994.4629 | 1083.861 | 1108.255 |
| ENSMUSG00000031596 | 0.202473635 | 0.07418 | 0.179688 | 743.4742 | 686.2979 | 807.404 | 740.9936 | 627.9717 | 707.2978 | 545.7877 | 711.2846 |
| ENSMUSG00000031297 | 0.863369996 | 4.63E-06 | 0.000126 | 127.7689 | 151.1775 | 118.4192 | 146.646 | 67.28268 | 63.69032 | 88.7146 | 78.30656 |
| ENSMUSG00000022756 | -0.235082494 | 0.331574 | 0.499409 | 150.9082 | 133.1802 | 186.324 | 123.3552 | 145.2916 | 126.2633 | 189.965 | 238.1824 |
| ENSMUSG00000040010 | -0.044733732 | 0.572721 | 0.715262 | 808.8677 | 746.289 | 819.8256 | 722.8785 | 785.9397 | 806.7441 | 765.6456 | 839.6203 |

| | | | | | | | | | | | |
|---|---|---|---|---|---|---|---|---|---|---|---|
| ENSMUSG00000031904 | -0.168384071 | 0.0931 | 0.209847 | 1073.46 | 1071.441 | 1209.864 | 1083.455 | 1294.948 | 1053.684 | 1336.505 | 1302.934 |
| ENSMUSG00000033106 | -0.02506277 | 0.798726 | 0.876883 | 1207.265 | 1352.199 | 1190.817 | 1295.66 | 1390.509 | 1401.187 | 1207.29 | 1133.27 |
| ENSMUSG00000000958 | -0.737487383 | 0.003997 | 0.021824 | 320.9313 | 463.1311 | 445.5214 | 452.8773 | 910.754 | 434.6585 | 536.1448 | 923.3648 |
| ENSMUSG00000022180 | -0.865765938 | 0.015853 | 0.059945 | 512.0817 | 263.9607 | 835.5596 | 339.8737 | 841.5211 | 547.5133 | 816.7529 | 1352.963 |
| ENSMUSG00000030492 | -2.223035781 | 8.86E-08 | 5.13E-06 | 64.38748 | 25.19625 | 101.029 | 70.73513 | 334.4632 | 159.7845 | 254.5723 | 477.4525 |
| ENSMUSG00000030495 | 0.866537673 | 0.106667 | 0.230871 | 28.16952 | 70.78947 | 48.03018 | 43.13117 | 20.47734 | 53.63396 | 8.678602 | 21.75182 |
| ENSMUSG00000027737 | -0.224560633 | 0.472786 | 0.632209 | 50.30272 | 31.19536 | 35.60859 | 31.91707 | 38.02934 | 54.75133 | 30.85725 | 51.11678 |
| ENSMUSG00000039710 | -5.842807431 | 1.29E-06 | 4.58E-05 | 0 | 0 | 10.76539 | 3.450494 | 175.52 | 52.51658 | 246.858 | 361.0802 |
| ENSMUSG00000041052 | -5.026631181 | 8.38E-09 | 7.40E-07 | 1.006054 | 4.799286 | 19.87456 | 6.038364 | 209.6489 | 79.33356 | 267.1081 | 490.5036 |
| ENSMUSG00000069072 | 1.227200148 | 0.01099 | 0.045808 | 55.33299 | 23.99643 | 22.35888 | 65.55939 | 20.47734 | 24.58223 | 12.53576 | 14.13868 |
| ENSMUSG00000054640 | 0.043598486 | 0.811238 | 0.885092 | 744.4802 | 757.0874 | 843.0126 | 637.4788 | 739.1344 | 487.1751 | 790.7171 | 875.5108 |
| ENSMUSG00000030376 | 0.189255755 | 0.217297 | 0.375825 | 122.7386 | 167.975 | 147.403 | 156.1349 | 135.5405 | 127.3806 | 131.1433 | 126.1606 |
| ENSMUSG00000079055 | 0.273903413 | 0.347789 | 0.516479 | 42.25428 | 27.5959 | 47.20208 | 48.30692 | 31.20356 | 40.22547 | 29.89296 | 36.9781 |
| ENSMUSG00000032754 | 0.412909493 | 5.44E-05 | 0.000845 | 702.2259 | 747.4888 | 713.8279 | 787.5752 | 596.7681 | 573.2129 | 575.6806 | 469.8393 |
| ENSMUSG00000028854 | 0.131020322 | 0.174486 | 0.322656 | 809.8738 | 712.694 | 854.606 | 714.2523 | 720.6073 | 652.5465 | 707.7882 | 743.9123 |
| ENSMUSG00000026062 | -0.551014086 | 0.090707 | 0.206346 | 248.4954 | 172.7743 | 289.8373 | 200.9913 | 264.2552 | 174.3104 | 352.9298 | 545.9707 |
| ENSMUSG00000036123 | -3.338832408 | 2.16E-06 | 6.84E-05 | 15.09082 | 3.599465 | 50.5145 | 11.21411 | 190.1467 | 41.34284 | 290.251 | 299.0875 |
| ENSMUSG00000020733 | -0.80028617 | 1.46E-07 | 7.71E-06 | 1469.845 | 1534.572 | 1902.989 | 1505.278 | 2607.448 | 2214.635 | 2862.01 | 3485.729 |
| ENSMUSG00000002504 | -0.785032401 | 4.45E-05 | 0.000729 | 1229.398 | 1131.432 | 1688.509 | 1335.341 | 2120.867 | 1643.657 | 2629.617 | 2887.554 |
| ENSMUSG00000026065 | 0.530065781 | 0.057305 | 0.149911 | 97.58727 | 50.3925 | 106.8258 | 84.5371 | 57.53157 | 56.98608 | 50.14304 | 71.78101 |
| ENSMUSG00000014786 | 0.611667747 | 7.91E-05 | 0.001131 | 301.8163 | 437.9349 | 377.6166 | 360.5766 | 241.8276 | 283.813 | 214.0722 | 227.3065 |
| ENSMUSG00000060681 | -0.070986337 | 0.262887 | 0.427212 | 808.8677 | 832.6761 | 827.2785 | 828.1186 | 897.1024 | 857.0259 | 840.8602 | 867.8977 |
| ENSMUSG00000037341 | -0.691894676 | 0.076591 | 0.183719 | 29.17558 | 19.19714 | 32.29616 | 12.07673 | 34.1289 | 27.93435 | 43.39301 | 44.59123 |
| ENSMUSG00000039463 | 0.103933877 | 0.208876 | 0.366165 | 1189.156 | 1232.217 | 1159.349 | 1186.97 | 1230.59 | 1115.139 | 998.0393 | 1090.854 |
| ENSMUSG00000031129 | -0.461580407 | 0.080537 | 0.189809 | 53.32088 | 93.58608 | 120.9036 | 85.39973 | 108.2374 | 112.8548 | 147.5362 | 118.5474 |
| ENSMUSG00000037994 | 0.759318661 | 0.004736 | 0.024759 | 207.2472 | 229.1659 | 112.6225 | 241.5346 | 119.9387 | 115.0895 | 141.7505 | 89.18247 |
| ENSMUSG00000021135 | -1.700370793 | 0.073673 | 0.178862 | 17.10292 | 1.199822 | 1.656213 | 7.763611 | 33.15378 | 6.704245 | 5.785735 | 44.59123 |
| ENSMUSG00000023073 | -4.577577047 | 3.35E-08 | 2.33E-06 | 2.012109 | 4.799286 | 2.48432 | 0.862623 | 31.20356 | 10.05637 | 91.60747 | 102.2336 |
| ENSMUSG00000032806 | -0.266196332 | 0.047322 | 0.130902 | 299.8042 | 293.9563 | 285.6968 | 299.3304 | 353.9654 | 298.3389 | 334.6083 | 430.6861 |
| ENSMUSG00000029219 | 1.270118792 | 0.059255 | 0.153404 | 49.29666 | 8.398751 | 18.21835 | 92.30071 | 20.47734 | 21.23011 | 17.3572 | 10.87591 |
| ENSMUSG00000058921 | -2.632876974 | 0.001816 | 0.01224 | 3.018163 | 1.199822 | 11.59349 | 1.725247 | 19.50223 | 8.938993 | 22.17865 | 60.9051 |
| ENSMUSG00000029321 | 1.234948999 | 0.021852 | 0.075006 | 77.46619 | 62.39072 | 36.43669 | 85.39973 | 12.67645 | 62.57295 | 22.17865 | 14.13868 |
| ENSMUSG00000031684 | 0.001372323 | 0.986283 | 0.992283 | 631.8021 | 693.4968 | 657.5167 | 630.5778 | 617.2455 | 632.4337 | 688.5025 | 669.9561 |
| ENSMUSG00000026177 | 0.390389881 | 0.230724 | 0.391167 | 53.32088 | 46.79304 | 81.98256 | 50.89479 | 44.85512 | 36.87335 | 64.60737 | 31.54014 |
| ENSMUSG00000023030 | -0.615793788 | 0.000315 | 0.003228 | 1137.847 | 927.462 | 1068.258 | 953.199 | 1632.336 | 1102.848 | 1550.577 | 1977.241 |
| ENSMUSG00000027202 | -2.418862845 | 0.030377 | 0.095285 | 567.4147 | 67.19001 | 5578.126 | 899.7163 | 7565.889 | 3592.358 | 12942.69 | 13935.3 |
| ENSMUSG00000024597 | 0.642559254 | 0.008216 | 0.037001 | 7048.417 | 8055.602 | 5841.464 | 7535.016 | 3912.147 | 7451.768 | 3517.727 | 3362.832 |
| ENSMUSG00000031766 | -2.81066857 | 3.23E-06 | 9.59E-05 | 160.9687 | 73.18911 | 341.1799 | 161.3106 | 992.6633 | 454.7713 | 1540.934 | 2184.97 |
| ENSMUSG00000017765 | 0.089169141 | 0.376402 | 0.544682 | 1011.085 | 1125.433 | 1197.442 | 1133.487 | 1129.179 | 992.2282 | 931.5033 | 1148.496 |
| ENSMUSG00000017740 | 0.100368664 | 0.369372 | 0.538266 | 262.5802 | 283.1579 | 287.353 | 253.6113 | 253.5289 | 263.7003 | 230.4651 | 266.4598 |
| ENSMUSG00000027130 | 0.382076441 | 0.008534 | 0.038027 | 1387.349 | 1262.212 | 1079.023 | 1446.62 | 945.858 | 1235.816 | 936.3248 | 853.759 |
| ENSMUSG00000017756 | -0.283410581 | 0.002935 | 0.017499 | 896.3944 | 820.6779 | 899.3238 | 781.5369 | 1071.647 | 949.768 | 972.9678 | 1141.971 |
| ENSMUSG00000035506 | 0.17010098 | 0.571354 | 0.714306 | 38.23007 | 39.59411 | 37.2648 | 46.58167 | 33.15378 | 44.69496 | 23.14294 | 43.50364 |
| ENSMUSG00000037344 | 0.017608838 | 0.836948 | 0.901367 | 488.9424 | 445.1338 | 493.5516 | 475.3055 | 442.7005 | 469.2971 | 488.8946 | 481.8029 |
| ENSMUSG00000029700 | -3.211759482 | 1.12E-08 | 9.52E-07 | 5.030272 | 5.999108 | 16.56213 | 18.97772 | 172.5947 | 36.87335 | 91.60747 | 137.0365 |
| ENSMUSG00000001095 | -2.568567286 | 1.66E-07 | 8.51E-06 | 9.054489 | 10.79839 | 24.01509 | 13.80198 | 102.3867 | 35.75597 | 69.42882 | 138.1241 |
| ENSMUSG00000018459 | -2.898627159 | 8.10E-06 | 0.000195 | 17.10292 | 9.598572 | 57.96746 | 30.19182 | 317.8863 | 52.51658 | 101.2504 | 390.4452 |
| ENSMUSG00000029843 | 0.191211658 | 0.766983 | 0.855469 | 18.10898 | 13.19804 | 7.45296 | 6.038364 | 7.80089 | 10.05637 | 3.857157 | 17.40146 |
| ENSMUSG00000020805 | 0.944498709 | 0.086744 | 0.199942 | 12.07265 | 25.19625 | 15.73403 | 11.21411 | 5.850668 | 12.29112 | 10.60718 | 4.350364 |
| ENSMUSG00000059336 | -1.265651492 | 0.015705 | 0.059687 | 81.4904 | 19.19714 | 171.4181 | 40.5433 | 227.2009 | 191.071 | 144.6434 | 191.416 |
| ENSMUSG00000024552 | -1.56598976 | 0.000823 | 0.006715 | 88.53278 | 99.58519 | 384.2415 | 108.6906 | 378.3432 | 311.7474 | 743.4669 | 586.2116 |
| ENSMUSG00000025557 | -1.079661193 | 0.090925 | 0.206612 | 12.07265 | 2.399643 | 19.04645 | 6.038364 | 14.62667 | 13.40849 | 23.14294 | 33.71532 |
| ENSMUSG00000022899 | -0.094595712 | 0.738062 | 0.836054 | 772.6497 | 914.264 | 635.9859 | 522.7498 | 418.3228 | 625.7295 | 947.8962 | 1045.175 |
| ENSMUSG00000024737 | -0.205923799 | 0.516686 | 0.670791 | 38.23007 | 41.99375 | 52.17072 | 43.9938 | 53.63112 | 26.81698 | 57.85735 | 65.25547 |
| ENSMUSG00000029416 | -0.060166294 | 0.649641 | 0.773185 | 284.7134 | 334.7502 | 330.4145 | 257.9244 | 303.2596 | 282.6956 | 327.8583 | 343.6788 |
| ENSMUSG00000032902 | 0.287684442 | 0.034526 | 0.104995 | 2896.43 | 4165.78 | 3042.464 | 3346.979 | 3163.261 | 2790.083 | 2618.045 | 2444.905 |
| ENSMUSG00000033965 | 0.662477025 | 3.94E-05 | 0.000666 | 907.461 | 1262.212 | 1252.097 | 1203.36 | 685.5032 | 935.2421 | 648.9666 | 653.6422 |
| ENSMUSG00000025161 | 0.758619556 | 2.16E-06 | 6.84E-05 | 608.6629 | 403.14 | 427.303 | 540.8649 | 313.0107 | 275.9914 | 311.4654 | 269.7226 |
| ENSMUSG00000027896 | -0.440994896 | 0.022521 | 0.076682 | 295.78 | 320.3523 | 281.5563 | 364.8897 | 429.049 | 353.0902 | 346.1798 | 586.2116 |
| ENSMUSG00000045775 | -1.618666938 | 6.24E-06 | 0.00016 | 172.0353 | 172.7743 | 390.8663 | 203.5791 | 645.5237 | 357.5597 | 764.6813 | 1120.219 |
| ENSMUSG00000041920 | -0.238484412 | 0.043238 | 0.123138 | 342.0585 | 377.9438 | 372.648 | 341.5989 | 456.3521 | 435.7759 | 351.9655 | 448.0875 |
| ENSMUSG00000020102 | -0.601962203 | 0.043311 | 0.123203 | 462.785 | 442.7341 | 808.2321 | 477.8934 | 627.9717 | 492.762 | 1015.396 | 1192 |
| ENSMUSG00000032988 | 0.438991186 | 0.323971 | 0.491966 | 23.13925 | 47.99286 | 35.60859 | 26.74133 | 16.57689 | 13.40849 | 39.53586 | 28.27737 |
| ENSMUSG00000037762 | -0.631524525 | 0.005357 | 0.026941 | 187.1261 | 250.7627 | 209.511 | 155.2722 | 305.2098 | 214.5358 | 312.4297 | 408.9342 |

| | | | | | | | | | | | |
|---|---|---|---|---|---|---|---|---|---|---|---|
| ENSMUSG00000019838 | -0.115940742 | 0.579092 | 0.720342 | 391.3551 | 284.3577 | 355.2577 | 354.5383 | 345.1894 | 280.4609 | 344.2512 | 532.9196 |
| ENSMUSG00000040938 | 0.119283446 | 0.315859 | 0.484252 | 187.1261 | 183.5727 | 197.0894 | 203.5791 | 182.3458 | 179.8972 | 178.3935 | 170.7518 |
| ENSMUSG00000009378 | -2.493031591 | 0.000333 | 0.003357 | 133.8052 | 32.39518 | 345.3205 | 127.6683 | 902.9531 | 404.4894 | 1020.218 | 1274.657 |
| ENSMUSG00000044367 | -0.551222222 | 0.00102 | 0.00796 | 316.9071 | 279.5584 | 328.7583 | 302.7808 | 446.601 | 345.2686 | 427.1801 | 581.8612 |
| ENSMUSG00000026220 | -0.576721812 | 0.223032 | 0.382554 | 14.08476 | 5.999108 | 23.18699 | 12.93935 | 18.52711 | 21.23011 | 17.3572 | 28.27737 |
| ENSMUSG00000021335 | -3.610677438 | 1.12E-08 | 9.52E-07 | 52.31483 | 17.99732 | 119.2474 | 56.07053 | 894.1771 | 313.9821 | 600.7521 | 1200.701 |
| ENSMUSG00000036110 | -2.184053654 | 0.003095 | 0.018205 | 3.018163 | 2.399643 | 19.04645 | 2.58787 | 28.27823 | 16.76061 | 32.78583 | 47.85401 |
| ENSMUSG00000036083 | -4.009541891 | 7.91E-12 | 1.87E-09 | 13.07871 | 4.799286 | 34.78048 | 15.52722 | 338.3636 | 89.38993 | 177.4292 | 506.8174 |
| ENSMUSG00000021336 | -1.009108673 | 0.000973 | 0.007672 | 65.39353 | 38.39429 | 76.18581 | 42.26855 | 144.3165 | 82.68568 | 92.57176 | 129.4233 |
| ENSMUSG00000049624 | -0.294331875 | 0.015217 | 0.058302 | 617.7174 | 704.2952 | 688.1566 | 654.7312 | 857.1228 | 668.1897 | 797.4671 | 944.0291 |
| ENSMUSG00000030500 | -0.219323555 | 0.613332 | 0.747221 | 8.048435 | 15.59768 | 14.07781 | 11.21411 | 14.62667 | 14.52586 | 13.50005 | 14.13868 |
| ENSMUSG00000070570 | 0.030583611 | 0.796806 | 0.875296 | 287.7315 | 307.1543 | 326.274 | 326.9343 | 287.6578 | 354.2076 | 280.6081 | 302.3503 |
| ENSMUSG00000019935 | -0.256202037 | 0.850384 | 0.910207 | 7.04238 | 4.799286 | 427.303 | 18.11509 | 122.864 | 45.81234 | 175.5006 | 202.2919 |
| ENSMUSG00000023393 | 0.589514073 | 0.161502 | 0.305367 | 43.26034 | 27.5959 | 28.98373 | 32.77969 | 20.47734 | 27.93435 | 30.85725 | 8.700729 |
| ENSMUSG00000036330 | -2.827499857 | 3.07E-05 | 0.000546 | 81.4904 | 16.7975 | 39.74912 | 97.47645 | 556.7886 | 130.7328 | 336.5369 | 650.3795 |
| ENSMUSG00000025094 | 1.027606431 | 0.01887 | 0.067676 | 293.7679 | 128.3809 | 90.26362 | 393.3563 | 88.73513 | 159.7845 | 107.0361 | 89.18247 |
| ENSMUSG00000037455 | 0.429255911 | 7.84E-06 | 0.000191 | 767.6195 | 753.4879 | 689.8128 | 819.4923 | 543.137 | 610.0863 | 515.8947 | 582.9488 |
| ENSMUSG00000001436 | -0.993473969 | 3.27E-06 | 9.68E-05 | 486.9303 | 401.9402 | 569.7374 | 493.4206 | 961.4597 | 619.0252 | 1015.396 | 1293.146 |
| ENSMUSG00000040918 | -0.104153767 | 0.317325 | 0.485577 | 351.113 | 337.1499 | 387.5539 | 346.7746 | 358.841 | 379.9072 | 423.3229 | 368.6934 |
| ENSMUSG00000038496 | -0.31452414 | 0.144455 | 0.282595 | 70.4238 | 65.99018 | 78.67013 | 66.42201 | 100.4365 | 62.57295 | 94.50034 | 92.44524 |
| ENSMUSG00000027397 | -0.226714176 | 0.004025 | 0.021935 | 1096.599 | 958.6574 | 1172.599 | 1067.928 | 1277.396 | 1252.576 | 1250.683 | 1252.905 |
| ENSMUSG00000037656 | -0.084939778 | 0.292496 | 0.459782 | 2089.575 | 2189.674 | 1983.315 | 1987.485 | 2270.059 | 1962.109 | 2144.579 | 2368.773 |
| ENSMUSG00000023829 | -3.163770175 | 8.10E-06 | 0.000195 | 60.36326 | 10.79839 | 107.6539 | 47.44429 | 612.3699 | 203.3621 | 372.2156 | 845.0583 |
| ENSMUSG00000040966 | -5.174584012 | 1.37E-09 | 1.71E-07 | 9.054489 | 4.799286 | 45.54586 | 18.11509 | 883.4508 | 151.9629 | 500.4661 | 1282.27 |
| ENSMUSG00000023828 | 0.185533683 | 0.550091 | 0.698052 | 49.29666 | 87.58697 | 91.91984 | 61.24627 | 85.8098 | 70.39457 | 44.3573 | 54.37955 |
| ENSMUSG00000020334 | -1.065579167 | 0.026728 | 0.087216 | 34.20585 | 41.99375 | 55.48314 | 16.38985 | 77.03379 | 33.52122 | 66.53595 | 132.6861 |
| ENSMUSG00000018900 | -0.361100222 | 0.02318 | 0.078469 | 929.5942 | 796.6815 | 872.8244 | 971.3141 | 1180.86 | 883.8429 | 1050.111 | 1472.598 |
| ENSMUSG00000024650 | -4.559778162 | 1.03E-06 | 3.82E-05 | 60.36326 | 9.598572 | 227.7293 | 56.07053 | 2274.935 | 611.2036 | 1394.362 | 4074.116 |
| ENSMUSG00000067144 | -5.674314347 | 1.68E-05 | 0.00034 | 0 | 1.199822 | 0 | 0.862623 | 15.60178 | 4.469496 | 15.42863 | 63.08028 |
| ENSMUSG00000063796 | -1.854957495 | 2.94E-06 | 8.88E-05 | 148.896 | 91.18644 | 279.0719 | 163.8985 | 772.2882 | 309.5126 | 421.3944 | 972.3064 |
| ENSMUSG00000061742 | -2.675125951 | 0.003258 | 0.018873 | 37.22401 | 10.79839 | 173.9024 | 46.58167 | 366.6419 | 121.7938 | 317.2511 | 912.4889 |
| ENSMUSG00000074028 | -4.309560554 | 3.26E-05 | 0.00057 | 3.018163 | 3.599465 | 43.88965 | 6.038364 | 190.1467 | 67.04245 | 272.8938 | 599.2627 |
| ENSMUSG00000033147 | 0.103085244 | 0.468316 | 0.628589 | 291.7558 | 283.1579 | 308.8838 | 304.5061 | 255.4792 | 343.0338 | 275.7867 | 233.8321 |
| ENSMUSG00000022199 | 0.228524485 | 0.004496 | 0.023786 | 2583.548 | 2905.968 | 2658.222 | 2897.552 | 2507.986 | 2500.683 | 2231.365 | 2186.058 |
| ENSMUSG00000000154 | -1.335082452 | 0.001083 | 0.008295 | 78.47224 | 76.78858 | 136.6376 | 64.69676 | 239.8774 | 90.5073 | 190.9293 | 379.5693 |
| ENSMUSG00000024757 | -4.972972177 | 4.41E-05 | 0.000725 | 14.08476 | 0 | 78.67013 | 22.42821 | 980.962 | 203.3621 | 743.4669 | 1699.905 |
| ENSMUSG00000063652 | 0.265407877 | 0.032941 | 0.101158 | 353.1251 | 369.545 | 372.648 | 416.6471 | 316.9112 | 363.1466 | 310.5011 | 268.635 |
| ENSMUSG00000022366 | -6.283077507 | 8.78E-09 | 7.60E-07 | 0 | 3.599465 | 0 | 0 | 48.75557 | 20.11273 | 38.57157 | 130.5109 |
| ENSMUSG00000038267 | -0.80463903 | 0.001065 | 0.008197 | 355.1372 | 293.9563 | 494.3797 | 287.2536 | 636.7477 | 395.5504 | 669.2167 | 799.3794 |
| ENSMUSG00000053303 | -4.571905625 | 1.75E-05 | 0.000354 | 2.012109 | 0 | 1.656213 | 0 | 12.67645 | 5.58687 | 37.60728 | 33.71532 |
| ENSMUSG00000067656 | -2.741472047 | 1.71E-06 | 5.78E-05 | 6.036326 | 4.799286 | 18.21835 | 13.80198 | 44.85512 | 32.40385 | 83.89316 | 130.5109 |
| ENSMUSG00000063590 | -3.156670398 | 3.22E-06 | 9.58E-05 | 1.006054 | 5.999108 | 3.312427 | 7.763611 | 42.9049 | 20.11273 | 22.17865 | 76.13138 |
| ENSMUSG00000075044 | -4.036426996 | 3.01E-19 | 3.47E-16 | 10.06054 | 4.799286 | 19.04645 | 10.35148 | 174.5449 | 92.74205 | 178.3935 | 295.8248 |
| ENSMUSG00000052562 | -3.233211126 | 1.11E-12 | 3.30E-10 | 7.04238 | 11.99822 | 7.45296 | 4.313117 | 69.2329 | 60.3382 | 44.3573 | 109.8467 |
| ENSMUSG00000024354 | -2.629234026 | 1.80E-05 | 0.000363 | 151.9142 | 53.99197 | 332.0708 | 144.0581 | 1004.365 | 453.6539 | 1006.718 | 1759.722 |
| ENSMUSG00000027340 | 0.445402942 | 0.008345 | 0.037423 | 1668.038 | 1100.236 | 991.2436 | 1135.213 | 896.1273 | 1010.106 | 849.5388 | 838.5327 |
| ENSMUSG00000026205 | -1.702891512 | 7.21E-07 | 2.85E-05 | 48.29061 | 35.99465 | 66.24853 | 46.58167 | 155.0427 | 79.33356 | 179.3578 | 230.5693 |
| ENSMUSG00000029847 | 0.472360022 | 0.154773 | 0.296408 | 27.16347 | 34.79482 | 28.98373 | 26.74133 | 28.27823 | 20.11273 | 16.39292 | 19.57664 |
| ENSMUSG00000034452 | 0.069482526 | 0.611411 | 0.746062 | 277.671 | 320.3523 | 305.5713 | 320.0333 | 305.2098 | 324.0385 | 232.3937 | 305.6131 |
| ENSMUSG00000037996 | 0.360529607 | 0.435782 | 0.599895 | 26.15741 | 10.79839 | 24.01509 | 31.91707 | 14.62667 | 29.05173 | 15.42863 | 14.13868 |
| ENSMUSG00000063873 | 0.790075432 | 0.008667 | 0.038451 | 471.8395 | 746.289 | 546.5504 | 476.1682 | 274.0063 | 557.5697 | 254.5723 | 209.9051 |
| ENSMUSG00000041771 | 0.06192904 | 0.860658 | 0.916562 | 19.11503 | 23.99643 | 21.53077 | 33.64232 | 22.42756 | 26.81698 | 18.32149 | 27.18978 |
| ENSMUSG00000035183 | 0.109108139 | 0.575291 | 0.717501 | 76.46013 | 121.182 | 93.57605 | 94.02596 | 86.78491 | 90.5073 | 93.53605 | 84.8321 |
| ENSMUSG00000003528 | -0.583101661 | 0.000298 | 0.003083 | 1120.745 | 1035.446 | 1291.018 | 1067.065 | 1704.495 | 1231.346 | 1726.078 | 2102.314 |
| ENSMUSG00000061904 | -0.501650339 | 0.000512 | 0.004669 | 13666.24 | 10788.8 | 13608.28 | 12135.39 | 17902.07 | 13357.09 | 19129.57 | 20685.98 |
| ENSMUSG00000031633 | -0.514832662 | 2.71E-05 | 0.000495 | 10064.57 | 11094.75 | 11321.87 | 11183.91 | 16148.82 | 12197.26 | 15743.95 | 18299.81 |
| ENSMUSG00000016319 | -1.285689171 | 2.48E-07 | 1.20E-05 | 4903.509 | 5234.821 | 7362.696 | 4583.119 | 13273.22 | 7408.19 | 15104.63 | 18056.19 |
| ENSMUSG00000025792 | -1.217626256 | 1.06E-05 | 0.000238 | 893.3763 | 782.2836 | 1283.565 | 873.8376 | 2289.561 | 1202.295 | 2049.114 | 3375.883 |
| ENSMUSG00000014606 | -0.537461199 | 0.000254 | 0.002745 | 1918.546 | 2072.092 | 2382.463 | 1952.117 | 2943.861 | 2293.969 | 3199.511 | 3646.693 |
| ENSMUSG00000027010 | -0.28201544 | 0.056072 | 0.147526 | 1548.318 | 1183.024 | 1427.656 | 1457.834 | 1688.893 | 1295.037 | 1931.471 | 1915.248 |
| ENSMUSG00000015112 | -0.987453102 | 4.82E-06 | 0.000131 | 775.6679 | 703.0954 | 858.7466 | 640.9293 | 1486.07 | 888.3124 | 1600.72 | 1930.474 |
| ENSMUSG00000031105 | 0.236495449 | 0.051085 | 0.138101 | 273.6468 | 278.3586 | 236.8385 | 264.8254 | 224.2756 | 249.1744 | 213.1079 | 206.6423 |
| ENSMUSG00000031482 | -0.57291101 | 1.95E-05 | 0.000385 | 481.9 | 487.1275 | 536.6131 | 530.5134 | 789.8402 | 611.2036 | 720.324 | 909.2261 |
| ENSMUSG00000071253 | -0.524471984 | 0.000108 | 0.001424 | 588.5418 | 507.5245 | 656.6886 | 615.9132 | 840.5459 | 689.4198 | 908.3604 | 972.3064 |

| | | | | | | | | | | | |
|---|---|---|---|---|---|---|---|---|---|---|---|
| ENSMUSG00000022404 | -0.224337736 | 0.010375 | 0.043785 | 2164.023 | 1949.71 | 1927.832 | 2160.872 | 2452.405 | 2101.781 | 2538.973 | 2488.408 |
| ENSMUSG00000004902 | 0.047795798 | 0.917704 | 0.951364 | 19.11503 | 25.19625 | 14.07781 | 18.97772 | 26.32801 | 25.6996 | 12.53576 | 9.78832 |
| ENSMUSG00000020744 | -0.343649152 | 0.020263 | 0.071112 | 641.8627 | 697.0963 | 647.5794 | 631.4404 | 885.4011 | 664.8376 | 729.9669 | 1040.825 |
| ENSMUSG00000032602 | -0.219044356 | 0.206881 | 0.363479 | 1196.199 | 1264.612 | 1273.628 | 1218.887 | 1394.409 | 1082.735 | 1295.04 | 1993.554 |
| ENSMUSG00000035472 | -0.538532725 | 0.061362 | 0.157418 | 65.39353 | 35.99465 | 40.57722 | 48.30692 | 69.2329 | 52.51658 | 63.64308 | 91.35765 |
| ENSMUSG00000019082 | 0.248490833 | 0.015327 | 0.05858 | 525.1604 | 567.5156 | 501.8326 | 522.7498 | 502.1823 | 398.9026 | 433.9301 | 443.7372 |
| ENSMUSG00000046329 | 0.031056194 | 0.697373 | 0.80906 | 1270.647 | 1112.235 | 1255.41 | 1193.008 | 1130.154 | 1124.078 | 1197.647 | 1280.095 |
| ENSMUSG00000040322 | 0.284635484 | 0.007106 | 0.033261 | 1357.167 | 1169.826 | 1289.362 | 1434.543 | 1126.254 | 1148.661 | 1119.54 | 917.9269 |
| ENSMUSG00000026819 | -0.306365924 | 0.056154 | 0.147653 | 486.9303 | 421.1374 | 388.382 | 365.7524 | 548.0126 | 393.3157 | 506.2518 | 605.7882 |
| ENSMUSG00000045100 | -0.031908503 | 0.812551 | 0.886 | 349.1009 | 325.1516 | 399.1474 | 356.2635 | 417.3476 | 298.3389 | 378.0014 | 368.6934 |
| ENSMUSG00000023912 | 0.683734941 | 0.002296 | 0.014489 | 746.4923 | 697.0963 | 505.145 | 880.7386 | 437.825 | 593.3256 | 419.4658 | 311.0511 |
| ENSMUSG00000040414 | -0.087207414 | 0.352692 | 0.521546 | 995.9938 | 916.6637 | 900.98 | 871.2497 | 1041.419 | 937.4769 | 1059.754 | 872.2481 |
| ENSMUSG00000021265 | -0.44431449 | 0.039707 | 0.116046 | 89.53884 | 121.182 | 122.5598 | 139.745 | 156.0178 | 186.6015 | 184.1792 | 117.4598 |
| ENSMUSG00000022003 | -0.753413966 | 0.007975 | 0.036144 | 297.7921 | 265.1606 | 419.8501 | 276.0395 | 537.2863 | 267.0524 | 606.5379 | 712.3722 |
| ENSMUSG00000069041 | -0.905469868 | | | 22.1332 | 19.19714 | 11.59349 | 12.93935 | 0.975111 | 119.559 | 2.892867 | 0 |
| ENSMUSG00000022299 | -0.171760321 | 0.104984 | 0.228462 | 278.6771 | 290.3568 | 321.3054 | 295.8799 | 346.1645 | 344.1512 | 302.7868 | 345.854 |
| ENSMUSG00000028982 | -0.545764576 | 0.02422 | 0.081141 | 377.2704 | 269.9598 | 381.7572 | 357.1261 | 391.9947 | 463.7102 | 778.1814 | 390.4452 |
| ENSMUSG00000040740 | -2.726239084 | 5.37E-07 | 2.23E-05 | 16.09687 | 11.99822 | 18.21835 | 13.80198 | 130.6649 | 25.6996 | 66.53595 | 176.1898 |
| ENSMUSG00000018740 | -0.631538379 | 0.001463 | 0.0104 | 154.9324 | 154.777 | 163.137 | 144.0581 | 213.5494 | 169.8409 | 301.8225 | 269.7226 |
| ENSMUSG00000032449 | 0.291167592 | 0.009933 | 0.042377 | 2894.418 | 3193.925 | 2935.638 | 3216.723 | 2381.222 | 3041.492 | 2392.401 | 2189.321 |
| ENSMUSG00000034248 | 0.290667316 | 0.028982 | 0.092256 | 1286.744 | 916.6637 | 907.6049 | 1043.774 | 911.7291 | 889.4298 | 781.0742 | 814.6057 |
| ENSMUSG00000032519 | -0.282716057 | 0.027199 | 0.088176 | 532.2028 | 459.5316 | 490.2391 | 603.8364 | 666.001 | 619.0252 | 708.7525 | 544.8831 |
| ENSMUSG00000018677 | -0.620762052 | 0.000178 | 0.002074 | 2956.794 | 2346.851 | 3024.245 | 2574.069 | 4324.619 | 2974.45 | 4329.658 | 5136.693 |
| ENSMUSG00000054099 | 0.074953541 | 0.588143 | 0.728361 | 423.5489 | 431.9358 | 380.9291 | 470.1298 | 377.3681 | 498.3488 | 390.5371 | 354.5547 |
| ENSMUSG00000002346 | -0.459993402 | 0.091529 | 0.207539 | 147.89 | 183.5727 | 129.1846 | 137.1571 | 210.624 | 129.6154 | 170.6792 | 309.9635 |
| ENSMUSG00000037636 | -0.029773063 | 0.943882 | 0.967579 | 32.19374 | 20.39697 | 22.35888 | 14.6646 | 19.50223 | 16.76061 | 19.28578 | 35.89051 |
| ENSMUSG00000050144 | -0.293551312 | 0.03695 | 0.1105 | 763.5953 | 671.9001 | 740.3273 | 695.2745 | 987.7878 | 677.1287 | 835.0744 | 1019.073 |
| ENSMUSG00000024818 | -0.857583124 | 0.000346 | 0.003454 | 202.2169 | 171.5745 | 270.7909 | 189.7772 | 380.2934 | 244.7049 | 378.9656 | 510.0802 |
| ENSMUSG00000024259 | 0.032850958 | 0.704615 | 0.814096 | 1496.003 | 1664.152 | 1455.811 | 1584.639 | 1462.667 | 1705.113 | 1434.862 | 1457.372 |
| ENSMUSG00000048856 | -0.063579802 | 0.690696 | 0.804105 | 185.114 | 153.5772 | 148.2311 | 144.0581 | 168.6943 | 188.8362 | 162.0006 | 139.2117 |
| ENSMUSG00000021509 | 0.104702334 | 0.597634 | 0.736409 | 223.3441 | 172.7743 | 227.7293 | 195.8155 | 158.9431 | 197.7752 | 251.6795 | 154.4379 |
| ENSMUSG00000045973 | -0.088504811 | 0.099198 | 0.219818 | 2884.358 | 2880.772 | 2728.611 | 2904.453 | 3017.969 | 3057.136 | 2890.939 | 3152.927 |
| ENSMUSG00000044348 | 0.27998579 | 0.051964 | 0.1397 | 422.5428 | 380.3434 | 362.7107 | 483.9318 | 319.8365 | 393.3157 | 353.8941 | 292.562 |
| ENSMUSG00000046959 | -2.012515508 | 7.77E-07 | 3.03E-05 | 85.51462 | 47.99286 | 104.3414 | 62.97152 | 322.7618 | 120.6764 | 256.5009 | 516.6058 |
| ENSMUSG00000034320 | -0.247517663 | 0.112158 | 0.238643 | 663.9959 | 574.7145 | 857.0904 | 548.6285 | 845.4215 | 712.8847 | 829.2887 | 753.7006 |
| ENSMUSG00000020651 | -1.649642411 | 0.000861 | 0.006955 | 31.18769 | 27.5959 | 43.88965 | 39.68068 | 102.3867 | 29.05173 | 110.8933 | 205.5547 |
| ENSMUSG00000023259 | -0.309842487 | 0.050063 | 0.136196 | 208.2533 | 181.1731 | 177.2148 | 163.0358 | 230.1263 | 182.132 | 226.608 | 264.2846 |
| ENSMUSG00000040569 | -0.522749197 | 0.028639 | 0.091448 | 256.5439 | 230.3657 | 221.9326 | 263.1002 | 244.7529 | 536.3396 | 315.3226 | 301.2627 |
| ENSMUSG00000036196 | 0.50840589 | 0.123954 | 0.253674 | 66.39959 | 68.38983 | 57.13936 | 62.97152 | 34.1289 | 77.09881 | 34.71441 | 33.71532 |
| ENSMUSG00000040441 | 0.770725005 | 0.003776 | 0.020951 | 121.7326 | 93.58608 | 96.06037 | 130.2561 | 69.2329 | 39.10809 | 64.60737 | 85.9197 |
| ENSMUSG00000039908 | 0.075599149 | 0.57486 | 0.717108 | 380.2885 | 278.3586 | 298.9465 | 354.5383 | 300.3343 | 292.752 | 337.5012 | 315.4014 |
| ENSMUSG00000031808 | 0.365800917 | 3.38E-07 | 1.52E-05 | 2354.167 | 2054.094 | 2084.344 | 2127.23 | 1603.083 | 1772.155 | 1642.184 | 1673.803 |
| ENSMUSG00000027359 | -3.949421069 | 5.19E-07 | 2.17E-05 | 125.7568 | 28.79572 | 423.9906 | 170.7995 | 2991.641 | 1130.783 | 2665.295 | 4798.452 |
| ENSMUSG00000027932 | -0.125195497 | 0.477761 | 0.63706 | 557.3541 | 428.3363 | 362.7107 | 398.5321 | 488.5308 | 392.1983 | 444.5373 | 578.5985 |
| ENSMUSG00000059316 | -0.525091554 | 0.000823 | 0.006715 | 730.3955 | 770.2854 | 896.0114 | 845.371 | 1093.1 | 884.9603 | 1237.183 | 1451.934 |
| ENSMUSG00000024600 | 0.387652748 | 0.014634 | 0.056634 | 637.8385 | 782.2836 | 626.0486 | 803.9651 | 610.4197 | 604.4994 | 553.502 | 408.9342 |
| ENSMUSG00000025726 | -0.811674875 | 0.165568 | 0.311015 | 19.11503 | 5.999108 | 4.96864 | 10.35148 | 19.50223 | 8.938993 | 17.3572 | 25.0146 |
| ENSMUSG00000027219 | -0.380377493 | 0.377674 | 0.545752 | 12.07265 | 26.39607 | 13.24971 | 15.52722 | 21.45245 | 25.6996 | 13.50005 | 26.10219 |
| ENSMUSG00000023942 | -0.161089038 | 0.255729 | 0.419644 | 1262.598 | 1396.592 | 1657.869 | 1312.913 | 1564.079 | 1231.346 | 1733.792 | 1766.248 |
| ENSMUSG00000024891 | -0.056003821 | 0.721169 | 0.825522 | 260.5681 | 196.7707 | 260.0255 | 213.9306 | 258.4045 | 197.7752 | 249.7509 | 263.197 |
| ENSMUSG00000020100 | -0.21877757 | 0.115043 | 0.242689 | 512.0817 | 555.5174 | 585.4714 | 552.079 | 625.0463 | 536.3396 | 598.8236 | 806.9926 |
| ENSMUSG00000050822 | 0.557901017 | 0.026214 | 0.08598 | 177.0656 | 89.98662 | 101.029 | 131.9814 | 92.63557 | 89.38993 | 83.89316 | 73.95619 |
| ENSMUSG00000037434 | -0.405019238 | 0.000481 | 0.00445 | 565.4025 | 571.1151 | 621.08 | 609.8748 | 692.329 | 709.5326 | 810.9672 | 924.4524 |
| ENSMUSG00000028836 | 0.261027976 | 0.507987 | 0.663699 | 270.6286 | 91.18644 | 115.1068 | 104.3774 | 136.5156 | 78.21619 | 97.39321 | 172.927 |
| ENSMUSG00000029151 | 2.927377171 | 4.72E-29 | 1.74E-25 | 286.7255 | 202.7698 | 213.6515 | 248.4356 | 27.30312 | 45.81234 | 33.75012 | 18.48905 |
| ENSMUSG00000005802 | -0.082132436 | 0.368248 | 0.536797 | 941.6669 | 916.6637 | 956.4632 | 1044.637 | 1003.39 | 1119.609 | 1054.932 | 910.3137 |
| ENSMUSG00000021629 | -0.142326968 | 0.013912 | 0.054502 | 1902.449 | 1952.11 | 1824.319 | 2017.676 | 2097.464 | 2144.241 | 2184.115 | 2066.423 |
| ENSMUSG00000024069 | -0.127159494 | 0.335452 | 0.503669 | 577.4752 | 517.1231 | 479.4737 | 530.5134 | 629.9219 | 458.1234 | 566.0377 | 642.7663 |
| ENSMUSG00000054414 | 0.074690877 | 0.234316 | 0.395152 | 993.9817 | 1096.637 | 1063.289 | 1047.225 | 1008.265 | 993.3456 | 986.4678 | 999.4962 |
| ENSMUSG00000029221 | -0.153840595 | 0.037324 | 0.11116 | 2231.429 | 2016.9 | 1972.55 | 2149.658 | 2172.548 | 2267.152 | 2399.151 | 2474.27 |
| ENSMUSG00000026614 | 1.316142293 | 0.002327 | 0.014615 | 62.37537 | 76.78858 | 55.48314 | 44.85642 | 12.67645 | 46.92971 | 17.3572 | 19.57664 |
| ENSMUSG00000066150 | -0.46112672 | 0.000236 | 0.002591 | 1535.239 | 1327.003 | 1494.732 | 1517.355 | 2117.942 | 1581.084 | 2079.007 | 2310.043 |
| ENSMUSG00000066152 | -0.319528507 | 0.009787 | 0.041905 | 506.0453 | 483.5281 | 501.0045 | 487.3823 | 584.0917 | 519.579 | 743.4669 | 619.9269 |
| ENSMUSG00000037771 | 0.269809577 | 0.321386 | 0.489397 | 39.23612 | 44.3934 | 46.37397 | 38.81806 | 42.9049 | 37.99072 | 32.78583 | 26.10219 |

| | | | | | | | | | | | |
|---|---|---|---|---|---|---|---|---|---|---|---|
| ENSMUSG00000027822 | -0.327832947 | 0.000982 | 0.007744 | 601.6205 | 663.5013 | 606.1741 | 598.6607 | 744.0099 | 700.5936 | 766.6099 | 886.3867 |
| ENSMUSG00000021490 | -3.04464571 | 8.27E-05 | 0.001166 | 535.2209 | 105.5843 | 1284.393 | 628.8525 | 6439.635 | 1949.818 | 3478.191 | 9211.897 |
| ENSMUSG00000029188 | -2.867176583 | 3.06E-07 | 1.39E-05 | 2.012109 | 8.398751 | 24.01509 | 10.35148 | 69.2329 | 51.39921 | 91.60747 | 119.635 |
| ENSMUSG00000006469 | -3.852462647 | 5.41E-05 | 0.000842 | 1.006054 | 1.199822 | 4.96864 | 0 | 49.73068 | 12.29112 | 8.678602 | 35.89051 |
| ENSMUSG00000028293 | -0.044585711 | 0.488945 | 0.646694 | 1038.248 | 1061.842 | 1070.742 | 1060.164 | 1064.822 | 1127.43 | 1033.718 | 1140.883 |
| ENSMUSG00000031156 | 0.049745751 | 0.642256 | 0.768658 | 603.6326 | 549.5183 | 544.8942 | 498.5964 | 506.0828 | 473.7666 | 574.7163 | 565.5474 |
| ENSMUSG00000027957 | -0.187760517 | 0.087507 | 0.201124 | 1018.127 | 824.2774 | 897.6676 | 873.8376 | 877.6002 | 1062.623 | 1055.897 | 1122.394 |
| ENSMUSG00000033272 | -0.384132017 | 0.000968 | 0.007642 | 1927.6 | 2153.68 | 2488.46 | 2116.878 | 3019.92 | 2373.303 | 2839.832 | 3105.073 |
| ENSMUSG00000022664 | 0.021871614 | 0.846194 | 0.907479 | 556.3481 | 562.7163 | 467.0521 | 485.657 | 506.0828 | 451.4191 | 515.8947 | 563.3722 |
| ENSMUSG00000020873 | -0.325066964 | 3.73E-05 | 0.000636 | 1014.103 | 982.6538 | 1094.757 | 1038.599 | 1213.038 | 1238.05 | 1318.183 | 1409.518 |
| ENSMUSG00000037089 | -0.302981742 | 0.000706 | 0.005989 | 1416.525 | 1625.758 | 1674.432 | 1524.256 | 2009.704 | 1710.7 | 2032.722 | 1944.613 |
| ENSMUSG00000021432 | 0.067703171 | 0.507623 | 0.663318 | 982.9151 | 1021.048 | 1046.727 | 1037.736 | 952.6837 | 1155.365 | 885.2175 | 910.3137 |
| ENSMUSG00000018999 | -0.193012028 | 0.220515 | 0.379329 | 1454.755 | 1437.386 | 1743.164 | 1468.185 | 1597.232 | 1390.013 | 1699.078 | 2292.642 |
| ENSMUSG00000049922 | -0.240514178 | 0.137491 | 0.272589 | 355.1372 | 335.95 | 344.4924 | 297.6051 | 379.3183 | 302.8084 | 396.3228 | 495.9415 |
| ENSMUSG00000017664 | -0.296237196 | 0.001519 | 0.010649 | 780.6982 | 724.6922 | 732.0463 | 730.6421 | 881.5006 | 839.1479 | 879.4317 | 1045.175 |
| ENSMUSG00000028521 | -0.028710432 | 0.765122 | 0.854012 | 545.2815 | 501.5254 | 461.2554 | 546.9033 | 488.5308 | 547.5133 | 525.5376 | 535.0948 |
| ENSMUSG00000033114 | 0.018634818 | 0.889821 | 0.933475 | 200.2048 | 199.1704 | 218.6202 | 193.2277 | 195.9974 | 173.193 | 201.5364 | 230.5693 |
| ENSMUSG00000050473 | 2.563481718 | 0.000272 | 0.002874 | 61.36932 | 8.398751 | 15.73403 | 30.19182 | 3.900445 | 4.469496 | 5.785735 | 5.437955 |
| ENSMUSG00000019731 | -0.357375802 | 0.000155 | 0.001873 | 811.8859 | 770.2854 | 814.0288 | 756.5208 | 1053.12 | 921.8336 | 1132.075 | 929.8904 |
| ENSMUSG00000042202 | 0.268235421 | 0.05637 | 0.148073 | 1399.422 | 1340.201 | 1199.098 | 1209.398 | 946.8331 | 1358.727 | 1059.754 | 909.2261 |
| ENSMUSG00000060181 | -0.038331991 | 0.702591 | 0.812521 | 502.0211 | 506.3247 | 585.4714 | 590.8971 | 547.0374 | 606.7341 | 527.4662 | 567.7225 |
| ENSMUSG00000048807 | -0.160925976 | 0.326083 | 0.493883 | 222.338 | 188.372 | 261.6817 | 203.5791 | 240.8525 | 198.8926 | 276.751 | 264.2846 |
| ENSMUSG00000038602 | 0.327796474 | 0.065704 | 0.165029 | 210.2654 | 153.5772 | 167.2775 | 164.7611 | 109.2125 | 153.0803 | 139.8219 | 153.3503 |
| ENSMUSG00000042195 | -0.618554061 | 0.001239 | 0.009173 | 430.5913 | 353.9474 | 570.5655 | 472.7177 | 665.0259 | 538.5743 | 721.2883 | 885.2991 |
| ENSMUSG00000057060 | -0.16278687 | 0.618544 | 0.751043 | 48.29061 | 32.39518 | 51.34261 | 62.10889 | 55.58134 | 40.22547 | 42.42872 | 80.48174 |
| ENSMUSG00000026342 | -0.122652429 | 0.236433 | 0.397487 | 1482.924 | 1298.207 | 1651.245 | 1480.262 | 1612.834 | 1410.126 | 1703.899 | 1714.044 |
| ENSMUSG00000029175 | -0.240042333 | 0.014047 | 0.054885 | 503.0272 | 473.9295 | 566.4249 | 522.7498 | 618.2206 | 555.3349 | 610.395 | 660.1678 |
| ENSMUSG00000044026 | -0.250909844 | 0.240748 | 0.403015 | 843.0736 | 719.8929 | 1397.016 | 791.8884 | 1111.627 | 912.8946 | 1277.683 | 1164.81 |
| ENSMUSG00000070287 | 0.297994193 | 0.248748 | 0.412183 | 98.59333 | 57.59143 | 91.91984 | 82.81186 | 48.75557 | 73.74669 | 65.57166 | 82.65692 |
| ENSMUSG00000020261 | 0.151505312 | 0.190857 | 0.343486 | 561.3783 | 491.9268 | 539.9255 | 622.8142 | 497.3068 | 560.9218 | 442.6087 | 498.1167 |
| ENSMUSG00000020264 | 0.787567172 | 0.148953 | 0.288738 | 30.18163 | 32.39518 | 11.59349 | 26.74133 | 12.67645 | 25.6996 | 14.46434 | 5.437955 |
| ENSMUSG00000043885 | 0.079013635 | 0.296556 | 0.464315 | 1197.205 | 1126.632 | 1175.083 | 1294.798 | 1139.905 | 1182.182 | 1161.968 | 1057.139 |
| ENSMUSG00000024036 | -0.375344471 | 0.150816 | 0.290973 | 95.57516 | 88.78679 | 101.029 | 87.12497 | 117.9885 | 77.09881 | 176.4649 | 110.9343 |
| ENSMUSG00000032122 | 0.252267652 | 0.318276 | 0.486549 | 117.7084 | 62.39072 | 81.15445 | 68.14726 | 72.15824 | 74.86406 | 64.60737 | 65.25547 |
| ENSMUSG00000029924 | 0.040393693 | 0.569257 | 0.713058 | 1996.012 | 2216.07 | 1976.691 | 2049.593 | 2068.211 | 1898.419 | 1918.935 | 2120.803 |
| ENSMUSG00000032114 | -1.702396406 | 1.52E-12 | 4.24E-10 | 579.4873 | 373.1445 | 592.9243 | 598.6607 | 1766.902 | 1050.332 | 1815.756 | 2350.284 |
| ENSMUSG00000023169 | 0.177365437 | 0.00101 | 0.007894 | 3140.902 | 2915.566 | 2903.342 | 2991.578 | 2531.389 | 2685.05 | 2666.26 | 2687.438 |
| ENSMUSG00000022462 | 0.365686723 | 1.62E-06 | 5.53E-05 | 8515.244 | 8414.348 | 7226.058 | 8278.598 | 6328.472 | 6675.193 | 6270.772 | 5894.744 |
| ENSMUSG00000010064 | -1.086796083 | 0.000177 | 0.002063 | 293.7679 | 256.7618 | 342.8361 | 324.3464 | 617.2455 | 332.9775 | 604.6093 | 1033.212 |
| ENSMUSG00000022464 | 0.265948754 | 0.025683 | 0.084625 | 1891.382 | 1989.304 | 2529.038 | 2035.791 | 1527.024 | 1810.146 | 1769.471 | 1920.686 |
| ENSMUSG00000031170 | 0.970996254 | 0.025188 | 0.083461 | 17.10292 | 51.59233 | 44.71776 | 46.58167 | 22.42756 | 15.64324 | 27.96439 | 15.22628 |
| ENSMUSG00000044712 | 0.266983661 | 0.016645 | 0.061903 | 388.337 | 429.5361 | 403.2879 | 445.9763 | 349.0898 | 395.5504 | 323.0369 | 318.6642 |
| ENSMUSG00000036534 | 0.021607921 | 0.81371 | 0.886897 | 571.4389 | 500.3256 | 514.2542 | 487.3823 | 537.2863 | 490.5272 | 523.609 | 489.416 |
| ENSMUSG00000047789 | 0.392114129 | 0.011169 | 0.046405 | 993.9817 | 740.2899 | 640.9545 | 873.8376 | 625.0463 | 703.9457 | 591.1093 | 555.759 |
| ENSMUSG00000061306 | -0.343128 | 0.003105 | 0.018253 | 3055.387 | 3231.119 | 3361.285 | 3143.4 | 4071.09 | 3226.976 | 4242.872 | 4684.255 |
| ENSMUSG00000061171 | 0.685854883 | 0.243946 | 0.406525 | 42.25428 | 9.598572 | 15.73403 | 34.50494 | 23.40267 | 8.938993 | 21.21436 | 9.78832 |
| ENSMUSG00000052310 | -0.199712769 | 0.059172 | 0.153209 | 2608.699 | 2595.214 | 3074.76 | 2824.229 | 3314.403 | 2686.167 | 3218.797 | 3534.671 |
| ENSMUSG00000072572 | 0.910815906 | 0.005121 | 0.026168 | 40.24217 | 35.99465 | 35.60859 | 44.85642 | 27.30312 | 24.58223 | 17.3572 | 14.13868 |
| ENSMUSG00000046822 | -0.026590946 | 0.816738 | 0.88878 | 733.4136 | 749.8885 | 780.9046 | 756.5208 | 713.7815 | 668.1897 | 771.4313 | 924.4524 |
| ENSMUSG00000063354 | 0.214075874 | 0.585303 | 0.725623 | 22.1332 | 26.39607 | 28.98373 | 18.97772 | 22.42756 | 31.28647 | 14.46434 | 15.22628 |
| ENSMUSG00000039878 | -2.658234437 | 3.41E-06 | 0.0001 | 116.7023 | 35.99465 | 181.3554 | 123.3552 | 752.7859 | 321.8037 | 609.4308 | 1208.314 |
| ENSMUSG00000024270 | 0.222866348 | 0.006233 | 0.030311 | 1607.675 | 1712.145 | 1544.419 | 1709.72 | 1416.837 | 1545.328 | 1389.541 | 1280.095 |
| ENSMUSG00000024327 | -0.517139706 | 0.001532 | 0.010697 | 702.2259 | 667.1008 | 757.7176 | 668.5332 | 1026.792 | 700.5936 | 1058.79 | 1214.839 |
| ENSMUSG00000053897 | -0.398232274 | 0.063228 | 0.160933 | 744.4802 | 904.6654 | 918.3703 | 1022.209 | 1004.365 | 782.1619 | 1338.433 | 1606.372 |
| ENSMUSG00000048833 | -0.118099711 | 0.233325 | 0.393913 | 1307.871 | 1227.417 | 1387.907 | 1340.517 | 1395.384 | 1271.572 | 1391.469 | 1657.489 |
| ENSMUSG00000025986 | -0.064343401 | 0.459041 | 0.620528 | 1675.081 | 1977.306 | 1604.871 | 1653.649 | 1707.42 | 1877.188 | 1839.864 | 1795.613 |
| ENSMUSG00000041654 | -0.381311425 | 0.047264 | 0.130781 | 346.0827 | 417.5379 | 403.2879 | 314.8576 | 475.8543 | 337.447 | 497.5732 | 617.7517 |
| ENSMUSG00000002105 | -0.05343208 | 0.649482 | 0.773046 | 836.0312 | 867.471 | 934.1043 | 931.6334 | 915.6295 | 749.758 | 1005.754 | 1033.212 |
| ENSMUSG00000022094 | 0.178193666 | 0.109815 | 0.235476 | 530.1906 | 455.9322 | 453.8024 | 549.4912 | 485.6054 | 426.8369 | 434.8944 | 411.1094 |
| ENSMUSG00000025993 | 0.451487548 | 0.000826 | 0.006728 | 3917.576 | 5684.754 | 5250.196 | 4720.276 | 3946.275 | 3881.758 | 3091.511 | 3393.284 |
| ENSMUSG00000013275 | 0.177205847 | 0.00672 | 0.032025 | 993.9817 | 938.2604 | 958.9475 | 915.2435 | 839.5708 | 824.6221 | 833.1458 | 868.9853 |
| ENSMUSG00000034591 | 0.32183465 | 0.123774 | 0.253453 | 74.44802 | 65.99018 | 71.21717 | 62.97152 | 54.60623 | 62.57295 | 56.89306 | 45.67883 |
| ENSMUSG00000030089 | 0.184831102 | 0.110042 | 0.235853 | 233.4046 | 223.1668 | 248.432 | 246.7103 | 190.1467 | 217.8879 | 213.1079 | 218.6058 |
| ENSMUSG00000027075 | -0.056832771 | 0.684231 | 0.798746 | 516.1059 | 488.3274 | 403.2879 | 498.5964 | 510.9583 | 449.1844 | 590.145 | 429.5985 |

| | | | | | | | | | | | |
|---|---|---|---|---|---|---|---|---|---|---|---|
| ENSMUSG00000038178 | -1.437725 | 1.42E-06 | 4.95E-05 | 277.671 | 231.5656 | 370.1637 | 256.1992 | 795.6908 | 372.0856 | 767.5742 | 1143.058 |
| ENSMUSG00000027074 | 0.100432241 | 0.439248 | 0.602643 | 2110.702 | 1960.508 | 1758.07 | 2306.655 | 1821.508 | 1945.348 | 2234.258 | 1586.795 |
| ENSMUSG00000028412 | 0.090708499 | 0.173302 | 0.321099 | 2817.958 | 2945.562 | 3150.946 | 2895.827 | 2672.78 | 2918.581 | 2869.725 | 2631.97 |
| ENSMUSG00000057193 | 0.22079019 | 0.015023 | 0.057753 | 3125.811 | 3009.152 | 3323.192 | 3095.956 | 2748.839 | 2297.321 | 2775.224 | 2951.722 |
| ENSMUSG00000039865 | 0.150425695 | 0.302579 | 0.47086 | 449.7063 | 391.1418 | 548.2066 | 398.5321 | 363.7165 | 391.0809 | 396.3228 | 462.2262 |
| ENSMUSG00000007034 | -1.370332491 | 2.21E-05 | 0.000425 | 159.9626 | 94.7859 | 196.2613 | 142.3329 | 405.6463 | 208.949 | 330.7512 | 591.6496 |
| ENSMUSG00000028360 | -0.553857707 | 0.109485 | 0.234988 | 23.13925 | 14.39786 | 24.8432 | 31.91707 | 33.15378 | 45.81234 | 30.85725 | 30.45255 |
| ENSMUSG00000039838 | 1.865465914 | 0.009796 | 0.04193 | 23.13925 | 21.59679 | 5.796746 | 35.36756 | 8.776002 | 7.821619 | 5.785735 | 1.087591 |
| ENSMUSG00000026435 | -0.24678013 | 0.277674 | 0.444018 | 456.7487 | 425.9366 | 403.2879 | 425.2734 | 405.6463 | 402.2547 | 791.6814 | 429.5985 |
| ENSMUSG00000079020 | -0.213323432 | 0.063621 | 0.161622 | 894.3823 | 785.8831 | 1058.32 | 951.4737 | 1006.315 | 1046.98 | 1026.004 | 1205.051 |
| ENSMUSG00000020829 | -0.62033744 | 4.11E-05 | 0.000689 | 296.786 | 259.1615 | 254.2287 | 244.1224 | 368.5921 | 331.8601 | 432.9658 | 486.1532 |
| ENSMUSG00000029650 | -0.233951681 | 0.374543 | 0.543269 | 54.32694 | 79.18822 | 62.108 | 56.93315 | 51.6809 | 65.92507 | 94.50034 | 83.74451 |
| ENSMUSG00000010122 | -2.90255208 | 1.10E-05 | 0.000246 | 98.59333 | 35.99465 | 278.2438 | 122.4925 | 1127.229 | 414.5458 | 858.2174 | 1608.547 |
| ENSMUSG00000081534 | -0.33308255 | 0.007108 | 0.033261 | 1798.825 | 1702.547 | 2104.219 | 1915.887 | 2458.256 | 1892.832 | 2462.795 | 2663.511 |
| ENSMUSG00000027953 | 0.210043588 | 0.044495 | 0.125603 | 1147.908 | 1240.615 | 932.4481 | 1164.542 | 972.186 | 1021.28 | 948.8605 | 932.0656 |
| ENSMUSG00000035699 | -2.966333336 | 5.18E-05 | 0.000814 | 5.030272 | 8.398751 | 19.04645 | 5.175741 | 117.0134 | 14.52586 | 30.85725 | 133.7737 |
| ENSMUSG00000053862 | -3.788001849 | 6.97E-11 | 1.34E-08 | 10.06054 | 2.399643 | 20.70267 | 8.626235 | 158.9431 | 49.16446 | 133.0719 | 246.8832 |
| ENSMUSG00000022560 | -0.526053809 | 0.007551 | 0.03472 | 303.8284 | 341.9491 | 336.2113 | 331.2474 | 441.7254 | 335.2122 | 456.1088 | 657.9926 |
| ENSMUSG00000027463 | -0.899372924 | 0.009247 | 0.040236 | 171.0292 | 127.1811 | 262.5098 | 168.2116 | 283.7574 | 203.3621 | 277.7153 | 597.0875 |
| ENSMUSG00000030237 | 1.192736343 | 0.001313 | 0.009599 | 528.1785 | 551.9179 | 401.6317 | 527.9256 | 163.8187 | 446.9496 | 150.4291 | 118.5474 |
| ENSMUSG00000079262 | -4.638955917 | 3.34E-05 | 0.000581 | 11.0666 | 0 | 30.63995 | 6.900988 | 254.5041 | 54.75133 | 284.4653 | 625.3649 |
| ENSMUSG00000030235 | 0.178979098 | 0.783383 | 0.866794 | 4.024217 | 17.99732 | 15.73403 | 20.70296 | 9.751113 | 25.6996 | 6.750024 | 9.78832 |
| ENSMUSG00000032548 | -0.950726671 | 0.001027 | 0.007993 | 292.7618 | 203.9697 | 421.5063 | 271.7264 | 474.8792 | 345.2686 | 722.2526 | 760.2262 |
| ENSMUSG00000030737 | 0.326224131 | 0.107423 | 0.231883 | 511.0756 | 394.7413 | 348.6329 | 421.8229 | 312.0356 | 298.3389 | 460.9302 | 264.2846 |
| ENSMUSG00000025790 | 0.329164854 | 2.06E-06 | 6.57E-05 | 1182.114 | 1123.033 | 1053.352 | 1117.96 | 876.6251 | 858.1433 | 921.8604 | 903.7882 |
| ENSMUSG00000038963 | -1.844700665 | 9.48E-06 | 0.000218 | 65.39353 | 47.99286 | 139.1219 | 36.23019 | 227.2009 | 167.6061 | 258.4295 | 387.1824 |
| ENSMUSG00000040693 | 0.617967836 | 0.013861 | 0.054385 | 1389.361 | 1622.159 | 1019.399 | 1269.782 | 785.9397 | 1301.741 | 836.0387 | 529.6569 |
| ENSMUSG00000025938 | 0.846388894 | 0.001291 | 0.009465 | 131.7931 | 112.7832 | 91.91984 | 123.3552 | 52.65601 | 96.09417 | 51.10733 | 56.55474 |
