## Supplementary material for "Lysine-specific demethylase 1a is obligatory for gene regulation during kidney development": Suppl Table-6

**Supplementary table-6**

| Chr | Start | End | CpGs | KWT | KKO | Diff | p_val1 | p_val2 | adj.pval1 | adj.pval2 |
|---|---|---|---|---|---|---|---|---|---|---|
| chr2 | 152686872 | 152686971 | 28 | 0.29996 | 0.64698 | -0.34702 | 4.73E-14 | 3.52E-19 | 5.23E-10 | 6.46E-15 |
| chr7 | 128688375 | 128688695 | 20 | 0.74935 | 0.4197 | 0.32965 | 5.71E-14 | 3.36E-14 | 5.23E-10 | 3.08E-10 |
| chr9 | 33728226 | 33728363 | 39 | 0.24405 | 0.4256 | -0.18155 | 1.66E-13 | 9.23E-08 | 1.02E-09 | 9.96E-05 |
| chr2 | 33644081 | 33644308 | 37 | 0.21828 | 0.39223 | -0.17395 | 3.12E-13 | 5.59E-09 | 1.43E-09 | 1.03E-05 |
| chr7 | 60004710 | 60005042 | 26 | 0.58819 | 0.45025 | 0.137942 | 9.03E-13 | 2.44E-09 | 3.31E-09 | 5.38E-06 |
| chr8 | 122865152 | 122865364 | 25 | 0.50646 | 0.71142 | -0.20496 | 7.30E-12 | 8.81E-08 | 2.23E-08 | 9.96E-05 |
| chr5 | 135455604 | 135455850 | 18 | 0.47406 | 0.64567 | -0.17161 | 3.82E-11 | 4.16E-08 | 1.00E-07 | 5.45E-05 |
| chr13 | 119488375 | 119488592 | 40 | 0.55081 | 0.73324 | -0.18243 | 4.95E-11 | 8.04E-13 | 1.13E-07 | 3.69E-09 |
| chr2 | 168268886 | 168269092 | 23 | 0.52954 | 0.82587 | -0.29633 | 7.01E-11 | 1.26E-06 | 1.30E-07 | 0.000681 |
| chr6 | 48629431 | 48629990 | 86 | 0.039994 | 0.31475 | -0.27476 | 7.09E-11 | 1.06E-13 | 1.30E-07 | 6.48E-10 |
| chr7 | 128687931 | 128688028 | 14 | 0.63561 | 0.15043 | 0.485179 | 1.33E-10 | 4.47E-11 | 2.22E-07 | 1.64E-07 |
| chr17 | 6492663 | 6493053 | 32 | 0.27702 | 0.44456 | -0.16755 | 2.19E-10 | 7.05E-06 | 3.34E-07 | 0.002346 |
| chr1 | 172244696 | 172245076 | 30 | 0.34113 | 0.66995 | -0.32882 | 6.29E-10 | 7.15E-06 | 8.41E-07 | 0.002346 |
| chr5 | 143254827 | 143254916 | 15 | 0.57123 | 0.1039 | 0.467333 | 6.42E-10 | 2.74E-07 | 8.41E-07 | 0.000218 |
| chr7 | 141269106 | 141269346 | 27 | 0.63152 | 0.84113 | -0.20961 | 1.37E-09 | 3.15E-06 | 1.68E-06 | 0.001363 |
| chrX | 169997079 | 169997215 | 28 | 0.4798 | 0.68138 | -0.20157 | 1.65E-09 | 7.85E-10 | 1.90E-06 | 2.40E-06 |
| chr15 | 99819377 | 99819888 | 20 | 0.3867 | 0.1926 | 0.1941 | 2.90E-09 | 6.18E-07 | 2.69E-06 | 0.00042 |
| chr19 | 44367194 | 44367391 | 12 | 0.32258 | 0.50296 | -0.18038 | 2.88E-09 | 1.77E-09 | 2.69E-06 | 4.65E-06 |
| chr3 | 129256579 | 129256692 | 15 | 0.41537 | 0.7193 | -0.30393 | 2.55E-09 | 2.66E-07 | 2.69E-06 | 0.000218 |
| chr5 | 33759845 | 33760261 | 25 | 0.83062 | 0.93672 | -0.1061 | 2.93E-09 | 4.62E-06 | 2.69E-06 | 0.001882 |
| chr15 | 12920047 | 12920305 | 23 | 0.48465 | 0.73487 | -0.25022 | 3.24E-09 | 2.11E-06 | 2.70E-06 | 0.001048 |
| chr9 | 22188138 | 22188437 | 19 | 0.003947 | 0.11326 | -0.10932 | 3.19E-09 | 2.64E-09 | 2.70E-06 | 5.38E-06 |
| chr10 | 80312555 | 80312777 | 25 | 0.23552 | 0.46128 | -0.22576 | 5.04E-09 | 1.12E-06 | 4.02E-06 | 0.000624 |
| chr19 | 5654012 | 5654126 | 21 | 0.042762 | 0.19748 | -0.15471 | 5.45E-09 | 8.05E-07 | 4.17E-06 | 0.000476 |
| chr8 | 11279402 | 11279717 | 21 | 0.33357 | 0.56288 | -0.22931 | 7.51E-09 | 5.87E-06 | 5.51E-06 | 0.002197 |
| chr11 | 103333174 | 103333373 | 21 | 0.33362 | 0.16319 | 0.170429 | 8.35E-09 | 2.84E-05 | 5.89E-06 | 0.006197 |
| chr10 | 13090743 | 13090975 | 20 | 0.38003 | 0.53423 | -0.1542 | 1.03E-08 | 1.39E-08 | 6.82E-06 | 1.96E-05 |
| chr8 | 108956791 | 108956964 | 23 | 0.54735 | 0.72713 | -0.17978 | 1.04E-08 | 1.53E-07 | 6.82E-06 | 0.000141 |
| chr7 | 44461885 | 44462061 | 29 | 0.43178 | 0.68212 | -0.25035 | 1.47E-08 | 3.49E-05 | 9.29E-06 | 0.006484 |
| chr1 | 152730713 | 152730822 | 16 | 0.59869 | 0.91256 | -0.31388 | 1.84E-08 | 4.28E-07 | 1.13E-05 | 0.000314 |
| chr1 | 136468112 | 136468486 | 28 | 0.3145 | 0.4998 | -0.1853 | 2.83E-08 | 1.44E-05 | 1.64E-05 | 0.004028 |
| chr15 | 85267942 | 85268205 | 29 | 0.57122 | 0.77436 | -0.20314 | 2.87E-08 | 3.46E-05 | 1.64E-05 | 0.006484 |
| chr5 | 44143103 | 44143229 | 23 | 0.8442 | 0.95674 | -0.11254 | 3.23E-08 | 2.32E-05 | 1.79E-05 | 0.005436 |
| chr13 | 98366918 | 98367148 | 21 | 0.12431 | 0.3184 | -0.1941 | 5.09E-08 | 1.66E-05 | 2.71E-05 | 0.00422 |
| chr6 | 130811103 | 130811225 | 11 | 0.89014 | 0.58382 | 0.306318 | 5.16E-08 | 1.48E-06 | 2.71E-05 | 0.000774 |
| chr10 | 130470552 | 130470656 | 24 | 0.006771 | 0.12696 | -0.12019 | 6.38E-08 | 1.18E-07 | 3.25E-05 | 0.00012 |
| chr2 | 174297148 | 174297353 | 20 | 0.3273 | 0.46273 | -0.13543 | 6.73E-08 | 3.44E-07 | 3.34E-05 | 0.000263 |
| chr14 | 42543317 | 42543423 | 23 | 0.05487 | 0.15961 | -0.10474 | 1.14E-07 | 9.15E-06 | 5.52E-05 | 0.002798 |
| chr8 | 22789262 | 22789377 | 11 | 0.64095 | 0.82477 | -0.18382 | 1.28E-07 | 6.99E-07 | 6.03E-05 | 0.000428 |
| chr13 | 41232246 | 41232532 | 22 | 0.58561 | 0.81593 | -0.23032 | 1.46E-07 | 0.000645 | 6.53E-05 | 0.031559 |
| chr16 | 89988367 | 89988438 | 11 | 0.59745 | 0.84291 | -0.24546 | 1.46E-07 | 7.15E-06 | 6.53E-05 | 0.002346 |
| chr4 | 103419258 | 103419431 | 21 | 0.72869 | 0.89205 | -0.16336 | 1.83E-07 | 0.000621 | 8.00E-05 | 0.031028 |
| chr1 | 36269584 | 36269756 | 35 | 0.27287 | 0.40817 | -0.1353 | 2.03E-07 | 0.000201 | 8.64E-05 | 0.01516 |
| chr2 | 174285292 | 174285580 | 25 | 0.33564 | 0.5119 | -0.17626 | 2.09E-07 | 1.68E-05 | 8.72E-05 | 0.00422 |
| chr13 | 119488678 | 119488750 | 12 | 0.23004 | 0.54 | -0.30996 | 2.27E-07 | 6.48E-06 | 9.14E-05 | 0.00233 |
| chr16 | 4648114 | 4648625 | 30 | 0.51315 | 0.69863 | -0.18548 | 2.34E-07 | 3.53E-05 | 9.14E-05 | 0.006484 |
| chr7 | 63891626 | 63891720 | 24 | 0.837 | 0.66954 | 0.167458 | 2.30E-07 | 5.69E-05 | 9.14E-05 | 0.008929 |
| chr12 | 109539771 | 109540297 | 24 | 0.65994 | 0.49333 | 0.166604 | 2.91E-07 | 1.91E-05 | 0.000105 | 0.004546 |
| chr15 | 101053043 | 101053112 | 18 | 0.11222 | 0.32511 | -0.21289 | 2.99E-07 | 0.000129 | 0.000105 | 0.013041 |

| | | | | | | | | | | |
|---|---|---|---|---|---|---|---|---|---|---|
| chr16 | 56029620 | 56029697 | 9 | 0.357 | 0.74761 | -0.39061 | 2.97E-07 | 2.13E-07 | 0.000105 | 0.000186 |
| chr6 | 30738246 | 30738290 | 9 | 0.40594 | 0.18978 | 0.216167 | 2.97E-07 | 5.40E-07 | 0.000105 | 0.000381 |
| chr9 | 61405675 | 61405702 | 13 | 0.35146 | 0.1425 | 0.208962 | 2.99E-07 | 7.14E-05 | 0.000105 | 0.009945 |
| chr4 | 127312648 | 127312793 | 17 | 0.49829 | 0.71115 | -0.21285 | 3.58E-07 | 6.15E-05 | 0.000124 | 0.009327 |
| chr8 | 3452062 | 3452122 | 15 | 0.9782 | 0.83227 | 0.145933 | 4.62E-07 | 1.29E-05 | 0.000157 | 0.003706 |
| chr2 | 174284194 | 174284690 | 18 | 0.28975 | 0.51319 | -0.22344 | 5.09E-07 | 1.15E-08 | 0.00017 | 1.91E-05 |
| chr13 | 4942996 | 4943101 | 11 | 0.48145 | 0.696 | -0.21455 | 5.74E-07 | 3.40E-06 | 0.000179 | 0.001417 |
| chr3 | 131466340 | 131466409 | 10 | 0.5803 | 0.27595 | 0.30435 | 5.61E-07 | 1.64E-05 | 0.000179 | 0.00422 |
| chr3 | 89238689 | 89238823 | 13 | 0.30577 | 0.65519 | -0.34942 | 5.85E-07 | 0.000394 | 0.000179 | 0.023015 |
| chr5 | 129602152 | 129602277 | 9 | 0.78572 | 0.3565 | 0.429222 | 5.77E-07 | 3.19E-06 | 0.000179 | 0.001363 |
| chr8 | 108956525 | 108956589 | 18 | 0.51686 | 0.729 | -0.21214 | 5.72E-07 | 4.55E-05 | 0.000179 | 0.007728 |
| chr6 | 64065695 | 64065767 | 12 | 0.98567 | 0.88192 | 0.10375 | 6.04E-07 | 1.25E-05 | 0.000182 | 0.003706 |
| chr19 | 40621800 | 40621926 | 22 | 0.29939 | 0.49389 | -0.1945 | 6.29E-07 | 0.000945 | 0.000186 | 0.031559 |
| chr3 | 54811275 | 54811390 | 24 | 0.36696 | 0.57563 | -0.20867 | 6.51E-07 | 0.000208 | 0.00019 | 0.015468 |
| chr11 | 120347182 | 120347299 | 6 | 0.22292 | 0.062833 | 0.160083 | 7.40E-07 | 6.94E-05 | 0.000206 | 0.009945 |
| chr2 | 165867054 | 165867075 | 6 | 0.51692 | 0.25833 | 0.258583 | 7.40E-07 | 2.82E-05 | 0.000206 | 0.006197 |
| chr7 | 24082142 | 24082203 | 6 | 0.42275 | 0.78942 | -0.36667 | 7.40E-07 | 2.89E-06 | 0.000206 | 0.001357 |
| chr11 | 22972854 | 22972990 | 24 | 0.42292 | 0.53785 | -0.11494 | 7.86E-07 | 1.29E-08 | 0.000215 | 1.96E-05 |
| chr14 | 54297138 | 54297448 | 28 | 0.7212 | 0.86104 | -0.13984 | 9.04E-07 | 4.74E-05 | 0.000244 | 0.007834 |
| chr11 | 120012167 | 120012231 | 19 | 0.49268 | 0.25332 | 0.239368 | 1.05E-06 | 3.11E-05 | 0.000263 | 0.006475 |
| chr15 | 89068154 | 89068331 | 19 | 0.27171 | 0.46118 | -0.18947 | 9.93E-07 | 0.000144 | 0.000263 | 0.01319 |
| chr18 | 64531369 | 64531501 | 21 | 0.183 | 0.33619 | -0.15319 | 1.04E-06 | 2.81E-05 | 0.000263 | 0.006197 |
| chr2 | 34671469 | 34671754 | 12 | 0.25688 | 0.078667 | 0.178208 | 1.02E-06 | 6.45E-06 | 0.000263 | 0.00233 |
| chr5 | 149002494 | 149002736 | 20 | 0.20572 | 0.33792 | -0.1322 | 1.02E-06 | 0.000314 | 0.000263 | 0.021124 |
| chr12 | 85963992 | 85964015 | 10 | 0.8376 | 0.42385 | 0.41375 | 1.29E-06 | 7.08E-05 | 0.000319 | 0.009945 |
| chr17 | 84187422 | 84187567 | 13 | 0.047538 | 0.23915 | -0.19162 | 1.36E-06 | 0.000228 | 0.000331 | 0.016263 |
| chr16 | 18976495 | 18976773 | 12 | 0.59113 | 0.86846 | -0.27733 | 1.40E-06 | 9.17E-05 | 0.000335 | 0.011255 |
| chr8 | 119896141 | 119896226 | 8 | 0.27737 | 0.049 | 0.228375 | 1.41E-06 | 1.39E-07 | 0.000335 | 0.000135 |
| chr17 | 78232661 | 78232786 | 6 | 0.53767 | 0.86392 | -0.32625 | 1.48E-06 | 0.00018 | 0.000336 | 0.015013 |
| chr4 | 59914551 | 59914581 | 6 | 0.064333 | 0.35017 | -0.28583 | 1.48E-06 | 6.97E-05 | 0.000336 | 0.009945 |
| chr4 | 84189414 | 84189508 | 13 | 0.26658 | 0.055423 | 0.211154 | 1.49E-06 | 7.11E-05 | 0.000336 | 0.009945 |
| chr6 | 88200244 | 88200364 | 16 | 0.19106 | 0.061313 | 0.12975 | 1.43E-06 | 0.00025 | 0.000336 | 0.017624 |
| chr5 | 103365471 | 103365630 | 9 | 0.71339 | 0.83833 | -0.12494 | 1.52E-06 | 7.45E-06 | 0.000339 | 0.002346 |
| chr5 | 75835690 | 75835796 | 12 | 0.085417 | 0.274 | -0.18858 | 1.55E-06 | 9.11E-05 | 0.000343 | 0.011255 |
| chr10 | 4038711 | 4038980 | 12 | 0.35392 | 0.25217 | 0.10175 | 1.72E-06 | 9.08E-05 | 0.000376 | 0.011255 |
| chr3 | 35357263 | 35357415 | 13 | 0.53742 | 0.73562 | -0.19819 | 1.78E-06 | 0.000226 | 0.000376 | 0.016181 |
| chr6 | 137689862 | 137690015 | 9 | 0.949 | 0.61361 | 0.335389 | 1.78E-06 | 0.000167 | 0.000376 | 0.014451 |
| chr8 | 105728018 | 105728185 | 16 | 0.71784 | 0.85328 | -0.13544 | 1.75E-06 | 0.000245 | 0.000376 | 0.017351 |
| chr10 | 119169119 | 119169670 | 39 | 0.63722 | 0.52676 | 0.110462 | 2.00E-06 | 0.000732 | 0.000418 | 0.031559 |
| chr5 | 125068469 | 125068610 | 10 | 0.3825 | 0.0604 | 0.3221 | 2.20E-06 | 1.52E-06 | 0.000454 | 0.000774 |
| chr11 | 99350723 | 99350976 | 21 | 0.69902 | 0.86274 | -0.16371 | 2.42E-06 | 7.00E-05 | 0.000489 | 0.009945 |
| chr5 | 124951525 | 124951625 | 9 | 0.58278 | 0.16644 | 0.416333 | 2.43E-06 | 8.01E-05 | 0.000489 | 0.010671 |
| chr17 | 39848495 | 39848522 | 9 | 0.34906 | 0.473 | -0.12394 | 2.83E-06 | 3.19E-06 | 0.000565 | 0.001363 |
| chr18 | 78260622 | 78260878 | 20 | 0.3715 | 0.60547 | -0.23398 | 2.92E-06 | 1.51E-05 | 0.000575 | 0.004083 |
| chr15 | 102243531 | 102243669 | 11 | 0.41545 | 0.19623 | 0.219227 | 3.00E-06 | 0.001297 | 0.000579 | 0.033685 |
| chr15 | 80051494 | 80051765 | 16 | 0.34391 | 0.55484 | -0.21094 | 2.97E-06 | 9.13E-05 | 0.000579 | 0.011255 |
| chr11 | 118487259 | 118487604 | 13 | 0.92992 | 0.64335 | 0.286577 | 3.06E-06 | 2.79E-06 | 0.000585 | 0.001348 |
| chr19 | 26776175 | 26776259 | 14 | 0.85307 | 0.54714 | 0.305929 | 3.26E-06 | 9.71E-05 | 0.000616 | 0.011723 |
| chr3 | 88365924 | 88366165 | 29 | 0.41716 | 0.61943 | -0.20228 | 3.33E-06 | 0.000146 | 0.000622 | 0.013255 |
| chr5 | 133842187 | 133842473 | 11 | 0.24486 | 0.42545 | -0.18059 | 3.36E-06 | 0.000402 | 0.000622 | 0.023015 |
| chr11 | 22971951 | 22972120 | 18 | 0.61217 | 0.44267 | 0.1695 | 3.48E-06 | 3.14E-06 | 0.000639 | 0.001363 |

| | | | | | | | | | | |
|---|---|---|---|---|---|---|---|---|---|---|
| chr6 | 136720600 | 136720688 | 8 | 0.88644 | 0.65037 | 0.236063 | 3.56E-06 | 3.90E-05 | 0.000646 | 0.006751 |
| chr17 | 67715644 | 67715871 | 23 | 0.45461 | 0.30567 | 0.148935 | 3.78E-06 | 0.000128 | 0.000681 | 0.013041 |
| chr12 | 56718415 | 56718465 | 14 | 0.14368 | 0.2785 | -0.13482 | 4.13E-06 | 0.000496 | 0.000736 | 0.025613 |
| chr11 | 106033856 | 106034035 | 10 | 0.55025 | 0.78045 | -0.2302 | 4.26E-06 | 0.000141 | 0.000751 | 0.013041 |
| chr1 | 151250936 | 151251141 | 13 | 0.084077 | 0.21419 | -0.13012 | 4.36E-06 | 3.88E-05 | 0.000759 | 0.006751 |
| chr4 | 140661772 | 140661841 | 16 | 0.56912 | 0.31606 | 0.253062 | 4.39E-06 | 0.000202 | 0.000759 | 0.015164 |
| chr15 | 83366932 | 83366999 | 9 | 0.32994 | 0.1065 | 0.223444 | 4.48E-06 | 7.45E-06 | 0.000769 | 0.002346 |
| chr6 | 22010464 | 22010722 | 13 | 0.83854 | 0.708 | 0.130538 | 4.76E-06 | 0.000684 | 0.000793 | 0.031559 |
| chr7 | 45638775 | 45638866 | 11 | 0.16236 | 0.029682 | 0.132682 | 4.71E-06 | 3.02E-05 | 0.000793 | 0.006362 |
| chr11 | 22972602 | 22972758 | 24 | 0.33798 | 0.14721 | 0.190771 | 4.82E-06 | 6.13E-05 | 0.000795 | 0.009327 |
| chr4 | 152999604 | 152999714 | 19 | 0.33429 | 0.54326 | -0.20897 | 4.85E-06 | 0.001292 | 0.000795 | 0.033605 |
| chr11 | 116807223 | 116807372 | 8 | 0.38862 | 0.19425 | 0.194375 | 5.11E-06 | 1.60E-05 | 0.000829 | 0.00422 |
| chr18 | 30267969 | 30268177 | 27 | 0.74104 | 0.85885 | -0.11782 | 5.44E-06 | 0.00031 | 0.000876 | 0.020918 |
| chrY | 90784112 | 90784147 | 19 | 0.58597 | 0.74605 | -0.16008 | 5.63E-06 | 0.000224 | 0.000898 | 0.016087 |
| chr12 | 97921263 | 97921408 | 12 | 0.46975 | 0.75433 | -0.28458 | 5.72E-06 | 9.20E-05 | 0.000905 | 0.011255 |
| chr2 | 174299898 | 174300052 | 11 | 0.45627 | 0.28109 | 0.175182 | 5.89E-06 | 6.99E-07 | 0.000924 | 0.000428 |
| chr15 | 78254803 | 78254907 | 20 | 0.10658 | 0.25143 | -0.14485 | 6.11E-06 | 0.000387 | 0.000934 | 0.023015 |
| chr8 | 34380760 | 34380903 | 8 | 0.47138 | 0.26281 | 0.208563 | 6.11E-06 | 0.000187 | 0.000934 | 0.015013 |
| chrX | 44871848 | 44871940 | 16 | 0.86947 | 0.97269 | -0.10322 | 6.04E-06 | 5.16E-05 | 0.000934 | 0.008303 |
| chr2 | 174287131 | 174287234 | 7 | 0.42471 | 0.66671 | -0.242 | 6.70E-06 | 6.48E-07 | 0.001016 | 0.000425 |
| chr10 | 81332726 | 81332911 | 24 | 0.28125 | 0.51771 | -0.23646 | 6.82E-06 | 0.000279 | 0.001026 | 0.019312 |
| chr2 | 32095970 | 32096095 | 15 | 0.8542 | 0.593 | 0.2612 | 6.97E-06 | 0.000567 | 0.001032 | 0.028497 |
| chr8 | 106884275 | 106884391 | 12 | 0.60779 | 0.78146 | -0.17367 | 6.95E-06 | 0.000303 | 0.001032 | 0.020494 |
| chr4 | 152108410 | 152108491 | 10 | 0.16375 | 0.00715 | 0.1566 | 7.11E-06 | 7.54E-06 | 0.001044 | 0.002346 |
| chr4 | 152888362 | 152888502 | 8 | 0.015562 | 0.2625 | -0.24694 | 7.29E-06 | 3.74E-05 | 0.001062 | 0.006668 |
| chr5 | 26754145 | 26754342 | 13 | 0.296 | 0.10946 | 0.186538 | 7.99E-06 | 0.001745 | 0.001154 | 0.042567 |
| chr2 | 173042914 | 173043127 | 11 | 0.2755 | 0.46045 | -0.18496 | 8.20E-06 | 6.63E-06 | 0.001176 | 0.002337 |
| chr18 | 65506977 | 65507216 | 7 | 0.87571 | 0.53414 | 0.341571 | 8.31E-06 | 4.99E-06 | 0.001176 | 0.001946 |
| chr9 | 123157013 | 123157082 | 22 | 0.20259 | 0.34032 | -0.13773 | 8.33E-06 | 0.000949 | 0.001176 | 0.031559 |
| chr10 | 85473827 | 85473857 | 5 | 0.5222 | 0.3456 | 0.1766 | 1.08E-05 | 0.000133 | 0.001211 | 0.013041 |
| chr11 | 102666266 | 102666301 | 5 | 0.5345 | 0.2592 | 0.2753 | 1.08E-05 | 0.000433 | 0.001211 | 0.023757 |
| chr11 | 116806890 | 116806914 | 5 | 0.5433 | 0.288 | 0.2553 | 1.08E-05 | 5.10E-05 | 0.001211 | 0.008278 |
| chr11 | 44555965 | 44556204 | 5 | 0.743 | 0.9835 | -0.2405 | 1.08E-05 | 0.000145 | 0.001211 | 0.01319 |
| chr11 | 77876612 | 77876650 | 5 | 0.2998 | 0.6748 | -0.375 | 1.08E-05 | 0.000138 | 0.001211 | 0.013041 |
| chr12 | 111347303 | 111347480 | 5 | 0.815 | 0.3704 | 0.4446 | 1.08E-05 | 0.000132 | 0.001211 | 0.013041 |
| chr12 | 26456603 | 26456639 | 5 | 0.2286 | 0.0497 | 0.1789 | 1.08E-05 | 0.000952 | 0.001211 | 0.031559 |
| chr12 | 48340380 | 48340527 | 5 | 0.7036 | 0.9742 | -0.2706 | 1.08E-05 | 2.34E-05 | 0.001211 | 0.005436 |
| chr12 | 76405145 | 76405179 | 5 | 0.1459 | 0.0217 | 0.1242 | 1.08E-05 | 0.00013 | 0.001211 | 0.013041 |
| chr14 | 117945342 | 117945548 | 5 | 0.3625 | 0.7586 | -0.3961 | 1.08E-05 | 1.91E-05 | 0.001211 | 0.004546 |
| chr14 | 54530868 | 54530882 | 5 | 0.8582 | 0.9801 | -0.1219 | 1.08E-05 | 0.000136 | 0.001211 | 0.013041 |
| chr15 | 83117424 | 83117518 | 5 | 0.7042 | 0.9413 | -0.2371 | 1.08E-05 | 0.000134 | 0.001211 | 0.013041 |
| chr16 | 31892763 | 31892784 | 5 | 0.9033 | 0.6495 | 0.2538 | 1.08E-05 | 0.000131 | 0.001211 | 0.013041 |
| chr16 | 95399146 | 95399166 | 5 | 0.4864 | 0.2127 | 0.2737 | 1.08E-05 | 1.47E-05 | 0.001211 | 0.004028 |
| chr19 | 47451658 | 47451766 | 5 | 0.6578 | 0.3376 | 0.3202 | 1.08E-05 | 0.000126 | 0.001211 | 0.013041 |
| chr2 | 167808854 | 167808907 | 5 | 0.9019 | 0.6888 | 0.2131 | 1.08E-05 | 0.000452 | 0.001211 | 0.023757 |
| chr2 | 174328397 | 174328466 | 7 | 0.73736 | 0.27807 | 0.459286 | 1.03E-05 | 4.96E-06 | 0.001211 | 0.001946 |
| chr2 | 99551167 | 99551186 | 5 | 0.8604 | 0.9652 | -0.1048 | 1.08E-05 | 1.47E-05 | 0.001211 | 0.004028 |
| chr3 | 51308473 | 51308566 | 5 | 0.8139 | 0.5828 | 0.2311 | 1.08E-05 | 4.61E-05 | 0.001211 | 0.007766 |
| chr4 | 125415605 | 125415633 | 5 | 0.6239 | 0.3316 | 0.2923 | 1.08E-05 | 0.000168 | 0.001211 | 0.014451 |
| chr4 | 131173739 | 131173779 | 5 | 0.3739 | 0.2376 | 0.1363 | 1.08E-05 | 0.001031 | 0.001211 | 0.031559 |
| chr5 | 100988365 | 100988721 | 8 | 0.77462 | 0.60081 | 0.173812 | 8.70E-06 | 8.61E-05 | 0.001211 | 0.01097 |

| | | | | | | | | | | |
|---|---|---|---|---|---|---|---|---|---|---|
| chr5 | 134765626 | 134765664 | 5 | 0.8649 | 0.6175 | 0.2474 | 1.08E-05 | 0.000138 | 0.001211 | 0.013041 |
| chr6 | 48629110 | 48629431 | 39 | 0.066244 | 0.21623 | -0.14999 | 9.71E-06 | 1.81E-05 | 0.001211 | 0.004428 |
| chr7 | 48964626 | 48964767 | 5 | 0.8819 | 0.756 | 0.1259 | 1.08E-05 | 4.71E-05 | 0.001211 | 0.007834 |
| chr7 | 73209165 | 73209600 | 21 | 0.18693 | 0.079 | 0.107929 | 9.11E-06 | 0.000833 | 0.001211 | 0.031559 |
| chr8 | 119633112 | 119633164 | 5 | 0.37 | 0.9 | -0.53 | 1.08E-05 | 4.81E-05 | 0.001211 | 0.00787 |
| chr8 | 3994254 | 3994493 | 19 | 0.38187 | 0.6795 | -0.29763 | 1.06E-05 | 0.001274 | 0.001211 | 0.033527 |
| chr8 | 70745165 | 70745441 | 26 | 0.43467 | 0.6351 | -0.20042 | 9.54E-06 | 0.000647 | 0.001211 | 0.031559 |
| chr8 | 84706851 | 84706888 | 5 | 0.0552 | 0.3189 | -0.2637 | 1.08E-05 | 0.00013 | 0.001211 | 0.013041 |
| chr9 | 122281899 | 122282046 | 5 | 0.9214 | 0.7788 | 0.1426 | 1.08E-05 | 4.36E-05 | 0.001211 | 0.007483 |
| chr9 | 89066524 | 89066570 | 5 | 0.191 | 0.3766 | -0.1856 | 1.08E-05 | 1.29E-05 | 0.001211 | 0.003706 |
| chr15 | 101930704 | 101930775 | 9 | 0.42817 | 0.20256 | 0.225611 | 1.09E-05 | 0.000671 | 0.001216 | 0.031559 |
| chr2 | 169754519 | 169754672 | 11 | 0.6315 | 0.77018 | -0.13868 | 1.14E-05 | 0.0004 | 0.001253 | 0.023015 |
| chr5 | 117087015 | 117087651 | 23 | 0.35687 | 0.58728 | -0.23041 | 1.14E-05 | 0.000422 | 0.001253 | 0.023445 |
| chr15 | 87068407 | 87068516 | 8 | 0.28856 | 0.039375 | 0.249188 | 1.23E-05 | 1.01E-06 | 0.001345 | 0.000579 |
| chr16 | 4718667 | 4718754 | 13 | 0.084462 | 0.21946 | -0.135 | 1.33E-05 | 0.001164 | 0.00144 | 0.032048 |
| chr11 | 100330449 | 100330601 | 19 | 0.27218 | 0.42697 | -0.15479 | 1.34E-05 | 0.0013 | 0.00145 | 0.033685 |
| chr12 | 85475616 | 85475662 | 6 | 0.34858 | 0.117 | 0.231583 | 1.41E-05 | 0.00103 | 0.001494 | 0.031559 |
| chr18 | 25478406 | 25478494 | 19 | 0.68289 | 0.81963 | -0.13674 | 1.41E-05 | 5.68E-05 | 0.001494 | 0.008929 |
| chr14 | 31669067 | 31669435 | 25 | 0.75618 | 0.85638 | -0.1002 | 1.45E-05 | 9.60E-05 | 0.001528 | 0.011657 |
| chr14 | 32201800 | 32201869 | 9 | 0.575 | 0.20317 | 0.371833 | 1.46E-05 | 0.00034 | 0.001531 | 0.022256 |
| chr10 | 80132136 | 80132204 | 10 | 0.15165 | 0.39595 | -0.2443 | 1.50E-05 | 0.000984 | 0.001568 | 0.031559 |
| chr10 | 121247097 | 121247183 | 14 | 0.81539 | 0.93257 | -0.11718 | 1.52E-05 | 0.000826 | 0.001573 | 0.031559 |
| chr10 | 90050032 | 90050100 | 7 | 0.673 | 0.87757 | -0.20457 | 1.57E-05 | 8.18E-05 | 0.001598 | 0.010671 |
| chr15 | 86082646 | 86082833 | 11 | 0.43768 | 0.68327 | -0.24559 | 1.57E-05 | 0.000401 | 0.001598 | 0.023015 |
| chr17 | 12596553 | 12596620 | 11 | 0.94395 | 0.72236 | 0.221591 | 1.57E-05 | 0.000114 | 0.001598 | 0.013041 |
| chr11 | 20035390 | 20035684 | 14 | 0.73771 | 0.47543 | 0.262286 | 1.63E-05 | 0.000799 | 0.001649 | 0.031559 |
| chr17 | 71494589 | 71494657 | 12 | 0.67662 | 0.85029 | -0.17367 | 1.64E-05 | 0.002728 | 0.001649 | 0.049584 |
| chr19 | 36815604 | 36815649 | 10 | 0.27465 | 0.1021 | 0.17255 | 1.70E-05 | 7.00E-05 | 0.001705 | 0.009945 |
| chr4 | 152041990 | 152042346 | 13 | 0.64819 | 0.76019 | -0.112 | 1.85E-05 | 0.001883 | 0.001843 | 0.045024 |
| chr1 | 180762623 | 180762920 | 10 | 0.93285 | 0.73185 | 0.201 | 1.92E-05 | 0.000978 | 0.001861 | 0.031559 |
| chr1 | 34121603 | 34121724 | 10 | 0.3831 | 0.19655 | 0.18655 | 2.17E-05 | 0.000521 | 0.001861 | 0.026686 |
| chr1 | 65523934 | 65523973 | 5 | 0.9776 | 0.8315 | 0.1461 | 2.17E-05 | 0.000427 | 0.001861 | 0.023618 |
| chr10 | 86816914 | 86817003 | 5 | 0.7719 | 0.3787 | 0.3932 | 2.17E-05 | 0.000137 | 0.001861 | 0.013041 |
| chr11 | 101879131 | 101879174 | 5 | 0.799 | 0.5486 | 0.2504 | 2.17E-05 | 0.001337 | 0.001861 | 0.034162 |
| chr11 | 104800171 | 104800269 | 5 | 0.5992 | 0.7459 | -0.1467 | 2.17E-05 | 0.002465 | 0.001861 | 0.049423 |
| chr11 | 106818449 | 106818634 | 11 | 0.52632 | 0.34073 | 0.185591 | 1.94E-05 | 0.000733 | 0.001861 | 0.031559 |
| chr11 | 97923686 | 97923777 | 5 | 0.374 | 0.6011 | -0.2271 | 2.17E-05 | 0.000358 | 0.001861 | 0.023015 |
| chr14 | 41723784 | 41723800 | 5 | 0.6027 | 0.868 | -0.2653 | 2.17E-05 | 0.000387 | 0.001861 | 0.023015 |
| chr14 | 70310144 | 70310382 | 12 | 0.73946 | 0.48142 | 0.258042 | 1.97E-05 | 9.12E-05 | 0.001861 | 0.011255 |
| chr15 | 101052739 | 101052778 | 19 | 0.62711 | 0.3945 | 0.232605 | 1.96E-05 | 0.000358 | 0.001861 | 0.023015 |
| chr16 | 31931415 | 31931593 | 19 | 0.58966 | 0.36426 | 0.225395 | 1.96E-05 | 0.000529 | 0.001861 | 0.02698 |
| chr17 | 28208259 | 28208375 | 5 | 0.679 | 0.4144 | 0.2646 | 2.17E-05 | 0.000602 | 0.001861 | 0.030193 |
| chr17 | 72656227 | 72656416 | 10 | 0.5793 | 0.7629 | -0.1836 | 2.17E-05 | 0.000961 | 0.001861 | 0.031559 |
| chr17 | 78283602 | 78283718 | 10 | 0.3479 | 0.5053 | -0.1574 | 1.92E-05 | 6.80E-05 | 0.001861 | 0.009945 |
| chr18 | 61042075 | 61042150 | 5 | 0.9093 | 0.7146 | 0.1947 | 2.17E-05 | 0.000136 | 0.001861 | 0.013041 |
| chr19 | 45575187 | 45575283 | 9 | 0.76439 | 0.53117 | 0.233222 | 1.94E-05 | 0.002408 | 0.001861 | 0.049423 |
| chr2 | 157705460 | 157705469 | 5 | 0.4871 | 0.7238 | -0.2367 | 2.17E-05 | 0.001135 | 0.001861 | 0.03198 |
| chr2 | 173266506 | 173266551 | 5 | 0.4418 | 0.7979 | -0.3561 | 2.17E-05 | 0.000394 | 0.001861 | 0.023015 |
| chr5 | 119680690 | 119681086 | 29 | 0.20709 | 0.35897 | -0.15188 | 2.02E-05 | 0.001497 | 0.001861 | 0.037778 |
| chr5 | 136219103 | 136219140 | 5 | 0.8031 | 0.9219 | -0.1188 | 2.17E-05 | 0.001028 | 0.001861 | 0.031559 |
| chr5 | 146737977 | 146738078 | 5 | 0.9832 | 0.7693 | 0.2139 | 2.17E-05 | 0.001046 | 0.001861 | 0.031559 |

| | | | | | | | | | | |
|---|---|---|---|---|---|---|---|---|---|---|
| chr5 | 34252529 | 34252577 | 5 | 0.3251 | 0.1011 | 0.224 | 2.17E-05 | 0.000152 | 0.001861 | 0.0137 |
| chr6 | 85125789 | 85125853 | 5 | 0.0732 | 0.3 | -0.2268 | 2.17E-05 | 0.000126 | 0.001861 | 0.013041 |
| chr8 | 111161634 | 111161748 | 5 | 0.7924 | 0.9502 | -0.1578 | 2.17E-05 | 0.00042 | 0.001861 | 0.023411 |
| chr8 | 24245795 | 24245944 | 5 | 0.7605 | 0.9038 | -0.1433 | 2.17E-05 | 0.001235 | 0.001861 | 0.033033 |
| chr8 | 25986614 | 25986703 | 5 | 0.7392 | 0.9567 | -0.2175 | 2.17E-05 | 0.00105 | 0.001861 | 0.031559 |
| chr11 | 52077259 | 52077381 | 9 | 0.89061 | 0.72167 | 0.168944 | 2.24E-05 | 0.00034 | 0.001862 | 0.022256 |
| chr11 | 76324099 | 76324131 | 9 | 0.87539 | 0.57917 | 0.296222 | 2.24E-05 | 0.001289 | 0.001862 | 0.033582 |
| chr15 | 97323504 | 97323768 | 39 | 0.52915 | 0.71554 | -0.18639 | 2.24E-05 | 0.000548 | 0.001862 | 0.027747 |
| chr4 | 147220155 | 147220216 | 9 | 0.74544 | 0.93606 | -0.19061 | 2.24E-05 | 8.01E-05 | 0.001862 | 0.010671 |
| chr4 | 32386408 | 32386544 | 9 | 0.43806 | 0.64428 | -0.20622 | 2.24E-05 | 0.00034 | 0.001862 | 0.022256 |
| chr6 | 52179988 | 52180107 | 6 | 0.005167 | 0.23167 | -0.2265 | 2.22E-05 | 2.66E-05 | 0.001862 | 0.006033 |
| chr7 | 37293573 | 37293862 | 6 | 0.77683 | 0.92308 | -0.14625 | 2.22E-05 | 0.000178 | 0.001862 | 0.014922 |
| chr3 | 128087315 | 128087674 | 17 | 0.14026 | 0.28312 | -0.14285 | 2.32E-05 | 0.00174 | 0.001917 | 0.042497 |
| chr13 | 66007674 | 66007797 | 28 | 0.23573 | 0.39234 | -0.15661 | 2.33E-05 | 0.002148 | 0.001918 | 0.04812 |
| chr4 | 149744558 | 149744765 | 7 | 0.98564 | 0.81436 | 0.171286 | 2.37E-05 | 0.000473 | 0.001938 | 0.024595 |
| chr7 | 139576663 | 139576779 | 11 | 0.27559 | 0.43623 | -0.16064 | 2.39E-05 | 0.001307 | 0.001939 | 0.033685 |
| chr15 | 77736870 | 77737249 | 10 | 0.5438 | 0.32695 | 0.21685 | 2.45E-05 | 0.000141 | 0.001976 | 0.013041 |
| chr2 | 37786535 | 37786728 | 46 | 0.31367 | 0.19712 | 0.116554 | 2.49E-05 | 0.001009 | 0.001999 | 0.031559 |
| chr9 | 118814354 | 118814617 | 20 | 0.17545 | 0.075425 | 0.100025 | 2.50E-05 | 0.000901 | 0.002004 | 0.031559 |
| chr4 | 150114680 | 150114693 | 9 | 0.81228 | 0.94067 | -0.12839 | 2.58E-05 | 0.001289 | 0.002055 | 0.033582 |
| chr3 | 108940310 | 108940486 | 11 | 0.49205 | 0.20005 | 0.292 | 2.65E-05 | 1.27E-05 | 0.002096 | 0.003706 |
| chr4 | 139556149 | 139556202 | 11 | 0.60523 | 0.75686 | -0.15164 | 2.65E-05 | 0.001308 | 0.002096 | 0.033685 |
| chr15 | 59057462 | 59057664 | 13 | 0.62919 | 0.77096 | -0.14177 | 2.78E-05 | 0.000129 | 0.002168 | 0.013041 |
| chr17 | 83593194 | 83593421 | 13 | 0.68573 | 0.45985 | 0.225885 | 2.78E-05 | 0.000377 | 0.002168 | 0.023015 |
| chr11 | 44546381 | 44546712 | 16 | 0.64922 | 0.78819 | -0.13897 | 2.80E-05 | 0.000549 | 0.002175 | 0.027747 |
| chr15 | 99294541 | 99294583 | 7 | 0.37279 | 0.18979 | 0.183 | 2.90E-05 | 3.82E-05 | 0.002244 | 0.006735 |
| chr19 | 15328536 | 15328692 | 11 | 0.73486 | 0.87714 | -0.14227 | 2.94E-05 | 0.000406 | 0.002247 | 0.023015 |
| chr3 | 128003089 | 128003280 | 11 | 0.23577 | 0.36895 | -0.13318 | 2.94E-05 | 0.002272 | 0.002247 | 0.049423 |
| chr3 | 32591111 | 32591217 | 11 | 0.096909 | 0.28782 | -0.19091 | 2.94E-05 | 0.000407 | 0.002247 | 0.023015 |
| chr17 | 43391507 | 43391717 | 9 | 0.72972 | 0.94267 | -0.21294 | 2.96E-05 | 0.000671 | 0.002255 | 0.031559 |
| chr15 | 73809823 | 73809892 | 10 | 0.5151 | 0.1718 | 0.3433 | 3.10E-05 | 7.54E-06 | 0.002343 | 0.002346 |
| chr2 | 164200040 | 164200105 | 10 | 0.24925 | 0.5188 | -0.26955 | 3.10E-05 | 3.44E-05 | 0.002343 | 0.006484 |
| chr15 | 101053132 | 101053300 | 22 | 0.18382 | 0.33641 | -0.15259 | 3.12E-05 | 0.000942 | 0.002348 | 0.031559 |
| chr11 | 24977759 | 24978086 | 15 | 0.38383 | 0.57467 | -0.19083 | 3.26E-05 | 0.001521 | 0.002422 | 0.038296 |
| chr13 | 45568236 | 45568312 | 11 | 0.60164 | 0.33709 | 0.264545 | 3.26E-05 | 5.98E-05 | 0.002422 | 0.009216 |
| chr14 | 74974149 | 74974384 | 11 | 0.49836 | 0.67914 | -0.18077 | 3.26E-05 | 0.001309 | 0.002422 | 0.033685 |
| chr10 | 92298787 | 92298870 | 6 | 0.98058 | 0.6725 | 0.308083 | 3.33E-05 | 0.000177 | 0.002442 | 0.014922 |
| chr17 | 34672728 | 34672746 | 6 | 0.80542 | 0.50167 | 0.30375 | 3.33E-05 | 0.001085 | 0.002442 | 0.031559 |
| chr2 | 80267189 | 80267259 | 6 | 0.34742 | 0.55342 | -0.206 | 3.33E-05 | 0.000215 | 0.002442 | 0.015643 |
| chr10 | 14130597 | 14130853 | 26 | 0.59017 | 0.69813 | -0.10796 | 3.36E-05 | 0.00032 | 0.002454 | 0.021402 |
| chr13 | 24471859 | 24472170 | 14 | 0.58454 | 0.75446 | -0.16993 | 3.39E-05 | 0.000813 | 0.002464 | 0.031559 |
| chr1 | 180936747 | 180936786 | 7 | 0.008857 | 0.119 | -0.11014 | 3.55E-05 | 8.48E-05 | 0.00255 | 0.010937 |
| chr11 | 98419018 | 98419109 | 7 | 0.4465 | 0.66886 | -0.22236 | 3.55E-05 | 0.0002 | 0.00255 | 0.01516 |
| chr8 | 87719713 | 87719759 | 7 | 0.63486 | 0.48086 | 0.154 | 3.55E-05 | 8.20E-05 | 0.00255 | 0.010671 |
| chr10 | 80132024 | 80132115 | 17 | 0.1255 | 0.23971 | -0.11421 | 3.57E-05 | 0.000289 | 0.00256 | 0.019842 |
| chr13 | 45568367 | 45568439 | 12 | 0.47417 | 0.72937 | -0.25521 | 3.72E-05 | 0.000955 | 0.002659 | 0.031559 |
| chr1 | 132115184 | 132115401 | 5 | 0.821 | 0.9653 | -0.1443 | 4.33E-05 | 0.000172 | 0.00272 | 0.014645 |
| chr1 | 170109699 | 170109984 | 12 | 0.44413 | 0.63008 | -0.18596 | 4.07E-05 | 0.000545 | 0.00272 | 0.027711 |
| chr10 | 59835854 | 59835934 | 5 | 0.257 | 0.5626 | -0.3056 | 4.33E-05 | 0.002788 | 0.00272 | 0.049718 |
| chr10 | 60839043 | 60839106 | 5 | 0.4279 | 0.684 | -0.2561 | 4.33E-05 | 0.001137 | 0.00272 | 0.03198 |
| chr10 | 68572763 | 68572785 | 5 | 0.8406 | 0.9896 | -0.149 | 4.33E-05 | 0.001033 | 0.00272 | 0.031559 |

| | | | | | | | | | | |
|---|---|---|---|---|---|---|---|---|---|---|
| chr10 | 94095804 | 94095957 | 11 | 0.62764 | 0.78327 | -0.15564 | 4.00E-05 | 3.02E-05 | 0.00272 | 0.006362 |
| chr11 | 109770230 | 109770409 | 14 | 0.73904 | 0.60771 | 0.131321 | 3.90E-05 | 0.000274 | 0.00272 | 0.019204 |
| chr11 | 70520193 | 70520198 | 5 | 0.114 | 0.2507 | -0.1367 | 4.33E-05 | 0.000951 | 0.00272 | 0.031559 |
| chr14 | 17885048 | 17885075 | 5 | 0.4164 | 0.7257 | -0.3093 | 4.33E-05 | 0.000951 | 0.00272 | 0.031559 |
| chr14 | 66045177 | 66045287 | 5 | 0.9227 | 0.7531 | 0.1696 | 4.33E-05 | 0.000381 | 0.00272 | 0.023015 |
| chr15 | 100636242 | 100636290 | 5 | 0.0135 | 0.1635 | -0.15 | 4.33E-05 | 0.00096 | 0.00272 | 0.031559 |
| chr17 | 28962963 | 28963030 | 5 | 0.8553 | 0.9685 | -0.1132 | 4.33E-05 | 0.000974 | 0.00272 | 0.031559 |
| chr17 | 36866912 | 36866983 | 5 | 0.0552 | 0.2374 | -0.1822 | 4.33E-05 | 0.001113 | 0.00272 | 0.031602 |
| chr17 | 65548627 | 65548643 | 5 | 0.8362 | 0.6411 | 0.1951 | 4.33E-05 | 0.000128 | 0.00272 | 0.013041 |
| chr18 | 38181098 | 38181239 | 5 | 0.9767 | 0.8038 | 0.1729 | 4.33E-05 | 0.00253 | 0.00272 | 0.049423 |
| chr18 | 53469655 | 53469668 | 5 | 0.175 | 0.0596 | 0.1154 | 4.33E-05 | 0.000373 | 0.00272 | 0.023015 |
| chr19 | 3814968 | 3815024 | 5 | 0.7404 | 0.9502 | -0.2098 | 4.33E-05 | 0.002496 | 0.00272 | 0.049423 |
| chr19 | 55511497 | 55511586 | 10 | 0.7341 | 0.9185 | -0.1844 | 3.93E-05 | 1.63E-05 | 0.00272 | 0.00422 |
| chr2 | 121301514 | 121301692 | 16 | 0.2775 | 0.47194 | -0.19444 | 4.22E-05 | 0.000641 | 0.00272 | 0.031559 |
| chr3 | 27022050 | 27022101 | 5 | 0.9862 | 0.8164 | 0.1698 | 4.33E-05 | 0.001151 | 0.00272 | 0.032048 |
| chr5 | 124123429 | 124123525 | 5 | 0.8467 | 0.5913 | 0.2554 | 4.33E-05 | 0.000419 | 0.00272 | 0.023411 |
| chr6 | 120950983 | 120951099 | 5 | 0.7925 | 0.9451 | -0.1526 | 4.33E-05 | 0.00099 | 0.00272 | 0.031559 |
| chr6 | 51332965 | 51333024 | 5 | 0.4126 | 0.2035 | 0.2091 | 4.33E-05 | 0.001211 | 0.00272 | 0.032629 |
| chr6 | 54485701 | 54485726 | 5 | 0.1822 | 0.0125 | 0.1697 | 4.33E-05 | 0.000997 | 0.00272 | 0.031559 |
| chr6 | 72603790 | 72603930 | 7 | 0.91064 | 0.72357 | 0.187071 | 4.33E-05 | 0.000193 | 0.00272 | 0.015013 |
| chr7 | 54910273 | 54910384 | 5 | 0.3549 | 0.4911 | -0.1362 | 4.33E-05 | 0.000989 | 0.00272 | 0.031559 |
| chr8 | 70596879 | 70597003 | 5 | 0.6447 | 0.9312 | -0.2865 | 4.33E-05 | 0.000125 | 0.00272 | 0.013041 |
| chr9 | 102183861 | 102184095 | 23 | 0.59185 | 0.71133 | -0.11948 | 4.28E-05 | 0.001376 | 0.00272 | 0.034917 |
| chr9 | 119485409 | 119485505 | 5 | 0.8015 | 0.9583 | -0.1568 | 4.33E-05 | 0.001006 | 0.00272 | 0.031559 |
| chr9 | 44710845 | 44711027 | 5 | 0.7601 | 0.9415 | -0.1814 | 4.33E-05 | 0.001033 | 0.00272 | 0.031559 |
| chr9 | 61229108 | 61229342 | 5 | 0.9727 | 0.7686 | 0.2041 | 4.33E-05 | 0.000986 | 0.00272 | 0.031559 |
| chr9 | 78177250 | 78177372 | 5 | 0.6083 | 0.4057 | 0.2026 | 4.33E-05 | 0.002729 | 0.00272 | 0.049584 |
| chrX | 42053397 | 42053659 | 7 | 0.96314 | 0.69557 | 0.267571 | 4.33E-05 | 0.000444 | 0.00272 | 0.023757 |
| chr2 | 174329010 | 174329254 | 33 | 0.33311 | 0.19811 | 0.135 | 4.37E-05 | 3.46E-05 | 0.002727 | 0.006484 |
| chr14 | 118043092 | 118043220 | 10 | 0.72165 | 0.91615 | -0.1945 | 4.42E-05 | 0.001789 | 0.002741 | 0.042972 |
| chr4 | 100335756 | 100335829 | 11 | 0.55341 | 0.39109 | 0.162318 | 4.42E-05 | 0.000212 | 0.002741 | 0.015641 |
| chr18 | 56761385 | 56761658 | 12 | 0.83487 | 0.64133 | 0.193542 | 4.45E-05 | 0.000526 | 0.00275 | 0.026869 |
| chr8 | 108956972 | 108957015 | 8 | 0.66775 | 0.80319 | -0.13544 | 4.69E-05 | 0.000399 | 0.002889 | 0.023015 |
| chr4 | 132113799 | 132113866 | 12 | 0.48746 | 0.6985 | -0.21104 | 4.86E-05 | 0.00089 | 0.002964 | 0.031559 |
| chr12 | 59143343 | 59143453 | 6 | 0.98058 | 0.87892 | 0.101667 | 4.96E-05 | 0.001059 | 0.002983 | 0.031559 |
| chr19 | 5503367 | 5503552 | 10 | 0.33995 | 0.5261 | -0.18615 | 4.96E-05 | 0.000983 | 0.002983 | 0.031559 |
| chr6 | 115000675 | 115000943 | 6 | 0.4965 | 0.81367 | -0.31717 | 4.96E-05 | 0.000183 | 0.002983 | 0.015013 |
| chr12 | 30904795 | 30905131 | 9 | 0.98611 | 0.87094 | 0.115167 | 5.13E-05 | 3.73E-05 | 0.003064 | 0.006668 |
| chr8 | 123421384 | 123421474 | 9 | 0.93822 | 0.72122 | 0.217 | 5.13E-05 | 0.000671 | 0.003064 | 0.031559 |
| chr4 | 135337853 | 135337972 | 14 | 0.91914 | 0.80389 | 0.11525 | 5.18E-05 | 0.000163 | 0.003083 | 0.014451 |
| chr1 | 91130878 | 91130974 | 7 | 0.93257 | 0.72307 | 0.2095 | 5.27E-05 | 0.000448 | 0.003107 | 0.023757 |
| chr15 | 79500244 | 79500253 | 7 | 0.85157 | 0.673 | 0.178571 | 5.27E-05 | 0.000204 | 0.003107 | 0.0153 |
| chr6 | 135679995 | 135680180 | 13 | 0.3775 | 0.52238 | -0.14489 | 5.24E-05 | 0.001926 | 0.003107 | 0.045635 |
| chrX | 169986687 | 169986940 | 22 | 0.62305 | 0.7782 | -0.15516 | 5.36E-05 | 0.000911 | 0.003144 | 0.031559 |
| chr12 | 43959986 | 43960348 | 11 | 0.96423 | 0.862 | 0.102227 | 5.41E-05 | 0.000116 | 0.003159 | 0.013041 |
| chr8 | 114764443 | 114764513 | 8 | 0.76587 | 0.92006 | -0.15419 | 5.51E-05 | 0.000189 | 0.003191 | 0.015013 |
| chr9 | 108462281 | 108462407 | 14 | 0.38168 | 0.52839 | -0.14671 | 5.55E-05 | 2.99E-05 | 0.003202 | 0.006362 |
| chr2 | 28326273 | 28326503 | 12 | 0.44767 | 0.69421 | -0.24654 | 5.80E-05 | 0.001552 | 0.00332 | 0.038883 |
| chr12 | 99645064 | 99645143 | 9 | 0.97178 | 0.79972 | 0.172056 | 5.87E-05 | 8.01E-05 | 0.003332 | 0.010671 |
| chr10 | 88336764 | 88336934 | 11 | 0.44423 | 0.70795 | -0.26373 | 5.97E-05 | 0.000735 | 0.003383 | 0.031559 |
| chr9 | 18779479 | 18779631 | 20 | 0.80467 | 0.63535 | 0.169325 | 6.01E-05 | 0.001986 | 0.00339 | 0.046791 |

| | | | | | | | | | | |
|---|---|---|---|---|---|---|---|---|---|---|
| chr12 | 82190552 | 82190858 | 13 | 0.60208 | 0.84542 | -0.24335 | 6.12E-05 | 0.001902 | 0.003446 | 0.045305 |
| chr2 | 66091512 | 66091847 | 16 | 0.5635 | 0.39244 | 0.171063 | 6.30E-05 | 0.00106 | 0.003534 | 0.031559 |
| chr6 | 89835496 | 89835590 | 17 | 0.65141 | 0.83606 | -0.18465 | 6.37E-05 | 0.000727 | 0.003564 | 0.031559 |
| chr17 | 84571671 | 84571750 | 7 | 0.60814 | 0.85479 | -0.24664 | 6.40E-05 | 3.32E-05 | 0.00357 | 0.006484 |
| chr4 | 132181894 | 132182046 | 8 | 0.30825 | 0.15188 | 0.156375 | 6.47E-05 | 0.000404 | 0.003585 | 0.023015 |
| chr5 | 112269392 | 112269473 | 8 | 0.050875 | 0.33281 | -0.28194 | 6.47E-05 | 8.53E-05 | 0.003585 | 0.010937 |
| chr13 | 73117734 | 73117943 | 13 | 0.43715 | 0.21558 | 0.221577 | 6.62E-05 | 0.001923 | 0.003656 | 0.045635 |
| chr5 | 127290878 | 127290963 | 9 | 0.793 | 0.58122 | 0.211778 | 6.71E-05 | 0.000167 | 0.003683 | 0.014451 |
| chr1 | 136789269 | 136789350 | 12 | 0.57629 | 0.8115 | -0.23521 | 6.90E-05 | 0.000951 | 0.003707 | 0.031559 |
| chr1 | 180464281 | 180464344 | 5 | 0.8547 | 0.487 | 0.3677 | 7.58E-05 | 0.002735 | 0.003707 | 0.049584 |
| chr10 | 43162493 | 43162576 | 6 | 0.40675 | 0.11317 | 0.293583 | 7.17E-05 | 0.00045 | 0.003707 | 0.023757 |
| chr10 | 78718407 | 78718462 | 5 | 0.5376 | 0.7773 | -0.2397 | 7.58E-05 | 0.00043 | 0.003707 | 0.02369 |
| chr11 | 94663958 | 94664166 | 10 | 0.96365 | 0.80845 | 0.1552 | 7.00E-05 | 0.001784 | 0.003707 | 0.042972 |
| chr12 | 112675068 | 112675091 | 5 | 0.0154 | 0.1217 | -0.1063 | 7.58E-05 | 0.000964 | 0.003707 | 0.031559 |
| chr16 | 66312006 | 66312067 | 10 | 0.9868 | 0.85895 | 0.12785 | 7.00E-05 | 7.02E-05 | 0.003707 | 0.009945 |
| chr17 | 32905512 | 32905607 | 10 | 0.2375 | 0.0331 | 0.2044 | 7.00E-05 | 3.44E-05 | 0.003707 | 0.006484 |
| chr17 | 6324767 | 6324875 | 19 | 0.54382 | 0.7115 | -0.16768 | 7.55E-05 | 0.000859 | 0.003707 | 0.031559 |
| chr17 | 88141062 | 88141077 | 5 | 0.61 | 0.861 | -0.251 | 7.58E-05 | 0.002535 | 0.003707 | 0.049423 |
| chr19 | 36712710 | 36712774 | 5 | 0.9067 | 0.6613 | 0.2454 | 7.58E-05 | 0.000131 | 0.003707 | 0.013041 |
| chr2 | 171873131 | 171873181 | 5 | 0.1134 | 0.352 | -0.2386 | 7.58E-05 | 0.00098 | 0.003707 | 0.031559 |
| chr2 | 36102716 | 36102802 | 5 | 0.1082 | 0.0059 | 0.1023 | 7.58E-05 | 0.000367 | 0.003707 | 0.023015 |
| chr2 | 52928282 | 52928486 | 5 | 0.6555 | 0.4186 | 0.2369 | 7.58E-05 | 0.000369 | 0.003707 | 0.023015 |
| chr2 | 80444532 | 80444582 | 5 | 0.7059 | 0.413 | 0.2929 | 7.58E-05 | 0.00111 | 0.003707 | 0.031572 |
| chr3 | 29964054 | 29964131 | 5 | 0.955 | 0.6763 | 0.2787 | 7.58E-05 | 0.001041 | 0.003707 | 0.031559 |
| chr4 | 126240876 | 126240955 | 5 | 0.5708 | 0.8101 | -0.2393 | 7.58E-05 | 0.002597 | 0.003707 | 0.049453 |
| chr4 | 136681024 | 136681078 | 5 | 0.678 | 0.3646 | 0.3134 | 7.58E-05 | 0.001023 | 0.003707 | 0.031559 |
| chr4 | 136778893 | 136779356 | 5 | 0.8111 | 0.3714 | 0.4397 | 7.58E-05 | 0.000368 | 0.003707 | 0.023015 |
| chr4 | 139578463 | 139578633 | 5 | 0.8737 | 0.6688 | 0.2049 | 7.58E-05 | 0.00277 | 0.003707 | 0.04971 |
| chr4 | 4269066 | 4269120 | 5 | 0.6167 | 0.95 | -0.3333 | 7.58E-05 | 0.002541 | 0.003707 | 0.049423 |
| chr5 | 113285284 | 113285303 | 5 | 0.978 | 0.845 | 0.133 | 7.58E-05 | 0.000126 | 0.003707 | 0.013041 |
| chr6 | 136822074 | 136822127 | 5 | 0.7744 | 0.9625 | -0.1881 | 7.58E-05 | 0.000974 | 0.003707 | 0.031559 |
| chr6 | 59353435 | 59353457 | 5 | 0.7818 | 0.9451 | -0.1633 | 7.58E-05 | 0.002404 | 0.003707 | 0.049423 |
| chr7 | 13007004 | 13007078 | 5 | 0.3437 | 0.6925 | -0.3488 | 7.58E-05 | 0.002541 | 0.003707 | 0.049423 |
| chr7 | 143154509 | 143154540 | 5 | 0.4545 | 0.1441 | 0.3104 | 7.58E-05 | 0.002591 | 0.003707 | 0.049443 |
| chr7 | 49525409 | 49525442 | 5 | 0.0458 | 0.3454 | -0.2996 | 7.58E-05 | 0.002471 | 0.003707 | 0.049423 |
| chr8 | 47652524 | 47652572 | 5 | 0.5434 | 0.6903 | -0.1469 | 7.58E-05 | 0.001008 | 0.003707 | 0.031559 |
| chr8 | 67035803 | 67035825 | 5 | 0.8546 | 0.9903 | -0.1357 | 7.58E-05 | 0.000139 | 0.003707 | 0.013041 |
| chr8 | 71682085 | 71682172 | 12 | 0.17067 | 0.35692 | -0.18625 | 6.90E-05 | 0.001621 | 0.003707 | 0.040407 |
| chr9 | 24952680 | 24952705 | 5 | 0.4456 | 0.0972 | 0.3484 | 7.58E-05 | 0.001009 | 0.003707 | 0.031559 |
| chr9 | 38005306 | 38005387 | 5 | 0.835 | 0.9385 | -0.1035 | 7.58E-05 | 0.001 | 0.003707 | 0.031559 |
| chr9 | 43231206 | 43231224 | 5 | 0.6279 | 0.3766 | 0.2513 | 7.58E-05 | 0.00046 | 0.003707 | 0.024027 |
| chr9 | 7873208 | 7873245 | 5 | 0.1937 | 0.0563 | 0.1374 | 7.58E-05 | 0.002402 | 0.003707 | 0.049423 |
| chr10 | 43004374 | 43004510 | 9 | 0.53456 | 0.69961 | -0.16506 | 7.66E-05 | 8.01E-05 | 0.003717 | 0.010671 |
| chr13 | 56197219 | 56197262 | 9 | 0.7355 | 0.52856 | 0.206944 | 7.66E-05 | 0.002408 | 0.003717 | 0.049423 |
| chr18 | 34863194 | 34863215 | 9 | 0.33306 | 0.66783 | -0.33478 | 7.66E-05 | 0.000671 | 0.003717 | 0.031559 |
| chr10 | 93276243 | 93276435 | 13 | 0.37519 | 0.19942 | 0.175769 | 7.72E-05 | 0.001915 | 0.00372 | 0.045565 |
| chr13 | 53121735 | 53121814 | 7 | 0.30479 | 0.76336 | -0.45857 | 7.77E-05 | 0.000477 | 0.00372 | 0.024706 |
| chr16 | 13135563 | 13135795 | 7 | 0.79879 | 0.68243 | 0.116357 | 7.77E-05 | 0.000956 | 0.00372 | 0.031559 |
| chr5 | 146946721 | 146946838 | 7 | 0.76986 | 0.91121 | -0.14136 | 7.77E-05 | 3.33E-05 | 0.00372 | 0.006484 |
| chr7 | 100288949 | 100288982 | 7 | 0.27086 | 0.11479 | 0.156071 | 7.77E-05 | 0.002018 | 0.00372 | 0.046791 |
| chr15 | 76666994 | 76667076 | 14 | 0.032607 | 0.14 | -0.10739 | 7.84E-05 | 5.21E-05 | 0.003736 | 0.008314 |

| | | | | | | | | | | |
|---|---|---|---|---|---|---|---|---|---|---|
| chr7 | 35747489 | 35747786 | 17 | 0.30103 | 0.49274 | -0.19171 | 7.83E-05 | 0.000293 | 0.003736 | 0.020007 |
| chr11 | 51177119 | 51177370 | 15 | 0.60817 | 0.7532 | -0.14503 | 7.90E-05 | 0.000197 | 0.003754 | 0.015125 |
| chr2 | 119749539 | 119749854 | 11 | 0.66955 | 0.86932 | -0.19977 | 8.03E-05 | 0.002266 | 0.003798 | 0.049423 |
| chr4 | 141464880 | 141464978 | 11 | 0.84573 | 0.69605 | 0.149682 | 8.03E-05 | 0.002207 | 0.003798 | 0.049155 |
| chr13 | 112188893 | 112189260 | 9 | 0.30889 | 0.47528 | -0.16639 | 8.74E-05 | 0.000167 | 0.00411 | 0.014451 |
| chr2 | 167175031 | 167175543 | 11 | 0.79186 | 0.89414 | -0.10227 | 8.86E-05 | 0.000116 | 0.004134 | 0.013041 |
| chr8 | 121089192 | 121089527 | 11 | 0.025818 | 0.15836 | -0.13255 | 8.86E-05 | 0.000218 | 0.004134 | 0.015765 |
| chr5 | 119667953 | 119668244 | 15 | 0.175 | 0.33963 | -0.16463 | 8.93E-05 | 0.000551 | 0.004149 | 0.027747 |
| chr10 | 121173818 | 121173922 | 7 | 0.87021 | 0.46157 | 0.408643 | 9.40E-05 | 3.38E-05 | 0.00429 | 0.006484 |
| chr12 | 18853408 | 18853526 | 20 | 0.16415 | 0.29255 | -0.1284 | 9.36E-05 | 0.001942 | 0.00429 | 0.045914 |
| chr13 | 44614423 | 44614579 | 7 | 0.67986 | 0.52129 | 0.158571 | 9.40E-05 | 0.000961 | 0.00429 | 0.031559 |
| chr17 | 49528650 | 49529003 | 7 | 0.54314 | 0.30793 | 0.235214 | 9.40E-05 | 8.19E-05 | 0.00429 | 0.010671 |
| chrX | 84545451 | 84545525 | 7 | 0.79529 | 0.93936 | -0.14407 | 9.40E-05 | 3.53E-05 | 0.00429 | 0.006484 |
| chr4 | 41522061 | 41522108 | 11 | 0.54941 | 0.73164 | -0.18223 | 9.76E-05 | 0.000218 | 0.004432 | 0.015765 |
| chr6 | 90232101 | 90232312 | 9 | 0.765 | 0.48306 | 0.281944 | 9.96E-05 | 0.00034 | 0.004512 | 0.022256 |
| chr2 | 127075029 | 127075268 | 6 | 0.97825 | 0.86542 | 0.112833 | 0.000103 | 0.000469 | 0.004611 | 0.024436 |
| chr4 | 154727103 | 154727237 | 6 | 0.411 | 0.125 | 0.286 | 0.000103 | 0.000178 | 0.004611 | 0.014922 |
| chr8 | 114793548 | 114793571 | 6 | 0.65208 | 0.49 | 0.162083 | 0.000103 | 0.001085 | 0.004611 | 0.031559 |
| chr9 | 64088924 | 64089073 | 6 | 0.53183 | 0.8095 | -0.27767 | 0.000103 | 0.001103 | 0.004611 | 0.031572 |
| chr13 | 97892658 | 97892950 | 13 | 0.34535 | 0.20377 | 0.141577 | 0.000105 | 0.000399 | 0.004665 | 0.023015 |
| chr7 | 37146671 | 37146847 | 13 | 0.79819 | 0.69423 | 0.103962 | 0.000105 | 0.001811 | 0.004665 | 0.043429 |
| chr12 | 105468671 | 105468827 | 16 | 0.47253 | 0.31594 | 0.156594 | 0.00011 | 0.001583 | 0.00487 | 0.039562 |
| chr4 | 129206951 | 129207072 | 10 | 0.74905 | 0.91515 | -0.1661 | 0.00011 | 0.001773 | 0.00487 | 0.042972 |
| chr7 | 24382007 | 24382235 | 14 | 0.76836 | 0.89961 | -0.13125 | 0.00011 | 0.001357 | 0.00487 | 0.034467 |
| chr18 | 76532828 | 76532935 | 7 | 0.97479 | 0.83371 | 0.141071 | 0.000114 | 0.000963 | 0.004914 | 0.031559 |
| chr19 | 4447802 | 4447848 | 9 | 0.80522 | 0.92156 | -0.11633 | 0.000113 | 0.002408 | 0.004914 | 0.049423 |
| chr4 | 132113629 | 132113679 | 7 | 0.11907 | 0.50886 | -0.38979 | 0.000114 | 0.000985 | 0.004914 | 0.031559 |
| chr4 | 134345937 | 134346034 | 7 | 0.97207 | 0.78164 | 0.190429 | 0.000114 | 3.41E-05 | 0.004914 | 0.006484 |
| chr4 | 141631329 | 141631457 | 9 | 0.97856 | 0.87433 | 0.104222 | 0.000113 | 3.73E-05 | 0.004914 | 0.006668 |
| chrX | 151743151 | 151743270 | 7 | 0.62186 | 0.79779 | -0.17593 | 0.000114 | 0.000199 | 0.004914 | 0.01516 |
| chr1 | 153159701 | 153159753 | 5 | 0.4224 | 0.6497 | -0.2273 | 0.00013 | 0.00013 | 0.004956 | 0.013041 |
| chr1 | 39655091 | 39655127 | 5 | 0.7259 | 0.829 | -0.1031 | 0.00013 | 0.001209 | 0.004956 | 0.032629 |
| chr1 | 40924912 | 40925028 | 5 | 0.1434 | 0.2917 | -0.1483 | 0.00013 | 0.001165 | 0.004956 | 0.032048 |
| chr10 | 79819458 | 79819504 | 5 | 0.8436 | 0.9758 | -0.1322 | 0.00013 | 0.000127 | 0.004956 | 0.013041 |
| chr11 | 114062658 | 114062714 | 5 | 0.824 | 0.9291 | -0.1051 | 0.00013 | 0.002442 | 0.004956 | 0.049423 |
| chr11 | 119267652 | 119267732 | 5 | 0.3978 | 0.2 | 0.1978 | 0.00013 | 0.001001 | 0.004956 | 0.031559 |
| chr11 | 88625650 | 88625782 | 5 | 0.5289 | 0.1796 | 0.3493 | 0.00013 | 0.002742 | 0.004956 | 0.049584 |
| chr12 | 43929074 | 43929115 | 5 | 0.8856 | 0.9968 | -0.1112 | 0.00013 | 0.000427 | 0.004956 | 0.023618 |
| chr14 | 66811876 | 66811902 | 5 | 0.601 | 0.8119 | -0.2109 | 0.00013 | 0.001201 | 0.004956 | 0.032601 |
| chr15 | 98496005 | 98496029 | 5 | 0.3678 | 0.5724 | -0.2046 | 0.00013 | 0.002593 | 0.004956 | 0.049443 |
| chr17 | 27434756 | 27434971 | 11 | 0.66195 | 0.88368 | -0.22173 | 0.00013 | 0.000713 | 0.004956 | 0.031559 |
| chr17 | 3313910 | 3313947 | 5 | 0.6418 | 0.323 | 0.3188 | 0.00013 | 0.002448 | 0.004956 | 0.049423 |
| chr17 | 35218712 | 35218932 | 5 | 0.9377 | 0.7632 | 0.1745 | 0.00013 | 0.002519 | 0.004956 | 0.049423 |
| chr2 | 120039010 | 120039096 | 8 | 0.7915 | 0.974 | -0.1825 | 0.000121 | 0.0004 | 0.004956 | 0.023015 |
| chr2 | 122644051 | 122644095 | 5 | 0.6492 | 0.9089 | -0.2597 | 0.00013 | 0.000131 | 0.004956 | 0.013041 |
| chr2 | 131159717 | 131159937 | 18 | 0.006556 | 0.15147 | -0.14492 | 0.000117 | 0.000792 | 0.004956 | 0.031559 |
| chr2 | 150385882 | 150385982 | 12 | 0.55558 | 0.36908 | 0.1865 | 0.000125 | 2.53E-05 | 0.004956 | 0.005798 |
| chr2 | 44104489 | 44104789 | 5 | 0.685 | 0.55 | 0.135 | 0.00013 | 0.000406 | 0.004956 | 0.023015 |
| chr2 | 72437004 | 72437049 | 5 | 0.585 | 0.8406 | -0.2556 | 0.00013 | 0.000967 | 0.004956 | 0.031559 |
| chr3 | 137827574 | 137827584 | 5 | 0.5138 | 0.3291 | 0.1847 | 0.00013 | 0.000999 | 0.004956 | 0.031559 |
| chr4 | 118167196 | 118167407 | 5 | 0.6735 | 0.8786 | -0.2051 | 0.00013 | 0.00276 | 0.004956 | 0.049584 |

| | | | | | | | | | | |
|---|---|---|---|---|---|---|---|---|---|---|
| chr4 | 118390743 | 118390823 | 5 | 0.8046 | 0.9857 | -0.1811 | 0.00013 | 0.000133 | 0.004956 | 0.013041 |
| chr4 | 135727816 | 135727912 | 5 | 0.6389 | 0.4269 | 0.212 | 0.00013 | 0.002479 | 0.004956 | 0.049423 |
| chr4 | 149107930 | 149107944 | 5 | 0.666 | 0.8011 | -0.1351 | 0.00013 | 0.000184 | 0.004956 | 0.015013 |
| chr4 | 58184415 | 58184424 | 5 | 0.4812 | 0.6964 | -0.2152 | 0.00013 | 0.002467 | 0.004956 | 0.049423 |
| chr4 | 62510268 | 62510448 | 8 | 0.92737 | 0.80275 | 0.124625 | 0.000121 | 0.00166 | 0.004956 | 0.040881 |
| chr5 | 125109877 | 125110246 | 13 | 0.17338 | 0.4205 | -0.24712 | 0.000121 | 0.000668 | 0.004956 | 0.031559 |
| chr5 | 72211019 | 72211091 | 5 | 0.5905 | 0.7726 | -0.1821 | 0.00013 | 0.000969 | 0.004956 | 0.031559 |
| chr6 | 109546580 | 109546739 | 5 | 0.7391 | 0.3106 | 0.4285 | 0.00013 | 0.000361 | 0.004956 | 0.023015 |
| chr6 | 135636598 | 135636619 | 5 | 0.8928 | 0.6461 | 0.2467 | 0.00013 | 0.001338 | 0.004956 | 0.034162 |
| chr6 | 143500309 | 143500402 | 5 | 0.4841 | 0.7699 | -0.2858 | 0.00013 | 0.00014 | 0.004956 | 0.013041 |
| chr7 | 20311986 | 20312105 | 5 | 0.6014 | 0.8783 | -0.2769 | 0.00013 | 0.001399 | 0.004956 | 0.035387 |
| chr7 | 49525711 | 49525795 | 5 | 0.0736 | 0.4883 | -0.4147 | 0.00013 | 0.000379 | 0.004956 | 0.023015 |
| chr7 | 6116855 | 6116932 | 8 | 0.46863 | 0.76706 | -0.29844 | 0.000121 | 0.000192 | 0.004956 | 0.015013 |
| chr7 | 83745742 | 83745755 | 5 | 0.555 | 0.3904 | 0.1646 | 0.00013 | 0.00256 | 0.004956 | 0.049423 |
| chr8 | 119632993 | 119633091 | 13 | 0.42727 | 0.61323 | -0.18596 | 0.000121 | 0.001151 | 0.004956 | 0.032048 |
| chr8 | 120562451 | 120562514 | 5 | 0.8575 | 0.5789 | 0.2786 | 0.00013 | 0.002449 | 0.004956 | 0.049423 |
| chr8 | 120693899 | 120694102 | 16 | 0.26056 | 0.13694 | 0.123625 | 0.000116 | 0.000153 | 0.004956 | 0.013705 |
| chr8 | 87832638 | 87832688 | 12 | 0.60758 | 0.36596 | 0.241625 | 0.000125 | 0.000948 | 0.004956 | 0.031559 |
| chrX | 115855306 | 115855647 | 5 | 0.8077 | 0.9477 | -0.14 | 0.00013 | 0.000957 | 0.004956 | 0.031559 |
| chr10 | 117478192 | 117478304 | 10 | 0.24065 | 0.3927 | -0.15205 | 0.000137 | 0.001787 | 0.005112 | 0.042972 |
| chr10 | 81561091 | 81561325 | 16 | 0.12525 | 0.021531 | 0.103719 | 0.000137 | 0.000661 | 0.005112 | 0.031559 |
| chr11 | 107897533 | 107897593 | 7 | 0.67479 | 0.87929 | -0.2045 | 0.000137 | 0.000994 | 0.005112 | 0.031559 |
| chr17 | 55306919 | 55307092 | 10 | 0.87 | 0.56 | 0.31 | 0.000137 | 0.001787 | 0.005112 | 0.042972 |
| chr6 | 124778789 | 124778846 | 7 | 0.62814 | 0.8785 | -0.25036 | 0.000137 | 0.000445 | 0.005112 | 0.023757 |
| chr6 | 138421568 | 138421603 | 7 | 0.021571 | 0.13893 | -0.11736 | 0.000137 | 0.000454 | 0.005112 | 0.02377 |
| chr8 | 20557811 | 20557880 | 10 | 0.15015 | 0.0355 | 0.11465 | 0.000137 | 7.08E-05 | 0.005112 | 0.009945 |
| chr9 | 77319159 | 77319250 | 7 | 0.94821 | 0.74557 | 0.202643 | 0.000137 | 0.000445 | 0.005112 | 0.023757 |
| chr2 | 129947879 | 129948038 | 13 | 0.51138 | 0.34315 | 0.168231 | 0.000141 | 0.00179 | 0.005211 | 0.042972 |
| chr2 | 127141515 | 127142030 | 22 | 0.78839 | 0.68486 | 0.103523 | 0.000142 | 0.002425 | 0.005224 | 0.049423 |
| chr15 | 91503138 | 91503249 | 11 | 0.50545 | 0.35318 | 0.152273 | 0.000143 | 0.002221 | 0.005265 | 0.049256 |
| chr8 | 80168839 | 80168954 | 11 | 0.20841 | 0.36964 | -0.16123 | 0.000143 | 0.00127 | 0.005265 | 0.033515 |
| chr5 | 115373459 | 115373540 | 6 | 0.39 | 0.12292 | 0.267083 | 0.000144 | 0.001038 | 0.005281 | 0.031559 |
| chr8 | 115001039 | 115001173 | 6 | 0.08925 | 0.23767 | -0.14842 | 0.000144 | 0.000441 | 0.005281 | 0.023757 |
| chr8 | 108956634 | 108956790 | 15 | 0.47173 | 0.6785 | -0.20677 | 0.000145 | 0.001522 | 0.005296 | 0.038296 |
| chr11 | 92694597 | 92694717 | 9 | 0.89794 | 0.73311 | 0.164833 | 0.000147 | 0.000167 | 0.005339 | 0.014451 |
| chr16 | 91514744 | 91514805 | 12 | 0.89483 | 0.76642 | 0.128417 | 0.000148 | 0.000519 | 0.005363 | 0.026686 |
| chr3 | 95622274 | 95622496 | 12 | 0.97804 | 0.85083 | 0.127208 | 0.000148 | 0.002727 | 0.005363 | 0.049584 |
| chr2 | 163376080 | 163376129 | 11 | 0.32709 | 0.16186 | 0.165227 | 0.000157 | 0.002249 | 0.00565 | 0.049423 |
| chr7 | 62463810 | 62463836 | 11 | 0.15886 | 0.34041 | -0.18155 | 0.000157 | 0.001303 | 0.00565 | 0.033685 |
| chr11 | 86955186 | 86955262 | 8 | 0.3735 | 0.15225 | 0.22125 | 0.000164 | 0.000849 | 0.005779 | 0.031559 |
| chr13 | 55855596 | 55855894 | 7 | 0.79014 | 0.46743 | 0.322714 | 0.000165 | 0.001005 | 0.005779 | 0.031559 |
| chr16 | 43974094 | 43974130 | 8 | 0.67756 | 0.90356 | -0.226 | 0.000164 | 0.000823 | 0.005779 | 0.031559 |
| chr19 | 55839267 | 55839373 | 7 | 0.41779 | 0.10171 | 0.316071 | 0.000165 | 0.000192 | 0.005779 | 0.015013 |
| chr2 | 122319531 | 122319617 | 7 | 0.20864 | 0.625 | -0.41636 | 0.000165 | 0.002014 | 0.005779 | 0.046791 |
| chr3 | 89211212 | 89211472 | 8 | 0.91563 | 0.80606 | 0.109563 | 0.000164 | 0.000187 | 0.005779 | 0.015013 |
| chr8 | 13953739 | 13953864 | 13 | 0.81038 | 0.91981 | -0.10942 | 0.000163 | 0.000402 | 0.005779 | 0.023015 |
| chrX | 169994046 | 169994084 | 7 | 0.14914 | 0.40271 | -0.25357 | 0.000165 | 0.002018 | 0.005779 | 0.046791 |
| chr8 | 119826180 | 119826542 | 9 | 0.84 | 0.65611 | 0.183889 | 0.000167 | 0.000671 | 0.005831 | 0.031559 |
| chr10 | 71330598 | 71330891 | 10 | 0.53965 | 0.76275 | -0.2231 | 0.00017 | 0.000264 | 0.005927 | 0.018571 |
| chr3 | 86547274 | 86547317 | 11 | 0.078091 | 0.20077 | -0.12268 | 0.000173 | 5.95E-05 | 0.006019 | 0.009216 |
| chr1 | 119443813 | 119444027 | 5 | 0.8445 | 0.9512 | -0.1067 | 0.000206 | 0.000981 | 0.006155 | 0.031559 |

| | | | | | | | | | | |
|---|---|---|---|---|---|---|---|---|---|---|
| chr1 | 152242156 | 152242223 | 5 | 0.8898 | 0.698 | 0.1918 | 0.000206 | 0.002504 | 0.006155 | 0.049423 |
| chr1 | 174293823 | 174293903 | 5 | 0.9143 | 0.7954 | 0.1189 | 0.000206 | 0.002665 | 0.006155 | 0.04951 |
| chr1 | 180358650 | 180358716 | 5 | 0.985 | 0.5063 | 0.4787 | 0.000206 | 0.000993 | 0.006155 | 0.031559 |
| chr1 | 180617932 | 180617945 | 5 | 0.9932 | 0.618 | 0.3752 | 0.000206 | 0.001042 | 0.006155 | 0.031559 |
| chr1 | 89974113 | 89974212 | 5 | 0.8105 | 0.5164 | 0.2941 | 0.000206 | 0.000992 | 0.006155 | 0.031559 |
| chr1 | 91313193 | 91313206 | 6 | 0.60325 | 0.32 | 0.28325 | 0.000201 | 0.000434 | 0.006155 | 0.023757 |
| chr10 | 38957285 | 38957357 | 5 | 0.9818 | 0.8646 | 0.1172 | 0.000206 | 0.000134 | 0.006155 | 0.013041 |
| chr11 | 113800235 | 113800253 | 5 | 0.9698 | 0.8551 | 0.1147 | 0.000206 | 0.00259 | 0.006155 | 0.049443 |
| chr11 | 113877709 | 113877783 | 5 | 0.5276 | 0.8548 | -0.3272 | 0.000206 | 0.002612 | 0.006155 | 0.049453 |
| chr11 | 79806402 | 79806423 | 5 | 0.4745 | 0.252 | 0.2225 | 0.000206 | 0.002461 | 0.006155 | 0.049423 |
| chr11 | 84481652 | 84481801 | 5 | 0.4971 | 0.2847 | 0.2124 | 0.000206 | 0.001056 | 0.006155 | 0.031559 |
| chr11 | 90035704 | 90035833 | 13 | 0.34142 | 0.22115 | 0.120269 | 0.000189 | 0.000687 | 0.006155 | 0.031559 |
| chr13 | 41334716 | 41334881 | 5 | 0.6148 | 0.8263 | -0.2115 | 0.000206 | 0.001279 | 0.006155 | 0.033582 |
| chr13 | 51573826 | 51573898 | 5 | 0.1744 | 0.4413 | -0.2669 | 0.000206 | 0.000128 | 0.006155 | 0.013041 |
| chr13 | 51702688 | 51702699 | 5 | 0.7729 | 0.9209 | -0.148 | 0.000206 | 0.001203 | 0.006155 | 0.032605 |
| chr13 | 53130906 | 53130944 | 5 | 0.8776 | 0.4346 | 0.443 | 0.000206 | 0.000128 | 0.006155 | 0.013041 |
| chr13 | 94025206 | 94025217 | 5 | 0.9751 | 0.8655 | 0.1096 | 0.000206 | 0.000982 | 0.006155 | 0.031559 |
| chr15 | 31345548 | 31345591 | 6 | 0.87883 | 0.71617 | 0.162667 | 0.000201 | 0.000448 | 0.006155 | 0.023757 |
| chr15 | 84430578 | 84430618 | 5 | 0.6895 | 0.4829 | 0.2066 | 0.000206 | 0.001211 | 0.006155 | 0.032629 |
| chr16 | 91856044 | 91856058 | 5 | 0.5508 | 0.7806 | -0.2298 | 0.000206 | 0.000986 | 0.006155 | 0.031559 |
| chr17 | 28860968 | 28861015 | 5 | 0.8195 | 0.5059 | 0.3136 | 0.000206 | 0.002573 | 0.006155 | 0.049423 |
| chr17 | 46916713 | 46916946 | 10 | 0.9722 | 0.86535 | 0.10685 | 0.000189 | 0.00179 | 0.006155 | 0.042972 |
| chr17 | 67912093 | 67912157 | 5 | 0.5208 | 0.655 | -0.1342 | 0.000206 | 0.001074 | 0.006155 | 0.031559 |
| chr18 | 30271945 | 30272006 | 5 | 0.8423 | 0.9498 | -0.1075 | 0.000206 | 0.001025 | 0.006155 | 0.031559 |
| chr19 | 5328099 | 5328192 | 5 | 0.7516 | 0.3958 | 0.3558 | 0.000206 | 0.002748 | 0.006155 | 0.049584 |
| chr19 | 57007350 | 57007463 | 5 | 0.509 | 0.8251 | -0.3161 | 0.000206 | 0.001105 | 0.006155 | 0.031572 |
| chr19 | 7096403 | 7096419 | 5 | 0.1458 | 0.5723 | -0.4265 | 0.000206 | 0.00096 | 0.006155 | 0.031559 |
| chr2 | 12175500 | 12175827 | 5 | 0.9285 | 0.7371 | 0.1914 | 0.000206 | 0.002438 | 0.006155 | 0.049423 |
| chr2 | 154603460 | 154603505 | 5 | 0.3133 | 0.0143 | 0.299 | 0.000206 | 0.001067 | 0.006155 | 0.031559 |
| chr2 | 174024658 | 174024700 | 5 | 0.6124 | 0.7976 | -0.1852 | 0.000206 | 0.00101 | 0.006155 | 0.031559 |
| chr2 | 174296278 | 174296544 | 21 | 0.45557 | 0.62148 | -0.16591 | 0.000184 | 0.000415 | 0.006155 | 0.023358 |
| chr2 | 180659442 | 180659533 | 10 | 0.083 | 0.29105 | -0.20805 | 0.000189 | 3.34E-05 | 0.006155 | 0.006484 |
| chr2 | 32776434 | 32776450 | 5 | 0.1528 | 0.0059 | 0.1469 | 0.000206 | 0.001005 | 0.006155 | 0.031559 |
| chr2 | 74675059 | 74675149 | 5 | 0.3692 | 0.1276 | 0.2416 | 0.000206 | 0.00115 | 0.006155 | 0.032048 |
| chr2 | 90233210 | 90233293 | 5 | 0.8743 | 0.9898 | -0.1155 | 0.000206 | 0.000132 | 0.006155 | 0.013041 |
| chr4 | 41522197 | 41522334 | 5 | 0.5525 | 0.2137 | 0.3388 | 0.000206 | 0.002781 | 0.006155 | 0.049718 |
| chr4_. | 1845 | 1948 | 6 | 0.04675 | 0.204 | -0.15725 | 0.000201 | 0.002384 | 0.006155 | 0.049423 |
| chr5 | 102710510 | 102710634 | 5 | 0.4513 | 0.6381 | -0.1868 | 0.000206 | 0.002432 | 0.006155 | 0.049423 |
| chr5 | 106816100 | 106816135 | 5 | 0.7093 | 0.3312 | 0.3781 | 0.000206 | 0.001157 | 0.006155 | 0.032048 |
| chr5 | 121968692 | 121969133 | 12 | 0.82175 | 0.60212 | 0.219625 | 0.00019 | 0.000955 | 0.006155 | 0.031559 |
| chr5 | 121982081 | 121982096 | 5 | 0.2478 | 0.05 | 0.1978 | 0.000206 | 0.002434 | 0.006155 | 0.049423 |
| chr5 | 125180805 | 125180853 | 5 | 0.1765 | 0.4833 | -0.3068 | 0.000206 | 0.000366 | 0.006155 | 0.023015 |
| chr5 | 125420933 | 125421143 | 11 | 0.90309 | 0.78727 | 0.115818 | 0.00019 | 0.000392 | 0.006155 | 0.023015 |
| chr5 | 129166249 | 129166337 | 5 | 0.2055 | 0.6665 | -0.461 | 0.000206 | 0.002569 | 0.006155 | 0.049423 |
| chr5 | 93097744 | 93097869 | 5 | 0.4663 | 0.2767 | 0.1896 | 0.000206 | 0.002447 | 0.006155 | 0.049423 |
| chr6 | 115792599 | 115792685 | 5 | 0.5988 | 0.8583 | -0.2595 | 0.000206 | 0.001109 | 0.006155 | 0.031572 |
| chr6 | 35251990 | 35252126 | 8 | 0.77931 | 0.59463 | 0.184688 | 0.000191 | 0.000188 | 0.006155 | 0.015013 |
| chr7 | 119740347 | 119740398 | 5 | 0.3172 | 0.0782 | 0.239 | 0.000206 | 0.002429 | 0.006155 | 0.049423 |
| chr7 | 142335269 | 142335410 | 5 | 0.8866 | 0.6664 | 0.2202 | 0.000206 | 0.001047 | 0.006155 | 0.031559 |
| chr7 | 25628054 | 25628157 | 5 | 0.443 | 0.1625 | 0.2805 | 0.000206 | 0.002547 | 0.006155 | 0.049423 |
| chr7 | 46038500 | 46038749 | 13 | 0.59488 | 0.81712 | -0.22223 | 0.000203 | 0.000362 | 0.006155 | 0.023015 |

| | | | | | | | | | | |
|---|---|---|---|---|---|---|---|---|---|---|
| chr8 | 115358121 | 115358270 | 7 | 0.51671 | 0.27664 | 0.240071 | 0.000198 | 0.00097 | 0.006155 | 0.031559 |
| chr8 | 117377622 | 117377716 | 5 | 0.9549 | 0.8473 | 0.1076 | 0.000206 | 0.000967 | 0.006155 | 0.031559 |
| chr8 | 120992986 | 120993201 | 5 | 0.6126 | 0.8732 | -0.2606 | 0.000206 | 0.002426 | 0.006155 | 0.049423 |
| chr8 | 20557742 | 20557780 | 8 | 0.15381 | 0.030125 | 0.123687 | 0.000191 | 0.000826 | 0.006155 | 0.031559 |
| chr9 | 100339546 | 100339714 | 5 | 0.7331 | 0.8456 | -0.1125 | 0.000206 | 0.002667 | 0.006155 | 0.04951 |
| chrX | 12297697 | 12297775 | 5 | 0.9166 | 0.4429 | 0.4737 | 0.000206 | 0.000965 | 0.006155 | 0.031559 |
| chrX | 53054873 | 53054990 | 5 | 0.7252 | 0.5457 | 0.1795 | 0.000206 | 0.001054 | 0.006155 | 0.031559 |
| chr5 | 111175089 | 111175256 | 10 | 0.1228 | 0.24425 | -0.12145 | 0.000211 | 0.001786 | 0.006263 | 0.042972 |
| chr8 | 120454103 | 120454166 | 10 | 0.9645 | 0.85015 | 0.11435 | 0.000211 | 0.000985 | 0.006263 | 0.031559 |
| chr1 | 190007159 | 190007347 | 14 | 0.46196 | 0.33929 | 0.122679 | 0.000213 | 0.001262 | 0.006304 | 0.033445 |
| chr2 | 156362246 | 156362391 | 14 | 0.75475 | 0.89161 | -0.13686 | 0.000213 | 0.001234 | 0.006304 | 0.033033 |
| chr8 | 106406010 | 106406029 | 9 | 0.61528 | 0.41328 | 0.202 | 0.000214 | 0.002408 | 0.006336 | 0.049423 |
| chr11 | 58195417 | 58195688 | 15 | 0.86493 | 0.69013 | 0.1748 | 0.000219 | 0.000949 | 0.006457 | 0.031559 |
| chr3 | 138464634 | 138464994 | 8 | 0.81838 | 0.41006 | 0.408313 | 0.000221 | 0.000401 | 0.006502 | 0.023015 |
| chr8 | 70598130 | 70598359 | 14 | 0.15886 | 0.038536 | 0.120321 | 0.000227 | 0.000292 | 0.006648 | 0.019993 |
| chr1 | 53978258 | 53978371 | 7 | 0.6945 | 0.42064 | 0.273857 | 0.000237 | 0.000979 | 0.006871 | 0.031559 |
| chr10 | 34128407 | 34128603 | 7 | 0.80614 | 0.575 | 0.231143 | 0.000237 | 0.002025 | 0.006871 | 0.046791 |
| chr3 | 114066286 | 114066399 | 7 | 0.86036 | 0.56386 | 0.2965 | 0.000237 | 0.000195 | 0.006871 | 0.015044 |
| chr4 | 132113872 | 132113905 | 7 | 0.74186 | 0.87543 | -0.13357 | 0.000237 | 0.000957 | 0.006871 | 0.031559 |
| chr9 | 96413249 | 96413434 | 7 | 0.87093 | 0.74186 | 0.129071 | 0.000237 | 0.002067 | 0.006871 | 0.046827 |
| chr2 | 31808521 | 31808638 | 9 | 0.088167 | 0.34906 | -0.26089 | 0.000242 | 0.000167 | 0.006983 | 0.014451 |
| chr5 | 90925708 | 90925880 | 12 | 0.74917 | 0.89958 | -0.15042 | 0.000242 | 0.002653 | 0.006983 | 0.04949 |
| chr11 | 88803657 | 88803782 | 19 | 0.50208 | 0.63679 | -0.13471 | 0.000245 | 0.001938 | 0.00705 | 0.045869 |
| chr18 | 41465202 | 41465335 | 8 | 0.56419 | 0.77038 | -0.20619 | 0.000256 | 0.000192 | 0.007314 | 0.015013 |
| chrX | 103694789 | 103694861 | 8 | 0.30444 | 0.11687 | 0.187562 | 0.000256 | 0.001648 | 0.007314 | 0.040781 |
| chr2 | 164393205 | 164393348 | 10 | 0.40345 | 0.77595 | -0.3725 | 0.00026 | 0.001788 | 0.007396 | 0.042972 |
| chrX | 169997443 | 169997539 | 13 | 0.4635 | 0.68258 | -0.21908 | 0.000271 | 7.16E-05 | 0.007648 | 0.009945 |
| chr15 | 86030544 | 86030594 | 6 | 0.764 | 0.93467 | -0.17067 | 0.000274 | 0.001091 | 0.007687 | 0.031572 |
| chr1 | 134096766 | 134096841 | 5 | 0.4328 | 0.1723 | 0.2605 | 0.000325 | 0.000134 | 0.007986 | 0.013041 |
| chr10 | 117885171 | 117885223 | 5 | 0.335 | 0.6618 | -0.3268 | 0.000325 | 0.00243 | 0.007986 | 0.049423 |
| chr11 | 116698080 | 116698158 | 5 | 0.9297 | 0.7624 | 0.1673 | 0.000325 | 0.002722 | 0.007986 | 0.049584 |
| chr11 | 119405767 | 119406057 | 5 | 0.9808 | 0.7965 | 0.1843 | 0.000325 | 0.0004 | 0.007986 | 0.023015 |
| chr11 | 90002311 | 90002357 | 5 | 0.092 | 0.3628 | -0.2708 | 0.000325 | 0.000451 | 0.007986 | 0.023757 |
| chr11 | 98108347 | 98108365 | 5 | 0.6916 | 0.8422 | -0.1506 | 0.000325 | 0.001097 | 0.007986 | 0.031572 |
| chr11 | 98914174 | 98914199 | 5 | 0.8316 | 0.6749 | 0.1567 | 0.000325 | 0.000997 | 0.007986 | 0.031559 |
| chr12 | 18387289 | 18387325 | 5 | 0.5438 | 0.0755 | 0.4683 | 0.000325 | 0.000971 | 0.007986 | 0.031559 |
| chr12 | 76993305 | 76993333 | 5 | 0.9314 | 0.7641 | 0.1673 | 0.000325 | 0.001043 | 0.007986 | 0.031559 |
| chr12 | 77009479 | 77009505 | 5 | 0.2054 | 0.4224 | -0.217 | 0.000325 | 0.001045 | 0.007986 | 0.031559 |
| chr13 | 29956796 | 29956893 | 5 | 0.9775 | 0.8095 | 0.168 | 0.000325 | 0.000449 | 0.007986 | 0.023757 |
| chr13 | 95778688 | 95779007 | 13 | 0.12369 | 0.31415 | -0.19046 | 0.00029 | 0.001002 | 0.007986 | 0.031559 |
| chr14 | 115964817 | 115964863 | 5 | 0.8553 | 0.6509 | 0.2044 | 0.000325 | 0.001159 | 0.007986 | 0.032048 |
| chr15 | 98206509 | 98206763 | 5 | 0.9223 | 0.692 | 0.2303 | 0.000325 | 0.000387 | 0.007986 | 0.023015 |
| chr17 | 12451380 | 12451436 | 5 | 0.5074 | 0.2953 | 0.2121 | 0.000325 | 0.000993 | 0.007986 | 0.031559 |
| chr17 | 43455046 | 43455088 | 5 | 0.8875 | 0.587 | 0.3005 | 0.000325 | 0.000136 | 0.007986 | 0.013041 |
| chr17 | 48647639 | 48647680 | 5 | 0.4615 | 0.6723 | -0.2108 | 0.000325 | 0.002686 | 0.007986 | 0.049584 |
| chr2 | 170838866 | 170838996 | 5 | 0.513 | 0.6754 | -0.1624 | 0.000325 | 0.00124 | 0.007986 | 0.033118 |
| chr2 | 174295488 | 174295632 | 9 | 0.15783 | 0.38144 | -0.22361 | 0.00031 | 0.001289 | 0.007986 | 0.033582 |
| chr2 | 177001068 | 177001281 | 5 | 0.4888 | 0.2488 | 0.24 | 0.000325 | 0.000967 | 0.007986 | 0.031559 |
| chr2 | 31204218 | 31204267 | 5 | 0.7949 | 0.6073 | 0.1876 | 0.000325 | 0.00112 | 0.007986 | 0.031662 |
| chr2 | 76869362 | 76869385 | 5 | 0.3756 | 0.4913 | -0.1157 | 0.000325 | 0.002429 | 0.007986 | 0.049423 |
| chr3 | 87800622 | 87800652 | 11 | 0.73486 | 0.86273 | -0.12786 | 0.000301 | 0.000738 | 0.007986 | 0.031559 |

| | | | | | | | | | | |
|---|---|---|---|---|---|---|---|---|---|---|
| chr3 | 95172720 | 95172731 | 5 | 0.6054 | 0.3255 | 0.2799 | 0.000325 | 0.000968 | 0.007986 | 0.031559 |
| chr4 | 135074259 | 135074365 | 5 | 0.7857 | 0.9919 | -0.2062 | 0.000325 | 0.001065 | 0.007986 | 0.031559 |
| chr4 | 154905108 | 154905148 | 5 | 0.2399 | 0.0291 | 0.2108 | 0.000325 | 0.000382 | 0.007986 | 0.023015 |
| chr5 | 102100537 | 102100564 | 5 | 0.6594 | 0.4959 | 0.1635 | 0.000325 | 0.000374 | 0.007986 | 0.023015 |
| chr5 | 122063650 | 122063680 | 5 | 0.8847 | 0.6919 | 0.1928 | 0.000325 | 0.001185 | 0.007986 | 0.032253 |
| chr5 | 143364645 | 143364750 | 5 | 0.142 | 0.0165 | 0.1255 | 0.000325 | 0.00041 | 0.007986 | 0.023134 |
| chr5 | 55653080 | 55653179 | 5 | 0.6275 | 0.798 | -0.1705 | 0.000325 | 0.001187 | 0.007986 | 0.032253 |
| chr6 | 114610159 | 114610199 | 5 | 0.742 | 0.9298 | -0.1878 | 0.000325 | 0.000376 | 0.007986 | 0.023015 |
| chr6 | 3052414 | 3052443 | 5 | 0.9936 | 0.8346 | 0.159 | 0.000325 | 0.002532 | 0.007986 | 0.049423 |
| chr7 | 141269051 | 141269086 | 5 | 0.4199 | 0.8164 | -0.3965 | 0.000325 | 0.001024 | 0.007986 | 0.031559 |
| chr7 | 6132459 | 6132487 | 5 | 0.6914 | 0.9212 | -0.2298 | 0.000325 | 0.000144 | 0.007986 | 0.01319 |
| chr8 | 114622190 | 114622201 | 5 | 0.5027 | 0.3888 | 0.1139 | 0.000325 | 0.000959 | 0.007986 | 0.031559 |
| chr8 | 121753233 | 121753297 | 5 | 0.5623 | 0.91 | -0.3477 | 0.000325 | 0.000967 | 0.007986 | 0.031559 |
| chr8 | 13050689 | 13050746 | 5 | 0.7714 | 0.9458 | -0.1744 | 0.000325 | 0.002455 | 0.007986 | 0.049423 |
| chr8 | 53457012 | 53457079 | 5 | 0.9857 | 0.7842 | 0.2015 | 0.000325 | 0.001116 | 0.007986 | 0.031641 |
| chr8 | 71709676 | 71709701 | 5 | 0.1669 | 0.3632 | -0.1963 | 0.000325 | 0.002628 | 0.007986 | 0.049456 |
| chr8 | 92194269 | 92194303 | 5 | 0.8761 | 0.6534 | 0.2227 | 0.000325 | 0.000982 | 0.007986 | 0.031559 |
| chr9 | 102301809 | 102301853 | 5 | 0.99 | 0.8734 | 0.1166 | 0.000325 | 0.000971 | 0.007986 | 0.031559 |
| chr2 | 37793392 | 37793475 | 18 | 0.54842 | 0.32978 | 0.218639 | 0.000327 | 0.002803 | 0.008027 | 0.049913 |
| chr14 | 100005605 | 100005724 | 7 | 0.84321 | 0.52671 | 0.3165 | 0.000338 | 0.002082 | 0.00817 | 0.046915 |
| chr16 | 98100701 | 98100800 | 7 | 0.71121 | 0.81729 | -0.10607 | 0.000338 | 0.001 | 0.00817 | 0.031559 |
| chr17 | 56627958 | 56628031 | 7 | 0.11793 | 0.24564 | -0.12771 | 0.000338 | 0.000214 | 0.00817 | 0.015641 |
| chr2 | 34897376 | 34897475 | 7 | 0.39793 | 0.64971 | -0.25179 | 0.000338 | 0.000444 | 0.00817 | 0.023757 |
| chr3 | 29813426 | 29813499 | 7 | 0.55471 | 0.27807 | 0.276643 | 0.000338 | 0.002026 | 0.00817 | 0.046791 |
| chr4 | 139724954 | 139725038 | 7 | 0.23043 | 0.46321 | -0.23279 | 0.000338 | 0.002026 | 0.00817 | 0.046791 |
| chr4 | 57093383 | 57093744 | 13 | 0.13877 | 0.42777 | -0.289 | 0.000334 | 0.000675 | 0.00817 | 0.031559 |
| chr7 | 144764922 | 144765125 | 7 | 0.92879 | 0.69636 | 0.232429 | 0.000338 | 0.002064 | 0.00817 | 0.046827 |
| chr12 | 54287088 | 54287144 | 8 | 0.69831 | 0.41338 | 0.284937 | 0.000343 | 0.000829 | 0.008225 | 0.031559 |
| chr16 | 22765622 | 22765773 | 8 | 0.5765 | 0.73237 | -0.15588 | 0.000343 | 0.00166 | 0.008225 | 0.040881 |
| chr4 | 12140877 | 12140907 | 8 | 0.19762 | 0.06775 | 0.129875 | 0.000343 | 0.000828 | 0.008225 | 0.031559 |
| chrX | 134475411 | 134475521 | 9 | 0.88806 | 0.62194 | 0.266111 | 0.00035 | 0.002408 | 0.008338 | 0.049423 |
| chr2 | 157561447 | 157561611 | 11 | 0.45482 | 0.61555 | -0.16073 | 0.00036 | 0.000404 | 0.008531 | 0.023015 |
| chr1 | 18658703 | 18658760 | 6 | 0.78283 | 0.96667 | -0.18383 | 0.000371 | 0.000187 | 0.008743 | 0.015013 |
| chr14 | 47823585 | 47823813 | 6 | 0.58058 | 0.35833 | 0.22225 | 0.000371 | 0.002528 | 0.008743 | 0.049423 |
| chr4 | 148467948 | 148468073 | 6 | 0.78633 | 0.95042 | -0.16408 | 0.000371 | 0.002608 | 0.008743 | 0.049453 |
| chr8 | 40896743 | 40896918 | 14 | 0.60071 | 0.49325 | 0.107464 | 0.000377 | 0.001924 | 0.008846 | 0.045635 |
| chr14 | 32304596 | 32305326 | 13 | 0.63515 | 0.8395 | -0.20435 | 0.000385 | 0.001885 | 0.009022 | 0.045024 |
| chr12 | 83370473 | 83370624 | 18 | 0.79169 | 0.68675 | 0.104944 | 0.000389 | 0.000776 | 0.009107 | 0.031559 |
| chr9 | 99319492 | 99319644 | 9 | 0.37217 | 0.23622 | 0.135944 | 0.000395 | 0.002408 | 0.009191 | 0.049423 |
| chr14 | 66170597 | 66170775 | 7 | 0.96064 | 0.85243 | 0.108214 | 0.000403 | 0.000959 | 0.009269 | 0.031559 |
| chr16 | 62711211 | 62711295 | 7 | 0.98207 | 0.879 | 0.103071 | 0.000403 | 0.002031 | 0.009269 | 0.046791 |
| chr5 | 111944488 | 111944559 | 7 | 0.034714 | 0.33279 | -0.29807 | 0.000403 | 8.12E-05 | 0.009269 | 0.010671 |
| chr8 | 33881792 | 33881926 | 7 | 0.47193 | 0.70157 | -0.22964 | 0.000403 | 0.000194 | 0.009269 | 0.015013 |
| chr8 | 88664841 | 88665110 | 7 | 0.82043 | 0.54143 | 0.279 | 0.000403 | 0.002113 | 0.009269 | 0.047506 |
| chr9 | 19160058 | 19160147 | 7 | 0.81714 | 0.929 | -0.11186 | 0.000403 | 0.002057 | 0.009269 | 0.046827 |
| chr10 | 81846724 | 81846919 | 18 | 0.31419 | 0.45289 | -0.13869 | 0.000406 | 0.001538 | 0.009314 | 0.038656 |
| chr1 | 187279174 | 187279349 | 5 | 0.8035 | 0.6413 | 0.1622 | 0.000487 | 0.002646 | 0.009702 | 0.049456 |
| chr1 | 89878641 | 89878696 | 5 | 0.8885 | 0.6174 | 0.2711 | 0.000487 | 0.001044 | 0.009702 | 0.031559 |
| chr1 | 89891726 | 89891775 | 5 | 0.8848 | 0.9871 | -0.1023 | 0.000487 | 0.00097 | 0.009702 | 0.031559 |
| chr11 | 102885980 | 102886033 | 5 | 0.7148 | 0.9209 | -0.2061 | 0.000487 | 0.000419 | 0.009702 | 0.023411 |
| chr11 | 78216273 | 78216379 | 5 | 0.833 | 0.9385 | -0.1055 | 0.000487 | 0.002643 | 0.009702 | 0.049456 |

| | | | | | | | | | | |
|---|---|---|---|---|---|---|---|---|---|---|
| chr11 | 80827972 | 80828163 | 5 | 0.7975 | 0.9733 | -0.1758 | 0.000487 | 0.001108 | 0.009702 | 0.031572 |
| chr11 | 97718861 | 97718893 | 5 | 0.3774 | 0.6572 | -0.2798 | 0.000487 | 0.001171 | 0.009702 | 0.032101 |
| chr12 | 112277805 | 112277903 | 11 | 0.62882 | 0.41582 | 0.213 | 0.00043 | 0.001253 | 0.009702 | 0.033371 |
| chr13 | 119488037 | 119488169 | 31 | 0.027484 | 0.19873 | -0.17124 | 0.000477 | 1.71E-05 | 0.009702 | 0.004237 |
| chr13 | 43289452 | 43289543 | 12 | 0.88925 | 0.75679 | 0.132458 | 0.000456 | 0.000942 | 0.009702 | 0.031559 |
| chr14 | 65890425 | 65890506 | 5 | 0.4039 | 0.0775 | 0.3264 | 0.000487 | 0.001355 | 0.009702 | 0.034467 |
| chr15 | 46984641 | 46984712 | 7 | 0.831 | 0.9535 | -0.1225 | 0.000479 | 0.002036 | 0.009702 | 0.046791 |
| chr15 | 74529869 | 74529920 | 5 | 0.2537 | 0.0716 | 0.1821 | 0.000487 | 0.000128 | 0.009702 | 0.013041 |
| chr15 | 76157750 | 76157782 | 5 | 0.9203 | 0.8134 | 0.1069 | 0.000487 | 0.001107 | 0.009702 | 0.031572 |
| chr15 | 86081210 | 86081240 | 5 | 0.774 | 0.5099 | 0.2641 | 0.000487 | 0.002474 | 0.009702 | 0.049423 |
| chr15 | 89067744 | 89067766 | 5 | 0.1205 | 0.3436 | -0.2231 | 0.000487 | 0.002617 | 0.009702 | 0.049456 |
| chr16 | 18460233 | 18460258 | 5 | 0.2653 | 0.0937 | 0.1716 | 0.000487 | 0.001032 | 0.009702 | 0.031559 |
| chr16 | 24015783 | 24015805 | 5 | 0.5567 | 0.4246 | 0.1321 | 0.000487 | 0.002483 | 0.009702 | 0.049423 |
| chr17 | 40580579 | 40580675 | 7 | 0.83443 | 0.64714 | 0.187286 | 0.000479 | 0.000969 | 0.009702 | 0.031559 |
| chr17 | 48461813 | 48461959 | 5 | 0.6439 | 0.8078 | -0.1639 | 0.000487 | 0.002623 | 0.009702 | 0.049456 |
| chr17 | 79335080 | 79335544 | 9 | 0.92506 | 0.79406 | 0.131 | 0.000445 | 0.002408 | 0.009702 | 0.049423 |
| chr17 | 81201122 | 81201371 | 11 | 0.17427 | 0.31723 | -0.14296 | 0.00043 | 0.002207 | 0.009702 | 0.049155 |
| chr17 | 84116608 | 84116678 | 5 | 0.8661 | 0.981 | -0.1149 | 0.000487 | 0.002478 | 0.009702 | 0.049423 |
| chr18 | 23618023 | 23618096 | 5 | 0.019 | 0.1679 | -0.1489 | 0.000487 | 0.001035 | 0.009702 | 0.031559 |
| chr18 | 80630233 | 80630308 | 5 | 0.6739 | 0.8785 | -0.2046 | 0.000487 | 0.001168 | 0.009702 | 0.032083 |
| chr18 | 85763459 | 85763484 | 5 | 0.9667 | 0.7366 | 0.2301 | 0.000487 | 0.000135 | 0.009702 | 0.013041 |
| chr19 | 44301244 | 44301281 | 5 | 0.4223 | 0.6854 | -0.2631 | 0.000487 | 0.002484 | 0.009702 | 0.049423 |
| chr19 | 47896896 | 47897000 | 7 | 0.57007 | 0.81679 | -0.24671 | 0.000479 | 0.000981 | 0.009702 | 0.031559 |
| chr2 | 115081886 | 115082067 | 5 | 0.4676 | 0.2839 | 0.1837 | 0.000487 | 0.000953 | 0.009702 | 0.031559 |
| chr2 | 168107204 | 168107246 | 7 | 0.55721 | 0.81107 | -0.25386 | 0.000479 | 0.002092 | 0.009702 | 0.047096 |
| chr2 | 169833497 | 169833770 | 5 | 0.2303 | 0.4462 | -0.2159 | 0.000487 | 0.002496 | 0.009702 | 0.049423 |
| chr2 | 25313307 | 25313382 | 5 | 0.4745 | 0.2293 | 0.2452 | 0.000487 | 0.00017 | 0.009702 | 0.01455 |
| chr2 | 35267357 | 35267530 | 5 | 0.7813 | 0.548 | 0.2333 | 0.000487 | 0.002579 | 0.009702 | 0.049443 |
| chr3 | 128667082 | 128667234 | 5 | 0.8516 | 0.7276 | 0.124 | 0.000487 | 0.002513 | 0.009702 | 0.049423 |
| chr3 | 89688705 | 89688864 | 5 | 0.0261 | 0.162 | -0.1359 | 0.000487 | 0.001214 | 0.009702 | 0.032651 |
| chr4 | 123116434 | 123116511 | 17 | 0.038941 | 0.17615 | -0.13721 | 0.000452 | 0.000287 | 0.009702 | 0.019758 |
| chr4 | 123116966 | 123117002 | 9 | 0.20389 | 0.386 | -0.18211 | 0.000445 | 0.000167 | 0.009702 | 0.014451 |
| chr4 | 126735970 | 126736023 | 7 | 0.144 | 0.024 | 0.12 | 0.000479 | 0.002056 | 0.009702 | 0.046827 |
| chr4 | 130263075 | 130263149 | 5 | 0.58 | 0.7299 | -0.1499 | 0.000487 | 0.00275 | 0.009702 | 0.049584 |
| chr4 | 141053973 | 141053986 | 5 | 0.1236 | 0.2977 | -0.1741 | 0.000487 | 0.000983 | 0.009702 | 0.031559 |
| chr4 | 141762100 | 141762284 | 5 | 0.7186 | 0.9024 | -0.1838 | 0.000487 | 0.00242 | 0.009702 | 0.049423 |
| chr4 | 156141429 | 156141634 | 5 | 0.9909 | 0.8862 | 0.1047 | 0.000487 | 0.001097 | 0.009702 | 0.031572 |
| chr4 | 70087298 | 70087344 | 9 | 0.81811 | 0.9555 | -0.13739 | 0.000445 | 0.002408 | 0.009702 | 0.049423 |
| chr4 | 71434371 | 71434415 | 5 | 0.7882 | 0.9522 | -0.164 | 0.000487 | 0.00102 | 0.009702 | 0.031559 |
| chr4 | 86316356 | 86316486 | 5 | 0.7086 | 0.8537 | -0.1451 | 0.000487 | 0.001041 | 0.009702 | 0.031559 |
| chr5 | 104926776 | 104926845 | 5 | 0.8665 | 0.985 | -0.1185 | 0.000487 | 0.002561 | 0.009702 | 0.049423 |
| chr5 | 112292496 | 112292535 | 5 | 0.8231 | 0.5951 | 0.228 | 0.000487 | 0.002611 | 0.009702 | 0.049453 |
| chr5 | 143758074 | 143758100 | 5 | 0.226 | 0.4057 | -0.1797 | 0.000487 | 0.001228 | 0.009702 | 0.032928 |
| chr5 | 65863588 | 65863652 | 5 | 0.7802 | 0.4639 | 0.3163 | 0.000487 | 0.001006 | 0.009702 | 0.031559 |
| chr6 | 113171746 | 113171809 | 5 | 0.1053 | 0.2965 | -0.1912 | 0.000487 | 0.000355 | 0.009702 | 0.023015 |
| chr7 | 101003608 | 101003746 | 5 | 0.4952 | 0.6714 | -0.1762 | 0.000487 | 0.002647 | 0.009702 | 0.049456 |
| chr8 | 121753681 | 121753733 | 11 | 0.73914 | 0.8425 | -0.10336 | 0.00043 | 0.001311 | 0.009702 | 0.033685 |
| chr8 | 14054297 | 14054466 | 5 | 0.7534 | 0.9356 | -0.1822 | 0.000487 | 0.002452 | 0.009702 | 0.049423 |
| chr8 | 26205175 | 26205337 | 11 | 0.65691 | 0.53173 | 0.125182 | 0.00047 | 0.001313 | 0.009702 | 0.033691 |
| chr9 | 116126470 | 116126606 | 5 | 0.5566 | 0.1899 | 0.3667 | 0.000487 | 0.000139 | 0.009702 | 0.013041 |
| chr9 | 119034866 | 119034903 | 9 | 0.92156 | 0.76222 | 0.159333 | 0.000445 | 0.002408 | 0.009702 | 0.049423 |

| | | | | | | | | | | |
|---|---|---|---|---|---|---|---|---|---|---|
| chr9 | 44381407 | 44381455 | 5 | 0.8082 | 0.9798 | -0.1716 | 0.000487 | 0.002755 | 0.009702 | 0.049584 |
| chr9 | 61618550 | 61618640 | 8 | 0.17156 | 0.0375 | 0.134063 | 0.000456 | 0.000401 | 0.009702 | 0.023015 |
| chr9 | 70269666 | 70269801 | 5 | 0.62 | 0.862 | -0.242 | 0.000487 | 0.000139 | 0.009702 | 0.013041 |
| chrX | 37422416 | 37422490 | 5 | 0.8078 | 0.9607 | -0.1529 | 0.000487 | 0.001039 | 0.009702 | 0.031559 |
| chr12 | 89187277 | 89187334 | 9 | 0.671 | 0.87067 | -0.19967 | 0.000501 | 0.002408 | 0.009849 | 0.049423 |
| chr13 | 40263428 | 40263483 | 9 | 0.22372 | 0.32511 | -0.10139 | 0.000501 | 0.00034 | 0.009849 | 0.022256 |
| chr1 | 101336743 | 101336833 | 10 | 0.8377 | 0.98825 | -0.15055 | 0.000535 | 0.001788 | 0.010381 | 0.042972 |
| chr1 | 189755224 | 189755260 | 10 | 0.71765 | 0.56005 | 0.1576 | 0.000535 | 0.000983 | 0.010381 | 0.031559 |
| chr3 | 152763487 | 152763771 | 10 | 0.0551 | 0.211 | -0.1559 | 0.000535 | 0.00014 | 0.010381 | 0.013041 |
| chr1 | 132265927 | 132266010 | 9 | 0.72494 | 0.47456 | 0.250389 | 0.000563 | 0.002408 | 0.010869 | 0.049423 |
| chr18 | 76751198 | 76751335 | 7 | 0.58436 | 0.83929 | -0.25493 | 0.000569 | 0.002068 | 0.010903 | 0.046827 |
| chr8 | 83588449 | 83588593 | 7 | 0.85593 | 0.65657 | 0.199357 | 0.000569 | 0.000971 | 0.010903 | 0.031559 |
| chr4 | 133275700 | 133275857 | 8 | 0.76138 | 0.91125 | -0.14988 | 0.000604 | 0.00084 | 0.011535 | 0.031559 |
| chr1 | 155529700 | 155529708 | 5 | 0.9093 | 0.7073 | 0.202 | 0.000725 | 0.002542 | 0.011681 | 0.049423 |
| chr1 | 165172346 | 165172427 | 5 | 0.6409 | 0.7866 | -0.1457 | 0.000725 | 0.00241 | 0.011681 | 0.049423 |
| chr1 | 179831808 | 179831986 | 11 | 0.703 | 0.81986 | -0.11686 | 0.000725 | 0.002208 | 0.011681 | 0.049155 |
| chr1 | 185026179 | 185026236 | 5 | 0.6784 | 0.8653 | -0.1869 | 0.000725 | 0.000129 | 0.011681 | 0.013041 |
| chr10 | 107642970 | 107643098 | 6 | 0.8445 | 0.65775 | 0.18675 | 0.000656 | 0.001053 | 0.011681 | 0.031559 |
| chr10 | 23399045 | 23399122 | 5 | 0.6198 | 0.8642 | -0.2444 | 0.000725 | 0.002478 | 0.011681 | 0.049423 |
| chr10 | 41486000 | 41486048 | 5 | 0.3309 | 0.747 | -0.4161 | 0.000725 | 0.001011 | 0.011681 | 0.031559 |
| chr10 | 76525109 | 76525208 | 5 | 0.9417 | 0.6855 | 0.2562 | 0.000725 | 0.001045 | 0.011681 | 0.031559 |
| chr10 | 79729572 | 79729625 | 9 | 0.74394 | 0.57089 | 0.173056 | 0.000711 | 0.001289 | 0.011681 | 0.033582 |
| chr11 | 107777395 | 107777725 | 8 | 0.26831 | 0.143 | 0.125313 | 0.000694 | 0.000854 | 0.011681 | 0.031559 |
| chr11 | 64701365 | 64701475 | 5 | 0.7025 | 0.5238 | 0.1787 | 0.000725 | 0.001178 | 0.011681 | 0.032247 |
| chr11 | 72402361 | 72402446 | 5 | 0.285 | 0.5039 | -0.2189 | 0.000725 | 0.001547 | 0.011681 | 0.038826 |
| chr11 | 96345988 | 96346015 | 5 | 0.2476 | 0.1223 | 0.1253 | 0.000725 | 0.000963 | 0.011681 | 0.031559 |
| chr12 | 101905130 | 101905209 | 5 | 0.7903 | 0.9247 | -0.1344 | 0.000725 | 0.002478 | 0.011681 | 0.049423 |
| chr12 | 111824830 | 111824940 | 5 | 0.96 | 0.8278 | 0.1322 | 0.000725 | 0.000358 | 0.011681 | 0.023015 |
| chr12 | 112329963 | 112329999 | 6 | 0.50125 | 0.27325 | 0.228 | 0.000656 | 0.002415 | 0.011681 | 0.049423 |
| chr13 | 42288907 | 42289376 | 5 | 0.6979 | 0.5519 | 0.146 | 0.000725 | 0.00015 | 0.011681 | 0.013574 |
| chr13 | 45242852 | 45242879 | 5 | 0.3444 | 0.5525 | -0.2081 | 0.000725 | 0.002642 | 0.011681 | 0.049456 |
| chr14 | 59557603 | 59557850 | 5 | 0.6687 | 0.7735 | -0.1048 | 0.000725 | 0.002724 | 0.011681 | 0.049584 |
| chr14 | 7728572 | 7728584 | 5 | 0.9521 | 0.8401 | 0.112 | 0.000725 | 0.000975 | 0.011681 | 0.031559 |
| chr15 | 78672104 | 78672177 | 12 | 0.30762 | 0.44942 | -0.14179 | 0.000668 | 0.000956 | 0.011681 | 0.031559 |
| chr15 | 97863721 | 97863864 | 9 | 0.6155 | 0.82511 | -0.20961 | 0.000633 | 0.001289 | 0.011681 | 0.033582 |
| chr16 | 33576647 | 33576760 | 6 | 0.13092 | 0.59375 | -0.46283 | 0.000656 | 0.001076 | 0.011681 | 0.031559 |
| chr17 | 32415905 | 32415941 | 11 | 0.28591 | 0.47345 | -0.18755 | 0.000725 | 0.002279 | 0.011681 | 0.049423 |
| chr17 | 34934357 | 34934506 | 5 | 0.5479 | 0.8353 | -0.2874 | 0.000725 | 0.000393 | 0.011681 | 0.023015 |
| chr18 | 57287967 | 57288016 | 5 | 0.7751 | 0.9549 | -0.1798 | 0.000725 | 0.001058 | 0.011681 | 0.031559 |
| chr18 | 76087353 | 76087445 | 5 | 0.529 | 0.3169 | 0.2121 | 0.000725 | 0.000383 | 0.011681 | 0.023015 |
| chr19 | 10712405 | 10712614 | 5 | 0.9479 | 0.716 | 0.2319 | 0.000725 | 0.001027 | 0.011681 | 0.031559 |
| chr2 | 152687141 | 152687289 | 22 | 0.26595 | 0.41359 | -0.14764 | 0.000725 | 0.00012 | 0.011681 | 0.013041 |
| chr2 | 164404211 | 164404255 | 5 | 0.0151 | 0.1462 | -0.1311 | 0.000725 | 0.001014 | 0.011681 | 0.031559 |
| chr2 | 171000120 | 171000291 | 7 | 0.77579 | 0.91257 | -0.13679 | 0.000674 | 0.000201 | 0.011681 | 0.01516 |
| chr2 | 173047108 | 173047178 | 5 | 0.7602 | 0.5445 | 0.2157 | 0.000725 | 0.001064 | 0.011681 | 0.031559 |
| chr2 | 31750647 | 31750713 | 11 | 0.79818 | 0.61632 | 0.181864 | 0.000665 | 0.002282 | 0.011681 | 0.049423 |
| chr2 | 3195400 | 3195496 | 5 | 0.6875 | 0.857 | -0.1695 | 0.000725 | 0.000966 | 0.011681 | 0.031559 |
| chr2 | 33487402 | 33487518 | 5 | 0.7768 | 0.4181 | 0.3587 | 0.000725 | 0.001152 | 0.011681 | 0.032048 |
| chr2 | 50373074 | 50373152 | 5 | 0.696 | 0.5521 | 0.1439 | 0.000725 | 0.000953 | 0.011681 | 0.031559 |
| chr2 | 86650949 | 86650988 | 5 | 0.9509 | 0.8023 | 0.1486 | 0.000725 | 0.002559 | 0.011681 | 0.049423 |
| chr3 | 89427293 | 89427322 | 5 | 0.7168 | 0.8435 | -0.1267 | 0.000725 | 0.002646 | 0.011681 | 0.049456 |

| | | | | | | | | | | |
|---|---|---|---|---|---|---|---|---|---|---|
| chr4 | 148952144 | 148952186 | 5 | 0.164 | 0.0214 | 0.1426 | 0.000725 | 0.00099 | 0.011681 | 0.031559 |
| chr4 | 34616181 | 34616247 | 5 | 0.6827 | 0.4784 | 0.2043 | 0.000725 | 0.002539 | 0.011681 | 0.049423 |
| chr4 | 45630782 | 45630824 | 5 | 0.5741 | 0.2623 | 0.3118 | 0.000725 | 0.001013 | 0.011681 | 0.031559 |
| chr4 | 63171156 | 63171298 | 7 | 0.21129 | 0.46457 | -0.25329 | 0.000674 | 0.000959 | 0.011681 | 0.031559 |
| chr5 | 107040838 | 107040937 | 14 | 0.26368 | 0.38018 | -0.1165 | 0.000615 | 0.002184 | 0.011681 | 0.048853 |
| chr5 | 109971976 | 109972018 | 5 | 0.9429 | 0.7996 | 0.1433 | 0.000725 | 0.00052 | 0.011681 | 0.026686 |
| chr5 | 111958306 | 111958359 | 5 | 0.3434 | 0.1231 | 0.2203 | 0.000725 | 0.000448 | 0.011681 | 0.023757 |
| chr5 | 128581676 | 128581686 | 5 | 0.4031 | 0.2612 | 0.1419 | 0.000725 | 0.002707 | 0.011681 | 0.049584 |
| chr5 | 63951599 | 63951841 | 5 | 0.5698 | 0.3647 | 0.2051 | 0.000725 | 0.001138 | 0.011681 | 0.03198 |
| chr5 | 64024858 | 64024929 | 5 | 0.987 | 0.846 | 0.141 | 0.000725 | 0.002498 | 0.011681 | 0.049423 |
| chr6 | 134227837 | 134227846 | 5 | 0.9548 | 0.8026 | 0.1522 | 0.000725 | 0.002416 | 0.011681 | 0.049423 |
| chr6 | 90474176 | 90474196 | 5 | 0.9841 | 0.8555 | 0.1286 | 0.000725 | 0.000963 | 0.011681 | 0.031559 |
| chr7 | 125424869 | 125424926 | 5 | 0.8304 | 0.9366 | -0.1062 | 0.000725 | 0.000965 | 0.011681 | 0.031559 |
| chr7 | 29432728 | 29432970 | 5 | 0.7471 | 0.361 | 0.3861 | 0.000725 | 0.002611 | 0.011681 | 0.049453 |
| chr7 | 44462076 | 44462094 | 5 | 0.2198 | 0.5512 | -0.3314 | 0.000725 | 0.00104 | 0.011681 | 0.031559 |
| chr7 | 68183269 | 68183406 | 9 | 0.7435 | 0.93 | -0.1865 | 0.000711 | 0.000671 | 0.011681 | 0.031559 |
| chr8 | 111765370 | 111765449 | 5 | 0.9824 | 0.8654 | 0.117 | 0.000725 | 0.001001 | 0.011681 | 0.031559 |
| chr8 | 67054194 | 67054230 | 5 | 0.8627 | 0.9655 | -0.1028 | 0.000725 | 0.000972 | 0.011681 | 0.031559 |
| chr8 | 69399536 | 69399580 | 13 | 0.061538 | 0.20738 | -0.14585 | 0.000664 | 0.000402 | 0.011681 | 0.023015 |
| chr9 | 56619653 | 56619971 | 14 | 0.94468 | 0.83854 | 0.106143 | 0.000615 | 0.001351 | 0.011681 | 0.034413 |
| chrX | 7739223 | 7739398 | 5 | 0.1985 | 0.4375 | -0.239 | 0.000725 | 0.002427 | 0.011681 | 0.049423 |
| chr19 | 56089874 | 56090088 | 16 | 0.61581 | 0.48469 | 0.131125 | 0.000751 | 0.000976 | 0.012086 | 0.031559 |
| chr2 | 174286075 | 174286376 | 33 | 0.63762 | 0.53686 | 0.100758 | 0.00077 | 0.000769 | 0.012382 | 0.031559 |
| chr1 | 174254784 | 174254872 | 7 | 0.75993 | 0.91643 | -0.1565 | 0.000796 | 0.000194 | 0.012545 | 0.015013 |
| chr10 | 60808052 | 60808098 | 7 | 0.1785 | 0.39793 | -0.21943 | 0.000796 | 0.002079 | 0.012545 | 0.046907 |
| chr13 | 81137087 | 81137169 | 7 | 0.63921 | 0.39221 | 0.247 | 0.000796 | 0.000445 | 0.012545 | 0.023757 |
| chr17 | 34957988 | 34958044 | 7 | 0.068857 | 0.28564 | -0.21679 | 0.000796 | 0.000959 | 0.012545 | 0.031559 |
| chr3 | 119911622 | 119911675 | 8 | 0.19431 | 0.0835 | 0.110813 | 0.000796 | 0.001648 | 0.012545 | 0.040781 |
| chr5 | 107040979 | 107041041 | 7 | 0.34107 | 0.59364 | -0.25257 | 0.000796 | 0.00097 | 0.012545 | 0.031559 |
| chr7 | 16472338 | 16472396 | 10 | 0.0561 | 0.2345 | -0.1784 | 0.000796 | 0.001787 | 0.012545 | 0.042972 |
| chr8 | 35589888 | 35589947 | 7 | 0.59207 | 0.79721 | -0.20514 | 0.000796 | 0.002062 | 0.012545 | 0.046827 |
| chr2 | 87526007 | 87526043 | 13 | 0.20454 | 0.34688 | -0.14235 | 0.000811 | 0.001163 | 0.012732 | 0.032048 |
| chr19 | 5690856 | 5691502 | 21 | 0.502 | 0.6666 | -0.1646 | 0.00082 | 0.00275 | 0.012873 | 0.049584 |
| chr9 | 62882187 | 62882312 | 6 | 0.75233 | 0.46417 | 0.288167 | 0.000858 | 0.000175 | 0.013272 | 0.014823 |
| chr11 | 106795767 | 106795936 | 13 | 0.29508 | 0.41208 | -0.117 | 0.000866 | 0.000386 | 0.013359 | 0.023015 |
| chr10 | 118004155 | 118004290 | 9 | 0.095167 | 0.28339 | -0.18822 | 0.000894 | 0.001289 | 0.013683 | 0.033582 |
| chr3 | 37063546 | 37063662 | 12 | 0.59046 | 0.39946 | 0.191 | 0.000901 | 0.000939 | 0.013772 | 0.031559 |
| chr1 | 180870442 | 180870477 | 7 | 0.67386 | 0.4405 | 0.233357 | 0.000939 | 0.00101 | 0.01414 | 0.031559 |
| chr11 | 75182516 | 75182895 | 14 | 0.50925 | 0.71357 | -0.20432 | 0.000932 | 0.00118 | 0.01414 | 0.032247 |
| chr11 | 98690954 | 98691019 | 7 | 0.805 | 0.91779 | -0.11279 | 0.000939 | 0.000965 | 0.01414 | 0.031559 |
| chr3 | 123013541 | 123013681 | 7 | 0.74457 | 0.87536 | -0.13079 | 0.000939 | 0.002073 | 0.01414 | 0.046827 |
| chr4 | 150140012 | 150140061 | 7 | 0.74664 | 0.92329 | -0.17664 | 0.000939 | 0.00203 | 0.01414 | 0.046791 |
| chr6 | 32527941 | 32528022 | 7 | 0.82643 | 0.68029 | 0.146143 | 0.000939 | 0.002135 | 0.01414 | 0.047869 |
| chr6 | 34160394 | 34160495 | 7 | 0.405 | 0.6945 | -0.2895 | 0.000939 | 0.002022 | 0.01414 | 0.046791 |
| chr6 | 35288837 | 35289188 | 7 | 0.53036 | 0.36421 | 0.166143 | 0.000939 | 0.002022 | 0.01414 | 0.046791 |
| chr7 | 89657798 | 89657871 | 7 | 0.31279 | 0.55207 | -0.23929 | 0.000939 | 0.002058 | 0.01414 | 0.046827 |
| chr1 | 119428101 | 119428132 | 5 | 0.6108 | 0.7577 | -0.1469 | 0.00105 | 0.002467 | 0.014268 | 0.049423 |
| chr1 | 171251309 | 171251364 | 5 | 0.1317 | 0.3505 | -0.2188 | 0.00105 | 0.002556 | 0.014268 | 0.049423 |
| chr1 | 89102726 | 89102863 | 5 | 0.4276 | 0.662 | -0.2344 | 0.00105 | 0.002573 | 0.014268 | 0.049423 |
| chr10 | 95576950 | 95576995 | 5 | 0.4953 | 0.3049 | 0.1904 | 0.00105 | 0.002524 | 0.014268 | 0.049423 |
| chr11 | 101083352 | 101083409 | 5 | 0.2014 | 0.0397 | 0.1617 | 0.00105 | 0.001078 | 0.014268 | 0.031559 |

| | | | | | | | | | | |
|---|---|---|---|---|---|---|---|---|---|---|
| chr11 | 53237265 | 53237395 | 11 | 0.54395 | 0.69168 | -0.14773 | 0.001016 | 0.002205 | 0.014268 | 0.049155 |
| chr11 | 77773447 | 77773653 | 5 | 0.9881 | 0.881 | 0.1071 | 0.00105 | 0.000996 | 0.014268 | 0.031559 |
| chr11 | 78531316 | 78531343 | 5 | 0.24 | 0.5237 | -0.2837 | 0.00105 | 0.000361 | 0.014268 | 0.023015 |
| chr15 | 100401307 | 100401369 | 8 | 0.65988 | 0.86975 | -0.20988 | 0.001042 | 0.001675 | 0.014268 | 0.041055 |
| chr16 | 47733297 | 47733357 | 5 | 0.9923 | 0.8782 | 0.1141 | 0.00105 | 0.002501 | 0.014268 | 0.049423 |
| chr17 | 24728323 | 24728533 | 5 | 0.7122 | 0.8153 | -0.1031 | 0.00105 | 0.002453 | 0.014268 | 0.049423 |
| chr19 | 25568491 | 25568537 | 11 | 0.25173 | 0.40518 | -0.15346 | 0.001016 | 0.002268 | 0.014268 | 0.049423 |
| chr19 | 4262439 | 4262529 | 5 | 0.6074 | 0.9043 | -0.2969 | 0.00105 | 0.001024 | 0.014268 | 0.031559 |
| chr2 | 173203090 | 173203199 | 5 | 0.5584 | 0.7633 | -0.2049 | 0.00105 | 0.001118 | 0.014268 | 0.031641 |
| chr2 | 74700798 | 74700852 | 5 | 0.1877 | 0.0696 | 0.1181 | 0.00105 | 0.000972 | 0.014268 | 0.031559 |
| chr3 | 95140430 | 95140455 | 5 | 0.7118 | 0.5297 | 0.1821 | 0.00105 | 0.002482 | 0.014268 | 0.049423 |
| chr4 | 108308511 | 108308542 | 5 | 0.8419 | 0.7292 | 0.1127 | 0.00105 | 0.001265 | 0.014268 | 0.033445 |
| chr4 | 129360053 | 129360131 | 8 | 0.86656 | 0.66706 | 0.1995 | 0.001042 | 0.001645 | 0.014268 | 0.040781 |
| chr4 | 151719279 | 151719298 | 5 | 0.18 | 0.0552 | 0.1248 | 0.00105 | 0.000953 | 0.014268 | 0.031559 |
| chr4 | 45271067 | 45271078 | 5 | 0.9435 | 0.7938 | 0.1497 | 0.00105 | 0.002619 | 0.014268 | 0.049456 |
| chr5 | 106334459 | 106334553 | 5 | 0.4612 | 0.2704 | 0.1908 | 0.00105 | 0.000141 | 0.014268 | 0.013041 |
| chr5 | 111869202 | 111869275 | 5 | 0.2115 | 0.393 | -0.1815 | 0.00105 | 0.000981 | 0.014268 | 0.031559 |
| chr5 | 138275782 | 138275818 | 5 | 0.2146 | 0.0394 | 0.1752 | 0.00105 | 0.000972 | 0.014268 | 0.031559 |
| chr5 | 81794538 | 81794577 | 11 | 0.82286 | 0.97118 | -0.14832 | 0.001016 | 0.002282 | 0.014268 | 0.049423 |
| chr6 | 108087502 | 108087552 | 5 | 0.7152 | 0.9445 | -0.2293 | 0.00105 | 0.000996 | 0.014268 | 0.031559 |
| chr7 | 4927664 | 4927821 | 5 | 0.4684 | 0.193 | 0.2754 | 0.00105 | 0.002469 | 0.014268 | 0.049423 |
| chr8 | 105259137 | 105259162 | 5 | 0.1356 | 0.0282 | 0.1074 | 0.00105 | 0.002461 | 0.014268 | 0.049423 |
| chr8 | 123165531 | 123165563 | 5 | 0.96 | 0.7652 | 0.1948 | 0.00105 | 0.000135 | 0.014268 | 0.013041 |
| chr9 | 43754696 | 43754788 | 5 | 0.1963 | 0.4132 | -0.2169 | 0.00105 | 0.001051 | 0.014268 | 0.031559 |
| chr9 | 53720850 | 53720997 | 5 | 0.7364 | 0.8496 | -0.1132 | 0.00105 | 0.000982 | 0.014268 | 0.031559 |
| chr16 | 33576965 | 33577052 | 7 | 0.030643 | 0.39886 | -0.36821 | 0.001105 | 0.00096 | 0.014692 | 0.031559 |
| chr2 | 112504768 | 112505037 | 6 | 0.97033 | 0.85475 | 0.115583 | 0.001115 | 0.002475 | 0.014692 | 0.049423 |
| chr2 | 34106439 | 34106577 | 6 | 0.2955 | 0.094583 | 0.200917 | 0.001115 | 0.001081 | 0.014692 | 0.031559 |
| chr2 | 5021507 | 5021690 | 6 | 0.98808 | 0.877 | 0.111083 | 0.001115 | 0.001046 | 0.014692 | 0.031559 |
| chr4 | 125652212 | 125652225 | 7 | 0.885 | 0.71893 | 0.166071 | 0.001105 | 0.000194 | 0.014692 | 0.015013 |
| chr5 | 44287319 | 44287564 | 11 | 0.79955 | 0.6495 | 0.150045 | 0.001104 | 0.000738 | 0.014692 | 0.031559 |
| chrX | 73176955 | 73177048 | 7 | 0.69929 | 0.94129 | -0.242 | 0.001105 | 0.000212 | 0.014692 | 0.015641 |
| chr1 | 136789357 | 136789505 | 13 | 0.40685 | 0.576 | -0.16915 | 0.001124 | 0.000373 | 0.014711 | 0.023015 |
| chr10 | 70168697 | 70168956 | 8 | 0.33656 | 0.50712 | -0.17056 | 0.00119 | 0.00083 | 0.015472 | 0.031559 |
| chr12 | 18387474 | 18387567 | 8 | 0.3305 | 0.048688 | 0.281812 | 0.00119 | 0.000822 | 0.015472 | 0.031559 |
| chr1 | 57790179 | 57790320 | 9 | 0.90306 | 0.76089 | 0.142167 | 0.00125 | 0.002408 | 0.01613 | 0.049423 |
| chr10 | 81347050 | 81347108 | 9 | 0.14028 | 0.34878 | -0.2085 | 0.00125 | 0.002408 | 0.01613 | 0.049423 |
| chr9 | 47250405 | 47250644 | 9 | 0.94528 | 0.84433 | 0.100944 | 0.00125 | 0.002408 | 0.01613 | 0.049423 |
| chr15 | 72809890 | 72810097 | 33 | 0.43868 | 0.55533 | -0.11665 | 0.001252 | 5.68E-06 | 0.01614 | 0.002172 |
| chr1 | 82441647 | 82441764 | 5 | 0.8354 | 0.9357 | -0.1003 | 0.001505 | 0.000981 | 0.01707 | 0.031559 |
| chr10 | 110600859 | 110600965 | 5 | 0.6153 | 0.8471 | -0.2318 | 0.001505 | 0.001146 | 0.01707 | 0.032048 |
| chr10 | 82953380 | 82953414 | 5 | 0.7617 | 0.9243 | -0.1626 | 0.001505 | 0.002504 | 0.01707 | 0.049423 |
| chr11 | 109517188 | 109517258 | 5 | 0.5295 | 0.7344 | -0.2049 | 0.001505 | 0.002423 | 0.01707 | 0.049423 |
| chr11 | 32190472 | 32190740 | 6 | 0.97025 | 0.81825 | 0.152 | 0.001433 | 0.001039 | 0.01707 | 0.031559 |
| chr11 | 87729472 | 87729523 | 11 | 0.16036 | 0.32618 | -0.16582 | 0.001412 | 0.002217 | 0.01707 | 0.049256 |
| chr12 | 21925151 | 21925511 | 5 | 0.9788 | 0.8087 | 0.1701 | 0.001505 | 0.000978 | 0.01707 | 0.031559 |
| chr12 | 84881624 | 84881645 | 5 | 0.314 | 0.5709 | -0.2569 | 0.001505 | 0.002658 | 0.01707 | 0.049503 |
| chr13 | 104319078 | 104319173 | 5 | 0.8727 | 0.9856 | -0.1129 | 0.001505 | 0.001146 | 0.01707 | 0.032048 |
| chr15 | 27997655 | 27997766 | 5 | 0.6037 | 0.4111 | 0.1926 | 0.001505 | 0.000957 | 0.01707 | 0.031559 |
| chr15 | 55396343 | 55396426 | 5 | 0.922 | 0.7187 | 0.2033 | 0.001505 | 0.00096 | 0.01707 | 0.031559 |
| chr16 | 38975625 | 38975788 | 5 | 0.6231 | 0.3751 | 0.248 | 0.001505 | 0.00257 | 0.01707 | 0.049423 |

| | | | | | | | | | | |
|---|---|---|---|---|---|---|---|---|---|---|
| chr17 | 88405314 | 88405345 | 5 | 0.45 | 0.2395 | 0.2105 | 0.001505 | 0.002593 | 0.01707 | 0.049443 |
| chr18 | 61910834 | 61910851 | 5 | 0.0734 | 0.2256 | -0.1522 | 0.001505 | 0.000978 | 0.01707 | 0.031559 |
| chr18 | 62968499 | 62968723 | 5 | 0.4081 | 0.7468 | -0.3387 | 0.001505 | 0.002572 | 0.01707 | 0.049423 |
| chr18 | 69125901 | 69125944 | 5 | 0.9591 | 0.7743 | 0.1848 | 0.001505 | 0.002718 | 0.01707 | 0.049584 |
| chr2 | 127978734 | 127978818 | 5 | 0.9153 | 0.7788 | 0.1365 | 0.001505 | 0.002772 | 0.01707 | 0.04971 |
| chr2 | 168120349 | 168120511 | 5 | 0.8636 | 0.9692 | -0.1056 | 0.001505 | 0.002585 | 0.01707 | 0.049443 |
| chr2 | 169158653 | 169158704 | 5 | 0.0071 | 0.1428 | -0.1357 | 0.001505 | 0.001086 | 0.01707 | 0.031559 |
| chr2 | 34129256 | 34129500 | 5 | 0.8602 | 0.4601 | 0.4001 | 0.001505 | 0.000991 | 0.01707 | 0.031559 |
| chr2 | 74675449 | 74675511 | 5 | 0.3037 | 0.0594 | 0.2443 | 0.001505 | 0.001265 | 0.01707 | 0.033445 |
| chr4 | 106623081 | 106623099 | 5 | 0.2984 | 0.5718 | -0.2734 | 0.001505 | 0.00278 | 0.01707 | 0.049718 |
| chr5 | 103890025 | 103890056 | 6 | 0.52233 | 0.31483 | 0.2075 | 0.001433 | 0.002378 | 0.01707 | 0.049423 |
| chr5 | 121965533 | 121965548 | 5 | 0.025 | 0.1797 | -0.1547 | 0.001505 | 0.001047 | 0.01707 | 0.031559 |
| chr5 | 128697205 | 128697260 | 10 | 0.4133 | 0.23635 | 0.17695 | 0.001413 | 0.000277 | 0.01707 | 0.019235 |
| chr5 | 24364982 | 24365163 | 10 | 0.7556 | 0.8967 | -0.1411 | 0.001413 | 0.001786 | 0.01707 | 0.042972 |
| chr5 | 41750438 | 41750492 | 5 | 0.7016 | 0.916 | -0.2144 | 0.001505 | 0.000969 | 0.01707 | 0.031559 |
| chr8 | 108419720 | 108419816 | 5 | 0.1695 | 0.3449 | -0.1754 | 0.001505 | 0.002648 | 0.01707 | 0.049456 |
| chr9 | 116383844 | 116383956 | 5 | 0.1977 | 0.0059 | 0.1918 | 0.001505 | 0.002584 | 0.01707 | 0.049443 |
| chr9 | 60492677 | 60492703 | 8 | 0.1775 | 0.3425 | -0.165 | 0.001357 | 0.000187 | 0.01707 | 0.015013 |
| chrX | 164706690 | 164706702 | 5 | 0.8594 | 0.9676 | -0.1082 | 0.001505 | 0.000956 | 0.01707 | 0.031559 |
| chrX | 27261857 | 27261884 | 5 | 0.8635 | 0.9757 | -0.1122 | 0.001505 | 0.002494 | 0.01707 | 0.049423 |
| chrX | 72499599 | 72499705 | 5 | 0.887 | 0.7303 | 0.1567 | 0.001505 | 0.000982 | 0.01707 | 0.031559 |
| chr17 | 46504048 | 46504191 | 7 | 0.63279 | 0.34871 | 0.284071 | 0.001522 | 0.002129 | 0.017132 | 0.0478 |
| chr9 | 42901558 | 42901658 | 7 | 0.96064 | 0.84393 | 0.116714 | 0.001522 | 0.002041 | 0.017132 | 0.046791 |
| chr2 | 168262940 | 168263242 | 11 | 0.52727 | 0.76577 | -0.2385 | 0.001531 | 0.001311 | 0.017205 | 0.033685 |
| chr4 | 156064885 | 156065179 | 8 | 0.67631 | 0.87431 | -0.198 | 0.001546 | 0.000825 | 0.017274 | 0.031559 |
| chr5 | 124743538 | 124743662 | 8 | 0.9125 | 0.72744 | 0.185063 | 0.001546 | 0.000192 | 0.017274 | 0.015013 |
| chr8 | 121480591 | 121480801 | 13 | 0.63265 | 0.39292 | 0.239731 | 0.001545 | 0.001896 | 0.017274 | 0.045225 |
| chr8 | 20297492 | 20297516 | 8 | 0.02775 | 0.22919 | -0.20144 | 0.001546 | 0.000822 | 0.017274 | 0.031559 |
| chr12 | 106291142 | 106291406 | 9 | 0.73767 | 0.51628 | 0.221389 | 0.001735 | 0.00034 | 0.019114 | 0.022256 |
| chr8 | 115462821 | 115462939 | 9 | 0.33706 | 0.15889 | 0.178167 | 0.001735 | 0.002408 | 0.019114 | 0.049423 |
| chr3 | 148569095 | 148569218 | 8 | 0.76525 | 0.89006 | -0.12481 | 0.001759 | 0.000835 | 0.019206 | 0.031559 |
| chr11 | 88803551 | 88803642 | 7 | 0.24 | 0.40079 | -0.16079 | 0.001781 | 0.001036 | 0.01929 | 0.031559 |
| chr2 | 87635044 | 87635331 | 7 | 0.7915 | 0.93779 | -0.14629 | 0.001781 | 0.000965 | 0.01929 | 0.031559 |
| chr4 | 118238631 | 118238756 | 7 | 0.50643 | 0.74821 | -0.24179 | 0.001781 | 0.002039 | 0.01929 | 0.046791 |
| chr1 | 162858763 | 162858803 | 5 | 0.7748 | 0.6389 | 0.1359 | 0.002089 | 0.00263 | 0.019583 | 0.049456 |
| chr1 | 91504439 | 91504577 | 5 | 0.8579 | 0.6557 | 0.2022 | 0.002089 | 0.002789 | 0.019583 | 0.049718 |
| chr11 | 113114220 | 113114266 | 5 | 0.7012 | 0.8096 | -0.1084 | 0.002089 | 0.00268 | 0.019583 | 0.049584 |
| chr11 | 120016089 | 120016282 | 5 | 0.6658 | 0.8084 | -0.1426 | 0.002089 | 0.000981 | 0.019583 | 0.031559 |
| chr11 | 6010289 | 6010326 | 5 | 0.8163 | 0.985 | -0.1687 | 0.002089 | 0.002705 | 0.019583 | 0.049584 |
| chr11 | 96045256 | 96045306 | 5 | 0.6957 | 0.9268 | -0.2311 | 0.002089 | 0.000991 | 0.019583 | 0.031559 |
| chr12 | 4807699 | 4807939 | 5 | 0.9203 | 0.7571 | 0.1632 | 0.002089 | 0.002558 | 0.019583 | 0.049423 |
| chr13 | 20676546 | 20676611 | 5 | 0.8225 | 0.98 | -0.1575 | 0.002089 | 0.001008 | 0.019583 | 0.031559 |
| chr14 | 122179355 | 122179472 | 7 | 0.90129 | 0.64286 | 0.258429 | 0.002081 | 0.002039 | 0.019583 | 0.046791 |
| chr14 | 4018867 | 4018879 | 5 | 0.1335 | 0.0073 | 0.1262 | 0.002089 | 0.000978 | 0.019583 | 0.031559 |
| chr14 | 68355692 | 68355761 | 5 | 0.6586 | 0.8308 | -0.1722 | 0.002089 | 0.001003 | 0.019583 | 0.031559 |
| chr14 | 69859345 | 69859404 | 5 | 0.6878 | 0.8171 | -0.1293 | 0.002089 | 0.00271 | 0.019583 | 0.049584 |
| chr15 | 78544248 | 78544322 | 7 | 0.31921 | 0.61971 | -0.3005 | 0.002081 | 0.000982 | 0.019583 | 0.031559 |
| chr15 | 99243270 | 99243310 | 5 | 0.9512 | 0.731 | 0.2202 | 0.002089 | 0.000966 | 0.019583 | 0.031559 |
| chr16 | 7224495 | 7224526 | 5 | 0.738 | 0.9866 | -0.2486 | 0.002089 | 0.001082 | 0.019583 | 0.031559 |
| chr16 | 91766077 | 91766138 | 5 | 0.9852 | 0.7078 | 0.2774 | 0.002089 | 0.002655 | 0.019583 | 0.04949 |
| chr17 | 46520824 | 46520852 | 6 | 0.42333 | 0.56517 | -0.14183 | 0.00183 | 0.001105 | 0.019583 | 0.031572 |

| | | | | | | | | | | |
|---|---|---|---|---|---|---|---|---|---|---|
| chr18 | 57708629 | 57708799 | 5 | 0.9487 | 0.7769 | 0.1718 | 0.002089 | 0.002783 | 0.019583 | 0.049718 |
| chr18 | 65563944 | 65564064 | 5 | 0.5386 | 0.3327 | 0.2059 | 0.002089 | 0.002477 | 0.019583 | 0.049423 |
| chr19 | 10603714 | 10603847 | 5 | 0.4192 | 0.6726 | -0.2534 | 0.002089 | 0.000981 | 0.019583 | 0.031559 |
| chr19 | 56826215 | 56826349 | 28 | 0.49098 | 0.38027 | 0.110714 | 0.001963 | 0.001676 | 0.019583 | 0.041055 |
| chr2 | 174300611 | 174300980 | 35 | 0.4576 | 0.29424 | 0.163357 | 0.002071 | 7.47E-05 | 0.019583 | 0.010298 |
| chr2 | 25440666 | 25440686 | 5 | 0.97 | 0.8174 | 0.1526 | 0.002089 | 0.000966 | 0.019583 | 0.031559 |
| chr2 | 68366178 | 68366485 | 5 | 0.2459 | 0.127 | 0.1189 | 0.002089 | 0.001024 | 0.019583 | 0.031559 |
| chr2 | 68399578 | 68399851 | 7 | 0.8945 | 0.66036 | 0.234143 | 0.002081 | 0.002071 | 0.019583 | 0.046827 |
| chr2 | 9869404 | 9869425 | 5 | 0.8193 | 0.6426 | 0.1767 | 0.002089 | 0.000996 | 0.019583 | 0.031559 |
| chr4 | 107442380 | 107442505 | 7 | 0.2455 | 0.12986 | 0.115643 | 0.002081 | 0.002035 | 0.019583 | 0.046791 |
| chr4 | 140425160 | 140425189 | 5 | 0.1282 | 0.3127 | -0.1845 | 0.002089 | 0.001181 | 0.019583 | 0.032247 |
| chr4 | 62510174 | 62510229 | 5 | 0.8824 | 0.7596 | 0.1228 | 0.002089 | 0.00108 | 0.019583 | 0.031559 |
| chr5 | 112957123 | 112957199 | 5 | 0.3014 | 0.5942 | -0.2928 | 0.002089 | 0.002486 | 0.019583 | 0.049423 |
| chr5 | 113899304 | 113899459 | 5 | 0.975 | 0.8691 | 0.1059 | 0.002089 | 0.000987 | 0.019583 | 0.031559 |
| chr5 | 116954407 | 116954779 | 7 | 0.87864 | 0.71707 | 0.161571 | 0.002081 | 0.002037 | 0.019583 | 0.046791 |
| chr7 | 106436069 | 106436108 | 5 | 0.8513 | 0.9673 | -0.116 | 0.002089 | 0.001051 | 0.019583 | 0.031559 |
| chr7 | 3240802 | 3240853 | 5 | 0.7092 | 0.8881 | -0.1789 | 0.002089 | 0.002535 | 0.019583 | 0.049423 |
| chr7 | 39354067 | 39354142 | 8 | 0.97744 | 0.85163 | 0.125812 | 0.001998 | 0.001651 | 0.019583 | 0.040781 |
| chr9 | 115152431 | 115152461 | 5 | 0.7786 | 0.9842 | -0.2056 | 0.002089 | 0.001006 | 0.019583 | 0.031559 |
| chr9 | 98806637 | 98806726 | 5 | 0.8447 | 0.9525 | -0.1078 | 0.002089 | 0.001072 | 0.019583 | 0.031559 |
| chrX | 8662607 | 8662632 | 5 | 0.863 | 0.9726 | -0.1096 | 0.002089 | 0.001042 | 0.019583 | 0.031559 |
| chr8 | 67793067 | 67793172 | 11 | 0.67886 | 0.789 | -0.11014 | 0.002106 | 0.000728 | 0.019677 | 0.031559 |
| chr13 | 64210320 | 64210602 | 13 | 0.89685 | 0.77038 | 0.126462 | 0.002108 | 0.001559 | 0.019678 | 0.039018 |
| chr12 | 20815294 | 20815442 | 9 | 0.068444 | 0.25578 | -0.18733 | 0.002148 | 0.002408 | 0.019963 | 0.049423 |
| chr7 | 143465353 | 143465472 | 13 | 0.48977 | 0.3515 | 0.138269 | 0.002241 | 0.001848 | 0.020782 | 0.044244 |
| chr1 | 168425927 | 168425976 | 8 | 0.47531 | 0.23981 | 0.2355 | 0.002267 | 0.000189 | 0.02091 | 0.015013 |
| chr12 | 95248572 | 95248766 | 8 | 0.88962 | 0.72881 | 0.160812 | 0.002267 | 0.001647 | 0.02091 | 0.040781 |
| chr14 | 124415255 | 124415393 | 6 | 0.87025 | 0.98067 | -0.11042 | 0.002316 | 0.002379 | 0.02113 | 0.049423 |
| chr8 | 10307952 | 10308006 | 6 | 0.089833 | 0.2535 | -0.16367 | 0.002316 | 0.001087 | 0.02113 | 0.031559 |
| chr16 | 22573057 | 22573080 | 7 | 0.094357 | 0.19486 | -0.1005 | 0.002425 | 0.002021 | 0.021795 | 0.046791 |
| chr4 | 119208123 | 119208239 | 7 | 0.20664 | 0.38007 | -0.17343 | 0.002425 | 0.000445 | 0.021795 | 0.023757 |
| chr4 | 134601770 | 134601904 | 7 | 0.39207 | 0.67764 | -0.28557 | 0.002425 | 0.002067 | 0.021795 | 0.046827 |
| chr9 | 78821817 | 78821867 | 12 | 0.13513 | 0.034333 | 0.100792 | 0.002437 | 0.002737 | 0.02189 | 0.049584 |
| chr10 | 40319251 | 40319316 | 5 | 0.9899 | 0.8841 | 0.1058 | 0.00288 | 0.001165 | 0.022946 | 0.032048 |
| chr10 | 47805526 | 47805612 | 5 | 0.7909 | 0.9634 | -0.1725 | 0.00288 | 0.002501 | 0.022946 | 0.049423 |
| chr11 | 12025931 | 12026173 | 52 | 0.3827 | 0.52904 | -0.14634 | 0.002848 | 6.58E-08 | 0.022946 | 8.05E-05 |
| chr11 | 32811506 | 32811551 | 5 | 0.7743 | 0.6346 | 0.1397 | 0.00288 | 0.001164 | 0.022946 | 0.032048 |
| chr12 | 111266078 | 111266239 | 5 | 0.9055 | 0.6544 | 0.2511 | 0.00288 | 0.002606 | 0.022946 | 0.049453 |
| chr13 | 100611681 | 100611870 | 5 | 0.8782 | 0.9924 | -0.1142 | 0.00288 | 0.002433 | 0.022946 | 0.049423 |
| chr13 | 67217926 | 67217991 | 5 | 0.7539 | 0.8713 | -0.1174 | 0.00288 | 0.001073 | 0.022946 | 0.031559 |
| chr14 | 43029723 | 43029910 | 5 | 0.5673 | 0.7665 | -0.1992 | 0.00288 | 0.001003 | 0.022946 | 0.031559 |
| chr15 | 34509914 | 34509982 | 5 | 0.5888 | 0.7642 | -0.1754 | 0.00288 | 0.000978 | 0.022946 | 0.031559 |
| chr15 | 85017297 | 85017555 | 5 | 0.0174 | 0.1318 | -0.1144 | 0.00288 | 0.00096 | 0.022946 | 0.031559 |
| chr17 | 26024528 | 26024636 | 5 | 0.7999 | 0.9539 | -0.154 | 0.00288 | 0.000954 | 0.022946 | 0.031559 |
| chr17 | 86013884 | 86013955 | 5 | 0.3611 | 0.6735 | -0.3124 | 0.00288 | 0.002428 | 0.022946 | 0.049423 |
| chr18 | 74452055 | 74452159 | 5 | 0.3906 | 0.6266 | -0.236 | 0.00288 | 0.002543 | 0.022946 | 0.049423 |
| chr2 | 130654707 | 130654876 | 5 | 0.8669 | 0.9922 | -0.1253 | 0.00288 | 0.002455 | 0.022946 | 0.049423 |
| chr2 | 167768373 | 167768392 | 5 | 0.627 | 0.4082 | 0.2188 | 0.00288 | 0.001014 | 0.022946 | 0.031559 |
| chr2 | 83324361 | 83324386 | 5 | 0.869 | 0.9868 | -0.1178 | 0.00288 | 0.002759 | 0.022946 | 0.049584 |
| chr4 | 13366767 | 13366828 | 5 | 0.9781 | 0.8669 | 0.1112 | 0.00288 | 0.002713 | 0.022946 | 0.049584 |
| chr5 | 101666943 | 101666961 | 5 | 0.0347 | 0.174 | -0.1393 | 0.00288 | 0.002559 | 0.022946 | 0.049423 |

| | | | | | | | | | | |
|---|---|---|---|---|---|---|---|---|---|---|
| chr5 | 109973996 | 109974020 | 5 | 0.592 | 0.437 | 0.155 | 0.00288 | 0.001633 | 0.022946 | 0.040636 |
| chr5 | 117907502 | 117907697 | 5 | 0.5161 | 0.2995 | 0.2166 | 0.00288 | 0.002407 | 0.022946 | 0.049423 |
| chr5 | 124089924 | 124089961 | 5 | 0.9418 | 0.76 | 0.1818 | 0.00288 | 0.001068 | 0.022946 | 0.031559 |
| chr5 | 65348252 | 65348297 | 7 | 0.32629 | 0.49721 | -0.17093 | 0.002821 | 0.000208 | 0.022946 | 0.015468 |
| chr6 | 22752694 | 22752720 | 5 | 0.7683 | 0.9702 | -0.2019 | 0.00288 | 0.00098 | 0.022946 | 0.031559 |
| chr6 | 5829654 | 5829771 | 5 | 0.6397 | 0.5216 | 0.1181 | 0.00288 | 0.002735 | 0.022946 | 0.049584 |
| chr7 | 46394693 | 46394866 | 5 | 0.4313 | 0.6757 | -0.2444 | 0.00288 | 0.00108 | 0.022946 | 0.031559 |
| chr8 | 104548423 | 104548507 | 5 | 0.5004 | 0.7219 | -0.2215 | 0.00288 | 0.002721 | 0.022946 | 0.049584 |
| chr9 | 31845885 | 31845932 | 5 | 0.9727 | 0.7557 | 0.217 | 0.00288 | 0.002411 | 0.022946 | 0.049423 |
| chr9 | 97001860 | 97001868 | 5 | 0.9009 | 0.7664 | 0.1345 | 0.00288 | 0.002709 | 0.022946 | 0.049584 |
| chr2 | 167897846 | 167897879 | 6 | 0.55167 | 0.36092 | 0.19075 | 0.002914 | 0.001397 | 0.023031 | 0.035387 |
| chr13 | 28823161 | 28823336 | 9 | 0.89306 | 0.73228 | 0.160778 | 0.002939 | 0.000671 | 0.02316 | 0.031559 |
| chr1 | 75492742 | 75492946 | 7 | 0.5775 | 0.74893 | -0.17143 | 0.003275 | 0.000957 | 0.025308 | 0.031559 |
| chr2 | 157557853 | 157557940 | 7 | 0.69907 | 0.80871 | -0.10964 | 0.003275 | 0.002038 | 0.025308 | 0.046791 |
| chr3 | 67449800 | 67449839 | 8 | 0.76769 | 0.90725 | -0.13956 | 0.003285 | 0.001664 | 0.025308 | 0.040925 |
| chr5 | 124725102 | 124725146 | 8 | 0.12594 | 0.021313 | 0.104625 | 0.003285 | 0.001652 | 0.025308 | 0.040781 |
| chr1 | 182368852 | 182369210 | 11 | 0.24745 | 0.132 | 0.115455 | 0.003345 | 0.000674 | 0.025731 | 0.031559 |
| chr1 | 34732053 | 34732145 | 5 | 0.0775 | 0.3959 | -0.3184 | 0.003886 | 0.001274 | 0.026355 | 0.033527 |
| chr10 | 8549222 | 8549363 | 7 | 0.95207 | 0.85043 | 0.101643 | 0.003795 | 0.000962 | 0.026355 | 0.031559 |
| chr11 | 117790191 | 117790273 | 7 | 0.80514 | 0.61257 | 0.192571 | 0.003795 | 0.000277 | 0.026355 | 0.019235 |
| chr11 | 82820816 | 82820847 | 5 | 0.7148 | 0.4903 | 0.2245 | 0.003886 | 0.002568 | 0.026355 | 0.049423 |
| chr12 | 108107567 | 108107639 | 5 | 0.9417 | 0.7608 | 0.1809 | 0.003886 | 0.00104 | 0.026355 | 0.031559 |
| chr12 | 19954233 | 19954256 | 5 | 0.052 | 0.2047 | -0.1527 | 0.003886 | 0.000999 | 0.026355 | 0.031559 |
| chr13 | 101210330 | 101210367 | 7 | 0.86229 | 0.97307 | -0.11079 | 0.003795 | 0.000962 | 0.026355 | 0.031559 |
| chr14 | 49300676 | 49300900 | 5 | 0.8961 | 0.7537 | 0.1424 | 0.003886 | 0.001611 | 0.026355 | 0.040217 |
| chr14 | 67614318 | 67614449 | 9 | 0.47361 | 0.74417 | -0.27056 | 0.003606 | 0.002408 | 0.026355 | 0.049423 |
| chr2 | 165195607 | 165195640 | 7 | 0.113 | 0.21636 | -0.10336 | 0.003795 | 0.002017 | 0.026355 | 0.046791 |
| chr2 | 32328145 | 32328216 | 5 | 0.9418 | 0.8294 | 0.1124 | 0.003886 | 0.000984 | 0.026355 | 0.031559 |
| chr4 | 118053647 | 118053947 | 5 | 0.856 | 0.9816 | -0.1256 | 0.003886 | 0.001251 | 0.026355 | 0.033352 |
| chr4 | 153439739 | 153439803 | 5 | 0.7411 | 0.8748 | -0.1337 | 0.003886 | 0.001 | 0.026355 | 0.031559 |
| chr4 | 47294031 | 47294136 | 5 | 0.3318 | 0.5776 | -0.2458 | 0.003886 | 0.002479 | 0.026355 | 0.049423 |
| chr5 | 3064536 | 3064600 | 6 | 0.823 | 0.951 | -0.128 | 0.003637 | 0.001031 | 0.026355 | 0.031559 |
| chr5 | 83588207 | 83588333 | 5 | 0.79 | 0.9605 | -0.1705 | 0.003886 | 0.001002 | 0.026355 | 0.031559 |
| chr5 | 91686793 | 91686854 | 7 | 0.76964 | 0.89264 | -0.123 | 0.003795 | 0.002034 | 0.026355 | 0.046791 |
| chr7 | 43808258 | 43808293 | 5 | 0.9204 | 0.7512 | 0.1692 | 0.003886 | 0.000998 | 0.026355 | 0.031559 |
| chr7 | 49400895 | 49401027 | 5 | 0.6982 | 0.5869 | 0.1113 | 0.003886 | 0.001101 | 0.026355 | 0.031572 |
| chr8 | 114512553 | 114512588 | 5 | 0.394 | 0.2296 | 0.1644 | 0.003886 | 0.001044 | 0.026355 | 0.031559 |
| chr9 | 121982632 | 121982807 | 5 | 0.6823 | 0.8863 | -0.204 | 0.003886 | 0.000983 | 0.026355 | 0.031559 |
| chr5 | 74126406 | 74126486 | 11 | 0.60177 | 0.71836 | -0.11659 | 0.004188 | 0.001304 | 0.028068 | 0.033685 |
| chr11 | 35475780 | 35475884 | 7 | 0.5905 | 0.46679 | 0.123714 | 0.004389 | 0.002039 | 0.029058 | 0.046791 |
| chr13 | 51937791 | 51938148 | 7 | 0.45764 | 0.17621 | 0.281429 | 0.004389 | 0.002036 | 0.029058 | 0.046791 |
| chr17 | 12769950 | 12769991 | 7 | 0.30386 | 0.48321 | -0.17936 | 0.004389 | 0.002024 | 0.029058 | 0.046791 |
| chr19 | 8770994 | 8771061 | 7 | 0.022 | 0.384 | -0.362 | 0.004389 | 0.002065 | 0.029058 | 0.046827 |
| chr4 | 40509542 | 40509835 | 7 | 0.87 | 0.68493 | 0.185071 | 0.004389 | 0.000964 | 0.029058 | 0.031559 |
| chr1 | 64927507 | 64927718 | 5 | 0.7735 | 0.8829 | -0.1094 | 0.005196 | 0.002439 | 0.030002 | 0.049423 |
| chr10 | 122822568 | 122822601 | 5 | 0.8865 | 0.7384 | 0.1481 | 0.005196 | 0.002555 | 0.030002 | 0.049423 |
| chr10 | 62356273 | 62356384 | 5 | 0.9268 | 0.6921 | 0.2347 | 0.005196 | 0.002759 | 0.030002 | 0.049584 |
| chr12 | 103314306 | 103314522 | 5 | 0.3164 | 0.558 | -0.2416 | 0.005196 | 0.001093 | 0.030002 | 0.031572 |
| chr13 | 49369508 | 49369535 | 5 | 0.9669 | 0.8575 | 0.1094 | 0.005196 | 0.001264 | 0.030002 | 0.033445 |
| chr15 | 73521585 | 73521668 | 5 | 0.5583 | 0.7977 | -0.2394 | 0.005196 | 0.002738 | 0.030002 | 0.049584 |
| chr16 | 94267880 | 94267935 | 9 | 0.81622 | 0.92667 | -0.11044 | 0.004865 | 0.002408 | 0.030002 | 0.049423 |

| | | | | | | | | | | |
|---|---|---|---|---|---|---|---|---|---|---|
| chr17 | 52227274 | 52227446 | 5 | 0.9559 | 0.8352 | 0.1207 | 0.005196 | 0.002663 | 0.030002 | 0.04951 |
| chr18 | 34085174 | 34085556 | 5 | 0.8585 | 0.9662 | -0.1077 | 0.005196 | 0.001053 | 0.030002 | 0.031559 |
| chr19 | 41088781 | 41088824 | 5 | 0.3769 | 0.6916 | -0.3147 | 0.005196 | 0.002669 | 0.030002 | 0.04951 |
| chr19 | 42760554 | 42760658 | 5 | 0.3732 | 0.603 | -0.2298 | 0.005196 | 0.001012 | 0.030002 | 0.031559 |
| chr2 | 105948539 | 105948563 | 5 | 0.9875 | 0.8356 | 0.1519 | 0.005196 | 0.002638 | 0.030002 | 0.049456 |
| chr2 | 29845828 | 29845841 | 5 | 0.3358 | 0.1241 | 0.2117 | 0.005196 | 0.002472 | 0.030002 | 0.049423 |
| chr3 | 145648338 | 145648710 | 20 | 0.11583 | 0.22043 | -0.1046 | 0.004669 | 0.000747 | 0.030002 | 0.031559 |
| chr5 | 118234199 | 118234275 | 5 | 0.9699 | 0.8666 | 0.1033 | 0.005196 | 0.001012 | 0.030002 | 0.031559 |
| chr5 | 147950230 | 147950265 | 5 | 0.7211 | 0.915 | -0.1939 | 0.005196 | 0.001221 | 0.030002 | 0.03278 |
| chr7 | 112122349 | 112122389 | 5 | 0.7845 | 0.9143 | -0.1298 | 0.005196 | 0.002702 | 0.030002 | 0.049584 |
| chr7 | 128495230 | 128495288 | 5 | 0.9363 | 0.7202 | 0.2161 | 0.005196 | 0.001183 | 0.030002 | 0.032253 |
| chr7 | 25763992 | 25764246 | 9 | 0.37133 | 0.53894 | -0.16761 | 0.004865 | 0.002408 | 0.030002 | 0.049423 |
| chr8 | 36122703 | 36122742 | 5 | 0.7263 | 0.859 | -0.1327 | 0.005196 | 0.002753 | 0.030002 | 0.049584 |
| chr8 | 90886557 | 90886637 | 12 | 0.43392 | 0.53758 | -0.10367 | 0.005043 | 0.002731 | 0.030002 | 0.049584 |
| chr13 | 30986018 | 30986043 | 8 | 0.21475 | 0.38631 | -0.17156 | 0.005287 | 0.000837 | 0.030319 | 0.031559 |
| chr7 | 126851864 | 126851998 | 6 | 0.68567 | 0.89725 | -0.21158 | 0.00556 | 0.002374 | 0.031516 | 0.049423 |
| chr7 | 68294330 | 68294434 | 6 | 0.28692 | 0.542 | -0.25508 | 0.00556 | 0.001072 | 0.031516 | 0.031559 |
| chrX | 11665458 | 11665546 | 7 | 0.48857 | 0.20693 | 0.281643 | 0.005836 | 0.002218 | 0.032559 | 0.049256 |
| chr1 | 180453845 | 180453962 | 5 | 0.8569 | 0.5348 | 0.3221 | 0.006842 | 0.001404 | 0.03344 | 0.035475 |
| chr10 | 108454832 | 108454859 | 5 | 0.3128 | 0.4188 | -0.106 | 0.006842 | 0.002477 | 0.03344 | 0.049423 |
| chr12 | 70216217 | 70216462 | 7 | 0.70329 | 0.5115 | 0.191786 | 0.00671 | 0.001071 | 0.03344 | 0.031559 |
| chr17 | 33912158 | 33912193 | 5 | 0.5377 | 0.6664 | -0.1287 | 0.006842 | 0.002523 | 0.03344 | 0.049423 |
| chr18 | 43174504 | 43174550 | 5 | 0.4056 | 0.0765 | 0.3291 | 0.006842 | 0.002438 | 0.03344 | 0.049423 |
| chr2 | 138278542 | 138278600 | 5 | 0.084 | 0.2033 | -0.1193 | 0.006842 | 0.000953 | 0.03344 | 0.031559 |
| chr2 | 60717646 | 60717675 | 5 | 0.2644 | 0.3674 | -0.103 | 0.006842 | 0.002475 | 0.03344 | 0.049423 |
| chr3 | 14711353 | 14711559 | 5 | 0.8066 | 0.9339 | -0.1273 | 0.006842 | 0.002438 | 0.03344 | 0.049423 |
| chr4 | 43937602 | 43937617 | 5 | 0.7927 | 0.9327 | -0.14 | 0.006842 | 0.001125 | 0.03344 | 0.031758 |
| chr8 | 119632799 | 119632873 | 5 | 0.9178 | 0.8074 | 0.1104 | 0.006842 | 0.002414 | 0.03344 | 0.049423 |
| chr8 | 92445497 | 92445583 | 5 | 0.2812 | 0.4514 | -0.1702 | 0.006842 | 0.001257 | 0.03344 | 0.033415 |
| chrX | 168928543 | 168928591 | 5 | 0.8175 | 0.9622 | -0.1447 | 0.006842 | 0.001208 | 0.03344 | 0.032629 |
| chr5 | 99734745 | 99734829 | 9 | 0.47794 | 0.33756 | 0.140389 | 0.007869 | 0.002408 | 0.037483 | 0.049423 |
| chr6 | 136487141 | 136487225 | 9 | 0.23017 | 0.51806 | -0.28789 | 0.007869 | 0.000671 | 0.037483 | 0.031559 |
| chr12 | 85963753 | 85963963 | 5 | 0.3461 | 0.118 | 0.2281 | 0.008931 | 0.002757 | 0.038028 | 0.049584 |
| chr14 | 117786043 | 117786082 | 5 | 0.4118 | 0.5635 | -0.1517 | 0.008931 | 0.001015 | 0.038028 | 0.031559 |
| chr16 | 18154611 | 18154675 | 5 | 0.094 | 0.2622 | -0.1682 | 0.008931 | 0.002478 | 0.038028 | 0.049423 |
| chr19 | 37686655 | 37686768 | 5 | 0.2353 | 0.0845 | 0.1508 | 0.008931 | 0.002479 | 0.038028 | 0.049423 |
| chr3 | 27455582 | 27455704 | 5 | 0.3733 | 0.7336 | -0.3603 | 0.008931 | 0.002604 | 0.038028 | 0.049453 |
| chr5 | 76634256 | 76634347 | 5 | 0.3339 | 0.6204 | -0.2865 | 0.008931 | 0.001035 | 0.038028 | 0.031559 |
| chr6 | 58906654 | 58906769 | 13 | 0.37742 | 0.50862 | -0.13119 | 0.008869 | 0.00023 | 0.038028 | 0.016384 |
| chr7 | 18469547 | 18469606 | 6 | 0.9505 | 0.78733 | 0.163167 | 0.008293 | 0.001317 | 0.038028 | 0.033749 |
| chr7 | 63544348 | 63544386 | 5 | 0.3848 | 0.27 | 0.1148 | 0.008931 | 0.000951 | 0.038028 | 0.031559 |
| chr8 | 44687168 | 44687187 | 5 | 0.8028 | 0.9159 | -0.1131 | 0.008931 | 0.002548 | 0.038028 | 0.049423 |
| chr8 | 99422762 | 99422838 | 5 | 0.481 | 0.6279 | -0.1469 | 0.008931 | 0.001339 | 0.038028 | 0.034162 |
| chrX | 16305941 | 16306022 | 5 | 0.2326 | 0.4329 | -0.2003 | 0.008931 | 0.001045 | 0.038028 | 0.031559 |
| chr16 | 44531184 | 44531427 | 10 | 0.28555 | 0.18355 | 0.102 | 0.009409 | 0.00167 | 0.039817 | 0.041008 |
| chr13 | 56197404 | 56197475 | 5 | 0.4621 | 0.3092 | 0.1529 | 0.011496 | 0.002689 | 0.043114 | 0.049584 |
| chr14 | 10180145 | 10180355 | 7 | 0.85636 | 0.97714 | -0.12079 | 0.0115 | 0.002067 | 0.043114 | 0.046827 |
| chr14 | 77590998 | 77591056 | 5 | 0.5522 | 0.4051 | 0.1471 | 0.011496 | 0.00246 | 0.043114 | 0.049423 |
| chr15 | 101231784 | 101231801 | 5 | 0.4017 | 0.6088 | -0.2071 | 0.011496 | 0.002449 | 0.043114 | 0.049423 |
| chrX | 150657159 | 150657425 | 25 | 0.06804 | 0.23286 | -0.16482 | 0.013073 | 0.000214 | 0.047548 | 0.015641 |
| chr15 | 97754744 | 97754778 | 5 | 0.4157 | 0.1484 | 0.2673 | 0.01469 | 0.002644 | 0.049013 | 0.049456 |

| | | | | | | | | | | |
|---|---|---|---|---|---|---|---|---|---|---|
| chr3 | 57845260 | 57845333 | 5 | 0.6842 | 0.8802 | -0.196 | 0.01469 | 0.000979 | 0.049013 | 0.031559 |
| chrX | 103067417 | 103067590 | 32 | 0.062422 | 0.16697 | -0.10455 | 0.013931 | 0.000302 | 0.049013 | 0.020494 |
